# Spatial profiling and neurovascular communication in the developing and adolescent cortex following prenatal alcohol exposure

**DOI:** 10.64898/2026.09.01.748632

**Authors:** Monica A. Long, Marissa R. Westenskow, Gabriela Perales, Sylvia F. Burris, Gurdeep Singh, Amy S. Gardiner

**Affiliations:** Department of Cell Biology and Physiology, University of New Mexico School of Medicine, Albuquerque, New Mexico, United States; Department of Systems Biology, George Mason University, Manassas, Virginia, United States; Department of Ophthalmology and Visual Sciences, University of New Mexico School of Medicine, Albuquerque, New Mexico, United States; University of New Mexico Comprehensive Cancer Center, Albuquerque, New Mexico, United States

## Abstract

Fetal alcohol spectrum disorders (FASD) constitute a wide range of developmental, cognitive, and behavioral impairments caused by prenatal alcohol exposure (PAE). Although neuronal and vascular consequences of PAE have been studied, how alcohol affects the cerebrovasculature within the framework of the neurovascular unit (NVU) across development remains poorly understood. At minimum, the NVU comprises neurons, astrocyte endfeet, and endothelial cells (ECs), which coordinate to maintain brain homeostasis. Here, we used NanoString’s Digital Spatial Profiling platform to characterize spatial transcriptomic data from neurons, astrocytes, and ECs from PAE and saccharin (SAC) control cortices at embryonic day 18 (E18) and postnatal day 28 (P28). Differentially expressed genes were then used for Ingenuity Pathway Analysis (IPA) to identify altered biological pathways and perform comparison analyses across developmental time points, while CellChat was used to infer cell–cell communication networks. We uncovered thousands of differentially expressed genes and numerous altered pathways and biological processes in PAE cortices across development. Both IPA and CellChat analyses implicated dysregulation of vascular and extracellular matrix (ECM) remodeling, cell adhesion, and neuroinflammatory signaling. CellChat further predicted the loss of several key bidirectional relationships and altered ligand-receptor interactions among neurovascular cell types at E18 and P28. Overall, these findings identify PAE-associated alterations in neurovascular gene expression and intercellular signaling across development, providing potential mechanisms by which PAE may disrupt neurodevelopment.

## INTRODUCTION

Alcohol exposure during pregnancy can produce teratogenic effects on fetal development, as ethanol readily crosses the placental barrier, impacting vital organs such as the brain^1^. Prenatal alcohol exposure (PAE) can lead to an array of physical, developmental, behavioral and cognitive impairments in children that can persist into adulthood^1,2^. These deficits are classified as fetal alcohol spectrum disorders (FASD), with adverse effects varying across individuals. It is often difficult to identify children affected by PAE because many do not present the physical characteristics typically associated with exposure, such as craniofacial dysmorphologies^2,3^.

Despite this, prevalence of FASD in the U.S. is high, estimated to be between 1%-5% of first-grade children^4^. FASD is one of the most common and underdiagnosed developmental disabilities, necessitating a greater understanding of its molecular and cellular impacts in the brain for early intervention and therapeutic efforts aimed at reducing long-term impairments^1,2^. Recent studies have demonstrated PAE-mediated alterations to gene expression in the brain at both genome-wide and single-cell levels^5,6^. Altered gene expression has been associated with pathways involved in alcohol metabolism, erythrocyte differentiation, angiogenesis, and embryonic development^6–8^. These pathways are critical for fetal brain development, and their disruption may contribute to cerebral growth deficits and potential neurovascular unit (NVU) dysfunction. The NVU, composed of neurons, astrocyte endfeet, pericytes, and endothelial cells (ECs), forms the blood-brain barrier (BBB), which is essential for maintaining brain homeostasis^9,10^. Understanding the molecular mechanisms underlying FASD remains an important area for further research^6,9,11^.

Although current sequencing and transcriptomic methods are powerful for analyzing gene expression and protein profiles, they are limited in their ability to capture tissue heterogeneity and perform high-level multiplexing^12^. Digital spatial profiling (DSP) is an emerging technology that addresses these limitations. Using NanoString’s GeoMx® Digital Spatial Profiler, formalin fixed and paraffin-embedded (FFPE) tissue samples are incubated with cell-specific antibodies and RNA probes linked to photocleavable oligonucleotides. Regions of interest (ROI) are then selectively exposed to UV light, releasing oligos for downstream quantification. This approach enables multiplexed transcriptomic analysis across distinct cell populations, allowing for improved characterization of heterogeneous tissues^13^. Applying DSP to the study of PAE-induced gene expression changes in the developing brain may provide valuable insights into the mechanisms underlying FASD as well as inform future therapeutic strategies.

While characterization of gene expression profiles provides important insights into molecular alterations due to PAE, transcriptomic data can also be leveraged to investigate intercellular communication networks. Cell-cell signaling among ECs, neurons, and astrocytes is critical for NVU function, regulating vascular tone, stability, and growth throughout development and into adulthood. As a teratogen, alcohol may disrupt molecular targets and signaling pathways that are essential for proper brain development^9,14^. Therefore, in addition to identifying differentially expressed genes, we sought to characterize ligand-receptor signaling interactions within the developing cortex to gain further insight into cellular communication dynamics associated with PAE.

Here we present the first spatial transcriptomic characterization of the NVU in the developing cortex using a mouse model of PAE at embryonic day 18 (E18) and postnatal day 28 (P28). Differential expression profiles were generated for ECs, neurons, and astrocytes (P28 only) to identify cell type-specific transcriptional alterations associated with PAE. In addition, inferred ligand-receptor signaling networks were examined to explore potential changes in intercellular communication within the NVU. Together, these analyses provide new insight into the molecular landscape and intercellular communication networks associated with PAE and identify candidate mechanisms that may contribute to FASD-associated pathogenesis. RESULTS

### Spatial transcriptomic and data analysis workflow

To understand gene expression signatures in the NVU of PAE and SAC mice, tissues were analyzed using the GeoMx® Digital Spatial Profiler (NanoString Technologies). An established voluntary PAE drinking paradigm was used^15,16^, where the PAE dams received access to saccharine-sweetened ethanol and the control animals received water with saccharine (SAC). PAE and SAC E18 and P28 brains were harvested, sliced, and formalin-fixed paraffin-embedded (FFPE) (Fig. 1A). The DSP workflow has five steps: 1) FFPE samples were incubated with fluorescently labeled cell-specific antibodies and RNA-specific probes with photocleavable oligos, 2) ROIs with sufficient numbers of NVU cell-types were selected, 3) Sequential UV light was applied to the ROIs, releasing the fluorescently labeled photocleavable oligos attached to transcripts, 4) Oligos were collected and dispensed into 96-well plates, 5) Transcripts were quantified using the nCounter Digital Analyzer (Fig. 1B). Data underwent quality control (QC) assessment, normalization, and differential gene expression analyses using NanoString pipelines before downstream analyses were performed using Qiagen’s Ingenuity Pathway Analysis (IPA) and CellChat (Fig. 1C).

**Figure 1.**
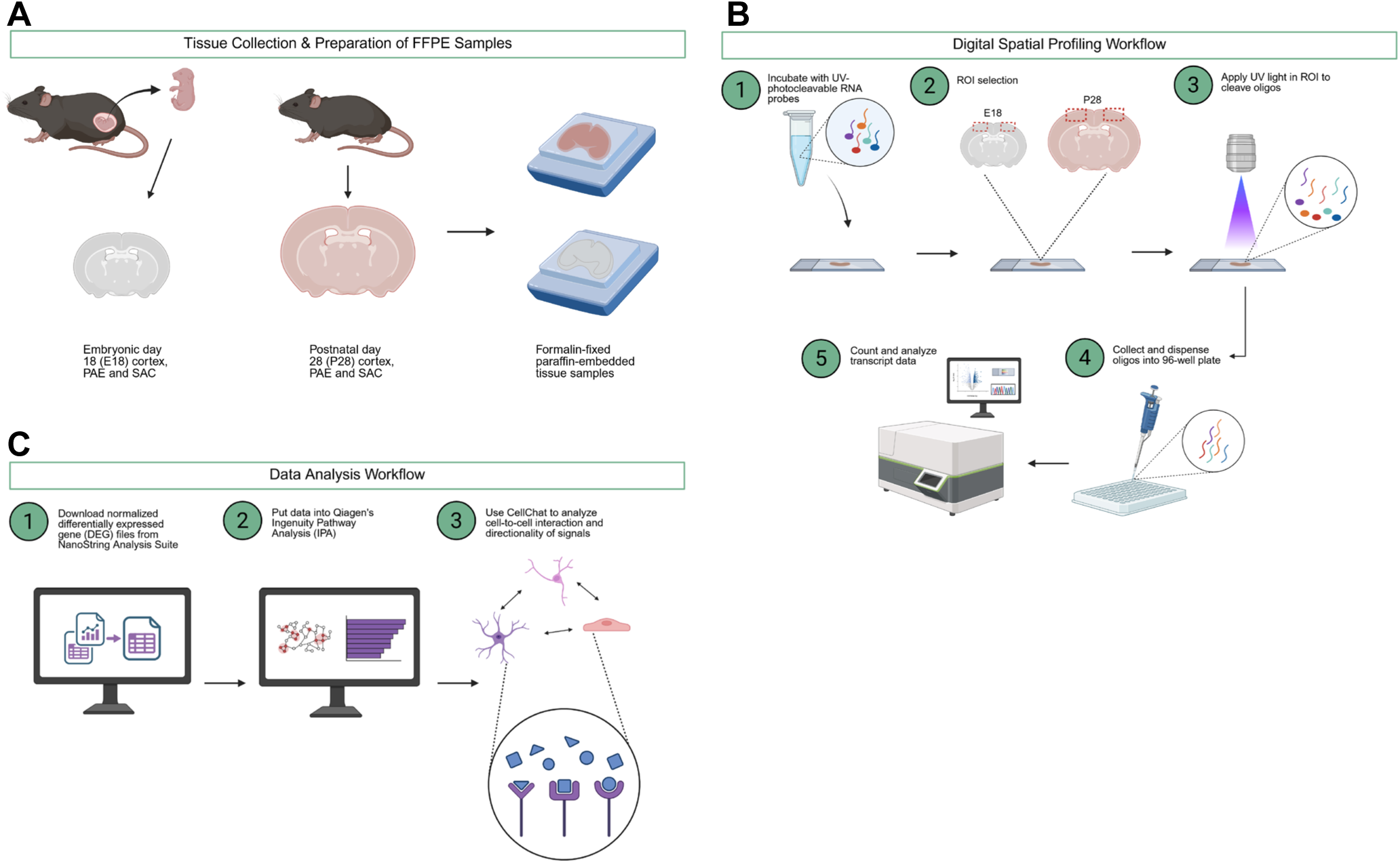
Experimental workflow for digital spatial profiling and data analysis. **a)** Schematic of tissue collection of embryonic day 18 (E18) and postnatal day 28 (P28) brains from prenatal alcohol exposed (PAE) and saccharin control (SAC) mice. Tissues were formalin-fixed and paraffin-embedded (FFPE) before being sent to NanoString for processing and digital spatial profiling. **b)** FFPE samples were incubated with UV-photocleavable cell type-specific and RNA probes. ROI’s in the somatosensory cortex of E18 and P28 tissues were selected for analysis. UV light was applied to the ROI’s and fluorescently barcoded oligos were cleaved from the probes. Oligos were collected and dispensed into 96-well plates and counted using the NanoString nCounter® system. Transcript-level data were analyzed using NanoString’s in-house software. **c)** Differential gene expression datasets were downloaded from NanoString’s Analysis Suite and used for Qiagen’s Ingenuity Pathway Analysis (IPA) and cell-cell interaction analysis via the CellChat package in R.

### Gene expression, canonical pathways, and biological processes are altered by PAE at E18 and P28 in cortical endothelial cells

A large number of differentially expressed genes (DEGs) was identified via Digital Spatial Profiling between PAE and SAC conditions in ECs at E18 and P28, 1127 and 472 genes respectively, FC>1.2, p<0.05 (Supp. Tables 1 and 2). At E18, upregulated genes included the repressive Roundabout guidance receptor genes 1 and 2 (Robo1 and Robo2), SAC3 domain-containing protein 1 (Sac3d1), and Protein O-linked mannose N-acetylglucosaminyltransferase 2 (Pomgnt2). Downregulated genes included Distal-less homeobox 2 (Dlx2), Transforming growth factor-β activated kinase 1 binding protein 1 (Tab1), Tensin 1 (Tns1), Tensin 2 (Tns2), Nuclear receptor subfamily 2 group F member 1 (Nr2f1), and Zinc finger protein 24 (Zfp24) (Fig. 2A). Our findings regarding Robo genes, which are associated with inhibited angiogenesis in ECs^17^, align with previous results indicating reduced cerebral microvascular angiogenesis as a result of PAE^15,18^. The downregulation of Tns1, Nr2f1, and Zfp24, which promote angiogenesis^19–21^, also aligns with these previous findings. At P28, upregulated genes included Forkhead box C1 (Foxc1), Wnt family member 5A (Wnt5a), v-Rel avian reticuloendotheliosis viral oncogene homolog A (Rela), and Transforming growth factor alpha (Tgfa). Downregulated genes included Eukaryotic translation initiation factor 2D (Eif2d), Zfp24, and Nr2f1 (Fig. 2B). The upregulation of Foxc1, Wnt5a, and Tgfa, which promote angiogenesis and EC proliferation^22–24^, at P28 is an interesting contrast from gene expression patterns observed at E18 in ECs. The upregulation of Rela suggests vascular inflammation at P28 as a result of PAE^25^. Additionally, while significant differences emerged in the profile of differentially expressed genes between E18 and P28, the downregulation of Nr2f1 and Zfp24 in PAE ECs persisted.

**Figure 2.**
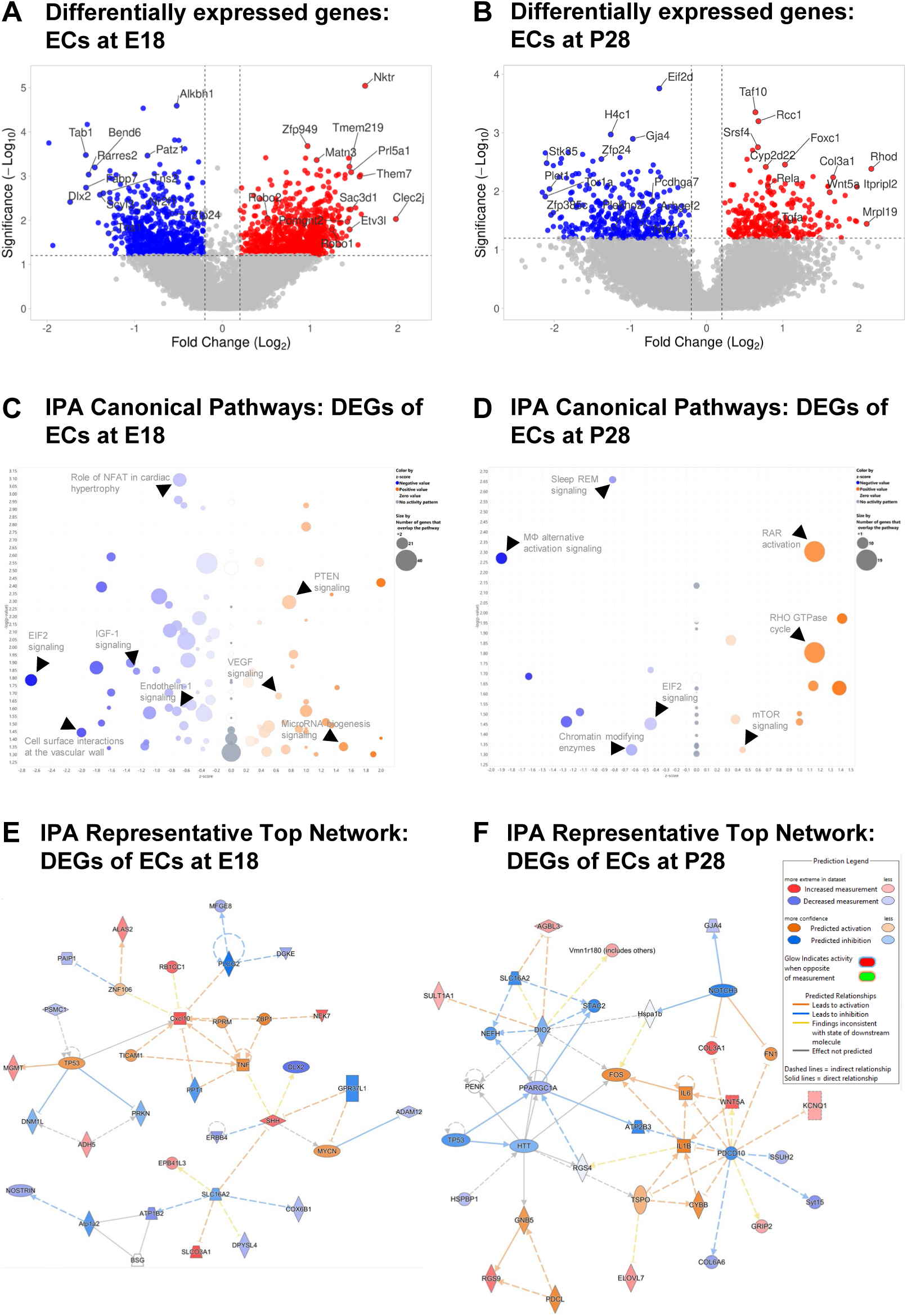
Differential gene expression and pathway analysis in endothelial cells at E18 and P28. **a)** Volcano plot of DEGs in ECs at E18 between PAE samples and SAC controls (n=6-8, fold change >1.2 and p<0.05). **b)** Volcano plot of DEGs in ECs at P28 between PAE samples and SAC controls (n=3, fold change >1.2 and p<0.05). **c)** IPA bubble volcano chart depicting canonical pathways predicted to be activated/inhibited in ECs at E18 between PAE and SAC. **d)** IPA bubble volcano chart depicting canonical pathways predicted to be activated/inhibited in ECs at P28 between PAE and SAC. **e)** IPA top molecular network consisting of DEGs and other genes predicted to be activated/inhibited in ECs at E18 associated with neurodevelopmental disease, organismal injury and disease, and abnormalities. **f)** IPA top molecular network consisting of DEGs and other genes predicted to be activated/inhibited in ECs at P28 associated with neurological disease, organismal injury and abnormalities, and psychological disorders. Legend on right shows the relationship between colors and gene expression or predicted activation/inhibition.

Ingenuity Pathway Analysis (IPA) of DEGs between PAE and SAC in ECs at E18 revealed many canonical pathways predicted to be upregulated or downregulated. Some upregulated pathways of interest in ECs at E18 included PTEN signaling, VEGF signaling, and MicroRNA biogenesis signaling (Fig. 2C). The upregulation of both PTEN signaling, which inhibits angiogenesis^26^, and VEGF signaling, which promotes vascular growth^27^, indicates complicated, multi-pronged mechanisms regulating altered angiogenesis in the brain during PAE. Some downregulated pathways of interest included the Role of NFAT in cardiac hypertrophy, EIF2 signaling, IGF-1 signaling, Endothelin-1 signaling, and Cell surface interactions at the vascular wall (Fig. 2C). Alterations to EIF2 signaling indicate perturbations in inflammation due to PAE^28^, downregulation of IGF-1 signaling has implications for impaired vascular barrier function^29^, and a decrease in Endothelin-1 signaling indicates alterations to vasoreactivity as a result of PAE at E18^30^. A subset of additional canonical pathways predicted to have increased or decreased activation, such as ROBO SLIT signaling and FGF signaling respectively, as well as the DEGs associated with them, are depicted in Table 1. All canonical pathways, and associated DEGs, predicted to be impacted by PAE in ECs at E18 are listed in Supp. Table 3. By P28, there were significant changes to canonical pathways predicted to be upregulated and downregulated between PAE and SAC samples. At P28, upregulated pathways included RAR activation, RHO GTPase cycle, and mTOR signaling (Fig. 2D). These pathways promote vascular barrier integrity, endothelial cell adhesion, and endothelial cell proliferation, respectively^31–33^, demonstrating an interesting contrast between active pathways in cortical ECs following PAE in the embryonic and adolescent brain. Downregulated pathways included Sleep REM signaling, Macrophage alternative activation signaling, EIF2 signaling, and Chromatin modifying enzymes (Fig. 2D). Interestingly, these results align with existing findings indicating disturbances to sleep following PAE^34^. While EIF2 signaling remained decreased in the P28 PAE samples, other pathways recovered to be more similar to SAC (Role of NFAT), and other pathways involving DEGs emerged (RAR activation). A subset of additional canonical pathways predicted to have increased or decreased activation, such as IL-6 signaling and TX/RXR activation respectively, as well as the DEGs associated with them, are depicted in Table 2. All canonical pathways, and associated DEGs, predicted to be impacted by PAE in ECs at P28 are listed in Supp. Table 4.

We also identified gene interaction networks associated with various biological processes using the DEGs between PAE and SAC ECs at both E18 and P28. One of the top networks that connected Dlx2, Sonic hedgehog (Shh), Erb-b2 receptor tyrosine kinase 4 (Erbb4), MYCN proto-oncogene (Mycn), ADAM metallopeptidase domain 12 (Adam12), Milk fat globule epidermal growth factor 8 (Mfge8), Alcohol dehydrogenase 5 (Adh5), and Tumor protein p53 (TP53) was identified from the E18 gene expression data and was associated with Organismal injury and abnormalities, Organismal survival, and Neurological disease (Fig. 2E). Another network connecting Wnt5a, Interleukin-1 beta (IL-1B), Cytochrome b-245 beta chain (Cybb), interleukin-6 (IL-6), and Fos proto-oncogene (Fos) was identified from the P28 gene expression data and was associated with Neurological disease, Organismal injury and abnormalities, and Psychological disorders (Fig. 2F). Additionally, IPA comparison analysis revealed developmental differences in the predicted activity of Endothelin-1, IGF-1, and VEGF signaling in ECs at E18 and P28 (Supp. Figs. 1-3). At E18, many pathway-associated molecules showed significant changes and contributed to directional pathway predictions, whereas at P28, many showed no significant changes, and the pathways lacked clear predicted activation or inhibition. Categorical heat maps depicting diseases and biological functions, such as Neurological disease and Nervous system development and function, associated with the DEGs in ECs at E18 and P28 are presented in Supp. Fig. 4.

### Gene expression, canonical pathways, and biological processes are altered by PAE at E18 and P28 in neurons

Digital Spatial Profiling also identified a large number of DEGs between PAE and SAC conditions in neurons at E18 and P28, 1787 and 517 genes respectively, FC>1.2, p<0.05 (Supp. Tables 5 and 6). At E18, Centrosomal protein 170 (Cep170), SATB homeobox 2 (Satb2), G protein subunit alpha Q (Gnaq), Calmodulin 1 (Calm1), Calcium voltage-gated channel subunit alpha1 A (Cacna1a), Calcium voltage-gated channel auxiliary subunit gamma 8 (Cacng8), and Neurotrophin 3 (Nft3) were upregulated, and Nr2f1, Cytohesin (Cyth1), Unc-5 netrin receptor D (Unc5d), MET proto-oncogene (Met), Forkhead box O1 (Foxo1), Nuclear receptor subfamily 4 group A member 1 (Nr4a1), and Ephrin A (EphA) and Ephrin B (EphB) family genes were downregulated (Fig. 3A). The upregulation of Cep170, Calm1, and Nft3 indicates alterations to neuronal migration and morphology^35–37^, and upregulation of Satb2 and Cacna1a indicates alterations to neurogenesis^38,39^. Gnaq upregulation may occur in response to elevated reactive oxygen species generated by the metabolism of alcohol in PAE conditions, and alterations to Cacng8 expression are related to behavioral phenotypes associated with PAE^40,41^. Downregulation of Cyth1, Unc5d, and Nr4a1 suggests impairments to axonal development^42–44^. Additionally, Nr2f1 and Unc5d downregulation could indicate impairment to neurogenesis and neuronal migration^44,45^. Neuronal growth and morphology and synapse maturation could be impacted by Met downregulation, and downregulation of Foxo1 could indicate alterations to energy handling in neurons as a result of PAE at E18^46–48^. By P28, differences in the profile of DEGs between PAE and SAC neurons could be observed compared to E18. Ten-eleven translocase 3 (Tet3), Potassium calcium-activated channel subfamily N member 3 (Kcnn3), tubulin beta 6 class V (Tubb6), lecithin-cholesterol acyltransferase (Lcat), and Claudin 11 (Cldn11) were upregulated, and Fos, Nr4a1, Activity-regulated cytoskeleton-associated protein (Arc), BTG anti-proliferation factor 2 (Btg2), RAD50 double strand break repair protein (Rad50), Cluster of differentiation 109 (Cd109), and Dual specificity phosphatase 1 (Dusp1) were included in the downregulated genes (Fig. 3B). The upregulation of Tet3 is particularly interesting in light of our previous findings characterizing the role of TET family-induced DNA demethylation in brain microvascular endothelial cells following PAE^49^. The overexpression of Kcnn3, also known as SK3, could contribute to memory-related symptoms following PAE^50^. Tubb6 activity has been linked to neuroprotection^51^. Lcat is produced in a variety of neural cell types and is involved in brain lipid metabolism^52^. Cldn11 is included in tight junctions, and in the brain, it is most associated with myelination^53^. Loss of Rad50 has been linked to disrupted embryonic gene expression programs through alterations to histone modification patterns, so its downregulation could be significant in general gene expression patterns in neurons following PAE^54^. Arc is also associated with neurodevelopmental disorders, which suggests that its alteration may contribute to neurodevelopmental impairments in PAE^55^. Both Btg2 and Dusp1 are associated with cell identity, and Dusp1 is linked to neuroprotection^56,57^. Cd109 contributes to neurite outgrowth^58^, and Fos is an established marker of neural activity; its downregulation could signal decreases in neuronal activation following PAE^59^.

**Figure 3.**
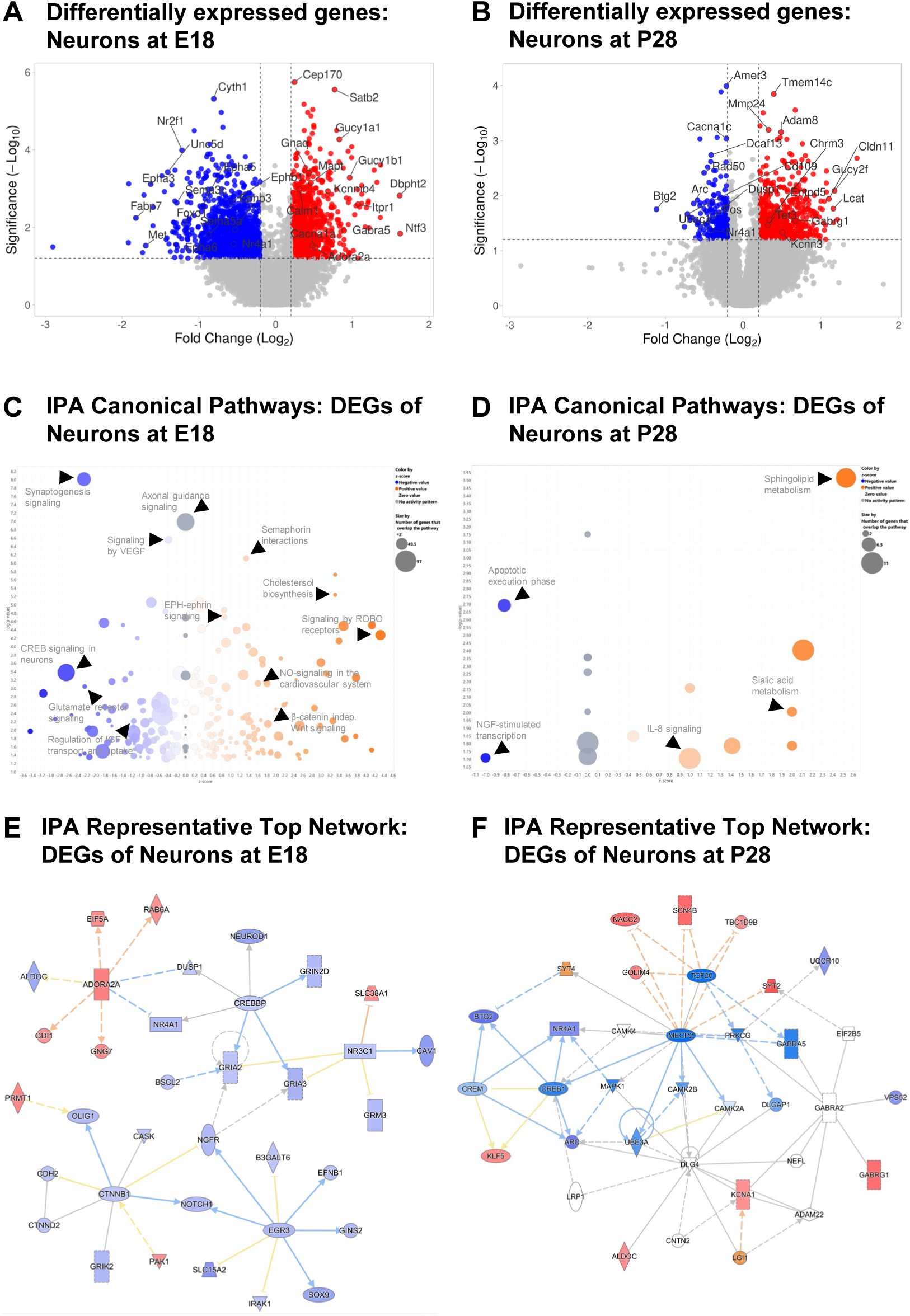
Differential gene expression and pathway analysis in neurons at E18 and P28. a) Volcano plot of DEGs in neurons at E18 between PAE samples and SAC controls (n=6-8, fold change >1.2 and p<0.05). b) Volcano plot of DEGs in neurons at P28 between PAE samples and SAC controls (n=3, fold change >1.2 and p<0.05). c) IPA bubble volcano chart depicting canonical pathways predicted to be activated/inhibited in neurons at E18 between PAE and SAC. d) IPA bubble volcano chart depicting canonical pathways predicted to be activated/inhibited in neurons at P28 between PAE and SAC. e) IPA top molecular network consisting of DEGs and other genes predicted to be activated/inhibited in neurons at E18 associated with cell morphology, cellular assembly and organization, and cellular development. f) IPA top molecular network consisting of DEGs and other genes predicted to be activated/inhibited in neurons at P28 associated with behavior, cell development, and cellular growth and proliferation.

Additionally, IPA analysis of the differential gene expression data revealed canonical pathways predicted to be upregulated and downregulated in the PAE neurons compared to SAC controls. Pathways predicted to be upregulated at E18 included Signaling by ROBO receptors, Semaphorin interactions, Cholesterol biosynthesis, EPH-ephrin signaling, NO-signaling in the cardiovascular system, and β-catenin-independent Wnt signaling (Fig. 3C). Some of these identified upregulated pathways have interesting implications for alterations to neuronal function at E18 in PAE in light of extensive evidence of neurological impairment resulting from PAE. Neural circuit formation is promoted by semaphorin interactions and Wnt signaling^60,61^. Axon guidance is promoted by Ephrin signaling, ROBO receptor signaling, and Wnt signaling^60,62,63^. Ephrin signaling also promotes neuronal migration^62^, and Wnt signaling promotes synapse function^60^. Some downregulated pathways of interest at E18 included Synaptogenesis signaling, VEGF signaling, CREB signaling in neurons, Glutamate receptor signaling, Axonal guidance signaling, and Regulation of IGF transport and uptake (Fig. 3C). The downregulation of VEGF signaling in neurons implies a decrease in promotion of vascular development by neurons^64^. Downregulated CREB signaling indicates impairments to neuronal plasticity and neuroprotection^65^. Decreased glutamate receptor signaling indicates an overall decrease in excitatory neuronal activity^66^. In addition to the identification of downregulated axonal guidance signaling, IGF transport and regulation have diverse roles that are potentially disrupted with its downregulation including axon growth, synaptogenesis, and myelination^67^. A subset of additional canonical pathways predicted to have increased or decreased activation, including decreased Wnt/β-catenin signaling, as well as the DEGs associated with them, are depicted in Table 3. All canonical pathways, and associated DEGs, predicted to be impacted by PAE in neurons at E18 are listed in Supp. Table 7. At P28, upregulated canonical pathways included Sphingolipid metabolism, Sialic acid metabolism, and IL-8 signaling. Downregulated pathways included the Apoptotic execution phase and NGF-stimulated transcription (Fig. 3D). Some sphingolipid metabolism contributes to neurodegeneration^68^, and sialic acid metabolism contributes to many facets of neuronal development and function including neuronal sprouting and plasticity, axon myelination, myelin stability, and neuronal connection remodeling^69^. Upregulated IL-8 signaling indicates inflammatory signaling in the brain as a result of PAE^70^. Downregulation of apoptotic execution implies alterations to neuronal survival following PAE^71^, and downregulation of NGF-mediated transcription has widespread implications for global gene expression in neurons at P28 following PAE^72^. A subset of additional canonical pathways predicted to have increased or decreased activation, including Neurovascular coupling signaling and Centrosomal KIAA0586 signaling (primary cilia signaling) respectively, as well as the DEGs associated with them, are depicted in Table 4. All canonical pathways, and associated DEGs, predicted to be impacted by PAE in neurons at P28 are listed in Supp. Table 8.

We identified interaction networks associated with biological processes using the DEGs between PAE and SAC neurons at both E18 and P28. For E18, one relevant gene interaction network connected CREB binding protein (Crebbp), Glutamate ionotropic receptor AMPA type subunit 2 (Gria2) and 3 (Gria3), NGF receptor (Ngfr), and Catenin beta 1 (Ctnnb1) and was associated with Cell morphology, Cellular assembly and organization, and Cellular development (Fig. 3E). At P28, one relevant gene interaction network connected Arc, Btg2, Nr4a1, Ubiquitin protein ligase E3A (Ube3A), Nucleus accumbens-associated protein 2 (Nacc2), Sodium voltage-gated channel beta subunit 4 (Scn4B), Synaptotagmin 2 (Syt2), Potassium voltage-gated channel subfamily A member 1 (Kcna1) and Gamma-aminobutyric acid type A receptor subunit gamma 1 (Gabrg1) and was associated with Behavior, Cellular development, and Cellular growth and proliferation (Fig. 3F). IPA comparison analysis revealed developmental differences in the predicted activity of CREB, Ephrin, and Netrin signaling in neurons at E18 and P28 (Supp. Figs. 5-7). Unlike in ECs, many pathway-associated molecules showed significant changes and contributed to directional pathway predictions at both timepoints. Notably, Netrin signaling at P28 showed a stark difference from its E18 counterpart, with most components lacking clear predicted activation or inhibition (Supp. Fig. 7). Categorical heat maps depicting diseases and biological functions, again including Neurological disease and Nervous system development and function, associated with the DEGs in neurons at E18 and P28 are presented in Supp. Fig. 8.

### Gene expression, canonical pathways, and biological processes are altered by PAE at P28 in astrocytes

We also assessed astrocytes at P28 via Digital Spatial Profiling and identified 508 DEGs between PAE and SAC samples (Supp. Table 9). Upregulated genes included Ependymin related 1 (Epdr1), PYD and CARD domain containing (Pycard), Mitochondrial ribosomal protein S3 (Mrps3), Solute carrier family 5 member 5 (Slc5a5), Killer cell lectin-like receptor family I member 2 (Klri2), and Centromere protein K (Cenpk), and downregulated genes included Myristoylated alanine-rich C-kinase substrate (Marcks), Cytochrome c oxidase copper chaperone COX11 (Cox11), Chymase 2, mast cell (Cma2), DEAD-box helicase 27 (Ddx27), and Ubinuclein 1 (Ubn1) (Fig. 4A). The identified upregulated genes indicate widespread alterations to various basic cellular functions and metabolism in astrocytes following PAE with the upregulation of Pycard especially indicating increased inflammation^73–75^. Likewise, the significantly downregulated genes are associated with a wide array of basic cellular functions as opposed to astrocyte-specific functions. One exception is Marcks, which is related to the regulation of astrocyte migration^76^.

**Figure 4.**
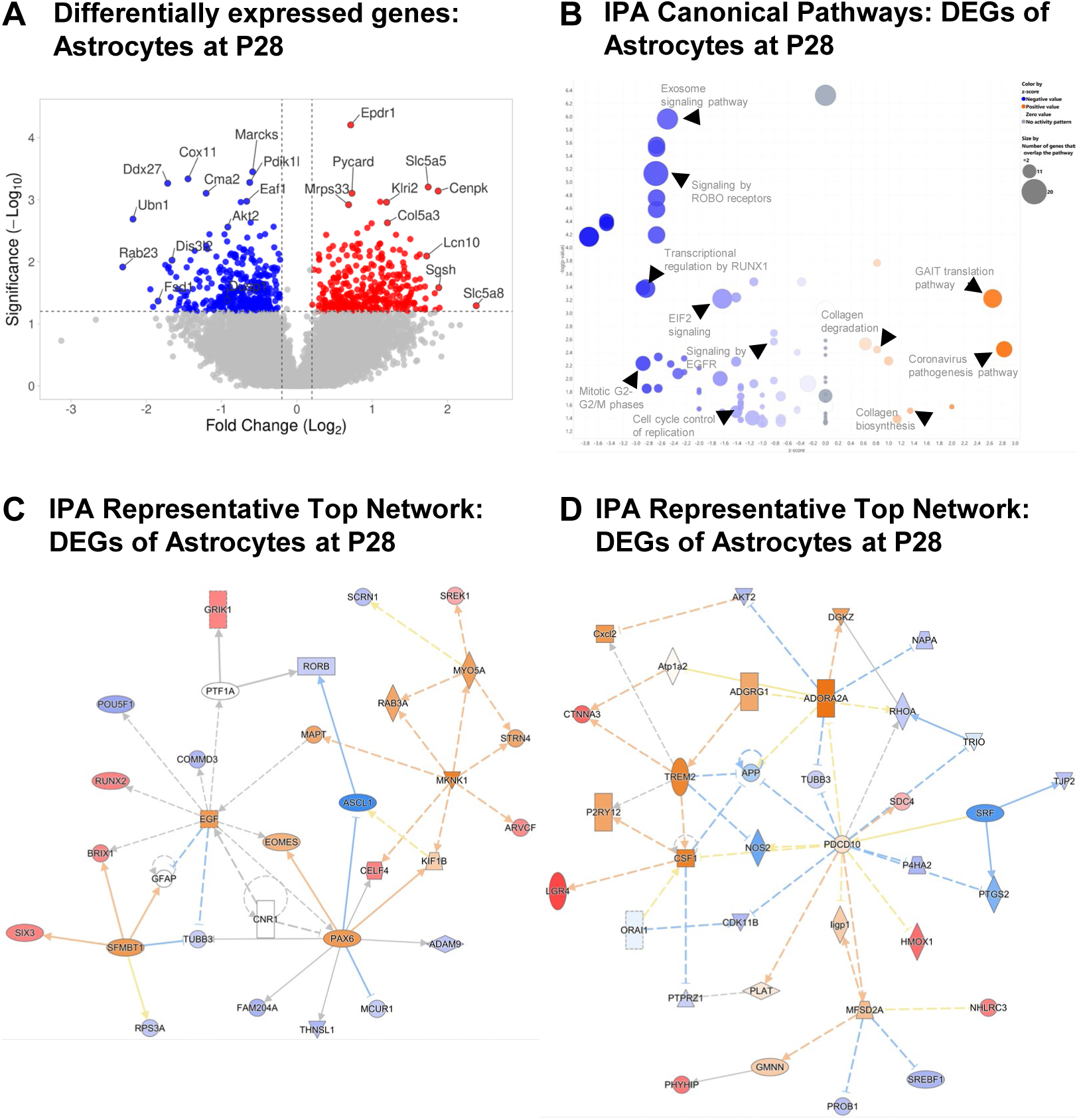
Differential gene expression and pathway analysis in astrocytes at P28. **a)** Volcano plot of DEGs in astrocytes at P28 between PAE samples and SAC controls (n=3, fold change >1.2 and p<0.05). **b)** IPA bubble volcano chart depicting canonical pathways predicted to be activated/inhibited in astrocytes at P28 between PAE and SAC. **c)** IPA top molecular network consisting of DEGs and other genes predicted to be activated/inhibited in astrocytes at P28 associated with cell development, nervous system development and function, and tissue development. **d)** IPA top molecular network consisting of DEGs and other genes predicted to be activated/inhibited in astrocytes at P28 associated with cellular movement, hematological system development and function, and immune cell trafficking.

IPA analysis of the DEGs identified canonical pathways predicted to be upregulated and downregulated resulting from PAE in astrocytes at P28. Upregulated pathways included the GAIT translation pathway and both Collagen degradation and biosynthesis, and downregulated pathways included the Exosome signaling pathway, Signaling by ROBO receptors, Transcriptional regulation by RUNX1, EIF2 signaling, Signaling by EGFR, and Mitotic G2-G2/M phases (Fig. 4B). Elevated GAIT translation indicates elevated inflammation^77^, and alterations to extracellular matrix composition related to collagen degradation could have an effect on astrocyte activation following neurological injury such as PAE^78^. Downregulated ROBO receptor signaling could represent interference in neuron-derived signaling to astrocytes resulting in alterations to astrocyte migration^79^. RUNX1-mediated transcriptional regulation and EIF2 signaling in astrocytes are both associated with inflammation, especially related to neurological injury^80–82^. In cortical astrocytes specifically, loss of EGFR signaling promotes apoptosis and is deleterious for surrounding neurons^83,84^. A subset of additional canonical pathways predicted to have increased or decreased activation, such as Integrin cell surface interactions and Apoptosis signaling respectively, as well as the DEGs associated with them, are depicted in Table 5. All canonical pathways, and associated DEGs, predicted to be impacted by PAE in astrocytes at P28 are listed in Supp. Table 10.

Gene interaction networks were also identified in the IPA analysis of DEGs between PAE and SAC astrocytes at P28. One network connecting Epidermal growth factor (Egf), Scm-like with four MBT domains 1 (Sfmbt1), SIX homeobox 3 (Six3), POU class 5 homeobox 1 (Pou5f1), CUGBP Elav-like family member 4 (Celf4), ADAM metallopeptidase domain 9 (Adam9), RAR-related orphan receptor B (Rorb), and Armadillo repeat gene deleted in velo-cardio-facial syndrome (Arvcf) was associated with Cellular development, Nervous system development and function, and Tissue development (Fig. 4C). Another network connecting AKT serine/threonine kinase 2 (Akt2), Triggering receptor expressed on myeloid cells 2 (Trem2), Ras homolog family member A (Rhoa), Tubulin beta 3 class III (Tubb3), Prolyl 4-hydroxylase subunit alpha 2 (P4ha2), and Cyclin dependent kinase 11B (Cdk11b) was associated with Cellular movement, Hematological system development and function, and Immune cell trafficking (Fig. 4D).

### CellChat analysis reveals altered neurovascular communication in PAE cortices

Cell-to-cell communication networks were inferred using CellChat v2.2.0^85^. This tool uses ligand-receptor interaction predictions from spatial data to measure autocrine and paracrine signaling as well as bidirectionality. Across all cell types (ECs, neurons, and astrocytes) and two developmental timepoints (E18 and P28), CellChat identified 1,970 predicted interactions spanning 142 signaling pathways with varying probabilities across the two conditions (PAE vs SAC) at each timepoint with some cell types serving predominantly as interaction producers (Supp. Tables 11-14).

### Communication signals and pathways are altered between endothelial cells and neurons at E18

At E18, ECs and neurons formed an active communication network in both SAC and PAE samples. The cellular relationship with the fewest interactions across SAC and PAE was EC to EC, and PAE had little effect on the level of overall autocrine ligand-receptor interactions between ECs (Fig. 5A-B), but some specific alterations to interactions were observed. The interaction between midkine (Mdk) ligand and syndecan-4 (Sdc4) receptor was increased, and the same was true for the interactions between Mdk and nucleolin (Ncl) and Mdk and low-density lipoprotein receptor-related protein 1 (Lrp1) (Supp. Fig. 9). The interaction between ephrin-B2 (Efnb2) and ephrin type-B receptor 3 (Ephb3) was increased with PAE as well. Notably, the interaction between pleiotropin (Ptn) and syndecan-2 (Sdc2) was decreased (Supp. Fig. 9). In both SAC and PAE samples, the cellular relationship with the most interactions was neuron to neuron, and PAE also had little effect on the level of overall autocrine ligand-receptor interactions between neurons (Fig. 5A-B), but some specific interactions were altered. The interaction between Ptn and Sdc2 was increased with PAE, and the interactions between Mdk and Sdc4, Mdk and Ncl, Mdk and Lrp1, Efnb2 and Ephb3, and Cd99 and Cd99 were decreased (Supp. Fig. 9). The z-scored normalized signaling changes in ECs at E18 under PAE conditions in contrast to SAC show strongly decreased incoming/outgoing signals (ΔZ≤2) for various factors involved in regulating blood vessel growth/repair and barrier integrity, such as EGF, somatostatin (SST), fibroblast growth factor (FGF) and PECAM2 (Fig. 6A). Interestingly, ECs displayed strongly increased outgoing (ΔZ≥2) rather than incoming signals for synaptic optimization and structural stabilization associated paracrine signals such as reelin (RELN) and neurexin (NRXN), thus suggesting enhanced endothelial involvement in neuroprotective paracrine communication under PAE relative to SAC (Fig. 6A). Signaling changes in neurons at E18 show decreased incoming/outgoing signals for genes regulating axon guidance and neural circuit remodeling such as junction adhesion molecules (JAM), semaphorin 5 (SEMA5), and natriuretic peptide receptor 1 (NRP1) as well as increased outgoing signals for neuromodulators such as neurotensin (NT) and secreted Ly6/uPAR-related protein-1 (SLURP) (Fig. 6C).

**Figure 5.**
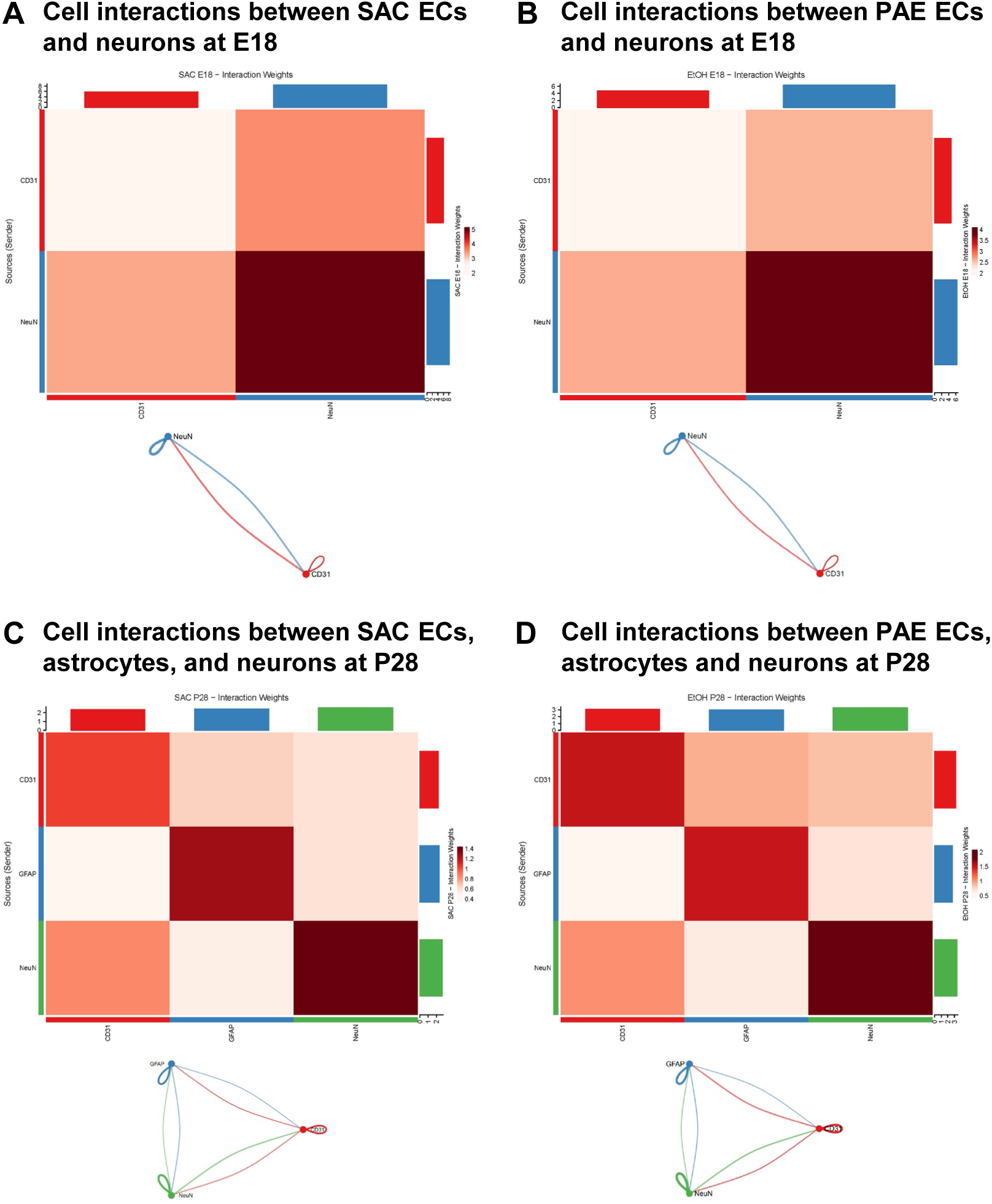
Neurovascular cell-cell interactions in SAC and PAE at E18 and P28. Heatmaps and corresponding circle plots show the aggregated communication interaction weights between **a)** SAC ECs and neurons at E18, **b)** PAE ECs and neurons at E18, **c)** SAC ECs, astrocytes, and neurons at P28, and **d)** PAE ECs, astrocytes, and neurons at P28. Heatmap rows represent the sending cell type and columns represent the receiving cell type. Color intensity reflects interaction weight. Bar plots along the top and right margins show the total outgoing (top) and incoming (right) interaction weight summed across all partners for each cell type, color-coded as follows: CD31 (red), NeuN (blue in a,c and green in c,d), and GFAP (blue in c,d). Circle plot nodes represent cell type and curved lines represent the directional signaling between cell types. Small loops indicate autocrine signaling while line thickness reflects communication strength.

**Figure 6.**
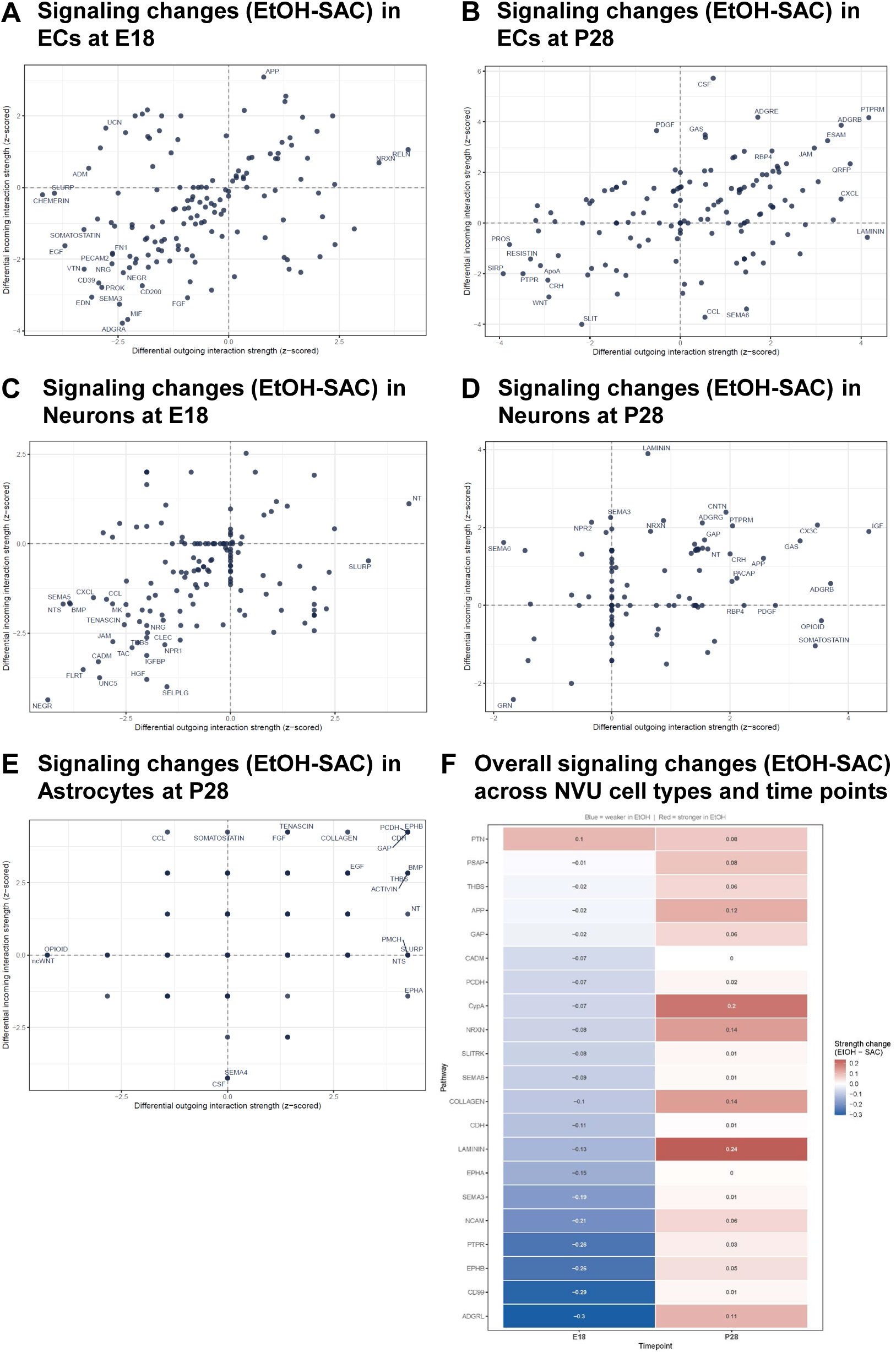
PAE alters neurovascular signaling across development. Scatter plots show the differential incoming (y-axis) and outgoing (x-axis) z-scored interaction strengths for **a-b)** E18 and P28 EC signaling pathways, **c-d)** E18 and P28 neuronal signaling pathways, and **e)** P28 astrocytic signaling pathways. **f)** Two-phase heatmap showing the change in pathway-level communication strength between PAE and SAC conditions (EtOH-SAC) at E18 and P28. Each row represents a signaling pathway, and each column is a timepoint. Chamber color and the numeric value reflect the direction and magnitude of change – blue indicates weaker signaling in PAE cortices relative to SAC cortices.

Overall, at E18, CD99 exhibited the highest communication activity, followed by cyclophilin A (CypA) (Table 6). CD99 autocrine signaling was strongest in neuronal cells for both conditions and also demonstrated substantial bidirectional communication between neurons and ECs. CD99-CD99 ligand-receptor pairings’ interaction probabilities were decreased in PAE (Supp. Fig. 9). Comparison of neuron-endothelial signaling between groups revealed an overall reduction in communication in PAE samples (Fig. 5A-B). Nine of the ten most altered signaling pathways showed decreased interaction strength relative to SAC controls (Table 6). PTN was the only pathway that exhibited increased signaling in PAE cortices (prediction score change = +0.10; Table 6).

### Communication signals and pathways are altered between endothelial cells, neurons and astrocytes at P28

At timepoint P28, astrocytes joined active neurovascular communication networks to further drive changes in both SAC and PAE samples. Autocrine ligand-receptor signaling increased across every cell type at P28, with EC-EC (Δ = +0.45) and neuron-neuron (Δ = +0.38) showing the largest increases in total interaction weight, followed by astrocyte-astrocyte (Δ = +0.19; Fig. 6C-D). In astrocytes, some specific alterations to interactions were observed such as an increase in the ligand-receptor pairing Ptn-Sdc2 and decreases in Ptn-Ncl, as well as interactions between contactin-2 (Cntn2) and L1 cell adhesion molecule (L1cam), and neurofascin (Nfasc) and contactin-1 + contactin-associated protein 1 (Cntn1+Cntnap1) (Supp. Fig. 10). Similar to E18, the cellular relationship with the most interactions at P28 was neuron to neuron, and PAE also had little effect on the level of overall autocrine ligand-receptor interactions between neurons (Fig. 5C-D), but some specific interactions were altered. Of note, PAE neurons gained interactions of Cntn2-L1cam and Nfasc-Cntn1+Cntnap1 with the help of astrocyte-to-neuron ligand-receptor pairings. In addition, neurons displayed increased fibronectin leucine-rich transmembrane protein 3 (Flrt3) and latrophilin-2 (Adgrl2), teneurin transmembrane protein 2 (Tenm2)+Flrt3 and Adgrl2, and both Tenm2-Adgrl2 and Tenm3-Adgrl2 interactions (Supp. Fig. 10). Remarkably, these four ligand-receptor interactions are decreased in PAE EC-EC signaling. Fifteen ligand-receptor pairings were lost in PAE relative to SAC for neuronal autocrine signaling, including Reln and very low-density lipoprotein receptor (Vldlr), Efnb3-Epha4, Efnb1-Ephb3, progranulin (Grn) and sortilin 1 (Sort1), and laminin subunit beta 1 (Lamb1) and synaptic vesicle glycoprotein 2A (Sv2a), indicating that neuronal autocrine signaling was not uniformly amplified at P28 despite the overall increase in total interaction weight. The signaling changes in ECs at P28 show decreased incoming/outgoing signals of factors involved in angiogenesis and neuroinflammation, such as slit guidance ligand (SLIT), WNT, resistin, and apolipoprotein A (ApoA) (Fig. 6B). Meanwhile, ECs displayed strongly increased incoming/outgoing (ΔZ≥2) signaling for structural stability and barrier integrity signals, such as JAM, adhesion G protein-coupled receptor B (ADGRB), and endothelial cell adhesion molecule (ESAM). Neurons at P28 show many increased incoming/outgoing signals; of note are ADGRB, laminin, NT, and NRXN involved in cellular structure and synaptic organization signaling. There are few decreased incoming/outgoing signals, but most interestingly is GRN, which is involved in cell survival and neuroprotective pathways (Fig. 6D). Astrocyte signaling changes at P28 show increased incoming/outgoing communications involved in vascular repair/remodeling and structural organization, such as collagen, FGF, EGF, EPHB, NT, and tenascin (Fig. 6E).

CypA remained dominant in communication activity, taking the top spot (prediction score change = +0.20) with NRXN as second (prediction score change = +0.14; Table 7). Interestingly, all top 10 interaction pathways at P28 showed an increase in interaction strength (Table 7; Fig. 6F), potentially indicating an adaptive mechanism for PAE. ADGRL showed the greatest magnitude of change from being suppressed at E18 (prediction score change = −0.30) and amplified at P28 (prediction score change = +0.11) (Fig. 6F). Overall, PAE altered neurovascular signaling across development from a mostly suppressive signaling pattern with the strongest magnitude of change being pathways associated with ADGRL, CD99, PTPR to an amplified pattern, including laminin, CypA, collagen, and NRXN (Fig. 6F).

## DISCUSSION

Using spatial genomics and IPA, we detected and analyzed differential gene expression and predicted alterations to canonical pathways and gene interaction networks in ECs, neurons, and astrocytes in intact tissue slices from E18 and P28 mouse brain from PAE and SAC control conditions. Additionally, we used CellChat analysis to characterize ligand-receptor interactions between brain ECs and neurons at E18 and P28 in PAE and SAC control conditions.

We identified hundreds of DEGs in ECs as a result of PAE at both developmental timepoints. The biological processes related to the functional activity of cortical ECs are diverse, and as such, the predicted impact of PAE-induced differential gene expression is also diverse. Some of the processes most represented in the documented functions of the identified DEGs and differentially regulated canonical pathways included angiogenesis, vascular barrier function, and EC proliferation. These findings align with existing research describing the impact of PAE on vascular development and integrity in the brain^15^. We also identified upregulated and downregulated genes, canonical pathways, and biological processes resulting from PAE in neurons at E18 and P28. These alterations are most notably documented to influence neuronal migration and morphology, especially axon development and guidance. In our analysis of alterations to gene expression, canonical pathways, and biological processes in astrocytes at P28 following PAE, differential regulation of genes and processes related to cell migration, both for astrocytes and surrounding neurons, was also a frequent finding.

As our studies were focused on ECs, neurons, and astrocytes and the interactions between these cell types, many of the implications of our findings in PAE are directly related to vascular development and neurovascular interactions. The brain vascular impacts of the observed alterations to intercellular interactions have been described, and virtually all observed changes to gene expression and canonical pathways in ECs could contribute to the alterations to brain vascular structure and function documented in existing literature regarding animal models of PAE and human studies of severe FASDs. However, our diverse findings regarding differential gene expression and regulation of canonical pathways as a result of PAE also have more widespread implications for the molecular mechanisms driving the development of symptoms and features of FASDs.

One of the most prominent features of FASDs is a deficit in learning and memory. Several alterations to gene expression and signaling pathways as a result of PAE that we detected in these studies have documented associations with learning and memory. We detected a downregulation of CREB signaling in neurons at E18, and loss of CREB has been shown to induce defects in memory^65^. Fos, which is downregulated in neurons as a result of PAE as well, also facilitates and serves as a marker for learning and memory^59^.

Other significant cellular and anatomical features of FASDs are related to neuronal migration, a process significantly represented in the function of differentially regulated genes and pathways identified in our studies. Both early and late-generated neurons have delayed migration in embryonic development as a result of PAE in a rat model^86^, and this delayed migration is thought to result in heterotopic clusters of neurons and other neural cells, potentially including astrocytes. Neuroglial heterotopias have also been observed in autopsy studies of severe cases of fetal alcohol syndrome (FAS), the most severe form of FASD^87–89^. The differential expression of Cep170, Calm1, Nft3, Nr2f1, Unc5d in neurons and the downregulation of Marcks expression and ROBO signaling in astrocytes as observed in our studies could contribute to the development of neuroglial heterotopias by disrupting neuron and astrocyte migration.

A very consistent process of brain development related to the differentially regulated genes and canonical pathways identified in our studies is axon development. Axon development, especially in the corpus callosum, is prominently altered by PAE, resulting in notable reductions in white matter in the brain^90^. The alterations to the expression of Cyth1, Unc5d, and Nr4a1 as well as Ephrin signaling, ROBO receptor signaling, Wnt signaling, axonal guidance, and IGF transport and regulation in neurons as well as alterations to intercellular interactions involving Mdk, Ptn, and syndecans that we observed as a result of PAE could contribute to thinning of the corpus callosum, one of the most prevalent neuroanatomical features of FASDs, through disruptions to axon development.

Animal models of PAE reveal significant alterations to the landscape of neuroinflammation in the brain of exposed subjects compared to controls. These alterations occur primarily through cytokine dysregulation and the activation of glial cells, particularly microglia^91,92^. Additionally, these alterations can result in molecular abnormalities in the brain that can persist into adulthood. The elevated neuroinflammation in the brain in PAE animals is implicated in deficits in memory and cognition as well as psychiatric conditions, which occur commonly alongside FASDs^93^. We identified alterations to gene expression and canonical pathways in all three examined cell types that could be related to an increase in inflammation as a result of PAE. Elevation of RelA and EIF2 signaling in ECs, elevated IL-8 signaling in neurons, elevated Pycard expression, GAIT signaling, and RUNX1-mediated transcription in astrocytes, and alterations to ligand-receptor interactions related to Mdk signaling could all contribute to driving neurocognitive effects of PAE through elevated neuroinflammation.

It is interesting to note that the predicted impact of the upregulation and downregulation of the identified genes, pathways, and biological processes were not always consistent and did not always correlate with observed biological features of PAE. For example, VEGF signaling, which should promote vascular development, is increased in PAE samples at E18 while previous research has thoroughly characterized deficits in brain vascular development as a result of PAE. In neurons, various components of neuron migration and morphological and functional development are promoted by Ephrin signaling, ROBO receptor signaling, and Wnt signaling, which we show in these studies are upregulated in PAE. As noted, impairments to these processes are documented in models of PAE. Therefore, as previously speculated regarding EphB/Ephrin-B signaling, there is the possibility that some of the alterations to these genes and pathways could be the result of mechanisms operating in opposition to mechanisms driving the cellular, anatomical, and physiological symptoms of PAE. This may be particularly true for contradictory mechanisms occurring at the P28 timepoint; mechanisms may begin to counteract alterations to typical cellular function that occurred during alcohol exposure once the exposure has halted. We do not aim to speculate that specific alterations to gene expression in ECs, neurons, and astrocytes represent compensatory mechanisms against specific features of PAE in our analysis of these studies, but our data present several avenues for future research.

In addition to analyzing our spatial transcriptomic data with IPA, we used CellChat to investigate how PAE alters neurovascular communication across development. We identified over 1,970 predicted cell-cell interactions, including 142 signaling pathways across developmental timepoints and neurovascular cell types, revealing substantial signaling alterations following PAE. The primary alterations in ligand-receptor pairings and signaling pathways at E18 and P28 are associated with cell adhesion, ECM and structural remodeling, neurovascular patterning, neurotrophic support, and synaptic remodeling^94–100^. The reprogramming of cellular communications under PAE conditions may contribute to some of the neuronal and vascular deficits seen in FASD. Many of the altered receptor-ligand pairings have not yet been implicated in FASD and PAE studies, highlighting our study as a novel interpretation into PAE-mediated neurovascular communication.

In late embryonic development, many PAE-induced communication alterations are tied to Mdk, Ptn and Ephrin-B signaling networks. Mdk is a small, heparin-binding neurotrophic growth factor involved in proliferation, survival, migration, neuroinflammation, angiogenesis, and repair^101,102^. Since PAE is known to disrupt neurotrophic signaling and vascular development, there is potential indication that Mdk signaling may play a role in the PAE-mediated dysfunction of these biological processes^33,103,104^. Increased EC autocrine signaling through Mdk-Sdc4, Mdk-Ncl, and Mdk-Lrp1 suggests activation of pathways involved in stress response, vascular remodeling and cell survival^102,105,106^. Of note, the increase in Mdk-Lrp1 is of interest because Lrp1 serves as a receptor for Apolipoprotein E (APOE), a pathway associated with neurobehavioral deficits in FASDs^107^.

EphB/Ephrin-B signaling is classically associated with angiogenic sprouting and vascular remodeling^108^ and its upregulation in our data may show an adaptive mechanistic response rather than enhanced vascular function. Eph-ephrin signaling regulates multiple aspects of vascular development through tightly controlled signaling, and disruption of this normal balance can produce aberrant vessel patterning and compromised vascular integrity^109,110^. Therefore, the increased Efnb2-Ephb3 signaling observed in PAE ECs may represent a maladaptive response to developmental stress rather than successful angiogenesis. This interpretation aligns with ours and others findings of reduced BMVEC angiogenesis following alcohol exposure and altered cerebrovascular orientation^15,33,104,111^. Overall, the increased Mdk- and Eph-mediated signaling observed in ECs may reflect activation of stress-responsive pathways associated with abnormal vessel architecture, neuroinflammation and compensatory angiogenesis^101,102,109,110^.

Interestingly, E18 EC and neuronal autocrine signaling exhibited opposing responses to PAE. Neurons displayed reduced Mdk and Eph signaling but increased Ptn-Sdc2 signaling. Ptn is a neurotrophic growth factor that promotes neuronal survival, synaptic development, and neurite extension, suggesting that increases in Ptn-Sdc2 signaling may reflect an adaptive response to PAE-induced neurodevelopmental stress^103,112^. Syndecans are transmembrane heparan sulfate proteoglycans and despite not extensively investigated in neurons in FASD, they function as key co-receptors for neurotrophic factors, including Mdk and Ptn, and are critical regulators of axonal growth and synaptic plasticity^113^. Together, these findings suggest that at E18, ECs and neurons respond differently to PAE, and that PAE may be causing a separation between vascular and neuronal development rather than uniform suppression.

On a global scale, the overall neurovascular communication between neurons and ECs was largely suppressed at E18. Nine of the ten altered signaling pathways showed substantial decreases in interaction, with the exception of PTN. CD99 displayed the highest communication activity overall and decreased following PAE. CD99 is a transmembrane glycoprotein important for early development, playing a role in cell adhesion, migration, differentiation, inflammation, and apoptosis^114^. Suppression of this pathway may contribute to impaired neurovascular interactions during late embryonic development.

Unlike the broad suppression seen at E18, communication networks at P28 demonstrated widespread amplification. All ten of the most altered pathways were increased following PAE, with CypA displaying the highest communication activity. CypA is a peptidyl-prolyl cis-trans isomerase (PPIase) and is important for protein folding, inflammatory responses, intracellular trafficking, and participates in the pathophysiology of conditions such as neurodegeneration and autoimmune diseases^115^. This shift in enhanced signaling during adolescence may suggest activation of adaptive neurovascular mechanisms following PAE.

At P28, the most notable autocrine changes occurred within astrocytes, related to Cntn2, Ptn, Nfasc signaling pathways. Both Cntn2 and L1cam are cell adhesion molecules critical for brain development, axonal guidance, and myelination^116,117^. While Cntn2 expression has not been extensively characterized in astrocytes, they are recognized as active participants in adhesion-mediated signaling that regulates synapse development, neuronal connectivity, and neural circuit organization^118–120^. Therefore, the predicted decrease in Cntn2-L1cam signaling within PAE astrocyte communication networks may reflect disrupted adhesion-mediated interactions that support neuronal connectivity and circuit maintenance.

Similarly, Nfasc, a member of the L1 family of cell adhesion molecules, is a critical regulator of axon-glial interactions and paranodal organization^121^. Its decreased interaction with Cntn1+Cntnap1 in PAE could indicate disruption of adhesion pathways involved in paranode formation and maintenance^122^. Together, the increase in Ptn-Sdc2 signaling and reduction in adhesion-associated signaling suggest that astrocytes may shift from structural support and connectivity-related functions towards adaptive neurotrophic communication following PAE.

In contrast, neuronal autocrine signaling displayed increases in adhesion and synapse associated communications, with no notable decreases in ligand-receptor pairings. The increases in synaptic adhesion molecule interactions Tenm2/3, Flrt3, and Adgrl2 could suggest ongoing neural circuit remodeling or adaptations in response to earlier developmental disruptions^123^. Notably, astrocytes and neurons demonstrated opposing signaling responses, similar to those observed between ECs and neurons at E18. This mirroring effect across developmental timepoints may suggest that PAE induces cell type specific adaptive responses that persist into adolescence, potentially contributing to long-term alterations in neural circuit organization.

In addition to modulating individual ligand-receptor interactions, PAE modified the signaling roles of neurovascular cell populations. At E18, ECs showed decreased incoming/outgoing signaling of SST, EGF, FGF, and PECAM2. SST is a peptide hormone that regulates cellular growth by inhibiting EGF- and FGF-mediated signaling^124^. Therefore, its decreased incoming/outgoing signaling impacts EGF and FGF pathways and may reflect the altered cerebrovascular structures seen in PAE^18^. ECs received increased RELN and NRXN incoming/outgoing signals, which are both involved in promoting angiogenesis in ECs^125,126^. Because RELN signaling is essential for proper development, its disruption by PAE may contribute to the altered cell-cell communication observed in our study^127^. The simultaneous activation of both growth-promoting and growth-suppressive pathways may represent the attempt of PAE ECs to maintain signaling homeostasis. Neurons at this timepoint demonstrated decreased incoming/outgoing signals associated with cell adhesion and axonal guidance pathways such as JAM, SEMA5, and NRP1^128,129^. Neurons experienced increased outgoing signals that support neuronal maturation, survival, neuroimmune responses, and synaptic development, such as NT and SLURP1^130,131^. These patterns at E18 highlight the complex landscape of neurovascular crosstalk and how PAE modulates these communications.

PAE ECs at P28 have increased incoming/outgoing signals, including JAM and ESAM, which are collectively important for cell adhesion, tissue architecture, and barrier integrity^128,132^. Moreover, incoming/outgoing signaling through the brain-specific angiogenesis inhibitor (BAI1/ADGRB) pathway was increased in PAE ECs. Given the established role of ADGRB signaling in regulating angiogenesis^133^, this increase may represent an adaptive response to alcohol-induced disruptions carried into adolescence. Neurons at P28 also exhibited increased BAI1/ADGRB incoming/outgoing signaling, potentially strengthening this pro-angiogenic pathway across the neurovasculature. Notably, neurons showed relatively few decreases in incoming/outgoing interactions, suggesting that neuronal communication remains broadly preserved and may be particularly responsive to signals from surrounding neurovascular cell types. Meanwhile, increased astrocytic collagen, EGF, FGF, EPHB, and tenascin incoming/outgoing signaling may suggest reactive astrocyte activation^85,134–137^. Alcohol increases the expression of pro-inflammatory cytokines and profoundly affects astrocyte activation, referred to as astrogliosis; these increased outward signals may indicate PAE’s ability to affect the developing CNS^138^.

Our studies have some practical limitations; most notably, our brain samples are not separated by sex. Additionally, our *in vivo* model represents only one pattern and dosage of maternal alcohol consumption, and therefore, may not be widely applicable to human FASDs since human drinking patterns and dosages are highly variable. Finally, our analysis is limited to one section of the brain; other differential gene expression patterns and intercellular interactions may be observable in other locations in the brain.

Despite these limitations, future studies can use our moderate model of PAE^139^ to examine the cortical microvascular architecture of adolescent mice to determine how the observed transcriptional alterations correspond to structural changes within the cerebrovasculature. Additionally, because of the critical role of the neurovascular unit in maintaining blood-brain barrier (BBB) integrity, it will also be important to investigate the effects of PAE on vascular permeability in both our embryonic and adolescent models. Evaluating BBB function alongside vascular morphology will provide novel insights into the long-term effects of PAE on cerebrovascular function during adolescence.

Together, these findings provide a molecular framework for understanding how PAE disrupts the developing neurovascular unit and what transcriptional and cell-cell communication mechanisms may be of interest. DEG and CellChat analyses identified a broad range of PAE-associated changes. Among recurring themes, pathways related to cell adhesion and tissue remodeling, vascular patterning and neuroinflammation were consistent with prior studies^1,104,138,140^. These pathways represent only a subset of the alterations detected, and the broader set of transcriptional and intercellular signaling changes provides additional mechanisms for future investigation.

## MATERIALS AND METHODS

### PAE mouse model

We utilized a previously established limited access paradigm^15,16,141^. In summary, C57BL/6J female mice had access to a 10% (w/v) ethanol (EtOH) in 0.066% (w/v) saccharin (SAC) drinking mixture or a 0.066% (w/v) SAC solution for 4hrs daily prior to breeding. C57BL/6J female mice continued this drinking pattern throughout gestation and were weaned off alcohol postpartum. Embryonic brains were collected on day 18 (E18) prior to birth. For adolescent mice experiments, pups were weaned between P20-P24, and brain tissue was collected on postnatal day 28 (P28) for further analysis. All animal experiment protocols were approved by the University of New Mexico Institutional Animal Care and Use Committee (IACUC).

### FFPE and sample preparation

Samples were prepared according to the GeoMx® NGS RNA slide preparation protocol. SAC and PAE whole brains were collected at E18 and P28. Samples were fixed in 4% PFA, cryopreserved, and frozen in TissueTek OCT. 5 µm coronal sections from the mid-cortex were washed with dH_2_O and dehydrated in increasing concentrations of EtOH (30%, 50%, and 70%). Once dehydrated, samples were treated with Hemo-De and embedded with Paraplast Plus. Coronal sections were mounted to SuperFrost Plus slides (ThermoFisher) and left on a slide warmer overnight (O/N) at 35°C per NanoString’s guidelines. Following sample preparation, slides were shipped to NanoString Technologies.

### Digital spatial profiling (DSP)

Tissue slides were incubated O/N with fluorescently labeled cell-specific antibodies (anti-CD31/PECAM for endothelial cells; anti-NeuN for neurons; and anti-GFAP for astrocytes), to enable identification of regions of interest (ROIs), and with the UV-photocleavable GeoMx® Mouse Whole Transcriptome Atlas (WTA v1.0) RNA probe mix. ROIs with high expression of anti-CD31/PECAM, anti-NeuN, and anti-GFAP fluorescent labeling were then selected and exposed to UV light. UV-photocleavable probes were collected using a microcapillary tube, dispensed into a 96-well plate, and quantified using the nCounter Analysis System. Quality control, normalization, and differential gene expression analyses were performed by NanoString using the NanoString Analysis Suite. Data was normalized to the third quartile (Q3). Differential gene expression was performed between PAE and SAC samples for each cell type within each timepoint, using linear mixed modeling (LMM) with Benjamini-Hochberg correction (FDR-adjusted p<0.05), and genes meeting these significance thresholds and absolute fold change (FC)≥1.2 were included for downstream analyses. Volcano plots of DEGs generated from DSP data were uploaded into the VolcaNoseR interactive web application^142^ based on the following parameters: FC threshold –1.2 to 1.2 (|log_2_FC|≥0.2) and significance threshold p<0.05 (-log_10_>1.2), using the Manhattan distance criterion.

### Ingenuity Pathway Analysis (IPA) software

Differential gene expression data generated from DSP were analyzed using Ingenuity Pathway Analysis (IPA; Qiagen, Redwood City, CA, USA). Excel files of DEGs for each cell type within each timepoint were downloaded from the NanoString Analysis Suite. DEG lists, including gene identifiers, fold change values, and corresponding statistical significance metrics, were uploaded into IPA’s Core Expression Data Analysis. The following core analyses were performed: 1) Canonical Pathway Analysis to identify significantly enriched biological pathways based on the IPA Knowledge Base. Significance was determined using right-tailed Fisher’s exact test (p<0.05), and pathway activation states were inferred using z-scores where available; 2) Network Analysis to identify top molecular interactions and predicted relationships; 3) Disease and Biological Function Analysis to identify biological functions and disease processes associated with the observed gene expression changes. Separate comparison analyses were conducted for each developmental stage (E18 and P28) to assess time-dependent effects of EtOH exposure. Where appropriate, comparative analyses were performed within IPA to identify shared and distinct pathway perturbations between developmental timepoints.

### NanoString data preprocessing

The raw gene expression data was exported as Q3-normalized counts from the NanoString DSPDA software in .xlsx format. Probe assignments and initial counting of hybridization events for the Mouse Whole Transcriptome Atlas (WTA v1.0) probe panel targeting the mouse reference genome was performed using the NanoString GeoMx® platform. Data was imported into R v4.4.2 via the readxl, resulting in a full-expression matrix of 14,129 genes × 46 segments. Each segment, derived from one of the regions of interest (ROIs), was annotated by cell type (based on the marker used for segmentation: CD31, NeuN, or GFAP), treatment condition (SAC or PAE), and time point (E18 or P28). For each cell type, four unique treatment condition-timepoint combination based pseudobulk gene expression matrices were constructed resulting in following segment subsets: SAC E18 (n=12 segments), PAE E18 (n=16 segments), SAC P28 (n=9 segments) and PAE P28 (n=9 segments). Each ROI segment was treated as a single observation in its condition group. Processed matrices were written to .rds for downstream CellChat analysis.

### Cell-cell communication analysis

Cell-cell communication analysis was performed in R v4.4.2 using CellChat v2.2.0^85^. Statistical analysis was conducted using the mouse signaling database (n=945 ligand-receptor pairs, n=142 pathways, derived from 877 signaling genes). As GeoMx® DSP generates bulk expression profiles per segment rather than single-cell data, the standard CellChat workflow was adapted using a pseudobulk approach where each cell type’s aggregated expression profile per condition served as input. Three standard pseudobulk preprocessing steps were modified as follows:

1. identifyOverExpressedGenes was skipped and all 877 signaling genes were used,
2. identifyOverExpressedInteractions was bypassed, instead ligand-receptor pair matching was performed using case-insensitive string matching, recovering 945 pairs, and
3. filterCommunication was also skipped. The communication probabilities were calculated using the computeCommunProb function with truncatedMean (trim = 0.1, distance.use = FALSE) for each cell type to define the incoming and outgoing interaction strength at E18 and P28 timepoints and for both PAE and SAC conditions. For both E18 and P28 timepoints, the cell type-specific differential incoming vs outgoing pathway strength was contrasted for PAE vs SAC condition by subtracting SAC z-score normalized ligand-receptor pair signaling communication probabilities from the PAE induced z-score normalized communication probabilities. This normalization was used to account for the effect arising from the potential slide-level technical variabilities. Scatterplots were used for accessing the differential pathway-sender-receiver dynamics changes. Pathway-sender-receiver combinations with mean probabilities below 0.01 were excluded prior to normalization. Code for analyses can be found at https://github.com/singhepicurelab/Nanostring_Endothelial_Neuron_CellChat/

## REFERENCES

1. Chung, D. D. et al. Toxic and Teratogenic Effects of Prenatal Alcohol Exposure on Fetal Development, Adolescence, and Adulthood. Int. J. Mol. Sci. 22, 8785 (2021).

2. Caputo, C., Wood, E. & Jabbour, L. Impact of fetal alcohol exposure on body systems: A systematic review. Birth Defects Res. Part C Embryo Today Rev. 108, 174–180 (2016).

3. Gardiner, A. S. et al. Alcohol Use During Pregnancy is Associated with Specific Alterations in MicroRNA Levels in Maternal Serum. Alcohol. Clin. Exp. Res. 40, 826–837 (2016).

4. May, P. A. et al. Prevalence of Fetal Alcohol Spectrum Disorders in 4 US Communities. JAMA 319, 474–482 (2018).

5. Frey, S. et al. Prenatal Alcohol Exposure Is Associated With Adverse Cognitive Effects and Distinct Whole-Genome DNA Methylation Patterns in Primary School Children. Front. Behav. Neurosci. 12, (2018).

6. Salem, N. A., Mahnke, A. H., Konganti, K., Hillhouse, A. E. & Miranda, R. C. Cell-type and fetal-sex-specific targets of prenatal alcohol exposure in developing mouse cerebral cortex. iScience 24, (2021).

7. Deyssenroth, M. A. et al. Prenatal alcohol exposure is associated with changes in placental gene co-expression networks. Sci. Rep. 14, 2687 (2024).

8. Sambo, D. & Goldman, D. Genetic Influences on Fetal Alcohol Spectrum Disorder. Genes 14, 195 (2023).

9. Paredes, I., Himmels, P. & Ruiz de Almodóvar, C. Neurovascular Communication during CNS Development. Dev. Cell 45, 10–32 (2018).

10. Iadecola, C. The Neurovascular Unit Coming of Age: A Journey through Neurovascular Coupling in Health and Disease. Neuron 96, 17–42 (2017).

11. Zhou, Y., Song, H. & Ming, G. Genetics of human brain development. Nat. Rev. Genet. 25, 26–45 (2024).

12. Van, T. M. & Blank, C. U. A user’s perspective on GeoMxTM digital spatial profiling. Immuno-Oncol. Technol. 1, 11–18 (2019).

13. Hernandez, S. et al. Challenges and Opportunities for Immunoprofiling Using a Spatial High-Plex Technology: The NanoString GeoMx® Digital Spatial Profiler. Front. Oncol. 12, (2022).

14. Saha, P. S. & Mayhan, W. G. Prenatal exposure to alcohol: mechanisms of cerebral vascular damage and lifelong consequences. Adv. Drug Alcohol Res. 2, 10818 (2022).

15. Perales, G. et al. MicroRNA-150-5p is upregulated in the brain microvasculature during prenatal alcohol exposure and inhibits the angiogenic factor Vezf1. Alcohol. Clin. Exp. Res. 46, 1953–1966 (2022).

16. Brady, M. L., Allan, A. M. & Caldwell, K. K. A Limited Access Mouse Model of Prenatal Alcohol Exposure that Produces Long-Lasting Deficits in Hippocampal-Dependent Learning and Memory. Alcohol. Clin. Exp. Res. 36, 457–466 (2012).

17. Park, K. W. et al. Robo4 is a vascular-specific receptor that inhibits endothelial migration. Dev. Biol. 261, 251–267 (2003).

18. Jégou, S. et al. Prenatal Alcohol Exposure Affects Vasculature Development in the Neonatal Brain. Ann. Neurol. 72, 952–960 (2012).

19. Jia, D., Huang, L., Bischoff, J. & Moses, M. A. The endogenous zinc finger transcription factor, ZNF24, modulates the angiogenic potential of human microvascular endothelial cells. FASEB J. 29, 1371–1382 (2015).

20. Shih, Y.-P., Sun, P., Wang, A. & Lo, S. H. Tensin1 positively regulates RhoA activity through its interaction with DLC1. Biochim. Biophys. Acta BBA - Mol. Cell Res. 1853, 3258–3265 (2015).

21. Yang, J., Sun, W. & Cui, G. Roles of the NR2F Family in the Development, Disease, and Cancer of the Lung. J. Dev. Biol. 12, 24 (2024).

22. Bhakuni, T. et al. FOXC1 regulates endothelial CD98 (LAT1/4F2hc) expression in retinal angiogenesis and blood-retina barrier formation. Nat. Commun. 15, 4097 (2024).

23. Leker, R. R. et al. Transforming growth factor alpha induces angiogenesis and neurogenesis following stroke. Neuroscience 163, 233–243 (2009).

24. Masckauchán, T. N. H. et al. Wnt5a Signaling Induces Proliferation and Survival of Endothelial Cells In Vitro and Expression of MMP-1 and Tie-2. Mol. Biol. Cell 17, 5163–5172 (2006).

25. Bijli, K. M., Fazal, F. & Rahman, A. Regulation of Rela/p65 and Endothelial Cell Inflammation by Proline-Rich Tyrosine Kinase 2. Am. J. Respir. Cell Mol. Biol. 47, 660–668 (2012).

26. Huang, J. & Kontos, C. D. PTEN modulates vascular endothelial growth factor-mediated signaling and angiogenic effects. J. Biol. Chem. 277, 10760–10766 (2002).

27. Ferrara, N. Role of vascular endothelial growth factor in the regulation of angiogenesis. Kidney Int. 56, 794–814 (1999).

28. Shrestha, N. et al. Eukaryotic Initiation Factor 2 (eIF2) Signaling Regulates Proinflammatory Cytokine Expression and Bacterial Invasion. J. Biol. Chem. 287, 28738–28744 (2012).

29. Drozd, M. et al. Endothelial insulin-like growth factor-1 signalling regulates vascular barrier function and atherogenesis. Cardiovasc. Res. 121, 1108–1120 (2025).

30. Houde, M., Desbiens, L. & D’Orléans-Juste, P. Endothelin-1: Biosynthesis, Signaling and Vasoreactivity. Adv. Pharmacol. 77, 143–175 (2016).

31. Stebbins, M. J. et al. Activation of RARα, RARγ, or RXRα increases barrier tightness in human induced pluripotent stem cell-derived brain endothelial cells. Biotechnol. J. 13, 10.1002/biot.201700093 (2018).

32. van Buul, J. D., Geerts, D. & Huveneers, S. Rho GAPs and GEFs. Cell Adhes. Migr. 8, 108–124 (2014).

33. Wang, S. et al. The mTOR/AP-1/VEGF signaling pathway regulates vascular endothelial cell growth. Oncotarget 7, 53269–53276 (2016).

34. Inkelis, S. M. & Thomas, J. D. SLEEP IN INFANTS AND CHILDREN WITH PRENATAL ALCOHOL EXPOSURE. Alcohol. Clin. Exp. Res. 10.1111/acer.13803 (2018) doi:10.1111/acer.13803.

35. Kasakura, N., Murata, Y., Suzuki, K. & Segi-Nishida, E. Role of endogenous NT-3 in neuronal activity and neurogenesis in the hippocampal dentate gyrus. Neurosci. Res. 218, 104923 (2025).

36. Kobayashi, H. et al. Calm1 signaling pathway is essential for the migration of mouse precerebellar neurons. Development 142, 375–384 (2015).

37. Liao, Y.-C. et al. CEP170 as a novel molecular link between centrosomal function and cerebral cortical development. J. Biomed. Sci. 33, 33 (2026).

38. Musante, I. et al. CACNA1A loss-of-function affects neurogenesis in human iPSC-derived neural models. Cell. Mol. Life Sci. CMLS 82, 234 (2025).

39. Zhang, Q. et al. Satb2 regulates the development of dopaminergic neurons in the arcuate nucleus by Dlx1. Cell Death Dis. 12, 879 (2021).

40. Bai, W.-J. et al. Deficiency of transmembrane AMPA receptor regulatory protein γ-8 leads to attention-deficit hyperactivity disorder-like behavior in mice. Zool. Res. 43, 851–870 (2022).

41. Jia, N. et al. Protective role and related mechanism of Gnaq in neural cells damaged by oxidative stress. Acta Biochim. Biophys. Sin. 49, 428–434 (2017).

42. Huang, M. et al. Nr4a1 regulates cell-specific transcriptional programs in inhibitory GABAergic interneurons. Neuron 112, 2031–2044.e7 (2024).

43. Ito, A., Fukaya, M., Okamoto, H. & Sakagami, H. Physiological and Pathological Roles of the Cytohesin Family in Neurons. Int. J. Mol. Sci. 23, 5087 (2022).

44. Takemoto, M. et al. Laminar and areal expression of unc5d and its role in cortical cell survival. Cereb. Cortex 21, 1925–1934 (2011).

45. Tocco, C., Bertacchi, M. & Studer, M. Structural and Functional Aspects of the Neurodevelopmental Gene NR2F1: From Animal Models to Human Pathology. Front. Mol. Neurosci. 14, 767965 (2021).

46. Doan, K. V. et al. FoxO1 in dopaminergic neurons regulates energy homeostasis and targets tyrosine hydroxylase. Nat. Commun. 7, 12733 (2016).

47. Peng, Y. et al. The autism associated MET receptor tyrosine kinase engages early neuronal growth mechanism and controls glutamatergic circuits development in the forebrain. Mol. Psychiatry 21, 925–935 (2016).

48. Santo, E. E. & Paik, J. FOXO in Neural Cells and Diseases of the Nervous System. Curr. Top. Dev. Biol. 127, 105–118 (2018).

49. Westenskow, M. R., Amdor, A. G., Gutierrez, R., Perales, G. & Gardiner, A. S. Prenatal alcohol exposure-mediated Tet1 upregulation promotes DNA demethylation and elevated transcription at the miR-150 promoter. Am. J. Physiol. Cell Physiol. 331, C1–C12 (2026).

50. Blank, T., Nijholt, I., Kye, M.-J., Radulovic, J. & Spiess, J. Small-conductance, Ca2+-activated K+ channel SK3 generates age-related memory and LTP deficits. Nat. Neurosci. 6, 911–912 (2003).

51. Zhang, L., Bai, W., Peng, Y., Lin, Y. & Tian, M. Human umbilical cord mesenchymal stem cell-derived exosomes provide neuroprotection in traumatic brain injury through the lncRNA TUBB6/Nrf2 pathway. Brain Res. 1824, 148689 (2024).

52. Hirsch-Reinshagen, V. et al. LCAT synthesized by primary astrocytes esterifies cholesterol on glia-derived lipoproteins. J. Lipid Res. 50, 885–893 (2009).

53. Gjervan, S. C., Ozgoren, O. K., Gow, A., Stockler-Ipsiroglu, S. & Pouladi, M. A. Claudin-11 in health and disease: implications for myelin disorders, hearing, and fertility. Front. Cell. Neurosci. 17, 1344090 (2024).

54. Boutet, T. et al. Developmental stage dominates cell-type identity and reveals a chromatin regulatory function for Rad50 in Drosophila. Nucleic Acids Res. 54, gkag294 (2026).

55. Pastuzyn, E. D. et al. The Neuronal Gene Arc Encodes a Repurposed Retrotransposon Gag Protein that Mediates Intercellular RNA Transfer. Cell 172, 275–288.e18 (2018).

56. el-Ghissassi, F. et al. BTG2(TIS21/PC3) induces neuronal differentiation and prevents apoptosis of terminally differentiated PC12 cells. Oncogene 21, 6772–6778 (2002).

57. Pérez-Sen, R. et al. Dual-Specificity Phosphatase Regulation in Neurons and Glial Cells. Int. J. Mol. Sci. 20, 1999 (2019).

58. Hosen, S., Ikeda-Yorifuji, I. & Yamashita, T. Asporin and CD109, expressed in the injured neonatal spinal cord, attenuate axonal re-growth *in vitro*. Neurosci. Lett. 833, 137832 (2024).

59. Chung, L. A Brief Introduction to the Transduction of Neural Activity into Fos Signal. Dev. Reprod. 19, 61–67 (2015).

60. Rosso, S. B. & Inestrosa, N. C. WNT signaling in neuronal maturation and synaptogenesis. Front. Cell. Neurosci. 7, 103 (2013).

61. Yoshida, Y. Semaphorin Signaling in Vertebrate Neural Circuit Assembly. Front. Mol. Neurosci. 5, 71 (2012).

62. Cramer, K. S. & Miko, I. J. Eph-ephrin signaling in nervous system development. F1000Research 5, F1000 Faculty Rev-413 (2016).

63. Tong, M., Jun, T., Nie, Y., Hao, J. & Fan, D. The Role of the Slit/Robo Signaling Pathway. J. Cancer 10, 2694–2705 (2019).

64. Rosenstein, J. M., Krum, J. M. & Ruhrberg, C. VEGF in the nervous system. Organogenesis 6, 107–114 (2010).

65. Sakamoto, K., Karelina, K. & Obrietan, K. CREB: a multifaceted regulator of neuronal plasticity and protection. J. Neurochem. 116, 1–9 (2011).

66. Willard, S. S. & Koochekpour, S. Glutamate, Glutamate Receptors, and Downstream Signaling Pathways. Int. J. Biol. Sci. 9, 948–959 (2013).

67. O’Kusky, J. & Ye, P. Neurodevelopmental effects of insulin-like growth factor signaling. Front. Neuroendocrinol. 33, 230–251 (2012).

68. Alaamery, M. et al. Role of Sphingolipid Metabolism in Neurodegeneration. J. Neurochem. 158, 25–35 (2021).

69. Rawal, P. & Zhao, L. Sialometabolism in Brain Health and Alzheimer’s Disease. Front. Neurosci. 15, 648617 (2021).

70. Bickel, M. The role of interleukin-8 in inflammation and mechanisms of regulation. J. Periodontol. 64, 456–460 (1993).

71. Hollville, E., Romero, S. E. & Deshmukh, M. Apoptotic Cell Death Regulation in Neurons. FEBS J. 286, 3276–3298 (2019).

72. Koch, M. et al. NGF stimulation alters the transcriptome and surface TrkB expression in axons of dorsal root ganglion neurons. Neurobiol. Pain 18, 100194 (2025).

73. Cataldo, L. R. et al. The human batokine EPDR1 regulates β-cell metabolism and function. Mol. Metab. 66, 101629 (2022).

74. Chen, X. et al. The cell cycle gene centromere protein K (CENPK) contributes to the malignant progression and prognosis of prostate cancer. Transl. Cancer Res. 11, (2022).

75. Liu, W. et al. The Inflammatory Gene PYCARD of the Entorhinal Cortex as an Early Diagnostic Target for Alzheimer’s Disease. Biomedicines 11, 194 (2023).

76. Lee, S. et al. Lipocalin-type prostaglandin D2 synthase protein regulates glial cell migration and morphology through myristoylated alanine-rich C-kinase substrate: prostaglandin D2-independent effects. J. Biol. Chem. 287, 9414–9428 (2012).

77. Mukhopadhyay, R., Jia, J., Arif, A., Ray, P. S. & Fox, P. L. The GAIT system: a gatekeeper of inflammatory gene expression. Trends Biochem. Sci. 34, 324–331 (2009).

78. Hu, Y. et al. Matrix stiffness changes affect astrocyte phenotype in an in vitro injury model. NPG Asia Mater. 13, 35 (2021).

79. Kaneko, N. et al. New Neurons Clear the Path of Astrocytic Processes for Their Rapid Migration in the Adult Brain. Neuron 67, 213–223 (2010).

80. Huang, M. et al. RUNX1 Induces Central Neuropathic Pain by Activating Microglia and Triggering the Inflammatory Response in Spinal Cord Injury. Inflammation 48, 4443–4460 (2025).

81. Ianevski, A. et al. Early transcriptional responses reveal cell type-specific vulnerability and neuroprotective mechanisms in the neonatal ischemic hippocampus. Acta Neuropathol. Commun. 13, 147 (2025).

82. Wang, J. et al. The EIF2α-PERK Signaling Pathway Mediates Manganese Exposure-Induced A1-Type Astrocytes Activation via Endoplasmic Reticulum Stress. Toxics 13, 910 (2025).

83. Stevens, H. E. et al. Neonatal loss of FGFR2 in astroglial cells affects locomotion, sociability, working memory, and glia-neuron interactions in mice. Transl. Psychiatry 13, 89 (2023).

84. Wagner, B. et al. Neuronal survival depends on EGFR signaling in cortical but not midbrain astrocytes. EMBO J. 25, 752–762 (2006).

85. Jin, S., Plikus, M. V. & Nie, Q. CellChat for systematic analysis of cell–cell communication from single-cell transcriptomics. Nat. Protoc. 20, 180–219 (2025).

86. Miller, M. W. Migration of cortical neurons is altered by gestational exposure to ethanol. Alcohol. Clin. Exp. Res. 17, 304–314 (1993).

87. Clarren, S. K., Alvord, E. C., Sumi, S. M., Streissguth, A. P. & Smith, D. W. Brain malformations related to prenatal exposure to ethanol. J. Pediatr. 92, 64–67 (1978).

88. Jones, K. L. & Smith, D. W. Recognition of the fetal alcohol syndrome in early infancy. Lancet 302, 999–1001 (1973).

89. Wisniewski, K., Dambska, M., Sher, J. H. & Qazi, Q. A clinical neuropathological study of the fetal alcohol syndrome. Neuropediatrics 14, 197–201 (1983).

90. Mathews, E., Dewees, K., Diaz, D. & Favero, C. White matter abnormalities in fetal alcohol spectrum disorders: Focus on axon growth and guidance. Exp. Biol. Med. 246, 812–821 (2021).

91. Padilla-Valdez, M. M., Díaz-Iñiguez, M. I., Ortuño-Sahagún, D. & Rojas-Mayorquín, A. E. Neuroinflammation in fetal alcohol spectrum disorders and related novel therapeutic approaches. Biochim. Biophys. Acta BBA - Mol. Basis Dis. 1870, 166854 (2024).

92. Terasaki, L. S. & Schwarz, J. M. Effects of Moderate Prenatal Alcohol Exposure during Early Gestation in Rats on Inflammation across the Maternal-Fetal-Immune Interface and Later-Life Immune Function in the Offspring. J. Neuroimmune Pharmacol. Off. J. Soc. NeuroImmune Pharmacol. 11, 680–692 (2016).

93. Noor, S. & Milligan, E. D. Lifelong Impacts of Moderate Prenatal Alcohol Exposure on Neuroimmune Function. Front. Immunol. 9, 1107 (2018).

94. Dalva, M. B., McClelland, A. C. & Kayser, M. S. Cell adhesion molecules: signalling functions at the synapse. Nat. Rev. Neurosci. 8, 206–220 (2007).

95. Gomez, A. M., Traunmüller, L. & Scheiffele, P. Neurexins: molecular codes for shaping neuronal synapses. Nat. Rev. Neurosci. 22, 137–151 (2021).

96. Hudson, N. & Campbell, M. Tight Junctions of the Neurovascular Unit. Front. Mol. Neurosci. 14, 752781 (2021).

97. Sijilmassi, O. Collagen IV and laminin-1 as key macromolecules in ocular structure and pathology: A review. Int. J. Biol. Macromol. 334, 149013 (2025).

98. Sosa, L. J. et al. The physiological role of the amyloid precursor protein as an adhesion molecule in the developing nervous system. J. Neurochem. 143, 11–29 (2017).

99. Wang, X. Pleiotrophin: Activity and Mechanism. Adv. Clin. Chem. 98, 51–89 (2020).

100. Wettschureck, N., Strilic, B. & Offermanns, S. Passing the Vascular Barrier: Endothelial Signaling Processes Controlling Extravasation. Physiol. Rev. 99, 1467–1525 (2019).

101. Neumaier, E. E., Rothhammer, V. & Linnerbauer, M. The role of midkine in health and disease. Front. Immunol. 14, 1310094 (2023).

102. Weckbach, L. T., Preissner, K. T. & Deindl, E. The Role of Midkine in Arteriogenesis, Involving Mechanosensing, Endothelial Cell Proliferation, and Vasodilation. Int. J. Mol. Sci. 19, 2559 (2018).

103. Boschen, K. E. & Klintsova, A. Y. Neurotrophins in the Brain: Interaction With Alcohol Exposure During Development. Vitam. Horm. 104, 197–242 (2017).

104. Jégou, S. et al. Prenatal alcohol exposure affects vasculature development in the neonatal brain. Ann. Neurol. 72, 952–960 (2012).

105. Nair, H. B. et al. Midkine (MDK) as a central regulator of the tumor microenvironment: From developmental cytokine to therapeutic target. Cancer Lett. 641, 218258 (2026).

106. Yýldýrým, B., Kulak, K. & Bilir, A. Midkine (MDK) in cancer and drug resistance: from inflammation to therapy. Discov. Oncol. 16, 1062 (2025).

107. Hwang, H. M. et al. Reduction of APOE accounts for neurobehavioral deficits in fetal alcohol spectrum disorders. Mol. Psychiatry 29, 3364–3380 (2024).

108. Salvucci, O. & Tosato, G. Essential Roles of EphB Receptors and EphrinB Ligands in Endothelial Cell Function and Angiogenesis. Adv. Cancer Res. 114, 21–57 (2012).

109. Rudno-Rudziñska, J. et al. A review on Eph/ephrin, angiogenesis and lymphangiogenesis in gastric, colorectal and pancreatic cancers. Chin. J. Cancer Res. 29, 303–312 (2017).

110. Stewen, J. et al. Eph-ephrin signaling couples endothelial cell sorting and arterial specification. Nat. Commun. 15, 2539 (2024).

111. Siqueira, M. et al. Ethanol Alters DNMT1/3a/3b Expression Profile, Promotes Persistent DNA Hypomethylation in Human Brain Endothelial Cells and Impairs Late Cortical Angiogenesis. J. Neurochem. 170, e70375 (2026).

112. Yanagisawa, H., Komuta, Y., Kawano, H., Toyoda, M. & Sango, K. Pleiotrophin induces neurite outgrowth and up-regulates growth-associated protein (GAP)-43 mRNA through the ALK/GSK3β/β-catenin signaling in developing mouse neurons. Neurosci. Res. 66, 111–116 (2010).

113. Edwards, T. J. & Hammarlund, M. Syndecan Promotes Axon Regeneration by Stabilizing Growth Cone Migration. Cell Rep. 8, 272–283 (2014).

114. Pasello, M., Manara, M. C. & Scotlandi, K. CD99 at the crossroads of physiology and pathology. J. Cell Commun. Signal. 12, 55–68 (2018).

115. Jurkova, K., Navratilova, H., Musilek, K. & Benek, O. Human Cyclophilins—An Emerging Class of Drug Targets. Med. Res. Rev. 46, 475–512 (2026).

116. Chatterjee, M., Schild, D. & Teunissen, C. E. Contactins in the central nervous system: role in health and disease. Neural Regen. Res. 14, 206–216 (2019).

117. Maness, P. F. & Schachner, M. Neural recognition molecules of the immunoglobulin superfamily: signaling transducers of axon guidance and neuronal migration. Nat. Neurosci. 10, 19–26 (2007).

118. Dallérac, G., Zapata, J. & Rouach, N. Versatile control of synaptic circuits by astrocytes: where, when and how? Nat. Rev. Neurosci. 19, 729–743 (2018).

119. Dewa, K. & Arimura, N. Neuronal and astrocytic protein connections and associated adhesion molecules. Neurosci. Res. 187, 14–20 (2023).

120. Tan, C. X. & Eroglu, C. Cell adhesion molecules regulating astrocyte-neuron interactions. Curr. Opin. Neurobiol. 69, 170–177 (2021).

121. Herron, L. R., Hill, M., Davey, F. & Gunn-Moore, F. J. The intracellular interactions of the L1 family of cell adhesion molecules. Biochem. J. 419, 519–531 (2009).

122. Sherman, D. L. et al. Neurofascins Are Required to Establish Axonal Domains for Saltatory Conduction. Neuron 48, 737–742 (2005).

123. Liakath-Ali, K., Refaee, R. & Südhof, T. C. Cartography of teneurin and latrophilin expression reveals spatiotemporal axis heterogeneity in the mouse hippocampus during development. PLOS Biol. 22, e3002599 (2024).

124. Chalabi, M. et al. Somatostatin analogs: does pharmacology impact antitumor efficacy? Trends Endocrinol. Metab. 25, 115–127 (2014).

125. Feng, Z. et al. Reelin promotes cerebral angiogenesis via mTOR/HIF-1α-mediated transcriptional upregulation of Netrin-4. Tissue Cell 103, 103727 (2026).

126. Rissone, A. et al. The Synaptic Proteins β-Neurexin and Neuroligin Synergize With Extracellular Matrix-Binding Vascular Endothelial Growth Factor A During Zebrafish Vascular Development. Arterioscler. Thromb. Vasc. Biol. 32, 1563–1572 (2012).

127. Wang, D., Howell, B. W. & Olson, E. C. Maternal Ethanol Exposure Acutely Elevates Src Family Kinase Activity in the Fetal Cortex. Mol. Neurobiol. 58, 5210–5223 (2021).

128. Ebnet, K. Junctional Adhesion Molecules (JAMs): Cell Adhesion Receptors With Pleiotropic Functions in Cell Physiology and Development. Physiol. Rev. 97, 1529–1554 (2017).

129. Gu, C. et al. Neuropilin-1 Conveys Semaphorin and VEGF Signaling during Neural and Cardiovascular Development. Dev. Cell 5, 45–57 (2003).

130. Lyukmanova, E. N. et al. Human Secreted Ly-6/uPAR Related Protein-1 (SLURP-1) Is a Selective Allosteric Antagonist of α7 Nicotinic Acetylcholine Receptor. PLoS ONE 11, e0149733 (2016).

131. St-Gelais, F., Jomphe, C. & Trudeau, L.-É. The role of neurotensin in central nervous system pathophysiology: What is the evidence? J. Psychiatry Neurosci. 31, 229–245 (2006).

132. Ishida, T. et al. Targeted Disruption of Endothelial Cell-selective Adhesion Molecule Inhibits Angiogenic Processes *in Vitro* and *in Vivo* *. J. Biol. Chem. 278, 34598–34604 (2003).

133. Li, A. et al. Functional diversity of BAI1 (ADGRB1): From angiostasis to synaptic remodeling and disease therapeutics. iScience 29, 114656 (2026).

134. Hara, M. et al. Interaction of reactive astrocytes with type I collagen induces astrocytic scar formation through the integrin–N-cadherin pathway after spinal cord injury. Nat. Med. 23, 818–828 (2017).

135. Kang, W. et al. Astrocyte activation is suppressed in both normal and injured brain by FGF signaling. Proc. Natl. Acad. Sci. U. S. A. 111, E2987–E2995 (2014).

136. Liu, B., Chen, H., Johns, T. G. & Neufeld, A. H. Epidermal Growth Factor Receptor Activation: An Upstream Signal for Transition of Quiescent Astrocytes into Reactive Astrocytes after Neural Injury. J. Neurosci. 26, 7532–7540 (2006).

137. Tyzack, G. E. et al. A neuroprotective astrocyte state is induced by neuronal signal EphB1 but fails in ALS models. Nat. Commun. 8, 1164 (2017).

138. Kane, C. J. M. & Drew, P. D. Neuroinflammatory Contribution of Microglia and Astrocytes in Fetal Alcohol Spectrum Disorders. J. Neurosci. Res. 99, 1973–1985 (2021).

139. Myrick, A. et al. Maternal Alcohol Drinking Patterns Predict Offspring Neurobehavioral Outcomes. Neuropharmacology 257, 110044 (2024).

140. Licheri, V. & Brigman, J. L. Altering Cell-Cell Interaction in Prenatal Alcohol Exposure Models: Insight on Cell-Adhesion Molecules During Brain Development. Front. Mol. Neurosci. 14, (2021).

141. Westenskow, M. R., Amdor, A. G., Gutierrez, R., Perales, G. & Gardiner, A. S. Prenatal alcohol exposure-mediated Tet1 upregulation promotes DNA demethylation and elevated transcription at the miR-150 promoter. Am. J. Physiol.-Cell Physiol. 331, C1–C12 (2026).

142. Goedhart, J. & Luijsterburg, M. S. VolcaNoseR is a web app for creating, exploring, labeling and sharing volcano plots. Sci. Rep. 10, 20560 (2020).

