## Supplemental Files for "Spatial profiling and neurovascular communication in the developing and adolescent cortex following prenatal alcohol exposure"

**A IPA Comparison Analysis: Endothelin-1 signaling in ECs at E18**

**B IPA Comparison Analysis: Endothelin-1 signaling in ECs at P28**

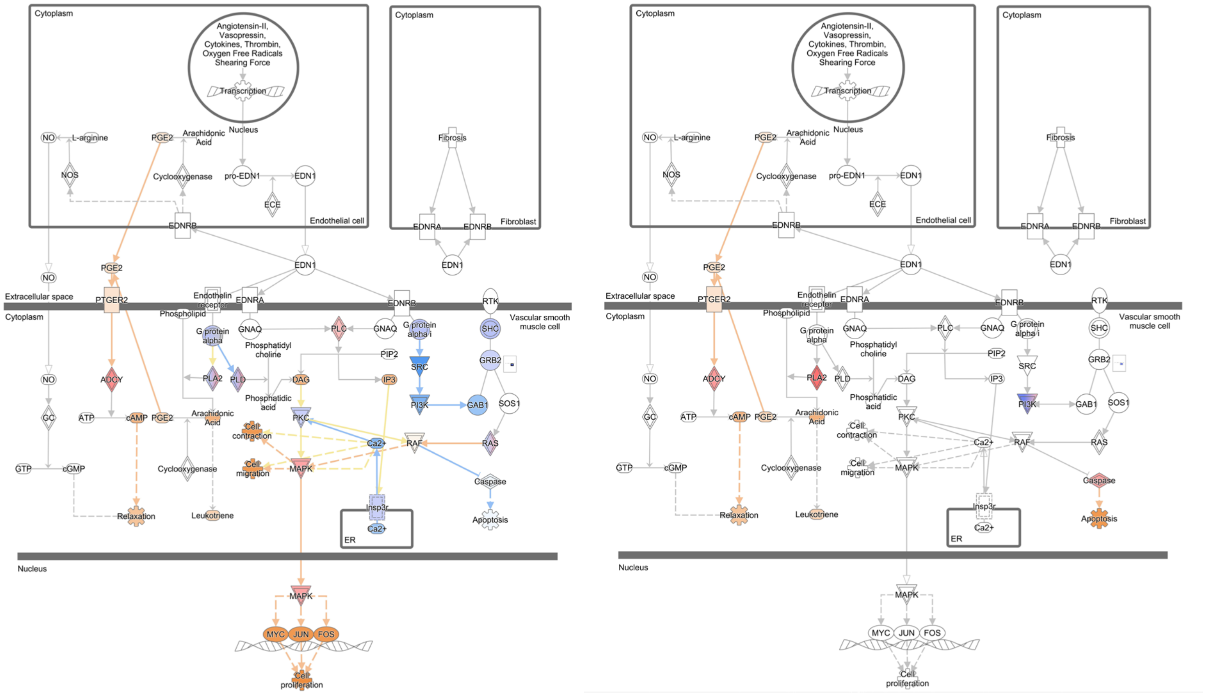

**Supplemental Figure 1. IPA comparison analysis of endothelin-1 signaling in endothelial cells at E18 and P28.** Graphical depiction of predicted endothelin-1 signaling pathways in ECs at **a)** E18 and **b)** P28. Node colors represent measured changes, with red indicating increased measurements, light blue indicating decreased measurements, orange indicating predicted activation, and dark blue indicating predicted inhibition. Predicted relationships are encoded by line color and style: orange lines indicate activation and blue lines indicate inhibition, solid lines indicate direct relationships, and dashed lines indicate indirect relationships.

**A** IPA Comparison Analysis: IGF-1 signaling in ECs at E18

**B** IPA Comparison Analysis: IGF-1 signaling in ECs at P28

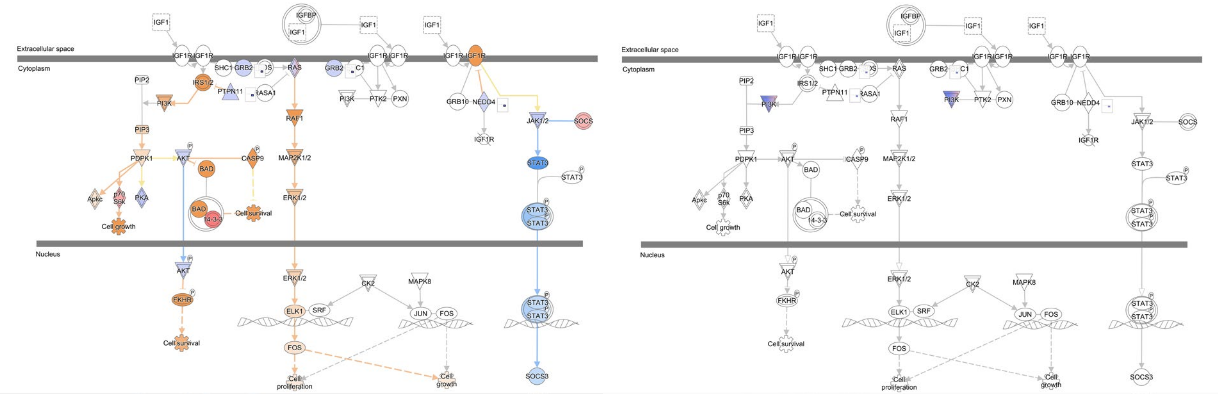

**Supplemental Figure 2. IPA comparison analysis of IGF-1 signaling in endothelial cells at E18 and P28.** Graphical depiction of predicted IGF-1 signaling pathways in ECs at **a)** E18 and **b)** P28. Node colors represent measured changes, with red indicating increased measurements, light blue indicating decreased measurements, orange indicating predicted activation, and dark blue indicating predicted inhibition. Predicted relationships are encoded by line color and style: orange lines indicate activation, and blue lines indicate inhibition, solid lines indicate direct relationships, and dashed lines indicate indirect relationships.

**A** **IPA Comparison Analysis: VEGF signaling in ECs at E18**

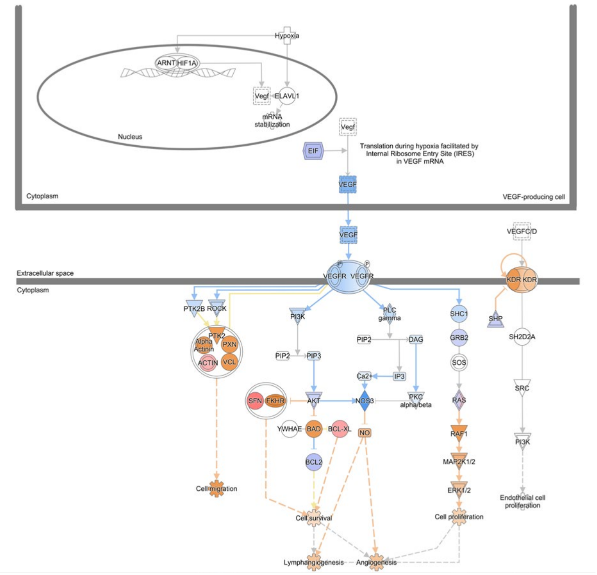

**B** **IPA Comparison Analysis: VEGF signaling in ECs at P28**

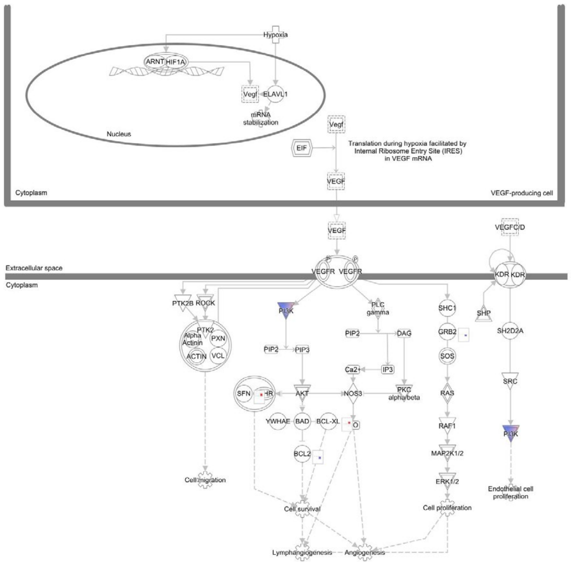

**Supplemental Figure 3. IPA comparison analysis of VEGF signaling in endothelial cells at E18 and P28.** Graphical depiction of predicted VEGF signaling pathways in ECs at **a)** E18 and **b)** P28. Node colors represent measured changes, with red indicating increased measurements, light blue indicating decreased measurements, orange indicating predicted activation, and dark blue indicating predicted inhibition. Predicted relationships are encoded by line color and style: orange lines indicate activation and blue lines indicate inhibition, solid lines indicate direct relationships, and dashed lines indicate indirect relationships.

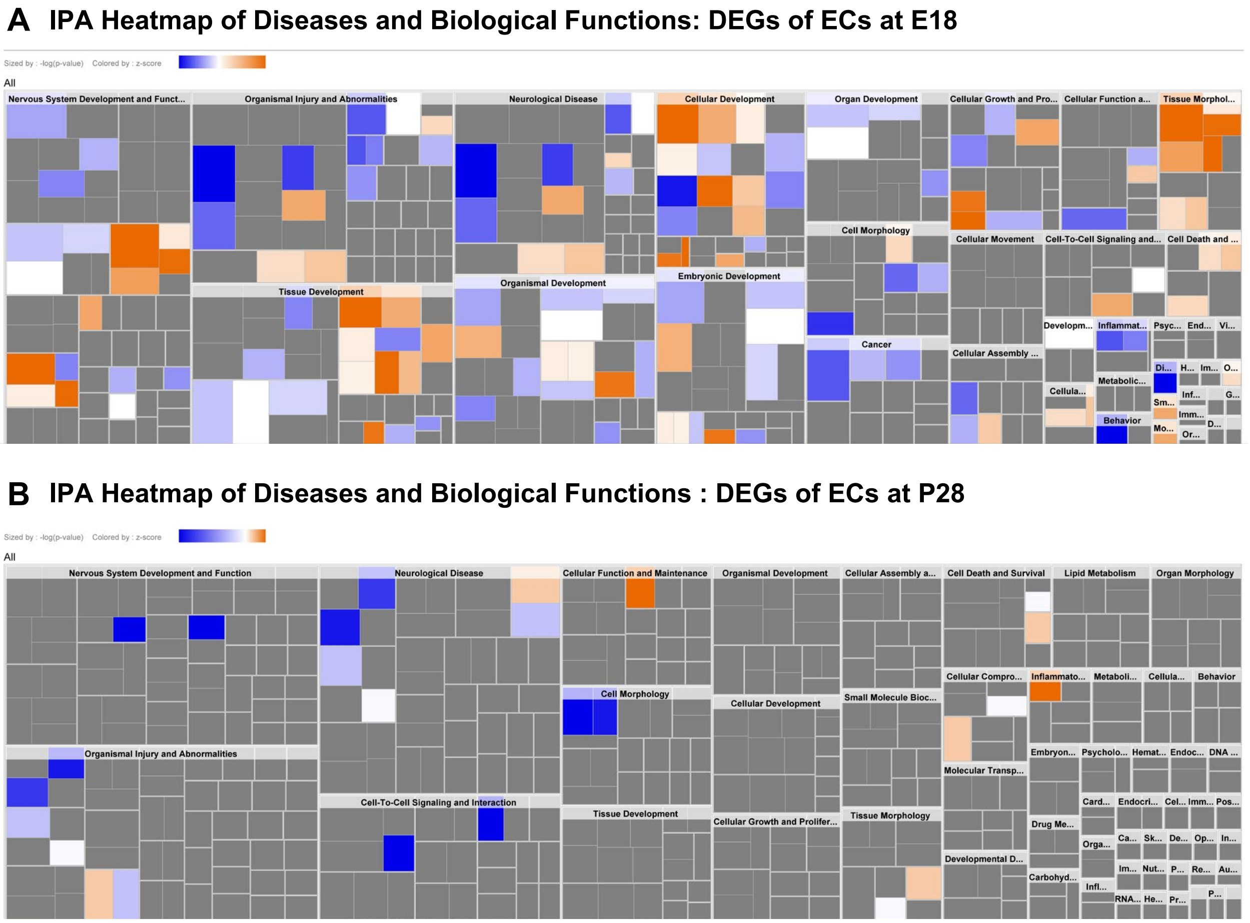

**Supplemental Figure 4. Diseases and biological functions associated with differentially expressed genes in endothelial cells at E18 and P28.** Categorical heatmap diseases and biological functions associated with DEGs in ECs at **a)** E18 and **b)** P28. Sections are sized by -log(p-value) and colored by z-score, with blue indicating decreased expression and orange displaying increased expression in PAE compared to SAC.

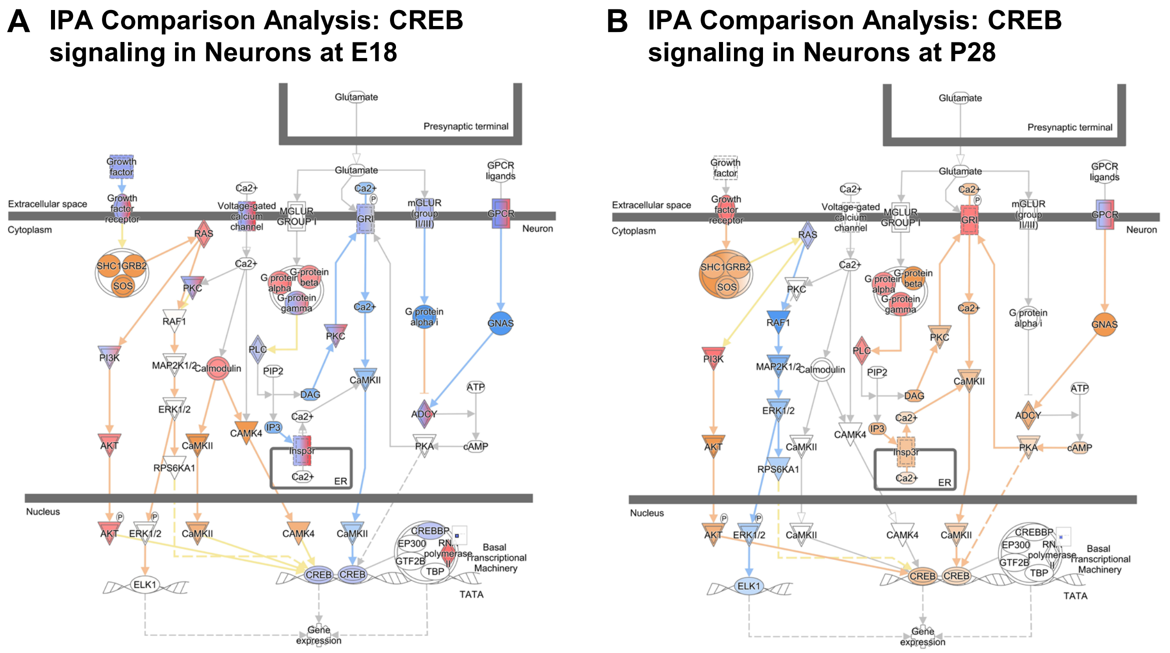

**Supplemental Figure 5. IPA comparison analysis of neuronal CREB signaling at E18 and P28.** Graphical depiction of predicted CREB signaling pathways in neurons at **a)** E18 and **b)** P28. Node colors represent measured changes, with red indicating increased measurements, light blue indicating decreased measurements, orange indicating predicted activation, and dark blue indicating predicted inhibition. Predicted relationships are encoded by line color and style: orange lines indicate activation and blue lines indicate inhibition, solid lines indicate direct relationships, and dashed lines indicate indirect relationships.

**A IPA Comparison Analysis: Ephrin signaling in Neurons at E18**

**B IPA Comparison Analysis: Ephrin signaling in Neurons at P28**

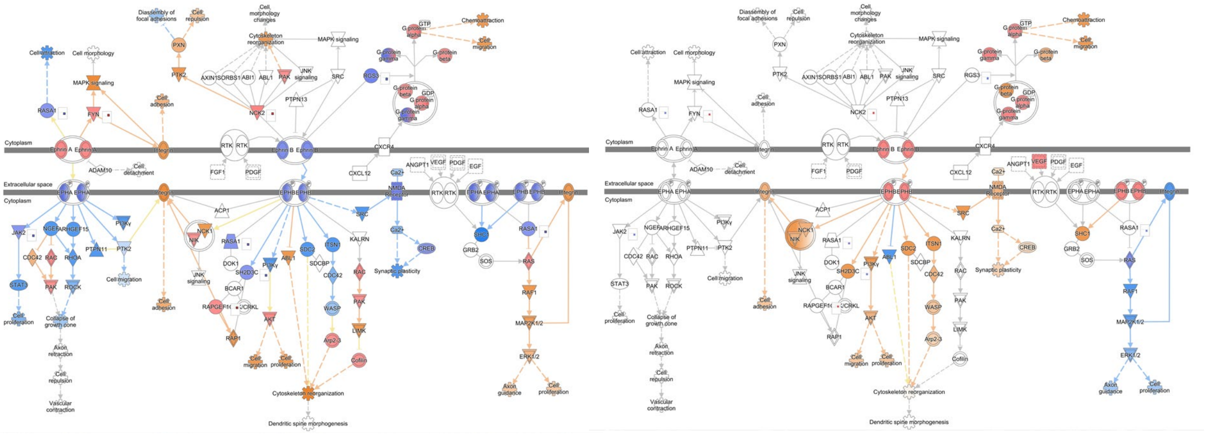

**Supplemental Figure 6. IPA comparison analysis of neuronal ephrin signaling at E18 and P28.** Graphical depiction of predicted ephrin signaling pathways in neurons at **a)** E18 and **b)** P28. Node colors represent measured changes, with red indicating increased measurements, light blue indicating decreased measurements, orange indicating predicted activation, and dark blue indicating predicted inhibition. Predicted relationships are encoded by line color and style: orange lines indicate activation and blue lines indicate inhibition, solid lines indicate direct relationships, and dashed lines indicate indirect relationships.

**A** **IPA Comparison Analysis: Netrin signaling in Neurons at E18**

**B** **IPA Comparison Analysis: Netrin signaling in Neurons at P28**

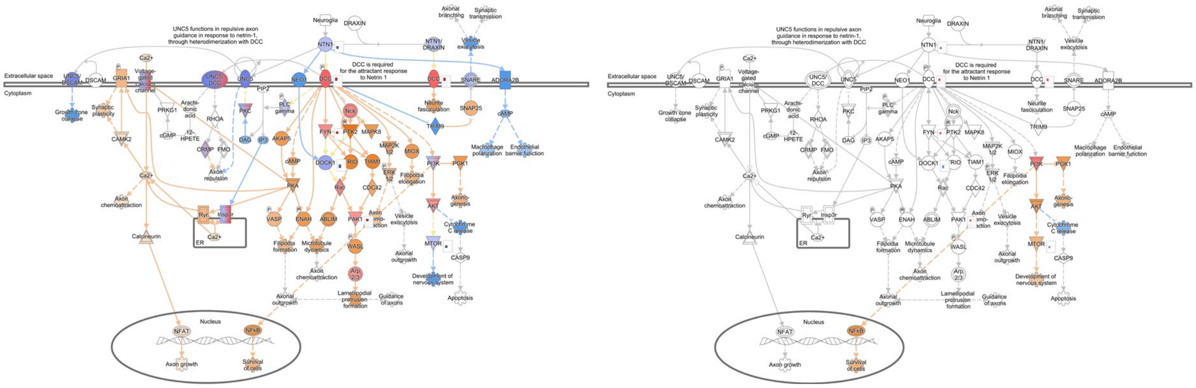

**Supplemental Figure 7. IPA comparison analysis of neuronal netrin signaling at E18 and P28.** Graphical depiction of predicted netrin signaling pathways in neurons at **a)** E18 and **b)** P28. Node colors represent measured changes, with red indicating increased measurements, light blue indicating decreased measurements, orange indicating predicted activation, and dark blue indicating predicted inhibition. Predicted relationships are encoded by line color and style: orange lines indicate activation and blue lines indicate inhibition, solid lines indicate direct relationships, and dashed lines indicate indirect relationships.

**A** IPA Heatmap of Diseases and Biological Functions: DEGs of Neurons at E18

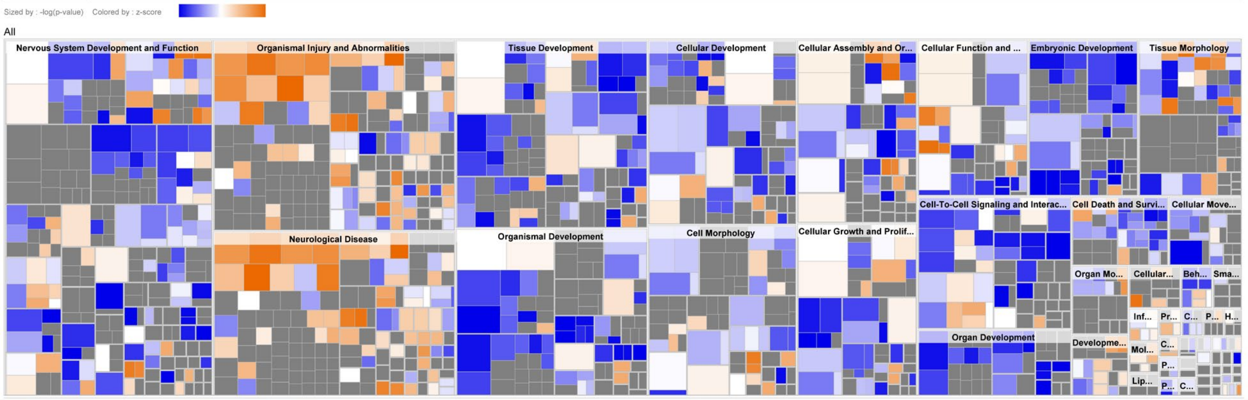

**B** IPA Heatmap of Diseases and Biological Functions : DEGs of Neurons at P28

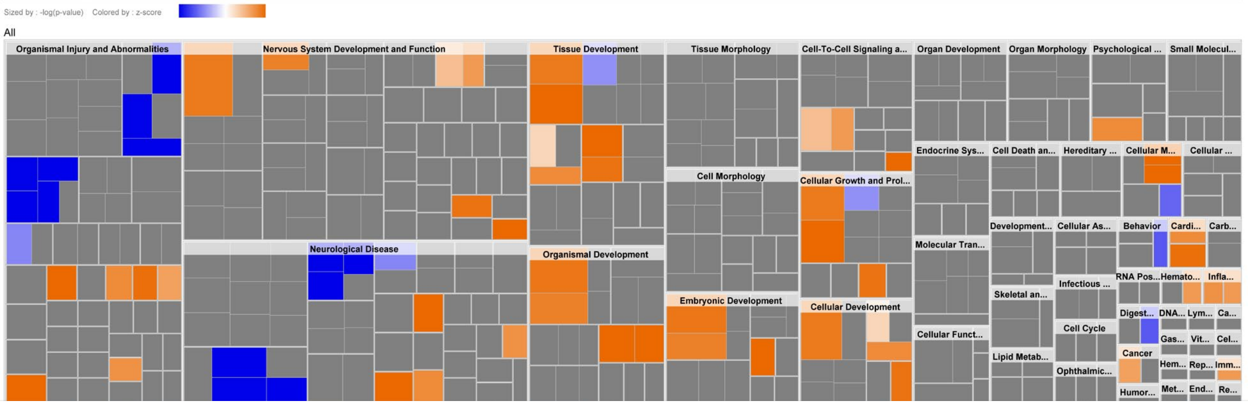

**Supplemental Figure 8. Diseases and biological functions associated with differentially expressed genes in neurons at E18 and P28.** Categorical heatmap of diseases and biological functions associated with DEGs in neurons at **a)** E18 and **b)** P28. Sections are sized by -log(p-value) and colored by z-score, with blue indicating decreased expression and orange displaying increased expression in PAE compared to SAC.

### Cell Chat: Ligand-receptor pair interaction between cells at E18 in SAC and PAE

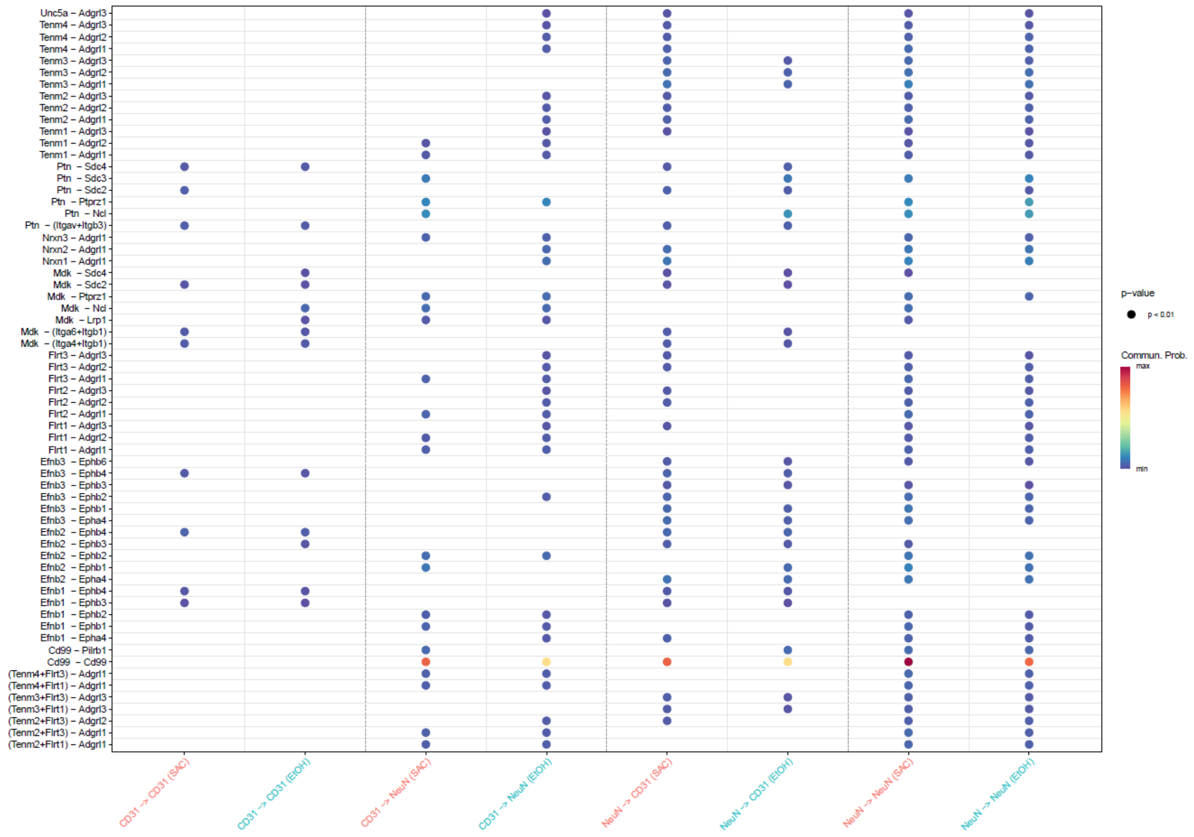

**Supplemental Figure 9. CellChat ligand-receptor pair interaction between neurovascular cells at E18 in SAC and PAE.** Bubble plot of the top ligand-receptor pairing pathways altered at E18 in ECs and neurons. Each row represents a ligand-receptor pair and each column represents a sending and receiving cell type direction, colored for each condition: SAC (red) and EtOH/PAE (green). Dot size indicates statistical significance (filled dot = p < 0.01). Dot color reflects communication probability, ranging from minimum (dark blue) to maximum (red).

**Cell Chat: Ligand-receptor pair interaction between cells at P28 in SAC and PAE**

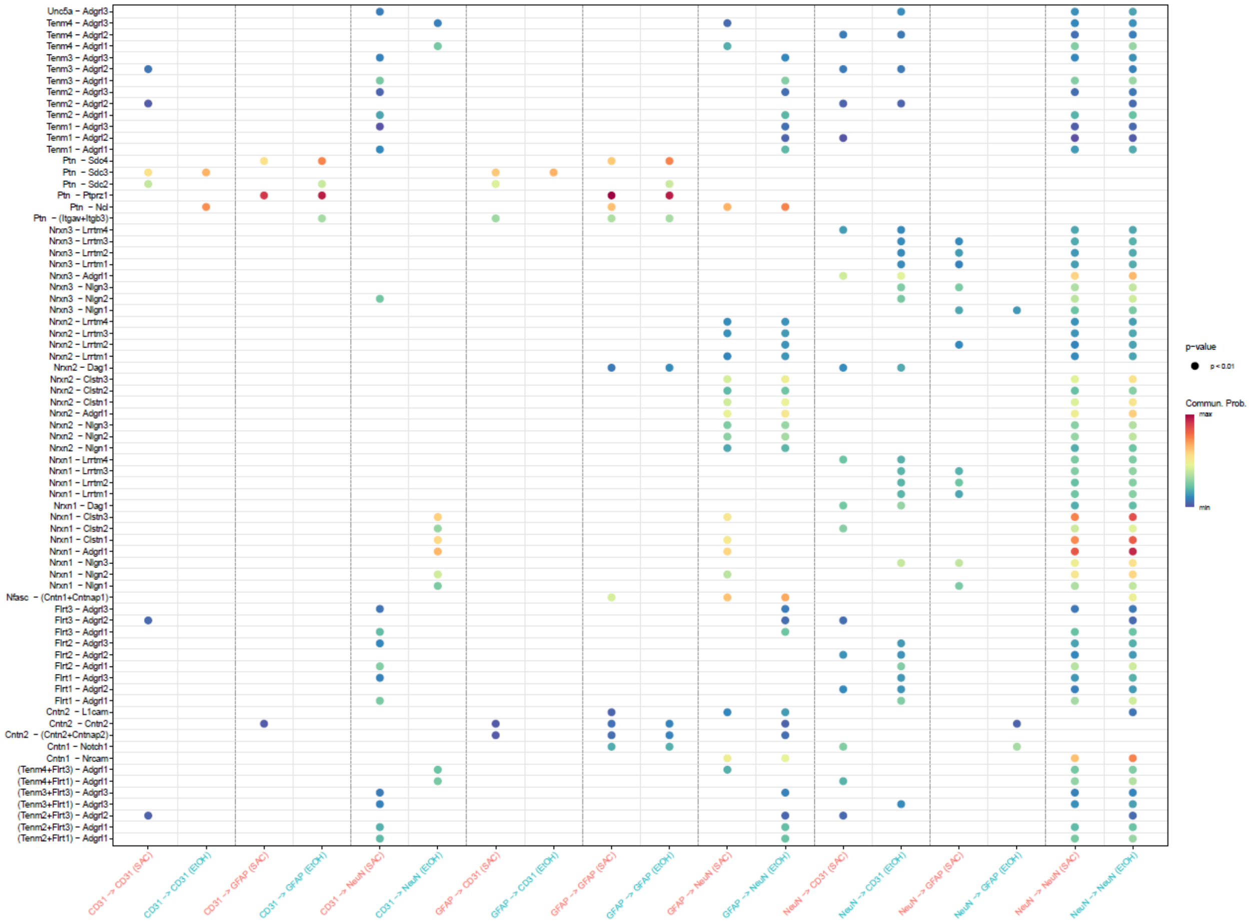

**Supplemental Figure 10. CellChat ligand-receptor pair interaction between neurovascular cells at P28 in SAC and PAE.** Bubble plot of the top ligand-receptor pairing pathways altered in ECs, neurons, and astrocytes. Each row represents a ligand-receptor pair and each column represents a sending and receiving cell type direction, colored for condition: SAC (red) and EtOH/PAE (green). Dot size indicates statistical significance (filled dot = p < 0.01). Dot color reflects communication probability, ranging from minimum (dark blue) to maximum (red).

**Supplementary Table 1: Differentially expressed genes between PAE and SAC in ECs at E18**

**Increased expression (1.2 FC, p<0.05)**

| Target nam | Log2 FC | Pvalue | Adjusted p | -log10 pval | -log10 adjusted pvalue |
|---|---|---|---|---|---|
| Clec2j | 1.977459 | 0.009353 | 0.466943 | 2.029030 | 0.330736 |
| Nktr | 1.626196 | 0.000009 | 0.128328 | 5.041760 | 0.891680 |
| Gm3233 | 1.582530 | 0.007249 | 0.459495 | 2.139734 | 0.337720 |
| Gm17455 | 1.574024 | 0.000938 | 0.346132 | 3.027738 | 0.460759 |
| Them7 | 1.563518 | 0.001030 | 0.346132 | 2.987002 | 0.460759 |
| S100a8 | 1.545703 | 0.036158 | 0.560743 | 1.441799 | 0.251236 |
| D17H6S56E | 1.519000 | 0.005226 | 0.447186 | 2.281800 | 0.349512 |
| Prl5a1 | 1.467743 | 0.000817 | 0.346132 | 3.087548 | 0.460759 |
| Sac3d1 | 1.460370 | 0.005680 | 0.450937 | 2.245639 | 0.345884 |
| Etv3l | 1.450157 | 0.016130 | 0.508849 | 1.792356 | 0.293411 |
| Tmem219 | 1.447305 | 0.000612 | 0.346132 | 3.213553 | 0.460759 |
| Rslcan18 | 1.441932 | 0.008890 | 0.459495 | 2.051076 | 0.337720 |
| Fhl2 | 1.439944 | 0.005666 | 0.450937 | 2.246745 | 0.345884 |
| Il6st | 1.438450 | 0.000393 | 0.346132 | 3.405057 | 0.460759 |
| Pomgnt2 | 1.424358 | 0.010748 | 0.477528 | 1.968665 | 0.321001 |
| Pxk | 1.412944 | 0.022872 | 0.512726 | 1.640690 | 0.290115 |
| Rnf212 | 1.398277 | 0.000930 | 0.346132 | 3.031373 | 0.460759 |
| Slc51b | 1.395426 | 0.001128 | 0.346132 | 2.947703 | 0.460759 |
| Pxt1 | 1.363972 | 0.019119 | 0.508849 | 1.718534 | 0.293411 |
| Isoc2a | 1.362265 | 0.025540 | 0.518442 | 1.592771 | 0.285299 |
| Cmtm8 | 1.358610 | 0.005451 | 0.447186 | 2.263510 | 0.349512 |
| Dzip1 | 1.354307 | 0.011230 | 0.477528 | 1.949602 | 0.321001 |
| 2310039H0 | 1.338466 | 0.011791 | 0.477528 | 1.928448 | 0.321001 |
| Sfn | 1.332530 | 0.011183 | 0.477528 | 1.951444 | 0.321001 |
| Hc | 1.332518 | 0.035866 | 0.559928 | 1.445312 | 0.251868 |
| Cfp | 1.332237 | 0.007949 | 0.459495 | 2.099713 | 0.337720 |
| Ptrhd1 | 1.325157 | 0.008251 | 0.459495 | 2.083489 | 0.337720 |
| Olfr297 | 1.323731 | 0.000913 | 0.346132 | 3.039456 | 0.460759 |
| Tspyl2 | 1.320076 | 0.004563 | 0.435548 | 2.340783 | 0.360964 |
| Galm | 1.318358 | 0.006292 | 0.454055 | 2.201195 | 0.342892 |
| Ccdc57 | 1.315236 | 0.008726 | 0.459495 | 2.059199 | 0.337720 |
| Nupl2 | 1.310610 | 0.027063 | 0.523236 | 1.567623 | 0.281303 |
| Cyba | 1.309571 | 0.023759 | 0.512726 | 1.624180 | 0.290115 |
| C030006K1 | 1.299650 | 0.021864 | 0.508849 | 1.660280 | 0.293411 |
| Olfr16 | 1.299358 | 0.000973 | 0.346132 | 3.011797 | 0.460759 |
| Col1a2 | 1.285490 | 0.005832 | 0.450937 | 2.234167 | 0.345884 |
| Slco3a1 | 1.283490 | 0.002291 | 0.385644 | 2.640020 | 0.413814 |

**Decreased expression (1.2 FC, p<0.05)**

| Target nam | Log2 FC | Pvalue | Adjusted p | -log10 pval | -log10 adjusted pvalue |
|---|---|---|---|---|---|
| Gm21828 | -1.979136 | 0.000179 | 0.346132 | 3.746475 | 0.460759 |
| Gm21783 | -1.933811 | 0.037388 | 0.566151 | 1.427267 | 0.247068 |
| Dlx2 | -1.736669 | 0.003847 | 0.408654 | 2.414903 | 0.388644 |
| Tab1 | -1.557298 | 0.000336 | 0.346132 | 3.474036 | 0.460759 |
| Fabp7 | -1.554786 | 0.001818 | 0.367785 | 2.740417 | 0.434406 |
| Gm31035 | -1.541530 | 0.000068 | 0.240122 | 4.167589 | 0.619569 |
| Rarres2 | -1.524927 | 0.000925 | 0.346132 | 3.033769 | 0.460759 |
| Bend6 | -1.454848 | 0.000638 | 0.346132 | 3.195186 | 0.460759 |
| Srsx | -1.415751 | 0.011362 | 0.477528 | 1.944555 | 0.321001 |
| Scyl3 | -1.396286 | 0.002656 | 0.399584 | 2.575774 | 0.398392 |
| Tns2 | -1.358964 | 0.002442 | 0.396032 | 2.612329 | 0.402270 |
| Sox9 | -1.321808 | 0.013031 | 0.489076 | 1.885018 | 0.310624 |
| Tns1 | -1.298557 | 0.019648 | 0.508849 | 1.706688 | 0.293411 |
| Btnl1 | -1.286100 | 0.001109 | 0.346132 | 2.954925 | 0.460759 |
| Ddx56 | -1.285140 | 0.002347 | 0.385644 | 2.629396 | 0.413814 |
| Cnbd2 | -1.282675 | 0.003218 | 0.402288 | 2.492378 | 0.395463 |
| Mylip | -1.281242 | 0.000762 | 0.346132 | 3.118267 | 0.460759 |
| Fmc1 | -1.272100 | 0.004074 | 0.422417 | 2.389979 | 0.374259 |
| Gm20815 | -1.265674 | 0.038085 | 0.566795 | 1.419241 | 0.246574 |
| Bcr | -1.255946 | 0.001151 | 0.346132 | 2.938742 | 0.460759 |
| Adm | -1.242422 | 0.011715 | 0.477528 | 1.931241 | 0.321001 |
| Tm2d1 | -1.233964 | 0.007705 | 0.459495 | 2.113223 | 0.337720 |
| Rapgefl1 | -1.230845 | 0.026038 | 0.518654 | 1.584400 | 0.285122 |
| Gm10256 | -1.227140 | 0.008677 | 0.459495 | 2.061609 | 0.337720 |
| Dtna | -1.219119 | 0.015594 | 0.508849 | 1.807054 | 0.293411 |
| Lipo2 | -1.216671 | 0.011356 | 0.477528 | 1.944783 | 0.321001 |
| Klk1b21 | -1.212723 | 0.024330 | 0.513619 | 1.613855 | 0.289359 |
| Vwa5b2 | -1.211059 | 0.000939 | 0.346132 | 3.027315 | 0.460759 |
| Hinfp | -1.209897 | 0.045299 | 0.579023 | 1.343911 | 0.237304 |
| Gm51592 | -1.201472 | 0.003259 | 0.402288 | 2.486903 | 0.395463 |
| Zfp637 | -1.194683 | 0.002650 | 0.399584 | 2.576754 | 0.398392 |
| Atp1b2 | -1.192604 | 0.005096 | 0.444459 | 2.292734 | 0.352168 |
| 4930467E2 | -1.188674 | 0.007625 | 0.459495 | 2.117746 | 0.337720 |
| Gm40469 | -1.179645 | 0.017298 | 0.508849 | 1.762002 | 0.293411 |
| Nr2e1 | -1.176221 | 0.005473 | 0.447186 | 2.261802 | 0.349512 |
| Cwf19l1 | -1.163084 | 0.001542 | 0.367785 | 2.812048 | 0.434406 |
| 9530068E0 | -1.154138 | 0.023859 | 0.512726 | 1.622355 | 0.290115 |

| | | | | | |
|---|---|---|---|---|---|
| Lep | 1.273277 | 0.017457 | 0.508849 | 1.758023 | 0.293411 |
| Ehd1 | 1.270426 | 0.010100 | 0.477528 | 1.995667 | 0.321001 |
| Il34 | 1.268197 | 0.025689 | 0.518654 | 1.590256 | 0.285122 |
| Cxcl10 | 1.261638 | 0.021946 | 0.508849 | 1.658637 | 0.293411 |
| Tmem252 | 1.258449 | 0.007431 | 0.459495 | 2.128976 | 0.337720 |
| Poc1a | 1.253743 | 0.009652 | 0.472983 | 2.015376 | 0.325154 |
| Wdpcp | 1.252036 | 0.009255 | 0.463691 | 2.033603 | 0.333772 |
| Olfr194 | 1.252036 | 0.013931 | 0.495749 | 1.856014 | 0.304738 |
| Cstdc5 | 1.247759 | 0.022346 | 0.510028 | 1.650796 | 0.292406 |
| Robo1 | 1.247405 | 0.016016 | 0.508849 | 1.795437 | 0.293411 |
| Rhbdd1 | 1.238972 | 0.020676 | 0.508849 | 1.684537 | 0.293411 |
| Rbfox3 | 1.235959 | 0.038102 | 0.566795 | 1.419048 | 0.246574 |
| Parm1 | 1.234128 | 0.005332 | 0.447186 | 2.273080 | 0.349512 |
| Ccn4 | 1.231610 | 0.003165 | 0.402288 | 2.499670 | 0.395463 |
| Ccl11 | 1.228759 | 0.007945 | 0.459495 | 2.099886 | 0.337720 |
| Trmt9b | 1.227956 | 0.039391 | 0.570881 | 1.404605 | 0.243454 |
| Tmcc1 | 1.221622 | 0.002800 | 0.399584 | 2.552838 | 0.398392 |
| Lpgat1 | 1.216306 | 0.001842 | 0.367785 | 2.734769 | 0.434406 |
| Siae | 1.214599 | 0.016909 | 0.508849 | 1.771880 | 0.293411 |
| Gtf2h2 | 1.213679 | 0.029849 | 0.536762 | 1.525075 | 0.270218 |
| Gemin6 | 1.211184 | 0.003714 | 0.408654 | 2.430206 | 0.388644 |
| Defa34 | 1.207518 | 0.000776 | 0.346132 | 3.110095 | 0.460759 |
| Fkbp14 | 1.207518 | 0.001372 | 0.362707 | 2.862670 | 0.440445 |
| St6galnac2 | 1.200157 | 0.020537 | 0.508849 | 1.687454 | 0.293411 |
| Atf1 | 1.198731 | 0.002467 | 0.396032 | 2.607868 | 0.402270 |
| Dbr1 | 1.198424 | 0.026545 | 0.522318 | 1.576021 | 0.282065 |
| Rcan1 | 1.194784 | 0.002776 | 0.399584 | 2.556624 | 0.398392 |
| Prdx4 | 1.187317 | 0.008249 | 0.459495 | 2.083577 | 0.337720 |
| Rce1 | 1.187092 | 0.037546 | 0.566795 | 1.425434 | 0.246574 |
| Spopl | 1.185678 | 0.007309 | 0.459495 | 2.136159 | 0.337720 |
| Gpr149 | 1.183145 | 0.007554 | 0.459495 | 2.121815 | 0.337720 |
| Nek7 | 1.181332 | 0.013847 | 0.495749 | 1.858632 | 0.304738 |
| Atp6v1c1 | 1.179956 | 0.023308 | 0.512726 | 1.632486 | 0.290115 |
| C130074G1 | 1.176706 | 0.006840 | 0.454055 | 2.164929 | 0.342892 |
| Begain | 1.176346 | 0.005975 | 0.451422 | 2.223657 | 0.345417 |
| Scgb1b30 | 1.167277 | 0.003506 | 0.407603 | 2.455139 | 0.389763 |
| Adamts15 | 1.167277 | 0.015841 | 0.508849 | 1.800224 | 0.293411 |
| Setd4 | 1.165570 | 0.018296 | 0.508849 | 1.737654 | 0.293411 |
| Tmed3 | 1.161904 | 0.010200 | 0.477528 | 1.991380 | 0.321001 |
| Fer1l6 | 1.157346 | 0.015894 | 0.508849 | 1.798771 | 0.293411 |
| Mypop | 1.157064 | 0.024508 | 0.513619 | 1.610693 | 0.289359 |

| | | | | | |
|---|---|---|---|---|---|
| Phgdh | -1.146047 | 0.032513 | 0.548023 | 1.487944 | 0.261201 |
| Vps35l | -1.138295 | 0.007858 | 0.459495 | 2.104667 | 0.337720 |
| Cd38 | -1.133594 | 0.009233 | 0.463691 | 2.034639 | 0.333772 |
| Zswim5 | -1.119177 | 0.019822 | 0.508849 | 1.702863 | 0.293411 |
| Cacng7 | -1.113591 | 0.010177 | 0.477528 | 1.992396 | 0.321001 |
| Tigd3 | -1.111393 | 0.011711 | 0.477528 | 1.931409 | 0.321001 |
| Prdm2 | -1.108928 | 0.010104 | 0.477528 | 1.995521 | 0.321001 |
| Ide | -1.101743 | 0.009850 | 0.476580 | 2.006562 | 0.321864 |
| Echs1 | -1.095283 | 0.014571 | 0.498513 | 1.836514 | 0.302323 |
| Krt2 | -1.090152 | 0.026049 | 0.518654 | 1.584203 | 0.285122 |
| Map3k20 | -1.086579 | 0.003513 | 0.407603 | 2.454284 | 0.389763 |
| Papolg | -1.082112 | 0.016889 | 0.508849 | 1.772395 | 0.293411 |
| Nr4a2 | -1.080962 | 0.040342 | 0.570893 | 1.394240 | 0.243445 |
| Prickle2 | -1.076361 | 0.008733 | 0.459495 | 2.058857 | 0.337720 |
| Zfp623 | -1.075406 | 0.029938 | 0.536762 | 1.523773 | 0.270218 |
| Lemd2 | -1.072923 | 0.042560 | 0.574283 | 1.371002 | 0.240874 |
| Nhp2 | -1.070033 | 0.003205 | 0.402288 | 2.494158 | 0.395463 |
| Itga6 | -1.069445 | 0.015112 | 0.508340 | 1.820677 | 0.293846 |
| Mbnl1 | -1.069433 | 0.021239 | 0.508849 | 1.672868 | 0.293411 |
| Doc2b | -1.068979 | 0.049871 | 0.592038 | 1.302153 | 0.227650 |
| Zfp978 | -1.067604 | 0.001891 | 0.367785 | 2.723286 | 0.434406 |
| Wipf2 | -1.064776 | 0.011861 | 0.477528 | 1.925890 | 0.321001 |
| Nfu1 | -1.062165 | 0.011171 | 0.477528 | 1.951927 | 0.321001 |
| 5730409E0 | -1.056991 | 0.046500 | 0.583451 | 1.332550 | 0.233995 |
| Serpinb3d | -1.055566 | 0.017167 | 0.508849 | 1.765301 | 0.293411 |
| Trafd1 | -1.054394 | 0.037740 | 0.566795 | 1.423197 | 0.246574 |
| Erbb4 | -1.052483 | 0.002175 | 0.385644 | 2.662469 | 0.413814 |
| Impact | -1.052433 | 0.028621 | 0.528456 | 1.543309 | 0.276991 |
| Cacng2 | -1.051430 | 0.040450 | 0.570922 | 1.393084 | 0.243423 |
| Nlrp1a | -1.050658 | 0.008847 | 0.459495 | 2.053200 | 0.337720 |
| Nt5dc2 | -1.049294 | 0.038972 | 0.569974 | 1.409249 | 0.244145 |
| Arl4c | -1.047899 | 0.039634 | 0.570881 | 1.401933 | 0.243454 |
| Rsbn1 | -1.044538 | 0.003824 | 0.408654 | 2.417449 | 0.388644 |
| Kcnc2 | -1.042220 | 0.013855 | 0.495749 | 1.858389 | 0.304738 |
| 4930402H2 | -1.041762 | 0.024164 | 0.513619 | 1.616823 | 0.289359 |
| Inafm2 | -1.041405 | 0.005410 | 0.447186 | 2.266834 | 0.349512 |
| Kifbp | -1.037178 | 0.037308 | 0.566146 | 1.428203 | 0.247072 |
| Lrig2 | -1.036615 | 0.019830 | 0.508849 | 1.702668 | 0.293411 |
| Ccdc18 | -1.034325 | 0.008804 | 0.459495 | 2.055334 | 0.337720 |
| Ttc7b | -1.031658 | 0.006584 | 0.454055 | 2.181490 | 0.342892 |
| Tmem19 | -1.025631 | 0.016679 | 0.508849 | 1.777831 | 0.293411 |

| | | | | | |
|---|---|---|---|---|---|
| Flt3l | 1.155117 | 0.011759 | 0.477528 | 1.929618 | 0.321001 |
| Prn | 1.154835 | 0.011085 | 0.477528 | 1.955273 | 0.321001 |
| Smim41 | 1.154543 | 0.026937 | 0.523236 | 1.569649 | 0.281303 |
| B3gat2 | 1.151691 | 0.049130 | 0.592038 | 1.308657 | 0.227650 |
| Sppl2a | 1.151410 | 0.001710 | 0.367785 | 2.767099 | 0.434406 |
| Nos1ap | 1.149984 | 0.008860 | 0.459495 | 2.052544 | 0.337720 |
| Dhcr24 | 1.149024 | 0.026007 | 0.518654 | 1.584906 | 0.285122 |
| Lrtm2 | 1.148277 | 0.008807 | 0.459495 | 2.055163 | 0.337720 |
| Ocstamp | 1.146851 | 0.014608 | 0.498513 | 1.835404 | 0.302323 |
| Sat1 | 1.145851 | 0.013623 | 0.495208 | 1.865712 | 0.305213 |
| AB124611 | 1.144622 | 0.036039 | 0.560124 | 1.443232 | 0.251716 |
| Cenpl | 1.144465 | 0.016011 | 0.508849 | 1.795592 | 0.293411 |
| Abra | 1.144330 | 0.020038 | 0.508849 | 1.698139 | 0.293411 |
| Rnf128 | 1.138956 | 0.006846 | 0.454055 | 2.164593 | 0.342892 |
| Abca6 | 1.134398 | 0.038269 | 0.566795 | 1.417151 | 0.246574 |
| Fkbp2 | 1.131048 | 0.003759 | 0.408654 | 2.424953 | 0.388644 |
| Gimap5 | 1.127318 | 0.030300 | 0.537364 | 1.518562 | 0.269731 |
| 2610528A1 | 1.127036 | 0.039719 | 0.570881 | 1.401001 | 0.243454 |
| Tm6sf1 | 1.125892 | 0.047055 | 0.585976 | 1.327391 | 0.232120 |
| Mfap4 | 1.125611 | 0.029512 | 0.533867 | 1.529997 | 0.272567 |
| Fam20a | 1.124650 | 0.016091 | 0.508849 | 1.793415 | 0.293411 |
| Ntn5 | 1.122771 | 0.017412 | 0.508849 | 1.759147 | 0.293411 |
| Ggcx | 1.121663 | 0.023711 | 0.512726 | 1.625054 | 0.290115 |
| Tspan13 | 1.118573 | 0.000983 | 0.346132 | 3.007394 | 0.460759 |
| Hesx1 | 1.117105 | 0.015350 | 0.508849 | 1.813898 | 0.293411 |
| Olfr322 | 1.116823 | 0.000713 | 0.346132 | 3.147045 | 0.460759 |
| Adcy5 | 1.115398 | 0.005544 | 0.450117 | 2.256206 | 0.346674 |
| Rhox2f | 1.115398 | 0.036522 | 0.562320 | 1.437450 | 0.250017 |
| Ebi3 | 1.114876 | 0.022628 | 0.512726 | 1.645360 | 0.290115 |
| Slc26a2 | 1.113972 | 0.017904 | 0.508849 | 1.747047 | 0.293411 |
| Dhx35 | 1.113972 | 0.003768 | 0.408654 | 2.423944 | 0.388644 |
| Atp9b | 1.113690 | 0.022576 | 0.512726 | 1.646358 | 0.290115 |
| Tlnrd1 | 1.111450 | 0.045978 | 0.583451 | 1.337447 | 0.233995 |
| Peak1 | 1.110490 | 0.010611 | 0.477528 | 1.974248 | 0.321001 |
| Traip | 1.106610 | 0.006427 | 0.454055 | 2.191970 | 0.342892 |
| Olfr503 | 1.106610 | 0.041193 | 0.572249 | 1.385175 | 0.242415 |
| Olfr799 | 1.106610 | 0.027985 | 0.527961 | 1.553074 | 0.277398 |
| Chd9 | 1.106271 | 0.008931 | 0.459495 | 2.049118 | 0.337720 |
| Zfp691 | 1.102663 | 0.019825 | 0.508849 | 1.702786 | 0.293411 |
| Cox6b2 | 1.101530 | 0.030526 | 0.537364 | 1.515327 | 0.269731 |
| Echdc3 | 1.101530 | 0.033171 | 0.548500 | 1.479235 | 0.260823 |

| | | | | | |
|---|---|---|---|---|---|
| Cngb1 | -1.024112 | 0.004862 | 0.443121 | 2.313226 | 0.353477 |
| Kiss1r | -1.014714 | 0.035703 | 0.559928 | 1.447301 | 0.251868 |
| Cyb5d2 | -1.010766 | 0.024268 | 0.513619 | 1.614974 | 0.289359 |
| Gm5640 | -1.007591 | 0.044314 | 0.574283 | 1.353455 | 0.240874 |
| Ackr3 | -1.004714 | 0.018013 | 0.508849 | 1.744403 | 0.293411 |
| Mon1a | -1.004647 | 0.019368 | 0.508849 | 1.712916 | 0.293411 |
| Tmed7 | -1.002781 | 0.002272 | 0.385644 | 2.643602 | 0.413814 |
| Gcdh | -1.001911 | 0.025462 | 0.518442 | 1.594107 | 0.285299 |
| Kpna2 | -1.000113 | 0.031238 | 0.537978 | 1.505322 | 0.269235 |
| Etnk2 | -0.999739 | 0.029678 | 0.534804 | 1.527570 | 0.271805 |
| Arhgef4 | -0.998778 | 0.000911 | 0.346132 | 3.040708 | 0.460759 |
| Ube2r2 | -0.996620 | 0.020210 | 0.508849 | 1.694434 | 0.293411 |
| Pdcd1lg2 | -0.996606 | 0.027703 | 0.526055 | 1.557476 | 0.278969 |
| Trim11 | -0.996606 | 0.039140 | 0.570881 | 1.407384 | 0.243454 |
| Pts | -0.996606 | 0.020355 | 0.508849 | 1.691338 | 0.293411 |
| Afap1 | -0.996338 | 0.017283 | 0.508849 | 1.762382 | 0.293411 |
| Lrpap1 | -0.993687 | 0.016375 | 0.508849 | 1.785821 | 0.293411 |
| Zic1 | -0.992921 | 0.014356 | 0.498513 | 1.842978 | 0.302323 |
| Eif2b2 | -0.991008 | 0.020880 | 0.508849 | 1.680267 | 0.293411 |
| Cisd1 | -0.990777 | 0.040106 | 0.570881 | 1.396795 | 0.243454 |
| Igsf9b | -0.988565 | 0.010150 | 0.477528 | 1.993537 | 0.321001 |
| Vcan | -0.986699 | 0.049701 | 0.592038 | 1.303639 | 0.227650 |
| Ttc38 | -0.986393 | 0.006444 | 0.454055 | 2.190839 | 0.342892 |
| Sall1 | -0.984686 | 0.006155 | 0.454055 | 2.210751 | 0.342892 |
| Ctnnal1 | -0.984618 | 0.008801 | 0.459495 | 2.055478 | 0.337720 |
| Ndufaf2 | -0.984618 | 0.048436 | 0.589415 | 1.314827 | 0.229579 |
| Trim9 | -0.983871 | 0.006568 | 0.454055 | 2.182569 | 0.342892 |
| Aadacl2fm1 | -0.983327 | 0.011458 | 0.477528 | 1.940882 | 0.321001 |
| Nlrp4e | -0.982048 | 0.020459 | 0.508849 | 1.689111 | 0.293411 |
| 4930453N2 | -0.980273 | 0.004308 | 0.422624 | 2.365764 | 0.374046 |
| Ier3ip1 | -0.978659 | 0.024560 | 0.513619 | 1.609770 | 0.289359 |
| Zfp280d | -0.976898 | 0.012722 | 0.486498 | 1.895428 | 0.312919 |
| Kcns1 | -0.975219 | 0.011534 | 0.477528 | 1.938030 | 0.321001 |
| Lce1b | -0.974711 | 0.012858 | 0.487219 | 1.890818 | 0.312276 |
| L3mbtl3 | -0.973684 | 0.014548 | 0.498513 | 1.837208 | 0.302323 |
| Olfr1313 | -0.972232 | 0.018306 | 0.508849 | 1.737416 | 0.293411 |
| Nr2f1 | -0.970935 | 0.006545 | 0.454055 | 2.184097 | 0.342892 |
| Cd200r1 | -0.970060 | 0.028485 | 0.528131 | 1.545384 | 0.277259 |
| Elp5 | -0.967470 | 0.017943 | 0.508849 | 1.746107 | 0.293411 |
| Gm52436 | -0.966690 | 0.001692 | 0.367785 | 2.771577 | 0.434406 |
| Tob1 | -0.965924 | 0.017124 | 0.508849 | 1.766400 | 0.293411 |

| | | | | | |
|---|---|---|---|---|---|
| Gnb1l | 1.099812 | 0.011402 | 0.477528 | 1.943023 | 0.321001 |
| Pbld2 | 1.096397 | 0.035039 | 0.555332 | 1.455454 | 0.255447 |
| Rb1cc1 | 1.095864 | 0.004183 | 0.422417 | 2.378480 | 0.374259 |
| Slc25a10 | 1.094904 | 0.008749 | 0.459495 | 2.058059 | 0.337720 |
| Ddx43 | 1.092450 | 0.011759 | 0.477528 | 1.929626 | 0.321001 |
| Lmln | 1.092450 | 0.009663 | 0.472983 | 2.014874 | 0.325154 |
| Actr6 | 1.091024 | 0.040258 | 0.570881 | 1.395152 | 0.243454 |
| Poc5 | 1.089317 | 0.044743 | 0.575184 | 1.349276 | 0.240193 |
| Cdca4 | 1.089249 | 0.014175 | 0.498168 | 1.848479 | 0.302624 |
| Dact2 | 1.086184 | 0.008480 | 0.459495 | 2.071602 | 0.337720 |
| Colca2 | 1.085370 | 0.016598 | 0.508849 | 1.779937 | 0.293411 |
| Olfr332 | 1.085370 | 0.021946 | 0.508849 | 1.658653 | 0.293411 |
| Cd177 | 1.084409 | 0.016906 | 0.508849 | 1.771952 | 0.293411 |
| Calm4 | 1.082530 | 0.012984 | 0.489076 | 1.886582 | 0.310624 |
| Egfl6 | 1.082237 | 0.004081 | 0.422417 | 2.389218 | 0.374259 |
| Herc6 | 1.082237 | 0.005813 | 0.450937 | 2.235595 | 0.345884 |
| Galnt9 | 1.082237 | 0.039794 | 0.570881 | 1.400179 | 0.243454 |
| Abcc12 | 1.081422 | 0.047748 | 0.587289 | 1.321049 | 0.231148 |
| Ube2d2b | 1.080931 | 0.012244 | 0.481842 | 1.912082 | 0.317096 |
| Pla2g12b | 1.080811 | 0.017110 | 0.508849 | 1.766757 | 0.293411 |
| Gask1b | 1.078582 | 0.006428 | 0.454055 | 2.191894 | 0.342892 |
| B3galnt1 | 1.076864 | 0.004229 | 0.422417 | 2.373779 | 0.374259 |
| Zfp831 | 1.075157 | 0.000932 | 0.346132 | 3.030816 | 0.460759 |
| Ctxn2 | 1.075157 | 0.016053 | 0.508849 | 1.794433 | 0.293411 |
| Matn3 | 1.073731 | 0.000436 | 0.346132 | 3.360633 | 0.460759 |
| B4galnt2 | 1.073731 | 0.024291 | 0.513619 | 1.614547 | 0.289359 |
| Alyref2 | 1.071956 | 0.048040 | 0.587289 | 1.318398 | 0.231148 |
| Ell3 | 1.069784 | 0.022425 | 0.510999 | 1.649269 | 0.291580 |
| Khk | 1.068358 | 0.018276 | 0.508849 | 1.738120 | 0.293411 |
| Frmpd1 | 1.068358 | 0.021121 | 0.508849 | 1.675276 | 0.293411 |
| Pml | 1.066651 | 0.027073 | 0.523236 | 1.567466 | 0.281303 |
| Noxa1 | 1.061289 | 0.047264 | 0.585976 | 1.325470 | 0.232120 |
| Syt10 | 1.060996 | 0.005065 | 0.444428 | 2.295453 | 0.352199 |
| Tent4b | 1.059571 | 0.041506 | 0.572650 | 1.381891 | 0.242111 |
| Mfsd4b5 | 1.059571 | 0.044278 | 0.574283 | 1.353812 | 0.240874 |
| Spg20 | 1.058892 | 0.041567 | 0.572694 | 1.381251 | 0.242077 |
| Pld4 | 1.058438 | 0.029039 | 0.529730 | 1.537014 | 0.275945 |
| Adam29 | 1.054731 | 0.020204 | 0.508849 | 1.694571 | 0.293411 |
| Rorb | 1.053959 | 0.012774 | 0.486498 | 1.893667 | 0.312919 |
| Shh | 1.053916 | 0.021467 | 0.508849 | 1.668219 | 0.293411 |
| 4930595D1 | 1.053916 | 0.017520 | 0.508849 | 1.756467 | 0.293411 |

| | | | | | |
|---|---|---|---|---|---|
| Fam207a | -0.965152 | 0.035861 | 0.559928 | 1.445381 | 0.251868 |
| Dhx40 | -0.963464 | 0.038518 | 0.566795 | 1.414339 | 0.246574 |
| Serpina3b | -0.962723 | 0.006231 | 0.454055 | 2.205473 | 0.342892 |
| Rgs4 | -0.958216 | 0.012044 | 0.477528 | 1.919221 | 0.321001 |
| Dnah7b | -0.957883 | 0.016787 | 0.508849 | 1.775038 | 0.293411 |
| Pramel46 | -0.952767 | 0.011467 | 0.477528 | 1.940537 | 0.321001 |
| Crat | -0.951110 | 0.044466 | 0.574283 | 1.351970 | 0.240874 |
| Zfp804a | -0.951060 | 0.023444 | 0.512726 | 1.629967 | 0.290115 |
| Ftsj3 | -0.950338 | 0.037257 | 0.566132 | 1.428786 | 0.247083 |
| Sall4 | -0.948563 | 0.036146 | 0.560743 | 1.441944 | 0.251236 |
| Cfap46 | -0.944726 | 0.047956 | 0.587289 | 1.319161 | 0.231148 |
| Ypel1 | -0.943737 | 0.022758 | 0.512726 | 1.642867 | 0.290115 |
| Ptdss1 | -0.943723 | 0.030904 | 0.537693 | 1.509991 | 0.269466 |
| Bcl2 | -0.943258 | 0.038486 | 0.566795 | 1.414699 | 0.246574 |
| Cep70 | -0.942554 | 0.042984 | 0.574283 | 1.366696 | 0.240874 |
| Trit1 | -0.941244 | 0.012768 | 0.486498 | 1.893874 | 0.312919 |
| Map1a | -0.939832 | 0.017854 | 0.508849 | 1.748273 | 0.293411 |
| Hmox1 | -0.939818 | 0.044107 | 0.574283 | 1.355488 | 0.240874 |
| Cdkl3 | -0.937953 | 0.038471 | 0.566795 | 1.414869 | 0.246574 |
| Bicd2 | -0.937249 | 0.015586 | 0.508849 | 1.807274 | 0.293411 |
| Mtmr1 | -0.933376 | 0.025646 | 0.518654 | 1.590979 | 0.285122 |
| Reg1 | -0.931380 | 0.010628 | 0.477528 | 1.973551 | 0.321001 |
| Mindy4 | -0.929912 | 0.036538 | 0.562320 | 1.437255 | 0.250017 |
| Pheta1 | -0.929912 | 0.021667 | 0.508849 | 1.664208 | 0.293411 |
| Kif24 | -0.929605 | 0.022813 | 0.512726 | 1.641824 | 0.290115 |
| Fam163b | -0.928791 | 0.039696 | 0.570881 | 1.401255 | 0.243454 |
| Gusb | -0.927433 | 0.012863 | 0.487219 | 1.890648 | 0.312276 |
| Usp24 | -0.927433 | 0.014251 | 0.498513 | 1.846165 | 0.302323 |
| Iars2 | -0.923088 | 0.002344 | 0.385644 | 2.630114 | 0.413814 |
| Fbxo28 | -0.921871 | 0.012775 | 0.486498 | 1.893626 | 0.312919 |
| Pds5b | -0.921231 | 0.040287 | 0.570881 | 1.394840 | 0.243454 |
| Tafa3 | -0.917220 | 0.010387 | 0.477528 | 1.983506 | 0.321001 |
| Mamstr | -0.917220 | 0.016335 | 0.508849 | 1.786888 | 0.293411 |
| Gm35498 | -0.910140 | 0.008805 | 0.459495 | 2.055274 | 0.337720 |
| Slc20a1 | -0.910140 | 0.018556 | 0.508849 | 1.731514 | 0.293411 |
| Taf3 | -0.908829 | 0.036671 | 0.562521 | 1.435682 | 0.249861 |
| Fam124a | -0.908170 | 0.025725 | 0.518654 | 1.589641 | 0.285122 |
| Shc3 | -0.907249 | 0.024067 | 0.512870 | 1.618581 | 0.289993 |
| Rnf122 | -0.905232 | 0.020807 | 0.508849 | 1.681786 | 0.293411 |
| Cul7 | -0.904070 | 0.005006 | 0.444428 | 2.300505 | 0.352199 |
| C1qtnf4 | -0.903017 | 0.002872 | 0.402288 | 2.541797 | 0.395463 |

| | | | | | |
|---|---|---|---|---|---|
| Cd79b | 1.053916 | 0.038769 | 0.568184 | 1.411514 | 0.245511 |
| Gm51965 | 1.053579 | 0.027784 | 0.526769 | 1.556202 | 0.278380 |
| Mus81 | 1.052490 | 0.006839 | 0.454055 | 2.165020 | 0.342892 |
| Gsg1l2 | 1.051879 | 0.022152 | 0.508849 | 1.654595 | 0.293411 |
| Opcml | 1.051462 | 0.025293 | 0.518442 | 1.597005 | 0.285299 |
| Reln | 1.051076 | 0.004396 | 0.428364 | 2.356900 | 0.368187 |
| Caskin1 | 1.051076 | 0.007784 | 0.459495 | 2.108813 | 0.337720 |
| Abcc1 | 1.050783 | 0.033972 | 0.550478 | 1.468875 | 0.259260 |
| Zbtb6 | 1.047932 | 0.049421 | 0.592038 | 1.306088 | 0.227650 |
| Mfsd4a | 1.044277 | 0.018836 | 0.508849 | 1.725018 | 0.293411 |
| Msh3 | 1.044277 | 0.022302 | 0.509834 | 1.651664 | 0.292571 |
| Lgals9 | 1.043827 | 0.019856 | 0.508849 | 1.702103 | 0.293411 |
| Prdm1 | 1.043317 | 0.039242 | 0.570881 | 1.406249 | 0.243454 |
| Ripor2 | 1.042277 | 0.016974 | 0.508849 | 1.770207 | 0.293411 |
| Wfs1 | 1.040570 | 0.016399 | 0.508849 | 1.785179 | 0.293411 |
| Serping1 | 1.040570 | 0.033564 | 0.550478 | 1.474128 | 0.259260 |
| Txlna | 1.040570 | 0.009593 | 0.472983 | 2.018029 | 0.325154 |
| 1700013F0 | 1.036904 | 0.010303 | 0.477528 | 1.987048 | 0.321001 |
| Cnpy2 | 1.034879 | 0.033194 | 0.548500 | 1.478938 | 0.260823 |
| Mtfp1 | 1.034518 | 0.031428 | 0.539514 | 1.502678 | 0.267997 |
| Utp11 | 1.033771 | 0.020270 | 0.508849 | 1.693146 | 0.293411 |
| Olfr458 | 1.033490 | 0.013114 | 0.489076 | 1.882255 | 0.310624 |
| 1110017D1 | 1.033490 | 0.026861 | 0.523236 | 1.570885 | 0.281303 |
| Lrrc40 | 1.032289 | 0.011535 | 0.477528 | 1.937997 | 0.321001 |
| Slc15a4 | 1.032064 | 0.043505 | 0.574283 | 1.361466 | 0.240874 |
| Fcf1 | 1.030582 | 0.048727 | 0.590861 | 1.312234 | 0.228515 |
| Tmem39a | 1.030357 | 0.013581 | 0.495208 | 1.867077 | 0.305213 |
| Adgrd1 | 1.028117 | 0.010693 | 0.477528 | 1.970886 | 0.321001 |
| Arpin | 1.028117 | 0.037180 | 0.566132 | 1.429686 | 0.247083 |
| Ckmt2 | 1.026703 | 0.034171 | 0.550478 | 1.466341 | 0.259260 |
| Tmem104 | 1.026691 | 0.019010 | 0.508849 | 1.721027 | 0.293411 |
| Cck | 1.025266 | 0.018081 | 0.508849 | 1.742779 | 0.293411 |
| Hemk1 | 1.023851 | 0.001880 | 0.367785 | 2.725753 | 0.434406 |
| Prkg2 | 1.023277 | 0.032764 | 0.548023 | 1.484599 | 0.261201 |
| Aldh5a1 | 1.022755 | 0.013253 | 0.491435 | 1.877690 | 0.308534 |
| Myocos | 1.021851 | 0.030438 | 0.537364 | 1.516581 | 0.269731 |
| Trp53bp2 | 1.021329 | 0.021430 | 0.508849 | 1.668987 | 0.293411 |
| Rit2 | 1.021329 | 0.005860 | 0.450937 | 2.232115 | 0.345884 |
| Sec11a | 1.020477 | 0.024597 | 0.513619 | 1.609124 | 0.289359 |
| Itgam | 1.019622 | 0.021280 | 0.508849 | 1.672030 | 0.293411 |
| Rad54l | 1.019330 | 0.013856 | 0.495749 | 1.858348 | 0.304738 |

| | | | | | |
|---|---|---|---|---|---|
| Gm52433 | -0.902908 | 0.000029 | 0.138592 | 4.531221 | 0.858262 |
| Zfand3 | -0.902285 | 0.021782 | 0.508849 | 1.661896 | 0.293411 |
| Prickle1 | -0.901310 | 0.031772 | 0.541568 | 1.497960 | 0.266347 |
| Pak4 | -0.899927 | 0.014789 | 0.502249 | 1.830068 | 0.299081 |
| Rab3c | -0.894520 | 0.044414 | 0.574283 | 1.352484 | 0.240874 |
| Fam219a | -0.893525 | 0.036479 | 0.562320 | 1.437955 | 0.250017 |
| Tomm70a | -0.892804 | 0.016264 | 0.508849 | 1.788770 | 0.293411 |
| Gnai1 | -0.887033 | 0.015741 | 0.508849 | 1.802955 | 0.293411 |
| Paip2 | -0.884599 | 0.038435 | 0.566795 | 1.415277 | 0.246574 |
| Frmd4b | -0.884507 | 0.023036 | 0.512726 | 1.637600 | 0.290115 |
| Mms22l | -0.880872 | 0.035969 | 0.559928 | 1.444072 | 0.251868 |
| Cyp4f16 | -0.880461 | 0.002752 | 0.399584 | 2.560281 | 0.398392 |
| Zfp740 | -0.879923 | 0.008910 | 0.459495 | 2.050104 | 0.337720 |
| Dimt1 | -0.878750 | 0.008875 | 0.459495 | 2.051850 | 0.337720 |
| Olfr1445 | -0.877726 | 0.001640 | 0.367785 | 2.785164 | 0.434406 |
| Tcf15 | -0.874739 | 0.024577 | 0.513619 | 1.609471 | 0.289359 |
| Nostrin | -0.866561 | 0.039307 | 0.570881 | 1.405527 | 0.243454 |
| Pex13 | -0.863125 | 0.021939 | 0.508849 | 1.658783 | 0.293411 |
| Sh3rf1 | -0.862250 | 0.024843 | 0.513889 | 1.604791 | 0.289131 |
| Dcun1d4 | -0.859748 | 0.032097 | 0.543729 | 1.493532 | 0.264618 |
| Adig | -0.859618 | 0.010611 | 0.477528 | 1.974261 | 0.321001 |
| Smyd4 | -0.859618 | 0.007719 | 0.459495 | 2.112454 | 0.337720 |
| Smim22 | -0.858989 | 0.034782 | 0.552758 | 1.458644 | 0.257465 |
| Dgke | -0.858260 | 0.005058 | 0.444428 | 2.295994 | 0.352199 |
| Patz1 | -0.855084 | 0.000345 | 0.346132 | 3.462073 | 0.460759 |
| Usf3 | -0.854313 | 0.010490 | 0.477528 | 1.979220 | 0.321001 |
| Dnal4 | -0.854313 | 0.018618 | 0.508849 | 1.730077 | 0.293411 |
| Eif1b | -0.854252 | 0.001989 | 0.369738 | 2.701373 | 0.432105 |
| Bhlhe40 | -0.851187 | 0.035631 | 0.559928 | 1.448176 | 0.251868 |
| Npnt | -0.849405 | 0.023959 | 0.512726 | 1.620539 | 0.290115 |
| Cnot7 | -0.846818 | 0.003973 | 0.418918 | 2.400847 | 0.377871 |
| Sh3bp4 | -0.846536 | 0.017870 | 0.508849 | 1.747886 | 0.293411 |
| Ppp1r12a | -0.843573 | 0.027815 | 0.526769 | 1.555722 | 0.278380 |
| Syde1 | -0.840924 | 0.008325 | 0.459495 | 2.079627 | 0.337720 |
| Hps3 | -0.840924 | 0.017468 | 0.508849 | 1.757751 | 0.293411 |
| Ankfy1 | -0.839175 | 0.004943 | 0.444428 | 2.306038 | 0.352199 |
| Chst14 | -0.838794 | 0.038239 | 0.566795 | 1.417491 | 0.246574 |
| Chst15 | -0.837723 | 0.035062 | 0.555332 | 1.455163 | 0.255447 |
| 1110004F1 | -0.837154 | 0.025519 | 0.518442 | 1.593140 | 0.285299 |
| Tpst1 | -0.835923 | 0.041796 | 0.572694 | 1.378861 | 0.242077 |
| Magoh | -0.835265 | 0.046534 | 0.583451 | 1.332226 | 0.233995 |

| | | | | | |
|---|---|---|---|---|---|
| Vps37c | 1.017904 | 0.028839 | 0.528456 | 1.540017 | 0.276991 |
| Lca5 | 1.013289 | 0.040468 | 0.570922 | 1.392888 | 0.243423 |
| Rpgrip1 | 1.013064 | 0.024373 | 0.513619 | 1.613083 | 0.289359 |
| Tbr1 | 1.012976 | 0.015294 | 0.508849 | 1.815483 | 0.293411 |
| Med8 | 1.012222 | 0.021004 | 0.508849 | 1.677706 | 0.293411 |
| Gins2 | 1.011638 | 0.020272 | 0.508849 | 1.693102 | 0.293411 |
| Rnf113a1 | 1.009409 | 0.012700 | 0.486498 | 1.896185 | 0.312919 |
| Klhl22 | 1.007984 | 0.001949 | 0.368901 | 2.710255 | 0.433090 |
| Slc25a44 | 1.007691 | 0.022223 | 0.508849 | 1.653206 | 0.293411 |
| Zfp865 | 1.002180 | 0.030586 | 0.537364 | 1.514477 | 0.269731 |
| Olfr1450 | 1.002036 | 0.049069 | 0.592007 | 1.309197 | 0.227673 |
| Bmp3 | 1.002036 | 0.025757 | 0.518654 | 1.589111 | 0.285122 |
| 1700037C1 | 1.002036 | 0.043055 | 0.574283 | 1.365973 | 0.240874 |
| 5730522E0 | 1.002036 | 0.026475 | 0.522318 | 1.577164 | 0.282065 |
| Tas2r109 | 1.000611 | 0.034079 | 0.550478 | 1.467516 | 0.259260 |
| 1700129C0 | 0.999128 | 0.007787 | 0.459495 | 2.108607 | 0.337720 |
| Olfr684 | 0.998903 | 0.033006 | 0.548023 | 1.481408 | 0.261201 |
| Dnajc2 | 0.997353 | 0.019224 | 0.508849 | 1.716149 | 0.293411 |
| Tmem86a | 0.996663 | 0.014501 | 0.498513 | 1.838590 | 0.302323 |
| Lss | 0.995249 | 0.047924 | 0.587289 | 1.319444 | 0.231148 |
| Fcor | 0.995238 | 0.029267 | 0.532148 | 1.533627 | 0.273968 |
| Serpinb11 | 0.993812 | 0.039594 | 0.570881 | 1.402375 | 0.243454 |
| Borcs7 | 0.993530 | 0.042509 | 0.574283 | 1.371523 | 0.240874 |
| Mfsd4b2 | 0.991823 | 0.013671 | 0.495245 | 1.864196 | 0.305180 |
| Tac2 | 0.990398 | 0.019275 | 0.508849 | 1.715001 | 0.293411 |
| Pygm | 0.988690 | 0.020984 | 0.508849 | 1.678119 | 0.293411 |
| Wdr59 | 0.987208 | 0.043694 | 0.574283 | 1.359583 | 0.240874 |
| Apba3 | 0.985025 | 0.018402 | 0.508849 | 1.735126 | 0.293411 |
| Col19a1 | 0.981610 | 0.033760 | 0.550478 | 1.471599 | 0.259260 |
| Ifit2 | 0.981610 | 0.039456 | 0.570881 | 1.403884 | 0.243454 |
| Ankrd49 | 0.981610 | 0.047433 | 0.585976 | 1.323920 | 0.232120 |
| Pemt | 0.977663 | 0.002032 | 0.372917 | 2.691977 | 0.428387 |
| Kcnk13 | 0.977663 | 0.007553 | 0.459495 | 2.121891 | 0.337720 |
| Slc39a14 | 0.971397 | 0.006123 | 0.454055 | 2.213035 | 0.342892 |
| Zfp949 | 0.970583 | 0.000210 | 0.346132 | 3.677859 | 0.460759 |
| Dnah5 | 0.970583 | 0.000861 | 0.346132 | 3.064989 | 0.460759 |
| Olfr1044 | 0.969157 | 0.026517 | 0.522318 | 1.576479 | 0.282065 |
| Dcc | 0.968514 | 0.020154 | 0.508849 | 1.695630 | 0.293411 |
| Zdhhc23 | 0.968197 | 0.047275 | 0.585976 | 1.325373 | 0.232120 |
| Slitrk5 | 0.966024 | 0.043222 | 0.574283 | 1.364292 | 0.240874 |
| Tnpo1 | 0.962212 | 0.017574 | 0.508849 | 1.755133 | 0.293411 |

| | | | | | |
|---|---|---|---|---|---|
| Rbm26 | -0.834498 | 0.023688 | 0.512726 | 1.625477 | 0.290115 |
| Fcgr3 | -0.833072 | 0.020434 | 0.508849 | 1.689638 | 0.293411 |
| Gng3 | -0.828404 | 0.004506 | 0.433081 | 2.346194 | 0.363431 |
| Ube2n | -0.823563 | 0.009422 | 0.468720 | 2.025849 | 0.329086 |
| Pde7b | -0.823563 | 0.030075 | 0.537364 | 1.521794 | 0.269731 |
| Mgme1 | -0.823233 | 0.025468 | 0.518442 | 1.594000 | 0.285299 |
| Hmgcll1 | -0.821899 | 0.020405 | 0.508849 | 1.690259 | 0.293411 |
| Mfap2 | -0.819233 | 0.026416 | 0.522318 | 1.578138 | 0.282065 |
| Prkaca | -0.816991 | 0.033951 | 0.550478 | 1.469152 | 0.259260 |
| Pnpla3 | -0.816593 | 0.003552 | 0.407941 | 2.449578 | 0.389403 |
| Washc2 | -0.816593 | 0.014280 | 0.498513 | 1.845267 | 0.302323 |
| Ube2e1 | -0.815854 | 0.019858 | 0.508849 | 1.702057 | 0.293411 |
| Cenpj | -0.815590 | 0.020483 | 0.508849 | 1.688600 | 0.293411 |
| 4930430F0 | -0.814421 | 0.009957 | 0.477528 | 2.001868 | 0.321001 |
| Nob1 | -0.811259 | 0.042734 | 0.574283 | 1.369225 | 0.240874 |
| Slc31a1 | -0.809527 | 0.041973 | 0.573444 | 1.377031 | 0.241509 |
| Plekho1 | -0.808449 | 0.013548 | 0.495208 | 1.868130 | 0.305213 |
| Atp8a1 | -0.807084 | 0.030290 | 0.537364 | 1.518695 | 0.269731 |
| Wsb1 | -0.807048 | 0.026951 | 0.523236 | 1.569424 | 0.281303 |
| Dbi | -0.806059 | 0.014370 | 0.498513 | 1.842536 | 0.302323 |
| Fnip2 | -0.803859 | 0.006917 | 0.454560 | 2.160051 | 0.342409 |
| Yipf6 | -0.802697 | 0.023929 | 0.512726 | 1.621075 | 0.290115 |
| Armh3 | -0.798511 | 0.001274 | 0.362707 | 2.894691 | 0.440445 |
| Fra10ac1 | -0.789698 | 0.023838 | 0.512726 | 1.622724 | 0.290115 |
| Hmgcs2 | -0.789044 | 0.021133 | 0.508849 | 1.675046 | 0.293411 |
| Enkur | -0.788273 | 0.013400 | 0.494300 | 1.872883 | 0.306009 |
| Polr1c | -0.786455 | 0.033478 | 0.550478 | 1.475236 | 0.259260 |
| Tbl1x | -0.785751 | 0.001348 | 0.362707 | 2.870339 | 0.440445 |
| Msh2 | -0.782137 | 0.002644 | 0.399584 | 2.577659 | 0.398392 |
| Defb50 | -0.781658 | 0.044232 | 0.574283 | 1.354268 | 0.240874 |
| Atad3a | -0.781499 | 0.024718 | 0.513619 | 1.606994 | 0.289359 |
| Nfyc | -0.780051 | 0.024068 | 0.512870 | 1.618560 | 0.289993 |
| Gm960 | -0.778060 | 0.032796 | 0.548023 | 1.484177 | 0.261201 |
| Rpl7l1 | -0.776501 | 0.000884 | 0.346132 | 3.053592 | 0.460759 |
| Nceh1 | -0.774112 | 0.004231 | 0.422417 | 2.373542 | 0.374259 |
| Slurp2 | -0.763899 | 0.040706 | 0.572249 | 1.390342 | 0.242415 |
| Fam227a | -0.762939 | 0.005417 | 0.447186 | 2.266237 | 0.349512 |
| Cdkn2b | -0.762192 | 0.030274 | 0.537364 | 1.518925 | 0.269731 |
| Spx | -0.762124 | 0.003118 | 0.402288 | 2.506071 | 0.395463 |
| Sfi1 | -0.761974 | 0.011879 | 0.477528 | 1.925205 | 0.321001 |
| Ube2w | -0.761684 | 0.032049 | 0.543564 | 1.494185 | 0.264750 |

| | | | | | |
|---|---|---|---|---|---|
| Olfr1209 | 0.961184 | 0.028464 | 0.528131 | 1.545706 | 0.277259 |
| Osr1 | 0.958944 | 0.023429 | 0.512726 | 1.630251 | 0.290115 |
| Kif20a | 0.958944 | 0.023975 | 0.512726 | 1.620242 | 0.290115 |
| Rasgrf1 | 0.957984 | 0.036787 | 0.563078 | 1.434310 | 0.249431 |
| Kdm8 | 0.956422 | 0.044190 | 0.574283 | 1.354678 | 0.240874 |
| Tyk2 | 0.954329 | 0.023116 | 0.512726 | 1.636082 | 0.290115 |
| Arhgap22 | 0.953582 | 0.045328 | 0.579023 | 1.343630 | 0.237304 |
| Olfr710 | 0.953289 | 0.022213 | 0.508849 | 1.653387 | 0.293411 |
| Ccm2l | 0.952206 | 0.007082 | 0.456158 | 2.149817 | 0.340885 |
| Gm40556 | 0.950157 | 0.021275 | 0.508849 | 1.672131 | 0.293411 |
| Wipf1 | 0.950157 | 0.023208 | 0.512726 | 1.634367 | 0.290115 |
| Rnf114 | 0.949597 | 0.009596 | 0.472983 | 2.017904 | 0.325154 |
| Pla2g4b | 0.948731 | 0.042867 | 0.574283 | 1.367874 | 0.240874 |
| Lztfl1 | 0.947909 | 0.000393 | 0.346132 | 3.405228 | 0.460759 |
| Als2cl | 0.947305 | 0.030889 | 0.537693 | 1.510198 | 0.269466 |
| Mri1 | 0.946956 | 0.033067 | 0.548023 | 1.480612 | 0.261201 |
| Nemp2 | 0.939944 | 0.003299 | 0.402288 | 2.481601 | 0.395463 |
| Pi16 | 0.939944 | 0.007619 | 0.459495 | 2.118110 | 0.337720 |
| Rab5if | 0.939557 | 0.000749 | 0.346132 | 3.125631 | 0.460759 |
| Ccdc74a | 0.938518 | 0.049316 | 0.592038 | 1.307015 | 0.227650 |
| Rttn | 0.936289 | 0.038379 | 0.566795 | 1.415909 | 0.246574 |
| Rnf112 | 0.935996 | 0.008779 | 0.459495 | 2.056569 | 0.337720 |
| Bag4 | 0.931370 | 0.042317 | 0.573809 | 1.373481 | 0.241233 |
| Mysm1 | 0.930005 | 0.012494 | 0.486498 | 1.903295 | 0.312919 |
| Shprh | 0.928251 | 0.010762 | 0.477528 | 1.968109 | 0.321001 |
| Sct | 0.926076 | 0.035944 | 0.559928 | 1.444378 | 0.251868 |
| Thsd7b | 0.925783 | 0.015678 | 0.508849 | 1.804718 | 0.293411 |
| Zpld2 | 0.925783 | 0.031049 | 0.537739 | 1.507949 | 0.269429 |
| Susd6 | 0.924953 | 0.005801 | 0.450937 | 2.236472 | 0.345884 |
| Pkd1l3 | 0.924358 | 0.041813 | 0.572694 | 1.378684 | 0.242077 |
| Tmem147 | 0.923543 | 0.046050 | 0.583451 | 1.336767 | 0.233995 |
| Dsc2 | 0.922583 | 0.032569 | 0.548023 | 1.487192 | 0.261201 |
| Gabrp | 0.920410 | 0.011019 | 0.477528 | 1.957849 | 0.321001 |
| Zfand1 | 0.918703 | 0.047820 | 0.587289 | 1.320390 | 0.231148 |
| Epop | 0.918703 | 0.013120 | 0.489076 | 1.882065 | 0.310624 |
| Lpar6 | 0.917559 | 0.032883 | 0.548023 | 1.483032 | 0.261201 |
| Fv1 | 0.917277 | 0.042137 | 0.573516 | 1.375337 | 0.241454 |
| Ltbp2 | 0.917277 | 0.046107 | 0.583451 | 1.336233 | 0.233995 |
| Agfg2 | 0.911623 | 0.003847 | 0.408654 | 2.414873 | 0.388644 |
| Frzb | 0.908490 | 0.015168 | 0.508849 | 1.819078 | 0.293411 |
| Clybl | 0.908490 | 0.017706 | 0.508849 | 1.751875 | 0.293411 |

| | | | | | |
|---|---|---|---|---|---|
| Sin3a | -0.760687 | 0.048882 | 0.591266 | 1.310855 | 0.228217 |
| Slc16a1 | -0.760258 | 0.028793 | 0.528456 | 1.540709 | 0.276991 |
| Npat | -0.758337 | 0.044487 | 0.574283 | 1.351763 | 0.240874 |
| Mchr1 | -0.758177 | 0.045941 | 0.583451 | 1.337800 | 0.233995 |
| Mfge8 | -0.757591 | 0.041167 | 0.572249 | 1.385449 | 0.242415 |
| Add3 | -0.757284 | 0.023219 | 0.512726 | 1.634156 | 0.290115 |
| Lztr1 | -0.755418 | 0.025309 | 0.518442 | 1.596732 | 0.285299 |
| Mcc | -0.754647 | 0.025465 | 0.518442 | 1.594049 | 0.285299 |
| Plpp2 | -0.744831 | 0.048744 | 0.590861 | 1.312076 | 0.228515 |
| Zc3h11a | -0.743124 | 0.031683 | 0.541568 | 1.499170 | 0.266347 |
| Srcap | -0.742196 | 0.034378 | 0.551303 | 1.463714 | 0.258610 |
| Jak1 | -0.740078 | 0.022192 | 0.508849 | 1.653808 | 0.293411 |
| Yeats4 | -0.737471 | 0.033576 | 0.550478 | 1.473971 | 0.259260 |
| Tdp2 | -0.736044 | 0.026472 | 0.522318 | 1.577210 | 0.282065 |
| Prss8 | -0.734618 | 0.007360 | 0.459495 | 2.133135 | 0.337720 |
| Gramd1b | -0.733656 | 0.033015 | 0.548023 | 1.481283 | 0.261201 |
| Ikzf2 | -0.732445 | 0.035838 | 0.559928 | 1.445655 | 0.251868 |
| Gm10265 | -0.729883 | 0.001433 | 0.362707 | 2.843603 | 0.440445 |
| Leo1 | -0.727140 | 0.019319 | 0.508849 | 1.714009 | 0.293411 |
| D8Ertd738 | -0.725709 | 0.003388 | 0.402288 | 2.469996 | 0.395463 |
| Mrpl49 | -0.724469 | 0.032710 | 0.548023 | 1.485320 | 0.261201 |
| Rfesd | -0.724405 | 0.006279 | 0.454055 | 2.202079 | 0.342892 |
| Dzank1 | -0.724405 | 0.002779 | 0.399584 | 2.556077 | 0.398392 |
| Mettl2 | -0.722936 | 0.018727 | 0.508849 | 1.727527 | 0.293411 |
| Inhbb | -0.722936 | 0.021530 | 0.508849 | 1.666959 | 0.293411 |
| Itpa | -0.722827 | 0.011353 | 0.477528 | 1.944897 | 0.321001 |
| Stxbp6 | -0.721272 | 0.018443 | 0.508849 | 1.734158 | 0.293411 |
| Ap1g2 | -0.720457 | 0.008907 | 0.459495 | 2.050292 | 0.337720 |
| Myog | -0.720457 | 0.048295 | 0.588712 | 1.316094 | 0.230097 |
| Hgf | -0.720457 | 0.001900 | 0.367785 | 2.721164 | 0.434406 |
| Dcps | -0.716431 | 0.039678 | 0.570881 | 1.401446 | 0.243454 |
| Speer4b | -0.716353 | 0.016809 | 0.508849 | 1.774461 | 0.293411 |
| Mrpl51 | -0.713067 | 0.038264 | 0.566795 | 1.417212 | 0.246574 |
| Gm52416 | -0.712783 | 0.000546 | 0.346132 | 3.262664 | 0.460759 |
| Ostf1 | -0.707149 | 0.021934 | 0.508849 | 1.658887 | 0.293411 |
| Itga8 | -0.707112 | 0.013311 | 0.492293 | 1.875794 | 0.307777 |
| Slc35e4 | -0.707112 | 0.031816 | 0.541568 | 1.497350 | 0.266347 |
| Kdm5a | -0.700307 | 0.010553 | 0.477528 | 1.976616 | 0.321001 |
| Gm3893 | -0.699457 | 0.040368 | 0.570893 | 1.393961 | 0.243445 |
| Mfsd6l | -0.697859 | 0.026900 | 0.523236 | 1.570243 | 0.281303 |
| Lbh | -0.696962 | 0.049817 | 0.592038 | 1.302620 | 0.227650 |

| | | | | | |
|---|---|---|---|---|---|
| Cfap65 | 0.908490 | 0.007490 | 0.459495 | 2.125490 | 0.337720 |
| Morc2b | 0.907064 | 0.011645 | 0.477528 | 1.933855 | 0.321001 |
| Gng7 | 0.905571 | 0.049934 | 0.592038 | 1.301604 | 0.227650 |
| Tex10 | 0.905357 | 0.035834 | 0.559928 | 1.445709 | 0.251868 |
| F8a | 0.905357 | 0.045495 | 0.580102 | 1.342036 | 0.236496 |
| Amt | 0.903117 | 0.018949 | 0.508849 | 1.722412 | 0.293411 |
| Zdhhc13 | 0.903117 | 0.047050 | 0.585976 | 1.327441 | 0.232120 |
| Klk6 | 0.899635 | 0.044406 | 0.574283 | 1.352561 | 0.240874 |
| Zbtb26 | 0.897755 | 0.013999 | 0.495749 | 1.853903 | 0.304738 |
| Inava | 0.896851 | 0.030619 | 0.537364 | 1.514016 | 0.269731 |
| Zbtb37 | 0.895076 | 0.021977 | 0.508849 | 1.658032 | 0.293411 |
| Gabrg3 | 0.894622 | 0.041406 | 0.572650 | 1.382939 | 0.242111 |
| Acss2 | 0.894330 | 0.019930 | 0.508849 | 1.700488 | 0.293411 |
| Ecpas | 0.893932 | 0.035986 | 0.559928 | 1.443863 | 0.251868 |
| Nr4a1 | 0.892904 | 0.019422 | 0.508849 | 1.711717 | 0.293411 |
| Alox5ap | 0.892904 | 0.019883 | 0.508849 | 1.701516 | 0.293411 |
| Lrrtm2 | 0.892904 | 0.035695 | 0.559928 | 1.447387 | 0.251868 |
| 1700020D( | 0.890382 | 0.047420 | 0.585976 | 1.324038 | 0.232120 |
| Fstl1 | 0.888818 | 0.009154 | 0.463691 | 2.038386 | 0.333772 |
| Pard6b | 0.888064 | 0.048137 | 0.587289 | 1.317520 | 0.231148 |
| Tshz2 | 0.887531 | 0.014963 | 0.504512 | 1.824995 | 0.297129 |
| Pycr1 | 0.885824 | 0.018896 | 0.508849 | 1.723641 | 0.293411 |
| Olfr10 | 0.884680 | 0.032932 | 0.548023 | 1.482384 | 0.261201 |
| 4930444P1 | 0.884117 | 0.040258 | 0.570881 | 1.395151 | 0.243454 |
| Abcb7 | 0.882691 | 0.026713 | 0.523236 | 1.573277 | 0.281303 |
| Knstrn | 0.882342 | 0.005260 | 0.447186 | 2.279008 | 0.349512 |
| Kcnmb4 | 0.880718 | 0.028066 | 0.527961 | 1.551817 | 0.277398 |
| Kdelr2 | 0.880711 | 0.026643 | 0.522920 | 1.574414 | 0.281565 |
| Keap1 | 0.879036 | 0.004813 | 0.443121 | 2.317614 | 0.353477 |
| Crb3 | 0.878743 | 0.002328 | 0.385644 | 2.632980 | 0.413814 |
| Cp | 0.878743 | 0.044512 | 0.574283 | 1.351521 | 0.240874 |
| Olfm3 | 0.877036 | 0.010664 | 0.477528 | 1.972068 | 0.321001 |
| Synb | 0.877036 | 0.012589 | 0.486498 | 1.900000 | 0.312919 |
| Plekha7 | 0.877036 | 0.011940 | 0.477528 | 1.923009 | 0.321001 |
| Apmap | 0.876796 | 0.004251 | 0.422417 | 2.371459 | 0.374259 |
| Prpf38a | 0.876650 | 0.021284 | 0.508849 | 1.671938 | 0.293411 |
| Aspdh | 0.875611 | 0.020574 | 0.508849 | 1.686692 | 0.293411 |
| Nf2 | 0.871115 | 0.018145 | 0.508849 | 1.741231 | 0.293411 |
| Nipa2 | 0.871052 | 0.030115 | 0.537364 | 1.521215 | 0.269731 |
| Rpa2 | 0.870249 | 0.040965 | 0.572249 | 1.387582 | 0.242415 |
| Limd2 | 0.868767 | 0.030854 | 0.537693 | 1.510687 | 0.269466 |

| | | | | | |
|---|---|---|---|---|---|
| Zwilch | -0.696899 | 0.020846 | 0.508849 | 1.680973 | 0.293411 |
| Olfr1126 | -0.694726 | 0.022174 | 0.508849 | 1.654147 | 0.293411 |
| Zfp14 | -0.693258 | 0.001537 | 0.367785 | 2.813380 | 0.434406 |
| Prtg | -0.693258 | 0.024709 | 0.513619 | 1.607139 | 0.289359 |
| Mrpl39 | -0.692443 | 0.023074 | 0.512726 | 1.636886 | 0.290115 |
| Pigyl | -0.690779 | 0.046048 | 0.583451 | 1.336789 | 0.233995 |
| Acadsb | -0.690002 | 0.017524 | 0.508849 | 1.756369 | 0.293411 |
| Dnajc6 | -0.685871 | 0.046855 | 0.585550 | 1.329243 | 0.232436 |
| Gm6958 | -0.685233 | 0.001792 | 0.367785 | 2.746758 | 0.434406 |
| Slc6a15 | -0.684189 | 0.005937 | 0.450937 | 2.226452 | 0.345884 |
| Agpat5 | -0.683510 | 0.023438 | 0.512726 | 1.630073 | 0.290115 |
| Kmt5a | -0.682403 | 0.017526 | 0.508849 | 1.756325 | 0.293411 |
| Gm46156 | -0.679819 | 0.045321 | 0.579023 | 1.343699 | 0.237304 |
| Tas2r123 | -0.677433 | 0.028197 | 0.527961 | 1.549795 | 0.277398 |
| Irf4 | -0.677433 | 0.034718 | 0.552431 | 1.459440 | 0.257722 |
| Slc8a2 | -0.676221 | 0.048710 | 0.590861 | 1.312385 | 0.228515 |
| Cbln1 | -0.675658 | 0.021448 | 0.508849 | 1.668612 | 0.293411 |
| Pam | -0.675459 | 0.018773 | 0.508849 | 1.726455 | 0.293411 |
| Spata7 | -0.674843 | 0.002606 | 0.399584 | 2.583975 | 0.398392 |
| Olfr299 | -0.672482 | 0.040273 | 0.570881 | 1.394987 | 0.243454 |
| Shmt2 | -0.671792 | 0.003166 | 0.402288 | 2.499503 | 0.395463 |
| Dpysl4 | -0.667428 | 0.041327 | 0.572650 | 1.383770 | 0.242111 |
| Bri3bp | -0.664301 | 0.037731 | 0.566795 | 1.423302 | 0.246574 |
| Erdr1 | -0.663435 | 0.005769 | 0.450937 | 2.238902 | 0.345884 |
| Gtf2h1 | -0.663256 | 0.043968 | 0.574283 | 1.356864 | 0.240874 |
| Dab2ip | -0.662084 | 0.012400 | 0.485483 | 1.906579 | 0.313826 |
| Hmgn5 | -0.661407 | 0.006602 | 0.454055 | 2.180356 | 0.342892 |
| Kansl2 | -0.660205 | 0.000880 | 0.346132 | 3.055305 | 0.460759 |
| Nhsl1 | -0.656150 | 0.017262 | 0.508849 | 1.762910 | 0.293411 |
| Grem2 | -0.655232 | 0.039868 | 0.570881 | 1.399378 | 0.243454 |
| B4galt2 | -0.651548 | 0.044083 | 0.574283 | 1.355732 | 0.240874 |
| Gstm7 | -0.649256 | 0.031301 | 0.537978 | 1.504444 | 0.269235 |
| Pkib | -0.649112 | 0.048135 | 0.587289 | 1.317542 | 0.231148 |
| Setd5 | -0.648546 | 0.020391 | 0.508849 | 1.690558 | 0.293411 |
| Idh3a | -0.644579 | 0.005366 | 0.447186 | 2.270332 | 0.349512 |
| Fam43b | -0.643807 | 0.014479 | 0.498513 | 1.839258 | 0.302323 |
| Cldn17 | -0.643807 | 0.002191 | 0.385644 | 2.659369 | 0.413814 |
| Ifitm10 | -0.641072 | 0.047807 | 0.587289 | 1.320509 | 0.231148 |
| Faf2 | -0.637482 | 0.033652 | 0.550478 | 1.472988 | 0.259260 |
| Dnmt1 | -0.636204 | 0.007061 | 0.456158 | 2.151145 | 0.340885 |
| Cmtr1 | -0.634524 | 0.008458 | 0.459495 | 2.072757 | 0.337720 |

| | | | | | |
|---|---|---|---|---|---|
| Pgap3 | 0.866823 | 0.019644 | 0.508849 | 1.706776 | 0.293411 |
| Sash3 | 0.866823 | 0.020424 | 0.508849 | 1.689856 | 0.293411 |
| Thoc6 | 0.866755 | 0.049003 | 0.591726 | 1.309774 | 0.227880 |
| Slc16a9 | 0.865023 | 0.001463 | 0.362707 | 2.834650 | 0.440445 |
| 2410004B1 | 0.863904 | 0.044531 | 0.574283 | 1.351333 | 0.240874 |
| Endou | 0.861450 | 0.047516 | 0.586293 | 1.323161 | 0.231886 |
| Pabpc6 | 0.859743 | 0.040980 | 0.572249 | 1.387425 | 0.242415 |
| Coro1a | 0.857531 | 0.027469 | 0.525245 | 1.561162 | 0.279638 |
| Slc30a8 | 0.856610 | 0.011997 | 0.477528 | 1.920940 | 0.321001 |
| Bmp6 | 0.852663 | 0.020280 | 0.508849 | 1.692930 | 0.293411 |
| Olfr169 | 0.852663 | 0.021536 | 0.508849 | 1.666826 | 0.293411 |
| 2310002L0 | 0.852663 | 0.036201 | 0.560796 | 1.441281 | 0.251195 |
| Pcdhgb7 | 0.851871 | 0.006680 | 0.454055 | 2.175249 | 0.342892 |
| Mmgt2 | 0.849530 | 0.046163 | 0.583451 | 1.335708 | 0.233995 |
| Il1f10 | 0.844157 | 0.002523 | 0.399584 | 2.598037 | 0.398392 |
| ccdc198 | 0.842450 | 0.039198 | 0.570881 | 1.406732 | 0.243454 |
| Kcnip4 | 0.842450 | 0.024455 | 0.513619 | 1.611635 | 0.289359 |
| Ciao3 | 0.839542 | 0.028224 | 0.527961 | 1.549382 | 0.277398 |
| Snapc2 | 0.837944 | 0.049846 | 0.592038 | 1.302367 | 0.227650 |
| Ankrd39 | 0.837077 | 0.040164 | 0.570881 | 1.396167 | 0.243454 |
| Depdc1a | 0.835370 | 0.023385 | 0.512726 | 1.631070 | 0.290115 |
| Mkrn2 | 0.833408 | 0.005474 | 0.447186 | 2.261668 | 0.349512 |
| Ube3c | 0.831465 | 0.027597 | 0.525245 | 1.559142 | 0.279638 |
| 2310079G1 | 0.831422 | 0.041644 | 0.572694 | 1.380449 | 0.242077 |
| Zfp575 | 0.830051 | 0.030839 | 0.537693 | 1.510895 | 0.269466 |
| Grm4 | 0.828571 | 0.009709 | 0.472983 | 2.012837 | 0.325154 |
| Timm21 | 0.826864 | 0.019027 | 0.508849 | 1.720640 | 0.293411 |
| Trim32 | 0.825731 | 0.008151 | 0.459495 | 2.088805 | 0.337720 |
| Slc24a3 | 0.825157 | 0.024695 | 0.513619 | 1.607388 | 0.289359 |
| Lbx2 | 0.822305 | 0.034149 | 0.550478 | 1.466619 | 0.259260 |
| Cdc45 | 0.818358 | 0.015361 | 0.508849 | 1.813585 | 0.293411 |
| Frrs1 | 0.818076 | 0.028741 | 0.528456 | 1.541503 | 0.276991 |
| Gtf3c5 | 0.816932 | 0.018202 | 0.508849 | 1.739888 | 0.293411 |
| Klf5 | 0.814944 | 0.040208 | 0.570881 | 1.395683 | 0.243454 |
| F830045P1 | 0.814944 | 0.023588 | 0.512726 | 1.627317 | 0.290115 |
| Tm2d3 | 0.814944 | 0.048481 | 0.589453 | 1.314426 | 0.229551 |
| Mical2 | 0.813450 | 0.040258 | 0.570881 | 1.395146 | 0.243454 |
| Chil6 | 0.810996 | 0.043409 | 0.574283 | 1.362423 | 0.240874 |
| Rrp9 | 0.810036 | 0.035449 | 0.559576 | 1.450398 | 0.252141 |
| Prl3d1 | 0.809571 | 0.046342 | 0.583451 | 1.334022 | 0.233995 |
| Mzt2 | 0.808903 | 0.037373 | 0.566151 | 1.427437 | 0.247068 |
| Fgf13 | -0.634298 | 0.028634 | 0.528456 | 1.543124 | 0.276991 |
| Lin54 | -0.633551 | 0.021673 | 0.508849 | 1.664083 | 0.293411 |
| Ccny | -0.632629 | 0.028289 | 0.527961 | 1.548384 | 0.277398 |
| Dmac1 | -0.632341 | 0.026065 | 0.518654 | 1.583945 | 0.285122 |
| Tstd2 | -0.630566 | 0.032799 | 0.548023 | 1.484139 | 0.261201 |
| Ppp3cc | -0.630044 | 0.007923 | 0.459495 | 2.101125 | 0.337720 |
| Chrnd | -0.623778 | 0.034538 | 0.551979 | 1.461706 | 0.258077 |
| Galnt10 | -0.621606 | 0.031904 | 0.541763 | 1.496148 | 0.266191 |
| D430019H1 | -0.619877 | 0.030537 | 0.537364 | 1.515180 | 0.269731 |
| Atg9a | -0.617470 | 0.047204 | 0.585976 | 1.326020 | 0.232120 |
| Sox2 | -0.612619 | 0.040491 | 0.570922 | 1.392636 | 0.243423 |
| Ppp1cc | -0.611827 | 0.006437 | 0.454055 | 2.191289 | 0.342892 |
| Zp3r | -0.611393 | 0.007497 | 0.459495 | 2.125126 | 0.337720 |
| Cttnbp2nl | -0.611111 | 0.003668 | 0.408654 | 2.435606 | 0.388644 |
| Abca7 | -0.609618 | 0.019546 | 0.508849 | 1.708951 | 0.293411 |
| Ccnd1 | -0.608717 | 0.042684 | 0.574283 | 1.369734 | 0.240874 |
| Nyap1 | -0.605868 | 0.038040 | 0.566795 | 1.419764 | 0.246574 |
| Mex3d | -0.597725 | 0.019468 | 0.508849 | 1.710674 | 0.293411 |
| Mxd4 | -0.596554 | 0.039515 | 0.570881 | 1.403233 | 0.243454 |
| Atg12 | -0.593769 | 0.043995 | 0.574283 | 1.356601 | 0.240874 |
| Htr2a | -0.592325 | 0.042234 | 0.573736 | 1.374336 | 0.241288 |
| Gm52520 | -0.586814 | 0.012570 | 0.486498 | 1.900650 | 0.312919 |
| Atp5o | -0.586808 | 0.011296 | 0.477528 | 1.947065 | 0.321001 |
| Klhl28 | -0.584847 | 0.019422 | 0.508849 | 1.711702 | 0.293411 |
| Gch1 | -0.577767 | 0.015649 | 0.508849 | 1.805525 | 0.293411 |
| Nr2f6 | -0.577578 | 0.020518 | 0.508849 | 1.687864 | 0.293411 |
| Fgfr1 | -0.576006 | 0.034506 | 0.551979 | 1.462104 | 0.258077 |
| Zfp24 | -0.573538 | 0.007046 | 0.456158 | 2.152083 | 0.340885 |
| Jpt2 | -0.570551 | 0.034041 | 0.550478 | 1.467997 | 0.259260 |
| Ptpn11 | -0.568965 | 0.003357 | 0.402288 | 2.474070 | 0.395463 |
| Fam71e1 | -0.567297 | 0.042740 | 0.574283 | 1.369161 | 0.240874 |
| Ddc | -0.567137 | 0.022661 | 0.512726 | 1.644724 | 0.290115 |
| Pank2 | -0.566577 | 0.036532 | 0.562320 | 1.437330 | 0.250017 |
| Col6a2 | -0.565779 | 0.034340 | 0.551303 | 1.464194 | 0.258610 |
| Atp5c1 | -0.565190 | 0.044551 | 0.574283 | 1.351144 | 0.240874 |
| Marchf2 | -0.563375 | 0.041773 | 0.572694 | 1.379100 | 0.242077 |
| Nedd4l | -0.562854 | 0.022068 | 0.508849 | 1.656240 | 0.293411 |
| Ttbk2 | -0.562297 | 0.004609 | 0.437009 | 2.336404 | 0.359510 |
| Cox6b1 | -0.560034 | 0.026873 | 0.523236 | 1.570681 | 0.281303 |
| Glg1 | -0.559528 | 0.010893 | 0.477528 | 1.962847 | 0.321001 |
| Gm52720 | -0.559436 | 0.043470 | 0.574283 | 1.361809 | 0.240874 |

| | | | | | |
|---|---|---|---|---|---|
| Nr2f2 | 0.808842 | 0.030784 | 0.537693 | 1.511680 | 0.269466 |
| Bcl2l1 | 0.806800 | 0.003230 | 0.402288 | 2.490840 | 0.395463 |
| Col13a1 | 0.799358 | 0.043494 | 0.574283 | 1.361575 | 0.240874 |
| Ncs1 | 0.797671 | 0.021844 | 0.508849 | 1.660662 | 0.293411 |
| Taf1a | 0.791756 | 0.031160 | 0.537739 | 1.506404 | 0.269429 |
| Ccdc91 | 0.789077 | 0.044104 | 0.574283 | 1.355523 | 0.240874 |
| Alas2 | 0.788376 | 0.041719 | 0.572694 | 1.379667 | 0.242077 |
| Pcsk1 | 0.788330 | 0.046279 | 0.583451 | 1.334615 | 0.233995 |
| Gabrb2 | 0.787662 | 0.005875 | 0.450937 | 2.231000 | 0.345884 |
| Mbd4 | 0.787370 | 0.012009 | 0.477528 | 1.920488 | 0.321001 |
| Klhl8 | 0.786623 | 0.042686 | 0.574283 | 1.369714 | 0.240874 |
| Mapk7 | 0.786380 | 0.048999 | 0.591726 | 1.309816 | 0.227880 |
| Smndc1 | 0.785490 | 0.038234 | 0.566795 | 1.417550 | 0.246574 |
| Dedd | 0.785471 | 0.044514 | 0.574283 | 1.351501 | 0.240874 |
| Cd164l2 | 0.783490 | 0.024758 | 0.513619 | 1.606292 | 0.289359 |
| Gm46901 | 0.781715 | 0.021238 | 0.508849 | 1.672878 | 0.293411 |
| Lgals8 | 0.779543 | 0.043540 | 0.574283 | 1.361116 | 0.240874 |
| D17H6S53E | 0.779543 | 0.008042 | 0.459495 | 2.094643 | 0.337720 |
| Bmpr2 | 0.778578 | 0.012068 | 0.477528 | 1.918369 | 0.321001 |
| Pnoc | 0.778117 | 0.046420 | 0.583451 | 1.333297 | 0.233995 |
| Camk2d | 0.778117 | 0.011118 | 0.477528 | 1.953968 | 0.321001 |
| 2010300C0 | 0.778117 | 0.025338 | 0.518442 | 1.596235 | 0.285299 |
| Stx16 | 0.776212 | 0.030445 | 0.537364 | 1.516482 | 0.269731 |
| Piwil1 | 0.775595 | 0.046875 | 0.585550 | 1.329054 | 0.232436 |
| Thoc5 | 0.773617 | 0.017018 | 0.508849 | 1.769094 | 0.293411 |
| Arid3b | 0.773277 | 0.020628 | 0.508849 | 1.685549 | 0.293411 |
| Aldh3a2 | 0.773209 | 0.004449 | 0.430491 | 2.351764 | 0.366036 |
| Rps6kb1 | 0.772426 | 0.010144 | 0.477528 | 1.993792 | 0.321001 |
| Ccpg1 | 0.771851 | 0.040799 | 0.572249 | 1.389351 | 0.242415 |
| Cenpw | 0.769904 | 0.003222 | 0.402288 | 2.491849 | 0.395463 |
| Olfr750 | 0.769330 | 0.009972 | 0.477528 | 2.001217 | 0.321001 |
| Calu | 0.768484 | 0.001459 | 0.362707 | 2.835829 | 0.440445 |
| Olfr446 | 0.767904 | 0.041432 | 0.572650 | 1.382662 | 0.242111 |
| Rpf2 | 0.766943 | 0.037979 | 0.566795 | 1.420451 | 0.246574 |
| Ube2q1 | 0.763582 | 0.010440 | 0.477528 | 1.981309 | 0.321001 |
| Zfp689 | 0.763289 | 0.033865 | 0.550478 | 1.470244 | 0.259260 |
| Pfkl | 0.763287 | 0.017848 | 0.508849 | 1.748408 | 0.293411 |
| Klhl20 | 0.762531 | 0.008522 | 0.459495 | 2.069476 | 0.337720 |
| G6pc2 | 0.762249 | 0.013025 | 0.489076 | 1.885228 | 0.310624 |
| C87436 | 0.762249 | 0.033059 | 0.548023 | 1.480715 | 0.261201 |
| Pex11b | 0.761105 | 0.038499 | 0.566795 | 1.414550 | 0.246574 |

| | | | | | |
|---|---|---|---|---|---|
| Mtg1 | -0.559262 | 0.002983 | 0.402288 | 2.525284 | 0.395463 |
| Ntrk2 | -0.557334 | 0.035962 | 0.559928 | 1.444157 | 0.251868 |
| Apba2 | -0.556206 | 0.033437 | 0.550478 | 1.475770 | 0.259260 |
| Cyp2c55 | -0.554605 | 0.021071 | 0.508849 | 1.676320 | 0.293411 |
| Ros1 | -0.553791 | 0.023730 | 0.512726 | 1.624708 | 0.290115 |
| Cdk16 | -0.553144 | 0.031130 | 0.537739 | 1.506822 | 0.269429 |
| 4833420G1 | -0.548740 | 0.003767 | 0.408654 | 2.423960 | 0.388644 |
| Gm36327 | -0.548639 | 0.003225 | 0.402288 | 2.491476 | 0.395463 |
| Ccdc8 | -0.548486 | 0.023717 | 0.512726 | 1.624937 | 0.290115 |
| N4bp2 | -0.548045 | 0.017702 | 0.508849 | 1.751967 | 0.293411 |
| Rragc | -0.547832 | 0.000903 | 0.346132 | 3.044529 | 0.460759 |
| Paip1 | -0.545817 | 0.041001 | 0.572249 | 1.387205 | 0.242415 |
| Gm52432 | -0.545121 | 0.000153 | 0.346132 | 3.815116 | 0.460759 |
| Qser1 | -0.539363 | 0.037778 | 0.566795 | 1.422758 | 0.246574 |
| Cd68 | -0.538273 | 0.022984 | 0.512726 | 1.638576 | 0.290115 |
| Trip11 | -0.536100 | 0.033999 | 0.550478 | 1.468536 | 0.259260 |
| Prpf8 | -0.535805 | 0.021139 | 0.508849 | 1.674912 | 0.293411 |
| Nap1l5 | -0.535751 | 0.049840 | 0.592038 | 1.302422 | 0.227650 |
| Plekhj1 | -0.535295 | 0.029620 | 0.534804 | 1.528419 | 0.271805 |
| Vps16 | -0.532082 | 0.006959 | 0.455172 | 2.157452 | 0.341825 |
| Prkd3 | -0.531153 | 0.042762 | 0.574283 | 1.368941 | 0.240874 |
| Ckb | -0.528342 | 0.039992 | 0.570881 | 1.398029 | 0.243454 |
| Klf3 | -0.527199 | 0.043656 | 0.574283 | 1.359952 | 0.240874 |
| Nop53 | -0.525864 | 0.049465 | 0.592038 | 1.305704 | 0.227650 |
| Ccar1 | -0.525523 | 0.011571 | 0.477528 | 1.936630 | 0.321001 |
| Rab35 | -0.524133 | 0.047896 | 0.587289 | 1.319704 | 0.231148 |
| Tha1 | -0.524112 | 0.001060 | 0.346132 | 2.974553 | 0.460759 |
| Alkbh1 | -0.520979 | 0.000026 | 0.138592 | 4.589003 | 0.858262 |
| Rbbp8 | -0.520979 | 0.018496 | 0.508849 | 1.732918 | 0.293411 |
| Sdf2 | -0.518807 | 0.006218 | 0.454055 | 2.206381 | 0.342892 |
| Adam12 | -0.512192 | 0.028806 | 0.528456 | 1.540517 | 0.276991 |
| Tmem181a | -0.511073 | 0.020061 | 0.508849 | 1.697646 | 0.293411 |
| Ramp2 | -0.511068 | 0.025839 | 0.518654 | 1.587718 | 0.285122 |
| Hps5 | -0.510766 | 0.018555 | 0.508849 | 1.731549 | 0.293411 |
| Itpr1 | -0.508486 | 0.044714 | 0.575184 | 1.349552 | 0.240193 |
| Insr | -0.506359 | 0.036957 | 0.564471 | 1.432298 | 0.248359 |
| Rtf1 | -0.504881 | 0.045183 | 0.578926 | 1.345030 | 0.237377 |
| Gtf3c1 | -0.504385 | 0.001365 | 0.362707 | 2.864927 | 0.440445 |
| Lyn | -0.503032 | 0.034260 | 0.551279 | 1.465215 | 0.258628 |
| Rap1a | -0.499111 | 0.014039 | 0.495749 | 1.852650 | 0.304738 |
| Lrr1 | -0.498830 | 0.000809 | 0.346132 | 3.091939 | 0.460759 |

| | | | | | |
|---|---|---|---|---|---|
| Rdh13 | 0.759117 | 0.042038 | 0.573444 | 1.376360 | 0.241509 |
| Cep120 | 0.755493 | 0.031146 | 0.537739 | 1.506591 | 0.269429 |
| Kcns2 | 0.752036 | 0.007766 | 0.459495 | 2.109806 | 0.337720 |
| Tmem9b | 0.752036 | 0.040061 | 0.570881 | 1.397281 | 0.243454 |
| Ccdc148 | 0.750611 | 0.039327 | 0.570881 | 1.405314 | 0.243454 |
| Arfgef1 | 0.746999 | 0.024806 | 0.513865 | 1.605447 | 0.289151 |
| Tada3 | 0.745703 | 0.009223 | 0.463691 | 2.035130 | 0.333772 |
| Timm8a1 | 0.740679 | 0.008402 | 0.459495 | 2.075629 | 0.337720 |
| Grn | 0.739433 | 0.004757 | 0.443121 | 2.322675 | 0.353477 |
| Manbal | 0.739397 | 0.034722 | 0.552431 | 1.459389 | 0.257722 |
| Rras | 0.738637 | 0.039890 | 0.570881 | 1.399135 | 0.243454 |
| Nsun3 | 0.736450 | 0.049351 | 0.592038 | 1.306705 | 0.227650 |
| Dis3 | 0.735236 | 0.014731 | 0.501497 | 1.831764 | 0.299732 |
| Iqcb1 | 0.735168 | 0.043594 | 0.574283 | 1.360578 | 0.240874 |
| Slc25a26 | 0.735025 | 0.001399 | 0.362707 | 2.854172 | 0.440445 |
| Nars2 | 0.734743 | 0.005759 | 0.450937 | 2.239647 | 0.345884 |
| Lypd1 | 0.732503 | 0.036413 | 0.562320 | 1.438743 | 0.250017 |
| Pign | 0.732446 | 0.004205 | 0.422417 | 2.376248 | 0.374259 |
| Mmp3 | 0.731610 | 0.028063 | 0.527961 | 1.551860 | 0.277398 |
| Wdr4 | 0.731610 | 0.028446 | 0.528131 | 1.545979 | 0.277259 |
| Pianp | 0.729783 | 0.028955 | 0.529730 | 1.538280 | 0.275945 |
| Pes1 | 0.729300 | 0.010711 | 0.477528 | 1.970180 | 0.321001 |
| Dip2c | 0.728272 | 0.018232 | 0.508849 | 1.739177 | 0.293411 |
| Ggt6 | 0.727663 | 0.000826 | 0.346132 | 3.083055 | 0.460759 |
| Cldn8 | 0.727663 | 0.049498 | 0.592038 | 1.305414 | 0.227650 |
| Cx3cr1 | 0.720583 | 0.003356 | 0.402288 | 2.474124 | 0.395463 |
| Psmd13 | 0.719833 | 0.001651 | 0.367785 | 2.782299 | 0.434406 |
| Prss54 | 0.717450 | 0.024930 | 0.514918 | 1.603286 | 0.288262 |
| Atmin | 0.714599 | 0.002335 | 0.385644 | 2.631713 | 0.413814 |
| Spink10 | 0.713502 | 0.014445 | 0.498513 | 1.840273 | 0.302323 |
| Slc24a4 | 0.713502 | 0.014938 | 0.504512 | 1.825715 | 0.297129 |
| Pradc1 | 0.712077 | 0.018941 | 0.508849 | 1.722605 | 0.293411 |
| Tmem8b | 0.711863 | 0.043212 | 0.574283 | 1.364394 | 0.240874 |
| Gsdma | 0.701514 | 0.026316 | 0.521440 | 1.579787 | 0.282796 |
| Srsf4 | 0.701121 | 0.038643 | 0.566918 | 1.412934 | 0.246480 |
| Pcdh15 | 0.700157 | 0.046332 | 0.583451 | 1.334118 | 0.233995 |
| Nmrk2 | 0.700157 | 0.023940 | 0.512726 | 1.620869 | 0.290115 |
| Nae1 | 0.698956 | 0.027996 | 0.527961 | 1.552897 | 0.277398 |
| Trip6 | 0.694784 | 0.007103 | 0.456158 | 2.148543 | 0.340885 |
| Hira | 0.691876 | 0.044144 | 0.574283 | 1.355133 | 0.240874 |
| Styx | 0.690450 | 0.041035 | 0.572249 | 1.386851 | 0.242415 |

| | | | | | |
|---|---|---|---|---|---|
| Gm52523 | -0.497440 | 0.000156 | 0.346132 | 3.807261 | 0.460759 |
| Sema6d | -0.494971 | 0.038576 | 0.566795 | 1.413682 | 0.246574 |
| Spcs2 | -0.490549 | 0.038880 | 0.569222 | 1.410272 | 0.244718 |
| Nsg2 | -0.488290 | 0.003156 | 0.402288 | 2.500829 | 0.395463 |
| Med11 | -0.486607 | 0.012702 | 0.486498 | 1.896121 | 0.312919 |
| Hmgn1 | -0.482102 | 0.034594 | 0.552250 | 1.461001 | 0.257864 |
| Rnf138 | -0.481060 | 0.042926 | 0.574283 | 1.367283 | 0.240874 |
| Eloa | -0.480580 | 0.011166 | 0.477528 | 1.952100 | 0.321001 |
| Cbfb | -0.480447 | 0.011850 | 0.477528 | 1.926287 | 0.321001 |
| Dysf | -0.479458 | 0.028687 | 0.528456 | 1.542308 | 0.276991 |
| Nup35 | -0.479020 | 0.032864 | 0.548023 | 1.483284 | 0.261201 |
| Kbtbd11 | -0.475365 | 0.030582 | 0.537364 | 1.514528 | 0.269731 |
| Mrto4 | -0.474968 | 0.005193 | 0.447186 | 2.284594 | 0.349512 |
| Gid8 | -0.474909 | 0.019231 | 0.508849 | 1.715993 | 0.293411 |
| Pfkfb2 | -0.471365 | 0.043871 | 0.574283 | 1.357823 | 0.240874 |
| Parp10 | -0.468750 | 0.003370 | 0.402288 | 2.472312 | 0.395463 |
| Dok4 | -0.466991 | 0.034699 | 0.552431 | 1.459684 | 0.257722 |
| Nes | -0.463328 | 0.006698 | 0.454055 | 2.174056 | 0.342892 |
| Tbpl1 | -0.457137 | 0.038292 | 0.566795 | 1.416894 | 0.246574 |
| Dcaf5 | -0.456949 | 0.027026 | 0.523236 | 1.568224 | 0.281303 |
| Fam57b | -0.455873 | 0.008588 | 0.459495 | 2.066107 | 0.337720 |
| Ndufs4 | -0.455675 | 0.044290 | 0.574283 | 1.353696 | 0.240874 |
| Kat6b | -0.453955 | 0.030099 | 0.537364 | 1.521454 | 0.269731 |
| Ptx4 | -0.453164 | 0.036742 | 0.563010 | 1.434834 | 0.249484 |
| Ppih | -0.452128 | 0.017249 | 0.508849 | 1.763245 | 0.293411 |
| Tigar | -0.446342 | 0.011974 | 0.477528 | 1.921772 | 0.321001 |
| Rab11b | -0.445141 | 0.014064 | 0.495749 | 1.851880 | 0.304738 |
| Grb2 | -0.444603 | 0.027425 | 0.525245 | 1.561851 | 0.279638 |
| Gm10406 | -0.442091 | 0.020031 | 0.508849 | 1.698299 | 0.293411 |
| Hmgb1 | -0.437332 | 0.022938 | 0.512726 | 1.639451 | 0.290115 |
| Gm37416 | -0.428972 | 0.000242 | 0.346132 | 3.616393 | 0.460759 |
| Ap3s1 | -0.426260 | 0.025519 | 0.518442 | 1.593135 | 0.285299 |
| Dhdds | -0.425397 | 0.022787 | 0.512726 | 1.642308 | 0.290115 |
| Naxe | -0.416223 | 0.021793 | 0.508849 | 1.661691 | 0.293411 |
| Sephs1 | -0.415507 | 0.049797 | 0.592038 | 1.302794 | 0.227650 |
| Akt1 | -0.412574 | 0.027063 | 0.523236 | 1.567626 | 0.281303 |
| Arl2bp | -0.411871 | 0.029058 | 0.529730 | 1.536740 | 0.275945 |
| Psmc1 | -0.409010 | 0.015991 | 0.508849 | 1.796112 | 0.293411 |
| Tecpr1 | -0.407967 | 0.042321 | 0.573809 | 1.373446 | 0.241233 |
| H4c8 | -0.404334 | 0.016538 | 0.508849 | 1.781510 | 0.293411 |
| Fis1 | -0.402908 | 0.046218 | 0.583451 | 1.335192 | 0.233995 |

| | | | | | |
|---|---|---|---|---|---|
| Ctns | 0.689411 | 0.036628 | 0.562486 | 1.436181 | 0.249888 |
| Krt80 | 0.685996 | 0.046559 | 0.583451 | 1.331995 | 0.233995 |
| Rimbp2 | 0.683145 | 0.026948 | 0.523236 | 1.569475 | 0.281303 |
| Myt1 | 0.680248 | 0.003738 | 0.408654 | 2.427368 | 0.388644 |
| Tmem101 | 0.679731 | 0.040592 | 0.571770 | 1.391559 | 0.242779 |
| 4930444G2 | 0.678916 | 0.021342 | 0.508849 | 1.670768 | 0.293411 |
| Zcchc7 | 0.675304 | 0.020492 | 0.508849 | 1.688412 | 0.293411 |
| Rufy1 | 0.675261 | 0.004788 | 0.443121 | 2.319837 | 0.353477 |
| Fez2 | 0.673671 | 0.011751 | 0.477528 | 1.929910 | 0.321001 |
| Lpl | 0.668703 | 0.038081 | 0.566795 | 1.419295 | 0.246574 |
| Iffo1 | 0.667277 | 0.019820 | 0.508849 | 1.702888 | 0.293411 |
| Eif4e | 0.663368 | 0.029411 | 0.533396 | 1.531494 | 0.272951 |
| St3gal4 | 0.661277 | 0.015758 | 0.508849 | 1.802502 | 0.293411 |
| Wdr3 | 0.660197 | 0.043987 | 0.574283 | 1.356676 | 0.240874 |
| Cenpv | 0.658760 | 0.017332 | 0.508849 | 1.761159 | 0.293411 |
| Trmt10b | 0.655639 | 0.006761 | 0.454055 | 2.170013 | 0.342892 |
| Gltpd2 | 0.654543 | 0.027567 | 0.525245 | 1.559613 | 0.279638 |
| Olfr854 | 0.653117 | 0.043445 | 0.574283 | 1.362059 | 0.240874 |
| Insm2 | 0.651342 | 0.012405 | 0.485483 | 1.906400 | 0.313826 |
| Lman1 | 0.649200 | 0.013199 | 0.490741 | 1.879445 | 0.309148 |
| Olfr1432 | 0.646851 | 0.028290 | 0.527961 | 1.548366 | 0.277398 |
| Galnt14 | 0.644330 | 0.015807 | 0.508849 | 1.801138 | 0.293411 |
| Bap1 | 0.642904 | 0.046368 | 0.583451 | 1.333785 | 0.233995 |
| Ift172 | 0.641197 | 0.027532 | 0.525245 | 1.560164 | 0.279638 |
| Ctbp2 | 0.639996 | 0.014608 | 0.498513 | 1.835411 | 0.302323 |
| Mgmt | 0.637542 | 0.014071 | 0.495749 | 1.851675 | 0.304738 |
| Cdh5 | 0.636739 | 0.005476 | 0.447186 | 2.261547 | 0.349512 |
| Pkn3 | 0.635824 | 0.028370 | 0.528084 | 1.547136 | 0.277297 |
| Rab11fip4 | 0.634451 | 0.037842 | 0.566795 | 1.422024 | 0.246574 |
| Pcmt1 | 0.631806 | 0.014564 | 0.498513 | 1.836726 | 0.302323 |
| Smarcal1 | 0.631674 | 0.016622 | 0.508849 | 1.779313 | 0.293411 |
| Slc39a9 | 0.630169 | 0.019579 | 0.508849 | 1.708200 | 0.293411 |
| Epb41l3 | 0.626661 | 0.047263 | 0.585976 | 1.325482 | 0.232120 |
| Crip3 | 0.622455 | 0.024555 | 0.513619 | 1.609855 | 0.289359 |
| Polr2g | 0.621997 | 0.025194 | 0.518442 | 1.598700 | 0.285299 |
| Klf6 | 0.620793 | 0.033027 | 0.548023 | 1.481132 | 0.261201 |
| Olfr202 | 0.616823 | 0.023735 | 0.512726 | 1.624602 | 0.290115 |
| Ece2 | 0.616823 | 0.045542 | 0.580182 | 1.341585 | 0.236435 |
| C1d | 0.615890 | 0.004196 | 0.422417 | 2.377187 | 0.374259 |
| Apol7c | 0.615330 | 0.031083 | 0.537739 | 1.507471 | 0.269429 |
| Ergic2 | 0.613450 | 0.009202 | 0.463691 | 2.036118 | 0.333772 |

| | | | | | |
|---|---|---|---|---|---|
| Nudc | -0.402345 | 0.003520 | 0.407603 | 2.453484 | 0.389763 |
| Rpl36 | -0.399382 | 0.001586 | 0.367785 | 2.799811 | 0.434406 |
| Armt1 | -0.399112 | 0.047660 | 0.587289 | 1.321848 | 0.231148 |
| Gm4724 | -0.398538 | 0.040072 | 0.570881 | 1.397158 | 0.243454 |
| Nnat | -0.398062 | 0.027271 | 0.525245 | 1.564299 | 0.279638 |
| Akr1a1 | -0.392395 | 0.009706 | 0.472983 | 2.012965 | 0.325154 |
| Fancd2 | -0.392032 | 0.046183 | 0.583451 | 1.335513 | 0.233995 |
| Sdc3 | -0.390761 | 0.047407 | 0.585976 | 1.324158 | 0.232120 |
| 1810010H2 | -0.390325 | 0.001777 | 0.367785 | 2.750432 | 0.434406 |
| Coa5 | -0.387840 | 0.020822 | 0.508849 | 1.681479 | 0.293411 |
| Sin3b | -0.387705 | 0.007967 | 0.459495 | 2.098720 | 0.337720 |
| Rnf220 | -0.387018 | 0.003017 | 0.402288 | 2.520405 | 0.395463 |
| Nrsn1 | -0.384513 | 0.008944 | 0.459495 | 2.048468 | 0.337720 |
| Pard6g | -0.382952 | 0.038594 | 0.566795 | 1.413479 | 0.246574 |
| Svbp | -0.374413 | 0.006917 | 0.454560 | 2.160097 | 0.342409 |
| Atp1b3 | -0.372771 | 0.040929 | 0.572249 | 1.387969 | 0.242415 |
| Gm52795 | -0.372141 | 0.022549 | 0.512726 | 1.646881 | 0.290115 |
| Rpl7 | -0.367007 | 0.001125 | 0.346132 | 2.948676 | 0.460759 |
| Cystm1 | -0.365486 | 0.023266 | 0.512726 | 1.633272 | 0.290115 |
| Snrnp200 | -0.360107 | 0.044541 | 0.574283 | 1.351242 | 0.240874 |
| Fgd6 | -0.359539 | 0.016418 | 0.508849 | 1.784675 | 0.293411 |
| Ago1 | -0.358421 | 0.046586 | 0.583451 | 1.331746 | 0.233995 |
| Plin3 | -0.358113 | 0.031606 | 0.541256 | 1.500224 | 0.266598 |
| Fam117b | -0.355638 | 0.016017 | 0.508849 | 1.795429 | 0.293411 |
| Taok2 | -0.354763 | 0.002766 | 0.399584 | 2.558210 | 0.398392 |
| Cox7b | -0.353046 | 0.014899 | 0.504512 | 1.826830 | 0.297129 |
| Srp14 | -0.350373 | 0.004276 | 0.422417 | 2.369004 | 0.374259 |
| Opalin | -0.349405 | 0.045198 | 0.578926 | 1.344882 | 0.237377 |
| Atrn | -0.346418 | 0.001116 | 0.346132 | 2.952480 | 0.460759 |
| Olfr382 | -0.345525 | 0.010553 | 0.477528 | 1.976636 | 0.321001 |
| Scrn1 | -0.341801 | 0.042611 | 0.574283 | 1.370477 | 0.240874 |
| Kif5b | -0.341595 | 0.037748 | 0.566795 | 1.423103 | 0.246574 |
| Matr3 | -0.336346 | 0.028326 | 0.527961 | 1.547810 | 0.277398 |
| Csnk1a1 | -0.335835 | 0.035164 | 0.555708 | 1.453896 | 0.255153 |
| Ehd4 | -0.333776 | 0.006474 | 0.454055 | 2.188840 | 0.342892 |
| Nedd4 | -0.326671 | 0.025759 | 0.518654 | 1.589067 | 0.285122 |
| Commd3 | -0.321952 | 0.024753 | 0.513619 | 1.606368 | 0.289359 |
| Snrpd3 | -0.320614 | 0.020803 | 0.508849 | 1.681872 | 0.293411 |
| Sumo2 | -0.319309 | 0.012663 | 0.486498 | 1.897463 | 0.312919 |
| Gm52800 | -0.318929 | 0.008165 | 0.459495 | 2.088024 | 0.337720 |
| Septin8 | -0.315280 | 0.027595 | 0.525245 | 1.559172 | 0.279638 |

| | | | | | |
|---|---|---|---|---|---|
| Ipp | 0.613157 | 0.003586 | 0.408554 | 2.445410 | 0.388751 |
| Glt8d2 | 0.612876 | 0.048002 | 0.587289 | 1.318738 | 0.231148 |
| Saxo2 | 0.612876 | 0.013923 | 0.495749 | 1.856265 | 0.304738 |
| Eif2ak3 | 0.610340 | 0.003229 | 0.402288 | 2.490920 | 0.395463 |
| Ggn | 0.604370 | 0.005044 | 0.444428 | 2.297199 | 0.352199 |
| Cep290 | 0.602663 | 0.016528 | 0.508849 | 1.781788 | 0.293411 |
| Elmod2 | 0.601635 | 0.047449 | 0.585976 | 1.323775 | 0.232120 |
| Neurod6 | 0.601495 | 0.029663 | 0.534804 | 1.527782 | 0.271805 |
| Pinlyp | 0.600888 | 0.018029 | 0.508849 | 1.744030 | 0.293411 |
| Pramef8 | 0.599008 | 0.023479 | 0.512726 | 1.629312 | 0.290115 |
| Kcnj6 | 0.598715 | 0.039641 | 0.570881 | 1.401856 | 0.243454 |
| Vamp7 | 0.596397 | 0.021110 | 0.508849 | 1.675505 | 0.293411 |
| Hapln2 | 0.595583 | 0.040879 | 0.572249 | 1.388504 | 0.242415 |
| Gigyf2 | 0.594436 | 0.009844 | 0.476580 | 2.006823 | 0.321864 |
| Sema7a | 0.593743 | 0.008494 | 0.459495 | 2.070909 | 0.337720 |
| Acta1 | 0.592450 | 0.011735 | 0.477528 | 1.930528 | 0.321001 |
| Ammecr1 | 0.591260 | 0.030618 | 0.537364 | 1.514029 | 0.269731 |
| Cdk12 | 0.586972 | 0.011500 | 0.477528 | 1.939289 | 0.321001 |
| Sec14l1 | 0.584825 | 0.006019 | 0.452298 | 2.220498 | 0.344575 |
| Get1 | 0.576478 | 0.037212 | 0.566132 | 1.429312 | 0.247083 |
| Tmem203 | 0.575157 | 0.048087 | 0.587289 | 1.317976 | 0.231148 |
| Rpp25l | 0.572237 | 0.043055 | 0.574283 | 1.365973 | 0.240874 |
| Id4 | 0.569784 | 0.013062 | 0.489076 | 1.883988 | 0.310624 |
| Dctn3 | 0.562585 | 0.001595 | 0.367785 | 2.797169 | 0.434406 |
| Mtr | 0.551008 | 0.010632 | 0.477528 | 1.973368 | 0.321001 |
| Phf23 | 0.547526 | 0.041333 | 0.572650 | 1.383705 | 0.242111 |
| Golm1 | 0.544277 | 0.026225 | 0.520379 | 1.581281 | 0.283680 |
| Ndufs8 | 0.544158 | 0.001230 | 0.362167 | 2.909930 | 0.441091 |
| Miip | 0.540570 | 0.029867 | 0.536762 | 1.524803 | 0.270218 |
| Rab27a | 0.540570 | 0.041833 | 0.572694 | 1.378478 | 0.242077 |
| Ccdc151 | 0.538330 | 0.015631 | 0.508849 | 1.806024 | 0.293411 |
| Ndufs3 | 0.536994 | 0.021424 | 0.508849 | 1.669100 | 0.293411 |
| Rpa1 | 0.536904 | 0.042091 | 0.573444 | 1.375811 | 0.241509 |
| Socs2 | 0.530272 | 0.020304 | 0.508849 | 1.692418 | 0.293411 |
| Sco2 | 0.529824 | 0.042618 | 0.574283 | 1.370407 | 0.240874 |
| Gja6 | 0.529543 | 0.021035 | 0.508849 | 1.677056 | 0.293411 |
| Fam104a | 0.528793 | 0.025219 | 0.518442 | 1.598267 | 0.285299 |
| Ppan | 0.524224 | 0.042018 | 0.573444 | 1.376566 | 0.241509 |
| Tmem208 | 0.517904 | 0.030511 | 0.537364 | 1.515540 | 0.269731 |
| Robo2 | 0.516520 | 0.009063 | 0.463691 | 2.042714 | 0.333772 |
| Cox17 | 0.513976 | 0.019322 | 0.508849 | 1.713952 | 0.293411 |

| | | | | | |
|---|---|---|---|---|---|
| St8sia4 | -0.314887 | 0.006812 | 0.454055 | 2.166742 | 0.342892 |
| Gm51591 | -0.313880 | 0.042187 | 0.573650 | 1.374818 | 0.241353 |
| Agbl5 | -0.310474 | 0.034126 | 0.550478 | 1.466915 | 0.259260 |
| Rtn1 | -0.309597 | 0.031146 | 0.537739 | 1.506600 | 0.269429 |
| Rpl37 | -0.301075 | 0.033566 | 0.550478 | 1.474097 | 0.259260 |
| Sacm1l | -0.296270 | 0.018316 | 0.508849 | 1.737167 | 0.293411 |
| Gnao1 | -0.295482 | 0.010686 | 0.477528 | 1.971170 | 0.321001 |
| Rpl18a | -0.291572 | 0.020347 | 0.508849 | 1.691502 | 0.293411 |
| 4930503L1 | -0.289724 | 0.021277 | 0.508849 | 1.672099 | 0.293411 |
| Rpl22 | -0.277075 | 0.013496 | 0.495208 | 1.869793 | 0.305213 |
| Hnrnpd | -0.275597 | 0.029361 | 0.533183 | 1.532224 | 0.273123 |
| Ybx1 | -0.275534 | 0.011998 | 0.477528 | 1.920902 | 0.321001 |
| Septin9 | -0.271637 | 0.000479 | 0.346132 | 3.319852 | 0.460759 |
| Prelid1 | -0.269500 | 0.026171 | 0.520042 | 1.582173 | 0.283962 |
| Tubb2b | -0.262395 | 0.043805 | 0.574283 | 1.358476 | 0.240874 |
| Srrm2 | -0.258433 | 0.029229 | 0.532148 | 1.534183 | 0.273968 |

| | | | | | |
|---|---|---|---|---|---|
| Rtf2 | 0.513748 | 0.028760 | 0.528456 | 1.541214 | 0.276991 |
| Vwa1 | 0.512521 | 0.035047 | 0.555332 | 1.455351 | 0.255447 |
| Slc45a1 | 0.512249 | 0.031173 | 0.537739 | 1.506226 | 0.269429 |
| Prr18 | 0.512249 | 0.020329 | 0.508849 | 1.691892 | 0.293411 |
| Fank1 | 0.511289 | 0.011814 | 0.477528 | 1.927617 | 0.321001 |
| Eml2 | 0.510810 | 0.023916 | 0.512726 | 1.621305 | 0.290115 |
| Atg16l1 | 0.509293 | 0.031508 | 0.540218 | 1.501584 | 0.267431 |
| Zbtb21 | 0.509117 | 0.044003 | 0.574283 | 1.356521 | 0.240874 |
| Akr1e1 | 0.509049 | 0.033892 | 0.550478 | 1.469906 | 0.259260 |
| Pak7 | 0.508742 | 0.008737 | 0.459495 | 2.058622 | 0.337720 |
| Rfx3 | 0.505743 | 0.023866 | 0.512726 | 1.622224 | 0.290115 |
| Stx12 | 0.505687 | 0.041672 | 0.572694 | 1.380152 | 0.242077 |
| Pcdhga8 | 0.501968 | 0.031903 | 0.541763 | 1.496167 | 0.266191 |
| Emc2 | 0.501797 | 0.042082 | 0.573444 | 1.375905 | 0.241509 |
| Stam2 | 0.497840 | 0.042984 | 0.574283 | 1.366698 | 0.240874 |
| Heatr1 | 0.493463 | 0.000385 | 0.346132 | 3.414828 | 0.460759 |
| Ache | 0.487876 | 0.039719 | 0.570881 | 1.401004 | 0.243454 |
| Adh5 | 0.487874 | 0.037163 | 0.566132 | 1.429894 | 0.247083 |
| Dmxl2 | 0.486945 | 0.030151 | 0.537364 | 1.520703 | 0.269731 |
| Fxr1 | 0.479062 | 0.022098 | 0.508849 | 1.655654 | 0.293411 |
| Hcfc1r1 | 0.477950 | 0.032977 | 0.548023 | 1.481782 | 0.261201 |
| Pate10 | 0.477663 | 0.006791 | 0.454055 | 2.168065 | 0.342892 |
| Olfr1277 | 0.477663 | 0.047334 | 0.585976 | 1.324825 | 0.232120 |
| Atxn7l3b | 0.476251 | 0.032537 | 0.548023 | 1.487627 | 0.261201 |
| Lima1 | 0.473104 | 0.023697 | 0.512726 | 1.625302 | 0.290115 |
| Zfp939 | 0.473037 | 0.024616 | 0.513619 | 1.608778 | 0.289359 |
| Rhob | 0.471897 | 0.010487 | 0.477528 | 1.979342 | 0.321001 |
| Ttc27 | 0.470583 | 0.033088 | 0.548023 | 1.480333 | 0.261201 |
| Mdh1 | 0.468034 | 0.030206 | 0.537364 | 1.519906 | 0.269731 |
| Zkscan6 | 0.463570 | 0.046446 | 0.583451 | 1.333055 | 0.233995 |
| Pou6f1 | 0.463060 | 0.013873 | 0.495749 | 1.857827 | 0.304738 |
| H4c1 | 0.454997 | 0.013635 | 0.495208 | 1.865344 | 0.305213 |
| Kras | 0.448826 | 0.047178 | 0.585976 | 1.326262 | 0.232120 |
| Pgrmc1 | 0.447020 | 0.037267 | 0.566132 | 1.428680 | 0.247083 |
| Ddah2 | 0.445343 | 0.028197 | 0.527961 | 1.549793 | 0.277398 |
| Otud1 | 0.444784 | 0.036394 | 0.562320 | 1.438970 | 0.250017 |
| Aimp1 | 0.444572 | 0.027623 | 0.525245 | 1.558730 | 0.279638 |
| Rarb | 0.443823 | 0.033009 | 0.548023 | 1.481374 | 0.261201 |
| Ssx2ip | 0.443358 | 0.047206 | 0.585976 | 1.326005 | 0.232120 |
| Ddx4 | 0.439944 | 0.027347 | 0.525245 | 1.563083 | 0.279638 |
| Arhgap35 | 0.434874 | 0.003283 | 0.402288 | 2.483719 | 0.395463 |

| | | | | | |
|---|---|---|---|---|---|
| Pgrmc2 | 0.434571 | 0.023074 | 0.512726 | 1.636886 | 0.290115 |
| Cep295nl | 0.428916 | 0.025971 | 0.518654 | 1.585507 | 0.285122 |
| Cox7c | 0.425458 | 0.025952 | 0.518654 | 1.585837 | 0.285122 |
| Ttc4 | 0.425261 | 0.049679 | 0.592038 | 1.303829 | 0.227650 |
| Map1lc3a | 0.421729 | 0.029923 | 0.536762 | 1.524001 | 0.270218 |
| Hdac2 | 0.413958 | 0.021383 | 0.508849 | 1.669925 | 0.293411 |
| Ppip5k2 | 0.408176 | 0.001958 | 0.368901 | 2.708110 | 0.433090 |
| Cct8 | 0.406864 | 0.023558 | 0.512726 | 1.627864 | 0.290115 |
| Phyhipl | 0.405103 | 0.011272 | 0.477528 | 1.948012 | 0.321001 |
| Daam1 | 0.404370 | 0.004851 | 0.443121 | 2.314159 | 0.353477 |
| Ppp1r10 | 0.399865 | 0.021948 | 0.508849 | 1.658607 | 0.293411 |
| Zfp704 | 0.399005 | 0.016140 | 0.508849 | 1.792090 | 0.293411 |
| Satb2 | 0.395461 | 0.041500 | 0.572650 | 1.381949 | 0.242111 |
| Zfp532 | 0.395239 | 0.041180 | 0.572249 | 1.385316 | 0.242415 |
| Scarf2 | 0.394330 | 0.028115 | 0.527961 | 1.551061 | 0.277398 |
| Xylt2 | 0.394330 | 0.040045 | 0.570881 | 1.397450 | 0.243454 |
| Pdss1 | 0.390664 | 0.043380 | 0.574283 | 1.362714 | 0.240874 |
| Wnt4 | 0.388956 | 0.033833 | 0.550478 | 1.470657 | 0.259260 |
| 1110065P2 | 0.386452 | 0.027364 | 0.525245 | 1.562815 | 0.279638 |
| Lsm7 | 0.383213 | 0.020005 | 0.508849 | 1.698856 | 0.293411 |
| Rnf7 | 0.383100 | 0.021151 | 0.508849 | 1.674676 | 0.293411 |
| Sf3b1 | 0.376204 | 0.007898 | 0.459495 | 2.102493 | 0.337720 |
| Gls | 0.375747 | 0.034491 | 0.551979 | 1.462288 | 0.258077 |
| Smc3 | 0.371114 | 0.005019 | 0.444428 | 2.299418 | 0.352199 |
| Rims1 | 0.367153 | 0.029485 | 0.533867 | 1.530404 | 0.272567 |
| Ski | 0.365677 | 0.017737 | 0.508849 | 1.751130 | 0.293411 |
| Elovl1 | 0.364907 | 0.044674 | 0.575184 | 1.349947 | 0.240193 |
| Wdr74 | 0.363623 | 0.007893 | 0.459495 | 2.102767 | 0.337720 |
| Cct5 | 0.362386 | 0.027558 | 0.525245 | 1.559752 | 0.279638 |
| Iqcm | 0.356610 | 0.002753 | 0.399584 | 2.560182 | 0.398392 |
| Pdia3 | 0.351123 | 0.045718 | 0.581899 | 1.339910 | 0.235153 |
| Pik3ap1 | 0.347290 | 0.037732 | 0.566795 | 1.423292 | 0.246574 |
| Ddx47 | 0.345061 | 0.021785 | 0.508849 | 1.661841 | 0.293411 |
| Pfdn5 | 0.339920 | 0.012100 | 0.477528 | 1.917198 | 0.321001 |
| Hars | 0.338341 | 0.049999 | 0.592038 | 1.301043 | 0.227650 |
| Msl2 | 0.336886 | 0.048764 | 0.590861 | 1.311897 | 0.228515 |
| Hspg2 | 0.333501 | 0.036922 | 0.564471 | 1.432717 | 0.248359 |
| Tmem18 | 0.331422 | 0.022047 | 0.508849 | 1.656651 | 0.293411 |
| Ift80 | 0.325157 | 0.025870 | 0.518654 | 1.587210 | 0.285122 |
| Fstl4 | 0.321209 | 0.015305 | 0.508849 | 1.815167 | 0.293411 |
| Ppfia1 | 0.316944 | 0.031793 | 0.541568 | 1.497672 | 0.266347 |

| | | | | | |
|---|---|---|---|---|---|
| Parp6 | 0.312850 | 0.045098 | 0.578698 | 1.345841 | 0.237548 |
| Fbxo45 | 0.312437 | 0.047709 | 0.587289 | 1.321398 | 0.231148 |
| Med17 | 0.311918 | 0.018094 | 0.508849 | 1.742463 | 0.293411 |
| Bdh1 | 0.307304 | 0.049877 | 0.592038 | 1.302102 | 0.227650 |
| Epm2aip1 | 0.298707 | 0.046625 | 0.583451 | 1.331382 | 0.233995 |
| Hsp90aa1 | 0.285317 | 0.037547 | 0.566795 | 1.425425 | 0.246574 |
| Mta1 | 0.283541 | 0.033622 | 0.550478 | 1.473379 | 0.259260 |
| Rbm15b | 0.279458 | 0.048406 | 0.589415 | 1.315096 | 0.229579 |
| Cbx2 | 0.277302 | 0.021578 | 0.508849 | 1.665990 | 0.293411 |
| Olfr328 | 0.272462 | 0.021165 | 0.508849 | 1.674373 | 0.293411 |
| Luc7l3 | 0.263221 | 0.019312 | 0.508849 | 1.714171 | 0.293411 |
| Sltm | 0.262540 | 0.033429 | 0.550478 | 1.475873 | 0.259260 |

**Supplementary Table 2: Differentially expressed genes between PAE and SAC in ECs at P28**

**Increased expression (1.2 FC, p<0.05)**

| Target nam | Log2 FC | Pvalue | Adjusted p | -log10 pval | -log10 adjusted pvalue |
|---|---|---|---|---|---|
| Rhod | 2.161809 | 0.004133 | 0.994385 | 2.383708 | 0.002445 |
| Mrpl19 | 2.101225 | 0.035937 | 0.994385 | 1.444462 | 0.002445 |
| Itpripl2 | 1.971704 | 0.008355 | 0.994385 | 2.078042 | 0.002445 |
| Ublcp1 | 1.966821 | 0.032714 | 0.994385 | 1.485268 | 0.002445 |
| 2610528J1 | 1.953323 | 0.031915 | 0.994385 | 1.496010 | 0.002445 |
| Cracr2b | 1.875201 | 0.029685 | 0.994385 | 1.527468 | 0.002445 |
| Gpatch1 | 1.818560 | 0.011808 | 0.994385 | 1.927818 | 0.002445 |
| Gm21062 | 1.736856 | 0.039678 | 0.994385 | 1.401449 | 0.002445 |
| Dbx1 | 1.699845 | 0.032140 | 0.994385 | 1.492961 | 0.002445 |
| Asb1 | 1.680214 | 0.015005 | 0.994385 | 1.823769 | 0.002445 |
| Col3a1 | 1.660742 | 0.005774 | 0.994385 | 2.238547 | 0.002445 |
| Tprn | 1.647035 | 0.038072 | 0.994385 | 1.419390 | 0.002445 |
| Wnt5a | 1.615999 | 0.010427 | 0.994385 | 1.981853 | 0.002445 |
| Pcdha10 | 1.592536 | 0.008257 | 0.994385 | 2.083176 | 0.002445 |
| Ctu1 | 1.592536 | 0.028307 | 0.994385 | 1.548101 | 0.002445 |
| Serpinf2 | 1.592536 | 0.036127 | 0.994385 | 1.442167 | 0.002445 |
| Asb10 | 1.592536 | 0.041027 | 0.994385 | 1.386934 | 0.002445 |
| Tk1 | 1.592536 | 0.008257 | 0.994385 | 2.083176 | 0.002445 |
| Fundc1 | 1.495142 | 0.023048 | 0.994385 | 1.637374 | 0.002445 |
| Tmco2 | 1.485226 | 0.026257 | 0.994385 | 1.580760 | 0.002445 |
| F2 | 1.485226 | 0.045553 | 0.994385 | 1.341486 | 0.002445 |
| Tbc1d4 | 1.458022 | 0.025535 | 0.994385 | 1.592863 | 0.002445 |
| Plaat3 | 1.454190 | 0.049131 | 0.994385 | 1.308640 | 0.002445 |
| Gm5464 | 1.421011 | 0.037774 | 0.994385 | 1.422805 | 0.002445 |
| Neil1 | 1.397548 | 0.005612 | 0.994385 | 2.250851 | 0.002445 |
| Islr2 | 1.383891 | 0.016504 | 0.994385 | 1.782403 | 0.002445 |
| Slfn5 | 1.380059 | 0.041474 | 0.994385 | 1.382227 | 0.002445 |
| Rgs9 | 1.366512 | 0.030551 | 0.994385 | 1.514971 | 0.002445 |
| Clint1 | 1.346881 | 0.039306 | 0.994385 | 1.405544 | 0.002445 |
| Xpo4 | 1.346881 | 0.018967 | 0.994385 | 1.722006 | 0.002445 |
| Trpm2 | 1.346881 | 0.022768 | 0.994385 | 1.642681 | 0.002445 |
| Mus81 | 1.346881 | 0.013919 | 0.994385 | 1.856388 | 0.002445 |
| Ttc4 | 1.290239 | 0.017564 | 0.994385 | 1.755388 | 0.002445 |
| Bend5 | 1.290239 | 0.028246 | 0.994385 | 1.549043 | 0.002445 |
| Polb | 1.281574 | 0.037619 | 0.994385 | 1.424596 | 0.002445 |
| Klk6 | 1.278834 | 0.030523 | 0.994385 | 1.515375 | 0.002445 |
| Cdhr2 | 1.259203 | 0.007281 | 0.994385 | 2.137814 | 0.002445 |

**Decreased expression (1.2 FC, p<0.05)**

| Target nam | Log2 FC | Pvalue | Adjusted p | -log10 pval | -log10 adjusted pvalue |
|---|---|---|---|---|---|
| S100a9 | -2.157860 | 0.010419 | 0.994385 | 1.982191 | 0.002445 |
| Chrna4 | -2.117225 | 0.002208 | 0.994385 | 2.655988 | 0.002445 |
| Tor1a | -2.117225 | 0.012335 | 0.994385 | 1.908854 | 0.002445 |
| Stk35 | -2.099736 | 0.003310 | 0.994385 | 2.480153 | 0.002445 |
| Olfr1138 | -2.086189 | 0.046574 | 0.994385 | 1.331859 | 0.002445 |
| Plet1 | -2.062726 | 0.009150 | 0.994385 | 2.038576 | 0.002445 |
| Zfp385c | -2.043094 | 0.025423 | 0.994385 | 1.594769 | 0.002445 |
| Klk5 | -2.012058 | 0.039986 | 0.994385 | 1.398088 | 0.002445 |
| Slc1a6 | -2.012058 | 0.023154 | 0.994385 | 1.635380 | 0.002445 |
| Clmn | -2.012058 | 0.003638 | 0.994385 | 2.439190 | 0.002445 |
| Pik3cg | -1.937036 | 0.005724 | 0.994385 | 2.242291 | 0.002445 |
| Mroh4 | -1.924380 | 0.028764 | 0.994385 | 1.541157 | 0.002445 |
| Atic | -1.919547 | 0.040683 | 0.994385 | 1.390588 | 0.002445 |
| Lrp3 | -1.919547 | 0.003303 | 0.994385 | 2.481150 | 0.002445 |
| Pgr | -1.904749 | 0.027556 | 0.994385 | 1.559782 | 0.002445 |
| Plcxd1 | -1.904749 | 0.002763 | 0.994385 | 2.558567 | 0.002445 |
| Olfr355 | -1.873712 | 0.048621 | 0.994385 | 1.313177 | 0.002445 |
| Ttc33 | -1.848107 | 0.043337 | 0.994385 | 1.363144 | 0.002445 |
| Sec14l5 | -1.848107 | 0.036050 | 0.994385 | 1.443091 | 0.002445 |
| Cilp2 | -1.840533 | 0.005930 | 0.994385 | 2.226946 | 0.002445 |
| Fdxacb1 | -1.817070 | 0.002129 | 0.994385 | 2.671816 | 0.002445 |
| Tbl1xr1 | -1.798690 | 0.049909 | 0.994385 | 1.301822 | 0.002445 |
| Rps6ka1 | -1.797439 | 0.041062 | 0.994385 | 1.386559 | 0.002445 |
| Naaladl2 | -1.783892 | 0.010702 | 0.994385 | 1.970542 | 0.002445 |
| Rab33a | -1.783892 | 0.013913 | 0.994385 | 1.856589 | 0.002445 |
| Zfp82 | -1.766403 | 0.010350 | 0.994385 | 1.985078 | 0.002445 |
| Trappc5 | -1.766403 | 0.038312 | 0.994385 | 1.416661 | 0.002445 |
| Hs3st3b1 | -1.766403 | 0.049600 | 0.994385 | 1.304521 | 0.002445 |
| Obsl1 | -1.766403 | 0.006954 | 0.994385 | 2.157750 | 0.002445 |
| Oprl1 | -1.760429 | 0.005362 | 0.994385 | 2.270644 | 0.002445 |
| Syt15 | -1.719677 | 0.041871 | 0.994385 | 1.378085 | 0.002445 |
| Znfx1 | -1.717217 | 0.005010 | 0.994385 | 2.300162 | 0.002445 |
| Dcaf12l1 | -1.709761 | 0.017506 | 0.994385 | 1.756803 | 0.002445 |
| Ptrhd1 | -1.709761 | 0.018762 | 0.994385 | 1.726727 | 0.002445 |
| Hdhd3 | -1.709761 | 0.044480 | 0.994385 | 1.351836 | 0.002445 |
| Ccdc177 | -1.678725 | 0.004987 | 0.994385 | 2.302186 | 0.002445 |
| Ces1h | -1.678725 | 0.017115 | 0.994385 | 1.766629 | 0.002445 |

| | | | | | |
|---|---|---|---|---|---|
| 4930402H2 | 1.249487 | 0.049003 | 0.994385 | 1.309778 | 0.002445 |
| Cln5 | 1.248395 | 0.008276 | 0.994385 | 2.082158 | 0.002445 |
| Anpep | 1.226024 | 0.014269 | 0.994385 | 1.845592 | 0.002445 |
| Rhcg | 1.202561 | 0.049835 | 0.994385 | 1.302465 | 0.002445 |
| Sh2b3 | 1.190386 | 0.007408 | 0.994385 | 2.130291 | 0.002445 |
| Tmcc1 | 1.185072 | 0.013653 | 0.994385 | 1.864783 | 0.002445 |
| Fam187a | 1.171524 | 0.042278 | 0.994385 | 1.373882 | 0.002445 |
| Msi1 | 1.151893 | 0.032736 | 0.994385 | 1.484975 | 0.002445 |
| Olfr65 | 1.151893 | 0.048805 | 0.994385 | 1.311532 | 0.002445 |
| Gm52215 | 1.151893 | 0.030742 | 0.994385 | 1.512261 | 0.002445 |
| Zmym4 | 1.147291 | 0.048462 | 0.994385 | 1.314599 | 0.002445 |
| Zfp7 | 1.138346 | 0.010900 | 0.994385 | 1.962588 | 0.002445 |
| Emc8 | 1.134404 | 0.015649 | 0.994385 | 1.805515 | 0.002445 |
| Trpc3 | 1.134404 | 0.016247 | 0.994385 | 1.789215 | 0.002445 |
| Snapc1 | 1.134404 | 0.046846 | 0.994385 | 1.329330 | 0.002445 |
| Irf5 | 1.120857 | 0.014061 | 0.994385 | 1.851979 | 0.002445 |
| Tmem181a | 1.118715 | 0.019096 | 0.994385 | 1.719065 | 0.002445 |
| Glmp | 1.081753 | 0.016085 | 0.994385 | 1.793567 | 0.002445 |
| Zfp367 | 1.077762 | 0.033184 | 0.994385 | 1.479078 | 0.002445 |
| Ccdc117 | 1.068047 | 0.048750 | 0.994385 | 1.312024 | 0.002445 |
| Nln | 1.064215 | 0.016786 | 0.994385 | 1.775057 | 0.002445 |
| Septin2 | 1.049186 | 0.041550 | 0.994385 | 1.381426 | 0.002445 |
| Map10 | 1.044584 | 0.031579 | 0.994385 | 1.500600 | 0.002445 |
| Zfp689 | 1.044584 | 0.031579 | 0.994385 | 1.500600 | 0.002445 |
| Plk4 | 1.033179 | 0.027736 | 0.994385 | 1.556960 | 0.002445 |
| Elovl7 | 1.031036 | 0.007475 | 0.994385 | 2.126388 | 0.002445 |
| Foxc1 | 1.027904 | 0.003495 | 0.994385 | 2.456532 | 0.002445 |
| Mtfr1l | 1.017489 | 0.037875 | 0.994385 | 1.421651 | 0.002445 |
| Ctag2l1 | 1.013547 | 0.044151 | 0.994385 | 1.355064 | 0.002445 |
| Cavin1 | 1.013547 | 0.040963 | 0.994385 | 1.387610 | 0.002445 |
| Ins1 | 1.013547 | 0.044277 | 0.994385 | 1.353824 | 0.002445 |
| Tmem59 | 1.007063 | 0.009736 | 0.994385 | 2.011615 | 0.002445 |
| Timd2 | 1.000000 | 0.045975 | 0.994385 | 1.337482 | 0.002445 |
| Rwdd3 | 0.980369 | 0.008685 | 0.994385 | 2.061215 | 0.002445 |
| 4930524J0 | 0.980369 | 0.034372 | 0.994385 | 1.463790 | 0.002445 |
| Adcy6 | 0.952072 | 0.047803 | 0.994385 | 1.320549 | 0.002445 |
| Nsmce3 | 0.946990 | 0.031942 | 0.994385 | 1.495637 | 0.002445 |
| Kctd11 | 0.945500 | 0.048617 | 0.994385 | 1.313210 | 0.002445 |
| Psme1 | 0.944730 | 0.041861 | 0.994385 | 1.378195 | 0.002445 |
| Dennd6b | 0.934534 | 0.042166 | 0.994385 | 1.375039 | 0.002445 |
| Agbl3 | 0.925869 | 0.044053 | 0.994385 | 1.356027 | 0.002445 |

| | | | | | |
|---|---|---|---|---|---|
| Slc7a6os | -1.667917 | 0.008223 | 0.994385 | 2.084972 | 0.002445 |
| Pex7 | -1.665177 | 0.043530 | 0.994385 | 1.361210 | 0.002445 |
| Tmem87a | -1.628057 | 0.037065 | 0.994385 | 1.431032 | 0.002445 |
| Exosc10 | -1.628057 | 0.002338 | 0.994385 | 2.631179 | 0.002445 |
| Ighmbp2 | -1.622083 | 0.034791 | 0.994385 | 1.458529 | 0.002445 |
| Papss1 | -1.622083 | 0.017218 | 0.994385 | 1.764026 | 0.002445 |
| Stum | -1.588904 | 0.009501 | 0.994385 | 2.022249 | 0.002445 |
| Atg3 | -1.577389 | 0.036465 | 0.994385 | 1.438122 | 0.002445 |
| C1rb | -1.571415 | 0.007789 | 0.994385 | 2.108508 | 0.002445 |
| Olfr1463 | -1.557868 | 0.004426 | 0.994385 | 2.353982 | 0.002445 |
| Tigd5 | -1.557868 | 0.045470 | 0.994385 | 1.342274 | 0.002445 |
| Sass6 | -1.540379 | 0.009050 | 0.994385 | 2.043363 | 0.002445 |
| Spsb3 | -1.526831 | 0.005307 | 0.994385 | 2.275123 | 0.002445 |
| Gm38666 | -1.524689 | 0.040673 | 0.994385 | 1.390691 | 0.002445 |
| Ankrd26 | -1.514774 | 0.021591 | 0.994385 | 1.665732 | 0.002445 |
| Col6a6 | -1.507200 | 0.018537 | 0.994385 | 1.731952 | 0.002445 |
| Nanos2 | -1.507200 | 0.027588 | 0.994385 | 1.559285 | 0.002445 |
| Erg28 | -1.507200 | 0.038797 | 0.994385 | 1.411205 | 0.002445 |
| Catsperg1 | -1.493653 | 0.003054 | 0.994385 | 2.515176 | 0.002445 |
| Cln6 | -1.483737 | 0.036855 | 0.994385 | 1.433501 | 0.002445 |
| Ago4 | -1.483737 | 0.027626 | 0.994385 | 1.558680 | 0.002445 |
| Zg16 | -1.483737 | 0.045237 | 0.994385 | 1.344509 | 0.002445 |
| Ddx21 | -1.483737 | 0.005140 | 0.994385 | 2.289034 | 0.002445 |
| F10 | -1.483737 | 0.011665 | 0.994385 | 1.933112 | 0.002445 |
| Osr2 | -1.483737 | 0.020881 | 0.994385 | 1.680246 | 0.002445 |
| Ppargc1a | -1.470190 | 0.033290 | 0.994385 | 1.477687 | 0.002445 |
| Resf1 | -1.456532 | 0.029287 | 0.994385 | 1.533329 | 0.002445 |
| Tbc1d30 | -1.450558 | 0.044053 | 0.994385 | 1.356028 | 0.002445 |
| Nr4a1 | -1.447426 | 0.015522 | 0.994385 | 1.809043 | 0.002445 |
| Zscan18 | -1.433069 | 0.015057 | 0.994385 | 1.822256 | 0.002445 |
| Ifrd2 | -1.433069 | 0.044052 | 0.994385 | 1.356034 | 0.002445 |
| Kat6b | -1.427095 | 0.014521 | 0.994385 | 1.838012 | 0.002445 |
| Zdhhc16 | -1.419522 | 0.007406 | 0.994385 | 2.130388 | 0.002445 |
| Ssmem1 | -1.419522 | 0.007406 | 0.994385 | 2.130388 | 0.002445 |
| Calb2 | -1.419522 | 0.025188 | 0.994385 | 1.598806 | 0.002445 |
| BC052040 | -1.419522 | 0.025188 | 0.994385 | 1.598806 | 0.002445 |
| Zbed5 | -1.419522 | 0.049478 | 0.994385 | 1.305585 | 0.002445 |
| Rgp1 | -1.414920 | 0.035757 | 0.994385 | 1.446637 | 0.002445 |
| Zkscan4 | -1.408715 | 0.027122 | 0.994385 | 1.566679 | 0.002445 |
| Sema4g | -1.408715 | 0.039752 | 0.994385 | 1.400639 | 0.002445 |
| Gal3st4 | -1.402033 | 0.017703 | 0.994385 | 1.751964 | 0.002445 |

| | | | | | |
|---|---|---|---|---|---|
| Cast | 0.923727 | 0.023479 | 0.994385 | 1.629317 | 0.002445 |
| Amotl1 | 0.916154 | 0.047519 | 0.994385 | 1.323137 | 0.002445 |
| Crim1 | 0.915062 | 0.034168 | 0.994385 | 1.466374 | 0.002445 |
| Sf3a3 | 0.915062 | 0.042683 | 0.994385 | 1.369744 | 0.002445 |
| Dynlt1a | 0.906238 | 0.038655 | 0.994385 | 1.412799 | 0.002445 |
| Tgfa | 0.906238 | 0.044269 | 0.994385 | 1.353902 | 0.002445 |
| Ddx46 | 0.902406 | 0.027619 | 0.994385 | 1.558789 | 0.002445 |
| Kcnq3 | 0.902175 | 0.024405 | 0.994385 | 1.612525 | 0.002445 |
| Tbc1d19 | 0.892691 | 0.017112 | 0.994385 | 1.766696 | 0.002445 |
| Rgl2 | 0.890604 | 0.006618 | 0.994385 | 2.179287 | 0.002445 |
| Arhgap31 | 0.885358 | 0.017021 | 0.994385 | 1.769027 | 0.002445 |
| Isyna1 | 0.881369 | 0.010386 | 0.994385 | 1.983542 | 0.002445 |
| Ptdss1 | 0.864394 | 0.039608 | 0.994385 | 1.402220 | 0.002445 |
| Rbm17 | 0.861654 | 0.025533 | 0.994385 | 1.592898 | 0.002445 |
| Spen | 0.860403 | 0.016092 | 0.994385 | 1.793392 | 0.002445 |
| Ssbp1 | 0.857052 | 0.047945 | 0.994385 | 1.319253 | 0.002445 |
| Tprkb | 0.851738 | 0.011416 | 0.994385 | 1.942477 | 0.002445 |
| Slc9a3r2 | 0.847719 | 0.026649 | 0.994385 | 1.574313 | 0.002445 |
| Tgm2 | 0.833358 | 0.009905 | 0.994385 | 2.004149 | 0.002445 |
| Emsy | 0.832107 | 0.029285 | 0.994385 | 1.533358 | 0.002445 |
| Dicer1 | 0.832107 | 0.037559 | 0.994385 | 1.425289 | 0.002445 |
| Mfge8 | 0.829495 | 0.010577 | 0.994385 | 1.975640 | 0.002445 |
| Srebf1 | 0.828476 | 0.048312 | 0.994385 | 1.315941 | 0.002445 |
| Casp6 | 0.818560 | 0.024337 | 0.994385 | 1.613729 | 0.002445 |
| Ndst2 | 0.818560 | 0.005674 | 0.994385 | 2.246122 | 0.002445 |
| Vmn1r113 | 0.816242 | 0.042770 | 0.994385 | 1.368858 | 0.002445 |
| A530032D1 | 0.815582 | 0.005970 | 0.994385 | 2.224044 | 0.002445 |
| Anxa7 | 0.811104 | 0.006301 | 0.994385 | 2.200624 | 0.002445 |
| Rela | 0.805012 | 0.008335 | 0.994385 | 2.079072 | 0.002445 |
| B4galt5 | 0.805012 | 0.010135 | 0.994385 | 1.994184 | 0.002445 |
| Mettl7a3 | 0.805012 | 0.027021 | 0.994385 | 1.568294 | 0.002445 |
| Mapre1 | 0.802553 | 0.025177 | 0.994385 | 1.599001 | 0.002445 |
| Ash1l | 0.800411 | 0.007615 | 0.994385 | 2.118338 | 0.002445 |
| Vps37a | 0.798929 | 0.034863 | 0.994385 | 1.457641 | 0.002445 |
| Anks1 | 0.798929 | 0.049289 | 0.994385 | 1.307248 | 0.002445 |
| Plin2 | 0.782690 | 0.047363 | 0.994385 | 1.324558 | 0.002445 |
| Cyp2d22 | 0.776716 | 0.003821 | 0.994385 | 2.417855 | 0.002445 |
| Rpap3 | 0.767892 | 0.036501 | 0.994385 | 1.437694 | 0.002445 |
| Rhobtb2 | 0.754345 | 0.047451 | 0.994385 | 1.323756 | 0.002445 |
| Grip2 | 0.754345 | 0.019386 | 0.994385 | 1.712506 | 0.002445 |
| Rccd1 | 0.754345 | 0.031793 | 0.994385 | 1.497662 | 0.002445 |

| | | | | | |
|---|---|---|---|---|---|
| Stard5 | -1.396059 | 0.032714 | 0.994385 | 1.485260 | 0.002445 |
| Faap100 | -1.396059 | 0.016168 | 0.994385 | 1.791353 | 0.002445 |
| Nhs | -1.391226 | 0.032934 | 0.994385 | 1.482360 | 0.002445 |
| Cblb | -1.386343 | 0.022883 | 0.994385 | 1.640478 | 0.002445 |
| Zkscan6 | -1.385252 | 0.041053 | 0.994385 | 1.386660 | 0.002445 |
| Cnih3 | -1.383884 | 0.016285 | 0.994385 | 1.788209 | 0.002445 |
| Zfp24 | -1.381742 | 0.002808 | 0.994385 | 2.551616 | 0.002445 |
| Thrsp | -1.377678 | 0.030752 | 0.994385 | 1.512121 | 0.002445 |
| Msln | -1.376428 | 0.021475 | 0.994385 | 1.668070 | 0.002445 |
| Brca2 | -1.345391 | 0.033240 | 0.994385 | 1.478336 | 0.002445 |
| Cel | -1.345391 | 0.035467 | 0.994385 | 1.450178 | 0.002445 |
| Olfr1362 | -1.331844 | 0.045184 | 0.994385 | 1.345019 | 0.002445 |
| Tob2 | -1.331844 | 0.049807 | 0.994385 | 1.302709 | 0.002445 |
| Fam20b | -1.325760 | 0.015203 | 0.994385 | 1.818084 | 0.002445 |
| E130311K1 | -1.312213 | 0.008607 | 0.994385 | 2.065161 | 0.002445 |
| Vmn1r89 | -1.312213 | 0.027139 | 0.994385 | 1.566404 | 0.002445 |
| Sema3f | -1.308861 | 0.027377 | 0.994385 | 1.562621 | 0.002445 |
| 2310002L0 | -1.298665 | 0.007980 | 0.994385 | 2.098022 | 0.002445 |
| Mavs | -1.294724 | 0.043860 | 0.994385 | 1.357930 | 0.002445 |
| Lcn10 | -1.288750 | 0.023845 | 0.994385 | 1.622600 | 0.002445 |
| D630045J1 | -1.288750 | 0.025533 | 0.994385 | 1.592898 | 0.002445 |
| Mbip | -1.263027 | 0.018839 | 0.994385 | 1.724935 | 0.002445 |
| H4c1 | -1.257713 | 0.001071 | 0.994385 | 2.970194 | 0.002445 |
| Acmsd | -1.255571 | 0.012020 | 0.994385 | 1.920103 | 0.002445 |
| Celf4 | -1.243396 | 0.044159 | 0.994385 | 1.354977 | 0.002445 |
| Klhl30 | -1.238082 | 0.009692 | 0.994385 | 2.013606 | 0.002445 |
| Kcna4 | -1.238082 | 0.029699 | 0.994385 | 1.527265 | 0.002445 |
| Pex26 | -1.224535 | 0.038728 | 0.994385 | 1.411979 | 0.002445 |
| Ssu2 | -1.224535 | 0.038728 | 0.994385 | 1.411979 | 0.002445 |
| Plekho2 | -1.212359 | 0.042503 | 0.994385 | 1.371579 | 0.002445 |
| Csk | -1.210594 | 0.008796 | 0.994385 | 2.055705 | 0.002445 |
| Josd1 | -1.210594 | 0.029314 | 0.994385 | 1.532932 | 0.002445 |
| Micall2 | -1.201071 | 0.024564 | 0.994385 | 1.609706 | 0.002445 |
| Ttll5 | -1.201071 | 0.028676 | 0.994385 | 1.542488 | 0.002445 |
| Eef2k | -1.196238 | 0.024925 | 0.994385 | 1.603369 | 0.002445 |
| Ercc5 | -1.191356 | 0.044854 | 0.994385 | 1.348203 | 0.002445 |
| Diaph3 | -1.181440 | 0.045789 | 0.994385 | 1.339235 | 0.002445 |
| Taf13 | -1.168751 | 0.031257 | 0.994385 | 1.505051 | 0.002445 |
| Abcd2 | -1.160319 | 0.040171 | 0.994385 | 1.396083 | 0.002445 |
| Wsb1 | -1.150404 | 0.033577 | 0.994385 | 1.473963 | 0.002445 |
| Trpm4 | -1.150404 | 0.019034 | 0.994385 | 1.720466 | 0.002445 |

| | | | | | |
|---|---|---|---|---|---|
| Kcnq1 | 0.754345 | 0.036762 | 0.994385 | 1.434596 | 0.002445 |
| Vim | 0.744655 | 0.026205 | 0.994385 | 1.581610 | 0.002445 |
| Hnrnpk | 0.739706 | 0.049367 | 0.994385 | 1.306560 | 0.002445 |
| Atf5 | 0.730882 | 0.017204 | 0.994385 | 1.764366 | 0.002445 |
| Il18 | 0.730882 | 0.041788 | 0.994385 | 1.378951 | 0.002445 |
| Eif4e | 0.730073 | 0.013040 | 0.994385 | 1.884735 | 0.002445 |
| Hyou1 | 0.722058 | 0.042811 | 0.994385 | 1.368444 | 0.002445 |
| Tln1 | 0.718706 | 0.028286 | 0.994385 | 1.548430 | 0.002445 |
| Arl6ip6 | 0.712501 | 0.021020 | 0.994385 | 1.677368 | 0.002445 |
| Elf2 | 0.711250 | 0.018641 | 0.994385 | 1.729522 | 0.002445 |
| Wwtr1 | 0.707419 | 0.006993 | 0.994385 | 2.155313 | 0.002445 |
| Tmem130 | 0.703908 | 0.043438 | 0.994385 | 1.362131 | 0.002445 |
| Pip4p2 | 0.697703 | 0.044471 | 0.994385 | 1.351925 | 0.002445 |
| Sec61g | 0.696380 | 0.026554 | 0.994385 | 1.575862 | 0.002445 |
| Lama5 | 0.695243 | 0.026048 | 0.994385 | 1.584224 | 0.002445 |
| H2aw | 0.690130 | 0.041311 | 0.994385 | 1.383937 | 0.002445 |
| Npdc1 | 0.688229 | 0.043558 | 0.994385 | 1.360936 | 0.002445 |
| Phkg2 | 0.680214 | 0.028293 | 0.994385 | 1.548316 | 0.002445 |
| Asb7 | 0.680214 | 0.028683 | 0.994385 | 1.542381 | 0.002445 |
| Rcc1 | 0.680214 | 0.000634 | 0.994385 | 3.197791 | 0.002445 |
| Smim12 | 0.680214 | 0.046064 | 0.994385 | 1.336636 | 0.002445 |
| Srsf4 | 0.672872 | 0.001764 | 0.994385 | 2.753472 | 0.002445 |
| Lamtor5 | 0.662065 | 0.038278 | 0.994385 | 1.417048 | 0.002445 |
| Armcx3 | 0.661353 | 0.017182 | 0.994385 | 1.764916 | 0.002445 |
| Pnpla6 | 0.657843 | 0.033341 | 0.994385 | 1.477022 | 0.002445 |
| Sult1a1 | 0.651869 | 0.016921 | 0.994385 | 1.771563 | 0.002445 |
| Rnf44 | 0.651060 | 0.042972 | 0.994385 | 1.366815 | 0.002445 |
| Gid8 | 0.647927 | 0.010961 | 0.994385 | 1.960165 | 0.002445 |
| Gpatch2 | 0.647035 | 0.037729 | 0.994385 | 1.423325 | 0.002445 |
| C130074G1 | 0.641722 | 0.027881 | 0.994385 | 1.554693 | 0.002445 |
| Taf10 | 0.640666 | 0.000447 | 0.994385 | 3.349727 | 0.002445 |
| Ptp4a3 | 0.633034 | 0.039971 | 0.994385 | 1.398253 | 0.002445 |
| Rps27 | 0.627772 | 0.049737 | 0.994385 | 1.303316 | 0.002445 |
| Kansl2 | 0.625337 | 0.032342 | 0.994385 | 1.490227 | 0.002445 |
| Fgf10 | 0.603941 | 0.001985 | 0.994385 | 2.702236 | 0.002445 |
| 9930021J0 | 0.600109 | 0.040002 | 0.994385 | 1.397913 | 0.002445 |
| Naca | 0.599595 | 0.014373 | 0.994385 | 1.842452 | 0.002445 |
| Gpr12 | 0.592536 | 0.036022 | 0.994385 | 1.443437 | 0.002445 |
| Prdm4 | 0.592536 | 0.033900 | 0.994385 | 1.469804 | 0.002445 |
| Zfp799 | 0.592536 | 0.035845 | 0.994385 | 1.445572 | 0.002445 |
| Oprm1 | 0.592536 | 0.033527 | 0.994385 | 1.474600 | 0.002445 |

| | | | | | |
|---|---|---|---|---|---|
| Nfkbid | -1.150404 | 0.023306 | 0.994385 | 1.632525 | 0.002445 |
| Olfr1350 | -1.150404 | 0.045949 | 0.994385 | 1.337723 | 0.002445 |
| Fv1 | -1.150404 | 0.045949 | 0.994385 | 1.337723 | 0.002445 |
| Vps13d | -1.139597 | 0.019215 | 0.994385 | 1.716366 | 0.002445 |
| Amacr | -1.139597 | 0.012959 | 0.994385 | 1.887412 | 0.002445 |
| Msantd3 | -1.136856 | 0.002656 | 0.994385 | 2.575765 | 0.002445 |
| Lrrc3 | -1.117225 | 0.018206 | 0.994385 | 1.739776 | 0.002445 |
| Ccdc121 | -1.117225 | 0.028521 | 0.994385 | 1.544839 | 0.002445 |
| Tbr1 | -1.113393 | 0.020536 | 0.994385 | 1.687480 | 0.002445 |
| Vps35 | -1.107588 | 0.007507 | 0.994385 | 2.124553 | 0.002445 |
| Junb | -1.100210 | 0.019121 | 0.994385 | 1.718483 | 0.002445 |
| Csf2ra | -1.099736 | 0.015609 | 0.994385 | 1.806634 | 0.002445 |
| Cdyl | -1.093762 | 0.005172 | 0.994385 | 2.286352 | 0.002445 |
| Ccbe1 | -1.086189 | 0.049537 | 0.994385 | 1.305068 | 0.002445 |
| Bbs7 | -1.086189 | 0.049537 | 0.994385 | 1.305068 | 0.002445 |
| Cngb1 | -1.086189 | 0.049537 | 0.994385 | 1.305068 | 0.002445 |
| Nol4l | -1.086189 | 0.042988 | 0.994385 | 1.366651 | 0.002445 |
| Tnks2 | -1.075382 | 0.032413 | 0.994385 | 1.489276 | 0.002445 |
| Perp | -1.053010 | 0.030551 | 0.994385 | 1.514971 | 0.002445 |
| Mex3d | -1.047085 | 0.007842 | 0.994385 | 2.105549 | 0.002445 |
| Aknad1 | -1.043094 | 0.030493 | 0.994385 | 1.515793 | 0.002445 |
| Nr2e3 | -1.043094 | 0.027068 | 0.994385 | 1.567542 | 0.002445 |
| 1700010B0 | -1.043094 | 0.044968 | 0.994385 | 1.347099 | 0.002445 |
| Hspbp1 | -1.035521 | 0.023614 | 0.994385 | 1.626827 | 0.002445 |
| Eepd1 | -1.029547 | 0.034983 | 0.994385 | 1.456140 | 0.002445 |
| Arhgef40 | -1.029547 | 0.037417 | 0.994385 | 1.426934 | 0.002445 |
| Paxip1 | -1.027252 | 0.026329 | 0.994385 | 1.579558 | 0.002445 |
| Ppm1e | -1.024714 | 0.006783 | 0.994385 | 2.168591 | 0.002445 |
| Pnpo | -1.012058 | 0.008688 | 0.994385 | 2.061060 | 0.002445 |
| Snrpa | -1.001251 | 0.004713 | 0.994385 | 2.326718 | 0.002445 |
| Atrx | -1.001251 | 0.024567 | 0.994385 | 1.609651 | 0.002445 |
| Tomm34 | -0.999883 | 0.022441 | 0.994385 | 1.648949 | 0.002445 |
| Srsx | -0.993909 | 0.040965 | 0.994385 | 1.387590 | 0.002445 |
| Mep1a | -0.986453 | 0.025854 | 0.994385 | 1.587467 | 0.002445 |
| Wdr75 | -0.978879 | 0.031825 | 0.994385 | 1.497233 | 0.002445 |
| Brd7 | -0.975047 | 0.011360 | 0.994385 | 1.944609 | 0.002445 |
| Zfp36 | -0.970214 | 0.009341 | 0.994385 | 2.029606 | 0.002445 |
| Exosc4 | -0.970214 | 0.049301 | 0.994385 | 1.307141 | 0.002445 |
| Gja4 | -0.968072 | 0.001269 | 0.994385 | 2.896680 | 0.002445 |
| Cdh2 | -0.964939 | 0.019797 | 0.994385 | 1.703391 | 0.002445 |
| Six3 | -0.955416 | 0.035747 | 0.994385 | 1.446761 | 0.002445 |

| | | | | | |
|---|---|---|---|---|---|
| Tspan3 | 0.587065 | 0.023411 | 0.994385 | 1.630585 | 0.002445 |
| Camta2 | 0.568914 | 0.004417 | 0.994385 | 2.354910 | 0.002445 |
| Lxn | 0.566699 | 0.041342 | 0.994385 | 1.383609 | 0.002445 |
| Slc25a36 | 0.559357 | 0.023962 | 0.994385 | 1.620475 | 0.002445 |
| Olfr972 | 0.549442 | 0.026768 | 0.994385 | 1.572392 | 0.002445 |
| Nhsl1 | 0.549442 | 0.042591 | 0.994385 | 1.370677 | 0.002445 |
| P2rx3 | 0.541868 | 0.021475 | 0.994385 | 1.668070 | 0.002445 |
| Slc22a8 | 0.540274 | 0.046666 | 0.994385 | 1.331001 | 0.002445 |
| Cog4 | 0.535894 | 0.030193 | 0.994385 | 1.520095 | 0.002445 |
| Clk1 | 0.529924 | 0.002832 | 0.994385 | 2.547893 | 0.002445 |
| Gm52416 | 0.526253 | 0.011690 | 0.994385 | 1.932202 | 0.002445 |
| Fryl | 0.518405 | 0.023309 | 0.994385 | 1.632468 | 0.002445 |
| Ddrgk1 | 0.518405 | 0.041932 | 0.994385 | 1.377455 | 0.002445 |
| Tmem222 | 0.518405 | 0.027085 | 0.994385 | 1.567270 | 0.002445 |
| Pde7b | 0.516263 | 0.032306 | 0.994385 | 1.490712 | 0.002445 |
| Ly6c1 | 0.511958 | 0.013804 | 0.994385 | 1.859990 | 0.002445 |
| Cdc42ep4 | 0.508921 | 0.039062 | 0.994385 | 1.408248 | 0.002445 |
| Babam1 | 0.504858 | 0.046452 | 0.994385 | 1.332995 | 0.002445 |
| Snapin | 0.502716 | 0.033373 | 0.994385 | 1.476609 | 0.002445 |
| Fbxl17 | 0.488162 | 0.023248 | 0.994385 | 1.633609 | 0.002445 |
| Timm50 | 0.485226 | 0.023270 | 0.994385 | 1.633212 | 0.002445 |
| Pds5a | 0.482073 | 0.023911 | 0.994385 | 1.621406 | 0.002445 |
| Ube2d2a | 0.479021 | 0.041343 | 0.994385 | 1.383595 | 0.002445 |
| Adipor1 | 0.477092 | 0.030641 | 0.994385 | 1.513693 | 0.002445 |
| Slc25a44 | 0.476561 | 0.024053 | 0.994385 | 1.618828 | 0.002445 |
| Srsf11 | 0.452048 | 0.019900 | 0.994385 | 1.701153 | 0.002445 |
| Zfp384 | 0.443224 | 0.048685 | 0.994385 | 1.312607 | 0.002445 |
| Mpnd | 0.438583 | 0.017864 | 0.994385 | 1.748017 | 0.002445 |
| Rpl27 | 0.434464 | 0.044237 | 0.994385 | 1.354215 | 0.002445 |
| Ctdnep1 | 0.432263 | 0.023449 | 0.994385 | 1.629877 | 0.002445 |
| Mt1 | 0.427279 | 0.048039 | 0.994385 | 1.318407 | 0.002445 |
| Sparcl1 | 0.427243 | 0.032001 | 0.994385 | 1.494831 | 0.002445 |
| Serpini1 | 0.426289 | 0.041014 | 0.994385 | 1.387072 | 0.002445 |
| Cst3 | 0.426096 | 0.035291 | 0.994385 | 1.452340 | 0.002445 |
| Syt11 | 0.424293 | 0.034884 | 0.994385 | 1.457370 | 0.002445 |
| Mcf2l | 0.420205 | 0.044618 | 0.994385 | 1.350493 | 0.002445 |
| Slc49a4 | 0.419920 | 0.023966 | 0.994385 | 1.620409 | 0.002445 |
| Srsf3 | 0.398519 | 0.041002 | 0.994385 | 1.387197 | 0.002445 |
| Ndfip2 | 0.387440 | 0.020873 | 0.994385 | 1.680425 | 0.002445 |
| Gm52800 | 0.385857 | 0.022035 | 0.994385 | 1.656883 | 0.002445 |
| Mfn2 | 0.383192 | 0.031195 | 0.994385 | 1.505920 | 0.002445 |

| | | | | | |
|---|---|---|---|---|---|
| Vps39 | -0.955416 | 0.035224 | 0.994385 | 1.453167 | 0.002445 |
| Pcdhb13 | -0.955416 | 0.038952 | 0.994385 | 1.409467 | 0.002445 |
| Mpv17l | -0.955416 | 0.038952 | 0.994385 | 1.409467 | 0.002445 |
| Gm7827 | -0.955416 | 0.038952 | 0.994385 | 1.409467 | 0.002445 |
| Tcf19 | -0.947843 | 0.033844 | 0.994385 | 1.470521 | 0.002445 |
| Klhl9 | -0.945500 | 0.023377 | 0.994385 | 1.631212 | 0.002445 |
| Mrps2 | -0.937036 | 0.040216 | 0.994385 | 1.395599 | 0.002445 |
| Taf9b | -0.937036 | 0.026042 | 0.994385 | 1.584317 | 0.002445 |
| Pdzd2 | -0.930749 | 0.032491 | 0.994385 | 1.488235 | 0.002445 |
| Tspyl3 | -0.929694 | 0.006191 | 0.994385 | 2.208205 | 0.002445 |
| Anxa11 | -0.927929 | 0.031335 | 0.994385 | 1.503973 | 0.002445 |
| Tmem126b | -0.922238 | 0.033896 | 0.994385 | 1.469852 | 0.002445 |
| Fam53a | -0.922238 | 0.045546 | 0.994385 | 1.341551 | 0.002445 |
| Chchd4 | -0.922238 | 0.028969 | 0.994385 | 1.538070 | 0.002445 |
| Trabd2b | -0.919547 | 0.038162 | 0.994385 | 1.418374 | 0.002445 |
| Sgk1 | -0.908077 | 0.032117 | 0.994385 | 1.493268 | 0.002445 |
| Fam160b2 | -0.904749 | 0.047325 | 0.994385 | 1.324911 | 0.002445 |
| Pdcd7 | -0.904749 | 0.046903 | 0.994385 | 1.328796 | 0.002445 |
| Car15 | -0.904749 | 0.041224 | 0.994385 | 1.384848 | 0.002445 |
| Spag11b | -0.904749 | 0.004592 | 0.994385 | 2.338029 | 0.002445 |
| Smim20 | -0.891201 | 0.032961 | 0.994385 | 1.481998 | 0.002445 |
| Hpf1 | -0.891201 | 0.015616 | 0.994385 | 1.806440 | 0.002445 |
| Slc5a11 | -0.882536 | 0.049877 | 0.994385 | 1.302102 | 0.002445 |
| Khk | -0.873712 | 0.011446 | 0.994385 | 1.941333 | 0.002445 |
| Nr2f1 | -0.867738 | 0.035305 | 0.994385 | 1.452163 | 0.002445 |
| Mmachc | -0.848107 | 0.015825 | 0.994385 | 1.800653 | 0.002445 |
| Atcay | -0.848107 | 0.012343 | 0.994385 | 1.908595 | 0.002445 |
| Sall2 | -0.834560 | 0.017648 | 0.994385 | 1.753297 | 0.002445 |
| Acbd4 | -0.834560 | 0.031793 | 0.994385 | 1.497662 | 0.002445 |
| Lrrc4 | -0.825124 | 0.033925 | 0.994385 | 1.469483 | 0.002445 |
| Arhgef2 | -0.809393 | 0.027484 | 0.994385 | 1.560920 | 0.002445 |
| Nudcd2 | -0.806263 | 0.024972 | 0.994385 | 1.602541 | 0.002445 |
| G2e3 | -0.801430 | 0.040271 | 0.994385 | 1.395003 | 0.002445 |
| Ap1s2 | -0.797439 | 0.030170 | 0.994385 | 1.520419 | 0.002445 |
| Grm2 | -0.791465 | 0.022110 | 0.994385 | 1.655416 | 0.002445 |
| Mrpl23 | -0.788774 | 0.010706 | 0.994385 | 1.970359 | 0.002445 |
| Klf16 | -0.783892 | 0.024721 | 0.994385 | 1.606926 | 0.002445 |
| Kirrel3 | -0.783892 | 0.046660 | 0.994385 | 1.331058 | 0.002445 |
| Egflam | -0.779290 | 0.026100 | 0.994385 | 1.583363 | 0.002445 |
| Rnf169 | -0.775227 | 0.043806 | 0.994385 | 1.358462 | 0.002445 |
| Sf3b3 | -0.766403 | 0.023215 | 0.994385 | 1.634235 | 0.002445 |

| | | | | | |
|---|---|---|---|---|---|
| Scarb2 | 0.370061 | 0.037417 | 0.994385 | 1.426934 | 0.002445 |
| Lpgat1 | 0.357607 | 0.040732 | 0.994385 | 1.390065 | 0.002445 |
| Rps15 | 0.354355 | 0.007798 | 0.994385 | 2.107999 | 0.002445 |
| Abcb1a | 0.351790 | 0.021113 | 0.994385 | 1.675451 | 0.002445 |
| Atg4b | 0.327249 | 0.007730 | 0.994385 | 2.111805 | 0.002445 |
| Esyt2 | 0.308203 | 0.014544 | 0.994385 | 1.837316 | 0.002445 |
| Pik3r2 | 0.292862 | 0.045625 | 0.994385 | 1.340798 | 0.002445 |
| 2210016L2 | 0.286610 | 0.022809 | 0.994385 | 1.641888 | 0.002445 |

| | | | | | |
|---|---|---|---|---|---|
| Apmap | -0.766403 | 0.048459 | 0.994385 | 1.314628 | 0.002445 |
| Zdhhc2 | -0.766403 | 0.048459 | 0.994385 | 1.314628 | 0.002445 |
| Krt83 | -0.766403 | 0.020255 | 0.994385 | 1.693468 | 0.002445 |
| Zfp975 | -0.765311 | 0.011079 | 0.994385 | 1.955494 | 0.002445 |
| Ap3m2 | -0.755596 | 0.049737 | 0.994385 | 1.303321 | 0.002445 |
| AI837181 | -0.750713 | 0.033917 | 0.994385 | 1.469583 | 0.002445 |
| Zdhhc21 | -0.742940 | 0.011349 | 0.994385 | 1.945030 | 0.002445 |
| Jmjd4 | -0.742048 | 0.042087 | 0.994385 | 1.375855 | 0.002445 |
| Lrrc49 | -0.733224 | 0.004533 | 0.994385 | 2.343641 | 0.002445 |
| Pcdhga7 | -0.719677 | 0.015033 | 0.994385 | 1.822961 | 0.002445 |
| Man2c1 | -0.718585 | 0.034798 | 0.994385 | 1.458445 | 0.002445 |
| Anapc7 | -0.709761 | 0.028631 | 0.994385 | 1.543164 | 0.002445 |
| Imp4 | -0.709761 | 0.042832 | 0.994385 | 1.368233 | 0.002445 |
| Pamr1 | -0.703787 | 0.014321 | 0.994385 | 1.844017 | 0.002445 |
| Faah | -0.701096 | 0.003539 | 0.994385 | 2.451096 | 0.002445 |
| Nav2 | -0.696214 | 0.043017 | 0.994385 | 1.366358 | 0.002445 |
| Hspe1 | -0.678725 | 0.045560 | 0.994385 | 1.341420 | 0.002445 |
| Efcab2 | -0.667917 | 0.026396 | 0.994385 | 1.578469 | 0.002445 |
| Ralgps2 | -0.663035 | 0.032153 | 0.994385 | 1.492779 | 0.002445 |
| Creb3 | -0.660575 | 0.045399 | 0.994385 | 1.342955 | 0.002445 |
| Olfr1415 | -0.653119 | 0.035792 | 0.994385 | 1.446214 | 0.002445 |
| Flywch1 | -0.648975 | 0.011391 | 0.994385 | 1.943451 | 0.002445 |
| Mrpl21 | -0.645546 | 0.012579 | 0.994385 | 1.900371 | 0.002445 |
| Jakmip3 | -0.645546 | 0.037365 | 0.994385 | 1.427532 | 0.002445 |
| Lsm7 | -0.645546 | 0.047746 | 0.994385 | 1.321059 | 0.002445 |
| Ddx51 | -0.636881 | 0.024709 | 0.994385 | 1.607151 | 0.002445 |
| Mrps23 | -0.635630 | 0.048234 | 0.994385 | 1.316647 | 0.002445 |
| Tm2d1 | -0.634739 | 0.004687 | 0.994385 | 2.329107 | 0.002445 |
| Gm13889 | -0.634739 | 0.046148 | 0.994385 | 1.335843 | 0.002445 |
| Neurl4 | -0.633088 | 0.022103 | 0.994385 | 1.655548 | 0.002445 |
| Mri1 | -0.622083 | 0.028591 | 0.994385 | 1.543775 | 0.002445 |
| Eif2d | -0.622083 | 0.000175 | 0.994385 | 3.757168 | 0.002445 |
| Katnb1 | -0.617250 | 0.017643 | 0.994385 | 1.753425 | 0.002445 |
| Atl1 | -0.612367 | 0.020547 | 0.994385 | 1.687253 | 0.002445 |
| Mboat7 | -0.604594 | 0.045208 | 0.994385 | 1.344789 | 0.002445 |
| Chrm3 | -0.602452 | 0.005676 | 0.994385 | 2.245980 | 0.002445 |
| Ddx19a | -0.588904 | 0.033520 | 0.994385 | 1.474691 | 0.002445 |
| Pacsin2 | -0.582420 | 0.034066 | 0.994385 | 1.467682 | 0.002445 |
| Camkk1 | -0.571415 | 0.032779 | 0.994385 | 1.484409 | 0.002445 |
| Chek1 | -0.565441 | 0.034079 | 0.994385 | 1.467515 | 0.002445 |
| Kif26a | -0.559240 | 0.010319 | 0.994385 | 1.986346 | 0.002445 |

| | | | | | |
|---|---|---|---|---|---|
| Pxmp4 | -0.540379 | 0.045597 | 0.994385 | 1.341062 | 0.002445 |
| Edil3 | -0.534405 | 0.043975 | 0.994385 | 1.356796 | 0.002445 |
| Lzts2 | -0.514774 | 0.018662 | 0.994385 | 1.729043 | 0.002445 |
| Cdr2 | -0.514774 | 0.048135 | 0.994385 | 1.317542 | 0.002445 |
| Anp32e | -0.505114 | 0.044511 | 0.994385 | 1.351537 | 0.002445 |
| Dppa2 | -0.483737 | 0.009670 | 0.994385 | 2.014568 | 0.002445 |
| Kat7 | -0.476559 | 0.035993 | 0.994385 | 1.443777 | 0.002445 |
| Meaf6 | -0.466644 | 0.028205 | 0.994385 | 1.549674 | 0.002445 |
| Ryr2 | -0.460755 | 0.022587 | 0.994385 | 1.646135 | 0.002445 |
| Tor1aip2 | -0.460755 | 0.034126 | 0.994385 | 1.466916 | 0.002445 |
| Spout1 | -0.445957 | 0.018246 | 0.994385 | 1.738824 | 0.002445 |
| Snd1 | -0.414920 | 0.044079 | 0.994385 | 1.355772 | 0.002445 |
| Ccndbp1 | -0.410318 | 0.021833 | 0.994385 | 1.660880 | 0.002445 |
| Rab11fip3 | -0.398799 | 0.019511 | 0.994385 | 1.709716 | 0.002445 |
| Grik2 | -0.376428 | 0.008660 | 0.994385 | 2.062457 | 0.002445 |
| Nans | -0.346103 | 0.025147 | 0.994385 | 1.599521 | 0.002445 |
| Zgpat | -0.343249 | 0.038418 | 0.994385 | 1.415467 | 0.002445 |
| Nudcd3 | -0.324869 | 0.017648 | 0.994385 | 1.753297 | 0.002445 |
| Flrt3 | -0.319786 | 0.034061 | 0.994385 | 1.467745 | 0.002445 |
| Plxnc1 | -0.288750 | 0.021475 | 0.994385 | 1.668070 | 0.002445 |
| Ces1g | -0.286607 | 0.048617 | 0.994385 | 1.313210 | 0.002445 |

**Supplementary Table 3: IPA Canonical Pathways of differentially expressed genes in ECs at E18**

| Ingenuity Canonical Pathways | -log(p-valı | Ratio | z-score | Molecules |
|---|---|---|---|---|
| Role of NFAT in Cardiac Hypertrophy | 3.090 | 0.114 | -0.688 | ADCY5,AKT1,CACNG2,CACNG7,Calm4,CAMK2D,CSNK1A1,GNAI1,GNB1L,GNG3,GNG7,GRB2,HDAC2,IL6ST,ITPR1,KRAS,PDIA3,PPP3CC,PRKACA,PRKD3,RAP1A,RCAN1,RRAS,SHC3,SLC8A2 |
| α-Adrenergic Signaling | 2.960 | 0.140 | -0.707 | ADCY5,Calm4,GNAI1,GNAO1,GNB1L,GNG3,GNG7,ITPR1,KRAS,PRKACA,PRKD3,PYGM,RAP1A,RRAS,SLC8A2 |
| CNTF Signaling | 2.940 | 0.179 | 0.000 | AKT1,GRB2,IL6ST,JAK1,KRAS,PTPN11,RAP1A,RPS6KB1,RRAS,TYK2 |
| GM-CSF Signaling | 2.920 | 0.167 | -0.632 | AKT1,BCL2L1,CAMK2D,CCND1,GRB2,KRAS,LYN,PPP3CC,PTPN11,RAP1A,RRAS |
| Complex IV assembly | 2.920 | 0.191 | 1.000 | COA5,COX17,COX6B1,COX6B2,COX7B,COX7C,RAB5IF,SCO2,TIMM21 |
| Signaling by CSF3 (G-CSF) | 2.910 | 0.233 | -0.378 | GRB2,JAK1,KRAS,LYN,PTPN11,RNF7,TYK2 |
| Adipogenesis pathway | 2.780 | 0.127 | 1.000 | AKT1,BMPR2,CTBP2,FGFR1,GTF2H1,Gtf2h2,HDAC2,KAT6B,KLF3,KLF5,LEP,LPL,NR2F2,RPS6KB1,SIN3A,SIN3B,SOX9 |
| Metabolism of cofactors | 2.620 | 0.240 | 0.000 | AKT1,GCH1,HSP90AA1,PDSS1,PRKG2,PTS |
| G Beta Gamma Signaling | 2.590 | 0.125 | -1.604 | AKT1,CACNG2,CACNG7,GNAI1,GNAO1,GNB1L,GNG3,GNG7,GRB2,ITPR1,KCNJ6,KRAS,PRKACA,PRKD3,RAP1A,RRAS |
| CCR3 Signaling in Eosinophils | 2.550 | 0.124 | 0.378 | Calm4,GNAI1,GNAO1,GNB1L,GNG3,GNG7,ITPR1,KRAS,PAK4,PLA2G12B,PLA2G4B,PPP1CC,PPP1R12A,PRKD3,RAP1A,RRAS |
| Serotonin Receptor Signaling | 2.550 | 0.090 | -0.324 | ADCY5,AKT1,ARFGEF1,BCL2,CACNG2,CACNG7,Calm4,CAMK2D,CD38,DDC,G6PC2,GCH1,GNAI1,GNAO1,GNB1L,GNG3,GNG7,HMOX1,HTR2A,KCNJ6,KRAS,LEP,LYN,PAK4,PDIA3,PLA2G12B,PLA2G4B,PLD4,PNPLA3,PPP1CC,PPP1R12A,PRKACA,PRKD3,PTS,RAB27A,RAP1A,RASGRF1,RHOB,RPS6KB1,RRAS |
| Sperm Motility | 2.510 | 0.103 | 0.000 | Calm4,CNGB1,ERBB4,FGFR1,FLT3LG,GNAI1,GNAO1,GNB1L,GNG3,GNG7,INSR,ITPR1,JAK1,LYN,NTRK2,PDIA3,PEAK1,PLA2G12B,PLA2G4B,PNPLA3,PRKACA,PRKD3,PRKG2,ROS1,TYK2 |
| GABAergic Receptor Signaling Pathway (Enhanced) | 2.420 | 0.120 | 2.000 | ADCY5,ALDH5A1,CACNG2,CACNG7,GABRB2,GABRG3,GABRP,GLS,GNAI1,GNAO1,GNB1L,GNG3,GNG7,ITPR1,KCNJ6,PRKACA |
| Role of NFAT in Regulation of the Immune Response | 2.390 | 0.109 | -1.732 | AKT1,Calm4,CD79B,CSNK1A1,Fcgr3,GNAI1,GNAO1,GNB1L,GNG3,GNG7,GRB2,ITPR1,KPNA2,KRAS,LYN,PPP3CC,RAP1A,RCAN1,RRAS,TNPO1 |
| Synthesis, secretion, and deacylation of Ghrelin | 2.340 | 0.250 | 1.342 | ACHE,LEP,PCSK1,SEC11A,SPCS2 |
| Interleukin-12 family signaling | 2.340 | 0.250 | 1.342 | EBI3,IL6ST,JAK1,TYK2,VAMP7 |
| Myelination Signaling Pathway | 2.330 | 0.095 | -0.962 | AKT1,ARHGAP35,BMP3,BMP6,BMPR2,Calm4,EIF4E,ERBB4,FGFR1,GNB1L,GRB2,HDAC2,ID4,ITGA6,Klk1b1 (includes others),KRAS,LYN,NTRK2,PPP3CC,PRKACA,PTPN11,RAB35,RAP1A,RPS6KB1,RRAS,SHH,SLC16A1,SOX2,WNT4,ZNF24 |
| Cytoprotection by HMOX1 | 2.330 | 0.158 | -0.333 | ABCC1,CHD9,COX6B1,COX6B2,COX7B,COX7C,HMOX1,SIN3A,SIN3B |
| Phospholipase C Signaling | 2.290 | 0.098 | 0.775 | ADCY5,ARHGEF4,Calm4,CD79B,GNB1L,GNG3,GNG7,GRB2,HDAC2,HMOX1,ITGA6,ITGA8,ITGAM,ITPR1,KRAS,LYN,PLA2G12B,PLA2G4B,PLD4,PPP1CC,PPP1R12A,PPP3CC,PRKD3,RAP1A,RHOB,RRAS |
| Interleukin-4 and Interleukin-13 signaling | 2.270 | 0.127 | -0.832 | AKT1,BCL2,BCL2L1,CCND1,COL1A2,HGF,HMOX1,HSP90AA1,IRF4,ITGAM,JAK1,SOX2,TYK2 |
| Mouse Embryonic Stem Cell Pluripotency | 2.270 | 0.127 | -0.832 | AKT1,BMPR2,GRB2,ID4,IL6ST,JAK1,KRAS,PTPN11,RAP1A,RRAS,SOX2,TAB1,TYK2 |
| STAT5 Activation | 2.260 | 0.429 | #NUM! | FLT3LG,GRB2,PTPN11 |
| Insulin Secretion Signaling Pathway | 2.190 | 0.096 | -0.426 | ADCY5,CACNG2,CACNG7,CAMK2D,CCK,EIF2AK3,EIF2B2,EIF4E,ELP5,INSR,ITPR1,JAK1,LYN,PAIP1,PCSK1,PDIA3,PRKACA,PRKD3,RAP1A,RPS6KB1,SEC11A,SPCS2,SRP14,STX16,TYK2,VPS37C |
| Mismatch Repair | 2.180 | 0.286 | 1.000 | MSH2,MSH3,RPA1,RPA2 |
| Oncostatin M Signaling | 2.160 | 0.175 | 0.378 | GRB2,IL6ST,JAK1,KRAS,RAP1A,RRAS,TYK2 |
| Oxytocin in Brain Signaling Pathway | 2.140 | 0.106 | -0.728 | AKT1,CACNG2,CACNG7,Calm4,GNAI1,GNAO1,GNB1L,GNG3,GNG7,ITPR1,KRAS,MAPK7,NLRP1,PLA2G12B,PLA2G4B,PNPLA3,PRKD3,RAP1A,RRAS |
| Signaling by CSF1 (M-CSF) in myeloid cells | 2.130 | 0.194 | 0.000 | GRB2,IL34,KRAS,LYN,PTPN11,THOC5 |
| Oxytocin Signaling Pathway | 2.110 | 0.095 | -0.816 | AKT1,CACNG2,CACNG7,Calm4,GNAI1,GNAO1,GNB1L,GNG3,GNG7,GRB2,ITPR1,KRAS,LEP,LPL,MAPK7,PLA2G12B,PLA2G4B,PPP1CC,PPP1R12A,PPP3CC,PRKACA,PRKD3,PRKG2,RAP1A,RRAS,SHC3 |
| Role of NANOG in Mammalian Embryonic Stem Cell Pluripotenc | 2.100 | 0.118 | -0.632 | AKT1,BMP3,BMP6,BMPR2,GRB2,IL6ST,JAK1,KRAS,RAP1A,RRAS,SALL4,SOX2,TYK2,WNT4 |
| Human Embryonic Stem Cell Pluripotency | 2.090 | 0.103 | -0.229 | AKT1,BMP3,BMP6,BMPR2,FGFR1,GRB2,HSP90AA1,ID4,JAK1,Klk1b1 (includes others),KRAS,NFYC,NTRK2,PRKD3,RAP1A,RRAS,SALL4,SOX2,TYK2,WNT4 |
| TP53 Regulates Metabolic Genes | 2.050 | 0.129 | -0.302 | AGO1,AKT1,COX6B1,COX6B2,COX7B,COX7C,GLS,RRAGC,SCO2,SFN,TIGAR |
| Estrogen Receptor Signaling | 2.040 | 0.087 | -0.600 | ADCY5,AKT1,BCL2,CACNG2,CACNG7,CCND1,CTBP2,EIF2B2,EIF4E,GNAI1,GNAO1,GNB1L,GNG3,GNG7,GRB2,HNRNPD,HSP90AA1,JAK1,KRAS,LEP,MED17,NDUFS3,NDUFS4,NDUFS8,PDIA3,PPP1CC,PPP1R12A,PRKACA,PRKD3,RAP1A,RPS6KB1,RRAS,SHC3,TYK2 |
| Tetrahydrobiopterin Biosynthesis I | 2.030 | 0.667 | #NUM! | GCH1,PTS |
| Tetrahydrobiopterin Biosynthesis II | 2.030 | 0.667 | #NUM! | GCH1,PTS |
| Regulation of RUNX1 Expression and Activity | 1.990 | 0.208 | -0.447 | AGO1,CBFB,CCND1,PML,PTPN11 |
| Role of JAK2 in Hormone-like Cytokine Signaling | 1.960 | 0.148 | 0.000 | BCL2,BCL2L1,CCND1,JAK1,LEP,PTPN11,SOCS2,TYK2 |
| PTEN Signaling | 1.950 | 0.107 | 0.832 | AKT1,BCL2,BCL2L1,BMPR2,CCND1,FGFR1,GRB2,INSR,ITGA6,ITGA8,ITGAM,KRAS,NTRK2,RAP1A,RPS6KB1,RRAS |
| Insulin Receptor Signaling | 1.950 | 0.113 | -0.277 | AKT1,EIF2B2,EIF4E,GRB2,INSR,JAK1,KRAS,PPP1CC,PPP1R10,PRKACA,PTPN11,RAP1A,RPS6KB1,RRAS |
| Glutaminergic Receptor Signaling Pathway (Enhanced) | 1.910 | 0.090 | -0.577 | ADCY5,AKT1,CACNG2,CACNG7,Calm4,CAMK2D,DGKE,EIF4E,GABRB2,GABRG3,GABRP,GLS,GNAI1,GNAO1,GNB1L,GRM4,HMOX1,ITPR1,PDIA3,PLA2G12B,PLA2G4B,PLD4,PNPLA3,PPP3CC,PRKACA,PRKD3,RPS6KB1,STX16 |
| Androgen Signaling | 1.900 | 0.104 | -1.342 | CACNG2,CACNG7,Calm4,CCND1,GNAI1,GNAO1,GNB1L,GNG3,GNG7,GTF2H1,Gtf2h2,HSP90AA1,ITPR1,POLR2G,PRKACA,PRKD3,TAF3 |
| Nuclear RNA decay | 1.870 | 0.171 | 0.816 | C1D,DIS3,PAPOLG,RBM26,TENT4B,ZCCHC7 |
| CLEAR Signaling Pathway | 1.870 | 0.092 | -1.800 | AKT1,ATP6V1C1,BMP6,BMPR2,CTNS,EIF2AK3,FGFR1,GNB1L,GUSB,HGF,HPS3,HPS5,INSR,ITPR1,KRAS,MAPK7,NTRK2,PML,PPP3CC,PRKD3,RAP1A,RRAGC,RRAS,SFN,WDR59 |
| Neuregulin Signaling | 1.850 | 0.113 | -1.000 | AKT1,ERBB4,GRB2,HSP90AA1,ITGA6,ITGA8,ITGAM,KRAS,PRKD3,PTPN11,RAP1A,RPS6KB1,RRAS |
| Docosahexaenoic Acid (DHA) Signaling | 1.840 | 0.094 | 0.000 | ADCY5,AKT1,BCL2,BCL2L1,BMP6,CCND1,EIF2AK3,FABP7,GNB1L,HGF,HMOX1,ITPR1,NTRK2,PDIA3,PLA2G12B,PLA2G4B,PNPLA3,PRKACA,PRKD3,RPS6KB1,STXBP6,SYT10 |
| IGF-1 Signaling | 1.840 | 0.117 | -1.265 | AKT1,GRB2,JAK1,KRAS,NEDD4,PRKACA,PTPN11,RAP1A,RPS6KB1,RRAS,SFN,SOCS2 |
| Warburg Effect Signaling Pathway | 1.840 | 0.109 | 0.277 | AKT1,BMP6,GLS,GNB1L,HGF,KEAP1,KRAS,MDH1,PFKFB2,PFKL,PHGDH,PRDX4,SLC16A1,TIGAR |
| EIF2 Signaling | 1.780 | 0.093 | -2.673 | ACTA1,AKT1,BCL2,CCND1,EIF2AK3,EIF2B2,EIF4E,ELP5,GRB2,INSR,KRAS,PAIP1,PPP1CC,RAP1A,RPL18A,RPL22,RPL36,RPL37,RPL7,RPL7L1,RRAS,VPS37C |
| Ketone body metabolism | 1.780 | 0.300 | #NUM! | BDH1,HMGCLL1,HMGCS2 |
| NoRC negatively regulates rRNA expression | 1.780 | 0.138 | -0.707 | DNMT1,GTF2H1,H4C1,HDAC2,POLR1C,SIN3A,SIN3B,TAF1A |
| Cilium Assembly | 1.770 | 0.097 | 0.229 | CCT5,CCT8,CEP290,CEP70,CNGB1,CPAP,DCTN3,HSP90AA1,IFT172,IFT80,IQCB1,KIF24,LZTFL1,MCHR1,PRKACA,SFI1,TNPO1,TRIP11,TTBK2 |
| Formation of the ureteric bud | 1.770 | 0.222 | #NUM! | ITGA8,NPNT,ROBO2,SALL1 |
| Synaptic Long Term Potentiation | 1.760 | 0.110 | -0.302 | Calm4,CAMK2D,GRM4,ITPR1,KRAS,PDIA3,PPP1CC,PPP1R10,PPP3CC,PRKACA,PRKD3,RAP1A,RRAS |
| JAK/STAT Signaling | 1.740 | 0.122 | -0.632 | AKT1,BCL2L1,GRB2,JAK1,KRAS,PTPN11,RAP1A,RRAS,SOCS2,TYK2 |
| rRNA modification in the nucleus and cytosol | 1.740 | 0.136 | 1.414 | DDX47,DIMT1,FCF1,HEATR1,NHP2,RRP9,UTP11,WDR3 |
| FLT3 Signaling | 1.710 | 0.158 | 0.000 | AKT1,FLT3LG,GRB2,KRAS,PTPN11,SOCS2 |
| Nicotinate metabolism | 1.700 | 0.179 | -0.447 | CD38,NAXE,NMRK2,PARP10,PARP6 |
| Hereditary Breast Cancer Signaling | 1.700 | 0.105 | -1.604 | AKT1,BMP6,CCND1,FANCD2,FGF13,HDAC2,HGF,KRAS,MSH2,POLR2G,RAP1A,RPA1,RRAS,SFN |
| Endocannabinoid Neuronal Synapse Pathway | 1.700 | 0.103 | -0.832 | ADCY5,CACNG2,CACNG7,GNAI1,GNAO1,GNB1L,GNG3,GNG7,ITPR1,KCNJ6,MAPK7,PDIA3,PPP3CC,PRKACA,RIMS1 |
| VEGF Signaling | 1.680 | 0.115 | 0.632 | ACTA1,AKT1,BCL2,BCL2L1,EIF2B2,GRB2,KRAS,PTPN11,RAP1A,RRAS,SFN |
| Ketogenesis | 1.660 | 0.273 | #NUM! | BDH1,HMGCLL1,HMGCS2 |
| GNRH Signaling | 1.650 | 0.096 | -0.277 | ADCY5,CACNG2,CACNG7,Calm4,CAMK2D,GNAI1,GNB1L,GNG3,GNG7,GRB2,ITPR1,KRAS,MAPK7,PAK4,PRKACA,PRKD3,RAP1A,RRAS |
| ERK5 Signaling | 1.640 | 0.123 | 1.000 | AKT1,IL6ST,KRAS,MAPK7,PTPN11,RAP1A,RPS6KB1,RRAS,SFN |
| Pancreatic Adenocarcinoma Signaling | 1.630 | 0.106 | -0.632 | AKT1,BCL2,BCL2L1,CCND1,CDKN2B,GRB2,HDAC2,HMOX1,JAK1,KRAS,PLD4,SIN3A,TYK2 |
| Thrombin Signaling | 1.620 | 0.092 | -0.535 | ADCY5,AKT1,ARHGEF4,CAMK2D,GNAI1,GNAO1,GNB1L,GNG3,GNG7,GRB2,ITPR1,KRAS,PDIA3,PPP1CC,PPP1R12A,PRKD3,RAP1A,RHOB,RPS6KB1,RRAS |
| FcγRIIB Signaling in B Lymphocytes | 1.610 | 0.116 | -0.378 | AKT1,CACNG2,CACNG7,CD79B,GRB2,ITPR1,KRAS,LYN,RAP1A,RRAS |
| RET signaling | 1.600 | 0.150 | -1.633 | DOK4,GRB2,MAPK7,PRKACA,PTPN11,SHC3 |
| STAT3 Pathway | 1.600 | 0.102 | -0.832 | BCL2,BMP6,BMPR2,FGFR1,HGF,IL6ST,INSR,KRAS,MAP3K20,NTRK2,RAP1A,RRAS,SOCS2,TYK2 |
| Cardiac Hypertrophy Signaling | 1.580 | 0.089 | 1.000 | ADCY5,AKT1,CACNG2,CACNG7,Calm4,EIF2B2,EIF4E,GNAI1,GNAO1,GNB1L,GNG3,GNG7,GRB2,KRAS,PDIA3,PPP3CC,PRKACA,RAP1A,RHOB,RPS6KB1,RRAS,TAB1 |
| Endothelin-1 Signaling | 1.580 | 0.096 | -0.728 | ADCY5,GNAI1,GNAO1,GRB2,HMOX1,ITPR1,KRAS,MAPK7,PDIA3,PLA2G12B,PLA2G4B,PLD4,PNPLA3,PRKD3,RAP1A,RRAS,SHC3 |
| Opioid Signaling Pathway | 1.570 | 0.087 | -1.091 | ADCY5,AKT1,CACNG2,CACNG7,Calm4,CAMK2D,GNAI1,GNAO1,GNB1L,GNG3,GNG7,ITPR1,KCNJ6,KRAS,LYN,MAPK7,PNOC,PPP3CC,PRKACA,PRKD3,RAP1A,RGS4,RPS6KB1,RRAS |
| Class I MHC mediated antigen processing and presentation | 1.560 | 0.083 | -0.365 | CUL7,CYBA,HERC6,HMGB1,KEAP1,KLHL20,KLHL22,LRR1,MYLIP,NEDD4,NEDD4L,PDIA3,PSMC1,PSMD13,RNF114,RNF138,RNF220,RNF7,S100A8,SH3RF1,TRAIP,TRIM11,TRIM32,TRIM9,UBE2N,UBE2Q1,UBE2R2,UBE2W,UBE3C,WSB1 |
| Endocannabinoid Developing Neuron Pathway | 1.550 | 0.103 | -0.302 | ADCY5,AKT1,CCND1,GNAI1,GNAO1,GNB1L,GNG3,GNG7,KRAS,MAPK7,PRKACA,RAP1A,RRAS |
| Creatine-phosphate Biosynthesis | 1.540 | 0.400 | #NUM! | CKB,CKMT2 |
| Superpathway of Serine and Glycine Biosynthesis I | 1.540 | 0.400 | #NUM! | PHGDH,SHMT2 |
| PI3K/AKT Signaling | 1.530 | 0.093 | 0.000 | AKT1,BCL2,BCL2L1,CCND1,EIF4E,GRB2,HSP90AA1,IL6ST,ITGA6,ITGA8,ITGAM,JAK1,KRAS,RAP1A,RPS6KB1,RRAS,SFN,TYK2 |
| Gαi Signaling | 1.510 | 0.099 | 1.265 | ADCY5,GNAI1,GNAO1,GNB1L,GNG3,GNG7,GRB2,GRM4,KRAS,PRKACA,RAP1A,RGS4,RRAS,SHC3 |
| NUR77 Signaling in T Lymphocytes | 1.510 | 0.117 | 0.707 | BCL2,Calm4,HDAC2,MAPK7,NR4A1,PPP3CC,PRKD3,SIN3A,SIN3B |
| Cell surface interactions at the vascular wall | 1.500 | 0.104 | -1.732 | ATP1B2,ATP1B3,COL1A2,GLG1,GRB2,ITGA6,ITGAM,KRAS,LYN,PTPN11,SDC3,SLC16A1 |
| Actin Nucleation by ARP-WASP Complex | 1.490 | 0.111 | 1.342 | GRB2,ITGA6,ITGA8,ITGAM,KRAS,PPP1R12A,RAP1A,RHOB,RRAS,WIPF1 |
| Dopamine-DARPP32 Feedback in cAMP Signaling | 1.490 | 0.095 | -0.632 | ADCY5,ATF1,CACNG2,CACNG7,Calm4,CSNK1A1,GNAI1,ITPR1,KCNJ6,PDIA3,PPP1CC,PPP1R10,PPP3CC,PRKACA,PRKD3,PRKG2 |
| Colorectal Cancer Metastasis Signaling | 1.480 | 0.087 | 0.500 | ADCY5,AKT1,BCL2L1,CCND1,DCC,GNAI1,GNAO1,GNB1L,GNG3,GNG7,GRB2,IL6ST,JAK1,KRAS,MSH2,MSH3,PRKACA,RAP1A,RHOB,RRAS,TYK2,WNT4 |
| Folate Signaling Pathway | 1.480 | 0.130 | 0.378 | BCL2,IL6ST,JAK1,MTR,SHMT2,SOX2,TYK2 |
| Dopamine Receptor Signaling | 1.470 | 0.121 | 0.447 | ADCY5,DDC,GCH1,NCS1,PPP1CC,PPP1R10,PRKACA,PTS |
| Other interleukin signaling | 1.470 | 0.182 | 1.000 | IL34,JAK1,TXLNA,TYK2 |
| Semaphorin Neuronal Repulsive Signaling Pathway | 1.460 | 0.098 | 0.905 | AKT1,DPYSL4,ITGA6,ITGA8,ITGAM,PAK4,PPP1CC,PPP1R12A,PRKACA,PRKG2,RRAS,SEMA6D,SMC3,VCAN |
| POU5F1 (OCT4), SOX2, NANOG activate genes related to prolife | 1.460 | 0.231 | #NUM! | SALL1,SALL4,SOX2 |
| GABA Receptor Signaling | 1.460 | 0.100 | #NUM! | ADCY5,ALDH5A1,CACNG2,CACNG7,GABRB2,GABRG3,GABRP,GNAI1,GNAO1,GNB1L,GNG3,GNG7,ITPR1 |
| IL-15 Signaling | 1.450 | 0.114 | 1.000 | AKT1,BCL2L1,GNB1L,JAK1,KRAS,RAP1A,RPS6KB1,RRAS,TYK2 |
| Systemic Lupus Erythematosus in B Cell Signaling Pathway | 1.440 | 0.087 | 0.447 | AKT1,BCL2,BCL2L1,Calm4,CCND1,CD79B,GRB2,IFIT2,IL6ST,JAK1,KRAS,LEP,LYN,PIK3AP1,PPP3CC,PRKD3,PTPN11,RAP1A,RRAS,SHC3,TYK2 |
| Acetylcholine Receptor Signaling Pathway | 1.440 | 0.092 | -2.000 | ACHE,ADCY5,AKT1,BCL2,CACNG2,CACNG7,Calm4,CAMK2D,CHRND,GNAI1,GNAO1,HMOX1,ITPR1,KEAP1,PDIA3,PRKACA,PRKD3 |
| High laminar flow shear stress activates signaling by PIEZO1 and | 1.410 | 0.118 | -0.707 | ADCY5,ADM,AKT1,CDH5,GNG3,GNG7,PRKACA,RAMP2 |
| RAF-independent MAPK1/3 activation | 1.410 | 0.174 | 0.000 | IL6ST,JAK1,PTPN11,TYK2 |
| RAS processing | 1.410 | 0.174 | 2.000 | BCL2L1,KRAS,PRKG2,RCE1 |
| Circadian Rhythm Signaling | 1.400 | 0.085 | #NUM! | ADCY5,BHLHE40,CACNG2,CACNG7,CAMK2D,EIF4E,GNAI1,GNB1L,GNG3,GNG7,ITPR1,Klk1b1 (includes others),KRAS,LYN,NTRK2,PDIA3,PRKACA,PRKD3,PRKG2,RAP1A,RORB,RRAS |
| p70S6K Signaling | 1.390 | 0.098 | -0.577 | AKT1,CD79B,GNAI1,GRB2,JAK1,KRAS,LYN,PDIA3,PRKD3,RAP1A,RPS6KB1,RRAS,SFN |
| Signaling by NTRK1 (TRKA) | 1.390 | 0.111 | -0.333 | ATF1,DNAL4,GRB2,KRAS,MAPK7,NTRK2,RAP1A,RIT2,SHC3 |
| FLT3 Signaling in Hematopoietic Progenitor Cells | 1.390 | 0.111 | 0.333 | AKT1,EIF4E,FLT3LG,GRB2,KRAS,PTPN11,RAP1A,RPS6KB1,RRAS |
| IL-8 Signaling | 1.390 | 0.089 | 0.258 | AKT1,BCL2,BCL2L1,CCND1,GNAI1,GNAO1,GNB1L,GNG3,GNG7,HMOX1,ITGAM,KRAS,PLD4,PRKD3,RAP1A,RHOB,RPS6KB1,RRAS |
| Non-Small Cell Lung Cancer Signaling | 1.380 | 0.106 | -1.134 | AKT1,CCND1,GRB2,HDAC2,ITPR1,KRAS,RAP1A,RARB,RRAS,SIN3A |

| | | | | |
|---|---|---|---|---|
| Nucleotide Excision Repair Pathway | 1.370 | 0.147 | #NUM! | GTF2H1,Gtf2h2,POLR2G,RPA1,RPA2 |
| Melatonin Signaling | 1.370 | 0.116 | -1.134 | Calm4,CAMK2D,GNAI1,GNAO1,PDIA3,PRKACA,PRKD3,RORB |
| Protein Sorting Signaling Pathway | 1.370 | 0.091 | 0.000 | ADCY5,AP1G2,AP3S1,ARFGEF1,HMOX1,KDELR2,LMAN1,LYN,PLD4,PLPP2,PRKACA,RAB11B,SACM1L,TMED3,TMED7,WASHC2C |
| CXCR4 Signaling | 1.350 | 0.093 | -1.155 | ADCY5,AKT1,GNAI1,GNAO1,GNB1L,GNG3,GNG7,ITPR1,KRAS,LYN,PAK4,PRKD3,RAP1A,RHOB,RRAS |
| MicroRNA Biogenesis Signaling Pathway | 1.350 | 0.091 | 1.500 | AGO1,AKT1,BMP6,DBR1,DIS3,HGF,HMOX1,HNRNPD,HSP90AA1,KRAS,NUP35,PIWIL1,POLR2G,RAP1A,RPS6KB1,RRAS |
| Incretin synthesis, secretion, and inactivation | 1.350 | 0.167 | 1.000 | LEP,PCSK1,SEC11A,SPCS2 |
| Interleukin-6 family signaling | 1.350 | 0.167 | 0.000 | IL6ST,JAK1,PTPN11,TYK2 |
| M-decay: degradation of maternal mRNAs by maternally stored | 1.340 | 0.130 | -1.633 | CNOT7,EIF4E,KDM5A,KPNA2,PAIP1,PAIP2 |
| Signaling by ERBB4 | 1.330 | 0.121 | -0.378 | ERBB4,GABRB2,GABRG3,GRB2,KRAS,MXD4,NEDD4 |
| Cilia Biogenesis Signaling Pathway | 1.330 | 0.089 | 0.728 | BMP6,Calm4,CEP120,CEP290,CPAP,EHD1,GNAI1,GNAO1,GNB1L,GNG3,GNG7,HGF,KIF24,SFN,SSX2IP,TTBK2,TXLNA |
| Major pathway of rRNA processing in the nucleolus and cytosol | 1.320 | 0.087 | 0.471 | C1D,DDX47,DIS3,FCF1,FTSJ3,HEATR1,NOB1,PES1,RPL18A,RPL22,RPL36,RPL37,RPL7,RPL7L1,RRP9,TEX10,UTP11,WDR3 |
| Axonal Guidance Signaling | 1.310 | 0.076 | #NUM! | ADAM12,ADAM29,ADAMTS15,AKT1,BMP3,BMP6,DCC,EIF4E,GNAI1,GNAO1,GNB1L,GNG3,GNG7,GRB2,ITGA6,ITGA8,ITGAM,KRAS,MYSM1,NTN5,NTRK2,PAK4,PDIA3,PPP3CC,PRKACA,PRKD3,PTPN11,RAP1A,ROBO1,ROBO2,RRAS,SEMA6D,SEMA7A,SHH,WIPF1,WNT4 |
| Pancreatic Secretion Signaling Pathway | 1.300 | 0.086 | 0.000 | ADCY5,AKT1,ARHGEF4,CCK,CD38,GNB1L,ITPR1,LPL,PAK4,PDIA3,PPP1CC,PPP1R10,PRKACA,PRKD3,RAP1A,RPS6KB1,SCT,STX16,STXBP6 |
| GABA receptor activation | 1.300 | 0.119 | 1.134 | ADCY5,GABRB2,GABRG3,GNAI1,GNG3,GNG7,KCNJ6 |
| Post-translational protein phosphorylation | 1.300 | 0.103 | 1.897 | CALU,CP,FAM20A,FSTL1,GOLM1,MATN3,MFGE8,VCAN,VWA1,WFS1 |
| Gαq Signaling | 1.290 | 0.091 | -0.632 | AKT1,Calm4,GNAI1,GNAO1,GNB1L,GNG3,GNG7,HMOX1,HTR2A,ITPR1,PLD4,PPP3CC,PRKD3,RGS4,RHOB |
| Cardiac β-adrenergic Signaling | 1.290 | 0.091 | -0.816 | ADCY5,CACNG2,CACNG7,GNAI1,GNAO1,GNB1L,GNG3,GNG7,PDE7B,PKIB,PPP1CC,PPP1R10,PRKACA,SLC8A2,TDP2 |
| IL-4 Signaling | 1.290 | 0.091 | 0.258 | AKT1,BCL2L1,COL13A1,COL19A1,COL1A2,COL6A2,GNAI1,GNB1L,GRB2,IRF4,JAK1,KRAS,RAP1A,RPS6KB1,RRAS |
| Glioma Signaling | 1.290 | 0.097 | -0.378 | AKT1,Calm4,CAMK2D,CCND1,CDKN2B,GRB2,HDAC2,KRAS,PRKD3,RAP1A,RRAS,SIN3A |
| Insulin processing | 1.290 | 0.160 | 1.000 | KIF5B,PCSK1,RAB27A,SLC30A8 |
| Signaling by NTRK2 (TRKB) | 1.290 | 0.160 | -1.000 | GRB2,KRAS,NTRK2,PTPN11 |
| Phospholipases | 1.270 | 0.117 | 0.816 | HMOX1,LPL,PDIA3,PLA2G12B,PLA2G4B,PLD4,PNPLA3 |
| Systemic Lupus Erythematosus Signaling | 1.260 | 0.090 | #NUM! | AKT1,C5,CD79B,Fcgr3,GRB2,KRAS,LSM7,LYN,PPIH,PRPF38A,PRPF8,RAP1A,RRAS,SNRNP200,SNRPD3 |
| Extra-nuclear estrogen signaling | 1.250 | 0.110 | -0.707 | AKT1,BCL2,CCND1,GNAI1,GNG3,GNG7,HSP90AA1,KRAS |
| BMP signaling pathway | 1.250 | 0.105 | 0.333 | BMP3,BMP6,BMPR2,GRB2,KRAS,PRKACA,RAP1A,RRAS,TAB1 |
| Glycoaminoglycan-protein Linkage Region Biosynthesis | 1.250 | 0.286 | #NUM! | B3GAT2,XYLT2 |
| Lanosterol Biosynthesis | 1.240 | 1.000 | #NUM! | LSS |
| PI3K Signaling in B Lymphocytes | 1.240 | 0.093 | 0.000 | AKT1,ATF1,Calm4,CAMK2D,CD79B,ITPR1,KRAS,LYN,PDIA3,PIK3AP1,PPP3CC,RAP1A,RRAS |
| Apelin Endothelial Signaling Pathway | 1.240 | 0.093 | -1.000 | ADCY5,AKT1,Calm4,GNAI1,GNAO1,GNB1L,GNG3,GNG7,KRAS,PRKD3,RAP1A,RPS6KB1,RRAS |
| Ethanol Degradation II | 1.240 | 0.154 | #NUM! | ACSS2,ADH5,AKR1A1,Aldh3a2 |
| Sleep NREM Signaling Pathway | 1.240 | 0.097 | 0.302 | ADCY5,CAMK2D,GABRB2,GABRG3,GABRP,GNAI1,KRAS,PRKACA,RAP1A,RRAS,SFN |
| IL-2 Signaling | 1.240 | 0.115 | -1.134 | AKT1,GRB2,JAK1,KRAS,PTPN11,RAP1A,RRAS |
| Apelin Muscle Signaling Pathway | 1.230 | 0.122 | #NUM! | AKT1,GNAI1,GNAO1,GNB1L,GNG3,GNG7 |
| Opioid Signalling | 1.200 | 0.102 | -1.000 | ADCY5,CAMK2D,GNAI1,GNG3,GNG7,ITPR1,KPNA2,PPP3CC,PRKACA |
| Regulation of Cellular Mechanics by Calpain Protease | 1.200 | 0.102 | 0.447 | CCND1,CNGB1,GRB2,ITGA6,ITGA8,ITGAM,KRAS,RAP1A,RRAS |
| Neuroinflammation Signaling Pathway | 1.200 | 0.081 | 0.000 | AKT1,BCL2,BMPR2,CD200R1,CX3CR1,GABRB2,GABRG3,GABRP,GLS,HMGB1,HMOX1,IDE,IL34,JAK1,KCNJ6,Klk1b1 (includes others),MAPK7,MFGE8,PLA2G12B,PLA2G4B,PPP3CC,TYK2 |
| Role of MAPK Signaling in the Pathogenesis of Influenza | 1.200 | 0.107 | #NUM! | AKT1,BCL2,KRAS,PLA2G12B,PLA2G4B,PNPLA3,RAP1A,RRAS |
| Chemokine Signaling | 1.200 | 0.107 | 0.816 | Calm4,CAMK2D,GNAI1,KRAS,PPP1CC,PPP1R12A,RAP1A,RRAS |
| GPCR-Mediated Nutrient Sensing in Enteroendocrine Cells | 1.190 | 0.096 | -0.302 | ADCY5,CACNG2,CACNG7,CCK,GNAI1,GNG3,GNG7,ITPR1,PDIA3,PRKACA,PRKD3 |
| IL-10 Signaling | 1.190 | 0.093 | 0.000 | AKT1,BCL2,BCL2L1,BHLHE40,CCN4,CCND1,HMOX1,IL6ST,JAK1,PRDM1,TAB1,TYK2 |
| fMLP Signaling in Neutrophils | 1.190 | 0.093 | -0.707 | Calm4,GNAI1,GNAO1,GNB1L,GNG3,GNG7,ITPR1,KRAS,PPP3CC,PRKD3,RAP1A,RRAS |
| Transport of inorganic cations/anions and amino acids/oligopep | 1.180 | 0.098 | 1.265 | CTNS,SLC15A4,SLC20A1,SLC24A3,SLC24A4,SLC25A10,SLC25A26,SLC26A2,SLC6A15,SLC8A2 |
| CDK5 Signaling | 1.180 | 0.098 | -0.707 | ADCY5,ITGA6,KRAS,MAPK7,NTRK2,PPP1CC,PPP1R10,PRKACA,RAP1A,RRAS |
| ERB2-ERBB3 Signaling | 1.170 | 0.111 | -0.378 | AKT1,CCND1,GRB2,KRAS,RAP1A,RRAS,TYK2 |
| Thyroid Cancer Signaling | 1.170 | 0.105 | -1.134 | AKT1,CCND1,INSR,Klk1b1 (includes others),KRAS,NTRK2,RAP1A,RRAS |
| G Protein Signaling Mediated by Tubby | 1.170 | 0.105 | -1.414 | GNAI1,GNAO1,GNB1L,GNG3,GNG7,GRB2,INSR,MCHR1 |
| HGF Signaling | 1.170 | 0.092 | -1.667 | AKT1,CCND1,GRB2,HGF,ITGA6,ITGA8,ITGAM,KRAS,PRKD3,PTPN11,RAP1A,RRAS |
| P2Y Purinergic Receptor Signaling Pathway | 1.170 | 0.092 | 0.632 | ADCY5,AKT1,GNAI1,GNB1L,GNG3,GNG7,KRAS,PDIA3,PRKACA,PRKD3,RAP1A,RRAS |
| CSDE1 Signaling Pathway | 1.160 | 0.118 | -1.000 | AGO1,AKT1,FABP7,HNRNPD,PAIP1,PIWIL1 |
| Metabolism of nitric oxide: NOS3 activation and regulation | 1.160 | 0.176 | #NUM! | AKT1,HSP90AA1,NOSTRIN |
| Gastrin-CREB signalling pathway via PKC and MAPK | 1.160 | 0.176 | #NUM! | GRB2,KRAS,MAPK7 |
| Mechanical load activates signaling by PIEZO1 and integrins in o | 1.160 | 0.176 | #NUM! | AKT1,CACNG7,HSPG2 |
| Mismatch Repair in Eukaryotes | 1.160 | 0.176 | #NUM! | MSH2,MSH3,RPA1 |
| Isoleucine Degradation I | 1.160 | 0.176 | #NUM! | ACADSB,ECHDC3,ECHS1 |
| DAG and IP3 signaling | 1.160 | 0.128 | -0.447 | ADCY5,CAMK2D,ITPR1,KPNA2,PRKACA |
| Mitophagy | 1.160 | 0.128 | -1.342 | ATG12,ATG9A,MAP1LC3A,TOMM70,UBE2N |
| Synaptic Long Term Depression | 1.160 | 0.086 | -0.500 | CACNG2,CACNG7,GNAI1,GNAO1,GRM4,ITPR1,KRAS,LYN,PDIA3,PLA2G12B,PLA2G4B,PNPLA3,PRKD3,PRKG2,RAP1A,RRAS |
| Ferroptosis Signaling Pathway | 1.150 | 0.094 | 2.111 | BAP1,EIF2AK3,FANCD2,GCH1,HMOX1,KEAP1,KRAS,RAP1A,RRAS,SAT1,SLC39A14 |
| RNA Polymerase II Transcription | 1.150 | 0.090 | -0.832 | ELOA,GTF2H1,Gtf2h2,LEO1,MAGOH,POLR2G,RTF1,SNRPD3,SRSF4,TAF3,THOC5,THOC6,ZC3H11A |
| Semaphorin interactions | 1.140 | 0.109 | 1.134 | ARHGAP35,DPYSL4,HSP90AA1,RHOB,RRAS,SEMA6D,SEMA7A |
| RNA Polymerase I Transcription | 1.140 | 0.109 | 1.134 | GTF2H1,Gtf2h2,H4C1,HDAC2,MTA1,POLR1C,TAF1A |
| IL-3 Signaling | 1.140 | 0.104 | -1.414 | AKT1,GRB2,JAK1,KRAS,PPP3CC,PRKD3,RAP1A,RRAS |
| Creatine metabolism | 1.140 | 0.250 | #NUM! | CKB,CKMT2 |
| Sucrose Degradation V (Mammalian) | 1.140 | 0.250 | #NUM! | GALM,KHK |
| MTOR signalling | 1.140 | 0.143 | 0.000 | AKT1,EIF4E,RPS6KB1,RRAGC |
| RHO GTPases activate PKNs | 1.140 | 0.143 | 1.000 | PKN3,PPP1R12A,RHOB,SFN |
| NLR signaling pathways | 1.130 | 0.115 | -1.633 | BCL2,BCL2L1,HMOX1,NLRP1,TAB1,UBE2N |
| Acute Myeloid Leukemia Signaling | 1.120 | 0.099 | -0.333 | AKT1,CCND1,FLT3LG,GRB2,KRAS,PML,RAP1A,RPS6KB1,RRAS |
| Signaling by Rho Family GTPases | 1.120 | 0.080 | -0.905 | ACTA1,ARHGEF4,CDH5,GNAI1,GNAO1,GNB1L,GNG3,GNG7,ITGA6,ITGA8,ITGAM,MAP3K20,NEDD4,PAK4,PARD6B,PARD6G,PPP1R12A,RHOB,SEPTIN8,SEPTIN9,WIPF1 |
| Renal Cell Carcinoma Signaling | 1.120 | 0.103 | -1.414 | AKT1,GRB2,HGF,KRAS,PAK4,PTPN11,RAP1A,RRAS |
| Neurotrophin/TRK Signaling | 1.120 | 0.103 | -0.816 | AKT1,GRB2,Klk1b1 (includes others),KRAS,NTRK2,PTPN11,RAP1A,RRAS |
| CCR5 Signaling in Macrophages | 1.110 | 0.095 | #NUM! | CACNG2,CACNG7,Calm4,GNAI1,GNAO1,GNB1L,GNG3,GNG7,ITPR1,PRKD3 |
| Adrenomedullin signaling pathway | 1.110 | 0.084 | -0.258 | ADCY5,ADM,AKT1,BCL2,Calm4,GRB2,ITPR1,KRAS,MAPK7,PDIA3,PRKACA,PRKG2,RAMP2,RAP1A,RRAS,SHC3 |
| Processing of Capped Intron-Containing Pre-mRNA | 1.100 | 0.079 | 0.426 | CCAR1,DHX35,EIF4E,HNRNPD,LSM7,LUC7L3,MAGOH,NUP35,POLR2G,PPIH,PRPF38A,PRPF8,RNF113A,SF3B1,SMNDC1,SNRNP200,SNRPD3,SRSF4,THOC5,THOC6,YBX1,ZC3H11A |
| Formation of the nephric duct | 1.100 | 0.167 | #NUM! | ID4,NPNT,OSR1 |
| Intrinsic Pathway for Apoptosis | 1.090 | 0.113 | 0.000 | AKT1,BCL2,BCL2L1,PPP3CC,SFN,TP53BP2 |
| SPINK1 General Cancer Pathway | 1.090 | 0.113 | 0.000 | AKT1,JAK1,KRAS,RAP1A,RRAS,TYK2 |
| Signaling by BMP | 1.090 | 0.138 | 1.000 | BMPR2,FSTL1,GREM2,SKI |
| ER-Mitochondrial Communication Signaling Pathway | 1.090 | 0.101 | 1.134 | AKT1,EIF2AK3,FIS1,PEMT,PML,PRELID1,PTDSS1,TOMM70 |
| ERBB4 Signaling | 1.090 | 0.106 | -1.134 | AKT1,ERBB4,GRB2,KRAS,PRKD3,RAP1A,RRAS |
| Generic Transcription Pathway | 1.080 | 0.083 | 1.000 | HDAC2,MED17,MED8,NR2E1,NR2F1,NR2F6,NR4A1,NR4A2,RARB,RORB,ZFP14,ZNF18,ZNF689,ZNF691,ZNF704,ZNF740 |
| ERBB Signaling | 1.080 | 0.097 | -1.000 | AKT1,ERBB4,GRB2,KRAS,PAK4,PRKD3,RAP1A,RPS6KB1,RRAS |
| HER-2 Signaling in Breast Cancer | 1.070 | 0.081 | -0.728 | AKT1,CCND1,COX17,COX6B1,COX6B2,COX7B,COX7C,GNB1L,GRB2,KRAS,LYN,NUP35,PARD6B,PARD6G,PRKD3,RAP1A,RPS6KB1,RRAS |
| Xenobiotic Metabolism General Signaling Pathway | 1.070 | 0.091 | 0.000 | AKT1,EIF2AK3,Gstm7,HMOX1,KEAP1,KRAS,MAPK7,PRKD3,RAP1A,RRAS,SLC51B |
| Eicosanoid Signaling | 1.070 | 0.079 | -0.218 | ABCC1,ADCY5,AKT1,ALOX5AP,AP1G2,CCND1,GNAI1,GNAO1,GNG3,GNG7,KCNMB4,KRAS,PDIA3,PLA2G12B,PLA2G4B,PNPLA3,PRKACA,PRKD3,RAP1A,RRAS,SOX9 |
| TP53 Regulates Transcription of DNA Repair Genes | 1.060 | 0.104 | -1.134 | AP1G2,CDK12,ELOA,FANCD2,GTF2H1,MSH2,POLR2G |
| Signaling by ALK | 1.050 | 0.133 | -1.000 | DNMT1,HDAC2,PRDM1,SIN3A |
| Glycosaminoglycan metabolism | 1.050 | 0.090 | -0.302 | B3GAT2,B4GALT2,CHST14,CHST15,GUSB,HSPG2,SDC3,SLC26A2,ST3GAL4,VCAN,XYLT2 |
| Folate Transformations I | 1.050 | 0.222 | #NUM! | MTR,SHMT2 |
| VEGF Family Ligand-Receptor Interactions | 1.040 | 0.099 | 0.000 | AKT1,GRB2,KRAS,PLA2G12B,PLA2G4B,PRKD3,RAP1A,RRAS |
| Class I peroxisomal membrane protein import | 1.040 | 0.158 | #NUM! | FIS1,PEX11B,PEX13 |
| Neurexins and neuroligins | 1.030 | 0.109 | 1.633 | APBA2,APBA3,BEGAIN,EPB41L3,LRRTM2,SYT10 |
| Calcium-induced T Lymphocyte Apoptosis | 1.030 | 0.109 | -1.000 | Calm4,HDAC2,ITPR1,NR4A1,PPP3CC,PRKD3 |
| p53 Signaling | 1.030 | 0.095 | 0.447 | AKT1,BCL2,BCL2L1,CCND1,PML,SCO2,SFN,TIGAR,TP53BP2 |
| TGF-β Signaling | 1.030 | 0.095 | -1.134 | BCL2,BMPR2,GRB2,INHBB,KRAS,RAP1A,RRAS,SKI,TAB1 |
| Protein Kinase A Signaling | 1.030 | 0.074 | -0.229 | ADCY5,ADD3,ATF1,Calm4,CAMK2D,CNGB1,DCC,EBI3,GNAI1,GNB1L,GNG3,GNG7,ITPR1,KDELR2,PDE7B,PDIA3,PPP1CC,PPP1R10,PPP3CC,PRKACA,PRKD3,PTPN11,PYGM,RAP1A,SFN,SHH,TDP2,TNS2 |
| Transcriptional regulation of white adipocyte differentiation | 1.020 | 0.098 | 0.707 | CHD9,KLF5,LEP,LPL,MED11,MED17,MED8,NR2F2 |
| E3 ubiquitin ligases ubiquitinate target proteins | 1.010 | 0.116 | -1.342 | LEO1,PEX13,RTF1,SHPRH,UBE2N |
| Mitochondrial protein degradation | 1.010 | 0.094 | -1.667 | ACADSB,BDH1,HMGCS2,IARS2,IDH3A,NDUFS3,PRELID1,PRKACA,SHMT2 |
| Regulation of Insulin-like Growth Factor (IGF) transport and upta | 1.010 | 0.091 | 1.897 | CALU,CP,FAM20A,FSTL1,GOLM1,MATN3,MFGE8,VCAN,VWA1,WFS1 |
| ESR-mediated signaling | 1.010 | 0.091 | -0.632 | AGO1,BCL2,CBFB,CCND1,ERBB4,H4C1,HSP90AA1,KPNA2,POLR2G,SMC3 |
| TEC Kinase Signaling | 1.010 | 0.082 | -0.378 | ACTA1,GNAI1,GNAO1,GNB1L,GNG3,GNG7,ITGA6,ITGA8,ITGAM,JAK1,LYN,PAK4,PRKD3,RHOB,TYK2 |

| | | | | |
|---|---|---|---|---|
| Deadenylation-dependent mRNA decay | 1.000 | 0.107 | 0.000 | CNOT7,DCPS,DIS3,EIF4E,LSM7,PAIP1 |
| RHO GTPase cycle | 1.000 | 0.072 | 0.707 | ADD3,ANKFY1,ARHGAP22,ARHGAP35,ARHGEF4,BCR,CKB,CYBA,DAAM1,DDX4,FAF2,GRB2,HSP90AA1,LMAN1,MTMR1,MTR,NDUFS3,NOXA1,NUDC,PAK4,PARD6B,PEAK1,PGRMC2,PKN3,RHOB,SH3RF1,SHMT2,SLITRK5,STAM2,SYDE1,WIPF1,WIPF2 |
| Epithelial Membrane Protein Signaling Pathway | 0.998 | 0.083 | -1.069 | AKT1,BMPR2,FGFR1,INSR,ITGA6,ITGA8,ITGAM,KRAS,LYN,NTRK2,RAP1A,RRAS,SOX9,TAB1 |
| Melanocyte Development and Pigmentation Signaling | 0.990 | 0.093 | -0.333 | ADCY5,BCL2,GRB2,KRAS,PRKACA,PTPN11,RAP1A,RPS6KB1,RRAS |
| Fc Epsilon RI Signaling | 0.990 | 0.090 | -0.632 | AKT1,GRB2,KRAS,LYN,PLA2G12B,PLA2G4B,PRKD3,PTPN11,RAP1A,RRAS |
| Prostate Cancer Signaling | 0.990 | 0.090 | -0.632 | AKT1,BCL2,CCND1,GRB2,HDAC2,HSP90AA1,KRAS,RAP1A,RRAS,SIN3A |
| Valine Degradation I | 0.988 | 0.150 | #NUM! | ACADSB,ECHDC3,ECHS1 |
| Orexin Signaling Pathway | 0.987 | 0.079 | -0.728 | ADCY5,AKT1,ATP6V1C1,CACNG2,CACNG7,Calm4,GNB1L,GNG3,GNG7,ITPR1,KCNJ6,PDIA3,PLA2G12B,PLA2G4B,PRKACA,PRKD3,PTPN11,SLC8A2 |
| Signaling by Insulin receptor | 0.981 | 0.114 | -0.447 | ATP6V1C1,GRB2,IDE,INSR,KRAS |
| Iron homeostasis signaling pathway | 0.975 | 0.087 | #NUM! | ALAS2,ATP6V1C1,BMP3,BMP6,BMPR2,CIAO3,CP,HMOX1,JAK1,SLC39A14,TYK2 |
| Neurovascular Coupling Signaling Pathway | 0.974 | 0.079 | 1.807 | CACNG2,CACNG7,Calm4,GABRB2,GABRG3,GABRP,ITPR1,KCNJ6,KCNMB4,PLA2G12B,PLA2G4B,PNPLA3,PPP1CC,PPP1R12A,PRKACA,PRKG2,SLC16A1 |
| Triglyceride metabolism | 0.969 | 0.125 | -2.000 | FABP7,PLIN3,PPP1CC,PRKACA |
| IL-7 Signaling Pathway | 0.959 | 0.099 | -2.646 | AKT1,BCL2,CCND1,GRB2,HGF,JAK1,LYN |
| Dolichol and Dolichyl Phosphate Biosynthesis | 0.955 | 0.500 | #NUM! | DHDDS |
| All-trans-decaprenyl Diphosphate Biosynthesis | 0.955 | 0.500 | #NUM! | PDSS1 |
| S-methyl-5-thio-α-D-ribose 1-phosphate Degradation | 0.955 | 0.500 | #NUM! | MRI1 |
| Formaldehyde Oxidation II (Glutathione-dependent) | 0.955 | 0.500 | #NUM! | ADH5 |
| Glycine Biosynthesis I | 0.955 | 0.500 | #NUM! | SHMT2 |
| Glutamine Degradation I | 0.955 | 0.500 | #NUM! | GLS |
| Cancer Drug Resistance by Drug Efflux | 0.950 | 0.103 | 0.000 | ABCC1,AKT1,KRAS,RAP1A,RRAS,YBX1 |
| Branched-chain amino acid catabolism | 0.939 | 0.143 | #NUM! | ACADSB,ECHS1,SLC25A44 |
| Interleukin-20 family signaling | 0.939 | 0.143 | #NUM! | JAK1,PTPN11,TYK2 |
| Co-inhibition by CTLA4 | 0.939 | 0.143 | #NUM! | AKT1,LYN,PTPN11 |
| Role of JAK family kinases in IL-6-type Cytokine Signaling | 0.935 | 0.097 | 1.134 | BCL2L1,CCND1,HSP90AA1,IL6ST,JAK1,PTPN11,TYK2 |
| Regulation of the Epithelial Mesenchymal Transition by Growth I | 0.935 | 0.080 | -1.387 | AKT1,FGF13,FGFR1,GRB2,HGF,JAK1,KRAS,PARD6B,PARD6G,PTPN11,RAP1A,RRAS,SHC3,TAB1,TYK2 |
| Ephrin Receptor Signaling | 0.934 | 0.079 | -0.832 | AKT1,GNAI1,GNAO1,GNB1L,GNG3,GNG7,GRB2,ITGA6,ITGA8,ITGAM,KRAS,PAK4,PTPN11,RAP1A,RRAS,WIPF1 |
| Cardiac Hypertrophy Signaling (Enhanced) | 0.929 | 0.070 | -0.192 | ADCY5,AKT1,BMPR2,CACNG2,CACNG7,Calm4,CAMK2D,EIF2B2,EIF4E,FGF13,FGFR1,GNAI1,GNB1L,GNG3,GNG7,HDAC2,IL6ST,ITGA6,ITGA8,ITGAM,ITPR1,KRAS,LEP,MAP3K20,MAPK7,PDE7B,PDIA3,PPP3CC,PRKACA,PRKD3,RAP1A,RCAN1,RPS6KB1,RRAS,TDP2,WNT4 |
| Role of OCT4 in Mammalian Embryonic Stem Cell Pluripotency | 0.919 | 0.109 | #NUM! | NR2F1,NR2F2,NR2F6,SALL4,SOX2 |
| TRIM21 Intracellular Antibody Signaling Pathway | 0.915 | 0.087 | -0.632 | IRF4,JAK1,MAP1LC3A,PRDM1,PSMC1,PSMD13,TAB1,TYK2,UBE2N,UBE2W |
| Hypoxia Signaling in the Cardiovascular System | 0.912 | 0.096 | #NUM! | AKT1,HSP90AA1,Ube2e1,UBE2N,UBE2Q1,UBE2R2,UBE2W |
| NRF2-mediated Oxidative Stress Response | 0.907 | 0.078 | 0.000 | ABCC1,ACTA1,AKR1A1,AKT1,Cyp2c55,DNAJC6,EIF2AK3,Gstm7,HMOX1,HSP90AA1,KEAP1,KRAS,MAPK7,PRKD3,RAP1A,RRAS |
| T Cell Exhaustion Signaling Pathway | 0.899 | 0.083 | 1.000 | AKT1,BMPR2,IRF4,JAK1,KRAS,LGALS9,PDCD1LG2,PRDM1,PTPN11,RAP1A,RRAS,TYK2 |
| Endometrial Cancer Signaling | 0.897 | 0.100 | -0.816 | AKT1,CCND1,GRB2,KRAS,RAP1A,RRAS |
| IL-15 Production | 0.897 | 0.086 | -1.265 | ERBB4,FGFR1,FLT3LG,INSR,JAK1,LYN,NTRK2,PEAK1,ROS1,TYK2 |
| GPVI-mediated activation cascade | 0.897 | 0.118 | 0.000 | COL1A2,LYN,PTPN11,RHOB |
| Peptide hormone biosynthesis | 0.895 | 0.182 | #NUM! | INHBB,PCSK1 |
| Signaling by Leptin | 0.895 | 0.182 | #NUM! | LEP,PTPN11 |
| NFE2L2 regulating tumorigenic genes | 0.895 | 0.182 | #NUM! | BCL2,BCL2L1 |
| Metabolism of porphyrins | 0.893 | 0.136 | #NUM! | ABCC1,ALAS2,HMOX1 |
| XBP1(S) activates chaperone genes | 0.890 | 0.106 | 1.342 | CUL7,FKBP14,PLA2G4B,TSPYL2,WFS1 |
| G alpha (z) signalling events | 0.890 | 0.106 | -0.447 | ADCY5,GNAI1,GNG3,GNG7,RGS4 |
| Leptin Signaling in Obesity | 0.889 | 0.095 | -1.000 | ADCY5,AKT1,GRB2,LEP,PDIA3,PRKACA,PTPN11 |
| Fcgamma receptor (FCGR) dependent phagocytosis | 0.888 | 0.091 | 0.000 | GRB2,HSP90AA1,ITPR1,LYN,NF2,PLD4,WIPF1,WIPF2 |
| ERK/MAPK Signaling | 0.881 | 0.078 | 0.302 | ATF1,EIF4E,GRB2,ITGA6,ITGA8,ITGAM,KRAS,PAK4,PLA2G12B,PLA2G4B,PPP1CC,PPP1R10,PPP1R12A,PRKACA,RAP1A,RRAS |
| Tumor Microenvironment Pathway | 0.881 | 0.081 | -0.832 | AKT1,BCL2,CCND1,COL1A2,FGF13,HGF,KRAS,LEP,LGALS9,PDCD1LG2,RAP1A,RRAS,SLC16A1 |
| SUMOylation of chromatin organization proteins | 0.873 | 0.098 | 0.816 | CBX2,H4C1,HDAC2,NUP35,SATB2,SUMO2 |
| Egg-Sperm Fusion Signaling Pathway | 0.872 | 0.087 | -0.333 | AKT1,CAMK2D,ENKUR,ITPR1,PRKACA,RAB27A,RAP1A,STX16,SYT10 |
| Corticotropin Releasing Hormone Signaling | 0.868 | 0.082 | 0.000 | ADCY5,CACNG2,CACNG7,Calm4,GNAI1,GNAO1,ITPR1,NR4A1,PRKACA,PRKD3,RAP1A,SHH |
| Antiproliferative Role of Somatostatin Receptor 2 | 0.867 | 0.093 | 0.447 | GNB1L,GNG3,GNG7,KRAS,PTPN11,RAP1A,RRAS |
| HIF1α Signaling | 0.867 | 0.078 | -0.258 | ADM,AKT1,BMP6,CAMK2D,EIF4E,HGF,HMOX1,HSP90AA1,KRAS,PPP3CC,PRKD3,RAP1A,RPS6KB1,RRAS,SAT1 |
| Transcriptional Regulatory Network in Embryonic Stem Cells | 0.866 | 0.080 | 0.277 | AKT1,BMPR2,FGFR1,HESX1,IL6ST,JAK1,KRAS,RAP1A,RRAS,SALL4,SOX2,TYK2,WNT4 |
| Autism Signaling Pathway | 0.866 | 0.074 | 0.229 | AKT1,Aldh3a2,ALDH5A1,CACNG2,CACNG7,CAMK2D,EIF4E,GNB1L,GRM4,HGF,KRAS,LEP,NTRK2,PRKACA,RAP1A,RARB,RELN,RPS6KB1,RRAS,SHH,WNT4 |
| Interferon alpha/beta signaling | 0.862 | 0.104 | -0.447 | IFIT2,IRF4,JAK1,PTPN11,TYK2 |
| Dilated Cardiomyopathy Signaling Pathway | 0.853 | 0.081 | 1.134 | ACTA1,ADCY5,AKT1,BCL2,CACNG2,CACNG7,CAMK2D,DSC2,ITPR1,KDM5A,PRKACA,PTPN11 |
| DYRK1A Signaling Pathway | 0.852 | 0.080 | 1.387 | AKT1,CCND1,FGFR1,IL6ST,KRAS,LIN54,MAPK7,POLR2G,PPP3CC,RAP1A,RCAN1,RRAS,SFN |
| IL-22 Signaling | 0.850 | 0.130 | #NUM! | AKT1,JAK1,TYK2 |
| Transcriptional Regulation by MECP2 | 0.848 | 0.097 | -0.816 | AGO1,CAMK2D,HDAC2,PRKACA,SIN3A,SOX2 |
| Fcγ Receptor-mediated Phagocytosis in Macrophages and Mono | 0.848 | 0.089 | -0.707 | ACTA1,AKT1,HMOX1,LYN,PLD4,PRKD3,RAB11B,RPS6KB1 |
| Angiopoietin Signaling | 0.845 | 0.092 | -1.134 | AKT1,GRB2,KRAS,PAK4,PTPN11,RAP1A,RRAS |
| White Adipose Tissue Browning Pathway | 0.841 | 0.082 | 0.333 | ADCY5,CACNG2,CACNG7,CTBP2,FGFR1,ITPR1,LEP,PRKACA,PRKG2,RARB,SLC16A1 |
| PTEN Regulation | 0.838 | 0.081 | -0.577 | AGO1,AKT1,CBX2,HDAC2,MTA1,NEDD4,NR2E1,PML,PSMC1,PSMD13,RRAGC,SALL4 |
| Lysosome Positioning Signaling Pathway | 0.838 | 0.081 | 0.000 | ACTA1,AKT1,BMPR2,BORCS7,DCTN3,FGFR1,GNB1L,INSR,KIF5B,NTRK2,PPP3CC,SEPTIN9 |
| RUNX1 regulates megakaryocyte differentiation and platelet fun | 0.835 | 0.102 | -1.342 | AGO1,CBFB,H4C1,SIN3A,SIN3B |
| Melanoma Signaling | 0.835 | 0.102 | -0.447 | AKT1,CCND1,KRAS,RAP1A,RRAS |
| Signal regulatory protein family interactions | 0.832 | 0.167 | #NUM! | GRB2,PTPN11 |
| Transcriptional regulation of testis differentiation | 0.832 | 0.167 | #NUM! | SOX9,WNT4 |
| Z-decay: degradation of maternal mRNAs by zygotically expresse | 0.832 | 0.167 | #NUM! | EIF4E,PAIP1 |
| Assembly of RNA Polymerase I Complex | 0.832 | 0.167 | #NUM! | POLR1C,TAF1A |
| Renin-Angiotensin Signaling | 0.829 | 0.083 | -1.667 | ADCY5,GRB2,ITPR1,KRAS,PAK4,PRKACA,PRKD3,RAP1A,RRAS,SHC3 |
| GP6 Signaling Pathway | 0.829 | 0.083 | -1.000 | AKT1,Calm4,COL13A1,COL19A1,COL1A2,COL6A2,GRB2,ITPR1,LYN,PRKD3 |
| B Cell Receptor Signaling | 0.825 | 0.078 | 0.277 | AKT1,BCL2L1,Calm4,CAMK2D,CD79B,GRB2,KRAS,LYN,PIK3AP1,PPP3CC,PTPN11,RAP1A,RPS6KB1,RRAS |
| Thrombopoietin Signaling | 0.825 | 0.095 | -0.816 | GRB2,KRAS,PRKD3,PTPN11,RAP1A,RRAS |
| Immunogenic Cell Death Signaling Pathway | 0.824 | 0.091 | 1.134 | ATG12,ATG16L1,BCL2,EIF2AK3,HMGB1,HSP90AA1,PDIA3 |
| Acute Phase Response Signaling | 0.824 | 0.079 | 0.000 | AKT1,C5,CP,GRB2,HMOX1,IL6ST,KRAS,PTPN11,RAP1A,RRAS,SERPING1,SOCS2,TAB1 |
| Autophagy | 0.819 | 0.076 | 0.500 | AKT1,ATG12,ATG16L1,ATG9A,BCL2,BMP6,Calm4,EIF2AK3,GNB1L,HGF,INSR,MAP1LC3A,PPP3CC,PRKACA,RB1CC1,VPS16 |
| Paxillin Signaling | 0.819 | 0.085 | 0.000 | ACTA1,GRB2,ITGA6,ITGA8,ITGAM,KRAS,PAK4,RAP1A,RRAS |
| Integrin signaling | 0.810 | 0.125 | #NUM! | AKT1,GRB2,RAP1A |
| Miscellaneous transport and binding events | 0.810 | 0.125 | #NUM! | ADD3,CTNS,NIPA2 |
| RHO GTPases Activate NADPH Oxidases | 0.810 | 0.125 | #NUM! | CYBA,NOXA1,S100A8 |
| Signaling by ERBB2 | 0.808 | 0.100 | -0.447 | AKT1,ERBB4,GRB2,HSP90AA1,KRAS |
| Transcriptional activity of SMAD2/SMAD3:SMAD4 heterotrimer | 0.808 | 0.100 | -1.342 | CDKN2B,COL1A2,NEDD4L,SKI,YBX1 |
| Transcriptional regulation by RUNX2 | 0.808 | 0.100 | -1.342 | AKT1,CBFB,CCND1,SATB2,SOX9 |
| Regulation of Actin-based Motility by Rho | 0.801 | 0.084 | 1.342 | ACTA1,ITGA6,ITGA8,ITGAM,PAK4,PPP1CC,PPP1R12A,RHOB,WIPF1 |
| Fanconi Anemia Pathway | 0.799 | 0.108 | 1.000 | FANCD2,MUS81,RPA1,RPA2 |
| GAIT Translation Signaling Pathway | 0.795 | 0.075 | 1.000 | AIMP1,CP,EIF2B2,EIF4E,ELP5,GNB1L,PAIP1,RPL18A,RPL22,RPL36,RPL37,RPL7,RPL7L1,RPS6KB1,S100A8,VPS37C |
| NADH Repair | 0.791 | 0.333 | #NUM! | NAXE |
| Serine Biosynthesis | 0.791 | 0.333 | #NUM! | PHGDH |
| L-cysteine Degradation III | 0.791 | 0.333 | #NUM! | CISD1 |
| Methionine Salvage II (Mammalian) | 0.791 | 0.333 | #NUM! | MTR |
| 4-aminobutyrate Degradation I | 0.791 | 0.333 | #NUM! | ALDH5A1 |
| Synaptogenesis Signaling Pathway | 0.790 | 0.072 | 1.091 | ADCY5,AKT1,Calm4,CAMK2D,CDH5,GRB2,GRM4,ITPR1,KRAS,LRRTM2,LYN,NTRK2,PRKACA,RAP1A,RASGRF1,RELN,RPS6KB1,RRAS,SHC3,STX16,STXBP6,SYT10 |
| Signaling by MET | 0.784 | 0.089 | -0.378 | COL1A2,GRB2,HGF,KRAS,PTPN11,RAP1A,STAM2 |
| Glioblastoma Multiforme Signaling | 0.783 | 0.077 | 0.000 | AKT1,CCND1,GRB2,ITPR1,KRAS,NF2,PDIA3,RAP1A,RHOB,RPS6KB1,RRAS,SHC3,WNT4 |
| Interleukin-3, Interleukin-5 and GM-CSF signaling | 0.783 | 0.098 | -2.236 | GRB2,JAK1,LYN,PRKACA,PTPN11 |
| Gap Junction Signaling | 0.781 | 0.071 | 0.894 | ACTA1,ADCY5,AKT1,Calm4,CCND1,CSNK1A1,GNAI1,GRB2,HTR2A,ITPR1,KRAS,LPAR6,MAPK7,PDIA3,PLA2G12B,PLA2G4B,PPP3CC,PRKACA,PRKD3,PRKG2,RAP1A,RRAS |
| Mitochondrial Dysfunction | 0.779 | 0.071 | 0.229 | ATP1B2,ATP1B3,BCL2,CACNG2,CACNG7,Calm4,CAMK2D,COX17,COX6B1,COX6B2,COX7B,COX7C,CP,FIS1,Gstm7,ITPR1,MAPK7,NDUFS3,NDUFS4,NDUFS8,PPP3CC,PRKACA,TOMM70 |
| Cytosolic iron-sulfur cluster assembly | 0.775 | 0.154 | #NUM! | ABCB7,CIAO3 |
| Platelet Adhesion to exposed collagen | 0.775 | 0.154 | #NUM! | COL1A2,LYN |
| Glutamate and glutamine metabolism | 0.775 | 0.154 | #NUM! | GLS,PYCR1 |
| Replacement of protamines by nucleosomes in the male pronucl | 0.775 | 0.154 | #NUM! | H4C1,HIRA |
| Co-inhibition by PD-1 | 0.775 | 0.154 | #NUM! | PDCD1LG2,PTPN11 |
| Cholesterol Biosynthesis I | 0.775 | 0.154 | #NUM! | DHCR24,LSS |

| | | | | |
|---|---|---|---|---|
| Cholesterol Biosynthesis II (via 24,25-dihydrolanosterol) | 0.775 | 0.154 | #NUM! | DHCR24,LSS |
| Cholesterol Biosynthesis III (via Desmosterol) | 0.775 | 0.154 | #NUM! | DHCR24,LSS |
| Signaling by Erythropoietin | 0.772 | 0.120 | #NUM! | GRB2,KRAS,LYN |
| Ovarian Cancer Signaling | 0.767 | 0.078 | 0.000 | AKT1,BCL2,CCND1,HDAC2,KRAS,MSH2,PRKACA,RAP1A,RPS6KB1,RRAS,SIN3A,WNT4 |
| Epithelial Adherens Junction Signaling | 0.767 | 0.078 | -0.577 | AKT1,ARHGAP35,BMPR2,FGFR1,HGF,KRAS,NF2,PAK4,RAP1A,RRAS,SFN,TNS1 |
| Regulation of eIF4 and p70S6K Signaling | 0.766 | 0.075 | 0.000 | AKT1,EIF2B2,EIF4E,ELP5,GRB2,ITGA6,ITGA8,ITGAM,KRAS,PAIP1,PAIP2,RAP1A,RPS6KB1,RRAS,VPS37C |
| Gαs Signaling | 0.766 | 0.081 | 0.447 | ADCY5,ADD3,CNGB1,GNAI1,GNAO1,GNB1L,GNG3,GNG7,PRKACA,RAP1A |
| RAC Signaling | 0.766 | 0.079 | 0.000 | ITGA6,ITGA8,ITGAM,KRAS,PAK4,PARD6B,PARD6G,RAP1A,RPS6KB1,RRAS,SH3RF1 |
| Macroautophagy | 0.759 | 0.091 | 0.000 | ATG12,ATG16L1,ATG9A,MAP1LC3A,RB1CC1,RRAGC |
| Interferon gamma signaling | 0.759 | 0.091 | -0.816 | CAMK2D,IRF4,JAK1,PML,PTPN11,YBX1 |
| Role of JAK1 and JAK3 in γc Cytokine Signaling | 0.759 | 0.091 | -0.816 | GRB2,JAK1,KRAS,PTPN11,RAP1A,RRAS |
| Glutamate Receptor Signaling | 0.759 | 0.091 | #NUM! | Calm4,GLS,GNB1L,GNG3,GNG7,GRM4 |
| DNA Methylation and Transcriptional Repression Signaling | 0.756 | 0.084 | 0.378 | CDK12,CDK16,DNMT1,HDAC2,MBD4,MTA1,SIN3A,TCF15 |
| Prolactin Signaling | 0.756 | 0.084 | -1.134 | GRB2,KCNMB4,KRAS,PRKD3,PTPN11,RAP1A,RRAS,SOCS2 |
| XIST Epigenetic Regulation Signaling Pathway | 0.751 | 0.080 | 0.632 | BCL2,CBX2,CMTR1,HDAC2,MATR3,MBD4,MTA1,RBM15B,SOX2,SOX9 |
| Integrin cell surface interactions | 0.745 | 0.086 | 0.378 | COL13A1,COL1A2,COL6A2,HSPG2,ITGA6,ITGA8,ITGAM |
| NF-κB Signaling | 0.745 | 0.076 | -1.732 | AKT1,BMPR2,FGFR1,HDAC2,INSR,KRAS,NTRK2,PRKACA,RAP1A,RRAS,TAB1,TDP2,UBE2N |
| MyD88:MAL(TIRAP) cascade initiated on plasma membrane | 0.741 | 0.103 | -1.000 | HMGB1,S100A8,TAB1,UBE2N |
| IL-1 Signaling | 0.739 | 0.083 | #NUM! | ADCY5,GNAI1,GNAO1,GNB1L,GNG3,GNG7,PRKACA,TAB1 |
| UVA-Induced MAPK Signaling | 0.739 | 0.083 | 1.000 | BCL2L1,KRAS,PARP10,PARP6,PDIA3,RAP1A,RPS6KB1,RRAS |
| CDP-diacylglycerol Biosynthesis I | 0.736 | 0.115 | #NUM! | AGPAT5,LPGAT1,PNPLA3 |
| Cholecystokinin/Gastrin-mediated Signaling | 0.736 | 0.081 | 0.333 | CCK,GRB2,ITPR1,KRAS,MAPK7,PRKD3,RAP1A,RHOB,RRAS |
| SAPK/JNK Signaling | 0.736 | 0.081 | -0.378 | GNB1L,GNG3,GNG7,GRB2,KRAS,MAP3K20,RAP1A,RRAS,TAB1 |
| Cardiac conduction | 0.736 | 0.079 | -1.265 | ATP1B2,ATP1B3,CACNG7,CAMK2D,FGF13,ITPR1,KCNIP4,KCNK13,PRKACA,SLC8A2 |
| Signaling by EGFR | 0.735 | 0.094 | -0.447 | ADAM12,GRB2,KRAS,PTPN11,STAM2 |
| Signaling by PTK6 | 0.735 | 0.094 | -0.447 | AKT1,ARHGAP35,CCND1,ERBB4,KRAS |
| Erythropoietin Signaling Pathway | 0.728 | 0.076 | -0.905 | AKT1,BCL2L1,CCND1,GRB2,ITPR1,KRAS,LEP,PRKD3,RAP1A,RPS6KB1,RRAS,SHC3 |
| ISGylation Signaling Pathway | 0.727 | 0.085 | 1.134 | CCND1,IRF4,JAK1,KRAS,NEDD4,TYK2,UBE2N |
| Regulation of the Epithelial-Mesenchymal Transition Pathway | 0.725 | 0.075 | #NUM! | AKT1,FGF13,FGFR1,GRB2,HGF,JAK1,KRAS,PARD6B,PARD6G,PTPN11,RAP1A,RRAS,TYK2,WNT4 |
| Regulation of TP53 Activity through Association with Co-factors | 0.724 | 0.143 | #NUM! | AKT1,TP53BP2 |
| Mitotic Telophase/Cytokinesis | 0.724 | 0.143 | #NUM! | PDS5B,SMC3 |
| Interleukin-15 signaling | 0.724 | 0.143 | #NUM! | GRB2,JAK1 |
| Assembly of RNA Polymerase III Complex | 0.724 | 0.143 | #NUM! | GTF3C1,GTF3C5 |
| Choline Biosynthesis III | 0.724 | 0.143 | #NUM! | HMOX1,PLD4 |
| Small Cell Lung Cancer Signaling | 0.722 | 0.083 | -0.447 | AKT1,BCL2,BCL2L1,CCND1,CDKN2B,HDAC2,RARB,SIN3A |
| Regulation of lipid metabolism by PPARalpha | 0.721 | 0.080 | -0.333 | CHD9,FHL2,HMGCS2,MED11,MED17,MED8,NFYC,SIN3A,SIN3B |
| O-linked glycosylation | 0.721 | 0.080 | 2.333 | ADAMTS15,CFP,GALNT10,GALNT14,GALNT9,POMGNT2,ST3GAL4,ST6GALNAC2,THSD7B |
| ISG15 antiviral mechanism | 0.717 | 0.088 | -1.633 | EIF4E,JAK1,KPNA2,NEDD4,NUP35,UBE2N |
| Agrin Interactions at Neuromuscular Junction | 0.717 | 0.088 | 0.000 | ACTA1,ERBB4,KRAS,PAK4,RAP1A,RRAS |
| Vasopressin regulates renal water homeostasis via Aquaporins | 0.714 | 0.100 | 0.000 | ADCY5,GNG3,GNG7,PRKACA |
| Meiotic recombination | 0.714 | 0.100 | 1.000 | H4C1,RBBP8,RPA1,RPA2 |
| Mechanisms of Viral Exit from Host Cells | 0.714 | 0.100 | #NUM! | ACTA1,NEDD4,PRKD3,VPS37C |
| Amino acids regulate mTORC1 | 0.712 | 0.093 | -0.447 | ATP6V1C1,FNIP2,RRAGC,SH3BP4,WDR59 |
| Interleukin-7 signaling | 0.703 | 0.111 | #NUM! | HGF,JAK1,SOCS2 |
| Syndecan interactions | 0.703 | 0.111 | #NUM! | COL1A2,ITGA6,SDC3 |
| Noradrenaline and Adrenaline Degradation | 0.703 | 0.111 | #NUM! | ADH5,AKR1A1,Aldh3a2 |
| Superpathway of Cholesterol Biosynthesis | 0.703 | 0.111 | #NUM! | DHCR24,HMGCS2,LSS |
| Interferon Signaling | 0.703 | 0.111 | #NUM! | BCL2,JAK1,TYK2 |
| Chronic Myeloid Leukemia Signaling | 0.699 | 0.071 | 0.000 | AKT1,BCL2,BCL2L1,Calm4,CCND1,CTBP2,GNB1L,GRB2,HDAC2,HSP90AA1,KRAS,PDIA3,PPP3CC,PTPN11,RAP1A,RRAS,SHH,SIN3A,WNT4 |
| Centrosomal KIAA0586 Signaling Pathway | 0.693 | 0.078 | 0.000 | CEP120,CEP290,FGF13,FGFR1,PRKACA,PSMC1,PSMD13,SHH,SMC3,VCAN |
| PEDF Signaling | 0.691 | 0.083 | -0.378 | AKT1,ARHGAP22,BCL2,BCL2L1,KRAS,RAP1A,RRAS |
| FGF Signaling | 0.691 | 0.083 | -2.646 | AKT1,FGF13,FGFR1,GRB2,HGF,ITPR1,PTPN11 |
| Nitric Oxide Signaling in the Cardiovascular System | 0.690 | 0.079 | -0.816 | AKT1,CACNG2,CACNG7,Calm4,HSP90AA1,ITPR1,PRKACA,PRKD3,PRKG2 |
| EGF Signaling | 0.690 | 0.091 | -1.342 | AKT1,GRB2,ITPR1,JAK1,RPS6KB1 |
| Elastic fibre formation | 0.688 | 0.098 | 0.000 | ITGA8,LTBP2,MFAP2,MFAP4 |
| RNA Polymerase III Transcription | 0.688 | 0.098 | 0.000 | GTF3C1,GTF3C5,POLR1C,SNAPC2 |
| DNA damage-induced 14-3-3σ Signaling | 0.688 | 0.098 | 1.000 | AKT1,BCL2,BCL2L1,SFN |
| eNOS Signaling | 0.684 | 0.076 | -0.816 | ADCY5,AKT1,Calm4,CHRND,CNGB1,HSP90AA1,ITPR1,LPAR6,NOSTRIN,PRKACA,PRKD3 |
| Mitotic Prometaphase | 0.680 | 0.073 | -0.535 | CENPL,CEP290,CEP70,CPAP,DCTN3,HSP90AA1,NEK7,NUDC,PDS5B,PPP1CC,PRKACA,SFI1,SMC3,ZWILCH |
| Role of Tissue Factor in Cancer | 0.680 | 0.073 | 0.535 | AKT1,BCL2,BCL2L1,BMPR2,EIF4E,HGF,ITGA6,KRAS,LYN,MAPK7,PTPN11,RAP1A,RPS6KB1,RRAS |
| Heme Degradation | 0.678 | 0.250 | #NUM! | HMOX1 |
| Spermine and Spermidine Degradation I | 0.678 | 0.250 | #NUM! | SAT1 |
| Catecholamine Biosynthesis | 0.678 | 0.250 | #NUM! | DDC |
| D-glucuronate Degradation I | 0.678 | 0.250 | #NUM! | AKR1A1 |
| Proline Biosynthesis I | 0.678 | 0.250 | #NUM! | PYCR1 |
| dTMP De Novo Biosynthesis | 0.678 | 0.250 | #NUM! | SHMT2 |
| Rap1 signalling | 0.678 | 0.133 | #NUM! | PRKACA,RAP1A |
| PPAR Signaling | 0.674 | 0.080 | 0.378 | GRB2,HSP90AA1,INSR,KRAS,NR2F1,RAP1A,RRAS,TAB1 |
| PIP3 activates AKT signaling | 0.671 | 0.075 | -1.508 | AKT1,ERBB4,FGFR1,FLT3LG,GRB2,HGF,INSR,NR4A1,NTRK2,PIK3AP1,PTPN11 |
| Sulfur amino acid metabolism | 0.671 | 0.107 | #NUM! | MRI1,MTR,SLC25A10 |
| Tryptophan Degradation X (Mammalian, via Tryptamine) | 0.671 | 0.107 | #NUM! | AKR1A1,Aldh3a2,DDC |
| Phosphatidylglycerol Biosynthesis II (Non-plastidic) | 0.671 | 0.107 | #NUM! | AGPAT5,LPGAT1,PNPLA3 |
| SLAMF Signaling Pathway | 0.666 | 0.076 | 0.000 | AKT1,BCL2,CD79B,GRB2,JAK1,KRAS,PTPN11,RAP1A,RRAS,TYK2 |
| Xenobiotic Metabolism PXR Signaling Pathway | 0.666 | 0.076 | -0.378 | Aldh3a2,ALDH5A1,CAMK2D,CHST15,Gstm7,HSP90AA1,PPP1CC,PPP1R10,PRKACA,PRKD3 |
| Signaling by SCF-KIT | 0.663 | 0.095 | -1.000 | GRB2,KRAS,LYN,PTPN11 |
| Exocyst Signaling Pathway | 0.658 | 0.079 | 0.000 | AKT1,CD79B,INSR,PARD6B,PARD6G,PRKD3,RAB11B,STX16 |
| Platelet homeostasis | 0.657 | 0.081 | -0.378 | GNG3,GNG7,ITPR1,KCNMB4,PRKG2,PTPN11,SLC8A2 |
| Sirtuin Signaling Pathway | 0.653 | 0.069 | -0.258 | ACSS2,AKT1,ATG12,ATG16L1,ATG9A,GLS,HMGCS2,MAP1LC3A,MAPK7,NDUFS3,NDUFS4,NDUFS8,NEDD4,POLR1C,RARB,RBBP8,RRP9,TIMM8A,TOMM70 |
| Polyamine Regulation in Colon Cancer | 0.647 | 0.088 | 0.447 | AKT1,GNB1L,KRAS,MTA1,SAT1 |
| Proteasomal PSMD10 Signaling Pathway | 0.647 | 0.077 | -0.378 | AKT1,BMP6,CCND1,HGF,KRAS,PSMC1,PSMD13,RAP1A,RRAS |
| Potassium Channels | 0.643 | 0.078 | 0.707 | GNG3,GNG7,KCNC2,KCNJ6,KCNK13,KCNMB4,KCNS1,KCNS2 |
| Oxidative Phosphorylation | 0.643 | 0.078 | 0.707 | COX17,COX6B1,COX6B2,COX7B,COX7C,NDUFS3,NDUFS4,NDUFS8 |
| Hepatitis B Chronic Liver Pathogenesis Signaling Pathway | 0.642 | 0.073 | -1.155 | AKT1,AP1G2,DNMT1,HMGB1,IL34,JAK1,KRAS,MSL2,PRKD3,RAP1A,RRAS,TYK2 |
| ERCC6 (CSB) and EHMT2 (G9a) positively regulate rRNA expressi | 0.641 | 0.103 | #NUM! | H4C1,HDAC2,MTA1 |
| mRNA Capping | 0.641 | 0.103 | #NUM! | GTF2H1,Gtf2h2,POLR2G |
| Respiratory electron transport | 0.641 | 0.081 | 0.378 | COX6B1,COX6B2,COX7B,COX7C,NDUFS3,NDUFS4,NDUFS8 |
| PI3K Cascade | 0.639 | 0.093 | -1.000 | FGFR1,FLT3LG,GRB2,PTPN11 |
| Collagen chain trimerization | 0.639 | 0.093 | 1.000 | COL13A1,COL19A1,COL1A2,COL6A2 |
| Transcriptional regulation of granulopoiesis | 0.639 | 0.093 | 1.000 | CBFB,H4C1,KLF5,PML |
| Transcriptional regulation of pluripotent stem cells | 0.635 | 0.125 | #NUM! | SALL4,SOX2 |
| RAN Signaling | 0.635 | 0.125 | #NUM! | KPNA2,TNPO1 |
| Extracellular matrix organization | 0.628 | 0.078 | -0.707 | ADAM12,COL1A2,COL6A2,HSPG2,ITGA6,ITGA8,MATN3,VCAN |
| PPARα/RXRα Activation | 0.627 | 0.072 | 0.302 | ADCY5,BMPR2,Cyp2c55,GRB2,HSP90AA1,INSR,KRAS,LPL,NR2F1,PDIA3,PRKACA,RAP1A,RRAS |
| Triacylglycerol Biosynthesis | 0.627 | 0.086 | 0.000 | AGPAT5,ELOVL1,LPGAT1,PLPP2,PNPLA3 |
| NER (Nucleotide Excision Repair, Enhanced Pathway) | 0.625 | 0.080 | 0.447 | GTF2H1,Gtf2h2,HMGN1,POLR2G,RPA1,RPA2,UBE2N |
| GPCR-Mediated Integration of Enteroendocrine Signaling Exemp | 0.624 | 0.082 | -0.816 | ADCY5,CCK,GNAI1,ITPR1,PDIA3,PRKACA |
| Relaxin Signaling | 0.622 | 0.073 | -0.447 | ADCY5,AKT1,GNAI1,GNAO1,GNB1L,GNG3,GNG7,PDE7B,PRKACA,RAP1A,TDP2 |
| B-WICH complex positively regulates rRNA expression | 0.616 | 0.091 | 1.000 | H4C1,POLR1C,SF3B1,TAF1A |
| RHO GTPases Activate WASPs and WAVEs | 0.616 | 0.091 | -1.000 | GRB2,LYN,WIPF1,WIPF2 |
| Systemic Lupus Erythematosus in T Cell Signaling Pathway | 0.615 | 0.071 | -0.535 | AKT1,DNMT1,GNAI1,GNB1L,GRB2,IRF4,ITPR1,KRAS,LEP,PPP3CC,RAP1A,RHOB,RPS6KB1,RRAS |
| RHOGDI Signaling | 0.615 | 0.070 | -1.134 | ACTA1,ARHGAP35,ARHGEF4,CDH5,GNAI1,GNAO1,GNB1L,GNG3,GNG7,ITGA6,ITGA8,ITGAM,PAK4,PPP1R12A,RHOB |
| Ephrin A Signaling | 0.614 | 0.074 | 0.632 | ACHE,AKT1,ARHGAP35,BMP6,HGF,LYN,PRKACA,PRKG2,PTPN11,RAP1A |

| | | | | |
|---|---|---|---|---|
| Regulation of TP53 Activity through Acetylation | 0.613 | 0.100 | #NUM! | AKT1,HDAC2,PML |
| Complement System | 0.613 | 0.100 | #NUM! | C5,ITGAM,SERPING1 |
| Role of Osteoclasts in Rheumatoid Arthritis Signaling Pathway | 0.609 | 0.068 | -0.775 | ADAM12,ADAM29,AKT1,BCL2,Calm4,CCND1,COL13A1,COL19A1,COL1A2,COL6A2,Fcgr3,FRZB,GRB2,KRAS,PPP3CC,RAP1A,RHOB,RRAS,SHC3 |
| UFMylation Signaling Pathway | 0.608 | 0.085 | -0.447 | AKT1,CCND1,EIF4E,ELP5,PAIP1 |
| WNT/β-catenin Signaling | 0.608 | 0.072 | -1.265 | AKT1,BMPR2,CCND1,CDH5,CSNK1A1,FRZB,GNAO1,RARB,SOX2,SOX9,TAB1,WNT4 |
| NAD Signaling Pathway | 0.602 | 0.074 | -0.333 | AKT1,BMP6,CD38,EIF2AK3,HGF,HMGCS2,NMRK2,PARP10,PARP6,POLR2G |
| Role of MAPK Signaling in Promoting the Pathogenesis of Influer | 0.600 | 0.076 | 1.414 | ATP6V1C1,BCL2,KRAS,PLA2G12B,PLA2G4B,PNPLA3,RAP1A,RRAS |
| Signaling by Type 1 Insulin-like Growth Factor 1 Receptor (IGF1R | 0.597 | 0.118 | #NUM! | GRB2,KRAS |
| SUMOylation of DNA methylation proteins | 0.597 | 0.118 | #NUM! | CBX2,DNMT1 |
| Regulation of TLR by endogenous ligand | 0.597 | 0.118 | #NUM! | HMGB1,S100A8 |
| Specification of the neural plate border | 0.597 | 0.118 | #NUM! | SOX2,ZIC1 |
| Formation of xylulose-5-phosphate | 0.594 | 0.200 | #NUM! | AKR1A1 |
| Reelin signalling pathway | 0.594 | 0.200 | #NUM! | RELN |
| Formation of lateral plate mesoderm | 0.594 | 0.200 | #NUM! | SHH |
| Proline Biosynthesis II (from Arginine) | 0.594 | 0.200 | #NUM! | PYCR1 |
| Serotonin and Melatonin Biosynthesis | 0.594 | 0.200 | #NUM! | DDC |
| Tetrapyrrole Biosynthesis II | 0.594 | 0.200 | #NUM! | ALAS2 |
| NAD Salvage Pathway III | 0.594 | 0.200 | #NUM! | NMRK2 |
| Glycine Cleavage Complex | 0.594 | 0.200 | #NUM! | AMT |
| Glutathione Redox Reactions II | 0.594 | 0.200 | #NUM! | PDIA3 |
| Galactose Degradation I (Leloir Pathway) | 0.594 | 0.200 | #NUM! | GALM |
| Glutamate Degradation III (via 4-aminobutyrate) | 0.594 | 0.200 | #NUM! | ALDH5A1 |
| Folate Polyglutamylation | 0.594 | 0.200 | #NUM! | SHMT2 |
| Acetate Conversion to Acetyl-CoA | 0.594 | 0.200 | #NUM! | ACSS2 |
| Glycerophospholipid biosynthesis | 0.594 | 0.074 | 0.333 | ACHE,AGPAT5,ETNK2,LPGAT1,PEMT,PLA2G4B,PLD4,PNPLA3,PTDSS1 |
| NGF Signaling | 0.594 | 0.074 | 0.000 | AKT1,GRB2,KRAS,MAPK7,PTPN11,RAP1A,RPS6KB1,RRAS,SHC3 |
| Integration of energy metabolism | 0.586 | 0.076 | -1.414 | ADCY5,GNAI1,GNG3,GNG7,ITPR1,KCNC2,PRKACA,RAP1A |
| Beta-catenin independent WNT signaling | 0.576 | 0.069 | 0.535 | AGO1,DAAM1,GNAO1,GNG3,GNG7,ITPR1,KRAS,PRICKLE1,PRKG2,PSMC1,PSMD13,RPS6KB1,TYK2,WNT4 |
| Netrin Signaling | 0.575 | 0.071 | -0.905 | AKT1,CACNG2,CACNG7,CAMK2D,DCC,DPYSL4,ITPR1,PPP3CC,PRKACA,PRKD3,STX16,TRIM9 |
| SUMOylation of DNA damage response and repair proteins | 0.574 | 0.079 | 0.816 | CBX2,NUP35,PML,RPA1,SMC3,SUMO2 |
| Transport of bile salts and organic acids, metal ions and amine c | 0.574 | 0.079 | 0.000 | CP,SLC16A1,SLC30A8,SLC31A1,SLC39A14,SLC6A15 |
| Activation of NMDA receptors and postsynaptic events | 0.574 | 0.079 | 0.000 | CAMK2D,ERBB4,KPNA2,KRAS,PRKACA,RASGRF1 |
| GDNF Family Ligand-Receptor Interactions | 0.574 | 0.079 | -0.447 | DOK4,GRB2,ITPR1,KRAS,RAP1A,RRAS |
| Macropinocytosis Signaling | 0.574 | 0.079 | -0.816 | ANKFY1,HGF,KRAS,PRKD3,RAP1A,RRAS |
| PXR/RXR Activation | 0.573 | 0.087 | -1.000 | AKT1,HMGCS2,INSR,PRKACA |
| nNOS Signaling in Neurons | 0.573 | 0.087 | #NUM! | Calm4,NOS1AP,PPP3CC,PRKD3 |
| MITF-M-dependent gene expression | 0.573 | 0.075 | -0.707 | AGO1,ATP6V1C1,BCL2,CCND1,IRF4,RAB27A,SIN3A,SOX2 |
| Class B/2 (Secretin family receptors) | 0.565 | 0.076 | 0.378 | ADM,GNG3,GNG7,RAMP2,SCT,SHH,WNT4 |
| DNA methylation | 0.561 | 0.111 | #NUM! | DNMT1,H4C1 |
| GABA synthesis, release, reuptake and degradation | 0.561 | 0.111 | #NUM! | ALDH5A1,RIMS1 |
| Putrescine Degradation III | 0.561 | 0.111 | #NUM! | Aldh3a2,SAT1 |
| G-protein beta:gamma signalling | 0.560 | 0.094 | #NUM! | AKT1,GNG3,GNG7 |
| Sialic acid metabolism | 0.560 | 0.094 | #NUM! | ST3GAL4,ST6GALNAC2,ST8SIA4 |
| NF-κB Activation by Viruses | 0.558 | 0.078 | -0.816 | AKT1,ITGA6,KRAS,PRKD3,RAP1A,RRAS |
| Integrin to Cytoskeleton Signaling Pathway | 0.557 | 0.069 | 0.000 | ACTA1,AKT1,GRB2,ITGA6,ITGA8,ITGAM,KRAS,PAK4,PPP1CC,PPP1R12A,RAP1A,RHOB,RRAS,WIPF1 |
| Ribonucleotide Reductase Signaling Pathway | 0.554 | 0.071 | -1.265 | AKT1,BCL2,CCND1,EIF4E,GNB1L,HGF,KRAS,NFYC,PARP10,PARP6,YBX1 |
| NCAM signaling for neurite out-growth | 0.553 | 0.081 | -1.342 | COL6A2,FGFR1,GRB2,KRAS,ST8SIA4 |
| iNOS Signaling | 0.552 | 0.085 | #NUM! | Calm4,JAK1,TAB1,TYK2 |
| Gene Silencing by RNA | 0.551 | 0.075 | 1.134 | AGO1,DDX4,H4C1,HSP90AA1,NUP35,PIWIL1,POLR2G |
| Neuroprotective Role of THOP1 in Alzheimer's Disease | 0.551 | 0.075 | #NUM! | ENDOU,IDE,Klk1b1 (includes others),PNOC,PRKACA,PRSS8,RELN |
| Neuropeptides in Migraine Signaling Pathway | 0.543 | 0.070 | 1.508 | ADCY5,CAMK2D,GNAI1,GNG3,GNG7,HTR2A,KCNMB4,PDIA3,PRKACA,PRKG2,RAMP2 |
| Unfolded Protein Response (UPR) | 0.536 | 0.091 | #NUM! | EIF2AK3,EIF2B2,VPS37C |
| Signal amplification | 0.536 | 0.091 | #NUM! | GNAI1,GNG3,GNG7 |
| Exosome Signaling Pathway | 0.533 | 0.070 | -0.632 | C1D,DIS3,POLR2G,RPL18A,RPL22,RPL36,RPL37,RPL7,RPL7L1,TENT4B,ZCCHC7 |
| Assembly of RNA Polymerase II Complex | 0.533 | 0.083 | #NUM! | GTF2H1,Gtf2h2,POLR2G,TAF3 |
| nNOS Signaling in Skeletal Muscle Cells | 0.533 | 0.083 | #NUM! | CACNG2,CACNG7,Calm4,ITPR1 |
| Dermatan Sulfate Biosynthesis | 0.533 | 0.083 | 0.000 | B3GAT2,CHST14,CHST15,XYLT2 |
| SUMOylation of transcription factors | 0.528 | 0.105 | #NUM! | MTA1,SUMO2 |
| RHO GTPases activate CIT | 0.528 | 0.105 | #NUM! | PPP1R12A,RHOB |
| RHO GTPases Activate ROCKs | 0.528 | 0.105 | #NUM! | PPP1R12A,RHOB |
| FOXO-mediated transcription | 0.528 | 0.105 | #NUM! | AKT1,SFN |
| Inflammasome pathway | 0.528 | 0.105 | #NUM! | NEK7,NLRP1 |
| Biosynthesis of Lipoxins (LX) | 0.527 | 0.167 | #NUM! | ALOX5AP |
| Fructose metabolism | 0.527 | 0.167 | #NUM! | KHK |
| Galactose catabolism | 0.527 | 0.167 | #NUM! | GALM |
| Epithelial-Mesenchymal Transition (EMT) during gastrulation | 0.527 | 0.167 | #NUM! | FGFR1 |
| Protein repair | 0.527 | 0.167 | #NUM! | PCMT1 |
| SLC15A4:TASL-dependent IRF5 activation | 0.527 | 0.167 | #NUM! | SLC15A4 |
| Role of IL-17A in Psoriasis | 0.527 | 0.167 | #NUM! | S100A8 |
| Selenocysteine Biosynthesis II (Archaea and Eukaryotes) | 0.527 | 0.167 | #NUM! | SEPHS1 |
| Aspartate Degradation II | 0.527 | 0.167 | #NUM! | MDH1 |
| Regulation of IL-2 Expression in Activated and Anergic T Lympho | 0.523 | 0.074 | 0.000 | Calm4,GRB2,KRAS,PPP3CC,RAP1A,RRAS,TOB1 |
| Apelin Cardiomyocyte Signaling Pathway | 0.523 | 0.074 | -1.134 | AKT1,GNAI1,ITPR1,MAPK7,PDIA3,PRKD3,SLC8A2 |
| CREB Signaling in Neurons | 0.523 | 0.063 | -0.928 | ACKR3,ADCY5,ADGRD1,AKT1,BMP6,BMPR2,CACNG2,CACNG7,Calm4,CAMK2D,CX3CR1,FGFR1,GNAI1,GNAO1,GNB1L,GNG3,GNG7,GPR149,GRB2,GRM4,HGF,HTR2A,INSR,ITPR1,KISS1R,KRAS,LPAR6,MCHR1,NTRK2,PDIA3,POLR2G,PRKACA,PRKD3,RAP1A,RRAS,SHC3 |
| Cell Cycle Control of Chromosomal Replication | 0.520 | 0.078 | 1.342 | CDC45,CDK12,CDK16,RPA1,RPA2 |
| DNA Double Strand Break Response | 0.514 | 0.082 | 0.000 | BAP1,H4C1,KPNA2,UBE2N |
| Activation of the pre-replicative complex | 0.513 | 0.088 | #NUM! | CDC45,RPA1,RPA2 |
| Transcriptional Regulation by NPAS4 | 0.513 | 0.088 | #NUM! | AGO1,RBFOX3,SYT10 |
| Retinol Biosynthesis | 0.513 | 0.088 | #NUM! | LPL,PNPLA3,RDH13 |
| Oxidative Ethanol Degradation III | 0.513 | 0.088 | #NUM! | ACSS2,Aldh3a2,Cyp2c55 |
| DHCR24 Signaling Pathway | 0.510 | 0.070 | -0.333 | AKT1,DHCR24,HMOX1,KLF5,KRAS,PRKD3,RAP1A,RRAS,SOX9 |
| Response of EIF2AK4 (GCN2) to amino acid deficiency | 0.509 | 0.071 | -2.121 | EIF2B2,IMPACT,RPL18A,RPL22,RPL36,RPL37,RPL7,VPS37C |
| Myocardin Signaling Pathway | 0.509 | 0.071 | 0.000 | AKT1,BMP3,BMP6,BMPR2,HGF,MYOG,SALL1,SALL4 |
| Receptor-type tyrosine-protein phosphatases | 0.498 | 0.100 | #NUM! | PPFIA1,SLITRK5 |
| TCA Cycle II (Eukaryotic) | 0.498 | 0.100 | #NUM! | IDH3A,MDH1 |
| Signaling by the B Cell Receptor (BCR) | 0.497 | 0.071 | 0.000 | CD79B,GRB2,ITPR1,KRAS,LYN,PIK3AP1,PSMC1,PSMD13 |
| Cellular response to heat stress | 0.497 | 0.072 | 1.890 | BAG4,CAMK2D,DNAJC2,HSP90AA1,NUP35,RPA1,RPA2 |
| GPER1 signaling | 0.497 | 0.072 | 0.378 | ADCY5,GNAI1,GNG3,GNG7,PRKACA,RPS6KB1,TYK2 |
| Signaling by FGFR1 | 0.495 | 0.080 | -1.000 | FGFR1,GRB2,KRAS,PTPN11 |
| UVB-Induced MAPK Signaling | 0.495 | 0.080 | 0.000 | AKT1,EIF4E,PRKD3,RPS6KB1 |
| Pulmonary Fibrosis Idiopathic Signaling Pathway | 0.489 | 0.064 | 1.147 | ACTA1,AKT1,BCL2,BMPR2,CCN4,CCND1,COL13A1,COL19A1,COL1A2,COL6A2,FGFR1,GRB2,IL6ST,KRAS,PEX13,RAP1A,RPS6KB1,RRAS,TYK2,WNT4 |
| Complex I biogenesis | 0.488 | 0.076 | -0.447 | DMAC1,NDUFAF2,NDUFS3,NDUFS4,NDUFS8 |
| Hepatic Fibrosis Signaling Pathway | 0.486 | 0.063 | 0.471 | AKT1,BCL2,BMPR2,CACNG2,CACNG7,Calm4,CCND1,COL1A2,CSNK1A1,FGFR1,GNAI1,ITGA6,ITGA8,ITGAM,JAK1,KRAS,LEP,PRKACA,PRKD3,RAP1A,RHOB,RPS6KB1,RRAS,SHH,WNT4 |
| PAK Signaling | 0.486 | 0.070 | -0.447 | GRB2,ITGA6,ITGA8,ITGAM,KRAS,PAK4,RAP1A,RRAS |
| Mitochondrial Division Signaling Pathway | 0.483 | 0.068 | -0.632 | CACNG2,CACNG7,CAMK2D,FIS1,HGF,ITPR1,KRAS,PPP3CC,PRKACA,RAP1A,RRAS |
| Granulocyte Adhesion and Diapedesis | 0.480 | 0.068 | #NUM! | ACKR3,C5,Ccl11,CDH5,CLDN8,Cxcl10,GLG1,GNAI1,ITGAM,SDC3 |
| UVC-Induced MAPK Signaling | 0.478 | 0.078 | 0.000 | KRAS,PRKD3,RAP1A,RRAS |
| Osteoarthritis Pathway | 0.477 | 0.066 | 0.000 | BMPR2,C1QTNF4,CCN4,FGFR1,FRZB,HMGB1,ITGA6,ITGA8,ITGAM,LEP,MATN3,RARRES2,S100A8,SOX9 |
| SNARE Signaling Pathway | 0.477 | 0.069 | 1.414 | ADCY5,Calm4,CAMK2D,PRKACA,RAB27A,RIMS1,STX16,STXBP6,SYT10 |
| ID1 Signaling Pathway | 0.473 | 0.066 | -0.832 | AKT1,BCL2,BCL2L1,BHLHE40,BMPR2,CCND1,CHRND,FGFR1,KRAS,LYN,RAP1A,RRAS,SHH |
| Neuropathic Pain Signaling in Dorsal Horn Neurons | 0.473 | 0.071 | -0.378 | CAMK2D,GRM4,ITPR1,NTRK2,PDIA3,PRKACA,PRKD3 |
| Apelin Adipocyte Signaling Pathway | 0.472 | 0.072 | 0.000 | ADCY5,CP,GNAI1,Gstm7,MAPK7,PRKACA |

| | | | | |
|---|---|---|---|---|
| Formation of intermediate mesoderm | 0.472 | 0.143 | #NUM! | OSR1 |
| Arginine Degradation VI (Arginase 2 Pathway) | 0.472 | 0.143 | #NUM! | PYCR1 |
| Inositol Pyrophosphates Biosynthesis | 0.472 | 0.143 | #NUM! | PPIP5K2 |
| WNT/SHH Axonal Guidance Signaling Pathway | 0.470 | 0.068 | 0.632 | ADCY5,CAMK2D,ITPR1,LYN,PARD6B,PARD6G,PRKACA,SFN,SHH,WNT4 |
| Formation of the dystrophin-glycoprotein complex (DGC) | 0.470 | 0.083 | #NUM! | ACTA1,DTNA,HSPG2 |
| SIRT1 negatively regulates rRNA expression | 0.469 | 0.095 | #NUM! | H4C1,TAF1A |
| RHO GTPases activate PAKs | 0.469 | 0.095 | #NUM! | NF2,PPP1R12A |
| Insertion of tail-anchored proteins into the endoplasmic reticulu | 0.469 | 0.095 | #NUM! | GET1,HMOX1 |
| Chaperone Mediated Autophagy | 0.469 | 0.095 | #NUM! | HSP90AA1,PLIN3 |
| Adrenergic Receptor Signaling Pathway (Enhanced) | 0.469 | 0.066 | -1.732 | ADCY5,ATP6V1C1,CACNG2,CACNG7,DDC,GNAI1,GNAO1,ITPR1,LEP,PDIA3,PRKACA,PRKD3 |
| Cyclophilin Signaling Pathway | 0.466 | 0.065 | 0.258 | AKT1,BCL2L1,CCND1,IL6ST,JAK1,PPP3CC,RBBP8,SF3B1,SNRNP200,SOX2,TIMM8A,TNPO1,TOMM70,TYK2,WNT4 |
| Natural Killer Cell Signaling | 0.465 | 0.067 | 0.000 | AKT1,COL1A2,Fcgr3,GRB2,KRAS,PAK4,PTPN11,RAP1A,RRAS,TYK2,WIPF1 |
| Spliceosomal Cycle | 0.461 | 0.077 | #NUM! | MAGOH,RNF113A,SF3B1,SNRNP200 |
| Neutrophil Extracellular Trap Signaling Pathway | 0.454 | 0.063 | -0.655 | AKT1,C5,COL13A1,COL19A1,COL1A2,COL6A2,HMGB1,ITGAM,ITPR1,LYN,NDUFS3,NDUFS4,NDUFS8,PDIA3,PLA2G12B,PLA2G4B,PNPLA3,PPP3CC,PRKD3,TIMM8A,TOMM70 |
| MyD88-independent TLR4 cascade | 0.450 | 0.081 | #NUM! | PTPN11,TAB1,UBE2N |
| Nucleosome assembly | 0.450 | 0.081 | #NUM! | CENPL,CENPW,H4C1 |
| S Phase | 0.449 | 0.069 | -0.378 | AKT1,CCND1,LIN54,PDS5B,PSMC1,PSMD13,SMC3 |
| PDGF Signaling | 0.446 | 0.071 | 0.000 | GRB2,JAK1,KRAS,RAP1A,RRAS,TYK2 |
| Growth hormone receptor signaling | 0.443 | 0.091 | #NUM! | LYN,SOCS2 |
| Glutathione Redox Reactions I | 0.443 | 0.091 | #NUM! | CP,Gstm7 |
| Ethanol Degradation IV | 0.443 | 0.091 | #NUM! | ACSS2,Aldh3a2 |
| TP53 Regulates Transcription of Cell Death Genes | 0.431 | 0.079 | #NUM! | PRELID1,TMEM219,TP53BP2 |
| Resolution of Abasic Sites (AP sites) | 0.431 | 0.079 | #NUM! | MBD4,RPA1,RPA2 |
| Serotonin Degradation | 0.431 | 0.079 | #NUM! | ADH5,AKR1A1,Aldh3a2 |
| Wound Healing Signaling Pathway | 0.430 | 0.064 | 1.069 | BMPR2,COL13A1,COL19A1,COL1A2,COL6A2,GRB2,ITGA6,JAK1,KRAS,LEP,RAP1A,RRAS,SHC3,TYK2 |
| Signaling by TGF-beta Receptor Complex | 0.429 | 0.074 | -1.000 | ITGA8,LTBP2,NEDD4L,PPP1CC |
| Sleep REM Signaling Pathway | 0.429 | 0.074 | 0.000 | EIF4E,NR4A2,PRKACA,RPS6KB1 |
| Necroptosis Signaling Pathway | 0.427 | 0.066 | -0.333 | CAMK2D,JAK1,PLA2G12B,PLA2G4B,PPP3CC,SLC25A10,TAB1,TIMM8A,TOMM70 |
| MSP-RON Signaling in Cancer Cells Pathway | 0.427 | 0.066 | 0.000 | AKT1,CCND1,GRB2,ITGA6,Klk1b1 (includes others),KRAS,RAP1A,RRAS,SFN |
| HDR through Homologous Recombination (HRR) or Single Stranc | 0.427 | 0.068 | 0.378 | H4C1,MUS81,RBBP8,RPA1,RPA2,SUMO2,UBE2N |
| Bladder Cancer Signaling | 0.427 | 0.068 | -0.447 | CCND1,FGF13,HDAC2,KRAS,RAP1A,RRAS,SIN3A |
| Neurotransmitter clearance | 0.425 | 0.125 | #NUM! | ACHE |
| DNA Damage Reversal | 0.425 | 0.125 | #NUM! | MGMT |
| NFE2L2 regulating MDR associated enzymes | 0.425 | 0.125 | #NUM! | ABCC1 |
| Embryonic Stem Cell Differentiation into Cardiac Lineages | 0.425 | 0.125 | #NUM! | SOX2 |
| Phosphatidylethanolamine Biosynthesis II | 0.425 | 0.125 | #NUM! | ETNK2 |
| Ketolysis | 0.425 | 0.125 | #NUM! | BDH1 |
| ID3 Signaling Pathway | 0.421 | 0.067 | 0.000 | AKT1,BMP6,BMPR2,CD79B,KRAS,RAP1A,RRAS,SOX2 |
| RHOA Signaling | 0.421 | 0.067 | -0.378 | ACTA1,ARHGAP35,LPAR6,NEDD4,PPP1CC,PPP1R12A,SEPTIN8,SEPTIN9 |
| Cellular response to mitochondrial stress | 0.419 | 0.087 | #NUM! | EIF2B2,VPS37C |
| Role of JAK1, JAK2 and TYK2 in Interferon Signaling | 0.419 | 0.087 | #NUM! | JAK1,TYK2 |
| SRP-dependent cotranslational protein targeting to membrane | 0.417 | 0.066 | -2.333 | RPL18A,RPL22,RPL36,RPL37,RPL7,RPL7L1,SEC11A,SPCS2,SRP14 |
| Telomerase Signaling | 0.416 | 0.067 | 0.000 | AKT1,GRB2,HDAC2,HSP90AA1,KRAS,RAP1A,RRAS |
| Eukaryotic Translation Initiation | 0.415 | 0.065 | -1.265 | CP,EIF2B2,EIF4E,RPL18A,RPL22,RPL36,RPL37,RPL7,RPL7L1,VPS37C |
| Molecular Mechanisms of Cancer | 0.414 | 0.060 | 0.000 | ACKR3,ADCY5,ADGRD1,AKT1,ARHGEF4,BCL2,BCL2L1,BMP3,BMP6,BMPR2,CAMK2D,CCND1,CDKN2B,CX3CR1,FANCD2,GNAI1,GNAO1,GNB1L,GNG3,GNG7,GPR149,GRB2,GRM4,HDAC2,HGF,HTR2A,IL6ST,ITGA6,ITGA8,ITGAM,JAK1,KISS1R,KRAS,LPAR6,MCHR1,PAK4,PRKACA,PRKD3,PTPN11,RAP1A,RASGRF1,RHOB,RRAS,SHC3,SHH,SIN3A,TAB1,TYK2 |
| Neurotransmitter release cycle | 0.413 | 0.077 | #NUM! | GLS,PPFIA1,RIMS1 |
| Nonhomologous End-Joining (NHEJ) | 0.413 | 0.077 | #NUM! | H4C1,TDP2,UBE2N |
| Retinoid metabolism and transport | 0.413 | 0.077 | #NUM! | HSPG2,LPL,SDC3 |
| Retinoic acid Mediated Apoptosis Signaling | 0.413 | 0.077 | #NUM! | PARP10,PARP6,RARB |
| CTLA4 Signaling in Cytotoxic T Lymphocytes | 0.412 | 0.064 | -0.905 | AKT1,AP1G2,BCL2L1,GRB2,HMOX1,KRAS,LYN,PLD4,PTPN11,RAP1A,RRAS |
| ROBO SLIT Signaling Pathway | 0.411 | 0.066 | 1.414 | AKT1,DCC,LYN,NEDD4,PAK4,PRKACA,ROBO1,ROBO2 |
| IL-6 Signaling | 0.411 | 0.066 | -0.707 | AKT1,GRB2,IL6ST,KRAS,PTPN11,RAP1A,RRAS,TAB1 |
| Aryl Hydrocarbon Receptor Signaling | 0.408 | 0.065 | -1.342 | Aldh3a2,ALDH5A1,CCND1,Cyp2c55,Gstm7,HSP90AA1,NR2F1,RARB,TRIP11 |
| AMPK Signaling | 0.407 | 0.063 | 0.000 | AKT1,CCND1,CHRND,GNAI1,GNAO1,GNB1L,GNG3,GNG7,INSR,LEP,PFKFB2,PFKL,PRKACA,RAB27A,RPS6KB1 |
| Xenobiotic Metabolism Signaling | 0.407 | 0.063 | #NUM! | Aldh3a2,ALDH5A1,CAMK2D,CHST15,EIF2AK3,Gstm7,HMOX1,HSP90AA1,KEAP1,KRAS,MAPK7,MGMT,PRKD3,RAP1A,RRAS |
| Microautophagy Signaling Pathway | 0.406 | 0.065 | 0.816 | COL13A1,COL19A1,COL1A2,COL6A2,MAP1LC3A,PSMC1,PSMD13,STAM2,VPS16,VPS37C |
| 14-3-3-mediated Signaling | 0.402 | 0.066 | 0.000 | AKT1,GRB2,KRAS,PDIA3,PRKD3,RAP1A,RRAS,SFN |
| Neddylation | 0.400 | 0.063 | -0.258 | CCDC8,COMMD3,CUL7,DCAF5,DCUN1D4,KEAP1,KLHL20,KLHL22,LRR1,NAE1,PSMC1,PSMD13,RNF7,SOCS2,WSB1 |
| Collagen degradation | 0.399 | 0.071 | 1.000 | COL13A1,COL19A1,COL1A2,COL6A2 |
| TNF signaling | 0.399 | 0.071 | 1.000 | BAG4,OTUD1,SPPL2A,TAB1 |
| Vitamin K Signaling Pathway | 0.399 | 0.071 | -1.000 | BCL2,CALU,GGCX,PRKACA |
| CGRP in Inflammation and Nociception Signaling Pathway | 0.399 | 0.065 | -1.000 | ADCY5,AKT1,IDE,Klk1b1 (includes others),PDIA3,PPP3CC,PRKACA,PRKD3,RAMP2 |
| Cell junction organization | 0.399 | 0.067 | 0.816 | CDH5,CLDN8,CRB3,ITGA6,PARD6B,PARD6G |
| Inhibition of ARE-Mediated mRNA Degradation Pathway | 0.397 | 0.064 | 0.000 | AGO1,AKT1,CNOT7,DIS3,MAPK7,PAPOLG,PRKACA,PSMC1,PSMD13,SFN |
| Gustation Pathway | 0.396 | 0.064 | 0.905 | ADCY5,CACNG2,CACNG7,GABRB2,GABRG3,GABRP,ITPR1,LPAR6,LPL,PKD1L3,PRKACA |
| MyD88 cascade initiated on plasma membrane | 0.396 | 0.083 | #NUM! | TAB1,UBE2N |
| Chromatin modifications during the maternal to zygotic transitic | 0.396 | 0.083 | #NUM! | H4C1,KDM5A |
| Activation of gene expression by SREBF (SREBP) | 0.395 | 0.075 | #NUM! | CHD9,LSS,NFYC |
| Signaling by FGFR3 | 0.395 | 0.075 | #NUM! | GRB2,KRAS,PTPN11 |
| Signaling by FGFR4 | 0.395 | 0.075 | #NUM! | GRB2,KRAS,PTPN11 |
| Signaling by VEGF | 0.395 | 0.066 | 0.378 | AKT1,CDH5,CYBA,HSP90AA1,ITPR1,KRAS,PRKACA |
| Caveolar-mediated Endocytosis Signaling | 0.392 | 0.069 | -0.447 | ACTA1,INSR,ITGA6,ITGA8,ITGAM |
| Ephrin B Signaling | 0.392 | 0.069 | #NUM! | GNAI1,GNAO1,GNB1L,GNG3,GNG7 |
| Neutrophil degranulation | 0.391 | 0.061 | 0.600 | ATP8A1,CCT8,CD68,CEP290,CFP,COMMD3,CYBA,CYSTM1,FAF2,GRN,GUSB,HMGB1,HSP90AA1,ITGAM,OSTF1,PFKL,PGRMC1,PRDX4,PSMD13,RAB27A,RAP1A,S100A8,SLC15A4,SRP14,VPS35L |
| Regulation of mRNA stability by proteins that bind AU-rich elem | 0.388 | 0.067 | 0.000 | AKT1,DIS3,HNRNPD,PSMC1,PSMD13,TNPO1 |
| POU5F1 (OCT4), SOX2, NANOG repress genes related to differer | 0.386 | 0.111 | #NUM! | SOX2 |
| GP1b-IX-V activation signalling | 0.386 | 0.111 | #NUM! | COL1A2 |
| RHO GTPases Activate Rhotekin and Rhophilins | 0.386 | 0.111 | #NUM! | RHOB |
| RUNX1 and FOXP3 control the development of regulatory T lymp | 0.386 | 0.111 | #NUM! | CBFB |
| Interleukin-9 signaling | 0.386 | 0.111 | #NUM! | JAK1 |
| Lipophagy | 0.386 | 0.111 | #NUM! | PLIN3 |
| Malate-aspartate shuttle | 0.386 | 0.111 | #NUM! | MDH1 |
| Leucine Degradation I | 0.386 | 0.111 | #NUM! | HMGCLL1 |
| Heme Biosynthesis II | 0.386 | 0.111 | #NUM! | ALAS2 |
| CDC42 Signaling | 0.382 | 0.064 | #NUM! | ITGA6,ITGA8,ITGAM,PAK4,PARD6B,PARD6G,PPP1CC,PPP1R12A,WIPF1 |
| Calcium Signaling | 0.380 | 0.062 | -2.333 | ACTA1,CACNG2,CACNG7,Calm4,CAMK2D,CHRND,HDAC2,ITPR1,PPP3CC,PRKACA,RAP1A,RCAN1,SLC8A2 |
| trans-Golgi Network Vesicle Budding | 0.380 | 0.068 | -1.342 | AP1G2,AP3S1,DNAJC6,VAMP7,YIPF6 |
| Interleukin-2 family signaling | 0.379 | 0.073 | #NUM! | GRB2,JAK1,LGALS9 |
| PD-1, PD-L1 cancer immunotherapy pathway | 0.377 | 0.066 | -0.447 | AKT1,BCL2L1,JAK1,PDCD1LG2,PTPN11,TYK2 |
| Nucleotide Excision Repair | 0.375 | 0.065 | 0.378 | GTF2H1,Gtf2h2,POLR2G,RPA1,RPA2,SUMO2,UBE2N |
| BMAL1:CLOCK,NPAS2 activates circadian expression | 0.374 | 0.080 | #NUM! | BHLHE40,CHD9 |
| Cholesterol biosynthesis | 0.374 | 0.080 | #NUM! | DHCR24,LSS |
| Glycogen metabolism | 0.374 | 0.080 | #NUM! | AKR1E2,PYGM |
| Signaling by NTRK3 (TRKC) | 0.374 | 0.080 | #NUM! | GRB2,KRAS |
| Signaling by PDGF | 0.372 | 0.069 | -1.000 | COL6A2,GRB2,KRAS,PTPN11 |
| Semaphorin Signaling in Neurons | 0.372 | 0.069 | #NUM! | DPYSL4,PAK4,RHOB,SEMA7A |
| HIPPO signaling | 0.368 | 0.067 | #NUM! | NF2,PPP1CC,PPP1R10,SFN,TP53BP2 |
| Atherosclerosis Signaling | 0.365 | 0.064 | 1.134 | COL1A2,GLG1,LPL,PLA2G12B,PLA2G4B,PNPLA3,S100A8 |
| TAK1-dependent IKK and NF-kappa-B activation | 0.363 | 0.071 | #NUM! | HMGB1,TAB1,UBE2N |
| Co-inhibition by BTLA | 0.359 | 0.068 | 0.000 | GRB2,PTPN11,RPS6KB1,TYK2 |
| Leukocyte Extravasation Signaling | 0.358 | 0.062 | -0.632 | ACTA1,ARHGAP35,CDH5,CLDN8,CYBA,GNAI1,ITGAM,PRKD3,PTPN11,RAP1A,WIPF1 |
| Estrogen-Dependent Breast Cancer Signaling | 0.357 | 0.066 | -0.447 | AKT1,CCND1,KRAS,RAP1A,RRAS |

| | | | | |
|---|---|---|---|---|
| ATF4 activates genes in response to endoplasmic reticulum stres | 0.354 | 0.077 | #NUM! | DIS3,NFYC |
| Processing and activation of SUMO | 0.352 | 0.100 | #NUM! | SUMO2 |
| Platelet Aggregation (Plug Formation) | 0.352 | 0.100 | #NUM! | COL1A2 |
| ATF6 (ATF6-alpha) activates chaperone genes | 0.352 | 0.100 | #NUM! | NFYC |
| Caspase activation via Dependence Receptors in the absence of l | 0.352 | 0.100 | #NUM! | DCC |
| RHOBTB3 ATPase cycle | 0.352 | 0.100 | #NUM! | PLIN3 |
| Formation of the posterior neural plate | 0.352 | 0.100 | #NUM! | SOX2 |
| Protein lipoylation | 0.352 | 0.100 | #NUM! | NFU1 |
| Glycogen Degradation II | 0.352 | 0.100 | #NUM! | PYGM |
| Citrulline Biosynthesis | 0.352 | 0.100 | #NUM! | GLS |
| Glycine Betaine Degradation | 0.352 | 0.100 | #NUM! | SHMT2 |
| Production of Nitric Oxide and Reactive Oxygen Species in Macro | 0.350 | 0.062 | 0.333 | AKT1,CYBA,JAK1,PPP1CC,PPP1R10,PPP1R12A,PRKD3,RAP1A,RHOB,S100A8,TYK2 |
| HEY1 Signaling Pathway | 0.348 | 0.062 | -0.333 | AKT1,BMP3,BMP6,BMPR2,ERBB4,GRB2,MYOG,NR2F2,SIN3A |
| SUMOylation of transcription cofactors | 0.348 | 0.070 | #NUM! | CBX2,SIN3A,SUMO2 |
| BER (Base Excision Repair) Pathway | 0.348 | 0.070 | #NUM! | HMGB1,MBD4,RPA1 |
| cAMP-mediated signaling | 0.347 | 0.061 | -0.277 | ADCY5,Calm4,CAMK2D,CNGB1,GNAI1,GNAO1,GRM4,PDE7B,PKIB,PPP3CC,PRKACA,RAP1A,RGS4,TDP2 |
| UBE2C in Cell Cycle Progression Signaling Pathway | 0.346 | 0.064 | 0.816 | AKT1,GNB1L,KEAP1,KRAS,PRDM2,RPS6KB1 |
| Telomere Maintenance | 0.346 | 0.065 | 1.342 | H4C1,NHP2,POLR2G,RPA1,RPA2 |
| ABC-family proteins mediated transport | 0.338 | 0.063 | 0.378 | ABCA7,ABCB7,ABCC1,EIF2B2,PSMC1,PSMD13,VPS37C |
| CDX Gastrointestinal Cancer Signaling Pathway | 0.336 | 0.061 | 0.905 | AKT1,BMP3,BMP6,CCND1,JAK1,KLF5,LEP,SALL4,SOX2,TYK2,WNT4 |
| Kinetochore Metaphase Signaling Pathway | 0.336 | 0.063 | #NUM! | CENPL,CENPW,PPP1CC,PPP1R10,SMC3,ZWILCH |
| Effects of PIP2 hydrolysis | 0.336 | 0.074 | #NUM! | DGKE,ITPR1 |
| PRC2 methylates histones and DNA | 0.336 | 0.074 | #NUM! | DNMT1,H4C1 |
| Senescence Pathway | 0.334 | 0.060 | 0.832 | AKT1,BHLHE40,BMPR2,CACNG2,CACNG7,Calm4,CBX2,CCND1,CDKN2B,DHCR24,EIF4E,KRAS,MAPK7,PML,PPP3CC,RAP1A,RRAS |
| Complement cascade | 0.334 | 0.068 | #NUM! | C5,CFP,SERPING1 |
| Netrin-1 signaling | 0.334 | 0.068 | #NUM! | DCC,PTPN11,ROBO1 |
| Tuberculosis Active Signaling Pathway | 0.331 | 0.060 | -0.905 | AKT1,ATP6V1C1,BCL2,CORO1A,CYBA,EIF2AK3,ITGAM,JAK1,NLRP1,PPP3CC,RPS6KB1,TYK2 |
| Virus Entry via Endocytic Pathways | 0.329 | 0.062 | -0.378 | ACTA1,AP1G2,AP3S1,KRAS,PRKD3,RAP1A,RRAS |
| Actin Cytoskeleton Signaling | 0.323 | 0.060 | -0.333 | ACTA1,ARHGAP35,ARHGEF4,FGF13,GRB2,ITGA6,ITGA8,ITGAM,KRAS,PAK4,PPP1CC,PPP1R12A,RAP1A,RRAS |
| Mitotic G1 phase and G1/S transition | 0.323 | 0.061 | -1.414 | AKT1,CCND1,CDC45,CDKN2B,LIN54,LYN,PSMC1,PSMD13 |
| Pulmonary Healing Signaling Pathway | 0.323 | 0.060 | -0.905 | ACKR3,BMPR2,CCND1,FGFR1,GRB2,KRAS,LYN,PRKD3,RAP1A,RRAS,WNT4 |
| Keratinization | 0.322 | 0.065 | 0.000 | DSC2,KRT2,KRT80,PRSS8 |
| Transcriptional and post-translational regulation of MITF-M expr | 0.322 | 0.065 | 2.000 | AIMP1,SOX2,SOX9,ZIC1 |
| RHO GTPases activate KTN1 | 0.322 | 0.091 | #NUM! | KIF5B |
| Advanced glycosylation endproduct receptor signaling | 0.322 | 0.091 | #NUM! | HMGB1 |
| Alpha-protein kinase 1 signaling pathway | 0.322 | 0.091 | #NUM! | TAB1 |
| Serine metabolism | 0.322 | 0.091 | #NUM! | PHGDH |
| Formation of the anterior neural plate | 0.322 | 0.091 | #NUM! | SOX2 |
| NR1H2 and NR1H3-mediated signaling | 0.320 | 0.067 | #NUM! | AGO1,ARL4C,MYLIP |
| Inhibition of Matrix Metalloproteases | 0.318 | 0.071 | #NUM! | ADAM12,HSPG2 |
| mRNA 3 Prime End Processing Signaling Pathway | 0.318 | 0.062 | -1.342 | CNOT7,DIS3,MBNL1,PAPOLG,PPP1CC,PPP1R10 |
| C-type lectin receptors (CLRs) | 0.315 | 0.061 | -1.414 | ITPR1,KRAS,LYN,PRKACA,PSMC1,PSMD13,TAB1,UBE2N |
| Role of BRCA1 in DNA Damage Response | 0.315 | 0.063 | #NUM! | ATF1,FANCD2,MSH2,RBBP8,RPA1 |
| NF1 RAS Signaling Pathway | 0.314 | 0.058 | 0.557 | ACKR3,ADCY5,ADGRD1,AKT1,BMP6,BMPR2,CX3CR1,FGFR1,GNB1L,GNG3,GNG7,GPR149,GRB2,GRM4,HGF,HTR2A,INSR,KISS1R,KRAS,LPAR6,MCHR1,NTRK2,PRKACA,PRKD3,PTPN11,RAP1A,RNF7,RRAS,SFN |
| RAF/MAP kinase cascade | 0.314 | 0.059 | -1.291 | CAMK2D,DAB2IP,ERBB4,FGFR1,FLT3LG,GRB2,HGF,JAK1,KRAS,PPP1CC,PSMC1,PSMD13,RAP1A,RASGRF1,SHC3 |
| IL-27 Signaling Pathway | 0.312 | 0.061 | -0.378 | AKT1,C5,EBI3,IL6ST,JAK1,LPAR6,TYK2 |
| RAR Activation | 0.311 | 0.058 | -0.426 | ADCY5,AKT1,BMPR2,CAMK2D,CDKN2B,CNGB1,COL1A2,GTF2H1,Gtf2h2,LEP,NR2F1,NR2F2,NR2F6,PDE7B,PML,PRKACA,RARB,RDH13,RHOB,RORB,SALL4,SOX9,TDP2 |
| Mitochondrial protein import | 0.311 | 0.064 | 1.000 | COX17,TIMM21,TIMM8A,TOMM70 |
| RAB geranylgeranylation | 0.311 | 0.064 | -1.000 | RAB11B,RAB27A,RAB35,RAB3C |
| Th17 Activation Pathway | 0.311 | 0.064 | -1.000 | HSP90AA1,IRF4,JAK1,TYK2 |
| Parkinson's Signaling Pathway | 0.311 | 0.059 | -0.832 | ACTA1,Aldh3a2,ALDH5A1,ATP6V1C1,CACNG2,CACNG7,Calm4,CAMK2D,DDC,EIF2AK3,Fcgr3,ITPR1,LEP,NDUFS3,NDUFS4,NDUFS8 |
| Heme signaling | 0.307 | 0.065 | #NUM! | CHD9,HMOX1,PGRMC2 |
| Chondroitin Sulfate Biosynthesis | 0.307 | 0.065 | #NUM! | B3GAT2,CHST15,XYLT2 |
| SUMOylation of intracellular receptors | 0.301 | 0.069 | #NUM! | NR4A2,SUMO2 |
| Protein ubiquitination | 0.301 | 0.069 | #NUM! | UBE2R2,UBE2W |
| FOXO-mediated transcription of oxidative stress, metabolic and | 0.301 | 0.069 | #NUM! | HDAC2,SIN3A |
| Response of EIF2AK1 (HRI) to heme deficiency | 0.301 | 0.069 | #NUM! | EIF2B2,VPS37C |
| Sonic Hedgehog Signaling | 0.301 | 0.069 | #NUM! | PRKACA,SHH |
| Phase II - Conjugation of compounds | 0.300 | 0.063 | 0.000 | AKR1A1,MTR,SLC26A2,TPST1 |
| Huntington's Disease Signaling | 0.300 | 0.058 | -0.816 | AKT1,BCL2L1,GLS,GNB1L,GNG3,GNG7,GRB2,HDAC2,ITPR1,POLR2G,PRKD3,PSMC1,PSMD13,SHC3,SIN3A,STX16 |
| Stress Granule Signaling Pathway | 0.300 | 0.059 | 0.832 | AKT1,EIF2AK3,EIF2B2,EIF4E,ELP5,GNB1L,HDAC2,HSP90AA1,INSR,KRAS,PAIP1,RPS6KB1,VPS37C,ZFAND1 |
| Fc epsilon receptor (FCERI) signaling | 0.300 | 0.060 | -1.414 | GRB2,ITPR1,KRAS,LYN,PSMC1,PSMD13,TAB1,UBE2N |
| G alpha (s) signalling events | 0.300 | 0.060 | -0.707 | ADCY5,ADM,GNAI1,GNG3,GNG7,PDE7B,RAMP2,SCT |
| Sumoylation Pathway | 0.300 | 0.061 | -2.000 | CTBP2,HDAC2,PML,RHOB,RPA1,SUMO2 |
| Glucocorticoid Receptor Signaling | 0.298 | 0.058 | #NUM! | AKT1,BCL2,BCL2L1,GRB2,GTF2H1,Gtf2h2,HMGB1,HSP90AA1,IL6ST,JAK1,KRAS,KRT2,KRT80,NDUFS3,NDUFS4,NDUFS8,PLA2G12B,PLA2G4B,POLR2G,PPP3CC,PRKACA,RAP1A,RARB,RRAS,TAB1,TAF3,TYK2 |
| Presynaptic depolarization and calcium channel opening | 0.295 | 0.083 | #NUM! | CACNG2 |
| Vitamin D (calciferol) metabolism | 0.295 | 0.083 | #NUM! | SUMO2 |
| HDR through MMEJ (alt-NHEJ) | 0.295 | 0.083 | #NUM! | RBBP8 |
| Lysine catabolism | 0.295 | 0.083 | #NUM! | GCDH |
| Specification of primordial germ cells | 0.295 | 0.083 | #NUM! | PRDM1 |
| Signaling by LTK | 0.295 | 0.083 | #NUM! | GRB2 |
| Leukotriene Biosynthesis | 0.295 | 0.083 | #NUM! | ALOX5AP |
| Glycogen Degradation III | 0.295 | 0.083 | #NUM! | PYGM |
| Mevalonate Pathway I | 0.295 | 0.083 | #NUM! | HMGCS2 |
| Glucose metabolism | 0.295 | 0.061 | -0.447 | G6PC2,NUP35,PFKFB2,PFKL,PRKACA |
| SUMOylation of RNA binding proteins | 0.294 | 0.064 | #NUM! | CBX2,NUP35,SUMO2 |
| PKCθ Signaling in T Lymphocytes | 0.290 | 0.059 | 0.000 | CACNG2,CACNG7,CAMK2D,GRB2,ITPR1,KRAS,PPP3CC,RAP1A,RRAS |
| DDX58/IFIH1-mediated induction of interferon-alpha/beta | 0.290 | 0.062 | -1.000 | ATG12,HMGB1,HSP90AA1,TOMM70 |
| Signaling by ROBO receptors | 0.289 | 0.058 | -1.604 | DCC,MAGOH,PAK4,PRKACA,PSMC1,PSMD13,ROBO1,ROBO2,RPL18A,RPL22,RPL36,RPL37,RPL7,RPL7L1 |
| Cyclins and Cell Cycle Regulation | 0.286 | 0.060 | #NUM! | CCND1,CDKN2B,HDAC2,MYT1,SIN3A |
| DAP12 interactions | 0.286 | 0.067 | #NUM! | GRB2,KRAS |
| Activation of kainate receptors upon glutamate binding | 0.286 | 0.067 | #NUM! | GNG3,GNG7 |
| Endosomal Sorting Complex Required For Transport (ESCRT) | 0.286 | 0.067 | #NUM! | STAM2,VPS37C |
| Role of Macrophages, Fibroblasts and Endothelial Cells in Rheum | 0.283 | 0.058 | 0.500 | AKT1,C5,Calm4,CAMK2D,CCND1,CSNK1A1,DAAM1,FRZB,GNAO1,IL6ST,KRAS,PDIA3,PPP3CC,PRKD3,RAP1A,RRAS,WNT4 |
| MYC Mediated Apoptosis Signaling | 0.282 | 0.063 | #NUM! | BCL2,BCL2L1,PRKACA |
| Collagen biosynthesis and modifying enzymes | 0.279 | 0.061 | 1.000 | COL13A1,COL19A1,COL1A2,COL6A2 |
| Deubiquitination | 0.279 | 0.058 | 1.069 | BAP1,H4C1,IDE,KEAP1,MYSM1,PSMC1,PSMD13,RCE1,RNF128,STAM2,TAB1,TADA3,TOMM70,USP24 |
| FXR/RXR Activation | 0.277 | 0.058 | 0.000 | AKT1,EIF2AK3,G6PC2,Gstm7,HMGB1,HMOX1,LEP,LPL,SLC51B |
| WNK Renal Signaling Pathway | 0.275 | 0.059 | -0.447 | AKT1,Calm4,INSR,NEDD4,OSR1,PRKD3 |
| Acetylcholine binding and downstream events | 0.272 | 0.077 | #NUM! | CHRND |
| Olfactory Signaling Pathway | 0.272 | 0.077 | #NUM! | CNGB1 |
| SUMOylation of immune response proteins | 0.272 | 0.077 | #NUM! | SUMO2 |
| Formation of axial mesoderm | 0.272 | 0.077 | #NUM! | SHH |
| Nephron development | 0.272 | 0.077 | #NUM! | WNT4 |
| Glutamate binding, activation of AMPA receptors and synaptic p | 0.271 | 0.065 | #NUM! | CACNG2,CAMK2D |
| MyD88 dependent cascade initiated on endosome | 0.271 | 0.065 | #NUM! | TAB1,UBE2N |
| Transcriptional regulation of brown and beige adipocyte differer | 0.271 | 0.065 | #NUM! | HDAC2,MTA1 |
| Ribotoxic Stress Signaling Pathway | 0.271 | 0.065 | #NUM! | GIGYF2,MAP3K20 |
| TP53 Regulates Transcription of Cell Cycle Genes | 0.271 | 0.061 | #NUM! | CNOT7,CPAP,SFN |
| Co-stimulation by CD28 | 0.271 | 0.061 | #NUM! | AKT1,GRB2,LYN |
| Amyloid Processing | 0.271 | 0.061 | #NUM! | AKT1,CSNK1A1,PRKACA |
| Cell Cycle: G1/S Checkpoint Regulation | 0.270 | 0.060 | #NUM! | CCND1,CDKN2B,HDAC2,SIN3A |

| | | | | |
|---|---|---|---|---|
| Xenobiotic Metabolism AHR Signaling Pathway | 0.270 | 0.060 | #NUM! | Aldh3a2,ALDH5A1,Gstm7,HSP90AA1 |
| ABRA Signaling Pathway | 0.269 | 0.059 | -0.447 | ABRA,ACTA1,CKMT2,MYOG,PPP3CC |
| LPS-stimulated MAPK Signaling | 0.269 | 0.059 | 0.000 | ATF1,KRAS,PRKD3,RAP1A,RRAS |
| ICOS-ICOSL Signaling in T Helper Cells | 0.267 | 0.058 | -1.000 | AKT1,Calm4,CAMK2D,GRB2,ITPR1,PPP3CC |
| Antioxidant Action of Vitamin C | 0.267 | 0.058 | -0.816 | HMOX1,PDIA3,PLA2G12B,PLA2G4B,PLD4,PNPLA3 |
| MSP-RON Signaling in Macrophages Pathway | 0.267 | 0.058 | 0.447 | GRB2,ITGAM,Klk1b1 (includes others),KRAS,RAP1A,RRAS |
| Bone Mineralization Signaling Pathway | 0.265 | 0.058 | -0.707 | BMPR2,COL1A2,FGF13,FGFR1,NR4A2,PDIA3,SLC20A1,WNT4 |
| Basal Cell Carcinoma Signaling | 0.260 | 0.059 | #NUM! | BMP3,BMP6,SHH,WNT4 |
| Toll Like Receptor 3 (TLR3) Cascade | 0.257 | 0.063 | #NUM! | TAB1,UBE2N |
| Thrombin signalling through proteinase activated receptors (PAF | 0.257 | 0.063 | #NUM! | GNG3,GNG7 |
| Late endosomal microautophagy | 0.257 | 0.063 | #NUM! | PLIN3,VPS37C |
| Inhibition of Angiogenesis by TSP1 | 0.257 | 0.063 | #NUM! | AKT1,HSPG2 |
| Plasma lipoprotein assembly, remodeling, and clearance | 0.251 | 0.058 | -1.000 | LPL,MYLIP,NCEH1,PRKACA |
| Sensory processing of sound by inner hair cells of the cochlea | 0.251 | 0.058 | 2.000 | CTBP2,EPB41L3,PCDH15,RIPOR2 |
| Prolactin receptor signaling | 0.251 | 0.071 | #NUM! | PTPN11 |
| LGI-ADAM interactions | 0.251 | 0.071 | #NUM! | CACNG2 |
| Phosphorylation and nuclear translocation of the CRY:PER:kinase | 0.251 | 0.071 | #NUM! | PPP1CC |
| DNA Double-Strand Break Repair by Homologous Recombination | 0.251 | 0.071 | #NUM! | RPA1 |
| FAT10 Cancer Signaling Pathway | 0.249 | 0.059 | #NUM! | ACKR3,AKT1,BMPR2 |
| Arachidonate metabolism | 0.244 | 0.061 | #NUM! | ABCC1,ALOX5AP |
| MAPK targets/ Nuclear events mediated by MAP kinases | 0.244 | 0.061 | #NUM! | ATF1,MAPK7 |
| MIF-mediated Glucocorticoid Regulation | 0.244 | 0.061 | #NUM! | PLA2G12B,PLA2G4B |
| Pre-NOTCH Expression and Processing | 0.242 | 0.057 | 0.000 | AGO1,CCND1,H4C1,ST3GAL4 |
| Signaling by FGFR2 | 0.242 | 0.057 | 0.000 | GRB2,KRAS,POLR2G,PTPN11 |
| Glioma Invasiveness Signaling | 0.242 | 0.057 | 1.000 | KRAS,RAP1A,RHOB,RRAS |
| Metabolism of non-coding RNA | 0.239 | 0.058 | #NUM! | GEMIN6,NUP35,SNRPD3 |
| Oncogene Induced Senescence | 0.232 | 0.059 | #NUM! | AGO1,CDKN2B |
| SUMOylation of SUMOylation proteins | 0.232 | 0.059 | #NUM! | NUP35,SUMO2 |
| Fatty Acid β-oxidation I | 0.232 | 0.059 | #NUM! | ECHDC3,ECHS1 |
| Superpathway of Methionine Degradation | 0.232 | 0.059 | #NUM! | MGMT,MTR |
| Aspirin ADME | 0.232 | 0.067 | #NUM! | SLC16A1 |
| Germ layer formation at gastrulation | 0.232 | 0.067 | #NUM! | SOX2 |
| Paracetamol ADME | 0.232 | 0.067 | #NUM! | ABCC1 |
| Glutaryl-CoA Degradation | 0.232 | 0.067 | #NUM! | GCDH |
| Histamine Degradation | 0.232 | 0.067 | #NUM! | Aldh3a2 |
| Mitochondrial Fatty Acid Beta-Oxidation | 0.220 | 0.057 | #NUM! | DBI,ECHS1 |
| Expression of BMAL (ARNTL), CLOCK, and NPAS2 | 0.220 | 0.057 | #NUM! | CHD9,RORB |
| IL-9 Signaling | 0.220 | 0.057 | #NUM! | JAK1,SOCS2 |
| Intrinsic Prothrombin Activation Pathway | 0.220 | 0.057 | #NUM! | COL1A2,Klk1b1 (includes others) |
| Dermatan Sulfate Biosynthesis (Late Stages) | 0.220 | 0.057 | #NUM! | CHST14,CHST15 |
| Metabolism of folate and pterines | 0.215 | 0.063 | #NUM! | SHMT2 |
| FOXO-mediated transcription of cell death genes | 0.215 | 0.063 | #NUM! | NFYC |
| Mitochondrial unfolded protein response (UPRmt) | 0.215 | 0.063 | #NUM! | AKT1 |
| Modulation of host responses by IFN-stimulated genes | 0.215 | 0.063 | #NUM! | EIF2AK3 |
| Superpathway of Citrulline Metabolism | 0.215 | 0.063 | #NUM! | GLS |
| Superpathway of Geranylgeranyldiphosphate Biosynthesis I (via | 0.215 | 0.063 | #NUM! | HMGCS2 |
| NFE2L2 regulating anti-oxidant/detoxification enzymes | 0.199 | 0.059 | #NUM! | HMOX1 |
| Pyroptosis Signaling Pathway | 0.000 | 0.056 | #NUM! | HMGB1,NEK7,NLRP1,PRKACA |
| IL-13 Signaling Pathway | 0.000 | 0.056 | -0.447 | AKT1,BCL2L1,JAK1,LYN,TYK2 |
| Oxytocin in Spinal Neurons Signaling Pathway | 0.000 | 0.030 | #NUM! | PRKG2 |
| Macrophage Classical Activation Signaling Pathway | 0.000 | 0.021 | #NUM! | JAK1,LEP,TYK2 |
| Multiple Sclerosis Signaling Pathway | 0.000 | 0.039 | 0.816 | C5,IRF4,LEP,NLRP1,PARP10,PARP6,SLC8A2 |
| Pathogen Induced Cytokine Storm Signaling Pathway | 0.000 | 0.053 | 1.155 | BHLHE40,C5,Ccl11,COL13A1,COL19A1,COL1A2,COL6A2,Cxcl10,IL6ST,JAK1,LEP,MAPK7,NLRP1,PRDM1,RAB27A |
| Role of Chondrocytes in Rheumatoid Arthritis Signaling Pathway | 0.000 | 0.024 | #NUM! | GNB1L,IL6ST,PPP3CC |
| Role of Osteoblasts in Rheumatoid Arthritis Signaling Pathway | 0.000 | 0.052 | 0.905 | AKT1,BCL2,BMP3,BMPR2,CSNK1A1,FRZB,IL6ST,JAK1,LEP,TYK2,WNT4 |
| Macrophage Alternative Activation Signaling Pathway | 0.000 | 0.046 | 0.378 | GNB1L,GRB2,IRF4,JAK1,LPL,RPS6KB1,TYK2 |
| S100 Family Signaling Pathway | 0.000 | 0.045 | -0.756 | ABCC1,ACKR3,ADGRD1,AKT1,BCL2,CACNG2,CACNG7,Calm4,CAMK2D,CX3CR1,Fcgr3,FGFR1,GPR149,GRM4,HTR2A,ITPR1,JAK1,KISS1R,Klk1b1 (includes others),LPAR6,MAPK7,MCHR1,NTRK2,PDIA3,PLA2G12B,PLA2G4B,PRKACA,PRKD3,S100A8,TYK2,WNT4 |
| Amyotrophic Lateral Sclerosis Signaling | 0.000 | 0.036 | #NUM! | BCL2,BCL2L1,CACNG2,CACNG7 |
| Chaperone Mediated Autophagy Signaling Pathway | 0.000 | 0.056 | 1.732 | AKT1,ATP6V1C1,CCND1,EIF2AK3,HDAC2,HMGB1,HSP90AA1,PLIN3,PPP3CC,RAB11B,RCAN1,SOX2 |
| IL-33 Signaling Pathway | 0.000 | 0.029 | 0.000 | AKT1,BCL2L1,PRKACA,RPS6KB1 |
| NOD1/2 Signaling Pathway | 0.000 | 0.027 | 2.000 | ATG16L1,HSP90AA1,LEP,SLC15A4 |
| Activin Inhibin Signaling Pathway | 0.000 | 0.027 | 0.447 | AKT1,COL1A2,INHBB,MAPK7,SKI |
| Cachexia Signaling Pathway | 0.000 | 0.057 | 1.414 | ADCY5,AKT1,BCL2,BMPR2,EIF2AK3,EIF2B2,HMGB1,HSP90AA1,INHBB,INSR,LEP,LPL,MYOG,PHGDH,PRKACA,PRKD3,PSMC1,PSMD13,VPS37C |
| NFKBIE Signaling Pathway | 0.000 | 0.018 | #NUM! | DYSF |
| Sertoli Cell-Germ Cell Junction Signaling Pathway (Enhanced) | 0.000 | 0.044 | -1.265 | ACTA1,AKT1,CLDN8,ITGA6,KRAS,PRKACA,PRKG2,RAP1A,RRAS,TAB1 |
| LPS/IL-1 Mediated Inhibition of RXR Function | 0.000 | 0.044 | #NUM! | Aldh3a2,ALDH5A1,CHST15,CRAT,Cyp2c55,FABP7,Gstm7,HMGCS2,MGMT |
| G alpha (i) signalling events | 0.000 | 0.051 | 0.302 | ACKR3,ADCY5,C5,CX3CR1,GNAI1,GNG3,GNG7,GRM4,MCHR1,PNOC,RGS4 |
| Meiotic synapsis | 0.000 | 0.047 | #NUM! | H4C1,SMC3 |
| Cellular response to hypoxia | 0.000 | 0.026 | #NUM! | PSMC1,PSMD13 |
| Formation of Fibrin Clot (Clotting Cascade) | 0.000 | 0.035 | #NUM! | SERPING1 |
| Degradation of the extracellular matrix | 0.000 | 0.015 | #NUM! | HSPG2 |
| Translocation of SLC2A4 (GLUT4) to the plasma membrane | 0.000 | 0.031 | #NUM! | AKT1,SFN |
| Antimicrobial peptides | 0.000 | 0.037 | #NUM! | S100A8 |
| PI Metabolism | 0.000 | 0.037 | #NUM! | MTMR1,RUFY1,SACM1L |
| Signaling by Activin | 0.000 | 0.053 | #NUM! | INHBB |
| Eukaryotic Translation Elongation | 0.000 | 0.051 | -2.449 | RPL18A,RPL22,RPL36,RPL37,RPL7,RPL7L1 |
| Mitochondrial biogenesis | 0.000 | 0.033 | #NUM! | ACSS2,CHD9 |
| Gamma carboxylation, hypusinylation, hydroxylation, and arylsu | 0.000 | 0.054 | #NUM! | GGCX,KDM8,TPST1 |
| Toll-like Receptor Cascades | 0.000 | 0.035 | #NUM! | ITGAM |
| Regulation of Apoptosis | 0.000 | 0.037 | #NUM! | PSMC1,PSMD13 |
| Cytosolic sensors of pathogen-associated DNA | 0.000 | 0.032 | #NUM! | POLR1C,TRIM32 |
| Regulation of beta-cell development | 0.000 | 0.024 | #NUM! | AKT1 |
| p75 NTR receptor-mediated signalling | 0.000 | 0.027 | 1.000 | ARHGEF4,HDAC2,RPS6KB1,TYK2 |
| Bile acid and bile salt metabolism | 0.000 | 0.026 | #NUM! | SLC51B |
| Degradation of beta-catenin by the destruction complex | 0.000 | 0.044 | 0.000 | CSNK1A1,CTBP2,PSMC1,PSMD13 |
| Metabolism of water-soluble vitamins and cofactors | 0.000 | 0.043 | #NUM! | ABCC1,MTR,PANK2 |
| Signaling by NOTCH1 | 0.000 | 0.013 | #NUM! | HDAC2 |
| Immunoregulatory interactions between a Lymphoid and a non- | 0.000 | 0.037 | #NUM! | CD200R1,COL1A2,PIANP |
| TCF dependent signaling in response to WNT | 0.000 | 0.048 | -1.000 | AKT1,CSNK1A1,H4C1,LEO1,PSMC1,PSMD13,SOX2,SOX9,WNT4 |
| Assembly of collagen fibrils and other multimeric structures | 0.000 | 0.054 | #NUM! | COL1A2,COL6A2,ITGA6 |
| TCR signaling | 0.000 | 0.027 | #NUM! | PSMC1,PSMD13,UBE2N |
| COPII-mediated vesicle transport | 0.000 | 0.014 | #NUM! | LMAN1 |
| Asparagine N-linked glycosylation | 0.000 | 0.055 | 0.378 | B4GALNT2,B4GALT2,DHDDS,LMAN1,PDIA3,PSMC1,ST3GAL4 |
| Metabolism of amine-derived hormones | 0.000 | 0.056 | #NUM! | DDC |
| Phase I - Functionalization of compounds | 0.000 | 0.026 | #NUM! | ACSS2,ADH5 |
| MHC class II antigen presentation | 0.000 | 0.019 | #NUM! | DCTN3,KIF5B |
| Abacavir ADME | 0.000 | 0.033 | #NUM! | RPS6KB1,TYK2 |
| Binding and Uptake of Ligands by Scavenger Receptors | 0.000 | 0.054 | #NUM! | COL1A2,HSP90AA1 |
| Visual phototransduction | 0.000 | 0.018 | #NUM! | CNGB1 |
| Selenoamino acid metabolism | 0.000 | 0.049 | -1.890 | AIMP1,RPL18A,RPL22,RPL36,RPL37,RPL7,RPL7L1 |
| Mitotic Metaphase and Anaphase | 0.000 | 0.049 | -0.905 | CENPL,LEMD2,NUDC,NUP35,PDS5B,PPP1CC,PSMC1,PSMD13,SMC3,TNPO1,ZWILCH |
| Oxidative Stress Induced Senescence | 0.000 | 0.051 | 1.000 | AGO1,CBX2,CDKN2B,H4C1 |

| | | | | |
|---|---|---|---|---|
| Senescence-Associated Secretory Phenotype (SASP) | 0.000 | 0.048 | #NUM! | CDKN2B,H4C1,MAPK7 |
| DNA Damage/Telomere Stress Induced Senescence | 0.000 | 0.048 | #NUM! | H4C1,HIRA |
| EPH-Ephrin signaling | 0.000 | 0.011 | #NUM! | LYN |
| SUMOylation of ubiquitinylation proteins | 0.000 | 0.053 | #NUM! | NUP35,PML |
| WNT ligand biogenesis and trafficking | 0.000 | 0.039 | #NUM! | WNT4 |
| Detoxification of Reactive Oxygen Species | 0.000 | 0.029 | #NUM! | CYBA |
| HSP90 chaperone cycle for steroid hormone receptors in the pre | 0.000 | 0.043 | #NUM! | DCTN3,HSP90AA1 |
| Metabolism of polyamines | 0.000 | 0.050 | #NUM! | PSMC1,PSMD13,SAT1 |
| Class A/1 (Rhodopsin-like receptors) | 0.000 | 0.035 | 0.632 | ACKR3,C5,CCK,CX3CR1,HTR2A,KISS1R,LPAR6,MCHR1,PNOC,TAC3 |
| L1CAM interactions | 0.000 | 0.009 | #NUM! | FGFR1 |
| Energy dependent regulation of mTOR by LKB1-AMPK | 0.000 | 0.035 | #NUM! | RRAGC |
| Glyoxylate metabolism and glycine degradation | 0.000 | 0.056 | #NUM! | AMT |
| Striated Muscle Contraction | 0.000 | 0.028 | #NUM! | ACTA1 |
| Peroxisomal lipid metabolism | 0.000 | 0.039 | #NUM! | CRAT |
| Protein folding | 0.000 | 0.055 | 1.342 | CCT5,CCT8,GNG3,GNG7,PFDN5 |
| G alpha (q) signalling events | 0.000 | 0.055 | -0.333 | CCK,GNG3,GNG7,HTR2A,KISS1R,LPAR6,MCHR1,RGS4,TAC3 |
| G alpha (12/13) signalling events | 0.000 | 0.051 | 0.000 | ARHGEF4,GNG3,GNG7,RHOB |
| Class C/3 (Metabotropic glutamate/pheromone receptors) | 0.000 | 0.046 | #NUM! | GRM4 |
| Transport of vitamins, nucleosides, and related molecules | 0.000 | 0.053 | #NUM! | ARL2BP,SLCO3A1 |
| Sphingolipid metabolism | 0.000 | 0.040 | 0.000 | ABCC1,B3GALNT1,PLPP2,PRKD3 |
| Smooth Muscle Contraction | 0.000 | 0.026 | #NUM! | DYSF |
| Interleukin-1 family signaling | 0.000 | 0.040 | -1.342 | PSMC1,PSMD13,PTPN11,TAB1,UBE2N |
| MAP kinase activation | 0.000 | 0.051 | #NUM! | TAB1,UBE2N |
| Mitotic G2-G2/M phases | 0.000 | 0.057 | -0.905 | CEP290,CEP70,CPAP,DCTN3,HSP90AA1,LIN54,PPP1R12A,PRKACA,PSMC1,PSMD13,SFI1 |
| Regulation of mitotic cell cycle | 0.000 | 0.022 | #NUM! | PSMC1,PSMD13 |
| SUMOylation of DNA replication proteins | 0.000 | 0.044 | #NUM! | NUP35,SUMO2 |
| RIPK1-mediated regulated necrosis | 0.000 | 0.033 | #NUM! | HSP90AA1 |
| Myogenesis | 0.000 | 0.035 | #NUM! | MYOG |
| Hedgehog ligand biogenesis | 0.000 | 0.046 | #NUM! | PSMC1,PSMD13,SHH |
| Signaling by Retinoic Acid | 0.000 | 0.054 | #NUM! | RARB,RDH13 |
| Mitochondrial translation | 0.000 | 0.028 | #NUM! | MRPL39,MRPL49,MRPL51 |
| Hedgehog 'off' state | 0.000 | 0.055 | 0.000 | ADCY5,CSNK1A1,IFT172,PRKACA,PSMC1,PSMD13 |
| Activation of anterior HOX genes in hindbrain during early embr | 0.000 | 0.040 | #NUM! | H4C1,POLR2G,RARB |
| Pyroptosis | 0.000 | 0.044 | #NUM! | HMGB1 |
| Activated PKN1 stimulates transcription of AR regulated genes K | 0.000 | 0.056 | #NUM! | H4C1 |
| Hedgehog 'on' state | 0.000 | 0.057 | 1.414 | CSNK1A1,DZIP1,PSMC1,PSMD13,RPS6KB1,SHH,SPOPL,TYK2 |
| RHO GTPases Activate Formins | 0.000 | 0.046 | 0.000 | CENPL,DAAM1,NUDC,PPP1CC,RHOB,ZWILCH |
| TNFR2 non-canonical NF-kB pathway | 0.000 | 0.029 | #NUM! | PSMC1,PSMD13 |
| MAPK6/MAPK4 signaling | 0.000 | 0.044 | -1.000 | AGO1,PRKACA,PSMC1,PSMD13 |
| Cargo concentration in the ER | 0.000 | 0.031 | #NUM! | LMAN1 |
| Interleukin-10 signaling | 0.000 | 0.056 | #NUM! | JAK1,TYK2 |
| Regulation of TP53 Activity through Phosphorylation | 0.000 | 0.044 | 0.000 | RBBP8,RPA1,RPA2,TAF3 |
| Regulation of TP53 Activity through Methylation | 0.000 | 0.053 | #NUM! | KMT5A |
| Regulation of TP53 Expression and Degradation | 0.000 | 0.054 | #NUM! | AKT1,PRDM1 |
| RNA polymerase II transcribes snRNA genes | 0.000 | 0.041 | #NUM! | ELL3,POLR2G,SNAPC2 |
| COPI-mediated anterograde transport | 0.000 | 0.044 | 1.000 | DCTN3,KDELR2,TMED3,TMED7 |
| Intra-Golgi and retrograde Golgi-to-ER traffic | 0.000 | 0.048 | -0.333 | BICD2,DCTN3,KDELR2,KIF5B,PLIN3,STX16,TMED3,TMED7,TRIP11 |
| Mitotic Prophase | 0.000 | 0.053 | -0.447 | H4C1,KMT5A,LEMD2,NEK7,NUP35 |
| DNA Replication Pre-Initiation | 0.000 | 0.031 | #NUM! | H4C1,PSMC1,PSMD13 |
| Synthesis of DNA | 0.000 | 0.049 | 1.633 | CDC45,GINS2,PSMC1,PSMD13,RPA1,RPA2 |
| Cell Cycle Checkpoints | 0.000 | 0.050 | 0.277 | CDC45,CENPL,H4C1,NUDC,PPP1CC,PSMC1,PSMD13,RBBP8,RPA1,RPA2,SFN,UBE2N,ZWILCH |
| Eukaryotic Translation Termination | 0.000 | 0.051 | -2.449 | RPL18A,RPL22,RPL36,RPL37,RPL7,RPL7L1 |
| DNA Damage Bypass | 0.000 | 0.042 | #NUM! | RPA1,RPA2 |
| Processing of Capped Intronless Pre-mRNA | 0.000 | 0.035 | #NUM! | SNRPD3 |
| Fatty acyl-CoA biosynthesis | 0.000 | 0.028 | #NUM! | ELOVL1 |
| Apoptotic execution phase | 0.000 | 0.020 | #NUM! | HMGB1 |
| Response to elevated platelet cytosolic Ca2+ | 0.000 | 0.033 | 0.000 | CALU,HGF,RARRES2,SERPING1 |
| Transcriptional Regulation by VENTX | 0.000 | 0.053 | #NUM! | AGO1,CCND1 |
| TBC/RABGAPs | 0.000 | 0.047 | #NUM! | RAB11B,RAB35 |
| Cargo recognition for clathrin-mediated endocytosis | 0.000 | 0.029 | #NUM! | GRB2,STAM2,VAMP7 |
| Clathrin-mediated endocytosis | 0.000 | 0.032 | 0.000 | DNAJC6,GRB2,STAM2,VAMP7 |
| Transcriptional regulation by the AP-2 (TFAP2) family of transcri | 0.000 | 0.029 | #NUM! | YEATS4 |
| RAB GEFs exchange GTP for GDP on RABs | 0.000 | 0.056 | -0.447 | AKT1,ALS2CL,MON1A,RAB27A,RAB35 |
| Transcriptional regulation by RUNX1 | 0.000 | 0.035 | 0.447 | CBFB,CBX2,H4C1,PSMC1,PSMD13 |
| Transcriptional regulation by RUNX3 | 0.000 | 0.052 | -0.447 | CBFB,CCND1,KRAS,PSMC1,PSMD13 |
| Regulation of RUNX2 expression and activity | 0.000 | 0.042 | #NUM! | CBFB,PSMC1,PSMD13 |
| Transcriptional Regulation by E2F6 | 0.000 | 0.029 | #NUM! | RBBP8 |
| Carboxyterminal post-translational modifications of tubulin | 0.000 | 0.053 | #NUM! | AGBL5,SVBP |
| Nucleotide catabolism | 0.000 | 0.031 | #NUM! | ITPA |
| Signaling by NOTCH3 | 0.000 | 0.041 | #NUM! | FABP7,YBX1 |
| Signaling by NOTCH4 | 0.000 | 0.036 | 0.447 | AKT1,PSMC1,PSMD13,RPS6KB1,TYK2 |
| NGF-stimulated transcription | 0.000 | 0.051 | #NUM! | ATF1,ID4 |
| Peroxisomal protein import | 0.000 | 0.051 | #NUM! | CRAT,IDE,PEX13 |
| Iron uptake and transport | 0.000 | 0.052 | #NUM! | ATP6V1C1,CP,HMOX1 |
| Nonsense-Mediated Decay (NMD) | 0.000 | 0.054 | -2.449 | MAGOH,RPL18A,RPL22,RPL36,RPL37,RPL7 |
| Assembly and cell surface presentation of NMDA receptors | 0.000 | 0.028 | #NUM! | CAMK2D |
| EGR2 and SOX10-mediated initiation of Schwann cell myelinatio | 0.000 | 0.035 | #NUM! | HDAC2 |
| Aggrephagy | 0.000 | 0.056 | #NUM! | HSP90AA1,UBE2N |
| Sensory processing of sound by outer hair cells of the cochlea | 0.000 | 0.055 | #NUM! | EPB41L3,PCDH15,RIPOR2 |
| Sensory perception of taste | 0.000 | 0.033 | #NUM! | GRM4 |
| KEAP1-NFE2L2 pathway | 0.000 | 0.048 | 1.134 | AKT1,EIF2AK3,KEAP1,PSMC1,PSMD13,RPS6KB1,TYK2 |
| Amyloid fiber formation | 0.000 | 0.048 | #NUM! | H4C1,HSPG2,MFGE8 |
| Formation of paraxial mesoderm | 0.000 | 0.050 | #NUM! | FGFR1 |
| Kinesins | 0.000 | 0.020 | #NUM! | KIF5B |
| Factors involved in megakaryocyte development and platelet pr | 0.000 | 0.051 | 0.000 | EHD1,HDAC2,PRKACA,SIN3A |
| Ion channel transport | 0.000 | 0.041 | -0.378 | ATP1B2,ATP1B3,ATP6V1C1,ATP8A1,ATP9B,CAMK2D,NEDD4L |
| Formation of WDR5-containing histone-modifying complexes | 0.000 | 0.048 | #NUM! | KANSL2,TADA3 |
| Hepatic Cholestasis | 0.000 | 0.046 | 0.707 | ABCC1,ADCY5,AKT1,Calm4,INSR,LEP,PPP3CC,PRKACA,PRKD3 |
| LXR/RXR Activation | 0.000 | 0.038 | #NUM! | ECHS1,LPL,MYLIP,S100A8 |
| Hepatic Fibrosis / Hepatic Stellate Cell Activation | 0.000 | 0.049 | #NUM! | BCL2,COL13A1,COL19A1,COL1A2,COL6A2,FGFR1,HGF,KLF6,LEP |
| CGAS-STING Signaling Pathway | 0.000 | 0.052 | 2.449 | ABCC1,ATG16L1,ATP6V1C1,EIF2AK3,LEP,MAP1LC3A |
| NAFLD Signaling Pathway | 0.000 | 0.049 | 1.897 | AKT1,EIF2AK3,G6PC2,GNB1L,INSR,JAK1,LEP,PNPLA3,SOCS2,TYK2 |
| VDR/RXR Activation | 0.000 | 0.028 | #NUM! | COL13A1,PRKD3 |
| BBSome Signaling Pathway | 0.000 | 0.051 | 1.091 | ACKR3,ADGRD1,CBX2,CCT5,CCT8,CEP290,CX3CR1,DZIP1,GPR149,GRM4,HTR2A,IFT172,IFT80,INSR,IQCB1,KISS1R,LEP,LPAR6,LZTFL1,MCHR1,NTRK2,PRKACA,SSX2IP |
| Cohesin Chromatin Regulation Pathway | 0.000 | 0.043 | 1.414 | H4C1,MED11,MED17,MED8,PCDH15,POLR2G,SMC3,SOX2,ZBTB21 |
| Histone Modification Signaling Pathway | 0.000 | 0.046 | -0.302 | BAP1,HDAC2,KAT6B,KDM5A,KDM8,KMT5A,L3MBTL3,PHF23,PRDM2,SFN,SMNDC1,TAF3 |
| Nuclear Cytoskeleton Signaling Pathway | 0.000 | 0.044 | -0.707 | ACTA1,CDH5,DCTN3,ITGA6,ITGA8,ITGAM,KIF5B,LYN,NUP35 |
| Hematoma Resolution Signaling Pathway | 0.000 | 0.040 | -0.333 | C5,CX3CR1,HMGB1,HMOX1,ITGAM,LEP,NDUFS3,NDUFS4,NDUFS8 |
| Glycation Signaling Pathway | 0.000 | 0.051 | 0.302 | AKT1,EIF2AK3,HMGB1,JAK1,KRAS,LEP,MAPK7,PFKL,PRKD3,RAP1A,RRAS |
| Ribosomal Quality Control Signaling Pathway | 0.000 | 0.056 | -2.111 | AGO1,DIS3,N4BP2,PSMC1,PSMD13,RPL18A,RPL22,RPL36,RPL37,RPL7,RPL7L1 |
| Sheddase Signaling Pathway | 0.000 | 0.023 | -2.000 | COL1A2,ERBB4,GNAI1,HGF |
| Inositol phosphate metabolism | 0.000 | 0.043 | #NUM! | PLD4,PPIP5K2 |

| | | | | |
|---|---|---|---|---|
| Post-translational modification: synthesis of GPI-anchored prote | 0.000 | 0.040 | #NUM! | LYPD1,OPCML,PIGN |
| Pyruvate metabolism | 0.000 | 0.021 | #NUM! | GID8 |
| Citric acid cycle (TCA cycle) | 0.000 | 0.048 | #NUM! | IDH3A |
| Ribavirin ADME | 0.000 | 0.048 | #NUM! | ITPA,RPS6KB1,TYK2 |
| Regulation of CDH11 Expression and Function | 0.000 | 0.033 | #NUM! | AGO1 |
| Somitogenesis | 0.000 | 0.036 | #NUM! | PSMC1,PSMD13 |
| PKR-mediated signaling | 0.000 | 0.037 | #NUM! | EIF2B2,SUMO2,VPS37C |
| Signaling by TGFBR3 | 0.000 | 0.041 | #NUM! | AGO1,MYOG |
| Regulation of endogenous retroelements | 0.000 | 0.056 | 1.000 | H4C1,HDAC2,MTA1,SUMO2 |
| Irritable Bowel Syndrome Signaling Pathway | 0.000 | 0.023 | 0.447 | ATG16L1,LEP,RHOB,TAB1,WNT4 |
| Lung Ionic Balance Signaling Pathway | 0.000 | 0.040 | 0.688 | ACKR3,ADGRD1,ATP1B2,ATP1B3,CNGB1,CX3CR1,GPR149,GRM4,HTR2A,KCNMB4,KISS1R,LPAR6,LYN,MCHR1,NEDD4,PDIA3,PRKACA,PRKD3,PRSS8 |
| NAP1L1 Transcription Regulation Signaling Pathway | 0.000 | 0.027 | #NUM! | AKT1,CCND1 |
| Cerebral Malformation Signaling Pathway | 0.000 | 0.055 | 0.000 | AKT1,BMP6,CDH5,CDKN2B,MAPK7,RAP1A,TLNRD1 |
| Preeclampsia Signaling Pathway | 0.000 | 0.055 | 0.000 | AKT1,CCND1,GNB1L,HMOX1,JAK1,KCNMB4,PKN3,TYK2 |
| Tuberculosis Latent Signaling Pathway | 0.000 | 0.036 | #NUM! | ADCY5,GNAI1,PRKACA |
| Zn Homeostasis Signaling Pathway | 0.000 | 0.041 | 0.408 | ACKR3,ADCY5,ADGRD1,AKT1,BCR,BMP3,BMP6,CX3CR1,CYBA,GPR149,GRM4,HTR2A,INSR,ITPR1,KISS1R,LPAR6,LYN,MCHR1,PDE7B,PPP3CC,PRKACA,PRKD3,SLC30A8,SLC39A14,TDP2 |
| Meiosis Initiation Signaling Pathway | 0.000 | 0.042 | 1.000 | HDAC2,RPA1,RPA2,SMC3 |
| Membrane Repair Signaling Pathway | 0.000 | 0.039 | 1.000 | DYSF,STX16,SYT10,TSPAN13 |
| Chromatin modifying enzymes | 0.000 | 0.055 | -0.577 | CCND1,ELP5,H4C1,HDAC2,KANSL2,KAT6B,KDM5A,KMT5A,MSL2,MTA1,TADA3,YEATS4 |
| NIK-->noncanonical NF-kB signaling | 0.000 | 0.033 | #NUM! | PSMC1,PSMD13 |
| Epigenetic regulation of gene expression by MLL3 and MLL4 com | 0.000 | 0.035 | #NUM! | H4C1,LPL,MED17 |
| Proteasome assembly | 0.000 | 0.039 | #NUM! | PSMC1,PSMD13 |
| Phosphorylated BMAL1:CLOCK (ARNTL:CLOCK) activates express | 0.000 | 0.046 | #NUM! | BHLHE40 |
| Degradation of CRY and PER proteins | 0.000 | 0.032 | #NUM! | PSMC1,PSMD13 |
| RORA,B,C and NR1D1 (REV-ERBA) regulate gene expression | 0.000 | 0.040 | #NUM! | CHD9 |
| Thioredoxin Signaling Pathway | 0.000 | 0.029 | #NUM! | AKT1,C5 |
| Ceramide Signaling | 0.000 | 0.057 | 1.342 | AKT1,BCL2,KRAS,RAP1A,RRAS |
| Tight Junction Signaling | 0.000 | 0.035 | #NUM! | ACTA1,AKT1,CLDN8,CRB3,PRKACA,STX16 |
| TR/RXR Activation | 0.000 | 0.025 | #NUM! | AKT1,CCND1,RPS6KB1 |
| Activation of IRF by Cytosolic Pattern Recognition Receptors | 0.000 | 0.020 | #NUM! | IFIT2 |
| Clathrin-mediated Endocytosis Signaling | 0.000 | 0.046 | #NUM! | ACTA1,AP1G2,AP3S1,FGF13,GRB2,PPP3CC,RAB11B,S100A8,SH3BP4 |
| IL-12 Signaling and Production in Macrophages | 0.000 | 0.052 | 0.707 | AKT1,Calm4,COL1A2,EBI3,Fcgr3,IL6ST,ITGAM,PRKD3,S100A8,TYK2 |
| Role of Pattern Recognition Receptors in Recognition of Bacteria | 0.000 | 0.023 | #NUM! | C5,LEP,PRKD3 |
| Role of PKR in Interferon Induction and Antiviral Response | 0.000 | 0.043 | -1.000 | HMGB1,HSP90AA1,JAK1,NLRP1,TYK2 |
| TREM1 Signaling | 0.000 | 0.049 | #NUM! | AKT1,GRB2,NLRP1 |
| MIF Regulation of Innate Immunity | 0.000 | 0.049 | #NUM! | PLA2G12B,PLA2G4B |
| CD40 Signaling | 0.000 | 0.015 | #NUM! | ATF1 |
| Cytotoxic T Lymphocyte-mediated Apoptosis of Target Cells | 0.000 | 0.032 | #NUM! | BCL2 |
| CD27 Signaling in Lymphocytes | 0.000 | 0.018 | #NUM! | BCL2L1 |
| Lymphotoxin β Receptor Signaling | 0.000 | 0.037 | #NUM! | AKT1,BCL2L1 |
| IL-17 Signaling | 0.000 | 0.049 | 0.378 | AKT1,HSP90AA1,JAK1,KRAS,LEP,RAP1A,RRAS |
| Airway Pathology in Chronic Obstructive Pulmonary Disease | 0.000 | 0.022 | #NUM! | FGF13,LEP |
| Induction of Apoptosis by HIV1 | 0.000 | 0.048 | #NUM! | BCL2,BCL2L1,SLC25A10 |
| T Helper Cell Differentiation | 0.000 | 0.015 | #NUM! | IL6ST |
| CD28 Signaling in T Helper Cells | 0.000 | 0.053 | -1.342 | AKT1,Calm4,GRB2,ITPR1,PPP3CC,PTPN11 |
| Dendritic Cell Maturation | 0.000 | 0.038 | 0.447 | AKT1,COL1A2,Fcgr3,LEP,PDIA3,TAB1 |
| Reelin Signaling in Neurons | 0.000 | 0.043 | -0.816 | AKT1,ARHGEF4,CAMK2D,LYN,RAP1A,RELN |
| Cellular Effects of Sildenafil (Viagra) | 0.000 | 0.048 | -0.756 | ACKR3,ACTA1,ADCY5,ADGRD1,AKT1,BCL2,BMPR2,CACNG2,CACNG7,Calm4,CNGB1,CX3CR1,GNAI1,GNG3,GNG7,GPR149,GRM4,HSP90AA1,HTR2A,KISS1R,KRAS,LPAR6,MAPK7,MCHR1,PPP1CC,PPP1R10,PPP1R12A,PPP3CC,PRKACA,PRKG2 |
| Factors Promoting Cardiogenesis in Vertebrates | 0.000 | 0.054 | 1.414 | BMP3,BMP6,BMPR2,CAMK2D,CCND1,PDIA3,PRKD3,WNT4 |
| Cardiomyocyte Differentiation via BMP Receptors | 0.000 | 0.042 | #NUM! | BMPR2 |
| Lipid Antigen Presentation by CD1 | 0.000 | 0.053 | #NUM! | PDIA3 |
| Cell Cycle Regulation by BTG Family Proteins | 0.000 | 0.054 | #NUM! | CCND1,CNOT7 |
| Mitotic Roles of Polo-Like Kinase | 0.000 | 0.031 | #NUM! | HSP90AA1,SMC3 |
| HMGB1 Signaling | 0.000 | 0.054 | -0.378 | AKT1,HMGB1,KAT6B,KRAS,LEP,RAP1A,RHOB,RRAS |
| Maturity Onset Diabetes of Young (MODY) Signaling | 0.000 | 0.044 | #NUM! | CACNG2,CACNG7,INSR |
| Role of CHK Proteins in Cell Cycle Checkpoint Control | 0.000 | 0.035 | #NUM! | ATMIN,RPA1 |
| ATM Signaling | 0.000 | 0.052 | #NUM! | ATF1,FANCD2,PPP1CC,RBBP8,SMC3 |
| Germ Cell-Sertoli Cell Junction Signaling | 0.000 | 0.056 | #NUM! | ACTA1,AKT1,ITGA6,KEAP1,KRAS,PAK4,RAP1A,RHOB,RRAS |
| Aldosterone Signaling in Epithelial Cells | 0.000 | 0.054 | 0.447 | DNAJC2,DNAJC6,HSP90AA1,ITPR1,KCNMB4,KRAS,NEDD4,PDIA3,PRKD3 |
| Growth Hormone Signaling | 0.000 | 0.045 | #NUM! | PRKD3,RPS6KB1,SOCS2 |
| Type I Diabetes Mellitus Signaling | 0.000 | 0.029 | #NUM! | BCL2,JAK1,SOCS2 |
| Type II Diabetes Mellitus Signaling | 0.000 | 0.047 | #NUM! | AKT1,CACNG2,CACNG7,INSR,ITPR1,PRKD3,SOCS2 |
| Gα12/13 Signaling | 0.000 | 0.055 | 0.378 | AKT1,CDH5,KRAS,LPAR6,MAPK7,RAP1A,RRAS |
| mTOR Signaling | 0.000 | 0.054 | 0.577 | AKT1,EIF4E,GNB1L,HMOX1,INSR,KRAS,PLD4,PRKD3,RAP1A,RHOB,RPS6KB1,RRAS |
| Communication between Innate and Adaptive Immune Cells | 0.000 | 0.015 | #NUM! | CD79B |
| Sphingosine-1-phosphate Signaling | 0.000 | 0.046 | -0.447 | ADCY5,AKT1,GNAI1,PDIA3,RHOB |
| Crosstalk between Dendritic Cells and Natural Killer Cells | 0.000 | 0.032 | #NUM! | ACTA1,CAMK2D |
| ILK Signaling | 0.000 | 0.026 | 0.000 | ACTA1,AKT1,CCND1,PPP1R12A,RHOB |
| FAK Signaling | 0.000 | 0.046 | 0.378 | ACKR3,ADGRD1,AKT1,ARHGAP35,BMPR2,CCND1,CDH5,COL1A2,CX3CR1,GPR149,GRB2,GRM4,HTR2A,IL6ST,ITGA6,ITGA8,ITGAM,KISS1R,KRAS,LPAR6,MAPK7,MCHR1,PAK4,RAP1A,RB1CC1,RRAS,SOCS2,WIPF1 |
| Role of Osteoblasts, Osteoclasts and Chondrocytes in Rheumato | 0.000 | 0.047 | #NUM! | AKT1,BCL2,BMP3,BMP6,BMPR2,Calm4,CSNK1A1,FRZB,PPP3CC,WNT4 |
| Altered T Cell and B Cell Signaling in Rheumatoid Arthritis | 0.000 | 0.013 | #NUM! | CD79B |
| B Cell Development | 0.000 | 0.037 | #NUM! | CD79B |
| Breast Cancer Regulation by Stathmin1 | 0.000 | 0.052 | -0.200 | ACKR3,ADGRD1,ARHGEF4,BMP6,CAMK2D,CCND1,CX3CR1,GNB1L,GNG3,GNG7,GPR149,GRB2,GRM4,HGF,HTR2A,KISS1R,KRAS,LPAR6,MCHR1,PPP1CC,PPP1R10,PPP1R12A,PRKACA,PRKD3,RAP1A,RB1CC1,RPS6KB1,RRAS,SHC3 |
| RANK Signaling in Osteoclasts | 0.000 | 0.034 | #NUM! | AKT1,Calm4,PPP3CC |
| Granzyme A Signaling | 0.000 | 0.045 | #NUM! | NDUFS3,NDUFS4,NDUFS8 |
| Role of WNT/GSK-3β Signaling in the Pathogenesis of Influenza | 0.000 | 0.033 | #NUM! | CSNK1A1,WNT4 |
| TWEAK Signaling | 0.000 | 0.028 | #NUM! | BAG4 |
| TNFR1 Signaling | 0.000 | 0.020 | #NUM! | PAK4 |
| Role of PI3K/AKT Signaling in the Pathogenesis of Influenza | 0.000 | 0.042 | #NUM! | AKT1,GNAI1 |
| Role of Hypercytokinemia/hyperchemokinemia in the Pathogene | 0.000 | 0.055 | #NUM! | IFIT2,JAK1,TYK2 |
| Antiproliferative Role of TOB in T Cell Signaling | 0.000 | 0.029 | #NUM! | TOB1 |
| MSP-RON Signaling Pathway | 0.000 | 0.056 | #NUM! | ACTA1,ITGAM,Klk1b1 (includes others) |
| OX40 Signaling Pathway | 0.000 | 0.044 | #NUM! | BCL2,BCL2L1 |
| IL-17A Signaling in Airway Cells | 0.000 | 0.050 | #NUM! | AKT1,JAK1,TYK2 |
| IL-17A Signaling in Fibroblasts | 0.000 | 0.028 | #NUM! | BCL2,COL1A2 |
| Sertoli Cell-Sertoli Cell Junction Signaling | 0.000 | 0.052 | 0.000 | ACTA1,AKT1,CDH5,CLDN8,GRB2,ITGA6,KEAP1,KRAS,MAP3K20,PAK4,RAP1A,RRAS |
| Estrogen-mediated S-phase Entry | 0.000 | 0.039 | #NUM! | CCND1 |
| GADD45 Signaling | 0.000 | 0.017 | #NUM! | CCND1 |
| Heparan Sulfate Biosynthesis | 0.000 | 0.050 | #NUM! | B3GAT2,CHST15,XYLT2 |
| Retinoate Biosynthesis I | 0.000 | 0.032 | #NUM! | RDH13 |
| Dopamine Degradation | 0.000 | 0.050 | #NUM! | Aldh3a2 |
| Heparan Sulfate Biosynthesis (Late Stages) | 0.000 | 0.019 | #NUM! | CHST15 |
| Ubiquinol-10 Biosynthesis (Eukaryotic) | 0.000 | 0.042 | #NUM! | MICAL2 |
| tRNA Splicing | 0.000 | 0.044 | #NUM! | PDE7B,TDP2 |
| The Visual Cycle | 0.000 | 0.053 | #NUM! | RDH13 |
| D-myo-inositol-5-phosphate Metabolism | 0.000 | 0.046 | 0.000 | G6PC2,PPFIA1,PPP1CC,PPP1R10,PPP1R12A,PPP3CC,PTPN11,SACM1L,STYX |
| NAD Salvage Pathway II | 0.000 | 0.048 | #NUM! | NMRK2 |
| Pyridoxal 5'-phosphate Salvage Pathway | 0.000 | 0.030 | #NUM! | CSNK1A1,MAPK7 |
| Triacylglycerol Degradation | 0.000 | 0.051 | #NUM! | LPL,PNPLA3 |
| Glutathione-mediated Detoxification | 0.000 | 0.048 | #NUM! | Gstm7 |
| tRNA Charging | 0.000 | 0.053 | #NUM! | IARS2,NARS2 |
| D-myo-inositol (1,4,5,6)-Tetrakisphosphate Biosynthesis | 0.000 | 0.050 | 0.000 | G6PC2,PPFIA1,PPP1CC,PPP1R10,PPP1R12A,PPP3CC,PTPN11,SACM1L,STYX |

| | | | | |
|---|---|---|---|---|
| Superpathway of Inositol Phosphate Compounds | 0.000 | 0.044 | 0.378 | G6PC2,PPFIA1,PPIP5K2,PPP1CC,PPP1R10,PPP1R12A,PPP3CC,PTPN11,SACM1L,STYX |
| D-myo-inositol (3,4,5,6)-tetrakisphosphate Biosynthesis | 0.000 | 0.050 | 0.000 | G6PC2,PPFIA1,PPP1CC,PPP1R10,PPP1R12A,PPP3CC,PTPN11,SACM1L,STYX |
| 3-phosphoinositide Degradation | 0.000 | 0.053 | -0.378 | G6PC2,MTMR1,PPFIA1,PPP1CC,PPP1R10,PPP1R12A,PPP3CC,PTPN11,SACM1L,STYX |
| 3-phosphoinositide Biosynthesis | 0.000 | 0.044 | 0.000 | G6PC2,PPFIA1,PPP1CC,PPP1R10,PPP1R12A,PPP3CC,PTPN11,SACM1L,STYX |
| Tryptophan Degradation III (Eukaryotic) | 0.000 | 0.046 | #NUM! | GCDH |
| Salvage Pathways of Pyrimidine Ribonucleotides | 0.000 | 0.022 | #NUM! | CSNK1A1,MAPK7 |
| Glycolysis I | 0.000 | 0.044 | #NUM! | PFKL |
| Gluconeogenesis I | 0.000 | 0.044 | #NUM! | MDH1 |
| Methylglyoxal Degradation III | 0.000 | 0.053 | #NUM! | AKR1A1 |
| Chondroitin Sulfate Biosynthesis (Late Stages) | 0.000 | 0.026 | #NUM! | CHST15 |
| Histidine Degradation VI | 0.000 | 0.050 | #NUM! | MICAL2 |
| Stearate Biosynthesis I (Animals) | 0.000 | 0.050 | #NUM! | DHCR24,ELOVL1,LPGAT1 |
| Pregnenolone Biosynthesis | 0.000 | 0.053 | #NUM! | MICAL2 |
| Fatty Acid α-oxidation | 0.000 | 0.056 | #NUM! | Aldh3a2 |
| Methionine Degradation I (to Homocysteine) | 0.000 | 0.053 | #NUM! | MGMT |
| Remodeling of Epithelial Adherens Junctions | 0.000 | 0.032 | #NUM! | ACTA1,HGF |
| Cysteine Biosynthesis III (mammalia) | 0.000 | 0.048 | #NUM! | MGMT |
| Agranulocyte Adhesion and Diapedesis | 0.000 | 0.054 | #NUM! | ACKR3,ACTA1,C5,Ccl11,CDH5,CLDN8,Cxcl10,GLG1,GNAI1 |
| PCP (Planar Cell Polarity) Pathway | 0.000 | 0.051 | #NUM! | DAAM1,PRICKLE1,WNT4 |
| Unfolded protein response | 0.000 | 0.034 | #NUM! | BCL2,DNAJC6,EIF2AK3 |
| WNT/Ca2+ Signaling Pathway | 0.000 | 0.015 | #NUM! | PDIA3 |
| Toll-like Receptor Signaling | 0.000 | 0.014 | #NUM! | TAB1 |
| Protein Ubiquitination Pathway | 0.000 | 0.054 | -0.577 | BAP1,DNAJC2,DNAJC6,HSP90AA1,NEDD4,NEDD4L,PSMC1,PSMD13,Ube2e1,UBE2N,UBE2Q1,UBE2R2,UBE2W,USP24 |
| Endoplasmic Reticulum Stress Pathway | 0.000 | 0.050 | #NUM! | EIF2AK3 |
| Cell Cycle: G2/M DNA Damage Checkpoint Regulation | 0.000 | 0.020 | #NUM! | SFN |
| Antigen Presentation Pathway | 0.000 | 0.040 | #NUM! | PDIA3 |
| Phototransduction Pathway | 0.000 | 0.041 | #NUM! | CNGB1,PRKACA |
| Death Receptor Signaling | 0.000 | 0.043 | 1.000 | ACTA1,BCL2,PARP10,PARP6 |
| p38 MAPK Signaling | 0.000 | 0.055 | 0.000 | ATF1,HMGN1,PLA2G12B,PLA2G4B,RPS6KB1,TAB1 |
| Apoptosis Signaling | 0.000 | 0.050 | -0.447 | BCL2,BCL2L1,KRAS,RAP1A,RRAS |
| T Cell Receptor Signaling | 0.000 | 0.056 | 0.333 | AKT1,Calm4,GNB1L,GRB2,KRAS,PPP3CC,PTPN11,RAP1A,RRAS,SHC3 |
| G-Protein Coupled Receptor Signaling | 0.000 | 0.048 | 0.180 | ACKR3,ADCY5,ADGRD1,AKT1,Calm4,CAMK2D,CCND1,CNGB1,CX3CR1,GNAI1,GNAO1,GNB1L,GNG3,GNG7,GPR149,GRB2,GRM4,HTR2A,KISS1R,KRAS,LPAR6,MAPK7,MCHR1,PAK4,PDE7B,PPP3CC,PRKACA,RAP1A,RGS4,RRAS,SHC3,TDP2 |
| Phagosome Formation | 0.000 | 0.050 | -0.365 | ACKR3,ADGRD1,AKT1,AP1G2,CX3CR1,Fcgr3,GPR149,GRB2,GRM4,HMOX1,HTR2A,ITGA6,ITGA8,ITGAM,ITPR1,KISS1R,KRAS,LPAR6,LYN,MAPK7,MCHR1,PAK4,PLA2G12B,PLA2G4B,PLD4,PNPLA3,PRKD3,RAP1A,RPS6KB1,RRAS,WIPF1 |
| Phagosome Maturation | 0.000 | 0.028 | #NUM! | ATP6V1C1,STX16,VPS16,VPS37C |
| Th1 and Th2 Activation Pathway | 0.000 | 0.034 | #NUM! | BMPR2,GRB2,JAK1,LGALS9,TYK2 |
| Th1 Pathway | 0.000 | 0.039 | 0.000 | GRB2,JAK1,LGALS9,TYK2 |
| Th2 Pathway | 0.000 | 0.034 | -1.000 | BMPR2,GRB2,JAK1,TYK2 |
| SPINK1 Pancreatic Cancer Pathway | 0.000 | 0.022 | #NUM! | Klk1b1 (includes others) |
| Endocannabinoid Cancer Inhibition Pathway | 0.000 | 0.050 | -0.378 | ADCY5,AKT1,CCND1,GNAI1,GNAO1,GNB1L,PRKACA |
| Apelin Pancreas Signaling Pathway | 0.000 | 0.046 | #NUM! | EIF2AK3,PRKACA |
| Apelin Liver Signaling Pathway | 0.000 | 0.053 | #NUM! | COL1A2 |
| Apelin Cardiac Fibroblast Signaling Pathway | 0.000 | 0.044 | #NUM! | AKT1 |
| BAG2 Signaling Pathway | 0.000 | 0.037 | #NUM! | HSP90AA1,PSMC1,PSMD13 |
| FAT10 Signaling Pathway | 0.000 | 0.036 | #NUM! | PSMC1,PSMD13 |
| PFKFB4 Signaling Pathway | 0.000 | 0.021 | #NUM! | PRKACA |
| IL-23 Signaling Pathway | 0.000 | 0.048 | #NUM! | AKT1,TYK2 |
| HOTAIR Regulatory Pathway | 0.000 | 0.035 | 1.342 | AGO1,AKT1,COL1A2,HDAC2,WNT4 |
| BEX2 Signaling Pathway | 0.000 | 0.051 | #NUM! | AKT1,BCL2,CCND1,Klk1b1 (includes others) |
| Xenobiotic Metabolism CAR Signaling Pathway | 0.000 | 0.049 | 1.342 | ABCC1,Aldh3a2,ALDH5A1,CHST15,Gstm7,HSP90AA1,PRKD3 |
| Regulation of the Epithelial Mesenchymal Transition in Developr | 0.000 | 0.024 | #NUM! | SHH,WNT4 |
| Coronavirus Pathogenesis Pathway | 0.000 | 0.049 | 0.333 | BCL2,CCND1,EIF2AK3,EIF4E,HDAC2,JAK1,SIN3A,TNPO1,TOMM70,TYK2 |
| Coronavirus Replication Pathway | 0.000 | 0.047 | #NUM! | IFITM10,RPL18A,RPL22,RPL36,RPL37,RPL7,RPL7L1 |
| Role of MAPK Signaling in Inhibiting the Pathogenesis of Influenz | 0.000 | 0.044 | #NUM! | PLA2G12B,PLA2G4B,PNPLA3 |

**Supplementary Table 4: IPA Canonical Pathways of differentially expressed genes in ECs at P28**

| **Ingenuity Canonical Pathways** | **-log(p-valu** | **Ratio** | **z-score** | **Molecules** |
|---|---|---|---|---|
| Sleep REM Signaling Pathway | 2.660 | 0.111 | -0.816 | CHRM3,CREB3,EIF4E,IL18,Mt1,RPS6KA1 |
| RAR Activation | 2.300 | 0.048 | 1.147 | ADCY6,BRD7,CNGB1,COL3A1,CREB3,CSK,DBX1,FGF10,IL18,NR2F1,PDE7B,PIK3CG,PIK3R2,PPARGC1A,RAB11FIP3,RELA,RHOBTB2,RHOD,TGM2 |
| Macrophage Alternative Activation Signaling Pathway | 2.270 | 0.065 | -1.897 | CREB3,IL18,IRF5,NFKBID,PIK3CG,PIK3R2,PPARGC1A,RELA,RPS6KA1,SREBF1 |
| Role of IL-17A in Arthritis | 2.130 | 0.102 | #NUM! | NFKBID,PIK3CG,PIK3R2,RELA,RPS6KA1 |
| HOTAIR Regulatory Pathway | 1.970 | 0.062 | 1.414 | AGO4,ATG3,COL3A1,LAMTOR5,PIK3CG,PIK3R2,RELA,VIM,WNT5A |
| Intrinsic Prothrombin Activation Pathway | 1.960 | 0.114 | 0.000 | COL3A1,F10,F2,KLK5 |
| Class I peroxisomal membrane protein import | 1.950 | 0.158 | #NUM! | ABCD2,PEX26,PXMP4 |
| Interleukin-1 processing | 1.920 | 0.286 | #NUM! | IL18,RELA |
| p75 NTR receptor-mediated signalling | 1.860 | 0.060 | 0.333 | ARHGEF2,ARHGEF40,MCF2L,PIK3CG,PIK3R2,PLK4,PRDM4,RELA,STK35 |
| RHO GTPase cycle | 1.800 | 0.043 | 1.147 | ARHGAP31,ARHGEF2,ARHGEF40,ARMCX3,CAVIN1,CDC42EP4,CSK,DDRGK1,DIAPH3,HSPE1,MCF2L,NHS,PIK3R2,RHOBTB2,RHOD,SPEN,TMEM59,TMEM87A,VIM |
| Transcriptional Regulation by MECP2 | 1.720 | 0.081 | -0.447 | AGO4,OPRM1,SGK1,TBL1XR1,TRPC3 |
| Epigenetic regulation of gene expression by MLL3 and MLL4 con | 1.690 | 0.070 | -1.633 | H4C1,PAXIP1,PLIN2,PPARGC1A,TBL1XR1,THRSP |
| NAD Signaling Pathway | 1.680 | 0.059 | 0.000 | PIK3CG,PIK3R2,PPARGC1A,RYR2,SREBF1,TGFA,TNKS2,TRPM2 |
| Acute Phase Response Signaling | 1.640 | 0.055 | 1.134 | C1ra/C1rb,F2,HNRNPK,IL18,NFKBID,PIK3CG,PIK3R2,RELA,SERPINF2 |
| Processing of Capped Intron-Containing Pre-mRNA | 1.630 | 0.047 | 1.387 | DDX46,EIF4E,GPATCH1,HNRNPK,LSM7,PDCD7,RBM17,SF3A3,SF3B3,SNRPA,SRSF11,SRSF3,SRSF4 |
| RORA,B,C and NR1D1 (REV-ERBA) regulate gene expression | 1.630 | 0.120 | #NUM! | PPARGC1A,SREBF1,TBL1XR1 |
| 2-amino-3-carboxymuconate Semialdehyde Degradation to Glut | 1.600 | 1.000 | #NUM! | ACMSD |
| NR1H2 and NR1H3-mediated signaling | 1.590 | 0.089 | 0.000 | AGO4,EEPD1,SREBF1,TBL1XR1 |
| Co-stimulation by ICOS | 1.530 | 0.182 | #NUM! | PIK3CG,PIK3R2 |
| TR/RXR Activation | 1.510 | 0.058 | -1.134 | F10,PIK3CG,PIK3R2,PPARGC1A,SREBF1,TBL1XR1,THRSP |
| Role of PI3K/AKT Signaling in the Pathogenesis of Influenza | 1.500 | 0.083 | 1.000 | NFKBID,PIK3CG,PIK3R2,RELA |
| MTOR signalling | 1.500 | 0.107 | #NUM! | EEF2K,EIF4E,LAMTOR5 |
| HMGB1 Signaling | 1.470 | 0.054 | 0.378 | IL18,KAT6B,KAT7,PIK3CG,PIK3R2,RELA,RHOBTB2,RHOD |
| Major pathway of rRNA processing in the nucleolus and cytosol | 1.460 | 0.049 | -1.265 | DDX21,EXOSC10,EXOSC4,IMP4,MRPS2,MRPS23,RPL27,RPS15,RPS27,WDR75 |
| mRNA 3 Prime End Processing Signaling Pathway | 1.460 | 0.062 | 1.000 | DICER1,EXOSC10,EXOSC4,HNRNPK,SNRPA,SRSF3 |
| Coagulation System | 1.460 | 0.103 | #NUM! | F10,F2,SERPINF2 |
| EIF2 Signaling | 1.450 | 0.047 | -0.447 | ATF5,EIF4E,MRPS2,MRPS23,PIK3CG,PIK3R2,RPL27,RPS15,RPS27,SREBF1,VPS37A |
| Extrinsic Prothrombin Activation Pathway | 1.390 | 0.154 | #NUM! | F10,F2 |
| Late endosomal microautophagy | 1.350 | 0.094 | #NUM! | PLIN2,VIM,VPS37A |
| Role of RIG1-like Receptors in Antiviral Innate Immunity | 1.350 | 0.094 | #NUM! | MAVS,NFKBID,RELA |
| Lymphotoxin β Receptor Signaling | 1.340 | 0.074 | #NUM! | NFKBID,PIK3CG,PIK3R2,RELA |
| DNA Double-Strand Break Repair by Homologous Recombinatio | 1.330 | 0.143 | #NUM! | ATRX,BRCA2 |
| ER-Mitochondrial Communication Signaling Pathway | 1.320 | 0.063 | 0.447 | IL18,MFN2,PTDSS1,TOMM34,VPS13D |
| Chromatin modifying enzymes | 1.320 | 0.046 | -0.632 | ASH1L,H4C1,KANSL2,KAT6B,KAT7,MBIP,MEAF6,RELA,TAF10,TBL1XR1 |
| S-methyl-5-thio-α-D-ribose 1-phosphate Degradation | 1.310 | 0.500 | #NUM! | MRI1 |
| Sulfate Activation for Sulfonation | 1.310 | 0.500 | #NUM! | PAPSS1 |
| Anandamide Degradation | 1.310 | 0.500 | #NUM! | FAAH |
| Role of BRCA1 in DNA Damage Response | 1.300 | 0.063 | #NUM! | BABAM1,BRCA2,BRD7,CHEK1,FAAP100 |
| Gamma carboxylation, hypusinylation, hydroxylation, and arylsu | 1.290 | 0.071 | 0.000 | F10,F2,JMJD4,RCCD1 |
| Adipogenesis pathway | 1.290 | 0.052 | -1.134 | KAT6B,KAT7,PAXIP1,RPS6KA1,SREBF1,TBL1XR1,WNT5A |
| FLT3 Signaling in Hematopoietic Progenitor Cells | 1.280 | 0.062 | -0.447 | CREB3,EIF4E,PIK3CG,PIK3R2,RPS6KA1 |
| mTOR Signaling | 1.280 | 0.045 | 0.447 | EIF4E,MRPS2,MRPS23,PIK3CG,PIK3R2,RHOBTB2,RHOD,RPS15,RPS27,RPS6KA1 |
| MSP-RON Signaling in Cancer Cells Pathway | 1.260 | 0.052 | 0.378 | CREB3,ELF2,KLK5,PIK3CG,PIK3R2,RELA,VIM |
| Gαq Signaling | 1.260 | 0.049 | 0.000 | CHRM3,CSK,NFKBID,PIK3CG,PIK3R2,RELA,RHOBTB2,RHOD |
| IL-9 Signaling | 1.250 | 0.086 | #NUM! | PIK3CG,PIK3R2,RELA |
| Ribosomal Quality Control Signaling Pathway | 1.250 | 0.046 | 0.816 | AGO4,EXOSC10,EXOSC4,MRPS2,MRPS23,PSME1,RPL27,RPS15,RPS27 |
| Sphingosine-1-phosphate Signaling | 1.240 | 0.055 | -0.447 | ADCY6,CASP6,PIK3CG,PIK3R2,RHOBTB2,RHOD |
| IL-33 Signaling Pathway | 1.240 | 0.051 | -0.378 | CASP6,CREB3,IL18,NFKBID,PIK3CG,PIK3R2,RPS6KA1 |
| Mitochondrial unfolded protein response (UPRmt) | 1.230 | 0.125 | #NUM! | ATF5,HSPE1 |
| Peroxisomal protein import | 1.220 | 0.068 | -1.000 | AMACR,PEX26,PEX7,UBE2D2 |
| Co-inhibition by BTLA | 1.220 | 0.068 | 0.000 | PIK3CG,PIK3R2,PLK4,STK35 |
| TWEAK Signaling | 1.220 | 0.083 | #NUM! | CASP6,NFKBID,RELA |
| IL-17A Signaling in Airway Cells | 1.200 | 0.067 | 1.000 | NFKBID,PIK3CG,PIK3R2,RELA |
| Role of IL-17F in Allergic Inflammatory Airway Diseases | 1.190 | 0.081 | #NUM! | CREB3,RELA,RPS6KA1 |
| Cyclophilin Signaling Pathway | 1.190 | 0.043 | 0.632 | CREB3,F2,NFKBID,PIK3CG,PIK3R2,SF3A3,SF3B3,TIMM50,TOMM34,WNT5A |

| | | | | |
|---|---|---|---|---|
| Abacavir ADME | 1.180 | 0.066 | 0.000 | PIK3CG,PIK3R2,PLK4,STK35 |
| Signaling by Rho Family GTPases | 1.180 | 0.042 | 0.632 | ARHGEF2,CDC42EP4,CDH2,DIAPH3,PIK3CG,PIK3R2,RELA,RHOBTB2,RHOD,SEPTIN2,VIM |
| Transcriptional Regulation by VENTX | 1.160 | 0.079 | #NUM! | AGO4,ANAPC7,RELA |
| Carboxyterminal post-translational modifications of tubulin | 1.160 | 0.079 | #NUM! | AGBL3,LRRC49,TTLL5 |
| Ribavirin ADME | 1.160 | 0.065 | 0.000 | PIK3CG,PIK3R2,PLK4,STK35 |
| Senescence-Associated Secretory Phenotype (SASP) | 1.140 | 0.064 | -1.000 | ANAPC7,H4C1,RELA,RPS6KA1 |
| Mitophagy | 1.140 | 0.077 | #NUM! | FUNDC1,MFN2,UBE2D2 |
| Nonhomologous End-Joining (NHEJ) | 1.140 | 0.077 | #NUM! | BABAM1,H4C1,PAXIP1 |
| Triacylglycerol Degradation | 1.140 | 0.077 | #NUM! | CEL,FAAH,NDST2 |
| Ciprofloxacin ADME | 1.140 | 0.333 | #NUM! | SLC22A8 |
| Inosine-5'-phosphate Biosynthesis II | 1.140 | 0.333 | #NUM! | ATIC |
| MicroRNA Biogenesis Signaling Pathway | 1.130 | 0.046 | 0.378 | AGO4,DICER1,EXOSC10,EXOSC4,PIK3CG,PIK3R2,SRSF3,TGFA |
| Stress Granule Signaling Pathway | 1.110 | 0.042 | 0.333 | EIF4E,MRPS2,MRPS23,MSI1,PIK3CG,PIK3R2,RPS15,RPS27,RPS6KA1,VPS37A |
| GP6 Signaling Pathway | 1.090 | 0.050 | 0.816 | COL3A1,COL6A6,LAMA5,PIK3CG,PIK3R2,TLN1 |
| CD40 Signaling | 1.080 | 0.061 | 1.000 | NFKBID,PIK3CG,PIK3R2,RELA |
| Signaling by SCF-KIT | 1.060 | 0.071 | #NUM! | CHEK1,PIK3R2,SH2B3 |
| Formation of WDR5-containing histone-modifying complexes | 1.060 | 0.071 | #NUM! | KANSL2,MBIP,PAXIP1 |
| IL-23 Signaling Pathway | 1.060 | 0.071 | #NUM! | PIK3CG,PIK3R2,RELA |
| GAIT Translation Signaling Pathway | 1.060 | 0.042 | -1.342 | EIF4E,IL18,MRPS2,MRPS23,RPL27,RPS15,RPS27,RPS6KA1,VPS37A |
| Signaling by Hippo | 1.060 | 0.100 | #NUM! | AMOTL1,WWTR1 |
| p53 Signaling | 1.050 | 0.053 | 0.447 | CASP6,CHEK1,PERP,PIK3CG,PIK3R2 |
| Pulmonary Fibrosis Idiopathic Signaling Pathway | 1.040 | 0.039 | 1.155 | CDH2,COL3A1,COL6A6,F2,PIK3CG,PIK3R2,RELA,RPS6KA1,TGFA,VIM,WNT5A,WWTR1 |
| Eukaryotic Translation Initiation | 1.040 | 0.046 | 1.134 | EIF4E,MRPS2,MRPS23,RPL27,RPS15,RPS27,VPS37A |
| Thrombin Signaling | 1.020 | 0.042 | 0.378 | ADCY6,ARHGEF2,ARHGEF40,F2,PIK3CG,PIK3R2,RELA,RHOBTB2,RHOD |
| GPER1 signaling | 1.020 | 0.052 | 0.447 | ADCY6,PIK3CG,PIK3R2,PLK4,STK35 |
| Melanocyte Development and Pigmentation Signaling | 1.020 | 0.052 | -0.447 | ADCY6,CREB3,PIK3CG,PIK3R2,RPS6KA1 |
| Chaperone Mediated Autophagy | 1.020 | 0.095 | #NUM! | PLIN2,VIM |
| Lysosomal oligosaccharide catabolism | 1.020 | 0.250 | #NUM! | MAN2C1 |
| Ribonucleotide Reductase Signaling Pathway | 1.010 | 0.045 | -1.134 | BRD7,CHEK1,CREB3,EIF4E,PIK3CG,PIK3R2,TNKS2 |
| Apelin Pancreas Signaling Pathway | 1.010 | 0.068 | #NUM! | PIK3CG,PIK3R2,RELA |
| Glioma Invasiveness Signaling | 1.010 | 0.057 | #NUM! | PIK3CG,PIK3R2,RHOBTB2,RHOD |
| Exosome Signaling Pathway | 0.993 | 0.044 | -0.378 | EXOSC10,EXOSC4,MRPS2,MRPS23,RPL27,RPS15,RPS27 |
| Dendritic Cell Maturation | 0.993 | 0.044 | 0.378 | COL3A1,CREB3,IL18,NFKBID,PIK3CG,PIK3R2,RELA |
| Gα12/13 Signaling | 0.992 | 0.047 | 1.342 | CDH2,F2,NFKBID,PIK3CG,PIK3R2,RELA |
| PPAR Signaling | 0.977 | 0.050 | -1.000 | IL18,NFKBID,NR2F1,PPARGC1A,RELA |
| ILK Signaling | 0.972 | 0.042 | 0.816 | CREB3,NACA,PIK3CG,PIK3R2,RELA,RHOBTB2,RHOD,VIM |
| Generic Transcription Pathway | 0.962 | 0.042 | -2.121 | NR2E3,NR2F1,NR4A1,PGR,TBL1XR1,ZKSCAN4,ZNF18,ZNF689 |
| Nephrin family interactions | 0.952 | 0.087 | #NUM! | KIRREL3,PIK3R2 |
| Surfactant metabolism | 0.952 | 0.087 | #NUM! | Csf2ra,ZDHHC2 |
| Interleukin-4 and Interleukin-13 signaling | 0.950 | 0.049 | 0.447 | IL18,JUNB,LAMA5,OPRM1,VIM |
| HDR through Homologous Recombination (HRR) or Single Stranc | 0.937 | 0.049 | -0.447 | BABAM1,BRCA2,CHEK1,H4C1,MUS81 |
| Cargo recognition for clathrin-mediated endocytosis | 0.937 | 0.049 | 1.342 | SCARB2,SYT11,TGFA,TOR1A,WNT5A |
| ICOS-ICOSL Signaling in T Helper Cells | 0.937 | 0.049 | 1.000 | CSK,NFKBID,PIK3CG,PIK3R2,RELA |
| MSP-RON Signaling in Macrophages Pathway | 0.937 | 0.049 | -1.342 | CREB3,KLK5,PIK3CG,PIK3R2,RELA |
| Hereditary Breast Cancer Signaling | 0.934 | 0.045 | 0.816 | BRCA2,CHEK1,FAAP100,PIK3CG,PIK3R2,TGFA |
| Hepatitis B Chronic Liver Pathogenesis Signaling Pathway | 0.930 | 0.043 | -0.378 | AP1S2,CAMKK1,CREB3,IL18,MAVS,PIK3CG,PIK3R2 |
| Assembly of RNA Polymerase II Complex | 0.929 | 0.063 | #NUM! | TAF10,TAF13,TAF9B |
| CMP-N-acetylneuraminate Biosynthesis I (Eukaryotes) | 0.925 | 0.200 | #NUM! | NANS |
| Myo-inositol Biosynthesis | 0.925 | 0.200 | #NUM! | ISYNA1 |
| Integrin signaling | 0.921 | 0.083 | #NUM! | CSK,TLN1 |
| IL-4 Signaling | 0.920 | 0.042 | -0.378 | COL3A1,COL6A6,CREB3,PIK3CG,PIK3R2,RPS6KA1,TGM2 |
| Angiopoietin Signaling | 0.913 | 0.053 | #NUM! | NFKBID,PIK3CG,PIK3R2,RELA |
| Estrogen-Dependent Breast Cancer Signaling | 0.913 | 0.053 | 0.000 | CREB3,PIK3CG,PIK3R2,RELA |
| cAMP-mediated signaling | 0.900 | 0.039 | -0.333 | ADCY6,CHRM3,CNGB1,CREB3,GRM2,OPRL1,OPRM1,PDE7B,RPS6KA1 |
| Mitochondrial translation | 0.898 | 0.047 | -1.342 | MRPL19,MRPL21,MRPL23,MRPS2,MRPS23 |
| NF-κB Activation by Viruses | 0.898 | 0.052 | 0.000 | NFKBID,PIK3CG,PIK3R2,RELA |
| TNFR1 Signaling | 0.890 | 0.060 | #NUM! | CASP6,NFKBID,RELA |
| UVB-Induced MAPK Signaling | 0.890 | 0.060 | #NUM! | EIF4E,PIK3CG,PIK3R2 |
| SRP-dependent cotranslational protein targeting to membrane | 0.890 | 0.044 | 0.816 | MRPS2,MRPS23,RPL27,RPS15,RPS27,SEC61G |

| | | | | |
|---|---|---|---|---|
| Regulation of eIF4 and p70S6K Signaling | 0.889 | 0.040 | #NUM! | EIF4E,MRPS2,MRPS23,PIK3CG,PIK3R2,RPS15,RPS27,VPS37A |
| MITF-M-dependent gene expression | 0.886 | 0.047 | 0.447 | AGO4,CDH2,DICER1,EDIL3,PPARGC1A |
| Neurotrophin/TRK Signaling | 0.883 | 0.051 | -1.000 | CREB3,PIK3CG,PIK3R2,RPS6KA1 |
| Activation of IRF by Cytosolic Pattern Recognition Receptors | 0.871 | 0.059 | #NUM! | MAVS,NFKBID,RELA |
| WNT ligand biogenesis and trafficking | 0.864 | 0.077 | #NUM! | VPS35,WNT5A |
| CDP-diacylglycerol Biosynthesis I | 0.864 | 0.077 | #NUM! | LPGAT1,MBOAT7 |
| Endocannabinoid Cancer Inhibition Pathway | 0.858 | 0.043 | 0.000 | ADCY6,CASP6,CREB3,PIK3CG,PIK3R2,VIM |
| Insulin Secretion Signaling Pathway | 0.853 | 0.037 | 0.000 | ADCY6,CHRM3,CREB3,EIF4E,KCNA4,PIK3CG,PIK3R2,RYR2,SEC61G,VPS37A |
| Spliceosomal Cycle | 0.853 | 0.058 | #NUM! | DDX46,SF3A3,SF3B3 |
| Fructose metabolism | 0.851 | 0.167 | #NUM! | KHK |
| SLC15A4:TASL-dependent IRF5 activation | 0.851 | 0.167 | #NUM! | IRF5 |
| Role of IL-17A in Psoriasis | 0.851 | 0.167 | #NUM! | S100A9 |
| Salvage Pathways of Pyrimidine Deoxyribonucleotides | 0.851 | 0.167 | #NUM! | TK1 |
| TCR signaling | 0.850 | 0.046 | 1.342 | CSK,PIK3R2,PSME1,RELA,UBE2D2 |
| Regulation of Insulin-like Growth Factor (IGF) transport and upta | 0.850 | 0.046 | 0.447 | CDH2,F2,MFGE8,MSLN,SPARCL1 |
| Prostate Cancer Signaling | 0.838 | 0.045 | 0.447 | CREB3,NFKBID,PIK3CG,PIK3R2,RELA |
| Coronavirus Pathogenesis Pathway | 0.838 | 0.039 | -0.816 | EIF4E,MAVS,MRPS2,MRPS23,NFKBID,RELA,RPS15,RPS27 |
| Selenoamino acid metabolism | 0.837 | 0.042 | 0.000 | MRPS2,MRPS23,PAPSS1,RPL27,RPS15,RPS27 |
| SPINK1 General Cancer Pathway | 0.835 | 0.057 | #NUM! | Mt1,PIK3CG,PIK3R2 |
| Neddylation | 0.829 | 0.038 | 0.333 | ASB1,ASB10,ASB7,KLHL9,OBSL1,PSME1,SPSB3,UBE2D2,WSB1 |
| Transcriptional regulation of white adipocyte differentiation | 0.827 | 0.049 | -1.000 | PPARGC1A,RELA,SREBF1,TBL1XR1 |
| MSP-RON Signaling Pathway | 0.818 | 0.056 | #NUM! | KLK5,PIK3CG,PIK3R2 |
| CD28 Signaling in T Helper Cells | 0.815 | 0.044 | 1.000 | CSK,NFKBID,PIK3CG,PIK3R2,RELA |
| Phosphatidylglycerol Biosynthesis II (Non-plastidic) | 0.812 | 0.071 | #NUM! | LPGAT1,MBOAT7 |
| Role of p14/p19ARF in Tumor Suppression | 0.812 | 0.071 | #NUM! | PIK3CG,PIK3R2 |
| RNA Polymerase II Transcription | 0.808 | 0.041 | 0.816 | SRSF11,SRSF3,SRSF4,TAF10,TAF13,TAF9B |
| Deubiquitination | 0.807 | 0.037 | -0.333 | BABAM1,H4C1,JOSD1,MAVS,POLB,PSME1,TAF10,TAF9B,TNKS2 |
| PEDF Signaling | 0.800 | 0.048 | #NUM! | NFKBID,PIK3CG,PIK3R2,RELA |
| FGF Signaling | 0.800 | 0.048 | 0.000 | CREB3,FGF10,PIK3CG,PIK3R2 |
| PIP3 activates AKT signaling | 0.798 | 0.041 | 0.000 | FGF10,NR4A1,PIK3CG,PIK3R2,SGK1,TGFA |
| KEAP1-NFE2L2 pathway | 0.798 | 0.041 | 0.816 | FBXL17,PIK3CG,PIK3R2,PLK4,PSME1,STK35 |
| Autophagy | 0.797 | 0.038 | -0.707 | ATG3,ATG4B,CREB3,FUNDC1,PIK3CG,PIK3R2,TGFA,VPS39 |
| Role of Osteoclasts in Rheumatoid Arthritis Signaling Pathway | 0.790 | 0.036 | 0.000 | COL3A1,COL6A6,CREB3,IL18,NFKBID,PIK3CG,PIK3R2,RELA,RHOBTB2,RHOD |
| Formation of intermediate mesoderm | 0.790 | 0.143 | #NUM! | FOXC1 |
| Formation of Fibrin Clot (Clotting Cascade) | 0.788 | 0.069 | #NUM! | F10,F2 |
| FOXO-mediated transcription of oxidative stress, metabolic and | 0.788 | 0.069 | #NUM! | PPARGC1A,SREBF1 |
| Response of EIF2AK1 (HRI) to heme deficiency | 0.788 | 0.069 | #NUM! | ATF5,VPS37A |
| LPS-stimulated MAPK Signaling | 0.787 | 0.047 | 0.000 | NFKBID,PIK3CG,PIK3R2,RELA |
| Deadenylation-dependent mRNA decay | 0.785 | 0.054 | #NUM! | EIF4E,EXOSC4,LSM7 |
| Vitamin K Signaling Pathway | 0.785 | 0.054 | #NUM! | F10,F2,MAVS |
| CNTF Signaling | 0.785 | 0.054 | #NUM! | PIK3CG,PIK3R2,RPS6KA1 |
| WNT/SHH Axonal Guidance Signaling Pathway | 0.779 | 0.041 | 0.000 | ADCY6,PIK3CG,PIK3R2,SEMA3F,TRPC3,WNT5A |
| Eukaryotic Translation Termination | 0.772 | 0.043 | 0.447 | MRPS2,MRPS23,RPL27,RPS15,RPS27 |
| Chaperone Mediated Autophagy Signaling Pathway | 0.766 | 0.037 | -1.414 | CHEK1,HSPBP1,IL18,PIK3CG,PIK3R2,PLIN2,PSME1,RELA |
| Regulation of TP53 Activity through Acetylation | 0.765 | 0.067 | #NUM! | BRD7,MEAF6 |
| Eukaryotic Translation Elongation | 0.761 | 0.042 | 0.447 | MRPS2,MRPS23,RPL27,RPS15,RPS27 |
| Relaxin Signaling | 0.760 | 0.040 | 1.342 | ADCY6,NFKBID,PDE7B,PIK3CG,PIK3R2,RELA |
| Signaling by PDGF | 0.753 | 0.052 | #NUM! | COL3A1,COL6A6,PIK3R2 |
| Triacylglycerol Biosynthesis | 0.753 | 0.052 | #NUM! | CLN5,LPGAT1,MBOAT7 |
| TNFR2 Signaling | 0.742 | 0.065 | #NUM! | NFKBID,RELA |
| Acetylcholine Receptor Signaling Pathway | 0.741 | 0.038 | -1.134 | ADCY6,CASP6,CHRM3,CHRNA4,CREB3,PIK3CG,PIK3R2 |
| RANK Signaling in Osteoclasts | 0.738 | 0.045 | #NUM! | NFKBID,PIK3CG,PIK3R2,RELA |
| Miro GTPase Cycle | 0.737 | 0.125 | #NUM! | MFN2 |
| Regulation of NFE2L2 gene expression | 0.737 | 0.125 | #NUM! | RELA |
| Glycosphingolipid transport | 0.737 | 0.125 | #NUM! | ESYT2 |
| Sucrose Degradation V (Mammalian) | 0.737 | 0.125 | #NUM! | KHK |
| Wound Healing Signaling Pathway | 0.737 | 0.037 | 1.414 | COL3A1,COL6A6,F2,IL18,LAMA5,NFKBID,TGFA,VIM |
| NGF Signaling | 0.731 | 0.041 | -0.447 | CREB3,PIK3CG,PIK3R2,RELA,RPS6KA1 |
| IL-6 Signaling | 0.731 | 0.041 | 0.447 | IL18,NFKBID,PIK3CG,PIK3R2,RELA |

| | | | | |
|---|---|---|---|---|
| Regulation of the Epithelial Mesenchymal Transition by Growth I | 0.725 | 0.037 | 0.816 | CDH2,FGF10,PIK3CG,PIK3R2,RELA,VIM,WWTR1 |
| Stearate Biosynthesis I (Animals) | 0.723 | 0.050 | #NUM! | CLN5,LPGAT1,MBOAT7 |
| Cargo concentration in the ER | 0.721 | 0.063 | #NUM! | CNIH3,TGFA |
| HER-2 Signaling in Breast Cancer | 0.715 | 0.036 | 0.707 | CASP6,ELF2,NFKBID,PGR,PIK3CG,PIK3R2,RELA,TGFA |
| Regulation of TP53 Activity through Phosphorylation | 0.714 | 0.044 | -1.000 | CHEK1,TAF10,TAF13,TAF9B |
| Acute Myeloid Leukemia Signaling | 0.714 | 0.044 | #NUM! | Csf2ra,PIK3CG,PIK3R2,RELA |
| PD-1, PD-L1 cancer immunotherapy pathway | 0.714 | 0.044 | -1.000 | CBLB,CSK,PIK3CG,PIK3R2 |
| Pancreatic Adenocarcinoma Signaling | 0.711 | 0.041 | 1.000 | BRCA2,PIK3CG,PIK3R2,RELA,TGFA |
| Mitochondrial biogenesis | 0.709 | 0.049 | #NUM! | PPARGC1A,SSBP1,TBL1XR1 |
| Adrenomedullin signaling pathway | 0.702 | 0.037 | 1.134 | ADCY6,CSK,IL18,KCNQ3,PIK3CG,PIK3R2,RELA |
| Gαs Signaling | 0.701 | 0.040 | -1.000 | ADCY6,CHRM3,CNGB1,CREB3,RYR2 |
| Unfolded Protein Response (UPR) | 0.701 | 0.061 | #NUM! | CREB3,VPS37A |
| MIF-mediated Glucocorticoid Regulation | 0.701 | 0.061 | #NUM! | NFKBID,RELA |
| Erythropoietin Signaling Pathway | 0.699 | 0.038 | 0.816 | IL18,NFKBID,PIK3CG,PIK3R2,RELA,TRPC3 |
| Death Receptor Signaling | 0.692 | 0.043 | 1.000 | CASP6,NFKBID,RELA,TNKS2 |
| Lipophagy | 0.691 | 0.111 | #NUM! | PLIN2 |
| Zygotic genome activation (ZGA) | 0.691 | 0.111 | #NUM! | DPPA2 |
| γ-glutamyl Cycle | 0.691 | 0.111 | #NUM! | ANPEP |
| Mineralocorticoid Biosynthesis | 0.691 | 0.111 | #NUM! | Cyp2d22 |
| Role of Tissue Factor in Cancer | 0.687 | 0.037 | 0.378 | EIF4E,F10,F2,PIK3CG,PIK3R2,RPS6KA1,TGFA |
| GPVI-mediated activation cascade | 0.681 | 0.059 | #NUM! | PIK3CG,PIK3R2 |
| 4-1BB Signaling in T Lymphocytes | 0.681 | 0.059 | #NUM! | NFKBID,RELA |
| Crosstalk between Dendritic Cells and Natural Killer Cells | 0.681 | 0.048 | #NUM! | IL18,RELA,TLN1 |
| UBE2C in Cell Cycle Progression Signaling Pathway | 0.681 | 0.043 | -1.000 | ANAPC7,PIK3CG,PIK3R2,RPS6KA1 |
| Clathrin-mediated endocytosis | 0.673 | 0.039 | 1.342 | PACSIN2,SCARB2,SYT11,TGFA,WNT5A |
| Germ Cell-Sertoli Cell Junction Signaling | 0.666 | 0.037 | #NUM! | CDH2,CLINT1,PIK3CG,PIK3R2,RHOBTB2,RHOD |
| Warburg Effect Signaling Pathway | 0.664 | 0.039 | 1.342 | PIK3CG,PIK3R2,SREBF1,TGFA,WWTR1 |
| Nuclear RNA decay | 0.663 | 0.057 | #NUM! | EXOSC10,EXOSC4 |
| Expression of BMAL (ARNTL), CLOCK, and NPAS2 | 0.663 | 0.057 | #NUM! | PPARGC1A,TBL1XR1 |
| Dermatan Sulfate Biosynthesis (Late Stages) | 0.663 | 0.057 | #NUM! | NDST2,SULT1A1 |
| UVA-Induced MAPK Signaling | 0.660 | 0.042 | #NUM! | PIK3CG,PIK3R2,RPS6KA1,TNKS2 |
| Transcriptional Regulatory Network in Embryonic Stem Cells | 0.658 | 0.037 | 0.816 | CDYL,FOXC1,PIK3CG,PIK3R2,SIX3,WNT5A |
| Role of Pattern Recognition Receptors in Recognition of Bacteria | 0.655 | 0.039 | 0.000 | IL18,MAVS,PIK3CG,PIK3R2,RELA |
| DDX58/IFIH1-mediated induction of interferon-alpha/beta | 0.655 | 0.046 | #NUM! | MAVS,RELA,UBE2D2 |
| WNT/Ca2+ Signaling Pathway | 0.655 | 0.046 | #NUM! | CREB3,RELA,WNT5A |
| Processing and activation of SUMO | 0.651 | 0.100 | #NUM! | RWDD3 |
| Platelet Aggregation (Plug Formation) | 0.651 | 0.100 | #NUM! | F2 |
| Glucocorticoid Biosynthesis | 0.651 | 0.100 | #NUM! | Cyp2d22 |
| Post-translational protein phosphorylation | 0.649 | 0.041 | 0.000 | CDH2,MFGE8,MSLN,SPARCL1 |
| Small Cell Lung Cancer Signaling | 0.649 | 0.041 | #NUM! | NFKBID,PIK3CG,PIK3R2,RELA |
| P2Y Purinergic Receptor Signaling Pathway | 0.647 | 0.039 | 0.447 | ADCY6,CREB3,PIK3CG,PIK3R2,RELA |
| Macroautophagy | 0.642 | 0.046 | #NUM! | ATG3,ATG4B,LAMTOR5 |
| Pyridoxal 5'-phosphate Salvage Pathway | 0.642 | 0.046 | #NUM! | FAM20B,PNPO,SGK1 |
| Glutamate Receptor Signaling | 0.642 | 0.046 | #NUM! | GRIK2,GRM2,SLC1A6 |
| Docosahexaenoic Acid (DHA) Signaling | 0.641 | 0.034 | 0.378 | ADCY6,CREB3,PIK3CG,PIK3R2,PLAAT3,RELA,SYT11,TGFA |
| Growth Hormone Signaling | 0.630 | 0.045 | #NUM! | PIK3CG,PIK3R2,RPS6KA1 |
| Neuropathic Pain Signaling in Dorsal Horn Neurons | 0.629 | 0.040 | 0.000 | GRM2,KCNQ3,PIK3CG,PIK3R2 |
| Binding and Uptake of Ligands by Scavenger Receptors | 0.627 | 0.054 | #NUM! | COL3A1,HYOU1 |
| Fanconi Anemia Pathway | 0.627 | 0.054 | #NUM! | FAAP100,MUS81 |
| High laminar flow shear stress activates signaling by PIEZO1 and | 0.617 | 0.044 | #NUM! | ADCY6,PIK3R2,TLN1 |
| Beta-catenin independent WNT signaling | 0.616 | 0.035 | 0.378 | AGO4,PIK3CG,PIK3R2,PLK4,PSME1,STK35,WNT5A |
| Dissolution of Fibrin Clot | 0.614 | 0.091 | #NUM! | SERPINF2 |
| Bile Acid Biosynthesis, Neutral Pathway | 0.614 | 0.091 | #NUM! | AMACR |
| Purine Nucleotides De Novo Biosynthesis II | 0.614 | 0.091 | #NUM! | ATIC |
| Bile acid and bile salt metabolism | 0.610 | 0.053 | #NUM! | AMACR,STARD5 |
| TP53 Regulates Transcription of Cell Death Genes | 0.610 | 0.053 | #NUM! | CASP6,PERP |
| Resolution of Abasic Sites (AP sites) | 0.610 | 0.053 | #NUM! | NEIL1,POLB |
| FLT3 Signaling | 0.610 | 0.053 | #NUM! | CSK,SH2B3 |
| Chondroitin Sulfate Biosynthesis (Late Stages) | 0.610 | 0.053 | #NUM! | NDST2,SULT1A1 |

| | | | | |
|---|---|---|---|---|
| Apoptosis Signaling | 0.609 | 0.040 | 0.000 | CASP6,NFKBID,RELA,RPS6KA1 |
| AMPK Signaling | 0.603 | 0.034 | -1.342 | BRD7,CHRNA4,CREB3,EEF2K,PIK3CG,PIK3R2,PPARGC1A,PPM1E |
| NAFLD Signaling Pathway | 0.603 | 0.034 | 1.134 | CASP6,IL18,MBOAT7,PIK3CG,PIK3R2,PPARGC1A,SREBF1 |
| DAG and IP3 signaling | 0.594 | 0.051 | #NUM! | ADCY6,CAMKK1 |
| NGF-stimulated transcription | 0.594 | 0.051 | #NUM! | JUNB,SGK1 |
| Signaling by ROBO receptors | 0.591 | 0.033 | 0.707 | FLRT3,MRPS2,MRPS23,MSI1,PSME1,RPL27,RPS15,RPS27 |
| NF-κB Signaling | 0.591 | 0.035 | 1.633 | IL18,NFKBID,PIK3CG,PIK3R2,RELA,TGFA |
| Egg-Sperm Fusion Signaling Pathway | 0.591 | 0.039 | 1.000 | PIK3CG,PIK3R2,SYT11,TRPC3 |
| Antioxidant Action of Vitamin C | 0.591 | 0.039 | #NUM! | Csf2ra,NFKBID,PLAAT3,RELA |
| ERK/MAPK Signaling | 0.590 | 0.034 | 0.378 | CREB3,EIF4E,ELF2,PIK3CG,PIK3R2,RPS6KA1,TLN1 |
| Sheddase Signaling Pathway | 0.584 | 0.035 | 1.633 | CDH2,ELF2,PIK3CG,PIK3R2,RELA,TGFA |
| COPII-mediated vesicle transport | 0.583 | 0.042 | #NUM! | CNIH3,TGFA,TRAPPC5 |
| HDR through MMEJ (alt-NHEJ) | 0.582 | 0.083 | #NUM! | BRCA2 |
| Z-decay: degradation of maternal mRNAs by zygotically expresse | 0.582 | 0.083 | #NUM! | EIF4E |
| Signaling by LTK | 0.582 | 0.083 | #NUM! | PIK3R2 |
| Signaling by NOTCH4 | 0.580 | 0.036 | 0.447 | PIK3CG,PIK3R2,PLK4,PSME1,STK35 |
| Activation of gene expression by SREBF (SREBP) | 0.578 | 0.050 | #NUM! | SREBF1,TBL1XR1 |
| Meiotic recombination | 0.578 | 0.050 | #NUM! | BRCA2,H4C1 |
| Cohesin Chromatin Regulation Pathway | 0.577 | 0.034 | -0.447 | ANAPC7,ATRX,CDHR2,CHEK1,H4C1,PCDHA10,PDS5A |
| Reelin Signaling in Neurons | 0.572 | 0.036 | -1.000 | ARHGEF2,ARHGEF40,CDH2,PIK3CG,PIK3R2 |
| PI3K Signaling in B Lymphocytes | 0.565 | 0.036 | 0.447 | ATF5,NFKBID,PIK3CG,PIK3R2,RELA |
| LXR/RXR Activation | 0.563 | 0.038 | #NUM! | IL18,RELA,SERPINF2,SREBF1 |
| Paxillin Signaling | 0.563 | 0.038 | 1.000 | CSK,PIK3CG,PIK3R2,TLN1 |
| MIF Regulation of Innate Immunity | 0.563 | 0.049 | #NUM! | NFKBID,RELA |
| DNA damage-induced 14-3-3σ Signaling | 0.563 | 0.049 | #NUM! | CASP6,CHEK1 |
| Protein Sorting Signaling Pathway | 0.563 | 0.034 | 0.000 | ADCY6,AP1S2,AP3M2,CLINT1,SCARB2,VPS35 |
| Extra-nuclear estrogen signaling | 0.561 | 0.041 | #NUM! | PIK3R2,TGFA,ZDHHC21 |
| ERK5 Signaling | 0.561 | 0.041 | #NUM! | CREB3,RPS6KA1,SGK1 |
| Hypoxia Signaling in the Cardiovascular System | 0.561 | 0.041 | #NUM! | CREB3,NFKBID,UBE2D2 |
| Hedgehog 'on' state | 0.557 | 0.036 | 0.447 | PIK3CG,PIK3R2,PLK4,PSME1,STK35 |
| Acetylcholine binding and downstream events | 0.552 | 0.077 | #NUM! | CHRNA4 |
| Olfactory Signaling Pathway | 0.552 | 0.077 | #NUM! | CNGB1 |
| SUMOylation of immune response proteins | 0.552 | 0.077 | #NUM! | RELA |
| Nucleotide biosynthesis | 0.552 | 0.077 | #NUM! | ATIC |
| Replacement of protamines by nucleosomes in the male pronuc | 0.552 | 0.077 | #NUM! | H4C1 |
| Co-inhibition by PD-1 | 0.552 | 0.077 | #NUM! | CSK |
| trans-Golgi Network Vesicle Budding | 0.550 | 0.041 | #NUM! | AP1S2,CLINT1,SNAPIN |
| Leptin Signaling in Obesity | 0.550 | 0.041 | #NUM! | ADCY6,PIK3CG,PIK3R2 |
| April Mediated Signaling | 0.549 | 0.048 | #NUM! | NFKBID,RELA |
| Atherosclerosis Signaling | 0.538 | 0.037 | #NUM! | COL3A1,IL18,PLAAT3,RELA |
| Production of Nitric Oxide and Reactive Oxygen Species in Macr | 0.537 | 0.034 | 0.447 | NFKBID,PIK3CG,PIK3R2,RELA,RHOBTB2,RHOD |
| PI3K Cascade | 0.535 | 0.047 | #NUM! | FGF10,PIK3R2 |
| Collagen chain trimerization | 0.535 | 0.047 | #NUM! | COL3A1,COL6A6 |
| B Cell Activating Factor Signaling | 0.535 | 0.047 | #NUM! | NFKBID,RELA |
| BER (Base Excision Repair) Pathway | 0.535 | 0.047 | #NUM! | NEIL1,POLB |
| B Cell Receptor Signaling | 0.530 | 0.033 | 0.000 | CREB3,CSK,NFKBID,PIK3CG,PIK3R2,RELA |
| Transport of bile salts and organic acids, metal ions and amine c | 0.529 | 0.040 | #NUM! | RHCG,SLC22A8,SLC5A11 |
| Activation of NMDA receptors and postsynaptic events | 0.529 | 0.040 | #NUM! | CAMKK1,PPM1E,RPS6KA1 |
| ESR-mediated signaling | 0.529 | 0.036 | -1.000 | AGO4,H4C1,PGR,TGFA |
| p38 MAPK Signaling | 0.529 | 0.036 | #NUM! | CREB3,EEF2K,IL18,RPS6KA1 |
| eNOS Signaling | 0.528 | 0.035 | 0.000 | ADCY6,CHRNA4,CNGB1,PIK3CG,PIK3R2 |
| Carnitine shuttle | 0.525 | 0.071 | #NUM! | THRSP |
| YAP1- and WWTR1 (TAZ)-stimulated gene expression | 0.525 | 0.071 | #NUM! | WWTR1 |
| Mitotic Telophase/Cytokinesis | 0.525 | 0.071 | #NUM! | PDS5A |
| Tryptophan catabolism | 0.525 | 0.071 | #NUM! | ACMSD |
| Digestion | 0.525 | 0.071 | #NUM! | CEL |
| PPARα/RXRα Activation | 0.524 | 0.033 | 0.000 | ADCY6,ADIPOR1,NFKBID,NR2F1,PPARGC1A,RELA |
| Class I MHC mediated antigen processing and presentation | 0.524 | 0.030 | 0.302 | ANAPC7,ASB1,ASB10,ASB7,CBLB,KLHL9,PSME1,S100A9,SEC61G,UBE2D2,WSB1 |
| B-WICH complex positively regulates rRNA expression | 0.521 | 0.046 | #NUM! | DDX21,H4C1 |

| | | | | |
|---|---|---|---|---|
| Regulation of lipid metabolism by PPARalpha | 0.513 | 0.036 | 0.000 | PLIN2,PPARGC1A,SREBF1,TBL1XR1 |
| Response of EIF2AK4 (GCN2) to amino acid deficiency | 0.513 | 0.036 | 2.000 | RPL27,RPS15,RPS27,VPS37A |
| Myocardin Signaling Pathway | 0.513 | 0.036 | 0.000 | PIK3CG,PIK3R2,RELA,TGFA |
| Renal Cell Carcinoma Signaling | 0.510 | 0.039 | #NUM! | PIK3CG,PIK3R2,TGFA |
| OX40 Signaling Pathway | 0.508 | 0.044 | #NUM! | NFKBID,RELA |
| NOD1/2 Signaling Pathway | 0.507 | 0.034 | 1.342 | IL18,IRF5,MAVS,NFKBID,RELA |
| Type II Diabetes Mellitus Signaling | 0.507 | 0.034 | 0.000 | ADIPOR1,NFKBID,PIK3CG,PIK3R2,RELA |
| Virus Entry via Endocytic Pathways | 0.505 | 0.035 | -1.000 | AP1S2,AP3M2,PIK3CG,PIK3R2 |
| Pancreatic Secretion Signaling Pathway | 0.501 | 0.032 | -0.378 | ADCY6,ARHGEF2,CHRM3,PIK3CG,PIK3R2,RYR2,TRPC3 |
| Regulation of cholesterol biosynthesis by SREBP (SREBF) | 0.500 | 0.067 | #NUM! | SREBF1 |
| Metabolism of Angiotensinogen to Angiotensins | 0.500 | 0.067 | #NUM! | ANPEP |
| Paracetamol ADME | 0.500 | 0.067 | #NUM! | SULT1A1 |
| Telomere Extension by Telomerase | 0.500 | 0.067 | #NUM! | TNKS2 |
| G alpha (12/13) signalling events | 0.500 | 0.038 | #NUM! | ARHGEF2,ARHGEF40,MCF2L |
| IL-15 Signaling | 0.500 | 0.038 | #NUM! | PIK3CG,PIK3R2,RELA |
| BEX2 Signaling Pathway | 0.500 | 0.038 | #NUM! | CDH2,PLXNC1,RELA |
| Coronavirus Replication Pathway | 0.500 | 0.034 | #NUM! | MRPS2,MRPS23,RPL27,RPS15,RPS27 |
| Heme signaling | 0.495 | 0.044 | #NUM! | PPARGC1A,TBL1XR1 |
| M-decay: degradation of maternal mRNAs by maternally stored | 0.495 | 0.044 | #NUM! | DICER1,EIF4E |
| PXR/RXR Activation | 0.495 | 0.044 | #NUM! | PPARGC1A,RELA |
| Role of OCT4 in Mammalian Embryonic Stem Cell Pluripotency | 0.495 | 0.044 | #NUM! | CASP6,NR2F1 |
| Chondroitin Sulfate Biosynthesis | 0.495 | 0.044 | #NUM! | NDST2,SULT1A1 |
| IL-27 Signaling Pathway | 0.490 | 0.035 | 0.000 | PIK3CG,PIK3R2,PPARGC1A,RELA |
| XBP1(S) activates chaperone genes | 0.483 | 0.043 | #NUM! | HYOU1,TLN1 |
| iNOS Signaling | 0.483 | 0.043 | #NUM! | NFKBID,RELA |
| Role of PKR in Interferon Induction and Antiviral Response | 0.482 | 0.035 | 1.000 | IL18,MAVS,NFKBID,RELA |
| IL-15 Production | 0.482 | 0.035 | #NUM! | CLK1,Csf2ra,CSK,RELA |
| PI Metabolism | 0.482 | 0.037 | #NUM! | PIK3CG,PIK3R2,PNPLA6 |
| PKR-mediated signaling | 0.482 | 0.037 | #NUM! | FAAP100,MAVS,VPS37A |
| Regulation of the Epithelial-Mesenchymal Transition Pathway | 0.481 | 0.032 | #NUM! | CDH2,FGF10,PIK3CG,PIK3R2,RELA,WNT5A |
| RAN Signaling | 0.478 | 0.063 | #NUM! | RCC1 |
| Proteasomal PSMD10 Signaling Pathway | 0.475 | 0.034 | 1.000 | PIK3CG,PIK3R2,RELA,TGFA |
| Histone Modification Signaling Pathway | 0.473 | 0.030 | -1.414 | ASH1L,ATRX,BRD7,CDYL,KAT6B,KAT7,SND1,TGM2 |
| JAK/STAT Signaling | 0.473 | 0.037 | #NUM! | PIK3CG,PIK3R2,RELA |
| Dermatan Sulfate Biosynthesis | 0.471 | 0.042 | #NUM! | NDST2,SULT1A1 |
| Protein Kinase A Signaling | 0.465 | 0.029 | -0.378 | ADCY6,ANAPC7,CNGB1,CREB3,NFKBID,PDE7B,PHKG2,PTP4A3,RELA,RYR2,TIMM50 |
| Microautophagy Signaling Pathway | 0.461 | 0.032 | #NUM! | COL3A1,COL6A6,PSME1,VPS37A,VPS39 |
| DNA Double Strand Break Response | 0.459 | 0.041 | #NUM! | BABAM1,H4C1 |
| RUNX1 regulates megakaryocyte differentiation and platelet fur | 0.459 | 0.041 | #NUM! | AGO4,H4C1 |
| Signaling by TGFBR3 | 0.459 | 0.041 | #NUM! | AGO4,KLF16 |
| Co-stimulation by CD28 | 0.459 | 0.041 | #NUM! | PIK3CG,PIK3R2 |
| Melanoma Signaling | 0.459 | 0.041 | #NUM! | PIK3CG,PIK3R2 |
| Phototransduction Pathway | 0.459 | 0.041 | #NUM! | CNGB1,RGS9 |
| Metabolism of nitric oxide: NOS3 activation and regulation | 0.456 | 0.059 | #NUM! | ZDHHC21 |
| Signaling by Type 1 Insulin-like Growth Factor 1 Receptor (IGF1R | 0.456 | 0.059 | #NUM! | PIK3R2 |
| Regulation of TLR by endogenous ligand | 0.456 | 0.059 | #NUM! | S100A9 |
| Gastrin-CREB signalling pathway via PKC and MAPK | 0.456 | 0.059 | #NUM! | RPS6KA1 |
| Thyroid Hormone Metabolism II (via Conjugation and/or Degrad | 0.456 | 0.059 | #NUM! | SULT1A1 |
| Inhibition of ARE-Mediated mRNA Degradation Pathway | 0.455 | 0.032 | 2.000 | AGO4,EXOSC10,EXOSC4,PSME1,ZFP36 |
| Eicosanoid Signaling | 0.454 | 0.030 | 0.000 | ADCY6,AP1S2,CREB3,P2RX3,PIK3CG,PIK3R2,PLAAT3,RPS6KA1 |
| Renin-Angiotensin Signaling | 0.453 | 0.033 | #NUM! | ADCY6,PIK3CG,PIK3R2,RELA |
| Transcriptional activity of SMAD2/SMAD3:SMAD4 heterotrimer | 0.448 | 0.040 | #NUM! | JUNB,WWTR1 |
| Signaling by FGFR1 | 0.448 | 0.040 | #NUM! | FGF10,FLRT3 |
| Apoptotic execution phase | 0.448 | 0.040 | #NUM! | CASP6,VIM |
| Cell Cycle: G2/M DNA Damage Checkpoint Regulation | 0.448 | 0.040 | #NUM! | CHEK1,RPS6KA1 |
| Glycerophospholipid biosynthesis | 0.446 | 0.033 | 1.000 | LPGAT1,MBOAT7,PLAAT3,PTDSS1 |
| Sertoli Cell-Sertoli Cell Junction Signaling | 0.443 | 0.030 | -1.134 | CDH2,CLINT1,CREB3,LAMA5,PIK3CG,PIK3R2,RPS6KA1 |
| Clathrin-mediated Endocytosis Signaling | 0.442 | 0.031 | #NUM! | AP1S2,AP3M2,F2,FGF10,PIK3CG,PIK3R2 |
| 14-3-3-mediated Signaling | 0.439 | 0.033 | 0.000 | PIK3CG,PIK3R2,RPS6KA1,VIM |

| | | | | |
|---|---|---|---|---|
| Gap Junction Signaling | 0.439 | 0.029 | 0.000 | ADCY6,CDH2,CREB3,GJA4,GRIK2,KCNQ3,PIK3CG,PIK3R2,RELA |
| CSDE1 Signaling Pathway | 0.437 | 0.039 | #NUM! | AGO4,VIM |
| FAT10 Cancer Signaling Pathway | 0.437 | 0.039 | #NUM! | NFKBID,RELA |
| Cilium Assembly | 0.437 | 0.031 | 0.000 | BBS7,CNGB1,MAPRE1,PLK4,RAB11FIP3,SEPTIN2 |
| DNA methylation | 0.437 | 0.056 | #NUM! | H4C1 |
| Activated PKN1 stimulates transcription of AR regulated genes K | 0.437 | 0.056 | #NUM! | H4C1 |
| Formation of the ureteric bud | 0.437 | 0.056 | #NUM! | FOXC1 |
| ID1 Signaling Pathway | 0.432 | 0.031 | 0.447 | CHRNA4,PIK3CG,PIK3R2,S100A9,TGM2,VIM |
| Actin Cytoskeleton Signaling | 0.429 | 0.030 | 0.000 | CSK,DIAPH3,F2,FGF10,PIK3CG,PIK3R2,TLN1 |
| Systemic Lupus Erythematosus in T Cell Signaling Pathway | 0.427 | 0.030 | 0.816 | CASP6,CREB3,PIK3CG,PIK3R2,RHOBTB2,RHOD |
| Insulin Receptor Signaling | 0.426 | 0.032 | 0.000 | EIF4E,PIK3CG,PIK3R2,SGK1 |
| Tumor Microenvironment Pathway | 0.425 | 0.031 | 1.000 | COL3A1,FGF10,PIK3CG,PIK3R2,RELA |
| Opioid Signalling | 0.423 | 0.034 | #NUM! | ADCY6,CAMKK1,OPRM1 |
| Ceramide Signaling | 0.423 | 0.034 | #NUM! | PIK3CG,PIK3R2,RELA |
| Regulation of Cellular Mechanics by Calpain Protease | 0.423 | 0.034 | #NUM! | CAST,CNGB1,TLN1 |
| Huntington's Disease Signaling | 0.423 | 0.029 | -1.342 | CASP6,CREB3,PIK3CG,PIK3R2,PSME1,SGK1,TAF9B,TGM2 |
| Glutaminergic Receptor Signaling Pathway (Enhanced) | 0.422 | 0.029 | -0.707 | ADCY6,CREB3,EIF4E,GRIK2,GRM2,PIK3CG,PIK3R2,PLAAT3,SLC1A6 |
| CXCR4 Signaling | 0.420 | 0.031 | 0.000 | ADCY6,PIK3CG,PIK3R2,RHOBTB2,RHOD |
| Pregnenolone Biosynthesis | 0.418 | 0.053 | #NUM! | Cyp2d22 |
| Methionine Degradation I (to Homocysteine) | 0.418 | 0.053 | #NUM! | SPOUT1 |
| Inflammasome pathway | 0.418 | 0.053 | #NUM! | IL18 |
| Signaling by EGFR | 0.416 | 0.038 | #NUM! | CSK,TGFA |
| Heparan Sulfate Biosynthesis (Late Stages) | 0.416 | 0.038 | #NUM! | NDST2,SULT1A1 |
| RAB GEFs exchange GTP for GDP on RABs | 0.415 | 0.034 | #NUM! | DENND6B,RGP1,TRAPPC5 |
| Opioid Signaling Pathway | 0.414 | 0.029 | -2.121 | ADCY6,CREB3,OPRL1,OPRM1,PIK3CG,RGS9,RPS6KA1,RYR2 |
| Endocannabinoid Developing Neuron Pathway | 0.413 | 0.032 | -1.000 | ADCY6,CREB3,PIK3CG,PIK3R2 |
| Androgen Signaling | 0.408 | 0.031 | #NUM! | KAT7,RELA,TAF10,TAF13,TAF9B |
| Regulation of mRNA stability by proteins that bind AU-rich elem | 0.408 | 0.033 | #NUM! | EXOSC4,PSME1,ZFP36 |
| Fcγ Receptor-mediated Phagocytosis in Macrophages and Monc | 0.408 | 0.033 | #NUM! | PIK3CG,PIK3R2,TLN1 |
| Cerebral Malformation Signaling Pathway | 0.407 | 0.032 | 1.000 | F2,IL18,PIK3CG,PIK3R2 |
| IL-8 Signaling | 0.406 | 0.030 | 1.000 | NFKBID,PIK3CG,PIK3R2,RELA,RHOBTB2,RHOD |
| Natural Killer Cell Signaling | 0.403 | 0.030 | 0.447 | COL3A1,IL18,PIK3CG,PIK3R2,RELA |
| Systemic Lupus Erythematosus in B Cell Signaling Pathway | 0.401 | 0.029 | 1.134 | CSK,IL18,IRF5,MAVS,PIK3CG,PIK3R2,RELA |
| Dopamine Degradation | 0.401 | 0.050 | #NUM! | SULT1A1 |
| Histidine Degradation VI | 0.401 | 0.050 | #NUM! | Cyp2d22 |
| DHCR24 Signaling Pathway | 0.400 | 0.031 | 0.000 | PIK3CG,PIK3R2,SERPINF2,SREBF1 |
| Aldosterone Signaling in Epithelial Cells | 0.398 | 0.030 | #NUM! | HSPE1,PIK3CG,PIK3R2,SASS6,SGK1 |
| EGF Signaling | 0.397 | 0.036 | #NUM! | PIK3CG,PIK3R2 |
| LPS/IL-1 Mediated Inhibition of RXR Function | 0.397 | 0.029 | #NUM! | Cyp2d22,IL18,NDST2,PPARGC1A,SREBF1,SULT1A1 |
| Integrin to Cytoskeleton Signaling Pathway | 0.397 | 0.029 | 1.000 | PIK3CG,PIK3R2,RHOBTB2,RHOD,TLN1,TSPAN3 |
| IL-10 Signaling | 0.394 | 0.031 | -1.000 | CREB3,IL18,NFKBID,RELA |
| Centrosomal KIAA0586 Signaling Pathway | 0.394 | 0.031 | 0.000 | FGF10,PIK3CG,PIK3R2,PSME1 |
| fMLP Signaling in Neutrophils | 0.394 | 0.031 | 1.000 | NFKBID,PIK3CG,PIK3R2,RELA |
| Systemic Lupus Erythematosus Signaling | 0.392 | 0.030 | #NUM! | IL18,LSM7,PIK3CG,PIK3R2,SNRPA |
| NFKBIE Signaling Pathway | 0.387 | 0.036 | #NUM! | NFKBID,RELA |
| Collagen degradation | 0.387 | 0.036 | #NUM! | COL3A1,COL6A6 |
| Assembly of collagen fibrils and other multimeric structures | 0.387 | 0.036 | #NUM! | COL3A1,COL6A6 |
| CD27 Signaling in Lymphocytes | 0.387 | 0.036 | #NUM! | NFKBID,RELA |
| Epithelial Membrane Protein Signaling Pathway | 0.387 | 0.030 | 0.447 | ANPEP,JUNB,PIK3CG,PIK3R2,WWTR1 |
| Glioblastoma Multiforme Signaling | 0.387 | 0.030 | 1.000 | PIK3CG,PIK3R2,RHOBTB2,RHOD,WNT5A |
| Gene Silencing by RNA | 0.386 | 0.032 | #NUM! | AGO4,DICER1,H4C1 |
| Airway Pathology in Chronic Obstructive Pulmonary Disease | 0.386 | 0.032 | #NUM! | FGF10,IL18,LCN10 |
| ERBB Signaling | 0.386 | 0.032 | #NUM! | PIK3CG,PIK3R2,TGFA |
| Fertilization | 0.385 | 0.048 | #NUM! | CATSPERG |
| SIRT1 negatively regulates rRNA expression | 0.385 | 0.048 | #NUM! | H4C1 |
| Branched-chain amino acid catabolism | 0.385 | 0.048 | #NUM! | SLC25A44 |
| Insertion of tail-anchored proteins into the endoplasmic reticulu | 0.385 | 0.048 | #NUM! | SEC61G |
| IL-17A Signaling in Gastric Cells | 0.385 | 0.048 | #NUM! | RELA |
| Glutathione-mediated Detoxification | 0.385 | 0.048 | #NUM! | ANPEP |

| | | | | |
|---|---|---|---|---|
| Cysteine Biosynthesis III (mammalia) | 0.385 | 0.048 | #NUM! | SPOUT1 |
| RHO GTPases Activate Formins | 0.382 | 0.031 | 1.000 | DIAPH3,MAPRE1,RHOD,RPS27 |
| Autism Signaling Pathway | 0.381 | 0.028 | 0.707 | CAMKK1,EIF4E,GRM2,IL18,PIK3CG,PIK3R2,RPS6KA1,WNT5A |
| Non-Small Cell Lung Cancer Signaling | 0.379 | 0.032 | #NUM! | PIK3CG,PIK3R2,TGFA |
| Visual phototransduction | 0.378 | 0.035 | #NUM! | CNGB1,RGS9 |
| Cardiac Hypertrophy Signaling | 0.375 | 0.028 | 0.816 | ADCY6,EIF4E,PIK3CG,PIK3R2,RHOBTB2,RHOD,RPS6KA1 |
| DNA Methylation and Transcriptional Repression Signaling | 0.372 | 0.032 | #NUM! | ERCC5,RELA,TCF19 |
| p70S6K Signaling | 0.371 | 0.030 | 1.000 | EEF2K,F2,PIK3CG,PIK3R2 |
| Class C/3 (Metabotropic glutamate/pheromone receptors) | 0.370 | 0.046 | #NUM! | GRM2 |
| Tryptophan Degradation III (Eukaryotic) | 0.370 | 0.046 | #NUM! | ACMSD |
| Semaphorin Signaling in Neurons | 0.369 | 0.035 | #NUM! | RHOBTB2,RHOD |
| Cancer Drug Resistance by Drug Efflux | 0.369 | 0.035 | #NUM! | PIK3CG,PIK3R2 |
| Serotonin Receptor Signaling | 0.368 | 0.027 | 0.302 | ADCY6,CREB3,KCNQ3,PIK3CG,PIK3R2,PLAAT3,PPARGC1A,RHOBTB2,RHOD,RYR2,TGM2,TRPC3 |
| Ion channel transport | 0.367 | 0.029 | -0.447 | RYR2,SGK1,TRPC3,TRPM2,TRPM4 |
| Fc epsilon receptor (FCERI) signaling | 0.365 | 0.030 | 2.000 | PIK3R2,PSME1,RELA,UBE2D2 |
| DNA Replication Pre-Initiation | 0.365 | 0.031 | #NUM! | ANAPC7,H4C1,PSME1 |
| IL-1 Signaling | 0.365 | 0.031 | #NUM! | ADCY6,NFKBID,RELA |
| Gustation Pathway | 0.362 | 0.029 | -0.447 | ADCY6,CHRM3,KCNQ3,P2RX3,TRPM4 |
| UFMylation Signaling Pathway | 0.361 | 0.034 | #NUM! | DDRGK1,EIF4E |
| rRNA modification in the nucleus and cytosol | 0.361 | 0.034 | #NUM! | IMP4,WDR75 |
| PCP (Planar Cell Polarity) Pathway | 0.361 | 0.034 | #NUM! | JUNB,WNT5A |
| Role of Osteoblasts, Osteoclasts and Chondrocytes in Rheumato | 0.360 | 0.028 | #NUM! | IL18,NFKBID,PIK3CG,PIK3R2,RELA,WNT5A |
| Pyroptosis | 0.355 | 0.044 | #NUM! | IL18 |
| Nucleotide salvage | 0.355 | 0.044 | #NUM! | TK1 |
| Cellular response to mitochondrial stress | 0.355 | 0.044 | #NUM! | VPS37A |
| Tumoricidal Function of Hepatic Natural Killer Cells | 0.355 | 0.044 | #NUM! | CASP6 |
| Role of JAK1, JAK2 and TYK2 in Interferon Signaling | 0.355 | 0.044 | #NUM! | RELA |
| Endometrial Cancer Signaling | 0.352 | 0.033 | #NUM! | PIK3CG,PIK3R2 |
| Heparan Sulfate Biosynthesis | 0.352 | 0.033 | #NUM! | NDST2,SULT1A1 |
| Colorectal Cancer Metastasis Signaling | 0.351 | 0.028 | 1.633 | ADCY6,PIK3CG,PIK3R2,RELA,RHOBTB2,RHOD,WNT5A |
| RAF/MAP kinase cascade | 0.347 | 0.028 | 1.890 | CSK,FGF10,PIK3R2,PSME1,RGL2,TGFA,TLN1 |
| Sumoylation Pathway | 0.346 | 0.030 | #NUM! | RCC1,RHOBTB2,RHOD |
| TREM1 Signaling | 0.344 | 0.033 | #NUM! | IL18,RELA |
| IL-2 Signaling | 0.344 | 0.033 | #NUM! | PIK3CG,PIK3R2 |
| Aryl Hydrocarbon Receptor Signaling | 0.343 | 0.029 | 0.000 | CHEK1,NR2F1,RELA,TGM2 |
| RHO GTPases Activate NADPH Oxidases | 0.342 | 0.042 | #NUM! | S100A9 |
| Regulation of RUNX1 Expression and Activity | 0.342 | 0.042 | #NUM! | AGO4 |
| Chromatin modifications during the maternal to zygotic transitic | 0.342 | 0.042 | #NUM! | H4C1 |
| Ubiquinol-10 Biosynthesis (Eukaryotic) | 0.342 | 0.042 | #NUM! | Cyp2d22 |
| CGRP in Inflammation and Nociception Signaling Pathway | 0.338 | 0.029 | 0.000 | ADCY6,CREB3,PIK3CG,PIK3R2 |
| RAC Signaling | 0.338 | 0.029 | 1.000 | MCF2L,PIK3CG,PIK3R2,RELA |
| Endothelin-1 Signaling | 0.338 | 0.028 | 0.000 | ADCY6,CASP6,PIK3CG,PIK3R2,PLAAT3 |
| NCAM signaling for neurite out-growth | 0.336 | 0.032 | #NUM! | COL3A1,COL6A6 |
| Keratinization | 0.336 | 0.032 | #NUM! | KLK5,PERP |
| Amyloid fiber formation | 0.336 | 0.032 | #NUM! | H4C1,MFGE8 |
| T Cell Receptor Signaling | 0.333 | 0.028 | 1.000 | CSK,NFKBID,PIK3CG,PIK3R2,RELA |
| Apelin Endothelial Signaling Pathway | 0.333 | 0.029 | 0.000 | ADCY6,PIK3CG,PIK3R2,RELA |
| BMAL1:CLOCK,NPAS2 activates circadian expression | 0.329 | 0.040 | #NUM! | TBL1XR1 |
| Cholesterol biosynthesis | 0.329 | 0.040 | #NUM! | SREBF1 |
| Glycogen metabolism | 0.329 | 0.040 | #NUM! | PHKG2 |
| Signaling by Erythropoietin | 0.329 | 0.040 | #NUM! | PIK3CG |
| Signaling by NTRK3 (TRKC) | 0.329 | 0.040 | #NUM! | CASP6 |
| Oxytocin in Brain Signaling Pathway | 0.329 | 0.028 | -1.000 | CAMKK1,CREB3,PIK3CG,PIK3R2,PLAAT3 |
| Mitochondrial protein import | 0.329 | 0.032 | #NUM! | CHCHD4,TIMM50 |
| Thrombopoietin Signaling | 0.329 | 0.032 | #NUM! | PIK3CG,PIK3R2 |
| Induction of Apoptosis by HIV1 | 0.329 | 0.032 | #NUM! | NFKBID,RELA |
| ERB2-ERBB3 Signaling | 0.329 | 0.032 | #NUM! | PIK3CG,PIK3R2 |
| Gαi Signaling | 0.328 | 0.028 | -1.000 | ADCY6,GRM2,OPRL1,OPRM1 |
| WNK Renal Signaling Pathway | 0.327 | 0.029 | #NUM! | PIK3CG,PIK3R2,SGK1 |

| | | | | |
|---|---|---|---|---|
| Potassium Channels | 0.327 | 0.029 | #NUM! | KCNA4,KCNQ1,KCNQ3 |
| Membrane Repair Signaling Pathway | 0.327 | 0.029 | #NUM! | CAVIN1,SYT11,TSPAN3 |
| Th1 Pathway | 0.327 | 0.029 | #NUM! | IL18,PIK3CG,PIK3R2 |
| Irritable Bowel Syndrome Signaling Pathway | 0.326 | 0.027 | 0.816 | IL18,PIK3CG,PIK3R2,RHOBTB2,RHOD,WNT5A |
| CDX Gastrointestinal Cancer Signaling Pathway | 0.324 | 0.028 | -1.342 | IL18,PIK3CG,PIK3R2,RELA,WNT5A |
| IL-17 Signaling | 0.323 | 0.028 | 1.000 | IL18,PIK3CG,PIK3R2,RELA |
| Extracellular matrix organization | 0.321 | 0.029 | #NUM! | COL3A1,COL6A6,LAMA5 |
| Phase II - Conjugation of compounds | 0.321 | 0.031 | #NUM! | PAPSS1,SULT1A1 |
| Semaphorin interactions | 0.321 | 0.031 | #NUM! | PLXNC1,TLN1 |
| RNA Polymerase I Transcription | 0.321 | 0.031 | #NUM! | CAVIN1,H4C1 |
| TEC Kinase Signaling | 0.320 | 0.028 | 1.000 | PIK3CG,PIK3R2,RELA,RHOBTB2,RHOD |
| ATF4 activates genes in response to endoplasmic reticulum stres | 0.317 | 0.039 | #NUM! | EXOSC4 |
| Peroxisomal lipid metabolism | 0.317 | 0.039 | #NUM! | AMACR |
| Melatonin Degradation I | 0.317 | 0.039 | #NUM! | SULT1A1 |
| Cell Cycle Checkpoints | 0.317 | 0.027 | 0.378 | ANAPC7,BABAM1,CHEK1,H4C1,MAPRE1,PSME1,RPS27 |
| Telomerase Signaling | 0.316 | 0.029 | #NUM! | ELF2,PIK3CG,PIK3R2 |
| Mitotic Roles of Polo-Like Kinase | 0.314 | 0.031 | #NUM! | ANAPC7,PLK4 |
| Macrophage Classical Activation Signaling Pathway | 0.313 | 0.028 | 2.000 | IL18,IRF5,NFKBID,RELA |
| G-Protein Coupled Receptor Signaling | 0.308 | 0.026 | -0.728 | ADCY6,CHRM3,CNGB1,CREB3,GPR12,GRM2,KCNQ3,NFKBID,OPRL1,OPRM1,PDE7B,PIK3CG,PIK3R2,RELA,RPS6KA1,SIX3,WWTR1 |
| Collagen biosynthesis and modifying enzymes | 0.307 | 0.030 | #NUM! | COL3A1,COL6A6 |
| Role of JAK1 and JAK3 in γc Cytokine Signaling | 0.307 | 0.030 | #NUM! | PIK3CG,PIK3R2 |
| ERBB4 Signaling | 0.307 | 0.030 | #NUM! | PIK3CG,PIK3R2 |
| GM-CSF Signaling | 0.307 | 0.030 | #NUM! | PIK3CG,PIK3R2 |
| Mitotic Metaphase and Anaphase | 0.306 | 0.027 | 1.633 | ANAPC7,MAPRE1,PDS5A,PSME1,RCC1,RPS27 |
| Phospholipase C Signaling | 0.306 | 0.026 | -0.447 | ADCY6,ARHGEF2,CREB3,RELA,RHOBTB2,RHOD,TGM2 |
| Effects of PIP2 hydrolysis | 0.305 | 0.037 | #NUM! | TRPC3 |
| Interleukin-7 signaling | 0.305 | 0.037 | #NUM! | PIK3R2 |
| Antimicrobial peptides | 0.305 | 0.037 | #NUM! | S100A9 |
| PRC2 methylates histones and DNA | 0.305 | 0.037 | #NUM! | H4C1 |
| Syndecan interactions | 0.305 | 0.037 | #NUM! | COL3A1 |
| Role of Cytokines in Mediating Communication between Immun | 0.305 | 0.037 | #NUM! | IL18 |
| Interferon Signaling | 0.305 | 0.037 | #NUM! | RELA |
| Integration of energy metabolism | 0.304 | 0.028 | #NUM! | ADCY6,ADIPOR1,CHRM3 |
| Preeclampsia Signaling Pathway | 0.303 | 0.027 | 0.000 | IL18,JUNB,PIK3CG,PIK3R2 |
| TP53 Regulates Transcription of DNA Repair Genes | 0.300 | 0.030 | #NUM! | AP1S2,CHEK1 |
| Synaptogenesis Signaling Pathway | 0.299 | 0.026 | 0.000 | ADCY6,CDH2,CREB3,GRM2,PIK3CG,PIK3R2,SYT11,TLN1 |
| Sulfur amino acid metabolism | 0.294 | 0.036 | #NUM! | MRI1 |
| Estrogen Receptor Signaling | 0.294 | 0.026 | 0.000 | ADCY6,CREB3,EIF4E,PGR,PIK3CG,PIK3R2,PPARGC1A,RELA,TBL1XR1,ZDHHC21 |
| Degradation of the extracellular matrix | 0.293 | 0.029 | #NUM! | CAST,LAMA5 |
| PTEN Regulation | 0.290 | 0.027 | 1.000 | AGO4,LAMTOR5,PSME1,TNKS2 |
| Lysosome Positioning Signaling Pathway | 0.290 | 0.027 | 1.000 | ANXA11,PIK3CG,PIK3R2,SNAPIN |
| Role of MAPK Signaling in Inhibiting the Pathogenesis of Influenz | 0.286 | 0.029 | #NUM! | NFKBID,PLAAT3 |
| Energy dependent regulation of mTOR by LKB1-AMPK | 0.284 | 0.035 | #NUM! | LAMTOR5 |
| SUMOylation of intracellular receptors | 0.284 | 0.035 | #NUM! | PGR |
| ERCC6 (CSB) and EHMT2 (G9a) positively regulate rRNA expressi | 0.284 | 0.035 | #NUM! | H4C1 |
| Myogenesis | 0.284 | 0.035 | #NUM! | CDH2 |
| Protein ubiquitination | 0.284 | 0.035 | #NUM! | UBE2D2 |
| EGR2 and SOX10-mediated initiation of Schwann cell myelinatio | 0.284 | 0.035 | #NUM! | WWTR1 |
| Cilia Biogenesis Signaling Pathway | 0.283 | 0.026 | -1.342 | ANAPC7,BBS7,PLK4,SASS6,TGFA |
| Neuroinflammation Signaling Pathway | 0.282 | 0.026 | -0.378 | CALB2,CREB3,IL18,MFGE8,PIK3CG,PIK3R2,RELA |
| Pre-NOTCH Expression and Processing | 0.280 | 0.029 | #NUM! | AGO4,H4C1 |
| Metabolism of water-soluble vitamins and cofactors | 0.280 | 0.029 | #NUM! | MMACHC,PNPO |
| HIF1α Signaling | 0.279 | 0.026 | 1.342 | EIF4E,PIK3CG,PIK3R2,TGFA,VIM |
| Nonsense-Mediated Decay (NMD) | 0.278 | 0.027 | #NUM! | RPL27,RPS15,RPS27 |
| Cholecystokinin/Gastrin-mediated Signaling | 0.278 | 0.027 | #NUM! | IL18,RHOBTB2,RHOD |
| SAPK/JNK Signaling | 0.278 | 0.027 | #NUM! | HNRNPK,PIK3CG,PIK3R2 |
| Hepatic Fibrosis Signaling Pathway | 0.277 | 0.025 | 2.530 | COL3A1,CREB3,IL18,NFKBID,PIK3CG,PIK3R2,RELA,RHOBTB2,RHOD,WNT5A |
| Sirtuin Signaling Pathway | 0.276 | 0.026 | -1.342 | ATG3,ATG4B,PPARGC1A,RELA,SREBF1,TIMM50,TOMM34 |
| IL-12 Signaling and Production in Macrophages | 0.275 | 0.026 | 1.342 | IL18,IRF5,PIK3CG,PIK3R2,RELA |

| | | | | |
|---|---|---|---|---|
| Signaling by ALK | 0.274 | 0.033 | #NUM! | PIK3R2 |
| DAP12 interactions | 0.274 | 0.033 | #NUM! | PIK3R2 |
| Activation of kainate receptors upon glutamate binding | 0.274 | 0.033 | #NUM! | GRIK2 |
| Endosomal Sorting Complex Required For Transport (ESCRT) | 0.274 | 0.033 | #NUM! | VPS37A |
| Signaling by CSF3 (G-CSF) | 0.274 | 0.033 | #NUM! | UBE2D2 |
| Sensory perception of taste | 0.274 | 0.033 | #NUM! | TRPM4 |
| Regulation of CDH11 Expression and Function | 0.274 | 0.033 | #NUM! | AGO4 |
| Superpathway of Melatonin Degradation | 0.274 | 0.033 | #NUM! | SULT1A1 |
| IL-17A Signaling in Fibroblasts | 0.273 | 0.028 | #NUM! | NFKBID,RELA |
| IL-7 Signaling Pathway | 0.273 | 0.028 | #NUM! | PIK3CG,PIK3R2 |
| Amyotrophic Lateral Sclerosis Signaling | 0.273 | 0.027 | #NUM! | GRIK2,PIK3CG,PIK3R2 |
| ABC-family proteins mediated transport | 0.273 | 0.027 | #NUM! | ABCD2,PSME1,VPS37A |
| PKCθ Signaling in T Lymphocytes | 0.272 | 0.026 | 0.000 | NFKBID,PIK3CG,PIK3R2,RELA |
| PI3K/AKT Signaling | 0.272 | 0.026 | -0.447 | EIF4E,NFKBID,PIK3CG,PIK3R2,RELA |
| Myelination Signaling Pathway | 0.269 | 0.025 | 1.134 | CREB3,EIF4E,LAMA5,PIK3CG,PIK3R2,SREBF1,WNT5A,ZNF24 |
| Ovarian Cancer Signaling | 0.268 | 0.026 | #NUM! | BRCA2,PIK3CG,PIK3R2,WNT5A |
| Regulation of RUNX2 expression and activity | 0.267 | 0.028 | #NUM! | PPARGC1A,PSME1 |
| Toll-like Receptor Signaling | 0.267 | 0.028 | #NUM! | IL18,RELA |
| Glutamate binding, activation of AMPA receptors and synaptic p | 0.264 | 0.032 | #NUM! | GRIP2 |
| Transcriptional regulation of brown and beige adipocyte differer | 0.264 | 0.032 | #NUM! | PPARGC1A |
| Turbulent (oscillatory, disturbed) flow shear stress activates sign | 0.264 | 0.032 | #NUM! | RELA |
| SWI/SNF chromatin remodelers | 0.264 | 0.032 | #NUM! | BRD7 |
| Ribotoxic Stress Signaling Pathway | 0.264 | 0.032 | #NUM! | ZFP36 |
| Cytotoxic T Lymphocyte-mediated Apoptosis of Target Cells | 0.264 | 0.032 | #NUM! | CASP6 |
| FXR/RXR Activation | 0.264 | 0.026 | -1.000 | IL18,Mt1,PPARGC1A,SREBF1 |
| Nitric Oxide Signaling in the Cardiovascular System | 0.263 | 0.026 | #NUM! | PIK3CG,PIK3R2,RYR2 |
| Xenobiotic Metabolism Signaling | 0.257 | 0.025 | #NUM! | NDST2,PIK3CG,PIK3R2,PPARGC1A,RELA,SULT1A1 |
| Neuropeptides in Migraine Signaling Pathway | 0.256 | 0.026 | 1.000 | ADCY6,CREB3,IL18,P2RX3 |
| RNA polymerase II transcribes snRNA genes | 0.256 | 0.027 | #NUM! | SNAPC1,TAF13 |
| NAP1L1 Transcription Regulation Signaling Pathway | 0.256 | 0.027 | #NUM! | PIK3CG,PIK3R2 |
| Toll Like Receptor 3 (TLR3) Cascade | 0.255 | 0.031 | #NUM! | UBE2D2 |
| G-protein beta:gamma signalling | 0.255 | 0.031 | #NUM! | PIK3CG |
| Thrombin signalling through proteinase activated receptors (PAF | 0.255 | 0.031 | #NUM! | F2 |
| Sialic acid metabolism | 0.255 | 0.031 | #NUM! | NANS |
| Post-translational modification: synthesis of GPI-anchored prote | 0.250 | 0.027 | #NUM! | MSLN,PLET1 |
| Antiproliferative Role of Somatostatin Receptor 2 | 0.250 | 0.027 | #NUM! | PIK3CG,PIK3R2 |
| Oxytocin in Spinal Neurons Signaling Pathway | 0.247 | 0.030 | #NUM! | CAMKK1 |
| Arachidonate metabolism | 0.247 | 0.030 | #NUM! | FAAH |
| MAPK targets/ Nuclear events mediated by MAP kinases | 0.247 | 0.030 | #NUM! | RPS6KA1 |
| Synaptic Long Term Potentiation | 0.245 | 0.025 | #NUM! | CREB3,GRM2,RPS6KA1 |
| Cellular response to hypoxia | 0.244 | 0.026 | #NUM! | PSME1,UBE2D2 |
| Activation of anterior HOX genes in hindbrain during early embr | 0.244 | 0.026 | #NUM! | H4C1,PAXIP1 |
| Thyroid Cancer Signaling | 0.244 | 0.026 | #NUM! | PIK3CG,PIK3R2 |
| GDNF Family Ligand-Receptor Interactions | 0.244 | 0.026 | #NUM! | PIK3CG,PIK3R2 |
| Macropinocytosis Signaling | 0.244 | 0.026 | #NUM! | PIK3CG,PIK3R2 |
| Role of NANOG in Mammalian Embryonic Stem Cell Pluripotency | 0.241 | 0.025 | #NUM! | PIK3CG,PIK3R2,WNT5A |
| Telomere Maintenance | 0.239 | 0.026 | #NUM! | ATRX,H4C1 |
| IL-3 Signaling | 0.239 | 0.026 | #NUM! | PIK3CG,PIK3R2 |
| Oncogene Induced Senescence | 0.239 | 0.029 | #NUM! | AGO4 |
| Transcriptional Regulation by E2F6 | 0.239 | 0.029 | #NUM! | CHEK1 |
| Transcriptional Regulation by NPAS4 | 0.239 | 0.029 | #NUM! | AGO4 |
| Retinol Biosynthesis | 0.239 | 0.029 | #NUM! | CEL |
| Oxidative Ethanol Degradation III | 0.239 | 0.029 | #NUM! | Cyp2d22 |
| Superpathway of Methionine Degradation | 0.239 | 0.029 | #NUM! | SPOUT1 |
| Nucleotide Excision Repair Pathway | 0.239 | 0.029 | #NUM! | ERCC5 |
| RHOA Signaling | 0.237 | 0.025 | #NUM! | CDC42EP4,SEMA3F,SEPTIN2 |
| Oxidative Stress Induced Senescence | 0.234 | 0.026 | #NUM! | AGO4,H4C1 |
| Factors involved in megakaryocyte development and platelet pr | 0.234 | 0.026 | #NUM! | MFN2,SH2B3 |
| Transcriptional regulation by the AP-2 (TFAP2) family of transcri | 0.231 | 0.029 | #NUM! | TGFA |

| | | | | |
|---|---|---|---|---|
| Graft-versus-Host Disease Signaling | 0.231 | 0.029 | #NUM! | IL18 |
| Antiproliferative Role of TOB in T Cell Signaling | 0.231 | 0.029 | #NUM! | RPS6KA1 |
| Signaling by MET | 0.229 | 0.025 | #NUM! | COL3A1,LAMA5 |
| Altered T Cell and B Cell Signaling in Rheumatoid Arthritis | 0.229 | 0.025 | #NUM! | IL18,RELA |
| Striated Muscle Contraction | 0.223 | 0.028 | #NUM! | VIM |
| Interleukin-10 signaling | 0.223 | 0.028 | #NUM! | IL18 |
| Fatty acyl-CoA biosynthesis | 0.223 | 0.028 | #NUM! | ELOVL7 |
| Aggrephagy | 0.223 | 0.028 | #NUM! | VIM |
| Formation of the dystrophin-glycoprotein complex (DGC) | 0.223 | 0.028 | #NUM! | LAMA5 |
| MyD88-independent TLR4 cascade | 0.216 | 0.027 | #NUM! | UBE2D2 |
| Regulation of TP53 Expression and Degradation | 0.216 | 0.027 | #NUM! | SGK1 |
| Nucleosome assembly | 0.216 | 0.027 | #NUM! | H4C1 |
| Serotonin Degradation | 0.209 | 0.026 | #NUM! | SULT1A1 |
| Neurotransmitter release cycle | 0.202 | 0.026 | #NUM! | SLC1A6 |
| Gap junction trafficking and regulation | 0.202 | 0.026 | #NUM! | GJA4 |
| MyD88:MAL(TIRAP) cascade initiated on plasma membrane | 0.202 | 0.026 | #NUM! | S100A9 |
| Smooth Muscle Contraction | 0.202 | 0.026 | #NUM! | TLN1 |
| Retinoic acid Mediated Apoptosis Signaling | 0.202 | 0.026 | #NUM! | TNKS2 |
| Vasopressin regulates renal water homeostasis via Aquaporins | 0.196 | 0.025 | #NUM! | ADCY6 |
| RET signaling | 0.196 | 0.025 | #NUM! | PIK3R2 |
| Dilated Cardiomyopathy Signaling Pathway | 0.000 | 0.014 | #NUM! | ADCY6,RYR2 |
| Neurovascular Coupling Signaling Pathway | 0.000 | 0.019 | #NUM! | CHRM3,PLAAT3,RYR2,TRPC3 |
| Oxytocin Signaling Pathway | 0.000 | 0.022 | 0.000 | CAMKK1,CREB3,MFN2,PIK3CG,PIK3R2,RELA |
| Pulmonary Healing Signaling Pathway | 0.000 | 0.016 | #NUM! | FGF10,WNT5A,WWTR1 |
| Pyroptosis Signaling Pathway | 0.000 | 0.014 | #NUM! | IL18 |
| CLEAR Signaling Pathway | 0.000 | 0.022 | -1.633 | CREB3,GLMP,LAMTOR5,PPARGC1A,TGFA,VPS35 |
| IL-13 Signaling Pathway | 0.000 | 0.023 | #NUM! | PIK3CG,PIK3R2 |
| SNARE Signaling Pathway | 0.000 | 0.023 | #NUM! | ADCY6,SYT11,Syt15 |
| Multiple Sclerosis Signaling Pathway | 0.000 | 0.011 | #NUM! | IL18,TNKS2 |
| Pathogen Induced Cytokine Storm Signaling Pathway | 0.000 | 0.018 | 0.000 | COL3A1,COL6A6,Csf2ra,IL18,RYR2 |
| Role of Chondrocytes in Rheumatoid Arthritis Signaling Pathway | 0.000 | 0.008 | #NUM! | IL18 |
| Role of Osteoblasts in Rheumatoid Arthritis Signaling Pathway | 0.000 | 0.019 | 0.000 | IL18,PIK3CG,PIK3R2,WNT5A |
| Glucocorticoid Receptor Signaling | 0.000 | 0.021 | #NUM! | BRD7,NFKBID,PGR,PIK3CG,PIK3R2,RELA,SGK1,TAF10,TAF13,TAF9B |
| Neutrophil Extracellular Trap Signaling Pathway | 0.000 | 0.024 | -0.378 | CASP6,COL3A1,COL6A6,PIK3CG,PIK3R2,PLAAT3,TIMM50,TOMM34 |
| S100 Family Signaling Pathway | 0.000 | 0.021 | -0.277 | CHRM3,CREB3,GPR12,GRM2,IL18,OPRL1,OPRM1,PIK3CG,PIK3R2,RYR2,S100A9,SERPINF2,TGFA,WNT5A |
| Neuregulin Signaling | 0.000 | 0.017 | #NUM! | PIK3R2,TGFA |
| Circadian Rhythm Signaling | 0.000 | 0.016 | #NUM! | ADCY6,CREB3,EIF4E,RYR2 |
| Leukocyte Extravasation Signaling | 0.000 | 0.017 | #NUM! | EDIL3,PIK3CG,PIK3R2 |
| Axonal Guidance Signaling | 0.000 | 0.021 | #NUM! | ANPEP,EIF4E,MEP1A,PIK3CG,PIK3R2,PLXNC1,RHOD,SEMA3F,SEMA4G,WNT5A |
| Fc Epsilon RI Signaling | 0.000 | 0.018 | #NUM! | PIK3CG,PIK3R2 |
| Synaptic Long Term Depression | 0.000 | 0.016 | #NUM! | GRM2,PLAAT3,RYR2 |
| ABRA Signaling Pathway | 0.000 | 0.024 | #NUM! | JUNB,PPARGC1A |
| Activin Inhibin Signaling Pathway | 0.000 | 0.016 | #NUM! | IL18,PIK3CG,PIK3R2 |
| Adrenergic Receptor Signaling Pathway (Enhanced) | 0.000 | 0.011 | #NUM! | ADCY6,IL18 |
| GABAergic Receptor Signaling Pathway (Enhanced) | 0.000 | 0.008 | #NUM! | ADCY6 |
| Cachexia Signaling Pathway | 0.000 | 0.024 | 1.134 | ADCY6,CASP6,IL18,MFN2,PIK3CG,PIK3R2,PSME1,VPS37A |
| ISGylation Signaling Pathway | 0.000 | 0.024 | #NUM! | IRF5,RPS15 |
| Orexin Signaling Pathway | 0.000 | 0.022 | 0.447 | ADCY6,PIK3CG,PIK3R2,PPARGC1A,TRPC3 |
| Sertoli Cell-Germ Cell Junction Signaling Pathway (Enhanced) | 0.000 | 0.022 | 0.447 | CDH2,LAMA5,PIK3CG,PIK3R2,RELA |
| NRF2-mediated Oxidative Stress Response | 0.000 | 0.020 | #NUM! | Cyp2d22,JUNB,PIK3CG,PIK3R2 |
| Signaling by Insulin receptor | 0.000 | 0.023 | #NUM! | PIK3R2 |
| G alpha (i) signalling events | 0.000 | 0.023 | 0.447 | ADCY6,GRM2,OPRL1,OPRM1,RGS9 |
| ISG15 antiviral mechanism | 0.000 | 0.015 | #NUM! | EIF4E |
| Meiotic synapsis | 0.000 | 0.023 | #NUM! | H4C1 |
| Signaling by ERBB4 | 0.000 | 0.017 | #NUM! | PGR |
| Translocation of SLC2A4 (GLUT4) to the plasma membrane | 0.000 | 0.016 | #NUM! | TBC1D4 |
| Glycosaminoglycan metabolism | 0.000 | 0.025 | #NUM! | B4GALT5,NDST2,PAPSS1 |
| Complement cascade | 0.000 | 0.023 | #NUM! | F2 |
| NLR signaling pathways | 0.000 | 0.019 | #NUM! | RELA |

| | | | | |
|---|---|---|---|---|
| Regulation of Apoptosis | 0.000 | 0.019 | #NUM! | PSME1 |
| Cytosolic sensors of pathogen-associated DNA | 0.000 | 0.016 | #NUM! | RELA |
| Regulation of beta-cell development | 0.000 | 0.024 | #NUM! | FGF10 |
| Signaling by NTRK1 (TRKA) | 0.000 | 0.025 | #NUM! | PIK3R2,RPS6KA1 |
| Signaling by VEGF | 0.000 | 0.009 | #NUM! | PIK3R2 |
| Degradation of beta-catenin by the destruction complex | 0.000 | 0.011 | #NUM! | PSME1 |
| Signaling by NOTCH1 | 0.000 | 0.013 | #NUM! | TBL1XR1 |
| Immunoregulatory interactions between a Lymphoid and a non- | 0.000 | 0.025 | #NUM! | COL3A1,NPDC1 |
| TCF dependent signaling in response to WNT | 0.000 | 0.021 | 0.000 | H4C1,PSME1,TNKS2,WNT5A |
| Cell surface interactions at the vascular wall | 0.000 | 0.017 | #NUM! | F2,PIK3R2 |
| Fcgamma receptor (FCGR) dependent phagocytosis | 0.000 | 0.011 | #NUM! | PIK3R2 |
| Asparagine N-linked glycosylation | 0.000 | 0.008 | #NUM! | B4GALT5 |
| MHC class II antigen presentation | 0.000 | 0.019 | #NUM! | AP1S2,KIF26A |
| Integrin cell surface interactions | 0.000 | 0.025 | #NUM! | COL3A1,COL6A6 |
| DNA Damage/Telomere Stress Induced Senescence | 0.000 | 0.024 | #NUM! | H4C1 |
| SUMOylation of DNA damage response and repair proteins | 0.000 | 0.013 | #NUM! | NSMCE3 |
| HSP90 chaperone cycle for steroid hormone receptors in the pre | 0.000 | 0.021 | #NUM! | PGR |
| Metabolism of polyamines | 0.000 | 0.017 | #NUM! | PSME1 |
| Class A/1 (Rhodopsin-like receptors) | 0.000 | 0.017 | 0.447 | CHRM3,F2,NLN,OPRL1,OPRM1 |
| Class B/2 (Secretin family receptors) | 0.000 | 0.011 | #NUM! | WNT5A |
| Netrin-1 signaling | 0.000 | 0.023 | #NUM! | TRPC3 |
| L1CAM interactions | 0.000 | 0.018 | #NUM! | KCNQ3,RPS6KA1 |
| SUMOylation of transcription cofactors | 0.000 | 0.023 | #NUM! | PPARGC1A |
| Protein folding | 0.000 | 0.011 | #NUM! | RGS9 |
| G alpha (q) signalling events | 0.000 | 0.018 | #NUM! | CHRM3,F2,PIK3R2 |
| Platelet homeostasis | 0.000 | 0.023 | #NUM! | P2RX3,TRPC3 |
| G alpha (s) signalling events | 0.000 | 0.015 | #NUM! | ADCY6,PDE7B |
| G alpha (z) signalling events | 0.000 | 0.021 | #NUM! | ADCY6 |
| Transport of inorganic cations/anions and amino acids/oligopept | 0.000 | 0.010 | #NUM! | SLC1A6 |
| NoRC negatively regulates rRNA expression | 0.000 | 0.017 | #NUM! | H4C1 |
| Sphingolipid metabolism | 0.000 | 0.010 | #NUM! | B4GALT5 |
| TAK1-dependent IKK and NF-kappa-B activation | 0.000 | 0.024 | #NUM! | RELA |
| Interleukin-1 family signaling | 0.000 | 0.024 | #NUM! | IL18,PSME1,RELA |
| Cell junction organization | 0.000 | 0.011 | #NUM! | CDH2 |
| Interleukin-2 family signaling | 0.000 | 0.024 | #NUM! | PIK3R2 |
| Mitotic G2-G2/M phases | 0.000 | 0.016 | #NUM! | MAPRE1,PLK4,PSME1 |
| Regulation of mitotic cell cycle | 0.000 | 0.022 | #NUM! | ANAPC7,PSME1 |
| Mitotic G1 phase and G1/S transition | 0.000 | 0.015 | #NUM! | PSME1,TK1 |
| SUMOylation of chromatin organization proteins | 0.000 | 0.016 | #NUM! | H4C1 |
| SUMOylation of RNA binding proteins | 0.000 | 0.021 | #NUM! | HNRNPK |
| Interleukin-3, Interleukin-5 and GM-CSF signaling | 0.000 | 0.020 | #NUM! | PIK3R2 |
| O-linked glycosylation | 0.000 | 0.009 | #NUM! | B4GALT5 |
| Hedgehog ligand biogenesis | 0.000 | 0.015 | #NUM! | PSME1 |
| Cardiac conduction | 0.000 | 0.024 | #NUM! | KCNQ1,RYR2,WWTR1 |
| Hedgehog 'off' state | 0.000 | 0.018 | #NUM! | ADCY6,PSME1 |
| C-type lectin receptors (CLRs) | 0.000 | 0.023 | #NUM! | PSME1,RELA,UBE2D2 |
| TP53 Regulates Metabolic Genes | 0.000 | 0.024 | #NUM! | AGO4,LAMTOR5 |
| Signaling by FGFR2 | 0.000 | 0.014 | #NUM! | FGF10 |
| TNFR2 non-canonical NF-kB pathway | 0.000 | 0.015 | #NUM! | PSME1 |
| MAPK6/MAPK4 signaling | 0.000 | 0.022 | #NUM! | AGO4,PSME1 |
| Nucleotide Excision Repair | 0.000 | 0.009 | #NUM! | ERCC5 |
| Neutrophil degranulation | 0.000 | 0.010 | 0.000 | ANPEP,PLEKHO2,S100A9,TRPM2 |
| COPI-mediated anterograde transport | 0.000 | 0.011 | #NUM! | COG4 |
| Intra-Golgi and retrograde Golgi-to-ER traffic | 0.000 | 0.016 | #NUM! | COG4,KIF26A,RGP1 |
| Mitotic Prophase | 0.000 | 0.021 | #NUM! | CTDNEP1,H4C1 |
| Mitotic Prometaphase | 0.000 | 0.021 | 2.000 | MAPRE1,PDS5A,PLK4,RPS27 |
| Synthesis of DNA | 0.000 | 0.016 | #NUM! | ANAPC7,PSME1 |
| S Phase | 0.000 | 0.020 | #NUM! | PDS5A,PSME1 |
| RNA Polymerase III Transcription | 0.000 | 0.024 | #NUM! | SNAPC1 |

| | | | | |
|---|---|---|---|---|
| TNF signaling | 0.000 | 0.018 | #NUM! | UBE2D2 |
| Response to elevated platelet cytosolic Ca2+ | 0.000 | 0.016 | #NUM! | SERPINF2,TLN1 |
| Interferon gamma signaling | 0.000 | 0.015 | #NUM! | IRF5 |
| TBC/RABGAPs | 0.000 | 0.023 | #NUM! | RAB33A |
| E3 ubiquitin ligases ubiquitinate target proteins | 0.000 | 0.023 | #NUM! | UBE2D2 |
| RAB geranylgeranylation | 0.000 | 0.016 | #NUM! | RAB33A |
| Transcriptional regulation by RUNX1 | 0.000 | 0.021 | #NUM! | ELF2,H4C1,PSME1 |
| Transcriptional regulation by RUNX3 | 0.000 | 0.021 | #NUM! | PSME1,WWTR1 |
| Transcriptional regulation by RUNX2 | 0.000 | 0.020 | #NUM! | WWTR1 |
| Interferon alpha/beta signaling | 0.000 | 0.021 | #NUM! | IRF5 |
| Transcriptional regulation of granulopoiesis | 0.000 | 0.023 | #NUM! | H4C1 |
| Amino acids regulate mTORC1 | 0.000 | 0.019 | #NUM! | LAMTOR5 |
| Sensory processing of sound by inner hair cells of the cochlea | 0.000 | 0.015 | #NUM! | TPRN |
| Sensory processing of sound by outer hair cells of the cochlea | 0.000 | 0.018 | #NUM! | TPRN |
| Cytoprotection by HMOX1 | 0.000 | 0.018 | #NUM! | TBL1XR1 |
| GABA receptor activation | 0.000 | 0.017 | #NUM! | ADCY6 |
| Kinesins | 0.000 | 0.020 | #NUM! | KIF26A |
| Signaling by the B Cell Receptor (BCR) | 0.000 | 0.018 | #NUM! | PSME1,RELA |
| Hepatic Cholestasis | 0.000 | 0.020 | 1.000 | ADCY6,IL18,NFKBID,RELA |
| Hepatic Fibrosis / Hepatic Stellate Cell Activation | 0.000 | 0.022 | #NUM! | COL3A1,COL6A6,RELA,TGFA |
| Mitochondrial Dysfunction | 0.000 | 0.019 | 0.816 | CREB3,MFN2,PIK3CG,PIK3R2,PPARGC1A,TOMM34 |
| CGAS-STING Signaling Pathway | 0.000 | 0.017 | #NUM! | IL18,RELA |
| ROBO SLIT Signaling Pathway | 0.000 | 0.008 | #NUM! | CDH2 |
| BBSome Signaling Pathway | 0.000 | 0.013 | -0.816 | BBS7,CHRM3,GPR12,GRM2,OPRL1,OPRM1 |
| Folate Signaling Pathway | 0.000 | 0.019 | #NUM! | ATIC |
| HEY1 Signaling Pathway | 0.000 | 0.014 | #NUM! | PIK3CG,PIK3R2 |
| Sleep NREM Signaling Pathway | 0.000 | 0.018 | #NUM! | ADCY6,CREB3 |
| Nuclear Cytoskeleton Signaling Pathway | 0.000 | 0.020 | -1.000 | CDH2,CTDNEP1,KIF26A,TOR1A |
| Hematoma Resolution Signaling Pathway | 0.000 | 0.018 | #NUM! | ADIPOR1,F2,IL18,VPS35 |
| Glycation Signaling Pathway | 0.000 | 0.019 | 1.000 | IL18,PIK3CG,PIK3R2,RELA |
| Inositol phosphate metabolism | 0.000 | 0.021 | #NUM! | ISYNA1 |
| Pyruvate metabolism | 0.000 | 0.021 | #NUM! | GID8 |
| Somitogenesis | 0.000 | 0.018 | #NUM! | PSME1 |
| Mitochondrial protein degradation | 0.000 | 0.021 | #NUM! | MRPS2,SSBP1 |
| Transcriptional and post-translational regulation of MITF-M expı | 0.000 | 0.016 | #NUM! | RPS6KA1 |
| Complex I biogenesis | 0.000 | 0.015 | #NUM! | TMEM126B |
| Regulation of endogenous retroelements | 0.000 | 0.014 | #NUM! | H4C1 |
| Complex IV assembly | 0.000 | 0.021 | #NUM! | SMIM20 |
| Lung Ionic Balance Signaling Pathway | 0.000 | 0.017 | 1.414 | ADCY6,CHRM3,CNGB1,GPR12,GRM2,KCNQ1,OPRL1,OPRM1 |
| TRIM21 Intracellular Antibody Signaling Pathway | 0.000 | 0.017 | #NUM! | IRF5,PSME1 |
| Bone Mineralization Signaling Pathway | 0.000 | 0.022 | #NUM! | COL3A1,FGF10,WNT5A |
| ID3 Signaling Pathway | 0.000 | 0.017 | #NUM! | ANAPC7,SREBF1 |
| Tuberculosis Active Signaling Pathway | 0.000 | 0.010 | #NUM! | PIK3CG,PIK3R2 |
| Tuberculosis Latent Signaling Pathway | 0.000 | 0.024 | #NUM! | ADCY6,PLIN2 |
| Zn Homeostasis Signaling Pathway | 0.000 | 0.021 | -0.832 | ADCY6,CHRM3,CREB3,GPR12,GRM2,IL18,Mt1,OPRL1,OPRM1,PDE7B,PIK3CG,PIK3R2,RYR2 |
| DYRK1A Signaling Pathway | 0.000 | 0.012 | #NUM! | CREB3,RNF169 |
| Meiosis Initiation Signaling Pathway | 0.000 | 0.010 | #NUM! | BRCA2 |
| NF1 RAS Signaling Pathway | 0.000 | 0.020 | -0.632 | ADCY6,CHRM3,Csf2ra,GPR12,GRM2,OPRL1,OPRM1,PIK3CG,PIK3R2,TGFA |
| NIK-->noncanonical NF-kB signaling | 0.000 | 0.016 | #NUM! | PSME1 |
| Proteasome assembly | 0.000 | 0.020 | #NUM! | PSME1 |
| Degradation of CRY and PER proteins | 0.000 | 0.016 | #NUM! | PSME1 |
| SLAMF Signaling Pathway | 0.000 | 0.015 | #NUM! | PIK3CG,PIK3R2 |
| Exocyst Signaling Pathway | 0.000 | 0.020 | #NUM! | ARHGEF2,CDH2 |
| XIST Epigenetic Regulation Signaling Pathway | 0.000 | 0.024 | #NUM! | HNRNPK,IL18,SPEN |
| Tight Junction Signaling | 0.000 | 0.012 | #NUM! | ARHGEF2,RELA |
| Regulation of Actin-based Motility by Rho | 0.000 | 0.019 | #NUM! | RHOBTB2,RHOD |
| α-Adrenergic Signaling | 0.000 | 0.019 | #NUM! | ADCY6,PHKG2 |
| Caveolar-mediated Endocytosis Signaling | 0.000 | 0.014 | #NUM! | CAVIN1 |
| Role of NFAT in Regulation of the Immune Response | 0.000 | 0.022 | #NUM! | NFKBID,PIK3CG,PIK3R2,RELA |

| | | | | |
|---|---|---|---|---|
| FcγRIIB Signaling in B Lymphocytes | 0.000 | 0.023 | #NUM! | PIK3CG,PIK3R2 |
| Calcium-induced T Lymphocyte Apoptosis | 0.000 | 0.018 | #NUM! | NR4A1 |
| CTLA4 Signaling in Cytotoxic T Lymphocytes | 0.000 | 0.023 | -1.000 | AP1S2,CBLB,PIK3CG,PIK3R2 |
| T Helper Cell Differentiation | 0.000 | 0.015 | #NUM! | IL18 |
| CCR3 Signaling in Eosinophils | 0.000 | 0.016 | #NUM! | PIK3CG,PIK3R2 |
| Cellular Effects of Sildenafil (Viagra) | 0.000 | 0.019 | 0.000 | ADCY6,CHRM3,CNGB1,CREB3,GPR12,GRM2,KCNQ3,OPRL1,OPRM1,PIK3CG,PIK3R2,PPARGC1A |
| Factors Promoting Cardiogenesis in Vertebrates | 0.000 | 0.014 | #NUM! | CREB3,WNT5A |
| CDK5 Signaling | 0.000 | 0.020 | #NUM! | ADCY6,LAMA5 |
| Molecular Mechanisms of Cancer | 0.000 | 0.023 | 0.471 | ADCY6,ARHGEF2,CASP6,CHEK1,CHRM3,F2,GPR12,GRM2,NFKBID,OPRL1,OPRM1,PIK3CG,PIK3R2,RELA,RHOBTB2,RHOD,TGFA,WNT5A |
| Corticotropin Releasing Hormone Signaling | 0.000 | 0.020 | #NUM! | ADCY6,CREB3,NR4A1 |
| HGF Signaling | 0.000 | 0.023 | #NUM! | ELF2,PIK3CG,PIK3R2 |
| Maturity Onset Diabetes of Young (MODY) Signaling | 0.000 | 0.015 | #NUM! | CEL |
| Role of CHK Proteins in Cell Cycle Checkpoint Control | 0.000 | 0.018 | #NUM! | CHEK1 |
| GNRH Signaling | 0.000 | 0.016 | #NUM! | ADCY6,CREB3,RELA |
| Human Embryonic Stem Cell Pluripotency | 0.000 | 0.015 | #NUM! | PIK3CG,PIK3R2,WNT5A |
| ATM Signaling | 0.000 | 0.021 | #NUM! | CHEK1,CREB3 |
| Prolactin Signaling | 0.000 | 0.021 | #NUM! | PIK3CG,PIK3R2 |
| CREB Signaling in Neurons | 0.000 | 0.021 | 0.000 | ADCY6,CHRM3,CREB3,GPR12,GRIK2,GRM2,OPRL1,OPRM1,PIK3CG,PIK3R2,RPS6KA1,TGFA |
| Type I Diabetes Mellitus Signaling | 0.000 | 0.019 | #NUM! | NFKBID,RELA |
| Basal Cell Carcinoma Signaling | 0.000 | 0.015 | #NUM! | WNT5A |
| Glioma Signaling | 0.000 | 0.024 | #NUM! | PIK3CG,PIK3R2,TGFA |
| Bladder Cancer Signaling | 0.000 | 0.010 | #NUM! | FGF10 |
| Chronic Myeloid Leukemia Signaling | 0.000 | 0.019 | 0.447 | PIK3CG,PIK3R2,RCC1,RELA,WNT5A |
| G Beta Gamma Signaling | 0.000 | 0.008 | #NUM! | PIK3CG |
| Communication between Innate and Adaptive Immune Cells | 0.000 | 0.015 | #NUM! | IL18 |
| CDC42 Signaling | 0.000 | 0.007 | #NUM! | DIAPH3 |
| FAK Signaling | 0.000 | 0.017 | 0.000 | CHRM3,CSK,ELF2,GPR12,GRM2,OPRL1,OPRM1,PIK3CG,PIK3R2,SPARCL1 |
| PAK Signaling | 0.000 | 0.018 | #NUM! | PIK3CG,PIK3R2 |
| Role of Macrophages, Fibroblasts and Endothelial Cells in Rheum | 0.000 | 0.024 | 0.378 | CREB3,IL18,NFKBID,PIK3CG,PIK3R2,RELA,WNT5A |
| Role of NFAT in Cardiac Hypertrophy | 0.000 | 0.014 | #NUM! | ADCY6,PIK3CG,PIK3R2 |
| Breast Cancer Regulation by Stathmin1 | 0.000 | 0.018 | -0.632 | ARHGEF2,CHRM3,CREB3,GPR12,GRM2,OPRL1,OPRM1,PIK3CG,PIK3R2,TGFA |
| Neuroprotective Role of THOP1 in Alzheimer's Disease | 0.000 | 0.011 | #NUM! | KLK5 |
| Regulation of IL-2 Expression in Activated and Anergic T Lympho | 0.000 | 0.021 | #NUM! | NFKBID,RELA |
| Role of WNT/GSK-3β Signaling in the Pathogenesis of Influenza | 0.000 | 0.017 | #NUM! | WNT5A |
| NUR77 Signaling in T Lymphocytes | 0.000 | 0.013 | #NUM! | NR4A1 |
| Role of MAPK Signaling in the Pathogenesis of Influenza | 0.000 | 0.013 | #NUM! | PLAAT3 |
| Role of Hypercytokinemia/hyperchemokinemia in the Pathogene | 0.000 | 0.018 | #NUM! | IL18 |
| Role of JAK2 in Hormone-like Cytokine Signaling | 0.000 | 0.019 | #NUM! | SH2B3 |
| Role of JAK family kinases in IL-6-type Cytokine Signaling | 0.000 | 0.014 | #NUM! | JUNB |
| Actin Nucleation by ARP-WASP Complex | 0.000 | 0.022 | #NUM! | RHOBTB2,RHOD |
| Dopamine-DARPP32 Feedback in cAMP Signaling | 0.000 | 0.018 | #NUM! | ADCY6,CAMKK1,CREB3 |
| RHOGDI Signaling | 0.000 | 0.019 | #NUM! | ARHGEF2,CDH2,RHOBTB2,RHOD |
| Mouse Embryonic Stem Cell Pluripotency | 0.000 | 0.020 | #NUM! | PIK3CG,PIK3R2 |
| nNOS Signaling in Skeletal Muscle Cells | 0.000 | 0.021 | #NUM! | RYR2 |
| VEGF Family Ligand-Receptor Interactions | 0.000 | 0.025 | #NUM! | PIK3CG,PIK3R2 |
| Ephrin A Signaling | 0.000 | 0.022 | #NUM! | PIK3CG,PIK3R2,TGFA |
| Ephrin B Signaling | 0.000 | 0.014 | #NUM! | HNRNPK |
| GADD45 Signaling | 0.000 | 0.017 | #NUM! | RELA |
| Netrin Signaling | 0.000 | 0.018 | #NUM! | PIK3CG,PIK3R2,RYR2 |
| tRNA Splicing | 0.000 | 0.022 | #NUM! | PDE7B |
| D-myo-inositol-5-phosphate Metabolism | 0.000 | 0.010 | #NUM! | PIP4P2,UBLCP1 |
| Phospholipases | 0.000 | 0.017 | #NUM! | PLAAT3 |
| D-myo-inositol (1,4,5,6)-Tetrakisphosphate Biosynthesis | 0.000 | 0.006 | #NUM! | UBLCP1 |
| Superpathway of Inositol Phosphate Compounds | 0.000 | 0.018 | 1.000 | PIK3CG,PIK3R2,PIP4P2,UBLCP1 |
| D-myo-inositol (3,4,5,6)-tetrakisphosphate Biosynthesis | 0.000 | 0.006 | #NUM! | UBLCP1 |
| 3-phosphoinositide Degradation | 0.000 | 0.011 | #NUM! | PIP4P2,UBLCP1 |
| 3-phosphoinositide Biosynthesis | 0.000 | 0.015 | #NUM! | PIK3CG,PIK3R2,UBLCP1 |
| Salvage Pathways of Pyrimidine Ribonucleotides | 0.000 | 0.022 | #NUM! | FAM20B,SGK1 |
| Epithelial Adherens Junction Signaling | 0.000 | 0.006 | #NUM! | CDH2 |

| | | | | |
|---|---|---|---|---|
| Remodeling of Epithelial Adherens Junctions | 0.000 | 0.016 | #NUM! | MAPRE1 |
| Agranulocyte Adhesion and Diapedesis | 0.000 | 0.006 | #NUM! | IL18 |
| Granulocyte Adhesion and Diapedesis | 0.000 | 0.007 | #NUM! | IL18 |
| Sperm Motility | 0.000 | 0.021 | #NUM! | CLK1,CNGB1,Csf2ra,CSK,PLAAT3 |
| STAT3 Pathway | 0.000 | 0.007 | #NUM! | TGFA |
| HIPPO signaling | 0.000 | 0.013 | #NUM! | WWTR1 |
| Unfolded protein response | 0.000 | 0.011 | #NUM! | SREBF1 |
| Parkinson's Signaling Pathway | 0.000 | 0.007 | #NUM! | IL18,PPARGC1A |
| PTEN Signaling | 0.000 | 0.020 | #NUM! | PIK3CG,PIK3R2,RELA |
| Cardiac β-adrenergic Signaling | 0.000 | 0.018 | #NUM! | ADCY6,PDE7B,RYR2 |
| Protein Ubiquitination Pathway | 0.000 | 0.015 | 0.000 | HSPE1,PSME1,SASS6,UBE2D2 |
| GABA Receptor Signaling | 0.000 | 0.015 | #NUM! | ADCY6,KCNQ3 |
| WNT/β-catenin Signaling | 0.000 | 0.012 | #NUM! | CDH2,WNT5A |
| Chemokine Signaling | 0.000 | 0.013 | #NUM! | PIK3CG |
| IGF-1 Signaling | 0.000 | 0.019 | #NUM! | PIK3CG,PIK3R2 |
| Dopamine Receptor Signaling | 0.000 | 0.015 | #NUM! | ADCY6 |
| VEGF Signaling | 0.000 | 0.021 | #NUM! | PIK3CG,PIK3R2 |
| PDGF Signaling | 0.000 | 0.024 | #NUM! | PIK3CG,PIK3R2 |
| BMP signaling pathway | 0.000 | 0.012 | #NUM! | RELA |
| GPCR-Mediated Integration of Enteroendocrine Signaling Exemp | 0.000 | 0.014 | #NUM! | ADCY6 |
| GPCR-Mediated Nutrient Sensing in Enteroendocrine Cells | 0.000 | 0.009 | #NUM! | ADCY6 |
| Phagosome Formation | 0.000 | 0.016 | -0.333 | AP1S2,CHRM3,GPR12,GRM2,OPRL1,OPRM1,PIK3CG,PIK3R2,PLAAT3,TLN1 |
| Phagosome Maturation | 0.000 | 0.021 | #NUM! | Dynlt1b (includes others),VPS37A,VPS39 |
| Th1 and Th2 Activation Pathway | 0.000 | 0.020 | #NUM! | IL18,PIK3CG,PIK3R2 |
| Th2 Pathway | 0.000 | 0.017 | #NUM! | PIK3CG,PIK3R2 |
| Osteoarthritis Pathway | 0.000 | 0.024 | 2.000 | CASP6,CREB3,PPARGC1A,RELA,S100A9 |
| Th17 Activation Pathway | 0.000 | 0.016 | #NUM! | RELA |
| SPINK1 Pancreatic Cancer Pathway | 0.000 | 0.022 | #NUM! | KLK5 |
| NER (Nucleotide Excision Repair, Enhanced Pathway) | 0.000 | 0.011 | #NUM! | ERCC5 |
| Endocannabinoid Neuronal Synapse Pathway | 0.000 | 0.014 | #NUM! | ADCY6,FAAH |
| Apelin Adipocyte Signaling Pathway | 0.000 | 0.024 | #NUM! | ADCY6,PPARGC1A |
| Apelin Cardiomyocyte Signaling Pathway | 0.000 | 0.021 | #NUM! | PIK3CG,PIK3R2 |
| Apelin Muscle Signaling Pathway | 0.000 | 0.020 | #NUM! | PPARGC1A |
| BAG2 Signaling Pathway | 0.000 | 0.025 | #NUM! | PSME1,RELA |
| Cardiac Hypertrophy Signaling (Enhanced) | 0.000 | 0.021 | 1.508 | ADCY6,DIAPH3,EIF4E,FGF10,IL18,PDE7B,PIK3CG,PIK3R2,RELA,RYR2,WNT5A |
| FAT10 Signaling Pathway | 0.000 | 0.018 | #NUM! | PSME1 |
| PFKFB4 Signaling Pathway | 0.000 | 0.021 | #NUM! | CREB3 |
| T Cell Exhaustion Signaling Pathway | 0.000 | 0.014 | #NUM! | PIK3CG,PIK3R2 |
| Senescence Pathway | 0.000 | 0.021 | 0.447 | ANAPC7,CHEK1,EIF4E,ELF2,PIK3CG,PIK3R2 |
| White Adipose Tissue Browning Pathway | 0.000 | 0.022 | #NUM! | ADCY6,CREB3,PPARGC1A |
| Necroptosis Signaling Pathway | 0.000 | 0.015 | #NUM! | TIMM50,TOMM34 |
| Xenobiotic Metabolism General Signaling Pathway | 0.000 | 0.017 | #NUM! | PIK3CG,PIK3R2 |
| Xenobiotic Metabolism AHR Signaling Pathway | 0.000 | 0.015 | #NUM! | RELA |
| Xenobiotic Metabolism CAR Signaling Pathway | 0.000 | 0.021 | #NUM! | NDST2,PPARGC1A,SULT1A1 |
| Xenobiotic Metabolism PXR Signaling Pathway | 0.000 | 0.023 | #NUM! | NDST2,PPARGC1A,SULT1A1 |
| Semaphorin Neuronal Repulsive Signaling Pathway | 0.000 | 0.021 | #NUM! | PIK3CG,PIK3R2,SEMA3F |
| Regulation of the Epithelial Mesenchymal Transition in Developr | 0.000 | 0.024 | #NUM! | RELA,WNT5A |
| Kinetochore Metaphase Signaling Pathway | 0.000 | 0.011 | #NUM! | ANAPC7 |
| Calcium Signaling | 0.000 | 0.024 | -2.000 | CAMKK1,CHRNA4,CREB3,RYR2,TRPC3 |
| Ephrin Receptor Signaling | 0.000 | 0.010 | #NUM! | CREB3,PIK3CG |
| Ferroptosis Signaling Pathway | 0.000 | 0.017 | #NUM! | PHKG2,WWTR1 |
| Role of MAPK Signaling in Promoting the Pathogenesis of Influer | 0.000 | 0.019 | #NUM! | NFKBID,PLAAT3 |

**Supplementary Table 5: Differentially expressed genes between PAE and SAC in Neurons at E18**

**Increased expression (1.2 FC, p<0.05)**

| Target nam | Log2 FC | Pvalue | Adjusted p | -log10 pval | -log10 adjusted pvalue |
|---|---|---|---|---|---|
| Ntf3 | 1.621543 | 0.014606 | 0.211638 | 1.835483 | 0.674407 |
| Dbpht2 | 1.615996 | 0.001527 | 0.105319 | 2.816105 | 0.977493 |
| Ccbe1 | 1.368355 | 0.000248 | 0.071776 | 3.604991 | 1.144022 |
| Ptprt | 1.363805 | 0.005604 | 0.159383 | 2.251468 | 0.797558 |
| 1700086L1 | 1.320441 | 0.000692 | 0.081349 | 3.160132 | 1.089648 |
| Clec2j | 1.283594 | 0.000337 | 0.072211 | 3.472375 | 1.141399 |
| Rasgrf2 | 1.218581 | 0.010458 | 0.191032 | 1.980539 | 0.718893 |
| S100a10 | 1.207932 | 0.002105 | 0.116540 | 2.676721 | 0.933524 |
| Kcnh5 | 1.202346 | 0.002404 | 0.119712 | 2.619150 | 0.921864 |
| Ptpro | 1.183417 | 0.003148 | 0.127797 | 2.501981 | 0.893479 |
| Nog | 1.168097 | 0.009790 | 0.189070 | 2.009196 | 0.723377 |
| Masp1 | 1.121444 | 0.001405 | 0.105319 | 2.852189 | 0.977493 |
| Arhgap28 | 1.121298 | 0.000831 | 0.083704 | 3.080260 | 1.077252 |
| Hs6st2 | 1.120917 | 0.002421 | 0.119712 | 2.616005 | 0.921864 |
| Lhfp | 1.093838 | 0.003006 | 0.127513 | 2.521973 | 0.894446 |
| Itpr1 | 1.072700 | 0.002541 | 0.120865 | 2.594933 | 0.917700 |
| Dkk3 | 1.067612 | 0.001073 | 0.093200 | 2.969351 | 1.030586 |
| Kif26b | 1.046128 | 0.002803 | 0.126635 | 2.552364 | 0.897446 |
| Tmtc1 | 1.030695 | 0.004713 | 0.149991 | 2.326740 | 0.823934 |
| Inhba | 1.011176 | 0.040005 | 0.295412 | 1.397885 | 0.529572 |
| Aff2 | 0.991212 | 0.000084 | 0.053674 | 4.077894 | 1.270236 |
| Mast4 | 0.989510 | 0.000160 | 0.062244 | 3.795856 | 1.205900 |
| Shc3 | 0.967924 | 0.046100 | 0.312375 | 1.336299 | 0.505324 |
| Gucy1b1 | 0.965336 | 0.000528 | 0.073869 | 3.277669 | 1.131538 |
| Fer1l6 | 0.962906 | 0.001853 | 0.113004 | 2.732015 | 0.946906 |
| Fat1 | 0.952014 | 0.043845 | 0.305662 | 1.358076 | 0.514758 |
| Klhl5 | 0.949070 | 0.002312 | 0.119065 | 2.636057 | 0.924217 |
| Tgfbr1 | 0.947711 | 0.008471 | 0.179457 | 2.072072 | 0.746039 |
| Slc16a9 | 0.944337 | 0.000111 | 0.061208 | 3.956505 | 1.213194 |
| Adgrl2 | 0.920880 | 0.001752 | 0.111471 | 2.756566 | 0.952839 |
| Syndig1 | 0.909173 | 0.037633 | 0.289746 | 1.424427 | 0.537983 |
| Gabra5 | 0.892260 | 0.012246 | 0.200942 | 1.912007 | 0.696929 |
| Opcml | 0.891165 | 0.020679 | 0.231686 | 1.684466 | 0.635100 |
| Trmt9b | 0.886338 | 0.028680 | 0.262087 | 1.542426 | 0.581554 |
| Phldb2 | 0.885741 | 0.011270 | 0.196420 | 1.948077 | 0.706814 |
| Csrnp3 | 0.881999 | 0.000298 | 0.072211 | 3.526146 | 1.141399 |
| Scarb1 | 0.850137 | 0.008258 | 0.178185 | 2.083109 | 0.749130 |

**Decreased expression (1.2 FC, p<0.05)**

| Target nam | Log2 FC | Pvalue | Adjusted p | -log10 pval | -log10 adjusted pvalue |
|---|---|---|---|---|---|
| Gm21783 | -2.902291 | 0.032053 | 0.270354 | 1.494133 | 0.568068 |
| Gm20792 | -1.917714 | 0.024865 | 0.246706 | 1.604405 | 0.607820 |
| Gm20815 | -1.913673 | 0.000757 | 0.082956 | 3.120645 | 1.081154 |
| Fabp7 | -1.819254 | 0.005747 | 0.161414 | 2.240573 | 0.792060 |
| Eif2s3y | -1.732460 | 0.044752 | 0.307668 | 1.349186 | 0.511917 |
| Srsx | -1.714478 | 0.001473 | 0.105319 | 2.831917 | 0.977493 |
| Met | -1.691354 | 0.029605 | 0.265389 | 1.528641 | 0.576117 |
| Epha3 | -1.627954 | 0.000775 | 0.083581 | 3.110699 | 1.077890 |
| H3c2 | -1.596150 | 0.009152 | 0.184984 | 2.038468 | 0.732865 |
| Tafa1 | -1.591453 | 0.003045 | 0.127513 | 2.516342 | 0.894446 |
| Gm17027 | -1.481546 | 0.000473 | 0.072211 | 3.325180 | 1.141399 |
| Rasgef1b | -1.447751 | 0.001202 | 0.101706 | 2.920017 | 0.992653 |
| Ddx3y | -1.426943 | 0.020633 | 0.231349 | 1.685443 | 0.635733 |
| Olfr895 | -1.413839 | 0.018902 | 0.224790 | 1.723487 | 0.648223 |
| Unc5d | -1.404553 | 0.000378 | 0.072211 | 3.422341 | 1.141399 |
| Acbd4 | -1.388387 | 0.026857 | 0.257241 | 1.570939 | 0.589659 |
| Abcg1 | -1.370667 | 0.042330 | 0.300976 | 1.373352 | 0.521469 |
| Pcp4 | -1.350829 | 0.000810 | 0.083704 | 3.091466 | 1.077252 |
| Mt3 | -1.341685 | 0.013091 | 0.206018 | 1.883042 | 0.686094 |
| Nr4a2 | -1.329676 | 0.007410 | 0.172342 | 2.130169 | 0.763610 |
| Gm21828 | -1.319707 | 0.007613 | 0.173360 | 2.118447 | 0.761051 |
| H1f5 | -1.318333 | 0.013470 | 0.207605 | 1.870644 | 0.682762 |
| Sly | -1.311699 | 0.002106 | 0.116540 | 2.676496 | 0.933524 |
| Vcan | -1.310170 | 0.000337 | 0.072211 | 3.471828 | 1.141399 |
| 4930555G( | -1.301951 | 0.009634 | 0.187913 | 2.016203 | 0.726043 |
| Tmem198 | -1.300760 | 0.027569 | 0.257555 | 1.559579 | 0.589130 |
| S100a1 | -1.298358 | 0.013964 | 0.209134 | 1.854977 | 0.679575 |
| H3f4 | -1.293664 | 0.007790 | 0.174684 | 2.108487 | 0.757747 |
| Epha6 | -1.291498 | 0.027977 | 0.259189 | 1.553194 | 0.586383 |
| Twist1 | -1.289964 | 0.040911 | 0.297907 | 1.388164 | 0.525919 |
| Sema3c | -1.288244 | 0.002342 | 0.119065 | 2.630502 | 0.924217 |
| Pdzrn3 | -1.286169 | 0.010773 | 0.192565 | 1.967679 | 0.715424 |
| Susd3 | -1.280591 | 0.032600 | 0.272531 | 1.486778 | 0.564584 |
| Fn1 | -1.271077 | 0.010006 | 0.190219 | 1.999752 | 0.720746 |
| Agbl4 | -1.254826 | 0.017168 | 0.219296 | 1.765271 | 0.658969 |
| Adgra2 | -1.221903 | 0.019747 | 0.227182 | 1.704509 | 0.643626 |
| Nr2f1 | -1.218642 | 0.000104 | 0.060933 | 3.985020 | 1.215151 |

| | | | | | |
|---|---|---|---|---|---|
| Pkd1l3 | 0.848910 | 0.012918 | 0.205756 | 1.888805 | 0.686648 |
| 5430427O1 | 0.837080 | 0.025863 | 0.251093 | 1.587317 | 0.600165 |
| Defb28 | 0.833271 | 0.009372 | 0.186226 | 2.028170 | 0.729959 |
| Cep126 | 0.829438 | 0.016858 | 0.219296 | 1.773200 | 0.658969 |
| Gucy1a1 | 0.829224 | 0.000054 | 0.044707 | 4.269257 | 1.349625 |
| Sec14l1 | 0.826388 | 0.000207 | 0.068135 | 3.684263 | 1.166630 |
| Dcc | 0.821847 | 0.006341 | 0.167451 | 2.197840 | 0.776113 |
| Frg2f4 | 0.818650 | 0.007813 | 0.174840 | 2.107176 | 0.757358 |
| Zfp423 | 0.810660 | 0.003999 | 0.140345 | 2.398073 | 0.852804 |
| Slco3a1 | 0.801821 | 0.006309 | 0.167451 | 2.200012 | 0.776113 |
| Nlgn1 | 0.794622 | 0.000032 | 0.032340 | 4.501067 | 1.490255 |
| Speer2 | 0.793731 | 0.001056 | 0.093200 | 2.976321 | 1.030586 |
| Pth1r | 0.773116 | 0.042695 | 0.302115 | 1.369621 | 0.519827 |
| Satb2 | 0.768709 | 0.000003 | 0.019808 | 5.552221 | 1.703170 |
| Manba | 0.762758 | 0.002992 | 0.127513 | 2.524094 | 0.894446 |
| Cdkl1 | 0.750004 | 0.016943 | 0.219296 | 1.771012 | 0.658969 |
| Taar2 | 0.745021 | 0.023435 | 0.241644 | 1.630131 | 0.616823 |
| Tyro3 | 0.736646 | 0.048914 | 0.319909 | 1.310567 | 0.494973 |
| Lrrc4c | 0.736470 | 0.003128 | 0.127788 | 2.504718 | 0.893510 |
| Mical2 | 0.734280 | 0.028994 | 0.262944 | 1.537696 | 0.580137 |
| Mycn | 0.732485 | 0.010580 | 0.191142 | 1.975518 | 0.718644 |
| Mdga1 | 0.732038 | 0.028326 | 0.260695 | 1.547810 | 0.583868 |
| Gm52799 | 0.731877 | 0.002549 | 0.120865 | 2.593564 | 0.917700 |
| Prkch | 0.730945 | 0.012886 | 0.205688 | 1.889871 | 0.686791 |
| Ptn | 0.729589 | 0.044225 | 0.306545 | 1.354332 | 0.513506 |
| Dtx4 | 0.728965 | 0.003048 | 0.127513 | 2.515936 | 0.894446 |
| Bdkrb1 | 0.728191 | 0.001422 | 0.105319 | 2.847249 | 0.977493 |
| Taar6 | 0.728145 | 0.004774 | 0.149991 | 2.321137 | 0.823934 |
| Vgf | 0.723199 | 0.026317 | 0.253968 | 1.579761 | 0.595221 |
| Neurod6 | 0.719964 | 0.005940 | 0.163948 | 2.226189 | 0.785294 |
| Cplx1 | 0.719884 | 0.003080 | 0.127513 | 2.511462 | 0.894446 |
| Ube2d2b | 0.719028 | 0.005132 | 0.151647 | 2.289690 | 0.819166 |
| Ube2ql1 | 0.717398 | 0.000125 | 0.061208 | 3.904480 | 1.213194 |
| Srm | 0.714696 | 0.000388 | 0.072211 | 3.410957 | 1.141399 |
| Ddx39 | 0.712381 | 0.000592 | 0.077963 | 3.227914 | 1.108113 |
| Zeb2 | 0.710734 | 0.023144 | 0.240948 | 1.635559 | 0.618077 |
| Sema7a | 0.703262 | 0.008131 | 0.177271 | 2.089868 | 0.751362 |
| Clstn2 | 0.699162 | 0.000207 | 0.068135 | 3.683243 | 1.166630 |
| Arid5b | 0.695386 | 0.025489 | 0.249150 | 1.593651 | 0.603540 |
| Mfsd4b5 | 0.694673 | 0.043194 | 0.303453 | 1.364578 | 0.517909 |
| Polr2l | 0.693759 | 0.000376 | 0.072211 | 3.424653 | 1.141399 |

| | | | | | |
|---|---|---|---|---|---|
| Pclaf | -1.214701 | 0.040605 | 0.297207 | 1.391424 | 0.526941 |
| Foxo1 | -1.197393 | 0.007955 | 0.175399 | 2.099358 | 0.755974 |
| Ovol2 | -1.183275 | 0.012008 | 0.199121 | 1.920524 | 0.700883 |
| B3galt5 | -1.182190 | 0.006956 | 0.171400 | 2.157662 | 0.765990 |
| Prl7c1 | -1.171017 | 0.011362 | 0.196420 | 1.944532 | 0.706814 |
| Slc15a2 | -1.170878 | 0.014234 | 0.210194 | 1.846660 | 0.677379 |
| Ehd4 | -1.169813 | 0.009998 | 0.190219 | 2.000091 | 0.720746 |
| Meis2 | -1.165126 | 0.032730 | 0.272973 | 1.485048 | 0.563881 |
| Cntn3 | -1.157220 | 0.010855 | 0.193151 | 1.964362 | 0.714102 |
| Ccnd2 | -1.154602 | 0.013916 | 0.209073 | 1.856496 | 0.679702 |
| Usp35 | -1.152443 | 0.010374 | 0.191032 | 1.984070 | 0.718893 |
| Prex1 | -1.151793 | 0.000552 | 0.074779 | 3.258354 | 1.126219 |
| Gm5798 | -1.151269 | 0.002230 | 0.119016 | 2.651775 | 0.924396 |
| Evi2a | -1.150627 | 0.020046 | 0.227717 | 1.697979 | 0.642605 |
| Fam205a1 | -1.147423 | 0.017920 | 0.220548 | 1.746668 | 0.656497 |
| Cd209d | -1.145323 | 0.020924 | 0.232953 | 1.679351 | 0.632732 |
| 1700049E1 | -1.134037 | 0.006898 | 0.171274 | 2.161277 | 0.766308 |
| Negr1 | -1.131623 | 0.008356 | 0.178880 | 2.077976 | 0.747439 |
| Foxp2 | -1.126649 | 0.005995 | 0.164635 | 2.222220 | 0.783478 |
| Gm52229 | -1.124223 | 0.040828 | 0.297844 | 1.389038 | 0.526011 |
| Msi2 | -1.124201 | 0.000380 | 0.072211 | 3.420091 | 1.141399 |
| Vmn1r125 | -1.124011 | 0.015548 | 0.214097 | 1.808322 | 0.669389 |
| Gda | -1.123547 | 0.001420 | 0.105319 | 2.847829 | 0.977493 |
| H1f1 | -1.123521 | 0.006116 | 0.165529 | 2.213536 | 0.781126 |
| Gm31035 | -1.117062 | 0.001427 | 0.105319 | 2.845681 | 0.977493 |
| C3ar1 | -1.110278 | 0.000712 | 0.081349 | 3.147611 | 1.089648 |
| Hmgb2 | -1.106181 | 0.002339 | 0.119065 | 2.630959 | 0.924217 |
| Cav1 | -1.102003 | 0.001580 | 0.105319 | 2.801238 | 0.977493 |
| Sstr2 | -1.090698 | 0.011385 | 0.196420 | 1.943680 | 0.706814 |
| Acr | -1.088804 | 0.014023 | 0.209206 | 1.853171 | 0.679425 |
| Olfr1426 | -1.059929 | 0.000032 | 0.032340 | 4.494208 | 1.490255 |
| Dscc1 | -1.059864 | 0.004958 | 0.150630 | 2.304718 | 0.822090 |
| Tekt2 | -1.053479 | 0.016850 | 0.219296 | 1.773403 | 0.658969 |
| A830018L1 | -1.049705 | 0.021784 | 0.237026 | 1.661867 | 0.625204 |
| Rpl32l | -1.030498 | 0.039803 | 0.294835 | 1.400089 | 0.530420 |
| Zbtb45 | -1.027396 | 0.040385 | 0.296642 | 1.393780 | 0.527768 |
| Mfge8 | -1.025456 | 0.028414 | 0.260823 | 1.546468 | 0.583653 |
| Neurod1 | -1.009110 | 0.010025 | 0.190219 | 1.998905 | 0.720746 |
| Rrm2 | -1.004783 | 0.009643 | 0.187913 | 2.015786 | 0.726043 |
| Mid1 | -1.003450 | 0.000399 | 0.072211 | 3.399492 | 1.141399 |
| Tle4 | -1.001003 | 0.000489 | 0.072211 | 3.310532 | 1.141399 |

| | | | | | |
|---|---|---|---|---|---|
| Ly75 | 0.693200 | 0.024013 | 0.244245 | 1.619553 | 0.612175 |
| Wdr60 | 0.690588 | 0.038239 | 0.290930 | 1.417497 | 0.536211 |
| Gnal | 0.690297 | 0.034199 | 0.278963 | 1.465986 | 0.554454 |
| Mvk | 0.688700 | 0.018599 | 0.222795 | 1.730521 | 0.652095 |
| Eps8 | 0.687173 | 0.023007 | 0.240762 | 1.638136 | 0.618413 |
| Rtn4rl2 | 0.684842 | 0.000442 | 0.072211 | 3.354136 | 1.141399 |
| Hsph1 | 0.684780 | 0.000528 | 0.073869 | 3.277297 | 1.131538 |
| Hps4 | 0.683892 | 0.001852 | 0.113004 | 2.732442 | 0.946906 |
| Kcnmb4 | 0.681743 | 0.001171 | 0.099650 | 2.931495 | 1.001523 |
| Pou3f1 | 0.681684 | 0.012223 | 0.200792 | 1.912836 | 0.697254 |
| Prss27 | 0.678188 | 0.014825 | 0.212593 | 1.829012 | 0.672451 |
| Slc26a5 | 0.677367 | 0.019630 | 0.226239 | 1.707084 | 0.645432 |
| Ybx3 | 0.675257 | 0.004636 | 0.148870 | 2.333860 | 0.827194 |
| Eif2s3x | 0.675018 | 0.001468 | 0.105319 | 2.833182 | 0.977493 |
| Adamts20 | 0.674709 | 0.023713 | 0.242939 | 1.625018 | 0.614502 |
| Olfr394 | 0.663215 | 0.006775 | 0.170567 | 2.169120 | 0.768105 |
| Chst8 | 0.661899 | 0.043678 | 0.305208 | 1.359735 | 0.515404 |
| Psg17 | 0.661350 | 0.024927 | 0.246706 | 1.603324 | 0.607820 |
| Gm1110 | 0.661123 | 0.018256 | 0.221774 | 1.738590 | 0.654089 |
| Lancl3 | 0.656516 | 0.029703 | 0.265780 | 1.527201 | 0.575477 |
| Vcpkmt | 0.655885 | 0.017872 | 0.220523 | 1.747822 | 0.656547 |
| Sprr1a | 0.650411 | 0.034145 | 0.278681 | 1.466676 | 0.554892 |
| Slc45a1 | 0.644170 | 0.011280 | 0.196420 | 1.947707 | 0.706814 |
| Tubb6 | 0.638881 | 0.005656 | 0.159819 | 2.247481 | 0.796371 |
| Fbn2 | 0.633832 | 0.006493 | 0.168913 | 2.187535 | 0.772338 |
| Gcat | 0.631934 | 0.006340 | 0.167451 | 2.197903 | 0.776113 |
| Arrdc2 | 0.628225 | 0.035073 | 0.283146 | 1.455032 | 0.547990 |
| Hmgcs1 | 0.628097 | 0.001564 | 0.105319 | 2.805852 | 0.977493 |
| Mtrr | 0.624663 | 0.007367 | 0.172342 | 2.132702 | 0.763610 |
| Dexi | 0.624644 | 0.039906 | 0.295181 | 1.398959 | 0.529912 |
| Mkrn3 | 0.623097 | 0.021275 | 0.234459 | 1.672126 | 0.629933 |
| Abracl | 0.622406 | 0.007215 | 0.172254 | 2.141753 | 0.763830 |
| Ajap1 | 0.616953 | 0.042134 | 0.300932 | 1.375372 | 0.521532 |
| Rhou | 0.614580 | 0.005198 | 0.152675 | 2.284168 | 0.816232 |
| Camkk2 | 0.614233 | 0.000641 | 0.079395 | 3.193384 | 1.100208 |
| Akr1b3 | 0.611458 | 0.014984 | 0.212593 | 1.824368 | 0.672451 |
| Rbfox3 | 0.604693 | 0.008733 | 0.182254 | 2.058815 | 0.739323 |
| Nuak1 | 0.604592 | 0.004524 | 0.147613 | 2.344467 | 0.830874 |
| Fasn | 0.604221 | 0.000631 | 0.079395 | 3.200304 | 1.100208 |
| Tspan5 | 0.602922 | 0.000697 | 0.081349 | 3.157057 | 1.089648 |
| Nkiras1 | 0.598126 | 0.008964 | 0.183815 | 2.047479 | 0.735618 |

| | | | | | |
|---|---|---|---|---|---|
| Cpne8 | -0.994306 | 0.042779 | 0.302156 | 1.368772 | 0.519769 |
| Trf | -0.991123 | 0.034628 | 0.280840 | 1.460573 | 0.551541 |
| Zfp691 | -0.989549 | 0.034242 | 0.279155 | 1.465436 | 0.554154 |
| Gm52606 | -0.988308 | 0.002826 | 0.126635 | 2.548822 | 0.897446 |
| Gm10256 | -0.985736 | 0.043930 | 0.305662 | 1.357235 | 0.514758 |
| Cdo1 | -0.980763 | 0.016998 | 0.219296 | 1.769596 | 0.658969 |
| Spata25 | -0.971187 | 0.016283 | 0.216823 | 1.788260 | 0.663895 |
| Smad6 | -0.970232 | 0.027415 | 0.257555 | 1.562015 | 0.589130 |
| 4930467E2 | -0.967383 | 0.003070 | 0.127513 | 2.512837 | 0.894446 |
| Gm10354 | -0.967091 | 0.000310 | 0.072211 | 3.508851 | 1.141399 |
| Fst | -0.966816 | 0.003981 | 0.140345 | 2.400023 | 0.852804 |
| H1f3 | -0.966547 | 0.000698 | 0.081349 | 3.155971 | 1.089648 |
| Cox16 | -0.965215 | 0.023889 | 0.243505 | 1.621810 | 0.613493 |
| Wnt7b | -0.962283 | 0.014204 | 0.210194 | 1.847599 | 0.677379 |
| Igf2 | -0.961788 | 0.046286 | 0.312437 | 1.334549 | 0.505237 |
| Dock1 | -0.958919 | 0.038724 | 0.292408 | 1.412018 | 0.534011 |
| Spag1 | -0.957168 | 0.020243 | 0.229164 | 1.693720 | 0.639854 |
| C2cd4d | -0.955616 | 0.022676 | 0.239850 | 1.644441 | 0.620060 |
| Psg16 | -0.953357 | 0.043878 | 0.305662 | 1.357752 | 0.514758 |
| Fgd5 | -0.952124 | 0.033115 | 0.274397 | 1.479978 | 0.561621 |
| Fgf21 | -0.948555 | 0.015895 | 0.216107 | 1.798734 | 0.665330 |
| Zfand4 | -0.947857 | 0.023750 | 0.243142 | 1.624342 | 0.614141 |
| Gm10334 | -0.945493 | 0.023333 | 0.241144 | 1.632035 | 0.617723 |
| Arhgef17 | -0.945477 | 0.030162 | 0.266162 | 1.520544 | 0.574854 |
| Epha5 | -0.943253 | 0.002735 | 0.125045 | 2.563057 | 0.902935 |
| Zfp786 | -0.942049 | 0.003027 | 0.127513 | 2.519044 | 0.894446 |
| H1f4 | -0.939344 | 0.002920 | 0.127176 | 2.534606 | 0.895594 |
| Tifab | -0.937328 | 0.023313 | 0.241119 | 1.632399 | 0.617769 |
| Gm52795 | -0.934244 | 0.000556 | 0.074779 | 3.255110 | 1.126219 |
| Hgf | -0.933369 | 0.013925 | 0.209073 | 1.856193 | 0.679702 |
| Angptl3 | -0.933245 | 0.021113 | 0.233694 | 1.675460 | 0.631352 |
| Metrn | -0.929854 | 0.001506 | 0.105319 | 2.822203 | 0.977493 |
| Sox9 | -0.929545 | 0.013894 | 0.209052 | 1.857161 | 0.679746 |
| Zswim7 | -0.929407 | 0.008862 | 0.182536 | 2.052482 | 0.738651 |
| Amy2a1 | -0.926980 | 0.000541 | 0.074779 | 3.267199 | 1.126219 |
| Igfbp3 | -0.925824 | 0.023223 | 0.241019 | 1.634073 | 0.617949 |
| Endou | -0.918696 | 0.013620 | 0.207695 | 1.865810 | 0.682575 |
| Cldn11 | -0.918457 | 0.030332 | 0.266662 | 1.518103 | 0.574038 |
| Sorl1 | -0.917840 | 0.001905 | 0.113600 | 2.720004 | 0.944621 |
| Adcy8 | -0.917103 | 0.010781 | 0.192565 | 1.967328 | 0.715424 |
| Plpp4 | -0.916445 | 0.027147 | 0.257555 | 1.566273 | 0.589130 |

| | | | | | |
|---|---|---|---|---|---|
| Cbx8 | 0.597489 | 0.000625 | 0.079395 | 3.204205 | 1.100208 |
| Gtf2h3 | 0.596687 | 0.020345 | 0.229578 | 1.691545 | 0.639069 |
| Gatm | 0.596255 | 0.019182 | 0.225156 | 1.717098 | 0.647516 |
| Bach2 | 0.594036 | 0.001256 | 0.103771 | 2.901009 | 0.983924 |
| Fam49b | 0.593613 | 0.020629 | 0.231349 | 1.685522 | 0.635733 |
| Bean1 | 0.593092 | 0.019081 | 0.225156 | 1.719404 | 0.647516 |
| Snx7 | 0.591793 | 0.004819 | 0.150323 | 2.317029 | 0.822975 |
| Tspyl5 | 0.589444 | 0.010318 | 0.191032 | 1.986404 | 0.718893 |
| Lrrtm1 | 0.589055 | 0.005653 | 0.159819 | 2.247720 | 0.796371 |
| Epb41l3 | 0.585944 | 0.003800 | 0.138880 | 2.420220 | 0.857359 |
| Limch1 | 0.580386 | 0.000095 | 0.058239 | 4.023142 | 1.234789 |
| Setd7 | 0.578164 | 0.007044 | 0.171400 | 2.152197 | 0.765990 |
| Foxp1 | 0.577956 | 0.009876 | 0.189569 | 2.005436 | 0.722234 |
| Atp13a3 | 0.576849 | 0.017976 | 0.220548 | 1.745303 | 0.656497 |
| Insig1 | 0.576390 | 0.002434 | 0.119712 | 2.613721 | 0.921864 |
| Wipf3 | 0.574348 | 0.003859 | 0.139603 | 2.413500 | 0.855104 |
| Shisa9 | 0.571508 | 0.007046 | 0.171400 | 2.152039 | 0.765990 |
| Nfix | 0.569977 | 0.021633 | 0.236690 | 1.664880 | 0.625819 |
| Sec22c | 0.566774 | 0.002127 | 0.116540 | 2.672203 | 0.933524 |
| Dync1i1 | 0.566374 | 0.001283 | 0.103883 | 2.891759 | 0.983457 |
| Nsun7 | 0.565128 | 0.020114 | 0.228252 | 1.696497 | 0.641585 |
| Tmem25 | 0.564761 | 0.045045 | 0.308481 | 1.346352 | 0.510771 |
| Slitrk4 | 0.559707 | 0.002960 | 0.127513 | 2.528692 | 0.894446 |
| Gm5820 | 0.559598 | 0.025875 | 0.251093 | 1.587117 | 0.600165 |
| Maff | 0.559003 | 0.043926 | 0.305662 | 1.357281 | 0.514758 |
| St3gal1 | 0.556509 | 0.029811 | 0.265832 | 1.525622 | 0.575392 |
| Mif | 0.555511 | 0.000596 | 0.077963 | 3.224770 | 1.108113 |
| Cenpk | 0.553901 | 0.028001 | 0.259241 | 1.552822 | 0.586296 |
| Asah1 | 0.552137 | 0.021645 | 0.236690 | 1.664637 | 0.625819 |
| Klrg2 | 0.550812 | 0.001899 | 0.113600 | 2.721463 | 0.944621 |
| Kcnq5 | 0.549115 | 0.008274 | 0.178185 | 2.082307 | 0.749130 |
| Gpr12 | 0.548842 | 0.000420 | 0.072211 | 3.377007 | 1.141399 |
| Homer2 | 0.547859 | 0.041932 | 0.300399 | 1.377456 | 0.522301 |
| Aprt | 0.547286 | 0.010551 | 0.191032 | 1.976700 | 0.718893 |
| Acaca | 0.542592 | 0.004770 | 0.149991 | 2.321503 | 0.823934 |
| Ydjc | 0.540506 | 0.004799 | 0.150323 | 2.318849 | 0.822975 |
| Pgls | 0.535934 | 0.022269 | 0.238730 | 1.652300 | 0.622092 |
| Kdm8 | 0.534967 | 0.021081 | 0.233694 | 1.676100 | 0.631352 |
| Prr7 | 0.534790 | 0.007930 | 0.175320 | 2.100747 | 0.756167 |
| Parm1 | 0.533154 | 0.048834 | 0.319852 | 1.311282 | 0.495051 |
| Acat2 | 0.531579 | 0.007094 | 0.171400 | 2.149133 | 0.765990 |

| | | | | | |
|---|---|---|---|---|---|
| Rnd2 | -0.915849 | 0.002935 | 0.127176 | 2.532457 | 0.895594 |
| Xrcc2 | -0.914370 | 0.009240 | 0.185437 | 2.034312 | 0.731804 |
| Washc2 | -0.913287 | 0.027294 | 0.257555 | 1.563926 | 0.589130 |
| Ahnak | -0.912393 | 0.013124 | 0.206018 | 1.881948 | 0.686094 |
| Rprm | -0.908382 | 0.010017 | 0.190219 | 1.999270 | 0.720746 |
| Gm52436 | -0.908320 | 0.000401 | 0.072211 | 3.397201 | 1.141399 |
| Gm10654 | -0.908078 | 0.029358 | 0.264080 | 1.532269 | 0.578264 |
| Uty | -0.907026 | 0.012453 | 0.202927 | 1.904722 | 0.692661 |
| Rassf4 | -0.905562 | 0.040042 | 0.295412 | 1.397484 | 0.529572 |
| Chd1l | -0.905485 | 0.000501 | 0.072211 | 3.300254 | 1.141399 |
| Pla2g2f | -0.904744 | 0.018952 | 0.225156 | 1.722344 | 0.647516 |
| Synb | -0.901414 | 0.048943 | 0.319909 | 1.310305 | 0.494973 |
| Wdyhv1 | -0.900664 | 0.000483 | 0.072211 | 3.316135 | 1.141399 |
| Rgs4 | -0.898789 | 0.028815 | 0.262944 | 1.540374 | 0.580137 |
| Arsb | -0.898300 | 0.013187 | 0.206640 | 1.879848 | 0.684785 |
| Abca1 | -0.897873 | 0.000152 | 0.062244 | 3.817581 | 1.205900 |
| Gnpda1 | -0.893583 | 0.010699 | 0.192257 | 1.970666 | 0.716119 |
| Cpvl | -0.893537 | 0.008210 | 0.178185 | 2.085670 | 0.749130 |
| Dock2 | -0.893452 | 0.019625 | 0.226239 | 1.707192 | 0.645432 |
| Olfr316 | -0.892254 | 0.038977 | 0.292754 | 1.409189 | 0.533497 |
| Tfpi | -0.891708 | 0.013989 | 0.209134 | 1.854223 | 0.679575 |
| Tcerg1l | -0.885066 | 0.038942 | 0.292754 | 1.409583 | 0.533497 |
| Dtl | -0.882846 | 0.017822 | 0.220365 | 1.749051 | 0.656857 |
| Gm52433 | -0.881961 | 0.000130 | 0.061208 | 3.886154 | 1.213194 |
| Rai14 | -0.880558 | 0.001353 | 0.105319 | 2.868588 | 0.977493 |
| Ppargc1a | -0.880279 | 0.035813 | 0.285697 | 1.445957 | 0.544095 |
| Aldoc | -0.880017 | 0.001049 | 0.093200 | 2.979325 | 1.030586 |
| Prss53 | -0.879219 | 0.041446 | 0.298749 | 1.382517 | 0.524693 |
| Cox4i2 | -0.876575 | 0.000356 | 0.072211 | 3.448926 | 1.141399 |
| Zfp780b | -0.876165 | 0.042425 | 0.301045 | 1.372377 | 0.521368 |
| Syce2 | -0.875916 | 0.012650 | 0.204016 | 1.897913 | 0.690337 |
| 1700020D( | -0.873516 | 0.042133 | 0.300932 | 1.375382 | 0.521532 |
| Sema5a | -0.872453 | 0.015525 | 0.214097 | 1.808962 | 0.669389 |
| Myt1 | -0.871759 | 0.002079 | 0.116540 | 2.682172 | 0.933524 |
| Arsk | -0.871340 | 0.004908 | 0.150369 | 2.309095 | 0.822841 |
| Taar8c | -0.870587 | 0.038308 | 0.290930 | 1.416710 | 0.536211 |
| Scel | -0.869136 | 0.029365 | 0.264080 | 1.532169 | 0.578264 |
| Dclk3 | -0.868548 | 0.042411 | 0.301045 | 1.372522 | 0.521368 |
| Tmem176b | -0.868008 | 0.023406 | 0.241553 | 1.630664 | 0.616987 |
| Mroh2b | -0.867679 | 0.042704 | 0.302115 | 1.369530 | 0.519827 |
| Cdk1 | -0.865212 | 0.018015 | 0.220548 | 1.744372 | 0.656497 |

| | | | | | |
|---|---|---|---|---|---|
| Dbt | 0.530468 | 0.006014 | 0.164635 | 2.220851 | 0.783478 |
| Lrrc75a | 0.529760 | 0.003286 | 0.129601 | 2.483284 | 0.887393 |
| Rps3a1 | 0.529455 | 0.000971 | 0.090863 | 3.012715 | 1.041611 |
| Acsl4 | 0.526540 | 0.002657 | 0.123993 | 2.575675 | 0.906602 |
| Mtss2 | 0.524603 | 0.019253 | 0.225156 | 1.715500 | 0.647516 |
| Mef2c | 0.524405 | 0.000295 | 0.072211 | 3.530197 | 1.141399 |
| Ripor3 | 0.523797 | 0.025166 | 0.247931 | 1.599184 | 0.605669 |
| Rasl10b | 0.521640 | 0.015779 | 0.215179 | 1.801921 | 0.667200 |
| Gng4 | 0.521526 | 0.007584 | 0.173360 | 2.120130 | 0.761051 |
| Msmo1 | 0.521249 | 0.047310 | 0.315038 | 1.325045 | 0.501637 |
| Bex4 | 0.518724 | 0.002376 | 0.119091 | 2.624126 | 0.924121 |
| Emc10 | 0.518470 | 0.000342 | 0.072211 | 3.466585 | 1.141399 |
| Dok5 | 0.518320 | 0.000978 | 0.090892 | 3.009711 | 1.041474 |
| Gskip | 0.517868 | 0.003130 | 0.127788 | 2.504459 | 0.893510 |
| Col9a3 | 0.517380 | 0.009562 | 0.187913 | 2.019433 | 0.726043 |
| Atox1 | 0.517284 | 0.008441 | 0.179457 | 2.073628 | 0.746039 |
| Adora2a | 0.516963 | 0.033162 | 0.274468 | 1.479355 | 0.561508 |
| Tmem185a | 0.515820 | 0.028402 | 0.260823 | 1.546652 | 0.583653 |
| Zfp428 | 0.515465 | 0.000199 | 0.068135 | 3.701587 | 1.166630 |
| Tle3 | 0.515358 | 0.015462 | 0.213716 | 1.810736 | 0.670162 |
| Pno1 | 0.513117 | 0.017828 | 0.220365 | 1.748892 | 0.656857 |
| Usp12 | 0.512372 | 0.016201 | 0.216756 | 1.790445 | 0.664028 |
| Masp2 | 0.511087 | 0.024572 | 0.245689 | 1.609552 | 0.609614 |
| Nap1l5 | 0.511059 | 0.045811 | 0.311164 | 1.339028 | 0.507010 |
| Ppp1r16b | 0.510958 | 0.016345 | 0.216922 | 1.786607 | 0.663696 |
| Polr1d | 0.508579 | 0.004691 | 0.149991 | 2.328756 | 0.823934 |
| Gipr | 0.508275 | 0.024549 | 0.245630 | 1.609964 | 0.609718 |
| Rfxap | 0.507712 | 0.000323 | 0.072211 | 3.491423 | 1.141399 |
| Fhod3 | 0.505172 | 0.012448 | 0.202927 | 1.904902 | 0.692661 |
| St6galnac4 | 0.504691 | 0.018457 | 0.222746 | 1.733842 | 0.652190 |
| Adgrb2 | 0.503460 | 0.007534 | 0.173075 | 2.122972 | 0.761766 |
| Slc37a4 | 0.502254 | 0.039210 | 0.293481 | 1.406606 | 0.532421 |
| Stk17b | 0.501606 | 0.039814 | 0.294835 | 1.399963 | 0.530420 |
| Vamp7 | 0.498106 | 0.040966 | 0.297907 | 1.387581 | 0.525919 |
| Ppp1r14b | 0.497572 | 0.000026 | 0.032340 | 4.580837 | 1.490255 |
| Nsdhl | 0.496920 | 0.002788 | 0.126634 | 2.554771 | 0.897450 |
| Slc39a10 | 0.496129 | 0.000310 | 0.072211 | 3.508306 | 1.141399 |
| Gabrb3 | 0.495841 | 0.000023 | 0.032340 | 4.632840 | 1.490255 |
| Myo1b | 0.494173 | 0.004301 | 0.144008 | 2.366383 | 0.841615 |
| Rars | 0.492187 | 0.003324 | 0.130068 | 2.478404 | 0.885830 |
| Pank4 | 0.491411 | 0.016915 | 0.219296 | 1.771729 | 0.658969 |

| | | | | | |
|---|---|---|---|---|---|
| D630023F1 | -0.861444 | 0.008836 | 0.182536 | 2.053751 | 0.738651 |
| Cacng3 | -0.860431 | 0.006071 | 0.164944 | 2.216740 | 0.782663 |
| Syn2 | -0.858333 | 0.000072 | 0.049254 | 4.142320 | 1.307557 |
| Fam163b | -0.857515 | 0.007136 | 0.171607 | 2.146553 | 0.765465 |
| Bmp8b | -0.857335 | 0.046570 | 0.313170 | 1.331891 | 0.504220 |
| Mcm6 | -0.854916 | 0.016690 | 0.219296 | 1.777551 | 0.658969 |
| Celsr1 | -0.853627 | 0.001553 | 0.105319 | 2.808837 | 0.977493 |
| Ntrk2 | -0.852742 | 0.007632 | 0.173360 | 2.117341 | 0.761051 |
| Pgap1 | -0.852550 | 0.000128 | 0.061208 | 3.891593 | 1.213194 |
| Smtn | -0.852424 | 0.031939 | 0.269885 | 1.495674 | 0.568821 |
| Cgrrf1 | -0.852373 | 0.040479 | 0.296930 | 1.392770 | 0.527346 |
| Akap13 | -0.849479 | 0.007424 | 0.172342 | 2.129341 | 0.763610 |
| Prc1 | -0.848823 | 0.029732 | 0.265780 | 1.526778 | 0.575477 |
| St6galnac5 | -0.848566 | 0.027020 | 0.257278 | 1.568321 | 0.589597 |
| Sfxn5 | -0.848454 | 0.047051 | 0.314447 | 1.327428 | 0.502452 |
| Fxyd2 | -0.848423 | 0.015055 | 0.212593 | 1.822312 | 0.672451 |
| Cntn6 | -0.844274 | 0.027428 | 0.257555 | 1.561800 | 0.589130 |
| Dscaml1 | -0.844162 | 0.004703 | 0.149991 | 2.327633 | 0.823934 |
| Hsf4 | -0.842805 | 0.048459 | 0.319066 | 1.314628 | 0.496119 |
| Ednrb | -0.835392 | 0.001032 | 0.093200 | 2.986197 | 1.030586 |
| Pnp | -0.832821 | 0.011301 | 0.196420 | 1.946889 | 0.706814 |
| Astn2 | -0.829600 | 0.025187 | 0.247931 | 1.598831 | 0.605669 |
| Med9 | -0.828017 | 0.008895 | 0.182648 | 2.050877 | 0.738385 |
| Hap1 | -0.827893 | 0.028530 | 0.261227 | 1.544696 | 0.582981 |
| Fbxw22 | -0.827846 | 0.018480 | 0.222772 | 1.733293 | 0.652140 |
| Zfp503 | -0.826778 | 0.017354 | 0.219296 | 1.760608 | 0.658969 |
| Sema3a | -0.825731 | 0.040823 | 0.297844 | 1.389098 | 0.526011 |
| 4430402I1 | -0.825459 | 0.005358 | 0.156393 | 2.271019 | 0.805784 |
| Plxna2 | -0.822727 | 0.000178 | 0.064517 | 3.749341 | 1.190325 |
| Serpinb6d | -0.822440 | 0.037448 | 0.289265 | 1.426569 | 0.538704 |
| Tmem176a | -0.821950 | 0.015772 | 0.215179 | 1.802107 | 0.667200 |
| Themis2 | -0.821679 | 0.003814 | 0.138894 | 2.418567 | 0.857318 |
| Mcf2 | -0.821238 | 0.006173 | 0.165622 | 2.209489 | 0.780882 |
| Scgb2b10 | -0.817807 | 0.008404 | 0.179457 | 2.075502 | 0.746039 |
| Lfng | -0.816208 | 0.045747 | 0.311164 | 1.339637 | 0.507010 |
| Rbm41 | -0.815781 | 0.003251 | 0.129398 | 2.487925 | 0.888073 |
| Slc26a9 | -0.815466 | 0.017074 | 0.219296 | 1.767657 | 0.658969 |
| Cartpt | -0.813644 | 0.011589 | 0.196420 | 1.935972 | 0.706814 |
| Klhl1 | -0.808739 | 0.036772 | 0.287676 | 1.434487 | 0.541097 |
| H3c1 | -0.808570 | 0.010275 | 0.191032 | 1.988231 | 0.718893 |
| Mcm5 | -0.807873 | 0.015370 | 0.213513 | 1.813334 | 0.670575 |

| | | | | | |
|---|---|---|---|---|---|
| Pitx2 | 0.489292 | 0.002352 | 0.119065 | 2.628586 | 0.924217 |
| Aoah | 0.487874 | 0.026873 | 0.257241 | 1.570687 | 0.589659 |
| Hsp90aa1 | 0.486802 | 0.000495 | 0.072211 | 3.305090 | 1.141399 |
| Sertad4 | 0.486724 | 0.004845 | 0.150369 | 2.314706 | 0.822841 |
| Ola1 | 0.486250 | 0.001117 | 0.096207 | 2.952030 | 1.016794 |
| Rhox1 | 0.486015 | 0.048461 | 0.319066 | 1.314604 | 0.496119 |
| Nrg1 | 0.485024 | 0.015975 | 0.216107 | 1.796569 | 0.665330 |
| Abcb7 | 0.484064 | 0.003569 | 0.136286 | 2.447429 | 0.865550 |
| Add2 | 0.483955 | 0.000009 | 0.022016 | 5.037621 | 1.657271 |
| Zfp672 | 0.483191 | 0.027112 | 0.257419 | 1.566837 | 0.589359 |
| Elovl4 | 0.482082 | 0.001345 | 0.105319 | 2.871408 | 0.977493 |
| Imp3 | 0.481078 | 0.044389 | 0.306701 | 1.352728 | 0.513284 |
| Armcx1 | 0.480350 | 0.004178 | 0.143187 | 2.379040 | 0.844095 |
| Usp43 | 0.478944 | 0.009431 | 0.186376 | 2.025447 | 0.729610 |
| Crip2 | 0.477977 | 0.000038 | 0.035890 | 4.419010 | 1.445021 |
| Cacna1a | 0.477787 | 0.029782 | 0.265832 | 1.526051 | 0.575392 |
| Sh3pxd2b | 0.477408 | 0.049237 | 0.320122 | 1.307706 | 0.494685 |
| Nt5dc3 | 0.476204 | 0.017171 | 0.219296 | 1.765193 | 0.658969 |
| Nop9 | 0.476011 | 0.007102 | 0.171400 | 2.148625 | 0.765990 |
| Coro1a | 0.475457 | 0.024944 | 0.246706 | 1.603029 | 0.607820 |
| Gas7 | 0.471917 | 0.017153 | 0.219296 | 1.765672 | 0.658969 |
| Lgi3 | 0.471536 | 0.048705 | 0.319307 | 1.312426 | 0.495792 |
| Ptges2 | 0.470109 | 0.005833 | 0.162877 | 2.234071 | 0.788141 |
| Clec2l | 0.468460 | 0.018634 | 0.222795 | 1.729688 | 0.652095 |
| Tmem151a | 0.468072 | 0.029833 | 0.265832 | 1.525303 | 0.575392 |
| Atp2b2 | 0.467445 | 0.009335 | 0.186226 | 2.029872 | 0.729959 |
| Cry1 | 0.466744 | 0.010298 | 0.191032 | 1.987265 | 0.718893 |
| Klf13 | 0.466029 | 0.007604 | 0.173360 | 2.118933 | 0.761051 |
| Chchd3 | 0.464843 | 0.005542 | 0.159133 | 2.256356 | 0.798240 |
| Slc35a1 | 0.464787 | 0.008041 | 0.176091 | 2.094670 | 0.754262 |
| Vps35 | 0.464776 | 0.001954 | 0.115118 | 2.709081 | 0.938859 |
| Gjb1 | 0.462711 | 0.045032 | 0.308481 | 1.346483 | 0.510771 |
| Gpr68 | 0.462296 | 0.032643 | 0.272728 | 1.486207 | 0.564270 |
| Psmb5 | 0.461761 | 0.007192 | 0.172254 | 2.143136 | 0.763830 |
| Acot7 | 0.461532 | 0.001656 | 0.106852 | 2.780856 | 0.971219 |
| Tkt | 0.460908 | 0.000013 | 0.023766 | 4.871034 | 1.624043 |
| Hspa12a | 0.460879 | 0.001828 | 0.113004 | 2.738054 | 0.946906 |
| Mapt | 0.460721 | 0.000713 | 0.081349 | 3.146673 | 1.089648 |
| Lamb1 | 0.458364 | 0.003657 | 0.137687 | 2.436847 | 0.861108 |
| Gars | 0.457541 | 0.003087 | 0.127513 | 2.510501 | 0.894446 |
| Gale | 0.457354 | 0.013338 | 0.207521 | 1.874907 | 0.682937 |

| | | | | | |
|---|---|---|---|---|---|
| Rasgrf1 | -0.807649 | 0.038089 | 0.290716 | 1.419206 | 0.536531 |
| Prdm16 | -0.807128 | 0.025233 | 0.248003 | 1.598032 | 0.605544 |
| Chp2 | -0.805787 | 0.013520 | 0.207605 | 1.869009 | 0.682762 |
| C1qtnf1 | -0.805128 | 0.001982 | 0.116056 | 2.702853 | 0.935334 |
| Tenm3 | -0.804760 | 0.025297 | 0.248021 | 1.596929 | 0.605512 |
| Cyth1 | -0.804698 | 0.000005 | 0.022016 | 5.314273 | 1.657271 |
| Fn3k | -0.804164 | 0.006158 | 0.165622 | 2.210561 | 0.780882 |
| Abhd6 | -0.803613 | 0.007481 | 0.172342 | 2.126039 | 0.763610 |
| Spink8 | -0.801143 | 0.039458 | 0.294582 | 1.403868 | 0.530794 |
| Depdc1b | -0.800275 | 0.032811 | 0.273158 | 1.483985 | 0.563586 |
| Zeb1 | -0.799407 | 0.023677 | 0.242939 | 1.625667 | 0.614502 |
| Gm14548 | -0.798949 | 0.020398 | 0.229626 | 1.690415 | 0.638978 |
| Galnt18 | -0.797557 | 0.035238 | 0.283672 | 1.452987 | 0.547183 |
| Ankrd44 | -0.795465 | 0.005000 | 0.150941 | 2.301028 | 0.821193 |
| Olfr1350 | -0.794757 | 0.033130 | 0.274397 | 1.479774 | 0.561621 |
| Ambp | -0.794736 | 0.013208 | 0.206640 | 1.879178 | 0.684785 |
| Kirrel3 | -0.794732 | 0.027573 | 0.257555 | 1.559515 | 0.589130 |
| Vwa3a | -0.792099 | 0.027260 | 0.257555 | 1.564478 | 0.589130 |
| Cdk6 | -0.789500 | 0.000453 | 0.072211 | 3.344207 | 1.141399 |
| Vim | -0.789194 | 0.010479 | 0.191032 | 1.979698 | 0.718893 |
| Acaa1b | -0.788623 | 0.004060 | 0.141411 | 2.391519 | 0.849518 |
| Clybl | -0.788084 | 0.040408 | 0.296642 | 1.393533 | 0.527768 |
| Fcrls | -0.787814 | 0.007871 | 0.174840 | 2.103982 | 0.757358 |
| Gm34826 | -0.787153 | 0.021382 | 0.234825 | 1.669960 | 0.629256 |
| Nudt17 | -0.786861 | 0.005870 | 0.163579 | 2.231345 | 0.786273 |
| Lrrtm4 | -0.786499 | 0.019940 | 0.227717 | 1.700279 | 0.642605 |
| Flp Recomb | -0.786289 | 0.029861 | 0.265832 | 1.524896 | 0.575392 |
| Gsdmc4 | -0.785667 | 0.009311 | 0.186131 | 2.030991 | 0.730181 |
| Cdh24 | -0.783436 | 0.032148 | 0.270512 | 1.492844 | 0.567814 |
| Plekhf2 | -0.783199 | 0.018990 | 0.225156 | 1.721477 | 0.647516 |
| Cadm1 | -0.782507 | 0.046695 | 0.313687 | 1.330726 | 0.503503 |
| Cfap45 | -0.782343 | 0.032542 | 0.272441 | 1.487559 | 0.564727 |
| Tmpo | -0.781613 | 0.010810 | 0.192722 | 1.966185 | 0.715069 |
| Actl10 | -0.781507 | 0.005135 | 0.151647 | 2.289475 | 0.819166 |
| Eml3 | -0.780796 | 0.003005 | 0.127513 | 2.522089 | 0.894446 |
| Cdh4 | -0.779657 | 0.028306 | 0.260695 | 1.548128 | 0.583868 |
| Cdca8 | -0.777731 | 0.036305 | 0.287007 | 1.440039 | 0.542108 |
| Samd5 | -0.777010 | 0.048623 | 0.319066 | 1.313156 | 0.496119 |
| Cabp7 | -0.776998 | 0.000250 | 0.071776 | 3.601634 | 1.144022 |
| Tubgcp2 | -0.774348 | 0.015997 | 0.216107 | 1.795967 | 0.665330 |
| Tead2 | -0.773777 | 0.008271 | 0.178185 | 2.082428 | 0.749130 |

| | | | | | |
|---|---|---|---|---|---|
| Aamp | 0.456706 | 0.002145 | 0.116540 | 2.668632 | 0.933524 |
| Nck2 | 0.456454 | 0.000353 | 0.072211 | 3.451611 | 1.141399 |
| Ddx1 | 0.455914 | 0.002549 | 0.120865 | 2.593623 | 0.917700 |
| Tmcc2 | 0.454953 | 0.015475 | 0.213716 | 1.810367 | 0.670162 |
| Rpl12 | 0.454844 | 0.000161 | 0.062244 | 3.793942 | 1.205900 |
| Mpp6 | 0.454537 | 0.000942 | 0.088766 | 3.025744 | 1.051755 |
| Rpl8 | 0.454361 | 0.006779 | 0.170567 | 2.168856 | 0.768105 |
| Nrbp2 | 0.453804 | 0.021116 | 0.233694 | 1.675382 | 0.631352 |
| Nelfa | 0.453772 | 0.025288 | 0.248021 | 1.597085 | 0.605512 |
| Rab9b | 0.453318 | 0.025824 | 0.251093 | 1.587969 | 0.600165 |
| Snap47 | 0.452925 | 0.005107 | 0.151584 | 2.291820 | 0.819346 |
| Lzic | 0.452387 | 0.042159 | 0.300932 | 1.375112 | 0.521532 |
| Znrf2 | 0.449942 | 0.001054 | 0.093200 | 2.977049 | 1.030586 |
| Bbs7 | 0.449741 | 0.011502 | 0.196420 | 1.939220 | 0.706814 |
| Igsf3 | 0.447616 | 0.021392 | 0.234825 | 1.669758 | 0.629256 |
| Erg28 | 0.446696 | 0.029002 | 0.262944 | 1.537577 | 0.580137 |
| Enc1 | 0.446267 | 0.000808 | 0.083704 | 3.092510 | 1.077252 |
| Lrpprc | 0.444659 | 0.007863 | 0.174840 | 2.104402 | 0.757358 |
| Fam169a | 0.443983 | 0.013290 | 0.207235 | 1.876489 | 0.683536 |
| Rnf41 | 0.443426 | 0.007109 | 0.171400 | 2.148173 | 0.765990 |
| Pycrl | 0.443388 | 0.027194 | 0.257555 | 1.565534 | 0.589130 |
| Sqle | 0.443224 | 0.013745 | 0.207695 | 1.861844 | 0.682575 |
| Canx | 0.442663 | 0.002043 | 0.116473 | 2.689762 | 0.933775 |
| Idh3a | 0.440951 | 0.010695 | 0.192257 | 1.970806 | 0.716119 |
| Pcdhb17 | 0.439977 | 0.030150 | 0.266162 | 1.520711 | 0.574854 |
| Fam20a | 0.439906 | 0.021725 | 0.237026 | 1.663048 | 0.625204 |
| Olfr1272 | 0.439339 | 0.042897 | 0.302421 | 1.367570 | 0.519387 |
| Ran | 0.438925 | 0.000073 | 0.049254 | 4.135418 | 1.307557 |
| Rab39b | 0.438327 | 0.004268 | 0.144008 | 2.369823 | 0.841615 |
| Nop58 | 0.438085 | 0.001324 | 0.105062 | 2.878216 | 0.978555 |
| Ptms | 0.437505 | 0.000051 | 0.044707 | 4.290769 | 1.349625 |
| Ccdc83 | 0.436289 | 0.040643 | 0.297207 | 1.391015 | 0.526941 |
| Gnaq | 0.435943 | 0.000396 | 0.072211 | 3.402377 | 1.141399 |
| Zdhhc3 | 0.435935 | 0.014129 | 0.209793 | 1.849883 | 0.678210 |
| Wdr88 | 0.435257 | 0.032143 | 0.270512 | 1.492912 | 0.567814 |
| Angptl6 | 0.433543 | 0.011525 | 0.196420 | 1.938375 | 0.706814 |
| Atxn7l3b | 0.432811 | 0.000067 | 0.049254 | 4.171724 | 1.307557 |
| Degs2 | 0.432788 | 0.014317 | 0.210194 | 1.844153 | 0.677379 |
| Il6st | 0.432771 | 0.010081 | 0.190660 | 1.996501 | 0.719741 |
| Wdr91 | 0.432018 | 0.017231 | 0.219296 | 1.763701 | 0.658969 |
| Gng7 | 0.429670 | 0.014453 | 0.210511 | 1.840035 | 0.676726 |

| | | | | | |
|---|---|---|---|---|---|
| Insm1 | -0.773119 | 0.044584 | 0.307142 | 1.350820 | 0.512661 |
| Myo6 | -0.773005 | 0.030290 | 0.266662 | 1.518701 | 0.574038 |
| Garem2 | -0.772622 | 0.035858 | 0.285833 | 1.445413 | 0.543888 |
| Rpa2 | -0.771318 | 0.026913 | 0.257262 | 1.570030 | 0.589624 |
| Tmem132b | -0.770843 | 0.006188 | 0.165622 | 2.208479 | 0.780882 |
| Odf3l1 | -0.770182 | 0.048065 | 0.317739 | 1.318168 | 0.497930 |
| Gpr156 | -0.768325 | 0.030460 | 0.267006 | 1.516271 | 0.573479 |
| Syne4 | -0.767952 | 0.031050 | 0.268555 | 1.507938 | 0.570967 |
| Zhx2 | -0.765720 | 0.003705 | 0.138127 | 2.431163 | 0.859722 |
| Nxph3 | -0.765336 | 0.016194 | 0.216756 | 1.790649 | 0.664028 |
| Pramel46 | -0.765286 | 0.011458 | 0.196420 | 1.940874 | 0.706814 |
| Thsd7b | -0.762303 | 0.008098 | 0.176827 | 2.091629 | 0.752452 |
| Sesn1 | -0.761796 | 0.010157 | 0.190810 | 1.993248 | 0.719400 |
| Nrgn | -0.761705 | 0.012899 | 0.205688 | 1.889437 | 0.686791 |
| Mecom | -0.760886 | 0.032960 | 0.273830 | 1.482015 | 0.562519 |
| Sphkap | -0.759113 | 0.032720 | 0.272973 | 1.485192 | 0.563881 |
| Kif11 | -0.758918 | 0.011923 | 0.198399 | 1.923633 | 0.702460 |
| Pskh1 | -0.758485 | 0.040278 | 0.296642 | 1.394933 | 0.527768 |
| Mapkapk2 | -0.757439 | 0.016983 | 0.219296 | 1.769987 | 0.658969 |
| Nlrp4e | -0.755797 | 0.000470 | 0.072211 | 3.328244 | 1.141399 |
| Nmur1 | -0.755408 | 0.012722 | 0.204944 | 1.895445 | 0.688364 |
| Lcor | -0.754733 | 0.000455 | 0.072211 | 3.342365 | 1.141399 |
| Prkcb | -0.754056 | 0.011225 | 0.196420 | 1.949821 | 0.706814 |
| 1700016D0 | -0.754047 | 0.047205 | 0.314730 | 1.326011 | 0.502061 |
| Zfp830 | -0.751890 | 0.000627 | 0.079395 | 3.203005 | 1.100208 |
| Postn | -0.751822 | 0.003745 | 0.138446 | 2.426495 | 0.858721 |
| 1700001C1 | -0.750276 | 0.017543 | 0.219316 | 1.755887 | 0.658929 |
| Mchr1 | -0.749748 | 0.003189 | 0.128007 | 2.496306 | 0.892768 |
| Vsig2 | -0.748815 | 0.014911 | 0.212593 | 1.826492 | 0.672451 |
| Smim15 | -0.746737 | 0.027448 | 0.257555 | 1.561497 | 0.589130 |
| Mfng | -0.746374 | 0.022303 | 0.238730 | 1.651632 | 0.622092 |
| Etv4 | -0.745592 | 0.026821 | 0.257241 | 1.571530 | 0.589659 |
| Ncaph | -0.745416 | 0.017518 | 0.219316 | 1.756515 | 0.658929 |
| Sec24d | -0.745262 | 0.000300 | 0.072211 | 3.523049 | 1.141399 |
| H2bc3 | -0.744813 | 0.004032 | 0.141015 | 2.394434 | 0.850734 |
| Gabrp | -0.742977 | 0.031000 | 0.268555 | 1.508642 | 0.570967 |
| Dpy19l1 | -0.741249 | 0.048608 | 0.319066 | 1.313288 | 0.496119 |
| Cct6b | -0.741201 | 0.010224 | 0.190810 | 1.990384 | 0.719400 |
| Fut2 | -0.740123 | 0.001818 | 0.113004 | 2.740427 | 0.946906 |
| Adgrg1 | -0.734487 | 0.010481 | 0.191032 | 1.979613 | 0.718893 |
| Slc26a8 | -0.733234 | 0.039309 | 0.293769 | 1.405514 | 0.531994 |

| | | | | | |
|---|---|---|---|---|---|
| Prmt3 | 0.427359 | 0.002006 | 0.116056 | 2.697731 | 0.935334 |
| Sub1 | 0.427093 | 0.000445 | 0.072211 | 3.352055 | 1.141399 |
| Ubc | 0.426934 | 0.047516 | 0.315909 | 1.323160 | 0.500439 |
| Sgsm2 | 0.426174 | 0.001565 | 0.105319 | 2.805437 | 0.977493 |
| Ipo4 | 0.425728 | 0.001300 | 0.103883 | 2.885973 | 0.983457 |
| Dctpp1 | 0.424643 | 0.014753 | 0.212593 | 1.831111 | 0.672451 |
| Crmp1 | 0.423978 | 0.000451 | 0.072211 | 3.345742 | 1.141399 |
| Kcns3 | 0.423958 | 0.035751 | 0.285359 | 1.446716 | 0.544609 |
| Armc6 | 0.422685 | 0.011167 | 0.196224 | 1.952072 | 0.707247 |
| Nme2 | 0.422683 | 0.001406 | 0.105319 | 2.852080 | 0.977493 |
| Nefm | 0.422424 | 0.027048 | 0.257278 | 1.567868 | 0.589597 |
| Scd1 | 0.422276 | 0.022455 | 0.238892 | 1.648682 | 0.621799 |
| Ubxn1 | 0.421524 | 0.003553 | 0.136286 | 2.449447 | 0.865550 |
| Iars | 0.421040 | 0.048590 | 0.319066 | 1.313452 | 0.496119 |
| Rpl3 | 0.420795 | 0.000067 | 0.049254 | 4.171040 | 1.307557 |
| Wdr5 | 0.420779 | 0.045679 | 0.311014 | 1.340283 | 0.507220 |
| Polr3g | 0.419867 | 0.046357 | 0.312502 | 1.333883 | 0.505148 |
| Bex3 | 0.419582 | 0.000548 | 0.074779 | 3.260878 | 1.126219 |
| Banf1 | 0.419335 | 0.015475 | 0.213716 | 1.810368 | 0.670162 |
| Josd1 | 0.418730 | 0.016526 | 0.218821 | 1.781828 | 0.659911 |
| Map1b | 0.418700 | 0.000011 | 0.022016 | 4.963293 | 1.657271 |
| Gprin3 | 0.418228 | 0.013726 | 0.207695 | 1.862458 | 0.682575 |
| Mafg | 0.417908 | 0.027018 | 0.257278 | 1.568340 | 0.589597 |
| Emc4 | 0.417620 | 0.033393 | 0.275731 | 1.476339 | 0.559515 |
| Fdps | 0.417211 | 0.015341 | 0.213387 | 1.814144 | 0.670832 |
| Olfr1510 | 0.417025 | 0.020261 | 0.229187 | 1.693329 | 0.639811 |
| Yae1d1 | 0.416884 | 0.002453 | 0.119712 | 2.610271 | 0.921864 |
| Mvd | 0.416816 | 0.042326 | 0.300976 | 1.373395 | 0.521469 |
| Ndufb2 | 0.416153 | 0.002443 | 0.119712 | 2.611999 | 0.921864 |
| Impdh1 | 0.415684 | 0.005536 | 0.159133 | 2.256778 | 0.798240 |
| Rnf183 | 0.415288 | 0.022616 | 0.239679 | 1.645589 | 0.620370 |
| Pa2g4 | 0.415127 | 0.049084 | 0.319909 | 1.309060 | 0.494973 |
| Pabpc1 | 0.414965 | 0.004499 | 0.147489 | 2.346842 | 0.831239 |
| Macrod2 | 0.414222 | 0.039513 | 0.294582 | 1.403265 | 0.530794 |
| Taf7 | 0.414172 | 0.037340 | 0.289062 | 1.427826 | 0.539008 |
| Mien1 | 0.413996 | 0.000835 | 0.083704 | 3.078114 | 1.077252 |
| Syt16 | 0.412496 | 0.003852 | 0.139603 | 2.414358 | 0.855104 |
| Mpped1 | 0.412138 | 0.000120 | 0.061208 | 3.921139 | 1.213194 |
| Cyp20a1 | 0.411620 | 0.041132 | 0.298312 | 1.385820 | 0.525329 |
| Mast3 | 0.410538 | 0.024169 | 0.244729 | 1.616741 | 0.611314 |
| Cnih2 | 0.410106 | 0.000938 | 0.088766 | 3.027840 | 1.051755 |

| | | | | | |
|---|---|---|---|---|---|
| Mif4gd | -0.732736 | 0.011138 | 0.196183 | 1.953186 | 0.707340 |
| Dppa5a | -0.732582 | 0.006785 | 0.170567 | 2.168449 | 0.768105 |
| Vmn2r114 | -0.730983 | 0.009417 | 0.186376 | 2.026104 | 0.729610 |
| Gm51592 | -0.730774 | 0.002476 | 0.120197 | 2.606294 | 0.920106 |
| 4931406B1 | -0.730553 | 0.012944 | 0.205815 | 1.887922 | 0.686522 |
| Wwox | -0.729840 | 0.037905 | 0.290550 | 1.421300 | 0.536779 |
| Prrt4 | -0.729810 | 0.012745 | 0.204978 | 1.894660 | 0.688293 |
| Coro6 | -0.729620 | 0.009085 | 0.184416 | 2.041672 | 0.734200 |
| Phyhip | -0.729265 | 0.028053 | 0.259399 | 1.552028 | 0.586032 |
| Timd2 | -0.728554 | 0.019184 | 0.225156 | 1.717053 | 0.647516 |
| Gng3 | -0.727251 | 0.025891 | 0.251093 | 1.586857 | 0.600165 |
| Ccdc137 | -0.726866 | 0.031482 | 0.269421 | 1.501937 | 0.569568 |
| Etv1 | -0.726838 | 0.024999 | 0.246854 | 1.602082 | 0.607561 |
| Grrp1 | -0.726169 | 0.035278 | 0.283694 | 1.452492 | 0.547150 |
| Ikzf2 | -0.725794 | 0.020761 | 0.232418 | 1.682752 | 0.633731 |
| D430042O( | -0.723850 | 0.019246 | 0.225156 | 1.715668 | 0.647516 |
| Raet1e | -0.721477 | 0.023025 | 0.240762 | 1.637803 | 0.618413 |
| Eef1akmt4 | -0.721352 | 0.045497 | 0.310440 | 1.342017 | 0.508022 |
| Adgrb1 | -0.721271 | 0.039507 | 0.294582 | 1.403321 | 0.530794 |
| Tas2r125 | -0.720912 | 0.013668 | 0.207695 | 1.864304 | 0.682575 |
| Ina | -0.717487 | 0.028616 | 0.261843 | 1.543385 | 0.581959 |
| Sox1 | -0.717046 | 0.001856 | 0.113004 | 2.731499 | 0.946906 |
| Rex2 | -0.712561 | 0.024302 | 0.244729 | 1.614355 | 0.611314 |
| Zfp978 | -0.712071 | 0.000513 | 0.073220 | 3.289815 | 1.135370 |
| Oxr1 | -0.712009 | 0.019482 | 0.226162 | 1.710370 | 0.645580 |
| Nab2 | -0.711654 | 0.006865 | 0.171274 | 2.163378 | 0.766308 |
| Kcnk12 | -0.711582 | 0.011491 | 0.196420 | 1.939637 | 0.706814 |
| Fgfbp3 | -0.710518 | 0.002297 | 0.119065 | 2.638750 | 0.924217 |
| Dner | -0.710413 | 0.011551 | 0.196420 | 1.937389 | 0.706814 |
| Chd7 | -0.710372 | 0.004003 | 0.140345 | 2.397580 | 0.852804 |
| Rps6ka2 | -0.710325 | 0.006247 | 0.166531 | 2.204309 | 0.778504 |
| Gm10662 | -0.709089 | 0.032496 | 0.272419 | 1.488166 | 0.564763 |
| Pdk2 | -0.708721 | 0.024355 | 0.244729 | 1.613410 | 0.611314 |
| Gm3095 | -0.708699 | 0.007142 | 0.171607 | 2.146169 | 0.765465 |
| Cyp2d12 | -0.707967 | 0.000011 | 0.022016 | 4.962254 | 1.657271 |
| Glcci1 | -0.707403 | 0.005414 | 0.157361 | 2.266507 | 0.803104 |
| Grik3 | -0.706648 | 0.048977 | 0.319909 | 1.310009 | 0.494973 |
| Pde1a | -0.703902 | 0.000500 | 0.072211 | 3.301387 | 1.141399 |
| Shd | -0.703779 | 0.001021 | 0.093200 | 2.990773 | 1.030586 |
| Stk38l | -0.703425 | 0.013458 | 0.207605 | 1.871019 | 0.682762 |
| Rbmx2 | -0.702516 | 0.012565 | 0.203366 | 1.900846 | 0.691721 |

| | | | | | |
|---|---|---|---|---|---|
| Mia3 | 0.409522 | 0.015334 | 0.213387 | 1.814346 | 0.670832 |
| Dpysl2 | 0.409432 | 0.003076 | 0.127513 | 2.512030 | 0.894446 |
| Lrfn5 | 0.407835 | 0.012063 | 0.199793 | 1.918551 | 0.699419 |
| Abcb10 | 0.407821 | 0.013115 | 0.206018 | 1.882230 | 0.686094 |
| Taf1d | 0.407223 | 0.001648 | 0.106793 | 2.783080 | 0.971455 |
| Mrps21 | 0.407193 | 0.009070 | 0.184382 | 2.042378 | 0.734282 |
| Mrps36 | 0.406462 | 0.009978 | 0.190219 | 2.000938 | 0.720746 |
| Dhcr24 | 0.406401 | 0.017565 | 0.219316 | 1.755343 | 0.658929 |
| Rnf182 | 0.406309 | 0.011797 | 0.198182 | 1.928220 | 0.702935 |
| Smc3 | 0.406298 | 0.008301 | 0.178232 | 2.080868 | 0.749013 |
| Hibadh | 0.404888 | 0.045192 | 0.309040 | 1.344935 | 0.509985 |
| Fabp5 | 0.404886 | 0.007838 | 0.174840 | 2.105815 | 0.757358 |
| Peli1 | 0.404834 | 0.001004 | 0.092748 | 2.998086 | 1.032696 |
| Vps28 | 0.403027 | 0.003173 | 0.127926 | 2.498535 | 0.893039 |
| Spred2 | 0.402593 | 0.015734 | 0.215179 | 1.803156 | 0.667200 |
| Prmt1 | 0.400269 | 0.001829 | 0.113004 | 2.737830 | 0.946906 |
| Rtl8a | 0.400184 | 0.018085 | 0.221033 | 1.742692 | 0.655543 |
| Kmt5a | 0.400118 | 0.013563 | 0.207605 | 1.867641 | 0.682762 |
| Pik3r2 | 0.397984 | 0.003998 | 0.140345 | 2.398179 | 0.852804 |
| Arl2bp | 0.396851 | 0.004298 | 0.144008 | 2.366760 | 0.841615 |
| Amelx | 0.395609 | 0.049924 | 0.322361 | 1.301691 | 0.491658 |
| Trmt11 | 0.395350 | 0.031132 | 0.268555 | 1.506797 | 0.570967 |
| Cisd3 | 0.393890 | 0.027033 | 0.257278 | 1.568098 | 0.589597 |
| Nup210 | 0.393087 | 0.044632 | 0.307142 | 1.350352 | 0.512661 |
| Rangap1 | 0.392914 | 0.002932 | 0.127176 | 2.532901 | 0.895594 |
| Fam135b | 0.392790 | 0.048109 | 0.317739 | 1.317774 | 0.497930 |
| B4galt5 | 0.392416 | 0.023561 | 0.242612 | 1.627815 | 0.615088 |
| Churc1 | 0.391638 | 0.024051 | 0.244275 | 1.618875 | 0.612121 |
| Smdt1 | 0.391426 | 0.016031 | 0.216107 | 1.795050 | 0.665330 |
| Prkaa2 | 0.391067 | 0.048570 | 0.319066 | 1.313631 | 0.496119 |
| Fam131b | 0.390669 | 0.034360 | 0.279468 | 1.463949 | 0.553668 |
| Cluh | 0.390592 | 0.001553 | 0.105319 | 2.808871 | 0.977493 |
| Rac3 | 0.390046 | 0.006646 | 0.169635 | 2.177455 | 0.770483 |
| Jph3 | 0.389831 | 0.003535 | 0.136074 | 2.451640 | 0.866226 |
| Rbsn | 0.389733 | 0.002090 | 0.116540 | 2.679779 | 0.933524 |
| Eif4a1 | 0.389261 | 0.000288 | 0.072211 | 3.540230 | 1.141399 |
| Thoc7 | 0.389170 | 0.018244 | 0.221774 | 1.738879 | 0.654089 |
| Dnajc19 | 0.388812 | 0.012793 | 0.204978 | 1.893040 | 0.688293 |
| Eif5a | 0.386477 | 0.003973 | 0.140345 | 2.400907 | 0.852804 |
| Yipf3 | 0.385318 | 0.037071 | 0.288088 | 1.430962 | 0.540475 |
| Rnpep | 0.384805 | 0.010425 | 0.191032 | 1.981911 | 0.718893 |

| | | | | | |
|---|---|---|---|---|---|
| Kirrel | -0.701795 | 0.048533 | 0.319066 | 1.313960 | 0.496119 |
| Pcp4l1 | -0.701516 | 0.003724 | 0.138446 | 2.428938 | 0.858721 |
| Lipo2 | -0.701509 | 0.033594 | 0.276331 | 1.473737 | 0.558570 |
| Cdca3 | -0.699854 | 0.008666 | 0.181647 | 2.062193 | 0.740772 |
| Elmo1 | -0.699607 | 0.001357 | 0.105319 | 2.867304 | 0.977493 |
| Top2a | -0.697461 | 0.002545 | 0.120865 | 2.594298 | 0.917700 |
| Opn4 | -0.697454 | 0.024046 | 0.244275 | 1.618958 | 0.612121 |
| Epha7 | -0.697415 | 0.020037 | 0.227717 | 1.698157 | 0.642605 |
| Phox2b | -0.696764 | 0.014884 | 0.212593 | 1.827274 | 0.672451 |
| Nfkbid | -0.696461 | 0.010914 | 0.193278 | 1.962032 | 0.713818 |
| 4833420G1 | -0.695818 | 0.000236 | 0.071776 | 3.627847 | 1.144022 |
| Pim1 | -0.695176 | 0.021242 | 0.234277 | 1.672802 | 0.630271 |
| Prickle2 | -0.693744 | 0.013986 | 0.209134 | 1.854310 | 0.679575 |
| Gria3 | -0.693702 | 0.010917 | 0.193278 | 1.961895 | 0.713818 |
| Nab1 | -0.693481 | 0.011518 | 0.196420 | 1.938641 | 0.706814 |
| Lrrc7 | -0.692616 | 0.004873 | 0.150369 | 2.312202 | 0.822841 |
| Nubpl | -0.692061 | 0.041995 | 0.300557 | 1.376807 | 0.522073 |
| Borcs6 | -0.692054 | 0.031821 | 0.269885 | 1.497280 | 0.568821 |
| Zfp784 | -0.691948 | 0.023644 | 0.242939 | 1.626274 | 0.614502 |
| Hdac9 | -0.690868 | 0.011303 | 0.196420 | 1.946797 | 0.706814 |
| Erbb4 | -0.690127 | 0.003971 | 0.140345 | 2.401091 | 0.852804 |
| Olfr136 | -0.689658 | 0.040925 | 0.297907 | 1.388008 | 0.525919 |
| Tex30 | -0.689496 | 0.005884 | 0.163629 | 2.230357 | 0.786140 |
| 3110040M | -0.688740 | 0.002112 | 0.116540 | 2.675309 | 0.933524 |
| Gm3893 | -0.686614 | 0.019279 | 0.225156 | 1.714908 | 0.647516 |
| Gm52432 | -0.686208 | 0.000026 | 0.032340 | 4.579258 | 1.490255 |
| Bfar | -0.685746 | 0.006034 | 0.164635 | 2.219359 | 0.783478 |
| Tbc1d23 | -0.685434 | 0.004518 | 0.147613 | 2.345075 | 0.830874 |
| Lsamp | -0.684686 | 0.031971 | 0.269982 | 1.495250 | 0.568666 |
| Cyp2j6 | -0.684496 | 0.014121 | 0.209793 | 1.850121 | 0.678210 |
| Sema6d | -0.684337 | 0.044394 | 0.306701 | 1.352679 | 0.513284 |
| Mad2l1 | -0.683915 | 0.029034 | 0.262944 | 1.537093 | 0.580137 |
| Tfdp2 | -0.682059 | 0.007664 | 0.173761 | 2.115517 | 0.760048 |
| Mdk | -0.677812 | 0.008978 | 0.183835 | 2.046803 | 0.735572 |
| Echdc3 | -0.676695 | 0.030090 | 0.266125 | 1.521572 | 0.574915 |
| Acnat1 | -0.674350 | 0.016352 | 0.216922 | 1.786427 | 0.663696 |
| H1f0 | -0.674073 | 0.004621 | 0.148870 | 2.335241 | 0.827194 |
| Pde6d | -0.673584 | 0.003872 | 0.139603 | 2.412116 | 0.855104 |
| A530032D1 | -0.672721 | 0.007767 | 0.174684 | 2.109754 | 0.757747 |
| C87414 | -0.672151 | 0.013462 | 0.207605 | 1.870877 | 0.682762 |
| Lbh | -0.671911 | 0.008876 | 0.182536 | 2.051775 | 0.738651 |

| | | | | | |
|---|---|---|---|---|---|
| Fbll1 | 0.384407 | 0.020904 | 0.232953 | 1.679773 | 0.632732 |
| Dnajc15 | 0.383841 | 0.005956 | 0.164020 | 2.225066 | 0.785102 |
| Pcbp1 | 0.383800 | 0.001539 | 0.105319 | 2.812816 | 0.977493 |
| Rab33a | 0.383482 | 0.013742 | 0.207695 | 1.861965 | 0.682575 |
| Strap | 0.382646 | 0.000360 | 0.072211 | 3.443794 | 1.141399 |
| Kctd13 | 0.382426 | 0.022563 | 0.239339 | 1.646606 | 0.620986 |
| Cyp51 | 0.379915 | 0.041020 | 0.297963 | 1.386999 | 0.525838 |
| Pomp | 0.379533 | 0.005913 | 0.163948 | 2.228170 | 0.785294 |
| Slc25a1 | 0.378402 | 0.004969 | 0.150636 | 2.303767 | 0.822073 |
| Rnasek | 0.378283 | 0.011282 | 0.196420 | 1.947627 | 0.706814 |
| Ptp4a1 | 0.378062 | 0.011199 | 0.196420 | 1.950832 | 0.706814 |
| Sh3bgrl3 | 0.377327 | 0.003062 | 0.127513 | 2.514045 | 0.894446 |
| Plaa | 0.377308 | 0.004438 | 0.146848 | 2.352785 | 0.833132 |
| Efna3 | 0.377038 | 0.015936 | 0.216107 | 1.797618 | 0.665330 |
| Gpd1l | 0.376944 | 0.001897 | 0.113600 | 2.721832 | 0.944621 |
| Mak16 | 0.376812 | 0.019974 | 0.227717 | 1.699525 | 0.642605 |
| Fyn | 0.376115 | 0.010737 | 0.192257 | 1.969122 | 0.716119 |
| E130308A1 | 0.374940 | 0.048113 | 0.317739 | 1.317737 | 0.497930 |
| Psmc5 | 0.373956 | 0.000342 | 0.072211 | 3.465922 | 1.141399 |
| Mrpl58 | 0.373891 | 0.006016 | 0.164635 | 2.220706 | 0.783478 |
| Mbtps2 | 0.373337 | 0.005933 | 0.163948 | 2.226745 | 0.785294 |
| Kdm4b | 0.373037 | 0.006703 | 0.169993 | 2.173704 | 0.769568 |
| Cox8a | 0.372866 | 0.030720 | 0.267913 | 1.512572 | 0.572006 |
| Dnlz | 0.372848 | 0.029499 | 0.264946 | 1.530194 | 0.576842 |
| Hsp90ab1 | 0.372672 | 0.000032 | 0.032340 | 4.499732 | 1.490255 |
| Morc4 | 0.372175 | 0.036665 | 0.287676 | 1.435744 | 0.541097 |
| Gpm6a | 0.371957 | 0.015992 | 0.216107 | 1.796103 | 0.665330 |
| Amd1 | 0.371824 | 0.012170 | 0.200164 | 1.914701 | 0.698614 |
| Pdia6 | 0.371473 | 0.001553 | 0.105319 | 2.808920 | 0.977493 |
| Dst | 0.371320 | 0.012138 | 0.199874 | 1.915838 | 0.699245 |
| Hcfc1 | 0.370972 | 0.001065 | 0.093200 | 2.972483 | 1.030586 |
| Cycs | 0.370564 | 0.000906 | 0.088302 | 3.042740 | 1.054028 |
| Sdhd | 0.370314 | 0.006895 | 0.171274 | 2.161455 | 0.766308 |
| Hdac2 | 0.369848 | 0.000007 | 0.022016 | 5.172449 | 1.657271 |
| Gm51965 | 0.369482 | 0.038817 | 0.292481 | 1.410982 | 0.533902 |
| Ndufa10 | 0.369246 | 0.018011 | 0.220548 | 1.744471 | 0.656497 |
| Ndufa13 | 0.366556 | 0.004435 | 0.146848 | 2.353105 | 0.833132 |
| Pcbp4 | 0.365208 | 0.015019 | 0.212593 | 1.823350 | 0.672451 |
| Txnrd1 | 0.365022 | 0.013065 | 0.206018 | 1.883893 | 0.686094 |
| Cltb | 0.365008 | 0.009858 | 0.189569 | 2.006198 | 0.722234 |
| Trappc2 | 0.364534 | 0.037301 | 0.288916 | 1.428285 | 0.539229 |

| | | | | | |
|---|---|---|---|---|---|
| Haus4 | -0.671675 | 0.034099 | 0.278466 | 1.467262 | 0.555228 |
| Smim22 | -0.671615 | 0.027247 | 0.257555 | 1.564682 | 0.589130 |
| Gpr137c | -0.671419 | 0.027523 | 0.257555 | 1.560299 | 0.589130 |
| Gm34733 | -0.671181 | 0.002053 | 0.116473 | 2.687656 | 0.933775 |
| Arhgef26 | -0.671111 | 0.002292 | 0.119065 | 2.639812 | 0.924217 |
| B4galnt2 | -0.668072 | 0.002332 | 0.119065 | 2.632345 | 0.924217 |
| Gm34758 | -0.668028 | 0.039219 | 0.293481 | 1.406499 | 0.532421 |
| Nnt | -0.663632 | 0.008524 | 0.179475 | 2.069353 | 0.745995 |
| Gpr146 | -0.663628 | 0.044822 | 0.307700 | 1.348508 | 0.511872 |
| Frmpd4 | -0.662346 | 0.026618 | 0.256150 | 1.574819 | 0.591506 |
| Nudt16 | -0.660217 | 0.037162 | 0.288631 | 1.429905 | 0.539657 |
| Naa16 | -0.659337 | 0.024702 | 0.245956 | 1.607263 | 0.609143 |
| Ntng2 | -0.658736 | 0.036497 | 0.287257 | 1.437745 | 0.541729 |
| Plppr4 | -0.658551 | 0.006712 | 0.169993 | 2.173155 | 0.769568 |
| Mcm3 | -0.658036 | 0.016513 | 0.218821 | 1.782167 | 0.659911 |
| Sorbs2 | -0.657131 | 0.016919 | 0.219296 | 1.771617 | 0.658969 |
| Clpx | -0.655779 | 0.002529 | 0.120865 | 2.597078 | 0.917700 |
| Tcf19 | -0.655599 | 0.026817 | 0.257241 | 1.571589 | 0.589659 |
| Gad2 | -0.653861 | 0.029130 | 0.263311 | 1.535653 | 0.579531 |
| Vmn1r101 | -0.653032 | 0.011920 | 0.198399 | 1.923738 | 0.702460 |
| Rasl11a | -0.651642 | 0.015228 | 0.213005 | 1.817370 | 0.671611 |
| Gad1 | -0.651097 | 0.001508 | 0.105319 | 2.821463 | 0.977493 |
| Vamp3 | -0.650460 | 0.004068 | 0.141411 | 2.390584 | 0.849518 |
| Slc6a1 | -0.650400 | 0.001644 | 0.106793 | 2.784130 | 0.971455 |
| Pigb | -0.650141 | 0.044001 | 0.305817 | 1.356540 | 0.514539 |
| H2ac10 | -0.649352 | 0.002838 | 0.126635 | 2.547004 | 0.897446 |
| Tnrc6b | -0.647144 | 0.009551 | 0.187913 | 2.019962 | 0.726043 |
| Dchs2 | -0.646116 | 0.030152 | 0.266162 | 1.520684 | 0.574854 |
| Tesmin | -0.645307 | 0.022682 | 0.239850 | 1.644317 | 0.620060 |
| Sfi1 | -0.644581 | 0.000367 | 0.072211 | 3.435502 | 1.141399 |
| 5730522E0 | -0.644282 | 0.020195 | 0.228979 | 1.694767 | 0.640204 |
| Cnmd | -0.643252 | 0.014261 | 0.210194 | 1.845840 | 0.677379 |
| Adgrb3 | -0.642495 | 0.011151 | 0.196183 | 1.952705 | 0.707340 |
| Cux2 | -0.642246 | 0.037821 | 0.290144 | 1.422270 | 0.537386 |
| 2010300CC | -0.641493 | 0.035648 | 0.284863 | 1.447962 | 0.545364 |
| Rogdi | -0.640096 | 0.010456 | 0.191032 | 1.980625 | 0.718893 |
| Slco4c1 | -0.639996 | 0.017574 | 0.219316 | 1.755123 | 0.658929 |
| Rbm12b1 | -0.639548 | 0.034610 | 0.280840 | 1.460800 | 0.551541 |
| Prickle1 | -0.639110 | 0.017377 | 0.219316 | 1.760013 | 0.658929 |
| S1pr1 | -0.638722 | 0.017936 | 0.220548 | 1.746266 | 0.656497 |
| Dll3 | -0.637716 | 0.028343 | 0.260695 | 1.547557 | 0.583868 |

| | | | | | |
|---|---|---|---|---|---|
| Akt1 | 0.364322 | 0.004862 | 0.150369 | 2.313162 | 0.822841 |
| Dpysl5 | 0.363638 | 0.001046 | 0.093200 | 2.980536 | 1.030586 |
| Hspe1 | 0.363252 | 0.003302 | 0.129601 | 2.481171 | 0.887393 |
| Kif3b | 0.363045 | 0.047004 | 0.314279 | 1.327865 | 0.502684 |
| Slc30a7 | 0.362861 | 0.024312 | 0.244729 | 1.614173 | 0.611314 |
| Rab21 | 0.361849 | 0.000937 | 0.088766 | 3.028058 | 1.051755 |
| Zfp282 | 0.361414 | 0.011954 | 0.198696 | 1.922473 | 0.701811 |
| Eef1b2 | 0.359757 | 0.009606 | 0.187913 | 2.017436 | 0.726043 |
| Capns1 | 0.359446 | 0.000640 | 0.079395 | 3.193962 | 1.100208 |
| Sfr1 | 0.358897 | 0.015955 | 0.216107 | 1.797101 | 0.665330 |
| Ift57 | 0.358785 | 0.023820 | 0.243163 | 1.623062 | 0.614103 |
| Arpc5l | 0.358482 | 0.011656 | 0.196638 | 1.933447 | 0.706334 |
| Rsl24d1 | 0.358032 | 0.017220 | 0.219296 | 1.763964 | 0.658969 |
| Spry2 | 0.357855 | 0.012795 | 0.204978 | 1.892953 | 0.688293 |
| Praf2 | 0.357812 | 0.004775 | 0.149991 | 2.321050 | 0.823934 |
| Abr | 0.357650 | 0.016686 | 0.219296 | 1.777661 | 0.658969 |
| Trp53i11 | 0.357139 | 0.010200 | 0.190810 | 1.991401 | 0.719400 |
| Cyfip2 | 0.356806 | 0.002284 | 0.119065 | 2.641210 | 0.924217 |
| Ppp1r1a | 0.356030 | 0.010212 | 0.190810 | 1.990908 | 0.719400 |
| Sms | 0.355733 | 0.012865 | 0.205688 | 1.890607 | 0.686791 |
| Mturn | 0.355465 | 0.001530 | 0.105319 | 2.815259 | 0.977493 |
| Snx27 | 0.354575 | 0.002592 | 0.122077 | 2.586326 | 0.913366 |
| Atp6v1e1 | 0.354290 | 0.038815 | 0.292481 | 1.411000 | 0.533902 |
| Fkbp10 | 0.353848 | 0.007684 | 0.173761 | 2.114398 | 0.760048 |
| Myl12b | 0.353621 | 0.002080 | 0.116540 | 2.682033 | 0.933524 |
| Plxna4 | 0.353551 | 0.014387 | 0.210194 | 1.842033 | 0.677379 |
| Eef1g | 0.353457 | 0.000664 | 0.080349 | 3.177868 | 1.095018 |
| Rae1 | 0.352304 | 0.018000 | 0.220548 | 1.744733 | 0.656497 |
| Copb2 | 0.351845 | 0.006989 | 0.171400 | 2.155614 | 0.765990 |
| Park7 | 0.351517 | 0.007412 | 0.172342 | 2.130051 | 0.763610 |
| Etf1 | 0.351127 | 0.013674 | 0.207695 | 1.864095 | 0.682575 |
| 2310061I0 | 0.349489 | 0.001555 | 0.105319 | 2.808332 | 0.977493 |
| Cbll1 | 0.349462 | 0.027274 | 0.257555 | 1.564250 | 0.589130 |
| Rab18 | 0.349125 | 0.000496 | 0.072211 | 3.304573 | 1.141399 |
| Fscn1 | 0.348976 | 0.001578 | 0.105319 | 2.801996 | 0.977493 |
| Emc6 | 0.348709 | 0.016065 | 0.216116 | 1.794119 | 0.665312 |
| Spata7 | 0.347748 | 0.036324 | 0.287007 | 1.439809 | 0.542108 |
| Clptm1l | 0.347450 | 0.016087 | 0.216116 | 1.793538 | 0.665312 |
| Mdh1 | 0.347295 | 0.001835 | 0.113004 | 2.736436 | 0.946906 |
| Zbtb37 | 0.346035 | 0.022110 | 0.238286 | 1.655404 | 0.622901 |
| Scaf1 | 0.345879 | 0.002457 | 0.119712 | 2.609546 | 0.921864 |

| | | | | | |
|---|---|---|---|---|---|
| Txndc16 | -0.636339 | 0.006576 | 0.169325 | 2.182027 | 0.771279 |
| H4c8 | -0.636264 | 0.018328 | 0.222266 | 1.736881 | 0.653126 |
| Ypel4 | -0.636208 | 0.008412 | 0.179457 | 2.075125 | 0.746039 |
| Ifitm2 | -0.635973 | 0.009731 | 0.188600 | 2.011832 | 0.724459 |
| Thy1 | -0.635691 | 0.029931 | 0.265965 | 1.523877 | 0.575176 |
| Cyp11a1 | -0.635652 | 0.036281 | 0.287007 | 1.440317 | 0.542108 |
| Prtn3 | -0.634807 | 0.026223 | 0.253421 | 1.581318 | 0.596158 |
| Aamdc | -0.634239 | 0.036144 | 0.286874 | 1.441969 | 0.542309 |
| Nipbl | -0.633864 | 0.000743 | 0.082167 | 3.129176 | 1.085300 |
| Nanp | -0.633467 | 0.000475 | 0.072211 | 3.322948 | 1.141399 |
| Hs3st4 | -0.632270 | 0.030763 | 0.267955 | 1.511967 | 0.571937 |
| Fastkd1 | -0.630803 | 0.007868 | 0.174840 | 2.104132 | 0.757358 |
| C330018D2 | -0.630446 | 0.047930 | 0.317423 | 1.319390 | 0.498361 |
| Fzd1 | -0.629836 | 0.002776 | 0.126536 | 2.556507 | 0.897788 |
| Mertk | -0.629758 | 0.024774 | 0.246307 | 1.606010 | 0.608524 |
| Ube2e3 | -0.628211 | 0.007083 | 0.171400 | 2.149760 | 0.765990 |
| Clasrp | -0.627888 | 0.004467 | 0.147296 | 2.350019 | 0.831808 |
| Snrnp48 | -0.627876 | 0.007363 | 0.172342 | 2.132922 | 0.763610 |
| Micu1 | -0.626860 | 0.014958 | 0.212593 | 1.825131 | 0.672451 |
| Zfp59 | -0.626749 | 0.027542 | 0.257555 | 1.560004 | 0.589130 |
| Gm31714 | -0.626383 | 0.023787 | 0.243163 | 1.623661 | 0.614103 |
| Fxyd3 | -0.626126 | 0.030054 | 0.266039 | 1.522102 | 0.575054 |
| Thbs3 | -0.625555 | 0.028210 | 0.260151 | 1.549597 | 0.584775 |
| Pir | -0.624919 | 0.009040 | 0.184321 | 2.043853 | 0.734426 |
| Rad54b | -0.624325 | 0.017069 | 0.219296 | 1.767784 | 0.658969 |
| Crygb | -0.624154 | 0.017687 | 0.219906 | 1.752351 | 0.657762 |
| Nr2f2 | -0.623842 | 0.011568 | 0.196420 | 1.936724 | 0.706814 |
| Zp3 | -0.623464 | 0.026894 | 0.257247 | 1.570349 | 0.589649 |
| Cldn20 | -0.622973 | 0.024833 | 0.246706 | 1.604968 | 0.607820 |
| Katnal2 | -0.622838 | 0.024070 | 0.244297 | 1.618522 | 0.612081 |
| Adamts10 | -0.621858 | 0.003622 | 0.136867 | 2.441032 | 0.863701 |
| Kcnj8 | -0.621562 | 0.031940 | 0.269885 | 1.495665 | 0.568821 |
| Dstyk | -0.621290 | 0.014668 | 0.212103 | 1.833640 | 0.673454 |
| Abraxas1 | -0.619619 | 0.035223 | 0.283672 | 1.453171 | 0.547183 |
| Pbx4 | -0.619396 | 0.045109 | 0.308768 | 1.345738 | 0.510367 |
| Zscan25 | -0.618148 | 0.025485 | 0.249150 | 1.593718 | 0.603540 |
| Ano1 | -0.617819 | 0.002278 | 0.119065 | 2.642469 | 0.924217 |
| Sparcl1 | -0.617799 | 0.033433 | 0.275743 | 1.475818 | 0.559495 |
| Ccdc181 | -0.616427 | 0.039074 | 0.293169 | 1.408112 | 0.532882 |
| Dnajc21 | -0.613352 | 0.002323 | 0.119065 | 2.633866 | 0.924217 |
| Amotl2 | -0.612677 | 0.000441 | 0.072211 | 3.355577 | 1.141399 |

| | | | | | |
|---|---|---|---|---|---|
| Rpl41 | 0.345494 | 0.002326 | 0.119065 | 2.633299 | 0.924217 |
| Capzb | 0.345329 | 0.002930 | 0.127176 | 2.533155 | 0.895594 |
| Slc4a7 | 0.343725 | 0.017811 | 0.220365 | 1.749314 | 0.656857 |
| Ankrd46 | 0.342946 | 0.006604 | 0.169325 | 2.180208 | 0.771279 |
| Gnaz | 0.342780 | 0.011828 | 0.198399 | 1.927096 | 0.702460 |
| Zbtb43 | 0.342204 | 0.006510 | 0.168913 | 2.186437 | 0.772338 |
| Cox5a | 0.341749 | 0.001569 | 0.105319 | 2.804468 | 0.977493 |
| Rap2a | 0.341081 | 0.012631 | 0.203941 | 1.898569 | 0.690496 |
| Stmn2 | 0.340638 | 0.004286 | 0.144008 | 2.367947 | 0.841615 |
| 6330403K0 | 0.340087 | 0.030883 | 0.268288 | 1.510277 | 0.571399 |
| Ythdf2 | 0.339942 | 0.001887 | 0.113600 | 2.724261 | 0.944621 |
| Pip4k2b | 0.338895 | 0.003340 | 0.130212 | 2.476277 | 0.885349 |
| Atp5b | 0.338567 | 0.019320 | 0.225209 | 1.713993 | 0.647415 |
| Jkamp | 0.337839 | 0.009043 | 0.184321 | 2.043671 | 0.734426 |
| Hyou1 | 0.337408 | 0.042200 | 0.300932 | 1.374684 | 0.521532 |
| Trp53inp2 | 0.337381 | 0.011997 | 0.199121 | 1.920917 | 0.700883 |
| Smcr8 | 0.336732 | 0.024392 | 0.244752 | 1.612752 | 0.611274 |
| Zfp451 | 0.336004 | 0.000655 | 0.080349 | 3.183677 | 1.095018 |
| Cct3 | 0.335594 | 0.014070 | 0.209678 | 1.851719 | 0.678447 |
| Fbxw4 | 0.335065 | 0.046859 | 0.314198 | 1.329212 | 0.502796 |
| Cct7 | 0.334453 | 0.007016 | 0.171400 | 2.153910 | 0.765990 |
| Wsb2 | 0.333813 | 0.012543 | 0.203366 | 1.901613 | 0.691721 |
| Oxct1 | 0.333677 | 0.009613 | 0.187913 | 2.017151 | 0.726043 |
| Dedd | 0.333643 | 0.031753 | 0.269762 | 1.498211 | 0.569020 |
| Rnf4 | 0.332777 | 0.004896 | 0.150369 | 2.310194 | 0.822841 |
| Snn | 0.331760 | 0.007466 | 0.172342 | 2.126908 | 0.763610 |
| Abcc8 | 0.331541 | 0.010900 | 0.193278 | 1.962558 | 0.713818 |
| Fem1b | 0.330958 | 0.001594 | 0.105718 | 2.797553 | 0.975852 |
| Rps19 | 0.330750 | 0.004196 | 0.143187 | 2.377175 | 0.844095 |
| Klhl2 | 0.329594 | 0.007624 | 0.173360 | 2.117834 | 0.761051 |
| Dis3 | 0.329177 | 0.005283 | 0.154865 | 2.277080 | 0.810046 |
| Clns1a | 0.329151 | 0.003623 | 0.136867 | 2.440910 | 0.863701 |
| Tmem35a | 0.328552 | 0.009865 | 0.189569 | 2.005909 | 0.722234 |
| Ndufa4 | 0.328007 | 0.008573 | 0.180246 | 2.066845 | 0.744134 |
| Cox6a1 | 0.327645 | 0.000712 | 0.081349 | 3.147412 | 1.089648 |
| Nomo1 | 0.327404 | 0.046900 | 0.314198 | 1.328823 | 0.502796 |
| Rps26 | 0.327092 | 0.014661 | 0.212103 | 1.833850 | 0.673454 |
| Mpp3 | 0.326914 | 0.015621 | 0.214689 | 1.806278 | 0.668191 |
| Hmgcr | 0.326371 | 0.009268 | 0.185733 | 2.033003 | 0.731112 |
| Shank2 | 0.325786 | 0.015193 | 0.212934 | 1.818366 | 0.671755 |
| Bola2 | 0.325047 | 0.017509 | 0.219316 | 1.756743 | 0.658929 |

| | | | | | |
|---|---|---|---|---|---|
| Shroom3 | -0.611938 | 0.035354 | 0.283990 | 1.451563 | 0.546697 |
| Phgdh | -0.611759 | 0.001999 | 0.116056 | 2.699096 | 0.935334 |
| Gm52416 | -0.611656 | 0.000336 | 0.072211 | 3.473931 | 1.141399 |
| Ccdc90b | -0.611183 | 0.016596 | 0.219296 | 1.780006 | 0.658969 |
| Gm52520 | -0.610552 | 0.000153 | 0.062244 | 3.814299 | 1.205900 |
| Pcdh8 | -0.609919 | 0.034513 | 0.280226 | 1.462023 | 0.552492 |
| Gcnt2 | -0.609822 | 0.008790 | 0.182527 | 2.056029 | 0.738673 |
| Olfr1415 | -0.609606 | 0.047191 | 0.314730 | 1.326140 | 0.502061 |
| Afap1 | -0.609257 | 0.024935 | 0.246706 | 1.603192 | 0.607820 |
| Tle2 | -0.607371 | 0.021009 | 0.233526 | 1.677599 | 0.631664 |
| Rab13 | -0.607255 | 0.000243 | 0.071776 | 3.614706 | 1.144022 |
| Tmem219 | -0.606950 | 0.027339 | 0.257555 | 1.563224 | 0.589130 |
| Olfr1034 | -0.606933 | 0.043540 | 0.304845 | 1.361108 | 0.515921 |
| Nnat | -0.606638 | 0.002724 | 0.125045 | 2.564827 | 0.902935 |
| Hspb8 | -0.606010 | 0.038884 | 0.292607 | 1.410231 | 0.533716 |
| H2bc11 | -0.605905 | 0.004938 | 0.150369 | 2.306415 | 0.822841 |
| Grik2 | -0.605534 | 0.004918 | 0.150369 | 2.308208 | 0.822841 |
| Rexo5 | -0.605463 | 0.040354 | 0.296642 | 1.394116 | 0.527768 |
| Cep95 | -0.605451 | 0.036751 | 0.287676 | 1.434726 | 0.541097 |
| Edil3 | -0.605307 | 0.031620 | 0.269725 | 1.500035 | 0.569079 |
| Egr3 | -0.604851 | 0.028873 | 0.262944 | 1.539505 | 0.580137 |
| Dennd4c | -0.604796 | 0.018412 | 0.222669 | 1.734893 | 0.652340 |
| Tmem38b | -0.602173 | 0.011863 | 0.198399 | 1.925797 | 0.702460 |
| Slc4a10 | -0.602040 | 0.010560 | 0.191032 | 1.976323 | 0.718893 |
| Zscan20 | -0.602011 | 0.034014 | 0.278160 | 1.468341 | 0.555706 |
| Edn3 | -0.601917 | 0.019287 | 0.225156 | 1.714746 | 0.647516 |
| Gm6958 | -0.601509 | 0.009115 | 0.184767 | 2.040224 | 0.733376 |
| Kcnh8 | -0.600752 | 0.039556 | 0.294597 | 1.402785 | 0.530772 |
| Gm10413 | -0.599050 | 0.000428 | 0.072211 | 3.368394 | 1.141399 |
| Dbf4 | -0.598561 | 0.022322 | 0.238730 | 1.651270 | 0.622092 |
| Il1rapl1 | -0.595514 | 0.030457 | 0.267006 | 1.516316 | 0.573479 |
| Tnks1bp1 | -0.594886 | 0.021697 | 0.237026 | 1.663606 | 0.625204 |
| Fcer1g | -0.594742 | 0.006857 | 0.171274 | 2.163864 | 0.766308 |
| Tshz1 | -0.594623 | 0.003804 | 0.138880 | 2.419729 | 0.857359 |
| Zcchc9 | -0.594567 | 0.024450 | 0.245063 | 1.611723 | 0.610723 |
| Zfp217 | -0.594073 | 0.047831 | 0.317257 | 1.320290 | 0.498589 |
| Cdk5rap2 | -0.593424 | 0.029006 | 0.262944 | 1.537517 | 0.580137 |
| H2-T10 | -0.592653 | 0.021793 | 0.237026 | 1.661675 | 0.625204 |
| Kcnc2 | -0.592639 | 0.015275 | 0.213247 | 1.816018 | 0.671118 |
| Arhgef40 | -0.592574 | 0.014941 | 0.212593 | 1.825620 | 0.672451 |
| Acadl | -0.591897 | 0.004483 | 0.147296 | 2.348421 | 0.831808 |

| | | | | | |
|---|---|---|---|---|---|
| Chchd1 | 0.324890 | 0.013629 | 0.207695 | 1.865535 | 0.682575 |
| Slc1a4 | 0.323365 | 0.022565 | 0.239339 | 1.646562 | 0.620986 |
| Pcca | 0.322883 | 0.028634 | 0.261843 | 1.543111 | 0.581959 |
| Rack1 | 0.322538 | 0.004719 | 0.149991 | 2.326181 | 0.823934 |
| Gm52481 | 0.322489 | 0.004636 | 0.148870 | 2.333822 | 0.827194 |
| Slc38a1 | 0.322334 | 0.003178 | 0.127926 | 2.497813 | 0.893039 |
| Lgals7 | 0.321739 | 0.022528 | 0.239339 | 1.647280 | 0.620986 |
| Gpr27 | 0.321353 | 0.029320 | 0.264012 | 1.532833 | 0.578376 |
| Hk1 | 0.320803 | 0.008752 | 0.182368 | 2.057902 | 0.739051 |
| Tub | 0.320800 | 0.018193 | 0.221383 | 1.740104 | 0.654856 |
| Pik3cd | 0.320695 | 0.016797 | 0.219296 | 1.774773 | 0.658969 |
| Epn1 | 0.320421 | 0.011569 | 0.196420 | 1.936698 | 0.706814 |
| Fdft1 | 0.320177 | 0.014214 | 0.210194 | 1.847275 | 0.677379 |
| C1qbp | 0.319156 | 0.023614 | 0.242939 | 1.626833 | 0.614502 |
| Hypk | 0.319104 | 0.027388 | 0.257555 | 1.562444 | 0.589130 |
| Rpl10a | 0.319060 | 0.004986 | 0.150827 | 2.302285 | 0.821521 |
| Zfp689 | 0.319021 | 0.037035 | 0.287962 | 1.431390 | 0.540664 |
| Ppp1ca | 0.318414 | 0.002047 | 0.116473 | 2.688965 | 0.933775 |
| Ndufb5 | 0.318333 | 0.020051 | 0.227717 | 1.697866 | 0.642605 |
| Hras | 0.318225 | 0.015030 | 0.212593 | 1.823050 | 0.672451 |
| Rpl36a | 0.317179 | 0.002360 | 0.119065 | 2.627140 | 0.924217 |
| Dcun1d5 | 0.316676 | 0.011415 | 0.196420 | 1.942520 | 0.706814 |
| Eid2b | 0.316598 | 0.040071 | 0.295473 | 1.397167 | 0.529482 |
| Tecr | 0.316511 | 0.007009 | 0.171400 | 2.154346 | 0.765990 |
| Kcnk13 | 0.315247 | 0.002426 | 0.119712 | 2.615064 | 0.921864 |
| Dynll1 | 0.314974 | 0.002154 | 0.116597 | 2.666753 | 0.933312 |
| Bex2 | 0.314569 | 0.008005 | 0.175894 | 2.096619 | 0.754749 |
| Glrx3 | 0.313768 | 0.016705 | 0.219296 | 1.777153 | 0.658969 |
| Dyrk2 | 0.313394 | 0.022060 | 0.238274 | 1.656397 | 0.622924 |
| Ndufc2 | 0.313274 | 0.016930 | 0.219296 | 1.771341 | 0.658969 |
| Comt | 0.313213 | 0.036393 | 0.287007 | 1.438983 | 0.542108 |
| Zmat2 | 0.312435 | 0.007300 | 0.172342 | 2.136653 | 0.763610 |
| Mdh2 | 0.312208 | 0.003694 | 0.138071 | 2.432487 | 0.859898 |
| Mtch2 | 0.311870 | 0.006915 | 0.171392 | 2.160214 | 0.766008 |
| Eif3i | 0.311410 | 0.003974 | 0.140345 | 2.400749 | 0.852804 |
| Rrp1 | 0.311202 | 0.000801 | 0.083704 | 3.096119 | 1.077252 |
| Hcfc1r1 | 0.310706 | 0.016093 | 0.216116 | 1.793358 | 0.665312 |
| Slc25a3 | 0.310451 | 0.002232 | 0.119016 | 2.651231 | 0.924396 |
| 5031439G( | 0.310078 | 0.010307 | 0.191032 | 1.986884 | 0.718893 |
| Atp5a1 | 0.309910 | 0.001236 | 0.103771 | 2.907892 | 0.983924 |
| Atp5g1 | 0.309353 | 0.004074 | 0.141411 | 2.390004 | 0.849518 |

| | | | | | |
|---|---|---|---|---|---|
| Gins2 | -0.591570 | 0.017580 | 0.219316 | 1.754988 | 0.658929 |
| Fezf2 | -0.591243 | 0.022756 | 0.240226 | 1.642900 | 0.619381 |
| Magi1 | -0.590687 | 0.001725 | 0.110743 | 2.763263 | 0.955684 |
| Yipf5 | -0.589826 | 0.031142 | 0.268555 | 1.506648 | 0.570967 |
| Fbln1 | -0.588934 | 0.003991 | 0.140345 | 2.398969 | 0.852804 |
| Zbtb14 | -0.586704 | 0.018678 | 0.222828 | 1.728673 | 0.652030 |
| Cryab | -0.585815 | 0.011391 | 0.196420 | 1.943442 | 0.706814 |
| Cct8l1 | -0.584858 | 0.007031 | 0.171400 | 2.152999 | 0.765990 |
| Cttnbp2 | -0.584699 | 0.001956 | 0.115118 | 2.708728 | 0.938859 |
| Olfr180 | -0.583839 | 0.032918 | 0.273731 | 1.482564 | 0.562676 |
| Btg1 | -0.582961 | 0.018536 | 0.222795 | 1.731983 | 0.652095 |
| Ephb1 | -0.582853 | 0.002272 | 0.119065 | 2.643537 | 0.924217 |
| Tmem163 | -0.581807 | 0.022113 | 0.238286 | 1.655343 | 0.622901 |
| Ccser1 | -0.581400 | 0.046191 | 0.312437 | 1.335440 | 0.505237 |
| Spon1 | -0.580678 | 0.035491 | 0.284410 | 1.449883 | 0.546055 |
| Abcd2 | -0.579902 | 0.006365 | 0.167780 | 2.196176 | 0.775260 |
| Bcor | -0.579629 | 0.008798 | 0.182527 | 2.055607 | 0.738673 |
| Ppfibp1 | -0.578017 | 0.036276 | 0.287007 | 1.440385 | 0.542108 |
| Gpsm2 | -0.577542 | 0.045256 | 0.309326 | 1.344323 | 0.509583 |
| Nptx2 | -0.577325 | 0.020610 | 0.231349 | 1.685931 | 0.635733 |
| Wtip | -0.577283 | 0.026099 | 0.252551 | 1.583379 | 0.597651 |
| Cobl | -0.576891 | 0.017937 | 0.220548 | 1.746247 | 0.656497 |
| Sox2 | -0.576550 | 0.025118 | 0.247812 | 1.600015 | 0.605878 |
| Herc4 | -0.575979 | 0.003750 | 0.138446 | 2.425980 | 0.858721 |
| Eomes | -0.575746 | 0.023240 | 0.241019 | 1.633755 | 0.617949 |
| Tigd2 | -0.575725 | 0.024388 | 0.244752 | 1.612816 | 0.611274 |
| Sirpa | -0.575214 | 0.016765 | 0.219296 | 1.775602 | 0.658969 |
| Vcl | -0.574367 | 0.019856 | 0.227717 | 1.702100 | 0.642605 |
| Syn3 | -0.573552 | 0.001151 | 0.098565 | 2.938873 | 1.006277 |
| Gm20765 | -0.573083 | 0.031045 | 0.268555 | 1.508006 | 0.570967 |
| Crtap | -0.572927 | 0.000204 | 0.068135 | 3.689661 | 1.166630 |
| Atm | -0.572392 | 0.007204 | 0.172254 | 2.142419 | 0.763830 |
| Abhd10 | -0.572325 | 0.005618 | 0.159383 | 2.250423 | 0.797558 |
| Zc3h3 | -0.572227 | 0.022916 | 0.240762 | 1.639860 | 0.618413 |
| Intu | -0.571908 | 0.016961 | 0.219296 | 1.770554 | 0.658969 |
| Rad51ap1 | -0.570798 | 0.036985 | 0.287933 | 1.431975 | 0.540709 |
| Gm16434 | -0.569706 | 0.010200 | 0.190810 | 1.991397 | 0.719400 |
| Trpt1 | -0.566415 | 0.007297 | 0.172342 | 2.136880 | 0.763610 |
| Mgst1 | -0.565681 | 0.008299 | 0.178232 | 2.080973 | 0.749013 |
| Itpr2 | -0.564324 | 0.037262 | 0.288916 | 1.428737 | 0.539229 |
| Nim1k | -0.563971 | 0.014479 | 0.210662 | 1.839275 | 0.676414 |

| | | | | | |
|---|---|---|---|---|---|
| Slirp | 0.309130 | 0.017601 | 0.219316 | 1.754472 | 0.658929 |
| Ckap4 | 0.308941 | 0.017693 | 0.219906 | 1.752206 | 0.657762 |
| Slc25a25 | 0.308857 | 0.019300 | 0.225156 | 1.714453 | 0.647516 |
| Rab6a | 0.308792 | 0.027578 | 0.257555 | 1.559445 | 0.589130 |
| B4galt2 | 0.308290 | 0.027952 | 0.259121 | 1.553592 | 0.586497 |
| Rpl15 | 0.307488 | 0.000892 | 0.087553 | 3.049445 | 1.057727 |
| Ranbp1 | 0.307469 | 0.015775 | 0.215179 | 1.802025 | 0.667200 |
| Calm1 | 0.307392 | 0.001051 | 0.093200 | 2.978451 | 1.030586 |
| Fbxo9 | 0.307382 | 0.006832 | 0.171274 | 2.165457 | 0.766308 |
| Pkm | 0.307009 | 0.033700 | 0.276653 | 1.472365 | 0.558065 |
| Rnf208 | 0.306610 | 0.009706 | 0.188600 | 2.012941 | 0.724459 |
| Khsrp | 0.306493 | 0.002096 | 0.116540 | 2.678680 | 0.933524 |
| Babam1 | 0.305747 | 0.004595 | 0.148870 | 2.337758 | 0.827194 |
| Psmd13 | 0.305618 | 0.014261 | 0.210194 | 1.845860 | 0.677379 |
| Zmynd10 | 0.305068 | 0.047487 | 0.315909 | 1.323428 | 0.500439 |
| Adgrl3 | 0.305041 | 0.044789 | 0.307700 | 1.348826 | 0.511872 |
| Actb | 0.304565 | 0.009711 | 0.188600 | 2.012747 | 0.724459 |
| Cat | 0.304307 | 0.016245 | 0.216823 | 1.789272 | 0.663895 |
| Ptp4a3 | 0.304112 | 0.001572 | 0.105319 | 2.803620 | 0.977493 |
| Eef2 | 0.303669 | 0.007958 | 0.175399 | 2.099196 | 0.755974 |
| Laptm4a | 0.303519 | 0.014344 | 0.210194 | 1.843325 | 0.677379 |
| Rapgef1 | 0.301463 | 0.001392 | 0.105319 | 2.856243 | 0.977493 |
| Atp5g3 | 0.300343 | 0.015093 | 0.212813 | 1.821215 | 0.672002 |
| Rpsa | 0.300134 | 0.013270 | 0.207235 | 1.877114 | 0.683536 |
| Sart3 | 0.300054 | 0.011415 | 0.196420 | 1.942511 | 0.706814 |
| Sbf1 | 0.299525 | 0.014803 | 0.212593 | 1.829649 | 0.672451 |
| Hsp90b1 | 0.298772 | 0.010731 | 0.192257 | 1.969362 | 0.716119 |
| Dpysl3 | 0.298401 | 0.000495 | 0.072211 | 3.305593 | 1.141399 |
| Mbd3 | 0.298355 | 0.016134 | 0.216258 | 1.792268 | 0.665028 |
| Srgap2 | 0.298352 | 0.002921 | 0.127176 | 2.534444 | 0.895594 |
| Cdk16 | 0.297552 | 0.018171 | 0.221383 | 1.740621 | 0.654856 |
| Skp1a | 0.297481 | 0.001809 | 0.113004 | 2.742560 | 0.946906 |
| Lamtor4 | 0.297234 | 0.048270 | 0.318230 | 1.316321 | 0.497260 |
| Mpped2 | 0.296718 | 0.030849 | 0.268203 | 1.510763 | 0.571536 |
| Mapre2 | 0.296452 | 0.000888 | 0.087553 | 3.051679 | 1.057727 |
| Pym1 | 0.295757 | 0.016077 | 0.216116 | 1.793794 | 0.665312 |
| Exosc9 | 0.295544 | 0.006089 | 0.165124 | 2.215432 | 0.782189 |
| Rbx1 | 0.295495 | 0.004384 | 0.146069 | 2.358158 | 0.835443 |
| Lrrc75b | 0.295318 | 0.025500 | 0.249150 | 1.593452 | 0.603540 |
| Lrp12 | 0.294382 | 0.041162 | 0.298381 | 1.385498 | 0.525230 |
| Wnk2 | 0.293957 | 0.004535 | 0.147616 | 2.343458 | 0.830867 |

| | | | | | |
|---|---|---|---|---|---|
| Sirt4 | -0.563542 | 0.038063 | 0.290716 | 1.419492 | 0.536531 |
| Igsf9b | -0.563299 | 0.015547 | 0.214097 | 1.808346 | 0.669389 |
| Gpr88 | -0.563289 | 0.023711 | 0.242939 | 1.625052 | 0.614502 |
| Brca2 | -0.563096 | 0.017341 | 0.219296 | 1.760930 | 0.658969 |
| Apol7c | -0.563058 | 0.012951 | 0.205815 | 1.887701 | 0.686522 |
| Map4k2 | -0.563010 | 0.041275 | 0.298605 | 1.384317 | 0.524902 |
| Fbxl4 | -0.562807 | 0.009944 | 0.190219 | 2.002450 | 0.720746 |
| Fbln2 | -0.562793 | 0.024274 | 0.244729 | 1.614862 | 0.611314 |
| Nhlh1 | -0.562616 | 0.033490 | 0.276046 | 1.475088 | 0.559018 |
| Muc1 | -0.562380 | 0.024931 | 0.246706 | 1.603267 | 0.607820 |
| Rbms3 | -0.562189 | 0.046251 | 0.312437 | 1.334878 | 0.505237 |
| Nphp1 | -0.561810 | 0.003753 | 0.138446 | 2.425603 | 0.858721 |
| Sag | -0.560878 | 0.029001 | 0.262944 | 1.537586 | 0.580137 |
| Cox7a1 | -0.560681 | 0.022631 | 0.239679 | 1.645295 | 0.620370 |
| Mcpt8 | -0.559936 | 0.028416 | 0.260823 | 1.546432 | 0.583653 |
| Cited2 | -0.559567 | 0.031712 | 0.269762 | 1.498778 | 0.569020 |
| Csrp2 | -0.558797 | 0.004737 | 0.149991 | 2.324473 | 0.823934 |
| Gm32584 | -0.557235 | 0.032273 | 0.271077 | 1.491163 | 0.566908 |
| Flywch1 | -0.557221 | 0.040214 | 0.296372 | 1.395621 | 0.528163 |
| Ocstamp | -0.556844 | 0.010132 | 0.190810 | 1.994296 | 0.719400 |
| Ngfr | -0.556446 | 0.038660 | 0.292118 | 1.412733 | 0.534442 |
| Grin2d | -0.556446 | 0.039056 | 0.293169 | 1.408309 | 0.532882 |
| Rmdn1 | -0.556258 | 0.011842 | 0.198399 | 1.926586 | 0.702460 |
| Mepce | -0.555876 | 0.001638 | 0.106793 | 2.785761 | 0.971455 |
| Serpini1 | -0.555333 | 0.003873 | 0.139603 | 2.411899 | 0.855104 |
| Gnptg | -0.554945 | 0.006436 | 0.168913 | 2.191381 | 0.772338 |
| Xab2 | -0.554362 | 0.015345 | 0.213387 | 1.814019 | 0.670832 |
| Slc12a6 | -0.553504 | 0.021961 | 0.237931 | 1.658353 | 0.623550 |
| Olig1 | -0.552365 | 0.000264 | 0.071776 | 3.578100 | 1.144022 |
| Olfr642 | -0.551116 | 0.041690 | 0.300051 | 1.379966 | 0.522805 |
| Elk4 | -0.551105 | 0.004939 | 0.150369 | 2.306404 | 0.822841 |
| Pcdhb21 | -0.550763 | 0.048963 | 0.319909 | 1.310135 | 0.494973 |
| Cplx3 | -0.549815 | 0.034735 | 0.281221 | 1.459237 | 0.550952 |
| Rab30 | -0.548704 | 0.027942 | 0.259121 | 1.553741 | 0.586497 |
| Gm45927 | -0.548534 | 0.002358 | 0.119065 | 2.627390 | 0.924217 |
| Kctd18 | -0.548459 | 0.023243 | 0.241019 | 1.633700 | 0.617949 |
| Nek11 | -0.548365 | 0.026235 | 0.253421 | 1.581114 | 0.596158 |
| Plpp1 | -0.547312 | 0.010387 | 0.191032 | 1.983505 | 0.718893 |
| Nr4a1 | -0.547116 | 0.026952 | 0.257278 | 1.569417 | 0.589597 |
| Nusap1 | -0.546971 | 0.036408 | 0.287007 | 1.438805 | 0.542108 |
| Vstm5 | -0.546481 | 0.012126 | 0.199874 | 1.916285 | 0.699245 |

| | | | | | | | | | | | |
|---|---|---|---|---|---|---|---|---|---|---|---|
| Srsf2 | 0.293039 | 0.039817 | 0.294835 | 1.399927 | 0.530420 | Notch1 | -0.546432 | 0.043058 | 0.303009 | 1.365947 | 0.518544 |
| Mrps35 | 0.292241 | 0.013590 | 0.207695 | 1.866785 | 0.682575 | Adpgk | -0.546001 | 0.005055 | 0.150961 | 2.296252 | 0.821135 |
| Eif3c | 0.292153 | 0.010038 | 0.190219 | 1.998373 | 0.720746 | Dll1 | -0.545301 | 0.038563 | 0.291859 | 1.413828 | 0.534826 |
| Ndufa3 | 0.291501 | 0.014768 | 0.212593 | 1.830665 | 0.672451 | Osr2 | -0.545246 | 0.003346 | 0.130212 | 2.475523 | 0.885349 |
| Rps3 | 0.291149 | 0.032563 | 0.272441 | 1.487279 | 0.564727 | Pik3ip1 | -0.545063 | 0.002897 | 0.127176 | 2.538011 | 0.895594 |
| Idh1 | 0.291061 | 0.014931 | 0.212593 | 1.825905 | 0.672451 | Csdc2 | -0.544743 | 0.019382 | 0.225739 | 1.712612 | 0.646393 |
| Hira | 0.290910 | 0.000447 | 0.072211 | 3.349307 | 1.141399 | Fchsd1 | -0.544588 | 0.002004 | 0.116056 | 2.698018 | 0.935334 |
| Eid2 | 0.290869 | 0.035563 | 0.284673 | 1.449006 | 0.545654 | Kctd12 | -0.543906 | 0.036866 | 0.287676 | 1.433374 | 0.541097 |
| Tkfc | 0.289144 | 0.036723 | 0.287676 | 1.435064 | 0.541097 | Uspl1 | -0.542756 | 0.032081 | 0.270429 | 1.493754 | 0.567947 |
| Pum3 | 0.287457 | 0.032389 | 0.271726 | 1.489607 | 0.565868 | Erdr1 | -0.542426 | 0.001575 | 0.105319 | 2.802662 | 0.977493 |
| Eif4e | 0.287401 | 0.005513 | 0.159133 | 2.258600 | 0.798240 | Rgs3 | -0.542349 | 0.024686 | 0.245956 | 1.607549 | 0.609143 |
| Tk2 | 0.287231 | 0.011576 | 0.196420 | 1.936452 | 0.706814 | Dop1a | -0.542079 | 0.039241 | 0.293486 | 1.406261 | 0.532412 |
| Swi5 | 0.287209 | 0.007365 | 0.172342 | 2.132846 | 0.763610 | Sstr3 | -0.542006 | 0.004000 | 0.140345 | 2.397921 | 0.852804 |
| Ncs1 | 0.287068 | 0.013204 | 0.206640 | 1.879287 | 0.684785 | Gca | -0.541552 | 0.028431 | 0.260823 | 1.546213 | 0.583653 |
| Bri3bp | 0.286804 | 0.018556 | 0.222795 | 1.731509 | 0.652095 | Adam26b | -0.541333 | 0.016960 | 0.219296 | 1.770579 | 0.658969 |
| Elavl2 | 0.286525 | 0.013604 | 0.207695 | 1.866319 | 0.682575 | Gm46894 | -0.539349 | 0.005065 | 0.150961 | 2.295437 | 0.821135 |
| Aqr | 0.285257 | 0.037288 | 0.288916 | 1.428435 | 0.539229 | Zhx1 | -0.539226 | 0.004192 | 0.143187 | 2.377558 | 0.844095 |
| Gsto1 | 0.285011 | 0.024257 | 0.244729 | 1.615166 | 0.611314 | Ubn1 | -0.537876 | 0.017233 | 0.219296 | 1.763647 | 0.658969 |
| Faxc | 0.284748 | 0.000121 | 0.061208 | 3.916225 | 1.213194 | Mybl1 | -0.537778 | 0.003605 | 0.136867 | 2.443058 | 0.863701 |
| Eif3e | 0.284520 | 0.002589 | 0.122077 | 2.586905 | 0.913366 | Mapk9 | -0.537426 | 0.026789 | 0.257241 | 1.572037 | 0.589659 |
| Agfg1 | 0.284473 | 0.013779 | 0.207753 | 1.860795 | 0.682454 | Olfr1097 | -0.537265 | 0.023274 | 0.241019 | 1.633134 | 0.617949 |
| Tmem167b | 0.284295 | 0.049814 | 0.322310 | 1.302649 | 0.491727 | Chl1 | -0.536770 | 0.001366 | 0.105319 | 2.864644 | 0.977493 |
| Dgcr6 | 0.284114 | 0.021770 | 0.237026 | 1.662143 | 0.625204 | Bnip5 | -0.536643 | 0.007083 | 0.171400 | 2.149807 | 0.765990 |
| Sema4g | 0.284108 | 0.022422 | 0.238845 | 1.649323 | 0.621884 | Slc51a | -0.536037 | 0.046482 | 0.313077 | 1.332713 | 0.504348 |
| Abcc10 | 0.283658 | 0.049729 | 0.322254 | 1.303392 | 0.491801 | Nlrp3 | -0.535697 | 0.020796 | 0.232506 | 1.682020 | 0.633566 |
| Atp6ap2 | 0.283302 | 0.028325 | 0.260695 | 1.547832 | 0.583868 | Mrtfa | -0.535289 | 0.000639 | 0.079395 | 3.194703 | 1.100208 |
| Gm8909 | 0.283148 | 0.042216 | 0.300932 | 1.374527 | 0.521532 | Pde2a | -0.534336 | 0.008814 | 0.182536 | 2.054817 | 0.738651 |
| Jph4 | 0.283002 | 0.003911 | 0.140345 | 2.407691 | 0.852804 | Csprs | -0.534068 | 0.005564 | 0.159361 | 2.254631 | 0.797618 |
| Rpl18 | 0.282836 | 0.006227 | 0.166308 | 2.205713 | 0.779088 | Cbfa2t3 | -0.533697 | 0.018678 | 0.222828 | 1.728663 | 0.652030 |
| Tmsb4x | 0.282703 | 0.006597 | 0.169325 | 2.180676 | 0.771279 | Ang2 | -0.533420 | 0.046362 | 0.312502 | 1.333837 | 0.505148 |
| Cacng8 | 0.282042 | 0.033134 | 0.274397 | 1.479723 | 0.561621 | Dyrk3 | -0.532960 | 0.021221 | 0.234277 | 1.673236 | 0.630271 |
| Med11 | 0.281719 | 0.027854 | 0.258722 | 1.555118 | 0.587167 | Zbtb5 | -0.532480 | 0.043420 | 0.304589 | 1.362310 | 0.516286 |
| Hspb11 | 0.280220 | 0.023094 | 0.240948 | 1.636506 | 0.618077 | Pdcd6ip | -0.532369 | 0.000261 | 0.071776 | 3.583606 | 1.144022 |
| Hnrnpa1 | 0.280104 | 0.009181 | 0.185052 | 2.037096 | 0.732705 | Plxnb1 | -0.532259 | 0.019556 | 0.226239 | 1.708726 | 0.645432 |
| Pdzd11 | 0.280040 | 0.024458 | 0.245063 | 1.611585 | 0.610723 | Preb | -0.532137 | 0.034486 | 0.280226 | 1.462363 | 0.552492 |
| Cct2 | 0.279817 | 0.007483 | 0.172342 | 2.125908 | 0.763610 | Pgghg | -0.530022 | 0.015776 | 0.215179 | 1.802006 | 0.667200 |
| Trim37 | 0.279622 | 0.019180 | 0.225156 | 1.717145 | 0.647516 | Pcdhgb1 | -0.529723 | 0.000832 | 0.083704 | 3.079822 | 1.077252 |
| Pfn1 | 0.279552 | 0.001075 | 0.093200 | 2.968479 | 1.030586 | Ephb3 | -0.529336 | 0.011366 | 0.196420 | 1.944387 | 0.706814 |
| 1700030K0 | 0.278923 | 0.034022 | 0.278160 | 1.468243 | 0.555706 | Skida1 | -0.529292 | 0.017064 | 0.219296 | 1.767918 | 0.658969 |
| Nop10 | 0.278864 | 0.036721 | 0.287676 | 1.435086 | 0.541097 | P4ha2 | -0.529281 | 0.035704 | 0.285144 | 1.447288 | 0.544935 |
| Hif1a | 0.278797 | 0.029101 | 0.263209 | 1.536099 | 0.579699 | Hecw2 | -0.529267 | 0.031567 | 0.269588 | 1.500773 | 0.569299 |

| | | | | | |
|---|---|---|---|---|---|
| Pak1 | 0.278487 | 0.023961 | 0.243896 | 1.620486 | 0.612795 |
| Irgq | 0.278384 | 0.044263 | 0.306545 | 1.353956 | 0.513506 |
| Cops5 | 0.278236 | 0.017179 | 0.219296 | 1.765008 | 0.658969 |
| Polr1c | 0.277698 | 0.049090 | 0.319909 | 1.309005 | 0.494973 |
| Cnih4 | 0.277120 | 0.011115 | 0.196038 | 1.954108 | 0.707660 |
| Zfp787 | 0.277002 | 0.014940 | 0.212593 | 1.825657 | 0.672451 |
| Mmp24 | 0.276672 | 0.016833 | 0.219296 | 1.773847 | 0.658969 |
| Apbb2 | 0.276639 | 0.040836 | 0.297844 | 1.388962 | 0.526011 |
| Psmb4 | 0.275444 | 0.013461 | 0.207605 | 1.870937 | 0.682762 |
| Rplp1 | 0.274692 | 0.004120 | 0.141979 | 2.385072 | 0.847776 |
| Hacd3 | 0.274660 | 0.029846 | 0.265832 | 1.525120 | 0.575392 |
| Vdac3 | 0.274313 | 0.004777 | 0.149991 | 2.320802 | 0.823934 |
| Cul2 | 0.274110 | 0.022162 | 0.238286 | 1.654386 | 0.622901 |
| Cfl1 | 0.273836 | 0.000258 | 0.071776 | 3.588125 | 1.144022 |
| Ppib | 0.273657 | 0.004279 | 0.144008 | 2.368617 | 0.841615 |
| Parp1 | 0.272822 | 0.000167 | 0.062244 | 3.778450 | 1.205900 |
| Gnb2 | 0.272190 | 0.007087 | 0.171400 | 2.149546 | 0.765990 |
| Slc25a22 | 0.271567 | 0.046820 | 0.314184 | 1.329566 | 0.502815 |
| Ccdc71l | 0.271546 | 0.000829 | 0.083704 | 3.081235 | 1.077252 |
| Mrpl51 | 0.271259 | 0.043833 | 0.305662 | 1.358199 | 0.514758 |
| Calr | 0.271070 | 0.013500 | 0.207605 | 1.869656 | 0.682762 |
| Grcc10 | 0.270182 | 0.001343 | 0.105319 | 2.872003 | 0.977493 |
| Rnf187 | 0.269573 | 0.045888 | 0.311386 | 1.338301 | 0.506701 |
| Osbp | 0.268705 | 0.036832 | 0.287676 | 1.433772 | 0.541097 |
| Ywhaz | 0.268270 | 0.025198 | 0.247931 | 1.598629 | 0.605669 |
| Myl6 | 0.267816 | 0.000744 | 0.082167 | 3.128171 | 1.085300 |
| Rnf220 | 0.266469 | 0.013549 | 0.207605 | 1.868082 | 0.682762 |
| Arhgdia | 0.266446 | 0.031402 | 0.269038 | 1.503047 | 0.570187 |
| Cct6a | 0.266402 | 0.012339 | 0.202233 | 1.908722 | 0.694149 |
| Gpr85 | 0.265961 | 0.025384 | 0.248697 | 1.595444 | 0.604329 |
| Lsm7 | 0.265703 | 0.020226 | 0.229158 | 1.694080 | 0.639866 |
| Map4k4 | 0.265322 | 0.000134 | 0.061211 | 3.871892 | 1.213173 |
| Faim | 0.264730 | 0.043549 | 0.304845 | 1.361024 | 0.515921 |
| Ndufs7 | 0.264254 | 0.037781 | 0.290144 | 1.422722 | 0.537386 |
| Tmem14c | 0.263708 | 0.031539 | 0.269588 | 1.501152 | 0.569299 |
| Psmb2 | 0.262530 | 0.036905 | 0.287676 | 1.432912 | 0.541097 |
| Agpat1 | 0.262223 | 0.037401 | 0.289226 | 1.427121 | 0.538762 |
| Arpc3 | 0.262221 | 0.007418 | 0.172342 | 2.129696 | 0.763610 |
| Gdi1 | 0.261996 | 0.023286 | 0.241019 | 1.632897 | 0.617949 |
| Psmd6 | 0.261753 | 0.036832 | 0.287676 | 1.433776 | 0.541097 |
| Gna12 | 0.261694 | 0.016108 | 0.216116 | 1.792965 | 0.665312 |

| | | | | | |
|---|---|---|---|---|---|
| Pcdha5 | -0.529216 | 0.007906 | 0.175076 | 2.102033 | 0.756773 |
| Trim23 | -0.528909 | 0.019628 | 0.226239 | 1.707120 | 0.645432 |
| Dpysl4 | -0.528646 | 0.028992 | 0.262944 | 1.537716 | 0.580137 |
| Fbxo6 | -0.528226 | 0.022785 | 0.240226 | 1.642356 | 0.619381 |
| Nudt15 | -0.526439 | 0.006190 | 0.165622 | 2.208329 | 0.780882 |
| Coq10b | -0.525974 | 0.006516 | 0.168913 | 2.186022 | 0.772338 |
| Gm10310 | -0.525407 | 0.040637 | 0.297207 | 1.391074 | 0.526941 |
| Exph5 | -0.524949 | 0.014194 | 0.210194 | 1.847906 | 0.677379 |
| Parp8 | -0.523095 | 0.019629 | 0.226239 | 1.707108 | 0.645432 |
| Cyb561 | -0.522935 | 0.038665 | 0.292118 | 1.412681 | 0.534442 |
| Rab23 | -0.520865 | 0.003139 | 0.127788 | 2.503261 | 0.893510 |
| Lin28b | -0.520578 | 0.031211 | 0.268609 | 1.505698 | 0.570879 |
| Mcm2 | -0.520174 | 0.014434 | 0.210441 | 1.840627 | 0.676870 |
| Mcmdc2 | -0.520007 | 0.034698 | 0.281086 | 1.459695 | 0.551161 |
| Grm3 | -0.519894 | 0.030669 | 0.267795 | 1.513299 | 0.572197 |
| Cpne2 | -0.519806 | 0.038361 | 0.290930 | 1.416115 | 0.536211 |
| Tmc5 | -0.519477 | 0.026623 | 0.256150 | 1.574738 | 0.591506 |
| Hic1 | -0.519324 | 0.017543 | 0.219316 | 1.755896 | 0.658929 |
| Rbm12b2 | -0.518710 | 0.037786 | 0.290144 | 1.422674 | 0.537386 |
| Rgma | -0.517996 | 0.042398 | 0.301045 | 1.372659 | 0.521368 |
| Pde1b | -0.517318 | 0.041313 | 0.298605 | 1.383913 | 0.524902 |
| Lrp8 | -0.516291 | 0.010417 | 0.191032 | 1.982247 | 0.718893 |
| Fbxl20 | -0.515230 | 0.036644 | 0.287676 | 1.435999 | 0.541097 |
| Gper1 | -0.514646 | 0.024675 | 0.245956 | 1.607747 | 0.609143 |
| Six1 | -0.514599 | 0.047756 | 0.317072 | 1.320975 | 0.498842 |
| Gdf3 | -0.514196 | 0.046938 | 0.314232 | 1.328478 | 0.502750 |
| Cfap97d2 | -0.514042 | 0.006468 | 0.168913 | 2.189261 | 0.772338 |
| Xrcc6 | -0.513451 | 0.047080 | 0.314475 | 1.327168 | 0.502414 |
| Fgfr3 | -0.512388 | 0.013667 | 0.207695 | 1.864315 | 0.682575 |
| Fbxo46 | -0.510639 | 0.012122 | 0.199874 | 1.916410 | 0.699245 |
| Stil | -0.508081 | 0.043433 | 0.304589 | 1.362180 | 0.516286 |
| Pam | -0.507955 | 0.020309 | 0.229546 | 1.692301 | 0.639130 |
| S100a9 | -0.506952 | 0.014596 | 0.211638 | 1.835759 | 0.674407 |
| Slc17a7 | -0.505950 | 0.047146 | 0.314636 | 1.326552 | 0.502192 |
| Pwwp3a | -0.505139 | 0.016906 | 0.219296 | 1.771955 | 0.658969 |
| Rasl11b | -0.504115 | 0.039188 | 0.293481 | 1.406852 | 0.532421 |
| Atmin | -0.503125 | 0.020556 | 0.231224 | 1.687058 | 0.635967 |
| Gins4 | -0.502264 | 0.011638 | 0.196638 | 1.934124 | 0.706334 |
| Cdc20 | -0.501874 | 0.013775 | 0.207753 | 1.860896 | 0.682454 |
| Rgs17 | -0.501611 | 0.007490 | 0.172342 | 2.125522 | 0.763610 |
| Dbi | -0.500974 | 0.006874 | 0.171274 | 2.162801 | 0.766308 |

| | | | | | |
|---|---|---|---|---|---|
| Ahcyl1 | 0.261613 | 0.019633 | 0.226239 | 1.707023 | 0.645432 |
| Micos10 | 0.261539 | 0.000921 | 0.088539 | 3.035972 | 1.052867 |
| Fkbp1a | 0.261483 | 0.018129 | 0.221314 | 1.741632 | 0.654991 |
| Rps10 | 0.261435 | 0.008708 | 0.181987 | 2.060095 | 0.739961 |
| Arfip2 | 0.261292 | 0.003679 | 0.137871 | 2.434265 | 0.860526 |
| Erh | 0.261151 | 0.003080 | 0.127513 | 2.511411 | 0.894446 |
| Ank | 0.261074 | 0.013378 | 0.207605 | 1.873615 | 0.682762 |
| Nudc | 0.259684 | 0.040023 | 0.295412 | 1.397685 | 0.529572 |
| Tuba1a | 0.259460 | 0.040747 | 0.297814 | 1.389904 | 0.526055 |
| Ap2m1 | 0.258539 | 0.000167 | 0.062244 | 3.776197 | 1.205900 |
| Arpc5 | 0.258203 | 0.041761 | 0.300125 | 1.379224 | 0.522697 |
| Eif3a | 0.257425 | 0.002141 | 0.116540 | 2.669438 | 0.933524 |

| | | | | | |
|---|---|---|---|---|---|
| Skp2 | -0.500731 | 0.015193 | 0.212934 | 1.818369 | 0.671755 |
| Apaf1 | -0.500665 | 0.000427 | 0.072211 | 3.369421 | 1.141399 |
| Dyrk1b | -0.499953 | 0.034321 | 0.279468 | 1.464436 | 0.553668 |
| 1500009L1 | -0.499037 | 0.032510 | 0.272419 | 1.487986 | 0.564763 |
| Tmem126a | -0.498825 | 0.010181 | 0.190810 | 1.992228 | 0.719400 |
| Cog7 | -0.498412 | 0.042221 | 0.300932 | 1.374472 | 0.521532 |
| Tbc1d9b | -0.498315 | 0.048932 | 0.319909 | 1.310403 | 0.494973 |
| Jak3 | -0.498267 | 0.021028 | 0.233557 | 1.677201 | 0.631607 |
| Numa1 | -0.498224 | 0.017337 | 0.219296 | 1.761032 | 0.658969 |
| Dbp | -0.497738 | 0.021432 | 0.234904 | 1.668938 | 0.629110 |
| Sh3bp4 | -0.497370 | 0.009229 | 0.185437 | 2.034851 | 0.731804 |
| Ccrl2 | -0.497369 | 0.033623 | 0.276331 | 1.473369 | 0.558570 |
| Lhx2 | -0.496956 | 0.010436 | 0.191032 | 1.981486 | 0.718893 |
| Nemp2 | -0.496696 | 0.031901 | 0.269885 | 1.496199 | 0.568821 |
| Trmt10b | -0.496627 | 0.027764 | 0.258396 | 1.556524 | 0.587715 |
| Foxn3 | -0.495374 | 0.036916 | 0.287676 | 1.432780 | 0.541097 |
| Axl | -0.494834 | 0.010541 | 0.191032 | 1.977120 | 0.718893 |
| Mup13 | -0.494812 | 0.017993 | 0.220548 | 1.744888 | 0.656497 |
| Vmn1r104 | -0.493514 | 0.023681 | 0.242939 | 1.625605 | 0.614502 |
| Mroh8 | -0.493208 | 0.009054 | 0.184321 | 2.043147 | 0.734426 |
| Gon4l | -0.493201 | 0.027690 | 0.258218 | 1.557682 | 0.588014 |
| Mrps31 | -0.491361 | 0.038059 | 0.290716 | 1.419545 | 0.536531 |
| Aldh16a1 | -0.490114 | 0.047899 | 0.317423 | 1.319672 | 0.498361 |
| Ccnb2 | -0.490040 | 0.034898 | 0.282221 | 1.457198 | 0.549411 |
| Cldn34b1 | -0.489938 | 0.025812 | 0.251093 | 1.588175 | 0.600165 |
| Appl2 | -0.489465 | 0.026875 | 0.257241 | 1.570654 | 0.589659 |
| Pcdhgb4 | -0.489317 | 0.022229 | 0.238730 | 1.653084 | 0.622092 |
| Itm2b | -0.489130 | 0.023404 | 0.241553 | 1.630711 | 0.616987 |
| Zcchc12 | -0.487656 | 0.043132 | 0.303170 | 1.365199 | 0.518314 |
| Elavl4 | -0.487031 | 0.002928 | 0.127176 | 2.533443 | 0.895594 |
| Gm10265 | -0.486370 | 0.007076 | 0.171400 | 2.150238 | 0.765990 |
| Dppa2 | -0.485322 | 0.002451 | 0.119712 | 2.610709 | 0.921864 |
| Tmub2 | -0.484887 | 0.041256 | 0.298605 | 1.384516 | 0.524902 |
| Hipk2 | -0.484867 | 0.017871 | 0.220523 | 1.747843 | 0.656547 |
| Apoc2 | -0.484456 | 0.048029 | 0.317677 | 1.318493 | 0.498014 |
| Inha | -0.484367 | 0.028101 | 0.259655 | 1.551277 | 0.585603 |
| Usp29 | -0.483786 | 0.023138 | 0.240948 | 1.635670 | 0.618077 |
| Tmem135 | -0.483394 | 0.048658 | 0.319145 | 1.312847 | 0.496012 |
| Olfr1335 | -0.482656 | 0.025243 | 0.248003 | 1.597865 | 0.605544 |
| Lypd1 | -0.481850 | 0.026992 | 0.257278 | 1.568757 | 0.589597 |
| Wasf1 | -0.481697 | 0.022997 | 0.240762 | 1.638336 | 0.618413 |

| | | | | | |
|---|---|---|---|---|---|
| Catsper2 | -0.481514 | 0.027329 | 0.257555 | 1.563369 | 0.589130 |
| Ubxn2a | -0.481054 | 0.013704 | 0.207695 | 1.863166 | 0.682575 |
| Neurog3 | -0.480853 | 0.038778 | 0.292481 | 1.411417 | 0.533902 |
| Gm46934 | -0.480562 | 0.003239 | 0.129398 | 2.489589 | 0.888073 |
| Pcdhga12 | -0.480530 | 0.005524 | 0.159133 | 2.257744 | 0.798240 |
| Sema4b | -0.480502 | 0.004429 | 0.146848 | 2.353727 | 0.833132 |
| Amph | -0.480253 | 0.001243 | 0.103771 | 2.905563 | 0.983924 |
| Qser1 | -0.480018 | 0.007860 | 0.174840 | 2.104562 | 0.757358 |
| Iffo2 | -0.479140 | 0.044964 | 0.308227 | 1.347132 | 0.511129 |
| Pigv | -0.478996 | 0.040002 | 0.295412 | 1.397914 | 0.529572 |
| Tcp11l2 | -0.478980 | 0.033197 | 0.274595 | 1.478900 | 0.561307 |
| Tusc2 | -0.478501 | 0.015247 | 0.213071 | 1.816806 | 0.671476 |
| Gm52720 | -0.477962 | 0.002850 | 0.126635 | 2.545099 | 0.897446 |
| Leng9 | -0.476736 | 0.026634 | 0.256150 | 1.574565 | 0.591506 |
| Mrps27 | -0.476390 | 0.001553 | 0.105319 | 2.808933 | 0.977493 |
| Dll4 | -0.476331 | 0.035228 | 0.283672 | 1.453116 | 0.547183 |
| Sptlc2 | -0.475774 | 0.017303 | 0.219296 | 1.761868 | 0.658969 |
| Pgap3 | -0.475152 | 0.030026 | 0.265965 | 1.522496 | 0.575176 |
| Wbp1 | -0.474983 | 0.008770 | 0.182470 | 2.057020 | 0.738809 |
| Usp24 | -0.474257 | 0.015063 | 0.212593 | 1.822098 | 0.672451 |
| Scgb1b10 | -0.473740 | 0.037489 | 0.289265 | 1.426096 | 0.538704 |
| Slc16a12 | -0.473576 | 0.044885 | 0.307985 | 1.347895 | 0.511470 |
| Etnppl | -0.472937 | 0.011610 | 0.196433 | 1.935181 | 0.706786 |
| Specc1l | -0.472802 | 0.037717 | 0.290075 | 1.423461 | 0.537489 |
| Aldh3a2 | -0.471529 | 0.018282 | 0.221895 | 1.737980 | 0.653852 |
| Gm9507 | -0.468248 | 0.028845 | 0.262944 | 1.539937 | 0.580137 |
| Hepacam | -0.467883 | 0.013490 | 0.207605 | 1.870003 | 0.682762 |
| Dock9 | -0.467398 | 0.011773 | 0.198014 | 1.929105 | 0.703303 |
| Rragb | -0.467121 | 0.010475 | 0.191032 | 1.979845 | 0.718893 |
| Gm39557 | -0.466948 | 0.045844 | 0.311239 | 1.338715 | 0.506906 |
| Pea15a | -0.466641 | 0.003368 | 0.130736 | 2.472584 | 0.883604 |
| Gdf10 | -0.466402 | 0.005045 | 0.150961 | 2.297108 | 0.821135 |
| Adamts4 | -0.465753 | 0.035628 | 0.284863 | 1.448207 | 0.545364 |
| P4htm | -0.465316 | 0.042795 | 0.302156 | 1.368602 | 0.519769 |
| Rps6ka5 | -0.465287 | 0.037292 | 0.288916 | 1.428381 | 0.539229 |
| Ice1 | -0.464998 | 0.018178 | 0.221383 | 1.740466 | 0.654856 |
| Odf2l | -0.464666 | 0.046194 | 0.312437 | 1.335417 | 0.505237 |
| Lgi1 | -0.464653 | 0.035871 | 0.285833 | 1.445260 | 0.543888 |
| Anks1 | -0.463436 | 0.003618 | 0.136867 | 2.441508 | 0.863701 |
| Slit1 | -0.463083 | 0.038198 | 0.290930 | 1.417957 | 0.536211 |
| Kdm2a | -0.462590 | 0.001301 | 0.103883 | 2.885565 | 0.983457 |

| | | | | | |
|---|---|---|---|---|---|
| Rgsl1 | -0.462413 | 0.007571 | 0.173360 | 2.120831 | 0.761051 |
| Slc25a27 | -0.461876 | 0.020600 | 0.231349 | 1.686127 | 0.635733 |
| Plch2 | -0.461710 | 0.009816 | 0.189205 | 2.008045 | 0.723068 |
| Ccnl2 | -0.460331 | 0.031320 | 0.268838 | 1.504182 | 0.570509 |
| Rapgef4 | -0.459382 | 0.019276 | 0.225156 | 1.714989 | 0.647516 |
| H2az2 | -0.459306 | 0.019192 | 0.225156 | 1.716889 | 0.647516 |
| Dnajc16 | -0.458252 | 0.047758 | 0.317072 | 1.320951 | 0.498842 |
| Rad51c | -0.457878 | 0.033288 | 0.275187 | 1.477711 | 0.560373 |
| Fam135a | -0.457687 | 0.006339 | 0.167451 | 2.197957 | 0.776113 |
| Ttc21b | -0.456985 | 0.036449 | 0.287039 | 1.438317 | 0.542059 |
| Gm29758 | -0.456636 | 0.019200 | 0.225156 | 1.716705 | 0.647516 |
| Cdip1 | -0.456556 | 0.039477 | 0.294582 | 1.403654 | 0.530794 |
| Pank2 | -0.455931 | 0.007416 | 0.172342 | 2.129833 | 0.763610 |
| Dusp2 | -0.455755 | 0.029601 | 0.265389 | 1.528699 | 0.576117 |
| Pex26 | -0.455494 | 0.016278 | 0.216823 | 1.788393 | 0.663895 |
| Smg8 | -0.455107 | 0.031062 | 0.268555 | 1.507777 | 0.570967 |
| Tmem161a | -0.454763 | 0.008514 | 0.179475 | 2.069891 | 0.745995 |
| Zfp143 | -0.454543 | 0.037923 | 0.290550 | 1.421099 | 0.536779 |
| Nub1 | -0.454465 | 0.006176 | 0.165622 | 2.209313 | 0.780882 |
| Ankzf1 | -0.453635 | 0.030330 | 0.266662 | 1.518127 | 0.574038 |
| Zfp397 | -0.453011 | 0.029931 | 0.265965 | 1.523879 | 0.575176 |
| Zcrb1 | -0.452582 | 0.035378 | 0.283990 | 1.451265 | 0.546697 |
| Btnl4 | -0.452182 | 0.036405 | 0.287007 | 1.438841 | 0.542108 |
| Rcn1 | -0.452094 | 0.009796 | 0.189070 | 2.008946 | 0.723377 |
| Hmgxb3 | -0.451656 | 0.027624 | 0.257776 | 1.558713 | 0.588758 |
| Chd9 | -0.451533 | 0.002730 | 0.125045 | 2.563881 | 0.902935 |
| Nxt1 | -0.451464 | 0.042985 | 0.302738 | 1.366683 | 0.518933 |
| Med16 | -0.451460 | 0.035281 | 0.283694 | 1.452459 | 0.547150 |
| Prom1 | -0.451146 | 0.044600 | 0.307142 | 1.350664 | 0.512661 |
| Dennd6b | -0.450298 | 0.018612 | 0.222795 | 1.730204 | 0.652095 |
| Cdk12 | -0.449170 | 0.022972 | 0.240762 | 1.638803 | 0.618413 |
| H2bc8\|H2b | -0.448266 | 0.032046 | 0.270354 | 1.494231 | 0.568068 |
| Aurkc | -0.447156 | 0.016029 | 0.216107 | 1.795097 | 0.665330 |
| Slc10a4l | -0.446036 | 0.010272 | 0.191032 | 1.988337 | 0.718893 |
| Arhgap12 | -0.445271 | 0.012512 | 0.203366 | 1.902688 | 0.691721 |
| Prss41 | -0.445143 | 0.002874 | 0.127176 | 2.541456 | 0.895594 |
| Wsb1 | -0.445102 | 0.004093 | 0.141729 | 2.387961 | 0.848541 |
| Ikzf4 | -0.444583 | 0.004747 | 0.149991 | 2.323595 | 0.823934 |
| Idnk | -0.444081 | 0.032570 | 0.272441 | 1.487178 | 0.564727 |
| Golga1 | -0.441023 | 0.013059 | 0.206018 | 1.884098 | 0.686094 |
| Qk | -0.440958 | 0.020816 | 0.232506 | 1.681592 | 0.633566 |

| | | | | | |
|---|---|---|---|---|---|
| Gm3642 | -0.440779 | 0.003463 | 0.133683 | 2.460525 | 0.873925 |
| Bub1 | -0.439377 | 0.011451 | 0.196420 | 1.941170 | 0.706814 |
| Cnot1 | -0.438829 | 0.012562 | 0.203366 | 1.900955 | 0.691721 |
| Golim4 | -0.438449 | 0.041130 | 0.298312 | 1.385845 | 0.525329 |
| Ipo8 | -0.437539 | 0.000714 | 0.081349 | 3.146307 | 1.089648 |
| Pou2f2 | -0.437461 | 0.044627 | 0.307142 | 1.350407 | 0.512661 |
| Rasgrp2 | -0.436907 | 0.033022 | 0.273948 | 1.481198 | 0.562332 |
| Fam214a | -0.436355 | 0.005769 | 0.161716 | 2.238898 | 0.791248 |
| Zfp341 | -0.435039 | 0.004190 | 0.143187 | 2.377817 | 0.844095 |
| Gabrg3 | -0.434581 | 0.015836 | 0.215539 | 1.800358 | 0.666475 |
| Hdac7 | -0.433485 | 0.005062 | 0.150961 | 2.295691 | 0.821135 |
| Armcx6 | -0.433281 | 0.042947 | 0.302621 | 1.367067 | 0.519101 |
| Serpinb1c | -0.432924 | 0.026542 | 0.255934 | 1.576059 | 0.591872 |
| Ccdc66 | -0.432613 | 0.045561 | 0.310659 | 1.341406 | 0.507715 |
| Ctnna1 | -0.432603 | 0.036714 | 0.287676 | 1.435169 | 0.541097 |
| Glrx2 | -0.432107 | 0.002032 | 0.116473 | 2.691970 | 0.933775 |
| Rapgef2 | -0.431880 | 0.009019 | 0.184321 | 2.044847 | 0.734426 |
| St8sia4 | -0.429795 | 0.025200 | 0.247931 | 1.598596 | 0.605669 |
| Rad1 | -0.428942 | 0.041442 | 0.298749 | 1.382556 | 0.524693 |
| A430033K( | -0.428684 | 0.041339 | 0.298605 | 1.383641 | 0.524902 |
| Dnal1 | -0.426917 | 0.019611 | 0.226239 | 1.707498 | 0.645432 |
| Vps4b | -0.426456 | 0.017713 | 0.219906 | 1.751701 | 0.657762 |
| Flrt2 | -0.425879 | 0.036915 | 0.287676 | 1.432802 | 0.541097 |
| Ptgis | -0.425845 | 0.005046 | 0.150961 | 2.297089 | 0.821135 |
| Ddx23 | -0.425400 | 0.015163 | 0.212934 | 1.819204 | 0.671755 |
| Armcx4 | -0.424099 | 0.009732 | 0.188600 | 2.011812 | 0.724459 |
| Glrx | -0.423925 | 0.046975 | 0.314232 | 1.328137 | 0.502750 |
| St7 | -0.423713 | 0.032754 | 0.273008 | 1.484736 | 0.563825 |
| Zfp654 | -0.423349 | 0.018140 | 0.221314 | 1.741364 | 0.654991 |
| Cdr2l | -0.423346 | 0.009314 | 0.186131 | 2.030842 | 0.730181 |
| Paxbp1 | -0.423302 | 0.015817 | 0.215495 | 1.800864 | 0.666562 |
| Paqr8 | -0.422382 | 0.003272 | 0.129601 | 2.485176 | 0.887393 |
| Cd200r2 | -0.422307 | 0.016729 | 0.219296 | 1.776523 | 0.658969 |
| Cfap69 | -0.420604 | 0.023010 | 0.240762 | 1.638082 | 0.618413 |
| Tubg1 | -0.420473 | 0.004930 | 0.150369 | 2.307194 | 0.822841 |
| Atoh7 | -0.420260 | 0.012877 | 0.205688 | 1.890190 | 0.686791 |
| Ska2 | -0.419959 | 0.015166 | 0.212934 | 1.819125 | 0.671755 |
| Mphosph9 | -0.419828 | 0.018558 | 0.222795 | 1.731480 | 0.652095 |
| Zgpat | -0.418881 | 0.030475 | 0.267006 | 1.516057 | 0.573479 |
| Pex3 | -0.417847 | 0.000921 | 0.088539 | 3.035630 | 1.052867 |
| Zfp318 | -0.417402 | 0.044412 | 0.306701 | 1.352499 | 0.513284 |

| | | | | | |
|---|---|---|---|---|---|
| Sdr39u1 | -0.417086 | 0.027807 | 0.258504 | 1.555843 | 0.587532 |
| Iqce | -0.416435 | 0.005141 | 0.151647 | 2.288911 | 0.819166 |
| Ndrg3 | -0.416115 | 0.008033 | 0.176091 | 2.095140 | 0.754262 |
| Sox21 | -0.413915 | 0.019481 | 0.226162 | 1.710393 | 0.645580 |
| Cdk18 | -0.413638 | 0.009140 | 0.184984 | 2.039074 | 0.732865 |
| Pigs | -0.413406 | 0.010127 | 0.190810 | 1.994540 | 0.719400 |
| Nrxn3 | -0.413110 | 0.018605 | 0.222795 | 1.730367 | 0.652095 |
| Pomt1 | -0.412922 | 0.015910 | 0.216107 | 1.798317 | 0.665330 |
| Oas1c | -0.412827 | 0.036816 | 0.287676 | 1.433958 | 0.541097 |
| Syt9 | -0.412093 | 0.035413 | 0.284109 | 1.450837 | 0.546516 |
| Bhlhe22 | -0.411975 | 0.014500 | 0.210750 | 1.838647 | 0.676232 |
| Lpin1 | -0.411426 | 0.017179 | 0.219296 | 1.764995 | 0.658969 |
| Gpt2 | -0.411378 | 0.038895 | 0.292607 | 1.410101 | 0.533716 |
| Gm52856 | -0.410982 | 0.001833 | 0.113004 | 2.736786 | 0.946906 |
| Zfp740 | -0.409910 | 0.012101 | 0.199874 | 1.917185 | 0.699245 |
| Mfap4 | -0.409456 | 0.020344 | 0.229578 | 1.691559 | 0.639069 |
| Enoph1 | -0.409217 | 0.000858 | 0.085339 | 3.066646 | 1.068854 |
| Nup93 | -0.409063 | 0.001775 | 0.112438 | 2.750863 | 0.949087 |
| Zfp90 | -0.409056 | 0.004619 | 0.148870 | 2.335452 | 0.827194 |
| Atp1a1 | -0.408221 | 0.042283 | 0.300976 | 1.373833 | 0.521469 |
| Adamts3 | -0.408110 | 0.040521 | 0.297086 | 1.392317 | 0.527118 |
| Kifc1 | -0.408014 | 0.028480 | 0.260941 | 1.545454 | 0.583458 |
| Grhpr | -0.407707 | 0.002511 | 0.120865 | 2.600174 | 0.917700 |
| Kcnn3 | -0.407702 | 0.030009 | 0.265965 | 1.522745 | 0.575176 |
| Pofut1 | -0.407450 | 0.049669 | 0.322039 | 1.303915 | 0.492092 |
| Ints1 | -0.407018 | 0.000824 | 0.083704 | 3.084027 | 1.077252 |
| Gm5640 | -0.406587 | 0.003122 | 0.127788 | 2.505618 | 0.893510 |
| Vps45 | -0.405798 | 0.039320 | 0.293769 | 1.405383 | 0.531994 |
| Oxgr1 | -0.404468 | 0.012378 | 0.202522 | 1.907366 | 0.693528 |
| Ntn1 | -0.403429 | 0.029935 | 0.265965 | 1.523824 | 0.575176 |
| Slc49a4 | -0.403100 | 0.006441 | 0.168913 | 2.191035 | 0.772338 |
| Pcdhgc4 | -0.401779 | 0.041370 | 0.298605 | 1.383318 | 0.524902 |
| Sap130 | -0.401532 | 0.038079 | 0.290716 | 1.419310 | 0.536531 |
| Sepsecs | -0.401493 | 0.014368 | 0.210194 | 1.842603 | 0.677379 |
| Ezh2 | -0.400728 | 0.002659 | 0.123993 | 2.575240 | 0.906602 |
| Wnk1 | -0.399503 | 0.018690 | 0.222828 | 1.728392 | 0.652030 |
| Vwa8 | -0.398992 | 0.045789 | 0.311164 | 1.339240 | 0.507010 |
| Mrps22 | -0.397505 | 0.020006 | 0.227717 | 1.698848 | 0.642605 |
| Gm8094 | -0.396997 | 0.010545 | 0.191032 | 1.976974 | 0.718893 |
| Efnb1 | -0.396790 | 0.008481 | 0.179457 | 2.071570 | 0.746039 |
| Zfp458 | -0.395053 | 0.017699 | 0.219906 | 1.752043 | 0.657762 |

| | | | | | |
|---|---|---|---|---|---|
| Osbpl6 | -0.393932 | 0.031700 | 0.269762 | 1.498941 | 0.569020 |
| Socs2 | -0.393758 | 0.005584 | 0.159361 | 2.253093 | 0.797618 |
| Hid1 | -0.393606 | 0.027342 | 0.257555 | 1.563175 | 0.589130 |
| Zfp57 | -0.393013 | 0.003993 | 0.140345 | 2.398709 | 0.852804 |
| Gm36327 | -0.392956 | 0.001390 | 0.105319 | 2.857009 | 0.977493 |
| Vti1a | -0.392802 | 0.011905 | 0.198399 | 1.924282 | 0.702460 |
| Wapl | -0.392765 | 0.000153 | 0.062244 | 3.814320 | 1.205900 |
| Gse1 | -0.392290 | 0.007288 | 0.172342 | 2.137412 | 0.763610 |
| Cbfa2t2 | -0.392243 | 0.045657 | 0.311011 | 1.340496 | 0.507224 |
| Brd8 | -0.392150 | 0.007788 | 0.174684 | 2.108600 | 0.757747 |
| Gm46156 | -0.391865 | 0.043668 | 0.305208 | 1.359834 | 0.515404 |
| Epb41l2 | -0.391650 | 0.046282 | 0.312437 | 1.334590 | 0.505237 |
| Prune2 | -0.391615 | 0.008241 | 0.178185 | 2.084013 | 0.749130 |
| Prpf39 | -0.391194 | 0.021553 | 0.236051 | 1.666486 | 0.626995 |
| Lrr1 | -0.390824 | 0.011595 | 0.196420 | 1.935728 | 0.706814 |
| Ppp1r16a | -0.389219 | 0.024477 | 0.245080 | 1.611245 | 0.610692 |
| Pkdcc | -0.389059 | 0.005446 | 0.157673 | 2.263902 | 0.802241 |
| Dck | -0.388755 | 0.037444 | 0.289265 | 1.426618 | 0.538704 |
| Kdm4d | -0.388560 | 0.035564 | 0.284673 | 1.448990 | 0.545654 |
| Tmem18 | -0.388152 | 0.048271 | 0.318230 | 1.316318 | 0.497260 |
| Ethe1 | -0.387478 | 0.031868 | 0.269885 | 1.496649 | 0.568821 |
| Lrig2 | -0.387470 | 0.007024 | 0.171400 | 2.153406 | 0.765990 |
| Lrrc8a | -0.387359 | 0.007982 | 0.175662 | 2.097867 | 0.755321 |
| Lrp1 | -0.386979 | 0.028464 | 0.260941 | 1.545710 | 0.583458 |
| D830030K2 | -0.386663 | 0.016767 | 0.219296 | 1.775535 | 0.658969 |
| Nfic | -0.386375 | 0.008461 | 0.179457 | 2.072565 | 0.746039 |
| Des | -0.386330 | 0.031905 | 0.269885 | 1.496137 | 0.568821 |
| Lima1 | -0.385907 | 0.016761 | 0.219296 | 1.775690 | 0.658969 |
| Cask | -0.384831 | 0.013881 | 0.209052 | 1.857584 | 0.679746 |
| Sptb | -0.384542 | 0.016969 | 0.219296 | 1.770331 | 0.658969 |
| Ttc26 | -0.383739 | 0.030461 | 0.267006 | 1.516257 | 0.573479 |
| Dbndd2 | -0.383730 | 0.022863 | 0.240694 | 1.640863 | 0.618535 |
| St3gal5 | -0.383708 | 0.010529 | 0.191032 | 1.977612 | 0.718893 |
| Mdfi | -0.382627 | 0.013124 | 0.206018 | 1.881932 | 0.686094 |
| Gxylt1 | -0.381993 | 0.011673 | 0.196638 | 1.932828 | 0.706334 |
| Pogz | -0.380896 | 0.001541 | 0.105319 | 2.812262 | 0.977493 |
| Slf2 | -0.380349 | 0.038881 | 0.292607 | 1.410258 | 0.533716 |
| Pot1a | -0.380087 | 0.030814 | 0.268068 | 1.511249 | 0.571754 |
| Ezh1 | -0.379396 | 0.029291 | 0.264012 | 1.533271 | 0.578376 |
| Cpped1 | -0.379263 | 0.019417 | 0.225777 | 1.711824 | 0.646319 |
| Gm52800 | -0.378965 | 0.008876 | 0.182536 | 2.051807 | 0.738651 |

| | | | | | |
|---|---|---|---|---|---|
| Vmn1r113 | -0.378615 | 0.019066 | 0.225156 | 1.719744 | 0.647516 |
| Zfp109 | -0.378575 | 0.042239 | 0.300932 | 1.374290 | 0.521532 |
| Zfp275 | -0.378126 | 0.013964 | 0.209134 | 1.854988 | 0.679575 |
| Coq7 | -0.377869 | 0.002377 | 0.119091 | 2.623953 | 0.924121 |
| Gm19470 | -0.377515 | 0.017600 | 0.219316 | 1.754483 | 0.658929 |
| Dnajc3 | -0.376221 | 0.014575 | 0.211623 | 1.836405 | 0.674437 |
| Bloc1s1 | -0.375940 | 0.019879 | 0.227717 | 1.701610 | 0.642605 |
| Rft1 | -0.375366 | 0.034049 | 0.278223 | 1.467893 | 0.555607 |
| Pi4kb | -0.374298 | 0.039741 | 0.294835 | 1.400763 | 0.530420 |
| Chd2 | -0.373679 | 0.007726 | 0.174090 | 2.112040 | 0.759227 |
| Psmb10 | -0.373523 | 0.006660 | 0.169635 | 2.176500 | 0.770483 |
| Pcdhac1 | -0.373178 | 0.036990 | 0.287933 | 1.431913 | 0.540709 |
| Tnfaip3 | -0.373057 | 0.049056 | 0.319909 | 1.309312 | 0.494973 |
| Wipf2 | -0.372040 | 0.048129 | 0.317739 | 1.317596 | 0.497930 |
| Hjurp | -0.370428 | 0.030551 | 0.267386 | 1.514981 | 0.572862 |
| Etfdh | -0.369989 | 0.037187 | 0.288670 | 1.429608 | 0.539599 |
| Kctd12b | -0.369974 | 0.046903 | 0.314198 | 1.328800 | 0.502796 |
| Zfp939 | -0.369716 | 0.031154 | 0.268555 | 1.506487 | 0.570967 |
| Paf1 | -0.369605 | 0.035621 | 0.284863 | 1.448290 | 0.545364 |
| Ms4a4c | -0.369541 | 0.026965 | 0.257278 | 1.569199 | 0.589597 |
| Emx1 | -0.369458 | 0.035949 | 0.286211 | 1.444315 | 0.543314 |
| Med18 | -0.368891 | 0.007479 | 0.172342 | 2.126135 | 0.763610 |
| Cmss1 | -0.368027 | 0.024209 | 0.244729 | 1.616020 | 0.611314 |
| Rsrp1 | -0.367477 | 0.006732 | 0.170138 | 2.171868 | 0.769199 |
| Safb2 | -0.366910 | 0.017566 | 0.219316 | 1.755327 | 0.658929 |
| Ptar1 | -0.366679 | 0.036843 | 0.287676 | 1.433641 | 0.541097 |
| Cyp4a29 | -0.366022 | 0.049159 | 0.319909 | 1.308396 | 0.494973 |
| Nav1 | -0.365986 | 0.030306 | 0.266662 | 1.518467 | 0.574038 |
| Diras2 | -0.365762 | 0.013352 | 0.207521 | 1.874454 | 0.682937 |
| Tmem123 | -0.364811 | 0.018424 | 0.222669 | 1.734606 | 0.652340 |
| Cdca4 | -0.364132 | 0.015020 | 0.212593 | 1.823320 | 0.672451 |
| Sbno1 | -0.363803 | 0.010892 | 0.193278 | 1.962885 | 0.713818 |
| Prl3d1 | -0.363588 | 0.028924 | 0.262944 | 1.538749 | 0.580137 |
| Rnase4 | -0.363587 | 0.002846 | 0.126635 | 2.545831 | 0.897446 |
| Meltf | -0.361690 | 0.042669 | 0.302115 | 1.369889 | 0.519827 |
| Svbp | -0.361176 | 0.011381 | 0.196420 | 1.943822 | 0.706814 |
| Myl12a | -0.360930 | 0.037011 | 0.287933 | 1.431674 | 0.540709 |
| Zfp995 | -0.360141 | 0.028055 | 0.259399 | 1.551989 | 0.586032 |
| Zfp513 | -0.359825 | 0.019854 | 0.227717 | 1.702143 | 0.642605 |
| Brd3os | -0.359619 | 0.031032 | 0.268555 | 1.508189 | 0.570967 |
| Cdc42bpa | -0.359391 | 0.009619 | 0.187913 | 2.016884 | 0.726043 |

| | | | | | |
|---|---|---|---|---|---|
| Cenpj | -0.359106 | 0.045756 | 0.311164 | 1.339552 | 0.507010 |
| Zscan18 | -0.358790 | 0.031219 | 0.268609 | 1.505587 | 0.570879 |
| Os9 | -0.358600 | 0.046492 | 0.313077 | 1.332623 | 0.504348 |
| Erbin | -0.358356 | 0.015580 | 0.214332 | 1.807423 | 0.668913 |
| Lsm1 | -0.357564 | 0.013091 | 0.206018 | 1.883018 | 0.686094 |
| Bbs4 | -0.357447 | 0.011486 | 0.196420 | 1.939844 | 0.706814 |
| Trnau1ap | -0.357191 | 0.048546 | 0.319066 | 1.313848 | 0.496119 |
| Rassf1 | -0.357026 | 0.021346 | 0.234691 | 1.670681 | 0.629503 |
| Slc25a19 | -0.356745 | 0.008693 | 0.181951 | 2.060821 | 0.740045 |
| Ddit4 | -0.355735 | 0.008194 | 0.178185 | 2.086495 | 0.749130 |
| Bscl2 | -0.355480 | 0.022434 | 0.238845 | 1.649094 | 0.621884 |
| Gria2 | -0.355106 | 0.021002 | 0.233526 | 1.677745 | 0.631664 |
| Gm35498 | -0.355014 | 0.041743 | 0.300125 | 1.379421 | 0.522697 |
| Wars | -0.354882 | 0.009214 | 0.185437 | 2.035541 | 0.731804 |
| Nmral1 | -0.353497 | 0.035980 | 0.286211 | 1.443934 | 0.543314 |
| Irak1 | -0.353380 | 0.032137 | 0.270512 | 1.492995 | 0.567814 |
| Mettl4 | -0.353321 | 0.023450 | 0.241644 | 1.629866 | 0.616823 |
| Ctnnd2 | -0.353129 | 0.014335 | 0.210194 | 1.843588 | 0.677379 |
| Adra2c | -0.351916 | 0.003297 | 0.129601 | 2.481931 | 0.887393 |
| C2cd3 | -0.351591 | 0.001301 | 0.103883 | 2.885644 | 0.983457 |
| Fbxl6 | -0.351533 | 0.019936 | 0.227717 | 1.700362 | 0.642605 |
| Hip1 | -0.351189 | 0.019075 | 0.225156 | 1.719543 | 0.647516 |
| Ulk2 | -0.350618 | 0.018829 | 0.224105 | 1.725178 | 0.649548 |
| Sh3bgrl | -0.350443 | 0.009387 | 0.186273 | 2.027451 | 0.729850 |
| Zfyve19 | -0.349738 | 0.024953 | 0.246706 | 1.602869 | 0.607820 |
| Kpna2 | -0.349049 | 0.032907 | 0.273731 | 1.482712 | 0.562676 |
| Carmil3 | -0.347580 | 0.044259 | 0.306545 | 1.354002 | 0.513506 |
| Pcmtd2 | -0.346268 | 0.001906 | 0.113600 | 2.719953 | 0.944621 |
| Mgrn1 | -0.345934 | 0.025741 | 0.250981 | 1.589371 | 0.600359 |
| Rasal2 | -0.345777 | 0.021758 | 0.237026 | 1.662390 | 0.625204 |
| Olfr1311 | -0.345624 | 0.017783 | 0.220365 | 1.749990 | 0.656857 |
| Crat | -0.345349 | 0.039553 | 0.294597 | 1.402821 | 0.530772 |
| Rpia | -0.344863 | 0.015655 | 0.214947 | 1.805333 | 0.667668 |
| Mtor | -0.344842 | 0.027582 | 0.257555 | 1.559372 | 0.589130 |
| Ddc | -0.343680 | 0.016550 | 0.218930 | 1.781205 | 0.659696 |
| Slc33a1 | -0.342732 | 0.021832 | 0.237201 | 1.660901 | 0.624883 |
| Akr1c18 | -0.342057 | 0.013550 | 0.207605 | 1.868076 | 0.682762 |
| Pxdn | -0.341492 | 0.036596 | 0.287676 | 1.436571 | 0.541097 |
| Fip1l1 | -0.341388 | 0.048257 | 0.318230 | 1.316443 | 0.497260 |
| Gm4724 | -0.341246 | 0.000290 | 0.072211 | 3.537031 | 1.141399 |
| Gpr155 | -0.341238 | 0.039959 | 0.295412 | 1.398387 | 0.529572 |

| | | | | | |
|---|---|---|---|---|---|
| Lrch2 | -0.341181 | 0.012781 | 0.204978 | 1.893433 | 0.688293 |
| Cspp1 | -0.340974 | 0.031736 | 0.269762 | 1.498454 | 0.569020 |
| Cep97 | -0.340033 | 0.027461 | 0.257555 | 1.561282 | 0.589130 |
| Dppa3 | -0.339972 | 0.025713 | 0.250883 | 1.589842 | 0.600530 |
| Mkrn1 | -0.339845 | 0.019023 | 0.225156 | 1.720714 | 0.647516 |
| B3galt6 | -0.339408 | 0.022421 | 0.238845 | 1.649355 | 0.621884 |
| Surf6 | -0.339307 | 0.015409 | 0.213579 | 1.812228 | 0.670442 |
| Slk | -0.339206 | 0.014713 | 0.212541 | 1.832299 | 0.672557 |
| Cdh2 | -0.339137 | 0.001288 | 0.103883 | 2.890162 | 0.983457 |
| Chrna4 | -0.338255 | 0.028936 | 0.262944 | 1.538565 | 0.580137 |
| Marchf6 | -0.337995 | 0.005740 | 0.161414 | 2.241115 | 0.792060 |
| Adcy1 | -0.337972 | 0.039601 | 0.294619 | 1.402295 | 0.530739 |
| Htatip2 | -0.337515 | 0.040861 | 0.297878 | 1.388688 | 0.525961 |
| Tmem243 | -0.337308 | 0.037597 | 0.289624 | 1.424846 | 0.538165 |
| Zfp9 | -0.337038 | 0.047314 | 0.315038 | 1.325011 | 0.501637 |
| Lmf2 | -0.336807 | 0.018537 | 0.222795 | 1.731949 | 0.652095 |
| Ddx19a | -0.336734 | 0.009358 | 0.186226 | 2.028822 | 0.729959 |
| Ccl27b | -0.336510 | 0.015420 | 0.213579 | 1.811922 | 0.670442 |
| Zfp821 | -0.335823 | 0.004820 | 0.150323 | 2.316957 | 0.822975 |
| Slc35e4 | -0.335635 | 0.041765 | 0.300125 | 1.379192 | 0.522697 |
| Senp3 | -0.334811 | 0.029305 | 0.264012 | 1.533060 | 0.578376 |
| Jcad | -0.334445 | 0.043088 | 0.303009 | 1.365645 | 0.518544 |
| Tmem223 | -0.333854 | 0.010512 | 0.191032 | 1.978334 | 0.718893 |
| Mre11a | -0.333718 | 0.019976 | 0.227717 | 1.699497 | 0.642605 |
| Fancc | -0.333044 | 0.044803 | 0.307700 | 1.348692 | 0.511872 |
| Wdr48 | -0.332288 | 0.040983 | 0.297907 | 1.387397 | 0.525919 |
| Tmem270 | -0.331716 | 0.026243 | 0.253421 | 1.580994 | 0.596158 |
| Atxn7l1 | -0.331238 | 0.030945 | 0.268541 | 1.509415 | 0.570989 |
| Armt1 | -0.330994 | 0.045293 | 0.309429 | 1.343969 | 0.509439 |
| Tpst1 | -0.330293 | 0.018400 | 0.222669 | 1.735187 | 0.652340 |
| Nbeal1 | -0.329730 | 0.017455 | 0.219316 | 1.758076 | 0.658929 |
| Ncaph2 | -0.329201 | 0.008238 | 0.178185 | 2.084175 | 0.749130 |
| Ints14 | -0.328953 | 0.017315 | 0.219296 | 1.761589 | 0.658969 |
| Atp6v1g2 | -0.328278 | 0.049110 | 0.319909 | 1.308834 | 0.494973 |
| Cers6 | -0.328245 | 0.030597 | 0.267495 | 1.514324 | 0.572685 |
| 2700062CC | -0.327961 | 0.010817 | 0.192722 | 1.965876 | 0.715069 |
| Zfp646 | -0.327584 | 0.000258 | 0.071776 | 3.588617 | 1.144022 |
| Atg13 | -0.327567 | 0.038044 | 0.290716 | 1.419715 | 0.536531 |
| H2ac18 | -0.326898 | 0.036360 | 0.287007 | 1.439373 | 0.542108 |
| Slc35c2 | -0.325154 | 0.009452 | 0.186501 | 2.024486 | 0.729318 |
| Prkcd | -0.324301 | 0.042045 | 0.300611 | 1.376289 | 0.521996 |

| | | | | | |
|---|---|---|---|---|---|
| Gpn1 | -0.322949 | 0.002676 | 0.124096 | 2.572591 | 0.906240 |
| Pik3ca | -0.322782 | 0.043448 | 0.304589 | 1.362032 | 0.516286 |
| Plch1 | -0.322573 | 0.019404 | 0.225777 | 1.712106 | 0.646319 |
| Nr3c1 | -0.322439 | 0.031651 | 0.269725 | 1.499612 | 0.569079 |
| Man2b1 | -0.322375 | 0.043463 | 0.304589 | 1.361876 | 0.516286 |
| Grhl1 | -0.322113 | 0.022312 | 0.238730 | 1.651461 | 0.622092 |
| Cetn3 | -0.321407 | 0.006848 | 0.171274 | 2.164461 | 0.766308 |
| 4932438A1 | -0.318348 | 0.014287 | 0.210194 | 1.845059 | 0.677379 |
| Idh3b | -0.318016 | 0.044154 | 0.306387 | 1.355032 | 0.513729 |
| Snrk | -0.317662 | 0.015328 | 0.213387 | 1.814516 | 0.670832 |
| Zfp617 | -0.316585 | 0.001475 | 0.105319 | 2.831111 | 0.977493 |
| Arih2 | -0.316470 | 0.029261 | 0.264012 | 1.533708 | 0.578376 |
| Ip6k1 | -0.316378 | 0.019828 | 0.227717 | 1.702711 | 0.642605 |
| Defb39 | -0.315731 | 0.041089 | 0.298307 | 1.386274 | 0.525337 |
| Mau2 | -0.315394 | 0.026039 | 0.252148 | 1.584371 | 0.598345 |
| Mrap2 | -0.315046 | 0.049503 | 0.321112 | 1.305366 | 0.493343 |
| Greb1l | -0.313824 | 0.037539 | 0.289496 | 1.425512 | 0.538357 |
| Gm37416 | -0.312984 | 0.030692 | 0.267826 | 1.512981 | 0.572148 |
| Phf20l1 | -0.312189 | 0.042792 | 0.302156 | 1.368638 | 0.519769 |
| Snx11 | -0.309760 | 0.012385 | 0.202522 | 1.907095 | 0.693528 |
| Cep85l | -0.308961 | 0.046959 | 0.314232 | 1.328277 | 0.502750 |
| Castor2 | -0.308647 | 0.032727 | 0.272973 | 1.485092 | 0.563881 |
| Sh2d3c | -0.308273 | 0.041489 | 0.298907 | 1.382067 | 0.524464 |
| Chd5 | -0.307754 | 0.049893 | 0.322310 | 1.301959 | 0.491727 |
| Gpr19 | -0.307738 | 0.043914 | 0.305662 | 1.357398 | 0.514758 |
| Usp54 | -0.307171 | 0.049801 | 0.322310 | 1.302764 | 0.491727 |
| Atl1 | -0.306313 | 0.043069 | 0.303009 | 1.365832 | 0.518544 |
| Rnasel | -0.304504 | 0.017198 | 0.219296 | 1.764532 | 0.658969 |
| Pcdhgc3 | -0.304285 | 0.018086 | 0.221033 | 1.742666 | 0.655543 |
| Rasa1 | -0.304193 | 0.032219 | 0.270945 | 1.491890 | 0.567118 |
| Mbtps1 | -0.304039 | 0.049888 | 0.322310 | 1.302004 | 0.491727 |
| Dusp1 | -0.303680 | 0.041861 | 0.300125 | 1.378189 | 0.522697 |
| Ctdspl2 | -0.303652 | 0.042484 | 0.301310 | 1.371777 | 0.520986 |
| Zfp446 | -0.303602 | 0.038229 | 0.290930 | 1.417604 | 0.536211 |
| 4933434E2 | -0.303436 | 0.010696 | 0.192257 | 1.970792 | 0.716119 |
| Sprtn | -0.302881 | 0.043565 | 0.304845 | 1.360866 | 0.515921 |
| Zdhhc20 | -0.302133 | 0.044742 | 0.307668 | 1.349283 | 0.511917 |
| Speer4b | -0.301986 | 0.029210 | 0.263862 | 1.534468 | 0.578624 |
| Ddr1 | -0.301483 | 0.048776 | 0.319624 | 1.311793 | 0.495361 |
| Nsg2 | -0.301286 | 0.008843 | 0.182536 | 2.053412 | 0.738651 |
| Rab34 | -0.301267 | 0.027710 | 0.258240 | 1.557358 | 0.587976 |

| | | | | | |
|---|---|---|---|---|---|
| Armc1 | -0.300979 | 0.049487 | 0.321112 | 1.305511 | 0.493343 |
| Ppp2r2b | -0.300802 | 0.019952 | 0.227717 | 1.700020 | 0.642605 |
| Rmi2 | -0.300146 | 0.036424 | 0.287007 | 1.438608 | 0.542108 |
| Wdr82 | -0.299538 | 0.022270 | 0.238730 | 1.652280 | 0.622092 |
| Slc5a6 | -0.298987 | 0.013095 | 0.206018 | 1.882885 | 0.686094 |
| N4bp2 | -0.297371 | 0.008348 | 0.178880 | 2.078404 | 0.747439 |
| Sh2b1 | -0.296485 | 0.049139 | 0.319909 | 1.308570 | 0.494973 |
| Ift140 | -0.295797 | 0.007351 | 0.172342 | 2.133625 | 0.763610 |
| Gm52523 | -0.295284 | 0.027083 | 0.257318 | 1.567300 | 0.589530 |
| Glmn | -0.294767 | 0.000729 | 0.081735 | 3.137304 | 1.087593 |
| Kat6b | -0.294454 | 0.029721 | 0.265780 | 1.526939 | 0.575477 |
| Ctnnb1 | -0.293416 | 0.030006 | 0.265965 | 1.522797 | 0.575176 |
| Wdr1 | -0.293410 | 0.006664 | 0.169635 | 2.176271 | 0.770483 |
| Azi2 | -0.292755 | 0.021334 | 0.234691 | 1.670920 | 0.629503 |
| Mcf2l | -0.292654 | 0.041266 | 0.298605 | 1.384405 | 0.524902 |
| Gm52006 | -0.291871 | 0.030652 | 0.267795 | 1.513539 | 0.572197 |
| Fbxl5 | -0.291485 | 0.020845 | 0.232618 | 1.681003 | 0.633356 |
| Lasp1 | -0.291451 | 0.029461 | 0.264772 | 1.530756 | 0.577128 |
| Olfr1042 | -0.290807 | 0.043941 | 0.305662 | 1.357129 | 0.514758 |
| Zfp808 | -0.290214 | 0.006714 | 0.169993 | 2.173015 | 0.769568 |
| Zc4h2 | -0.289980 | 0.038506 | 0.291849 | 1.414477 | 0.534842 |
| Prpf38a | -0.287358 | 0.031321 | 0.268838 | 1.504159 | 0.570509 |
| Jak2 | -0.287225 | 0.048583 | 0.319066 | 1.313513 | 0.496119 |
| Gstm7 | -0.286463 | 0.021154 | 0.233851 | 1.674610 | 0.631060 |
| Trdmt1 | -0.286345 | 0.040629 | 0.297207 | 1.391159 | 0.526941 |
| Sufu | -0.285137 | 0.003776 | 0.138880 | 2.422992 | 0.857359 |
| Plcg1 | -0.283005 | 0.002808 | 0.126635 | 2.551528 | 0.897446 |
| Paip1 | -0.282774 | 0.031832 | 0.269885 | 1.497139 | 0.568821 |
| Limd2 | -0.282601 | 0.021737 | 0.237026 | 1.662803 | 0.625204 |
| Srsf11 | -0.282382 | 0.021860 | 0.237201 | 1.660353 | 0.624883 |
| Ndrg4 | -0.282145 | 0.017540 | 0.219316 | 1.755973 | 0.658929 |
| H4c17 | -0.279408 | 0.041860 | 0.300125 | 1.378196 | 0.522697 |
| Tmed4 | -0.279289 | 0.039161 | 0.293481 | 1.407146 | 0.532421 |
| Tnpo2 | -0.278354 | 0.013117 | 0.206018 | 1.882178 | 0.686094 |
| Tia1 | -0.278306 | 0.005357 | 0.156393 | 2.271042 | 0.805784 |
| Fli1 | -0.275475 | 0.047100 | 0.314475 | 1.326979 | 0.502414 |
| Nbea | -0.274219 | 0.039597 | 0.294619 | 1.402334 | 0.530739 |
| Septin10 | -0.272763 | 0.010152 | 0.190810 | 1.993449 | 0.719400 |
| Clca3b | -0.272581 | 0.014926 | 0.212593 | 1.826065 | 0.672451 |
| Rab31 | -0.271908 | 0.033927 | 0.277997 | 1.469457 | 0.555960 |
| Usf1 | -0.270889 | 0.011435 | 0.196420 | 1.941764 | 0.706814 |

| | | | | | |
|---|---|---|---|---|---|
| Ubac1 | -0.269886 | 0.031211 | 0.268609 | 1.505698 | 0.570879 |
| Crebbp | -0.268958 | 0.041871 | 0.300125 | 1.378091 | 0.522697 |
| Haus5 | -0.268673 | 0.046188 | 0.312437 | 1.335469 | 0.505237 |
| Setx | -0.268167 | 0.023786 | 0.243163 | 1.623678 | 0.614103 |
| Vmn1r117 | -0.267930 | 0.018462 | 0.222746 | 1.733714 | 0.652190 |
| Map3k11 | -0.267918 | 0.015672 | 0.214962 | 1.804881 | 0.667637 |
| Golgb1 | -0.267260 | 0.038663 | 0.292118 | 1.412708 | 0.534442 |
| Acadsb | -0.267120 | 0.026007 | 0.252004 | 1.584915 | 0.598592 |
| Dusp12 | -0.266898 | 0.005424 | 0.157361 | 2.265656 | 0.803104 |
| Zc3h14 | -0.266714 | 0.000578 | 0.076982 | 3.238387 | 1.113612 |
| Ndel1 | -0.265693 | 0.032272 | 0.271077 | 1.491172 | 0.566908 |
| Ythdf1 | -0.263481 | 0.005062 | 0.150961 | 2.295640 | 0.821135 |
| Ncstn | -0.261069 | 0.042742 | 0.302156 | 1.369148 | 0.519769 |
| Gm51591 | -0.260927 | 0.002710 | 0.125045 | 2.567101 | 0.902935 |
| B230307C2 | -0.259908 | 0.021898 | 0.237437 | 1.659587 | 0.624451 |
| Scai | -0.259031 | 0.048612 | 0.319066 | 1.313260 | 0.496119 |
| Sapcd1 | -0.258498 | 0.029742 | 0.265780 | 1.526626 | 0.575477 |
| Prrc2c | -0.258436 | 0.005098 | 0.151584 | 2.292577 | 0.819346 |
| Mapk7 | -0.257974 | 0.043271 | 0.303842 | 1.363805 | 0.517352 |
| Olfr906 | -0.257851 | 0.042693 | 0.302115 | 1.369647 | 0.519827 |

**Supplementary Table 6:Differentially expressed genes between PAE and SAC in Neurons at P28**

**Increased expression (1.2 FC, p<0.05)**

| Target nam | Log2 FC | Pvalue | Adjusted p | -log10 pval | -log10 adjusted pvalue |
|---|---|---|---|---|---|
| Alas2 | 1.464161 | 0.002094 | 0.862404 | 2.679034 | 0.064289 |
| Ros1 | 1.212300 | 0.027228 | 0.862404 | 1.564991 | 0.064289 |
| Gucy2f | 1.175750 | 0.008208 | 0.862404 | 2.085755 | 0.064289 |
| Slc51a | 1.161633 | 0.031196 | 0.862404 | 1.505897 | 0.064289 |
| Lcat | 1.159656 | 0.017292 | 0.862404 | 1.762148 | 0.064289 |
| Cldn11 | 1.102909 | 0.011515 | 0.862404 | 1.938731 | 0.064289 |
| Aknad1 | 1.070746 | 0.003594 | 0.862404 | 2.444417 | 0.064289 |
| Vcan | 1.070603 | 0.008667 | 0.862404 | 2.062144 | 0.064289 |
| Eif2ak2 | 1.069631 | 0.005947 | 0.862404 | 2.225731 | 0.064289 |
| Gfap | 1.057903 | 0.034372 | 0.862404 | 1.463797 | 0.064289 |
| Cd163 | 1.045297 | 0.042703 | 0.862404 | 1.369537 | 0.064289 |
| Olfr222 | 1.044241 | 0.012762 | 0.862404 | 1.894071 | 0.064289 |
| Apod | 1.033132 | 0.031531 | 0.862404 | 1.501267 | 0.064289 |
| Nde1 | 1.019598 | 0.017893 | 0.862404 | 1.747306 | 0.064289 |
| Mfsd4b5 | 1.001228 | 0.012653 | 0.862404 | 1.897808 | 0.064289 |
| Dab2 | 0.990186 | 0.017001 | 0.862404 | 1.769517 | 0.064289 |
| Bpi | 0.979266 | 0.024683 | 0.862404 | 1.607610 | 0.064289 |
| Tubb6 | 0.948378 | 0.021111 | 0.862404 | 1.675498 | 0.064289 |
| Otx1 | 0.945827 | 0.041773 | 0.862404 | 1.379108 | 0.064289 |
| Hhex | 0.937769 | 0.023642 | 0.862404 | 1.626312 | 0.064289 |
| Npsr1 | 0.929913 | 0.045005 | 0.862404 | 1.346743 | 0.064289 |
| Irx3 | 0.921947 | 0.022088 | 0.862404 | 1.655837 | 0.064289 |
| H2ac1 | 0.917459 | 0.020994 | 0.862404 | 1.677897 | 0.064289 |
| Sprr2d | 0.915373 | 0.009885 | 0.862404 | 2.005029 | 0.064289 |
| Cyp2j12 | 0.906941 | 0.019437 | 0.862404 | 1.711361 | 0.064289 |
| Tyw1 | 0.904968 | 0.033115 | 0.862404 | 1.479982 | 0.064289 |
| Syt2 | 0.901226 | 0.004086 | 0.862404 | 2.388709 | 0.064289 |
| Gm45935 | 0.896303 | 0.023778 | 0.862404 | 1.623827 | 0.064289 |
| Trpv6 | 0.892514 | 0.047671 | 0.862404 | 1.321747 | 0.064289 |
| Myo3b | 0.887795 | 0.039317 | 0.862404 | 1.405421 | 0.064289 |
| Gabrp | 0.881322 | 0.019162 | 0.862404 | 1.717551 | 0.064289 |
| Mag | 0.863243 | 0.018872 | 0.862404 | 1.724172 | 0.064289 |
| Otp | 0.858360 | 0.040001 | 0.862404 | 1.397928 | 0.064289 |
| St6galnac2 | 0.858176 | 0.049521 | 0.862404 | 1.305215 | 0.064289 |
| Lep | 0.851362 | 0.022363 | 0.862404 | 1.650464 | 0.064289 |
| Gm7102 | 0.847270 | 0.046079 | 0.862404 | 1.336499 | 0.064289 |
| Vangl1 | 0.842815 | 0.012240 | 0.862404 | 1.912203 | 0.064289 |

**Decreased expression (1.2 FC, p<0.05)**

| Target nam | Log2 FC | Pvalue | Adjusted p | -log10 pval | -log10 adjusted pvalue |
|---|---|---|---|---|---|
| Btg2 | -1.115168 | 0.017804 | 0.862404 | 1.749483 | 0.064289 |
| Ubac1 | -0.755235 | 0.037163 | 0.862404 | 1.429891 | 0.064289 |
| Cd109 | -0.715350 | 0.021586 | 0.862404 | 1.665834 | 0.064289 |
| Clspn | -0.689672 | 0.024894 | 0.862404 | 1.603908 | 0.064289 |
| Tbx1 | -0.680354 | 0.027921 | 0.862404 | 1.554062 | 0.064289 |
| Igfbp2 | -0.656955 | 0.025204 | 0.862404 | 1.598526 | 0.064289 |
| Arc | -0.635887 | 0.013960 | 0.862404 | 1.855127 | 0.064289 |
| Rfx4 | -0.610861 | 0.028473 | 0.862404 | 1.545569 | 0.064289 |
| Gm51415 | -0.592091 | 0.033512 | 0.862404 | 1.474804 | 0.064289 |
| Dhx37 | -0.587398 | 0.012027 | 0.862404 | 1.919840 | 0.064289 |
| Sfmbt1 | -0.579695 | 0.031458 | 0.862404 | 1.502268 | 0.064289 |
| Ier2 | -0.576682 | 0.049662 | 0.862404 | 1.303975 | 0.064289 |
| Phkb | -0.576416 | 0.030567 | 0.862404 | 1.514743 | 0.064289 |
| Raph1 | -0.556298 | 0.015270 | 0.862404 | 1.816173 | 0.064289 |
| 1700037C1 | -0.555965 | 0.000932 | 0.862404 | 3.030677 | 0.064289 |
| Gpr88 | -0.553222 | 0.043162 | 0.862404 | 1.364897 | 0.064289 |
| L1td1 | -0.553016 | 0.013554 | 0.862404 | 1.867941 | 0.064289 |
| Dusp1 | -0.552265 | 0.035982 | 0.862404 | 1.443918 | 0.064289 |
| Slc12a4 | -0.548989 | 0.021096 | 0.862404 | 1.675807 | 0.064289 |
| Elk3 | -0.508154 | 0.041652 | 0.862404 | 1.380361 | 0.064289 |
| Cntnap5c | -0.503836 | 0.017441 | 0.862404 | 1.758417 | 0.064289 |
| Rad50 | -0.502490 | 0.003876 | 0.862404 | 2.411662 | 0.064289 |
| Stag2 | -0.495570 | 0.013537 | 0.862404 | 1.868464 | 0.064289 |
| Zc3h3 | -0.493430 | 0.048942 | 0.862404 | 1.310321 | 0.064289 |
| C2cd3 | -0.486630 | 0.032944 | 0.862404 | 1.482222 | 0.064289 |
| Pex2 | -0.476976 | 0.019001 | 0.862404 | 1.721216 | 0.064289 |
| Fos | -0.470462 | 0.034433 | 0.862404 | 1.463021 | 0.064289 |
| H1f1 | -0.463630 | 0.020856 | 0.862404 | 1.680765 | 0.064289 |
| Dync2h1 | -0.461815 | 0.012480 | 0.862404 | 1.903787 | 0.064289 |
| Melk | -0.457486 | 0.003025 | 0.862404 | 2.519214 | 0.064289 |
| Tek | -0.449053 | 0.031632 | 0.862404 | 1.499876 | 0.064289 |
| Echdc2 | -0.442305 | 0.040675 | 0.862404 | 1.390674 | 0.064289 |
| Bmp1 | -0.438897 | 0.047375 | 0.862404 | 1.324449 | 0.064289 |
| Arntl | -0.438873 | 0.005797 | 0.862404 | 2.236760 | 0.064289 |
| Usp40 | -0.436476 | 0.027048 | 0.862404 | 1.567870 | 0.064289 |
| Cebpzos | -0.427120 | 0.018182 | 0.862404 | 1.740347 | 0.064289 |
| Kank3 | -0.426993 | 0.038985 | 0.862404 | 1.409106 | 0.064289 |

| | | | | | |
|---|---|---|---|---|---|
| Rsph4a | 0.842006 | 0.014303 | 0.862404 | 1.844582 | 0.064289 |
| Cxcr3 | 0.841011 | 0.017360 | 0.862404 | 1.760459 | 0.064289 |
| Tpx2 | 0.837237 | 0.035618 | 0.862404 | 1.448327 | 0.064289 |
| Prr27 | 0.836519 | 0.002369 | 0.862404 | 2.625525 | 0.064289 |
| Tent5c | 0.829781 | 0.017220 | 0.862404 | 1.763976 | 0.064289 |
| Flt3 | 0.817606 | 0.010852 | 0.862404 | 1.964495 | 0.064289 |
| 2200002D( | 0.813779 | 0.017708 | 0.862404 | 1.751830 | 0.064289 |
| Slc15a3 | 0.812292 | 0.025656 | 0.862404 | 1.590803 | 0.064289 |
| Dnah9 | 0.807594 | 0.031705 | 0.862404 | 1.498878 | 0.064289 |
| Slurp2 | 0.803508 | 0.036533 | 0.862404 | 1.437317 | 0.064289 |
| Psg27 | 0.800490 | 0.001880 | 0.862404 | 2.725894 | 0.064289 |
| Ms4a10 | 0.799715 | 0.011766 | 0.862404 | 1.929367 | 0.064289 |
| Cbln1 | 0.795382 | 0.048192 | 0.862404 | 1.317027 | 0.064289 |
| Ubash3b | 0.793651 | 0.005006 | 0.862404 | 2.300540 | 0.064289 |
| Gdf9 | 0.791171 | 0.017796 | 0.862404 | 1.749674 | 0.064289 |
| Mitf | 0.790656 | 0.020320 | 0.862404 | 1.692080 | 0.064289 |
| Prdm6 | 0.788854 | 0.008240 | 0.862404 | 2.084069 | 0.064289 |
| Lhx4 | 0.788452 | 0.005340 | 0.862404 | 2.272430 | 0.064289 |
| Chil6 | 0.785359 | 0.008216 | 0.862404 | 2.085331 | 0.064289 |
| Trf | 0.783416 | 0.035269 | 0.862404 | 1.452605 | 0.064289 |
| Clec2j | 0.776658 | 0.012631 | 0.862404 | 1.898576 | 0.064289 |
| Cd82 | 0.773380 | 0.014576 | 0.862404 | 1.836355 | 0.064289 |
| Lcp1 | 0.769130 | 0.047400 | 0.862404 | 1.324221 | 0.064289 |
| Pgr | 0.768618 | 0.024809 | 0.862404 | 1.605387 | 0.064289 |
| Ankrd29 | 0.764231 | 0.037816 | 0.862404 | 1.422322 | 0.064289 |
| Casq2 | 0.762365 | 0.001145 | 0.862404 | 2.941126 | 0.064289 |
| Armcx6 | 0.762020 | 0.029904 | 0.862404 | 1.524272 | 0.064289 |
| Olfr1126 | 0.760906 | 0.005253 | 0.862404 | 2.279570 | 0.064289 |
| Scml2 | 0.759280 | 0.016853 | 0.862404 | 1.773319 | 0.064289 |
| Ptx4 | 0.758647 | 0.020610 | 0.862404 | 1.685929 | 0.064289 |
| Nacc2 | 0.755413 | 0.002216 | 0.862404 | 2.654358 | 0.064289 |
| Lilr4b | 0.747219 | 0.033768 | 0.862404 | 1.471496 | 0.064289 |
| Fgf5 | 0.744214 | 0.040701 | 0.862404 | 1.390396 | 0.064289 |
| Mtcp1 | 0.742842 | 0.039284 | 0.862404 | 1.405783 | 0.064289 |
| Clmp | 0.742793 | 0.003053 | 0.862404 | 2.515297 | 0.064289 |
| Fzd6 | 0.738479 | 0.047340 | 0.862404 | 1.324773 | 0.064289 |
| Gdnf | 0.734292 | 0.035298 | 0.862404 | 1.452244 | 0.064289 |
| Dsc1 | 0.733640 | 0.019838 | 0.862404 | 1.702512 | 0.064289 |
| Olfr1085 | 0.731638 | 0.010842 | 0.862404 | 1.964898 | 0.064289 |
| Etd | 0.727172 | 0.007718 | 0.862404 | 2.112502 | 0.064289 |
| Olfr644 | 0.718923 | 0.044680 | 0.862404 | 1.349882 | 0.064289 |

| | | | | | |
|---|---|---|---|---|---|
| Cfap52 | -0.426444 | 0.015205 | 0.862404 | 1.818010 | 0.064289 |
| Adh5 | -0.416279 | 0.002404 | 0.862404 | 2.619109 | 0.064289 |
| Ccdc112 | -0.414128 | 0.042553 | 0.862404 | 1.371071 | 0.064289 |
| Vldlr | -0.413218 | 0.017732 | 0.862404 | 1.751237 | 0.064289 |
| Dcaf13 | -0.411191 | 0.001824 | 0.862404 | 2.739079 | 0.064289 |
| Nr4a1 | -0.410623 | 0.042409 | 0.862404 | 1.372545 | 0.064289 |
| Dkc1 | -0.405317 | 0.013393 | 0.862404 | 1.873109 | 0.064289 |
| Fosb | -0.400238 | 0.044227 | 0.862404 | 1.354312 | 0.064289 |
| Fst | -0.399571 | 0.013970 | 0.862404 | 1.854813 | 0.064289 |
| Cutc | -0.398385 | 0.012386 | 0.862404 | 1.907065 | 0.064289 |
| Dipk2a | -0.396943 | 0.027895 | 0.862404 | 1.554467 | 0.064289 |
| Acbd5 | -0.396866 | 0.023002 | 0.862404 | 1.638227 | 0.064289 |
| Tprkb | -0.391930 | 0.006290 | 0.862404 | 2.201336 | 0.064289 |
| Rras2 | -0.390940 | 0.020350 | 0.862404 | 1.691432 | 0.064289 |
| Tle1 | -0.388465 | 0.029678 | 0.862404 | 1.527569 | 0.064289 |
| Anapc15 | -0.387347 | 0.029660 | 0.862404 | 1.527829 | 0.064289 |
| Rab18 | -0.386970 | 0.013825 | 0.862404 | 1.859342 | 0.064289 |
| Fam122a | -0.380385 | 0.009820 | 0.862404 | 2.007876 | 0.064289 |
| Ubap2l | -0.377953 | 0.003507 | 0.862404 | 2.455109 | 0.064289 |
| Gal3st2 | -0.373871 | 0.045324 | 0.862404 | 1.343673 | 0.064289 |
| Ncoa6 | -0.371812 | 0.006459 | 0.862404 | 2.189807 | 0.064289 |
| Shq1 | -0.370021 | 0.009950 | 0.862404 | 2.002198 | 0.064289 |
| Mss51 | -0.367513 | 0.043572 | 0.862404 | 1.360793 | 0.064289 |
| Mettl8 | -0.363013 | 0.044418 | 0.862404 | 1.352443 | 0.064289 |
| Spaca6 | -0.360630 | 0.017266 | 0.862404 | 1.762809 | 0.064289 |
| Pmpcb | -0.358750 | 0.020062 | 0.862404 | 1.697633 | 0.064289 |
| Atg13 | -0.357997 | 0.005241 | 0.862404 | 2.280571 | 0.064289 |
| Anapc10 | -0.356133 | 0.045293 | 0.862404 | 1.343971 | 0.064289 |
| Atf7ip | -0.354049 | 0.003521 | 0.862404 | 2.453288 | 0.064289 |
| Fam57a | -0.351250 | 0.030116 | 0.862404 | 1.521197 | 0.064289 |
| Nt5c2 | -0.349989 | 0.026058 | 0.862404 | 1.584057 | 0.064289 |
| Epc2 | -0.348641 | 0.045981 | 0.862404 | 1.337422 | 0.064289 |
| Prpsap2 | -0.348569 | 0.040542 | 0.862404 | 1.392094 | 0.064289 |
| Hirip3 | -0.343160 | 0.033913 | 0.862404 | 1.469636 | 0.064289 |
| Rsl1d1 | -0.337624 | 0.049619 | 0.862404 | 1.304353 | 0.064289 |
| Fam91a1 | -0.337111 | 0.030084 | 0.862404 | 1.521659 | 0.064289 |
| Fyco1 | -0.335703 | 0.045918 | 0.862404 | 1.338015 | 0.064289 |
| Lrrc57 | -0.335662 | 0.015495 | 0.862404 | 1.809812 | 0.064289 |
| Ivns1abp | -0.335454 | 0.000879 | 0.862404 | 3.055972 | 0.064289 |
| Trerf1 | -0.330239 | 0.037796 | 0.862404 | 1.422557 | 0.064289 |
| Tbl1x | -0.330005 | 0.024944 | 0.862404 | 1.603037 | 0.064289 |

| | | | | | |
|---|---|---|---|---|---|
| Scn4b | 0.718510 | 0.013867 | 0.862404 | 1.858009 | 0.064289 |
| Zap70 | 0.717158 | 0.006446 | 0.862404 | 2.190691 | 0.064289 |
| Gjb1 | 0.715638 | 0.016575 | 0.862404 | 1.780547 | 0.064289 |
| Adamts7 | 0.714899 | 0.029718 | 0.862404 | 1.526982 | 0.064289 |
| Stmnd1 | 0.714616 | 0.034763 | 0.862404 | 1.458878 | 0.064289 |
| Pvalb | 0.714220 | 0.003838 | 0.862404 | 2.415867 | 0.064289 |
| Nusap1 | 0.713134 | 0.021600 | 0.862404 | 1.665556 | 0.064289 |
| Dmrta2 | 0.709816 | 0.008227 | 0.862404 | 2.084776 | 0.064289 |
| Fads6 | 0.708612 | 0.013531 | 0.862404 | 1.868683 | 0.064289 |
| Arx | 0.704983 | 0.021090 | 0.862404 | 1.675913 | 0.064289 |
| 4930564D( | 0.704480 | 0.043597 | 0.862404 | 1.360548 | 0.064289 |
| Faap24 | 0.702428 | 0.047121 | 0.862404 | 1.326789 | 0.064289 |
| Dpep2 | 0.695787 | 0.016491 | 0.862404 | 1.782746 | 0.064289 |
| Slc12a7 | 0.695164 | 0.032806 | 0.862404 | 1.484052 | 0.064289 |
| P2ry14 | 0.694454 | 0.028717 | 0.862404 | 1.541866 | 0.064289 |
| Krtap22-2 | 0.693854 | 0.029816 | 0.862404 | 1.525551 | 0.064289 |
| Slc32a1 | 0.688654 | 0.008558 | 0.862404 | 2.067610 | 0.064289 |
| Prkra | 0.687806 | 0.011172 | 0.862404 | 1.951852 | 0.064289 |
| Sowahd | 0.687445 | 0.040687 | 0.862404 | 1.390547 | 0.064289 |
| Shc4 | 0.687235 | 0.011740 | 0.862404 | 1.930314 | 0.064289 |
| Cpa1 | 0.685939 | 0.025157 | 0.862404 | 1.599342 | 0.064289 |
| Rassf10 | 0.682443 | 0.007286 | 0.862404 | 2.137535 | 0.064289 |
| Ano5 | 0.681912 | 0.035584 | 0.862404 | 1.448741 | 0.064289 |
| Dhrs3 | 0.681075 | 0.008009 | 0.862404 | 2.096426 | 0.064289 |
| Gimap4 | 0.679174 | 0.006452 | 0.862404 | 2.190282 | 0.064289 |
| Itm2a | 0.676617 | 0.022173 | 0.862404 | 1.654181 | 0.064289 |
| Bcan | 0.673043 | 0.027759 | 0.862404 | 1.556604 | 0.064289 |
| Ide | 0.670673 | 0.014757 | 0.862404 | 1.830990 | 0.064289 |
| Olfr1408 | 0.669922 | 0.000280 | 0.862404 | 3.553436 | 0.064289 |
| Gabrg1 | 0.669064 | 0.028490 | 0.862404 | 1.545301 | 0.064289 |
| Fv1 | 0.668390 | 0.002431 | 0.862404 | 2.614270 | 0.064289 |
| Spry3 | 0.665951 | 0.044229 | 0.862404 | 1.354294 | 0.064289 |
| Adgrg3 | 0.662089 | 0.021313 | 0.862404 | 1.671358 | 0.064289 |
| Fgf11 | 0.660664 | 0.008730 | 0.862404 | 2.058985 | 0.064289 |
| Prl5a1 | 0.655797 | 0.049893 | 0.862404 | 1.301956 | 0.064289 |
| Gpr25 | 0.650733 | 0.007779 | 0.862404 | 2.109085 | 0.064289 |
| Sema5b | 0.649368 | 0.029952 | 0.862404 | 1.523571 | 0.064289 |
| Lyplal1 | 0.647222 | 0.031039 | 0.862404 | 1.508092 | 0.064289 |
| Thap2 | 0.646969 | 0.024743 | 0.862404 | 1.606543 | 0.064289 |
| F2 | 0.646023 | 0.011689 | 0.862404 | 1.932210 | 0.064289 |
| Slfnl1 | 0.643305 | 0.030607 | 0.862404 | 1.514183 | 0.064289 |

| | | | | | |
|---|---|---|---|---|---|
| Fbh1 | -0.326590 | 0.048122 | 0.862404 | 1.317653 | 0.064289 |
| Srsf4 | -0.325805 | 0.048225 | 0.862404 | 1.316732 | 0.064289 |
| Actn1 | -0.320195 | 0.040159 | 0.862404 | 1.396221 | 0.064289 |
| Zscan18 | -0.319941 | 0.036324 | 0.862404 | 1.439801 | 0.064289 |
| Uqcr10 | -0.319302 | 0.009229 | 0.862404 | 2.034830 | 0.064289 |
| Vps52 | -0.314887 | 0.030422 | 0.862404 | 1.516812 | 0.064289 |
| Asnsd1 | -0.314014 | 0.035036 | 0.862404 | 1.455491 | 0.064289 |
| Rex1bd | -0.314013 | 0.038006 | 0.862404 | 1.420153 | 0.064289 |
| Ankrd24 | -0.312288 | 0.028449 | 0.862404 | 1.545939 | 0.064289 |
| Iglon5 | -0.312157 | 0.032743 | 0.862404 | 1.484888 | 0.064289 |
| Selenoo | -0.310303 | 0.022759 | 0.862404 | 1.642838 | 0.064289 |
| Samd8 | -0.308686 | 0.019945 | 0.862404 | 1.700167 | 0.064289 |
| H1f4 | -0.308448 | 0.025206 | 0.862404 | 1.598501 | 0.064289 |
| Zfp160 | -0.305010 | 0.044176 | 0.862404 | 1.354812 | 0.064289 |
| Pced1a | -0.304710 | 0.025574 | 0.862404 | 1.592202 | 0.064289 |
| Prkab2 | -0.302125 | 0.014543 | 0.862404 | 1.837356 | 0.064289 |
| Pias1 | -0.301950 | 0.047122 | 0.862404 | 1.326775 | 0.064289 |
| Usp20 | -0.301210 | 0.039329 | 0.862404 | 1.405287 | 0.064289 |
| Esd | -0.300918 | 0.011220 | 0.862404 | 1.949989 | 0.064289 |
| Ube2e2 | -0.297031 | 0.032038 | 0.862404 | 1.494337 | 0.064289 |
| Dcaf6 | -0.294910 | 0.045498 | 0.862404 | 1.342009 | 0.064289 |
| Pde6d | -0.293908 | 0.027978 | 0.862404 | 1.553183 | 0.064289 |
| Stx6 | -0.288049 | 0.004512 | 0.862404 | 2.345586 | 0.064289 |
| H2bc8\|H2b | -0.287856 | 0.034790 | 0.862404 | 1.458541 | 0.064289 |
| Rab5c | -0.287745 | 0.049439 | 0.862404 | 1.305933 | 0.064289 |
| Cdk17 | -0.286687 | 0.042076 | 0.862404 | 1.375971 | 0.064289 |
| Tbl1xr1 | -0.285432 | 0.028551 | 0.862404 | 1.544383 | 0.064289 |
| Tufm | -0.285369 | 0.035549 | 0.862404 | 1.449176 | 0.064289 |
| Armc8 | -0.285138 | 0.046244 | 0.862404 | 1.334943 | 0.064289 |
| Rnf40 | -0.284986 | 0.010727 | 0.862404 | 1.969534 | 0.064289 |
| Dbpht2 | -0.283477 | 0.000130 | 0.672561 | 3.886430 | 0.172268 |
| Sptan1 | -0.275667 | 0.036086 | 0.862404 | 1.442665 | 0.064289 |
| Apopt1 | -0.270790 | 0.025606 | 0.862404 | 1.591652 | 0.064289 |
| Nr2c2 | -0.270106 | 0.021950 | 0.862404 | 1.658560 | 0.064289 |
| Mnat1 | -0.269551 | 0.027612 | 0.862404 | 1.558907 | 0.064289 |
| Bccip | -0.268637 | 0.025135 | 0.862404 | 1.599717 | 0.064289 |
| Psmb6 | -0.266850 | 0.011188 | 0.862404 | 1.951233 | 0.064289 |
| Arid1a | -0.266708 | 0.047212 | 0.862404 | 1.325951 | 0.064289 |
| Dync1li2 | -0.263411 | 0.041989 | 0.862404 | 1.376861 | 0.064289 |
| Msantd4 | -0.262954 | 0.048916 | 0.862404 | 1.310545 | 0.064289 |
| Supt5 | -0.260142 | 0.013317 | 0.862404 | 1.875583 | 0.064289 |

| | | | | | |
|---|---|---|---|---|---|
| Cmtm5 | 0.643085 | 0.022964 | 0.862404 | 1.638957 | 0.064289 |
| Nufip1 | 0.641276 | 0.020918 | 0.862404 | 1.679489 | 0.064289 |
| Spink10 | 0.636741 | 0.008334 | 0.862404 | 2.079150 | 0.064289 |
| Rnaseh1 | 0.636521 | 0.012428 | 0.862404 | 1.905583 | 0.064289 |
| Mrgpra9 | 0.630621 | 0.021769 | 0.862404 | 1.662158 | 0.064289 |
| Mid2 | 0.630502 | 0.025211 | 0.862404 | 1.598410 | 0.064289 |
| Gm46487 | 0.629961 | 0.004548 | 0.862404 | 2.342167 | 0.064289 |
| Ctrb1 | 0.629006 | 0.035941 | 0.862404 | 1.444414 | 0.064289 |
| Gipr | 0.628988 | 0.002451 | 0.862404 | 2.610610 | 0.064289 |
| Cybc1 | 0.628824 | 0.043711 | 0.862404 | 1.359406 | 0.064289 |
| Tacc3 | 0.626942 | 0.011791 | 0.862404 | 1.928446 | 0.064289 |
| Slc22a8 | 0.626506 | 0.018768 | 0.862404 | 1.726587 | 0.064289 |
| Sema6d | 0.625870 | 0.035407 | 0.862404 | 1.450909 | 0.064289 |
| Rnd2 | 0.624903 | 0.001820 | 0.862404 | 2.739955 | 0.064289 |
| Stk3 | 0.624577 | 0.013695 | 0.862404 | 1.863439 | 0.064289 |
| Mmrn2 | 0.615163 | 0.048331 | 0.862404 | 1.315771 | 0.064289 |
| Siah3 | 0.614209 | 0.021711 | 0.862404 | 1.663319 | 0.064289 |
| Gm10334 | 0.613356 | 0.044371 | 0.862404 | 1.352898 | 0.064289 |
| Psg26 | 0.612422 | 0.041479 | 0.862404 | 1.382176 | 0.064289 |
| Alkbh2 | 0.607763 | 0.013222 | 0.862404 | 1.878689 | 0.064289 |
| Lrrd1 | 0.605503 | 0.023799 | 0.862404 | 1.623446 | 0.064289 |
| Mt2 | 0.605324 | 0.039276 | 0.862404 | 1.405868 | 0.064289 |
| Gdf10 | 0.604436 | 0.022708 | 0.862404 | 1.643815 | 0.064289 |
| Fam78a | 0.604129 | 0.047894 | 0.862404 | 1.319715 | 0.064289 |
| 1110032F0 | 0.603427 | 0.006192 | 0.862404 | 2.208163 | 0.064289 |
| Fndc3b | 0.601992 | 0.035103 | 0.862404 | 1.454652 | 0.064289 |
| Lmln | 0.600806 | 0.001713 | 0.862404 | 2.766196 | 0.064289 |
| Ghr | 0.597673 | 0.016195 | 0.862404 | 1.790623 | 0.064289 |
| Col1a2 | 0.597159 | 0.041133 | 0.862404 | 1.385804 | 0.064289 |
| Pik3c2b | 0.596590 | 0.012275 | 0.862404 | 1.910967 | 0.064289 |
| Srd5a3 | 0.596346 | 0.040280 | 0.862404 | 1.394907 | 0.064289 |
| Dysf | 0.596104 | 0.045290 | 0.862404 | 1.344002 | 0.064289 |
| Olfr799 | 0.591468 | 0.039566 | 0.862404 | 1.402682 | 0.064289 |
| Esp18 | 0.585296 | 0.034479 | 0.862404 | 1.462450 | 0.064289 |
| 4921507P0 | 0.584126 | 0.022619 | 0.862404 | 1.645531 | 0.064289 |
| Olfr1105 | 0.577756 | 0.042124 | 0.862404 | 1.375474 | 0.064289 |
| Nox4 | 0.575461 | 0.047492 | 0.862404 | 1.323381 | 0.064289 |
| Lce1b | 0.573839 | 0.038136 | 0.862404 | 1.418666 | 0.064289 |
| Rhbdl2 | 0.572656 | 0.034449 | 0.862404 | 1.462821 | 0.064289 |
| Tpst1 | 0.572267 | 0.036504 | 0.862404 | 1.437661 | 0.064289 |
| Spata6 | 0.571278 | 0.007971 | 0.862404 | 2.098488 | 0.064289 |
| Pard6a | -0.259205 | 0.047801 | 0.862404 | 1.320559 | 0.064289 |
| Trafd1 | -0.259065 | 0.035644 | 0.862404 | 1.448013 | 0.064289 |
| Smarcb1 | -0.257911 | 0.030925 | 0.862404 | 1.509686 | 0.064289 |

| | | | | | |
|---|---|---|---|---|---|
| Rbp4 | 0.570234 | 0.003071 | 0.862404 | 2.512690 | 0.064289 |
| F12 | 0.569770 | 0.016202 | 0.862404 | 1.790443 | 0.064289 |
| Myo5c | 0.569375 | 0.017059 | 0.862404 | 1.768053 | 0.064289 |
| Ucp2 | 0.565746 | 0.043039 | 0.862404 | 1.366136 | 0.064289 |
| Mmp23 | 0.565467 | 0.008448 | 0.862404 | 2.073271 | 0.064289 |
| Klhl33 | 0.561755 | 0.029089 | 0.862404 | 1.536266 | 0.064289 |
| Selenop | 0.561301 | 0.000948 | 0.862404 | 3.023221 | 0.064289 |
| Smok3c | 0.560663 | 0.032985 | 0.862404 | 1.481687 | 0.064289 |
| Fam13a | 0.556753 | 0.028211 | 0.862404 | 1.549579 | 0.064289 |
| Slc27a5 | 0.555910 | 0.035419 | 0.862404 | 1.450758 | 0.064289 |
| Ifi27 | 0.555549 | 0.016214 | 0.862404 | 1.790106 | 0.064289 |
| Armc2 | 0.554730 | 0.023989 | 0.862404 | 1.619988 | 0.064289 |
| Gpr173 | 0.554462 | 0.047018 | 0.862404 | 1.327733 | 0.064289 |
| Ccdc18 | 0.551886 | 0.029464 | 0.862404 | 1.530707 | 0.064289 |
| Rad51c | 0.546981 | 0.021079 | 0.862404 | 1.676152 | 0.064289 |
| Sptssb | 0.544838 | 0.019611 | 0.862404 | 1.707491 | 0.064289 |
| Utp14b | 0.543374 | 0.044666 | 0.862404 | 1.350022 | 0.064289 |
| Plpp1 | 0.540487 | 0.029883 | 0.862404 | 1.524578 | 0.064289 |
| Dhfr | 0.539955 | 0.002602 | 0.862404 | 2.584709 | 0.064289 |
| Slx4 | 0.535474 | 0.036672 | 0.862404 | 1.435663 | 0.064289 |
| Trmt5 | 0.533656 | 0.035823 | 0.862404 | 1.445844 | 0.064289 |
| Zkscan4 | 0.533458 | 0.031868 | 0.862404 | 1.496643 | 0.064289 |
| Gtf2h4 | 0.530907 | 0.023717 | 0.862404 | 1.624938 | 0.064289 |
| Vmn1r8 | 0.529393 | 0.007309 | 0.862404 | 2.136152 | 0.064289 |
| Irf8 | 0.529234 | 0.045347 | 0.862404 | 1.343453 | 0.064289 |
| Rhbdf2 | 0.529134 | 0.002372 | 0.862404 | 2.624818 | 0.064289 |
| Grik1 | 0.526859 | 0.030055 | 0.862404 | 1.522078 | 0.064289 |
| Trim25 | 0.526281 | 0.016076 | 0.862404 | 1.793818 | 0.064289 |
| Myo1e | 0.525885 | 0.032228 | 0.862404 | 1.491767 | 0.064289 |
| Cdx2 | 0.523262 | 0.019093 | 0.862404 | 1.719130 | 0.064289 |
| Aqp4 | 0.522171 | 0.043245 | 0.862404 | 1.364066 | 0.064289 |
| Xrcc5 | 0.521468 | 0.007018 | 0.862404 | 2.153783 | 0.064289 |
| Mdm1 | 0.520048 | 0.014090 | 0.862404 | 1.851086 | 0.064289 |
| Simc1 | 0.516541 | 0.036308 | 0.862404 | 1.439995 | 0.064289 |
| Mis12 | 0.515067 | 0.032346 | 0.862404 | 1.490186 | 0.064289 |
| Spin2c | 0.514177 | 0.028835 | 0.862404 | 1.540084 | 0.064289 |
| Enox2 | 0.512270 | 0.005098 | 0.862404 | 2.292635 | 0.064289 |
| Gm6358 | 0.509161 | 0.020234 | 0.862404 | 1.693928 | 0.064289 |
| Fat1 | 0.507210 | 0.002306 | 0.862404 | 2.637120 | 0.064289 |
| Nt5dc1 | 0.506085 | 0.045584 | 0.862404 | 1.341188 | 0.064289 |
| Htr1f | 0.504739 | 0.021096 | 0.862404 | 1.675798 | 0.064289 |

| | | | | | |
|---|---|---|---|---|---|
| Kcnn3 | 0.503675 | 0.046636 | 0.862404 | 1.331280 | 0.064289 |
| Dpy19l4 | 0.503540 | 0.022655 | 0.862404 | 1.644844 | 0.064289 |
| Klrg2 | 0.502788 | 0.020519 | 0.862404 | 1.687852 | 0.064289 |
| Abca14 | 0.501484 | 0.023402 | 0.862404 | 1.630739 | 0.064289 |
| tTA/rtTA | 0.501449 | 0.012187 | 0.862404 | 1.914107 | 0.064289 |
| Adgra3 | 0.494516 | 0.042258 | 0.862404 | 1.374094 | 0.064289 |
| Ebi3 | 0.488992 | 0.048033 | 0.862404 | 1.318457 | 0.064289 |
| Adam8 | 0.488471 | 0.000702 | 0.862404 | 3.153705 | 0.064289 |
| Col14a1 | 0.484646 | 0.039770 | 0.862404 | 1.400445 | 0.064289 |
| Gm5414 | 0.484412 | 0.048321 | 0.862404 | 1.315864 | 0.064289 |
| Aff2 | 0.483836 | 0.044605 | 0.862404 | 1.350620 | 0.064289 |
| Ripply1 | 0.483301 | 0.027408 | 0.862404 | 1.562115 | 0.064289 |
| Golim4 | 0.481616 | 0.047509 | 0.862404 | 1.323223 | 0.064289 |
| Bdkrb1 | 0.478925 | 0.006960 | 0.862404 | 2.157399 | 0.064289 |
| Zhx2 | 0.476881 | 0.038746 | 0.862404 | 1.411772 | 0.064289 |
| Speer2 | 0.476006 | 0.000875 | 0.862404 | 3.058052 | 0.064289 |
| Hsd17b7 | 0.474362 | 0.006972 | 0.862404 | 2.156638 | 0.064289 |
| Gsn | 0.472689 | 0.030620 | 0.862404 | 1.513999 | 0.064289 |
| Gm8909 | 0.470290 | 0.022068 | 0.862404 | 1.656238 | 0.064289 |
| Zkscan14 | 0.466969 | 0.033510 | 0.862404 | 1.474828 | 0.064289 |
| Neu2 | 0.465632 | 0.005090 | 0.862404 | 2.293280 | 0.064289 |
| Ccdc166 | 0.462324 | 0.029654 | 0.862404 | 1.527911 | 0.064289 |
| Gm40755 | 0.460244 | 0.040560 | 0.862404 | 1.391899 | 0.064289 |
| Gng5 | 0.459688 | 0.046669 | 0.862404 | 1.330970 | 0.064289 |
| Zscan12 | 0.456051 | 0.026463 | 0.862404 | 1.577357 | 0.064289 |
| Ssc5d | 0.455976 | 0.030264 | 0.862404 | 1.519071 | 0.064289 |
| Zfp217 | 0.455649 | 0.025029 | 0.862404 | 1.601553 | 0.064289 |
| Mybphl | 0.450516 | 0.025638 | 0.862404 | 1.591123 | 0.064289 |
| Ube2d2b | 0.450394 | 0.030723 | 0.862404 | 1.512538 | 0.064289 |
| Entpd5 | 0.446562 | 0.018415 | 0.862404 | 1.734817 | 0.064289 |
| Tfb1m | 0.445560 | 0.031955 | 0.862404 | 1.495457 | 0.064289 |
| Mlkl | 0.442926 | 0.034491 | 0.862404 | 1.462295 | 0.064289 |
| Ficd | 0.442158 | 0.043340 | 0.862404 | 1.363114 | 0.064289 |
| Zfp7 | 0.441898 | 0.044765 | 0.862404 | 1.349061 | 0.064289 |
| Arl14epl | 0.441021 | 0.009516 | 0.862404 | 2.021556 | 0.064289 |
| Saysd1 | 0.440728 | 0.016225 | 0.862404 | 1.789809 | 0.064289 |
| Prom1 | 0.438023 | 0.022176 | 0.862404 | 1.654121 | 0.064289 |
| Sec14l5 | 0.436143 | 0.000958 | 0.862404 | 3.018418 | 0.064289 |
| Rfc4 | 0.435782 | 0.040270 | 0.862404 | 1.395020 | 0.064289 |
| Lig3 | 0.432932 | 0.027658 | 0.862404 | 1.558177 | 0.064289 |
| Lgi3 | 0.431527 | 0.015297 | 0.862404 | 1.815387 | 0.064289 |

| | | | | | |
|---|---|---|---|---|---|
| Otos | 0.428370 | 0.025038 | 0.862404 | 1.601397 | 0.064289 |
| F3 | 0.426850 | 0.008506 | 0.862404 | 2.070292 | 0.064289 |
| Olfr429 | 0.424777 | 0.025394 | 0.862404 | 1.595264 | 0.064289 |
| Slfn9 | 0.423175 | 0.027435 | 0.862404 | 1.561699 | 0.064289 |
| Efnb1 | 0.422827 | 0.003873 | 0.862404 | 2.411992 | 0.064289 |
| Shroom1 | 0.422317 | 0.027303 | 0.862404 | 1.563789 | 0.064289 |
| Sparc | 0.421651 | 0.047219 | 0.862404 | 1.325881 | 0.064289 |
| Endod1 | 0.418762 | 0.047636 | 0.862404 | 1.322068 | 0.064289 |
| Mthfd1 | 0.415980 | 0.036911 | 0.862404 | 1.432841 | 0.064289 |
| Cox15 | 0.415583 | 0.023362 | 0.862404 | 1.631494 | 0.064289 |
| Gm19426 | 0.415439 | 0.020143 | 0.862404 | 1.695879 | 0.064289 |
| Nptx2 | 0.413833 | 0.021564 | 0.862404 | 1.666263 | 0.064289 |
| Msmo1 | 0.413310 | 0.031115 | 0.862404 | 1.507037 | 0.064289 |
| Prl2c3 | 0.408595 | 0.035105 | 0.862404 | 1.454628 | 0.064289 |
| Eepd1 | 0.408461 | 0.033911 | 0.862404 | 1.469664 | 0.064289 |
| Zcchc24 | 0.407288 | 0.024991 | 0.862404 | 1.602224 | 0.064289 |
| Gpr1 | 0.407288 | 0.037546 | 0.862404 | 1.425432 | 0.064289 |
| Sox1 | 0.407207 | 0.007858 | 0.862404 | 2.104707 | 0.064289 |
| Bmx | 0.407049 | 0.047972 | 0.862404 | 1.319009 | 0.064289 |
| Cdk5rap1 | 0.406538 | 0.029109 | 0.862404 | 1.535967 | 0.064289 |
| Rlbp1 | 0.403079 | 0.021358 | 0.862404 | 1.670440 | 0.064289 |
| Rarb | 0.401715 | 0.018429 | 0.862404 | 1.734491 | 0.064289 |
| Fbn2 | 0.400148 | 0.034976 | 0.862404 | 1.456232 | 0.064289 |
| Neu3 | 0.399993 | 0.046301 | 0.862404 | 1.334409 | 0.064289 |
| Nfic | 0.395448 | 0.020181 | 0.862404 | 1.695066 | 0.064289 |
| Chad | 0.395305 | 0.046112 | 0.862404 | 1.336189 | 0.064289 |
| Tmem14c | 0.393506 | 0.000143 | 0.672561 | 3.845228 | 0.172268 |
| Vegfb | 0.392768 | 0.006628 | 0.862404 | 2.178621 | 0.064289 |
| Serpinb3c | 0.392292 | 0.025204 | 0.862404 | 1.598524 | 0.064289 |
| Tmem165 | 0.392258 | 0.021489 | 0.862404 | 1.667787 | 0.064289 |
| Mrpl4 | 0.389925 | 0.044116 | 0.862404 | 1.355407 | 0.064289 |
| Cers2 | 0.389158 | 0.032874 | 0.862404 | 1.483146 | 0.064289 |
| Mbp | 0.389011 | 0.005713 | 0.862404 | 2.243127 | 0.064289 |
| P2ry10b | 0.386601 | 0.037053 | 0.862404 | 1.431175 | 0.064289 |
| Car5b | 0.385589 | 0.033549 | 0.862404 | 1.474324 | 0.064289 |
| Sspn | 0.383825 | 0.015636 | 0.862404 | 1.805873 | 0.064289 |
| Zfp74 | 0.382862 | 0.041310 | 0.862404 | 1.383948 | 0.064289 |
| Nkain1 | 0.382646 | 0.043965 | 0.862404 | 1.356892 | 0.064289 |
| Tnfaip1 | 0.382420 | 0.049472 | 0.862404 | 1.305637 | 0.064289 |
| T2 | 0.379230 | 0.020267 | 0.862404 | 1.693217 | 0.064289 |
| Timm17b | 0.376605 | 0.026847 | 0.862404 | 1.571097 | 0.064289 |

| | | | | | |
|---|---|---|---|---|---|
| Panx2 | 0.375533 | 0.022505 | 0.862404 | 1.647726 | 0.064289 |
| Sft2d3 | 0.375110 | 0.039489 | 0.862404 | 1.403529 | 0.064289 |
| Psmg1 | 0.374257 | 0.033553 | 0.862404 | 1.474273 | 0.064289 |
| Usb1 | 0.373714 | 0.049171 | 0.862404 | 1.308292 | 0.064289 |
| Lrfn4 | 0.371541 | 0.016007 | 0.862404 | 1.795690 | 0.064289 |
| Mad1l1 | 0.368115 | 0.044229 | 0.862404 | 1.354297 | 0.064289 |
| Trim13 | 0.367800 | 0.022527 | 0.862404 | 1.647300 | 0.064289 |
| Nek6 | 0.365498 | 0.047107 | 0.862404 | 1.326913 | 0.064289 |
| Pnkp | 0.365287 | 0.042843 | 0.862404 | 1.368122 | 0.064289 |
| Pex10 | 0.364727 | 0.016391 | 0.862404 | 1.785397 | 0.064289 |
| Gnal | 0.363412 | 0.030991 | 0.862404 | 1.508769 | 0.064289 |
| Gm35410 | 0.361319 | 0.047172 | 0.862404 | 1.326314 | 0.064289 |
| Ppm1l | 0.360394 | 0.006107 | 0.862404 | 2.214186 | 0.064289 |
| Ftl1 | 0.356533 | 0.044734 | 0.862404 | 1.349362 | 0.064289 |
| 1810030O( | 0.355484 | 0.014700 | 0.862404 | 1.832674 | 0.064289 |
| Pcnt | 0.353739 | 0.006130 | 0.862404 | 2.212512 | 0.064289 |
| Insig1 | 0.351561 | 0.009444 | 0.862404 | 2.024842 | 0.064289 |
| Klf5 | 0.349455 | 0.029735 | 0.862404 | 1.526736 | 0.064289 |
| Eqtn | 0.347861 | 0.036353 | 0.862404 | 1.439458 | 0.064289 |
| Rpl36a | 0.347386 | 0.042695 | 0.862404 | 1.369622 | 0.064289 |
| Tjap1 | 0.347336 | 0.027278 | 0.862404 | 1.564187 | 0.064289 |
| Spry4 | 0.346227 | 0.041169 | 0.862404 | 1.385431 | 0.064289 |
| Pdrg1 | 0.345877 | 0.021212 | 0.862404 | 1.673411 | 0.064289 |
| Slc27a4 | 0.345697 | 0.022409 | 0.862404 | 1.649579 | 0.064289 |
| Chrm3 | 0.338659 | 0.027654 | 0.862404 | 1.558243 | 0.064289 |
| Hsd17b12 | 0.337944 | 0.035996 | 0.862404 | 1.443751 | 0.064289 |
| Cisd2 | 0.337746 | 0.025749 | 0.862404 | 1.589243 | 0.064289 |
| Spag4 | 0.336250 | 0.037325 | 0.862404 | 1.427999 | 0.064289 |
| Slc9a1 | 0.333693 | 0.015874 | 0.862404 | 1.799304 | 0.064289 |
| Cep72 | 0.329525 | 0.013383 | 0.862404 | 1.873453 | 0.064289 |
| Nudt5 | 0.328275 | 0.045752 | 0.862404 | 1.339586 | 0.064289 |
| Mmp24 | 0.327946 | 0.000638 | 0.862404 | 3.194890 | 0.064289 |
| Gna12 | 0.325587 | 0.041396 | 0.862404 | 1.383046 | 0.064289 |
| Lamp1 | 0.324499 | 0.028888 | 0.862404 | 1.539277 | 0.064289 |
| Ppp1r1b | 0.324275 | 0.019743 | 0.862404 | 1.704592 | 0.064289 |
| Cers5 | 0.322743 | 0.010980 | 0.862404 | 1.959392 | 0.064289 |
| Tssc4 | 0.322702 | 0.046389 | 0.862404 | 1.333588 | 0.064289 |
| Mknk1 | 0.319841 | 0.044521 | 0.862404 | 1.351437 | 0.064289 |
| Tmx2 | 0.318577 | 0.043139 | 0.862404 | 1.365134 | 0.064289 |
| Tet3 | 0.318064 | 0.034731 | 0.862404 | 1.459287 | 0.064289 |
| Trmt9b | 0.317740 | 0.025267 | 0.862404 | 1.597452 | 0.064289 |

| | | | | | |
|---|---|---|---|---|---|
| Wdpcp | 0.317581 | 0.005396 | 0.862404 | 2.267893 | 0.064289 |
| Aldoc | 0.316632 | 0.043437 | 0.862404 | 1.362139 | 0.064289 |
| Tbc1d9b | 0.315940 | 0.028819 | 0.862404 | 1.540326 | 0.064289 |
| Kcnip3 | 0.315406 | 0.026701 | 0.862404 | 1.573465 | 0.064289 |
| Creg1 | 0.314826 | 0.032332 | 0.862404 | 1.490365 | 0.064289 |
| Wdr45 | 0.314726 | 0.036299 | 0.862404 | 1.440100 | 0.064289 |
| Mroh1 | 0.314601 | 0.007971 | 0.862404 | 2.098467 | 0.064289 |
| Dennd4b | 0.312755 | 0.046667 | 0.862404 | 1.330987 | 0.064289 |
| Gdpd1 | 0.311531 | 0.009781 | 0.862404 | 2.009604 | 0.064289 |
| Nus1 | 0.310997 | 0.026903 | 0.862404 | 1.570192 | 0.064289 |
| Smpd1 | 0.310841 | 0.002061 | 0.862404 | 2.685821 | 0.064289 |
| Nfkbiz | 0.308659 | 0.030407 | 0.862404 | 1.517021 | 0.064289 |
| Zswim1 | 0.308066 | 0.016030 | 0.862404 | 1.795068 | 0.064289 |
| Rhcg | 0.307434 | 0.033453 | 0.862404 | 1.475569 | 0.064289 |
| Arntl2 | 0.306695 | 0.029109 | 0.862404 | 1.535965 | 0.064289 |
| Arl6ip4 | 0.305432 | 0.035537 | 0.862404 | 1.449320 | 0.064289 |
| Gm35486 | 0.305162 | 0.036604 | 0.862404 | 1.436470 | 0.064289 |
| Epas1 | 0.303978 | 0.023210 | 0.862404 | 1.634317 | 0.064289 |
| Ccdc107 | 0.303549 | 0.014810 | 0.862404 | 1.829448 | 0.064289 |
| Ptgs2 | 0.303530 | 0.004045 | 0.862404 | 2.393041 | 0.064289 |
| Rpgr | 0.302219 | 0.005525 | 0.862404 | 2.257682 | 0.064289 |
| Tnpo1 | 0.301174 | 0.028673 | 0.862404 | 1.542534 | 0.064289 |
| Fdx1 | 0.299435 | 0.014293 | 0.862404 | 1.844885 | 0.064289 |
| Asrgl1 | 0.298286 | 0.026192 | 0.862404 | 1.581827 | 0.064289 |
| Stox2 | 0.297937 | 0.049062 | 0.862404 | 1.309252 | 0.064289 |
| S100a1 | 0.297841 | 0.009495 | 0.862404 | 2.022514 | 0.064289 |
| Sh2b3 | 0.297689 | 0.025775 | 0.862404 | 1.588800 | 0.064289 |
| Elovl7 | 0.297566 | 0.029899 | 0.862404 | 1.524338 | 0.064289 |
| 4933403O( | 0.296944 | 0.002862 | 0.862404 | 2.543329 | 0.064289 |
| Prpf40a | 0.296499 | 0.014688 | 0.862404 | 1.833049 | 0.064289 |
| Bmf | 0.295769 | 0.023203 | 0.862404 | 1.634460 | 0.064289 |
| Rabif | 0.295358 | 0.027533 | 0.862404 | 1.560154 | 0.064289 |
| Ddost | 0.295286 | 0.030355 | 0.862404 | 1.517764 | 0.064289 |
| Nppc | 0.294351 | 0.022463 | 0.862404 | 1.648525 | 0.064289 |
| Papolg | 0.293658 | 0.010124 | 0.862404 | 1.994662 | 0.064289 |
| Srrm4 | 0.292974 | 0.025957 | 0.862404 | 1.585747 | 0.064289 |
| Septin1 | 0.290341 | 0.006234 | 0.862404 | 2.205257 | 0.064289 |
| Mrpl54 | 0.288961 | 0.028915 | 0.862404 | 1.538882 | 0.064289 |
| Klhl32 | 0.288854 | 0.040883 | 0.862404 | 1.388460 | 0.064289 |
| Fdft1 | 0.286816 | 0.047675 | 0.862404 | 1.321711 | 0.064289 |
| Chst2 | 0.285951 | 0.046166 | 0.862404 | 1.335676 | 0.064289 |

| | | | | | |
|---|---|---|---|---|---|
| Zfp397 | 0.285900 | 0.012867 | 0.862404 | 1.890534 | 0.064289 |
| Rrs1 | 0.285297 | 0.044219 | 0.862404 | 1.354387 | 0.064289 |
| Kcna1 | 0.284667 | 0.017862 | 0.862404 | 1.748082 | 0.064289 |
| Manf | 0.284111 | 0.011102 | 0.862404 | 1.954595 | 0.064289 |
| Gm46382 | 0.282905 | 0.006447 | 0.862404 | 2.190622 | 0.064289 |
| Rnf165 | 0.282197 | 0.036887 | 0.862404 | 1.433129 | 0.064289 |
| Kcnc1 | 0.279449 | 0.047451 | 0.862404 | 1.323757 | 0.064289 |
| Haghl | 0.279315 | 0.021144 | 0.862404 | 1.674803 | 0.064289 |
| Abracl | 0.277991 | 0.035771 | 0.862404 | 1.446465 | 0.064289 |
| Glul | 0.276570 | 0.020324 | 0.862404 | 1.691990 | 0.064289 |
| Depdc5 | 0.276393 | 0.005588 | 0.862404 | 2.252739 | 0.064289 |
| Lrrfip1 | 0.276019 | 0.046521 | 0.862404 | 1.332355 | 0.064289 |
| Brpf1 | 0.275882 | 0.036027 | 0.862404 | 1.443374 | 0.064289 |
| 2610002M | 0.273826 | 0.036239 | 0.862404 | 1.440829 | 0.064289 |
| Retreg3 | 0.273376 | 0.046518 | 0.862404 | 1.332383 | 0.064289 |
| Smim4 | 0.272655 | 0.030367 | 0.862404 | 1.517602 | 0.064289 |
| Cep170b | 0.270632 | 0.017905 | 0.862404 | 1.747031 | 0.064289 |
| Ep400 | 0.270595 | 0.026749 | 0.862404 | 1.572700 | 0.064289 |
| Coch | 0.269495 | 0.046250 | 0.862404 | 1.334890 | 0.064289 |
| Kif5a | 0.267877 | 0.025307 | 0.862404 | 1.596759 | 0.064289 |
| Malrd1 | 0.267568 | 0.036401 | 0.862404 | 1.438889 | 0.064289 |
| Frs2 | 0.266463 | 0.030827 | 0.862404 | 1.511066 | 0.064289 |
| St8sia5 | 0.266207 | 0.012058 | 0.862404 | 1.918719 | 0.064289 |
| Kdm4c | 0.263484 | 0.021509 | 0.862404 | 1.667372 | 0.064289 |
| Chga | 0.262463 | 0.043771 | 0.862404 | 1.358810 | 0.064289 |
| Qsox2 | 0.258153 | 0.003007 | 0.862404 | 2.521815 | 0.064289 |
| Usf1 | 0.258152 | 0.014221 | 0.862404 | 1.847077 | 0.064289 |
| Bivm | 0.257988 | 0.017687 | 0.862404 | 1.752341 | 0.064289 |
| Mfsd9 | 0.257400 | 0.000314 | 0.862404 | 3.502870 | 0.064289 |
| Mcf2l | 0.257263 | 0.013398 | 0.862404 | 1.872973 | 0.064289 |

**Supplementary Table 7: IPA Canonical Pathways of differentially expressed genes in Neurons at E18**

| Ingenuity Canonical Pathways | -log(p-va | Ratio | z-score | Molecules |
|---|---|---|---|---|
| Synaptogenesis Signaling Pathway | 8.010 | 0.199 | -2.252 | [illegible] |
| Axonal Guidance Signaling | 6.980 | 0.169 | #NUM! | [illegible] |
| Signaling by VEGF | 6.550 | 0.264 | -0.378 | [illegible] |
| Semaphorin interactions | 6.110 | 0.312 | 1.342 | CFL1,CRMP1,DPYSL2,DPYSL3,DPYSL4,DPYSL5,FYN,HSP90AA1,HSP90AB1,MET,MYL12B,MYL6,PAK1,PLXNA2,PLXNA4,PLXNB1,SEMA3A,SEMA5A,SEMA6D,SEMA7A |
| Superpathway of Cholesterol Biosynthesis | 5.720 | 0.444 | 3.317 | Acat2/Acat3,CYP51A1,DHCR24,FDFT1,FDPS,HMGCR,HMGCS1,MSMO1,MVD,MVK,NSDHL,SQLE |
| Cholesterol biosynthesis | 5.230 | 0.440 | 3.317 | CYP51A1,DHCR24,FDFT1,FDPS,HMGCR,HMGCS1,MSMO1,MVD,MVK,NSDHL,SQLE |
| Stress Granule Signaling Pathway | 5.050 | 0.184 | -0.762 | [illegible] |
| Estrogen Receptor Signaling | 5.000 | 0.162 | 0.309 | [illegible] |
| Fcgamma receptor (FCGR) dependent phagocytosis | 4.870 | 0.250 | 0.853 | ACTB,AHCYL1,ARPC3,ARPC5,CFL1,CYFIP2,DOCK1,ELMO1,FYN,HSP90AA1,HSP90AB1,ITPR1,ITPR2,PAK1,PIK3CA,PIK3R2,PLCG1,PLPP4,PRKCD,WASF1,WIPF2,WIPF3 |
| Activation of NMDA receptors and postsynaptic events | 4.840 | 0.263 | -0.447 | ADCY1,ADCY8,CALM1,CAMKK2,ERBB4,GRIA2,GRIA3,GRIN2D,HRAS,KPNA2,LRRC7,MAPT,NRG1,NRGN,PRKAA2,RASGRF1,RASGRF2,RPS6KA2,TUBA1A,TUBB6 |
| EPH-Ephrin signaling | 4.700 | 0.244 | 0.853 | ACTB,AP2M1,ARPC3,ARPC5,CFL1,CLTB,EFNA3,EFNB1,EPHA3,EPHA5,EPHA6,EPHA7,EPHB1,EPHB3,FYN,HRAS,MYL12B,MYL6,NCK2,NCSTN,PAK1,RASA1 |
| GABA Receptor Signaling | 4.690 | 0.215 | #NUM! | [illegible] |
| DAG and IP3 signaling | 4.510 | 0.333 | -1.387 | ADCY1,ADCY8,AHCYL1,CALM1,CAMKK2,ITPR1,ITPR2,KPNA2,NBEA,PDE1A,PDE1B,PLCG1,PRKCD |
| Eukaryotic Translation Initiation | 4.500 | 0.201 | 4.131 | [illegible] |
| EIF2 Signaling | 4.490 | 0.178 | 3.500 | [illegible] |
| CXCR4 Signaling | 4.460 | 0.198 | 1.342 | [illegible] |
| Netrin Signaling | 4.420 | 0.194 | 0.354 | [illegible] |
| Coronavirus Replication Pathway | 4.380 | 0.201 | 1.134 | [illegible] |
| Signaling by ROBO receptors | 4.270 | 0.174 | 4.320 | [illegible] |
| Semaphorin Signaling in Neurons | 4.260 | 0.276 | #NUM! | CFL1,CRMP1,DPYSL2,DPYSL3,DPYSL4,DPYSL5,FYN,MET,PAK1,PLXNA2,PLXNB1,RAC3,RHOU,RND2,SEMA3A,SEMA7A |
| G alpha (z) signalling events | 4.200 | 0.298 | -1.069 | ADCY1,ADCY8,ADRA2C,GNAZ,GNB2,GNG3,GNG4,GNG7,PRKCB,PRKCD,PRKCH,RGS17,RGS4,RGSL1 |
| Eukaryotic Translation Elongation | 4.130 | 0.212 | 3.400 | [illegible] |
| mTOR Signaling | 4.090 | 0.176 | 0.894 | [illegible] |
| Ephrin B Signaling | 4.010 | 0.247 | 1.387 | [illegible] |
| High laminar flow shear stress activates signaling by PIEZO1 | 3.900 | 0.250 | 0.728 | [illegible] |
| NRF2-mediated Oxidative Stress Response | 3.860 | 0.176 | 1.500 | [illegible] |
| Activation of gene expression by SREBF (SREBP) | 3.710 | 0.300 | 2.111 | [illegible] |
| Apelin Endothelial Signaling Pathway | 3.680 | 0.193 | 1.698 | [illegible] |
| Ephrin Receptor Signaling | 3.610 | 0.173 | 3.000 | [illegible] |
| Integration of energy metabolism | 3.570 | 0.208 | 1.706 | [illegible] |
| Granzyme A Signaling | 3.460 | 0.239 | 0.277 | [illegible] |
| Opioid Signalling | 3.390 | 0.216 | 0.229 | [illegible] |
| CREB Signaling in Neurons | 3.370 | 0.136 | -2.646 | [illegible] |
| RHO GTPase cycle | 3.350 | 0.143 | -0.126 | [illegible] |
| Signaling by NOTCH1 | 3.350 | 0.227 | -1.213 | [illegible] |
| Eukaryotic Translation Termination | 3.350 | 0.197 | 3.128 | [illegible] |
| Huntington's Disease Signaling | 3.310 | 0.157 | 1.279 | [illegible] |
| Circadian Rhythm Signaling | 3.300 | 0.159 | #NUM! | [illegible] |
| G Protein Signaling Mediated by Tubby | 3.280 | 0.224 | 0.258 | [illegible] |
| GABAergic Receptor Signaling Pathway (Enhanced) | 3.280 | 0.188 | 0.200 | [illegible] |
| Ribosomal Quality Control Signaling Pathway | 3.250 | 0.169 | 3.780 | [illegible] |
| CCR5 Signaling in Macrophages | 3.210 | 0.200 | -1.342 | [illegible] |
| Role of NFAT in Cardiac Hypertrophy | 3.210 | 0.164 | 0.600 | [illegible] |
| Cholesterol Biosynthesis I | 3.200 | 0.462 | 2.449 | [illegible] |
| Cholesterol Biosynthesis II (via 24,25-dihydrolanosterol) | 3.200 | 0.462 | 2.449 | [illegible] |
| Cholesterol Biosynthesis III (via Desmosterol) | 3.200 | 0.462 | 2.449 | [illegible] |
| Semaphorin Neuronal Repulsive Signaling Pathway | 3.160 | 0.182 | -1.633 | [illegible] |
| G Beta Gamma Signaling | 3.150 | 0.188 | 1.789 | [illegible] |
| Nitric Oxide Signaling in the Cardiovascular System | 3.110 | 0.193 | 1.606 | [illegible] |
| Cardiac Hypertrophy Signaling | 3.110 | 0.158 | 0.200 | [illegible] |
| fMLP Signaling in Neutrophils | 3.100 | 0.186 | 1.807 | [illegible] |
| α-Adrenergic Signaling | 3.100 | 0.196 | 0.333 | [illegible] |
| Glutamate Receptor Signaling | 3.040 | 0.227 | -2.236 | [illegible] |
| Thrombin Signaling | 3.020 | 0.161 | 0.894 | [illegible] |
| Orexin Signaling Pathway | 2.920 | 0.158 | 0.507 | [illegible] |
| Signaling by ALK | 2.910 | 0.300 | 0.333 | [illegible] |
| GAIT Translation Signaling Pathway | 2.870 | 0.160 | -3.153 | [illegible] |
| Remodeling of Epithelial Adherens Junctions | 2.840 | 0.226 | 0.378 | [illegible] |
| Aldosterone Signaling in Epithelial Cells | 2.830 | 0.169 | -0.577 | [illegible] |
| Signaling by ERBB2 | 2.750 | 0.240 | 0.577 | [illegible] |
| TP53 Regulates Metabolic Genes | 2.710 | 0.200 | 0.728 | [illegible] |
| GPCR-Mediated Nutrient Sensing in Enteroendocrine Cells | 2.690 | 0.183 | -0.943 | [illegible] |
| Endocannabinoid Neuronal Synapse Pathway | 2.680 | 0.171 | -0.243 | [illegible] |
| Epithelial Adherens Junction Signaling | 2.670 | 0.169 | -1.000 | [illegible] |
| Reelin Signaling in Neurons | 2.640 | 0.173 | 0.000 | [illegible] |
| G-Protein Coupled Receptor Signaling | 2.640 | 0.127 | -0.556 | [illegible] |
| Superpathway of Geranylgeranyldiphosphate Biosynthesis I ( | 2.640 | 0.375 | 2.236 | [illegible] |
| RAF/MAP kinase cascade | 2.620 | 0.150 | -0.649 | [illegible] |
| Respiratory electron transport | 2.600 | 0.195 | 2.183 | [illegible] |
| PPARα/RXRα Activation | 2.560 | 0.160 | -1.342 | [illegible] |
| Synaptic Long Term Potentiation | 2.540 | 0.178 | -1.789 | [illegible] |
| Endocannabinoid Developing Neuron Pathway | 2.530 | 0.175 | 0.728 | [illegible] |
| Selenoamino acid metabolism | 2.520 | 0.169 | 2.858 | [illegible] |
| Mevalonate Pathway I | 2.500 | 0.417 | 2.000 | [illegible] |
| Cohesin Chromatin Regulation Pathway | 2.480 | 0.154 | -1.633 | [illegible] |
| Opioid Signaling Pathway | 2.470 | 0.145 | 0.186 | [illegible] |
| HIF1α Signaling | 2.460 | 0.156 | 0.192 | [illegible] |
| Post-translational protein phosphorylation | 2.460 | 0.186 | -0.943 | [illegible] |
| Acetylcholine Receptor Signaling Pathway | 2.420 | 0.157 | -0.378 | [illegible] |
| CCR3 Signaling in Eosinophils | 2.400 | 0.171 | 1.000 | [illegible] |
| EGR2 and SOX10-mediated initiation of Schwann cell myelina | 2.390 | 0.276 | 1.414 | [illegible] |
| Somitogenesis | 2.380 | 0.218 | 0.577 | [illegible] |
| Transcriptional Regulation by MECP2 | 2.380 | 0.210 | -1.941 | [illegible] |
| 14-3-3-mediated Signaling | 2.370 | 0.172 | -0.500 | [illegible] |
| P2Y Purinergic Receptor Signaling Pathway | 2.360 | 0.169 | -0.243 | [illegible] |
| Molecular Mechanisms of Cancer | 2.360 | 0.121 | -0.513 | [illegible] |
| DNA Damage/Telomere Stress Induced Senescence | 2.350 | 0.238 | -2.530 | [illegible] |
| Apelin Muscle Signaling Pathway | 2.330 | 0.224 | #NUM! | [illegible] |
| Oxytocin in Brain Signaling Pathway | 2.310 | 0.156 | 1.347 | [illegible] |
| Renal Cell Carcinoma Signaling | 2.290 | 0.192 | 1.265 | [illegible] |
| ERBB Signaling | 2.280 | 0.183 | 0.728 | [illegible] |
| Glioma Signaling | 2.280 | 0.169 | 0.000 | [illegible] |
| Cell Cycle Control of Chromosomal Replication | 2.260 | 0.203 | -2.887 | [illegible] |
| UVB-Induced MAPK Signaling | 2.260 | 0.220 | -0.302 | [illegible] |
| Hedgehog 'off' state | 2.250 | 0.174 | 0.688 | [illegible] |
| G alpha (12/13) signalling events | 2.230 | 0.190 | -0.775 | [illegible] |
| Binding and Uptake of Ligands by Scavenger Receptors | 2.230 | 0.243 | 1.667 | [illegible] |
| Interleukin-4 and Interleukin-13 signaling | 2.220 | 0.176 | -1.698 | [illegible] |
| Exosome Signaling Pathway | 2.220 | 0.158 | 3.266 | [illegible] |
| Regulation of Insulin-like Growth Factor (IGF) transport and u | 2.210 | 0.173 | -1.147 | [illegible] |
| Regulation of eIF4 and p70S6K Signaling | 2.200 | 0.150 | 1.414 | [illegible] |
| Apelin Cardiomyocyte Signaling Pathway | 2.190 | 0.179 | 0.000 | [illegible] |
| Insulin Secretion Signaling Pathway | 2.170 | 0.141 | -0.898 | [illegible] |
| ERK5 Signaling | 2.160 | 0.192 | 0.535 | [illegible] |
| Beta-catenin independent WNT signaling | 2.140 | 0.149 | 1.826 | [illegible] |
| ERBB4 Signaling | 2.140 | 0.197 | 0.577 | [illegible] |
| Renin-Angiotensin Signaling | 2.120 | 0.167 | 0.500 | [illegible] |
| Gap Junction Signaling | 2.100 | 0.136 | 2.530 | [illegible] |
| eNOS Signaling | 2.100 | 0.159 | 0.500 | [illegible] |
| Formation of paraxial mesoderm | 2.090 | 0.300 | -1.633 | [illegible] |
| SRP-dependent cotranslational protein targeting to membra | 2.090 | 0.161 | 2.985 | [illegible] |
| Growth Hormone Signaling | 2.080 | 0.194 | -1.155 | [illegible] |
| Regulation of mRNA stability by proteins that bind AU-rich e | 2.060 | 0.178 | 2.500 | [illegible] |
| Fcγ Receptor-mediated Phagocytosis in Macrophages and M | 2.060 | 0.178 | 1.500 | [illegible] |
| Epoxysqualene Biosynthesis | 2.060 | 1.000 | #NUM! | FDFT1,SQLE |
| Spermine Biosynthesis | 2.060 | 1.000 | #NUM! | AMD1,SMS |
| Ascorbate Recycling (Cytosolic) | 2.060 | 1.000 | #NUM! | GLRX,GSTO1 |
| Spermidine Biosynthesis I | 2.060 | 1.000 | #NUM! | AMD1,SRM |
| Glutamate Dependent Acid Resistance | 2.060 | 1.000 | #NUM! | GAD1,GAD2 |
| Post-translational modification: synthesis of GPI-anchored p | 2.050 | 0.187 | -2.138 | [illegible] |
| Regulation of cholesterol biosynthesis by SREBP (SREBF) | 2.020 | 0.333 | 0.447 | [illegible] |
| Neuropathic Pain Signaling in Dorsal Horn Neurons | 2.010 | 0.172 | -2.183 | [illegible] |
| Cellular response to hypoxia | 2.000 | 0.184 | 1.604 | [illegible] |
| Formation of the posterior neural plate | 2.000 | 0.400 | 0.000 | [illegible] |
| Docosahexaenoic Acid (DHA) Signaling | 1.990 | 0.142 | -0.522 | [illegible] |
| Citric acid cycle (TCA cycle) | 1.980 | 0.286 | 0.816 | [illegible] |
| IL-15 Production | 1.960 | 0.164 | -3.441 | [illegible] |
| Protein Kinase A Signaling | 1.960 | 0.130 | -2.058 | [illegible] |
| Cilia Biogenesis Signaling Pathway | 1.960 | 0.147 | -0.756 | [illegible] |
| NF1 RAS Signaling Pathway | 1.940 | 0.124 | -1.152 | [illegible] |
| Hereditary Breast Cancer Signaling | 1.940 | 0.158 | -0.447 | [illegible] |
| Neurexins and neuroligins | 1.940 | 0.200 | -0.302 | [illegible] |
| Relaxin Signaling | 1.920 | 0.153 | 1.265 | [illegible] |
| Factors involved in megakaryocyte development and platele | 1.910 | 0.179 | -0.535 | [illegible] |
| Fc epsilon receptor (FCERI) signaling | 1.900 | 0.157 | 1.528 | [illegible] |
| Platelet homeostasis | 1.890 | 0.174 | 0.775 | [illegible] |
| Myelination Signaling Pathway | 1.890 | 0.132 | 0.000 | [illegible] |
| Non-Small Cell Lung Cancer Signaling | 1.890 | 0.170 | 0.632 | [illegible] |
| Sphingosine-1-phosphate Signaling | 1.880 | 0.164 | 0.535 | [illegible] |
| Arsenate Detoxification I (Glutaredoxin) | 1.880 | 0.500 | #NUM! | [illegible] |
| Zymosterol Biosynthesis | 1.880 | 0.500 | #NUM! | [illegible] |
| CNTF Signaling | 1.880 | 0.196 | 0.302 | [illegible] |
| Ephrin A Signaling | 1.870 | 0.156 | -0.894 | [illegible] |
| Oxytocin Signaling Pathway | 1.850 | 0.135 | 2.058 | [illegible] |
| Pancreatic Secretion Signaling Pathway | 1.850 | 0.140 | -0.186 | [illegible] |
| Nonsense-Mediated Decay (NMD) | 1.850 | 0.162 | 3.771 | [illegible] |
| Prostate Cancer Signaling | 1.850 | 0.162 | 1.414 | [illegible] |
| Prolactin Signaling | 1.840 | 0.168 | 0.577 | [illegible] |
| Formation of the anterior neural plate | 1.840 | 0.364 | 0.000 | [illegible] |
| Regulation of RUNX2 expression and activity | 1.830 | 0.181 | 1.387 | [illegible] |
| Cilium Assembly | 1.820 | 0.143 | -1.134 | [illegible] |
| Cell Cycle: G2/M DNA Damage Checkpoint Regulation | 1.800 | 0.200 | 1.000 | [illegible] |
| UVA-Induced MAPK Signaling | 1.800 | 0.167 | -0.632 | [illegible] |
| Glutaminergic Receptor Signaling Pathway (Enhanced) | 1.800 | 0.131 | -1.121 | [illegible] |
| Aggrephagy | 1.800 | 0.222 | 2.121 | [illegible] |
| RAF-independent MAPK1/3 activation | 1.780 | 0.261 | -1.633 | [illegible] |
| Nephrin family interactions | 1.780 | 0.261 | 0.000 | [illegible] |
| Major pathway of rRNA processing in the nucleolus and cyto | 1.780 | 0.141 | 3.528 | [illegible] |
| KEAP1-NFE2L2 pathway | 1.780 | 0.151 | 1.706 | [illegible] |
| Signaling by NTRK1 (TRKA) | 1.770 | 0.173 | 0.000 | [illegible] |
| Signaling by NOTCH4 | 1.770 | 0.152 | 1.528 | [illegible] |
| Androgen Signaling | 1.750 | 0.146 | -0.707 | [illegible] |
| Pyridoxal 5'-phosphate Salvage Pathway | 1.750 | 0.182 | -1.508 | [illegible] |
| Activation of kainate receptors upon glutamate binding | 1.740 | 0.233 | 0.378 | [illegible] |
| Leptin Signaling in Obesity | 1.730 | 0.176 | 0.816 | [illegible] |
| Netrin-1 signaling | 1.730 | 0.205 | -1.667 | [illegible] |
| Notch Signaling | 1.730 | 0.216 | -1.134 | [illegible] |
| Gαq Signaling | 1.720 | 0.145 | 0.775 | [illegible] |
| Cardiac β-adrenergic Signaling | 1.720 | 0.145 | -1.265 | [illegible] |
| Type II Diabetes Mellitus Signaling | 1.720 | 0.149 | -0.832 | [illegible] |
| Cell Cycle Checkpoints | 1.710 | 0.134 | -0.845 | [illegible] |
| Cell Cycle: G1/S Checkpoint Regulation | 1.700 | 0.179 | 0.000 | [illegible] |
| RHO GTPases activate IQGAPs | 1.690 | 0.250 | 0.816 | [illegible] |
| Z-decay: degradation of maternal mRNAs by zygotically expr | 1.690 | 0.333 | 1.000 | [illegible] |
| Mitotic Prometaphase | 1.690 | 0.141 | -1.732 | [illegible] |
| C-type lectin receptors (CLRs) | 1.690 | 0.152 | 1.342 | [illegible] |
| Antiproliferative Role of Somatostatin Receptor 2 | 1.690 | 0.173 | 1.134 | [illegible] |
| Cyclins and Cell Cycle Regulation | 1.680 | 0.169 | -0.378 | [illegible] |
| Protein folding | 1.680 | 0.165 | 2.324 | [illegible] |
| Gαs Signaling | 1.680 | 0.153 | -1.265 | [illegible] |
| Gαi Signaling | 1.670 | 0.149 | 0.632 | [illegible] |
| Glutamate binding, activation of AMPA receptors and synap | 1.660 | 0.226 | -1.134 | [illegible] |
| IL-8 Signaling | 1.660 | 0.139 | 0.447 | [illegible] |
| Macropinocytosis Signaling | 1.640 | 0.171 | -0.277 | [illegible] |
| Signaling by PTK6 | 1.640 | 0.189 | 0.000 | [illegible] |
| Glioblastoma Multiforme Signaling | 1.630 | 0.143 | -0.894 | [illegible] |
| p75 NTR receptor-mediated signalling | 1.630 | 0.146 | -0.426 | [illegible] |

| | | | | |
|---|---|---|---|---|
| Human Embryonic Stem Cell Pluripotency | 1.610 | 0.138 | -0.192 | AKT1,BMP8B,CTNNB1,FGFR3,FOXO1,FZD1,HRAS,HSP90AA1,HSP90AB1,HSP90B1,INHBA,JAK2,JAK3,NOG,NTF3,NTRK2,PIK3CA,PIK3CD,PIK3R2,PRKCB,PRKCD,PRKCH,RAP2A,SMAD6,SOX2,TGFBR1,WNT7B |
| Adenine and Adenosine Salvage I | 1.610 | 0.667 | #NUM! | APRT,PNP |
| Signaling by NTRK2 (TRKB) | 1.610 | 0.240 | 0.000 | FYN,HRAS,NTF3,NTRK2,PIK3CA,PLCG1 |
| Signaling by Erythropoietin | 1.610 | 0.240 | 0.000 | HRAS,JAK2,PIK3CA,PIK3CD,PLCG1,RAPGEF1 |
| IL-3 Signaling | 1.600 | 0.169 | 1.387 | AKT1,FOXO1,HRAS,JAK2,PAK1,PIK3CA,PIK3CD,PIK3R2,PRKCB,PRKCD,PRKCH,RAP2A,RAPGEF1 |
| Endothelin-1 Signaling | 1.600 | 0.140 | 0.200 | ADCY1,ADCY8,EDNRB,GNA12,GNAL,GNAQ,GNAZ,GUCY1A1,GUCY1B1,HRAS,ITPR1,ITPR2,MAPK7,MAPK9,PIK3CA,PIK3CD,PIK3R2,PLCG1,PLCH1,PLCH2,PRKCB,PRKCD,PRKCH,RAP2A,SHC3 |
| Sialic acid metabolism | 1.590 | 0.219 | -0.378 | NANP,SLC35A1,ST3GAL1,ST3GAL5,ST6GALNAC4,ST6GALNAC5,ST8SIA4 |
| Germ Cell-Sertoli Cell Junction Signaling | 1.590 | 0.143 | #NUM! | ACTB,AKT1,CDH2,CFL1,CTNNA1,CTNNB1,EPN1,HRAS,MAP3K11,MAPK9,PAK1,PIK3CA,PIK3CD,PIK3R2,RAC3,RAP2A,RHOU,RND2,TGFBR1,TUBA1A,TUBB6,TUBG1,VCL |
| CDK5 Signaling | 1.570 | 0.157 | 0.535 | ADCY1,ADCY8,CACNA1A,GNAL,GUCY1A1,GUCY1B1,HRAS,LAMB1,MAPK7,MAPK9,MAPT,NGFR,NTRK2,PPP1CA,PPP2R2B,RAP2A |
| Cytosolic sensors of pathogen-associated DNA | 1.570 | 0.177 | 0.905 | CREBBP,CTNNB1,DTX4,MRE11,NKIRAS1,POLR1C,POLR1D,POLR2L,POLR3G,UBC,XRCC6 |
| GNRH Signaling | 1.570 | 0.138 | -0.500 | ADCY1,ADCY8,CACNA1A,CACNG3,CACNG8,CALM1,CATSPER2,CREBBP,GNAQ,GNB2,GNG3,GNG4,GNG7,GUCY1A1,GUCY1B1,HRAS,ITPR1,ITPR2,MAP3K11,MAPK7,MAPK9,PAK1,PRKCB,PRKCD,PRKCH,RAP2A |
| Chronic Myeloid Leukemia Signaling | 1.560 | 0.130 | -1.029 | AKT1,CALM1,CCND2,CDK6,CTNNB1,DYRK2,FZD1,HDAC2,HDAC7,HDAC9,HNRNPA1,HRAS,HSP90AA1,HSP90AB1,HSP90B1,JAK2,LIN28B,MECOM,MSI2,MTOR,NOTCH1,PA2G4,PIK3CA,PIK3CD,PIK3R2,PLCG1,PLCH1,PLCH2,PRKAA2,RAC3,RAN,RAP2A,SKP2,TGFBR1,WNT7B |
| Chaperone Mediated Autophagy Signaling Pathway | 1.560 | 0.135 | -0.186 | AKT1,ATP6AP2,ATP6V1E1,ATP6V1G2,CREBBP,EIF4A1,EPS8,FCER1G,HDAC2,HIF1A,HSP90AA1,HSP90AB1,HSP90B1,IDH1,MAPT,MMP24,MTOR,NLRP3,PARK7,PEA15,PIK3CA,PIK3CD,PIK3R2,PRKAA2,PSMB10,PSMB2,PSMB4,PSMB5,SOX2 |
| Complex IV assembly | 1.560 | 0.191 | -0.333 | COQ10B,COX16,COX4I2,COX5A,COX6A1,COX7A1,COX8A,COXFA4,TMEM223 |
| DHCR24 Signaling Pathway | 1.550 | 0.148 | 0.688 | AKT1,AMBP,APOC2,CYP51A1,DHCR24,HMGCR,HRAS,MAPT,NLGN1,PIK3CA,PIK3CD,PIK3R2,PRKCB,PRKCD,PRKCH,RAP2A,SCARB1,SOX9,TF |
| Warburg Effect Signaling Pathway | 1.550 | 0.148 | 1.414 | ACACA,AKT1,FASN,GPT2,HGF,HIF1A,MDH1,MDH2,MTOR,NNT,PHGDH,PIK3CA,PIK3CD,PIK3R2,PKM,PRKAA2,Scd1,SIRT4,TKT |
| Calcium-induced T Lymphocyte Apoptosis | 1.530 | 0.182 | -1.134 | CALM1,FCER1G,HDAC2,ITPR1,ITPR2,NR4A1,PLCG1,PRKCB,PRKCD,PRKCH |
| EGF Signaling | 1.530 | 0.182 | 0.632 | AKT1,HRAS,ITPR1,ITPR2,MTOR,PIK3CA,PIK3CD,PIK3R2,PLCG1,RASA1 |
| RAB geranylgeranylation | 1.520 | 0.175 | 0.302 | RAB13,RAB18,RAB21,RAB23,RAB30,RAB31,RAB33A,RAB34,RAB39B,RAB6A,RAB9B |
| Amyotrophic Lateral Sclerosis Signaling | 1.520 | 0.152 | -0.832 | APAF1,CACNA1A,CACNG3,CACNG8,CAPNS1,CAT,CYCS,GRIA2,GRIA3,GRIK2,GRIK3,GRIN2D,NEFM,PAK1,PIK3CA,PIK3CD,PIK3R2 |
| ABC-family proteins mediated transport | 1.520 | 0.152 | 1.698 | ABCB10,ABCB7,ABCC10,ABCD2,ABCG1,EIF2S3,OS9,PEX3,PSMB10,PSMB2,PSMB4,PSMB5,PSMC5,PSMD13,PSMD6,UBC,VPS28 |
| Response of EIF2AK4 (GCN2) to amino acid deficiency | 1.520 | 0.152 | 4.123 | EIF2S3,RPL10A,RPL12,RPL15,RPL18,RPL3,RPL36A,RPL41,RPL8,RPLP1,RPS10,RPS19,RPS26,RPS3,RPS3A,RPSA,VPS28 |
| Xenobiotic Metabolism General Signaling Pathway | 1.500 | 0.149 | -0.500 | AKT1,CREBBP,Gstm7,GSTO1,HRAS,MAP3K11,MAPK7,MAPK9,MGST1,PIK3CA,PIK3CD,PIK3R2,PRKCB,PRKCD,PRKCH,RAP2A,RBX1,SLC51A |
| NFE2L2 regulates pentose phosphate pathway genes | 1.500 | 0.375 | #NUM! | CREBBP,MAFG,TKT |
| Signaling by Rho Family GTPases | 1.490 | 0.129 | -0.209 | ACTB,ARFIP2,ARHGEF17,ARPC3,ARPC5,ARPC5L,CDH2,CDH24,CDH4,CFL1,DES,GNA12,GNAL,GNAQ,GNAZ,GNB2,GNG3,GNG4,GNG7,MAP3K11,MAPK7,MAPK9,MYL12B,MYL6,PAK1,PIK3CA,PIK3CD,PIK3R2,PIP4K2B,RAC3,RHOU,RND2,SEPTIN10,VIM,WASF1 |
| TEC Kinase Signaling | 1.490 | 0.137 | -0.500 | ACTB,FCER1G,FYN,GNA12,GNAL,GNAQ,GNAZ,GNB2,GNG3,GNG4,GNG7,JAK2,JAK3,MAPK9,PAK1,PIK3CA,PIK3CD,PIK3R2,PLCG1,PRKCB,PRKCD,PRKCH,RAC3,RHOU,RND2 |
| IL-1 Signaling | 1.490 | 0.156 | #NUM! | ADCY1,ADCY8,GNA12,GNAL,GNAQ,GNAZ,GNB2,GNG3,GNG4,GNG7,GUCY1A1,GUCY1B1,IRAK1,MAPK9,NFKBID |
| Corticotropin Releasing Hormone Signaling | 1.490 | 0.143 | -0.775 | ADCY1,ADCY8,ARPC5,CACNA1A,CACNG3,CACNG8,CALM1,CATSPER2,CREBBP,GAD1,GNAQ,GUCY1A1,GUCY1B1,ITPR1,ITPR2,MEF2C,NR4A1,PLCG1,PRKCB,PRKCD,PRKCH |
| Breast Cancer Regulation by Stathmin1 | 1.490 | 0.117 | -1.838 | ADGRA2,ADGRB1,ADGRB2,ADGRB3,ADGRG1,ADGRL3,ADORA2A,ADRA2C,ARHGEF17,AURKC,BDKRB1,C3AR1,CCND2,CCRL2,CDK1,CDK6,CELSR1,CREBBP,EDNRB,FZD1,GIPR,GNAQ,GNB2,GNG3,GNG4,GNG7,GPER1,GPR12,GPR137C,GPR146,GPR155,GPR156,GPR19,GPR27,GPR68,GPR85,GPR88,GRM3,HGF,HIF1A,HRAS,MCHR1,MTOR,NMUR1,OXGR1,PAK1,PIK3CA,PIK3CD,PIK3R2,PPP1CA,PPP1R14B,PPP2R2B,PRKCB,PRKCD,PRKCH,PTH1R,RAC3,RAP2A,S1PR1,SHC3,SSTR2,SSTR3,TUBA1A,TUBB6,TUBG1 |
| Deadenylation-dependent mRNA decay | 1.480 | 0.179 | 0.632 | CNOT1,DIS3,EIF4A1,EIF4E,EXOSC9,LSM1,LSM7,PABPC1,PAIP1,TNKS1BP1 |
| Ribonucleotide Reductase Signaling Pathway | 1.480 | 0.141 | 0.000 | ACTB,AKT1,ATM,CDK1,CDK6,CREBBP,CTNNB1,EIF4E,FBXL5,HGF,HIF1A,MAPK9,MET,MTOR,PARP1,PARP8,PIK3CA,PIK3CD,PIK3R2,RBX1,RRM2,SKP2 |
| Signaling by the B Cell Receptor (BCR) | 1.480 | 0.150 | 2.668 | AHCYL1,CALM1,FKBP1A,FYN,HRAS,ITPR1,ITPR2,PIK3CD,PRKCB,PSMB10,PSMB2,PSMB4,PSMB5,PSMC5,PSMD13,PSMD6,UBC |
| Unfolded protein response | 1.480 | 0.159 | 1.890 | CALR,CANX,DNAJC15,DNAJC16,DNAJC19,DNAJC21,DNAJC3,HSP90B1,HSPH1,INSIG1,MBTPS1,MBTPS2,OS9,PDIA6 |
| TCA Cycle II (Eukaryotic) | 1.480 | 0.250 | 1.342 | IDH3A,IDH3B,MDH1,MDH2,SDHD |
| Endoplasmic Reticulum Stress Pathway | 1.480 | 0.250 | #NUM! | CALR,DNAJC3,HSP90B1,MBTPS1,MBTPS2 |
| CGRP in Inflammation and Nociception Signaling Pathway | 1.470 | 0.144 | -1.342 | ADCY1,ADCY8,AKT1,CREBBP,CTNNB1,GNAQ,GRIN2D,GUCY1A1,GUCY1B1,PIK3CA,PIK3CD,PIK3R2,PLCG1,PLCH1,PLCH2,PRKCB,PRKCD,PRKCH,RAPGEF4,WNT7B |
| GPER1 signaling | 1.460 | 0.155 | 0.258 | ADCY1,ADCY8,CDKL1,FN1,GNAZ,GNB2,GNG3,GNG4,GNG7,GPER1,PI4KB,PIK3CA,PIK3CD,PIK3R2,STK17B |
| Effects of PIP2 hydrolysis | 1.460 | 0.222 | -0.816 | ABHD6,ITPR1,ITPR2,PRKCD,PRKCH,RASGRP2 |
| Prolactin receptor signaling | 1.450 | 0.286 | -1.000 | JAK2,RBX1,SH2B1,SKP2 |
| LGI-ADAM interactions | 1.450 | 0.286 | 0.000 | CACNG3,CACNG8,LGI1,LGI3 |
| Mitotic Telophase/Cytokinesis | 1.450 | 0.286 | -1.000 | MAU2,NIPBL,SMC3,WAPL |
| DNA Double-Strand Break Repair by Non-Homologous End Jc | 1.450 | 0.286 | -1.000 | ATM,MRE11,PARP1,XRCC6 |
| RAB GEFs exchange GTP for GDP on RABs | 1.450 | 0.157 | 1.604 | AKT1,DENND4C,DENND6B,GDI1,HPS4,RAB13,RAB18,RAB21,RAB31,RAB39B,RAB6A,RAB9B,SBF1,TRAPPC2 |
| Endocannabinoid Cancer Inhibition Pathway | 1.440 | 0.143 | 1.886 | ADCY1,ADCY8,AKT1,CAMKK2,CCND2,CREBBP,CTNNB1,GNAL,GNAQ,GUCY1A1,GUCY1B1,HIF1A,MTOR,PIK3CA,PIK3CD,PIK3R2,PRKAA2,SPTLC2,TWIST1,VIM |
| Cytoprotection by HMOX1 | 1.440 | 0.175 | 0.000 | CHD9,COX4I2,COX5A,COX6A1,COX7A1,COX8A,COXFA4,CREBBP,CYCS,NLRP3 |
| Extra-nuclear estrogen signaling | 1.430 | 0.164 | 1.732 | AKT1,CALM1,CAV1,GNB2,GNG3,GNG4,GNG7,HRAS,HSP90AA1,PIK3CA,PIK3R2,PRMT1 |
| Mitotic Roles of Polo-Like Kinase | 1.430 | 0.169 | -2.236 | CCNB2,CDC20,CDK1,HSP90AA1,HSP90B1,KIF11,PPP2R2B,PRC1,SLK,SMC3 |
| MITF-M-dependent gene expression | 1.400 | 0.150 | 0.500 | ACTB,ASAH1,ATP6AP2,ATP6V1E1,CDH2,CTNNB1,EDIL3,MCM2,MCM5,MET,PPARGC1A,PXDN,SOX2,TNRC6B,USF1,ZEB1 |
| Regulation of Actin-based Motility by Rho | 1.400 | 0.150 | 2.496 | ACTB,ARHGDIA,ARPC3,ARPC5,ARPC5L,CFL1,MYL12B,MYL6,PAK1,PFN1,PIP4K2B,PPP1CA,RAC3,RHOU,RND2,WASF1 |
| Apoptotic execution phase | 1.400 | 0.180 | -2.333 | CTNNB1,H1-0,H1-1,H1-4,H1-5,HMGB2,MAPT,PRKCD,VIM |
| Integrin to Cytoskeleton Signaling Pathway | 1.400 | 0.132 | 2.041 | ACTB,AKT1,ARPC3,ARPC5,ARPC5L,CAPNS1,CAV1,DOCK1,FYN,HRAS,MAP3K11,MYL12B,NCK2,PAK1,PFN1,PIK3CA,PIK3CD,PIK3R2,PLCG1,PPP1CA,RAC3,RAP2A,RAPGEF1,RHOU,RND2,TSPAN5,VCL |
| Chaperone Mediated Autophagy | 1.390 | 0.238 | 1.342 | HSP90AA1,HSP90AB1,PARK7,UBC,VIM |
| Mitochondrial Dysfunction | 1.390 | 0.123 | -0.333 | ACADL,ATP1A1,CACNA1A,CACNG3,CACNG8,CALM1,CAPNS1,CATSPER2,COX4I2,COX5A,COX6A1,COX7A1,COX8A,COXFA4,CREBBP,CYCS,Gstm7,HAP1,ITPR1,ITPR2,MAPK7,MAPK9,MAPT,MGST1,MTOR,NCSTN,NDUFA10,NDUFA13,NDUFA3,NDUFB2,NDUFB5,NDUFS7,PARK7,PIK3CA,PIK3CD,PIK3R2,PPARGC1A,PRKAA2,SDHD,VDAC3 |
| MTOR signalling | 1.380 | 0.214 | 0.816 | AKT1,EIF4E,FKBP1A,LAMTOR4,MTOR,RRAGB |
| Degradation of beta-catenin by the destruction complex | 1.380 | 0.154 | 1.069 | CTNNB1,PSMB10,PSMB2,PSMB4,PSMB5,PSMC5,PSMD13,PSMD6,RBX1,SKP2,TLE2,TLE3,TLE4,UBC |
| Dopamine-DARPP32 Feedback in cAMP Signaling | 1.370 | 0.136 | -1.147 | ADCY1,ADCY8,CACNA1A,CACNG3,CACNG8,CALM1,CAMKK2,CREBBP,GNAQ,GRIN2D,GUCY1A1,GUCY1B1,ITPR1,ITPR2,KCNJ8,PLCG1,PLCH1,PLCH2,PPP1CA,PPP2R2B,PRKCB,PRKCD,PRKCH |
| RHOGDI Signaling | 1.370 | 0.131 | 0.000 | ACTB,ARHGAP12,ARHGDIA,ARHGEF17,ARPC3,ARPC5,ARPC5L,CDH2,CDH24,CDH4,CFL1,CREBBP,GNA12,GNAL,GNAQ,GNAZ,GNB2,GNG3,GNG4,GNG7,MYL12B,MYL6,PAK1,PIP4K2B,RAC3,RHOU,RND2,WASF1 |
| Synaptic Long Term Depression | 1.370 | 0.134 | 0.626 | CACNA1A,CACNG3,CACNG8,CATSPER2,GAD1,GNA12,GNAL,GNAQ,GNAZ,GRIA2,GRIA3,GRM3,GUCY1A1,GUCY1B1,HRAS,ITPR1,ITPR2,PLCG1,PLCH1,PLCH2,PPP2R2B,PRKCB,PRKCD,PRKCH,RAP2A |
| Apelin Adipocyte Signaling Pathway | 1.360 | 0.157 | 0.333 | ADCY1,ADCY8,CAT,Gstm7,GUCY1A1,GUCY1B1,HIF1A,MAPK7,MAPK9,MGST1,PPARGC1A,PRKAA2,RAC3 |
| Malate-aspartate shuttle | 1.350 | 0.333 | #NUM! | MDH1,MDH2,SLC25A22 |
| Interleukin-3, Interleukin-5 and GM-CSF signaling | 1.350 | 0.176 | 1.000 | FYN,JAK2,JAK3,PIK3CA,PIK3CD,PIK3R2,RAPGEF1,UBC,YWHAZ |
| Telomere Extension by Telomerase | 1.350 | 0.267 | -1.000 | HNRNPA1,MRE11,POT1,XRCC6 |
| Production of Nitric Oxide and Reactive Oxygen Species in M | 1.350 | 0.134 | -2.132 | AKT1,APOC2,CAT,CREBBP,JAK2,JAK3,MAP3K11,MAPK9,NFKBID,NGFR,PIK3CA,PIK3CD,PIK3R2,PLCG1,PPP1CA,PPP1R14B,PPP2R2B,PRKCB,PRKCD,PRKCH,RAC3,RHOU,RND2,SIRPA |
| GABA receptor activation | 1.350 | 0.169 | 0.632 | ADCY1,ADCY8,GABRA5,GABRB3,GABRG3,GNAL,GNB2,GNG3,GNG4,GNG7 |
| Lysosomal oligosaccharide catabolism | 1.340 | 0.500 | #NUM! | MAN2B1,MANBA |
| Mitotic Metaphase and Anaphase | 1.330 | 0.129 | 0.928 | BANF1,BUB1,CCNB2,CDC20,CDCA8,CDK1,CENPK,DYNC1I1,DYNLL1,MAD2L1,NDEL1,NUDC,NUP93,PSMB10,PSMB2,PSMB4,PSMB5,PSMC5,PSMD13,PSMD6,RAN,RANGAP1,SKA2,SMC3,TMPO,TUBA1A,TUBB6,UBC,WAPL |
| FGF Signaling | 1.330 | 0.155 | -0.832 | AKT1,CREBBP,FGFR3,HGF,HRAS,ITPR1,MAPKAPK2,MET,PIK3CA,PIK3CD,PIK3R2,PLCG1,RPS6KA5 |
| Colorectal Cancer Metastasis Signaling | 1.330 | 0.126 | -0.447 | ADCY1,ADCY8,AKT1,CTNNB1,DCC,FZD1,GNA12,GNAL,GNAQ,GNAZ,GNB2,GNG3,GNG4,GNG7,GUCY1A1,GUCY1B1,HRAS,IL6ST,JAK2,JAK3,LRP1,MAPK9,MMP24,PIK3CA,PIK3CD,PIK3R2,RAC3,RAP2A,RHOU,RND2,TGFBR1,WNT7B |
| FOXO-mediated transcription of oxidative stress, metabolic a | 1.320 | 0.207 | 0.000 | CAT,FOXO1,HDAC2,NR3C1,PLXNA4,PPARGC1A |
| RHO GTPases Activate WASPs and WAVEs | 1.310 | 0.182 | 1.414 | ACTB,ARPC3,ARPC5,CYFIP2,FYN,WASF1,WIPF2,WIPF3 |
| Metabolism of polyamines | 1.300 | 0.167 | 2.530 | AMD1,PSMB10,PSMB2,PSMB4,PSMB5,PSMC5,PSMD13,PSMD6,SMS,SRM |
| Adrenergic Receptor Signaling Pathway (Enhanced) | 1.300 | 0.133 | -0.408 | ADCY1,ADCY8,ADRA2C,ATP6AP2,ATP6V1E1,ATP6V1G2,CACNA1A,CACNG3,CACNG8,CATSPER2,COMT,DDC,GNAL,GNAQ,GUCY1A1,GUCY1B1,ITPR1,ITPR2,PLCG1,PLCH1,PLCH2,PRKCB,PRKCD,PRKCH |
| CLEAR Signaling Pathway | 1.300 | 0.125 | -0.686 | AKT1,ASAH1,ATP6AP2,ATP6V1E1,ATP6V1G2,BLOC1S1,CREBBP,DDIT4,FGFR3,GNPTG,HGF,HIF1A,HRAS,ITPR1,ITPR2,LAMTOR4,MAPK7,MAPK9,MID1,MTOR,NGFR,NTRK2,PLCG1,PPARGC1A,PPP2R2B,PRKAA2,PRKCB,PRKCD,PRKCH,RAP2A,RRAGB,TGFBR1,VPS35,YWHAZ |
| Gα12/13 Signaling | 1.290 | 0.141 | 2.000 | AKT1,CDH2,CDH24,CDH4,CTNNB1,GNA12,HRAS,MAPK7,MAPK9,MEF2C,MYL12B,MYL6,NFKBID,PIK3CA,PIK3CD,PIK3R2,RAP2A,RASA1 |
| ILK Signaling | 1.280 | 0.131 | 0.426 | ACTB,AKT1,CFL1,CREBBP,CTNNB1,DOCK1,FN1,HIF1A,MAPK9,MTOR,MUC1,MYL6,NCK2,PIK3CA,PIK3CD,PIK3R2,PPP1R14B,PPP2R2B,RAC3,RHOU,RND2,RPS6KA5,TMSB4X,VCL,VIM |
| SAPK/JNK Signaling | 1.280 | 0.144 | 0.832 | FCER1G,GNA12,GNB2,GNG3,GNG4,GNG7,HRAS,MAP3K11,MAP4K2,MAP4K4,MAPK9,PIK3CA,PIK3CD,PIK3R2,RAC3,RAP2A |
| NUR77 Signaling in T Lymphocytes | 1.280 | 0.156 | -0.905 | APAF1,CALM1,CYCS,FCER1G,HDAC2,HDAC7,HDAC9,MAPK7,NR4A1,PRKCB,PRKCD,PRKCH |
| Hepatitis B Chronic Liver Pathogenesis Signaling Pathway | 1.280 | 0.134 | 0.894 | AKT1,CAMKK2,CDC20,CREBBP,CTNNB1,ETV4,HIF1A,HRAS,MAD2L1,MAPK9,NLRP3,PIK3CA,PIK3CD,PIK3R2,PRKCB,PRKCD,PRKCH,RAP2A,RASSF1,TRIM37,VDAC3,ZHX2 |
| Mitotic Prophase | 1.270 | 0.149 | -1.604 | BANF1,CCNB2,CDK1,H2AZ2,H3C1,KMT5A,LPIN1,NCAPH2,NUMA1,NUP210,NUP93,PRKCB,RAE1,TMPO |
| Phospholipase C Signaling | 1.270 | 0.125 | 0.229 | ADCY1 ADCY8 AHNAK ARHGEF17 CALM1 CREBBP FCER1G FYN GNAQ GNB2 GNG3 GNG4 GNG7 GUCY1A1 GUCY1B1 HDAC2 HDAC7 HDAC9 HRAS ITPR1 ITPR2 MEF2C MYL12B MYL6 PLCG1 PPP1CA PRKCB PRKCD PRKCH RAC3 RAP2A RHOU RND2 |
| SPINK1 General Cancer Pathway | 1.260 | 0.170 | 1.000 | AKT1,HRAS,JAK2,JAK3,MT3,PIK3CA,PIK3CD,PIK3R2,RAP2A |
| FcγRIIB Signaling in B Lymphocytes | 1.260 | 0.151 | 1.134 | AKT1,CACNA1A,CACNG3,CACNG8,CATSPER2,HRAS,ITPR1,ITPR2,MAPK9,PIK3CA,PIK3CD,PIK3R2,RAP2A |
| Cellular Effects of Sildenafil (Viagra) | 1.260 | 0.113 | 1.698 | ACTB,ADCY1,ADCY8,ADGRA2,ADGRB1,ADGRB2,ADGRB3,ADGRG1,ADGRL3,ADORA2A,ADRA2C,AKT1,BDKRB1,C3AR1,CACNA1A,CACNG3,CACNG8,CALM1,CAPNS1,CCRL2,CELSR1,CREBBP,EDNRB,FZD1,GIPR,GNAQ,GNB2,GNG3,GNG4,GNG7,GPER1,GPR12,GPR137C,GPR146,GPR155,GPR156,GPR19,GPR27,GPR68,GPR85,GPR88,GRM3,GUCY1A1,GUCY1B1,HIF1A,HSP90AA1,HSP90AB1,HSP90B1,KCNN3,MAPK7,MAPK9,MCHR1,MEF2C,MTOR,MYL12B,MYL6,NMUR1,OXGR1,PDE6D,PIK3CA,PIK3CD,PIK3R2,PPARGC1A,PPP1CA,PPP2R2B,PRKAA2,PTH1R,S1PR1,SSTR2,SSTR3,TGFBR1 |
| RAN Signaling | 1.260 | 0.250 | #NUM! | KPNA2,RAN,RANBP1,RANGAP1 |
| Granzyme B Signaling | 1.260 | 0.250 | 0.000 | APAF1,CYCS,NUMA1,PARP1 |
| Regulation of CDH11 Expression and Function | 1.260 | 0.200 | -0.816 | BHLHE22,CDH24,CTNNA1,CTNNB1,TNRC6B,ZEB2 |
| Senescence Pathway | 1.250 | 0.123 | 0.180 | AKT1,ATM,CACNA1A,CACNG3,CACNG8,CALM1,CAPNS1,CAT,CBX8,CCNB2,CCND2,CDK1,CDK6,CREBBP,DHCR24,EIF4E,EZH2,HIPK2,HRAS,ITPR2,MAPK7,MAPKAPK2,MRE11,MTOR,PARP1,PDK2,PIK3CA,PIK3CD,PIK3R2,PPP1CA,PPP2R2B,RAP2A,RPS6KA5,SMAD6,TGFBR1 |
| Neurotrophin/TRK Signaling | 1.240 | 0.154 | 0.302 | AKT1,CREBBP,HRAS,NGFR,NTF3,NTRK2,PIK3CA,PIK3CD,PIK3R2,PLCG1,RAP2A,SPRY2 |
| Erythropoietin Signaling Pathway | 1.230 | 0.134 | 0.447 | AKT1,APAF1,CCND2,CTNNB1,HIF1A,HRAS,ITPR1,ITPR2,JAK2,MTOR,NFKBID,PIK3CA,PIK3CD,PIK3R2,PLCG1,PRKCB,PRKCD,PRKCH,RAC3,RAP2A,SHC3 |
| Pentose Phosphate Pathway | 1.230 | 0.300 | #NUM! | PGLS,RPIA,TKT |
| Transport of vitamins, nucleosides, and related molecules | 1.230 | 0.184 | 0.378 | ARL2BP,PDZD11,SLC33A1,SLC35A1,SLC5A6,SLCO3A1,SLCO4C1 |
| Degradation of CRY and PER proteins | 1.220 | 0.161 | 2.530 | CRY1,PSMB10,PSMB2,PSMB4,PSMB5,PSMC5,PSMD13,PSMD6,RBX1,UBC |
| CD28 Signaling in T Helper Cells | 1.220 | 0.142 | 1.387 | AKT1,ARPC3,ARPC5,ARPC5L,CALM1,FCER1G,FYN,ITPR1,ITPR2,MAPK9,NFKBID,PAK1,PIK3CA,PIK3CD,PIK3R2,PLCG1 |
| M-decay: degradation of maternal mRNAs by maternally sto | 1.220 | 0.174 | 0.000 | CNOT1,EIF4A1,EIF4E,KDM4B,KPNA2,PABPC1,PAIP1,TNKS1BP1 |
| Synthesis of DNA | 1.220 | 0.139 | -0.243 | GINS2,GINS4,MCM2,MCM3,MCM5,MCM6,PSMB10,PSMB2,PSMB4,PSMB5,PSMC5,PSMD13,PSMD6,RBX1,RPA2,SKP2,UBC |
| RHO GTPases Activate Formins | 1.210 | 0.137 | 0.471 | ACTB,BUB1,CDC20,CDCA8,CENPK,DYNC1I1,DYNLL1,MAD2L1,MRTFA,NDEL1,NUDC,PFN1,RANGAP1,SCAI,SKA2,SRGAP2,TUBA1A,TUBB6 |
| Xenobiotic Metabolism PXR Signaling Pathway | 1.210 | 0.137 | -1.000 | ALDH16A1,Aldh3a2,CAT,CITED2,CREBBP,Gstm7,GSTO1,HS3ST4,HS6ST2,HSP90AA1,HSP90AB1,HSP90B1,MGST1,PPARGC1A,PPP1CA,PRKCB,PRKCD,PRKCH |
| DNA Replication Pre-Initiation | 1.210 | 0.146 | 0.000 | H2AZ2,H3C1,MCM2,MCM3,MCM5,MCM6,PSMB10,PSMB2,PSMB4,PSMB5,PSMC5,PSMD13,PSMD6,UBC |
| PTEN Regulation | 1.210 | 0.134 | 1.342 | AKT1,CBX8,EZH2,HDAC2,HDAC7,LAMTOR4,MBD3,MECOM,MKRN1,MTOR,PSMB10,PSMB2,PSMB4,PSMB5,PSMC5,PSMD13,PSMD6,RRAGB,TNRC6B,UBC |
| Neutrophil degranulation | 1.210 | 0.117 | 0.289 | ACAA1,ALDOC,APAF1,APRT,ARPC5,ARSB,ASAH1,ATP6AP2,C3AR1,CAT,CCT2,CKAP4,CPPED1,DNAJC3,DOCK2,DYNLL1,EEF2,FABP5,FCER1G,GCA,HSP90AA1,HSP90AB1,IDH1,IMPDH1,LRRC7,MAN2B1,MANBA,MGST1,MIF,NCSTN,NDUFC2,PA2G4,PKM,PNP,PRKCD,PRTN3,PSMD13,PSMD6,PTGES2,RAB18,RAB31,RAB6A,RAB9B,S100A9,SIRPA,SLCO4C1,VCL,XRCC6 |
| Transcriptional regulation of brown and beige adipocyte diffe | 1.200 | 0.194 | 0.000 | COX7A1,HDAC2,MBD3,PPARGC1A,PRDM16,ZNF423 |
| Protein Ubiquitination Pathway | 1.190 | 0.123 | 0.928 | CDC20,CRYAB,CUL2,DNAJC15,DNAJC16,DNAJC19,DNAJC21,DNAJC3,HSP90AA1,HSP90AB1,HSP90B1,HSPA12A,HSPB8,HSPE1,HSPH1,PSMB10,PSMB2,PSMB4,PSMB5,PSMC5,PSMD13,PSMD6,RBX1,SKP2,UBC,UBE2E3,USP12,USP24,USP29,USP35,USP43,USP54 |
| Sertoli Cell-Sertoli Cell Junction Signaling | 1.190 | 0.125 | -0.756 | ACTB,AKT1,CDH2,CDH24,CDH4,CLDN11,CLDN20,CREBBP,CTNNA1,CTNNB1,EPN1,HRAS,IL1RAPL1,LAMB1,MAP3K11,MAPK9,NGFR,PAK1,PIK3CA,PIK3CD,PIK3R2,RAP2A,SPTB,TGFBR1,TUBA1A,TUBB6,TUBG1,VCL,YBX3 |
| Thrombopoietin Signaling | 1.180 | 0.159 | 0.000 | HRAS,JAK2,PIK3CA,PIK3CD,PIK3R2,PLCG1,PRKCB,PRKCD,PRKCH,RAP2A |
| ERB2-ERBB3 Signaling | 1.180 | 0.159 | 1.265 | AKT1,ETV4,FOXO1,HRAS,JAK3,NRG1,PIK3CA,PIK3CD,PIK3R2,RAP2A |
| Cellular response to heat stress | 1.180 | 0.144 | -0.535 | ATM,CREBBP,CRYAB,HSP90AA1,HSP90AB1,HSPA12A,HSPB8,HSPH1,MAPKAPK2,MTOR,NUP210,NUP93,RAE1,RPA2 |
| Small Cell Lung Cancer Signaling | 1.180 | 0.144 | 0.000 | AKT1,APAF1,CCND2,CDK6,CYCS,HDAC2,HDAC7,HDAC9,NFKBID,PA2G4,PIK3CA,PIK3CD,PIK3R2,SKP2 |
| Gap junction trafficking and regulation | 1.180 | 0.179 | 1.890 | ACTB,AP2M1,CLTB,GJB1,MYO6,TUBA1A,TUBB6 |
| Smooth Muscle Contraction | 1.180 | 0.179 | 1.890 | CALM1,GUCY1A1,GUCY1B1,MYL12B,MYL6,PAK1,VCL |
| Metabolism of nitric oxide: NOS3 activation and regulation | 1.170 | 0.235 | 1.000 | AKT1,CALM1,CAV1,HSP90AA1 |
| Signaling by Type 1 Insulin-like Growth Factor 1 Receptor (IG | 1.170 | 0.235 | 0.000 | HRAS,IGF2,PIK3CA,PIK3R2 |
| Protein methylation | 1.170 | 0.235 | 2.000 | CALM1,EEF2,PRMT3,VCPKMT |
| FOXO-mediated transcription of cell cycle genes | 1.170 | 0.235 | -1.000 | BTG1,CAV1,FOXO1,PCBP4 |
| HSP90 chaperone cycle for steroid hormone receptors in the | 1.170 | 0.170 | 2.121 | CAPZB,DYNC1I1,DYNLL1,HSP90AA1,HSP90AB1,NR3C1,TUBA1A,TUBB6 |
| Neuregulin Signaling | 1.170 | 0.139 | 0.000 | AKT1,ERBB4,ERBIN,HRAS,HSP90AA1,HSP90AB1,HSP90B1,MTOR,NRG1,PIK3R2,PLCG1,PRKCB,PRKCD,PRKCH,RAP2A,RNF41 |
| p70S6K Signaling | 1.160 | 0.135 | 0.728 | AKT1,EEF2,GNAQ,HRAS,MAPT,MTOR,PIK3CA,PIK3CD,PIK3R2,PLCG1,PLCH1,PLCH2,PPP2R2B,PRKCB,PRKCD,PRKCH,RAP2A,YWHAZ |
| ID1 Signaling Pathway | 1.150 | 0.127 | 0.209 | AKT1,BHLHE22,CAV1,CDC20,CHRNA4,CREBBP,CTNNB1,FGFR3,FN1,FYN,HIF1A,HRAS,IGF2,KIF11,NGFR,PARK7,PIK3CA,PIK3CD,PIK3R2,PLXNB1,RAP2A,S100A9,TGFBR1,VIM,WDR48 |
| Glutamate Degradation III (via 4-aminobutyrate) | 1.140 | 0.400 | #NUM! | GAD1,GAD2 |
| G-protein beta:gamma signalling | 1.140 | 0.188 | 1.633 | AKT1,GNB2,GNG3,GNG4,GNG7,PAK1 |
| Thrombin signalling through proteinase activated receptors ( | 1.140 | 0.188 | 1.633 | GNA12,GNAQ,GNB2,GNG3,GNG4,GNG7 |
| Inhibition of Angiogenesis by TSP1 | 1.140 | 0.188 | 1.000 | AKT1,FYN,GUCY1A1,GUCY1B1,MAPK9,TGFBR1 |
| Adipogenesis pathway | 1.130 | 0.134 | 0.471 | AKT1,BSCL2,CTNNB1,EZH2,FGFR3,FOXO1,FZD1,GTF2H3,HDAC2,HDAC7,HDAC9,HIF1A,KAT6B,LPIN1,NR2F2,SAP130,SOX9,ZNF423 |
| White Adipose Tissue Browning Pathway | 1.130 | 0.134 | -0.258 | ADCY1,ADCY8,CACNA1A,CACNG3,CACNG8,CAMKK2,CATSPER2,CREBBP,FCER1G,FGFR3,GUCY1A1,GUCY1B1,ITPR1,ITPR2,PPARGC1A,PRDM16,PRKAA2,VGF |
| Gamma carboxylation, hypusinylation, hydroxylation, and ar | 1.130 | 0.161 | 0.333 | ARSB,ARSK,EEF2,EIF5A,ETF1,FN3K,KDM8,RPL8,TPST1 |
| RET signaling | 1.130 | 0.175 | 0.378 | DOK5,MAPK7,PIK3CA,PIK3CD,PIK3R2,PLCG1,SHC3 |
| Meiotic recombination | 1.130 | 0.175 | -2.646 | ATM,BRCA2,H2AZ2,H3C1,MRE11,RAD51C,RPA2 |
| DNA Damage Bypass | 1.130 | 0.167 | -0.707 | DTL,PCLAF,RBX1,RPA2,SPRTN,UBC,USP43,WDR48 |
| GPCR-Mediated Integration of Enteroendocrine Signaling Exe | 1.120 | 0.151 | 0.333 | ADCY1,ADCY8,GIPR,GNAQ,GUCY1A1,GUCY1B1,ITPR1,ITPR2,PLCG1,PLCH1,PLCH2 |
| Sumoylation Pathway | 1.120 | 0.141 | 0.000 | ARHGDIA,CREBBP,HDAC2,MAPK9,NR3C1,RAC3,RAN,RANGAP1,RHOU,RND2,RNF4,SENP3,ZEB1,ZNF217 |
| Co-stimulation by ICOS | 1.120 | 0.273 | #NUM! | PIK3CA,PIK3CD,PIK3R2 |
| Metabolism of cofactors | 1.110 | 0.200 | 1.342 | AKT1,CALM1,COQ7,HSP90AA1,IDH1 |
| Signaling by NTRK3 (TRKC) | 1.110 | 0.200 | 0.447 | HRAS,NTF3,PIK3CA,PLCG1,PTPRO |
| MAPK6/MAPK4 signaling | 1.100 | 0.143 | 0.832 | CDK1,ETV4,FOXO1,PAK1,PSMB10,PSMB2,PSMB4,PSMB5,PSMC5,PSMD13,PSMD6,TNRC6B,UBC |
| GABA synthesis, release, reuptake and degradation | 1.100 | 0.222 | -1.000 | CPLX1,GAD1,GAD2,SLC6A1 |
| Oxytocin in Spinal Neurons Signaling Pathway | 1.090 | 0.182 | 0.816 | ABCC8,CAMKK2,GAD1,GUCY1A1,GUCY1B1,KCNJ8 |
| Signal amplification | 1.090 | 0.182 | 1.633 | AAMP,GNAQ,GNB2,GNG3,GNG4,GNG7 |
| MAPK targets/ Nuclear events mediated by MAP kinases | 1.090 | 0.182 | -1.633 | MAPK7,MAPK9,MAPKAPK2,MEF2C,RPS6KA2,RPS6KA5 |
| NAD Signaling Pathway | 1.090 | 0.132 | -1.500 | ACADL,AKT1,H1-0,H1-1,H1-4,H1-5,H1f3,HGF,HIF1A,PARP1,PARP8,PIK3CA,PIK3CD,PIK3R2,POLR2L,PPARGC1A,PRKAA2,SIRT4 |
| DNA Double Strand Break Response | 1.090 | 0.163 | 0.000 | ABRAXAS1,ATM,BABAM1,KDM4B,KPNA2,MRE11,UBC,UBXN1 |
| Regulation of beta-cell development | 1.080 | 0.171 | -1.890 | AKT1,CREBBP,FOXO1,INSM1,NEUROD1,NEUROG3,NOTCH1 |
| RNA Polymerase III Transcription | 1.080 | 0.171 | 1.134 | NFIC,NFIX,POLR1C,POLR1D,POLR2L,POLR3G,ZNF143 |
| GM-CSF Signaling | 1.070 | 0.152 | 1.000 | AKT1,HRAS,JAK2,PIK3CA,PIK3CD,PIK3R2,PIM1,PRKCB,RACK1,RAP2A |
| Eicosanoid Signaling | 1.070 | 0.120 | 0.000 | ADCY1,ADCY8,AKT1,CREBBP,CTNNB1,GNAL,GNAQ,GNB2,GNG3,GNG4,GNG7,GUCY1A1,GUCY1B1,HRAS,KCNJ8,KCNMB4,MAPKAPK2,PIK3CA,PIK3CD,PIK3R2,PLCG1,PLCH1,PLCH2,PRKCB,PRKCD,PRKCH,PTGES2,PTGIS,RAP2A,RAPGEF4,RPS6KA2,SOX9 |
| Cachexia Signaling Pathway | 1.070 | 0.117 | 0.324 | ADCY1,ADCY8,AKT1,CAPNS1,DDIT4,EIF2S3,FOXO1,GUCY1A1,GUCY1B1,HSP90AA1,HSP90AB1,HSP90B1,IGFBP3,IL1RAPL1,INHBA,MAPK9,MEF2C,MTOR,NGFR,NR3C1,PHGDH,PIK3CA,PIK3CD,PIK3R2,PRKAA2,PRKCB,PRKCD,PRKCH,PSMB10,PSMB2,PSMB4,PSMB5,PSMC5,PSMD13,PSMD6,PTH1R,S1PR1,TGFBR1,VPS28 |
| Role of Tissue Factor in Cancer | 1.070 | 0.125 | 0.408 | AKT1,CFL1,EIF4E,FYN,GNA12,GNAQ,HGF,HRAS,JAK2,MAPK7,MAPK9,MET,MTOR,NGFR,PDIA6,PIK3CA,PIK3CD,PIK3R2,RAP2A,RPS6KA2,RPS6KA5,TFPI,TGFBR1,ZEB1 |
| ESR-mediated signaling | 1.060 | 0.136 | -0.258 | CREBBP,ERBB4,H2AZ2,H3C1,HSP90AA1,HSP90AB1,KDM4B,KPNA2,POLR2L,PRMT1,SMC3,TLE3,TNRC6B,USF1,ZNF217 |
| Role of NFAT in Regulation of the Immune Response | 1.060 | 0.126 | 2.138 | AKT1,CALM1,FCER1G,FYN,GNA12,GNAL,GNAQ,GNAZ,GNB2,GNG3,GNG4,GNG7,HRAS,ITPR1,ITPR2,KPNA2,MEF2C,NFKBID,PIK3CA,PIK3CD,PIK3R2,PLCG1,RAP2A |
| Signaling by ERBB4 | 1.060 | 0.155 | -0.333 | ERBB4,GABRB3,GABRG3,HRAS,NCSTN,NRG1,PIK3CA,UBC,WWOX |
| Signaling by PDGF | 1.060 | 0.155 | 0.333 | COL9A3,HRAS,NCK2,PIK3CA,PIK3R2,PLCG1,RAPGEF1,RASA1,THBS3 |
| NoRC negatively regulates rRNA expression | 1.060 | 0.155 | 1.000 | GTF2H3,H2AZ2,H3C1,HDAC2,POLR1C,POLR1D,POLR2L,SAP130,TAF1D |
| D-myo-inositol (1,4,5)-Trisphosphate Biosynthesis | 1.050 | 0.192 | -1.342 | PI4KB,PIP4K2B,PLCG1,PLCH1,PLCH2 |
| Neddylation | 1.050 | 0.121 | 1.890 | COPS5,CUL2,DCUN1D5,DTL,FBXL20,FBXL4,FBXL5,FBXO9,FBXW4,FEM1B,HIF1A,KLHL2,KLHL5,LRR1,NUB1,PSMB10,PSMB2,PSMB4,PSMB5,PSMC5,PSMD13,PSMD6,RBX1,SKP2,SOCS2,UBC,WDR5,WSB1,WSB2 |
| WNK Renal Signaling Pathway | 1.040 | 0.137 | -0.302 | AKT1,CALM1,CUL2,LRRC8A,PIK3CA,PIK3CD,PIK3R2,PRKCB,PRKCD,PRKCH,RBX1,RPS6KA5,SLC12A6,WNK1 |
| Potassium Channels | 1.040 | 0.137 | 1.069 | ABCC8,GNB2,GNG3,GNG4,GNG7,KCNC2,KCNH5,KCNH8,KCNJ8,KCNK13,KCNMB4,KCNN3,KCNQ5,KCNS3 |
| Oxidative Phosphorylation | 1.040 | 0.137 | 2.309 | COX4I2,COX5A,COX6A1,COX7A1,COX8A,COXFA4,CYCS,NDUFA10,NDUFA13,NDUFA3,NDUFB2,NDUFB5,NDUFS7,SDHD |
| Centrosomal KIAA0586 Signaling Pathway | 1.040 | 0.132 | 1.213 | C2CD3,FGFR3,KIF3B,PIK3CA,PIK3CD,PIK3R2,PSMB10,PSMB2,PSMB4,PSMB5,PSMC5,PSMD13,PSMD6,RBX1,SKP2,SMC3,VCAN |
| Activation of the pre-replicative complex | 1.040 | 0.176 | -2.449 | DBF4,MCM2,MCM3,MCM5,MCM6,RPA2 |
| Superpathway of Methionine Degradation | 1.040 | 0.176 | 1.342 | AHCYL1,CDO1,EEF1AKMT4,PCCA,PRMT1,PRMT3 |
| Cholecystokinin/Gastrin-mediated Signaling | 1.040 | 0.135 | 0.535 | GNA12,GNAQ,HRAS,ITPR1,ITPR2,MAPK7,MAPK9,MEF2C,PRKCB,PRKCD,PRKCH,RAC3,RAP2A,RHOU,RND2 |
| UDP-N-acetyl-D-galactosamine Biosynthesis I | 1.030 | 1.000 | #NUM! | GALE |
| Mitotic G2-G2/M phases | 1.030 | 0.124 | 0.408 | CCNB2,CDK1,CDK5RAP2,CPAP,DYNLL1,HAUS4,HAUS5,HJURP,HSP90AA1,HSP90AB1,PSMB10,PSMB2,PSMB4,PSMB5,PSMC5,PSMD13,PSMD6,RBX1,SFI1,TUBA1A,TUBB6,TUBG1,TUBGCP2,UBC |
| Thyroid Cancer Signaling | 1.030 | 0.145 | 0.905 | AKT1,CTNNB1,FOXO1,HRAS,MTOR,NTF3,NTRK2,PIK3CA,PIK3CD,PIK3R2,RAP2A |
| GDNF Family Ligand-Receptor Interactions | 1.030 | 0.145 | 0.707 | DOK5,HRAS,ITPR1,ITPR2,MAPK9,PIK3CA,PIK3CD,PIK3R2,PLCG1,RAP2A,RASA1 |
| HDR through MMEJ (alt-NHEJ) | 1.030 | 0.250 | #NUM! | BRCA2,MRE11,PARP1 |
| Assembly of RNA Polymerase I Complex | 1.030 | 0.250 | #NUM! | POLR1C,POLR1D,TAF1D |
| UDP-N-acetyl-D-galactosamine Biosynthesis II | 1.030 | 0.250 | #NUM! | GALE,GNPDA1,HK1 |
| RHO GTPases Activate ROCKs | 1.020 | 0.211 | 2.000 | CFL1,MYL12B,MYL6,PAK1 |
| FOXO-mediated transcription | 1.020 | 0.211 | 0.000 | AKT1,CREBBP,FOXO1,YWHAZ |
| Lipid Antigen Presentation by CD1 | 1.020 | 0.211 | 1.000 | AP2M1,CALR,CANX,FCER1G |
| Methionine Degradation I (to Homocysteine) | 1.020 | 0.211 | #NUM! | AHCYL1,EEF1AKMT4,PRMT1,PRMT3 |
| UFMylation Signaling Pathway | 1.020 | 0.153 | -1.000 | ABCA1,ABCG1,AKT1,ATM,EIF4A1,EIF4E,MRE11,PABPC1,PAIP1 |
| RAC Signaling | 1.020 | 0.129 | 1.698 | ARFIP2,ARPC3,ARPC5,ARPC5L,CFL1,CYFIP2,ELK4,HRAS,MAP3K11,MCF2L,PAK1,PIK3CA,PIK3CD,PIK3R2,PIP4K2B,RAC3,RAP2A,WASF1 |
| HGF Signaling | 1.010 | 0.131 | -0.243 | AKT1,DOCK1,HGF,HRAS,MAP3K11,MAPK9,MET,PAK1,PIK3CA,PIK3CD,PIK3R2,PLCG1,PRKCB,PRKCD,PRKCH,RAP2A,RAPGEF1 |
| ROBO SLIT Signaling Pathway | 1.010 | 0.132 | -0.775 | AKT1,ARPC3,ARPC5,ARPC5L,CDH2,CTNNB1,DCC,FYN,MYL12B,NCK2,NRXN3,NTN1,PAK1,PRDM16,SLIT1,SRGAP2 |
| NGF Signaling | 1.010 | 0.132 | -1.291 | AKT1,CREBBP,HRAS,MAP3K11,MAPK7,MAPK9,NGFR,PIK3CA,PIK3CD,PIK3R2,PLCG1,PRKCD,RAP2A,RPS6KA2,RPS6KA5,SHC3 |
| Egg-Sperm Fusion Signaling Pathway | 1.010 | 0.136 | 0.000 | ACR,AKT1,CCNB2,CDK1,ITPR1,ITPR2,PIK3CA,PIK3CD,PIK3R2,SYT16,SYT9,VAMP3,VTI1A,ZP3 |
| CSDE1 Signaling Pathway | 1.010 | 0.157 | 0.000 | AKT1,APAF1,CTNNB1,FABP7,FN1,PABPC1,PAIP1,VIM |
| Proteasome assembly | 1.010 | 0.157 | 2.121 | POMP,PSMB10,PSMB2,PSMB4,PSMB5,PSMC5,PSMD13,PSMD6 |
| Interleukin-7 signaling | 0.995 | 0.185 | -1.342 | H3C1,HGF,JAK3,PIK3R2,SOCS2 |
| Pentose Phosphate Pathway (Non-oxidative Branch) | 0.994 | 0.333 | #NUM! | RPIA,TKT |
| Aspartate Degradation II | 0.994 | 0.333 | #NUM! | MDH1,MDH2 |
| Mitotic G1 phase and G1/S transition | 0.991 | 0.130 | 0.243 | AKT1,CCND2,CDK1,CDK6,JAK2,PSMB10,PSMB2,PSMB4,PSMB5,PSMC5,PSMD13,PSMD6,RRM2,SKP2,TFDP2,TOP2A,UBC |
| p53 Signaling | 0.986 | 0.137 | -2.111 | ADGRB1,AKT1,APAF1,ATM,CCND2,CTNNB1,HDAC9,HIF1A,HIPK2,PIK3CA,PIK3CD,PIK3R2,RPRM |
| Virus Entry via Endocytic Pathways | 0.986 | 0.133 | 1.291 | ACTB,AP2M1,CAV1,CLTB,FYN,HRAS,PIK3CA,PIK3CD,PIK3R2,PLCG1,PRKCB,PRKCD,PRKCH,RAC3,RAP2A |
| Telomerase Signaling | 0.985 | 0.135 | 2.530 | AKT1,HDAC2,HDAC7,HDAC9,HRAS,HSP90AA1,HSP90AB1,HSP90B1,PIK3CA,PIK3CD,PIK3R2,POT1,PPP2R2B,RAP2A |
| Plasma lipoprotein assembly, remodeling, and clearance | 0.973 | 0.145 | -0.632 | ABCA1,ABCG1,ANGPTL3,AP2M1,APOC2,LMF2,MBTPS1,MBTPS2,SCARB1,UBC |
| Oxidative Stress Induced Senescence | 0.966 | 0.141 | -0.905 | CBX8,CDK6,EZH2,H2AZ2,H3C1,MAP4K4,MAPK9,MAPKAPK2,TFDP2,TNRC6B,UBC |
| Sertoli Cell-Germ Cell Junction Signaling Pathway (Enhanced) | 0.963 | 0.119 | 1.732 | ACTB,AKT1,ARPC3,ARPC5,ARPC5L,CDH2,CLDN11,CLDN20,CTNNA1,CTNNB1,EPS8,GUCY1A1,GUCY1B1,HRAS,IL1RAPL1,LAMB1,MAP3K11,MAPK9,NGFR,PIK3CA,PIK3CD,PIK3R2,PPP2R2B,PRKAA2,RAC3,RAP2A,TGFBR1 |
| Sirtuin Signaling Pathway | 0.962 | 0.117 | -2.600 | ABCA1,ACADL,AKT1,ATG13,COXFA4,FOXO1,H1-0,H1-1,H1-4,H1-5,H1f3,HIF1A,MAPK7,MTOR,MYCN,NDUFA10,NDUFA13,NDUFA3,NDUFB2,NDUFB5,NDUFS7,PARP1,POLR1C,POLR1D,PPARGC1A,PRKAA2,SDHD,SIRT4,TUBA1A,VDAC3,XRCC6,ZBTB14 |
| VEGF Signaling | 0.960 | 0.135 | 2.111 | ACTB,AKT1,EIF2S3,FOXO1,HIF1A,HRAS,PIK3CA,PIK3CD,PIK3R2,PLCG1,PRKCB,RAP2A,VCL |

| | | | | |
|---|---|---|---|---|
| Interleukin-12 family signaling | 0.959 | 0.200 | 1.000 | CANX,IL6ST,JAK2,VAMP7 |
| Valine Degradation I | 0.959 | 0.200 | #NUM! | ACADSB,DBT,ECHDC3,HIBADH |
| B-WICH complex positively regulates rRNA expression | 0.953 | 0.159 | 1.134 | ACTB,H2AZ2,H3C1,POLR1C,POLR1D,POLR2L,TAF1D |
| Mitochondrial biogenesis | 0.952 | 0.148 | 0.333 | CALM1,CHD9,CREBBP,CYCS,HCFC1,MEF2C,PPARGC1A,PRKAA2,SIRT4 |
| Fatty acyl-CoA biosynthesis | 0.943 | 0.167 | 2.449 | ACACA,ACSL4,ELOVL4,FASN,HACD3,TECR |
| Assembly and cell surface presentation of NMDA receptors | 0.943 | 0.167 | -0.816 | CASK,GRIN2D,LRRC7,NBEA,TUBA1A,TUBB6 |
| Pre-NOTCH Expression and Processing | 0.942 | 0.143 | -2.530 | CREBBP,H2AZ2,H3C1,LFNG,MFNG,NOTCH1,POFUT1,RAB6A,TFDP2,TNRC6B |
| Metabolism of water-soluble vitamins and cofactors | 0.942 | 0.143 | 1.265 | ACACA,FASN,GSTO1,MTRR,PANK2,PANK4,PCCA,PDZD11,SLC25A19,SLC5A6 |
| Pentose phosphate pathway | 0.942 | 0.231 | #NUM! | PGLS,RPIA,TKT |
| Mitochondrial calcium ion transport | 0.942 | 0.231 | #NUM! | MICU1,SMDT1,VDAC3 |
| Class I MHC mediated antigen processing and presentation | 0.939 | 0.113 | 0.156 | ARIH2,CALR,CANX,CDC20,CUL2,FBXL20,FBXL4,FBXL5,FBXO9,FBXW4,GLMN,HECW2,HERC4,KLHL2,KLHL5,LRR1,MGRN1,MKRN1,PSMB10,PSMB2,PSMB4,PSMB5,PSMC5,PSMD13,PSMD6,RBX1,RNF182,RNF220,RNF4,RNF41,S100A1,S100A9,SEC24D,SKP2,TRIM37,UBAC1,UBC,UBE2E3,VAMP3,WSB1,ZNRF2 |
| Melanocyte Development and Pigmentation Signaling | 0.934 | 0.134 | -0.905 | ADCY1,ADCY8,CREBBP,GUCY1A1,GUCY1B1,HRAS,PIK3CA,PIK3CD,PIK3R2,PLCG1,RAP2A,RPS6KA2,RPS6KA5 |
| Xenobiotic Metabolism CAR Signaling Pathway | 0.930 | 0.126 | 0.000 | ALDH16A1,Aldh3a2,CITED2,CREBBP,Gstm7,GSTO1,HS3ST4,HS6ST2,HSP90AA1,HSP90AB1,HSP90B1,MGST1,PPARGC1A,PPP2R2B,PRKCB,PRKCD,PRKCH,RACK1 |
| Adrenomedullin signaling pathway | 0.929 | 0.121 | 0.218 | ADCY1,ADCY8,AKT1,CALM1,GAD1,GNAQ,GUCY1A1,GUCY1B1,HIF1A,HRAS,ITPR1,ITPR2,KCNN3,MAPK7,MAPK9,PIK3CA,PIK3CD,PIK3R2,PLCG1,PLCH1,PLCH2,RAP2A,SHC3 |
| Transcriptional Regulatory Network in Embryonic Stem Cells | 0.924 | 0.123 | 0.000 | AKT1,CTNNB1,EOMES,FGFR3,FZD1,GDF3,HRAS,IGF2,IL6ST,INHBA,JAK2,JAK3,NOG,PIK3CA,PIK3CD,PIK3R2,RAP2A,SOX2,TGFBR1,WNT7B |
| XIST Epigenetic Regulation Signaling Pathway | 0.919 | 0.128 | 0.000 | AXL,CBX8,CHD5,EZH2,H2AZ2,HDAC2,HIF1A,MBD3,MUC1,NOTCH1,NR3C1,SOX2,SOX9,TRDMT1,ZEB1,ZEB2 |
| AMPK Signaling | 0.908 | 0.117 | -0.500 | ACACA,ACTB,ADRA2C,AKT1,CAMKK2,CHRNA4,CREBBP,EEF2,FASN,FOXO1,GNA12,GNAL,GNAQ,GNAZ,GNB2,GNG3,GNG4,GNG7,HMGCR,MTOR,PIK3CA,PIK3CD,PIK3R2,PPARGC1A,PPP2R2B,PRKAA2,RAB6A,RAB8B |
| IL-13 Signaling Pathway | 0.907 | 0.135 | -1.155 | AKT1,ANO1,CTNNB1,DUSP1,FYN,JAK2,MTOR,PIK3CA,PIK3CD,PIK3R2,POSTN,PRKCD |
| Phagosome Formation | 0.907 | 0.107 | -1.364 | ADGRA2,ADGRB1,ADGRB2,ADGRB3,ADGRG1,ADGRL3,ADORA2A,ADRA2C,AKT1,ARFIP2,ARPC3,ARPC5,ARPC5L,BDKRB1,C3AR1,CCRL2,CELSR1,CFL1,DOCK1,EDNRB,ELMO1,FCER1G,FN1,FYN,FZD1,GIPR,GPER1,GPR12,GPR137C,GPR146,GPR155,GPR156,GPR19,GPR27,GPR68,GPR85,GPR88,GRM3,HRAS,ITPR1,ITPR2,MAPK7,MCHR1,MTOR,MYL12B,MYL6,NMUR1,OXGR1,PAK1,PIK3CA,PIK3CD,PIK3R2,PIP4K2B,PLCG1,PRKCB,PRKCD,PRKCH,PTH1R,RAC3,RAP2A,RAPGEF1,RAPGEF4,S1PR1,SAP130,SSTR2,SSTR3,WASF1 |
| RHO GTPases activate PAKs | 0.899 | 0.190 | 2.000 | CALM1,MYL12B,MYL6,PAK1 |
| Cysteine Biosynthesis III (mammalia) | 0.899 | 0.190 | #NUM! | AHCYL1,EEF1AKMT4,PRMT1,PRMT3 |
| Vitamin-C Transport | 0.899 | 0.190 | #NUM! | GLRX,GSTO1,LRRC8A,TXNRD1 |
| Regulation of Apoptosis | 0.897 | 0.148 | 2.121 | PSMB10,PSMB2,PSMB4,PSMB5,PSMC5,PSMD13,PSMD6,UBC |
| Amino acids regulate mTORC1 | 0.897 | 0.148 | -1.414 | ATP6V1E1,ATP6V1G2,CASTOR2,LAMTOR4,MTOR,RRAGB,SESN1,SH3BP4 |
| Lymphotoxin β Receptor Signaling | 0.897 | 0.148 | 0.816 | AKT1,APAF1,CREBBP,CYCS,NFKBID,PIK3CA,PIK3CD,PIK3R2 |
| Myogenesis | 0.893 | 0.172 | -1.342 | CDH2,CDH4,CTNNA1,CTNNB1,MEF2C |
| Proteasomal PSMD10 Signaling Pathway | 0.892 | 0.128 | 0.000 | AKT1,CDK6,CTNNB1,ETV1,HGF,HIF1A,HRAS,PIK3CA,PIK3CD,PIK3R2,PSMC5,PSMD13,PSMD6,RAP2A,TWIST1 |
| HEY1 Signaling Pathway | 0.889 | 0.124 | 0.000 | AKT1,BMP8B,DLL1,DLL3,DLL4,ERBB4,HIF1A,MEF2C,MMP24,NCSTN,NOTCH1,NR2F2,PIK3CA,PIK3CD,PIK3R2,PRKAA2,SMAD6,TGFBR1 |
| Th17 Activation Pathway | 0.888 | 0.143 | -2.333 | FCER1G,HIF1A,HSP90AA1,HSP90AB1,HSP90B1,IRAK1,JAK2,JAK3,MTOR |
| Regulation of mitotic cell cycle | 0.881 | 0.133 | 0.577 | CDC20,CDK1,MAD2L1,PSMB10,PSMB2,PSMB4,PSMB5,PSMC5,PSMD13,PSMD6,SKP2,UBC |
| FLT3 Signaling in Hematopoietic Progenitor Cells | 0.881 | 0.136 | 0.302 | AKT1,CREBBP,EIF4E,HRAS,MTOR,PIK3CA,PIK3CD,PIK3R2,RAP2A,RPS6KA2,RPS6KA5 |
| Regulation of CDH19 Expression and Function | 0.875 | 0.286 | #NUM! | CTNNA1,CTNNB1 |
| Ceramide Biosynthesis | 0.875 | 0.286 | #NUM! | DEGS2,SPTLC2 |
| PIP3 activates AKT signaling | 0.869 | 0.123 | -0.943 | AKT1,ERBB4,FGFR3,FOXO1,FYN,HGF,IRAK1,MET,MKRN1,MTOR,NR4A1,NRG1,NTF3,NTRK2,PIK3CA,PIK3CD,PIK3R2,PIP4K2B |
| Preeclampsia Signaling Pathway | 0.869 | 0.123 | 0.000 | AKT1,AMOTL2,COMT,CTNNB1,FZD1,GPER1,HIF1A,JAK2,JAK3,KCNMB4,MAPK9,MTOR,PIK3CA,PIK3CD,PIK3R2,PPP2R2B,PRKAA2,TWIST1 |
| Zn Homeostasis Signaling Pathway | 0.868 | 0.107 | -1.250 | ADCY1,ADCY8,ADGRA2,ADGRB1,ADGRB2,ADGRB3,ADGRG1,ADGRL3,ADORA2A,ADRA2C,AKT1,BDKRB1,BMP8B,C3AR1,CCRL2,CELSR1,CREBBP,EDNRB,FCER1G,FZD1,GIPR,GNAQ,GPER1,GPR12,GPR137C,GPR146,GPR155,GPR156,GPR19,GPR27,GPR68,GPR85,GPR88,GRIA2,GRIA3,GRIN2D,GRM3,GUCY1A1,GUCY1B1,ITPR1,ITPR2,MCHR1,MMP24,MPPED2,MT3,NMUR1,OXGR1,PARP1,PDE1A,PDE1B,PDE2A,PDE6D,PIK3CA,PIK3CD,PIK3R2,PRKCB,PRKCD,PRKCH,PTH1R,S1PR1,SIRT4,SLC30A7,SLC39A10,SMAD6,SSTR2,SSTR3 |
| DNA Double-Strand Break Repair by Homologous Recombina | 0.867 | 0.214 | #NUM! | ATM,BRCA2,MRE11 |
| FAT10 Signaling Pathway | 0.864 | 0.145 | #NUM! | NUB1,PSMB10,PSMB2,PSMB4,PSMB5,PSMC5,PSMD13,PSMD6 |
| PPAR Signaling | 0.859 | 0.130 | -1.732 | CITED2,CREBBP,HRAS,HSP90AA1,HSP90AB1,HSP90B1,IL1RAPL1,MAP4K4,NFKBID,NGFR,NR2F1,PPARGC1A,RAP2A |
| FLT3 Signaling | 0.859 | 0.158 | 0.816 | AKT1,FYN,HRAS,PIK3CA,SOCS2,UBC |
| Protein Sorting Signaling Pathway | 0.858 | 0.120 | 0.000 | ADCY1,ADCY8,CLTB,COPB2,FYN,GNAQ,GOLGA1,GUCY1A1,GUCY1B1,MIA3,PI4KB,PLPP1,PREB,SEC24D,SNX11,SNX27,SNX7,TBC1D23,TMED4,VPS35,WASHC2C |
| Translocation of SLC2A4 (GLUT4) to the plasma membrane | 0.857 | 0.141 | 2.333 | ACTB.AKT1.CALM1.KIF3B.PRKAA2.RAB13.TUBA1A.TUBB6.YWHAZ |
| RNA Polymerase I Transcription | 0.857 | 0.141 | 1.667 | GTF2H3,H2AZ2,H3C1,HDAC2,MBD3,POLR1C,POLR1D,POLR2L,TAF1D |
| Salvage Pathways of Pyrimidine Ribonucleotides | 0.856 | 0.132 | -1.508 | ADPGK,CDK1,CDK18,CDK6,IRAK1,MAPK7,MAPK9,PAK1,PIM1,PRKAA2,PRKCD,PRKCH |
| Hypoxia Signaling in the Cardiovascular System | 0.854 | 0.137 | 0.816 | AKT1,ATM,COPS5,CREBBP,HIF1A,HSP90AA1,HSP90AB1,HSP90B1,NFKBID,UBE2E3 |
| JAK/STAT Signaling | 0.854 | 0.134 | 0.905 | AKT1,GNAQ,HRAS,JAK2,JAK3,MTOR,PIK3CA,PIK3CD,PIK3R2,RAP2A,SOCS2 |
| Parkinson's Signaling Pathway | 0.849 | 0.114 | -3.138 | ACTB,ALDH16A1,Aldh3a2,ATP6AP2,ATP6V1E1,ATP6V1G2,CACNA1A,CACNG3,CACNG8,CALM1,CATSPER2,COXFA4,DDC,GRIA2,GRIA3,GRIN2D,ITPR1,ITPR2,NDUFA10,NDUFA13,NDUFA3,NDUFB2,NDUFB5,NDUFS7,NLRP3,PARK7,PPARGC1A,TUBA1A,TUBB6,TUBG1,UBC |
| DAP12 interactions | 0.846 | 0.167 | 0.447 | FYN,HRAS,PIK3CA,PIK3R2,PLCG1 |
| MicroRNA Biogenesis Signaling Pathway | 0.840 | 0.119 | 2.400 | AKT1,ATM,DDX1,DIS3,EXOSC9,HGF,HNRNPA1,HRAS,HSP90AA1,HSP90AB1,HSP90B1,KHSRP,MTOR,NUP93,PIK3CA,PIK3CD,PIK3R2,POLR2L,RAE1,RAN,RAP2A |
| Histone Modification Signaling Pathway | 0.839 | 0.114 | -1.512 | ATM,BUB1,CREBBP,EZH1,EZH2,FOXO1,H1-0,H1-1,H1-4,H1-5,H1f3,H2AZ2,HDAC2,KAT6B,KDM2A,KDM4B,KDM4D,KDM8,KMT5A,MAPK9,MORC4,PIM1,PRDM16,PRKCB,PRMT1,RPS6KA5,SETD7,WDR5,XRCC6,YWHAZ |
| Nuclear Cytoskeleton Signaling Pathway | 0.836 | 0.117 | -0.626 | ACTB,ATM,CAPZB,CDH2,CDH24,CDH4,CTNNB1,DES,DYNC1I1,DYNLL1,FN1,FYN,KIF11,KIF26B,KIF3B,MRTFA,NUP93,RAE1,RBX1,SKP2,SYNE4,TUBA1A,TUBB6,XRCC6 |
| S Phase | 0.836 | 0.129 | 1.387 | AKT1,PSMB10,PSMB2,PSMB4,PSMB5,PSMC5,PSMD13,PSMD6,SKP2,SMC3,TFDP2,UBC,WAPL |
| WNT/SHH Axonal Guidance Signaling Pathway | 0.831 | 0.122 | -0.471 | ADCY1,ADCY8,CTNNB1,FYN,FZD1,GUCY1A1,GUCY1B1,ITPR1,ITPR2,MAPK9,PIK3CA,PIK3CD,PIK3R2,PLXNA2,SEMA3A,SEMA3C,WNT7B,YWHAZ |
| NAP1L1 Transcription Regulation Signaling Pathway | 0.826 | 0.135 | 1.000 | AKT1.CTNNB1.MTOR.PIK3CA.PIK3CD.PIK3R2.SCARB1.TUBA1A.TUBB6.TUBG1 |
| RHOA Signaling | 0.826 | 0.125 | 1.941 | ACTB,ARHGAP12,ARPC3,ARPC5,ARPC5L,CFL1,GNA12,MYL12B,MYL6,PFN1,PIP4K2B,PPP1CA,RAPGEF2,SEPTIN10,WASF1 |
| GP6 Signaling Pathway | 0.826 | 0.125 | 0.832 | AKT1,CALM1,COL9A3,FCER1G,FYN,ITPR1,LAMB1,PIK3CA,PIK3CD,PIK3R2,PLCG1,PRKCB,PRKCD,PRKCH,RASGRP2 |
| ERK/MAPK Signaling | 0.820 | 0.117 | 0.626 | CREBBP,DOCK1,DUSP1,DUSP2,EIF4E,FYN,HRAS,MYCN,PAK1,PIK3CA,PIK3CD,PIK3R2,PLCG1,PPP1CA,PPP1R14B,PPP2R2B,PRKCB,PRKCD,RAC3,RAP2A,RAPGEF1,RAPGEF4,RPS6KA5,YWHAZ |
| Coronavirus Pathogenesis Pathway | 0.820 | 0.117 | -1.500 | EIF4E,HDAC2,HDAC7,HDAC9,HIF1A,KGD4,MAPK9,MRPS22,MRPS27,MRPS31,MRPS35,NFKBID,NLRP3,PA2G4,PTGES2,RACK1,RPS10,RPS19,RPS26,RPS3,RPS3A,RPSA,TGFBR1,TNPO2 |
| Fc Epsilon RI Signaling | 0.819 | 0.126 | 0.535 | AKT1,FCER1G,FYN,HRAS,MAPK9,PIK3CA,PIK3CD,PIK3R2,PLCG1,PRKCB,PRKCD,PRKCH,RAC3,RAP2A |
| TCF dependent signaling in response to WNT | 0.816 | 0.118 | 0.000 | AKT1,CAV1,CREBBP,CTNNB1,FZD1,H2AZ2,H3C1,PSMB10,PSMB2,PSMB4,PSMB5,PSMC5,PSMD13,PSMD6,RBX1,SOX2,SOX9,TLE2,TLE3,TLE4,UBC,YWHAZ |
| Regulation of the Epithelial Mesenchymal Transition by Grov | 0.816 | 0.118 | -1.528 | AKT1,CDH2,FGFR3,FOXO1,HGF,HRAS,JAK2,JAK3,MAPK9,MET,MTOR,NGFR,PIK3CA,PIK3CD,PIK3R2,RAP2A,SHC3,TGFBR1,TWIST1,VIM,ZEB1,ZEB2 |
| Cardiac Hypertrophy Signaling (Enhanced) | 0.815 | 0.107 | -1.769 | ADCY1,ADCY8,ADRA2C,AKAP13,AKT1,CACNA1A,CACNG3,CACNG8,CALM1,CTNNB1,EDNRB,EIF4E,FGFR3,FZD1,GNA12,GNAQ,GNB2,GNG3,GNG4,GNG7,GUCY1A1,GUCY1B1,HDAC2,HDAC7,HDAC9,HRAS,IL6ST,ITPR1,ITPR2,JAK2,MAP3K11,MAPK7,MAPK9,MAPKAPK2,MEF2C,MPPED2,MTOR,NGFR,PDE1A,PDE1B,PDE2A,PDE6D,PIK3CA,PIK3CD,PIK3R2,PLCG1,PLCH1,PLCH2,PRKCB,PRKCD,PRKCH,RAP2A,RPS6KA5,TGFBR1,WNT7B |
| BBSome Signaling Pathway | 0.813 | 0.108 | -2.236 | ADGRA2,ADGRB1,ADGRB2,ADGRB3,ADGRG1,ADGRL3,ADORA2A,ADRA2C,BBS4,BBS7,BDKRB1,C3AR1,CBX8,CCDC66,CCRL2,CCT2,CCT3,CCT6A,CCT6B,CCT7,CELSR1,EDNRB,EZH2,FZD1,GIPR,GPER1,GPR12,GPR137C,GPR146,GPR155,GPR156,GPR19,GPR27,GPR68,GPR85,GPR88,GRM3,IFT140,IFT57,MCHR1,NMUR1,NTRK2,OXGR1,PRKCD,PTH1R,S1PR1,SSTR2,SSTR3,TTC21B |
| Mouse Embryonic Stem Cell Pluripotency | 0.813 | 0.127 | -0.277 | AKT1,CREBBP,CTNNB1,FZD1,HRAS,IL6ST,JAK2,JAK3,PIK3CA,PIK3CD,PIK3R2,RAP2A,SOX2 |
| nNOS Signaling in Skeletal Muscle Cells | 0.806 | 0.146 | 0.000 | CACNA1A,CACNG3,CACNG8,CALM1,CATSPER2,ITPR1,ITPR2 |
| FAK Signaling | 0.802 | 0.106 | -1.890 | ADGRA2,ADGRB1,ADGRB2,ADGRB3,ADGRG1,ADGRL3,ADORA2A,ADRA2C,AKT1,ARPC3,ARPC5,ARPC5L,BDKRB1,C3AR1,CAPNS1,CCRL2,CELSR1,CTNNB1,DOCK1,EDNRB,EFNA3,EFNB1,ETV4,FCER1G,FYN,FZD1,GIPR,GPER1,GPR12,GPR137C,GPR146,GPR155,GPR156,GPR19,GPR27,GPR68,GPR85,GPR88,GRM3,HRAS,IL6ST,MAPK7,MAPK9,MCHR1,MET,MTOR,NCK2,NMUR1,OXGR1,PAK1,PIK3CA,PIK3CD,PIK3R2,PLCG1,PTH1R,RAP2A,RASA1,S1PR1,SOCS2,SPARCL1,SSTR2,SSTR3,TGFBR1,USF1 |
| Turbulent (oscillatory, disturbed) flow shear stress activates | 0.802 | 0.161 | -0.447 | CAPNS1,FN1,GNAQ,NLRP3,VCL |
| Regulation of the Epithelial-Mesenchymal Transition Pathwa | 0.800 | 0.117 | #NUM! | AKT1,CDH2,CTNNB1,FGFR3,FZD1,HGF,HIF1A,HRAS,JAK2,JAK3,MET,NCSTN,NOTCH1,PIK3CA,PIK3CD,PIK3R2,RAP2A,TGFBR1,TWIST1,WNT7B,ZEB1,ZEB2 |
| Rap1 signalling | 0.799 | 0.200 | #NUM! | RAPGEF4,RASGRP2,YWHAZ |
| Germ layer formation at gastrulation | 0.799 | 0.200 | #NUM! | CTNNB1,EOMES,SOX2 |
| Hedgehog ligand biogenesis | 0.798 | 0.136 | 1.667 | OS9,PSMB10,PSMB2,PSMB4,PSMB5,PSMC5,PSMD13,PSMD6,UBC |
| Complex I biogenesis | 0.798 | 0.136 | 1.667 | NDUFA10,NDUFA13,NDUFA3,NDUFB2,NDUFB5,NDUFC2,NDUFS7,NUBPL,TMEM126A |
| Dopamine Receptor Signaling | 0.798 | 0.136 | -1.000 | ADCY1,ADCY8,COMT,DDC,GUCY1A1,GUCY1B1,NCS1,PPP1CA,PPP2R2B |
| SNARE Signaling Pathway | 0.794 | 0.122 | 0.832 | ADCY1,ADCY8,CALM1,CPLX1,CPLX3,GUCY1A1,GUCY1B1,MYL6,RAB6A,RAB9B,SYN2,SYN3,SYT16,SYT9,VAMP3,VTI1A |
| Nucleotide salvage | 0.792 | 0.174 | 0.000 | APRT,DCK,PNP,TK2 |
| RAS processing | 0.792 | 0.174 | 1.000 | CALM1,HRAS,PDE6D,UBC |
| HDR through Homologous Recombination (HRR) or Single Str | 0.791 | 0.126 | -1.941 | ABRAXAS1,ATM,BABAM1,BRCA2,MRE11,RAD1,RAD51AP1,RAD51C,RMI2,RNF4,RPA2,UBC,XRCC2 |
| Hedgehog 'on' state | 0.784 | 0.121 | 1.698 | CDKL1,IQCE,PI4KB,PIK3CA,PIK3CD,PIK3R2,PSMB10,PSMB2,PSMB4,PSMB5,PSMC5,PSMD13,PSMD6,RBX1,STK17B,SUFU,UBC |
| Vasopressin regulates renal water homeostasis via Aquapori | 0.782 | 0.150 | 0.000 | ADCY1,ADCY8,GNB2,GNG3,GNG4,GNG7 |
| Signaling by FGFR3 | 0.782 | 0.150 | 0.000 | FGFR3,HRAS,PIK3CA,PLCG1,SPRY2,UBC |
| Mechanisms of Viral Exit from Host Cells | 0.782 | 0.150 | #NUM! | ACTB,PDCD6IP,PRKCB,PRKCD,PRKCH,VPS28 |
| LPS-stimulated MAPK Signaling | 0.778 | 0.129 | 0.302 | HRAS,MAPK9,NFKBID,PAK1,PIK3CA,PIK3CD,PIK3R2,PRKCB,PRKCD,PRKCH,RAP2A |
| PDGF Signaling | 0.778 | 0.129 | 0.302 | CAV1,HRAS,JAK2,JAK3,PIK3CA,PIK3CD,PIK3R2,PLCG1,PRKCB,RAP2A,RASA1 |
| Regulation of NFE2L2 gene expression | 0.777 | 0.250 | #NUM! | CREBBP,NOTCH1 |
| Sucrose Degradation V (Mammalian) | 0.777 | 0.250 | #NUM! | ALDOC,TKFC |
| Ketolysis | 0.777 | 0.250 | #NUM! | Acat2/Acat3,OXCT1 |
| Sleep NREM Signaling Pathway | 0.777 | 0.124 | 0.000 | ADCY1,ADCY8,ADORA2A,CREBBP,GABRA5,GABRB3,GABRG3,GABRP,GUCY1A1,GUCY1B1,HRAS,PARP1,RAP2A,YWHAZ |
| Angiopoietin Signaling | 0.774 | 0.132 | 1.890 | AKT1,FOXO1,HRAS,NFKBID,PAK1,PIK3CA,PIK3CD,PIK3R2,RAP2A,RASA1 |
| TP53 Regulates Transcription of Cell Cycle Genes | 0.773 | 0.143 | -1.134 | CDK1,CNOT1,CPAP,PCBP4,PRMT1,TFDP2,TNKS1BP1 |
| Signaling by NOTCH3 | 0.773 | 0.143 | -1.890 | CREBBP,DLL1,DLL4,FABP7,NCSTN,NOTCH1,UBC |
| Co-stimulation by CD28 | 0.773 | 0.143 | 1.134 | AKT1,FYN,MTOR,PAK1,PIK3CA,PIK3CD,PIK3R2 |
| Xenobiotic Metabolism AHR Signaling Pathway | 0.771 | 0.134 | -1.890 | ALDH16A1,Aldh3a2,CREBBP,Gstm7,GSTO1,HSP90AA1,HSP90AB1,HSP90B1,MGST1 |
| Cancer Drug Resistance by Drug Efflux | 0.770 | 0.138 | 1.414 | ABCC10,AKT1,FOXO1,HRAS,PIK3CA,PIK3CD,PIK3R2,RAP2A |
| Xenobiotic Metabolism Signaling | 0.764 | 0.113 | #NUM! | ALDH16A1,Aldh3a2,CAT,CITED2,CREBBP,Gstm7,GSTO1,HRAS,HS3ST4,HS6ST2,HSP90AA1,HSP90AB1,HSP90B1,MAP3K11,MAPK7,MAPK9,MGST1,PIK3CA,PIK3CD,PIK3R2,PPARGC1A,PPP2R2B,PRKCB,PRKCD,PRKCH,RAP2A,RBX1 |
| NF-κB Signaling | 0.762 | 0.117 | 0.000 | AKT1,AZI2,CREBBP,FCER1G,FGFR3,HDAC2,HRAS,IRAK1,MAP4K4,NFKBID,NGFR,NTRK2,PELI1,PIK3CA,PIK3CD,PIK3R2,PRKCB,RAP2A,TGFBR1,TNFAIP3 |
| Triglyceride metabolism | 0.761 | 0.156 | -0.447 | CAV1,FABP5,FABP7,LPIN1,PPP1CA |
| Alanine metabolism | 0.750 | 0.500 | #NUM! | GPT2 |
| L-DOPA Degradation | 0.750 | 0.500 | #NUM! | COMT |
| Alanine Degradation III | 0.750 | 0.500 | #NUM! | GPT2 |
| Alanine Biosynthesis II | 0.750 | 0.500 | #NUM! | GPT2 |
| Xanthine and Xanthosine Salvage | 0.750 | 0.500 | #NUM! | PNP |
| Fatty Acid Biosynthesis Initiation II | 0.750 | 0.500 | #NUM! | FASN |
| Glycine Degradation (Creatine Biosynthesis) | 0.750 | 0.500 | #NUM! | GATM |
| Threonine Degradation II | 0.750 | 0.500 | #NUM! | GCAT |
| NF-κB Activation by Viruses | 0.748 | 0.130 | 0.632 | AKT1,HRAS,NFKBID,PIK3CA,PIK3CD,PIK3R2,PRKCB,PRKCD,PRKCH,RAP2A |
| DNA damage-induced 14-3-3σ Signaling | 0.747 | 0.146 | -0.816 | AKT1,APAF1,ATM,CCNB2,CDK1,RAD1 |
| Agrin Interactions at Neuromuscular Junction | 0.744 | 0.132 | 1.414 | ACTB,ERBB4,HRAS,LAMB1,MAPK9,NRG1,PAK1,RAC3,RAP2A |
| Integrin signaling | 0.744 | 0.167 | -1.000 | AKT1,FN1,RAPGEF4,RASGRP2 |
| S100 Family Signaling Pathway | 0.743 | 0.104 | -2.668 | ADGRA2,ADGRB1,ADGRB2,ADGRB3,ADGRG1,ADGRL3,ADORA2A,ADRA2C,AHNAK,AKT1,BDKRB1,C3AR1,CACNA1A,CACNG3,CACNG8,CALM1,CATSPER2,CCRL2,CELSR1,COPS5,CREBBP,CTNNB1,EDNRB,ETV4,FGFR3,FZD1,GIPR,GPER1,GPR12,GPR137C,GPR146,GPR155,GPR156,GPR19,GPR27,GPR68,GPR85,GPR88,GRM3,HIF1A,IRAK1,ITPR1,ITPR2,MAPK7,MAPK9,MCHR1,MMP24,NMUR1,NR3C1,NTF3,NTRK2,OXGR1,PIK3CA,PIK3CD,PIK3R2,PLCG1,PLCH1,PLCH2,PRKCB,PRKCD,PRKCH,PTH1R,RAC3,S100A1,S100A10,S100A9,S1PR1,SMAD6,SSTR2,SSTR3,WNT7B |
| Signaling by FGFR1 | 0.742 | 0.140 | 0.378 | FLRT2,HRAS,PIK3CA,PLCG1,SPRED2,SPRY2,UBC |
| Transcriptional regulation by RUNX3 | 0.739 | 0.125 | 0.577 | CREBBP,CTNNB1,NOTCH1,PSMB10,PSMB2,PSMB4,PSMB5,PSMC5,PSMD13,PSMD6,TEAD2,UBC |
| Meiosis Initiation Signaling Pathway | 0.739 | 0.125 | -1.732 | BRCA2,FBXL5,HDAC2,MCMDC2,PRC1,RBX1,RPA2,SKP2,SMC3,SYCE2,TESMIN,WDR5 |
| FOXO-mediated transcription of cell death genes | 0.738 | 0.188 | #NUM! | CITED2,CREBBP,FOXO1 |
| Mitochondrial unfolded protein response (UPRmt) | 0.738 | 0.188 | #NUM! | AKT1,CAT,HSPE1 |
| DYRK1A Signaling Pathway | 0.735 | 0.117 | 0.688 | AKT1,CCND2,CREBBP,CUL2,FGFR3,FOXO1,HRAS,IL6ST,MAPK7,MAPK9,MAPT,MET,NOTCH1,POLR2L,RAP2A,RBX1,SPRY2,TFDP2,YWHAZ |
| Pulmonary Fibrosis Idiopathic Signaling Pathway | 0.728 | 0.109 | -1.219 | ACTB,AKT1,AXL,CAV1,CDH2,COL9A3,CREBBP,CTNNB1,EPHB3,EZH1,EZH2,FGFR3,FN1,FOXO1,FZD1,GNA12,HRAS,IL6ST,JAK2,MAPK9,MMP24,MRTFA,MTOR,MUC1,NOTCH1,PIK3CA,PIK3CD,PIK3R2,RAP2A,RPS6KA2,RPS6KA5,TGFBR1,VIM,WNT7B |
| LXR/RXR Activation | 0.727 | 0.123 | 0.378 | ABCA1,ABCG1,ACACA,AMBP,APOC2,CYP51A1,FASN,FDFT1,HMGCR,IL1RAPL1,NGFR,Scd1,TF |
| Signaling by NOTCH2 | 0.721 | 0.152 | -1.342 | DLL1,DLL4,MDK,NCSTN,UBC |
| Endometrial Cancer Signaling | 0.713 | 0.133 | 1.414 | AKT1,CTNNA1,CTNNB1,HRAS,PIK3CA,PIK3CD,PIK3R2,RAP2A |
| IL-17A Signaling in Airway Cells | 0.713 | 0.133 | 0.000 | AKT1,JAK2,JAK3,MAPK9,NFKBID,PIK3CA,PIK3CD,PIK3R2 |
| IL-23 Signaling Pathway | 0.713 | 0.143 | 0.816 | AKT1,HIF1A,JAK2,PIK3CA,PIK3CD,PIK3R2 |
| T Cell Exhaustion Signaling Pathway | 0.712 | 0.117 | -1.387 | AKT1,EOMES,FCER1G,FOXO1,FOXP1,HRAS,JAK2,JAK3,MAPK9,MTOR,PIK3CA,PIK3CD,PIK3R2,PLCG1,PPP2R2B,RAP2A,TGFBR1 |
| LPS/IL-1 Mediated Inhibition of RXR Function | 0.704 | 0.113 | -0.632 | ABCA1,ABCG1,ACAA1,ACSL4,ALDH16A1,Aldh3a2,APOC2,CAT,CRAT,FABP5,FABP7,Gstm7,GSTO1,HMGCS1,HS3ST4,HS6ST2,IL1RAPL1,IRAK1,MAPK9,MGST1,NGFR,PPARGC1A,SCARB1 |
| Neurovascular Coupling Signaling Pathway | 0.701 | 0.112 | 0.000 | ADORA2A,CACNA1A,CACNG3,CACNG8,CALM1,CATSPER2,GABRA5,GABRB3,GABRG3,GABRP,GAD1,GNAQ,GRIA2,GRIA3,GRIN2D,GUCY1A1,GUCY1B1,ITPR1,ITPR2,KCNJ8,KCNMB4,KCNN3,MAPT,PPP1CA |
| IL-15 Signaling | 0.700 | 0.127 | 0.632 | AKT1,HRAS,JAK2,JAK3,MTOR,PIK3CA,PIK3CD,PIK3R2,RAC3,RAP2A |
| Cardiogenesis | 0.699 | 0.160 | 0.000 | CTNNB1,EOMES,MEF2C,WDR5 |
| RAR Activation | 0.697 | 0.107 | 0.309 | ACTB,ADCY1,ADCY8,AKT1,CITED2,CREBBP,DUSP1,FABP5,GNAQ,GTF2H3,GUCY1A1,GUCY1B1,HIF1A,IGFBP3,JAK2,LAMB1,MAPK9,MEIS2,MPPED2,NR2F1,NR2F2,PARP1,PDE1A,PDE1B,PDE2A,PDE6D,PIK3CA,PIK3CD,PIK3R2,PITX2,PPARGC1A,PRDM16,PRKCD,PRMT1,PSMC5,RAC3,RHOU,RND2,RPS6KA2,RPS6KA5,SOX9,TGFBR1 |
| POU5F1 (OCT4), SOX2, NANOG repress genes related to diffe | 0.694 | 0.222 | #NUM! | EOMES,SOX2 |
| Zygotic genome activation (ZGA) | 0.694 | 0.222 | #NUM! | CREBBP,DPPA2 |
| Signaling by FGFR2 | 0.693 | 0.129 | 0.333 | FGFBP3,HNRNPA1,HRAS,PIK3CA,PLCG1,POLR2L,SPRY2,TIA1,UBC |
| NIK-->noncanonical NF-kB signaling | 0.686 | 0.131 | 2.121 | PSMB10,PSMB2,PSMB4,PSMB5,PSMC5,PSMD13,PSMD6,UBC |
| Gastrin-CREB signalling pathway via PKC and MAPK | 0.683 | 0.176 | #NUM! | HRAS,MAPK7,RPS6KA2 |
| Isoleucine Degradation I | 0.683 | 0.176 | #NUM! | ACADSB,Acat2/Acat3,ECHDC3 |
| SUMOylation of transcription cofactors | 0.681 | 0.140 | -0.816 | CBX8,CREBBP,HIPK2,MRTFA,PARK7,PPARGC1A |
| Sphingolipid metabolism | 0.676 | 0.121 | -1.155 | ARSB,ARSK,ASAH1,B4GALT5,CERS6,DEGS2,FUT2,OSBP,PLPP1,SPTLC2,ST3GAL5,ST6GALNAC5 |
| Neuropeptides in Migraine Signaling Pathway | 0.674 | 0.115 | -1.414 | ABCC8,ADCY1,ADCY8,CREBBP,GAD1,GNAQ,GNB2,GNG3,GNG4,GNG7,GUCY1A1,GUCY1B1,KCNJ8,KCNMB4,PLCG1,PLCH1,PLCH2,RAPGEF4 |
| WNT/β-catenin Signaling | 0.671 | 0.114 | -0.258 | AKT1,APPL2,CDH2,CREBBP,CTNNB1,DKK3,FZD1,GNAQ,LRP1,PPP2R2B,SOX1,SOX2,SOX21,SOX9,TGFBR1,TLE3,TLE4,UBC,WNT7B |
| IL-7 Signaling Pathway | 0.668 | 0.127 | 0.000 | AKT1,FOXO1,FYN,HGF,JAK3,MET,PIK3CA,PIK3CD,PIK3R2 |
| Intra-Golgi and retrograde Golgi-to-ER traffic | 0.665 | 0.112 | 0.655 | ARFIP2,CAPZB,COG7,COPB2,CYTH1,DYNC1I1,DYNLL1,GOLGA1,GOLIM4,KIF11,KIF26B,KIF3B,RAB18,RAB30,RAB6A,RAB9B,TUBA1A,TUBB6,VAMP3,VPS45,VTI1A |
| Actin Nucleation by ARP-WASP Complex | 0.665 | 0.122 | 1.667 | ARPC3,ARPC5,ARPC5L,GNA12,HRAS,NCK2,RAC3,RAP2A,RHOU,RND2,WASF1 |
| Role of NANOG in Mammalian Embryonic Stem Cell Pluripot | 0.660 | 0.118 | 0.302 | AKT1,BMP8B,CTNNB1,FZD1,HRAS,IL6ST,JAK2,JAK3,PIK3CA,PIK3CD,PIK3R2,RAP2A,SOX2,WNT7B |
| VEGF Family Ligand-Receptor Interactions | 0.655 | 0.123 | 0.632 | AKT1,HRAS,PIK3CA,PIK3CD,PIK3R2,PLCG1,PRKCB,PRKCD,PRKCH,RAP2A |
| BAG2 Signaling Pathway | 0.655 | 0.123 | #NUM! | HSP90AA1,MAPKAPK2,MAPT,PSMB10,PSMB2,PSMB4,PSMB5,PSMC5,PSMD13,PSMD6 |
| Intrinsic Pathway for Apoptosis | 0.654 | 0.132 | 1.134 | AKT1,APAF1,C1QBP,CYCS,DYNLL1,TFDP2,YWHAZ |
| Signaling by EGFR | 0.654 | 0.132 | 1.134 | AAMP,EPN1,HRAS,PIK3CA,PLCG1,SPRY2,UBC |
| Leukocyte Extravasation Signaling | 0.653 | 0.112 | -0.243 | ACTB,ARHGAP12,CLDN11,CLDN20,CTNNA1,CTNNB1,EDIL3,MAPK9,MMP24,MYL6,PIK3CA,PIK3CD,PIK3R2,PLCG1,PRKCB,PRKCD,PRKCH,RAPGEF4,THY1,VCL |
| IL-9 Signaling | 0.649 | 0.143 | 0.000 | JAK3,PIK3CA,PIK3CD,PIK3R2,SOCS2 |
| Lysosome Positioning Signaling Pathway | 0.646 | 0.114 | -1.213 | ACTB,AKT1,BLOC1S1,BORCS6,CAPZB,DYNC1I1,DYNLL1,FGFR3,MTOR,NDEL1,NGFR,NTRK2,PIK3CA,PIK3CD,PIK3R2,RAB34,TGFBR1 |
| Regulation of TP53 Activity through Phosphorylation | 0.645 | 0.121 | -0.302 | ATM,DYRK2,HIPK2,MRE11,NUAK1,PRKAA2,RAD1,RMI2,RPA2,TAF7,UBC |
| Purine Nucleotides Degradation II (Aerobic) | 0.633 | 0.167 | #NUM! | GDA,IMPDH1,PNP |
| Sleep REM Signaling Pathway | 0.627 | 0.130 | -1.134 | CFL1,CREBBP,EIF4E,NGFR,NR4A2,RPS6KA2,RPS6KA5 |
| IL-6 Signaling | 0.625 | 0.116 | 0.000 | AKT1,HRAS,IL1RAPL1,IL6ST,JAK2,MAP4K4,MAPK9,MAPKAPK2,NFKBID,NGFR,PIK3CA,PIK3CD,PIK3R2,RAP2A |
| ATF6 (ATF6-alpha) activates chaperone genes | 0.623 | 0.200 | #NUM! | CALR,HSP90B1 |
| Prostanoid Biosynthesis | 0.623 | 0.200 | #NUM! | PTGES2,PTGIS |
| Oleate Biosynthesis II (Animals) | 0.623 | 0.200 | #NUM! | PTPRT,Scd1 |
| SUMOylation of DNA replication proteins | 0.620 | 0.133 | 0.000 | CDCA8,NUP210,NUP93,RAE1,RANGAP1,TOP2A |
| NR1H2 and NR1H3-mediated signaling | 0.620 | 0.133 | -0.816 | ABCA1,ABCG1,ANGPTL3,APOC2,FASN,TNRC6B |
| tRNA Splicing | 0.620 | 0.133 | -2.236 | MPPED2,PDE1A,PDE1B,PDE2A,PDE6D,TRPT1 |
| Antimicrobial peptides | 0.618 | 0.148 | 0.000 | ATOX1,PDZD11,PRTN3,S100A9 |
| Interconversion of nucleotide di- and triphosphates | 0.618 | 0.148 | 0.000 | DCTPP1,GLRX,RRM2,TXNRD1 |
| Serotonin Receptor Signaling | 0.617 | 0.104 | 0.905 | ADCY1,ADCY8,ADRA2C,AKT1,CACNA1A,CACNG3,CACNG8,CALM1,CAV1,COPS5,CREBBP,DDC,FOXO1,FYN,GNA12,GNAL,GNAQ,GNB2,GNG3,GNG4,GNG7,GUCY1A1,GUCY1B1,HIF1A,HRAS,JAK2,KCNN3,MAPT,MTOR,PAK1,PIK3CA,PIK3CD,PIK3R2,PLCG1,PLCH1,PLCH2,PPARGC1A,PPP1CA,PRKCB,PRKCD,PRKCH,RAC3,RAP2A,RASGRF1,RHOU,RND2 |
| Regulation of lipid metabolism by PPARalpha | 0.612 | 0.116 | -0.832 | ABCA1,CHD9,CREBBP,FDFT1,GRHL1,HMGCR,HMGCS1,MED11,MED16,MED18,MED9,PPARGC1A,TXNRD1 |
| O-linked glycosylation | 0.612 | 0.116 | -1.387 | ADAMTS10,ADAMTS20,ADAMTS3,ADAMTS4,B4GALT5,GALNT18,MUC1,POMT1,SEMA5A,SPON1,ST3GAL1,ST6GALNAC4,THSD7B |
| Myocardin Signaling Pathway | 0.612 | 0.116 | -0.277 | ABCA1,AKT1,BMP8B,CDK1,HGF,MEF2C,MRTFA,PIK3CA,PIK3CD,PIK3R2,PIM1,TEAD2,TGFBR1 |
| Gene Silencing by RNA | 0.605 | 0.118 | -0.302 | H2AZ2,H3C1,HSP90AA1,IPO8,MYBL1,NUP210,NUP93,POLR2L,RAE1,RAN,TNRC6B |
| Neuroinflammation Signaling Pathway | 0.601 | 0.107 | -1.528 | AKT1.CREBBP.CTNNB1.FZD1.GABRA5.GABRB3.GABRG3.GABRP.GAD1.GAD2.GJB1.GRIN2D.IRAK1.JAK2.JAK3.MAPK7.MAPK9.MAPT.MFGE8.NCSTN.NLRP3.NTF3.PIK3CA.PIK3CD.PIK3R2.PLCG1.RAC3.SLC6A1.TGFBR1 |
| IGF-1 Signaling | 0.599 | 0.117 | 1.667 | AKT1,FOXO1,HRAS,IGFBP3,JAK2,PIK3CA,PIK3CD,PIK3R2,RAP2A,RASA1,SOCS2,YWHAZ |
| Metallothioneins bind metals | 0.594 | 0.333 | #NUM! | MT3 |
| SUMOylation of nuclear envelope proteins | 0.594 | 0.333 | #NUM! | RANGAP1 |
| NFE2L2 regulating TCA cycle genes | 0.594 | 0.333 | #NUM! | IDH1 |
| Diphthamide Biosynthesis | 0.594 | 0.333 | #NUM! | EEF2 |
| Serine Biosynthesis | 0.594 | 0.333 | #NUM! | PHGDH |
| Guanine and Guanosine Salvage I | 0.594 | 0.333 | #NUM! | PNP |
| Palmitate Biosynthesis I (Animals) | 0.594 | 0.333 | #NUM! | FASN |
| Hypusine Biosynthesis | 0.594 | 0.333 | #NUM! | EIF5A |
| Biotin-carboxyl Carrier Protein Assembly | 0.594 | 0.333 | #NUM! | ACACA |
| Trans, trans-farnesyl Diphosphate Biosynthesis | 0.594 | 0.333 | #NUM! | FDPS |
| N-acetylglucosamine Degradation I | 0.594 | 0.333 | #NUM! | GNPDA1 |
| nNOS Signaling in Neurons | 0.592 | 0.130 | #NUM! | CALM1,CAPNS1,GRIN2D,PRKCB,PRKCD,PRKCH |
| Role of Chondrocytes in Rheumatoid Arthritis Signaling Path | 0.591 | 0.114 | -0.535 | ADAMTS4,CAMKK2,FN1,IL1RAPL1,IL6ST,IRAK1,JAK2,LRP1,MAPK9,MMP24,MTOR,NGFR,NLRP3,PRKAA2 |
| Pancreatic Adenocarcinoma Signaling | 0.591 | 0.114 | -0.707 | AKT1,BRCA2,HDAC2,HDAC7,HDAC9,JAK2,JAK3,MAPK9,NOTCH1,PA2G4,PIK3CA,PIK3CD,PIK3R2,TGFBR1 |
| Signaling by Activin | 0.587 | 0.158 | #NUM! | FST,INHA,INHBA |
| RHO GTPases activate CIT | 0.587 | 0.158 | #NUM! | MYL12B,MYL6,PRC1 |
| Regulation of TP53 Activity through Methylation | 0.587 | 0.158 | #NUM! | ATM,KMT5A,UBC |
| Class I peroxisomal membrane protein import | 0.587 | 0.158 | #NUM! | ABCD2,PEX26,PEX3 |
| Pregnenolone Biosynthesis | 0.587 | 0.158 | #NUM! | COQ7,CYP11A1,MICAL2 |
| UBE2C in Cell Cycle Progression Signaling Pathway | 0.586 | 0.117 | 0.000 | AKT1,CCNB2,CDC20,CDK1,MAD2L1,MTOR,NOTCH1,PIK3CA,PIK3CD,PIK3R2,UBC |
| PKCθ Signaling in T Lymphocytes | 0.585 | 0.111 | 0.302 | CACNA1A,CACNG3,CACNG8,CATSPER2,FCER1G,FYN,HRAS,ITPR1,ITPR2,MAP3K11,NFKBID,PIK3CA,PIK3CD,PIK3R2,PLCG1,RAC3,RAP2A |
| Regulation of TP53 Expression and Degradation | 0.585 | 0.135 | -0.447 | AKT1,ATM,CDK1,MTOR,UBC |
| Mitochondrial Division Signaling Pathway | 0.584 | 0.110 | -0.943 | CACNA1A,CACNG3,CACNG8,CATSPER2,CCNB2,CDK1,GNAQ,HGF,HRAS,ITPR1,ITPR2,MET,NGFR,PPP2R2B,PRKAA2,PRKCD,RAP2A,S1PR1 |

| | | | | |
|---|---|---|---|---|
| Osteoarthritis Pathway | 0.584 | 0.108 | -0.218 | ADAMTS4,ANKH,CNMD,CREBBP,CTNNA1,CTNNB1,DDIT4,FGFR3,FN1,FZD1,HIF1A,IL1RAPL1,LRP1,MEF2C,MTOR,NOTCH1,PPARGC1A,PRKAA2,PTH1R,S100A9,SMAD6,SOX9,TGFBR1 |
| Gustation Pathway | 0.584 | 0.110 | -0.243 | ABCC8,ADCY1,ADCY8,CACNA1A,CACNG3,CACNG8,CATSPER2,GABRA5,GABRB3,GABRG3,GABRP,GUCY1A1,GUCY1B1,ITPR1,ITPR2,KCNJ8,KCNN3,LRRC8A,PKD1L3 |
| Sulfur amino acid metabolism | 0.581 | 0.143 | -1.000 | CDO1,ENOPH1,ETHE1,MTRR |
| RHO GTPases activate PKNs | 0.581 | 0.143 | 2.000 | MYL12B,MYL6,PAK1,YWHAZ |
| Role of p14/p19ARF in Tumor Suppression | 0.581 | 0.143 | 0.000 | PIK3CA,PIK3CD,PIK3R2,SENP3 |
| PAK Signaling | 0.578 | 0.114 | 0.905 | CFL1,EPHA3,EPHB3,HRAS,MAPK9,MYL12B,MYL6,NCK2,PAK1,PIK3CA,PIK3CD,PIK3R2,RAP2A |
| Role of Macrophages, Fibroblasts and Endothelial Cells in Rh | 0.571 | 0.105 | -1.671 | ADAMTS4,AKT1,CALM1,CREBBP,CTNNB1,DKK3,FN1,FZD1,GNAQ,HRAS,IL1RAPL1,IL6ST,IRAK1,JAK2,LRP1,MAPK9,MAPKAPK2,MIF,NFKBID,NGFR,PIK3CA,PIK3CD,PIK3R2,PLCG1,PLCH1,PLCH2,PRKCB,PRKCD,PRKCH,RAP2A,WNT7B |
| G alpha (q) signalling events | 0.570 | 0.110 | -0.943 | BDKRB1,EDN3,EDNRB,GNAQ,GNB2,GNG3,GNG4,GNG7,GPR68,MCHR1,NMUR1,OPN4,PIK3CA,PIK3R2,RGS17,RGS3,RGS4,RGSL1 |
| DNA Methylation and Transcriptional Repression Signaling | 0.567 | 0.116 | -1.000 | CDK1,CDK12,CDK16,CDK18,CDK6,CHD5,HDAC2,MBD3,SAP130,TCF19,ZEB1 |
| CD40 Signaling | 0.566 | 0.121 | 0.000 | JAK3,MAPK9,MAPKAPK2,NFKBID,PIK3CA,PIK3CD,PIK3R2,TNFAIP3 |
| iNOS Signaling | 0.565 | 0.128 | -2.236 | CALM1,CREBBP,IRAK1,JAK2,JAK3,NFKBID |
| Peptide hormone biosynthesis | 0.562 | 0.182 | #NUM! | INHA,INHBA |
| Signaling by Leptin | 0.562 | 0.182 | #NUM! | JAK2,SH2B1 |
| Pexophagy | 0.562 | 0.182 | #NUM! | ATM,UBC |
| NFE2L2 regulating tumorigenic genes | 0.562 | 0.182 | #NUM! | CREBBP,NOTCH1 |
| Phosphorylation of CLOCK, acetylation of BMAL1 (ARNTL) at | 0.562 | 0.182 | #NUM! | CREBBP,CRY1 |
| Ketogenesis | 0.562 | 0.182 | #NUM! | Acat2/Acat3,HMGCS1 |
| IL-27 Signaling Pathway | 0.561 | 0.113 | 0.832 | AKT1,ATG13,IL1RAPL1,IL6ST,JAK2,JAK3,MTOR,NGFR,NLRP3,PIK3CA,PIK3CD,PIK3R2,PPARGC1A |
| Acute Phase Response Signaling | 0.557 | 0.109 | 0.243 | AKT1,AMBP,FN1,HRAS,IL6ST,IRAK1,JAK2,MAPK9,MTOR,NFKBID,NGFR,NR3C1,PIK3CA,PIK3CD,PIK3R2,RAP2A,SOCS2,TF |
| TP53 Regulates Transcription of Cell Death Genes | 0.555 | 0.132 | -2.236 | APAF1,ATM,CREBBP,IGFBP3,TMEM219 |
| BMP signaling pathway | 0.553 | 0.116 | 0.000 | BMP8B,CREBBP,FST,HRAS,MAPK9,NOG,PITX2,RAP2A,SMAD6,ZNF423 |
| Signaling by BMP | 0.547 | 0.138 | 0.000 | INHA,INHBA,NOG,SMAD6 |
| Energy dependent regulation of mTOR by LKB1-AMPK | 0.547 | 0.138 | 0.000 | LAMTOR4,MTOR,PRKAA2,RRAGB |
| ERCC6 (CSB) and EHMT2 (G9a) positively regulate rRNA expre | 0.547 | 0.138 | 0.000 | H2AZ2,H3C1,HDAC2,MBD3 |
| Sonic Hedgehog Signaling | 0.547 | 0.138 | #NUM! | CCNB2,CDK1,DYRK1B,SUFU |
| Paxillin Signaling | 0.547 | 0.113 | 1.155 | ACTB,ARFIP2,DOCK1,HRAS,MAPK9,NCK2,PAK1,PIK3CA,PIK3CD,PIK3R2,RAP2A,VCL |
| Receptor-type tyrosine-protein phosphatases | 0.545 | 0.150 | #NUM! | IL1RAPL1,PPFIBP1,SLITRK4 |
| Histidine Degradation VI | 0.545 | 0.150 | #NUM! | COQ7,CYP11A1,MICAL2 |
| Iron homeostasis signaling pathway | 0.544 | 0.111 | #NUM! | ABCB10,ATP6AP2,ATP6V1E1,ATP6V1G2,BMP8B,FBXL5,HIF1A,JAK2,JAK3,LRP1,PCBP1,RBX1,SKP2,TF |
| Inhibition of ARE-Mediated mRNA Degradation Pathway | 0.543 | 0.109 | 0.000 | AKT1,CNOT1,DIS3,EXOSC9,MAPK7,MAPKAPK2,NGFR,PPP2R2B,PSMB10,PSMB2,PSMB4,PSMB5,PSMC5,PSMD13,PSMD6,TIA1,YWHAZ |
| Triacylglycerol Biosynthesis | 0.530 | 0.121 | -0.816 | AGPAT1,DBT,LPIN1,PLPP1,PLPP4,PLPPR4,SIRT4 |
| Factors Promoting Cardiogenesis in Vertebrates | 0.529 | 0.109 | -2.324 | BMP8B,CREBBP,CTNNB1,FZD1,LRP1,MAPK9,MEF2C,NOG,PLCG1,PLCH1,PLCH2,PRKCB,PRKCD,PRKCH,TGFBR1,WNT7B |
| Clathrin-mediated endocytosis | 0.529 | 0.110 | 1.069 | ACTB,AGFG1,AMPH,AP2M1,ARPC3,ARPC5,CLTB,EPN1,HIP1,SYT9,TF,UBC,VAMP3,VAMP7 |
| Cerebral Malformation Signaling Pathway | 0.529 | 0.110 | 1.387 | AKT1,CDK1,CTNNA1,CTNNB1,FOXO1,MAPK7,MAPK9,MEF2C,MTOR,MYL12B,PIK3CA,PIK3CD,PIK3R2,PPP2R2B |
| Neurotransmitter release cycle | 0.527 | 0.128 | -1.342 | CASK,CPLX1,SLC17A7,SYN2,SYN3 |
| MyD88:MAL(TIRAP) cascade initiated on plasma membrane | 0.527 | 0.128 | -0.447 | IRAK1,PELI1,S100A1,S100A9,UBC |
| Nonhomologous End-Joining (NHEJ) | 0.527 | 0.128 | -1.342 | ABRAXAS1,ATM,BABAM1,MRE11,XRCC6 |
| Chromatin modifying enzymes | 0.525 | 0.106 | -0.209 | ACTB,ARID5B,BRD8,CREBBP,EZH2,H2AZ2,H3C1,HCFC1,HDAC2,JAK2,KAT6B,KDM2A,KDM4B,KDM4D,KMT5A,MBD3,MECOM,PRDM16,PRMT1,PRMT3,SAP130,SETD7,WDR5 |
| ISG15 antiviral mechanism | 0.523 | 0.118 | 0.707 | EIF4A1,EIF4E,KPNA2,NUP210,NUP93,PLCG1,RAE1,UBC |
| TNFR2 non-canonical NF-kB pathway | 0.523 | 0.118 | 2.121 | PSMB10,PSMB2,PSMB4,PSMB5,PSMC5,PSMD13,PSMD6,UBC |
| Maturity Onset Diabetes of Young (MODY) Signaling | 0.523 | 0.118 | #NUM! | ABCC8,APOC2,CACNA1A,CACNG3,CACNG8,CREBBP,KCNJ8,NEUROD1 |
| Ceramide Signaling | 0.517 | 0.114 | -0.632 | AKT1,CYCS,HRAS,NGFR,PIK3CA,PIK3CD,PIK3R2,PPP2R2B,RAP2A,S1PR1 |
| Superpathway of Inositol Phosphate Compounds | 0.516 | 0.105 | -0.894 | ATP1A1,DUSP1,DUSP12,DUSP2,IP6K1,NUDT15,PANK4,PI4KB,PIK3CA,PIK3CD,PIK3R2,PIP4K2B,PLCG1,PLCH1,PLCH2,PLPPR4,PPP1CA,PPP1R14B,PPP1R16B,PPP1R1A,PPP2R2B,PTPRO,RASA1,SIRPA |
| RIPK1-mediated regulated necrosis | 0.515 | 0.133 | 1.000 | HSP90AA1,PDCD6IP,PELI1,UBC |
| Regulation of TP53 Activity through Acetylation | 0.515 | 0.133 | 2.000 | AKT1,HDAC2,MBD3,PIP4K2B |
| Complement System | 0.515 | 0.133 | #NUM! | C1QBP,C3AR1,MASP1,MASP2 |
| Dilated Cardiomyopathy Signaling Pathway | 0.515 | 0.108 | -1.890 | ACTB,ADCY1,ADCY8,AKT1,CACNA1A,CACNG3,CACNG8,CATSPER2,DES,GUCY1A1,GUCY1B1,ITPR1,ITPR2,MTOR,MYL6,PDE2A |
| Signaling by TGFBR3 | 0.515 | 0.122 | 0.000 | INHA,INHBA,MYCN,NCSTN,TGFBR1,TNRC6B |
| Melanoma Signaling | 0.515 | 0.122 | 1.633 | AKT1,HRAS,PIK3CA,PIK3CD,PIK3R2,RAP2A |
| Role of IL-17A in Arthritis | 0.515 | 0.122 | #NUM! | MAPK9,MAPKAPK2,NFKBID,PIK3CA,PIK3CD,PIK3R2 |
| Aryl Hydrocarbon Receptor Signaling | 0.514 | 0.109 | -1.667 | ALDH16A1,Aldh3a2,APAF1,ATM,CCND2,CDK6,Gstm7,GSTO1,HSP90AA1,HSP90AB1,HSP90B1,MGST1,NFIC,NFIX,NR2F1 |
| Nucleotide Excision Repair | 0.514 | 0.111 | 1.155 | ACTB,AQR,CHD1L,COPS5,GTF2H3,PARP1,POLR2L,RBX1,RPA2,UBC,XAB2,ZNF830 |
| Calcium Signaling | 0.512 | 0.105 | 0.000 | ATP2B2,CACNA1A,CACNG3,CACNG8,CALM1,CALR,CAMKK2,CATSPER2,CHRNA4,CREBBP,GRIA2,GRIA3,GRIN2D,HDAC2,HDAC7,HDAC9,ITPR1,ITPR2,MEF2C,MICU1,MYL6,RAP2A |
| Specification of primordial germ cells | 0.509 | 0.167 | #NUM! | CBFA2T2,EOMES |
| Signaling by LTK | 0.509 | 0.167 | #NUM! | PIK3CA,PIK3R2 |
| Guanosine Nucleotides Degradation III | 0.509 | 0.167 | #NUM! | GDA,PNP |
| PCP (Planar Cell Polarity) Pathway | 0.508 | 0.119 | -0.816 | CELSR1,EFNB1,FZD1,MAPK9,PFN1,PRICKLE1,WNT7B |
| Fertilization | 0.506 | 0.143 | #NUM! | ACR,CATSPER2,ZP3 |
| SIRT1 negatively regulates rRNA expression | 0.506 | 0.143 | #NUM! | H2AZ2,H3C1,TAF1D |
| Branched-chain amino acid catabolism | 0.506 | 0.143 | #NUM! | ACADSB,DBT,HIBADH |
| Glutathione-mediated Detoxification | 0.506 | 0.143 | #NUM! | Gstm7,GSTO1,MGST1 |
| Melatonin Signaling | 0.503 | 0.116 | -1.134 | CALM1,GNAQ,PLCG1,PLCH1,PLCH2,PRKCB,PRKCD,PRKCH |
| BEX2 Signaling Pathway | 0.501 | 0.114 | 0.000 | AKT1,BEX1,CDH2,CTNNB1,HIF1A,MAPK9,MTOR,NGFR,PPP2R2B |
| Signaling by FGFR4 | 0.500 | 0.125 | 0.447 | HRAS,PIK3CA,PLCG1,SPRY2,UBC |
| Oncostatin M Signaling | 0.500 | 0.125 | 0.447 | HRAS,IL6ST,JAK2,JAK3,RAP2A |
| Cell junction organization | 0.499 | 0.112 | -1.897 | ACTB,CADM1,CDH2,CDH24,CDH4,CLDN11,CLDN20,CTNNA1,CTNNB1,DST |
| Transcriptional regulation by RUNX2 | 0.492 | 0.120 | -0.816 | AKT1,CDK1,SATB2,SMAD6,SOX9,TWIST1 |
| TNFR1 Signaling | 0.492 | 0.120 | -0.447 | APAF1,CYCS,MAP4K2,NFKBID,PAK1,TNFAIP3 |
| Neurotransmitter uptake and metabolism In glial cells | 0.489 | 0.250 | #NUM! | SLC38A1 |
| Uracil Degradation II (Reductive) | 0.489 | 0.250 | #NUM! | DPYSL2 |
| Acyl-CoA Hydrolysis | 0.489 | 0.250 | #NUM! | ACOT7 |
| Pentose Phosphate Pathway (Oxidative Branch) | 0.489 | 0.250 | #NUM! | PGLS |
| Branched-chain α-keto acid Dehydrogenase Complex | 0.489 | 0.250 | #NUM! | DBT |
| Catecholamine Biosynthesis | 0.489 | 0.250 | #NUM! | DDC |
| Eumelanin Biosynthesis | 0.489 | 0.250 | #NUM! | MIF |
| Methylmalonyl Pathway | 0.489 | 0.250 | #NUM! | PCCA |
| Taurine Biosynthesis | 0.489 | 0.250 | #NUM! | CDO1 |
| Thymine Degradation | 0.489 | 0.250 | #NUM! | DPYSL2 |
| L-cysteine Degradation I | 0.489 | 0.250 | #NUM! | CDO1 |
| N-acetylglucosamine Degradation II | 0.489 | 0.250 | #NUM! | GNPDA1 |
| PI3K Signaling in B Lymphocytes | 0.487 | 0.107 | 0.000 | AKT1,CALM1,FYN,HRAS,ITPR1,ITPR2,NFKBID,PIK3CA,PIK3CD,PIK3R2,PLCG1,PLCH1,PLCH2,PRKCB,RAP2A |
| Stearate Biosynthesis I (Animals) | 0.487 | 0.117 | 2.236 | ACOT7,ACSL4,DBT,DHCR24,FASN,SIRT4,TECR |
| Signaling by CSF1 (M-CSF) in myeloid cells | 0.485 | 0.129 | 1.000 | FYN,PIK3CA,PIK3CD,UBC |
| Glioma Invasiveness Signaling | 0.484 | 0.114 | 1.134 | HRAS,PIK3CA,PIK3CD,PIK3R2,RAC3,RAP2A,RHOU,RND2 |
| TCR signaling | 0.483 | 0.109 | 1.732 | PAK1,PIK3CA,PIK3R2,PLCG1,PSMB10,PSMB2,PSMB4,PSMB5,PSMC5,PSMD13,PSMD6,UBC |
| HER-2 Signaling in Breast Cancer | 0.482 | 0.104 | 0.655 | AKT1,COX4I2,COX5A,COX6A1,COX7A1,COX8A,FCER1G,FOXO1,FYN,HRAS,MTOR,NFKBID,NRG1,NUP93,PIK3CA,PIK3CD,PIK3R2,PLCG1,PRKCB,PRKCD,PRKCH,RAE1,RAP2A |
| Elastic fibre formation | 0.475 | 0.122 | -1.342 | FBLN1,FBLN2,FBN2,FN1,MFAP4 |
| Interleukin-2 family signaling | 0.475 | 0.122 | -0.447 | JAK2,JAK3,PIK3CA,PIK3CD,PIK3R2 |
| SLAMF Signaling Pathway | 0.472 | 0.107 | 0.535 | AKT1,AP2M1,FCER1G,FYN,HRAS,JAK2,JAK3,MTOR,PIK3CA,PIK3CD,PIK3R2,PLCG1,RAP2A,RASA1 |
| Growth hormone receptor signaling | 0.471 | 0.136 | #NUM! | JAK2,SH2B1,SOCS2 |
| Phosphorylated BMAL1:CLOCK (ARNTL:CLOCK) activates expi | 0.471 | 0.136 | #NUM! | CREBBP,CRY1,DBP |
| UVC-Induced MAPK Signaling | 0.469 | 0.118 | 0.000 | HRAS MAPK9 PRKCB PRKCD PRKCH RAP2A |
| FAT10 Cancer Signaling Pathway | 0.469 | 0.118 | 0.447 | AKT1,CTNNB1,MAD2L1,NFKBID,NGFR,TGFBR1 |
| IL-2 Signaling | 0.467 | 0.115 | 1.134 | AKT1,HRAS,JAK3,PIK3CA,PIK3CD,PIK3R2,RAP2A |
| Acute Myeloid Leukemia Signaling | 0.466 | 0.110 | 1.134 | AKT1,HRAS,IDH1,IDH3B,MTOR,PIK3CA,PIK3CD,PIK3R2,PIM1,RAP2A |
| IL-17A Signaling in Fibroblasts | 0.465 | 0.113 | -1.414 | FN1,HIF1A,JAK2,MAPK9,NFKBID,P4HA2,PRKCB,THY1 |
| Synthesis of Prostaglandins (PG) and Thromboxanes (TX) | 0.461 | 0.154 | #NUM! | PTGES2,PTGIS |
| Platelet Adhesion to exposed collagen | 0.461 | 0.154 | #NUM! | FCER1G,FYN |
| Formation of axial mesoderm | 0.461 | 0.154 | #NUM! | CTNNB1,TEAD2 |
| Formation of definitive endoderm | 0.461 | 0.154 | #NUM! | CTNNB1,EOMES |
| Nephron development | 0.461 | 0.154 | #NUM! | DLL1,LFNG |
| Urate Biosynthesis/Inosine 5'-phosphate Degradation | 0.461 | 0.154 | #NUM! | IMPDH1,PNP |
| Cargo concentration in the ER | 0.457 | 0.125 | 0.000 | CNIH2,MIA3,PREB,SEC24D |
| Nucleotide catabolism | 0.457 | 0.125 | -2.000 | GDA,NUDT15,NUDT16,PNP |
| Late endosomal microautophagy | 0.457 | 0.125 | 1.000 | PARK7,UBC,VIM,VPS28 |
| L1CAM interactions | 0.454 | 0.107 | 0.577 | ACTB,AP2M1,CHL1,CNTN6,DPYSL2,LAMB1,PAK1,RPS6KA2,RPS6KA5,SPTB,TUBA1A,TUBB6 |
| Membrane Repair Signaling Pathway | 0.452 | 0.108 | -0.905 | AHNAK,CAPNS1,CAV1,PDCD6IP,S100A10,SYT16,SYT9,TSPAN5,VAMP3,VPS4B,VTI1A |
| Signaling by SCF-KIT | 0.451 | 0.119 | 0.447 | FYN,HRAS,JAK2,PIK3CA,PIK3R2 |
| Glucose metabolism | 0.448 | 0.110 | 0.333 | ADPGK,ALDOC,GNPDA1,HK1,NUP210,NUP93,PKM,RAE1,SLC37A4 |
| NLR signaling pathways | 0.448 | 0.115 | 0.000 | AAMP,HSP90AB1,IRAK1,NLRP3,TNFAIP3,UBC |
| Amyloid fiber formation | 0.447 | 0.113 | -1.134 | H3C1,ITM2B,MFGE8,NCSTN,SORL1,TSPAN5,UBC |
| Ribavirin ADME | 0.447 | 0.113 | 0.378 | CDKL1,PI4KB,PIK3CA,PIK3CD,PIK3R2,PNP,STK17B |
| Transcriptional and post-translational regulation of MITF-M ( | 0.447 | 0.113 | -0.378 | CREBBP,CTNNB1,EDN3,EDNRB,SOX2,SOX9,YWHAZ |
| Macrophage Alternative Activation Signaling Pathway | 0.439 | 0.104 | -2.000 | ABCA1,ADORA2A,CREBBP,DUSP1,FCER1G,JAK2,JAK3,MTOR,NFKBID,NR3C1,PIK3CA,PIK3CD,PIK3R2,PPARGC1A,RPS6KA5,SCARB1 |
| Surfactant metabolism | 0.438 | 0.130 | #NUM! | ADORA2A,ADRA2C,CKAP4 |
| Gluconeogenesis I | 0.438 | 0.130 | #NUM! | ALDOC,MDH1,MDH2 |
| G alpha (i) signalling events | 0.438 | 0.102 | -2.558 | ADCY1,ADCY8,ADRA2C,BDKRB1,C3AR1,GNAZ,GNB2,GNG3,GNG4,GNG7,GPER1,GPSM2,GRM3,MCHR1,NMUR1,OXGR1,RGS17,RGS3,RGS4,RGSL1,SSTR2,SSTR3 |
| Type I Diabetes Mellitus Signaling | 0.437 | 0.107 | -1.667 | APAF1,CYCS,FCER1G,GAD1,GAD2,IRAK1,JAK2,MAPK9,NFKBID,NGFR,SOCS2 |
| Phagosome Maturation | 0.436 | 0.104 | #NUM! | ATP6AP2,ATP6V1E1,ATP6V1G2,CALR,CANX,DYNC1I1,DYNLL1,RAC3,TUBA1A,TUBB6,TUBG1,VAMP3,VPS28,VPS45,VTI1A |
| G alpha (s) signalling events | 0.433 | 0.104 | 0.535 | ADCY1,ADCY8,ADORA2A,GIPR,GNAZ,GNB2,GNG3,GNG4,GNG7,GPR27,PDE1A,PDE1B,PDE2A,PTH1R |
| Unfolded Protein Response (UPR) | 0.430 | 0.121 | 1.000 | EIF2S3,MBTPS1,MBTPS2,VPS28 |
| Insulin Receptor Signaling | 0.429 | 0.105 | 1.941 | AKT1,EIF4E,FOXO1,FYN,HRAS,JAK2,MTOR,PIK3CA,PIK3CD,PIK3R2,PPP1CA,RAP2A,RAPGEF1 |
| Carnitine shuttle | 0.419 | 0.143 | #NUM! | ACACA,PRKAA2 |
| YAP1- and WWTR1 (TAZ)-stimulated gene expression | 0.419 | 0.143 | #NUM! | HIPK2,TEAD2 |
| The CRY:PER:kinase complex represses transactivation by the | 0.419 | 0.143 | #NUM! | CREBBP,CRY1 |
| Phosphorylation and nuclear translocation of the CRY:PER:ki | 0.419 | 0.143 | #NUM! | CRY1,PPP1CA |
| Interleukin-1 family signaling | 0.416 | 0.104 | 1.387 | IL1RAPL1,IRAK1,PELI1,PSMB10,PSMB2,PSMB4,PSMB5,PSMC5,PSMD13,PSMD6,RBX1,SKP2,UBC |
| Reelin signalling pathway | 0.412 | 0.200 | #NUM! | FYN |
| NFE2L2 regulating inflammation associated genes | 0.412 | 0.200 | #NUM! | CREBBP |
| NFE2L2 regulating ER-stress associated genes | 0.412 | 0.200 | #NUM! | CREBBP |
| Creatine-phosphate Biosynthesis | 0.412 | 0.200 | #NUM! | MAP4K4 |
| Superpathway of Serine and Glycine Biosynthesis I | 0.412 | 0.200 | #NUM! | PHGDH |
| Serotonin and Melatonin Biosynthesis | 0.412 | 0.200 | #NUM! | DDC |
| CMP-N-acetylneuraminate Biosynthesis I (Eukaryotes) | 0.412 | 0.200 | #NUM! | NANP |
| Glutathione Redox Reactions II | 0.412 | 0.200 | #NUM! | GLRX |
| 2-oxobutanoate Degradation I | 0.412 | 0.200 | #NUM! | PCCA |
| Galactose Degradation I (Leloir Pathway) | 0.412 | 0.200 | #NUM! | GALE |
| MyD88 cascade initiated on plasma membrane | 0.408 | 0.125 | #NUM! | IRAK1,PELI1,UBC |
| Miscellaneous transport and binding events | 0.408 | 0.125 | #NUM! | ADD2,ANKH,LRRC8A |
| RHO GTPases Activate NADPH Oxidases | 0.408 | 0.125 | #NUM! | PRKCB,PRKCD,S100A9 |
| Regulation of RUNX1 Expression and Activity | 0.408 | 0.125 | #NUM! | CCND2,CDK6,TNRC6B |
| Chromatin modifications during the maternal to zygotic tran | 0.408 | 0.125 | #NUM! | DPPA3,H2AZ2,H3C1 |
| Ubiquinol-10 Biosynthesis (Eukaryotic) | 0.408 | 0.125 | #NUM! | COQ7,CYP11A1,MICAL2 |
| MSP-RON Signaling in Cancer Cells Pathway | 0.408 | 0.103 | 0.000 | AKT1,CREBBP,CTNNB1,HIF1A,HRAS,MET,MTOR,PIK3CA,PIK3CD,PIK3R2,RAP2A,RPS6KA2,VIM,YWHAZ |
| MSP-RON Signaling Pathway | 0.408 | 0.111 | #NUM! | ACTB,JAK2,PIK3CA,PIK3CD,PIK3R2,RPS6KA2 |
| Signaling by Insulin receptor | 0.406 | 0.114 | 0.447 | ATP6V1E1,ATP6V1G2,HRAS,PIK3CA,PIK3R2 |
| Apelin Pancreas Signaling Pathway | 0.406 | 0.114 | 1.342 | MAPK9,PIK3CA,PIK3CD,PIK3R2,PRKAA2 |
| GPVI-mediated activation cascade | 0.406 | 0.118 | 0.000 | FCER1G,FYN,PIK3CA,PIK3R2 |
| Oncogene Induced Senescence | 0.406 | 0.118 | 0.000 | CDK6,TFDP2,TNRC6B,UBC |
| Transcriptional Regulation by E2F6 | 0.406 | 0.118 | 1.000 | APAF1,EZH2,RRM2,TFDP2 |
| Transcriptional Regulation by NPAS4 | 0.406 | 0.118 | 0.000 | CREBBP,NR3C1,RBFOX3,TNRC6B |
| TGF-β Signaling | 0.405 | 0.105 | 0.333 | CREBBP,HRAS,INHA,INHBA,MAPK9,PITX2,RAP2A,SMAD6,TGFBR1,ZNF423 |
| Kinetochore Metaphase Signaling Pathway | 0.405 | 0.105 | -1.134 | BUB1,CDC20,CDCA8,CDK1,CENPK,H2AZ2,MAD2L1,PPP1CA,SKA2,SMC3 |
| ABRA Signaling Pathway | 0.401 | 0.106 | -1.667 | ACTB CFL1 MAPK9 MEF2C MRTFA PPARGC1A PPP1CA SMTN VCL |
| Tuberculosis Active Signaling Pathway | 0.400 | 0.101 | 1.147 | AKT1,ATP6AP2,ATP6V1E1,ATP6V1G2,C3AR1,CORO1A,EZH2,FCER1G,FOXO1,IL1RAPL1,MAP4K2,MAPK9,MKRN1,MTOR,NGFR,NLRP3,PIK3CA,PIK3CD,PIK3R2,RACK1 |
| Lung Ionic Balance Signaling Pathway | 0.398 | 0.098 | -1.938 | ADGRA2,ADGRB1,ADGRB2,ADGRB3,ADGRG1,ADGRL3,ADORA2A,ADRA2C,ATP1A1,BDKRB1,C3AR1,CCRL2,CELSR1,Clca3b,EDNRB,FYN,FZD1,GIPR,GPER1,GPR12,GPR137C,GPR146,GPR155,GPR156,GPR19,GPR27,GPR68,GPR85,GPR88,GRM3,KCNMB4,MCHR1,NMUR1,OXGR1,PLCG1,PLCH1,PLCH2,PPP2R2B,PRKAA2,PRKCB,PRKCD,PRKCH,PTH1R,S1PR1,SSTR2,SSTR3 |
| Chemokine Signaling | 0.397 | 0.107 | 0.000 | CALM1,CFL1,GNAQ,HRAS,PLCG1,PPP1CA,PRKCB,RAP2A |
| STAT3 Pathway | 0.396 | 0.102 | -1.155 | FGFR3,HGF,HRAS,IL6ST,JAK2,MAP3K11,MAPK9,NDUFA13,NGFR,NTRK2,PIM1,RAP2A,SOCS2,TGFBR1 |
| Cyclophilin Signaling Pathway | 0.394 | 0.100 | 0.209 | ABCA1,AKT1,CALR,CREBBP,CTNNB1,CYCS,HIF1A,HNRNPA1,IL6ST,JAK2,JAK3,LRP1,MAPK9,MRE11,NFKBID,PDIA6,PIK3CA,PIK3CD,PIK3R2,PPIB,SOX2,VDAC3,WNT7B |
| cAMP-mediated signaling | 0.394 | 0.100 | -1.886 | ADCY1,ADCY8,ADORA2A,ADRA2C,AKAP13,CALM1,CREBBP,DUSP1,GNAL,GPER1,GRM3,GUCY1A1,GUCY1B1,MPPED2,PDE1A,PDE1B,PDE2A,PDE6D,PTH1R,RAPGEF4,RGS4,S1PR1,SSTR3 |
| DDX58/IFIH1-mediated induction of interferon-alpha/beta | 0.393 | 0.108 | 1.134 | CREBBP,HSP90AA1,HSP90AB1,NKIRAS1,TKFC,TNFAIP3,UBC |
| Asparagine N-linked glycosylation | 0.392 | 0.102 | 1.387 | B4GALNT2,B4GALT2,B4GALT5,CALR,CANX,CHST8,HK1,MARCHF6,MVD,OS9,RFT1,UBC,UBXN1 |
| Sensory processing of sound by outer hair cells of the cochle | 0.389 | 0.109 | 1.633 | ACTB,ATP2B2,CASK,EPB41L3,EPS8,SLC26A5 |
| Ferroptosis Signaling Pathway | 0.387 | 0.103 | 1.265 | ABCA1,ACACA,ACSL4,FDFT1,H2AZ2,HMGCR,HRAS,PRKAA2,RAP2A,SLC38A1,TF,TXNRD1 |
| Glutaryl-CoA Degradation | 0.382 | 0.133 | #NUM! | Acat2/Acat3,PARK7 |
| Detoxification of Reactive Oxygen Species | 0.382 | 0.114 | 2.000 | ATOX1,CAT,CYCS,TXNRD1 |
| Mitochondrial Fatty Acid Beta-Oxidation | 0.382 | 0.114 | 0.000 | ACADL,ACOT7,DBI,PCCA |
| Transcriptional regulation by the AP-2 (TFAP2) family of tran | 0.382 | 0.114 | -1.000 | CITED2,CREBBP,PITX2,WWOX |
| Nuclear RNA decay | 0.382 | 0.114 | 0.000 | DIS3,EXOSC9,WDR82,ZC3H3 |
| Expression of BMAL (ARNTL), CLOCK, and NPAS2 | 0.382 | 0.114 | -1.000 | CHD9,CREBBP,MEF2C,PPARGC1A |
| Antiproliferative Role of TOB in T Cell Signaling | 0.382 | 0.114 | #NUM! | FCER1G,PABPC1,SKP2,TGFBR1 |
| Deubiquitination | 0.381 | 0.099 | 1.225 | ABRAXAS1,ACTB,BABAM1,CDC20,CDK1,HCFC1,HIF1A,JOSD1,NLRP3,PSMB10,PSMB2,PSMB4,PSMB5,PSMC5,PSMD13,PSMD6,SKP2,TGFBR1,TNFAIP3,UBC,USP12,USP24,VDAC3,WDR48 |
| RORA,B,C and NR1D1 (REV-ERBA) regulate gene expression | 0.380 | 0.120 | #NUM! | CHD9,CREBBP,PPARGC1A |
| Antigen Presentation Pathway | 0.380 | 0.120 | #NUM! | CALR,CANX,PSMB5 |
| Role of JAK1 and JAK3 in γc Cytokine Signaling | 0.377 | 0.106 | 0.378 | HRAS,JAK2,JAK3,PIK3CA,PIK3CD,PIK3R2,RAP2A |
| Generic Transcription Pathway | 0.367 | 0.099 | -2.982 | CREBBP,HDAC2,HDAC7,HDAC9,MED16,NOTCH1,NR2F1,NR3C1,NR4A1,NR4A2,ZFP90,ZNF25,ZNF282,ZNF446,ZNF689,ZNF691,ZNF740,ZNF786,ZSCAN25 |
| PXR/RXR Activation | 0.366 | 0.109 | 1.000 | AKT1,FOXO1,NR3C1,PPARGC1A,Scd1 |
| Role of OCT4 in Mammalian Embryonic Stem Cell Pluripoten | 0.366 | 0.109 | #NUM! | AQR,NR2F1,NR2F2,PARP1,SOX2 |
| 3-phosphoinositide Biosynthesis | 0.363 | 0.099 | 0.000 | ATP1A1 DUSP1 DUSP12 DUSP2 NUDT15 PANK4 PI4KB PIK3CA PIK3CD PIK3R2 PIP4K2B PLPPR4 PPP1CA PPP1R14B PPP1R16B PPP1R1A PPP2R2B PTPRO RASA1 SIRPA |
| Actin Cytoskeleton Signaling | 0.360 | 0.098 | 1.606 | ACTB,ARPC3,ARPC5,ARPC5L,CFL1,CYFIP2,DOCK1,FN1,GNA12,HRAS,MYL12B,MYL6,PAK1,PFN1,PIK3CA,PIK3CD,PIK3R2,PPP1CA,RAC3,RAP2A,TMSB4X,VCL,WASF1 |
| Glycation Signaling Pathway | 0.359 | 0.098 | 0.218 | AKT1,FN3K,HIF1A,HK1,HRAS,IRAK1,JAK2,MAPK7,MAPK9,MAPT,MTOR,PIK3CA,PIK3CD,PIK3R2,PRKCB,PRKCD,PRKCH,RAC3,RAP2A,SCARB1,SETD7 |
| Sheddase Signaling Pathway | 0.358 | 0.099 | 0.243 | CCRL2,CDH2,DDIT4,ERBB4,FN1,HGF,HIF1A,IGF2,IGFBP3,MAPK9,MET,MMP24,MTOR,PIK3CA,PIK3CD,PIK3R2,POSTN |
| Role of CHK Proteins in Cell Cycle Checkpoint Control | 0.354 | 0.105 | -1.342 | ATM,ATMIN,CDK1,MRE11,PPP2R2B,RAD1 |
| Polyamine Regulation in Colon Cancer | 0.354 | 0.105 | 0.447 | AKT1,CTNNB1,EIF5A,MTOR,SMS,SRM |
| ATF4 activates genes in response to endoplasmic reticulum s | 0.354 | 0.115 | #NUM! | DIS3,EXOSC9,KHSRP |
| Fructose metabolism | 0.352 | 0.167 | #NUM! | TKFC |
| Galactose catabolism | 0.352 | 0.167 | #NUM! | GALE |
| OAS antiviral response | 0.352 | 0.167 | #NUM! | RNASEL |
| Epithelial-Mesenchymal Transition (EMT) during gastrulation | 0.352 | 0.167 | #NUM! | EOMES |
| Role of Lipids/Lipid Rafts in the Pathogenesis of Influenza | 0.352 | 0.167 | #NUM! | FDPS |

| | | | | |
|---|---|---|---|---|
| Role of IL-17A in Psoriasis | 0.352 | 0.167 | #NUM! | S100A9 |
| Trehalose Degradation II (Trehalase) | 0.352 | 0.167 | #NUM! | HK1 |
| Selenocysteine Biosynthesis II (Archaea and Eukaryotes) | 0.352 | 0.167 | #NUM! | SEPSECS |
| Ceramide Degradation | 0.352 | 0.167 | #NUM! | ASAH1 |
| Salvage Pathways of Pyrimidine Deoxyribonucleotides | 0.352 | 0.167 | #NUM! | TK2 |
| PI3K/AKT Signaling | 0.349 | 0.098 | 0.728 | AKT1,CTNNB1,EIF4E,FOXO1,HRAS,HSP90AA1,HSP90AB1,HSP90B1,IL6ST,JAK2,JAK3,MTOR,NFKBID,PIK3CA,PIK3CD,PIK3R2,PPP2R2B,RAP2A,YWHAZ |
| Transcriptional regulation of pluripotent stem cells | 0.349 | 0.125 | #NUM! | FOXP1,SOX2 |
| Processing of SMDT1 | 0.349 | 0.125 | #NUM! | MICU1,SMDT1 |
| SUMOylation of RNA binding proteins | 0.347 | 0.106 | 1.342 | CBX8,NOP58,NUP210,NUP93,RAE1 |
| Inositol phosphate metabolism | 0.347 | 0.106 | -1.342 | CALM1,IP6K1,PLCG1,PLCH1,PLCH2 |
| Thioredoxin Signaling Pathway | 0.345 | 0.103 | 1.134 | AKT1,HIF1A,MAPK9,MIF,PRKAA2,RRM2,TXNRD1 |
| Transcriptional regulation by RUNX1 | 0.341 | 0.099 | 0.535 | CBX8,CREBBP,H2AZ2,H3C1,HIPK2,PRKCB,PSMB10,PSMB2,PSMB4,PSMB5,PSMC5,PSMD13,PSMD6,UBC |
| Fanconi Anemia Pathway | 0.339 | 0.108 | -1.000 | FANCC,RPA2,UBC,WDR48 |
| Signaling by MET | 0.338 | 0.101 | 0.000 | FN1,HGF,HRAS,LAMB1,MET,PIK3CA,RAPGEF1,UBC |
| Iron uptake and transport | 0.338 | 0.103 | 0.000 | ATP6V1E1,ATP6V1G2,FBXL5,GLRX3,TF,UBC |
| PRC2 methylates histones and DNA | 0.330 | 0.111 | #NUM! | EZH2,H2AZ2,H3C1 |
| Role of PI3K/AKT Signaling in the Pathogenesis of Influenza | 0.329 | 0.104 | 1.342 | AKT1,NFKBID,PIK3CA,PIK3CD,PIK3R2 |
| Response to elevated platelet cytosolic Ca2+ | 0.329 | 0.098 | -0.577 | CALM1,CFL1,FN1,HGF,IGF2,OLA1,PFN1,PRKCB,TF,TMSB4X,VCL,WDR1 |
| Ovarian Cancer Signaling | 0.327 | 0.097 | 1.134 | AKT1,BRCA2,CTNNB1,FZD1,HDAC2,HDAC7,HDAC9,HRAS,MTOR,PA2G4,PIK3CA,PIK3CD,PIK3R2,RAP2A,WNT7B |
| Exocyst Signaling Pathway | 0.327 | 0.099 | 0.632 | AKT1,ARPC3,ARPC5,ARPC5L,CDH2,PRKCB,PRKCD,PRKCH,VAMP3,VTI1A |
| Apoptosis Signaling | 0.327 | 0.099 | -1.000 | APAF1,CAPNS1,CDK1,CYCS,HRAS,MAP4K4,NFKBID,PARP1,PLCG1,RAP2A |
| Natural Killer Cell Signaling | 0.324 | 0.097 | 0.000 | AKT1,CFL1,FCER1G,FYN,HRAS,JAK2,JAK3,MAP3K11,MTOR,PAK1,PIK3CA,PIK3CD,PIK3R2,PLCG1,RAC3,RAP2A |
| Peroxisomal protein import | 0.322 | 0.102 | 0.000 | ACAA1,CAT,CRAT,IDH1,PEX26,UBC |
| Co-inhibition by BTLA | 0.322 | 0.102 | 0.816 | CDKL1,PI4KB,PIK3CA,PIK3CD,PIK3R2,STK17B |
| Carboxyterminal post-translational modifications of tubulin | 0.320 | 0.105 | 0.000 | AGBL4,SVBP,TUBA1A,TUBB6 |
| COPI-mediated anterograde transport | 0.320 | 0.099 | 1.000 | CAPZB,COG7,COPB2,DYNC1I1,DYNLL1,GOLGB1,SPTB,TUBA1A,TUBB6 |
| PD-1, PD-L1 cancer immunotherapy pathway | 0.320 | 0.099 | 0.333 | AKT1,JAK2,JAK3,NGFR,PIK3CA,PIK3CD,PIK3R2,PLCG1,SKP2 |
| Regulation of TLR by endogenous ligand | 0.319 | 0.118 | #NUM! | S100A1,S100A9 |
| Azathioprine ADME | 0.319 | 0.118 | #NUM! | IMPDH1,NUDT15 |
| NFE2L2 regulating anti-oxidant/detoxification enzymes | 0.319 | 0.118 | #NUM! | CREBBP,TXNRD1 |
| Specification of the neural plate border | 0.319 | 0.118 | #NUM! | CTNNB1,SOX2 |
| FXR/RXR Activation | 0.318 | 0.097 | 1.069 | AKT1,APOC2,CREBBP,FASN,FOXO1,Gstm7,GSTO1,MAFG,MAPK9,MGST1,NLRP3,PPARGC1A,PRKAA2,SCARB1,SLC51A |
| Transport of inorganic cations/anions and amino acids/oligo | 0.315 | 0.098 | 0.632 | CALM1,SLC12A6,SLC17A7,SLC1A4,SLC25A1,SLC25A22,SLC26A9,SLC38A1,SLC4A10,SLC4A7 |
| RUNX1 regulates megakaryocyte differentiation and platelet | 0.313 | 0.102 | -0.447 | H2AZ2,H3C1,PRMT1,TNRC6B,WDR5 |
| Kinesins | 0.313 | 0.102 | 1.342 | KIF11,KIF26B,KIF3B,TUBA1A,TUBB6 |
| Phototransduction Pathway | 0.313 | 0.102 | -1.342 | GUCY1A1,GUCY1B1,OPN4,PDE6D,SAG |
| PKR-mediated signaling | 0.312 | 0.099 | 0.707 | CDK1,DNAJC3,EIF2S3,FANCC,MAPT,TUBA1A,TUBB6,VPS28 |
| HOTAIR Regulatory Pathway | 0.312 | 0.097 | -1.069 | AKT1,CREBBP,CTNNB1,EZH2,HDAC2,MET,MMP24,MRTFA,PIK3CA,PIK3CD,PIK3R2,TWIST1,VIM,WNT7B |
| Autism Signaling Pathway | 0.311 | 0.095 | -1.177 | AKT1,ALDH16A1,Aldh3a2,CACNA1A,CACNG3,CACNG8,CAMKK2,CREBBP,CTNNB1,DBT,EIF4E,FZD1,GRIN2D,GRM3,HGF,HRAS,IGF2,MET,MTOR,NLGN1,NTRK2,PCCA,PIK3CA,PIK3CD,PIK3R2,RAP2A,WNT7B |
| Aryl hydrocarbon receptor signalling | 0.304 | 0.143 | #NUM! | HSP90AB1 |
| TYSND1 cleaves peroxisomal proteins | 0.304 | 0.143 | #NUM! | ACAA1 |
| Mitochondrial Uncoupling | 0.304 | 0.143 | #NUM! | SLC25A27 |
| Thioredoxin Pathway | 0.304 | 0.143 | #NUM! | TXNRD1 |
| Acetyl-CoA Biosynthesis I (Pyruvate Dehydrogenase Complex | 0.304 | 0.143 | #NUM! | DBT |
| Inositol Pyrophosphates Biosynthesis | 0.304 | 0.143 | #NUM! | IP6K1 |
| Adenine and Adenosine Salvage III | 0.304 | 0.143 | #NUM! | PNP |
| Purine Ribonucleosides Degradation to Ribose-1-phosphate | 0.304 | 0.143 | #NUM! | PNP |
| Glycoaminoglycan-protein Linkage Region Biosynthesis | 0.304 | 0.143 | #NUM! | B3GALT6 |
| Cargo recognition for clathrin-mediated endocytosis | 0.304 | 0.097 | 1.265 | AGFG1,AP2M1,CLTB,COPS5,EPN1,SYT9,TF,UBC,VAMP3,VAMP7 |
| ICOS-ICOSL Signaling in T Helper Cells | 0.304 | 0.097 | 0.447 | AKT1,CALM1,FCER1G,ITPR1,ITPR2,NFKBID,PIK3CA,PIK3CD,PIK3R2,PLCG1 |
| Bladder Cancer Signaling | 0.304 | 0.097 | 0.000 | FGFR3,HDAC2,HDAC7,HDAC9,HRAS,MMP24,PA2G4,RAP2A,RASSF1,RPS6KA5 |
| MAP kinase activation | 0.301 | 0.103 | -1.000 | IRAK1,MAPK9,MAPKAPK2,UBC |
| NGF-stimulated transcription | 0.301 | 0.103 | -1.000 | EGR3,NAB1,NAB2,VGF |
| Retinoid metabolism and transport | 0.301 | 0.103 | -1.000 | APOC2,LRP1,LRP12,LRP8 |
| Retinoic acid Mediated Apoptosis Signaling | 0.301 | 0.103 | 0.000 | APAF1,CYCS,PARP1,PARP8 |
| Transcriptional regulation of white adipocyte differentiation | 0.299 | 0.098 | -1.414 | CHD9,CREBBP,MED11,MED16,MED18,MED9,NR2F2,PPARGC1A |
| Epithelial Membrane Protein Signaling Pathway | 0.297 | 0.095 | 1.000 | AKT1,FGFR3,FYN,HIF1A,HRAS,MTOR,NGFR,NTRK2,PIK3CA,PIK3CD,PIK3R2,RAP2A,SMAD6,SOX9,TGFBR1,TWIST1 |
| Abacavir ADME | 0.293 | 0.098 | 0.816 | CDKL1,PI4KB,PIK3CA,PIK3CD,PIK3R2,STK17B |
| SUMOylation of chromatin organization proteins | 0.293 | 0.098 | 1.633 | CBX8,HDAC2,NUP210,NUP93,RAE1,SATB2 |
| Glyoxylate metabolism and glycine degradation | 0.292 | 0.111 | #NUM! | GRHPR,KGD4 |
| DNA methylation | 0.292 | 0.111 | #NUM! | H2AZ2,H3C1 |
| Activated PKN1 stimulates transcription of AR regulated gen | 0.292 | 0.111 | #NUM! | H2AZ2,H3C1 |
| Formation of the nephric duct | 0.292 | 0.111 | #NUM! | CTNNB1,MECOM |
| Dendritic Cell Maturation | 0.291 | 0.095 | -1.069 | AKT1,CREBBP,FCER1G,FSCN1,JAK2,LY75,MAPK9,NFKBID,NGFR,PIK3CA,PIK3CD,PIK3R2,PLCG1,PLCH1,PLCH2 |
| Cell surface interactions at the vascular wall | 0.291 | 0.096 | -0.905 | CAV1,FCER1G,FN1,FYN,HRAS,MERTK,MIF,PIK3CA,PIK3R2,PLCG1,SIRPA |
| Cardiac conduction | 0.288 | 0.095 | 0.000 | AHCYL1,ATP1A1,ATP2B2,CACNG8,CALM1,FXYD2,FXYD3,HIPK2,ITPR1,ITPR2,KCNK12,KCNK13 |
| Hepatic Fibrosis Signaling Pathway | 0.288 | 0.094 | 0.000 | AKT1 CACNA1A CACNG3 CACNG8 CALM1 CREBBP CTNNB1 EZH2 FOXO1 FZD1 HIF1A HRAS IL1RAPL1 IRAK1 JAK2 LRP1 MAPK9 MTOR MYL12B MYL6 NFKBID NGFR PIK3CA PIK3CD PIK3R2 PLCG1 PRKCB PRKCD PRKCH RAC3 RAP2A RHOU RND2 SDHD TF TGFBR1 WNT7B |
| Regulation of the Epithelial Mesenchymal Transition in Devel | 0.287 | 0.096 | -1.414 | CTNNB1,FZD1,HIF1A,NCSTN,NOTCH1,SUFU,TWIST1,WNT7B |
| Formation of Fibrin Clot (Clotting Cascade) | 0.287 | 0.103 | #NUM! | C1QBP,PRTN3,TFPI |
| B Cell Receptor Signaling | 0.287 | 0.094 | 0.000 | AKT1,CALM1,CFL1,CREBBP,FOXO1,HRAS,MAP3K11,MAPK9,MEF2C,MTOR,NFKBID,PIK3CA,PIK3CD,PIK3R2,POU2F2,PRKCB,RAP2A |
| MHC class II antigen presentation | 0.283 | 0.095 | 1.897 | AP2M1,CANX,CAPZB,DYNC1I1,DYNLL1,KIF11,KIF3B,SEC24D,TUBA1A,TUBB6 |
| NCAM signaling for neurite out-growth | 0.279 | 0.097 | 0.000 | COL9A3,FYN,HRAS,RPS6KA5,SPTB,ST8SIA4 |
| IL-33 Signaling Pathway | 0.277 | 0.094 | -1.155 | AKT1,CREBBP,H2AZ2,IRAK1,JAK2,LPIN1,MAPK9,MAPKAPK2,MTOR,NFKBID,PIK3CA,PIK3CD,PIK3R2 |
| PEDF Signaling | 0.275 | 0.095 | 1.134 | AKT1,HRAS,NFKBID,PIK3CA,PIK3CD,PIK3R2,RAP2A,ZEB1 |
| Regulation of IL-2 Expression in Activated and Anergic T Lym | 0.274 | 0.095 | -0.707 | CALM1,FCER1G,FYN,HRAS,MAPK9,NFKBID,PLCG1,RAP2A,TGFBR1 |
| Bone Mineralization Signaling Pathway | 0.268 | 0.094 | -1.387 | ANKH,CTNNB1,DBP,FGFR3,FZD1,GNAQ,IGF2,NR4A2,PLCG1,PLCH1,PLCH2,PTH1R,WNT7B |
| Endosomal Sorting Complex Required For Transport (ESCRT) | 0.268 | 0.100 | #NUM! | UBC,VPS28,VPS4B |
| Apelin Liver Signaling Pathway | 0.267 | 0.105 | #NUM! | EDN3,MAPK9 |
| Spliceosomal Cycle | 0.267 | 0.096 | -1.000 | AQR,DDX23,RBMX2,XAB2,ZNF830 |
| Senescence-Associated Secretory Phenotype (SASP) | 0.266 | 0.095 | -1.633 | CDK6,H2AZ2,H3C1,MAPK7,RPS6KA2,UBC |
| Induction of Apoptosis by HIV1 | 0.266 | 0.095 | 0.000 | APAF1,CYCS,MAPK9,NFKBID,NGFR,SLC25A3 |
| Neurotransmitter clearance | 0.265 | 0.125 | #NUM! | COMT |
| Neurotransmitter receptors and postsynaptic signal transmi | 0.265 | 0.125 | #NUM! | NBEA |
| Creatine metabolism | 0.265 | 0.125 | #NUM! | GATM |
| NFE2L2 regulating MDR associated enzymes | 0.265 | 0.125 | #NUM! | CREBBP |
| Embryonic Stem Cell Differentiation into Cardiac Lineages | 0.265 | 0.125 | #NUM! | SOX2 |
| Sphingosine and Sphingosine-1-phosphate Metabolism | 0.265 | 0.125 | #NUM! | ASAH1 |
| Superoxide Radicals Degradation | 0.265 | 0.125 | #NUM! | CAT |
| Mitochondrial protein degradation | 0.263 | 0.094 | 1.667 | ACADSB,CLPX,COX5A,DBT,IDH3A,MDH2,NDUFA13,OXCT1,SMDT1 |
| HIPPO signaling | 0.254 | 0.093 | 0.000 | PPP1CA,PPP2R2B,RASSF1,RBX1,SKP2,TEAD2,YWHAZ |
| Phase II - Conjugation of compounds | 0.254 | 0.094 | 0.000 | ABHD10,COMT,GSTO1,MGST1,MTRR,TPST1 |
| Formation of WDR5-containing histone-modifying complexes | 0.252 | 0.095 | 0.000 | HCFC1,PHF20L1,WDR5,WDR82 |
| MyD88 dependent cascade initiated on endosome | 0.250 | 0.097 | #NUM! | IRAK1,PELI1,UBC |
| Cytotoxic T Lymphocyte-mediated Apoptosis of Target Cells | 0.250 | 0.097 | #NUM! | APAF1,CYCS,FCER1G |
| Dopamine Degradation | 0.245 | 0.100 | #NUM! | Aldh3a2,COMT |
| GP1b-IX-V activation signalling | 0.232 | 0.111 | #NUM! | YWHAZ |
| Urea cycle | 0.232 | 0.111 | #NUM! | NMRAL1 |
| Interleukin-9 signaling | 0.232 | 0.111 | #NUM! | JAK3 |
| Lipophagy | 0.232 | 0.111 | #NUM! | PRKAA2 |
| GDP-glucose Biosynthesis | 0.232 | 0.111 | #NUM! | HK1 |
| Mineralocorticoid Biosynthesis | 0.232 | 0.111 | #NUM! | NSDHL |
| Synaptic adhesion-like molecules | 0.225 | 0.095 | #NUM! | GRIA3,GRIN2D |
| Interleukin-20 family signaling | 0.225 | 0.095 | #NUM! | JAK2,JAK3 |
| Co-inhibition by CTLA4 | 0.225 | 0.095 | #NUM! | AKT1,FYN |
| Platelet Aggregation (Plug Formation) | 0.204 | 0.100 | #NUM! | ADRA2C |
| Caspase activation via Dependence Receptors in the absence | 0.204 | 0.100 | #NUM! | DCC |
| Ketone body metabolism | 0.204 | 0.100 | #NUM! | OXCT1 |
| RHOBTB3 ATPase cycle | 0.204 | 0.100 | #NUM! | RAB9B |
| Protein lipoylation | 0.204 | 0.100 | #NUM! | DBT |
| Glucose and Glucose-1-phosphate Degradation | 0.204 | 0.100 | #NUM! | HK1 |
| Calcium Transport I | 0.204 | 0.100 | #NUM! | ATP2B2 |
| NAD Phosphorylation and Dephosphorylation | 0.204 | 0.100 | #NUM! | NNT |
| Glucocorticoid Biosynthesis | 0.204 | 0.100 | #NUM! | NSDHL |
| Pulmonary Healing Signaling Pathway | 0.000 | 0.087 | -0.500 | CFL1,CTNNB1,FYN,FZD1,HIF1A,HRAS,MAPK9,MMP24,NGFR,NOTCH1,PRKAA2,PRKCB,PRKCD,PRKCH,RAP2A,WNT7B |
| Pyroptosis Signaling Pathway | 0.000 | 0.028 | #NUM! | NGFR,NLRP3 |
| Wound Healing Signaling Pathway | 0.000 | 0.059 | -0.277 | COL9A3,FN1,HRAS,IL1RAPL1,JAK2,LAMB1,MAPK9,NFKBID,NGFR,RAP2A,SHC3,TGFBR1,VIM |
| CDX Gastrointestinal Cancer Signaling Pathway | 0.000 | 0.077 | -0.535 | AKT1,BMP8B,CTNNB1,DLL1,EFNB1,FZD1,JAK2,JAK3,PIK3CA,PIK3CD,PIK3R2,RAC3,SOX2,WNT7B |
| Immunogenic Cell Death Signaling Pathway | 0.000 | 0.091 | 0.378 | CALR,HSP90AA1,HSP90AB1,HSP90B1,LRP1,NGFR,NLRP3 |
| Macrophage Classical Activation Signaling Pathway | 0.000 | 0.056 | 0.000 | EZH2,HIF1A,IRAK1,JAK2,MERTK,NFKBID,PKM,SDHD |
| Multiple Sclerosis Signaling Pathway | 0.000 | 0.044 | 0.000 | CAPNS1,DUSP1,EZH2,GRIN2D,MASP1,NLRP3,PARP1,PARP8 |
| Pathogen Induced Cytokine Storm Signaling Pathway | 0.000 | 0.035 | -1.265 | COL9A3,EOMES,HMGCR,IL1RAPL1,IL6ST,JAK2,MAPK7,MAPK9,NGFR,NLRP3 |
| Role of Osteoblasts in Rheumatoid Arthritis Signaling Pathwa | 0.000 | 0.062 | 0.277 | AKT1,CTNNB1,FZD1,IL6ST,JAK2,JAK3,LRP1,MMP24,PIK3CA,PIK3CD,PIK3R2,SMAD6,WNT7B |
| Role of Osteoclasts in Rheumatoid Arthritis Signaling Pathwa | 0.000 | 0.082 | 0.655 | Adam26a/Adam26b,AKT1,CALM1,COL9A3,CREBBP,CTNNA1,CTNNB1,FOXO1,HRAS,IL1RAPL1,MAPK9,MMP24,NFKBID,NGFR,PIK3CA,PIK3CD,PIK3R2,PLCG1,RAC3,RAP2A,RHOU,RND2,SHC3 |
| Glucocorticoid Receptor Signaling | 0.000 | 0.073 | #NUM! | ACTB,AKT1,CAV1,COXFA4,CREBBP,DUSP1,GTF2H3,HRAS,HSP90AA1,HSP90AB1,HSP90B1,IL6ST,JAK2,JAK3,MAPK9,NDUFA10,NDUFA13,NDUFA3,NDUFB2,NDUFB5,NDUFS7,NFKBID,NR3C1,PIK3CA,PIK3CD,PIK3R2,POLR2L,PRKAA2,RAP2A,RPS6KA5,SDHD,SLC37A4,TAF7,TGFBR1 |
| IL-10 Signaling | 0.000 | 0.093 | -0.577 | AKT1,CREBBP,DDIT4,DUSP1,HIF1A,IL1RAPL1,IL6ST,MAP4K4,MTOR,NFKBID,PKM,PRKAA2 |
| Neutrophil Extracellular Trap Signaling Pathway | 0.000 | 0.093 | 0.378 | AKT1,COL9A3,COXFA4,DOCK1,FYN,HIF1A,ITPR1,ITPR2,KCNN3,MAPK9,MTOR,NDUFA10,NDUFA13,NDUFA3,NDUFB2,NDUFB5,NDUFS7,NLRP3,PIK3CA,PIK3CD,PIK3R2,PLCG1,PLCH1,PLCH2,PRKCB,PRKCD,PRKCH,PRTN3,RAC3,SDHD,VDAC3 |
| NOD1/2 Signaling Pathway | 0.000 | 0.068 | 1.265 | ERBIN,HSP90AA1,HSP90AB1,HSP90B1,MAP4K4,MAPK9,MAPKAPK2,NFKBID,NLRP3,TNFAIP3 |
| Activin Inhibin Signaling Pathway | 0.000 | 0.081 | -1.807 | AKT1,CCND2,CREBBP,CTNNB1,FST,IL1RAPL1,INHA,INHBA,MAPK7,MAPK9,NEUROG3,NGFR,PIK3CA,PIK3CD,PIK3R2 |
| ISGylation Signaling Pathway | 0.000 | 0.037 | #NUM! | CREBBP,HIF1A,MAPK9 |
| Microautophagy Signaling Pathway | 0.000 | 0.084 | 1.155 | COL9A3,PRKAA2,PSMB10,PSMB2,PSMB4,PSMB5,PSMC5,PSMD13,PSMD6,SEC24D,VPS28,VPS45,VPS4B |
| NFKBIE Signaling Pathway | 0.000 | 0.071 | 0.000 | ANKRD44,FCER1G,NFKBID,NGFR |
| PI3K Cascade | 0.000 | 0.070 | #NUM! | FGFR3,PIK3CA,PIK3R2 |
| Presynaptic depolarization and calcium channel opening | 0.000 | 0.083 | #NUM! | CACNA1A |
| Meiotic synapsis | 0.000 | 0.093 | -1.000 | H2AZ2,POT1,SMC3,SYCE2 |
| Mitochondrial protein import | 0.000 | 0.032 | #NUM! | CHCHD3,DNAJC19 |
| BMAL1:CLOCK,NPAS2 activates circadian expression | 0.000 | 0.080 | #NUM! | CHD9,CREBBP |
| Collagen degradation | 0.000 | 0.018 | #NUM! | COL9A3 |
| Degradation of the extracellular matrix | 0.000 | 0.088 | 0.000 | ADAMTS4,CAPNS1,FBN2,FN1,LAMB1,NCSTN |
| Extracellular matrix organization | 0.000 | 0.049 | -0.447 | COL9A3,DDR1,FN1,LAMB1,VCAN |
| Glycerophospholipid biosynthesis | 0.000 | 0.033 | 0.000 | AGPAT1,ETNPPL,GPD1L,LPIN1 |
| PI Metabolism | 0.000 | 0.074 | 0.816 | PI4KB,PIK3CA,PIK3CD,PIK3R2,PIP4K2B,SBF1 |
| Telomere Maintenance | 0.000 | 0.091 | -1.134 | DSCC1,H2AZ2,H3C1,NOP10,POLR2L,POT1,RPA2 |
| Activation of Matrix Metalloproteinases | 0.000 | 0.053 | #NUM! | MMP24 |
| Glycosaminoglycan metabolism | 0.000 | 0.066 | 0.000 | ARSB,B3GALT6,B4GALT2,B4GALT5,HS3ST4,HS6ST2,ST3GAL1,VCAN |
| Macroautophagy | 0.000 | 0.091 | 0.000 | ATG13,DYNLL1,LAMTOR4,MTOR,PRKAA2,RRAGB |
| Collagen biosynthesis and modifying enzymes | 0.000 | 0.076 | -0.447 | ADAMTS3,COL9A3,CRTAP,P4HA2,PPIB |
| Toll-like Receptor Cascades | 0.000 | 0.035 | #NUM! | RBSN |
| MyD88-independent TLR4 cascade | 0.000 | 0.027 | #NUM! | UBC |
| Complement cascade | 0.000 | 0.068 | #NUM! | C3AR1,MASP1,MASP2 |
| Trafficking and processing of endosomal TLR | 0.000 | 0.083 | #NUM! | HSP90B1 |
| Toll Like Receptor 3 (TLR3) Cascade | 0.000 | 0.031 | #NUM! | UBC |
| Signaling by TGF-beta Receptor Complex | 0.000 | 0.093 | 2.236 | FKBP1A,PPP1CA,STRAP,TGFBR1,UBC |
| Acetylcholine binding and downstream events | 0.000 | 0.077 | #NUM! | CHRNA4 |
| Bile acid and bile salt metabolism | 0.000 | 0.079 | #NUM! | OSBP,OSBPL6,SLC51A |
| Metabolism of non-coding RNA | 0.000 | 0.077 | 1.000 | CLNS1A,NUP210,NUP93,RAE1 |
| Metabolism of steroid hormones | 0.000 | 0.029 | #NUM! | CYP11A1 |
| Nicotinate metabolism | 0.000 | 0.036 | #NUM! | PARP8 |
| Immunoregulatory interactions between a Lymphoid and a n | 0.000 | 0.012 | #NUM! | FCER1G |
| trans-Golgi Network Vesicle Budding | 0.000 | 0.054 | 0.000 | BLOC1S1,CLTB,GOLGB1,VAMP7 |
| Assembly of collagen fibrils and other multimeric structures | 0.000 | 0.054 | #NUM! | COL9A3,DST,PXDN |
| Metabolism of Angiotensinogen to Angiotensins | 0.000 | 0.067 | #NUM! | ATP6AP2 |
| Signaling by Hippo | 0.000 | 0.050 | #NUM! | AMOTL2 |
| RNA Polymerase II Transcription | 0.000 | 0.062 | 1.000 | FIP1L1,GTF2H3,NELFA,PAF1,POLR2L,SRSF11,SRSF2,TAF7,THOC7 |
| COPII-mediated vesicle transport | 0.000 | 0.085 | 0.000 | ANKRD44,CNIH2,PREB,SEC22C,SEC24D,TRAPPC2 |
| Metabolism of amine-derived hormones | 0.000 | 0.056 | #NUM! | DDC |
| Phase I - Functionalization of compounds | 0.000 | 0.051 | 0.000 | CYP11A1,CYP20A1,CYP51A1,PTGIS |
| Arachidonate metabolism | 0.000 | 0.030 | #NUM! | MAPKAPK2 |
| Integrin cell surface interactions | 0.000 | 0.025 | #NUM! | COL9A3,FN1 |
| Transcriptional activity of SMAD2/SMAD3:SMAD4 heterotrir | 0.000 | 0.060 | #NUM! | PARP1,TFDP2,UBC |
| Visual phototransduction | 0.000 | 0.035 | #NUM! | CALM1,SAG |
| Cytosolic iron-sulfur cluster assembly | 0.000 | 0.077 | #NUM! | ABCB7 |
| POU5F1 (OCT4), SOX2, NANOG activate genes related to pro | 0.000 | 0.077 | #NUM! | SOX2 |
| Syndecan interactions | 0.000 | 0.074 | #NUM! | CASK,FN1 |
| SUMOylation of DNA damage response and repair proteins | 0.000 | 0.092 | 1.134 | CBX8,HDAC7,NUP210,NUP93,PARP1,RAE1,SMC3 |
| SUMOylation of transcription factors | 0.000 | 0.053 | #NUM! | HIC1 |
| SUMOylation of ubiquitinylation proteins | 0.000 | 0.079 | #NUM! | NUP210,NUP93,RAE1 |
| WNT ligand biogenesis and trafficking | 0.000 | 0.077 | #NUM! | VPS35,WNT7B |
| Class A/1 (Rhodopsin-like receptors) | 0.000 | 0.059 | -2.668 | ADORA2A,ADRA2C,BDKRB1,C3AR1,CCRL2,EDN3,EDNRB,GPER1,GPR68,MCHR1,NMUR1,OPN4,OXGR1,PLPPR4,S1PR1,SSTR2,SSTR3 |
| Class B/2 (Secretin family receptors) | 0.000 | 0.087 | 0.707 | FZD1,GIPR,GNB2,GNG3,GNG4,GNG7,PTH1R,WNT7B |
| XBP1(S) activates chaperone genes | 0.000 | 0.085 | 0.000 | DNAJC3,HYOU1,PDIA6,PREB |
| Olfactory Signaling Pathway | 0.000 | 0.077 | #NUM! | GNAL |
| Striated Muscle Contraction | 0.000 | 0.056 | #NUM! | DES,VIM |
| Peroxisomal lipid metabolism | 0.000 | 0.077 | #NUM! | ACAA1,CRAT |
| Signal regulatory protein family interactions | 0.000 | 0.083 | #NUM! | SIRPA |
| Incretin synthesis, secretion, and inactivation | 0.000 | 0.042 | #NUM! | CTNNB1 |
| SUMOylation of SUMOylation proteins | 0.000 | 0.088 | #NUM! | NUP210,NUP93,RAE1 |
| SUMOylation of intracellular receptors | 0.000 | 0.069 | #NUM! | NR3C1,NR4A2 |
| Class C/3 (Metabotropic glutamate/pheromone receptors) | 0.000 | 0.046 | #NUM! | GRM3 |
| Transport of bile salts and organic acids, metal ions and amir | 0.000 | 0.026 | #NUM! | SLC39A10,SLC6A1 |
| TAK1-dependent IKK and NF-kappa-B activation | 0.000 | 0.071 | #NUM! | IRAK1,NKIRAS1,UBC |

| | | | | |
|---|---|---|---|---|
| Other interleukin signaling | 0.000 | 0.046 | #NUM! | PRTN3 |
| SUMOylation of DNA methylation proteins | 0.000 | 0.059 | #NUM! | CBX8 |
| Mitophagy | 0.000 | 0.051 | #NUM! | UBC,VDAC3 |
| Mismatch Repair | 0.000 | 0.071 | #NUM! | RPA2 |
| Signaling by Retinoic Acid | 0.000 | 0.054 | #NUM! | FABP5,PDK2 |
| Mitochondrial translation | 0.000 | 0.085 | 1.000 | CHCHD1,KGD4,MRPL51,MRPL58,MRPS21,MRPS22,MRPS27,MRPS31,MRPS35 |
| Activation of anterior HOX genes in hindbrain during early er | 0.000 | 0.079 | -0.816 | CREBBP,EZH2,H2AZ2,H3C1,POLR2L,WDR5 |
| Pyroptosis | 0.000 | 0.044 | #NUM! | CYCS |
| Interleukin-6 family signaling | 0.000 | 0.083 | #NUM! | IL6ST,JAK2 |
| TP53 Regulates Transcription of DNA Repair Genes | 0.000 | 0.090 | 0.000 | ATM,CDK12,FANCC,GTF2H3,NELFA,POLR2L |
| Regulation of TP53 Activity through Association with Co-fact | 0.000 | 0.071 | #NUM! | AKT1 |
| Keratinization | 0.000 | 0.016 | #NUM! | CAPNS1 |
| RNA polymerase II transcribes snRNA genes | 0.000 | 0.081 | -1.633 | ICE1,INTS1,INTS14,POLR2L,POU2F2,ZNF143 |
| mRNA Capping | 0.000 | 0.069 | #NUM! | GTF2H3,POLR2L |
| Processing of Capped Intron-Containing Pre-mRNA | 0.000 | 0.079 | 0.000 | AQR,DDX23,EIF4E,FIP1L1,HNRNPA1,LSM7,NUP210,NUP93,NXT1,PCBP1,POLR2L,PRPF38A,RAE1,RBMX2,SNRNP48,SRSF11,SRSF2,THOC7,XAB2,ZCRB1,ZMAT2,ZNF830 |
| Resolution of Abasic Sites (AP sites) | 0.000 | 0.053 | #NUM! | PARP1,RPA2 |
| Processing of Capped Intronless Pre-mRNA | 0.000 | 0.035 | #NUM! | FIP1L1 |
| Dissolution of Fibrin Clot | 0.000 | 0.091 | #NUM! | S100A10 |
| TNF signaling | 0.000 | 0.071 | 0.000 | MAPKAPK2,RACK1,TNFAIP3,UBC |
| Nucleosome assembly | 0.000 | 0.081 | #NUM! | CENPK,H2AZ2,HJURP |
| Interferon gamma signaling | 0.000 | 0.046 | #NUM! | JAK2,MID1,PRKCD |
| Protein ubiquitination | 0.000 | 0.069 | #NUM! | UBC,UBE2E3 |
| Transcriptional Regulation by VENTX | 0.000 | 0.053 | #NUM! | CTNNB1,TNRC6B |
| TBC/RABGAPs | 0.000 | 0.047 | #NUM! | RAB33A,RAB6A |
| E3 ubiquitin ligases ubiquitinate target proteins | 0.000 | 0.047 | #NUM! | PAF1,UBC |
| Digestion | 0.000 | 0.071 | #NUM! | PIR |
| Collagen chain trimerization | 0.000 | 0.023 | #NUM! | COL9A3 |
| Cristae formation | 0.000 | 0.063 | #NUM! | CHCHD3,MICOS10 |
| Gene and protein expression by JAK-STAT signaling after IL-1 | 0.000 | 0.086 | #NUM! | CFL1,GSTO1,MIF |
| Glycogen metabolism | 0.000 | 0.080 | #NUM! | CALM1,UBC |
| Interleukin-15 signaling | 0.000 | 0.071 | #NUM! | JAK3 |
| Interferon alpha/beta signaling | 0.000 | 0.042 | #NUM! | RNASEL,UBC |
| Transcriptional regulation of granulopoiesis | 0.000 | 0.093 | -1.000 | FLI1,H2AZ2,H3C1,TFDP2 |
| Alpha-protein kinase 1 signaling pathway | 0.000 | 0.091 | #NUM! | UBC |
| Response of EIF2AK1 (HRI) to heme deficiency | 0.000 | 0.069 | #NUM! | EIF2S3,VPS28 |
| Sensory processing of sound by inner hair cells of the cochle | 0.000 | 0.087 | 1.633 | ACTB,ATP2B2,CAPZB,CASK,EPB41L3,EPS8 |
| Signaling by CSF3 (G-CSF) | 0.000 | 0.067 | #NUM! | JAK2,UBC |
| Transcriptional regulation of testis differentiation | 0.000 | 0.083 | #NUM! | SOX9 |
| Heme signaling | 0.000 | 0.087 | 1.000 | CHD9,CREBBP,MEF2C,PPARGC1A |
| Expression and translocation of olfactory receptors | 0.000 | 0.024 | #NUM! | LHX2 |
| Serine metabolism | 0.000 | 0.091 | #NUM! | PHGDH |
| Ion channel transport | 0.000 | 0.070 | 0.000 | ANO1,ATP1A1,ATP2B2,ATP6V1E1,ATP6V1G2,CALM1,FXYD2,FXYD3,PDZD11,UBC,WNK1,WNK2 |
| Coagulation System | 0.000 | 0.069 | #NUM! | BDKRB1,TFPI |
| Hepatic Cholestasis | 0.000 | 0.087 | -2.500 | ADCY1,ADCY8,AKT1,CALM1,FOXO1,GPER1,GUCY1A1,GUCY1B1,IL1RAPL1,IRAK1,MAP4K2,MAPK9,NFKBID,NGFR,PRKCB,PRKCD,PRKCH |
| Hepatic Fibrosis / Hepatic Stellate Cell Activation | 0.000 | 0.065 | #NUM! | COL9A3,EDNRB,FN1,HGF,IGF2,IGFBP3,IL1RAPL1,LHX2,MET,MYL6,NGFR,TGFBR1 |
| CGAS-STING Signaling Pathway | 0.000 | 0.043 | -0.447 | ATP6AP2,ATP6V1E1,ATP6V1G2,LRRC8A,NLRP3 |
| NAFLD Signaling Pathway | 0.000 | 0.093 | -2.065 | ACACA,AKT1,FASN,FOXO1,HMGCR,JAK2,JAK3,LPIN1,MAPK9,MTOR,NGFR,NLRP3,PIK3CA,PIK3CD,PIK3R2,PPARGC1A,PPP2R2B,PRKAA2,SOCS2 |
| VDR/RXR Activation | 0.000 | 0.083 | #NUM! | FOXO1,IGFBP3,PRKCB,PRKCD,PRKCH,PSMC5 |
| Folate Signaling Pathway | 0.000 | 0.074 | 0.000 | IL6ST,JAK2,JAK3,SOX2 |
| Hematoma Resolution Signaling Pathway | 0.000 | 0.090 | 1.213 | AXL,CAT,COXFA4,DUSP1,GRIN2D,LRP1,MAPK9,MERTK,MMP24,NDUFA10,NDUFA13,NDUFA3,NDUFB2,NDUFB5,NDUFS7,PRKAA2,S1PR1,SIRPA,TUBA1A,VPS35 |
| mRNA 3 Prime End Processing Signaling Pathway | 0.000 | 0.093 | 0.378 | CNOT1,DIS3,EXOSC9,FIP1L1,PABPC1,PCBP1,PCBP4,PPP1CA,WNK1 |
| rRNA modification in the nucleus and cytosol | 0.000 | 0.068 | 2.000 | IMP3,NOP10,NOP58,PNO1 |
| Pyruvate metabolism | 0.000 | 0.085 | 0.000 | PDK2,PKM,SIRT4,UBC |
| Nucleotide biosynthesis | 0.000 | 0.077 | #NUM! | IMPDH1 |
| Replacement of protamines by nucleosomes in the male pro | 0.000 | 0.077 | #NUM! | HIRA |
| Formation of the ureteric bud | 0.000 | 0.056 | #NUM! | SIX1 |
| Mitochondrial RNA degradation | 0.000 | 0.091 | #NUM! | LRPPRC,SLIRP |
| Cellular response to mitochondrial stress | 0.000 | 0.087 | #NUM! | EIF2S3,VPS28 |
| Maturation of TCA enzymes and regulation of TCA cycle | 0.000 | 0.053 | #NUM! | SDHD |
| Regulation of endogenous retroelements | 0.000 | 0.070 | 0.447 | ACTB,H2AZ2,H3C1,HDAC2,MBD3 |
| Irritable Bowel Syndrome Signaling Pathway | 0.000 | 0.059 | 1.941 | IL1RAPL1,LAMB1,MMP24,MYL12B,MYL6,PIK3CA,PIK3CD,PIK3R2,RAC3,RHOU,RND2,UBC,WNT7B |
| TRIM21 Intracellular Antibody Signaling Pathway | 0.000 | 0.087 | 1.897 | CTNNB1,JAK2,PSMB10,PSMB2,PSMB4,PSMB5,PSMC5,PSMD13,PSMD6,UBC |
| ID3 Signaling Pathway | 0.000 | 0.067 | 0.707 | AKT1,FCER1G,HRAS,RAC3,RAP2A,SMAD6,SOX2,TGFBR1 |
| Tuberculosis Latent Signaling Pathway | 0.000 | 0.060 | 0.447 | ADCY1,ADCY8,GUCY1A1,GUCY1B1,NGFR |
| ER-Mitochondrial Communication Signaling Pathway | 0.000 | 0.089 | -0.816 | AKT1,CDIP1,CHCHD3,MICOS10,NCSTN,NLRP3,NR3C1 |
| Epigenetic regulation of gene expression by MLL3 and MLL4 | 0.000 | 0.081 | -1.890 | CREBBP,H2AZ2,H3C1,LPIN1,MED16,PPARGC1A,WDR5 |
| Mechanical load activates signaling by PIEZO1 and integrins i | 0.000 | 0.059 | #NUM! | AKT1 |
| Formation of the dystrophin-glycoprotein complex (DGC) | 0.000 | 0.056 | #NUM! | ACTB,LAMB1 |
| SWI/SNF chromatin remodelers | 0.000 | 0.032 | #NUM! | ACTB |
| Ribotoxic Stress Signaling Pathway | 0.000 | 0.065 | #NUM! | MAPK9,RNASEL |
| Tight Junction Signaling | 0.000 | 0.093 | #NUM! | ACTB,AKT1,CASK,CLDN11,CLDN20,CTNNA1,CTNNB1,MYL6,NGFR,PPP2R2B,RAB13,TGFBR1,VAMP3,VCL,VTI1A,YBX3 |
| TR/RXR Activation | 0.000 | 0.092 | 0.905 | ACACA,AKT1,FASN,HIF1A,MTOR,NRGN,PIK3CA,PIK3CD,PIK3R2,PPARGC1A,SCARB1 |
| Role of BRCA1 in DNA Damage Response | 0.000 | 0.088 | #NUM! | ABRAXAS1,ACTB,ATM,BABAM1,BRCA2,FANCC,MRE11 |
| Activation of IRF by Cytosolic Pattern Recognition Receptors | 0.000 | 0.078 | 0.000 | CREBBP,MAPK9,NFKBID,PPIB |
| Caveolar-mediated Endocytosis Signaling | 0.000 | 0.069 | #NUM! | ACTB,CAV1,COPB2,DYRK3,FYN |
| Clathrin-mediated Endocytosis Signaling | 0.000 | 0.092 | #NUM! | ACTB,AMPH,AP2M1,APOC2,ARPC3,ARPC5,ARPC5L,CLTB,EPN1,HIP1,MET,MYO6,PIK3CA,PIK3CD,PIK3R2,SH3BP4,TF,UBC |
| IL-12 Signaling and Production in Macrophages | 0.000 | 0.083 | -1.291 | AKT1,APOC2,CALM1,CAMKK2,IL6ST,JAK2,MTOR,PIK3CA,PIK3CD,PIK3R2,POU2F2,PRKAA2,PRKCB,PRKCD,PRKCH,XRCC6 |
| Role of RIG1-like Receptors in Antiviral Innate Immunity | 0.000 | 0.063 | #NUM! | CREBBP,NFKBID |
| Role of Pattern Recognition Receptors in Recognition of Bact | 0.000 | 0.078 | -0.707 | C3AR1,MAPK9,NLRP3,PIK3CA,PIK3CD,PIK3R2,PRKCB,PRKCD,PRKCH,RNASEL |
| Role of PKR in Interferon Induction and Antiviral Response | 0.000 | 0.086 | -0.333 | APAF1,CYCS,DNAJC3,FANCC,HSP90AA1,HSP90AB1,HSP90B1,MAPK9,NFKBID,NLRP3 |
| TREM1 Signaling | 0.000 | 0.082 | -1.342 | AKT1,IRAK1,JAK2,NLRP3,PLCG1 |
| MIF Regulation of Innate Immunity | 0.000 | 0.073 | #NUM! | MAPK9,MIF,NFKBID |
| IL-22 Signaling | 0.000 | 0.087 | #NUM! | AKT1,MAPK9 |
| CD27 Signaling in Lymphocytes | 0.000 | 0.089 | -1.000 | APAF1,CYCS,MAP3K11,MAPK9,NFKBID |
| IL-17 Signaling | 0.000 | 0.078 | 1.508 | AKT1,HRAS,HSP90AA1,HSP90AB1,HSP90B1,JAK2,MAPK9,PIK3CA,PIK3CD,PIK3R2,RAP2A |
| Airway Pathology in Chronic Obstructive Pulmonary Disease | 0.000 | 0.022 | #NUM! | AMBP,PRTN3 |
| 4-1BB Signaling in T Lymphocytes | 0.000 | 0.059 | #NUM! | MAPK9,NFKBID |
| CTLA4 Signaling in Cytotoxic T Lymphocytes | 0.000 | 0.082 | 0.000 | AKT1,AP2M1,CLTB,FCER1G,FYN,HRAS,MAPK9,PIK3CA,PIK3CD,PIK3R2,PLCG1,PPP2R2B,RAC3,RAP2A |
| B Cell Activating Factor Signaling | 0.000 | 0.047 | #NUM! | MAPK9,NFKBID |
| T Helper Cell Differentiation | 0.000 | 0.061 | #NUM! | FCER1G,IL6ST,NGFR,TGFBR1 |
| Cardiomyocyte Differentiation via BMP Receptors | 0.000 | 0.083 | #NUM! | MEF2C,SMAD6 |
| Cell Cycle Regulation by BTG Family Proteins | 0.000 | 0.081 | #NUM! | BTG1,PPP2R2B,PRMT1 |
| HMGB1 Signaling | 0.000 | 0.081 | 0.302 | AKT1,HRAS,KAT6B,MAPK9,NGFR,PIK3CA,PIK3CD,PIK3R2,RAC3,RAP2A,RHOU,RND2 |
| ATM Signaling | 0.000 | 0.093 | 0.707 | ATM,CCNB2,CDK1,CREBBP,MAPK9,MRE11,PPP2R2B,SMC3,ZEB1 |
| Basal Cell Carcinoma Signaling | 0.000 | 0.074 | -2.000 | BMP8B,CTNNB1,FZD1,SUFU,WNT7B |
| Primary Immunodeficiency Signaling | 0.000 | 0.051 | #NUM! | JAK3,RFXAP |
| Allograft Rejection Signaling | 0.000 | 0.025 | #NUM! | FCER1G |
| Autoimmune Thyroid Disease Signaling | 0.000 | 0.026 | #NUM! | FCER1G |
| Graft-versus-Host Disease Signaling | 0.000 | 0.029 | #NUM! | FCER1G |
| MYC Mediated Apoptosis Signaling | 0.000 | 0.083 | -1.000 | APAF1,CREBBP,CYCS,NGFR |
| Communication between Innate and Adaptive Immune Cells | 0.000 | 0.015 | #NUM! | FCER1G |
| Crosstalk between Dendritic Cells and Natural Killer Cells | 0.000 | 0.032 | #NUM! | ACTB,FSCN1 |
| Systemic Lupus Erythematosus Signaling | 0.000 | 0.090 | #NUM! | AKT1,FCER1G,HRAS,LSM1,LSM7,MTOR,PIK3CA,PIK3CD,PIK3R2,PLCG1,PRPF38A,PRPF39,RAP2A,SNRNP48,ZCRB1 |
| CDC42 Signaling | 0.000 | 0.092 | 0.632 | ARPC3,ARPC5,ARPC5L,CDC42BPA,CFL1,FCER1G,MAP3K11,MAPK9,MYL12B,MYL6,PAK1,PPP1CA,RASA1 |
| April Mediated Signaling | 0.000 | 0.048 | #NUM! | MAPK9,NFKBID |
| Role of Osteoblasts, Osteoclasts and Chondrocytes in Rheum | 0.000 | 0.085 | #NUM! | ADAMTS4,AKT1,BMP8B,CALM1,CTNNB1,DKK3,FOXO1,FZD1,IL1RAPL1,LRP1,MAPK9,NFKBID,NGFR,PIK3CA,PIK3CD,PIK3R2,SMAD6,WNT7B |
| Altered T Cell and B Cell Signaling in Rheumatoid Arthritis | 0.000 | 0.025 | #NUM! | FCER1G,PRTN3 |
| Atherosclerosis Signaling | 0.000 | 0.009 | #NUM! | APOC2 |
| Estrogen-Dependent Breast Cancer Signaling | 0.000 | 0.092 | 1.134 | AKT1,CREBBP,HRAS,PIK3CA,PIK3CD,PIK3R2,RAP2A |
| RANK Signaling in Osteoclasts | 0.000 | 0.090 | 0.000 | AKT1,CALM1,MAP3K11,MAPK9,NFKBID,PIK3CA,PIK3CD,PIK3R2 |
| Neuroprotective Role of THOP1 in Alzheimer's Disease | 0.000 | 0.065 | -1.000 | ACR,ENDOU,MAPT,MASP1,PRSS53,PRTN3 |
| Role of WNT/GSK-3β Signaling in the Pathogenesis of Influen | 0.000 | 0.050 | #NUM! | CTNNB1,FZD1,WNT7B |
| TWEAK Signaling | 0.000 | 0.083 | #NUM! | APAF1,CYCS,NFKBID |
| Role of MAPK Signaling in the Pathogenesis of Influenza | 0.000 | 0.053 | #NUM! | AKT1,HRAS,MAPK9,RAP2A |
| TNFR2 Signaling | 0.000 | 0.065 | #NUM! | NFKBID,TNFAIP3 |
| Role of Hypercytokinemia/hyperchemokinemia in the Pathoi | 0.000 | 0.036 | #NUM! | NLRP3,S1PR1 |
| OX40 Signaling Pathway | 0.000 | 0.067 | #NUM! | FCER1G,MAPK9,NFKBID |
| Extrinsic Prothrombin Activation Pathway | 0.000 | 0.077 | #NUM! | TFPI |
| MIF-mediated Glucocorticoid Regulation | 0.000 | 0.091 | #NUM! | MIF,NFKBID,NR3C1 |
| Tumoricidal Function of Hepatic Natural Killer Cells | 0.000 | 0.087 | #NUM! | APAF1,CYCS |
| Assembly of RNA Polymerase II Complex | 0.000 | 0.063 | #NUM! | GTF2H3,POLR2L,TAF7 |
| IL-17A Signaling in Gastric Cells | 0.000 | 0.048 | #NUM! | MAPK9 |
| Role of IL-17F in Allergic Inflammatory Airway Diseases | 0.000 | 0.081 | #NUM! | CREBBP,RPS6KA2,RPS6KA5 |
| Role of JAK1, JAK2 and TYK2 in Interferon Signaling | 0.000 | 0.044 | #NUM! | JAK2 |
| Role of JAK2 in Hormone-like Cytokine Signaling | 0.000 | 0.093 | -2.000 | CCND2,JAK2,SH2B1,SIRPA,SOCS2 |
| Role of JAK family kinases in IL-6-type Cytokine Signaling | 0.000 | 0.083 | 0.816 | HSP90AA1,HSP90AB1,HSP90B1,IL6ST,JAK2,MAPK9 |
| Hematopoiesis from Pluripotent Stem Cells | 0.000 | 0.032 | #NUM! | FCER1G |
| Estrogen-mediated S-phase Entry | 0.000 | 0.077 | #NUM! | CDK1,SKP2 |
| GADD45 Signaling | 0.000 | 0.086 | 0.447 | ATM,CCND2,CDK1,MAPK9,TGFBR1 |
| Heparan Sulfate Biosynthesis | 0.000 | 0.050 | #NUM! | B3GALT6,HS3ST4,HS6ST2 |
| Fatty Acid Activation | 0.000 | 0.067 | #NUM! | ACSL4 |
| Bupropion Degradation | 0.000 | 0.083 | #NUM! | CYP51A1 |
| Nicotine Degradation III | 0.000 | 0.037 | #NUM! | CYP51A1 |
| Chondroitin Sulfate Degradation (Metazoa) | 0.000 | 0.067 | #NUM! | ARSB |
| Heparan Sulfate Biosynthesis (Late Stages) | 0.000 | 0.038 | #NUM! | HS3ST4,HS6ST2 |
| Pyrimidine Ribonucleotides De Novo Biosynthesis | 0.000 | 0.053 | #NUM! | NUDT15,NUDT16 |
| Androgen Biosynthesis | 0.000 | 0.083 | #NUM! | NSDHL |
| Glutathione Redox Reactions I | 0.000 | 0.091 | #NUM! | Gstm7,MGST1 |
| CDP-diacylglycerol Biosynthesis I | 0.000 | 0.039 | #NUM! | AGPAT1 |
| D-myo-inositol-5-phosphate Metabolism | 0.000 | 0.093 | -1.069 | ATP1A1,DUSP1,DUSP12,DUSP2,NUDT15,PANK4,PLCG1,PLCH1,PLCH2,PLPPR4,PPP1CA,PPP1R14B,PPP1R16B,PPP1R1A,PPP2R2B,PTPRO,RASA1,SIRPA |
| Melatonin Degradation I | 0.000 | 0.039 | #NUM! | CYP51A1 |
| γ-linolenate Biosynthesis II (Animals) | 0.000 | 0.053 | #NUM! | ACSL4 |
| Triacylglycerol Degradation | 0.000 | 0.026 | #NUM! | ABHD6 |
| Phospholipases | 0.000 | 0.050 | #NUM! | PLCG1,PLCH1,PLCH2 |
| Estrogen Biosynthesis | 0.000 | 0.037 | #NUM! | CYP51A1 |
| Tryptophan Degradation X (Mammalian, via Tryptamine) | 0.000 | 0.071 | #NUM! | Aldh3a2,DDC |
| Chondroitin Sulfate Biosynthesis | 0.000 | 0.065 | #NUM! | B3GALT6,HS3ST4,HS6ST2 |
| Dermatan Sulfate Biosynthesis | 0.000 | 0.063 | #NUM! | B3GALT6,HS3ST4,HS6ST2 |
| Dermatan Sulfate Biosynthesis (Late Stages) | 0.000 | 0.057 | #NUM! | HS3ST4,HS6ST2 |
| Phosphatidylglycerol Biosynthesis II (Non-plastidic) | 0.000 | 0.036 | #NUM! | AGPAT1 |
| D-myo-inositol (1,4,5,6)-Tetrakisphosphate Biosynthesis | 0.000 | 0.084 | -0.302 | ATP1A1,DUSP1,DUSP12,DUSP2,NUDT15,PANK4,PLPPR4,PPP1CA,PPP1R14B,PPP1R16B,PPP1R1A,PPP2R2B,PTPRO,RASA1,SIRPA |
| Nicotine Degradation II | 0.000 | 0.029 | #NUM! | CYP51A1 |
| Mitochondrial L-carnitine Shuttle Pathway | 0.000 | 0.053 | #NUM! | ACSL4 |
| Serotonin Degradation | 0.000 | 0.026 | #NUM! | Aldh3a2 |
| D-myo-inositol (3,4,5,6)-tetrakisphosphate Biosynthesis | 0.000 | 0.084 | -0.302 | ATP1A1,DUSP1,DUSP12,DUSP2,NUDT15,PANK4,PLPPR4,PPP1CA,PPP1R14B,PPP1R16B,PPP1R1A,PPP2R2B,PTPRO,RASA1,SIRPA |
| 3-phosphoinositide Degradation | 0.000 | 0.079 | -0.302 | ATP1A1,DUSP1,DUSP12,DUSP2,NUDT15,PANK4,PLPPR4,PPP1CA,PPP1R14B,PPP1R16B,PPP1R1A,PPP2R2B,PTPRO,RASA1,SIRPA |
| Colanic Acid Building Blocks Biosynthesis | 0.000 | 0.071 | #NUM! | GALE |
| Tryptophan Degradation III (Eukaryotic) | 0.000 | 0.091 | #NUM! | Acat2/Acat3,PARK7 |
| Pyrimidine Ribonucleotides Interconversion | 0.000 | 0.057 | #NUM! | NUDT15,NUDT16 |
| Purine Nucleotides De Novo Biosynthesis II | 0.000 | 0.091 | #NUM! | IMPDH1 |
| Glycolysis I | 0.000 | 0.087 | #NUM! | ALDOC,PKM |
| Histamine Degradation | 0.000 | 0.067 | #NUM! | Aldh3a2 |
| Chondroitin Sulfate Biosynthesis (Late Stages) | 0.000 | 0.053 | #NUM! | HS3ST4,HS6ST2 |
| Adenosine Nucleotides Degradation II | 0.000 | 0.067 | #NUM! | PNP |
| Superpathway of Melatonin Degradation | 0.000 | 0.033 | #NUM! | CYP51A1 |
| Noradrenaline and Adrenaline Degradation | 0.000 | 0.074 | #NUM! | Aldh3a2,COMT |
| Acetone Degradation I (to Methylglyoxal) | 0.000 | 0.046 | #NUM! | CYP51A1 |
| Putrescine Degradation III | 0.000 | 0.056 | #NUM! | Aldh3a2 |
| Fatty Acid α-oxidation | 0.000 | 0.056 | #NUM! | Aldh3a2 |
| Oxidative Ethanol Degradation III | 0.000 | 0.029 | #NUM! | Aldh3a2 |
| Ethanol Degradation IV | 0.000 | 0.091 | #NUM! | Aldh3a2,CAT |
| Ethanol Degradation II | 0.000 | 0.039 | #NUM! | Aldh3a2 |
| Fatty Acid β-oxidation I | 0.000 | 0.088 | #NUM! | ACAA1,ACSL4,ECHDC3 |
| Antioxidant Action of Vitamin C | 0.000 | 0.087 | 2.449 | GLRX,GSTO1,JAK2,MAPK9,NFKBID,PLCG1,PLCH1,PLCH2,TXNRD1 |
| Inhibition of Matrix Metalloproteases | 0.000 | 0.071 | #NUM! | LRP1,MMP24 |
| Regulation of Cellular Mechanics by Calpain Protease | 0.000 | 0.080 | 2.000 | CAPNS1,CCND2,CDK1,CDK6,HRAS,RAP2A,VCL |
| Agranulocyte Adhesion and Diapedesis | 0.000 | 0.042 | #NUM! | ACTB,CCRL2,CLDN11,CLDN20,FN1,MMP24,MYL6 |
| Granulocyte Adhesion and Diapedesis | 0.000 | 0.048 | #NUM! | CCRL2,CLDN11,CLDN20,IL1RAPL1,MMP24,NGFR,THY1 |
| BER (Base Excision Repair) Pathway | 0.000 | 0.023 | #NUM! | PARP1 |
| WNT/Ca2+ Signaling Pathway | 0.000 | 0.092 | -2.449 | CREBBP,FZD1,PDE6D,PLCG1,PLCH1,PLCH2 |
| Toll-like Receptor Signaling | 0.000 | 0.056 | #NUM! | IRAK1,MAP4K4,TNFAIP3,UBC |
| Nucleotide Excision Repair Pathway | 0.000 | 0.088 | #NUM! | GTF2H3,POLR2L,RPA2 |
| PTEN Signaling | 0.000 | 0.087 | 0.000 | AKT1,FGFR3,FOXO1,HRAS,MAGI1,NGFR,NTRK2,PIK3CA,PIK3CD,PIK3R2,RAC3,RAP2A,TGFBR1 |
| Interferon Signaling | 0.000 | 0.037 | #NUM! | JAK2 |
| Amyloid Processing | 0.000 | 0.082 | #NUM! | AKT1,CAPNS1,MAPT,NCSTN |
| IL-4 Signaling | 0.000 | 0.079 | 0.277 | AKT1,ANO1,COL9A3,CREBBP,FCER1G,HRAS,JAK3,MTOR,NR3C1,PIK3CA,PIK3CD,PIK3R2,RAP2A |
| Death Receptor Signaling | 0.000 | 0.075 | 1.134 | ACTB,APAF1,CYCS,MAP4K4,NFKBID,PARP1,PARP8 |
| p38 MAPK Signaling | 0.000 | 0.091 | -0.707 | CREBBP,DUSP1,IL1RAPL1,IRAK1,MAPKAPK2,MAPT,MEF2C,RPS6KA2,RPS6KA5,TGFBR1 |
| T Cell Receptor Signaling | 0.000 | 0.084 | 0.277 | AKT1,CALM1,CTNNB1,FCER1G,FYN,HRAS,MTOR,NFKBID,PIK3CA,PIK3CD,PIK3R2,PLCG1,RAP2A,RASA1,SHC3 |
| Autophagy | 0.000 | 0.090 | -1.147 | AKT1,ATG13,ATM,CALM1,CAMKK2,CREBBP,FOXO1,HGF,HIF1A,MAPK9,MTOR,NGFR,PIK3CA,PIK3CD,PIK3R2,PPP2R2B,PRKAA2,SESN1,VPS45 |

| | | | | |
|---|---|---|---|---|
| Inflammasome pathway | 0.000 | 0.053 | #NUM! | NLRP3 |
| Th1 and Th2 Activation Pathway | 0.000 | 0.074 | #NUM! | DLL1,DLL4,JAK2,JAK3,NCSTN,NOTCH1,PIK3CA,PIK3CD,PIK3R2,S1PR1,TGFBR1 |
| Th1 Pathway | 0.000 | 0.088 | -1.667 | DLL1,DLL4,JAK2,JAK3,NCSTN,NOTCH1,PIK3CA,PIK3CD,PIK3R2 |
| Th2 Pathway | 0.000 | 0.085 | -1.265 | DLL1,JAK2,JAK3,NCSTN,NOTCH1,PIK3CA,PIK3CD,PIK3R2,S1PR1,TGFBR1 |
| SPINK1 Pancreatic Cancer Pathway | 0.000 | 0.022 | #NUM! | TGFBR1 |
| NER (Nucleotide Excision Repair, Enhanced Pathway) | 0.000 | 0.080 | -0.447 | COPS5,GTF2H3,POLR2L,RBX1,RPA2,TOP2A,XAB2 |
| Apelin Cardiac Fibroblast Signaling Pathway | 0.000 | 0.087 | #NUM! | AKT1,PRKAA2 |
| PFKFB4 Signaling Pathway | 0.000 | 0.083 | 0.000 | CREBBP,HK1,RPS6KA5,TKT |
| Systemic Lupus Erythematosus in T Cell Signaling Pathway | 0.000 | 0.081 | 0.000 | AKT1,CREBBP,FCER1G,GNA12,HRAS,ITPR1,MTOR,PIK3CA,PIK3CD,PIK3R2,PLCG1,PPP2R2B,RAC3,RAP2A,RHOU,RND2 |
| Systemic Lupus Erythematosus in B Cell Signaling Pathway | 0.000 | 0.091 | 0.655 | AKT1,CALM1,CCND2,CTNNB1,FOXO1,FYN,HRAS,IL6ST,IRAK1,JAK2,MAP4K4,MTOR,PIK3CA,PIK3CD,PIK3R2,PRKCB,PRKCD,PRKCH,RAC3,RAP2A,RASGRP2,SHC3 |
| Necroptosis Signaling Pathway | 0.000 | 0.081 | 0.302 | AXL,CAPNS1,FKBP1A,MERTK,NGFR,NLRP3,PELI1,SLC25A3,TYRO3,UBC,VDAC3 |
| Tumor Microenvironment Pathway | 0.000 | 0.087 | 1.069 | AKT1,FN1,FOXO1,HGF,HIF1A,HRAS,IGF2,JAK2,MMP24,PIK3CA,PIK3CD,PIK3R2,RAP2A,SLC1A4 |
| MSP-RON Signaling in Macrophages Pathway | 0.000 | 0.068 | 1.134 | CREBBP,HRAS,JAK2,PIK3CA,PIK3CD,PIK3R2,RAP2A |
| Role of MAPK Signaling in Promoting the Pathogenesis of Inf | 0.000 | 0.067 | 0.816 | ATP6AP2,ATP6V1E1,ATP6V1G2,HRAS,MAPK9,NFKBID,RAP2A |
| Role of MAPK Signaling in Inhibiting the Pathogenesis of Influ | 0.000 | 0.029 | #NUM! | MAPK9,NFKBID |

**Supplementary Table 8: IPA Canonical Pathways of differentially expressed genes in Neurons at P28**

| Ingenuity Canonical Pathways | -log(p-valı | Ratio | z-score | Molecules |
|---|---|---|---|---|
| Sphingolipid metabolism | 3.510 | 0.101 | 2.530 | CERS2,CERS5,NEU2,NEU3,PLPP1,PPM1L,SAMD8,SMPD1,SPTSSB,ST8SIA5 |
| Formaldehyde Oxidation II (Glutathione-dependent) | 3.150 | 1.000 | #NUM! | ADH5,ESD |
| Apoptotic execution phase | 2.690 | 0.120 | -0.816 | BMX,CLSPN,GSN,H1-1,H1-4,SPTAN1 |
| Tumor Microenvironment Pathway | 2.400 | 0.068 | 2.111 | COL1A2,FGF11,FGF5,FOS,LEP,MMP23B,MMP24,PIK3C2B,PTGS2,RRAS2,VEGFB |
| Extrinsic Prothrombin Activation Pathway | 2.360 | 0.231 | #NUM! | F12,F2,F3 |
| Cholesterol Biosynthesis I | 2.360 | 0.231 | #NUM! | FDFT1,HSD17B7,MSMO1 |
| Cholesterol Biosynthesis II (via 24,25-dihydrolanosterol) | 2.360 | 0.231 | #NUM! | FDFT1,HSD17B7,MSMO1 |
| Cholesterol Biosynthesis III (via Desmosterol) | 2.360 | 0.231 | #NUM! | FDFT1,HSD17B7,MSMO1 |
| DNA Double-Strand Break Repair by Non-Homologous End Joini | 2.260 | 0.214 | #NUM! | LIG3,RAD50,XRCC5 |
| Coagulation System | 2.160 | 0.138 | 1.000 | BDKRB1,F12,F2,F3 |
| Zymosterol Biosynthesis | 2.000 | 0.333 | #NUM! | HSD17B7,MSMO1 |
| Sialic acid metabolism | 2.000 | 0.125 | 2.000 | NEU2,NEU3,ST6GALNAC2,ST8SIA5 |
| Role of JAK2 in Hormone-like Cytokine Signaling | 1.850 | 0.093 | 0.447 | FOS,GHR,LEP,SH2B3,VEGFB |
| Clathrin-mediated Endocytosis Signaling | 1.810 | 0.056 | #NUM! | APOD,DAB2,F2,FGF11,FGF5,MYO1E,PIK3C2B,RAB5C,RBP4,TF,VEGFB |
| Binding and Uptake of Ligands by Scavenger Receptors | 1.790 | 0.108 | 2.000 | COL1A2,FTL,SPARC,SSC5D |
| Response to elevated platelet cytosolic Ca2+ | 1.780 | 0.066 | 1.414 | ACTN1,CD109,ENDOD1,MANF,SELENOP,SPARC,TF,VEGFB |
| Relaxin Signaling | 1.720 | 0.060 | #NUM! | FICD,FOS,GDPD1,GNA12,GNAL,GNG5,GUCY2F,PDE6D,PIK3C2B |
| NGF-stimulated transcription | 1.710 | 0.103 | -1.000 | ARC,F3,FOS,FOSB |
| IL-8 Signaling | 1.700 | 0.055 | 1.000 | FOS,GNA12,GNAL,GNG5,NOX4,PIK3C2B,PTGS2,RND2,RRAS2,TEK,VEGFB |
| POU5F1 (OCT4), SOX2, NANOG repress genes related to differe | 1.650 | 0.222 | #NUM! | CDX2,HHEX |
| CDX Gastrointestinal Cancer Signaling Pathway | 1.630 | 0.055 | -0.632 | BMP1,CDX2,EFNB1,FOS,FZD6,KLF5,LEP,NOX4,PIK3C2B,PTGS2 |
| DNA Damage/Telomere Stress Induced Senescence | 1.600 | 0.095 | -1.000 | EP400,H1-1,H1-4,RAD50 |
| Glutamine Biosynthesis I | 1.570 | 1.000 | #NUM! | GLUL |
| Platelet Aggregation (Plug Formation) | 1.560 | 0.200 | #NUM! | COL1A2,F2 |
| Cholesterol biosynthesis | 1.550 | 0.120 | #NUM! | FDFT1,HSD17B7,MSMO1 |
| Apelin Endothelial Signaling Pathway | 1.470 | 0.057 | -0.816 | FOS,GNA12,GNAL,GNG5,PIK3C2B,PRKAB2,RRAS2,TEK |
| Superpathway of Cholesterol Biosynthesis | 1.470 | 0.111 | #NUM! | FDFT1,HSD17B7,MSMO1 |
| IL-12 Signaling and Production in Macrophages | 1.460 | 0.052 | 1.265 | APOD,COL1A2,CXCR3,EBI3,FOS,IRF8,PIK3C2B,PRKAB2,RBP4,XRCC5 |
| Colorectal Cancer Metastasis Signaling | 1.410 | 0.047 | 1.667 | FOS,FZD6,GNA12,GNAL,GNG5,MMP23B,MMP24,PIK3C2B,PTGS2,RND2,RRAS2,VEGFB |
| HDR through MMEJ (alt-NHEJ) | 1.410 | 0.167 | #NUM! | LIG3,RAD50 |
| Apelin Muscle Signaling Pathway | 1.390 | 0.082 | #NUM! | GNA12,GNAL,GNG5,PRKAB2 |
| Formation of Fibrin Clot (Clotting Cascade) | 1.390 | 0.103 | #NUM! | F12,F2,F3 |
| mRNA Capping | 1.390 | 0.103 | #NUM! | GTF2H4,MNAT1,SUPT5H |
| EGR2 and SOX10-mediated initiation of Schwann cell myelinatio | 1.390 | 0.103 | #NUM! | GJB1,MAG,MBP |
| TR/RXR Activation | 1.380 | 0.058 | 1.890 | MBP,NCOA6,PIK3C2B,SYT2,TBL1XR1,UCP2,VEGFB |
| Role of Chondrocytes in Rheumatoid Arthritis Signaling Pathwa | 1.330 | 0.057 | 1.134 | FOS,MLKL,MMP23B,MMP24,PRKAB2,PTGS2,VEGFB |
| Estrogen-Dependent Breast Cancer Signaling | 1.280 | 0.066 | #NUM! | FOS,HSD17B12,HSD17B7,PIK3C2B,RRAS2 |
| GDNF Family Ligand-Receptor Interactions | 1.280 | 0.066 | 0.000 | FOS,FRS2,GDNF,PIK3C2B,RRAS2 |
| Asparagine Degradation I | 1.280 | 0.500 | #NUM! | ASRGL1 |
| Epoxysqualene Biosynthesis | 1.280 | 0.500 | #NUM! | FDFT1 |
| Thrombin signalling through proteinase activated receptors (PA | 1.280 | 0.094 | #NUM! | F2,GNA12,GNG5 |
| Nucleotide catabolism | 1.280 | 0.094 | #NUM! | ENTPD5,NT5C2,NUDT5 |
| DHCR24 Signaling Pathway | 1.260 | 0.055 | 1.890 | APOD,KLF5,LCAT,PIK3C2B,RBP4,RRAS2,TF |
| Sleep REM Signaling Pathway | 1.260 | 0.074 | 0.000 | ARC,CHRM3,FOS,MKNK1 |
| Bladder Cancer Signaling | 1.240 | 0.058 | #NUM! | FGF11,FGF5,MMP23B,MMP24,RRAS2,VEGFB |
| MSP-RON Signaling in Macrophages Pathway | 1.240 | 0.058 | -0.816 | F12,FOS,NFKBIZ,PIK3C2B,PTGS2,RRAS2 |
| Centrosomal KIAA0586 Signaling Pathway | 1.240 | 0.054 | -2.646 | BCAN,C2CD3,FGF11,FGF5,PIK3C2B,PSMB6,VCAN |
| Telomere Extension by Telomerase | 1.230 | 0.133 | #NUM! | RAD50,XRCC5 |
| Fatty Acid Activation | 1.230 | 0.133 | #NUM! | SLC27A4,SLC27A5 |
| GABA Receptor Signaling | 1.230 | 0.054 | #NUM! | GABRG1,GABRP,GNA12,GNAL,GNG5,KCNN3,SLC32A1 |
| Metabolism of steroid hormones | 1.210 | 0.088 | #NUM! | FDX1,HSD17B12,SRD5A3 |
| Role of BRCA1 in DNA Damage Response | 1.210 | 0.063 | #NUM! | ARID1A,FAAP24,RAD50,RFC4,SMARCB1 |
| RAR Activation | 1.200 | 0.041 | 0.000 | ARID1A,COL1A2,DHRS3,DUSP1,FICD,FOS,GDPD1,GTF2H4,LEP,MNAT1,PDE6D,PIK3C2B,RARB,RBP4,RND2,SMARCB1 |
| LXR/RXR Activation | 1.190 | 0.057 | 1.342 | APOD,FDFT1,LCAT,PTGS2,RBP4,TF |
| PKR-mediated signaling | 1.190 | 0.062 | 2.236 | EIF2AK2,FAAP24,PRKRA,TRIM25,TUBB6 |
| Visual phototransduction | 1.190 | 0.070 | 2.000 | DHRS3,GUCY2F,RBP4,RLBP1 |
| Role of CHK Proteins in Cell Cycle Checkpoint Control | 1.190 | 0.070 | #NUM! | CLSPN,PPM1L,RAD50,RFC4 |
| Hereditary Breast Cancer Signaling | 1.190 | 0.053 | -0.378 | FAAP24,PIK3C2B,RAD50,RAD51C,RFC4,RRAS2,VEGFB |
| Intrinsic Prothrombin Activation Pathway | 1.180 | 0.086 | #NUM! | COL1A2,F12,F2 |
| Metabolism of folate and pterines | 1.180 | 0.125 | #NUM! | DHFR,MTHFD1 |
| Aggrephagy | 1.150 | 0.083 | #NUM! | DYNC1LI2,PCNT,TUBB6 |

| | | | | |
|---|---|---|---|---|
| Mitotic Prometaphase | 1.140 | 0.047 | 1.667 | CEP72,DYNC1LI2,MAD1L1,MIS12,NDE1,NEK6,PCNT,STAG2,TUBB6 |
| ESR-mediated signaling | 1.130 | 0.055 | 0.000 | FOS,FOSB,PGR,STAG2,USF1,ZNF217 |
| Stearate Biosynthesis I (Animals) | 1.120 | 0.067 | #NUM! | HSD17B12,SLC27A4,SLC27A5,SRD5A3 |
| Hematoma Resolution Signaling Pathway | 1.120 | 0.045 | -1.414 | DUSP1,F2,FOS,FTL,LEP,MMP23B,MMP24,NOX4,PRKAB2,UCP2 |
| ABRA Signaling Pathway | 1.120 | 0.059 | -0.447 | ACTN1,FOS,FOSB,GSN,VEGFB |
| Metallothioneins bind metals | 1.110 | 0.333 | #NUM! | MT2A |
| Ciprofloxacin ADME | 1.110 | 0.333 | #NUM! | SLC22A8 |
| Methylglyoxal Degradation I | 1.110 | 0.333 | #NUM! | HAGHL |
| Amyotrophic Lateral Sclerosis Signaling | 1.100 | 0.054 | 1.342 | GDNF,GLUL,GRIK1,PIK3C2B,RAB5C,VEGFB |
| Mitochondrial biogenesis | 1.100 | 0.066 | -1.000 | NCOA6,PRKAB2,TBL1XR1,TFB1M |
| Mitotic Metaphase and Anaphase | 1.100 | 0.044 | 0.000 | ANAPC10,ANAPC15,DYNC1LI2,MAD1L1,MIS12,NDE1,PSMB6,STAG2,TNPO1,TUBB6 |
| Resolution of Abasic Sites (AP sites) | 1.100 | 0.079 | #NUM! | LIG3,PNKP,RFC4 |
| Axonal Guidance Signaling | 1.080 | 0.038 | #NUM! | ADAM8,ADAMTS7,BMP1,EFNB1,FZD6,GNA12,GNAL,GNG5,MAG,MKNK1,MMP23B,MMP24,PIK3C2B,PMPCB,RRAS2,SEMA6D,TUBB6,VEGFB |
| Gap junction trafficking and regulation | 1.070 | 0.077 | #NUM! | DAB2,GJB1,TUBB6 |
| Ceramide Signaling | 1.070 | 0.057 | -1.000 | FOS,PIK3C2B,PPM1L,RRAS2,SMPD1 |
| IL-27 Signaling Pathway | 1.060 | 0.052 | 1.633 | ATG13,CXCR3,EBI3,IRF8,PIK3C2B,PTGS2 |
| Sheddase Signaling Pathway | 1.050 | 0.047 | 0.707 | ADAM8,COL1A2,ELK3,FOS,IGFBP2,MMP23B,MMP24,PIK3C2B |
| SUMOylation of transcription factors | 1.050 | 0.105 | #NUM! | MITF,PIAS1 |
| Class I peroxisomal membrane protein import | 1.050 | 0.105 | #NUM! | ACBD5,PEX2 |
| The Visual Cycle | 1.050 | 0.105 | #NUM! | DHRS3,RLBP1 |
| γ-linolenate Biosynthesis II (Animals) | 1.050 | 0.105 | #NUM! | SLC27A4,SLC27A5 |
| Mitochondrial L-carnitine Shuttle Pathway | 1.050 | 0.105 | #NUM! | SLC27A4,SLC27A5 |
| Activation of gene expression by SREBF (SREBP) | 1.050 | 0.075 | #NUM! | FDFT1,NCOA6,TBL1XR1 |
| Signaling by FGFR3 | 1.050 | 0.075 | #NUM! | FGF5,FRS2,MKNK1 |
| Oncostatin M Signaling | 1.050 | 0.075 | #NUM! | EPAS1,MT2A,RRAS2 |
| Ferroptosis Signaling Pathway | 1.030 | 0.051 | 0.000 | FDFT1,FTL,NOX4,PRKAB2,RRAS2,TF |
| Signaling by Rho Family GTPases | 1.020 | 0.042 | 1.414 | FOS,GFAP,GNA12,GNAL,GNG5,NOX4,PARD6A,PIK3C2B,RND2,SEPTIN1,SLC9A1 |
| Mitotic Roles of Polo-Like Kinase | 1.020 | 0.062 | #NUM! | ANAPC10,ANAPC15,PPM1L,STAG2 |
| Actin Cytoskeleton Signaling | 1.010 | 0.043 | 0.378 | ACTN1,F2,FGF11,FGF5,GNA12,GSN,MYO5C,PIK3C2B,RRAS2,SLC9A1 |
| Interferon gamma signaling | 1.000 | 0.061 | 1.000 | IRF8,MT2A,PIAS1,TRIM25 |
| Neurotransmitter uptake and metabolism In glial cells | 0.990 | 0.250 | #NUM! | GLUL |
| Pathogenesis of Multiple Sclerosis | 0.990 | 0.250 | #NUM! | CXCR3 |
| dTMP De Novo Biosynthesis | 0.990 | 0.250 | #NUM! | DHFR |
| Airway Pathology in Chronic Obstructive Pulmonary Disease | 0.989 | 0.054 | #NUM! | APOD,FGF11,FGF5,LEP,RBP4 |
| TP53 Regulates Transcription of DNA Repair Genes | 0.986 | 0.060 | -1.000 | FOS,GTF2H4,MNAT1,SUPT5H |
| Zn Homeostasis Signaling Pathway | 0.986 | 0.036 | 2.400 | ADGRA3,ADGRG3,BDKRB1,BMP1,CHRM3,FICD,FZD6,GDPD1,GIPR,GPR173,GPR25,GPR88,HTR1F,MMP23B,MMP24,MT2A,NOX4,P2RY14,PDE6D,PIK3C2B,SLC12A4,SLC9A1 |
| Endothelin-1 Signaling | 0.983 | 0.045 | 0.707 | FOS,GNA12,GNAL,GUCY2F,LCAT,PIK3C2B,PTGS2,RRAS2 |
| PI3K Cascade | 0.974 | 0.070 | #NUM! | FGF5,FLT3,FRS2 |
| E3 ubiquitin ligases ubiquitinate target proteins | 0.974 | 0.070 | #NUM! | PEX10,PEX2,RNF40 |
| BER (Base Excision Repair) Pathway | 0.974 | 0.070 | #NUM! | LIG3,PNKP,RFC4 |
| Chaperone Mediated Autophagy | 0.972 | 0.095 | #NUM! | GFAP,PCNT |
| Degradation of the extracellular matrix | 0.968 | 0.059 | 1.000 | ADAM8,BCAN,BMP1,FBN2 |
| Thioredoxin Signaling Pathway | 0.968 | 0.059 | -1.000 | F3,FOS,PRKAB2,VEGFB |
| Metabolism of porphyrins | 0.938 | 0.091 | #NUM! | ALAS2,COX15 |
| Oxytocin Signaling Pathway | 0.934 | 0.040 | 0.632 | FOS,GNA12,GNAL,GNG5,GUCY2F,LEP,MYO5C,PIK3C2B,PRKAB2,PTGS2,RRAS2 |
| Role of NFAT in Regulation of the Immune Response | 0.933 | 0.044 | 0.816 | FOS,GNA12,GNAL,GNG5,PIK3C2B,RRAS2,TNPO1,ZAP70 |
| NR1H2 and NR1H3-mediated signaling | 0.929 | 0.067 | #NUM! | APOD,EEPD1,TBL1XR1 |
| tRNA Splicing | 0.929 | 0.067 | #NUM! | FICD,GDPD1,PDE6D |
| G-Protein Coupled Receptor Signaling | 0.924 | 0.035 | 1.706 | ADGRA3,ADGRG3,BDKRB1,CHRM3,DUSP1,FICD,FOS,FZD6,GDPD1,GIPR,GNA12,GNAL,GNG5,GPR173,GPR25,GPR88,HTR1F,KCNN3,MKNK1,P2RY14,PDE6D,PIK3C2B,RRAS2 |
| Cellular Effects of Sildenafil (Viagra) | 0.918 | 0.035 | -0.655 | ADGRA3,ADGRG3,BDKRB1,CHRM3,FZD6,GIPR,GNG5,GPR173,GPR25,GPR88,GUCY2F,HTR1F,KCNN3,MYO5C,NPPC,P2RY14,PDE6D,PIK3C2B,PPM1L,PRKAB2,SLC9A1,VEGFB |
| Nephrin family interactions | 0.906 | 0.087 | #NUM! | ACTN1,SPTAN1 |
| Sumoylation Pathway | 0.904 | 0.051 | -1.000 | FOS,PIAS1,RFC4,RND2,ZNF217 |
| Transmission across Electrical Synapses | 0.899 | 0.200 | #NUM! | PANX2 |
| Reelin signalling pathway | 0.899 | 0.200 | #NUM! | VLDLR |
| Tetrahydrofolate Salvage from 5,10-methenyltetrahydrofolate | 0.899 | 0.200 | #NUM! | MTHFD1 |
| Tetrapyrrole Biosynthesis II | 0.899 | 0.200 | #NUM! | ALAS2 |
| Folate Polyglutamylation | 0.899 | 0.200 | #NUM! | MTHFD1 |
| HSP90 chaperone cycle for steroid hormone receptors in the pr | 0.888 | 0.064 | #NUM! | DYNC1LI2,PGR,TUBB6 |
| Ephrin B Signaling | 0.886 | 0.055 | #NUM! | EFNB1,GNA12,GNAL,GNG5 |
| fMLP Signaling in Neutrophils | 0.884 | 0.047 | #NUM! | GNA12,GNAL,GNG5,NOX4,PIK3C2B,RRAS2 |
| Incretin synthesis, secretion, and inactivation | 0.875 | 0.083 | #NUM! | CDX2,LEP |
| WNK Renal Signaling Pathway | 0.865 | 0.049 | 1.000 | ANAPC10,NPPC,PIK3C2B,SLC12A4,SLC12A7 |
| Transport of inorganic cations/anions and amino acids/oligopep | 0.865 | 0.049 | 1.342 | SLC12A4,SLC12A7,SLC15A3,SLC32A1,SLC9A1 |
| Membrane Repair Signaling Pathway | 0.865 | 0.049 | 2.236 | ANO5,CD82,DYSF,SMPD1,SYT2 |

| | | | | |
|---|---|---|---|---|
| CDK5 Signaling | 0.865 | 0.049 | 0.447 | FOSB,GNAL,PPM1L,PPP1R1B,RRAS2 |
| Neutrophil degranulation | 0.858 | 0.036 | 1.807 | ADAM8,ADGRG3,ALDOC,ARMC8,BPI,CREG1,DDOST,DSC1,FTL,GSN,LAMP1,RAB18,RAB5C,SPTAN1,XRCC5 |
| ILK Signaling | 0.858 | 0.042 | 0.378 | ACTN1,FOS,MYO5C,PIK3C2B,PPM1L,PTGS2,RND2,VEGFB |
| Antiproliferative Role of Somatostatin Receptor 2 | 0.855 | 0.053 | #NUM! | GNG5,GUCY2F,PIK3C2B,RRAS2 |
| HDR through Homologous Recombination (HRR) or Single Stran | 0.852 | 0.049 | 0.447 | CLSPN,RAD50,RAD51C,RFC4,SLX4 |
| Generic Transcription Pathway | 0.849 | 0.042 | 0.707 | NR2C2,NR4A1,PGR,RARB,TBL1XR1,ZKSCAN4,ZNF394,ZNF569 |
| HIF1α Signaling | 0.849 | 0.042 | 2.121 | MKNK1,MMP23B,MMP24,NOX4,PIK3C2B,RRAS2,TF,VEGFB |
| Role of Tissue Factor in Cancer | 0.849 | 0.042 | 1.414 | F2,F3,FGF5,FOS,GNA12,PIK3C2B,PTGS2,RRAS2 |
| Melanoma Signaling | 0.849 | 0.061 | #NUM! | MITF,PIK3C2B,RRAS2 |
| CXCR4 Signaling | 0.847 | 0.043 | #NUM! | FOS,GNA12,GNAL,GNG5,PIK3C2B,RND2,RRAS2 |
| BMAL1:CLOCK,NPAS2 activates circadian expression | 0.846 | 0.080 | #NUM! | NCOA6,TBL1XR1 |
| RORA,B,C and NR1D1 (REV-ERBA) regulate gene expression | 0.846 | 0.080 | #NUM! | NCOA6,TBL1XR1 |
| Thyroid Cancer Signaling | 0.840 | 0.053 | 0.000 | FOS,GDNF,PIK3C2B,RRAS2 |
| Telomerase Signaling | 0.840 | 0.048 | -1.000 | DKC1,ELK3,PIK3C2B,PPM1L,RRAS2 |
| GABAergic Receptor Signaling Pathway (Enhanced) | 0.840 | 0.045 | 2.236 | GABRG1,GABRP,GLUL,GNAL,GNG5,SLC32A1 |
| Signaling by FGFR1 | 0.830 | 0.060 | #NUM! | FGF5,FRS2,MKNK1 |
| Adipogenesis pathway | 0.829 | 0.045 | 0.816 | FZD6,GTF2H4,KLF5,LEP,MNAT1,TBL1XR1 |
| rRNA modification in the mitochondrion | 0.825 | 0.167 | #NUM! | TFB1M |
| Sphingomyelin Metabolism | 0.825 | 0.167 | #NUM! | SMPD1 |
| Cardiac β-adrenergic Signaling | 0.818 | 0.042 | 0.000 | FICD,GDPD1,GNA12,GNAL,GNG5,PDE6D,PPM1L |
| Cilium Assembly | 0.814 | 0.041 | 0.707 | C2CD3,CEP72,DYNC2H1,NDE1,PCNT,PDE6D,TNPO1,TUBB6 |
| UVC-Induced MAPK Signaling | 0.812 | 0.059 | #NUM! | FOS,RRAS2,SMPD1 |
| Neurotrophin/TRK Signaling | 0.811 | 0.051 | #NUM! | FOS,FRS2,PIK3C2B,RRAS2 |
| MSP-RON Signaling in Cancer Cells Pathway | 0.808 | 0.044 | 0.000 | ELK3,F12,FOS,PIK3C2B,RRAS2,VEGFB |
| α-Adrenergic Signaling | 0.804 | 0.047 | #NUM! | GNA12,GNAL,GNG5,PHKB,RRAS2 |
| Antimicrobial peptides | 0.793 | 0.074 | #NUM! | BPI,CHGA |
| Syndecan interactions | 0.793 | 0.074 | #NUM! | ACTN1,COL1A2 |
| Estrogen Biosynthesis | 0.793 | 0.074 | #NUM! | HSD17B12,HSD17B7 |
| Nucleotide Excision Repair | 0.792 | 0.046 | 0.447 | GTF2H4,LIG3,MNAT1,PIAS1,RFC4 |
| Cell Cycle Checkpoints | 0.788 | 0.038 | -0.632 | ANAPC10,ANAPC15,CLSPN,DYNC1LI2,MAD1L1,MIS12,NDE1,PSMB6,RAD50,RFC4 |
| Atherosclerosis Signaling | 0.780 | 0.046 | 2.236 | APOD,COL1A2,F3,LCAT,RBP4 |
| SPINK1 General Cancer Pathway | 0.777 | 0.057 | #NUM! | MT2A,PIK3C2B,RRAS2 |
| VEGF Family Ligand-Receptor Interactions | 0.770 | 0.049 | 0.000 | FOS,PIK3C2B,RRAS2,VEGFB |
| Inhibition of Matrix Metalloproteases | 0.768 | 0.071 | #NUM! | MMP23B,MMP24 |
| STAT5 Activation | 0.764 | 0.143 | #NUM! | FLT3 |
| Mitochondrial Uncoupling | 0.764 | 0.143 | #NUM! | UCP2 |
| Ceramide Biosynthesis | 0.764 | 0.143 | #NUM! | SPTSSB |
| Cholecystokinin/Gastrin-mediated Signaling | 0.758 | 0.045 | 0.447 | FOS,GNA12,PTGS2,RND2,RRAS2 |
| Gαi Signaling | 0.757 | 0.043 | #NUM! | GNA12,GNAL,GNG5,HTR1F,P2RY14,RRAS2 |
| Transcriptional regulation of white adipocyte differentiation | 0.757 | 0.049 | -1.000 | KLF5,LEP,NCOA6,TBL1XR1 |
| JAK/STAT Signaling | 0.757 | 0.049 | 0.000 | FOS,PIAS1,PIK3C2B,RRAS2 |
| IL-17 Signaling | 0.748 | 0.042 | 0.816 | FOS,LEP,PIK3C2B,PTGS2,RRAS2,VEGFB |
| SUMOylation of intracellular receptors | 0.744 | 0.069 | #NUM! | PGR,PIAS1 |
| Sensory processing of sound by outer hair cells of the cochlea | 0.744 | 0.055 | #NUM! | GSN,MYO3B,SPTAN1 |
| FGF Signaling | 0.731 | 0.048 | 2.000 | FGF11,FGF5,FRS2,PIK3C2B |
| Phagosome Maturation | 0.729 | 0.042 | #NUM! | DYNC1LI2,DYNC2H1,LAMP1,NOX4,RAB5C,TUBB6 |
| Assembly of collagen fibrils and other multimeric structures | 0.728 | 0.054 | #NUM! | BMP1,COL14A1,COL1A2 |
| AMPK Signaling | 0.724 | 0.038 | 0.000 | ARID1A,GNA12,GNAL,GNG5,LEP,PIK3C2B,PPM1L,PRKAB2,SMARCB1 |
| Activation of kainate receptors upon glutamate binding | 0.722 | 0.067 | #NUM! | GNG5,GRIK1 |
| HEY1 Signaling Pathway | 0.720 | 0.041 | 0.816 | BMP1,MMP23B,MMP24,PIK3C2B,PRKAB2,VEGFB |
| PDGF Signaling | 0.719 | 0.047 | 0.000 | EIF2AK2,FOS,PIK3C2B,RRAS2 |
| Neuroinflammation Signaling Pathway | 0.719 | 0.037 | 0.447 | FOS,GABRG1,GABRP,GDNF,GJB1,GLUL,IDE,NOX4,PIK3C2B,PTGS2 |
| DNA Damage Reversal | 0.711 | 0.125 | #NUM! | ALKBH2 |
| Sucrose Degradation V (Mammalian) | 0.711 | 0.125 | #NUM! | ALDOC |
| Histidine Degradation III | 0.711 | 0.125 | #NUM! | MTHFD1 |
| Preeclampsia Signaling Pathway | 0.710 | 0.041 | -1.633 | FOS,FZD6,PIK3C2B,PRKAB2,PTGS2,VEGFB |
| Leukocyte Extravasation Signaling | 0.704 | 0.039 | 1.342 | ACTN1,BMX,CLDN11,MMP23B,MMP24,NOX4,PIK3C2B |
| Sperm Motility | 0.703 | 0.037 | #NUM! | BMX,GNA12,GNAL,GNG5,LCAT,NPPC,ROS1,TEK,ZAP70 |
| SWI/SNF chromatin remodelers | 0.700 | 0.065 | #NUM! | ARID1A,SMARCB1 |
| Signaling by ERBB4 | 0.697 | 0.052 | #NUM! | GFAP,PGR,SPARC |
| Production of Nitric Oxide and Reactive Oxygen Species in Maci | 0.696 | 0.039 | 1.134 | APOD,FOS,IRF8,PIK3C2B,PPM1L,RBP4,RND2 |
| Lysosome Positioning Signaling Pathway | 0.684 | 0.040 | 1.633 | DYNC1LI2,DYNC2H1,FYCO1,GHR,KIF5A,PIK3C2B |
| NER (Nucleotide Excision Repair, Enhanced Pathway) | 0.683 | 0.046 | 1.000 | GTF2H4,LIG3,MNAT1,RFC4 |
| Peroxisomal protein import | 0.683 | 0.051 | #NUM! | IDE,PEX10,PEX2 |

| | | | | |
|---|---|---|---|---|
| rRNA modification in the nucleus and cytosol | 0.683 | 0.051 | #NUM! | DCAF13,DHX37,DKC1 |
| PCP (Planar Cell Polarity) Pathway | 0.683 | 0.051 | #NUM! | EFNB1,FZD6,VANGL1 |
| Protein Kinase A Signaling | 0.682 | 0.035 | 0.302 | ANAPC10,DUSP1,EBI3,FICD,GDPD1,GNG5,H1-1,H1-4,PDE6D,PHKB,PPP1R1B,PTGS2,UBASH3B |
| Role of NANOG in Mammalian Embryonic Stem Cell Pluripotenc | 0.675 | 0.042 | #NUM! | BMP1,CDX2,FZD6,PIK3C2B,RRAS2 |
| Role of Osteoclasts in Rheumatoid Arthritis Signaling Pathway | 0.674 | 0.036 | 1.667 | ADAM8,COL1A2,FOS,GSN,MITF,MMP23B,MMP24,PIK3C2B,RND2,RRAS2 |
| Neurovascular Coupling Signaling Pathway | 0.672 | 0.037 | 2.828 | CHRM3,ENTPD5,GABRG1,GABRP,GUCY2F,KCNN3,LCAT,PTGS2 |
| TEC Kinase Signaling | 0.672 | 0.039 | 1.000 | BMX,FOS,GNA12,GNAL,GNG5,PIK3C2B,RND2 |
| RANK Signaling in Osteoclasts | 0.671 | 0.045 | 1.000 | FOS,GSN,MITF,PIK3C2B |
| GP1b-IX-V activation signalling | 0.666 | 0.111 | #NUM! | COL1A2 |
| Lipophagy | 0.666 | 0.111 | #NUM! | PRKAB2 |
| Heme Biosynthesis II | 0.666 | 0.111 | #NUM! | ALAS2 |
| Folate Transformations I | 0.666 | 0.111 | #NUM! | MTHFD1 |
| Arachidonate metabolism | 0.659 | 0.061 | #NUM! | DPEP2,PTGS2 |
| Abacavir ADME | 0.654 | 0.049 | #NUM! | NT5C2,PIK3C2B,STK3 |
| IL-2 Signaling | 0.654 | 0.049 | #NUM! | FOS,PIK3C2B,RRAS2 |
| Regulation of TP53 Activity through Phosphorylation | 0.649 | 0.044 | 0.000 | PRKAB2,RAD50,RFC4,TPX2 |
| 14-3-3-mediated Signaling | 0.646 | 0.041 | #NUM! | FOS,GFAP,PIK3C2B,RRAS2,TUBB6 |
| Ribavirin ADME | 0.641 | 0.048 | #NUM! | NT5C2,PIK3C2B,STK3 |
| Remodeling of Epithelial Adherens Junctions | 0.641 | 0.048 | #NUM! | ACTN1,RAB5C,TUBB6 |
| Transcriptional Regulation by NPAS4 | 0.640 | 0.059 | #NUM! | FOS,KCNIP3 |
| Retinol Biosynthesis | 0.640 | 0.059 | #NUM! | DHRS3,ESD |
| Fatty Acid β-oxidation I | 0.640 | 0.059 | #NUM! | SLC27A4,SLC27A5 |
| Nucleotide Excision Repair Pathway | 0.640 | 0.059 | #NUM! | GTF2H4,MNAT1 |
| Intra-Golgi and retrograde Golgi-to-ER traffic | 0.634 | 0.037 | -0.378 | DYNC1LI2,GOLIM4,KIF5A,RAB18,STX6,TUBB6,VPS52 |
| Regulation of the Epithelial Mesenchymal Transition by Growth | 0.634 | 0.037 | -0.447 | FGF11,FGF5,FOS,FRS2,PARD6A,PIK3C2B,RRAS2 |
| FXR/RXR Activation | 0.633 | 0.039 | -0.816 | LEP,PRKAB2,PTGS2,SLC27A5,SLC51A,VLDLR |
| Molecular Mechanisms of Cancer | 0.631 | 0.031 | 3.266 | ADGRA3,ADGRG3,BDKRB1,BMP1,CHRM3,F2,FOS,FZD6,GHR,GIPR,GNA12,GNAL,GNG5,GPR173,GPR25,GPR88,HTR1F,MMP23B,MMP24,P2RY14,PIK3C2B,RABIF,RAD50,RND2,RRAS2 |
| Senescence-Associated Secretory Phenotype (SASP) | 0.628 | 0.048 | #NUM! | ANAPC10,ANAPC15,FOS |
| Thrombopoietin Signaling | 0.628 | 0.048 | #NUM! | FOS,PIK3C2B,RRAS2 |
| Regulation of the Epithelial-Mesenchymal Transition Pathway | 0.627 | 0.037 | #NUM! | FGF11,FGF5,FRS2,FZD6,PARD6A,PIK3C2B,RRAS2 |
| Passive transport by Aquaporins | 0.626 | 0.100 | #NUM! | AQP4 |
| Phenylalanine and tyrosine metabolism | 0.626 | 0.100 | #NUM! | ASRGL1 |
| Formation of the posterior neural plate | 0.626 | 0.100 | #NUM! | SOX1 |
| Protein lipoylation | 0.626 | 0.100 | #NUM! | FDX1 |
| Prostanoid Biosynthesis | 0.626 | 0.100 | #NUM! | PTGS2 |
| Oleate Biosynthesis II (Animals) | 0.626 | 0.100 | #NUM! | FADS6 |
| Nuclear RNA decay | 0.622 | 0.057 | #NUM! | PAPOLG,ZC3H3 |
| Expression of BMAL (ARNTL), CLOCK, and NPAS2 | 0.622 | 0.057 | #NUM! | NCOA6,TBL1XR1 |
| Pyrimidine Ribonucleotides Interconversion | 0.622 | 0.057 | #NUM! | ENTPD5,NUDT5 |
| Estrogen Receptor Signaling | 0.620 | 0.033 | 1.732 | FOS,GNA12,GNAL,GNG5,LEP,MMP23B,MMP24,PGR,PIK3C2B,PRKAB2,RRAS2,TBL1XR1,VEGFB |
| HER-2 Signaling in Breast Cancer | 0.617 | 0.036 | -0.707 | COX15,ELK3,FOS,PARD6A,PGR,PIK3C2B,PTGS2,RRAS2 |
| Cardiac conduction | 0.610 | 0.040 | 2.236 | CASQ2,FGF11,KCNIP3,NPPC,SCN4B |
| Iron homeostasis signaling pathway | 0.610 | 0.040 | #NUM! | ALAS2,BMP1,EPAS1,FTL,TF |
| Kinetochore Metaphase Signaling Pathway | 0.607 | 0.042 | 1.000 | ANAPC10,MAD1L1,MIS12,STAG2 |
| Cilia Biogenesis Signaling Pathway | 0.605 | 0.037 | -1.134 | ANAPC10,GNA12,GNAL,GNG5,NDE1,PCNT,TUBB6 |
| Fatty acyl-CoA biosynthesis | 0.604 | 0.056 | #NUM! | ELOVL7,HSD17B12 |
| Cerebral Malformation Signaling Pathway | 0.601 | 0.039 | 2.000 | F2,PIK3C2B,PPM1L,PTGS2,VEGFB |
| IL-1 Signaling | 0.597 | 0.042 | #NUM! | FOS,GNA12,GNAL,GNG5 |
| UVA-Induced MAPK Signaling | 0.597 | 0.042 | #NUM! | FOS,PIK3C2B,RRAS2,SMPD1 |
| VEGF Signaling | 0.597 | 0.042 | 0.000 | ACTN1,PIK3C2B,RRAS2,VEGFB |
| Signaling by Leptin | 0.590 | 0.091 | #NUM! | LEP |
| RHO GTPases activate KTN1 | 0.590 | 0.091 | #NUM! | KIF5A |
| Advanced glycosylation endproduct receptor signaling | 0.590 | 0.091 | #NUM! | DDOST |
| Pexophagy | 0.590 | 0.091 | #NUM! | EPAS1 |
| Formation of the anterior neural plate | 0.590 | 0.091 | #NUM! | SOX1 |
| Bile Acid Biosynthesis, Neutral Pathway | 0.590 | 0.091 | #NUM! | SLC27A5 |
| Macroautophagy | 0.590 | 0.046 | #NUM! | ATG13,PRKAB2,WDR45 |
| Collagen biosynthesis and modifying enzymes | 0.590 | 0.046 | #NUM! | BMP1,COL14A1,COL1A2 |
| CD40 Signaling | 0.590 | 0.046 | #NUM! | FOS,PIK3C2B,PTGS2 |
| Glutamate Receptor Signaling | 0.590 | 0.046 | #NUM! | GLUL,GNG5,GRIK1 |
| Signaling by Retinoic Acid | 0.587 | 0.054 | #NUM! | DHRS3,RARB |
| Fanconi Anemia Pathway | 0.587 | 0.054 | #NUM! | FAAP24,SLX4 |
| Cell Cycle Regulation by BTG Family Proteins | 0.587 | 0.054 | #NUM! | BTG2,PPM1L |
| Germ Cell-Sertoli Cell Junction Signaling | 0.586 | 0.037 | #NUM! | ACTN1,GSN,PIK3C2B,RND2,RRAS2,TUBB6 |

| | | | | |
|---|---|---|---|---|
| Mitotic G2-G2/M phases | 0.584 | 0.036 | 1.134 | CEP72,MNAT1,NDE1,PCNT,PSMB6,TPX2,TUBB6 |
| CCR3 Signaling in Eosinophils | 0.584 | 0.039 | #NUM! | GNA12,GNAL,GNG5,PIK3C2B,RRAS2 |
| FAK Signaling | 0.581 | 0.031 | 2.524 | ADGRA3,ADGRG3,BDKRB1,CHRM3,COL1A2,EFNB1,ELK3,FOS,FZD6,GHR,GIPR,GPR173,GPR25,GPR88,HTR1F,P2RY14,PIK3C2B,RRAS2,USF1 |
| Transcriptional Regulatory Network in Embryonic Stem Cells | 0.579 | 0.037 | -0.816 | CDX2,FZD6,OTX1,PIK3C2B,RFX4,RRAS2 |
| Growth Hormone Signaling | 0.578 | 0.045 | #NUM! | FOS,GHR,PIK3C2B |
| Granzyme A Signaling | 0.578 | 0.045 | #NUM! | H1-1,H1-4,MBP |
| Glucocorticoid Receptor Signaling | 0.577 | 0.032 | #NUM! | ARID1A,DCAF6,DUSP1,FOS,GHR,GTF2H4,MNAT1,PGR,PIK3C2B,PRKAB2,PTGS2,RARB,RRAS2,SMARCB1,TFB1M |
| HGF Signaling | 0.575 | 0.039 | #NUM! | ELK3,FOS,PIK3C2B,PTGS2,RRAS2 |
| Bile acid and bile salt metabolism | 0.571 | 0.053 | #NUM! | SLC27A5,SLC51A |
| Transport of vitamins, nucleosides, and related molecules | 0.571 | 0.053 | #NUM! | APOD,SLC27A4 |
| Transcriptional Regulation by VENTX | 0.571 | 0.053 | #NUM! | ANAPC10,ANAPC15 |
| FLT3 Signaling | 0.571 | 0.053 | #NUM! | FLT3,SH2B3 |
| Pyrimidine Ribonucleotides De Novo Biosynthesis | 0.571 | 0.053 | #NUM! | ENTPD5,NUDT5 |
| CREB Signaling in Neurons | 0.570 | 0.031 | 2.840 | ADGRA3,ADGRG3,BDKRB1,CHRM3,FZD6,GHR,GIPR,GNA12,GNAL,GNG5,GPR173,GPR25,GPR88,GRIK1,HTR1F,P2RY14,PIK3C2B,RRAS2 |
| RHO GTPases Activate Formins | 0.567 | 0.038 | 1.342 | DYNC1LI2,MAD1L1,MIS12,NDE1,TUBB6 |
| cAMP-mediated signaling | 0.561 | 0.035 | 2.121 | CHRM3,DUSP1,FICD,GDPD1,GNAL,HTR1F,P2RY14,PDE6D |
| Systemic Lupus Erythematosus in T Cell Signaling Pathway | 0.558 | 0.035 | 2.646 | FOS,GNA12,LEP,PIK3C2B,PPM1L,RND2,RRAS2 |
| Reversible hydration of carbon dioxide | 0.558 | 0.083 | #NUM! | CA5B |
| Androgen Biosynthesis | 0.558 | 0.083 | #NUM! | SRD5A3 |
| Leukotriene Biosynthesis | 0.558 | 0.083 | #NUM! | DPEP2 |
| Guanosine Nucleotides Degradation III | 0.558 | 0.083 | #NUM! | NT5C2 |
| Acute Phase Response Signaling | 0.557 | 0.036 | -1.000 | F2,FOS,FTL,RBP4,RRAS2,TF |
| Gαq Signaling | 0.557 | 0.036 | #NUM! | CHRM3,GNA12,GNAL,GNG5,PIK3C2B,RND2 |
| Nonhomologous End-Joining (NHEJ) | 0.555 | 0.051 | #NUM! | RAD50,XRCC5 |
| Plasma lipoprotein assembly, remodeling, and clearance | 0.555 | 0.044 | #NUM! | BMP1,LCAT,VLDLR |
| Role of MAPK Signaling in Inhibiting the Pathogenesis of Influer | 0.555 | 0.044 | #NUM! | EIF2AK2,LCAT,PTGS2 |
| Signaling by FGFR2 | 0.543 | 0.043 | #NUM! | FGF5,FRS2,MKNK1 |
| Glioma Invasiveness Signaling | 0.543 | 0.043 | #NUM! | PIK3C2B,RND2,RRAS2 |
| Potassium Channels | 0.540 | 0.039 | 2.000 | GNG5,KCNA1,KCNC1,KCNN3 |
| Vasopressin regulates renal water homeostasis via Aquaporins | 0.540 | 0.050 | #NUM! | AQP4,GNG5 |
| Signaling by FGFR4 | 0.540 | 0.050 | #NUM! | FRS2,MKNK1 |
| RET signaling | 0.540 | 0.050 | #NUM! | FRS2,GDNF |
| Meiotic recombination | 0.540 | 0.050 | #NUM! | RAD50,RAD51C |
| Regulation of endogenous retroelements | 0.532 | 0.042 | #NUM! | ARID1A,ATF7IP,SMARCB1 |
| IL-17A Signaling in Fibroblasts | 0.532 | 0.042 | #NUM! | COL1A2,FOS,NFKBIZ |
| Extracellular matrix organization | 0.531 | 0.039 | 2.000 | BCAN,COL1A2,SPARC,VCAN |
| IGF-1 Signaling | 0.531 | 0.039 | #NUM! | FOS,IGFBP2,PIK3C2B,RRAS2 |
| Mitochondrial iron-sulfur cluster biogenesis | 0.529 | 0.077 | #NUM! | FDX1 |
| Synthesis of Prostaglandins (PG) and Thromboxanes (TX) | 0.529 | 0.077 | #NUM! | PTGS2 |
| Olfactory Signaling Pathway | 0.529 | 0.077 | #NUM! | GNAL |
| SUMOylation of immune response proteins | 0.529 | 0.077 | #NUM! | EIF2AK2 |
| Platelet Adhesion to exposed collagen | 0.529 | 0.077 | #NUM! | COL1A2 |
| Glutamate and glutamine metabolism | 0.529 | 0.077 | #NUM! | GLUL |
| Urate Biosynthesis/Inosine 5'-phosphate Degradation | 0.529 | 0.077 | #NUM! | NT5C2 |
| MIF Regulation of Innate Immunity | 0.525 | 0.049 | #NUM! | FOS,PTGS2 |
| Role of JAK family kinases in IL-6-type Cytokine Signaling | 0.522 | 0.042 | #NUM! | FOS,PIAS1,VEGFB |
| NAFLD Signaling Pathway | 0.520 | 0.034 | 1.134 | FOS,LEP,NOX4,PIK3C2B,PPM1L,PRKAB2,PTGS2 |
| CLEAR Signaling Pathway | 0.516 | 0.033 | 1.667 | DEPDC5,GHR,KCNIP3,LAMP1,MITF,PPM1L,PRKAB2,RRAS2,WDR45 |
| CCR5 Signaling in Macrophages | 0.513 | 0.038 | #NUM! | FOS,GNA12,GNAL,GNG5 |
| ERK5 Signaling | 0.511 | 0.041 | #NUM! | FOS,GNA12,RRAS2 |
| ERK/MAPK Signaling | 0.508 | 0.034 | -0.378 | DUSP1,ELK3,FOS,MKNK1,PIK3C2B,PPM1L,RRAS2 |
| Breast Cancer Regulation by Stathmin1 | 0.506 | 0.031 | 2.500 | ADGRA3,ADGRG3,BDKRB1,CHRM3,FZD6,GIPR,GNG5,GPR173,GPR25,GPR88,HTR1F,P2RY14,PIK3C2B,PPM1L,RRAS2,TUBB6,VEGFB |
| Bone Mineralization Signaling Pathway | 0.505 | 0.036 | 2.236 | COL1A2,FGF11,FGF5,FZD6,TRPV6 |
| RAC Signaling | 0.505 | 0.036 | 0.447 | MCF2L,NOX4,PARD6A,PIK3C2B,RRAS2 |
| Tight Junction Signaling | 0.502 | 0.035 | #NUM! | CLDN11,FOS,MYO5C,PARD6A,PPM1L,SPTAN1 |
| Gustation Pathway | 0.502 | 0.035 | 1.633 | CHRM3,GABRG1,GABRP,KCNN3,P2RY14,SCN4B |
| Prolactin receptor signaling | 0.502 | 0.071 | #NUM! | GHR |
| Carnitine shuttle | 0.502 | 0.071 | #NUM! | PRKAB2 |
| LGI-ADAM interactions | 0.502 | 0.071 | #NUM! | LGI3 |
| Mitotic Telophase/Cytokinesis | 0.502 | 0.071 | #NUM! | STAG2 |
| DNA Double-Strand Break Repair by Homologous Recombinatic | 0.502 | 0.071 | #NUM! | RAD50 |
| NAP1L1 Transcription Regulation Signaling Pathway | 0.501 | 0.041 | #NUM! | PIK3C2B,RASSF10,TUBB6 |
| Pulmonary Fibrosis Idiopathic Signaling Pathway | 0.499 | 0.032 | 1.897 | COL1A2,F2,FOS,FZD6,GNA12,MMP23B,MMP24,NOX4,PIK3C2B,RRAS2 |
| Endocannabinoid Cancer Inhibition Pathway | 0.498 | 0.036 | -0.447 | GNAL,PIK3C2B,PRKAB2,SMPD1,VEGFB |

| | | | | |
|---|---|---|---|---|
| Collagen chain trimerization | 0.497 | 0.047 | #NUM! | COL14A1,COL1A2 |
| MITF-M-dependent gene expression | 0.497 | 0.037 | 0.000 | ARID1A,MITF,SMARCB1,USF1 |
| Role of MAPK Signaling in the Pathogenesis of Influenza | 0.491 | 0.040 | #NUM! | LCAT,PTGS2,RRAS2 |
| BBSome Signaling Pathway | 0.491 | 0.031 | 3.207 | ADGRA3,ADGRG3,BDKRB1,CEP72,CHRM3,FZD6,GIPR,GPR173,GPR25,GPR88,HTR1F,LEP,P2RY14,RAB5C |
| Role of Osteoblasts in Rheumatoid Arthritis Signaling Pathway | 0.485 | 0.033 | 1.134 | FZD6,LEP,MMP23B,MMP24,PIK3C2B,PTGS2,VEGFB |
| Apelin Pancreas Signaling Pathway | 0.484 | 0.046 | #NUM! | PIK3C2B,PRKAB2 |
| Angiopoietin Signaling | 0.481 | 0.040 | #NUM! | PIK3C2B,RRAS2,TEK |
| G Protein Signaling Mediated by Tubby | 0.481 | 0.040 | #NUM! | GNA12,GNAL,GNG5 |
| Regulation of cholesterol biosynthesis by SREBP (SREBF) | 0.477 | 0.067 | #NUM! | INSIG1 |
| Adenosine Nucleotides Degradation II | 0.477 | 0.067 | #NUM! | NT5C2 |
| Regulation of Insulin-like Growth Factor (IGF) transport and upt | 0.472 | 0.036 | 1.000 | F2,IGFBP2,TF,VCAN |
| Sphingosine-1-phosphate Signaling | 0.472 | 0.036 | 2.000 | GNA12,PIK3C2B,RND2,SMPD1 |
| Telomere Maintenance | 0.472 | 0.039 | #NUM! | DKC1,RFC4,SHQ1 |
| NF-κB Activation by Viruses | 0.472 | 0.039 | #NUM! | EIF2AK2,PIK3C2B,RRAS2 |
| IL-3 Signaling | 0.472 | 0.039 | #NUM! | FOS,PIK3C2B,RRAS2 |
| Osteoarthritis Pathway | 0.468 | 0.033 | 2.646 | EPAS1,FZD6,LEP,PRKAB2,PTGS2,RBP4,VEGFB |
| SAPK/JNK Signaling | 0.465 | 0.036 | #NUM! | GNA12,GNG5,PIK3C2B,RRAS2 |
| eNOS Signaling | 0.464 | 0.035 | #NUM! | AQP4,BDKRB1,PIK3C2B,PRKAB2,VEGFB |
| T Cell Exhaustion Signaling Pathway | 0.464 | 0.035 | #NUM! | FOS,PIK3C2B,PPM1L,RRAS2,ZAP70 |
| Renal Cell Carcinoma Signaling | 0.463 | 0.039 | #NUM! | FOS,PIK3C2B,RRAS2 |
| Glycation Signaling Pathway | 0.462 | 0.033 | 0.378 | DDOST,FOS,LEP,NOX4,PIK3C2B,RRAS2,VEGFB |
| Oxytocin in Brain Signaling Pathway | 0.459 | 0.033 | 1.342 | GNAL,GNG5,LCAT,PIK3C2B,PTGS2,RRAS2 |
| Heme signaling | 0.459 | 0.044 | #NUM! | NCOA6,TBL1XR1 |
| O-linked glycosylation | 0.457 | 0.036 | 2.000 | ADAMTS7,MMRN2,SEMA5B,ST6GALNAC2 |
| Chaperone Mediated Autophagy Signaling Pathway | 0.457 | 0.033 | -1.134 | FLT3,GFAP,MMP23B,MMP24,PIK3C2B,PRKAB2,PSMB6 |
| NF1 RAS Signaling Pathway | 0.456 | 0.030 | 2.840 | ADGRA3,ADGRG3,BDKRB1,CHRM3,FZD6,GHR,GIPR,GNG5,GPR173,GPR25,GPR88,HTR1F,P2RY14,PIK3C2B,RRAS2 |
| Transcriptional regulation of pluripotent stem cells | 0.455 | 0.063 | #NUM! | EPAS1 |
| Processing of SMDT1 | 0.455 | 0.063 | #NUM! | PMPCB |
| Modulation of host responses by IFN-stimulated genes | 0.455 | 0.063 | #NUM! | TRIM25 |
| RAN Signaling | 0.455 | 0.063 | #NUM! | TNPO1 |
| G alpha (12/13) signalling events | 0.453 | 0.038 | #NUM! | GNA12,GNG5,MCF2L |
| BEX2 Signaling Pathway | 0.453 | 0.038 | #NUM! | PPM1L,SMARCB1,VEGFB |
| Sleep NREM Signaling Pathway | 0.450 | 0.035 | 2.000 | GABRG1,GABRP,RRAS2,XRCC5 |
| Thrombin Signaling | 0.446 | 0.032 | 1.342 | F2,GNA12,GNAL,GNG5,PIK3C2B,RND2,RRAS2 |
| Type II Diabetes Mellitus Signaling | 0.444 | 0.034 | #NUM! | PIK3C2B,PRKAB2,SLC27A4,SLC27A5,SMPD1 |
| Pulmonary Healing Signaling Pathway | 0.442 | 0.033 | 1.633 | FZD6,MMP23B,MMP24,PRKAB2,RRAS2,VEGFB |
| Chromatin modifying enzymes | 0.441 | 0.032 | -0.378 | ARID1A,ATF7IP,BRPF1,EP400,KDM4C,SMARCB1,TBL1XR1 |
| HMGB1 Signaling | 0.438 | 0.034 | 0.000 | FOS,LEP,PIK3C2B,RND2,RRAS2 |
| RAF/MAP kinase cascade | 0.437 | 0.032 | 0.000 | DUSP1,FGF5,FLT3,FRS2,GDNF,PSMB6,SPTAN1,TEK |
| PI Metabolism | 0.436 | 0.037 | #NUM! | BMX,GDPD1,PIK3C2B |
| FLT3 Signaling in Hematopoietic Progenitor Cells | 0.436 | 0.037 | #NUM! | FLT3,PIK3C2B,RRAS2 |
| DNA Damage Bypass | 0.435 | 0.042 | #NUM! | RFC4,TRIM25 |
| Interferon alpha/beta signaling | 0.435 | 0.042 | #NUM! | EIF2AK2,IRF8 |
| Assembly of RNA Polymerase II Complex | 0.435 | 0.042 | #NUM! | GTF2H4,MNAT1 |
| Cell surface interactions at the vascular wall | 0.435 | 0.035 | 1.000 | COL1A2,F2,MAG,TEK |
| Regulation of TLR by endogenous ligand | 0.434 | 0.059 | #NUM! | S100A1 |
| Mismatch Repair in Eukaryotes | 0.434 | 0.059 | #NUM! | RFC4 |
| Activin Inhibin Signaling Pathway | 0.430 | 0.032 | 0.816 | COL1A2,FOS,FOSB,FST,PIK3C2B,PTGS2 |
| IL-15 Production | 0.428 | 0.035 | 1.000 | BMX,ROS1,TEK,ZAP70 |
| ISGylation Signaling Pathway | 0.427 | 0.037 | #NUM! | EIF2AK2,IRF8,TRIM25 |
| Kinesins | 0.424 | 0.041 | #NUM! | KIF5A,TUBB6 |
| Role of IL-17A in Arthritis | 0.424 | 0.041 | #NUM! | PIK3C2B,PTGS2 |
| Phototransduction Pathway | 0.424 | 0.041 | #NUM! | GUCY2F,PDE6D |
| Lung Ionic Balance Signaling Pathway | 0.421 | 0.030 | 1.604 | ADGRA3,ADGRG3,BDKRB1,CHRM3,FZD6,GIPR,GPR173,GPR25,GPR88,HTR1F,NOX4,P2RY14,PPM1L,PRKAB2 |
| mTOR Signaling | 0.421 | 0.032 | 0.816 | ATG13,PIK3C2B,PPM1L,PRKAB2,RND2,RRAS2,VEGFB |
| Synaptic Long Term Depression | 0.419 | 0.032 | 0.816 | GNA12,GNAL,GUCY2F,LCAT,PPM1L,RRAS2 |
| Activated PKN1 stimulates transcription of AR regulated genes | 0.414 | 0.056 | #NUM! | KDM4C |
| GABA synthesis, release, reuptake and degradation | 0.414 | 0.056 | #NUM! | SLC32A1 |
| Formation of the ureteric bud | 0.414 | 0.056 | #NUM! | GDNF |
| Purine Nucleotides Degradation II (Aerobic) | 0.414 | 0.056 | #NUM! | NT5C2 |
| Fatty Acid α-oxidation | 0.414 | 0.056 | #NUM! | PTGS2 |
| UVB-Induced MAPK Signaling | 0.413 | 0.040 | #NUM! | FOS,PIK3C2B |
| PEDF Signaling | 0.411 | 0.036 | #NUM! | GDNF,PIK3C2B,RRAS2 |
| Ovarian Cancer Signaling | 0.407 | 0.033 | #NUM! | FZD6,PIK3C2B,PTGS2,RRAS2,VEGFB |

| | | | | |
|---|---|---|---|---|
| LPS-stimulated MAPK Signaling | 0.403 | 0.035 | #NUM! | FOS,PIK3C2B,RRAS2 |
| Proteasome assembly | 0.403 | 0.039 | #NUM! | PSMB6,PSMG1 |
| Sertoli Cell-Germ Cell Junction Signaling Pathway (Enhanced) | 0.401 | 0.031 | -1.134 | CLDN11,FOS,FOSB,PIK3C2B,PPM1L,PRKAB2,RRAS2 |
| ID3 Signaling Pathway | 0.401 | 0.033 | #NUM! | ANAPC10,ANAPC15,NOX4,RRAS2 |
| RHO GTPase cycle | 0.401 | 0.029 | 0.277 | ACBD5,ACTN1,FAM13A,FAM91A1,FRS2,MCF2L,PARD6A,RND2,RRAS2,SPTAN1,TNFAIP1,VANGL1,ZAP70 |
| Signaling by Activin | 0.396 | 0.053 | #NUM! | FST |
| Activation of Matrix Metalloproteinases | 0.396 | 0.053 | #NUM! | MMP24 |
| Methylglyoxal Degradation III | 0.396 | 0.053 | #NUM! | HSD17B12 |
| Pregnenolone Biosynthesis | 0.396 | 0.053 | #NUM! | FDX1 |
| Apelin Liver Signaling Pathway | 0.396 | 0.053 | #NUM! | COL1A2 |
| Histone Modification Signaling Pathway | 0.395 | 0.030 | 0.000 | BRPF1,H1-1,H1-4,KDM4C,RNF40,SFMBT1,VEGFB,XRCC5 |
| BMP signaling pathway | 0.395 | 0.035 | #NUM! | BMP1,FST,RRAS2 |
| Xenobiotic Metabolism General Signaling Pathway | 0.394 | 0.033 | #NUM! | FTL,PIK3C2B,RRAS2,SLC51A |
| Glycosaminoglycan metabolism | 0.388 | 0.033 | 2.000 | BCAN,CHST2,SLC9A1,VCAN |
| Synthesis of DNA | 0.388 | 0.033 | -1.000 | ANAPC10,ANAPC15,PSMB6,RFC4 |
| Opioid Signalling | 0.380 | 0.034 | #NUM! | GNAL,GNG5,PPP1R1B |
| Signaling by Hippo | 0.379 | 0.050 | #NUM! | STK3 |
| Synthesis, secretion, and deacylation of Ghrelin | 0.379 | 0.050 | #NUM! | LEP |
| Interleukin-12 family signaling | 0.379 | 0.050 | #NUM! | EBI3 |
| Eicosanoid Signaling | 0.378 | 0.030 | 1.633 | DPEP2,GNAL,GNG5,LCAT,PIK3C2B,PTGS2,RRAS2,VEGFB |
| Gαs Signaling | 0.375 | 0.032 | #NUM! | CHRM3,GNA12,GNAL,GNG5 |
| Sertoli Cell-Sertoli Cell Junction Signaling | 0.373 | 0.030 | 0.378 | ACTN1,CLDN11,PIK3C2B,RRAS2,SPTAN1,TJAP1,TUBB6 |
| Folate Signaling Pathway | 0.373 | 0.037 | #NUM! | DHFR,MTHFD1 |
| MSP-RON Signaling Pathway | 0.373 | 0.037 | #NUM! | F12,PIK3C2B |
| Cell junction organization | 0.372 | 0.034 | #NUM! | ACTN1,CLDN11,PARD6A |
| RAB GEFs exchange GTP for GDP on RABs | 0.372 | 0.034 | #NUM! | DENND4B,RAB18,RAB5C |
| Regulation of mitotic cell cycle | 0.365 | 0.033 | #NUM! | ANAPC10,ANAPC15,PSMB6 |
| Actin Nucleation by ARP-WASP Complex | 0.365 | 0.033 | #NUM! | GNA12,RND2,RRAS2 |
| Calcium-induced T Lymphocyte Apoptosis | 0.364 | 0.036 | #NUM! | NR4A1,ZAP70 |
| EGF Signaling | 0.364 | 0.036 | #NUM! | FOS,PIK3C2B |
| Synaptic adhesion-like molecules | 0.364 | 0.048 | #NUM! | LRFN4 |
| IL-17A Signaling in Gastric Cells | 0.364 | 0.048 | #NUM! | FOS |
| NAD Salvage Pathway II | 0.364 | 0.048 | #NUM! | NT5C2 |
| Endocannabinoid Developing Neuron Pathway | 0.363 | 0.032 | #NUM! | GNAL,GNG5,PIK3C2B,RRAS2 |
| COPI-mediated anterograde transport | 0.358 | 0.033 | #NUM! | DYNC1LI2,SPTAN1,TUBB6 |
| Acute Myeloid Leukemia Signaling | 0.358 | 0.033 | #NUM! | FLT3,PIK3C2B,RRAS2 |
| Mitochondrial Division Signaling Pathway | 0.358 | 0.031 | 0.447 | MLKL,PPM1L,PRKAB2,RRAS2,VEGFB |
| Asparagine N-linked glycosylation | 0.357 | 0.032 | 2.000 | DDOST,NUS1,SRD5A3,TRIM13 |
| Clathrin-mediated endocytosis | 0.357 | 0.032 | 1.000 | DAB2,RAB5C,SYT2,TF |
| Collagen degradation | 0.355 | 0.036 | #NUM! | COL14A1,COL1A2 |
| Gamma carboxylation, hypusinylation, hydroxylation, and aryls | 0.355 | 0.036 | #NUM! | F2,TPST1 |
| CNTF Signaling | 0.355 | 0.036 | #NUM! | PIK3C2B,RRAS2 |
| G alpha (q) signalling events | 0.352 | 0.031 | 2.236 | BDKRB1,CHRM3,F2,GNG5,NPSR1 |
| Androgen Signaling | 0.352 | 0.031 | #NUM! | GNA12,GNAL,GNG5,GTF2H4,MNAT1 |
| Gα12/13 Signaling | 0.351 | 0.031 | 1.000 | F2,GNA12,PIK3C2B,RRAS2 |
| G Beta Gamma Signaling | 0.351 | 0.031 | #NUM! | GNA12,GNAL,GNG5,RRAS2 |
| Class B/2 (Secretin family receptors) | 0.351 | 0.033 | #NUM! | FZD6,GIPR,GNG5 |
| Sirtuin Signaling Pathway | 0.350 | 0.029 | -0.378 | ATG13,EPAS1,H1-1,H1-4,RARB,TIMM17B,UCP2,XRCC5 |
| Growth hormone receptor signaling | 0.349 | 0.046 | #NUM! | GHR |
| Acetone Degradation I (to Methylglyoxal) | 0.349 | 0.046 | #NUM! | HSD17B12 |
| Cytoprotection by HMOX1 | 0.346 | 0.035 | #NUM! | NCOA6,TBL1XR1 |
| Glutaminergic Receptor Signaling Pathway (Enhanced) | 0.345 | 0.029 | 1.667 | GABRG1,GABRP,GLUL,GNAL,GRIK1,LCAT,MKNK1,PIK3C2B,SCN4B |
| Beta-catenin independent WNT signaling | 0.345 | 0.030 | 0.816 | FZD6,GNG5,PARD6A,PIK3C2B,PSMB6,STK3 |
| Ephrin Receptor Signaling | 0.345 | 0.030 | #NUM! | EFNB1,GNA12,GNAL,GNG5,RRAS2,VEGFB |
| ERBB Signaling | 0.344 | 0.032 | #NUM! | FOS,PIK3C2B,RRAS2 |
| S100 Family Signaling Pathway | 0.340 | 0.028 | 3.441 | ADGRA3,ADGRG3,BDKRB1,CHRM3,FOS,FZD6,GIPR,GPR173,GPR25,GPR88,HTR1F,MMP23B,MMP24,NOX4,P2RY14,PIK3C2B,S100A1,TRPV6,VEGFB |
| P2Y Purinergic Receptor Signaling Pathway | 0.340 | 0.031 | #NUM! | FOS,GNG5,PIK3C2B,RRAS2 |
| UBE2C in Cell Cycle Progression Signaling Pathway | 0.338 | 0.032 | #NUM! | ANAPC10,ANAPC15,PIK3C2B |
| Non-Small Cell Lung Cancer Signaling | 0.338 | 0.032 | #NUM! | PIK3C2B,RARB,RRAS2 |
| Agranulocyte Adhesion and Diapedesis | 0.337 | 0.030 | #NUM! | CLDN11,CXCR3,MMP23B,MMP24,MYO5C |
| WNT/β-catenin Signaling | 0.337 | 0.030 | -1.000 | FZD6,PPM1L,RARB,SOX1,Tle1 |
| NoRC negatively regulates rRNA expression | 0.337 | 0.035 | #NUM! | GTF2H4,MNAT1 |
| Iron uptake and transport | 0.337 | 0.035 | #NUM! | FTL,TF |
| Cancer Drug Resistance by Drug Efflux | 0.337 | 0.035 | #NUM! | PTGS2,RRAS2 |

| | | | | |
|---|---|---|---|---|
| RAF-independent MAPK1/3 activation | 0.335 | 0.044 | #NUM! | DUSP1 |
| RAS processing | 0.335 | 0.044 | #NUM! | PDE6D |
| Glycolysis I | 0.335 | 0.044 | #NUM! | ALDOC |
| Gluconeogenesis I | 0.335 | 0.044 | #NUM! | ALDOC |
| Apelin Cardiac Fibroblast Signaling Pathway | 0.335 | 0.044 | #NUM! | PRKAB2 |
| SLAMF Signaling Pathway | 0.334 | 0.031 | 0.000 | FOS,PIK3C2B,RRAS2,ZAP70 |
| Prolactin Signaling | 0.331 | 0.032 | #NUM! | FOS,PIK3C2B,RRAS2 |
| Regulation of IL-2 Expression in Activated and Anergic T Lymph | 0.331 | 0.032 | #NUM! | FOS,RRAS2,ZAP70 |
| GABA receptor activation | 0.329 | 0.034 | #NUM! | GNAL,GNG5 |
| Co-inhibition by BTLA | 0.329 | 0.034 | #NUM! | PIK3C2B,STK3 |
| DNA Replication Pre-Initiation | 0.325 | 0.031 | #NUM! | ANAPC10,ANAPC15,PSMB6 |
| p70S6K Signaling | 0.324 | 0.030 | 0.000 | F2,PIK3C2B,PPM1L,RRAS2 |
| RHO GTPases activate IQGAPs | 0.321 | 0.042 | #NUM! | TUBB6 |
| Chromatin modifications during the maternal to zygotic transiti | 0.321 | 0.042 | #NUM! | TET3 |
| Endometrial Cancer Signaling | 0.321 | 0.033 | #NUM! | PIK3C2B,RRAS2 |
| GPER1 signaling | 0.319 | 0.031 | #NUM! | GNG5,PIK3C2B,STK3 |
| Melanocyte Development and Pigmentation Signaling | 0.319 | 0.031 | #NUM! | MITF,PIK3C2B,RRAS2 |
| Small Cell Lung Cancer Signaling | 0.319 | 0.031 | #NUM! | PIK3C2B,PTGS2,RARB |
| CTLA4 Signaling in Cytotoxic T Lymphocytes | 0.318 | 0.029 | 0.447 | FOS,PIK3C2B,PPM1L,RRAS2,ZAP70 |
| NF-κB Signaling | 0.318 | 0.029 | 1.342 | EIF2AK2,GHR,PIK3C2B,RRAS2,ZAP70 |
| G alpha (s) signalling events | 0.318 | 0.030 | 2.000 | GIPR,GNG5,GPR25,NPSR1 |
| Hepatic Fibrosis Signaling Pathway | 0.315 | 0.028 | 1.667 | COL1A2,FOS,FTL,FZD6,LEP,NOX4,PIK3C2B,RND2,RRAS2,TF,VEGFB |
| Ion channel transport | 0.314 | 0.029 | 1.342 | ANO5,CASQ2,CUTC,MLKL,TRPV6 |
| Senescence Pathway | 0.309 | 0.028 | 0.000 | ANAPC10,ANAPC15,ELK3,EP400,PIK3C2B,PPM1L,RAD50,RRAS2 |
| Insulin processing | 0.309 | 0.040 | #NUM! | KIF5A |
| Metabolism of cofactors | 0.309 | 0.040 | #NUM! | DHFR |
| Glycogen metabolism | 0.309 | 0.040 | #NUM! | PHKB |
| Signaling by NTRK2 (TRKB) | 0.309 | 0.040 | #NUM! | FRS2 |
| Cardiogenesis | 0.309 | 0.040 | #NUM! | TBX1 |
| Complex III assembly | 0.309 | 0.040 | #NUM! | UQCR10 |
| Antigen Presentation Pathway | 0.309 | 0.040 | #NUM! | PSMB6 |
| NAD Signaling Pathway | 0.308 | 0.029 | -1.000 | H1-1,H1-4,PIK3C2B,PRKAB2 |
| Necroptosis Signaling Pathway | 0.308 | 0.029 | 2.000 | EIF2AK2,GLUL,MLKL,TIMM17B |
| Neuropathic Pain Signaling in Dorsal Horn Neurons | 0.307 | 0.030 | #NUM! | FOS,KCNN3,PIK3C2B |
| Autophagy | 0.306 | 0.028 | -1.342 | ATG13,FOS,LAMP1,PIK3C2B,PPM1L,PRKAB2 |
| Cytosolic sensors of pathogen-associated DNA | 0.306 | 0.032 | #NUM! | LRRFIP1,XRCC5 |
| NCAM signaling for neurite out-growth | 0.306 | 0.032 | #NUM! | GDNF,SPTAN1 |
| Transcriptional Regulation by MECP2 | 0.306 | 0.032 | #NUM! | PVALB,TBL1XR1 |
| PPAR Signaling | 0.301 | 0.030 | #NUM! | FOS,PTGS2,RRAS2 |
| Cardiac Hypertrophy Signaling (Enhanced) | 0.299 | 0.027 | 2.309 | FGF11,FGF5,FICD,FZD6,GDPD1,GHR,GNA12,GNG5,LEP,MKNK1,PDE6D,PIK3C2B,PTGS2,RRAS2 |
| Mitochondrial protein import | 0.298 | 0.032 | #NUM! | PMPCB,TIMM17B |
| RAB geranylgeranylation | 0.298 | 0.032 | #NUM! | RAB18,RAB5C |
| ERB2-ERBB3 Signaling | 0.298 | 0.032 | #NUM! | PIK3C2B,RRAS2 |
| Aryl Hydrocarbon Receptor Signaling | 0.298 | 0.029 | 0.000 | DHFR,FOS,NFIC,RARB |
| Signaling by NOTCH4 | 0.298 | 0.029 | 1.000 | PIK3C2B,PSMB6,STK3,TACC3 |
| Peroxisomal lipid metabolism | 0.297 | 0.039 | #NUM! | ACBD5 |
| Ethanol Degradation II | 0.297 | 0.039 | #NUM! | ADH5 |
| S Phase | 0.295 | 0.030 | #NUM! | MNAT1,PSMB6,STAG2 |
| Translocation of SLC2A4 (GLUT4) to the plasma membrane | 0.291 | 0.031 | #NUM! | PRKAB2,TUBB6 |
| Phase II - Conjugation of compounds | 0.291 | 0.031 | #NUM! | ESD,TPST1 |
| RNA Polymerase I Transcription | 0.291 | 0.031 | #NUM! | GTF2H4,MNAT1 |
| Cell Cycle Control of Chromosomal Replication | 0.291 | 0.031 | #NUM! | CDK17,RFC4 |
| Interleukin-4 and Interleukin-13 signaling | 0.289 | 0.029 | #NUM! | COL1A2,FOS,PTGS2 |
| Mouse Embryonic Stem Cell Pluripotency | 0.289 | 0.029 | #NUM! | FZD6,PIK3C2B,RRAS2 |
| Noradrenaline and Adrenaline Degradation | 0.286 | 0.037 | #NUM! | ADH5 |
| Interferon Signaling | 0.286 | 0.037 | #NUM! | PIAS1 |
| WNT/Ca2+ Signaling Pathway | 0.284 | 0.031 | #NUM! | FZD6,PDE6D |
| Cargo recognition for clathrin-mediated endocytosis | 0.284 | 0.029 | #NUM! | DAB2,SYT2,TF |
| Egg-Sperm Fusion Signaling Pathway | 0.284 | 0.029 | #NUM! | PIK3C2B,SPACA6,SYT2 |
| Transcriptional regulation by RUNX1 | 0.279 | 0.028 | -2.000 | ARID1A,MNAT1,PSMB6,SMARCB1 |
| Role of JAK1 and JAK3 in γc Cytokine Signaling | 0.277 | 0.030 | #NUM! | PIK3C2B,RRAS2 |
| ERBB4 Signaling | 0.277 | 0.030 | #NUM! | PIK3C2B,RRAS2 |
| Dopamine Receptor Signaling | 0.277 | 0.030 | #NUM! | PPM1L,PPP1R1B |
| GM-CSF Signaling | 0.277 | 0.030 | #NUM! | PIK3C2B,RRAS2 |

| | | | | |
|---|---|---|---|---|
| Wound Healing Signaling Pathway | 0.276 | 0.027 | 0.816 | COL1A2,F2,FOS,LEP,RRAS2,VEGFB |
| PPARα/RXRα Activation | 0.275 | 0.028 | 0.000 | GHR,NCOA6,NR2C2,PRKAB2,RRAS2 |
| Role of p14/p19ARF in Tumor Suppression | 0.275 | 0.036 | #NUM! | PIK3C2B |
| Semaphorin Neuronal Repulsive Signaling Pathway | 0.275 | 0.028 | 1.000 | BCAN,PIK3C2B,SEMA6D,VCAN |
| MHC class II antigen presentation | 0.273 | 0.029 | #NUM! | DYNC1LI2,KIF5A,TUBB6 |
| Role of MAPK Signaling in Promoting the Pathogenesis of Influe | 0.273 | 0.029 | #NUM! | LCAT,PTGS2,RRAS2 |
| Irritable Bowel Syndrome Signaling Pathway | 0.272 | 0.027 | 2.449 | LEP,MLKL,MMP23B,MMP24,PIK3C2B,RND2 |
| Integration of energy metabolism | 0.268 | 0.028 | #NUM! | CHRM3,GNG5,PRKAB2 |
| Mitochondrial translation | 0.268 | 0.028 | #NUM! | MRPL4,MRPL54,TUFM |
| Paxillin Signaling | 0.268 | 0.028 | #NUM! | ACTN1,PIK3C2B,RRAS2 |
| RNA Polymerase II Transcription | 0.266 | 0.028 | -1.000 | GTF2H4,MNAT1,SRSF4,SUPT5H |
| HOTAIR Regulatory Pathway | 0.266 | 0.028 | 2.000 | COL1A2,MMP23B,MMP24,PIK3C2B |
| Toll-like Receptor Cascades | 0.265 | 0.035 | #NUM! | BPI |
| Energy dependent regulation of mTOR by LKB1-AMPK | 0.265 | 0.035 | #NUM! | PRKAB2 |
| ISG15 antiviral mechanism | 0.264 | 0.029 | #NUM! | EIF2AK2,TRIM25 |
| Maturity Onset Diabetes of Young (MODY) Signaling | 0.264 | 0.029 | #NUM! | APOD,UCP2 |
| Basal Cell Carcinoma Signaling | 0.264 | 0.029 | #NUM! | BMP1,FZD6 |
| Hepatic Fibrosis / Hepatic Stellate Cell Activation | 0.263 | 0.027 | #NUM! | COL1A2,CXCR3,LEP,MYO5C,VEGFB |
| PIP3 activates AKT signaling | 0.261 | 0.027 | 1.000 | FGF5,FLT3,FRS2,NR4A1 |
| Sensory processing of sound by inner hair cells of the cochlea | 0.258 | 0.029 | #NUM! | MYO3B,SPTAN1 |
| Corticotropin Releasing Hormone Signaling | 0.257 | 0.027 | #NUM! | FOS,GUCY2F,NR4A1,PTGS2 |
| Granulocyte Adhesion and Diapedesis | 0.257 | 0.027 | #NUM! | CLDN11,CXCR3,MMP23B,MMP24 |
| Signaling by ALK | 0.255 | 0.033 | #NUM! | FRS2 |
| RIPK1-mediated regulated necrosis | 0.255 | 0.033 | #NUM! | MLKL |
| Regulation of TP53 Activity through Acetylation | 0.255 | 0.033 | #NUM! | BRPF1 |
| Sensory perception of taste | 0.255 | 0.033 | #NUM! | SCN4B |
| Hedgehog 'off' state | 0.253 | 0.028 | #NUM! | DYNC2H1,PSMB6,TUBB6 |
| WNT/SHH Axonal Guidance Signaling Pathway | 0.253 | 0.027 | 1.000 | FZD6,PARD6A,PIK3C2B,TRPV6 |
| TNFR2 Signaling | 0.246 | 0.032 | #NUM! | FOS |
| Retinoate Biosynthesis I | 0.246 | 0.032 | #NUM! | DHRS3 |
| Toll-like Receptor Signaling | 0.240 | 0.028 | #NUM! | EIF2AK2,FOS |
| L1CAM interactions | 0.239 | 0.027 | #NUM! | SCN4B,SPTAN1,TUBB6 |
| Regulation of lipid metabolism by PPARalpha | 0.239 | 0.027 | #NUM! | FDFT1,NCOA6,TBL1XR1 |
| Myocardin Signaling Pathway | 0.239 | 0.027 | #NUM! | BMP1,PIK3C2B,SPRY4 |
| G-protein beta:gamma signalling | 0.237 | 0.031 | #NUM! | GNG5 |
| Late endosomal microautophagy | 0.237 | 0.031 | #NUM! | PCNT |
| Role of RIG1-like Receptors in Antiviral Innate Immunity | 0.237 | 0.031 | #NUM! | TRIM25 |
| Extra-nuclear estrogen signaling | 0.234 | 0.027 | #NUM! | FOS,GNG5 |
| Oxytocin in Spinal Neurons Signaling Pathway | 0.229 | 0.030 | #NUM! | GUCY2F |
| Signal amplification | 0.229 | 0.030 | #NUM! | GNG5 |
| MAPK targets/ Nuclear events mediated by MAP kinases | 0.229 | 0.030 | #NUM! | FOS |
| MIF-mediated Glucocorticoid Regulation | 0.229 | 0.030 | #NUM! | PTGS2 |
| trans-Golgi Network Vesicle Budding | 0.229 | 0.027 | #NUM! | FTL,RAB5C |
| Leptin Signaling in Obesity | 0.229 | 0.027 | #NUM! | LEP,PIK3C2B |
| Signaling by NOTCH1 | 0.223 | 0.027 | #NUM! | TBL1XR1,Tle1 |
| HIPPO signaling | 0.223 | 0.027 | #NUM! | PPM1L,STK3 |
| Chemokine Signaling | 0.223 | 0.027 | #NUM! | FOS,RRAS2 |
| GPVI-mediated activation cascade | 0.221 | 0.029 | #NUM! | COL1A2 |
| Detoxification of Reactive Oxygen Species | 0.213 | 0.029 | #NUM! | NOX4 |
| Gene and protein expression by JAK-STAT signaling after IL-12 s | 0.213 | 0.029 | #NUM! | LCP1 |
| IL-9 Signaling | 0.213 | 0.029 | #NUM! | PIK3C2B |
| Dermatan Sulfate Biosynthesis (Late Stages) | 0.213 | 0.029 | #NUM! | CHST2 |
| Interleukin-10 signaling | 0.206 | 0.028 | #NUM! | PTGS2 |
| Assembly and cell surface presentation of NMDA receptors | 0.206 | 0.028 | #NUM! | TUBB6 |
| Formation of the dystrophin-glycoprotein complex (DGC) | 0.206 | 0.028 | #NUM! | SSPN |
| Notch Signaling | 0.199 | 0.027 | #NUM! | MAG |
| Dilated Cardiomyopathy Signaling Pathway | 0.000 | 0.007 | #NUM! | MYO5C |
| CSDE1 Signaling Pathway | 0.000 | 0.020 | #NUM! | FOS |
| ID1 Signaling Pathway | 0.000 | 0.015 | #NUM! | PIK3C2B,RRAS2,VEGFB |
| IL-13 Signaling Pathway | 0.000 | 0.023 | #NUM! | DUSP1,PIK3C2B |
| SNARE Signaling Pathway | 0.000 | 0.015 | #NUM! | MYO5C,SYT2 |
| Immunogenic Cell Death Signaling Pathway | 0.000 | 0.013 | #NUM! | MLKL |
| Macrophage Classical Activation Signaling Pathway | 0.000 | 0.021 | #NUM! | BPI,IRF8,LEP |
| MicroRNA Biogenesis Signaling Pathway | 0.000 | 0.017 | #NUM! | PIK3C2B,PRKRA,RRAS2 |

| | | | | |
|---|---|---|---|---|
| Multiple Sclerosis Signaling Pathway | 0.000 | 0.017 | #NUM! | DUSP1,LEP,MBP |
| Pathogen Induced Cytokine Storm Signaling Pathway | 0.000 | 0.025 | 1.633 | COL1A2,CXCR3,FOS,FTL,LEP,MLKL,VEGFB |
| Ribonucleotide Reductase Signaling Pathway | 0.000 | 0.026 | -1.000 | ARID1A,FOS,PIK3C2B,SMARCB1 |
| Natural Killer Cell Signaling | 0.000 | 0.024 | 0.000 | COL1A2,PIK3C2B,RRAS2,ZAP70 |
| IL-10 Signaling | 0.000 | 0.023 | #NUM! | DUSP1,FOS,PRKAB2 |
| Macrophage Alternative Activation Signaling Pathway | 0.000 | 0.026 | 1.000 | DUSP1,EPAS1,FOS,PIK3C2B |
| Neutrophil Extracellular Trap Signaling Pathway | 0.000 | 0.024 | 2.121 | BPI,COL1A2,KCNN3,LCAT,MLKL,NOX4,PIK3C2B,TIMM17B |
| Neuregulin Signaling | 0.000 | 0.009 | #NUM! | RRAS2 |
| Circadian Rhythm Signaling | 0.000 | 0.019 | #NUM! | GNG5,GUCY2F,MKNK1,PRKAB2,RRAS2 |
| Synaptic Long Term Potentiation | 0.000 | 0.008 | #NUM! | RRAS2 |
| Fc Epsilon RI Signaling | 0.000 | 0.018 | #NUM! | PIK3C2B,RRAS2 |
| Huntington's Disease Signaling | 0.000 | 0.011 | #NUM! | GNG5,PIK3C2B,PSMB6 |
| IL-33 Signaling Pathway | 0.000 | 0.022 | #NUM! | FOS,PIK3C2B,PTGS2 |
| Myelination Signaling Pathway | 0.000 | 0.022 | 1.134 | BMP1,FOS,FZD6,MAG,MBP,PIK3C2B,RRAS2 |
| NOD1/2 Signaling Pathway | 0.000 | 0.020 | #NUM! | FOS,LEP,SLC15A3 |
| Acetylcholine Receptor Signaling Pathway | 0.000 | 0.016 | #NUM! | CHRM3,GNAL,PIK3C2B |
| Adrenergic Receptor Signaling Pathway (Enhanced) | 0.000 | 0.011 | #NUM! | GNAL,LEP |
| Cachexia Signaling Pathway | 0.000 | 0.018 | 0.000 | EIF2AK2,LEP,NOX4,PIK3C2B,PRKAB2,PSMB6 |
| Microautophagy Signaling Pathway | 0.000 | 0.019 | #NUM! | COL1A2,PRKAB2,PSMB6 |
| NFKBIE Signaling Pathway | 0.000 | 0.018 | #NUM! | DYSF |
| Orexin Signaling Pathway | 0.000 | 0.013 | #NUM! | GNG5,PIK3C2B,PRKAB2 |
| NRF2-mediated Oxidative Stress Response | 0.000 | 0.020 | #NUM! | FOS,FTL,PIK3C2B,RRAS2 |
| LPS/IL-1 Mediated Inhibition of RXR Function | 0.000 | 0.015 | #NUM! | CHST2,SLC27A4,SLC27A5 |
| p53 Signaling | 0.000 | 0.021 | #NUM! | PIAS1,PIK3C2B |
| Intrinsic Pathway for Apoptosis | 0.000 | 0.019 | #NUM! | BMF |
| Signaling by Insulin receptor | 0.000 | 0.023 | #NUM! | IDE |
| G alpha (i) signalling events | 0.000 | 0.023 | 2.236 | BDKRB1,CXCR3,GNG5,HTR1F,P2RY14 |
| Meiotic synapsis | 0.000 | 0.023 | #NUM! | STAG2 |
| Cellular response to hypoxia | 0.000 | 0.026 | #NUM! | EPAS1,PSMB6 |
| Signaling by SCF-KIT | 0.000 | 0.024 | #NUM! | SH2B3 |
| Elastic fibre formation | 0.000 | 0.024 | #NUM! | FBN2 |
| Eukaryotic Translation Elongation | 0.000 | 0.008 | #NUM! | RPL36A |
| MyD88:MAL(TIRAP) cascade initiated on plasma membrane | 0.000 | 0.026 | #NUM! | S100A1 |
| Complement cascade | 0.000 | 0.023 | #NUM! | F2 |
| DDX58/IFIH1-mediated induction of interferon-alpha/beta | 0.000 | 0.015 | #NUM! | TRIM25 |
| Regulation of Apoptosis | 0.000 | 0.019 | #NUM! | PSMB6 |
| Signaling by TGF-beta Receptor Complex | 0.000 | 0.019 | #NUM! | PARD6A |
| SRP-dependent cotranslational protein targeting to membrane | 0.000 | 0.015 | #NUM! | DDOST,RPL36A |
| Signaling by NTRK1 (TRKA) | 0.000 | 0.012 | #NUM! | FRS2 |
| p75 NTR receptor-mediated signalling | 0.000 | 0.027 | 2.000 | MAG,MCF2L,PIK3C2B,STK3 |
| Signaling by VEGF | 0.000 | 0.009 | #NUM! | VEGFB |
| Degradation of beta-catenin by the destruction complex | 0.000 | 0.022 | #NUM! | PSMB6,Tle1 |
| Immunoregulatory interactions between a Lymphoid and a nor | 0.000 | 0.012 | #NUM! | COL1A2 |
| TCF dependent signaling in response to WNT | 0.000 | 0.016 | #NUM! | FZD6,PSMB6,Tle1 |
| TCR signaling | 0.000 | 0.018 | #NUM! | PSMB6,ZAP70 |
| Gene Silencing by RNA | 0.000 | 0.011 | #NUM! | PRKRA |
| Phase I - Functionalization of compounds | 0.000 | 0.026 | #NUM! | ADH5,FDX1 |
| Integrin cell surface interactions | 0.000 | 0.012 | #NUM! | COL1A2 |
| Transcriptional activity of SMAD2/SMAD3:SMAD4 heterotrimer | 0.000 | 0.020 | #NUM! | COL1A2 |
| Selenoamino acid metabolism | 0.000 | 0.007 | #NUM! | RPL36A |
| Fc epsilon receptor (FCERI) signaling | 0.000 | 0.015 | #NUM! | FOS,PSMB6 |
| Oxidative Stress Induced Senescence | 0.000 | 0.013 | #NUM! | FOS |
| EPH-Ephrin signaling | 0.000 | 0.011 | #NUM! | EFNB1 |
| SUMOylation of DNA damage response and repair proteins | 0.000 | 0.026 | #NUM! | PIAS1,STAG2 |
| SUMOylation of ubiquitinylation proteins | 0.000 | 0.026 | #NUM! | PIAS1 |
| Metabolism of polyamines | 0.000 | 0.017 | #NUM! | PSMB6 |
| Class A/1 (Rhodopsin-like receptors) | 0.000 | 0.024 | 2.646 | BDKRB1,CHRM3,CXCR3,F2,HTR1F,NPSR1,P2RY14 |
| Semaphorin interactions | 0.000 | 0.016 | #NUM! | SEMA6D |
| Signaling by ROBO receptors | 0.000 | 0.012 | #NUM! | LHX4,PSMB6,RPL36A |
| ABC-family proteins mediated transport | 0.000 | 0.009 | #NUM! | PSMB6 |
| SUMOylation of transcription cofactors | 0.000 | 0.023 | #NUM! | PIAS1 |
| Protein folding | 0.000 | 0.022 | #NUM! | GNG5,TUBB6 |
| Platelet homeostasis | 0.000 | 0.012 | #NUM! | GNG5 |
| G alpha (z) signalling events | 0.000 | 0.021 | #NUM! | GNG5 |

| | | | | |
|---|---|---|---|---|
| Transport of bile salts and organic acids, metal ions and amine c | 0.000 | 0.026 | #NUM! | RHCG,SLC22A8 |
| Activation of NMDA receptors and postsynaptic events | 0.000 | 0.026 | #NUM! | PRKAB2,TUBB6 |
| Smooth Muscle Contraction | 0.000 | 0.026 | #NUM! | DYSF |
| Interleukin-1 family signaling | 0.000 | 0.008 | #NUM! | PSMB6 |
| Regulation of mRNA stability by proteins that bind AU-rich elem | 0.000 | 0.022 | #NUM! | PSMB6,TNPO1 |
| Mitotic G1 phase and G1/S transition | 0.000 | 0.023 | #NUM! | DHFR,MNAT1,PSMB6 |
| SUMOylation of chromatin organization proteins | 0.000 | 0.016 | #NUM! | PIAS1 |
| Hedgehog ligand biogenesis | 0.000 | 0.015 | #NUM! | PSMB6 |
| Activation of anterior HOX genes in hindbrain during early emb | 0.000 | 0.026 | #NUM! | NCOA6,RARB |
| C-type lectin receptors (CLRs) | 0.000 | 0.008 | #NUM! | PSMB6 |
| TP53 Regulates Metabolic Genes | 0.000 | 0.012 | #NUM! | PRKAB2 |
| Hedgehog 'on' state | 0.000 | 0.021 | #NUM! | PIK3C2B,PSMB6,STK3 |
| TNFR2 non-canonical NF-kB pathway | 0.000 | 0.015 | #NUM! | PSMB6 |
| MAPK6/MAPK4 signaling | 0.000 | 0.011 | #NUM! | PSMB6 |
| Deubiquitination | 0.000 | 0.021 | -0.447 | CLSPN,IDE,PSMB6,TRIM25,USP20 |
| DNA Double Strand Break Response | 0.000 | 0.020 | #NUM! | RAD50 |
| Major pathway of rRNA processing in the nucleolus and cytosol | 0.000 | 0.015 | #NUM! | DCAF13,DHX37,RPL36A |
| TP53 Regulates Transcription of Cell Cycle Genes | 0.000 | 0.020 | #NUM! | BTG2 |
| Neurexins and neuroligins | 0.000 | 0.018 | #NUM! | SYT2 |
| Keratinization | 0.000 | 0.016 | #NUM! | DSC1 |
| Signaling by MET | 0.000 | 0.013 | #NUM! | COL1A2 |
| PTEN Regulation | 0.000 | 0.007 | #NUM! | PSMB6 |
| RNA polymerase II transcribes snRNA genes | 0.000 | 0.014 | #NUM! | SUPT5H |
| Mitotic Prophase | 0.000 | 0.011 | #NUM! | NEK6 |
| Glucose metabolism | 0.000 | 0.012 | #NUM! | ALDOC |
| Processing of Capped Intron-Containing Pre-mRNA | 0.000 | 0.007 | #NUM! | PRPF40A,SRSF4 |
| Eukaryotic Translation Initiation | 0.000 | 0.006 | #NUM! | RPL36A |
| Eukaryotic Translation Termination | 0.000 | 0.009 | #NUM! | RPL36A |
| RNA Polymerase III Transcription | 0.000 | 0.024 | #NUM! | NFIC |
| Signaling by PTK6 | 0.000 | 0.019 | #NUM! | EPAS1 |
| TBC/RABGAPs | 0.000 | 0.023 | #NUM! | RAB5C |
| Transcriptional regulation by RUNX3 | 0.000 | 0.010 | #NUM! | PSMB6 |
| Regulation of RUNX2 expression and activity | 0.000 | 0.014 | #NUM! | PSMB6 |
| Neddylation | 0.000 | 0.017 | -1.000 | DCAF13,DCAF6,EPAS1,PSMB6 |
| Carboxyterminal post-translational modifications of tubulin | 0.000 | 0.026 | #NUM! | TUBB6 |
| Post-translational protein phosphorylation | 0.000 | 0.021 | #NUM! | TF,VCAN |
| Signaling by NOTCH3 | 0.000 | 0.020 | #NUM! | TACC3 |
| Nonsense-Mediated Decay (NMD) | 0.000 | 0.009 | #NUM! | RPL36A |
| Transcriptional regulation of granulopoiesis | 0.000 | 0.023 | #NUM! | KLF5 |
| Response of EIF2AK4 (GCN2) to amino acid deficiency | 0.000 | 0.009 | #NUM! | RPL36A |
| Amino acids regulate mTORC1 | 0.000 | 0.019 | #NUM! | DEPDC5 |
| KEAP1-NFE2L2 pathway | 0.000 | 0.021 | #NUM! | PIK3C2B,PSMB6,STK3 |
| Retinoid metabolism and transport | 0.000 | 0.026 | #NUM! | RBP4 |
| Amyloid fiber formation | 0.000 | 0.016 | #NUM! | GSN |
| Class I MHC mediated antigen processing and presentation | 0.000 | 0.014 | -1.342 | ANAPC10,PSMB6,S100A1,UBAC1,UBE2E2 |
| Factors involved in megakaryocyte development and platelet p | 0.000 | 0.026 | #NUM! | RAD51C,SH2B3 |
| Signaling by the B Cell Receptor (BCR) | 0.000 | 0.009 | #NUM! | PSMB6 |
| Formation of WDR5-containing histone-modifying complexes | 0.000 | 0.024 | #NUM! | NCOA6 |
| Hepatic Cholestasis | 0.000 | 0.005 | #NUM! | LEP |
| Mitochondrial Dysfunction | 0.000 | 0.015 | 0.447 | COX15,PIK3C2B,PRKAB2,UCP2,UQCR10 |
| Autism Signaling Pathway | 0.000 | 0.018 | 1.000 | FZD6,LEP,PIK3C2B,RARB,RRAS2 |
| CGAS-STING Signaling Pathway | 0.000 | 0.009 | #NUM! | LEP |
| Pancreatic Secretion Signaling Pathway | 0.000 | 0.023 | 2.236 | CA5B,CHRM3,GNA12,PIK3C2B,SLC9A1 |
| Protein Sorting Signaling Pathway | 0.000 | 0.006 | #NUM! | PLPP1 |
| ROBO SLIT Signaling Pathway | 0.000 | 0.017 | #NUM! | FOS,VEGFB |
| VDR/RXR Activation | 0.000 | 0.014 | #NUM! | TRPV6 |
| UFMylation Signaling Pathway | 0.000 | 0.017 | #NUM! | RAD50 |
| Cohesin Chromatin Regulation Pathway | 0.000 | 0.019 | 0.000 | ANAPC10,FAT1,PPM1L,STAG2 |
| Nuclear Cytoskeleton Signaling Pathway | 0.000 | 0.024 | 1.342 | DHFR,DYNC1LI2,KIF5A,TUBB6,XRCC5 |
| mRNA 3 Prime End Processing Signaling Pathway | 0.000 | 0.021 | #NUM! | PAPOLG,SUPT5H |
| Ribosomal Quality Control Signaling Pathway | 0.000 | 0.010 | #NUM! | PSMB6,RPL36A |
| Post-translational modification: synthesis of GPI-anchored prot | 0.000 | 0.013 | #NUM! | CD109 |
| Pyruvate metabolism | 0.000 | 0.021 | #NUM! | ARMC8 |
| M-decay: degradation of maternal mRNAs by maternally storec | 0.000 | 0.022 | #NUM! | KDM4C |

| | | | | |
|---|---|---|---|---|
| Somitogenesis | 0.000 | 0.018 | #NUM! | PSMB6 |
| Transcriptional and post-translational regulation of MITF-M exp | 0.000 | 0.016 | #NUM! | MITF |
| Respiratory electron transport | 0.000 | 0.012 | #NUM! | UQCR10 |
| Complex IV assembly | 0.000 | 0.021 | #NUM! | COX15 |
| Cyclophilin Signaling Pathway | 0.000 | 0.026 | 1.633 | F2,NOX4,PIK3C2B,RAD50,TIMM17B,TNPO1 |
| Hepatitis B Chronic Liver Pathogenesis Signaling Pathway | 0.000 | 0.018 | #NUM! | PIK3C2B,RRAS2,ZHX2 |
| TRIM21 Intracellular Antibody Signaling Pathway | 0.000 | 0.026 | #NUM! | FOS,IRF8,PSMB6 |
| Tuberculosis Active Signaling Pathway | 0.000 | 0.020 | 1.000 | FOS,NFKBIZ,PIK3C2B,PTGS2 |
| Tuberculosis Latent Signaling Pathway | 0.000 | 0.012 | #NUM! | VEGFB |
| Epithelial Membrane Protein Signaling Pathway | 0.000 | 0.024 | 1.000 | GHR,NOX4,PIK3C2B,RRAS2 |
| Proteasomal PSMD10 Signaling Pathway | 0.000 | 0.026 | #NUM! | PIK3C2B,RRAS2,VEGFB |
| Warburg Effect Signaling Pathway | 0.000 | 0.023 | #NUM! | PIK3C2B,PRKAB2,VEGFB |
| DYRK1A Signaling Pathway | 0.000 | 0.018 | #NUM! | EPAS1,RRAS2,VEGFB |
| GAIT Translation Signaling Pathway | 0.000 | 0.014 | #NUM! | APOD,RBP4,RPL36A |
| Meiosis Initiation Signaling Pathway | 0.000 | 0.010 | #NUM! | STAG2 |
| Stress Granule Signaling Pathway | 0.000 | 0.017 | 0.000 | EIF2AK2,PIK3C2B,PTGS2,UBAP2L |
| NIK-->noncanonical NF-kB signaling | 0.000 | 0.016 | #NUM! | PSMB6 |
| Epigenetic regulation of gene expression by MLL3 and MLL4 co | 0.000 | 0.023 | #NUM! | NCOA6,TBL1XR1 |
| High laminar flow shear stress activates signaling by PIEZO1 an | 0.000 | 0.015 | #NUM! | GNG5 |
| Degradation of CRY and PER proteins | 0.000 | 0.016 | #NUM! | PSMB6 |
| Vitamin K Signaling Pathway | 0.000 | 0.018 | #NUM! | F2 |
| CGRP in Inflammation and Nociception Signaling Pathway | 0.000 | 0.022 | #NUM! | IDE,PIK3C2B,VEGFB |
| Neuropeptides in Migraine Signaling Pathway | 0.000 | 0.026 | -1.000 | FOS,GNG5,GUCY2F,HTR1F |
| Exocyst Signaling Pathway | 0.000 | 0.010 | #NUM! | PARD6A |
| Exosome Signaling Pathway | 0.000 | 0.006 | #NUM! | RPL36A |
| Regulation of Actin-based Motility by Rho | 0.000 | 0.019 | #NUM! | GSN,RND2 |
| Erythropoietin Signaling Pathway | 0.000 | 0.026 | -1.000 | FOS,LEP,PIK3C2B,RRAS2 |
| Caveolar-mediated Endocytosis Signaling | 0.000 | 0.014 | #NUM! | RAB5C |
| Role of Pattern Recognition Receptors in Recognition of Bacteri | 0.000 | 0.023 | #NUM! | EIF2AK2,LEP,PIK3C2B |
| Role of PKR in Interferon Induction and Antiviral Response | 0.000 | 0.026 | #NUM! | EIF2AK2,FOS,PRKRA |
| FcγRIIB Signaling in B Lymphocytes | 0.000 | 0.023 | #NUM! | PIK3C2B,RRAS2 |
| CD27 Signaling in Lymphocytes | 0.000 | 0.018 | #NUM! | FOS |
| Lymphotoxin β Receptor Signaling | 0.000 | 0.019 | #NUM! | PIK3C2B |
| B Cell Activating Factor Signaling | 0.000 | 0.023 | #NUM! | FOS |
| CD28 Signaling in T Helper Cells | 0.000 | 0.027 | #NUM! | FOS,PIK3C2B,ZAP70 |
| IL-15 Signaling | 0.000 | 0.025 | #NUM! | PIK3C2B,RRAS2 |
| Virus Entry via Endocytic Pathways | 0.000 | 0.018 | #NUM! | PIK3C2B,RRAS2 |
| Dendritic Cell Maturation | 0.000 | 0.025 | 2.000 | COL1A2,IRF8,LEP,PIK3C2B |
| Reelin Signaling in Neurons | 0.000 | 0.014 | #NUM! | PIK3C2B,VLDLR |
| Factors Promoting Cardiogenesis in Vertebrates | 0.000 | 0.020 | #NUM! | BMP1,FZD6,NOX4 |
| Agrin Interactions at Neuromuscular Junction | 0.000 | 0.015 | #NUM! | RRAS2 |
| Renin-Angiotensin Signaling | 0.000 | 0.025 | #NUM! | FOS,PIK3C2B,RRAS2 |
| Docosahexaenoic Acid (DHA) Signaling | 0.000 | 0.022 | 1.342 | LCAT,PIK3C2B,PRKAB2,PTGS2,SYT2 |
| Semaphorin Signaling in Neurons | 0.000 | 0.017 | #NUM! | RND2 |
| Cardiac Hypertrophy Signaling | 0.000 | 0.024 | #NUM! | GNA12,GNAL,GNG5,PIK3C2B,RND2,RRAS2 |
| ICOS-ICOSL Signaling in T Helper Cells | 0.000 | 0.019 | #NUM! | PIK3C2B,ZAP70 |
| Polyamine Regulation in Colon Cancer | 0.000 | 0.018 | #NUM! | FOS |
| GNRH Signaling | 0.000 | 0.016 | #NUM! | FOS,GNG5,RRAS2 |
| Human Embryonic Stem Cell Pluripotency | 0.000 | 0.021 | 0.000 | BMP1,FZD6,PIK3C2B,RRAS2 |
| DNA Methylation and Transcriptional Repression Signaling | 0.000 | 0.021 | #NUM! | CDK17,TET3 |
| ATM Signaling | 0.000 | 0.021 | #NUM! | PPM1L,RAD50 |
| Role of OCT4 in Mammalian Embryonic Stem Cell Pluripotency | 0.000 | 0.022 | #NUM! | CDX2 |
| Aldosterone Signaling in Epithelial Cells | 0.000 | 0.018 | #NUM! | DUSP1,PIK3C2B,SLC9A1 |
| Prostate Cancer Signaling | 0.000 | 0.018 | #NUM! | PIK3C2B,RRAS2 |
| Type I Diabetes Mellitus Signaling | 0.000 | 0.010 | #NUM! | PIAS1 |
| Primary Immunodeficiency Signaling | 0.000 | 0.026 | #NUM! | ZAP70 |
| Glioma Signaling | 0.000 | 0.016 | #NUM! | PIK3C2B,RRAS2 |
| Chronic Myeloid Leukemia Signaling | 0.000 | 0.026 | 0.378 | FOS,FZD6,IRF8,LIG3,PIK3C2B,PRKAB2,RRAS2 |
| Pancreatic Adenocarcinoma Signaling | 0.000 | 0.024 | #NUM! | PIK3C2B,PTGS2,VEGFB |
| Systemic Lupus Erythematosus Signaling | 0.000 | 0.024 | #NUM! | FOS,PIK3C2B,PRPF40A,RRAS2 |
| CDC42 Signaling | 0.000 | 0.014 | #NUM! | FOS,PARD6A |
| EIF2 Signaling | 0.000 | 0.021 | 0.447 | EIF2AK2,NOX4,PIK3C2B,RPL36A,RRAS2 |
| April Mediated Signaling | 0.000 | 0.024 | #NUM! | FOS |
| Retinoic acid Mediated Apoptosis Signaling | 0.000 | 0.026 | #NUM! | RARB |

| | | | | |
|---|---|---|---|---|
| PAK Signaling | 0.000 | 0.018 | #NUM! | PIK3C2B,RRAS2 |
| RHOA Signaling | 0.000 | 0.017 | #NUM! | GNA12,SEPTIN1 |
| Role of Osteoblasts, Osteoclasts and Chondrocytes in Rheumat | 0.000 | 0.024 | #NUM! | BMP1,FOS,FZD6,GSN,PIK3C2B |
| Phospholipase C Signaling | 0.000 | 0.015 | #NUM! | GNG5,RND2,RRAS2,ZAP70 |
| Role of Macrophages, Fibroblasts and Endothelial Cells in Rheu | 0.000 | 0.017 | 0.447 | FOS,FZD6,PIK3C2B,RRAS2,VEGFB |
| Regulation of eIF4 and p70S6K Signaling | 0.000 | 0.020 | #NUM! | MKNK1,PIK3C2B,PPM1L,RRAS2 |
| Role of NFAT in Cardiac Hypertrophy | 0.000 | 0.014 | #NUM! | GNG5,PIK3C2B,RRAS2 |
| Glioblastoma Multiforme Signaling | 0.000 | 0.024 | 1.000 | FZD6,PIK3C2B,RND2,RRAS2 |
| Neuroprotective Role of THOP1 in Alzheimer's Disease | 0.000 | 0.022 | #NUM! | Ctrb1,IDE |
| Role of WNT/GSK-3β Signaling in the Pathogenesis of Influenza | 0.000 | 0.017 | #NUM! | FZD6 |
| NUR77 Signaling in T Lymphocytes | 0.000 | 0.013 | #NUM! | NR4A1 |
| PKCθ Signaling in T Lymphocytes | 0.000 | 0.026 | 0.000 | FOS,PIK3C2B,RRAS2,ZAP70 |
| TNFR1 Signaling | 0.000 | 0.020 | #NUM! | FOS |
| Role of PI3K/AKT Signaling in the Pathogenesis of Influenza | 0.000 | 0.021 | #NUM! | PIK3C2B |
| Role of Hypercytokinemia/hyperchemokinemia in the Pathoger | 0.000 | 0.018 | #NUM! | EIF2AK2 |
| PI3K Signaling in B Lymphocytes | 0.000 | 0.014 | #NUM! | FOS,RRAS2 |
| Cyclins and Cell Cycle Regulation | 0.000 | 0.012 | #NUM! | PPM1L |
| IL-17A Signaling in Airway Cells | 0.000 | 0.017 | #NUM! | PIK3C2B |
| Dopamine-DARPP32 Feedback in cAMP Signaling | 0.000 | 0.012 | #NUM! | PPM1L,PPP1R1B |
| NGF Signaling | 0.000 | 0.025 | #NUM! | PIK3C2B,RRAS2,SMPD1 |
| RHOGDI Signaling | 0.000 | 0.023 | #NUM! | GNA12,GNAL,GNG5,MYO5C,RND2 |
| Gap Junction Signaling | 0.000 | 0.026 | 0.000 | GJB1,GRIK1,GUCY2F,KCNN3,P2RY14,PIK3C2B,RRAS2,TUBB6 |
| iNOS Signaling | 0.000 | 0.021 | #NUM! | FOS |
| Ephrin A Signaling | 0.000 | 0.015 | #NUM! | EPAS1,PIK3C2B |
| Netrin Signaling | 0.000 | 0.006 | #NUM! | PIK3C2B |
| Heparan Sulfate Biosynthesis | 0.000 | 0.017 | #NUM! | CHST2 |
| Heparan Sulfate Biosynthesis (Late Stages) | 0.000 | 0.019 | #NUM! | CHST2 |
| D-myo-inositol-5-phosphate Metabolism | 0.000 | 0.021 | #NUM! | DUSP1,NUDT5,PPM1L,PPP1R1B |
| Pyridoxal 5'-phosphate Salvage Pathway | 0.000 | 0.015 | #NUM! | EIF2AK2 |
| Phospholipases | 0.000 | 0.017 | #NUM! | LCAT |
| Chondroitin Sulfate Biosynthesis | 0.000 | 0.022 | #NUM! | CHST2 |
| Dermatan Sulfate Biosynthesis | 0.000 | 0.021 | #NUM! | CHST2 |
| D-myo-inositol (1,4,5,6)-Tetrakisphosphate Biosynthesis | 0.000 | 0.022 | #NUM! | DUSP1,NUDT5,PPM1L,PPP1R1B |
| Superpathway of Inositol Phosphate Compounds | 0.000 | 0.022 | 1.000 | DUSP1,NUDT5,PIK3C2B,PPM1L,PPP1R1B |
| Serotonin Degradation | 0.000 | 0.026 | #NUM! | ADH5 |
| D-myo-inositol (3,4,5,6)-tetrakisphosphate Biosynthesis | 0.000 | 0.022 | #NUM! | DUSP1,NUDT5,PPM1L,PPP1R1B |
| 3-phosphoinositide Degradation | 0.000 | 0.021 | #NUM! | DUSP1,NUDT5,PPM1L,PPP1R1B |
| 3-phosphoinositide Biosynthesis | 0.000 | 0.025 | 1.000 | DUSP1,NUDT5,PIK3C2B,PPM1L,PPP1R1B |
| Triacylglycerol Biosynthesis | 0.000 | 0.017 | #NUM! | PLPP1 |
| Salvage Pathways of Pyrimidine Ribonucleotides | 0.000 | 0.011 | #NUM! | EIF2AK2 |
| Chondroitin Sulfate Biosynthesis (Late Stages) | 0.000 | 0.026 | #NUM! | CHST2 |
| Antioxidant Action of Vitamin C | 0.000 | 0.010 | #NUM! | LCAT |
| Epithelial Adherens Junction Signaling | 0.000 | 0.026 | 0.000 | PPM1L,PRKAB2,RRAS2,STK3 |
| Regulation of Cellular Mechanics by Calpain Protease | 0.000 | 0.023 | #NUM! | ACTN1,RRAS2 |
| STAT3 Pathway | 0.000 | 0.015 | #NUM! | GHR,RRAS2 |
| Oxidative Phosphorylation | 0.000 | 0.020 | #NUM! | COX15,UQCR10 |
| Unfolded protein response | 0.000 | 0.023 | #NUM! | CD82,INSIG1 |
| Parkinson's Signaling Pathway | 0.000 | 0.007 | #NUM! | LEP,TUBB6 |
| PI3K/AKT Signaling | 0.000 | 0.021 | #NUM! | GHR,PPM1L,PTGS2,RRAS2 |
| PTEN Signaling | 0.000 | 0.013 | #NUM! | GHR,RRAS2 |
| Nitric Oxide Signaling in the Cardiovascular System | 0.000 | 0.026 | #NUM! | GUCY2F,PIK3C2B,VEGFB |
| Protein Ubiquitination Pathway | 0.000 | 0.023 | -2.000 | ANAPC10,CFAP52,PSMB6,UBE2E2,USP20,USP40 |
| Xenobiotic Metabolism Signaling | 0.000 | 0.025 | #NUM! | CHST2,ESD,FTL,PIK3C2B,PPM1L,RRAS2 |
| IL-4 Signaling | 0.000 | 0.018 | #NUM! | COL1A2,PIK3C2B,RRAS2 |
| B Cell Receptor Signaling | 0.000 | 0.011 | #NUM! | PIK3C2B,RRAS2 |
| Serotonin Receptor Signaling | 0.000 | 0.025 | 1.265 | GNA12,GNAL,GNG5,KCNN3,LCAT,LEP,PIK3C2B,PTGS2,RND2,RRAS2,TRPV6 |
| Insulin Receptor Signaling | 0.000 | 0.016 | #NUM! | PIK3C2B,RRAS2 |
| Integrin to Cytoskeleton Signaling Pathway | 0.000 | 0.025 | 0.000 | ACTN1,GSN,PIK3C2B,RND2,RRAS2 |
| Death Receptor Signaling | 0.000 | 0.011 | #NUM! | SPTAN1 |
| p38 MAPK Signaling | 0.000 | 0.018 | #NUM! | DUSP1,MKNK1 |
| TGF-β Signaling | 0.000 | 0.021 | #NUM! | FOS,RRAS2 |
| Apoptosis Signaling | 0.000 | 0.020 | #NUM! | RRAS2,SPTAN1 |
| Hypoxia Signaling in the Cardiovascular System | 0.000 | 0.014 | #NUM! | UBE2E2 |
| T Cell Receptor Signaling | 0.000 | 0.022 | 0.000 | FOS,PIK3C2B,RRAS2,ZAP70 |

| | | | | |
|---|---|---|---|---|
| IL-6 Signaling | 0.000 | 0.025 | #NUM! | FOS,PIK3C2B,RRAS2 |
| GPCR-Mediated Integration of Enteroendocrine Signaling Exem | 0.000 | 0.014 | #NUM! | GIPR |
| GPCR-Mediated Nutrient Sensing in Enteroendocrine Cells | 0.000 | 0.009 | #NUM! | GNG5 |
| Phagosome Formation | 0.000 | 0.024 | 2.840 | ADGRA3,ADGRG3,BDKRB1,CHRM3,FZD6,GIPR,GPR173,GPR25,GPR88,HTR1F,LCAT,MYO5C,P2RY14,PIK3C2B,RRAS2 |
| Macropinocytosis Signaling | 0.000 | 0.026 | #NUM! | PIK3C2B,RRAS2 |
| PD-1, PD-L1 cancer immunotherapy pathway | 0.000 | 0.022 | #NUM! | PIK3C2B,ZAP70 |
| Th1 and Th2 Activation Pathway | 0.000 | 0.013 | #NUM! | CXCR3,PIK3C2B |
| Th1 Pathway | 0.000 | 0.020 | #NUM! | CXCR3,PIK3C2B |
| Th2 Pathway | 0.000 | 0.008 | #NUM! | PIK3C2B |
| GP6 Signaling Pathway | 0.000 | 0.017 | #NUM! | COL1A2,PIK3C2B |
| IL-7 Signaling Pathway | 0.000 | 0.014 | #NUM! | PIK3C2B |
| Opioid Signaling Pathway | 0.000 | 0.022 | #NUM! | FOS,FOSB,GNA12,GNAL,GNG5,RRAS2 |
| Adrenomedullin signaling pathway | 0.000 | 0.026 | 0.447 | FOS,GUCY2F,KCNN3,PIK3C2B,RRAS2 |
| SPINK1 Pancreatic Cancer Pathway | 0.000 | 0.022 | #NUM! | Ctrb1 |
| Endocannabinoid Neuronal Synapse Pathway | 0.000 | 0.021 | #NUM! | GNAL,GNG5,PTGS2 |
| Apelin Adipocyte Signaling Pathway | 0.000 | 0.024 | #NUM! | NOX4,PRKAB2 |
| Apelin Cardiomyocyte Signaling Pathway | 0.000 | 0.021 | #NUM! | PIK3C2B,SLC9A1 |
| BAG2 Signaling Pathway | 0.000 | 0.012 | #NUM! | PSMB6 |
| FAT10 Signaling Pathway | 0.000 | 0.018 | #NUM! | PSMB6 |
| Synaptogenesis Signaling Pathway | 0.000 | 0.020 | 0.000 | EFNB1,PIK3C2B,RAB5C,RRAS2,SYT2,VLDLR |
| IL-23 Signaling Pathway | 0.000 | 0.024 | #NUM! | PIK3C2B |
| Systemic Lupus Erythematosus in B Cell Signaling Pathway | 0.000 | 0.017 | 0.000 | FOS,LEP,PIK3C2B,RRAS2 |
| White Adipose Tissue Browning Pathway | 0.000 | 0.022 | #NUM! | LEP,PRKAB2,RARB |
| Inhibition of ARE-Mediated mRNA Degradation Pathway | 0.000 | 0.019 | #NUM! | PAPOLG,PPM1L,PSMB6 |
| Xenobiotic Metabolism CAR Signaling Pathway | 0.000 | 0.014 | #NUM! | CHST2,PPM1L |
| Xenobiotic Metabolism PXR Signaling Pathway | 0.000 | 0.015 | #NUM! | CHST2,ESD |
| Insulin Secretion Signaling Pathway | 0.000 | 0.015 | 2.000 | CHRM3,GHR,GIPR,PIK3C2B |
| Regulation of the Epithelial Mesenchymal Transition in Develop | 0.000 | 0.012 | #NUM! | FZD6 |
| Coronavirus Pathogenesis Pathway | 0.000 | 0.019 | -1.000 | FOS,PTGS2,TNPO1,TRIM25 |
| Coronavirus Replication Pathway | 0.000 | 0.013 | #NUM! | RPL36A,TUBB6 |
| Calcium Signaling | 0.000 | 0.019 | #NUM! | CASQ2,GRIK1,MYO5C,TRPV6 |

**Supplementary Table 9:Differentially expressed genes between PAE and SAC in Astrocytes at P28**

**Increased expression (1.2 FC, p<0.05)**

| Target nam | Log2 FC | Pvalue | Adjusted p | -log10 pval | -log10 adjusted pvalue |
|---|---|---|---|---|---|
| Sgsh | 1.883353 | 0.026089 | 0.992917 | 1.583536 | 0.003087 |
| Cenpk | 1.875779 | 0.000728 | 0.992917 | 3.138035 | 0.003087 |
| Gm11549 | 1.829945 | 0.032111 | 0.992917 | 1.493351 | 0.003087 |
| Slc5a5 | 1.742267 | 0.000626 | 0.992917 | 3.203199 | 0.003087 |
| Lcn10 | 1.722635 | 0.008106 | 0.992917 | 2.091216 | 0.003087 |
| Olfr1167 | 1.707996 | 0.039078 | 0.992917 | 1.408064 | 0.003087 |
| Frmd7 | 1.699172 | 0.024999 | 0.992917 | 1.602081 | 0.003087 |
| Nlrc5 | 1.634957 | 0.018905 | 0.992917 | 1.723434 | 0.003087 |
| Phf5a | 1.634957 | 0.025689 | 0.992917 | 1.590255 | 0.003087 |
| Washc3 | 1.631125 | 0.007595 | 0.992917 | 2.119495 | 0.003087 |
| Lgr4 | 1.591863 | 0.008324 | 0.992917 | 2.079663 | 0.003087 |
| C2cd2 | 1.584289 | 0.026960 | 0.992917 | 1.569279 | 0.003087 |
| Cacna1g | 1.570742 | 0.019631 | 0.992917 | 1.707066 | 0.003087 |
| Spata19 | 1.560826 | 0.046023 | 0.992917 | 1.337024 | 0.003087 |
| Fancf | 1.547279 | 0.045892 | 0.992917 | 1.338266 | 0.003087 |
| Btd | 1.547279 | 0.048253 | 0.992917 | 1.316473 | 0.003087 |
| Smim3 | 1.527648 | 0.028040 | 0.992917 | 1.552221 | 0.003087 |
| Cib4 | 1.514100 | 0.031753 | 0.992917 | 1.498213 | 0.003087 |
| Ubiad1 | 1.504185 | 0.034728 | 0.992917 | 1.459319 | 0.003087 |
| Msrb2 | 1.504185 | 0.004316 | 0.992917 | 2.364891 | 0.003087 |
| Sptssb | 1.487946 | 0.040043 | 0.992917 | 1.397475 | 0.003087 |
| Entpd6 | 1.485804 | 0.036476 | 0.992917 | 1.437996 | 0.003087 |
| Msmp | 1.483064 | 0.045230 | 0.992917 | 1.344574 | 0.003087 |
| Kcnc4 | 1.473148 | 0.017710 | 0.992917 | 1.751789 | 0.003087 |
| Piwil4 | 1.473148 | 0.016052 | 0.992917 | 1.794475 | 0.003087 |
| Lmbr1l | 1.473148 | 0.024426 | 0.992917 | 1.612145 | 0.003087 |
| Entpd7 | 1.468315 | 0.041943 | 0.992917 | 1.377341 | 0.003087 |
| Scd4 | 1.454999 | 0.024102 | 0.992917 | 1.617952 | 0.003087 |
| Zbtb26 | 1.453517 | 0.044090 | 0.992917 | 1.355660 | 0.003087 |
| Il10rb | 1.449685 | 0.038392 | 0.992917 | 1.415755 | 0.003087 |
| Wdr55 | 1.439970 | 0.046671 | 0.992917 | 1.330952 | 0.003087 |
| Oprd1 | 1.430054 | 0.007363 | 0.992917 | 2.132952 | 0.003087 |
| Spg21 | 1.426030 | 0.014095 | 0.992917 | 1.850926 | 0.003087 |
| Chac1 | 1.422481 | 0.011982 | 0.992917 | 1.921487 | 0.003087 |
| Crygb | 1.422481 | 0.048383 | 0.992917 | 1.315310 | 0.003087 |
| Pacrgl | 1.416507 | 0.019130 | 0.992917 | 1.718292 | 0.003087 |
| Kif24 | 1.408933 | 0.010279 | 0.992917 | 1.988057 | 0.003087 |

**Decreased expression (1.2 FC, p<0.05)**

| Target nam | Log2 FC | Pvalue | Adjusted p | -log10 pval | -log10 adjusted pvalue |
|---|---|---|---|---|---|
| Rab23 | -2.312886 | 0.012146 | 0.992917 | 1.915559 | 0.003087 |
| Ubn1 | -2.177280 | 0.002058 | 0.992917 | 2.686603 | 0.003087 |
| H1f5 | -1.946654 | 0.027814 | 0.992917 | 1.555731 | 0.003087 |
| Fsd1 | -1.843947 | 0.043041 | 0.992917 | 1.366118 | 0.003087 |
| Zfp626 | -1.747444 | 0.011253 | 0.992917 | 1.948732 | 0.003087 |
| Ddx27 | -1.715748 | 0.000546 | 0.992917 | 3.262942 | 0.003087 |
| Csmd2 | -1.710434 | 0.012899 | 0.992917 | 1.889438 | 0.003087 |
| Vps33b | -1.700719 | 0.012109 | 0.992917 | 1.916898 | 0.003087 |
| H3c1 | -1.677614 | 0.037118 | 0.992917 | 1.430412 | 0.003087 |
| Tdrkh | -1.677256 | 0.014831 | 0.992917 | 1.828818 | 0.003087 |
| Dis3l2 | -1.658280 | 0.009440 | 0.992917 | 2.025023 | 0.003087 |
| Fdxacb1 | -1.603125 | 0.017616 | 0.992917 | 1.754091 | 0.003087 |
| Brca2 | -1.583493 | 0.027582 | 0.992917 | 1.559376 | 0.003087 |
| Noc4l | -1.562373 | 0.045279 | 0.992917 | 1.344099 | 0.003087 |
| Ier3ip1 | -1.552457 | 0.040014 | 0.992917 | 1.397787 | 0.003087 |
| Igflr1 | -1.534308 | 0.029877 | 0.992917 | 1.524658 | 0.003087 |
| H4c14 | -1.495815 | 0.035504 | 0.992917 | 1.449720 | 0.003087 |
| Nacad | -1.472553 | 0.035642 | 0.992917 | 1.448040 | 0.003087 |
| Ccdc30 | -1.464779 | 0.048192 | 0.992917 | 1.317025 | 0.003087 |
| Pdyn | -1.462637 | 0.012279 | 0.992917 | 1.910829 | 0.003087 |
| 1700022I1 | -1.455063 | 0.030678 | 0.992917 | 1.513177 | 0.003087 |
| Cox11 | -1.446398 | 0.000463 | 0.992917 | 3.333955 | 0.003087 |
| Cep192 | -1.445148 | 0.044791 | 0.992917 | 1.348807 | 0.003087 |
| Gm11563 | -1.414111 | 0.009424 | 0.992917 | 2.025780 | 0.003087 |
| Fam13c | -1.381092 | 0.027491 | 0.992917 | 1.560813 | 0.003087 |
| Tlcd5 | -1.357469 | 0.006666 | 0.992917 | 2.176147 | 0.003087 |
| Pcdhgb8 | -1.357469 | 0.048104 | 0.992917 | 1.317818 | 0.003087 |
| Gpatch2 | -1.331747 | 0.019212 | 0.992917 | 1.716428 | 0.003087 |
| Pus10 | -1.293254 | 0.025112 | 0.992917 | 1.600122 | 0.003087 |
| Ddx10 | -1.293254 | 0.020532 | 0.992917 | 1.687559 | 0.003087 |
| Pou5f1 | -1.293254 | 0.014910 | 0.992917 | 1.826514 | 0.003087 |
| C1galt1c1 | -1.264958 | 0.013889 | 0.992917 | 1.857344 | 0.003087 |
| Ggt6 | -1.248278 | 0.047660 | 0.992917 | 1.321844 | 0.003087 |
| Fbxo8 | -1.236613 | 0.045120 | 0.992917 | 1.345631 | 0.003087 |
| Trappc6b | -1.208317 | 0.014422 | 0.992917 | 1.840975 | 0.003087 |
| Esyt1 | -1.205576 | 0.005461 | 0.992917 | 2.262749 | 0.003087 |
| Cma2 | -1.205576 | 0.000789 | 0.992917 | 3.103184 | 0.003087 |

| | | | | | |
|---|---|---|---|---|---|
| Nectin2 | 1.404331 | 0.047369 | 0.992917 | 1.324502 | 0.003087 |
| Hells | 1.398126 | 0.025192 | 0.992917 | 1.598745 | 0.003087 |
| Smyd4 | 1.396875 | 0.043877 | 0.992917 | 1.357765 | 0.003087 |
| Stac2 | 1.365839 | 0.048273 | 0.992917 | 1.316295 | 0.003087 |
| Oosp2 | 1.365839 | 0.006460 | 0.992917 | 2.189773 | 0.003087 |
| Reck | 1.365839 | 0.016033 | 0.992917 | 1.794992 | 0.003087 |
| Gpr63 | 1.365839 | 0.016788 | 0.992917 | 1.774992 | 0.003087 |
| Dedd2 | 1.365839 | 0.027368 | 0.992917 | 1.562750 | 0.003087 |
| Asic4 | 1.365839 | 0.027368 | 0.992917 | 1.562750 | 0.003087 |
| Prl2a1 | 1.365839 | 0.036053 | 0.992917 | 1.443063 | 0.003087 |
| Map3k6 | 1.334802 | 0.031818 | 0.992917 | 1.497324 | 0.003087 |
| Ndufaf8 | 1.334802 | 0.045951 | 0.992917 | 1.337708 | 0.003087 |
| Clec14a | 1.334802 | 0.015963 | 0.992917 | 1.796878 | 0.003087 |
| Gda | 1.330200 | 0.018730 | 0.992917 | 1.727471 | 0.003087 |
| Ctnna3 | 1.315171 | 0.003673 | 0.992917 | 2.434994 | 0.003087 |
| Tnfsfm13 | 1.310448 | 0.024606 | 0.992917 | 1.608967 | 0.003087 |
| Tgfbr3l | 1.309197 | 0.043210 | 0.992917 | 1.364421 | 0.003087 |
| Zfp786 | 1.309197 | 0.027848 | 0.992917 | 1.555200 | 0.003087 |
| Tmem106c | 1.288076 | 0.048467 | 0.992917 | 1.314557 | 0.003087 |
| Rfwd3 | 1.278161 | 0.006408 | 0.992917 | 2.193268 | 0.003087 |
| Rcvrn | 1.278161 | 0.035193 | 0.992917 | 1.453538 | 0.003087 |
| Dusp14 | 1.278161 | 0.016271 | 0.992917 | 1.788592 | 0.003087 |
| Ereg | 1.268445 | 0.021084 | 0.992917 | 1.676047 | 0.003087 |
| Cdk10 | 1.260672 | 0.021680 | 0.992917 | 1.663942 | 0.003087 |
| Nsmaf | 1.255178 | 0.011773 | 0.992917 | 1.929121 | 0.003087 |
| Gpr52 | 1.241040 | 0.046251 | 0.992917 | 1.334875 | 0.003087 |
| Hmox1 | 1.240149 | 0.025081 | 0.992917 | 1.600653 | 0.003087 |
| 1700066B1 | 1.227493 | 0.010452 | 0.992917 | 1.980787 | 0.003087 |
| Zcchc8 | 1.227493 | 0.040538 | 0.992917 | 1.392140 | 0.003087 |
| Zfp398 | 1.227493 | 0.047310 | 0.992917 | 1.325048 | 0.003087 |
| Acvrl1 | 1.216686 | 0.011518 | 0.992917 | 1.938632 | 0.003087 |
| Ccdc38 | 1.213946 | 0.016592 | 0.992917 | 1.780090 | 0.003087 |
| Cyp17a1 | 1.201888 | 0.034559 | 0.992917 | 1.461442 | 0.003087 |
| Col5a3 | 1.201888 | 0.002359 | 0.992917 | 2.627243 | 0.003087 |
| Exoc6 | 1.194314 | 0.017950 | 0.992917 | 1.745931 | 0.003087 |
| Klri2 | 1.190483 | 0.001097 | 0.992917 | 2.959614 | 0.003087 |
| Gdf9 | 1.190483 | 0.031615 | 0.992917 | 1.500100 | 0.003087 |
| Spata5 | 1.176825 | 0.041253 | 0.992917 | 1.384540 | 0.003087 |
| Fcgr2b | 1.176825 | 0.041801 | 0.992917 | 1.378810 | 0.003087 |
| Nanog | 1.157304 | 0.045446 | 0.992917 | 1.342507 | 0.003087 |
| Gm32168 | 1.147589 | 0.022247 | 0.992917 | 1.652726 | 0.003087 |

| | | | | | |
|---|---|---|---|---|---|
| Pcdhb19 | -1.205576 | 0.042157 | 0.992917 | 1.375125 | 0.003087 |
| Swap70 | -1.191059 | 0.026477 | 0.992917 | 1.577133 | 0.003087 |
| Psg21 | -1.187427 | 0.006231 | 0.992917 | 2.205475 | 0.003087 |
| Mrpl11 | -1.172398 | 0.018489 | 0.992917 | 1.733089 | 0.003087 |
| Ube2r2 | -1.162482 | 0.003608 | 0.992917 | 2.442751 | 0.003087 |
| H1f1 | -1.135962 | 0.042635 | 0.992917 | 1.370231 | 0.003087 |
| Ankrd39 | -1.131446 | 0.022406 | 0.992917 | 1.649634 | 0.003087 |
| Dpp6 | -1.089602 | 0.045783 | 0.992917 | 1.339295 | 0.003087 |
| Tom1l1 | -1.087460 | 0.049675 | 0.992917 | 1.303858 | 0.003087 |
| Lsm14b | -1.069971 | 0.008268 | 0.992917 | 2.082601 | 0.003087 |
| Tax1bp3 | -1.057315 | 0.044770 | 0.992917 | 1.349009 | 0.003087 |
| H4c17 | -1.042067 | 0.046469 | 0.992917 | 1.332832 | 0.003087 |
| Cntn6 | -1.041625 | 0.047043 | 0.992917 | 1.327509 | 0.003087 |
| Igf2bp3 | -1.034052 | 0.036630 | 0.992917 | 1.436157 | 0.003087 |
| Siva1 | -1.034052 | 0.013326 | 0.992917 | 1.875294 | 0.003087 |
| Fam204a | -1.033852 | 0.040524 | 0.992917 | 1.392287 | 0.003087 |
| Slc46a1 | -1.024136 | 0.045045 | 0.992917 | 1.346356 | 0.003087 |
| Srebf1 | -1.024136 | 0.014982 | 0.992917 | 1.824434 | 0.003087 |
| Ap1m1 | -1.014499 | 0.012088 | 0.992917 | 1.917630 | 0.003087 |
| Lyst | -1.010589 | 0.039250 | 0.992917 | 1.406163 | 0.003087 |
| Alg11 | -1.005987 | 0.025691 | 0.992917 | 1.590222 | 0.003087 |
| Frg2f10 | -1.005987 | 0.037368 | 0.992917 | 1.427501 | 0.003087 |
| Pkn1 | -1.004664 | 0.025424 | 0.992917 | 1.594749 | 0.003087 |
| Mcm4 | -0.993100 | 0.043410 | 0.992917 | 1.362407 | 0.003087 |
| P4ha2 | -0.990958 | 0.025734 | 0.992917 | 1.589497 | 0.003087 |
| Lcp2 | -0.988498 | 0.039528 | 0.992917 | 1.403093 | 0.003087 |
| 4921524J1 | -0.982293 | 0.036125 | 0.992917 | 1.442188 | 0.003087 |
| Uchl3 | -0.977410 | 0.047373 | 0.992917 | 1.324468 | 0.003087 |
| Arrb2 | -0.974950 | 0.019725 | 0.992917 | 1.704978 | 0.003087 |
| Cdk1 | -0.973468 | 0.013500 | 0.992917 | 1.869671 | 0.003087 |
| Olfm1 | -0.971485 | 0.038802 | 0.992917 | 1.411146 | 0.003087 |
| Plekha3 | -0.961010 | 0.003611 | 0.992917 | 2.442363 | 0.003087 |
| Med7 | -0.959921 | 0.028458 | 0.992917 | 1.545791 | 0.003087 |
| Commd3 | -0.957461 | 0.018599 | 0.992917 | 1.730511 | 0.003087 |
| Fbxo6 | -0.951256 | 0.032503 | 0.992917 | 1.488075 | 0.003087 |
| Rbbp4 | -0.946270 | 0.010041 | 0.992917 | 1.998205 | 0.003087 |
| Cdk11b | -0.938363 | 0.032189 | 0.992917 | 1.492287 | 0.003087 |
| Thnsl1 | -0.936458 | 0.014741 | 0.992917 | 1.831487 | 0.003087 |
| Dnah7b | -0.936458 | 0.044859 | 0.992917 | 1.348148 | 0.003087 |
| Mdm1 | -0.936458 | 0.044859 | 0.992917 | 1.348148 | 0.003087 |
| Scyl3 | -0.926743 | 0.004706 | 0.992917 | 2.327346 | 0.003087 |

| | | | | | |
|---|---|---|---|---|---|
| Rab9b | 1.147589 | 0.022247 | 0.992917 | 1.652726 | 0.003087 |
| Rlf | 1.146501 | 0.039691 | 0.992917 | 1.401307 | 0.003087 |
| Zbtb1 | 1.139815 | 0.017837 | 0.992917 | 1.748676 | 0.003087 |
| Tek | 1.139815 | 0.029180 | 0.992917 | 1.534916 | 0.003087 |
| Dap | 1.137673 | 0.013024 | 0.992917 | 1.885249 | 0.003087 |
| Phyhip | 1.134026 | 0.008492 | 0.992917 | 2.070989 | 0.003087 |
| Prkcd | 1.130331 | 0.041265 | 0.992917 | 1.384422 | 0.003087 |
| Ccdc158 | 1.126268 | 0.007347 | 0.992917 | 2.133865 | 0.003087 |
| Epop | 1.125875 | 0.009189 | 0.992917 | 2.036742 | 0.003087 |
| Cul4a | 1.120184 | 0.012795 | 0.992917 | 1.892962 | 0.003087 |
| Wdr70 | 1.120184 | 0.015386 | 0.992917 | 1.812864 | 0.003087 |
| Gm46913 | 1.116552 | 0.008033 | 0.992917 | 2.095095 | 0.003087 |
| Tesk1 | 1.115582 | 0.025570 | 0.992917 | 1.592274 | 0.003087 |
| Epg5 | 1.106636 | 0.030606 | 0.992917 | 1.514187 | 0.003087 |
| Olfr618 | 1.106636 | 0.001081 | 0.992917 | 2.966164 | 0.003087 |
| Ap1s3 | 1.106636 | 0.005094 | 0.992917 | 2.292910 | 0.003087 |
| Apip | 1.106636 | 0.023098 | 0.992917 | 1.636418 | 0.003087 |
| Pcdhb20 | 1.096721 | 0.005201 | 0.992917 | 2.283916 | 0.003087 |
| Relb | 1.089147 | 0.027178 | 0.992917 | 1.565789 | 0.003087 |
| Fstl1 | 1.083173 | 0.008649 | 0.992917 | 2.063011 | 0.003087 |
| Lmtk3 | 1.080482 | 0.024871 | 0.992917 | 1.604300 | 0.003087 |
| H6pd | 1.073458 | 0.038727 | 0.992917 | 1.411988 | 0.003087 |
| Pnmal1 | 1.073458 | 0.041563 | 0.992917 | 1.381295 | 0.003087 |
| Ptprj | 1.070998 | 0.005937 | 0.992917 | 2.226398 | 0.003087 |
| Tinagl1 | 1.069516 | 0.027044 | 0.992917 | 1.567924 | 0.003087 |
| Rsl24d1 | 1.065684 | 0.039500 | 0.992917 | 1.403403 | 0.003087 |
| Ntng1 | 1.049995 | 0.044757 | 0.992917 | 1.349144 | 0.003087 |
| Tcp11l1 | 1.042421 | 0.031916 | 0.992917 | 1.495992 | 0.003087 |
| Tsc2 | 1.035360 | 0.030266 | 0.992917 | 1.519047 | 0.003087 |
| Malsu1 | 1.033987 | 0.032211 | 0.992917 | 1.491997 | 0.003087 |
| Nhlrc3 | 1.032506 | 0.039067 | 0.992917 | 1.408186 | 0.003087 |
| Gm36375 | 1.032506 | 0.021460 | 0.992917 | 1.668379 | 0.003087 |
| Runx2 | 1.032506 | 0.036493 | 0.992917 | 1.437791 | 0.003087 |
| Clec12a | 1.032506 | 0.042575 | 0.992917 | 1.370843 | 0.003087 |
| Celf4 | 1.024932 | 0.029895 | 0.992917 | 1.524408 | 0.003087 |
| Six3 | 1.018958 | 0.011483 | 0.992917 | 1.939949 | 0.003087 |
| Mkks | 1.004209 | 0.039094 | 0.992917 | 1.407895 | 0.003087 |
| 4931406B1 | 0.999327 | 0.037758 | 0.992917 | 1.422988 | 0.003087 |
| Grik1 | 0.995495 | 0.026724 | 0.992917 | 1.573099 | 0.003087 |
| Mfhas1 | 0.991553 | 0.016241 | 0.992917 | 1.789386 | 0.003087 |
| Shisa7 | 0.990662 | 0.002409 | 0.992917 | 2.618152 | 0.003087 |

| | | | | | |
|---|---|---|---|---|---|
| Dusp1 | -0.925651 | 0.041460 | 0.992917 | 1.382366 | 0.003087 |
| Naa25 | -0.922911 | 0.026292 | 0.992917 | 1.580173 | 0.003087 |
| Faxc | -0.920818 | 0.022553 | 0.992917 | 1.646800 | 0.003087 |
| Akt2 | -0.918078 | 0.002756 | 0.992917 | 2.559664 | 0.003087 |
| Gm20172 | -0.916827 | 0.034285 | 0.992917 | 1.464896 | 0.003087 |
| Sra1 | -0.914945 | 0.037394 | 0.992917 | 1.427202 | 0.003087 |
| Ankrd45 | -0.895706 | 0.019419 | 0.992917 | 1.711783 | 0.003087 |
| Yrdc | -0.894614 | 0.039113 | 0.992917 | 1.407683 | 0.003087 |
| Prpf8 | -0.893941 | 0.036691 | 0.992917 | 1.435438 | 0.003087 |
| Zcrb1 | -0.893246 | 0.024966 | 0.992917 | 1.602650 | 0.003087 |
| Pcna | -0.891104 | 0.013848 | 0.992917 | 1.858610 | 0.003087 |
| Sall2 | -0.885790 | 0.042758 | 0.992917 | 1.368983 | 0.003087 |
| Scn3a | -0.885790 | 0.007774 | 0.992917 | 2.109376 | 0.003087 |
| Aamp | -0.882813 | 0.017511 | 0.992917 | 1.756682 | 0.003087 |
| Mmp3 | -0.879816 | 0.039371 | 0.992917 | 1.404822 | 0.003087 |
| Ulk2 | -0.872243 | 0.007331 | 0.992917 | 2.134844 | 0.003087 |
| Umad1 | -0.872243 | 0.031785 | 0.992917 | 1.497783 | 0.003087 |
| Nt5c3 | -0.870101 | 0.028814 | 0.992917 | 1.540402 | 0.003087 |
| Astn2 | -0.857494 | 0.015744 | 0.992917 | 1.802895 | 0.003087 |
| Mmp15 | -0.852458 | 0.045585 | 0.992917 | 1.341181 | 0.003087 |
| Gpkow | -0.848010 | 0.031957 | 0.992917 | 1.495427 | 0.003087 |
| Drg2 | -0.843947 | 0.031175 | 0.992917 | 1.506200 | 0.003087 |
| Plppr3 | -0.843947 | 0.038174 | 0.992917 | 1.418229 | 0.003087 |
| Penk | -0.839064 | 0.048804 | 0.992917 | 1.311545 | 0.003087 |
| Pkd2l1 | -0.835123 | 0.041049 | 0.992917 | 1.386697 | 0.003087 |
| H1f4 | -0.833566 | 0.032806 | 0.992917 | 1.484051 | 0.003087 |
| Mettl11b | -0.831291 | 0.007507 | 0.992917 | 2.124524 | 0.003087 |
| Pbrm1 | -0.829149 | 0.007739 | 0.992917 | 2.111328 | 0.003087 |
| Ppan | -0.829149 | 0.008367 | 0.992917 | 2.077433 | 0.003087 |
| Gm3233 | -0.817743 | 0.037661 | 0.992917 | 1.424111 | 0.003087 |
| Dtx4 | -0.815601 | 0.030376 | 0.992917 | 1.517466 | 0.003087 |
| Raly | -0.812468 | 0.007750 | 0.992917 | 2.110713 | 0.003087 |
| Rps15a | -0.809387 | 0.008629 | 0.992917 | 2.064053 | 0.003087 |
| Zbtb43 | -0.798112 | 0.029499 | 0.992917 | 1.530192 | 0.003087 |
| Amz2 | -0.795970 | 0.015141 | 0.992917 | 1.819847 | 0.003087 |
| Pigu | -0.787305 | 0.015902 | 0.992917 | 1.798551 | 0.003087 |
| Ldah | -0.785937 | 0.010449 | 0.992917 | 1.980936 | 0.003087 |
| Taar6 | -0.784565 | 0.028981 | 0.992917 | 1.537884 | 0.003087 |
| Scrn1 | -0.769560 | 0.046886 | 0.992917 | 1.328954 | 0.003087 |
| Hpca | -0.768904 | 0.016106 | 0.992917 | 1.793009 | 0.003087 |
| Siah1a | -0.763683 | 0.040771 | 0.992917 | 1.389647 | 0.003087 |

| | | | | | |
|---|---|---|---|---|---|
| Ptpro | 0.990662 | 0.010862 | 0.992917 | 1.964098 | 0.003087 |
| Bri3bp | 0.987529 | 0.040353 | 0.992917 | 1.394129 | 0.003087 |
| Slc26a4 | 0.985780 | 0.024879 | 0.992917 | 1.604171 | 0.003087 |
| Inhbb | 0.985780 | 0.039380 | 0.992917 | 1.404721 | 0.003087 |
| Slit2 | 0.981838 | 0.043093 | 0.992917 | 1.365593 | 0.003087 |
| Brix1 | 0.968290 | 0.006742 | 0.992917 | 2.171237 | 0.003087 |
| Kti12 | 0.968290 | 0.018654 | 0.992917 | 1.729221 | 0.003087 |
| Nudt15 | 0.968290 | 0.045058 | 0.992917 | 1.346227 | 0.003087 |
| Septin1 | 0.968290 | 0.039725 | 0.992917 | 1.400936 | 0.003087 |
| Tasor2 | 0.963689 | 0.008027 | 0.992917 | 2.095446 | 0.003087 |
| Olfr123 | 0.962317 | 0.030285 | 0.992917 | 1.518771 | 0.003087 |
| Scube1 | 0.944827 | 0.012983 | 0.992917 | 1.886619 | 0.003087 |
| Rhox2f | 0.935112 | 0.002839 | 0.992917 | 2.546868 | 0.003087 |
| Arvcf | 0.934020 | 0.037334 | 0.992917 | 1.427890 | 0.003087 |
| Orai3 | 0.925196 | 0.044171 | 0.992917 | 1.354859 | 0.003087 |
| Calcr | 0.911649 | 0.045252 | 0.992917 | 1.344366 | 0.003087 |
| Rab11fip1 | 0.907707 | 0.033960 | 0.992917 | 1.469029 | 0.003087 |
| Fkbp9 | 0.907424 | 0.012826 | 0.992917 | 1.891921 | 0.003087 |
| Rspo2 | 0.906816 | 0.034492 | 0.992917 | 1.462278 | 0.003087 |
| Spry1 | 0.901733 | 0.043466 | 0.992917 | 1.361850 | 0.003087 |
| Fancg | 0.894160 | 0.048395 | 0.992917 | 1.315196 | 0.003087 |
| Purg | 0.885365 | 0.017997 | 0.992917 | 1.744802 | 0.003087 |
| Dennd2c | 0.880612 | 0.013787 | 0.992917 | 1.860535 | 0.003087 |
| Slco3a1 | 0.878519 | 0.024399 | 0.992917 | 1.612625 | 0.003087 |
| Fam78a | 0.870897 | 0.022808 | 0.992917 | 1.641921 | 0.003087 |
| Aftph | 0.867186 | 0.027775 | 0.992917 | 1.556348 | 0.003087 |
| Slc38a5 | 0.860981 | 0.029735 | 0.992917 | 1.526728 | 0.003087 |
| Atl3 | 0.858521 | 0.037445 | 0.992917 | 1.426602 | 0.003087 |
| Col23a1 | 0.837518 | 0.008452 | 0.992917 | 2.073020 | 0.003087 |
| Lypd4 | 0.837518 | 0.005124 | 0.992917 | 2.290399 | 0.003087 |
| 1700020D( | 0.837518 | 0.049623 | 0.992917 | 1.304316 | 0.003087 |
| Mrps36 | 0.829945 | 0.035772 | 0.992917 | 1.446459 | 0.003087 |
| Nell1 | 0.811403 | 0.006672 | 0.992917 | 2.175742 | 0.003087 |
| 2900026A( | 0.806482 | 0.005157 | 0.992917 | 2.287631 | 0.003087 |
| Ctbp2 | 0.806482 | 0.046815 | 0.992917 | 1.329615 | 0.003087 |
| S100a11 | 0.804339 | 0.037319 | 0.992917 | 1.428071 | 0.003087 |
| Rgs17 | 0.801880 | 0.028101 | 0.992917 | 1.551278 | 0.003087 |
| Mme | 0.800398 | 0.003472 | 0.992917 | 2.459360 | 0.003087 |
| Bambi | 0.796766 | 0.036071 | 0.992917 | 1.442842 | 0.003087 |
| Dnajc1 | 0.794306 | 0.006360 | 0.992917 | 2.196541 | 0.003087 |
| Slc2a9 | 0.786850 | 0.011533 | 0.992917 | 1.938069 | 0.003087 |

| | | | | | |
|---|---|---|---|---|---|
| Prob1 | -0.762474 | 0.005883 | 0.992917 | 2.230370 | 0.003087 |
| 1600012H( | -0.761582 | 0.027639 | 0.992917 | 1.558477 | 0.003087 |
| Nt5c2 | -0.757360 | 0.029887 | 0.992917 | 1.524521 | 0.003087 |
| Ddit4 | -0.747757 | 0.025626 | 0.992917 | 1.591318 | 0.003087 |
| Tm9sf3 | -0.747444 | 0.043741 | 0.992917 | 1.359110 | 0.003087 |
| Heatr5b | -0.745302 | 0.037275 | 0.992917 | 1.428581 | 0.003087 |
| Dctpp1 | -0.745302 | 0.001093 | 0.992917 | 2.961432 | 0.003087 |
| Coq5 | -0.742843 | 0.029683 | 0.992917 | 1.527498 | 0.003087 |
| Wdr88 | -0.741471 | 0.030282 | 0.992917 | 1.518819 | 0.003087 |
| Olfr1446 | -0.741471 | 0.019280 | 0.992917 | 1.714901 | 0.003087 |
| Snap29 | -0.736637 | 0.015319 | 0.992917 | 1.814760 | 0.003087 |
| Gm42372 | -0.736637 | 0.037760 | 0.992917 | 1.422973 | 0.003087 |
| Hmgn1 | -0.734551 | 0.045557 | 0.992917 | 1.341441 | 0.003087 |
| H2aw | -0.725530 | 0.049694 | 0.992917 | 1.303696 | 0.003087 |
| Cct3 | -0.724547 | 0.010914 | 0.992917 | 1.961997 | 0.003087 |
| Tomm7 | -0.724182 | 0.031997 | 0.992917 | 1.494892 | 0.003087 |
| Mybbp1a | -0.721839 | 0.045507 | 0.992917 | 1.341918 | 0.003087 |
| Tjp2 | -0.720691 | 0.042911 | 0.992917 | 1.367429 | 0.003087 |
| Phldb1 | -0.720094 | 0.009040 | 0.992917 | 2.043817 | 0.003087 |
| Napa | -0.713125 | 0.044998 | 0.992917 | 1.346807 | 0.003087 |
| Lrrc8c | -0.710434 | 0.006499 | 0.992917 | 2.187174 | 0.003087 |
| Adam9 | -0.710434 | 0.026005 | 0.992917 | 1.584944 | 0.003087 |
| Gba | -0.708139 | 0.008376 | 0.992917 | 2.076939 | 0.003087 |
| H2ax | -0.708139 | 0.018541 | 0.992917 | 1.731877 | 0.003087 |
| Slc35f1 | -0.703515 | 0.027240 | 0.992917 | 1.564795 | 0.003087 |
| Atp2b4 | -0.703459 | 0.034517 | 0.992917 | 1.461961 | 0.003087 |
| Pcdhga12 | -0.702208 | 0.006318 | 0.992917 | 2.199447 | 0.003087 |
| Cuedc2 | -0.700326 | 0.037588 | 0.992917 | 1.424951 | 0.003087 |
| Impa1 | -0.687452 | 0.018418 | 0.992917 | 1.734751 | 0.003087 |
| Stat2 | -0.677256 | 0.017454 | 0.992917 | 1.758094 | 0.003087 |
| Eaf1 | -0.667340 | 0.001062 | 0.992917 | 2.973905 | 0.003087 |
| Rimklb | -0.667340 | 0.049320 | 0.992917 | 1.306975 | 0.003087 |
| Sppl3 | -0.665159 | 0.041854 | 0.992917 | 1.378260 | 0.003087 |
| Kansl2 | -0.663513 | 0.006991 | 0.992917 | 2.155474 | 0.003087 |
| Actn2 | -0.659766 | 0.004083 | 0.992917 | 2.388972 | 0.003087 |
| Psma7 | -0.658379 | 0.021016 | 0.992917 | 1.677453 | 0.003087 |
| Rps3a1 | -0.650185 | 0.038843 | 0.992917 | 1.410684 | 0.003087 |
| Ripor1 | -0.649851 | 0.040914 | 0.992917 | 1.388133 | 0.003087 |
| Rbp4 | -0.644077 | 0.047288 | 0.992917 | 1.325250 | 0.003087 |
| Phc1 | -0.630810 | 0.047015 | 0.992917 | 1.327763 | 0.003087 |
| Pdik1l | -0.626588 | 0.000528 | 0.992917 | 3.277463 | 0.003087 |

| | | | | | |
|---|---|---|---|---|---|
| 1700012A( | 0.785959 | 0.019844 | 0.992917 | 1.702380 | 0.003087 |
| Cyp20a1 | 0.773303 | 0.007948 | 0.992917 | 2.099753 | 0.003087 |
| Calcb | 0.763387 | 0.044674 | 0.992917 | 1.349942 | 0.003087 |
| Sugp1 | 0.749840 | 0.039088 | 0.992917 | 1.407959 | 0.003087 |
| Cphx1 | 0.749840 | 0.040140 | 0.992917 | 1.396424 | 0.003087 |
| Noc2l | 0.745007 | 0.006010 | 0.992917 | 2.221142 | 0.003087 |
| Ces1b | 0.736183 | 0.043764 | 0.992917 | 1.358884 | 0.003087 |
| Pcdhb18 | 0.732351 | 0.037278 | 0.992917 | 1.428546 | 0.003087 |
| Sc5d | 0.732091 | 0.005539 | 0.992917 | 2.256576 | 0.003087 |
| Pycard | 0.730209 | 0.000789 | 0.992917 | 3.103184 | 0.003087 |
| Slirp | 0.725173 | 0.047930 | 0.992917 | 1.319392 | 0.003087 |
| Gpr3 | 0.722635 | 0.031458 | 0.992917 | 1.502271 | 0.003087 |
| Epdr1 | 0.713528 | 0.000063 | 0.886668 | 4.202320 | 0.052239 |
| Col6a1 | 0.709088 | 0.014509 | 0.992917 | 1.838357 | 0.003087 |
| Ncoa4 | 0.708888 | 0.035572 | 0.992917 | 1.448894 | 0.003087 |
| Mtus2 | 0.699172 | 0.032182 | 0.992917 | 1.492393 | 0.003087 |
| Orm2 | 0.699172 | 0.049214 | 0.992917 | 1.307913 | 0.003087 |
| Ly6c1 | 0.698935 | 0.013521 | 0.992917 | 1.868997 | 0.003087 |
| Stk4 | 0.690507 | 0.024637 | 0.992917 | 1.608412 | 0.003087 |
| Col6a2 | 0.689256 | 0.033792 | 0.992917 | 1.471188 | 0.003087 |
| Mrps33 | 0.686041 | 0.001208 | 0.992917 | 2.917977 | 0.003087 |
| Flrt1 | 0.685625 | 0.034522 | 0.992917 | 1.461910 | 0.003087 |
| Tmem143 | 0.679541 | 0.012199 | 0.992917 | 1.913677 | 0.003087 |
| Csrnp1 | 0.665994 | 0.033448 | 0.992917 | 1.475624 | 0.003087 |
| Tax1bp1 | 0.662347 | 0.007894 | 0.992917 | 2.102722 | 0.003087 |
| Fam149a | 0.662162 | 0.044530 | 0.992917 | 1.351343 | 0.003087 |
| Slc2a8 | 0.661319 | 0.022023 | 0.992917 | 1.657119 | 0.003087 |
| Fgf2 | 0.652446 | 0.043539 | 0.992917 | 1.361124 | 0.003087 |
| Ppp1r12b | 0.646521 | 0.017080 | 0.992917 | 1.767514 | 0.003087 |
| Sidt1 | 0.642531 | 0.022680 | 0.992917 | 1.644362 | 0.003087 |
| Lratd2 | 0.642531 | 0.036395 | 0.992917 | 1.438955 | 0.003087 |
| Fbxo11 | 0.638506 | 0.017080 | 0.992917 | 1.767513 | 0.003087 |
| P2rx6 | 0.634957 | 0.049669 | 0.992917 | 1.303919 | 0.003087 |
| Gas2 | 0.634957 | 0.005411 | 0.992917 | 2.266710 | 0.003087 |
| Prdm11 | 0.634957 | 0.014414 | 0.992917 | 1.841230 | 0.003087 |
| Cpne2 | 0.621690 | 0.046811 | 0.992917 | 1.329652 | 0.003087 |
| Odf2l | 0.617468 | 0.022598 | 0.992917 | 1.645922 | 0.003087 |
| Sco2 | 0.601779 | 0.006381 | 0.992917 | 2.195130 | 0.003087 |
| Myh4 | 0.601779 | 0.047250 | 0.992917 | 1.325598 | 0.003087 |
| Paxbp1 | 0.600687 | 0.041452 | 0.992917 | 1.382450 | 0.003087 |
| Plekhg1 | 0.586827 | 0.027841 | 0.992917 | 1.555320 | 0.003087 |

| | | | | | |
|---|---|---|---|---|---|
| 2810004N2 | -0.624128 | 0.045182 | 0.992917 | 1.345034 | 0.003087 |
| Nfat5 | -0.624128 | 0.031463 | 0.992917 | 1.502194 | 0.003087 |
| Zbtb20 | -0.619882 | 0.049832 | 0.992917 | 1.302488 | 0.003087 |
| Kmt5b | -0.614130 | 0.002336 | 0.992917 | 2.631607 | 0.003087 |
| Smarce1 | -0.610698 | 0.027196 | 0.992917 | 1.565494 | 0.003087 |
| Ak3 | -0.607116 | 0.006340 | 0.992917 | 2.197915 | 0.003087 |
| Taf15 | -0.605007 | 0.045974 | 0.992917 | 1.337489 | 0.003087 |
| Dazap1 | -0.603125 | 0.021184 | 0.992917 | 1.673996 | 0.003087 |
| Olfr1106 | -0.603125 | 0.042597 | 0.992917 | 1.370625 | 0.003087 |
| Rpl12 | -0.602302 | 0.009073 | 0.992917 | 2.042269 | 0.003087 |
| Gas6 | -0.597151 | 0.007711 | 0.992917 | 2.112904 | 0.003087 |
| Gm36327 | -0.595855 | 0.048743 | 0.992917 | 1.312088 | 0.003087 |
| Nisch | -0.589185 | 0.046552 | 0.992917 | 1.332059 | 0.003087 |
| Marcks | -0.588118 | 0.000357 | 0.992917 | 3.447883 | 0.003087 |
| Msi2 | -0.582658 | 0.027867 | 0.992917 | 1.554912 | 0.003087 |
| Ina | -0.578458 | 0.047303 | 0.992917 | 1.325115 | 0.003087 |
| Dad1 | -0.578260 | 0.009490 | 0.992917 | 2.022744 | 0.003087 |
| Phc2 | -0.576151 | 0.020307 | 0.992917 | 1.692349 | 0.003087 |
| Psma4 | -0.573970 | 0.043327 | 0.992917 | 1.363238 | 0.003087 |
| Cfap20 | -0.568181 | 0.038653 | 0.992917 | 1.412821 | 0.003087 |
| Kank1 | -0.562172 | 0.038982 | 0.992917 | 1.409136 | 0.003087 |
| 1500009C( | -0.561876 | 0.028290 | 0.992917 | 1.548373 | 0.003087 |
| Ppm1m | -0.557771 | 0.049951 | 0.992917 | 1.301456 | 0.003087 |
| Swi5 | -0.557418 | 0.010435 | 0.992917 | 1.981505 | 0.003087 |
| Rps27a | -0.556855 | 0.017199 | 0.992917 | 1.764503 | 0.003087 |
| Akr1a1 | -0.555115 | 0.021617 | 0.992917 | 1.665213 | 0.003087 |
| Hdgfl3 | -0.554400 | 0.024401 | 0.992917 | 1.612600 | 0.003087 |
| Hsp90aa1 | -0.549630 | 0.026050 | 0.992917 | 1.584196 | 0.003087 |
| Zcchc12 | -0.546483 | 0.030301 | 0.992917 | 1.518541 | 0.003087 |
| Sash1 | -0.543927 | 0.025066 | 0.992917 | 1.600916 | 0.003087 |
| Dtnb | -0.541650 | 0.043825 | 0.992917 | 1.358277 | 0.003087 |
| Abcd1 | -0.541650 | 0.043825 | 0.992917 | 1.358277 | 0.003087 |
| Phb2 | -0.530245 | 0.048471 | 0.992917 | 1.314522 | 0.003087 |
| H2az1 | -0.528464 | 0.025154 | 0.992917 | 1.599389 | 0.003087 |
| H2ac18 | -0.527944 | 0.015399 | 0.992917 | 1.812499 | 0.003087 |
| Pwp1 | -0.518996 | 0.042594 | 0.992917 | 1.370650 | 0.003087 |
| Cfap61 | -0.515447 | 0.012790 | 0.992917 | 1.893127 | 0.003087 |
| Fam102a | -0.511878 | 0.049583 | 0.992917 | 1.304666 | 0.003087 |
| Zfp101 | -0.509363 | 0.021403 | 0.992917 | 1.669533 | 0.003087 |
| Mcur1 | -0.509363 | 0.029398 | 0.992917 | 1.531689 | 0.003087 |
| Rhoa | -0.508512 | 0.031581 | 0.992917 | 1.500569 | 0.003087 |

| | | | | | |
|---|---|---|---|---|---|
| Rmc1 | 0.579207 | 0.036445 | 0.992917 | 1.438357 | 0.003087 |
| Polr3gl | 0.579207 | 0.037295 | 0.992917 | 1.428353 | 0.003087 |
| Fat4 | 0.574374 | 0.040175 | 0.992917 | 1.396048 | 0.003087 |
| Nucb1 | 0.573714 | 0.044979 | 0.992917 | 1.346994 | 0.003087 |
| Slc12a6 | 0.565660 | 0.040179 | 0.992917 | 1.395997 | 0.003087 |
| Mcee | 0.558684 | 0.029575 | 0.992917 | 1.529071 | 0.003087 |
| Ppm1g | 0.554852 | 0.017792 | 0.992917 | 1.749770 | 0.003087 |
| Slc27a4 | 0.554852 | 0.041955 | 0.992917 | 1.377213 | 0.003087 |
| Dtwd2 | 0.554852 | 0.009321 | 0.992917 | 2.030516 | 0.003087 |
| Rab6b | 0.551386 | 0.040626 | 0.992917 | 1.391192 | 0.003087 |
| Fam189b | 0.550019 | 0.015062 | 0.992917 | 1.822130 | 0.003087 |
| Fbxl17 | 0.547362 | 0.017979 | 0.992917 | 1.745246 | 0.003087 |
| Tmem63a | 0.539128 | 0.044076 | 0.992917 | 1.355793 | 0.003087 |
| Snx1 | 0.530502 | 0.018425 | 0.992917 | 1.734598 | 0.003087 |
| Pex13 | 0.528539 | 0.028890 | 0.992917 | 1.539253 | 0.003087 |
| Chp2 | 0.527648 | 0.021137 | 0.992917 | 1.674947 | 0.003087 |
| Slc10a4l | 0.527648 | 0.033299 | 0.992917 | 1.477575 | 0.003087 |
| Tmem200b | 0.514100 | 0.035567 | 0.992917 | 1.448955 | 0.003087 |
| Esf1 | 0.504185 | 0.028875 | 0.992917 | 1.539476 | 0.003087 |
| Ppp1r14c | 0.494469 | 0.037012 | 0.992917 | 1.431661 | 0.003087 |
| Grik5 | 0.490896 | 0.034659 | 0.992917 | 1.460184 | 0.003087 |
| Vac14 | 0.490637 | 0.017992 | 0.992917 | 1.744921 | 0.003087 |
| Amn1 | 0.485804 | 0.044062 | 0.992917 | 1.355938 | 0.003087 |
| Enkd1 | 0.485804 | 0.018260 | 0.992917 | 1.738499 | 0.003087 |
| S100a4 | 0.477139 | 0.005576 | 0.992917 | 2.253692 | 0.003087 |
| Plekhb2 | 0.477139 | 0.010899 | 0.992917 | 1.962620 | 0.003087 |
| Srek1 | 0.471897 | 0.030730 | 0.992917 | 1.512441 | 0.003087 |
| Hmg20a | 0.465692 | 0.047415 | 0.992917 | 1.324082 | 0.003087 |
| Cap2 | 0.462341 | 0.024300 | 0.992917 | 1.614387 | 0.003087 |
| Fgf9 | 0.449685 | 0.048505 | 0.992917 | 1.314212 | 0.003087 |
| Qsox1 | 0.447543 | 0.021495 | 0.992917 | 1.667654 | 0.003087 |
| Plekhm1 | 0.439970 | 0.045837 | 0.992917 | 1.338787 | 0.003087 |
| Sccpdh | 0.430404 | 0.006250 | 0.992917 | 2.204096 | 0.003087 |
| Ell2 | 0.425172 | 0.008226 | 0.992917 | 2.084820 | 0.003087 |
| Mrps21 | 0.387537 | 0.003683 | 0.992917 | 2.433768 | 0.003087 |
| Sec14l2 | 0.386842 | 0.048470 | 0.992917 | 1.314526 | 0.003087 |
| Trabd2b | 0.385470 | 0.028058 | 0.992917 | 1.551946 | 0.003087 |
| Galnt2 | 0.374663 | 0.045665 | 0.992917 | 1.340414 | 0.003087 |
| Cops9 | 0.370672 | 0.041072 | 0.992917 | 1.386453 | 0.003087 |
| Tmem151b | 0.364588 | 0.041351 | 0.992917 | 1.383519 | 0.003087 |
| D830031N | 0.360525 | 0.040916 | 0.992917 | 1.388104 | 0.003087 |

| | | | | | |
|---|---|---|---|---|---|
| Cgrrf1 | -0.505531 | 0.041577 | 0.992917 | 1.381146 | 0.003087 |
| Mthfs | -0.501789 | 0.038155 | 0.992917 | 1.418448 | 0.003087 |
| Vmn1r113 | -0.500604 | 0.021397 | 0.992917 | 1.669643 | 0.003087 |
| Smarcb1 | -0.499878 | 0.016627 | 0.992917 | 1.779193 | 0.003087 |
| E2f6 | -0.493355 | 0.009510 | 0.992917 | 2.021838 | 0.003087 |
| Stip1 | -0.493355 | 0.039232 | 0.992917 | 1.406357 | 0.003087 |
| Rpl3 | -0.491200 | 0.003861 | 0.992917 | 2.413270 | 0.003087 |
| Spred2 | -0.490780 | 0.037146 | 0.992917 | 1.430084 | 0.003087 |
| Nol7 | -0.488896 | 0.029587 | 0.992917 | 1.528892 | 0.003087 |
| Tubb5 | -0.485106 | 0.016337 | 0.992917 | 1.786838 | 0.003087 |
| Spry2 | -0.483630 | 0.024179 | 0.992917 | 1.616566 | 0.003087 |
| Sbk1 | -0.478604 | 0.044574 | 0.992917 | 1.350916 | 0.003087 |
| Rbm7 | -0.477666 | 0.017072 | 0.992917 | 1.767722 | 0.003087 |
| Rpl27 | -0.473793 | 0.027424 | 0.992917 | 1.561867 | 0.003087 |
| Ptma | -0.473212 | 0.042179 | 0.992917 | 1.374908 | 0.003087 |
| Cdh12 | -0.464779 | 0.022614 | 0.992917 | 1.645622 | 0.003087 |
| Rnf122 | -0.447290 | 0.032035 | 0.992917 | 1.494373 | 0.003087 |
| Mrpl46 | -0.445148 | 0.042581 | 0.992917 | 1.370783 | 0.003087 |
| Ip6k1 | -0.443973 | 0.022865 | 0.992917 | 1.640825 | 0.003087 |
| Pcbp1 | -0.439540 | 0.013254 | 0.992917 | 1.877644 | 0.003087 |
| Khsrp | -0.436856 | 0.015767 | 0.992917 | 1.802252 | 0.003087 |
| Itgb8 | -0.433426 | 0.035584 | 0.992917 | 1.448740 | 0.003087 |
| Rorb | -0.431600 | 0.028730 | 0.992917 | 1.541667 | 0.003087 |
| Ppp1r14b | -0.429642 | 0.029161 | 0.992917 | 1.535203 | 0.003087 |
| Csprs | -0.426286 | 0.045316 | 0.992917 | 1.343747 | 0.003087 |
| Tubb3 | -0.422430 | 0.003592 | 0.992917 | 2.444675 | 0.003087 |
| Sfpq | -0.417094 | 0.015658 | 0.992917 | 1.805252 | 0.003087 |
| Tbl1x | -0.416851 | 0.042689 | 0.992917 | 1.369679 | 0.003087 |
| Rpl41 | -0.412138 | 0.011696 | 0.992917 | 1.931949 | 0.003087 |
| Tectb | -0.408137 | 0.022908 | 0.992917 | 1.640015 | 0.003087 |
| Atp5b | -0.403186 | 0.047985 | 0.992917 | 1.318896 | 0.003087 |
| Rps4x | -0.393812 | 0.017122 | 0.992917 | 1.766453 | 0.003087 |
| Rpl15 | -0.393042 | 0.030841 | 0.992917 | 1.510878 | 0.003087 |
| Nrf1 | -0.390648 | 0.033049 | 0.992917 | 1.480843 | 0.003087 |
| Atp5pb | -0.389818 | 0.027613 | 0.992917 | 1.558883 | 0.003087 |
| Dnaja2 | -0.380050 | 0.039903 | 0.992917 | 1.398993 | 0.003087 |
| Cdk5rap1 | -0.377101 | 0.041956 | 0.992917 | 1.377205 | 0.003087 |
| Exoc6b | -0.377101 | 0.044476 | 0.992917 | 1.351874 | 0.003087 |
| Capza2 | -0.375831 | 0.016570 | 0.992917 | 1.780677 | 0.003087 |
| Ptprz1 | -0.373294 | 0.046180 | 0.992917 | 1.335551 | 0.003087 |
| Phlpp1 | -0.360577 | 0.026815 | 0.992917 | 1.571621 | 0.003087 |

| | | | | | |
|---|---|---|---|---|---|
| Wasf2 | 0.354472 | 0.026198 | 0.992917 | 1.581729 | 0.003087 |
| Cstdc5 | 0.346208 | 0.032582 | 0.992917 | 1.487025 | 0.003087 |
| Ube2e1 | 0.345116 | 0.031785 | 0.992917 | 1.497783 | 0.003087 |
| Pura | 0.334665 | 0.046230 | 0.992917 | 1.335077 | 0.003087 |
| Sdc4 | 0.309616 | 0.023163 | 0.992917 | 1.635213 | 0.003087 |
| Hcfc1 | 0.297022 | 0.008781 | 0.992917 | 2.056453 | 0.003087 |
| Mcl1 | 0.296427 | 0.038785 | 0.992917 | 1.411335 | 0.003087 |
| Spock2 | 0.294484 | 0.009304 | 0.992917 | 2.031342 | 0.003087 |
| Prr13 | 0.288968 | 0.007593 | 0.992917 | 2.119558 | 0.003087 |
| Drosha | 0.287499 | 0.013916 | 0.992917 | 1.856486 | 0.003087 |

| | | | | | |
|---|---|---|---|---|---|
| Capn10 | -0.343922 | 0.032582 | 0.992917 | 1.487025 | 0.003087 |
| Tmsb4x | -0.334691 | 0.012665 | 0.992917 | 1.897390 | 0.003087 |
| Gnai2 | -0.314851 | 0.013198 | 0.992917 | 1.879494 | 0.003087 |
| Polr2h | -0.311833 | 0.016407 | 0.992917 | 1.784981 | 0.003087 |
| Rps24 | -0.310136 | 0.002727 | 0.992917 | 2.564394 | 0.003087 |
| 2610507B1 | -0.302297 | 0.005855 | 0.992917 | 2.232490 | 0.003087 |
| Ppp2r2c | -0.302102 | 0.012298 | 0.992917 | 1.910173 | 0.003087 |
| Skp1a | -0.290884 | 0.020322 | 0.992917 | 1.692027 | 0.003087 |
| Zeb2 | -0.287941 | 0.042816 | 0.992917 | 1.368395 | 0.003087 |
| Ubb | -0.280582 | 0.028217 | 0.992917 | 1.549490 | 0.003087 |
| Rpl23a | -0.275851 | 0.033933 | 0.992917 | 1.469381 | 0.003087 |
| Commd10 | -0.273623 | 0.023415 | 0.992917 | 1.630508 | 0.003087 |
| Rps29 | -0.270256 | 0.013999 | 0.992917 | 1.853894 | 0.003087 |
| Rilpl1 | -0.258984 | 0.012820 | 0.992917 | 1.892115 | 0.003087 |

**Supplementary Table 10: IPA Canonical Pathways of differentially expressed genes in Astrocytes at P28**

| **Ingenuity Canonical Pathways** | **-log(p-valu** | **Ratio** | **z-score** | **Molecules** |
|---|---|---|---|---|
| Coronavirus Replication Pathway | 6.320 | 0.114 | #NUM! | KGD4,MRPS33,RPL12,RPL15,RPL23A,RPL27,RPL3,RPL41,RPS15A,RPS24,RPS27A,RPS29,RPS3A,RPS4X,TUBB,TUBB3,ZCRB1 |
| Exosome Signaling Pathway | 5.960 | 0.108 | -2.500 | KGD4,MRPS33,POLR2H,RBM7,RPL12,RPL15,RPL23A,RPL27,RPL3,RPL41,RPS15A,RPS24,RPS27A,RPS29,RPS3A,RPS4X,ZCCHC8 |
| Eukaryotic Translation Termination | 5.560 | 0.120 | -2.673 | KGD4,MRPS33,RPL12,RPL15,RPL23A,RPL27,RPL3,RPL41,RPS15A,RPS24,RPS27A,RPS29,RPS3A,RPS4X |
| Eukaryotic Translation Elongation | 5.510 | 0.119 | -2.673 | KGD4,MRPS33,RPL12,RPL15,RPL23A,RPL27,RPL3,RPL41,RPS15A,RPS24,RPS27A,RPS29,RPS3A,RPS4X |
| Signaling by ROBO receptors | 5.130 | 0.083 | -2.683 | CAP2,KGD4,MRPS33,PSMA4,PSMA7,RHOA,RPL12,RPL15,RPL23A,RPL27,RPL3,RPL41,RPS15A,RPS24,RPS27A,RPS29,RPS3A,RPS4X,SLIT2,UBB |
| SRP-dependent cotranslational protein targeting to membrane | 4.750 | 0.102 | -2.673 | KGD4,MRPS33,RPL12,RPL15,RPL23A,RPL27,RPL3,RPL41,RPS15A,RPS24,RPS27A,RPS29,RPS3A,RPS4X |
| Selenoamino acid metabolism | 4.580 | 0.099 | -2.673 | KGD4,MRPS33,RPL12,RPL15,RPL23A,RPL27,RPL3,RPL41,RPS15A,RPS24,RPS27A,RPS29,RPS3A,RPS4X |
| Nonsense-Mediated Decay (NMD) | 4.400 | 0.108 | -3.464 | RPL12,RPL15,RPL23A,RPL27,RPL3,RPL41,RPS15A,RPS24,RPS27A,RPS29,RPS3A,RPS4X |
| Response of EIF2AK4 (GCN2) to amino acid deficiency | 4.360 | 0.107 | -3.464 | RPL12,RPL15,RPL23A,RPL27,RPL3,RPL41,RPS15A,RPS24,RPS27A,RPS29,RPS3A,RPS4X |
| Eukaryotic Translation Initiation | 4.190 | 0.091 | -2.673 | KGD4,MRPS33,RPL12,RPL15,RPL23A,RPL27,RPL3,RPL41,RPS15A,RPS24,RPS27A,RPS29,RPS3A,RPS4X |
| Ribosomal Quality Control Signaling Pathway | 4.160 | 0.082 | -3.742 | KGD4,MRPS33,PSMA4,PSMA7,RPL12,RPL15,RPL23A,RPL27,RPL3,RPL41,RPS15A,RPS24,RPS27A,RPS29,RPS3A,RPS4X |
| Nucleotide catabolism | 3.760 | 0.188 | 0.816 | ENTPD6,ENTPD7,GDA,NT5C2,NT5C3A,NUDT15 |
| Signaling by ERBB2 | 3.470 | 0.140 | -1.134 | AKT2,EREG,HSP90AA1,PRKCD,RHOA,RPS27A,UBB |
| Signaling by FGFR1 | 3.470 | 0.140 | -0.378 | FGF2,FGF9,FLRT1,RPS27A,SPRED2,SPRY2,UBB |
| Transcriptional regulation by RUNX1 | 3.380 | 0.085 | -2.887 | H2AX,H3C1,PBRM1,PHC1,PHC2,PSMA4,PSMA7,RPS27A,RUNX2,SMARCB1,SMARCE1,UBB |
| Major pathway of rRNA processing in the nucleolus and cytosol | 3.370 | 0.073 | -2.840 | KGD4,MRPS33,NOC4L,RPL12,RPL15,RPL23A,RPL27,RPL3,RPL41,RPS15A,RPS24,RPS27A,RPS29,RPS3A,RPS4X |
| Regulation of endogenous retroelements | 3.240 | 0.113 | -1.414 | H2AX,H3C1,PIWIL4,RBBP4,RBM7,SMARCB1,SMARCE1,ZCCHC8 |
| GAIT Translation Signaling Pathway | 3.220 | 0.070 | 2.646 | KGD4,MRPS33,RBP4,RPL12,RPL15,RPL23A,RPL27,RPL3,RPL41,RPS15A,RPS24,RPS27A,RPS29,RPS3A,RPS4X |
| EIF2 Signaling | 3.220 | 0.068 | -1.633 | AKT2,KGD4,MRPS33,RPL12,RPL15,RPL23A,RPL27,RPL3,RPL41,RPS15A,RPS24,RPS27A,RPS29,RPS3A,RPS4X,SREBF1 |
| mTOR Signaling | 3.030 | 0.068 | 0.000 | AKT2,DDIT4,HMOX1,KGD4,MRPS33,PPP2R2C,PRKCD,RHOA,RPS15A,RPS24,RPS27A,RPS29,RPS3A,RPS4X,TSC2 |
| Apoptotic execution phase | 2.690 | 0.120 | -0.816 | GAS2,H1-1,H1-4,H1-5,PRKCD,TJP2 |
| Formation of the anterior neural plate | 2.580 | 0.273 | #NUM! | NANOG,POU5F1,ZEB2 |
| Signaling by EGFR | 2.560 | 0.113 | -0.816 | AAMP,EREG,RPS27A,SPRY1,SPRY2,UBB |
| Hereditary Breast Cancer Signaling | 2.530 | 0.075 | 0.632 | AKT2,BRCA2,CDK1,FANCF,FANCG,FGF2,H2AX,POLR2H,RPS27A,UBB |
| Guanosine Nucleotides Degradation III | 2.460 | 0.250 | #NUM! | GDA,NT5C2,NT5C3A |
| Coronavirus Pathogenesis Pathway | 2.450 | 0.063 | 2.828 | ADAM9,E2F6,KGD4,MRPS33,Pycard,RELB,RPS15A,RPS24,RPS27A,RPS29,RPS3A,RPS4X,STAT2 |
| Collagen degradation | 2.440 | 0.107 | 0.816 | ADAM9,COL23A1,COL5A3,COL6A1,COL6A2,MMP15 |
| Signaling by FGFR3 | 2.400 | 0.125 | -0.447 | FGF2,FGF9,RPS27A,SPRY2,UBB |
| Signaling by FGFR4 | 2.400 | 0.125 | -0.447 | FGF2,FGF9,RPS27A,SPRY2,UBB |
| POU5F1 (OCT4), SOX2, NANOG activate genes related to prolife | 2.360 | 0.231 | #NUM! | FGF2,NANOG,POU5F1 |
| Oxidative Stress Induced Senescence | 2.330 | 0.090 | -2.646 | H2AX,H3C1,PHC1,PHC2,RBBP4,RPS27A,UBB |
| DNA Damage/Telomere Stress Induced Senescence | 2.310 | 0.119 | -2.236 | H1-1,H1-4,H1-5,H2AX,UBN1 |
| Role of BRCA1 in DNA Damage Response | 2.270 | 0.088 | 1.000 | BRCA2,E2F6,FANCF,FANCG,PBRM1,SMARCB1,SMARCE1 |
| Exocyst Signaling Pathway | 2.260 | 0.079 | -1.414 | AKT2,CDH12,EXOC6,EXOC6B,NAPA,PRKCD,RHOA,WASF2 |
| Mitotic G2-G2/M phases | 2.230 | 0.062 | -2.887 | CDK1,CDK11B,CEP192,HSP90AA1,PPP1R12B,PSMA4,PSMA7,RBBP4,RPS27A,TUBB,TUBB3,UBB |
| Amyloid fiber formation | 2.220 | 0.097 | -2.449 | H2AX,H2AZ1,H3C1,RPS27A,SIAH1,UBB |
| Protein ubiquitination | 2.160 | 0.138 | -2.000 | RPS27A,UBB,UBE2R2,UCHL3 |
| HSP90 chaperone cycle for steroid hormone receptors in the pr | 2.100 | 0.106 | -2.236 | CAPZA2,DNAJA2,HSP90AA1,STIP1,TUBB3 |
| Mitotic G1 phase and G1/S transition | 2.070 | 0.069 | -2.333 | AKT2,CDK1,E2F6,PCNA,PSMA4,PSMA7,RBBP4,RPS27A,UBB |
| Purine Nucleotides Degradation II (Aerobic) | 2.020 | 0.176 | #NUM! | GDA,NT5C2,NT5C3A |
| Cohesin Chromatin Regulation Pathway | 2.000 | 0.058 | -1.667 | CDK1,H2AX,H2AZ1,H3C1,MED7,NANOG,PCDHB14,PCDHGA12,POLR2H,POU5F1,PPP2R2C,SRA1 |
| Signaling by FGFR2 | 1.970 | 0.086 | -0.816 | FGF2,FGF9,POLR2H,RPS27A,SPRY2,UBB |
| Activated PKN1 stimulates transcription of AR regulated genes I | 1.940 | 0.167 | #NUM! | H2AX,H3C1,PKN1 |
| Stress Granule Signaling Pathway | 1.920 | 0.054 | -0.277 | AKT2,HSP90AA1,KGD4,MRPS33,Pycard,RPS15A,RPS24,RPS27A,RPS29,RPS3A,RPS4X,TSC2,ULK2 |
| Gene Silencing by RNA | 1.920 | 0.075 | -1.134 | DROSHA,H2AX,H3C1,HSP90AA1,PIWIL4,POLR2H,TDRKH |
| Signaling by PTK6 | 1.880 | 0.094 | -1.342 | EREG,RHOA,RPS27A,SFPQ,UBB |
| DNA Replication Pre-Initiation | 1.850 | 0.073 | -2.646 | H2AX,H3C1,MCM4,PSMA4,PSMA7,RPS27A,UBB |
| Signaling by TGF-beta Receptor Complex | 1.850 | 0.093 | -1.342 | BAMBI,ITGB8,RHOA,RPS27A,UBB |
| Maternal to Zygotic Transition Signaling Pathway | 1.850 | 0.067 | -2.828 | CDK1,DIS3L2,PBRM1,POU5F1,PSMA4,PSMA7,SMARCB1,SMARCE1 |
| Aggrephagy | 1.830 | 0.111 | -2.000 | HSP90AA1,RPS27A,TUBB3,UBB |
| Fanconi Anemia Pathway | 1.790 | 0.108 | 0.000 | FANCF,FANCG,RPS27A,UBB |
| Regulation of TP53 Expression and Degradation | 1.790 | 0.108 | -2.000 | AKT2,CDK1,RPS27A,UBB |

| | | | | |
|---|---|---|---|---|
| Chaperone Mediated Autophagy | 1.760 | 0.143 | #NUM! | HSP90AA1,RPS27A,UBB |
| Embryonic Stem Cell Differentiation into Cardiac Lineages | 1.750 | 0.250 | #NUM! | NANOG,POU5F1 |
| FLT3 Signaling | 1.750 | 0.105 | -1.000 | AKT2,PTPRJ,RPS27A,UBB |
| Regulation of eIF4 and p70S6K Signaling | 1.730 | 0.055 | #NUM! | AKT2,ITGB8,KGD4,MRPS33,PPP2R2C,RPS15A,RPS24,RPS27A,RPS29,RPS3A,RPS4X |
| XIST Epigenetic Regulation Signaling Pathway | 1.730 | 0.064 | -0.707 | H2AX,H2AZ1,NANOG,PHC1,PHC2,POU5F1,RBBP4,ZEB2 |
| BEX2 Signaling Pathway | 1.730 | 0.076 | -1.342 | AKT2,PPP2R2C,RELB,SMARCB1,SWAP70,TSC2 |
| POU5F1 (OCT4), SOX2, NANOG repress genes related to differe | 1.650 | 0.222 | #NUM! | NANOG,POU5F1 |
| RHO GTPases Activate Rhotekin and Rhophilins | 1.650 | 0.222 | #NUM! | RHOA,TAX1BP3 |
| NIK-->noncanonical NF-kB signaling | 1.640 | 0.082 | -1.342 | PSMA4,PSMA7,RELB,RPS27A,UBB |
| Cytosolic sensors of pathogen-associated DNA | 1.610 | 0.081 | -1.342 | DTX4,POLR2H,POLR3GL,RPS27A,UBB |
| AMPK Signaling | 1.580 | 0.050 | 0.000 | AK3,AKT2,GNAI2,PBRM1,PHLPP1,PPM1G,PPM1M,PPP2R2C,RAB9B,SMARCB1,SMARCE1,TSC2 |
| E3 ubiquitin ligases ubiquitinate target proteins | 1.570 | 0.093 | -1.000 | PCNA,PEX13,RPS27A,UBB |
| Collagen chain trimerization | 1.570 | 0.093 | 2.000 | COL23A1,COL5A3,COL6A1,COL6A2 |
| Cell Cycle Control of Chromosomal Replication | 1.560 | 0.078 | -1.342 | CDK1,CDK10,CDK11B,MCM4,PCNA |
| B-WICH complex positively regulates rRNA expression | 1.540 | 0.091 | -2.000 | H2AX,H3C1,MYBBP1A,POLR2H |
| DDX58/IFIH1-mediated induction of interferon-alpha/beta | 1.530 | 0.077 | -0.447 | HSP90AA1,NLRC5,RPS27A,TAX1BP1,UBB |
| Collagen biosynthesis and modifying enzymes | 1.510 | 0.076 | 1.342 | COL23A1,COL5A3,COL6A1,COL6A2,P4HA2 |
| Pyridoxal 5'-phosphate Salvage Pathway | 1.510 | 0.076 | -0.447 | AKT2,CDK1,MAP3K6,PKN1,PRKCD |
| ILK Signaling | 1.490 | 0.052 | -1.414 | ACTN2,AKT2,ITGB8,MYH4,PPP1R14B,PPP2R2C,RELB,RHOA,TESK1,TMSB4X |
| Regulation of mRNA stability by proteins that bind AU-rich elem | 1.480 | 0.067 | -1.633 | KHSRP,PRKCD,PSMA4,PSMA7,RPS27A,UBB |
| Alpha-protein kinase 1 signaling pathway | 1.480 | 0.182 | #NUM! | RPS27A,UBB |
| Pexophagy | 1.480 | 0.182 | #NUM! | RPS27A,UBB |
| PRC2 methylates histones and DNA | 1.470 | 0.111 | #NUM! | H2AX,H3C1,RBBP4 |
| Syndecan interactions | 1.470 | 0.111 | #NUM! | COL5A3,FGF2,SDC4 |
| RHO GTPases activate PKNs | 1.430 | 0.107 | #NUM! | PKN1,PPP1R12B,RHOA |
| DNA Damage Bypass | 1.410 | 0.083 | -1.000 | CUL4A,PCNA,RPS27A,UBB |
| Specification of primordial germ cells | 1.410 | 0.167 | #NUM! | NANOG,POU5F1 |
| RAF/MAP kinase cascade | 1.400 | 0.047 | -1.155 | ACTN2,ARRB2,DUSP1,EREG,FGF2,FGF9,PSMA4,PSMA7,RPS27A,SPRED2,TEK,UBB |
| ERCC6 (CSB) and EHMT2 (G9a) positively regulate rRNA express | 1.390 | 0.103 | #NUM! | H2AX,H3C1,RBBP4 |
| GP6 Signaling Pathway | 1.380 | 0.058 | 1.134 | AKT2,COL23A1,COL5A3,COL6A1,COL6A2,LCP2,PRKCD |
| KEAP1-NFE2L2 pathway | 1.380 | 0.055 | -0.707 | AKT2,FBXL17,PRKCD,PSMA4,PSMA7,RPS27A,STK4,UBB |
| Regulation of RUNX2 expression and activity | 1.370 | 0.069 | -1.342 | PSMA4,PSMA7,RPS27A,RUNX2,UBB |
| RIPK1-mediated regulated necrosis | 1.350 | 0.100 | #NUM! | HSP90AA1,RPS27A,UBB |
| Urate Biosynthesis/Inosine 5'-phosphate Degradation | 1.340 | 0.154 | #NUM! | NT5C2,NT5C3A |
| PTEN Regulation | 1.340 | 0.054 | -0.707 | AKT2,PHC1,PHC2,PSMA4,PSMA7,RBBP4,RPS27A,UBB |
| MicroRNA Biogenesis Signaling Pathway | 1.330 | 0.051 | -1.000 | AKT2,DNAJA2,DROSHA,FGF2,HMOX1,HSP90AA1,KHSRP,PIWIL4,POLR2H |
| CSDE1 Signaling Pathway | 1.330 | 0.078 | #NUM! | AKT2,CDK11B,PIWIL4,SDC4 |
| SWI/SNF chromatin remodelers | 1.310 | 0.097 | #NUM! | PBRM1,SMARCB1,SMARCE1 |
| NLR signaling pathways | 1.310 | 0.077 | -1.000 | AAMP,HMOX1,RPS27A,UBB |
| Production of Nitric Oxide and Reactive Oxygen Species in Macr | 1.300 | 0.050 | 0.000 | AKT2,MAP3K6,PPP1R14B,PPP1R14C,PPP2R2C,PRKCD,RBP4,RELB,RHOA |
| Adenosine Nucleotides Degradation II | 1.280 | 0.143 | #NUM! | NT5C2,NT5C3A |
| S-methyl-5-thio-α-D-ribose 1-phosphate Degradation | 1.280 | 0.500 | #NUM! | APIP |
| S Phase | 1.280 | 0.059 | -1.633 | AKT2,PSMA4,PSMA7,RBBP4,RPS27A,UBB |
| D-myo-inositol (1,4,5,6)-Tetrakisphosphate Biosynthesis | 1.270 | 0.050 | 0.447 | DUSP1,DUSP14,NUDT15,PPP1R12B,PPP1R14B,PPP1R14C,PPP2R2C,PTPRJ,PTPRO |
| D-myo-inositol (3,4,5,6)-tetrakisphosphate Biosynthesis | 1.270 | 0.050 | 0.447 | DUSP1,DUSP14,NUDT15,PPP1R12B,PPP1R14B,PPP1R14C,PPP2R2C,PTPRJ,PTPRO |
| Interleukin-4 and Interleukin-13 signaling | 1.260 | 0.059 | 1.633 | FGF2,HMOX1,HSP90AA1,MCL1,NANOG,OPRD1 |
| DHCR24 Signaling Pathway | 1.260 | 0.055 | 1.134 | AKT2,CYP17A1,HMOX1,PRKCD,RBP4,SC5D,SREBF1 |
| Regulation of Apoptosis | 1.260 | 0.074 | -2.000 | PSMA4,PSMA7,RPS27A,UBB |
| IL-10 Signaling | 1.240 | 0.054 | -0.816 | AKT2,DDIT4,DUSP1,Fcgr2b,HMOX1,IL10RB,RELB |
| Germ layer formation at gastrulation | 1.230 | 0.133 | #NUM! | NANOG,POU5F1 |
| Transcriptional Regulation by E2F6 | 1.210 | 0.088 | #NUM! | E2F6,PHC1,RBBP4 |
| TNF signaling | 1.210 | 0.071 | 0.000 | NSMAF,RPS27A,TAX1BP1,UBB |
| TCF dependent signaling in response to WNT | 1.200 | 0.048 | -1.667 | AKT2,H2AX,H3C1,LGR4,PSMA4,PSMA7,RPS27A,RSPO2,UBB |
| Signaling by VEGF | 1.190 | 0.057 | -0.816 | AAMP,AKT2,HSP90AA1,PRKCD,RHOA,WASF2 |
| Integrin cell surface interactions | 1.190 | 0.062 | 1.342 | COL23A1,COL5A3,COL6A1,COL6A2,ITGB8 |

| | | | | |
|---|---|---|---|---|
| Role of CHK Proteins in Cell Cycle Checkpoint Control | 1.190 | 0.070 | #NUM! | CDK1,E2F6,PCNA,PPP2R2C |
| Pyrimidine Ribonucleotides Interconversion | 1.180 | 0.086 | #NUM! | ENTPD6,ENTPD7,NUDT15 |
| Metabolism of folate and pterines | 1.180 | 0.125 | #NUM! | MTHFS,SLC46A1 |
| Transcriptional regulation of pluripotent stem cells | 1.180 | 0.125 | #NUM! | NANOG,POU5F1 |
| ISGylation Signaling Pathway | 1.170 | 0.061 | 0.447 | IL10RB,NFAT5,PCNA,S100A4,STAT2 |
| Iron uptake and transport | 1.160 | 0.069 | -1.000 | HMOX1,RPS27A,SLC46A1,UBB |
| Role of Osteoclasts in Rheumatoid Arthritis Signaling Pathway | 1.150 | 0.043 | 0.333 | ADAM9,AKT2,CALCR,COL23A1,COL5A3,COL6A1,COL6A2,CTNNA3,Fcgr2b,LCP2,MMP15,RHOA |
| NAD Signaling Pathway | 1.150 | 0.052 | -1.633 | AKT2,FGF2,H1-1,H1-4,H1-5,POLR2H,SREBF1 |
| 3-phosphoinositide Degradation | 1.140 | 0.047 | 0.447 | DUSP1,DUSP14,NUDT15,PPP1R12B,PPP1R14B,PPP1R14C,PPP2R2C,PTPRJ,PTPRO |
| Androgen Signaling | 1.140 | 0.049 | #NUM! | CACNA1G,GNAI2,HSP90AA1,NCOA4,POLR2H,PRKCD,RELB,TAF15 |
| TCR signaling | 1.130 | 0.055 | -1.633 | LCP2,PSMA4,PSMA7,PTPRJ,RPS27A,UBB |
| PI3K/AKT Signaling | 1.130 | 0.047 | -1.000 | AKT2,HSP90AA1,IL10RB,ITGB8,MCL1,NANOG,PPP2R2C,RELB,TSC2 |
| SUMOylation of DNA methylation proteins | 1.130 | 0.118 | #NUM! | PHC1,PHC2 |
| Nucleosome assembly | 1.130 | 0.081 | #NUM! | CENPK,H2AX,RBBP4 |
| Clathrin-mediated Endocytosis Signaling | 1.110 | 0.046 | #NUM! | AP1M1,AP1S3,ARRB2,FGF2,FGF9,ITGB8,RBP4,RPS27A,UBB |
| Metabolism of vitamin K | 1.110 | 0.333 | #NUM! | UBIAD1 |
| Myocardin Signaling Pathway | 1.100 | 0.054 | -0.816 | AKT2,CDK1,FGF2,RHOA,RUNX2,SPRY1 |
| Cilium Assembly | 1.100 | 0.046 | -1.000 | CCT3,CDK1,CEP192,EXOC6,HSP90AA1,KIF24,MKKS,TUBB,TUBB3 |
| D-myo-inositol-5-phosphate Metabolism | 1.100 | 0.046 | 0.447 | DUSP1,DUSP14,NUDT15,PPP1R12B,PPP1R14B,PPP1R14C,PPP2R2C,PTPRJ,PTPRO |
| Pyrimidine Ribonucleotides De Novo Biosynthesis | 1.100 | 0.079 | #NUM! | ENTPD6,ENTPD7,NUDT15 |
| Cellular hexose transport | 1.090 | 0.111 | #NUM! | SLC2A8,SLC2A9 |
| DNA methylation | 1.090 | 0.111 | #NUM! | H2AX,H3C1 |
| NCAM signaling for neurite out-growth | 1.080 | 0.065 | 2.000 | CACNA1G,COL5A3,COL6A1,COL6A2 |
| Degradation of CRY and PER proteins | 1.080 | 0.065 | -2.000 | PSMA4,PSMA7,RPS27A,UBB |
| Mitophagy | 1.070 | 0.077 | #NUM! | RPS27A,TOMM7,UBB |
| Neutrophil degranulation | 1.070 | 0.039 | 0.500 | AP1M1,C6orf120,CLEC12A,COMMD3,HSP90AA1,MME,NHLRC3,PRKCD,PTPRJ,QSOX1,RAB9B,RHOA,S100A11,SNAP29,TMEM63A,TUBB |
| NER (Nucleotide Excision Repair, Enhanced Pathway) | 1.070 | 0.057 | #NUM! | CUL4A,HMGN1,PCNA,POLR2H,RBBP4 |
| Senescence-Associated Secretory Phenotype (SASP) | 1.060 | 0.064 | -2.000 | H2AX,H3C1,RPS27A,UBB |
| RAB GEFs exchange GTP for GDP on RABs | 1.050 | 0.056 | 0.447 | AKT2,DENND2C,RAB6B,RAB9B,TRAPPC6B |
| RHO GTPases activate CIT | 1.050 | 0.105 | #NUM! | PPP1R12B,RHOA |
| RHO GTPases Activate ROCKs | 1.050 | 0.105 | #NUM! | PPP1R12B,RHOA |
| Regulation of TP53 Activity through Methylation | 1.050 | 0.105 | #NUM! | RPS27A,UBB |
| Class I peroxisomal membrane protein import | 1.050 | 0.105 | #NUM! | ABCD1,PEX13 |
| Meiotic recombination | 1.050 | 0.075 | #NUM! | BRCA2,H2AX,H3C1 |
| Superpathway of Inositol Phosphate Compounds | 1.040 | 0.043 | 0.000 | DUSP1,DUSP14,IP6K1,NUDT15,PPP1R12B,PPP1R14B,PPP1R14C,PPP2R2C,PTPRJ,PTPRO |
| RNA Polymerase I Transcription | 1.040 | 0.063 | -2.000 | H2AX,H3C1,POLR2H,RBBP4 |
| Tight Junction Signaling | 1.040 | 0.046 | #NUM! | AKT2,MYH4,NAPA,NECTIN2,PPP2R2C,RELB,RHOA,TJP2 |
| Regulation of mitotic cell cycle | 1.040 | 0.056 | -2.236 | CDK1,PSMA4,PSMA7,RPS27A,UBB |
| Degradation of beta-catenin by the destruction complex | 1.020 | 0.055 | -1.342 | CTBP2,PSMA4,PSMA7,RPS27A,UBB |
| MAPK6/MAPK4 signaling | 1.020 | 0.055 | -2.236 | CDK1,PSMA4,PSMA7,RPS27A,UBB |
| Salvage Pathways of Pyrimidine Ribonucleotides | 1.020 | 0.055 | -0.447 | AKT2,CDK1,MAP3K6,PKN1,PRKCD |
| 3-phosphoinositide Biosynthesis | 1.010 | 0.044 | 0.447 | DUSP1,DUSP14,NUDT15,PPP1R12B,PPP1R14B,PPP1R14C,PPP2R2C,PTPRJ,PTPRO |
| Signaling by Hippo | 1.010 | 0.100 | #NUM! | STK4,TJP2 |
| Hedgehog ligand biogenesis | 1.000 | 0.061 | -2.000 | PSMA4,PSMA7,RPS27A,UBB |
| TAK1-dependent IKK and NF-kappa-B activation | 0.997 | 0.071 | #NUM! | NLRC5,RPS27A,UBB |
| Pentose Phosphate Pathway (Oxidative Branch) | 0.990 | 0.250 | #NUM! | H6PD |
| Heme Degradation | 0.990 | 0.250 | #NUM! | HMOX1 |
| Methylmalonyl Pathway | 0.990 | 0.250 | #NUM! | MCEE |
| D-glucuronate Degradation I | 0.990 | 0.250 | #NUM! | AKR1A1 |
| PI3K Cascade | 0.974 | 0.070 | #NUM! | AKT2,FGF2,FGF9 |
| SIRT1 negatively regulates rRNA expression | 0.972 | 0.095 | #NUM! | H2AX,H3C1 |
| Interleukin-20 family signaling | 0.972 | 0.095 | #NUM! | IL10RB,STAT2 |
| NAD Salvage Pathway II | 0.972 | 0.095 | #NUM! | NT5C2,NT5C3A |
| TNFR2 non-canonical NF-kB pathway | 0.969 | 0.059 | -2.000 | PSMA4,PSMA7,RPS27A,UBB |
| Synthesis of DNA | 0.969 | 0.049 | -2.449 | MCM4,PCNA,PSMA4,PSMA7,RPS27A,UBB |

| | | | | |
|---|---|---|---|---|
| DNA Methylation and Transcriptional Repression Signaling | 0.960 | 0.053 | -1.342 | CDK1,CDK10,CDK11B,E2F6,RBBP4 |
| Other interleukin signaling | 0.938 | 0.091 | #NUM! | IL10RB,PTPRZ1 |
| Huntington's Disease Signaling | 0.935 | 0.040 | -1.342 | AKT2,ATP5PB,CAPN10,NAPA,PENK,POLR2H,PRKCD,PSMA4,PSMA7,RPS27A,UBB |
| Osteoarthritis Pathway | 0.933 | 0.042 | 1.633 | ACVRL1,DDIT4,FGF2,ITGB8,MYBBP1A,RBP4,RELB,RUNX2,SDC4 |
| Post-translational protein phosphorylation | 0.932 | 0.052 | 0.447 | FSTL1,GAS6,NUCB1,PENK,QSOX1 |
| Hepatic Fibrosis / Hepatic Stellate Cell Activation | 0.924 | 0.044 | #NUM! | BAMBI,COL23A1,COL5A3,COL6A1,COL6A2,FGF2,MYH4,RELB |
| Microautophagy Signaling Pathway | 0.919 | 0.045 | #NUM! | COL23A1,COL5A3,COL6A1,COL6A2,PSMA4,PSMA7,VPS33B |
| Chaperone Mediated Autophagy Signaling Pathway | 0.916 | 0.042 | 0.707 | HSP90AA1,MCL1,MMP15,PHLPP1,POU5F1,PSMA4,PSMA7,Pycard,STIP1 |
| Role of OCT4 in Mammalian Embryonic Stem Cell Pluripotency | 0.909 | 0.065 | #NUM! | NANOG,PHC1,POU5F1 |
| RAF-independent MAPK1/3 activation | 0.906 | 0.087 | #NUM! | CDK1,DUSP1 |
| RAS processing | 0.906 | 0.087 | #NUM! | RPS27A,UBB |
| IL-22 Signaling | 0.906 | 0.087 | #NUM! | AKT2,IL10RB |
| Role of JAK1, JAK2 and TYK2 in Interferon Signaling | 0.906 | 0.087 | #NUM! | RELB,STAT2 |
| Formation of xylulose-5-phosphate | 0.899 | 0.200 | #NUM! | AKR1A1 |
| Myo-inositol Biosynthesis | 0.899 | 0.200 | #NUM! | IMPA1 |
| 2-oxobutanoate Degradation I | 0.899 | 0.200 | #NUM! | MCEE |
| Pulmonary Fibrosis Idiopathic Signaling Pathway | 0.892 | 0.039 | 0.577 | AKT2,COL23A1,COL5A3,COL6A1,COL6A2,FGF2,FGF9,MMP15,PEX13,RBBP4,RELB,RHOA |
| Chromatin modifying enzymes | 0.889 | 0.041 | -2.333 | H2AX,H3C1,HCFC1,KANSL2,KMT5B,PBRM1,RBBP4,SMARCB1,SMARCE1 |
| G alpha (z) signalling events | 0.888 | 0.064 | #NUM! | GNAI2,PRKCD,RGS17 |
| Extra-nuclear estrogen signaling | 0.886 | 0.055 | -1.000 | AKT2,EREG,GNAI2,HSP90AA1 |
| Apoptosis Signaling | 0.878 | 0.050 | -1.000 | CAPN10,CDK1,GAS2,MCL1,RELB |
| MyD88 cascade initiated on plasma membrane | 0.876 | 0.083 | #NUM! | RPS27A,UBB |
| Chromatin modifications during the maternal to zygotic transiti | 0.876 | 0.083 | #NUM! | H2AX,H3C1 |
| NAP1L1 Transcription Regulation Signaling Pathway | 0.871 | 0.054 | -1.000 | AKT2,MCL1,TUBB,TUBB3 |
| Interferon alpha/beta signaling | 0.868 | 0.063 | #NUM! | RPS27A,STAT2,UBB |
| Role of PI3K/AKT Signaling in the Pathogenesis of Influenza | 0.868 | 0.063 | #NUM! | AKT2,GNAI2,RELB |
| Signaling by NOTCH1 | 0.855 | 0.053 | -2.000 | ARRB2,DTX4,RPS27A,UBB |
| Post-translational modification: synthesis of GPI-anchored prot | 0.855 | 0.053 | 0.000 | NTNG1,PIGU,RECK,TECTB |
| HIPPO signaling | 0.855 | 0.053 | #NUM! | FAT4,PPP2R2C,STK4,TJP2 |
| HDR through Homologous Recombination (HRR) or Single Stran | 0.853 | 0.049 | -2.236 | BRCA2,H2AX,PCNA,RPS27A,UBB |
| Cargo recognition for clathrin-mediated endocytosis | 0.853 | 0.049 | -0.447 | ARRB2,EREG,RPS27A,SLC2A8,UBB |
| ICOS-ICOSL Signaling in T Helper Cells | 0.853 | 0.049 | #NUM! | AKT2,LCP2,NFAT5,PLEKHA3,RELB |
| C-type lectin receptors (CLRs) | 0.851 | 0.046 | -0.816 | PRKCD,PSMA4,PSMA7,RELB,RPS27A,UBB |
| DNA Double Strand Break Response | 0.849 | 0.061 | #NUM! | H2AX,RPS27A,UBB |
| Signaling by TGFBR3 | 0.849 | 0.061 | #NUM! | ARRB2,FGF2,HELLS |
| Apelin Muscle Signaling Pathway | 0.849 | 0.061 | #NUM! | AKT2,GNAI2,NRF1 |
| Cholesterol biosynthesis | 0.847 | 0.080 | #NUM! | SC5D,SREBF1 |
| Metabolism of cofactors | 0.847 | 0.080 | #NUM! | COQ5,HSP90AA1 |
| Glycogen metabolism | 0.847 | 0.080 | #NUM! | RPS27A,UBB |
| Cellular response to hypoxia | 0.841 | 0.053 | -2.000 | PSMA4,PSMA7,RPS27A,UBB |
| Activation of anterior HOX genes in hindbrain during early emb | 0.841 | 0.053 | -2.000 | H2AX,H3C1,POLR2H,RBBP4 |
| Transcriptional regulation by RUNX2 | 0.830 | 0.060 | #NUM! | AKT2,CDK1,RUNX2 |
| Gαq Signaling | 0.828 | 0.043 | 0.447 | AKT2,CALCR,GNAI2,HMOX1,PRKCD,RELB,RHOA |
| Telomere Maintenance | 0.826 | 0.052 | -2.000 | H2AX,H3C1,PCNA,POLR2H |
| Protein repair | 0.825 | 0.167 | #NUM! | MSRB2 |
| Estrogen-mediated S-phase Entry | 0.819 | 0.077 | #NUM! | CDK1,E2F6 |
| Natural Killer Cell Signaling | 0.818 | 0.042 | -0.816 | AKT2,COL5A3,LCP2,MAP3K6,NECTIN2,NFAT5,RELB |
| Mitochondrial translation | 0.816 | 0.047 | 0.447 | KGD4,MRPL11,MRPL46,MRPS21,MRPS33 |
| FAT10 Cancer Signaling Pathway | 0.812 | 0.059 | #NUM! | AKT2,PCNA,RELB |
| Signaling by MET | 0.798 | 0.051 | 0.000 | COL5A3,PTPRJ,RPS27A,UBB |
| ER-Mitochondrial Communication Signaling Pathway | 0.798 | 0.051 | 0.000 | AKT2,ESYT1,Pycard,TOMM7 |
| Estrogen Receptor Signaling | 0.796 | 0.036 | -0.577 | AKT2,ATP5PB,CTBP2,GNAI2,HSP90AA1,MMP15,NRF1,PCNA,PPP1R12B,PRKCD,RELB,RHOA,RUNX2,SRA1 |
| Nucleotide Excision Repair | 0.792 | 0.046 | -1.342 | CUL4A,PCNA,POLR2H,RPS27A,UBB |
| Hedgehog 'off' state | 0.781 | 0.046 | -2.236 | PSMA4,PSMA7,RPS27A,TUBB3,UBB |
| PKR-mediated signaling | 0.771 | 0.049 | 0.000 | CDK1,FANCF,FANCG,TUBB3 |

| | | | | |
|---|---|---|---|---|
| BAG2 Signaling Pathway | 0.771 | 0.049 | #NUM! | HSP90AA1,PSMA4,PSMA7,RELB |
| Regulation of Insulin-like Growth Factor (IGF) transport and upt | 0.770 | 0.046 | 0.447 | FSTL1,GAS6,NUCB1,PENK,QSOX1 |
| ESR-mediated signaling | 0.770 | 0.046 | -2.236 | H2AX,H3C1,HSP90AA1,KANK1,POLR2H |
| MTOR signalling | 0.768 | 0.071 | #NUM! | AKT2,TSC2 |
| Inhibition of Matrix Metalloproteases | 0.768 | 0.071 | #NUM! | MMP15,RECK |
| Phospholipase C Signaling | 0.767 | 0.038 | -1.633 | Fcgr2b,HMOX1,ITGB8,LCP2,MARCKS,NFAT5,PPP1R12B,PRKCD,RELB,RHOA |
| Formation of intermediate mesoderm | 0.764 | 0.143 | #NUM! | FGF2 |
| Ceramide Biosynthesis | 0.764 | 0.143 | #NUM! | SPTSSB |
| Inositol Pyrophosphates Biosynthesis | 0.764 | 0.143 | #NUM! | IP6K1 |
| CTLA4 Signaling in Cytotoxic T Lymphocytes | 0.764 | 0.041 | 1.134 | AKT2,AP1M1,AP1S3,HMOX1,LCP2,NFAT5,PPP2R2C |
| Hedgehog 'on' state | 0.758 | 0.043 | -1.633 | ARRB2,PSMA4,PSMA7,RPS27A,STK4,UBB |
| Sheddase Signaling Pathway | 0.755 | 0.041 | -0.378 | CDH12,DDIT4,E2F6,GNAI2,ITGB8,MMP15,RELB |
| Actin Cytoskeleton Signaling | 0.753 | 0.038 | -0.447 | ACTN2,FGF2,FGF9,ITGB8,MYH4,PPP1R12B,RHOA,TMSB4X,WASF2 |
| NRF2-mediated Oxidative Stress Response | 0.748 | 0.039 | -0.447 | AKR1A1,DNAJA2,DNAJC1,HMOX1,HSP90AA1,PRKCD,STIP1,UBB |
| ABC-family proteins mediated transport | 0.747 | 0.045 | -2.236 | ABCD1,PSMA4,PSMA7,RPS27A,UBB |
| Germ Cell Differentiation Signaling Pathway | 0.747 | 0.045 | -0.447 | ACVRL1,FGF9,INHBB,NANOG,POU5F1 |
| Signaling by BMP | 0.745 | 0.069 | #NUM! | ACVRL1,FSTL1 |
| Sonic Hedgehog Signaling | 0.745 | 0.069 | #NUM! | ARRB2,CDK1 |
| PEDF Signaling | 0.732 | 0.048 | #NUM! | AKT2,RELB,RHOA,WASF2 |
| Assembly of collagen fibrils and other multimeric structures | 0.728 | 0.054 | #NUM! | COL5A3,COL6A1,COL6A2 |
| CD27 Signaling in Lymphocytes | 0.728 | 0.054 | #NUM! | MAP3K6,RELB,SIVA1 |
| Activation of kainate receptors upon glutamate binding | 0.722 | 0.067 | #NUM! | GRIK1,GRIK5 |
| Regulation of TP53 Activity through Acetylation | 0.722 | 0.067 | #NUM! | AKT2,RBBP4 |
| Endosomal Sorting Complex Required For Transport (ESCRT) | 0.722 | 0.067 | #NUM! | RPS27A,UBB |
| Signaling by CSF3 (G-CSF) | 0.722 | 0.067 | #NUM! | RPS27A,UBB |
| TP53 Regulates Metabolic Genes | 0.719 | 0.047 | 0.000 | AKT2,DDIT4,SCO2,TSC2 |
| Neddylation | 0.717 | 0.038 | -1.667 | COMMD10,COMMD3,CUL4A,FBXO11,PSMA4,PSMA7,RPS27A,UBB,UCHL3 |
| Neuregulin Signaling | 0.715 | 0.044 | #NUM! | AKT2,EREG,HSP90AA1,ITGB8,PRKCD |
| TRIM21 Intracellular Antibody Signaling Pathway | 0.715 | 0.044 | -2.236 | PSMA4,PSMA7,RPS27A,STAT2,UBB |
| Glycosphingolipid transport | 0.712 | 0.125 | #NUM! | ESYT1 |
| PIP3 activates AKT signaling | 0.711 | 0.041 | 0.816 | AKT2,EREG,FGF2,FGF9,PHLPP1,TSC2 |
| Epigenetic regulation of gene expression by MLL3 and MLL4 cor | 0.707 | 0.047 | -1.000 | H2AX,H3C1,MED7,NCOA4 |
| Signaling by CSF1 (M-CSF) in myeloid cells | 0.700 | 0.065 | #NUM! | RPS27A,UBB |
| MyD88 dependent cascade initiated on endosome | 0.700 | 0.065 | #NUM! | RPS27A,UBB |
| Signaling by ERBB4 | 0.698 | 0.052 | #NUM! | EREG,RPS27A,UBB |
| Signaling by PDGF | 0.698 | 0.052 | #NUM! | COL5A3,COL6A1,COL6A2 |
| NoRC negatively regulates rRNA expression | 0.698 | 0.052 | #NUM! | H2AX,H3C1,POLR2H |
| GADD45 Signaling | 0.698 | 0.052 | #NUM! | CDK1,PCNA,RELB |
| Deubiquitination | 0.696 | 0.037 | -2.333 | ARRB2,CDK1,HCFC1,PSMA4,PSMA7,RHOA,RPS27A,UBB,UCHL3 |
| Ferroptosis Signaling Pathway | 0.695 | 0.043 | 2.236 | CHAC1,H2AX,H2AZ1,HMOX1,NCOA4 |
| Autophagy | 0.695 | 0.038 | 0.000 | AKT2,EPG5,FGF2,PLEKHM1,PPP2R2C,SNAP29,TSC2,VPS33B |
| Type II Diabetes Mellitus Signaling | 0.693 | 0.041 | #NUM! | AKT2,CACNA1G,NSMAF,PRKCD,RELB,SLC27A4 |
| Ceramide Signaling | 0.683 | 0.046 | #NUM! | AKT2,NSMAF,PPP2R2C,RELB |
| Regulation of Cellular Mechanics by Calpain Protease | 0.683 | 0.046 | #NUM! | ACTN2,CAPN10,CDK1,ITGB8 |
| Peroxisomal protein import | 0.683 | 0.051 | #NUM! | PEX13,RPS27A,UBB |
| Toll Like Receptor 3 (TLR3) Cascade | 0.679 | 0.063 | #NUM! | RPS27A,UBB |
| G-protein beta:gamma signalling | 0.679 | 0.063 | #NUM! | AKT2,RHOA |
| Late endosomal microautophagy | 0.679 | 0.063 | #NUM! | RPS27A,UBB |
| Protein Kinase A Signaling | 0.678 | 0.034 | -1.155 | DUSP1,GNAI2,H1-1,H1-4,H1-5,MYH4,NFAT5,PRKCD,PTPRJ,PTPRO,PTPRZ1,RELB,RHOA |
| IL-13 Signaling Pathway | 0.672 | 0.045 | -1.000 | AKT2,DUSP1,PRKCD,RHOA |
| Folate Transformations I | 0.666 | 0.111 | #NUM! | MTHFS |
| ID3 Signaling Pathway | 0.665 | 0.042 | -0.447 | ACVRL1,AKT2,POU5F1,RUNX2,SREBF1 |
| TR/RXR Activation | 0.665 | 0.042 | -0.447 | AKT2,NCOA4,NRF1,PCNA,SREBF1 |
| Fcγ Receptor-mediated Phagocytosis in Macrophages and Mon | 0.661 | 0.044 | 0.000 | AKT2,HMOX1,LCP2,PRKCD |
| Signaling by NOTCH2 | 0.659 | 0.061 | #NUM! | RPS27A,UBB |

| | | | | |
|---|---|---|---|---|
| Signal amplification | 0.659 | 0.061 | #NUM! | AAMP,GNAI2 |
| NGF Signaling | 0.656 | 0.041 | 0.000 | AKT2,MAP3K6,PRKCD,RELB,RHOA |
| TREM1 Signaling | 0.655 | 0.049 | #NUM! | AKT2,NLRC5,RELB |
| Regulation of TP53 Activity through Phosphorylation | 0.649 | 0.044 | -1.000 | NOC2L,RPS27A,TAF15,UBB |
| Response to elevated platelet cytosolic Ca2+ | 0.646 | 0.041 | -0.447 | ACTN2,GAS6,QSOX1,SCCPDH,TMSB4X |
| Epithelial Adherens Junction Signaling | 0.642 | 0.039 | 1.633 | AKT2,MYH4,NECTIN2,PPP2R2C,RHOA,STK4 |
| Remodeling of Epithelial Adherens Junctions | 0.641 | 0.048 | #NUM! | ACTN2,TUBB,TUBB3 |
| GPVI-mediated activation cascade | 0.640 | 0.059 | #NUM! | LCP2,RHOA |
| Oncogene Induced Senescence | 0.640 | 0.059 | #NUM! | RPS27A,UBB |
| 14-3-3-mediated Signaling | 0.637 | 0.041 | #NUM! | AKT2,PRKCD,TSC2,TUBB,TUBB3 |
| Pancreatic Adenocarcinoma Signaling | 0.637 | 0.041 | #NUM! | AKT2,BRCA2,E2F6,HMOX1,RELB |
| Intra-Golgi and retrograde Golgi-to-ER traffic | 0.635 | 0.037 | -0.378 | CAPZA2,GALNT2,NAPA,RAB6B,RAB9B,SNAP29,TUBB3 |
| Regulation of the Epithelial Mesenchymal Transition by Growth | 0.635 | 0.037 | -1.000 | AKT2,CDH12,FGF2,FGF9,RELB,RHOA,ZEB2 |
| RAB geranylgeranylation | 0.628 | 0.048 | #NUM! | RAB23,RAB6B,RAB9B |
| Th17 Activation Pathway | 0.628 | 0.048 | #NUM! | HSP90AA1,NFAT5,RELB |
| Airway Pathology in Chronic Obstructive Pulmonary Disease | 0.628 | 0.043 | #NUM! | FGF2,FGF9,LCN10,RBP4 |
| Regulation of the Epithelial-Mesenchymal Transition Pathway | 0.627 | 0.037 | #NUM! | AKT2,CDH12,FGF2,FGF9,RELB,RHOA,ZEB2 |
| RHOBTB3 ATPase cycle | 0.626 | 0.100 | #NUM! | RAB9B |
| Formation of the posterior neural plate | 0.626 | 0.100 | #NUM! | ZEB2 |
| Calcium Transport I | 0.626 | 0.100 | #NUM! | ATP2B4 |
| Pentose Phosphate Pathway | 0.626 | 0.100 | #NUM! | H6PD |
| Glucocorticoid Biosynthesis | 0.626 | 0.100 | #NUM! | CYP17A1 |
| Ribonucleotide Reductase Signaling Pathway | 0.626 | 0.039 | -1.633 | AKT2,CDK1,E2F6,PBRM1,SMARCB1,SMARCE1 |
| Nuclear RNA decay | 0.622 | 0.057 | #NUM! | RBM7,ZCCHC8 |
| UBE2C in Cell Cycle Progression Signaling Pathway | 0.617 | 0.043 | #NUM! | AKT2,CDK1,RPS27A,UBB |
| Translocation of SLC2A4 (GLUT4) to the plasma membrane | 0.615 | 0.047 | #NUM! | AKT2,EXOC6,TUBB3 |
| Kinetochore Metaphase Signaling Pathway | 0.607 | 0.042 | #NUM! | CDK1,CENPK,H2AX,H2AZ1 |
| Cilia Biogenesis Signaling Pathway | 0.606 | 0.037 | -1.134 | CDK1,CEP192,FGF2,GNAI2,KIF24,TUBB,TUBB3 |
| Assembly and cell surface presentation of NMDA receptors | 0.604 | 0.056 | #NUM! | ACTN2,TUBB3 |
| Mitotic Roles of Polo-Like Kinase | 0.602 | 0.046 | #NUM! | CDK1,HSP90AA1,PPP2R2C |
| Clathrin-mediated endocytosis | 0.601 | 0.039 | -0.447 | ARRB2,EREG,RPS27A,SLC2A8,UBB |
| HIF1α Signaling | 0.599 | 0.037 | 0.378 | AKT2,FGF2,HMOX1,HSP90AA1,MMP15,PRKCD,SLC2A8 |
| Transcriptional regulation by RUNX3 | 0.597 | 0.042 | -2.000 | PSMA4,PSMA7,RPS27A,UBB |
| IL-12 Signaling and Production in Macrophages | 0.592 | 0.036 | 0.816 | AKT2,Fcgr2b,IL10RB,NFAT5,PRKCD,RBP4,RELB |
| Peptide hormone biosynthesis | 0.590 | 0.091 | #NUM! | INHBB |
| RHO GTPases activate KTN1 | 0.590 | 0.091 | #NUM! | RHOA |
| Advanced glycosylation endproduct receptor signaling | 0.590 | 0.091 | #NUM! | CAPZA2 |
| Protein Ubiquitination Pathway | 0.588 | 0.035 | -1.890 | DNAJC1,HSP90AA1,PSMA4,PSMA7,RPS27A,UBB,Ube2e1,UBE2R2,UCHL3 |
| MyD88-independent TLR4 cascade | 0.587 | 0.054 | #NUM! | RPS27A,UBB |
| Cell Cycle Regulation by BTG Family Proteins | 0.587 | 0.054 | #NUM! | E2F6,PPP2R2C |
| Germ Cell-Sertoli Cell Junction Signaling | 0.587 | 0.037 | #NUM! | ACTN2,MAP3K6,NECTIN2,RHOA,TUBB,TUBB3 |
| Human Embryonic Stem Cell Pluripotency | 0.578 | 0.036 | 0.378 | ACVRL1,AKT2,FGF2,HSP90AA1,NANOG,POU5F1,PRKCD |
| TP53 Regulates Transcription of DNA Repair Genes | 0.578 | 0.045 | #NUM! | AP1M1,AP1S3,POLR2H |
| Granzyme A Signaling | 0.578 | 0.045 | #NUM! | H1-1,H1-4,H1-5 |
| Transport of vitamins, nucleosides, and related molecules | 0.571 | 0.053 | #NUM! | SLC27A4,SLCO3A1 |
| Degradation of the extracellular matrix | 0.566 | 0.044 | #NUM! | CAPN10,MMP15,SCUBE1 |
| Hepatitis B Chronic Liver Pathogenesis Signaling Pathway | 0.565 | 0.037 | 0.447 | AKT2,AP1M1,AP1S3,PRKCD,Pycard,STAT2 |
| HDR through MMEJ (alt-NHEJ) | 0.558 | 0.083 | #NUM! | BRCA2 |
| Transcriptional regulation of testis differentiation | 0.558 | 0.083 | #NUM! | FGF9 |
| Z-decay: degradation of maternal mRNAs by zygotically express | 0.558 | 0.083 | #NUM! | DIS3L2 |
| Dolichyl-diphosphooligosaccharide Biosynthesis | 0.558 | 0.083 | #NUM! | ALG11 |
| IL-4 Signaling | 0.557 | 0.036 | 0.816 | AKT2,COL23A1,COL5A3,COL6A1,COL6A2,GNAI2 |
| Neutrophil Extracellular Trap Signaling Pathway | 0.555 | 0.033 | -1.265 | AKT2,ATP5PB,COL23A1,COL5A3,COL6A1,COL6A2,MCL1,ORAI3,PRKCD,Pycard,TOMM7 |
| Sertoli Cell-Sertoli Cell Junction Signaling | 0.555 | 0.035 | -1.414 | ACTN2,AKT2,CDH12,MAP3K6,NECTIN2,TJP2,TUBB,TUBB3 |
| MyD88:MAL(TIRAP) cascade initiated on plasma membrane | 0.555 | 0.051 | #NUM! | RPS27A,UBB |

| | | | | |
|---|---|---|---|---|
| MAP kinase activation | 0.555 | 0.051 | #NUM! | RPS27A,UBB |
| Retinoid metabolism and transport | 0.555 | 0.051 | #NUM! | RBP4,SDC4 |
| Melatonin Signaling | 0.544 | 0.043 | #NUM! | GNAI2,PRKCD,RORB |
| Fc epsilon receptor (FCERI) signaling | 0.544 | 0.037 | -2.236 | LCP2,PSMA4,PSMA7,RPS27A,UBB |
| Adipogenesis pathway | 0.544 | 0.037 | 0.447 | ACVRL1,CTBP2,FGF2,RBBP4,SREBF1 |
| Systemic Lupus Erythematosus Signaling | 0.543 | 0.036 | #NUM! | AKT2,Fcgr2b,LSM14B,NFAT5,PRPF8,ZCRB1 |
| WNT/β-catenin Signaling | 0.543 | 0.036 | #NUM! | AKT2,CDH12,POU5F1,PPP2R2C,RPS27A,UBB |
| WNK Renal Signaling Pathway | 0.540 | 0.039 | -1.000 | AKT2,LRRC8C,PRKCD,SLC12A6 |
| Transport of inorganic cations/anions and amino acids/oligopep | 0.540 | 0.039 | 2.000 | SLC12A6,SLC26A4,SLC38A5,SLC5A5 |
| Activation of gene expression by SREBF (SREBP) | 0.540 | 0.050 | #NUM! | SC5D,SREBF1 |
| Beta-catenin independent WNT signaling | 0.533 | 0.035 | -1.890 | ARRB2,PSMA4,PSMA7,RHOA,RPS27A,STK4,UBB |
| IL-8 Signaling | 0.533 | 0.035 | -0.378 | AKT2,ARRB2,GNAI2,HMOX1,PRKCD,RHOA,TEK |
| Extracellular matrix organization | 0.531 | 0.039 | 2.000 | COL5A3,COL6A1,COL6A2,FGF2 |
| Bladder Cancer Signaling | 0.531 | 0.039 | #NUM! | E2F6,FGF2,FGF9,MMP15 |
| Glutamate and glutamine metabolism | 0.529 | 0.077 | #NUM! | RIMKLB |
| Replacement of protamines by nucleosomes in the male pronu | 0.529 | 0.077 | #NUM! | H2AX |
| Cholesterol Biosynthesis I | 0.529 | 0.077 | #NUM! | SC5D |
| Cholesterol Biosynthesis II (via 24,25-dihydrolanosterol) | 0.529 | 0.077 | #NUM! | SC5D |
| Cholesterol Biosynthesis III (via Desmosterol) | 0.529 | 0.077 | #NUM! | SC5D |
| RNA Polymerase III Transcription | 0.525 | 0.049 | #NUM! | POLR2H,POLR3GL |
| DNA damage-induced 14-3-3σ Signaling | 0.525 | 0.049 | #NUM! | AKT2,CDK1 |
| Netrin Signaling | 0.523 | 0.035 | 0.000 | AKT2,CACNA1G,NAPA,NFAT5,PRKCD,RHOA |
| Toll-like Receptor Signaling | 0.522 | 0.042 | #NUM! | RELB,RPS27A,UBB |
| Pulmonary Hypertension Signaling Pathway | 0.517 | 0.032 | 0.302 | ACVRL1,AKT2,HMOX1,NFAT5,PCNA,POU5F1,PPP1R12B,PRKCD,RUNX2,S100A4,TEK |
| Nuclear Cytoskeleton Signaling Pathway | 0.515 | 0.034 | -1.890 | CAPZA2,CDH12,H2AX,ITGB8,RHOA,RUNX2,TUBB3 |
| MHC class II antigen presentation | 0.514 | 0.038 | -1.000 | AP1M1,AP1S3,CAPZA2,TUBB3 |
| Signaling by NOTCH4 | 0.513 | 0.036 | -1.342 | PSMA4,PSMA7,RPS27A,STK4,UBB |
| Hypoxia Signaling in the Cardiovascular System | 0.512 | 0.041 | #NUM! | HSP90AA1,Ube2e1,UBE2R2 |
| Oxytocin Signaling Pathway | 0.511 | 0.033 | -0.333 | AKT2,ARRB2,ATP2B4,GNAI2,MYH4,PPP1R12B,PRKCD,RELB,RHOA |
| Formation of WDR5-containing histone-modifying complexes | 0.511 | 0.048 | #NUM! | HCFC1,KANSL2 |
| April Mediated Signaling | 0.511 | 0.048 | #NUM! | NFAT5,RELB |
| IL-23 Signaling Pathway | 0.511 | 0.048 | #NUM! | AKT2,RELB |
| Gap Junction Signaling | 0.509 | 0.032 | -1.000 | AKT2,CACNA1G,GNAI2,GRIK1,PRKCD,RELB,RUNX2,TJP2,TUBB,TUBB3 |
| Breast Cancer Regulation by Stathmin1 | 0.507 | 0.031 | 2.183 | CALCR,CDK1,E2F6,FGF2,GPR3,GPR52,GPR63,LGR4,OPRD1,PPP1R14B,PPP1R14C,PPP2R2C,PRKCD,RHOA,SIVA1,TUBB,TUBB3 |
| YAP1- and WWTR1 (TAZ)-stimulated gene expression | 0.502 | 0.071 | #NUM! | RUNX2 |
| Mismatch Repair | 0.502 | 0.071 | #NUM! | PCNA |
| Regulation of TP53 Activity through Association with Co-factors | 0.502 | 0.071 | #NUM! | AKT2 |
| DNA Double-Strand Break Repair by Homologous Recombinatic | 0.502 | 0.071 | #NUM! | BRCA2 |
| Choline Biosynthesis III | 0.502 | 0.071 | #NUM! | HMOX1 |
| trans-Golgi Network Vesicle Budding | 0.501 | 0.041 | #NUM! | AP1M1,AP1S3,NAPA |
| Apelin Endothelial Signaling Pathway | 0.499 | 0.036 | 0.000 | AKT2,GNAI2,PRKCD,RELB,TEK |
| SUMOylation of transcription cofactors | 0.497 | 0.047 | #NUM! | PHC1,PHC2 |
| TBC/RABGAPs | 0.497 | 0.047 | #NUM! | RAB6B,TSC2 |
| Transcriptional regulation of granulopoiesis | 0.497 | 0.047 | #NUM! | H2AX,H3C1 |
| B Cell Activating Factor Signaling | 0.497 | 0.047 | #NUM! | NFAT5,RELB |
| Chemokine Signaling | 0.491 | 0.040 | #NUM! | GNAI2,PPP1R12B,RHOA |
| Protein Sorting Signaling Pathway | 0.490 | 0.034 | 1.633 | AP1M1,AP1S3,HMOX1,RAB11FIP1,SNX1,WASHC3 |
| Glutaminergic Receptor Signaling Pathway (Enhanced) | 0.490 | 0.032 | 0.632 | AKT2,CACNA1G,GNAI2,GRIK1,GRIK5,HMOX1,NAPA,PRKCD,SCN3A,SLC38A5 |
| Processing of Capped Intron-Containing Pre-mRNA | 0.486 | 0.032 | -1.000 | GPKOW,PCBP1,PHF5A,POLR2H,PRPF8,RBM7,SUGP1,WDR70,ZCRB1 |
| Netrin-1 signaling | 0.484 | 0.046 | #NUM! | SIAH1,SLIT2 |
| Angiopoietin Signaling | 0.482 | 0.040 | #NUM! | AKT2,RELB,TEK |
| Macropinocytosis Signaling | 0.482 | 0.040 | #NUM! | ITGB8,PRKCD,RHOA |
| Regulation of cholesterol biosynthesis by SREBP (SREBF) | 0.478 | 0.067 | #NUM! | SREBF1 |
| Metabolism of Angiotensinogen to Angiotensins | 0.478 | 0.067 | #NUM! | MME |
| Fatty Acid Activation | 0.478 | 0.067 | #NUM! | SLC27A4 |

| | | | | |
|---|---|---|---|---|
| Androgen Biosynthesis | 0.478 | 0.067 | #NUM! | CYP17A1 |
| NF-κB Activation by Viruses | 0.472 | 0.039 | #NUM! | AKT2,PRKCD,RELB |
| Leukocyte Extravasation Signaling | 0.472 | 0.034 | -0.816 | ACTN2,CTNNA3,GNAI2,MMP15,PRKCD,RHOA |
| Opioid Signaling Pathway | 0.471 | 0.032 | -0.707 | AKT2,ARRB2,CACNA1G,GNAI2,OPRD1,PDYN,PENK,PRKCD,RGS17 |
| Prostate Cancer Signaling | 0.465 | 0.036 | -1.000 | AKT2,E2F6,HSP90AA1,RELB |
| HEY1 Signaling Pathway | 0.464 | 0.035 | -1.342 | ACVRL1,AKT2,E2F6,MMP15,RUNX2 |
| T Cell Exhaustion Signaling Pathway | 0.464 | 0.035 | -0.447 | AKT2,IL10RB,NFAT5,PPP2R2C,STAT2 |
| RHOGDI Signaling | 0.463 | 0.033 | 1.000 | CDH12,GNAI2,ITGB8,MYH4,PPP1R12B,RHOA,WASF2 |
| Factors involved in megakaryocyte development and platelet pı | 0.463 | 0.039 | #NUM! | AK3,CAPZA2,H3C1 |
| Renal Cell Carcinoma Signaling | 0.463 | 0.039 | #NUM! | AKT2,RPS27A,UBB |
| Heme signaling | 0.459 | 0.044 | #NUM! | HMOX1,SLC46A1 |
| nNOS Signaling in Neurons | 0.459 | 0.044 | #NUM! | CAPN10,PRKCD |
| Processing of SMDT1 | 0.455 | 0.063 | #NUM! | PHB2 |
| Mitochondrial unfolded protein response (UPRmt) | 0.455 | 0.063 | #NUM! | NRF1 |
| D-myo-inositol (1,4,5)-trisphosphate Degradation | 0.455 | 0.063 | #NUM! | IMPA1 |
| Signaling by the B Cell Receptor (BCR) | 0.450 | 0.035 | -2.000 | PSMA4,PSMA7,RPS27A,UBB |
| CD28 Signaling in T Helper Cells | 0.450 | 0.035 | #NUM! | AKT2,LCP2,NFAT5,RELB |
| Virus Entry via Endocytic Pathways | 0.450 | 0.035 | 0.000 | AP1M1,AP1S3,ITGB8,PRKCD |
| TEC Kinase Signaling | 0.448 | 0.033 | #NUM! | GNAI2,ITGB8,PRKCD,RELB,RHOA,STAT2 |
| SUMOylation of RNA binding proteins | 0.447 | 0.043 | #NUM! | PHC1,PHC2 |
| Inositol phosphate metabolism | 0.447 | 0.043 | #NUM! | IMPA1,IP6K1 |
| Pyruvate metabolism | 0.447 | 0.043 | #NUM! | RPS27A,UBB |
| Complex IV assembly | 0.447 | 0.043 | #NUM! | COX11,SCO2 |
| RAR Activation | 0.445 | 0.031 | -1.732 | AKT2,DUSP1,PBRM1,POU5F1,PRKCD,RBP4,RELB,RHOA,RORB,SMARCB1,SMARCE1,SRA1 |
| Role of NFAT in Regulation of the Immune Response | 0.442 | 0.033 | -2.000 | AKT2,Fcgr2b,GNAI2,LCP2,NFAT5,RELB |
| Assembly of RNA Polymerase II Complex | 0.435 | 0.042 | #NUM! | POLR2H,TAF15 |
| IL-27 Signaling Pathway | 0.435 | 0.035 | #NUM! | AKT2,MCL1,Pycard,RELB |
| Metabolism of nitric oxide: NOS3 activation and regulation | 0.434 | 0.059 | #NUM! | HSP90AA1 |
| Azathioprine ADME | 0.434 | 0.059 | #NUM! | NUDT15 |
| NFE2L2 regulating anti-oxidant/detoxification enzymes | 0.434 | 0.059 | #NUM! | HMOX1 |
| Specification of the neural plate border | 0.434 | 0.059 | #NUM! | POU5F1 |
| Mismatch Repair in Eukaryotes | 0.434 | 0.059 | #NUM! | PCNA |
| Activin Inhibin Signaling Pathway | 0.431 | 0.032 | -1.633 | AKT2,BAMBI,INHBB,NANOG,POU5F1,RHOA |
| JAK/STAT Signaling | 0.428 | 0.037 | #NUM! | AKT2,RELB,STAT2 |
| Acetylcholine Receptor Signaling Pathway | 0.425 | 0.032 | 0.000 | ADAM9,AKT2,CACNA1G,GNAI2,HMOX1,PRKCD |
| TP53 Regulates Transcription of Cell Cycle Genes | 0.424 | 0.041 | #NUM! | CDK1,PCNA |
| RUNX1 regulates megakaryocyte differentiation and platelet fuı | 0.424 | 0.041 | #NUM! | H2AX,H3C1 |
| Signaling by NOTCH3 | 0.424 | 0.041 | #NUM! | RPS27A,UBB |
| Amyloid Processing | 0.424 | 0.041 | #NUM! | AKT2,CAPN10 |
| Proteasomal PSMD10 Signaling Pathway | 0.421 | 0.034 | -1.000 | AKT2,E2F6,FGF2,POU5F1 |
| Cyclins and Cell Cycle Regulation | 0.419 | 0.036 | #NUM! | CDK1,E2F6,PPP2R2C |
| Metabolism of amine-derived hormones | 0.415 | 0.056 | #NUM! | SLC5A5 |
| Glyoxylate metabolism and glycine degradation | 0.415 | 0.056 | #NUM! | KGD4 |
| Formation of the nephric duct | 0.415 | 0.056 | #NUM! | FGF2 |
| Formation of the ureteric bud | 0.415 | 0.056 | #NUM! | SLIT2 |
| PKCθ Signaling in T Lymphocytes | 0.414 | 0.033 | #NUM! | CACNA1G,LCP2,MAP3K6,NFAT5,RELB |
| Transcriptional activity of SMAD2/SMAD3:SMAD4 heterotrimer | 0.413 | 0.040 | #NUM! | RPS27A,UBB |
| FGF Signaling | 0.411 | 0.036 | #NUM! | AKT2,FGF2,FGF9 |
| Role of NANOG in Mammalian Embryonic Stem Cell Pluripotenc | 0.408 | 0.034 | #NUM! | ACVRL1,AKT2,NANOG,POU5F1 |
| Macrophage Alternative Activation Signaling Pathway | 0.408 | 0.033 | -0.447 | AKT2,DUSP1,IL10RB,RELB,SREBF1 |
| Mitotic Metaphase and Anaphase | 0.406 | 0.031 | -1.890 | CDK1,CENPK,PSMA4,PSMA7,RPS27A,TUBB3,UBB |
| Interleukin-3, Interleukin-5 and GM-CSF signaling | 0.403 | 0.039 | #NUM! | RPS27A,UBB |
| Proteasome assembly | 0.403 | 0.039 | #NUM! | PSMA4,PSMA7 |
| Activation of IRF by Cytosolic Pattern Recognition Receptors | 0.403 | 0.039 | #NUM! | RELB,STAT2 |
| RHOA Signaling | 0.401 | 0.033 | -1.000 | PKN1,PPP1R12B,RHOA,SEPTIN1 |

| | | | | |
|---|---|---|---|---|
| Cell Cycle Checkpoints | 0.400 | 0.031 | -2.121 | CDK1,CENPK,H2AX,MCM4,PSMA4,PSMA7,RPS27A,UBB |
| Signaling by Activin | 0.396 | 0.053 | #NUM! | INHBB |
| Activation of Matrix Metalloproteinases | 0.396 | 0.053 | #NUM! | MMP15 |
| FOXO-mediated transcription | 0.396 | 0.053 | #NUM! | AKT2 |
| Formation of ATP by chemiosmotic coupling | 0.396 | 0.053 | #NUM! | ATP5PB |
| γ-linolenate Biosynthesis II (Animals) | 0.396 | 0.053 | #NUM! | SLC27A4 |
| Mitochondrial L-carnitine Shuttle Pathway | 0.396 | 0.053 | #NUM! | SLC27A4 |
| Methylglyoxal Degradation III | 0.396 | 0.053 | #NUM! | AKR1A1 |
| Inflammasome pathway | 0.396 | 0.053 | #NUM! | Pycard |
| Signaling by Rho Family GTPases | 0.396 | 0.030 | -1.342 | CDH12,GNAI2,ITGB8,PKN1,PPP1R12B,RELB,RHOA,SEPTIN1 |
| BMP signaling pathway | 0.395 | 0.035 | #NUM! | ACVRL1,RELB,RUNX2 |
| Xenobiotic Metabolism General Signaling Pathway | 0.394 | 0.033 | 1.000 | AKT2,HMOX1,MAP3K6,PRKCD |
| Mitotic Prometaphase | 0.393 | 0.031 | -1.633 | CDK1,CENPK,CEP192,HSP90AA1,TUBB,TUBB3 |
| Intrinsic Pathway for Apoptosis | 0.383 | 0.038 | #NUM! | AKT2,APIP |
| Opioid Signalling | 0.380 | 0.034 | #NUM! | GNAI2,PDYN,PRKCD |
| Fcgamma receptor (FCGR) dependent phagocytosis | 0.380 | 0.034 | #NUM! | HSP90AA1,PRKCD,WASF2 |
| Unfolded protein response | 0.380 | 0.034 | #NUM! | DNAJA2,DNAJC1,SREBF1 |
| Dendritic Cell Maturation | 0.379 | 0.031 | 1.000 | AKT2,COL5A3,Fcgr2b,RELB,STAT2 |
| Folate Signaling Pathway | 0.373 | 0.037 | #NUM! | POU5F1,SLC46A1 |
| Lymphotoxin β Receptor Signaling | 0.373 | 0.037 | #NUM! | AKT2,RELB |
| Cell junction organization | 0.373 | 0.034 | #NUM! | CDH12,NECTIN2,TESK1 |
| RANK Signaling in Osteoclasts | 0.373 | 0.034 | #NUM! | AKT2,MAP3K6,RELB |
| Interleukin-1 family signaling | 0.369 | 0.032 | -2.000 | PSMA4,PSMA7,RPS27A,UBB |
| Tumor Microenvironment Pathway | 0.368 | 0.031 | 0.000 | AKT2,FGF2,FGF9,MMP15,RELB |
| Chronic Myeloid Leukemia Signaling | 0.366 | 0.030 | -1.414 | AKT2,CTBP2,E2F6,HSP90AA1,MCL1,MSI2,NFAT5,RELB |
| Actin Nucleation by ARP-WASP Complex | 0.365 | 0.033 | #NUM! | ITGB8,PPP1R12B,RHOA |
| Somitogenesis | 0.364 | 0.036 | #NUM! | PSMA4,PSMA7 |
| Role of Hypercytokinemia/hyperchemokinemia in the Pathoger | 0.364 | 0.036 | #NUM! | Pycard,STAT2 |
| FAT10 Signaling Pathway | 0.364 | 0.036 | #NUM! | PSMA4,PSMA7 |
| RHO GTPases activate PAKs | 0.364 | 0.048 | #NUM! | PPP1R12B |
| Insertion of tail-anchored proteins into the endoplasmic reticulu | 0.364 | 0.048 | #NUM! | HMOX1 |
| Citric acid cycle (TCA cycle) | 0.364 | 0.048 | #NUM! | KGD4 |
| Co-inhibition by CTLA4 | 0.364 | 0.048 | #NUM! | AKT2 |
| Vitamin-C Transport | 0.364 | 0.048 | #NUM! | LRRC8C |
| Transcriptional Regulatory Network in Embryonic Stem Cells | 0.363 | 0.031 | -0.447 | AKT2,FGF2,NANOG,POU5F1,SIX3 |
| Protein folding | 0.358 | 0.033 | #NUM! | CCT3,FKBP9,TUBB3 |
| COPI-mediated anterograde transport | 0.358 | 0.033 | #NUM! | CAPZA2,NAPA,TUBB3 |
| Mitochondrial Division Signaling Pathway | 0.358 | 0.031 | -0.447 | CACNA1G,CDK1,PPP2R2C,PRKCD,RHOA |
| DYRK1A Signaling Pathway | 0.358 | 0.031 | -1.342 | AKT2,MME,POLR2H,RBBP4,SPRY2 |
| Asparagine N-linked glycosylation | 0.357 | 0.032 | -2.000 | ALG11,DAD1,RPS27A,UBB |
| Cerebral Malformation Signaling Pathway | 0.357 | 0.032 | #NUM! | AKT2,CDK1,PPP2R2C,RHOA |
| Gamma carboxylation, hypusinylation, hydroxylation, and arylsı | 0.355 | 0.036 | #NUM! | DRG2,GAS6 |
| Vitamin K Signaling Pathway | 0.355 | 0.036 | #NUM! | GAS6,UBIAD1 |
| CLEAR Signaling Pathway | 0.354 | 0.029 | 0.707 | AKT2,DDIT4,FGF2,PPP2R2C,PRKCD,SGSH,STK4,TSC2 |
| Gα12/13 Signaling | 0.352 | 0.031 | #NUM! | AKT2,CDH12,RELB,RHOA |
| G Beta Gamma Signaling | 0.352 | 0.031 | 0.000 | AKT2,CACNA1G,GNAI2,PRKCD |
| Metabolism of porphyrins | 0.349 | 0.046 | #NUM! | HMOX1 |
| Mitochondrial RNA degradation | 0.349 | 0.046 | #NUM! | SLIRP |
| Superpathway of D-myo-inositol (1,4,5)-trisphosphate Metaboli | 0.349 | 0.046 | #NUM! | IMPA1 |
| Acute Phase Response Signaling | 0.348 | 0.030 | #NUM! | AKT2,HMOX1,Orm2/Orm3,RBP4,RELB |
| Visual phototransduction | 0.346 | 0.035 | #NUM! | RBP4,RCVRN |
| Polyamine Regulation in Colon Cancer | 0.346 | 0.035 | #NUM! | AKT2,RHOA |
| Centrosomal KIAA0586 Signaling Pathway | 0.346 | 0.031 | -2.000 | FGF2,FGF9,PSMA4,PSMA7 |
| fMLP Signaling in Neutrophils | 0.346 | 0.031 | #NUM! | GNAI2,NFAT5,PRKCD,RELB |
| CCR3 Signaling in Eosinophils | 0.346 | 0.031 | #NUM! | GNAI2,PPP1R12B,PRKCD,RHOA |

| | | | | |
|---|---|---|---|---|
| HGF Signaling | 0.340 | 0.031 | #NUM! | AKT2,ITGB8,MAP3K6,PRKCD |
| GABA Receptor Signaling | 0.340 | 0.031 | #NUM! | CACNA1G,GNAI2,RPS27A,UBB |
| Xenobiotic Metabolism Signaling | 0.340 | 0.029 | #NUM! | HMOX1,HSP90AA1,MAP3K6,PPP2R2C,PRKCD,RELB,SRA1 |
| Mitotic Prophase | 0.338 | 0.032 | #NUM! | CDK1,H2AX,H3C1 |
| Non-Small Cell Lung Cancer Signaling | 0.338 | 0.032 | #NUM! | AKT2,E2F6,STK4 |
| Integrin to Cytoskeleton Signaling Pathway | 0.336 | 0.029 | -2.236 | ACTN2,AKT2,CAPN10,ITGB8,PPP1R12B,RHOA |
| Systemic Lupus Erythematosus in B Cell Signaling Pathway | 0.336 | 0.029 | -0.447 | AKT2,Fcgr2b,MCL1,NFAT5,PRKCD,RELB,STAT2 |
| SNARE Signaling Pathway | 0.335 | 0.031 | #NUM! | MYH4,NAPA,RAB9B,SNAP29 |
| SLAMF Signaling Pathway | 0.335 | 0.031 | -1.000 | AKT2,LCP2,NFAT5,PHB2 |
| P2Y Purinergic Receptor Signaling Pathway | 0.335 | 0.031 | #NUM! | AKT2,GNAI2,PRKCD,RELB |
| Nephrin family interactions | 0.335 | 0.044 | #NUM! | ACTN2 |
| Cellular response to mitochondrial stress | 0.335 | 0.044 | #NUM! | PHB2 |
| Apelin Cardiac Fibroblast Signaling Pathway | 0.335 | 0.044 | #NUM! | AKT2 |
| p53 Signaling | 0.332 | 0.032 | #NUM! | AKT2,PCNA,SCO2 |
| PCP (Planar Cell Polarity) Pathway | 0.329 | 0.034 | #NUM! | LGR4,RHOA |
| Glioblastoma Multiforme Signaling | 0.328 | 0.030 | -1.000 | AKT2,E2F6,PRKCD,RHOA,TSC2 |
| Myelination Signaling Pathway | 0.328 | 0.028 | -1.667 | ACVRL1,AKT2,CNTN6,FGF2,ITGB8,RHOA,SREBF1,TSC2,ZEB2 |
| ERK/MAPK Signaling | 0.328 | 0.029 | 1.342 | DUSP1,ITGB8,PPP1R14B,PPP1R14C,PPP2R2C,PRKCD |
| Apelin Cardiomyocyte Signaling Pathway | 0.325 | 0.031 | #NUM! | AKT2,GNAI2,PRKCD |
| Miscellaneous transport and binding events | 0.321 | 0.042 | #NUM! | LRRC8C |
| RHO GTPases activate IQGAPs | 0.321 | 0.042 | #NUM! | TUBB3 |
| RHO GTPases Activate NADPH Oxidases | 0.321 | 0.042 | #NUM! | PRKCD |
| Cardiomyocyte Differentiation via BMP Receptors | 0.321 | 0.042 | #NUM! | ACVRL1 |
| Metabolism of polyamines | 0.321 | 0.033 | #NUM! | PSMA4,PSMA7 |
| Role of WNT/GSK-3β Signaling in the Pathogenesis of Influenza | 0.321 | 0.033 | #NUM! | NCOA4,SIAH1 |
| IL-17A Signaling in Airway Cells | 0.321 | 0.033 | #NUM! | AKT2,RELB |
| Cellular response to heat stress | 0.319 | 0.031 | #NUM! | DEDD2,HSP90AA1,UBB |
| ATM Signaling | 0.319 | 0.031 | #NUM! | CDK1,H2AX,PPP2R2C |
| Small Cell Lung Cancer Signaling | 0.319 | 0.031 | #NUM! | AKT2,E2F6,RELB |
| p70S6K Signaling | 0.319 | 0.030 | 0.000 | AKT2,GNAI2,PPP2R2C,PRKCD |
| Ephrin A Signaling | 0.313 | 0.030 | 0.000 | AKT2,FGF2,RHOA,TSC2 |
| Mitochondrial biogenesis | 0.313 | 0.033 | #NUM! | HCFC1,NRF1 |
| Abacavir ADME | 0.313 | 0.033 | #NUM! | NT5C2,STK4 |
| SUMOylation of chromatin organization proteins | 0.313 | 0.033 | #NUM! | PHC1,PHC2 |
| mRNA 3 Prime End Processing Signaling Pathway | 0.313 | 0.031 | #NUM! | DIS3L2,E2F6,PCBP1 |
| Gustation Pathway | 0.310 | 0.029 | -0.447 | CACNA1G,LRRC8C,P2RX6,PKD2L1,SCN3A |
| Pathogen Induced Cytokine Storm Signaling Pathway | 0.310 | 0.028 | 2.646 | COL23A1,COL5A3,COL6A1,COL6A2,LYST,NLRC5,Pycard,SLC2A8 |
| Insulin processing | 0.309 | 0.040 | #NUM! | EXOC6 |
| Signaling by NTRK3 (TRKC) | 0.309 | 0.040 | #NUM! | PTPRO |
| RORA,B,C and NR1D1 (REV-ERBA) regulate gene expression | 0.309 | 0.040 | #NUM! | SREBF1 |
| Antigen Presentation Pathway | 0.309 | 0.040 | #NUM! | NLRC5 |
| Necroptosis Signaling Pathway | 0.308 | 0.029 | #NUM! | CAPN10,Pycard,STAT2,TOMM7 |
| Sumoylation Pathway | 0.307 | 0.030 | #NUM! | CTBP2,PCNA,RHOA |
| Ribavirin ADME | 0.306 | 0.032 | #NUM! | NT5C2,STK4 |
| Axonal Guidance Signaling | 0.304 | 0.027 | #NUM! | ADAM9,AKT2,GNAI2,ITGB8,MME,MMP15,NFAT5,NTNG1,PRKCD,RHOA,SLIT2,TUBB,TUBB3 |
| Phagosome Formation | 0.304 | 0.027 | 2.000 | AKT2,AP1M1,AP1S3,CALCR,Fcgr2b,GPR3,GPR52,GPR63,HMOX1,ITGB8,LGR4,MARCKS,MYH4,OPRD1,PRKCD,RHOA,WASF2 |
| PPAR Signaling | 0.301 | 0.030 | #NUM! | HSP90AA1,RELB,SRA1 |
| Crosstalk between Dendritic Cells and Natural Killer Cells | 0.298 | 0.032 | #NUM! | NECTIN2,RELB |
| ATF4 activates genes in response to endoplasmic reticulum stre | 0.297 | 0.039 | #NUM! | KHSRP |
| Peroxisomal lipid metabolism | 0.297 | 0.039 | #NUM! | ABCD1 |
| Ethanol Degradation II | 0.297 | 0.039 | #NUM! | AKR1A1 |
| Phase II - Conjugation of compounds | 0.291 | 0.031 | #NUM! | AKR1A1,CHAC1 |
| Semaphorin interactions | 0.291 | 0.031 | #NUM! | HSP90AA1,RHOA |
| Membrane Repair Signaling Pathway | 0.290 | 0.029 | #NUM! | CAPN10,MYH4,NAPA |
| Mouse Embryonic Stem Cell Pluripotency | 0.290 | 0.029 | #NUM! | AKT2,NANOG,POU5F1 |

| | | | | |
|---|---|---|---|---|
| Endocannabinoid Cancer Inhibition Pathway | 0.289 | 0.029 | 1.000 | AKT2,GNAI2,NSMAF,RHOA |
| Effects of PIP2 hydrolysis | 0.286 | 0.037 | #NUM! | PRKCD |
| Interleukin-7 signaling | 0.286 | 0.037 | #NUM! | H3C1 |
| Interconversion of nucleotide di- and triphosphates | 0.286 | 0.037 | #NUM! | DCTPP1 |
| Noradrenaline and Adrenaline Degradation | 0.286 | 0.037 | #NUM! | AKR1A1 |
| Superpathway of Cholesterol Biosynthesis | 0.286 | 0.037 | #NUM! | SC5D |
| Interferon Signaling | 0.286 | 0.037 | #NUM! | STAT2 |
| Egg-Sperm Fusion Signaling Pathway | 0.284 | 0.029 | #NUM! | AKT2,CDK1,NAPA |
| CDC42 Signaling | 0.284 | 0.028 | #NUM! | EXOC6,EXOC6B,ITGB8,PPP1R12B |
| PI3K Signaling in B Lymphocytes | 0.284 | 0.028 | #NUM! | AKT2,NFAT5,PLEKHA3,RELB |
| Thrombin Signaling | 0.280 | 0.028 | -1.000 | AKT2,GNAI2,PPP1R12B,PRKCD,RELB,RHOA |
| B Cell Receptor Signaling | 0.280 | 0.028 | #NUM! | AKT2,Fcgr2b,MAP3K6,NFAT5,RELB |
| Telomerase Signaling | 0.279 | 0.029 | #NUM! | AKT2,HSP90AA1,PPP2R2C |
| Antioxidant Action of Vitamin C | 0.279 | 0.029 | #NUM! | HMOX1,RELB,SLC2A8 |
| WNT/Ca2+ Signaling Pathway | 0.277 | 0.030 | #NUM! | NFAT5,RELB |
| Glutamate Receptor Signaling | 0.277 | 0.030 | #NUM! | GRIK1,GRIK5 |
| Sulfur amino acid metabolism | 0.275 | 0.036 | #NUM! | APIP |
| Semaphorin Neuronal Repulsive Signaling Pathway | 0.275 | 0.028 | #NUM! | AKT2,ITGB8,PPP1R12B,RHOA |
| CCR5 Signaling in Macrophages | 0.273 | 0.029 | #NUM! | CACNA1G,GNAI2,PRKCD |
| Cachexia Signaling Pathway | 0.273 | 0.027 | -1.000 | AKT2,CAPN10,DDIT4,HSP90AA1,INHBB,PRKCD,PSMA4,PSMA7,STAT2 |
| Multiple Sclerosis Signaling Pathway | 0.272 | 0.028 | 0.000 | CAPN10,CTNNA3,DUSP1,Pycard,RBBP4 |
| Xenobiotic Metabolism AHR Signaling Pathway | 0.271 | 0.030 | #NUM! | HSP90AA1,RELB |
| Phagosome Maturation | 0.270 | 0.028 | #NUM! | NAPA,TUBB,TUBB3,VPS33B |
| Xenobiotic Metabolism CAR Signaling Pathway | 0.270 | 0.028 | -2.000 | HSP90AA1,PPP2R2C,PRKCD,SRA1 |
| LXR/RXR Activation | 0.268 | 0.028 | #NUM! | RBP4,RELB,SREBF1 |
| HOTAIR Regulatory Pathway | 0.266 | 0.028 | #NUM! | AKT2,MMP15,RBBP4,RELB |
| HER-2 Signaling in Breast Cancer | 0.265 | 0.027 | 0.000 | AKT2,COX11,ITGB8,PRKCD,RELB,TSC2 |
| Energy dependent regulation of mTOR by LKB1-AMPK | 0.265 | 0.035 | #NUM! | TSC2 |
| mRNA Capping | 0.265 | 0.035 | #NUM! | POLR2H |
| FOXO-mediated transcription of oxidative stress, metabolic and | 0.265 | 0.035 | #NUM! | SREBF1 |
| Response of EIF2AK1 (HRI) to heme deficiency | 0.265 | 0.035 | #NUM! | CHAC1 |
| ISG15 antiviral mechanism | 0.264 | 0.029 | #NUM! | RPS27A,UBB |
| MITF-M-dependent gene expression | 0.263 | 0.028 | #NUM! | AKT2,SMARCB1,SMARCE1 |
| Regulation of Actin-based Motility by Rho | 0.263 | 0.028 | #NUM! | ITGB8,PPP1R12B,RHOA |
| Preeclampsia Signaling Pathway | 0.262 | 0.027 | -1.000 | AKT2,HMOX1,NOC2L,RHOA |
| Plasma lipoprotein assembly, remodeling, and clearance | 0.258 | 0.029 | #NUM! | RPS27A,UBB |
| Sensory processing of sound by inner hair cells of the cochlea | 0.258 | 0.029 | #NUM! | CAPZA2,CTBP2 |
| Signaling by ALK | 0.255 | 0.033 | #NUM! | PTPRZ1 |
| DAP12 interactions | 0.255 | 0.033 | #NUM! | LCP2 |
| Sensory perception of taste | 0.255 | 0.033 | #NUM! | SCN3A |
| Regulation of CDH11 Expression and Function | 0.255 | 0.033 | #NUM! | ZEB2 |
| Tryptophan Degradation X (Mammalian, via Tryptamine) | 0.255 | 0.033 | #NUM! | AKR1A1 |
| Atherosclerosis Signaling | 0.253 | 0.028 | #NUM! | COL5A3,RBP4,RELB |
| RNA m6A Methylation Signaling Pathway | 0.252 | 0.027 | -1.342 | AKT2,IGF2BP3,POU5F1,RELB,SPRED2 |
| Pre-NOTCH Expression and Processing | 0.252 | 0.029 | #NUM! | H2AX,H3C1 |
| Sphingosine-1-phosphate Signaling | 0.248 | 0.027 | #NUM! | AKT2,GNAI2,RHOA |
| Transcriptional regulation of brown and beige adipocyte differe | 0.246 | 0.032 | #NUM! | RBBP4 |
| TNFR2 Signaling | 0.246 | 0.032 | #NUM! | RELB |
| Pyroptosis Signaling Pathway | 0.246 | 0.028 | #NUM! | Pycard,RHOA |
| COPII-mediated vesicle transport | 0.246 | 0.028 | #NUM! | NAPA,TRAPPC6B |
| IL-17A Signaling in Fibroblasts | 0.246 | 0.028 | #NUM! | P4HA2,RELB |
| IL-7 Signaling Pathway | 0.246 | 0.028 | #NUM! | AKT2,MCL1 |
| Fc Epsilon RI Signaling | 0.244 | 0.027 | #NUM! | AKT2,LCP2,PRKCD |
| VDR/RXR Activation | 0.240 | 0.028 | #NUM! | PRKCD,RUNX2 |
| Role of JAK family kinases in IL-6-type Cytokine Signaling | 0.240 | 0.028 | #NUM! | HSP90AA1,MCL1 |

| | | | | |
|---|---|---|---|---|
| Amyotrophic Lateral Sclerosis Signaling | 0.239 | 0.027 | #NUM! | CAPN10,GRIK1,GRIK5 |
| L1CAM interactions | 0.239 | 0.027 | #NUM! | CNTN6,SCN3A,TUBB3 |
| Regulation of lipid metabolism by PPARalpha | 0.239 | 0.027 | #NUM! | MED7,NRF1,SREBF1 |
| Thrombin signalling through proteinase activated receptors (PA | 0.238 | 0.031 | #NUM! | ARRB2 |
| Cristae formation | 0.238 | 0.031 | #NUM! | ATP5PB |
| Role of RIG1-like Receptors in Antiviral Innate Immunity | 0.238 | 0.031 | #NUM! | RELB |
| Inhibition of Angiogenesis by TSP1 | 0.238 | 0.031 | #NUM! | AKT2 |
| Ephrin B Signaling | 0.235 | 0.027 | #NUM! | GNAI2,RHOA |
| MIF-mediated Glucocorticoid Regulation | 0.229 | 0.030 | #NUM! | RELB |
| RNA polymerase II transcribes snRNA genes | 0.229 | 0.027 | #NUM! | ELL2,POLR2H |
| Metabolism of steroid hormones | 0.221 | 0.029 | #NUM! | CYP17A1 |
| Activation of the pre-replicative complex | 0.221 | 0.029 | #NUM! | MCM4 |
| 4-1BB Signaling in T Lymphocytes | 0.221 | 0.029 | #NUM! | RELB |
| Fatty Acid β-oxidation I | 0.221 | 0.029 | #NUM! | SLC27A4 |
| Superpathway of Methionine Degradation | 0.221 | 0.029 | #NUM! | MCEE |
| Nucleotide Excision Repair Pathway | 0.221 | 0.029 | #NUM! | POLR2H |
| Mitochondrial Fatty Acid Beta-Oxidation | 0.214 | 0.029 | #NUM! | MCEE |
| Expression of BMAL (ARNTL), CLOCK, and NPAS2 | 0.214 | 0.029 | #NUM! | RORB |
| IL-9 Signaling | 0.214 | 0.029 | #NUM! | RELB |
| Intrinsic Prothrombin Activation Pathway | 0.214 | 0.029 | #NUM! | COL5A3 |
| Striated Muscle Contraction | 0.206 | 0.028 | #NUM! | ACTN2 |
| Interleukin-10 signaling | 0.206 | 0.028 | #NUM! | IL10RB |
| Formation of the dystrophin-glycoprotein complex (DGC) | 0.206 | 0.028 | #NUM! | DTNB |
| TWEAK Signaling | 0.206 | 0.028 | #NUM! | RELB |
| Binding and Uptake of Ligands by Scavenger Receptors | 0.199 | 0.027 | #NUM! | HSP90AA1 |
| Role of IL-17F in Allergic Inflammatory Airway Diseases | 0.199 | 0.027 | #NUM! | RELB |
| Notch Signaling | 0.199 | 0.027 | #NUM! | DTX4 |
| Dilated Cardiomyopathy Signaling Pathway | 0.000 | 0.020 | #NUM! | AKT2,CACNA1G,MYH4 |
| Neurovascular Coupling Signaling Pathway | 0.000 | 0.023 | 1.342 | CACNA1G,ENTPD6,ENTPD7,PPP1R12B,RHOA |
| Oxytocin in Brain Signaling Pathway | 0.000 | 0.022 | 0.000 | AKT2,CACNA1G,GNAI2,PRKCD |
| Pulmonary Healing Signaling Pathway | 0.000 | 0.011 | #NUM! | MMP15,PRKCD |
| Wound Healing Signaling Pathway | 0.000 | 0.023 | 2.236 | COL23A1,COL5A3,COL6A1,COL6A2,FGF2 |
| ID1 Signaling Pathway | 0.000 | 0.025 | 0.447 | ACVRL1,AKT2,FGF2,NANOG,POU5F1 |
| CDX Gastrointestinal Cancer Signaling Pathway | 0.000 | 0.011 | #NUM! | AKT2,RELB |
| Immunogenic Cell Death Signaling Pathway | 0.000 | 0.026 | #NUM! | HSP90AA1,Pycard |
| Macrophage Classical Activation Signaling Pathway | 0.000 | 0.014 | #NUM! | RBBP4,STAT2 |
| Role of Chondrocytes in Rheumatoid Arthritis Signaling Pathway | 0.000 | 0.024 | #NUM! | MMP15,Pycard,RUNX2 |
| Role of Osteoblasts in Rheumatoid Arthritis Signaling Pathway | 0.000 | 0.024 | -1.000 | ACVRL1,AKT2,MMP15,RUNX2,STAT2 |
| Glucocorticoid Receptor Signaling | 0.000 | 0.026 | #NUM! | AKT2,ATP5PB,DUSP1,HSP90AA1,IL10RB,NFAT5,PBRM1,POLR2H,SMARCB1,SMARCE1,SRA1,TAF15 |
| S100 Family Signaling Pathway | 0.000 | 0.023 | 2.138 | AKT2,CACNA1G,CALCR,EREG,Fcgr2b,GPR3,GPR52,GPR63,IL10RB,LGR4,MMP15,OPRD1,PRKCD,RHOA,S100A11,S100A4 |
| Circadian Rhythm Signaling | 0.000 | 0.015 | #NUM! | CACNA1G,GNAI2,PRKCD,RORB |
| Synaptic Long Term Potentiation | 0.000 | 0.008 | #NUM! | PRKCD |
| Synaptic Long Term Depression | 0.000 | 0.021 | 1.000 | CACNA1G,GNAI2,PPP2R2C,PRKCD |
| IL-33 Signaling Pathway | 0.000 | 0.022 | #NUM! | AKT2,H2AX,H2AZ1 |
| NOD1/2 Signaling Pathway | 0.000 | 0.014 | #NUM! | HSP90AA1,Pycard |
| ABRA Signaling Pathway | 0.000 | 0.012 | #NUM! | RHOA |
| Adrenergic Receptor Signaling Pathway (Enhanced) | 0.000 | 0.017 | #NUM! | CACNA1G,GNAI2,PRKCD |
| GABAergic Receptor Signaling Pathway (Enhanced) | 0.000 | 0.015 | #NUM! | CACNA1G,GNAI2 |
| NFKBIE Signaling Pathway | 0.000 | 0.018 | #NUM! | RELB |
| Orexin Signaling Pathway | 0.000 | 0.013 | #NUM! | AKT2,CACNA1G,PRKCD |
| Sertoli Cell-Germ Cell Junction Signaling Pathway (Enhanced) | 0.000 | 0.018 | 0.000 | AKT2,MAP3K6,NECTIN2,PPP2R2C |
| PPARα/RXRα Activation | 0.000 | 0.011 | #NUM! | HSP90AA1,RELB |
| Aryl Hydrocarbon Receptor Signaling | 0.000 | 0.015 | #NUM! | HSP90AA1,RELB |
| LPS/IL-1 Mediated Inhibition of RXR Function | 0.000 | 0.010 | #NUM! | SLC27A4,SREBF1 |
| G alpha (i) signalling events | 0.000 | 0.023 | -0.447 | GNAI2,OPRD1,PDYN,PENK,RGS17 |

| | | | | |
|---|---|---|---|---|
| Meiotic synapsis | 0.000 | 0.023 | #NUM! | H2AX |
| Mitochondrial protein import | 0.000 | 0.016 | #NUM! | TOMM7 |
| Potassium Channels | 0.000 | 0.010 | #NUM! | KCNC4 |
| PI Metabolism | 0.000 | 0.025 | #NUM! | PLEKHA3,VAC14 |
| DAG and IP3 signaling | 0.000 | 0.026 | #NUM! | PRKCD |
| Elastic fibre formation | 0.000 | 0.024 | #NUM! | ITGB8 |
| Gap junction trafficking and regulation | 0.000 | 0.026 | #NUM! | TUBB3 |
| Glycosaminoglycan metabolism | 0.000 | 0.016 | #NUM! | SDC4,SGSH |
| Macroautophagy | 0.000 | 0.015 | #NUM! | TSC2 |
| Integration of energy metabolism | 0.000 | 0.019 | #NUM! | GNAI2,MARCKS |
| Regulation of beta-cell development | 0.000 | 0.024 | #NUM! | AKT2 |
| Signaling by NTRK1 (TRKA) | 0.000 | 0.012 | #NUM! | RHOA |
| p75 NTR receptor-mediated signalling | 0.000 | 0.027 | -1.000 | RHOA,RPS27A,STK4,UBB |
| Metabolism of water-soluble vitamins and cofactors | 0.000 | 0.014 | #NUM! | BTD |
| Immunoregulatory interactions between a Lymphoid and a non | 0.000 | 0.012 | #NUM! | NECTIN2 |
| Cell surface interactions at the vascular wall | 0.000 | 0.026 | #NUM! | GAS6,SDC4,TEK |
| Generic Transcription Pathway | 0.000 | 0.016 | #NUM! | MED7,RORB,ZNF786 |
| RNA Polymerase II Transcription | 0.000 | 0.021 | #NUM! | EAF1,POLR2H,TAF15 |
| Phase I - Functionalization of compounds | 0.000 | 0.026 | #NUM! | CYP17A1,CYP20A1 |
| EPH-Ephrin signaling | 0.000 | 0.011 | #NUM! | RHOA |
| SUMOylation of DNA damage response and repair proteins | 0.000 | 0.026 | #NUM! | PHC1,PHC2 |
| Class A/1 (Rhodopsin-like receptors) | 0.000 | 0.014 | -1.000 | OPRD1,PDYN,PENK,PLPPR3 |
| Class B/2 (Secretin family receptors) | 0.000 | 0.011 | #NUM! | CALCR |
| Transcriptional regulation of white adipocyte differentiation | 0.000 | 0.024 | #NUM! | MED7,SREBF1 |
| G alpha (q) signalling events | 0.000 | 0.006 | #NUM! | RGS17 |
| G alpha (12/13) signalling events | 0.000 | 0.013 | #NUM! | RHOA |
| Platelet homeostasis | 0.000 | 0.023 | #NUM! | ATP2B4,P2RX6 |
| G alpha (s) signalling events | 0.000 | 0.022 | #NUM! | ARRB2,CALCR,GNAI2 |
| Sphingolipid metabolism | 0.000 | 0.010 | #NUM! | SPTSSB |
| Activation of NMDA receptors and postsynaptic events | 0.000 | 0.026 | #NUM! | ACTN2,TUBB3 |
| Smooth Muscle Contraction | 0.000 | 0.026 | #NUM! | CACNA1G |
| SUMOylation of DNA replication proteins | 0.000 | 0.022 | #NUM! | PCNA |
| O-linked glycosylation | 0.000 | 0.018 | #NUM! | C1GALT1C1,GALNT2 |
| Cardiac conduction | 0.000 | 0.024 | #NUM! | ATP2B4,MME,SCN3A |
| RHO GTPases Activate WASPs and WAVEs | 0.000 | 0.023 | #NUM! | WASF2 |
| RHO GTPases Activate Formins | 0.000 | 0.023 | #NUM! | CENPK,RHOA,TUBB3 |
| Nonhomologous End-Joining (NHEJ) | 0.000 | 0.026 | #NUM! | H2AX |
| Resolution of Abasic Sites (AP sites) | 0.000 | 0.026 | #NUM! | PCNA |
| Interferon gamma signaling | 0.000 | 0.015 | #NUM! | PRKCD |
| Carboxyterminal post-translational modifications of tubulin | 0.000 | 0.026 | #NUM! | TUBB3 |
| RHO GTPase cycle | 0.000 | 0.025 | -2.111 | ESYT1,HSP90AA1,MSI2,NISCH,PKN1,PLEKHG1,RHOA,STIP1,SWAP70,TJP2,WASF2 |
| NR1H2 and NR1H3-mediated signaling | 0.000 | 0.022 | #NUM! | SREBF1 |
| GPER1 signaling | 0.000 | 0.021 | #NUM! | GNAI2,STK4 |
| Cytoprotection by HMOX1 | 0.000 | 0.018 | #NUM! | HMOX1 |
| GABA receptor activation | 0.000 | 0.017 | #NUM! | GNAI2 |
| Class I MHC mediated antigen processing and presentation | 0.000 | 0.019 | -1.890 | FBXO11,PSMA4,PSMA7,RPS27A,SIAH1,UBB,UBE2R2 |
| Kinesins | 0.000 | 0.020 | #NUM! | TUBB3 |
| Ion channel transport | 0.000 | 0.023 | -1.000 | ASIC4,ATP2B4,RPS27A,UBB |
| Hepatic Cholestasis | 0.000 | 0.020 | 2.000 | AKT2,PRKCD,RELB,TJP2 |
| Mitochondrial Dysfunction | 0.000 | 0.019 | 0.816 | ATP5PB,CACNA1G,CAPN10,COX11,NRF1,TOMM7 |
| Autism Signaling Pathway | 0.000 | 0.007 | #NUM! | AKT2,TSC2 |
| CGAS-STING Signaling Pathway | 0.000 | 0.009 | #NUM! | Pycard |
| NAFLD Signaling Pathway | 0.000 | 0.020 | #NUM! | AKT2,PPP2R2C,Pycard,SREBF1 |
| Pancreatic Secretion Signaling Pathway | 0.000 | 0.023 | -1.342 | AKT2,ATP2B4,NAPA,PRKCD,RHOA |
| ROBO SLIT Signaling Pathway | 0.000 | 0.025 | #NUM! | AKT2,RHOA,SLIT2 |

| | | | | |
|---|---|---|---|---|
| FXR/RXR Activation | 0.000 | 0.019 | #NUM! | AKT2,HMOX1,SREBF1 |
| BBSome Signaling Pathway | 0.000 | 0.024 | 2.530 | ARRB2,CALCR,CCT3,GPR3,GPR52,GPR63,LGR4,MKKS,OPRD1,PRKCD,RHOA |
| Sleep NREM Signaling Pathway | 0.000 | 0.009 | #NUM! | GNAI2 |
| UFMylation Signaling Pathway | 0.000 | 0.017 | #NUM! | AKT2 |
| Histone Modification Signaling Pathway | 0.000 | 0.027 | -2.646 | H1-1,H1-4,H1-5,H2AX,H2AZ1,KMT5B,PBRM1 |
| Hematoma Resolution Signaling Pathway | 0.000 | 0.018 | 1.000 | DUSP1,HMOX1,MMP15,RHOA |
| Glycation Signaling Pathway | 0.000 | 0.009 | #NUM! | AKT2,PRKCD |
| rRNA modification in the nucleus and cytosol | 0.000 | 0.017 | #NUM! | NOC4L |
| M-decay: degradation of maternal mRNAs by maternally stored | 0.000 | 0.022 | #NUM! | ELL2 |
| Complex I biogenesis | 0.000 | 0.015 | #NUM! | NDUFAF8 |
| Cyclophilin Signaling Pathway | 0.000 | 0.017 | -2.000 | AKT2,NFAT5,RHOA,TOMM7 |
| Irritable Bowel Syndrome Signaling Pathway | 0.000 | 0.014 | #NUM! | IL10RB,MMP15,RHOA |
| Lung Ionic Balance Signaling Pathway | 0.000 | 0.019 | 3.000 | CALCR,GPR3,GPR52,GPR63,LGR4,OPRD1,PPP2R2C,PRKCD,RHOA |
| Bone Mineralization Signaling Pathway | 0.000 | 0.021 | #NUM! | FGF2,FGF9,RUNX2 |
| Tuberculosis Active Signaling Pathway | 0.000 | 0.015 | #NUM! | AKT2,IL10RB,Pycard |
| Tuberculosis Latent Signaling Pathway | 0.000 | 0.012 | #NUM! | GNAI2 |
| Epithelial Membrane Protein Signaling Pathway | 0.000 | 0.024 | -2.000 | AKT2,ITGB8,RHOA,RUNX2 |
| Warburg Effect Signaling Pathway | 0.000 | 0.023 | #NUM! | AKT2,FGF2,SREBF1 |
| Zn Homeostasis Signaling Pathway | 0.000 | 0.018 | 2.111 | AKT2,CALCR,GPR3,GPR52,GPR63,LGR4,MMP15,NFAT5,OPRD1,PRKCD,S100A4 |
| Meiosis Initiation Signaling Pathway | 0.000 | 0.021 | #NUM! | BRCA2,E2F6 |
| NF1 RAS Signaling Pathway | 0.000 | 0.020 | 1.265 | AKT2,CALCR,FGF2,GPR3,GPR52,GPR63,LGR4,OPRD1,PRKCD,RHOA |
| Co-stimulation by CD28 | 0.000 | 0.020 | #NUM! | AKT2 |
| Co-inhibition by BTLA | 0.000 | 0.017 | #NUM! | STK4 |
| Lysosome Positioning Signaling Pathway | 0.000 | 0.020 | #NUM! | AKT2,CAPZA2,PLEKHM1 |
| Thioredoxin Signaling Pathway | 0.000 | 0.015 | #NUM! | AKT2 |
| CGRP in Inflammation and Nociception Signaling Pathway | 0.000 | 0.022 | #NUM! | AKT2,PRKCD,RUNX2 |
| Neuropeptides in Migraine Signaling Pathway | 0.000 | 0.006 | #NUM! | GNAI2 |
| Cereblon and Thalidomide Signaling Pathway | 0.000 | 0.024 | 0.000 | CUL4A,DDIT4,E2F6,Pycard |
| Ovarian Folliculogenesis Signaling Pathway | 0.000 | 0.027 | -0.816 | AKT2,GDF9,HMOX1,Pycard,STK4,TSC2 |
| S-Adenosylmethionine Regulatory Signaling Pathway | 0.000 | 0.024 | #NUM! | IGF2BP3,PPP2R2C |
| PXR/RXR Activation | 0.000 | 0.022 | #NUM! | AKT2 |
| Erythropoietin Signaling Pathway | 0.000 | 0.019 | #NUM! | AKT2,PRKCD,RELB |
| α-Adrenergic Signaling | 0.000 | 0.019 | #NUM! | GNAI2,PRKCD |
| Caveolar-mediated Endocytosis Signaling | 0.000 | 0.014 | #NUM! | ITGB8 |
| Role of Pattern Recognition Receptors in Recognition of Bacteri | 0.000 | 0.016 | #NUM! | PRKCD,RELB |
| Role of PKR in Interferon Induction and Antiviral Response | 0.000 | 0.026 | #NUM! | HSP90AA1,RELB,STAT2 |
| MIF Regulation of Innate Immunity | 0.000 | 0.024 | #NUM! | RELB |
| FcγRIIB Signaling in B Lymphocytes | 0.000 | 0.012 | #NUM! | CACNA1G |
| LPS-stimulated MAPK Signaling | 0.000 | 0.024 | #NUM! | PRKCD,RELB |
| CD40 Signaling | 0.000 | 0.015 | #NUM! | RELB |
| Calcium-induced T Lymphocyte Apoptosis | 0.000 | 0.018 | #NUM! | PRKCD |
| IL-3 Signaling | 0.000 | 0.026 | #NUM! | AKT2,PRKCD |
| IL-17 Signaling | 0.000 | 0.014 | #NUM! | AKT2,HSP90AA1 |
| CXCR4 Signaling | 0.000 | 0.025 | -1.000 | AKT2,GNAI2,PRKCD,RHOA |
| Thrombopoietin Signaling | 0.000 | 0.016 | #NUM! | PRKCD |
| Induction of Apoptosis by HIV1 | 0.000 | 0.016 | #NUM! | RELB |
| IL-15 Production | 0.000 | 0.026 | #NUM! | LMTK3,RELB,TEK |
| T Helper Cell Differentiation | 0.000 | 0.015 | #NUM! | IL10RB |
| IL-15 Signaling | 0.000 | 0.025 | #NUM! | AKT2,RELB |
| Mechanisms of Viral Exit from Host Cells | 0.000 | 0.025 | #NUM! | PRKCD |
| Reelin Signaling in Neurons | 0.000 | 0.014 | #NUM! | AKT2,RHOA |
| Cellular Effects of Sildenafil (Viagra) | 0.000 | 0.022 | -1.604 | AKT2,CALCR,CAPN10,GNAI2,GPR3,GPR52,GPR63,HSP90AA1,LGR4,MYH4,NFAT5,OPRD1,PPP1R12B,PPP2R2C |
| Neuropathic Pain Signaling in Dorsal Horn Neurons | 0.000 | 0.010 | #NUM! | PRKCD |
| Endothelin-1 Signaling | 0.000 | 0.017 | #NUM! | GNAI2,HMOX1,PRKCD |
| Relaxin Signaling | 0.000 | 0.020 | #NUM! | AKT2,GNAI2,RELB |

| | | | | |
|---|---|---|---|---|
| Factors Promoting Cardiogenesis in Vertebrates | 0.000 | 0.020 | #NUM! | ACVRL1,PRKCD,RHOA |
| Renin-Angiotensin Signaling | 0.000 | 0.017 | #NUM! | PRKCD,RELB |
| Docosahexaenoic Acid (DHA) Signaling | 0.000 | 0.017 | 1.000 | AKT2,FGF2,HMOX1,PRKCD |
| Semaphorin Signaling in Neurons | 0.000 | 0.017 | #NUM! | RHOA |
| Cardiac Hypertrophy Signaling | 0.000 | 0.012 | #NUM! | GNAI2,MAP3K6,RHOA |
| CDK5 Signaling | 0.000 | 0.010 | #NUM! | PPP2R2C |
| Molecular Mechanisms of Cancer | 0.000 | 0.021 | 1.213 | ACVRL1,AKT2,CALCR,E2F6,FGF2,GNAI2,GPR3,GPR52,GPR63,IL10RB,ITGB8,LGR4,MMP15,OPRD1,PRKCD,RELB,RHOA |
| Corticotropin Releasing Hormone Signaling | 0.000 | 0.020 | #NUM! | CACNA1G,GNAI2,PRKCD |
| HMGB1 Signaling | 0.000 | 0.020 | #NUM! | AKT2,RELB,RHOA |
| FLT3 Signaling in Hematopoietic Progenitor Cells | 0.000 | 0.025 | #NUM! | AKT2,STAT2 |
| GNRH Signaling | 0.000 | 0.027 | #NUM! | CACNA1G,GNAI2,MAP3K6,PRKCD,RELB |
| Cholecystokinin/Gastrin-mediated Signaling | 0.000 | 0.018 | #NUM! | PRKCD,RHOA |
| Aldosterone Signaling in Epithelial Cells | 0.000 | 0.024 | #NUM! | DNAJC1,DUSP1,HSP90AA1,PRKCD |
| Growth Hormone Signaling | 0.000 | 0.015 | #NUM! | PRKCD |
| Prolactin Signaling | 0.000 | 0.011 | #NUM! | PRKCD |
| CREB Signaling in Neurons | 0.000 | 0.024 | 3.051 | AKT2,CACNA1G,CALCR,FGF2,GNAI2,GPR3,GPR52,GPR63,GRIK1,GRIK5,LGR4,OPRD1,POLR2H,PRKCD |
| Melanoma Signaling | 0.000 | 0.020 | #NUM! | AKT2 |
| Type I Diabetes Mellitus Signaling | 0.000 | 0.010 | #NUM! | RELB |
| Endometrial Cancer Signaling | 0.000 | 0.017 | #NUM! | AKT2 |
| Glioma Signaling | 0.000 | 0.024 | #NUM! | AKT2,E2F6,PRKCD |
| Acute Myeloid Leukemia Signaling | 0.000 | 0.022 | #NUM! | AKT2,RELB |
| Thyroid Cancer Signaling | 0.000 | 0.013 | #NUM! | AKT2 |
| Colorectal Cancer Metastasis Signaling | 0.000 | 0.024 | -1.000 | AKT2,GNAI2,MMP15,RELB,RHOA,SIAH1 |
| MYC Mediated Apoptosis Signaling | 0.000 | 0.021 | #NUM! | MCL1 |
| G Protein Signaling Mediated by Tubby | 0.000 | 0.026 | #NUM! | GNAI2,LCP2 |
| FAK Signaling | 0.000 | 0.020 | 2.309 | AKT2,CALCR,CAPN10,GPR3,GPR52,GPR63,IL10RB,ITGB8,LGR4,NANOG,OPRD1,RHOA |
| PAK Signaling | 0.000 | 0.009 | #NUM! | ITGB8 |
| RAC Signaling | 0.000 | 0.022 | #NUM! | ITGB8,RELB,RHOA |
| Role of Osteoblasts, Osteoclasts and Chondrocytes in Rheumato | 0.000 | 0.024 | #NUM! | ACVRL1,AKT2,CALCR,NFAT5,RUNX2 |
| Ovarian Cancer Signaling | 0.000 | 0.026 | #NUM! | AKT2,BRCA2,E2F6,FGF9 |
| Altered T Cell and B Cell Signaling in Rheumatoid Arthritis | 0.000 | 0.013 | #NUM! | RELB |
| Role of Macrophages, Fibroblasts and Endothelial Cells in Rheur | 0.000 | 0.017 | -0.447 | AKT2,FGF2,NFAT5,PRKCD,RHOA |
| Estrogen-Dependent Breast Cancer Signaling | 0.000 | 0.026 | #NUM! | AKT2,RELB |
| Leptin Signaling in Obesity | 0.000 | 0.013 | #NUM! | AKT2 |
| Role of NFAT in Cardiac Hypertrophy | 0.000 | 0.018 | 1.000 | AKT2,CACNA1G,GNAI2,PRKCD |
| Glioma Invasiveness Signaling | 0.000 | 0.014 | #NUM! | RHOA |
| IL-1 Signaling | 0.000 | 0.021 | #NUM! | GNAI2,RELB |
| Neuroprotective Role of THOP1 in Alzheimer's Disease | 0.000 | 0.022 | #NUM! | MME,PDYN |
| Regulation of IL-2 Expression in Activated and Anergic T Lympho | 0.000 | 0.021 | #NUM! | NFAT5,RELB |
| NUR77 Signaling in T Lymphocytes | 0.000 | 0.026 | #NUM! | PCNA,PRKCD |
| TNFR1 Signaling | 0.000 | 0.020 | #NUM! | RELB |
| Role of MAPK Signaling in the Pathogenesis of Influenza | 0.000 | 0.013 | #NUM! | AKT2 |
| OX40 Signaling Pathway | 0.000 | 0.022 | #NUM! | RELB |
| Role of Tissue Factor in Cancer | 0.000 | 0.010 | #NUM! | AKT2,ARRB2 |
| Spliceosomal Cycle | 0.000 | 0.019 | #NUM! | GPKOW |
| Role of IL-17A in Arthritis | 0.000 | 0.020 | #NUM! | RELB |
| Dopamine-DARPP32 Feedback in cAMP Signaling | 0.000 | 0.018 | #NUM! | GNAI2,PPP2R2C,PRKCD |
| Paxillin Signaling | 0.000 | 0.019 | #NUM! | ACTN2,ITGB8 |
| eNOS Signaling | 0.000 | 0.021 | #NUM! | AKT2,HSP90AA1,PRKCD |
| iNOS Signaling | 0.000 | 0.021 | #NUM! | RELB |
| nNOS Signaling in Skeletal Muscle Cells | 0.000 | 0.021 | #NUM! | CACNA1G |
| VEGF Family Ligand-Receptor Interactions | 0.000 | 0.025 | #NUM! | AKT2,PRKCD |
| ERBB Signaling | 0.000 | 0.022 | #NUM! | EREG,PRKCD |
| ERBB4 Signaling | 0.000 | 0.015 | #NUM! | PRKCD |
| Phospholipases | 0.000 | 0.016 | #NUM! | HMOX1 |

| | | | | |
|---|---|---|---|---|
| Serotonin Degradation | 0.000 | 0.026 | #NUM! | AKR1A1 |
| Triacylglycerol Biosynthesis | 0.000 | 0.017 | #NUM! | PLPPR3 |
| Stearate Biosynthesis I (Animals) | 0.000 | 0.017 | #NUM! | SLC27A4 |
| Gαi Signaling | 0.000 | 0.014 | #NUM! | GNAI2,OPRD1 |
| Gαs Signaling | 0.000 | 0.008 | #NUM! | GNAI2 |
| Agranulocyte Adhesion and Diapedesis | 0.000 | 0.024 | #NUM! | GNAI2,MMP15,MYH4,SDC4 |
| Granulocyte Adhesion and Diapedesis | 0.000 | 0.020 | #NUM! | GNAI2,MMP15,SDC4 |
| Sperm Motility | 0.000 | 0.021 | #NUM! | CACNA1G,GNAI2,LMTK3,PRKCD,TEK |
| UVB-Induced MAPK Signaling | 0.000 | 0.020 | #NUM! | PRKCD |
| UVC-Induced MAPK Signaling | 0.000 | 0.020 | #NUM! | PRKCD |
| STAT3 Pathway | 0.000 | 0.015 | #NUM! | FGF2,IL10RB |
| Oxidative Phosphorylation | 0.000 | 0.020 | #NUM! | ATP5PB,COX11 |
| BER (Base Excision Repair) Pathway | 0.000 | 0.023 | #NUM! | PCNA |
| Parkinson's Signaling Pathway | 0.000 | 0.026 | 2.236 | CACNA1G,Fcgr2b,Pycard,RPS27A,TUBB,TUBB3,UBB |
| EGF Signaling | 0.000 | 0.018 | #NUM! | AKT2 |
| Cell Cycle: G1/S Checkpoint Regulation | 0.000 | 0.015 | #NUM! | E2F6 |
| PTEN Signaling | 0.000 | 0.020 | #NUM! | AKT2,ITGB8,RELB |
| Nitric Oxide Signaling in the Cardiovascular System | 0.000 | 0.026 | #NUM! | AKT2,HSP90AA1,PRKCD |
| Cardiac β-adrenergic Signaling | 0.000 | 0.012 | #NUM! | GNAI2,PPP2R2C |
| IL-2 Signaling | 0.000 | 0.016 | #NUM! | AKT2 |
| Cell Cycle: G2/M DNA Damage Checkpoint Regulation | 0.000 | 0.020 | #NUM! | CDK1 |
| Serotonin Receptor Signaling | 0.000 | 0.020 | -1.000 | AKT2,ARRB2,CACNA1G,GNAI2,HMOX1,ORAI3,PPP1R12B,PRKCD,RHOA |
| Insulin Receptor Signaling | 0.000 | 0.016 | #NUM! | AKT2,TSC2 |
| Phototransduction Pathway | 0.000 | 0.020 | #NUM! | RCVRN |
| Neurotrophin/TRK Signaling | 0.000 | 0.026 | #NUM! | SPRY1,SPRY2 |
| Death Receptor Signaling | 0.000 | 0.022 | #NUM! | GAS2,RELB |
| IGF-1 Signaling | 0.000 | 0.010 | #NUM! | AKT2 |
| Dopamine Receptor Signaling | 0.000 | 0.015 | #NUM! | PPP2R2C |
| cAMP-mediated signaling | 0.000 | 0.013 | #NUM! | DUSP1,GNAI2,OPRD1 |
| p38 MAPK Signaling | 0.000 | 0.018 | #NUM! | DUSP1,HMGN1 |
| TGF-β Signaling | 0.000 | 0.021 | #NUM! | INHBB,RUNX2 |
| NF-κB Signaling | 0.000 | 0.012 | #NUM! | AKT2,RELB |
| VEGF Signaling | 0.000 | 0.021 | #NUM! | ACTN2,AKT2 |
| T Cell Receptor Signaling | 0.000 | 0.022 | -1.000 | AKT2,LCP2,NFAT5,RELB |
| G-Protein Coupled Receptor Signaling | 0.000 | 0.021 | 1.604 | AKT2,ARRB2,CALCR,DUSP1,GNAI2,GPR3,GPR52,GPR63,LGR4,MAP3K6,NFAT5,OPRD1,RELB,SIX3 |
| IL-6 Signaling | 0.000 | 0.025 | #NUM! | AKT2,MCL1,RELB |
| GPCR-Mediated Integration of Enteroendocrine Signaling Exem | 0.000 | 0.014 | #NUM! | GNAI2 |
| GPCR-Mediated Nutrient Sensing in Enteroendocrine Cells | 0.000 | 0.026 | #NUM! | CACNA1G,GNAI2,PRKCD |
| PD-1, PD-L1 cancer immunotherapy pathway | 0.000 | 0.022 | #NUM! | AKT2,LCP2 |
| Cancer Drug Resistance by Drug Efflux | 0.000 | 0.017 | #NUM! | AKT2 |
| Th1 and Th2 Activation Pathway | 0.000 | 0.007 | #NUM! | IL10RB |
| Th1 Pathway | 0.000 | 0.010 | #NUM! | IL10RB |
| Sirtuin Signaling Pathway | 0.000 | 0.026 | -1.000 | ATP5PB,H1-1,H1-4,H1-5,RELB,SREBF1,TOMM7 |
| Neuroinflammation Signaling Pathway | 0.000 | 0.018 | #NUM! | AKT2,HMOX1,NFAT5,Pycard,RELB |
| Adrenomedullin signaling pathway | 0.000 | 0.011 | #NUM! | AKT2,RELB |
| Iron homeostasis signaling pathway | 0.000 | 0.024 | #NUM! | HMOX1,PCBP1,SLC46A1 |
| SPINK1 General Cancer Pathway | 0.000 | 0.019 | #NUM! | AKT2 |
| Endocannabinoid Neuronal Synapse Pathway | 0.000 | 0.014 | #NUM! | CACNA1G,GNAI2 |
| Endocannabinoid Developing Neuron Pathway | 0.000 | 0.024 | #NUM! | AKT2,GNAI2,RHOA |
| Apelin Pancreas Signaling Pathway | 0.000 | 0.023 | #NUM! | RELB |
| Apelin Adipocyte Signaling Pathway | 0.000 | 0.012 | #NUM! | GNAI2 |
| Cardiac Hypertrophy Signaling (Enhanced) | 0.000 | 0.025 | -0.302 | AKT2,FGF2,FGF9,GNAI2,IL10RB,ITGB8,MAP3K6,NFAT5,PKN1,PRKCD,RELB,RHOA,TSC2 |
| PFKFB4 Signaling Pathway | 0.000 | 0.021 | #NUM! | FGF2 |
| Synaptogenesis Signaling Pathway | 0.000 | 0.020 | 0.000 | AKT2,CDH12,MARCKS,NAPA,PRKCD,RHOA |
| Systemic Lupus Erythematosus in T Cell Signaling Pathway | 0.000 | 0.020 | -2.000 | AKT2,GNAI2,PPP2R2C,RHOA |

| | | | | |
|---|---|---|---|---|
| Senescence Pathway | 0.000 | 0.021 | -0.816 | AKT2,CAPN10,CDK1,E2F6,NFAT5,PPP2R2C |
| White Adipose Tissue Browning Pathway | 0.000 | 0.022 | #NUM! | CACNA1G,CTBP2,NRF1 |
| Inhibition of ARE-Mediated mRNA Degradation Pathway | 0.000 | 0.026 | #NUM! | AKT2,PPP2R2C,PSMA4,PSMA7 |
| Hepatic Fibrosis Signaling Pathway | 0.000 | 0.020 | -0.816 | AKT2,BAMBI,FGF2,GNAI2,ITGB8,PRKCD,RELB,RHOA |
| Xenobiotic Metabolism PXR Signaling Pathway | 0.000 | 0.023 | #NUM! | HSP90AA1,PRKCD,SRA1 |
| Insulin Secretion Signaling Pathway | 0.000 | 0.019 | 0.447 | CACNA1G,NAPA,PRKCD,SLC2A8,STAT2 |
| Regulation of the Epithelial Mesenchymal Transition in Develop | 0.000 | 0.024 | #NUM! | RELB,S100A4 |
| MSP-RON Signaling in Cancer Cells Pathway | 0.000 | 0.015 | #NUM! | AKT2,RELB |
| MSP-RON Signaling in Macrophages Pathway | 0.000 | 0.010 | #NUM! | RELB |
| Calcium Signaling | 0.000 | 0.024 | 0.000 | ATP2B4,CACNA1G,GRIK1,MYH4,NFAT5 |
| GM-CSF Signaling | 0.000 | 0.015 | #NUM! | AKT2 |
| Eicosanoid Signaling | 0.000 | 0.026 | -1.134 | AKT2,AP1M1,AP1S3,ARRB2,GNAI2,PRKCD,RHOA |
| Ephrin Receptor Signaling | 0.000 | 0.020 | #NUM! | AKT2,GNAI2,ITGB8,RHOA |

**Supplementary Table 11: Cell Chat Interactions at E18 in SAC**

| source | target | ligand | receptor | prob | pval | interaction_name | interaction_name_2 | pathway_name | annotation | evidence |
|---|---|---|---|---|---|---|---|---|---|---|
| CD31 | CD31 | Tgfb1 | ACVR1B_TGFbR2 | 0.001611 | 0 | TGFB1_ACVR1B_TGFBR2 | Tgfb1 - (Acvr1b+Tgfbr2) | TGFb | Secreted Signaling | PMID: 27449815 |
| CD31 | NeuN | Tgfb1 | ACVR1B_TGFbR2 | 0.00122 | 0 | TGFB1_ACVR1B_TGFBR2 | Tgfb1 - (Acvr1b+Tgfbr2) | TGFb | Secreted Signaling | PMID: 27449815 |
| CD31 | CD31 | Tgfb1 | ACVR1C_TGFbR2 | 0.000816 | 0 | TGFB1_ACVR1C_TGFBR2 | Tgfb1 - (Acvr1c+Tgfbr2) | TGFb | Secreted Signaling | PMID: 27449815 |
| CD31 | NeuN | Tgfb1 | ACVR1C_TGFbR2 | 0.000491 | 0 | TGFB1_ACVR1C_TGFBR2 | Tgfb1 - (Acvr1c+Tgfbr2) | TGFb | Secreted Signaling | PMID: 27449815 |
| CD31 | CD31 | Tgfb3 | ACVR1B_TGFbR2 | 0.00099 | 0 | TGFB3_ACVR1B_TGFBR2 | Tgfb3 - (Acvr1b+Tgfbr2) | TGFb | Secreted Signaling | PMID: 27449815 |
| NeuN | CD31 | Tgfb3 | ACVR1B_TGFbR2 | 0.000995 | 0 | TGFB3_ACVR1B_TGFBR2 | Tgfb3 - (Acvr1b+Tgfbr2) | TGFb | Secreted Signaling | PMID: 27449815 |
| CD31 | CD31 | Tgfb3 | ACVR1C_TGFbR2 | 0.000501 | 0 | TGFB3_ACVR1C_TGFBR2 | Tgfb3 - (Acvr1c+Tgfbr2) | TGFb | Secreted Signaling | PMID: 27449815 |
| NeuN | CD31 | Tgfb3 | ACVR1C_TGFbR2 | 0.000503 | 0 | TGFB3_ACVR1C_TGFBR2 | Tgfb3 - (Acvr1c+Tgfbr2) | TGFb | Secreted Signaling | PMID: 27449815 |
| CD31 | NeuN | Bmp2 | BMPR1A_ACVR2A | 0.001286 | 0 | BMP2_BMPR1A_ACVR2A | Bmp2 - (Bmpr1a+Acvr2a) | BMP | Secreted Signaling | KEGG: mmu04350; PMID:26893264 |
| NeuN | NeuN | Bmp2 | BMPR1A_ACVR2A | 0.00096 | 0 | BMP2_BMPR1A_ACVR2A | Bmp2 - (Bmpr1a+Acvr2a) | BMP | Secreted Signaling | KEGG: mmu04350; PMID:26893264 |
| CD31 | CD31 | Bmp2 | BMPR1A_ACVR2B | 0.000739 | 0 | BMP2_BMPR1A_ACVR2B | Bmp2 - (Bmpr1a+Acvr2b) | BMP | Secreted Signaling | KEGG: mmu04350; PMID:26893264 |
| CD31 | NeuN | Bmp2 | BMPR1A_ACVR2B | 0.000892 | 0 | BMP2_BMPR1A_ACVR2B | Bmp2 - (Bmpr1a+Acvr2b) | BMP | Secreted Signaling | KEGG: mmu04350; PMID:26893264 |
| CD31 | CD31 | Bmp2 | BMPR1A_BMPR2 | 0.00122 | 0 | BMP2_BMPR1A_BMPR2 | Bmp2 - (Bmpr1a+Bmpr2) | BMP | Secreted Signaling | KEGG: mmu04350; PMID:26893264 |
| CD31 | NeuN | Bmp2 | BMPR1A_BMPR2 | 0.001428 | 0 | BMP2_BMPR1A_BMPR2 | Bmp2 - (Bmpr1a+Bmpr2) | BMP | Secreted Signaling | KEGG: mmu04350; PMID:26893264 |
| CD31 | NeuN | Bmp2 | BMPR1B_ACVR2A | 0.000836 | 0 | BMP2_BMPR1B_ACVR2A | Bmp2 - (Bmpr1b+Acvr2a) | BMP | Secreted Signaling | KEGG: mmu04350; PMID:26893264 |
| NeuN | NeuN | Bmp2 | BMPR1B_ACVR2A | 0.000623 | 0 | BMP2_BMPR1B_ACVR2A | Bmp2 - (Bmpr1b+Acvr2a) | BMP | Secreted Signaling | KEGG: mmu04350; PMID:26893264 |
| CD31 | CD31 | Bmp2 | BMPR1B_ACVR2B | 0.000567 | 0 | BMP2_BMPR1B_ACVR2B | Bmp2 - (Bmpr1b+Acvr2b) | BMP | Secreted Signaling | KEGG: mmu04350; PMID:26893264 |
| CD31 | NeuN | Bmp2 | BMPR1B_ACVR2B | 0.000579 | 0 | BMP2_BMPR1B_ACVR2B | Bmp2 - (Bmpr1b+Acvr2b) | BMP | Secreted Signaling | KEGG: mmu04350; PMID:26893264 |
| CD31 | CD31 | Bmp2 | BMPR1B_BMPR2 | 0.000938 | 0 | BMP2_BMPR1B_BMPR2 | Bmp2 - (Bmpr1b+Bmpr2) | BMP | Secreted Signaling | KEGG: mmu04350; PMID:26893264 |
| CD31 | NeuN | Bmp2 | BMPR1B_BMPR2 | 0.000928 | 0 | BMP2_BMPR1B_BMPR2 | Bmp2 - (Bmpr1b+Bmpr2) | BMP | Secreted Signaling | KEGG: mmu04350; PMID:26893264 |
| CD31 | CD31 | Gdf6 | BMPR1A_ACVR2A | 0.000792 | 0 | GDF6_BMPR1A_ACVR2A | Gdf6 - (Bmpr1a+Acvr2a) | BMP | Secreted Signaling | KEGG: mmu04350; PMID:26893264 |
| CD31 | NeuN | Gdf6 | BMPR1A_ACVR2A | 0.001323 | 0 | GDF6_BMPR1A_ACVR2A | Gdf6 - (Bmpr1a+Acvr2a) | BMP | Secreted Signaling | KEGG: mmu04350; PMID:26893264 |
| CD31 | CD31 | Gdf6 | BMPR1A_ACVR2B | 0.00076 | 0 | GDF6_BMPR1A_ACVR2B | Gdf6 - (Bmpr1a+Acvr2b) | BMP | Secreted Signaling | KEGG: mmu04350; PMID:26893264 |
| CD31 | NeuN | Gdf6 | BMPR1A_ACVR2B | 0.000917 | 0 | GDF6_BMPR1A_ACVR2B | Gdf6 - (Bmpr1a+Acvr2b) | BMP | Secreted Signaling | KEGG: mmu04350; PMID:26893264 |
| CD31 | CD31 | Gdf6 | BMPR1A_BMPR2 | 0.001255 | 0 | GDF6_BMPR1A_BMPR2 | Gdf6 - (Bmpr1a+Bmpr2) | BMP | Secreted Signaling | KEGG: mmu04350; PMID:26893264 |
| CD31 | NeuN | Gdf6 | BMPR1A_BMPR2 | 0.001469 | 0 | GDF6_BMPR1A_BMPR2 | Gdf6 - (Bmpr1a+Bmpr2) | BMP | Secreted Signaling | KEGG: mmu04350; PMID:26893264 |
| CD31 | CD31 | Gdf6 | BMPR1B_ACVR2A | 0.000608 | 0 | GDF6_BMPR1B_ACVR2A | Gdf6 - (Bmpr1b+Acvr2a) | BMP | Secreted Signaling | KEGG: mmu04350; PMID:26893264 |
| CD31 | NeuN | Gdf6 | BMPR1B_ACVR2A | 0.000859 | 0 | GDF6_BMPR1B_ACVR2A | Gdf6 - (Bmpr1b+Acvr2a) | BMP | Secreted Signaling | KEGG: mmu04350; PMID:26893264 |
| CD31 | CD31 | Gdf6 | BMPR1B_ACVR2B | 0.000584 | 0 | GDF6_BMPR1B_ACVR2B | Gdf6 - (Bmpr1b+Acvr2b) | BMP | Secreted Signaling | KEGG: mmu04350; PMID:26893264 |
| CD31 | NeuN | Gdf6 | BMPR1B_ACVR2B | 0.000596 | 0 | GDF6_BMPR1B_ACVR2B | Gdf6 - (Bmpr1b+Acvr2b) | BMP | Secreted Signaling | KEGG: mmu04350; PMID:26893264 |
| CD31 | CD31 | Gdf6 | BMPR1B_BMPR2 | 0.000964 | 0 | GDF6_BMPR1B_BMPR2 | Gdf6 - (Bmpr1b+Bmpr2) | BMP | Secreted Signaling | KEGG: mmu04350; PMID:26893264 |
| CD31 | NeuN | Gdf6 | BMPR1B_BMPR2 | 0.000954 | 0 | GDF6_BMPR1B_BMPR2 | Gdf6 - (Bmpr1b+Bmpr2) | BMP | Secreted Signaling | KEGG: mmu04350; PMID:26893264 |
| CD31 | CD31 | Bmp5 | ACVR1_ACVR2A | 0.000734 | 0 | BMP5_ACVR1_ACVR2A | Bmp5 - (Acvr1+Acvr2a) | BMP | Secreted Signaling | KEGG: mmu04350; PMID:26893264 |
| CD31 | NeuN | Bmp5 | ACVR1_ACVR2A | 0.001645 | 0 | BMP5_ACVR1_ACVR2A | Bmp5 - (Acvr1+Acvr2a) | BMP | Secreted Signaling | KEGG: mmu04350; PMID:26893264 |
| CD31 | CD31 | Bmp5 | ACVR1_ACVR2B | 0.000704 | 0 | BMP5_ACVR1_ACVR2B | Bmp5 - (Acvr1+Acvr2b) | BMP | Secreted Signaling | KEGG: mmu04350; PMID:26893264 |
| CD31 | NeuN | Bmp5 | ACVR1_ACVR2B | 0.001141 | 0 | BMP5_ACVR1_ACVR2B | Bmp5 - (Acvr1+Acvr2b) | BMP | Secreted Signaling | KEGG: mmu04350; PMID:26893264 |
| CD31 | CD31 | Bmp5 | ACVR1_BMPR2 | 0.001162 | 0 | BMP5_ACVR1_BMPR2 | Bmp5 - (Acvr1+Bmpr2) | BMP | Secreted Signaling | KEGG: mmu04350; PMID:26893264 |
| CD31 | NeuN | Bmp5 | ACVR1_BMPR2 | 0.001826 | 0 | BMP5_ACVR1_BMPR2 | Bmp5 - (Acvr1+Bmpr2) | BMP | Secreted Signaling | KEGG: mmu04350; PMID:26893264 |
| CD31 | CD31 | Bmp5 | BMPR1A_ACVR2A | 0.000799 | 0 | BMP5_BMPR1A_ACVR2A | Bmp5 - (Bmpr1a+Acvr2a) | BMP | Secreted Signaling | KEGG: mmu04350; PMID:26893264 |
| CD31 | NeuN | Bmp5 | BMPR1A_ACVR2A | 0.001335 | 0 | BMP5_BMPR1A_ACVR2A | Bmp5 - (Bmpr1a+Acvr2a) | BMP | Secreted Signaling | KEGG: mmu04350; PMID:26893264 |
| CD31 | CD31 | Bmp5 | BMPR1A_ACVR2B | 0.000767 | 0 | BMP5_BMPR1A_ACVR2B | Bmp5 - (Bmpr1a+Acvr2b) | BMP | Secreted Signaling | KEGG: mmu04350; PMID:26893264 |
| CD31 | NeuN | Bmp5 | BMPR1A_ACVR2B | 0.000926 | 0 | BMP5_BMPR1A_ACVR2B | Bmp5 - (Bmpr1a+Acvr2b) | BMP | Secreted Signaling | KEGG: mmu04350; PMID:26893264 |
| CD31 | CD31 | Bmp5 | BMPR1A_BMPR2 | 0.001266 | 0 | BMP5_BMPR1A_BMPR2 | Bmp5 - (Bmpr1a+Bmpr2) | BMP | Secreted Signaling | KEGG: mmu04350; PMID:26893264 |
| CD31 | NeuN | Bmp5 | BMPR1A_BMPR2 | 0.001482 | 0 | BMP5_BMPR1A_BMPR2 | Bmp5 - (Bmpr1a+Bmpr2) | BMP | Secreted Signaling | KEGG: mmu04350; PMID:26893264 |
| CD31 | CD31 | Bmp5 | BMPR1B_ACVR2A | 0.000614 | 0 | BMP5_BMPR1B_ACVR2A | Bmp5 - (Bmpr1b+Acvr2a) | BMP | Secreted Signaling | KEGG: mmu04350; PMID:26893264 |
| CD31 | NeuN | Bmp5 | BMPR1B_ACVR2A | 0.000867 | 0 | BMP5_BMPR1B_ACVR2A | Bmp5 - (Bmpr1b+Acvr2a) | BMP | Secreted Signaling | KEGG: mmu04350; PMID:26893264 |
| CD31 | CD31 | Bmp5 | BMPR1B_ACVR2B | 0.000589 | 0 | BMP5_BMPR1B_ACVR2B | Bmp5 - (Bmpr1b+Acvr2b) | BMP | Secreted Signaling | KEGG: mmu04350; PMID:26893264 |
| CD31 | NeuN | Bmp5 | BMPR1B_ACVR2B | 0.000601 | 0 | BMP5_BMPR1B_ACVR2B | Bmp5 - (Bmpr1b+Acvr2b) | BMP | Secreted Signaling | KEGG: mmu04350; PMID:26893264 |
| CD31 | CD31 | Bmp5 | BMPR1B_BMPR2 | 0.000973 | 0 | BMP5_BMPR1B_BMPR2 | Bmp5 - (Bmpr1b+Bmpr2) | BMP | Secreted Signaling | KEGG: mmu04350; PMID:26893264 |
| CD31 | NeuN | Bmp5 | BMPR1B_BMPR2 | 0.000963 | 0 | BMP5_BMPR1B_BMPR2 | Bmp5 - (Bmpr1b+Bmpr2) | BMP | Secreted Signaling | KEGG: mmu04350; PMID:26893264 |
| CD31 | NeuN | Bmp6 | ACVR1_ACVR2A | 0.00105 | 0 | BMP6_ACVR1_ACVR2A | Bmp6 - (Acvr1+Acvr2a) | BMP | Secreted Signaling | KEGG: mmu04350; PMID:26893264 |
| NeuN | NeuN | Bmp6 | ACVR1_ACVR2A | 0.001233 | 0 | BMP6_ACVR1_ACVR2A | Bmp6 - (Acvr1+Acvr2a) | BMP | Secreted Signaling | KEGG: mmu04350; PMID:26893264 |
| CD31 | NeuN | Bmp6 | ACVR1_ACVR2B | 0.000728 | 0 | BMP6_ACVR1_ACVR2B | Bmp6 - (Acvr1+Acvr2b) | BMP | Secreted Signaling | KEGG: mmu04350; PMID:26893264 |
| NeuN | NeuN | Bmp6 | ACVR1_ACVR2B | 0.000855 | 0 | BMP6_ACVR1_ACVR2B | Bmp6 - (Acvr1+Acvr2b) | BMP | Secreted Signaling | KEGG: mmu04350; PMID:26893264 |
| CD31 | NeuN | Bmp6 | ACVR1_BMPR2 | 0.001166 | 0 | BMP6_ACVR1_BMPR2 | Bmp6 - (Acvr1+Bmpr2) | BMP | Secreted Signaling | KEGG: mmu04350; PMID:26893264 |
| NeuN | NeuN | Bmp6 | ACVR1_BMPR2 | 0.001369 | 0 | BMP6_ACVR1_BMPR2 | Bmp6 - (Acvr1+Bmpr2) | BMP | Secreted Signaling | KEGG: mmu04350; PMID:26893264 |

| | | | | | | | | | | |
|---|---|---|---|---|---|---|---|---|---|---|
| CD31 | NeuN | Bmp6 | BMPR1A_ACVR2A | 0.000852 | 0 | BMP6_BMPR1A_ACVR2A | Bmp6 - (Bmpr1a+Acvr2a) | BMP | Secreted Signaling | KEGG: mmu04350; PMID:26893264 |
| NeuN | NeuN | Bmp6 | BMPR1A_ACVR2A | 0.001001 | 0 | BMP6_BMPR1A_ACVR2A | Bmp6 - (Bmpr1a+Acvr2a) | BMP | Secreted Signaling | KEGG: mmu04350; PMID:26893264 |
| CD31 | NeuN | Bmp6 | BMPR1A_ACVR2B | 0.000591 | 0 | BMP6_BMPR1A_ACVR2B | Bmp6 - (Bmpr1a+Acvr2b) | BMP | Secreted Signaling | KEGG: mmu04350; PMID:26893264 |
| NeuN | NeuN | Bmp6 | BMPR1A_ACVR2B | 0.000694 | 0 | BMP6_BMPR1A_ACVR2B | Bmp6 - (Bmpr1a+Acvr2b) | BMP | Secreted Signaling | KEGG: mmu04350; PMID:26893264 |
| NeuN | CD31 | Bmp6 | BMPR1A_BMPR2 | 0.000949 | 0 | BMP6_BMPR1A_BMPR2 | Bmp6 - (Bmpr1a+Bmpr2) | BMP | Secreted Signaling | KEGG: mmu04350; PMID:26893264 |
| NeuN | NeuN | Bmp6 | BMPR1A_BMPR2 | 0.001111 | 0 | BMP6_BMPR1A_BMPR2 | Bmp6 - (Bmpr1a+Bmpr2) | BMP | Secreted Signaling | KEGG: mmu04350; PMID:26893264 |
| CD31 | NeuN | Bmp6 | BMPR1B_ACVR2A | 0.000553 | 0 | BMP6_BMPR1B_ACVR2A | Bmp6 - (Bmpr1b+Acvr2a) | BMP | Secreted Signaling | KEGG: mmu04350; PMID:26893264 |
| NeuN | NeuN | Bmp6 | BMPR1B_ACVR2A | 0.00065 | 0 | BMP6_BMPR1B_ACVR2A | Bmp6 - (Bmpr1b+Acvr2a) | BMP | Secreted Signaling | KEGG: mmu04350; PMID:26893264 |
| NeuN | CD31 | Bmp6 | BMPR1B_ACVR2B | 0.000441 | 0 | BMP6_BMPR1B_ACVR2B | Bmp6 - (Bmpr1b+Acvr2b) | BMP | Secreted Signaling | KEGG: mmu04350; PMID:26893264 |
| NeuN | NeuN | Bmp6 | BMPR1B_ACVR2B | 0.000451 | 0 | BMP6_BMPR1B_ACVR2B | Bmp6 - (Bmpr1b+Acvr2b) | BMP | Secreted Signaling | KEGG: mmu04350; PMID:26893264 |
| NeuN | CD31 | Bmp6 | BMPR1B_BMPR2 | 0.000729 | 0 | BMP6_BMPR1B_BMPR2 | Bmp6 - (Bmpr1b+Bmpr2) | BMP | Secreted Signaling | KEGG: mmu04350; PMID:26893264 |
| NeuN | NeuN | Bmp6 | BMPR1B_BMPR2 | 0.000722 | 0 | BMP6_BMPR1B_BMPR2 | Bmp6 - (Bmpr1b+Bmpr2) | BMP | Secreted Signaling | KEGG: mmu04350; PMID:26893264 |
| CD31 | NeuN | Bmp8b | ACVR1_ACVR2A | 0.001319 | 0 | BMP8B_ACVR1_ACVR2A | Bmp8b - (Acvr1+Acvr2a) | BMP | Secreted Signaling | KEGG: mmu04350; PMID:26893264 |
| NeuN | NeuN | Bmp8b | ACVR1_ACVR2A | 0.00101 | 0 | BMP8B_ACVR1_ACVR2A | Bmp8b - (Acvr1+Acvr2a) | BMP | Secreted Signaling | KEGG: mmu04350; PMID:26893264 |
| CD31 | NeuN | Bmp8b | ACVR1_ACVR2B | 0.000915 | 0 | BMP8B_ACVR1_ACVR2B | Bmp8b - (Acvr1+Acvr2b) | BMP | Secreted Signaling | KEGG: mmu04350; PMID:26893264 |
| NeuN | NeuN | Bmp8b | ACVR1_ACVR2B | 0.000701 | 0 | BMP8B_ACVR1_ACVR2B | Bmp8b - (Acvr1+Acvr2b) | BMP | Secreted Signaling | KEGG: mmu04350; PMID:26893264 |
| CD31 | NeuN | Bmp8b | ACVR1_BMPR2 | 0.001465 | 0 | BMP8B_ACVR1_BMPR2 | Bmp8b - (Acvr1+Bmpr2) | BMP | Secreted Signaling | KEGG: mmu04350; PMID:26893264 |
| NeuN | NeuN | Bmp8b | ACVR1_BMPR2 | 0.001122 | 0 | BMP8B_ACVR1_BMPR2 | Bmp8b - (Acvr1+Bmpr2) | BMP | Secreted Signaling | KEGG: mmu04350; PMID:26893264 |
| CD31 | NeuN | Bmp8b | BMPR1A_ACVR2A | 0.001071 | 0 | BMP8B_BMPR1A_ACVR2A | Bmp8b - (Bmpr1a+Acvr2a) | BMP | Secreted Signaling | KEGG: mmu04350; PMID:26893264 |
| NeuN | NeuN | Bmp8b | BMPR1A_ACVR2A | 0.00082 | 0 | BMP8B_BMPR1A_ACVR2A | Bmp8b - (Bmpr1a+Acvr2a) | BMP | Secreted Signaling | KEGG: mmu04350; PMID:26893264 |
| CD31 | CD31 | Bmp8b | BMPR1A_ACVR2B | 0.000615 | 0 | BMP8B_BMPR1A_ACVR2B | Bmp8b - (Bmpr1a+Acvr2b) | BMP | Secreted Signaling | KEGG: mmu04350; PMID:26893264 |
| CD31 | NeuN | Bmp8b | BMPR1A_ACVR2B | 0.000742 | 0 | BMP8B_BMPR1A_ACVR2B | Bmp8b - (Bmpr1a+Acvr2b) | BMP | Secreted Signaling | KEGG: mmu04350; PMID:26893264 |
| CD31 | CD31 | Bmp8b | BMPR1A_BMPR2 | 0.001016 | 0 | BMP8B_BMPR1A_BMPR2 | Bmp8b - (Bmpr1a+Bmpr2) | BMP | Secreted Signaling | KEGG: mmu04350; PMID:26893264 |
| CD31 | NeuN | Bmp8b | BMPR1A_BMPR2 | 0.001189 | 0 | BMP8B_BMPR1A_BMPR2 | Bmp8b - (Bmpr1a+Bmpr2) | BMP | Secreted Signaling | KEGG: mmu04350; PMID:26893264 |
| CD31 | NeuN | Bmp8b | BMPR1B_ACVR2A | 0.000695 | 0 | BMP8B_BMPR1B_ACVR2A | Bmp8b - (Bmpr1b+Acvr2a) | BMP | Secreted Signaling | KEGG: mmu04350; PMID:26893264 |
| NeuN | NeuN | Bmp8b | BMPR1B_ACVR2A | 0.000533 | 0 | BMP8B_BMPR1B_ACVR2A | Bmp8b - (Bmpr1b+Acvr2a) | BMP | Secreted Signaling | KEGG: mmu04350; PMID:26893264 |
| CD31 | CD31 | Bmp8b | BMPR1B_ACVR2B | 0.000472 | 0 | BMP8B_BMPR1B_ACVR2B | Bmp8b - (Bmpr1b+Acvr2b) | BMP | Secreted Signaling | KEGG: mmu04350; PMID:26893264 |
| CD31 | NeuN | Bmp8b | BMPR1B_ACVR2B | 0.000482 | 0 | BMP8B_BMPR1B_ACVR2B | Bmp8b - (Bmpr1b+Acvr2b) | BMP | Secreted Signaling | KEGG: mmu04350; PMID:26893264 |
| CD31 | CD31 | Bmp8b | BMPR1B_BMPR2 | 0.00078 | 0 | BMP8B_BMPR1B_BMPR2 | Bmp8b - (Bmpr1b+Bmpr2) | BMP | Secreted Signaling | KEGG: mmu04350; PMID:26893264 |
| CD31 | NeuN | Bmp8b | BMPR1B_BMPR2 | 0.000772 | 0 | BMP8B_BMPR1B_BMPR2 | Bmp8b - (Bmpr1b+Bmpr2) | BMP | Secreted Signaling | KEGG: mmu04350; PMID:26893264 |
| CD31 | CD31 | Gdf9 | ACVR1B_BMPR2 | 0.002715 | 0 | GDF9_ACVR1B_BMPR2 | Gdf9 - (Acvr1b+Bmpr2) | GDF | Secreted Signaling | KEGG: mmu04350 |
| CD31 | NeuN | Gdf9 | ACVR1B_BMPR2 | 0.004017 | 0 | GDF9_ACVR1B_BMPR2 | Gdf9 - (Acvr1b+Bmpr2) | GDF | Secreted Signaling | KEGG: mmu04350 |
| CD31 | NeuN | Gdf10 | ACVR1B_ACVR2A | 0.002561 | 0 | GDF10_ACVR1B_ACVR2A | Gdf10 - (Acvr1b+Acvr2a) | GDF | Secreted Signaling | KEGG: mmu04350 |
| NeuN | NeuN | Gdf10 | ACVR1B_ACVR2A | 0.003144 | 0 | GDF10_ACVR1B_ACVR2A | Gdf10 - (Acvr1b+Acvr2a) | GDF | Secreted Signaling | KEGG: mmu04350 |
| CD31 | NeuN | Gdf11 | TGFBR1_ACVR2A | 0.002272 | 0 | GDF11_TGFBR1_ACVR2A | Gdf11 - (Tgfbr1+Acvr2a) | GDF | Secreted Signaling | KEGG: mmu04350 |
| NeuN | NeuN | Gdf11 | TGFBR1_ACVR2A | 0.002326 | 0 | GDF11_TGFBR1_ACVR2A | Gdf11 - (Tgfbr1+Acvr2a) | GDF | Secreted Signaling | KEGG: mmu04350 |
| CD31 | NeuN | Gdf11 | TGFBR1_ACVR2B | 0.001575 | 0 | GDF11_TGFBR1_ACVR2B | Gdf11 - (Tgfbr1+Acvr2b) | GDF | Secreted Signaling | KEGG: mmu04350 |
| NeuN | NeuN | Gdf11 | TGFBR1_ACVR2B | 0.001613 | 0 | GDF11_TGFBR1_ACVR2B | Gdf11 - (Tgfbr1+Acvr2b) | GDF | Secreted Signaling | KEGG: mmu04350 |
| CD31 | NeuN | Gdf11 | ACVR1B_ACVR2B | 0.002384 | 0 | GDF11_ACVR1B_ACVR2B | Gdf11 - (Acvr1b+Acvr2b) | GDF | Secreted Signaling | KEGG: mmu04350 |
| NeuN | NeuN | Gdf11 | ACVR1B_ACVR2B | 0.002441 | 0 | GDF11_ACVR1B_ACVR2B | Gdf11 - (Acvr1b+Acvr2b) | GDF | Secreted Signaling | KEGG: mmu04350 |
| CD31 | CD31 | Gdnf | GFRA1_RET | 0.000606 | 0 | GDNF_GFRA1 | Gdnf - Gfra1 | GDNF | Secreted Signaling | PMID: 11544105 |
| NeuN | CD31 | Gdnf | GFRA1_RET | 0.000577 | 0 | GDNF_GFRA1 | Gdnf - Gfra1 | GDNF | Secreted Signaling | PMID: 11544105 |
| CD31 | CD31 | Nrtn | GFRA2_RET | 0.000531 | 0 | NRTN_GFRA2 | Nrtn - Gfra2 | GDNF | Secreted Signaling | PMID: 11544105 |
| CD31 | NeuN | Nrtn | GFRA2_RET | 0.000414 | 0 | NRTN_GFRA2 | Nrtn - Gfra2 | GDNF | Secreted Signaling | PMID: 11544105 |
| CD31 | NeuN | Inhba | ACVR1B_ACVR2A | 0.00205 | 0 | INHBA_ACVR1B_ACVR2A | Inhba - (Acvr1b+Acvr2a) | ACTIVIN | Secreted Signaling | KEGG: mmu04350 |
| NeuN | NeuN | Inhba | ACVR1B_ACVR2A | 0.002806 | 0 | INHBA_ACVR1B_ACVR2A | Inhba - (Acvr1b+Acvr2a) | ACTIVIN | Secreted Signaling | KEGG: mmu04350 |
| CD31 | NeuN | Inhba | ACVR1B_ACVR2B | 0.001422 | 0 | INHBA_ACVR1B_ACVR2B | Inhba - (Acvr1b+Acvr2b) | ACTIVIN | Secreted Signaling | KEGG: mmu04350 |
| NeuN | NeuN | Inhba | ACVR1B_ACVR2B | 0.001947 | 0 | INHBA_ACVR1B_ACVR2B | Inhba - (Acvr1b+Acvr2b) | ACTIVIN | Secreted Signaling | KEGG: mmu04350 |
| CD31 | NeuN | Inhbb | ACVR1B_ACVR2A | 0.003672 | 0 | INHBB_ACVR1B_ACVR2A | Inhbb - (Acvr1b+Acvr2a) | ACTIVIN | Secreted Signaling | KEGG: mmu04350 |
| NeuN | NeuN | Inhbb | ACVR1B_ACVR2A | 0.002992 | 0 | INHBB_ACVR1B_ACVR2A | Inhbb - (Acvr1b+Acvr2a) | ACTIVIN | Secreted Signaling | KEGG: mmu04350 |
| CD31 | NeuN | Inhbb | ACVR1B_ACVR2B | 0.002548 | 0 | INHBB_ACVR1B_ACVR2B | Inhbb - (Acvr1b+Acvr2b) | ACTIVIN | Secreted Signaling | KEGG: mmu04350 |
| NeuN | NeuN | Inhbb | ACVR1B_ACVR2B | 0.002076 | 0 | INHBB_ACVR1B_ACVR2B | Inhbb - (Acvr1b+Acvr2b) | ACTIVIN | Secreted Signaling | KEGG: mmu04350 |
| CD31 | NeuN | Inhbb | ACVR1C_ACVR2A | 0.001479 | 0 | INHBB_ACVR1C_ACVR2A | Inhbb - (Acvr1c+Acvr2a) | ACTIVIN | Secreted Signaling | KEGG: mmu04350 |
| NeuN | NeuN | Inhbb | ACVR1C_ACVR2A | 0.001204 | 0 | INHBB_ACVR1C_ACVR2A | Inhbb - (Acvr1c+Acvr2a) | ACTIVIN | Secreted Signaling | KEGG: mmu04350 |
| CD31 | NeuN | Inhbb | ACVR1C_ACVR2B | 0.001025 | 0 | INHBB_ACVR1C_ACVR2B | Inhbb - (Acvr1c+Acvr2b) | ACTIVIN | Secreted Signaling | KEGG: mmu04350 |
| NeuN | NeuN | Inhbb | ACVR1C_ACVR2B | 0.000835 | 0 | INHBB_ACVR1C_ACVR2B | Inhbb - (Acvr1c+Acvr2b) | ACTIVIN | Secreted Signaling | KEGG: mmu04350 |
| CD31 | CD31 | Wnt10b | FZD1_LRP5 | 0.000275 | 0 | WNT10B_FZD1_LRP5 | Wnt10b - (Fzd1+Lrp5) | WNT | Secreted Signaling | KEGG: mmu04310; PMID: 23209147 |
| NeuN | CD31 | Wnt10b | FZD1_LRP5 | 0.000337 | 0 | WNT10B_FZD1_LRP5 | Wnt10b - (Fzd1+Lrp5) | WNT | Secreted Signaling | KEGG: mmu04310; PMID: 23209147 |
| NeuN | CD31 | Wnt10b | FZD2_LRP5 | 0.000268 | 0 | WNT10B_FZD2_LRP5 | Wnt10b - (Fzd2+Lrp5) | WNT | Secreted Signaling | KEGG: mmu04310; PMID: 23209147 |

| | | | | | | | | | | |
|---|---|---|---|---|---|---|---|---|---|---|
| NeuN | NeuN | Wnt10b | FZD2_LRP5 | 0.000222 | 0 | WNT10B_FZD2_LRP5 | Wnt10b - (Fzd2+Lrp5) | WNT | Secreted Signaling | KEGG: mmu04310; PMID: 23209147 |
| NeuN | CD31 | Wnt10b | FZD3_LRP5 | 0.000457 | 0 | WNT10B_FZD3_LRP5 | Wnt10b - (Fzd3+Lrp5) | WNT | Secreted Signaling | KEGG: mmu04310; PMID: 23209147 |
| NeuN | NeuN | Wnt10b | FZD3_LRP5 | 0.000547 | 0 | WNT10B_FZD3_LRP5 | Wnt10b - (Fzd3+Lrp5) | WNT | Secreted Signaling | KEGG: mmu04310; PMID: 23209147 |
| CD31 | CD31 | Wnt10b | FZD4_LRP5 | 0.000174 | 0 | WNT10B_FZD4_LRP5 | Wnt10b - (Fzd4+Lrp5) | WNT | Secreted Signaling | KEGG: mmu04310; PMID: 23209147 |
| NeuN | CD31 | Wnt10b | FZD4_LRP5 | 0.000213 | 0 | WNT10B_FZD4_LRP5 | Wnt10b - (Fzd4+Lrp5) | WNT | Secreted Signaling | KEGG: mmu04310; PMID: 23209147 |
| NeuN | CD31 | Wnt10b | FZD5_LRP5 | 0.000219 | 0 | WNT10B_FZD5_LRP5 | Wnt10b - (Fzd5+Lrp5) | WNT | Secreted Signaling | KEGG: mmu04310; PMID: 23209147 |
| NeuN | NeuN | Wnt10b | FZD5_LRP5 | 0.000207 | 0 | WNT10B_FZD5_LRP5 | Wnt10b - (Fzd5+Lrp5) | WNT | Secreted Signaling | KEGG: mmu04310; PMID: 23209147 |
| CD31 | CD31 | Wnt10b | FZD6_LRP5 | 0.000383 | 0 | WNT10B_FZD6_LRP5 | Wnt10b - (Fzd6+Lrp5) | WNT | Secreted Signaling | KEGG: mmu04310; PMID: 23209147 |
| NeuN | CD31 | Wnt10b | FZD6_LRP5 | 0.000469 | 0 | WNT10B_FZD6_LRP5 | Wnt10b - (Fzd6+Lrp5) | WNT | Secreted Signaling | KEGG: mmu04310; PMID: 23209147 |
| NeuN | CD31 | Wnt10b | FZD8_LRP5 | 0.000237 | 0 | WNT10B_FZD8_LRP5 | Wnt10b - (Fzd8+Lrp5) | WNT | Secreted Signaling | KEGG: mmu04310; PMID: 23209147 |
| NeuN | NeuN | Wnt10b | FZD8_LRP5 | 0.000211 | 0 | WNT10B_FZD8_LRP5 | Wnt10b - (Fzd8+Lrp5) | WNT | Secreted Signaling | KEGG: mmu04310; PMID: 23209147 |
| NeuN | CD31 | Wnt10b | FZD9_LRP5 | 0.000197 | 0 | WNT10B_FZD9_LRP5 | Wnt10b - (Fzd9+Lrp5) | WNT | Secreted Signaling | KEGG: mmu04310; PMID: 23209147 |
| NeuN | NeuN | Wnt10b | FZD9_LRP5 | 0.000213 | 0 | WNT10B_FZD9_LRP5 | Wnt10b - (Fzd9+Lrp5) | WNT | Secreted Signaling | KEGG: mmu04310; PMID: 23209147 |
| CD31 | CD31 | Wnt4 | FZD1_LRP5 | 0.000371 | 0 | WNT4_FZD1_LRP5 | Wnt4 - (Fzd1+Lrp5) | WNT | Secreted Signaling | KEGG: mmu04310; PMID: 23209147 |
| NeuN | CD31 | Wnt4 | FZD1_LRP5 | 0.000341 | 0 | WNT4_FZD1_LRP5 | Wnt4 - (Fzd1+Lrp5) | WNT | Secreted Signaling | KEGG: mmu04310; PMID: 23209147 |
| CD31 | CD31 | Wnt4 | FZD2_LRP5 | 0.000295 | 0 | WNT4_FZD2_LRP5 | Wnt4 - (Fzd2+Lrp5) | WNT | Secreted Signaling | KEGG: mmu04310; PMID: 23209147 |
| NeuN | CD31 | Wnt4 | FZD2_LRP5 | 0.000271 | 0 | WNT4_FZD2_LRP5 | Wnt4 - (Fzd2+Lrp5) | WNT | Secreted Signaling | KEGG: mmu04310; PMID: 23209147 |
| CD31 | NeuN | Wnt4 | FZD3_LRP5 | 0.000603 | 0 | WNT4_FZD3_LRP5 | Wnt4 - (Fzd3+Lrp5) | WNT | Secreted Signaling | KEGG: mmu04310; PMID: 23209147 |
| NeuN | NeuN | Wnt4 | FZD3_LRP5 | 0.000554 | 0 | WNT4_FZD3_LRP5 | Wnt4 - (Fzd3+Lrp5) | WNT | Secreted Signaling | KEGG: mmu04310; PMID: 23209147 |
| CD31 | CD31 | Wnt4 | FZD4_LRP5 | 0.000235 | 0 | WNT4_FZD4_LRP5 | Wnt4 - (Fzd4+Lrp5) | WNT | Secreted Signaling | KEGG: mmu04310; PMID: 23209147 |
| NeuN | CD31 | Wnt4 | FZD4_LRP5 | 0.000215 | 0 | WNT4_FZD4_LRP5 | Wnt4 - (Fzd4+Lrp5) | WNT | Secreted Signaling | KEGG: mmu04310; PMID: 23209147 |
| CD31 | CD31 | Wnt4 | FZD5_LRP5 | 0.000242 | 0 | WNT4_FZD5_LRP5 | Wnt4 - (Fzd5+Lrp5) | WNT | Secreted Signaling | KEGG: mmu04310; PMID: 23209147 |
| CD31 | NeuN | Wnt4 | FZD5_LRP5 | 0.000228 | 0 | WNT4_FZD5_LRP5 | Wnt4 - (Fzd5+Lrp5) | WNT | Secreted Signaling | KEGG: mmu04310; PMID: 23209147 |
| CD31 | CD31 | Wnt4 | FZD6_LRP5 | 0.000517 | 0 | WNT4_FZD6_LRP5 | Wnt4 - (Fzd6+Lrp5) | WNT | Secreted Signaling | KEGG: mmu04310; PMID: 23209147 |
| NeuN | CD31 | Wnt4 | FZD6_LRP5 | 0.000475 | 0 | WNT4_FZD6_LRP5 | Wnt4 - (Fzd6+Lrp5) | WNT | Secreted Signaling | KEGG: mmu04310; PMID: 23209147 |
| CD31 | CD31 | Wnt4 | FZD8_LRP5 | 0.000261 | 0 | WNT4_FZD8_LRP5 | Wnt4 - (Fzd8+Lrp5) | WNT | Secreted Signaling | KEGG: mmu04310; PMID: 23209147 |
| NeuN | CD31 | Wnt4 | FZD8_LRP5 | 0.00024 | 0 | WNT4_FZD8_LRP5 | Wnt4 - (Fzd8+Lrp5) | WNT | Secreted Signaling | KEGG: mmu04310; PMID: 23209147 |
| CD31 | CD31 | Wnt4 | FZD9_LRP5 | 0.000217 | 0 | WNT4_FZD9_LRP5 | Wnt4 - (Fzd9+Lrp5) | WNT | Secreted Signaling | KEGG: mmu04310; PMID: 23209147 |
| CD31 | NeuN | Wnt4 | FZD9_LRP5 | 0.000235 | 0 | WNT4_FZD9_LRP5 | Wnt4 - (Fzd9+Lrp5) | WNT | Secreted Signaling | KEGG: mmu04310; PMID: 23209147 |
| CD31 | CD31 | Wnt7a | FZD1_LRP5 | 0.00057 | 0 | WNT7A_FZD1_LRP5 | Wnt7a - (Fzd1+Lrp5) | WNT | Secreted Signaling | KEGG: mmu04310; PMID: 23209147 |
| CD31 | NeuN | Wnt7a | FZD1_LRP5 | 0.000457 | 0 | WNT7A_FZD1_LRP5 | Wnt7a - (Fzd1+Lrp5) | WNT | Secreted Signaling | KEGG: mmu04310; PMID: 23209147 |
| CD31 | CD31 | Wnt7a | FZD2_LRP5 | 0.000453 | 0 | WNT7A_FZD2_LRP5 | Wnt7a - (Fzd2+Lrp5) | WNT | Secreted Signaling | KEGG: mmu04310; PMID: 23209147 |
| CD31 | NeuN | Wnt7a | FZD2_LRP5 | 0.000376 | 0 | WNT7A_FZD2_LRP5 | Wnt7a - (Fzd2+Lrp5) | WNT | Secreted Signaling | KEGG: mmu04310; PMID: 23209147 |
| CD31 | CD31 | Wnt7a | FZD3_LRP5 | 0.000775 | 0 | WNT7A_FZD3_LRP5 | Wnt7a - (Fzd3+Lrp5) | WNT | Secreted Signaling | KEGG: mmu04310; PMID: 23209147 |
| CD31 | NeuN | Wnt7a | FZD3_LRP5 | 0.000926 | 0 | WNT7A_FZD3_LRP5 | Wnt7a - (Fzd3+Lrp5) | WNT | Secreted Signaling | KEGG: mmu04310; PMID: 23209147 |
| CD31 | CD31 | Wnt7a | FZD4_LRP5 | 0.00036 | 0 | WNT7A_FZD4_LRP5 | Wnt7a - (Fzd4+Lrp5) | WNT | Secreted Signaling | KEGG: mmu04310; PMID: 23209147 |
| CD31 | NeuN | Wnt7a | FZD4_LRP5 | 0.000284 | 0 | WNT7A_FZD4_LRP5 | Wnt7a - (Fzd4+Lrp5) | WNT | Secreted Signaling | KEGG: mmu04310; PMID: 23209147 |
| CD31 | CD31 | Wnt7a | FZD5_LRP5 | 0.000371 | 0 | WNT7A_FZD5_LRP5 | Wnt7a - (Fzd5+Lrp5) | WNT | Secreted Signaling | KEGG: mmu04310; PMID: 23209147 |
| CD31 | NeuN | Wnt7a | FZD5_LRP5 | 0.00035 | 0 | WNT7A_FZD5_LRP5 | Wnt7a - (Fzd5+Lrp5) | WNT | Secreted Signaling | KEGG: mmu04310; PMID: 23209147 |
| CD31 | CD31 | Wnt7a | FZD6_LRP5 | 0.000794 | 0 | WNT7A_FZD6_LRP5 | Wnt7a - (Fzd6+Lrp5) | WNT | Secreted Signaling | KEGG: mmu04310; PMID: 23209147 |
| NeuN | CD31 | Wnt7a | FZD6_LRP5 | 0.000557 | 0 | WNT7A_FZD6_LRP5 | Wnt7a - (Fzd6+Lrp5) | WNT | Secreted Signaling | KEGG: mmu04310; PMID: 23209147 |
| CD31 | CD31 | Wnt7a | FZD8_LRP5 | 0.000401 | 0 | WNT7A_FZD8_LRP5 | Wnt7a - (Fzd8+Lrp5) | WNT | Secreted Signaling | KEGG: mmu04310; PMID: 23209147 |
| CD31 | NeuN | Wnt7a | FZD8_LRP5 | 0.000358 | 0 | WNT7A_FZD8_LRP5 | Wnt7a - (Fzd8+Lrp5) | WNT | Secreted Signaling | KEGG: mmu04310; PMID: 23209147 |
| CD31 | CD31 | Wnt7a | FZD9_LRP5 | 0.000333 | 0 | WNT7A_FZD9_LRP5 | Wnt7a - (Fzd9+Lrp5) | WNT | Secreted Signaling | KEGG: mmu04310; PMID: 23209147 |
| CD31 | NeuN | Wnt7a | FZD9_LRP5 | 0.000361 | 0 | WNT7A_FZD9_LRP5 | Wnt7a - (Fzd9+Lrp5) | WNT | Secreted Signaling | KEGG: mmu04310; PMID: 23209147 |
| NeuN | CD31 | Wnt7b | FZD1_LRP5 | 0.001765 | 0 | WNT7B_FZD1_LRP5 | Wnt7b - (Fzd1+Lrp5) | WNT | Secreted Signaling | KEGG: mmu04310; PMID: 23209147 |
| NeuN | NeuN | Wnt7b | FZD1_LRP5 | 0.001413 | 0 | WNT7B_FZD1_LRP5 | Wnt7b - (Fzd1+Lrp5) | WNT | Secreted Signaling | KEGG: mmu04310; PMID: 23209147 |
| NeuN | CD31 | Wnt7b | FZD2_LRP5 | 0.001403 | 0 | WNT7B_FZD2_LRP5 | Wnt7b - (Fzd2+Lrp5) | WNT | Secreted Signaling | KEGG: mmu04310; PMID: 23209147 |
| NeuN | NeuN | Wnt7b | FZD2_LRP5 | 0.001166 | 0 | WNT7B_FZD2_LRP5 | Wnt7b - (Fzd2+Lrp5) | WNT | Secreted Signaling | KEGG: mmu04310; PMID: 23209147 |
| NeuN | CD31 | Wnt7b | FZD3_LRP5 | 0.002395 | 0 | WNT7B_FZD3_LRP5 | Wnt7b - (Fzd3+Lrp5) | WNT | Secreted Signaling | KEGG: mmu04310; PMID: 23209147 |
| NeuN | NeuN | Wnt7b | FZD3_LRP5 | 0.002863 | 0 | WNT7B_FZD3_LRP5 | Wnt7b - (Fzd3+Lrp5) | WNT | Secreted Signaling | KEGG: mmu04310; PMID: 23209147 |
| NeuN | CD31 | Wnt7b | FZD4_LRP5 | 0.001115 | 0 | WNT7B_FZD4_LRP5 | Wnt7b - (Fzd4+Lrp5) | WNT | Secreted Signaling | KEGG: mmu04310; PMID: 23209147 |
| NeuN | NeuN | Wnt7b | FZD4_LRP5 | 0.000878 | 0 | WNT7B_FZD4_LRP5 | Wnt7b - (Fzd4+Lrp5) | WNT | Secreted Signaling | KEGG: mmu04310; PMID: 23209147 |
| NeuN | CD31 | Wnt7b | FZD5_LRP5 | 0.001148 | 0 | WNT7B_FZD5_LRP5 | Wnt7b - (Fzd5+Lrp5) | WNT | Secreted Signaling | KEGG: mmu04310; PMID: 23209147 |
| NeuN | NeuN | Wnt7b | FZD5_LRP5 | 0.001084 | 0 | WNT7B_FZD5_LRP5 | Wnt7b - (Fzd5+Lrp5) | WNT | Secreted Signaling | KEGG: mmu04310; PMID: 23209147 |
| CD31 | CD31 | Wnt7b | FZD6_LRP5 | 0.001258 | 0 | WNT7B_FZD6_LRP5 | Wnt7b - (Fzd6+Lrp5) | WNT | Secreted Signaling | KEGG: mmu04310; PMID: 23209147 |
| NeuN | CD31 | Wnt7b | FZD6_LRP5 | 0.002454 | 0 | WNT7B_FZD6_LRP5 | Wnt7b - (Fzd6+Lrp5) | WNT | Secreted Signaling | KEGG: mmu04310; PMID: 23209147 |
| NeuN | CD31 | Wnt7b | FZD8_LRP5 | 0.001243 | 0 | WNT7B_FZD8_LRP5 | Wnt7b - (Fzd8+Lrp5) | WNT | Secreted Signaling | KEGG: mmu04310; PMID: 23209147 |
| NeuN | NeuN | Wnt7b | FZD8_LRP5 | 0.001108 | 0 | WNT7B_FZD8_LRP5 | Wnt7b - (Fzd8+Lrp5) | WNT | Secreted Signaling | KEGG: mmu04310; PMID: 23209147 |

| | | | | | | | | | | |
|---|---|---|---|---|---|---|---|---|---|---|
| NeuN | CD31 | Wnt7b | FZD9_LRP5 | 0.001031 | 0 | WNT7B_FZD9_LRP5 | Wnt7b - (Fzd9+Lrp5) | WNT | Secreted Signaling | KEGG: mmu04310; PMID: 23209147 |
| NeuN | NeuN | Wnt7b | FZD9_LRP5 | 0.001118 | 0 | WNT7B_FZD9_LRP5 | Wnt7b - (Fzd9+Lrp5) | WNT | Secreted Signaling | KEGG: mmu04310; PMID: 23209147 |
| CD31 | CD31 | Wnt9a | FZD1_LRP5 | 0.000288 | 0 | WNT9A_FZD1_LRP5 | Wnt9a - (Fzd1+Lrp5) | WNT | Secreted Signaling | KEGG: mmu04310; PMID: 23209147 |
| NeuN | CD31 | Wnt9a | FZD1_LRP5 | 0.00024 | 0 | WNT9A_FZD1_LRP5 | Wnt9a - (Fzd1+Lrp5) | WNT | Secreted Signaling | KEGG: mmu04310; PMID: 23209147 |
| CD31 | CD31 | Wnt9a | FZD2_LRP5 | 0.000229 | 0 | WNT9A_FZD2_LRP5 | Wnt9a - (Fzd2+Lrp5) | WNT | Secreted Signaling | KEGG: mmu04310; PMID: 23209147 |
| NeuN | CD31 | Wnt9a | FZD2_LRP5 | 0.000191 | 0 | WNT9A_FZD2_LRP5 | Wnt9a - (Fzd2+Lrp5) | WNT | Secreted Signaling | KEGG: mmu04310; PMID: 23209147 |
| CD31 | CD31 | Wnt9a | FZD3_LRP5 | 0.000392 | 0 | WNT9A_FZD3_LRP5 | Wnt9a - (Fzd3+Lrp5) | WNT | Secreted Signaling | KEGG: mmu04310; PMID: 23209147 |
| CD31 | NeuN | Wnt9a | FZD3_LRP5 | 0.000468 | 0 | WNT9A_FZD3_LRP5 | Wnt9a - (Fzd3+Lrp5) | WNT | Secreted Signaling | KEGG: mmu04310; PMID: 23209147 |
| CD31 | CD31 | Wnt9a | FZD4_LRP5 | 0.000182 | 0 | WNT9A_FZD4_LRP5 | Wnt9a - (Fzd4+Lrp5) | WNT | Secreted Signaling | KEGG: mmu04310; PMID: 23209147 |
| NeuN | CD31 | Wnt9a | FZD4_LRP5 | 0.000152 | 0 | WNT9A_FZD4_LRP5 | Wnt9a - (Fzd4+Lrp5) | WNT | Secreted Signaling | KEGG: mmu04310; PMID: 23209147 |
| CD31 | CD31 | Wnt9a | FZD5_LRP5 | 0.000187 | 0 | WNT9A_FZD5_LRP5 | Wnt9a - (Fzd5+Lrp5) | WNT | Secreted Signaling | KEGG: mmu04310; PMID: 23209147 |
| CD31 | NeuN | Wnt9a | FZD5_LRP5 | 0.000177 | 0 | WNT9A_FZD5_LRP5 | Wnt9a - (Fzd5+Lrp5) | WNT | Secreted Signaling | KEGG: mmu04310; PMID: 23209147 |
| CD31 | CD31 | Wnt9a | FZD6_LRP5 | 0.000401 | 0 | WNT9A_FZD6_LRP5 | Wnt9a - (Fzd6+Lrp5) | WNT | Secreted Signaling | KEGG: mmu04310; PMID: 23209147 |
| NeuN | CD31 | Wnt9a | FZD6_LRP5 | 0.000334 | 0 | WNT9A_FZD6_LRP5 | Wnt9a - (Fzd6+Lrp5) | WNT | Secreted Signaling | KEGG: mmu04310; PMID: 23209147 |
| CD31 | CD31 | Wnt9a | FZD8_LRP5 | 0.000203 | 0 | WNT9A_FZD8_LRP5 | Wnt9a - (Fzd8+Lrp5) | WNT | Secreted Signaling | KEGG: mmu04310; PMID: 23209147 |
| CD31 | NeuN | Wnt9a | FZD8_LRP5 | 0.000181 | 0 | WNT9A_FZD8_LRP5 | Wnt9a - (Fzd8+Lrp5) | WNT | Secreted Signaling | KEGG: mmu04310; PMID: 23209147 |
| CD31 | CD31 | Wnt9a | FZD9_LRP5 | 0.000168 | 0 | WNT9A_FZD9_LRP5 | Wnt9a - (Fzd9+Lrp5) | WNT | Secreted Signaling | KEGG: mmu04310; PMID: 23209147 |
| CD31 | NeuN | Wnt9a | FZD9_LRP5 | 0.000182 | 0 | WNT9A_FZD9_LRP5 | Wnt9a - (Fzd9+Lrp5) | WNT | Secreted Signaling | KEGG: mmu04310; PMID: 23209147 |
| NeuN | CD31 | Wnt10b | FZD1_LRP6 | 0.000416 | 0 | WNT10B_FZD1_LRP6 | Wnt10b - (Fzd1+Lrp6) | WNT | Secreted Signaling | KEGG: mmu04310; PMID: 23209147 |
| NeuN | NeuN | Wnt10b | FZD1_LRP6 | 0.000424 | 0 | WNT10B_FZD1_LRP6 | Wnt10b - (Fzd1+Lrp6) | WNT | Secreted Signaling | KEGG: mmu04310; PMID: 23209147 |
| NeuN | CD31 | Wnt10b | FZD2_LRP6 | 0.000331 | 0 | WNT10B_FZD2_LRP6 | Wnt10b - (Fzd2+Lrp6) | WNT | Secreted Signaling | KEGG: mmu04310; PMID: 23209147 |
| NeuN | NeuN | Wnt10b | FZD2_LRP6 | 0.000349 | 0 | WNT10B_FZD2_LRP6 | Wnt10b - (Fzd2+Lrp6) | WNT | Secreted Signaling | KEGG: mmu04310; PMID: 23209147 |
| CD31 | NeuN | Wnt10b | FZD3_LRP6 | 0.000703 | 0 | WNT10B_FZD3_LRP6 | Wnt10b - (Fzd3+Lrp6) | WNT | Secreted Signaling | KEGG: mmu04310; PMID: 23209147 |
| NeuN | NeuN | Wnt10b | FZD3_LRP6 | 0.00086 | 0 | WNT10B_FZD3_LRP6 | Wnt10b - (Fzd3+Lrp6) | WNT | Secreted Signaling | KEGG: mmu04310; PMID: 23209147 |
| NeuN | CD31 | Wnt10b | FZD4_LRP6 | 0.000263 | 0 | WNT10B_FZD4_LRP6 | Wnt10b - (Fzd4+Lrp6) | WNT | Secreted Signaling | KEGG: mmu04310; PMID: 23209147 |
| NeuN | NeuN | Wnt10b | FZD4_LRP6 | 0.000263 | 0 | WNT10B_FZD4_LRP6 | Wnt10b - (Fzd4+Lrp6) | WNT | Secreted Signaling | KEGG: mmu04310; PMID: 23209147 |
| NeuN | CD31 | Wnt10b | FZD5_LRP6 | 0.00027 | 0 | WNT10B_FZD5_LRP6 | Wnt10b - (Fzd5+Lrp6) | WNT | Secreted Signaling | KEGG: mmu04310; PMID: 23209147 |
| NeuN | NeuN | Wnt10b | FZD5_LRP6 | 0.000325 | 0 | WNT10B_FZD5_LRP6 | Wnt10b - (Fzd5+Lrp6) | WNT | Secreted Signaling | KEGG: mmu04310; PMID: 23209147 |
| CD31 | CD31 | Wnt10b | FZD6_LRP6 | 0.000473 | 0 | WNT10B_FZD6_LRP6 | Wnt10b - (Fzd6+Lrp6) | WNT | Secreted Signaling | KEGG: mmu04310; PMID: 23209147 |
| NeuN | CD31 | Wnt10b | FZD6_LRP6 | 0.000579 | 0 | WNT10B_FZD6_LRP6 | Wnt10b - (Fzd6+Lrp6) | WNT | Secreted Signaling | KEGG: mmu04310; PMID: 23209147 |
| NeuN | CD31 | Wnt10b | FZD8_LRP6 | 0.000293 | 0 | WNT10B_FZD8_LRP6 | Wnt10b - (Fzd8+Lrp6) | WNT | Secreted Signaling | KEGG: mmu04310; PMID: 23209147 |
| NeuN | NeuN | Wnt10b | FZD8_LRP6 | 0.000332 | 0 | WNT10B_FZD8_LRP6 | Wnt10b - (Fzd8+Lrp6) | WNT | Secreted Signaling | KEGG: mmu04310; PMID: 23209147 |
| CD31 | NeuN | Wnt10b | FZD9_LRP6 | 0.000274 | 0 | WNT10B_FZD9_LRP6 | Wnt10b - (Fzd9+Lrp6) | WNT | Secreted Signaling | KEGG: mmu04310; PMID: 23209147 |
| NeuN | NeuN | Wnt10b | FZD9_LRP6 | 0.000335 | 0 | WNT10B_FZD9_LRP6 | Wnt10b - (Fzd9+Lrp6) | WNT | Secreted Signaling | KEGG: mmu04310; PMID: 23209147 |
| CD31 | CD31 | Wnt4 | FZD1_LRP6 | 0.000459 | 0 | WNT4_FZD1_LRP6 | Wnt4 - (Fzd1+Lrp6) | WNT | Secreted Signaling | KEGG: mmu04310; PMID: 23209147 |
| CD31 | NeuN | Wnt4 | FZD1_LRP6 | 0.000467 | 0 | WNT4_FZD1_LRP6 | Wnt4 - (Fzd1+Lrp6) | WNT | Secreted Signaling | KEGG: mmu04310; PMID: 23209147 |
| CD31 | CD31 | Wnt4 | FZD2_LRP6 | 0.000365 | 0 | WNT4_FZD2_LRP6 | Wnt4 - (Fzd2+Lrp6) | WNT | Secreted Signaling | KEGG: mmu04310; PMID: 23209147 |
| CD31 | NeuN | Wnt4 | FZD2_LRP6 | 0.000385 | 0 | WNT4_FZD2_LRP6 | Wnt4 - (Fzd2+Lrp6) | WNT | Secreted Signaling | KEGG: mmu04310; PMID: 23209147 |
| CD31 | NeuN | Wnt4 | FZD3_LRP6 | 0.000948 | 0 | WNT4_FZD3_LRP6 | Wnt4 - (Fzd3+Lrp6) | WNT | Secreted Signaling | KEGG: mmu04310; PMID: 23209147 |
| NeuN | NeuN | Wnt4 | FZD3_LRP6 | 0.00087 | 0 | WNT4_FZD3_LRP6 | Wnt4 - (Fzd3+Lrp6) | WNT | Secreted Signaling | KEGG: mmu04310; PMID: 23209147 |
| CD31 | CD31 | Wnt4 | FZD4_LRP6 | 0.00029 | 0 | WNT4_FZD4_LRP6 | Wnt4 - (Fzd4+Lrp6) | WNT | Secreted Signaling | KEGG: mmu04310; PMID: 23209147 |
| CD31 | NeuN | Wnt4 | FZD4_LRP6 | 0.00029 | 0 | WNT4_FZD4_LRP6 | Wnt4 - (Fzd4+Lrp6) | WNT | Secreted Signaling | KEGG: mmu04310; PMID: 23209147 |
| CD31 | NeuN | Wnt4 | FZD5_LRP6 | 0.000359 | 0 | WNT4_FZD5_LRP6 | Wnt4 - (Fzd5+Lrp6) | WNT | Secreted Signaling | KEGG: mmu04310; PMID: 23209147 |
| NeuN | NeuN | Wnt4 | FZD5_LRP6 | 0.000329 | 0 | WNT4_FZD5_LRP6 | Wnt4 - (Fzd5+Lrp6) | WNT | Secreted Signaling | KEGG: mmu04310; PMID: 23209147 |
| CD31 | CD31 | Wnt4 | FZD6_LRP6 | 0.000639 | 0 | WNT4_FZD6_LRP6 | Wnt4 - (Fzd6+Lrp6) | WNT | Secreted Signaling | KEGG: mmu04310; PMID: 23209147 |
| NeuN | CD31 | Wnt4 | FZD6_LRP6 | 0.000586 | 0 | WNT4_FZD6_LRP6 | Wnt4 - (Fzd6+Lrp6) | WNT | Secreted Signaling | KEGG: mmu04310; PMID: 23209147 |
| CD31 | NeuN | Wnt4 | FZD8_LRP6 | 0.000366 | 0 | WNT4_FZD8_LRP6 | Wnt4 - (Fzd8+Lrp6) | WNT | Secreted Signaling | KEGG: mmu04310; PMID: 23209147 |
| NeuN | NeuN | Wnt4 | FZD8_LRP6 | 0.000336 | 0 | WNT4_FZD8_LRP6 | Wnt4 - (Fzd8+Lrp6) | WNT | Secreted Signaling | KEGG: mmu04310; PMID: 23209147 |
| CD31 | NeuN | Wnt4 | FZD9_LRP6 | 0.00037 | 0 | WNT4_FZD9_LRP6 | Wnt4 - (Fzd9+Lrp6) | WNT | Secreted Signaling | KEGG: mmu04310; PMID: 23209147 |
| NeuN | NeuN | Wnt4 | FZD9_LRP6 | 0.000339 | 0 | WNT4_FZD9_LRP6 | Wnt4 - (Fzd9+Lrp6) | WNT | Secreted Signaling | KEGG: mmu04310; PMID: 23209147 |
| CD31 | CD31 | Wnt7a | FZD1_LRP6 | 0.000704 | 0 | WNT7A_FZD1_LRP6 | Wnt7a - (Fzd1+Lrp6) | WNT | Secreted Signaling | KEGG: mmu04310; PMID: 23209147 |
| CD31 | NeuN | Wnt7a | FZD1_LRP6 | 0.000718 | 0 | WNT7A_FZD1_LRP6 | Wnt7a - (Fzd1+Lrp6) | WNT | Secreted Signaling | KEGG: mmu04310; PMID: 23209147 |
| CD31 | CD31 | Wnt7a | FZD2_LRP6 | 0.00056 | 0 | WNT7A_FZD2_LRP6 | Wnt7a - (Fzd2+Lrp6) | WNT | Secreted Signaling | KEGG: mmu04310; PMID: 23209147 |
| CD31 | NeuN | Wnt7a | FZD2_LRP6 | 0.000592 | 0 | WNT7A_FZD2_LRP6 | Wnt7a - (Fzd2+Lrp6) | WNT | Secreted Signaling | KEGG: mmu04310; PMID: 23209147 |
| CD31 | NeuN | Wnt7a | FZD3_LRP6 | 0.001456 | 0 | WNT7A_FZD3_LRP6 | Wnt7a - (Fzd3+Lrp6) | WNT | Secreted Signaling | KEGG: mmu04310; PMID: 23209147 |
| NeuN | NeuN | Wnt7a | FZD3_LRP6 | 0.001021 | 0 | WNT7A_FZD3_LRP6 | Wnt7a - (Fzd3+Lrp6) | WNT | Secreted Signaling | KEGG: mmu04310; PMID: 23209147 |
| CD31 | CD31 | Wnt7a | FZD4_LRP6 | 0.000445 | 0 | WNT7A_FZD4_LRP6 | Wnt7a - (Fzd4+Lrp6) | WNT | Secreted Signaling | KEGG: mmu04310; PMID: 23209147 |
| CD31 | NeuN | Wnt7a | FZD4_LRP6 | 0.000446 | 0 | WNT7A_FZD4_LRP6 | Wnt7a - (Fzd4+Lrp6) | WNT | Secreted Signaling | KEGG: mmu04310; PMID: 23209147 |
| CD31 | CD31 | Wnt7a | FZD5_LRP6 | 0.000458 | 0 | WNT7A_FZD5_LRP6 | Wnt7a - (Fzd5+Lrp6) | WNT | Secreted Signaling | KEGG: mmu04310; PMID: 23209147 |

| | | | | | | | | | | |
|---|---|---|---|---|---|---|---|---|---|---|
| CD31 | NeuN | Wnt7a | FZD5_LRP6 | 0.000551 | 0 | WNT7A_FZD5_LRP6 | Wnt7a - (Fzd5+Lrp6) | WNT | Secreted Signaling | KEGG: mmu04310; PMID: 23209147 |
| CD31 | CD31 | Wnt7a | FZD6_LRP6 | 0.00098 | 0 | WNT7A_FZD6_LRP6 | Wnt7a - (Fzd6+Lrp6) | WNT | Secreted Signaling | KEGG: mmu04310; PMID: 23209147 |
| NeuN | CD31 | Wnt7a | FZD6_LRP6 | 0.000687 | 0 | WNT7A_FZD6_LRP6 | Wnt7a - (Fzd6+Lrp6) | WNT | Secreted Signaling | KEGG: mmu04310; PMID: 23209147 |
| CD31 | CD31 | Wnt7a | FZD8_LRP6 | 0.000496 | 0 | WNT7A_FZD8_LRP6 | Wnt7a - (Fzd8+Lrp6) | WNT | Secreted Signaling | KEGG: mmu04310; PMID: 23209147 |
| CD31 | NeuN | Wnt7a | FZD8_LRP6 | 0.000562 | 0 | WNT7A_FZD8_LRP6 | Wnt7a - (Fzd8+Lrp6) | WNT | Secreted Signaling | KEGG: mmu04310; PMID: 23209147 |
| CD31 | CD31 | Wnt7a | FZD9_LRP6 | 0.000411 | 0 | WNT7A_FZD9_LRP6 | Wnt7a - (Fzd9+Lrp6) | WNT | Secreted Signaling | KEGG: mmu04310; PMID: 23209147 |
| CD31 | NeuN | Wnt7a | FZD9_LRP6 | 0.000568 | 0 | WNT7A_FZD9_LRP6 | Wnt7a - (Fzd9+Lrp6) | WNT | Secreted Signaling | KEGG: mmu04310; PMID: 23209147 |
| NeuN | CD31 | Wnt7b | FZD1_LRP6 | 0.002178 | 0 | WNT7B_FZD1_LRP6 | Wnt7b - (Fzd1+Lrp6) | WNT | Secreted Signaling | KEGG: mmu04310; PMID: 23209147 |
| NeuN | NeuN | Wnt7b | FZD1_LRP6 | 0.00222 | 0 | WNT7B_FZD1_LRP6 | Wnt7b - (Fzd1+Lrp6) | WNT | Secreted Signaling | KEGG: mmu04310; PMID: 23209147 |
| NeuN | CD31 | Wnt7b | FZD2_LRP6 | 0.001732 | 0 | WNT7B_FZD2_LRP6 | Wnt7b - (Fzd2+Lrp6) | WNT | Secreted Signaling | KEGG: mmu04310; PMID: 23209147 |
| NeuN | NeuN | Wnt7b | FZD2_LRP6 | 0.001831 | 0 | WNT7B_FZD2_LRP6 | Wnt7b - (Fzd2+Lrp6) | WNT | Secreted Signaling | KEGG: mmu04310; PMID: 23209147 |
| NeuN | CD31 | Wnt7b | FZD3_LRP6 | 0.002954 | 0 | WNT7B_FZD3_LRP6 | Wnt7b - (Fzd3+Lrp6) | WNT | Secreted Signaling | KEGG: mmu04310; PMID: 23209147 |
| NeuN | NeuN | Wnt7b | FZD3_LRP6 | 0.004491 | 0 | WNT7B_FZD3_LRP6 | Wnt7b - (Fzd3+Lrp6) | WNT | Secreted Signaling | KEGG: mmu04310; PMID: 23209147 |
| NeuN | CD31 | Wnt7b | FZD4_LRP6 | 0.001377 | 0 | WNT7B_FZD4_LRP6 | Wnt7b - (Fzd4+Lrp6) | WNT | Secreted Signaling | KEGG: mmu04310; PMID: 23209147 |
| NeuN | NeuN | Wnt7b | FZD4_LRP6 | 0.00138 | 0 | WNT7B_FZD4_LRP6 | Wnt7b - (Fzd4+Lrp6) | WNT | Secreted Signaling | KEGG: mmu04310; PMID: 23209147 |
| NeuN | CD31 | Wnt7b | FZD5_LRP6 | 0.001418 | 0 | WNT7B_FZD5_LRP6 | Wnt7b - (Fzd5+Lrp6) | WNT | Secreted Signaling | KEGG: mmu04310; PMID: 23209147 |
| NeuN | NeuN | Wnt7b | FZD5_LRP6 | 0.001704 | 0 | WNT7B_FZD5_LRP6 | Wnt7b - (Fzd5+Lrp6) | WNT | Secreted Signaling | KEGG: mmu04310; PMID: 23209147 |
| CD31 | CD31 | Wnt7b | FZD6_LRP6 | 0.001553 | 0 | WNT7B_FZD6_LRP6 | Wnt7b - (Fzd6+Lrp6) | WNT | Secreted Signaling | KEGG: mmu04310; PMID: 23209147 |
| NeuN | CD31 | Wnt7b | FZD6_LRP6 | 0.003028 | 0 | WNT7B_FZD6_LRP6 | Wnt7b - (Fzd6+Lrp6) | WNT | Secreted Signaling | KEGG: mmu04310; PMID: 23209147 |
| NeuN | CD31 | Wnt7b | FZD8_LRP6 | 0.001534 | 0 | WNT7B_FZD8_LRP6 | Wnt7b - (Fzd8+Lrp6) | WNT | Secreted Signaling | KEGG: mmu04310; PMID: 23209147 |
| NeuN | NeuN | Wnt7b | FZD8_LRP6 | 0.00174 | 0 | WNT7B_FZD8_LRP6 | Wnt7b - (Fzd8+Lrp6) | WNT | Secreted Signaling | KEGG: mmu04310; PMID: 23209147 |
| NeuN | CD31 | Wnt7b | FZD9_LRP6 | 0.001273 | 0 | WNT7B_FZD9_LRP6 | Wnt7b - (Fzd9+Lrp6) | WNT | Secreted Signaling | KEGG: mmu04310; PMID: 23209147 |
| NeuN | NeuN | Wnt7b | FZD9_LRP6 | 0.001757 | 0 | WNT7B_FZD9_LRP6 | Wnt7b - (Fzd9+Lrp6) | WNT | Secreted Signaling | KEGG: mmu04310; PMID: 23209147 |
| CD31 | CD31 | Wnt9a | FZD1_LRP6 | 0.000356 | 0 | WNT9A_FZD1_LRP6 | Wnt9a - (Fzd1+Lrp6) | WNT | Secreted Signaling | KEGG: mmu04310; PMID: 23209147 |
| CD31 | NeuN | Wnt9a | FZD1_LRP6 | 0.000363 | 0 | WNT9A_FZD1_LRP6 | Wnt9a - (Fzd1+Lrp6) | WNT | Secreted Signaling | KEGG: mmu04310; PMID: 23209147 |
| CD31 | CD31 | Wnt9a | FZD2_LRP6 | 0.000283 | 0 | WNT9A_FZD2_LRP6 | Wnt9a - (Fzd2+Lrp6) | WNT | Secreted Signaling | KEGG: mmu04310; PMID: 23209147 |
| CD31 | NeuN | Wnt9a | FZD2_LRP6 | 0.000299 | 0 | WNT9A_FZD2_LRP6 | Wnt9a - (Fzd2+Lrp6) | WNT | Secreted Signaling | KEGG: mmu04310; PMID: 23209147 |
| CD31 | NeuN | Wnt9a | FZD3_LRP6 | 0.000736 | 0 | WNT9A_FZD3_LRP6 | Wnt9a - (Fzd3+Lrp6) | WNT | Secreted Signaling | KEGG: mmu04310; PMID: 23209147 |
| NeuN | NeuN | Wnt9a | FZD3_LRP6 | 0.000613 | 0 | WNT9A_FZD3_LRP6 | Wnt9a - (Fzd3+Lrp6) | WNT | Secreted Signaling | KEGG: mmu04310; PMID: 23209147 |
| CD31 | CD31 | Wnt9a | FZD4_LRP6 | 0.000225 | 0 | WNT9A_FZD4_LRP6 | Wnt9a - (Fzd4+Lrp6) | WNT | Secreted Signaling | KEGG: mmu04310; PMID: 23209147 |
| CD31 | NeuN | Wnt9a | FZD4_LRP6 | 0.000225 | 0 | WNT9A_FZD4_LRP6 | Wnt9a - (Fzd4+Lrp6) | WNT | Secreted Signaling | KEGG: mmu04310; PMID: 23209147 |
| CD31 | NeuN | Wnt9a | FZD5_LRP6 | 0.000278 | 0 | WNT9A_FZD5_LRP6 | Wnt9a - (Fzd5+Lrp6) | WNT | Secreted Signaling | KEGG: mmu04310; PMID: 23209147 |
| NeuN | NeuN | Wnt9a | FZD5_LRP6 | 0.000232 | 0 | WNT9A_FZD5_LRP6 | Wnt9a - (Fzd5+Lrp6) | WNT | Secreted Signaling | KEGG: mmu04310; PMID: 23209147 |
| CD31 | CD31 | Wnt9a | FZD6_LRP6 | 0.000496 | 0 | WNT9A_FZD6_LRP6 | Wnt9a - (Fzd6+Lrp6) | WNT | Secreted Signaling | KEGG: mmu04310; PMID: 23209147 |
| NeuN | CD31 | Wnt9a | FZD6_LRP6 | 0.000413 | 0 | WNT9A_FZD6_LRP6 | Wnt9a - (Fzd6+Lrp6) | WNT | Secreted Signaling | KEGG: mmu04310; PMID: 23209147 |
| CD31 | CD31 | Wnt9a | FZD8_LRP6 | 0.00025 | 0 | WNT9A_FZD8_LRP6 | Wnt9a - (Fzd8+Lrp6) | WNT | Secreted Signaling | KEGG: mmu04310; PMID: 23209147 |
| CD31 | NeuN | Wnt9a | FZD8_LRP6 | 0.000284 | 0 | WNT9A_FZD8_LRP6 | Wnt9a - (Fzd8+Lrp6) | WNT | Secreted Signaling | KEGG: mmu04310; PMID: 23209147 |
| CD31 | NeuN | Wnt9a | FZD9_LRP6 | 0.000287 | 0 | WNT9A_FZD9_LRP6 | Wnt9a - (Fzd9+Lrp6) | WNT | Secreted Signaling | KEGG: mmu04310; PMID: 23209147 |
| NeuN | NeuN | Wnt9a | FZD9_LRP6 | 0.000239 | 0 | WNT9A_FZD9_LRP6 | Wnt9a - (Fzd9+Lrp6) | WNT | Secreted Signaling | KEGG: mmu04310; PMID: 23209147 |
| CD31 | CD31 | Wnt5a | Fzd1 | 0.001706 | 0 | WNT5A_FZD1 | Wnt5a - Fzd1 | ncWNT | Secreted Signaling | KEGG: mmu04310 |
| CD31 | NeuN | Wnt5a | Fzd1 | 0.001192 | 0 | WNT5A_FZD1 | Wnt5a - Fzd1 | ncWNT | Secreted Signaling | KEGG: mmu04310 |
| CD31 | CD31 | Wnt5a | Fzd2 | 0.001078 | 0 | WNT5A_FZD2 | Wnt5a - Fzd2 | ncWNT | Secreted Signaling | KEGG: mmu04310 |
| CD31 | NeuN | Wnt5a | Fzd2 | 0.00081 | 0 | WNT5A_FZD2 | Wnt5a - Fzd2 | ncWNT | Secreted Signaling | KEGG: mmu04310 |
| CD31 | CD31 | Wnt5a | Fzd3 | 0.003145 | 0 | WNT5A_FZD3 | Wnt5a - Fzd3 | ncWNT | Secreted Signaling | KEGG: mmu04310 |
| CD31 | NeuN | Wnt5a | Fzd3 | 0.004896 | 0 | WNT5A_FZD3 | Wnt5a - Fzd3 | ncWNT | Secreted Signaling | KEGG: mmu04310 |
| CD31 | CD31 | Wnt5a | Fzd4 | 0.00068 | 0 | WNT5A_FZD4 | Wnt5a - Fzd4 | ncWNT | Secreted Signaling | KEGG: mmu04310 |
| CD31 | NeuN | Wnt5a | Fzd4 | 0.00046 | 0 | WNT5A_FZD4 | Wnt5a - Fzd4 | ncWNT | Secreted Signaling | KEGG: mmu04310 |
| CD31 | CD31 | Wnt5a | Fzd5 | 0.000722 | 0 | WNT5A_FZD5 | Wnt5a - Fzd5 | ncWNT | Secreted Signaling | KEGG: mmu04310 |
| CD31 | NeuN | Wnt5a | Fzd5 | 0.000701 | 0 | WNT5A_FZD5 | Wnt5a - Fzd5 | ncWNT | Secreted Signaling | KEGG: mmu04310 |
| CD31 | CD31 | Wnt5a | Fzd6 | 0.003304 | 0 | WNT5A_FZD6 | Wnt5a - Fzd6 | ncWNT | Secreted Signaling | KEGG: mmu04310 |
| NeuN | CD31 | Wnt5a | Fzd6 | 0.001934 | 0 | WNT5A_FZD6 | Wnt5a - Fzd6 | ncWNT | Secreted Signaling | KEGG: mmu04310 |
| CD31 | CD31 | Wnt5a | Fzd8 | 0.000845 | 0 | WNT5A_FZD8 | Wnt5a - Fzd8 | ncWNT | Secreted Signaling | KEGG: mmu04310 |
| CD31 | NeuN | Wnt5a | Fzd8 | 0.000732 | 0 | WNT5A_FZD8 | Wnt5a - Fzd8 | ncWNT | Secreted Signaling | KEGG: mmu04310 |
| CD31 | CD31 | Wnt5a | Fzd9 | 0.000581 | 0 | WNT5A_FZD9 | Wnt5a - Fzd9 | ncWNT | Secreted Signaling | KEGG: mmu04310 |
| CD31 | NeuN | Wnt5a | Fzd9 | 0.000746 | 0 | WNT5A_FZD9 | Wnt5a - Fzd9 | ncWNT | Secreted Signaling | KEGG: mmu04310 |
| CD31 | CD31 | Wnt5a | Mcam | 0.003517 | 0 | WNT5A_MCAM | Wnt5a - Mcam | ncWNT | Secreted Signaling | PMID: 24335906 |
| NeuN | CD31 | Wnt5a | Mcam | 0.002059 | 0 | WNT5A_MCAM | Wnt5a - Mcam | ncWNT | Secreted Signaling | PMID: 24335906 |
| CD31 | CD31 | Hbegf | Erbb4 | 0.003127 | 0 | HBEGF_ERBB4 | Hbegf - Erbb4 | EGF | Secreted Signaling | KEGG: mmu04012 |
| CD31 | NeuN | Hbegf | Erbb4 | 0.003473 | 0 | HBEGF_ERBB4 | Hbegf - Erbb4 | EGF | Secreted Signaling | KEGG: mmu04012 |

| | | | | | | | | | | |
|---|---|---|---|---|---|---|---|---|---|---|
| CD31 | CD31 | Hbegf | ERBB2_ERBB4 | 0.002154 | 0 | HBEGF_ERBB2_ERBB4 | Hbegf - (Erbb2+Erbb4) | EGF | Secreted Signaling | KEGG: mmu04012 |
| CD31 | NeuN | Hbegf | ERBB2_ERBB4 | 0.001753 | 0 | HBEGF_ERBB2_ERBB4 | Hbegf - (Erbb2+Erbb4) | EGF | Secreted Signaling | KEGG: mmu04012 |
| CD31 | CD31 | Ereg | Erbb4 | 0.001303 | 0 | EREG_ERBB4 | Ereg - Erbb4 | EGF | Secreted Signaling | KEGG: mmu04012 |
| CD31 | NeuN | Ereg | Erbb4 | 0.001448 | 0 | EREG_ERBB4 | Ereg - Erbb4 | EGF | Secreted Signaling | KEGG: mmu04012 |
| CD31 | CD31 | Ereg | ERBB2_ERBB4 | 0.000897 | 0 | EREG_ERBB2_ERBB4 | Ereg - (Erbb2+Erbb4) | EGF | Secreted Signaling | KEGG: mmu04012 |
| NeuN | CD31 | Ereg | ERBB2_ERBB4 | 0.000806 | 0 | EREG_ERBB2_ERBB4 | Ereg - (Erbb2+Erbb4) | EGF | Secreted Signaling | KEGG: mmu04012 |
| CD31 | CD31 | Nrg1 | Erbb3 | 0.000679 | 0 | NRG1_ERBB3 | Nrg1 - Erbb3 | NRG | Secreted Signaling | KEGG: mmu04012 |
| NeuN | CD31 | Nrg1 | Erbb3 | 0.000562 | 0 | NRG1_ERBB3 | Nrg1 - Erbb3 | NRG | Secreted Signaling | KEGG: mmu04012 |
| CD31 | CD31 | Nrg1 | ERBB2_ERBB3 | 0.000929 | 0 | NRG1_ERBB2_ERBB3 | Nrg1 - (Erbb2+Erbb3) | NRG | Secreted Signaling | KEGG: mmu04012 |
| NeuN | CD31 | Nrg1 | ERBB2_ERBB3 | 0.000769 | 0 | NRG1_ERBB2_ERBB3 | Nrg1 - (Erbb2+Erbb3) | NRG | Secreted Signaling | KEGG: mmu04012 |
| CD31 | CD31 | Nrg1 | Erbb4 | 0.002683 | 0 | NRG1_ERBB4 | Nrg1 - Erbb4 | NRG | Secreted Signaling | KEGG: mmu04012 |
| CD31 | NeuN | Nrg1 | Erbb4 | 0.00298 | 0 | NRG1_ERBB4 | Nrg1 - Erbb4 | NRG | Secreted Signaling | KEGG: mmu04012 |
| CD31 | CD31 | Nrg1 | ERBB2_ERBB4 | 0.001848 | 0 | NRG1_ERBB2_ERBB4 | Nrg1 - (Erbb2+Erbb4) | NRG | Secreted Signaling | KEGG: mmu04012 |
| NeuN | CD31 | Nrg1 | ERBB2_ERBB4 | 0.001529 | 0 | NRG1_ERBB2_ERBB4 | Nrg1 - (Erbb2+Erbb4) | NRG | Secreted Signaling | KEGG: mmu04012 |
| NeuN | CD31 | Nrg2 | Erbb3 | 0.000531 | 0 | NRG2_ERBB3 | Nrg2 - Erbb3 | NRG | Secreted Signaling | KEGG: mmu04012 |
| NeuN | NeuN | Nrg2 | Erbb3 | 0.000429 | 0 | NRG2_ERBB3 | Nrg2 - Erbb3 | NRG | Secreted Signaling | KEGG: mmu04012 |
| CD31 | CD31 | Nrg2 | ERBB2_ERBB3 | 0.000577 | 0 | NRG2_ERBB2_ERBB3 | Nrg2 - (Erbb2+Erbb3) | NRG | Secreted Signaling | KEGG: mmu04012 |
| NeuN | CD31 | Nrg2 | ERBB2_ERBB3 | 0.000727 | 0 | NRG2_ERBB2_ERBB3 | Nrg2 - (Erbb2+Erbb3) | NRG | Secreted Signaling | KEGG: mmu04012 |
| NeuN | CD31 | Nrg2 | Erbb4 | 0.002099 | 0 | NRG2_ERBB4 | Nrg2 - Erbb4 | NRG | Secreted Signaling | KEGG: mmu04012 |
| NeuN | NeuN | Nrg2 | Erbb4 | 0.002331 | 0 | NRG2_ERBB4 | Nrg2 - Erbb4 | NRG | Secreted Signaling | KEGG: mmu04012 |
| NeuN | CD31 | Nrg2 | ERBB2_ERBB4 | 0.001445 | 0 | NRG2_ERBB2_ERBB4 | Nrg2 - (Erbb2+Erbb4) | NRG | Secreted Signaling | KEGG: mmu04012 |
| NeuN | NeuN | Nrg2 | ERBB2_ERBB4 | 0.001176 | 0 | NRG2_ERBB2_ERBB4 | Nrg2 - (Erbb2+Erbb4) | NRG | Secreted Signaling | KEGG: mmu04012 |
| NeuN | CD31 | Nrg3 | Erbb4 | 0.008906 | 0 | NRG3_ERBB4 | Nrg3 - Erbb4 | NRG | Secreted Signaling | KEGG: mmu04012 |
| NeuN | NeuN | Nrg3 | Erbb4 | 0.009886 | 0 | NRG3_ERBB4 | Nrg3 - Erbb4 | NRG | Secreted Signaling | KEGG: mmu04012 |
| NeuN | CD31 | Nrg3 | ERBB2_ERBB4 | 0.006146 | 0 | NRG3_ERBB2_ERBB4 | Nrg3 - (Erbb2+Erbb4) | NRG | Secreted Signaling | KEGG: mmu04012 |
| NeuN | NeuN | Nrg3 | ERBB2_ERBB4 | 0.005007 | 0 | NRG3_ERBB2_ERBB4 | Nrg3 - (Erbb2+Erbb4) | NRG | Secreted Signaling | KEGG: mmu04012 |
| CD31 | CD31 | Nrg1 | ITGAV_ITGB3 | 0.00164 | 0 | NRG1_ITGAV_ITGB3 | Nrg1 - (Itgav+Itgb3) | NRG | Secreted Signaling | PMID: 20682778 |
| NeuN | CD31 | Nrg1 | ITGAV_ITGB3 | 0.001358 | 0 | NRG1_ITGAV_ITGB3 | Nrg1 - (Itgav+Itgb3) | NRG | Secreted Signaling | PMID: 20682778 |
| CD31 | CD31 | Fgf2 | Fgfr1 | 0.001281 | 0 | FGF2_FGFR1 | Fgf2 - Fgfr1 | FGF | Secreted Signaling | PMC: 4393358 |
| NeuN | CD31 | Fgf2 | Fgfr1 | 0.00101 | 0 | FGF2_FGFR1 | Fgf2 - Fgfr1 | FGF | Secreted Signaling | PMC: 4393358 |
| CD31 | CD31 | Fgf2 | Fgfr2 | 0.000985 | 0 | FGF2_FGFR2 | Fgf2 - Fgfr2 | FGF | Secreted Signaling | PMC: 4393358 |
| CD31 | NeuN | Fgf2 | Fgfr2 | 0.000781 | 0 | FGF2_FGFR2 | Fgf2 - Fgfr2 | FGF | Secreted Signaling | PMC: 4393358 |
| CD31 | CD31 | Fgf2 | Fgfr3 | 0.00073 | 0 | FGF2_FGFR3 | Fgf2 - Fgfr3 | FGF | Secreted Signaling | PMC: 4393358 |
| CD31 | NeuN | Fgf2 | Fgfr3 | 0.000734 | 0 | FGF2_FGFR3 | Fgf2 - Fgfr3 | FGF | Secreted Signaling | PMC: 4393358 |
| CD31 | CD31 | Fgf5 | Fgfr1 | 0.001342 | 0 | FGF5_FGFR1 | Fgf5 - Fgfr1 | FGF | Secreted Signaling | PMC: 4393358 |
| CD31 | NeuN | Fgf5 | Fgfr1 | 0.000983 | 0 | FGF5_FGFR1 | Fgf5 - Fgfr1 | FGF | Secreted Signaling | PMC: 4393358 |
| CD31 | CD31 | Fgf5 | Fgfr2 | 0.001031 | 0 | FGF5_FGFR2 | Fgf5 - Fgfr2 | FGF | Secreted Signaling | PMC: 4393358 |
| CD31 | NeuN | Fgf5 | Fgfr2 | 0.000817 | 0 | FGF5_FGFR2 | Fgf5 - Fgfr2 | FGF | Secreted Signaling | PMC: 4393358 |
| CD31 | CD31 | Fgf5 | Fgfr3 | 0.000765 | 0 | FGF5_FGFR3 | Fgf5 - Fgfr3 | FGF | Secreted Signaling | PMC: 4393358 |
| CD31 | NeuN | Fgf5 | Fgfr3 | 0.000769 | 0 | FGF5_FGFR3 | Fgf5 - Fgfr3 | FGF | Secreted Signaling | PMC: 4393358 |
| CD31 | CD31 | Fgf3 | Fgfr1 | 0.001372 | 0 | FGF3_FGFR1 | Fgf3 - Fgfr1 | FGF | Secreted Signaling | PMC: 4393358 |
| CD31 | NeuN | Fgf3 | Fgfr1 | 0.001006 | 0 | FGF3_FGFR1 | Fgf3 - Fgfr1 | FGF | Secreted Signaling | PMC: 4393358 |
| CD31 | CD31 | Fgf3 | Fgfr2 | 0.001055 | 0 | FGF3_FGFR2 | Fgf3 - Fgfr2 | FGF | Secreted Signaling | PMC: 4393358 |
| CD31 | NeuN | Fgf3 | Fgfr2 | 0.000836 | 0 | FGF3_FGFR2 | Fgf3 - Fgfr2 | FGF | Secreted Signaling | PMC: 4393358 |
| CD31 | CD31 | Fgf10 | Fgfr1 | 0.001993 | 0 | FGF10_FGFR1 | Fgf10 - Fgfr1 | FGF | Secreted Signaling | PMC: 4393358 |
| CD31 | NeuN | Fgf10 | Fgfr1 | 0.00146 | 0 | FGF10_FGFR1 | Fgf10 - Fgfr1 | FGF | Secreted Signaling | PMC: 4393358 |
| CD31 | CD31 | Fgf10 | Fgfr2 | 0.001532 | 0 | FGF10_FGFR2 | Fgf10 - Fgfr2 | FGF | Secreted Signaling | PMC: 4393358 |
| CD31 | NeuN | Fgf10 | Fgfr2 | 0.001214 | 0 | FGF10_FGFR2 | Fgf10 - Fgfr2 | FGF | Secreted Signaling | PMC: 4393358 |
| CD31 | CD31 | Fgf22 | Fgfr2 | 0.000903 | 0 | FGF22_FGFR2 | Fgf22 - Fgfr2 | FGF | Secreted Signaling | PMC: 4393358 |
| NeuN | CD31 | Fgf22 | Fgfr2 | 0.000717 | 0 | FGF22_FGFR2 | Fgf22 - Fgfr2 | FGF | Secreted Signaling | PMC: 4393358 |
| CD31 | CD31 | Fgf18 | Fgfr1 | 0.001051 | 0 | FGF18_FGFR1 | Fgf18 - Fgfr1 | FGF | Secreted Signaling | PMC: 4393358 |
| NeuN | CD31 | Fgf18 | Fgfr1 | 0.001013 | 0 | FGF18_FGFR1 | Fgf18 - Fgfr1 | FGF | Secreted Signaling | PMC: 4393358 |
| CD31 | CD31 | Fgf18 | Fgfr2 | 0.000808 | 0 | FGF18_FGFR2 | Fgf18 - Fgfr2 | FGF | Secreted Signaling | PMC: 4393358 |
| NeuN | CD31 | Fgf18 | Fgfr2 | 0.000778 | 0 | FGF18_FGFR2 | Fgf18 - Fgfr2 | FGF | Secreted Signaling | PMC: 4393358 |
| CD31 | CD31 | Fgf18 | Fgfr3 | 0.000599 | 0 | FGF18_FGFR3 | Fgf18 - Fgfr3 | FGF | Secreted Signaling | PMC: 4393358 |
| CD31 | NeuN | Fgf18 | Fgfr3 | 0.000602 | 0 | FGF18_FGFR3 | Fgf18 - Fgfr3 | FGF | Secreted Signaling | PMC: 4393358 |
| CD31 | CD31 | Fgf9 | Fgfr1 | 0.001181 | 0 | FGF9_FGFR1 | Fgf9 - Fgfr1 | FGF | Secreted Signaling | PMC: 4393358 |
| NeuN | CD31 | Fgf9 | Fgfr1 | 0.001035 | 0 | FGF9_FGFR1 | Fgf9 - Fgfr1 | FGF | Secreted Signaling | PMC: 4393358 |
| CD31 | CD31 | Fgf9 | Fgfr2 | 0.000908 | 0 | FGF9_FGFR2 | Fgf9 - Fgfr2 | FGF | Secreted Signaling | PMC: 4393358 |

| | | | | | | | | | |
|---|---|---|---|---|---|---|---|---|---|
| NeuN | CD31 | Fgf9 | Fgfr2 | 0.000795 | 0 | FGF9_FGFR2 | Fgf9 - Fgfr2 | FGF | Secreted Signaling | PMC: 4393358 |
| CD31 | CD31 | Fgf9 | Fgfr3 | 0.000673 | 0 | FGF9_FGFR3 | Fgf9 - Fgfr3 | FGF | Secreted Signaling | PMC: 4393358 |
| CD31 | NeuN | Fgf9 | Fgfr3 | 0.000676 | 0 | FGF9_FGFR3 | Fgf9 - Fgfr3 | FGF | Secreted Signaling | PMC: 4393358 |
| CD31 | CD31 | Fgf15 | Fgfr1 | 0.00113 | 0 | FGF15_FGFR1 | Fgf15 - Fgfr1 | FGF | Secreted Signaling | PMC: 4393358 |
| NeuN | CD31 | Fgf15 | Fgfr1 | 0.001181 | 0 | FGF15_FGFR1 | Fgf15 - Fgfr1 | FGF | Secreted Signaling | PMC: 4393358 |
| CD31 | CD31 | Fgf15 | Fgfr2 | 0.000869 | 0 | FGF15_FGFR2 | Fgf15 - Fgfr2 | FGF | Secreted Signaling | PMC: 4393358 |
| NeuN | CD31 | Fgf15 | Fgfr2 | 0.000907 | 0 | FGF15_FGFR2 | Fgf15 - Fgfr2 | FGF | Secreted Signaling | PMC: 4393358 |
| NeuN | CD31 | Fgf15 | Fgfr3 | 0.000673 | 0 | FGF15_FGFR3 | Fgf15 - Fgfr3 | FGF | Secreted Signaling | PMC: 4393358 |
| NeuN | NeuN | Fgf15 | Fgfr3 | 0.000673 | 0 | FGF15_FGFR3 | Fgf15 - Fgfr3 | FGF | Secreted Signaling | PMC: 4393358 |
| CD31 | CD31 | Fgf21 | Fgfr1 | 0.001235 | 0 | FGF21_FGFR1 | Fgf21 - Fgfr1 | FGF | Secreted Signaling | PMC: 4393358 |
| NeuN | CD31 | Fgf21 | Fgfr1 | 0.001184 | 0 | FGF21_FGFR1 | Fgf21 - Fgfr1 | FGF | Secreted Signaling | PMC: 4393358 |
| CD31 | CD31 | Fgf21 | Fgfr3 | 0.000703 | 0 | FGF21_FGFR3 | Fgf21 - Fgfr3 | FGF | Secreted Signaling | PMC: 4393358 |
| CD31 | NeuN | Fgf21 | Fgfr3 | 0.000704 | 0 | FGF21_FGFR3 | Fgf21 - Fgfr3 | FGF | Secreted Signaling | PMC: 4393358 |
| NeuN | CD31 | Pdgfa | Pdgfra | 0.000823 | 0 | PDGFA_PDGFRA | Pdgfa - Pdgfra | PDGF | Secreted Signaling | PMID: 15207812 |
| NeuN | NeuN | Pdgfa | Pdgfra | 0.000583 | 0 | PDGFA_PDGFRA | Pdgfa - Pdgfra | PDGF | Secreted Signaling | PMID: 15207812 |
| CD31 | CD31 | Pdgfa | Pdgfrb | 0.000947 | 0 | PDGFA_PDGFRB | Pdgfa - Pdgfrb | PDGF | Secreted Signaling | PMID: 15207812 |
| NeuN | CD31 | Pdgfa | Pdgfrb | 0.001658 | 0 | PDGFA_PDGFRB | Pdgfa - Pdgfrb | PDGF | Secreted Signaling | PMID: 15207812 |
| CD31 | CD31 | Pdgfb | Pdgfra | 0.001849 | 0 | PDGFB_PDGFRA | Pdgfb - Pdgfra | PDGF | Secreted Signaling | PMID: 15207812 |
| CD31 | NeuN | Pdgfb | Pdgfra | 0.001313 | 0 | PDGFB_PDGFRA | Pdgfb - Pdgfra | PDGF | Secreted Signaling | PMID: 15207812 |
| CD31 | CD31 | Pdgfb | Pdgfrb | 0.003713 | 0 | PDGFB_PDGFRB | Pdgfb - Pdgfrb | PDGF | Secreted Signaling | PMID: 15207812 |
| CD31 | NeuN | Pdgfb | Pdgfrb | 0.000879 | 0 | PDGFB_PDGFRB | Pdgfb - Pdgfrb | PDGF | Secreted Signaling | PMID: 15207812 |
| NeuN | CD31 | Pdgfc | Pdgfra | 0.000668 | 0 | PDGFC_PDGFRA | Pdgfc - Pdgfra | PDGF | Secreted Signaling | PMID: 15207812 |
| NeuN | NeuN | Pdgfc | Pdgfra | 0.000473 | 0 | PDGFC_PDGFRA | Pdgfc - Pdgfra | PDGF | Secreted Signaling | PMID: 15207812 |
| CD31 | CD31 | Pdgfd | Pdgfrb | 0.001352 | 0 | PDGFD_PDGFRB | Pdgfd - Pdgfrb | PDGF | Secreted Signaling | PMID: 15207812 |
| NeuN | CD31 | Pdgfd | Pdgfrb | 0.000843 | 0 | PDGFD_PDGFRB | Pdgfd - Pdgfrb | PDGF | Secreted Signaling | PMID: 15207812 |
| CD31 | CD31 | Vegfa | Flt1 | 0.011216 | 0 | VEGFA_VEGFR1 | Vegfa - Vegfr1 | VEGF | Secreted Signaling | KEGG: mmu04370; PMID: 16633338 |
| NeuN | CD31 | Vegfa | Flt1 | 0.015384 | 0 | VEGFA_VEGFR1 | Vegfa - Vegfr1 | VEGF | Secreted Signaling | KEGG: mmu04370; PMID: 16633338 |
| CD31 | CD31 | Vegfa | Kdr | 0.008511 | 0 | VEGFA_VEGFR2 | Vegfa - Vegfr2 | VEGF | Secreted Signaling | KEGG: mmu04370; PMID: 16633338 |
| NeuN | CD31 | Vegfa | Kdr | 0.011687 | 0 | VEGFA_VEGFR2 | Vegfa - Vegfr2 | VEGF | Secreted Signaling | KEGG: mmu04370; PMID: 16633338 |
| CD31 | CD31 | Vegfb | Flt1 | 0.009807 | 0 | VEGFB_VEGFR1 | Vegfb - Vegfr1 | VEGF | Secreted Signaling | KEGG: mmu04370; PMID: 16633338 |
| NeuN | CD31 | Vegfb | Flt1 | 0.010268 | 0 | VEGFB_VEGFR1 | Vegfb - Vegfr1 | VEGF | Secreted Signaling | KEGG: mmu04370; PMID: 16633338 |
| CD31 | CD31 | Pgf | Flt1 | 0.009171 | 0 | PGF_VEGFR1 | Pgf - Vegfr1 | VEGF | Secreted Signaling | KEGG: mmu04370; PMID: 16633338 |
| NeuN | CD31 | Pgf | Flt1 | 0.006255 | 0 | PGF_VEGFR1 | Pgf - Vegfr1 | VEGF | Secreted Signaling | KEGG: mmu04370; PMID: 16633338 |
| CD31 | CD31 | Vegfa | FLT1_KDR | 0.010194 | 0 | VEGFA_VEGFR1R2 | Vegfa - Vegfr1r2 | VEGF | Secreted Signaling | PMID: 16633338 |
| NeuN | CD31 | Vegfa | FLT1_KDR | 0.013988 | 0 | VEGFA_VEGFR1R2 | Vegfa - Vegfr1r2 | VEGF | Secreted Signaling | PMID: 16633338 |
| CD31 | CD31 | Igf1 | Igf1r | 0.000921 | 0 | IGF1_IGF1R | Igf1 - Igf1r | IGF | Secreted Signaling | PMID: 14604834 |
| CD31 | NeuN | Igf1 | Igf1r | 0.000779 | 0 | IGF1_IGF1R | Igf1 - Igf1r | IGF | Secreted Signaling | PMID: 14604834 |
| CD31 | CD31 | Igf2 | Igf1r | 0.010745 | 0 | IGF2_IGF1R | Igf2 - Igf1r | IGF | Secreted Signaling | PMID: 14604834 |
| CD31 | NeuN | Igf2 | Igf1r | 0.009134 | 0 | IGF2_IGF1R | Igf2 - Igf1r | IGF | Secreted Signaling | PMID: 14604834 |
| CD31 | CD31 | Igf2 | Igf2r | 0.004444 | 0 | IGF2_IGF2R | Igf2 - Igf2r | IGF | Secreted Signaling | PMID: 14604834 |
| CD31 | NeuN | Igf2 | Igf2r | 0.003827 | 0 | IGF2_IGF2R | Igf2 - Igf2r | IGF | Secreted Signaling | PMID: 14604834 |
| CD31 | CD31 | Igf1 | ITGAV_ITGB3 | 0.000269 | 0 | IGF1_ITGAV_ITGB3 | Igf1 - (Itgav+Itgb3) | IGF | Secreted Signaling | PMID: 28190785; PMID: 28873464 |
| NeuN | CD31 | Igf1 | ITGAV_ITGB3 | 0.000223 | 0 | IGF1_ITGAV_ITGB3 | Igf1 - (Itgav+Itgb3) | IGF | Secreted Signaling | PMID: 28190785; PMID: 28873464 |
| CD31 | CD31 | Igf2 | ITGAV_ITGB3 | 0.0032 | 0 | IGF2_ITGAV_ITGB3 | Igf2 - (Itgav+Itgb3) | IGF | Secreted Signaling | PMID: 28190785; PMID: 28873464 |
| CD31 | NeuN | Igf2 | ITGAV_ITGB3 | 0.002012 | 0 | IGF2_ITGAV_ITGB3 | Igf2 - (Itgav+Itgb3) | IGF | Secreted Signaling | PMID: 28190785; PMID: 28873464 |
| CD31 | CD31 | Ins1 | Insr | 0.003796 | 0 | INS1_INSSR | Ins1 - Inssr | INSULIN | Secreted Signaling | KEGG: mmu04910; PMID: 24633815 |
| NeuN | CD31 | Ins1 | Insr | 0.003672 | 0 | INS1_INSSR | Ins1 - Inssr | INSULIN | Secreted Signaling | KEGG: mmu04910; PMID: 24633815 |
| CD31 | CD31 | Apln | Aplnr | 0.010903 | 0 | APLN_APLNR | Apln - Aplnr | APELIN | Secreted Signaling | KEGG: mmu04371 |
| NeuN | CD31 | Apln | Aplnr | 0.001975 | 0 | APLN_APLNR | Apln - Aplnr | APELIN | Secreted Signaling | KEGG: mmu04371 |
| CD31 | CD31 | Ihh | PTCH1_SMO | 0.001478 | 0 | IHH_PTCH1 | Ihh - Ptch1 | HH | Secreted Signaling | KEGG: mmu04340; PMID: 25701396 |
| NeuN | CD31 | Ihh | PTCH1_SMO | 0.001238 | 0 | IHH_PTCH1 | Ihh - Ptch1 | HH | Secreted Signaling | KEGG: mmu04340; PMID: 25701396 |
| CD31 | CD31 | Shh | PTCH1_SMO | 0.000828 | 0 | SHH_PTCH1 | Shh - Ptch1 | HH | Secreted Signaling | KEGG: mmu04340; PMID: 25701396 |
| NeuN | CD31 | Shh | PTCH1_SMO | 0.000839 | 0 | SHH_PTCH1 | Shh - Ptch1 | HH | Secreted Signaling | KEGG: mmu04340; PMID: 25701396 |
| CD31 | CD31 | Ihh | PTCH2_SMO | 0.00116 | 0 | IHH_PTCH2 | Ihh - Ptch2 | HH | Secreted Signaling | KEGG: mmu04340; PMID: 25701396 |
| NeuN | CD31 | Ihh | PTCH2_SMO | 0.000972 | 0 | IHH_PTCH2 | Ihh - Ptch2 | HH | Secreted Signaling | KEGG: mmu04340; PMID: 25701396 |
| CD31 | CD31 | Shh | PTCH2_SMO | 0.000649 | 0 | SHH_PTCH2 | Shh - Ptch2 | HH | Secreted Signaling | KEGG: mmu04340; PMID: 25701396 |
| NeuN | CD31 | Shh | PTCH2_SMO | 0.000658 | 0 | SHH_PTCH2 | Shh - Ptch2 | HH | Secreted Signaling | KEGG: mmu04340; PMID: 25701396 |
| CD31 | CD31 | Ccl8 | Ccr1 | 0.000646 | 0 | CCL8_CCR1 | Ccl8 - Ccr1 | CCL | Secreted Signaling | KEGG: mmu04060 |
| NeuN | CD31 | Ccl8 | Ccr1 | 0.000534 | 0 | CCL8_CCR1 | Ccl8 - Ccr1 | CCL | Secreted Signaling | KEGG: mmu04060 |

| | | | | | | | | | | |
|---|---|---|---|---|---|---|---|---|---|---|
| CD31 | CD31 | Ccl6 | Ccr1 | 0.000742 | 0 | CCL6_CCR1 | Ccl6 - Ccr1 | CCL | Secreted Signaling | KEGG: mmu04060 |
| NeuN | CD31 | Ccl6 | Ccr1 | 0.000635 | 0 | CCL6_CCR1 | Ccl6 - Ccr1 | CCL | Secreted Signaling | KEGG: mmu04060 |
| CD31 | CD31 | Ccl8 | Ccr1l1 | 0.000604 | 0 | CCL8_CCR1L1 | Ccl8 - Ccr1l1 | CCL | Secreted Signaling | KEGG: mmu04060 |
| NeuN | CD31 | Ccl8 | Ccr1l1 | 0.000499 | 0 | CCL8_CCR1L1 | Ccl8 - Ccr1l1 | CCL | Secreted Signaling | KEGG: mmu04060 |
| CD31 | CD31 | Ccl6 | Ccr1l1 | 0.000693 | 0 | CCL6_CCR1L1 | Ccl6 - Ccr1l1 | CCL | Secreted Signaling | KEGG: mmu04060 |
| NeuN | CD31 | Ccl6 | Ccr1l1 | 0.000593 | 0 | CCL6_CCR1L1 | Ccl6 - Ccr1l1 | CCL | Secreted Signaling | KEGG: mmu04060 |
| CD31 | CD31 | Ccl6 | Ccr3 | 0.000685 | 0 | CCL6_CCR3 | Ccl6 - Ccr3 | CCL | Secreted Signaling | KEGG: mmu04060 |
| NeuN | CD31 | Ccl6 | Ccr3 | 0.000586 | 0 | CCL6_CCR3 | Ccl6 - Ccr3 | CCL | Secreted Signaling | KEGG: mmu04060 |
| CD31 | CD31 | Ccl11 | Ccr3 | 0.000491 | 0 | CCL11_CCR3 | Ccl11 - Ccr3 | CCL | Secreted Signaling | KEGG: mmu04060 |
| NeuN | CD31 | Ccl11 | Ccr3 | 0.000544 | 0 | CCL11_CCR3 | Ccl11 - Ccr3 | CCL | Secreted Signaling | KEGG: mmu04060 |
| NeuN | CD31 | Ccl27a | Ccr3 | 0.008027 | 0 | CCL27A_CCR3 | Ccl27a - Ccr3 | CCL | Secreted Signaling | KEGG: mmu04060 |
| NeuN | NeuN | Ccl27a | Ccr3 | 0.006061 | 0 | CCL27A_CCR3 | Ccl27a - Ccr3 | CCL | Secreted Signaling | KEGG: mmu04060 |
| NeuN | CD31 | Ccl27b | Ccr3 | 0.001775 | 0 | CCL27B_CCR3 | Ccl27b - Ccr3 | CCL | Secreted Signaling | KEGG: mmu04060 |
| NeuN | NeuN | Ccl27b | Ccr3 | 0.001338 | 0 | CCL27B_CCR3 | Ccl27b - Ccr3 | CCL | Secreted Signaling | KEGG: mmu04060 |
| CD31 | CD31 | Ccl19 | Ccr7 | 0.001412 | 0 | CCL19_CCR7 | Ccl19 - Ccr7 | CCL | Secreted Signaling | KEGG: mmu04060 |
| NeuN | CD31 | Ccl19 | Ccr7 | 0.0012 | 0 | CCL19_CCR7 | Ccl19 - Ccr7 | CCL | Secreted Signaling | KEGG: mmu04060 |
| CD31 | CD31 | Ccl21a | Ccr7 | 0.000845 | 0 | CCL21A_CCR7 | Ccl21a - Ccr7 | CCL | Secreted Signaling | KEGG: mmu04060 |
| NeuN | CD31 | Ccl21a | Ccr7 | 0.000939 | 0 | CCL21A_CCR7 | Ccl21a - Ccr7 | CCL | Secreted Signaling | KEGG: mmu04060 |
| CD31 | CD31 | Ccl25 | Ccr9 | 0.001092 | 0 | CCL25_CCR9 | Ccl25 - Ccr9 | CCL | Secreted Signaling | KEGG: mmu04060 |
| CD31 | NeuN | Ccl25 | Ccr9 | 0.000817 | 0 | CCL25_CCR9 | Ccl25 - Ccr9 | CCL | Secreted Signaling | KEGG: mmu04060 |
| CD31 | CD31 | Ccl4 | Ackr2 | 0.001484 | 0 | CCL4_ACKR2 | Ccl4 - Ackr2 | CCL | Secreted Signaling | PMID: 26740381 |
| NeuN | CD31 | Ccl4 | Ackr2 | 0.001378 | 0 | CCL4_ACKR2 | Ccl4 - Ackr2 | CCL | Secreted Signaling | PMID: 26740381 |
| CD31 | NeuN | Ccl8 | Ackr1 | 0.001134 | 0 | CCL8_ACKR1 | Ccl8 - Ackr1 | CCL | Secreted Signaling | PMID: 26740381 |
| NeuN | NeuN | Ccl8 | Ackr1 | 0.000937 | 0 | CCL8_ACKR1 | Ccl8 - Ackr1 | CCL | Secreted Signaling | PMID: 26740381 |
| CD31 | NeuN | Ccl11 | Ackr1 | 0.000931 | 0 | CCL11_ACKR1 | Ccl11 - Ackr1 | CCL | Secreted Signaling | PMID: 26740381 |
| NeuN | NeuN | Ccl11 | Ackr1 | 0.001033 | 0 | CCL11_ACKR1 | Ccl11 - Ackr1 | CCL | Secreted Signaling | PMID: 26740381 |
| CD31 | CD31 | Ccl11 | Ackr2 | 0.000943 | 0 | CCL11_ACKR2 | Ccl11 - Ackr2 | CCL | Secreted Signaling | PMID: 26740381 |
| NeuN | CD31 | Ccl11 | Ackr2 | 0.001046 | 0 | CCL11_ACKR2 | Ccl11 - Ackr2 | CCL | Secreted Signaling | PMID: 26740381 |
| CD31 | CD31 | Pf4 | Cxcr3 | 0.000827 | 0 | CXCL4_CXCR3 | Cxcl4 - Cxcr3 | CXCL | Secreted Signaling | KEGG: mmu04060 |
| CD31 | NeuN | Pf4 | Cxcr3 | 0.000741 | 0 | CXCL4_CXCR3 | Cxcl4 - Cxcr3 | CXCL | Secreted Signaling | KEGG: mmu04060 |
| CD31 | CD31 | Cxcl10 | Cxcr3 | 0.00035 | 0 | CXCL10_CXCR3 | Cxcl10 - Cxcr3 | CXCL | Secreted Signaling | KEGG: mmu04060 |
| NeuN | CD31 | Cxcl10 | Cxcr3 | 0.000365 | 0 | CXCL10_CXCR3 | Cxcl10 - Cxcr3 | CXCL | Secreted Signaling | KEGG: mmu04060 |
| CD31 | CD31 | Cxcl12 | Cxcr4 | 0.012738 | 0 | CXCL12_CXCR4 | Cxcl12 - Cxcr4 | CXCL | Secreted Signaling | KEGG: mmu04060 |
| CD31 | NeuN | Cxcl12 | Cxcr4 | 0.004339 | 0 | CXCL12_CXCR4 | Cxcl12 - Cxcr4 | CXCL | Secreted Signaling | KEGG: mmu04060 |
| CD31 | CD31 | Cxcl12 | Ackr3 | 0.005984 | 0 | CXCL12_ACKR3 | Cxcl12 - Ackr3 | CXCL | Secreted Signaling | KEGG: mmu04060 |
| CD31 | NeuN | Cxcl12 | Ackr3 | 0.002376 | 0 | CXCL12_ACKR3 | Cxcl12 - Ackr3 | CXCL | Secreted Signaling | KEGG: mmu04060 |
| CD31 | NeuN | Cxcl1 | Ackr1 | 0.001231 | 0 | CXCL1_ACKR1 | Cxcl1 - Ackr1 | CXCL | Secreted Signaling | KEGG: mmu04060 |
| NeuN | NeuN | Cxcl1 | Ackr1 | 0.00094 | 0 | CXCL1_ACKR1 | Cxcl1 - Ackr1 | CXCL | Secreted Signaling | KEGG: mmu04060 |
| CD31 | NeuN | Cxcl10 | Ackr1 | 0.000633 | 0 | CXCL10_ACKR1 | Cxcl10 - Ackr1 | CXCL | Secreted Signaling | KEGG: mmu04060 |
| NeuN | NeuN | Cxcl10 | Ackr1 | 0.00066 | 0 | CXCL10_ACKR1 | Cxcl10 - Ackr1 | CXCL | Secreted Signaling | KEGG: mmu04060 |
| CD31 | CD31 | Mif | Ackr3 | 0.01614 | 0 | MIF_ACKR3 | Mif - Ackr3 | MIF | Secreted Signaling | PMID: 26175090 |
| NeuN | CD31 | Mif | Ackr3 | 0.022327 | 0 | MIF_ACKR3 | Mif - Ackr3 | MIF | Secreted Signaling | PMID: 26175090 |
| CD31 | NeuN | Cx3cl1 | Cx3cr1 | 0.000528 | 0 | CX3CL1_CX3CR1 | Cx3cl1 - Cx3cr1 | CX3C | Secreted Signaling | KEGG: mmu04060 |
| NeuN | NeuN | Cx3cl1 | Cx3cr1 | 0.000559 | 0 | CX3CL1_CX3CR1 | Cx3cl1 - Cx3cr1 | CX3C | Secreted Signaling | KEGG: mmu04060 |
| CD31 | CD31 | Clcf1 | CNTFR_LIFR | 0.000834 | 0 | CLCF1_CNTFR_LIFR | Clcf1 - (Cntfr+Lifr) | LIFR | Secreted Signaling | KEGG: mmu04060 |
| CD31 | NeuN | Clcf1 | CNTFR_LIFR | 0.001061 | 0 | CLCF1_CNTFR_LIFR | Clcf1 - (Cntfr+Lifr) | LIFR | Secreted Signaling | KEGG: mmu04060 |
| CD31 | NeuN | Ctf1 | LIFR_IL6ST | 0.000903 | 0 | CTF1_LIFR_IL6ST | Ctf1 - (Lifr+Il6st) | LIFR | Secreted Signaling | KEGG: mmu04060 |
| NeuN | NeuN | Ctf1 | LIFR_IL6ST | 0.000806 | 0 | CTF1_LIFR_IL6ST | Ctf1 - (Lifr+Il6st) | LIFR | Secreted Signaling | KEGG: mmu04060 |
| CD31 | CD31 | Il34 | Csf1r | 0.000707 | 0 | IL34_CSF1R | Il34 - Csf1r | CSF | Secreted Signaling | KEGG: mmu04060 |
| NeuN | CD31 | Il34 | Csf1r | 0.000522 | 0 | IL34_CSF1R | Il34 - Csf1r | CSF | Secreted Signaling | KEGG: mmu04060 |
| CD31 | CD31 | Csf1 | Csf1r | 0.001184 | 0 | CSF1_CSF1R | Csf1 - Csf1r | CSF | Secreted Signaling | KEGG: mmu04060 |
| CD31 | NeuN | Csf1 | Csf1r | 0.000543 | 0 | CSF1_CSF1R | Csf1 - Csf1r | CSF | Secreted Signaling | KEGG: mmu04060 |
| CD31 | CD31 | Tnfsf10 | Tnfrsf10b | 0.00066 | 0 | TNFSF10_TNFRSF10B | Tnfsf10 - Tnfrsf10b | TRAIL | Secreted Signaling | KEGG: mmu04060 |
| NeuN | CD31 | Tnfsf10 | Tnfrsf10b | 0.000458 | 0 | TNFSF10_TNFRSF10B | Tnfsf10 - Tnfrsf10b | TRAIL | Secreted Signaling | KEGG: mmu04060 |
| CD31 | CD31 | Retn | Cap1 | 0.003443 | 0 | RETN_CAP1 | Retn - Cap1 | RESISTIN | Secreted Signaling | PMID: 30809105 |
| CD31 | NeuN | Retn | Cap1 | 0.002777 | 0 | RETN_CAP1 | Retn - Cap1 | RESISTIN | Secreted Signaling | PMID: 30809105 |
| CD31 | CD31 | Nampt | ITGA5_ITGB1 | 0.020489 | 0 | NAMPT_ITGA5_ITGB1 | Nampt - (Itga5+Itgb1) | VISFATIN | Secreted Signaling | PMID: 28490838 |
| CD31 | NeuN | Nampt | ITGA5_ITGB1 | 0.006504 | 0 | NAMPT_ITGA5_ITGB1 | Nampt - (Itga5+Itgb1) | VISFATIN | Secreted Signaling | PMID: 28490838 |
| CD31 | CD31 | Angptl4 | ITGA5_ITGB1 | 0.006168 | 0 | ANGPTL4_ITGA5_ITGB1 | Angptl4 - (Itga5+Itgb1) | ANGPTL | Secreted Signaling | PMID: 30049845 |

| | | | | | | | | | | |
|---|---|---|---|---|---|---|---|---|---|---|
| NeuN | CD31 | Angptl4 | ITGA5_ITGB1 | 0.00241 | 0 | ANGPTL4_ITGA5_ITGB1 | Angptl4 - (Itga5+Itgb1) | ANGPTL | Secreted Signaling | PMID: 30049845 |
| CD31 | CD31 | Angptl4 | ITGAV_ITGB3 | 0.002237 | 0 | ANGPTL4_ITGAV_ITGB3 | Angptl4 - (Itgav+Itgb3) | ANGPTL | Secreted Signaling | PMID: 30049845 |
| CD31 | NeuN | Angptl4 | ITGAV_ITGB3 | 0.001324 | 0 | ANGPTL4_ITGAV_ITGB3 | Angptl4 - (Itgav+Itgb3) | ANGPTL | Secreted Signaling | PMID: 30049845 |
| CD31 | CD31 | Angptl4 | Cdh5 | 0.007667 | 0 | ANGPTL4_CDH5 | Angptl4 - Cdh5 | ANGPTL | Secreted Signaling | PMID: 30049845 |
| NeuN | CD31 | Angptl4 | Cdh5 | 0.002999 | 0 | ANGPTL4_CDH5 | Angptl4 - Cdh5 | ANGPTL | Secreted Signaling | PMID: 30049845 |
| CD31 | CD31 | Angptl4 | Cdh11 | 0.00516 | 0 | ANGPTL4_CDH11 | Angptl4 - Cdh11 | ANGPTL | Secreted Signaling | PMID: 30049845 |
| CD31 | NeuN | Angptl4 | Cdh11 | 0.004137 | 0 | ANGPTL4_CDH11 | Angptl4 - Cdh11 | ANGPTL | Secreted Signaling | PMID: 30049845 |
| CD31 | CD31 | Angptl4 | Sdc2 | 0.00271 | 0 | ANGPTL4_SDC2 | Angptl4 - Sdc2 | ANGPTL | Secreted Signaling | PMID: 29017031 |
| CD31 | NeuN | Angptl4 | Sdc2 | 0.00112 | 0 | ANGPTL4_SDC2 | Angptl4 - Sdc2 | ANGPTL | Secreted Signaling | PMID: 29017031 |
| CD31 | CD31 | Angptl4 | Sdc3 | 0.010479 | 0 | ANGPTL4_SDC3 | Angptl4 - Sdc3 | ANGPTL | Secreted Signaling | PMID: 29017031 |
| CD31 | NeuN | Angptl4 | Sdc3 | 0.01237 | 0 | ANGPTL4_SDC3 | Angptl4 - Sdc3 | ANGPTL | Secreted Signaling | PMID: 29017031 |
| CD31 | CD31 | Angptl4 | Sdc4 | 0.001216 | 0 | ANGPTL4_SDC4 | Angptl4 - Sdc4 | ANGPTL | Secreted Signaling | PMID: 29017031 |
| CD31 | NeuN | Angptl4 | Sdc4 | 0.001148 | 0 | ANGPTL4_SDC4 | Angptl4 - Sdc4 | ANGPTL | Secreted Signaling | PMID: 29017031 |
| CD31 | CD31 | Angptl3 | ITGAV_ITGB3 | 0.001301 | 0 | ANGPTL3_ITGAV_ITGB3 | Angptl3 - (Itgav+Itgb3) | ANGPTL | Secreted Signaling | PMID: 11877390 |
| NeuN | CD31 | Angptl3 | ITGAV_ITGB3 | 0.00083 | 0 | ANGPTL3_ITGAV_ITGB3 | Angptl3 - (Itgav+Itgb3) | ANGPTL | Secreted Signaling | PMID: 11877390 |
| CD31 | CD31 | Mdk | Sdc2 | 0.00355 | 0 | MDK_SDC2 | Mdk - Sdc2 | MK | Secreted Signaling | PMID: 28356350 |
| NeuN | CD31 | Mdk | Sdc2 | 0.004031 | 0 | MDK_SDC2 | Mdk - Sdc2 | MK | Secreted Signaling | PMID: 28356350 |
| NeuN | CD31 | Mdk | Sdc4 | 0.001811 | 0 | MDK_SDC4 | Mdk - Sdc4 | MK | Secreted Signaling | PMID: 28356350 |
| NeuN | NeuN | Mdk | Sdc4 | 0.001709 | 0 | MDK_SDC4 | Mdk - Sdc4 | MK | Secreted Signaling | PMID: 28356350 |
| CD31 | NeuN | Mdk | Ptprz1 | 0.024759 | 0 | MDK_PTPRZ1 | Mdk - Ptprz1 | MK | Secreted Signaling | PMID: 28356350 |
| NeuN | NeuN | Mdk | Ptprz1 | 0.028032 | 0 | MDK_PTPRZ1 | Mdk - Ptprz1 | MK | Secreted Signaling | PMID: 28356350 |
| CD31 | CD31 | Mdk | ITGA4_ITGB1 | 0.008059 | 0 | MDK_ITGA4_ITGB1 | Mdk - (Itga4+Itgb1) | MK | Secreted Signaling | PMID: 28356350 |
| NeuN | CD31 | Mdk | ITGA4_ITGB1 | 0.009145 | 0 | MDK_ITGA4_ITGB1 | Mdk - (Itga4+Itgb1) | MK | Secreted Signaling | PMID: 28356350 |
| CD31 | CD31 | Mdk | ITGA6_ITGB1 | 0.007691 | 0 | MDK_ITGA6_ITGB1 | Mdk - (Itga6+Itgb1) | MK | Secreted Signaling | PMID: 28356350 |
| NeuN | CD31 | Mdk | ITGA6_ITGB1 | 0.008728 | 0 | MDK_ITGA6_ITGB1 | Mdk - (Itga6+Itgb1) | MK | Secreted Signaling | PMID: 28356350 |
| CD31 | NeuN | Mdk | Lrp1 | 0.006756 | 0 | MDK_LRP1 | Mdk - Lrp1 | MK | Secreted Signaling | PMID: 28356350 |
| NeuN | NeuN | Mdk | Lrp1 | 0.007668 | 0 | MDK_LRP1 | Mdk - Lrp1 | MK | Secreted Signaling | PMID: 28356350 |
| CD31 | NeuN | Mdk | Ncl | 0.028593 | 0 | MDK_NCL | Mdk - Ncl | MK | Secreted Signaling | PMID: 28356350 |
| NeuN | NeuN | Mdk | Ncl | 0.032357 | 0 | MDK_NCL | Mdk - Ncl | MK | Secreted Signaling | PMID: 28356350 |
| CD31 | NeuN | Ptn | Ptprz1 | 0.08332 | 0 | PTN_PTPRZ1 | Ptn - Ptprz1 | PTN | Secreted Signaling | PMID: 28356350; PMID: 25620911 |
| NeuN | NeuN | Ptn | Ptprz1 | 0.087228 | 0 | PTN_PTPRZ1 | Ptn - Ptprz1 | PTN | Secreted Signaling | PMID: 28356350; PMID: 25620911 |
| CD31 | CD31 | Ptn | Sdc2 | 0.012595 | 0 | PTN_SDC2 | Ptn - Sdc2 | PTN | Secreted Signaling | PMID: 28356350; PMID: 25620911 |
| NeuN | CD31 | Ptn | Sdc2 | 0.013234 | 0 | PTN_SDC2 | Ptn - Sdc2 | PTN | Secreted Signaling | PMID: 28356350; PMID: 25620911 |
| CD31 | NeuN | Ptn | Sdc3 | 0.05553 | 0 | PTN_SDC3 | Ptn - Sdc3 | PTN | Secreted Signaling | PMID: 28356350; PMID: 25620911 |
| NeuN | NeuN | Ptn | Sdc3 | 0.058218 | 0 | PTN_SDC3 | Ptn - Sdc3 | PTN | Secreted Signaling | PMID: 28356350; PMID: 25620911 |
| CD31 | CD31 | Ptn | Sdc4 | 0.005685 | 0 | PTN_SDC4 | Ptn - Sdc4 | PTN | Secreted Signaling | PMID: 28356350; PMID: 25620911 |
| NeuN | CD31 | Ptn | Sdc4 | 0.005975 | 0 | PTN_SDC4 | Ptn - Sdc4 | PTN | Secreted Signaling | PMID: 28356350; PMID: 25620911 |
| CD31 | CD31 | Ptn | ITGAV_ITGB3 | 0.010417 | 0 | PTN_ITGAV_ITGB3 | Ptn - (Itgav+Itgb3) | PTN | Secreted Signaling | PMID: 28356350; PMID: 25620911 |
| NeuN | CD31 | Ptn | ITGAV_ITGB3 | 0.010947 | 0 | PTN_ITGAV_ITGB3 | Ptn - (Itgav+Itgb3) | PTN | Secreted Signaling | PMID: 28356350; PMID: 25620911 |
| CD31 | NeuN | Ptn | Ncl | 0.095338 | 0 | PTN_NCL | Ptn - Ncl | PTN | Secreted Signaling | PMID: 28356350; PMID: 25620911 |
| NeuN | NeuN | Ptn | Ncl | 0.099749 | 0 | PTN_NCL | Ptn - Ncl | PTN | Secreted Signaling | PMID: 28356350; PMID: 25620911 |
| CD31 | CD31 | Postn | ITGAV_ITGB3 | 0.001986 | 0 | POSTN_ITGAV_ITGB3 | Postn - (Itgav+Itgb3) | PERIOSTIN | Secreted Signaling | PMID: 26721738 |
| CD31 | NeuN | Postn | ITGAV_ITGB3 | 0.001175 | 0 | POSTN_ITGAV_ITGB3 | Postn - (Itgav+Itgb3) | PERIOSTIN | Secreted Signaling | PMID: 26721738 |
| CD31 | CD31 | Postn | ITGAV_ITGB5 | 0.001763 | 0 | POSTN_ITGAV_ITGB5 | Postn - (Itgav+Itgb5) | PERIOSTIN | Secreted Signaling | PMID: 26721738 |
| CD31 | NeuN | Postn | ITGAV_ITGB5 | 0.000974 | 0 | POSTN_ITGAV_ITGB5 | Postn - (Itgav+Itgb5) | PERIOSTIN | Secreted Signaling | PMID: 26721738 |
| CD31 | CD31 | Nmb | Nmbr | 0.000592 | 0 | NMB_NMBR | Nmb - Nmbr | GPR | Secreted Signaling | KEGG: mmu04080 |
| NeuN | CD31 | Nmb | Nmbr | 0.000483 | 0 | NMB_NMBR | Nmb - Nmbr | GPR | Secreted Signaling | KEGG: mmu04080 |
| CD31 | CD31 | Cck | Cckbr | 0.000576 | 0 | CCK_CCKBR | Cck - Cckbr | CCK | Secreted Signaling | KEGG: mmu04080 |
| NeuN | CD31 | Cck | Cckbr | 0.000987 | 0 | CCK_CCKBR | Cck - Cckbr | CCK | Secreted Signaling | KEGG: mmu04080 |
| CD31 | CD31 | Edn1 | Ednra | 0.003469 | 0 | EDN1_EDNRA | Edn1 - Ednra | EDN | Secreted Signaling | KEGG: mmu04080 |
| NeuN | CD31 | Edn1 | Ednra | 0.001752 | 0 | EDN1_EDNRA | Edn1 - Ednra | EDN | Secreted Signaling | KEGG: mmu04080 |
| CD31 | CD31 | Edn3 | Ednra | 0.003125 | 0 | EDN3_EDNRA | Edn3 - Ednra | EDN | Secreted Signaling | KEGG: mmu04080 |
| NeuN | CD31 | Edn3 | Ednra | 0.001811 | 0 | EDN3_EDNRA | Edn3 - Ednra | EDN | Secreted Signaling | KEGG: mmu04080 |
| CD31 | CD31 | Edn1 | Ednrb | 0.001072 | 0 | EDN1_EDNRB | Edn1 - Ednrb | EDN | Secreted Signaling | KEGG: mmu04080 |
| CD31 | NeuN | Edn1 | Ednrb | 0.000799 | 0 | EDN1_EDNRB | Edn1 - Ednrb | EDN | Secreted Signaling | KEGG: mmu04080 |
| CD31 | CD31 | Edn3 | Ednrb | 0.000966 | 0 | EDN3_EDNRB | Edn3 - Ednrb | EDN | Secreted Signaling | KEGG: mmu04080 |
| CD31 | NeuN | Edn3 | Ednrb | 0.000719 | 0 | EDN3_EDNRB | Edn3 - Ednrb | EDN | Secreted Signaling | KEGG: mmu04080 |
| CD31 | CD31 | Nms | Nmur1 | 0.000749 | 0 | NMS_NMUR1 | Nms - Nmur1 | NMU | Secreted Signaling | PMID: 29288685 |
| CD31 | NeuN | Nms | Nmur1 | 0.000845 | 0 | NMS_NMUR1 | Nms - Nmur1 | NMU | Secreted Signaling | PMID: 29288685 |

| | | | | | | | | | | |
|---|---|---|---|---|---|---|---|---|---|---|
| CD31 | NeuN | Npy | Npy1r | 0.003506 | 0 | NPY_NPY1R | Npy - Npy1r | NPY | Secreted Signaling | KEGG: mmu04080 |
| NeuN | NeuN | Npy | Npy1r | 0.004185 | 0 | NPY_NPY1R | Npy - Npy1r | NPY | Secreted Signaling | KEGG: mmu04080 |
| CD31 | CD31 | Npy | Npy2r | 0.003878 | 0 | NPY_NPY2R | Npy - Npy2r | NPY | Secreted Signaling | KEGG: mmu04080 |
| NeuN | CD31 | Npy | Npy2r | 0.004628 | 0 | NPY_NPY2R | Npy - Npy2r | NPY | Secreted Signaling | KEGG: mmu04080 |
| CD31 | CD31 | Npy | Npy5r | 0.001689 | 0 | NPY_NPY5R | Npy - Npy5r | NPY | Secreted Signaling | KEGG: mmu04080 |
| NeuN | CD31 | Npy | Npy5r | 0.002017 | 0 | NPY_NPY5R | Npy - Npy5r | NPY | Secreted Signaling | KEGG: mmu04080 |
| CD31 | CD31 | Npy | Npy6r | 0.001797 | 0 | NPY_NPY6R | Npy - Npy6r | NPY | Secreted Signaling | KEGG: mmu04080 |
| NeuN | CD31 | Npy | Npy6r | 0.002145 | 0 | NPY_NPY6R | Npy - Npy6r | NPY | Secreted Signaling | KEGG: mmu04080 |
| CD31 | CD31 | Npb | Npbwr1 | 0.000585 | 0 | NPB_NPBWR1 | Npb - Npbwr1 | NPW-B | Secreted Signaling | KEGG: mmu04080 |
| NeuN | CD31 | Npb | Npbwr1 | 0.000553 | 0 | NPB_NPBWR1 | Npb - Npbwr1 | NPW-B | Secreted Signaling | KEGG: mmu04080 |
| CD31 | NeuN | Nts | Ntsr1 | 0.002398 | 0 | NTS_NTSR1 | Nts - Ntsr1 | NTS | Secreted Signaling | KEGG: mmu04080 |
| NeuN | NeuN | Nts | Ntsr1 | 0.002452 | 0 | NTS_NTSR1 | Nts - Ntsr1 | NTS | Secreted Signaling | KEGG: mmu04080 |
| CD31 | CD31 | Nts | Ntsr2 | 0.000392 | 0 | NTS_NTSR2 | Nts - Ntsr2 | NTS | Secreted Signaling | KEGG: mmu04080 |
| NeuN | CD31 | Nts | Ntsr2 | 0.000401 | 0 | NTS_NTSR2 | Nts - Ntsr2 | NTS | Secreted Signaling | KEGG: mmu04080 |
| CD31 | CD31 | Pdyn | Oprd1 | 0.000339 | 0 | PDYN_OPRD1 | Pdyn - Oprd1 | OPIOID | Secreted Signaling | KEGG: mmu04080 |
| CD31 | NeuN | Pdyn | Oprd1 | 0.00039 | 0 | PDYN_OPRD1 | Pdyn - Oprd1 | OPIOID | Secreted Signaling | KEGG: mmu04080 |
| CD31 | CD31 | Pdyn | Oprk1 | 0.000544 | 0 | PDYN_OPRK1 | Pdyn - Oprk1 | OPIOID | Secreted Signaling | KEGG: mmu04080 |
| CD31 | NeuN | Pdyn | Oprk1 | 0.000439 | 0 | PDYN_OPRK1 | Pdyn - Oprk1 | OPIOID | Secreted Signaling | KEGG: mmu04080 |
| CD31 | NeuN | Pdyn | Oprl1 | 0.001038 | 0 | PDYN_OPRL1 | Pdyn - Oprl1 | OPIOID | Secreted Signaling | KEGG: mmu04080 |
| NeuN | NeuN | Pdyn | Oprl1 | 0.000817 | 0 | PDYN_OPRL1 | Pdyn - Oprl1 | OPIOID | Secreted Signaling | KEGG: mmu04080 |
| CD31 | CD31 | Pdyn | Oprm1 | 0.000464 | 0 | PDYN_OPRM1 | Pdyn - Oprm1 | OPIOID | Secreted Signaling | KEGG: mmu04080 |
| CD31 | NeuN | Pdyn | Oprm1 | 0.000435 | 0 | PDYN_OPRM1 | Pdyn - Oprm1 | OPIOID | Secreted Signaling | KEGG: mmu04080 |
| CD31 | CD31 | Penk | Oprd1 | 0.000554 | 0 | PENK_OPRD1 | Penk - Oprd1 | OPIOID | Secreted Signaling | KEGG: mmu04080 |
| CD31 | NeuN | Penk | Oprd1 | 0.000638 | 0 | PENK_OPRD1 | Penk - Oprd1 | OPIOID | Secreted Signaling | KEGG: mmu04080 |
| CD31 | CD31 | Penk | Oprk1 | 0.00089 | 0 | PENK_OPRK1 | Penk - Oprk1 | OPIOID | Secreted Signaling | KEGG: mmu04080 |
| CD31 | NeuN | Penk | Oprk1 | 0.000718 | 0 | PENK_OPRK1 | Penk - Oprk1 | OPIOID | Secreted Signaling | KEGG: mmu04080 |
| CD31 | CD31 | Penk | Oprl1 | 0.001058 | 0 | PENK_OPRL1 | Penk - Oprl1 | OPIOID | Secreted Signaling | KEGG: mmu04080 |
| CD31 | NeuN | Penk | Oprl1 | 0.001697 | 0 | PENK_OPRL1 | Penk - Oprl1 | OPIOID | Secreted Signaling | KEGG: mmu04080 |
| CD31 | CD31 | Penk | Oprm1 | 0.000759 | 0 | PENK_OPRM1 | Penk - Oprm1 | OPIOID | Secreted Signaling | KEGG: mmu04080 |
| CD31 | NeuN | Penk | Oprm1 | 0.000711 | 0 | PENK_OPRM1 | Penk - Oprm1 | OPIOID | Secreted Signaling | KEGG: mmu04080 |
| CD31 | CD31 | Oxt | Oxtr | 0.000702 | 0 | OXT_OXTR | Oxt - Oxtr | OXT | Secreted Signaling | KEGG: mmu04080 |
| NeuN | CD31 | Oxt | Oxtr | 0.000638 | 0 | OXT_OXTR | Oxt - Oxtr | OXT | Secreted Signaling | KEGG: mmu04080 |
| CD31 | CD31 | Sst | Sstr1 | 0.000853 | 0 | SST_SSTR1 | Sst - Sstr1 | SOMATOSTATIN | Secreted Signaling | KEGG: mmu04080 |
| NeuN | CD31 | Sst | Sstr1 | 0.000832 | 0 | SST_SSTR1 | Sst - Sstr1 | SOMATOSTATIN | Secreted Signaling | KEGG: mmu04080 |
| CD31 | NeuN | Sst | Sstr2 | 0.001341 | 0 | SST_SSTR2 | Sst - Sstr2 | SOMATOSTATIN | Secreted Signaling | KEGG: mmu04080 |
| NeuN | NeuN | Sst | Sstr2 | 0.001307 | 0 | SST_SSTR2 | Sst - Sstr2 | SOMATOSTATIN | Secreted Signaling | KEGG: mmu04080 |
| CD31 | CD31 | Sst | Sstr3 | 0.000509 | 0 | SST_SSTR3 | Sst - Sstr3 | SOMATOSTATIN | Secreted Signaling | KEGG: mmu04080 |
| CD31 | NeuN | Sst | Sstr3 | 0.000512 | 0 | SST_SSTR3 | Sst - Sstr3 | SOMATOSTATIN | Secreted Signaling | KEGG: mmu04080 |
| CD31 | CD31 | Sst | Sstr4 | 0.000546 | 0 | SST_SSTR4 | Sst - Sstr4 | SOMATOSTATIN | Secreted Signaling | KEGG: mmu04080 |
| NeuN | CD31 | Sst | Sstr4 | 0.000532 | 0 | SST_SSTR4 | Sst - Sstr4 | SOMATOSTATIN | Secreted Signaling | KEGG: mmu04080 |
| CD31 | CD31 | Cort | Sstr1 | 0.001632 | 0 | CORT_SSTR1 | Cort - Sstr1 | SOMATOSTATIN | Secreted Signaling | KEGG: mmu04080 |
| NeuN | CD31 | Cort | Sstr1 | 0.001357 | 0 | CORT_SSTR1 | Cort - Sstr1 | SOMATOSTATIN | Secreted Signaling | KEGG: mmu04080 |
| CD31 | NeuN | Cort | Sstr2 | 0.002563 | 0 | CORT_SSTR2 | Cort - Sstr2 | SOMATOSTATIN | Secreted Signaling | KEGG: mmu04080 |
| NeuN | NeuN | Cort | Sstr2 | 0.002132 | 0 | CORT_SSTR2 | Cort - Sstr2 | SOMATOSTATIN | Secreted Signaling | KEGG: mmu04080 |
| CD31 | CD31 | Cort | Sstr3 | 0.000974 | 0 | CORT_SSTR3 | Cort - Sstr3 | SOMATOSTATIN | Secreted Signaling | KEGG: mmu04080 |
| CD31 | NeuN | Cort | Sstr3 | 0.000979 | 0 | CORT_SSTR3 | Cort - Sstr3 | SOMATOSTATIN | Secreted Signaling | KEGG: mmu04080 |
| CD31 | CD31 | Cort | Sstr4 | 0.001044 | 0 | CORT_SSTR4 | Cort - Sstr4 | SOMATOSTATIN | Secreted Signaling | KEGG: mmu04080 |
| NeuN | CD31 | Cort | Sstr4 | 0.000868 | 0 | CORT_SSTR4 | Cort - Sstr4 | SOMATOSTATIN | Secreted Signaling | KEGG: mmu04080 |
| CD31 | CD31 | Tac1 | Tacr1 | 0.000576 | 0 | TAC1_TACR1 | Tac1 - Tacr1 | TAC | Secreted Signaling | KEGG: mmu04080 |
| CD31 | NeuN | Tac1 | Tacr1 | 0.000628 | 0 | TAC1_TACR1 | Tac1 - Tacr1 | TAC | Secreted Signaling | KEGG: mmu04080 |
| NeuN | CD31 | Tac2 | Tacr3 | 0.000419 | 0 | TAC2_TACR3 | Tac2 - Tacr3 | TAC | Secreted Signaling | KEGG: mmu04080 |
| NeuN | NeuN | Tac2 | Tacr3 | 0.000364 | 0 | TAC2_TACR3 | Tac2 - Tacr3 | TAC | Secreted Signaling | KEGG: mmu04080 |
| CD31 | CD31 | Ctsg | F2r | 0.001143 | 0 | CTSG_F2R | Ctsg - F2r | PARs | Secreted Signaling | KEGG: mmu04080 |
| NeuN | CD31 | Ctsg | F2r | 0.001254 | 0 | CTSG_F2R | Ctsg - F2r | PARs | Secreted Signaling | KEGG: mmu04080 |
| CD31 | CD31 | F2 | F2r | 0.002212 | 0 | F2_F2R | F2 - F2r | PARs | Secreted Signaling | KEGG: mmu04080 |
| NeuN | CD31 | F2 | F2r | 0.001607 | 0 | F2_F2R | F2 - F2r | PARs | Secreted Signaling | KEGG: mmu04080 |
| CD31 | CD31 | Ctsg | F2rl2 | 0.000429 | 0 | CTSG_F2RL2 | Ctsg - F2rl2 | PARs | Secreted Signaling | KEGG: mmu04080 |
| NeuN | CD31 | Ctsg | F2rl2 | 0.00047 | 0 | CTSG_F2RL2 | Ctsg - F2rl2 | PARs | Secreted Signaling | KEGG: mmu04080 |
| CD31 | CD31 | F2 | F2rl2 | 0.00083 | 0 | F2_F2RL2 | F2 - F2rl2 | PARs | Secreted Signaling | KEGG: mmu04080 |

| | | | | | | | | | |
|---|---|---|---|---|---|---|---|---|---|
| NeuN | CD31 | F2 | F2rl2 | 0.000603 | 0 F2_F2RL2 | F2 - F2rl2 | PARs | Secreted Signaling | KEGG: mmu04080 |
| CD31 | CD31 | Ctsg | Pard3 | 0.000818 | 0 CTSG_PARD3 | Ctsg - Pard3 | PARs | Secreted Signaling | KEGG: mmu04080 |
| NeuN | CD31 | Ctsg | Pard3 | 0.000898 | 0 CTSG_PARD3 | Ctsg - Pard3 | PARs | Secreted Signaling | KEGG: mmu04080 |
| CD31 | CD31 | F2 | Pard3 | 0.001584 | 0 F2_PARD3 | F2 - Pard3 | PARs | Secreted Signaling | KEGG: mmu04080 |
| NeuN | CD31 | F2 | Pard3 | 0.001151 | 0 F2_PARD3 | F2 - Pard3 | PARs | Secreted Signaling | KEGG: mmu04080 |
| CD31 | CD31 | Plg | Pard3 | 0.00157 | 0 PLG_PARD3 | Plg - Pard3 | PARs | Secreted Signaling | KEGG: mmu04080 |
| CD31 | NeuN | Plg | Pard3 | 0.001034 | 0 PLG_PARD3 | Plg - Pard3 | PARs | Secreted Signaling | KEGG: mmu04080 |
| CD31 | CD31 | Pmch | Mchr1 | 0.000926 | 0 PMCH_MCHR1 | Pmch - Mchr1 | PMCH | Secreted Signaling | KEGG: mmu04080 |
| NeuN | CD31 | Pmch | Mchr1 | 0.000547 | 0 PMCH_MCHR1 | Pmch - Mchr1 | PMCH | Secreted Signaling | KEGG: mmu04080 |
| CD31 | CD31 | Prok2 | Prokr1 | 0.001186 | 0 PROK2_PROKR1 | Prok2 - Prokr1 | PROK | Secreted Signaling | PMID: 24496457 |
| NeuN | CD31 | Prok2 | Prokr1 | 0.001014 | 0 PROK2_PROKR1 | Prok2 - Prokr1 | PROK | Secreted Signaling | PMID: 24496457 |
| CD31 | CD31 | Prok2 | Prokr2 | 0.001639 | 0 PROK2_PROKR2 | Prok2 - Prokr2 | PROK | Secreted Signaling | PMID: 24496457 |
| CD31 | NeuN | Prok2 | Prokr2 | 0.001588 | 0 PROK2_PROKR2 | Prok2 - Prokr2 | PROK | Secreted Signaling | PMID: 24496457 |
| CD31 | NeuN | Adcyap1 | Adcyap1r1 | 0.001629 | 0 ADCYAP1_ADCYAP1R1 | Adcyap1 - Adcyap1r1 | PACAP | Secreted Signaling | PMID: 21350482 |
| NeuN | NeuN | Adcyap1 | Adcyap1r1 | 0.001421 | 0 ADCYAP1_ADCYAP1R1 | Adcyap1 - Adcyap1r1 | PACAP | Secreted Signaling | PMID: 21350482 |
| CD31 | CD31 | Vip | Vipr1 | 0.000417 | 0 VIP_VIPR1 | Vip - Vipr1 | VIP | Secreted Signaling | PMID: 10655512 |
| NeuN | CD31 | Vip | Vipr1 | 0.000325 | 0 VIP_VIPR1 | Vip - Vipr1 | VIP | Secreted Signaling | PMID: 10655512 |
| NeuN | CD31 | Nppa | Npr1 | 0.003149 | 0 NPPA_NPR1 | Nppa - Npr1 | NPR1 | Secreted Signaling | KEGG: mmu04080; PMID:29934640 |
| NeuN | NeuN | Nppa | Npr1 | 0.002163 | 0 NPPA_NPR1 | Nppa - Npr1 | NPR1 | Secreted Signaling | KEGG: mmu04080; PMID:29934640 |
| CD31 | CD31 | Nppb | Npr1 | 0.000899 | 0 NPPB_NPR1 | Nppb - Npr1 | NPR1 | Secreted Signaling | KEGG: mmu04080; PMID:29934640 |
| CD31 | NeuN | Nppb | Npr1 | 0.000617 | 0 NPPB_NPR1 | Nppb - Npr1 | NPR1 | Secreted Signaling | KEGG: mmu04080; PMID:29934640 |
| CD31 | NeuN | Nppc | Npr2 | 0.002096 | 0 NPPC_NPR2 | Nppc - Npr2 | NPR2 | Secreted Signaling | KEGG: mmu04080; PMID:29934640 |
| NeuN | NeuN | Nppc | Npr2 | 0.001363 | 0 NPPC_NPR2 | Nppc - Npr2 | NPR2 | Secreted Signaling | KEGG: mmu04080; PMID:29934640 |
| CD31 | CD31 | Kitl | Kit | 0.005279 | 0 KITL_KIT | Kitl - Kit | KIT | Secreted Signaling | KEGG: mmu04080 |
| CD31 | NeuN | Kitl | Kit | 0.005218 | 0 KITL_KIT | Kitl - Kit | KIT | Secreted Signaling | KEGG: mmu04080 |
| CD31 | CD31 | Gip | Gipr | 0.001613 | 0 GIP_GIPR | Gip - Gipr | GIPR | Secreted Signaling | KEGG: mmu04080 |
| CD31 | NeuN | Gip | Gipr | 0.002488 | 0 GIP_GIPR | Gip - Gipr | GIPR | Secreted Signaling | KEGG: mmu04080 |
| CD31 | NeuN | Bdnf | Ntrk2 | 0.006328 | 0 BDNF_NTRK2 | Bdnf - Ntrk2 | NT | Secreted Signaling | PMID: 27908981 |
| NeuN | NeuN | Bdnf | Ntrk2 | 0.003725 | 0 BDNF_NTRK2 | Bdnf - Ntrk2 | NT | Secreted Signaling | PMID: 27908981 |
| CD31 | CD31 | Flt3l | Flt3 | 0.000525 | 0 FLT3L_FLT3 | Flt3l - Flt3 | FLT3 | Secreted Signaling | PMID: 25992210 |
| NeuN | CD31 | Flt3l | Flt3 | 0.000494 | 0 FLT3L_FLT3 | Flt3l - Flt3 | FLT3 | Secreted Signaling | PMID: 25992210 |
| CD31 | NeuN | Hgf | Met | 0.00293 | 0 HGF_MET | Hgf - Met | HGF | Secreted Signaling | PMID: 28475121 |
| NeuN | NeuN | Hgf | Met | 0.002053 | 0 HGF_MET | Hgf - Met | HGF | Secreted Signaling | PMID: 28475121 |
| CD31 | CD31 | Ntf3 | Ntrk1 | 0.000599 | 0 NTF3_NTRK1 | Ntf3 - Ntrk1 | NT | Secreted Signaling | PMID: 27908981 |
| CD31 | NeuN | Ntf3 | Ntrk1 | 0.000411 | 0 NTF3_NTRK1 | Ntf3 - Ntrk1 | NT | Secreted Signaling | PMID: 27908981 |
| CD31 | NeuN | Ntf3 | Ntrk2 | 0.007886 | 0 NTF3_NTRK2 | Ntf3 - Ntrk2 | NT | Secreted Signaling | PMID: 27908981 |
| NeuN | NeuN | Ntf3 | Ntrk2 | 0.003588 | 0 NTF3_NTRK2 | Ntf3 - Ntrk2 | NT | Secreted Signaling | PMID: 27908981 |
| CD31 | NeuN | Ntf3 | Ntrk3 | 0.011528 | 0 NTF3_NTRK3 | Ntf3 - Ntrk3 | NT | Secreted Signaling | PMID: 27908981 |
| NeuN | NeuN | Ntf3 | Ntrk3 | 0.005256 | 0 NTF3_NTRK3 | Ntf3 - Ntrk3 | NT | Secreted Signaling | PMID: 27908981 |
| NeuN | CD31 | Sema3a | NRP1_PLXNA1 | 0.005336 | 0 SEMA3A_NRP1_PLXNA1 | Sema3a - (Nrp1+Plxna1) | SEMA3 | Secreted Signaling | PMID: 27533782 |
| NeuN | NeuN | Sema3a | NRP1_PLXNA1 | 0.006451 | 0 SEMA3A_NRP1_PLXNA1 | Sema3a - (Nrp1+Plxna1) | SEMA3 | Secreted Signaling | PMID: 27533782 |
| NeuN | CD31 | Sema3a | NRP1_PLXNA2 | 0.005577 | 0 SEMA3A_NRP1_PLXNA2 | Sema3a - (Nrp1+Plxna2) | SEMA3 | Secreted Signaling | PMID: 27533782 |
| NeuN | NeuN | Sema3a | NRP1_PLXNA2 | 0.008055 | 0 SEMA3A_NRP1_PLXNA2 | Sema3a - (Nrp1+Plxna2) | SEMA3 | Secreted Signaling | PMID: 27533782 |
| NeuN | CD31 | Sema3a | NRP1_PLXNA3 | 0.004711 | 0 SEMA3A_NRP1_PLXNA3 | Sema3a - (Nrp1+Plxna3) | SEMA3 | Secreted Signaling | PMID: 27533782 |
| NeuN | NeuN | Sema3a | NRP1_PLXNA3 | 0.006742 | 0 SEMA3A_NRP1_PLXNA3 | Sema3a - (Nrp1+Plxna3) | SEMA3 | Secreted Signaling | PMID: 27533782 |
| NeuN | CD31 | Sema3a | NRP1_PLXNA4 | 0.008825 | 0 SEMA3A_NRP1_PLXNA4 | Sema3a - (Nrp1+Plxna4) | SEMA3 | Secreted Signaling | PMID: 27533782 |
| NeuN | NeuN | Sema3a | NRP1_PLXNA4 | 0.011689 | 0 SEMA3A_NRP1_PLXNA4 | Sema3a - (Nrp1+Plxna4) | SEMA3 | Secreted Signaling | PMID: 27533782 |
| CD31 | CD31 | Sema3b | NRP1_PLXNA1 | 0.002506 | 0 SEMA3B_NRP1_PLXNA1 | Sema3b - (Nrp1+Plxna1) | SEMA3 | Secreted Signaling | PMID: 27533782 |
| CD31 | NeuN | Sema3b | NRP1_PLXNA1 | 0.003032 | 0 SEMA3B_NRP1_PLXNA1 | Sema3b - (Nrp1+Plxna1) | SEMA3 | Secreted Signaling | PMID: 27533782 |
| CD31 | CD31 | Sema3b | NRP1_PLXNA2 | 0.00262 | 0 SEMA3B_NRP1_PLXNA2 | Sema3b - (Nrp1+Plxna2) | SEMA3 | Secreted Signaling | PMID: 27533782 |
| CD31 | NeuN | Sema3b | NRP1_PLXNA2 | 0.003789 | 0 SEMA3B_NRP1_PLXNA2 | Sema3b - (Nrp1+Plxna2) | SEMA3 | Secreted Signaling | PMID: 27533782 |
| CD31 | CD31 | Sema3b | NRP1_PLXNA3 | 0.002212 | 0 SEMA3B_NRP1_PLXNA3 | Sema3b - (Nrp1+Plxna3) | SEMA3 | Secreted Signaling | PMID: 27533782 |
| CD31 | NeuN | Sema3b | NRP1_PLXNA3 | 0.003169 | 0 SEMA3B_NRP1_PLXNA3 | Sema3b - (Nrp1+Plxna3) | SEMA3 | Secreted Signaling | PMID: 27533782 |
| CD31 | CD31 | Sema3b | NRP1_PLXNA4 | 0.004153 | 0 SEMA3B_NRP1_PLXNA4 | Sema3b - (Nrp1+Plxna4) | SEMA3 | Secreted Signaling | PMID: 27533782 |
| CD31 | NeuN | Sema3b | NRP1_PLXNA4 | 0.005509 | 0 SEMA3B_NRP1_PLXNA4 | Sema3b - (Nrp1+Plxna4) | SEMA3 | Secreted Signaling | PMID: 27533782 |
| NeuN | CD31 | Sema3c | NRP1_PLXNA1 | 0.006276 | 0 SEMA3C_NRP1_PLXNA1 | Sema3c - (Nrp1+Plxna1) | SEMA3 | Secreted Signaling | PMID: 27533782 |
| NeuN | NeuN | Sema3c | NRP1_PLXNA1 | 0.007586 | 0 SEMA3C_NRP1_PLXNA1 | Sema3c - (Nrp1+Plxna1) | SEMA3 | Secreted Signaling | PMID: 27533782 |
| NeuN | CD31 | Sema3c | NRP1_PLXNA2 | 0.006559 | 0 SEMA3C_NRP1_PLXNA2 | Sema3c - (Nrp1+Plxna2) | SEMA3 | Secreted Signaling | PMID: 27533782 |
| NeuN | NeuN | Sema3c | NRP1_PLXNA2 | 0.009469 | 0 SEMA3C_NRP1_PLXNA2 | Sema3c - (Nrp1+Plxna2) | SEMA3 | Secreted Signaling | PMID: 27533782 |

| | | | | | | | | | | |
|---|---|---|---|---|---|---|---|---|---|---|
| NeuN | CD31 | Sema3c | NRP1_PLXNA3 | 0.005541 | 0 | SEMA3C_NRP1_PLXNA3 | Sema3c - (Nrp1+Plxna3) | SEMA3 | Secreted Signaling | PMID: 27533782 |
| NeuN | NeuN | Sema3c | NRP1_PLXNA3 | 0.007927 | 0 | SEMA3C_NRP1_PLXNA3 | Sema3c - (Nrp1+Plxna3) | SEMA3 | Secreted Signaling | PMID: 27533782 |
| NeuN | CD31 | Sema3c | NRP1_PLXNA4 | 0.010373 | 0 | SEMA3C_NRP1_PLXNA4 | Sema3c - (Nrp1+Plxna4) | SEMA3 | Secreted Signaling | PMID: 27533782 |
| NeuN | NeuN | Sema3c | NRP1_PLXNA4 | 0.013733 | 0 | SEMA3C_NRP1_PLXNA4 | Sema3c - (Nrp1+Plxna4) | SEMA3 | Secreted Signaling | PMID: 27533782 |
| CD31 | CD31 | Sema3d | NRP1_PLXNA1 | 0.002237 | 0 | SEMA3D_NRP1_PLXNA1 | Sema3d - (Nrp1+Plxna1) | SEMA3 | Secreted Signaling | PMID: 27533782 |
| CD31 | NeuN | Sema3d | NRP1_PLXNA1 | 0.002707 | 0 | SEMA3D_NRP1_PLXNA1 | Sema3d - (Nrp1+Plxna1) | SEMA3 | Secreted Signaling | PMID: 27533782 |
| CD31 | CD31 | Sema3d | NRP1_PLXNA2 | 0.002338 | 0 | SEMA3D_NRP1_PLXNA2 | Sema3d - (Nrp1+Plxna2) | SEMA3 | Secreted Signaling | PMID: 27533782 |
| CD31 | NeuN | Sema3d | NRP1_PLXNA2 | 0.003382 | 0 | SEMA3D_NRP1_PLXNA2 | Sema3d - (Nrp1+Plxna2) | SEMA3 | Secreted Signaling | PMID: 27533782 |
| CD31 | CD31 | Sema3d | NRP1_PLXNA3 | 0.001974 | 0 | SEMA3D_NRP1_PLXNA3 | Sema3d - (Nrp1+Plxna3) | SEMA3 | Secreted Signaling | PMID: 27533782 |
| CD31 | NeuN | Sema3d | NRP1_PLXNA3 | 0.002829 | 0 | SEMA3D_NRP1_PLXNA3 | Sema3d - (Nrp1+Plxna3) | SEMA3 | Secreted Signaling | PMID: 27533782 |
| CD31 | CD31 | Sema3d | NRP1_PLXNA4 | 0.003708 | 0 | SEMA3D_NRP1_PLXNA4 | Sema3d - (Nrp1+Plxna4) | SEMA3 | Secreted Signaling | PMID: 27533782 |
| CD31 | NeuN | Sema3d | NRP1_PLXNA4 | 0.004919 | 0 | SEMA3D_NRP1_PLXNA4 | Sema3d - (Nrp1+Plxna4) | SEMA3 | Secreted Signaling | PMID: 27533782 |
| CD31 | CD31 | Sema3b | NRP2_PLXNA1 | 0.001586 | 0 | SEMA3B_NRP2_PLXNA1 | Sema3b - (Nrp2+Plxna1) | SEMA3 | Secreted Signaling | PMID: 27533782 |
| CD31 | NeuN | Sema3b | NRP2_PLXNA1 | 0.001843 | 0 | SEMA3B_NRP2_PLXNA1 | Sema3b - (Nrp2+Plxna1) | SEMA3 | Secreted Signaling | PMID: 27533782 |
| CD31 | CD31 | Sema3b | NRP2_PLXNA2 | 0.001658 | 0 | SEMA3B_NRP2_PLXNA2 | Sema3b - (Nrp2+Plxna2) | SEMA3 | Secreted Signaling | PMID: 27533782 |
| CD31 | NeuN | Sema3b | NRP2_PLXNA2 | 0.002304 | 0 | SEMA3B_NRP2_PLXNA2 | Sema3b - (Nrp2+Plxna2) | SEMA3 | Secreted Signaling | PMID: 27533782 |
| CD31 | CD31 | Sema3b | NRP2_PLXNA3 | 0.0014 | 0 | SEMA3B_NRP2_PLXNA3 | Sema3b - (Nrp2+Plxna3) | SEMA3 | Secreted Signaling | PMID: 27533782 |
| CD31 | NeuN | Sema3b | NRP2_PLXNA3 | 0.001927 | 0 | SEMA3B_NRP2_PLXNA3 | Sema3b - (Nrp2+Plxna3) | SEMA3 | Secreted Signaling | PMID: 27533782 |
| CD31 | CD31 | Sema3b | NRP2_PLXNA4 | 0.00263 | 0 | SEMA3B_NRP2_PLXNA4 | Sema3b - (Nrp2+Plxna4) | SEMA3 | Secreted Signaling | PMID: 27533782 |
| CD31 | NeuN | Sema3b | NRP2_PLXNA4 | 0.003353 | 0 | SEMA3B_NRP2_PLXNA4 | Sema3b - (Nrp2+Plxna4) | SEMA3 | Secreted Signaling | PMID: 27533782 |
| NeuN | CD31 | Sema3c | NRP2_PLXNA1 | 0.003977 | 0 | SEMA3C_NRP2_PLXNA1 | Sema3c - (Nrp2+Plxna1) | SEMA3 | Secreted Signaling | PMID: 27533782 |
| NeuN | NeuN | Sema3c | NRP2_PLXNA1 | 0.004621 | 0 | SEMA3C_NRP2_PLXNA1 | Sema3c - (Nrp2+Plxna1) | SEMA3 | Secreted Signaling | PMID: 27533782 |
| NeuN | CD31 | Sema3c | NRP2_PLXNA2 | 0.004157 | 0 | SEMA3C_NRP2_PLXNA2 | Sema3c - (Nrp2+Plxna2) | SEMA3 | Secreted Signaling | PMID: 27533782 |
| NeuN | NeuN | Sema3c | NRP2_PLXNA2 | 0.005772 | 0 | SEMA3C_NRP2_PLXNA2 | Sema3c - (Nrp2+Plxna2) | SEMA3 | Secreted Signaling | PMID: 27533782 |
| NeuN | CD31 | Sema3c | NRP2_PLXNA3 | 0.003511 | 0 | SEMA3C_NRP2_PLXNA3 | Sema3c - (Nrp2+Plxna3) | SEMA3 | Secreted Signaling | PMID: 27533782 |
| NeuN | NeuN | Sema3c | NRP2_PLXNA3 | 0.004829 | 0 | SEMA3C_NRP2_PLXNA3 | Sema3c - (Nrp2+Plxna3) | SEMA3 | Secreted Signaling | PMID: 27533782 |
| NeuN | CD31 | Sema3c | NRP2_PLXNA4 | 0.006583 | 0 | SEMA3C_NRP2_PLXNA4 | Sema3c - (Nrp2+Plxna4) | SEMA3 | Secreted Signaling | PMID: 27533782 |
| NeuN | NeuN | Sema3c | NRP2_PLXNA4 | 0.008385 | 0 | SEMA3C_NRP2_PLXNA4 | Sema3c - (Nrp2+Plxna4) | SEMA3 | Secreted Signaling | PMID: 27533782 |
| CD31 | CD31 | Sema3d | NRP2_PLXNA1 | 0.001416 | 0 | SEMA3D_NRP2_PLXNA1 | Sema3d - (Nrp2+Plxna1) | SEMA3 | Secreted Signaling | PMID: 27533782 |
| CD31 | NeuN | Sema3d | NRP2_PLXNA1 | 0.001645 | 0 | SEMA3D_NRP2_PLXNA1 | Sema3d - (Nrp2+Plxna1) | SEMA3 | Secreted Signaling | PMID: 27533782 |
| CD31 | CD31 | Sema3d | NRP2_PLXNA2 | 0.00148 | 0 | SEMA3D_NRP2_PLXNA2 | Sema3d - (Nrp2+Plxna2) | SEMA3 | Secreted Signaling | PMID: 27533782 |
| CD31 | NeuN | Sema3d | NRP2_PLXNA2 | 0.002057 | 0 | SEMA3D_NRP2_PLXNA2 | Sema3d - (Nrp2+Plxna2) | SEMA3 | Secreted Signaling | PMID: 27533782 |
| CD31 | CD31 | Sema3d | NRP2_PLXNA3 | 0.001249 | 0 | SEMA3D_NRP2_PLXNA3 | Sema3d - (Nrp2+Plxna3) | SEMA3 | Secreted Signaling | PMID: 27533782 |
| CD31 | NeuN | Sema3d | NRP2_PLXNA3 | 0.00172 | 0 | SEMA3D_NRP2_PLXNA3 | Sema3d - (Nrp2+Plxna3) | SEMA3 | Secreted Signaling | PMID: 27533782 |
| CD31 | CD31 | Sema3d | NRP2_PLXNA4 | 0.002347 | 0 | SEMA3D_NRP2_PLXNA4 | Sema3d - (Nrp2+Plxna4) | SEMA3 | Secreted Signaling | PMID: 27533782 |
| CD31 | NeuN | Sema3d | NRP2_PLXNA4 | 0.002993 | 0 | SEMA3D_NRP2_PLXNA4 | Sema3d - (Nrp2+Plxna4) | SEMA3 | Secreted Signaling | PMID: 27533782 |
| NeuN | CD31 | Sema3f | NRP2_PLXNA1 | 0.006777 | 0 | SEMA3F_NRP2_PLXNA1 | Sema3f - (Nrp2+Plxna1) | SEMA3 | Secreted Signaling | PMID: 27533782 |
| NeuN | NeuN | Sema3f | NRP2_PLXNA1 | 0.007871 | 0 | SEMA3F_NRP2_PLXNA1 | Sema3f - (Nrp2+Plxna1) | SEMA3 | Secreted Signaling | PMID: 27533782 |
| NeuN | CD31 | Sema3f | NRP2_PLXNA2 | 0.007083 | 0 | SEMA3F_NRP2_PLXNA2 | Sema3f - (Nrp2+Plxna2) | SEMA3 | Secreted Signaling | PMID: 27533782 |
| NeuN | NeuN | Sema3f | NRP2_PLXNA2 | 0.009824 | 0 | SEMA3F_NRP2_PLXNA2 | Sema3f - (Nrp2+Plxna2) | SEMA3 | Secreted Signaling | PMID: 27533782 |
| NeuN | CD31 | Sema3f | NRP2_PLXNA3 | 0.005985 | 0 | SEMA3F_NRP2_PLXNA3 | Sema3f - (Nrp2+Plxna3) | SEMA3 | Secreted Signaling | PMID: 27533782 |
| NeuN | NeuN | Sema3f | NRP2_PLXNA3 | 0.008224 | 0 | SEMA3F_NRP2_PLXNA3 | Sema3f - (Nrp2+Plxna3) | SEMA3 | Secreted Signaling | PMID: 27533782 |
| NeuN | CD31 | Sema3f | NRP2_PLXNA4 | 0.011198 | 0 | SEMA3F_NRP2_PLXNA4 | Sema3f - (Nrp2+Plxna4) | SEMA3 | Secreted Signaling | PMID: 27533782 |
| NeuN | NeuN | Sema3f | NRP2_PLXNA4 | 0.014245 | 0 | SEMA3F_NRP2_PLXNA4 | Sema3f - (Nrp2+Plxna4) | SEMA3 | Secreted Signaling | PMID: 27533782 |
| CD31 | CD31 | Sema3g | NRP2_PLXNA1 | 0.001697 | 0 | SEMA3G_NRP2_PLXNA1 | Sema3g - (Nrp2+Plxna1) | SEMA3 | Secreted Signaling | PMID: 27533782 |
| CD31 | NeuN | Sema3g | NRP2_PLXNA1 | 0.001973 | 0 | SEMA3G_NRP2_PLXNA1 | Sema3g - (Nrp2+Plxna1) | SEMA3 | Secreted Signaling | PMID: 27533782 |
| CD31 | CD31 | Sema3g | NRP2_PLXNA2 | 0.001774 | 0 | SEMA3G_NRP2_PLXNA2 | Sema3g - (Nrp2+Plxna2) | SEMA3 | Secreted Signaling | PMID: 27533782 |
| CD31 | NeuN | Sema3g | NRP2_PLXNA2 | 0.002466 | 0 | SEMA3G_NRP2_PLXNA2 | Sema3g - (Nrp2+Plxna2) | SEMA3 | Secreted Signaling | PMID: 27533782 |
| CD31 | CD31 | Sema3g | NRP2_PLXNA3 | 0.001498 | 0 | SEMA3G_NRP2_PLXNA3 | Sema3g - (Nrp2+Plxna3) | SEMA3 | Secreted Signaling | PMID: 27533782 |
| CD31 | NeuN | Sema3g | NRP2_PLXNA3 | 0.002062 | 0 | SEMA3G_NRP2_PLXNA3 | Sema3g - (Nrp2+Plxna3) | SEMA3 | Secreted Signaling | PMID: 27533782 |
| CD31 | CD31 | Sema3g | NRP2_PLXNA4 | 0.002814 | 0 | SEMA3G_NRP2_PLXNA4 | Sema3g - (Nrp2+Plxna4) | SEMA3 | Secreted Signaling | PMID: 27533782 |
| CD31 | NeuN | Sema3g | NRP2_PLXNA4 | 0.003588 | 0 | SEMA3G_NRP2_PLXNA4 | Sema3g - (Nrp2+Plxna4) | SEMA3 | Secreted Signaling | PMID: 27533782 |
| NeuN | CD31 | Sema3c | NRP1_NRP2 | 0.006693 | 0 | SEMA3C_NRP1_NRP2 | Sema3c - (Nrp1+Nrp2) | SEMA3 | Secreted Signaling | PMID: 27533782 |
| NeuN | NeuN | Sema3c | NRP1_NRP2 | 0.003423 | 0 | SEMA3C_NRP1_NRP2 | Sema3c - (Nrp1+Nrp2) | SEMA3 | Secreted Signaling | PMID: 27533782 |
| CD31 | CD31 | Sema3c | Plxnd1 | 0.006131 | 0 | SEMA3C_PLXND1 | Sema3c - Plxnd1 | SEMA3 | Secreted Signaling | PMID: 27533782 |
| NeuN | CD31 | Sema3c | Plxnd1 | 0.012648 | 0 | SEMA3C_PLXND1 | Sema3c - Plxnd1 | SEMA3 | Secreted Signaling | PMID: 27533782 |
| CD31 | CD31 | Sema3e | Plxnd1 | 0.004066 | 0 | SEMA3E_PLXND1 | Sema3e - Plxnd1 | SEMA3 | Secreted Signaling | PMID: 22325954 |
| NeuN | CD31 | Sema3e | Plxnd1 | 0.003311 | 0 | SEMA3E_PLXND1 | Sema3e - Plxnd1 | SEMA3 | Secreted Signaling | PMID: 22325954 |
| CD31 | CD31 | Crh | Crhr1 | 0.000543 | 0 | CRH_CRHR1 | Crh - Crhr1 | CRH | Secreted Signaling | KEGG: mmu04080 |

| | | | | | | | | | | |
|---|---|---|---|---|---|---|---|---|---|---|
| CD31 | NeuN | Crh | Crhr1 | 0.000623 | 0 | CRH_CRHR1 | Crh - Crhr1 | CRH | Secreted Signaling | KEGG: mmu04080 |
| CD31 | CD31 | Crh | Crhr2 | 0.000358 | 0 | CRH_CRHR2 | Crh - Crhr2 | CRH | Secreted Signaling | KEGG: mmu04080 |
| CD31 | NeuN | Crh | Crhr2 | 0.000351 | 0 | CRH_CRHR2 | Crh - Crhr2 | CRH | Secreted Signaling | KEGG: mmu04080 |
| NeuN | CD31 | Enho | Gpr19 | 0.002371 | 0 | ENHO_GPR19 | Enho - Gpr19 | ENHO | Secreted Signaling | PMID: 28476646 |
| NeuN | NeuN | Enho | Gpr19 | 0.003527 | 0 | ENHO_GPR19 | Enho - Gpr19 | ENHO | Secreted Signaling | PMID: 28476646 |
| CD31 | CD31 | Gas6 | Axl | 0.000722 | 0 | GAS6_AXL | Gas6 - Axl | GAS | Secreted Signaling | PMID: 27801848 |
| NeuN | CD31 | Gas6 | Axl | 0.001299 | 0 | GAS6_AXL | Gas6 - Axl | GAS | Secreted Signaling | PMID: 27801848 |
| NeuN | CD31 | Gas6 | Mertk | 0.000737 | 0 | GAS6_MERTK | Gas6 - Mertk | GAS | Secreted Signaling | PMID: 27801848 |
| NeuN | NeuN | Gas6 | Mertk | 0.000557 | 0 | GAS6_MERTK | Gas6 - Mertk | GAS | Secreted Signaling | PMID: 27801848 |
| NeuN | CD31 | Gas6 | Tyro3 | 0.000593 | 0 | GAS6_TYRO3 | Gas6 - Tyro3 | GAS | Secreted Signaling | PMID: 27801848 |
| NeuN | NeuN | Gas6 | Tyro3 | 0.000454 | 0 | GAS6_TYRO3 | Gas6 - Tyro3 | GAS | Secreted Signaling | PMID: 27801848 |
| CD31 | NeuN | Grn | Sort1 | 0.005063 | 0 | GRN_SORT1 | Grn - Sort1 | GRN | Secreted Signaling | PMID: 29555433 |
| NeuN | NeuN | Grn | Sort1 | 0.004334 | 0 | GRN_SORT1 | Grn - Sort1 | GRN | Secreted Signaling | PMID: 29555433 |
| CD31 | CD31 | Insl5 | Rxfp4 | 0.001088 | 0 | INSL5_RXFP4 | Insl5 - Rxfp4 | RLN | Secreted Signaling | KEGG: mmu04080 |
| NeuN | CD31 | Insl5 | Rxfp4 | 0.000734 | 0 | INSL5_RXFP4 | Insl5 - Rxfp4 | RLN | Secreted Signaling | KEGG: mmu04080 |
| CD31 | CD31 | Insl5 | Rxfp3 | 0.000882 | 0 | INSL5_RXFP3 | Insl5 - Rxfp3 | RLN | Secreted Signaling | KEGG: mmu04080 |
| NeuN | CD31 | Insl5 | Rxfp3 | 0.000596 | 0 | INSL5_RXFP3 | Insl5 - Rxfp3 | RLN | Secreted Signaling | KEGG: mmu04080 |
| CD31 | CD31 | Insl5 | Rxfp2 | 0.000874 | 0 | INSL5_RXFP2 | Insl5 - Rxfp2 | RLN | Secreted Signaling | KEGG: mmu04080 |
| NeuN | CD31 | Insl5 | Rxfp2 | 0.00059 | 0 | INSL5_RXFP2 | Insl5 - Rxfp2 | RLN | Secreted Signaling | KEGG: mmu04080 |
| CD31 | CD31 | Insl5 | Rxfp1 | 0.000923 | 0 | INSL5_RXFP1 | Insl5 - Rxfp1 | RLN | Secreted Signaling | KEGG: mmu04080 |
| CD31 | NeuN | Insl5 | Rxfp1 | 0.000835 | 0 | INSL5_RXFP1 | Insl5 - Rxfp1 | RLN | Secreted Signaling | KEGG: mmu04080 |
| CD31 | CD31 | Lgals9 | Ighm | 0.000877 | 0 | LGALS9_IGHM | Lgals9 - Ighm | GALECTIN | Secreted Signaling | PMID: 30120235 |
| CD31 | NeuN | Lgals9 | Ighm | 0.000687 | 0 | LGALS9_IGHM | Lgals9 - Ighm | GALECTIN | Secreted Signaling | PMID: 30120235 |
| CD31 | CD31 | Pmch | Mertk | 0.000518 | 0 | PMCH_MERTK | Pmch - Mertk | PMCH | Secreted Signaling | PMID: 30501104 |
| CD31 | NeuN | Pmch | Mertk | 0.000391 | 0 | PMCH_MERTK | Pmch - Mertk | PMCH | Secreted Signaling | PMID: 30501104 |
| CD31 | CD31 | Pros1 | Axl | 0.001344 | 0 | PROS1_AXL | Pros1 - Axl | PROS | Secreted Signaling | PMID: 29531161 |
| CD31 | NeuN | Pros1 | Axl | 0.000621 | 0 | PROS1_AXL | Pros1 - Axl | PROS | Secreted Signaling | PMID: 29531161 |
| CD31 | CD31 | Pros1 | Tyro3 | 0.000614 | 0 | PROS1_TYRO3 | Pros1 - Tyro3 | PROS | Secreted Signaling | PMID: 30501104 |
| CD31 | NeuN | Pros1 | Tyro3 | 0.00047 | 0 | PROS1_TYRO3 | Pros1 - Tyro3 | PROS | Secreted Signaling | PMID: 30501104 |
| NeuN | CD31 | Psap | Gpr37l1 | 0.006082 | 0 | PSAP_GPR37L1 | Psap - Gpr37l1 | PSAP | Secreted Signaling | PMID: 23690594 |
| NeuN | NeuN | Psap | Gpr37l1 | 0.005334 | 0 | PSAP_GPR37L1 | Psap - Gpr37l1 | PSAP | Secreted Signaling | PMID: 23690594 |
| NeuN | CD31 | Psap | Gpr37 | 0.007795 | 0 | PSAP_GPR37 | Psap - Gpr37 | PSAP | Secreted Signaling | PMID: 23690594 |
| NeuN | NeuN | Psap | Gpr37 | 0.00672 | 0 | PSAP_GPR37 | Psap - Gpr37 | PSAP | Secreted Signaling | PMID: 23690594 |
| CD31 | CD31 | Qrfp | Qrfpr | 0.000676 | 0 | QRFP_QRFPR | Qrfp - Qrfpr | QRFP | Secreted Signaling | PMID: 25386115 |
| CD31 | NeuN | Qrfp | Qrfpr | 0.000653 | 0 | QRFP_QRFPR | Qrfp - Qrfpr | QRFP | Secreted Signaling | PMID: 25386115 |
| CD31 | CD31 | Rarres2 | Cmklr1 | 0.001431 | 0 | RARRES2_CMKLR1 | Rarres2 - Cmklr1 | CHEMERIN | Secreted Signaling | PMID: 24671882 |
| CD31 | NeuN | Rarres2 | Cmklr1 | 0.000682 | 0 | RARRES2_CMKLR1 | Rarres2 - Cmklr1 | CHEMERIN | Secreted Signaling | PMID: 24671882 |
| CD31 | CD31 | Rln3 | Rxfp3 | 0.000788 | 0 | RLN3_RXFP3 | Rln3 - Rxfp3 | RLN | Secreted Signaling | KEGG: mmu04080 |
| NeuN | CD31 | Rln3 | Rxfp3 | 0.000561 | 0 | RLN3_RXFP3 | Rln3 - Rxfp3 | RLN | Secreted Signaling | KEGG: mmu04080 |
| CD31 | CD31 | Rln3 | Rxfp1 | 0.000825 | 0 | RLN3_RXFP1 | Rln3 - Rxfp1 | RLN | Secreted Signaling | KEGG: mmu04080 |
| CD31 | NeuN | Rln3 | Rxfp1 | 0.000746 | 0 | RLN3_RXFP1 | Rln3 - Rxfp1 | RLN | Secreted Signaling | KEGG: mmu04080 |
| CD31 | CD31 | Rln3 | Rxfp4 | 0.000971 | 0 | RLN3_RXFP4 | Rln3 - Rxfp4 | RLN | Secreted Signaling | KEGG: mmu04080 |
| NeuN | CD31 | Rln3 | Rxfp4 | 0.000691 | 0 | RLN3_RXFP4 | Rln3 - Rxfp4 | RLN | Secreted Signaling | KEGG: mmu04080 |
| CD31 | CD31 | Rln3 | Rxfp2 | 0.000781 | 0 | RLN3_RXFP2 | Rln3 - Rxfp2 | RLN | Secreted Signaling | KEGG: mmu04080 |
| NeuN | CD31 | Rln3 | Rxfp2 | 0.000556 | 0 | RLN3_RXFP2 | Rln3 - Rxfp2 | RLN | Secreted Signaling | KEGG: mmu04080 |
| CD31 | CD31 | Slurp2 | Chrnb2 | 0.00087 | 0 | SLURP2_CHRNB2 | Slurp2 - Chrnb2 | SLURP | Secreted Signaling | KEGG: mmu04080 |
| CD31 | NeuN | Slurp2 | Chrnb2 | 0.001246 | 0 | SLURP2_CHRNB2 | Slurp2 - Chrnb2 | SLURP | Secreted Signaling | KEGG: mmu04080 |
| CD31 | CD31 | Slurp2 | Chrnb3 | 0.000554 | 0 | SLURP2_CHRNB3 | Slurp2 - Chrnb3 | SLURP | Secreted Signaling | KEGG: mmu04080 |
| CD31 | NeuN | Slurp2 | Chrnb3 | 0.000457 | 0 | SLURP2_CHRNB3 | Slurp2 - Chrnb3 | SLURP | Secreted Signaling | KEGG: mmu04080 |
| CD31 | CD31 | Slurp2 | Chrnb4 | 0.000719 | 0 | SLURP2_CHRNB4 | Slurp2 - Chrnb4 | SLURP | Secreted Signaling | KEGG: mmu04080 |
| CD31 | NeuN | Slurp2 | Chrnb4 | 0.000591 | 0 | SLURP2_CHRNB4 | Slurp2 - Chrnb4 | SLURP | Secreted Signaling | KEGG: mmu04080 |
| CD31 | CD31 | Slurp2 | Chrna1 | 0.000448 | 0 | SLURP2_CHRNA1 | Slurp2 - Chrna1 | SLURP | Secreted Signaling | KEGG: mmu04080 |
| CD31 | NeuN | Slurp2 | Chrna1 | 0.000474 | 0 | SLURP2_CHRNA1 | Slurp2 - Chrna1 | SLURP | Secreted Signaling | KEGG: mmu04080 |
| CD31 | CD31 | Slurp2 | Chrna4 | 0.00138 | 0 | SLURP2_CHRNA4 | Slurp2 - Chrna4 | SLURP | Secreted Signaling | KEGG: mmu04080 |
| CD31 | NeuN | Slurp2 | Chrna4 | 0.000913 | 0 | SLURP2_CHRNA4 | Slurp2 - Chrna4 | SLURP | Secreted Signaling | KEGG: mmu04080 |
| CD31 | CD31 | Slurp2 | Chrna5 | 0.000709 | 0 | SLURP2_CHRNA5 | Slurp2 - Chrna5 | SLURP | Secreted Signaling | KEGG: mmu04080 |
| NeuN | CD31 | Slurp2 | Chrna5 | 0.000451 | 0 | SLURP2_CHRNA5 | Slurp2 - Chrna5 | SLURP | Secreted Signaling | KEGG: mmu04080 |
| CD31 | NeuN | Slurp2 | Chrna7 | 0.0012 | 0 | SLURP2_CHRNA7 | Slurp2 - Chrna7 | SLURP | Secreted Signaling | KEGG: mmu04080 |
| NeuN | NeuN | Slurp2 | Chrna7 | 0.000763 | 0 | SLURP2_CHRNA7 | Slurp2 - Chrna7 | SLURP | Secreted Signaling | KEGG: mmu04080 |

| | | | | | | | | | | |
|---|---|---|---|---|---|---|---|---|---|---|
| CD31 | CD31 | Slurp2 | Chrna9 | 0.000751 | 0 | SLURP2_CHRNA9 | Slurp2 - Chrna9 | SLURP | Secreted Signaling | KEGG: mmu04080 |
| CD31 | NeuN | Slurp2 | Chrna9 | 0.000503 | 0 | SLURP2_CHRNA9 | Slurp2 - Chrna9 | SLURP | Secreted Signaling | KEGG: mmu04080 |
| CD31 | CD31 | Slurp2 | Chrnd | 0.000684 | 0 | SLURP2_CHRND | Slurp2 - Chrnd | SLURP | Secreted Signaling | KEGG: mmu04080 |
| CD31 | NeuN | Slurp2 | Chrnd | 0.000517 | 0 | SLURP2_CHRND | Slurp2 - Chrnd | SLURP | Secreted Signaling | KEGG: mmu04080 |
| CD31 | CD31 | Ucn | Crhr1 | 0.001487 | 0 | UCN_CRHR1 | Ucn - Crhr1 | UCN | Secreted Signaling | KEGG: mmu04080 |
| CD31 | NeuN | Ucn | Crhr1 | 0.001707 | 0 | UCN_CRHR1 | Ucn - Crhr1 | UCN | Secreted Signaling | KEGG: mmu04080 |
| CD31 | CD31 | Ucn | Crhr2 | 0.00098 | 0 | UCN_CRHR2 | Ucn - Crhr2 | UCN | Secreted Signaling | KEGG: mmu04080 |
| CD31 | NeuN | Ucn | Crhr2 | 0.000963 | 0 | UCN_CRHR2 | Ucn - Crhr2 | UCN | Secreted Signaling | KEGG: mmu04080 |
| CD31 | CD31 | Klk6 | F2r | 0.001778 | 0 | KLK6_F2R | Klk6 - F2r | KLK | Secreted Signaling | PMID: 35102281 |
| NeuN | CD31 | Klk6 | F2r | 0.001947 | 0 | KLK6_F2R | Klk6 - F2r | KLK | Secreted Signaling | PMID: 35102281 |
| CD31 | CD31 | Klk8 | F2r | 0.002958 | 0 | KLK8_F2R | Klk8 - F2r | KLK | Secreted Signaling | PMID: 34552064 |
| CD31 | NeuN | Klk8 | F2r | 0.00193 | 0 | KLK8_F2R | Klk8 - F2r | KLK | Secreted Signaling | PMID: 34552064 |
| CD31 | CD31 | Adm | CALCR_RAMP1 | 0.002828 | 0 | ADM_CALCR_RAMP1 | Adm - (Calcr+Ramp1) | ADM | Secreted Signaling | PMID: 33462236 |
| CD31 | NeuN | Adm | CALCR_RAMP1 | 0.002083 | 0 | ADM_CALCR_RAMP1 | Adm - (Calcr+Ramp1) | ADM | Secreted Signaling | PMID: 33462236 |
| CD31 | CD31 | Adm | CALCR_RAMP2 | 0.006561 | 0 | ADM_CALCR_RAMP2 | Adm - (Calcr+Ramp2) | ADM | Secreted Signaling | PMID: 33462236 |
| CD31 | NeuN | Adm | CALCR_RAMP2 | 0.002484 | 0 | ADM_CALCR_RAMP2 | Adm - (Calcr+Ramp2) | ADM | Secreted Signaling | PMID: 33462236 |
| CD31 | CD31 | Adm | CALCR_RAMP3 | 0.002579 | 0 | ADM_CALCR_RAMP3 | Adm - (Calcr+Ramp3) | ADM | Secreted Signaling | PMID: 33462236 |
| CD31 | NeuN | Adm | CALCR_RAMP3 | 0.002555 | 0 | ADM_CALCR_RAMP3 | Adm - (Calcr+Ramp3) | ADM | Secreted Signaling | PMID: 33462236 |
| CD31 | CD31 | Calca | CALCR_RAMP1 | 0.000763 | 0 | CALCA_CALCR_RAMP1 | Calca - (Calcr+Ramp1) | CALC | Secreted Signaling | PMID: 30741933;PMID: 18599553 |
| CD31 | NeuN | Calca | CALCR_RAMP1 | 0.000562 | 0 | CALCA_CALCR_RAMP1 | Calca - (Calcr+Ramp1) | CALC | Secreted Signaling | PMID: 30741933;PMID: 18599553 |
| CD31 | CD31 | Calca | CALCR_RAMP2 | 0.001774 | 0 | CALCA_CALCR_RAMP2 | Calca - (Calcr+Ramp2) | CALC | Secreted Signaling | PMID: 30741933;PMID: 18599553 |
| NeuN | CD31 | Calca | CALCR_RAMP2 | 0.001292 | 0 | CALCA_CALCR_RAMP2 | Calca - (Calcr+Ramp2) | CALC | Secreted Signaling | PMID: 30741933;PMID: 18599553 |
| CD31 | CD31 | Calca | CALCR_RAMP3 | 0.000695 | 0 | CALCA_CALCR_RAMP3 | Calca - (Calcr+Ramp3) | CALC | Secreted Signaling | PMID: 30741933;PMID: 18599553 |
| CD31 | NeuN | Calca | CALCR_RAMP3 | 0.000689 | 0 | CALCA_CALCR_RAMP3 | Calca - (Calcr+Ramp3) | CALC | Secreted Signaling | PMID: 30741933;PMID: 18599553 |
| CD31 | CD31 | Calcb | CALCR_RAMP1 | 0.000774 | 0 | CALCB_CALCR_RAMP1 | Calcb - (Calcr+Ramp1) | CALC | Secreted Signaling | PMID: 30741933;PMID: 18599553 |
| CD31 | NeuN | Calcb | CALCR_RAMP1 | 0.00057 | 0 | CALCB_CALCR_RAMP1 | Calcb - (Calcr+Ramp1) | CALC | Secreted Signaling | PMID: 30741933;PMID: 18599553 |
| CD31 | CD31 | Calcb | CALCR_RAMP2 | 0.0018 | 0 | CALCB_CALCR_RAMP2 | Calcb - (Calcr+Ramp2) | CALC | Secreted Signaling | PMID: 30741933;PMID: 18599553 |
| NeuN | CD31 | Calcb | CALCR_RAMP2 | 0.001306 | 0 | CALCB_CALCR_RAMP2 | Calcb - (Calcr+Ramp2) | CALC | Secreted Signaling | PMID: 30741933;PMID: 18599553 |
| CD31 | CD31 | Calcb | CALCR_RAMP3 | 0.000705 | 0 | CALCB_CALCR_RAMP3 | Calcb - (Calcr+Ramp3) | CALC | Secreted Signaling | PMID: 30741933;PMID: 18599553 |
| CD31 | NeuN | Calcb | CALCR_RAMP3 | 0.000699 | 0 | CALCB_CALCR_RAMP3 | Calcb - (Calcr+Ramp3) | CALC | Secreted Signaling | PMID: 30741933;PMID: 18599553 |
| CD31 | CD31 | Ctsg | Fpr1 | 0.000318 | 0 | CTSG_FPR1 | Ctsg - Fpr1 | CTSG | Secreted Signaling | PMID: 26516201 |
| NeuN | CD31 | Ctsg | Fpr1 | 0.00035 | 0 | CTSG_FPR1 | Ctsg - Fpr1 | CTSG | Secreted Signaling | PMID: 26516201 |
| CD31 | CD31 | Grp | Nmbr | 0.000649 | 0 | GRP_NMBR | Grp - Nmbr | GRP | Secreted Signaling | PMID:25209280;PMID:29133874 |
| NeuN | CD31 | Grp | Nmbr | 0.000658 | 0 | GRP_NMBR | Grp - Nmbr | GRP | Secreted Signaling | PMID:25209280;PMID:29133874 |
| CD31 | NeuN | Ptprs | Ntrk3 | 0.095621 | 0 | PTPRS_NTRK3 | Ptprs - Ntrk3 | PTPR | Secreted Signaling | PMID:21262467;PMID:25385546; |
| NeuN | NeuN | Ptprs | Ntrk3 | 0.19348 | 0 | PTPRS_NTRK3 | Ptprs - Ntrk3 | PTPR | Secreted Signaling | PMID:21262467;PMID:25385546; |
| CD31 | NeuN | Slitrk1 | Ptprd | 0.016221 | 0 | SLITRK1_PTPRD | Slitrk1 - Ptprd | SLITRK | Secreted Signaling | PMID: 30648269;PMID: 30822649 |
| NeuN | NeuN | Slitrk1 | Ptprd | 0.030365 | 0 | SLITRK1_PTPRD | Slitrk1 - Ptprd | SLITRK | Secreted Signaling | PMID: 30648269;PMID: 30822649 |
| CD31 | NeuN | Slitrk1 | Ptprs | 0.026143 | 0 | SLITRK1_PTPRS | Slitrk1 - Ptprs | SLITRK | Secreted Signaling | PMID: 30648269;PMID: 30822649 |
| NeuN | NeuN | Slitrk1 | Ptprs | 0.048512 | 0 | SLITRK1_PTPRS | Slitrk1 - Ptprs | SLITRK | Secreted Signaling | PMID: 30648269;PMID: 30822649 |
| CD31 | NeuN | Slitrk2 | Ptprd | 0.010837 | 0 | SLITRK2_PTPRD | Slitrk2 - Ptprd | SLITRK | Secreted Signaling | PMID: 30648269;PMID: 30822649 |
| NeuN | NeuN | Slitrk2 | Ptprd | 0.010436 | 0 | SLITRK2_PTPRD | Slitrk2 - Ptprd | SLITRK | Secreted Signaling | PMID: 30648269;PMID: 30822649 |
| CD31 | NeuN | Slitrk2 | Ptprs | 0.017525 | 0 | SLITRK2_PTPRS | Slitrk2 - Ptprs | SLITRK | Secreted Signaling | PMID: 30648269;PMID: 30822649 |
| NeuN | NeuN | Slitrk2 | Ptprs | 0.01688 | 0 | SLITRK2_PTPRS | Slitrk2 - Ptprs | SLITRK | Secreted Signaling | PMID: 30648269;PMID: 30822649 |
| CD31 | NeuN | Slitrk3 | Ptprd | 0.008826 | 0 | SLITRK3_PTPRD | Slitrk3 - Ptprd | SLITRK | Secreted Signaling | PMID: 30648269;PMID: 30822649 |
| NeuN | NeuN | Slitrk3 | Ptprd | 0.013371 | 0 | SLITRK3_PTPRD | Slitrk3 - Ptprd | SLITRK | Secreted Signaling | PMID: 30648269;PMID: 30822649 |
| CD31 | NeuN | Slitrk4 | Ptprs | 0.015444 | 0 | SLITRK4_PTPRS | Slitrk4 - Ptprs | SLITRK | Secreted Signaling | PMID: 30648269;PMID: 30822649 |
| NeuN | NeuN | Slitrk4 | Ptprs | 0.0089 | 0 | SLITRK4_PTPRS | Slitrk4 - Ptprs | SLITRK | Secreted Signaling | PMID: 30648269;PMID: 30822649 |
| CD31 | NeuN | Slitrk5 | Ptprs | 0.012701 | 0 | SLITRK5_PTPRS | Slitrk5 - Ptprs | SLITRK | Secreted Signaling | PMID: 30648269;PMID: 30822649 |
| NeuN | NeuN | Slitrk5 | Ptprs | 0.021237 | 0 | SLITRK5_PTPRS | Slitrk5 - Ptprs | SLITRK | Secreted Signaling | PMID: 30648269;PMID: 30822649 |
| CD31 | CD31 | Bdnf | Sort1 | 0.001576 | 0 | BDNF_SORT1 | Bdnf - Sort1 | NT | Secreted Signaling | uniprot |
| CD31 | NeuN | Bdnf | Sort1 | 0.002151 | 0 | BDNF_SORT1 | Bdnf - Sort1 | NT | Secreted Signaling | uniprot |
| CD31 | CD31 | Bdnf | Ngfr | 0.000475 | 0 | BDNF_NGFR | Bdnf - Ngfr | NT | Secreted Signaling | uniprot |
| CD31 | NeuN | Bdnf | Ngfr | 0.000673 | 0 | BDNF_NGFR | Bdnf - Ngfr | NT | Secreted Signaling | uniprot |
| CD31 | NeuN | Igfbp3 | Tmem219 | 0.005278 | 0 | IGFBP3_TMEM219 | Igfbp3 - Tmem219 | IGFBP | Secreted Signaling | PMID: 35115561;PMID:20353938 |
| NeuN | NeuN | Igfbp3 | Tmem219 | 0.004836 | 0 | IGFBP3_TMEM219 | Igfbp3 - Tmem219 | IGFBP | Secreted Signaling | PMID: 35115561;PMID:20353938 |
| CD31 | CD31 | Lgals9 | P4hb | 0.003995 | 0 | LGALS9_P4HB | Lgals9 - P4hb | GALECTIN | Secreted Signaling | PMID:21670307;uniprot |
| CD31 | NeuN | Lgals9 | P4hb | 0.00366 | 0 | LGALS9_P4HB | Lgals9 - P4hb | GALECTIN | Secreted Signaling | PMID:21670307;uniprot |
| CD31 | NeuN | Nts | Sort1 | 0.001988 | 0 | NTS_SORT1 | Nts - Sort1 | NTS | Secreted Signaling | PMID: 34038712 |

| | | | | | | | | | | |
|---|---|---|---|---|---|---|---|---|---|---|
| NeuN | NeuN | Nts | Sort1 | 0.002033 | 0 | NTS_SORT1 | Nts - Sort1 | NTS | Secreted Signaling | PMID: 34038712 |
| CD31 | CD31 | Ntf3 | Ngfr | 0.000593 | 0 | NTF3_NGFR | Ntf3 - Ngfr | NT | Secreted Signaling | uniprot |
| CD31 | NeuN | Ntf3 | Ngfr | 0.00084 | 0 | NTF3_NGFR | Ntf3 - Ngfr | NT | Secreted Signaling | uniprot |
| CD31 | CD31 | Rbp4 | Stra6 | 0.000591 | 0 | RBP4_STRA6 | Rbp4 - Stra6 | RBP4 | Secreted Signaling | PMID: 33199363 |
| NeuN | CD31 | Rbp4 | Stra6 | 0.000487 | 0 | RBP4_STRA6 | Rbp4 - Stra6 | RBP4 | Secreted Signaling | PMID: 33199363 |
| CD31 | NeuN | Slit1 | Robo1 | 0.010846 | 0 | SLIT1_ROBO1 | Slit1 - Robo1 | SLIT | Secreted Signaling | PMID:10102268 |
| NeuN | NeuN | Slit1 | Robo1 | 0.013365 | 0 | SLIT1_ROBO1 | Slit1 - Robo1 | SLIT | Secreted Signaling | PMID:10102268 |
| CD31 | NeuN | Slit2 | Robo1 | 0.006956 | 0 | SLIT2_ROBO1 | Slit2 - Robo1 | SLIT | Secreted Signaling | PMID:10102268 |
| NeuN | NeuN | Slit2 | Robo1 | 0.00509 | 0 | SLIT2_ROBO1 | Slit2 - Robo1 | SLIT | Secreted Signaling | PMID:10102268 |
| CD31 | NeuN | Slit2 | Robo2 | 0.010383 | 0 | SLIT2_ROBO2 | Slit2 - Robo2 | SLIT | Secreted Signaling | PMID:10102268 |
| NeuN | NeuN | Slit2 | Robo2 | 0.007605 | 0 | SLIT2_ROBO2 | Slit2 - Robo2 | SLIT | Secreted Signaling | PMID:10102268 |
| NeuN | CD31 | Tub | Mertk | 0.001805 | 0 | TUB_MERTK | Tub - Mertk | TULP | Secreted Signaling | PMID: 20978475 |
| NeuN | NeuN | Tub | Mertk | 0.001363 | 0 | TUB_MERTK | Tub - Mertk | TULP | Secreted Signaling | PMID: 20978475 |
| CD31 | CD31 | Pros1 | Mertk | 0.000763 | 0 | PROS1_MERTK | Pros1 - Mertk | PROS | Secreted Signaling | PMID: 34631419 |
| CD31 | NeuN | Pros1 | Mertk | 0.000576 | 0 | PROS1_MERTK | Pros1 - Mertk | PROS | Secreted Signaling | PMID: 34631419 |
| CD31 | CD31 | Col1a1 | ITGA1_ITGB1 | 0.002913 | 0 | COL1A1_ITGA1_ITGB1 | Col1a1 - (Itga1+Itgb1) | COLLAGEN | ECM-Receptor | KEGG: mmu04512 |
| NeuN | CD31 | Col1a1 | ITGA1_ITGB1 | 0.002227 | 0 | COL1A1_ITGA1_ITGB1 | Col1a1 - (Itga1+Itgb1) | COLLAGEN | ECM-Receptor | KEGG: mmu04512 |
| CD31 | CD31 | Col1a2 | ITGA1_ITGB1 | 0.00249 | 0 | COL1A2_ITGA1_ITGB1 | Col1a2 - (Itga1+Itgb1) | COLLAGEN | ECM-Receptor | KEGG: mmu04512 |
| NeuN | CD31 | Col1a2 | ITGA1_ITGB1 | 0.002481 | 0 | COL1A2_ITGA1_ITGB1 | Col1a2 - (Itga1+Itgb1) | COLLAGEN | ECM-Receptor | KEGG: mmu04512 |
| CD31 | CD31 | Col4a1 | ITGA1_ITGB1 | 0.076591 | 0 | COL4A1_ITGA1_ITGB1 | Col4a1 - (Itga1+Itgb1) | COLLAGEN | ECM-Receptor | KEGG: mmu04512 |
| CD31 | NeuN | Col4a1 | ITGA1_ITGB1 | 0.019763 | 0 | COL4A1_ITGA1_ITGB1 | Col4a1 - (Itga1+Itgb1) | COLLAGEN | ECM-Receptor | KEGG: mmu04512 |
| CD31 | CD31 | Col4a2 | ITGA1_ITGB1 | 0.061182 | 0 | COL4A2_ITGA1_ITGB1 | Col4a2 - (Itga1+Itgb1) | COLLAGEN | ECM-Receptor | KEGG: mmu04512 |
| CD31 | NeuN | Col4a2 | ITGA1_ITGB1 | 0.015594 | 0 | COL4A2_ITGA1_ITGB1 | Col4a2 - (Itga1+Itgb1) | COLLAGEN | ECM-Receptor | KEGG: mmu04512 |
| CD31 | CD31 | Col6a1 | ITGA1_ITGB1 | 0.003453 | 0 | COL6A1_ITGA1_ITGB1 | Col6a1 - (Itga1+Itgb1) | COLLAGEN | ECM-Receptor | KEGG: mmu04512 |
| NeuN | CD31 | Col6a1 | ITGA1_ITGB1 | 0.001968 | 0 | COL6A1_ITGA1_ITGB1 | Col6a1 - (Itga1+Itgb1) | COLLAGEN | ECM-Receptor | KEGG: mmu04512 |
| CD31 | CD31 | Col6a2 | ITGA1_ITGB1 | 0.0046 | 0 | COL6A2_ITGA1_ITGB1 | Col6a2 - (Itga1+Itgb1) | COLLAGEN | ECM-Receptor | KEGG: mmu04512 |
| NeuN | CD31 | Col6a2 | ITGA1_ITGB1 | 0.002855 | 0 | COL6A2_ITGA1_ITGB1 | Col6a2 - (Itga1+Itgb1) | COLLAGEN | ECM-Receptor | KEGG: mmu04512 |
| CD31 | CD31 | Col6a4 | ITGA1_ITGB1 | 0.002682 | 0 | COL6A4_ITGA1_ITGB1 | Col6a4 - (Itga1+Itgb1) | COLLAGEN | ECM-Receptor | KEGG: mmu04512 |
| NeuN | CD31 | Col6a4 | ITGA1_ITGB1 | 0.002949 | 0 | COL6A4_ITGA1_ITGB1 | Col6a4 - (Itga1+Itgb1) | COLLAGEN | ECM-Receptor | KEGG: mmu04512 |
| CD31 | CD31 | Col6a6 | ITGA1_ITGB1 | 0.002259 | 0 | COL6A6_ITGA1_ITGB1 | Col6a6 - (Itga1+Itgb1) | COLLAGEN | ECM-Receptor | KEGG: mmu04512 |
| NeuN | CD31 | Col6a6 | ITGA1_ITGB1 | 0.002317 | 0 | COL6A6_ITGA1_ITGB1 | Col6a6 - (Itga1+Itgb1) | COLLAGEN | ECM-Receptor | KEGG: mmu04512 |
| CD31 | CD31 | Col9a2 | ITGA1_ITGB1 | 0.003414 | 0 | COL9A2_ITGA1_ITGB1 | Col9a2 - (Itga1+Itgb1) | COLLAGEN | ECM-Receptor | KEGG: mmu04512 |
| NeuN | CD31 | Col9a2 | ITGA1_ITGB1 | 0.004137 | 0 | COL9A2_ITGA1_ITGB1 | Col9a2 - (Itga1+Itgb1) | COLLAGEN | ECM-Receptor | KEGG: mmu04512 |
| CD31 | CD31 | Col9a3 | ITGA1_ITGB1 | 0.004338 | 0 | COL9A3_ITGA1_ITGB1 | Col9a3 - (Itga1+Itgb1) | COLLAGEN | ECM-Receptor | KEGG: mmu04512 |
| NeuN | CD31 | Col9a3 | ITGA1_ITGB1 | 0.006021 | 0 | COL9A3_ITGA1_ITGB1 | Col9a3 - (Itga1+Itgb1) | COLLAGEN | ECM-Receptor | KEGG: mmu04512 |
| CD31 | CD31 | Fn1 | ITGA3_ITGB1 | 0.011508 | 0 | FN1_ITGA3_ITGB1 | Fn1 - (Itga3+Itgb1) | FN1 | ECM-Receptor | KEGG: mmu04512 |
| CD31 | NeuN | Fn1 | ITGA3_ITGB1 | 0.006764 | 0 | FN1_ITGA3_ITGB1 | Fn1 - (Itga3+Itgb1) | FN1 | ECM-Receptor | KEGG: mmu04512 |
| CD31 | CD31 | Fn1 | ITGA4_ITGB1 | 0.016123 | 0 | FN1_ITGA4_ITGB1 | Fn1 - (Itga4+Itgb1) | FN1 | ECM-Receptor | KEGG: mmu04512 |
| CD31 | NeuN | Fn1 | ITGA4_ITGB1 | 0.004993 | 0 | FN1_ITGA4_ITGB1 | Fn1 - (Itga4+Itgb1) | FN1 | ECM-Receptor | KEGG: mmu04512 |
| CD31 | CD31 | Fn1 | ITGA5_ITGB1 | 0.016149 | 0 | FN1_ITGA5_ITGB1 | Fn1 - (Itga5+Itgb1) | FN1 | ECM-Receptor | KEGG: mmu04512 |
| CD31 | NeuN | Fn1 | ITGA5_ITGB1 | 0.00511 | 0 | FN1_ITGA5_ITGB1 | Fn1 - (Itga5+Itgb1) | FN1 | ECM-Receptor | KEGG: mmu04512 |
| CD31 | CD31 | Fn1 | ITGA8_ITGB1 | 0.009876 | 0 | FN1_ITGA8_ITGB1 | Fn1 - (Itga8+Itgb1) | FN1 | ECM-Receptor | KEGG: mmu04512 |
| CD31 | NeuN | Fn1 | ITGA8_ITGB1 | 0.005018 | 0 | FN1_ITGA8_ITGB1 | Fn1 - (Itga8+Itgb1) | FN1 | ECM-Receptor | KEGG: mmu04512 |
| CD31 | CD31 | Lama1 | ITGA1_ITGB1 | 0.00406 | 0 | LAMA1_ITGA1_ITGB1 | Lama1 - (Itga1+Itgb1) | LAMININ | ECM-Receptor | KEGG: mmu04512 |
| NeuN | CD31 | Lama1 | ITGA1_ITGB1 | 0.002852 | 0 | LAMA1_ITGA1_ITGB1 | Lama1 - (Itga1+Itgb1) | LAMININ | ECM-Receptor | KEGG: mmu04512 |
| CD31 | CD31 | Lama4 | ITGA1_ITGB1 | 0.023963 | 0 | LAMA4_ITGA1_ITGB1 | Lama4 - (Itga1+Itgb1) | LAMININ | ECM-Receptor | KEGG: mmu04512 |
| CD31 | NeuN | Lama4 | ITGA1_ITGB1 | 0.005932 | 0 | LAMA4_ITGA1_ITGB1 | Lama4 - (Itga1+Itgb1) | LAMININ | ECM-Receptor | KEGG: mmu04512 |
| CD31 | CD31 | Lama5 | ITGA1_ITGB1 | 0.00653 | 0 | LAMA5_ITGA1_ITGB1 | Lama5 - (Itga1+Itgb1) | LAMININ | ECM-Receptor | KEGG: mmu04512 |
| NeuN | CD31 | Lama5 | ITGA1_ITGB1 | 0.002613 | 0 | LAMA5_ITGA1_ITGB1 | Lama5 - (Itga1+Itgb1) | LAMININ | ECM-Receptor | KEGG: mmu04512 |
| CD31 | CD31 | Lamb1 | ITGA1_ITGB1 | 0.022544 | 0 | LAMB1_ITGA1_ITGB1 | Lamb1 - (Itga1+Itgb1) | LAMININ | ECM-Receptor | KEGG: mmu04512 |
| CD31 | NeuN | Lamb1 | ITGA1_ITGB1 | 0.005575 | 0 | LAMB1_ITGA1_ITGB1 | Lamb1 - (Itga1+Itgb1) | LAMININ | ECM-Receptor | KEGG: mmu04512 |
| CD31 | CD31 | Lamb2 | ITGA1_ITGB1 | 0.006088 | 0 | LAMB2_ITGA1_ITGB1 | Lamb2 - (Itga1+Itgb1) | LAMININ | ECM-Receptor | KEGG: mmu04512 |
| NeuN | CD31 | Lamb2 | ITGA1_ITGB1 | 0.002655 | 0 | LAMB2_ITGA1_ITGB1 | Lamb2 - (Itga1+Itgb1) | LAMININ | ECM-Receptor | KEGG: mmu04512 |
| CD31 | CD31 | Lamc1 | ITGA1_ITGB1 | 0.011745 | 0 | LAMC1_ITGA1_ITGB1 | Lamc1 - (Itga1+Itgb1) | LAMININ | ECM-Receptor | KEGG: mmu04512 |
| CD31 | NeuN | Lamc1 | ITGA1_ITGB1 | 0.002881 | 0 | LAMC1_ITGA1_ITGB1 | Lamc1 - (Itga1+Itgb1) | LAMININ | ECM-Receptor | KEGG: mmu04512 |
| CD31 | CD31 | Lamc2 | ITGA1_ITGB1 | 0.004023 | 0 | LAMC2_ITGA1_ITGB1 | Lamc2 - (Itga1+Itgb1) | LAMININ | ECM-Receptor | KEGG: mmu04512 |
| NeuN | CD31 | Lamc2 | ITGA1_ITGB1 | 0.002417 | 0 | LAMC2_ITGA1_ITGB1 | Lamc2 - (Itga1+Itgb1) | LAMININ | ECM-Receptor | KEGG: mmu04512 |
| CD31 | CD31 | Lamc3 | ITGA1_ITGB1 | 0.007372 | 0 | LAMC3_ITGA1_ITGB1 | Lamc3 - (Itga1+Itgb1) | LAMININ | ECM-Receptor | KEGG: mmu04512 |
| NeuN | CD31 | Lamc3 | ITGA1_ITGB1 | 0.003199 | 0 | LAMC3_ITGA1_ITGB1 | Lamc3 - (Itga1+Itgb1) | LAMININ | ECM-Receptor | KEGG: mmu04512 |

| | | | | | | | | | | |
|---|---|---|---|---|---|---|---|---|---|---|
| CD31 | CD31 | Lama1 | ITGA3_ITGB1 | 0.002287 | 0 | LAMA1_ITGA3_ITGB1 | Lama1 - (Itga3+Itgb1) | LAMININ | ECM-Receptor | KEGG: mmu04512 |
| NeuN | CD31 | Lama1 | ITGA3_ITGB1 | 0.001605 | 0 | LAMA1_ITGA3_ITGB1 | Lama1 - (Itga3+Itgb1) | LAMININ | ECM-Receptor | KEGG: mmu04512 |
| CD31 | CD31 | Lama4 | ITGA3_ITGB1 | 0.013616 | 0 | LAMA4_ITGA3_ITGB1 | Lama4 - (Itga3+Itgb1) | LAMININ | ECM-Receptor | KEGG: mmu04512 |
| CD31 | NeuN | Lama4 | ITGA3_ITGB1 | 0.008009 | 0 | LAMA4_ITGA3_ITGB1 | Lama4 - (Itga3+Itgb1) | LAMININ | ECM-Receptor | KEGG: mmu04512 |
| CD31 | CD31 | Lama5 | ITGA3_ITGB1 | 0.003682 | 0 | LAMA5_ITGA3_ITGB1 | Lama5 - (Itga3+Itgb1) | LAMININ | ECM-Receptor | KEGG: mmu04512 |
| CD31 | NeuN | Lama5 | ITGA3_ITGB1 | 0.002157 | 0 | LAMA5_ITGA3_ITGB1 | Lama5 - (Itga3+Itgb1) | LAMININ | ECM-Receptor | KEGG: mmu04512 |
| CD31 | CD31 | Lamb1 | ITGA3_ITGB1 | 0.012801 | 0 | LAMB1_ITGA3_ITGB1 | Lamb1 - (Itga3+Itgb1) | LAMININ | ECM-Receptor | KEGG: mmu04512 |
| CD31 | NeuN | Lamb1 | ITGA3_ITGB1 | 0.007528 | 0 | LAMB1_ITGA3_ITGB1 | Lamb1 - (Itga3+Itgb1) | LAMININ | ECM-Receptor | KEGG: mmu04512 |
| CD31 | CD31 | Lamb2 | ITGA3_ITGB1 | 0.003432 | 0 | LAMB2_ITGA3_ITGB1 | Lamb2 - (Itga3+Itgb1) | LAMININ | ECM-Receptor | KEGG: mmu04512 |
| CD31 | NeuN | Lamb2 | ITGA3_ITGB1 | 0.00201 | 0 | LAMB2_ITGA3_ITGB1 | Lamb2 - (Itga3+Itgb1) | LAMININ | ECM-Receptor | KEGG: mmu04512 |
| CD31 | CD31 | Lamc1 | ITGA3_ITGB1 | 0.006638 | 0 | LAMC1_ITGA3_ITGB1 | Lamc1 - (Itga3+Itgb1) | LAMININ | ECM-Receptor | KEGG: mmu04512 |
| CD31 | NeuN | Lamc1 | ITGA3_ITGB1 | 0.003893 | 0 | LAMC1_ITGA3_ITGB1 | Lamc1 - (Itga3+Itgb1) | LAMININ | ECM-Receptor | KEGG: mmu04512 |
| CD31 | CD31 | Lamc2 | ITGA3_ITGB1 | 0.002266 | 0 | LAMC2_ITGA3_ITGB1 | Lamc2 - (Itga3+Itgb1) | LAMININ | ECM-Receptor | KEGG: mmu04512 |
| NeuN | CD31 | Lamc2 | ITGA3_ITGB1 | 0.00136 | 0 | LAMC2_ITGA3_ITGB1 | Lamc2 - (Itga3+Itgb1) | LAMININ | ECM-Receptor | KEGG: mmu04512 |
| CD31 | CD31 | Lamc3 | ITGA3_ITGB1 | 0.004158 | 0 | LAMC3_ITGA3_ITGB1 | Lamc3 - (Itga3+Itgb1) | LAMININ | ECM-Receptor | KEGG: mmu04512 |
| CD31 | NeuN | Lamc3 | ITGA3_ITGB1 | 0.002437 | 0 | LAMC3_ITGA3_ITGB1 | Lamc3 - (Itga3+Itgb1) | LAMININ | ECM-Receptor | KEGG: mmu04512 |
| NeuN | CD31 | Reln | ITGA3_ITGB1 | 0.003613 | 0 | RELN_ITGA3_ITGB1 | Reln - (Itga3+Itgb1) | RELN | ECM-Receptor | KEGG: mmu04512 |
| NeuN | NeuN | Reln | ITGA3_ITGB1 | 0.002116 | 0 | RELN_ITGA3_ITGB1 | Reln - (Itga3+Itgb1) | RELN | ECM-Receptor | KEGG: mmu04512 |
| CD31 | CD31 | Thbs1 | ITGA3_ITGB1 | 0.002681 | 0 | THBS1_ITGA3_ITGB1 | Thbs1 - (Itga3+Itgb1) | THBS | ECM-Receptor | KEGG: mmu04512 |
| NeuN | CD31 | Thbs1 | ITGA3_ITGB1 | 0.002197 | 0 | THBS1_ITGA3_ITGB1 | Thbs1 - (Itga3+Itgb1) | THBS | ECM-Receptor | KEGG: mmu04512 |
| NeuN | CD31 | Thbs3 | ITGA3_ITGB1 | 0.005792 | 0 | THBS3_ITGA3_ITGB1 | Thbs3 - (Itga3+Itgb1) | THBS | ECM-Receptor | KEGG: mmu04512 |
| NeuN | NeuN | Thbs3 | ITGA3_ITGB1 | 0.003396 | 0 | THBS3_ITGA3_ITGB1 | Thbs3 - (Itga3+Itgb1) | THBS | ECM-Receptor | KEGG: mmu04512 |
| CD31 | CD31 | Thbs4 | ITGA3_ITGB1 | 0.00151 | 0 | THBS4_ITGA3_ITGB1 | Thbs4 - (Itga3+Itgb1) | THBS | ECM-Receptor | KEGG: mmu04512 |
| NeuN | CD31 | Thbs4 | ITGA3_ITGB1 | 0.001581 | 0 | THBS4_ITGA3_ITGB1 | Thbs4 - (Itga3+Itgb1) | THBS | ECM-Receptor | KEGG: mmu04512 |
| CD31 | CD31 | Col1a1 | ITGA3_ITGB1 | 0.00164 | 0 | COL1A1_ITGA3_ITGB1 | Col1a1 - (Itga3+Itgb1) | COLLAGEN | ECM-Receptor | KEGG: mmu04512 |
| NeuN | CD31 | Col1a1 | ITGA3_ITGB1 | 0.001253 | 0 | COL1A1_ITGA3_ITGB1 | Col1a1 - (Itga3+Itgb1) | COLLAGEN | ECM-Receptor | KEGG: mmu04512 |
| CD31 | CD31 | Col1a2 | ITGA3_ITGB1 | 0.001402 | 0 | COL1A2_ITGA3_ITGB1 | Col1a2 - (Itga3+Itgb1) | COLLAGEN | ECM-Receptor | KEGG: mmu04512 |
| NeuN | CD31 | Col1a2 | ITGA3_ITGB1 | 0.001396 | 0 | COL1A2_ITGA3_ITGB1 | Col1a2 - (Itga3+Itgb1) | COLLAGEN | ECM-Receptor | KEGG: mmu04512 |
| CD31 | CD31 | Col4a1 | ITGA3_ITGB1 | 0.044556 | 0 | COL4A1_ITGA3_ITGB1 | Col4a1 - (Itga3+Itgb1) | COLLAGEN | ECM-Receptor | KEGG: mmu04512 |
| CD31 | NeuN | Col4a1 | ITGA3_ITGB1 | 0.026553 | 0 | COL4A1_ITGA3_ITGB1 | Col4a1 - (Itga3+Itgb1) | COLLAGEN | ECM-Receptor | KEGG: mmu04512 |
| CD31 | CD31 | Col4a2 | ITGA3_ITGB1 | 0.035345 | 0 | COL4A2_ITGA3_ITGB1 | Col4a2 - (Itga3+Itgb1) | COLLAGEN | ECM-Receptor | KEGG: mmu04512 |
| CD31 | NeuN | Col4a2 | ITGA3_ITGB1 | 0.020982 | 0 | COL4A2_ITGA3_ITGB1 | Col4a2 - (Itga3+Itgb1) | COLLAGEN | ECM-Receptor | KEGG: mmu04512 |
| CD31 | CD31 | Col6a1 | ITGA3_ITGB1 | 0.001944 | 0 | COL6A1_ITGA3_ITGB1 | Col6a1 - (Itga3+Itgb1) | COLLAGEN | ECM-Receptor | KEGG: mmu04512 |
| CD31 | NeuN | Col6a1 | ITGA3_ITGB1 | 0.001138 | 0 | COL6A1_ITGA3_ITGB1 | Col6a1 - (Itga3+Itgb1) | COLLAGEN | ECM-Receptor | KEGG: mmu04512 |
| CD31 | CD31 | Col6a2 | ITGA3_ITGB1 | 0.002592 | 0 | COL6A2_ITGA3_ITGB1 | Col6a2 - (Itga3+Itgb1) | COLLAGEN | ECM-Receptor | KEGG: mmu04512 |
| NeuN | CD31 | Col6a2 | ITGA3_ITGB1 | 0.001607 | 0 | COL6A2_ITGA3_ITGB1 | Col6a2 - (Itga3+Itgb1) | COLLAGEN | ECM-Receptor | KEGG: mmu04512 |
| CD31 | CD31 | Col6a4 | ITGA3_ITGB1 | 0.00151 | 0 | COL6A4_ITGA3_ITGB1 | Col6a4 - (Itga3+Itgb1) | COLLAGEN | ECM-Receptor | KEGG: mmu04512 |
| NeuN | CD31 | Col6a4 | ITGA3_ITGB1 | 0.00166 | 0 | COL6A4_ITGA3_ITGB1 | Col6a4 - (Itga3+Itgb1) | COLLAGEN | ECM-Receptor | KEGG: mmu04512 |
| CD31 | CD31 | Col6a6 | ITGA3_ITGB1 | 0.001272 | 0 | COL6A6_ITGA3_ITGB1 | Col6a6 - (Itga3+Itgb1) | COLLAGEN | ECM-Receptor | KEGG: mmu04512 |
| NeuN | CD31 | Col6a6 | ITGA3_ITGB1 | 0.001304 | 0 | COL6A6_ITGA3_ITGB1 | Col6a6 - (Itga3+Itgb1) | COLLAGEN | ECM-Receptor | KEGG: mmu04512 |
| CD31 | CD31 | Col9a2 | ITGA3_ITGB1 | 0.001922 | 0 | COL9A2_ITGA3_ITGB1 | Col9a2 - (Itga3+Itgb1) | COLLAGEN | ECM-Receptor | KEGG: mmu04512 |
| NeuN | CD31 | Col9a2 | ITGA3_ITGB1 | 0.00233 | 0 | COL9A2_ITGA3_ITGB1 | Col9a2 - (Itga3+Itgb1) | COLLAGEN | ECM-Receptor | KEGG: mmu04512 |
| CD31 | CD31 | Col9a3 | ITGA3_ITGB1 | 0.002443 | 0 | COL9A3_ITGA3_ITGB1 | Col9a3 - (Itga3+Itgb1) | COLLAGEN | ECM-Receptor | KEGG: mmu04512 |
| NeuN | CD31 | Col9a3 | ITGA3_ITGB1 | 0.003394 | 0 | COL9A3_ITGA3_ITGB1 | Col9a3 - (Itga3+Itgb1) | COLLAGEN | ECM-Receptor | KEGG: mmu04512 |
| CD31 | CD31 | Lama1 | ITGA6_ITGB1 | 0.003069 | 0 | LAMA1_ITGA6_ITGB1 | Lama1 - (Itga6+Itgb1) | LAMININ | ECM-Receptor | KEGG: mmu04512 |
| NeuN | CD31 | Lama1 | ITGA6_ITGB1 | 0.002155 | 0 | LAMA1_ITGA6_ITGB1 | Lama1 - (Itga6+Itgb1) | LAMININ | ECM-Receptor | KEGG: mmu04512 |
| CD31 | CD31 | Lama4 | ITGA6_ITGB1 | 0.018199 | 0 | LAMA4_ITGA6_ITGB1 | Lama4 - (Itga6+Itgb1) | LAMININ | ECM-Receptor | KEGG: mmu04512 |
| CD31 | NeuN | Lama4 | ITGA6_ITGB1 | 0.00748 | 0 | LAMA4_ITGA6_ITGB1 | Lama4 - (Itga6+Itgb1) | LAMININ | ECM-Receptor | KEGG: mmu04512 |
| CD31 | CD31 | Lama5 | ITGA6_ITGB1 | 0.004938 | 0 | LAMA5_ITGA6_ITGB1 | Lama5 - (Itga6+Itgb1) | LAMININ | ECM-Receptor | KEGG: mmu04512 |
| CD31 | NeuN | Lama5 | ITGA6_ITGB1 | 0.002014 | 0 | LAMA5_ITGA6_ITGB1 | Lama5 - (Itga6+Itgb1) | LAMININ | ECM-Receptor | KEGG: mmu04512 |
| CD31 | CD31 | Lamb1 | ITGA6_ITGB1 | 0.017116 | 0 | LAMB1_ITGA6_ITGB1 | Lamb1 - (Itga6+Itgb1) | LAMININ | ECM-Receptor | KEGG: mmu04512 |
| CD31 | NeuN | Lamb1 | ITGA6_ITGB1 | 0.00703 | 0 | LAMB1_ITGA6_ITGB1 | Lamb1 - (Itga6+Itgb1) | LAMININ | ECM-Receptor | KEGG: mmu04512 |
| CD31 | CD31 | Lamb2 | ITGA6_ITGB1 | 0.004603 | 0 | LAMB2_ITGA6_ITGB1 | Lamb2 - (Itga6+Itgb1) | LAMININ | ECM-Receptor | KEGG: mmu04512 |
| NeuN | CD31 | Lamb2 | ITGA6_ITGB1 | 0.002006 | 0 | LAMB2_ITGA6_ITGB1 | Lamb2 - (Itga6+Itgb1) | LAMININ | ECM-Receptor | KEGG: mmu04512 |
| CD31 | CD31 | Lamc1 | ITGA6_ITGB1 | 0.008893 | 0 | LAMC1_ITGA6_ITGB1 | Lamc1 - (Itga6+Itgb1) | LAMININ | ECM-Receptor | KEGG: mmu04512 |
| CD31 | NeuN | Lamc1 | ITGA6_ITGB1 | 0.003635 | 0 | LAMC1_ITGA6_ITGB1 | Lamc1 - (Itga6+Itgb1) | LAMININ | ECM-Receptor | KEGG: mmu04512 |
| CD31 | CD31 | Lamc2 | ITGA6_ITGB1 | 0.00304 | 0 | LAMC2_ITGA6_ITGB1 | Lamc2 - (Itga6+Itgb1) | LAMININ | ECM-Receptor | KEGG: mmu04512 |
| NeuN | CD31 | Lamc2 | ITGA6_ITGB1 | 0.001826 | 0 | LAMC2_ITGA6_ITGB1 | Lamc2 - (Itga6+Itgb1) | LAMININ | ECM-Receptor | KEGG: mmu04512 |
| CD31 | CD31 | Lamc3 | ITGA6_ITGB1 | 0.005576 | 0 | LAMC3_ITGA6_ITGB1 | Lamc3 - (Itga6+Itgb1) | LAMININ | ECM-Receptor | KEGG: mmu04512 |

| | | | | | | | | | | |
|---|---|---|---|---|---|---|---|---|---|---|
| NeuN | CD31 | Lamc3 | ITGA6_ITGB1 | 0.002417 | 0 | LAMC3_ITGA6_ITGB1 | Lamc3 - (Itga6+Itgb1) | LAMININ | ECM-Receptor | KEGG: mmu04512 |
| CD31 | CD31 | Lama1 | ITGA7_ITGB1 | 0.002104 | 0 | LAMA1_ITGA7_ITGB1 | Lama1 - (Itga7+Itgb1) | LAMININ | ECM-Receptor | KEGG: mmu04512 |
| NeuN | CD31 | Lama1 | ITGA7_ITGB1 | 0.001477 | 0 | LAMA1_ITGA7_ITGB1 | Lama1 - (Itga7+Itgb1) | LAMININ | ECM-Receptor | KEGG: mmu04512 |
| CD31 | CD31 | Lama4 | ITGA7_ITGB1 | 0.012537 | 0 | LAMA4_ITGA7_ITGB1 | Lama4 - (Itga7+Itgb1) | LAMININ | ECM-Receptor | KEGG: mmu04512 |
| CD31 | NeuN | Lama4 | ITGA7_ITGB1 | 0.005838 | 0 | LAMA4_ITGA7_ITGB1 | Lama4 - (Itga7+Itgb1) | LAMININ | ECM-Receptor | KEGG: mmu04512 |
| CD31 | CD31 | Lama5 | ITGA7_ITGB1 | 0.003388 | 0 | LAMA5_ITGA7_ITGB1 | Lama5 - (Itga7+Itgb1) | LAMININ | ECM-Receptor | KEGG: mmu04512 |
| CD31 | NeuN | Lama5 | ITGA7_ITGB1 | 0.00157 | 0 | LAMA5_ITGA7_ITGB1 | Lama5 - (Itga7+Itgb1) | LAMININ | ECM-Receptor | KEGG: mmu04512 |
| CD31 | CD31 | Lamb1 | ITGA7_ITGB1 | 0.011787 | 0 | LAMB1_ITGA7_ITGB1 | Lamb1 - (Itga7+Itgb1) | LAMININ | ECM-Receptor | KEGG: mmu04512 |
| CD31 | NeuN | Lamb1 | ITGA7_ITGB1 | 0.005486 | 0 | LAMB1_ITGA7_ITGB1 | Lamb1 - (Itga7+Itgb1) | LAMININ | ECM-Receptor | KEGG: mmu04512 |
| CD31 | CD31 | Lamb2 | ITGA7_ITGB1 | 0.003157 | 0 | LAMB2_ITGA7_ITGB1 | Lamb2 - (Itga7+Itgb1) | LAMININ | ECM-Receptor | KEGG: mmu04512 |
| CD31 | NeuN | Lamb2 | ITGA7_ITGB1 | 0.001463 | 0 | LAMB2_ITGA7_ITGB1 | Lamb2 - (Itga7+Itgb1) | LAMININ | ECM-Receptor | KEGG: mmu04512 |
| CD31 | CD31 | Lamc1 | ITGA7_ITGB1 | 0.006108 | 0 | LAMC1_ITGA7_ITGB1 | Lamc1 - (Itga7+Itgb1) | LAMININ | ECM-Receptor | KEGG: mmu04512 |
| CD31 | NeuN | Lamc1 | ITGA7_ITGB1 | 0.002834 | 0 | LAMC1_ITGA7_ITGB1 | Lamc1 - (Itga7+Itgb1) | LAMININ | ECM-Receptor | KEGG: mmu04512 |
| CD31 | CD31 | Lamc2 | ITGA7_ITGB1 | 0.002084 | 0 | LAMC2_ITGA7_ITGB1 | Lamc2 - (Itga7+Itgb1) | LAMININ | ECM-Receptor | KEGG: mmu04512 |
| NeuN | CD31 | Lamc2 | ITGA7_ITGB1 | 0.001251 | 0 | LAMC2_ITGA7_ITGB1 | Lamc2 - (Itga7+Itgb1) | LAMININ | ECM-Receptor | KEGG: mmu04512 |
| CD31 | CD31 | Lamc3 | ITGA7_ITGB1 | 0.003826 | 0 | LAMC3_ITGA7_ITGB1 | Lamc3 - (Itga7+Itgb1) | LAMININ | ECM-Receptor | KEGG: mmu04512 |
| CD31 | NeuN | Lamc3 | ITGA7_ITGB1 | 0.001773 | 0 | LAMC3_ITGA7_ITGB1 | Lamc3 - (Itga7+Itgb1) | LAMININ | ECM-Receptor | KEGG: mmu04512 |
| CD31 | CD31 | Vtn | ITGA8_ITGB1 | 0.005928 | 0 | VTN_ITGA8_ITGB1 | Vtn - (Itga8+Itgb1) | VTN | ECM-Receptor | KEGG: mmu04512 |
| CD31 | NeuN | Vtn | ITGA8_ITGB1 | 0.003006 | 0 | VTN_ITGA8_ITGB1 | Vtn - (Itga8+Itgb1) | VTN | ECM-Receptor | KEGG: mmu04512 |
| CD31 | CD31 | Tnc | ITGA8_ITGB1 | 0.002536 | 0 | TNC_ITGA8_ITGB1 | Tnc - (Itga8+Itgb1) | TENASCIN | ECM-Receptor | KEGG: mmu04512 |
| NeuN | CD31 | Tnc | ITGA8_ITGB1 | 0.001496 | 0 | TNC_ITGA8_ITGB1 | Tnc - (Itga8+Itgb1) | TENASCIN | ECM-Receptor | KEGG: mmu04512 |
| CD31 | CD31 | Tnr | ITGA8_ITGB1 | 0.000847 | 0 | TNR_ITGA8_ITGB1 | Tnr - (Itga8+Itgb1) | TENASCIN | ECM-Receptor | KEGG: mmu04512 |
| NeuN | CD31 | Tnr | ITGA8_ITGB1 | 0.000978 | 0 | TNR_ITGA8_ITGB1 | Tnr - (Itga8+Itgb1) | TENASCIN | ECM-Receptor | KEGG: mmu04512 |
| CD31 | CD31 | Col1a1 | ITGA11_ITGB1 | 0.001398 | 0 | COL1A1_ITGA11_ITGB1 | Col1a1 - (Itga11+Itgb1) | COLLAGEN | ECM-Receptor | KEGG: mmu04512 |
| NeuN | CD31 | Col1a1 | ITGA11_ITGB1 | 0.001068 | 0 | COL1A1_ITGA11_ITGB1 | Col1a1 - (Itga11+Itgb1) | COLLAGEN | ECM-Receptor | KEGG: mmu04512 |
| CD31 | CD31 | Col1a2 | ITGA11_ITGB1 | 0.001194 | 0 | COL1A2_ITGA11_ITGB1 | Col1a2 - (Itga11+Itgb1) | COLLAGEN | ECM-Receptor | KEGG: mmu04512 |
| NeuN | CD31 | Col1a2 | ITGA11_ITGB1 | 0.00119 | 0 | COL1A2_ITGA11_ITGB1 | Col1a2 - (Itga11+Itgb1) | COLLAGEN | ECM-Receptor | KEGG: mmu04512 |
| CD31 | CD31 | Col4a1 | ITGA11_ITGB1 | 0.038215 | 0 | COL4A1_ITGA11_ITGB1 | Col4a1 - (Itga11+Itgb1) | COLLAGEN | ECM-Receptor | KEGG: mmu04512 |
| CD31 | NeuN | Col4a1 | ITGA11_ITGB1 | 0.021952 | 0 | COL4A1_ITGA11_ITGB1 | Col4a1 - (Itga11+Itgb1) | COLLAGEN | ECM-Receptor | KEGG: mmu04512 |
| CD31 | CD31 | Col4a2 | ITGA11_ITGB1 | 0.030273 | 0 | COL4A2_ITGA11_ITGB1 | Col4a2 - (Itga11+Itgb1) | COLLAGEN | ECM-Receptor | KEGG: mmu04512 |
| CD31 | NeuN | Col4a2 | ITGA11_ITGB1 | 0.017329 | 0 | COL4A2_ITGA11_ITGB1 | Col4a2 - (Itga11+Itgb1) | COLLAGEN | ECM-Receptor | KEGG: mmu04512 |
| CD31 | CD31 | Col6a1 | ITGA11_ITGB1 | 0.001657 | 0 | COL6A1_ITGA11_ITGB1 | Col6a1 - (Itga11+Itgb1) | COLLAGEN | ECM-Receptor | KEGG: mmu04512 |
| NeuN | CD31 | Col6a1 | ITGA11_ITGB1 | 0.000944 | 0 | COL6A1_ITGA11_ITGB1 | Col6a1 - (Itga11+Itgb1) | COLLAGEN | ECM-Receptor | KEGG: mmu04512 |
| CD31 | CD31 | Col6a2 | ITGA11_ITGB1 | 0.002209 | 0 | COL6A2_ITGA11_ITGB1 | Col6a2 - (Itga11+Itgb1) | COLLAGEN | ECM-Receptor | KEGG: mmu04512 |
| NeuN | CD31 | Col6a2 | ITGA11_ITGB1 | 0.00137 | 0 | COL6A2_ITGA11_ITGB1 | Col6a2 - (Itga11+Itgb1) | COLLAGEN | ECM-Receptor | KEGG: mmu04512 |
| CD31 | CD31 | Col6a4 | ITGA11_ITGB1 | 0.001286 | 0 | COL6A4_ITGA11_ITGB1 | Col6a4 - (Itga11+Itgb1) | COLLAGEN | ECM-Receptor | KEGG: mmu04512 |
| NeuN | CD31 | Col6a4 | ITGA11_ITGB1 | 0.001415 | 0 | COL6A4_ITGA11_ITGB1 | Col6a4 - (Itga11+Itgb1) | COLLAGEN | ECM-Receptor | KEGG: mmu04512 |
| CD31 | CD31 | Col6a6 | ITGA11_ITGB1 | 0.001084 | 0 | COL6A6_ITGA11_ITGB1 | Col6a6 - (Itga11+Itgb1) | COLLAGEN | ECM-Receptor | KEGG: mmu04512 |
| NeuN | CD31 | Col6a6 | ITGA11_ITGB1 | 0.001111 | 0 | COL6A6_ITGA11_ITGB1 | Col6a6 - (Itga11+Itgb1) | COLLAGEN | ECM-Receptor | KEGG: mmu04512 |
| CD31 | CD31 | Col9a2 | ITGA11_ITGB1 | 0.001638 | 0 | COL9A2_ITGA11_ITGB1 | Col9a2 - (Itga11+Itgb1) | COLLAGEN | ECM-Receptor | KEGG: mmu04512 |
| NeuN | CD31 | Col9a2 | ITGA11_ITGB1 | 0.001986 | 0 | COL9A2_ITGA11_ITGB1 | Col9a2 - (Itga11+Itgb1) | COLLAGEN | ECM-Receptor | KEGG: mmu04512 |
| CD31 | CD31 | Col9a3 | ITGA11_ITGB1 | 0.002083 | 0 | COL9A3_ITGA11_ITGB1 | Col9a3 - (Itga11+Itgb1) | COLLAGEN | ECM-Receptor | KEGG: mmu04512 |
| NeuN | CD31 | Col9a3 | ITGA11_ITGB1 | 0.002893 | 0 | COL9A3_ITGA11_ITGB1 | Col9a3 - (Itga11+Itgb1) | COLLAGEN | ECM-Receptor | KEGG: mmu04512 |
| CD31 | CD31 | Fn1 | ITGAV_ITGB1 | 0.014562 | 0 | FN1_ITGAV_ITGB1 | Fn1 - (Itgav+Itgb1) | FN1 | ECM-Receptor | KEGG: mmu04512 |
| CD31 | NeuN | Fn1 | ITGAV_ITGB1 | 0.005839 | 0 | FN1_ITGAV_ITGB1 | Fn1 - (Itgav+Itgb1) | FN1 | ECM-Receptor | KEGG: mmu04512 |
| CD31 | CD31 | Vtn | ITGAV_ITGB1 | 0.008758 | 0 | VTN_ITGAV_ITGB1 | Vtn - (Itgav+Itgb1) | VTN | ECM-Receptor | KEGG: mmu04512 |
| CD31 | NeuN | Vtn | ITGAV_ITGB1 | 0.003499 | 0 | VTN_ITGAV_ITGB1 | Vtn - (Itgav+Itgb1) | VTN | ECM-Receptor | KEGG: mmu04512 |
| CD31 | CD31 | Fn1 | ITGA4_ITGB7 | 0.007435 | 0 | FN1_ITGA4_ITGB7 | Fn1 - (Itga4+Itgb7) | FN1 | ECM-Receptor | KEGG: mmu04512 |
| CD31 | NeuN | Fn1 | ITGA4_ITGB7 | 0.003215 | 0 | FN1_ITGA4_ITGB7 | Fn1 - (Itga4+Itgb7) | FN1 | ECM-Receptor | KEGG: mmu04512 |
| CD31 | CD31 | Fn1 | ITGAV_ITGB3 | 0.005896 | 0 | FN1_ITGAV_ITGB3 | Fn1 - (Itgav+Itgb3) | FN1 | ECM-Receptor | KEGG: mmu04512 |
| CD31 | NeuN | Fn1 | ITGAV_ITGB3 | 0.003494 | 0 | FN1_ITGAV_ITGB3 | Fn1 - (Itgav+Itgb3) | FN1 | ECM-Receptor | KEGG: mmu04512 |
| CD31 | CD31 | Vtn | ITGAV_ITGB3 | 0.003533 | 0 | VTN_ITGAV_ITGB3 | Vtn - (Itgav+Itgb3) | VTN | ECM-Receptor | KEGG: mmu04512 |
| CD31 | NeuN | Vtn | ITGAV_ITGB3 | 0.002092 | 0 | VTN_ITGAV_ITGB3 | Vtn - (Itgav+Itgb3) | VTN | ECM-Receptor | KEGG: mmu04512 |
| CD31 | CD31 | Vwf | ITGAV_ITGB3 | 0.003692 | 0 | VWF_ITGAV_ITGB3 | Vwf - (Itgav+Itgb3) | VWF | ECM-Receptor | KEGG: mmu04512 |
| CD31 | NeuN | Vwf | ITGAV_ITGB3 | 0.002186 | 0 | VWF_ITGAV_ITGB3 | Vwf - (Itgav+Itgb3) | VWF | ECM-Receptor | KEGG: mmu04512 |
| CD31 | CD31 | Tnc | ITGAV_ITGB3 | 0.00151 | 0 | TNC_ITGAV_ITGB3 | Tnc - (Itgav+Itgb3) | TENASCIN | ECM-Receptor | KEGG: mmu04512 |
| CD31 | NeuN | Tnc | ITGAV_ITGB3 | 0.000893 | 0 | TNC_ITGAV_ITGB3 | Tnc - (Itgav+Itgb3) | TENASCIN | ECM-Receptor | KEGG: mmu04512 |
| CD31 | CD31 | Tnr | ITGAV_ITGB3 | 0.000504 | 0 | TNR_ITGAV_ITGB3 | Tnr - (Itgav+Itgb3) | TENASCIN | ECM-Receptor | KEGG: mmu04512 |
| NeuN | CD31 | Tnr | ITGAV_ITGB3 | 0.000582 | 0 | TNR_ITGAV_ITGB3 | Tnr - (Itgav+Itgb3) | TENASCIN | ECM-Receptor | KEGG: mmu04512 |

| | | | | | | | | | | |
|---|---|---|---|---|---|---|---|---|---|---|
| CD31 | CD31 | Thbs1 | ITGAV_ITGB3 | 0.001368 | 0 | THBS1_ITGAV_ITGB3 | Thbs1 - (Itgav+Itgb3) | THBS | ECM-Receptor | KEGG: mmu04512 |
| NeuN | CD31 | Thbs1 | ITGAV_ITGB3 | 0.001121 | 0 | THBS1_ITGAV_ITGB3 | Thbs1 - (Itgav+Itgb3) | THBS | ECM-Receptor | KEGG: mmu04512 |
| NeuN | CD31 | Thbs3 | ITGAV_ITGB3 | 0.002959 | 0 | THBS3_ITGAV_ITGB3 | Thbs3 - (Itgav+Itgb3) | THBS | ECM-Receptor | KEGG: mmu04512 |
| NeuN | NeuN | Thbs3 | ITGAV_ITGB3 | 0.001751 | 0 | THBS3_ITGAV_ITGB3 | Thbs3 - (Itgav+Itgb3) | THBS | ECM-Receptor | KEGG: mmu04512 |
| CD31 | CD31 | Thbs4 | ITGAV_ITGB3 | 0.00077 | 0 | THBS4_ITGAV_ITGB3 | Thbs4 - (Itgav+Itgb3) | THBS | ECM-Receptor | KEGG: mmu04512 |
| NeuN | CD31 | Thbs4 | ITGAV_ITGB3 | 0.000806 | 0 | THBS4_ITGAV_ITGB3 | Thbs4 - (Itgav+Itgb3) | THBS | ECM-Receptor | KEGG: mmu04512 |
| CD31 | CD31 | Fn1 | ITGA2B_ITGB3 | 0.004063 | 0 | FN1_ITGA2B_ITGB3 | Fn1 - (Itga2b+Itgb3) | FN1 | ECM-Receptor | KEGG: mmu04512 |
| CD31 | NeuN | Fn1 | ITGA2B_ITGB3 | 0.002804 | 0 | FN1_ITGA2B_ITGB3 | Fn1 - (Itga2b+Itgb3) | FN1 | ECM-Receptor | KEGG: mmu04512 |
| CD31 | CD31 | Vtn | ITGA2B_ITGB3 | 0.002433 | 0 | VTN_ITGA2B_ITGB3 | Vtn - (Itga2b+Itgb3) | VTN | ECM-Receptor | KEGG: mmu04512 |
| CD31 | NeuN | Vtn | ITGA2B_ITGB3 | 0.001678 | 0 | VTN_ITGA2B_ITGB3 | Vtn - (Itga2b+Itgb3) | VTN | ECM-Receptor | KEGG: mmu04512 |
| CD31 | CD31 | Vwf | ITGA2B_ITGB3 | 0.002543 | 0 | VWF_ITGA2B_ITGB3 | Vwf - (Itga2b+Itgb3) | VWF | ECM-Receptor | KEGG: mmu04512 |
| CD31 | NeuN | Vwf | ITGA2B_ITGB3 | 0.001754 | 0 | VWF_ITGA2B_ITGB3 | Vwf - (Itga2b+Itgb3) | VWF | ECM-Receptor | KEGG: mmu04512 |
| CD31 | CD31 | Tnc | ITGA2B_ITGB3 | 0.001039 | 0 | TNC_ITGA2B_ITGB3 | Tnc - (Itga2b+Itgb3) | TENASCIN | ECM-Receptor | KEGG: mmu04512 |
| CD31 | NeuN | Tnc | ITGA2B_ITGB3 | 0.000716 | 0 | TNC_ITGA2B_ITGB3 | Tnc - (Itga2b+Itgb3) | TENASCIN | ECM-Receptor | KEGG: mmu04512 |
| CD31 | CD31 | Tnr | ITGA2B_ITGB3 | 0.000347 | 0 | TNR_ITGA2B_ITGB3 | Tnr - (Itga2b+Itgb3) | TENASCIN | ECM-Receptor | KEGG: mmu04512 |
| NeuN | CD31 | Tnr | ITGA2B_ITGB3 | 0.0004 | 0 | TNR_ITGA2B_ITGB3 | Tnr - (Itga2b+Itgb3) | TENASCIN | ECM-Receptor | KEGG: mmu04512 |
| CD31 | CD31 | Vtn | ITGAV_ITGB5 | 0.003138 | 0 | VTN_ITGAV_ITGB5 | Vtn - (Itgav+Itgb5) | VTN | ECM-Receptor | KEGG: mmu04512 |
| CD31 | NeuN | Vtn | ITGAV_ITGB5 | 0.001735 | 0 | VTN_ITGAV_ITGB5 | Vtn - (Itgav+Itgb5) | VTN | ECM-Receptor | KEGG: mmu04512 |
| CD31 | CD31 | Vtn | ITGAV_ITGB8 | 0.003572 | 0 | VTN_ITGAV_ITGB8 | Vtn - (Itgav+Itgb8) | VTN | ECM-Receptor | KEGG: mmu04512 |
| CD31 | NeuN | Vtn | ITGAV_ITGB8 | 0.00251 | 0 | VTN_ITGAV_ITGB8 | Vtn - (Itgav+Itgb8) | VTN | ECM-Receptor | KEGG: mmu04512 |
| CD31 | CD31 | Fn1 | ITGAV_ITGB8 | 0.00596 | 0 | FN1_ITGAV_ITGB8 | Fn1 - (Itgav+Itgb8) | FN1 | ECM-Receptor | KEGG: mmu04512 |
| CD31 | NeuN | Fn1 | ITGAV_ITGB8 | 0.004191 | 0 | FN1_ITGAV_ITGB8 | Fn1 - (Itgav+Itgb8) | FN1 | ECM-Receptor | KEGG: mmu04512 |
| CD31 | CD31 | Lama1 | ITGAV_ITGB8 | 0.001179 | 0 | LAMA1_ITGAV_ITGB8 | Lama1 - (Itgav+Itgb8) | LAMININ | ECM-Receptor | KEGG: mmu04512 |
| CD31 | NeuN | Lama1 | ITGAV_ITGB8 | 0.000828 | 0 | LAMA1_ITGAV_ITGB8 | Lama1 - (Itgav+Itgb8) | LAMININ | ECM-Receptor | KEGG: mmu04512 |
| CD31 | CD31 | Lama4 | ITGAV_ITGB8 | 0.007059 | 0 | LAMA4_ITGAV_ITGB8 | Lama4 - (Itgav+Itgb8) | LAMININ | ECM-Receptor | KEGG: mmu04512 |
| CD31 | NeuN | Lama4 | ITGAV_ITGB8 | 0.004966 | 0 | LAMA4_ITGAV_ITGB8 | Lama4 - (Itgav+Itgb8) | LAMININ | ECM-Receptor | KEGG: mmu04512 |
| CD31 | CD31 | Lama5 | ITGAV_ITGB8 | 0.0019 | 0 | LAMA5_ITGAV_ITGB8 | Lama5 - (Itgav+Itgb8) | LAMININ | ECM-Receptor | KEGG: mmu04512 |
| CD31 | NeuN | Lama5 | ITGAV_ITGB8 | 0.001334 | 0 | LAMA5_ITGAV_ITGB8 | Lama5 - (Itgav+Itgb8) | LAMININ | ECM-Receptor | KEGG: mmu04512 |
| CD31 | CD31 | Lamb1 | ITGAV_ITGB8 | 0.006634 | 0 | LAMB1_ITGAV_ITGB8 | Lamb1 - (Itgav+Itgb8) | LAMININ | ECM-Receptor | KEGG: mmu04512 |
| CD31 | NeuN | Lamb1 | ITGAV_ITGB8 | 0.004666 | 0 | LAMB1_ITGAV_ITGB8 | Lamb1 - (Itgav+Itgb8) | LAMININ | ECM-Receptor | KEGG: mmu04512 |
| CD31 | CD31 | Lamb2 | ITGAV_ITGB8 | 0.00177 | 0 | LAMB2_ITGAV_ITGB8 | Lamb2 - (Itgav+Itgb8) | LAMININ | ECM-Receptor | KEGG: mmu04512 |
| CD31 | NeuN | Lamb2 | ITGAV_ITGB8 | 0.001243 | 0 | LAMB2_ITGAV_ITGB8 | Lamb2 - (Itgav+Itgb8) | LAMININ | ECM-Receptor | KEGG: mmu04512 |
| CD31 | CD31 | Lamc1 | ITGAV_ITGB8 | 0.003429 | 0 | LAMC1_ITGAV_ITGB8 | Lamc1 - (Itgav+Itgb8) | LAMININ | ECM-Receptor | KEGG: mmu04512 |
| CD31 | NeuN | Lamc1 | ITGAV_ITGB8 | 0.00241 | 0 | LAMC1_ITGAV_ITGB8 | Lamc1 - (Itgav+Itgb8) | LAMININ | ECM-Receptor | KEGG: mmu04512 |
| CD31 | CD31 | Lamc2 | ITGAV_ITGB8 | 0.001168 | 0 | LAMC2_ITGAV_ITGB8 | Lamc2 - (Itgav+Itgb8) | LAMININ | ECM-Receptor | KEGG: mmu04512 |
| CD31 | NeuN | Lamc2 | ITGAV_ITGB8 | 0.00082 | 0 | LAMC2_ITGAV_ITGB8 | Lamc2 - (Itgav+Itgb8) | LAMININ | ECM-Receptor | KEGG: mmu04512 |
| CD31 | CD31 | Lamc3 | ITGAV_ITGB8 | 0.002146 | 0 | LAMC3_ITGAV_ITGB8 | Lamc3 - (Itgav+Itgb8) | LAMININ | ECM-Receptor | KEGG: mmu04512 |
| CD31 | NeuN | Lamc3 | ITGAV_ITGB8 | 0.001507 | 0 | LAMC3_ITGAV_ITGB8 | Lamc3 - (Itgav+Itgb8) | LAMININ | ECM-Receptor | KEGG: mmu04512 |
| CD31 | CD31 | Col1a1 | ITGAV_ITGB8 | 0.000845 | 0 | COL1A1_ITGAV_ITGB8 | Col1a1 - (Itgav+Itgb8) | COLLAGEN | ECM-Receptor | KEGG: mmu04512 |
| NeuN | CD31 | Col1a1 | ITGAV_ITGB8 | 0.000646 | 0 | COL1A1_ITGAV_ITGB8 | Col1a1 - (Itgav+Itgb8) | COLLAGEN | ECM-Receptor | KEGG: mmu04512 |
| CD31 | CD31 | Col1a2 | ITGAV_ITGB8 | 0.000722 | 0 | COL1A2_ITGAV_ITGB8 | Col1a2 - (Itgav+Itgb8) | COLLAGEN | ECM-Receptor | KEGG: mmu04512 |
| NeuN | CD31 | Col1a2 | ITGAV_ITGB8 | 0.00072 | 0 | COL1A2_ITGAV_ITGB8 | Col1a2 - (Itgav+Itgb8) | COLLAGEN | ECM-Receptor | KEGG: mmu04512 |
| CD31 | CD31 | Col4a1 | ITGAV_ITGB8 | 0.023453 | 0 | COL4A1_ITGAV_ITGB8 | Col4a1 - (Itgav+Itgb8) | COLLAGEN | ECM-Receptor | KEGG: mmu04512 |
| CD31 | NeuN | Col4a1 | ITGAV_ITGB8 | 0.01658 | 0 | COL4A1_ITGAV_ITGB8 | Col4a1 - (Itgav+Itgb8) | COLLAGEN | ECM-Receptor | KEGG: mmu04512 |
| CD31 | CD31 | Col4a2 | ITGAV_ITGB8 | 0.01852 | 0 | COL4A2_ITGAV_ITGB8 | Col4a2 - (Itgav+Itgb8) | COLLAGEN | ECM-Receptor | KEGG: mmu04512 |
| CD31 | NeuN | Col4a2 | ITGAV_ITGB8 | 0.013074 | 0 | COL4A2_ITGAV_ITGB8 | Col4a2 - (Itgav+Itgb8) | COLLAGEN | ECM-Receptor | KEGG: mmu04512 |
| CD31 | CD31 | Col6a1 | ITGAV_ITGB8 | 0.001002 | 0 | COL6A1_ITGAV_ITGB8 | Col6a1 - (Itgav+Itgb8) | COLLAGEN | ECM-Receptor | KEGG: mmu04512 |
| CD31 | NeuN | Col6a1 | ITGAV_ITGB8 | 0.000704 | 0 | COL6A1_ITGAV_ITGB8 | Col6a1 - (Itgav+Itgb8) | COLLAGEN | ECM-Receptor | KEGG: mmu04512 |
| CD31 | CD31 | Col6a2 | ITGAV_ITGB8 | 0.001336 | 0 | COL6A2_ITGAV_ITGB8 | Col6a2 - (Itgav+Itgb8) | COLLAGEN | ECM-Receptor | KEGG: mmu04512 |
| CD31 | NeuN | Col6a2 | ITGAV_ITGB8 | 0.000939 | 0 | COL6A2_ITGAV_ITGB8 | Col6a2 - (Itgav+Itgb8) | COLLAGEN | ECM-Receptor | KEGG: mmu04512 |
| CD31 | CD31 | Col6a4 | ITGAV_ITGB8 | 0.000778 | 0 | COL6A4_ITGAV_ITGB8 | Col6a4 - (Itgav+Itgb8) | COLLAGEN | ECM-Receptor | KEGG: mmu04512 |
| NeuN | CD31 | Col6a4 | ITGAV_ITGB8 | 0.000856 | 0 | COL6A4_ITGAV_ITGB8 | Col6a4 - (Itgav+Itgb8) | COLLAGEN | ECM-Receptor | KEGG: mmu04512 |
| CD31 | CD31 | Col6a6 | ITGAV_ITGB8 | 0.000655 | 0 | COL6A6_ITGAV_ITGB8 | Col6a6 - (Itgav+Itgb8) | COLLAGEN | ECM-Receptor | KEGG: mmu04512 |
| NeuN | CD31 | Col6a6 | ITGAV_ITGB8 | 0.000672 | 0 | COL6A6_ITGAV_ITGB8 | Col6a6 - (Itgav+Itgb8) | COLLAGEN | ECM-Receptor | KEGG: mmu04512 |
| CD31 | CD31 | Col9a2 | ITGAV_ITGB8 | 0.000991 | 0 | COL9A2_ITGAV_ITGB8 | Col9a2 - (Itgav+Itgb8) | COLLAGEN | ECM-Receptor | KEGG: mmu04512 |
| NeuN | CD31 | Col9a2 | ITGAV_ITGB8 | 0.001201 | 0 | COL9A2_ITGAV_ITGB8 | Col9a2 - (Itgav+Itgb8) | COLLAGEN | ECM-Receptor | KEGG: mmu04512 |
| CD31 | CD31 | Col9a3 | ITGAV_ITGB8 | 0.00126 | 0 | COL9A3_ITGAV_ITGB8 | Col9a3 - (Itgav+Itgb8) | COLLAGEN | ECM-Receptor | KEGG: mmu04512 |
| NeuN | CD31 | Col9a3 | ITGAV_ITGB8 | 0.001751 | 0 | COL9A3_ITGAV_ITGB8 | Col9a3 - (Itgav+Itgb8) | COLLAGEN | ECM-Receptor | KEGG: mmu04512 |
| CD31 | CD31 | Col1a1 | Sdc4 | 0.000454 | 0 | COL1A1_SDC4 | Col1a1 - Sdc4 | COLLAGEN | ECM-Receptor | KEGG: mmu04512 |

| | | | | | | | | | | |
|---|---|---|---|---|---|---|---|---|---|---|
| CD31 | NeuN | Col1a1 | Sdc4 | 0.000429 | 0 | COL1A1_SDC4 | Col1a1 - Sdc4 | COLLAGEN | ECM-Receptor | KEGG: mmu04512 |
| CD31 | CD31 | Col1a2 | Sdc4 | 0.000388 | 0 | COL1A2_SDC4 | Col1a2 - Sdc4 | COLLAGEN | ECM-Receptor | KEGG: mmu04512 |
| NeuN | CD31 | Col1a2 | Sdc4 | 0.000387 | 0 | COL1A2_SDC4 | Col1a2 - Sdc4 | COLLAGEN | ECM-Receptor | KEGG: mmu04512 |
| CD31 | CD31 | Col4a1 | Sdc4 | 0.012738 | 0 | COL4A1_SDC4 | Col4a1 - Sdc4 | COLLAGEN | ECM-Receptor | KEGG: mmu04512 |
| CD31 | NeuN | Col4a1 | Sdc4 | 0.012031 | 0 | COL4A1_SDC4 | Col4a1 - Sdc4 | COLLAGEN | ECM-Receptor | KEGG: mmu04512 |
| CD31 | CD31 | Col4a2 | Sdc4 | 0.010035 | 0 | COL4A2_SDC4 | Col4a2 - Sdc4 | COLLAGEN | ECM-Receptor | KEGG: mmu04512 |
| CD31 | NeuN | Col4a2 | Sdc4 | 0.009477 | 0 | COL4A2_SDC4 | Col4a2 - Sdc4 | COLLAGEN | ECM-Receptor | KEGG: mmu04512 |
| CD31 | CD31 | Col6a1 | Sdc4 | 0.000539 | 0 | COL6A1_SDC4 | Col6a1 - Sdc4 | COLLAGEN | ECM-Receptor | KEGG: mmu04512 |
| CD31 | NeuN | Col6a1 | Sdc4 | 0.000508 | 0 | COL6A1_SDC4 | Col6a1 - Sdc4 | COLLAGEN | ECM-Receptor | KEGG: mmu04512 |
| CD31 | CD31 | Col6a2 | Sdc4 | 0.000718 | 0 | COL6A2_SDC4 | Col6a2 - Sdc4 | COLLAGEN | ECM-Receptor | KEGG: mmu04512 |
| CD31 | NeuN | Col6a2 | Sdc4 | 0.000678 | 0 | COL6A2_SDC4 | Col6a2 - Sdc4 | COLLAGEN | ECM-Receptor | KEGG: mmu04512 |
| NeuN | CD31 | Col6a4 | Sdc4 | 0.00046 | 0 | COL6A4_SDC4 | Col6a4 - Sdc4 | COLLAGEN | ECM-Receptor | KEGG: mmu04512 |
| NeuN | NeuN | Col6a4 | Sdc4 | 0.000434 | 0 | COL6A4_SDC4 | Col6a4 - Sdc4 | COLLAGEN | ECM-Receptor | KEGG: mmu04512 |
| CD31 | CD31 | Col6a6 | Sdc4 | 0.000352 | 0 | COL6A6_SDC4 | Col6a6 - Sdc4 | COLLAGEN | ECM-Receptor | KEGG: mmu04512 |
| NeuN | CD31 | Col6a6 | Sdc4 | 0.000361 | 0 | COL6A6_SDC4 | Col6a6 - Sdc4 | COLLAGEN | ECM-Receptor | KEGG: mmu04512 |
| NeuN | CD31 | Col9a2 | Sdc4 | 0.000646 | 0 | COL9A2_SDC4 | Col9a2 - Sdc4 | COLLAGEN | ECM-Receptor | KEGG: mmu04512 |
| NeuN | NeuN | Col9a2 | Sdc4 | 0.00061 | 0 | COL9A2_SDC4 | Col9a2 - Sdc4 | COLLAGEN | ECM-Receptor | KEGG: mmu04512 |
| NeuN | CD31 | Col9a3 | Sdc4 | 0.000941 | 0 | COL9A3_SDC4 | Col9a3 - Sdc4 | COLLAGEN | ECM-Receptor | KEGG: mmu04512 |
| NeuN | NeuN | Col9a3 | Sdc4 | 0.000888 | 0 | COL9A3_SDC4 | Col9a3 - Sdc4 | COLLAGEN | ECM-Receptor | KEGG: mmu04512 |
| CD31 | CD31 | Fn1 | Sdc4 | 0.003211 | 0 | FN1_SDC4 | Fn1 - Sdc4 | FN1 | ECM-Receptor | KEGG: mmu04512 |
| CD31 | NeuN | Fn1 | Sdc4 | 0.003031 | 0 | FN1_SDC4 | Fn1 - Sdc4 | FN1 | ECM-Receptor | KEGG: mmu04512 |
| CD31 | CD31 | Tnc | Sdc4 | 0.00082 | 0 | TNC_SDC4 | Tnc - Sdc4 | TENASCIN | ECM-Receptor | KEGG: mmu04512 |
| CD31 | NeuN | Tnc | Sdc4 | 0.000774 | 0 | TNC_SDC4 | Tnc - Sdc4 | TENASCIN | ECM-Receptor | KEGG: mmu04512 |
| NeuN | CD31 | Tnr | Sdc4 | 0.000316 | 0 | TNR_SDC4 | Tnr - Sdc4 | TENASCIN | ECM-Receptor | KEGG: mmu04512 |
| NeuN | NeuN | Tnr | Sdc4 | 0.000298 | 0 | TNR_SDC4 | Tnr - Sdc4 | TENASCIN | ECM-Receptor | KEGG: mmu04512 |
| CD31 | CD31 | Thbs1 | Sdc4 | 0.000743 | 0 | THBS1_SDC4 | Thbs1 - Sdc4 | THBS | ECM-Receptor | KEGG: mmu04512 |
| CD31 | NeuN | Thbs1 | Sdc4 | 0.000702 | 0 | THBS1_SDC4 | Thbs1 - Sdc4 | THBS | ECM-Receptor | KEGG: mmu04512 |
| NeuN | CD31 | Thbs3 | Sdc4 | 0.001609 | 0 | THBS3_SDC4 | Thbs3 - Sdc4 | THBS | ECM-Receptor | KEGG: mmu04512 |
| NeuN | NeuN | Thbs3 | Sdc4 | 0.001519 | 0 | THBS3_SDC4 | Thbs3 - Sdc4 | THBS | ECM-Receptor | KEGG: mmu04512 |
| CD31 | CD31 | Thbs4 | Sdc4 | 0.000418 | 0 | THBS4_SDC4 | Thbs4 - Sdc4 | THBS | ECM-Receptor | KEGG: mmu04512 |
| NeuN | CD31 | Thbs4 | Sdc4 | 0.000438 | 0 | THBS4_SDC4 | Thbs4 - Sdc4 | THBS | ECM-Receptor | KEGG: mmu04512 |
| CD31 | NeuN | Lama1 | Sv2a | 0.003331 | 0 | LAMA1_SV2A | Lama1 - Sv2a | LAMININ | ECM-Receptor | KEGG: mmu04512 |
| NeuN | NeuN | Lama1 | Sv2a | 0.002339 | 0 | LAMA1_SV2A | Lama1 - Sv2a | LAMININ | ECM-Receptor | KEGG: mmu04512 |
| CD31 | CD31 | Lama4 | Sv2a | 0.008838 | 0 | LAMA4_SV2A | Lama4 - Sv2a | LAMININ | ECM-Receptor | KEGG: mmu04512 |
| CD31 | NeuN | Lama4 | Sv2a | 0.019726 | 0 | LAMA4_SV2A | Lama4 - Sv2a | LAMININ | ECM-Receptor | KEGG: mmu04512 |
| CD31 | CD31 | Lama5 | Sv2a | 0.002382 | 0 | LAMA5_SV2A | Lama5 - Sv2a | LAMININ | ECM-Receptor | KEGG: mmu04512 |
| CD31 | NeuN | Lama5 | Sv2a | 0.005359 | 0 | LAMA5_SV2A | Lama5 - Sv2a | LAMININ | ECM-Receptor | KEGG: mmu04512 |
| CD31 | CD31 | Lamb1 | Sv2a | 0.008307 | 0 | LAMB1_SV2A | Lamb1 - Sv2a | LAMININ | ECM-Receptor | KEGG: mmu04512 |
| CD31 | NeuN | Lamb1 | Sv2a | 0.018554 | 0 | LAMB1_SV2A | Lamb1 - Sv2a | LAMININ | ECM-Receptor | KEGG: mmu04512 |
| CD31 | CD31 | Lamb2 | Sv2a | 0.00222 | 0 | LAMB2_SV2A | Lamb2 - Sv2a | LAMININ | ECM-Receptor | KEGG: mmu04512 |
| CD31 | NeuN | Lamb2 | Sv2a | 0.004995 | 0 | LAMB2_SV2A | Lamb2 - Sv2a | LAMININ | ECM-Receptor | KEGG: mmu04512 |
| CD31 | CD31 | Lamc1 | Sv2a | 0.004298 | 0 | LAMC1_SV2A | Lamc1 - Sv2a | LAMININ | ECM-Receptor | KEGG: mmu04512 |
| CD31 | NeuN | Lamc1 | Sv2a | 0.009647 | 0 | LAMC1_SV2A | Lamc1 - Sv2a | LAMININ | ECM-Receptor | KEGG: mmu04512 |
| CD31 | NeuN | Lamc2 | Sv2a | 0.003299 | 0 | LAMC2_SV2A | Lamc2 - Sv2a | LAMININ | ECM-Receptor | KEGG: mmu04512 |
| NeuN | NeuN | Lamc2 | Sv2a | 0.001982 | 0 | LAMC2_SV2A | Lamc2 - Sv2a | LAMININ | ECM-Receptor | KEGG: mmu04512 |
| CD31 | CD31 | Lamc3 | Sv2a | 0.00269 | 0 | LAMC3_SV2A | Lamc3 - Sv2a | LAMININ | ECM-Receptor | KEGG: mmu04512 |
| CD31 | NeuN | Lamc3 | Sv2a | 0.006051 | 0 | LAMC3_SV2A | Lamc3 - Sv2a | LAMININ | ECM-Receptor | KEGG: mmu04512 |
| CD31 | NeuN | Lama1 | Sv2b | 0.001412 | 0 | LAMA1_SV2B | Lama1 - Sv2b | LAMININ | ECM-Receptor | KEGG: mmu04512 |
| NeuN | NeuN | Lama1 | Sv2b | 0.000991 | 0 | LAMA1_SV2B | Lama1 - Sv2b | LAMININ | ECM-Receptor | KEGG: mmu04512 |
| CD31 | CD31 | Lama4 | Sv2b | 0.005712 | 0 | LAMA4_SV2B | Lama4 - Sv2b | LAMININ | ECM-Receptor | KEGG: mmu04512 |
| CD31 | NeuN | Lama4 | Sv2b | 0.008442 | 0 | LAMA4_SV2B | Lama4 - Sv2b | LAMININ | ECM-Receptor | KEGG: mmu04512 |
| CD31 | CD31 | Lama5 | Sv2b | 0.001536 | 0 | LAMA5_SV2B | Lama5 - Sv2b | LAMININ | ECM-Receptor | KEGG: mmu04512 |
| CD31 | NeuN | Lama5 | Sv2b | 0.002274 | 0 | LAMA5_SV2B | Lama5 - Sv2b | LAMININ | ECM-Receptor | KEGG: mmu04512 |
| CD31 | CD31 | Lamb1 | Sv2b | 0.005368 | 0 | LAMB1_SV2B | Lamb1 - Sv2b | LAMININ | ECM-Receptor | KEGG: mmu04512 |
| CD31 | NeuN | Lamb1 | Sv2b | 0.007935 | 0 | LAMB1_SV2B | Lamb1 - Sv2b | LAMININ | ECM-Receptor | KEGG: mmu04512 |
| CD31 | CD31 | Lamb2 | Sv2b | 0.001431 | 0 | LAMB2_SV2B | Lamb2 - Sv2b | LAMININ | ECM-Receptor | KEGG: mmu04512 |
| CD31 | NeuN | Lamb2 | Sv2b | 0.00212 | 0 | LAMB2_SV2B | Lamb2 - Sv2b | LAMININ | ECM-Receptor | KEGG: mmu04512 |
| CD31 | CD31 | Lamc1 | Sv2b | 0.002773 | 0 | LAMC1_SV2B | Lamc1 - Sv2b | LAMININ | ECM-Receptor | KEGG: mmu04512 |
| CD31 | NeuN | Lamc1 | Sv2b | 0.004105 | 0 | LAMC1_SV2B | Lamc1 - Sv2b | LAMININ | ECM-Receptor | KEGG: mmu04512 |

| | | | | | | | | | | |
|---|---|---|---|---|---|---|---|---|---|---|
| CD31 | CD31 | Lamc2 | Sv2b | 0.000944 | 0 | LAMC2_SV2B | Lamc2 - Sv2b | LAMININ | ECM-Receptor | KEGG: mmu04512 |
| CD31 | NeuN | Lamc2 | Sv2b | 0.001399 | 0 | LAMC2_SV2B | Lamc2 - Sv2b | LAMININ | ECM-Receptor | KEGG: mmu04512 |
| CD31 | CD31 | Lamc3 | Sv2b | 0.001735 | 0 | LAMC3_SV2B | Lamc3 - Sv2b | LAMININ | ECM-Receptor | KEGG: mmu04512 |
| CD31 | NeuN | Lamc3 | Sv2b | 0.002569 | 0 | LAMC3_SV2B | Lamc3 - Sv2b | LAMININ | ECM-Receptor | KEGG: mmu04512 |
| CD31 | CD31 | Lama1 | Sv2c | 0.001126 | 0 | LAMA1_SV2C | Lama1 - Sv2c | LAMININ | ECM-Receptor | KEGG: mmu04512 |
| CD31 | NeuN | Lama1 | Sv2c | 0.000915 | 0 | LAMA1_SV2C | Lama1 - Sv2c | LAMININ | ECM-Receptor | KEGG: mmu04512 |
| CD31 | CD31 | Lama4 | Sv2c | 0.006744 | 0 | LAMA4_SV2C | Lama4 - Sv2c | LAMININ | ECM-Receptor | KEGG: mmu04512 |
| CD31 | NeuN | Lama4 | Sv2c | 0.005486 | 0 | LAMA4_SV2C | Lama4 - Sv2c | LAMININ | ECM-Receptor | KEGG: mmu04512 |
| CD31 | CD31 | Lama5 | Sv2c | 0.001815 | 0 | LAMA5_SV2C | Lama5 - Sv2c | LAMININ | ECM-Receptor | KEGG: mmu04512 |
| CD31 | NeuN | Lama5 | Sv2c | 0.001475 | 0 | LAMA5_SV2C | Lama5 - Sv2c | LAMININ | ECM-Receptor | KEGG: mmu04512 |
| CD31 | CD31 | Lamb1 | Sv2c | 0.006338 | 0 | LAMB1_SV2C | Lamb1 - Sv2c | LAMININ | ECM-Receptor | KEGG: mmu04512 |
| CD31 | NeuN | Lamb1 | Sv2c | 0.005155 | 0 | LAMB1_SV2C | Lamb1 - Sv2c | LAMININ | ECM-Receptor | KEGG: mmu04512 |
| CD31 | CD31 | Lamb2 | Sv2c | 0.001691 | 0 | LAMB2_SV2C | Lamb2 - Sv2c | LAMININ | ECM-Receptor | KEGG: mmu04512 |
| CD31 | NeuN | Lamb2 | Sv2c | 0.001374 | 0 | LAMB2_SV2C | Lamb2 - Sv2c | LAMININ | ECM-Receptor | KEGG: mmu04512 |
| CD31 | CD31 | Lamc1 | Sv2c | 0.003276 | 0 | LAMC1_SV2C | Lamc1 - Sv2c | LAMININ | ECM-Receptor | KEGG: mmu04512 |
| CD31 | NeuN | Lamc1 | Sv2c | 0.002663 | 0 | LAMC1_SV2C | Lamc1 - Sv2c | LAMININ | ECM-Receptor | KEGG: mmu04512 |
| CD31 | CD31 | Lamc2 | Sv2c | 0.001116 | 0 | LAMC2_SV2C | Lamc2 - Sv2c | LAMININ | ECM-Receptor | KEGG: mmu04512 |
| CD31 | NeuN | Lamc2 | Sv2c | 0.000907 | 0 | LAMC2_SV2C | Lamc2 - Sv2c | LAMININ | ECM-Receptor | KEGG: mmu04512 |
| CD31 | CD31 | Lamc3 | Sv2c | 0.00205 | 0 | LAMC3_SV2C | Lamc3 - Sv2c | LAMININ | ECM-Receptor | KEGG: mmu04512 |
| CD31 | NeuN | Lamc3 | Sv2c | 0.001666 | 0 | LAMC3_SV2C | Lamc3 - Sv2c | LAMININ | ECM-Receptor | KEGG: mmu04512 |
| CD31 | CD31 | Thbs1 | Cd36 | 0.001037 | 0 | THBS1_CD36 | Thbs1 - Cd36 | THBS | ECM-Receptor | KEGG: mmu04512 |
| NeuN | CD31 | Thbs1 | Cd36 | 0.00085 | 0 | THBS1_CD36 | Thbs1 - Cd36 | THBS | ECM-Receptor | KEGG: mmu04512 |
| CD31 | CD31 | Thbs3 | Cd36 | 0.001257 | 0 | THBS3_CD36 | Thbs3 - Cd36 | THBS | ECM-Receptor | KEGG: mmu04512 |
| NeuN | CD31 | Thbs3 | Cd36 | 0.002244 | 0 | THBS3_CD36 | Thbs3 - Cd36 | THBS | ECM-Receptor | KEGG: mmu04512 |
| CD31 | CD31 | Thbs4 | Cd36 | 0.000583 | 0 | THBS4_CD36 | Thbs4 - Cd36 | THBS | ECM-Receptor | KEGG: mmu04512 |
| NeuN | CD31 | Thbs4 | Cd36 | 0.000611 | 0 | THBS4_CD36 | Thbs4 - Cd36 | THBS | ECM-Receptor | KEGG: mmu04512 |
| CD31 | NeuN | Thbs1 | Cd47 | 0.001786 | 0 | THBS1_CD47 | Thbs1 - Cd47 | THBS | ECM-Receptor | KEGG: mmu04512 |
| NeuN | NeuN | Thbs1 | Cd47 | 0.001463 | 0 | THBS1_CD47 | Thbs1 - Cd47 | THBS | ECM-Receptor | KEGG: mmu04512 |
| CD31 | NeuN | Thbs3 | Cd47 | 0.002165 | 0 | THBS3_CD47 | Thbs3 - Cd47 | THBS | ECM-Receptor | KEGG: mmu04512 |
| NeuN | NeuN | Thbs3 | Cd47 | 0.003862 | 0 | THBS3_CD47 | Thbs3 - Cd47 | THBS | ECM-Receptor | KEGG: mmu04512 |
| CD31 | NeuN | Thbs4 | Cd47 | 0.001005 | 0 | THBS4_CD47 | Thbs4 - Cd47 | THBS | ECM-Receptor | KEGG: mmu04512 |
| NeuN | NeuN | Thbs4 | Cd47 | 0.001053 | 0 | THBS4_CD47 | Thbs4 - Cd47 | THBS | ECM-Receptor | KEGG: mmu04512 |
| CD31 | CD31 | Agrn | Dag1 | 0.00772 | 0 | AGRN_DAG1 | Agrn - Dag1 | AGRN | ECM-Receptor | KEGG: mmu04512 |
| CD31 | NeuN | Agrn | Dag1 | 0.007391 | 0 | AGRN_DAG1 | Agrn - Dag1 | AGRN | ECM-Receptor | KEGG: mmu04512 |
| CD31 | CD31 | Hspg2 | Dag1 | 0.002337 | 0 | HSPG2_DAG1 | Hspg2 - Dag1 | HSPG | ECM-Receptor | KEGG: mmu04512 |
| CD31 | NeuN | Hspg2 | Dag1 | 0.002237 | 0 | HSPG2_DAG1 | Hspg2 - Dag1 | HSPG | ECM-Receptor | KEGG: mmu04512 |
| CD31 | CD31 | Lama1 | Dag1 | 0.001037 | 0 | LAMA1_DAG1 | Lama1 - Dag1 | LAMININ | ECM-Receptor | KEGG: mmu04512 |
| CD31 | NeuN | Lama1 | Dag1 | 0.000992 | 0 | LAMA1_DAG1 | Lama1 - Dag1 | LAMININ | ECM-Receptor | KEGG: mmu04512 |
| CD31 | CD31 | Lama4 | Dag1 | 0.006211 | 0 | LAMA4_DAG1 | Lama4 - Dag1 | LAMININ | ECM-Receptor | KEGG: mmu04512 |
| CD31 | NeuN | Lama4 | Dag1 | 0.005947 | 0 | LAMA4_DAG1 | Lama4 - Dag1 | LAMININ | ECM-Receptor | KEGG: mmu04512 |
| CD31 | CD31 | Lama5 | Dag1 | 0.001671 | 0 | LAMA5_DAG1 | Lama5 - Dag1 | LAMININ | ECM-Receptor | KEGG: mmu04512 |
| CD31 | NeuN | Lama5 | Dag1 | 0.001599 | 0 | LAMA5_DAG1 | Lama5 - Dag1 | LAMININ | ECM-Receptor | KEGG: mmu04512 |
| CD31 | CD31 | Lamb1 | Dag1 | 0.005837 | 0 | LAMB1_DAG1 | Lamb1 - Dag1 | LAMININ | ECM-Receptor | KEGG: mmu04512 |
| CD31 | NeuN | Lamb1 | Dag1 | 0.005588 | 0 | LAMB1_DAG1 | Lamb1 - Dag1 | LAMININ | ECM-Receptor | KEGG: mmu04512 |
| CD31 | CD31 | Lamb2 | Dag1 | 0.001557 | 0 | LAMB2_DAG1 | Lamb2 - Dag1 | LAMININ | ECM-Receptor | KEGG: mmu04512 |
| CD31 | NeuN | Lamb2 | Dag1 | 0.00149 | 0 | LAMB2_DAG1 | Lamb2 - Dag1 | LAMININ | ECM-Receptor | KEGG: mmu04512 |
| CD31 | CD31 | Lamc1 | Dag1 | 0.003016 | 0 | LAMC1_DAG1 | Lamc1 - Dag1 | LAMININ | ECM-Receptor | KEGG: mmu04512 |
| CD31 | NeuN | Lamc1 | Dag1 | 0.002887 | 0 | LAMC1_DAG1 | Lamc1 - Dag1 | LAMININ | ECM-Receptor | KEGG: mmu04512 |
| CD31 | CD31 | Lamc2 | Dag1 | 0.001027 | 0 | LAMC2_DAG1 | Lamc2 - Dag1 | LAMININ | ECM-Receptor | KEGG: mmu04512 |
| CD31 | NeuN | Lamc2 | Dag1 | 0.000983 | 0 | LAMC2_DAG1 | Lamc2 - Dag1 | LAMININ | ECM-Receptor | KEGG: mmu04512 |
| CD31 | CD31 | Lamc3 | Dag1 | 0.001887 | 0 | LAMC3_DAG1 | Lamc3 - Dag1 | LAMININ | ECM-Receptor | KEGG: mmu04512 |
| CD31 | NeuN | Lamc3 | Dag1 | 0.001806 | 0 | LAMC3_DAG1 | Lamc3 - Dag1 | LAMININ | ECM-Receptor | KEGG: mmu04512 |
| NeuN | CD31 | Reln | Lrp8 | 0.008715 | 0 | RELN_LRP8 | Reln - Lrp8 | RELN | ECM-Receptor | PMID: 14578885 |
| NeuN | NeuN | Reln | Lrp8 | 0.00442 | 0 | RELN_LRP8 | Reln - Lrp8 | RELN | ECM-Receptor | PMID: 14578885 |
| NeuN | CD31 | Reln | Vldlr | 0.001526 | 0 | RELN_VLDLR | Reln - Vldlr | RELN | ECM-Receptor | PMID: 14578885 |
| NeuN | NeuN | Reln | Vldlr | 0.003333 | 0 | RELN_VLDLR | Reln - Vldlr | RELN | ECM-Receptor | PMID: 14578885 |
| CD31 | NeuN | Pnoc | Oprl1 | 0.000736 | 0 | PNOC_OPRL1 | Pnoc - Oprl1 | OPIOID | Non-protein Signaling | PMID: 29934561;PMID: 34724150 |
| NeuN | NeuN | Pnoc | Oprl1 | 0.000798 | 0 | PNOC_OPRL1 | Pnoc - Oprl1 | OPIOID | Non-protein Signaling | PMID: 29934561;PMID: 34724150 |
| CD31 | CD31 | Ppia | Bsg | 0.538073 | 0 | PPIA_BSG | Ppia - Bsg | CypA | Non-protein Signaling | PMID: 36012604 |

| | | | | | | | | | | |
|---|---|---|---|---|---|---|---|---|---|---|
| NeuN | CD31 | Ppia | Bsg | 0.590644 | 0 | PPIA_BSG | Ppia - Bsg | CypA | Non-protein Signaling | PMID: 36012604 |
| NeuN | CD31 | Cadm1 | Cadm1 | 0.017754 | 0 | CADM1_CADM1 | Cadm1 - Cadm1 | CADM | Cell-Cell Contact | PMID: 24503895 |
| CD31 | NeuN | Cadm1 | Cadm1 | 0.017754 | 0 | CADM1_CADM1 | Cadm1 - Cadm1 | CADM | Cell-Cell Contact | PMID: 24503895 |
| NeuN | NeuN | Cadm1 | Cadm1 | 0.045834 | 0 | CADM1_CADM1 | Cadm1 - Cadm1 | CADM | Cell-Cell Contact | PMID: 24503895 |
| CD31 | CD31 | Cd200 | Cd200r1 | 0.003135 | 0 | CD200_CD200R1 | Cd200 - Cd200r1 | CD200 | Cell-Cell Contact | PMID: 15187158 |
| NeuN | CD31 | Cd200 | Cd200r1 | 0.005744 | 0 | CD200_CD200R1 | Cd200 - Cd200r1 | CD200 | Cell-Cell Contact | PMID: 15187158 |
| NeuN | CD31 | Cd200 | Cd200r2 | 0.005449 | 0 | CD200_CD200R2 | Cd200 - Cd200r2 | CD200 | Cell-Cell Contact | KEGG: mmu04514 |
| NeuN | NeuN | Cd200 | Cd200r2 | 0.005719 | 0 | CD200_CD200R2 | Cd200 - Cd200r2 | CD200 | Cell-Cell Contact | KEGG: mmu04514 |
| NeuN | CD31 | Cd200 | Cd200r3 | 0.004559 | 0 | CD200_CD200R3 | Cd200 - Cd200r3 | CD200 | Cell-Cell Contact | PMID: 22020332 |
| NeuN | NeuN | Cd200 | Cd200r3 | 0.003634 | 0 | CD200_CD200R3 | Cd200 - Cd200r3 | CD200 | Cell-Cell Contact | PMID: 22020332 |
| CD31 | CD31 | Entpd1 | Adora1 | 0.001408 | 0 | ENTPD1_ADORA1 | Entpd1 - Adora1 | CD39 | Cell-Cell Contact | PMID: 21677139 |
| CD31 | NeuN | Entpd1 | Adora1 | 0.002492 | 0 | ENTPD1_ADORA1 | Entpd1 - Adora1 | CD39 | Cell-Cell Contact | PMID: 21677139 |
| CD31 | CD31 | Entpd1 | Adora2a | 0.005649 | 0 | ENTPD1_ADORA2A | Entpd1 - Adora2a | CD39 | Cell-Cell Contact | PMID: 21677139 |
| NeuN | CD31 | Entpd1 | Adora2a | 0.00196 | 0 | ENTPD1_ADORA2A | Entpd1 - Adora2a | CD39 | Cell-Cell Contact | PMID: 21677139 |
| CD31 | CD31 | Entpd1 | Adora2b | 0.00123 | 0 | ENTPD1_ADORA2B | Entpd1 - Adora2b | CD39 | Cell-Cell Contact | PMID: 21677139 |
| CD31 | NeuN | Entpd1 | Adora2b | 0.000929 | 0 | ENTPD1_ADORA2B | Entpd1 - Adora2b | CD39 | Cell-Cell Contact | PMID: 21677139 |
| CD31 | CD31 | Cd46 | Jag1 | 0.001446 | 0 | CD46_JAG1 | Cd46 - Jag1 | CD46 | Cell-Cell Contact | PMID: 23086448 |
| NeuN | CD31 | Cd46 | Jag1 | 0.001209 | 0 | CD46_JAG1 | Cd46 - Jag1 | CD46 | Cell-Cell Contact | PMID: 23086448 |
| CD31 | CD31 | Cd6 | Alcam | 0.000957 | 0 | CD6_ALCAM | Cd6 - Alcam | CD6 | Cell-Cell Contact | PMID: 23602662 |
| CD31 | NeuN | Cd6 | Alcam | 0.001021 | 0 | CD6_ALCAM | Cd6 - Alcam | CD6 | Cell-Cell Contact | PMID: 23602662 |
| CD31 | CD31 | Cd96 | Pvr | 0.000788 | 0 | CD96_PVR | Cd96 - Pvr | CD96 | Cell-Cell Contact | PMID: 15034010 |
| CD31 | NeuN | Cd96 | Pvr | 0.000557 | 0 | CD96_PVR | Cd96 - Pvr | CD96 | Cell-Cell Contact | PMID: 15034010 |
| NeuN | CD31 | Cd99 | Cd99 | 0.608457 | 0 | CD99_CD99 | Cd99 - Cd99 | CD99 | Cell-Cell Contact | KEGG: mmu04514 |
| CD31 | NeuN | Cd99 | Cd99 | 0.608457 | 0 | CD99_CD99 | Cd99 - Cd99 | CD99 | Cell-Cell Contact | KEGG: mmu04514 |
| NeuN | NeuN | Cd99 | Cd99 | 0.666667 | 0 | CD99_CD99 | Cd99 - Cd99 | CD99 | Cell-Cell Contact | KEGG: mmu04514 |
| CD31 | CD31 | Cdh4 | Cdh4 | 0.002183 | 0 | CDH4_CDH4 | Cdh4 - Cdh4 | CDH | Cell-Cell Contact | KEGG: mmu04514 |
| NeuN | CD31 | Cdh4 | Cdh4 | 0.002118 | 0 | CDH4_CDH4 | Cdh4 - Cdh4 | CDH | Cell-Cell Contact | KEGG: mmu04514 |
| CD31 | NeuN | Cdh4 | Cdh4 | 0.002118 | 0 | CDH4_CDH4 | Cdh4 - Cdh4 | CDH | Cell-Cell Contact | KEGG: mmu04514 |
| NeuN | CD31 | Cdh2 | Cdh2 | 0.054476 | 0 | CDH2_CDH2 | Cdh2 - Cdh2 | CDH | Cell-Cell Contact | KEGG: mmu04514 |
| CD31 | NeuN | Cdh2 | Cdh2 | 0.054476 | 0 | CDH2_CDH2 | Cdh2 - Cdh2 | CDH | Cell-Cell Contact | KEGG: mmu04514 |
| NeuN | NeuN | Cdh2 | Cdh2 | 0.092885 | 0 | CDH2_CDH2 | Cdh2 - Cdh2 | CDH | Cell-Cell Contact | KEGG: mmu04514 |
| CD31 | CD31 | Cdh5 | Cdh5 | 0.017963 | 0 | CDH5_CDH5 | Cdh5 - Cdh5 | CDH5 | Cell-Cell Contact | KEGG: mmu04514 |
| NeuN | CD31 | Cdh5 | Cdh5 | 0.002422 | 0 | CDH5_CDH5 | Cdh5 - Cdh5 | CDH5 | Cell-Cell Contact | KEGG: mmu04514 |
| CD31 | NeuN | Cdh5 | Cdh5 | 0.002422 | 0 | CDH5_CDH5 | Cdh5 - Cdh5 | CDH5 | Cell-Cell Contact | KEGG: mmu04514 |
| NeuN | CD31 | Cldn11 | Cldn11 | 0.000176 | 0 | CLDN11_CLDN11 | Cldn11 - Cldn11 | CLDN | Cell-Cell Contact | KEGG: mmu04514 |
| CD31 | NeuN | Cldn11 | Cldn11 | 0.000176 | 0 | CLDN11_CLDN11 | Cldn11 - Cldn11 | CLDN | Cell-Cell Contact | KEGG: mmu04514 |
| NeuN | NeuN | Cldn11 | Cldn11 | 0.000188 | 0 | CLDN11_CLDN11 | Cldn11 - Cldn11 | CLDN | Cell-Cell Contact | KEGG: mmu04514 |
| CD31 | NeuN | Clec2f | Klrb1f | 0.00069 | 0 | CLEC2F_KLRB1F | Clec2f - Klrb1f | CLEC | Cell-Cell Contact | PMID: 24223577 |
| NeuN | NeuN | Clec2f | Klrb1f | 0.000585 | 0 | CLEC2F_KLRB1F | Clec2f - Klrb1f | CLEC | Cell-Cell Contact | PMID: 24223577 |
| NeuN | CD31 | Nfasc | CNTN1_CNTNAP1 | 0.005072 | 0 | NFASC_CNTN1_CNTNAP1 | Nfasc - (Cntn1+Cntnap1) | CNTN | Cell-Cell Contact | KEGG: mmu04514 |
| NeuN | NeuN | Nfasc | CNTN1_CNTNAP1 | 0.006144 | 0 | NFASC_CNTN1_CNTNAP1 | Nfasc - (Cntn1+Cntnap1) | CNTN | Cell-Cell Contact | KEGG: mmu04514 |
| NeuN | CD31 | Cntn2 | CNTN2_CNTNAP2 | 0.010984 | 0 | CNTN2_CNTN2_CNTNAP2 | Cntn2 - (Cntn2+Cntnap2) | CNTN | Cell-Cell Contact | KEGG: mmu04514 |
| NeuN | NeuN | Cntn2 | CNTN2_CNTNAP2 | 0.01246 | 0 | CNTN2_CNTN2_CNTNAP2 | Cntn2 - (Cntn2+Cntnap2) | CNTN | Cell-Cell Contact | KEGG: mmu04514 |
| CD31 | NeuN | Cntn2 | L1cam | 0.034178 | 0 | CNTN2_L1CAM | Cntn2 - L1cam | CNTN | Cell-Cell Contact | KEGG: mmu04514 |
| NeuN | NeuN | Cntn2 | L1cam | 0.063817 | 0 | CNTN2_L1CAM | Cntn2 - L1cam | CNTN | Cell-Cell Contact | KEGG: mmu04514 |
| NeuN | CD31 | Cntn2 | Cntn2 | 0.017161 | 0 | CNTN2_CNTN2 | Cntn2 - Cntn2 | CNTN | Cell-Cell Contact | KEGG: mmu04514 |
| CD31 | NeuN | Cntn2 | Cntn2 | 0.017161 | 0 | CNTN2_CNTN2 | Cntn2 - Cntn2 | CNTN | Cell-Cell Contact | KEGG: mmu04514 |
| NeuN | NeuN | Cntn2 | Cntn2 | 0.03254 | 0 | CNTN2_CNTN2 | Cntn2 - Cntn2 | CNTN | Cell-Cell Contact | KEGG: mmu04514 |
| CD31 | NeuN | Cntn1 | Nrcam | 0.010408 | 0 | CNTN1_NRCAM | Cntn1 - Nrcam | CNTN | Cell-Cell Contact | KEGG: mmu04514 |
| NeuN | NeuN | Cntn1 | Nrcam | 0.024925 | 0 | CNTN1_NRCAM | Cntn1 - Nrcam | CNTN | Cell-Cell Contact | KEGG: mmu04514 |
| CD31 | CD31 | Cspg4 | ITGA3_ITGB1 | 0.004089 | 0 | CSPG4_ITGA2_ITGB1 | Cspg4 - (Itga2+Itgb1) | CSPG4 | Cell-Cell Contact | PMID: 30213051 |
| CD31 | NeuN | Cspg4 | ITGA3_ITGB1 | 0.002396 | 0 | CSPG4_ITGA2_ITGB1 | Cspg4 - (Itga2+Itgb1) | CSPG4 | Cell-Cell Contact | PMID: 30213051 |
| CD31 | CD31 | Efna1 | Epha3 | 0.003456 | 0 | EFNA1_EPHA3 | Efna1 - Epha3 | EPHA | Cell-Cell Contact | PMID: 15114347 |
| CD31 | NeuN | Efna1 | Epha3 | 0.0064 | 0 | EFNA1_EPHA3 | Efna1 - Epha3 | EPHA | Cell-Cell Contact | PMID: 15114347 |
| CD31 | CD31 | Efna1 | Epha4 | 0.00849 | 0 | EFNA1_EPHA4 | Efna1 - Epha4 | EPHA | Cell-Cell Contact | PMID: 15114347 |
| CD31 | NeuN | Efna1 | Epha4 | 0.010392 | 0 | EFNA1_EPHA4 | Efna1 - Epha4 | EPHA | Cell-Cell Contact | PMID: 15114347 |
| CD31 | CD31 | Efna1 | Epha5 | 0.009516 | 0 | EFNA1_EPHA5 | Efna1 - Epha5 | EPHA | Cell-Cell Contact | PMID: 15114347 |
| CD31 | NeuN | Efna1 | Epha5 | 0.01471 | 0 | EFNA1_EPHA5 | Efna1 - Epha5 | EPHA | Cell-Cell Contact | PMID: 15114347 |
| CD31 | CD31 | Efna1 | Epha7 | 0.003471 | 0 | EFNA1_EPHA7 | Efna1 - Epha7 | EPHA | Cell-Cell Contact | PMID: 15114347 |

| | | | | | | | | | | |
|---|---|---|---|---|---|---|---|---|---|---|
| CD31 | NeuN | Efna1 | Epha7 | 0.009973 | 0 | EFNA1_EPHA7 | Efna1 - Epha7 | EPHA | Cell-Cell Contact | PMID: 15114347 |
| CD31 | CD31 | Efna1 | Epha8 | 0.001681 | 0 | EFNA1_EPHA8 | Efna1 - Epha8 | EPHA | Cell-Cell Contact | PMID: 15114347 |
| CD31 | NeuN | Efna1 | Epha8 | 0.001668 | 0 | EFNA1_EPHA8 | Efna1 - Epha8 | EPHA | Cell-Cell Contact | PMID: 15114347 |
| CD31 | NeuN | Efna2 | Epha3 | 0.005066 | 0 | EFNA2_EPHA3 | Efna2 - Epha3 | EPHA | Cell-Cell Contact | PMID: 15114347 |
| NeuN | NeuN | Efna2 | Epha3 | 0.007491 | 0 | EFNA2_EPHA3 | Efna2 - Epha3 | EPHA | Cell-Cell Contact | PMID: 15114347 |
| NeuN | CD31 | Efna2 | Epha4 | 0.009933 | 0 | EFNA2_EPHA4 | Efna2 - Epha4 | EPHA | Cell-Cell Contact | PMID: 15114347 |
| NeuN | NeuN | Efna2 | Epha4 | 0.012155 | 0 | EFNA2_EPHA4 | Efna2 - Epha4 | EPHA | Cell-Cell Contact | PMID: 15114347 |
| CD31 | NeuN | Efna2 | Epha5 | 0.011665 | 0 | EFNA2_EPHA5 | Efna2 - Epha5 | EPHA | Cell-Cell Contact | PMID: 15114347 |
| NeuN | NeuN | Efna2 | Epha5 | 0.017193 | 0 | EFNA2_EPHA5 | Efna2 - Epha5 | EPHA | Cell-Cell Contact | PMID: 15114347 |
| CD31 | NeuN | Efna2 | Epha7 | 0.007901 | 0 | EFNA2_EPHA7 | Efna2 - Epha7 | EPHA | Cell-Cell Contact | PMID: 15114347 |
| NeuN | NeuN | Efna2 | Epha7 | 0.011666 | 0 | EFNA2_EPHA7 | Efna2 - Epha7 | EPHA | Cell-Cell Contact | PMID: 15114347 |
| NeuN | CD31 | Efna2 | Epha8 | 0.001969 | 0 | EFNA2_EPHA8 | Efna2 - Epha8 | EPHA | Cell-Cell Contact | PMID: 15114347 |
| NeuN | NeuN | Efna2 | Epha8 | 0.001954 | 0 | EFNA2_EPHA8 | Efna2 - Epha8 | EPHA | Cell-Cell Contact | PMID: 15114347 |
| NeuN | CD31 | Efna3 | Epha3 | 0.004328 | 0 | EFNA3_EPHA3 | Efna3 - Epha3 | EPHA | Cell-Cell Contact | PMID: 15114347 |
| NeuN | NeuN | Efna3 | Epha3 | 0.008009 | 0 | EFNA3_EPHA3 | Efna3 - Epha3 | EPHA | Cell-Cell Contact | PMID: 15114347 |
| NeuN | CD31 | Efna3 | Epha4 | 0.010618 | 0 | EFNA3_EPHA4 | Efna3 - Epha4 | EPHA | Cell-Cell Contact | PMID: 15114347 |
| NeuN | NeuN | Efna3 | Epha4 | 0.012991 | 0 | EFNA3_EPHA4 | Efna3 - Epha4 | EPHA | Cell-Cell Contact | PMID: 15114347 |
| NeuN | CD31 | Efna3 | Epha5 | 0.011899 | 0 | EFNA3_EPHA5 | Efna3 - Epha5 | EPHA | Cell-Cell Contact | PMID: 15114347 |
| NeuN | NeuN | Efna3 | Epha5 | 0.01837 | 0 | EFNA3_EPHA5 | Efna3 - Epha5 | EPHA | Cell-Cell Contact | PMID: 15114347 |
| CD31 | NeuN | Efna3 | Epha7 | 0.005614 | 0 | EFNA3_EPHA7 | Efna3 - Epha7 | EPHA | Cell-Cell Contact | PMID: 15114347 |
| NeuN | NeuN | Efna3 | Epha7 | 0.012469 | 0 | EFNA3_EPHA7 | Efna3 - Epha7 | EPHA | Cell-Cell Contact | PMID: 15114347 |
| NeuN | CD31 | Efna3 | Epha8 | 0.002106 | 0 | EFNA3_EPHA8 | Efna3 - Epha8 | EPHA | Cell-Cell Contact | PMID: 15114347 |
| NeuN | NeuN | Efna3 | Epha8 | 0.00209 | 0 | EFNA3_EPHA8 | Efna3 - Epha8 | EPHA | Cell-Cell Contact | PMID: 15114347 |
| CD31 | NeuN | Efna4 | Epha3 | 0.00468 | 0 | EFNA4_EPHA3 | Efna4 - Epha3 | EPHA | Cell-Cell Contact | PMID: 15114347 |
| NeuN | NeuN | Efna4 | Epha3 | 0.006125 | 0 | EFNA4_EPHA3 | Efna4 - Epha3 | EPHA | Cell-Cell Contact | PMID: 15114347 |
| NeuN | CD31 | Efna4 | Epha4 | 0.008126 | 0 | EFNA4_EPHA4 | Efna4 - Epha4 | EPHA | Cell-Cell Contact | PMID: 15114347 |
| NeuN | NeuN | Efna4 | Epha4 | 0.009947 | 0 | EFNA4_EPHA4 | Efna4 - Epha4 | EPHA | Cell-Cell Contact | PMID: 15114347 |
| CD31 | NeuN | Efna4 | Epha5 | 0.010782 | 0 | EFNA4_EPHA5 | Efna4 - Epha5 | EPHA | Cell-Cell Contact | PMID: 15114347 |
| NeuN | NeuN | Efna4 | Epha5 | 0.014084 | 0 | EFNA4_EPHA5 | Efna4 - Epha5 | EPHA | Cell-Cell Contact | PMID: 15114347 |
| CD31 | NeuN | Efna4 | Epha7 | 0.0073 | 0 | EFNA4_EPHA7 | Efna4 - Epha7 | EPHA | Cell-Cell Contact | PMID: 15114347 |
| NeuN | NeuN | Efna4 | Epha7 | 0.009546 | 0 | EFNA4_EPHA7 | Efna4 - Epha7 | EPHA | Cell-Cell Contact | PMID: 15114347 |
| NeuN | CD31 | Efna4 | Epha8 | 0.001609 | 0 | EFNA4_EPHA8 | Efna4 - Epha8 | EPHA | Cell-Cell Contact | PMID: 15114347 |
| NeuN | NeuN | Efna4 | Epha8 | 0.001596 | 0 | EFNA4_EPHA8 | Efna4 - Epha8 | EPHA | Cell-Cell Contact | PMID: 15114347 |
| CD31 | NeuN | Efna5 | Epha3 | 0.001987 | 0 | EFNA5_EPHA3 | Efna5 - Epha3 | EPHA | Cell-Cell Contact | PMID: 15114347 |
| NeuN | NeuN | Efna5 | Epha3 | 0.003173 | 0 | EFNA5_EPHA3 | Efna5 - Epha3 | EPHA | Cell-Cell Contact | PMID: 15114347 |
| NeuN | CD31 | Efna5 | Epha4 | 0.004214 | 0 | EFNA5_EPHA4 | Efna5 - Epha4 | EPHA | Cell-Cell Contact | PMID: 15114347 |
| NeuN | NeuN | Efna5 | Epha4 | 0.005163 | 0 | EFNA5_EPHA4 | Efna5 - Epha4 | EPHA | Cell-Cell Contact | PMID: 15114347 |
| NeuN | CD31 | Efna5 | Epha5 | 0.004726 | 0 | EFNA5_EPHA5 | Efna5 - Epha5 | EPHA | Cell-Cell Contact | PMID: 15114347 |
| NeuN | NeuN | Efna5 | Epha5 | 0.007325 | 0 | EFNA5_EPHA5 | Efna5 - Epha5 | EPHA | Cell-Cell Contact | PMID: 15114347 |
| CD31 | NeuN | Efna5 | Epha7 | 0.003104 | 0 | EFNA5_EPHA7 | Efna5 - Epha7 | EPHA | Cell-Cell Contact | PMID: 15114347 |
| NeuN | NeuN | Efna5 | Epha7 | 0.004954 | 0 | EFNA5_EPHA7 | Efna5 - Epha7 | EPHA | Cell-Cell Contact | PMID: 15114347 |
| NeuN | CD31 | Efna5 | Epha8 | 0.000832 | 0 | EFNA5_EPHA8 | Efna5 - Epha8 | EPHA | Cell-Cell Contact | PMID: 15114347 |
| NeuN | NeuN | Efna5 | Epha8 | 0.000825 | 0 | EFNA5_EPHA8 | Efna5 - Epha8 | EPHA | Cell-Cell Contact | PMID: 15114347 |
| CD31 | NeuN | Efna5 | Ephb2 | 0.002954 | 0 | EFNA5_EPHB2 | Efna5 - Ephb2 | EPHA | Cell-Cell Contact | PMID:15107857; PMID: 15114347 |
| NeuN | NeuN | Efna5 | Ephb2 | 0.004714 | 0 | EFNA5_EPHB2 | Efna5 - Ephb2 | EPHA | Cell-Cell Contact | PMID:15107857; PMID: 15114347 |
| NeuN | CD31 | Efnb1 | Epha4 | 0.012499 | 0 | EFNB1_EPHA4 | Efnb1 - Epha4 | EPHB | Cell-Cell Contact | PMID: 15114347 |
| NeuN | NeuN | Efnb1 | Epha4 | 0.015286 | 0 | EFNB1_EPHA4 | Efnb1 - Epha4 | EPHB | Cell-Cell Contact | PMID: 15114347 |
| CD31 | NeuN | Efnb1 | Ephb1 | 0.016899 | 0 | EFNB1_EPHB1 | Efnb1 - Ephb1 | EPHB | Cell-Cell Contact | PMID: 15114347 |
| NeuN | NeuN | Efnb1 | Ephb1 | 0.021856 | 0 | EFNB1_EPHB1 | Efnb1 - Ephb1 | EPHB | Cell-Cell Contact | PMID: 15114347 |
| CD31 | NeuN | Efnb1 | Ephb2 | 0.01078 | 0 | EFNB1_EPHB2 | Efnb1 - Ephb2 | EPHB | Cell-Cell Contact | PMID: 15114347 |
| NeuN | NeuN | Efnb1 | Ephb2 | 0.013968 | 0 | EFNB1_EPHB2 | Efnb1 - Ephb2 | EPHB | Cell-Cell Contact | PMID: 15114347 |
| CD31 | CD31 | Efnb1 | Ephb3 | 0.002184 | 0 | EFNB1_EPHB3 | Efnb1 - Ephb3 | EPHB | Cell-Cell Contact | PMID: 15114347 |
| NeuN | CD31 | Efnb1 | Ephb3 | 0.002837 | 0 | EFNB1_EPHB3 | Efnb1 - Ephb3 | EPHB | Cell-Cell Contact | PMID: 15114347 |
| CD31 | CD31 | Efnb1 | Ephb4 | 0.005631 | 0 | EFNB1_EPHB4 | Efnb1 - Ephb4 | EPHB | Cell-Cell Contact | PMID: 15114347 |
| NeuN | CD31 | Efnb1 | Ephb4 | 0.007307 | 0 | EFNB1_EPHB4 | Efnb1 - Ephb4 | EPHB | Cell-Cell Contact | PMID: 15114347 |
| NeuN | CD31 | Efnb2 | Epha4 | 0.041295 | 0 | EFNB2_EPHA4 | Efnb2 - Epha4 | EPHB | Cell-Cell Contact | PMID: 15114347 |
| NeuN | NeuN | Efnb2 | Epha4 | 0.050176 | 0 | EFNB2_EPHA4 | Efnb2 - Epha4 | EPHB | Cell-Cell Contact | PMID: 15114347 |
| CD31 | NeuN | Efnb2 | Ephb1 | 0.046352 | 0 | EFNB2_EPHB1 | Efnb2 - Ephb1 | EPHB | Cell-Cell Contact | PMID: 15114347 |
| NeuN | NeuN | Efnb2 | Ephb1 | 0.070667 | 0 | EFNB2_EPHB1 | Efnb2 - Ephb1 | EPHB | Cell-Cell Contact | PMID: 15114347 |

| | | | | | | | | | | |
|---|---|---|---|---|---|---|---|---|---|---|
| CD31 | NeuN | Efnb2 | Ephb2 | 0.029893 | 0 | EFNB2_EPHB2 | Efnb2 - Ephb2 | EPHB | Cell-Cell Contact | PMID: 15114347 |
| NeuN | NeuN | Efnb2 | Ephb2 | 0.04599 | 0 | EFNB2_EPHB2 | Efnb2 - Ephb2 | EPHB | Cell-Cell Contact | PMID: 15114347 |
| NeuN | CD31 | Efnb2 | Ephb3 | 0.009589 | 0 | EFNB2_EPHB3 | Efnb2 - Ephb3 | EPHB | Cell-Cell Contact | PMID: 15114347 |
| NeuN | NeuN | Efnb2 | Ephb3 | 0.006851 | 0 | EFNB2_EPHB3 | Efnb2 - Ephb3 | EPHB | Cell-Cell Contact | PMID: 15114347 |
| CD31 | CD31 | Efnb2 | Ephb4 | 0.01576 | 0 | EFNB2_EPHB4 | Efnb2 - Ephb4 | EPHB | Cell-Cell Contact | PMID: 15114347 |
| NeuN | CD31 | Efnb2 | Ephb4 | 0.024439 | 0 | EFNB2_EPHB4 | Efnb2 - Ephb4 | EPHB | Cell-Cell Contact | PMID: 15114347 |
| NeuN | CD31 | Efnb3 | Epha4 | 0.026464 | 0 | EFNB3_EPHA4 | Efnb3 - Epha4 | EPHB | Cell-Cell Contact | PMID: 15114347 |
| NeuN | NeuN | Efnb3 | Epha4 | 0.032263 | 0 | EFNB3_EPHA4 | Efnb3 - Epha4 | EPHB | Cell-Cell Contact | PMID: 15114347 |
| NeuN | CD31 | Efnb3 | Ephb1 | 0.021657 | 0 | EFNB3_EPHB1 | Efnb3 - Ephb1 | EPHB | Cell-Cell Contact | PMID: 15114347 |
| NeuN | NeuN | Efnb3 | Ephb1 | 0.045791 | 0 | EFNB3_EPHB1 | Efnb3 - Ephb1 | EPHB | Cell-Cell Contact | PMID: 15114347 |
| NeuN | CD31 | Efnb3 | Ephb2 | 0.017954 | 0 | EFNB3_EPHB2 | Efnb3 - Ephb2 | EPHB | Cell-Cell Contact | PMID: 15114347 |
| NeuN | NeuN | Efnb3 | Ephb2 | 0.029525 | 0 | EFNB3_EPHB2 | Efnb3 - Ephb2 | EPHB | Cell-Cell Contact | PMID: 15114347 |
| NeuN | CD31 | Efnb3 | Ephb3 | 0.006073 | 0 | EFNB3_EPHB3 | Efnb3 - Ephb3 | EPHB | Cell-Cell Contact | PMID: 15114347 |
| NeuN | NeuN | Efnb3 | Ephb3 | 0.004334 | 0 | EFNB3_EPHB3 | Efnb3 - Ephb3 | EPHB | Cell-Cell Contact | PMID: 15114347 |
| CD31 | CD31 | Efnb3 | Ephb4 | 0.006968 | 0 | EFNB3_EPHB4 | Efnb3 - Ephb4 | EPHB | Cell-Cell Contact | PMID: 15114347 |
| NeuN | CD31 | Efnb3 | Ephb4 | 0.015564 | 0 | EFNB3_EPHB4 | Efnb3 - Ephb4 | EPHB | Cell-Cell Contact | PMID: 15114347 |
| NeuN | CD31 | Efnb3 | Ephb6 | 0.008912 | 0 | EFNB3_EPHB6 | Efnb3 - Ephb6 | EPHB | Cell-Cell Contact | PMID: 15114347 |
| NeuN | NeuN | Efnb3 | Ephb6 | 0.00632 | 0 | EFNB3_EPHB6 | Efnb3 - Ephb6 | EPHB | Cell-Cell Contact | PMID: 15114347 |
| CD31 | CD31 | Esam | Esam | 0.019117 | 0 | ESAM_ESAM | Esam - Esam | ESAM | Cell-Cell Contact | KEGG: mmu04514; PMID: 11279107 |
| NeuN | CD31 | Esam | Esam | 0.002678 | 0 | ESAM_ESAM | Esam - Esam | ESAM | Cell-Cell Contact | KEGG: mmu04514; PMID: 11279107 |
| CD31 | NeuN | Esam | Esam | 0.002678 | 0 | ESAM_ESAM | Esam - Esam | ESAM | Cell-Cell Contact | KEGG: mmu04514; PMID: 11279107 |
| CD31 | CD31 | Icam5 | Cd209a | 0.000472 | 0 | ICAM5_CD209A | Icam5 - Cd209a | ICAM | Cell-Cell Contact | PMID: 15795245 |
| CD31 | NeuN | Icam5 | Cd209a | 0.000341 | 0 | ICAM5_CD209A | Icam5 - Cd209a | ICAM | Cell-Cell Contact | PMID: 15795245 |
| CD31 | CD31 | Jam2 | ITGA3_ITGB1 | 0.002578 | 0 | JAM2_ITGA3_ITGB1 | Jam2 - (Itga3+Itgb1) | JAM | Cell-Cell Contact | PMID: 12070135 |
| CD31 | NeuN | Jam2 | ITGA3_ITGB1 | 0.001509 | 0 | JAM2_ITGA3_ITGB1 | Jam2 - (Itga3+Itgb1) | JAM | Cell-Cell Contact | PMID: 12070135 |
| CD31 | CD31 | Jam2 | ITGAV_ITGB1 | 0.00327 | 0 | JAM2_ITGAV_ITGB1 | Jam2 - (Itgav+Itgb1) | JAM | Cell-Cell Contact | KEGG: mmu04514 |
| NeuN | CD31 | Jam2 | ITGAV_ITGB1 | 0.001355 | 0 | JAM2_ITGAV_ITGB1 | Jam2 - (Itgav+Itgb1) | JAM | Cell-Cell Contact | KEGG: mmu04514 |
| CD31 | CD31 | Jam2 | Jam3 | 0.001464 | 0 | JAM2_JAM3 | Jam2 - Jam3 | JAM | Cell-Cell Contact | PMC1237098; PMID: 11590146 |
| CD31 | NeuN | Jam2 | Jam3 | 0.001512 | 0 | JAM2_JAM3 | Jam2 - Jam3 | JAM | Cell-Cell Contact | PMC1237098; PMID: 11590146 |
| CD31 | CD31 | F11r | F11r | 0.002328 | 0 | F11R_F11R | F11r - F11r | JAM | Cell-Cell Contact | KEGG: mmu04514 |
| NeuN | CD31 | F11r | F11r | 0.001039 | 0 | F11R_F11R | F11r - F11r | JAM | Cell-Cell Contact | KEGG: mmu04514 |
| CD31 | NeuN | F11r | F11r | 0.001039 | 0 | F11R_F11R | F11r - F11r | JAM | Cell-Cell Contact | KEGG: mmu04514 |
| CD31 | CD31 | Jam2 | F11r | 0.001618 | 0 | JAM2_F11R | Jam2 - F11r | JAM | Cell-Cell Contact | KEGG: mmu04514 |
| CD31 | NeuN | Jam2 | F11r | 0.000721 | 0 | JAM2_F11R | Jam2 - F11r | JAM | Cell-Cell Contact | KEGG: mmu04514 |
| CD31 | CD31 | Jam2 | Jam2 | 0.001124 | 0 | JAM2_JAM2 | Jam2 - Jam2 | JAM | Cell-Cell Contact | KEGG: mmu04514 |
| NeuN | CD31 | Jam2 | Jam2 | 0.000465 | 0 | JAM2_JAM2 | Jam2 - Jam2 | JAM | Cell-Cell Contact | KEGG: mmu04514 |
| CD31 | NeuN | Jam2 | Jam2 | 0.000465 | 0 | JAM2_JAM2 | Jam2 - Jam2 | JAM | Cell-Cell Contact | KEGG: mmu04514 |
| CD31 | CD31 | Jam3 | F11r | 0.002108 | 0 | JAM3_F11R | Jam3 - F11r | JAM | Cell-Cell Contact | KEGG: mmu04514 |
| NeuN | CD31 | Jam3 | F11r | 0.002176 | 0 | JAM3_F11R | Jam3 - F11r | JAM | Cell-Cell Contact | KEGG: mmu04514 |
| NeuN | CD31 | Jam3 | Jam3 | 0.00197 | 0 | JAM3_JAM3 | Jam3 - Jam3 | JAM | Cell-Cell Contact | KEGG: mmu04514 |
| CD31 | NeuN | Jam3 | Jam3 | 0.00197 | 0 | JAM3_JAM3 | Jam3 - Jam3 | JAM | Cell-Cell Contact | KEGG: mmu04514 |
| NeuN | NeuN | Jam3 | Jam3 | 0.002033 | 0 | JAM3_JAM3 | Jam3 - Jam3 | JAM | Cell-Cell Contact | KEGG: mmu04514 |
| NeuN | CD31 | L1cam | ITGAV_ITGB3 | 0.014381 | 0 | L1CAM_ITGAV_ITGB3 | L1cam - (Itgav+Itgb3) | L1CAM | Cell-Cell Contact | PMID: 9396761 |
| NeuN | NeuN | L1cam | ITGAV_ITGB3 | 0.008551 | 0 | L1CAM_ITGAV_ITGB3 | L1cam - (Itgav+Itgb3) | L1CAM | Cell-Cell Contact | PMID: 9396761 |
| NeuN | CD31 | L1cam | ITGA4_ITGB7 | 0.018094 | 0 | L1CAM_ITGA4_ITGB7 | L1cam - (Itga4+Itgb7) | L1CAM | Cell-Cell Contact | PMID: 8557754 |
| NeuN | NeuN | L1cam | ITGA4_ITGB7 | 0.007872 | 0 | L1CAM_ITGA4_ITGB7 | L1cam - (Itga4+Itgb7) | L1CAM | Cell-Cell Contact | PMID: 8557754 |
| NeuN | CD31 | L1cam | L1cam | 0.04315 | 0 | L1CAM_L1CAM | L1cam - L1cam | L1CAM | Cell-Cell Contact | PMID: 18701456 |
| CD31 | NeuN | L1cam | L1cam | 0.04315 | 0 | L1CAM_L1CAM | L1cam - L1cam | L1CAM | Cell-Cell Contact | PMID: 18701456 |
| NeuN | NeuN | L1cam | L1cam | 0.121386 | 0 | L1CAM_L1CAM | L1cam - L1cam | L1CAM | Cell-Cell Contact | PMID: 18701456 |
| CD31 | CD31 | Mag | Mag | 0.000211 | 0 | MAG_MAG | Mag - Mag | MAG | Cell-Cell Contact | KEGG: mmu04514 |
| NeuN | CD31 | Mag | Mag | 0.000184 | 0 | MAG_MAG | Mag - Mag | MAG | Cell-Cell Contact | KEGG: mmu04514 |
| CD31 | NeuN | Mag | Mag | 0.000184 | 0 | MAG_MAG | Mag - Mag | MAG | Cell-Cell Contact | KEGG: mmu04514 |
| CD31 | CD31 | Mpzl1 | Mpzl1 | 0.022943 | 0 | MPZL1_MPZL1 | Mpzl1 - Mpzl1 | MPZ | Cell-Cell Contact | KEGG: mmu04514 |
| NeuN | CD31 | Mpzl1 | Mpzl1 | 0.010568 | 0 | MPZL1_MPZL1 | Mpzl1 - Mpzl1 | MPZ | Cell-Cell Contact | KEGG: mmu04514 |
| CD31 | NeuN | Mpzl1 | Mpzl1 | 0.010568 | 0 | MPZL1_MPZL1 | Mpzl1 - Mpzl1 | MPZ | Cell-Cell Contact | KEGG: mmu04514 |
| NeuN | CD31 | Ncam1 | Fgfr1 | 0.03642 | 0 | NCAM1_FGFR1 | Ncam1 - Fgfr1 | NCAM | Cell-Cell Contact | PMID: 12791257 |
| NeuN | NeuN | Ncam1 | Fgfr1 | 0.026942 | 0 | NCAM1_FGFR1 | Ncam1 - Fgfr1 | NCAM | Cell-Cell Contact | PMID: 12791257 |
| NeuN | CD31 | Ncam1 | Ncam1 | 0.173738 | 0 | NCAM1_NCAM1 | Ncam1 - Ncam1 | NCAM | Cell-Cell Contact | KEGG: mmu04514 |
| CD31 | NeuN | Ncam1 | Ncam1 | 0.173738 | 0 | NCAM1_NCAM1 | Ncam1 - Ncam1 | NCAM | Cell-Cell Contact | KEGG: mmu04514 |

| | | | | | | | | | | |
|---|---|---|---|---|---|---|---|---|---|---|
| NeuN | NeuN | Ncam1 | Ncam1 | 0.330971 | 0 | NCAM1_NCAM1 | Ncam1 - Ncam1 | NCAM | Cell-Cell Contact | KEGG: mmu04514 |
| NeuN | CD31 | Ncam1 | Ncam2 | 0.018206 | 0 | NCAM1_NCAM2 | Ncam1 - Ncam2 | NCAM | Cell-Cell Contact | KEGG: mmu04514 |
| NeuN | NeuN | Ncam1 | Ncam2 | 0.0271 | 0 | NCAM1_NCAM2 | Ncam1 - Ncam2 | NCAM | Cell-Cell Contact | KEGG: mmu04514 |
| CD31 | NeuN | Ncam1 | L1cam | 0.100006 | 0 | NCAM1_L1CAM | Ncam1 - L1cam | NCAM | Cell-Cell Contact | KEGG: mmu04514 |
| NeuN | NeuN | Ncam1 | L1cam | 0.207249 | 0 | NCAM1_L1CAM | Ncam1 - L1cam | NCAM | Cell-Cell Contact | KEGG: mmu04514 |
| CD31 | NeuN | Ncam2 | L1cam | 0.009704 | 0 | NCAM2_L1CAM | Ncam2 - L1cam | NCAM | Cell-Cell Contact | KEGG: mmu04514 |
| NeuN | NeuN | Ncam2 | L1cam | 0.014506 | 0 | NCAM2_L1CAM | Ncam2 - L1cam | NCAM | Cell-Cell Contact | KEGG: mmu04514 |
| CD31 | CD31 | Nectin1 | Cd96 | 0.001121 | 0 | NECTIN1_CD96 | Nectin1 - Cd96 | NECTIN | Cell-Cell Contact | PMID: 23027581 |
| NeuN | CD31 | Nectin1 | Cd96 | 0.00171 | 0 | NECTIN1_CD96 | Nectin1 - Cd96 | NECTIN | Cell-Cell Contact | PMID: 23027581 |
| CD31 | CD31 | Nectin3 | Nectin2 | 0.002399 | 0 | NECTIN3_NECTIN2 | Nectin3 - Nectin2 | NECTIN | Cell-Cell Contact | PMID: 23027581 |
| NeuN | CD31 | Nectin3 | Nectin2 | 0.003135 | 0 | NECTIN3_NECTIN2 | Nectin3 - Nectin2 | NECTIN | Cell-Cell Contact | PMID: 23027581 |
| CD31 | CD31 | Nectin3 | Pvr | 0.001223 | 0 | NECTIN3_PVR | Nectin3 - Pvr | NECTIN | Cell-Cell Contact | PMID: 23027581 |
| NeuN | CD31 | Nectin3 | Pvr | 0.001599 | 0 | NECTIN3_PVR | Nectin3 - Pvr | NECTIN | Cell-Cell Contact | PMID: 23027581 |
| NeuN | CD31 | Nectin1 | Nectin3 | 0.002653 | 0 | NECTIN1_NECTIN3 | Nectin1 - Nectin3 | NECTIN | Cell-Cell Contact | KEGG: mmu04514 |
| NeuN | NeuN | Nectin1 | Nectin3 | 0.003468 | 0 | NECTIN1_NECTIN3 | Nectin1 - Nectin3 | NECTIN | Cell-Cell Contact | KEGG: mmu04514 |
| NeuN | CD31 | Negr1 | Negr1 | 0.005057 | 0 | NEGR1_NEGR1 | Negr1 - Negr1 | NEGR | Cell-Cell Contact | KEGG: mmu04514 |
| CD31 | NeuN | Negr1 | Negr1 | 0.005057 | 0 | NEGR1_NEGR1 | Negr1 - Negr1 | NEGR | Cell-Cell Contact | KEGG: mmu04514 |
| NeuN | NeuN | Negr1 | Negr1 | 0.00909 | 0 | NEGR1_NEGR1 | Negr1 - Negr1 | NEGR | Cell-Cell Contact | KEGG: mmu04514 |
| NeuN | CD31 | Lrrc4c | Ntng1 | 0.001116 | 0 | LRRC4C_NTNG1 | Lrrc4c - Ntng1 | NGL | Cell-Cell Contact | KEGG: mmu04514 |
| NeuN | NeuN | Lrrc4c | Ntng1 | 0.001199 | 0 | LRRC4C_NTNG1 | Lrrc4c - Ntng1 | NGL | Cell-Cell Contact | KEGG: mmu04514 |
| CD31 | NeuN | Lrrc4 | Ntng2 | 0.006548 | 0 | LRRC4_NTNG2 | Lrrc4 - Ntng2 | NGL | Cell-Cell Contact | KEGG: mmu04514 |
| NeuN | NeuN | Lrrc4 | Ntng2 | 0.006257 | 0 | LRRC4_NTNG2 | Lrrc4 - Ntng2 | NGL | Cell-Cell Contact | KEGG: mmu04514 |
| NeuN | CD31 | Lrrc4b | Ptprf | 0.016346 | 0 | LRRC4B_PTPRF | Lrrc4b - Ptprf | NGL | Cell-Cell Contact | KEGG: mmu04514 |
| NeuN | NeuN | Lrrc4b | Ptprf | 0.030692 | 0 | LRRC4B_PTPRF | Lrrc4b - Ptprf | NGL | Cell-Cell Contact | KEGG: mmu04514 |
| CD31 | CD31 | Dlk1 | Notch1 | 0.005419 | 0 | DLK1_NOTCH1 | Dlk1 - Notch1 | NOTCH | Cell-Cell Contact | PMID: 22353464 |
| NeuN | CD31 | Dlk1 | Notch1 | 0.002294 | 0 | DLK1_NOTCH1 | Dlk1 - Notch1 | NOTCH | Cell-Cell Contact | PMID: 22353464 |
| CD31 | CD31 | Dlk1 | Notch3 | 0.006448 | 0 | DLK1_NOTCH3 | Dlk1 - Notch3 | NOTCH | Cell-Cell Contact | PMID: 22353464 |
| NeuN | CD31 | Dlk1 | Notch3 | 0.002731 | 0 | DLK1_NOTCH3 | Dlk1 - Notch3 | NOTCH | Cell-Cell Contact | PMID: 22353464 |
| CD31 | CD31 | Dlk1 | Notch4 | 0.003952 | 0 | DLK1_NOTCH4 | Dlk1 - Notch4 | NOTCH | Cell-Cell Contact | PMID: 22353464 |
| NeuN | CD31 | Dlk1 | Notch4 | 0.001672 | 0 | DLK1_NOTCH4 | Dlk1 - Notch4 | NOTCH | Cell-Cell Contact | PMID: 22353464 |
| CD31 | CD31 | Dll1 | Notch1 | 0.003899 | 0 | DLL1_NOTCH1 | Dll1 - Notch1 | NOTCH | Cell-Cell Contact | PMID: 22353464 |
| NeuN | CD31 | Dll1 | Notch1 | 0.003233 | 0 | DLL1_NOTCH1 | Dll1 - Notch1 | NOTCH | Cell-Cell Contact | PMID: 22353464 |
| CD31 | CD31 | Dll1 | Notch3 | 0.00464 | 0 | DLL1_NOTCH3 | Dll1 - Notch3 | NOTCH | Cell-Cell Contact | PMID: 22353464 |
| NeuN | CD31 | Dll1 | Notch3 | 0.003848 | 0 | DLL1_NOTCH3 | Dll1 - Notch3 | NOTCH | Cell-Cell Contact | PMID: 22353464 |
| CD31 | CD31 | Dll1 | Notch4 | 0.002842 | 0 | DLL1_NOTCH4 | Dll1 - Notch4 | NOTCH | Cell-Cell Contact | PMID: 22353464 |
| NeuN | CD31 | Dll1 | Notch4 | 0.002356 | 0 | DLL1_NOTCH4 | Dll1 - Notch4 | NOTCH | Cell-Cell Contact | PMID: 22353464 |
| CD31 | CD31 | Dll3 | Notch1 | 0.001973 | 0 | DLL3_NOTCH1 | Dll3 - Notch1 | NOTCH | Cell-Cell Contact | PMID: 22353464 |
| NeuN | CD31 | Dll3 | Notch1 | 0.002041 | 0 | DLL3_NOTCH1 | Dll3 - Notch1 | NOTCH | Cell-Cell Contact | PMID: 22353464 |
| CD31 | CD31 | Dll3 | Notch3 | 0.002349 | 0 | DLL3_NOTCH3 | Dll3 - Notch3 | NOTCH | Cell-Cell Contact | PMID: 22353464 |
| NeuN | CD31 | Dll3 | Notch3 | 0.00243 | 0 | DLL3_NOTCH3 | Dll3 - Notch3 | NOTCH | Cell-Cell Contact | PMID: 22353464 |
| CD31 | CD31 | Dll3 | Notch4 | 0.001438 | 0 | DLL3_NOTCH4 | Dll3 - Notch4 | NOTCH | Cell-Cell Contact | PMID: 22353464 |
| NeuN | CD31 | Dll3 | Notch4 | 0.001487 | 0 | DLL3_NOTCH4 | Dll3 - Notch4 | NOTCH | Cell-Cell Contact | PMID: 22353464 |
| CD31 | CD31 | Dll4 | Notch1 | 0.007023 | 0 | DLL4_NOTCH1 | Dll4 - Notch1 | NOTCH | Cell-Cell Contact | PMID: 22353464 |
| CD31 | NeuN | Dll4 | Notch1 | 0.002184 | 0 | DLL4_NOTCH1 | Dll4 - Notch1 | NOTCH | Cell-Cell Contact | PMID: 22353464 |
| CD31 | CD31 | Dll4 | Notch3 | 0.008353 | 0 | DLL4_NOTCH3 | Dll4 - Notch3 | NOTCH | Cell-Cell Contact | PMID: 22353464 |
| CD31 | NeuN | Dll4 | Notch3 | 0.002222 | 0 | DLL4_NOTCH3 | Dll4 - Notch3 | NOTCH | Cell-Cell Contact | PMID: 22353464 |
| CD31 | CD31 | Dll4 | Notch4 | 0.005124 | 0 | DLL4_NOTCH4 | Dll4 - Notch4 | NOTCH | Cell-Cell Contact | PMID: 22353464 |
| CD31 | NeuN | Dll4 | Notch4 | 0.001256 | 0 | DLL4_NOTCH4 | Dll4 - Notch4 | NOTCH | Cell-Cell Contact | PMID: 22353464 |
| CD31 | CD31 | Jag1 | Notch1 | 0.003285 | 0 | JAG1_NOTCH1 | Jag1 - Notch1 | NOTCH | Cell-Cell Contact | PMID: 22353464 |
| NeuN | CD31 | Jag1 | Notch1 | 0.001269 | 0 | JAG1_NOTCH1 | Jag1 - Notch1 | NOTCH | Cell-Cell Contact | PMID: 22353464 |
| CD31 | CD31 | Jag1 | Notch3 | 0.00391 | 0 | JAG1_NOTCH3 | Jag1 - Notch3 | NOTCH | Cell-Cell Contact | PMID: 22353464 |
| NeuN | CD31 | Jag1 | Notch3 | 0.001511 | 0 | JAG1_NOTCH3 | Jag1 - Notch3 | NOTCH | Cell-Cell Contact | PMID: 22353464 |
| CD31 | CD31 | Jag1 | Notch4 | 0.002394 | 0 | JAG1_NOTCH4 | Jag1 - Notch4 | NOTCH | Cell-Cell Contact | PMID: 22353464 |
| NeuN | CD31 | Jag1 | Notch4 | 0.000924 | 0 | JAG1_NOTCH4 | Jag1 - Notch4 | NOTCH | Cell-Cell Contact | PMID: 22353464 |
| CD31 | CD31 | Jag2 | Notch1 | 0.003824 | 0 | JAG2_NOTCH1 | Jag2 - Notch1 | NOTCH | Cell-Cell Contact | PMID: 22353464 |
| NeuN | CD31 | Jag2 | Notch1 | 0.002665 | 0 | JAG2_NOTCH1 | Jag2 - Notch1 | NOTCH | Cell-Cell Contact | PMID: 22353464 |
| CD31 | CD31 | Jag2 | Notch3 | 0.004551 | 0 | JAG2_NOTCH3 | Jag2 - Notch3 | NOTCH | Cell-Cell Contact | PMID: 22353464 |
| NeuN | CD31 | Jag2 | Notch3 | 0.003172 | 0 | JAG2_NOTCH3 | Jag2 - Notch3 | NOTCH | Cell-Cell Contact | PMID: 22353464 |
| CD31 | CD31 | Jag2 | Notch4 | 0.002787 | 0 | JAG2_NOTCH4 | Jag2 - Notch4 | NOTCH | Cell-Cell Contact | PMID: 22353464 |

| | | | | | | | | | | |
|---|---|---|---|---|---|---|---|---|---|---|
| NeuN | CD31 | Jag2 | Notch4 | 0.001942 | 0 | JAG2_NOTCH4 | Jag2 - Notch4 | NOTCH | Cell-Cell Contact | PMID: 22353464 |
| NeuN | CD31 | Nrxn1 | Nlgn1 | 0.008379 | 0 | NRXN1_NLGN1 | Nrxn1 - Nlgn1 | NRXN | Cell-Cell Contact | KEGG: mmu04514 |
| NeuN | NeuN | Nrxn1 | Nlgn1 | 0.018192 | 0 | NRXN1_NLGN1 | Nrxn1 - Nlgn1 | NRXN | Cell-Cell Contact | KEGG: mmu04514 |
| NeuN | CD31 | Nrxn1 | Nlgn2 | 0.02096 | 0 | NRXN1_NLGN2 | Nrxn1 - Nlgn2 | NRXN | Cell-Cell Contact | KEGG: mmu04514 |
| NeuN | NeuN | Nrxn1 | Nlgn2 | 0.039525 | 0 | NRXN1_NLGN2 | Nrxn1 - Nlgn2 | NRXN | Cell-Cell Contact | KEGG: mmu04514 |
| NeuN | CD31 | Nrxn1 | Nlgn3 | 0.02049 | 0 | NRXN1_NLGN3 | Nrxn1 - Nlgn3 | NRXN | Cell-Cell Contact | KEGG: mmu04514 |
| NeuN | NeuN | Nrxn1 | Nlgn3 | 0.033895 | 0 | NRXN1_NLGN3 | Nrxn1 - Nlgn3 | NRXN | Cell-Cell Contact | KEGG: mmu04514 |
| CD31 | NeuN | Nrxn2 | Nlgn1 | 0.005779 | 0 | NRXN2_NLGN1 | Nrxn2 - Nlgn1 | NRXN | Cell-Cell Contact | KEGG: mmu04514 |
| NeuN | NeuN | Nrxn2 | Nlgn1 | 0.012334 | 0 | NRXN2_NLGN1 | Nrxn2 - Nlgn1 | NRXN | Cell-Cell Contact | KEGG: mmu04514 |
| NeuN | CD31 | Nrxn2 | Nlgn2 | 0.014223 | 0 | NRXN2_NLGN2 | Nrxn2 - Nlgn2 | NRXN | Cell-Cell Contact | KEGG: mmu04514 |
| NeuN | NeuN | Nrxn2 | Nlgn2 | 0.026986 | 0 | NRXN2_NLGN2 | Nrxn2 - Nlgn2 | NRXN | Cell-Cell Contact | KEGG: mmu04514 |
| NeuN | CD31 | Nrxn2 | Nlgn3 | 0.013903 | 0 | NRXN2_NLGN3 | Nrxn2 - Nlgn3 | NRXN | Cell-Cell Contact | KEGG: mmu04514 |
| NeuN | NeuN | Nrxn2 | Nlgn3 | 0.023099 | 0 | NRXN2_NLGN3 | Nrxn2 - Nlgn3 | NRXN | Cell-Cell Contact | KEGG: mmu04514 |
| CD31 | NeuN | Nrxn3 | Nlgn1 | 0.003618 | 0 | NRXN3_NLGN1 | Nrxn3 - Nlgn1 | NRXN | Cell-Cell Contact | KEGG: mmu04514 |
| NeuN | NeuN | Nrxn3 | Nlgn1 | 0.004661 | 0 | NRXN3_NLGN1 | Nrxn3 - Nlgn1 | NRXN | Cell-Cell Contact | KEGG: mmu04514 |
| CD31 | NeuN | Nrxn3 | Nlgn2 | 0.008 | 0 | NRXN3_NLGN2 | Nrxn3 - Nlgn2 | NRXN | Cell-Cell Contact | KEGG: mmu04514 |
| NeuN | NeuN | Nrxn3 | Nlgn2 | 0.010294 | 0 | NRXN3_NLGN2 | Nrxn3 - Nlgn2 | NRXN | Cell-Cell Contact | KEGG: mmu04514 |
| CD31 | NeuN | Nrxn3 | Nlgn3 | 0.006828 | 0 | NRXN3_NLGN3 | Nrxn3 - Nlgn3 | NRXN | Cell-Cell Contact | KEGG: mmu04514 |
| NeuN | NeuN | Nrxn3 | Nlgn3 | 0.00879 | 0 | NRXN3_NLGN3 | Nrxn3 - Nlgn3 | NRXN | Cell-Cell Contact | KEGG: mmu04514 |
| CD31 | CD31 | Ocln | Ocln | 0.002024 | 0 | OCLN_OCLN | Ocln - Ocln | OCLN | Cell-Cell Contact | KEGG: mmu04514 |
| NeuN | CD31 | Ocln | Ocln | 0.000662 | 0 | OCLN_OCLN | Ocln - Ocln | OCLN | Cell-Cell Contact | KEGG: mmu04514 |
| CD31 | NeuN | Ocln | Ocln | 0.000662 | 0 | OCLN_OCLN | Ocln - Ocln | OCLN | Cell-Cell Contact | KEGG: mmu04514 |
| CD31 | CD31 | Pecam1 | Pecam1 | 0.0073 | 0 | PECAM1_PECAM1 | Pecam1 - Pecam1 | PECAM1 | Cell-Cell Contact | KEGG: mmu04514 |
| NeuN | CD31 | Pecam1 | Pecam1 | 0.001108 | 0 | PECAM1_PECAM1 | Pecam1 - Pecam1 | PECAM1 | Cell-Cell Contact | KEGG: mmu04514 |
| CD31 | NeuN | Pecam1 | Pecam1 | 0.001108 | 0 | PECAM1_PECAM1 | Pecam1 - Pecam1 | PECAM1 | Cell-Cell Contact | KEGG: mmu04514 |
| CD31 | CD31 | Ptprm | Ptprm | 0.01683 | 0 | PTPRM_PTPRM | Ptprm - Ptprm | PTPRM | Cell-Cell Contact | KEGG: mmu04514 |
| NeuN | CD31 | Ptprm | Ptprm | 0.005497 | 0 | PTPRM_PTPRM | Ptprm - Ptprm | PTPRM | Cell-Cell Contact | KEGG: mmu04514 |
| CD31 | NeuN | Ptprm | Ptprm | 0.005497 | 0 | PTPRM_PTPRM | Ptprm - Ptprm | PTPRM | Cell-Cell Contact | KEGG: mmu04514 |
| CD31 | CD31 | Sele | Glg1 | 0.001303 | 0 | SELE_GLG1 | Sele - Glg1 | SELE | Cell-Cell Contact | PMID: 11404363 |
| NeuN | CD31 | Sele | Glg1 | 0.001536 | 0 | SELE_GLG1 | Sele - Glg1 | SELE | Cell-Cell Contact | PMID: 11404363 |
| CD31 | CD31 | Sell | Cd34 | 0.006958 | 0 | SELL_CD34 | Sell - Cd34 | SELL | Cell-Cell Contact | PMID: 7692600; PMID: 8977216 |
| NeuN | CD31 | Sell | Cd34 | 0.003965 | 0 | SELL_CD34 | Sell - Cd34 | SELL | Cell-Cell Contact | PMID: 7692600; PMID: 8977216 |
| CD31 | CD31 | Sell | Podxl | 0.005494 | 0 | SELL_PODXL | Sell - Podxl | SELL | Cell-Cell Contact | PMID: 22814396 |
| NeuN | CD31 | Sell | Podxl | 0.003129 | 0 | SELL_PODXL | Sell - Podxl | SELL | Cell-Cell Contact | PMID: 22814396 |
| CD31 | NeuN | Selplg | Sele | 0.000356 | 0 | SELPLG_SELE | Selplg - Sele | SELPLG | Cell-Cell Contact | PMC4571854 |
| NeuN | NeuN | Selplg | Sele | 0.000355 | 0 | SELPLG_SELE | Selplg - Sele | SELPLG | Cell-Cell Contact | PMC4571854 |
| CD31 | CD31 | Selplg | Sell | 0.000764 | 0 | SELPLG_SELL | Selplg - Sell | SELPLG | Cell-Cell Contact | PMC2431087; PMID: 8892633,PMID: 8896607 |
| NeuN | CD31 | Selplg | Sell | 0.000762 | 0 | SELPLG_SELL | Selplg - Sell | SELPLG | Cell-Cell Contact | PMC2431087; PMID: 8892633,PMID: 8896607 |
| NeuN | CD31 | Sema4a | NRP1_PLXNA1 | 0.003696 | 0 | SEMA4A_NRP1_PLXNA1 | Sema4a - (Nrp1+Plxna1) | SEMA4 | Cell-Cell Contact | PMID: 27533782 |
| NeuN | NeuN | Sema4a | NRP1_PLXNA1 | 0.00447 | 0 | SEMA4A_NRP1_PLXNA1 | Sema4a - (Nrp1+Plxna1) | SEMA4 | Cell-Cell Contact | PMID: 27533782 |
| NeuN | CD31 | Sema4a | NRP1_PLXNA2 | 0.003863 | 0 | SEMA4A_NRP1_PLXNA2 | Sema4a - (Nrp1+Plxna2) | SEMA4 | Cell-Cell Contact | PMID: 27533782 |
| NeuN | NeuN | Sema4a | NRP1_PLXNA2 | 0.005584 | 0 | SEMA4A_NRP1_PLXNA2 | Sema4a - (Nrp1+Plxna2) | SEMA4 | Cell-Cell Contact | PMID: 27533782 |
| NeuN | CD31 | Sema4a | NRP1_PLXNA3 | 0.003262 | 0 | SEMA4A_NRP1_PLXNA3 | Sema4a - (Nrp1+Plxna3) | SEMA4 | Cell-Cell Contact | PMID: 27533782 |
| NeuN | NeuN | Sema4a | NRP1_PLXNA3 | 0.004671 | 0 | SEMA4A_NRP1_PLXNA3 | Sema4a - (Nrp1+Plxna3) | SEMA4 | Cell-Cell Contact | PMID: 27533782 |
| NeuN | CD31 | Sema4a | NRP1_PLXNA4 | 0.006119 | 0 | SEMA4A_NRP1_PLXNA4 | Sema4a - (Nrp1+Plxna4) | SEMA4 | Cell-Cell Contact | PMID: 27533782 |
| NeuN | NeuN | Sema4a | NRP1_PLXNA4 | 0.008112 | 0 | SEMA4A_NRP1_PLXNA4 | Sema4a - (Nrp1+Plxna4) | SEMA4 | Cell-Cell Contact | PMID: 27533782 |
| CD31 | NeuN | Sema4a | Plxnb1 | 0.001724 | 0 | SEMA4A_PLXNB1 | Sema4a - Plxnb1 | SEMA4 | Cell-Cell Contact | PMID: 27533782 |
| NeuN | NeuN | Sema4a | Plxnb1 | 0.002803 | 0 | SEMA4A_PLXNB1 | Sema4a - Plxnb1 | SEMA4 | Cell-Cell Contact | PMID: 27533782 |
| NeuN | CD31 | Sema4a | Plxnb2 | 0.004002 | 0 | SEMA4A_PLXNB2 | Sema4a - Plxnb2 | SEMA4 | Cell-Cell Contact | PMID: 27533782 |
| NeuN | NeuN | Sema4a | Plxnb2 | 0.005499 | 0 | SEMA4A_PLXNB2 | Sema4a - Plxnb2 | SEMA4 | Cell-Cell Contact | PMID: 27533782 |
| NeuN | CD31 | Sema4a | Plxnb3 | 0.001004 | 0 | SEMA4A_PLXNB3 | Sema4a - Plxnb3 | SEMA4 | Cell-Cell Contact | PMID: 27533782 |
| NeuN | NeuN | Sema4a | Plxnb3 | 0.00064 | 0 | SEMA4A_PLXNB3 | Sema4a - Plxnb3 | SEMA4 | Cell-Cell Contact | PMID: 27533782 |
| CD31 | NeuN | Sema4d | Plxnb1 | 0.002173 | 0 | SEMA4D_PLXNB1 | Sema4d - Plxnb1 | SEMA4 | Cell-Cell Contact | PMID: 27533782 |
| NeuN | NeuN | Sema4d | Plxnb1 | 0.002391 | 0 | SEMA4D_PLXNB1 | Sema4d - Plxnb1 | SEMA4 | Cell-Cell Contact | PMID: 27533782 |
| CD31 | NeuN | Sema4d | Plxnb2 | 0.004265 | 0 | SEMA4D_PLXNB2 | Sema4d - Plxnb2 | SEMA4 | Cell-Cell Contact | PMID: 27533782 |
| NeuN | NeuN | Sema4d | Plxnb2 | 0.004692 | 0 | SEMA4D_PLXNB2 | Sema4d - Plxnb2 | SEMA4 | Cell-Cell Contact | PMID: 27533782 |
| CD31 | CD31 | Sema4d | Plxnb3 | 0.000778 | 0 | SEMA4D_PLXNB3 | Sema4d - Plxnb3 | SEMA4 | Cell-Cell Contact | PMID: 27533782 |
| NeuN | CD31 | Sema4d | Plxnb3 | 0.000856 | 0 | SEMA4D_PLXNB3 | Sema4d - Plxnb3 | SEMA4 | Cell-Cell Contact | PMID: 27533782 |
| CD31 | NeuN | Sema4c | Plxnb2 | 0.010647 | 0 | SEMA4C_PLXNB2 | Sema4c - Plxnb2 | SEMA4 | Cell-Cell Contact | PMID: 27533782 |

| | | | | | | | | | | |
|---|---|---|---|---|---|---|---|---|---|---|
| NeuN | NeuN | Sema4c | Plxnb2 | 0.013121 | 0 | SEMA4C_PLXNB2 | Sema4c - Plxnb2 | SEMA4 | Cell-Cell Contact | PMID: 27533782 |
| NeuN | CD31 | Sema4g | Plxnb2 | 0.00581 | 0 | SEMA4G_PLXNB2 | Sema4g - Plxnb2 | SEMA4 | Cell-Cell Contact | PMID: 27533782 |
| NeuN | NeuN | Sema4g | Plxnb2 | 0.007977 | 0 | SEMA4G_PLXNB2 | Sema4g - Plxnb2 | SEMA4 | Cell-Cell Contact | PMID: 27533782 |
| NeuN | CD31 | Sema4a | Timd2 | 0.00083 | 0 | SEMA4A_TIMD2 | Sema4a - Timd2 | SEMA4 | Cell-Cell Contact | PMID: 22325954 |
| NeuN | NeuN | Sema4a | Timd2 | 0.001049 | 0 | SEMA4A_TIMD2 | Sema4a - Timd2 | SEMA4 | Cell-Cell Contact | PMID: 22325954 |
| CD31 | NeuN | Sema5a | Plxna1 | 0.002929 | 0 | SEMA5A_PLXNA1 | Sema5a - Plxna1 | SEMA5 | Cell-Cell Contact | PMID: 27533782 |
| NeuN | NeuN | Sema5a | Plxna1 | 0.004306 | 0 | SEMA5A_PLXNA1 | Sema5a - Plxna1 | SEMA5 | Cell-Cell Contact | PMID: 27533782 |
| CD31 | NeuN | Sema5b | Plxna1 | 0.002885 | 0 | SEMA5B_PLXNA1 | Sema5b - Plxna1 | SEMA5 | Cell-Cell Contact | PMID: 27533782 |
| NeuN | NeuN | Sema5b | Plxna1 | 0.002987 | 0 | SEMA5B_PLXNA1 | Sema5b - Plxna1 | SEMA5 | Cell-Cell Contact | PMID: 27533782 |
| CD31 | NeuN | Sema5a | Plxna3 | 0.0032 | 0 | SEMA5A_PLXNA3 | Sema5a - Plxna3 | SEMA5 | Cell-Cell Contact | PMID: 27533782 |
| NeuN | NeuN | Sema5a | Plxna3 | 0.004703 | 0 | SEMA5A_PLXNA3 | Sema5a - Plxna3 | SEMA5 | Cell-Cell Contact | PMID: 27533782 |
| CD31 | NeuN | Sema5b | Plxna3 | 0.003152 | 0 | SEMA5B_PLXNA3 | Sema5b - Plxna3 | SEMA5 | Cell-Cell Contact | PMID: 27533782 |
| NeuN | NeuN | Sema5b | Plxna3 | 0.003263 | 0 | SEMA5B_PLXNA3 | Sema5b - Plxna3 | SEMA5 | Cell-Cell Contact | PMID: 27533782 |
| CD31 | CD31 | Sema5a | Plxnb3 | 0.00059 | 0 | SEMA5A_PLXNB3 | Sema5a - Plxnb3 | SEMA5 | Cell-Cell Contact | PMID: 27533782 |
| NeuN | CD31 | Sema5a | Plxnb3 | 0.000868 | 0 | SEMA5A_PLXNB3 | Sema5a - Plxnb3 | SEMA5 | Cell-Cell Contact | PMID: 27533782 |
| CD31 | CD31 | Sema6d | PLXNA1_KDR | 0.033756 | 0 | SEMA6D_PLXNA1_KDR | Sema6d - (Plxna1+Kdr) | SEMA6 | Cell-Cell Contact | PMID: 22325954 |
| CD31 | NeuN | Sema6d | PLXNA1_KDR | 0.012166 | 0 | SEMA6D_PLXNA1_KDR | Sema6d - (Plxna1+Kdr) | SEMA6 | Cell-Cell Contact | PMID: 22325954 |
| CD31 | CD31 | Sema6d | PLXNA1_TREM2 | 0.009289 | 0 | SEMA6D_PLXNA1_TREM2 | Sema6d - (Plxna1+Trem2) | SEMA6 | Cell-Cell Contact | PMID: 22325954 |
| CD31 | NeuN | Sema6d | PLXNA1_TREM2 | 0.008312 | 0 | SEMA6D_PLXNA1_TREM2 | Sema6d - (Plxna1+Trem2) | SEMA6 | Cell-Cell Contact | PMID: 22325954 |
| CD31 | CD31 | Sema6d | Plxna1 | 0.018948 | 0 | SEMA6D_PLXNA1 | Sema6d - Plxna1 | SEMA6 | Cell-Cell Contact | PMID: 27533782 |
| CD31 | NeuN | Sema6d | Plxna1 | 0.030864 | 0 | SEMA6D_PLXNA1 | Sema6d - Plxna1 | SEMA6 | Cell-Cell Contact | PMID: 27533782 |
| CD31 | NeuN | Sema6a | Plxna2 | 0.018208 | 0 | SEMA6A_PLXNA2 | Sema6a - Plxna2 | SEMA6 | Cell-Cell Contact | PMID: 27533782 |
| NeuN | NeuN | Sema6a | Plxna2 | 0.008316 | 0 | SEMA6A_PLXNA2 | Sema6a - Plxna2 | SEMA6 | Cell-Cell Contact | PMID: 27533782 |
| CD31 | NeuN | Sema6b | Plxna2 | 0.007323 | 0 | SEMA6B_PLXNA2 | Sema6b - Plxna2 | SEMA6 | Cell-Cell Contact | PMID: 27533782 |
| NeuN | NeuN | Sema6b | Plxna2 | 0.011674 | 0 | SEMA6B_PLXNA2 | Sema6b - Plxna2 | SEMA6 | Cell-Cell Contact | PMID: 27533782 |
| CD31 | CD31 | Sema6a | Plxna4 | 0.019426 | 0 | SEMA6A_PLXNA4 | Sema6a - Plxna4 | SEMA6 | Cell-Cell Contact | PMID: 27533782 |
| CD31 | NeuN | Sema6a | Plxna4 | 0.037857 | 0 | SEMA6A_PLXNA4 | Sema6a - Plxna4 | SEMA6 | Cell-Cell Contact | PMID: 27533782 |
| CD31 | NeuN | Sema6b | Plxna4 | 0.01541 | 0 | SEMA6B_PLXNA4 | Sema6b - Plxna4 | SEMA6 | Cell-Cell Contact | PMID: 27533782 |
| NeuN | NeuN | Sema6b | Plxna4 | 0.024446 | 0 | SEMA6B_PLXNA4 | Sema6b - Plxna4 | SEMA6 | Cell-Cell Contact | PMID: 27533782 |
| CD31 | CD31 | Sema7a | ITGB1_ITGA1 | 0.012758 | 0 | SEMA7A_ITGB1_ITGA1 | Sema7a - (Itgb1+Itga1) | SEMA7 | Cell-Cell Contact | PMID: 22325954 |
| NeuN | CD31 | Sema7a | ITGB1_ITGA1 | 0.016807 | 0 | SEMA7A_ITGB1_ITGA1 | Sema7a - (Itgb1+Itga1) | SEMA7 | Cell-Cell Contact | PMID: 22325954 |
| NeuN | CD31 | Sema7a | Plxnc1 | 0.003053 | 0 | SEMA7A_PLXNC1 | Sema7a - Plxnc1 | SEMA7 | Cell-Cell Contact | PMID: 27533782 |
| NeuN | NeuN | Sema7a | Plxnc1 | 0.002616 | 0 | SEMA7A_PLXNC1 | Sema7a - Plxnc1 | SEMA7 | Cell-Cell Contact | PMID: 27533782 |
| CD31 | CD31 | Thy1 | ITGAV_ITGB3 | 0.00139 | 0 | THY1_ITGAV_ITGB3 | Thy1 - (Itgav+Itgb3) | THY1 | Cell-Cell Contact | PMID: 29212879; PMID: 18346467 |
| NeuN | CD31 | Thy1 | ITGAV_ITGB3 | 0.001908 | 0 | THY1_ITGAV_ITGB3 | Thy1 - (Itgav+Itgb3) | THY1 | Cell-Cell Contact | PMID: 29212879; PMID: 18346467 |
| NeuN | CD31 | Cdh3 | Cdh3 | 0.000636 | 0 | CDH3_CDH3 | Cdh3 - Cdh3 | CDH | Cell-Cell Contact | KEGG: hsa04514 |
| CD31 | NeuN | Cdh3 | Cdh3 | 0.000636 | 0 | CDH3_CDH3 | Cdh3 - Cdh3 | CDH | Cell-Cell Contact | KEGG: hsa04514 |
| NeuN | NeuN | Cdh3 | Cdh3 | 0.000744 | 0 | CDH3_CDH3 | Cdh3 - Cdh3 | CDH | Cell-Cell Contact | KEGG: hsa04514 |
| CD31 | CD31 | Cdh12 | ITGA1_ITGB1 | 0.003699 | 0 | CDH12_ITGA1_ITGB1 | Cdh12 - ITGA1_ITGB1 | CDH | Cell-Cell Contact | PMID: 34385456 |
| NeuN | CD31 | Cdh12 | ITGA1_ITGB1 | 0.009884 | 0 | CDH12_ITGA1_ITGB1 | Cdh12 - ITGA1_ITGB1 | CDH | Cell-Cell Contact | PMID: 34385456 |
| CD31 | CD31 | Gja1 | Gja1 | 0.006145 | 0 | GJA1_GJA1 | Gja1 - Gja1 | GAP | Cell-Cell Contact | PMID: 32532966;PMID: 15094344 |
| NeuN | CD31 | Gja1 | Gja1 | 0.001553 | 0 | GJA1_GJA1 | Gja1 - Gja1 | GAP | Cell-Cell Contact | PMID: 32532966;PMID: 15094344 |
| CD31 | NeuN | Gja1 | Gja1 | 0.001553 | 0 | GJA1_GJA1 | Gja1 - Gja1 | GAP | Cell-Cell Contact | PMID: 32532966;PMID: 15094344 |
| CD31 | CD31 | Gja1 | Gja6 | 0.001541 | 0 | GJA1_GJA6 | Gja1 - Gja6 | GAP | Cell-Cell Contact | PMID: 32532966;PMID: 15094344 |
| CD31 | NeuN | Gja1 | Gja6 | 0.003032 | 0 | GJA1_GJA6 | Gja1 - Gja6 | GAP | Cell-Cell Contact | PMID: 32532966;PMID: 15094344 |
| NeuN | CD31 | Gja6 | Gja6 | 0.000759 | 0 | GJA6_GJA6 | Gja6 - Gja6 | GAP | Cell-Cell Contact | PMID: 32532966;PMID: 15094344 |
| CD31 | NeuN | Gja6 | Gja6 | 0.000759 | 0 | GJA6_GJA6 | Gja6 - Gja6 | GAP | Cell-Cell Contact | PMID: 32532966;PMID: 15094344 |
| NeuN | NeuN | Gja6 | Gja6 | 0.001494 | 0 | GJA6_GJA6 | Gja6 - Gja6 | GAP | Cell-Cell Contact | PMID: 32532966;PMID: 15094344 |
| CD31 | CD31 | Gjc1 | Gja1 | 0.008442 | 0 | GJC1_GJA1 | Gjc1 - Gja1 | GAP | Cell-Cell Contact | PMID: 32532966;PMID: 15094344 |
| NeuN | CD31 | Gjc1 | Gja1 | 0.006513 | 0 | GJC1_GJA1 | Gjc1 - Gja1 | GAP | Cell-Cell Contact | PMID: 32532966;PMID: 15094344 |
| CD31 | NeuN | Gjc1 | Gja6 | 0.00417 | 0 | GJC1_GJA6 | Gjc1 - Gja6 | GAP | Cell-Cell Contact | PMID: 32532966;PMID: 15094344 |
| NeuN | NeuN | Gjc1 | Gja6 | 0.003214 | 0 | GJC1_GJA6 | Gjc1 - Gja6 | GAP | Cell-Cell Contact | PMID: 32532966;PMID: 15094344 |
| CD31 | CD31 | Gjc1 | Gjc1 | 0.011587 | 0 | GJC1_GJC1 | Gjc1 - Gjc1 | GAP | Cell-Cell Contact | PMID: 32532966;PMID: 15094344 |
| NeuN | CD31 | Gjc1 | Gjc1 | 0.008946 | 0 | GJC1_GJC1 | Gjc1 - Gjc1 | GAP | Cell-Cell Contact | PMID: 32532966;PMID: 15094344 |
| CD31 | NeuN | Gjc1 | Gjc1 | 0.008946 | 0 | GJC1_GJC1 | Gjc1 - Gjc1 | GAP | Cell-Cell Contact | PMID: 32532966;PMID: 15094344 |
| CD31 | CD31 | Cldn1 | Cldn1 | 0.001465 | 0 | CLDN1_CLDN1 | Cldn1 - Cldn1 | CLDN | Cell-Cell Contact | PMID: 31942240 |
| NeuN | CD31 | Cldn1 | Cldn1 | 0.000948 | 0 | CLDN1_CLDN1 | Cldn1 - Cldn1 | CLDN | Cell-Cell Contact | PMID: 31942240 |
| CD31 | NeuN | Cldn1 | Cldn1 | 0.000948 | 0 | CLDN1_CLDN1 | Cldn1 - Cldn1 | CLDN | Cell-Cell Contact | PMID: 31942240 |
| CD31 | CD31 | Cldn3 | Cldn3 | 0.000822 | 0 | CLDN3_CLDN3 | Cldn3 - Cldn3 | CLDN | Cell-Cell Contact | PMID: 31942240 |

| | | | | | | | | | | |
|---|---|---|---|---|---|---|---|---|---|---|
| NeuN | CD31 | Cldn3 | Cldn3 | 0.000722 | 0 | CLDN3_CLDN3 | Cldn3 - Cldn3 | CLDN | Cell-Cell Contact | PMID: 31942240 |
| CD31 | NeuN | Cldn3 | Cldn3 | 0.000722 | 0 | CLDN3_CLDN3 | Cldn3 - Cldn3 | CLDN | Cell-Cell Contact | PMID: 31942240 |
| CD31 | CD31 | Pcdha9 | Pcdha9 | 0.000709 | 0 | PCDHA9_PCDHA9 | Pcdha9 - Pcdha9 | PCDH | Cell-Cell Contact | PMID: 35253643; PMID: 32694982 |
| NeuN | CD31 | Pcdha9 | Pcdha9 | 0.000684 | 0 | PCDHA9_PCDHA9 | Pcdha9 - Pcdha9 | PCDH | Cell-Cell Contact | PMID: 35253643; PMID: 32694982 |
| CD31 | NeuN | Pcdha9 | Pcdha9 | 0.000684 | 0 | PCDHA9_PCDHA9 | Pcdha9 - Pcdha9 | PCDH | Cell-Cell Contact | PMID: 35253643; PMID: 32694982 |
| NeuN | CD31 | Pcdha11 | Pcdha11 | 0.001563 | 0 | PCDHA11_PCDHA11 | Pcdha11 - Pcdha11 | PCDH | Cell-Cell Contact | PMID: 35253643; PMID: 32694982 |
| CD31 | NeuN | Pcdha11 | Pcdha11 | 0.001563 | 0 | PCDHA11_PCDHA11 | Pcdha11 - Pcdha11 | PCDH | Cell-Cell Contact | PMID: 35253643; PMID: 32694982 |
| NeuN | NeuN | Pcdha11 | Pcdha11 | 0.003258 | 0 | PCDHA11_PCDHA11 | Pcdha11 - Pcdha11 | PCDH | Cell-Cell Contact | PMID: 35253643; PMID: 32694982 |
| NeuN | CD31 | Pcdha10 | Pcdha10 | 0.000487 | 0 | PCDHA10_PCDHA10 | Pcdha10 - Pcdha10 | PCDH | Cell-Cell Contact | PMID: 35253643; PMID: 32694982 |
| CD31 | NeuN | Pcdha10 | Pcdha10 | 0.000487 | 0 | PCDHA10_PCDHA10 | Pcdha10 - Pcdha10 | PCDH | Cell-Cell Contact | PMID: 35253643; PMID: 32694982 |
| NeuN | NeuN | Pcdha10 | Pcdha10 | 0.000508 | 0 | PCDHA10_PCDHA10 | Pcdha10 - Pcdha10 | PCDH | Cell-Cell Contact | PMID: 35253643; PMID: 32694982 |
| NeuN | CD31 | Pcdha12 | Pcdha12 | 0.001169 | 0 | PCDHA12_PCDHA12 | Pcdha12 - Pcdha12 | PCDH | Cell-Cell Contact | PMID: 35253643; PMID: 32694982 |
| CD31 | NeuN | Pcdha12 | Pcdha12 | 0.001169 | 0 | PCDHA12_PCDHA12 | Pcdha12 - Pcdha12 | PCDH | Cell-Cell Contact | PMID: 35253643; PMID: 32694982 |
| NeuN | NeuN | Pcdha12 | Pcdha12 | 0.002009 | 0 | PCDHA12_PCDHA12 | Pcdha12 - Pcdha12 | PCDH | Cell-Cell Contact | PMID: 35253643; PMID: 32694982 |
| CD31 | CD31 | Pcdha2 | Pcdha2 | 0.001139 | 0 | PCDHA2_PCDHA2 | Pcdha2 - Pcdha2 | PCDH | Cell-Cell Contact | PMID: 35253643; PMID: 32694982 |
| NeuN | CD31 | Pcdha2 | Pcdha2 | 0.000962 | 0 | PCDHA2_PCDHA2 | Pcdha2 - Pcdha2 | PCDH | Cell-Cell Contact | PMID: 35253643; PMID: 32694982 |
| CD31 | NeuN | Pcdha2 | Pcdha2 | 0.000962 | 0 | PCDHA2_PCDHA2 | Pcdha2 - Pcdha2 | PCDH | Cell-Cell Contact | PMID: 35253643; PMID: 32694982 |
| NeuN | CD31 | Pcdha3 | Pcdha3 | 0.000986 | 0 | PCDHA3_PCDHA3 | Pcdha3 - Pcdha3 | PCDH | Cell-Cell Contact | PMID: 35253643; PMID: 32694982 |
| CD31 | NeuN | Pcdha3 | Pcdha3 | 0.000986 | 0 | PCDHA3_PCDHA3 | Pcdha3 - Pcdha3 | PCDH | Cell-Cell Contact | PMID: 35253643; PMID: 32694982 |
| NeuN | NeuN | Pcdha3 | Pcdha3 | 0.001206 | 0 | PCDHA3_PCDHA3 | Pcdha3 - Pcdha3 | PCDH | Cell-Cell Contact | PMID: 35253643; PMID: 32694982 |
| NeuN | CD31 | Pcdha5 | Pcdha5 | 0.00115 | 0 | PCDHA5_PCDHA5 | Pcdha5 - Pcdha5 | PCDH | Cell-Cell Contact | PMID: 35253643; PMID: 32694982 |
| CD31 | NeuN | Pcdha5 | Pcdha5 | 0.00115 | 0 | PCDHA5_PCDHA5 | Pcdha5 - Pcdha5 | PCDH | Cell-Cell Contact | PMID: 35253643; PMID: 32694982 |
| NeuN | NeuN | Pcdha5 | Pcdha5 | 0.002355 | 0 | PCDHA5_PCDHA5 | Pcdha5 - Pcdha5 | PCDH | Cell-Cell Contact | PMID: 35253643; PMID: 32694982 |
| NeuN | CD31 | Pcdha8 | Pcdha8 | 0.000605 | 0 | PCDHA8_PCDHA8 | Pcdha8 - Pcdha8 | PCDH | Cell-Cell Contact | PMID: 35253643; PMID: 32694982 |
| CD31 | NeuN | Pcdha8 | Pcdha8 | 0.000605 | 0 | PCDHA8_PCDHA8 | Pcdha8 - Pcdha8 | PCDH | Cell-Cell Contact | PMID: 35253643; PMID: 32694982 |
| NeuN | NeuN | Pcdha8 | Pcdha8 | 0.000674 | 0 | PCDHA8_PCDHA8 | Pcdha8 - Pcdha8 | PCDH | Cell-Cell Contact | PMID: 35253643; PMID: 32694982 |
| NeuN | CD31 | Pcdhac1 | Pcdhac1 | 0.000784 | 0 | PCDHAC1_PCDHAC1 | Pcdhac1 - Pcdhac1 | PCDH | Cell-Cell Contact | PMID: 35253643; PMID: 32694982 |
| CD31 | NeuN | Pcdhac1 | Pcdhac1 | 0.000784 | 0 | PCDHAC1_PCDHAC1 | Pcdhac1 - Pcdhac1 | PCDH | Cell-Cell Contact | PMID: 35253643; PMID: 32694982 |
| NeuN | NeuN | Pcdhac1 | Pcdhac1 | 0.001352 | 0 | PCDHAC1_PCDHAC1 | Pcdhac1 - Pcdhac1 | PCDH | Cell-Cell Contact | PMID: 35253643; PMID: 32694982 |
| NeuN | CD31 | Pcdhac2 | Pcdhac2 | 0.000967 | 0 | PCDHAC2_PCDHAC2 | Pcdhac2 - Pcdhac2 | PCDH | Cell-Cell Contact | PMID: 35253643; PMID: 32694982 |
| CD31 | NeuN | Pcdhac2 | Pcdhac2 | 0.000967 | 0 | PCDHAC2_PCDHAC2 | Pcdhac2 - Pcdhac2 | PCDH | Cell-Cell Contact | PMID: 35253643; PMID: 32694982 |
| NeuN | NeuN | Pcdhac2 | Pcdhac2 | 0.001463 | 0 | PCDHAC2_PCDHAC2 | Pcdhac2 - Pcdhac2 | PCDH | Cell-Cell Contact | PMID: 35253643; PMID: 32694982 |
| CD31 | CD31 | Pcdhb18 | Pcdhb18 | 0.001046 | 0 | PCDHB18_PCDHB18 | Pcdhb18 - Pcdhb18 | PCDH | Cell-Cell Contact | PMID: 35253643; PMID: 32694982 |
| NeuN | CD31 | Pcdhb18 | Pcdhb18 | 0.000733 | 0 | PCDHB18_PCDHB18 | Pcdhb18 - Pcdhb18 | PCDH | Cell-Cell Contact | PMID: 35253643; PMID: 32694982 |
| CD31 | NeuN | Pcdhb18 | Pcdhb18 | 0.000733 | 0 | PCDHB18_PCDHB18 | Pcdhb18 - Pcdhb18 | PCDH | Cell-Cell Contact | PMID: 35253643; PMID: 32694982 |
| NeuN | CD31 | Pcdhb19 | Pcdhb19 | 0.000608 | 0 | PCDHB19_PCDHB19 | Pcdhb19 - Pcdhb19 | PCDH | Cell-Cell Contact | PMID: 35253643; PMID: 32694982 |
| CD31 | NeuN | Pcdhb19 | Pcdhb19 | 0.000608 | 0 | PCDHB19_PCDHB19 | Pcdhb19 - Pcdhb19 | PCDH | Cell-Cell Contact | PMID: 35253643; PMID: 32694982 |
| NeuN | NeuN | Pcdhb19 | Pcdhb19 | 0.000961 | 0 | PCDHB19_PCDHB19 | Pcdhb19 - Pcdhb19 | PCDH | Cell-Cell Contact | PMID: 35253643; PMID: 32694982 |
| NeuN | CD31 | Pcdhb19 | Pcdhb21 | 0.000818 | 0 | PCDHB19_PCDHB21 | Pcdhb19 - Pcdhb21 | PCDH | Cell-Cell Contact | PMID: 35253643; PMID: 32694982 |
| NeuN | NeuN | Pcdhb19 | Pcdhb21 | 0.000914 | 0 | PCDHB19_PCDHB21 | Pcdhb19 - Pcdhb21 | PCDH | Cell-Cell Contact | PMID: 35253643; PMID: 32694982 |
| NeuN | CD31 | Pcdhb21 | Pcdhb21 | 0.000778 | 0 | PCDHB21_PCDHB21 | Pcdhb21 - Pcdhb21 | PCDH | Cell-Cell Contact | PMID: 35253643; PMID: 32694982 |
| CD31 | NeuN | Pcdhb21 | Pcdhb21 | 0.000778 | 0 | PCDHB21_PCDHB21 | Pcdhb21 - Pcdhb21 | PCDH | Cell-Cell Contact | PMID: 35253643; PMID: 32694982 |
| NeuN | NeuN | Pcdhb21 | Pcdhb21 | 0.000869 | 0 | PCDHB21_PCDHB21 | Pcdhb21 - Pcdhb21 | PCDH | Cell-Cell Contact | PMID: 35253643; PMID: 32694982 |
| CD31 | CD31 | Pcdhb16 | Pcdhb16 | 0.000743 | 0 | PCDHB16_PCDHB16 | Pcdhb16 - Pcdhb16 | PCDH | Cell-Cell Contact | PMID: 35253643; PMID: 32694982 |
| NeuN | CD31 | Pcdhb16 | Pcdhb16 | 0.000608 | 0 | PCDHB16_PCDHB16 | Pcdhb16 - Pcdhb16 | PCDH | Cell-Cell Contact | PMID: 35253643; PMID: 32694982 |
| CD31 | NeuN | Pcdhb16 | Pcdhb16 | 0.000608 | 0 | PCDHB16_PCDHB16 | Pcdhb16 - Pcdhb16 | PCDH | Cell-Cell Contact | PMID: 35253643; PMID: 32694982 |
| NeuN | CD31 | Pcdhb20 | Pcdhb20 | 0.001248 | 0 | PCDHB20_PCDHB20 | Pcdhb20 - Pcdhb20 | PCDH | Cell-Cell Contact | PMID: 35253643; PMID: 32694982 |
| CD31 | NeuN | Pcdhb20 | Pcdhb20 | 0.001248 | 0 | PCDHB20_PCDHB20 | Pcdhb20 - Pcdhb20 | PCDH | Cell-Cell Contact | PMID: 35253643; PMID: 32694982 |
| NeuN | NeuN | Pcdhb20 | Pcdhb20 | 0.001539 | 0 | PCDHB20_PCDHB20 | Pcdhb20 - Pcdhb20 | PCDH | Cell-Cell Contact | PMID: 35253643; PMID: 32694982 |
| NeuN | CD31 | Pcdhb22 | Pcdhb22 | 0.000562 | 0 | PCDHB22_PCDHB22 | Pcdhb22 - Pcdhb22 | PCDH | Cell-Cell Contact | PMID: 35253643; PMID: 32694982 |
| CD31 | NeuN | Pcdhb22 | Pcdhb22 | 0.000562 | 0 | PCDHB22_PCDHB22 | Pcdhb22 - Pcdhb22 | PCDH | Cell-Cell Contact | PMID: 35253643; PMID: 32694982 |
| NeuN | NeuN | Pcdhb22 | Pcdhb22 | 0.000798 | 0 | PCDHB22_PCDHB22 | Pcdhb22 - Pcdhb22 | PCDH | Cell-Cell Contact | PMID: 35253643; PMID: 32694982 |
| NeuN | CD31 | Pcdhb17 | Pcdhb17 | 0.000754 | 0 | PCDHB17_PCDHB17 | Pcdhb17 - Pcdhb17 | PCDH | Cell-Cell Contact | PMID: 35253643; PMID: 32694982 |
| CD31 | NeuN | Pcdhb17 | Pcdhb17 | 0.000754 | 0 | PCDHB17_PCDHB17 | Pcdhb17 - Pcdhb17 | PCDH | Cell-Cell Contact | PMID: 35253643; PMID: 32694982 |
| NeuN | NeuN | Pcdhb17 | Pcdhb17 | 0.000881 | 0 | PCDHB17_PCDHB17 | Pcdhb17 - Pcdhb17 | PCDH | Cell-Cell Contact | PMID: 35253643; PMID: 32694982 |
| CD31 | CD31 | Pcdhb2 | Pcdhb2 | 0.000653 | 0 | PCDHB2_PCDHB2 | Pcdhb2 - Pcdhb2 | PCDH | Cell-Cell Contact | PMID: 35253643; PMID: 32694982 |
| NeuN | CD31 | Pcdhb2 | Pcdhb2 | 0.000552 | 0 | PCDHB2_PCDHB2 | Pcdhb2 - Pcdhb2 | PCDH | Cell-Cell Contact | PMID: 35253643; PMID: 32694982 |
| CD31 | NeuN | Pcdhb2 | Pcdhb2 | 0.000552 | 0 | PCDHB2_PCDHB2 | Pcdhb2 - Pcdhb2 | PCDH | Cell-Cell Contact | PMID: 35253643; PMID: 32694982 |
| CD31 | CD31 | Pcdhb3 | Pcdhb3 | 0.001158 | 0 | PCDHB3_PCDHB3 | Pcdhb3 - Pcdhb3 | PCDH | Cell-Cell Contact | PMID: 35253643; PMID: 32694982 |

| | | | | | | | | | | |
|---|---|---|---|---|---|---|---|---|---|---|
| NeuN | CD31 | Pcdhb3 | Pcdhb3 | 0.000593 | 0 | PCDHB3_PCDHB3 | Pcdhb3 - Pcdhb3 | PCDH | Cell-Cell Contact | PMID: 35253643; PMID: 32694982 |
| CD31 | NeuN | Pcdhb3 | Pcdhb3 | 0.000593 | 0 | PCDHB3_PCDHB3 | Pcdhb3 - Pcdhb3 | PCDH | Cell-Cell Contact | PMID: 35253643; PMID: 32694982 |
| NeuN | CD31 | Pcdhb7 | Pcdhb7 | 0.000541 | 0 | PCDHB7_PCDHB7 | Pcdhb7 - Pcdhb7 | PCDH | Cell-Cell Contact | PMID: 35253643; PMID: 32694982 |
| CD31 | NeuN | Pcdhb7 | Pcdhb7 | 0.000541 | 0 | PCDHB7_PCDHB7 | Pcdhb7 - Pcdhb7 | PCDH | Cell-Cell Contact | PMID: 35253643; PMID: 32694982 |
| NeuN | NeuN | Pcdhb7 | Pcdhb7 | 0.000605 | 0 | PCDHB7_PCDHB7 | Pcdhb7 - Pcdhb7 | PCDH | Cell-Cell Contact | PMID: 35253643; PMID: 32694982 |
| CD31 | CD31 | Pcdhb7 | Pcdhb5 | 0.000581 | 0 | PCDHB7_PCDHB5 | Pcdhb7 - Pcdhb5 | PCDH | Cell-Cell Contact | PMID: 35253643; PMID: 32694982 |
| NeuN | CD31 | Pcdhb7 | Pcdhb5 | 0.00065 | 0 | PCDHB7_PCDHB5 | Pcdhb7 - Pcdhb5 | PCDH | Cell-Cell Contact | PMID: 35253643; PMID: 32694982 |
| CD31 | NeuN | Pcdhb7 | Pcdhb9 | 0.000497 | 0 | PCDHB7_PCDHB9 | Pcdhb7 - Pcdhb9 | PCDH | Cell-Cell Contact | PMID: 35253643; PMID: 32694982 |
| NeuN | NeuN | Pcdhb7 | Pcdhb9 | 0.000556 | 0 | PCDHB7_PCDHB9 | Pcdhb7 - Pcdhb9 | PCDH | Cell-Cell Contact | PMID: 35253643; PMID: 32694982 |
| CD31 | CD31 | Pcdhb5 | Pcdhb5 | 0.000697 | 0 | PCDHB5_PCDHB5 | Pcdhb5 - Pcdhb5 | PCDH | Cell-Cell Contact | PMID: 35253643; PMID: 32694982 |
| NeuN | CD31 | Pcdhb5 | Pcdhb5 | 0.000555 | 0 | PCDHB5_PCDHB5 | Pcdhb5 - Pcdhb5 | PCDH | Cell-Cell Contact | PMID: 35253643; PMID: 32694982 |
| CD31 | NeuN | Pcdhb5 | Pcdhb5 | 0.000555 | 0 | PCDHB5_PCDHB5 | Pcdhb5 - Pcdhb5 | PCDH | Cell-Cell Contact | PMID: 35253643; PMID: 32694982 |
| CD31 | NeuN | Pcdhb5 | Pcdhb9 | 0.000597 | 0 | PCDHB5_PCDHB9 | Pcdhb5 - Pcdhb9 | PCDH | Cell-Cell Contact | PMID: 35253643; PMID: 32694982 |
| NeuN | NeuN | Pcdhb5 | Pcdhb9 | 0.000475 | 0 | PCDHB5_PCDHB9 | Pcdhb5 - Pcdhb9 | PCDH | Cell-Cell Contact | PMID: 35253643; PMID: 32694982 |
| NeuN | CD31 | Pcdhb9 | Pcdhb9 | 0.000405 | 0 | PCDHB9_PCDHB9 | Pcdhb9 - Pcdhb9 | PCDH | Cell-Cell Contact | PMID: 35253643; PMID: 32694982 |
| CD31 | NeuN | Pcdhb9 | Pcdhb9 | 0.000405 | 0 | PCDHB9_PCDHB9 | Pcdhb9 - Pcdhb9 | PCDH | Cell-Cell Contact | PMID: 35253643; PMID: 32694982 |
| NeuN | NeuN | Pcdhb9 | Pcdhb9 | 0.000511 | 0 | PCDHB9_PCDHB9 | Pcdhb9 - Pcdhb9 | PCDH | Cell-Cell Contact | PMID: 35253643; PMID: 32694982 |
| CD31 | CD31 | Pcdhb12 | Pcdhb12 | 0.000575 | 0 | PCDHB12_PCDHB12 | Pcdhb12 - Pcdhb12 | PCDH | Cell-Cell Contact | PMID: 35253643; PMID: 32694982 |
| NeuN | CD31 | Pcdhb12 | Pcdhb12 | 0.000465 | 0 | PCDHB12_PCDHB12 | Pcdhb12 - Pcdhb12 | PCDH | Cell-Cell Contact | PMID: 35253643; PMID: 32694982 |
| CD31 | NeuN | Pcdhb12 | Pcdhb12 | 0.000465 | 0 | PCDHB12_PCDHB12 | Pcdhb12 - Pcdhb12 | PCDH | Cell-Cell Contact | PMID: 35253643; PMID: 32694982 |
| CD31 | CD31 | Pcdhb12 | Pcdhb10 | 0.000525 | 0 | PCDHB12_PCDHB10 | Pcdhb12 - Pcdhb10 | PCDH | Cell-Cell Contact | PMID: 35253643; PMID: 32694982 |
| NeuN | CD31 | Pcdhb12 | Pcdhb10 | 0.000424 | 0 | PCDHB12_PCDHB10 | Pcdhb12 - Pcdhb10 | PCDH | Cell-Cell Contact | PMID: 35253643; PMID: 32694982 |
| CD31 | CD31 | Pcdhb12 | Pcdhb11 | 0.000494 | 0 | PCDHB12_PCDHB11 | Pcdhb12 - Pcdhb11 | PCDH | Cell-Cell Contact | PMID: 35253643; PMID: 32694982 |
| CD31 | NeuN | Pcdhb12 | Pcdhb11 | 0.000505 | 0 | PCDHB12_PCDHB11 | Pcdhb12 - Pcdhb11 | PCDH | Cell-Cell Contact | PMID: 35253643; PMID: 32694982 |
| CD31 | CD31 | Pcdhb12 | Pcdhb8 | 0.000429 | 0 | PCDHB12_PCDHB8 | Pcdhb12 - Pcdhb8 | PCDH | Cell-Cell Contact | PMID: 35253643; PMID: 32694982 |
| CD31 | NeuN | Pcdhb12 | Pcdhb8 | 0.000364 | 0 | PCDHB12_PCDHB8 | Pcdhb12 - Pcdhb8 | PCDH | Cell-Cell Contact | PMID: 35253643; PMID: 32694982 |
| CD31 | CD31 | Pcdhb12 | Pcdhb6 | 0.000552 | 0 | PCDHB12_PCDHB6 | Pcdhb12 - Pcdhb6 | PCDH | Cell-Cell Contact | PMID: 35253643; PMID: 32694982 |
| CD31 | NeuN | Pcdhb12 | Pcdhb6 | 0.00052 | 0 | PCDHB12_PCDHB6 | Pcdhb12 - Pcdhb6 | PCDH | Cell-Cell Contact | PMID: 35253643; PMID: 32694982 |
| CD31 | CD31 | Pcdhb10 | Pcdhb10 | 0.000479 | 0 | PCDHB10_PCDHB10 | Pcdhb10 - Pcdhb10 | PCDH | Cell-Cell Contact | PMID: 35253643; PMID: 32694982 |
| NeuN | CD31 | Pcdhb10 | Pcdhb10 | 0.000301 | 0 | PCDHB10_PCDHB10 | Pcdhb10 - Pcdhb10 | PCDH | Cell-Cell Contact | PMID: 35253643; PMID: 32694982 |
| CD31 | NeuN | Pcdhb10 | Pcdhb10 | 0.000301 | 0 | PCDHB10_PCDHB10 | Pcdhb10 - Pcdhb10 | PCDH | Cell-Cell Contact | PMID: 35253643; PMID: 32694982 |
| CD31 | CD31 | Pcdhb10 | Pcdhb11 | 0.000451 | 0 | PCDHB10_PCDHB11 | Pcdhb10 - Pcdhb11 | PCDH | Cell-Cell Contact | PMID: 35253643; PMID: 32694982 |
| CD31 | NeuN | Pcdhb10 | Pcdhb11 | 0.00046 | 0 | PCDHB10_PCDHB11 | Pcdhb10 - Pcdhb11 | PCDH | Cell-Cell Contact | PMID: 35253643; PMID: 32694982 |
| CD31 | CD31 | Pcdhb10 | Pcdhb8 | 0.000391 | 0 | PCDHB10_PCDHB8 | Pcdhb10 - Pcdhb8 | PCDH | Cell-Cell Contact | PMID: 35253643; PMID: 32694982 |
| CD31 | NeuN | Pcdhb10 | Pcdhb8 | 0.000333 | 0 | PCDHB10_PCDHB8 | Pcdhb10 - Pcdhb8 | PCDH | Cell-Cell Contact | PMID: 35253643; PMID: 32694982 |
| CD31 | CD31 | Pcdhb10 | Pcdhb6 | 0.000503 | 0 | PCDHB10_PCDHB6 | Pcdhb10 - Pcdhb6 | PCDH | Cell-Cell Contact | PMID: 35253643; PMID: 32694982 |
| CD31 | NeuN | Pcdhb10 | Pcdhb6 | 0.000474 | 0 | PCDHB10_PCDHB6 | Pcdhb10 - Pcdhb6 | PCDH | Cell-Cell Contact | PMID: 35253643; PMID: 32694982 |
| NeuN | CD31 | Pcdhb11 | Pcdhb11 | 0.000434 | 0 | PCDHB11_PCDHB11 | Pcdhb11 - Pcdhb11 | PCDH | Cell-Cell Contact | PMID: 35253643; PMID: 32694982 |
| CD31 | NeuN | Pcdhb11 | Pcdhb11 | 0.000434 | 0 | PCDHB11_PCDHB11 | Pcdhb11 - Pcdhb11 | PCDH | Cell-Cell Contact | PMID: 35253643; PMID: 32694982 |
| NeuN | NeuN | Pcdhb11 | Pcdhb11 | 0.000443 | 0 | PCDHB11_PCDHB11 | Pcdhb11 - Pcdhb11 | PCDH | Cell-Cell Contact | PMID: 35253643; PMID: 32694982 |
| CD31 | CD31 | Pcdhb11 | Pcdhb8 | 0.000368 | 0 | PCDHB11_PCDHB8 | Pcdhb11 - Pcdhb8 | PCDH | Cell-Cell Contact | PMID: 35253643; PMID: 32694982 |
| NeuN | CD31 | Pcdhb11 | Pcdhb8 | 0.000376 | 0 | PCDHB11_PCDHB8 | Pcdhb11 - Pcdhb8 | PCDH | Cell-Cell Contact | PMID: 35253643; PMID: 32694982 |
| CD31 | CD31 | Pcdhb11 | Pcdhb6 | 0.000474 | 0 | PCDHB11_PCDHB6 | Pcdhb11 - Pcdhb6 | PCDH | Cell-Cell Contact | PMID: 35253643; PMID: 32694982 |
| NeuN | CD31 | Pcdhb11 | Pcdhb6 | 0.000484 | 0 | PCDHB11_PCDHB6 | Pcdhb11 - Pcdhb6 | PCDH | Cell-Cell Contact | PMID: 35253643; PMID: 32694982 |
| CD31 | CD31 | Pcdhb8 | Pcdhb8 | 0.00032 | 0 | PCDHB8_PCDHB8 | Pcdhb8 - Pcdhb8 | PCDH | Cell-Cell Contact | PMID: 35253643; PMID: 32694982 |
| NeuN | CD31 | Pcdhb8 | Pcdhb8 | 0.000272 | 0 | PCDHB8_PCDHB8 | Pcdhb8 - Pcdhb8 | PCDH | Cell-Cell Contact | PMID: 35253643; PMID: 32694982 |
| CD31 | NeuN | Pcdhb8 | Pcdhb8 | 0.000272 | 0 | PCDHB8_PCDHB8 | Pcdhb8 - Pcdhb8 | PCDH | Cell-Cell Contact | PMID: 35253643; PMID: 32694982 |
| CD31 | CD31 | Pcdhb8 | Pcdhb6 | 0.000411 | 0 | PCDHB8_PCDHB6 | Pcdhb8 - Pcdhb6 | PCDH | Cell-Cell Contact | PMID: 35253643; PMID: 32694982 |
| CD31 | NeuN | Pcdhb8 | Pcdhb6 | 0.000388 | 0 | PCDHB8_PCDHB6 | Pcdhb8 - Pcdhb6 | PCDH | Cell-Cell Contact | PMID: 35253643; PMID: 32694982 |
| CD31 | CD31 | Pcdhb6 | Pcdhb6 | 0.000529 | 0 | PCDHB6_PCDHB6 | Pcdhb6 - Pcdhb6 | PCDH | Cell-Cell Contact | PMID: 35253643; PMID: 32694982 |
| NeuN | CD31 | Pcdhb6 | Pcdhb6 | 0.000499 | 0 | PCDHB6_PCDHB6 | Pcdhb6 - Pcdhb6 | PCDH | Cell-Cell Contact | PMID: 35253643; PMID: 32694982 |
| CD31 | NeuN | Pcdhb6 | Pcdhb6 | 0.000499 | 0 | PCDHB6_PCDHB6 | Pcdhb6 - Pcdhb6 | PCDH | Cell-Cell Contact | PMID: 35253643; PMID: 32694982 |
| CD31 | CD31 | Pcdhb13 | Pcdhb13 | 0.001225 | 0 | PCDHB13_PCDHB13 | Pcdhb13 - Pcdhb13 | PCDH | Cell-Cell Contact | PMID: 35253643; PMID: 32694982 |
| NeuN | CD31 | Pcdhb13 | Pcdhb13 | 0.001069 | 0 | PCDHB13_PCDHB13 | Pcdhb13 - Pcdhb13 | PCDH | Cell-Cell Contact | PMID: 35253643; PMID: 32694982 |
| CD31 | NeuN | Pcdhb13 | Pcdhb13 | 0.001069 | 0 | PCDHB13_PCDHB13 | Pcdhb13 - Pcdhb13 | PCDH | Cell-Cell Contact | PMID: 35253643; PMID: 32694982 |
| NeuN | CD31 | Pcdhb15 | Pcdhb15 | 0.000262 | 0 | PCDHB15_PCDHB15 | Pcdhb15 - Pcdhb15 | PCDH | Cell-Cell Contact | PMID: 35253643; PMID: 32694982 |
| CD31 | NeuN | Pcdhb15 | Pcdhb15 | 0.000262 | 0 | PCDHB15_PCDHB15 | Pcdhb15 - Pcdhb15 | PCDH | Cell-Cell Contact | PMID: 35253643; PMID: 32694982 |
| NeuN | NeuN | Pcdhb15 | Pcdhb15 | 0.000325 | 0 | PCDHB15_PCDHB15 | Pcdhb15 - Pcdhb15 | PCDH | Cell-Cell Contact | PMID: 35253643; PMID: 32694982 |
| CD31 | CD31 | Pcdhga1 | Pcdhga1 | 0.00143 | 0 | PCDHGA1_PCDHGA1 | Pcdhga1 - Pcdhga1 | PCDH | Cell-Cell Contact | PMID: 35253643; PMID: 32694982 |

| | | | | | | | | | | |
|---|---|---|---|---|---|---|---|---|---|---|
| NeuN | CD31 | Pcdhga1 | Pcdhga1 | 0.00104 | 0 | PCDHGA1_PCDHGA1 | Pcdhga1 - Pcdhga1 | PCDH | Cell-Cell Contact | PMID: 35253643; PMID: 32694982 |
| CD31 | NeuN | Pcdhga1 | Pcdhga1 | 0.00104 | 0 | PCDHGA1_PCDHGA1 | Pcdhga1 - Pcdhga1 | PCDH | Cell-Cell Contact | PMID: 35253643; PMID: 32694982 |
| CD31 | CD31 | Pcdhga10 | Pcdhga10 | 0.00077 | 0 | PCDHGA10_PCDHGA10 | Pcdhga10 - Pcdhga10 | PCDH | Cell-Cell Contact | PMID: 35253643; PMID: 32694982 |
| NeuN | CD31 | Pcdhga10 | Pcdhga10 | 0.000512 | 0 | PCDHGA10_PCDHGA10 | Pcdhga10 - Pcdhga10 | PCDH | Cell-Cell Contact | PMID: 35253643; PMID: 32694982 |
| CD31 | NeuN | Pcdhga10 | Pcdhga10 | 0.000512 | 0 | PCDHGA10_PCDHGA10 | Pcdhga10 - Pcdhga10 | PCDH | Cell-Cell Contact | PMID: 35253643; PMID: 32694982 |
| NeuN | CD31 | Pcdhga11 | Pcdhga11 | 0.002279 | 0 | PCDHGA11_PCDHGA11 | Pcdhga11 - Pcdhga11 | PCDH | Cell-Cell Contact | PMID: 35253643; PMID: 32694982 |
| CD31 | NeuN | Pcdhga11 | Pcdhga11 | 0.002279 | 0 | PCDHGA11_PCDHGA11 | Pcdhga11 - Pcdhga11 | PCDH | Cell-Cell Contact | PMID: 35253643; PMID: 32694982 |
| NeuN | NeuN | Pcdhga11 | Pcdhga11 | 0.002876 | 0 | PCDHGA11_PCDHGA11 | Pcdhga11 - Pcdhga11 | PCDH | Cell-Cell Contact | PMID: 35253643; PMID: 32694982 |
| NeuN | CD31 | Pcdhga12 | Pcdhga12 | 0.001891 | 0 | PCDHGA12_PCDHGA12 | Pcdhga12 - Pcdhga12 | PCDH | Cell-Cell Contact | PMID: 35253643; PMID: 32694982 |
| CD31 | NeuN | Pcdhga12 | Pcdhga12 | 0.001891 | 0 | PCDHGA12_PCDHGA12 | Pcdhga12 - Pcdhga12 | PCDH | Cell-Cell Contact | PMID: 35253643; PMID: 32694982 |
| NeuN | NeuN | Pcdhga12 | Pcdhga12 | 0.002027 | 0 | PCDHGA12_PCDHGA12 | Pcdhga12 - Pcdhga12 | PCDH | Cell-Cell Contact | PMID: 35253643; PMID: 32694982 |
| NeuN | CD31 | Pcdhga2 | Pcdhga2 | 0.000731 | 0 | PCDHGA2_PCDHGA2 | Pcdhga2 - Pcdhga2 | PCDH | Cell-Cell Contact | PMID: 35253643; PMID: 32694982 |
| CD31 | NeuN | Pcdhga2 | Pcdhga2 | 0.000731 | 0 | PCDHGA2_PCDHGA2 | Pcdhga2 - Pcdhga2 | PCDH | Cell-Cell Contact | PMID: 35253643; PMID: 32694982 |
| NeuN | NeuN | Pcdhga2 | Pcdhga2 | 0.000769 | 0 | PCDHGA2_PCDHGA2 | Pcdhga2 - Pcdhga2 | PCDH | Cell-Cell Contact | PMID: 35253643; PMID: 32694982 |
| NeuN | CD31 | Pcdhga3 | Pcdhga3 | 0.000472 | 0 | PCDHGA3_PCDHGA3 | Pcdhga3 - Pcdhga3 | PCDH | Cell-Cell Contact | PMID: 35253643; PMID: 32694982 |
| CD31 | NeuN | Pcdhga3 | Pcdhga3 | 0.000472 | 0 | PCDHGA3_PCDHGA3 | Pcdhga3 - Pcdhga3 | PCDH | Cell-Cell Contact | PMID: 35253643; PMID: 32694982 |
| NeuN | NeuN | Pcdhga3 | Pcdhga3 | 0.000599 | 0 | PCDHGA3_PCDHGA3 | Pcdhga3 - Pcdhga3 | PCDH | Cell-Cell Contact | PMID: 35253643; PMID: 32694982 |
| NeuN | CD31 | Pcdhga4 | Pcdhga4 | 0.000963 | 0 | PCDHGA4_PCDHGA4 | Pcdhga4 - Pcdhga4 | PCDH | Cell-Cell Contact | PMID: 35253643; PMID: 32694982 |
| CD31 | NeuN | Pcdhga4 | Pcdhga4 | 0.000963 | 0 | PCDHGA4_PCDHGA4 | Pcdhga4 - Pcdhga4 | PCDH | Cell-Cell Contact | PMID: 35253643; PMID: 32694982 |
| NeuN | NeuN | Pcdhga4 | Pcdhga4 | 0.001225 | 0 | PCDHGA4_PCDHGA4 | Pcdhga4 - Pcdhga4 | PCDH | Cell-Cell Contact | PMID: 35253643; PMID: 32694982 |
| NeuN | CD31 | Pcdhga5 | Pcdhga5 | 0.000875 | 0 | PCDHGA5_PCDHGA5 | Pcdhga5 - Pcdhga5 | PCDH | Cell-Cell Contact | PMID: 35253643; PMID: 32694982 |
| CD31 | NeuN | Pcdhga5 | Pcdhga5 | 0.000875 | 0 | PCDHGA5_PCDHGA5 | Pcdhga5 - Pcdhga5 | PCDH | Cell-Cell Contact | PMID: 35253643; PMID: 32694982 |
| NeuN | NeuN | Pcdhga5 | Pcdhga5 | 0.001188 | 0 | PCDHGA5_PCDHGA5 | Pcdhga5 - Pcdhga5 | PCDH | Cell-Cell Contact | PMID: 35253643; PMID: 32694982 |
| NeuN | CD31 | Pcdhga6 | Pcdhga6 | 0.001046 | 0 | PCDHGA6_PCDHGA6 | Pcdhga6 - Pcdhga6 | PCDH | Cell-Cell Contact | PMID: 35253643; PMID: 32694982 |
| CD31 | NeuN | Pcdhga6 | Pcdhga6 | 0.001046 | 0 | PCDHGA6_PCDHGA6 | Pcdhga6 - Pcdhga6 | PCDH | Cell-Cell Contact | PMID: 35253643; PMID: 32694982 |
| NeuN | NeuN | Pcdhga6 | Pcdhga6 | 0.001057 | 0 | PCDHGA6_PCDHGA6 | Pcdhga6 - Pcdhga6 | PCDH | Cell-Cell Contact | PMID: 35253643; PMID: 32694982 |
| CD31 | CD31 | Pcdhga7 | Pcdhga7 | 0.00112 | 0 | PCDHGA7_PCDHGA7 | Pcdhga7 - Pcdhga7 | PCDH | Cell-Cell Contact | PMID: 35253643; PMID: 32694982 |
| NeuN | CD31 | Pcdhga7 | Pcdhga7 | 0.0011 | 0 | PCDHGA7_PCDHGA7 | Pcdhga7 - Pcdhga7 | PCDH | Cell-Cell Contact | PMID: 35253643; PMID: 32694982 |
| CD31 | NeuN | Pcdhga7 | Pcdhga7 | 0.0011 | 0 | PCDHGA7_PCDHGA7 | Pcdhga7 - Pcdhga7 | PCDH | Cell-Cell Contact | PMID: 35253643; PMID: 32694982 |
| CD31 | CD31 | Pcdhga8 | Pcdhga8 | 0.000712 | 0 | PCDHGA8_PCDHGA8 | Pcdhga8 - Pcdhga8 | PCDH | Cell-Cell Contact | PMID: 35253643; PMID: 32694982 |
| NeuN | CD31 | Pcdhga8 | Pcdhga8 | 0.000632 | 0 | PCDHGA8_PCDHGA8 | Pcdhga8 - Pcdhga8 | PCDH | Cell-Cell Contact | PMID: 35253643; PMID: 32694982 |
| CD31 | NeuN | Pcdhga8 | Pcdhga8 | 0.000632 | 0 | PCDHGA8_PCDHGA8 | Pcdhga8 - Pcdhga8 | PCDH | Cell-Cell Contact | PMID: 35253643; PMID: 32694982 |
| CD31 | CD31 | Pcdhga9 | Pcdhga9 | 0.000653 | 0 | PCDHGA9_PCDHGA9 | Pcdhga9 - Pcdhga9 | PCDH | Cell-Cell Contact | PMID: 35253643; PMID: 32694982 |
| NeuN | CD31 | Pcdhga9 | Pcdhga9 | 0.000389 | 0 | PCDHGA9_PCDHGA9 | Pcdhga9 - Pcdhga9 | PCDH | Cell-Cell Contact | PMID: 35253643; PMID: 32694982 |
| CD31 | NeuN | Pcdhga9 | Pcdhga9 | 0.000389 | 0 | PCDHGA9_PCDHGA9 | Pcdhga9 - Pcdhga9 | PCDH | Cell-Cell Contact | PMID: 35253643; PMID: 32694982 |
| NeuN | CD31 | Pcdhgb1 | Pcdhgb1 | 0.001443 | 0 | PCDHGB1_PCDHGB1 | Pcdhgb1 - Pcdhgb1 | PCDH | Cell-Cell Contact | PMID: 35253643; PMID: 32694982 |
| CD31 | NeuN | Pcdhgb1 | Pcdhgb1 | 0.001443 | 0 | PCDHGB1_PCDHGB1 | Pcdhgb1 - Pcdhgb1 | PCDH | Cell-Cell Contact | PMID: 35253643; PMID: 32694982 |
| NeuN | NeuN | Pcdhgb1 | Pcdhgb1 | 0.00175 | 0 | PCDHGB1_PCDHGB1 | Pcdhgb1 - Pcdhgb1 | PCDH | Cell-Cell Contact | PMID: 35253643; PMID: 32694982 |
| CD31 | CD31 | Pcdhgb2 | Pcdhgb2 | 0.002666 | 0 | PCDHGB2_PCDHGB2 | Pcdhgb2 - Pcdhgb2 | PCDH | Cell-Cell Contact | PMID: 35253643; PMID: 32694982 |
| NeuN | CD31 | Pcdhgb2 | Pcdhgb2 | 0.002437 | 0 | PCDHGB2_PCDHGB2 | Pcdhgb2 - Pcdhgb2 | PCDH | Cell-Cell Contact | PMID: 35253643; PMID: 32694982 |
| CD31 | NeuN | Pcdhgb2 | Pcdhgb2 | 0.002437 | 0 | PCDHGB2_PCDHGB2 | Pcdhgb2 - Pcdhgb2 | PCDH | Cell-Cell Contact | PMID: 35253643; PMID: 32694982 |
| NeuN | CD31 | Pcdhgb8 | Pcdhgb8 | 0.000287 | 0 | PCDHGB8_PCDHGB8 | Pcdhgb8 - Pcdhgb8 | PCDH | Cell-Cell Contact | PMID: 35253643; PMID: 32694982 |
| CD31 | NeuN | Pcdhgb8 | Pcdhgb8 | 0.000287 | 0 | PCDHGB8_PCDHGB8 | Pcdhgb8 - Pcdhgb8 | PCDH | Cell-Cell Contact | PMID: 35253643; PMID: 32694982 |
| NeuN | NeuN | Pcdhgb8 | Pcdhgb8 | 0.000302 | 0 | PCDHGB8_PCDHGB8 | Pcdhgb8 - Pcdhgb8 | PCDH | Cell-Cell Contact | PMID: 35253643; PMID: 32694982 |
| NeuN | CD31 | Pcdhgb4 | Pcdhgb4 | 0.001378 | 0 | PCDHGB4_PCDHGB4 | Pcdhgb4 - Pcdhgb4 | PCDH | Cell-Cell Contact | PMID: 35253643; PMID: 32694982 |
| CD31 | NeuN | Pcdhgb4 | Pcdhgb4 | 0.001378 | 0 | PCDHGB4_PCDHGB4 | Pcdhgb4 - Pcdhgb4 | PCDH | Cell-Cell Contact | PMID: 35253643; PMID: 32694982 |
| NeuN | NeuN | Pcdhgb4 | Pcdhgb4 | 0.001565 | 0 | PCDHGB4_PCDHGB4 | Pcdhgb4 - Pcdhgb4 | PCDH | Cell-Cell Contact | PMID: 35253643; PMID: 32694982 |
| NeuN | CD31 | Pcdhgb5 | Pcdhgb5 | 0.000918 | 0 | PCDHGB5_PCDHGB5 | Pcdhgb5 - Pcdhgb5 | PCDH | Cell-Cell Contact | PMID: 35253643; PMID: 32694982 |
| CD31 | NeuN | Pcdhgb5 | Pcdhgb5 | 0.000918 | 0 | PCDHGB5_PCDHGB5 | Pcdhgb5 - Pcdhgb5 | PCDH | Cell-Cell Contact | PMID: 35253643; PMID: 32694982 |
| NeuN | NeuN | Pcdhgb5 | Pcdhgb5 | 0.001045 | 0 | PCDHGB5_PCDHGB5 | Pcdhgb5 - Pcdhgb5 | PCDH | Cell-Cell Contact | PMID: 35253643; PMID: 32694982 |
| NeuN | CD31 | Pcdhgb6 | Pcdhgb6 | 0.002405 | 0 | PCDHGB6_PCDHGB6 | Pcdhgb6 - Pcdhgb6 | PCDH | Cell-Cell Contact | PMID: 35253643; PMID: 32694982 |
| CD31 | NeuN | Pcdhgb6 | Pcdhgb6 | 0.002405 | 0 | PCDHGB6_PCDHGB6 | Pcdhgb6 - Pcdhgb6 | PCDH | Cell-Cell Contact | PMID: 35253643; PMID: 32694982 |
| NeuN | NeuN | Pcdhgb6 | Pcdhgb6 | 0.00291 | 0 | PCDHGB6_PCDHGB6 | Pcdhgb6 - Pcdhgb6 | PCDH | Cell-Cell Contact | PMID: 35253643; PMID: 32694982 |
| NeuN | CD31 | Pcdhgb7 | Pcdhgb7 | 0.005158 | 0 | PCDHGB7_PCDHGB7 | Pcdhgb7 - Pcdhgb7 | PCDH | Cell-Cell Contact | PMID: 35253643; PMID: 32694982 |
| CD31 | NeuN | Pcdhgb7 | Pcdhgb7 | 0.005158 | 0 | PCDHGB7_PCDHGB7 | Pcdhgb7 - Pcdhgb7 | PCDH | Cell-Cell Contact | PMID: 35253643; PMID: 32694982 |
| NeuN | NeuN | Pcdhgb7 | Pcdhgb7 | 0.009852 | 0 | PCDHGB7_PCDHGB7 | Pcdhgb7 - Pcdhgb7 | PCDH | Cell-Cell Contact | PMID: 35253643; PMID: 32694982 |
| NeuN | CD31 | Pcdhgc3 | Pcdhgc3 | 0.019006 | 0 | PCDHGC3_PCDHGC3 | Pcdhgc3 - Pcdhgc3 | PCDH | Cell-Cell Contact | PMID: 35253643; PMID: 32694982 |
| CD31 | NeuN | Pcdhgc3 | Pcdhgc3 | 0.019006 | 0 | PCDHGC3_PCDHGC3 | Pcdhgc3 - Pcdhgc3 | PCDH | Cell-Cell Contact | PMID: 35253643; PMID: 32694982 |
| NeuN | NeuN | Pcdhgc3 | Pcdhgc3 | 0.022907 | 0 | PCDHGC3_PCDHGC3 | Pcdhgc3 - Pcdhgc3 | PCDH | Cell-Cell Contact | PMID: 35253643; PMID: 32694982 |

| | | | | | | | | | |
|---|---|---|---|---|---|---|---|---|---|
| NeuN | CD31 | Pcdhgc4 | Pcdhgc4 | 0.001564 | 0 PCDHGC4_PCDHGC4 | Pcdhgc4 - Pcdhgc4 | PCDH | Cell-Cell Contact | PMID: 35253643; PMID: 32694982 |
| CD31 | NeuN | Pcdhgc4 | Pcdhgc4 | 0.001564 | 0 PCDHGC4_PCDHGC4 | Pcdhgc4 - Pcdhgc4 | PCDH | Cell-Cell Contact | PMID: 35253643; PMID: 32694982 |
| NeuN | NeuN | Pcdhgc4 | Pcdhgc4 | 0.002361 | 0 PCDHGC4_PCDHGC4 | Pcdhgc4 - Pcdhgc4 | PCDH | Cell-Cell Contact | PMID: 35253643; PMID: 32694982 |
| CD31 | CD31 | Pcdhgc5 | Pcdhgc5 | 0.000845 | 0 PCDHGC5_PCDHGC5 | Pcdhgc5 - Pcdhgc5 | PCDH | Cell-Cell Contact | PMID: 35253643; PMID: 32694982 |
| NeuN | CD31 | Pcdhgc5 | Pcdhgc5 | 0.000692 | 0 PCDHGC5_PCDHGC5 | Pcdhgc5 - Pcdhgc5 | PCDH | Cell-Cell Contact | PMID: 35253643; PMID: 32694982 |
| CD31 | NeuN | Pcdhgc5 | Pcdhgc5 | 0.000692 | 0 PCDHGC5_PCDHGC5 | Pcdhgc5 - Pcdhgc5 | PCDH | Cell-Cell Contact | PMID: 35253643; PMID: 32694982 |
| CD31 | CD31 | Sdc2 | Adgra2 | 0.002796 | 0 SDC2_ADGRA2 | Sdc2 - Adgra2 | ADGRA | Cell-Cell Contact | PMID: 31462748 |
| NeuN | CD31 | Sdc2 | Adgra2 | 0.001156 | 0 SDC2_ADGRA2 | Sdc2 - Adgra2 | ADGRA | Cell-Cell Contact | PMID: 31462748 |
| CD31 | CD31 | ITGAV_ITG | Adgra2 | 0.002308 | 0 ITGAV_ITGB3_ADGRA2 | (Itgav+Itgb3) - Adgra2 | ADGRA | Cell-Cell Contact | PMID: 31462748 |
| NeuN | CD31 | ITGAV_ITG | Adgra2 | 0.001366 | 0 ITGAV_ITGB3_ADGRA2 | (Itgav+Itgb3) - Adgra2 | ADGRA | Cell-Cell Contact | PMID: 31462748 |
| CD31 | NeuN | Tax1bp3 | Adgrb1 | 0.009065 | 0 TAX1BP3_ADGRB1 | Tax1bp3 - Adgrb1 | ADGRB | Cell-Cell Contact | PMID: 31462748 |
| NeuN | NeuN | Tax1bp3 | Adgrb1 | 0.004701 | 0 TAX1BP3_ADGRB1 | Tax1bp3 - Adgrb1 | ADGRB | Cell-Cell Contact | PMID: 31462748 |
| CD31 | NeuN | Rtn4r | Adgrb1 | 0.002127 | 0 RTN4R_ADGRB1 | Rtn4r - Adgrb1 | ADGRB | Cell-Cell Contact | PMID: 31462748 |
| NeuN | NeuN | Rtn4r | Adgrb1 | 0.003099 | 0 RTN4R_ADGRB1 | Rtn4r - Adgrb1 | ADGRB | Cell-Cell Contact | PMID: 31462748 |
| CD31 | NeuN | ITGAV_ITG | Adgrb1 | 0.001936 | 0 ITGAV_ITGB5_ADGRB1 | (Itgav+Itgb5) - Adgrb1 | ADGRB | Cell-Cell Contact | PMID: 31462748 |
| NeuN | NeuN | ITGAV_ITG | Adgrb1 | 0.00107 | 0 ITGAV_ITGB5_ADGRB1 | (Itgav+Itgb5) - Adgrb1 | ADGRB | Cell-Cell Contact | PMID: 31462748 |
| CD31 | CD31 | Tax1bp3 | Adgrb2 | 0.011958 | 0 TAX1BP3_ADGRB2 | Tax1bp3 - Adgrb2 | ADGRB | Cell-Cell Contact | PMID: 31462748 |
| CD31 | NeuN | Tax1bp3 | Adgrb2 | 0.021491 | 0 TAX1BP3_ADGRB2 | Tax1bp3 - Adgrb2 | ADGRB | Cell-Cell Contact | PMID: 31462748 |
| CD31 | NeuN | C1ql1 | Adgrb3 | 0.001998 | 0 C1QL1_ADGRB3 | C1ql1 - Adgrb3 | ADGRB | Cell-Cell Contact | PMID: 31462748 |
| NeuN | NeuN | C1ql1 | Adgrb3 | 0.001405 | 0 C1QL1_ADGRB3 | C1ql1 - Adgrb3 | ADGRB | Cell-Cell Contact | PMID: 31462748 |
| CD31 | NeuN | C1ql2 | Adgrb3 | 0.002254 | 0 C1QL2_ADGRB3 | C1ql2 - Adgrb3 | ADGRB | Cell-Cell Contact | PMID: 31462748 |
| NeuN | NeuN | C1ql2 | Adgrb3 | 0.00155 | 0 C1QL2_ADGRB3 | C1ql2 - Adgrb3 | ADGRB | Cell-Cell Contact | PMID: 31462748 |
| CD31 | NeuN | C1ql3 | Adgrb3 | 0.00197 | 0 C1QL3_ADGRB3 | C1ql3 - Adgrb3 | ADGRB | Cell-Cell Contact | PMID: 31462748 |
| NeuN | NeuN | C1ql3 | Adgrb3 | 0.001423 | 0 C1QL3_ADGRB3 | C1ql3 - Adgrb3 | ADGRB | Cell-Cell Contact | PMID: 31462748 |
| CD31 | CD31 | Thy1 | Adgre5 | 0.003593 | 0 THY1_ADGRE5 | Thy1 - Adgre5 | ADGRE | Cell-Cell Contact | PMID: 31462748 |
| NeuN | CD31 | Thy1 | Adgre5 | 0.004928 | 0 THY1_ADGRE5 | Thy1 - Adgre5 | ADGRE | Cell-Cell Contact | PMID: 31462748 |
| CD31 | CD31 | ITGAV_ITG | Adgre5 | 0.009849 | 0 ITGAV_ITGB1_ADGRE5 | (Itgav+Itgb1) - Adgre5 | ADGRE | Cell-Cell Contact | PMID: 31462748 |
| NeuN | CD31 | ITGAV_ITG | Adgre5 | 0.003938 | 0 ITGAV_ITGB1_ADGRE5 | (Itgav+Itgb1) - Adgre5 | ADGRE | Cell-Cell Contact | PMID: 31462748 |
| CD31 | CD31 | ITGAV_ITG | Adgre5 | 0.003976 | 0 ITGAV_ITGB3_ADGRE5 | (Itgav+Itgb3) - Adgre5 | ADGRE | Cell-Cell Contact | PMID: 31462748 |
| NeuN | CD31 | ITGAV_ITG | Adgre5 | 0.002354 | 0 ITGAV_ITGB3_ADGRE5 | (Itgav+Itgb3) - Adgre5 | ADGRE | Cell-Cell Contact | PMID: 31462748 |
| CD31 | CD31 | Cd55 | Adgre5 | 0.001877 | 0 CD55_ADGRE5 | Cd55 - Adgre5 | ADGRE | Cell-Cell Contact | PMID: 31462748 |
| NeuN | CD31 | Cd55 | Adgre5 | 0.001522 | 0 CD55_ADGRE5 | Cd55 - Adgre5 | ADGRE | Cell-Cell Contact | PMID: 31462748 |
| CD31 | CD31 | Lpar1 | Adgre5 | 0.003045 | 0 LPAR1_ADGRE5 | Lpar1 - Adgre5 | ADGRE | Cell-Cell Contact | PMID: 31462748 |
| NeuN | CD31 | Lpar1 | Adgre5 | 0.001747 | 0 LPAR1_ADGRE5 | Lpar1 - Adgre5 | ADGRE | Cell-Cell Contact | PMID: 31462748 |
| CD31 | CD31 | Lpar2 | Adgre5 | 0.002507 | 0 LPAR2_ADGRE5 | Lpar2 - Adgre5 | ADGRE | Cell-Cell Contact | PMID: 31462748 |
| NeuN | CD31 | Lpar2 | Adgre5 | 0.002483 | 0 LPAR2_ADGRE5 | Lpar2 - Adgre5 | ADGRE | Cell-Cell Contact | PMID: 31462748 |
| CD31 | CD31 | Col3a1 | Adgrg1 | 0.002707 | 0 COL3A1_ADGRG1 | Col3a1 - Adgrg1 | ADGRG | Cell-Cell Contact | PMID: 31462748;UniProtKB |
| CD31 | NeuN | Col3a1 | Adgrg1 | 0.004033 | 0 COL3A1_ADGRG1 | Col3a1 - Adgrg1 | ADGRG | Cell-Cell Contact | PMID: 31462748;UniProtKB |
| CD31 | NeuN | Tgm2 | Adgrg1 | 0.001752 | 0 TGM2_ADGRG1 | Tgm2 - Adgrg1 | ADGRG | Cell-Cell Contact | PMID: 31462748 |
| NeuN | NeuN | Tgm2 | Adgrg1 | 0.001432 | 0 TGM2_ADGRG1 | Tgm2 - Adgrg1 | ADGRG | Cell-Cell Contact | PMID: 31462748 |
| CD31 | CD31 | Col4a1 | Adgrg6 | 0.011739 | 0 COL4A1_ADGRG6 | Col4a1 - Adgrg6 | ADGRG | Cell-Cell Contact | PMID: 31462748 |
| CD31 | NeuN | Col4a1 | Adgrg6 | 0.012598 | 0 COL4A1_ADGRG6 | Col4a1 - Adgrg6 | ADGRG | Cell-Cell Contact | PMID: 31462748 |
| CD31 | CD31 | Col4a2 | Adgrg6 | 0.009246 | 0 COL4A2_ADGRG6 | Col4a2 - Adgrg6 | ADGRG | Cell-Cell Contact | PMID: 31462748 |
| CD31 | NeuN | Col4a2 | Adgrg6 | 0.009925 | 0 COL4A2_ADGRG6 | Col4a2 - Adgrg6 | ADGRG | Cell-Cell Contact | PMID: 31462748 |
| CD31 | CD31 | Prnp | Adgrg6 | 0.003264 | 0 PRNP_ADGRG6 | Prnp - Adgrg6 | ADGRG | Cell-Cell Contact | PMID: 31462748 |
| CD31 | NeuN | Prnp | Adgrg6 | 0.003506 | 0 PRNP_ADGRG6 | Prnp - Adgrg6 | ADGRG | Cell-Cell Contact | PMID: 31462748 |
| CD31 | NeuN | Flrt1 | Adgrl1 | 0.011909 | 0 FLRT1_ADGRL1 | Flrt1 - Adgrl1 | ADGRL | Cell-Cell Contact | PMID: 25728924;PMID: 34022279;PMID: 27091502;PMID: 35907405 |
| NeuN | NeuN | Flrt1 | Adgrl1 | 0.015375 | 0 FLRT1_ADGRL1 | Flrt1 - Adgrl1 | ADGRL | Cell-Cell Contact | PMID: 25728924;PMID: 34022279;PMID: 27091502;PMID: 35907405 |
| CD31 | NeuN | Flrt2 | Adgrl1 | 0.018126 | 0 FLRT2_ADGRL1 | Flrt2 - Adgrl1 | ADGRL | Cell-Cell Contact | PMID: 25728924;PMID: 34022279;PMID: 27091502;PMID: 35907405 |
| NeuN | NeuN | Flrt2 | Adgrl1 | 0.026775 | 0 FLRT2_ADGRL1 | Flrt2 - Adgrl1 | ADGRL | Cell-Cell Contact | PMID: 25728924;PMID: 34022279;PMID: 27091502;PMID: 35907405 |
| CD31 | NeuN | Flrt3 | Adgrl1 | 0.014988 | 0 FLRT3_ADGRL1 | Flrt3 - Adgrl1 | ADGRL | Cell-Cell Contact | PMID: 25728924;PMID: 34022279;PMID: 27091502;PMID: 35907405 |
| NeuN | NeuN | Flrt3 | Adgrl1 | 0.022718 | 0 FLRT3_ADGRL1 | Flrt3 - Adgrl1 | ADGRL | Cell-Cell Contact | PMID: 25728924;PMID: 34022279;PMID: 27091502;PMID: 35907405 |
| NeuN | CD31 | Nrxn1 | Adgrl1 | 0.043318 | 0 NRXN1_ADGRL1 | Nrxn1 - Adgrl1 | ADGRL | Cell-Cell Contact | PMID: 25728924;PMID: 34022279;PMID: 27091502;PMID: 35907405 |
| NeuN | NeuN | Nrxn1 | Adgrl1 | 0.076363 | 0 NRXN1_ADGRL1 | Nrxn1 - Adgrl1 | ADGRL | Cell-Cell Contact | PMID: 25728924;PMID: 34022279;PMID: 27091502;PMID: 35907405 |
| NeuN | CD31 | Nrxn2 | Adgrl1 | 0.029613 | 0 NRXN2_ADGRL1 | Nrxn2 - Adgrl1 | ADGRL | Cell-Cell Contact | PMID: 25728924;PMID: 34022279;PMID: 27091502;PMID: 35907405 |
| NeuN | NeuN | Nrxn2 | Adgrl1 | 0.05278 | 0 NRXN2_ADGRL1 | Nrxn2 - Adgrl1 | ADGRL | Cell-Cell Contact | PMID: 25728924;PMID: 34022279;PMID: 27091502;PMID: 35907405 |
| CD31 | NeuN | Nrxn3 | Adgrl1 | 0.015943 | 0 NRXN3_ADGRL1 | Nrxn3 - Adgrl1 | ADGRL | Cell-Cell Contact | PMID: 25728924;PMID: 34022279;PMID: 27091502;PMID: 35907405 |
| NeuN | NeuN | Nrxn3 | Adgrl1 | 0.020469 | 0 NRXN3_ADGRL1 | Nrxn3 - Adgrl1 | ADGRL | Cell-Cell Contact | PMID: 25728924;PMID: 34022279;PMID: 27091502;PMID: 35907405 |
| CD31 | NeuN | Tenm1 | Adgrl1 | 0.007177 | 0 TENM1_ADGRL1 | Tenm1 - Adgrl1 | ADGRL | Cell-Cell Contact | PMID: 25728924;PMID: 34022279;PMID: 27091502;PMID: 35907405 |

| | | | | | | | | | | |
|---|---|---|---|---|---|---|---|---|---|---|
| NeuN | NeuN | Tenm1 | Adgrl1 | 0.008652 | 0 | TENM1_ADGRL1 | Tenm1 - Adgrl1 | ADGRL | Cell-Cell Contact | PMID: 25728924;PMID: 34022279;PMID: 27091502;PMID: 35907405 |
| NeuN | CD31 | Tenm2 | Adgrl1 | 0.013935 | 0 | TENM2_ADGRL1 | Tenm2 - Adgrl1 | ADGRL | Cell-Cell Contact | PMID: 25728924;PMID: 34022279;PMID: 27091502;PMID: 35907405 |
| NeuN | NeuN | Tenm2 | Adgrl1 | 0.025154 | 0 | TENM2_ADGRL1 | Tenm2 - Adgrl1 | ADGRL | Cell-Cell Contact | PMID: 25728924;PMID: 34022279;PMID: 27091502;PMID: 35907405 |
| NeuN | CD31 | Tenm3 | Adgrl1 | 0.037521 | 0 | TENM3_ADGRL1 | Tenm3 - Adgrl1 | ADGRL | Cell-Cell Contact | PMID: 25728924;PMID: 34022279;PMID: 27091502;PMID: 35907405 |
| NeuN | NeuN | Tenm3 | Adgrl1 | 0.066452 | 0 | TENM3_ADGRL1 | Tenm3 - Adgrl1 | ADGRL | Cell-Cell Contact | PMID: 25728924;PMID: 34022279;PMID: 27091502;PMID: 35907405 |
| NeuN | CD31 | Tenm4 | Adgrl1 | 0.014377 | 0 | TENM4_ADGRL1 | Tenm4 - Adgrl1 | ADGRL | Cell-Cell Contact | PMID: 25728924;PMID: 34022279;PMID: 27091502;PMID: 35907405 |
| NeuN | NeuN | Tenm4 | Adgrl1 | 0.025944 | 0 | TENM4_ADGRL1 | Tenm4 - Adgrl1 | ADGRL | Cell-Cell Contact | PMID: 25728924;PMID: 34022279;PMID: 27091502;PMID: 35907405 |
| CD31 | NeuN | TENM2_FL | Adgrl1 | 0.012381 | 0 | TENM2_FLRT1_ADGRL1 | (Tenm2+Flrt1) - Adgrl1 | ADGRL | Cell-Cell Contact | PMID: 25728924;PMID: 34022279;PMID: 27091502;PMID: 35907405 |
| NeuN | NeuN | TENM2_FL | Adgrl1 | 0.019678 | 0 | TENM2_FLRT1_ADGRL1 | (Tenm2+Flrt1) - Adgrl1 | ADGRL | Cell-Cell Contact | PMID: 25728924;PMID: 34022279;PMID: 27091502;PMID: 35907405 |
| CD31 | NeuN | TENM2_FL | Adgrl1 | 0.01389 | 0 | TENM2_FLRT3_ADGRL1 | (Tenm2+Flrt3) - Adgrl1 | ADGRL | Cell-Cell Contact | PMID: 25728924;PMID: 34022279;PMID: 27091502;PMID: 35907405 |
| NeuN | NeuN | TENM2_FL | Adgrl1 | 0.023906 | 0 | TENM2_FLRT3_ADGRL1 | (Tenm2+Flrt3) - Adgrl1 | ADGRL | Cell-Cell Contact | PMID: 25728924;PMID: 34022279;PMID: 27091502;PMID: 35907405 |
| CD31 | NeuN | TENM4_FL | Adgrl1 | 0.012372 | 0 | TENM4_FLRT1_ADGRL1 | (Tenm4+Flrt1) - Adgrl1 | ADGRL | Cell-Cell Contact | PMID: 25728924;PMID: 34022279;PMID: 27091502;PMID: 35907405 |
| NeuN | NeuN | TENM4_FL | Adgrl1 | 0.019986 | 0 | TENM4_FLRT1_ADGRL1 | (Tenm4+Flrt1) - Adgrl1 | ADGRL | Cell-Cell Contact | PMID: 25728924;PMID: 34022279;PMID: 27091502;PMID: 35907405 |
| CD31 | NeuN | TENM4_FL | Adgrl1 | 0.01388 | 0 | TENM4_FLRT3_ADGRL1 | (Tenm4+Flrt3) - Adgrl1 | ADGRL | Cell-Cell Contact | PMID: 25728924;PMID: 34022279;PMID: 27091502;PMID: 35907405 |
| NeuN | NeuN | TENM4_FL | Adgrl1 | 0.024279 | 0 | TENM4_FLRT3_ADGRL1 | (Tenm4+Flrt3) - Adgrl1 | ADGRL | Cell-Cell Contact | PMID: 25728924;PMID: 34022279;PMID: 27091502;PMID: 35907405 |
| CD31 | NeuN | Flrt1 | Adgrl2 | 0.006076 | 0 | FLRT1_ADGRL2 | Flrt1 - Adgrl2 | ADGRL | Cell-Cell Contact | PMID: 25728924;PMID: 34022279;PMID: 27091502;PMID: 35907405 |
| NeuN | NeuN | Flrt1 | Adgrl2 | 0.007858 | 0 | FLRT1_ADGRL2 | Flrt1 - Adgrl2 | ADGRL | Cell-Cell Contact | PMID: 25728924;PMID: 34022279;PMID: 27091502;PMID: 35907405 |
| NeuN | CD31 | Flrt2 | Adgrl2 | 0.009804 | 0 | FLRT2_ADGRL2 | Flrt2 - Adgrl2 | ADGRL | Cell-Cell Contact | PMID: 25728924;PMID: 34022279;PMID: 27091502;PMID: 35907405 |
| NeuN | NeuN | Flrt2 | Adgrl2 | 0.013761 | 0 | FLRT2_ADGRL2 | Flrt2 - Adgrl2 | ADGRL | Cell-Cell Contact | PMID: 25728924;PMID: 34022279;PMID: 27091502;PMID: 35907405 |
| NeuN | CD31 | Flrt3 | Adgrl2 | 0.008297 | 0 | FLRT3_ADGRL2 | Flrt3 - Adgrl2 | ADGRL | Cell-Cell Contact | PMID: 25728924;PMID: 34022279;PMID: 27091502;PMID: 35907405 |
| NeuN | NeuN | Flrt3 | Adgrl2 | 0.011653 | 0 | FLRT3_ADGRL2 | Flrt3 - Adgrl2 | ADGRL | Cell-Cell Contact | PMID: 25728924;PMID: 34022279;PMID: 27091502;PMID: 35907405 |
| CD31 | NeuN | Tenm1 | Adgrl2 | 0.003653 | 0 | TENM1_ADGRL2 | Tenm1 - Adgrl2 | ADGRL | Cell-Cell Contact | PMID: 25728924;PMID: 34022279;PMID: 27091502;PMID: 35907405 |
| NeuN | NeuN | Tenm1 | Adgrl2 | 0.004407 | 0 | TENM1_ADGRL2 | Tenm1 - Adgrl2 | ADGRL | Cell-Cell Contact | PMID: 25728924;PMID: 34022279;PMID: 27091502;PMID: 35907405 |
| NeuN | CD31 | Tenm2 | Adgrl2 | 0.009201 | 0 | TENM2_ADGRL2 | Tenm2 - Adgrl2 | ADGRL | Cell-Cell Contact | PMID: 25728924;PMID: 34022279;PMID: 27091502;PMID: 35907405 |
| NeuN | NeuN | Tenm2 | Adgrl2 | 0.012918 | 0 | TENM2_ADGRL2 | Tenm2 - Adgrl2 | ADGRL | Cell-Cell Contact | PMID: 25728924;PMID: 34022279;PMID: 27091502;PMID: 35907405 |
| NeuN | CD31 | Tenm3 | Adgrl2 | 0.024977 | 0 | TENM3_ADGRL2 | Tenm3 - Adgrl2 | ADGRL | Cell-Cell Contact | PMID: 25728924;PMID: 34022279;PMID: 27091502;PMID: 35907405 |
| NeuN | NeuN | Tenm3 | Adgrl2 | 0.034844 | 0 | TENM3_ADGRL2 | Tenm3 - Adgrl2 | ADGRL | Cell-Cell Contact | PMID: 25728924;PMID: 34022279;PMID: 27091502;PMID: 35907405 |
| NeuN | CD31 | Tenm4 | Adgrl2 | 0.009494 | 0 | TENM4_ADGRL2 | Tenm4 - Adgrl2 | ADGRL | Cell-Cell Contact | PMID: 25728924;PMID: 34022279;PMID: 27091502;PMID: 35907405 |
| NeuN | NeuN | Tenm4 | Adgrl2 | 0.013329 | 0 | TENM4_ADGRL2 | Tenm4 - Adgrl2 | ADGRL | Cell-Cell Contact | PMID: 25728924;PMID: 34022279;PMID: 27091502;PMID: 35907405 |
| NeuN | CD31 | TENM2_FL | Adgrl2 | 0.008737 | 0 | TENM2_FLRT3_ADGRL2 | (Tenm2+Flrt3) - Adgrl2 | ADGRL | Cell-Cell Contact | PMID: 25728924;PMID: 34022279;PMID: 27091502;PMID: 35907405 |
| NeuN | NeuN | TENM2_FL | Adgrl2 | 0.012269 | 0 | TENM2_FLRT3_ADGRL2 | (Tenm2+Flrt3) - Adgrl2 | ADGRL | Cell-Cell Contact | PMID: 25728924;PMID: 34022279;PMID: 27091502;PMID: 35907405 |
| NeuN | CD31 | Flrt1 | Adgrl3 | 0.004235 | 0 | FLRT1_ADGRL3 | Flrt1 - Adgrl3 | ADGRL | Cell-Cell Contact | PMID: 25728924;PMID: 34022279;PMID: 27091502;PMID: 35907405 |
| NeuN | NeuN | Flrt1 | Adgrl3 | 0.004909 | 0 | FLRT1_ADGRL3 | Flrt1 - Adgrl3 | ADGRL | Cell-Cell Contact | PMID: 25728924;PMID: 34022279;PMID: 27091502;PMID: 35907405 |
| NeuN | CD31 | Flrt2 | Adgrl3 | 0.007438 | 0 | FLRT2_ADGRL3 | Flrt2 - Adgrl3 | ADGRL | Cell-Cell Contact | PMID: 25728924;PMID: 34022279;PMID: 27091502;PMID: 35907405 |
| NeuN | NeuN | Flrt2 | Adgrl3 | 0.008616 | 0 | FLRT2_ADGRL3 | Flrt2 - Adgrl3 | ADGRL | Cell-Cell Contact | PMID: 25728924;PMID: 34022279;PMID: 27091502;PMID: 35907405 |
| NeuN | CD31 | Flrt3 | Adgrl3 | 0.006292 | 0 | FLRT3_ADGRL3 | Flrt3 - Adgrl3 | ADGRL | Cell-Cell Contact | PMID: 25728924;PMID: 34022279;PMID: 27091502;PMID: 35907405 |
| NeuN | NeuN | Flrt3 | Adgrl3 | 0.00729 | 0 | FLRT3_ADGRL3 | Flrt3 - Adgrl3 | ADGRL | Cell-Cell Contact | PMID: 25728924;PMID: 34022279;PMID: 27091502;PMID: 35907405 |
| NeuN | CD31 | Tenm1 | Adgrl3 | 0.002371 | 0 | TENM1_ADGRL3 | Tenm1 - Adgrl3 | ADGRL | Cell-Cell Contact | PMID: 25728924;PMID: 34022279;PMID: 27091502;PMID: 35907405 |
| NeuN | NeuN | Tenm1 | Adgrl3 | 0.002749 | 0 | TENM1_ADGRL3 | Tenm1 - Adgrl3 | ADGRL | Cell-Cell Contact | PMID: 25728924;PMID: 34022279;PMID: 27091502;PMID: 35907405 |
| NeuN | CD31 | Tenm2 | Adgrl3 | 0.006979 | 0 | TENM2_ADGRL3 | Tenm2 - Adgrl3 | ADGRL | Cell-Cell Contact | PMID: 25728924;PMID: 34022279;PMID: 27091502;PMID: 35907405 |
| NeuN | NeuN | Tenm2 | Adgrl3 | 0.008086 | 0 | TENM2_ADGRL3 | Tenm2 - Adgrl3 | ADGRL | Cell-Cell Contact | PMID: 25728924;PMID: 34022279;PMID: 27091502;PMID: 35907405 |
| NeuN | CD31 | Tenm3 | Adgrl3 | 0.019019 | 0 | TENM3_ADGRL3 | Tenm3 - Adgrl3 | ADGRL | Cell-Cell Contact | PMID: 25728924;PMID: 34022279;PMID: 27091502;PMID: 35907405 |
| NeuN | NeuN | Tenm3 | Adgrl3 | 0.021992 | 0 | TENM3_ADGRL3 | Tenm3 - Adgrl3 | ADGRL | Cell-Cell Contact | PMID: 25728924;PMID: 34022279;PMID: 27091502;PMID: 35907405 |
| NeuN | CD31 | Tenm4 | Adgrl3 | 0.007202 | 0 | TENM4_ADGRL3 | Tenm4 - Adgrl3 | ADGRL | Cell-Cell Contact | PMID: 25728924;PMID: 34022279;PMID: 27091502;PMID: 35907405 |
| NeuN | NeuN | Tenm4 | Adgrl3 | 0.008344 | 0 | TENM4_ADGRL3 | Tenm4 - Adgrl3 | ADGRL | Cell-Cell Contact | PMID: 25728924;PMID: 34022279;PMID: 27091502;PMID: 35907405 |
| NeuN | CD31 | Unc5a | Adgrl3 | 0.00301 | 0 | UNC5A_ADGRL3 | Unc5a - Adgrl3 | ADGRL | Cell-Cell Contact | PMID: 25728924;PMID: 34022279;PMID: 27091502;PMID: 35907405 |
| NeuN | NeuN | Unc5a | Adgrl3 | 0.00349 | 0 | UNC5A_ADGRL3 | Unc5a - Adgrl3 | ADGRL | Cell-Cell Contact | PMID: 25728924;PMID: 34022279;PMID: 27091502;PMID: 35907405 |
| NeuN | CD31 | TENM3_FL | Adgrl3 | 0.008999 | 0 | TENM3_FLRT1_ADGRL3 | (Tenm3+Flrt1) - Adgrl3 | ADGRL | Cell-Cell Contact | PMID: 25728924;PMID: 34022279;PMID: 27091502;PMID: 35907405 |
| NeuN | NeuN | TENM3_FL | Adgrl3 | 0.010422 | 0 | TENM3_FLRT1_ADGRL3 | (Tenm3+Flrt1) - Adgrl3 | ADGRL | Cell-Cell Contact | PMID: 25728924;PMID: 34022279;PMID: 27091502;PMID: 35907405 |
| NeuN | CD31 | TENM3_FL | Adgrl3 | 0.010958 | 0 | TENM3_FLRT3_ADGRL3 | (Tenm3+Flrt3) - Adgrl3 | ADGRL | Cell-Cell Contact | PMID: 25728924;PMID: 34022279;PMID: 27091502;PMID: 35907405 |
| NeuN | NeuN | TENM3_FL | Adgrl3 | 0.012687 | 0 | TENM3_FLRT3_ADGRL3 | (Tenm3+Flrt3) - Adgrl3 | ADGRL | Cell-Cell Contact | PMID: 25728924;PMID: 34022279;PMID: 27091502;PMID: 35907405 |
| NeuN | CD31 | Flrt1 | Flrt1 | 0.001426 | 0 | FLRT1_FLRT1 | Flrt1 - Flrt1 | FLRT | Cell-Cell Contact | PMID: 27091502 |
| CD31 | NeuN | Flrt1 | Flrt1 | 0.001426 | 0 | FLRT1_FLRT1 | Flrt1 - Flrt1 | FLRT | Cell-Cell Contact | PMID: 27091502 |
| NeuN | NeuN | Flrt1 | Flrt1 | 0.001847 | 0 | FLRT1_FLRT1 | Flrt1 - Flrt1 | FLRT | Cell-Cell Contact | PMID: 27091502 |
| NeuN | CD31 | Flrt2 | Flrt2 | 0.003839 | 0 | FLRT2_FLRT2 | Flrt2 - Flrt2 | FLRT | Cell-Cell Contact | PMID: 27091502 |
| CD31 | NeuN | Flrt2 | Flrt2 | 0.003839 | 0 | FLRT2_FLRT2 | Flrt2 - Flrt2 | FLRT | Cell-Cell Contact | PMID: 27091502 |
| NeuN | NeuN | Flrt2 | Flrt2 | 0.00571 | 0 | FLRT2_FLRT2 | Flrt2 - Flrt2 | FLRT | Cell-Cell Contact | PMID: 27091502 |
| NeuN | CD31 | Flrt3 | Flrt3 | 0.002677 | 0 | FLRT3_FLRT3 | Flrt3 - Flrt3 | FLRT | Cell-Cell Contact | PMID: 27091502 |
| CD31 | NeuN | Flrt3 | Flrt3 | 0.002677 | 0 | FLRT3_FLRT3 | Flrt3 - Flrt3 | FLRT | Cell-Cell Contact | PMID: 27091502 |

| | | | | | | | | | | |
|---|---|---|---|---|---|---|---|---|---|---|
| NeuN | NeuN | Flrt3 | Flrt3 | 0.004083 | 0 | FLRT3_FLRT3 | Flrt3 - Flrt3 | FLRT | Cell-Cell Contact | PMID: 27091502 |
| CD31 | NeuN | Flrt1 | Unc5a | 0.001013 | 0 | FLRT1_UNC5A | Flrt1 - Unc5a | UNC5 | Cell-Cell Contact | PMID: 27091502 |
| NeuN | NeuN | Flrt1 | Unc5a | 0.001312 | 0 | FLRT1_UNC5A | Flrt1 - Unc5a | UNC5 | Cell-Cell Contact | PMID: 27091502 |
| CD31 | NeuN | Flrt2 | Unc5a | 0.00155 | 0 | FLRT2_UNC5A | Flrt2 - Unc5a | UNC5 | Cell-Cell Contact | PMID: 27091502 |
| NeuN | NeuN | Flrt2 | Unc5a | 0.002309 | 0 | FLRT2_UNC5A | Flrt2 - Unc5a | UNC5 | Cell-Cell Contact | PMID: 27091502 |
| CD31 | NeuN | Flrt3 | Unc5a | 0.001278 | 0 | FLRT3_UNC5A | Flrt3 - Unc5a | UNC5 | Cell-Cell Contact | PMID: 27091502 |
| NeuN | NeuN | Flrt3 | Unc5a | 0.001951 | 0 | FLRT3_UNC5A | Flrt3 - Unc5a | UNC5 | Cell-Cell Contact | PMID: 27091502 |
| CD31 | CD31 | Flrt1 | Unc5b | 0.001843 | 0 | FLRT1_UNC5B | Flrt1 - Unc5b | UNC5 | Cell-Cell Contact | PMID: 27091502 |
| NeuN | CD31 | Flrt1 | Unc5b | 0.002386 | 0 | FLRT1_UNC5B | Flrt1 - Unc5b | UNC5 | Cell-Cell Contact | PMID: 27091502 |
| CD31 | CD31 | Flrt2 | Unc5b | 0.002819 | 0 | FLRT2_UNC5B | Flrt2 - Unc5b | UNC5 | Cell-Cell Contact | PMID: 27091502 |
| NeuN | CD31 | Flrt2 | Unc5b | 0.004196 | 0 | FLRT2_UNC5B | Flrt2 - Unc5b | UNC5 | Cell-Cell Contact | PMID: 27091502 |
| CD31 | CD31 | Flrt3 | Unc5b | 0.002325 | 0 | FLRT3_UNC5B | Flrt3 - Unc5b | UNC5 | Cell-Cell Contact | PMID: 27091502 |
| NeuN | CD31 | Flrt3 | Unc5b | 0.003548 | 0 | FLRT3_UNC5B | Flrt3 - Unc5b | UNC5 | Cell-Cell Contact | PMID: 27091502 |
| NeuN | CD31 | Flrt1 | Unc5c | 0.000856 | 0 | FLRT1_UNC5C | Flrt1 - Unc5 | UNC5 | Cell-Cell Contact | PMID: 27091502 |
| NeuN | NeuN | Flrt1 | Unc5c | 0.001059 | 0 | FLRT1_UNC5C | Flrt1 - Unc5 | UNC5 | Cell-Cell Contact | PMID: 27091502 |
| NeuN | CD31 | Flrt2 | Unc5c | 0.001507 | 0 | FLRT2_UNC5C | Flrt2 - Unc5 | UNC5 | Cell-Cell Contact | PMID: 27091502 |
| NeuN | NeuN | Flrt2 | Unc5c | 0.001864 | 0 | FLRT2_UNC5C | Flrt2 - Unc5 | UNC5 | Cell-Cell Contact | PMID: 27091502 |
| NeuN | CD31 | Flrt3 | Unc5c | 0.001273 | 0 | FLRT3_UNC5C | Flrt3 - Unc5 | UNC5 | Cell-Cell Contact | PMID: 27091502 |
| NeuN | NeuN | Flrt3 | Unc5c | 0.001575 | 0 | FLRT3_UNC5C | Flrt3 - Unc5 | UNC5 | Cell-Cell Contact | PMID: 27091502 |
| CD31 | NeuN | Flrt1 | Unc5d | 0.002149 | 0 | FLRT1_UNC5D | Flrt1 - Unc5d | UNC5 | Cell-Cell Contact | PMID: 27091502 |
| NeuN | NeuN | Flrt1 | Unc5d | 0.002783 | 0 | FLRT1_UNC5D | Flrt1 - Unc5d | UNC5 | Cell-Cell Contact | PMID: 27091502 |
| NeuN | CD31 | Flrt2 | Unc5d | 0.003656 | 0 | FLRT2_UNC5D | Flrt2 - Unc5d | UNC5 | Cell-Cell Contact | PMID: 27091502 |
| NeuN | NeuN | Flrt2 | Unc5d | 0.004893 | 0 | FLRT2_UNC5D | Flrt2 - Unc5d | UNC5 | Cell-Cell Contact | PMID: 27091502 |
| NeuN | CD31 | Flrt3 | Unc5d | 0.003091 | 0 | FLRT3_UNC5D | Flrt3 - Unc5d | UNC5 | Cell-Cell Contact | PMID: 27091502 |
| NeuN | NeuN | Flrt3 | Unc5d | 0.004137 | 0 | FLRT3_UNC5D | Flrt3 - Unc5d | UNC5 | Cell-Cell Contact | PMID: 27091502 |
| NeuN | CD31 | Nrxn1 | Clstn1 | 0.008695 | 0 | NRXN1_CLSTN1 | Nrxn1 - Clstn1 | NRXN | Cell-Cell Contact | PMID: 32434929 |
| NeuN | NeuN | Nrxn1 | Clstn1 | 0.016238 | 0 | NRXN1_CLSTN1 | Nrxn1 - Clstn1 | NRXN | Cell-Cell Contact | PMID: 32434929 |
| NeuN | CD31 | Nrxn2 | Clstn1 | 0.005877 | 0 | NRXN2_CLSTN1 | Nrxn2 - Clstn1 | NRXN | Cell-Cell Contact | PMID: 32434929 |
| NeuN | NeuN | Nrxn2 | Clstn1 | 0.011002 | 0 | NRXN2_CLSTN1 | Nrxn2 - Clstn1 | NRXN | Cell-Cell Contact | PMID: 32434929 |
| NeuN | CD31 | Nrxn1 | Clstn2 | 0.025056 | 0 | NRXN1_CLSTN2 | Nrxn1 - Clstn2 | NRXN | Cell-Cell Contact | PMID: 32434929 |
| NeuN | NeuN | Nrxn1 | Clstn2 | 0.038575 | 0 | NRXN1_CLSTN2 | Nrxn1 - Clstn2 | NRXN | Cell-Cell Contact | PMID: 32434929 |
| NeuN | CD31 | Nrxn2 | Clstn2 | 0.017026 | 0 | NRXN2_CLSTN2 | Nrxn2 - Clstn2 | NRXN | Cell-Cell Contact | PMID: 32434929 |
| NeuN | NeuN | Nrxn2 | Clstn2 | 0.026329 | 0 | NRXN2_CLSTN2 | Nrxn2 - Clstn2 | NRXN | Cell-Cell Contact | PMID: 32434929 |
| NeuN | CD31 | Nrxn1 | Clstn3 | 0.007807 | 0 | NRXN1_CLSTN3 | Nrxn1 - Clstn3 | NRXN | Cell-Cell Contact | PMID: 32434929 |
| NeuN | NeuN | Nrxn1 | Clstn3 | 0.012871 | 0 | NRXN1_CLSTN3 | Nrxn1 - Clstn3 | NRXN | Cell-Cell Contact | PMID: 32434929 |
| NeuN | CD31 | Nrxn2 | Clstn3 | 0.005275 | 0 | NRXN2_CLSTN3 | Nrxn2 - Clstn3 | NRXN | Cell-Cell Contact | PMID: 32434929 |
| NeuN | NeuN | Nrxn2 | Clstn3 | 0.008711 | 0 | NRXN2_CLSTN3 | Nrxn2 - Clstn3 | NRXN | Cell-Cell Contact | PMID: 32434929 |
| NeuN | CD31 | Nrxn1 | Dag1 | 0.007883 | 0 | NRXN1_DAG1 | Nrxn1 - Dag1 | NRXN | Cell-Cell Contact | PMID: 21628410 |
| NeuN | NeuN | Nrxn1 | Dag1 | 0.007548 | 0 | NRXN1_DAG1 | Nrxn1 - Dag1 | NRXN | Cell-Cell Contact | PMID: 21628410 |
| NeuN | CD31 | Nrxn2 | Dag1 | 0.005327 | 0 | NRXN2_DAG1 | Nrxn2 - Dag1 | NRXN | Cell-Cell Contact | PMID: 21628410 |
| NeuN | NeuN | Nrxn2 | Dag1 | 0.005099 | 0 | NRXN2_DAG1 | Nrxn2 - Dag1 | NRXN | Cell-Cell Contact | PMID: 21628410 |
| CD31 | NeuN | Lrfn4 | Ptprd | 0.014696 | 0 | LRFN4_PTPRD | Lrfn4 - Ptprd | PTPR | Cell-Cell Contact | PMID: 29674953;PMID: 23916315 |
| NeuN | NeuN | Lrfn4 | Ptprd | 0.023296 | 0 | LRFN4_PTPRD | Lrfn4 - Ptprd | PTPR | Cell-Cell Contact | PMID: 29674953;PMID: 23916315 |
| CD31 | NeuN | Lrfn4 | Ptprf | 0.006876 | 0 | LRFN4_PTPRF | Lrfn4 - Ptprf | PTPR | Cell-Cell Contact | PMID: 29674953;PMID: 23916315 |
| NeuN | NeuN | Lrfn4 | Ptprf | 0.01095 | 0 | LRFN4_PTPRF | Lrfn4 - Ptprf | PTPR | Cell-Cell Contact | PMID: 29674953;PMID: 23916315 |
| CD31 | NeuN | Lrfn4 | Ptprs | 0.023707 | 0 | LRFN4_PTPRS | Lrfn4 - Ptprs | PTPR | Cell-Cell Contact | PMID: 29674953;PMID: 23916315 |
| NeuN | NeuN | Lrfn4 | Ptprs | 0.037381 | 0 | LRFN4_PTPRS | Lrfn4 - Ptprs | PTPR | Cell-Cell Contact | PMID: 29674953;PMID: 23916315 |
| CD31 | NeuN | Lrfn5 | Ptprd | 0.018774 | 0 | LRFN5_PTPRD | Lrfn5 - Ptprd | PTPR | Cell-Cell Contact | PMID: 29674953;PMID: 23916315 |
| NeuN | NeuN | Lrfn5 | Ptprd | 0.024285 | 0 | LRFN5_PTPRD | Lrfn5 - Ptprd | PTPR | Cell-Cell Contact | PMID: 29674953;PMID: 23916315 |
| CD31 | NeuN | Lrfn5 | Ptprf | 0.008803 | 0 | LRFN5_PTPRF | Lrfn5 - Ptprf | PTPR | Cell-Cell Contact | PMID: 29674953;PMID: 23916315 |
| NeuN | NeuN | Lrfn5 | Ptprf | 0.011421 | 0 | LRFN5_PTPRF | Lrfn5 - Ptprf | PTPR | Cell-Cell Contact | PMID: 29674953;PMID: 23916315 |
| CD31 | NeuN | Lrfn5 | Ptprs | 0.03021 | 0 | LRFN5_PTPRS | Lrfn5 - Ptprs | PTPR | Cell-Cell Contact | PMID: 29674953;PMID: 23916315 |
| NeuN | NeuN | Lrfn5 | Ptprs | 0.038944 | 0 | LRFN5_PTPRS | Lrfn5 - Ptprs | PTPR | Cell-Cell Contact | PMID: 29674953;PMID: 23916315 |
| CD31 | NeuN | Lrrc4b | Ptprd | 0.031545 | 0 | LRRC4B_PTPRD | Lrrc4b - Ptprd | PTPR | Cell-Cell Contact | PMID: 29674953;PMID: 23916315 |
| NeuN | NeuN | Lrrc4b | Ptprd | 0.06386 | 0 | LRRC4B_PTPRD | Lrrc4b - Ptprd | PTPR | Cell-Cell Contact | PMID: 29674953;PMID: 23916315 |
| CD31 | NeuN | Lrrc4b | Ptprs | 0.05036 | 0 | LRRC4B_PTPRS | Lrrc4b - Ptprs | PTPR | Cell-Cell Contact | PMID: 29674953;PMID: 23916315 |
| NeuN | NeuN | Lrrc4b | Ptprs | 0.09996 | 0 | LRRC4B_PTPRS | Lrrc4b - Ptprs | PTPR | Cell-Cell Contact | PMID: 29674953;PMID: 23916315 |
| CD31 | NeuN | Lrrc4c | Ptprf | 0.006263 | 0 | LRRC4C_PTPRF | Lrrc4c - Ptprf | PTPR | Cell-Cell Contact | PMID: 29674953;PMID: 23916315 |
| NeuN | NeuN | Lrrc4c | Ptprf | 0.009978 | 0 | LRRC4C_PTPRF | Lrrc4c - Ptprf | PTPR | Cell-Cell Contact | PMID: 29674953;PMID: 23916315 |

| | | | | | | | | | | |
|---|---|---|---|---|---|---|---|---|---|---|
| NeuN | CD31 | Nrxn1 | Lrrtm1 | 0.006031 | 0 | NRXN1_LRRTM1 | Nrxn1 - Lrrtm1 | NRXN | Cell-Cell Contact | PMID:20519524;PMID:20064387;PMID:20064388; |
| NeuN | NeuN | Nrxn1 | Lrrtm1 | 0.012164 | 0 | NRXN1_LRRTM1 | Nrxn1 - Lrrtm1 | NRXN | Cell-Cell Contact | PMID:20519524;PMID:20064387;PMID:20064388; |
| NeuN | CD31 | Nrxn2 | Lrrtm1 | 0.004073 | 0 | NRXN2_LRRTM1 | Nrxn2 - Lrrtm1 | NRXN | Cell-Cell Contact | PMID:20519524;PMID:20064387;PMID:20064388; |
| NeuN | NeuN | Nrxn2 | Lrrtm1 | 0.008231 | 0 | NRXN2_LRRTM1 | Nrxn2 - Lrrtm1 | NRXN | Cell-Cell Contact | PMID:20519524;PMID:20064387;PMID:20064388; |
| CD31 | NeuN | Nrxn3 | Lrrtm1 | 0.002407 | 0 | NRXN3_LRRTM1 | Nrxn3 - Lrrtm1 | NRXN | Cell-Cell Contact | PMID:20519524;PMID:20064387;PMID:20064388; |
| NeuN | NeuN | Nrxn3 | Lrrtm1 | 0.003103 | 0 | NRXN3_LRRTM1 | Nrxn3 - Lrrtm1 | NRXN | Cell-Cell Contact | PMID:20519524;PMID:20064387;PMID:20064388; |
| NeuN | CD31 | Nrxn1 | Lrrtm2 | 0.003299 | 0 | NRXN1_LRRTM2 | Nrxn1 - Lrrtm2 | NRXN | Cell-Cell Contact | PMID:20519524;PMID:20064387;PMID:20064388; |
| NeuN | NeuN | Nrxn1 | Lrrtm2 | 0.006626 | 0 | NRXN1_LRRTM2 | Nrxn1 - Lrrtm2 | NRXN | Cell-Cell Contact | PMID:20519524;PMID:20064387;PMID:20064388; |
| NeuN | CD31 | Nrxn2 | Lrrtm2 | 0.002226 | 0 | NRXN2_LRRTM2 | Nrxn2 - Lrrtm2 | NRXN | Cell-Cell Contact | PMID:20519524;PMID:20064387;PMID:20064388; |
| NeuN | NeuN | Nrxn2 | Lrrtm2 | 0.004475 | 0 | NRXN2_LRRTM2 | Nrxn2 - Lrrtm2 | NRXN | Cell-Cell Contact | PMID:20519524;PMID:20064387;PMID:20064388; |
| CD31 | NeuN | Nrxn3 | Lrrtm2 | 0.001305 | 0 | NRXN3_LRRTM2 | Nrxn3 - Lrrtm2 | NRXN | Cell-Cell Contact | PMID:20519524;PMID:20064387;PMID:20064388; |
| NeuN | NeuN | Nrxn3 | Lrrtm2 | 0.001683 | 0 | NRXN3_LRRTM2 | Nrxn3 - Lrrtm2 | NRXN | Cell-Cell Contact | PMID:20519524;PMID:20064387;PMID:20064388; |
| NeuN | CD31 | Nrxn1 | Lrrtm3 | 0.005497 | 0 | NRXN1_LRRTM3 | Nrxn1 - Lrrtm3 | NRXN | Cell-Cell Contact | PMID:20519524;PMID:20064387;PMID:20064388; |
| NeuN | NeuN | Nrxn1 | Lrrtm3 | 0.006324 | 0 | NRXN1_LRRTM3 | Nrxn1 - Lrrtm3 | NRXN | Cell-Cell Contact | PMID:20519524;PMID:20064387;PMID:20064388; |
| NeuN | CD31 | Nrxn2 | Lrrtm3 | 0.003711 | 0 | NRXN2_LRRTM3 | Nrxn2 - Lrrtm3 | NRXN | Cell-Cell Contact | PMID:20519524;PMID:20064387;PMID:20064388; |
| NeuN | NeuN | Nrxn2 | Lrrtm3 | 0.004271 | 0 | NRXN2_LRRTM3 | Nrxn2 - Lrrtm3 | NRXN | Cell-Cell Contact | PMID:20519524;PMID:20064387;PMID:20064388; |
| NeuN | CD31 | Nrxn3 | Lrrtm3 | 0.001395 | 0 | NRXN3_LRRTM3 | Nrxn3 - Lrrtm3 | NRXN | Cell-Cell Contact | PMID:20519524;PMID:20064387;PMID:20064388; |
| NeuN | NeuN | Nrxn3 | Lrrtm3 | 0.001606 | 0 | NRXN3_LRRTM3 | Nrxn3 - Lrrtm3 | NRXN | Cell-Cell Contact | PMID:20519524;PMID:20064387;PMID:20064388; |
| NeuN | CD31 | Nrxn1 | Lrrtm4 | 0.00734 | 0 | NRXN1_LRRTM4 | Nrxn1 - Lrrtm4 | NRXN | Cell-Cell Contact | PMID:20519524;PMID:20064387;PMID:20064388; |
| NeuN | NeuN | Nrxn1 | Lrrtm4 | 0.010226 | 0 | NRXN1_LRRTM4 | Nrxn1 - Lrrtm4 | NRXN | Cell-Cell Contact | PMID:20519524;PMID:20064387;PMID:20064388; |
| NeuN | CD31 | Nrxn2 | Lrrtm4 | 0.004959 | 0 | NRXN2_LRRTM4 | Nrxn2 - Lrrtm4 | NRXN | Cell-Cell Contact | PMID:20519524;PMID:20064387;PMID:20064388; |
| NeuN | NeuN | Nrxn2 | Lrrtm4 | 0.006915 | 0 | NRXN2_LRRTM4 | Nrxn2 - Lrrtm4 | NRXN | Cell-Cell Contact | PMID:20519524;PMID:20064387;PMID:20064388; |
| CD31 | NeuN | Nrxn3 | Lrrtm4 | 0.00202 | 0 | NRXN3_LRRTM4 | Nrxn3 - Lrrtm4 | NRXN | Cell-Cell Contact | PMID:20519524;PMID:20064387;PMID:20064388; |
| NeuN | NeuN | Nrxn3 | Lrrtm4 | 0.002604 | 0 | NRXN3_LRRTM4 | Nrxn3 - Lrrtm4 | NRXN | Cell-Cell Contact | PMID:20519524;PMID:20064387;PMID:20064388; |
| CD31 | NeuN | Lrrtm4 | Ptprs | 0.018081 | 0 | LRRTM4_PTPRS | Lrrtm4 - Ptprs | PTPR | Cell-Cell Contact | PMID:31995730 |
| NeuN | NeuN | Lrrtm4 | Ptprs | 0.025082 | 0 | LRRTM4_PTPRS | Lrrtm4 - Ptprs | PTPR | Cell-Cell Contact | PMID:31995730 |
| CD31 | NeuN | App | Sorl1 | 0.025119 | 0 | APP_SORL1 | App - Sorl1 | APP | Cell-Cell Contact | PMID: 24523320 |
| NeuN | NeuN | App | Sorl1 | 0.026339 | 0 | APP_SORL1 | App - Sorl1 | APP | Cell-Cell Contact | PMID: 24523320 |
| CD31 | CD31 | App | Tnfrsf21 | 0.007366 | 0 | APP_TNFRSF21 | App - Tnfrsf21 | APP | Cell-Cell Contact | PMID: 33075465 |
| NeuN | CD31 | App | Tnfrsf21 | 0.00773 | 0 | APP_TNFRSF21 | App - Tnfrsf21 | APP | Cell-Cell Contact | PMID: 33075465 |
| CD31 | CD31 | App | TREM2_TYROBP | 0.006418 | 0 | APP_TREM2_TYROBP | App - (Trem2+Tyrobp) | APP | Cell-Cell Contact | PMID:27477013 |
| NeuN | CD31 | App | TREM2_TYROBP | 0.006736 | 0 | APP_TREM2_TYROBP | App - (Trem2+Tyrobp) | APP | Cell-Cell Contact | PMID:27477013 |
| CD31 | CD31 | Apoa2 | TREM2_TYROBP | 0.000393 | 0 | APOA2_TREM2_TYROBP | Apoa2 - (Trem2+Tyrobp) | ApoA | Cell-Cell Contact | uniprot;PMID:27477017 |
| NeuN | CD31 | Apoa2 | TREM2_TYROBP | 0.000429 | 0 | APOA2_TREM2_TYROBP | Apoa2 - (Trem2+Tyrobp) | ApoA | Cell-Cell Contact | uniprot;PMID:27477017 |
| CD31 | CD31 | Apoe | TREM2_TYROBP | 0.000791 | 0 | APOE_TREM2_TYROBP | Apoe - (Trem2+Tyrobp) | ApoE | Cell-Cell Contact | uniprot;PMID:27477015 |
| CD31 | NeuN | Apoe | TREM2_TYROBP | 0.000521 | 0 | APOE_TREM2_TYROBP | Apoe - (Trem2+Tyrobp) | ApoE | Cell-Cell Contact | uniprot;PMID:27477015 |
| NeuN | CD31 | Cadm1 | Nectin3 | 0.008761 | 0 | CADM1_NECTIN3 | Cadm1 - Nectin3 | CADM | Cell-Cell Contact | uniprot |
| NeuN | NeuN | Cadm1 | Nectin3 | 0.011429 | 0 | CADM1_NECTIN3 | Cadm1 - Nectin3 | CADM | Cell-Cell Contact | uniprot |
| CD31 | NeuN | Cd99 | Pilrb1 | 0.024158 | 0 | CD99_PILRB1 | Cd99 - Pilrb1 | CD99 | Cell-Cell Contact | uniprot;PMID:18234675 |
| NeuN | NeuN | Cd99 | Pilrb1 | 0.030878 | 0 | CD99_PILRB1 | Cd99 - Pilrb1 | CD99 | Cell-Cell Contact | uniprot;PMID:18234675 |
| CD31 | CD31 | Ceacam2 | Cd209c | 0.000736 | 0 | CEACAM2_CD209C | Ceacam2 - Cd209c | CEACAM | Cell-Cell Contact | PMID:16282604;PMID:16246332 |
| CD31 | NeuN | Ceacam2 | Cd209c | 0.000593 | 0 | CEACAM2_CD209C | Ceacam2 - Cd209c | CEACAM | Cell-Cell Contact | PMID:16282604;PMID:16246332 |
| CD31 | CD31 | Ceacam2 | Cd209a | 0.000989 | 0 | CEACAM2_CD209A | Ceacam2 - Cd209a | CEACAM | Cell-Cell Contact | PMID:16282604;PMID:16246332 |
| CD31 | NeuN | Ceacam2 | Cd209a | 0.000715 | 0 | CEACAM2_CD209A | Ceacam2 - Cd209a | CEACAM | Cell-Cell Contact | PMID:16282604;PMID:16246332 |
| CD31 | CD31 | Ceacam2 | Cd209d | 0.000576 | 0 | CEACAM2_CD209D | Ceacam2 - Cd209d | CEACAM | Cell-Cell Contact | PMID:16282604;PMID:16246332 |
| CD31 | NeuN | Ceacam2 | Cd209d | 0.0006 | 0 | CEACAM2_CD209D | Ceacam2 - Cd209d | CEACAM | Cell-Cell Contact | PMID:16282604;PMID:16246332 |
| CD31 | CD31 | Ceacam2 | Cd209b | 0.001329 | 0 | CEACAM2_CD209B | Ceacam2 - Cd209b | CEACAM | Cell-Cell Contact | PMID:16282604;PMID:16246332 |
| NeuN | CD31 | Ceacam2 | Cd209b | 0.000936 | 0 | CEACAM2_CD209B | Ceacam2 - Cd209b | CEACAM | Cell-Cell Contact | PMID:16282604;PMID:16246332 |
| CD31 | CD31 | Ceacam2 | Ceacam2 | 0.000928 | 0 | CEACAM2_CEACAM2 | Ceacam2 - Ceacam2 | CEACAM | Cell-Cell Contact | PMID:21982860 |
| NeuN | CD31 | Ceacam2 | Ceacam2 | 0.000653 | 0 | CEACAM2_CEACAM2 | Ceacam2 - Ceacam2 | CEACAM | Cell-Cell Contact | PMID:21982860 |
| CD31 | NeuN | Ceacam2 | Ceacam2 | 0.000653 | 0 | CEACAM2_CEACAM2 | Ceacam2 - Ceacam2 | CEACAM | Cell-Cell Contact | PMID:21982860 |
| CD31 | CD31 | Ceacam2 | Sele | 0.000495 | 0 | CEACAM2_SELE | Ceacam2 - Sele | CEACAM | Cell-Cell Contact | PMID:1378450 |
| CD31 | NeuN | Ceacam2 | Sele | 0.000583 | 0 | CEACAM2_SELE | Ceacam2 - Sele | CEACAM | Cell-Cell Contact | PMID:1378450 |
| CD31 | CD31 | Ceacam2 | Cd1d1 | 0.001711 | 0 | CEACAM2_CD1D1 | Ceacam2 - Cd1d1 | CEACAM | Cell-Cell Contact | PMID:24104458 |
| NeuN | CD31 | Ceacam2 | Cd1d1 | 0.001205 | 0 | CEACAM2_CD1D1 | Ceacam2 - Cd1d1 | CEACAM | Cell-Cell Contact | PMID:24104458 |
| CD31 | CD31 | Cntn1 | Notch1 | 0.003041 | 0 | CNTN1_NOTCH1 | Cntn1 - Notch1 | CNTN | Cell-Cell Contact | PMID:14567914 |
| NeuN | CD31 | Cntn1 | Notch1 | 0.007359 | 0 | CNTN1_NOTCH1 | Cntn1 - Notch1 | CNTN | Cell-Cell Contact | PMID:14567914 |
| CD31 | CD31 | Clec2h | Klrb1f | 0.000831 | 0 | CLEC2H_KLRB1F | Klrb1f - Clec2h | CLEC | Cell-Cell Contact | PMID:24223577 |
| CD31 | NeuN | Clec2h | Klrb1f | 0.000988 | 0 | CLEC2H_KLRB1F | Klrb1f - Clec2h | CLEC | Cell-Cell Contact | PMID:24223577 |

| | | | | | | | | | | |
|---|---|---|---|---|---|---|---|---|---|---|
| CD31 | CD31 | Clec2d | Klrb1f | 0.001435 | 0 | CLEC2D_KLRB1F | Klrb1f - Clec2d | CLEC | Cell-Cell Contact | PMID:24223577 |
| CD31 | NeuN | Clec2d | Klrb1f | 0.001706 | 0 | CLEC2D_KLRB1F | Klrb1f - Clec2d | CLEC | Cell-Cell Contact | PMID:24223577 |
| CD31 | CD31 | Mmp2 | MMP14_ITGAV_ITGB3 | 0.001291 | 0 | MMP2_INTEGRIN_AVB3_COMP | Mmp2 - Integrin_avb3 | MMP | Cell-Cell Contact | PMID: 29986882 |
| NeuN | CD31 | Mmp2 | MMP14_ITGAV_ITGB3 | 0.000961 | 0 | MMP2_INTEGRIN_AVB3_COMP | Mmp2 - Integrin_avb3 | MMP | Cell-Cell Contact | PMID: 29986882 |
| CD31 | NeuN | Ntn1 | Dcc | 0.002097 | 0 | NTN1_DCC | Ntn1 - Dcc | Netrin | Cell-Cell Contact | PMID: 22685302 |
| NeuN | NeuN | Ntn1 | Dcc | 0.002221 | 0 | NTN1_DCC | Ntn1 - Dcc | Netrin | Cell-Cell Contact | PMID: 22685302 |
| CD31 | NeuN | Ntn1 | Dscam | 0.001321 | 0 | NTN1_DSCAM | Ntn1 - Dscam | Netrin | Cell-Cell Contact | PMID:18585357;PMID: 22685302 |
| NeuN | NeuN | Ntn1 | Dscam | 0.0014 | 0 | NTN1_DSCAM | Ntn1 - Dscam | Netrin | Cell-Cell Contact | PMID:18585357;PMID: 22685302 |
| CD31 | NeuN | Ntn1 | Unc5a | 0.000749 | 0 | NTN1_UNC5A | Ntn1 - Unc5a | Netrin | Cell-Cell Contact | PMID: 22685302 |
| NeuN | NeuN | Ntn1 | Unc5a | 0.000794 | 0 | NTN1_UNC5A | Ntn1 - Unc5a | Netrin | Cell-Cell Contact | PMID: 22685302 |
| CD31 | CD31 | Ntn1 | Unc5b | 0.001364 | 0 | NTN1_UNC5B | Ntn1 - Unc5b | Netrin | Cell-Cell Contact | PMID: 22685302 |
| NeuN | CD31 | Ntn1 | Unc5b | 0.001445 | 0 | NTN1_UNC5B | Ntn1 - Unc5b | Netrin | Cell-Cell Contact | PMID: 22685302 |
| CD31 | NeuN | Ntn1 | Unc5c | 0.000605 | 0 | NTN1_UNC5C | Ntn1 - Unc5c | Netrin | Cell-Cell Contact | PMID: 22685302 |
| NeuN | NeuN | Ntn1 | Unc5c | 0.000641 | 0 | NTN1_UNC5C | Ntn1 - Unc5c | Netrin | Cell-Cell Contact | PMID: 22685302 |
| CD31 | NeuN | Ntn4 | Ntrk2 | 0.005224 | 0 | NTN4_NTRK2 | Ntn4 - Ntrk2 | Netrin | Cell-Cell Contact | PMID: 34531526 |
| NeuN | NeuN | Ntn4 | Ntrk2 | 0.007878 | 0 | NTN4_NTRK2 | Ntn4 - Ntrk2 | Netrin | Cell-Cell Contact | PMID: 34531526 |
| CD31 | CD31 | Ntn4 | Unc5b | 0.001012 | 0 | NTN4_UNC5B | Ntn4 - Unc5b | Netrin | Cell-Cell Contact | PMID: 31914695 |
| NeuN | CD31 | Ntn4 | Unc5b | 0.001529 | 0 | NTN4_UNC5B | Ntn4 - Unc5b | Netrin | Cell-Cell Contact | PMID: 31914695 |
| NeuN | CD31 | Ntn4 | Unc5d | 0.001332 | 0 | NTN4_UNC5D | Ntn4 - Unc5d | Netrin | Cell-Cell Contact | uniprot |
| NeuN | NeuN | Ntn4 | Unc5d | 0.001784 | 0 | NTN4_UNC5D | Ntn4 - Unc5d | Netrin | Cell-Cell Contact | uniprot |
| CD31 | CD31 | Pecam1 | Cd177 | 0.001705 | 0 | PECAM1_CD177 | Pecam1 - Cd177 | PECAM1 | Cell-Cell Contact | PMID:17580308 |
| CD31 | NeuN | Pecam1 | Cd177 | 0.001331 | 0 | PECAM1_CD177 | Pecam1 - Cd177 | PECAM1 | Cell-Cell Contact | PMID:17580308 |
| CD31 | CD31 | Pecam1 | Cd38 | 0.003673 | 0 | PECAM1_CD38 | Pecam1 - Cd38 | PECAM2 | Cell-Cell Contact | PMID:26407101 |
| CD31 | NeuN | Pecam1 | Cd38 | 0.001114 | 0 | PECAM1_CD38 | Pecam1 - Cd38 | PECAM2 | Cell-Cell Contact | PMID:26407101 |
| CD31 | NeuN | Sirpb1a | Cd47 | 0.001555 | 0 | SIRPB1A_CD47 | Sirpb1a - Cd47 | SIRP | Cell-Cell Contact | PMID:15294972 |
| NeuN | NeuN | Sirpb1a | Cd47 | 0.00127 | 0 | SIRPB1A_CD47 | Sirpb1a - Cd47 | SIRP | Cell-Cell Contact | PMID:15294972 |

**Supplementary Table 12: Cell Chat Interactions at E18 in PAE**

| source | target | ligand | receptor | prob | pval | interaction_name | interaction_name_2 | pathway_name | annotation | evidence |
|---|---|---|---|---|---|---|---|---|---|---|
| CD31 | CD31 | Tgfb1 | ACVR1B_TGFbR2 | 0.001821 | 0 | TGFB1_ACVR1B_TGFBR2 | Tgfb1 - (Acvr1b+Tgfbr2) | TGFb | Secreted Signaling | PMID: 27449815 |
| CD31 | NeuN | Tgfb1 | ACVR1B_TGFbR2 | 0.001207 | 0 | TGFB1_ACVR1B_TGFBR2 | Tgfb1 - (Acvr1b+Tgfbr2) | TGFb | Secreted Signaling | PMID: 27449815 |
| CD31 | CD31 | Tgfb1 | ACVR1C_TGFbR2 | 0.001151 | 0 | TGFB1_ACVR1C_TGFBR2 | Tgfb1 - (Acvr1c+Tgfbr2) | TGFb | Secreted Signaling | PMID: 27449815 |
| CD31 | NeuN | Tgfb1 | ACVR1C_TGFbR2 | 0.000543 | 0 | TGFB1_ACVR1C_TGFBR2 | Tgfb1 - (Acvr1c+Tgfbr2) | TGFb | Secreted Signaling | PMID: 27449815 |
| CD31 | CD31 | Tgfb3 | ACVR1B_TGFbR2 | 0.000811 | 0 | TGFB3_ACVR1B_TGFBR2 | Tgfb3 - (Acvr1b+Tgfbr2) | TGFb | Secreted Signaling | PMID: 27449815 |
| CD31 | NeuN | Tgfb3 | ACVR1B_TGFbR2 | 0.000538 | 0 | TGFB3_ACVR1B_TGFBR2 | Tgfb3 - (Acvr1b+Tgfbr2) | TGFb | Secreted Signaling | PMID: 27449815 |
| CD31 | CD31 | Tgfb3 | ACVR1C_TGFbR2 | 0.000513 | 0 | TGFB3_ACVR1C_TGFBR2 | Tgfb3 - (Acvr1c+Tgfbr2) | TGFb | Secreted Signaling | PMID: 27449815 |
| NeuN | CD31 | Tgfb3 | ACVR1C_TGFbR2 | 0.000323 | 0 | TGFB3_ACVR1C_TGFBR2 | Tgfb3 - (Acvr1c+Tgfbr2) | TGFb | Secreted Signaling | PMID: 27449815 |
| CD31 | CD31 | Bmp2 | BMPR1A_ACVR2A | 0.000987 | 0 | BMP2_BMPR1A_ACVR2A | Bmp2 - (Bmpr1a+Acvr2a) | BMP | Secreted Signaling | KEGG: mmu04350; PMID:26893264 |
| CD31 | NeuN | Bmp2 | BMPR1A_ACVR2A | 0.001162 | 0 | BMP2_BMPR1A_ACVR2A | Bmp2 - (Bmpr1a+Acvr2a) | BMP | Secreted Signaling | KEGG: mmu04350; PMID:26893264 |
| CD31 | CD31 | Bmp2 | BMPR1A_ACVR2B | 0.000731 | 0 | BMP2_BMPR1A_ACVR2B | Bmp2 - (Bmpr1a+Acvr2b) | BMP | Secreted Signaling | KEGG: mmu04350; PMID:26893264 |
| CD31 | NeuN | Bmp2 | BMPR1A_ACVR2B | 0.000757 | 0 | BMP2_BMPR1A_ACVR2B | Bmp2 - (Bmpr1a+Acvr2b) | BMP | Secreted Signaling | KEGG: mmu04350; PMID:26893264 |
| CD31 | CD31 | Bmp2 | BMPR1A_BMPR2 | 0.001488 | 0 | BMP2_BMPR1A_BMPR2 | Bmp2 - (Bmpr1a+Bmpr2) | BMP | Secreted Signaling | KEGG: mmu04350; PMID:26893264 |
| CD31 | NeuN | Bmp2 | BMPR1A_BMPR2 | 0.001383 | 0 | BMP2_BMPR1A_BMPR2 | Bmp2 - (Bmpr1a+Bmpr2) | BMP | Secreted Signaling | KEGG: mmu04350; PMID:26893264 |
| CD31 | CD31 | Bmp2 | BMPR1B_ACVR2A | 0.000724 | 0 | BMP2_BMPR1B_ACVR2A | Bmp2 - (Bmpr1b+Acvr2a) | BMP | Secreted Signaling | KEGG: mmu04350; PMID:26893264 |
| CD31 | NeuN | Bmp2 | BMPR1B_ACVR2A | 0.000818 | 0 | BMP2_BMPR1B_ACVR2A | Bmp2 - (Bmpr1b+Acvr2a) | BMP | Secreted Signaling | KEGG: mmu04350; PMID:26893264 |
| CD31 | CD31 | Bmp2 | BMPR1B_ACVR2B | 0.000536 | 0 | BMP2_BMPR1B_ACVR2B | Bmp2 - (Bmpr1b+Acvr2b) | BMP | Secreted Signaling | KEGG: mmu04350; PMID:26893264 |
| CD31 | NeuN | Bmp2 | BMPR1B_ACVR2B | 0.000533 | 0 | BMP2_BMPR1B_ACVR2B | Bmp2 - (Bmpr1b+Acvr2b) | BMP | Secreted Signaling | KEGG: mmu04350; PMID:26893264 |
| CD31 | CD31 | Bmp2 | BMPR1B_BMPR2 | 0.001092 | 0 | BMP2_BMPR1B_BMPR2 | Bmp2 - (Bmpr1b+Bmpr2) | BMP | Secreted Signaling | KEGG: mmu04350; PMID:26893264 |
| CD31 | NeuN | Bmp2 | BMPR1B_BMPR2 | 0.000974 | 0 | BMP2_BMPR1B_BMPR2 | Bmp2 - (Bmpr1b+Bmpr2) | BMP | Secreted Signaling | KEGG: mmu04350; PMID:26893264 |
| CD31 | CD31 | Gdf6 | BMPR1A_ACVR2A | 0.000817 | 0 | GDF6_BMPR1A_ACVR2A | Gdf6 - (Bmpr1a+Acvr2a) | BMP | Secreted Signaling | KEGG: mmu04350; PMID:26893264 |
| CD31 | NeuN | Gdf6 | BMPR1A_ACVR2A | 0.000963 | 0 | GDF6_BMPR1A_ACVR2A | Gdf6 - (Bmpr1a+Acvr2a) | BMP | Secreted Signaling | KEGG: mmu04350; PMID:26893264 |
| CD31 | CD31 | Gdf6 | BMPR1A_ACVR2B | 0.000605 | 0 | GDF6_BMPR1A_ACVR2B | Gdf6 - (Bmpr1a+Acvr2b) | BMP | Secreted Signaling | KEGG: mmu04350; PMID:26893264 |
| CD31 | NeuN | Gdf6 | BMPR1A_ACVR2B | 0.000626 | 0 | GDF6_BMPR1A_ACVR2B | Gdf6 - (Bmpr1a+Acvr2b) | BMP | Secreted Signaling | KEGG: mmu04350; PMID:26893264 |
| CD31 | CD31 | Gdf6 | BMPR1A_BMPR2 | 0.001232 | 0 | GDF6_BMPR1A_BMPR2 | Gdf6 - (Bmpr1a+Bmpr2) | BMP | Secreted Signaling | KEGG: mmu04350; PMID:26893264 |
| CD31 | NeuN | Gdf6 | BMPR1A_BMPR2 | 0.001146 | 0 | GDF6_BMPR1A_BMPR2 | Gdf6 - (Bmpr1a+Bmpr2) | BMP | Secreted Signaling | KEGG: mmu04350; PMID:26893264 |
| CD31 | CD31 | Gdf6 | BMPR1B_ACVR2A | 0.0006 | 0 | GDF6_BMPR1B_ACVR2A | Gdf6 - (Bmpr1b+Acvr2a) | BMP | Secreted Signaling | KEGG: mmu04350; PMID:26893264 |
| CD31 | NeuN | Gdf6 | BMPR1B_ACVR2A | 0.000678 | 0 | GDF6_BMPR1B_ACVR2A | Gdf6 - (Bmpr1b+Acvr2a) | BMP | Secreted Signaling | KEGG: mmu04350; PMID:26893264 |
| CD31 | CD31 | Gdf6 | BMPR1B_ACVR2B | 0.000444 | 0 | GDF6_BMPR1B_ACVR2B | Gdf6 - (Bmpr1b+Acvr2b) | BMP | Secreted Signaling | KEGG: mmu04350; PMID:26893264 |
| CD31 | NeuN | Gdf6 | BMPR1B_ACVR2B | 0.000441 | 0 | GDF6_BMPR1B_ACVR2B | Gdf6 - (Bmpr1b+Acvr2b) | BMP | Secreted Signaling | KEGG: mmu04350; PMID:26893264 |
| CD31 | CD31 | Gdf6 | BMPR1B_BMPR2 | 0.000904 | 0 | GDF6_BMPR1B_BMPR2 | Gdf6 - (Bmpr1b+Bmpr2) | BMP | Secreted Signaling | KEGG: mmu04350; PMID:26893264 |
| CD31 | NeuN | Gdf6 | BMPR1B_BMPR2 | 0.000807 | 0 | GDF6_BMPR1B_BMPR2 | Gdf6 - (Bmpr1b+Bmpr2) | BMP | Secreted Signaling | KEGG: mmu04350; PMID:26893264 |
| CD31 | CD31 | Bmp5 | ACVR1_ACVR2A | 0.000833 | 0 | BMP5_ACVR1_ACVR2A | Bmp5 - (Acvr1+Acvr2a) | BMP | Secreted Signaling | KEGG: mmu04350; PMID:26893264 |
| CD31 | NeuN | Bmp5 | ACVR1_ACVR2A | 0.001689 | 0 | BMP5_ACVR1_ACVR2A | Bmp5 - (Acvr1+Acvr2a) | BMP | Secreted Signaling | KEGG: mmu04350; PMID:26893264 |
| CD31 | CD31 | Bmp5 | ACVR1_ACVR2B | 0.000616 | 0 | BMP5_ACVR1_ACVR2B | Bmp5 - (Acvr1+Acvr2b) | BMP | Secreted Signaling | KEGG: mmu04350; PMID:26893264 |
| CD31 | NeuN | Bmp5 | ACVR1_ACVR2B | 0.001099 | 0 | BMP5_ACVR1_ACVR2B | Bmp5 - (Acvr1+Acvr2b) | BMP | Secreted Signaling | KEGG: mmu04350; PMID:26893264 |
| CD31 | CD31 | Bmp5 | ACVR1_BMPR2 | 0.001255 | 0 | BMP5_ACVR1_BMPR2 | Bmp5 - (Acvr1+Bmpr2) | BMP | Secreted Signaling | KEGG: mmu04350; PMID:26893264 |
| CD31 | NeuN | Bmp5 | ACVR1_BMPR2 | 0.002009 | 0 | BMP5_ACVR1_BMPR2 | Bmp5 - (Acvr1+Bmpr2) | BMP | Secreted Signaling | KEGG: mmu04350; PMID:26893264 |
| CD31 | CD31 | Bmp5 | BMPR1A_ACVR2A | 0.000997 | 0 | BMP5_BMPR1A_ACVR2A | Bmp5 - (Bmpr1a+Acvr2a) | BMP | Secreted Signaling | KEGG: mmu04350; PMID:26893264 |
| CD31 | NeuN | Bmp5 | BMPR1A_ACVR2A | 0.001174 | 0 | BMP5_BMPR1A_ACVR2A | Bmp5 - (Bmpr1a+Acvr2a) | BMP | Secreted Signaling | KEGG: mmu04350; PMID:26893264 |
| CD31 | CD31 | Bmp5 | BMPR1A_ACVR2B | 0.000738 | 0 | BMP5_BMPR1A_ACVR2B | Bmp5 - (Bmpr1a+Acvr2b) | BMP | Secreted Signaling | KEGG: mmu04350; PMID:26893264 |
| CD31 | NeuN | Bmp5 | BMPR1A_ACVR2B | 0.000764 | 0 | BMP5_BMPR1A_ACVR2B | Bmp5 - (Bmpr1a+Acvr2b) | BMP | Secreted Signaling | KEGG: mmu04350; PMID:26893264 |
| CD31 | CD31 | Bmp5 | BMPR1A_BMPR2 | 0.001503 | 0 | BMP5_BMPR1A_BMPR2 | Bmp5 - (Bmpr1a+Bmpr2) | BMP | Secreted Signaling | KEGG: mmu04350; PMID:26893264 |
| CD31 | NeuN | Bmp5 | BMPR1A_BMPR2 | 0.001397 | 0 | BMP5_BMPR1A_BMPR2 | Bmp5 - (Bmpr1a+Bmpr2) | BMP | Secreted Signaling | KEGG: mmu04350; PMID:26893264 |
| CD31 | CD31 | Bmp5 | BMPR1B_ACVR2A | 0.000732 | 0 | BMP5_BMPR1B_ACVR2A | Bmp5 - (Bmpr1b+Acvr2a) | BMP | Secreted Signaling | KEGG: mmu04350; PMID:26893264 |
| CD31 | NeuN | Bmp5 | BMPR1B_ACVR2A | 0.000827 | 0 | BMP5_BMPR1B_ACVR2A | Bmp5 - (Bmpr1b+Acvr2a) | BMP | Secreted Signaling | KEGG: mmu04350; PMID:26893264 |
| CD31 | CD31 | Bmp5 | BMPR1B_ACVR2B | 0.000541 | 0 | BMP5_BMPR1B_ACVR2B | Bmp5 - (Bmpr1b+Acvr2b) | BMP | Secreted Signaling | KEGG: mmu04350; PMID:26893264 |
| CD31 | NeuN | Bmp5 | BMPR1B_ACVR2B | 0.000538 | 0 | BMP5_BMPR1B_ACVR2B | Bmp5 - (Bmpr1b+Acvr2b) | BMP | Secreted Signaling | KEGG: mmu04350; PMID:26893264 |
| CD31 | CD31 | Bmp5 | BMPR1B_BMPR2 | 0.001103 | 0 | BMP5_BMPR1B_BMPR2 | Bmp5 - (Bmpr1b+Bmpr2) | BMP | Secreted Signaling | KEGG: mmu04350; PMID:26893264 |
| CD31 | NeuN | Bmp5 | BMPR1B_BMPR2 | 0.000984 | 0 | BMP5_BMPR1B_BMPR2 | Bmp5 - (Bmpr1b+Bmpr2) | BMP | Secreted Signaling | KEGG: mmu04350; PMID:26893264 |
| CD31 | NeuN | Bmp6 | ACVR1_ACVR2A | 0.001611 | 0 | BMP6_ACVR1_ACVR2A | Bmp6 - (Acvr1+Acvr2a) | BMP | Secreted Signaling | KEGG: mmu04350; PMID:26893264 |
| NeuN | NeuN | Bmp6 | ACVR1_ACVR2A | 0.000969 | 0 | BMP6_ACVR1_ACVR2A | Bmp6 - (Acvr1+Acvr2a) | BMP | Secreted Signaling | KEGG: mmu04350; PMID:26893264 |
| CD31 | NeuN | Bmp6 | ACVR1_ACVR2B | 0.001049 | 0 | BMP6_ACVR1_ACVR2B | Bmp6 - (Acvr1+Acvr2b) | BMP | Secreted Signaling | KEGG: mmu04350; PMID:26893264 |
| NeuN | NeuN | Bmp6 | ACVR1_ACVR2B | 0.00063 | 0 | BMP6_ACVR1_ACVR2B | Bmp6 - (Acvr1+Acvr2b) | BMP | Secreted Signaling | KEGG: mmu04350; PMID:26893264 |
| CD31 | CD31 | Bmp6 | ACVR1_BMPR2 | 0.001197 | 0 | BMP6_ACVR1_BMPR2 | Bmp6 - (Acvr1+Bmpr2) | BMP | Secreted Signaling | KEGG: mmu04350; PMID:26893264 |
| CD31 | NeuN | Bmp6 | ACVR1_BMPR2 | 0.001917 | 0 | BMP6_ACVR1_BMPR2 | Bmp6 - (Acvr1+Bmpr2) | BMP | Secreted Signaling | KEGG: mmu04350; PMID:26893264 |
| CD31 | CD31 | Bmp6 | BMPR1A_ACVR2A | 0.000951 | 0 | BMP6_BMPR1A_ACVR2A | Bmp6 - (Bmpr1a+Acvr2a) | BMP | Secreted Signaling | KEGG: mmu04350; PMID:26893264 |

| | | | | | | | | | | |
|---|---|---|---|---|---|---|---|---|---|---|
| CD31 | NeuN | Bmp6 | BMPR1A_ACVR2A | 0.00112 | 0 | BMP6_BMPR1A_ACVR2A | Bmp6 - (Bmpr1a+Acvr2a) | BMP | Secreted Signaling | KEGG: mmu04350; PMID:26893264 |
| CD31 | CD31 | Bmp6 | BMPR1A_ACVR2B | 0.000704 | 0 | BMP6_BMPR1A_ACVR2B | Bmp6 - (Bmpr1a+Acvr2b) | BMP | Secreted Signaling | KEGG: mmu04350; PMID:26893264 |
| CD31 | NeuN | Bmp6 | BMPR1A_ACVR2B | 0.000729 | 0 | BMP6_BMPR1A_ACVR2B | Bmp6 - (Bmpr1a+Acvr2b) | BMP | Secreted Signaling | KEGG: mmu04350; PMID:26893264 |
| CD31 | CD31 | Bmp6 | BMPR1A_BMPR2 | 0.001433 | 0 | BMP6_BMPR1A_BMPR2 | Bmp6 - (Bmpr1a+Bmpr2) | BMP | Secreted Signaling | KEGG: mmu04350; PMID:26893264 |
| CD31 | NeuN | Bmp6 | BMPR1A_BMPR2 | 0.001333 | 0 | BMP6_BMPR1A_BMPR2 | Bmp6 - (Bmpr1a+Bmpr2) | BMP | Secreted Signaling | KEGG: mmu04350; PMID:26893264 |
| CD31 | CD31 | Bmp6 | BMPR1B_ACVR2A | 0.000698 | 0 | BMP6_BMPR1B_ACVR2A | Bmp6 - (Bmpr1b+Acvr2a) | BMP | Secreted Signaling | KEGG: mmu04350; PMID:26893264 |
| CD31 | NeuN | Bmp6 | BMPR1B_ACVR2A | 0.000788 | 0 | BMP6_BMPR1B_ACVR2A | Bmp6 - (Bmpr1b+Acvr2a) | BMP | Secreted Signaling | KEGG: mmu04350; PMID:26893264 |
| CD31 | CD31 | Bmp6 | BMPR1B_ACVR2B | 0.000516 | 0 | BMP6_BMPR1B_ACVR2B | Bmp6 - (Bmpr1b+Acvr2b) | BMP | Secreted Signaling | KEGG: mmu04350; PMID:26893264 |
| CD31 | NeuN | Bmp6 | BMPR1B_ACVR2B | 0.000513 | 0 | BMP6_BMPR1B_ACVR2B | Bmp6 - (Bmpr1b+Acvr2b) | BMP | Secreted Signaling | KEGG: mmu04350; PMID:26893264 |
| CD31 | CD31 | Bmp6 | BMPR1B_BMPR2 | 0.001052 | 0 | BMP6_BMPR1B_BMPR2 | Bmp6 - (Bmpr1b+Bmpr2) | BMP | Secreted Signaling | KEGG: mmu04350; PMID:26893264 |
| CD31 | NeuN | Bmp6 | BMPR1B_BMPR2 | 0.000938 | 0 | BMP6_BMPR1B_BMPR2 | Bmp6 - (Bmpr1b+Bmpr2) | BMP | Secreted Signaling | KEGG: mmu04350; PMID:26893264 |
| CD31 | CD31 | Bmp8b | ACVR1_ACVR2A | 0.000696 | 0 | BMP8B_ACVR1_ACVR2A | Bmp8b - (Acvr1+Acvr2a) | BMP | Secreted Signaling | KEGG: mmu04350; PMID:26893264 |
| CD31 | NeuN | Bmp8b | ACVR1_ACVR2A | 0.001411 | 0 | BMP8B_ACVR1_ACVR2A | Bmp8b - (Acvr1+Acvr2a) | BMP | Secreted Signaling | KEGG: mmu04350; PMID:26893264 |
| CD31 | CD31 | Bmp8b | ACVR1_ACVR2B | 0.000515 | 0 | BMP8B_ACVR1_ACVR2B | Bmp8b - (Acvr1+Acvr2b) | BMP | Secreted Signaling | KEGG: mmu04350; PMID:26893264 |
| CD31 | NeuN | Bmp8b | ACVR1_ACVR2B | 0.000918 | 0 | BMP8B_ACVR1_ACVR2B | Bmp8b - (Acvr1+Acvr2b) | BMP | Secreted Signaling | KEGG: mmu04350; PMID:26893264 |
| CD31 | CD31 | Bmp8b | ACVR1_BMPR2 | 0.001049 | 0 | BMP8B_ACVR1_BMPR2 | Bmp8b - (Acvr1+Bmpr2) | BMP | Secreted Signaling | KEGG: mmu04350; PMID:26893264 |
| CD31 | NeuN | Bmp8b | ACVR1_BMPR2 | 0.001679 | 0 | BMP8B_ACVR1_BMPR2 | Bmp8b - (Acvr1+Bmpr2) | BMP | Secreted Signaling | KEGG: mmu04350; PMID:26893264 |
| CD31 | CD31 | Bmp8b | BMPR1A_ACVR2A | 0.000833 | 0 | BMP8B_BMPR1A_ACVR2A | Bmp8b - (Bmpr1a+Acvr2a) | BMP | Secreted Signaling | KEGG: mmu04350; PMID:26893264 |
| CD31 | NeuN | Bmp8b | BMPR1A_ACVR2A | 0.000981 | 0 | BMP8B_BMPR1A_ACVR2A | Bmp8b - (Bmpr1a+Acvr2a) | BMP | Secreted Signaling | KEGG: mmu04350; PMID:26893264 |
| CD31 | CD31 | Bmp8b | BMPR1A_ACVR2B | 0.000616 | 0 | BMP8B_BMPR1A_ACVR2B | Bmp8b - (Bmpr1a+Acvr2b) | BMP | Secreted Signaling | KEGG: mmu04350; PMID:26893264 |
| CD31 | NeuN | Bmp8b | BMPR1A_ACVR2B | 0.000638 | 0 | BMP8B_BMPR1A_ACVR2B | Bmp8b - (Bmpr1a+Acvr2b) | BMP | Secreted Signaling | KEGG: mmu04350; PMID:26893264 |
| CD31 | CD31 | Bmp8b | BMPR1A_BMPR2 | 0.001255 | 0 | BMP8B_BMPR1A_BMPR2 | Bmp8b - (Bmpr1a+Bmpr2) | BMP | Secreted Signaling | KEGG: mmu04350; PMID:26893264 |
| CD31 | NeuN | Bmp8b | BMPR1A_BMPR2 | 0.001167 | 0 | BMP8B_BMPR1A_BMPR2 | Bmp8b - (Bmpr1a+Bmpr2) | BMP | Secreted Signaling | KEGG: mmu04350; PMID:26893264 |
| CD31 | CD31 | Bmp8b | BMPR1B_ACVR2A | 0.000611 | 0 | BMP8B_BMPR1B_ACVR2A | Bmp8b - (Bmpr1b+Acvr2a) | BMP | Secreted Signaling | KEGG: mmu04350; PMID:26893264 |
| CD31 | NeuN | Bmp8b | BMPR1B_ACVR2A | 0.00069 | 0 | BMP8B_BMPR1B_ACVR2A | Bmp8b - (Bmpr1b+Acvr2a) | BMP | Secreted Signaling | KEGG: mmu04350; PMID:26893264 |
| CD31 | CD31 | Bmp8b | BMPR1B_ACVR2B | 0.000452 | 0 | BMP8B_BMPR1B_ACVR2B | Bmp8b - (Bmpr1b+Acvr2b) | BMP | Secreted Signaling | KEGG: mmu04350; PMID:26893264 |
| CD31 | NeuN | Bmp8b | BMPR1B_ACVR2B | 0.000449 | 0 | BMP8B_BMPR1B_ACVR2B | Bmp8b - (Bmpr1b+Acvr2b) | BMP | Secreted Signaling | KEGG: mmu04350; PMID:26893264 |
| CD31 | CD31 | Bmp8b | BMPR1B_BMPR2 | 0.000921 | 0 | BMP8B_BMPR1B_BMPR2 | Bmp8b - (Bmpr1b+Bmpr2) | BMP | Secreted Signaling | KEGG: mmu04350; PMID:26893264 |
| CD31 | NeuN | Bmp8b | BMPR1B_BMPR2 | 0.000822 | 0 | BMP8B_BMPR1B_BMPR2 | Bmp8b - (Bmpr1b+Bmpr2) | BMP | Secreted Signaling | KEGG: mmu04350; PMID:26893264 |
| CD31 | CD31 | Gdf9 | ACVR1B_BMPR2 | 0.00212 | 0 | GDF9_ACVR1B_BMPR2 | Gdf9 - (Acvr1b+Bmpr2) | GDF | Secreted Signaling | KEGG: mmu04350 |
| CD31 | NeuN | Gdf9 | ACVR1B_BMPR2 | 0.002257 | 0 | GDF9_ACVR1B_BMPR2 | Gdf9 - (Acvr1b+Bmpr2) | GDF | Secreted Signaling | KEGG: mmu04350 |
| CD31 | NeuN | Gdf10 | ACVR1B_ACVR2A | 0.00177 | 0 | GDF10_ACVR1B_ACVR2A | Gdf10 - (Acvr1b+Acvr2a) | GDF | Secreted Signaling | KEGG: mmu04350 |
| NeuN | NeuN | Gdf10 | ACVR1B_ACVR2A | 0.001508 | 0 | GDF10_ACVR1B_ACVR2A | Gdf10 - (Acvr1b+Acvr2a) | GDF | Secreted Signaling | KEGG: mmu04350 |
| CD31 | NeuN | Gdf11 | TGFBR1_ACVR2A | 0.002983 | 0 | GDF11_TGFBR1_ACVR2A | Gdf11 - (Tgfbr1+Acvr2a) | GDF | Secreted Signaling | KEGG: mmu04350 |
| NeuN | NeuN | Gdf11 | TGFBR1_ACVR2A | 0.002503 | 0 | GDF11_TGFBR1_ACVR2A | Gdf11 - (Tgfbr1+Acvr2a) | GDF | Secreted Signaling | KEGG: mmu04350 |
| CD31 | NeuN | Gdf11 | TGFBR1_ACVR2B | 0.001942 | 0 | GDF11_TGFBR1_ACVR2B | Gdf11 - (Tgfbr1+Acvr2b) | GDF | Secreted Signaling | KEGG: mmu04350 |
| NeuN | NeuN | Gdf11 | TGFBR1_ACVR2B | 0.001629 | 0 | GDF11_TGFBR1_ACVR2B | Gdf11 - (Tgfbr1+Acvr2b) | GDF | Secreted Signaling | KEGG: mmu04350 |
| CD31 | CD31 | Gdf11 | ACVR1B_ACVR2B | 0.001582 | 0 | GDF11_ACVR1B_ACVR2B | Gdf11 - (Acvr1b+Acvr2b) | GDF | Secreted Signaling | KEGG: mmu04350 |
| CD31 | NeuN | Gdf11 | ACVR1B_ACVR2B | 0.001876 | 0 | GDF11_ACVR1B_ACVR2B | Gdf11 - (Acvr1b+Acvr2b) | GDF | Secreted Signaling | KEGG: mmu04350 |
| CD31 | CD31 | Gdnf | GFRA1_RET | 0.000639 | 0 | GDNF_GFRA1 | Gdnf - Gfra1 | GDNF | Secreted Signaling | PMID: 11544105 |
| NeuN | CD31 | Gdnf | GFRA1_RET | 0.000501 | 0 | GDNF_GFRA1 | Gdnf - Gfra1 | GDNF | Secreted Signaling | PMID: 11544105 |
| CD31 | CD31 | Nrtn | GFRA2_RET | 0.000528 | 0 | NRTN_GFRA2 | Nrtn - Gfra2 | GDNF | Secreted Signaling | PMID: 11544105 |
| NeuN | CD31 | Nrtn | GFRA2_RET | 0.000487 | 0 | NRTN_GFRA2 | Nrtn - Gfra2 | GDNF | Secreted Signaling | PMID: 11544105 |
| NeuN | CD31 | Inhba | ACVR1B_ACVR2A | 0.003142 | 0 | INHBA_ACVR1B_ACVR2A | Inhba - (Acvr1b+Acvr2a) | ACTIVIN | Secreted Signaling | KEGG: mmu04350 |
| NeuN | NeuN | Inhba | ACVR1B_ACVR2A | 0.004354 | 0 | INHBA_ACVR1B_ACVR2A | Inhba - (Acvr1b+Acvr2a) | ACTIVIN | Secreted Signaling | KEGG: mmu04350 |
| NeuN | CD31 | Inhba | ACVR1B_ACVR2B | 0.002326 | 0 | INHBA_ACVR1B_ACVR2B | Inhba - (Acvr1b+Acvr2b) | ACTIVIN | Secreted Signaling | KEGG: mmu04350 |
| NeuN | NeuN | Inhba | ACVR1B_ACVR2B | 0.002837 | 0 | INHBA_ACVR1B_ACVR2B | Inhba - (Acvr1b+Acvr2b) | ACTIVIN | Secreted Signaling | KEGG: mmu04350 |
| NeuN | CD31 | Inhbb | ACVR1B_ACVR2A | 0.002052 | 0 | INHBB_ACVR1B_ACVR2A | Inhbb - (Acvr1b+Acvr2a) | ACTIVIN | Secreted Signaling | KEGG: mmu04350 |
| NeuN | NeuN | Inhbb | ACVR1B_ACVR2A | 0.002845 | 0 | INHBB_ACVR1B_ACVR2A | Inhbb - (Acvr1b+Acvr2a) | ACTIVIN | Secreted Signaling | KEGG: mmu04350 |
| NeuN | CD31 | Inhbb | ACVR1B_ACVR2B | 0.001519 | 0 | INHBB_ACVR1B_ACVR2B | Inhbb - (Acvr1b+Acvr2b) | ACTIVIN | Secreted Signaling | KEGG: mmu04350 |
| NeuN | NeuN | Inhbb | ACVR1B_ACVR2B | 0.001852 | 0 | INHBB_ACVR1B_ACVR2B | Inhbb - (Acvr1b+Acvr2b) | ACTIVIN | Secreted Signaling | KEGG: mmu04350 |
| NeuN | CD31 | Inhbb | ACVR1C_ACVR2A | 0.001297 | 0 | INHBB_ACVR1C_ACVR2A | Inhbb - (Acvr1c+Acvr2a) | ACTIVIN | Secreted Signaling | KEGG: mmu04350 |
| NeuN | NeuN | Inhbb | ACVR1C_ACVR2A | 0.00128 | 0 | INHBB_ACVR1C_ACVR2A | Inhbb - (Acvr1c+Acvr2a) | ACTIVIN | Secreted Signaling | KEGG: mmu04350 |
| NeuN | CD31 | Inhbb | ACVR1C_ACVR2B | 0.00096 | 0 | INHBB_ACVR1C_ACVR2B | Inhbb - (Acvr1c+Acvr2b) | ACTIVIN | Secreted Signaling | KEGG: mmu04350 |
| NeuN | NeuN | Inhbb | ACVR1C_ACVR2B | 0.000833 | 0 | INHBB_ACVR1C_ACVR2B | Inhbb - (Acvr1c+Acvr2b) | ACTIVIN | Secreted Signaling | KEGG: mmu04350 |
| CD31 | CD31 | Wnt10b | FZD1_LRP5 | 0.000457 | 0 | WNT10B_FZD1_LRP5 | Wnt10b - (Fzd1+Lrp5) | WNT | Secreted Signaling | KEGG: mmu04310; PMID: 23209147 |
| CD31 | NeuN | Wnt10b | FZD1_LRP5 | 0.000315 | 0 | WNT10B_FZD1_LRP5 | Wnt10b - (Fzd1+Lrp5) | WNT | Secreted Signaling | KEGG: mmu04310; PMID: 23209147 |
| CD31 | CD31 | Wnt10b | FZD2_LRP5 | 0.000286 | 0 | WNT10B_FZD2_LRP5 | Wnt10b - (Fzd2+Lrp5) | WNT | Secreted Signaling | KEGG: mmu04310; PMID: 23209147 |
| CD31 | NeuN | Wnt10b | FZD2_LRP5 | 0.000324 | 0 | WNT10B_FZD2_LRP5 | Wnt10b - (Fzd2+Lrp5) | WNT | Secreted Signaling | KEGG: mmu04310; PMID: 23209147 |
| CD31 | CD31 | Wnt10b | FZD3_LRP5 | 0.000503 | 0 | WNT10B_FZD3_LRP5 | Wnt10b - (Fzd3+Lrp5) | WNT | Secreted Signaling | KEGG: mmu04310; PMID: 23209147 |

| | | | | | | | | | | |
|---|---|---|---|---|---|---|---|---|---|---|
| CD31 | NeuN | Wnt10b | FZD3_LRP5 | 0.000819 | 0 | WNT10B_FZD3_LRP5 | Wnt10b - (Fzd3+Lrp5) | WNT | Secreted Signaling | KEGG: mmu04310; PMID: 23209147 |
| CD31 | CD31 | Wnt10b | FZD4_LRP5 | 0.000247 | 0 | WNT10B_FZD4_LRP5 | Wnt10b - (Fzd4+Lrp5) | WNT | Secreted Signaling | KEGG: mmu04310; PMID: 23209147 |
| CD31 | NeuN | Wnt10b | FZD4_LRP5 | 0.000226 | 0 | WNT10B_FZD4_LRP5 | Wnt10b - (Fzd4+Lrp5) | WNT | Secreted Signaling | KEGG: mmu04310; PMID: 23209147 |
| CD31 | CD31 | Wnt10b | FZD5_LRP5 | 0.0003 | 0 | WNT10B_FZD5_LRP5 | Wnt10b - (Fzd5+Lrp5) | WNT | Secreted Signaling | KEGG: mmu04310; PMID: 23209147 |
| CD31 | NeuN | Wnt10b | FZD5_LRP5 | 0.00025 | 0 | WNT10B_FZD5_LRP5 | Wnt10b - (Fzd5+Lrp5) | WNT | Secreted Signaling | KEGG: mmu04310; PMID: 23209147 |
| CD31 | CD31 | Wnt10b | FZD6_LRP5 | 0.00061 | 0 | WNT10B_FZD6_LRP5 | Wnt10b - (Fzd6+Lrp5) | WNT | Secreted Signaling | KEGG: mmu04310; PMID: 23209147 |
| NeuN | CD31 | Wnt10b | FZD6_LRP5 | 0.000306 | 0 | WNT10B_FZD6_LRP5 | Wnt10b - (Fzd6+Lrp5) | WNT | Secreted Signaling | KEGG: mmu04310; PMID: 23209147 |
| CD31 | CD31 | Wnt10b | FZD8_LRP5 | 0.000338 | 0 | WNT10B_FZD8_LRP5 | Wnt10b - (Fzd8+Lrp5) | WNT | Secreted Signaling | KEGG: mmu04310; PMID: 23209147 |
| CD31 | NeuN | Wnt10b | FZD8_LRP5 | 0.00031 | 0 | WNT10B_FZD8_LRP5 | Wnt10b - (Fzd8+Lrp5) | WNT | Secreted Signaling | KEGG: mmu04310; PMID: 23209147 |
| CD31 | CD31 | Wnt10b | FZD9_LRP5 | 0.000313 | 0 | WNT10B_FZD9_LRP5 | Wnt10b - (Fzd9+Lrp5) | WNT | Secreted Signaling | KEGG: mmu04310; PMID: 23209147 |
| CD31 | NeuN | Wnt10b | FZD9_LRP5 | 0.000327 | 0 | WNT10B_FZD9_LRP5 | Wnt10b - (Fzd9+Lrp5) | WNT | Secreted Signaling | KEGG: mmu04310; PMID: 23209147 |
| CD31 | CD31 | Wnt4 | FZD1_LRP5 | 0.00039 | 0 | WNT4_FZD1_LRP5 | Wnt4 - (Fzd1+Lrp5) | WNT | Secreted Signaling | KEGG: mmu04310; PMID: 23209147 |
| NeuN | CD31 | Wnt4 | FZD1_LRP5 | 0.000314 | 0 | WNT4_FZD1_LRP5 | Wnt4 - (Fzd1+Lrp5) | WNT | Secreted Signaling | KEGG: mmu04310; PMID: 23209147 |
| CD31 | CD31 | Wnt4 | FZD2_LRP5 | 0.000244 | 0 | WNT4_FZD2_LRP5 | Wnt4 - (Fzd2+Lrp5) | WNT | Secreted Signaling | KEGG: mmu04310; PMID: 23209147 |
| CD31 | NeuN | Wnt4 | FZD2_LRP5 | 0.000276 | 0 | WNT4_FZD2_LRP5 | Wnt4 - (Fzd2+Lrp5) | WNT | Secreted Signaling | KEGG: mmu04310; PMID: 23209147 |
| CD31 | NeuN | Wnt4 | FZD3_LRP5 | 0.000698 | 0 | WNT4_FZD3_LRP5 | Wnt4 - (Fzd3+Lrp5) | WNT | Secreted Signaling | KEGG: mmu04310; PMID: 23209147 |
| NeuN | NeuN | Wnt4 | FZD3_LRP5 | 0.000563 | 0 | WNT4_FZD3_LRP5 | Wnt4 - (Fzd3+Lrp5) | WNT | Secreted Signaling | KEGG: mmu04310; PMID: 23209147 |
| CD31 | CD31 | Wnt4 | FZD4_LRP5 | 0.000211 | 0 | WNT4_FZD4_LRP5 | Wnt4 - (Fzd4+Lrp5) | WNT | Secreted Signaling | KEGG: mmu04310; PMID: 23209147 |
| CD31 | NeuN | Wnt4 | FZD4_LRP5 | 0.000192 | 0 | WNT4_FZD4_LRP5 | Wnt4 - (Fzd4+Lrp5) | WNT | Secreted Signaling | KEGG: mmu04310; PMID: 23209147 |
| CD31 | CD31 | Wnt4 | FZD5_LRP5 | 0.000255 | 0 | WNT4_FZD5_LRP5 | Wnt4 - (Fzd5+Lrp5) | WNT | Secreted Signaling | KEGG: mmu04310; PMID: 23209147 |
| CD31 | NeuN | Wnt4 | FZD5_LRP5 | 0.000213 | 0 | WNT4_FZD5_LRP5 | Wnt4 - (Fzd5+Lrp5) | WNT | Secreted Signaling | KEGG: mmu04310; PMID: 23209147 |
| CD31 | CD31 | Wnt4 | FZD6_LRP5 | 0.00052 | 0 | WNT4_FZD6_LRP5 | Wnt4 - (Fzd6+Lrp5) | WNT | Secreted Signaling | KEGG: mmu04310; PMID: 23209147 |
| NeuN | CD31 | Wnt4 | FZD6_LRP5 | 0.000419 | 0 | WNT4_FZD6_LRP5 | Wnt4 - (Fzd6+Lrp5) | WNT | Secreted Signaling | KEGG: mmu04310; PMID: 23209147 |
| CD31 | CD31 | Wnt4 | FZD8_LRP5 | 0.000288 | 0 | WNT4_FZD8_LRP5 | Wnt4 - (Fzd8+Lrp5) | WNT | Secreted Signaling | KEGG: mmu04310; PMID: 23209147 |
| CD31 | NeuN | Wnt4 | FZD8_LRP5 | 0.000265 | 0 | WNT4_FZD8_LRP5 | Wnt4 - (Fzd8+Lrp5) | WNT | Secreted Signaling | KEGG: mmu04310; PMID: 23209147 |
| CD31 | CD31 | Wnt4 | FZD9_LRP5 | 0.000267 | 0 | WNT4_FZD9_LRP5 | Wnt4 - (Fzd9+Lrp5) | WNT | Secreted Signaling | KEGG: mmu04310; PMID: 23209147 |
| CD31 | NeuN | Wnt4 | FZD9_LRP5 | 0.000278 | 0 | WNT4_FZD9_LRP5 | Wnt4 - (Fzd9+Lrp5) | WNT | Secreted Signaling | KEGG: mmu04310; PMID: 23209147 |
| CD31 | CD31 | Wnt7a | FZD1_LRP5 | 0.000341 | 0 | WNT7A_FZD1_LRP5 | Wnt7a - (Fzd1+Lrp5) | WNT | Secreted Signaling | KEGG: mmu04310; PMID: 23209147 |
| NeuN | CD31 | Wnt7a | FZD1_LRP5 | 0.000335 | 0 | WNT7A_FZD1_LRP5 | Wnt7a - (Fzd1+Lrp5) | WNT | Secreted Signaling | KEGG: mmu04310; PMID: 23209147 |
| CD31 | NeuN | Wnt7a | FZD2_LRP5 | 0.000242 | 0 | WNT7A_FZD2_LRP5 | Wnt7a - (Fzd2+Lrp5) | WNT | Secreted Signaling | KEGG: mmu04310; PMID: 23209147 |
| NeuN | NeuN | Wnt7a | FZD2_LRP5 | 0.000237 | 0 | WNT7A_FZD2_LRP5 | Wnt7a - (Fzd2+Lrp5) | WNT | Secreted Signaling | KEGG: mmu04310; PMID: 23209147 |
| CD31 | NeuN | Wnt7a | FZD3_LRP5 | 0.000612 | 0 | WNT7A_FZD3_LRP5 | Wnt7a - (Fzd3+Lrp5) | WNT | Secreted Signaling | KEGG: mmu04310; PMID: 23209147 |
| NeuN | NeuN | Wnt7a | FZD3_LRP5 | 0.0006 | 0 | WNT7A_FZD3_LRP5 | Wnt7a - (Fzd3+Lrp5) | WNT | Secreted Signaling | KEGG: mmu04310; PMID: 23209147 |
| CD31 | CD31 | Wnt7a | FZD4_LRP5 | 0.000185 | 0 | WNT7A_FZD4_LRP5 | Wnt7a - (Fzd4+Lrp5) | WNT | Secreted Signaling | KEGG: mmu04310; PMID: 23209147 |
| NeuN | CD31 | Wnt7a | FZD4_LRP5 | 0.000181 | 0 | WNT7A_FZD4_LRP5 | Wnt7a - (Fzd4+Lrp5) | WNT | Secreted Signaling | KEGG: mmu04310; PMID: 23209147 |
| CD31 | CD31 | Wnt7a | FZD5_LRP5 | 0.000224 | 0 | WNT7A_FZD5_LRP5 | Wnt7a - (Fzd5+Lrp5) | WNT | Secreted Signaling | KEGG: mmu04310; PMID: 23209147 |
| NeuN | CD31 | Wnt7a | FZD5_LRP5 | 0.000219 | 0 | WNT7A_FZD5_LRP5 | Wnt7a - (Fzd5+Lrp5) | WNT | Secreted Signaling | KEGG: mmu04310; PMID: 23209147 |
| CD31 | CD31 | Wnt7a | FZD6_LRP5 | 0.000456 | 0 | WNT7A_FZD6_LRP5 | Wnt7a - (Fzd6+Lrp5) | WNT | Secreted Signaling | KEGG: mmu04310; PMID: 23209147 |
| NeuN | CD31 | Wnt7a | FZD6_LRP5 | 0.000447 | 0 | WNT7A_FZD6_LRP5 | Wnt7a - (Fzd6+Lrp5) | WNT | Secreted Signaling | KEGG: mmu04310; PMID: 23209147 |
| CD31 | CD31 | Wnt7a | FZD8_LRP5 | 0.000252 | 0 | WNT7A_FZD8_LRP5 | Wnt7a - (Fzd8+Lrp5) | WNT | Secreted Signaling | KEGG: mmu04310; PMID: 23209147 |
| NeuN | CD31 | Wnt7a | FZD8_LRP5 | 0.000247 | 0 | WNT7A_FZD8_LRP5 | Wnt7a - (Fzd8+Lrp5) | WNT | Secreted Signaling | KEGG: mmu04310; PMID: 23209147 |
| CD31 | NeuN | Wnt7a | FZD9_LRP5 | 0.000244 | 0 | WNT7A_FZD9_LRP5 | Wnt7a - (Fzd9+Lrp5) | WNT | Secreted Signaling | KEGG: mmu04310; PMID: 23209147 |
| NeuN | NeuN | Wnt7a | FZD9_LRP5 | 0.000239 | 0 | WNT7A_FZD9_LRP5 | Wnt7a - (Fzd9+Lrp5) | WNT | Secreted Signaling | KEGG: mmu04310; PMID: 23209147 |
| NeuN | CD31 | Wnt7b | FZD1_LRP5 | 0.000756 | 0 | WNT7B_FZD1_LRP5 | Wnt7b - (Fzd1+Lrp5) | WNT | Secreted Signaling | KEGG: mmu04310; PMID: 23209147 |
| NeuN | NeuN | Wnt7b | FZD1_LRP5 | 0.00052 | 0 | WNT7B_FZD1_LRP5 | Wnt7b - (Fzd1+Lrp5) | WNT | Secreted Signaling | KEGG: mmu04310; PMID: 23209147 |
| NeuN | CD31 | Wnt7b | FZD2_LRP5 | 0.000473 | 0 | WNT7B_FZD2_LRP5 | Wnt7b - (Fzd2+Lrp5) | WNT | Secreted Signaling | KEGG: mmu04310; PMID: 23209147 |
| NeuN | NeuN | Wnt7b | FZD2_LRP5 | 0.000536 | 0 | WNT7B_FZD2_LRP5 | Wnt7b - (Fzd2+Lrp5) | WNT | Secreted Signaling | KEGG: mmu04310; PMID: 23209147 |
| CD31 | NeuN | Wnt7b | FZD3_LRP5 | 0.000901 | 0 | WNT7B_FZD3_LRP5 | Wnt7b - (Fzd3+Lrp5) | WNT | Secreted Signaling | KEGG: mmu04310; PMID: 23209147 |
| NeuN | NeuN | Wnt7b | FZD3_LRP5 | 0.001354 | 0 | WNT7B_FZD3_LRP5 | Wnt7b - (Fzd3+Lrp5) | WNT | Secreted Signaling | KEGG: mmu04310; PMID: 23209147 |
| NeuN | CD31 | Wnt7b | FZD4_LRP5 | 0.000409 | 0 | WNT7B_FZD4_LRP5 | Wnt7b - (Fzd4+Lrp5) | WNT | Secreted Signaling | KEGG: mmu04310; PMID: 23209147 |
| NeuN | NeuN | Wnt7b | FZD4_LRP5 | 0.000373 | 0 | WNT7B_FZD4_LRP5 | Wnt7b - (Fzd4+Lrp5) | WNT | Secreted Signaling | KEGG: mmu04310; PMID: 23209147 |
| NeuN | CD31 | Wnt7b | FZD5_LRP5 | 0.000495 | 0 | WNT7B_FZD5_LRP5 | Wnt7b - (Fzd5+Lrp5) | WNT | Secreted Signaling | KEGG: mmu04310; PMID: 23209147 |
| NeuN | NeuN | Wnt7b | FZD5_LRP5 | 0.000414 | 0 | WNT7B_FZD5_LRP5 | Wnt7b - (Fzd5+Lrp5) | WNT | Secreted Signaling | KEGG: mmu04310; PMID: 23209147 |
| CD31 | CD31 | Wnt7b | FZD6_LRP5 | 0.000671 | 0 | WNT7B_FZD6_LRP5 | Wnt7b - (Fzd6+Lrp5) | WNT | Secreted Signaling | KEGG: mmu04310; PMID: 23209147 |
| NeuN | CD31 | Wnt7b | FZD6_LRP5 | 0.001009 | 0 | WNT7B_FZD6_LRP5 | Wnt7b - (Fzd6+Lrp5) | WNT | Secreted Signaling | KEGG: mmu04310; PMID: 23209147 |
| NeuN | CD31 | Wnt7b | FZD8_LRP5 | 0.000559 | 0 | WNT7B_FZD8_LRP5 | Wnt7b - (Fzd8+Lrp5) | WNT | Secreted Signaling | KEGG: mmu04310; PMID: 23209147 |
| NeuN | NeuN | Wnt7b | FZD8_LRP5 | 0.000513 | 0 | WNT7B_FZD8_LRP5 | Wnt7b - (Fzd8+Lrp5) | WNT | Secreted Signaling | KEGG: mmu04310; PMID: 23209147 |
| NeuN | CD31 | Wnt7b | FZD9_LRP5 | 0.000518 | 0 | WNT7B_FZD9_LRP5 | Wnt7b - (Fzd9+Lrp5) | WNT | Secreted Signaling | KEGG: mmu04310; PMID: 23209147 |
| NeuN | NeuN | Wnt7b | FZD9_LRP5 | 0.00054 | 0 | WNT7B_FZD9_LRP5 | Wnt7b - (Fzd9+Lrp5) | WNT | Secreted Signaling | KEGG: mmu04310; PMID: 23209147 |
| CD31 | CD31 | Wnt9a | FZD1_LRP5 | 0.000259 | 0 | WNT9A_FZD1_LRP5 | Wnt9a - (Fzd1+Lrp5) | WNT | Secreted Signaling | KEGG: mmu04310; PMID: 23209147 |

| | | | | | | | | | | |
|---|---|---|---|---|---|---|---|---|---|---|
| NeuN | CD31 | Wnt9a | FZD1_LRP5 | 0.000193 | 0 | WNT9A_FZD1_LRP5 | Wnt9a - (Fzd1+Lrp5) | WNT | Secreted Signaling | KEGG: mmu04310; PMID: 23209147 |
| CD31 | CD31 | Wnt9a | FZD2_LRP5 | 0.000162 | 0 | WNT9A_FZD2_LRP5 | Wnt9a - (Fzd2+Lrp5) | WNT | Secreted Signaling | KEGG: mmu04310; PMID: 23209147 |
| CD31 | NeuN | Wnt9a | FZD2_LRP5 | 0.000184 | 0 | WNT9A_FZD2_LRP5 | Wnt9a - (Fzd2+Lrp5) | WNT | Secreted Signaling | KEGG: mmu04310; PMID: 23209147 |
| CD31 | NeuN | Wnt9a | FZD3_LRP5 | 0.000464 | 0 | WNT9A_FZD3_LRP5 | Wnt9a - (Fzd3+Lrp5) | WNT | Secreted Signaling | KEGG: mmu04310; PMID: 23209147 |
| NeuN | NeuN | Wnt9a | FZD3_LRP5 | 0.000346 | 0 | WNT9A_FZD3_LRP5 | Wnt9a - (Fzd3+Lrp5) | WNT | Secreted Signaling | KEGG: mmu04310; PMID: 23209147 |
| CD31 | CD31 | Wnt9a | FZD4_LRP5 | 0.00014 | 0 | WNT9A_FZD4_LRP5 | Wnt9a - (Fzd4+Lrp5) | WNT | Secreted Signaling | KEGG: mmu04310; PMID: 23209147 |
| CD31 | NeuN | Wnt9a | FZD4_LRP5 | 0.000128 | 0 | WNT9A_FZD4_LRP5 | Wnt9a - (Fzd4+Lrp5) | WNT | Secreted Signaling | KEGG: mmu04310; PMID: 23209147 |
| CD31 | CD31 | Wnt9a | FZD5_LRP5 | 0.00017 | 0 | WNT9A_FZD5_LRP5 | Wnt9a - (Fzd5+Lrp5) | WNT | Secreted Signaling | KEGG: mmu04310; PMID: 23209147 |
| CD31 | NeuN | Wnt9a | FZD5_LRP5 | 0.000142 | 0 | WNT9A_FZD5_LRP5 | Wnt9a - (Fzd5+Lrp5) | WNT | Secreted Signaling | KEGG: mmu04310; PMID: 23209147 |
| CD31 | CD31 | Wnt9a | FZD6_LRP5 | 0.000346 | 0 | WNT9A_FZD6_LRP5 | Wnt9a - (Fzd6+Lrp5) | WNT | Secreted Signaling | KEGG: mmu04310; PMID: 23209147 |
| NeuN | CD31 | Wnt9a | FZD6_LRP5 | 0.000257 | 0 | WNT9A_FZD6_LRP5 | Wnt9a - (Fzd6+Lrp5) | WNT | Secreted Signaling | KEGG: mmu04310; PMID: 23209147 |
| CD31 | CD31 | Wnt9a | FZD8_LRP5 | 0.000191 | 0 | WNT9A_FZD8_LRP5 | Wnt9a - (Fzd8+Lrp5) | WNT | Secreted Signaling | KEGG: mmu04310; PMID: 23209147 |
| CD31 | NeuN | Wnt9a | FZD8_LRP5 | 0.000176 | 0 | WNT9A_FZD8_LRP5 | Wnt9a - (Fzd8+Lrp5) | WNT | Secreted Signaling | KEGG: mmu04310; PMID: 23209147 |
| CD31 | CD31 | Wnt9a | FZD9_LRP5 | 0.000177 | 0 | WNT9A_FZD9_LRP5 | Wnt9a - (Fzd9+Lrp5) | WNT | Secreted Signaling | KEGG: mmu04310; PMID: 23209147 |
| CD31 | NeuN | Wnt9a | FZD9_LRP5 | 0.000185 | 0 | WNT9A_FZD9_LRP5 | Wnt9a - (Fzd9+Lrp5) | WNT | Secreted Signaling | KEGG: mmu04310; PMID: 23209147 |
| CD31 | CD31 | Wnt10b | FZD1_LRP6 | 0.000584 | 0 | WNT10B_FZD1_LRP6 | Wnt10b - (Fzd1+Lrp6) | WNT | Secreted Signaling | KEGG: mmu04310; PMID: 23209147 |
| CD31 | NeuN | Wnt10b | FZD1_LRP6 | 0.000425 | 0 | WNT10B_FZD1_LRP6 | Wnt10b - (Fzd1+Lrp6) | WNT | Secreted Signaling | KEGG: mmu04310; PMID: 23209147 |
| CD31 | CD31 | Wnt10b | FZD2_LRP6 | 0.000366 | 0 | WNT10B_FZD2_LRP6 | Wnt10b - (Fzd2+Lrp6) | WNT | Secreted Signaling | KEGG: mmu04310; PMID: 23209147 |
| CD31 | NeuN | Wnt10b | FZD2_LRP6 | 0.000438 | 0 | WNT10B_FZD2_LRP6 | Wnt10b - (Fzd2+Lrp6) | WNT | Secreted Signaling | KEGG: mmu04310; PMID: 23209147 |
| CD31 | CD31 | Wnt10b | FZD3_LRP6 | 0.000642 | 0 | WNT10B_FZD3_LRP6 | Wnt10b - (Fzd3+Lrp6) | WNT | Secreted Signaling | KEGG: mmu04310; PMID: 23209147 |
| CD31 | NeuN | Wnt10b | FZD3_LRP6 | 0.001107 | 0 | WNT10B_FZD3_LRP6 | Wnt10b - (Fzd3+Lrp6) | WNT | Secreted Signaling | KEGG: mmu04310; PMID: 23209147 |
| CD31 | CD31 | Wnt10b | FZD4_LRP6 | 0.000316 | 0 | WNT10B_FZD4_LRP6 | Wnt10b - (Fzd4+Lrp6) | WNT | Secreted Signaling | KEGG: mmu04310; PMID: 23209147 |
| CD31 | NeuN | Wnt10b | FZD4_LRP6 | 0.000305 | 0 | WNT10B_FZD4_LRP6 | Wnt10b - (Fzd4+Lrp6) | WNT | Secreted Signaling | KEGG: mmu04310; PMID: 23209147 |
| CD31 | CD31 | Wnt10b | FZD5_LRP6 | 0.000383 | 0 | WNT10B_FZD5_LRP6 | Wnt10b - (Fzd5+Lrp6) | WNT | Secreted Signaling | KEGG: mmu04310; PMID: 23209147 |
| CD31 | NeuN | Wnt10b | FZD5_LRP6 | 0.000339 | 0 | WNT10B_FZD5_LRP6 | Wnt10b - (Fzd5+Lrp6) | WNT | Secreted Signaling | KEGG: mmu04310; PMID: 23209147 |
| CD31 | CD31 | Wnt10b | FZD6_LRP6 | 0.000779 | 0 | WNT10B_FZD6_LRP6 | Wnt10b - (Fzd6+Lrp6) | WNT | Secreted Signaling | KEGG: mmu04310; PMID: 23209147 |
| NeuN | CD31 | Wnt10b | FZD6_LRP6 | 0.000391 | 0 | WNT10B_FZD6_LRP6 | Wnt10b - (Fzd6+Lrp6) | WNT | Secreted Signaling | KEGG: mmu04310; PMID: 23209147 |
| CD31 | CD31 | Wnt10b | FZD8_LRP6 | 0.000431 | 0 | WNT10B_FZD8_LRP6 | Wnt10b - (Fzd8+Lrp6) | WNT | Secreted Signaling | KEGG: mmu04310; PMID: 23209147 |
| CD31 | NeuN | Wnt10b | FZD8_LRP6 | 0.00042 | 0 | WNT10B_FZD8_LRP6 | Wnt10b - (Fzd8+Lrp6) | WNT | Secreted Signaling | KEGG: mmu04310; PMID: 23209147 |
| CD31 | CD31 | Wnt10b | FZD9_LRP6 | 0.0004 | 0 | WNT10B_FZD9_LRP6 | Wnt10b - (Fzd9+Lrp6) | WNT | Secreted Signaling | KEGG: mmu04310; PMID: 23209147 |
| CD31 | NeuN | Wnt10b | FZD9_LRP6 | 0.000442 | 0 | WNT10B_FZD9_LRP6 | Wnt10b - (Fzd9+Lrp6) | WNT | Secreted Signaling | KEGG: mmu04310; PMID: 23209147 |
| CD31 | CD31 | Wnt4 | FZD1_LRP6 | 0.000498 | 0 | WNT4_FZD1_LRP6 | Wnt4 - (Fzd1+Lrp6) | WNT | Secreted Signaling | KEGG: mmu04310; PMID: 23209147 |
| NeuN | CD31 | Wnt4 | FZD1_LRP6 | 0.000401 | 0 | WNT4_FZD1_LRP6 | Wnt4 - (Fzd1+Lrp6) | WNT | Secreted Signaling | KEGG: mmu04310; PMID: 23209147 |
| CD31 | CD31 | Wnt4 | FZD2_LRP6 | 0.000312 | 0 | WNT4_FZD2_LRP6 | Wnt4 - (Fzd2+Lrp6) | WNT | Secreted Signaling | KEGG: mmu04310; PMID: 23209147 |
| CD31 | NeuN | Wnt4 | FZD2_LRP6 | 0.000374 | 0 | WNT4_FZD2_LRP6 | Wnt4 - (Fzd2+Lrp6) | WNT | Secreted Signaling | KEGG: mmu04310; PMID: 23209147 |
| CD31 | NeuN | Wnt4 | FZD3_LRP6 | 0.000944 | 0 | WNT4_FZD3_LRP6 | Wnt4 - (Fzd3+Lrp6) | WNT | Secreted Signaling | KEGG: mmu04310; PMID: 23209147 |
| NeuN | NeuN | Wnt4 | FZD3_LRP6 | 0.000761 | 0 | WNT4_FZD3_LRP6 | Wnt4 - (Fzd3+Lrp6) | WNT | Secreted Signaling | KEGG: mmu04310; PMID: 23209147 |
| CD31 | CD31 | Wnt4 | FZD4_LRP6 | 0.000269 | 0 | WNT4_FZD4_LRP6 | Wnt4 - (Fzd4+Lrp6) | WNT | Secreted Signaling | KEGG: mmu04310; PMID: 23209147 |
| CD31 | NeuN | Wnt4 | FZD4_LRP6 | 0.00026 | 0 | WNT4_FZD4_LRP6 | Wnt4 - (Fzd4+Lrp6) | WNT | Secreted Signaling | KEGG: mmu04310; PMID: 23209147 |
| CD31 | CD31 | Wnt4 | FZD5_LRP6 | 0.000326 | 0 | WNT4_FZD5_LRP6 | Wnt4 - (Fzd5+Lrp6) | WNT | Secreted Signaling | KEGG: mmu04310; PMID: 23209147 |
| CD31 | NeuN | Wnt4 | FZD5_LRP6 | 0.000289 | 0 | WNT4_FZD5_LRP6 | Wnt4 - (Fzd5+Lrp6) | WNT | Secreted Signaling | KEGG: mmu04310; PMID: 23209147 |
| CD31 | CD31 | Wnt4 | FZD6_LRP6 | 0.000664 | 0 | WNT4_FZD6_LRP6 | Wnt4 - (Fzd6+Lrp6) | WNT | Secreted Signaling | KEGG: mmu04310; PMID: 23209147 |
| NeuN | CD31 | Wnt4 | FZD6_LRP6 | 0.000535 | 0 | WNT4_FZD6_LRP6 | Wnt4 - (Fzd6+Lrp6) | WNT | Secreted Signaling | KEGG: mmu04310; PMID: 23209147 |
| CD31 | CD31 | Wnt4 | FZD8_LRP6 | 0.000368 | 0 | WNT4_FZD8_LRP6 | Wnt4 - (Fzd8+Lrp6) | WNT | Secreted Signaling | KEGG: mmu04310; PMID: 23209147 |
| CD31 | NeuN | Wnt4 | FZD8_LRP6 | 0.000358 | 0 | WNT4_FZD8_LRP6 | Wnt4 - (Fzd8+Lrp6) | WNT | Secreted Signaling | KEGG: mmu04310; PMID: 23209147 |
| CD31 | CD31 | Wnt4 | FZD9_LRP6 | 0.000341 | 0 | WNT4_FZD9_LRP6 | Wnt4 - (Fzd9+Lrp6) | WNT | Secreted Signaling | KEGG: mmu04310; PMID: 23209147 |
| CD31 | NeuN | Wnt4 | FZD9_LRP6 | 0.000377 | 0 | WNT4_FZD9_LRP6 | Wnt4 - (Fzd9+Lrp6) | WNT | Secreted Signaling | KEGG: mmu04310; PMID: 23209147 |
| CD31 | CD31 | Wnt7a | FZD1_LRP6 | 0.000436 | 0 | WNT7A_FZD1_LRP6 | Wnt7a - (Fzd1+Lrp6) | WNT | Secreted Signaling | KEGG: mmu04310; PMID: 23209147 |
| NeuN | CD31 | Wnt7a | FZD1_LRP6 | 0.000427 | 0 | WNT7A_FZD1_LRP6 | Wnt7a - (Fzd1+Lrp6) | WNT | Secreted Signaling | KEGG: mmu04310; PMID: 23209147 |
| CD31 | NeuN | Wnt7a | FZD2_LRP6 | 0.000327 | 0 | WNT7A_FZD2_LRP6 | Wnt7a - (Fzd2+Lrp6) | WNT | Secreted Signaling | KEGG: mmu04310; PMID: 23209147 |
| NeuN | NeuN | Wnt7a | FZD2_LRP6 | 0.000321 | 0 | WNT7A_FZD2_LRP6 | Wnt7a - (Fzd2+Lrp6) | WNT | Secreted Signaling | KEGG: mmu04310; PMID: 23209147 |
| CD31 | NeuN | Wnt7a | FZD3_LRP6 | 0.000827 | 0 | WNT7A_FZD3_LRP6 | Wnt7a - (Fzd3+Lrp6) | WNT | Secreted Signaling | KEGG: mmu04310; PMID: 23209147 |
| NeuN | NeuN | Wnt7a | FZD3_LRP6 | 0.000811 | 0 | WNT7A_FZD3_LRP6 | Wnt7a - (Fzd3+Lrp6) | WNT | Secreted Signaling | KEGG: mmu04310; PMID: 23209147 |
| CD31 | CD31 | Wnt7a | FZD4_LRP6 | 0.000236 | 0 | WNT7A_FZD4_LRP6 | Wnt7a - (Fzd4+Lrp6) | WNT | Secreted Signaling | KEGG: mmu04310; PMID: 23209147 |
| NeuN | CD31 | Wnt7a | FZD4_LRP6 | 0.000231 | 0 | WNT7A_FZD4_LRP6 | Wnt7a - (Fzd4+Lrp6) | WNT | Secreted Signaling | KEGG: mmu04310; PMID: 23209147 |
| CD31 | CD31 | Wnt7a | FZD5_LRP6 | 0.000286 | 0 | WNT7A_FZD5_LRP6 | Wnt7a - (Fzd5+Lrp6) | WNT | Secreted Signaling | KEGG: mmu04310; PMID: 23209147 |
| NeuN | CD31 | Wnt7a | FZD5_LRP6 | 0.00028 | 0 | WNT7A_FZD5_LRP6 | Wnt7a - (Fzd5+Lrp6) | WNT | Secreted Signaling | KEGG: mmu04310; PMID: 23209147 |
| CD31 | CD31 | Wnt7a | FZD6_LRP6 | 0.000582 | 0 | WNT7A_FZD6_LRP6 | Wnt7a - (Fzd6+Lrp6) | WNT | Secreted Signaling | KEGG: mmu04310; PMID: 23209147 |
| NeuN | CD31 | Wnt7a | FZD6_LRP6 | 0.000571 | 0 | WNT7A_FZD6_LRP6 | Wnt7a - (Fzd6+Lrp6) | WNT | Secreted Signaling | KEGG: mmu04310; PMID: 23209147 |
| CD31 | CD31 | Wnt7a | FZD8_LRP6 | 0.000322 | 0 | WNT7A_FZD8_LRP6 | Wnt7a - (Fzd8+Lrp6) | WNT | Secreted Signaling | KEGG: mmu04310; PMID: 23209147 |

| | | | | | | | | | | |
|---|---|---|---|---|---|---|---|---|---|---|
| NeuN | CD31 | Wnt7a | FZD8_LRP6 | 0.000316 | 0 | WNT7A_FZD8_LRP6 | Wnt7a - (Fzd8+Lrp6) | WNT | Secreted Signaling | KEGG: mmu04310; PMID: 23209147 |
| CD31 | NeuN | Wnt7a | FZD9_LRP6 | 0.00033 | 0 | WNT7A_FZD9_LRP6 | Wnt7a - (Fzd9+Lrp6) | WNT | Secreted Signaling | KEGG: mmu04310; PMID: 23209147 |
| NeuN | NeuN | Wnt7a | FZD9_LRP6 | 0.000323 | 0 | WNT7A_FZD9_LRP6 | Wnt7a - (Fzd9+Lrp6) | WNT | Secreted Signaling | KEGG: mmu04310; PMID: 23209147 |
| NeuN | CD31 | Wnt7b | FZD1_LRP6 | 0.000965 | 0 | WNT7B_FZD1_LRP6 | Wnt7b - (Fzd1+Lrp6) | WNT | Secreted Signaling | KEGG: mmu04310; PMID: 23209147 |
| NeuN | NeuN | Wnt7b | FZD1_LRP6 | 0.000703 | 0 | WNT7B_FZD1_LRP6 | Wnt7b - (Fzd1+Lrp6) | WNT | Secreted Signaling | KEGG: mmu04310; PMID: 23209147 |
| NeuN | CD31 | Wnt7b | FZD2_LRP6 | 0.000605 | 0 | WNT7B_FZD2_LRP6 | Wnt7b - (Fzd2+Lrp6) | WNT | Secreted Signaling | KEGG: mmu04310; PMID: 23209147 |
| NeuN | NeuN | Wnt7b | FZD2_LRP6 | 0.000725 | 0 | WNT7B_FZD2_LRP6 | Wnt7b - (Fzd2+Lrp6) | WNT | Secreted Signaling | KEGG: mmu04310; PMID: 23209147 |
| CD31 | NeuN | Wnt7b | FZD3_LRP6 | 0.001218 | 0 | WNT7B_FZD3_LRP6 | Wnt7b - (Fzd3+Lrp6) | WNT | Secreted Signaling | KEGG: mmu04310; PMID: 23209147 |
| NeuN | NeuN | Wnt7b | FZD3_LRP6 | 0.00183 | 0 | WNT7B_FZD3_LRP6 | Wnt7b - (Fzd3+Lrp6) | WNT | Secreted Signaling | KEGG: mmu04310; PMID: 23209147 |
| NeuN | CD31 | Wnt7b | FZD4_LRP6 | 0.000522 | 0 | WNT7B_FZD4_LRP6 | Wnt7b - (Fzd4+Lrp6) | WNT | Secreted Signaling | KEGG: mmu04310; PMID: 23209147 |
| NeuN | NeuN | Wnt7b | FZD4_LRP6 | 0.000504 | 0 | WNT7B_FZD4_LRP6 | Wnt7b - (Fzd4+Lrp6) | WNT | Secreted Signaling | KEGG: mmu04310; PMID: 23209147 |
| NeuN | CD31 | Wnt7b | FZD5_LRP6 | 0.000633 | 0 | WNT7B_FZD5_LRP6 | Wnt7b - (Fzd5+Lrp6) | WNT | Secreted Signaling | KEGG: mmu04310; PMID: 23209147 |
| NeuN | NeuN | Wnt7b | FZD5_LRP6 | 0.00056 | 0 | WNT7B_FZD5_LRP6 | Wnt7b - (Fzd5+Lrp6) | WNT | Secreted Signaling | KEGG: mmu04310; PMID: 23209147 |
| CD31 | CD31 | Wnt7b | FZD6_LRP6 | 0.000857 | 0 | WNT7B_FZD6_LRP6 | Wnt7b - (Fzd6+Lrp6) | WNT | Secreted Signaling | KEGG: mmu04310; PMID: 23209147 |
| NeuN | CD31 | Wnt7b | FZD6_LRP6 | 0.001288 | 0 | WNT7B_FZD6_LRP6 | Wnt7b - (Fzd6+Lrp6) | WNT | Secreted Signaling | KEGG: mmu04310; PMID: 23209147 |
| NeuN | CD31 | Wnt7b | FZD8_LRP6 | 0.000714 | 0 | WNT7B_FZD8_LRP6 | Wnt7b - (Fzd8+Lrp6) | WNT | Secreted Signaling | KEGG: mmu04310; PMID: 23209147 |
| NeuN | NeuN | Wnt7b | FZD8_LRP6 | 0.000694 | 0 | WNT7B_FZD8_LRP6 | Wnt7b - (Fzd8+Lrp6) | WNT | Secreted Signaling | KEGG: mmu04310; PMID: 23209147 |
| NeuN | CD31 | Wnt7b | FZD9_LRP6 | 0.000662 | 0 | WNT7B_FZD9_LRP6 | Wnt7b - (Fzd9+Lrp6) | WNT | Secreted Signaling | KEGG: mmu04310; PMID: 23209147 |
| NeuN | NeuN | Wnt7b | FZD9_LRP6 | 0.000731 | 0 | WNT7B_FZD9_LRP6 | Wnt7b - (Fzd9+Lrp6) | WNT | Secreted Signaling | KEGG: mmu04310; PMID: 23209147 |
| CD31 | CD31 | Wnt9a | FZD1_LRP6 | 0.000331 | 0 | WNT9A_FZD1_LRP6 | Wnt9a - (Fzd1+Lrp6) | WNT | Secreted Signaling | KEGG: mmu04310; PMID: 23209147 |
| NeuN | CD31 | Wnt9a | FZD1_LRP6 | 0.000246 | 0 | WNT9A_FZD1_LRP6 | Wnt9a - (Fzd1+Lrp6) | WNT | Secreted Signaling | KEGG: mmu04310; PMID: 23209147 |
| CD31 | CD31 | Wnt9a | FZD2_LRP6 | 0.000207 | 0 | WNT9A_FZD2_LRP6 | Wnt9a - (Fzd2+Lrp6) | WNT | Secreted Signaling | KEGG: mmu04310; PMID: 23209147 |
| CD31 | NeuN | Wnt9a | FZD2_LRP6 | 0.000248 | 0 | WNT9A_FZD2_LRP6 | Wnt9a - (Fzd2+Lrp6) | WNT | Secreted Signaling | KEGG: mmu04310; PMID: 23209147 |
| CD31 | NeuN | Wnt9a | FZD3_LRP6 | 0.000627 | 0 | WNT9A_FZD3_LRP6 | Wnt9a - (Fzd3+Lrp6) | WNT | Secreted Signaling | KEGG: mmu04310; PMID: 23209147 |
| NeuN | NeuN | Wnt9a | FZD3_LRP6 | 0.000467 | 0 | WNT9A_FZD3_LRP6 | Wnt9a - (Fzd3+Lrp6) | WNT | Secreted Signaling | KEGG: mmu04310; PMID: 23209147 |
| CD31 | CD31 | Wnt9a | FZD4_LRP6 | 0.000179 | 0 | WNT9A_FZD4_LRP6 | Wnt9a - (Fzd4+Lrp6) | WNT | Secreted Signaling | KEGG: mmu04310; PMID: 23209147 |
| CD31 | NeuN | Wnt9a | FZD4_LRP6 | 0.000173 | 0 | WNT9A_FZD4_LRP6 | Wnt9a - (Fzd4+Lrp6) | WNT | Secreted Signaling | KEGG: mmu04310; PMID: 23209147 |
| CD31 | CD31 | Wnt9a | FZD5_LRP6 | 0.000217 | 0 | WNT9A_FZD5_LRP6 | Wnt9a - (Fzd5+Lrp6) | WNT | Secreted Signaling | KEGG: mmu04310; PMID: 23209147 |
| CD31 | NeuN | Wnt9a | FZD5_LRP6 | 0.000192 | 0 | WNT9A_FZD5_LRP6 | Wnt9a - (Fzd5+Lrp6) | WNT | Secreted Signaling | KEGG: mmu04310; PMID: 23209147 |
| CD31 | CD31 | Wnt9a | FZD6_LRP6 | 0.000442 | 0 | WNT9A_FZD6_LRP6 | Wnt9a - (Fzd6+Lrp6) | WNT | Secreted Signaling | KEGG: mmu04310; PMID: 23209147 |
| NeuN | CD31 | Wnt9a | FZD6_LRP6 | 0.000329 | 0 | WNT9A_FZD6_LRP6 | Wnt9a - (Fzd6+Lrp6) | WNT | Secreted Signaling | KEGG: mmu04310; PMID: 23209147 |
| CD31 | CD31 | Wnt9a | FZD8_LRP6 | 0.000244 | 0 | WNT9A_FZD8_LRP6 | Wnt9a - (Fzd8+Lrp6) | WNT | Secreted Signaling | KEGG: mmu04310; PMID: 23209147 |
| CD31 | NeuN | Wnt9a | FZD8_LRP6 | 0.000238 | 0 | WNT9A_FZD8_LRP6 | Wnt9a - (Fzd8+Lrp6) | WNT | Secreted Signaling | KEGG: mmu04310; PMID: 23209147 |
| CD31 | CD31 | Wnt9a | FZD9_LRP6 | 0.000227 | 0 | WNT9A_FZD9_LRP6 | Wnt9a - (Fzd9+Lrp6) | WNT | Secreted Signaling | KEGG: mmu04310; PMID: 23209147 |
| CD31 | NeuN | Wnt9a | FZD9_LRP6 | 0.00025 | 0 | WNT9A_FZD9_LRP6 | Wnt9a - (Fzd9+Lrp6) | WNT | Secreted Signaling | KEGG: mmu04310; PMID: 23209147 |
| CD31 | CD31 | Wnt5a | Fzd1 | 0.001172 | 0 | WNT5A_FZD1 | Wnt5a - Fzd1 | ncWNT | Secreted Signaling | KEGG: mmu04310 |
| NeuN | CD31 | Wnt5a | Fzd1 | 0.000567 | 0 | WNT5A_FZD1 | Wnt5a - Fzd1 | ncWNT | Secreted Signaling | KEGG: mmu04310 |
| CD31 | CD31 | Wnt5a | Fzd2 | 0.00046 | 0 | WNT5A_FZD2 | Wnt5a - Fzd2 | ncWNT | Secreted Signaling | KEGG: mmu04310 |
| CD31 | NeuN | Wnt5a | Fzd2 | 0.000531 | 0 | WNT5A_FZD2 | Wnt5a - Fzd2 | ncWNT | Secreted Signaling | KEGG: mmu04310 |
| CD31 | NeuN | Wnt5a | Fzd3 | 0.003385 | 0 | WNT5A_FZD3 | Wnt5a - Fzd3 | ncWNT | Secreted Signaling | KEGG: mmu04310 |
| NeuN | NeuN | Wnt5a | Fzd3 | 0.001641 | 0 | WNT5A_FZD3 | Wnt5a - Fzd3 | ncWNT | Secreted Signaling | KEGG: mmu04310 |
| CD31 | CD31 | Wnt5a | Fzd4 | 0.000343 | 0 | WNT5A_FZD4 | Wnt5a - Fzd4 | ncWNT | Secreted Signaling | KEGG: mmu04310 |
| CD31 | NeuN | Wnt5a | Fzd4 | 0.000257 | 0 | WNT5A_FZD4 | Wnt5a - Fzd4 | ncWNT | Secreted Signaling | KEGG: mmu04310 |
| CD31 | CD31 | Wnt5a | Fzd5 | 0.000503 | 0 | WNT5A_FZD5 | Wnt5a - Fzd5 | ncWNT | Secreted Signaling | KEGG: mmu04310 |
| CD31 | NeuN | Wnt5a | Fzd5 | 0.000317 | 0 | WNT5A_FZD5 | Wnt5a - Fzd5 | ncWNT | Secreted Signaling | KEGG: mmu04310 |
| CD31 | CD31 | Wnt5a | Fzd6 | 0.002089 | 0 | WNT5A_FZD6 | Wnt5a - Fzd6 | ncWNT | Secreted Signaling | KEGG: mmu04310 |
| NeuN | CD31 | Wnt5a | Fzd6 | 0.001012 | 0 | WNT5A_FZD6 | Wnt5a - Fzd6 | ncWNT | Secreted Signaling | KEGG: mmu04310 |
| CD31 | CD31 | Wnt5a | Fzd8 | 0.00064 | 0 | WNT5A_FZD8 | Wnt5a - Fzd8 | ncWNT | Secreted Signaling | KEGG: mmu04310 |
| CD31 | NeuN | Wnt5a | Fzd8 | 0.000487 | 0 | WNT5A_FZD8 | Wnt5a - Fzd8 | ncWNT | Secreted Signaling | KEGG: mmu04310 |
| CD31 | CD31 | Wnt5a | Fzd9 | 0.00055 | 0 | WNT5A_FZD9 | Wnt5a - Fzd9 | ncWNT | Secreted Signaling | KEGG: mmu04310 |
| CD31 | NeuN | Wnt5a | Fzd9 | 0.000539 | 0 | WNT5A_FZD9 | Wnt5a - Fzd9 | ncWNT | Secreted Signaling | KEGG: mmu04310 |
| CD31 | CD31 | Wnt5a | Mcam | 0.002315 | 0 | WNT5A_MCAM | Wnt5a - Mcam | ncWNT | Secreted Signaling | PMID: 24335906 |
| NeuN | CD31 | Wnt5a | Mcam | 0.001122 | 0 | WNT5A_MCAM | Wnt5a - Mcam | ncWNT | Secreted Signaling | PMID: 24335906 |
| CD31 | NeuN | Hbegf | Erbb4 | 0.001085 | 0 | HBEGF_ERBB4 | Hbegf - Erbb4 | EGF | Secreted Signaling | KEGG: mmu04012 |
| NeuN | NeuN | Hbegf | Erbb4 | 0.000924 | 0 | HBEGF_ERBB4 | Hbegf - Erbb4 | EGF | Secreted Signaling | KEGG: mmu04012 |
| CD31 | CD31 | Hbegf | ERBB2_ERBB4 | 0.000824 | 0 | HBEGF_ERBB2_ERBB4 | Hbegf - (Erbb2+Erbb4) | EGF | Secreted Signaling | KEGG: mmu04012 |
| NeuN | CD31 | Hbegf | ERBB2_ERBB4 | 0.000702 | 0 | HBEGF_ERBB2_ERBB4 | Hbegf - (Erbb2+Erbb4) | EGF | Secreted Signaling | KEGG: mmu04012 |
| CD31 | NeuN | Ereg | Erbb4 | 0.000586 | 0 | EREG_ERBB4 | Ereg - Erbb4 | EGF | Secreted Signaling | KEGG: mmu04012 |
| NeuN | NeuN | Ereg | Erbb4 | 0.000492 | 0 | EREG_ERBB4 | Ereg - Erbb4 | EGF | Secreted Signaling | KEGG: mmu04012 |
| CD31 | CD31 | Ereg | ERBB2_ERBB4 | 0.000445 | 0 | EREG_ERBB2_ERBB4 | Ereg - (Erbb2+Erbb4) | EGF | Secreted Signaling | KEGG: mmu04012 |

| | | | | | | | | | |
|---|---|---|---|---|---|---|---|---|---|
| NeuN | CD31 | Ereg | ERBB2_ERBB4 | 0.000373 | 0 | EREG_ERBB2_ERBB4 | Ereg - (Erbb2+Erbb4) | EGF | Secreted Signaling | KEGG: mmu04012 |
| CD31 | CD31 | Nrg1 | Erbb3 | 0.000595 | 0 | NRG1_ERBB3 | Nrg1 - Erbb3 | NRG | Secreted Signaling | KEGG: mmu04012 |
| NeuN | CD31 | Nrg1 | Erbb3 | 0.000883 | 0 | NRG1_ERBB3 | Nrg1 - Erbb3 | NRG | Secreted Signaling | KEGG: mmu04012 |
| CD31 | CD31 | Nrg1 | ERBB2_ERBB3 | 0.000733 | 0 | NRG1_ERBB2_ERBB3 | Nrg1 - (Erbb2+Erbb3) | NRG | Secreted Signaling | KEGG: mmu04012 |
| NeuN | CD31 | Nrg1 | ERBB2_ERBB3 | 0.001088 | 0 | NRG1_ERBB2_ERBB3 | Nrg1 - (Erbb2+Erbb3) | NRG | Secreted Signaling | KEGG: mmu04012 |
| NeuN | CD31 | Nrg1 | Erbb4 | 0.00109 | 0 | NRG1_ERBB4 | Nrg1 - Erbb4 | NRG | Secreted Signaling | KEGG: mmu04012 |
| NeuN | NeuN | Nrg1 | Erbb4 | 0.001591 | 0 | NRG1_ERBB4 | Nrg1 - Erbb4 | NRG | Secreted Signaling | KEGG: mmu04012 |
| NeuN | CD31 | Nrg1 | ERBB2_ERBB4 | 0.001208 | 0 | NRG1_ERBB2_ERBB4 | Nrg1 - (Erbb2+Erbb4) | NRG | Secreted Signaling | KEGG: mmu04012 |
| NeuN | NeuN | Nrg1 | ERBB2_ERBB4 | 0.000891 | 0 | NRG1_ERBB2_ERBB4 | Nrg1 - (Erbb2+Erbb4) | NRG | Secreted Signaling | KEGG: mmu04012 |
| CD31 | CD31 | Nrg2 | Erbb3 | 0.000482 | 0 | NRG2_ERBB3 | Nrg2 - Erbb3 | NRG | Secreted Signaling | KEGG: mmu04012 |
| NeuN | CD31 | Nrg2 | Erbb3 | 0.000582 | 0 | NRG2_ERBB3 | Nrg2 - Erbb3 | NRG | Secreted Signaling | KEGG: mmu04012 |
| CD31 | CD31 | Nrg2 | ERBB2_ERBB3 | 0.000594 | 0 | NRG2_ERBB2_ERBB3 | Nrg2 - (Erbb2+Erbb3) | NRG | Secreted Signaling | KEGG: mmu04012 |
| NeuN | CD31 | Nrg2 | ERBB2_ERBB3 | 0.000717 | 0 | NRG2_ERBB2_ERBB3 | Nrg2 - (Erbb2+Erbb3) | NRG | Secreted Signaling | KEGG: mmu04012 |
| CD31 | NeuN | Nrg2 | Erbb4 | 0.000869 | 0 | NRG2_ERBB4 | Nrg2 - Erbb4 | NRG | Secreted Signaling | KEGG: mmu04012 |
| NeuN | NeuN | Nrg2 | Erbb4 | 0.001049 | 0 | NRG2_ERBB4 | Nrg2 - Erbb4 | NRG | Secreted Signaling | KEGG: mmu04012 |
| CD31 | CD31 | Nrg2 | ERBB2_ERBB4 | 0.00066 | 0 | NRG2_ERBB2_ERBB4 | Nrg2 - (Erbb2+Erbb4) | NRG | Secreted Signaling | KEGG: mmu04012 |
| NeuN | CD31 | Nrg2 | ERBB2_ERBB4 | 0.000797 | 0 | NRG2_ERBB2_ERBB4 | Nrg2 - (Erbb2+Erbb4) | NRG | Secreted Signaling | KEGG: mmu04012 |
| NeuN | CD31 | Nrg3 | Erbb4 | 0.003525 | 0 | NRG3_ERBB4 | Nrg3 - Erbb4 | NRG | Secreted Signaling | KEGG: mmu04012 |
| NeuN | NeuN | Nrg3 | Erbb4 | 0.005141 | 0 | NRG3_ERBB4 | Nrg3 - Erbb4 | NRG | Secreted Signaling | KEGG: mmu04012 |
| NeuN | CD31 | Nrg3 | ERBB2_ERBB4 | 0.003908 | 0 | NRG3_ERBB2_ERBB4 | Nrg3 - (Erbb2+Erbb4) | NRG | Secreted Signaling | KEGG: mmu04012 |
| NeuN | NeuN | Nrg3 | ERBB2_ERBB4 | 0.002884 | 0 | NRG3_ERBB2_ERBB4 | Nrg3 - (Erbb2+Erbb4) | NRG | Secreted Signaling | KEGG: mmu04012 |
| CD31 | CD31 | Nrg1 | ITGAV_ITGB3 | 0.000903 | 0 | NRG1_ITGAV_ITGB3 | Nrg1 - (Itgav+Itgb3) | NRG | Secreted Signaling | PMID: 20682778 |
| NeuN | CD31 | Nrg1 | ITGAV_ITGB3 | 0.001341 | 0 | NRG1_ITGAV_ITGB3 | Nrg1 - (Itgav+Itgb3) | NRG | Secreted Signaling | PMID: 20682778 |
| CD31 | NeuN | Fgf2 | Fgfr1 | 0.001081 | 0 | FGF2_FGFR1 | Fgf2 - Fgfr1 | FGF | Secreted Signaling | PMC: 4393358 |
| NeuN | NeuN | Fgf2 | Fgfr1 | 0.000931 | 0 | FGF2_FGFR1 | Fgf2 - Fgfr1 | FGF | Secreted Signaling | PMC: 4393358 |
| CD31 | NeuN | Fgf2 | Fgfr2 | 0.001189 | 0 | FGF2_FGFR2 | Fgf2 - Fgfr2 | FGF | Secreted Signaling | PMC: 4393358 |
| NeuN | NeuN | Fgf2 | Fgfr2 | 0.001024 | 0 | FGF2_FGFR2 | Fgf2 - Fgfr2 | FGF | Secreted Signaling | PMC: 4393358 |
| CD31 | CD31 | Fgf2 | Fgfr3 | 0.000748 | 0 | FGF2_FGFR3 | Fgf2 - Fgfr3 | FGF | Secreted Signaling | PMC: 4393358 |
| NeuN | CD31 | Fgf2 | Fgfr3 | 0.000644 | 0 | FGF2_FGFR3 | Fgf2 - Fgfr3 | FGF | Secreted Signaling | PMC: 4393358 |
| CD31 | NeuN | Fgf5 | Fgfr1 | 0.000702 | 0 | FGF5_FGFR1 | Fgf5 - Fgfr1 | FGF | Secreted Signaling | PMC: 4393358 |
| NeuN | NeuN | Fgf5 | Fgfr1 | 0.000546 | 0 | FGF5_FGFR1 | Fgf5 - Fgfr1 | FGF | Secreted Signaling | PMC: 4393358 |
| CD31 | NeuN | Fgf5 | Fgfr2 | 0.000772 | 0 | FGF5_FGFR2 | Fgf5 - Fgfr2 | FGF | Secreted Signaling | PMC: 4393358 |
| NeuN | NeuN | Fgf5 | Fgfr2 | 0.000601 | 0 | FGF5_FGFR2 | Fgf5 - Fgfr2 | FGF | Secreted Signaling | PMC: 4393358 |
| CD31 | CD31 | Fgf5 | Fgfr3 | 0.000486 | 0 | FGF5_FGFR3 | Fgf5 - Fgfr3 | FGF | Secreted Signaling | PMC: 4393358 |
| NeuN | CD31 | Fgf5 | Fgfr3 | 0.000378 | 0 | FGF5_FGFR3 | Fgf5 - Fgfr3 | FGF | Secreted Signaling | PMC: 4393358 |
| CD31 | NeuN | Fgf3 | Fgfr1 | 0.000627 | 0 | FGF3_FGFR1 | Fgf3 - Fgfr1 | FGF | Secreted Signaling | PMC: 4393358 |
| NeuN | NeuN | Fgf3 | Fgfr1 | 0.000549 | 0 | FGF3_FGFR1 | Fgf3 - Fgfr1 | FGF | Secreted Signaling | PMC: 4393358 |
| CD31 | NeuN | Fgf3 | Fgfr2 | 0.000689 | 0 | FGF3_FGFR2 | Fgf3 - Fgfr2 | FGF | Secreted Signaling | PMC: 4393358 |
| NeuN | NeuN | Fgf3 | Fgfr2 | 0.000604 | 0 | FGF3_FGFR2 | Fgf3 - Fgfr2 | FGF | Secreted Signaling | PMC: 4393358 |
| CD31 | NeuN | Fgf10 | Fgfr1 | 0.000881 | 0 | FGF10_FGFR1 | Fgf10 - Fgfr1 | FGF | Secreted Signaling | PMC: 4393358 |
| NeuN | NeuN | Fgf10 | Fgfr1 | 0.000788 | 0 | FGF10_FGFR1 | Fgf10 - Fgfr1 | FGF | Secreted Signaling | PMC: 4393358 |
| CD31 | NeuN | Fgf10 | Fgfr2 | 0.000969 | 0 | FGF10_FGFR2 | Fgf10 - Fgfr2 | FGF | Secreted Signaling | PMC: 4393358 |
| NeuN | NeuN | Fgf10 | Fgfr2 | 0.000867 | 0 | FGF10_FGFR2 | Fgf10 - Fgfr2 | FGF | Secreted Signaling | PMC: 4393358 |
| CD31 | CD31 | Fgf22 | Fgfr2 | 0.000673 | 0 | FGF22_FGFR2 | Fgf22 - Fgfr2 | FGF | Secreted Signaling | PMC: 4393358 |
| CD31 | NeuN | Fgf22 | Fgfr2 | 0.001188 | 0 | FGF22_FGFR2 | Fgf22 - Fgfr2 | FGF | Secreted Signaling | PMC: 4393358 |
| CD31 | CD31 | Fgf18 | Fgfr1 | 0.000828 | 0 | FGF18_FGFR1 | Fgf18 - Fgfr1 | FGF | Secreted Signaling | PMC: 4393358 |
| CD31 | NeuN | Fgf18 | Fgfr1 | 0.001141 | 0 | FGF18_FGFR1 | Fgf18 - Fgfr1 | FGF | Secreted Signaling | PMC: 4393358 |
| CD31 | NeuN | Fgf18 | Fgfr2 | 0.001255 | 0 | FGF18_FGFR2 | Fgf18 - Fgfr2 | FGF | Secreted Signaling | PMC: 4393358 |
| NeuN | NeuN | Fgf18 | Fgfr2 | 0.000809 | 0 | FGF18_FGFR2 | Fgf18 - Fgfr2 | FGF | Secreted Signaling | PMC: 4393358 |
| CD31 | CD31 | Fgf18 | Fgfr3 | 0.00079 | 0 | FGF18_FGFR3 | Fgf18 - Fgfr3 | FGF | Secreted Signaling | PMC: 4393358 |
| NeuN | CD31 | Fgf18 | Fgfr3 | 0.000509 | 0 | FGF18_FGFR3 | Fgf18 - Fgfr3 | FGF | Secreted Signaling | PMC: 4393358 |
| CD31 | NeuN | Fgf9 | Fgfr1 | 0.000905 | 0 | FGF9_FGFR1 | Fgf9 - Fgfr1 | FGF | Secreted Signaling | PMC: 4393358 |
| NeuN | NeuN | Fgf9 | Fgfr1 | 0.000906 | 0 | FGF9_FGFR1 | Fgf9 - Fgfr1 | FGF | Secreted Signaling | PMC: 4393358 |
| CD31 | NeuN | Fgf9 | Fgfr2 | 0.000995 | 0 | FGF9_FGFR2 | Fgf9 - Fgfr2 | FGF | Secreted Signaling | PMC: 4393358 |
| NeuN | NeuN | Fgf9 | Fgfr2 | 0.000997 | 0 | FGF9_FGFR2 | Fgf9 - Fgfr2 | FGF | Secreted Signaling | PMC: 4393358 |
| CD31 | CD31 | Fgf9 | Fgfr3 | 0.000626 | 0 | FGF9_FGFR3 | Fgf9 - Fgfr3 | FGF | Secreted Signaling | PMC: 4393358 |
| NeuN | CD31 | Fgf9 | Fgfr3 | 0.000627 | 0 | FGF9_FGFR3 | Fgf9 - Fgfr3 | FGF | Secreted Signaling | PMC: 4393358 |
| CD31 | CD31 | Fgf15 | Fgfr1 | 0.000676 | 0 | FGF15_FGFR1 | Fgf15 - Fgfr1 | FGF | Secreted Signaling | PMC: 4393358 |
| CD31 | NeuN | Fgf15 | Fgfr1 | 0.000928 | 0 | FGF15_FGFR1 | Fgf15 - Fgfr1 | FGF | Secreted Signaling | PMC: 4393358 |
| CD31 | NeuN | Fgf15 | Fgfr2 | 0.001021 | 0 | FGF15_FGFR2 | Fgf15 - Fgfr2 | FGF | Secreted Signaling | PMC: 4393358 |

| | | | | | | | | | |
|---|---|---|---|---|---|---|---|---|---|
| NeuN | NeuN | Fgf15 | Fgfr2 | 0.000713 | 0 FGF15_FGFR2 | Fgf15 - Fgfr2 | FGF | Secreted Signaling | PMC: 4393358 |
| CD31 | CD31 | Fgf15 | Fgfr3 | 0.000645 | 0 FGF15_FGFR3 | Fgf15 - Fgfr3 | FGF | Secreted Signaling | PMC: 4393358 |
| NeuN | CD31 | Fgf15 | Fgfr3 | 0.000451 | 0 FGF15_FGFR3 | Fgf15 - Fgfr3 | FGF | Secreted Signaling | PMC: 4393358 |
| CD31 | CD31 | Fgf21 | Fgfr1 | 0.000589 | 0 FGF21_FGFR1 | Fgf21 - Fgfr1 | FGF | Secreted Signaling | PMC: 4393358 |
| CD31 | NeuN | Fgf21 | Fgfr1 | 0.000808 | 0 FGF21_FGFR1 | Fgf21 - Fgfr1 | FGF | Secreted Signaling | PMC: 4393358 |
| CD31 | CD31 | Fgf21 | Fgfr3 | 0.000562 | 0 FGF21_FGFR3 | Fgf21 - Fgfr3 | FGF | Secreted Signaling | PMC: 4393358 |
| CD31 | NeuN | Fgf21 | Fgfr3 | 0.000339 | 0 FGF21_FGFR3 | Fgf21 - Fgfr3 | FGF | Secreted Signaling | PMC: 4393358 |
| NeuN | CD31 | Pdgfa | Pdgfra | 0.000399 | 0 PDGFA_PDGFRA | Pdgfa - Pdgfra | PDGF | Secreted Signaling | PMID: 15207812 |
| NeuN | NeuN | Pdgfa | Pdgfra | 0.000366 | 0 PDGFA_PDGFRA | Pdgfa - Pdgfra | PDGF | Secreted Signaling | PMID: 15207812 |
| CD31 | CD31 | Pdgfa | Pdgfrb | 0.000718 | 0 PDGFA_PDGFRB | Pdgfa - Pdgfrb | PDGF | Secreted Signaling | PMID: 15207812 |
| NeuN | CD31 | Pdgfa | Pdgfrb | 0.001361 | 0 PDGFA_PDGFRB | Pdgfa - Pdgfrb | PDGF | Secreted Signaling | PMID: 15207812 |
| CD31 | CD31 | Pdgfb | Pdgfra | 0.000866 | 0 PDGFB_PDGFRA | Pdgfb - Pdgfra | PDGF | Secreted Signaling | PMID: 15207812 |
| CD31 | NeuN | Pdgfb | Pdgfra | 0.000795 | 0 PDGFB_PDGFRA | Pdgfb - Pdgfra | PDGF | Secreted Signaling | PMID: 15207812 |
| CD31 | CD31 | Pdgfb | Pdgfrb | 0.002944 | 0 PDGFB_PDGFRB | Pdgfb - Pdgfrb | PDGF | Secreted Signaling | PMID: 15207812 |
| CD31 | NeuN | Pdgfb | Pdgfrb | 0.000725 | 0 PDGFB_PDGFRB | Pdgfb - Pdgfrb | PDGF | Secreted Signaling | PMID: 15207812 |
| CD31 | CD31 | Pdgfc | Pdgfra | 0.000285 | 0 PDGFC_PDGFRA | Pdgfc - Pdgfra | PDGF | Secreted Signaling | PMID: 15207812 |
| NeuN | CD31 | Pdgfc | Pdgfra | 0.000297 | 0 PDGFC_PDGFRA | Pdgfc - Pdgfra | PDGF | Secreted Signaling | PMID: 15207812 |
| CD31 | CD31 | Pdgfd | Pdgfrb | 0.001095 | 0 PDGFD_PDGFRB | Pdgfd - Pdgfrb | PDGF | Secreted Signaling | PMID: 15207812 |
| NeuN | CD31 | Pdgfd | Pdgfrb | 0.000542 | 0 PDGFD_PDGFRB | Pdgfd - Pdgfrb | PDGF | Secreted Signaling | PMID: 15207812 |
| CD31 | CD31 | Vegfa | Flt1 | 0.003954 | 0 VEGFA_VEGFR1 | Vegfa - Vegfr1 | VEGF | Secreted Signaling | KEGG: mmu04370; PMID: 16633338 |
| NeuN | CD31 | Vegfa | Flt1 | 0.010769 | 0 VEGFA_VEGFR1 | Vegfa - Vegfr1 | VEGF | Secreted Signaling | KEGG: mmu04370; PMID: 16633338 |
| CD31 | CD31 | Vegfa | Kdr | 0.004186 | 0 VEGFA_VEGFR2 | Vegfa - Vegfr2 | VEGF | Secreted Signaling | KEGG: mmu04370; PMID: 16633338 |
| NeuN | CD31 | Vegfa | Kdr | 0.011397 | 0 VEGFA_VEGFR2 | Vegfa - Vegfr2 | VEGF | Secreted Signaling | KEGG: mmu04370; PMID: 16633338 |
| CD31 | CD31 | Vegfb | Flt1 | 0.005349 | 0 VEGFB_VEGFR1 | Vegfb - Vegfr1 | VEGF | Secreted Signaling | KEGG: mmu04370; PMID: 16633338 |
| NeuN | CD31 | Vegfb | Flt1 | 0.007351 | 0 VEGFB_VEGFR1 | Vegfb - Vegfr1 | VEGF | Secreted Signaling | KEGG: mmu04370; PMID: 16633338 |
| CD31 | CD31 | Pgf | Flt1 | 0.004448 | 0 PGF_VEGFR1 | Pgf - Vegfr1 | VEGF | Secreted Signaling | KEGG: mmu04370; PMID: 16633338 |
| NeuN | CD31 | Pgf | Flt1 | 0.0041 | 0 PGF_VEGFR1 | Pgf - Vegfr1 | VEGF | Secreted Signaling | KEGG: mmu04370; PMID: 16633338 |
| CD31 | CD31 | Vegfa | FLT1_KDR | 0.004226 | 0 VEGFA_VEGFR1R2 | Vegfa - Vegfr1r2 | VEGF | Secreted Signaling | PMID: 16633338 |
| NeuN | CD31 | Vegfa | FLT1_KDR | 0.011504 | 0 VEGFA_VEGFR1R2 | Vegfa - Vegfr1r2 | VEGF | Secreted Signaling | PMID: 16633338 |
| CD31 | CD31 | Igf1 | Igf1r | 0.001153 | 0 IGF1_IGF1R | Igf1 - Igf1r | IGF | Secreted Signaling | PMID: 14604834 |
| CD31 | NeuN | Igf1 | Igf1r | 0.000913 | 0 IGF1_IGF1R | Igf1 - Igf1r | IGF | Secreted Signaling | PMID: 14604834 |
| CD31 | CD31 | Igf2 | Igf1r | 0.008046 | 0 IGF2_IGF1R | Igf2 - Igf1r | IGF | Secreted Signaling | PMID: 14604834 |
| CD31 | NeuN | Igf2 | Igf1r | 0.006407 | 0 IGF2_IGF1R | Igf2 - Igf1r | IGF | Secreted Signaling | PMID: 14604834 |
| CD31 | CD31 | Igf2 | Igf2r | 0.003569 | 0 IGF2_IGF2R | Igf2 - Igf2r | IGF | Secreted Signaling | PMID: 14604834 |
| CD31 | NeuN | Igf2 | Igf2r | 0.003644 | 0 IGF2_IGF2R | Igf2 - Igf2r | IGF | Secreted Signaling | PMID: 14604834 |
| CD31 | CD31 | Igf1 | ITGAV_ITGB3 | 0.000325 | 0 IGF1_ITGAV_ITGB3 | Igf1 - (Itgav+Itgb3) | IGF | Secreted Signaling | PMID: 28190785; PMID: 28873464 |
| CD31 | NeuN | Igf1 | ITGAV_ITGB3 | 0.000199 | 0 IGF1_ITGAV_ITGB3 | Igf1 - (Itgav+Itgb3) | IGF | Secreted Signaling | PMID: 28190785; PMID: 28873464 |
| CD31 | CD31 | Igf2 | ITGAV_ITGB3 | 0.002299 | 0 IGF2_ITGAV_ITGB3 | Igf2 - (Itgav+Itgb3) | IGF | Secreted Signaling | PMID: 28190785; PMID: 28873464 |
| CD31 | NeuN | Igf2 | ITGAV_ITGB3 | 0.001409 | 0 IGF2_ITGAV_ITGB3 | Igf2 - (Itgav+Itgb3) | IGF | Secreted Signaling | PMID: 28190785; PMID: 28873464 |
| CD31 | CD31 | Ins1 | Insr | 0.003267 | 0 INS1_INSSR | Ins1 - Inssr | INSULIN | Secreted Signaling | KEGG: mmu04910; PMID: 24633815 |
| CD31 | NeuN | Ins1 | Insr | 0.00227 | 0 INS1_INSSR | Ins1 - Inssr | INSULIN | Secreted Signaling | KEGG: mmu04910; PMID: 24633815 |
| CD31 | CD31 | Apln | Aplnr | 0.004386 | 0 APLN_APLNR | Apln - Aplnr | APELIN | Secreted Signaling | KEGG: mmu04371 |
| NeuN | CD31 | Apln | Aplnr | 0.001263 | 0 APLN_APLNR | Apln - Aplnr | APELIN | Secreted Signaling | KEGG: mmu04371 |
| CD31 | CD31 | Ihh | PTCH1_SMO | 0.001084 | 0 IHH_PTCH1 | Ihh - Ptch1 | HH | Secreted Signaling | KEGG: mmu04340; PMID: 25701396 |
| NeuN | CD31 | Ihh | PTCH1_SMO | 0.001061 | 0 IHH_PTCH1 | Ihh - Ptch1 | HH | Secreted Signaling | KEGG: mmu04340; PMID: 25701396 |
| CD31 | CD31 | Shh | PTCH1_SMO | 0.001203 | 0 SHH_PTCH1 | Shh - Ptch1 | HH | Secreted Signaling | KEGG: mmu04340; PMID: 25701396 |
| CD31 | NeuN | Shh | PTCH1_SMO | 0.000739 | 0 SHH_PTCH1 | Shh - Ptch1 | HH | Secreted Signaling | KEGG: mmu04340; PMID: 25701396 |
| CD31 | CD31 | Ihh | PTCH2_SMO | 0.000985 | 0 IHH_PTCH2 | Ihh - Ptch2 | HH | Secreted Signaling | KEGG: mmu04340; PMID: 25701396 |
| NeuN | CD31 | Ihh | PTCH2_SMO | 0.000964 | 0 IHH_PTCH2 | Ihh - Ptch2 | HH | Secreted Signaling | KEGG: mmu04340; PMID: 25701396 |
| CD31 | CD31 | Shh | PTCH2_SMO | 0.001093 | 0 SHH_PTCH2 | Shh - Ptch2 | HH | Secreted Signaling | KEGG: mmu04340; PMID: 25701396 |
| CD31 | NeuN | Shh | PTCH2_SMO | 0.000657 | 0 SHH_PTCH2 | Shh - Ptch2 | HH | Secreted Signaling | KEGG: mmu04340; PMID: 25701396 |
| CD31 | CD31 | Ccl8 | Ccr1 | 0.000683 | 0 CCL8_CCR1 | Ccl8 - Ccr1 | CCL | Secreted Signaling | KEGG: mmu04060 |
| NeuN | CD31 | Ccl8 | Ccr1 | 0.000439 | 0 CCL8_CCR1 | Ccl8 - Ccr1 | CCL | Secreted Signaling | KEGG: mmu04060 |
| CD31 | CD31 | Ccl6 | Ccr1 | 0.001059 | 0 CCL6_CCR1 | Ccl6 - Ccr1 | CCL | Secreted Signaling | KEGG: mmu04060 |
| NeuN | CD31 | Ccl6 | Ccr1 | 0.000485 | 0 CCL6_CCR1 | Ccl6 - Ccr1 | CCL | Secreted Signaling | KEGG: mmu04060 |
| CD31 | CD31 | Ccl8 | Ccr1l1 | 0.000666 | 0 CCL8_CCR1L1 | Ccl8 - Ccr1l1 | CCL | Secreted Signaling | KEGG: mmu04060 |
| NeuN | CD31 | Ccl8 | Ccr1l1 | 0.000428 | 0 CCL8_CCR1L1 | Ccl8 - Ccr1l1 | CCL | Secreted Signaling | KEGG: mmu04060 |
| CD31 | CD31 | Ccl6 | Ccr1l1 | 0.001032 | 0 CCL6_CCR1L1 | Ccl6 - Ccr1l1 | CCL | Secreted Signaling | KEGG: mmu04060 |
| CD31 | NeuN | Ccl6 | Ccr1l1 | 0.000543 | 0 CCL6_CCR1L1 | Ccl6 - Ccr1l1 | CCL | Secreted Signaling | KEGG: mmu04060 |
| CD31 | CD31 | Ccl6 | Ccr3 | 0.000995 | 0 CCL6_CCR3 | Ccl6 - Ccr3 | CCL | Secreted Signaling | KEGG: mmu04060 |

| | | | | | | | | | | |
|---|---|---|---|---|---|---|---|---|---|---|
| CD31 | NeuN | Ccl6 | Ccr3 | 0.000491 | 0 | CCL6_CCR3 | Ccl6 - Ccr3 | CCL | Secreted Signaling | KEGG: mmu04060 |
| CD31 | CD31 | Ccl11 | Ccr3 | 0.001147 | 0 | CCL11_CCR3 | Ccl11 - Ccr3 | CCL | Secreted Signaling | KEGG: mmu04060 |
| CD31 | NeuN | Ccl11 | Ccr3 | 0.000566 | 0 | CCL11_CCR3 | Ccl11 - Ccr3 | CCL | Secreted Signaling | KEGG: mmu04060 |
| CD31 | CD31 | Ccl27a | Ccr3 | 0.003658 | 0 | CCL27A_CCR3 | Ccl27a - Ccr3 | CCL | Secreted Signaling | KEGG: mmu04060 |
| NeuN | CD31 | Ccl27a | Ccr3 | 0.006487 | 0 | CCL27A_CCR3 | Ccl27a - Ccr3 | CCL | Secreted Signaling | KEGG: mmu04060 |
| CD31 | CD31 | Ccl27b | Ccr3 | 0.000826 | 0 | CCL27B_CCR3 | Ccl27b - Ccr3 | CCL | Secreted Signaling | KEGG: mmu04060 |
| NeuN | CD31 | Ccl27b | Ccr3 | 0.001386 | 0 | CCL27B_CCR3 | Ccl27b - Ccr3 | CCL | Secreted Signaling | KEGG: mmu04060 |
| CD31 | CD31 | Ccl19 | Ccr7 | 0.000694 | 0 | CCL19_CCR7 | Ccl19 - Ccr7 | CCL | Secreted Signaling | KEGG: mmu04060 |
| NeuN | CD31 | Ccl19 | Ccr7 | 0.000616 | 0 | CCL19_CCR7 | Ccl19 - Ccr7 | CCL | Secreted Signaling | KEGG: mmu04060 |
| CD31 | CD31 | Ccl21a | Ccr7 | 0.000693 | 0 | CCL21A_CCR7 | Ccl21a - Ccr7 | CCL | Secreted Signaling | KEGG: mmu04060 |
| NeuN | CD31 | Ccl21a | Ccr7 | 0.0006 | 0 | CCL21A_CCR7 | Ccl21a - Ccr7 | CCL | Secreted Signaling | KEGG: mmu04060 |
| CD31 | CD31 | Ccl25 | Ccr9 | 0.00055 | 0 | CCL25_CCR9 | Ccl25 - Ccr9 | CCL | Secreted Signaling | KEGG: mmu04060 |
| NeuN | CD31 | Ccl25 | Ccr9 | 0.000416 | 0 | CCL25_CCR9 | Ccl25 - Ccr9 | CCL | Secreted Signaling | KEGG: mmu04060 |
| CD31 | CD31 | Ccl4 | Ackr2 | 0.000948 | 0 | CCL4_ACKR2 | Ccl4 - Ackr2 | CCL | Secreted Signaling | PMID: 26740381 |
| NeuN | CD31 | Ccl4 | Ackr2 | 0.000875 | 0 | CCL4_ACKR2 | Ccl4 - Ackr2 | CCL | Secreted Signaling | PMID: 26740381 |
| CD31 | NeuN | Ccl8 | Ackr1 | 0.000788 | 0 | CCL8_ACKR1 | Ccl8 - Ackr1 | CCL | Secreted Signaling | PMID: 26740381 |
| NeuN | NeuN | Ccl8 | Ackr1 | 0.000506 | 0 | CCL8_ACKR1 | Ccl8 - Ackr1 | CCL | Secreted Signaling | PMID: 26740381 |
| CD31 | CD31 | Ccl11 | Ackr1 | 0.000802 | 0 | CCL11_ACKR1 | Ccl11 - Ackr1 | CCL | Secreted Signaling | PMID: 26740381 |
| CD31 | NeuN | Ccl11 | Ackr1 | 0.001408 | 0 | CCL11_ACKR1 | Ccl11 - Ackr1 | CCL | Secreted Signaling | PMID: 26740381 |
| CD31 | CD31 | Ccl11 | Ackr2 | 0.001209 | 0 | CCL11_ACKR2 | Ccl11 - Ackr2 | CCL | Secreted Signaling | PMID: 26740381 |
| NeuN | CD31 | Ccl11 | Ackr2 | 0.000568 | 0 | CCL11_ACKR2 | Ccl11 - Ackr2 | CCL | Secreted Signaling | PMID: 26740381 |
| CD31 | CD31 | Pf4 | Cxcr3 | 0.001078 | 0 | CXCL4_CXCR3 | Cxcl4 - Cxcr3 | CXCL | Secreted Signaling | KEGG: mmu04060 |
| CD31 | NeuN | Pf4 | Cxcr3 | 0.000647 | 0 | CXCL4_CXCR3 | Cxcl4 - Cxcr3 | CXCL | Secreted Signaling | KEGG: mmu04060 |
| CD31 | CD31 | Cxcl10 | Cxcr3 | 0.000983 | 0 | CXCL10_CXCR3 | Cxcl10 - Cxcr3 | CXCL | Secreted Signaling | KEGG: mmu04060 |
| CD31 | NeuN | Cxcl10 | Cxcr3 | 0.00059 | 0 | CXCL10_CXCR3 | Cxcl10 - Cxcr3 | CXCL | Secreted Signaling | KEGG: mmu04060 |
| CD31 | CD31 | Cxcl12 | Cxcr4 | 0.011029 | 0 | CXCL12_CXCR4 | Cxcl12 - Cxcr4 | CXCL | Secreted Signaling | KEGG: mmu04060 |
| CD31 | NeuN | Cxcl12 | Cxcr4 | 0.002866 | 0 | CXCL12_CXCR4 | Cxcl12 - Cxcr4 | CXCL | Secreted Signaling | KEGG: mmu04060 |
| CD31 | CD31 | Cxcl12 | Ackr3 | 0.002804 | 0 | CXCL12_ACKR3 | Cxcl12 - Ackr3 | CXCL | Secreted Signaling | KEGG: mmu04060 |
| CD31 | NeuN | Cxcl12 | Ackr3 | 0.001996 | 0 | CXCL12_ACKR3 | Cxcl12 - Ackr3 | CXCL | Secreted Signaling | KEGG: mmu04060 |
| CD31 | NeuN | Cxcl1 | Ackr1 | 0.000935 | 0 | CXCL1_ACKR1 | Cxcl1 - Ackr1 | CXCL | Secreted Signaling | KEGG: mmu04060 |
| NeuN | NeuN | Cxcl1 | Ackr1 | 0.000636 | 0 | CXCL1_ACKR1 | Cxcl1 - Ackr1 | CXCL | Secreted Signaling | KEGG: mmu04060 |
| CD31 | CD31 | Cxcl10 | Ackr1 | 0.000666 | 0 | CXCL10_ACKR1 | Cxcl10 - Ackr1 | CXCL | Secreted Signaling | KEGG: mmu04060 |
| CD31 | NeuN | Cxcl10 | Ackr1 | 0.001169 | 0 | CXCL10_ACKR1 | Cxcl10 - Ackr1 | CXCL | Secreted Signaling | KEGG: mmu04060 |
| NeuN | CD31 | Mif | Ackr3 | 0.01249 | 0 | MIF_ACKR3 | Mif - Ackr3 | MIF | Secreted Signaling | PMID: 26175090 |
| NeuN | NeuN | Mif | Ackr3 | 0.008914 | 0 | MIF_ACKR3 | Mif - Ackr3 | MIF | Secreted Signaling | PMID: 26175090 |
| CD31 | CD31 | Cx3cl1 | Cx3cr1 | 0.00037 | 0 | CX3CL1_CX3CR1 | Cx3cl1 - Cx3cr1 | CX3C | Secreted Signaling | KEGG: mmu04060 |
| NeuN | CD31 | Cx3cl1 | Cx3cr1 | 0.00032 | 0 | CX3CL1_CX3CR1 | Cx3cl1 - Cx3cr1 | CX3C | Secreted Signaling | KEGG: mmu04060 |
| CD31 | CD31 | Clcf1 | CNTFR_LIFR | 0.000718 | 0 | CLCF1_CNTFR_LIFR | Clcf1 - (Cntfr+Lifr) | LIFR | Secreted Signaling | KEGG: mmu04060 |
| CD31 | NeuN | Clcf1 | CNTFR_LIFR | 0.000705 | 0 | CLCF1_CNTFR_LIFR | Clcf1 - (Cntfr+Lifr) | LIFR | Secreted Signaling | KEGG: mmu04060 |
| CD31 | CD31 | Ctf1 | LIFR_IL6ST | 0.00093 | 0 | CTF1_LIFR_IL6ST | Ctf1 - (Lifr+Il6st) | LIFR | Secreted Signaling | KEGG: mmu04060 |
| CD31 | NeuN | Ctf1 | LIFR_IL6ST | 0.000955 | 0 | CTF1_LIFR_IL6ST | Ctf1 - (Lifr+Il6st) | LIFR | Secreted Signaling | KEGG: mmu04060 |
| CD31 | CD31 | Il34 | Csf1r | 0.00138 | 0 | IL34_CSF1R | Il34 - Csf1r | CSF | Secreted Signaling | KEGG: mmu04060 |
| CD31 | NeuN | Il34 | Csf1r | 0.000489 | 0 | IL34_CSF1R | Il34 - Csf1r | CSF | Secreted Signaling | KEGG: mmu04060 |
| CD31 | CD31 | Csf1 | Csf1r | 0.000911 | 0 | CSF1_CSF1R | Csf1 - Csf1r | CSF | Secreted Signaling | KEGG: mmu04060 |
| NeuN | CD31 | Csf1 | Csf1r | 0.000387 | 0 | CSF1_CSF1R | Csf1 - Csf1r | CSF | Secreted Signaling | KEGG: mmu04060 |
| CD31 | CD31 | Tnfsf10 | Tnfrsf10b | 0.000748 | 0 | TNFSF10_TNFRSF10B | Tnfsf10 - Tnfrsf10b | TRAIL | Secreted Signaling | KEGG: mmu04060 |
| CD31 | NeuN | Tnfsf10 | Tnfrsf10b | 0.000532 | 0 | TNFSF10_TNFRSF10B | Tnfsf10 - Tnfrsf10b | TRAIL | Secreted Signaling | KEGG: mmu04060 |
| CD31 | CD31 | Retn | Cap1 | 0.003449 | 0 | RETN_CAP1 | Retn - Cap1 | RESISTIN | Secreted Signaling | PMID: 30809105 |
| CD31 | NeuN | Retn | Cap1 | 0.003405 | 0 | RETN_CAP1 | Retn - Cap1 | RESISTIN | Secreted Signaling | PMID: 30809105 |
| CD31 | CD31 | Nampt | ITGA5_ITGB1 | 0.013342 | 0 | NAMPT_ITGA5_ITGB1 | Nampt - (Itga5+Itgb1) | VISFATIN | Secreted Signaling | PMID: 28490838 |
| NeuN | CD31 | Nampt | ITGA5_ITGB1 | 0.004729 | 0 | NAMPT_ITGA5_ITGB1 | Nampt - (Itga5+Itgb1) | VISFATIN | Secreted Signaling | PMID: 28490838 |
| CD31 | CD31 | Angptl4 | ITGA5_ITGB1 | 0.005418 | 0 | ANGPTL4_ITGA5_ITGB1 | Angptl4 - (Itga5+Itgb1) | ANGPTL | Secreted Signaling | PMID: 30049845 |
| CD31 | NeuN | Angptl4 | ITGA5_ITGB1 | 0.00151 | 0 | ANGPTL4_ITGA5_ITGB1 | Angptl4 - (Itga5+Itgb1) | ANGPTL | Secreted Signaling | PMID: 30049845 |
| CD31 | CD31 | Angptl4 | ITGAV_ITGB3 | 0.001951 | 0 | ANGPTL4_ITGAV_ITGB3 | Angptl4 - (Itgav+Itgb3) | ANGPTL | Secreted Signaling | PMID: 30049845 |
| CD31 | NeuN | Angptl4 | ITGAV_ITGB3 | 0.001039 | 0 | ANGPTL4_ITGAV_ITGB3 | Angptl4 - (Itgav+Itgb3) | ANGPTL | Secreted Signaling | PMID: 30049845 |
| CD31 | CD31 | Angptl4 | Cdh5 | 0.010341 | 0 | ANGPTL4_CDH5 | Angptl4 - Cdh5 | ANGPTL | Secreted Signaling | PMID: 30049845 |
| NeuN | CD31 | Angptl4 | Cdh5 | 0.002788 | 0 | ANGPTL4_CDH5 | Angptl4 - Cdh5 | ANGPTL | Secreted Signaling | PMID: 30049845 |
| CD31 | CD31 | Angptl4 | Cdh11 | 0.004883 | 0 | ANGPTL4_CDH11 | Angptl4 - Cdh11 | ANGPTL | Secreted Signaling | PMID: 30049845 |
| CD31 | NeuN | Angptl4 | Cdh11 | 0.004334 | 0 | ANGPTL4_CDH11 | Angptl4 - Cdh11 | ANGPTL | Secreted Signaling | PMID: 30049845 |
| CD31 | CD31 | Angptl4 | Sdc2 | 0.001496 | 0 | ANGPTL4_SDC2 | Angptl4 - Sdc2 | ANGPTL | Secreted Signaling | PMID: 29017031 |

| | | | | | | | | | | |
|---|---|---|---|---|---|---|---|---|---|---|
| CD31 | NeuN | Angptl4 | Sdc2 | 0.000894 | 0 | ANGPTL4_SDC2 | Angptl4 - Sdc2 | ANGPTL | Secreted Signaling | PMID: 29017031 |
| CD31 | CD31 | Angptl4 | Sdc3 | 0.007024 | 0 | ANGPTL4_SDC3 | Angptl4 - Sdc3 | ANGPTL | Secreted Signaling | PMID: 29017031 |
| CD31 | NeuN | Angptl4 | Sdc3 | 0.011693 | 0 | ANGPTL4_SDC3 | Angptl4 - Sdc3 | ANGPTL | Secreted Signaling | PMID: 29017031 |
| CD31 | CD31 | Angptl4 | Sdc4 | 0.001795 | 0 | ANGPTL4_SDC4 | Angptl4 - Sdc4 | ANGPTL | Secreted Signaling | PMID: 29017031 |
| CD31 | NeuN | Angptl4 | Sdc4 | 0.000831 | 0 | ANGPTL4_SDC4 | Angptl4 - Sdc4 | ANGPTL | Secreted Signaling | PMID: 29017031 |
| CD31 | CD31 | Angptl3 | ITGAV_ITGB3 | 0.000698 | 0 | ANGPTL3_ITGAV_ITGB3 | Angptl3 - (Itgav+Itgb3) | ANGPTL | Secreted Signaling | PMID: 11877390 |
| CD31 | NeuN | Angptl3 | ITGAV_ITGB3 | 0.000371 | 0 | ANGPTL3_ITGAV_ITGB3 | Angptl3 - (Itgav+Itgb3) | ANGPTL | Secreted Signaling | PMID: 11877390 |
| CD31 | CD31 | Mdk | Sdc2 | 0.001868 | 0 | MDK_SDC2 | Mdk - Sdc2 | MK | Secreted Signaling | PMID: 28356350 |
| NeuN | CD31 | Mdk | Sdc2 | 0.001149 | 0 | MDK_SDC2 | Mdk - Sdc2 | MK | Secreted Signaling | PMID: 28356350 |
| CD31 | CD31 | Mdk | Sdc4 | 0.002242 | 0 | MDK_SDC4 | Mdk - Sdc4 | MK | Secreted Signaling | PMID: 28356350 |
| NeuN | CD31 | Mdk | Sdc4 | 0.00138 | 0 | MDK_SDC4 | Mdk - Sdc4 | MK | Secreted Signaling | PMID: 28356350 |
| CD31 | NeuN | Mdk | Ptprz1 | 0.027068 | 0 | MDK_PTPRZ1 | Mdk - Ptprz1 | MK | Secreted Signaling | PMID: 28356350 |
| NeuN | NeuN | Mdk | Ptprz1 | 0.016818 | 0 | MDK_PTPRZ1 | Mdk - Ptprz1 | MK | Secreted Signaling | PMID: 28356350 |
| CD31 | CD31 | Mdk | ITGA4_ITGB1 | 0.007137 | 0 | MDK_ITGA4_ITGB1 | Mdk - (Itga4+Itgb1) | MK | Secreted Signaling | PMID: 28356350 |
| NeuN | CD31 | Mdk | ITGA4_ITGB1 | 0.004401 | 0 | MDK_ITGA4_ITGB1 | Mdk - (Itga4+Itgb1) | MK | Secreted Signaling | PMID: 28356350 |
| CD31 | CD31 | Mdk | ITGA6_ITGB1 | 0.005098 | 0 | MDK_ITGA6_ITGB1 | Mdk - (Itga6+Itgb1) | MK | Secreted Signaling | PMID: 28356350 |
| NeuN | CD31 | Mdk | ITGA6_ITGB1 | 0.003141 | 0 | MDK_ITGA6_ITGB1 | Mdk - (Itga6+Itgb1) | MK | Secreted Signaling | PMID: 28356350 |
| CD31 | CD31 | Mdk | Lrp1 | 0.002891 | 0 | MDK_LRP1 | Mdk - Lrp1 | MK | Secreted Signaling | PMID: 28356350 |
| CD31 | NeuN | Mdk | Lrp1 | 0.004431 | 0 | MDK_LRP1 | Mdk - Lrp1 | MK | Secreted Signaling | PMID: 28356350 |
| CD31 | CD31 | Mdk | Ncl | 0.021606 | 0 | MDK_NCL | Mdk - Ncl | MK | Secreted Signaling | PMID: 28356350 |
| CD31 | NeuN | Mdk | Ncl | 0.025904 | 0 | MDK_NCL | Mdk - Ncl | MK | Secreted Signaling | PMID: 28356350 |
| CD31 | NeuN | Ptn | Ptprz1 | 0.080262 | 0 | PTN_PTPRZ1 | Ptn - Ptprz1 | PTN | Secreted Signaling | PMID: 28356350; PMID: 25620911 |
| NeuN | NeuN | Ptn | Ptprz1 | 0.136203 | 0 | PTN_PTPRZ1 | Ptn - Ptprz1 | PTN | Secreted Signaling | PMID: 28356350; PMID: 25620911 |
| NeuN | CD31 | Ptn | Sdc2 | 0.010494 | 0 | PTN_SDC2 | Ptn - Sdc2 | PTN | Secreted Signaling | PMID: 28356350; PMID: 25620911 |
| NeuN | NeuN | Ptn | Sdc2 | 0.006293 | 0 | PTN_SDC2 | Ptn - Sdc2 | PTN | Secreted Signaling | PMID: 28356350; PMID: 25620911 |
| NeuN | CD31 | Ptn | Sdc3 | 0.047689 | 0 | PTN_SDC3 | Ptn - Sdc3 | PTN | Secreted Signaling | PMID: 28356350; PMID: 25620911 |
| NeuN | NeuN | Ptn | Sdc3 | 0.077291 | 0 | PTN_SDC3 | Ptn - Sdc3 | PTN | Secreted Signaling | PMID: 28356350; PMID: 25620911 |
| CD31 | CD31 | Ptn | Sdc4 | 0.006998 | 0 | PTN_SDC4 | Ptn - Sdc4 | PTN | Secreted Signaling | PMID: 28356350; PMID: 25620911 |
| NeuN | CD31 | Ptn | Sdc4 | 0.012574 | 0 | PTN_SDC4 | Ptn - Sdc4 | PTN | Secreted Signaling | PMID: 28356350; PMID: 25620911 |
| CD31 | CD31 | Ptn | ITGAV_ITGB3 | 0.007602 | 0 | PTN_ITGAV_ITGB3 | Ptn - (Itgav+Itgb3) | PTN | Secreted Signaling | PMID: 28356350; PMID: 25620911 |
| NeuN | CD31 | Ptn | ITGAV_ITGB3 | 0.013651 | 0 | PTN_ITGAV_ITGB3 | Ptn - (Itgav+Itgb3) | PTN | Secreted Signaling | PMID: 28356350; PMID: 25620911 |
| NeuN | CD31 | Ptn | Ncl | 0.111236 | 0 | PTN_NCL | Ptn - Ncl | PTN | Secreted Signaling | PMID: 28356350; PMID: 25620911 |
| NeuN | NeuN | Ptn | Ncl | 0.130975 | 0 | PTN_NCL | Ptn - Ncl | PTN | Secreted Signaling | PMID: 28356350; PMID: 25620911 |
| CD31 | CD31 | Postn | ITGAV_ITGB3 | 0.000998 | 0 | POSTN_ITGAV_ITGB3 | Postn - (Itgav+Itgb3) | PERIOSTIN | Secreted Signaling | PMID: 26721738 |
| CD31 | NeuN | Postn | ITGAV_ITGB3 | 0.000531 | 0 | POSTN_ITGAV_ITGB3 | Postn - (Itgav+Itgb3) | PERIOSTIN | Secreted Signaling | PMID: 26721738 |
| CD31 | CD31 | Postn | ITGAV_ITGB5 | 0.000889 | 0 | POSTN_ITGAV_ITGB5 | Postn - (Itgav+Itgb5) | PERIOSTIN | Secreted Signaling | PMID: 26721738 |
| CD31 | NeuN | Postn | ITGAV_ITGB5 | 0.000451 | 0 | POSTN_ITGAV_ITGB5 | Postn - (Itgav+Itgb5) | PERIOSTIN | Secreted Signaling | PMID: 26721738 |
| CD31 | CD31 | Nmb | Nmbr | 0.000427 | 0 | NMB_NMBR | Nmb - Nmbr | GPR | Secreted Signaling | KEGG: mmu04080 |
| NeuN | CD31 | Nmb | Nmbr | 0.00038 | 0 | NMB_NMBR | Nmb - Nmbr | GPR | Secreted Signaling | KEGG: mmu04080 |
| CD31 | CD31 | Cck | Cckbr | 0.000732 | 0 | CCK_CCKBR | Cck - Cckbr | CCK | Secreted Signaling | KEGG: mmu04080 |
| NeuN | CD31 | Cck | Cckbr | 0.000945 | 0 | CCK_CCKBR | Cck - Cckbr | CCK | Secreted Signaling | KEGG: mmu04080 |
| CD31 | CD31 | Edn1 | Ednra | 0.001941 | 0 | EDN1_EDNRA | Edn1 - Ednra | EDN | Secreted Signaling | KEGG: mmu04080 |
| NeuN | CD31 | Edn1 | Ednra | 0.000685 | 0 | EDN1_EDNRA | Edn1 - Ednra | EDN | Secreted Signaling | KEGG: mmu04080 |
| CD31 | CD31 | Edn3 | Ednra | 0.001548 | 0 | EDN3_EDNRA | Edn3 - Ednra | EDN | Secreted Signaling | KEGG: mmu04080 |
| NeuN | CD31 | Edn3 | Ednra | 0.000718 | 0 | EDN3_EDNRA | Edn3 - Ednra | EDN | Secreted Signaling | KEGG: mmu04080 |
| CD31 | CD31 | Edn1 | Ednrb | 0.000656 | 0 | EDN1_EDNRB | Edn1 - Ednrb | EDN | Secreted Signaling | KEGG: mmu04080 |
| CD31 | NeuN | Edn1 | Ednrb | 0.000321 | 0 | EDN1_EDNRB | Edn1 - Ednrb | EDN | Secreted Signaling | KEGG: mmu04080 |
| CD31 | CD31 | Edn3 | Ednrb | 0.000524 | 0 | EDN3_EDNRB | Edn3 - Ednrb | EDN | Secreted Signaling | KEGG: mmu04080 |
| CD31 | NeuN | Edn3 | Ednrb | 0.000256 | 0 | EDN3_EDNRB | Edn3 - Ednrb | EDN | Secreted Signaling | KEGG: mmu04080 |
| CD31 | CD31 | Nms | Nmur1 | 0.000571 | 0 | NMS_NMUR1 | Nms - Nmur1 | NMU | Secreted Signaling | PMID: 29288685 |
| CD31 | NeuN | Nms | Nmur1 | 0.000416 | 0 | NMS_NMUR1 | Nms - Nmur1 | NMU | Secreted Signaling | PMID: 29288685 |
| CD31 | NeuN | Npy | Npy1r | 0.002195 | 0 | NPY_NPY1R | Npy - Npy1r | NPY | Secreted Signaling | KEGG: mmu04080 |
| NeuN | NeuN | Npy | Npy1r | 0.002828 | 0 | NPY_NPY1R | Npy - Npy1r | NPY | Secreted Signaling | KEGG: mmu04080 |
| NeuN | CD31 | Npy | Npy2r | 0.002955 | 0 | NPY_NPY2R | Npy - Npy2r | NPY | Secreted Signaling | KEGG: mmu04080 |
| NeuN | NeuN | Npy | Npy2r | 0.002474 | 0 | NPY_NPY2R | Npy - Npy2r | NPY | Secreted Signaling | KEGG: mmu04080 |
| CD31 | CD31 | Npy | Npy5r | 0.001432 | 0 | NPY_NPY5R | Npy - Npy5r | NPY | Secreted Signaling | KEGG: mmu04080 |
| NeuN | CD31 | Npy | Npy5r | 0.001845 | 0 | NPY_NPY5R | Npy - Npy5r | NPY | Secreted Signaling | KEGG: mmu04080 |
| CD31 | CD31 | Npy | Npy6r | 0.002001 | 0 | NPY_NPY6R | Npy - Npy6r | NPY | Secreted Signaling | KEGG: mmu04080 |
| NeuN | CD31 | Npy | Npy6r | 0.002579 | 0 | NPY_NPY6R | Npy - Npy6r | NPY | Secreted Signaling | KEGG: mmu04080 |
| CD31 | CD31 | Npb | Npbwr1 | 0.00055 | 0 | NPB_NPBWR1 | Npb - Npbwr1 | NPW-B | Secreted Signaling | KEGG: mmu04080 |

| | | | | | | | | | | |
|---|---|---|---|---|---|---|---|---|---|---|
| NeuN | CD31 | Npb | Npbwr1 | 0.000403 | 0 | NPB_NPBWR1 | Npb - Npbwr1 | NPW-B | Secreted Signaling | KEGG: mmu04080 |
| CD31 | CD31 | Nts | Ntsr1 | 0.001311 | 0 | NTS_NTSR1 | Nts - Ntsr1 | NTS | Secreted Signaling | KEGG: mmu04080 |
| CD31 | NeuN | Nts | Ntsr1 | 0.002518 | 0 | NTS_NTSR1 | Nts - Ntsr1 | NTS | Secreted Signaling | KEGG: mmu04080 |
| CD31 | CD31 | Nts | Ntsr2 | 0.000456 | 0 | NTS_NTSR2 | Nts - Ntsr2 | NTS | Secreted Signaling | KEGG: mmu04080 |
| CD31 | NeuN | Nts | Ntsr2 | 0.000331 | 0 | NTS_NTSR2 | Nts - Ntsr2 | NTS | Secreted Signaling | KEGG: mmu04080 |
| CD31 | CD31 | Pdyn | Oprd1 | 0.000449 | 0 | PDYN_OPRD1 | Pdyn - Oprd1 | OPIOID | Secreted Signaling | KEGG: mmu04080 |
| NeuN | CD31 | Pdyn | Oprd1 | 0.000346 | 0 | PDYN_OPRD1 | Pdyn - Oprd1 | OPIOID | Secreted Signaling | KEGG: mmu04080 |
| CD31 | CD31 | Pdyn | Oprk1 | 0.000367 | 0 | PDYN_OPRK1 | Pdyn - Oprk1 | OPIOID | Secreted Signaling | KEGG: mmu04080 |
| NeuN | CD31 | Pdyn | Oprk1 | 0.000283 | 0 | PDYN_OPRK1 | Pdyn - Oprk1 | OPIOID | Secreted Signaling | KEGG: mmu04080 |
| CD31 | CD31 | Pdyn | Oprl1 | 0.000621 | 0 | PDYN_OPRL1 | Pdyn - Oprl1 | OPIOID | Secreted Signaling | KEGG: mmu04080 |
| CD31 | NeuN | Pdyn | Oprl1 | 0.000781 | 0 | PDYN_OPRL1 | Pdyn - Oprl1 | OPIOID | Secreted Signaling | KEGG: mmu04080 |
| CD31 | CD31 | Pdyn | Oprm1 | 0.000422 | 0 | PDYN_OPRM1 | Pdyn - Oprm1 | OPIOID | Secreted Signaling | KEGG: mmu04080 |
| NeuN | CD31 | Pdyn | Oprm1 | 0.000326 | 0 | PDYN_OPRM1 | Pdyn - Oprm1 | OPIOID | Secreted Signaling | KEGG: mmu04080 |
| CD31 | CD31 | Penk | Oprd1 | 0.000772 | 0 | PENK_OPRD1 | Penk - Oprd1 | OPIOID | Secreted Signaling | KEGG: mmu04080 |
| CD31 | NeuN | Penk | Oprd1 | 0.000585 | 0 | PENK_OPRD1 | Penk - Oprd1 | OPIOID | Secreted Signaling | KEGG: mmu04080 |
| CD31 | CD31 | Penk | Oprk1 | 0.000631 | 0 | PENK_OPRK1 | Penk - Oprk1 | OPIOID | Secreted Signaling | KEGG: mmu04080 |
| CD31 | NeuN | Penk | Oprk1 | 0.000432 | 0 | PENK_OPRK1 | Penk - Oprk1 | OPIOID | Secreted Signaling | KEGG: mmu04080 |
| CD31 | CD31 | Penk | Oprl1 | 0.001068 | 0 | PENK_OPRL1 | Penk - Oprl1 | OPIOID | Secreted Signaling | KEGG: mmu04080 |
| CD31 | NeuN | Penk | Oprl1 | 0.001345 | 0 | PENK_OPRL1 | Penk - Oprl1 | OPIOID | Secreted Signaling | KEGG: mmu04080 |
| CD31 | CD31 | Penk | Oprm1 | 0.000727 | 0 | PENK_OPRM1 | Penk - Oprm1 | OPIOID | Secreted Signaling | KEGG: mmu04080 |
| CD31 | NeuN | Penk | Oprm1 | 0.00046 | 0 | PENK_OPRM1 | Penk - Oprm1 | OPIOID | Secreted Signaling | KEGG: mmu04080 |
| CD31 | CD31 | Oxt | Oxtr | 0.000799 | 0 | OXT_OXTR | Oxt - Oxtr | OXT | Secreted Signaling | KEGG: mmu04080 |
| NeuN | CD31 | Oxt | Oxtr | 0.000734 | 0 | OXT_OXTR | Oxt - Oxtr | OXT | Secreted Signaling | KEGG: mmu04080 |
| CD31 | CD31 | Sst | Sstr1 | 0.000391 | 0 | SST_SSTR1 | Sst - Sstr1 | SOMATOSTATIN | Secreted Signaling | KEGG: mmu04080 |
| NeuN | CD31 | Sst | Sstr1 | 0.000335 | 0 | SST_SSTR1 | Sst - Sstr1 | SOMATOSTATIN | Secreted Signaling | KEGG: mmu04080 |
| CD31 | CD31 | Sst | Sstr2 | 0.000407 | 0 | SST_SSTR2 | Sst - Sstr2 | SOMATOSTATIN | Secreted Signaling | KEGG: mmu04080 |
| CD31 | NeuN | Sst | Sstr2 | 0.000399 | 0 | SST_SSTR2 | Sst - Sstr2 | SOMATOSTATIN | Secreted Signaling | KEGG: mmu04080 |
| CD31 | CD31 | Sst | Sstr3 | 0.000381 | 0 | SST_SSTR3 | Sst - Sstr3 | SOMATOSTATIN | Secreted Signaling | KEGG: mmu04080 |
| NeuN | CD31 | Sst | Sstr3 | 0.000326 | 0 | SST_SSTR3 | Sst - Sstr3 | SOMATOSTATIN | Secreted Signaling | KEGG: mmu04080 |
| CD31 | CD31 | Sst | Sstr4 | 0.000316 | 0 | SST_SSTR4 | Sst - Sstr4 | SOMATOSTATIN | Secreted Signaling | KEGG: mmu04080 |
| NeuN | CD31 | Sst | Sstr4 | 0.00027 | 0 | SST_SSTR4 | Sst - Sstr4 | SOMATOSTATIN | Secreted Signaling | KEGG: mmu04080 |
| CD31 | CD31 | Cort | Sstr1 | 0.0006 | 0 | CORT_SSTR1 | Cort - Sstr1 | SOMATOSTATIN | Secreted Signaling | KEGG: mmu04080 |
| NeuN | CD31 | Cort | Sstr1 | 0.000668 | 0 | CORT_SSTR1 | Cort - Sstr1 | SOMATOSTATIN | Secreted Signaling | KEGG: mmu04080 |
| NeuN | CD31 | Cort | Sstr2 | 0.000694 | 0 | CORT_SSTR2 | Cort - Sstr2 | SOMATOSTATIN | Secreted Signaling | KEGG: mmu04080 |
| NeuN | NeuN | Cort | Sstr2 | 0.000681 | 0 | CORT_SSTR2 | Cort - Sstr2 | SOMATOSTATIN | Secreted Signaling | KEGG: mmu04080 |
| CD31 | CD31 | Cort | Sstr3 | 0.000585 | 0 | CORT_SSTR3 | Cort - Sstr3 | SOMATOSTATIN | Secreted Signaling | KEGG: mmu04080 |
| NeuN | CD31 | Cort | Sstr3 | 0.00065 | 0 | CORT_SSTR3 | Cort - Sstr3 | SOMATOSTATIN | Secreted Signaling | KEGG: mmu04080 |
| CD31 | CD31 | Cort | Sstr4 | 0.000484 | 0 | CORT_SSTR4 | Cort - Sstr4 | SOMATOSTATIN | Secreted Signaling | KEGG: mmu04080 |
| NeuN | CD31 | Cort | Sstr4 | 0.000539 | 0 | CORT_SSTR4 | Cort - Sstr4 | SOMATOSTATIN | Secreted Signaling | KEGG: mmu04080 |
| CD31 | CD31 | Tac1 | Tacr1 | 0.000346 | 0 | TAC1_TACR1 | Tac1 - Tacr1 | TAC | Secreted Signaling | KEGG: mmu04080 |
| NeuN | CD31 | Tac1 | Tacr1 | 0.000341 | 0 | TAC1_TACR1 | Tac1 - Tacr1 | TAC | Secreted Signaling | KEGG: mmu04080 |
| CD31 | CD31 | Tac2 | Tacr3 | 0.000406 | 0 | TAC2_TACR3 | Tac2 - Tacr3 | TAC | Secreted Signaling | KEGG: mmu04080 |
| CD31 | NeuN | Tac2 | Tacr3 | 0.000352 | 0 | TAC2_TACR3 | Tac2 - Tacr3 | TAC | Secreted Signaling | KEGG: mmu04080 |
| CD31 | CD31 | Ctsg | F2r | 0.00131 | 0 | CTSG_F2R | Ctsg - F2r | PARs | Secreted Signaling | KEGG: mmu04080 |
| CD31 | NeuN | Ctsg | F2r | 0.000768 | 0 | CTSG_F2R | Ctsg - F2r | PARs | Secreted Signaling | KEGG: mmu04080 |
| CD31 | CD31 | F2 | F2r | 0.001783 | 0 | F2_F2R | F2 - F2r | PARs | Secreted Signaling | KEGG: mmu04080 |
| CD31 | NeuN | F2 | F2r | 0.001045 | 0 | F2_F2R | F2 - F2r | PARs | Secreted Signaling | KEGG: mmu04080 |
| CD31 | CD31 | Ctsg | F2rl2 | 0.000505 | 0 | CTSG_F2RL2 | Ctsg - F2rl2 | PARs | Secreted Signaling | KEGG: mmu04080 |
| CD31 | NeuN | Ctsg | F2rl2 | 0.000365 | 0 | CTSG_F2RL2 | Ctsg - F2rl2 | PARs | Secreted Signaling | KEGG: mmu04080 |
| CD31 | CD31 | F2 | F2rl2 | 0.000688 | 0 | F2_F2RL2 | F2 - F2rl2 | PARs | Secreted Signaling | KEGG: mmu04080 |
| CD31 | NeuN | F2 | F2rl2 | 0.000497 | 0 | F2_F2RL2 | F2 - F2rl2 | PARs | Secreted Signaling | KEGG: mmu04080 |
| CD31 | CD31 | Ctsg | Pard3 | 0.001012 | 0 | CTSG_PARD3 | Ctsg - Pard3 | PARs | Secreted Signaling | KEGG: mmu04080 |
| CD31 | NeuN | Ctsg | Pard3 | 0.00066 | 0 | CTSG_PARD3 | Ctsg - Pard3 | PARs | Secreted Signaling | KEGG: mmu04080 |
| CD31 | CD31 | F2 | Pard3 | 0.001378 | 0 | F2_PARD3 | F2 - Pard3 | PARs | Secreted Signaling | KEGG: mmu04080 |
| CD31 | NeuN | F2 | Pard3 | 0.000898 | 0 | F2_PARD3 | F2 - Pard3 | PARs | Secreted Signaling | KEGG: mmu04080 |
| CD31 | CD31 | Plg | Pard3 | 0.000851 | 0 | PLG_PARD3 | Plg - Pard3 | PARs | Secreted Signaling | KEGG: mmu04080 |
| CD31 | NeuN | Plg | Pard3 | 0.000554 | 0 | PLG_PARD3 | Plg - Pard3 | PARs | Secreted Signaling | KEGG: mmu04080 |
| CD31 | CD31 | Pmch | Mchr1 | 0.000507 | 0 | PMCH_MCHR1 | Pmch - Mchr1 | PMCH | Secreted Signaling | KEGG: mmu04080 |
| CD31 | NeuN | Pmch | Mchr1 | 0.000271 | 0 | PMCH_MCHR1 | Pmch - Mchr1 | PMCH | Secreted Signaling | KEGG: mmu04080 |
| CD31 | CD31 | Prok2 | Prokr1 | 0.000522 | 0 | PROK2_PROKR1 | Prok2 - Prokr1 | PROK | Secreted Signaling | PMID: 24496457 |

| | | | | | | | | | | |
|---|---|---|---|---|---|---|---|---|---|---|
| NeuN | CD31 | Prok2 | Prokr1 | 0.000516 | 0 | PROK2_PROKR1 | Prok2 - Prokr1 | PROK | Secreted Signaling | PMID: 24496457 |
| CD31 | NeuN | Prok2 | Prokr2 | 0.000755 | 0 | PROK2_PROKR2 | Prok2 - Prokr2 | PROK | Secreted Signaling | PMID: 24496457 |
| NeuN | NeuN | Prok2 | Prokr2 | 0.000745 | 0 | PROK2_PROKR2 | Prok2 - Prokr2 | PROK | Secreted Signaling | PMID: 24496457 |
| CD31 | NeuN | Adcyap1 | Adcyap1r1 | 0.00121 | 0 | ADCYAP1_ADCYAP1R1 | Adcyap1 - Adcyap1r1 | PACAP | Secreted Signaling | PMID: 21350482 |
| NeuN | NeuN | Adcyap1 | Adcyap1r1 | 0.001153 | 0 | ADCYAP1_ADCYAP1R1 | Adcyap1 - Adcyap1r1 | PACAP | Secreted Signaling | PMID: 21350482 |
| CD31 | CD31 | Vip | Vipr1 | 0.000334 | 0 | VIP_VIPR1 | Vip - Vipr1 | VIP | Secreted Signaling | PMID: 10655512 |
| NeuN | CD31 | Vip | Vipr1 | 0.00019 | 0 | VIP_VIPR1 | Vip - Vipr1 | VIP | Secreted Signaling | PMID: 10655512 |
| CD31 | CD31 | Nppa | Npr1 | 0.002456 | 0 | NPPA_NPR1 | Nppa - Npr1 | NPR1 | Secreted Signaling | KEGG: mmu04080; PMID:29934640 |
| NeuN | CD31 | Nppa | Npr1 | 0.003662 | 0 | NPPA_NPR1 | Nppa - Npr1 | NPR1 | Secreted Signaling | KEGG: mmu04080; PMID:29934640 |
| CD31 | CD31 | Nppb | Npr1 | 0.000727 | 0 | NPPB_NPR1 | Nppb - Npr1 | NPR1 | Secreted Signaling | KEGG: mmu04080; PMID:29934640 |
| CD31 | NeuN | Nppb | Npr1 | 0.000368 | 0 | NPPB_NPR1 | Nppb - Npr1 | NPR1 | Secreted Signaling | KEGG: mmu04080; PMID:29934640 |
| CD31 | NeuN | Nppc | Npr2 | 0.001488 | 0 | NPPC_NPR2 | Nppc - Npr2 | NPR2 | Secreted Signaling | KEGG: mmu04080; PMID:29934640 |
| NeuN | NeuN | Nppc | Npr2 | 0.000997 | 0 | NPPC_NPR2 | Nppc - Npr2 | NPR2 | Secreted Signaling | KEGG: mmu04080; PMID:29934640 |
| CD31 | CD31 | Kitl | Kit | 0.004085 | 0 | KITL_KIT | Kitl - Kit | KIT | Secreted Signaling | KEGG: mmu04080 |
| CD31 | NeuN | Kitl | Kit | 0.003795 | 0 | KITL_KIT | Kitl - Kit | KIT | Secreted Signaling | KEGG: mmu04080 |
| CD31 | NeuN | Gip | Gipr | 0.002368 | 0 | GIP_GIPR | Gip - Gipr | GIPR | Secreted Signaling | KEGG: mmu04080 |
| NeuN | NeuN | Gip | Gipr | 0.001642 | 0 | GIP_GIPR | Gip - Gipr | GIPR | Secreted Signaling | KEGG: mmu04080 |
| CD31 | NeuN | Bdnf | Ntrk2 | 0.002481 | 0 | BDNF_NTRK2 | Bdnf - Ntrk2 | NT | Secreted Signaling | PMID: 27908981 |
| NeuN | NeuN | Bdnf | Ntrk2 | 0.001925 | 0 | BDNF_NTRK2 | Bdnf - Ntrk2 | NT | Secreted Signaling | PMID: 27908981 |
| CD31 | CD31 | Flt3l | Flt3 | 0.000806 | 0 | FLT3L_FLT3 | Flt3l - Flt3 | FLT3 | Secreted Signaling | PMID: 25992210 |
| CD31 | NeuN | Flt3l | Flt3 | 0.000506 | 0 | FLT3L_FLT3 | Flt3l - Flt3 | FLT3 | Secreted Signaling | PMID: 25992210 |
| CD31 | CD31 | Hgf | Met | 0.000454 | 0 | HGF_MET | Hgf - Met | HGF | Secreted Signaling | PMID: 28475121 |
| CD31 | NeuN | Hgf | Met | 0.00054 | 0 | HGF_MET | Hgf - Met | HGF | Secreted Signaling | PMID: 28475121 |
| CD31 | CD31 | Ntf3 | Ntrk1 | 0.00073 | 0 | NTF3_NTRK1 | Ntf3 - Ntrk1 | NT | Secreted Signaling | PMID: 27908981 |
| NeuN | CD31 | Ntf3 | Ntrk1 | 0.001372 | 0 | NTF3_NTRK1 | Ntf3 - Ntrk1 | NT | Secreted Signaling | PMID: 27908981 |
| CD31 | NeuN | Ntf3 | Ntrk2 | 0.003338 | 0 | NTF3_NTRK2 | Ntf3 - Ntrk2 | NT | Secreted Signaling | PMID: 27908981 |
| NeuN | NeuN | Ntf3 | Ntrk2 | 0.006259 | 0 | NTF3_NTRK2 | Ntf3 - Ntrk2 | NT | Secreted Signaling | PMID: 27908981 |
| CD31 | NeuN | Ntf3 | Ntrk3 | 0.00829 | 0 | NTF3_NTRK3 | Ntf3 - Ntrk3 | NT | Secreted Signaling | PMID: 27908981 |
| NeuN | NeuN | Ntf3 | Ntrk3 | 0.015476 | 0 | NTF3_NTRK3 | Ntf3 - Ntrk3 | NT | Secreted Signaling | PMID: 27908981 |
| NeuN | CD31 | Sema3a | NRP1_PLXNA1 | 0.002765 | 0 | SEMA3A_NRP1_PLXNA1 | Sema3a - (Nrp1+Plxna1) | SEMA3 | Secreted Signaling | PMID: 27533782 |
| NeuN | NeuN | Sema3a | NRP1_PLXNA1 | 0.002674 | 0 | SEMA3A_NRP1_PLXNA1 | Sema3a - (Nrp1+Plxna1) | SEMA3 | Secreted Signaling | PMID: 27533782 |
| NeuN | CD31 | Sema3a | NRP1_PLXNA2 | 0.002757 | 0 | SEMA3A_NRP1_PLXNA2 | Sema3a - (Nrp1+Plxna2) | SEMA3 | Secreted Signaling | PMID: 27533782 |
| NeuN | NeuN | Sema3a | NRP1_PLXNA2 | 0.002397 | 0 | SEMA3A_NRP1_PLXNA2 | Sema3a - (Nrp1+Plxna2) | SEMA3 | Secreted Signaling | PMID: 27533782 |
| CD31 | NeuN | Sema3a | NRP1_PLXNA3 | 0.002427 | 0 | SEMA3A_NRP1_PLXNA3 | Sema3a - (Nrp1+Plxna3) | SEMA3 | Secreted Signaling | PMID: 27533782 |
| NeuN | NeuN | Sema3a | NRP1_PLXNA3 | 0.002797 | 0 | SEMA3A_NRP1_PLXNA3 | Sema3a - (Nrp1+Plxna3) | SEMA3 | Secreted Signaling | PMID: 27533782 |
| CD31 | NeuN | Sema3a | NRP1_PLXNA4 | 0.004544 | 0 | SEMA3A_NRP1_PLXNA4 | Sema3a - (Nrp1+Plxna4) | SEMA3 | Secreted Signaling | PMID: 27533782 |
| NeuN | NeuN | Sema3a | NRP1_PLXNA4 | 0.005235 | 0 | SEMA3A_NRP1_PLXNA4 | Sema3a - (Nrp1+Plxna4) | SEMA3 | Secreted Signaling | PMID: 27533782 |
| CD31 | CD31 | Sema3b | NRP1_PLXNA1 | 0.001267 | 0 | SEMA3B_NRP1_PLXNA1 | Sema3b - (Nrp1+Plxna1) | SEMA3 | Secreted Signaling | PMID: 27533782 |
| CD31 | NeuN | Sema3b | NRP1_PLXNA1 | 0.001225 | 0 | SEMA3B_NRP1_PLXNA1 | Sema3b - (Nrp1+Plxna1) | SEMA3 | Secreted Signaling | PMID: 27533782 |
| CD31 | CD31 | Sema3b | NRP1_PLXNA2 | 0.001263 | 0 | SEMA3B_NRP1_PLXNA2 | Sema3b - (Nrp1+Plxna2) | SEMA3 | Secreted Signaling | PMID: 27533782 |
| NeuN | CD31 | Sema3b | NRP1_PLXNA2 | 0.001112 | 0 | SEMA3B_NRP1_PLXNA2 | Sema3b - (Nrp1+Plxna2) | SEMA3 | Secreted Signaling | PMID: 27533782 |
| CD31 | NeuN | Sema3b | NRP1_PLXNA3 | 0.001281 | 0 | SEMA3B_NRP1_PLXNA3 | Sema3b - (Nrp1+Plxna3) | SEMA3 | Secreted Signaling | PMID: 27533782 |
| NeuN | NeuN | Sema3b | NRP1_PLXNA3 | 0.001128 | 0 | SEMA3B_NRP1_PLXNA3 | Sema3b - (Nrp1+Plxna3) | SEMA3 | Secreted Signaling | PMID: 27533782 |
| CD31 | NeuN | Sema3b | NRP1_PLXNA4 | 0.002402 | 0 | SEMA3B_NRP1_PLXNA4 | Sema3b - (Nrp1+Plxna4) | SEMA3 | Secreted Signaling | PMID: 27533782 |
| NeuN | NeuN | Sema3b | NRP1_PLXNA4 | 0.002114 | 0 | SEMA3B_NRP1_PLXNA4 | Sema3b - (Nrp1+Plxna4) | SEMA3 | Secreted Signaling | PMID: 27533782 |
| NeuN | CD31 | Sema3c | NRP1_PLXNA1 | 0.002224 | 0 | SEMA3C_NRP1_PLXNA1 | Sema3c - (Nrp1+Plxna1) | SEMA3 | Secreted Signaling | PMID: 27533782 |
| NeuN | NeuN | Sema3c | NRP1_PLXNA1 | 0.00215 | 0 | SEMA3C_NRP1_PLXNA1 | Sema3c - (Nrp1+Plxna1) | SEMA3 | Secreted Signaling | PMID: 27533782 |
| CD31 | CD31 | Sema3c | NRP1_PLXNA2 | 0.00197 | 0 | SEMA3C_NRP1_PLXNA2 | Sema3c - (Nrp1+Plxna2) | SEMA3 | Secreted Signaling | PMID: 27533782 |
| NeuN | CD31 | Sema3c | NRP1_PLXNA2 | 0.002217 | 0 | SEMA3C_NRP1_PLXNA2 | Sema3c - (Nrp1+Plxna2) | SEMA3 | Secreted Signaling | PMID: 27533782 |
| CD31 | NeuN | Sema3c | NRP1_PLXNA3 | 0.001998 | 0 | SEMA3C_NRP1_PLXNA3 | Sema3c - (Nrp1+Plxna3) | SEMA3 | Secreted Signaling | PMID: 27533782 |
| NeuN | NeuN | Sema3c | NRP1_PLXNA3 | 0.002249 | 0 | SEMA3C_NRP1_PLXNA3 | Sema3c - (Nrp1+Plxna3) | SEMA3 | Secreted Signaling | PMID: 27533782 |
| CD31 | NeuN | Sema3c | NRP1_PLXNA4 | 0.003742 | 0 | SEMA3C_NRP1_PLXNA4 | Sema3c - (Nrp1+Plxna4) | SEMA3 | Secreted Signaling | PMID: 27533782 |
| NeuN | NeuN | Sema3c | NRP1_PLXNA4 | 0.004212 | 0 | SEMA3C_NRP1_PLXNA4 | Sema3c - (Nrp1+Plxna4) | SEMA3 | Secreted Signaling | PMID: 27533782 |
| CD31 | CD31 | Sema3d | NRP1_PLXNA1 | 0.001383 | 0 | SEMA3D_NRP1_PLXNA1 | Sema3d - (Nrp1+Plxna1) | SEMA3 | Secreted Signaling | PMID: 27533782 |
| CD31 | NeuN | Sema3d | NRP1_PLXNA1 | 0.001337 | 0 | SEMA3D_NRP1_PLXNA1 | Sema3d - (Nrp1+Plxna1) | SEMA3 | Secreted Signaling | PMID: 27533782 |
| CD31 | CD31 | Sema3d | NRP1_PLXNA2 | 0.001379 | 0 | SEMA3D_NRP1_PLXNA2 | Sema3d - (Nrp1+Plxna2) | SEMA3 | Secreted Signaling | PMID: 27533782 |
| NeuN | CD31 | Sema3d | NRP1_PLXNA2 | 0.001208 | 0 | SEMA3D_NRP1_PLXNA2 | Sema3d - (Nrp1+Plxna2) | SEMA3 | Secreted Signaling | PMID: 27533782 |
| CD31 | NeuN | Sema3d | NRP1_PLXNA3 | 0.001399 | 0 | SEMA3D_NRP1_PLXNA3 | Sema3d - (Nrp1+Plxna3) | SEMA3 | Secreted Signaling | PMID: 27533782 |
| NeuN | NeuN | Sema3d | NRP1_PLXNA3 | 0.001225 | 0 | SEMA3D_NRP1_PLXNA3 | Sema3d - (Nrp1+Plxna3) | SEMA3 | Secreted Signaling | PMID: 27533782 |
| CD31 | NeuN | Sema3d | NRP1_PLXNA4 | 0.002621 | 0 | SEMA3D_NRP1_PLXNA4 | Sema3d - (Nrp1+Plxna4) | SEMA3 | Secreted Signaling | PMID: 27533782 |

| | | | | | | | | | | |
|---|---|---|---|---|---|---|---|---|---|---|
| NeuN | NeuN | Sema3d | NRP1_PLXNA4 | 0.002297 | 0 | SEMA3D_NRP1_PLXNA4 | Sema3d - (Nrp1+Plxna4) | SEMA3 | Secreted Signaling | PMID: 27533782 |
| CD31 | CD31 | Sema3b | NRP2_PLXNA1 | 0.00083 | 0 | SEMA3B_NRP2_PLXNA1 | Sema3b - (Nrp2+Plxna1) | SEMA3 | Secreted Signaling | PMID: 27533782 |
| CD31 | NeuN | Sema3b | NRP2_PLXNA1 | 0.00081 | 0 | SEMA3B_NRP2_PLXNA1 | Sema3b - (Nrp2+Plxna1) | SEMA3 | Secreted Signaling | PMID: 27533782 |
| CD31 | CD31 | Sema3b | NRP2_PLXNA2 | 0.000828 | 0 | SEMA3B_NRP2_PLXNA2 | Sema3b - (Nrp2+Plxna2) | SEMA3 | Secreted Signaling | PMID: 27533782 |
| NeuN | CD31 | Sema3b | NRP2_PLXNA2 | 0.000729 | 0 | SEMA3B_NRP2_PLXNA2 | Sema3b - (Nrp2+Plxna2) | SEMA3 | Secreted Signaling | PMID: 27533782 |
| CD31 | NeuN | Sema3b | NRP2_PLXNA3 | 0.000848 | 0 | SEMA3B_NRP2_PLXNA3 | Sema3b - (Nrp2+Plxna3) | SEMA3 | Secreted Signaling | PMID: 27533782 |
| NeuN | NeuN | Sema3b | NRP2_PLXNA3 | 0.000746 | 0 | SEMA3B_NRP2_PLXNA3 | Sema3b - (Nrp2+Plxna3) | SEMA3 | Secreted Signaling | PMID: 27533782 |
| CD31 | NeuN | Sema3b | NRP2_PLXNA4 | 0.001589 | 0 | SEMA3B_NRP2_PLXNA4 | Sema3b - (Nrp2+Plxna4) | SEMA3 | Secreted Signaling | PMID: 27533782 |
| NeuN | NeuN | Sema3b | NRP2_PLXNA4 | 0.001399 | 0 | SEMA3B_NRP2_PLXNA4 | Sema3b - (Nrp2+Plxna4) | SEMA3 | Secreted Signaling | PMID: 27533782 |
| NeuN | CD31 | Sema3c | NRP2_PLXNA1 | 0.001458 | 0 | SEMA3C_NRP2_PLXNA1 | Sema3c - (Nrp2+Plxna1) | SEMA3 | Secreted Signaling | PMID: 27533782 |
| NeuN | NeuN | Sema3c | NRP2_PLXNA1 | 0.001423 | 0 | SEMA3C_NRP2_PLXNA1 | Sema3c - (Nrp2+Plxna1) | SEMA3 | Secreted Signaling | PMID: 27533782 |
| CD31 | CD31 | Sema3c | NRP2_PLXNA2 | 0.001291 | 0 | SEMA3C_NRP2_PLXNA2 | Sema3c - (Nrp2+Plxna2) | SEMA3 | Secreted Signaling | PMID: 27533782 |
| NeuN | CD31 | Sema3c | NRP2_PLXNA2 | 0.001453 | 0 | SEMA3C_NRP2_PLXNA2 | Sema3c - (Nrp2+Plxna2) | SEMA3 | Secreted Signaling | PMID: 27533782 |
| CD31 | NeuN | Sema3c | NRP2_PLXNA3 | 0.001322 | 0 | SEMA3C_NRP2_PLXNA3 | Sema3c - (Nrp2+Plxna3) | SEMA3 | Secreted Signaling | PMID: 27533782 |
| NeuN | NeuN | Sema3c | NRP2_PLXNA3 | 0.001488 | 0 | SEMA3C_NRP2_PLXNA3 | Sema3c - (Nrp2+Plxna3) | SEMA3 | Secreted Signaling | PMID: 27533782 |
| CD31 | NeuN | Sema3c | NRP2_PLXNA4 | 0.002477 | 0 | SEMA3C_NRP2_PLXNA4 | Sema3c - (Nrp2+Plxna4) | SEMA3 | Secreted Signaling | PMID: 27533782 |
| NeuN | NeuN | Sema3c | NRP2_PLXNA4 | 0.002789 | 0 | SEMA3C_NRP2_PLXNA4 | Sema3c - (Nrp2+Plxna4) | SEMA3 | Secreted Signaling | PMID: 27533782 |
| CD31 | CD31 | Sema3d | NRP2_PLXNA1 | 0.000906 | 0 | SEMA3D_NRP2_PLXNA1 | Sema3d - (Nrp2+Plxna1) | SEMA3 | Secreted Signaling | PMID: 27533782 |
| CD31 | NeuN | Sema3d | NRP2_PLXNA1 | 0.000884 | 0 | SEMA3D_NRP2_PLXNA1 | Sema3d - (Nrp2+Plxna1) | SEMA3 | Secreted Signaling | PMID: 27533782 |
| CD31 | CD31 | Sema3d | NRP2_PLXNA2 | 0.000903 | 0 | SEMA3D_NRP2_PLXNA2 | Sema3d - (Nrp2+Plxna2) | SEMA3 | Secreted Signaling | PMID: 27533782 |
| CD31 | NeuN | Sema3d | NRP2_PLXNA2 | 0.000793 | 0 | SEMA3D_NRP2_PLXNA2 | Sema3d - (Nrp2+Plxna2) | SEMA3 | Secreted Signaling | PMID: 27533782 |
| CD31 | NeuN | Sema3d | NRP2_PLXNA3 | 0.000925 | 0 | SEMA3D_NRP2_PLXNA3 | Sema3d - (Nrp2+Plxna3) | SEMA3 | Secreted Signaling | PMID: 27533782 |
| NeuN | NeuN | Sema3d | NRP2_PLXNA3 | 0.000811 | 0 | SEMA3D_NRP2_PLXNA3 | Sema3d - (Nrp2+Plxna3) | SEMA3 | Secreted Signaling | PMID: 27533782 |
| CD31 | NeuN | Sema3d | NRP2_PLXNA4 | 0.001735 | 0 | SEMA3D_NRP2_PLXNA4 | Sema3d - (Nrp2+Plxna4) | SEMA3 | Secreted Signaling | PMID: 27533782 |
| NeuN | NeuN | Sema3d | NRP2_PLXNA4 | 0.00152 | 0 | SEMA3D_NRP2_PLXNA4 | Sema3d - (Nrp2+Plxna4) | SEMA3 | Secreted Signaling | PMID: 27533782 |
| NeuN | CD31 | Sema3f | NRP2_PLXNA1 | 0.005606 | 0 | SEMA3F_NRP2_PLXNA1 | Sema3f - (Nrp2+Plxna1) | SEMA3 | Secreted Signaling | PMID: 27533782 |
| NeuN | NeuN | Sema3f | NRP2_PLXNA1 | 0.005473 | 0 | SEMA3F_NRP2_PLXNA1 | Sema3f - (Nrp2+Plxna1) | SEMA3 | Secreted Signaling | PMID: 27533782 |
| NeuN | CD31 | Sema3f | NRP2_PLXNA2 | 0.005589 | 0 | SEMA3F_NRP2_PLXNA2 | Sema3f - (Nrp2+Plxna2) | SEMA3 | Secreted Signaling | PMID: 27533782 |
| NeuN | NeuN | Sema3f | NRP2_PLXNA2 | 0.004908 | 0 | SEMA3F_NRP2_PLXNA2 | Sema3f - (Nrp2+Plxna2) | SEMA3 | Secreted Signaling | PMID: 27533782 |
| NeuN | CD31 | Sema3f | NRP2_PLXNA3 | 0.00451 | 0 | SEMA3F_NRP2_PLXNA3 | Sema3f - (Nrp2+Plxna3) | SEMA3 | Secreted Signaling | PMID: 27533782 |
| NeuN | NeuN | Sema3f | NRP2_PLXNA3 | 0.005724 | 0 | SEMA3F_NRP2_PLXNA3 | Sema3f - (Nrp2+Plxna3) | SEMA3 | Secreted Signaling | PMID: 27533782 |
| NeuN | CD31 | Sema3f | NRP2_PLXNA4 | 0.00747 | 0 | SEMA3F_NRP2_PLXNA4 | Sema3f - (Nrp2+Plxna4) | SEMA3 | Secreted Signaling | PMID: 27533782 |
| NeuN | NeuN | Sema3f | NRP2_PLXNA4 | 0.010685 | 0 | SEMA3F_NRP2_PLXNA4 | Sema3f - (Nrp2+Plxna4) | SEMA3 | Secreted Signaling | PMID: 27533782 |
| CD31 | CD31 | Sema3g | NRP2_PLXNA1 | 0.00211 | 0 | SEMA3G_NRP2_PLXNA1 | Sema3g - (Nrp2+Plxna1) | SEMA3 | Secreted Signaling | PMID: 27533782 |
| CD31 | NeuN | Sema3g | NRP2_PLXNA1 | 0.00206 | 0 | SEMA3G_NRP2_PLXNA1 | Sema3g - (Nrp2+Plxna1) | SEMA3 | Secreted Signaling | PMID: 27533782 |
| CD31 | CD31 | Sema3g | NRP2_PLXNA2 | 0.002104 | 0 | SEMA3G_NRP2_PLXNA2 | Sema3g - (Nrp2+Plxna2) | SEMA3 | Secreted Signaling | PMID: 27533782 |
| CD31 | NeuN | Sema3g | NRP2_PLXNA2 | 0.001846 | 0 | SEMA3G_NRP2_PLXNA2 | Sema3g - (Nrp2+Plxna2) | SEMA3 | Secreted Signaling | PMID: 27533782 |
| CD31 | CD31 | Sema3g | NRP2_PLXNA3 | 0.001696 | 0 | SEMA3G_NRP2_PLXNA3 | Sema3g - (Nrp2+Plxna3) | SEMA3 | Secreted Signaling | PMID: 27533782 |
| CD31 | NeuN | Sema3g | NRP2_PLXNA3 | 0.002154 | 0 | SEMA3G_NRP2_PLXNA3 | Sema3g - (Nrp2+Plxna3) | SEMA3 | Secreted Signaling | PMID: 27533782 |
| CD31 | CD31 | Sema3g | NRP2_PLXNA4 | 0.002815 | 0 | SEMA3G_NRP2_PLXNA4 | Sema3g - (Nrp2+Plxna4) | SEMA3 | Secreted Signaling | PMID: 27533782 |
| CD31 | NeuN | Sema3g | NRP2_PLXNA4 | 0.004034 | 0 | SEMA3G_NRP2_PLXNA4 | Sema3g - (Nrp2+Plxna4) | SEMA3 | Secreted Signaling | PMID: 27533782 |
| CD31 | CD31 | Sema3c | NRP1_NRP2 | 0.002034 | 0 | SEMA3C_NRP1_NRP2 | Sema3c - (Nrp1+Nrp2) | SEMA3 | Secreted Signaling | PMID: 27533782 |
| NeuN | CD31 | Sema3c | NRP1_NRP2 | 0.00229 | 0 | SEMA3C_NRP1_NRP2 | Sema3c - (Nrp1+Nrp2) | SEMA3 | Secreted Signaling | PMID: 27533782 |
| CD31 | CD31 | Sema3c | Plxnd1 | 0.004319 | 0 | SEMA3C_PLXND1 | Sema3c - Plxnd1 | SEMA3 | Secreted Signaling | PMID: 27533782 |
| NeuN | CD31 | Sema3c | Plxnd1 | 0.004861 | 0 | SEMA3C_PLXND1 | Sema3c - Plxnd1 | SEMA3 | Secreted Signaling | PMID: 27533782 |
| CD31 | CD31 | Sema3e | Plxnd1 | 0.003657 | 0 | SEMA3E_PLXND1 | Sema3e - Plxnd1 | SEMA3 | Secreted Signaling | PMID: 22325954 |
| NeuN | CD31 | Sema3e | Plxnd1 | 0.002474 | 0 | SEMA3E_PLXND1 | Sema3e - Plxnd1 | SEMA3 | Secreted Signaling | PMID: 22325954 |
| CD31 | CD31 | Crh | Crhr1 | 0.000347 | 0 | CRH_CRHR1 | Crh - Crhr1 | CRH | Secreted Signaling | KEGG: mmu04080 |
| CD31 | NeuN | Crh | Crhr1 | 0.000317 | 0 | CRH_CRHR1 | Crh - Crhr1 | CRH | Secreted Signaling | KEGG: mmu04080 |
| CD31 | CD31 | Crh | Crhr2 | 0.000288 | 0 | CRH_CRHR2 | Crh - Crhr2 | CRH | Secreted Signaling | KEGG: mmu04080 |
| NeuN | CD31 | Crh | Crhr2 | 0.000233 | 0 | CRH_CRHR2 | Crh - Crhr2 | CRH | Secreted Signaling | KEGG: mmu04080 |
| NeuN | CD31 | Enho | Gpr19 | 0.002227 | 0 | ENHO_GPR19 | Enho - Gpr19 | ENHO | Secreted Signaling | PMID: 28476646 |
| NeuN | NeuN | Enho | Gpr19 | 0.002201 | 0 | ENHO_GPR19 | Enho - Gpr19 | ENHO | Secreted Signaling | PMID: 28476646 |
| CD31 | CD31 | Gas6 | Axl | 0.00112 | 0 | GAS6_AXL | Gas6 - Axl | GAS | Secreted Signaling | PMID: 27801848 |
| NeuN | CD31 | Gas6 | Axl | 0.000893 | 0 | GAS6_AXL | Gas6 - Axl | GAS | Secreted Signaling | PMID: 27801848 |
| CD31 | CD31 | Gas6 | Mertk | 0.000449 | 0 | GAS6_MERTK | Gas6 - Mertk | GAS | Secreted Signaling | PMID: 27801848 |
| NeuN | CD31 | Gas6 | Mertk | 0.000358 | 0 | GAS6_MERTK | Gas6 - Mertk | GAS | Secreted Signaling | PMID: 27801848 |
| CD31 | CD31 | Gas6 | Tyro3 | 0.000478 | 0 | GAS6_TYRO3 | Gas6 - Tyro3 | GAS | Secreted Signaling | PMID: 27801848 |
| CD31 | NeuN | Gas6 | Tyro3 | 0.000533 | 0 | GAS6_TYRO3 | Gas6 - Tyro3 | GAS | Secreted Signaling | PMID: 27801848 |
| CD31 | CD31 | Grn | Sort1 | 0.004985 | 0 | GRN_SORT1 | Grn - Sort1 | GRN | Secreted Signaling | PMID: 29555433 |

| | | | | | | | | | | |
|---|---|---|---|---|---|---|---|---|---|---|
| CD31 | NeuN | Grn | Sort1 | 0.006129 | 0 | GRN_SORT1 | Grn - Sort1 | GRN | Secreted Signaling | PMID: 29555433 |
| CD31 | CD31 | Insl5 | Rxfp4 | 0.000622 | 0 | INSL5_RXFP4 | Insl5 - Rxfp4 | RLN | Secreted Signaling | KEGG: mmu04080 |
| NeuN | CD31 | Insl5 | Rxfp4 | 0.00047 | 0 | INSL5_RXFP4 | Insl5 - Rxfp4 | RLN | Secreted Signaling | KEGG: mmu04080 |
| CD31 | CD31 | Insl5 | Rxfp3 | 0.000466 | 0 | INSL5_RXFP3 | Insl5 - Rxfp3 | RLN | Secreted Signaling | KEGG: mmu04080 |
| CD31 | NeuN | Insl5 | Rxfp3 | 0.000555 | 0 | INSL5_RXFP3 | Insl5 - Rxfp3 | RLN | Secreted Signaling | KEGG: mmu04080 |
| CD31 | CD31 | Insl5 | Rxfp2 | 0.000414 | 0 | INSL5_RXFP2 | Insl5 - Rxfp2 | RLN | Secreted Signaling | KEGG: mmu04080 |
| CD31 | NeuN | Insl5 | Rxfp2 | 0.000487 | 0 | INSL5_RXFP2 | Insl5 - Rxfp2 | RLN | Secreted Signaling | KEGG: mmu04080 |
| CD31 | CD31 | Insl5 | Rxfp1 | 0.00046 | 0 | INSL5_RXFP1 | Insl5 - Rxfp1 | RLN | Secreted Signaling | KEGG: mmu04080 |
| CD31 | NeuN | Insl5 | Rxfp1 | 0.00039 | 0 | INSL5_RXFP1 | Insl5 - Rxfp1 | RLN | Secreted Signaling | KEGG: mmu04080 |
| CD31 | CD31 | Lgals9 | Ighm | 0.001214 | 0 | LGALS9_IGHM | Lgals9 - Ighm | GALECTIN | Secreted Signaling | PMID: 30120235 |
| CD31 | NeuN | Lgals9 | Ighm | 0.001266 | 0 | LGALS9_IGHM | Lgals9 - Ighm | GALECTIN | Secreted Signaling | PMID: 30120235 |
| CD31 | CD31 | Pmch | Mertk | 0.000407 | 0 | PMCH_MERTK | Pmch - Mertk | PMCH | Secreted Signaling | PMID: 30501104 |
| CD31 | NeuN | Pmch | Mertk | 0.00025 | 0 | PMCH_MERTK | Pmch - Mertk | PMCH | Secreted Signaling | PMID: 30501104 |
| CD31 | CD31 | Pros1 | Axl | 0.00136 | 0 | PROS1_AXL | Pros1 - Axl | PROS | Secreted Signaling | PMID: 29531161 |
| NeuN | CD31 | Pros1 | Axl | 0.000396 | 0 | PROS1_AXL | Pros1 - Axl | PROS | Secreted Signaling | PMID: 29531161 |
| CD31 | CD31 | Pros1 | Tyro3 | 0.000581 | 0 | PROS1_TYRO3 | Pros1 - Tyro3 | PROS | Secreted Signaling | PMID: 30501104 |
| CD31 | NeuN | Pros1 | Tyro3 | 0.000647 | 0 | PROS1_TYRO3 | Pros1 - Tyro3 | PROS | Secreted Signaling | PMID: 30501104 |
| CD31 | CD31 | Psap | Gpr37l1 | 0.003615 | 0 | PSAP_GPR37L1 | Psap - Gpr37l1 | PSAP | Secreted Signaling | PMID: 23690594 |
| NeuN | CD31 | Psap | Gpr37l1 | 0.005553 | 0 | PSAP_GPR37L1 | Psap - Gpr37l1 | PSAP | Secreted Signaling | PMID: 23690594 |
| NeuN | CD31 | Psap | Gpr37 | 0.005206 | 0 | PSAP_GPR37 | Psap - Gpr37 | PSAP | Secreted Signaling | PMID: 23690594 |
| NeuN | NeuN | Psap | Gpr37 | 0.003418 | 0 | PSAP_GPR37 | Psap - Gpr37 | PSAP | Secreted Signaling | PMID: 23690594 |
| CD31 | CD31 | Qrfp | Qrfpr | 0.000826 | 0 | QRFP_QRFPR | Qrfp - Qrfpr | QRFP | Secreted Signaling | PMID: 25386115 |
| CD31 | NeuN | Qrfp | Qrfpr | 0.000598 | 0 | QRFP_QRFPR | Qrfp - Qrfpr | QRFP | Secreted Signaling | PMID: 25386115 |
| CD31 | CD31 | Rarres2 | Cmklr1 | 0.000319 | 0 | RARRES2_CMKLR1 | Rarres2 - Cmklr1 | CHEMERIN | Secreted Signaling | PMID: 24671882 |
| NeuN | CD31 | Rarres2 | Cmklr1 | 0.000287 | 0 | RARRES2_CMKLR1 | Rarres2 - Cmklr1 | CHEMERIN | Secreted Signaling | PMID: 24671882 |
| CD31 | CD31 | Rln3 | Rxfp3 | 0.000597 | 0 | RLN3_RXFP3 | Rln3 - Rxfp3 | RLN | Secreted Signaling | KEGG: mmu04080 |
| CD31 | NeuN | Rln3 | Rxfp3 | 0.00071 | 0 | RLN3_RXFP3 | Rln3 - Rxfp3 | RLN | Secreted Signaling | KEGG: mmu04080 |
| CD31 | CD31 | Rln3 | Rxfp1 | 0.000589 | 0 | RLN3_RXFP1 | Rln3 - Rxfp1 | RLN | Secreted Signaling | KEGG: mmu04080 |
| CD31 | NeuN | Rln3 | Rxfp1 | 0.000499 | 0 | RLN3_RXFP1 | Rln3 - Rxfp1 | RLN | Secreted Signaling | KEGG: mmu04080 |
| CD31 | CD31 | Rln3 | Rxfp4 | 0.000796 | 0 | RLN3_RXFP4 | Rln3 - Rxfp4 | RLN | Secreted Signaling | KEGG: mmu04080 |
| NeuN | CD31 | Rln3 | Rxfp4 | 0.000536 | 0 | RLN3_RXFP4 | Rln3 - Rxfp4 | RLN | Secreted Signaling | KEGG: mmu04080 |
| CD31 | CD31 | Rln3 | Rxfp2 | 0.000529 | 0 | RLN3_RXFP2 | Rln3 - Rxfp2 | RLN | Secreted Signaling | KEGG: mmu04080 |
| CD31 | NeuN | Rln3 | Rxfp2 | 0.000623 | 0 | RLN3_RXFP2 | Rln3 - Rxfp2 | RLN | Secreted Signaling | KEGG: mmu04080 |
| NeuN | CD31 | Slurp2 | Chrnb2 | 0.000498 | 0 | SLURP2_CHRNB2 | Slurp2 - Chrnb2 | SLURP | Secreted Signaling | KEGG: mmu04080 |
| NeuN | NeuN | Slurp2 | Chrnb2 | 0.000688 | 0 | SLURP2_CHRNB2 | Slurp2 - Chrnb2 | SLURP | Secreted Signaling | KEGG: mmu04080 |
| CD31 | CD31 | Slurp2 | Chrnb3 | 0.000275 | 0 | SLURP2_CHRNB3 | Slurp2 - Chrnb3 | SLURP | Secreted Signaling | KEGG: mmu04080 |
| NeuN | CD31 | Slurp2 | Chrnb3 | 0.00042 | 0 | SLURP2_CHRNB3 | Slurp2 - Chrnb3 | SLURP | Secreted Signaling | KEGG: mmu04080 |
| NeuN | CD31 | Slurp2 | Chrnb4 | 0.0003 | 0 | SLURP2_CHRNB4 | Slurp2 - Chrnb4 | SLURP | Secreted Signaling | KEGG: mmu04080 |
| NeuN | NeuN | Slurp2 | Chrnb4 | 0.000302 | 0 | SLURP2_CHRNB4 | Slurp2 - Chrnb4 | SLURP | Secreted Signaling | KEGG: mmu04080 |
| CD31 | CD31 | Slurp2 | Chrna1 | 0.000295 | 0 | SLURP2_CHRNA1 | Slurp2 - Chrna1 | SLURP | Secreted Signaling | KEGG: mmu04080 |
| NeuN | CD31 | Slurp2 | Chrna1 | 0.000451 | 0 | SLURP2_CHRNA1 | Slurp2 - Chrna1 | SLURP | Secreted Signaling | KEGG: mmu04080 |
| NeuN | CD31 | Slurp2 | Chrna4 | 0.000373 | 0 | SLURP2_CHRNA4 | Slurp2 - Chrna4 | SLURP | Secreted Signaling | KEGG: mmu04080 |
| NeuN | NeuN | Slurp2 | Chrna4 | 0.000425 | 0 | SLURP2_CHRNA4 | Slurp2 - Chrna4 | SLURP | Secreted Signaling | KEGG: mmu04080 |
| NeuN | CD31 | Slurp2 | Chrna5 | 0.000379 | 0 | SLURP2_CHRNA5 | Slurp2 - Chrna5 | SLURP | Secreted Signaling | KEGG: mmu04080 |
| NeuN | NeuN | Slurp2 | Chrna5 | 0.000264 | 0 | SLURP2_CHRNA5 | Slurp2 - Chrna5 | SLURP | Secreted Signaling | KEGG: mmu04080 |
| CD31 | NeuN | Slurp2 | Chrna7 | 0.000491 | 0 | SLURP2_CHRNA7 | Slurp2 - Chrna7 | SLURP | Secreted Signaling | KEGG: mmu04080 |
| NeuN | NeuN | Slurp2 | Chrna7 | 0.000749 | 0 | SLURP2_CHRNA7 | Slurp2 - Chrna7 | SLURP | Secreted Signaling | KEGG: mmu04080 |
| CD31 | CD31 | Slurp2 | Chrna9 | 0.000283 | 0 | SLURP2_CHRNA9 | Slurp2 - Chrna9 | SLURP | Secreted Signaling | KEGG: mmu04080 |
| NeuN | CD31 | Slurp2 | Chrna9 | 0.000431 | 0 | SLURP2_CHRNA9 | Slurp2 - Chrna9 | SLURP | Secreted Signaling | KEGG: mmu04080 |
| NeuN | CD31 | Slurp2 | Chrnd | 0.000281 | 0 | SLURP2_CHRND | Slurp2 - Chrnd | SLURP | Secreted Signaling | KEGG: mmu04080 |
| NeuN | NeuN | Slurp2 | Chrnd | 0.000254 | 0 | SLURP2_CHRND | Slurp2 - Chrnd | SLURP | Secreted Signaling | KEGG: mmu04080 |
| CD31 | CD31 | Ucn | Crhr1 | 0.001122 | 0 | UCN_CRHR1 | Ucn - Crhr1 | UCN | Secreted Signaling | KEGG: mmu04080 |
| NeuN | CD31 | Ucn | Crhr1 | 0.001073 | 0 | UCN_CRHR1 | Ucn - Crhr1 | UCN | Secreted Signaling | KEGG: mmu04080 |
| CD31 | CD31 | Ucn | Crhr2 | 0.000932 | 0 | UCN_CRHR2 | Ucn - Crhr2 | UCN | Secreted Signaling | KEGG: mmu04080 |
| NeuN | CD31 | Ucn | Crhr2 | 0.000891 | 0 | UCN_CRHR2 | Ucn - Crhr2 | UCN | Secreted Signaling | KEGG: mmu04080 |
| CD31 | CD31 | Klk6 | F2r | 0.001911 | 0 | KLK6_F2R | Klk6 - F2r | KLK | Secreted Signaling | PMID: 35102281 |
| NeuN | CD31 | Klk6 | F2r | 0.001396 | 0 | KLK6_F2R | Klk6 - F2r | KLK | Secreted Signaling | PMID: 35102281 |
| CD31 | CD31 | Klk8 | F2r | 0.001663 | 0 | KLK8_F2R | Klk8 - F2r | KLK | Secreted Signaling | PMID: 34552064 |
| NeuN | CD31 | Klk8 | F2r | 0.000981 | 0 | KLK8_F2R | Klk8 - F2r | KLK | Secreted Signaling | PMID: 34552064 |
| CD31 | CD31 | Adm | CALCR_RAMP1 | 0.001015 | 0 | ADM_CALCR_RAMP1 | Adm - (Calcr+Ramp1) | ADM | Secreted Signaling | PMID: 33462236 |

| | | | | | | | | | | |
|---|---|---|---|---|---|---|---|---|---|---|
| CD31 | NeuN | Adm | CALCR_RAMP1 | 0.000675 | 0 | ADM_CALCR_RAMP1 | Adm - (Calcr+Ramp1) | ADM | Secreted Signaling | PMID: 33462236 |
| CD31 | CD31 | Adm | CALCR_RAMP2 | 0.002324 | 0 | ADM_CALCR_RAMP2 | Adm - (Calcr+Ramp2) | ADM | Secreted Signaling | PMID: 33462236 |
| NeuN | CD31 | Adm | CALCR_RAMP2 | 0.000745 | 0 | ADM_CALCR_RAMP2 | Adm - (Calcr+Ramp2) | ADM | Secreted Signaling | PMID: 33462236 |
| CD31 | CD31 | Adm | CALCR_RAMP3 | 0.001022 | 0 | ADM_CALCR_RAMP3 | Adm - (Calcr+Ramp3) | ADM | Secreted Signaling | PMID: 33462236 |
| CD31 | NeuN | Adm | CALCR_RAMP3 | 0.000686 | 0 | ADM_CALCR_RAMP3 | Adm - (Calcr+Ramp3) | ADM | Secreted Signaling | PMID: 33462236 |
| CD31 | CD31 | Calca | CALCR_RAMP1 | 0.000719 | 0 | CALCA_CALCR_RAMP1 | Calca - (Calcr+Ramp1) | CALC | Secreted Signaling | PMID: 30741933;PMID: 18599553 |
| CD31 | NeuN | Calca | CALCR_RAMP1 | 0.000478 | 0 | CALCA_CALCR_RAMP1 | Calca - (Calcr+Ramp1) | CALC | Secreted Signaling | PMID: 30741933;PMID: 18599553 |
| CD31 | CD31 | Calca | CALCR_RAMP2 | 0.001647 | 0 | CALCA_CALCR_RAMP2 | Calca - (Calcr+Ramp2) | CALC | Secreted Signaling | PMID: 30741933;PMID: 18599553 |
| NeuN | CD31 | Calca | CALCR_RAMP2 | 0.000731 | 0 | CALCA_CALCR_RAMP2 | Calca - (Calcr+Ramp2) | CALC | Secreted Signaling | PMID: 30741933;PMID: 18599553 |
| CD31 | CD31 | Calca | CALCR_RAMP3 | 0.000724 | 0 | CALCA_CALCR_RAMP3 | Calca - (Calcr+Ramp3) | CALC | Secreted Signaling | PMID: 30741933;PMID: 18599553 |
| CD31 | NeuN | Calca | CALCR_RAMP3 | 0.000486 | 0 | CALCA_CALCR_RAMP3 | Calca - (Calcr+Ramp3) | CALC | Secreted Signaling | PMID: 30741933;PMID: 18599553 |
| CD31 | CD31 | Calcb | CALCR_RAMP1 | 0.000591 | 0 | CALCB_CALCR_RAMP1 | Calcb - (Calcr+Ramp1) | CALC | Secreted Signaling | PMID: 30741933;PMID: 18599553 |
| NeuN | CD31 | Calcb | CALCR_RAMP1 | 0.000527 | 0 | CALCB_CALCR_RAMP1 | Calcb - (Calcr+Ramp1) | CALC | Secreted Signaling | PMID: 30741933;PMID: 18599553 |
| CD31 | CD31 | Calcb | CALCR_RAMP2 | 0.001354 | 0 | CALCB_CALCR_RAMP2 | Calcb - (Calcr+Ramp2) | CALC | Secreted Signaling | PMID: 30741933;PMID: 18599553 |
| NeuN | CD31 | Calcb | CALCR_RAMP2 | 0.001208 | 0 | CALCB_CALCR_RAMP2 | Calcb - (Calcr+Ramp2) | CALC | Secreted Signaling | PMID: 30741933;PMID: 18599553 |
| CD31 | CD31 | Calcb | CALCR_RAMP3 | 0.000595 | 0 | CALCB_CALCR_RAMP3 | Calcb - (Calcr+Ramp3) | CALC | Secreted Signaling | PMID: 30741933;PMID: 18599553 |
| NeuN | CD31 | Calcb | CALCR_RAMP3 | 0.000531 | 0 | CALCB_CALCR_RAMP3 | Calcb - (Calcr+Ramp3) | CALC | Secreted Signaling | PMID: 30741933;PMID: 18599553 |
| CD31 | CD31 | Ctsg | Fpr1 | 0.000631 | 0 | CTSG_FPR1 | Ctsg - Fpr1 | CTSG | Secreted Signaling | PMID: 26516201 |
| NeuN | CD31 | Ctsg | Fpr1 | 0.00036 | 0 | CTSG_FPR1 | Ctsg - Fpr1 | CTSG | Secreted Signaling | PMID: 26516201 |
| CD31 | CD31 | Grp | Nmbr | 0.00079 | 0 | GRP_NMBR | Grp - Nmbr | GRP | Secreted Signaling | PMID:25209280;PMID:29133874 |
| NeuN | CD31 | Grp | Nmbr | 0.000623 | 0 | GRP_NMBR | Grp - Nmbr | GRP | Secreted Signaling | PMID:25209280;PMID:29133874 |
| CD31 | NeuN | Ptprs | Ntrk3 | 0.072439 | 0 | PTPRS_NTRK3 | Ptprs - Ntrk3 | PTPR | Secreted Signaling | PMID:21262467;PMID:25385546; |
| NeuN | NeuN | Ptprs | Ntrk3 | 0.131284 | 0 | PTPRS_NTRK3 | Ptprs - Ntrk3 | PTPR | Secreted Signaling | PMID:21262467;PMID:25385546; |
| NeuN | CD31 | Slitrk1 | Ptprd | 0.011552 | 0 | SLITRK1_PTPRD | Slitrk1 - Ptprd | SLITRK | Secreted Signaling | PMID: 30648269;PMID: 30822649 |
| NeuN | NeuN | Slitrk1 | Ptprd | 0.021008 | 0 | SLITRK1_PTPRD | Slitrk1 - Ptprd | SLITRK | Secreted Signaling | PMID: 30648269;PMID: 30822649 |
| NeuN | CD31 | Slitrk1 | Ptprs | 0.018535 | 0 | SLITRK1_PTPRS | Slitrk1 - Ptprs | SLITRK | Secreted Signaling | PMID: 30648269;PMID: 30822649 |
| NeuN | NeuN | Slitrk1 | Ptprs | 0.035257 | 0 | SLITRK1_PTPRS | Slitrk1 - Ptprs | SLITRK | Secreted Signaling | PMID: 30648269;PMID: 30822649 |
| CD31 | NeuN | Slitrk2 | Ptprd | 0.004848 | 0 | SLITRK2_PTPRD | Slitrk2 - Ptprd | SLITRK | Secreted Signaling | PMID: 30648269;PMID: 30822649 |
| NeuN | NeuN | Slitrk2 | Ptprd | 0.00652 | 0 | SLITRK2_PTPRD | Slitrk2 - Ptprd | SLITRK | Secreted Signaling | PMID: 30648269;PMID: 30822649 |
| CD31 | NeuN | Slitrk2 | Ptprs | 0.008229 | 0 | SLITRK2_PTPRS | Slitrk2 - Ptprs | SLITRK | Secreted Signaling | PMID: 30648269;PMID: 30822649 |
| NeuN | NeuN | Slitrk2 | Ptprs | 0.011053 | 0 | SLITRK2_PTPRS | Slitrk2 - Ptprs | SLITRK | Secreted Signaling | PMID: 30648269;PMID: 30822649 |
| CD31 | NeuN | Slitrk3 | Ptprd | 0.007335 | 0 | SLITRK3_PTPRD | Slitrk3 - Ptprd | SLITRK | Secreted Signaling | PMID: 30648269;PMID: 30822649 |
| NeuN | NeuN | Slitrk3 | Ptprd | 0.009165 | 0 | SLITRK3_PTPRD | Slitrk3 - Ptprd | SLITRK | Secreted Signaling | PMID: 30648269;PMID: 30822649 |
| CD31 | NeuN | Slitrk4 | Ptprs | 0.00864 | 0 | SLITRK4_PTPRS | Slitrk4 - Ptprs | SLITRK | Secreted Signaling | PMID: 30648269;PMID: 30822649 |
| NeuN | NeuN | Slitrk4 | Ptprs | 0.008454 | 0 | SLITRK4_PTPRS | Slitrk4 - Ptprs | SLITRK | Secreted Signaling | PMID: 30648269;PMID: 30822649 |
| CD31 | NeuN | Slitrk5 | Ptprs | 0.014054 | 0 | SLITRK5_PTPRS | Slitrk5 - Ptprs | SLITRK | Secreted Signaling | PMID: 30648269;PMID: 30822649 |
| NeuN | NeuN | Slitrk5 | Ptprs | 0.012399 | 0 | SLITRK5_PTPRS | Slitrk5 - Ptprs | SLITRK | Secreted Signaling | PMID: 30648269;PMID: 30822649 |
| CD31 | CD31 | Bdnf | Sort1 | 0.001209 | 0 | BDNF_SORT1 | Bdnf - Sort1 | NT | Secreted Signaling | uniprot |
| CD31 | NeuN | Bdnf | Sort1 | 0.001487 | 0 | BDNF_SORT1 | Bdnf - Sort1 | NT | Secreted Signaling | uniprot |
| CD31 | CD31 | Bdnf | Ngfr | 0.000551 | 0 | BDNF_NGFR | Bdnf - Ngfr | NT | Secreted Signaling | uniprot |
| NeuN | CD31 | Bdnf | Ngfr | 0.000427 | 0 | BDNF_NGFR | Bdnf - Ngfr | NT | Secreted Signaling | uniprot |
| CD31 | CD31 | Igfbp3 | Tmem219 | 0.004769 | 0 | IGFBP3_TMEM219 | Igfbp3 - Tmem219 | IGFBP | Secreted Signaling | PMID: 35115561;PMID:20353938 |
| CD31 | NeuN | Igfbp3 | Tmem219 | 0.002803 | 0 | IGFBP3_TMEM219 | Igfbp3 - Tmem219 | IGFBP | Secreted Signaling | PMID: 35115561;PMID:20353938 |
| CD31 | CD31 | Lgals9 | P4hb | 0.005514 | 0 | LGALS9_P4HB | Lgals9 - P4hb | GALECTIN | Secreted Signaling | PMID:21670307;uniprot |
| CD31 | NeuN | Lgals9 | P4hb | 0.006559 | 0 | LGALS9_P4HB | Lgals9 - P4hb | GALECTIN | Secreted Signaling | PMID:21670307;uniprot |
| CD31 | CD31 | Nts | Sort1 | 0.001831 | 0 | NTS_SORT1 | Nts - Sort1 | NTS | Secreted Signaling | PMID: 34038712 |
| CD31 | NeuN | Nts | Sort1 | 0.002253 | 0 | NTS_SORT1 | Nts - Sort1 | NTS | Secreted Signaling | PMID: 34038712 |
| NeuN | CD31 | Ntf3 | Ngfr | 0.001393 | 0 | NTF3_NGFR | Ntf3 - Ngfr | NT | Secreted Signaling | uniprot |
| NeuN | NeuN | Ntf3 | Ngfr | 0.000842 | 0 | NTF3_NGFR | Ntf3 - Ngfr | NT | Secreted Signaling | uniprot |
| CD31 | CD31 | Rbp4 | Stra6 | 0.000502 | 0 | RBP4_STRA6 | Rbp4 - Stra6 | RBP4 | Secreted Signaling | PMID: 33199363 |
| NeuN | CD31 | Rbp4 | Stra6 | 0.000337 | 0 | RBP4_STRA6 | Rbp4 - Stra6 | RBP4 | Secreted Signaling | PMID: 33199363 |
| CD31 | NeuN | Slit1 | Robo1 | 0.006158 | 0 | SLIT1_ROBO1 | Slit1 - Robo1 | SLIT | Secreted Signaling | PMID:10102268 |
| NeuN | NeuN | Slit1 | Robo1 | 0.008016 | 0 | SLIT1_ROBO1 | Slit1 - Robo1 | SLIT | Secreted Signaling | PMID:10102268 |
| CD31 | NeuN | Slit2 | Robo1 | 0.003536 | 0 | SLIT2_ROBO1 | Slit2 - Robo1 | SLIT | Secreted Signaling | PMID:10102268 |
| NeuN | NeuN | Slit2 | Robo1 | 0.002395 | 0 | SLIT2_ROBO1 | Slit2 - Robo1 | SLIT | Secreted Signaling | PMID:10102268 |
| CD31 | NeuN | Slit2 | Robo2 | 0.004311 | 0 | SLIT2_ROBO2 | Slit2 - Robo2 | SLIT | Secreted Signaling | PMID:10102268 |
| NeuN | NeuN | Slit2 | Robo2 | 0.002921 | 0 | SLIT2_ROBO2 | Slit2 - Robo2 | SLIT | Secreted Signaling | PMID:10102268 |
| NeuN | CD31 | Tub | Mertk | 0.001332 | 0 | TUB_MERTK | Tub - Mertk | TULP | Secreted Signaling | PMID: 20978475 |
| NeuN | NeuN | Tub | Mertk | 0.000817 | 0 | TUB_MERTK | Tub - Mertk | TULP | Secreted Signaling | PMID: 20978475 |
| CD31 | CD31 | Pros1 | Mertk | 0.000546 | 0 | PROS1_MERTK | Pros1 - Mertk | PROS | Secreted Signaling | PMID: 34631419 |

| | | | | | | | | | | |
|---|---|---|---|---|---|---|---|---|---|---|
| CD31 | NeuN | Pros1 | Mertk | 0.000335 | 0 | PROS1_MERTK | Pros1 - Mertk | PROS | Secreted Signaling | PMID: 34631419 |
| CD31 | CD31 | Col1a1 | ITGA1_ITGB1 | 0.001972 | 0 | COL1A1_ITGA1_ITGB1 | Col1a1 - (Itga1+Itgb1) | COLLAGEN | ECM-Receptor | KEGG: mmu04512 |
| NeuN | CD31 | Col1a1 | ITGA1_ITGB1 | 0.001262 | 0 | COL1A1_ITGA1_ITGB1 | Col1a1 - (Itga1+Itgb1) | COLLAGEN | ECM-Receptor | KEGG: mmu04512 |
| CD31 | CD31 | Col1a2 | ITGA1_ITGB1 | 0.004185 | 0 | COL1A2_ITGA1_ITGB1 | Col1a2 - (Itga1+Itgb1) | COLLAGEN | ECM-Receptor | KEGG: mmu04512 |
| NeuN | CD31 | Col1a2 | ITGA1_ITGB1 | 0.001798 | 0 | COL1A2_ITGA1_ITGB1 | Col1a2 - (Itga1+Itgb1) | COLLAGEN | ECM-Receptor | KEGG: mmu04512 |
| CD31 | CD31 | Col4a1 | ITGA1_ITGB1 | 0.056358 | 0 | COL4A1_ITGA1_ITGB1 | Col4a1 - (Itga1+Itgb1) | COLLAGEN | ECM-Receptor | KEGG: mmu04512 |
| CD31 | NeuN | Col4a1 | ITGA1_ITGB1 | 0.016811 | 0 | COL4A1_ITGA1_ITGB1 | Col4a1 - (Itga1+Itgb1) | COLLAGEN | ECM-Receptor | KEGG: mmu04512 |
| CD31 | CD31 | Col4a2 | ITGA1_ITGB1 | 0.048701 | 0 | COL4A2_ITGA1_ITGB1 | Col4a2 - (Itga1+Itgb1) | COLLAGEN | ECM-Receptor | KEGG: mmu04512 |
| CD31 | NeuN | Col4a2 | ITGA1_ITGB1 | 0.014444 | 0 | COL4A2_ITGA1_ITGB1 | Col4a2 - (Itga1+Itgb1) | COLLAGEN | ECM-Receptor | KEGG: mmu04512 |
| CD31 | CD31 | Col6a1 | ITGA1_ITGB1 | 0.002107 | 0 | COL6A1_ITGA1_ITGB1 | Col6a1 - (Itga1+Itgb1) | COLLAGEN | ECM-Receptor | KEGG: mmu04512 |
| NeuN | CD31 | Col6a1 | ITGA1_ITGB1 | 0.00155 | 0 | COL6A1_ITGA1_ITGB1 | Col6a1 - (Itga1+Itgb1) | COLLAGEN | ECM-Receptor | KEGG: mmu04512 |
| CD31 | CD31 | Col6a2 | ITGA1_ITGB1 | 0.002095 | 0 | COL6A2_ITGA1_ITGB1 | Col6a2 - (Itga1+Itgb1) | COLLAGEN | ECM-Receptor | KEGG: mmu04512 |
| NeuN | CD31 | Col6a2 | ITGA1_ITGB1 | 0.001451 | 0 | COL6A2_ITGA1_ITGB1 | Col6a2 - (Itga1+Itgb1) | COLLAGEN | ECM-Receptor | KEGG: mmu04512 |
| CD31 | CD31 | Col6a4 | ITGA1_ITGB1 | 0.002012 | 0 | COL6A4_ITGA1_ITGB1 | Col6a4 - (Itga1+Itgb1) | COLLAGEN | ECM-Receptor | KEGG: mmu04512 |
| NeuN | CD31 | Col6a4 | ITGA1_ITGB1 | 0.00166 | 0 | COL6A4_ITGA1_ITGB1 | Col6a4 - (Itga1+Itgb1) | COLLAGEN | ECM-Receptor | KEGG: mmu04512 |
| CD31 | CD31 | Col6a6 | ITGA1_ITGB1 | 0.002511 | 0 | COL6A6_ITGA1_ITGB1 | Col6a6 - (Itga1+Itgb1) | COLLAGEN | ECM-Receptor | KEGG: mmu04512 |
| NeuN | CD31 | Col6a6 | ITGA1_ITGB1 | 0.001543 | 0 | COL6A6_ITGA1_ITGB1 | Col6a6 - (Itga1+Itgb1) | COLLAGEN | ECM-Receptor | KEGG: mmu04512 |
| CD31 | CD31 | Col9a2 | ITGA1_ITGB1 | 0.003213 | 0 | COL9A2_ITGA1_ITGB1 | Col9a2 - (Itga1+Itgb1) | COLLAGEN | ECM-Receptor | KEGG: mmu04512 |
| NeuN | CD31 | Col9a2 | ITGA1_ITGB1 | 0.002431 | 0 | COL9A2_ITGA1_ITGB1 | Col9a2 - (Itga1+Itgb1) | COLLAGEN | ECM-Receptor | KEGG: mmu04512 |
| CD31 | CD31 | Col9a3 | ITGA1_ITGB1 | 0.001895 | 0 | COL9A3_ITGA1_ITGB1 | Col9a3 - (Itga1+Itgb1) | COLLAGEN | ECM-Receptor | KEGG: mmu04512 |
| NeuN | CD31 | Col9a3 | ITGA1_ITGB1 | 0.005714 | 0 | COL9A3_ITGA1_ITGB1 | Col9a3 - (Itga1+Itgb1) | COLLAGEN | ECM-Receptor | KEGG: mmu04512 |
| CD31 | CD31 | Fn1 | ITGA3_ITGB1 | 0.008263 | 0 | FN1_ITGA3_ITGB1 | Fn1 - (Itga3+Itgb1) | FN1 | ECM-Receptor | KEGG: mmu04512 |
| CD31 | NeuN | Fn1 | ITGA3_ITGB1 | 0.004156 | 0 | FN1_ITGA3_ITGB1 | Fn1 - (Itga3+Itgb1) | FN1 | ECM-Receptor | KEGG: mmu04512 |
| CD31 | CD31 | Fn1 | ITGA4_ITGB1 | 0.01119 | 0 | FN1_ITGA4_ITGB1 | Fn1 - (Itga4+Itgb1) | FN1 | ECM-Receptor | KEGG: mmu04512 |
| CD31 | NeuN | Fn1 | ITGA4_ITGB1 | 0.003008 | 0 | FN1_ITGA4_ITGB1 | Fn1 - (Itga4+Itgb1) | FN1 | ECM-Receptor | KEGG: mmu04512 |
| CD31 | CD31 | Fn1 | ITGA5_ITGB1 | 0.0106 | 0 | FN1_ITGA5_ITGB1 | Fn1 - (Itga5+Itgb1) | FN1 | ECM-Receptor | KEGG: mmu04512 |
| CD31 | NeuN | Fn1 | ITGA5_ITGB1 | 0.002966 | 0 | FN1_ITGA5_ITGB1 | Fn1 - (Itga5+Itgb1) | FN1 | ECM-Receptor | KEGG: mmu04512 |
| CD31 | CD31 | Fn1 | ITGA8_ITGB1 | 0.005168 | 0 | FN1_ITGA8_ITGB1 | Fn1 - (Itga8+Itgb1) | FN1 | ECM-Receptor | KEGG: mmu04512 |
| CD31 | NeuN | Fn1 | ITGA8_ITGB1 | 0.003338 | 0 | FN1_ITGA8_ITGB1 | Fn1 - (Itga8+Itgb1) | FN1 | ECM-Receptor | KEGG: mmu04512 |
| CD31 | CD31 | Lama1 | ITGA1_ITGB1 | 0.003296 | 0 | LAMA1_ITGA1_ITGB1 | Lama1 - (Itga1+Itgb1) | LAMININ | ECM-Receptor | KEGG: mmu04512 |
| NeuN | CD31 | Lama1 | ITGA1_ITGB1 | 0.002211 | 0 | LAMA1_ITGA1_ITGB1 | Lama1 - (Itga1+Itgb1) | LAMININ | ECM-Receptor | KEGG: mmu04512 |
| CD31 | CD31 | Lama4 | ITGA1_ITGB1 | 0.015901 | 0 | LAMA4_ITGA1_ITGB1 | Lama4 - (Itga1+Itgb1) | LAMININ | ECM-Receptor | KEGG: mmu04512 |
| CD31 | NeuN | Lama4 | ITGA1_ITGB1 | 0.004605 | 0 | LAMA4_ITGA1_ITGB1 | Lama4 - (Itga1+Itgb1) | LAMININ | ECM-Receptor | KEGG: mmu04512 |
| CD31 | CD31 | Lama5 | ITGA1_ITGB1 | 0.003053 | 0 | LAMA5_ITGA1_ITGB1 | Lama5 - (Itga1+Itgb1) | LAMININ | ECM-Receptor | KEGG: mmu04512 |
| NeuN | CD31 | Lama5 | ITGA1_ITGB1 | 0.001753 | 0 | LAMA5_ITGA1_ITGB1 | Lama5 - (Itga1+Itgb1) | LAMININ | ECM-Receptor | KEGG: mmu04512 |
| CD31 | CD31 | Lamb1 | ITGA1_ITGB1 | 0.021203 | 0 | LAMB1_ITGA1_ITGB1 | Lamb1 - (Itga1+Itgb1) | LAMININ | ECM-Receptor | KEGG: mmu04512 |
| CD31 | NeuN | Lamb1 | ITGA1_ITGB1 | 0.006164 | 0 | LAMB1_ITGA1_ITGB1 | Lamb1 - (Itga1+Itgb1) | LAMININ | ECM-Receptor | KEGG: mmu04512 |
| CD31 | CD31 | Lamb2 | ITGA1_ITGB1 | 0.0072 | 0 | LAMB2_ITGA1_ITGB1 | Lamb2 - (Itga1+Itgb1) | LAMININ | ECM-Receptor | KEGG: mmu04512 |
| CD31 | NeuN | Lamb2 | ITGA1_ITGB1 | 0.002072 | 0 | LAMB2_ITGA1_ITGB1 | Lamb2 - (Itga1+Itgb1) | LAMININ | ECM-Receptor | KEGG: mmu04512 |
| CD31 | CD31 | Lamc1 | ITGA1_ITGB1 | 0.00732 | 0 | LAMC1_ITGA1_ITGB1 | Lamc1 - (Itga1+Itgb1) | LAMININ | ECM-Receptor | KEGG: mmu04512 |
| NeuN | CD31 | Lamc1 | ITGA1_ITGB1 | 0.0022 | 0 | LAMC1_ITGA1_ITGB1 | Lamc1 - (Itga1+Itgb1) | LAMININ | ECM-Receptor | KEGG: mmu04512 |
| CD31 | CD31 | Lamc2 | ITGA1_ITGB1 | 0.002456 | 0 | LAMC2_ITGA1_ITGB1 | Lamc2 - (Itga1+Itgb1) | LAMININ | ECM-Receptor | KEGG: mmu04512 |
| NeuN | CD31 | Lamc2 | ITGA1_ITGB1 | 0.001295 | 0 | LAMC2_ITGA1_ITGB1 | Lamc2 - (Itga1+Itgb1) | LAMININ | ECM-Receptor | KEGG: mmu04512 |
| CD31 | CD31 | Lamc3 | ITGA1_ITGB1 | 0.00374 | 0 | LAMC3_ITGA1_ITGB1 | Lamc3 - (Itga1+Itgb1) | LAMININ | ECM-Receptor | KEGG: mmu04512 |
| NeuN | CD31 | Lamc3 | ITGA1_ITGB1 | 0.001521 | 0 | LAMC3_ITGA1_ITGB1 | Lamc3 - (Itga1+Itgb1) | LAMININ | ECM-Receptor | KEGG: mmu04512 |
| CD31 | CD31 | Lama1 | ITGA3_ITGB1 | 0.002175 | 0 | LAMA1_ITGA3_ITGB1 | Lama1 - (Itga3+Itgb1) | LAMININ | ECM-Receptor | KEGG: mmu04512 |
| NeuN | CD31 | Lama1 | ITGA3_ITGB1 | 0.001459 | 0 | LAMA1_ITGA3_ITGB1 | Lama1 - (Itga3+Itgb1) | LAMININ | ECM-Receptor | KEGG: mmu04512 |
| CD31 | CD31 | Lama4 | ITGA3_ITGB1 | 0.010541 | 0 | LAMA4_ITGA3_ITGB1 | Lama4 - (Itga3+Itgb1) | LAMININ | ECM-Receptor | KEGG: mmu04512 |
| CD31 | NeuN | Lama4 | ITGA3_ITGB1 | 0.005308 | 0 | LAMA4_ITGA3_ITGB1 | Lama4 - (Itga3+Itgb1) | LAMININ | ECM-Receptor | KEGG: mmu04512 |
| CD31 | CD31 | Lama5 | ITGA3_ITGB1 | 0.002015 | 0 | LAMA5_ITGA3_ITGB1 | Lama5 - (Itga3+Itgb1) | LAMININ | ECM-Receptor | KEGG: mmu04512 |
| NeuN | CD31 | Lama5 | ITGA3_ITGB1 | 0.001156 | 0 | LAMA5_ITGA3_ITGB1 | Lama5 - (Itga3+Itgb1) | LAMININ | ECM-Receptor | KEGG: mmu04512 |
| CD31 | CD31 | Lamb1 | ITGA3_ITGB1 | 0.014081 | 0 | LAMB1_ITGA3_ITGB1 | Lamb1 - (Itga3+Itgb1) | LAMININ | ECM-Receptor | KEGG: mmu04512 |
| CD31 | NeuN | Lamb1 | ITGA3_ITGB1 | 0.007103 | 0 | LAMB1_ITGA3_ITGB1 | Lamb1 - (Itga3+Itgb1) | LAMININ | ECM-Receptor | KEGG: mmu04512 |
| CD31 | CD31 | Lamb2 | ITGA3_ITGB1 | 0.004759 | 0 | LAMB2_ITGA3_ITGB1 | Lamb2 - (Itga3+Itgb1) | LAMININ | ECM-Receptor | KEGG: mmu04512 |
| CD31 | NeuN | Lamb2 | ITGA3_ITGB1 | 0.002389 | 0 | LAMB2_ITGA3_ITGB1 | Lamb2 - (Itga3+Itgb1) | LAMININ | ECM-Receptor | KEGG: mmu04512 |
| CD31 | CD31 | Lamc1 | ITGA3_ITGB1 | 0.004838 | 0 | LAMC1_ITGA3_ITGB1 | Lamc1 - (Itga3+Itgb1) | LAMININ | ECM-Receptor | KEGG: mmu04512 |
| CD31 | NeuN | Lamc1 | ITGA3_ITGB1 | 0.002429 | 0 | LAMC1_ITGA3_ITGB1 | Lamc1 - (Itga3+Itgb1) | LAMININ | ECM-Receptor | KEGG: mmu04512 |
| CD31 | CD31 | Lamc2 | ITGA3_ITGB1 | 0.00162 | 0 | LAMC2_ITGA3_ITGB1 | Lamc2 - (Itga3+Itgb1) | LAMININ | ECM-Receptor | KEGG: mmu04512 |
| NeuN | CD31 | Lamc2 | ITGA3_ITGB1 | 0.000854 | 0 | LAMC2_ITGA3_ITGB1 | Lamc2 - (Itga3+Itgb1) | LAMININ | ECM-Receptor | KEGG: mmu04512 |
| CD31 | CD31 | Lamc3 | ITGA3_ITGB1 | 0.002469 | 0 | LAMC3_ITGA3_ITGB1 | Lamc3 - (Itga3+Itgb1) | LAMININ | ECM-Receptor | KEGG: mmu04512 |

| | | | | | | | | | | |
|---|---|---|---|---|---|---|---|---|---|---|
| CD31 | NeuN | Lamc3 | ITGA3_ITGB1 | 0.001238 | 0 | LAMC3_ITGA3_ITGB1 | Lamc3 - (Itga3+Itgb1) | LAMININ | ECM-Receptor | KEGG: mmu04512 |
| CD31 | CD31 | Reln | ITGA3_ITGB1 | 0.002372 | 0 | RELN_ITGA3_ITGB1 | Reln - (Itga3+Itgb1) | RELN | ECM-Receptor | KEGG: mmu04512 |
| NeuN | CD31 | Reln | ITGA3_ITGB1 | 0.002939 | 0 | RELN_ITGA3_ITGB1 | Reln - (Itga3+Itgb1) | RELN | ECM-Receptor | KEGG: mmu04512 |
| CD31 | CD31 | Thbs1 | ITGA3_ITGB1 | 0.002346 | 0 | THBS1_ITGA3_ITGB1 | Thbs1 - (Itga3+Itgb1) | THBS | ECM-Receptor | KEGG: mmu04512 |
| NeuN | CD31 | Thbs1 | ITGA3_ITGB1 | 0.002086 | 0 | THBS1_ITGA3_ITGB1 | Thbs1 - (Itga3+Itgb1) | THBS | ECM-Receptor | KEGG: mmu04512 |
| CD31 | CD31 | Thbs3 | ITGA3_ITGB1 | 0.001929 | 0 | THBS3_ITGA3_ITGB1 | Thbs3 - (Itga3+Itgb1) | THBS | ECM-Receptor | KEGG: mmu04512 |
| NeuN | CD31 | Thbs3 | ITGA3_ITGB1 | 0.003052 | 0 | THBS3_ITGA3_ITGB1 | Thbs3 - (Itga3+Itgb1) | THBS | ECM-Receptor | KEGG: mmu04512 |
| CD31 | CD31 | Thbs4 | ITGA3_ITGB1 | 0.00144 | 0 | THBS4_ITGA3_ITGB1 | Thbs4 - (Itga3+Itgb1) | THBS | ECM-Receptor | KEGG: mmu04512 |
| NeuN | CD31 | Thbs4 | ITGA3_ITGB1 | 0.001342 | 0 | THBS4_ITGA3_ITGB1 | Thbs4 - (Itga3+Itgb1) | THBS | ECM-Receptor | KEGG: mmu04512 |
| CD31 | CD31 | Col1a1 | ITGA3_ITGB1 | 0.001301 | 0 | COL1A1_ITGA3_ITGB1 | Col1a1 - (Itga3+Itgb1) | COLLAGEN | ECM-Receptor | KEGG: mmu04512 |
| NeuN | CD31 | Col1a1 | ITGA3_ITGB1 | 0.000832 | 0 | COL1A1_ITGA3_ITGB1 | Col1a1 - (Itga3+Itgb1) | COLLAGEN | ECM-Receptor | KEGG: mmu04512 |
| CD31 | CD31 | Col1a2 | ITGA3_ITGB1 | 0.002763 | 0 | COL1A2_ITGA3_ITGB1 | Col1a2 - (Itga3+Itgb1) | COLLAGEN | ECM-Receptor | KEGG: mmu04512 |
| CD31 | NeuN | Col1a2 | ITGA3_ITGB1 | 0.001386 | 0 | COL1A2_ITGA3_ITGB1 | Col1a2 - (Itga3+Itgb1) | COLLAGEN | ECM-Receptor | KEGG: mmu04512 |
| CD31 | CD31 | Col4a1 | ITGA3_ITGB1 | 0.037883 | 0 | COL4A1_ITGA3_ITGB1 | Col4a1 - (Itga3+Itgb1) | COLLAGEN | ECM-Receptor | KEGG: mmu04512 |
| CD31 | NeuN | Col4a1 | ITGA3_ITGB1 | 0.019343 | 0 | COL4A1_ITGA3_ITGB1 | Col4a1 - (Itga3+Itgb1) | COLLAGEN | ECM-Receptor | KEGG: mmu04512 |
| CD31 | CD31 | Col4a2 | ITGA3_ITGB1 | 0.032649 | 0 | COL4A2_ITGA3_ITGB1 | Col4a2 - (Itga3+Itgb1) | COLLAGEN | ECM-Receptor | KEGG: mmu04512 |
| CD31 | NeuN | Col4a2 | ITGA3_ITGB1 | 0.016626 | 0 | COL4A2_ITGA3_ITGB1 | Col4a2 - (Itga3+Itgb1) | COLLAGEN | ECM-Receptor | KEGG: mmu04512 |
| CD31 | CD31 | Col6a1 | ITGA3_ITGB1 | 0.00139 | 0 | COL6A1_ITGA3_ITGB1 | Col6a1 - (Itga3+Itgb1) | COLLAGEN | ECM-Receptor | KEGG: mmu04512 |
| NeuN | CD31 | Col6a1 | ITGA3_ITGB1 | 0.001022 | 0 | COL6A1_ITGA3_ITGB1 | Col6a1 - (Itga3+Itgb1) | COLLAGEN | ECM-Receptor | KEGG: mmu04512 |
| CD31 | CD31 | Col6a2 | ITGA3_ITGB1 | 0.001382 | 0 | COL6A2_ITGA3_ITGB1 | Col6a2 - (Itga3+Itgb1) | COLLAGEN | ECM-Receptor | KEGG: mmu04512 |
| NeuN | CD31 | Col6a2 | ITGA3_ITGB1 | 0.000957 | 0 | COL6A2_ITGA3_ITGB1 | Col6a2 - (Itga3+Itgb1) | COLLAGEN | ECM-Receptor | KEGG: mmu04512 |
| CD31 | CD31 | Col6a4 | ITGA3_ITGB1 | 0.001327 | 0 | COL6A4_ITGA3_ITGB1 | Col6a4 - (Itga3+Itgb1) | COLLAGEN | ECM-Receptor | KEGG: mmu04512 |
| NeuN | CD31 | Col6a4 | ITGA3_ITGB1 | 0.001095 | 0 | COL6A4_ITGA3_ITGB1 | Col6a4 - (Itga3+Itgb1) | COLLAGEN | ECM-Receptor | KEGG: mmu04512 |
| CD31 | CD31 | Col6a6 | ITGA3_ITGB1 | 0.001657 | 0 | COL6A6_ITGA3_ITGB1 | Col6a6 - (Itga3+Itgb1) | COLLAGEN | ECM-Receptor | KEGG: mmu04512 |
| NeuN | CD31 | Col6a6 | ITGA3_ITGB1 | 0.001018 | 0 | COL6A6_ITGA3_ITGB1 | Col6a6 - (Itga3+Itgb1) | COLLAGEN | ECM-Receptor | KEGG: mmu04512 |
| CD31 | CD31 | Col9a2 | ITGA3_ITGB1 | 0.002121 | 0 | COL9A2_ITGA3_ITGB1 | Col9a2 - (Itga3+Itgb1) | COLLAGEN | ECM-Receptor | KEGG: mmu04512 |
| NeuN | CD31 | Col9a2 | ITGA3_ITGB1 | 0.001604 | 0 | COL9A2_ITGA3_ITGB1 | Col9a2 - (Itga3+Itgb1) | COLLAGEN | ECM-Receptor | KEGG: mmu04512 |
| NeuN | CD31 | Col9a3 | ITGA3_ITGB1 | 0.003775 | 0 | COL9A3_ITGA3_ITGB1 | Col9a3 - (Itga3+Itgb1) | COLLAGEN | ECM-Receptor | KEGG: mmu04512 |
| NeuN | NeuN | Col9a3 | ITGA3_ITGB1 | 0.001894 | 0 | COL9A3_ITGA3_ITGB1 | Col9a3 - (Itga3+Itgb1) | COLLAGEN | ECM-Receptor | KEGG: mmu04512 |
| CD31 | CD31 | Lama1 | ITGA6_ITGB1 | 0.002106 | 0 | LAMA1_ITGA6_ITGB1 | Lama1 - (Itga6+Itgb1) | LAMININ | ECM-Receptor | KEGG: mmu04512 |
| NeuN | CD31 | Lama1 | ITGA6_ITGB1 | 0.001412 | 0 | LAMA1_ITGA6_ITGB1 | Lama1 - (Itga6+Itgb1) | LAMININ | ECM-Receptor | KEGG: mmu04512 |
| CD31 | CD31 | Lama4 | ITGA6_ITGB1 | 0.010208 | 0 | LAMA4_ITGA6_ITGB1 | Lama4 - (Itga6+Itgb1) | LAMININ | ECM-Receptor | KEGG: mmu04512 |
| CD31 | NeuN | Lama4 | ITGA6_ITGB1 | 0.005087 | 0 | LAMA4_ITGA6_ITGB1 | Lama4 - (Itga6+Itgb1) | LAMININ | ECM-Receptor | KEGG: mmu04512 |
| CD31 | CD31 | Lama5 | ITGA6_ITGB1 | 0.001951 | 0 | LAMA5_ITGA6_ITGB1 | Lama5 - (Itga6+Itgb1) | LAMININ | ECM-Receptor | KEGG: mmu04512 |
| NeuN | CD31 | Lama5 | ITGA6_ITGB1 | 0.001119 | 0 | LAMA5_ITGA6_ITGB1 | Lama5 - (Itga6+Itgb1) | LAMININ | ECM-Receptor | KEGG: mmu04512 |
| CD31 | CD31 | Lamb1 | ITGA6_ITGB1 | 0.013639 | 0 | LAMB1_ITGA6_ITGB1 | Lamb1 - (Itga6+Itgb1) | LAMININ | ECM-Receptor | KEGG: mmu04512 |
| CD31 | NeuN | Lamb1 | ITGA6_ITGB1 | 0.006808 | 0 | LAMB1_ITGA6_ITGB1 | Lamb1 - (Itga6+Itgb1) | LAMININ | ECM-Receptor | KEGG: mmu04512 |
| CD31 | CD31 | Lamb2 | ITGA6_ITGB1 | 0.004608 | 0 | LAMB2_ITGA6_ITGB1 | Lamb2 - (Itga6+Itgb1) | LAMININ | ECM-Receptor | KEGG: mmu04512 |
| CD31 | NeuN | Lamb2 | ITGA6_ITGB1 | 0.00229 | 0 | LAMB2_ITGA6_ITGB1 | Lamb2 - (Itga6+Itgb1) | LAMININ | ECM-Receptor | KEGG: mmu04512 |
| CD31 | CD31 | Lamc1 | ITGA6_ITGB1 | 0.004685 | 0 | LAMC1_ITGA6_ITGB1 | Lamc1 - (Itga6+Itgb1) | LAMININ | ECM-Receptor | KEGG: mmu04512 |
| CD31 | NeuN | Lamc1 | ITGA6_ITGB1 | 0.002328 | 0 | LAMC1_ITGA6_ITGB1 | Lamc1 - (Itga6+Itgb1) | LAMININ | ECM-Receptor | KEGG: mmu04512 |
| CD31 | CD31 | Lamc2 | ITGA6_ITGB1 | 0.001569 | 0 | LAMC2_ITGA6_ITGB1 | Lamc2 - (Itga6+Itgb1) | LAMININ | ECM-Receptor | KEGG: mmu04512 |
| NeuN | CD31 | Lamc2 | ITGA6_ITGB1 | 0.000827 | 0 | LAMC2_ITGA6_ITGB1 | Lamc2 - (Itga6+Itgb1) | LAMININ | ECM-Receptor | KEGG: mmu04512 |
| CD31 | CD31 | Lamc3 | ITGA6_ITGB1 | 0.002391 | 0 | LAMC3_ITGA6_ITGB1 | Lamc3 - (Itga6+Itgb1) | LAMININ | ECM-Receptor | KEGG: mmu04512 |
| CD31 | NeuN | Lamc3 | ITGA6_ITGB1 | 0.001187 | 0 | LAMC3_ITGA6_ITGB1 | Lamc3 - (Itga6+Itgb1) | LAMININ | ECM-Receptor | KEGG: mmu04512 |
| CD31 | CD31 | Lama1 | ITGA7_ITGB1 | 0.001858 | 0 | LAMA1_ITGA7_ITGB1 | Lama1 - (Itga7+Itgb1) | LAMININ | ECM-Receptor | KEGG: mmu04512 |
| NeuN | CD31 | Lama1 | ITGA7_ITGB1 | 0.001246 | 0 | LAMA1_ITGA7_ITGB1 | Lama1 - (Itga7+Itgb1) | LAMININ | ECM-Receptor | KEGG: mmu04512 |
| CD31 | CD31 | Lama4 | ITGA7_ITGB1 | 0.009015 | 0 | LAMA4_ITGA7_ITGB1 | Lama4 - (Itga7+Itgb1) | LAMININ | ECM-Receptor | KEGG: mmu04512 |
| CD31 | NeuN | Lama4 | ITGA7_ITGB1 | 0.004034 | 0 | LAMA4_ITGA7_ITGB1 | Lama4 - (Itga7+Itgb1) | LAMININ | ECM-Receptor | KEGG: mmu04512 |
| CD31 | CD31 | Lama5 | ITGA7_ITGB1 | 0.001721 | 0 | LAMA5_ITGA7_ITGB1 | Lama5 - (Itga7+Itgb1) | LAMININ | ECM-Receptor | KEGG: mmu04512 |
| NeuN | CD31 | Lama5 | ITGA7_ITGB1 | 0.000988 | 0 | LAMA5_ITGA7_ITGB1 | Lama5 - (Itga7+Itgb1) | LAMININ | ECM-Receptor | KEGG: mmu04512 |
| CD31 | CD31 | Lamb1 | ITGA7_ITGB1 | 0.012049 | 0 | LAMB1_ITGA7_ITGB1 | Lamb1 - (Itga7+Itgb1) | LAMININ | ECM-Receptor | KEGG: mmu04512 |
| CD31 | NeuN | Lamb1 | ITGA7_ITGB1 | 0.005401 | 0 | LAMB1_ITGA7_ITGB1 | Lamb1 - (Itga7+Itgb1) | LAMININ | ECM-Receptor | KEGG: mmu04512 |
| CD31 | CD31 | Lamb2 | ITGA7_ITGB1 | 0.004067 | 0 | LAMB2_ITGA7_ITGB1 | Lamb2 - (Itga7+Itgb1) | LAMININ | ECM-Receptor | KEGG: mmu04512 |
| CD31 | NeuN | Lamb2 | ITGA7_ITGB1 | 0.001815 | 0 | LAMB2_ITGA7_ITGB1 | Lamb2 - (Itga7+Itgb1) | LAMININ | ECM-Receptor | KEGG: mmu04512 |
| CD31 | CD31 | Lamc1 | ITGA7_ITGB1 | 0.004134 | 0 | LAMC1_ITGA7_ITGB1 | Lamc1 - (Itga7+Itgb1) | LAMININ | ECM-Receptor | KEGG: mmu04512 |
| CD31 | NeuN | Lamc1 | ITGA7_ITGB1 | 0.001845 | 0 | LAMC1_ITGA7_ITGB1 | Lamc1 - (Itga7+Itgb1) | LAMININ | ECM-Receptor | KEGG: mmu04512 |
| CD31 | CD31 | Lamc2 | ITGA7_ITGB1 | 0.001384 | 0 | LAMC2_ITGA7_ITGB1 | Lamc2 - (Itga7+Itgb1) | LAMININ | ECM-Receptor | KEGG: mmu04512 |
| NeuN | CD31 | Lamc2 | ITGA7_ITGB1 | 0.00073 | 0 | LAMC2_ITGA7_ITGB1 | Lamc2 - (Itga7+Itgb1) | LAMININ | ECM-Receptor | KEGG: mmu04512 |
| CD31 | CD31 | Lamc3 | ITGA7_ITGB1 | 0.002109 | 0 | LAMC3_ITGA7_ITGB1 | Lamc3 - (Itga7+Itgb1) | LAMININ | ECM-Receptor | KEGG: mmu04512 |

| | | | | | | | | | | |
|---|---|---|---|---|---|---|---|---|---|---|
| CD31 | NeuN | Lamc3 | ITGA7_ITGB1 | 0.00094 | 0 | LAMC3_ITGA7_ITGB1 | Lamc3 - (Itga7+Itgb1) | LAMININ | ECM-Receptor | KEGG: mmu04512 |
| CD31 | CD31 | Vtn | ITGA8_ITGB1 | 0.002621 | 0 | VTN_ITGA8_ITGB1 | Vtn - (Itga8+Itgb1) | VTN | ECM-Receptor | KEGG: mmu04512 |
| CD31 | NeuN | Vtn | ITGA8_ITGB1 | 0.001692 | 0 | VTN_ITGA8_ITGB1 | Vtn - (Itga8+Itgb1) | VTN | ECM-Receptor | KEGG: mmu04512 |
| CD31 | CD31 | Tnc | ITGA8_ITGB1 | 0.001182 | 0 | TNC_ITGA8_ITGB1 | Tnc - (Itga8+Itgb1) | TENASCIN | ECM-Receptor | KEGG: mmu04512 |
| CD31 | NeuN | Tnc | ITGA8_ITGB1 | 0.000762 | 0 | TNC_ITGA8_ITGB1 | Tnc - (Itga8+Itgb1) | TENASCIN | ECM-Receptor | KEGG: mmu04512 |
| CD31 | CD31 | Tnr | ITGA8_ITGB1 | 0.000784 | 0 | TNR_ITGA8_ITGB1 | Tnr - (Itga8+Itgb1) | TENASCIN | ECM-Receptor | KEGG: mmu04512 |
| NeuN | CD31 | Tnr | ITGA8_ITGB1 | 0.000942 | 0 | TNR_ITGA8_ITGB1 | Tnr - (Itga8+Itgb1) | TENASCIN | ECM-Receptor | KEGG: mmu04512 |
| CD31 | CD31 | Col1a1 | ITGA11_ITGB1 | 0.001138 | 0 | COL1A1_ITGA11_ITGB1 | Col1a1 - (Itga11+Itgb1) | COLLAGEN | ECM-Receptor | KEGG: mmu04512 |
| NeuN | CD31 | Col1a1 | ITGA11_ITGB1 | 0.000728 | 0 | COL1A1_ITGA11_ITGB1 | Col1a1 - (Itga11+Itgb1) | COLLAGEN | ECM-Receptor | KEGG: mmu04512 |
| CD31 | CD31 | Col1a2 | ITGA11_ITGB1 | 0.002416 | 0 | COL1A2_ITGA11_ITGB1 | Col1a2 - (Itga11+Itgb1) | COLLAGEN | ECM-Receptor | KEGG: mmu04512 |
| CD31 | NeuN | Col1a2 | ITGA11_ITGB1 | 0.001301 | 0 | COL1A2_ITGA11_ITGB1 | Col1a2 - (Itga11+Itgb1) | COLLAGEN | ECM-Receptor | KEGG: mmu04512 |
| CD31 | CD31 | Col4a1 | ITGA11_ITGB1 | 0.033276 | 0 | COL4A1_ITGA11_ITGB1 | Col4a1 - (Itga11+Itgb1) | COLLAGEN | ECM-Receptor | KEGG: mmu04512 |
| CD31 | NeuN | Col4a1 | ITGA11_ITGB1 | 0.018174 | 0 | COL4A1_ITGA11_ITGB1 | Col4a1 - (Itga11+Itgb1) | COLLAGEN | ECM-Receptor | KEGG: mmu04512 |
| CD31 | CD31 | Col4a2 | ITGA11_ITGB1 | 0.028659 | 0 | COL4A2_ITGA11_ITGB1 | Col4a2 - (Itga11+Itgb1) | COLLAGEN | ECM-Receptor | KEGG: mmu04512 |
| CD31 | NeuN | Col4a2 | ITGA11_ITGB1 | 0.015618 | 0 | COL4A2_ITGA11_ITGB1 | Col4a2 - (Itga11+Itgb1) | COLLAGEN | ECM-Receptor | KEGG: mmu04512 |
| CD31 | CD31 | Col6a1 | ITGA11_ITGB1 | 0.001216 | 0 | COL6A1_ITGA11_ITGB1 | Col6a1 - (Itga11+Itgb1) | COLLAGEN | ECM-Receptor | KEGG: mmu04512 |
| NeuN | CD31 | Col6a1 | ITGA11_ITGB1 | 0.000894 | 0 | COL6A1_ITGA11_ITGB1 | Col6a1 - (Itga11+Itgb1) | COLLAGEN | ECM-Receptor | KEGG: mmu04512 |
| CD31 | CD31 | Col6a2 | ITGA11_ITGB1 | 0.001209 | 0 | COL6A2_ITGA11_ITGB1 | Col6a2 - (Itga11+Itgb1) | COLLAGEN | ECM-Receptor | KEGG: mmu04512 |
| NeuN | CD31 | Col6a2 | ITGA11_ITGB1 | 0.000837 | 0 | COL6A2_ITGA11_ITGB1 | Col6a2 - (Itga11+Itgb1) | COLLAGEN | ECM-Receptor | KEGG: mmu04512 |
| CD31 | CD31 | Col6a4 | ITGA11_ITGB1 | 0.00116 | 0 | COL6A4_ITGA11_ITGB1 | Col6a4 - (Itga11+Itgb1) | COLLAGEN | ECM-Receptor | KEGG: mmu04512 |
| NeuN | CD31 | Col6a4 | ITGA11_ITGB1 | 0.000958 | 0 | COL6A4_ITGA11_ITGB1 | Col6a4 - (Itga11+Itgb1) | COLLAGEN | ECM-Receptor | KEGG: mmu04512 |
| CD31 | CD31 | Col6a6 | ITGA11_ITGB1 | 0.001449 | 0 | COL6A6_ITGA11_ITGB1 | Col6a6 - (Itga11+Itgb1) | COLLAGEN | ECM-Receptor | KEGG: mmu04512 |
| NeuN | CD31 | Col6a6 | ITGA11_ITGB1 | 0.00089 | 0 | COL6A6_ITGA11_ITGB1 | Col6a6 - (Itga11+Itgb1) | COLLAGEN | ECM-Receptor | KEGG: mmu04512 |
| CD31 | CD31 | Col9a2 | ITGA11_ITGB1 | 0.001854 | 0 | COL9A2_ITGA11_ITGB1 | Col9a2 - (Itga11+Itgb1) | COLLAGEN | ECM-Receptor | KEGG: mmu04512 |
| NeuN | CD31 | Col9a2 | ITGA11_ITGB1 | 0.001402 | 0 | COL9A2_ITGA11_ITGB1 | Col9a2 - (Itga11+Itgb1) | COLLAGEN | ECM-Receptor | KEGG: mmu04512 |
| NeuN | CD31 | Col9a3 | ITGA11_ITGB1 | 0.003301 | 0 | COL9A3_ITGA11_ITGB1 | Col9a3 - (Itga11+Itgb1) | COLLAGEN | ECM-Receptor | KEGG: mmu04512 |
| NeuN | NeuN | Col9a3 | ITGA11_ITGB1 | 0.001778 | 0 | COL9A3_ITGA11_ITGB1 | Col9a3 - (Itga11+Itgb1) | COLLAGEN | ECM-Receptor | KEGG: mmu04512 |
| CD31 | CD31 | Fn1 | ITGAV_ITGB1 | 0.00863 | 0 | FN1_ITGAV_ITGB1 | Fn1 - (Itgav+Itgb1) | FN1 | ECM-Receptor | KEGG: mmu04512 |
| CD31 | NeuN | Fn1 | ITGAV_ITGB1 | 0.003451 | 0 | FN1_ITGAV_ITGB1 | Fn1 - (Itgav+Itgb1) | FN1 | ECM-Receptor | KEGG: mmu04512 |
| CD31 | CD31 | Vtn | ITGAV_ITGB1 | 0.004385 | 0 | VTN_ITGAV_ITGB1 | Vtn - (Itgav+Itgb1) | VTN | ECM-Receptor | KEGG: mmu04512 |
| CD31 | NeuN | Vtn | ITGAV_ITGB1 | 0.001749 | 0 | VTN_ITGAV_ITGB1 | Vtn - (Itgav+Itgb1) | VTN | ECM-Receptor | KEGG: mmu04512 |
| CD31 | CD31 | Fn1 | ITGA4_ITGB7 | 0.004558 | 0 | FN1_ITGA4_ITGB7 | Fn1 - (Itga4+Itgb7) | FN1 | ECM-Receptor | KEGG: mmu04512 |
| CD31 | NeuN | Fn1 | ITGA4_ITGB7 | 0.002131 | 0 | FN1_ITGA4_ITGB7 | Fn1 - (Itga4+Itgb7) | FN1 | ECM-Receptor | KEGG: mmu04512 |
| CD31 | CD31 | Fn1 | ITGAV_ITGB3 | 0.00383 | 0 | FN1_ITGAV_ITGB3 | Fn1 - (Itgav+Itgb3) | FN1 | ECM-Receptor | KEGG: mmu04512 |
| CD31 | NeuN | Fn1 | ITGAV_ITGB3 | 0.002041 | 0 | FN1_ITGAV_ITGB3 | Fn1 - (Itgav+Itgb3) | FN1 | ECM-Receptor | KEGG: mmu04512 |
| CD31 | CD31 | Vtn | ITGAV_ITGB3 | 0.001941 | 0 | VTN_ITGAV_ITGB3 | Vtn - (Itgav+Itgb3) | VTN | ECM-Receptor | KEGG: mmu04512 |
| CD31 | NeuN | Vtn | ITGAV_ITGB3 | 0.001033 | 0 | VTN_ITGAV_ITGB3 | Vtn - (Itgav+Itgb3) | VTN | ECM-Receptor | KEGG: mmu04512 |
| CD31 | CD31 | Vwf | ITGAV_ITGB3 | 0.002989 | 0 | VWF_ITGAV_ITGB3 | Vwf - (Itgav+Itgb3) | VWF | ECM-Receptor | KEGG: mmu04512 |
| CD31 | NeuN | Vwf | ITGAV_ITGB3 | 0.001592 | 0 | VWF_ITGAV_ITGB3 | Vwf - (Itgav+Itgb3) | VWF | ECM-Receptor | KEGG: mmu04512 |
| CD31 | CD31 | Tnc | ITGAV_ITGB3 | 0.000875 | 0 | TNC_ITGAV_ITGB3 | Tnc - (Itgav+Itgb3) | TENASCIN | ECM-Receptor | KEGG: mmu04512 |
| CD31 | NeuN | Tnc | ITGAV_ITGB3 | 0.000466 | 0 | TNC_ITGAV_ITGB3 | Tnc - (Itgav+Itgb3) | TENASCIN | ECM-Receptor | KEGG: mmu04512 |
| CD31 | CD31 | Tnr | ITGAV_ITGB3 | 0.000581 | 0 | TNR_ITGAV_ITGB3 | Tnr - (Itgav+Itgb3) | TENASCIN | ECM-Receptor | KEGG: mmu04512 |
| NeuN | CD31 | Tnr | ITGAV_ITGB3 | 0.000698 | 0 | TNR_ITGAV_ITGB3 | Tnr - (Itgav+Itgb3) | TENASCIN | ECM-Receptor | KEGG: mmu04512 |
| CD31 | CD31 | Thbs1 | ITGAV_ITGB3 | 0.001084 | 0 | THBS1_ITGAV_ITGB3 | Thbs1 - (Itgav+Itgb3) | THBS | ECM-Receptor | KEGG: mmu04512 |
| NeuN | CD31 | Thbs1 | ITGAV_ITGB3 | 0.000964 | 0 | THBS1_ITGAV_ITGB3 | Thbs1 - (Itgav+Itgb3) | THBS | ECM-Receptor | KEGG: mmu04512 |
| CD31 | CD31 | Thbs3 | ITGAV_ITGB3 | 0.000891 | 0 | THBS3_ITGAV_ITGB3 | Thbs3 - (Itgav+Itgb3) | THBS | ECM-Receptor | KEGG: mmu04512 |
| NeuN | CD31 | Thbs3 | ITGAV_ITGB3 | 0.001411 | 0 | THBS3_ITGAV_ITGB3 | Thbs3 - (Itgav+Itgb3) | THBS | ECM-Receptor | KEGG: mmu04512 |
| CD31 | CD31 | Thbs4 | ITGAV_ITGB3 | 0.000665 | 0 | THBS4_ITGAV_ITGB3 | Thbs4 - (Itgav+Itgb3) | THBS | ECM-Receptor | KEGG: mmu04512 |
| NeuN | CD31 | Thbs4 | ITGAV_ITGB3 | 0.00062 | 0 | THBS4_ITGAV_ITGB3 | Thbs4 - (Itgav+Itgb3) | THBS | ECM-Receptor | KEGG: mmu04512 |
| CD31 | CD31 | Fn1 | ITGA2B_ITGB3 | 0.002504 | 0 | FN1_ITGA2B_ITGB3 | Fn1 - (Itga2b+Itgb3) | FN1 | ECM-Receptor | KEGG: mmu04512 |
| CD31 | NeuN | Fn1 | ITGA2B_ITGB3 | 0.001945 | 0 | FN1_ITGA2B_ITGB3 | Fn1 - (Itga2b+Itgb3) | FN1 | ECM-Receptor | KEGG: mmu04512 |
| CD31 | CD31 | Vtn | ITGA2B_ITGB3 | 0.001268 | 0 | VTN_ITGA2B_ITGB3 | Vtn - (Itga2b+Itgb3) | VTN | ECM-Receptor | KEGG: mmu04512 |
| CD31 | NeuN | Vtn | ITGA2B_ITGB3 | 0.000985 | 0 | VTN_ITGA2B_ITGB3 | Vtn - (Itga2b+Itgb3) | VTN | ECM-Receptor | KEGG: mmu04512 |
| CD31 | CD31 | Vwf | ITGA2B_ITGB3 | 0.001954 | 0 | VWF_ITGA2B_ITGB3 | Vwf - (Itga2b+Itgb3) | VWF | ECM-Receptor | KEGG: mmu04512 |
| CD31 | NeuN | Vwf | ITGA2B_ITGB3 | 0.001518 | 0 | VWF_ITGA2B_ITGB3 | Vwf - (Itga2b+Itgb3) | VWF | ECM-Receptor | KEGG: mmu04512 |
| CD31 | CD31 | Tnc | ITGA2B_ITGB3 | 0.000571 | 0 | TNC_ITGA2B_ITGB3 | Tnc - (Itga2b+Itgb3) | TENASCIN | ECM-Receptor | KEGG: mmu04512 |
| CD31 | NeuN | Tnc | ITGA2B_ITGB3 | 0.000444 | 0 | TNC_ITGA2B_ITGB3 | Tnc - (Itga2b+Itgb3) | TENASCIN | ECM-Receptor | KEGG: mmu04512 |
| CD31 | CD31 | Tnr | ITGA2B_ITGB3 | 0.000379 | 0 | TNR_ITGA2B_ITGB3 | Tnr - (Itga2b+Itgb3) | TENASCIN | ECM-Receptor | KEGG: mmu04512 |
| NeuN | CD31 | Tnr | ITGA2B_ITGB3 | 0.000456 | 0 | TNR_ITGA2B_ITGB3 | Tnr - (Itga2b+Itgb3) | TENASCIN | ECM-Receptor | KEGG: mmu04512 |
| CD31 | CD31 | Vtn | ITGAV_ITGB5 | 0.001729 | 0 | VTN_ITGAV_ITGB5 | Vtn - (Itgav+Itgb5) | VTN | ECM-Receptor | KEGG: mmu04512 |

| | | | | | | | | | | |
|---|---|---|---|---|---|---|---|---|---|---|
| CD31 | NeuN | Vtn | ITGAV_ITGB5 | 0.000878 | 0 | VTN_ITGAV_ITGB5 | Vtn - (Itgav+Itgb5) | VTN | ECM-Receptor | KEGG: mmu04512 |
| CD31 | CD31 | Vtn | ITGAV_ITGB8 | 0.002213 | 0 | VTN_ITGAV_ITGB8 | Vtn - (Itgav+Itgb8) | VTN | ECM-Receptor | KEGG: mmu04512 |
| CD31 | NeuN | Vtn | ITGAV_ITGB8 | 0.001318 | 0 | VTN_ITGAV_ITGB8 | Vtn - (Itgav+Itgb8) | VTN | ECM-Receptor | KEGG: mmu04512 |
| CD31 | CD31 | Fn1 | ITGAV_ITGB8 | 0.004364 | 0 | FN1_ITGAV_ITGB8 | Fn1 - (Itgav+Itgb8) | FN1 | ECM-Receptor | KEGG: mmu04512 |
| CD31 | NeuN | Fn1 | ITGAV_ITGB8 | 0.002601 | 0 | FN1_ITGAV_ITGB8 | Fn1 - (Itgav+Itgb8) | FN1 | ECM-Receptor | KEGG: mmu04512 |
| CD31 | CD31 | Lama1 | ITGAV_ITGB8 | 0.001145 | 0 | LAMA1_ITGAV_ITGB8 | Lama1 - (Itgav+Itgb8) | LAMININ | ECM-Receptor | KEGG: mmu04512 |
| NeuN | CD31 | Lama1 | ITGAV_ITGB8 | 0.000768 | 0 | LAMA1_ITGAV_ITGB8 | Lama1 - (Itgav+Itgb8) | LAMININ | ECM-Receptor | KEGG: mmu04512 |
| CD31 | CD31 | Lama4 | ITGAV_ITGB8 | 0.005572 | 0 | LAMA4_ITGAV_ITGB8 | Lama4 - (Itgav+Itgb8) | LAMININ | ECM-Receptor | KEGG: mmu04512 |
| CD31 | NeuN | Lama4 | ITGAV_ITGB8 | 0.003323 | 0 | LAMA4_ITGAV_ITGB8 | Lama4 - (Itgav+Itgb8) | LAMININ | ECM-Receptor | KEGG: mmu04512 |
| CD31 | CD31 | Lama5 | ITGAV_ITGB8 | 0.001061 | 0 | LAMA5_ITGAV_ITGB8 | Lama5 - (Itgav+Itgb8) | LAMININ | ECM-Receptor | KEGG: mmu04512 |
| CD31 | NeuN | Lama5 | ITGAV_ITGB8 | 0.000632 | 0 | LAMA5_ITGAV_ITGB8 | Lama5 - (Itgav+Itgb8) | LAMININ | ECM-Receptor | KEGG: mmu04512 |
| CD31 | CD31 | Lamb1 | ITGAV_ITGB8 | 0.007457 | 0 | LAMB1_ITGAV_ITGB8 | Lamb1 - (Itgav+Itgb8) | LAMININ | ECM-Receptor | KEGG: mmu04512 |
| CD31 | NeuN | Lamb1 | ITGAV_ITGB8 | 0.00445 | 0 | LAMB1_ITGAV_ITGB8 | Lamb1 - (Itgav+Itgb8) | LAMININ | ECM-Receptor | KEGG: mmu04512 |
| CD31 | CD31 | Lamb2 | ITGAV_ITGB8 | 0.002509 | 0 | LAMB2_ITGAV_ITGB8 | Lamb2 - (Itgav+Itgb8) | LAMININ | ECM-Receptor | KEGG: mmu04512 |
| CD31 | NeuN | Lamb2 | ITGAV_ITGB8 | 0.001494 | 0 | LAMB2_ITGAV_ITGB8 | Lamb2 - (Itgav+Itgb8) | LAMININ | ECM-Receptor | KEGG: mmu04512 |
| CD31 | CD31 | Lamc1 | ITGAV_ITGB8 | 0.002551 | 0 | LAMC1_ITGAV_ITGB8 | Lamc1 - (Itgav+Itgb8) | LAMININ | ECM-Receptor | KEGG: mmu04512 |
| CD31 | NeuN | Lamc1 | ITGAV_ITGB8 | 0.001519 | 0 | LAMC1_ITGAV_ITGB8 | Lamc1 - (Itgav+Itgb8) | LAMININ | ECM-Receptor | KEGG: mmu04512 |
| CD31 | CD31 | Lamc2 | ITGAV_ITGB8 | 0.000853 | 0 | LAMC2_ITGAV_ITGB8 | Lamc2 - (Itgav+Itgb8) | LAMININ | ECM-Receptor | KEGG: mmu04512 |
| CD31 | NeuN | Lamc2 | ITGAV_ITGB8 | 0.000508 | 0 | LAMC2_ITGAV_ITGB8 | Lamc2 - (Itgav+Itgb8) | LAMININ | ECM-Receptor | KEGG: mmu04512 |
| CD31 | CD31 | Lamc3 | ITGAV_ITGB8 | 0.0013 | 0 | LAMC3_ITGAV_ITGB8 | Lamc3 - (Itgav+Itgb8) | LAMININ | ECM-Receptor | KEGG: mmu04512 |
| CD31 | NeuN | Lamc3 | ITGAV_ITGB8 | 0.000774 | 0 | LAMC3_ITGAV_ITGB8 | Lamc3 - (Itgav+Itgb8) | LAMININ | ECM-Receptor | KEGG: mmu04512 |
| CD31 | CD31 | Col1a1 | ITGAV_ITGB8 | 0.000685 | 0 | COL1A1_ITGAV_ITGB8 | Col1a1 - (Itgav+Itgb8) | COLLAGEN | ECM-Receptor | KEGG: mmu04512 |
| NeuN | CD31 | Col1a1 | ITGAV_ITGB8 | 0.000438 | 0 | COL1A1_ITGAV_ITGB8 | Col1a1 - (Itgav+Itgb8) | COLLAGEN | ECM-Receptor | KEGG: mmu04512 |
| CD31 | CD31 | Col1a2 | ITGAV_ITGB8 | 0.001455 | 0 | COL1A2_ITGAV_ITGB8 | Col1a2 - (Itgav+Itgb8) | COLLAGEN | ECM-Receptor | KEGG: mmu04512 |
| CD31 | NeuN | Col1a2 | ITGAV_ITGB8 | 0.000866 | 0 | COL1A2_ITGAV_ITGB8 | Col1a2 - (Itgav+Itgb8) | COLLAGEN | ECM-Receptor | KEGG: mmu04512 |
| CD31 | CD31 | Col4a1 | ITGAV_ITGB8 | 0.020292 | 0 | COL4A1_ITGAV_ITGB8 | Col4a1 - (Itgav+Itgb8) | COLLAGEN | ECM-Receptor | KEGG: mmu04512 |
| CD31 | NeuN | Col4a1 | ITGAV_ITGB8 | 0.012174 | 0 | COL4A1_ITGAV_ITGB8 | Col4a1 - (Itgav+Itgb8) | COLLAGEN | ECM-Receptor | KEGG: mmu04512 |
| CD31 | CD31 | Col4a2 | ITGAV_ITGB8 | 0.017444 | 0 | COL4A2_ITGAV_ITGB8 | Col4a2 - (Itgav+Itgb8) | COLLAGEN | ECM-Receptor | KEGG: mmu04512 |
| CD31 | NeuN | Col4a2 | ITGAV_ITGB8 | 0.010453 | 0 | COL4A2_ITGAV_ITGB8 | Col4a2 - (Itgav+Itgb8) | COLLAGEN | ECM-Receptor | KEGG: mmu04512 |
| CD31 | CD31 | Col6a1 | ITGAV_ITGB8 | 0.000732 | 0 | COL6A1_ITGAV_ITGB8 | Col6a1 - (Itgav+Itgb8) | COLLAGEN | ECM-Receptor | KEGG: mmu04512 |
| NeuN | CD31 | Col6a1 | ITGAV_ITGB8 | 0.000538 | 0 | COL6A1_ITGAV_ITGB8 | Col6a1 - (Itgav+Itgb8) | COLLAGEN | ECM-Receptor | KEGG: mmu04512 |
| CD31 | CD31 | Col6a2 | ITGAV_ITGB8 | 0.000728 | 0 | COL6A2_ITGAV_ITGB8 | Col6a2 - (Itgav+Itgb8) | COLLAGEN | ECM-Receptor | KEGG: mmu04512 |
| NeuN | CD31 | Col6a2 | ITGAV_ITGB8 | 0.000504 | 0 | COL6A2_ITGAV_ITGB8 | Col6a2 - (Itgav+Itgb8) | COLLAGEN | ECM-Receptor | KEGG: mmu04512 |
| CD31 | CD31 | Col6a4 | ITGAV_ITGB8 | 0.000699 | 0 | COL6A4_ITGAV_ITGB8 | Col6a4 - (Itgav+Itgb8) | COLLAGEN | ECM-Receptor | KEGG: mmu04512 |
| NeuN | CD31 | Col6a4 | ITGAV_ITGB8 | 0.000576 | 0 | COL6A4_ITGAV_ITGB8 | Col6a4 - (Itgav+Itgb8) | COLLAGEN | ECM-Receptor | KEGG: mmu04512 |
| CD31 | CD31 | Col6a6 | ITGAV_ITGB8 | 0.000872 | 0 | COL6A6_ITGAV_ITGB8 | Col6a6 - (Itgav+Itgb8) | COLLAGEN | ECM-Receptor | KEGG: mmu04512 |
| NeuN | CD31 | Col6a6 | ITGAV_ITGB8 | 0.000535 | 0 | COL6A6_ITGAV_ITGB8 | Col6a6 - (Itgav+Itgb8) | COLLAGEN | ECM-Receptor | KEGG: mmu04512 |
| CD31 | CD31 | Col9a2 | ITGAV_ITGB8 | 0.001117 | 0 | COL9A2_ITGAV_ITGB8 | Col9a2 - (Itgav+Itgb8) | COLLAGEN | ECM-Receptor | KEGG: mmu04512 |
| NeuN | CD31 | Col9a2 | ITGAV_ITGB8 | 0.000844 | 0 | COL9A2_ITGAV_ITGB8 | Col9a2 - (Itgav+Itgb8) | COLLAGEN | ECM-Receptor | KEGG: mmu04512 |
| NeuN | CD31 | Col9a3 | ITGAV_ITGB8 | 0.001989 | 0 | COL9A3_ITGAV_ITGB8 | Col9a3 - (Itgav+Itgb8) | COLLAGEN | ECM-Receptor | KEGG: mmu04512 |
| NeuN | NeuN | Col9a3 | ITGAV_ITGB8 | 0.001184 | 0 | COL9A3_ITGAV_ITGB8 | Col9a3 - (Itgav+Itgb8) | COLLAGEN | ECM-Receptor | KEGG: mmu04512 |
| CD31 | CD31 | Col1a1 | Sdc4 | 0.000553 | 0 | COL1A1_SDC4 | Col1a1 - Sdc4 | COLLAGEN | ECM-Receptor | KEGG: mmu04512 |
| NeuN | CD31 | Col1a1 | Sdc4 | 0.000354 | 0 | COL1A1_SDC4 | Col1a1 - Sdc4 | COLLAGEN | ECM-Receptor | KEGG: mmu04512 |
| CD31 | CD31 | Col1a2 | Sdc4 | 0.001175 | 0 | COL1A2_SDC4 | Col1a2 - Sdc4 | COLLAGEN | ECM-Receptor | KEGG: mmu04512 |
| CD31 | NeuN | Col1a2 | Sdc4 | 0.000544 | 0 | COL1A2_SDC4 | Col1a2 - Sdc4 | COLLAGEN | ECM-Receptor | KEGG: mmu04512 |
| CD31 | CD31 | Col4a1 | Sdc4 | 0.016441 | 0 | COL4A1_SDC4 | Col4a1 - Sdc4 | COLLAGEN | ECM-Receptor | KEGG: mmu04512 |
| CD31 | NeuN | Col4a1 | Sdc4 | 0.00767 | 0 | COL4A1_SDC4 | Col4a1 - Sdc4 | COLLAGEN | ECM-Receptor | KEGG: mmu04512 |
| CD31 | CD31 | Col4a2 | Sdc4 | 0.014126 | 0 | COL4A2_SDC4 | Col4a2 - Sdc4 | COLLAGEN | ECM-Receptor | KEGG: mmu04512 |
| CD31 | NeuN | Col4a2 | Sdc4 | 0.006581 | 0 | COL4A2_SDC4 | Col4a2 - Sdc4 | COLLAGEN | ECM-Receptor | KEGG: mmu04512 |
| CD31 | CD31 | Col6a1 | Sdc4 | 0.000591 | 0 | COL6A1_SDC4 | Col6a1 - Sdc4 | COLLAGEN | ECM-Receptor | KEGG: mmu04512 |
| NeuN | CD31 | Col6a1 | Sdc4 | 0.000434 | 0 | COL6A1_SDC4 | Col6a1 - Sdc4 | COLLAGEN | ECM-Receptor | KEGG: mmu04512 |
| CD31 | CD31 | Col6a2 | Sdc4 | 0.000587 | 0 | COL6A2_SDC4 | Col6a2 - Sdc4 | COLLAGEN | ECM-Receptor | KEGG: mmu04512 |
| NeuN | CD31 | Col6a2 | Sdc4 | 0.000407 | 0 | COL6A2_SDC4 | Col6a2 - Sdc4 | COLLAGEN | ECM-Receptor | KEGG: mmu04512 |
| CD31 | CD31 | Col6a4 | Sdc4 | 0.000564 | 0 | COL6A4_SDC4 | Col6a4 - Sdc4 | COLLAGEN | ECM-Receptor | KEGG: mmu04512 |
| NeuN | CD31 | Col6a4 | Sdc4 | 0.000465 | 0 | COL6A4_SDC4 | Col6a4 - Sdc4 | COLLAGEN | ECM-Receptor | KEGG: mmu04512 |
| CD31 | CD31 | Col6a6 | Sdc4 | 0.000704 | 0 | COL6A6_SDC4 | Col6a6 - Sdc4 | COLLAGEN | ECM-Receptor | KEGG: mmu04512 |
| NeuN | CD31 | Col6a6 | Sdc4 | 0.000432 | 0 | COL6A6_SDC4 | Col6a6 - Sdc4 | COLLAGEN | ECM-Receptor | KEGG: mmu04512 |
| CD31 | CD31 | Col9a2 | Sdc4 | 0.000901 | 0 | COL9A2_SDC4 | Col9a2 - Sdc4 | COLLAGEN | ECM-Receptor | KEGG: mmu04512 |
| NeuN | CD31 | Col9a2 | Sdc4 | 0.000682 | 0 | COL9A2_SDC4 | Col9a2 - Sdc4 | COLLAGEN | ECM-Receptor | KEGG: mmu04512 |
| NeuN | CD31 | Col9a3 | Sdc4 | 0.001606 | 0 | COL9A3_SDC4 | Col9a3 - Sdc4 | COLLAGEN | ECM-Receptor | KEGG: mmu04512 |

| | | | | | | | | | | |
|---|---|---|---|---|---|---|---|---|---|---|
| NeuN | NeuN | Col9a3 | Sdc4 | 0.000743 | 0 | COL9A3_SDC4 | Col9a3 - Sdc4 | COLLAGEN | ECM-Receptor | KEGG: mmu04512 |
| CD31 | CD31 | Fn1 | Sdc4 | 0.003525 | 0 | FN1_SDC4 | Fn1 - Sdc4 | FN1 | ECM-Receptor | KEGG: mmu04512 |
| CD31 | NeuN | Fn1 | Sdc4 | 0.001633 | 0 | FN1_SDC4 | Fn1 - Sdc4 | FN1 | ECM-Receptor | KEGG: mmu04512 |
| CD31 | CD31 | Tnc | Sdc4 | 0.000805 | 0 | TNC_SDC4 | Tnc - Sdc4 | TENASCIN | ECM-Receptor | KEGG: mmu04512 |
| CD31 | NeuN | Tnc | Sdc4 | 0.000372 | 0 | TNC_SDC4 | Tnc - Sdc4 | TENASCIN | ECM-Receptor | KEGG: mmu04512 |
| CD31 | CD31 | Tnr | Sdc4 | 0.000534 | 0 | TNR_SDC4 | Tnr - Sdc4 | TENASCIN | ECM-Receptor | KEGG: mmu04512 |
| NeuN | CD31 | Tnr | Sdc4 | 0.000642 | 0 | TNR_SDC4 | Tnr - Sdc4 | TENASCIN | ECM-Receptor | KEGG: mmu04512 |
| CD31 | CD31 | Thbs1 | Sdc4 | 0.000997 | 0 | THBS1_SDC4 | Thbs1 - Sdc4 | THBS | ECM-Receptor | KEGG: mmu04512 |
| NeuN | CD31 | Thbs1 | Sdc4 | 0.000887 | 0 | THBS1_SDC4 | Thbs1 - Sdc4 | THBS | ECM-Receptor | KEGG: mmu04512 |
| CD31 | CD31 | Thbs3 | Sdc4 | 0.00082 | 0 | THBS3_SDC4 | Thbs3 - Sdc4 | THBS | ECM-Receptor | KEGG: mmu04512 |
| NeuN | CD31 | Thbs3 | Sdc4 | 0.001298 | 0 | THBS3_SDC4 | Thbs3 - Sdc4 | THBS | ECM-Receptor | KEGG: mmu04512 |
| CD31 | CD31 | Thbs4 | Sdc4 | 0.000612 | 0 | THBS4_SDC4 | Thbs4 - Sdc4 | THBS | ECM-Receptor | KEGG: mmu04512 |
| NeuN | CD31 | Thbs4 | Sdc4 | 0.00057 | 0 | THBS4_SDC4 | Thbs4 - Sdc4 | THBS | ECM-Receptor | KEGG: mmu04512 |
| CD31 | NeuN | Lama1 | Sv2a | 0.003026 | 0 | LAMA1_SV2A | Lama1 - Sv2a | LAMININ | ECM-Receptor | KEGG: mmu04512 |
| NeuN | NeuN | Lama1 | Sv2a | 0.00203 | 0 | LAMA1_SV2A | Lama1 - Sv2a | LAMININ | ECM-Receptor | KEGG: mmu04512 |
| CD31 | CD31 | Lama4 | Sv2a | 0.006784 | 0 | LAMA4_SV2A | Lama4 - Sv2a | LAMININ | ECM-Receptor | KEGG: mmu04512 |
| CD31 | NeuN | Lama4 | Sv2a | 0.014617 | 0 | LAMA4_SV2A | Lama4 - Sv2a | LAMININ | ECM-Receptor | KEGG: mmu04512 |
| CD31 | NeuN | Lama5 | Sv2a | 0.002804 | 0 | LAMA5_SV2A | Lama5 - Sv2a | LAMININ | ECM-Receptor | KEGG: mmu04512 |
| NeuN | NeuN | Lama5 | Sv2a | 0.001609 | 0 | LAMA5_SV2A | Lama5 - Sv2a | LAMININ | ECM-Receptor | KEGG: mmu04512 |
| CD31 | CD31 | Lamb1 | Sv2a | 0.009074 | 0 | LAMB1_SV2A | Lamb1 - Sv2a | LAMININ | ECM-Receptor | KEGG: mmu04512 |
| CD31 | NeuN | Lamb1 | Sv2a | 0.019499 | 0 | LAMB1_SV2A | Lamb1 - Sv2a | LAMININ | ECM-Receptor | KEGG: mmu04512 |
| CD31 | CD31 | Lamb2 | Sv2a | 0.003056 | 0 | LAMB2_SV2A | Lamb2 - Sv2a | LAMININ | ECM-Receptor | KEGG: mmu04512 |
| CD31 | NeuN | Lamb2 | Sv2a | 0.006614 | 0 | LAMB2_SV2A | Lamb2 - Sv2a | LAMININ | ECM-Receptor | KEGG: mmu04512 |
| CD31 | CD31 | Lamc1 | Sv2a | 0.003107 | 0 | LAMC1_SV2A | Lamc1 - Sv2a | LAMININ | ECM-Receptor | KEGG: mmu04512 |
| CD31 | NeuN | Lamc1 | Sv2a | 0.006724 | 0 | LAMC1_SV2A | Lamc1 - Sv2a | LAMININ | ECM-Receptor | KEGG: mmu04512 |
| CD31 | NeuN | Lamc2 | Sv2a | 0.002255 | 0 | LAMC2_SV2A | Lamc2 - Sv2a | LAMININ | ECM-Receptor | KEGG: mmu04512 |
| NeuN | NeuN | Lamc2 | Sv2a | 0.001189 | 0 | LAMC2_SV2A | Lamc2 - Sv2a | LAMININ | ECM-Receptor | KEGG: mmu04512 |
| CD31 | CD31 | Lamc3 | Sv2a | 0.001585 | 0 | LAMC3_SV2A | Lamc3 - Sv2a | LAMININ | ECM-Receptor | KEGG: mmu04512 |
| CD31 | NeuN | Lamc3 | Sv2a | 0.003435 | 0 | LAMC3_SV2A | Lamc3 - Sv2a | LAMININ | ECM-Receptor | KEGG: mmu04512 |
| CD31 | CD31 | Lama1 | Sv2b | 0.000798 | 0 | LAMA1_SV2B | Lama1 - Sv2b | LAMININ | ECM-Receptor | KEGG: mmu04512 |
| CD31 | NeuN | Lama1 | Sv2b | 0.001058 | 0 | LAMA1_SV2B | Lama1 - Sv2b | LAMININ | ECM-Receptor | KEGG: mmu04512 |
| CD31 | CD31 | Lama4 | Sv2b | 0.003887 | 0 | LAMA4_SV2B | Lama4 - Sv2b | LAMININ | ECM-Receptor | KEGG: mmu04512 |
| CD31 | NeuN | Lama4 | Sv2b | 0.005148 | 0 | LAMA4_SV2B | Lama4 - Sv2b | LAMININ | ECM-Receptor | KEGG: mmu04512 |
| CD31 | CD31 | Lama5 | Sv2b | 0.000739 | 0 | LAMA5_SV2B | Lama5 - Sv2b | LAMININ | ECM-Receptor | KEGG: mmu04512 |
| CD31 | NeuN | Lama5 | Sv2b | 0.00098 | 0 | LAMA5_SV2B | Lama5 - Sv2b | LAMININ | ECM-Receptor | KEGG: mmu04512 |
| CD31 | CD31 | Lamb1 | Sv2b | 0.005204 | 0 | LAMB1_SV2B | Lamb1 - Sv2b | LAMININ | ECM-Receptor | KEGG: mmu04512 |
| CD31 | NeuN | Lamb1 | Sv2b | 0.00689 | 0 | LAMB1_SV2B | Lamb1 - Sv2b | LAMININ | ECM-Receptor | KEGG: mmu04512 |
| CD31 | CD31 | Lamb2 | Sv2b | 0.001748 | 0 | LAMB2_SV2B | Lamb2 - Sv2b | LAMININ | ECM-Receptor | KEGG: mmu04512 |
| CD31 | NeuN | Lamb2 | Sv2b | 0.002317 | 0 | LAMB2_SV2B | Lamb2 - Sv2b | LAMININ | ECM-Receptor | KEGG: mmu04512 |
| CD31 | CD31 | Lamc1 | Sv2b | 0.001778 | 0 | LAMC1_SV2B | Lamc1 - Sv2b | LAMININ | ECM-Receptor | KEGG: mmu04512 |
| CD31 | NeuN | Lamc1 | Sv2b | 0.002356 | 0 | LAMC1_SV2B | Lamc1 - Sv2b | LAMININ | ECM-Receptor | KEGG: mmu04512 |
| CD31 | CD31 | Lamc2 | Sv2b | 0.000594 | 0 | LAMC2_SV2B | Lamc2 - Sv2b | LAMININ | ECM-Receptor | KEGG: mmu04512 |
| CD31 | NeuN | Lamc2 | Sv2b | 0.000788 | 0 | LAMC2_SV2B | Lamc2 - Sv2b | LAMININ | ECM-Receptor | KEGG: mmu04512 |
| CD31 | CD31 | Lamc3 | Sv2b | 0.000906 | 0 | LAMC3_SV2B | Lamc3 - Sv2b | LAMININ | ECM-Receptor | KEGG: mmu04512 |
| CD31 | NeuN | Lamc3 | Sv2b | 0.001201 | 0 | LAMC3_SV2B | Lamc3 - Sv2b | LAMININ | ECM-Receptor | KEGG: mmu04512 |
| CD31 | CD31 | Lama1 | Sv2c | 0.000667 | 0 | LAMA1_SV2C | Lama1 - Sv2c | LAMININ | ECM-Receptor | KEGG: mmu04512 |
| CD31 | NeuN | Lama1 | Sv2c | 0.00087 | 0 | LAMA1_SV2C | Lama1 - Sv2c | LAMININ | ECM-Receptor | KEGG: mmu04512 |
| CD31 | CD31 | Lama4 | Sv2c | 0.00325 | 0 | LAMA4_SV2C | Lama4 - Sv2c | LAMININ | ECM-Receptor | KEGG: mmu04512 |
| CD31 | NeuN | Lama4 | Sv2c | 0.004236 | 0 | LAMA4_SV2C | Lama4 - Sv2c | LAMININ | ECM-Receptor | KEGG: mmu04512 |
| CD31 | CD31 | Lama5 | Sv2c | 0.000618 | 0 | LAMA5_SV2C | Lama5 - Sv2c | LAMININ | ECM-Receptor | KEGG: mmu04512 |
| CD31 | NeuN | Lama5 | Sv2c | 0.000806 | 0 | LAMA5_SV2C | Lama5 - Sv2c | LAMININ | ECM-Receptor | KEGG: mmu04512 |
| CD31 | CD31 | Lamb1 | Sv2c | 0.004352 | 0 | LAMB1_SV2C | Lamb1 - Sv2c | LAMININ | ECM-Receptor | KEGG: mmu04512 |
| CD31 | NeuN | Lamb1 | Sv2c | 0.005671 | 0 | LAMB1_SV2C | Lamb1 - Sv2c | LAMININ | ECM-Receptor | KEGG: mmu04512 |
| CD31 | CD31 | Lamb2 | Sv2c | 0.001461 | 0 | LAMB2_SV2C | Lamb2 - Sv2c | LAMININ | ECM-Receptor | KEGG: mmu04512 |
| CD31 | NeuN | Lamb2 | Sv2c | 0.001906 | 0 | LAMB2_SV2C | Lamb2 - Sv2c | LAMININ | ECM-Receptor | KEGG: mmu04512 |
| CD31 | CD31 | Lamc1 | Sv2c | 0.001486 | 0 | LAMC1_SV2C | Lamc1 - Sv2c | LAMININ | ECM-Receptor | KEGG: mmu04512 |
| CD31 | NeuN | Lamc1 | Sv2c | 0.001938 | 0 | LAMC1_SV2C | Lamc1 - Sv2c | LAMININ | ECM-Receptor | KEGG: mmu04512 |
| CD31 | CD31 | Lamc2 | Sv2c | 0.000496 | 0 | LAMC2_SV2C | Lamc2 - Sv2c | LAMININ | ECM-Receptor | KEGG: mmu04512 |
| CD31 | NeuN | Lamc2 | Sv2c | 0.000648 | 0 | LAMC2_SV2C | Lamc2 - Sv2c | LAMININ | ECM-Receptor | KEGG: mmu04512 |
| CD31 | CD31 | Lamc3 | Sv2c | 0.000757 | 0 | LAMC3_SV2C | Lamc3 - Sv2c | LAMININ | ECM-Receptor | KEGG: mmu04512 |

| | | | | | | | | | | |
|---|---|---|---|---|---|---|---|---|---|---|
| CD31 | NeuN | Lamc3 | Sv2c | 0.000988 | 0 | LAMC3_SV2C | Lamc3 - Sv2c | LAMININ | ECM-Receptor | KEGG: mmu04512 |
| CD31 | CD31 | Thbs1 | Cd36 | 0.000667 | 0 | THBS1_CD36 | Thbs1 - Cd36 | THBS | ECM-Receptor | KEGG: mmu04512 |
| NeuN | CD31 | Thbs1 | Cd36 | 0.000593 | 0 | THBS1_CD36 | Thbs1 - Cd36 | THBS | ECM-Receptor | KEGG: mmu04512 |
| CD31 | CD31 | Thbs3 | Cd36 | 0.000549 | 0 | THBS3_CD36 | Thbs3 - Cd36 | THBS | ECM-Receptor | KEGG: mmu04512 |
| NeuN | CD31 | Thbs3 | Cd36 | 0.000869 | 0 | THBS3_CD36 | Thbs3 - Cd36 | THBS | ECM-Receptor | KEGG: mmu04512 |
| CD31 | CD31 | Thbs4 | Cd36 | 0.000409 | 0 | THBS4_CD36 | Thbs4 - Cd36 | THBS | ECM-Receptor | KEGG: mmu04512 |
| NeuN | CD31 | Thbs4 | Cd36 | 0.000382 | 0 | THBS4_CD36 | Thbs4 - Cd36 | THBS | ECM-Receptor | KEGG: mmu04512 |
| CD31 | NeuN | Thbs1 | Cd47 | 0.001177 | 0 | THBS1_CD47 | Thbs1 - Cd47 | THBS | ECM-Receptor | KEGG: mmu04512 |
| NeuN | NeuN | Thbs1 | Cd47 | 0.001047 | 0 | THBS1_CD47 | Thbs1 - Cd47 | THBS | ECM-Receptor | KEGG: mmu04512 |
| CD31 | NeuN | Thbs3 | Cd47 | 0.000968 | 0 | THBS3_CD47 | Thbs3 - Cd47 | THBS | ECM-Receptor | KEGG: mmu04512 |
| NeuN | NeuN | Thbs3 | Cd47 | 0.001533 | 0 | THBS3_CD47 | Thbs3 - Cd47 | THBS | ECM-Receptor | KEGG: mmu04512 |
| CD31 | NeuN | Thbs4 | Cd47 | 0.000722 | 0 | THBS4_CD47 | Thbs4 - Cd47 | THBS | ECM-Receptor | KEGG: mmu04512 |
| NeuN | NeuN | Thbs4 | Cd47 | 0.000673 | 0 | THBS4_CD47 | Thbs4 - Cd47 | THBS | ECM-Receptor | KEGG: mmu04512 |
| CD31 | CD31 | Agrn | Dag1 | 0.007855 | 0 | AGRN_DAG1 | Agrn - Dag1 | AGRN | ECM-Receptor | KEGG: mmu04512 |
| NeuN | CD31 | Agrn | Dag1 | 0.006062 | 0 | AGRN_DAG1 | Agrn - Dag1 | AGRN | ECM-Receptor | KEGG: mmu04512 |
| CD31 | CD31 | Hspg2 | Dag1 | 0.003648 | 0 | HSPG2_DAG1 | Hspg2 - Dag1 | HSPG | ECM-Receptor | KEGG: mmu04512 |
| CD31 | NeuN | Hspg2 | Dag1 | 0.002577 | 0 | HSPG2_DAG1 | Hspg2 - Dag1 | HSPG | ECM-Receptor | KEGG: mmu04512 |
| CD31 | CD31 | Lama1 | Dag1 | 0.001588 | 0 | LAMA1_DAG1 | Lama1 - Dag1 | LAMININ | ECM-Receptor | KEGG: mmu04512 |
| CD31 | NeuN | Lama1 | Dag1 | 0.001121 | 0 | LAMA1_DAG1 | Lama1 - Dag1 | LAMININ | ECM-Receptor | KEGG: mmu04512 |
| CD31 | CD31 | Lama4 | Dag1 | 0.007713 | 0 | LAMA4_DAG1 | Lama4 - Dag1 | LAMININ | ECM-Receptor | KEGG: mmu04512 |
| CD31 | NeuN | Lama4 | Dag1 | 0.005455 | 0 | LAMA4_DAG1 | Lama4 - Dag1 | LAMININ | ECM-Receptor | KEGG: mmu04512 |
| CD31 | CD31 | Lama5 | Dag1 | 0.001471 | 0 | LAMA5_DAG1 | Lama5 - Dag1 | LAMININ | ECM-Receptor | KEGG: mmu04512 |
| CD31 | NeuN | Lama5 | Dag1 | 0.001038 | 0 | LAMA5_DAG1 | Lama5 - Dag1 | LAMININ | ECM-Receptor | KEGG: mmu04512 |
| CD31 | CD31 | Lamb1 | Dag1 | 0.010313 | 0 | LAMB1_DAG1 | Lamb1 - Dag1 | LAMININ | ECM-Receptor | KEGG: mmu04512 |
| CD31 | NeuN | Lamb1 | Dag1 | 0.007299 | 0 | LAMB1_DAG1 | Lamb1 - Dag1 | LAMININ | ECM-Receptor | KEGG: mmu04512 |
| CD31 | CD31 | Lamb2 | Dag1 | 0.003477 | 0 | LAMB2_DAG1 | Lamb2 - Dag1 | LAMININ | ECM-Receptor | KEGG: mmu04512 |
| CD31 | NeuN | Lamb2 | Dag1 | 0.002456 | 0 | LAMB2_DAG1 | Lamb2 - Dag1 | LAMININ | ECM-Receptor | KEGG: mmu04512 |
| CD31 | CD31 | Lamc1 | Dag1 | 0.003535 | 0 | LAMC1_DAG1 | Lamc1 - Dag1 | LAMININ | ECM-Receptor | KEGG: mmu04512 |
| CD31 | NeuN | Lamc1 | Dag1 | 0.002497 | 0 | LAMC1_DAG1 | Lamc1 - Dag1 | LAMININ | ECM-Receptor | KEGG: mmu04512 |
| CD31 | CD31 | Lamc2 | Dag1 | 0.001183 | 0 | LAMC2_DAG1 | Lamc2 - Dag1 | LAMININ | ECM-Receptor | KEGG: mmu04512 |
| CD31 | NeuN | Lamc2 | Dag1 | 0.000835 | 0 | LAMC2_DAG1 | Lamc2 - Dag1 | LAMININ | ECM-Receptor | KEGG: mmu04512 |
| CD31 | CD31 | Lamc3 | Dag1 | 0.001803 | 0 | LAMC3_DAG1 | Lamc3 - Dag1 | LAMININ | ECM-Receptor | KEGG: mmu04512 |
| CD31 | NeuN | Lamc3 | Dag1 | 0.001273 | 0 | LAMC3_DAG1 | Lamc3 - Dag1 | LAMININ | ECM-Receptor | KEGG: mmu04512 |
| CD31 | CD31 | Reln | Lrp8 | 0.004291 | 0 | RELN_LRP8 | Reln - Lrp8 | RELN | ECM-Receptor | PMID: 14578885 |
| NeuN | CD31 | Reln | Lrp8 | 0.005315 | 0 | RELN_LRP8 | Reln - Lrp8 | RELN | ECM-Receptor | PMID: 14578885 |
| CD31 | NeuN | Reln | Vldlr | 0.001632 | 0 | RELN_VLDLR | Reln - Vldlr | RELN | ECM-Receptor | PMID: 14578885 |
| NeuN | NeuN | Reln | Vldlr | 0.002024 | 0 | RELN_VLDLR | Reln - Vldlr | RELN | ECM-Receptor | PMID: 14578885 |
| CD31 | CD31 | Pnoc | Oprl1 | 0.000814 | 0 | PNOC_OPRL1 | Pnoc - Oprl1 | OPIOID | Non-protein Signaling | PMID: 29934561;PMID: 34724150 |
| CD31 | NeuN | Pnoc | Oprl1 | 0.001024 | 0 | PNOC_OPRL1 | Pnoc - Oprl1 | OPIOID | Non-protein Signaling | PMID: 29934561;PMID: 34724150 |
| CD31 | CD31 | Ppia | Bsg | 0.485021 | 0 | PPIA_BSG | Ppia - Bsg | CypA | Non-protein Signaling | PMID: 36012604 |
| NeuN | CD31 | Ppia | Bsg | 0.569947 | 0 | PPIA_BSG | Ppia - Bsg | CypA | Non-protein Signaling | PMID: 36012604 |
| NeuN | CD31 | Cadm1 | Cadm1 | 0.005092 | 0 | CADM1_CADM1 | Cadm1 - Cadm1 | CADM | Cell-Cell Contact | PMID: 24503895 |
| CD31 | NeuN | Cadm1 | Cadm1 | 0.005092 | 0 | CADM1_CADM1 | Cadm1 - Cadm1 | CADM | Cell-Cell Contact | PMID: 24503895 |
| NeuN | NeuN | Cadm1 | Cadm1 | 0.014982 | 0 | CADM1_CADM1 | Cadm1 - Cadm1 | CADM | Cell-Cell Contact | PMID: 24503895 |
| NeuN | CD31 | Cd200 | Cd200r1 | 0.002141 | 0 | CD200_CD200R1 | Cd200 - Cd200r1 | CD200 | Cell-Cell Contact | PMID: 15187158 |
| NeuN | NeuN | Cd200 | Cd200r1 | 0.001725 | 0 | CD200_CD200R1 | Cd200 - Cd200r1 | CD200 | Cell-Cell Contact | PMID: 15187158 |
| NeuN | CD31 | Cd200 | Cd200r2 | 0.003582 | 0 | CD200_CD200R2 | Cd200 - Cd200r2 | CD200 | Cell-Cell Contact | KEGG: mmu04514 |
| NeuN | NeuN | Cd200 | Cd200r2 | 0.003313 | 0 | CD200_CD200R2 | Cd200 - Cd200r2 | CD200 | Cell-Cell Contact | KEGG: mmu04514 |
| CD31 | CD31 | Cd200 | Cd200r3 | 0.001926 | 0 | CD200_CD200R3 | Cd200 - Cd200r3 | CD200 | Cell-Cell Contact | PMID: 22020332 |
| NeuN | CD31 | Cd200 | Cd200r3 | 0.004326 | 0 | CD200_CD200R3 | Cd200 - Cd200r3 | CD200 | Cell-Cell Contact | PMID: 22020332 |
| CD31 | CD31 | Entpd1 | Adora1 | 0.001196 | 0 | ENTPD1_ADORA1 | Entpd1 - Adora1 | CD39 | Cell-Cell Contact | PMID: 21677139 |
| CD31 | NeuN | Entpd1 | Adora1 | 0.001635 | 0 | ENTPD1_ADORA1 | Entpd1 - Adora1 | CD39 | Cell-Cell Contact | PMID: 21677139 |
| CD31 | CD31 | Entpd1 | Adora2a | 0.003647 | 0 | ENTPD1_ADORA2A | Entpd1 - Adora2a | CD39 | Cell-Cell Contact | PMID: 21677139 |
| NeuN | CD31 | Entpd1 | Adora2a | 0.001443 | 0 | ENTPD1_ADORA2A | Entpd1 - Adora2a | CD39 | Cell-Cell Contact | PMID: 21677139 |
| CD31 | CD31 | Entpd1 | Adora2b | 0.000689 | 0 | ENTPD1_ADORA2B | Entpd1 - Adora2b | CD39 | Cell-Cell Contact | PMID: 21677139 |
| CD31 | NeuN | Entpd1 | Adora2b | 0.000455 | 0 | ENTPD1_ADORA2B | Entpd1 - Adora2b | CD39 | Cell-Cell Contact | PMID: 21677139 |
| CD31 | CD31 | Cd46 | Jag1 | 0.001084 | 0 | CD46_JAG1 | Cd46 - Jag1 | CD46 | Cell-Cell Contact | PMID: 23086448 |
| NeuN | CD31 | Cd46 | Jag1 | 0.000845 | 0 | CD46_JAG1 | Cd46 - Jag1 | CD46 | Cell-Cell Contact | PMID: 23086448 |
| CD31 | CD31 | Cd6 | Alcam | 0.000658 | 0 | CD6_ALCAM | Cd6 - Alcam | CD6 | Cell-Cell Contact | PMID: 23602662 |
| CD31 | NeuN | Cd6 | Alcam | 0.000702 | 0 | CD6_ALCAM | Cd6 - Alcam | CD6 | Cell-Cell Contact | PMID: 23602662 |

| | | | | | | | | | | |
|---|---|---|---|---|---|---|---|---|---|---|
| CD31 | CD31 | Cd96 | Pvr | 0.000596 | 0 | CD96_PVR | Cd96 - Pvr | CD96 | Cell-Cell Contact | PMID: 15034010 |
| CD31 | NeuN | Cd96 | Pvr | 0.000459 | 0 | CD96_PVR | Cd96 - Pvr | CD96 | Cell-Cell Contact | PMID: 15034010 |
| NeuN | CD31 | Cd99 | Cd99 | 0.497081 | 0 | CD99_CD99 | Cd99 - Cd99 | CD99 | Cell-Cell Contact | KEGG: mmu04514 |
| CD31 | NeuN | Cd99 | Cd99 | 0.497081 | 0 | CD99_CD99 | Cd99 - Cd99 | CD99 | Cell-Cell Contact | KEGG: mmu04514 |
| NeuN | NeuN | Cd99 | Cd99 | 0.602333 | 0 | CD99_CD99 | Cd99 - Cd99 | CD99 | Cell-Cell Contact | KEGG: mmu04514 |
| CD31 | CD31 | Cdh4 | Cdh4 | 0.001269 | 0 | CDH4_CDH4 | Cdh4 - Cdh4 | CDH | Cell-Cell Contact | KEGG: mmu04514 |
| NeuN | CD31 | Cdh4 | Cdh4 | 0.000843 | 0 | CDH4_CDH4 | Cdh4 - Cdh4 | CDH | Cell-Cell Contact | KEGG: mmu04514 |
| CD31 | NeuN | Cdh4 | Cdh4 | 0.000843 | 0 | CDH4_CDH4 | Cdh4 - Cdh4 | CDH | Cell-Cell Contact | KEGG: mmu04514 |
| NeuN | CD31 | Cdh2 | Cdh2 | 0.027077 | 0 | CDH2_CDH2 | Cdh2 - Cdh2 | CDH | Cell-Cell Contact | KEGG: mmu04514 |
| CD31 | NeuN | Cdh2 | Cdh2 | 0.027077 | 0 | CDH2_CDH2 | Cdh2 - Cdh2 | CDH | Cell-Cell Contact | KEGG: mmu04514 |
| NeuN | NeuN | Cdh2 | Cdh2 | 0.043502 | 0 | CDH2_CDH2 | Cdh2 - Cdh2 | CDH | Cell-Cell Contact | KEGG: mmu04514 |
| CD31 | CD31 | Cdh5 | Cdh5 | 0.030105 | 0 | CDH5_CDH5 | Cdh5 - Cdh5 | CDH5 | Cell-Cell Contact | KEGG: mmu04514 |
| NeuN | CD31 | Cdh5 | Cdh5 | 0.002485 | 0 | CDH5_CDH5 | Cdh5 - Cdh5 | CDH5 | Cell-Cell Contact | KEGG: mmu04514 |
| CD31 | NeuN | Cdh5 | Cdh5 | 0.002485 | 0 | CDH5_CDH5 | Cdh5 - Cdh5 | CDH5 | Cell-Cell Contact | KEGG: mmu04514 |
| CD31 | CD31 | Cldn11 | Cldn11 | 0.000309 | 0 | CLDN11_CLDN11 | Cldn11 - Cldn11 | CLDN | Cell-Cell Contact | KEGG: mmu04514 |
| NeuN | CD31 | Cldn11 | Cldn11 | 0.000116 | 0 | CLDN11_CLDN11 | Cldn11 - Cldn11 | CLDN | Cell-Cell Contact | KEGG: mmu04514 |
| CD31 | NeuN | Cldn11 | Cldn11 | 0.000116 | 0 | CLDN11_CLDN11 | Cldn11 - Cldn11 | CLDN | Cell-Cell Contact | KEGG: mmu04514 |
| CD31 | CD31 | Clec2f | Klrb1f | 0.00068 | 0 | CLEC2F_KLRB1F | Clec2f - Klrb1f | CLEC | Cell-Cell Contact | PMID: 24223577 |
| CD31 | NeuN | Clec2f | Klrb1f | 0.000694 | 0 | CLEC2F_KLRB1F | Clec2f - Klrb1f | CLEC | Cell-Cell Contact | PMID: 24223577 |
| NeuN | CD31 | Nfasc | CNTN1_CNTNAP1 | 0.003049 | 0 | NFASC_CNTN1_CNTNAP1 | Nfasc - (Cntn1+Cntnap1) | CNTN | Cell-Cell Contact | KEGG: mmu04514 |
| NeuN | NeuN | Nfasc | CNTN1_CNTNAP1 | 0.004157 | 0 | NFASC_CNTN1_CNTNAP1 | Nfasc - (Cntn1+Cntnap1) | CNTN | Cell-Cell Contact | KEGG: mmu04514 |
| NeuN | CD31 | Cntn2 | CNTN2_CNTNAP2 | 0.007668 | 0 | CNTN2_CNTN2_CNTNAP2 | Cntn2 - (Cntn2+Cntnap2) | CNTN | Cell-Cell Contact | KEGG: mmu04514 |
| NeuN | NeuN | Cntn2 | CNTN2_CNTNAP2 | 0.009496 | 0 | CNTN2_CNTN2_CNTNAP2 | Cntn2 - (Cntn2+Cntnap2) | CNTN | Cell-Cell Contact | KEGG: mmu04514 |
| CD31 | NeuN | Cntn2 | L1cam | 0.03356 | 0 | CNTN2_L1CAM | Cntn2 - L1cam | CNTN | Cell-Cell Contact | KEGG: mmu04514 |
| NeuN | NeuN | Cntn2 | L1cam | 0.050918 | 0 | CNTN2_L1CAM | Cntn2 - L1cam | CNTN | Cell-Cell Contact | KEGG: mmu04514 |
| NeuN | CD31 | Cntn2 | Cntn2 | 0.016024 | 0 | CNTN2_CNTN2 | Cntn2 - Cntn2 | CNTN | Cell-Cell Contact | KEGG: mmu04514 |
| CD31 | NeuN | Cntn2 | Cntn2 | 0.016024 | 0 | CNTN2_CNTN2 | Cntn2 - Cntn2 | CNTN | Cell-Cell Contact | KEGG: mmu04514 |
| NeuN | NeuN | Cntn2 | Cntn2 | 0.024543 | 0 | CNTN2_CNTN2 | Cntn2 - Cntn2 | CNTN | Cell-Cell Contact | KEGG: mmu04514 |
| CD31 | NeuN | Cntn1 | Nrcam | 0.009872 | 0 | CNTN1_NRCAM | Cntn1 - Nrcam | CNTN | Cell-Cell Contact | KEGG: mmu04514 |
| NeuN | NeuN | Cntn1 | Nrcam | 0.020046 | 0 | CNTN1_NRCAM | Cntn1 - Nrcam | CNTN | Cell-Cell Contact | KEGG: mmu04514 |
| CD31 | CD31 | Cspg4 | ITGA3_ITGB1 | 0.003577 | 0 | CSPG4_ITGA2_ITGB1 | Cspg4 - (Itga2+Itgb1) | CSPG4 | Cell-Cell Contact | PMID: 30213051 |
| CD31 | NeuN | Cspg4 | ITGA3_ITGB1 | 0.001795 | 0 | CSPG4_ITGA2_ITGB1 | Cspg4 - (Itga2+Itgb1) | CSPG4 | Cell-Cell Contact | PMID: 30213051 |
| CD31 | CD31 | Efna1 | Epha3 | 0.001793 | 0 | EFNA1_EPHA3 | Efna1 - Epha3 | EPHA | Cell-Cell Contact | PMID: 15114347 |
| CD31 | NeuN | Efna1 | Epha3 | 0.001437 | 0 | EFNA1_EPHA3 | Efna1 - Epha3 | EPHA | Cell-Cell Contact | PMID: 15114347 |
| CD31 | CD31 | Efna1 | Epha4 | 0.004223 | 0 | EFNA1_EPHA4 | Efna1 - Epha4 | EPHA | Cell-Cell Contact | PMID: 15114347 |
| CD31 | NeuN | Efna1 | Epha4 | 0.005958 | 0 | EFNA1_EPHA4 | Efna1 - Epha4 | EPHA | Cell-Cell Contact | PMID: 15114347 |
| CD31 | CD31 | Efna1 | Epha5 | 0.004194 | 0 | EFNA1_EPHA5 | Efna1 - Epha5 | EPHA | Cell-Cell Contact | PMID: 15114347 |
| CD31 | NeuN | Efna1 | Epha5 | 0.004735 | 0 | EFNA1_EPHA5 | Efna1 - Epha5 | EPHA | Cell-Cell Contact | PMID: 15114347 |
| CD31 | CD31 | Efna1 | Epha7 | 0.002161 | 0 | EFNA1_EPHA7 | Efna1 - Epha7 | EPHA | Cell-Cell Contact | PMID: 15114347 |
| CD31 | NeuN | Efna1 | Epha7 | 0.004012 | 0 | EFNA1_EPHA7 | Efna1 - Epha7 | EPHA | Cell-Cell Contact | PMID: 15114347 |
| CD31 | CD31 | Efna1 | Epha8 | 0.001258 | 0 | EFNA1_EPHA8 | Efna1 - Epha8 | EPHA | Cell-Cell Contact | PMID: 15114347 |
| CD31 | NeuN | Efna1 | Epha8 | 0.001121 | 0 | EFNA1_EPHA8 | Efna1 - Epha8 | EPHA | Cell-Cell Contact | PMID: 15114347 |
| NeuN | CD31 | Efna2 | Epha3 | 0.003213 | 0 | EFNA2_EPHA3 | Efna2 - Epha3 | EPHA | Cell-Cell Contact | PMID: 15114347 |
| NeuN | NeuN | Efna2 | Epha3 | 0.002576 | 0 | EFNA2_EPHA3 | Efna2 - Epha3 | EPHA | Cell-Cell Contact | PMID: 15114347 |
| NeuN | CD31 | Efna2 | Epha4 | 0.007553 | 0 | EFNA2_EPHA4 | Efna2 - Epha4 | EPHA | Cell-Cell Contact | PMID: 15114347 |
| NeuN | NeuN | Efna2 | Epha4 | 0.010642 | 0 | EFNA2_EPHA4 | Efna2 - Epha4 | EPHA | Cell-Cell Contact | PMID: 15114347 |
| NeuN | CD31 | Efna2 | Epha5 | 0.007502 | 0 | EFNA2_EPHA5 | Efna2 - Epha5 | EPHA | Cell-Cell Contact | PMID: 15114347 |
| NeuN | NeuN | Efna2 | Epha5 | 0.008466 | 0 | EFNA2_EPHA5 | Efna2 - Epha5 | EPHA | Cell-Cell Contact | PMID: 15114347 |
| CD31 | NeuN | Efna2 | Epha7 | 0.004503 | 0 | EFNA2_EPHA7 | Efna2 - Epha7 | EPHA | Cell-Cell Contact | PMID: 15114347 |
| NeuN | NeuN | Efna2 | Epha7 | 0.007178 | 0 | EFNA2_EPHA7 | Efna2 - Epha7 | EPHA | Cell-Cell Contact | PMID: 15114347 |
| NeuN | CD31 | Efna2 | Epha8 | 0.002256 | 0 | EFNA2_EPHA8 | Efna2 - Epha8 | EPHA | Cell-Cell Contact | PMID: 15114347 |
| NeuN | NeuN | Efna2 | Epha8 | 0.00201 | 0 | EFNA2_EPHA8 | Efna2 - Epha8 | EPHA | Cell-Cell Contact | PMID: 15114347 |
| NeuN | CD31 | Efna3 | Epha3 | 0.003449 | 0 | EFNA3_EPHA3 | Efna3 - Epha3 | EPHA | Cell-Cell Contact | PMID: 15114347 |
| NeuN | NeuN | Efna3 | Epha3 | 0.002765 | 0 | EFNA3_EPHA3 | Efna3 - Epha3 | EPHA | Cell-Cell Contact | PMID: 15114347 |
| NeuN | CD31 | Efna3 | Epha4 | 0.008106 | 0 | EFNA3_EPHA4 | Efna3 - Epha4 | EPHA | Cell-Cell Contact | PMID: 15114347 |
| NeuN | NeuN | Efna3 | Epha4 | 0.011417 | 0 | EFNA3_EPHA4 | Efna3 - Epha4 | EPHA | Cell-Cell Contact | PMID: 15114347 |
| NeuN | CD31 | Efna3 | Epha5 | 0.00805 | 0 | EFNA3_EPHA5 | Efna3 - Epha5 | EPHA | Cell-Cell Contact | PMID: 15114347 |
| NeuN | NeuN | Efna3 | Epha5 | 0.009084 | 0 | EFNA3_EPHA5 | Efna3 - Epha5 | EPHA | Cell-Cell Contact | PMID: 15114347 |
| NeuN | CD31 | Efna3 | Epha7 | 0.004155 | 0 | EFNA3_EPHA7 | Efna3 - Epha7 | EPHA | Cell-Cell Contact | PMID: 15114347 |
| NeuN | NeuN | Efna3 | Epha7 | 0.007703 | 0 | EFNA3_EPHA7 | Efna3 - Epha7 | EPHA | Cell-Cell Contact | PMID: 15114347 |

| | | | | | | | | | |
|---|---|---|---|---|---|---|---|---|---|
| NeuN | CD31 | Efna3 | Epha8 | 0.002422 | 0 | EFNA3_EPHA8 | Efna3 - Epha8 | EPHA | Cell-Cell Contact | PMID: 15114347 |
| NeuN | NeuN | Efna3 | Epha8 | 0.002158 | 0 | EFNA3_EPHA8 | Efna3 - Epha8 | EPHA | Cell-Cell Contact | PMID: 15114347 |
| CD31 | CD31 | Efna4 | Epha3 | 0.001932 | 0 | EFNA4_EPHA3 | Efna4 - Epha3 | EPHA | Cell-Cell Contact | PMID: 15114347 |
| NeuN | CD31 | Efna4 | Epha3 | 0.002044 | 0 | EFNA4_EPHA3 | Efna4 - Epha3 | EPHA | Cell-Cell Contact | PMID: 15114347 |
| CD31 | NeuN | Efna4 | Epha4 | 0.006418 | 0 | EFNA4_EPHA4 | Efna4 - Epha4 | EPHA | Cell-Cell Contact | PMID: 15114347 |
| NeuN | NeuN | Efna4 | Epha4 | 0.006788 | 0 | EFNA4_EPHA4 | Efna4 - Epha4 | EPHA | Cell-Cell Contact | PMID: 15114347 |
| CD31 | NeuN | Efna4 | Epha5 | 0.005101 | 0 | EFNA4_EPHA5 | Efna4 - Epha5 | EPHA | Cell-Cell Contact | PMID: 15114347 |
| NeuN | NeuN | Efna4 | Epha5 | 0.005396 | 0 | EFNA4_EPHA5 | Efna4 - Epha5 | EPHA | Cell-Cell Contact | PMID: 15114347 |
| CD31 | NeuN | Efna4 | Epha7 | 0.004323 | 0 | EFNA4_EPHA7 | Efna4 - Epha7 | EPHA | Cell-Cell Contact | PMID: 15114347 |
| NeuN | NeuN | Efna4 | Epha7 | 0.004573 | 0 | EFNA4_EPHA7 | Efna4 - Epha7 | EPHA | Cell-Cell Contact | PMID: 15114347 |
| CD31 | CD31 | Efna4 | Epha8 | 0.001356 | 0 | EFNA4_EPHA8 | Efna4 - Epha8 | EPHA | Cell-Cell Contact | PMID: 15114347 |
| NeuN | CD31 | Efna4 | Epha8 | 0.001435 | 0 | EFNA4_EPHA8 | Efna4 - Epha8 | EPHA | Cell-Cell Contact | PMID: 15114347 |
| NeuN | CD31 | Efna5 | Epha3 | 0.001307 | 0 | EFNA5_EPHA3 | Efna5 - Epha3 | EPHA | Cell-Cell Contact | PMID: 15114347 |
| NeuN | NeuN | Efna5 | Epha3 | 0.001047 | 0 | EFNA5_EPHA3 | Efna5 - Epha3 | EPHA | Cell-Cell Contact | PMID: 15114347 |
| NeuN | CD31 | Efna5 | Epha4 | 0.00308 | 0 | EFNA5_EPHA4 | Efna5 - Epha4 | EPHA | Cell-Cell Contact | PMID: 15114347 |
| NeuN | NeuN | Efna5 | Epha4 | 0.004347 | 0 | EFNA5_EPHA4 | Efna5 - Epha4 | EPHA | Cell-Cell Contact | PMID: 15114347 |
| NeuN | CD31 | Efna5 | Epha5 | 0.003059 | 0 | EFNA5_EPHA5 | Efna5 - Epha5 | EPHA | Cell-Cell Contact | PMID: 15114347 |
| NeuN | NeuN | Efna5 | Epha5 | 0.003454 | 0 | EFNA5_EPHA5 | Efna5 - Epha5 | EPHA | Cell-Cell Contact | PMID: 15114347 |
| CD31 | NeuN | Efna5 | Epha7 | 0.001805 | 0 | EFNA5_EPHA7 | Efna5 - Epha7 | EPHA | Cell-Cell Contact | PMID: 15114347 |
| NeuN | NeuN | Efna5 | Epha7 | 0.002926 | 0 | EFNA5_EPHA7 | Efna5 - Epha7 | EPHA | Cell-Cell Contact | PMID: 15114347 |
| NeuN | CD31 | Efna5 | Epha8 | 0.000917 | 0 | EFNA5_EPHA8 | Efna5 - Epha8 | EPHA | Cell-Cell Contact | PMID: 15114347 |
| NeuN | NeuN | Efna5 | Epha8 | 0.000817 | 0 | EFNA5_EPHA8 | Efna5 - Epha8 | EPHA | Cell-Cell Contact | PMID: 15114347 |
| CD31 | NeuN | Efna5 | Ephb2 | 0.002701 | 0 | EFNA5_EPHB2 | Efna5 - Ephb2 | EPHA | Cell-Cell Contact | PMID:15107857; PMID: 15114347 |
| NeuN | NeuN | Efna5 | Ephb2 | 0.004375 | 0 | EFNA5_EPHB2 | Efna5 - Ephb2 | EPHA | Cell-Cell Contact | PMID:15107857; PMID: 15114347 |
| CD31 | NeuN | Efnb1 | Epha4 | 0.006291 | 0 | EFNB1_EPHA4 | Efnb1 - Epha4 | EPHB | Cell-Cell Contact | PMID: 15114347 |
| NeuN | NeuN | Efnb1 | Epha4 | 0.008019 | 0 | EFNB1_EPHA4 | Efnb1 - Epha4 | EPHB | Cell-Cell Contact | PMID: 15114347 |
| CD31 | NeuN | Efnb1 | Ephb1 | 0.006514 | 0 | EFNB1_EPHB1 | Efnb1 - Ephb1 | EPHB | Cell-Cell Contact | PMID: 15114347 |
| NeuN | NeuN | Efnb1 | Ephb1 | 0.008303 | 0 | EFNB1_EPHB1 | Efnb1 - Ephb1 | EPHB | Cell-Cell Contact | PMID: 15114347 |
| CD31 | NeuN | Efnb1 | Ephb2 | 0.006331 | 0 | EFNB1_EPHB2 | Efnb1 - Ephb2 | EPHB | Cell-Cell Contact | PMID: 15114347 |
| NeuN | NeuN | Efnb1 | Ephb2 | 0.008071 | 0 | EFNB1_EPHB2 | Efnb1 - Ephb2 | EPHB | Cell-Cell Contact | PMID: 15114347 |
| CD31 | CD31 | Efnb1 | Ephb3 | 0.001029 | 0 | EFNB1_EPHB3 | Efnb1 - Ephb3 | EPHB | Cell-Cell Contact | PMID: 15114347 |
| NeuN | CD31 | Efnb1 | Ephb3 | 0.001314 | 0 | EFNB1_EPHB3 | Efnb1 - Ephb3 | EPHB | Cell-Cell Contact | PMID: 15114347 |
| CD31 | CD31 | Efnb1 | Ephb4 | 0.003457 | 0 | EFNB1_EPHB4 | Efnb1 - Ephb4 | EPHB | Cell-Cell Contact | PMID: 15114347 |
| NeuN | CD31 | Efnb1 | Ephb4 | 0.004411 | 0 | EFNB1_EPHB4 | Efnb1 - Ephb4 | EPHB | Cell-Cell Contact | PMID: 15114347 |
| NeuN | CD31 | Efnb2 | Epha4 | 0.027021 | 0 | EFNB2_EPHA4 | Efnb2 - Epha4 | EPHB | Cell-Cell Contact | PMID: 15114347 |
| NeuN | NeuN | Efnb2 | Epha4 | 0.037766 | 0 | EFNB2_EPHA4 | Efnb2 - Epha4 | EPHB | Cell-Cell Contact | PMID: 15114347 |
| NeuN | CD31 | Efnb2 | Ephb1 | 0.025496 | 0 | EFNB2_EPHB1 | Efnb2 - Ephb1 | EPHB | Cell-Cell Contact | PMID: 15114347 |
| NeuN | NeuN | Efnb2 | Ephb1 | 0.039061 | 0 | EFNB2_EPHB1 | Efnb2 - Ephb1 | EPHB | Cell-Cell Contact | PMID: 15114347 |
| CD31 | NeuN | Efnb2 | Ephb2 | 0.024552 | 0 | EFNB2_EPHB2 | Efnb2 - Ephb2 | EPHB | Cell-Cell Contact | PMID: 15114347 |
| NeuN | NeuN | Efnb2 | Ephb2 | 0.038001 | 0 | EFNB2_EPHB2 | Efnb2 - Ephb2 | EPHB | Cell-Cell Contact | PMID: 15114347 |
| CD31 | CD31 | Efnb2 | Ephb3 | 0.004053 | 0 | EFNB2_EPHB3 | Efnb2 - Ephb3 | EPHB | Cell-Cell Contact | PMID: 15114347 |
| NeuN | CD31 | Efnb2 | Ephb3 | 0.006346 | 0 | EFNB2_EPHB3 | Efnb2 - Ephb3 | EPHB | Cell-Cell Contact | PMID: 15114347 |
| CD31 | CD31 | Efnb2 | Ephb4 | 0.013521 | 0 | EFNB2_EPHB4 | Efnb2 - Ephb4 | EPHB | Cell-Cell Contact | PMID: 15114347 |
| NeuN | CD31 | Efnb2 | Ephb4 | 0.021058 | 0 | EFNB2_EPHB4 | Efnb2 - Ephb4 | EPHB | Cell-Cell Contact | PMID: 15114347 |
| NeuN | CD31 | Efnb3 | Epha4 | 0.013322 | 0 | EFNB3_EPHA4 | Efnb3 - Epha4 | EPHB | Cell-Cell Contact | PMID: 15114347 |
| NeuN | NeuN | Efnb3 | Epha4 | 0.018724 | 0 | EFNB3_EPHA4 | Efnb3 - Epha4 | EPHB | Cell-Cell Contact | PMID: 15114347 |
| NeuN | CD31 | Efnb3 | Ephb1 | 0.01256 | 0 | EFNB3_EPHB1 | Efnb3 - Ephb1 | EPHB | Cell-Cell Contact | PMID: 15114347 |
| NeuN | NeuN | Efnb3 | Ephb1 | 0.019379 | 0 | EFNB3_EPHB1 | Efnb3 - Ephb1 | EPHB | Cell-Cell Contact | PMID: 15114347 |
| CD31 | NeuN | Efnb3 | Ephb2 | 0.00934 | 0 | EFNB3_EPHB2 | Efnb3 - Ephb2 | EPHB | Cell-Cell Contact | PMID: 15114347 |
| NeuN | NeuN | Efnb3 | Ephb2 | 0.018843 | 0 | EFNB3_EPHB2 | Efnb3 - Ephb2 | EPHB | Cell-Cell Contact | PMID: 15114347 |
| NeuN | CD31 | Efnb3 | Ephb3 | 0.003095 | 0 | EFNB3_EPHB3 | Efnb3 - Ephb3 | EPHB | Cell-Cell Contact | PMID: 15114347 |
| NeuN | NeuN | Efnb3 | Ephb3 | 0.001937 | 0 | EFNB3_EPHB3 | Efnb3 - Ephb3 | EPHB | Cell-Cell Contact | PMID: 15114347 |
| CD31 | CD31 | Efnb3 | Ephb4 | 0.005108 | 0 | EFNB3_EPHB4 | Efnb3 - Ephb4 | EPHB | Cell-Cell Contact | PMID: 15114347 |
| NeuN | CD31 | Efnb3 | Ephb4 | 0.01035 | 0 | EFNB3_EPHB4 | Efnb3 - Ephb4 | EPHB | Cell-Cell Contact | PMID: 15114347 |
| NeuN | CD31 | Efnb3 | Ephb6 | 0.004088 | 0 | EFNB3_EPHB6 | Efnb3 - Ephb6 | EPHB | Cell-Cell Contact | PMID: 15114347 |
| NeuN | NeuN | Efnb3 | Ephb6 | 0.005288 | 0 | EFNB3_EPHB6 | Efnb3 - Ephb6 | EPHB | Cell-Cell Contact | PMID: 15114347 |
| CD31 | CD31 | Esam | Esam | 0.013287 | 0 | ESAM_ESAM | Esam - Esam | ESAM | Cell-Cell Contact | KEGG: mmu04514; PMID: 11279107 |
| NeuN | CD31 | Esam | Esam | 0.001813 | 0 | ESAM_ESAM | Esam - Esam | ESAM | Cell-Cell Contact | KEGG: mmu04514; PMID: 11279107 |
| CD31 | NeuN | Esam | Esam | 0.001813 | 0 | ESAM_ESAM | Esam - Esam | ESAM | Cell-Cell Contact | KEGG: mmu04514; PMID: 11279107 |
| CD31 | CD31 | Icam5 | Cd209a | 0.00037 | 0 | ICAM5_CD209A | Icam5 - Cd209a | ICAM | Cell-Cell Contact | PMID: 15795245 |

| | | | | | | | | | | |
|---|---|---|---|---|---|---|---|---|---|---|
| CD31 | NeuN | Icam5 | Cd209a | 0.000297 | 0 | ICAM5_CD209A | Icam5 - Cd209a | ICAM | Cell-Cell Contact | PMID: 15795245 |
| CD31 | CD31 | Jam2 | ITGA3_ITGB1 | 0.001953 | 0 | JAM2_ITGA3_ITGB1 | Jam2 - (Itga3+Itgb1) | JAM | Cell-Cell Contact | PMID: 12070135 |
| CD31 | NeuN | Jam2 | ITGA3_ITGB1 | 0.000979 | 0 | JAM2_ITGA3_ITGB1 | Jam2 - (Itga3+Itgb1) | JAM | Cell-Cell Contact | PMID: 12070135 |
| CD31 | CD31 | Jam2 | ITGAV_ITGB1 | 0.00204 | 0 | JAM2_ITGAV_ITGB1 | Jam2 - (Itgav+Itgb1) | JAM | Cell-Cell Contact | KEGG: mmu04514 |
| NeuN | CD31 | Jam2 | ITGAV_ITGB1 | 0.000827 | 0 | JAM2_ITGAV_ITGB1 | Jam2 - (Itgav+Itgb1) | JAM | Cell-Cell Contact | KEGG: mmu04514 |
| CD31 | CD31 | Jam2 | Jam3 | 0.000993 | 0 | JAM2_JAM3 | Jam2 - Jam3 | JAM | Cell-Cell Contact | PMC1237098; PMID: 11590146 |
| CD31 | NeuN | Jam2 | Jam3 | 0.000952 | 0 | JAM2_JAM3 | Jam2 - Jam3 | JAM | Cell-Cell Contact | PMC1237098; PMID: 11590146 |
| CD31 | CD31 | F11r | F11r | 0.001235 | 0 | F11R_F11R | F11r - F11r | JAM | Cell-Cell Contact | KEGG: mmu04514 |
| NeuN | CD31 | F11r | F11r | 0.000607 | 0 | F11R_F11R | F11r - F11r | JAM | Cell-Cell Contact | KEGG: mmu04514 |
| CD31 | NeuN | F11r | F11r | 0.000607 | 0 | F11R_F11R | F11r - F11r | JAM | Cell-Cell Contact | KEGG: mmu04514 |
| CD31 | CD31 | Jam2 | F11r | 0.000962 | 0 | JAM2_F11R | Jam2 - F11r | JAM | Cell-Cell Contact | KEGG: mmu04514 |
| CD31 | NeuN | Jam2 | F11r | 0.000473 | 0 | JAM2_F11R | Jam2 - F11r | JAM | Cell-Cell Contact | KEGG: mmu04514 |
| CD31 | CD31 | Jam2 | Jam2 | 0.00075 | 0 | JAM2_JAM2 | Jam2 - Jam2 | JAM | Cell-Cell Contact | KEGG: mmu04514 |
| NeuN | CD31 | Jam2 | Jam2 | 0.000304 | 0 | JAM2_JAM2 | Jam2 - Jam2 | JAM | Cell-Cell Contact | KEGG: mmu04514 |
| CD31 | NeuN | Jam2 | Jam2 | 0.000304 | 0 | JAM2_JAM2 | Jam2 - Jam2 | JAM | Cell-Cell Contact | KEGG: mmu04514 |
| CD31 | CD31 | Jam3 | F11r | 0.001275 | 0 | JAM3_F11R | Jam3 - F11r | JAM | Cell-Cell Contact | KEGG: mmu04514 |
| NeuN | CD31 | Jam3 | F11r | 0.001221 | 0 | JAM3_F11R | Jam3 - F11r | JAM | Cell-Cell Contact | KEGG: mmu04514 |
| CD31 | CD31 | Jam3 | Jam3 | 0.001315 | 0 | JAM3_JAM3 | Jam3 - Jam3 | JAM | Cell-Cell Contact | KEGG: mmu04514 |
| NeuN | CD31 | Jam3 | Jam3 | 0.00126 | 0 | JAM3_JAM3 | Jam3 - Jam3 | JAM | Cell-Cell Contact | KEGG: mmu04514 |
| CD31 | NeuN | Jam3 | Jam3 | 0.00126 | 0 | JAM3_JAM3 | Jam3 - Jam3 | JAM | Cell-Cell Contact | KEGG: mmu04514 |
| NeuN | CD31 | L1cam | ITGAV_ITGB3 | 0.011026 | 0 | L1CAM_ITGAV_ITGB3 | L1cam - (Itgav+Itgb3) | L1CAM | Cell-Cell Contact | PMID: 9396761 |
| NeuN | NeuN | L1cam | ITGAV_ITGB3 | 0.005895 | 0 | L1CAM_ITGAV_ITGB3 | L1cam - (Itgav+Itgb3) | L1CAM | Cell-Cell Contact | PMID: 9396761 |
| NeuN | CD31 | L1cam | ITGA4_ITGB7 | 0.013106 | 0 | L1CAM_ITGA4_ITGB7 | L1cam - (Itga4+Itgb7) | L1CAM | Cell-Cell Contact | PMID: 8557754 |
| NeuN | NeuN | L1cam | ITGA4_ITGB7 | 0.006156 | 0 | L1CAM_ITGA4_ITGB7 | L1cam - (Itga4+Itgb7) | L1CAM | Cell-Cell Contact | PMID: 8557754 |
| NeuN | CD31 | L1cam | L1cam | 0.034293 | 0 | L1CAM_L1CAM | L1cam - L1cam | L1CAM | Cell-Cell Contact | PMID: 18701456 |
| CD31 | NeuN | L1cam | L1cam | 0.034293 | 0 | L1CAM_L1CAM | L1cam - L1cam | L1CAM | Cell-Cell Contact | PMID: 18701456 |
| NeuN | NeuN | L1cam | L1cam | 0.102656 | 0 | L1CAM_L1CAM | L1cam - L1cam | L1CAM | Cell-Cell Contact | PMID: 18701456 |
| CD31 | CD31 | Mag | Mag | 0.000264 | 0 | MAG_MAG | Mag - Mag | MAG | Cell-Cell Contact | KEGG: mmu04514 |
| NeuN | CD31 | Mag | Mag | 0.000142 | 0 | MAG_MAG | Mag - Mag | MAG | Cell-Cell Contact | KEGG: mmu04514 |
| CD31 | NeuN | Mag | Mag | 0.000142 | 0 | MAG_MAG | Mag - Mag | MAG | Cell-Cell Contact | KEGG: mmu04514 |
| CD31 | CD31 | Mpzl1 | Mpzl1 | 0.017436 | 0 | MPZL1_MPZL1 | Mpzl1 - Mpzl1 | MPZ | Cell-Cell Contact | KEGG: mmu04514 |
| NeuN | CD31 | Mpzl1 | Mpzl1 | 0.007994 | 0 | MPZL1_MPZL1 | Mpzl1 - Mpzl1 | MPZ | Cell-Cell Contact | KEGG: mmu04514 |
| CD31 | NeuN | Mpzl1 | Mpzl1 | 0.007994 | 0 | MPZL1_MPZL1 | Mpzl1 - Mpzl1 | MPZ | Cell-Cell Contact | KEGG: mmu04514 |
| NeuN | CD31 | Ncam1 | Fgfr1 | 0.020127 | 0 | NCAM1_FGFR1 | Ncam1 - Fgfr1 | NCAM | Cell-Cell Contact | PMID: 12791257 |
| NeuN | NeuN | Ncam1 | Fgfr1 | 0.027536 | 0 | NCAM1_FGFR1 | Ncam1 - Fgfr1 | NCAM | Cell-Cell Contact | PMID: 12791257 |
| NeuN | CD31 | Ncam1 | Ncam1 | 0.128665 | 0 | NCAM1_NCAM1 | Ncam1 - Ncam1 | NCAM | Cell-Cell Contact | KEGG: mmu04514 |
| CD31 | NeuN | Ncam1 | Ncam1 | 0.128665 | 0 | NCAM1_NCAM1 | Ncam1 - Ncam1 | NCAM | Cell-Cell Contact | KEGG: mmu04514 |
| NeuN | NeuN | Ncam1 | Ncam1 | 0.30062 | 0 | NCAM1_NCAM1 | Ncam1 - Ncam1 | NCAM | Cell-Cell Contact | KEGG: mmu04514 |
| NeuN | CD31 | Ncam1 | Ncam2 | 0.015817 | 0 | NCAM1_NCAM2 | Ncam1 - Ncam2 | NCAM | Cell-Cell Contact | KEGG: mmu04514 |
| NeuN | NeuN | Ncam1 | Ncam2 | 0.019068 | 0 | NCAM1_NCAM2 | Ncam1 - Ncam2 | NCAM | Cell-Cell Contact | KEGG: mmu04514 |
| CD31 | NeuN | Ncam1 | L1cam | 0.070786 | 0 | NCAM1_L1CAM | Ncam1 - L1cam | NCAM | Cell-Cell Contact | KEGG: mmu04514 |
| NeuN | NeuN | Ncam1 | L1cam | 0.181502 | 0 | NCAM1_L1CAM | Ncam1 - L1cam | NCAM | Cell-Cell Contact | KEGG: mmu04514 |
| CD31 | NeuN | Ncam2 | L1cam | 0.008223 | 0 | NCAM2_L1CAM | Ncam2 - L1cam | NCAM | Cell-Cell Contact | KEGG: mmu04514 |
| NeuN | NeuN | Ncam2 | L1cam | 0.009929 | 0 | NCAM2_L1CAM | Ncam2 - L1cam | NCAM | Cell-Cell Contact | KEGG: mmu04514 |
| CD31 | CD31 | Nectin1 | Cd96 | 0.000895 | 0 | NECTIN1_CD96 | Nectin1 - Cd96 | NECTIN | Cell-Cell Contact | PMID: 23027581 |
| NeuN | CD31 | Nectin1 | Cd96 | 0.001059 | 0 | NECTIN1_CD96 | Nectin1 - Cd96 | NECTIN | Cell-Cell Contact | PMID: 23027581 |
| CD31 | CD31 | Nectin3 | Nectin2 | 0.002331 | 0 | NECTIN3_NECTIN2 | Nectin3 - Nectin2 | NECTIN | Cell-Cell Contact | PMID: 23027581 |
| NeuN | CD31 | Nectin3 | Nectin2 | 0.00241 | 0 | NECTIN3_NECTIN2 | Nectin3 - Nectin2 | NECTIN | Cell-Cell Contact | PMID: 23027581 |
| CD31 | CD31 | Nectin3 | Pvr | 0.001155 | 0 | NECTIN3_PVR | Nectin3 - Pvr | NECTIN | Cell-Cell Contact | PMID: 23027581 |
| NeuN | CD31 | Nectin3 | Pvr | 0.001193 | 0 | NECTIN3_PVR | Nectin3 - Pvr | NECTIN | Cell-Cell Contact | PMID: 23027581 |
| NeuN | CD31 | Nectin1 | Nectin3 | 0.002051 | 0 | NECTIN1_NECTIN3 | Nectin1 - Nectin3 | NECTIN | Cell-Cell Contact | KEGG: mmu04514 |
| NeuN | NeuN | Nectin1 | Nectin3 | 0.00212 | 0 | NECTIN1_NECTIN3 | Nectin1 - Nectin3 | NECTIN | Cell-Cell Contact | KEGG: mmu04514 |
| NeuN | CD31 | Negr1 | Negr1 | 0.001332 | 0 | NEGR1_NEGR1 | Negr1 - Negr1 | NEGR | Cell-Cell Contact | KEGG: mmu04514 |
| CD31 | NeuN | Negr1 | Negr1 | 0.001332 | 0 | NEGR1_NEGR1 | Negr1 - Negr1 | NEGR | Cell-Cell Contact | KEGG: mmu04514 |
| NeuN | NeuN | Negr1 | Negr1 | 0.001453 | 0 | NEGR1_NEGR1 | Negr1 - Negr1 | NEGR | Cell-Cell Contact | KEGG: mmu04514 |
| NeuN | CD31 | Lrrc4c | Ntng1 | 0.001523 | 0 | LRRC4C_NTNG1 | Lrrc4c - Ntng1 | NGL | Cell-Cell Contact | KEGG: mmu04514 |
| NeuN | NeuN | Lrrc4c | Ntng1 | 0.001026 | 0 | LRRC4C_NTNG1 | Lrrc4c - Ntng1 | NGL | Cell-Cell Contact | KEGG: mmu04514 |
| CD31 | NeuN | Lrrc4 | Ntng2 | 0.003671 | 0 | LRRC4_NTNG2 | Lrrc4 - Ntng2 | NGL | Cell-Cell Contact | KEGG: mmu04514 |
| NeuN | NeuN | Lrrc4 | Ntng2 | 0.00337 | 0 | LRRC4_NTNG2 | Lrrc4 - Ntng2 | NGL | Cell-Cell Contact | KEGG: mmu04514 |
| CD31 | NeuN | Lrrc4b | Ptprf | 0.01249 | 0 | LRRC4B_PTPRF | Lrrc4b - Ptprf | NGL | Cell-Cell Contact | KEGG: mmu04514 |

| | | | | | | | | | |
|---|---|---|---|---|---|---|---|---|---|
| NeuN | NeuN | Lrrc4b | Ptprf | 0.023039 | 0 | LRRC4B_PTPRF | Lrrc4b - Ptprf | NGL | Cell-Cell Contact | KEGG: mmu04514 |
| CD31 | CD31 | Dlk1 | Notch1 | 0.003695 | 0 | DLK1_NOTCH1 | Dlk1 - Notch1 | NOTCH | Cell-Cell Contact | PMID: 22353464 |
| CD31 | NeuN | Dlk1 | Notch1 | 0.000788 | 0 | DLK1_NOTCH1 | Dlk1 - Notch1 | NOTCH | Cell-Cell Contact | PMID: 22353464 |
| CD31 | CD31 | Dlk1 | Notch3 | 0.004792 | 0 | DLK1_NOTCH3 | Dlk1 - Notch3 | NOTCH | Cell-Cell Contact | PMID: 22353464 |
| CD31 | NeuN | Dlk1 | Notch3 | 0.001072 | 0 | DLK1_NOTCH3 | Dlk1 - Notch3 | NOTCH | Cell-Cell Contact | PMID: 22353464 |
| CD31 | CD31 | Dlk1 | Notch4 | 0.002815 | 0 | DLK1_NOTCH4 | Dlk1 - Notch4 | NOTCH | Cell-Cell Contact | PMID: 22353464 |
| CD31 | NeuN | Dlk1 | Notch4 | 0.000797 | 0 | DLK1_NOTCH4 | Dlk1 - Notch4 | NOTCH | Cell-Cell Contact | PMID: 22353464 |
| CD31 | CD31 | Dll1 | Notch1 | 0.001794 | 0 | DLL1_NOTCH1 | Dll1 - Notch1 | NOTCH | Cell-Cell Contact | PMID: 22353464 |
| NeuN | CD31 | Dll1 | Notch1 | 0.001713 | 0 | DLL1_NOTCH1 | Dll1 - Notch1 | NOTCH | Cell-Cell Contact | PMID: 22353464 |
| CD31 | CD31 | Dll1 | Notch3 | 0.002328 | 0 | DLL1_NOTCH3 | Dll1 - Notch3 | NOTCH | Cell-Cell Contact | PMID: 22353464 |
| NeuN | CD31 | Dll1 | Notch3 | 0.002223 | 0 | DLL1_NOTCH3 | Dll1 - Notch3 | NOTCH | Cell-Cell Contact | PMID: 22353464 |
| CD31 | CD31 | Dll1 | Notch4 | 0.001366 | 0 | DLL1_NOTCH4 | Dll1 - Notch4 | NOTCH | Cell-Cell Contact | PMID: 22353464 |
| NeuN | CD31 | Dll1 | Notch4 | 0.001305 | 0 | DLL1_NOTCH4 | Dll1 - Notch4 | NOTCH | Cell-Cell Contact | PMID: 22353464 |
| CD31 | CD31 | Dll3 | Notch1 | 0.001648 | 0 | DLL3_NOTCH1 | Dll3 - Notch1 | NOTCH | Cell-Cell Contact | PMID: 22353464 |
| NeuN | CD31 | Dll3 | Notch1 | 0.001037 | 0 | DLL3_NOTCH1 | Dll3 - Notch1 | NOTCH | Cell-Cell Contact | PMID: 22353464 |
| CD31 | CD31 | Dll3 | Notch3 | 0.002139 | 0 | DLL3_NOTCH3 | Dll3 - Notch3 | NOTCH | Cell-Cell Contact | PMID: 22353464 |
| NeuN | CD31 | Dll3 | Notch3 | 0.001346 | 0 | DLL3_NOTCH3 | Dll3 - Notch3 | NOTCH | Cell-Cell Contact | PMID: 22353464 |
| CD31 | CD31 | Dll3 | Notch4 | 0.001255 | 0 | DLL3_NOTCH4 | Dll3 - Notch4 | NOTCH | Cell-Cell Contact | PMID: 22353464 |
| NeuN | CD31 | Dll3 | Notch4 | 0.00079 | 0 | DLL3_NOTCH4 | Dll3 - Notch4 | NOTCH | Cell-Cell Contact | PMID: 22353464 |
| CD31 | CD31 | Dll4 | Notch1 | 0.004191 | 0 | DLL4_NOTCH1 | Dll4 - Notch1 | NOTCH | Cell-Cell Contact | PMID: 22353464 |
| CD31 | NeuN | Dll4 | Notch1 | 0.000895 | 0 | DLL4_NOTCH1 | Dll4 - Notch1 | NOTCH | Cell-Cell Contact | PMID: 22353464 |
| CD31 | CD31 | Dll4 | Notch3 | 0.005435 | 0 | DLL4_NOTCH3 | Dll4 - Notch3 | NOTCH | Cell-Cell Contact | PMID: 22353464 |
| CD31 | NeuN | Dll4 | Notch3 | 0.001216 | 0 | DLL4_NOTCH3 | Dll4 - Notch3 | NOTCH | Cell-Cell Contact | PMID: 22353464 |
| CD31 | CD31 | Dll4 | Notch4 | 0.003193 | 0 | DLL4_NOTCH4 | Dll4 - Notch4 | NOTCH | Cell-Cell Contact | PMID: 22353464 |
| CD31 | NeuN | Dll4 | Notch4 | 0.000904 | 0 | DLL4_NOTCH4 | Dll4 - Notch4 | NOTCH | Cell-Cell Contact | PMID: 22353464 |
| CD31 | CD31 | Jag1 | Notch1 | 0.002848 | 0 | JAG1_NOTCH1 | Jag1 - Notch1 | NOTCH | Cell-Cell Contact | PMID: 22353464 |
| NeuN | CD31 | Jag1 | Notch1 | 0.000734 | 0 | JAG1_NOTCH1 | Jag1 - Notch1 | NOTCH | Cell-Cell Contact | PMID: 22353464 |
| CD31 | CD31 | Jag1 | Notch3 | 0.003694 | 0 | JAG1_NOTCH3 | Jag1 - Notch3 | NOTCH | Cell-Cell Contact | PMID: 22353464 |
| NeuN | CD31 | Jag1 | Notch3 | 0.000953 | 0 | JAG1_NOTCH3 | Jag1 - Notch3 | NOTCH | Cell-Cell Contact | PMID: 22353464 |
| CD31 | CD31 | Jag1 | Notch4 | 0.002169 | 0 | JAG1_NOTCH4 | Jag1 - Notch4 | NOTCH | Cell-Cell Contact | PMID: 22353464 |
| CD31 | NeuN | Jag1 | Notch4 | 0.000614 | 0 | JAG1_NOTCH4 | Jag1 - Notch4 | NOTCH | Cell-Cell Contact | PMID: 22353464 |
| CD31 | CD31 | Jag2 | Notch1 | 0.003828 | 0 | JAG2_NOTCH1 | Jag2 - Notch1 | NOTCH | Cell-Cell Contact | PMID: 22353464 |
| NeuN | CD31 | Jag2 | Notch1 | 0.002268 | 0 | JAG2_NOTCH1 | Jag2 - Notch1 | NOTCH | Cell-Cell Contact | PMID: 22353464 |
| CD31 | CD31 | Jag2 | Notch3 | 0.004964 | 0 | JAG2_NOTCH3 | Jag2 - Notch3 | NOTCH | Cell-Cell Contact | PMID: 22353464 |
| NeuN | CD31 | Jag2 | Notch3 | 0.002942 | 0 | JAG2_NOTCH3 | Jag2 - Notch3 | NOTCH | Cell-Cell Contact | PMID: 22353464 |
| CD31 | CD31 | Jag2 | Notch4 | 0.002916 | 0 | JAG2_NOTCH4 | Jag2 - Notch4 | NOTCH | Cell-Cell Contact | PMID: 22353464 |
| NeuN | CD31 | Jag2 | Notch4 | 0.001727 | 0 | JAG2_NOTCH4 | Jag2 - Notch4 | NOTCH | Cell-Cell Contact | PMID: 22353464 |
| CD31 | NeuN | Nrxn1 | Nlgn1 | 0.010856 | 0 | NRXN1_NLGN1 | Nrxn1 - Nlgn1 | NRXN | Cell-Cell Contact | KEGG: mmu04514 |
| NeuN | NeuN | Nrxn1 | Nlgn1 | 0.022902 | 0 | NRXN1_NLGN1 | Nrxn1 - Nlgn1 | NRXN | Cell-Cell Contact | KEGG: mmu04514 |
| CD31 | NeuN | Nrxn1 | Nlgn2 | 0.014995 | 0 | NRXN1_NLGN2 | Nrxn1 - Nlgn2 | NRXN | Cell-Cell Contact | KEGG: mmu04514 |
| NeuN | NeuN | Nrxn1 | Nlgn2 | 0.031488 | 0 | NRXN1_NLGN2 | Nrxn1 - Nlgn2 | NRXN | Cell-Cell Contact | KEGG: mmu04514 |
| NeuN | CD31 | Nrxn1 | Nlgn3 | 0.015771 | 0 | NRXN1_NLGN3 | Nrxn1 - Nlgn3 | NRXN | Cell-Cell Contact | KEGG: mmu04514 |
| NeuN | NeuN | Nrxn1 | Nlgn3 | 0.022278 | 0 | NRXN1_NLGN3 | Nrxn1 - Nlgn3 | NRXN | Cell-Cell Contact | KEGG: mmu04514 |
| CD31 | NeuN | Nrxn2 | Nlgn1 | 0.007752 | 0 | NRXN2_NLGN1 | Nrxn2 - Nlgn1 | NRXN | Cell-Cell Contact | KEGG: mmu04514 |
| NeuN | NeuN | Nrxn2 | Nlgn1 | 0.01563 | 0 | NRXN2_NLGN1 | Nrxn2 - Nlgn1 | NRXN | Cell-Cell Contact | KEGG: mmu04514 |
| CD31 | NeuN | Nrxn2 | Nlgn2 | 0.01072 | 0 | NRXN2_NLGN2 | Nrxn2 - Nlgn2 | NRXN | Cell-Cell Contact | KEGG: mmu04514 |
| NeuN | NeuN | Nrxn2 | Nlgn2 | 0.02155 | 0 | NRXN2_NLGN2 | Nrxn2 - Nlgn2 | NRXN | Cell-Cell Contact | KEGG: mmu04514 |
| NeuN | CD31 | Nrxn2 | Nlgn3 | 0.010738 | 0 | NRXN2_NLGN3 | Nrxn2 - Nlgn3 | NRXN | Cell-Cell Contact | KEGG: mmu04514 |
| NeuN | NeuN | Nrxn2 | Nlgn3 | 0.015201 | 0 | NRXN2_NLGN3 | Nrxn2 - Nlgn3 | NRXN | Cell-Cell Contact | KEGG: mmu04514 |
| CD31 | NeuN | Nrxn3 | Nlgn1 | 0.004605 | 0 | NRXN3_NLGN1 | Nrxn3 - Nlgn1 | NRXN | Cell-Cell Contact | KEGG: mmu04514 |
| NeuN | NeuN | Nrxn3 | Nlgn1 | 0.004432 | 0 | NRXN3_NLGN1 | Nrxn3 - Nlgn1 | NRXN | Cell-Cell Contact | KEGG: mmu04514 |
| CD31 | NeuN | Nrxn3 | Nlgn2 | 0.006377 | 0 | NRXN3_NLGN2 | Nrxn3 - Nlgn2 | NRXN | Cell-Cell Contact | KEGG: mmu04514 |
| NeuN | NeuN | Nrxn3 | Nlgn2 | 0.006138 | 0 | NRXN3_NLGN2 | Nrxn3 - Nlgn2 | NRXN | Cell-Cell Contact | KEGG: mmu04514 |
| CD31 | NeuN | Nrxn3 | Nlgn3 | 0.004477 | 0 | NRXN3_NLGN3 | Nrxn3 - Nlgn3 | NRXN | Cell-Cell Contact | KEGG: mmu04514 |
| NeuN | NeuN | Nrxn3 | Nlgn3 | 0.004309 | 0 | NRXN3_NLGN3 | Nrxn3 - Nlgn3 | NRXN | Cell-Cell Contact | KEGG: mmu04514 |
| CD31 | CD31 | Ocln | Ocln | 0.001495 | 0 | OCLN_OCLN | Ocln - Ocln | OCLN | Cell-Cell Contact | KEGG: mmu04514 |
| NeuN | CD31 | Ocln | Ocln | 0.000448 | 0 | OCLN_OCLN | Ocln - Ocln | OCLN | Cell-Cell Contact | KEGG: mmu04514 |
| CD31 | NeuN | Ocln | Ocln | 0.000448 | 0 | OCLN_OCLN | Ocln - Ocln | OCLN | Cell-Cell Contact | KEGG: mmu04514 |
| CD31 | CD31 | Pecam1 | Pecam1 | 0.004407 | 0 | PECAM1_PECAM1 | Pecam1 - Pecam1 | PECAM1 | Cell-Cell Contact | KEGG: mmu04514 |
| NeuN | CD31 | Pecam1 | Pecam1 | 0.000697 | 0 | PECAM1_PECAM1 | Pecam1 - Pecam1 | PECAM1 | Cell-Cell Contact | KEGG: mmu04514 |

| | | | | | | | | | | |
|---|---|---|---|---|---|---|---|---|---|---|
| CD31 | NeuN | Pecam1 | Pecam1 | 0.000697 | 0 | PECAM1_PECAM1 | Pecam1 - Pecam1 | PECAM1 | Cell-Cell Contact | KEGG: mmu04514 |
| CD31 | CD31 | Ptprm | Ptprm | 0.023218 | 0 | PTPRM_PTPRM | Ptprm - Ptprm | PTPRM | Cell-Cell Contact | KEGG: mmu04514 |
| NeuN | CD31 | Ptprm | Ptprm | 0.005285 | 0 | PTPRM_PTPRM | Ptprm - Ptprm | PTPRM | Cell-Cell Contact | KEGG: mmu04514 |
| CD31 | NeuN | Ptprm | Ptprm | 0.005285 | 0 | PTPRM_PTPRM | Ptprm - Ptprm | PTPRM | Cell-Cell Contact | KEGG: mmu04514 |
| CD31 | CD31 | Sele | Glg1 | 0.001346 | 0 | SELE_GLG1 | Sele - Glg1 | SELE | Cell-Cell Contact | PMID: 11404363 |
| CD31 | NeuN | Sele | Glg1 | 0.001657 | 0 | SELE_GLG1 | Sele - Glg1 | SELE | Cell-Cell Contact | PMID: 11404363 |
| CD31 | CD31 | Sell | Cd34 | 0.004894 | 0 | SELL_CD34 | Sell - Cd34 | SELL | Cell-Cell Contact | PMID: 7692600; PMID: 8977216 |
| NeuN | CD31 | Sell | Cd34 | 0.002435 | 0 | SELL_CD34 | Sell - Cd34 | SELL | Cell-Cell Contact | PMID: 7692600; PMID: 8977216 |
| CD31 | CD31 | Sell | Podxl | 0.003829 | 0 | SELL_PODXL | Sell - Podxl | SELL | Cell-Cell Contact | PMID: 22814396 |
| NeuN | CD31 | Sell | Podxl | 0.001904 | 0 | SELL_PODXL | Sell - Podxl | SELL | Cell-Cell Contact | PMID: 22814396 |
| CD31 | CD31 | Selplg | Sele | 0.000629 | 0 | SELPLG_SELE | Selplg - Sele | SELPLG | Cell-Cell Contact | PMC4571854 |
| NeuN | CD31 | Selplg | Sele | 0.000487 | 0 | SELPLG_SELE | Selplg - Sele | SELPLG | Cell-Cell Contact | PMC4571854 |
| CD31 | CD31 | Selplg | Sell | 0.00068 | 0 | SELPLG_SELL | Selplg - Sell | SELPLG | Cell-Cell Contact | PMC2431087; PMID: 8892633,PMID: 8896607 |
| NeuN | CD31 | Selplg | Sell | 0.000526 | 0 | SELPLG_SELL | Selplg - Sell | SELPLG | Cell-Cell Contact | PMC2431087; PMID: 8892633,PMID: 8896607 |
| NeuN | CD31 | Sema4a | NRP1_PLXNA1 | 0.002952 | 0 | SEMA4A_NRP1_PLXNA1 | Sema4a - (Nrp1+Plxna1) | SEMA4 | Cell-Cell Contact | PMID: 27533782 |
| NeuN | NeuN | Sema4a | NRP1_PLXNA1 | 0.002855 | 0 | SEMA4A_NRP1_PLXNA1 | Sema4a - (Nrp1+Plxna1) | SEMA4 | Cell-Cell Contact | PMID: 27533782 |
| NeuN | CD31 | Sema4a | NRP1_PLXNA2 | 0.002943 | 0 | SEMA4A_NRP1_PLXNA2 | Sema4a - (Nrp1+Plxna2) | SEMA4 | Cell-Cell Contact | PMID: 27533782 |
| NeuN | NeuN | Sema4a | NRP1_PLXNA2 | 0.002559 | 0 | SEMA4A_NRP1_PLXNA2 | Sema4a - (Nrp1+Plxna2) | SEMA4 | Cell-Cell Contact | PMID: 27533782 |
| NeuN | CD31 | Sema4a | NRP1_PLXNA3 | 0.002374 | 0 | SEMA4A_NRP1_PLXNA3 | Sema4a - (Nrp1+Plxna3) | SEMA4 | Cell-Cell Contact | PMID: 27533782 |
| NeuN | NeuN | Sema4a | NRP1_PLXNA3 | 0.002986 | 0 | SEMA4A_NRP1_PLXNA3 | Sema4a - (Nrp1+Plxna3) | SEMA4 | Cell-Cell Contact | PMID: 27533782 |
| NeuN | CD31 | Sema4a | NRP1_PLXNA4 | 0.003937 | 0 | SEMA4A_NRP1_PLXNA4 | Sema4a - (Nrp1+Plxna4) | SEMA4 | Cell-Cell Contact | PMID: 27533782 |
| NeuN | NeuN | Sema4a | NRP1_PLXNA4 | 0.005587 | 0 | SEMA4A_NRP1_PLXNA4 | Sema4a - (Nrp1+Plxna4) | SEMA4 | Cell-Cell Contact | PMID: 27533782 |
| NeuN | CD31 | Sema4a | Plxnb1 | 0.001217 | 0 | SEMA4A_PLXNB1 | Sema4a - Plxnb1 | SEMA4 | Cell-Cell Contact | PMID: 27533782 |
| NeuN | NeuN | Sema4a | Plxnb1 | 0.00139 | 0 | SEMA4A_PLXNB1 | Sema4a - Plxnb1 | SEMA4 | Cell-Cell Contact | PMID: 27533782 |
| NeuN | CD31 | Sema4a | Plxnb2 | 0.00233 | 0 | SEMA4A_PLXNB2 | Sema4a - Plxnb2 | SEMA4 | Cell-Cell Contact | PMID: 27533782 |
| NeuN | NeuN | Sema4a | Plxnb2 | 0.003274 | 0 | SEMA4A_PLXNB2 | Sema4a - Plxnb2 | SEMA4 | Cell-Cell Contact | PMID: 27533782 |
| CD31 | CD31 | Sema4a | Plxnb3 | 0.000684 | 0 | SEMA4A_PLXNB3 | Sema4a - Plxnb3 | SEMA4 | Cell-Cell Contact | PMID: 27533782 |
| NeuN | CD31 | Sema4a | Plxnb3 | 0.001041 | 0 | SEMA4A_PLXNB3 | Sema4a - Plxnb3 | SEMA4 | Cell-Cell Contact | PMID: 27533782 |
| CD31 | NeuN | Sema4d | Plxnb1 | 0.000984 | 0 | SEMA4D_PLXNB1 | Sema4d - Plxnb1 | SEMA4 | Cell-Cell Contact | PMID: 27533782 |
| NeuN | NeuN | Sema4d | Plxnb1 | 0.000953 | 0 | SEMA4D_PLXNB1 | Sema4d - Plxnb1 | SEMA4 | Cell-Cell Contact | PMID: 27533782 |
| CD31 | NeuN | Sema4d | Plxnb2 | 0.00232 | 0 | SEMA4D_PLXNB2 | Sema4d - Plxnb2 | SEMA4 | Cell-Cell Contact | PMID: 27533782 |
| NeuN | NeuN | Sema4d | Plxnb2 | 0.002246 | 0 | SEMA4D_PLXNB2 | Sema4d - Plxnb2 | SEMA4 | Cell-Cell Contact | PMID: 27533782 |
| CD31 | CD31 | Sema4d | Plxnb3 | 0.000737 | 0 | SEMA4D_PLXNB3 | Sema4d - Plxnb3 | SEMA4 | Cell-Cell Contact | PMID: 27533782 |
| NeuN | CD31 | Sema4d | Plxnb3 | 0.000714 | 0 | SEMA4D_PLXNB3 | Sema4d - Plxnb3 | SEMA4 | Cell-Cell Contact | PMID: 27533782 |
| CD31 | NeuN | Sema4c | Plxnb2 | 0.006308 | 0 | SEMA4C_PLXNB2 | Sema4c - Plxnb2 | SEMA4 | Cell-Cell Contact | PMID: 27533782 |
| NeuN | NeuN | Sema4c | Plxnb2 | 0.007483 | 0 | SEMA4C_PLXNB2 | Sema4c - Plxnb2 | SEMA4 | Cell-Cell Contact | PMID: 27533782 |
| NeuN | CD31 | Sema4g | Plxnb2 | 0.004318 | 0 | SEMA4G_PLXNB2 | Sema4g - Plxnb2 | SEMA4 | Cell-Cell Contact | PMID: 27533782 |
| NeuN | NeuN | Sema4g | Plxnb2 | 0.006064 | 0 | SEMA4G_PLXNB2 | Sema4g - Plxnb2 | SEMA4 | Cell-Cell Contact | PMID: 27533782 |
| CD31 | CD31 | Sema4a | Timd2 | 0.000602 | 0 | SEMA4A_TIMD2 | Sema4a - Timd2 | SEMA4 | Cell-Cell Contact | PMID: 22325954 |
| NeuN | CD31 | Sema4a | Timd2 | 0.000916 | 0 | SEMA4A_TIMD2 | Sema4a - Timd2 | SEMA4 | Cell-Cell Contact | PMID: 22325954 |
| CD31 | CD31 | Sema5a | Plxna1 | 0.002847 | 0 | SEMA5A_PLXNA1 | Sema5a - Plxna1 | SEMA5 | Cell-Cell Contact | PMID: 27533782 |
| CD31 | NeuN | Sema5a | Plxna1 | 0.003906 | 0 | SEMA5A_PLXNA1 | Sema5a - Plxna1 | SEMA5 | Cell-Cell Contact | PMID: 27533782 |
| CD31 | CD31 | Sema5b | Plxna1 | 0.001687 | 0 | SEMA5B_PLXNA1 | Sema5b - Plxna1 | SEMA5 | Cell-Cell Contact | PMID: 27533782 |
| CD31 | NeuN | Sema5b | Plxna1 | 0.002316 | 0 | SEMA5B_PLXNA1 | Sema5b - Plxna1 | SEMA5 | Cell-Cell Contact | PMID: 27533782 |
| CD31 | NeuN | Sema5a | Plxna3 | 0.004273 | 0 | SEMA5A_PLXNA3 | Sema5a - Plxna3 | SEMA5 | Cell-Cell Contact | PMID: 27533782 |
| NeuN | NeuN | Sema5a | Plxna3 | 0.002015 | 0 | SEMA5A_PLXNA3 | Sema5a - Plxna3 | SEMA5 | Cell-Cell Contact | PMID: 27533782 |
| CD31 | NeuN | Sema5b | Plxna3 | 0.002534 | 0 | SEMA5B_PLXNA3 | Sema5b - Plxna3 | SEMA5 | Cell-Cell Contact | PMID: 27533782 |
| NeuN | NeuN | Sema5b | Plxna3 | 0.001743 | 0 | SEMA5B_PLXNA3 | Sema5b - Plxna3 | SEMA5 | Cell-Cell Contact | PMID: 27533782 |
| CD31 | CD31 | Sema5a | Plxnb3 | 0.001082 | 0 | SEMA5A_PLXNB3 | Sema5a - Plxnb3 | SEMA5 | Cell-Cell Contact | PMID: 27533782 |
| CD31 | NeuN | Sema5a | Plxnb3 | 0.000541 | 0 | SEMA5A_PLXNB3 | Sema5a - Plxnb3 | SEMA5 | Cell-Cell Contact | PMID: 27533782 |
| CD31 | CD31 | Sema6d | PLXNA1_KDR | 0.022699 | 0 | SEMA6D_PLXNA1_KDR | Sema6d - (Plxna1+Kdr) | SEMA6 | Cell-Cell Contact | PMID: 22325954 |
| CD31 | NeuN | Sema6d | PLXNA1_KDR | 0.006194 | 0 | SEMA6D_PLXNA1_KDR | Sema6d - (Plxna1+Kdr) | SEMA6 | Cell-Cell Contact | PMID: 22325954 |
| CD31 | CD31 | Sema6d | PLXNA1_TREM2 | 0.007164 | 0 | SEMA6D_PLXNA1_TREM2 | Sema6d - (Plxna1+Trem2) | SEMA6 | Cell-Cell Contact | PMID: 22325954 |
| CD31 | NeuN | Sema6d | PLXNA1_TREM2 | 0.004866 | 0 | SEMA6D_PLXNA1_TREM2 | Sema6d - (Plxna1+Trem2) | SEMA6 | Cell-Cell Contact | PMID: 22325954 |
| CD31 | CD31 | Sema6d | Plxna1 | 0.012975 | 0 | SEMA6D_PLXNA1 | Sema6d - Plxna1 | SEMA6 | Cell-Cell Contact | PMID: 27533782 |
| CD31 | NeuN | Sema6d | Plxna1 | 0.017734 | 0 | SEMA6D_PLXNA1 | Sema6d - Plxna1 | SEMA6 | Cell-Cell Contact | PMID: 27533782 |
| CD31 | CD31 | Sema6a | Plxna2 | 0.006248 | 0 | SEMA6A_PLXNA2 | Sema6a - Plxna2 | SEMA6 | Cell-Cell Contact | PMID: 27533782 |
| CD31 | NeuN | Sema6a | Plxna2 | 0.006928 | 0 | SEMA6A_PLXNA2 | Sema6a - Plxna2 | SEMA6 | Cell-Cell Contact | PMID: 27533782 |
| NeuN | CD31 | Sema6b | Plxna2 | 0.004119 | 0 | SEMA6B_PLXNA2 | Sema6b - Plxna2 | SEMA6 | Cell-Cell Contact | PMID: 27533782 |
| NeuN | NeuN | Sema6b | Plxna2 | 0.004568 | 0 | SEMA6B_PLXNA2 | Sema6b - Plxna2 | SEMA6 | Cell-Cell Contact | PMID: 27533782 |

| | | | | | | | | | | |
|---|---|---|---|---|---|---|---|---|---|---|
| CD31 | NeuN | Sema6a | Plxna4 | 0.032375 | 0 | SEMA6A_PLXNA4 | Sema6a - Plxna4 | SEMA6 | Cell-Cell Contact | PMID: 27533782 |
| NeuN | NeuN | Sema6a | Plxna4 | 0.012962 | 0 | SEMA6A_PLXNA4 | Sema6a - Plxna4 | SEMA6 | Cell-Cell Contact | PMID: 27533782 |
| CD31 | NeuN | Sema6b | Plxna4 | 0.009245 | 0 | SEMA6B_PLXNA4 | Sema6b - Plxna4 | SEMA6 | Cell-Cell Contact | PMID: 27533782 |
| NeuN | NeuN | Sema6b | Plxna4 | 0.021534 | 0 | SEMA6B_PLXNA4 | Sema6b - Plxna4 | SEMA6 | Cell-Cell Contact | PMID: 27533782 |
| CD31 | CD31 | Sema7a | ITGB1_ITGA1 | 0.013288 | 0 | SEMA7A_ITGB1_ITGA1 | Sema7a - (Itgb1+Itga1) | SEMA7 | Cell-Cell Contact | PMID: 22325954 |
| NeuN | CD31 | Sema7a | ITGB1_ITGA1 | 0.018359 | 0 | SEMA7A_ITGB1_ITGA1 | Sema7a - (Itgb1+Itga1) | SEMA7 | Cell-Cell Contact | PMID: 22325954 |
| NeuN | CD31 | Sema7a | Plxnc1 | 0.002397 | 0 | SEMA7A_PLXNC1 | Sema7a - Plxnc1 | SEMA7 | Cell-Cell Contact | PMID: 27533782 |
| NeuN | NeuN | Sema7a | Plxnc1 | 0.002915 | 0 | SEMA7A_PLXNC1 | Sema7a - Plxnc1 | SEMA7 | Cell-Cell Contact | PMID: 27533782 |
| CD31 | CD31 | Thy1 | ITGAV_ITGB3 | 0.000834 | 0 | THY1_ITGAV_ITGB3 | Thy1 - (Itgav+Itgb3) | THY1 | Cell-Cell Contact | PMID: 29212879; PMID: 18346467 |
| NeuN | CD31 | Thy1 | ITGAV_ITGB3 | 0.000914 | 0 | THY1_ITGAV_ITGB3 | Thy1 - (Itgav+Itgb3) | THY1 | Cell-Cell Contact | PMID: 29212879; PMID: 18346467 |
| CD31 | CD31 | Cdh3 | Cdh3 | 0.000512 | 0 | CDH3_CDH3 | Cdh3 - Cdh3 | CDH | Cell-Cell Contact | KEGG: hsa04514 |
| NeuN | CD31 | Cdh3 | Cdh3 | 0.000412 | 0 | CDH3_CDH3 | Cdh3 - Cdh3 | CDH | Cell-Cell Contact | KEGG: hsa04514 |
| CD31 | NeuN | Cdh3 | Cdh3 | 0.000412 | 0 | CDH3_CDH3 | Cdh3 - Cdh3 | CDH | Cell-Cell Contact | KEGG: hsa04514 |
| CD31 | CD31 | Cdh12 | ITGA1_ITGB1 | 0.003293 | 0 | CDH12_ITGA1_ITGB1 | Cdh12 - ITGA1_ITGB1 | CDH | Cell-Cell Contact | PMID: 34385456 |
| NeuN | CD31 | Cdh12 | ITGA1_ITGB1 | 0.00709 | 0 | CDH12_ITGA1_ITGB1 | Cdh12 - ITGA1_ITGB1 | CDH | Cell-Cell Contact | PMID: 34385456 |
| CD31 | CD31 | Gja1 | Gja1 | 0.002588 | 0 | GJA1_GJA1 | Gja1 - Gja1 | GAP | Cell-Cell Contact | PMID: 32532966;PMID: 15094344 |
| NeuN | CD31 | Gja1 | Gja1 | 0.000624 | 0 | GJA1_GJA1 | Gja1 - Gja1 | GAP | Cell-Cell Contact | PMID: 32532966;PMID: 15094344 |
| CD31 | NeuN | Gja1 | Gja1 | 0.000624 | 0 | GJA1_GJA1 | Gja1 - Gja1 | GAP | Cell-Cell Contact | PMID: 32532966;PMID: 15094344 |
| CD31 | CD31 | Gja1 | Gja6 | 0.001248 | 0 | GJA1_GJA6 | Gja1 - Gja6 | GAP | Cell-Cell Contact | PMID: 32532966;PMID: 15094344 |
| CD31 | NeuN | Gja1 | Gja6 | 0.001908 | 0 | GJA1_GJA6 | Gja1 - Gja6 | GAP | Cell-Cell Contact | PMID: 32532966;PMID: 15094344 |
| NeuN | CD31 | Gja6 | Gja6 | 0.00092 | 0 | GJA6_GJA6 | Gja6 - Gja6 | GAP | Cell-Cell Contact | PMID: 32532966;PMID: 15094344 |
| CD31 | NeuN | Gja6 | Gja6 | 0.00092 | 0 | GJA6_GJA6 | Gja6 - Gja6 | GAP | Cell-Cell Contact | PMID: 32532966;PMID: 15094344 |
| NeuN | NeuN | Gja6 | Gja6 | 0.001407 | 0 | GJA6_GJA6 | Gja6 - Gja6 | GAP | Cell-Cell Contact | PMID: 32532966;PMID: 15094344 |
| CD31 | CD31 | Gjc1 | Gja1 | 0.004537 | 0 | GJC1_GJA1 | Gjc1 - Gja1 | GAP | Cell-Cell Contact | PMID: 32532966;PMID: 15094344 |
| NeuN | CD31 | Gjc1 | Gja1 | 0.003439 | 0 | GJC1_GJA1 | Gjc1 - Gja1 | GAP | Cell-Cell Contact | PMID: 32532966;PMID: 15094344 |
| CD31 | NeuN | Gjc1 | Gja6 | 0.003346 | 0 | GJC1_GJA6 | Gjc1 - Gja6 | GAP | Cell-Cell Contact | PMID: 32532966;PMID: 15094344 |
| NeuN | NeuN | Gjc1 | Gja6 | 0.002536 | 0 | GJC1_GJA6 | Gjc1 - Gja6 | GAP | Cell-Cell Contact | PMID: 32532966;PMID: 15094344 |
| CD31 | CD31 | Gjc1 | Gjc1 | 0.00794 | 0 | GJC1_GJC1 | Gjc1 - Gjc1 | GAP | Cell-Cell Contact | PMID: 32532966;PMID: 15094344 |
| NeuN | CD31 | Gjc1 | Gjc1 | 0.006023 | 0 | GJC1_GJC1 | Gjc1 - Gjc1 | GAP | Cell-Cell Contact | PMID: 32532966;PMID: 15094344 |
| CD31 | NeuN | Gjc1 | Gjc1 | 0.006023 | 0 | GJC1_GJC1 | Gjc1 - Gjc1 | GAP | Cell-Cell Contact | PMID: 32532966;PMID: 15094344 |
| NeuN | CD31 | Cldn1 | Cldn1 | 0.000546 | 0 | CLDN1_CLDN1 | Cldn1 - Cldn1 | CLDN | Cell-Cell Contact | PMID: 31942240 |
| CD31 | NeuN | Cldn1 | Cldn1 | 0.000546 | 0 | CLDN1_CLDN1 | Cldn1 - Cldn1 | CLDN | Cell-Cell Contact | PMID: 31942240 |
| NeuN | NeuN | Cldn1 | Cldn1 | 0.000592 | 0 | CLDN1_CLDN1 | Cldn1 - Cldn1 | CLDN | Cell-Cell Contact | PMID: 31942240 |
| CD31 | CD31 | Cldn3 | Cldn3 | 0.000693 | 0 | CLDN3_CLDN3 | Cldn3 - Cldn3 | CLDN | Cell-Cell Contact | PMID: 31942240 |
| NeuN | CD31 | Cldn3 | Cldn3 | 0.000582 | 0 | CLDN3_CLDN3 | Cldn3 - Cldn3 | CLDN | Cell-Cell Contact | PMID: 31942240 |
| CD31 | NeuN | Cldn3 | Cldn3 | 0.000582 | 0 | CLDN3_CLDN3 | Cldn3 - Cldn3 | CLDN | Cell-Cell Contact | PMID: 31942240 |
| CD31 | CD31 | Pcdha9 | Pcdha9 | 0.000524 | 0 | PCDHA9_PCDHA9 | Pcdha9 - Pcdha9 | PCDH | Cell-Cell Contact | PMID: 35253643; PMID: 32694982 |
| NeuN | CD31 | Pcdha9 | Pcdha9 | 0.000418 | 0 | PCDHA9_PCDHA9 | Pcdha9 - Pcdha9 | PCDH | Cell-Cell Contact | PMID: 35253643; PMID: 32694982 |
| CD31 | NeuN | Pcdha9 | Pcdha9 | 0.000418 | 0 | PCDHA9_PCDHA9 | Pcdha9 - Pcdha9 | PCDH | Cell-Cell Contact | PMID: 35253643; PMID: 32694982 |
| NeuN | CD31 | Pcdha11 | Pcdha11 | 0.001184 | 0 | PCDHA11_PCDHA11 | Pcdha11 - Pcdha11 | PCDH | Cell-Cell Contact | PMID: 35253643; PMID: 32694982 |
| CD31 | NeuN | Pcdha11 | Pcdha11 | 0.001184 | 0 | PCDHA11_PCDHA11 | Pcdha11 - Pcdha11 | PCDH | Cell-Cell Contact | PMID: 35253643; PMID: 32694982 |
| NeuN | NeuN | Pcdha11 | Pcdha11 | 0.002233 | 0 | PCDHA11_PCDHA11 | Pcdha11 - Pcdha11 | PCDH | Cell-Cell Contact | PMID: 35253643; PMID: 32694982 |
| CD31 | CD31 | Pcdha10 | Pcdha10 | 0.000443 | 0 | PCDHA10_PCDHA10 | Pcdha10 - Pcdha10 | PCDH | Cell-Cell Contact | PMID: 35253643; PMID: 32694982 |
| NeuN | CD31 | Pcdha10 | Pcdha10 | 0.000407 | 0 | PCDHA10_PCDHA10 | Pcdha10 - Pcdha10 | PCDH | Cell-Cell Contact | PMID: 35253643; PMID: 32694982 |
| CD31 | NeuN | Pcdha10 | Pcdha10 | 0.000407 | 0 | PCDHA10_PCDHA10 | Pcdha10 - Pcdha10 | PCDH | Cell-Cell Contact | PMID: 35253643; PMID: 32694982 |
| NeuN | CD31 | Pcdha12 | Pcdha12 | 0.001255 | 0 | PCDHA12_PCDHA12 | Pcdha12 - Pcdha12 | PCDH | Cell-Cell Contact | PMID: 35253643; PMID: 32694982 |
| CD31 | NeuN | Pcdha12 | Pcdha12 | 0.001255 | 0 | PCDHA12_PCDHA12 | Pcdha12 - Pcdha12 | PCDH | Cell-Cell Contact | PMID: 35253643; PMID: 32694982 |
| NeuN | NeuN | Pcdha12 | Pcdha12 | 0.001315 | 0 | PCDHA12_PCDHA12 | Pcdha12 - Pcdha12 | PCDH | Cell-Cell Contact | PMID: 35253643; PMID: 32694982 |
| CD31 | CD31 | Pcdha2 | Pcdha2 | 0.000762 | 0 | PCDHA2_PCDHA2 | Pcdha2 - Pcdha2 | PCDH | Cell-Cell Contact | PMID: 35253643; PMID: 32694982 |
| NeuN | CD31 | Pcdha2 | Pcdha2 | 0.000535 | 0 | PCDHA2_PCDHA2 | Pcdha2 - Pcdha2 | PCDH | Cell-Cell Contact | PMID: 35253643; PMID: 32694982 |
| CD31 | NeuN | Pcdha2 | Pcdha2 | 0.000535 | 0 | PCDHA2_PCDHA2 | Pcdha2 - Pcdha2 | PCDH | Cell-Cell Contact | PMID: 35253643; PMID: 32694982 |
| CD31 | CD31 | Pcdha3 | Pcdha3 | 0.000968 | 0 | PCDHA3_PCDHA3 | Pcdha3 - Pcdha3 | PCDH | Cell-Cell Contact | PMID: 35253643; PMID: 32694982 |
| NeuN | CD31 | Pcdha3 | Pcdha3 | 0.000949 | 0 | PCDHA3_PCDHA3 | Pcdha3 - Pcdha3 | PCDH | Cell-Cell Contact | PMID: 35253643; PMID: 32694982 |
| CD31 | NeuN | Pcdha3 | Pcdha3 | 0.000949 | 0 | PCDHA3_PCDHA3 | Pcdha3 - Pcdha3 | PCDH | Cell-Cell Contact | PMID: 35253643; PMID: 32694982 |
| NeuN | CD31 | Pcdha5 | Pcdha5 | 0.000493 | 0 | PCDHA5_PCDHA5 | Pcdha5 - Pcdha5 | PCDH | Cell-Cell Contact | PMID: 35253643; PMID: 32694982 |
| CD31 | NeuN | Pcdha5 | Pcdha5 | 0.000493 | 0 | PCDHA5_PCDHA5 | Pcdha5 - Pcdha5 | PCDH | Cell-Cell Contact | PMID: 35253643; PMID: 32694982 |
| NeuN | NeuN | Pcdha5 | Pcdha5 | 0.000886 | 0 | PCDHA5_PCDHA5 | Pcdha5 - Pcdha5 | PCDH | Cell-Cell Contact | PMID: 35253643; PMID: 32694982 |
| CD31 | CD31 | Pcdha8 | Pcdha8 | 0.000457 | 0 | PCDHA8_PCDHA8 | Pcdha8 - Pcdha8 | PCDH | Cell-Cell Contact | PMID: 35253643; PMID: 32694982 |
| NeuN | CD31 | Pcdha8 | Pcdha8 | 0.000365 | 0 | PCDHA8_PCDHA8 | Pcdha8 - Pcdha8 | PCDH | Cell-Cell Contact | PMID: 35253643; PMID: 32694982 |
| CD31 | NeuN | Pcdha8 | Pcdha8 | 0.000365 | 0 | PCDHA8_PCDHA8 | Pcdha8 - Pcdha8 | PCDH | Cell-Cell Contact | PMID: 35253643; PMID: 32694982 |

| | | | | | | | | | | |
|---|---|---|---|---|---|---|---|---|---|---|
| NeuN | CD31 | Pcdhac1 | Pcdhac1 | 0.000569 | 0 | PCDHAC1_PCDHAC1 | Pcdhac1 - Pcdhac1 | PCDH | Cell-Cell Contact | PMID: 35253643; PMID: 32694982 |
| CD31 | NeuN | Pcdhac1 | Pcdhac1 | 0.000569 | 0 | PCDHAC1_PCDHAC1 | Pcdhac1 - Pcdhac1 | PCDH | Cell-Cell Contact | PMID: 35253643; PMID: 32694982 |
| NeuN | NeuN | Pcdhac1 | Pcdhac1 | 0.000598 | 0 | PCDHAC1_PCDHAC1 | Pcdhac1 - Pcdhac1 | PCDH | Cell-Cell Contact | PMID: 35253643; PMID: 32694982 |
| NeuN | CD31 | Pcdhac2 | Pcdhac2 | 0.000521 | 0 | PCDHAC2_PCDHAC2 | Pcdhac2 - Pcdhac2 | PCDH | Cell-Cell Contact | PMID: 35253643; PMID: 32694982 |
| CD31 | NeuN | Pcdhac2 | Pcdhac2 | 0.000521 | 0 | PCDHAC2_PCDHAC2 | Pcdhac2 - Pcdhac2 | PCDH | Cell-Cell Contact | PMID: 35253643; PMID: 32694982 |
| NeuN | NeuN | Pcdhac2 | Pcdhac2 | 0.001025 | 0 | PCDHAC2_PCDHAC2 | Pcdhac2 - Pcdhac2 | PCDH | Cell-Cell Contact | PMID: 35253643; PMID: 32694982 |
| CD31 | CD31 | Pcdhb18 | Pcdhb18 | 0.000424 | 0 | PCDHB18_PCDHB18 | Pcdhb18 - Pcdhb18 | PCDH | Cell-Cell Contact | PMID: 35253643; PMID: 32694982 |
| NeuN | CD31 | Pcdhb18 | Pcdhb18 | 0.00033 | 0 | PCDHB18_PCDHB18 | Pcdhb18 - Pcdhb18 | PCDH | Cell-Cell Contact | PMID: 35253643; PMID: 32694982 |
| CD31 | NeuN | Pcdhb18 | Pcdhb18 | 0.00033 | 0 | PCDHB18_PCDHB18 | Pcdhb18 - Pcdhb18 | PCDH | Cell-Cell Contact | PMID: 35253643; PMID: 32694982 |
| CD31 | CD31 | Pcdhb19 | Pcdhb19 | 0.000824 | 0 | PCDHB19_PCDHB19 | Pcdhb19 - Pcdhb19 | PCDH | Cell-Cell Contact | PMID: 35253643; PMID: 32694982 |
| NeuN | CD31 | Pcdhb19 | Pcdhb19 | 0.000687 | 0 | PCDHB19_PCDHB19 | Pcdhb19 - Pcdhb19 | PCDH | Cell-Cell Contact | PMID: 35253643; PMID: 32694982 |
| CD31 | NeuN | Pcdhb19 | Pcdhb19 | 0.000687 | 0 | PCDHB19_PCDHB19 | Pcdhb19 - Pcdhb19 | PCDH | Cell-Cell Contact | PMID: 35253643; PMID: 32694982 |
| CD31 | CD31 | Pcdhb19 | Pcdhb21 | 0.000631 | 0 | PCDHB19_PCDHB21 | Pcdhb19 - Pcdhb21 | PCDH | Cell-Cell Contact | PMID: 35253643; PMID: 32694982 |
| NeuN | CD31 | Pcdhb19 | Pcdhb21 | 0.000526 | 0 | PCDHB19_PCDHB21 | Pcdhb19 - Pcdhb21 | PCDH | Cell-Cell Contact | PMID: 35253643; PMID: 32694982 |
| CD31 | CD31 | Pcdhb21 | Pcdhb21 | 0.000483 | 0 | PCDHB21_PCDHB21 | Pcdhb21 - Pcdhb21 | PCDH | Cell-Cell Contact | PMID: 35253643; PMID: 32694982 |
| NeuN | CD31 | Pcdhb21 | Pcdhb21 | 0.000388 | 0 | PCDHB21_PCDHB21 | Pcdhb21 - Pcdhb21 | PCDH | Cell-Cell Contact | PMID: 35253643; PMID: 32694982 |
| CD31 | NeuN | Pcdhb21 | Pcdhb21 | 0.000388 | 0 | PCDHB21_PCDHB21 | Pcdhb21 - Pcdhb21 | PCDH | Cell-Cell Contact | PMID: 35253643; PMID: 32694982 |
| CD31 | CD31 | Pcdhb16 | Pcdhb16 | 0.000705 | 0 | PCDHB16_PCDHB16 | Pcdhb16 - Pcdhb16 | PCDH | Cell-Cell Contact | PMID: 35253643; PMID: 32694982 |
| NeuN | CD31 | Pcdhb16 | Pcdhb16 | 0.000436 | 0 | PCDHB16_PCDHB16 | Pcdhb16 - Pcdhb16 | PCDH | Cell-Cell Contact | PMID: 35253643; PMID: 32694982 |
| CD31 | NeuN | Pcdhb16 | Pcdhb16 | 0.000436 | 0 | PCDHB16_PCDHB16 | Pcdhb16 - Pcdhb16 | PCDH | Cell-Cell Contact | PMID: 35253643; PMID: 32694982 |
| CD31 | CD31 | Pcdhb20 | Pcdhb20 | 0.0015 | 0 | PCDHB20_PCDHB20 | Pcdhb20 - Pcdhb20 | PCDH | Cell-Cell Contact | PMID: 35253643; PMID: 32694982 |
| NeuN | CD31 | Pcdhb20 | Pcdhb20 | 0.001358 | 0 | PCDHB20_PCDHB20 | Pcdhb20 - Pcdhb20 | PCDH | Cell-Cell Contact | PMID: 35253643; PMID: 32694982 |
| CD31 | NeuN | Pcdhb20 | Pcdhb20 | 0.001358 | 0 | PCDHB20_PCDHB20 | Pcdhb20 - Pcdhb20 | PCDH | Cell-Cell Contact | PMID: 35253643; PMID: 32694982 |
| CD31 | CD31 | Pcdhb22 | Pcdhb22 | 0.001191 | 0 | PCDHB22_PCDHB22 | Pcdhb22 - Pcdhb22 | PCDH | Cell-Cell Contact | PMID: 35253643; PMID: 32694982 |
| NeuN | CD31 | Pcdhb22 | Pcdhb22 | 0.000986 | 0 | PCDHB22_PCDHB22 | Pcdhb22 - Pcdhb22 | PCDH | Cell-Cell Contact | PMID: 35253643; PMID: 32694982 |
| CD31 | NeuN | Pcdhb22 | Pcdhb22 | 0.000986 | 0 | PCDHB22_PCDHB22 | Pcdhb22 - Pcdhb22 | PCDH | Cell-Cell Contact | PMID: 35253643; PMID: 32694982 |
| NeuN | CD31 | Pcdhb17 | Pcdhb17 | 0.000788 | 0 | PCDHB17_PCDHB17 | Pcdhb17 - Pcdhb17 | PCDH | Cell-Cell Contact | PMID: 35253643; PMID: 32694982 |
| CD31 | NeuN | Pcdhb17 | Pcdhb17 | 0.000788 | 0 | PCDHB17_PCDHB17 | Pcdhb17 - Pcdhb17 | PCDH | Cell-Cell Contact | PMID: 35253643; PMID: 32694982 |
| NeuN | NeuN | Pcdhb17 | Pcdhb17 | 0.001139 | 0 | PCDHB17_PCDHB17 | Pcdhb17 - Pcdhb17 | PCDH | Cell-Cell Contact | PMID: 35253643; PMID: 32694982 |
| CD31 | CD31 | Pcdhb2 | Pcdhb2 | 0.000446 | 0 | PCDHB2_PCDHB2 | Pcdhb2 - Pcdhb2 | PCDH | Cell-Cell Contact | PMID: 35253643; PMID: 32694982 |
| NeuN | CD31 | Pcdhb2 | Pcdhb2 | 0.000431 | 0 | PCDHB2_PCDHB2 | Pcdhb2 - Pcdhb2 | PCDH | Cell-Cell Contact | PMID: 35253643; PMID: 32694982 |
| CD31 | NeuN | Pcdhb2 | Pcdhb2 | 0.000431 | 0 | PCDHB2_PCDHB2 | Pcdhb2 - Pcdhb2 | PCDH | Cell-Cell Contact | PMID: 35253643; PMID: 32694982 |
| CD31 | CD31 | Pcdhb3 | Pcdhb3 | 0.000532 | 0 | PCDHB3_PCDHB3 | Pcdhb3 - Pcdhb3 | PCDH | Cell-Cell Contact | PMID: 35253643; PMID: 32694982 |
| NeuN | CD31 | Pcdhb3 | Pcdhb3 | 0.000372 | 0 | PCDHB3_PCDHB3 | Pcdhb3 - Pcdhb3 | PCDH | Cell-Cell Contact | PMID: 35253643; PMID: 32694982 |
| CD31 | NeuN | Pcdhb3 | Pcdhb3 | 0.000372 | 0 | PCDHB3_PCDHB3 | Pcdhb3 - Pcdhb3 | PCDH | Cell-Cell Contact | PMID: 35253643; PMID: 32694982 |
| NeuN | CD31 | Pcdhb7 | Pcdhb7 | 0.00032 | 0 | PCDHB7_PCDHB7 | Pcdhb7 - Pcdhb7 | PCDH | Cell-Cell Contact | PMID: 35253643; PMID: 32694982 |
| CD31 | NeuN | Pcdhb7 | Pcdhb7 | 0.00032 | 0 | PCDHB7_PCDHB7 | Pcdhb7 - Pcdhb7 | PCDH | Cell-Cell Contact | PMID: 35253643; PMID: 32694982 |
| NeuN | NeuN | Pcdhb7 | Pcdhb7 | 0.000321 | 0 | PCDHB7_PCDHB7 | Pcdhb7 - Pcdhb7 | PCDH | Cell-Cell Contact | PMID: 35253643; PMID: 32694982 |
| CD31 | CD31 | Pcdhb7 | Pcdhb5 | 0.000454 | 0 | PCDHB7_PCDHB5 | Pcdhb7 - Pcdhb5 | PCDH | Cell-Cell Contact | PMID: 35253643; PMID: 32694982 |
| NeuN | CD31 | Pcdhb7 | Pcdhb5 | 0.000455 | 0 | PCDHB7_PCDHB5 | Pcdhb7 - Pcdhb5 | PCDH | Cell-Cell Contact | PMID: 35253643; PMID: 32694982 |
| CD31 | CD31 | Pcdhb7 | Pcdhb9 | 0.000329 | 0 | PCDHB7_PCDHB9 | Pcdhb7 - Pcdhb9 | PCDH | Cell-Cell Contact | PMID: 35253643; PMID: 32694982 |
| NeuN | CD31 | Pcdhb7 | Pcdhb9 | 0.00033 | 0 | PCDHB7_PCDHB9 | Pcdhb7 - Pcdhb9 | PCDH | Cell-Cell Contact | PMID: 35253643; PMID: 32694982 |
| CD31 | CD31 | Pcdhb5 | Pcdhb5 | 0.000646 | 0 | PCDHB5_PCDHB5 | Pcdhb5 - Pcdhb5 | PCDH | Cell-Cell Contact | PMID: 35253643; PMID: 32694982 |
| NeuN | CD31 | Pcdhb5 | Pcdhb5 | 0.00036 | 0 | PCDHB5_PCDHB5 | Pcdhb5 - Pcdhb5 | PCDH | Cell-Cell Contact | PMID: 35253643; PMID: 32694982 |
| CD31 | NeuN | Pcdhb5 | Pcdhb5 | 0.00036 | 0 | PCDHB5_PCDHB5 | Pcdhb5 - Pcdhb5 | PCDH | Cell-Cell Contact | PMID: 35253643; PMID: 32694982 |
| CD31 | CD31 | Pcdhb5 | Pcdhb9 | 0.000467 | 0 | PCDHB5_PCDHB9 | Pcdhb5 - Pcdhb9 | PCDH | Cell-Cell Contact | PMID: 35253643; PMID: 32694982 |
| CD31 | NeuN | Pcdhb5 | Pcdhb9 | 0.000404 | 0 | PCDHB5_PCDHB9 | Pcdhb5 - Pcdhb9 | PCDH | Cell-Cell Contact | PMID: 35253643; PMID: 32694982 |
| CD31 | CD31 | Pcdhb9 | Pcdhb9 | 0.000338 | 0 | PCDHB9_PCDHB9 | Pcdhb9 - Pcdhb9 | PCDH | Cell-Cell Contact | PMID: 35253643; PMID: 32694982 |
| NeuN | CD31 | Pcdhb9 | Pcdhb9 | 0.000292 | 0 | PCDHB9_PCDHB9 | Pcdhb9 - Pcdhb9 | PCDH | Cell-Cell Contact | PMID: 35253643; PMID: 32694982 |
| CD31 | NeuN | Pcdhb9 | Pcdhb9 | 0.000292 | 0 | PCDHB9_PCDHB9 | Pcdhb9 - Pcdhb9 | PCDH | Cell-Cell Contact | PMID: 35253643; PMID: 32694982 |
| CD31 | CD31 | Pcdhb12 | Pcdhb12 | 0.000671 | 0 | PCDHB12_PCDHB12 | Pcdhb12 - Pcdhb12 | PCDH | Cell-Cell Contact | PMID: 35253643; PMID: 32694982 |
| NeuN | CD31 | Pcdhb12 | Pcdhb12 | 0.000365 | 0 | PCDHB12_PCDHB12 | Pcdhb12 - Pcdhb12 | PCDH | Cell-Cell Contact | PMID: 35253643; PMID: 32694982 |
| CD31 | NeuN | Pcdhb12 | Pcdhb12 | 0.000365 | 0 | PCDHB12_PCDHB12 | Pcdhb12 - Pcdhb12 | PCDH | Cell-Cell Contact | PMID: 35253643; PMID: 32694982 |
| CD31 | CD31 | Pcdhb12 | Pcdhb10 | 0.000576 | 0 | PCDHB12_PCDHB10 | Pcdhb12 - Pcdhb10 | PCDH | Cell-Cell Contact | PMID: 35253643; PMID: 32694982 |
| CD31 | NeuN | Pcdhb12 | Pcdhb10 | 0.00034 | 0 | PCDHB12_PCDHB10 | Pcdhb12 - Pcdhb10 | PCDH | Cell-Cell Contact | PMID: 35253643; PMID: 32694982 |
| CD31 | CD31 | Pcdhb12 | Pcdhb11 | 0.000637 | 0 | PCDHB12_PCDHB11 | Pcdhb12 - Pcdhb11 | PCDH | Cell-Cell Contact | PMID: 35253643; PMID: 32694982 |
| CD31 | NeuN | Pcdhb12 | Pcdhb11 | 0.000416 | 0 | PCDHB12_PCDHB11 | Pcdhb12 - Pcdhb11 | PCDH | Cell-Cell Contact | PMID: 35253643; PMID: 32694982 |
| CD31 | CD31 | Pcdhb12 | Pcdhb8 | 0.000605 | 0 | PCDHB12_PCDHB8 | Pcdhb12 - Pcdhb8 | PCDH | Cell-Cell Contact | PMID: 35253643; PMID: 32694982 |
| CD31 | NeuN | Pcdhb12 | Pcdhb8 | 0.00033 | 0 | PCDHB12_PCDHB8 | Pcdhb12 - Pcdhb8 | PCDH | Cell-Cell Contact | PMID: 35253643; PMID: 32694982 |
| CD31 | CD31 | Pcdhb12 | Pcdhb6 | 0.0007 | 0 | PCDHB12_PCDHB6 | Pcdhb12 - Pcdhb6 | PCDH | Cell-Cell Contact | PMID: 35253643; PMID: 32694982 |

| | | | | | | | | | | |
|---|---|---|---|---|---|---|---|---|---|---|
| CD31 | NeuN | Pcdhb12 | Pcdhb6 | 0.00058 | 0 | PCDHB12_PCDHB6 | Pcdhb12 - Pcdhb6 | PCDH | Cell-Cell Contact | PMID: 35253643; PMID: 32694982 |
| CD31 | CD31 | Pcdhb10 | Pcdhb10 | 0.000494 | 0 | PCDHB10_PCDHB10 | Pcdhb10 - Pcdhb10 | PCDH | Cell-Cell Contact | PMID: 35253643; PMID: 32694982 |
| NeuN | CD31 | Pcdhb10 | Pcdhb10 | 0.000292 | 0 | PCDHB10_PCDHB10 | Pcdhb10 - Pcdhb10 | PCDH | Cell-Cell Contact | PMID: 35253643; PMID: 32694982 |
| CD31 | NeuN | Pcdhb10 | Pcdhb10 | 0.000292 | 0 | PCDHB10_PCDHB10 | Pcdhb10 - Pcdhb10 | PCDH | Cell-Cell Contact | PMID: 35253643; PMID: 32694982 |
| CD31 | CD31 | Pcdhb10 | Pcdhb11 | 0.000546 | 0 | PCDHB10_PCDHB11 | Pcdhb10 - Pcdhb11 | PCDH | Cell-Cell Contact | PMID: 35253643; PMID: 32694982 |
| CD31 | NeuN | Pcdhb10 | Pcdhb11 | 0.000357 | 0 | PCDHB10_PCDHB11 | Pcdhb10 - Pcdhb11 | PCDH | Cell-Cell Contact | PMID: 35253643; PMID: 32694982 |
| CD31 | CD31 | Pcdhb10 | Pcdhb8 | 0.000519 | 0 | PCDHB10_PCDHB8 | Pcdhb10 - Pcdhb8 | PCDH | Cell-Cell Contact | PMID: 35253643; PMID: 32694982 |
| NeuN | CD31 | Pcdhb10 | Pcdhb8 | 0.000307 | 0 | PCDHB10_PCDHB8 | Pcdhb10 - Pcdhb8 | PCDH | Cell-Cell Contact | PMID: 35253643; PMID: 32694982 |
| CD31 | CD31 | Pcdhb10 | Pcdhb6 | 0.0006 | 0 | PCDHB10_PCDHB6 | Pcdhb10 - Pcdhb6 | PCDH | Cell-Cell Contact | PMID: 35253643; PMID: 32694982 |
| CD31 | NeuN | Pcdhb10 | Pcdhb6 | 0.000498 | 0 | PCDHB10_PCDHB6 | Pcdhb10 - Pcdhb6 | PCDH | Cell-Cell Contact | PMID: 35253643; PMID: 32694982 |
| CD31 | CD31 | Pcdhb11 | Pcdhb11 | 0.000604 | 0 | PCDHB11_PCDHB11 | Pcdhb11 - Pcdhb11 | PCDH | Cell-Cell Contact | PMID: 35253643; PMID: 32694982 |
| NeuN | CD31 | Pcdhb11 | Pcdhb11 | 0.000395 | 0 | PCDHB11_PCDHB11 | Pcdhb11 - Pcdhb11 | PCDH | Cell-Cell Contact | PMID: 35253643; PMID: 32694982 |
| CD31 | NeuN | Pcdhb11 | Pcdhb11 | 0.000395 | 0 | PCDHB11_PCDHB11 | Pcdhb11 - Pcdhb11 | PCDH | Cell-Cell Contact | PMID: 35253643; PMID: 32694982 |
| CD31 | CD31 | Pcdhb11 | Pcdhb8 | 0.000574 | 0 | PCDHB11_PCDHB8 | Pcdhb11 - Pcdhb8 | PCDH | Cell-Cell Contact | PMID: 35253643; PMID: 32694982 |
| NeuN | CD31 | Pcdhb11 | Pcdhb8 | 0.000375 | 0 | PCDHB11_PCDHB8 | Pcdhb11 - Pcdhb8 | PCDH | Cell-Cell Contact | PMID: 35253643; PMID: 32694982 |
| CD31 | CD31 | Pcdhb11 | Pcdhb6 | 0.000664 | 0 | PCDHB11_PCDHB6 | Pcdhb11 - Pcdhb6 | PCDH | Cell-Cell Contact | PMID: 35253643; PMID: 32694982 |
| CD31 | NeuN | Pcdhb11 | Pcdhb6 | 0.00055 | 0 | PCDHB11_PCDHB6 | Pcdhb11 - Pcdhb6 | PCDH | Cell-Cell Contact | PMID: 35253643; PMID: 32694982 |
| CD31 | CD31 | Pcdhb8 | Pcdhb8 | 0.000545 | 0 | PCDHB8_PCDHB8 | Pcdhb8 - Pcdhb8 | PCDH | Cell-Cell Contact | PMID: 35253643; PMID: 32694982 |
| NeuN | CD31 | Pcdhb8 | Pcdhb8 | 0.000297 | 0 | PCDHB8_PCDHB8 | Pcdhb8 - Pcdhb8 | PCDH | Cell-Cell Contact | PMID: 35253643; PMID: 32694982 |
| CD31 | NeuN | Pcdhb8 | Pcdhb8 | 0.000297 | 0 | PCDHB8_PCDHB8 | Pcdhb8 - Pcdhb8 | PCDH | Cell-Cell Contact | PMID: 35253643; PMID: 32694982 |
| CD31 | CD31 | Pcdhb8 | Pcdhb6 | 0.00063 | 0 | PCDHB8_PCDHB6 | Pcdhb8 - Pcdhb6 | PCDH | Cell-Cell Contact | PMID: 35253643; PMID: 32694982 |
| CD31 | NeuN | Pcdhb8 | Pcdhb6 | 0.000522 | 0 | PCDHB8_PCDHB6 | Pcdhb8 - Pcdhb6 | PCDH | Cell-Cell Contact | PMID: 35253643; PMID: 32694982 |
| CD31 | CD31 | Pcdhb6 | Pcdhb6 | 0.000729 | 0 | PCDHB6_PCDHB6 | Pcdhb6 - Pcdhb6 | PCDH | Cell-Cell Contact | PMID: 35253643; PMID: 32694982 |
| NeuN | CD31 | Pcdhb6 | Pcdhb6 | 0.000605 | 0 | PCDHB6_PCDHB6 | Pcdhb6 - Pcdhb6 | PCDH | Cell-Cell Contact | PMID: 35253643; PMID: 32694982 |
| CD31 | NeuN | Pcdhb6 | Pcdhb6 | 0.000605 | 0 | PCDHB6_PCDHB6 | Pcdhb6 - Pcdhb6 | PCDH | Cell-Cell Contact | PMID: 35253643; PMID: 32694982 |
| NeuN | CD31 | Pcdhb13 | Pcdhb13 | 0.00054 | 0 | PCDHB13_PCDHB13 | Pcdhb13 - Pcdhb13 | PCDH | Cell-Cell Contact | PMID: 35253643; PMID: 32694982 |
| CD31 | NeuN | Pcdhb13 | Pcdhb13 | 0.00054 | 0 | PCDHB13_PCDHB13 | Pcdhb13 - Pcdhb13 | PCDH | Cell-Cell Contact | PMID: 35253643; PMID: 32694982 |
| NeuN | NeuN | Pcdhb13 | Pcdhb13 | 0.000654 | 0 | PCDHB13_PCDHB13 | Pcdhb13 - Pcdhb13 | PCDH | Cell-Cell Contact | PMID: 35253643; PMID: 32694982 |
| CD31 | CD31 | Pcdhb15 | Pcdhb15 | 0.000385 | 0 | PCDHB15_PCDHB15 | Pcdhb15 - Pcdhb15 | PCDH | Cell-Cell Contact | PMID: 35253643; PMID: 32694982 |
| NeuN | CD31 | Pcdhb15 | Pcdhb15 | 0.000315 | 0 | PCDHB15_PCDHB15 | Pcdhb15 - Pcdhb15 | PCDH | Cell-Cell Contact | PMID: 35253643; PMID: 32694982 |
| CD31 | NeuN | Pcdhb15 | Pcdhb15 | 0.000315 | 0 | PCDHB15_PCDHB15 | Pcdhb15 - Pcdhb15 | PCDH | Cell-Cell Contact | PMID: 35253643; PMID: 32694982 |
| CD31 | CD31 | Pcdhga1 | Pcdhga1 | 0.000449 | 0 | PCDHGA1_PCDHGA1 | Pcdhga1 - Pcdhga1 | PCDH | Cell-Cell Contact | PMID: 35253643; PMID: 32694982 |
| NeuN | CD31 | Pcdhga1 | Pcdhga1 | 0.000333 | 0 | PCDHGA1_PCDHGA1 | Pcdhga1 - Pcdhga1 | PCDH | Cell-Cell Contact | PMID: 35253643; PMID: 32694982 |
| CD31 | NeuN | Pcdhga1 | Pcdhga1 | 0.000333 | 0 | PCDHGA1_PCDHGA1 | Pcdhga1 - Pcdhga1 | PCDH | Cell-Cell Contact | PMID: 35253643; PMID: 32694982 |
| CD31 | CD31 | Pcdhga10 | Pcdhga10 | 0.000218 | 0 | PCDHGA10_PCDHGA10 | Pcdhga10 - Pcdhga10 | PCDH | Cell-Cell Contact | PMID: 35253643; PMID: 32694982 |
| NeuN | CD31 | Pcdhga10 | Pcdhga10 | 0.000211 | 0 | PCDHGA10_PCDHGA10 | Pcdhga10 - Pcdhga10 | PCDH | Cell-Cell Contact | PMID: 35253643; PMID: 32694982 |
| CD31 | NeuN | Pcdhga10 | Pcdhga10 | 0.000211 | 0 | PCDHGA10_PCDHGA10 | Pcdhga10 - Pcdhga10 | PCDH | Cell-Cell Contact | PMID: 35253643; PMID: 32694982 |
| NeuN | CD31 | Pcdhga11 | Pcdhga11 | 0.001099 | 0 | PCDHGA11_PCDHGA11 | Pcdhga11 - Pcdhga11 | PCDH | Cell-Cell Contact | PMID: 35253643; PMID: 32694982 |
| CD31 | NeuN | Pcdhga11 | Pcdhga11 | 0.001099 | 0 | PCDHGA11_PCDHGA11 | Pcdhga11 - Pcdhga11 | PCDH | Cell-Cell Contact | PMID: 35253643; PMID: 32694982 |
| NeuN | NeuN | Pcdhga11 | Pcdhga11 | 0.001833 | 0 | PCDHGA11_PCDHGA11 | Pcdhga11 - Pcdhga11 | PCDH | Cell-Cell Contact | PMID: 35253643; PMID: 32694982 |
| NeuN | CD31 | Pcdhga12 | Pcdhga12 | 0.000607 | 0 | PCDHGA12_PCDHGA12 | Pcdhga12 - Pcdhga12 | PCDH | Cell-Cell Contact | PMID: 35253643; PMID: 32694982 |
| CD31 | NeuN | Pcdhga12 | Pcdhga12 | 0.000607 | 0 | PCDHGA12_PCDHGA12 | Pcdhga12 - Pcdhga12 | PCDH | Cell-Cell Contact | PMID: 35253643; PMID: 32694982 |
| NeuN | NeuN | Pcdhga12 | Pcdhga12 | 0.000749 | 0 | PCDHGA12_PCDHGA12 | Pcdhga12 - Pcdhga12 | PCDH | Cell-Cell Contact | PMID: 35253643; PMID: 32694982 |
| CD31 | CD31 | Pcdhga2 | Pcdhga2 | 0.000498 | 0 | PCDHGA2_PCDHGA2 | Pcdhga2 - Pcdhga2 | PCDH | Cell-Cell Contact | PMID: 35253643; PMID: 32694982 |
| NeuN | CD31 | Pcdhga2 | Pcdhga2 | 0.000432 | 0 | PCDHGA2_PCDHGA2 | Pcdhga2 - Pcdhga2 | PCDH | Cell-Cell Contact | PMID: 35253643; PMID: 32694982 |
| CD31 | NeuN | Pcdhga2 | Pcdhga2 | 0.000432 | 0 | PCDHGA2_PCDHGA2 | Pcdhga2 - Pcdhga2 | PCDH | Cell-Cell Contact | PMID: 35253643; PMID: 32694982 |
| NeuN | CD31 | Pcdhga3 | Pcdhga3 | 0.000329 | 0 | PCDHGA3_PCDHGA3 | Pcdhga3 - Pcdhga3 | PCDH | Cell-Cell Contact | PMID: 35253643; PMID: 32694982 |
| CD31 | NeuN | Pcdhga3 | Pcdhga3 | 0.000329 | 0 | PCDHGA3_PCDHGA3 | Pcdhga3 - Pcdhga3 | PCDH | Cell-Cell Contact | PMID: 35253643; PMID: 32694982 |
| NeuN | NeuN | Pcdhga3 | Pcdhga3 | 0.000344 | 0 | PCDHGA3_PCDHGA3 | Pcdhga3 - Pcdhga3 | PCDH | Cell-Cell Contact | PMID: 35253643; PMID: 32694982 |
| NeuN | CD31 | Pcdhga4 | Pcdhga4 | 0.000778 | 0 | PCDHGA4_PCDHGA4 | Pcdhga4 - Pcdhga4 | PCDH | Cell-Cell Contact | PMID: 35253643; PMID: 32694982 |
| CD31 | NeuN | Pcdhga4 | Pcdhga4 | 0.000778 | 0 | PCDHGA4_PCDHGA4 | Pcdhga4 - Pcdhga4 | PCDH | Cell-Cell Contact | PMID: 35253643; PMID: 32694982 |
| NeuN | NeuN | Pcdhga4 | Pcdhga4 | 0.001043 | 0 | PCDHGA4_PCDHGA4 | Pcdhga4 - Pcdhga4 | PCDH | Cell-Cell Contact | PMID: 35253643; PMID: 32694982 |
| NeuN | CD31 | Pcdhga5 | Pcdhga5 | 0.000487 | 0 | PCDHGA5_PCDHGA5 | Pcdhga5 - Pcdhga5 | PCDH | Cell-Cell Contact | PMID: 35253643; PMID: 32694982 |
| CD31 | NeuN | Pcdhga5 | Pcdhga5 | 0.000487 | 0 | PCDHGA5_PCDHGA5 | Pcdhga5 - Pcdhga5 | PCDH | Cell-Cell Contact | PMID: 35253643; PMID: 32694982 |
| NeuN | NeuN | Pcdhga5 | Pcdhga5 | 0.000538 | 0 | PCDHGA5_PCDHGA5 | Pcdhga5 - Pcdhga5 | PCDH | Cell-Cell Contact | PMID: 35253643; PMID: 32694982 |
| CD31 | CD31 | Pcdhga6 | Pcdhga6 | 0.001118 | 0 | PCDHGA6_PCDHGA6 | Pcdhga6 - Pcdhga6 | PCDH | Cell-Cell Contact | PMID: 35253643; PMID: 32694982 |
| NeuN | CD31 | Pcdhga6 | Pcdhga6 | 0.000793 | 0 | PCDHGA6_PCDHGA6 | Pcdhga6 - Pcdhga6 | PCDH | Cell-Cell Contact | PMID: 35253643; PMID: 32694982 |
| CD31 | NeuN | Pcdhga6 | Pcdhga6 | 0.000793 | 0 | PCDHGA6_PCDHGA6 | Pcdhga6 - Pcdhga6 | PCDH | Cell-Cell Contact | PMID: 35253643; PMID: 32694982 |
| CD31 | CD31 | Pcdhga7 | Pcdhga7 | 0.000615 | 0 | PCDHGA7_PCDHGA7 | Pcdhga7 - Pcdhga7 | PCDH | Cell-Cell Contact | PMID: 35253643; PMID: 32694982 |
| NeuN | CD31 | Pcdhga7 | Pcdhga7 | 0.000536 | 0 | PCDHGA7_PCDHGA7 | Pcdhga7 - Pcdhga7 | PCDH | Cell-Cell Contact | PMID: 35253643; PMID: 32694982 |

| | | | | | | | | | | |
|---|---|---|---|---|---|---|---|---|---|---|
| CD31 | NeuN | Pcdhga7 | Pcdhga7 | 0.000536 | 0 | PCDHGA7_PCDHGA7 | Pcdhga7 - Pcdhga7 | PCDH | Cell-Cell Contact | PMID: 35253643; PMID: 32694982 |
| CD31 | CD31 | Pcdhga8 | Pcdhga8 | 0.000929 | 0 | PCDHGA8_PCDHGA8 | Pcdhga8 - Pcdhga8 | PCDH | Cell-Cell Contact | PMID: 35253643; PMID: 32694982 |
| NeuN | CD31 | Pcdhga8 | Pcdhga8 | 0.000569 | 0 | PCDHGA8_PCDHGA8 | Pcdhga8 - Pcdhga8 | PCDH | Cell-Cell Contact | PMID: 35253643; PMID: 32694982 |
| CD31 | NeuN | Pcdhga8 | Pcdhga8 | 0.000569 | 0 | PCDHGA8_PCDHGA8 | Pcdhga8 - Pcdhga8 | PCDH | Cell-Cell Contact | PMID: 35253643; PMID: 32694982 |
| CD31 | CD31 | Pcdhga9 | Pcdhga9 | 0.000452 | 0 | PCDHGA9_PCDHGA9 | Pcdhga9 - Pcdhga9 | PCDH | Cell-Cell Contact | PMID: 35253643; PMID: 32694982 |
| NeuN | CD31 | Pcdhga9 | Pcdhga9 | 0.000224 | 0 | PCDHGA9_PCDHGA9 | Pcdhga9 - Pcdhga9 | PCDH | Cell-Cell Contact | PMID: 35253643; PMID: 32694982 |
| CD31 | NeuN | Pcdhga9 | Pcdhga9 | 0.000224 | 0 | PCDHGA9_PCDHGA9 | Pcdhga9 - Pcdhga9 | PCDH | Cell-Cell Contact | PMID: 35253643; PMID: 32694982 |
| NeuN | CD31 | Pcdhgb1 | Pcdhgb1 | 0.000483 | 0 | PCDHGB1_PCDHGB1 | Pcdhgb1 - Pcdhgb1 | PCDH | Cell-Cell Contact | PMID: 35253643; PMID: 32694982 |
| CD31 | NeuN | Pcdhgb1 | Pcdhgb1 | 0.000483 | 0 | PCDHGB1_PCDHGB1 | Pcdhgb1 - Pcdhgb1 | PCDH | Cell-Cell Contact | PMID: 35253643; PMID: 32694982 |
| NeuN | NeuN | Pcdhgb1 | Pcdhgb1 | 0.000618 | 0 | PCDHGB1_PCDHGB1 | Pcdhgb1 - Pcdhgb1 | PCDH | Cell-Cell Contact | PMID: 35253643; PMID: 32694982 |
| CD31 | CD31 | Pcdhgb2 | Pcdhgb2 | 0.002867 | 0 | PCDHGB2_PCDHGB2 | Pcdhgb2 - Pcdhgb2 | PCDH | Cell-Cell Contact | PMID: 35253643; PMID: 32694982 |
| NeuN | CD31 | Pcdhgb2 | Pcdhgb2 | 0.00232 | 0 | PCDHGB2_PCDHGB2 | Pcdhgb2 - Pcdhgb2 | PCDH | Cell-Cell Contact | PMID: 35253643; PMID: 32694982 |
| CD31 | NeuN | Pcdhgb2 | Pcdhgb2 | 0.00232 | 0 | PCDHGB2_PCDHGB2 | Pcdhgb2 - Pcdhgb2 | PCDH | Cell-Cell Contact | PMID: 35253643; PMID: 32694982 |
| CD31 | CD31 | Pcdhgb8 | Pcdhgb8 | 0.000434 | 0 | PCDHGB8_PCDHGB8 | Pcdhgb8 - Pcdhgb8 | PCDH | Cell-Cell Contact | PMID: 35253643; PMID: 32694982 |
| NeuN | CD31 | Pcdhgb8 | Pcdhgb8 | 0.000286 | 0 | PCDHGB8_PCDHGB8 | Pcdhgb8 - Pcdhgb8 | PCDH | Cell-Cell Contact | PMID: 35253643; PMID: 32694982 |
| CD31 | NeuN | Pcdhgb8 | Pcdhgb8 | 0.000286 | 0 | PCDHGB8_PCDHGB8 | Pcdhgb8 - Pcdhgb8 | PCDH | Cell-Cell Contact | PMID: 35253643; PMID: 32694982 |
| CD31 | CD31 | Pcdhgb4 | Pcdhgb4 | 0.000883 | 0 | PCDHGB4_PCDHGB4 | Pcdhgb4 - Pcdhgb4 | PCDH | Cell-Cell Contact | PMID: 35253643; PMID: 32694982 |
| NeuN | CD31 | Pcdhgb4 | Pcdhgb4 | 0.000722 | 0 | PCDHGB4_PCDHGB4 | Pcdhgb4 - Pcdhgb4 | PCDH | Cell-Cell Contact | PMID: 35253643; PMID: 32694982 |
| CD31 | NeuN | Pcdhgb4 | Pcdhgb4 | 0.000722 | 0 | PCDHGB4_PCDHGB4 | Pcdhgb4 - Pcdhgb4 | PCDH | Cell-Cell Contact | PMID: 35253643; PMID: 32694982 |
| CD31 | CD31 | Pcdhgb5 | Pcdhgb5 | 0.000763 | 0 | PCDHGB5_PCDHGB5 | Pcdhgb5 - Pcdhgb5 | PCDH | Cell-Cell Contact | PMID: 35253643; PMID: 32694982 |
| NeuN | CD31 | Pcdhgb5 | Pcdhgb5 | 0.000593 | 0 | PCDHGB5_PCDHGB5 | Pcdhgb5 - Pcdhgb5 | PCDH | Cell-Cell Contact | PMID: 35253643; PMID: 32694982 |
| CD31 | NeuN | Pcdhgb5 | Pcdhgb5 | 0.000593 | 0 | PCDHGB5_PCDHGB5 | Pcdhgb5 - Pcdhgb5 | PCDH | Cell-Cell Contact | PMID: 35253643; PMID: 32694982 |
| NeuN | CD31 | Pcdhgb6 | Pcdhgb6 | 0.001346 | 0 | PCDHGB6_PCDHGB6 | Pcdhgb6 - Pcdhgb6 | PCDH | Cell-Cell Contact | PMID: 35253643; PMID: 32694982 |
| CD31 | NeuN | Pcdhgb6 | Pcdhgb6 | 0.001346 | 0 | PCDHGB6_PCDHGB6 | Pcdhgb6 - Pcdhgb6 | PCDH | Cell-Cell Contact | PMID: 35253643; PMID: 32694982 |
| NeuN | NeuN | Pcdhgb6 | Pcdhgb6 | 0.001395 | 0 | PCDHGB6_PCDHGB6 | Pcdhgb6 - Pcdhgb6 | PCDH | Cell-Cell Contact | PMID: 35253643; PMID: 32694982 |
| NeuN | CD31 | Pcdhgb7 | Pcdhgb7 | 0.00698 | 0 | PCDHGB7_PCDHGB7 | Pcdhgb7 - Pcdhgb7 | PCDH | Cell-Cell Contact | PMID: 35253643; PMID: 32694982 |
| CD31 | NeuN | Pcdhgb7 | Pcdhgb7 | 0.00698 | 0 | PCDHGB7_PCDHGB7 | Pcdhgb7 - Pcdhgb7 | PCDH | Cell-Cell Contact | PMID: 35253643; PMID: 32694982 |
| NeuN | NeuN | Pcdhgb7 | Pcdhgb7 | 0.00914 | 0 | PCDHGB7_PCDHGB7 | Pcdhgb7 - Pcdhgb7 | PCDH | Cell-Cell Contact | PMID: 35253643; PMID: 32694982 |
| NeuN | CD31 | Pcdhgc3 | Pcdhgc3 | 0.009735 | 0 | PCDHGC3_PCDHGC3 | Pcdhgc3 - Pcdhgc3 | PCDH | Cell-Cell Contact | PMID: 35253643; PMID: 32694982 |
| CD31 | NeuN | Pcdhgc3 | Pcdhgc3 | 0.009735 | 0 | PCDHGC3_PCDHGC3 | Pcdhgc3 - Pcdhgc3 | PCDH | Cell-Cell Contact | PMID: 35253643; PMID: 32694982 |
| NeuN | NeuN | Pcdhgc3 | Pcdhgc3 | 0.010967 | 0 | PCDHGC3_PCDHGC3 | Pcdhgc3 - Pcdhgc3 | PCDH | Cell-Cell Contact | PMID: 35253643; PMID: 32694982 |
| NeuN | CD31 | Pcdhgc4 | Pcdhgc4 | 0.000942 | 0 | PCDHGC4_PCDHGC4 | Pcdhgc4 - Pcdhgc4 | PCDH | Cell-Cell Contact | PMID: 35253643; PMID: 32694982 |
| CD31 | NeuN | Pcdhgc4 | Pcdhgc4 | 0.000942 | 0 | PCDHGC4_PCDHGC4 | Pcdhgc4 - Pcdhgc4 | PCDH | Cell-Cell Contact | PMID: 35253643; PMID: 32694982 |
| NeuN | NeuN | Pcdhgc4 | Pcdhgc4 | 0.001017 | 0 | PCDHGC4_PCDHGC4 | Pcdhgc4 - Pcdhgc4 | PCDH | Cell-Cell Contact | PMID: 35253643; PMID: 32694982 |
| CD31 | CD31 | Pcdhgc5 | Pcdhgc5 | 0.00058 | 0 | PCDHGC5_PCDHGC5 | Pcdhgc5 - Pcdhgc5 | PCDH | Cell-Cell Contact | PMID: 35253643; PMID: 32694982 |
| NeuN | CD31 | Pcdhgc5 | Pcdhgc5 | 0.000556 | 0 | PCDHGC5_PCDHGC5 | Pcdhgc5 - Pcdhgc5 | PCDH | Cell-Cell Contact | PMID: 35253643; PMID: 32694982 |
| CD31 | NeuN | Pcdhgc5 | Pcdhgc5 | 0.000556 | 0 | PCDHGC5_PCDHGC5 | Pcdhgc5 - Pcdhgc5 | PCDH | Cell-Cell Contact | PMID: 35253643; PMID: 32694982 |
| CD31 | CD31 | Sdc2 | Adgra2 | 0.000674 | 0 | SDC2_ADGRA2 | Sdc2 - Adgra2 | ADGRA | Cell-Cell Contact | PMID: 31462748 |
| NeuN | CD31 | Sdc2 | Adgra2 | 0.000403 | 0 | SDC2_ADGRA2 | Sdc2 - Adgra2 | ADGRA | Cell-Cell Contact | PMID: 31462748 |
| CD31 | CD31 | ITGAV_ITG | Adgra2 | 0.00088 | 0 | ITGAV_ITGB3_ADGRA2 | (Itgav+Itgb3) - Adgra2 | ADGRA | Cell-Cell Contact | PMID: 31462748 |
| NeuN | CD31 | ITGAV_ITG | Adgra2 | 0.000468 | 0 | ITGAV_ITGB3_ADGRA2 | (Itgav+Itgb3) - Adgra2 | ADGRA | Cell-Cell Contact | PMID: 31462748 |
| CD31 | CD31 | Tax1bp3 | Adgrb1 | 0.003119 | 0 | TAX1BP3_ADGRB1 | Tax1bp3 - Adgrb1 | ADGRB | Cell-Cell Contact | PMID: 31462748 |
| CD31 | NeuN | Tax1bp3 | Adgrb1 | 0.003817 | 0 | TAX1BP3_ADGRB1 | Tax1bp3 - Adgrb1 | ADGRB | Cell-Cell Contact | PMID: 31462748 |
| NeuN | CD31 | Rtn4r | Adgrb1 | 0.001289 | 0 | RTN4R_ADGRB1 | Rtn4r - Adgrb1 | ADGRB | Cell-Cell Contact | PMID: 31462748 |
| NeuN | NeuN | Rtn4r | Adgrb1 | 0.001579 | 0 | RTN4R_ADGRB1 | Rtn4r - Adgrb1 | ADGRB | Cell-Cell Contact | PMID: 31462748 |
| CD31 | CD31 | ITGAV_ITG | Adgrb1 | 0.000739 | 0 | ITGAV_ITGB5_ADGRB1 | (Itgav+Itgb5) - Adgrb1 | ADGRB | Cell-Cell Contact | PMID: 31462748 |
| CD31 | NeuN | ITGAV_ITG | Adgrb1 | 0.000904 | 0 | ITGAV_ITGB5_ADGRB1 | (Itgav+Itgb5) - Adgrb1 | ADGRB | Cell-Cell Contact | PMID: 31462748 |
| CD31 | NeuN | Tax1bp3 | Adgrb2 | 0.019736 | 0 | TAX1BP3_ADGRB2 | Tax1bp3 - Adgrb2 | ADGRB | Cell-Cell Contact | PMID: 31462748 |
| NeuN | NeuN | Tax1bp3 | Adgrb2 | 0.01134 | 0 | TAX1BP3_ADGRB2 | Tax1bp3 - Adgrb2 | ADGRB | Cell-Cell Contact | PMID: 31462748 |
| CD31 | CD31 | C1ql1 | Adgrb3 | 0.000734 | 0 | C1QL1_ADGRB3 | C1ql1 - Adgrb3 | ADGRB | Cell-Cell Contact | PMID: 31462748 |
| CD31 | NeuN | C1ql1 | Adgrb3 | 0.001041 | 0 | C1QL1_ADGRB3 | C1ql1 - Adgrb3 | ADGRB | Cell-Cell Contact | PMID: 31462748 |
| CD31 | NeuN | C1ql2 | Adgrb3 | 0.00086 | 0 | C1QL2_ADGRB3 | C1ql2 - Adgrb3 | ADGRB | Cell-Cell Contact | PMID: 31462748 |
| NeuN | NeuN | C1ql2 | Adgrb3 | 0.000847 | 0 | C1QL2_ADGRB3 | C1ql2 - Adgrb3 | ADGRB | Cell-Cell Contact | PMID: 31462748 |
| CD31 | CD31 | C1ql3 | Adgrb3 | 0.000919 | 0 | C1QL3_ADGRB3 | C1ql3 - Adgrb3 | ADGRB | Cell-Cell Contact | PMID: 31462748 |
| CD31 | NeuN | C1ql3 | Adgrb3 | 0.001302 | 0 | C1QL3_ADGRB3 | C1ql3 - Adgrb3 | ADGRB | Cell-Cell Contact | PMID: 31462748 |
| CD31 | CD31 | Thy1 | Adgre5 | 0.002215 | 0 | THY1_ADGRE5 | Thy1 - Adgre5 | ADGRE | Cell-Cell Contact | PMID: 31462748 |
| NeuN | CD31 | Thy1 | Adgre5 | 0.002427 | 0 | THY1_ADGRE5 | Thy1 - Adgre5 | ADGRE | Cell-Cell Contact | PMID: 31462748 |
| CD31 | CD31 | ITGAV_ITG | Adgre5 | 0.0065 | 0 | ITGAV_ITGB1_ADGRE5 | (Itgav+Itgb1) - Adgre5 | ADGRE | Cell-Cell Contact | PMID: 31462748 |
| NeuN | CD31 | ITGAV_ITG | Adgre5 | 0.002596 | 0 | ITGAV_ITGB1_ADGRE5 | (Itgav+Itgb1) - Adgre5 | ADGRE | Cell-Cell Contact | PMID: 31462748 |
| CD31 | CD31 | ITGAV_ITG | Adgre5 | 0.002881 | 0 | ITGAV_ITGB3_ADGRE5 | (Itgav+Itgb3) - Adgre5 | ADGRE | Cell-Cell Contact | PMID: 31462748 |

| | | | | | | | | | | |
|---|---|---|---|---|---|---|---|---|---|---|
| NeuN | CD31 | ITGAV_ITG | Adgre5 | 0.001534 | 0 | ITGAV_ITGB3_ADGRE5 | (Itgav+Itgb3) - Adgre5 | ADGRE | Cell-Cell Contact | PMID: 31462748 |
| CD31 | CD31 | Cd55 | Adgre5 | 0.00188 | 0 | CD55_ADGRE5 | Cd55 - Adgre5 | ADGRE | Cell-Cell Contact | PMID: 31462748 |
| NeuN | CD31 | Cd55 | Adgre5 | 0.000935 | 0 | CD55_ADGRE5 | Cd55 - Adgre5 | ADGRE | Cell-Cell Contact | PMID: 31462748 |
| CD31 | CD31 | Lpar1 | Adgre5 | 0.002036 | 0 | LPAR1_ADGRE5 | Lpar1 - Adgre5 | ADGRE | Cell-Cell Contact | PMID: 31462748 |
| NeuN | CD31 | Lpar1 | Adgre5 | 0.001533 | 0 | LPAR1_ADGRE5 | Lpar1 - Adgre5 | ADGRE | Cell-Cell Contact | PMID: 31462748 |
| CD31 | CD31 | Lpar2 | Adgre5 | 0.002262 | 0 | LPAR2_ADGRE5 | Lpar2 - Adgre5 | ADGRE | Cell-Cell Contact | PMID: 31462748 |
| NeuN | CD31 | Lpar2 | Adgre5 | 0.00192 | 0 | LPAR2_ADGRE5 | Lpar2 - Adgre5 | ADGRE | Cell-Cell Contact | PMID: 31462748 |
| CD31 | CD31 | Col3a1 | Adgrg1 | 0.001133 | 0 | COL3A1_ADGRG1 | Col3a1 - Adgrg1 | ADGRG | Cell-Cell Contact | PMID: 31462748;UniProtKB |
| CD31 | NeuN | Col3a1 | Adgrg1 | 0.001527 | 0 | COL3A1_ADGRG1 | Col3a1 - Adgrg1 | ADGRG | Cell-Cell Contact | PMID: 31462748;UniProtKB |
| CD31 | CD31 | Tgm2 | Adgrg1 | 0.000824 | 0 | TGM2_ADGRG1 | Tgm2 - Adgrg1 | ADGRG | Cell-Cell Contact | PMID: 31462748 |
| CD31 | NeuN | Tgm2 | Adgrg1 | 0.001111 | 0 | TGM2_ADGRG1 | Tgm2 - Adgrg1 | ADGRG | Cell-Cell Contact | PMID: 31462748 |
| CD31 | CD31 | Col4a1 | Adgrg6 | 0.009983 | 0 | COL4A1_ADGRG6 | Col4a1 - Adgrg6 | ADGRG | Cell-Cell Contact | PMID: 31462748 |
| CD31 | NeuN | Col4a1 | Adgrg6 | 0.009482 | 0 | COL4A1_ADGRG6 | Col4a1 - Adgrg6 | ADGRG | Cell-Cell Contact | PMID: 31462748 |
| CD31 | CD31 | Col4a2 | Adgrg6 | 0.008569 | 0 | COL4A2_ADGRG6 | Col4a2 - Adgrg6 | ADGRG | Cell-Cell Contact | PMID: 31462748 |
| CD31 | NeuN | Col4a2 | Adgrg6 | 0.008139 | 0 | COL4A2_ADGRG6 | Col4a2 - Adgrg6 | ADGRG | Cell-Cell Contact | PMID: 31462748 |
| CD31 | CD31 | Prnp | Adgrg6 | 0.002648 | 0 | PRNP_ADGRG6 | Prnp - Adgrg6 | ADGRG | Cell-Cell Contact | PMID: 31462748 |
| CD31 | NeuN | Prnp | Adgrg6 | 0.002515 | 0 | PRNP_ADGRG6 | Prnp - Adgrg6 | ADGRG | Cell-Cell Contact | PMID: 31462748 |
| CD31 | NeuN | Flrt1 | Adgrl1 | 0.011061 | 0 | FLRT1_ADGRL1 | Flrt1 - Adgrl1 | ADGRL | Cell-Cell Contact | PMID: 25728924;PMID: 34022279;PMID: 27091502;PMID: 35907405 |
| NeuN | NeuN | Flrt1 | Adgrl1 | 0.01205 | 0 | FLRT1_ADGRL1 | Flrt1 - Adgrl1 | ADGRL | Cell-Cell Contact | PMID: 25728924;PMID: 34022279;PMID: 27091502;PMID: 35907405 |
| CD31 | NeuN | Flrt2 | Adgrl1 | 0.008642 | 0 | FLRT2_ADGRL1 | Flrt2 - Adgrl1 | ADGRL | Cell-Cell Contact | PMID: 25728924;PMID: 34022279;PMID: 27091502;PMID: 35907405 |
| NeuN | NeuN | Flrt2 | Adgrl1 | 0.016492 | 0 | FLRT2_ADGRL1 | Flrt2 - Adgrl1 | ADGRL | Cell-Cell Contact | PMID: 25728924;PMID: 34022279;PMID: 27091502;PMID: 35907405 |
| CD31 | NeuN | Flrt3 | Adgrl1 | 0.008714 | 0 | FLRT3_ADGRL1 | Flrt3 - Adgrl1 | ADGRL | Cell-Cell Contact | PMID: 25728924;PMID: 34022279;PMID: 27091502;PMID: 35907405 |
| NeuN | NeuN | Flrt3 | Adgrl1 | 0.011091 | 0 | FLRT3_ADGRL1 | Flrt3 - Adgrl1 | ADGRL | Cell-Cell Contact | PMID: 25728924;PMID: 34022279;PMID: 27091502;PMID: 35907405 |
| CD31 | NeuN | Nrxn1 | Adgrl1 | 0.030586 | 0 | NRXN1_ADGRL1 | Nrxn1 - Adgrl1 | ADGRL | Cell-Cell Contact | PMID: 25728924;PMID: 34022279;PMID: 27091502;PMID: 35907405 |
| NeuN | NeuN | Nrxn1 | Adgrl1 | 0.063129 | 0 | NRXN1_ADGRL1 | Nrxn1 - Adgrl1 | ADGRL | Cell-Cell Contact | PMID: 25728924;PMID: 34022279;PMID: 27091502;PMID: 35907405 |
| CD31 | NeuN | Nrxn2 | Adgrl1 | 0.021966 | 0 | NRXN2_ADGRL1 | Nrxn2 - Adgrl1 | ADGRL | Cell-Cell Contact | PMID: 25728924;PMID: 34022279;PMID: 27091502;PMID: 35907405 |
| NeuN | NeuN | Nrxn2 | Adgrl1 | 0.043654 | 0 | NRXN2_ADGRL1 | Nrxn2 - Adgrl1 | ADGRL | Cell-Cell Contact | PMID: 25728924;PMID: 34022279;PMID: 27091502;PMID: 35907405 |
| CD31 | NeuN | Nrxn3 | Adgrl1 | 0.013126 | 0 | NRXN3_ADGRL1 | Nrxn3 - Adgrl1 | ADGRL | Cell-Cell Contact | PMID: 25728924;PMID: 34022279;PMID: 27091502;PMID: 35907405 |
| NeuN | NeuN | Nrxn3 | Adgrl1 | 0.012637 | 0 | NRXN3_ADGRL1 | Nrxn3 - Adgrl1 | ADGRL | Cell-Cell Contact | PMID: 25728924;PMID: 34022279;PMID: 27091502;PMID: 35907405 |
| CD31 | NeuN | Tenm1 | Adgrl1 | 0.006612 | 0 | TENM1_ADGRL1 | Tenm1 - Adgrl1 | ADGRL | Cell-Cell Contact | PMID: 25728924;PMID: 34022279;PMID: 27091502;PMID: 35907405 |
| NeuN | NeuN | Tenm1 | Adgrl1 | 0.006435 | 0 | TENM1_ADGRL1 | Tenm1 - Adgrl1 | ADGRL | Cell-Cell Contact | PMID: 25728924;PMID: 34022279;PMID: 27091502;PMID: 35907405 |
| CD31 | NeuN | Tenm2 | Adgrl1 | 0.009932 | 0 | TENM2_ADGRL1 | Tenm2 - Adgrl1 | ADGRL | Cell-Cell Contact | PMID: 25728924;PMID: 34022279;PMID: 27091502;PMID: 35907405 |
| NeuN | NeuN | Tenm2 | Adgrl1 | 0.015367 | 0 | TENM2_ADGRL1 | Tenm2 - Adgrl1 | ADGRL | Cell-Cell Contact | PMID: 25728924;PMID: 34022279;PMID: 27091502;PMID: 35907405 |
| NeuN | CD31 | Tenm3 | Adgrl1 | 0.016781 | 0 | TENM3_ADGRL1 | Tenm3 - Adgrl1 | ADGRL | Cell-Cell Contact | PMID: 25728924;PMID: 34022279;PMID: 27091502;PMID: 35907405 |
| NeuN | NeuN | Tenm3 | Adgrl1 | 0.035639 | 0 | TENM3_ADGRL1 | Tenm3 - Adgrl1 | ADGRL | Cell-Cell Contact | PMID: 25728924;PMID: 34022279;PMID: 27091502;PMID: 35907405 |
| CD31 | NeuN | Tenm4 | Adgrl1 | 0.009457 | 0 | TENM4_ADGRL1 | Tenm4 - Adgrl1 | ADGRL | Cell-Cell Contact | PMID: 25728924;PMID: 34022279;PMID: 27091502;PMID: 35907405 |
| NeuN | NeuN | Tenm4 | Adgrl1 | 0.016651 | 0 | TENM4_ADGRL1 | Tenm4 - Adgrl1 | ADGRL | Cell-Cell Contact | PMID: 25728924;PMID: 34022279;PMID: 27091502;PMID: 35907405 |
| CD31 | NeuN | TENM2_FLI | Adgrl1 | 0.010481 | 0 | TENM2_FLRT1_ADGRL1 | (Tenm2+Flrt1) - Adgrl1 | ADGRL | Cell-Cell Contact | PMID: 25728924;PMID: 34022279;PMID: 27091502;PMID: 35907405 |
| NeuN | NeuN | TENM2_FLI | Adgrl1 | 0.013609 | 0 | TENM2_FLRT1_ADGRL1 | (Tenm2+Flrt1) - Adgrl1 | ADGRL | Cell-Cell Contact | PMID: 25728924;PMID: 34022279;PMID: 27091502;PMID: 35907405 |
| CD31 | NeuN | TENM2_FLI | Adgrl1 | 0.009303 | 0 | TENM2_FLRT3_ADGRL1 | (Tenm2+Flrt3) - Adgrl1 | ADGRL | Cell-Cell Contact | PMID: 25728924;PMID: 34022279;PMID: 27091502;PMID: 35907405 |
| NeuN | NeuN | TENM2_FLI | Adgrl1 | 0.013057 | 0 | TENM2_FLRT3_ADGRL1 | (Tenm2+Flrt3) - Adgrl1 | ADGRL | Cell-Cell Contact | PMID: 25728924;PMID: 34022279;PMID: 27091502;PMID: 35907405 |
| CD31 | NeuN | TENM4_FLI | Adgrl1 | 0.010228 | 0 | TENM4_FLRT1_ADGRL1 | (Tenm4+Flrt1) - Adgrl1 | ADGRL | Cell-Cell Contact | PMID: 25728924;PMID: 34022279;PMID: 27091502;PMID: 35907405 |
| NeuN | NeuN | TENM4_FLI | Adgrl1 | 0.014167 | 0 | TENM4_FLRT1_ADGRL1 | (Tenm4+Flrt1) - Adgrl1 | ADGRL | Cell-Cell Contact | PMID: 25728924;PMID: 34022279;PMID: 27091502;PMID: 35907405 |
| CD31 | NeuN | TENM4_FLI | Adgrl1 | 0.009078 | 0 | TENM4_FLRT3_ADGRL1 | (Tenm4+Flrt3) - Adgrl1 | ADGRL | Cell-Cell Contact | PMID: 25728924;PMID: 34022279;PMID: 27091502;PMID: 35907405 |
| NeuN | NeuN | TENM4_FLI | Adgrl1 | 0.013593 | 0 | TENM4_FLRT3_ADGRL1 | (Tenm4+Flrt3) - Adgrl1 | ADGRL | Cell-Cell Contact | PMID: 25728924;PMID: 34022279;PMID: 27091502;PMID: 35907405 |
| CD31 | NeuN | Flrt1 | Adgrl2 | 0.009928 | 0 | FLRT1_ADGRL2 | Flrt1 - Adgrl2 | ADGRL | Cell-Cell Contact | PMID: 25728924;PMID: 34022279;PMID: 27091502;PMID: 35907405 |
| NeuN | NeuN | Flrt1 | Adgrl2 | 0.010817 | 0 | FLRT1_ADGRL2 | Flrt1 - Adgrl2 | ADGRL | Cell-Cell Contact | PMID: 25728924;PMID: 34022279;PMID: 27091502;PMID: 35907405 |
| CD31 | NeuN | Flrt2 | Adgrl2 | 0.007755 | 0 | FLRT2_ADGRL2 | Flrt2 - Adgrl2 | ADGRL | Cell-Cell Contact | PMID: 25728924;PMID: 34022279;PMID: 27091502;PMID: 35907405 |
| NeuN | NeuN | Flrt2 | Adgrl2 | 0.014812 | 0 | FLRT2_ADGRL2 | Flrt2 - Adgrl2 | ADGRL | Cell-Cell Contact | PMID: 25728924;PMID: 34022279;PMID: 27091502;PMID: 35907405 |
| CD31 | NeuN | Flrt3 | Adgrl2 | 0.00782 | 0 | FLRT3_ADGRL2 | Flrt3 - Adgrl2 | ADGRL | Cell-Cell Contact | PMID: 25728924;PMID: 34022279;PMID: 27091502;PMID: 35907405 |
| NeuN | NeuN | Flrt3 | Adgrl2 | 0.009955 | 0 | FLRT3_ADGRL2 | Flrt3 - Adgrl2 | ADGRL | Cell-Cell Contact | PMID: 25728924;PMID: 34022279;PMID: 27091502;PMID: 35907405 |
| CD31 | NeuN | Tenm1 | Adgrl2 | 0.005932 | 0 | TENM1_ADGRL2 | Tenm1 - Adgrl2 | ADGRL | Cell-Cell Contact | PMID: 25728924;PMID: 34022279;PMID: 27091502;PMID: 35907405 |
| NeuN | NeuN | Tenm1 | Adgrl2 | 0.005773 | 0 | TENM1_ADGRL2 | Tenm1 - Adgrl2 | ADGRL | Cell-Cell Contact | PMID: 25728924;PMID: 34022279;PMID: 27091502;PMID: 35907405 |
| CD31 | NeuN | Tenm2 | Adgrl2 | 0.008914 | 0 | TENM2_ADGRL2 | Tenm2 - Adgrl2 | ADGRL | Cell-Cell Contact | PMID: 25728924;PMID: 34022279;PMID: 27091502;PMID: 35907405 |
| NeuN | NeuN | Tenm2 | Adgrl2 | 0.013799 | 0 | TENM2_ADGRL2 | Tenm2 - Adgrl2 | ADGRL | Cell-Cell Contact | PMID: 25728924;PMID: 34022279;PMID: 27091502;PMID: 35907405 |
| NeuN | CD31 | Tenm3 | Adgrl2 | 0.014408 | 0 | TENM3_ADGRL2 | Tenm3 - Adgrl2 | ADGRL | Cell-Cell Contact | PMID: 25728924;PMID: 34022279;PMID: 27091502;PMID: 35907405 |
| NeuN | NeuN | Tenm3 | Adgrl2 | 0.032071 | 0 | TENM3_ADGRL2 | Tenm3 - Adgrl2 | ADGRL | Cell-Cell Contact | PMID: 25728924;PMID: 34022279;PMID: 27091502;PMID: 35907405 |
| CD31 | NeuN | Tenm4 | Adgrl2 | 0.008487 | 0 | TENM4_ADGRL2 | Tenm4 - Adgrl2 | ADGRL | Cell-Cell Contact | PMID: 25728924;PMID: 34022279;PMID: 27091502;PMID: 35907405 |
| NeuN | NeuN | Tenm4 | Adgrl2 | 0.014955 | 0 | TENM4_ADGRL2 | Tenm4 - Adgrl2 | ADGRL | Cell-Cell Contact | PMID: 25728924;PMID: 34022279;PMID: 27091502;PMID: 35907405 |
| CD31 | NeuN | TENM2_FLI | Adgrl2 | 0.008349 | 0 | TENM2_FLRT3_ADGRL2 | (Tenm2+Flrt3) - Adgrl2 | ADGRL | Cell-Cell Contact | PMID: 25728924;PMID: 34022279;PMID: 27091502;PMID: 35907405 |

| | | | | | | | | | | |
|---|---|---|---|---|---|---|---|---|---|---|
| NeuN | NeuN | TENM2_FL | Adgrl2 | 0.011722 | 0 | TENM2_FLRT3_ADGRL2 | (Tenm2+Flrt3) - Adgrl2 | ADGRL | Cell-Cell Contact | PMID: 25728924;PMID: 34022279;PMID: 27091502;PMID: 35907405 |
| CD31 | NeuN | Flrt1 | Adgrl3 | 0.00387 | 0 | FLRT1_ADGRL3 | Flrt1 - Adgrl3 | ADGRL | Cell-Cell Contact | PMID: 25728924;PMID: 34022279;PMID: 27091502;PMID: 35907405 |
| NeuN | NeuN | Flrt1 | Adgrl3 | 0.004219 | 0 | FLRT1_ADGRL3 | Flrt1 - Adgrl3 | ADGRL | Cell-Cell Contact | PMID: 25728924;PMID: 34022279;PMID: 27091502;PMID: 35907405 |
| CD31 | NeuN | Flrt2 | Adgrl3 | 0.003019 | 0 | FLRT2_ADGRL3 | Flrt2 - Adgrl3 | ADGRL | Cell-Cell Contact | PMID: 25728924;PMID: 34022279;PMID: 27091502;PMID: 35907405 |
| NeuN | NeuN | Flrt2 | Adgrl3 | 0.005791 | 0 | FLRT2_ADGRL3 | Flrt2 - Adgrl3 | ADGRL | Cell-Cell Contact | PMID: 25728924;PMID: 34022279;PMID: 27091502;PMID: 35907405 |
| CD31 | NeuN | Flrt3 | Adgrl3 | 0.003044 | 0 | FLRT3_ADGRL3 | Flrt3 - Adgrl3 | ADGRL | Cell-Cell Contact | PMID: 25728924;PMID: 34022279;PMID: 27091502;PMID: 35907405 |
| NeuN | NeuN | Flrt3 | Adgrl3 | 0.00388 | 0 | FLRT3_ADGRL3 | Flrt3 - Adgrl3 | ADGRL | Cell-Cell Contact | PMID: 25728924;PMID: 34022279;PMID: 27091502;PMID: 35907405 |
| CD31 | NeuN | Tenm1 | Adgrl3 | 0.002307 | 0 | TENM1_ADGRL3 | Tenm1 - Adgrl3 | ADGRL | Cell-Cell Contact | PMID: 25728924;PMID: 34022279;PMID: 27091502;PMID: 35907405 |
| NeuN | NeuN | Tenm1 | Adgrl3 | 0.002245 | 0 | TENM1_ADGRL3 | Tenm1 - Adgrl3 | ADGRL | Cell-Cell Contact | PMID: 25728924;PMID: 34022279;PMID: 27091502;PMID: 35907405 |
| CD31 | NeuN | Tenm2 | Adgrl3 | 0.003472 | 0 | TENM2_ADGRL3 | Tenm2 - Adgrl3 | ADGRL | Cell-Cell Contact | PMID: 25728924;PMID: 34022279;PMID: 27091502;PMID: 35907405 |
| NeuN | NeuN | Tenm2 | Adgrl3 | 0.005392 | 0 | TENM2_ADGRL3 | Tenm2 - Adgrl3 | ADGRL | Cell-Cell Contact | PMID: 25728924;PMID: 34022279;PMID: 27091502;PMID: 35907405 |
| NeuN | CD31 | Tenm3 | Adgrl3 | 0.006355 | 0 | TENM3_ADGRL3 | Tenm3 - Adgrl3 | ADGRL | Cell-Cell Contact | PMID: 25728924;PMID: 34022279;PMID: 27091502;PMID: 35907405 |
| NeuN | NeuN | Tenm3 | Adgrl3 | 0.012674 | 0 | TENM3_ADGRL3 | Tenm3 - Adgrl3 | ADGRL | Cell-Cell Contact | PMID: 25728924;PMID: 34022279;PMID: 27091502;PMID: 35907405 |
| CD31 | NeuN | Tenm4 | Adgrl3 | 0.003305 | 0 | TENM4_ADGRL3 | Tenm4 - Adgrl3 | ADGRL | Cell-Cell Contact | PMID: 25728924;PMID: 34022279;PMID: 27091502;PMID: 35907405 |
| NeuN | NeuN | Tenm4 | Adgrl3 | 0.005847 | 0 | TENM4_ADGRL3 | Tenm4 - Adgrl3 | ADGRL | Cell-Cell Contact | PMID: 25728924;PMID: 34022279;PMID: 27091502;PMID: 35907405 |
| CD31 | NeuN | Unc5a | Adgrl3 | 0.002365 | 0 | UNC5A_ADGRL3 | Unc5a - Adgrl3 | ADGRL | Cell-Cell Contact | PMID: 25728924;PMID: 34022279;PMID: 27091502;PMID: 35907405 |
| NeuN | NeuN | Unc5a | Adgrl3 | 0.002729 | 0 | UNC5A_ADGRL3 | Unc5a - Adgrl3 | ADGRL | Cell-Cell Contact | PMID: 25728924;PMID: 34022279;PMID: 27091502;PMID: 35907405 |
| NeuN | CD31 | TENM3_FL | Adgrl3 | 0.003661 | 0 | TENM3_FLRT1_ADGRL3 | (Tenm3+Flrt1) - Adgrl3 | ADGRL | Cell-Cell Contact | PMID: 25728924;PMID: 34022279;PMID: 27091502;PMID: 35907405 |
| NeuN | NeuN | TENM3_FL | Adgrl3 | 0.00732 | 0 | TENM3_FLRT1_ADGRL3 | (Tenm3+Flrt1) - Adgrl3 | ADGRL | Cell-Cell Contact | PMID: 25728924;PMID: 34022279;PMID: 27091502;PMID: 35907405 |
| NeuN | CD31 | TENM3_FL | Adgrl3 | 0.003511 | 0 | TENM3_FLRT3_ADGRL3 | (Tenm3+Flrt3) - Adgrl3 | ADGRL | Cell-Cell Contact | PMID: 25728924;PMID: 34022279;PMID: 27091502;PMID: 35907405 |
| NeuN | NeuN | TENM3_FL | Adgrl3 | 0.007022 | 0 | TENM3_FLRT3_ADGRL3 | (Tenm3+Flrt3) - Adgrl3 | ADGRL | Cell-Cell Contact | PMID: 25728924;PMID: 34022279;PMID: 27091502;PMID: 35907405 |
| NeuN | CD31 | Flrt1 | Flrt1 | 0.001138 | 0 | FLRT1_FLRT1 | Flrt1 - Flrt1 | FLRT | Cell-Cell Contact | PMID: 27091502 |
| CD31 | NeuN | Flrt1 | Flrt1 | 0.001138 | 0 | FLRT1_FLRT1 | Flrt1 - Flrt1 | FLRT | Cell-Cell Contact | PMID: 27091502 |
| NeuN | NeuN | Flrt1 | Flrt1 | 0.001241 | 0 | FLRT1_FLRT1 | Flrt1 - Flrt1 | FLRT | Cell-Cell Contact | PMID: 27091502 |
| NeuN | CD31 | Flrt2 | Flrt2 | 0.00122 | 0 | FLRT2_FLRT2 | Flrt2 - Flrt2 | FLRT | Cell-Cell Contact | PMID: 27091502 |
| CD31 | NeuN | Flrt2 | Flrt2 | 0.00122 | 0 | FLRT2_FLRT2 | Flrt2 - Flrt2 | FLRT | Cell-Cell Contact | PMID: 27091502 |
| NeuN | NeuN | Flrt2 | Flrt2 | 0.002344 | 0 | FLRT2_FLRT2 | Flrt2 - Flrt2 | FLRT | Cell-Cell Contact | PMID: 27091502 |
| NeuN | CD31 | Flrt3 | Flrt3 | 0.000823 | 0 | FLRT3_FLRT3 | Flrt3 - Flrt3 | FLRT | Cell-Cell Contact | PMID: 27091502 |
| CD31 | NeuN | Flrt3 | Flrt3 | 0.000823 | 0 | FLRT3_FLRT3 | Flrt3 - Flrt3 | FLRT | Cell-Cell Contact | PMID: 27091502 |
| NeuN | NeuN | Flrt3 | Flrt3 | 0.00105 | 0 | FLRT3_FLRT3 | Flrt3 - Flrt3 | FLRT | Cell-Cell Contact | PMID: 27091502 |
| CD31 | NeuN | Flrt1 | Unc5a | 0.000735 | 0 | FLRT1_UNC5A | Flrt1 - Unc5a | UNC5 | Cell-Cell Contact | PMID: 27091502 |
| NeuN | NeuN | Flrt1 | Unc5a | 0.000802 | 0 | FLRT1_UNC5A | Flrt1 - Unc5a | UNC5 | Cell-Cell Contact | PMID: 27091502 |
| NeuN | CD31 | Flrt2 | Unc5a | 0.000955 | 0 | FLRT2_UNC5A | Flrt2 - Unc5a | UNC5 | Cell-Cell Contact | PMID: 27091502 |
| NeuN | NeuN | Flrt2 | Unc5a | 0.001102 | 0 | FLRT2_UNC5A | Flrt2 - Unc5a | UNC5 | Cell-Cell Contact | PMID: 27091502 |
| NeuN | CD31 | Flrt3 | Unc5a | 0.000639 | 0 | FLRT3_UNC5A | Flrt3 - Unc5a | UNC5 | Cell-Cell Contact | PMID: 27091502 |
| NeuN | NeuN | Flrt3 | Unc5a | 0.000737 | 0 | FLRT3_UNC5A | Flrt3 - Unc5a | UNC5 | Cell-Cell Contact | PMID: 27091502 |
| CD31 | CD31 | Flrt1 | Unc5b | 0.001894 | 0 | FLRT1_UNC5B | Flrt1 - Unc5b | UNC5 | Cell-Cell Contact | PMID: 27091502 |
| NeuN | CD31 | Flrt1 | Unc5b | 0.002065 | 0 | FLRT1_UNC5B | Flrt1 - Unc5b | UNC5 | Cell-Cell Contact | PMID: 27091502 |
| CD31 | CD31 | Flrt2 | Unc5b | 0.001477 | 0 | FLRT2_UNC5B | Flrt2 - Unc5b | UNC5 | Cell-Cell Contact | PMID: 27091502 |
| NeuN | CD31 | Flrt2 | Unc5b | 0.002837 | 0 | FLRT2_UNC5B | Flrt2 - Unc5b | UNC5 | Cell-Cell Contact | PMID: 27091502 |
| CD31 | CD31 | Flrt3 | Unc5b | 0.001489 | 0 | FLRT3_UNC5B | Flrt3 - Unc5b | UNC5 | Cell-Cell Contact | PMID: 27091502 |
| NeuN | CD31 | Flrt3 | Unc5b | 0.001899 | 0 | FLRT3_UNC5B | Flrt3 - Unc5b | UNC5 | Cell-Cell Contact | PMID: 27091502 |
| CD31 | NeuN | Flrt1 | Unc5c | 0.00053 | 0 | FLRT1_UNC5C | Flrt1 - Unc5 | UNC5 | Cell-Cell Contact | PMID: 27091502 |
| NeuN | NeuN | Flrt1 | Unc5c | 0.000578 | 0 | FLRT1_UNC5C | Flrt1 - Unc5 | UNC5 | Cell-Cell Contact | PMID: 27091502 |
| NeuN | CD31 | Flrt2 | Unc5c | 0.000549 | 0 | FLRT2_UNC5C | Flrt2 - Unc5 | UNC5 | Cell-Cell Contact | PMID: 27091502 |
| NeuN | NeuN | Flrt2 | Unc5c | 0.000795 | 0 | FLRT2_UNC5C | Flrt2 - Unc5 | UNC5 | Cell-Cell Contact | PMID: 27091502 |
| CD31 | NeuN | Flrt3 | Unc5c | 0.000417 | 0 | FLRT3_UNC5C | Flrt3 - Unc5 | UNC5 | Cell-Cell Contact | PMID: 27091502 |
| NeuN | NeuN | Flrt3 | Unc5c | 0.000532 | 0 | FLRT3_UNC5C | Flrt3 - Unc5 | UNC5 | Cell-Cell Contact | PMID: 27091502 |
| CD31 | CD31 | Flrt1 | Unc5d | 0.000968 | 0 | FLRT1_UNC5D | Flrt1 - Unc5d | UNC5 | Cell-Cell Contact | PMID: 27091502 |
| NeuN | CD31 | Flrt1 | Unc5d | 0.001055 | 0 | FLRT1_UNC5D | Flrt1 - Unc5d | UNC5 | Cell-Cell Contact | PMID: 27091502 |
| NeuN | CD31 | Flrt2 | Unc5d | 0.00145 | 0 | FLRT2_UNC5D | Flrt2 - Unc5d | UNC5 | Cell-Cell Contact | PMID: 27091502 |
| NeuN | NeuN | Flrt2 | Unc5d | 0.001085 | 0 | FLRT2_UNC5D | Flrt2 - Unc5d | UNC5 | Cell-Cell Contact | PMID: 27091502 |
| CD31 | CD31 | Flrt3 | Unc5d | 0.000761 | 0 | FLRT3_UNC5D | Flrt3 - Unc5d | UNC5 | Cell-Cell Contact | PMID: 27091502 |
| NeuN | CD31 | Flrt3 | Unc5d | 0.000971 | 0 | FLRT3_UNC5D | Flrt3 - Unc5d | UNC5 | Cell-Cell Contact | PMID: 27091502 |
| NeuN | CD31 | Nrxn1 | Clstn1 | 0.007805 | 0 | NRXN1_CLSTN1 | Nrxn1 - Clstn1 | NRXN | Cell-Cell Contact | PMID: 32434929 |
| NeuN | NeuN | Nrxn1 | Clstn1 | 0.010775 | 0 | NRXN1_CLSTN1 | Nrxn1 - Clstn1 | NRXN | Cell-Cell Contact | PMID: 32434929 |
| NeuN | CD31 | Nrxn2 | Clstn1 | 0.005301 | 0 | NRXN2_CLSTN1 | Nrxn2 - Clstn1 | NRXN | Cell-Cell Contact | PMID: 32434929 |
| NeuN | NeuN | Nrxn2 | Clstn1 | 0.007325 | 0 | NRXN2_CLSTN1 | Nrxn2 - Clstn1 | NRXN | Cell-Cell Contact | PMID: 32434929 |
| CD31 | NeuN | Nrxn1 | Clstn2 | 0.021196 | 0 | NRXN1_CLSTN2 | Nrxn1 - Clstn2 | NRXN | Cell-Cell Contact | PMID: 32434929 |
| NeuN | NeuN | Nrxn1 | Clstn2 | 0.044203 | 0 | NRXN1_CLSTN2 | Nrxn1 - Clstn2 | NRXN | Cell-Cell Contact | PMID: 32434929 |

| | | | | | | | | | | |
|---|---|---|---|---|---|---|---|---|---|---|
| CD31 | NeuN | Nrxn2 | Clstn2 | 0.015181 | 0 | NRXN2_CLSTN2 | Nrxn2 - Clstn2 | NRXN | Cell-Cell Contact | PMID: 32434929 |
| NeuN | NeuN | Nrxn2 | Clstn2 | 0.030378 | 0 | NRXN2_CLSTN2 | Nrxn2 - Clstn2 | NRXN | Cell-Cell Contact | PMID: 32434929 |
| NeuN | CD31 | Nrxn1 | Clstn3 | 0.005688 | 0 | NRXN1_CLSTN3 | Nrxn1 - Clstn3 | NRXN | Cell-Cell Contact | PMID: 32434929 |
| NeuN | NeuN | Nrxn1 | Clstn3 | 0.009623 | 0 | NRXN1_CLSTN3 | Nrxn1 - Clstn3 | NRXN | Cell-Cell Contact | PMID: 32434929 |
| NeuN | CD31 | Nrxn2 | Clstn3 | 0.00386 | 0 | NRXN2_CLSTN3 | Nrxn2 - Clstn3 | NRXN | Cell-Cell Contact | PMID: 32434929 |
| NeuN | NeuN | Nrxn2 | Clstn3 | 0.006539 | 0 | NRXN2_CLSTN3 | Nrxn2 - Clstn3 | NRXN | Cell-Cell Contact | PMID: 32434929 |
| NeuN | CD31 | Nrxn1 | Dag1 | 0.01005 | 0 | NRXN1_DAG1 | Nrxn1 - Dag1 | NRXN | Cell-Cell Contact | PMID: 21628410 |
| NeuN | NeuN | Nrxn1 | Dag1 | 0.007112 | 0 | NRXN1_DAG1 | Nrxn1 - Dag1 | NRXN | Cell-Cell Contact | PMID: 21628410 |
| NeuN | CD31 | Nrxn2 | Dag1 | 0.00683 | 0 | NRXN2_DAG1 | Nrxn2 - Dag1 | NRXN | Cell-Cell Contact | PMID: 21628410 |
| NeuN | NeuN | Nrxn2 | Dag1 | 0.004829 | 0 | NRXN2_DAG1 | Nrxn2 - Dag1 | NRXN | Cell-Cell Contact | PMID: 21628410 |
| CD31 | NeuN | Lrfn4 | Ptprd | 0.010342 | 0 | LRFN4_PTPRD | Lrfn4 - Ptprd | PTPR | Cell-Cell Contact | PMID: 29674953;PMID: 23916315 |
| NeuN | NeuN | Lrfn4 | Ptprd | 0.015541 | 0 | LRFN4_PTPRD | Lrfn4 - Ptprd | PTPR | Cell-Cell Contact | PMID: 29674953;PMID: 23916315 |
| CD31 | NeuN | Lrfn4 | Ptprf | 0.005971 | 0 | LRFN4_PTPRF | Lrfn4 - Ptprf | PTPR | Cell-Cell Contact | PMID: 29674953;PMID: 23916315 |
| NeuN | NeuN | Lrfn4 | Ptprf | 0.008993 | 0 | LRFN4_PTPRF | Lrfn4 - Ptprf | PTPR | Cell-Cell Contact | PMID: 29674953;PMID: 23916315 |
| CD31 | NeuN | Lrfn4 | Ptprs | 0.017486 | 0 | LRFN4_PTPRS | Lrfn4 - Ptprs | PTPR | Cell-Cell Contact | PMID: 29674953;PMID: 23916315 |
| NeuN | NeuN | Lrfn4 | Ptprs | 0.02618 | 0 | LRFN4_PTPRS | Lrfn4 - Ptprs | PTPR | Cell-Cell Contact | PMID: 29674953;PMID: 23916315 |
| NeuN | CD31 | Lrfn5 | Ptprd | 0.010729 | 0 | LRFN5_PTPRD | Lrfn5 - Ptprd | PTPR | Cell-Cell Contact | PMID: 29674953;PMID: 23916315 |
| NeuN | NeuN | Lrfn5 | Ptprd | 0.019523 | 0 | LRFN5_PTPRD | Lrfn5 - Ptprd | PTPR | Cell-Cell Contact | PMID: 29674953;PMID: 23916315 |
| NeuN | CD31 | Lrfn5 | Ptprf | 0.005271 | 0 | LRFN5_PTPRF | Lrfn5 - Ptprf | PTPR | Cell-Cell Contact | PMID: 29674953;PMID: 23916315 |
| NeuN | NeuN | Lrfn5 | Ptprf | 0.011317 | 0 | LRFN5_PTPRF | Lrfn5 - Ptprf | PTPR | Cell-Cell Contact | PMID: 29674953;PMID: 23916315 |
| NeuN | CD31 | Lrfn5 | Ptprs | 0.017223 | 0 | LRFN5_PTPRS | Lrfn5 - Ptprs | PTPR | Cell-Cell Contact | PMID: 29674953;PMID: 23916315 |
| NeuN | NeuN | Lrfn5 | Ptprs | 0.032799 | 0 | LRFN5_PTPRS | Lrfn5 - Ptprs | PTPR | Cell-Cell Contact | PMID: 29674953;PMID: 23916315 |
| NeuN | CD31 | Lrrc4b | Ptprd | 0.021855 | 0 | LRRC4B_PTPRD | Lrrc4b - Ptprd | PTPR | Cell-Cell Contact | PMID: 29674953;PMID: 23916315 |
| NeuN | NeuN | Lrrc4b | Ptprd | 0.039406 | 0 | LRRC4B_PTPRD | Lrrc4b - Ptprd | PTPR | Cell-Cell Contact | PMID: 29674953;PMID: 23916315 |
| CD31 | NeuN | Lrrc4b | Ptprs | 0.036118 | 0 | LRRC4B_PTPRS | Lrrc4b - Ptprs | PTPR | Cell-Cell Contact | PMID: 29674953;PMID: 23916315 |
| NeuN | NeuN | Lrrc4b | Ptprs | 0.065302 | 0 | LRRC4B_PTPRS | Lrrc4b - Ptprs | PTPR | Cell-Cell Contact | PMID: 29674953;PMID: 23916315 |
| CD31 | NeuN | Lrrc4c | Ptprf | 0.006485 | 0 | LRRC4C_PTPRF | Lrrc4c - Ptprf | PTPR | Cell-Cell Contact | PMID: 29674953;PMID: 23916315 |
| NeuN | NeuN | Lrrc4c | Ptprf | 0.012427 | 0 | LRRC4C_PTPRF | Lrrc4c - Ptprf | PTPR | Cell-Cell Contact | PMID: 29674953;PMID: 23916315 |
| NeuN | CD31 | Nrxn1 | Lrrtm1 | 0.009053 | 0 | NRXN1_LRRTM1 | Nrxn1 - Lrrtm1 | NRXN | Cell-Cell Contact | PMID:20519524;PMID:20064387;PMID:20064388; |
| NeuN | NeuN | Nrxn1 | Lrrtm1 | 0.013597 | 0 | NRXN1_LRRTM1 | Nrxn1 - Lrrtm1 | NRXN | Cell-Cell Contact | PMID:20519524;PMID:20064387;PMID:20064388; |
| NeuN | CD31 | Nrxn2 | Lrrtm1 | 0.006151 | 0 | NRXN2_LRRTM1 | Nrxn2 - Lrrtm1 | NRXN | Cell-Cell Contact | PMID:20519524;PMID:20064387;PMID:20064388; |
| NeuN | NeuN | Nrxn2 | Lrrtm1 | 0.009252 | 0 | NRXN2_LRRTM1 | Nrxn2 - Lrrtm1 | NRXN | Cell-Cell Contact | PMID:20519524;PMID:20064387;PMID:20064388; |
| CD31 | NeuN | Nrxn3 | Lrrtm1 | 0.002714 | 0 | NRXN3_LRRTM1 | Nrxn3 - Lrrtm1 | NRXN | Cell-Cell Contact | PMID:20519524;PMID:20064387;PMID:20064388; |
| NeuN | NeuN | Nrxn3 | Lrrtm1 | 0.002612 | 0 | NRXN3_LRRTM1 | Nrxn3 - Lrrtm1 | NRXN | Cell-Cell Contact | PMID:20519524;PMID:20064387;PMID:20064388; |
| NeuN | CD31 | Nrxn1 | Lrrtm2 | 0.004574 | 0 | NRXN1_LRRTM2 | Nrxn1 - Lrrtm2 | NRXN | Cell-Cell Contact | PMID:20519524;PMID:20064387;PMID:20064388; |
| NeuN | NeuN | Nrxn1 | Lrrtm2 | 0.004533 | 0 | NRXN1_LRRTM2 | Nrxn1 - Lrrtm2 | NRXN | Cell-Cell Contact | PMID:20519524;PMID:20064387;PMID:20064388; |
| NeuN | CD31 | Nrxn2 | Lrrtm2 | 0.003103 | 0 | NRXN2_LRRTM2 | Nrxn2 - Lrrtm2 | NRXN | Cell-Cell Contact | PMID:20519524;PMID:20064387;PMID:20064388; |
| NeuN | NeuN | Nrxn2 | Lrrtm2 | 0.003076 | 0 | NRXN2_LRRTM2 | Nrxn2 - Lrrtm2 | NRXN | Cell-Cell Contact | PMID:20519524;PMID:20064387;PMID:20064388; |
| CD31 | CD31 | Nrxn3 | Lrrtm2 | 0.000906 | 0 | NRXN3_LRRTM2 | Nrxn3 - Lrrtm2 | NRXN | Cell-Cell Contact | PMID:20519524;PMID:20064387;PMID:20064388; |
| CD31 | NeuN | Nrxn3 | Lrrtm2 | 0.000898 | 0 | NRXN3_LRRTM2 | Nrxn3 - Lrrtm2 | NRXN | Cell-Cell Contact | PMID:20519524;PMID:20064387;PMID:20064388; |
| NeuN | CD31 | Nrxn1 | Lrrtm3 | 0.005395 | 0 | NRXN1_LRRTM3 | Nrxn1 - Lrrtm3 | NRXN | Cell-Cell Contact | PMID:20519524;PMID:20064387;PMID:20064388; |
| NeuN | NeuN | Nrxn1 | Lrrtm3 | 0.003996 | 0 | NRXN1_LRRTM3 | Nrxn1 - Lrrtm3 | NRXN | Cell-Cell Contact | PMID:20519524;PMID:20064387;PMID:20064388; |
| NeuN | CD31 | Nrxn2 | Lrrtm3 | 0.003661 | 0 | NRXN2_LRRTM3 | Nrxn2 - Lrrtm3 | NRXN | Cell-Cell Contact | PMID:20519524;PMID:20064387;PMID:20064388; |
| NeuN | NeuN | Nrxn2 | Lrrtm3 | 0.00271 | 0 | NRXN2_LRRTM3 | Nrxn2 - Lrrtm3 | NRXN | Cell-Cell Contact | PMID:20519524;PMID:20064387;PMID:20064388; |
| CD31 | CD31 | Nrxn3 | Lrrtm3 | 0.00107 | 0 | NRXN3_LRRTM3 | Nrxn3 - Lrrtm3 | NRXN | Cell-Cell Contact | PMID:20519524;PMID:20064387;PMID:20064388; |
| NeuN | CD31 | Nrxn3 | Lrrtm3 | 0.001029 | 0 | NRXN3_LRRTM3 | Nrxn3 - Lrrtm3 | NRXN | Cell-Cell Contact | PMID:20519524;PMID:20064387;PMID:20064388; |
| NeuN | CD31 | Nrxn1 | Lrrtm4 | 0.004208 | 0 | NRXN1_LRRTM4 | Nrxn1 - Lrrtm4 | NRXN | Cell-Cell Contact | PMID:20519524;PMID:20064387;PMID:20064388; |
| NeuN | NeuN | Nrxn1 | Lrrtm4 | 0.004571 | 0 | NRXN1_LRRTM4 | Nrxn1 - Lrrtm4 | NRXN | Cell-Cell Contact | PMID:20519524;PMID:20064387;PMID:20064388; |
| NeuN | CD31 | Nrxn2 | Lrrtm4 | 0.002855 | 0 | NRXN2_LRRTM4 | Nrxn2 - Lrrtm4 | NRXN | Cell-Cell Contact | PMID:20519524;PMID:20064387;PMID:20064388; |
| NeuN | NeuN | Nrxn2 | Lrrtm4 | 0.003101 | 0 | NRXN2_LRRTM4 | Nrxn2 - Lrrtm4 | NRXN | Cell-Cell Contact | PMID:20519524;PMID:20064387;PMID:20064388; |
| CD31 | NeuN | Nrxn3 | Lrrtm4 | 0.000906 | 0 | NRXN3_LRRTM4 | Nrxn3 - Lrrtm4 | NRXN | Cell-Cell Contact | PMID:20519524;PMID:20064387;PMID:20064388; |
| NeuN | NeuN | Nrxn3 | Lrrtm4 | 0.000871 | 0 | NRXN3_LRRTM4 | Nrxn3 - Lrrtm4 | NRXN | Cell-Cell Contact | PMID:20519524;PMID:20064387;PMID:20064388; |
| CD31 | NeuN | Lrrtm4 | Ptprs | 0.009191 | 0 | LRRTM4_PTPRS | Lrrtm4 - Ptprs | PTPR | Cell-Cell Contact | PMID:31995730 |
| NeuN | NeuN | Lrrtm4 | Ptprs | 0.009979 | 0 | LRRTM4_PTPRS | Lrrtm4 - Ptprs | PTPR | Cell-Cell Contact | PMID:31995730 |
| CD31 | CD31 | App | Sorl1 | 0.016465 | 0 | APP_SORL1 | App - Sorl1 | APP | Cell-Cell Contact | PMID: 24523320 |
| NeuN | CD31 | App | Sorl1 | 0.015944 | 0 | APP_SORL1 | App - Sorl1 | APP | Cell-Cell Contact | PMID: 24523320 |
| CD31 | CD31 | App | Tnfrsf21 | 0.006528 | 0 | APP_TNFRSF21 | App - Tnfrsf21 | APP | Cell-Cell Contact | PMID: 33075465 |
| NeuN | CD31 | App | Tnfrsf21 | 0.006319 | 0 | APP_TNFRSF21 | App - Tnfrsf21 | APP | Cell-Cell Contact | PMID: 33075465 |
| CD31 | CD31 | App | TREM2_TYROBP | 0.008225 | 0 | APP_TREM2_TYROBP | App - (Trem2+Tyrobp) | APP | Cell-Cell Contact | PMID:27477013 |
| NeuN | CD31 | App | TREM2_TYROBP | 0.007962 | 0 | APP_TREM2_TYROBP | App - (Trem2+Tyrobp) | APP | Cell-Cell Contact | PMID:27477013 |

| | | | | | | | | | | |
|---|---|---|---|---|---|---|---|---|---|---|
| CD31 | CD31 | Apoa2 | TREM2_TYROBP | 0.000742 | 0 | APOA2_TREM2_TYROBP | Apoa2 - (Trem2+Tyrobp) | ApoA | Cell-Cell Contact | uniprot;PMID:27477017 |
| NeuN | CD31 | Apoa2 | TREM2_TYROBP | 0.000389 | 0 | APOA2_TREM2_TYROBP | Apoa2 - (Trem2+Tyrobp) | ApoA | Cell-Cell Contact | uniprot;PMID:27477017 |
| CD31 | CD31 | Apoe | TREM2_TYROBP | 0.001308 | 0 | APOE_TREM2_TYROBP | Apoe - (Trem2+Tyrobp) | ApoE | Cell-Cell Contact | uniprot;PMID:27477015 |
| CD31 | NeuN | Apoe | TREM2_TYROBP | 0.000623 | 0 | APOE_TREM2_TYROBP | Apoe - (Trem2+Tyrobp) | ApoE | Cell-Cell Contact | uniprot;PMID:27477015 |
| NeuN | CD31 | Cadm1 | Nectin3 | 0.00509 | 0 | CADM1_NECTIN3 | Cadm1 - Nectin3 | CADM | Cell-Cell Contact | uniprot |
| NeuN | NeuN | Cadm1 | Nectin3 | 0.005261 | 0 | CADM1_NECTIN3 | Cadm1 - Nectin3 | CADM | Cell-Cell Contact | uniprot |
| NeuN | CD31 | Cd99 | Pilrb1 | 0.029157 | 0 | CD99_PILRB1 | Cd99 - Pilrb1 | CD99 | Cell-Cell Contact | uniprot;PMID:18234675 |
| NeuN | NeuN | Cd99 | Pilrb1 | 0.020997 | 0 | CD99_PILRB1 | Cd99 - Pilrb1 | CD99 | Cell-Cell Contact | uniprot;PMID:18234675 |
| CD31 | CD31 | Ceacam2 | Cd209c | 0.000608 | 0 | CEACAM2_CD209C | Ceacam2 - Cd209c | CEACAM | Cell-Cell Contact | PMID:16282604;PMID:16246332 |
| NeuN | CD31 | Ceacam2 | Cd209c | 0.000543 | 0 | CEACAM2_CD209C | Ceacam2 - Cd209c | CEACAM | Cell-Cell Contact | PMID:16282604;PMID:16246332 |
| CD31 | CD31 | Ceacam2 | Cd209a | 0.000452 | 0 | CEACAM2_CD209A | Ceacam2 - Cd209a | CEACAM | Cell-Cell Contact | PMID:16282604;PMID:16246332 |
| NeuN | CD31 | Ceacam2 | Cd209a | 0.000403 | 0 | CEACAM2_CD209A | Ceacam2 - Cd209a | CEACAM | Cell-Cell Contact | PMID:16282604;PMID:16246332 |
| CD31 | CD31 | Ceacam2 | Cd209d | 0.000409 | 0 | CEACAM2_CD209D | Ceacam2 - Cd209d | CEACAM | Cell-Cell Contact | PMID:16282604;PMID:16246332 |
| NeuN | CD31 | Ceacam2 | Cd209d | 0.000365 | 0 | CEACAM2_CD209D | Ceacam2 - Cd209d | CEACAM | Cell-Cell Contact | PMID:16282604;PMID:16246332 |
| CD31 | CD31 | Ceacam2 | Cd209b | 0.000486 | 0 | CEACAM2_CD209B | Ceacam2 - Cd209b | CEACAM | Cell-Cell Contact | PMID:16282604;PMID:16246332 |
| NeuN | CD31 | Ceacam2 | Cd209b | 0.000434 | 0 | CEACAM2_CD209B | Ceacam2 - Cd209b | CEACAM | Cell-Cell Contact | PMID:16282604;PMID:16246332 |
| CD31 | CD31 | Ceacam2 | Ceacam2 | 0.000336 | 0 | CEACAM2_CEACAM2 | Ceacam2 - Ceacam2 | CEACAM | Cell-Cell Contact | PMID:21982860 |
| NeuN | CD31 | Ceacam2 | Ceacam2 | 0.0003 | 0 | CEACAM2_CEACAM2 | Ceacam2 - Ceacam2 | CEACAM | Cell-Cell Contact | PMID:21982860 |
| CD31 | NeuN | Ceacam2 | Ceacam2 | 0.0003 | 0 | CEACAM2_CEACAM2 | Ceacam2 - Ceacam2 | CEACAM | Cell-Cell Contact | PMID:21982860 |
| CD31 | CD31 | Ceacam2 | Sele | 0.000505 | 0 | CEACAM2_SELE | Ceacam2 - Sele | CEACAM | Cell-Cell Contact | PMID:1378450 |
| NeuN | CD31 | Ceacam2 | Sele | 0.000451 | 0 | CEACAM2_SELE | Ceacam2 - Sele | CEACAM | Cell-Cell Contact | PMID:1378450 |
| CD31 | CD31 | Ceacam2 | Cd1d1 | 0.00076 | 0 | CEACAM2_CD1D1 | Ceacam2 - Cd1d1 | CEACAM | Cell-Cell Contact | PMID:24104458 |
| NeuN | CD31 | Ceacam2 | Cd1d1 | 0.000679 | 0 | CEACAM2_CD1D1 | Ceacam2 - Cd1d1 | CEACAM | Cell-Cell Contact | PMID:24104458 |
| CD31 | CD31 | Cntn1 | Notch1 | 0.00263 | 0 | CNTN1_NOTCH1 | Cntn1 - Notch1 | CNTN | Cell-Cell Contact | PMID:14567914 |
| NeuN | CD31 | Cntn1 | Notch1 | 0.005381 | 0 | CNTN1_NOTCH1 | Cntn1 - Notch1 | CNTN | Cell-Cell Contact | PMID:14567914 |
| CD31 | CD31 | Clec2h | Klrb1f | 0.000589 | 0 | CLEC2H_KLRB1F | Klrb1f - Clec2h | CLEC | Cell-Cell Contact | PMID:24223577 |
| CD31 | NeuN | Clec2h | Klrb1f | 0.000601 | 0 | CLEC2H_KLRB1F | Klrb1f - Clec2h | CLEC | Cell-Cell Contact | PMID:24223577 |
| CD31 | CD31 | Clec2d | Klrb1f | 0.001006 | 0 | CLEC2D_KLRB1F | Klrb1f - Clec2d | CLEC | Cell-Cell Contact | PMID:24223577 |
| CD31 | NeuN | Clec2d | Klrb1f | 0.001026 | 0 | CLEC2D_KLRB1F | Klrb1f - Clec2d | CLEC | Cell-Cell Contact | PMID:24223577 |
| CD31 | CD31 | Mmp2 | MMP14_ITGAV_ITGB3 | 0.001071 | 0 | MMP2_INTEGRIN_AVB3_COMP | Mmp2 - Integrin_avb3 | MMP | Cell-Cell Contact | PMID: 29986882 |
| NeuN | CD31 | Mmp2 | MMP14_ITGAV_ITGB3 | 0.000868 | 0 | MMP2_INTEGRIN_AVB3_COMP | Mmp2 - Integrin_avb3 | MMP | Cell-Cell Contact | PMID: 29986882 |
| CD31 | NeuN | Ntn1 | Dcc | 0.003406 | 0 | NTN1_DCC | Ntn1 - Dcc | Netrin | Cell-Cell Contact | PMID: 22685302 |
| NeuN | NeuN | Ntn1 | Dcc | 0.002144 | 0 | NTN1_DCC | Ntn1 - Dcc | Netrin | Cell-Cell Contact | PMID: 22685302 |
| CD31 | CD31 | Ntn1 | Dscam | 0.000712 | 0 | NTN1_DSCAM | Ntn1 - Dscam | Netrin | Cell-Cell Contact | PMID:18585357;PMID: 22685302 |
| CD31 | NeuN | Ntn1 | Dscam | 0.000902 | 0 | NTN1_DSCAM | Ntn1 - Dscam | Netrin | Cell-Cell Contact | PMID:18585357;PMID: 22685302 |
| CD31 | CD31 | Ntn1 | Unc5a | 0.000528 | 0 | NTN1_UNC5A | Ntn1 - Unc5a | Netrin | Cell-Cell Contact | PMID: 22685302 |
| CD31 | NeuN | Ntn1 | Unc5a | 0.00061 | 0 | NTN1_UNC5A | Ntn1 - Unc5a | Netrin | Cell-Cell Contact | PMID: 22685302 |
| CD31 | CD31 | Ntn1 | Unc5b | 0.001571 | 0 | NTN1_UNC5B | Ntn1 - Unc5b | Netrin | Cell-Cell Contact | PMID: 22685302 |
| NeuN | CD31 | Ntn1 | Unc5b | 0.000988 | 0 | NTN1_UNC5B | Ntn1 - Unc5b | Netrin | Cell-Cell Contact | PMID: 22685302 |
| CD31 | CD31 | Ntn1 | Unc5c | 0.000303 | 0 | NTN1_UNC5C | Ntn1 - Unc5c | Netrin | Cell-Cell Contact | PMID: 22685302 |
| CD31 | NeuN | Ntn1 | Unc5c | 0.00044 | 0 | NTN1_UNC5C | Ntn1 - Unc5c | Netrin | Cell-Cell Contact | PMID: 22685302 |
| CD31 | NeuN | Ntn4 | Ntrk2 | 0.002722 | 0 | NTN4_NTRK2 | Ntn4 - Ntrk2 | Netrin | Cell-Cell Contact | PMID: 34531526 |
| NeuN | NeuN | Ntn4 | Ntrk2 | 0.002867 | 0 | NTN4_NTRK2 | Ntn4 - Ntrk2 | Netrin | Cell-Cell Contact | PMID: 34531526 |
| CD31 | CD31 | Ntn4 | Unc5b | 0.00113 | 0 | NTN4_UNC5B | Ntn4 - Unc5b | Netrin | Cell-Cell Contact | PMID: 31914695 |
| NeuN | CD31 | Ntn4 | Unc5b | 0.00119 | 0 | NTN4_UNC5B | Ntn4 - Unc5b | Netrin | Cell-Cell Contact | PMID: 31914695 |
| CD31 | CD31 | Ntn4 | Unc5d | 0.000577 | 0 | NTN4_UNC5D | Ntn4 - Unc5d | Netrin | Cell-Cell Contact | uniprot |
| NeuN | CD31 | Ntn4 | Unc5d | 0.000608 | 0 | NTN4_UNC5D | Ntn4 - Unc5d | Netrin | Cell-Cell Contact | uniprot |
| CD31 | CD31 | Pecam1 | Cd177 | 0.002046 | 0 | PECAM1_CD177 | Pecam1 - Cd177 | PECAM1 | Cell-Cell Contact | PMID:17580308 |
| CD31 | NeuN | Pecam1 | Cd177 | 0.001121 | 0 | PECAM1_CD177 | Pecam1 - Cd177 | PECAM1 | Cell-Cell Contact | PMID:17580308 |
| CD31 | CD31 | Pecam1 | Cd38 | 0.001069 | 0 | PECAM1_CD38 | Pecam1 - Cd38 | PECAM2 | Cell-Cell Contact | PMID:26407101 |
| CD31 | NeuN | Pecam1 | Cd38 | 0.000699 | 0 | PECAM1_CD38 | Pecam1 - Cd38 | PECAM2 | Cell-Cell Contact | PMID:26407101 |
| CD31 | NeuN | Sirpb1a | Cd47 | 0.001011 | 0 | SIRPB1A_CD47 | Sirpb1a - Cd47 | SIRP | Cell-Cell Contact | PMID:15294972 |
| NeuN | NeuN | Sirpb1a | Cd47 | 0.000719 | 0 | SIRPB1A_CD47 | Sirpb1a - Cd47 | SIRP | Cell-Cell Contact | PMID:15294972 |

**Supplementary Table 13: Cell Chat Interactions at P28 in SAC**

| source | target | ligand | receptor | prob | pval | interaction_name | interaction_name_2 | pathway_name | annotation | evidence |
|---|---|---|---|---|---|---|---|---|---|---|
| CD31 | CD31 | Tgfb1 | ACVR1B_TGFbR2 | 0.00034 | 0 | TGFB1_ACVR1B_TGFBR2 | Tgfb1 - (Acvr1b+Tgfbr2) | TGFb | Secreted Signaling | PMID: 27449815 |
| CD31 | GFAP | Tgfb1 | ACVR1B_TGFbR2 | 0.000296 | 0 | TGFB1_ACVR1B_TGFBR2 | Tgfb1 - (Acvr1b+Tgfbr2) | TGFb | Secreted Signaling | PMID: 27449815 |
| CD31 | CD31 | Tgfb1 | ACVR1C_TGFbR2 | 0.000271 | 0 | TGFB1_ACVR1C_TGFBR2 | Tgfb1 - (Acvr1c+Tgfbr2) | TGFb | Secreted Signaling | PMID: 27449815 |
| GFAP | CD31 | Tgfb1 | ACVR1C_TGFbR2 | 0.000198 | 0 | TGFB1_ACVR1C_TGFBR2 | Tgfb1 - (Acvr1c+Tgfbr2) | TGFb | Secreted Signaling | PMID: 27449815 |
| CD31 | CD31 | Tgfb3 | ACVR1B_TGFbR2 | 0.000504 | 0 | TGFB3_ACVR1B_TGFBR2 | Tgfb3 - (Acvr1b+Tgfbr2) | TGFb | Secreted Signaling | PMID: 27449815 |
| CD31 | GFAP | Tgfb3 | ACVR1B_TGFbR2 | 0.000439 | 0 | TGFB3_ACVR1B_TGFBR2 | Tgfb3 - (Acvr1b+Tgfbr2) | TGFb | Secreted Signaling | PMID: 27449815 |
| CD31 | CD31 | Tgfb3 | ACVR1C_TGFbR2 | 0.000401 | 0 | TGFB3_ACVR1C_TGFBR2 | Tgfb3 - (Acvr1c+Tgfbr2) | TGFb | Secreted Signaling | PMID: 27449815 |
| CD31 | GFAP | Tgfb3 | ACVR1C_TGFbR2 | 0.000272 | 0 | TGFB3_ACVR1C_TGFBR2 | Tgfb3 - (Acvr1c+Tgfbr2) | TGFb | Secreted Signaling | PMID: 27449815 |
| GFAP | CD31 | Bmp2 | BMPR1A_ACVR2A | 0.000465 | 0 | BMP2_BMPR1A_ACVR2A | Bmp2 - (Bmpr1a+Acvr2a) | BMP | Secreted Signaling | KEGG: mmu04350; PMID:26893264 |
| GFAP | GFAP | Bmp2 | BMPR1A_ACVR2A | 0.00041 | 0 | BMP2_BMPR1A_ACVR2A | Bmp2 - (Bmpr1a+Acvr2a) | BMP | Secreted Signaling | KEGG: mmu04350; PMID:26893264 |
| GFAP | CD31 | Bmp2 | BMPR1A_ACVR2B | 0.000404 | 0 | BMP2_BMPR1A_ACVR2B | Bmp2 - (Bmpr1a+Acvr2b) | BMP | Secreted Signaling | KEGG: mmu04350; PMID:26893264 |
| GFAP | GFAP | Bmp2 | BMPR1A_ACVR2B | 0.000349 | 0 | BMP2_BMPR1A_ACVR2B | Bmp2 - (Bmpr1a+Acvr2b) | BMP | Secreted Signaling | KEGG: mmu04350; PMID:26893264 |
| GFAP | CD31 | Bmp2 | BMPR1A_BMPR2 | 0.000723 | 0 | BMP2_BMPR1A_BMPR2 | Bmp2 - (Bmpr1a+Bmpr2) | BMP | Secreted Signaling | KEGG: mmu04350; PMID:26893264 |
| GFAP | GFAP | Bmp2 | BMPR1A_BMPR2 | 0.000696 | 0 | BMP2_BMPR1A_BMPR2 | Bmp2 - (Bmpr1a+Bmpr2) | BMP | Secreted Signaling | KEGG: mmu04350; PMID:26893264 |
| GFAP | CD31 | Bmp2 | BMPR1B_ACVR2A | 0.000332 | 0 | BMP2_BMPR1B_ACVR2A | Bmp2 - (Bmpr1b+Acvr2a) | BMP | Secreted Signaling | KEGG: mmu04350; PMID:26893264 |
| GFAP | GFAP | Bmp2 | BMPR1B_ACVR2A | 0.000375 | 0 | BMP2_BMPR1B_ACVR2A | Bmp2 - (Bmpr1b+Acvr2a) | BMP | Secreted Signaling | KEGG: mmu04350; PMID:26893264 |
| GFAP | CD31 | Bmp2 | BMPR1B_ACVR2B | 0.000288 | 0 | BMP2_BMPR1B_ACVR2B | Bmp2 - (Bmpr1b+Acvr2b) | BMP | Secreted Signaling | KEGG: mmu04350; PMID:26893264 |
| GFAP | GFAP | Bmp2 | BMPR1B_ACVR2B | 0.000319 | 0 | BMP2_BMPR1B_ACVR2B | Bmp2 - (Bmpr1b+Acvr2b) | BMP | Secreted Signaling | KEGG: mmu04350; PMID:26893264 |
| CD31 | GFAP | Bmp2 | BMPR1B_BMPR2 | 0.000549 | 0 | BMP2_BMPR1B_BMPR2 | Bmp2 - (Bmpr1b+Bmpr2) | BMP | Secreted Signaling | KEGG: mmu04350; PMID:26893264 |
| GFAP | GFAP | Bmp2 | BMPR1B_BMPR2 | 0.000636 | 0 | BMP2_BMPR1B_BMPR2 | Bmp2 - (Bmpr1b+Bmpr2) | BMP | Secreted Signaling | KEGG: mmu04350; PMID:26893264 |
| CD31 | CD31 | Gdf6 | BMPR1A_ACVR2A | 0.000359 | 0 | GDF6_BMPR1A_ACVR2A | Gdf6 - (Bmpr1a+Acvr2a) | BMP | Secreted Signaling | KEGG: mmu04350; PMID:26893264 |
| GFAP | CD31 | Gdf6 | BMPR1A_ACVR2A | 0.000356 | 0 | GDF6_BMPR1A_ACVR2A | Gdf6 - (Bmpr1a+Acvr2a) | BMP | Secreted Signaling | KEGG: mmu04350; PMID:26893264 |
| CD31 | CD31 | Gdf6 | BMPR1A_ACVR2B | 0.000312 | 0 | GDF6_BMPR1A_ACVR2B | Gdf6 - (Bmpr1a+Acvr2b) | BMP | Secreted Signaling | KEGG: mmu04350; PMID:26893264 |
| GFAP | CD31 | Gdf6 | BMPR1A_ACVR2B | 0.000309 | 0 | GDF6_BMPR1A_ACVR2B | Gdf6 - (Bmpr1a+Acvr2b) | BMP | Secreted Signaling | KEGG: mmu04350; PMID:26893264 |
| CD31 | CD31 | Gdf6 | BMPR1A_BMPR2 | 0.000558 | 0 | GDF6_BMPR1A_BMPR2 | Gdf6 - (Bmpr1a+Bmpr2) | BMP | Secreted Signaling | KEGG: mmu04350; PMID:26893264 |
| GFAP | CD31 | Gdf6 | BMPR1A_BMPR2 | 0.000553 | 0 | GDF6_BMPR1A_BMPR2 | Gdf6 - (Bmpr1a+Bmpr2) | BMP | Secreted Signaling | KEGG: mmu04350; PMID:26893264 |
| CD31 | GFAP | Gdf6 | BMPR1B_ACVR2A | 0.00029 | 0 | GDF6_BMPR1B_ACVR2A | Gdf6 - (Bmpr1b+Acvr2a) | BMP | Secreted Signaling | KEGG: mmu04350; PMID:26893264 |
| GFAP | GFAP | Gdf6 | BMPR1B_ACVR2A | 0.000287 | 0 | GDF6_BMPR1B_ACVR2A | Gdf6 - (Bmpr1b+Acvr2a) | BMP | Secreted Signaling | KEGG: mmu04350; PMID:26893264 |
| CD31 | GFAP | Gdf6 | BMPR1B_ACVR2B | 0.000246 | 0 | GDF6_BMPR1B_ACVR2B | Gdf6 - (Bmpr1b+Acvr2b) | BMP | Secreted Signaling | KEGG: mmu04350; PMID:26893264 |
| GFAP | GFAP | Gdf6 | BMPR1B_ACVR2B | 0.000244 | 0 | GDF6_BMPR1B_ACVR2B | Gdf6 - (Bmpr1b+Acvr2b) | BMP | Secreted Signaling | KEGG: mmu04350; PMID:26893264 |
| CD31 | GFAP | Gdf6 | BMPR1B_BMPR2 | 0.000491 | 0 | GDF6_BMPR1B_BMPR2 | Gdf6 - (Bmpr1b+Bmpr2) | BMP | Secreted Signaling | KEGG: mmu04350; PMID:26893264 |
| GFAP | GFAP | Gdf6 | BMPR1B_BMPR2 | 0.000487 | 0 | GDF6_BMPR1B_BMPR2 | Gdf6 - (Bmpr1b+Bmpr2) | BMP | Secreted Signaling | KEGG: mmu04350; PMID:26893264 |
| CD31 | CD31 | Bmp5 | ACVR1_ACVR2A | 0.000392 | 0 | BMP5_ACVR1_ACVR2A | Bmp5 - (Acvr1+Acvr2a) | BMP | Secreted Signaling | KEGG: mmu04350; PMID:26893264 |
| CD31 | NeuN | Bmp5 | ACVR1_ACVR2A | 0.000318 | 0 | BMP5_ACVR1_ACVR2A | Bmp5 - (Acvr1+Acvr2a) | BMP | Secreted Signaling | KEGG: mmu04350; PMID:26893264 |
| CD31 | CD31 | Bmp5 | ACVR1_ACVR2B | 0.00034 | 0 | BMP5_ACVR1_ACVR2B | Bmp5 - (Acvr1+Acvr2b) | BMP | Secreted Signaling | KEGG: mmu04350; PMID:26893264 |
| GFAP | CD31 | Bmp5 | ACVR1_ACVR2B | 0.000256 | 0 | BMP5_ACVR1_ACVR2B | Bmp5 - (Acvr1+Acvr2b) | BMP | Secreted Signaling | KEGG: mmu04350; PMID:26893264 |
| CD31 | CD31 | Bmp5 | ACVR1_BMPR2 | 0.000609 | 0 | BMP5_ACVR1_BMPR2 | Bmp5 - (Acvr1+Bmpr2) | BMP | Secreted Signaling | KEGG: mmu04350; PMID:26893264 |
| GFAP | CD31 | Bmp5 | ACVR1_BMPR2 | 0.000458 | 0 | BMP5_ACVR1_BMPR2 | Bmp5 - (Acvr1+Bmpr2) | BMP | Secreted Signaling | KEGG: mmu04350; PMID:26893264 |
| CD31 | CD31 | Bmp5 | BMPR1A_ACVR2A | 0.000379 | 0 | BMP5_BMPR1A_ACVR2A | Bmp5 - (Bmpr1a+Acvr2a) | BMP | Secreted Signaling | KEGG: mmu04350; PMID:26893264 |
| CD31 | GFAP | Bmp5 | BMPR1A_ACVR2A | 0.000335 | 0 | BMP5_BMPR1A_ACVR2A | Bmp5 - (Bmpr1a+Acvr2a) | BMP | Secreted Signaling | KEGG: mmu04350; PMID:26893264 |
| CD31 | CD31 | Bmp5 | BMPR1A_ACVR2B | 0.000329 | 0 | BMP5_BMPR1A_ACVR2B | Bmp5 - (Bmpr1a+Acvr2b) | BMP | Secreted Signaling | KEGG: mmu04350; PMID:26893264 |
| CD31 | GFAP | Bmp5 | BMPR1A_ACVR2B | 0.000284 | 0 | BMP5_BMPR1A_ACVR2B | Bmp5 - (Bmpr1a+Acvr2b) | BMP | Secreted Signaling | KEGG: mmu04350; PMID:26893264 |
| CD31 | CD31 | Bmp5 | BMPR1A_BMPR2 | 0.00059 | 0 | BMP5_BMPR1A_BMPR2 | Bmp5 - (Bmpr1a+Bmpr2) | BMP | Secreted Signaling | KEGG: mmu04350; PMID:26893264 |
| CD31 | GFAP | Bmp5 | BMPR1A_BMPR2 | 0.000568 | 0 | BMP5_BMPR1A_BMPR2 | Bmp5 - (Bmpr1a+Bmpr2) | BMP | Secreted Signaling | KEGG: mmu04350; PMID:26893264 |
| CD31 | CD31 | Bmp5 | BMPR1B_ACVR2A | 0.000271 | 0 | BMP5_BMPR1B_ACVR2A | Bmp5 - (Bmpr1b+Acvr2a) | BMP | Secreted Signaling | KEGG: mmu04350; PMID:26893264 |
| CD31 | GFAP | Bmp5 | BMPR1B_ACVR2A | 0.000306 | 0 | BMP5_BMPR1B_ACVR2A | Bmp5 - (Bmpr1b+Acvr2a) | BMP | Secreted Signaling | KEGG: mmu04350; PMID:26893264 |
| CD31 | CD31 | Bmp5 | BMPR1B_ACVR2B | 0.000235 | 0 | BMP5_BMPR1B_ACVR2B | Bmp5 - (Bmpr1b+Acvr2b) | BMP | Secreted Signaling | KEGG: mmu04350; PMID:26893264 |
| CD31 | GFAP | Bmp5 | BMPR1B_ACVR2B | 0.00026 | 0 | BMP5_BMPR1B_ACVR2B | Bmp5 - (Bmpr1b+Acvr2b) | BMP | Secreted Signaling | KEGG: mmu04350; PMID:26893264 |
| CD31 | CD31 | Bmp5 | BMPR1B_BMPR2 | 0.000421 | 0 | BMP5_BMPR1B_BMPR2 | Bmp5 - (Bmpr1b+Bmpr2) | BMP | Secreted Signaling | KEGG: mmu04350; PMID:26893264 |
| CD31 | GFAP | Bmp5 | BMPR1B_BMPR2 | 0.000519 | 0 | BMP5_BMPR1B_BMPR2 | Bmp5 - (Bmpr1b+Bmpr2) | BMP | Secreted Signaling | KEGG: mmu04350; PMID:26893264 |
| CD31 | CD31 | Bmp6 | ACVR1_ACVR2A | 0.000522 | 0 | BMP6_ACVR1_ACVR2A | Bmp6 - (Acvr1+Acvr2a) | BMP | Secreted Signaling | KEGG: mmu04350; PMID:26893264 |
| CD31 | NeuN | Bmp6 | ACVR1_ACVR2A | 0.000425 | 0 | BMP6_ACVR1_ACVR2A | Bmp6 - (Acvr1+Acvr2a) | BMP | Secreted Signaling | KEGG: mmu04350; PMID:26893264 |
| CD31 | CD31 | Bmp6 | ACVR1_ACVR2B | 0.000453 | 0 | BMP6_ACVR1_ACVR2B | Bmp6 - (Acvr1+Acvr2b) | BMP | Secreted Signaling | KEGG: mmu04350; PMID:26893264 |
| GFAP | CD31 | Bmp6 | ACVR1_ACVR2B | 0.000322 | 0 | BMP6_ACVR1_ACVR2B | Bmp6 - (Acvr1+Acvr2b) | BMP | Secreted Signaling | KEGG: mmu04350; PMID:26893264 |
| CD31 | CD31 | Bmp6 | ACVR1_BMPR2 | 0.000812 | 0 | BMP6_ACVR1_BMPR2 | Bmp6 - (Acvr1+Bmpr2) | BMP | Secreted Signaling | KEGG: mmu04350; PMID:26893264 |
| GFAP | CD31 | Bmp6 | ACVR1_BMPR2 | 0.000576 | 0 | BMP6_ACVR1_BMPR2 | Bmp6 - (Acvr1+Bmpr2) | BMP | Secreted Signaling | KEGG: mmu04350; PMID:26893264 |

| | | | | | | | | | | |
|---|---|---|---|---|---|---|---|---|---|---|
| CD31 | CD31 | Bmp6 | BMPR1A_ACVR2A | 0.000505 | 0 | BMP6_BMPR1A_ACVR2A | Bmp6 - (Bmpr1a+Acvr2a) | BMP | Secreted Signaling | KEGG: mmu04350; PMID:26893264 |
| CD31 | GFAP | Bmp6 | BMPR1A_ACVR2A | 0.000446 | 0 | BMP6_BMPR1A_ACVR2A | Bmp6 - (Bmpr1a+Acvr2a) | BMP | Secreted Signaling | KEGG: mmu04350; PMID:26893264 |
| CD31 | CD31 | Bmp6 | BMPR1A_ACVR2B | 0.000439 | 0 | BMP6_BMPR1A_ACVR2B | Bmp6 - (Bmpr1a+Acvr2b) | BMP | Secreted Signaling | KEGG: mmu04350; PMID:26893264 |
| CD31 | GFAP | Bmp6 | BMPR1A_ACVR2B | 0.000379 | 0 | BMP6_BMPR1A_ACVR2B | Bmp6 - (Bmpr1a+Acvr2b) | BMP | Secreted Signaling | KEGG: mmu04350; PMID:26893264 |
| CD31 | CD31 | Bmp6 | BMPR1A_BMPR2 | 0.000786 | 0 | BMP6_BMPR1A_BMPR2 | Bmp6 - (Bmpr1a+Bmpr2) | BMP | Secreted Signaling | KEGG: mmu04350; PMID:26893264 |
| CD31 | GFAP | Bmp6 | BMPR1A_BMPR2 | 0.000757 | 0 | BMP6_BMPR1A_BMPR2 | Bmp6 - (Bmpr1a+Bmpr2) | BMP | Secreted Signaling | KEGG: mmu04350; PMID:26893264 |
| CD31 | CD31 | Bmp6 | BMPR1B_ACVR2A | 0.000361 | 0 | BMP6_BMPR1B_ACVR2A | Bmp6 - (Bmpr1b+Acvr2a) | BMP | Secreted Signaling | KEGG: mmu04350; PMID:26893264 |
| CD31 | GFAP | Bmp6 | BMPR1B_ACVR2A | 0.000408 | 0 | BMP6_BMPR1B_ACVR2A | Bmp6 - (Bmpr1b+Acvr2a) | BMP | Secreted Signaling | KEGG: mmu04350; PMID:26893264 |
| CD31 | CD31 | Bmp6 | BMPR1B_ACVR2B | 0.000313 | 0 | BMP6_BMPR1B_ACVR2B | Bmp6 - (Bmpr1b+Acvr2b) | BMP | Secreted Signaling | KEGG: mmu04350; PMID:26893264 |
| CD31 | GFAP | Bmp6 | BMPR1B_ACVR2B | 0.000347 | 0 | BMP6_BMPR1B_ACVR2B | Bmp6 - (Bmpr1b+Acvr2b) | BMP | Secreted Signaling | KEGG: mmu04350; PMID:26893264 |
| CD31 | CD31 | Bmp6 | BMPR1B_BMPR2 | 0.000561 | 0 | BMP6_BMPR1B_BMPR2 | Bmp6 - (Bmpr1b+Bmpr2) | BMP | Secreted Signaling | KEGG: mmu04350; PMID:26893264 |
| CD31 | GFAP | Bmp6 | BMPR1B_BMPR2 | 0.000691 | 0 | BMP6_BMPR1B_BMPR2 | Bmp6 - (Bmpr1b+Bmpr2) | BMP | Secreted Signaling | KEGG: mmu04350; PMID:26893264 |
| CD31 | CD31 | Bmp8b | ACVR1_ACVR2A | 0.000263 | 0 | BMP8B_ACVR1_ACVR2A | Bmp8b - (Acvr1+Acvr2a) | BMP | Secreted Signaling | KEGG: mmu04350; PMID:26893264 |
| GFAP | CD31 | Bmp8b | ACVR1_ACVR2A | 0.000319 | 0 | BMP8B_ACVR1_ACVR2A | Bmp8b - (Acvr1+Acvr2a) | BMP | Secreted Signaling | KEGG: mmu04350; PMID:26893264 |
| CD31 | CD31 | Bmp8b | ACVR1_ACVR2B | 0.000228 | 0 | BMP8B_ACVR1_ACVR2B | Bmp8b - (Acvr1+Acvr2b) | BMP | Secreted Signaling | KEGG: mmu04350; PMID:26893264 |
| GFAP | CD31 | Bmp8b | ACVR1_ACVR2B | 0.000277 | 0 | BMP8B_ACVR1_ACVR2B | Bmp8b - (Acvr1+Acvr2b) | BMP | Secreted Signaling | KEGG: mmu04350; PMID:26893264 |
| CD31 | CD31 | Bmp8b | ACVR1_BMPR2 | 0.000408 | 0 | BMP8B_ACVR1_BMPR2 | Bmp8b - (Acvr1+Bmpr2) | BMP | Secreted Signaling | KEGG: mmu04350; PMID:26893264 |
| GFAP | CD31 | Bmp8b | ACVR1_BMPR2 | 0.000495 | 0 | BMP8B_ACVR1_BMPR2 | Bmp8b - (Acvr1+Bmpr2) | BMP | Secreted Signaling | KEGG: mmu04350; PMID:26893264 |
| GFAP | CD31 | Bmp8b | BMPR1A_ACVR2A | 0.000308 | 0 | BMP8B_BMPR1A_ACVR2A | Bmp8b - (Bmpr1a+Acvr2a) | BMP | Secreted Signaling | KEGG: mmu04350; PMID:26893264 |
| GFAP | GFAP | Bmp8b | BMPR1A_ACVR2A | 0.000272 | 0 | BMP8B_BMPR1A_ACVR2A | Bmp8b - (Bmpr1a+Acvr2a) | BMP | Secreted Signaling | KEGG: mmu04350; PMID:26893264 |
| GFAP | CD31 | Bmp8b | BMPR1A_ACVR2B | 0.000268 | 0 | BMP8B_BMPR1A_ACVR2B | Bmp8b - (Bmpr1a+Acvr2b) | BMP | Secreted Signaling | KEGG: mmu04350; PMID:26893264 |
| GFAP | GFAP | Bmp8b | BMPR1A_ACVR2B | 0.000231 | 0 | BMP8B_BMPR1A_ACVR2B | Bmp8b - (Bmpr1a+Acvr2b) | BMP | Secreted Signaling | KEGG: mmu04350; PMID:26893264 |
| GFAP | CD31 | Bmp8b | BMPR1A_BMPR2 | 0.00048 | 0 | BMP8B_BMPR1A_BMPR2 | Bmp8b - (Bmpr1a+Bmpr2) | BMP | Secreted Signaling | KEGG: mmu04350; PMID:26893264 |
| GFAP | GFAP | Bmp8b | BMPR1A_BMPR2 | 0.000462 | 0 | BMP8B_BMPR1A_BMPR2 | Bmp8b - (Bmpr1a+Bmpr2) | BMP | Secreted Signaling | KEGG: mmu04350; PMID:26893264 |
| GFAP | CD31 | Bmp8b | BMPR1B_ACVR2A | 0.00022 | 0 | BMP8B_BMPR1B_ACVR2A | Bmp8b - (Bmpr1b+Acvr2a) | BMP | Secreted Signaling | KEGG: mmu04350; PMID:26893264 |
| GFAP | GFAP | Bmp8b | BMPR1B_ACVR2A | 0.000249 | 0 | BMP8B_BMPR1B_ACVR2A | Bmp8b - (Bmpr1b+Acvr2a) | BMP | Secreted Signaling | KEGG: mmu04350; PMID:26893264 |
| GFAP | CD31 | Bmp8b | BMPR1B_ACVR2B | 0.000191 | 0 | BMP8B_BMPR1B_ACVR2B | Bmp8b - (Bmpr1b+Acvr2b) | BMP | Secreted Signaling | KEGG: mmu04350; PMID:26893264 |
| GFAP | GFAP | Bmp8b | BMPR1B_ACVR2B | 0.000211 | 0 | BMP8B_BMPR1B_ACVR2B | Bmp8b - (Bmpr1b+Acvr2b) | BMP | Secreted Signaling | KEGG: mmu04350; PMID:26893264 |
| CD31 | GFAP | Bmp8b | BMPR1B_BMPR2 | 0.000348 | 0 | BMP8B_BMPR1B_BMPR2 | Bmp8b - (Bmpr1b+Bmpr2) | BMP | Secreted Signaling | KEGG: mmu04350; PMID:26893264 |
| GFAP | GFAP | Bmp8b | BMPR1B_BMPR2 | 0.000422 | 0 | BMP8B_BMPR1B_BMPR2 | Bmp8b - (Bmpr1b+Bmpr2) | BMP | Secreted Signaling | KEGG: mmu04350; PMID:26893264 |
| CD31 | CD31 | Gdf9 | ACVR1B_BMPR2 | 0.000639 | 0 | GDF9_ACVR1B_BMPR2 | Gdf9 - (Acvr1b+Bmpr2) | GDF | Secreted Signaling | KEGG: mmu04350 |
| CD31 | NeuN | Gdf9 | ACVR1B_BMPR2 | 0.001031 | 0 | GDF9_ACVR1B_BMPR2 | Gdf9 - (Acvr1b+Bmpr2) | GDF | Secreted Signaling | KEGG: mmu04350 |
| CD31 | CD31 | Gdf10 | ACVR1B_ACVR2A | 0.000539 | 0 | GDF10_ACVR1B_ACVR2A | Gdf10 - (Acvr1b+Acvr2a) | GDF | Secreted Signaling | KEGG: mmu04350 |
| CD31 | NeuN | Gdf10 | ACVR1B_ACVR2A | 0.001012 | 0 | GDF10_ACVR1B_ACVR2A | Gdf10 - (Acvr1b+Acvr2a) | GDF | Secreted Signaling | KEGG: mmu04350 |
| CD31 | CD31 | Gdf11 | TGFBR1_ACVR2A | 0.000482 | 0 | GDF11_TGFBR1_ACVR2A | Gdf11 - (Tgfbr1+Acvr2a) | GDF | Secreted Signaling | KEGG: mmu04350 |
| GFAP | CD31 | Gdf11 | TGFBR1_ACVR2A | 0.000607 | 0 | GDF11_TGFBR1_ACVR2A | Gdf11 - (Tgfbr1+Acvr2a) | GDF | Secreted Signaling | KEGG: mmu04350 |
| CD31 | CD31 | Gdf11 | TGFBR1_ACVR2B | 0.000419 | 0 | GDF11_TGFBR1_ACVR2B | Gdf11 - (Tgfbr1+Acvr2b) | GDF | Secreted Signaling | KEGG: mmu04350 |
| GFAP | CD31 | Gdf11 | TGFBR1_ACVR2B | 0.000527 | 0 | GDF11_TGFBR1_ACVR2B | Gdf11 - (Tgfbr1+Acvr2b) | GDF | Secreted Signaling | KEGG: mmu04350 |
| GFAP | GFAP | Gdf11 | ACVR1B_ACVR2B | 0.00045 | 0 | GDF11_ACVR1B_ACVR2B | Gdf11 - (Acvr1b+Acvr2b) | GDF | Secreted Signaling | KEGG: mmu04350 |
| GFAP | NeuN | Gdf11 | ACVR1B_ACVR2B | 0.00072 | 0 | GDF11_ACVR1B_ACVR2B | Gdf11 - (Acvr1b+Acvr2b) | GDF | Secreted Signaling | KEGG: mmu04350 |
| CD31 | CD31 | Gdnf | GFRA1_RET | 0.000468 | 0 | GDNF_GFRA1 | Gdnf - Gfra1 | GDNF | Secreted Signaling | PMID: 11544105 |
| GFAP | CD31 | Gdnf | GFRA1_RET | 0.000439 | 0 | GDNF_GFRA1 | Gdnf - Gfra1 | GDNF | Secreted Signaling | PMID: 11544105 |
| CD31 | CD31 | Nrtn | GFRA2_RET | 0.00031 | 0 | NRTN_GFRA2 | Nrtn - Gfra2 | GDNF | Secreted Signaling | PMID: 11544105 |
| GFAP | CD31 | Nrtn | GFRA2_RET | 0.000407 | 0 | NRTN_GFRA2 | Nrtn - Gfra2 | GDNF | Secreted Signaling | PMID: 11544105 |
| CD31 | CD31 | Inhba | ACVR1B_ACVR2A | 0.000435 | 0 | INHBA_ACVR1B_ACVR2A | Inhba - (Acvr1b+Acvr2a) | ACTIVIN | Secreted Signaling | KEGG: mmu04350 |
| CD31 | NeuN | Inhba | ACVR1B_ACVR2A | 0.000837 | 0 | INHBA_ACVR1B_ACVR2A | Inhba - (Acvr1b+Acvr2a) | ACTIVIN | Secreted Signaling | KEGG: mmu04350 |
| CD31 | CD31 | Inhba | ACVR1B_ACVR2B | 0.000378 | 0 | INHBA_ACVR1B_ACVR2B | Inhba - (Acvr1b+Acvr2b) | ACTIVIN | Secreted Signaling | KEGG: mmu04350 |
| CD31 | NeuN | Inhba | ACVR1B_ACVR2B | 0.000502 | 0 | INHBA_ACVR1B_ACVR2B | Inhba - (Acvr1b+Acvr2b) | ACTIVIN | Secreted Signaling | KEGG: mmu04350 |
| CD31 | CD31 | Inhbb | ACVR1B_ACVR2A | 0.000437 | 0 | INHBB_ACVR1B_ACVR2A | Inhbb - (Acvr1b+Acvr2a) | ACTIVIN | Secreted Signaling | KEGG: mmu04350 |
| CD31 | NeuN | Inhbb | ACVR1B_ACVR2A | 0.00084 | 0 | INHBB_ACVR1B_ACVR2A | Inhbb - (Acvr1b+Acvr2a) | ACTIVIN | Secreted Signaling | KEGG: mmu04350 |
| CD31 | CD31 | Inhbb | ACVR1B_ACVR2B | 0.00038 | 0 | INHBB_ACVR1B_ACVR2B | Inhbb - (Acvr1b+Acvr2b) | ACTIVIN | Secreted Signaling | KEGG: mmu04350 |
| CD31 | NeuN | Inhbb | ACVR1B_ACVR2B | 0.000504 | 0 | INHBB_ACVR1B_ACVR2B | Inhbb - (Acvr1b+Acvr2b) | ACTIVIN | Secreted Signaling | KEGG: mmu04350 |
| CD31 | CD31 | Inhbb | ACVR1C_ACVR2A | 0.000348 | 0 | INHBB_ACVR1C_ACVR2A | Inhbb - (Acvr1c+Acvr2a) | ACTIVIN | Secreted Signaling | KEGG: mmu04350 |
| CD31 | NeuN | Inhbb | ACVR1C_ACVR2A | 0.000285 | 0 | INHBB_ACVR1C_ACVR2A | Inhbb - (Acvr1c+Acvr2a) | ACTIVIN | Secreted Signaling | KEGG: mmu04350 |
| CD31 | CD31 | Inhbb | ACVR1C_ACVR2B | 0.000302 | 0 | INHBB_ACVR1C_ACVR2B | Inhbb - (Acvr1c+Acvr2b) | ACTIVIN | Secreted Signaling | KEGG: mmu04350 |
| GFAP | CD31 | Inhbb | ACVR1C_ACVR2B | 0.000237 | 0 | INHBB_ACVR1C_ACVR2B | Inhbb - (Acvr1c+Acvr2b) | ACTIVIN | Secreted Signaling | KEGG: mmu04350 |
| GFAP | CD31 | Wnt10b | FZD1_LRP5 | 0.000249 | 0 | WNT10B_FZD1_LRP5 | Wnt10b - (Fzd1+Lrp5) | WNT | Secreted Signaling | KEGG: mmu04310; PMID: 23209147 |
| GFAP | GFAP | Wnt10b | FZD1_LRP5 | 0.000266 | 0 | WNT10B_FZD1_LRP5 | Wnt10b - (Fzd1+Lrp5) | WNT | Secreted Signaling | KEGG: mmu04310; PMID: 23209147 |
| CD31 | GFAP | Wnt10b | FZD2_LRP5 | 0.000271 | 0 | WNT10B_FZD2_LRP5 | Wnt10b - (Fzd2+Lrp5) | WNT | Secreted Signaling | KEGG: mmu04310; PMID: 23209147 |

| | | | | | | | | | | |
|---|---|---|---|---|---|---|---|---|---|---|
| GFAP | GFAP | Wnt10b | FZD2_LRP5 | 0.000315 | 0 | WNT10B_FZD2_LRP5 | Wnt10b - (Fzd2+Lrp5) | WNT | Secreted Signaling | KEGG: mmu04310; PMID: 23209147 |
| GFAP | CD31 | Wnt10b | FZD3_LRP5 | 0.000329 | 0 | WNT10B_FZD3_LRP5 | Wnt10b - (Fzd3+Lrp5) | WNT | Secreted Signaling | KEGG: mmu04310; PMID: 23209147 |
| GFAP | GFAP | Wnt10b | FZD3_LRP5 | 0.00031 | 0 | WNT10B_FZD3_LRP5 | Wnt10b - (Fzd3+Lrp5) | WNT | Secreted Signaling | KEGG: mmu04310; PMID: 23209147 |
| CD31 | CD31 | Wnt10b | FZD4_LRP5 | 0.0003 | 0 | WNT10B_FZD4_LRP5 | Wnt10b - (Fzd4+Lrp5) | WNT | Secreted Signaling | KEGG: mmu04310; PMID: 23209147 |
| GFAP | CD31 | Wnt10b | FZD4_LRP5 | 0.00035 | 0 | WNT10B_FZD4_LRP5 | Wnt10b - (Fzd4+Lrp5) | WNT | Secreted Signaling | KEGG: mmu04310; PMID: 23209147 |
| CD31 | GFAP | Wnt10b | FZD5_LRP5 | 0.000228 | 0 | WNT10B_FZD5_LRP5 | Wnt10b - (Fzd5+Lrp5) | WNT | Secreted Signaling | KEGG: mmu04310; PMID: 23209147 |
| GFAP | GFAP | Wnt10b | FZD5_LRP5 | 0.000266 | 0 | WNT10B_FZD5_LRP5 | Wnt10b - (Fzd5+Lrp5) | WNT | Secreted Signaling | KEGG: mmu04310; PMID: 23209147 |
| CD31 | CD31 | Wnt10b | FZD6_LRP5 | 0.000398 | 0 | WNT10B_FZD6_LRP5 | Wnt10b - (Fzd6+Lrp5) | WNT | Secreted Signaling | KEGG: mmu04310; PMID: 23209147 |
| GFAP | CD31 | Wnt10b | FZD6_LRP5 | 0.000464 | 0 | WNT10B_FZD6_LRP5 | Wnt10b - (Fzd6+Lrp5) | WNT | Secreted Signaling | KEGG: mmu04310; PMID: 23209147 |
| GFAP | CD31 | Wnt10b | FZD8_LRP5 | 0.000267 | 0 | WNT10B_FZD8_LRP5 | Wnt10b - (Fzd8+Lrp5) | WNT | Secreted Signaling | KEGG: mmu04310; PMID: 23209147 |
| GFAP | GFAP | Wnt10b | FZD8_LRP5 | 0.000235 | 0 | WNT10B_FZD8_LRP5 | Wnt10b - (Fzd8+Lrp5) | WNT | Secreted Signaling | KEGG: mmu04310; PMID: 23209147 |
| CD31 | GFAP | Wnt10b | FZD9_LRP5 | 0.0002 | 0 | WNT10B_FZD9_LRP5 | Wnt10b - (Fzd9+Lrp5) | WNT | Secreted Signaling | KEGG: mmu04310; PMID: 23209147 |
| GFAP | GFAP | Wnt10b | FZD9_LRP5 | 0.000232 | 0 | WNT10B_FZD9_LRP5 | Wnt10b - (Fzd9+Lrp5) | WNT | Secreted Signaling | KEGG: mmu04310; PMID: 23209147 |
| CD31 | GFAP | Wnt4 | FZD1_LRP5 | 0.000304 | 0 | WNT4_FZD1_LRP5 | Wnt4 - (Fzd1+Lrp5) | WNT | Secreted Signaling | KEGG: mmu04310; PMID: 23209147 |
| GFAP | GFAP | Wnt4 | FZD1_LRP5 | 0.000286 | 0 | WNT4_FZD1_LRP5 | Wnt4 - (Fzd1+Lrp5) | WNT | Secreted Signaling | KEGG: mmu04310; PMID: 23209147 |
| CD31 | GFAP | Wnt4 | FZD2_LRP5 | 0.00036 | 0 | WNT4_FZD2_LRP5 | Wnt4 - (Fzd2+Lrp5) | WNT | Secreted Signaling | KEGG: mmu04310; PMID: 23209147 |
| GFAP | GFAP | Wnt4 | FZD2_LRP5 | 0.000339 | 0 | WNT4_FZD2_LRP5 | Wnt4 - (Fzd2+Lrp5) | WNT | Secreted Signaling | KEGG: mmu04310; PMID: 23209147 |
| CD31 | CD31 | Wnt4 | FZD3_LRP5 | 0.000377 | 0 | WNT4_FZD3_LRP5 | Wnt4 - (Fzd3+Lrp5) | WNT | Secreted Signaling | KEGG: mmu04310; PMID: 23209147 |
| NeuN | CD31 | Wnt4 | FZD3_LRP5 | 0.000357 | 0 | WNT4_FZD3_LRP5 | Wnt4 - (Fzd3+Lrp5) | WNT | Secreted Signaling | KEGG: mmu04310; PMID: 23209147 |
| CD31 | CD31 | Wnt4 | FZD4_LRP5 | 0.0004 | 0 | WNT4_FZD4_LRP5 | Wnt4 - (Fzd4+Lrp5) | WNT | Secreted Signaling | KEGG: mmu04310; PMID: 23209147 |
| NeuN | CD31 | Wnt4 | FZD4_LRP5 | 0.000379 | 0 | WNT4_FZD4_LRP5 | Wnt4 - (Fzd4+Lrp5) | WNT | Secreted Signaling | KEGG: mmu04310; PMID: 23209147 |
| CD31 | GFAP | Wnt4 | FZD5_LRP5 | 0.000304 | 0 | WNT4_FZD5_LRP5 | Wnt4 - (Fzd5+Lrp5) | WNT | Secreted Signaling | KEGG: mmu04310; PMID: 23209147 |
| GFAP | GFAP | Wnt4 | FZD5_LRP5 | 0.000286 | 0 | WNT4_FZD5_LRP5 | Wnt4 - (Fzd5+Lrp5) | WNT | Secreted Signaling | KEGG: mmu04310; PMID: 23209147 |
| CD31 | CD31 | Wnt4 | FZD6_LRP5 | 0.00053 | 0 | WNT4_FZD6_LRP5 | Wnt4 - (Fzd6+Lrp5) | WNT | Secreted Signaling | KEGG: mmu04310; PMID: 23209147 |
| NeuN | CD31 | Wnt4 | FZD6_LRP5 | 0.000502 | 0 | WNT4_FZD6_LRP5 | Wnt4 - (Fzd6+Lrp5) | WNT | Secreted Signaling | KEGG: mmu04310; PMID: 23209147 |
| CD31 | CD31 | Wnt4 | FZD8_LRP5 | 0.000306 | 0 | WNT4_FZD8_LRP5 | Wnt4 - (Fzd8+Lrp5) | WNT | Secreted Signaling | KEGG: mmu04310; PMID: 23209147 |
| NeuN | CD31 | Wnt4 | FZD8_LRP5 | 0.00029 | 0 | WNT4_FZD8_LRP5 | Wnt4 - (Fzd8+Lrp5) | WNT | Secreted Signaling | KEGG: mmu04310; PMID: 23209147 |
| CD31 | GFAP | Wnt4 | FZD9_LRP5 | 0.000266 | 0 | WNT4_FZD9_LRP5 | Wnt4 - (Fzd9+Lrp5) | WNT | Secreted Signaling | KEGG: mmu04310; PMID: 23209147 |
| GFAP | GFAP | Wnt4 | FZD9_LRP5 | 0.00025 | 0 | WNT4_FZD9_LRP5 | Wnt4 - (Fzd9+Lrp5) | WNT | Secreted Signaling | KEGG: mmu04310; PMID: 23209147 |
| GFAP | CD31 | Wnt7a | FZD1_LRP5 | 0.000358 | 0 | WNT7A_FZD1_LRP5 | Wnt7a - (Fzd1+Lrp5) | WNT | Secreted Signaling | KEGG: mmu04310; PMID: 23209147 |
| GFAP | GFAP | Wnt7a | FZD1_LRP5 | 0.000382 | 0 | WNT7A_FZD1_LRP5 | Wnt7a - (Fzd1+Lrp5) | WNT | Secreted Signaling | KEGG: mmu04310; PMID: 23209147 |
| GFAP | CD31 | Wnt7a | FZD2_LRP5 | 0.000296 | 0 | WNT7A_FZD2_LRP5 | Wnt7a - (Fzd2+Lrp5) | WNT | Secreted Signaling | KEGG: mmu04310; PMID: 23209147 |
| GFAP | GFAP | Wnt7a | FZD2_LRP5 | 0.000453 | 0 | WNT7A_FZD2_LRP5 | Wnt7a - (Fzd2+Lrp5) | WNT | Secreted Signaling | KEGG: mmu04310; PMID: 23209147 |
| GFAP | CD31 | Wnt7a | FZD3_LRP5 | 0.000473 | 0 | WNT7A_FZD3_LRP5 | Wnt7a - (Fzd3+Lrp5) | WNT | Secreted Signaling | KEGG: mmu04310; PMID: 23209147 |
| GFAP | GFAP | Wnt7a | FZD3_LRP5 | 0.000446 | 0 | WNT7A_FZD3_LRP5 | Wnt7a - (Fzd3+Lrp5) | WNT | Secreted Signaling | KEGG: mmu04310; PMID: 23209147 |
| GFAP | CD31 | Wnt7a | FZD4_LRP5 | 0.000502 | 0 | WNT7A_FZD4_LRP5 | Wnt7a - (Fzd4+Lrp5) | WNT | Secreted Signaling | KEGG: mmu04310; PMID: 23209147 |
| GFAP | GFAP | Wnt7a | FZD4_LRP5 | 0.000371 | 0 | WNT7A_FZD4_LRP5 | Wnt7a - (Fzd4+Lrp5) | WNT | Secreted Signaling | KEGG: mmu04310; PMID: 23209147 |
| GFAP | CD31 | Wnt7a | FZD5_LRP5 | 0.000302 | 0 | WNT7A_FZD5_LRP5 | Wnt7a - (Fzd5+Lrp5) | WNT | Secreted Signaling | KEGG: mmu04310; PMID: 23209147 |
| GFAP | GFAP | Wnt7a | FZD5_LRP5 | 0.000382 | 0 | WNT7A_FZD5_LRP5 | Wnt7a - (Fzd5+Lrp5) | WNT | Secreted Signaling | KEGG: mmu04310; PMID: 23209147 |
| CD31 | CD31 | Wnt7a | FZD6_LRP5 | 0.000409 | 0 | WNT7A_FZD6_LRP5 | Wnt7a - (Fzd6+Lrp5) | WNT | Secreted Signaling | KEGG: mmu04310; PMID: 23209147 |
| GFAP | CD31 | Wnt7a | FZD6_LRP5 | 0.000666 | 0 | WNT7A_FZD6_LRP5 | Wnt7a - (Fzd6+Lrp5) | WNT | Secreted Signaling | KEGG: mmu04310; PMID: 23209147 |
| GFAP | CD31 | Wnt7a | FZD8_LRP5 | 0.000384 | 0 | WNT7A_FZD8_LRP5 | Wnt7a - (Fzd8+Lrp5) | WNT | Secreted Signaling | KEGG: mmu04310; PMID: 23209147 |
| GFAP | GFAP | Wnt7a | FZD8_LRP5 | 0.000338 | 0 | WNT7A_FZD8_LRP5 | Wnt7a - (Fzd8+Lrp5) | WNT | Secreted Signaling | KEGG: mmu04310; PMID: 23209147 |
| GFAP | CD31 | Wnt7a | FZD9_LRP5 | 0.000228 | 0 | WNT7A_FZD9_LRP5 | Wnt7a - (Fzd9+Lrp5) | WNT | Secreted Signaling | KEGG: mmu04310; PMID: 23209147 |
| GFAP | GFAP | Wnt7a | FZD9_LRP5 | 0.000334 | 0 | WNT7A_FZD9_LRP5 | Wnt7a - (Fzd9+Lrp5) | WNT | Secreted Signaling | KEGG: mmu04310; PMID: 23209147 |
| CD31 | CD31 | Wnt7b | FZD1_LRP5 | 0.000309 | 0 | WNT7B_FZD1_LRP5 | Wnt7b - (Fzd1+Lrp5) | WNT | Secreted Signaling | KEGG: mmu04310; PMID: 23209147 |
| CD31 | GFAP | Wnt7b | FZD1_LRP5 | 0.00033 | 0 | WNT7B_FZD1_LRP5 | Wnt7b - (Fzd1+Lrp5) | WNT | Secreted Signaling | KEGG: mmu04310; PMID: 23209147 |
| CD31 | GFAP | Wnt7b | FZD2_LRP5 | 0.000391 | 0 | WNT7B_FZD2_LRP5 | Wnt7b - (Fzd2+Lrp5) | WNT | Secreted Signaling | KEGG: mmu04310; PMID: 23209147 |
| GFAP | GFAP | Wnt7b | FZD2_LRP5 | 0.00027 | 0 | WNT7B_FZD2_LRP5 | Wnt7b - (Fzd2+Lrp5) | WNT | Secreted Signaling | KEGG: mmu04310; PMID: 23209147 |
| CD31 | CD31 | Wnt7b | FZD3_LRP5 | 0.000409 | 0 | WNT7B_FZD3_LRP5 | Wnt7b - (Fzd3+Lrp5) | WNT | Secreted Signaling | KEGG: mmu04310; PMID: 23209147 |
| CD31 | GFAP | Wnt7b | FZD3_LRP5 | 0.000385 | 0 | WNT7B_FZD3_LRP5 | Wnt7b - (Fzd3+Lrp5) | WNT | Secreted Signaling | KEGG: mmu04310; PMID: 23209147 |
| CD31 | CD31 | Wnt7b | FZD4_LRP5 | 0.000434 | 0 | WNT7B_FZD4_LRP5 | Wnt7b - (Fzd4+Lrp5) | WNT | Secreted Signaling | KEGG: mmu04310; PMID: 23209147 |
| CD31 | GFAP | Wnt7b | FZD4_LRP5 | 0.00032 | 0 | WNT7B_FZD4_LRP5 | Wnt7b - (Fzd4+Lrp5) | WNT | Secreted Signaling | KEGG: mmu04310; PMID: 23209147 |
| CD31 | CD31 | Wnt7b | FZD5_LRP5 | 0.000261 | 0 | WNT7B_FZD5_LRP5 | Wnt7b - (Fzd5+Lrp5) | WNT | Secreted Signaling | KEGG: mmu04310; PMID: 23209147 |
| CD31 | GFAP | Wnt7b | FZD5_LRP5 | 0.00033 | 0 | WNT7B_FZD5_LRP5 | Wnt7b - (Fzd5+Lrp5) | WNT | Secreted Signaling | KEGG: mmu04310; PMID: 23209147 |
| CD31 | CD31 | Wnt7b | FZD6_LRP5 | 0.000575 | 0 | WNT7B_FZD6_LRP5 | Wnt7b - (Fzd6+Lrp5) | WNT | Secreted Signaling | KEGG: mmu04310; PMID: 23209147 |
| GFAP | CD31 | Wnt7b | FZD6_LRP5 | 0.000398 | 0 | WNT7B_FZD6_LRP5 | Wnt7b - (Fzd6+Lrp5) | WNT | Secreted Signaling | KEGG: mmu04310; PMID: 23209147 |
| CD31 | CD31 | Wnt7b | FZD8_LRP5 | 0.000332 | 0 | WNT7B_FZD8_LRP5 | Wnt7b - (Fzd8+Lrp5) | WNT | Secreted Signaling | KEGG: mmu04310; PMID: 23209147 |
| CD31 | GFAP | Wnt7b | FZD8_LRP5 | 0.000292 | 0 | WNT7B_FZD8_LRP5 | Wnt7b - (Fzd8+Lrp5) | WNT | Secreted Signaling | KEGG: mmu04310; PMID: 23209147 |

| | | | | | | | | | | |
|---|---|---|---|---|---|---|---|---|---|---|
| CD31 | GFAP | Wnt7b | FZD9_LRP5 | 0.000288 | 0 | WNT7B_FZD9_LRP5 | Wnt7b - (Fzd9+Lrp5) | WNT | Secreted Signaling | KEGG: mmu04310; PMID: 23209147 |
| GFAP | GFAP | Wnt7b | FZD9_LRP5 | 0.000199 | 0 | WNT7B_FZD9_LRP5 | Wnt7b - (Fzd9+Lrp5) | WNT | Secreted Signaling | KEGG: mmu04310; PMID: 23209147 |
| GFAP | CD31 | Wnt9a | FZD1_LRP5 | 0.000334 | 0 | WNT9A_FZD1_LRP5 | Wnt9a - (Fzd1+Lrp5) | WNT | Secreted Signaling | KEGG: mmu04310; PMID: 23209147 |
| GFAP | GFAP | Wnt9a | FZD1_LRP5 | 0.000356 | 0 | WNT9A_FZD1_LRP5 | Wnt9a - (Fzd1+Lrp5) | WNT | Secreted Signaling | KEGG: mmu04310; PMID: 23209147 |
| CD31 | GFAP | Wnt9a | FZD2_LRP5 | 0.000394 | 0 | WNT9A_FZD2_LRP5 | Wnt9a - (Fzd2+Lrp5) | WNT | Secreted Signaling | KEGG: mmu04310; PMID: 23209147 |
| GFAP | GFAP | Wnt9a | FZD2_LRP5 | 0.000422 | 0 | WNT9A_FZD2_LRP5 | Wnt9a - (Fzd2+Lrp5) | WNT | Secreted Signaling | KEGG: mmu04310; PMID: 23209147 |
| GFAP | CD31 | Wnt9a | FZD3_LRP5 | 0.000441 | 0 | WNT9A_FZD3_LRP5 | Wnt9a - (Fzd3+Lrp5) | WNT | Secreted Signaling | KEGG: mmu04310; PMID: 23209147 |
| GFAP | GFAP | Wnt9a | FZD3_LRP5 | 0.000416 | 0 | WNT9A_FZD3_LRP5 | Wnt9a - (Fzd3+Lrp5) | WNT | Secreted Signaling | KEGG: mmu04310; PMID: 23209147 |
| CD31 | CD31 | Wnt9a | FZD4_LRP5 | 0.000437 | 0 | WNT9A_FZD4_LRP5 | Wnt9a - (Fzd4+Lrp5) | WNT | Secreted Signaling | KEGG: mmu04310; PMID: 23209147 |
| GFAP | CD31 | Wnt9a | FZD4_LRP5 | 0.000468 | 0 | WNT9A_FZD4_LRP5 | Wnt9a - (Fzd4+Lrp5) | WNT | Secreted Signaling | KEGG: mmu04310; PMID: 23209147 |
| CD31 | GFAP | Wnt9a | FZD5_LRP5 | 0.000332 | 0 | WNT9A_FZD5_LRP5 | Wnt9a - (Fzd5+Lrp5) | WNT | Secreted Signaling | KEGG: mmu04310; PMID: 23209147 |
| GFAP | GFAP | Wnt9a | FZD5_LRP5 | 0.000356 | 0 | WNT9A_FZD5_LRP5 | Wnt9a - (Fzd5+Lrp5) | WNT | Secreted Signaling | KEGG: mmu04310; PMID: 23209147 |
| CD31 | CD31 | Wnt9a | FZD6_LRP5 | 0.00058 | 0 | WNT9A_FZD6_LRP5 | Wnt9a - (Fzd6+Lrp5) | WNT | Secreted Signaling | KEGG: mmu04310; PMID: 23209147 |
| GFAP | CD31 | Wnt9a | FZD6_LRP5 | 0.000621 | 0 | WNT9A_FZD6_LRP5 | Wnt9a - (Fzd6+Lrp5) | WNT | Secreted Signaling | KEGG: mmu04310; PMID: 23209147 |
| CD31 | CD31 | Wnt9a | FZD8_LRP5 | 0.000334 | 0 | WNT9A_FZD8_LRP5 | Wnt9a - (Fzd8+Lrp5) | WNT | Secreted Signaling | KEGG: mmu04310; PMID: 23209147 |
| GFAP | CD31 | Wnt9a | FZD8_LRP5 | 0.000358 | 0 | WNT9A_FZD8_LRP5 | Wnt9a - (Fzd8+Lrp5) | WNT | Secreted Signaling | KEGG: mmu04310; PMID: 23209147 |
| CD31 | GFAP | Wnt9a | FZD9_LRP5 | 0.000291 | 0 | WNT9A_FZD9_LRP5 | Wnt9a - (Fzd9+Lrp5) | WNT | Secreted Signaling | KEGG: mmu04310; PMID: 23209147 |
| GFAP | GFAP | Wnt9a | FZD9_LRP5 | 0.000311 | 0 | WNT9A_FZD9_LRP5 | Wnt9a - (Fzd9+Lrp5) | WNT | Secreted Signaling | KEGG: mmu04310; PMID: 23209147 |
| GFAP | CD31 | Wnt10b | FZD1_LRP6 | 0.000286 | 0 | WNT10B_FZD1_LRP6 | Wnt10b - (Fzd1+Lrp6) | WNT | Secreted Signaling | KEGG: mmu04310; PMID: 23209147 |
| GFAP | GFAP | Wnt10b | FZD1_LRP6 | 0.000279 | 0 | WNT10B_FZD1_LRP6 | Wnt10b - (Fzd1+Lrp6) | WNT | Secreted Signaling | KEGG: mmu04310; PMID: 23209147 |
| CD31 | GFAP | Wnt10b | FZD2_LRP6 | 0.000284 | 0 | WNT10B_FZD2_LRP6 | Wnt10b - (Fzd2+Lrp6) | WNT | Secreted Signaling | KEGG: mmu04310; PMID: 23209147 |
| GFAP | GFAP | Wnt10b | FZD2_LRP6 | 0.000331 | 0 | WNT10B_FZD2_LRP6 | Wnt10b - (Fzd2+Lrp6) | WNT | Secreted Signaling | KEGG: mmu04310; PMID: 23209147 |
| GFAP | CD31 | Wnt10b | FZD3_LRP6 | 0.000378 | 0 | WNT10B_FZD3_LRP6 | Wnt10b - (Fzd3+Lrp6) | WNT | Secreted Signaling | KEGG: mmu04310; PMID: 23209147 |
| GFAP | GFAP | Wnt10b | FZD3_LRP6 | 0.000326 | 0 | WNT10B_FZD3_LRP6 | Wnt10b - (Fzd3+Lrp6) | WNT | Secreted Signaling | KEGG: mmu04310; PMID: 23209147 |
| CD31 | CD31 | Wnt10b | FZD4_LRP6 | 0.000344 | 0 | WNT10B_FZD4_LRP6 | Wnt10b - (Fzd4+Lrp6) | WNT | Secreted Signaling | KEGG: mmu04310; PMID: 23209147 |
| GFAP | CD31 | Wnt10b | FZD4_LRP6 | 0.000401 | 0 | WNT10B_FZD4_LRP6 | Wnt10b - (Fzd4+Lrp6) | WNT | Secreted Signaling | KEGG: mmu04310; PMID: 23209147 |
| GFAP | CD31 | Wnt10b | FZD5_LRP6 | 0.000241 | 0 | WNT10B_FZD5_LRP6 | Wnt10b - (Fzd5+Lrp6) | WNT | Secreted Signaling | KEGG: mmu04310; PMID: 23209147 |
| GFAP | GFAP | Wnt10b | FZD5_LRP6 | 0.000279 | 0 | WNT10B_FZD5_LRP6 | Wnt10b - (Fzd5+Lrp6) | WNT | Secreted Signaling | KEGG: mmu04310; PMID: 23209147 |
| CD31 | CD31 | Wnt10b | FZD6_LRP6 | 0.000457 | 0 | WNT10B_FZD6_LRP6 | Wnt10b - (Fzd6+Lrp6) | WNT | Secreted Signaling | KEGG: mmu04310; PMID: 23209147 |
| GFAP | CD31 | Wnt10b | FZD6_LRP6 | 0.000532 | 0 | WNT10B_FZD6_LRP6 | Wnt10b - (Fzd6+Lrp6) | WNT | Secreted Signaling | KEGG: mmu04310; PMID: 23209147 |
| CD31 | CD31 | Wnt10b | FZD8_LRP6 | 0.000263 | 0 | WNT10B_FZD8_LRP6 | Wnt10b - (Fzd8+Lrp6) | WNT | Secreted Signaling | KEGG: mmu04310; PMID: 23209147 |
| GFAP | CD31 | Wnt10b | FZD8_LRP6 | 0.000307 | 0 | WNT10B_FZD8_LRP6 | Wnt10b - (Fzd8+Lrp6) | WNT | Secreted Signaling | KEGG: mmu04310; PMID: 23209147 |
| CD31 | GFAP | Wnt10b | FZD9_LRP6 | 0.000209 | 0 | WNT10B_FZD9_LRP6 | Wnt10b - (Fzd9+Lrp6) | WNT | Secreted Signaling | KEGG: mmu04310; PMID: 23209147 |
| GFAP | GFAP | Wnt10b | FZD9_LRP6 | 0.000244 | 0 | WNT10B_FZD9_LRP6 | Wnt10b - (Fzd9+Lrp6) | WNT | Secreted Signaling | KEGG: mmu04310; PMID: 23209147 |
| CD31 | CD31 | Wnt4 | FZD1_LRP6 | 0.000327 | 0 | WNT4_FZD1_LRP6 | Wnt4 - (Fzd1+Lrp6) | WNT | Secreted Signaling | KEGG: mmu04310; PMID: 23209147 |
| CD31 | GFAP | Wnt4 | FZD1_LRP6 | 0.000319 | 0 | WNT4_FZD1_LRP6 | Wnt4 - (Fzd1+Lrp6) | WNT | Secreted Signaling | KEGG: mmu04310; PMID: 23209147 |
| CD31 | GFAP | Wnt4 | FZD2_LRP6 | 0.000378 | 0 | WNT4_FZD2_LRP6 | Wnt4 - (Fzd2+Lrp6) | WNT | Secreted Signaling | KEGG: mmu04310; PMID: 23209147 |
| GFAP | GFAP | Wnt4 | FZD2_LRP6 | 0.000356 | 0 | WNT4_FZD2_LRP6 | Wnt4 - (Fzd2+Lrp6) | WNT | Secreted Signaling | KEGG: mmu04310; PMID: 23209147 |
| CD31 | CD31 | Wnt4 | FZD3_LRP6 | 0.000432 | 0 | WNT4_FZD3_LRP6 | Wnt4 - (Fzd3+Lrp6) | WNT | Secreted Signaling | KEGG: mmu04310; PMID: 23209147 |
| CD31 | NeuN | Wnt4 | FZD3_LRP6 | 0.000422 | 0 | WNT4_FZD3_LRP6 | Wnt4 - (Fzd3+Lrp6) | WNT | Secreted Signaling | KEGG: mmu04310; PMID: 23209147 |
| CD31 | CD31 | Wnt4 | FZD4_LRP6 | 0.000459 | 0 | WNT4_FZD4_LRP6 | Wnt4 - (Fzd4+Lrp6) | WNT | Secreted Signaling | KEGG: mmu04310; PMID: 23209147 |
| NeuN | CD31 | Wnt4 | FZD4_LRP6 | 0.000435 | 0 | WNT4_FZD4_LRP6 | Wnt4 - (Fzd4+Lrp6) | WNT | Secreted Signaling | KEGG: mmu04310; PMID: 23209147 |
| CD31 | GFAP | Wnt4 | FZD5_LRP6 | 0.000319 | 0 | WNT4_FZD5_LRP6 | Wnt4 - (Fzd5+Lrp6) | WNT | Secreted Signaling | KEGG: mmu04310; PMID: 23209147 |
| GFAP | GFAP | Wnt4 | FZD5_LRP6 | 0.0003 | 0 | WNT4_FZD5_LRP6 | Wnt4 - (Fzd5+Lrp6) | WNT | Secreted Signaling | KEGG: mmu04310; PMID: 23209147 |
| CD31 | CD31 | Wnt4 | FZD6_LRP6 | 0.000608 | 0 | WNT4_FZD6_LRP6 | Wnt4 - (Fzd6+Lrp6) | WNT | Secreted Signaling | KEGG: mmu04310; PMID: 23209147 |
| NeuN | CD31 | Wnt4 | FZD6_LRP6 | 0.000576 | 0 | WNT4_FZD6_LRP6 | Wnt4 - (Fzd6+Lrp6) | WNT | Secreted Signaling | KEGG: mmu04310; PMID: 23209147 |
| CD31 | CD31 | Wnt4 | FZD8_LRP6 | 0.000351 | 0 | WNT4_FZD8_LRP6 | Wnt4 - (Fzd8+Lrp6) | WNT | Secreted Signaling | KEGG: mmu04310; PMID: 23209147 |
| NeuN | CD31 | Wnt4 | FZD8_LRP6 | 0.000332 | 0 | WNT4_FZD8_LRP6 | Wnt4 - (Fzd8+Lrp6) | WNT | Secreted Signaling | KEGG: mmu04310; PMID: 23209147 |
| CD31 | GFAP | Wnt4 | FZD9_LRP6 | 0.000279 | 0 | WNT4_FZD9_LRP6 | Wnt4 - (Fzd9+Lrp6) | WNT | Secreted Signaling | KEGG: mmu04310; PMID: 23209147 |
| GFAP | GFAP | Wnt4 | FZD9_LRP6 | 0.000263 | 0 | WNT4_FZD9_LRP6 | Wnt4 - (Fzd9+Lrp6) | WNT | Secreted Signaling | KEGG: mmu04310; PMID: 23209147 |
| GFAP | CD31 | Wnt7a | FZD1_LRP6 | 0.000411 | 0 | WNT7A_FZD1_LRP6 | Wnt7a - (Fzd1+Lrp6) | WNT | Secreted Signaling | KEGG: mmu04310; PMID: 23209147 |
| GFAP | GFAP | Wnt7a | FZD1_LRP6 | 0.000401 | 0 | WNT7A_FZD1_LRP6 | Wnt7a - (Fzd1+Lrp6) | WNT | Secreted Signaling | KEGG: mmu04310; PMID: 23209147 |
| GFAP | CD31 | Wnt7a | FZD2_LRP6 | 0.000339 | 0 | WNT7A_FZD2_LRP6 | Wnt7a - (Fzd2+Lrp6) | WNT | Secreted Signaling | KEGG: mmu04310; PMID: 23209147 |
| GFAP | GFAP | Wnt7a | FZD2_LRP6 | 0.000475 | 0 | WNT7A_FZD2_LRP6 | Wnt7a - (Fzd2+Lrp6) | WNT | Secreted Signaling | KEGG: mmu04310; PMID: 23209147 |
| GFAP | CD31 | Wnt7a | FZD3_LRP6 | 0.000543 | 0 | WNT7A_FZD3_LRP6 | Wnt7a - (Fzd3+Lrp6) | WNT | Secreted Signaling | KEGG: mmu04310; PMID: 23209147 |
| GFAP | GFAP | Wnt7a | FZD3_LRP6 | 0.000468 | 0 | WNT7A_FZD3_LRP6 | Wnt7a - (Fzd3+Lrp6) | WNT | Secreted Signaling | KEGG: mmu04310; PMID: 23209147 |
| GFAP | CD31 | Wnt7a | FZD4_LRP6 | 0.000576 | 0 | WNT7A_FZD4_LRP6 | Wnt7a - (Fzd4+Lrp6) | WNT | Secreted Signaling | KEGG: mmu04310; PMID: 23209147 |
| GFAP | GFAP | Wnt7a | FZD4_LRP6 | 0.000389 | 0 | WNT7A_FZD4_LRP6 | Wnt7a - (Fzd4+Lrp6) | WNT | Secreted Signaling | KEGG: mmu04310; PMID: 23209147 |
| GFAP | CD31 | Wnt7a | FZD5_LRP6 | 0.000346 | 0 | WNT7A_FZD5_LRP6 | Wnt7a - (Fzd5+Lrp6) | WNT | Secreted Signaling | KEGG: mmu04310; PMID: 23209147 |

| | | | | | | | | | | |
|---|---|---|---|---|---|---|---|---|---|---|
| GFAP | GFAP | Wnt7a | FZD5_LRP6 | 0.000401 | 0 | WNT7A_FZD5_LRP6 | Wnt7a - (Fzd5+Lrp6) | WNT | Secreted Signaling | KEGG: mmu04310; PMID: 23209147 |
| CD31 | CD31 | Wnt7a | FZD6_LRP6 | 0.000469 | 0 | WNT7A_FZD6_LRP6 | Wnt7a - (Fzd6+Lrp6) | WNT | Secreted Signaling | KEGG: mmu04310; PMID: 23209147 |
| GFAP | CD31 | Wnt7a | FZD6_LRP6 | 0.000764 | 0 | WNT7A_FZD6_LRP6 | Wnt7a - (Fzd6+Lrp6) | WNT | Secreted Signaling | KEGG: mmu04310; PMID: 23209147 |
| GFAP | CD31 | Wnt7a | FZD8_LRP6 | 0.000441 | 0 | WNT7A_FZD8_LRP6 | Wnt7a - (Fzd8+Lrp6) | WNT | Secreted Signaling | KEGG: mmu04310; PMID: 23209147 |
| GFAP | GFAP | Wnt7a | FZD8_LRP6 | 0.000354 | 0 | WNT7A_FZD8_LRP6 | Wnt7a - (Fzd8+Lrp6) | WNT | Secreted Signaling | KEGG: mmu04310; PMID: 23209147 |
| GFAP | CD31 | Wnt7a | FZD9_LRP6 | 0.000261 | 0 | WNT7A_FZD9_LRP6 | Wnt7a - (Fzd9+Lrp6) | WNT | Secreted Signaling | KEGG: mmu04310; PMID: 23209147 |
| GFAP | GFAP | Wnt7a | FZD9_LRP6 | 0.00035 | 0 | WNT7A_FZD9_LRP6 | Wnt7a - (Fzd9+Lrp6) | WNT | Secreted Signaling | KEGG: mmu04310; PMID: 23209147 |
| CD31 | CD31 | Wnt7b | FZD1_LRP6 | 0.000355 | 0 | WNT7B_FZD1_LRP6 | Wnt7b - (Fzd1+Lrp6) | WNT | Secreted Signaling | KEGG: mmu04310; PMID: 23209147 |
| CD31 | GFAP | Wnt7b | FZD1_LRP6 | 0.000346 | 0 | WNT7B_FZD1_LRP6 | Wnt7b - (Fzd1+Lrp6) | WNT | Secreted Signaling | KEGG: mmu04310; PMID: 23209147 |
| CD31 | CD31 | Wnt7b | FZD2_LRP6 | 0.000293 | 0 | WNT7B_FZD2_LRP6 | Wnt7b - (Fzd2+Lrp6) | WNT | Secreted Signaling | KEGG: mmu04310; PMID: 23209147 |
| CD31 | GFAP | Wnt7b | FZD2_LRP6 | 0.000411 | 0 | WNT7B_FZD2_LRP6 | Wnt7b - (Fzd2+Lrp6) | WNT | Secreted Signaling | KEGG: mmu04310; PMID: 23209147 |
| CD31 | CD31 | Wnt7b | FZD3_LRP6 | 0.000469 | 0 | WNT7B_FZD3_LRP6 | Wnt7b - (Fzd3+Lrp6) | WNT | Secreted Signaling | KEGG: mmu04310; PMID: 23209147 |
| CD31 | NeuN | Wnt7b | FZD3_LRP6 | 0.000458 | 0 | WNT7B_FZD3_LRP6 | Wnt7b - (Fzd3+Lrp6) | WNT | Secreted Signaling | KEGG: mmu04310; PMID: 23209147 |
| CD31 | CD31 | Wnt7b | FZD4_LRP6 | 0.000498 | 0 | WNT7B_FZD4_LRP6 | Wnt7b - (Fzd4+Lrp6) | WNT | Secreted Signaling | KEGG: mmu04310; PMID: 23209147 |
| GFAP | CD31 | Wnt7b | FZD4_LRP6 | 0.000344 | 0 | WNT7B_FZD4_LRP6 | Wnt7b - (Fzd4+Lrp6) | WNT | Secreted Signaling | KEGG: mmu04310; PMID: 23209147 |
| CD31 | CD31 | Wnt7b | FZD5_LRP6 | 0.000299 | 0 | WNT7B_FZD5_LRP6 | Wnt7b - (Fzd5+Lrp6) | WNT | Secreted Signaling | KEGG: mmu04310; PMID: 23209147 |
| CD31 | GFAP | Wnt7b | FZD5_LRP6 | 0.000346 | 0 | WNT7B_FZD5_LRP6 | Wnt7b - (Fzd5+Lrp6) | WNT | Secreted Signaling | KEGG: mmu04310; PMID: 23209147 |
| CD31 | CD31 | Wnt7b | FZD6_LRP6 | 0.00066 | 0 | WNT7B_FZD6_LRP6 | Wnt7b - (Fzd6+Lrp6) | WNT | Secreted Signaling | KEGG: mmu04310; PMID: 23209147 |
| GFAP | CD31 | Wnt7b | FZD6_LRP6 | 0.000456 | 0 | WNT7B_FZD6_LRP6 | Wnt7b - (Fzd6+Lrp6) | WNT | Secreted Signaling | KEGG: mmu04310; PMID: 23209147 |
| CD31 | CD31 | Wnt7b | FZD8_LRP6 | 0.000381 | 0 | WNT7B_FZD8_LRP6 | Wnt7b - (Fzd8+Lrp6) | WNT | Secreted Signaling | KEGG: mmu04310; PMID: 23209147 |
| CD31 | GFAP | Wnt7b | FZD8_LRP6 | 0.000306 | 0 | WNT7B_FZD8_LRP6 | Wnt7b - (Fzd8+Lrp6) | WNT | Secreted Signaling | KEGG: mmu04310; PMID: 23209147 |
| CD31 | CD31 | Wnt7b | FZD9_LRP6 | 0.000226 | 0 | WNT7B_FZD9_LRP6 | Wnt7b - (Fzd9+Lrp6) | WNT | Secreted Signaling | KEGG: mmu04310; PMID: 23209147 |
| CD31 | GFAP | Wnt7b | FZD9_LRP6 | 0.000303 | 0 | WNT7B_FZD9_LRP6 | Wnt7b - (Fzd9+Lrp6) | WNT | Secreted Signaling | KEGG: mmu04310; PMID: 23209147 |
| GFAP | CD31 | Wnt9a | FZD1_LRP6 | 0.000383 | 0 | WNT9A_FZD1_LRP6 | Wnt9a - (Fzd1+Lrp6) | WNT | Secreted Signaling | KEGG: mmu04310; PMID: 23209147 |
| GFAP | GFAP | Wnt9a | FZD1_LRP6 | 0.000373 | 0 | WNT9A_FZD1_LRP6 | Wnt9a - (Fzd1+Lrp6) | WNT | Secreted Signaling | KEGG: mmu04310; PMID: 23209147 |
| CD31 | GFAP | Wnt9a | FZD2_LRP6 | 0.000414 | 0 | WNT9A_FZD2_LRP6 | Wnt9a - (Fzd2+Lrp6) | WNT | Secreted Signaling | KEGG: mmu04310; PMID: 23209147 |
| GFAP | GFAP | Wnt9a | FZD2_LRP6 | 0.000443 | 0 | WNT9A_FZD2_LRP6 | Wnt9a - (Fzd2+Lrp6) | WNT | Secreted Signaling | KEGG: mmu04310; PMID: 23209147 |
| CD31 | CD31 | Wnt9a | FZD3_LRP6 | 0.000472 | 0 | WNT9A_FZD3_LRP6 | Wnt9a - (Fzd3+Lrp6) | WNT | Secreted Signaling | KEGG: mmu04310; PMID: 23209147 |
| GFAP | CD31 | Wnt9a | FZD3_LRP6 | 0.000506 | 0 | WNT9A_FZD3_LRP6 | Wnt9a - (Fzd3+Lrp6) | WNT | Secreted Signaling | KEGG: mmu04310; PMID: 23209147 |
| CD31 | CD31 | Wnt9a | FZD4_LRP6 | 0.000501 | 0 | WNT9A_FZD4_LRP6 | Wnt9a - (Fzd4+Lrp6) | WNT | Secreted Signaling | KEGG: mmu04310; PMID: 23209147 |
| GFAP | CD31 | Wnt9a | FZD4_LRP6 | 0.000537 | 0 | WNT9A_FZD4_LRP6 | Wnt9a - (Fzd4+Lrp6) | WNT | Secreted Signaling | KEGG: mmu04310; PMID: 23209147 |
| CD31 | GFAP | Wnt9a | FZD5_LRP6 | 0.000349 | 0 | WNT9A_FZD5_LRP6 | Wnt9a - (Fzd5+Lrp6) | WNT | Secreted Signaling | KEGG: mmu04310; PMID: 23209147 |
| GFAP | GFAP | Wnt9a | FZD5_LRP6 | 0.000373 | 0 | WNT9A_FZD5_LRP6 | Wnt9a - (Fzd5+Lrp6) | WNT | Secreted Signaling | KEGG: mmu04310; PMID: 23209147 |
| CD31 | CD31 | Wnt9a | FZD6_LRP6 | 0.000665 | 0 | WNT9A_FZD6_LRP6 | Wnt9a - (Fzd6+Lrp6) | WNT | Secreted Signaling | KEGG: mmu04310; PMID: 23209147 |
| GFAP | CD31 | Wnt9a | FZD6_LRP6 | 0.000712 | 0 | WNT9A_FZD6_LRP6 | Wnt9a - (Fzd6+Lrp6) | WNT | Secreted Signaling | KEGG: mmu04310; PMID: 23209147 |
| CD31 | CD31 | Wnt9a | FZD8_LRP6 | 0.000384 | 0 | WNT9A_FZD8_LRP6 | Wnt9a - (Fzd8+Lrp6) | WNT | Secreted Signaling | KEGG: mmu04310; PMID: 23209147 |
| GFAP | CD31 | Wnt9a | FZD8_LRP6 | 0.000411 | 0 | WNT9A_FZD8_LRP6 | Wnt9a - (Fzd8+Lrp6) | WNT | Secreted Signaling | KEGG: mmu04310; PMID: 23209147 |
| CD31 | GFAP | Wnt9a | FZD9_LRP6 | 0.000305 | 0 | WNT9A_FZD9_LRP6 | Wnt9a - (Fzd9+Lrp6) | WNT | Secreted Signaling | KEGG: mmu04310; PMID: 23209147 |
| GFAP | GFAP | Wnt9a | FZD9_LRP6 | 0.000327 | 0 | WNT9A_FZD9_LRP6 | Wnt9a - (Fzd9+Lrp6) | WNT | Secreted Signaling | KEGG: mmu04310; PMID: 23209147 |
| GFAP | CD31 | Wnt5a | Fzd1 | 0.000502 | 0 | WNT5A_FZD1 | Wnt5a - Fzd1 | ncWNT | Secreted Signaling | KEGG: mmu04310 |
| GFAP | GFAP | Wnt5a | Fzd1 | 0.000528 | 0 | WNT5A_FZD1 | Wnt5a - Fzd1 | ncWNT | Secreted Signaling | KEGG: mmu04310 |
| CD31 | GFAP | Wnt5a | Fzd2 | 0.000388 | 0 | WNT5A_FZD2 | Wnt5a - Fzd2 | ncWNT | Secreted Signaling | KEGG: mmu04310 |
| GFAP | GFAP | Wnt5a | Fzd2 | 0.000743 | 0 | WNT5A_FZD2 | Wnt5a - Fzd2 | ncWNT | Secreted Signaling | KEGG: mmu04310 |
| GFAP | GFAP | Wnt5a | Fzd3 | 0.000721 | 0 | WNT5A_FZD3 | Wnt5a - Fzd3 | ncWNT | Secreted Signaling | KEGG: mmu04310 |
| GFAP | NeuN | Wnt5a | Fzd3 | 0.001225 | 0 | WNT5A_FZD3 | Wnt5a - Fzd3 | ncWNT | Secreted Signaling | KEGG: mmu04310 |
| CD31 | CD31 | Wnt5a | Fzd4 | 0.000516 | 0 | WNT5A_FZD4 | Wnt5a - Fzd4 | ncWNT | Secreted Signaling | KEGG: mmu04310 |
| GFAP | CD31 | Wnt5a | Fzd4 | 0.000989 | 0 | WNT5A_FZD4 | Wnt5a - Fzd4 | ncWNT | Secreted Signaling | KEGG: mmu04310 |
| GFAP | CD31 | Wnt5a | Fzd5 | 0.000357 | 0 | WNT5A_FZD5 | Wnt5a - Fzd5 | ncWNT | Secreted Signaling | KEGG: mmu04310 |
| GFAP | GFAP | Wnt5a | Fzd5 | 0.000527 | 0 | WNT5A_FZD5 | Wnt5a - Fzd5 | ncWNT | Secreted Signaling | KEGG: mmu04310 |
| CD31 | CD31 | Wnt5a | Fzd6 | 0.000907 | 0 | WNT5A_FZD6 | Wnt5a - Fzd6 | ncWNT | Secreted Signaling | KEGG: mmu04310 |
| GFAP | CD31 | Wnt5a | Fzd6 | 0.001738 | 0 | WNT5A_FZD6 | Wnt5a - Fzd6 | ncWNT | Secreted Signaling | KEGG: mmu04310 |
| GFAP | CD31 | Wnt5a | Fzd8 | 0.000579 | 0 | WNT5A_FZD8 | Wnt5a - Fzd8 | ncWNT | Secreted Signaling | KEGG: mmu04310 |
| GFAP | GFAP | Wnt5a | Fzd8 | 0.000413 | 0 | WNT5A_FZD8 | Wnt5a - Fzd8 | ncWNT | Secreted Signaling | KEGG: mmu04310 |
| CD31 | GFAP | Wnt5a | Fzd9 | 0.000211 | 0 | WNT5A_FZD9 | Wnt5a - Fzd9 | ncWNT | Secreted Signaling | KEGG: mmu04310 |
| GFAP | GFAP | Wnt5a | Fzd9 | 0.000404 | 0 | WNT5A_FZD9 | Wnt5a - Fzd9 | ncWNT | Secreted Signaling | KEGG: mmu04310 |
| GFAP | CD31 | Wnt5a | Mcam | 0.001171 | 0 | WNT5A_MCAM | Wnt5a - Mcam | ncWNT | Secreted Signaling | PMID: 24335906 |
| GFAP | GFAP | Wnt5a | Mcam | 0.001398 | 0 | WNT5A_MCAM | Wnt5a - Mcam | ncWNT | Secreted Signaling | PMID: 24335906 |
| CD31 | CD31 | Hbegf | Erbb4 | 0.000802 | 0 | HBEGF_ERBB4 | Hbegf - Erbb4 | EGF | Secreted Signaling | KEGG: mmu04012 |
| CD31 | GFAP | Hbegf | Erbb4 | 0.000945 | 0 | HBEGF_ERBB4 | Hbegf - Erbb4 | EGF | Secreted Signaling | KEGG: mmu04012 |

| | | | | | | | | | |
|---|---|---|---|---|---|---|---|---|---|
| CD31 | CD31 | Hbegf | ERBB2_ERBB4 | 0.000666 | 0 | HBEGF_ERBB2_ERBB4 | Hbegf - (Erbb2+Erbb4) | EGF | Secreted Signaling | KEGG: mmu04012 |
| CD31 | GFAP | Hbegf | ERBB2_ERBB4 | 0.000579 | 0 | HBEGF_ERBB2_ERBB4 | Hbegf - (Erbb2+Erbb4) | EGF | Secreted Signaling | KEGG: mmu04012 |
| CD31 | GFAP | Ereg | Erbb4 | 0.000335 | 0 | EREG_ERBB4 | Ereg - Erbb4 | EGF | Secreted Signaling | KEGG: mmu04012 |
| GFAP | GFAP | Ereg | Erbb4 | 0.000387 | 0 | EREG_ERBB4 | Ereg - Erbb4 | EGF | Secreted Signaling | KEGG: mmu04012 |
| GFAP | CD31 | Ereg | ERBB2_ERBB4 | 0.000272 | 0 | EREG_ERBB2_ERBB4 | Ereg - (Erbb2+Erbb4) | EGF | Secreted Signaling | KEGG: mmu04012 |
| GFAP | GFAP | Ereg | ERBB2_ERBB4 | 0.000237 | 0 | EREG_ERBB2_ERBB4 | Ereg - (Erbb2+Erbb4) | EGF | Secreted Signaling | KEGG: mmu04012 |
| GFAP | GFAP | Nrg1 | Erbb3 | 0.00078 | 0 | NRG1_ERBB3 | Nrg1 - Erbb3 | NRG | Secreted Signaling | KEGG: mmu04012 |
| NeuN | GFAP | Nrg1 | Erbb3 | 0.000961 | 0 | NRG1_ERBB3 | Nrg1 - Erbb3 | NRG | Secreted Signaling | KEGG: mmu04012 |
| GFAP | GFAP | Nrg1 | ERBB2_ERBB3 | 0.000422 | 0 | NRG1_ERBB2_ERBB3 | Nrg1 - (Erbb2+Erbb3) | NRG | Secreted Signaling | KEGG: mmu04012 |
| NeuN | GFAP | Nrg1 | ERBB2_ERBB3 | 0.00052 | 0 | NRG1_ERBB2_ERBB3 | Nrg1 - (Erbb2+Erbb3) | NRG | Secreted Signaling | KEGG: mmu04012 |
| GFAP | GFAP | Nrg1 | Erbb4 | 0.00061 | 0 | NRG1_ERBB4 | Nrg1 - Erbb4 | NRG | Secreted Signaling | KEGG: mmu04012 |
| NeuN | GFAP | Nrg1 | Erbb4 | 0.000751 | 0 | NRG1_ERBB4 | Nrg1 - Erbb4 | NRG | Secreted Signaling | KEGG: mmu04012 |
| CD31 | CD31 | Nrg1 | ERBB2_ERBB4 | 0.000434 | 0 | NRG1_ERBB2_ERBB4 | Nrg1 - (Erbb2+Erbb4) | NRG | Secreted Signaling | KEGG: mmu04012 |
| NeuN | CD31 | Nrg1 | ERBB2_ERBB4 | 0.000529 | 0 | NRG1_ERBB2_ERBB4 | Nrg1 - (Erbb2+Erbb4) | NRG | Secreted Signaling | KEGG: mmu04012 |
| GFAP | GFAP | Nrg2 | Erbb3 | 0.001129 | 0 | NRG2_ERBB3 | Nrg2 - Erbb3 | NRG | Secreted Signaling | KEGG: mmu04012 |
| NeuN | GFAP | Nrg2 | Erbb3 | 0.00067 | 0 | NRG2_ERBB3 | Nrg2 - Erbb3 | NRG | Secreted Signaling | KEGG: mmu04012 |
| GFAP | CD31 | Nrg2 | ERBB2_ERBB3 | 0.00047 | 0 | NRG2_ERBB2_ERBB3 | Nrg2 - (Erbb2+Erbb3) | NRG | Secreted Signaling | KEGG: mmu04012 |
| GFAP | GFAP | Nrg2 | ERBB2_ERBB3 | 0.000611 | 0 | NRG2_ERBB2_ERBB3 | Nrg2 - (Erbb2+Erbb3) | NRG | Secreted Signaling | KEGG: mmu04012 |
| GFAP | CD31 | Nrg2 | Erbb4 | 0.000749 | 0 | NRG2_ERBB4 | Nrg2 - Erbb4 | NRG | Secreted Signaling | KEGG: mmu04012 |
| GFAP | GFAP | Nrg2 | Erbb4 | 0.000883 | 0 | NRG2_ERBB4 | Nrg2 - Erbb4 | NRG | Secreted Signaling | KEGG: mmu04012 |
| GFAP | CD31 | Nrg2 | ERBB2_ERBB4 | 0.000622 | 0 | NRG2_ERBB2_ERBB4 | Nrg2 - (Erbb2+Erbb4) | NRG | Secreted Signaling | KEGG: mmu04012 |
| GFAP | GFAP | Nrg2 | ERBB2_ERBB4 | 0.000541 | 0 | NRG2_ERBB2_ERBB4 | Nrg2 - (Erbb2+Erbb4) | NRG | Secreted Signaling | KEGG: mmu04012 |
| NeuN | GFAP | Nrg3 | Erbb4 | 0.003203 | 0 | NRG3_ERBB4 | Nrg3 - Erbb4 | NRG | Secreted Signaling | KEGG: mmu04012 |
| NeuN | NeuN | Nrg3 | Erbb4 | 0.001781 | 0 | NRG3_ERBB4 | Nrg3 - Erbb4 | NRG | Secreted Signaling | KEGG: mmu04012 |
| NeuN | CD31 | Nrg3 | ERBB2_ERBB4 | 0.002257 | 0 | NRG3_ERBB2_ERBB4 | Nrg3 - (Erbb2+Erbb4) | NRG | Secreted Signaling | KEGG: mmu04012 |
| NeuN | NeuN | Nrg3 | ERBB2_ERBB4 | 0.00099 | 0 | NRG3_ERBB2_ERBB4 | Nrg3 - (Erbb2+Erbb4) | NRG | Secreted Signaling | KEGG: mmu04012 |
| GFAP | GFAP | Nrg1 | ITGAV_ITGB3 | 0.000548 | 0 | NRG1_ITGAV_ITGB3 | Nrg1 - (Itgav+Itgb3) | NRG | Secreted Signaling | PMID: 20682778 |
| NeuN | GFAP | Nrg1 | ITGAV_ITGB3 | 0.000674 | 0 | NRG1_ITGAV_ITGB3 | Nrg1 - (Itgav+Itgb3) | NRG | Secreted Signaling | PMID: 20682778 |
| CD31 | GFAP | Fgf2 | Fgfr1 | 0.000916 | 0 | FGF2_FGFR1 | Fgf2 - Fgfr1 | FGF | Secreted Signaling | PMC: 4393358 |
| GFAP | GFAP | Fgf2 | Fgfr1 | 0.000808 | 0 | FGF2_FGFR1 | Fgf2 - Fgfr1 | FGF | Secreted Signaling | PMC: 4393358 |
| CD31 | GFAP | Fgf2 | Fgfr2 | 0.001051 | 0 | FGF2_FGFR2 | Fgf2 - Fgfr2 | FGF | Secreted Signaling | PMC: 4393358 |
| GFAP | GFAP | Fgf2 | Fgfr2 | 0.000927 | 0 | FGF2_FGFR2 | Fgf2 - Fgfr2 | FGF | Secreted Signaling | PMC: 4393358 |
| CD31 | GFAP | Fgf2 | Fgfr3 | 0.001267 | 0 | FGF2_FGFR3 | Fgf2 - Fgfr3 | FGF | Secreted Signaling | PMC: 4393358 |
| GFAP | GFAP | Fgf2 | Fgfr3 | 0.001117 | 0 | FGF2_FGFR3 | Fgf2 - Fgfr3 | FGF | Secreted Signaling | PMC: 4393358 |
| CD31 | CD31 | Fgf5 | Fgfr1 | 0.000321 | 0 | FGF5_FGFR1 | Fgf5 - Fgfr1 | FGF | Secreted Signaling | PMC: 4393358 |
| CD31 | GFAP | Fgf5 | Fgfr1 | 0.000651 | 0 | FGF5_FGFR1 | Fgf5 - Fgfr1 | FGF | Secreted Signaling | PMC: 4393358 |
| CD31 | GFAP | Fgf5 | Fgfr2 | 0.000747 | 0 | FGF5_FGFR2 | Fgf5 - Fgfr2 | FGF | Secreted Signaling | PMC: 4393358 |
| GFAP | GFAP | Fgf5 | Fgfr2 | 0.000339 | 0 | FGF5_FGFR2 | Fgf5 - Fgfr2 | FGF | Secreted Signaling | PMC: 4393358 |
| CD31 | CD31 | Fgf5 | Fgfr3 | 0.000409 | 0 | FGF5_FGFR3 | Fgf5 - Fgfr3 | FGF | Secreted Signaling | PMC: 4393358 |
| CD31 | GFAP | Fgf5 | Fgfr3 | 0.0009 | 0 | FGF5_FGFR3 | Fgf5 - Fgfr3 | FGF | Secreted Signaling | PMC: 4393358 |
| CD31 | GFAP | Fgf3 | Fgfr1 | 0.000976 | 0 | FGF3_FGFR1 | Fgf3 - Fgfr1 | FGF | Secreted Signaling | PMC: 4393358 |
| GFAP | GFAP | Fgf3 | Fgfr1 | 0.000901 | 0 | FGF3_FGFR1 | Fgf3 - Fgfr1 | FGF | Secreted Signaling | PMC: 4393358 |
| CD31 | GFAP | Fgf3 | Fgfr2 | 0.00112 | 0 | FGF3_FGFR2 | Fgf3 - Fgfr2 | FGF | Secreted Signaling | PMC: 4393358 |
| GFAP | GFAP | Fgf3 | Fgfr2 | 0.001034 | 0 | FGF3_FGFR2 | Fgf3 - Fgfr2 | FGF | Secreted Signaling | PMC: 4393358 |
| CD31 | GFAP | Fgf10 | Fgfr1 | 0.000766 | 0 | FGF10_FGFR1 | Fgf10 - Fgfr1 | FGF | Secreted Signaling | PMC: 4393358 |
| GFAP | GFAP | Fgf10 | Fgfr1 | 0.000662 | 0 | FGF10_FGFR1 | Fgf10 - Fgfr1 | FGF | Secreted Signaling | PMC: 4393358 |
| CD31 | GFAP | Fgf10 | Fgfr2 | 0.000879 | 0 | FGF10_FGFR2 | Fgf10 - Fgfr2 | FGF | Secreted Signaling | PMC: 4393358 |
| GFAP | GFAP | Fgf10 | Fgfr2 | 0.00076 | 0 | FGF10_FGFR2 | Fgf10 - Fgfr2 | FGF | Secreted Signaling | PMC: 4393358 |
| CD31 | GFAP | Fgf22 | Fgfr2 | 0.000796 | 0 | FGF22_FGFR2 | Fgf22 - Fgfr2 | FGF | Secreted Signaling | PMC: 4393358 |
| GFAP | GFAP | Fgf22 | Fgfr2 | 0.001105 | 0 | FGF22_FGFR2 | Fgf22 - Fgfr2 | FGF | Secreted Signaling | PMC: 4393358 |
| CD31 | GFAP | Fgf18 | Fgfr1 | 0.00073 | 0 | FGF18_FGFR1 | Fgf18 - Fgfr1 | FGF | Secreted Signaling | PMC: 4393358 |
| GFAP | GFAP | Fgf18 | Fgfr1 | 0.000752 | 0 | FGF18_FGFR1 | Fgf18 - Fgfr1 | FGF | Secreted Signaling | PMC: 4393358 |
| CD31 | GFAP | Fgf18 | Fgfr2 | 0.000838 | 0 | FGF18_FGFR2 | Fgf18 - Fgfr2 | FGF | Secreted Signaling | PMC: 4393358 |
| GFAP | GFAP | Fgf18 | Fgfr2 | 0.000863 | 0 | FGF18_FGFR2 | Fgf18 - Fgfr2 | FGF | Secreted Signaling | PMC: 4393358 |
| CD31 | GFAP | Fgf18 | Fgfr3 | 0.00101 | 0 | FGF18_FGFR3 | Fgf18 - Fgfr3 | FGF | Secreted Signaling | PMC: 4393358 |
| GFAP | GFAP | Fgf18 | Fgfr3 | 0.00104 | 0 | FGF18_FGFR3 | Fgf18 - Fgfr3 | FGF | Secreted Signaling | PMC: 4393358 |
| GFAP | GFAP | Fgf9 | Fgfr1 | 0.000697 | 0 | FGF9_FGFR1 | Fgf9 - Fgfr1 | FGF | Secreted Signaling | PMC: 4393358 |
| NeuN | GFAP | Fgf9 | Fgfr1 | 0.000813 | 0 | FGF9_FGFR1 | Fgf9 - Fgfr1 | FGF | Secreted Signaling | PMC: 4393358 |
| GFAP | GFAP | Fgf9 | Fgfr2 | 0.0008 | 0 | FGF9_FGFR2 | Fgf9 - Fgfr2 | FGF | Secreted Signaling | PMC: 4393358 |

| | | | | | | | | | | |
|---|---|---|---|---|---|---|---|---|---|---|
| NeuN | GFAP | Fgf9 | Fgfr2 | 0.000933 | 0 | FGF9_FGFR2 | Fgf9 - Fgfr2 | FGF | Secreted Signaling | PMC: 4393358 |
| GFAP | GFAP | Fgf9 | Fgfr3 | 0.000964 | 0 | FGF9_FGFR3 | Fgf9 - Fgfr3 | FGF | Secreted Signaling | PMC: 4393358 |
| NeuN | GFAP | Fgf9 | Fgfr3 | 0.001124 | 0 | FGF9_FGFR3 | Fgf9 - Fgfr3 | FGF | Secreted Signaling | PMC: 4393358 |
| CD31 | GFAP | Fgf15 | Fgfr1 | 0.001013 | 0 | FGF15_FGFR1 | Fgf15 - Fgfr1 | FGF | Secreted Signaling | PMC: 4393358 |
| GFAP | GFAP | Fgf15 | Fgfr1 | 0.001071 | 0 | FGF15_FGFR1 | Fgf15 - Fgfr1 | FGF | Secreted Signaling | PMC: 4393358 |
| CD31 | GFAP | Fgf15 | Fgfr2 | 0.001162 | 0 | FGF15_FGFR2 | Fgf15 - Fgfr2 | FGF | Secreted Signaling | PMC: 4393358 |
| GFAP | GFAP | Fgf15 | Fgfr2 | 0.00123 | 0 | FGF15_FGFR2 | Fgf15 - Fgfr2 | FGF | Secreted Signaling | PMC: 4393358 |
| CD31 | GFAP | Fgf15 | Fgfr3 | 0.001401 | 0 | FGF15_FGFR3 | Fgf15 - Fgfr3 | FGF | Secreted Signaling | PMC: 4393358 |
| GFAP | GFAP | Fgf15 | Fgfr3 | 0.001482 | 0 | FGF15_FGFR3 | Fgf15 - Fgfr3 | FGF | Secreted Signaling | PMC: 4393358 |
| CD31 | GFAP | Fgf21 | Fgfr1 | 0.000949 | 0 | FGF21_FGFR1 | Fgf21 - Fgfr1 | FGF | Secreted Signaling | PMC: 4393358 |
| GFAP | GFAP | Fgf21 | Fgfr1 | 0.000809 | 0 | FGF21_FGFR1 | Fgf21 - Fgfr1 | FGF | Secreted Signaling | PMC: 4393358 |
| CD31 | GFAP | Fgf21 | Fgfr3 | 0.001312 | 0 | FGF21_FGFR3 | Fgf21 - Fgfr3 | FGF | Secreted Signaling | PMC: 4393358 |
| GFAP | GFAP | Fgf21 | Fgfr3 | 0.001119 | 0 | FGF21_FGFR3 | Fgf21 - Fgfr3 | FGF | Secreted Signaling | PMC: 4393358 |
| GFAP | GFAP | Pdgfa | Pdgfra | 0.000517 | 0 | PDGFA_PDGFRA | Pdgfa - Pdgfra | PDGF | Secreted Signaling | PMID: 15207812 |
| NeuN | GFAP | Pdgfa | Pdgfra | 0.000712 | 0 | PDGFA_PDGFRA | Pdgfa - Pdgfra | PDGF | Secreted Signaling | PMID: 15207812 |
| CD31 | GFAP | Pdgfa | Pdgfrb | 0.000592 | 0 | PDGFA_PDGFRB | Pdgfa - Pdgfrb | PDGF | Secreted Signaling | PMID: 15207812 |
| NeuN | CD31 | Pdgfa | Pdgfrb | 0.000789 | 0 | PDGFA_PDGFRB | Pdgfa - Pdgfrb | PDGF | Secreted Signaling | PMID: 15207812 |
| CD31 | CD31 | Pdgfb | Pdgfra | 0.00065 | 0 | PDGFB_PDGFRA | Pdgfb - Pdgfra | PDGF | Secreted Signaling | PMID: 15207812 |
| CD31 | GFAP | Pdgfb | Pdgfra | 0.001111 | 0 | PDGFB_PDGFRA | Pdgfb - Pdgfra | PDGF | Secreted Signaling | PMID: 15207812 |
| CD31 | CD31 | Pdgfb | Pdgfrb | 0.001229 | 0 | PDGFB_PDGFRB | Pdgfb - Pdgfrb | PDGF | Secreted Signaling | PMID: 15207812 |
| CD31 | GFAP | Pdgfb | Pdgfrb | 0.000667 | 0 | PDGFB_PDGFRB | Pdgfb - Pdgfrb | PDGF | Secreted Signaling | PMID: 15207812 |
| CD31 | GFAP | Pdgfc | Pdgfra | 0.000396 | 0 | PDGFC_PDGFRA | Pdgfc - Pdgfra | PDGF | Secreted Signaling | PMID: 15207812 |
| GFAP | GFAP | Pdgfc | Pdgfra | 0.000542 | 0 | PDGFC_PDGFRA | Pdgfc - Pdgfra | PDGF | Secreted Signaling | PMID: 15207812 |
| CD31 | CD31 | Pdgfd | Pdgfrb | 0.000785 | 0 | PDGFD_PDGFRB | Pdgfd - Pdgfrb | PDGF | Secreted Signaling | PMID: 15207812 |
| CD31 | GFAP | Pdgfd | Pdgfrb | 0.000425 | 0 | PDGFD_PDGFRB | Pdgfd - Pdgfrb | PDGF | Secreted Signaling | PMID: 15207812 |
| CD31 | CD31 | Vegfa | Flt1 | 0.002998 | 0 | VEGFA_VEGFR1 | Vegfa - Vegfr1 | VEGF | Secreted Signaling | KEGG: mmu04370; PMID: 16633338 |
| GFAP | CD31 | Vegfa | Flt1 | 0.011057 | 0 | VEGFA_VEGFR1 | Vegfa - Vegfr1 | VEGF | Secreted Signaling | KEGG: mmu04370; PMID: 16633338 |
| CD31 | CD31 | Vegfa | Kdr | 0.00111 | 0 | VEGFA_VEGFR2 | Vegfa - Vegfr2 | VEGF | Secreted Signaling | KEGG: mmu04370; PMID: 16633338 |
| GFAP | CD31 | Vegfa | Kdr | 0.004115 | 0 | VEGFA_VEGFR2 | Vegfa - Vegfr2 | VEGF | Secreted Signaling | KEGG: mmu04370; PMID: 16633338 |
| CD31 | CD31 | Vegfb | Flt1 | 0.004951 | 0 | VEGFB_VEGFR1 | Vegfb - Vegfr1 | VEGF | Secreted Signaling | KEGG: mmu04370; PMID: 16633338 |
| GFAP | CD31 | Vegfb | Flt1 | 0.006065 | 0 | VEGFB_VEGFR1 | Vegfb - Vegfr1 | VEGF | Secreted Signaling | KEGG: mmu04370; PMID: 16633338 |
| CD31 | CD31 | Pgf | Flt1 | 0.00384 | 0 | PGF_VEGFR1 | Pgf - Vegfr1 | VEGF | Secreted Signaling | KEGG: mmu04370; PMID: 16633338 |
| GFAP | CD31 | Pgf | Flt1 | 0.002105 | 0 | PGF_VEGFR1 | Pgf - Vegfr1 | VEGF | Secreted Signaling | KEGG: mmu04370; PMID: 16633338 |
| CD31 | CD31 | Vegfa | FLT1_KDR | 0.001879 | 0 | VEGFA_VEGFR1R2 | Vegfa - Vegfr1r2 | VEGF | Secreted Signaling | PMID: 16633338 |
| GFAP | CD31 | Vegfa | FLT1_KDR | 0.006949 | 0 | VEGFA_VEGFR1R2 | Vegfa - Vegfr1r2 | VEGF | Secreted Signaling | PMID: 16633338 |
| CD31 | CD31 | Igf1 | Igf1r | 0.0006 | 0 | IGF1_IGF1R | Igf1 - Igf1r | IGF | Secreted Signaling | PMID: 14604834 |
| NeuN | CD31 | Igf1 | Igf1r | 0.000366 | 0 | IGF1_IGF1R | Igf1 - Igf1r | IGF | Secreted Signaling | PMID: 14604834 |
| CD31 | CD31 | Igf2 | Igf1r | 0.003563 | 0 | IGF2_IGF1R | Igf2 - Igf1r | IGF | Secreted Signaling | PMID: 14604834 |
| GFAP | CD31 | Igf2 | Igf1r | 0.002369 | 0 | IGF2_IGF1R | Igf2 - Igf1r | IGF | Secreted Signaling | PMID: 14604834 |
| CD31 | CD31 | Igf2 | Igf2r | 0.000744 | 0 | IGF2_IGF2R | Igf2 - Igf2r | IGF | Secreted Signaling | PMID: 14604834 |
| CD31 | NeuN | Igf2 | Igf2r | 0.000827 | 0 | IGF2_IGF2R | Igf2 - Igf2r | IGF | Secreted Signaling | PMID: 14604834 |
| CD31 | CD31 | Igf1 | ITGAV_ITGB3 | 0.000173 | 0 | IGF1_ITGAV_ITGB3 | Igf1 - (Itgav+Itgb3) | IGF | Secreted Signaling | PMID: 28190785; PMID: 28873464 |
| CD31 | GFAP | Igf1 | ITGAV_ITGB3 | 0.0002 | 0 | IGF1_ITGAV_ITGB3 | Igf1 - (Itgav+Itgb3) | IGF | Secreted Signaling | PMID: 28190785; PMID: 28873464 |
| CD31 | CD31 | Igf2 | ITGAV_ITGB3 | 0.001028 | 0 | IGF2_ITGAV_ITGB3 | Igf2 - (Itgav+Itgb3) | IGF | Secreted Signaling | PMID: 28190785; PMID: 28873464 |
| CD31 | GFAP | Igf2 | ITGAV_ITGB3 | 0.00119 | 0 | IGF2_ITGAV_ITGB3 | Igf2 - (Itgav+Itgb3) | IGF | Secreted Signaling | PMID: 28190785; PMID: 28873464 |
| CD31 | CD31 | Ins1 | Insr | 0.000686 | 0 | INS1_INSSR | Ins1 - Inssr | INSULIN | Secreted Signaling | KEGG: mmu04910; PMID: 24633815 |
| GFAP | CD31 | Ins1 | Insr | 0.001089 | 0 | INS1_INSSR | Ins1 - Inssr | INSULIN | Secreted Signaling | KEGG: mmu04910; PMID: 24633815 |
| CD31 | CD31 | Apln | Aplnr | 0.000737 | 0 | APLN_APLNR | Apln - Aplnr | APELIN | Secreted Signaling | KEGG: mmu04371 |
| GFAP | CD31 | Apln | Aplnr | 0.000475 | 0 | APLN_APLNR | Apln - Aplnr | APELIN | Secreted Signaling | KEGG: mmu04371 |
| CD31 | CD31 | Ihh | PTCH1_SMO | 0.000498 | 0 | IHH_PTCH1 | Ihh - Ptch1 | HH | Secreted Signaling | KEGG: mmu04340; PMID: 25701396 |
| GFAP | CD31 | Ihh | PTCH1_SMO | 0.000616 | 0 | IHH_PTCH1 | Ihh - Ptch1 | HH | Secreted Signaling | KEGG: mmu04340; PMID: 25701396 |
| CD31 | CD31 | Shh | PTCH1_SMO | 0.00054 | 0 | SHH_PTCH1 | Shh - Ptch1 | HH | Secreted Signaling | KEGG: mmu04340; PMID: 25701396 |
| GFAP | CD31 | Shh | PTCH1_SMO | 0.000405 | 0 | SHH_PTCH1 | Shh - Ptch1 | HH | Secreted Signaling | KEGG: mmu04340; PMID: 25701396 |
| CD31 | CD31 | Ihh | PTCH2_SMO | 0.000433 | 0 | IHH_PTCH2 | Ihh - Ptch2 | HH | Secreted Signaling | KEGG: mmu04340; PMID: 25701396 |
| GFAP | CD31 | Ihh | PTCH2_SMO | 0.000536 | 0 | IHH_PTCH2 | Ihh - Ptch2 | HH | Secreted Signaling | KEGG: mmu04340; PMID: 25701396 |
| CD31 | CD31 | Shh | PTCH2_SMO | 0.00047 | 0 | SHH_PTCH2 | Shh - Ptch2 | HH | Secreted Signaling | KEGG: mmu04340; PMID: 25701396 |
| GFAP | CD31 | Shh | PTCH2_SMO | 0.000353 | 0 | SHH_PTCH2 | Shh - Ptch2 | HH | Secreted Signaling | KEGG: mmu04340; PMID: 25701396 |
| CD31 | CD31 | Ccl8 | Ccr1 | 0.000591 | 0 | CCL8_CCR1 | Ccl8 - Ccr1 | CCL | Secreted Signaling | KEGG: mmu04060 |
| CD31 | GFAP | Ccl8 | Ccr1 | 0.000483 | 0 | CCL8_CCR1 | Ccl8 - Ccr1 | CCL | Secreted Signaling | KEGG: mmu04060 |

| | | | | | | | | | | |
|---|---|---|---|---|---|---|---|---|---|---|
| CD31 | CD31 | Ccl6 | Ccr1 | 0.000225 | 0 | CCL6_CCR1 | Ccl6 - Ccr1 | CCL | Secreted Signaling | KEGG: mmu04060 |
| GFAP | CD31 | Ccl6 | Ccr1 | 0.000249 | 0 | CCL6_CCR1 | Ccl6 - Ccr1 | CCL | Secreted Signaling | KEGG: mmu04060 |
| CD31 | CD31 | Ccl8 | Ccr1l1 | 0.000318 | 0 | CCL8_CCR1L1 | Ccl8 - Ccr1l1 | CCL | Secreted Signaling | KEGG: mmu04060 |
| CD31 | GFAP | Ccl8 | Ccr1l1 | 0.000408 | 0 | CCL8_CCR1L1 | Ccl8 - Ccr1l1 | CCL | Secreted Signaling | KEGG: mmu04060 |
| CD31 | GFAP | Ccl6 | Ccr1l1 | 0.000155 | 0 | CCL6_CCR1L1 | Ccl6 - Ccr1l1 | CCL | Secreted Signaling | KEGG: mmu04060 |
| GFAP | GFAP | Ccl6 | Ccr1l1 | 0.000172 | 0 | CCL6_CCR1L1 | Ccl6 - Ccr1l1 | CCL | Secreted Signaling | KEGG: mmu04060 |
| CD31 | CD31 | Ccl6 | Ccr3 | 0.000161 | 0 | CCL6_CCR3 | Ccl6 - Ccr3 | CCL | Secreted Signaling | KEGG: mmu04060 |
| GFAP | CD31 | Ccl6 | Ccr3 | 0.000179 | 0 | CCL6_CCR3 | Ccl6 - Ccr3 | CCL | Secreted Signaling | KEGG: mmu04060 |
| CD31 | CD31 | Ccl11 | Ccr3 | 0.000267 | 0 | CCL11_CCR3 | Ccl11 - Ccr3 | CCL | Secreted Signaling | KEGG: mmu04060 |
| GFAP | CD31 | Ccl11 | Ccr3 | 0.000309 | 0 | CCL11_CCR3 | Ccl11 - Ccr3 | CCL | Secreted Signaling | KEGG: mmu04060 |
| NeuN | CD31 | Ccl27a | Ccr3 | 0.007097 | 0 | CCL27A_CCR3 | Ccl27a - Ccr3 | CCL | Secreted Signaling | KEGG: mmu04060 |
| NeuN | NeuN | Ccl27a | Ccr3 | 0.001975 | 0 | CCL27A_CCR3 | Ccl27a - Ccr3 | CCL | Secreted Signaling | KEGG: mmu04060 |
| CD31 | CD31 | Ccl27b | Ccr3 | 0.001187 | 0 | CCL27B_CCR3 | Ccl27b - Ccr3 | CCL | Secreted Signaling | KEGG: mmu04060 |
| NeuN | CD31 | Ccl27b | Ccr3 | 0.003928 | 0 | CCL27B_CCR3 | Ccl27b - Ccr3 | CCL | Secreted Signaling | KEGG: mmu04060 |
| CD31 | CD31 | Ccl19 | Ccr7 | 0.000736 | 0 | CCL19_CCR7 | Ccl19 - Ccr7 | CCL | Secreted Signaling | KEGG: mmu04060 |
| GFAP | CD31 | Ccl19 | Ccr7 | 0.000525 | 0 | CCL19_CCR7 | Ccl19 - Ccr7 | CCL | Secreted Signaling | KEGG: mmu04060 |
| CD31 | CD31 | Ccl21a | Ccr7 | 0.000543 | 0 | CCL21A_CCR7 | Ccl21a - Ccr7 | CCL | Secreted Signaling | KEGG: mmu04060 |
| NeuN | CD31 | Ccl21a | Ccr7 | 0.000608 | 0 | CCL21A_CCR7 | Ccl21a - Ccr7 | CCL | Secreted Signaling | KEGG: mmu04060 |
| CD31 | CD31 | Ccl25 | Ccr9 | 0.000246 | 0 | CCL25_CCR9 | Ccl25 - Ccr9 | CCL | Secreted Signaling | KEGG: mmu04060 |
| GFAP | CD31 | Ccl25 | Ccr9 | 0.000293 | 0 | CCL25_CCR9 | Ccl25 - Ccr9 | CCL | Secreted Signaling | KEGG: mmu04060 |
| CD31 | CD31 | Ccl4 | Ackr2 | 0.000467 | 0 | CCL4_ACKR2 | Ccl4 - Ackr2 | CCL | Secreted Signaling | PMID: 26740381 |
| GFAP | CD31 | Ccl4 | Ackr2 | 0.000398 | 0 | CCL4_ACKR2 | Ccl4 - Ackr2 | CCL | Secreted Signaling | PMID: 26740381 |
| CD31 | CD31 | Ccl8 | Ackr1 | 0.0006 | 0 | CCL8_ACKR1 | Ccl8 - Ackr1 | CCL | Secreted Signaling | PMID: 26740381 |
| CD31 | NeuN | Ccl8 | Ackr1 | 0.001176 | 0 | CCL8_ACKR1 | Ccl8 - Ackr1 | CCL | Secreted Signaling | PMID: 26740381 |
| CD31 | CD31 | Ccl11 | Ackr1 | 0.000379 | 0 | CCL11_ACKR1 | Ccl11 - Ackr1 | CCL | Secreted Signaling | PMID: 26740381 |
| GFAP | NeuN | Ccl11 | Ackr1 | 0.000859 | 0 | CCL11_ACKR1 | Ccl11 - Ackr1 | CCL | Secreted Signaling | PMID: 26740381 |
| CD31 | CD31 | Ccl11 | Ackr2 | 0.000305 | 0 | CCL11_ACKR2 | Ccl11 - Ackr2 | CCL | Secreted Signaling | PMID: 26740381 |
| GFAP | CD31 | Ccl11 | Ackr2 | 0.000353 | 0 | CCL11_ACKR2 | Ccl11 - Ackr2 | CCL | Secreted Signaling | PMID: 26740381 |
| CD31 | CD31 | Pf4 | Cxcr3 | 0.000463 | 0 | CXCL4_CXCR3 | Cxcl4 - Cxcr3 | CXCL | Secreted Signaling | KEGG: mmu04060 |
| CD31 | GFAP | Pf4 | Cxcr3 | 0.000467 | 0 | CXCL4_CXCR3 | Cxcl4 - Cxcr3 | CXCL | Secreted Signaling | KEGG: mmu04060 |
| GFAP | CD31 | Cxcl10 | Cxcr3 | 0.000407 | 0 | CXCL10_CXCR3 | Cxcl10 - Cxcr3 | CXCL | Secreted Signaling | KEGG: mmu04060 |
| GFAP | GFAP | Cxcl10 | Cxcr3 | 0.000411 | 0 | CXCL10_CXCR3 | Cxcl10 - Cxcr3 | CXCL | Secreted Signaling | KEGG: mmu04060 |
| CD31 | CD31 | Cxcl12 | Cxcr4 | 0.005689 | 0 | CXCL12_CXCR4 | Cxcl12 - Cxcr4 | CXCL | Secreted Signaling | KEGG: mmu04060 |
| CD31 | GFAP | Cxcl12 | Cxcr4 | 0.00774 | 0 | CXCL12_CXCR4 | Cxcl12 - Cxcr4 | CXCL | Secreted Signaling | KEGG: mmu04060 |
| CD31 | CD31 | Cxcl12 | Ackr3 | 0.005353 | 0 | CXCL12_ACKR3 | Cxcl12 - Ackr3 | CXCL | Secreted Signaling | KEGG: mmu04060 |
| CD31 | GFAP | Cxcl12 | Ackr3 | 0.003876 | 0 | CXCL12_ACKR3 | Cxcl12 - Ackr3 | CXCL | Secreted Signaling | KEGG: mmu04060 |
| CD31 | CD31 | Cxcl1 | Ackr1 | 0.000551 | 0 | CXCL1_ACKR1 | Cxcl1 - Ackr1 | CXCL | Secreted Signaling | KEGG: mmu04060 |
| CD31 | NeuN | Cxcl1 | Ackr1 | 0.001079 | 0 | CXCL1_ACKR1 | Cxcl1 - Ackr1 | CXCL | Secreted Signaling | KEGG: mmu04060 |
| GFAP | GFAP | Cxcl10 | Ackr1 | 0.00028 | 0 | CXCL10_ACKR1 | Cxcl10 - Ackr1 | CXCL | Secreted Signaling | KEGG: mmu04060 |
| GFAP | NeuN | Cxcl10 | Ackr1 | 0.000866 | 0 | CXCL10_ACKR1 | Cxcl10 - Ackr1 | CXCL | Secreted Signaling | KEGG: mmu04060 |
| CD31 | CD31 | Mif | Ackr3 | 0.002567 | 0 | MIF_ACKR3 | Mif - Ackr3 | MIF | Secreted Signaling | PMID: 26175090 |
| NeuN | CD31 | Mif | Ackr3 | 0.005359 | 0 | MIF_ACKR3 | Mif - Ackr3 | MIF | Secreted Signaling | PMID: 26175090 |
| CD31 | CD31 | Cx3cl1 | Cx3cr1 | 0.004558 | 0 | CX3CL1_CX3CR1 | Cx3cl1 - Cx3cr1 | CX3C | Secreted Signaling | KEGG: mmu04060 |
| NeuN | GFAP | Cx3cl1 | Cx3cr1 | 0.015135 | 0 | CX3CL1_CX3CR1 | Cx3cl1 - Cx3cr1 | CX3C | Secreted Signaling | KEGG: mmu04060 |
| CD31 | CD31 | Clcf1 | CNTFR_LIFR | 0.000753 | 0 | CLCF1_CNTFR_LIFR | Clcf1 - (Cntfr+Lifr) | LIFR | Secreted Signaling | KEGG: mmu04060 |
| CD31 | GFAP | Clcf1 | CNTFR_LIFR | 0.000879 | 0 | CLCF1_CNTFR_LIFR | Clcf1 - (Cntfr+Lifr) | LIFR | Secreted Signaling | KEGG: mmu04060 |
| CD31 | CD31 | Ctf1 | LIFR_IL6ST | 0.000548 | 0 | CTF1_LIFR_IL6ST | Ctf1 - (Lifr+Il6st) | LIFR | Secreted Signaling | KEGG: mmu04060 |
| CD31 | GFAP | Ctf1 | LIFR_IL6ST | 0.000407 | 0 | CTF1_LIFR_IL6ST | Ctf1 - (Lifr+Il6st) | LIFR | Secreted Signaling | KEGG: mmu04060 |
| GFAP | GFAP | Il34 | Csf1r | 0.001595 | 0 | IL34_CSF1R | Il34 - Csf1r | CSF | Secreted Signaling | KEGG: mmu04060 |
| NeuN | GFAP | Il34 | Csf1r | 0.002837 | 0 | IL34_CSF1R | Il34 - Csf1r | CSF | Secreted Signaling | KEGG: mmu04060 |
| CD31 | GFAP | Csf1 | Csf1r | 0.00092 | 0 | CSF1_CSF1R | Csf1 - Csf1r | CSF | Secreted Signaling | KEGG: mmu04060 |
| GFAP | GFAP | Csf1 | Csf1r | 0.000933 | 0 | CSF1_CSF1R | Csf1 - Csf1r | CSF | Secreted Signaling | KEGG: mmu04060 |
| CD31 | CD31 | Tnfsf10 | Tnfrsf10b | 0.000252 | 0 | TNFSF10_TNFRSF10B | Tnfsf10 - Tnfrsf10b | TRAIL | Secreted Signaling | KEGG: mmu04060 |
| CD31 | GFAP | Tnfsf10 | Tnfrsf10b | 0.000225 | 0 | TNFSF10_TNFRSF10B | Tnfsf10 - Tnfrsf10b | TRAIL | Secreted Signaling | KEGG: mmu04060 |
| CD31 | CD31 | Retn | Cap1 | 0.002302 | 0 | RETN_CAP1 | Retn - Cap1 | RESISTIN | Secreted Signaling | PMID: 30809105 |
| CD31 | NeuN | Retn | Cap1 | 0.002923 | 0 | RETN_CAP1 | Retn - Cap1 | RESISTIN | Secreted Signaling | PMID: 30809105 |
| CD31 | CD31 | Nampt | ITGA5_ITGB1 | 0.002864 | 0 | NAMPT_ITGA5_ITGB1 | Nampt - (Itga5+Itgb1) | VISFATIN | Secreted Signaling | PMID: 28490838 |
| CD31 | GFAP | Nampt | ITGA5_ITGB1 | 0.002266 | 0 | NAMPT_ITGA5_ITGB1 | Nampt - (Itga5+Itgb1) | VISFATIN | Secreted Signaling | PMID: 28490838 |
| CD31 | CD31 | Angptl4 | ITGA5_ITGB1 | 0.000876 | 0 | ANGPTL4_ITGA5_ITGB1 | Angptl4 - (Itga5+Itgb1) | ANGPTL | Secreted Signaling | PMID: 30049845 |

| | | | | | | | | | | |
|---|---|---|---|---|---|---|---|---|---|---|
| CD31 | GFAP | Angptl4 | ITGA5_ITGB1 | 0.000692 | 0 | ANGPTL4_ITGA5_ITGB1 | Angptl4 - (Itga5+Itgb1) | ANGPTL | Secreted Signaling | PMID: 30049845 |
| CD31 | CD31 | Angptl4 | ITGAV_ITGB3 | 0.000588 | 0 | ANGPTL4_ITGAV_ITGB3 | Angptl4 - (Itgav+Itgb3) | ANGPTL | Secreted Signaling | PMID: 30049845 |
| CD31 | GFAP | Angptl4 | ITGAV_ITGB3 | 0.000684 | 0 | ANGPTL4_ITGAV_ITGB3 | Angptl4 - (Itgav+Itgb3) | ANGPTL | Secreted Signaling | PMID: 30049845 |
| CD31 | CD31 | Angptl4 | Cdh5 | 0.002067 | 0 | ANGPTL4_CDH5 | Angptl4 - Cdh5 | ANGPTL | Secreted Signaling | PMID: 30049845 |
| GFAP | CD31 | Angptl4 | Cdh5 | 0.001145 | 0 | ANGPTL4_CDH5 | Angptl4 - Cdh5 | ANGPTL | Secreted Signaling | PMID: 30049845 |
| CD31 | CD31 | Angptl4 | Cdh11 | 0.001 | 0 | ANGPTL4_CDH11 | Angptl4 - Cdh11 | ANGPTL | Secreted Signaling | PMID: 30049845 |
| CD31 | NeuN | Angptl4 | Cdh11 | 0.001777 | 0 | ANGPTL4_CDH11 | Angptl4 - Cdh11 | ANGPTL | Secreted Signaling | PMID: 30049845 |
| CD31 | CD31 | Angptl4 | Sdc2 | 0.001011 | 0 | ANGPTL4_SDC2 | Angptl4 - Sdc2 | ANGPTL | Secreted Signaling | PMID: 29017031 |
| CD31 | GFAP | Angptl4 | Sdc2 | 0.000779 | 0 | ANGPTL4_SDC2 | Angptl4 - Sdc2 | ANGPTL | Secreted Signaling | PMID: 29017031 |
| CD31 | CD31 | Angptl4 | Sdc3 | 0.002002 | 0 | ANGPTL4_SDC3 | Angptl4 - Sdc3 | ANGPTL | Secreted Signaling | PMID: 29017031 |
| CD31 | NeuN | Angptl4 | Sdc3 | 0.001468 | 0 | ANGPTL4_SDC3 | Angptl4 - Sdc3 | ANGPTL | Secreted Signaling | PMID: 29017031 |
| CD31 | GFAP | Angptl4 | Sdc4 | 0.001986 | 0 | ANGPTL4_SDC4 | Angptl4 - Sdc4 | ANGPTL | Secreted Signaling | PMID: 29017031 |
| GFAP | GFAP | Angptl4 | Sdc4 | 0.0011 | 0 | ANGPTL4_SDC4 | Angptl4 - Sdc4 | ANGPTL | Secreted Signaling | PMID: 29017031 |
| CD31 | CD31 | Angptl3 | ITGAV_ITGB3 | 0.000367 | 0 | ANGPTL3_ITGAV_ITGB3 | Angptl3 - (Itgav+Itgb3) | ANGPTL | Secreted Signaling | PMID: 11877390 |
| CD31 | GFAP | Angptl3 | ITGAV_ITGB3 | 0.000427 | 0 | ANGPTL3_ITGAV_ITGB3 | Angptl3 - (Itgav+Itgb3) | ANGPTL | Secreted Signaling | PMID: 11877390 |
| GFAP | CD31 | Mdk | Sdc2 | 0.00164 | 0 | MDK_SDC2 | Mdk - Sdc2 | MK | Secreted Signaling | PMID: 28356350 |
| GFAP | GFAP | Mdk | Sdc2 | 0.001263 | 0 | MDK_SDC2 | Mdk - Sdc2 | MK | Secreted Signaling | PMID: 28356350 |
| CD31 | GFAP | Mdk | Sdc4 | 0.00231 | 0 | MDK_SDC4 | Mdk - Sdc4 | MK | Secreted Signaling | PMID: 28356350 |
| GFAP | GFAP | Mdk | Sdc4 | 0.003217 | 0 | MDK_SDC4 | Mdk - Sdc4 | MK | Secreted Signaling | PMID: 28356350 |
| CD31 | GFAP | Mdk | Ptprz1 | 0.005863 | 0 | MDK_PTPRZ1 | Mdk - Ptprz1 | MK | Secreted Signaling | PMID: 28356350 |
| GFAP | GFAP | Mdk | Ptprz1 | 0.008156 | 0 | MDK_PTPRZ1 | Mdk - Ptprz1 | MK | Secreted Signaling | PMID: 28356350 |
| GFAP | CD31 | Mdk | ITGA4_ITGB1 | 0.001489 | 0 | MDK_ITGA4_ITGB1 | Mdk - (Itga4+Itgb1) | MK | Secreted Signaling | PMID: 28356350 |
| GFAP | GFAP | Mdk | ITGA4_ITGB1 | 0.001171 | 0 | MDK_ITGA4_ITGB1 | Mdk - (Itga4+Itgb1) | MK | Secreted Signaling | PMID: 28356350 |
| CD31 | CD31 | Mdk | ITGA6_ITGB1 | 0.001793 | 0 | MDK_ITGA6_ITGB1 | Mdk - (Itga6+Itgb1) | MK | Secreted Signaling | PMID: 28356350 |
| GFAP | CD31 | Mdk | ITGA6_ITGB1 | 0.002498 | 0 | MDK_ITGA6_ITGB1 | Mdk - (Itga6+Itgb1) | MK | Secreted Signaling | PMID: 28356350 |
| GFAP | GFAP | Mdk | Lrp1 | 0.001229 | 0 | MDK_LRP1 | Mdk - Lrp1 | MK | Secreted Signaling | PMID: 28356350 |
| GFAP | NeuN | Mdk | Lrp1 | 0.001717 | 0 | MDK_LRP1 | Mdk - Lrp1 | MK | Secreted Signaling | PMID: 28356350 |
| GFAP | GFAP | Mdk | Ncl | 0.003418 | 0 | MDK_NCL | Mdk - Ncl | MK | Secreted Signaling | PMID: 28356350 |
| GFAP | NeuN | Mdk | Ncl | 0.00377 | 0 | MDK_NCL | Mdk - Ncl | MK | Secreted Signaling | PMID: 28356350 |
| CD31 | GFAP | Ptn | Ptprz1 | 0.07704 | 0 | PTN_PTPRZ1 | Ptn - Ptprz1 | PTN | Secreted Signaling | PMID: 28356350; PMID: 25620911 |
| GFAP | GFAP | Ptn | Ptprz1 | 0.092201 | 0 | PTN_PTPRZ1 | Ptn - Ptprz1 | PTN | Secreted Signaling | PMID: 28356350; PMID: 25620911 |
| CD31 | CD31 | Ptn | Sdc2 | 0.016398 | 0 | PTN_SDC2 | Ptn - Sdc2 | PTN | Secreted Signaling | PMID: 28356350; PMID: 25620911 |
| GFAP | CD31 | Ptn | Sdc2 | 0.019882 | 0 | PTN_SDC2 | Ptn - Sdc2 | PTN | Secreted Signaling | PMID: 28356350; PMID: 25620911 |
| CD31 | CD31 | Ptn | Sdc3 | 0.03198 | 0 | PTN_SDC3 | Ptn - Sdc3 | PTN | Secreted Signaling | PMID: 28356350; PMID: 25620911 |
| GFAP | CD31 | Ptn | Sdc3 | 0.038645 | 0 | PTN_SDC3 | Ptn - Sdc3 | PTN | Secreted Signaling | PMID: 28356350; PMID: 25620911 |
| CD31 | GFAP | Ptn | Sdc4 | 0.031724 | 0 | PTN_SDC4 | Ptn - Sdc4 | PTN | Secreted Signaling | PMID: 28356350; PMID: 25620911 |
| GFAP | GFAP | Ptn | Sdc4 | 0.038337 | 0 | PTN_SDC4 | Ptn - Sdc4 | PTN | Secreted Signaling | PMID: 28356350; PMID: 25620911 |
| GFAP | CD31 | Ptn | ITGAV_ITGB3 | 0.011641 | 0 | PTN_ITGAV_ITGB3 | Ptn - (Itgav+Itgb3) | PTN | Secreted Signaling | PMID: 28356350; PMID: 25620911 |
| GFAP | GFAP | Ptn | ITGAV_ITGB3 | 0.013523 | 0 | PTN_ITGAV_ITGB3 | Ptn - (Itgav+Itgb3) | PTN | Secreted Signaling | PMID: 28356350; PMID: 25620911 |
| GFAP | GFAP | Ptn | Ncl | 0.04064 | 0 | PTN_NCL | Ptn - Ncl | PTN | Secreted Signaling | PMID: 28356350; PMID: 25620911 |
| GFAP | NeuN | Ptn | Ncl | 0.044646 | 0 | PTN_NCL | Ptn - Ncl | PTN | Secreted Signaling | PMID: 28356350; PMID: 25620911 |
| CD31 | GFAP | Postn | ITGAV_ITGB3 | 0.000963 | 0 | POSTN_ITGAV_ITGB3 | Postn - (Itgav+Itgb3) | PERIOSTIN | Secreted Signaling | PMID: 26721738 |
| GFAP | GFAP | Postn | ITGAV_ITGB3 | 0.000868 | 0 | POSTN_ITGAV_ITGB3 | Postn - (Itgav+Itgb3) | PERIOSTIN | Secreted Signaling | PMID: 26721738 |
| CD31 | GFAP | Postn | ITGAV_ITGB5 | 0.001205 | 0 | POSTN_ITGAV_ITGB5 | Postn - (Itgav+Itgb5) | PERIOSTIN | Secreted Signaling | PMID: 26721738 |
| GFAP | GFAP | Postn | ITGAV_ITGB5 | 0.001086 | 0 | POSTN_ITGAV_ITGB5 | Postn - (Itgav+Itgb5) | PERIOSTIN | Secreted Signaling | PMID: 26721738 |
| CD31 | GFAP | Nmb | Nmbr | 0.000284 | 0 | NMB_NMBR | Nmb - Nmbr | GPR | Secreted Signaling | KEGG: mmu04080 |
| GFAP | GFAP | Nmb | Nmbr | 0.000537 | 0 | NMB_NMBR | Nmb - Nmbr | GPR | Secreted Signaling | KEGG: mmu04080 |
| NeuN | CD31 | Cck | Cckbr | 0.007789 | 0 | CCK_CCKBR | Cck - Cckbr | CCK | Secreted Signaling | KEGG: mmu04080 |
| NeuN | NeuN | Cck | Cckbr | 0.023805 | 0 | CCK_CCKBR | Cck - Cckbr | CCK | Secreted Signaling | KEGG: mmu04080 |
| CD31 | CD31 | Edn1 | Ednra | 0.001174 | 0 | EDN1_EDNRA | Edn1 - Ednra | EDN | Secreted Signaling | KEGG: mmu04080 |
| CD31 | GFAP | Edn1 | Ednra | 0.001414 | 0 | EDN1_EDNRA | Edn1 - Ednra | EDN | Secreted Signaling | KEGG: mmu04080 |
| CD31 | CD31 | Edn3 | Ednra | 0.000889 | 0 | EDN3_EDNRA | Edn3 - Ednra | EDN | Secreted Signaling | KEGG: mmu04080 |
| CD31 | GFAP | Edn3 | Ednra | 0.001071 | 0 | EDN3_EDNRA | Edn3 - Ednra | EDN | Secreted Signaling | KEGG: mmu04080 |
| CD31 | CD31 | Edn1 | Ednrb | 0.00158 | 0 | EDN1_EDNRB | Edn1 - Ednrb | EDN | Secreted Signaling | KEGG: mmu04080 |
| CD31 | GFAP | Edn1 | Ednrb | 0.004652 | 0 | EDN1_EDNRB | Edn1 - Ednrb | EDN | Secreted Signaling | KEGG: mmu04080 |
| CD31 | GFAP | Edn3 | Ednrb | 0.003525 | 0 | EDN3_EDNRB | Edn3 - Ednrb | EDN | Secreted Signaling | KEGG: mmu04080 |
| GFAP | GFAP | Edn3 | Ednrb | 0.00128 | 0 | EDN3_EDNRB | Edn3 - Ednrb | EDN | Secreted Signaling | KEGG: mmu04080 |
| CD31 | CD31 | Nms | Nmur1 | 0.000413 | 0 | NMS_NMUR1 | Nms - Nmur1 | NMU | Secreted Signaling | PMID: 29288685 |
| GFAP | CD31 | Nms | Nmur1 | 0.000288 | 0 | NMS_NMUR1 | Nms - Nmur1 | NMU | Secreted Signaling | PMID: 29288685 |

| | | | | | | | | | | |
|---|---|---|---|---|---|---|---|---|---|---|
| GFAP | NeuN | Npy | Npy1r | 0.000887 | 0 | NPY_NPY1R | Npy - Npy1r | NPY | Secreted Signaling | KEGG: mmu04080 |
| NeuN | NeuN | Npy | Npy1r | 0.001153 | 0 | NPY_NPY1R | Npy - Npy1r | NPY | Secreted Signaling | KEGG: mmu04080 |
| GFAP | GFAP | Npy | Npy2r | 0.000711 | 0 | NPY_NPY2R | Npy - Npy2r | NPY | Secreted Signaling | KEGG: mmu04080 |
| NeuN | GFAP | Npy | Npy2r | 0.000925 | 0 | NPY_NPY2R | Npy - Npy2r | NPY | Secreted Signaling | KEGG: mmu04080 |
| NeuN | CD31 | Npy | Npy5r | 0.000912 | 0 | NPY_NPY5R | Npy - Npy5r | NPY | Secreted Signaling | KEGG: mmu04080 |
| GFAP | GFAP | Npy | Npy5r | 0.000558 | 0 | NPY_NPY5R | Npy - Npy5r | NPY | Secreted Signaling | KEGG: mmu04080 |
| NeuN | CD31 | Npy | Npy6r | 0.000794 | 0 | NPY_NPY6R | Npy - Npy6r | NPY | Secreted Signaling | KEGG: mmu04080 |
| GFAP | GFAP | Npy | Npy6r | 0.000361 | 0 | NPY_NPY6R | Npy - Npy6r | NPY | Secreted Signaling | KEGG: mmu04080 |
| GFAP | CD31 | Npb | Npbwr1 | 0.000382 | 0 | NPB_NPBWR1 | Npb - Npbwr1 | NPW-B | Secreted Signaling | KEGG: mmu04080 |
| GFAP | GFAP | Npb | Npbwr1 | 0.000474 | 0 | NPB_NPBWR1 | Npb - Npbwr1 | NPW-B | Secreted Signaling | KEGG: mmu04080 |
| CD31 | CD31 | Nts | Ntsr1 | 0.000208 | 0 | NTS_NTSR1 | Nts - Ntsr1 | NTS | Secreted Signaling | KEGG: mmu04080 |
| CD31 | NeuN | Nts | Ntsr1 | 0.000199 | 0 | NTS_NTSR1 | Nts - Ntsr1 | NTS | Secreted Signaling | KEGG: mmu04080 |
| CD31 | GFAP | Nts | Ntsr2 | 0.00096 | 0 | NTS_NTSR2 | Nts - Ntsr2 | NTS | Secreted Signaling | KEGG: mmu04080 |
| GFAP | GFAP | Nts | Ntsr2 | 0.000658 | 0 | NTS_NTSR2 | Nts - Ntsr2 | NTS | Secreted Signaling | KEGG: mmu04080 |
| CD31 | CD31 | Pdyn | Oprd1 | 0.000409 | 0 | PDYN_OPRD1 | Pdyn - Oprd1 | OPIOID | Secreted Signaling | KEGG: mmu04080 |
| GFAP | CD31 | Pdyn | Oprd1 | 0.000591 | 0 | PDYN_OPRD1 | Pdyn - Oprd1 | OPIOID | Secreted Signaling | KEGG: mmu04080 |
| GFAP | CD31 | Pdyn | Oprk1 | 0.000512 | 0 | PDYN_OPRK1 | Pdyn - Oprk1 | OPIOID | Secreted Signaling | KEGG: mmu04080 |
| GFAP | GFAP | Pdyn | Oprk1 | 0.000484 | 0 | PDYN_OPRK1 | Pdyn - Oprk1 | OPIOID | Secreted Signaling | KEGG: mmu04080 |
| GFAP | NeuN | Pdyn | Oprl1 | 0.000912 | 0 | PDYN_OPRL1 | Pdyn - Oprl1 | OPIOID | Secreted Signaling | KEGG: mmu04080 |
| NeuN | NeuN | Pdyn | Oprl1 | 0.000655 | 0 | PDYN_OPRL1 | Pdyn - Oprl1 | OPIOID | Secreted Signaling | KEGG: mmu04080 |
| GFAP | CD31 | Pdyn | Oprm1 | 0.000478 | 0 | PDYN_OPRM1 | Pdyn - Oprm1 | OPIOID | Secreted Signaling | KEGG: mmu04080 |
| GFAP | GFAP | Pdyn | Oprm1 | 0.000437 | 0 | PDYN_OPRM1 | Pdyn - Oprm1 | OPIOID | Secreted Signaling | KEGG: mmu04080 |
| CD31 | CD31 | Penk | Oprd1 | 0.000599 | 0 | PENK_OPRD1 | Penk - Oprd1 | OPIOID | Secreted Signaling | KEGG: mmu04080 |
| NeuN | CD31 | Penk | Oprd1 | 0.000583 | 0 | PENK_OPRD1 | Penk - Oprd1 | OPIOID | Secreted Signaling | KEGG: mmu04080 |
| CD31 | CD31 | Penk | Oprk1 | 0.000519 | 0 | PENK_OPRK1 | Penk - Oprk1 | OPIOID | Secreted Signaling | KEGG: mmu04080 |
| NeuN | CD31 | Penk | Oprk1 | 0.000505 | 0 | PENK_OPRK1 | Penk - Oprk1 | OPIOID | Secreted Signaling | KEGG: mmu04080 |
| CD31 | NeuN | Penk | Oprl1 | 0.000924 | 0 | PENK_OPRL1 | Penk - Oprl1 | OPIOID | Secreted Signaling | KEGG: mmu04080 |
| NeuN | NeuN | Penk | Oprl1 | 0.000899 | 0 | PENK_OPRL1 | Penk - Oprl1 | OPIOID | Secreted Signaling | KEGG: mmu04080 |
| CD31 | CD31 | Penk | Oprm1 | 0.000485 | 0 | PENK_OPRM1 | Penk - Oprm1 | OPIOID | Secreted Signaling | KEGG: mmu04080 |
| NeuN | CD31 | Penk | Oprm1 | 0.000472 | 0 | PENK_OPRM1 | Penk - Oprm1 | OPIOID | Secreted Signaling | KEGG: mmu04080 |
| CD31 | GFAP | Oxt | Oxtr | 0.000393 | 0 | OXT_OXTR | Oxt - Oxtr | OXT | Secreted Signaling | KEGG: mmu04080 |
| GFAP | GFAP | Oxt | Oxtr | 0.000387 | 0 | OXT_OXTR | Oxt - Oxtr | OXT | Secreted Signaling | KEGG: mmu04080 |
| NeuN | CD31 | Sst | Sstr1 | 0.004727 | 0 | SST_SSTR1 | Sst - Sstr1 | SOMATOSTATIN | Secreted Signaling | KEGG: mmu04080 |
| NeuN | NeuN | Sst | Sstr1 | 0.00515 | 0 | SST_SSTR1 | Sst - Sstr1 | SOMATOSTATIN | Secreted Signaling | KEGG: mmu04080 |
| GFAP | NeuN | Sst | Sstr2 | 0.004129 | 0 | SST_SSTR2 | Sst - Sstr2 | SOMATOSTATIN | Secreted Signaling | KEGG: mmu04080 |
| NeuN | NeuN | Sst | Sstr2 | 0.00619 | 0 | SST_SSTR2 | Sst - Sstr2 | SOMATOSTATIN | Secreted Signaling | KEGG: mmu04080 |
| NeuN | GFAP | Sst | Sstr3 | 0.002871 | 0 | SST_SSTR3 | Sst - Sstr3 | SOMATOSTATIN | Secreted Signaling | KEGG: mmu04080 |
| NeuN | NeuN | Sst | Sstr3 | 0.003611 | 0 | SST_SSTR3 | Sst - Sstr3 | SOMATOSTATIN | Secreted Signaling | KEGG: mmu04080 |
| GFAP | NeuN | Sst | Sstr4 | 0.009678 | 0 | SST_SSTR4 | Sst - Sstr4 | SOMATOSTATIN | Secreted Signaling | KEGG: mmu04080 |
| NeuN | NeuN | Sst | Sstr4 | 0.014468 | 0 | SST_SSTR4 | Sst - Sstr4 | SOMATOSTATIN | Secreted Signaling | KEGG: mmu04080 |
| CD31 | CD31 | Cort | Sstr1 | 0.000752 | 0 | CORT_SSTR1 | Cort - Sstr1 | SOMATOSTATIN | Secreted Signaling | KEGG: mmu04080 |
| CD31 | NeuN | Cort | Sstr1 | 0.000819 | 0 | CORT_SSTR1 | Cort - Sstr1 | SOMATOSTATIN | Secreted Signaling | KEGG: mmu04080 |
| CD31 | CD31 | Cort | Sstr2 | 0.0006 | 0 | CORT_SSTR2 | Cort - Sstr2 | SOMATOSTATIN | Secreted Signaling | KEGG: mmu04080 |
| CD31 | NeuN | Cort | Sstr2 | 0.000986 | 0 | CORT_SSTR2 | Cort - Sstr2 | SOMATOSTATIN | Secreted Signaling | KEGG: mmu04080 |
| CD31 | CD31 | Cort | Sstr3 | 0.000447 | 0 | CORT_SSTR3 | Cort - Sstr3 | SOMATOSTATIN | Secreted Signaling | KEGG: mmu04080 |
| CD31 | NeuN | Cort | Sstr3 | 0.000574 | 0 | CORT_SSTR3 | Cort - Sstr3 | SOMATOSTATIN | Secreted Signaling | KEGG: mmu04080 |
| CD31 | NeuN | Cort | Sstr4 | 0.00232 | 0 | CORT_SSTR4 | Cort - Sstr4 | SOMATOSTATIN | Secreted Signaling | KEGG: mmu04080 |
| NeuN | NeuN | Cort | Sstr4 | 0.001226 | 0 | CORT_SSTR4 | Cort - Sstr4 | SOMATOSTATIN | Secreted Signaling | KEGG: mmu04080 |
| CD31 | GFAP | Tac1 | Tacr1 | 0.000518 | 0 | TAC1_TACR1 | Tac1 - Tacr1 | TAC | Secreted Signaling | KEGG: mmu04080 |
| GFAP | GFAP | Tac1 | Tacr1 | 0.000402 | 0 | TAC1_TACR1 | Tac1 - Tacr1 | TAC | Secreted Signaling | KEGG: mmu04080 |
| CD31 | GFAP | Tac2 | Tacr3 | 0.000303 | 0 | TAC2_TACR3 | Tac2 - Tacr3 | TAC | Secreted Signaling | KEGG: mmu04080 |
| GFAP | GFAP | Tac2 | Tacr3 | 0.000414 | 0 | TAC2_TACR3 | Tac2 - Tacr3 | TAC | Secreted Signaling | KEGG: mmu04080 |
| CD31 | CD31 | Ctsg | F2r | 0.000313 | 0 | CTSG_F2R | Ctsg - F2r | PARs | Secreted Signaling | KEGG: mmu04080 |
| GFAP | CD31 | Ctsg | F2r | 0.000302 | 0 | CTSG_F2R | Ctsg - F2r | PARs | Secreted Signaling | KEGG: mmu04080 |
| GFAP | CD31 | F2 | F2r | 0.000245 | 0 | F2_F2R | F2 - F2r | PARs | Secreted Signaling | KEGG: mmu04080 |
| GFAP | GFAP | F2 | F2r | 0.000177 | 0 | F2_F2R | F2 - F2r | PARs | Secreted Signaling | KEGG: mmu04080 |
| CD31 | CD31 | Ctsg | F2rl2 | 0.000312 | 0 | CTSG_F2RL2 | Ctsg - F2rl2 | PARs | Secreted Signaling | KEGG: mmu04080 |
| GFAP | CD31 | Ctsg | F2rl2 | 0.000301 | 0 | CTSG_F2RL2 | Ctsg - F2rl2 | PARs | Secreted Signaling | KEGG: mmu04080 |
| GFAP | CD31 | F2 | F2rl2 | 0.000245 | 0 | F2_F2RL2 | F2 - F2rl2 | PARs | Secreted Signaling | KEGG: mmu04080 |

| | | | | | | | | | | |
|---|---|---|---|---|---|---|---|---|---|---|
| GFAP | GFAP | F2 | F2rl2 | 0.000208 | 0 | F2_F2RL2 | F2 - F2rl2 | PARs | Secreted Signaling | KEGG: mmu04080 |
| CD31 | GFAP | Ctsg | Pard3 | 0.000404 | 0 | CTSG_PARD3 | Ctsg - Pard3 | PARs | Secreted Signaling | KEGG: mmu04080 |
| GFAP | GFAP | Ctsg | Pard3 | 0.00039 | 0 | CTSG_PARD3 | Ctsg - Pard3 | PARs | Secreted Signaling | KEGG: mmu04080 |
| GFAP | CD31 | F2 | Pard3 | 0.000291 | 0 | F2_PARD3 | F2 - Pard3 | PARs | Secreted Signaling | KEGG: mmu04080 |
| GFAP | GFAP | F2 | Pard3 | 0.000317 | 0 | F2_PARD3 | F2 - Pard3 | PARs | Secreted Signaling | KEGG: mmu04080 |
| CD31 | GFAP | Plg | Pard3 | 0.000473 | 0 | PLG_PARD3 | Plg - Pard3 | PARs | Secreted Signaling | KEGG: mmu04080 |
| GFAP | GFAP | Plg | Pard3 | 0.000488 | 0 | PLG_PARD3 | Plg - Pard3 | PARs | Secreted Signaling | KEGG: mmu04080 |
| CD31 | CD31 | Pmch | Mchr1 | 0.000406 | 0 | PMCH_MCHR1 | Pmch - Mchr1 | PMCH | Secreted Signaling | KEGG: mmu04080 |
| CD31 | GFAP | Pmch | Mchr1 | 0.000347 | 0 | PMCH_MCHR1 | Pmch - Mchr1 | PMCH | Secreted Signaling | KEGG: mmu04080 |
| CD31 | CD31 | Prok2 | Prokr1 | 0.000221 | 0 | PROK2_PROKR1 | Prok2 - Prokr1 | PROK | Secreted Signaling | PMID: 24496457 |
| GFAP | CD31 | Prok2 | Prokr1 | 0.000306 | 0 | PROK2_PROKR1 | Prok2 - Prokr1 | PROK | Secreted Signaling | PMID: 24496457 |
| GFAP | CD31 | Prok2 | Prokr2 | 0.000285 | 0 | PROK2_PROKR2 | Prok2 - Prokr2 | PROK | Secreted Signaling | PMID: 24496457 |
| GFAP | GFAP | Prok2 | Prokr2 | 0.000236 | 0 | PROK2_PROKR2 | Prok2 - Prokr2 | PROK | Secreted Signaling | PMID: 24496457 |
| NeuN | GFAP | Adcyap1 | Adcyap1r1 | 0.001673 | 0 | ADCYAP1_ADCYAP1R1 | Adcyap1 - Adcyap1r1 | PACAP | Secreted Signaling | PMID: 21350482 |
| NeuN | NeuN | Adcyap1 | Adcyap1r1 | 0.000761 | 0 | ADCYAP1_ADCYAP1R1 | Adcyap1 - Adcyap1r1 | PACAP | Secreted Signaling | PMID: 21350482 |
| CD31 | NeuN | Vip | Vipr1 | 0.000321 | 0 | VIP_VIPR1 | Vip - Vipr1 | VIP | Secreted Signaling | PMID: 10655512 |
| NeuN | NeuN | Vip | Vipr1 | 0.000196 | 0 | VIP_VIPR1 | Vip - Vipr1 | VIP | Secreted Signaling | PMID: 10655512 |
| CD31 | CD31 | Nppa | Npr1 | 0.000675 | 0 | NPPA_NPR1 | Nppa - Npr1 | NPR1 | Secreted Signaling | KEGG: mmu04080; PMID:29934640 |
| NeuN | CD31 | Nppa | Npr1 | 0.001209 | 0 | NPPA_NPR1 | Nppa - Npr1 | NPR1 | Secreted Signaling | KEGG: mmu04080; PMID:29934640 |
| CD31 | CD31 | Nppb | Npr1 | 0.000221 | 0 | NPPB_NPR1 | Nppb - Npr1 | NPR1 | Secreted Signaling | KEGG: mmu04080; PMID:29934640 |
| GFAP | CD31 | Nppb | Npr1 | 0.000401 | 0 | NPPB_NPR1 | Nppb - Npr1 | NPR1 | Secreted Signaling | KEGG: mmu04080; PMID:29934640 |
| NeuN | GFAP | Nppc | Npr2 | 0.000649 | 0 | NPPC_NPR2 | Nppc - Npr2 | NPR2 | Secreted Signaling | KEGG: mmu04080; PMID:29934640 |
| NeuN | NeuN | Nppc | Npr2 | 0.000622 | 0 | NPPC_NPR2 | Nppc - Npr2 | NPR2 | Secreted Signaling | KEGG: mmu04080; PMID:29934640 |
| CD31 | CD31 | Kitl | Kit | 0.000732 | 0 | KITL_KIT | Kitl - Kit | KIT | Secreted Signaling | KEGG: mmu04080 |
| CD31 | GFAP | Kitl | Kit | 0.001469 | 0 | KITL_KIT | Kitl - Kit | KIT | Secreted Signaling | KEGG: mmu04080 |
| CD31 | GFAP | Gip | Gipr | 0.000227 | 0 | GIP_GIPR | Gip - Gipr | GIPR | Secreted Signaling | KEGG: mmu04080 |
| GFAP | GFAP | Gip | Gipr | 0.000294 | 0 | GIP_GIPR | Gip - Gipr | GIPR | Secreted Signaling | KEGG: mmu04080 |
| NeuN | GFAP | Bdnf | Ntrk2 | 0.010261 | 0 | BDNF_NTRK2 | Bdnf - Ntrk2 | NT | Secreted Signaling | PMID: 27908981 |
| NeuN | NeuN | Bdnf | Ntrk2 | 0.008953 | 0 | BDNF_NTRK2 | Bdnf - Ntrk2 | NT | Secreted Signaling | PMID: 27908981 |
| CD31 | CD31 | Flt3l | Flt3 | 0.000455 | 0 | FLT3L_FLT3 | Flt3l - Flt3 | FLT3 | Secreted Signaling | PMID: 25992210 |
| GFAP | CD31 | Flt3l | Flt3 | 0.000481 | 0 | FLT3L_FLT3 | Flt3l - Flt3 | FLT3 | Secreted Signaling | PMID: 25992210 |
| CD31 | CD31 | Hgf | Met | 0.000436 | 0 | HGF_MET | Hgf - Met | HGF | Secreted Signaling | PMID: 28475121 |
| CD31 | NeuN | Hgf | Met | 0.000386 | 0 | HGF_MET | Hgf - Met | HGF | Secreted Signaling | PMID: 28475121 |
| CD31 | GFAP | Ntf3 | Ntrk1 | 0.000284 | 0 | NTF3_NTRK1 | Ntf3 - Ntrk1 | NT | Secreted Signaling | PMID: 27908981 |
| GFAP | GFAP | Ntf3 | Ntrk1 | 0.000255 | 0 | NTF3_NTRK1 | Ntf3 - Ntrk1 | NT | Secreted Signaling | PMID: 27908981 |
| CD31 | GFAP | Ntf3 | Ntrk2 | 0.005661 | 0 | NTF3_NTRK2 | Ntf3 - Ntrk2 | NT | Secreted Signaling | PMID: 27908981 |
| GFAP | GFAP | Ntf3 | Ntrk2 | 0.005094 | 0 | NTF3_NTRK2 | Ntf3 - Ntrk2 | NT | Secreted Signaling | PMID: 27908981 |
| CD31 | NeuN | Ntf3 | Ntrk3 | 0.002908 | 0 | NTF3_NTRK3 | Ntf3 - Ntrk3 | NT | Secreted Signaling | PMID: 27908981 |
| NeuN | NeuN | Ntf3 | Ntrk3 | 0.000952 | 0 | NTF3_NTRK3 | Ntf3 - Ntrk3 | NT | Secreted Signaling | PMID: 27908981 |
| NeuN | CD31 | Sema3a | NRP1_PLXNA1 | 0.000982 | 0 | SEMA3A_NRP1_PLXNA1 | Sema3a - (Nrp1+Plxna1) | SEMA3 | Secreted Signaling | PMID: 27533782 |
| NeuN | NeuN | Sema3a | NRP1_PLXNA1 | 0.000872 | 0 | SEMA3A_NRP1_PLXNA1 | Sema3a - (Nrp1+Plxna1) | SEMA3 | Secreted Signaling | PMID: 27533782 |
| NeuN | CD31 | Sema3a | NRP1_PLXNA2 | 0.00145 | 0 | SEMA3A_NRP1_PLXNA2 | Sema3a - (Nrp1+Plxna2) | SEMA3 | Secreted Signaling | PMID: 27533782 |
| NeuN | NeuN | Sema3a | NRP1_PLXNA2 | 0.001058 | 0 | SEMA3A_NRP1_PLXNA2 | Sema3a - (Nrp1+Plxna2) | SEMA3 | Secreted Signaling | PMID: 27533782 |
| NeuN | CD31 | Sema3a | NRP1_PLXNA3 | 0.000606 | 0 | SEMA3A_NRP1_PLXNA3 | Sema3a - (Nrp1+Plxna3) | SEMA3 | Secreted Signaling | PMID: 27533782 |
| NeuN | NeuN | Sema3a | NRP1_PLXNA3 | 0.000442 | 0 | SEMA3A_NRP1_PLXNA3 | Sema3a - (Nrp1+Plxna3) | SEMA3 | Secreted Signaling | PMID: 27533782 |
| NeuN | CD31 | Sema3a | NRP1_PLXNA4 | 0.001167 | 0 | SEMA3A_NRP1_PLXNA4 | Sema3a - (Nrp1+Plxna4) | SEMA3 | Secreted Signaling | PMID: 27533782 |
| NeuN | NeuN | Sema3a | NRP1_PLXNA4 | 0.00103 | 0 | SEMA3A_NRP1_PLXNA4 | Sema3a - (Nrp1+Plxna4) | SEMA3 | Secreted Signaling | PMID: 27533782 |
| CD31 | CD31 | Sema3b | NRP1_PLXNA1 | 0.000608 | 0 | SEMA3B_NRP1_PLXNA1 | Sema3b - (Nrp1+Plxna1) | SEMA3 | Secreted Signaling | PMID: 27533782 |
| GFAP | CD31 | Sema3b | NRP1_PLXNA1 | 0.000681 | 0 | SEMA3B_NRP1_PLXNA1 | Sema3b - (Nrp1+Plxna1) | SEMA3 | Secreted Signaling | PMID: 27533782 |
| CD31 | CD31 | Sema3b | NRP1_PLXNA2 | 0.000897 | 0 | SEMA3B_NRP1_PLXNA2 | Sema3b - (Nrp1+Plxna2) | SEMA3 | Secreted Signaling | PMID: 27533782 |
| GFAP | CD31 | Sema3b | NRP1_PLXNA2 | 0.001005 | 0 | SEMA3B_NRP1_PLXNA2 | Sema3b - (Nrp1+Plxna2) | SEMA3 | Secreted Signaling | PMID: 27533782 |
| CD31 | CD31 | Sema3b | NRP1_PLXNA3 | 0.000375 | 0 | SEMA3B_NRP1_PLXNA3 | Sema3b - (Nrp1+Plxna3) | SEMA3 | Secreted Signaling | PMID: 27533782 |
| GFAP | CD31 | Sema3b | NRP1_PLXNA3 | 0.00042 | 0 | SEMA3B_NRP1_PLXNA3 | Sema3b - (Nrp1+Plxna3) | SEMA3 | Secreted Signaling | PMID: 27533782 |
| CD31 | CD31 | Sema3b | NRP1_PLXNA4 | 0.000722 | 0 | SEMA3B_NRP1_PLXNA4 | Sema3b - (Nrp1+Plxna4) | SEMA3 | Secreted Signaling | PMID: 27533782 |
| GFAP | CD31 | Sema3b | NRP1_PLXNA4 | 0.000809 | 0 | SEMA3B_NRP1_PLXNA4 | Sema3b - (Nrp1+Plxna4) | SEMA3 | Secreted Signaling | PMID: 27533782 |
| CD31 | CD31 | Sema3c | NRP1_PLXNA1 | 0.001589 | 0 | SEMA3C_NRP1_PLXNA1 | Sema3c - (Nrp1+Plxna1) | SEMA3 | Secreted Signaling | PMID: 27533782 |
| CD31 | NeuN | Sema3c | NRP1_PLXNA1 | 0.001411 | 0 | SEMA3C_NRP1_PLXNA1 | Sema3c - (Nrp1+Plxna1) | SEMA3 | Secreted Signaling | PMID: 27533782 |
| CD31 | CD31 | Sema3c | NRP1_PLXNA2 | 0.002345 | 0 | SEMA3C_NRP1_PLXNA2 | Sema3c - (Nrp1+Plxna2) | SEMA3 | Secreted Signaling | PMID: 27533782 |
| CD31 | NeuN | Sema3c | NRP1_PLXNA2 | 0.001711 | 0 | SEMA3C_NRP1_PLXNA2 | Sema3c - (Nrp1+Plxna2) | SEMA3 | Secreted Signaling | PMID: 27533782 |

| | | | | | | | | | | |
|---|---|---|---|---|---|---|---|---|---|---|
| CD31 | CD31 | Sema3c | NRP1_PLXNA3 | 0.000981 | 0 | SEMA3C_NRP1_PLXNA3 | Sema3c - (Nrp1+Plxna3) | SEMA3 | Secreted Signaling | PMID: 27533782 |
| CD31 | GFAP | Sema3c | NRP1_PLXNA3 | 0.000791 | 0 | SEMA3C_NRP1_PLXNA3 | Sema3c - (Nrp1+Plxna3) | SEMA3 | Secreted Signaling | PMID: 27533782 |
| CD31 | CD31 | Sema3c | NRP1_PLXNA4 | 0.001889 | 0 | SEMA3C_NRP1_PLXNA4 | Sema3c - (Nrp1+Plxna4) | SEMA3 | Secreted Signaling | PMID: 27533782 |
| CD31 | NeuN | Sema3c | NRP1_PLXNA4 | 0.001666 | 0 | SEMA3C_NRP1_PLXNA4 | Sema3c - (Nrp1+Plxna4) | SEMA3 | Secreted Signaling | PMID: 27533782 |
| CD31 | CD31 | Sema3d | NRP1_PLXNA1 | 0.000658 | 0 | SEMA3D_NRP1_PLXNA1 | Sema3d - (Nrp1+Plxna1) | SEMA3 | Secreted Signaling | PMID: 27533782 |
| GFAP | CD31 | Sema3d | NRP1_PLXNA1 | 0.001025 | 0 | SEMA3D_NRP1_PLXNA1 | Sema3d - (Nrp1+Plxna1) | SEMA3 | Secreted Signaling | PMID: 27533782 |
| CD31 | CD31 | Sema3d | NRP1_PLXNA2 | 0.000972 | 0 | SEMA3D_NRP1_PLXNA2 | Sema3d - (Nrp1+Plxna2) | SEMA3 | Secreted Signaling | PMID: 27533782 |
| GFAP | CD31 | Sema3d | NRP1_PLXNA2 | 0.001512 | 0 | SEMA3D_NRP1_PLXNA2 | Sema3d - (Nrp1+Plxna2) | SEMA3 | Secreted Signaling | PMID: 27533782 |
| GFAP | CD31 | Sema3d | NRP1_PLXNA3 | 0.000632 | 0 | SEMA3D_NRP1_PLXNA3 | Sema3d - (Nrp1+Plxna3) | SEMA3 | Secreted Signaling | PMID: 27533782 |
| GFAP | GFAP | Sema3d | NRP1_PLXNA3 | 0.00051 | 0 | SEMA3D_NRP1_PLXNA3 | Sema3d - (Nrp1+Plxna3) | SEMA3 | Secreted Signaling | PMID: 27533782 |
| CD31 | CD31 | Sema3d | NRP1_PLXNA4 | 0.000782 | 0 | SEMA3D_NRP1_PLXNA4 | Sema3d - (Nrp1+Plxna4) | SEMA3 | Secreted Signaling | PMID: 27533782 |
| GFAP | CD31 | Sema3d | NRP1_PLXNA4 | 0.001218 | 0 | SEMA3D_NRP1_PLXNA4 | Sema3d - (Nrp1+Plxna4) | SEMA3 | Secreted Signaling | PMID: 27533782 |
| GFAP | CD31 | Sema3b | NRP2_PLXNA1 | 0.000537 | 0 | SEMA3B_NRP2_PLXNA1 | Sema3b - (Nrp2+Plxna1) | SEMA3 | Secreted Signaling | PMID: 27533782 |
| GFAP | GFAP | Sema3b | NRP2_PLXNA1 | 0.000596 | 0 | SEMA3B_NRP2_PLXNA1 | Sema3b - (Nrp2+Plxna1) | SEMA3 | Secreted Signaling | PMID: 27533782 |
| CD31 | CD31 | Sema3b | NRP2_PLXNA2 | 0.000707 | 0 | SEMA3B_NRP2_PLXNA2 | Sema3b - (Nrp2+Plxna2) | SEMA3 | Secreted Signaling | PMID: 27533782 |
| GFAP | CD31 | Sema3b | NRP2_PLXNA2 | 0.000792 | 0 | SEMA3B_NRP2_PLXNA2 | Sema3b - (Nrp2+Plxna2) | SEMA3 | Secreted Signaling | PMID: 27533782 |
| CD31 | GFAP | Sema3b | NRP2_PLXNA3 | 0.000472 | 0 | SEMA3B_NRP2_PLXNA3 | Sema3b - (Nrp2+Plxna3) | SEMA3 | Secreted Signaling | PMID: 27533782 |
| GFAP | GFAP | Sema3b | NRP2_PLXNA3 | 0.000529 | 0 | SEMA3B_NRP2_PLXNA3 | Sema3b - (Nrp2+Plxna3) | SEMA3 | Secreted Signaling | PMID: 27533782 |
| GFAP | CD31 | Sema3b | NRP2_PLXNA4 | 0.000638 | 0 | SEMA3B_NRP2_PLXNA4 | Sema3b - (Nrp2+Plxna4) | SEMA3 | Secreted Signaling | PMID: 27533782 |
| GFAP | GFAP | Sema3b | NRP2_PLXNA4 | 0.000683 | 0 | SEMA3B_NRP2_PLXNA4 | Sema3b - (Nrp2+Plxna4) | SEMA3 | Secreted Signaling | PMID: 27533782 |
| CD31 | CD31 | Sema3c | NRP2_PLXNA1 | 0.001253 | 0 | SEMA3C_NRP2_PLXNA1 | Sema3c - (Nrp2+Plxna1) | SEMA3 | Secreted Signaling | PMID: 27533782 |
| CD31 | GFAP | Sema3c | NRP2_PLXNA1 | 0.001392 | 0 | SEMA3C_NRP2_PLXNA1 | Sema3c - (Nrp2+Plxna1) | SEMA3 | Secreted Signaling | PMID: 27533782 |
| CD31 | CD31 | Sema3c | NRP2_PLXNA2 | 0.001849 | 0 | SEMA3C_NRP2_PLXNA2 | Sema3c - (Nrp2+Plxna2) | SEMA3 | Secreted Signaling | PMID: 27533782 |
| CD31 | GFAP | Sema3c | NRP2_PLXNA2 | 0.001327 | 0 | SEMA3C_NRP2_PLXNA2 | Sema3c - (Nrp2+Plxna2) | SEMA3 | Secreted Signaling | PMID: 27533782 |
| CD31 | CD31 | Sema3c | NRP2_PLXNA3 | 0.000773 | 0 | SEMA3C_NRP2_PLXNA3 | Sema3c - (Nrp2+Plxna3) | SEMA3 | Secreted Signaling | PMID: 27533782 |
| CD31 | GFAP | Sema3c | NRP2_PLXNA3 | 0.001236 | 0 | SEMA3C_NRP2_PLXNA3 | Sema3c - (Nrp2+Plxna3) | SEMA3 | Secreted Signaling | PMID: 27533782 |
| CD31 | CD31 | Sema3c | NRP2_PLXNA4 | 0.001489 | 0 | SEMA3C_NRP2_PLXNA4 | Sema3c - (Nrp2+Plxna4) | SEMA3 | Secreted Signaling | PMID: 27533782 |
| CD31 | GFAP | Sema3c | NRP2_PLXNA4 | 0.001594 | 0 | SEMA3C_NRP2_PLXNA4 | Sema3c - (Nrp2+Plxna4) | SEMA3 | Secreted Signaling | PMID: 27533782 |
| GFAP | CD31 | Sema3d | NRP2_PLXNA1 | 0.000808 | 0 | SEMA3D_NRP2_PLXNA1 | Sema3d - (Nrp2+Plxna1) | SEMA3 | Secreted Signaling | PMID: 27533782 |
| GFAP | GFAP | Sema3d | NRP2_PLXNA1 | 0.000897 | 0 | SEMA3D_NRP2_PLXNA1 | Sema3d - (Nrp2+Plxna1) | SEMA3 | Secreted Signaling | PMID: 27533782 |
| GFAP | CD31 | Sema3d | NRP2_PLXNA2 | 0.001192 | 0 | SEMA3D_NRP2_PLXNA2 | Sema3d - (Nrp2+Plxna2) | SEMA3 | Secreted Signaling | PMID: 27533782 |
| GFAP | GFAP | Sema3d | NRP2_PLXNA2 | 0.000856 | 0 | SEMA3D_NRP2_PLXNA2 | Sema3d - (Nrp2+Plxna2) | SEMA3 | Secreted Signaling | PMID: 27533782 |
| CD31 | GFAP | Sema3d | NRP2_PLXNA3 | 0.000512 | 0 | SEMA3D_NRP2_PLXNA3 | Sema3d - (Nrp2+Plxna3) | SEMA3 | Secreted Signaling | PMID: 27533782 |
| GFAP | GFAP | Sema3d | NRP2_PLXNA3 | 0.000797 | 0 | SEMA3D_NRP2_PLXNA3 | Sema3d - (Nrp2+Plxna3) | SEMA3 | Secreted Signaling | PMID: 27533782 |
| GFAP | CD31 | Sema3d | NRP2_PLXNA4 | 0.00096 | 0 | SEMA3D_NRP2_PLXNA4 | Sema3d - (Nrp2+Plxna4) | SEMA3 | Secreted Signaling | PMID: 27533782 |
| GFAP | GFAP | Sema3d | NRP2_PLXNA4 | 0.001028 | 0 | SEMA3D_NRP2_PLXNA4 | Sema3d - (Nrp2+Plxna4) | SEMA3 | Secreted Signaling | PMID: 27533782 |
| CD31 | CD31 | Sema3f | NRP2_PLXNA1 | 0.001008 | 0 | SEMA3F_NRP2_PLXNA1 | Sema3f - (Nrp2+Plxna1) | SEMA3 | Secreted Signaling | PMID: 27533782 |
| CD31 | GFAP | Sema3f | NRP2_PLXNA1 | 0.00112 | 0 | SEMA3F_NRP2_PLXNA1 | Sema3f - (Nrp2+Plxna1) | SEMA3 | Secreted Signaling | PMID: 27533782 |
| CD31 | CD31 | Sema3f | NRP2_PLXNA2 | 0.001488 | 0 | SEMA3F_NRP2_PLXNA2 | Sema3f - (Nrp2+Plxna2) | SEMA3 | Secreted Signaling | PMID: 27533782 |
| CD31 | GFAP | Sema3f | NRP2_PLXNA2 | 0.001068 | 0 | SEMA3F_NRP2_PLXNA2 | Sema3f - (Nrp2+Plxna2) | SEMA3 | Secreted Signaling | PMID: 27533782 |
| CD31 | CD31 | Sema3f | NRP2_PLXNA3 | 0.000622 | 0 | SEMA3F_NRP2_PLXNA3 | Sema3f - (Nrp2+Plxna3) | SEMA3 | Secreted Signaling | PMID: 27533782 |
| CD31 | GFAP | Sema3f | NRP2_PLXNA3 | 0.000995 | 0 | SEMA3F_NRP2_PLXNA3 | Sema3f - (Nrp2+Plxna3) | SEMA3 | Secreted Signaling | PMID: 27533782 |
| CD31 | CD31 | Sema3f | NRP2_PLXNA4 | 0.001198 | 0 | SEMA3F_NRP2_PLXNA4 | Sema3f - (Nrp2+Plxna4) | SEMA3 | Secreted Signaling | PMID: 27533782 |
| CD31 | GFAP | Sema3f | NRP2_PLXNA4 | 0.001283 | 0 | SEMA3F_NRP2_PLXNA4 | Sema3f - (Nrp2+Plxna4) | SEMA3 | Secreted Signaling | PMID: 27533782 |
| CD31 | CD31 | Sema3g | NRP2_PLXNA1 | 0.001423 | 0 | SEMA3G_NRP2_PLXNA1 | Sema3g - (Nrp2+Plxna1) | SEMA3 | Secreted Signaling | PMID: 27533782 |
| CD31 | GFAP | Sema3g | NRP2_PLXNA1 | 0.00158 | 0 | SEMA3G_NRP2_PLXNA1 | Sema3g - (Nrp2+Plxna1) | SEMA3 | Secreted Signaling | PMID: 27533782 |
| CD31 | CD31 | Sema3g | NRP2_PLXNA2 | 0.002099 | 0 | SEMA3G_NRP2_PLXNA2 | Sema3g - (Nrp2+Plxna2) | SEMA3 | Secreted Signaling | PMID: 27533782 |
| CD31 | GFAP | Sema3g | NRP2_PLXNA2 | 0.001507 | 0 | SEMA3G_NRP2_PLXNA2 | Sema3g - (Nrp2+Plxna2) | SEMA3 | Secreted Signaling | PMID: 27533782 |
| CD31 | CD31 | Sema3g | NRP2_PLXNA3 | 0.000878 | 0 | SEMA3G_NRP2_PLXNA3 | Sema3g - (Nrp2+Plxna3) | SEMA3 | Secreted Signaling | PMID: 27533782 |
| CD31 | GFAP | Sema3g | NRP2_PLXNA3 | 0.001403 | 0 | SEMA3G_NRP2_PLXNA3 | Sema3g - (Nrp2+Plxna3) | SEMA3 | Secreted Signaling | PMID: 27533782 |
| CD31 | CD31 | Sema3g | NRP2_PLXNA4 | 0.001691 | 0 | SEMA3G_NRP2_PLXNA4 | Sema3g - (Nrp2+Plxna4) | SEMA3 | Secreted Signaling | PMID: 27533782 |
| CD31 | GFAP | Sema3g | NRP2_PLXNA4 | 0.00181 | 0 | SEMA3G_NRP2_PLXNA4 | Sema3g - (Nrp2+Plxna4) | SEMA3 | Secreted Signaling | PMID: 27533782 |
| CD31 | CD31 | Sema3c | NRP1_NRP2 | 0.001942 | 0 | SEMA3C_NRP1_NRP2 | Sema3c - (Nrp1+Nrp2) | SEMA3 | Secreted Signaling | PMID: 27533782 |
| CD31 | GFAP | Sema3c | NRP1_NRP2 | 0.001273 | 0 | SEMA3C_NRP1_NRP2 | Sema3c - (Nrp1+Nrp2) | SEMA3 | Secreted Signaling | PMID: 27533782 |
| CD31 | CD31 | Sema3c | Plxnd1 | 0.00235 | 0 | SEMA3C_PLXND1 | Sema3c - Plxnd1 | SEMA3 | Secreted Signaling | PMID: 27533782 |
| GFAP | CD31 | Sema3c | Plxnd1 | 0.001105 | 0 | SEMA3C_PLXND1 | Sema3c - Plxnd1 | SEMA3 | Secreted Signaling | PMID: 27533782 |
| CD31 | CD31 | Sema3e | Plxnd1 | 0.001318 | 0 | SEMA3E_PLXND1 | Sema3e - Plxnd1 | SEMA3 | Secreted Signaling | PMID: 22325954 |
| NeuN | CD31 | Sema3e | Plxnd1 | 0.002157 | 0 | SEMA3E_PLXND1 | Sema3e - Plxnd1 | SEMA3 | Secreted Signaling | PMID: 22325954 |
| CD31 | CD31 | Crh | Crhr1 | 0.000694 | 0 | CRH_CRHR1 | Crh - Crhr1 | CRH | Secreted Signaling | KEGG: mmu04080 |

| | | | | | | | | | | |
|---|---|---|---|---|---|---|---|---|---|---|
| CD31 | NeuN | Crh | Crhr1 | 0.001101 | 0 | CRH_CRHR1 | Crh - Crhr1 | CRH | Secreted Signaling | KEGG: mmu04080 |
| CD31 | CD31 | Crh | Crhr2 | 0.000341 | 0 | CRH_CRHR2 | Crh - Crhr2 | CRH | Secreted Signaling | KEGG: mmu04080 |
| CD31 | GFAP | Crh | Crhr2 | 0.000363 | 0 | CRH_CRHR2 | Crh - Crhr2 | CRH | Secreted Signaling | KEGG: mmu04080 |
| GFAP | GFAP | Enho | Gpr19 | 0.001078 | 0 | ENHO_GPR19 | Enho - Gpr19 | ENHO | Secreted Signaling | PMID: 28476646 |
| GFAP | NeuN | Enho | Gpr19 | 0.001274 | 0 | ENHO_GPR19 | Enho - Gpr19 | ENHO | Secreted Signaling | PMID: 28476646 |
| CD31 | CD31 | Gas6 | Axl | 0.001935 | 0 | GAS6_AXL | Gas6 - Axl | GAS | Secreted Signaling | PMID: 27801848 |
| CD31 | GFAP | Gas6 | Axl | 0.002263 | 0 | GAS6_AXL | Gas6 - Axl | GAS | Secreted Signaling | PMID: 27801848 |
| CD31 | CD31 | Gas6 | Mertk | 0.002529 | 0 | GAS6_MERTK | Gas6 - Mertk | GAS | Secreted Signaling | PMID: 27801848 |
| CD31 | GFAP | Gas6 | Mertk | 0.002416 | 0 | GAS6_MERTK | Gas6 - Mertk | GAS | Secreted Signaling | PMID: 27801848 |
| CD31 | NeuN | Gas6 | Tyro3 | 0.001827 | 0 | GAS6_TYRO3 | Gas6 - Tyro3 | GAS | Secreted Signaling | PMID: 27801848 |
| NeuN | NeuN | Gas6 | Tyro3 | 0.001666 | 0 | GAS6_TYRO3 | Gas6 - Tyro3 | GAS | Secreted Signaling | PMID: 27801848 |
| CD31 | NeuN | Grn | Sort1 | 0.00653 | 0 | GRN_SORT1 | Grn - Sort1 | GRN | Secreted Signaling | PMID: 29555433 |
| NeuN | NeuN | Grn | Sort1 | 0.004014 | 0 | GRN_SORT1 | Grn - Sort1 | GRN | Secreted Signaling | PMID: 29555433 |
| GFAP | CD31 | Insl5 | Rxfp4 | 0.000195 | 0 | INSL5_RXFP4 | Insl5 - Rxfp4 | RLN | Secreted Signaling | KEGG: mmu04080 |
| GFAP | GFAP | Insl5 | Rxfp4 | 0.000205 | 0 | INSL5_RXFP4 | Insl5 - Rxfp4 | RLN | Secreted Signaling | KEGG: mmu04080 |
| GFAP | CD31 | Insl5 | Rxfp3 | 0.000395 | 0 | INSL5_RXFP3 | Insl5 - Rxfp3 | RLN | Secreted Signaling | KEGG: mmu04080 |
| GFAP | GFAP | Insl5 | Rxfp3 | 0.000271 | 0 | INSL5_RXFP3 | Insl5 - Rxfp3 | RLN | Secreted Signaling | KEGG: mmu04080 |
| GFAP | CD31 | Insl5 | Rxfp2 | 0.000272 | 0 | INSL5_RXFP2 | Insl5 - Rxfp2 | RLN | Secreted Signaling | KEGG: mmu04080 |
| GFAP | GFAP | Insl5 | Rxfp2 | 0.000331 | 0 | INSL5_RXFP2 | Insl5 - Rxfp2 | RLN | Secreted Signaling | KEGG: mmu04080 |
| GFAP | GFAP | Insl5 | Rxfp1 | 0.000364 | 0 | INSL5_RXFP1 | Insl5 - Rxfp1 | RLN | Secreted Signaling | KEGG: mmu04080 |
| GFAP | NeuN | Insl5 | Rxfp1 | 0.000343 | 0 | INSL5_RXFP1 | Insl5 - Rxfp1 | RLN | Secreted Signaling | KEGG: mmu04080 |
| CD31 | CD31 | Lgals9 | Ighm | 0.0007 | 0 | LGALS9_IGHM | Lgals9 - Ighm | GALECTIN | Secreted Signaling | PMID: 30120235 |
| CD31 | GFAP | Lgals9 | Ighm | 0.00051 | 0 | LGALS9_IGHM | Lgals9 - Ighm | GALECTIN | Secreted Signaling | PMID: 30120235 |
| CD31 | CD31 | Pmch | Mertk | 0.000649 | 0 | PMCH_MERTK | Pmch - Mertk | PMCH | Secreted Signaling | PMID: 30501104 |
| CD31 | GFAP | Pmch | Mertk | 0.00062 | 0 | PMCH_MERTK | Pmch - Mertk | PMCH | Secreted Signaling | PMID: 30501104 |
| CD31 | CD31 | Pros1 | Axl | 0.000984 | 0 | PROS1_AXL | Pros1 - Axl | PROS | Secreted Signaling | PMID: 29531161 |
| CD31 | GFAP | Pros1 | Axl | 0.001151 | 0 | PROS1_AXL | Pros1 - Axl | PROS | Secreted Signaling | PMID: 29531161 |
| CD31 | CD31 | Pros1 | Tyro3 | 0.000612 | 0 | PROS1_TYRO3 | Pros1 - Tyro3 | PROS | Secreted Signaling | PMID: 30501104 |
| CD31 | NeuN | Pros1 | Tyro3 | 0.000929 | 0 | PROS1_TYRO3 | Pros1 - Tyro3 | PROS | Secreted Signaling | PMID: 30501104 |
| GFAP | GFAP | Psap | Gpr37l1 | 0.119773 | 0 | PSAP_GPR37L1 | Psap - Gpr37l1 | PSAP | Secreted Signaling | PMID: 23690594 |
| NeuN | GFAP | Psap | Gpr37l1 | 0.168835 | 0 | PSAP_GPR37L1 | Psap - Gpr37l1 | PSAP | Secreted Signaling | PMID: 23690594 |
| GFAP | GFAP | Psap | Gpr37 | 0.04137 | 0 | PSAP_GPR37 | Psap - Gpr37 | PSAP | Secreted Signaling | PMID: 23690594 |
| NeuN | GFAP | Psap | Gpr37 | 0.060524 | 0 | PSAP_GPR37 | Psap - Gpr37 | PSAP | Secreted Signaling | PMID: 23690594 |
| CD31 | GFAP | Qrfp | Qrfpr | 0.00067 | 0 | QRFP_QRFPR | Qrfp - Qrfpr | QRFP | Secreted Signaling | PMID: 25386115 |
| GFAP | GFAP | Qrfp | Qrfpr | 0.000669 | 0 | QRFP_QRFPR | Qrfp - Qrfpr | QRFP | Secreted Signaling | PMID: 25386115 |
| CD31 | CD31 | Rarres2 | Cmklr1 | 0.000465 | 0 | RARRES2_CMKLR1 | Rarres2 - Cmklr1 | CHEMERIN | Secreted Signaling | PMID: 24671882 |
| GFAP | CD31 | Rarres2 | Cmklr1 | 0.000335 | 0 | RARRES2_CMKLR1 | Rarres2 - Cmklr1 | CHEMERIN | Secreted Signaling | PMID: 24671882 |
| CD31 | CD31 | Rln3 | Rxfp3 | 0.000366 | 0 | RLN3_RXFP3 | Rln3 - Rxfp3 | RLN | Secreted Signaling | KEGG: mmu04080 |
| CD31 | GFAP | Rln3 | Rxfp3 | 0.000251 | 0 | RLN3_RXFP3 | Rln3 - Rxfp3 | RLN | Secreted Signaling | KEGG: mmu04080 |
| CD31 | CD31 | Rln3 | Rxfp1 | 0.000256 | 0 | RLN3_RXFP1 | Rln3 - Rxfp1 | RLN | Secreted Signaling | KEGG: mmu04080 |
| CD31 | GFAP | Rln3 | Rxfp1 | 0.000338 | 0 | RLN3_RXFP1 | Rln3 - Rxfp1 | RLN | Secreted Signaling | KEGG: mmu04080 |
| CD31 | CD31 | Rln3 | Rxfp4 | 0.000181 | 0 | RLN3_RXFP4 | Rln3 - Rxfp4 | RLN | Secreted Signaling | KEGG: mmu04080 |
| CD31 | GFAP | Rln3 | Rxfp4 | 0.00019 | 0 | RLN3_RXFP4 | Rln3 - Rxfp4 | RLN | Secreted Signaling | KEGG: mmu04080 |
| CD31 | CD31 | Rln3 | Rxfp2 | 0.000252 | 0 | RLN3_RXFP2 | Rln3 - Rxfp2 | RLN | Secreted Signaling | KEGG: mmu04080 |
| CD31 | GFAP | Rln3 | Rxfp2 | 0.000307 | 0 | RLN3_RXFP2 | Rln3 - Rxfp2 | RLN | Secreted Signaling | KEGG: mmu04080 |
| CD31 | CD31 | Slurp2 | Chrnb2 | 0.000394 | 0 | SLURP2_CHRNB2 | Slurp2 - Chrnb2 | SLURP | Secreted Signaling | KEGG: mmu04080 |
| CD31 | NeuN | Slurp2 | Chrnb2 | 0.000412 | 0 | SLURP2_CHRNB2 | Slurp2 - Chrnb2 | SLURP | Secreted Signaling | KEGG: mmu04080 |
| CD31 | CD31 | Slurp2 | Chrnb3 | 0.000377 | 0 | SLURP2_CHRNB3 | Slurp2 - Chrnb3 | SLURP | Secreted Signaling | KEGG: mmu04080 |
| CD31 | GFAP | Slurp2 | Chrnb3 | 0.000284 | 0 | SLURP2_CHRNB3 | Slurp2 - Chrnb3 | SLURP | Secreted Signaling | KEGG: mmu04080 |
| CD31 | CD31 | Slurp2 | Chrnb4 | 0.000339 | 0 | SLURP2_CHRNB4 | Slurp2 - Chrnb4 | SLURP | Secreted Signaling | KEGG: mmu04080 |
| CD31 | GFAP | Slurp2 | Chrnb4 | 0.000296 | 0 | SLURP2_CHRNB4 | Slurp2 - Chrnb4 | SLURP | Secreted Signaling | KEGG: mmu04080 |
| CD31 | GFAP | Slurp2 | Chrna1 | 0.00046 | 0 | SLURP2_CHRNA1 | Slurp2 - Chrna1 | SLURP | Secreted Signaling | KEGG: mmu04080 |
| GFAP | GFAP | Slurp2 | Chrna1 | 0.000297 | 0 | SLURP2_CHRNA1 | Slurp2 - Chrna1 | SLURP | Secreted Signaling | KEGG: mmu04080 |
| CD31 | CD31 | Slurp2 | Chrna4 | 0.000737 | 0 | SLURP2_CHRNA4 | Slurp2 - Chrna4 | SLURP | Secreted Signaling | KEGG: mmu04080 |
| CD31 | NeuN | Slurp2 | Chrna4 | 0.000803 | 0 | SLURP2_CHRNA4 | Slurp2 - Chrna4 | SLURP | Secreted Signaling | KEGG: mmu04080 |
| CD31 | CD31 | Slurp2 | Chrna5 | 0.00023 | 0 | SLURP2_CHRNA5 | Slurp2 - Chrna5 | SLURP | Secreted Signaling | KEGG: mmu04080 |
| CD31 | GFAP | Slurp2 | Chrna5 | 0.000331 | 0 | SLURP2_CHRNA5 | Slurp2 - Chrna5 | SLURP | Secreted Signaling | KEGG: mmu04080 |
| CD31 | GFAP | Slurp2 | Chrna7 | 0.000373 | 0 | SLURP2_CHRNA7 | Slurp2 - Chrna7 | SLURP | Secreted Signaling | KEGG: mmu04080 |
| GFAP | GFAP | Slurp2 | Chrna7 | 0.000241 | 0 | SLURP2_CHRNA7 | Slurp2 - Chrna7 | SLURP | Secreted Signaling | KEGG: mmu04080 |

| | | | | | | | | | | |
|---|---|---|---|---|---|---|---|---|---|---|
| CD31 | CD31 | Slurp2 | Chrna9 | 0.00034 | 0 | SLURP2_CHRNA9 | Slurp2 - Chrna9 | SLURP | Secreted Signaling | KEGG: mmu04080 |
| CD31 | GFAP | Slurp2 | Chrna9 | 0.000302 | 0 | SLURP2_CHRNA9 | Slurp2 - Chrna9 | SLURP | Secreted Signaling | KEGG: mmu04080 |
| CD31 | CD31 | Slurp2 | Chrnd | 0.000306 | 0 | SLURP2_CHRND | Slurp2 - Chrnd | SLURP | Secreted Signaling | KEGG: mmu04080 |
| CD31 | GFAP | Slurp2 | Chrnd | 0.000225 | 0 | SLURP2_CHRND | Slurp2 - Chrnd | SLURP | Secreted Signaling | KEGG: mmu04080 |
| CD31 | NeuN | Ucn | Crhr1 | 0.001275 | 0 | UCN_CRHR1 | Ucn - Crhr1 | UCN | Secreted Signaling | KEGG: mmu04080 |
| NeuN | NeuN | Ucn | Crhr1 | 0.000816 | 0 | UCN_CRHR1 | Ucn - Crhr1 | UCN | Secreted Signaling | KEGG: mmu04080 |
| CD31 | CD31 | Ucn | Crhr2 | 0.000395 | 0 | UCN_CRHR2 | Ucn - Crhr2 | UCN | Secreted Signaling | KEGG: mmu04080 |
| CD31 | GFAP | Ucn | Crhr2 | 0.00042 | 0 | UCN_CRHR2 | Ucn - Crhr2 | UCN | Secreted Signaling | KEGG: mmu04080 |
| GFAP | CD31 | Klk6 | F2r | 0.000526 | 0 | KLK6_F2R | Klk6 - F2r | KLK | Secreted Signaling | PMID: 35102281 |
| GFAP | GFAP | Klk6 | F2r | 0.00038 | 0 | KLK6_F2R | Klk6 - F2r | KLK | Secreted Signaling | PMID: 35102281 |
| CD31 | CD31 | Klk8 | F2r | 0.000377 | 0 | KLK8_F2R | Klk8 - F2r | KLK | Secreted Signaling | PMID: 34552064 |
| GFAP | CD31 | Klk8 | F2r | 0.000428 | 0 | KLK8_F2R | Klk8 - F2r | KLK | Secreted Signaling | PMID: 34552064 |
| CD31 | GFAP | Adm | CALCR_RAMP1 | 0.000465 | 0 | ADM_CALCR_RAMP1 | Adm - (Calcr+Ramp1) | ADM | Secreted Signaling | PMID: 33462236 |
| GFAP | GFAP | Adm | CALCR_RAMP1 | 0.000355 | 0 | ADM_CALCR_RAMP1 | Adm - (Calcr+Ramp1) | ADM | Secreted Signaling | PMID: 33462236 |
| CD31 | CD31 | Adm | CALCR_RAMP2 | 0.000498 | 0 | ADM_CALCR_RAMP2 | Adm - (Calcr+Ramp2) | ADM | Secreted Signaling | PMID: 33462236 |
| CD31 | GFAP | Adm | CALCR_RAMP2 | 0.000548 | 0 | ADM_CALCR_RAMP2 | Adm - (Calcr+Ramp2) | ADM | Secreted Signaling | PMID: 33462236 |
| CD31 | GFAP | Adm | CALCR_RAMP3 | 0.00037 | 0 | ADM_CALCR_RAMP3 | Adm - (Calcr+Ramp3) | ADM | Secreted Signaling | PMID: 33462236 |
| GFAP | GFAP | Adm | CALCR_RAMP3 | 0.000282 | 0 | ADM_CALCR_RAMP3 | Adm - (Calcr+Ramp3) | ADM | Secreted Signaling | PMID: 33462236 |
| CD31 | GFAP | Calca | CALCR_RAMP1 | 0.000409 | 0 | CALCA_CALCR_RAMP1 | Calca - (Calcr+Ramp1) | CALC | Secreted Signaling | PMID: 30741933;PMID: 18599553 |
| GFAP | GFAP | Calca | CALCR_RAMP1 | 0.000345 | 0 | CALCA_CALCR_RAMP1 | Calca - (Calcr+Ramp1) | CALC | Secreted Signaling | PMID: 30741933;PMID: 18599553 |
| CD31 | CD31 | Calca | CALCR_RAMP2 | 0.000438 | 0 | CALCA_CALCR_RAMP2 | Calca - (Calcr+Ramp2) | CALC | Secreted Signaling | PMID: 30741933;PMID: 18599553 |
| CD31 | GFAP | Calca | CALCR_RAMP2 | 0.000482 | 0 | CALCA_CALCR_RAMP2 | Calca - (Calcr+Ramp2) | CALC | Secreted Signaling | PMID: 30741933;PMID: 18599553 |
| CD31 | GFAP | Calca | CALCR_RAMP3 | 0.000325 | 0 | CALCA_CALCR_RAMP3 | Calca - (Calcr+Ramp3) | CALC | Secreted Signaling | PMID: 30741933;PMID: 18599553 |
| GFAP | GFAP | Calca | CALCR_RAMP3 | 0.000274 | 0 | CALCA_CALCR_RAMP3 | Calca - (Calcr+Ramp3) | CALC | Secreted Signaling | PMID: 30741933;PMID: 18599553 |
| CD31 | GFAP | Calcb | CALCR_RAMP1 | 0.000347 | 0 | CALCB_CALCR_RAMP1 | Calcb - (Calcr+Ramp1) | CALC | Secreted Signaling | PMID: 30741933;PMID: 18599553 |
| GFAP | GFAP | Calcb | CALCR_RAMP1 | 0.000345 | 0 | CALCB_CALCR_RAMP1 | Calcb - (Calcr+Ramp1) | CALC | Secreted Signaling | PMID: 30741933;PMID: 18599553 |
| CD31 | GFAP | Calcb | CALCR_RAMP2 | 0.000409 | 0 | CALCB_CALCR_RAMP2 | Calcb - (Calcr+Ramp2) | CALC | Secreted Signaling | PMID: 30741933;PMID: 18599553 |
| GFAP | GFAP | Calcb | CALCR_RAMP2 | 0.000406 | 0 | CALCB_CALCR_RAMP2 | Calcb - (Calcr+Ramp2) | CALC | Secreted Signaling | PMID: 30741933;PMID: 18599553 |
| CD31 | GFAP | Calcb | CALCR_RAMP3 | 0.000276 | 0 | CALCB_CALCR_RAMP3 | Calcb - (Calcr+Ramp3) | CALC | Secreted Signaling | PMID: 30741933;PMID: 18599553 |
| GFAP | GFAP | Calcb | CALCR_RAMP3 | 0.000274 | 0 | CALCB_CALCR_RAMP3 | Calcb - (Calcr+Ramp3) | CALC | Secreted Signaling | PMID: 30741933;PMID: 18599553 |
| CD31 | CD31 | Ctsg | Fpr1 | 0.00036 | 0 | CTSG_FPR1 | Ctsg - Fpr1 | CTSG | Secreted Signaling | PMID: 26516201 |
| GFAP | CD31 | Ctsg | Fpr1 | 0.000347 | 0 | CTSG_FPR1 | Ctsg - Fpr1 | CTSG | Secreted Signaling | PMID: 26516201 |
| GFAP | GFAP | Grp | Nmbr | 0.000634 | 0 | GRP_NMBR | Grp - Nmbr | GRP | Secreted Signaling | PMID:25209280;PMID:29133874 |
| NeuN | GFAP | Grp | Nmbr | 0.000684 | 0 | GRP_NMBR | Grp - Nmbr | GRP | Secreted Signaling | PMID:25209280;PMID:29133874 |
| GFAP | NeuN | Ptprs | Ntrk3 | 0.01882 | 0 | PTPRS_NTRK3 | Ptprs - Ntrk3 | PTPR | Secreted Signaling | PMID:21262467;PMID:25385546; |
| NeuN | NeuN | Ptprs | Ntrk3 | 0.036312 | 0 | PTPRS_NTRK3 | Ptprs - Ntrk3 | PTPR | Secreted Signaling | PMID:21262467;PMID:25385546; |
| NeuN | GFAP | Slitrk1 | Ptprd | 0.009995 | 0 | SLITRK1_PTPRD | Slitrk1 - Ptprd | SLITRK | Secreted Signaling | PMID: 30648269;PMID: 30822649 |
| NeuN | NeuN | Slitrk1 | Ptprd | 0.010977 | 0 | SLITRK1_PTPRD | Slitrk1 - Ptprd | SLITRK | Secreted Signaling | PMID: 30648269;PMID: 30822649 |
| CD31 | NeuN | Slitrk1 | Ptprs | 0.008317 | 0 | SLITRK1_PTPRS | Slitrk1 - Ptprs | SLITRK | Secreted Signaling | PMID: 30648269;PMID: 30822649 |
| NeuN | NeuN | Slitrk1 | Ptprs | 0.015066 | 0 | SLITRK1_PTPRS | Slitrk1 - Ptprs | SLITRK | Secreted Signaling | PMID: 30648269;PMID: 30822649 |
| GFAP | GFAP | Slitrk2 | Ptprd | 0.004607 | 0 | SLITRK2_PTPRD | Slitrk2 - Ptprd | SLITRK | Secreted Signaling | PMID: 30648269;PMID: 30822649 |
| GFAP | NeuN | Slitrk2 | Ptprd | 0.005062 | 0 | SLITRK2_PTPRD | Slitrk2 - Ptprd | SLITRK | Secreted Signaling | PMID: 30648269;PMID: 30822649 |
| GFAP | GFAP | Slitrk2 | Ptprs | 0.003557 | 0 | SLITRK2_PTPRS | Slitrk2 - Ptprs | SLITRK | Secreted Signaling | PMID: 30648269;PMID: 30822649 |
| GFAP | NeuN | Slitrk2 | Ptprs | 0.006964 | 0 | SLITRK2_PTPRS | Slitrk2 - Ptprs | SLITRK | Secreted Signaling | PMID: 30648269;PMID: 30822649 |
| NeuN | GFAP | Slitrk3 | Ptprd | 0.005088 | 0 | SLITRK3_PTPRD | Slitrk3 - Ptprd | SLITRK | Secreted Signaling | PMID: 30648269;PMID: 30822649 |
| NeuN | NeuN | Slitrk3 | Ptprd | 0.005591 | 0 | SLITRK3_PTPRD | Slitrk3 - Ptprd | SLITRK | Secreted Signaling | PMID: 30648269;PMID: 30822649 |
| CD31 | NeuN | Slitrk4 | Ptprs | 0.005393 | 0 | SLITRK4_PTPRS | Slitrk4 - Ptprs | SLITRK | Secreted Signaling | PMID: 30648269;PMID: 30822649 |
| NeuN | NeuN | Slitrk4 | Ptprs | 0.005346 | 0 | SLITRK4_PTPRS | Slitrk4 - Ptprs | SLITRK | Secreted Signaling | PMID: 30648269;PMID: 30822649 |
| GFAP | NeuN | Slitrk5 | Ptprs | 0.003851 | 0 | SLITRK5_PTPRS | Slitrk5 - Ptprs | SLITRK | Secreted Signaling | PMID: 30648269;PMID: 30822649 |
| NeuN | NeuN | Slitrk5 | Ptprs | 0.00749 | 0 | SLITRK5_PTPRS | Slitrk5 - Ptprs | SLITRK | Secreted Signaling | PMID: 30648269;PMID: 30822649 |
| CD31 | NeuN | Bdnf | Sort1 | 0.002366 | 0 | BDNF_SORT1 | Bdnf - Sort1 | NT | Secreted Signaling | uniprot |
| NeuN | NeuN | Bdnf | Sort1 | 0.003668 | 0 | BDNF_SORT1 | Bdnf - Sort1 | NT | Secreted Signaling | uniprot |
| CD31 | CD31 | Bdnf | Ngfr | 0.000225 | 0 | BDNF_NGFR | Bdnf - Ngfr | NT | Secreted Signaling | uniprot |
| NeuN | CD31 | Bdnf | Ngfr | 0.00035 | 0 | BDNF_NGFR | Bdnf - Ngfr | NT | Secreted Signaling | uniprot |
| CD31 | CD31 | Igfbp3 | Tmem219 | 0.001623 | 0 | IGFBP3_TMEM219 | Igfbp3 - Tmem219 | IGFBP | Secreted Signaling | PMID: 35115561;PMID:20353938 |
| CD31 | NeuN | Igfbp3 | Tmem219 | 0.001819 | 0 | IGFBP3_TMEM219 | Igfbp3 - Tmem219 | IGFBP | Secreted Signaling | PMID: 35115561;PMID:20353938 |
| CD31 | CD31 | Lgals9 | P4hb | 0.001348 | 0 | LGALS9_P4HB | Lgals9 - P4hb | GALECTIN | Secreted Signaling | PMID:21670307;uniprot |
| CD31 | GFAP | Lgals9 | P4hb | 0.001643 | 0 | LGALS9_P4HB | Lgals9 - P4hb | GALECTIN | Secreted Signaling | PMID:21670307;uniprot |
| CD31 | CD31 | Nts | Sort1 | 0.001196 | 0 | NTS_SORT1 | Nts - Sort1 | NTS | Secreted Signaling | PMID: 34038712 |

| | | | | | | | | | | |
|---|---|---|---|---|---|---|---|---|---|---|
| CD31 | NeuN | Nts | Sort1 | 0.00214 | 0 | NTS_SORT1 | Nts - Sort1 | NTS | Secreted Signaling | PMID: 34038712 |
| CD31 | CD31 | Ntf3 | Ngfr | 0.000192 | 0 | NTF3_NGFR | Ntf3 - Ngfr | NT | Secreted Signaling | uniprot |
| GFAP | CD31 | Ntf3 | Ngfr | 0.000173 | 0 | NTF3_NGFR | Ntf3 - Ngfr | NT | Secreted Signaling | uniprot |
| CD31 | CD31 | Rbp4 | Stra6 | 0.000523 | 0 | RBP4_STRA6 | Rbp4 - Stra6 | RBP4 | Secreted Signaling | PMID: 33199363 |
| GFAP | CD31 | Rbp4 | Stra6 | 0.000763 | 0 | RBP4_STRA6 | Rbp4 - Stra6 | RBP4 | Secreted Signaling | PMID: 33199363 |
| NeuN | CD31 | Slit1 | Robo1 | 0.001503 | 0 | SLIT1_ROBO1 | Slit1 - Robo1 | SLIT | Secreted Signaling | PMID:10102268 |
| NeuN | NeuN | Slit1 | Robo1 | 0.001488 | 0 | SLIT1_ROBO1 | Slit1 - Robo1 | SLIT | Secreted Signaling | PMID:10102268 |
| CD31 | CD31 | Slit2 | Robo1 | 0.000614 | 0 | SLIT2_ROBO1 | Slit2 - Robo1 | SLIT | Secreted Signaling | PMID:10102268 |
| CD31 | NeuN | Slit2 | Robo1 | 0.000608 | 0 | SLIT2_ROBO1 | Slit2 - Robo1 | SLIT | Secreted Signaling | PMID:10102268 |
| CD31 | NeuN | Slit2 | Robo2 | 0.00172 | 0 | SLIT2_ROBO2 | Slit2 - Robo2 | SLIT | Secreted Signaling | PMID:10102268 |
| NeuN | NeuN | Slit2 | Robo2 | 0.00158 | 0 | SLIT2_ROBO2 | Slit2 - Robo2 | SLIT | Secreted Signaling | PMID:10102268 |
| CD31 | CD31 | Tub | Mertk | 0.001129 | 0 | TUB_MERTK | Tub - Mertk | TULP | Secreted Signaling | PMID: 20978475 |
| NeuN | CD31 | Tub | Mertk | 0.002707 | 0 | TUB_MERTK | Tub - Mertk | TULP | Secreted Signaling | PMID: 20978475 |
| CD31 | CD31 | Pros1 | Mertk | 0.001286 | 0 | PROS1_MERTK | Pros1 - Mertk | PROS | Secreted Signaling | PMID: 34631419 |
| CD31 | GFAP | Pros1 | Mertk | 0.001229 | 0 | PROS1_MERTK | Pros1 - Mertk | PROS | Secreted Signaling | PMID: 34631419 |
| CD31 | CD31 | Col1a1 | ITGA1_ITGB1 | 0.002691 | 0 | COL1A1_ITGA1_ITGB1 | Col1a1 - (Itga1+Itgb1) | COLLAGEN | ECM-Receptor | KEGG: mmu04512 |
| GFAP | CD31 | Col1a1 | ITGA1_ITGB1 | 0.001376 | 0 | COL1A1_ITGA1_ITGB1 | Col1a1 - (Itga1+Itgb1) | COLLAGEN | ECM-Receptor | KEGG: mmu04512 |
| CD31 | CD31 | Col1a2 | ITGA1_ITGB1 | 0.001452 | 0 | COL1A2_ITGA1_ITGB1 | Col1a2 - (Itga1+Itgb1) | COLLAGEN | ECM-Receptor | KEGG: mmu04512 |
| GFAP | CD31 | Col1a2 | ITGA1_ITGB1 | 0.000917 | 0 | COL1A2_ITGA1_ITGB1 | Col1a2 - (Itga1+Itgb1) | COLLAGEN | ECM-Receptor | KEGG: mmu04512 |
| CD31 | CD31 | Col4a1 | ITGA1_ITGB1 | 0.007743 | 0 | COL4A1_ITGA1_ITGB1 | Col4a1 - (Itga1+Itgb1) | COLLAGEN | ECM-Receptor | KEGG: mmu04512 |
| CD31 | GFAP | Col4a1 | ITGA1_ITGB1 | 0.003749 | 0 | COL4A1_ITGA1_ITGB1 | Col4a1 - (Itga1+Itgb1) | COLLAGEN | ECM-Receptor | KEGG: mmu04512 |
| CD31 | CD31 | Col4a2 | ITGA1_ITGB1 | 0.006606 | 0 | COL4A2_ITGA1_ITGB1 | Col4a2 - (Itga1+Itgb1) | COLLAGEN | ECM-Receptor | KEGG: mmu04512 |
| CD31 | GFAP | Col4a2 | ITGA1_ITGB1 | 0.003196 | 0 | COL4A2_ITGA1_ITGB1 | Col4a2 - (Itga1+Itgb1) | COLLAGEN | ECM-Receptor | KEGG: mmu04512 |
| CD31 | CD31 | Col6a1 | ITGA1_ITGB1 | 0.001231 | 0 | COL6A1_ITGA1_ITGB1 | Col6a1 - (Itga1+Itgb1) | COLLAGEN | ECM-Receptor | KEGG: mmu04512 |
| NeuN | CD31 | Col6a1 | ITGA1_ITGB1 | 0.003186 | 0 | COL6A1_ITGA1_ITGB1 | Col6a1 - (Itga1+Itgb1) | COLLAGEN | ECM-Receptor | KEGG: mmu04512 |
| CD31 | CD31 | Col6a2 | ITGA1_ITGB1 | 0.001505 | 0 | COL6A2_ITGA1_ITGB1 | Col6a2 - (Itga1+Itgb1) | COLLAGEN | ECM-Receptor | KEGG: mmu04512 |
| GFAP | CD31 | Col6a2 | ITGA1_ITGB1 | 0.001234 | 0 | COL6A2_ITGA1_ITGB1 | Col6a2 - (Itga1+Itgb1) | COLLAGEN | ECM-Receptor | KEGG: mmu04512 |
| CD31 | CD31 | Col6a4 | ITGA1_ITGB1 | 0.001021 | 0 | COL6A4_ITGA1_ITGB1 | Col6a4 - (Itga1+Itgb1) | COLLAGEN | ECM-Receptor | KEGG: mmu04512 |
| GFAP | CD31 | Col6a4 | ITGA1_ITGB1 | 0.001109 | 0 | COL6A4_ITGA1_ITGB1 | Col6a4 - (Itga1+Itgb1) | COLLAGEN | ECM-Receptor | KEGG: mmu04512 |
| CD31 | CD31 | Col6a6 | ITGA1_ITGB1 | 0.001767 | 0 | COL6A6_ITGA1_ITGB1 | Col6a6 - (Itga1+Itgb1) | COLLAGEN | ECM-Receptor | KEGG: mmu04512 |
| GFAP | CD31 | Col6a6 | ITGA1_ITGB1 | 0.00094 | 0 | COL6A6_ITGA1_ITGB1 | Col6a6 - (Itga1+Itgb1) | COLLAGEN | ECM-Receptor | KEGG: mmu04512 |
| CD31 | CD31 | Col9a2 | ITGA1_ITGB1 | 0.001001 | 0 | COL9A2_ITGA1_ITGB1 | Col9a2 - (Itga1+Itgb1) | COLLAGEN | ECM-Receptor | KEGG: mmu04512 |
| GFAP | CD31 | Col9a2 | ITGA1_ITGB1 | 0.001316 | 0 | COL9A2_ITGA1_ITGB1 | Col9a2 - (Itga1+Itgb1) | COLLAGEN | ECM-Receptor | KEGG: mmu04512 |
| GFAP | CD31 | Col9a3 | ITGA1_ITGB1 | 0.003342 | 0 | COL9A3_ITGA1_ITGB1 | Col9a3 - (Itga1+Itgb1) | COLLAGEN | ECM-Receptor | KEGG: mmu04512 |
| GFAP | GFAP | Col9a3 | ITGA1_ITGB1 | 0.001614 | 0 | COL9A3_ITGA1_ITGB1 | Col9a3 - (Itga1+Itgb1) | COLLAGEN | ECM-Receptor | KEGG: mmu04512 |
| CD31 | CD31 | Fn1 | ITGA3_ITGB1 | 0.006825 | 0 | FN1_ITGA3_ITGB1 | Fn1 - (Itga3+Itgb1) | FN1 | ECM-Receptor | KEGG: mmu04512 |
| CD31 | GFAP | Fn1 | ITGA3_ITGB1 | 0.003669 | 0 | FN1_ITGA3_ITGB1 | Fn1 - (Itga3+Itgb1) | FN1 | ECM-Receptor | KEGG: mmu04512 |
| CD31 | CD31 | Fn1 | ITGA4_ITGB1 | 0.006501 | 0 | FN1_ITGA4_ITGB1 | Fn1 - (Itga4+Itgb1) | FN1 | ECM-Receptor | KEGG: mmu04512 |
| CD31 | GFAP | Fn1 | ITGA4_ITGB1 | 0.005118 | 0 | FN1_ITGA4_ITGB1 | Fn1 - (Itga4+Itgb1) | FN1 | ECM-Receptor | KEGG: mmu04512 |
| CD31 | CD31 | Fn1 | ITGA5_ITGB1 | 0.0062 | 0 | FN1_ITGA5_ITGB1 | Fn1 - (Itga5+Itgb1) | FN1 | ECM-Receptor | KEGG: mmu04512 |
| CD31 | GFAP | Fn1 | ITGA5_ITGB1 | 0.004909 | 0 | FN1_ITGA5_ITGB1 | Fn1 - (Itga5+Itgb1) | FN1 | ECM-Receptor | KEGG: mmu04512 |
| CD31 | CD31 | Fn1 | ITGA8_ITGB1 | 0.007003 | 0 | FN1_ITGA8_ITGB1 | Fn1 - (Itga8+Itgb1) | FN1 | ECM-Receptor | KEGG: mmu04512 |
| CD31 | GFAP | Fn1 | ITGA8_ITGB1 | 0.004748 | 0 | FN1_ITGA8_ITGB1 | Fn1 - (Itga8+Itgb1) | FN1 | ECM-Receptor | KEGG: mmu04512 |
| CD31 | CD31 | Lama1 | ITGA1_ITGB1 | 0.001373 | 0 | LAMA1_ITGA1_ITGB1 | Lama1 - (Itga1+Itgb1) | LAMININ | ECM-Receptor | KEGG: mmu04512 |
| GFAP | CD31 | Lama1 | ITGA1_ITGB1 | 0.0009 | 0 | LAMA1_ITGA1_ITGB1 | Lama1 - (Itga1+Itgb1) | LAMININ | ECM-Receptor | KEGG: mmu04512 |
| CD31 | CD31 | Lama4 | ITGA1_ITGB1 | 0.002392 | 0 | LAMA4_ITGA1_ITGB1 | Lama4 - (Itga1+Itgb1) | LAMININ | ECM-Receptor | KEGG: mmu04512 |
| GFAP | CD31 | Lama4 | ITGA1_ITGB1 | 0.001691 | 0 | LAMA4_ITGA1_ITGB1 | Lama4 - (Itga1+Itgb1) | LAMININ | ECM-Receptor | KEGG: mmu04512 |
| CD31 | CD31 | Lama5 | ITGA1_ITGB1 | 0.002937 | 0 | LAMA5_ITGA1_ITGB1 | Lama5 - (Itga1+Itgb1) | LAMININ | ECM-Receptor | KEGG: mmu04512 |
| GFAP | CD31 | Lama5 | ITGA1_ITGB1 | 0.001822 | 0 | LAMA5_ITGA1_ITGB1 | Lama5 - (Itga1+Itgb1) | LAMININ | ECM-Receptor | KEGG: mmu04512 |
| CD31 | CD31 | Lamb1 | ITGA1_ITGB1 | 0.001731 | 0 | LAMB1_ITGA1_ITGB1 | Lamb1 - (Itga1+Itgb1) | LAMININ | ECM-Receptor | KEGG: mmu04512 |
| GFAP | CD31 | Lamb1 | ITGA1_ITGB1 | 0.000997 | 0 | LAMB1_ITGA1_ITGB1 | Lamb1 - (Itga1+Itgb1) | LAMININ | ECM-Receptor | KEGG: mmu04512 |
| CD31 | CD31 | Lamb2 | ITGA1_ITGB1 | 0.005758 | 0 | LAMB2_ITGA1_ITGB1 | Lamb2 - (Itga1+Itgb1) | LAMININ | ECM-Receptor | KEGG: mmu04512 |
| GFAP | CD31 | Lamb2 | ITGA1_ITGB1 | 0.003362 | 0 | LAMB2_ITGA1_ITGB1 | Lamb2 - (Itga1+Itgb1) | LAMININ | ECM-Receptor | KEGG: mmu04512 |
| CD31 | CD31 | Lamc1 | ITGA1_ITGB1 | 0.002858 | 0 | LAMC1_ITGA1_ITGB1 | Lamc1 - (Itga1+Itgb1) | LAMININ | ECM-Receptor | KEGG: mmu04512 |
| CD31 | GFAP | Lamc1 | ITGA1_ITGB1 | 0.00138 | 0 | LAMC1_ITGA1_ITGB1 | Lamc1 - (Itga1+Itgb1) | LAMININ | ECM-Receptor | KEGG: mmu04512 |
| CD31 | CD31 | Lamc2 | ITGA1_ITGB1 | 0.001021 | 0 | LAMC2_ITGA1_ITGB1 | Lamc2 - (Itga1+Itgb1) | LAMININ | ECM-Receptor | KEGG: mmu04512 |
| NeuN | CD31 | Lamc2 | ITGA1_ITGB1 | 0.000886 | 0 | LAMC2_ITGA1_ITGB1 | Lamc2 - (Itga1+Itgb1) | LAMININ | ECM-Receptor | KEGG: mmu04512 |
| CD31 | CD31 | Lamc3 | ITGA1_ITGB1 | 0.002156 | 0 | LAMC3_ITGA1_ITGB1 | Lamc3 - (Itga1+Itgb1) | LAMININ | ECM-Receptor | KEGG: mmu04512 |
| GFAP | CD31 | Lamc3 | ITGA1_ITGB1 | 0.001182 | 0 | LAMC3_ITGA1_ITGB1 | Lamc3 - (Itga1+Itgb1) | LAMININ | ECM-Receptor | KEGG: mmu04512 |

| | | | | | | | | | | |
|---|---|---|---|---|---|---|---|---|---|---|
| CD31 | CD31 | Lama1 | ITGA3_ITGB1 | 0.000893 | 0 | LAMA1_ITGA3_ITGB1 | Lama1 - (Itga3+Itgb1) | LAMININ | ECM-Receptor | KEGG: mmu04512 |
| GFAP | CD31 | Lama1 | ITGA3_ITGB1 | 0.000585 | 0 | LAMA1_ITGA3_ITGB1 | Lama1 - (Itga3+Itgb1) | LAMININ | ECM-Receptor | KEGG: mmu04512 |
| CD31 | CD31 | Lama4 | ITGA3_ITGB1 | 0.001555 | 0 | LAMA4_ITGA3_ITGB1 | Lama4 - (Itga3+Itgb1) | LAMININ | ECM-Receptor | KEGG: mmu04512 |
| GFAP | CD31 | Lama4 | ITGA3_ITGB1 | 0.001099 | 0 | LAMA4_ITGA3_ITGB1 | Lama4 - (Itga3+Itgb1) | LAMININ | ECM-Receptor | KEGG: mmu04512 |
| CD31 | CD31 | Lama5 | ITGA3_ITGB1 | 0.00191 | 0 | LAMA5_ITGA3_ITGB1 | Lama5 - (Itga3+Itgb1) | LAMININ | ECM-Receptor | KEGG: mmu04512 |
| GFAP | CD31 | Lama5 | ITGA3_ITGB1 | 0.001185 | 0 | LAMA5_ITGA3_ITGB1 | Lama5 - (Itga3+Itgb1) | LAMININ | ECM-Receptor | KEGG: mmu04512 |
| CD31 | CD31 | Lamb1 | ITGA3_ITGB1 | 0.001125 | 0 | LAMB1_ITGA3_ITGB1 | Lamb1 - (Itga3+Itgb1) | LAMININ | ECM-Receptor | KEGG: mmu04512 |
| GFAP | CD31 | Lamb1 | ITGA3_ITGB1 | 0.000648 | 0 | LAMB1_ITGA3_ITGB1 | Lamb1 - (Itga3+Itgb1) | LAMININ | ECM-Receptor | KEGG: mmu04512 |
| CD31 | CD31 | Lamb2 | ITGA3_ITGB1 | 0.003749 | 0 | LAMB2_ITGA3_ITGB1 | Lamb2 - (Itga3+Itgb1) | LAMININ | ECM-Receptor | KEGG: mmu04512 |
| GFAP | CD31 | Lamb2 | ITGA3_ITGB1 | 0.002187 | 0 | LAMB2_ITGA3_ITGB1 | Lamb2 - (Itga3+Itgb1) | LAMININ | ECM-Receptor | KEGG: mmu04512 |
| CD31 | CD31 | Lamc1 | ITGA3_ITGB1 | 0.001859 | 0 | LAMC1_ITGA3_ITGB1 | Lamc1 - (Itga3+Itgb1) | LAMININ | ECM-Receptor | KEGG: mmu04512 |
| CD31 | GFAP | Lamc1 | ITGA3_ITGB1 | 0.000997 | 0 | LAMC1_ITGA3_ITGB1 | Lamc1 - (Itga3+Itgb1) | LAMININ | ECM-Receptor | KEGG: mmu04512 |
| CD31 | CD31 | Lamc2 | ITGA3_ITGB1 | 0.000664 | 0 | LAMC2_ITGA3_ITGB1 | Lamc2 - (Itga3+Itgb1) | LAMININ | ECM-Receptor | KEGG: mmu04512 |
| NeuN | CD31 | Lamc2 | ITGA3_ITGB1 | 0.000576 | 0 | LAMC2_ITGA3_ITGB1 | Lamc2 - (Itga3+Itgb1) | LAMININ | ECM-Receptor | KEGG: mmu04512 |
| CD31 | CD31 | Lamc3 | ITGA3_ITGB1 | 0.001402 | 0 | LAMC3_ITGA3_ITGB1 | Lamc3 - (Itga3+Itgb1) | LAMININ | ECM-Receptor | KEGG: mmu04512 |
| GFAP | CD31 | Lamc3 | ITGA3_ITGB1 | 0.000768 | 0 | LAMC3_ITGA3_ITGB1 | Lamc3 - (Itga3+Itgb1) | LAMININ | ECM-Receptor | KEGG: mmu04512 |
| CD31 | CD31 | Reln | ITGA3_ITGB1 | 0.00065 | 0 | RELN_ITGA3_ITGB1 | Reln - (Itga3+Itgb1) | RELN | ECM-Receptor | KEGG: mmu04512 |
| NeuN | CD31 | Reln | ITGA3_ITGB1 | 0.00112 | 0 | RELN_ITGA3_ITGB1 | Reln - (Itga3+Itgb1) | RELN | ECM-Receptor | KEGG: mmu04512 |
| CD31 | CD31 | Thbs1 | ITGA3_ITGB1 | 0.000595 | 0 | THBS1_ITGA3_ITGB1 | Thbs1 - (Itga3+Itgb1) | THBS | ECM-Receptor | KEGG: mmu04512 |
| GFAP | CD31 | Thbs1 | ITGA3_ITGB1 | 0.000735 | 0 | THBS1_ITGA3_ITGB1 | Thbs1 - (Itga3+Itgb1) | THBS | ECM-Receptor | KEGG: mmu04512 |
| CD31 | CD31 | Thbs3 | ITGA3_ITGB1 | 0.000513 | 0 | THBS3_ITGA3_ITGB1 | Thbs3 - (Itga3+Itgb1) | THBS | ECM-Receptor | KEGG: mmu04512 |
| GFAP | CD31 | Thbs3 | ITGA3_ITGB1 | 0.000691 | 0 | THBS3_ITGA3_ITGB1 | Thbs3 - (Itga3+Itgb1) | THBS | ECM-Receptor | KEGG: mmu04512 |
| GFAP | CD31 | Thbs4 | ITGA3_ITGB1 | 0.009418 | 0 | THBS4_ITGA3_ITGB1 | Thbs4 - (Itga3+Itgb1) | THBS | ECM-Receptor | KEGG: mmu04512 |
| GFAP | GFAP | Thbs4 | ITGA3_ITGB1 | 0.005069 | 0 | THBS4_ITGA3_ITGB1 | Thbs4 - (Itga3+Itgb1) | THBS | ECM-Receptor | KEGG: mmu04512 |
| CD31 | CD31 | Col1a1 | ITGA3_ITGB1 | 0.00175 | 0 | COL1A1_ITGA3_ITGB1 | Col1a1 - (Itga3+Itgb1) | COLLAGEN | ECM-Receptor | KEGG: mmu04512 |
| CD31 | GFAP | Col1a1 | ITGA3_ITGB1 | 0.000938 | 0 | COL1A1_ITGA3_ITGB1 | Col1a1 - (Itga3+Itgb1) | COLLAGEN | ECM-Receptor | KEGG: mmu04512 |
| CD31 | CD31 | Col1a2 | ITGA3_ITGB1 | 0.000944 | 0 | COL1A2_ITGA3_ITGB1 | Col1a2 - (Itga3+Itgb1) | COLLAGEN | ECM-Receptor | KEGG: mmu04512 |
| GFAP | CD31 | Col1a2 | ITGA3_ITGB1 | 0.000596 | 0 | COL1A2_ITGA3_ITGB1 | Col1a2 - (Itga3+Itgb1) | COLLAGEN | ECM-Receptor | KEGG: mmu04512 |
| CD31 | CD31 | Col4a1 | ITGA3_ITGB1 | 0.005045 | 0 | COL4A1_ITGA3_ITGB1 | Col4a1 - (Itga3+Itgb1) | COLLAGEN | ECM-Receptor | KEGG: mmu04512 |
| CD31 | GFAP | Col4a1 | ITGA3_ITGB1 | 0.00271 | 0 | COL4A1_ITGA3_ITGB1 | Col4a1 - (Itga3+Itgb1) | COLLAGEN | ECM-Receptor | KEGG: mmu04512 |
| CD31 | CD31 | Col4a2 | ITGA3_ITGB1 | 0.004302 | 0 | COL4A2_ITGA3_ITGB1 | Col4a2 - (Itga3+Itgb1) | COLLAGEN | ECM-Receptor | KEGG: mmu04512 |
| CD31 | GFAP | Col4a2 | ITGA3_ITGB1 | 0.00231 | 0 | COL4A2_ITGA3_ITGB1 | Col4a2 - (Itga3+Itgb1) | COLLAGEN | ECM-Receptor | KEGG: mmu04512 |
| NeuN | CD31 | Col6a1 | ITGA3_ITGB1 | 0.002072 | 0 | COL6A1_ITGA3_ITGB1 | Col6a1 - (Itga3+Itgb1) | COLLAGEN | ECM-Receptor | KEGG: mmu04512 |
| NeuN | NeuN | Col6a1 | ITGA3_ITGB1 | 0.000805 | 0 | COL6A1_ITGA3_ITGB1 | Col6a1 - (Itga3+Itgb1) | COLLAGEN | ECM-Receptor | KEGG: mmu04512 |
| CD31 | CD31 | Col6a2 | ITGA3_ITGB1 | 0.000978 | 0 | COL6A2_ITGA3_ITGB1 | Col6a2 - (Itga3+Itgb1) | COLLAGEN | ECM-Receptor | KEGG: mmu04512 |
| GFAP | CD31 | Col6a2 | ITGA3_ITGB1 | 0.000802 | 0 | COL6A2_ITGA3_ITGB1 | Col6a2 - (Itga3+Itgb1) | COLLAGEN | ECM-Receptor | KEGG: mmu04512 |
| CD31 | CD31 | Col6a4 | ITGA3_ITGB1 | 0.000664 | 0 | COL6A4_ITGA3_ITGB1 | Col6a4 - (Itga3+Itgb1) | COLLAGEN | ECM-Receptor | KEGG: mmu04512 |
| GFAP | CD31 | Col6a4 | ITGA3_ITGB1 | 0.000721 | 0 | COL6A4_ITGA3_ITGB1 | Col6a4 - (Itga3+Itgb1) | COLLAGEN | ECM-Receptor | KEGG: mmu04512 |
| CD31 | CD31 | Col6a6 | ITGA3_ITGB1 | 0.001149 | 0 | COL6A6_ITGA3_ITGB1 | Col6a6 - (Itga3+Itgb1) | COLLAGEN | ECM-Receptor | KEGG: mmu04512 |
| CD31 | GFAP | Col6a6 | ITGA3_ITGB1 | 0.000616 | 0 | COL6A6_ITGA3_ITGB1 | Col6a6 - (Itga3+Itgb1) | COLLAGEN | ECM-Receptor | KEGG: mmu04512 |
| CD31 | CD31 | Col9a2 | ITGA3_ITGB1 | 0.00065 | 0 | COL9A2_ITGA3_ITGB1 | Col9a2 - (Itga3+Itgb1) | COLLAGEN | ECM-Receptor | KEGG: mmu04512 |
| GFAP | CD31 | Col9a2 | ITGA3_ITGB1 | 0.000855 | 0 | COL9A2_ITGA3_ITGB1 | Col9a2 - (Itga3+Itgb1) | COLLAGEN | ECM-Receptor | KEGG: mmu04512 |
| GFAP | CD31 | Col9a3 | ITGA3_ITGB1 | 0.002174 | 0 | COL9A3_ITGA3_ITGB1 | Col9a3 - (Itga3+Itgb1) | COLLAGEN | ECM-Receptor | KEGG: mmu04512 |
| GFAP | GFAP | Col9a3 | ITGA3_ITGB1 | 0.001166 | 0 | COL9A3_ITGA3_ITGB1 | Col9a3 - (Itga3+Itgb1) | COLLAGEN | ECM-Receptor | KEGG: mmu04512 |
| CD31 | CD31 | Lama1 | ITGA6_ITGB1 | 0.001427 | 0 | LAMA1_ITGA6_ITGB1 | Lama1 - (Itga6+Itgb1) | LAMININ | ECM-Receptor | KEGG: mmu04512 |
| GFAP | CD31 | Lama1 | ITGA6_ITGB1 | 0.000936 | 0 | LAMA1_ITGA6_ITGB1 | Lama1 - (Itga6+Itgb1) | LAMININ | ECM-Receptor | KEGG: mmu04512 |
| CD31 | CD31 | Lama4 | ITGA6_ITGB1 | 0.002485 | 0 | LAMA4_ITGA6_ITGB1 | Lama4 - (Itga6+Itgb1) | LAMININ | ECM-Receptor | KEGG: mmu04512 |
| GFAP | CD31 | Lama4 | ITGA6_ITGB1 | 0.001757 | 0 | LAMA4_ITGA6_ITGB1 | Lama4 - (Itga6+Itgb1) | LAMININ | ECM-Receptor | KEGG: mmu04512 |
| CD31 | CD31 | Lama5 | ITGA6_ITGB1 | 0.003052 | 0 | LAMA5_ITGA6_ITGB1 | Lama5 - (Itga6+Itgb1) | LAMININ | ECM-Receptor | KEGG: mmu04512 |
| GFAP | CD31 | Lama5 | ITGA6_ITGB1 | 0.001893 | 0 | LAMA5_ITGA6_ITGB1 | Lama5 - (Itga6+Itgb1) | LAMININ | ECM-Receptor | KEGG: mmu04512 |
| CD31 | CD31 | Lamb1 | ITGA6_ITGB1 | 0.001798 | 0 | LAMB1_ITGA6_ITGB1 | Lamb1 - (Itga6+Itgb1) | LAMININ | ECM-Receptor | KEGG: mmu04512 |
| GFAP | CD31 | Lamb1 | ITGA6_ITGB1 | 0.001036 | 0 | LAMB1_ITGA6_ITGB1 | Lamb1 - (Itga6+Itgb1) | LAMININ | ECM-Receptor | KEGG: mmu04512 |
| CD31 | CD31 | Lamb2 | ITGA6_ITGB1 | 0.005983 | 0 | LAMB2_ITGA6_ITGB1 | Lamb2 - (Itga6+Itgb1) | LAMININ | ECM-Receptor | KEGG: mmu04512 |
| GFAP | CD31 | Lamb2 | ITGA6_ITGB1 | 0.003493 | 0 | LAMB2_ITGA6_ITGB1 | Lamb2 - (Itga6+Itgb1) | LAMININ | ECM-Receptor | KEGG: mmu04512 |
| CD31 | CD31 | Lamc1 | ITGA6_ITGB1 | 0.00297 | 0 | LAMC1_ITGA6_ITGB1 | Lamc1 - (Itga6+Itgb1) | LAMININ | ECM-Receptor | KEGG: mmu04512 |
| CD31 | GFAP | Lamc1 | ITGA6_ITGB1 | 0.001464 | 0 | LAMC1_ITGA6_ITGB1 | Lamc1 - (Itga6+Itgb1) | LAMININ | ECM-Receptor | KEGG: mmu04512 |
| CD31 | CD31 | Lamc2 | ITGA6_ITGB1 | 0.001061 | 0 | LAMC2_ITGA6_ITGB1 | Lamc2 - (Itga6+Itgb1) | LAMININ | ECM-Receptor | KEGG: mmu04512 |
| NeuN | CD31 | Lamc2 | ITGA6_ITGB1 | 0.00092 | 0 | LAMC2_ITGA6_ITGB1 | Lamc2 - (Itga6+Itgb1) | LAMININ | ECM-Receptor | KEGG: mmu04512 |
| CD31 | CD31 | Lamc3 | ITGA6_ITGB1 | 0.00224 | 0 | LAMC3_ITGA6_ITGB1 | Lamc3 - (Itga6+Itgb1) | LAMININ | ECM-Receptor | KEGG: mmu04512 |

| | | | | | | | | | | |
|---|---|---|---|---|---|---|---|---|---|---|
| GFAP | CD31 | Lamc3 | ITGA6_ITGB1 | 0.001228 | 0 | LAMC3_ITGA6_ITGB1 | Lamc3 - (Itga6+Itgb1) | LAMININ | ECM-Receptor | KEGG: mmu04512 |
| CD31 | CD31 | Lama1 | ITGA7_ITGB1 | 0.000957 | 0 | LAMA1_ITGA7_ITGB1 | Lama1 - (Itga7+Itgb1) | LAMININ | ECM-Receptor | KEGG: mmu04512 |
| GFAP | CD31 | Lama1 | ITGA7_ITGB1 | 0.000627 | 0 | LAMA1_ITGA7_ITGB1 | Lama1 - (Itga7+Itgb1) | LAMININ | ECM-Receptor | KEGG: mmu04512 |
| CD31 | CD31 | Lama4 | ITGA7_ITGB1 | 0.001668 | 0 | LAMA4_ITGA7_ITGB1 | Lama4 - (Itga7+Itgb1) | LAMININ | ECM-Receptor | KEGG: mmu04512 |
| GFAP | CD31 | Lama4 | ITGA7_ITGB1 | 0.001179 | 0 | LAMA4_ITGA7_ITGB1 | Lama4 - (Itga7+Itgb1) | LAMININ | ECM-Receptor | KEGG: mmu04512 |
| CD31 | CD31 | Lama5 | ITGA7_ITGB1 | 0.002048 | 0 | LAMA5_ITGA7_ITGB1 | Lama5 - (Itga7+Itgb1) | LAMININ | ECM-Receptor | KEGG: mmu04512 |
| GFAP | CD31 | Lama5 | ITGA7_ITGB1 | 0.00127 | 0 | LAMA5_ITGA7_ITGB1 | Lama5 - (Itga7+Itgb1) | LAMININ | ECM-Receptor | KEGG: mmu04512 |
| CD31 | CD31 | Lamb1 | ITGA7_ITGB1 | 0.001206 | 0 | LAMB1_ITGA7_ITGB1 | Lamb1 - (Itga7+Itgb1) | LAMININ | ECM-Receptor | KEGG: mmu04512 |
| CD31 | GFAP | Lamb1 | ITGA7_ITGB1 | 0.000706 | 0 | LAMB1_ITGA7_ITGB1 | Lamb1 - (Itga7+Itgb1) | LAMININ | ECM-Receptor | KEGG: mmu04512 |
| CD31 | CD31 | Lamb2 | ITGA7_ITGB1 | 0.004019 | 0 | LAMB2_ITGA7_ITGB1 | Lamb2 - (Itga7+Itgb1) | LAMININ | ECM-Receptor | KEGG: mmu04512 |
| CD31 | GFAP | Lamb2 | ITGA7_ITGB1 | 0.002356 | 0 | LAMB2_ITGA7_ITGB1 | Lamb2 - (Itga7+Itgb1) | LAMININ | ECM-Receptor | KEGG: mmu04512 |
| CD31 | CD31 | Lamc1 | ITGA7_ITGB1 | 0.001993 | 0 | LAMC1_ITGA7_ITGB1 | Lamc1 - (Itga7+Itgb1) | LAMININ | ECM-Receptor | KEGG: mmu04512 |
| CD31 | GFAP | Lamc1 | ITGA7_ITGB1 | 0.001167 | 0 | LAMC1_ITGA7_ITGB1 | Lamc1 - (Itga7+Itgb1) | LAMININ | ECM-Receptor | KEGG: mmu04512 |
| CD31 | CD31 | Lamc2 | ITGA7_ITGB1 | 0.000712 | 0 | LAMC2_ITGA7_ITGB1 | Lamc2 - (Itga7+Itgb1) | LAMININ | ECM-Receptor | KEGG: mmu04512 |
| NeuN | CD31 | Lamc2 | ITGA7_ITGB1 | 0.000617 | 0 | LAMC2_ITGA7_ITGB1 | Lamc2 - (Itga7+Itgb1) | LAMININ | ECM-Receptor | KEGG: mmu04512 |
| CD31 | CD31 | Lamc3 | ITGA7_ITGB1 | 0.001503 | 0 | LAMC3_ITGA7_ITGB1 | Lamc3 - (Itga7+Itgb1) | LAMININ | ECM-Receptor | KEGG: mmu04512 |
| CD31 | GFAP | Lamc3 | ITGA7_ITGB1 | 0.00088 | 0 | LAMC3_ITGA7_ITGB1 | Lamc3 - (Itga7+Itgb1) | LAMININ | ECM-Receptor | KEGG: mmu04512 |
| CD31 | CD31 | Vtn | ITGA8_ITGB1 | 0.003494 | 0 | VTN_ITGA8_ITGB1 | Vtn - (Itga8+Itgb1) | VTN | ECM-Receptor | KEGG: mmu04512 |
| CD31 | GFAP | Vtn | ITGA8_ITGB1 | 0.002366 | 0 | VTN_ITGA8_ITGB1 | Vtn - (Itga8+Itgb1) | VTN | ECM-Receptor | KEGG: mmu04512 |
| GFAP | CD31 | Tnc | ITGA8_ITGB1 | 0.001128 | 0 | TNC_ITGA8_ITGB1 | Tnc - (Itga8+Itgb1) | TENASCIN | ECM-Receptor | KEGG: mmu04512 |
| GFAP | GFAP | Tnc | ITGA8_ITGB1 | 0.000763 | 0 | TNC_ITGA8_ITGB1 | Tnc - (Itga8+Itgb1) | TENASCIN | ECM-Receptor | KEGG: mmu04512 |
| GFAP | CD31 | Tnr | ITGA8_ITGB1 | 0.001793 | 0 | TNR_ITGA8_ITGB1 | Tnr - (Itga8+Itgb1) | TENASCIN | ECM-Receptor | KEGG: mmu04512 |
| GFAP | GFAP | Tnr | ITGA8_ITGB1 | 0.001213 | 0 | TNR_ITGA8_ITGB1 | Tnr - (Itga8+Itgb1) | TENASCIN | ECM-Receptor | KEGG: mmu04512 |
| CD31 | CD31 | Col1a1 | ITGA11_ITGB1 | 0.001626 | 0 | COL1A1_ITGA11_ITGB1 | Col1a1 - (Itga11+Itgb1) | COLLAGEN | ECM-Receptor | KEGG: mmu04512 |
| CD31 | GFAP | Col1a1 | ITGA11_ITGB1 | 0.001229 | 0 | COL1A1_ITGA11_ITGB1 | Col1a1 - (Itga11+Itgb1) | COLLAGEN | ECM-Receptor | KEGG: mmu04512 |
| CD31 | CD31 | Col1a2 | ITGA11_ITGB1 | 0.000877 | 0 | COL1A2_ITGA11_ITGB1 | Col1a2 - (Itga11+Itgb1) | COLLAGEN | ECM-Receptor | KEGG: mmu04512 |
| CD31 | GFAP | Col1a2 | ITGA11_ITGB1 | 0.000663 | 0 | COL1A2_ITGA11_ITGB1 | Col1a2 - (Itga11+Itgb1) | COLLAGEN | ECM-Receptor | KEGG: mmu04512 |
| CD31 | CD31 | Col4a1 | ITGA11_ITGB1 | 0.004689 | 0 | COL4A1_ITGA11_ITGB1 | Col4a1 - (Itga11+Itgb1) | COLLAGEN | ECM-Receptor | KEGG: mmu04512 |
| CD31 | GFAP | Col4a1 | ITGA11_ITGB1 | 0.003547 | 0 | COL4A1_ITGA11_ITGB1 | Col4a1 - (Itga11+Itgb1) | COLLAGEN | ECM-Receptor | KEGG: mmu04512 |
| CD31 | CD31 | Col4a2 | ITGA11_ITGB1 | 0.003999 | 0 | COL4A2_ITGA11_ITGB1 | Col4a2 - (Itga11+Itgb1) | COLLAGEN | ECM-Receptor | KEGG: mmu04512 |
| CD31 | GFAP | Col4a2 | ITGA11_ITGB1 | 0.003024 | 0 | COL4A2_ITGA11_ITGB1 | Col4a2 - (Itga11+Itgb1) | COLLAGEN | ECM-Receptor | KEGG: mmu04512 |
| CD31 | CD31 | Col6a1 | ITGA11_ITGB1 | 0.000744 | 0 | COL6A1_ITGA11_ITGB1 | Col6a1 - (Itga11+Itgb1) | COLLAGEN | ECM-Receptor | KEGG: mmu04512 |
| NeuN | CD31 | Col6a1 | ITGA11_ITGB1 | 0.001926 | 0 | COL6A1_ITGA11_ITGB1 | Col6a1 - (Itga11+Itgb1) | COLLAGEN | ECM-Receptor | KEGG: mmu04512 |
| CD31 | CD31 | Col6a2 | ITGA11_ITGB1 | 0.000909 | 0 | COL6A2_ITGA11_ITGB1 | Col6a2 - (Itga11+Itgb1) | COLLAGEN | ECM-Receptor | KEGG: mmu04512 |
| GFAP | CD31 | Col6a2 | ITGA11_ITGB1 | 0.000745 | 0 | COL6A2_ITGA11_ITGB1 | Col6a2 - (Itga11+Itgb1) | COLLAGEN | ECM-Receptor | KEGG: mmu04512 |
| CD31 | CD31 | Col6a4 | ITGA11_ITGB1 | 0.000617 | 0 | COL6A4_ITGA11_ITGB1 | Col6a4 - (Itga11+Itgb1) | COLLAGEN | ECM-Receptor | KEGG: mmu04512 |
| GFAP | CD31 | Col6a4 | ITGA11_ITGB1 | 0.00067 | 0 | COL6A4_ITGA11_ITGB1 | Col6a4 - (Itga11+Itgb1) | COLLAGEN | ECM-Receptor | KEGG: mmu04512 |
| CD31 | CD31 | Col6a6 | ITGA11_ITGB1 | 0.001068 | 0 | COL6A6_ITGA11_ITGB1 | Col6a6 - (Itga11+Itgb1) | COLLAGEN | ECM-Receptor | KEGG: mmu04512 |
| CD31 | GFAP | Col6a6 | ITGA11_ITGB1 | 0.000807 | 0 | COL6A6_ITGA11_ITGB1 | Col6a6 - (Itga11+Itgb1) | COLLAGEN | ECM-Receptor | KEGG: mmu04512 |
| CD31 | CD31 | Col9a2 | ITGA11_ITGB1 | 0.000604 | 0 | COL9A2_ITGA11_ITGB1 | Col9a2 - (Itga11+Itgb1) | COLLAGEN | ECM-Receptor | KEGG: mmu04512 |
| GFAP | CD31 | Col9a2 | ITGA11_ITGB1 | 0.000795 | 0 | COL9A2_ITGA11_ITGB1 | Col9a2 - (Itga11+Itgb1) | COLLAGEN | ECM-Receptor | KEGG: mmu04512 |
| GFAP | CD31 | Col9a3 | ITGA11_ITGB1 | 0.00202 | 0 | COL9A3_ITGA11_ITGB1 | Col9a3 - (Itga11+Itgb1) | COLLAGEN | ECM-Receptor | KEGG: mmu04512 |
| GFAP | GFAP | Col9a3 | ITGA11_ITGB1 | 0.001527 | 0 | COL9A3_ITGA11_ITGB1 | Col9a3 - (Itga11+Itgb1) | COLLAGEN | ECM-Receptor | KEGG: mmu04512 |
| CD31 | CD31 | Fn1 | ITGAV_ITGB1 | 0.008968 | 0 | FN1_ITGAV_ITGB1 | Fn1 - (Itgav+Itgb1) | FN1 | ECM-Receptor | KEGG: mmu04512 |
| CD31 | GFAP | Fn1 | ITGAV_ITGB1 | 0.007602 | 0 | FN1_ITGAV_ITGB1 | Fn1 - (Itgav+Itgb1) | FN1 | ECM-Receptor | KEGG: mmu04512 |
| CD31 | CD31 | Vtn | ITGAV_ITGB1 | 0.00448 | 0 | VTN_ITGAV_ITGB1 | Vtn - (Itgav+Itgb1) | VTN | ECM-Receptor | KEGG: mmu04512 |
| CD31 | GFAP | Vtn | ITGAV_ITGB1 | 0.003794 | 0 | VTN_ITGAV_ITGB1 | Vtn - (Itgav+Itgb1) | VTN | ECM-Receptor | KEGG: mmu04512 |
| CD31 | CD31 | Fn1 | ITGA4_ITGB7 | 0.002808 | 0 | FN1_ITGA4_ITGB7 | Fn1 - (Itga4+Itgb7) | FN1 | ECM-Receptor | KEGG: mmu04512 |
| CD31 | GFAP | Fn1 | ITGA4_ITGB7 | 0.002778 | 0 | FN1_ITGA4_ITGB7 | Fn1 - (Itga4+Itgb7) | FN1 | ECM-Receptor | KEGG: mmu04512 |
| CD31 | CD31 | Fn1 | ITGAV_ITGB3 | 0.004168 | 0 | FN1_ITGAV_ITGB3 | Fn1 - (Itgav+Itgb3) | FN1 | ECM-Receptor | KEGG: mmu04512 |
| CD31 | GFAP | Fn1 | ITGAV_ITGB3 | 0.004848 | 0 | FN1_ITGAV_ITGB3 | Fn1 - (Itgav+Itgb3) | FN1 | ECM-Receptor | KEGG: mmu04512 |
| CD31 | CD31 | Vtn | ITGAV_ITGB3 | 0.002077 | 0 | VTN_ITGAV_ITGB3 | Vtn - (Itgav+Itgb3) | VTN | ECM-Receptor | KEGG: mmu04512 |
| CD31 | GFAP | Vtn | ITGAV_ITGB3 | 0.002416 | 0 | VTN_ITGAV_ITGB3 | Vtn - (Itgav+Itgb3) | VTN | ECM-Receptor | KEGG: mmu04512 |
| CD31 | CD31 | Vwf | ITGAV_ITGB3 | 0.001404 | 0 | VWF_ITGAV_ITGB3 | Vwf - (Itgav+Itgb3) | VWF | ECM-Receptor | KEGG: mmu04512 |
| CD31 | GFAP | Vwf | ITGAV_ITGB3 | 0.001633 | 0 | VWF_ITGAV_ITGB3 | Vwf - (Itgav+Itgb3) | VWF | ECM-Receptor | KEGG: mmu04512 |
| GFAP | CD31 | Tnc | ITGAV_ITGB3 | 0.00067 | 0 | TNC_ITGAV_ITGB3 | Tnc - (Itgav+Itgb3) | TENASCIN | ECM-Receptor | KEGG: mmu04512 |
| GFAP | GFAP | Tnc | ITGAV_ITGB3 | 0.000779 | 0 | TNC_ITGAV_ITGB3 | Tnc - (Itgav+Itgb3) | TENASCIN | ECM-Receptor | KEGG: mmu04512 |
| GFAP | CD31 | Tnr | ITGAV_ITGB3 | 0.001065 | 0 | TNR_ITGAV_ITGB3 | Tnr - (Itgav+Itgb3) | TENASCIN | ECM-Receptor | KEGG: mmu04512 |
| GFAP | GFAP | Tnr | ITGAV_ITGB3 | 0.001239 | 0 | TNR_ITGAV_ITGB3 | Tnr - (Itgav+Itgb3) | TENASCIN | ECM-Receptor | KEGG: mmu04512 |

| | | | | | | | | | | |
|---|---|---|---|---|---|---|---|---|---|---|
| GFAP | CD31 | Thbs1 | ITGAV_ITGB3 | 0.000448 | 0 | THBS1_ITGAV_ITGB3 | Thbs1 - (Itgav+Itgb3) | THBS | ECM-Receptor | KEGG: mmu04512 |
| GFAP | GFAP | Thbs1 | ITGAV_ITGB3 | 0.000521 | 0 | THBS1_ITGAV_ITGB3 | Thbs1 - (Itgav+Itgb3) | THBS | ECM-Receptor | KEGG: mmu04512 |
| GFAP | CD31 | Thbs3 | ITGAV_ITGB3 | 0.000421 | 0 | THBS3_ITGAV_ITGB3 | Thbs3 - (Itgav+Itgb3) | THBS | ECM-Receptor | KEGG: mmu04512 |
| GFAP | GFAP | Thbs3 | ITGAV_ITGB3 | 0.00049 | 0 | THBS3_ITGAV_ITGB3 | Thbs3 - (Itgav+Itgb3) | THBS | ECM-Receptor | KEGG: mmu04512 |
| GFAP | CD31 | Thbs4 | ITGAV_ITGB3 | 0.005757 | 0 | THBS4_ITGAV_ITGB3 | Thbs4 - (Itgav+Itgb3) | THBS | ECM-Receptor | KEGG: mmu04512 |
| GFAP | GFAP | Thbs4 | ITGAV_ITGB3 | 0.006695 | 0 | THBS4_ITGAV_ITGB3 | Thbs4 - (Itgav+Itgb3) | THBS | ECM-Receptor | KEGG: mmu04512 |
| CD31 | CD31 | Fn1 | ITGA2B_ITGB3 | 0.002914 | 0 | FN1_ITGA2B_ITGB3 | Fn1 - (Itga2b+Itgb3) | FN1 | ECM-Receptor | KEGG: mmu04512 |
| CD31 | GFAP | Fn1 | ITGA2B_ITGB3 | 0.003127 | 0 | FN1_ITGA2B_ITGB3 | Fn1 - (Itga2b+Itgb3) | FN1 | ECM-Receptor | KEGG: mmu04512 |
| CD31 | CD31 | Vtn | ITGA2B_ITGB3 | 0.001451 | 0 | VTN_ITGA2B_ITGB3 | Vtn - (Itga2b+Itgb3) | VTN | ECM-Receptor | KEGG: mmu04512 |
| CD31 | GFAP | Vtn | ITGA2B_ITGB3 | 0.001558 | 0 | VTN_ITGA2B_ITGB3 | Vtn - (Itga2b+Itgb3) | VTN | ECM-Receptor | KEGG: mmu04512 |
| CD31 | CD31 | Vwf | ITGA2B_ITGB3 | 0.000981 | 0 | VWF_ITGA2B_ITGB3 | Vwf - (Itga2b+Itgb3) | VWF | ECM-Receptor | KEGG: mmu04512 |
| CD31 | GFAP | Vwf | ITGA2B_ITGB3 | 0.001052 | 0 | VWF_ITGA2B_ITGB3 | Vwf - (Itga2b+Itgb3) | VWF | ECM-Receptor | KEGG: mmu04512 |
| GFAP | CD31 | Tnc | ITGA2B_ITGB3 | 0.000468 | 0 | TNC_ITGA2B_ITGB3 | Tnc - (Itga2b+Itgb3) | TENASCIN | ECM-Receptor | KEGG: mmu04512 |
| GFAP | GFAP | Tnc | ITGA2B_ITGB3 | 0.000502 | 0 | TNC_ITGA2B_ITGB3 | Tnc - (Itga2b+Itgb3) | TENASCIN | ECM-Receptor | KEGG: mmu04512 |
| GFAP | CD31 | Tnr | ITGA2B_ITGB3 | 0.000744 | 0 | TNR_ITGA2B_ITGB3 | Tnr - (Itga2b+Itgb3) | TENASCIN | ECM-Receptor | KEGG: mmu04512 |
| GFAP | GFAP | Tnr | ITGA2B_ITGB3 | 0.000798 | 0 | TNR_ITGA2B_ITGB3 | Tnr - (Itga2b+Itgb3) | TENASCIN | ECM-Receptor | KEGG: mmu04512 |
| CD31 | CD31 | Vtn | ITGAV_ITGB5 | 0.002679 | 0 | VTN_ITGAV_ITGB5 | Vtn - (Itgav+Itgb5) | VTN | ECM-Receptor | KEGG: mmu04512 |
| CD31 | GFAP | Vtn | ITGAV_ITGB5 | 0.003023 | 0 | VTN_ITGAV_ITGB5 | Vtn - (Itgav+Itgb5) | VTN | ECM-Receptor | KEGG: mmu04512 |
| CD31 | CD31 | Vtn | ITGAV_ITGB8 | 0.00214 | 0 | VTN_ITGAV_ITGB8 | Vtn - (Itgav+Itgb8) | VTN | ECM-Receptor | KEGG: mmu04512 |
| CD31 | GFAP | Vtn | ITGAV_ITGB8 | 0.003732 | 0 | VTN_ITGAV_ITGB8 | Vtn - (Itgav+Itgb8) | VTN | ECM-Receptor | KEGG: mmu04512 |
| CD31 | CD31 | Fn1 | ITGAV_ITGB8 | 0.004294 | 0 | FN1_ITGAV_ITGB8 | Fn1 - (Itgav+Itgb8) | FN1 | ECM-Receptor | KEGG: mmu04512 |
| CD31 | GFAP | Fn1 | ITGAV_ITGB8 | 0.007478 | 0 | FN1_ITGAV_ITGB8 | Fn1 - (Itgav+Itgb8) | FN1 | ECM-Receptor | KEGG: mmu04512 |
| CD31 | GFAP | Lama1 | ITGAV_ITGB8 | 0.000979 | 0 | LAMA1_ITGAV_ITGB8 | Lama1 - (Itgav+Itgb8) | LAMININ | ECM-Receptor | KEGG: mmu04512 |
| GFAP | GFAP | Lama1 | ITGAV_ITGB8 | 0.000641 | 0 | LAMA1_ITGAV_ITGB8 | Lama1 - (Itgav+Itgb8) | LAMININ | ECM-Receptor | KEGG: mmu04512 |
| CD31 | GFAP | Lama4 | ITGAV_ITGB8 | 0.001705 | 0 | LAMA4_ITGAV_ITGB8 | Lama4 - (Itgav+Itgb8) | LAMININ | ECM-Receptor | KEGG: mmu04512 |
| GFAP | GFAP | Lama4 | ITGAV_ITGB8 | 0.001205 | 0 | LAMA4_ITGAV_ITGB8 | Lama4 - (Itgav+Itgb8) | LAMININ | ECM-Receptor | KEGG: mmu04512 |
| CD31 | GFAP | Lama5 | ITGAV_ITGB8 | 0.002094 | 0 | LAMA5_ITGAV_ITGB8 | Lama5 - (Itgav+Itgb8) | LAMININ | ECM-Receptor | KEGG: mmu04512 |
| GFAP | GFAP | Lama5 | ITGAV_ITGB8 | 0.001299 | 0 | LAMA5_ITGAV_ITGB8 | Lama5 - (Itgav+Itgb8) | LAMININ | ECM-Receptor | KEGG: mmu04512 |
| CD31 | GFAP | Lamb1 | ITGAV_ITGB8 | 0.001233 | 0 | LAMB1_ITGAV_ITGB8 | Lamb1 - (Itgav+Itgb8) | LAMININ | ECM-Receptor | KEGG: mmu04512 |
| GFAP | GFAP | Lamb1 | ITGAV_ITGB8 | 0.00071 | 0 | LAMB1_ITGAV_ITGB8 | Lamb1 - (Itgav+Itgb8) | LAMININ | ECM-Receptor | KEGG: mmu04512 |
| CD31 | GFAP | Lamb2 | ITGAV_ITGB8 | 0.004109 | 0 | LAMB2_ITGAV_ITGB8 | Lamb2 - (Itgav+Itgb8) | LAMININ | ECM-Receptor | KEGG: mmu04512 |
| GFAP | GFAP | Lamb2 | ITGAV_ITGB8 | 0.002397 | 0 | LAMB2_ITGAV_ITGB8 | Lamb2 - (Itgav+Itgb8) | LAMININ | ECM-Receptor | KEGG: mmu04512 |
| CD31 | CD31 | Lamc1 | ITGAV_ITGB8 | 0.001167 | 0 | LAMC1_ITGAV_ITGB8 | Lamc1 - (Itgav+Itgb8) | LAMININ | ECM-Receptor | KEGG: mmu04512 |
| CD31 | GFAP | Lamc1 | ITGAV_ITGB8 | 0.002038 | 0 | LAMC1_ITGAV_ITGB8 | Lamc1 - (Itgav+Itgb8) | LAMININ | ECM-Receptor | KEGG: mmu04512 |
| CD31 | GFAP | Lamc2 | ITGAV_ITGB8 | 0.000728 | 0 | LAMC2_ITGAV_ITGB8 | Lamc2 - (Itgav+Itgb8) | LAMININ | ECM-Receptor | KEGG: mmu04512 |
| GFAP | GFAP | Lamc2 | ITGAV_ITGB8 | 0.000574 | 0 | LAMC2_ITGAV_ITGB8 | Lamc2 - (Itgav+Itgb8) | LAMININ | ECM-Receptor | KEGG: mmu04512 |
| CD31 | CD31 | Lamc3 | ITGAV_ITGB8 | 0.00088 | 0 | LAMC3_ITGAV_ITGB8 | Lamc3 - (Itgav+Itgb8) | LAMININ | ECM-Receptor | KEGG: mmu04512 |
| CD31 | GFAP | Lamc3 | ITGAV_ITGB8 | 0.001537 | 0 | LAMC3_ITGAV_ITGB8 | Lamc3 - (Itgav+Itgb8) | LAMININ | ECM-Receptor | KEGG: mmu04512 |
| CD31 | CD31 | Col1a1 | ITGAV_ITGB8 | 0.001099 | 0 | COL1A1_ITGAV_ITGB8 | Col1a1 - (Itgav+Itgb8) | COLLAGEN | ECM-Receptor | KEGG: mmu04512 |
| CD31 | GFAP | Col1a1 | ITGAV_ITGB8 | 0.001918 | 0 | COL1A1_ITGAV_ITGB8 | Col1a1 - (Itgav+Itgb8) | COLLAGEN | ECM-Receptor | KEGG: mmu04512 |
| CD31 | GFAP | Col1a2 | ITGAV_ITGB8 | 0.001035 | 0 | COL1A2_ITGAV_ITGB8 | Col1a2 - (Itgav+Itgb8) | COLLAGEN | ECM-Receptor | KEGG: mmu04512 |
| GFAP | GFAP | Col1a2 | ITGAV_ITGB8 | 0.000654 | 0 | COL1A2_ITGAV_ITGB8 | Col1a2 - (Itgav+Itgb8) | COLLAGEN | ECM-Receptor | KEGG: mmu04512 |
| CD31 | CD31 | Col4a1 | ITGAV_ITGB8 | 0.003172 | 0 | COL4A1_ITGAV_ITGB8 | Col4a1 - (Itgav+Itgb8) | COLLAGEN | ECM-Receptor | KEGG: mmu04512 |
| CD31 | GFAP | Col4a1 | ITGAV_ITGB8 | 0.005529 | 0 | COL4A1_ITGAV_ITGB8 | Col4a1 - (Itgav+Itgb8) | COLLAGEN | ECM-Receptor | KEGG: mmu04512 |
| CD31 | CD31 | Col4a2 | ITGAV_ITGB8 | 0.002704 | 0 | COL4A2_ITGAV_ITGB8 | Col4a2 - (Itgav+Itgb8) | COLLAGEN | ECM-Receptor | KEGG: mmu04512 |
| CD31 | GFAP | Col4a2 | ITGAV_ITGB8 | 0.004715 | 0 | COL4A2_ITGAV_ITGB8 | Col4a2 - (Itgav+Itgb8) | COLLAGEN | ECM-Receptor | KEGG: mmu04512 |
| NeuN | GFAP | Col6a1 | ITGAV_ITGB8 | 0.002272 | 0 | COL6A1_ITGAV_ITGB8 | Col6a1 - (Itgav+Itgb8) | COLLAGEN | ECM-Receptor | KEGG: mmu04512 |
| NeuN | NeuN | Col6a1 | ITGAV_ITGB8 | 0.000965 | 0 | COL6A1_ITGAV_ITGB8 | Col6a1 - (Itgav+Itgb8) | COLLAGEN | ECM-Receptor | KEGG: mmu04512 |
| CD31 | GFAP | Col6a2 | ITGAV_ITGB8 | 0.001073 | 0 | COL6A2_ITGAV_ITGB8 | Col6a2 - (Itgav+Itgb8) | COLLAGEN | ECM-Receptor | KEGG: mmu04512 |
| GFAP | GFAP | Col6a2 | ITGAV_ITGB8 | 0.000879 | 0 | COL6A2_ITGAV_ITGB8 | Col6a2 - (Itgav+Itgb8) | COLLAGEN | ECM-Receptor | KEGG: mmu04512 |
| CD31 | GFAP | Col6a4 | ITGAV_ITGB8 | 0.000728 | 0 | COL6A4_ITGAV_ITGB8 | Col6a4 - (Itgav+Itgb8) | COLLAGEN | ECM-Receptor | KEGG: mmu04512 |
| GFAP | GFAP | Col6a4 | ITGAV_ITGB8 | 0.00079 | 0 | COL6A4_ITGAV_ITGB8 | Col6a4 - (Itgav+Itgb8) | COLLAGEN | ECM-Receptor | KEGG: mmu04512 |
| CD31 | CD31 | Col6a6 | ITGAV_ITGB8 | 0.000721 | 0 | COL6A6_ITGAV_ITGB8 | Col6a6 - (Itgav+Itgb8) | COLLAGEN | ECM-Receptor | KEGG: mmu04512 |
| CD31 | GFAP | Col6a6 | ITGAV_ITGB8 | 0.00126 | 0 | COL6A6_ITGAV_ITGB8 | Col6a6 - (Itgav+Itgb8) | COLLAGEN | ECM-Receptor | KEGG: mmu04512 |
| CD31 | GFAP | Col9a2 | ITGAV_ITGB8 | 0.000713 | 0 | COL9A2_ITGAV_ITGB8 | Col9a2 - (Itgav+Itgb8) | COLLAGEN | ECM-Receptor | KEGG: mmu04512 |
| GFAP | GFAP | Col9a2 | ITGAV_ITGB8 | 0.000938 | 0 | COL9A2_ITGAV_ITGB8 | Col9a2 - (Itgav+Itgb8) | COLLAGEN | ECM-Receptor | KEGG: mmu04512 |
| GFAP | CD31 | Col9a3 | ITGAV_ITGB8 | 0.001365 | 0 | COL9A3_ITGAV_ITGB8 | Col9a3 - (Itgav+Itgb8) | COLLAGEN | ECM-Receptor | KEGG: mmu04512 |
| GFAP | GFAP | Col9a3 | ITGAV_ITGB8 | 0.002383 | 0 | COL9A3_ITGAV_ITGB8 | Col9a3 - (Itgav+Itgb8) | COLLAGEN | ECM-Receptor | KEGG: mmu04512 |
| CD31 | GFAP | Col1a1 | Sdc4 | 0.0036 | 0 | COL1A1_SDC4 | Col1a1 - Sdc4 | COLLAGEN | ECM-Receptor | KEGG: mmu04512 |

| | | | | | | | | | | |
|---|---|---|---|---|---|---|---|---|---|---|
| GFAP | GFAP | Col1a1 | Sdc4 | 0.001842 | 0 | COL1A1_SDC4 | Col1a1 - Sdc4 | COLLAGEN | ECM-Receptor | KEGG: mmu04512 |
| CD31 | GFAP | Col1a2 | Sdc4 | 0.001944 | 0 | COL1A2_SDC4 | Col1a2 - Sdc4 | COLLAGEN | ECM-Receptor | KEGG: mmu04512 |
| GFAP | GFAP | Col1a2 | Sdc4 | 0.001228 | 0 | COL1A2_SDC4 | Col1a2 - Sdc4 | COLLAGEN | ECM-Receptor | KEGG: mmu04512 |
| CD31 | CD31 | Col4a1 | Sdc4 | 0.004127 | 0 | COL4A1_SDC4 | Col4a1 - Sdc4 | COLLAGEN | ECM-Receptor | KEGG: mmu04512 |
| CD31 | GFAP | Col4a1 | Sdc4 | 0.010344 | 0 | COL4A1_SDC4 | Col4a1 - Sdc4 | COLLAGEN | ECM-Receptor | KEGG: mmu04512 |
| CD31 | CD31 | Col4a2 | Sdc4 | 0.003519 | 0 | COL4A2_SDC4 | Col4a2 - Sdc4 | COLLAGEN | ECM-Receptor | KEGG: mmu04512 |
| CD31 | GFAP | Col4a2 | Sdc4 | 0.008828 | 0 | COL4A2_SDC4 | Col4a2 - Sdc4 | COLLAGEN | ECM-Receptor | KEGG: mmu04512 |
| GFAP | GFAP | Col6a1 | Sdc4 | 0.001483 | 0 | COL6A1_SDC4 | Col6a1 - Sdc4 | COLLAGEN | ECM-Receptor | KEGG: mmu04512 |
| NeuN | GFAP | Col6a1 | Sdc4 | 0.004263 | 0 | COL6A1_SDC4 | Col6a1 - Sdc4 | COLLAGEN | ECM-Receptor | KEGG: mmu04512 |
| CD31 | GFAP | Col6a2 | Sdc4 | 0.002015 | 0 | COL6A2_SDC4 | Col6a2 - Sdc4 | COLLAGEN | ECM-Receptor | KEGG: mmu04512 |
| GFAP | GFAP | Col6a2 | Sdc4 | 0.001652 | 0 | COL6A2_SDC4 | Col6a2 - Sdc4 | COLLAGEN | ECM-Receptor | KEGG: mmu04512 |
| CD31 | GFAP | Col6a4 | Sdc4 | 0.001367 | 0 | COL6A4_SDC4 | Col6a4 - Sdc4 | COLLAGEN | ECM-Receptor | KEGG: mmu04512 |
| GFAP | GFAP | Col6a4 | Sdc4 | 0.001485 | 0 | COL6A4_SDC4 | Col6a4 - Sdc4 | COLLAGEN | ECM-Receptor | KEGG: mmu04512 |
| CD31 | GFAP | Col6a6 | Sdc4 | 0.002366 | 0 | COL6A6_SDC4 | Col6a6 - Sdc4 | COLLAGEN | ECM-Receptor | KEGG: mmu04512 |
| GFAP | GFAP | Col6a6 | Sdc4 | 0.001258 | 0 | COL6A6_SDC4 | Col6a6 - Sdc4 | COLLAGEN | ECM-Receptor | KEGG: mmu04512 |
| CD31 | GFAP | Col9a2 | Sdc4 | 0.00134 | 0 | COL9A2_SDC4 | Col9a2 - Sdc4 | COLLAGEN | ECM-Receptor | KEGG: mmu04512 |
| GFAP | GFAP | Col9a2 | Sdc4 | 0.001762 | 0 | COL9A2_SDC4 | Col9a2 - Sdc4 | COLLAGEN | ECM-Receptor | KEGG: mmu04512 |
| GFAP | CD31 | Col9a3 | Sdc4 | 0.001777 | 0 | COL9A3_SDC4 | Col9a3 - Sdc4 | COLLAGEN | ECM-Receptor | KEGG: mmu04512 |
| GFAP | GFAP | Col9a3 | Sdc4 | 0.004471 | 0 | COL9A3_SDC4 | Col9a3 - Sdc4 | COLLAGEN | ECM-Receptor | KEGG: mmu04512 |
| CD31 | CD31 | Fn1 | Sdc4 | 0.005585 | 0 | FN1_SDC4 | Fn1 - Sdc4 | FN1 | ECM-Receptor | KEGG: mmu04512 |
| CD31 | GFAP | Fn1 | Sdc4 | 0.013968 | 0 | FN1_SDC4 | Fn1 - Sdc4 | FN1 | ECM-Receptor | KEGG: mmu04512 |
| CD31 | GFAP | Tnc | Sdc4 | 0.001178 | 0 | TNC_SDC4 | Tnc - Sdc4 | TENASCIN | ECM-Receptor | KEGG: mmu04512 |
| GFAP | GFAP | Tnc | Sdc4 | 0.002263 | 0 | TNC_SDC4 | Tnc - Sdc4 | TENASCIN | ECM-Receptor | KEGG: mmu04512 |
| GFAP | GFAP | Tnr | Sdc4 | 0.003594 | 0 | TNR_SDC4 | Tnr - Sdc4 | TENASCIN | ECM-Receptor | KEGG: mmu04512 |
| NeuN | GFAP | Tnr | Sdc4 | 0.002186 | 0 | TNR_SDC4 | Tnr - Sdc4 | TENASCIN | ECM-Receptor | KEGG: mmu04512 |
| CD31 | GFAP | Thbs1 | Sdc4 | 0.001227 | 0 | THBS1_SDC4 | Thbs1 - Sdc4 | THBS | ECM-Receptor | KEGG: mmu04512 |
| GFAP | GFAP | Thbs1 | Sdc4 | 0.001514 | 0 | THBS1_SDC4 | Thbs1 - Sdc4 | THBS | ECM-Receptor | KEGG: mmu04512 |
| CD31 | GFAP | Thbs3 | Sdc4 | 0.001058 | 0 | THBS3_SDC4 | Thbs3 - Sdc4 | THBS | ECM-Receptor | KEGG: mmu04512 |
| GFAP | GFAP | Thbs3 | Sdc4 | 0.001424 | 0 | THBS3_SDC4 | Thbs3 - Sdc4 | THBS | ECM-Receptor | KEGG: mmu04512 |
| GFAP | CD31 | Thbs4 | Sdc4 | 0.007711 | 0 | THBS4_SDC4 | Thbs4 - Sdc4 | THBS | ECM-Receptor | KEGG: mmu04512 |
| GFAP | GFAP | Thbs4 | Sdc4 | 0.019223 | 0 | THBS4_SDC4 | Thbs4 - Sdc4 | THBS | ECM-Receptor | KEGG: mmu04512 |
| CD31 | CD31 | Lama1 | Sv2a | 0.001707 | 0 | LAMA1_SV2A | Lama1 - Sv2a | LAMININ | ECM-Receptor | KEGG: mmu04512 |
| CD31 | NeuN | Lama1 | Sv2a | 0.00487 | 0 | LAMA1_SV2A | Lama1 - Sv2a | LAMININ | ECM-Receptor | KEGG: mmu04512 |
| CD31 | CD31 | Lama4 | Sv2a | 0.002973 | 0 | LAMA4_SV2A | Lama4 - Sv2a | LAMININ | ECM-Receptor | KEGG: mmu04512 |
| CD31 | NeuN | Lama4 | Sv2a | 0.00846 | 0 | LAMA4_SV2A | Lama4 - Sv2a | LAMININ | ECM-Receptor | KEGG: mmu04512 |
| CD31 | CD31 | Lama5 | Sv2a | 0.00365 | 0 | LAMA5_SV2A | Lama5 - Sv2a | LAMININ | ECM-Receptor | KEGG: mmu04512 |
| CD31 | NeuN | Lama5 | Sv2a | 0.010372 | 0 | LAMA5_SV2A | Lama5 - Sv2a | LAMININ | ECM-Receptor | KEGG: mmu04512 |
| CD31 | NeuN | Lamb1 | Sv2a | 0.006131 | 0 | LAMB1_SV2A | Lamb1 - Sv2a | LAMININ | ECM-Receptor | KEGG: mmu04512 |
| NeuN | NeuN | Lamb1 | Sv2a | 0.002254 | 0 | LAMB1_SV2A | Lamb1 - Sv2a | LAMININ | ECM-Receptor | KEGG: mmu04512 |
| CD31 | CD31 | Lamb2 | Sv2a | 0.007151 | 0 | LAMB2_SV2A | Lamb2 - Sv2a | LAMININ | ECM-Receptor | KEGG: mmu04512 |
| CD31 | NeuN | Lamb2 | Sv2a | 0.020194 | 0 | LAMB2_SV2A | Lamb2 - Sv2a | LAMININ | ECM-Receptor | KEGG: mmu04512 |
| CD31 | CD31 | Lamc1 | Sv2a | 0.003552 | 0 | LAMC1_SV2A | Lamc1 - Sv2a | LAMININ | ECM-Receptor | KEGG: mmu04512 |
| CD31 | NeuN | Lamc1 | Sv2a | 0.010098 | 0 | LAMC1_SV2A | Lamc1 - Sv2a | LAMININ | ECM-Receptor | KEGG: mmu04512 |
| CD31 | NeuN | Lamc2 | Sv2a | 0.003625 | 0 | LAMC2_SV2A | Lamc2 - Sv2a | LAMININ | ECM-Receptor | KEGG: mmu04512 |
| NeuN | NeuN | Lamc2 | Sv2a | 0.003145 | 0 | LAMC2_SV2A | Lamc2 - Sv2a | LAMININ | ECM-Receptor | KEGG: mmu04512 |
| CD31 | CD31 | Lamc3 | Sv2a | 0.00268 | 0 | LAMC3_SV2A | Lamc3 - Sv2a | LAMININ | ECM-Receptor | KEGG: mmu04512 |
| CD31 | NeuN | Lamc3 | Sv2a | 0.00763 | 0 | LAMC3_SV2A | Lamc3 - Sv2a | LAMININ | ECM-Receptor | KEGG: mmu04512 |
| CD31 | CD31 | Lama1 | Sv2b | 0.001541 | 0 | LAMA1_SV2B | Lama1 - Sv2b | LAMININ | ECM-Receptor | KEGG: mmu04512 |
| CD31 | NeuN | Lama1 | Sv2b | 0.00532 | 0 | LAMA1_SV2B | Lama1 - Sv2b | LAMININ | ECM-Receptor | KEGG: mmu04512 |
| CD31 | CD31 | Lama4 | Sv2b | 0.002684 | 0 | LAMA4_SV2B | Lama4 - Sv2b | LAMININ | ECM-Receptor | KEGG: mmu04512 |
| CD31 | NeuN | Lama4 | Sv2b | 0.00924 | 0 | LAMA4_SV2B | Lama4 - Sv2b | LAMININ | ECM-Receptor | KEGG: mmu04512 |
| CD31 | CD31 | Lama5 | Sv2b | 0.003295 | 0 | LAMA5_SV2B | Lama5 - Sv2b | LAMININ | ECM-Receptor | KEGG: mmu04512 |
| CD31 | NeuN | Lama5 | Sv2b | 0.011327 | 0 | LAMA5_SV2B | Lama5 - Sv2b | LAMININ | ECM-Receptor | KEGG: mmu04512 |
| CD31 | NeuN | Lamb1 | Sv2b | 0.006698 | 0 | LAMB1_SV2B | Lamb1 - Sv2b | LAMININ | ECM-Receptor | KEGG: mmu04512 |
| NeuN | NeuN | Lamb1 | Sv2b | 0.002464 | 0 | LAMB1_SV2B | Lamb1 - Sv2b | LAMININ | ECM-Receptor | KEGG: mmu04512 |
| CD31 | CD31 | Lamb2 | Sv2b | 0.006458 | 0 | LAMB2_SV2B | Lamb2 - Sv2b | LAMININ | ECM-Receptor | KEGG: mmu04512 |
| CD31 | NeuN | Lamb2 | Sv2b | 0.022031 | 0 | LAMB2_SV2B | Lamb2 - Sv2b | LAMININ | ECM-Receptor | KEGG: mmu04512 |
| CD31 | CD31 | Lamc1 | Sv2b | 0.003207 | 0 | LAMC1_SV2B | Lamc1 - Sv2b | LAMININ | ECM-Receptor | KEGG: mmu04512 |
| CD31 | NeuN | Lamc1 | Sv2b | 0.011027 | 0 | LAMC1_SV2B | Lamc1 - Sv2b | LAMININ | ECM-Receptor | KEGG: mmu04512 |

| | | | | | | | | | | |
|---|---|---|---|---|---|---|---|---|---|---|
| CD31 | NeuN | Lamc2 | Sv2b | 0.003961 | 0 | LAMC2_SV2B | Lamc2 - Sv2b | LAMININ | ECM-Receptor | KEGG: mmu04512 |
| NeuN | NeuN | Lamc2 | Sv2b | 0.003436 | 0 | LAMC2_SV2B | Lamc2 - Sv2b | LAMININ | ECM-Receptor | KEGG: mmu04512 |
| CD31 | CD31 | Lamc3 | Sv2b | 0.002419 | 0 | LAMC3_SV2B | Lamc3 - Sv2b | LAMININ | ECM-Receptor | KEGG: mmu04512 |
| CD31 | NeuN | Lamc3 | Sv2b | 0.008334 | 0 | LAMC3_SV2B | Lamc3 - Sv2b | LAMININ | ECM-Receptor | KEGG: mmu04512 |
| CD31 | GFAP | Lama1 | Sv2c | 0.000473 | 0 | LAMA1_SV2C | Lama1 - Sv2c | LAMININ | ECM-Receptor | KEGG: mmu04512 |
| GFAP | GFAP | Lama1 | Sv2c | 0.00031 | 0 | LAMA1_SV2C | Lama1 - Sv2c | LAMININ | ECM-Receptor | KEGG: mmu04512 |
| CD31 | GFAP | Lama4 | Sv2c | 0.000825 | 0 | LAMA4_SV2C | Lama4 - Sv2c | LAMININ | ECM-Receptor | KEGG: mmu04512 |
| GFAP | GFAP | Lama4 | Sv2c | 0.000583 | 0 | LAMA4_SV2C | Lama4 - Sv2c | LAMININ | ECM-Receptor | KEGG: mmu04512 |
| CD31 | GFAP | Lama5 | Sv2c | 0.001013 | 0 | LAMA5_SV2C | Lama5 - Sv2c | LAMININ | ECM-Receptor | KEGG: mmu04512 |
| GFAP | GFAP | Lama5 | Sv2c | 0.000628 | 0 | LAMA5_SV2C | Lama5 - Sv2c | LAMININ | ECM-Receptor | KEGG: mmu04512 |
| CD31 | CD31 | Lamb1 | Sv2c | 0.000361 | 0 | LAMB1_SV2C | Lamb1 - Sv2c | LAMININ | ECM-Receptor | KEGG: mmu04512 |
| CD31 | GFAP | Lamb1 | Sv2c | 0.000596 | 0 | LAMB1_SV2C | Lamb1 - Sv2c | LAMININ | ECM-Receptor | KEGG: mmu04512 |
| CD31 | CD31 | Lamb2 | Sv2c | 0.001204 | 0 | LAMB2_SV2C | Lamb2 - Sv2c | LAMININ | ECM-Receptor | KEGG: mmu04512 |
| CD31 | GFAP | Lamb2 | Sv2c | 0.00199 | 0 | LAMB2_SV2C | Lamb2 - Sv2c | LAMININ | ECM-Receptor | KEGG: mmu04512 |
| CD31 | CD31 | Lamc1 | Sv2c | 0.000596 | 0 | LAMC1_SV2C | Lamc1 - Sv2c | LAMININ | ECM-Receptor | KEGG: mmu04512 |
| CD31 | GFAP | Lamc1 | Sv2c | 0.000986 | 0 | LAMC1_SV2C | Lamc1 - Sv2c | LAMININ | ECM-Receptor | KEGG: mmu04512 |
| CD31 | GFAP | Lamc2 | Sv2c | 0.000352 | 0 | LAMC2_SV2C | Lamc2 - Sv2c | LAMININ | ECM-Receptor | KEGG: mmu04512 |
| GFAP | GFAP | Lamc2 | Sv2c | 0.000277 | 0 | LAMC2_SV2C | Lamc2 - Sv2c | LAMININ | ECM-Receptor | KEGG: mmu04512 |
| CD31 | CD31 | Lamc3 | Sv2c | 0.000449 | 0 | LAMC3_SV2C | Lamc3 - Sv2c | LAMININ | ECM-Receptor | KEGG: mmu04512 |
| CD31 | GFAP | Lamc3 | Sv2c | 0.000743 | 0 | LAMC3_SV2C | Lamc3 - Sv2c | LAMININ | ECM-Receptor | KEGG: mmu04512 |
| CD31 | GFAP | Thbs1 | Cd36 | 0.00021 | 0 | THBS1_CD36 | Thbs1 - Cd36 | THBS | ECM-Receptor | KEGG: mmu04512 |
| GFAP | GFAP | Thbs1 | Cd36 | 0.000259 | 0 | THBS1_CD36 | Thbs1 - Cd36 | THBS | ECM-Receptor | KEGG: mmu04512 |
| CD31 | GFAP | Thbs3 | Cd36 | 0.000181 | 0 | THBS3_CD36 | Thbs3 - Cd36 | THBS | ECM-Receptor | KEGG: mmu04512 |
| GFAP | GFAP | Thbs3 | Cd36 | 0.000243 | 0 | THBS3_CD36 | Thbs3 - Cd36 | THBS | ECM-Receptor | KEGG: mmu04512 |
| GFAP | CD31 | Thbs4 | Cd36 | 0.001903 | 0 | THBS4_CD36 | Thbs4 - Cd36 | THBS | ECM-Receptor | KEGG: mmu04512 |
| GFAP | GFAP | Thbs4 | Cd36 | 0.003333 | 0 | THBS4_CD36 | Thbs4 - Cd36 | THBS | ECM-Receptor | KEGG: mmu04512 |
| GFAP | GFAP | Thbs1 | Cd47 | 0.000437 | 0 | THBS1_CD47 | Thbs1 - Cd47 | THBS | ECM-Receptor | KEGG: mmu04512 |
| GFAP | NeuN | Thbs1 | Cd47 | 0.00084 | 0 | THBS1_CD47 | Thbs1 - Cd47 | THBS | ECM-Receptor | KEGG: mmu04512 |
| GFAP | NeuN | Thbs3 | Cd47 | 0.000791 | 0 | THBS3_CD47 | Thbs3 - Cd47 | THBS | ECM-Receptor | KEGG: mmu04512 |
| NeuN | NeuN | Thbs3 | Cd47 | 0.000585 | 0 | THBS3_CD47 | Thbs3 - Cd47 | THBS | ECM-Receptor | KEGG: mmu04512 |
| GFAP | GFAP | Thbs4 | Cd47 | 0.005619 | 0 | THBS4_CD47 | Thbs4 - Cd47 | THBS | ECM-Receptor | KEGG: mmu04512 |
| GFAP | NeuN | Thbs4 | Cd47 | 0.010757 | 0 | THBS4_CD47 | Thbs4 - Cd47 | THBS | ECM-Receptor | KEGG: mmu04512 |
| CD31 | CD31 | Agrn | Dag1 | 0.002744 | 0 | AGRN_DAG1 | Agrn - Dag1 | AGRN | ECM-Receptor | KEGG: mmu04512 |
| CD31 | GFAP | Agrn | Dag1 | 0.002082 | 0 | AGRN_DAG1 | Agrn - Dag1 | AGRN | ECM-Receptor | KEGG: mmu04512 |
| CD31 | CD31 | Hspg2 | Dag1 | 0.001033 | 0 | HSPG2_DAG1 | Hspg2 - Dag1 | HSPG | ECM-Receptor | KEGG: mmu04512 |
| CD31 | GFAP | Hspg2 | Dag1 | 0.000784 | 0 | HSPG2_DAG1 | Hspg2 - Dag1 | HSPG | ECM-Receptor | KEGG: mmu04512 |
| CD31 | CD31 | Lama1 | Dag1 | 0.000621 | 0 | LAMA1_DAG1 | Lama1 - Dag1 | LAMININ | ECM-Receptor | KEGG: mmu04512 |
| CD31 | GFAP | Lama1 | Dag1 | 0.000471 | 0 | LAMA1_DAG1 | Lama1 - Dag1 | LAMININ | ECM-Receptor | KEGG: mmu04512 |
| CD31 | CD31 | Lama4 | Dag1 | 0.001083 | 0 | LAMA4_DAG1 | Lama4 - Dag1 | LAMININ | ECM-Receptor | KEGG: mmu04512 |
| CD31 | GFAP | Lama4 | Dag1 | 0.000821 | 0 | LAMA4_DAG1 | Lama4 - Dag1 | LAMININ | ECM-Receptor | KEGG: mmu04512 |
| CD31 | CD31 | Lama5 | Dag1 | 0.00133 | 0 | LAMA5_DAG1 | Lama5 - Dag1 | LAMININ | ECM-Receptor | KEGG: mmu04512 |
| CD31 | GFAP | Lama5 | Dag1 | 0.001009 | 0 | LAMA5_DAG1 | Lama5 - Dag1 | LAMININ | ECM-Receptor | KEGG: mmu04512 |
| CD31 | CD31 | Lamb1 | Dag1 | 0.000783 | 0 | LAMB1_DAG1 | Lamb1 - Dag1 | LAMININ | ECM-Receptor | KEGG: mmu04512 |
| CD31 | GFAP | Lamb1 | Dag1 | 0.000594 | 0 | LAMB1_DAG1 | Lamb1 - Dag1 | LAMININ | ECM-Receptor | KEGG: mmu04512 |
| CD31 | CD31 | Lamb2 | Dag1 | 0.002611 | 0 | LAMB2_DAG1 | Lamb2 - Dag1 | LAMININ | ECM-Receptor | KEGG: mmu04512 |
| CD31 | GFAP | Lamb2 | Dag1 | 0.001982 | 0 | LAMB2_DAG1 | Lamb2 - Dag1 | LAMININ | ECM-Receptor | KEGG: mmu04512 |
| CD31 | CD31 | Lamc1 | Dag1 | 0.001294 | 0 | LAMC1_DAG1 | Lamc1 - Dag1 | LAMININ | ECM-Receptor | KEGG: mmu04512 |
| CD31 | GFAP | Lamc1 | Dag1 | 0.000982 | 0 | LAMC1_DAG1 | Lamc1 - Dag1 | LAMININ | ECM-Receptor | KEGG: mmu04512 |
| CD31 | CD31 | Lamc2 | Dag1 | 0.000462 | 0 | LAMC2_DAG1 | Lamc2 - Dag1 | LAMININ | ECM-Receptor | KEGG: mmu04512 |
| NeuN | CD31 | Lamc2 | Dag1 | 0.000401 | 0 | LAMC2_DAG1 | Lamc2 - Dag1 | LAMININ | ECM-Receptor | KEGG: mmu04512 |
| CD31 | CD31 | Lamc3 | Dag1 | 0.000976 | 0 | LAMC3_DAG1 | Lamc3 - Dag1 | LAMININ | ECM-Receptor | KEGG: mmu04512 |
| CD31 | GFAP | Lamc3 | Dag1 | 0.00074 | 0 | LAMC3_DAG1 | Lamc3 - Dag1 | LAMININ | ECM-Receptor | KEGG: mmu04512 |
| NeuN | CD31 | Reln | Lrp8 | 0.000848 | 0 | RELN_LRP8 | Reln - Lrp8 | RELN | ECM-Receptor | PMID: 14578885 |
| NeuN | NeuN | Reln | Lrp8 | 0.000521 | 0 | RELN_LRP8 | Reln - Lrp8 | RELN | ECM-Receptor | PMID: 14578885 |
| NeuN | GFAP | Reln | Vldlr | 0.000707 | 0 | RELN_VLDLR | Reln - Vldlr | RELN | ECM-Receptor | PMID: 14578885 |
| NeuN | NeuN | Reln | Vldlr | 0.000514 | 0 | RELN_VLDLR | Reln - Vldlr | RELN | ECM-Receptor | PMID: 14578885 |
| GFAP | GFAP | Pnoc | Oprl1 | 0.000374 | 0 | PNOC_OPRL1 | Pnoc - Oprl1 | OPIOID | Non-protein Signaling | PMID: 29934561;PMID: 34724150 |
| GFAP | NeuN | Pnoc | Oprl1 | 0.000608 | 0 | PNOC_OPRL1 | Pnoc - Oprl1 | OPIOID | Non-protein Signaling | PMID: 29934561;PMID: 34724150 |
| CD31 | CD31 | Ppia | Bsg | 0.33433 | 0 | PPIA_BSG | Ppia - Bsg | CypA | Non-protein Signaling | PMID: 36012604 |

| | | | | | | | | | | |
|---|---|---|---|---|---|---|---|---|---|---|
| NeuN | CD31 | Ppia | Bsg | 0.464019 | 0 | PPIA_BSG | Ppia - Bsg | CypA | Non-protein Signaling | PMID: 36012604 |
| NeuN | GFAP | Cadm1 | Cadm1 | 0.004812 | 0 | CADM1_CADM1 | Cadm1 - Cadm1 | CADM | Cell-Cell Contact | PMID: 24503895 |
| GFAP | NeuN | Cadm1 | Cadm1 | 0.004812 | 0 | CADM1_CADM1 | Cadm1 - Cadm1 | CADM | Cell-Cell Contact | PMID: 24503895 |
| NeuN | NeuN | Cadm1 | Cadm1 | 0.008091 | 0 | CADM1_CADM1 | Cadm1 - Cadm1 | CADM | Cell-Cell Contact | PMID: 24503895 |
| CD31 | CD31 | Cd200 | Cd200r1 | 0.000998 | 0 | CD200_CD200R1 | Cd200 - Cd200r1 | CD200 | Cell-Cell Contact | PMID: 15187158 |
| NeuN | GFAP | Cd200 | Cd200r1 | 0.001335 | 0 | CD200_CD200R1 | Cd200 - Cd200r1 | CD200 | Cell-Cell Contact | PMID: 15187158 |
| NeuN | CD31 | Cd200 | Cd200r2 | 0.001145 | 0 | CD200_CD200R2 | Cd200 - Cd200r2 | CD200 | Cell-Cell Contact | KEGG: mmu04514 |
| NeuN | NeuN | Cd200 | Cd200r2 | 0.001094 | 0 | CD200_CD200R2 | Cd200 - Cd200r2 | CD200 | Cell-Cell Contact | KEGG: mmu04514 |
| CD31 | CD31 | Cd200 | Cd200r3 | 0.000738 | 0 | CD200_CD200R3 | Cd200 - Cd200r3 | CD200 | Cell-Cell Contact | PMID: 22020332 |
| NeuN | GFAP | Cd200 | Cd200r3 | 0.001096 | 0 | CD200_CD200R3 | Cd200 - Cd200r3 | CD200 | Cell-Cell Contact | PMID: 22020332 |
| CD31 | CD31 | Entpd1 | Adora1 | 0.0031 | 0 | ENTPD1_ADORA1 | Entpd1 - Adora1 | CD39 | Cell-Cell Contact | PMID: 21677139 |
| CD31 | NeuN | Entpd1 | Adora1 | 0.005387 | 0 | ENTPD1_ADORA1 | Entpd1 - Adora1 | CD39 | Cell-Cell Contact | PMID: 21677139 |
| CD31 | CD31 | Entpd1 | Adora2a | 0.001314 | 0 | ENTPD1_ADORA2A | Entpd1 - Adora2a | CD39 | Cell-Cell Contact | PMID: 21677139 |
| CD31 | GFAP | Entpd1 | Adora2a | 0.001546 | 0 | ENTPD1_ADORA2A | Entpd1 - Adora2a | CD39 | Cell-Cell Contact | PMID: 21677139 |
| CD31 | CD31 | Entpd1 | Adora2b | 0.001245 | 0 | ENTPD1_ADORA2B | Entpd1 - Adora2b | CD39 | Cell-Cell Contact | PMID: 21677139 |
| CD31 | GFAP | Entpd1 | Adora2b | 0.001519 | 0 | ENTPD1_ADORA2B | Entpd1 - Adora2b | CD39 | Cell-Cell Contact | PMID: 21677139 |
| CD31 | CD31 | Cd46 | Jag1 | 0.0008 | 0 | CD46_JAG1 | Cd46 - Jag1 | CD46 | Cell-Cell Contact | PMID: 23086448 |
| GFAP | CD31 | Cd46 | Jag1 | 0.000632 | 0 | CD46_JAG1 | Cd46 - Jag1 | CD46 | Cell-Cell Contact | PMID: 23086448 |
| CD31 | GFAP | Cd6 | Alcam | 0.000757 | 0 | CD6_ALCAM | Cd6 - Alcam | CD6 | Cell-Cell Contact | PMID: 23602662 |
| GFAP | GFAP | Cd6 | Alcam | 0.000493 | 0 | CD6_ALCAM | Cd6 - Alcam | CD6 | Cell-Cell Contact | PMID: 23602662 |
| CD31 | CD31 | Cd96 | Pvr | 0.000456 | 0 | CD96_PVR | Cd96 - Pvr | CD96 | Cell-Cell Contact | PMID: 15034010 |
| CD31 | GFAP | Cd96 | Pvr | 0.000299 | 0 | CD96_PVR | Cd96 - Pvr | CD96 | Cell-Cell Contact | PMID: 15034010 |
| NeuN | CD31 | Cd99 | Cd99 | 0.0912 | 0 | CD99_CD99 | Cd99 - Cd99 | CD99 | Cell-Cell Contact | KEGG: mmu04514 |
| CD31 | NeuN | Cd99 | Cd99 | 0.0912 | 0 | CD99_CD99 | Cd99 - Cd99 | CD99 | Cell-Cell Contact | KEGG: mmu04514 |
| NeuN | NeuN | Cd99 | Cd99 | 0.143313 | 0 | CD99_CD99 | Cd99 - Cd99 | CD99 | Cell-Cell Contact | KEGG: mmu04514 |
| GFAP | CD31 | Cdh4 | Cdh4 | 0.000551 | 0 | CDH4_CDH4 | Cdh4 - Cdh4 | CDH | Cell-Cell Contact | KEGG: mmu04514 |
| CD31 | GFAP | Cdh4 | Cdh4 | 0.000551 | 0 | CDH4_CDH4 | Cdh4 - Cdh4 | CDH | Cell-Cell Contact | KEGG: mmu04514 |
| GFAP | GFAP | Cdh4 | Cdh4 | 0.000709 | 0 | CDH4_CDH4 | Cdh4 - Cdh4 | CDH | Cell-Cell Contact | KEGG: mmu04514 |
| GFAP | GFAP | Cdh2 | Cdh2 | 0.004083 | 0 | CDH2_CDH2 | Cdh2 - Cdh2 | CDH | Cell-Cell Contact | KEGG: mmu04514 |
| NeuN | GFAP | Cdh2 | Cdh2 | 0.003617 | 0 | CDH2_CDH2 | Cdh2 - Cdh2 | CDH | Cell-Cell Contact | KEGG: mmu04514 |
| GFAP | NeuN | Cdh2 | Cdh2 | 0.003617 | 0 | CDH2_CDH2 | Cdh2 - Cdh2 | CDH | Cell-Cell Contact | KEGG: mmu04514 |
| CD31 | CD31 | Cdh5 | Cdh5 | 0.009112 | 0 | CDH5_CDH5 | Cdh5 - Cdh5 | CDH5 | Cell-Cell Contact | KEGG: mmu04514 |
| GFAP | CD31 | Cdh5 | Cdh5 | 0.001802 | 0 | CDH5_CDH5 | Cdh5 - Cdh5 | CDH5 | Cell-Cell Contact | KEGG: mmu04514 |
| CD31 | GFAP | Cdh5 | Cdh5 | 0.001802 | 0 | CDH5_CDH5 | Cdh5 - Cdh5 | CDH5 | Cell-Cell Contact | KEGG: mmu04514 |
| GFAP | CD31 | Cldn11 | Cldn11 | 0.068206 | 0 | CLDN11_CLDN11 | Cldn11 - Cldn11 | CLDN | Cell-Cell Contact | KEGG: mmu04514 |
| CD31 | GFAP | Cldn11 | Cldn11 | 0.068206 | 0 | CLDN11_CLDN11 | Cldn11 - Cldn11 | CLDN | Cell-Cell Contact | KEGG: mmu04514 |
| GFAP | GFAP | Cldn11 | Cldn11 | 0.355014 | 0 | CLDN11_CLDN11 | Cldn11 - Cldn11 | CLDN | Cell-Cell Contact | KEGG: mmu04514 |
| CD31 | GFAP | Clec2f | Klrb1f | 0.000246 | 0 | CLEC2F_KLRB1F | Clec2f - Klrb1f | CLEC | Cell-Cell Contact | PMID: 24223577 |
| GFAP | GFAP | Clec2f | Klrb1f | 0.000238 | 0 | CLEC2F_KLRB1F | Clec2f - Klrb1f | CLEC | Cell-Cell Contact | PMID: 24223577 |
| GFAP | GFAP | Nfasc | CNTN1_CNTNAP1 | 0.01886 | 0 | NFASC_CNTN1_CNTNAP1 | Nfasc - (Cntn1+Cntnap1) | CNTN | Cell-Cell Contact | KEGG: mmu04514 |
| GFAP | NeuN | Nfasc | CNTN1_CNTNAP1 | 0.040203 | 0 | NFASC_CNTN1_CNTNAP1 | Nfasc - (Cntn1+Cntnap1) | CNTN | Cell-Cell Contact | KEGG: mmu04514 |
| GFAP | CD31 | Cntn2 | CNTN2_CNTNAP2 | 0.000861 | 0 | CNTN2_CNTN2_CNTNAP2 | Cntn2 - (Cntn2+Cntnap2) | CNTN | Cell-Cell Contact | KEGG: mmu04514 |
| GFAP | GFAP | Cntn2 | CNTN2_CNTNAP2 | 0.001426 | 0 | CNTN2_CNTN2_CNTNAP2 | Cntn2 - (Cntn2+Cntnap2) | CNTN | Cell-Cell Contact | KEGG: mmu04514 |
| GFAP | GFAP | Cntn2 | L1cam | 0.001138 | 0 | CNTN2_L1CAM | Cntn2 - L1cam | CNTN | Cell-Cell Contact | KEGG: mmu04514 |
| GFAP | NeuN | Cntn2 | L1cam | 0.002485 | 0 | CNTN2_L1CAM | Cntn2 - L1cam | CNTN | Cell-Cell Contact | KEGG: mmu04514 |
| GFAP | CD31 | Cntn2 | Cntn2 | 0.00089 | 0 | CNTN2_CNTN2 | Cntn2 - Cntn2 | CNTN | Cell-Cell Contact | KEGG: mmu04514 |
| CD31 | GFAP | Cntn2 | Cntn2 | 0.00089 | 0 | CNTN2_CNTN2 | Cntn2 - Cntn2 | CNTN | Cell-Cell Contact | KEGG: mmu04514 |
| GFAP | GFAP | Cntn2 | Cntn2 | 0.001563 | 0 | CNTN2_CNTN2 | Cntn2 - Cntn2 | CNTN | Cell-Cell Contact | KEGG: mmu04514 |
| GFAP | NeuN | Cntn1 | Nrcam | 0.025197 | 0 | CNTN1_NRCAM | Cntn1 - Nrcam | CNTN | Cell-Cell Contact | KEGG: mmu04514 |
| NeuN | NeuN | Cntn1 | Nrcam | 0.040965 | 0 | CNTN1_NRCAM | Cntn1 - Nrcam | CNTN | Cell-Cell Contact | KEGG: mmu04514 |
| CD31 | CD31 | Cspg4 | ITGA3_ITGB1 | 0.00133 | 0 | CSPG4_ITGA2_ITGB1 | Cspg4 - (Itga2+Itgb1) | CSPG4 | Cell-Cell Contact | PMID: 30213051 |
| GFAP | CD31 | Cspg4 | ITGA3_ITGB1 | 0.001239 | 0 | CSPG4_ITGA2_ITGB1 | Cspg4 - (Itga2+Itgb1) | CSPG4 | Cell-Cell Contact | PMID: 30213051 |
| CD31 | CD31 | Efna1 | Epha3 | 0.00067 | 0 | EFNA1_EPHA3 | Efna1 - Epha3 | EPHA | Cell-Cell Contact | PMID: 15114347 |
| GFAP | CD31 | Efna1 | Epha3 | 0.000508 | 0 | EFNA1_EPHA3 | Efna1 - Epha3 | EPHA | Cell-Cell Contact | PMID: 15114347 |
| CD31 | CD31 | Efna1 | Epha4 | 0.002291 | 0 | EFNA1_EPHA4 | Efna1 - Epha4 | EPHA | Cell-Cell Contact | PMID: 15114347 |
| CD31 | NeuN | Efna1 | Epha4 | 0.006353 | 0 | EFNA1_EPHA4 | Efna1 - Epha4 | EPHA | Cell-Cell Contact | PMID: 15114347 |
| GFAP | GFAP | Efna1 | Epha5 | 0.00095 | 0 | EFNA1_EPHA5 | Efna1 - Epha5 | EPHA | Cell-Cell Contact | PMID: 15114347 |
| CD31 | NeuN | Efna1 | Epha5 | 0.002227 | 0 | EFNA1_EPHA5 | Efna1 - Epha5 | EPHA | Cell-Cell Contact | PMID: 15114347 |
| CD31 | CD31 | Efna1 | Epha7 | 0.001057 | 0 | EFNA1_EPHA7 | Efna1 - Epha7 | EPHA | Cell-Cell Contact | PMID: 15114347 |

| | | | | | | | | | | |
|---|---|---|---|---|---|---|---|---|---|---|
| CD31 | NeuN | Efna1 | Epha7 | 0.00158 | 0 | EFNA1_EPHA7 | Efna1 - Epha7 | EPHA | Cell-Cell Contact | PMID: 15114347 |
| CD31 | CD31 | Efna1 | Epha8 | 0.000634 | 0 | EFNA1_EPHA8 | Efna1 - Epha8 | EPHA | Cell-Cell Contact | PMID: 15114347 |
| CD31 | GFAP | Efna1 | Epha8 | 0.000528 | 0 | EFNA1_EPHA8 | Efna1 - Epha8 | EPHA | Cell-Cell Contact | PMID: 15114347 |
| GFAP | CD31 | Efna2 | Epha3 | 0.000505 | 0 | EFNA2_EPHA3 | Efna2 - Epha3 | EPHA | Cell-Cell Contact | PMID: 15114347 |
| GFAP | GFAP | Efna2 | Epha3 | 0.00036 | 0 | EFNA2_EPHA3 | Efna2 - Epha3 | EPHA | Cell-Cell Contact | PMID: 15114347 |
| GFAP | NeuN | Efna2 | Epha4 | 0.004796 | 0 | EFNA2_EPHA4 | Efna2 - Epha4 | EPHA | Cell-Cell Contact | PMID: 15114347 |
| NeuN | NeuN | Efna2 | Epha4 | 0.002197 | 0 | EFNA2_EPHA4 | Efna2 - Epha4 | EPHA | Cell-Cell Contact | PMID: 15114347 |
| GFAP | GFAP | Efna2 | Epha5 | 0.000943 | 0 | EFNA2_EPHA5 | Efna2 - Epha5 | EPHA | Cell-Cell Contact | PMID: 15114347 |
| GFAP | NeuN | Efna2 | Epha5 | 0.001679 | 0 | EFNA2_EPHA5 | Efna2 - Epha5 | EPHA | Cell-Cell Contact | PMID: 15114347 |
| GFAP | GFAP | Efna2 | Epha7 | 0.000664 | 0 | EFNA2_EPHA7 | Efna2 - Epha7 | EPHA | Cell-Cell Contact | PMID: 15114347 |
| GFAP | NeuN | Efna2 | Epha7 | 0.001191 | 0 | EFNA2_EPHA7 | Efna2 - Epha7 | EPHA | Cell-Cell Contact | PMID: 15114347 |
| GFAP | CD31 | Efna2 | Epha8 | 0.000478 | 0 | EFNA2_EPHA8 | Efna2 - Epha8 | EPHA | Cell-Cell Contact | PMID: 15114347 |
| GFAP | GFAP | Efna2 | Epha8 | 0.000398 | 0 | EFNA2_EPHA8 | Efna2 - Epha8 | EPHA | Cell-Cell Contact | PMID: 15114347 |
| CD31 | CD31 | Efna3 | Epha3 | 0.000434 | 0 | EFNA3_EPHA3 | Efna3 - Epha3 | EPHA | Cell-Cell Contact | PMID: 15114347 |
| NeuN | CD31 | Efna3 | Epha3 | 0.000871 | 0 | EFNA3_EPHA3 | Efna3 - Epha3 | EPHA | Cell-Cell Contact | PMID: 15114347 |
| CD31 | NeuN | Efna3 | Epha4 | 0.004127 | 0 | EFNA3_EPHA4 | Efna3 - Epha4 | EPHA | Cell-Cell Contact | PMID: 15114347 |
| NeuN | NeuN | Efna3 | Epha4 | 0.008244 | 0 | EFNA3_EPHA4 | Efna3 - Epha4 | EPHA | Cell-Cell Contact | PMID: 15114347 |
| NeuN | GFAP | Efna3 | Epha5 | 0.001626 | 0 | EFNA3_EPHA5 | Efna3 - Epha5 | EPHA | Cell-Cell Contact | PMID: 15114347 |
| NeuN | NeuN | Efna3 | Epha5 | 0.002893 | 0 | EFNA3_EPHA5 | Efna3 - Epha5 | EPHA | Cell-Cell Contact | PMID: 15114347 |
| NeuN | CD31 | Efna3 | Epha7 | 0.001374 | 0 | EFNA3_EPHA7 | Efna3 - Epha7 | EPHA | Cell-Cell Contact | PMID: 15114347 |
| NeuN | NeuN | Efna3 | Epha7 | 0.002053 | 0 | EFNA3_EPHA7 | Efna3 - Epha7 | EPHA | Cell-Cell Contact | PMID: 15114347 |
| NeuN | CD31 | Efna3 | Epha8 | 0.000824 | 0 | EFNA3_EPHA8 | Efna3 - Epha8 | EPHA | Cell-Cell Contact | PMID: 15114347 |
| NeuN | NeuN | Efna3 | Epha8 | 0.00046 | 0 | EFNA3_EPHA8 | Efna3 - Epha8 | EPHA | Cell-Cell Contact | PMID: 15114347 |
| CD31 | CD31 | Efna4 | Epha3 | 0.00052 | 0 | EFNA4_EPHA3 | Efna4 - Epha3 | EPHA | Cell-Cell Contact | PMID: 15114347 |
| CD31 | GFAP | Efna4 | Epha3 | 0.000371 | 0 | EFNA4_EPHA3 | Efna4 - Epha3 | EPHA | Cell-Cell Contact | PMID: 15114347 |
| CD31 | CD31 | Efna4 | Epha4 | 0.001777 | 0 | EFNA4_EPHA4 | Efna4 - Epha4 | EPHA | Cell-Cell Contact | PMID: 15114347 |
| CD31 | NeuN | Efna4 | Epha4 | 0.004933 | 0 | EFNA4_EPHA4 | Efna4 - Epha4 | EPHA | Cell-Cell Contact | PMID: 15114347 |
| CD31 | CD31 | Efna4 | Epha5 | 0.000687 | 0 | EFNA4_EPHA5 | Efna4 - Epha5 | EPHA | Cell-Cell Contact | PMID: 15114347 |
| CD31 | NeuN | Efna4 | Epha5 | 0.001728 | 0 | EFNA4_EPHA5 | Efna4 - Epha5 | EPHA | Cell-Cell Contact | PMID: 15114347 |
| CD31 | CD31 | Efna4 | Epha7 | 0.00082 | 0 | EFNA4_EPHA7 | Efna4 - Epha7 | EPHA | Cell-Cell Contact | PMID: 15114347 |
| CD31 | NeuN | Efna4 | Epha7 | 0.001226 | 0 | EFNA4_EPHA7 | Efna4 - Epha7 | EPHA | Cell-Cell Contact | PMID: 15114347 |
| CD31 | CD31 | Efna4 | Epha8 | 0.000492 | 0 | EFNA4_EPHA8 | Efna4 - Epha8 | EPHA | Cell-Cell Contact | PMID: 15114347 |
| CD31 | GFAP | Efna4 | Epha8 | 0.00041 | 0 | EFNA4_EPHA8 | Efna4 - Epha8 | EPHA | Cell-Cell Contact | PMID: 15114347 |
| NeuN | CD31 | Efna5 | Epha3 | 0.000868 | 0 | EFNA5_EPHA3 | Efna5 - Epha3 | EPHA | Cell-Cell Contact | PMID: 15114347 |
| NeuN | NeuN | Efna5 | Epha3 | 0.000414 | 0 | EFNA5_EPHA3 | Efna5 - Epha3 | EPHA | Cell-Cell Contact | PMID: 15114347 |
| NeuN | CD31 | Efna5 | Epha4 | 0.002967 | 0 | EFNA5_EPHA4 | Efna5 - Epha4 | EPHA | Cell-Cell Contact | PMID: 15114347 |
| NeuN | NeuN | Efna5 | Epha4 | 0.008218 | 0 | EFNA5_EPHA4 | Efna5 - Epha4 | EPHA | Cell-Cell Contact | PMID: 15114347 |
| NeuN | GFAP | Efna5 | Epha5 | 0.001621 | 0 | EFNA5_EPHA5 | Efna5 - Epha5 | EPHA | Cell-Cell Contact | PMID: 15114347 |
| NeuN | NeuN | Efna5 | Epha5 | 0.002884 | 0 | EFNA5_EPHA5 | Efna5 - Epha5 | EPHA | Cell-Cell Contact | PMID: 15114347 |
| NeuN | CD31 | Efna5 | Epha7 | 0.001369 | 0 | EFNA5_EPHA7 | Efna5 - Epha7 | EPHA | Cell-Cell Contact | PMID: 15114347 |
| NeuN | NeuN | Efna5 | Epha7 | 0.002047 | 0 | EFNA5_EPHA7 | Efna5 - Epha7 | EPHA | Cell-Cell Contact | PMID: 15114347 |
| NeuN | CD31 | Efna5 | Epha8 | 0.000821 | 0 | EFNA5_EPHA8 | Efna5 - Epha8 | EPHA | Cell-Cell Contact | PMID: 15114347 |
| NeuN | NeuN | Efna5 | Epha8 | 0.000459 | 0 | EFNA5_EPHA8 | Efna5 - Epha8 | EPHA | Cell-Cell Contact | PMID: 15114347 |
| NeuN | CD31 | Efna5 | Ephb2 | 0.001448 | 0 | EFNA5_EPHB2 | Efna5 - Ephb2 | EPHA | Cell-Cell Contact | PMID:15107857; PMID: 15114347 |
| NeuN | NeuN | Efna5 | Ephb2 | 0.001734 | 0 | EFNA5_EPHB2 | Efna5 - Ephb2 | EPHA | Cell-Cell Contact | PMID:15107857; PMID: 15114347 |
| CD31 | NeuN | Efnb1 | Epha4 | 0.005242 | 0 | EFNB1_EPHA4 | Efnb1 - Epha4 | EPHB | Cell-Cell Contact | PMID: 15114347 |
| NeuN | NeuN | Efnb1 | Epha4 | 0.002974 | 0 | EFNB1_EPHA4 | Efnb1 - Epha4 | EPHB | Cell-Cell Contact | PMID: 15114347 |
| CD31 | GFAP | Efnb1 | Ephb1 | 0.002655 | 0 | EFNB1_EPHB1 | Efnb1 - Ephb1 | EPHB | Cell-Cell Contact | PMID: 15114347 |
| GFAP | GFAP | Efnb1 | Ephb1 | 0.0025 | 0 | EFNB1_EPHB1 | Efnb1 - Ephb1 | EPHB | Cell-Cell Contact | PMID: 15114347 |
| CD31 | CD31 | Efnb1 | Ephb2 | 0.000921 | 0 | EFNB1_EPHB2 | Efnb1 - Ephb2 | EPHB | Cell-Cell Contact | PMID: 15114347 |
| CD31 | NeuN | Efnb1 | Ephb2 | 0.001103 | 0 | EFNB1_EPHB2 | Efnb1 - Ephb2 | EPHB | Cell-Cell Contact | PMID: 15114347 |
| CD31 | NeuN | Efnb1 | Ephb3 | 0.000672 | 0 | EFNB1_EPHB3 | Efnb1 - Ephb3 | EPHB | Cell-Cell Contact | PMID: 15114347 |
| NeuN | NeuN | Efnb1 | Ephb3 | 0.000381 | 0 | EFNB1_EPHB3 | Efnb1 - Ephb3 | EPHB | Cell-Cell Contact | PMID: 15114347 |
| CD31 | CD31 | Efnb1 | Ephb4 | 0.001536 | 0 | EFNB1_EPHB4 | Efnb1 - Ephb4 | EPHB | Cell-Cell Contact | PMID: 15114347 |
| GFAP | CD31 | Efnb1 | Ephb4 | 0.001446 | 0 | EFNB1_EPHB4 | Efnb1 - Ephb4 | EPHB | Cell-Cell Contact | PMID: 15114347 |
| CD31 | NeuN | Efnb2 | Epha4 | 0.005777 | 0 | EFNB2_EPHA4 | Efnb2 - Epha4 | EPHB | Cell-Cell Contact | PMID: 15114347 |
| NeuN | NeuN | Efnb2 | Epha4 | 0.009952 | 0 | EFNB2_EPHA4 | Efnb2 - Epha4 | EPHB | Cell-Cell Contact | PMID: 15114347 |
| NeuN | GFAP | Efnb2 | Ephb1 | 0.005053 | 0 | EFNB2_EPHB1 | Efnb2 - Ephb1 | EPHB | Cell-Cell Contact | PMID: 15114347 |
| NeuN | NeuN | Efnb2 | Ephb1 | 0.002392 | 0 | EFNB2_EPHB1 | Efnb2 - Ephb1 | EPHB | Cell-Cell Contact | PMID: 15114347 |

| | | | | | | | | | | |
|---|---|---|---|---|---|---|---|---|---|---|
| NeuN | CD31 | Efnb2 | Ephb2 | 0.001756 | 0 | EFNB2_EPHB2 | Efnb2 - Ephb2 | EPHB | Cell-Cell Contact | PMID: 15114347 |
| NeuN | NeuN | Efnb2 | Ephb2 | 0.002103 | 0 | EFNB2_EPHB2 | Efnb2 - Ephb2 | EPHB | Cell-Cell Contact | PMID: 15114347 |
| NeuN | GFAP | Efnb2 | Ephb3 | 0.000769 | 0 | EFNB2_EPHB3 | Efnb2 - Ephb3 | EPHB | Cell-Cell Contact | PMID: 15114347 |
| NeuN | NeuN | Efnb2 | Ephb3 | 0.001281 | 0 | EFNB2_EPHB3 | Efnb2 - Ephb3 | EPHB | Cell-Cell Contact | PMID: 15114347 |
| CD31 | CD31 | Efnb2 | Ephb4 | 0.001694 | 0 | EFNB2_EPHB4 | Efnb2 - Ephb4 | EPHB | Cell-Cell Contact | PMID: 15114347 |
| NeuN | CD31 | Efnb2 | Ephb4 | 0.002926 | 0 | EFNB2_EPHB4 | Efnb2 - Ephb4 | EPHB | Cell-Cell Contact | PMID: 15114347 |
| GFAP | NeuN | Efnb3 | Epha4 | 0.023899 | 0 | EFNB3_EPHA4 | Efnb3 - Epha4 | EPHB | Cell-Cell Contact | PMID: 15114347 |
| NeuN | NeuN | Efnb3 | Epha4 | 0.006221 | 0 | EFNB3_EPHA4 | Efnb3 - Epha4 | EPHB | Cell-Cell Contact | PMID: 15114347 |
| GFAP | GFAP | Efnb3 | Ephb1 | 0.012218 | 0 | EFNB3_EPHB1 | Efnb3 - Ephb1 | EPHB | Cell-Cell Contact | PMID: 15114347 |
| GFAP | NeuN | Efnb3 | Ephb1 | 0.005805 | 0 | EFNB3_EPHB1 | Efnb3 - Ephb1 | EPHB | Cell-Cell Contact | PMID: 15114347 |
| GFAP | GFAP | Efnb3 | Ephb2 | 0.002763 | 0 | EFNB3_EPHB2 | Efnb3 - Ephb2 | EPHB | Cell-Cell Contact | PMID: 15114347 |
| GFAP | NeuN | Efnb3 | Ephb2 | 0.005106 | 0 | EFNB3_EPHB2 | Efnb3 - Ephb2 | EPHB | Cell-Cell Contact | PMID: 15114347 |
| GFAP | GFAP | Efnb3 | Ephb3 | 0.001871 | 0 | EFNB3_EPHB3 | Efnb3 - Ephb3 | EPHB | Cell-Cell Contact | PMID: 15114347 |
| GFAP | NeuN | Efnb3 | Ephb3 | 0.003115 | 0 | EFNB3_EPHB3 | Efnb3 - Ephb3 | EPHB | Cell-Cell Contact | PMID: 15114347 |
| GFAP | CD31 | Efnb3 | Ephb4 | 0.007097 | 0 | EFNB3_EPHB4 | Efnb3 - Ephb4 | EPHB | Cell-Cell Contact | PMID: 15114347 |
| GFAP | GFAP | Efnb3 | Ephb4 | 0.003025 | 0 | EFNB3_EPHB4 | Efnb3 - Ephb4 | EPHB | Cell-Cell Contact | PMID: 15114347 |
| GFAP | NeuN | Efnb3 | Ephb6 | 0.014958 | 0 | EFNB3_EPHB6 | Efnb3 - Ephb6 | EPHB | Cell-Cell Contact | PMID: 15114347 |
| NeuN | NeuN | Efnb3 | Ephb6 | 0.003868 | 0 | EFNB3_EPHB6 | Efnb3 - Ephb6 | EPHB | Cell-Cell Contact | PMID: 15114347 |
| CD31 | CD31 | Esam | Esam | 0.012185 | 0 | ESAM_ESAM | Esam - Esam | ESAM | Cell-Cell Contact | KEGG: mmu04514; PMID: 11279107 |
| GFAP | CD31 | Esam | Esam | 0.001517 | 0 | ESAM_ESAM | Esam - Esam | ESAM | Cell-Cell Contact | KEGG: mmu04514; PMID: 11279107 |
| CD31 | GFAP | Esam | Esam | 0.001517 | 0 | ESAM_ESAM | Esam - Esam | ESAM | Cell-Cell Contact | KEGG: mmu04514; PMID: 11279107 |
| NeuN | GFAP | Icam5 | Cd209a | 0.00096 | 0 | ICAM5_CD209A | Icam5 - Cd209a | ICAM | Cell-Cell Contact | PMID: 15795245 |
| NeuN | NeuN | Icam5 | Cd209a | 0.000296 | 0 | ICAM5_CD209A | Icam5 - Cd209a | ICAM | Cell-Cell Contact | PMID: 15795245 |
| CD31 | CD31 | Jam2 | ITGA3_ITGB1 | 0.000849 | 0 | JAM2_ITGA3_ITGB1 | Jam2 - (Itga3+Itgb1) | JAM | Cell-Cell Contact | PMID: 12070135 |
| GFAP | CD31 | Jam2 | ITGA3_ITGB1 | 0.000989 | 0 | JAM2_ITGA3_ITGB1 | Jam2 - (Itga3+Itgb1) | JAM | Cell-Cell Contact | PMID: 12070135 |
| CD31 | CD31 | Jam2 | ITGAV_ITGB1 | 0.001117 | 0 | JAM2_ITGAV_ITGB1 | Jam2 - (Itgav+Itgb1) | JAM | Cell-Cell Contact | KEGG: mmu04514 |
| GFAP | CD31 | Jam2 | ITGAV_ITGB1 | 0.001302 | 0 | JAM2_ITGAV_ITGB1 | Jam2 - (Itgav+Itgb1) | JAM | Cell-Cell Contact | KEGG: mmu04514 |
| CD31 | GFAP | Jam2 | Jam3 | 0.001694 | 0 | JAM2_JAM3 | Jam2 - Jam3 | JAM | Cell-Cell Contact | PMC1237098; PMID: 11590146 |
| GFAP | GFAP | Jam2 | Jam3 | 0.001973 | 0 | JAM2_JAM3 | Jam2 - Jam3 | JAM | Cell-Cell Contact | PMC1237098; PMID: 11590146 |
| CD31 | CD31 | F11r | F11r | 0.002642 | 0 | F11R_F11R | F11r - F11r | JAM | Cell-Cell Contact | KEGG: mmu04514 |
| GFAP | CD31 | F11r | F11r | 0.001085 | 0 | F11R_F11R | F11r - F11r | JAM | Cell-Cell Contact | KEGG: mmu04514 |
| CD31 | GFAP | F11r | F11r | 0.001085 | 0 | F11R_F11R | F11r - F11r | JAM | Cell-Cell Contact | KEGG: mmu04514 |
| CD31 | CD31 | Jam2 | F11r | 0.000977 | 0 | JAM2_F11R | Jam2 - F11r | JAM | Cell-Cell Contact | KEGG: mmu04514 |
| GFAP | CD31 | Jam2 | F11r | 0.001139 | 0 | JAM2_F11R | Jam2 - F11r | JAM | Cell-Cell Contact | KEGG: mmu04514 |
| GFAP | CD31 | Jam2 | Jam2 | 0.000421 | 0 | JAM2_JAM2 | Jam2 - Jam2 | JAM | Cell-Cell Contact | KEGG: mmu04514 |
| CD31 | GFAP | Jam2 | Jam2 | 0.000421 | 0 | JAM2_JAM2 | Jam2 - Jam2 | JAM | Cell-Cell Contact | KEGG: mmu04514 |
| GFAP | GFAP | Jam2 | Jam2 | 0.00049 | 0 | JAM2_JAM2 | Jam2 - Jam2 | JAM | Cell-Cell Contact | KEGG: mmu04514 |
| GFAP | CD31 | Jam3 | F11r | 0.004575 | 0 | JAM3_F11R | Jam3 - F11r | JAM | Cell-Cell Contact | KEGG: mmu04514 |
| GFAP | GFAP | Jam3 | F11r | 0.00188 | 0 | JAM3_F11R | Jam3 - F11r | JAM | Cell-Cell Contact | KEGG: mmu04514 |
| GFAP | GFAP | Jam3 | Jam3 | 0.00791 | 0 | JAM3_JAM3 | Jam3 - Jam3 | JAM | Cell-Cell Contact | KEGG: mmu04514 |
| NeuN | GFAP | Jam3 | Jam3 | 0.002693 | 0 | JAM3_JAM3 | Jam3 - Jam3 | JAM | Cell-Cell Contact | KEGG: mmu04514 |
| GFAP | NeuN | Jam3 | Jam3 | 0.002693 | 0 | JAM3_JAM3 | Jam3 - Jam3 | JAM | Cell-Cell Contact | KEGG: mmu04514 |
| NeuN | GFAP | L1cam | ITGAV_ITGB3 | 0.001991 | 0 | L1CAM_ITGAV_ITGB3 | L1cam - (Itgav+Itgb3) | L1CAM | Cell-Cell Contact | PMID: 9396761 |
| NeuN | NeuN | L1cam | ITGAV_ITGB3 | 0.001146 | 0 | L1CAM_ITGAV_ITGB3 | L1cam - (Itgav+Itgb3) | L1CAM | Cell-Cell Contact | PMID: 9396761 |
| NeuN | CD31 | L1cam | ITGA4_ITGB7 | 0.001152 | 0 | L1CAM_ITGA4_ITGB7 | L1cam - (Itga4+Itgb7) | L1CAM | Cell-Cell Contact | PMID: 8557754 |
| NeuN | NeuN | L1cam | ITGA4_ITGB7 | 0.000593 | 0 | L1CAM_ITGA4_ITGB7 | L1cam - (Itga4+Itgb7) | L1CAM | Cell-Cell Contact | PMID: 8557754 |
| NeuN | GFAP | L1cam | L1cam | 0.00181 | 0 | L1CAM_L1CAM | L1cam - L1cam | L1CAM | Cell-Cell Contact | PMID: 18701456 |
| GFAP | NeuN | L1cam | L1cam | 0.00181 | 0 | L1CAM_L1CAM | L1cam - L1cam | L1CAM | Cell-Cell Contact | PMID: 18701456 |
| NeuN | NeuN | L1cam | L1cam | 0.003948 | 0 | L1CAM_L1CAM | L1cam - L1cam | L1CAM | Cell-Cell Contact | PMID: 18701456 |
| GFAP | CD31 | Mag | Mag | 0.007685 | 0 | MAG_MAG | Mag - Mag | MAG | Cell-Cell Contact | KEGG: mmu04514 |
| CD31 | GFAP | Mag | Mag | 0.007685 | 0 | MAG_MAG | Mag - Mag | MAG | Cell-Cell Contact | KEGG: mmu04514 |
| GFAP | GFAP | Mag | Mag | 0.05949 | 0 | MAG_MAG | Mag - Mag | MAG | Cell-Cell Contact | KEGG: mmu04514 |
| GFAP | CD31 | Mpzl1 | Mpzl1 | 0.003274 | 0 | MPZL1_MPZL1 | Mpzl1 - Mpzl1 | MPZ | Cell-Cell Contact | KEGG: mmu04514 |
| CD31 | GFAP | Mpzl1 | Mpzl1 | 0.003274 | 0 | MPZL1_MPZL1 | Mpzl1 - Mpzl1 | MPZ | Cell-Cell Contact | KEGG: mmu04514 |
| GFAP | GFAP | Mpzl1 | Mpzl1 | 0.003284 | 0 | MPZL1_MPZL1 | Mpzl1 - Mpzl1 | MPZ | Cell-Cell Contact | KEGG: mmu04514 |
| GFAP | GFAP | Ncam1 | Fgfr1 | 0.008299 | 0 | NCAM1_FGFR1 | Ncam1 - Fgfr1 | NCAM | Cell-Cell Contact | PMID: 12791257 |
| NeuN | GFAP | Ncam1 | Fgfr1 | 0.007117 | 0 | NCAM1_FGFR1 | Ncam1 - Fgfr1 | NCAM | Cell-Cell Contact | PMID: 12791257 |
| GFAP | GFAP | Ncam1 | Ncam1 | 0.025408 | 0 | NCAM1_NCAM1 | Ncam1 - Ncam1 | NCAM | Cell-Cell Contact | KEGG: mmu04514 |
| NeuN | GFAP | Ncam1 | Ncam1 | 0.021842 | 0 | NCAM1_NCAM1 | Ncam1 - Ncam1 | NCAM | Cell-Cell Contact | KEGG: mmu04514 |

| | | | | | | | | | | |
|---|---|---|---|---|---|---|---|---|---|---|
| GFAP | NeuN | Ncam1 | Ncam1 | 0.021842 | 0 | NCAM1_NCAM1 | Ncam1 - Ncam1 | NCAM | Cell-Cell Contact | KEGG: mmu04514 |
| GFAP | NeuN | Ncam1 | Ncam2 | 0.010236 | 0 | NCAM1_NCAM2 | Ncam1 - Ncam2 | NCAM | Cell-Cell Contact | KEGG: mmu04514 |
| NeuN | NeuN | Ncam1 | Ncam2 | 0.00878 | 0 | NCAM1_NCAM2 | Ncam1 - Ncam2 | NCAM | Cell-Cell Contact | KEGG: mmu04514 |
| GFAP | NeuN | Ncam1 | L1cam | 0.010064 | 0 | NCAM1_L1CAM | Ncam1 - L1cam | NCAM | Cell-Cell Contact | KEGG: mmu04514 |
| NeuN | NeuN | Ncam1 | L1cam | 0.008632 | 0 | NCAM1_L1CAM | Ncam1 - L1cam | NCAM | Cell-Cell Contact | KEGG: mmu04514 |
| GFAP | NeuN | Ncam2 | L1cam | 0.002531 | 0 | NCAM2_L1CAM | Ncam2 - L1cam | NCAM | Cell-Cell Contact | KEGG: mmu04514 |
| NeuN | NeuN | Ncam2 | L1cam | 0.004017 | 0 | NCAM2_L1CAM | Ncam2 - L1cam | NCAM | Cell-Cell Contact | KEGG: mmu04514 |
| CD31 | CD31 | Nectin1 | Cd96 | 0.000582 | 0 | NECTIN1_CD96 | Nectin1 - Cd96 | NECTIN | Cell-Cell Contact | PMID: 23027581 |
| NeuN | CD31 | Nectin1 | Cd96 | 0.000784 | 0 | NECTIN1_CD96 | Nectin1 - Cd96 | NECTIN | Cell-Cell Contact | PMID: 23027581 |
| CD31 | CD31 | Nectin3 | Nectin2 | 0.000622 | 0 | NECTIN3_NECTIN2 | Nectin3 - Nectin2 | NECTIN | Cell-Cell Contact | PMID: 23027581 |
| GFAP | CD31 | Nectin3 | Nectin2 | 0.000546 | 0 | NECTIN3_NECTIN2 | Nectin3 - Nectin2 | NECTIN | Cell-Cell Contact | PMID: 23027581 |
| CD31 | CD31 | Nectin3 | Pvr | 0.000873 | 0 | NECTIN3_PVR | Nectin3 - Pvr | NECTIN | Cell-Cell Contact | PMID: 23027581 |
| GFAP | CD31 | Nectin3 | Pvr | 0.000767 | 0 | NECTIN3_PVR | Nectin3 - Pvr | NECTIN | Cell-Cell Contact | PMID: 23027581 |
| CD31 | CD31 | Nectin1 | Nectin3 | 0.001114 | 0 | NECTIN1_NECTIN3 | Nectin1 - Nectin3 | NECTIN | Cell-Cell Contact | KEGG: mmu04514 |
| NeuN | CD31 | Nectin1 | Nectin3 | 0.001501 | 0 | NECTIN1_NECTIN3 | Nectin1 - Nectin3 | NECTIN | Cell-Cell Contact | KEGG: mmu04514 |
| NeuN | CD31 | Negr1 | Negr1 | 0.003768 | 0 | NEGR1_NEGR1 | Negr1 - Negr1 | NEGR | Cell-Cell Contact | KEGG: mmu04514 |
| CD31 | NeuN | Negr1 | Negr1 | 0.003768 | 0 | NEGR1_NEGR1 | Negr1 - Negr1 | NEGR | Cell-Cell Contact | KEGG: mmu04514 |
| NeuN | NeuN | Negr1 | Negr1 | 0.008903 | 0 | NEGR1_NEGR1 | Negr1 - Negr1 | NEGR | Cell-Cell Contact | KEGG: mmu04514 |
| NeuN | CD31 | Lrrc4c | Ntng1 | 0.00109 | 0 | LRRC4C_NTNG1 | Lrrc4c - Ntng1 | NGL | Cell-Cell Contact | KEGG: mmu04514 |
| NeuN | NeuN | Lrrc4c | Ntng1 | 0.001286 | 0 | LRRC4C_NTNG1 | Lrrc4c - Ntng1 | NGL | Cell-Cell Contact | KEGG: mmu04514 |
| NeuN | CD31 | Lrrc4 | Ntng2 | 0.001181 | 0 | LRRC4_NTNG2 | Lrrc4 - Ntng2 | NGL | Cell-Cell Contact | KEGG: mmu04514 |
| NeuN | NeuN | Lrrc4 | Ntng2 | 0.001174 | 0 | LRRC4_NTNG2 | Lrrc4 - Ntng2 | NGL | Cell-Cell Contact | KEGG: mmu04514 |
| NeuN | GFAP | Lrrc4b | Ptprf | 0.00431 | 0 | LRRC4B_PTPRF | Lrrc4b - Ptprf | NGL | Cell-Cell Contact | KEGG: mmu04514 |
| NeuN | NeuN | Lrrc4b | Ptprf | 0.003346 | 0 | LRRC4B_PTPRF | Lrrc4b - Ptprf | NGL | Cell-Cell Contact | KEGG: mmu04514 |
| CD31 | CD31 | Dlk1 | Notch1 | 0.000852 | 0 | DLK1_NOTCH1 | Dlk1 - Notch1 | NOTCH | Cell-Cell Contact | PMID: 22353464 |
| CD31 | GFAP | Dlk1 | Notch1 | 0.000746 | 0 | DLK1_NOTCH1 | Dlk1 - Notch1 | NOTCH | Cell-Cell Contact | PMID: 22353464 |
| CD31 | CD31 | Dlk1 | Notch3 | 0.001469 | 0 | DLK1_NOTCH3 | Dlk1 - Notch3 | NOTCH | Cell-Cell Contact | PMID: 22353464 |
| GFAP | CD31 | Dlk1 | Notch3 | 0.001119 | 0 | DLK1_NOTCH3 | Dlk1 - Notch3 | NOTCH | Cell-Cell Contact | PMID: 22353464 |
| CD31 | CD31 | Dlk1 | Notch4 | 0.000726 | 0 | DLK1_NOTCH4 | Dlk1 - Notch4 | NOTCH | Cell-Cell Contact | PMID: 22353464 |
| GFAP | CD31 | Dlk1 | Notch4 | 0.000553 | 0 | DLK1_NOTCH4 | Dlk1 - Notch4 | NOTCH | Cell-Cell Contact | PMID: 22353464 |
| GFAP | CD31 | Dll1 | Notch1 | 0.001154 | 0 | DLL1_NOTCH1 | Dll1 - Notch1 | NOTCH | Cell-Cell Contact | PMID: 22353464 |
| GFAP | GFAP | Dll1 | Notch1 | 0.001011 | 0 | DLL1_NOTCH1 | Dll1 - Notch1 | NOTCH | Cell-Cell Contact | PMID: 22353464 |
| CD31 | CD31 | Dll1 | Notch3 | 0.00126 | 0 | DLL1_NOTCH3 | Dll1 - Notch3 | NOTCH | Cell-Cell Contact | PMID: 22353464 |
| GFAP | CD31 | Dll1 | Notch3 | 0.001991 | 0 | DLL1_NOTCH3 | Dll1 - Notch3 | NOTCH | Cell-Cell Contact | PMID: 22353464 |
| CD31 | CD31 | Dll1 | Notch4 | 0.000622 | 0 | DLL1_NOTCH4 | Dll1 - Notch4 | NOTCH | Cell-Cell Contact | PMID: 22353464 |
| GFAP | CD31 | Dll1 | Notch4 | 0.000983 | 0 | DLL1_NOTCH4 | Dll1 - Notch4 | NOTCH | Cell-Cell Contact | PMID: 22353464 |
| CD31 | CD31 | Dll3 | Notch1 | 0.000893 | 0 | DLL3_NOTCH1 | Dll3 - Notch1 | NOTCH | Cell-Cell Contact | PMID: 22353464 |
| CD31 | GFAP | Dll3 | Notch1 | 0.000782 | 0 | DLL3_NOTCH1 | Dll3 - Notch1 | NOTCH | Cell-Cell Contact | PMID: 22353464 |
| CD31 | CD31 | Dll3 | Notch3 | 0.00154 | 0 | DLL3_NOTCH3 | Dll3 - Notch3 | NOTCH | Cell-Cell Contact | PMID: 22353464 |
| GFAP | CD31 | Dll3 | Notch3 | 0.000966 | 0 | DLL3_NOTCH3 | Dll3 - Notch3 | NOTCH | Cell-Cell Contact | PMID: 22353464 |
| CD31 | CD31 | Dll3 | Notch4 | 0.000761 | 0 | DLL3_NOTCH4 | Dll3 - Notch4 | NOTCH | Cell-Cell Contact | PMID: 22353464 |
| GFAP | CD31 | Dll3 | Notch4 | 0.000477 | 0 | DLL3_NOTCH4 | Dll3 - Notch4 | NOTCH | Cell-Cell Contact | PMID: 22353464 |
| CD31 | CD31 | Dll4 | Notch1 | 0.001724 | 0 | DLL4_NOTCH1 | Dll4 - Notch1 | NOTCH | Cell-Cell Contact | PMID: 22353464 |
| CD31 | GFAP | Dll4 | Notch1 | 0.00151 | 0 | DLL4_NOTCH1 | Dll4 - Notch1 | NOTCH | Cell-Cell Contact | PMID: 22353464 |
| CD31 | CD31 | Dll4 | Notch3 | 0.002974 | 0 | DLL4_NOTCH3 | Dll4 - Notch3 | NOTCH | Cell-Cell Contact | PMID: 22353464 |
| CD31 | GFAP | Dll4 | Notch3 | 0.001185 | 0 | DLL4_NOTCH3 | Dll4 - Notch3 | NOTCH | Cell-Cell Contact | PMID: 22353464 |
| CD31 | CD31 | Dll4 | Notch4 | 0.00147 | 0 | DLL4_NOTCH4 | Dll4 - Notch4 | NOTCH | Cell-Cell Contact | PMID: 22353464 |
| GFAP | CD31 | Dll4 | Notch4 | 0.000521 | 0 | DLL4_NOTCH4 | Dll4 - Notch4 | NOTCH | Cell-Cell Contact | PMID: 22353464 |
| CD31 | CD31 | Jag1 | Notch1 | 0.001654 | 0 | JAG1_NOTCH1 | Jag1 - Notch1 | NOTCH | Cell-Cell Contact | PMID: 22353464 |
| CD31 | GFAP | Jag1 | Notch1 | 0.001449 | 0 | JAG1_NOTCH1 | Jag1 - Notch1 | NOTCH | Cell-Cell Contact | PMID: 22353464 |
| CD31 | CD31 | Jag1 | Notch3 | 0.002853 | 0 | JAG1_NOTCH3 | Jag1 - Notch3 | NOTCH | Cell-Cell Contact | PMID: 22353464 |
| CD31 | GFAP | Jag1 | Notch3 | 0.001137 | 0 | JAG1_NOTCH3 | Jag1 - Notch3 | NOTCH | Cell-Cell Contact | PMID: 22353464 |
| CD31 | CD31 | Jag1 | Notch4 | 0.00141 | 0 | JAG1_NOTCH4 | Jag1 - Notch4 | NOTCH | Cell-Cell Contact | PMID: 22353464 |
| GFAP | CD31 | Jag1 | Notch4 | 0.000517 | 0 | JAG1_NOTCH4 | Jag1 - Notch4 | NOTCH | Cell-Cell Contact | PMID: 22353464 |
| CD31 | CD31 | Jag2 | Notch1 | 0.001684 | 0 | JAG2_NOTCH1 | Jag2 - Notch1 | NOTCH | Cell-Cell Contact | PMID: 22353464 |
| CD31 | GFAP | Jag2 | Notch1 | 0.001474 | 0 | JAG2_NOTCH1 | Jag2 - Notch1 | NOTCH | Cell-Cell Contact | PMID: 22353464 |
| CD31 | CD31 | Jag2 | Notch3 | 0.002903 | 0 | JAG2_NOTCH3 | Jag2 - Notch3 | NOTCH | Cell-Cell Contact | PMID: 22353464 |
| NeuN | CD31 | Jag2 | Notch3 | 0.00177 | 0 | JAG2_NOTCH3 | Jag2 - Notch3 | NOTCH | Cell-Cell Contact | PMID: 22353464 |
| CD31 | CD31 | Jag2 | Notch4 | 0.001435 | 0 | JAG2_NOTCH4 | Jag2 - Notch4 | NOTCH | Cell-Cell Contact | PMID: 22353464 |

| | | | | | | | | | |
|---|---|---|---|---|---|---|---|---|---|
| NeuN | CD31 | Jag2 | Notch4 | 0.000874 | 0 JAG2_NOTCH4 | Jag2 - Notch4 | NOTCH | Cell-Cell Contact | PMID: 22353464 |
| NeuN | GFAP | Nrxn1 | Nlgn1 | 0.008275 | 0 NRXN1_NLGN1 | Nrxn1 - Nlgn1 | NRXN | Cell-Cell Contact | KEGG: mmu04514 |
| NeuN | NeuN | Nrxn1 | Nlgn1 | 0.013222 | 0 NRXN1_NLGN1 | Nrxn1 - Nlgn1 | NRXN | Cell-Cell Contact | KEGG: mmu04514 |
| GFAP | NeuN | Nrxn1 | Nlgn2 | 0.015054 | 0 NRXN1_NLGN2 | Nrxn1 - Nlgn2 | NRXN | Cell-Cell Contact | KEGG: mmu04514 |
| NeuN | NeuN | Nrxn1 | Nlgn2 | 0.029189 | 0 NRXN1_NLGN2 | Nrxn1 - Nlgn2 | NRXN | Cell-Cell Contact | KEGG: mmu04514 |
| NeuN | GFAP | Nrxn1 | Nlgn3 | 0.015921 | 0 NRXN1_NLGN3 | Nrxn1 - Nlgn3 | NRXN | Cell-Cell Contact | KEGG: mmu04514 |
| NeuN | NeuN | Nrxn1 | Nlgn3 | 0.025595 | 0 NRXN1_NLGN3 | Nrxn1 - Nlgn3 | NRXN | Cell-Cell Contact | KEGG: mmu04514 |
| GFAP | NeuN | Nrxn2 | Nlgn1 | 0.004357 | 0 NRXN2_NLGN1 | Nrxn2 - Nlgn1 | NRXN | Cell-Cell Contact | KEGG: mmu04514 |
| NeuN | NeuN | Nrxn2 | Nlgn1 | 0.004829 | 0 NRXN2_NLGN1 | Nrxn2 - Nlgn1 | NRXN | Cell-Cell Contact | KEGG: mmu04514 |
| GFAP | NeuN | Nrxn2 | Nlgn2 | 0.009724 | 0 NRXN2_NLGN2 | Nrxn2 - Nlgn2 | NRXN | Cell-Cell Contact | KEGG: mmu04514 |
| NeuN | NeuN | Nrxn2 | Nlgn2 | 0.010771 | 0 NRXN2_NLGN2 | Nrxn2 - Nlgn2 | NRXN | Cell-Cell Contact | KEGG: mmu04514 |
| GFAP | NeuN | Nrxn2 | Nlgn3 | 0.008506 | 0 NRXN2_NLGN3 | Nrxn2 - Nlgn3 | NRXN | Cell-Cell Contact | KEGG: mmu04514 |
| NeuN | NeuN | Nrxn2 | Nlgn3 | 0.009423 | 0 NRXN2_NLGN3 | Nrxn2 - Nlgn3 | NRXN | Cell-Cell Contact | KEGG: mmu04514 |
| NeuN | GFAP | Nrxn3 | Nlgn1 | 0.004269 | 0 NRXN3_NLGN1 | Nrxn3 - Nlgn1 | NRXN | Cell-Cell Contact | KEGG: mmu04514 |
| NeuN | NeuN | Nrxn3 | Nlgn1 | 0.006837 | 0 NRXN3_NLGN1 | Nrxn3 - Nlgn1 | NRXN | Cell-Cell Contact | KEGG: mmu04514 |
| CD31 | NeuN | Nrxn3 | Nlgn2 | 0.007397 | 0 NRXN3_NLGN2 | Nrxn3 - Nlgn2 | NRXN | Cell-Cell Contact | KEGG: mmu04514 |
| NeuN | NeuN | Nrxn3 | Nlgn2 | 0.015213 | 0 NRXN3_NLGN2 | Nrxn3 - Nlgn2 | NRXN | Cell-Cell Contact | KEGG: mmu04514 |
| NeuN | GFAP | Nrxn3 | Nlgn3 | 0.008244 | 0 NRXN3_NLGN3 | Nrxn3 - Nlgn3 | NRXN | Cell-Cell Contact | KEGG: mmu04514 |
| NeuN | NeuN | Nrxn3 | Nlgn3 | 0.013316 | 0 NRXN3_NLGN3 | Nrxn3 - Nlgn3 | NRXN | Cell-Cell Contact | KEGG: mmu04514 |
| CD31 | CD31 | Ocln | Ocln | 0.006489 | 0 OCLN_OCLN | Ocln - Ocln | OCLN | Cell-Cell Contact | KEGG: mmu04514 |
| GFAP | CD31 | Ocln | Ocln | 0.001249 | 0 OCLN_OCLN | Ocln - Ocln | OCLN | Cell-Cell Contact | KEGG: mmu04514 |
| CD31 | GFAP | Ocln | Ocln | 0.001249 | 0 OCLN_OCLN | Ocln - Ocln | OCLN | Cell-Cell Contact | KEGG: mmu04514 |
| CD31 | CD31 | Pecam1 | Pecam1 | 0.002353 | 0 PECAM1_PECAM1 | Pecam1 - Pecam1 | PECAM1 | Cell-Cell Contact | KEGG: mmu04514 |
| GFAP | CD31 | Pecam1 | Pecam1 | 0.000969 | 0 PECAM1_PECAM1 | Pecam1 - Pecam1 | PECAM1 | Cell-Cell Contact | KEGG: mmu04514 |
| CD31 | GFAP | Pecam1 | Pecam1 | 0.000969 | 0 PECAM1_PECAM1 | Pecam1 - Pecam1 | PECAM1 | Cell-Cell Contact | KEGG: mmu04514 |
| CD31 | CD31 | Ptprm | Ptprm | 0.016137 | 0 PTPRM_PTPRM | Ptprm - Ptprm | PTPRM | Cell-Cell Contact | KEGG: mmu04514 |
| NeuN | CD31 | Ptprm | Ptprm | 0.005268 | 0 PTPRM_PTPRM | Ptprm - Ptprm | PTPRM | Cell-Cell Contact | KEGG: mmu04514 |
| CD31 | NeuN | Ptprm | Ptprm | 0.005268 | 0 PTPRM_PTPRM | Ptprm - Ptprm | PTPRM | Cell-Cell Contact | KEGG: mmu04514 |
| GFAP | GFAP | Sele | Glg1 | 0.000447 | 0 SELE_GLG1 | Sele - Glg1 | SELE | Cell-Cell Contact | PMID: 11404363 |
| GFAP | NeuN | Sele | Glg1 | 0.0011 | 0 SELE_GLG1 | Sele - Glg1 | SELE | Cell-Cell Contact | PMID: 11404363 |
| CD31 | CD31 | Sell | Cd34 | 0.00157 | 0 SELL_CD34 | Sell - Cd34 | SELL | Cell-Cell Contact | PMID: 7692600; PMID: 8977216 |
| GFAP | CD31 | Sell | Cd34 | 0.001909 | 0 SELL_CD34 | Sell - Cd34 | SELL | Cell-Cell Contact | PMID: 7692600; PMID: 8977216 |
| CD31 | CD31 | Sell | Podxl | 0.001285 | 0 SELL_PODXL | Sell - Podxl | SELL | Cell-Cell Contact | PMID: 22814396 |
| GFAP | CD31 | Sell | Podxl | 0.001563 | 0 SELL_PODXL | Sell - Podxl | SELL | Cell-Cell Contact | PMID: 22814396 |
| CD31 | GFAP | Selplg | Sele | 0.000757 | 0 SELPLG_SELE | Selplg - Sele | SELPLG | Cell-Cell Contact | PMC4571854 |
| GFAP | GFAP | Selplg | Sele | 0.000836 | 0 SELPLG_SELE | Selplg - Sele | SELPLG | Cell-Cell Contact | PMC4571854 |
| CD31 | GFAP | Selplg | Sell | 0.000561 | 0 SELPLG_SELL | Selplg - Sell | SELPLG | Cell-Cell Contact | PMC2431087; PMID: 8892633,PMID: 8896607 |
| GFAP | GFAP | Selplg | Sell | 0.000619 | 0 SELPLG_SELL | Selplg - Sell | SELPLG | Cell-Cell Contact | PMC2431087; PMID: 8892633,PMID: 8896607 |
| NeuN | CD31 | Sema4a | NRP1_PLXNA1 | 0.001673 | 0 SEMA4A_NRP1_PLXNA1 | Sema4a - (Nrp1+Plxna1) | SEMA4 | Cell-Cell Contact | PMID: 27533782 |
| NeuN | NeuN | Sema4a | NRP1_PLXNA1 | 0.001485 | 0 SEMA4A_NRP1_PLXNA1 | Sema4a - (Nrp1+Plxna1) | SEMA4 | Cell-Cell Contact | PMID: 27533782 |
| CD31 | CD31 | Sema4a | NRP1_PLXNA2 | 0.00185 | 0 SEMA4A_NRP1_PLXNA2 | Sema4a - (Nrp1+Plxna2) | SEMA4 | Cell-Cell Contact | PMID: 27533782 |
| NeuN | CD31 | Sema4a | NRP1_PLXNA2 | 0.002468 | 0 SEMA4A_NRP1_PLXNA2 | Sema4a - (Nrp1+Plxna2) | SEMA4 | Cell-Cell Contact | PMID: 27533782 |
| CD31 | CD31 | Sema4a | NRP1_PLXNA3 | 0.000774 | 0 SEMA4A_NRP1_PLXNA3 | Sema4a - (Nrp1+Plxna3) | SEMA4 | Cell-Cell Contact | PMID: 27533782 |
| NeuN | CD31 | Sema4a | NRP1_PLXNA3 | 0.001033 | 0 SEMA4A_NRP1_PLXNA3 | Sema4a - (Nrp1+Plxna3) | SEMA4 | Cell-Cell Contact | PMID: 27533782 |
| NeuN | CD31 | Sema4a | NRP1_PLXNA4 | 0.001988 | 0 SEMA4A_NRP1_PLXNA4 | Sema4a - (Nrp1+Plxna4) | SEMA4 | Cell-Cell Contact | PMID: 27533782 |
| NeuN | NeuN | Sema4a | NRP1_PLXNA4 | 0.001754 | 0 SEMA4A_NRP1_PLXNA4 | Sema4a - (Nrp1+Plxna4) | SEMA4 | Cell-Cell Contact | PMID: 27533782 |
| GFAP | GFAP | Sema4a | Plxnb1 | 0.001433 | 0 SEMA4A_PLXNB1 | Sema4a - Plxnb1 | SEMA4 | Cell-Cell Contact | PMID: 27533782 |
| NeuN | GFAP | Sema4a | Plxnb1 | 0.002184 | 0 SEMA4A_PLXNB1 | Sema4a - Plxnb1 | SEMA4 | Cell-Cell Contact | PMID: 27533782 |
| GFAP | GFAP | Sema4a | Plxnb2 | 0.001576 | 0 SEMA4A_PLXNB2 | Sema4a - Plxnb2 | SEMA4 | Cell-Cell Contact | PMID: 27533782 |
| NeuN | GFAP | Sema4a | Plxnb2 | 0.002402 | 0 SEMA4A_PLXNB2 | Sema4a - Plxnb2 | SEMA4 | Cell-Cell Contact | PMID: 27533782 |
| GFAP | GFAP | Sema4a | Plxnb3 | 0.001051 | 0 SEMA4A_PLXNB3 | Sema4a - Plxnb3 | SEMA4 | Cell-Cell Contact | PMID: 27533782 |
| NeuN | GFAP | Sema4a | Plxnb3 | 0.001603 | 0 SEMA4A_PLXNB3 | Sema4a - Plxnb3 | SEMA4 | Cell-Cell Contact | PMID: 27533782 |
| GFAP | CD31 | Sema4d | Plxnb1 | 0.002941 | 0 SEMA4D_PLXNB1 | Sema4d - Plxnb1 | SEMA4 | Cell-Cell Contact | PMID: 27533782 |
| GFAP | GFAP | Sema4d | Plxnb1 | 0.00558 | 0 SEMA4D_PLXNB1 | Sema4d - Plxnb1 | SEMA4 | Cell-Cell Contact | PMID: 27533782 |
| GFAP | CD31 | Sema4d | Plxnb2 | 0.004111 | 0 SEMA4D_PLXNB2 | Sema4d - Plxnb2 | SEMA4 | Cell-Cell Contact | PMID: 27533782 |
| GFAP | GFAP | Sema4d | Plxnb2 | 0.006133 | 0 SEMA4D_PLXNB2 | Sema4d - Plxnb2 | SEMA4 | Cell-Cell Contact | PMID: 27533782 |
| GFAP | CD31 | Sema4d | Plxnb3 | 0.001858 | 0 SEMA4D_PLXNB3 | Sema4d - Plxnb3 | SEMA4 | Cell-Cell Contact | PMID: 27533782 |
| GFAP | GFAP | Sema4d | Plxnb3 | 0.004099 | 0 SEMA4D_PLXNB3 | Sema4d - Plxnb3 | SEMA4 | Cell-Cell Contact | PMID: 27533782 |
| GFAP | GFAP | Sema4c | Plxnb2 | 0.000996 | 0 SEMA4C_PLXNB2 | Sema4c - Plxnb2 | SEMA4 | Cell-Cell Contact | PMID: 27533782 |

| | | | | | | | | | | |
|---|---|---|---|---|---|---|---|---|---|---|
| NeuN | GFAP | Sema4c | Plxnb2 | 0.000915 | 0 | SEMA4C_PLXNB2 | Sema4c - Plxnb2 | SEMA4 | Cell-Cell Contact | PMID: 27533782 |
| CD31 | GFAP | Sema4g | Plxnb2 | 0.001263 | 0 | SEMA4G_PLXNB2 | Sema4g - Plxnb2 | SEMA4 | Cell-Cell Contact | PMID: 27533782 |
| GFAP | GFAP | Sema4g | Plxnb2 | 0.001072 | 0 | SEMA4G_PLXNB2 | Sema4g - Plxnb2 | SEMA4 | Cell-Cell Contact | PMID: 27533782 |
| GFAP | GFAP | Sema4a | Timd2 | 0.000632 | 0 | SEMA4A_TIMD2 | Sema4a - Timd2 | SEMA4 | Cell-Cell Contact | PMID: 22325954 |
| NeuN | GFAP | Sema4a | Timd2 | 0.000964 | 0 | SEMA4A_TIMD2 | Sema4a - Timd2 | SEMA4 | Cell-Cell Contact | PMID: 22325954 |
| GFAP | NeuN | Sema5a | Plxna1 | 0.00208 | 0 | SEMA5A_PLXNA1 | Sema5a - Plxna1 | SEMA5 | Cell-Cell Contact | PMID: 27533782 |
| NeuN | NeuN | Sema5a | Plxna1 | 0.001013 | 0 | SEMA5A_PLXNA1 | Sema5a - Plxna1 | SEMA5 | Cell-Cell Contact | PMID: 27533782 |
| GFAP | NeuN | Sema5b | Plxna1 | 0.001038 | 0 | SEMA5B_PLXNA1 | Sema5b - Plxna1 | SEMA5 | Cell-Cell Contact | PMID: 27533782 |
| NeuN | NeuN | Sema5b | Plxna1 | 0.000791 | 0 | SEMA5B_PLXNA1 | Sema5b - Plxna1 | SEMA5 | Cell-Cell Contact | PMID: 27533782 |
| GFAP | GFAP | Sema5a | Plxna3 | 0.00078 | 0 | SEMA5A_PLXNA3 | Sema5a - Plxna3 | SEMA5 | Cell-Cell Contact | PMID: 27533782 |
| GFAP | NeuN | Sema5a | Plxna3 | 0.000535 | 0 | SEMA5A_PLXNA3 | Sema5a - Plxna3 | SEMA5 | Cell-Cell Contact | PMID: 27533782 |
| GFAP | GFAP | Sema5b | Plxna3 | 0.000389 | 0 | SEMA5B_PLXNA3 | Sema5b - Plxna3 | SEMA5 | Cell-Cell Contact | PMID: 27533782 |
| NeuN | GFAP | Sema5b | Plxna3 | 0.000296 | 0 | SEMA5B_PLXNA3 | Sema5b - Plxna3 | SEMA5 | Cell-Cell Contact | PMID: 27533782 |
| CD31 | GFAP | Sema5a | Plxnb3 | 0.000971 | 0 | SEMA5A_PLXNB3 | Sema5a - Plxnb3 | SEMA5 | Cell-Cell Contact | PMID: 27533782 |
| GFAP | GFAP | Sema5a | Plxnb3 | 0.001852 | 0 | SEMA5A_PLXNB3 | Sema5a - Plxnb3 | SEMA5 | Cell-Cell Contact | PMID: 27533782 |
| CD31 | CD31 | Sema6d | PLXNA1_KDR | 0.001931 | 0 | SEMA6D_PLXNA1_KDR | Sema6d - (Plxna1+Kdr) | SEMA6 | Cell-Cell Contact | PMID: 22325954 |
| GFAP | CD31 | Sema6d | PLXNA1_KDR | 0.002168 | 0 | SEMA6D_PLXNA1_KDR | Sema6d - (Plxna1+Kdr) | SEMA6 | Cell-Cell Contact | PMID: 22325954 |
| GFAP | CD31 | Sema6d | PLXNA1_TREM2 | 0.001123 | 0 | SEMA6D_PLXNA1_TREM2 | Sema6d - (Plxna1+Trem2) | SEMA6 | Cell-Cell Contact | PMID: 22325954 |
| GFAP | GFAP | Sema6d | PLXNA1_TREM2 | 0.00122 | 0 | SEMA6D_PLXNA1_TREM2 | Sema6d - (Plxna1+Trem2) | SEMA6 | Cell-Cell Contact | PMID: 22325954 |
| CD31 | CD31 | Sema6d | Plxna1 | 0.001111 | 0 | SEMA6D_PLXNA1 | Sema6d - Plxna1 | SEMA6 | Cell-Cell Contact | PMID: 27533782 |
| GFAP | NeuN | Sema6d | Plxna1 | 0.00189 | 0 | SEMA6D_PLXNA1 | Sema6d - Plxna1 | SEMA6 | Cell-Cell Contact | PMID: 27533782 |
| CD31 | CD31 | Sema6a | Plxna2 | 0.002409 | 0 | SEMA6A_PLXNA2 | Sema6a - Plxna2 | SEMA6 | Cell-Cell Contact | PMID: 27533782 |
| GFAP | NeuN | Sema6a | Plxna2 | 0.005129 | 0 | SEMA6A_PLXNA2 | Sema6a - Plxna2 | SEMA6 | Cell-Cell Contact | PMID: 27533782 |
| NeuN | CD31 | Sema6b | Plxna2 | 0.003344 | 0 | SEMA6B_PLXNA2 | Sema6b - Plxna2 | SEMA6 | Cell-Cell Contact | PMID: 27533782 |
| NeuN | NeuN | Sema6b | Plxna2 | 0.003427 | 0 | SEMA6B_PLXNA2 | Sema6b - Plxna2 | SEMA6 | Cell-Cell Contact | PMID: 27533782 |
| GFAP | GFAP | Sema6a | Plxna4 | 0.002176 | 0 | SEMA6A_PLXNA4 | Sema6a - Plxna4 | SEMA6 | Cell-Cell Contact | PMID: 27533782 |
| GFAP | NeuN | Sema6a | Plxna4 | 0.004863 | 0 | SEMA6A_PLXNA4 | Sema6a - Plxna4 | SEMA6 | Cell-Cell Contact | PMID: 27533782 |
| NeuN | CD31 | Sema6b | Plxna4 | 0.00217 | 0 | SEMA6B_PLXNA4 | Sema6b - Plxna4 | SEMA6 | Cell-Cell Contact | PMID: 27533782 |
| NeuN | NeuN | Sema6b | Plxna4 | 0.003249 | 0 | SEMA6B_PLXNA4 | Sema6b - Plxna4 | SEMA6 | Cell-Cell Contact | PMID: 27533782 |
| CD31 | CD31 | Sema7a | ITGB1_ITGA1 | 0.011604 | 0 | SEMA7A_ITGB1_ITGA1 | Sema7a - (Itgb1+Itga1) | SEMA7 | Cell-Cell Contact | PMID: 22325954 |
| NeuN | CD31 | Sema7a | ITGB1_ITGA1 | 0.006917 | 0 | SEMA7A_ITGB1_ITGA1 | Sema7a - (Itgb1+Itga1) | SEMA7 | Cell-Cell Contact | PMID: 22325954 |
| CD31 | CD31 | Sema7a | Plxnc1 | 0.002335 | 0 | SEMA7A_PLXNC1 | Sema7a - Plxnc1 | SEMA7 | Cell-Cell Contact | PMID: 27533782 |
| CD31 | NeuN | Sema7a | Plxnc1 | 0.003237 | 0 | SEMA7A_PLXNC1 | Sema7a - Plxnc1 | SEMA7 | Cell-Cell Contact | PMID: 27533782 |
| NeuN | GFAP | Thy1 | ITGAV_ITGB3 | 0.02002 | 0 | THY1_ITGAV_ITGB3 | Thy1 - (Itgav+Itgb3) | THY1 | Cell-Cell Contact | PMID: 29212879; PMID: 18346467 |
| NeuN | NeuN | Thy1 | ITGAV_ITGB3 | 0.011617 | 0 | THY1_ITGAV_ITGB3 | Thy1 - (Itgav+Itgb3) | THY1 | Cell-Cell Contact | PMID: 29212879; PMID: 18346467 |
| CD31 | CD31 | Cdh3 | Cdh3 | 0.000402 | 0 | CDH3_CDH3 | Cdh3 - Cdh3 | CDH | Cell-Cell Contact | KEGG: hsa04514 |
| GFAP | CD31 | Cdh3 | Cdh3 | 0.00035 | 0 | CDH3_CDH3 | Cdh3 - Cdh3 | CDH | Cell-Cell Contact | KEGG: hsa04514 |
| CD31 | GFAP | Cdh3 | Cdh3 | 0.00035 | 0 | CDH3_CDH3 | Cdh3 - Cdh3 | CDH | Cell-Cell Contact | KEGG: hsa04514 |
| CD31 | CD31 | Cdh12 | ITGA1_ITGB1 | 0.002151 | 0 | CDH12_ITGA1_ITGB1 | Cdh12 - ITGA1_ITGB1 | CDH | Cell-Cell Contact | PMID: 34385456 |
| NeuN | CD31 | Cdh12 | ITGA1_ITGB1 | 0.003759 | 0 | CDH12_ITGA1_ITGB1 | Cdh12 - ITGA1_ITGB1 | CDH | Cell-Cell Contact | PMID: 34385456 |
| GFAP | CD31 | Gja1 | Gja1 | 0.02545 | 0 | GJA1_GJA1 | Gja1 - Gja1 | GAP | Cell-Cell Contact | PMID: 32532966;PMID: 15094344 |
| CD31 | GFAP | Gja1 | Gja1 | 0.02545 | 0 | GJA1_GJA1 | Gja1 - Gja1 | GAP | Cell-Cell Contact | PMID: 32532966;PMID: 15094344 |
| GFAP | GFAP | Gja1 | Gja1 | 0.091232 | 0 | GJA1_GJA1 | Gja1 - Gja1 | GAP | Cell-Cell Contact | PMID: 32532966;PMID: 15094344 |
| GFAP | CD31 | Gja1 | Gja6 | 0.00471 | 0 | GJA1_GJA6 | Gja1 - Gja6 | GAP | Cell-Cell Contact | PMID: 32532966;PMID: 15094344 |
| GFAP | GFAP | Gja1 | Gja6 | 0.003836 | 0 | GJA1_GJA6 | Gja1 - Gja6 | GAP | Cell-Cell Contact | PMID: 32532966;PMID: 15094344 |
| CD31 | CD31 | Gja6 | Gja6 | 0.000223 | 0 | GJA6_GJA6 | Gja6 - Gja6 | GAP | Cell-Cell Contact | PMID: 32532966;PMID: 15094344 |
| GFAP | CD31 | Gja6 | Gja6 | 0.000182 | 0 | GJA6_GJA6 | Gja6 - Gja6 | GAP | Cell-Cell Contact | PMID: 32532966;PMID: 15094344 |
| CD31 | GFAP | Gja6 | Gja6 | 0.000182 | 0 | GJA6_GJA6 | Gja6 - Gja6 | GAP | Cell-Cell Contact | PMID: 32532966;PMID: 15094344 |
| CD31 | GFAP | Gjc1 | Gja1 | 0.013125 | 0 | GJC1_GJA1 | Gjc1 - Gja1 | GAP | Cell-Cell Contact | PMID: 32532966;PMID: 15094344 |
| GFAP | GFAP | Gjc1 | Gja1 | 0.003806 | 0 | GJC1_GJA1 | Gjc1 - Gja1 | GAP | Cell-Cell Contact | PMID: 32532966;PMID: 15094344 |
| CD31 | CD31 | Gjc1 | Gja6 | 0.000627 | 0 | GJC1_GJA6 | Gjc1 - Gja6 | GAP | Cell-Cell Contact | PMID: 32532966;PMID: 15094344 |
| CD31 | GFAP | Gjc1 | Gja6 | 0.00051 | 0 | GJC1_GJA6 | Gjc1 - Gja6 | GAP | Cell-Cell Contact | PMID: 32532966;PMID: 15094344 |
| CD31 | CD31 | Gjc1 | Gjc1 | 0.001759 | 0 | GJC1_GJC1 | Gjc1 - Gjc1 | GAP | Cell-Cell Contact | PMID: 32532966;PMID: 15094344 |
| GFAP | CD31 | Gjc1 | Gjc1 | 0.000506 | 0 | GJC1_GJC1 | Gjc1 - Gjc1 | GAP | Cell-Cell Contact | PMID: 32532966;PMID: 15094344 |
| CD31 | GFAP | Gjc1 | Gjc1 | 0.000506 | 0 | GJC1_GJC1 | Gjc1 - Gjc1 | GAP | Cell-Cell Contact | PMID: 32532966;PMID: 15094344 |
| GFAP | CD31 | Cldn1 | Cldn1 | 0.000485 | 0 | CLDN1_CLDN1 | Cldn1 - Cldn1 | CLDN | Cell-Cell Contact | PMID: 31942240 |
| CD31 | GFAP | Cldn1 | Cldn1 | 0.000485 | 0 | CLDN1_CLDN1 | Cldn1 - Cldn1 | CLDN | Cell-Cell Contact | PMID: 31942240 |
| GFAP | GFAP | Cldn1 | Cldn1 | 0.000555 | 0 | CLDN1_CLDN1 | Cldn1 - Cldn1 | CLDN | Cell-Cell Contact | PMID: 31942240 |
| CD31 | CD31 | Cldn3 | Cldn3 | 0.00045 | 0 | CLDN3_CLDN3 | Cldn3 - Cldn3 | CLDN | Cell-Cell Contact | PMID: 31942240 |

| | | | | | | | | | | |
|---|---|---|---|---|---|---|---|---|---|---|
| GFAP | CD31 | Cldn3 | Cldn3 | 0.00016 | 0 | CLDN3_CLDN3 | Cldn3 - Cldn3 | CLDN | Cell-Cell Contact | PMID: 31942240 |
| CD31 | GFAP | Cldn3 | Cldn3 | 0.00016 | 0 | CLDN3_CLDN3 | Cldn3 - Cldn3 | CLDN | Cell-Cell Contact | PMID: 31942240 |
| CD31 | CD31 | Pcdha9 | Pcdha9 | 0.000666 | 0 | PCDHA9_PCDHA9 | Pcdha9 - Pcdha9 | PCDH | Cell-Cell Contact | PMID: 35253643; PMID: 32694982 |
| NeuN | CD31 | Pcdha9 | Pcdha9 | 0.000287 | 0 | PCDHA9_PCDHA9 | Pcdha9 - Pcdha9 | PCDH | Cell-Cell Contact | PMID: 35253643; PMID: 32694982 |
| CD31 | NeuN | Pcdha9 | Pcdha9 | 0.000287 | 0 | PCDHA9_PCDHA9 | Pcdha9 - Pcdha9 | PCDH | Cell-Cell Contact | PMID: 35253643; PMID: 32694982 |
| GFAP | CD31 | Pcdha11 | Pcdha11 | 0.000504 | 0 | PCDHA11_PCDHA11 | Pcdha11 - Pcdha11 | PCDH | Cell-Cell Contact | PMID: 35253643; PMID: 32694982 |
| CD31 | GFAP | Pcdha11 | Pcdha11 | 0.000504 | 0 | PCDHA11_PCDHA11 | Pcdha11 - Pcdha11 | PCDH | Cell-Cell Contact | PMID: 35253643; PMID: 32694982 |
| GFAP | GFAP | Pcdha11 | Pcdha11 | 0.000667 | 0 | PCDHA11_PCDHA11 | Pcdha11 - Pcdha11 | PCDH | Cell-Cell Contact | PMID: 35253643; PMID: 32694982 |
| NeuN | CD31 | Pcdha10 | Pcdha10 | 0.000129 | 0 | PCDHA10_PCDHA10 | Pcdha10 - Pcdha10 | PCDH | Cell-Cell Contact | PMID: 35253643; PMID: 32694982 |
| CD31 | NeuN | Pcdha10 | Pcdha10 | 0.000129 | 0 | PCDHA10_PCDHA10 | Pcdha10 - Pcdha10 | PCDH | Cell-Cell Contact | PMID: 35253643; PMID: 32694982 |
| NeuN | NeuN | Pcdha10 | Pcdha10 | 0.000181 | 0 | PCDHA10_PCDHA10 | Pcdha10 - Pcdha10 | PCDH | Cell-Cell Contact | PMID: 35253643; PMID: 32694982 |
| NeuN | CD31 | Pcdha12 | Pcdha12 | 0.000418 | 0 | PCDHA12_PCDHA12 | Pcdha12 - Pcdha12 | PCDH | Cell-Cell Contact | PMID: 35253643; PMID: 32694982 |
| CD31 | NeuN | Pcdha12 | Pcdha12 | 0.000418 | 0 | PCDHA12_PCDHA12 | Pcdha12 - Pcdha12 | PCDH | Cell-Cell Contact | PMID: 35253643; PMID: 32694982 |
| NeuN | NeuN | Pcdha12 | Pcdha12 | 0.00043 | 0 | PCDHA12_PCDHA12 | Pcdha12 - Pcdha12 | PCDH | Cell-Cell Contact | PMID: 35253643; PMID: 32694982 |
| GFAP | CD31 | Pcdha2 | Pcdha2 | 0.00036 | 0 | PCDHA2_PCDHA2 | Pcdha2 - Pcdha2 | PCDH | Cell-Cell Contact | PMID: 35253643; PMID: 32694982 |
| CD31 | GFAP | Pcdha2 | Pcdha2 | 0.00036 | 0 | PCDHA2_PCDHA2 | Pcdha2 - Pcdha2 | PCDH | Cell-Cell Contact | PMID: 35253643; PMID: 32694982 |
| GFAP | GFAP | Pcdha2 | Pcdha2 | 0.000452 | 0 | PCDHA2_PCDHA2 | Pcdha2 - Pcdha2 | PCDH | Cell-Cell Contact | PMID: 35253643; PMID: 32694982 |
| CD31 | CD31 | Pcdha3 | Pcdha3 | 0.000714 | 0 | PCDHA3_PCDHA3 | Pcdha3 - Pcdha3 | PCDH | Cell-Cell Contact | PMID: 35253643; PMID: 32694982 |
| NeuN | CD31 | Pcdha3 | Pcdha3 | 0.00037 | 0 | PCDHA3_PCDHA3 | Pcdha3 - Pcdha3 | PCDH | Cell-Cell Contact | PMID: 35253643; PMID: 32694982 |
| CD31 | NeuN | Pcdha3 | Pcdha3 | 0.00037 | 0 | PCDHA3_PCDHA3 | Pcdha3 - Pcdha3 | PCDH | Cell-Cell Contact | PMID: 35253643; PMID: 32694982 |
| GFAP | GFAP | Pcdha5 | Pcdha5 | 0.000315 | 0 | PCDHA5_PCDHA5 | Pcdha5 - Pcdha5 | PCDH | Cell-Cell Contact | PMID: 35253643; PMID: 32694982 |
| NeuN | GFAP | Pcdha5 | Pcdha5 | 0.000256 | 0 | PCDHA5_PCDHA5 | Pcdha5 - Pcdha5 | PCDH | Cell-Cell Contact | PMID: 35253643; PMID: 32694982 |
| GFAP | NeuN | Pcdha5 | Pcdha5 | 0.000256 | 0 | PCDHA5_PCDHA5 | Pcdha5 - Pcdha5 | PCDH | Cell-Cell Contact | PMID: 35253643; PMID: 32694982 |
| CD31 | CD31 | Pcdha8 | Pcdha8 | 0.000415 | 0 | PCDHA8_PCDHA8 | Pcdha8 - Pcdha8 | PCDH | Cell-Cell Contact | PMID: 35253643; PMID: 32694982 |
| GFAP | CD31 | Pcdha8 | Pcdha8 | 0.000243 | 0 | PCDHA8_PCDHA8 | Pcdha8 - Pcdha8 | PCDH | Cell-Cell Contact | PMID: 35253643; PMID: 32694982 |
| CD31 | GFAP | Pcdha8 | Pcdha8 | 0.000243 | 0 | PCDHA8_PCDHA8 | Pcdha8 - Pcdha8 | PCDH | Cell-Cell Contact | PMID: 35253643; PMID: 32694982 |
| GFAP | GFAP | Pcdhac1 | Pcdhac1 | 0.000518 | 0 | PCDHAC1_PCDHAC1 | Pcdhac1 - Pcdhac1 | PCDH | Cell-Cell Contact | PMID: 35253643; PMID: 32694982 |
| NeuN | GFAP | Pcdhac1 | Pcdhac1 | 0.000339 | 0 | PCDHAC1_PCDHAC1 | Pcdhac1 - Pcdhac1 | PCDH | Cell-Cell Contact | PMID: 35253643; PMID: 32694982 |
| GFAP | NeuN | Pcdhac1 | Pcdhac1 | 0.000339 | 0 | PCDHAC1_PCDHAC1 | Pcdhac1 - Pcdhac1 | PCDH | Cell-Cell Contact | PMID: 35253643; PMID: 32694982 |
| NeuN | CD31 | Pcdhac2 | Pcdhac2 | 0.001543 | 0 | PCDHAC2_PCDHAC2 | Pcdhac2 - Pcdhac2 | PCDH | Cell-Cell Contact | PMID: 35253643; PMID: 32694982 |
| CD31 | NeuN | Pcdhac2 | Pcdhac2 | 0.001543 | 0 | PCDHAC2_PCDHAC2 | Pcdhac2 - Pcdhac2 | PCDH | Cell-Cell Contact | PMID: 35253643; PMID: 32694982 |
| NeuN | NeuN | Pcdhac2 | Pcdhac2 | 0.001576 | 0 | PCDHAC2_PCDHAC2 | Pcdhac2 - Pcdhac2 | PCDH | Cell-Cell Contact | PMID: 35253643; PMID: 32694982 |
| CD31 | CD31 | Pcdhb18 | Pcdhb18 | 0.000556 | 0 | PCDHB18_PCDHB18 | Pcdhb18 - Pcdhb18 | PCDH | Cell-Cell Contact | PMID: 35253643; PMID: 32694982 |
| GFAP | CD31 | Pcdhb18 | Pcdhb18 | 0.000317 | 0 | PCDHB18_PCDHB18 | Pcdhb18 - Pcdhb18 | PCDH | Cell-Cell Contact | PMID: 35253643; PMID: 32694982 |
| CD31 | GFAP | Pcdhb18 | Pcdhb18 | 0.000317 | 0 | PCDHB18_PCDHB18 | Pcdhb18 - Pcdhb18 | PCDH | Cell-Cell Contact | PMID: 35253643; PMID: 32694982 |
| GFAP | CD31 | Pcdhb19 | Pcdhb19 | 0.000249 | 0 | PCDHB19_PCDHB19 | Pcdhb19 - Pcdhb19 | PCDH | Cell-Cell Contact | PMID: 35253643; PMID: 32694982 |
| CD31 | GFAP | Pcdhb19 | Pcdhb19 | 0.000249 | 0 | PCDHB19_PCDHB19 | Pcdhb19 - Pcdhb19 | PCDH | Cell-Cell Contact | PMID: 35253643; PMID: 32694982 |
| GFAP | GFAP | Pcdhb19 | Pcdhb19 | 0.000338 | 0 | PCDHB19_PCDHB19 | Pcdhb19 - Pcdhb19 | PCDH | Cell-Cell Contact | PMID: 35253643; PMID: 32694982 |
| GFAP | CD31 | Pcdhb19 | Pcdhb21 | 0.000279 | 0 | PCDHB19_PCDHB21 | Pcdhb19 - Pcdhb21 | PCDH | Cell-Cell Contact | PMID: 35253643; PMID: 32694982 |
| GFAP | GFAP | Pcdhb19 | Pcdhb21 | 0.000338 | 0 | PCDHB19_PCDHB21 | Pcdhb19 - Pcdhb21 | PCDH | Cell-Cell Contact | PMID: 35253643; PMID: 32694982 |
| GFAP | CD31 | Pcdhb21 | Pcdhb21 | 0.000279 | 0 | PCDHB21_PCDHB21 | Pcdhb21 - Pcdhb21 | PCDH | Cell-Cell Contact | PMID: 35253643; PMID: 32694982 |
| CD31 | GFAP | Pcdhb21 | Pcdhb21 | 0.000279 | 0 | PCDHB21_PCDHB21 | Pcdhb21 - Pcdhb21 | PCDH | Cell-Cell Contact | PMID: 35253643; PMID: 32694982 |
| GFAP | GFAP | Pcdhb21 | Pcdhb21 | 0.000338 | 0 | PCDHB21_PCDHB21 | Pcdhb21 - Pcdhb21 | PCDH | Cell-Cell Contact | PMID: 35253643; PMID: 32694982 |
| CD31 | CD31 | Pcdhb16 | Pcdhb16 | 0.000372 | 0 | PCDHB16_PCDHB16 | Pcdhb16 - Pcdhb16 | PCDH | Cell-Cell Contact | PMID: 35253643; PMID: 32694982 |
| GFAP | CD31 | Pcdhb16 | Pcdhb16 | 0.000211 | 0 | PCDHB16_PCDHB16 | Pcdhb16 - Pcdhb16 | PCDH | Cell-Cell Contact | PMID: 35253643; PMID: 32694982 |
| CD31 | GFAP | Pcdhb16 | Pcdhb16 | 0.000211 | 0 | PCDHB16_PCDHB16 | Pcdhb16 - Pcdhb16 | PCDH | Cell-Cell Contact | PMID: 35253643; PMID: 32694982 |
| CD31 | CD31 | Pcdhb20 | Pcdhb20 | 0.000384 | 0 | PCDHB20_PCDHB20 | Pcdhb20 - Pcdhb20 | PCDH | Cell-Cell Contact | PMID: 35253643; PMID: 32694982 |
| NeuN | CD31 | Pcdhb20 | Pcdhb20 | 0.000297 | 0 | PCDHB20_PCDHB20 | Pcdhb20 - Pcdhb20 | PCDH | Cell-Cell Contact | PMID: 35253643; PMID: 32694982 |
| CD31 | NeuN | Pcdhb20 | Pcdhb20 | 0.000297 | 0 | PCDHB20_PCDHB20 | Pcdhb20 - Pcdhb20 | PCDH | Cell-Cell Contact | PMID: 35253643; PMID: 32694982 |
| CD31 | CD31 | Pcdhb22 | Pcdhb22 | 0.000318 | 0 | PCDHB22_PCDHB22 | Pcdhb22 - Pcdhb22 | PCDH | Cell-Cell Contact | PMID: 35253643; PMID: 32694982 |
| NeuN | CD31 | Pcdhb22 | Pcdhb22 | 0.000215 | 0 | PCDHB22_PCDHB22 | Pcdhb22 - Pcdhb22 | PCDH | Cell-Cell Contact | PMID: 35253643; PMID: 32694982 |
| CD31 | NeuN | Pcdhb22 | Pcdhb22 | 0.000215 | 0 | PCDHB22_PCDHB22 | Pcdhb22 - Pcdhb22 | PCDH | Cell-Cell Contact | PMID: 35253643; PMID: 32694982 |
| CD31 | CD31 | Pcdhb17 | Pcdhb17 | 0.000476 | 0 | PCDHB17_PCDHB17 | Pcdhb17 - Pcdhb17 | PCDH | Cell-Cell Contact | PMID: 35253643; PMID: 32694982 |
| GFAP | CD31 | Pcdhb17 | Pcdhb17 | 0.000312 | 0 | PCDHB17_PCDHB17 | Pcdhb17 - Pcdhb17 | PCDH | Cell-Cell Contact | PMID: 35253643; PMID: 32694982 |
| CD31 | GFAP | Pcdhb17 | Pcdhb17 | 0.000312 | 0 | PCDHB17_PCDHB17 | Pcdhb17 - Pcdhb17 | PCDH | Cell-Cell Contact | PMID: 35253643; PMID: 32694982 |
| CD31 | CD31 | Pcdhb2 | Pcdhb2 | 0.000344 | 0 | PCDHB2_PCDHB2 | Pcdhb2 - Pcdhb2 | PCDH | Cell-Cell Contact | PMID: 35253643; PMID: 32694982 |
| GFAP | CD31 | Pcdhb2 | Pcdhb2 | 0.000263 | 0 | PCDHB2_PCDHB2 | Pcdhb2 - Pcdhb2 | PCDH | Cell-Cell Contact | PMID: 35253643; PMID: 32694982 |
| CD31 | GFAP | Pcdhb2 | Pcdhb2 | 0.000263 | 0 | PCDHB2_PCDHB2 | Pcdhb2 - Pcdhb2 | PCDH | Cell-Cell Contact | PMID: 35253643; PMID: 32694982 |
| GFAP | CD31 | Pcdhb3 | Pcdhb3 | 0.000176 | 0 | PCDHB3_PCDHB3 | Pcdhb3 - Pcdhb3 | PCDH | Cell-Cell Contact | PMID: 35253643; PMID: 32694982 |

| | | | | | | | | | |
|---|---|---|---|---|---|---|---|---|---|
| CD31 | GFAP | Pcdhb3 | Pcdhb3 | 0.000176 | 0 | PCDHB3_PCDHB3 | Pcdhb3 - Pcdhb3 | PCDH | Cell-Cell Contact | PMID: 35253643; PMID: 32694982 |
| GFAP | GFAP | Pcdhb3 | Pcdhb3 | 0.000236 | 0 | PCDHB3_PCDHB3 | Pcdhb3 - Pcdhb3 | PCDH | Cell-Cell Contact | PMID: 35253643; PMID: 32694982 |
| CD31 | CD31 | Pcdhb7 | Pcdhb7 | 0.000235 | 0 | PCDHB7_PCDHB7 | Pcdhb7 - Pcdhb7 | PCDH | Cell-Cell Contact | PMID: 35253643; PMID: 32694982 |
| GFAP | CD31 | Pcdhb7 | Pcdhb7 | 0.00016 | 0 | PCDHB7_PCDHB7 | Pcdhb7 - Pcdhb7 | PCDH | Cell-Cell Contact | PMID: 35253643; PMID: 32694982 |
| CD31 | GFAP | Pcdhb7 | Pcdhb7 | 0.00016 | 0 | PCDHB7_PCDHB7 | Pcdhb7 - Pcdhb7 | PCDH | Cell-Cell Contact | PMID: 35253643; PMID: 32694982 |
| CD31 | CD31 | Pcdhb7 | Pcdhb5 | 0.000236 | 0 | PCDHB7_PCDHB5 | Pcdhb7 - Pcdhb5 | PCDH | Cell-Cell Contact | PMID: 35253643; PMID: 32694982 |
| GFAP | CD31 | Pcdhb7 | Pcdhb5 | 0.000161 | 0 | PCDHB7_PCDHB5 | Pcdhb7 - Pcdhb5 | PCDH | Cell-Cell Contact | PMID: 35253643; PMID: 32694982 |
| CD31 | CD31 | Pcdhb7 | Pcdhb9 | 0.000402 | 0 | PCDHB7_PCDHB9 | Pcdhb7 - Pcdhb9 | PCDH | Cell-Cell Contact | PMID: 35253643; PMID: 32694982 |
| CD31 | GFAP | Pcdhb7 | Pcdhb9 | 0.000305 | 0 | PCDHB7_PCDHB9 | Pcdhb7 - Pcdhb9 | PCDH | Cell-Cell Contact | PMID: 35253643; PMID: 32694982 |
| CD31 | CD31 | Pcdhb5 | Pcdhb5 | 0.000237 | 0 | PCDHB5_PCDHB5 | Pcdhb5 - Pcdhb5 | PCDH | Cell-Cell Contact | PMID: 35253643; PMID: 32694982 |
| GFAP | CD31 | Pcdhb5 | Pcdhb5 | 0.000123 | 0 | PCDHB5_PCDHB5 | Pcdhb5 - Pcdhb5 | PCDH | Cell-Cell Contact | PMID: 35253643; PMID: 32694982 |
| CD31 | GFAP | Pcdhb5 | Pcdhb5 | 0.000123 | 0 | PCDHB5_PCDHB5 | Pcdhb5 - Pcdhb5 | PCDH | Cell-Cell Contact | PMID: 35253643; PMID: 32694982 |
| CD31 | CD31 | Pcdhb5 | Pcdhb9 | 0.000404 | 0 | PCDHB5_PCDHB9 | Pcdhb5 - Pcdhb9 | PCDH | Cell-Cell Contact | PMID: 35253643; PMID: 32694982 |
| CD31 | GFAP | Pcdhb5 | Pcdhb9 | 0.000306 | 0 | PCDHB5_PCDHB9 | Pcdhb5 - Pcdhb9 | PCDH | Cell-Cell Contact | PMID: 35253643; PMID: 32694982 |
| CD31 | CD31 | Pcdhb9 | Pcdhb9 | 0.00069 | 0 | PCDHB9_PCDHB9 | Pcdhb9 - Pcdhb9 | PCDH | Cell-Cell Contact | PMID: 35253643; PMID: 32694982 |
| GFAP | CD31 | Pcdhb9 | Pcdhb9 | 0.000522 | 0 | PCDHB9_PCDHB9 | Pcdhb9 - Pcdhb9 | PCDH | Cell-Cell Contact | PMID: 35253643; PMID: 32694982 |
| CD31 | GFAP | Pcdhb9 | Pcdhb9 | 0.000522 | 0 | PCDHB9_PCDHB9 | Pcdhb9 - Pcdhb9 | PCDH | Cell-Cell Contact | PMID: 35253643; PMID: 32694982 |
| CD31 | CD31 | Pcdhb12 | Pcdhb12 | 0.000387 | 0 | PCDHB12_PCDHB12 | Pcdhb12 - Pcdhb12 | PCDH | Cell-Cell Contact | PMID: 35253643; PMID: 32694982 |
| GFAP | CD31 | Pcdhb12 | Pcdhb12 | 0.00029 | 0 | PCDHB12_PCDHB12 | Pcdhb12 - Pcdhb12 | PCDH | Cell-Cell Contact | PMID: 35253643; PMID: 32694982 |
| CD31 | GFAP | Pcdhb12 | Pcdhb12 | 0.00029 | 0 | PCDHB12_PCDHB12 | Pcdhb12 - Pcdhb12 | PCDH | Cell-Cell Contact | PMID: 35253643; PMID: 32694982 |
| CD31 | CD31 | Pcdhb12 | Pcdhb10 | 0.000418 | 0 | PCDHB12_PCDHB10 | Pcdhb12 - Pcdhb10 | PCDH | Cell-Cell Contact | PMID: 35253643; PMID: 32694982 |
| CD31 | GFAP | Pcdhb12 | Pcdhb10 | 0.000442 | 0 | PCDHB12_PCDHB10 | Pcdhb12 - Pcdhb10 | PCDH | Cell-Cell Contact | PMID: 35253643; PMID: 32694982 |
| CD31 | CD31 | Pcdhb12 | Pcdhb11 | 0.000279 | 0 | PCDHB12_PCDHB11 | Pcdhb12 - Pcdhb11 | PCDH | Cell-Cell Contact | PMID: 35253643; PMID: 32694982 |
| CD31 | GFAP | Pcdhb12 | Pcdhb11 | 0.000264 | 0 | PCDHB12_PCDHB11 | Pcdhb12 - Pcdhb11 | PCDH | Cell-Cell Contact | PMID: 35253643; PMID: 32694982 |
| CD31 | CD31 | Pcdhb12 | Pcdhb8 | 0.000205 | 0 | PCDHB12_PCDHB8 | Pcdhb12 - Pcdhb8 | PCDH | Cell-Cell Contact | PMID: 35253643; PMID: 32694982 |
| CD31 | GFAP | Pcdhb12 | Pcdhb8 | 0.000235 | 0 | PCDHB12_PCDHB8 | Pcdhb12 - Pcdhb8 | PCDH | Cell-Cell Contact | PMID: 35253643; PMID: 32694982 |
| CD31 | CD31 | Pcdhb12 | Pcdhb6 | 0.00027 | 0 | PCDHB12_PCDHB6 | Pcdhb12 - Pcdhb6 | PCDH | Cell-Cell Contact | PMID: 35253643; PMID: 32694982 |
| CD31 | GFAP | Pcdhb12 | Pcdhb6 | 0.000303 | 0 | PCDHB12_PCDHB6 | Pcdhb12 - Pcdhb6 | PCDH | Cell-Cell Contact | PMID: 35253643; PMID: 32694982 |
| GFAP | CD31 | Pcdhb10 | Pcdhb10 | 0.000477 | 0 | PCDHB10_PCDHB10 | Pcdhb10 - Pcdhb10 | PCDH | Cell-Cell Contact | PMID: 35253643; PMID: 32694982 |
| CD31 | GFAP | Pcdhb10 | Pcdhb10 | 0.000477 | 0 | PCDHB10_PCDHB10 | Pcdhb10 - Pcdhb10 | PCDH | Cell-Cell Contact | PMID: 35253643; PMID: 32694982 |
| GFAP | GFAP | Pcdhb10 | Pcdhb10 | 0.000505 | 0 | PCDHB10_PCDHB10 | Pcdhb10 - Pcdhb10 | PCDH | Cell-Cell Contact | PMID: 35253643; PMID: 32694982 |
| GFAP | CD31 | Pcdhb10 | Pcdhb11 | 0.000318 | 0 | PCDHB10_PCDHB11 | Pcdhb10 - Pcdhb11 | PCDH | Cell-Cell Contact | PMID: 35253643; PMID: 32694982 |
| GFAP | GFAP | Pcdhb10 | Pcdhb11 | 0.000302 | 0 | PCDHB10_PCDHB11 | Pcdhb10 - Pcdhb11 | PCDH | Cell-Cell Contact | PMID: 35253643; PMID: 32694982 |
| CD31 | GFAP | Pcdhb10 | Pcdhb8 | 0.000253 | 0 | PCDHB10_PCDHB8 | Pcdhb10 - Pcdhb8 | PCDH | Cell-Cell Contact | PMID: 35253643; PMID: 32694982 |
| GFAP | GFAP | Pcdhb10 | Pcdhb8 | 0.000268 | 0 | PCDHB10_PCDHB8 | Pcdhb10 - Pcdhb8 | PCDH | Cell-Cell Contact | PMID: 35253643; PMID: 32694982 |
| CD31 | GFAP | Pcdhb10 | Pcdhb6 | 0.000326 | 0 | PCDHB10_PCDHB6 | Pcdhb10 - Pcdhb6 | PCDH | Cell-Cell Contact | PMID: 35253643; PMID: 32694982 |
| GFAP | GFAP | Pcdhb10 | Pcdhb6 | 0.000346 | 0 | PCDHB10_PCDHB6 | Pcdhb10 - Pcdhb6 | PCDH | Cell-Cell Contact | PMID: 35253643; PMID: 32694982 |
| CD31 | CD31 | Pcdhb11 | Pcdhb11 | 0.000201 | 0 | PCDHB11_PCDHB11 | Pcdhb11 - Pcdhb11 | PCDH | Cell-Cell Contact | PMID: 35253643; PMID: 32694982 |
| GFAP | CD31 | Pcdhb11 | Pcdhb11 | 0.00019 | 0 | PCDHB11_PCDHB11 | Pcdhb11 - Pcdhb11 | PCDH | Cell-Cell Contact | PMID: 35253643; PMID: 32694982 |
| CD31 | GFAP | Pcdhb11 | Pcdhb11 | 0.00019 | 0 | PCDHB11_PCDHB11 | Pcdhb11 - Pcdhb11 | PCDH | Cell-Cell Contact | PMID: 35253643; PMID: 32694982 |
| CD31 | GFAP | Pcdhb11 | Pcdhb8 | 0.000169 | 0 | PCDHB11_PCDHB8 | Pcdhb11 - Pcdhb8 | PCDH | Cell-Cell Contact | PMID: 35253643; PMID: 32694982 |
| GFAP | GFAP | Pcdhb11 | Pcdhb8 | 0.00016 | 0 | PCDHB11_PCDHB8 | Pcdhb11 - Pcdhb8 | PCDH | Cell-Cell Contact | PMID: 35253643; PMID: 32694982 |
| CD31 | GFAP | Pcdhb11 | Pcdhb6 | 0.000218 | 0 | PCDHB11_PCDHB6 | Pcdhb11 - Pcdhb6 | PCDH | Cell-Cell Contact | PMID: 35253643; PMID: 32694982 |
| GFAP | GFAP | Pcdhb11 | Pcdhb6 | 0.000207 | 0 | PCDHB11_PCDHB6 | Pcdhb11 - Pcdhb6 | PCDH | Cell-Cell Contact | PMID: 35253643; PMID: 32694982 |
| GFAP | CD31 | Pcdhb8 | Pcdhb8 | 0.000124 | 0 | PCDHB8_PCDHB8 | Pcdhb8 - Pcdhb8 | PCDH | Cell-Cell Contact | PMID: 35253643; PMID: 32694982 |
| CD31 | GFAP | Pcdhb8 | Pcdhb8 | 0.000124 | 0 | PCDHB8_PCDHB8 | Pcdhb8 - Pcdhb8 | PCDH | Cell-Cell Contact | PMID: 35253643; PMID: 32694982 |
| GFAP | GFAP | Pcdhb8 | Pcdhb8 | 0.000142 | 0 | PCDHB8_PCDHB8 | Pcdhb8 - Pcdhb8 | PCDH | Cell-Cell Contact | PMID: 35253643; PMID: 32694982 |
| GFAP | CD31 | Pcdhb8 | Pcdhb6 | 0.000164 | 0 | PCDHB8_PCDHB6 | Pcdhb8 - Pcdhb6 | PCDH | Cell-Cell Contact | PMID: 35253643; PMID: 32694982 |
| GFAP | GFAP | Pcdhb8 | Pcdhb6 | 0.000184 | 0 | PCDHB8_PCDHB6 | Pcdhb8 - Pcdhb6 | PCDH | Cell-Cell Contact | PMID: 35253643; PMID: 32694982 |
| GFAP | CD31 | Pcdhb6 | Pcdhb6 | 0.000211 | 0 | PCDHB6_PCDHB6 | Pcdhb6 - Pcdhb6 | PCDH | Cell-Cell Contact | PMID: 35253643; PMID: 32694982 |
| CD31 | GFAP | Pcdhb6 | Pcdhb6 | 0.000211 | 0 | PCDHB6_PCDHB6 | Pcdhb6 - Pcdhb6 | PCDH | Cell-Cell Contact | PMID: 35253643; PMID: 32694982 |
| GFAP | GFAP | Pcdhb6 | Pcdhb6 | 0.000237 | 0 | PCDHB6_PCDHB6 | Pcdhb6 - Pcdhb6 | PCDH | Cell-Cell Contact | PMID: 35253643; PMID: 32694982 |
| GFAP | CD31 | Pcdhb13 | Pcdhb13 | 0.000385 | 0 | PCDHB13_PCDHB13 | Pcdhb13 - Pcdhb13 | PCDH | Cell-Cell Contact | PMID: 35253643; PMID: 32694982 |
| CD31 | GFAP | Pcdhb13 | Pcdhb13 | 0.000385 | 0 | PCDHB13_PCDHB13 | Pcdhb13 - Pcdhb13 | PCDH | Cell-Cell Contact | PMID: 35253643; PMID: 32694982 |
| GFAP | GFAP | Pcdhb13 | Pcdhb13 | 0.000486 | 0 | PCDHB13_PCDHB13 | Pcdhb13 - Pcdhb13 | PCDH | Cell-Cell Contact | PMID: 35253643; PMID: 32694982 |
| CD31 | CD31 | Pcdhb15 | Pcdhb15 | 0.000321 | 0 | PCDHB15_PCDHB15 | Pcdhb15 - Pcdhb15 | PCDH | Cell-Cell Contact | PMID: 35253643; PMID: 32694982 |
| NeuN | CD31 | Pcdhb15 | Pcdhb15 | 0.000155 | 0 | PCDHB15_PCDHB15 | Pcdhb15 - Pcdhb15 | PCDH | Cell-Cell Contact | PMID: 35253643; PMID: 32694982 |
| CD31 | NeuN | Pcdhb15 | Pcdhb15 | 0.000155 | 0 | PCDHB15_PCDHB15 | Pcdhb15 - Pcdhb15 | PCDH | Cell-Cell Contact | PMID: 35253643; PMID: 32694982 |
| GFAP | CD31 | Pcdhga1 | Pcdhga1 | 0.000185 | 0 | PCDHGA1_PCDHGA1 | Pcdhga1 - Pcdhga1 | PCDH | Cell-Cell Contact | PMID: 35253643; PMID: 32694982 |

| | | | | | | | | | | |
|---|---|---|---|---|---|---|---|---|---|---|
| CD31 | GFAP | Pcdhga1 | Pcdhga1 | 0.000185 | 0 | PCDHGA1_PCDHGA1 | Pcdhga1 - Pcdhga1 | PCDH | Cell-Cell Contact | PMID: 35253643; PMID: 32694982 |
| GFAP | GFAP | Pcdhga1 | Pcdhga1 | 0.000292 | 0 | PCDHGA1_PCDHGA1 | Pcdhga1 - Pcdhga1 | PCDH | Cell-Cell Contact | PMID: 35253643; PMID: 32694982 |
| GFAP | CD31 | Pcdhga10 | Pcdhga10 | 0.000353 | 0 | PCDHGA10_PCDHGA10 | Pcdhga10 - Pcdhga10 | PCDH | Cell-Cell Contact | PMID: 35253643; PMID: 32694982 |
| CD31 | GFAP | Pcdhga10 | Pcdhga10 | 0.000353 | 0 | PCDHGA10_PCDHGA10 | Pcdhga10 - Pcdhga10 | PCDH | Cell-Cell Contact | PMID: 35253643; PMID: 32694982 |
| GFAP | GFAP | Pcdhga10 | Pcdhga10 | 0.000375 | 0 | PCDHGA10_PCDHGA10 | Pcdhga10 - Pcdhga10 | PCDH | Cell-Cell Contact | PMID: 35253643; PMID: 32694982 |
| GFAP | CD31 | Pcdhga11 | Pcdhga11 | 0.000688 | 0 | PCDHGA11_PCDHGA11 | Pcdhga11 - Pcdhga11 | PCDH | Cell-Cell Contact | PMID: 35253643; PMID: 32694982 |
| CD31 | GFAP | Pcdhga11 | Pcdhga11 | 0.000688 | 0 | PCDHGA11_PCDHGA11 | Pcdhga11 - Pcdhga11 | PCDH | Cell-Cell Contact | PMID: 35253643; PMID: 32694982 |
| GFAP | GFAP | Pcdhga11 | Pcdhga11 | 0.000707 | 0 | PCDHGA11_PCDHGA11 | Pcdhga11 - Pcdhga11 | PCDH | Cell-Cell Contact | PMID: 35253643; PMID: 32694982 |
| GFAP | CD31 | Pcdhga12 | Pcdhga12 | 0.000693 | 0 | PCDHGA12_PCDHGA12 | Pcdhga12 - Pcdhga12 | PCDH | Cell-Cell Contact | PMID: 35253643; PMID: 32694982 |
| CD31 | GFAP | Pcdhga12 | Pcdhga12 | 0.000693 | 0 | PCDHGA12_PCDHGA12 | Pcdhga12 - Pcdhga12 | PCDH | Cell-Cell Contact | PMID: 35253643; PMID: 32694982 |
| GFAP | GFAP | Pcdhga12 | Pcdhga12 | 0.000781 | 0 | PCDHGA12_PCDHGA12 | Pcdhga12 - Pcdhga12 | PCDH | Cell-Cell Contact | PMID: 35253643; PMID: 32694982 |
| CD31 | CD31 | Pcdhga2 | Pcdhga2 | 0.000706 | 0 | PCDHGA2_PCDHGA2 | Pcdhga2 - Pcdhga2 | PCDH | Cell-Cell Contact | PMID: 35253643; PMID: 32694982 |
| GFAP | CD31 | Pcdhga2 | Pcdhga2 | 0.0003 | 0 | PCDHGA2_PCDHGA2 | Pcdhga2 - Pcdhga2 | PCDH | Cell-Cell Contact | PMID: 35253643; PMID: 32694982 |
| CD31 | GFAP | Pcdhga2 | Pcdhga2 | 0.0003 | 0 | PCDHGA2_PCDHGA2 | Pcdhga2 - Pcdhga2 | PCDH | Cell-Cell Contact | PMID: 35253643; PMID: 32694982 |
| CD31 | CD31 | Pcdhga3 | Pcdhga3 | 0.000219 | 0 | PCDHGA3_PCDHGA3 | Pcdhga3 - Pcdhga3 | PCDH | Cell-Cell Contact | PMID: 35253643; PMID: 32694982 |
| GFAP | CD31 | Pcdhga3 | Pcdhga3 | 0.000202 | 0 | PCDHGA3_PCDHGA3 | Pcdhga3 - Pcdhga3 | PCDH | Cell-Cell Contact | PMID: 35253643; PMID: 32694982 |
| CD31 | GFAP | Pcdhga3 | Pcdhga3 | 0.000202 | 0 | PCDHGA3_PCDHGA3 | Pcdhga3 - Pcdhga3 | PCDH | Cell-Cell Contact | PMID: 35253643; PMID: 32694982 |
| CD31 | CD31 | Pcdhga4 | Pcdhga4 | 0.000658 | 0 | PCDHGA4_PCDHGA4 | Pcdhga4 - Pcdhga4 | PCDH | Cell-Cell Contact | PMID: 35253643; PMID: 32694982 |
| GFAP | CD31 | Pcdhga4 | Pcdhga4 | 0.000398 | 0 | PCDHGA4_PCDHGA4 | Pcdhga4 - Pcdhga4 | PCDH | Cell-Cell Contact | PMID: 35253643; PMID: 32694982 |
| CD31 | GFAP | Pcdhga4 | Pcdhga4 | 0.000398 | 0 | PCDHGA4_PCDHGA4 | Pcdhga4 - Pcdhga4 | PCDH | Cell-Cell Contact | PMID: 35253643; PMID: 32694982 |
| GFAP | CD31 | Pcdhga5 | Pcdhga5 | 0.0003 | 0 | PCDHGA5_PCDHGA5 | Pcdhga5 - Pcdhga5 | PCDH | Cell-Cell Contact | PMID: 35253643; PMID: 32694982 |
| CD31 | GFAP | Pcdhga5 | Pcdhga5 | 0.0003 | 0 | PCDHGA5_PCDHGA5 | Pcdhga5 - Pcdhga5 | PCDH | Cell-Cell Contact | PMID: 35253643; PMID: 32694982 |
| GFAP | GFAP | Pcdhga5 | Pcdhga5 | 0.000412 | 0 | PCDHGA5_PCDHGA5 | Pcdhga5 - Pcdhga5 | PCDH | Cell-Cell Contact | PMID: 35253643; PMID: 32694982 |
| CD31 | CD31 | Pcdhga6 | Pcdhga6 | 0.000292 | 0 | PCDHGA6_PCDHGA6 | Pcdhga6 - Pcdhga6 | PCDH | Cell-Cell Contact | PMID: 35253643; PMID: 32694982 |
| GFAP | CD31 | Pcdhga6 | Pcdhga6 | 0.000176 | 0 | PCDHGA6_PCDHGA6 | Pcdhga6 - Pcdhga6 | PCDH | Cell-Cell Contact | PMID: 35253643; PMID: 32694982 |
| CD31 | GFAP | Pcdhga6 | Pcdhga6 | 0.000176 | 0 | PCDHGA6_PCDHGA6 | Pcdhga6 - Pcdhga6 | PCDH | Cell-Cell Contact | PMID: 35253643; PMID: 32694982 |
| GFAP | CD31 | Pcdhga7 | Pcdhga7 | 0.000503 | 0 | PCDHGA7_PCDHGA7 | Pcdhga7 - Pcdhga7 | PCDH | Cell-Cell Contact | PMID: 35253643; PMID: 32694982 |
| CD31 | GFAP | Pcdhga7 | Pcdhga7 | 0.000503 | 0 | PCDHGA7_PCDHGA7 | Pcdhga7 - Pcdhga7 | PCDH | Cell-Cell Contact | PMID: 35253643; PMID: 32694982 |
| GFAP | GFAP | Pcdhga7 | Pcdhga7 | 0.000611 | 0 | PCDHGA7_PCDHGA7 | Pcdhga7 - Pcdhga7 | PCDH | Cell-Cell Contact | PMID: 35253643; PMID: 32694982 |
| CD31 | CD31 | Pcdhga8 | Pcdhga8 | 0.000154 | 0 | PCDHGA8_PCDHGA8 | Pcdhga8 - Pcdhga8 | PCDH | Cell-Cell Contact | PMID: 35253643; PMID: 32694982 |
| GFAP | CD31 | Pcdhga8 | Pcdhga8 | 0.000131 | 0 | PCDHGA8_PCDHGA8 | Pcdhga8 - Pcdhga8 | PCDH | Cell-Cell Contact | PMID: 35253643; PMID: 32694982 |
| CD31 | GFAP | Pcdhga8 | Pcdhga8 | 0.000131 | 0 | PCDHGA8_PCDHGA8 | Pcdhga8 - Pcdhga8 | PCDH | Cell-Cell Contact | PMID: 35253643; PMID: 32694982 |
| GFAP | CD31 | Pcdhga9 | Pcdhga9 | 0.000113 | 0 | PCDHGA9_PCDHGA9 | Pcdhga9 - Pcdhga9 | PCDH | Cell-Cell Contact | PMID: 35253643; PMID: 32694982 |
| CD31 | GFAP | Pcdhga9 | Pcdhga9 | 0.000113 | 0 | PCDHGA9_PCDHGA9 | Pcdhga9 - Pcdhga9 | PCDH | Cell-Cell Contact | PMID: 35253643; PMID: 32694982 |
| GFAP | GFAP | Pcdhga9 | Pcdhga9 | 0.000113 | 0 | PCDHGA9_PCDHGA9 | Pcdhga9 - Pcdhga9 | PCDH | Cell-Cell Contact | PMID: 35253643; PMID: 32694982 |
| CD31 | CD31 | Pcdhgb1 | Pcdhgb1 | 0.000623 | 0 | PCDHGB1_PCDHGB1 | Pcdhgb1 - Pcdhgb1 | PCDH | Cell-Cell Contact | PMID: 35253643; PMID: 32694982 |
| GFAP | CD31 | Pcdhgb1 | Pcdhgb1 | 0.000434 | 0 | PCDHGB1_PCDHGB1 | Pcdhgb1 - Pcdhgb1 | PCDH | Cell-Cell Contact | PMID: 35253643; PMID: 32694982 |
| CD31 | GFAP | Pcdhgb1 | Pcdhgb1 | 0.000434 | 0 | PCDHGB1_PCDHGB1 | Pcdhgb1 - Pcdhgb1 | PCDH | Cell-Cell Contact | PMID: 35253643; PMID: 32694982 |
| GFAP | CD31 | Pcdhgb2 | Pcdhgb2 | 0.000318 | 0 | PCDHGB2_PCDHGB2 | Pcdhgb2 - Pcdhgb2 | PCDH | Cell-Cell Contact | PMID: 35253643; PMID: 32694982 |
| CD31 | GFAP | Pcdhgb2 | Pcdhgb2 | 0.000318 | 0 | PCDHGB2_PCDHGB2 | Pcdhgb2 - Pcdhgb2 | PCDH | Cell-Cell Contact | PMID: 35253643; PMID: 32694982 |
| GFAP | GFAP | Pcdhgb2 | Pcdhgb2 | 0.000366 | 0 | PCDHGB2_PCDHGB2 | Pcdhgb2 - Pcdhgb2 | PCDH | Cell-Cell Contact | PMID: 35253643; PMID: 32694982 |
| GFAP | CD31 | Pcdhgb8 | Pcdhgb8 | 0.000139 | 0 | PCDHGB8_PCDHGB8 | Pcdhgb8 - Pcdhgb8 | PCDH | Cell-Cell Contact | PMID: 35253643; PMID: 32694982 |
| CD31 | GFAP | Pcdhgb8 | Pcdhgb8 | 0.000139 | 0 | PCDHGB8_PCDHGB8 | Pcdhgb8 - Pcdhgb8 | PCDH | Cell-Cell Contact | PMID: 35253643; PMID: 32694982 |
| GFAP | GFAP | Pcdhgb8 | Pcdhgb8 | 0.000354 | 0 | PCDHGB8_PCDHGB8 | Pcdhgb8 - Pcdhgb8 | PCDH | Cell-Cell Contact | PMID: 35253643; PMID: 32694982 |
| CD31 | CD31 | Pcdhgb4 | Pcdhgb4 | 0.000483 | 0 | PCDHGB4_PCDHGB4 | Pcdhgb4 - Pcdhgb4 | PCDH | Cell-Cell Contact | PMID: 35253643; PMID: 32694982 |
| GFAP | CD31 | Pcdhgb4 | Pcdhgb4 | 0.000454 | 0 | PCDHGB4_PCDHGB4 | Pcdhgb4 - Pcdhgb4 | PCDH | Cell-Cell Contact | PMID: 35253643; PMID: 32694982 |
| CD31 | GFAP | Pcdhgb4 | Pcdhgb4 | 0.000454 | 0 | PCDHGB4_PCDHGB4 | Pcdhgb4 - Pcdhgb4 | PCDH | Cell-Cell Contact | PMID: 35253643; PMID: 32694982 |
| CD31 | CD31 | Pcdhgb5 | Pcdhgb5 | 0.000538 | 0 | PCDHGB5_PCDHGB5 | Pcdhgb5 - Pcdhgb5 | PCDH | Cell-Cell Contact | PMID: 35253643; PMID: 32694982 |
| GFAP | CD31 | Pcdhgb5 | Pcdhgb5 | 0.000394 | 0 | PCDHGB5_PCDHGB5 | Pcdhgb5 - Pcdhgb5 | PCDH | Cell-Cell Contact | PMID: 35253643; PMID: 32694982 |
| CD31 | GFAP | Pcdhgb5 | Pcdhgb5 | 0.000394 | 0 | PCDHGB5_PCDHGB5 | Pcdhgb5 - Pcdhgb5 | PCDH | Cell-Cell Contact | PMID: 35253643; PMID: 32694982 |
| GFAP | CD31 | Pcdhgb6 | Pcdhgb6 | 0.000329 | 0 | PCDHGB6_PCDHGB6 | Pcdhgb6 - Pcdhgb6 | PCDH | Cell-Cell Contact | PMID: 35253643; PMID: 32694982 |
| CD31 | GFAP | Pcdhgb6 | Pcdhgb6 | 0.000329 | 0 | PCDHGB6_PCDHGB6 | Pcdhgb6 - Pcdhgb6 | PCDH | Cell-Cell Contact | PMID: 35253643; PMID: 32694982 |
| GFAP | GFAP | Pcdhgb6 | Pcdhgb6 | 0.000467 | 0 | PCDHGB6_PCDHGB6 | Pcdhgb6 - Pcdhgb6 | PCDH | Cell-Cell Contact | PMID: 35253643; PMID: 32694982 |
| GFAP | GFAP | Pcdhgb7 | Pcdhgb7 | 0.000812 | 0 | PCDHGB7_PCDHGB7 | Pcdhgb7 - Pcdhgb7 | PCDH | Cell-Cell Contact | PMID: 35253643; PMID: 32694982 |
| NeuN | GFAP | Pcdhgb7 | Pcdhgb7 | 0.000617 | 0 | PCDHGB7_PCDHGB7 | Pcdhgb7 - Pcdhgb7 | PCDH | Cell-Cell Contact | PMID: 35253643; PMID: 32694982 |
| GFAP | NeuN | Pcdhgb7 | Pcdhgb7 | 0.000617 | 0 | PCDHGB7_PCDHGB7 | Pcdhgb7 - Pcdhgb7 | PCDH | Cell-Cell Contact | PMID: 35253643; PMID: 32694982 |
| GFAP | CD31 | Pcdhgc3 | Pcdhgc3 | 0.006889 | 0 | PCDHGC3_PCDHGC3 | Pcdhgc3 - Pcdhgc3 | PCDH | Cell-Cell Contact | PMID: 35253643; PMID: 32694982 |
| CD31 | GFAP | Pcdhgc3 | Pcdhgc3 | 0.006889 | 0 | PCDHGC3_PCDHGC3 | Pcdhgc3 - Pcdhgc3 | PCDH | Cell-Cell Contact | PMID: 35253643; PMID: 32694982 |
| GFAP | GFAP | Pcdhgc3 | Pcdhgc3 | 0.011144 | 0 | PCDHGC3_PCDHGC3 | Pcdhgc3 - Pcdhgc3 | PCDH | Cell-Cell Contact | PMID: 35253643; PMID: 32694982 |

| | | | | | | | | | | |
|---|---|---|---|---|---|---|---|---|---|---|
| CD31 | CD31 | Pcdhgc4 | Pcdhgc4 | 0.000682 | 0 | PCDHGC4_PCDHGC4 | Pcdhgc4 - Pcdhgc4 | PCDH | Cell-Cell Contact | PMID: 35253643; PMID: 32694982 |
| GFAP | CD31 | Pcdhgc4 | Pcdhgc4 | 0.00061 | 0 | PCDHGC4_PCDHGC4 | Pcdhgc4 - Pcdhgc4 | PCDH | Cell-Cell Contact | PMID: 35253643; PMID: 32694982 |
| CD31 | GFAP | Pcdhgc4 | Pcdhgc4 | 0.00061 | 0 | PCDHGC4_PCDHGC4 | Pcdhgc4 - Pcdhgc4 | PCDH | Cell-Cell Contact | PMID: 35253643; PMID: 32694982 |
| NeuN | CD31 | Pcdhgc5 | Pcdhgc5 | 0.003869 | 0 | PCDHGC5_PCDHGC5 | Pcdhgc5 - Pcdhgc5 | PCDH | Cell-Cell Contact | PMID: 35253643; PMID: 32694982 |
| CD31 | NeuN | Pcdhgc5 | Pcdhgc5 | 0.003869 | 0 | PCDHGC5_PCDHGC5 | Pcdhgc5 - Pcdhgc5 | PCDH | Cell-Cell Contact | PMID: 35253643; PMID: 32694982 |
| NeuN | NeuN | Pcdhgc5 | Pcdhgc5 | 0.006477 | 0 | PCDHGC5_PCDHGC5 | Pcdhgc5 - Pcdhgc5 | PCDH | Cell-Cell Contact | PMID: 35253643; PMID: 32694982 |
| CD31 | CD31 | Sdc2 | Adgra2 | 0.001785 | 0 | SDC2_ADGRA2 | Sdc2 - Adgra2 | ADGRA | Cell-Cell Contact | PMID: 31462748 |
| GFAP | CD31 | Sdc2 | Adgra2 | 0.001376 | 0 | SDC2_ADGRA2 | Sdc2 - Adgra2 | ADGRA | Cell-Cell Contact | PMID: 31462748 |
| CD31 | CD31 | ITGAV_ITG | Adgra2 | 0.001037 | 0 | ITGAV_ITGB3_ADGRA2 | (Itgav+Itgb3) - Adgra2 | ADGRA | Cell-Cell Contact | PMID: 31462748 |
| GFAP | CD31 | ITGAV_ITG | Adgra2 | 0.001207 | 0 | ITGAV_ITGB3_ADGRA2 | (Itgav+Itgb3) - Adgra2 | ADGRA | Cell-Cell Contact | PMID: 31462748 |
| CD31 | CD31 | Tax1bp3 | Adgrb1 | 0.001359 | 0 | TAX1BP3_ADGRB1 | Tax1bp3 - Adgrb1 | ADGRB | Cell-Cell Contact | PMID: 31462748 |
| GFAP | NeuN | Tax1bp3 | Adgrb1 | 0.003796 | 0 | TAX1BP3_ADGRB1 | Tax1bp3 - Adgrb1 | ADGRB | Cell-Cell Contact | PMID: 31462748 |
| CD31 | NeuN | Rtn4r | Adgrb1 | 0.004276 | 0 | RTN4R_ADGRB1 | Rtn4r - Adgrb1 | ADGRB | Cell-Cell Contact | PMID: 31462748 |
| NeuN | NeuN | Rtn4r | Adgrb1 | 0.010222 | 0 | RTN4R_ADGRB1 | Rtn4r - Adgrb1 | ADGRB | Cell-Cell Contact | PMID: 31462748 |
| GFAP | NeuN | ITGAV_ITG | Adgrb1 | 0.004291 | 0 | ITGAV_ITGB5_ADGRB1 | (Itgav+Itgb5) - Adgrb1 | ADGRB | Cell-Cell Contact | PMID: 31462748 |
| NeuN | NeuN | ITGAV_ITG | Adgrb1 | 0.001723 | 0 | ITGAV_ITGB5_ADGRB1 | (Itgav+Itgb5) - Adgrb1 | ADGRB | Cell-Cell Contact | PMID: 31462748 |
| CD31 | CD31 | Tax1bp3 | Adgrb2 | 0.002652 | 0 | TAX1BP3_ADGRB2 | Tax1bp3 - Adgrb2 | ADGRB | Cell-Cell Contact | PMID: 31462748 |
| GFAP | NeuN | Tax1bp3 | Adgrb2 | 0.009728 | 0 | TAX1BP3_ADGRB2 | Tax1bp3 - Adgrb2 | ADGRB | Cell-Cell Contact | PMID: 31462748 |
| GFAP | GFAP | C1ql1 | Adgrb3 | 0.00088 | 0 | C1QL1_ADGRB3 | C1ql1 - Adgrb3 | ADGRB | Cell-Cell Contact | PMID: 31462748 |
| GFAP | NeuN | C1ql1 | Adgrb3 | 0.001953 | 0 | C1QL1_ADGRB3 | C1ql1 - Adgrb3 | ADGRB | Cell-Cell Contact | PMID: 31462748 |
| CD31 | NeuN | C1ql2 | Adgrb3 | 0.001934 | 0 | C1QL2_ADGRB3 | C1ql2 - Adgrb3 | ADGRB | Cell-Cell Contact | PMID: 31462748 |
| NeuN | NeuN | C1ql2 | Adgrb3 | 0.000906 | 0 | C1QL2_ADGRB3 | C1ql2 - Adgrb3 | ADGRB | Cell-Cell Contact | PMID: 31462748 |
| CD31 | NeuN | C1ql3 | Adgrb3 | 0.003881 | 0 | C1QL3_ADGRB3 | C1ql3 - Adgrb3 | ADGRB | Cell-Cell Contact | PMID: 31462748 |
| NeuN | NeuN | C1ql3 | Adgrb3 | 0.008359 | 0 | C1QL3_ADGRB3 | C1ql3 - Adgrb3 | ADGRB | Cell-Cell Contact | PMID: 31462748 |
| CD31 | CD31 | Thy1 | Adgre5 | 0.012647 | 0 | THY1_ADGRE5 | Thy1 - Adgre5 | ADGRE | Cell-Cell Contact | PMID: 31462748 |
| NeuN | CD31 | Thy1 | Adgre5 | 0.039424 | 0 | THY1_ADGRE5 | Thy1 - Adgre5 | ADGRE | Cell-Cell Contact | PMID: 31462748 |
| CD31 | CD31 | ITGAV_ITG | Adgre5 | 0.003733 | 0 | ITGAV_ITGB1_ADGRE5 | (Itgav+Itgb1) - Adgre5 | ADGRE | Cell-Cell Contact | PMID: 31462748 |
| GFAP | CD31 | ITGAV_ITG | Adgre5 | 0.003161 | 0 | ITGAV_ITGB1_ADGRE5 | (Itgav+Itgb1) - Adgre5 | ADGRE | Cell-Cell Contact | PMID: 31462748 |
| CD31 | CD31 | ITGAV_ITG | Adgre5 | 0.00173 | 0 | ITGAV_ITGB3_ADGRE5 | (Itgav+Itgb3) - Adgre5 | ADGRE | Cell-Cell Contact | PMID: 31462748 |
| GFAP | CD31 | ITGAV_ITG | Adgre5 | 0.002013 | 0 | ITGAV_ITGB3_ADGRE5 | (Itgav+Itgb3) - Adgre5 | ADGRE | Cell-Cell Contact | PMID: 31462748 |
| CD31 | CD31 | Cd55 | Adgre5 | 0.000882 | 0 | CD55_ADGRE5 | Cd55 - Adgre5 | ADGRE | Cell-Cell Contact | PMID: 31462748 |
| GFAP | CD31 | Cd55 | Adgre5 | 0.001427 | 0 | CD55_ADGRE5 | Cd55 - Adgre5 | ADGRE | Cell-Cell Contact | PMID: 31462748 |
| CD31 | CD31 | Lpar1 | Adgre5 | 0.001446 | 0 | LPAR1_ADGRE5 | Lpar1 - Adgre5 | ADGRE | Cell-Cell Contact | PMID: 31462748 |
| GFAP | CD31 | Lpar1 | Adgre5 | 0.004393 | 0 | LPAR1_ADGRE5 | Lpar1 - Adgre5 | ADGRE | Cell-Cell Contact | PMID: 31462748 |
| CD31 | CD31 | Lpar2 | Adgre5 | 0.001213 | 0 | LPAR2_ADGRE5 | Lpar2 - Adgre5 | ADGRE | Cell-Cell Contact | PMID: 31462748 |
| GFAP | CD31 | Lpar2 | Adgre5 | 0.00076 | 0 | LPAR2_ADGRE5 | Lpar2 - Adgre5 | ADGRE | Cell-Cell Contact | PMID: 31462748 |
| CD31 | GFAP | Col3a1 | Adgrg1 | 0.001742 | 0 | COL3A1_ADGRG1 | Col3a1 - Adgrg1 | ADGRG | Cell-Cell Contact | PMID: 31462748;UniProtKB |
| GFAP | GFAP | Col3a1 | Adgrg1 | 0.001622 | 0 | COL3A1_ADGRG1 | Col3a1 - Adgrg1 | ADGRG | Cell-Cell Contact | PMID: 31462748;UniProtKB |
| CD31 | GFAP | Tgm2 | Adgrg1 | 0.001677 | 0 | TGM2_ADGRG1 | Tgm2 - Adgrg1 | ADGRG | Cell-Cell Contact | PMID: 31462748 |
| GFAP | GFAP | Tgm2 | Adgrg1 | 0.001076 | 0 | TGM2_ADGRG1 | Tgm2 - Adgrg1 | ADGRG | Cell-Cell Contact | PMID: 31462748 |
| CD31 | CD31 | Col4a1 | Adgrg6 | 0.002937 | 0 | COL4A1_ADGRG6 | Col4a1 - Adgrg6 | ADGRG | Cell-Cell Contact | PMID: 31462748 |
| CD31 | GFAP | Col4a1 | Adgrg6 | 0.002605 | 0 | COL4A1_ADGRG6 | Col4a1 - Adgrg6 | ADGRG | Cell-Cell Contact | PMID: 31462748 |
| CD31 | CD31 | Col4a2 | Adgrg6 | 0.002504 | 0 | COL4A2_ADGRG6 | Col4a2 - Adgrg6 | ADGRG | Cell-Cell Contact | PMID: 31462748 |
| CD31 | GFAP | Col4a2 | Adgrg6 | 0.002221 | 0 | COL4A2_ADGRG6 | Col4a2 - Adgrg6 | ADGRG | Cell-Cell Contact | PMID: 31462748 |
| NeuN | CD31 | Prnp | Adgrg6 | 0.009977 | 0 | PRNP_ADGRG6 | Prnp - Adgrg6 | ADGRG | Cell-Cell Contact | PMID: 31462748 |
| GFAP | GFAP | Prnp | Adgrg6 | 0.005271 | 0 | PRNP_ADGRG6 | Prnp - Adgrg6 | ADGRG | Cell-Cell Contact | PMID: 31462748 |
| CD31 | NeuN | Flrt1 | Adgrl1 | 0.008062 | 0 | FLRT1_ADGRL1 | Flrt1 - Adgrl1 | ADGRL | Cell-Cell Contact | PMID: 25728924;PMID: 34022279;PMID: 27091502;PMID: 35907405 |
| NeuN | NeuN | Flrt1 | Adgrl1 | 0.012049 | 0 | FLRT1_ADGRL1 | Flrt1 - Adgrl1 | ADGRL | Cell-Cell Contact | PMID: 25728924;PMID: 34022279;PMID: 27091502;PMID: 35907405 |
| CD31 | NeuN | Flrt2 | Adgrl1 | 0.008855 | 0 | FLRT2_ADGRL1 | Flrt2 - Adgrl1 | ADGRL | Cell-Cell Contact | PMID: 25728924;PMID: 34022279;PMID: 27091502;PMID: 35907405 |
| NeuN | NeuN | Flrt2 | Adgrl1 | 0.014417 | 0 | FLRT2_ADGRL1 | Flrt2 - Adgrl1 | ADGRL | Cell-Cell Contact | PMID: 25728924;PMID: 34022279;PMID: 27091502;PMID: 35907405 |
| CD31 | NeuN | Flrt3 | Adgrl1 | 0.00604 | 0 | FLRT3_ADGRL1 | Flrt3 - Adgrl1 | ADGRL | Cell-Cell Contact | PMID: 25728924;PMID: 34022279;PMID: 27091502;PMID: 35907405 |
| NeuN | NeuN | Flrt3 | Adgrl1 | 0.006563 | 0 | FLRT3_ADGRL1 | Flrt3 - Adgrl1 | ADGRL | Cell-Cell Contact | PMID: 25728924;PMID: 34022279;PMID: 27091502;PMID: 35907405 |
| GFAP | NeuN | Nrxn1 | Adgrl1 | 0.035391 | 0 | NRXN1_ADGRL1 | Nrxn1 - Adgrl1 | ADGRL | Cell-Cell Contact | PMID: 25728924;PMID: 34022279;PMID: 27091502;PMID: 35907405 |
| NeuN | NeuN | Nrxn1 | Adgrl1 | 0.067315 | 0 | NRXN1_ADGRL1 | Nrxn1 - Adgrl1 | ADGRL | Cell-Cell Contact | PMID: 25728924;PMID: 34022279;PMID: 27091502;PMID: 35907405 |
| GFAP | NeuN | Nrxn2 | Adgrl1 | 0.023029 | 0 | NRXN2_ADGRL1 | Nrxn2 - Adgrl1 | ADGRL | Cell-Cell Contact | PMID: 25728924;PMID: 34022279;PMID: 27091502;PMID: 35907405 |
| NeuN | NeuN | Nrxn2 | Adgrl1 | 0.025471 | 0 | NRXN2_ADGRL1 | Nrxn2 - Adgrl1 | ADGRL | Cell-Cell Contact | PMID: 25728924;PMID: 34022279;PMID: 27091502;PMID: 35907405 |
| NeuN | CD31 | Nrxn3 | Adgrl1 | 0.018102 | 0 | NRXN3_ADGRL1 | Nrxn3 - Adgrl1 | ADGRL | Cell-Cell Contact | PMID: 25728924;PMID: 34022279;PMID: 27091502;PMID: 35907405 |
| NeuN | NeuN | Nrxn3 | Adgrl1 | 0.035756 | 0 | NRXN3_ADGRL1 | Nrxn3 - Adgrl1 | ADGRL | Cell-Cell Contact | PMID: 25728924;PMID: 34022279;PMID: 27091502;PMID: 35907405 |
| CD31 | NeuN | Tenm1 | Adgrl1 | 0.002528 | 0 | TENM1_ADGRL1 | Tenm1 - Adgrl1 | ADGRL | Cell-Cell Contact | PMID: 25728924;PMID: 34022279;PMID: 27091502;PMID: 35907405 |

| | | | | | | | | | | |
|---|---|---|---|---|---|---|---|---|---|---|
| NeuN | NeuN | Tenm1 | Adgrl1 | 0.003503 | 0 | TENM1_ADGRL1 | Tenm1 - Adgrl1 | ADGRL | Cell-Cell Contact | PMID: 25728924;PMID: 34022279;PMID: 27091502;PMID: 35907405 |
| CD31 | NeuN | Tenm2 | Adgrl1 | 0.004128 | 0 | TENM2_ADGRL1 | Tenm2 - Adgrl1 | ADGRL | Cell-Cell Contact | PMID: 25728924;PMID: 34022279;PMID: 27091502;PMID: 35907405 |
| NeuN | NeuN | Tenm2 | Adgrl1 | 0.005112 | 0 | TENM2_ADGRL1 | Tenm2 - Adgrl1 | ADGRL | Cell-Cell Contact | PMID: 25728924;PMID: 34022279;PMID: 27091502;PMID: 35907405 |
| CD31 | NeuN | Tenm3 | Adgrl1 | 0.007901 | 0 | TENM3_ADGRL1 | Tenm3 - Adgrl1 | ADGRL | Cell-Cell Contact | PMID: 25728924;PMID: 34022279;PMID: 27091502;PMID: 35907405 |
| NeuN | NeuN | Tenm3 | Adgrl1 | 0.008907 | 0 | TENM3_ADGRL1 | Tenm3 - Adgrl1 | ADGRL | Cell-Cell Contact | PMID: 25728924;PMID: 34022279;PMID: 27091502;PMID: 35907405 |
| GFAP | NeuN | Tenm4 | Adgrl1 | 0.004863 | 0 | TENM4_ADGRL1 | Tenm4 - Adgrl1 | ADGRL | Cell-Cell Contact | PMID: 25728924;PMID: 34022279;PMID: 27091502;PMID: 35907405 |
| NeuN | NeuN | Tenm4 | Adgrl1 | 0.008806 | 0 | TENM4_ADGRL1 | Tenm4 - Adgrl1 | ADGRL | Cell-Cell Contact | PMID: 25728924;PMID: 34022279;PMID: 27091502;PMID: 35907405 |
| CD31 | NeuN | TENM2_FL | Adgrl1 | 0.005771 | 0 | TENM2_FLRT1_ADGRL1 | (Tenm2+Flrt1) - Adgrl1 | ADGRL | Cell-Cell Contact | PMID: 25728924;PMID: 34022279;PMID: 27091502;PMID: 35907405 |
| NeuN | NeuN | TENM2_FL | Adgrl1 | 0.007854 | 0 | TENM2_FLRT1_ADGRL1 | (Tenm2+Flrt1) - Adgrl1 | ADGRL | Cell-Cell Contact | PMID: 25728924;PMID: 34022279;PMID: 27091502;PMID: 35907405 |
| CD31 | NeuN | TENM2_FL | Adgrl1 | 0.004994 | 0 | TENM2_FLRT3_ADGRL1 | (Tenm2+Flrt3) - Adgrl1 | ADGRL | Cell-Cell Contact | PMID: 25728924;PMID: 34022279;PMID: 27091502;PMID: 35907405 |
| NeuN | NeuN | TENM2_FL | Adgrl1 | 0.005793 | 0 | TENM2_FLRT3_ADGRL1 | (Tenm2+Flrt3) - Adgrl1 | ADGRL | Cell-Cell Contact | PMID: 25728924;PMID: 34022279;PMID: 27091502;PMID: 35907405 |
| NeuN | CD31 | TENM4_FL | Adgrl1 | 0.005149 | 0 | TENM4_FLRT1_ADGRL1 | (Tenm4+Flrt1) - Adgrl1 | ADGRL | Cell-Cell Contact | PMID: 25728924;PMID: 34022279;PMID: 27091502;PMID: 35907405 |
| NeuN | NeuN | TENM4_FL | Adgrl1 | 0.010302 | 0 | TENM4_FLRT1_ADGRL1 | (Tenm4+Flrt1) - Adgrl1 | ADGRL | Cell-Cell Contact | PMID: 25728924;PMID: 34022279;PMID: 27091502;PMID: 35907405 |
| GFAP | NeuN | TENM4_FL | Adgrl1 | 0.004969 | 0 | TENM4_FLRT3_ADGRL1 | (Tenm4+Flrt3) - Adgrl1 | ADGRL | Cell-Cell Contact | PMID: 25728924;PMID: 34022279;PMID: 27091502;PMID: 35907405 |
| NeuN | NeuN | TENM4_FL | Adgrl1 | 0.007603 | 0 | TENM4_FLRT3_ADGRL1 | (Tenm4+Flrt3) - Adgrl1 | ADGRL | Cell-Cell Contact | PMID: 25728924;PMID: 34022279;PMID: 27091502;PMID: 35907405 |
| NeuN | CD31 | Flrt1 | Adgrl2 | 0.002592 | 0 | FLRT1_ADGRL2 | Flrt1 - Adgrl2 | ADGRL | Cell-Cell Contact | PMID: 25728924;PMID: 34022279;PMID: 27091502;PMID: 35907405 |
| NeuN | NeuN | Flrt1 | Adgrl2 | 0.002082 | 0 | FLRT1_ADGRL2 | Flrt1 - Adgrl2 | ADGRL | Cell-Cell Contact | PMID: 25728924;PMID: 34022279;PMID: 27091502;PMID: 35907405 |
| NeuN | CD31 | Flrt2 | Adgrl2 | 0.003107 | 0 | FLRT2_ADGRL2 | Flrt2 - Adgrl2 | ADGRL | Cell-Cell Contact | PMID: 25728924;PMID: 34022279;PMID: 27091502;PMID: 35907405 |
| NeuN | NeuN | Flrt2 | Adgrl2 | 0.002496 | 0 | FLRT2_ADGRL2 | Flrt2 - Adgrl2 | ADGRL | Cell-Cell Contact | PMID: 25728924;PMID: 34022279;PMID: 27091502;PMID: 35907405 |
| CD31 | CD31 | Flrt3 | Adgrl2 | 0.001293 | 0 | FLRT3_ADGRL2 | Flrt3 - Adgrl2 | ADGRL | Cell-Cell Contact | PMID: 25728924;PMID: 34022279;PMID: 27091502;PMID: 35907405 |
| NeuN | CD31 | Flrt3 | Adgrl2 | 0.001406 | 0 | FLRT3_ADGRL2 | Flrt3 - Adgrl2 | ADGRL | Cell-Cell Contact | PMID: 25728924;PMID: 34022279;PMID: 27091502;PMID: 35907405 |
| NeuN | CD31 | Tenm1 | Adgrl2 | 0.000748 | 0 | TENM1_ADGRL2 | Tenm1 - Adgrl2 | ADGRL | Cell-Cell Contact | PMID: 25728924;PMID: 34022279;PMID: 27091502;PMID: 35907405 |
| NeuN | NeuN | Tenm1 | Adgrl2 | 0.000601 | 0 | TENM1_ADGRL2 | Tenm1 - Adgrl2 | ADGRL | Cell-Cell Contact | PMID: 25728924;PMID: 34022279;PMID: 27091502;PMID: 35907405 |
| CD31 | CD31 | Tenm2 | Adgrl2 | 0.000882 | 0 | TENM2_ADGRL2 | Tenm2 - Adgrl2 | ADGRL | Cell-Cell Contact | PMID: 25728924;PMID: 34022279;PMID: 27091502;PMID: 35907405 |
| NeuN | CD31 | Tenm2 | Adgrl2 | 0.001093 | 0 | TENM2_ADGRL2 | Tenm2 - Adgrl2 | ADGRL | Cell-Cell Contact | PMID: 25728924;PMID: 34022279;PMID: 27091502;PMID: 35907405 |
| CD31 | CD31 | Tenm3 | Adgrl2 | 0.001694 | 0 | TENM3_ADGRL2 | Tenm3 - Adgrl2 | ADGRL | Cell-Cell Contact | PMID: 25728924;PMID: 34022279;PMID: 27091502;PMID: 35907405 |
| NeuN | CD31 | Tenm3 | Adgrl2 | 0.001911 | 0 | TENM3_ADGRL2 | Tenm3 - Adgrl2 | ADGRL | Cell-Cell Contact | PMID: 25728924;PMID: 34022279;PMID: 27091502;PMID: 35907405 |
| NeuN | CD31 | Tenm4 | Adgrl2 | 0.001889 | 0 | TENM4_ADGRL2 | Tenm4 - Adgrl2 | ADGRL | Cell-Cell Contact | PMID: 25728924;PMID: 34022279;PMID: 27091502;PMID: 35907405 |
| NeuN | NeuN | Tenm4 | Adgrl2 | 0.001517 | 0 | TENM4_ADGRL2 | Tenm4 - Adgrl2 | ADGRL | Cell-Cell Contact | PMID: 25728924;PMID: 34022279;PMID: 27091502;PMID: 35907405 |
| CD31 | CD31 | TENM2_FL | Adgrl2 | 0.001068 | 0 | TENM2_FLRT3_ADGRL2 | (Tenm2+Flrt3) - Adgrl2 | ADGRL | Cell-Cell Contact | PMID: 25728924;PMID: 34022279;PMID: 27091502;PMID: 35907405 |
| NeuN | CD31 | TENM2_FL | Adgrl2 | 0.00124 | 0 | TENM2_FLRT3_ADGRL2 | (Tenm2+Flrt3) - Adgrl2 | ADGRL | Cell-Cell Contact | PMID: 25728924;PMID: 34022279;PMID: 27091502;PMID: 35907405 |
| CD31 | NeuN | Flrt1 | Adgrl3 | 0.002245 | 0 | FLRT1_ADGRL3 | Flrt1 - Adgrl3 | ADGRL | Cell-Cell Contact | PMID: 25728924;PMID: 34022279;PMID: 27091502;PMID: 35907405 |
| NeuN | NeuN | Flrt1 | Adgrl3 | 0.003365 | 0 | FLRT1_ADGRL3 | Flrt1 - Adgrl3 | ADGRL | Cell-Cell Contact | PMID: 25728924;PMID: 34022279;PMID: 27091502;PMID: 35907405 |
| CD31 | NeuN | Flrt2 | Adgrl3 | 0.002467 | 0 | FLRT2_ADGRL3 | Flrt2 - Adgrl3 | ADGRL | Cell-Cell Contact | PMID: 25728924;PMID: 34022279;PMID: 27091502;PMID: 35907405 |
| NeuN | NeuN | Flrt2 | Adgrl3 | 0.004034 | 0 | FLRT2_ADGRL3 | Flrt2 - Adgrl3 | ADGRL | Cell-Cell Contact | PMID: 25728924;PMID: 34022279;PMID: 27091502;PMID: 35907405 |
| CD31 | NeuN | Flrt3 | Adgrl3 | 0.001679 | 0 | FLRT3_ADGRL3 | Flrt3 - Adgrl3 | ADGRL | Cell-Cell Contact | PMID: 25728924;PMID: 34022279;PMID: 27091502;PMID: 35907405 |
| NeuN | NeuN | Flrt3 | Adgrl3 | 0.001826 | 0 | FLRT3_ADGRL3 | Flrt3 - Adgrl3 | ADGRL | Cell-Cell Contact | PMID: 25728924;PMID: 34022279;PMID: 27091502;PMID: 35907405 |
| CD31 | NeuN | Tenm1 | Adgrl3 | 0.000701 | 0 | TENM1_ADGRL3 | Tenm1 - Adgrl3 | ADGRL | Cell-Cell Contact | PMID: 25728924;PMID: 34022279;PMID: 27091502;PMID: 35907405 |
| NeuN | NeuN | Tenm1 | Adgrl3 | 0.000972 | 0 | TENM1_ADGRL3 | Tenm1 - Adgrl3 | ADGRL | Cell-Cell Contact | PMID: 25728924;PMID: 34022279;PMID: 27091502;PMID: 35907405 |
| CD31 | NeuN | Tenm2 | Adgrl3 | 0.001146 | 0 | TENM2_ADGRL3 | Tenm2 - Adgrl3 | ADGRL | Cell-Cell Contact | PMID: 25728924;PMID: 34022279;PMID: 27091502;PMID: 35907405 |
| NeuN | NeuN | Tenm2 | Adgrl3 | 0.001421 | 0 | TENM2_ADGRL3 | Tenm2 - Adgrl3 | ADGRL | Cell-Cell Contact | PMID: 25728924;PMID: 34022279;PMID: 27091502;PMID: 35907405 |
| CD31 | NeuN | Tenm3 | Adgrl3 | 0.0022 | 0 | TENM3_ADGRL3 | Tenm3 - Adgrl3 | ADGRL | Cell-Cell Contact | PMID: 25728924;PMID: 34022279;PMID: 27091502;PMID: 35907405 |
| NeuN | NeuN | Tenm3 | Adgrl3 | 0.002482 | 0 | TENM3_ADGRL3 | Tenm3 - Adgrl3 | ADGRL | Cell-Cell Contact | PMID: 25728924;PMID: 34022279;PMID: 27091502;PMID: 35907405 |
| GFAP | NeuN | Tenm4 | Adgrl3 | 0.001351 | 0 | TENM4_ADGRL3 | Tenm4 - Adgrl3 | ADGRL | Cell-Cell Contact | PMID: 25728924;PMID: 34022279;PMID: 27091502;PMID: 35907405 |
| NeuN | NeuN | Tenm4 | Adgrl3 | 0.002454 | 0 | TENM4_ADGRL3 | Tenm4 - Adgrl3 | ADGRL | Cell-Cell Contact | PMID: 25728924;PMID: 34022279;PMID: 27091502;PMID: 35907405 |
| CD31 | NeuN | Unc5a | Adgrl3 | 0.001998 | 0 | UNC5A_ADGRL3 | Unc5a - Adgrl3 | ADGRL | Cell-Cell Contact | PMID: 25728924;PMID: 34022279;PMID: 27091502;PMID: 35907405 |
| NeuN | NeuN | Unc5a | Adgrl3 | 0.002899 | 0 | UNC5A_ADGRL3 | Unc5a - Adgrl3 | ADGRL | Cell-Cell Contact | PMID: 25728924;PMID: 34022279;PMID: 27091502;PMID: 35907405 |
| CD31 | NeuN | TENM3_FL | Adgrl3 | 0.002223 | 0 | TENM3_FLRT1_ADGRL3 | (Tenm3+Flrt1) - Adgrl3 | ADGRL | Cell-Cell Contact | PMID: 25728924;PMID: 34022279;PMID: 27091502;PMID: 35907405 |
| NeuN | NeuN | TENM3_FL | Adgrl3 | 0.00289 | 0 | TENM3_FLRT1_ADGRL3 | (Tenm3+Flrt1) - Adgrl3 | ADGRL | Cell-Cell Contact | PMID: 25728924;PMID: 34022279;PMID: 27091502;PMID: 35907405 |
| CD31 | NeuN | TENM3_FL | Adgrl3 | 0.001922 | 0 | TENM3_FLRT3_ADGRL3 | (Tenm3+Flrt3) - Adgrl3 | ADGRL | Cell-Cell Contact | PMID: 25728924;PMID: 34022279;PMID: 27091502;PMID: 35907405 |
| NeuN | NeuN | TENM3_FL | Adgrl3 | 0.002129 | 0 | TENM3_FLRT3_ADGRL3 | (Tenm3+Flrt3) - Adgrl3 | ADGRL | Cell-Cell Contact | PMID: 25728924;PMID: 34022279;PMID: 27091502;PMID: 35907405 |
| NeuN | CD31 | Flrt1 | Flrt1 | 0.001096 | 0 | FLRT1_FLRT1 | Flrt1 - Flrt1 | FLRT | Cell-Cell Contact | PMID: 27091502 |
| CD31 | NeuN | Flrt1 | Flrt1 | 0.001096 | 0 | FLRT1_FLRT1 | Flrt1 - Flrt1 | FLRT | Cell-Cell Contact | PMID: 27091502 |
| NeuN | NeuN | Flrt1 | Flrt1 | 0.001644 | 0 | FLRT1_FLRT1 | Flrt1 - Flrt1 | FLRT | Cell-Cell Contact | PMID: 27091502 |
| NeuN | CD31 | Flrt2 | Flrt2 | 0.001444 | 0 | FLRT2_FLRT2 | Flrt2 - Flrt2 | FLRT | Cell-Cell Contact | PMID: 27091502 |
| CD31 | NeuN | Flrt2 | Flrt2 | 0.001444 | 0 | FLRT2_FLRT2 | Flrt2 - Flrt2 | FLRT | Cell-Cell Contact | PMID: 27091502 |
| NeuN | NeuN | Flrt2 | Flrt2 | 0.002363 | 0 | FLRT2_FLRT2 | Flrt2 - Flrt2 | FLRT | Cell-Cell Contact | PMID: 27091502 |
| NeuN | CD31 | Flrt3 | Flrt3 | 0.000444 | 0 | FLRT3_FLRT3 | Flrt3 - Flrt3 | FLRT | Cell-Cell Contact | PMID: 27091502 |
| CD31 | NeuN | Flrt3 | Flrt3 | 0.000444 | 0 | FLRT3_FLRT3 | Flrt3 - Flrt3 | FLRT | Cell-Cell Contact | PMID: 27091502 |

| | | | | | | | | | | |
|---|---|---|---|---|---|---|---|---|---|---|
| NeuN | NeuN | Flrt3 | Flrt3 | 0.000483 | 0 | FLRT3_FLRT3 | Flrt3 - Flrt3 | FLRT | Cell-Cell Contact | PMID: 27091502 |
| NeuN | CD31 | Flrt1 | Unc5a | 0.000975 | 0 | FLRT1_UNC5A | Flrt1 - Unc5a | UNC5 | Cell-Cell Contact | PMID: 27091502 |
| NeuN | NeuN | Flrt1 | Unc5a | 0.001416 | 0 | FLRT1_UNC5A | Flrt1 - Unc5a | UNC5 | Cell-Cell Contact | PMID: 27091502 |
| NeuN | CD31 | Flrt2 | Unc5a | 0.001169 | 0 | FLRT2_UNC5A | Flrt2 - Unc5a | UNC5 | Cell-Cell Contact | PMID: 27091502 |
| NeuN | NeuN | Flrt2 | Unc5a | 0.001697 | 0 | FLRT2_UNC5A | Flrt2 - Unc5a | UNC5 | Cell-Cell Contact | PMID: 27091502 |
| CD31 | NeuN | Flrt3 | Unc5a | 0.000706 | 0 | FLRT3_UNC5A | Flrt3 - Unc5a | UNC5 | Cell-Cell Contact | PMID: 27091502 |
| NeuN | NeuN | Flrt3 | Unc5a | 0.000767 | 0 | FLRT3_UNC5A | Flrt3 - Unc5a | UNC5 | Cell-Cell Contact | PMID: 27091502 |
| CD31 | CD31 | Flrt1 | Unc5b | 0.000555 | 0 | FLRT1_UNC5B | Flrt1 - Unc5b | UNC5 | Cell-Cell Contact | PMID: 27091502 |
| NeuN | CD31 | Flrt1 | Unc5b | 0.000832 | 0 | FLRT1_UNC5B | Flrt1 - Unc5b | UNC5 | Cell-Cell Contact | PMID: 27091502 |
| CD31 | CD31 | Flrt2 | Unc5b | 0.00061 | 0 | FLRT2_UNC5B | Flrt2 - Unc5b | UNC5 | Cell-Cell Contact | PMID: 27091502 |
| NeuN | CD31 | Flrt2 | Unc5b | 0.000998 | 0 | FLRT2_UNC5B | Flrt2 - Unc5b | UNC5 | Cell-Cell Contact | PMID: 27091502 |
| CD31 | CD31 | Flrt3 | Unc5b | 0.000415 | 0 | FLRT3_UNC5B | Flrt3 - Unc5b | UNC5 | Cell-Cell Contact | PMID: 27091502 |
| NeuN | CD31 | Flrt3 | Unc5b | 0.000451 | 0 | FLRT3_UNC5B | Flrt3 - Unc5b | UNC5 | Cell-Cell Contact | PMID: 27091502 |
| NeuN | GFAP | Flrt1 | Unc5c | 0.000668 | 0 | FLRT1_UNC5C | Flrt1 - Unc5 | UNC5 | Cell-Cell Contact | PMID: 27091502 |
| NeuN | NeuN | Flrt1 | Unc5c | 0.000454 | 0 | FLRT1_UNC5C | Flrt1 - Unc5 | UNC5 | Cell-Cell Contact | PMID: 27091502 |
| NeuN | GFAP | Flrt2 | Unc5c | 0.000801 | 0 | FLRT2_UNC5C | Flrt2 - Unc5 | UNC5 | Cell-Cell Contact | PMID: 27091502 |
| NeuN | NeuN | Flrt2 | Unc5c | 0.000545 | 0 | FLRT2_UNC5C | Flrt2 - Unc5 | UNC5 | Cell-Cell Contact | PMID: 27091502 |
| GFAP | GFAP | Flrt3 | Unc5c | 0.00028 | 0 | FLRT3_UNC5C | Flrt3 - Unc5 | UNC5 | Cell-Cell Contact | PMID: 27091502 |
| NeuN | GFAP | Flrt3 | Unc5c | 0.000362 | 0 | FLRT3_UNC5C | Flrt3 - Unc5 | UNC5 | Cell-Cell Contact | PMID: 27091502 |
| NeuN | GFAP | Flrt1 | Unc5d | 0.000594 | 0 | FLRT1_UNC5D | Flrt1 - Unc5d | UNC5 | Cell-Cell Contact | PMID: 27091502 |
| NeuN | NeuN | Flrt1 | Unc5d | 0.000604 | 0 | FLRT1_UNC5D | Flrt1 - Unc5d | UNC5 | Cell-Cell Contact | PMID: 27091502 |
| NeuN | GFAP | Flrt2 | Unc5d | 0.000712 | 0 | FLRT2_UNC5D | Flrt2 - Unc5d | UNC5 | Cell-Cell Contact | PMID: 27091502 |
| NeuN | NeuN | Flrt2 | Unc5d | 0.000725 | 0 | FLRT2_UNC5D | Flrt2 - Unc5d | UNC5 | Cell-Cell Contact | PMID: 27091502 |
| NeuN | GFAP | Flrt3 | Unc5d | 0.000322 | 0 | FLRT3_UNC5D | Flrt3 - Unc5d | UNC5 | Cell-Cell Contact | PMID: 27091502 |
| NeuN | NeuN | Flrt3 | Unc5d | 0.000327 | 0 | FLRT3_UNC5D | Flrt3 - Unc5d | UNC5 | Cell-Cell Contact | PMID: 27091502 |
| GFAP | NeuN | Nrxn1 | Clstn1 | 0.027942 | 0 | NRXN1_CLSTN1 | Nrxn1 - Clstn1 | NRXN | Cell-Cell Contact | PMID: 32434929 |
| NeuN | NeuN | Nrxn1 | Clstn1 | 0.053519 | 0 | NRXN1_CLSTN1 | Nrxn1 - Clstn1 | NRXN | Cell-Cell Contact | PMID: 32434929 |
| GFAP | NeuN | Nrxn2 | Clstn1 | 0.018133 | 0 | NRXN2_CLSTN1 | Nrxn2 - Clstn1 | NRXN | Cell-Cell Contact | PMID: 32434929 |
| NeuN | NeuN | Nrxn2 | Clstn1 | 0.020067 | 0 | NRXN2_CLSTN1 | Nrxn2 - Clstn1 | NRXN | Cell-Cell Contact | PMID: 32434929 |
| NeuN | CD31 | Nrxn1 | Clstn2 | 0.009275 | 0 | NRXN1_CLSTN2 | Nrxn1 - Clstn2 | NRXN | Cell-Cell Contact | PMID: 32434929 |
| NeuN | NeuN | Nrxn1 | Clstn2 | 0.017517 | 0 | NRXN1_CLSTN2 | Nrxn1 - Clstn2 | NRXN | Cell-Cell Contact | PMID: 32434929 |
| GFAP | NeuN | Nrxn2 | Clstn2 | 0.005789 | 0 | NRXN2_CLSTN2 | Nrxn2 - Clstn2 | NRXN | Cell-Cell Contact | PMID: 32434929 |
| NeuN | NeuN | Nrxn2 | Clstn2 | 0.006415 | 0 | NRXN2_CLSTN2 | Nrxn2 - Clstn2 | NRXN | Cell-Cell Contact | PMID: 32434929 |
| GFAP | NeuN | Nrxn1 | Clstn3 | 0.029014 | 0 | NRXN1_CLSTN3 | Nrxn1 - Clstn3 | NRXN | Cell-Cell Contact | PMID: 32434929 |
| NeuN | NeuN | Nrxn1 | Clstn3 | 0.055517 | 0 | NRXN1_CLSTN3 | Nrxn1 - Clstn3 | NRXN | Cell-Cell Contact | PMID: 32434929 |
| GFAP | NeuN | Nrxn2 | Clstn3 | 0.018835 | 0 | NRXN2_CLSTN3 | Nrxn2 - Clstn3 | NRXN | Cell-Cell Contact | PMID: 32434929 |
| NeuN | NeuN | Nrxn2 | Clstn3 | 0.020843 | 0 | NRXN2_CLSTN3 | Nrxn2 - Clstn3 | NRXN | Cell-Cell Contact | PMID: 32434929 |
| NeuN | CD31 | Nrxn1 | Dag1 | 0.007411 | 0 | NRXN1_DAG1 | Nrxn1 - Dag1 | NRXN | Cell-Cell Contact | PMID: 21628410 |
| NeuN | NeuN | Nrxn1 | Dag1 | 0.004744 | 0 | NRXN1_DAG1 | Nrxn1 - Dag1 | NRXN | Cell-Cell Contact | PMID: 21628410 |
| NeuN | CD31 | Nrxn2 | Dag1 | 0.002697 | 0 | NRXN2_DAG1 | Nrxn2 - Dag1 | NRXN | Cell-Cell Contact | PMID: 21628410 |
| GFAP | GFAP | Nrxn2 | Dag1 | 0.001846 | 0 | NRXN2_DAG1 | Nrxn2 - Dag1 | NRXN | Cell-Cell Contact | PMID: 21628410 |
| CD31 | NeuN | Lrfn4 | Ptprd | 0.004856 | 0 | LRFN4_PTPRD | Lrfn4 - Ptprd | PTPR | Cell-Cell Contact | PMID: 29674953;PMID: 23916315 |
| NeuN | NeuN | Lrfn4 | Ptprd | 0.004337 | 0 | LRFN4_PTPRD | Lrfn4 - Ptprd | PTPR | Cell-Cell Contact | PMID: 29674953;PMID: 23916315 |
| CD31 | CD31 | Lrfn4 | Ptprf | 0.001539 | 0 | LRFN4_PTPRF | Lrfn4 - Ptprf | PTPR | Cell-Cell Contact | PMID: 29674953;PMID: 23916315 |
| CD31 | GFAP | Lrfn4 | Ptprf | 0.001979 | 0 | LRFN4_PTPRF | Lrfn4 - Ptprf | PTPR | Cell-Cell Contact | PMID: 29674953;PMID: 23916315 |
| CD31 | NeuN | Lrfn4 | Ptprs | 0.006681 | 0 | LRFN4_PTPRS | Lrfn4 - Ptprs | PTPR | Cell-Cell Contact | PMID: 29674953;PMID: 23916315 |
| NeuN | NeuN | Lrfn4 | Ptprs | 0.005967 | 0 | LRFN4_PTPRS | Lrfn4 - Ptprs | PTPR | Cell-Cell Contact | PMID: 29674953;PMID: 23916315 |
| NeuN | GFAP | Lrfn5 | Ptprd | 0.004743 | 0 | LRFN5_PTPRD | Lrfn5 - Ptprd | PTPR | Cell-Cell Contact | PMID: 29674953;PMID: 23916315 |
| NeuN | NeuN | Lrfn5 | Ptprd | 0.005211 | 0 | LRFN5_PTPRD | Lrfn5 - Ptprd | PTPR | Cell-Cell Contact | PMID: 29674953;PMID: 23916315 |
| NeuN | GFAP | Lrfn5 | Ptprf | 0.002124 | 0 | LRFN5_PTPRF | Lrfn5 - Ptprf | PTPR | Cell-Cell Contact | PMID: 29674953;PMID: 23916315 |
| NeuN | NeuN | Lrfn5 | Ptprf | 0.001648 | 0 | LRFN5_PTPRF | Lrfn5 - Ptprf | PTPR | Cell-Cell Contact | PMID: 29674953;PMID: 23916315 |
| CD31 | NeuN | Lrfn5 | Ptprs | 0.006311 | 0 | LRFN5_PTPRS | Lrfn5 - Ptprs | PTPR | Cell-Cell Contact | PMID: 29674953;PMID: 23916315 |
| NeuN | NeuN | Lrfn5 | Ptprs | 0.007168 | 0 | LRFN5_PTPRS | Lrfn5 - Ptprs | PTPR | Cell-Cell Contact | PMID: 29674953;PMID: 23916315 |
| NeuN | GFAP | Lrrc4b | Ptprd | 0.009597 | 0 | LRRC4B_PTPRD | Lrrc4b - Ptprd | PTPR | Cell-Cell Contact | PMID: 29674953;PMID: 23916315 |
| NeuN | NeuN | Lrrc4b | Ptprd | 0.01054 | 0 | LRRC4B_PTPRD | Lrrc4b - Ptprd | PTPR | Cell-Cell Contact | PMID: 29674953;PMID: 23916315 |
| GFAP | NeuN | Lrrc4b | Ptprs | 0.010159 | 0 | LRRC4B_PTPRS | Lrrc4b - Ptprs | PTPR | Cell-Cell Contact | PMID: 29674953;PMID: 23916315 |
| NeuN | NeuN | Lrrc4b | Ptprs | 0.014469 | 0 | LRRC4B_PTPRS | Lrrc4b - Ptprs | PTPR | Cell-Cell Contact | PMID: 29674953;PMID: 23916315 |
| NeuN | GFAP | Lrrc4c | Ptprf | 0.004402 | 0 | LRRC4C_PTPRF | Lrrc4c - Ptprf | PTPR | Cell-Cell Contact | PMID: 29674953;PMID: 23916315 |
| NeuN | NeuN | Lrrc4c | Ptprf | 0.003418 | 0 | LRRC4C_PTPRF | Lrrc4c - Ptprf | PTPR | Cell-Cell Contact | PMID: 29674953;PMID: 23916315 |

| | | | | | | | | | | |
|---|---|---|---|---|---|---|---|---|---|---|
| NeuN | GFAP | Nrxn1 | Lrrtm1 | 0.004294 | 0 | NRXN1_LRRTM1 | Nrxn1 - Lrrtm1 | NRXN | Cell-Cell Contact | PMID:20519524;PMID:20064387;PMID:20064388; |
| NeuN | NeuN | Nrxn1 | Lrrtm1 | 0.007218 | 0 | NRXN1_LRRTM1 | Nrxn1 - Lrrtm1 | NRXN | Cell-Cell Contact | PMID:20519524;PMID:20064387;PMID:20064388; |
| GFAP | NeuN | Nrxn2 | Lrrtm1 | 0.002369 | 0 | NRXN2_LRRTM1 | Nrxn2 - Lrrtm1 | NRXN | Cell-Cell Contact | PMID:20519524;PMID:20064387;PMID:20064388; |
| NeuN | NeuN | Nrxn2 | Lrrtm1 | 0.002626 | 0 | NRXN2_LRRTM1 | Nrxn2 - Lrrtm1 | NRXN | Cell-Cell Contact | PMID:20519524;PMID:20064387;PMID:20064388; |
| NeuN | GFAP | Nrxn3 | Lrrtm1 | 0.002211 | 0 | NRXN3_LRRTM1 | Nrxn3 - Lrrtm1 | NRXN | Cell-Cell Contact | PMID:20519524;PMID:20064387;PMID:20064388; |
| NeuN | NeuN | Nrxn3 | Lrrtm1 | 0.003722 | 0 | NRXN3_LRRTM1 | Nrxn3 - Lrrtm1 | NRXN | Cell-Cell Contact | PMID:20519524;PMID:20064387;PMID:20064388; |
| NeuN | GFAP | Nrxn1 | Lrrtm2 | 0.007027 | 0 | NRXN1_LRRTM2 | Nrxn1 - Lrrtm2 | NRXN | Cell-Cell Contact | PMID:20519524;PMID:20064387;PMID:20064388; |
| NeuN | NeuN | Nrxn1 | Lrrtm2 | 0.006445 | 0 | NRXN1_LRRTM2 | Nrxn1 - Lrrtm2 | NRXN | Cell-Cell Contact | PMID:20519524;PMID:20064387;PMID:20064388; |
| NeuN | GFAP | Nrxn2 | Lrrtm2 | 0.002556 | 0 | NRXN2_LRRTM2 | Nrxn2 - Lrrtm2 | NRXN | Cell-Cell Contact | PMID:20519524;PMID:20064387;PMID:20064388; |
| NeuN | NeuN | Nrxn2 | Lrrtm2 | 0.002344 | 0 | NRXN2_LRRTM2 | Nrxn2 - Lrrtm2 | NRXN | Cell-Cell Contact | PMID:20519524;PMID:20064387;PMID:20064388; |
| NeuN | GFAP | Nrxn3 | Lrrtm2 | 0.003623 | 0 | NRXN3_LRRTM2 | Nrxn3 - Lrrtm2 | NRXN | Cell-Cell Contact | PMID:20519524;PMID:20064387;PMID:20064388; |
| NeuN | NeuN | Nrxn3 | Lrrtm2 | 0.003322 | 0 | NRXN3_LRRTM2 | Nrxn3 - Lrrtm2 | NRXN | Cell-Cell Contact | PMID:20519524;PMID:20064387;PMID:20064388; |
| NeuN | GFAP | Nrxn1 | Lrrtm3 | 0.005058 | 0 | NRXN1_LRRTM3 | Nrxn1 - Lrrtm3 | NRXN | Cell-Cell Contact | PMID:20519524;PMID:20064387;PMID:20064388; |
| NeuN | NeuN | Nrxn1 | Lrrtm3 | 0.008756 | 0 | NRXN1_LRRTM3 | Nrxn1 - Lrrtm3 | NRXN | Cell-Cell Contact | PMID:20519524;PMID:20064387;PMID:20064388; |
| GFAP | NeuN | Nrxn2 | Lrrtm3 | 0.002877 | 0 | NRXN2_LRRTM3 | Nrxn2 - Lrrtm3 | NRXN | Cell-Cell Contact | PMID:20519524;PMID:20064387;PMID:20064388; |
| NeuN | NeuN | Nrxn2 | Lrrtm3 | 0.003189 | 0 | NRXN2_LRRTM3 | Nrxn2 - Lrrtm3 | NRXN | Cell-Cell Contact | PMID:20519524;PMID:20064387;PMID:20064388; |
| NeuN | GFAP | Nrxn3 | Lrrtm3 | 0.002605 | 0 | NRXN3_LRRTM3 | Nrxn3 - Lrrtm3 | NRXN | Cell-Cell Contact | PMID:20519524;PMID:20064387;PMID:20064388; |
| NeuN | NeuN | Nrxn3 | Lrrtm3 | 0.004518 | 0 | NRXN3_LRRTM3 | Nrxn3 - Lrrtm3 | NRXN | Cell-Cell Contact | PMID:20519524;PMID:20064387;PMID:20064388; |
| NeuN | CD31 | Nrxn1 | Lrrtm4 | 0.007013 | 0 | NRXN1_LRRTM4 | Nrxn1 - Lrrtm4 | NRXN | Cell-Cell Contact | PMID:20519524;PMID:20064387;PMID:20064388; |
| NeuN | NeuN | Nrxn1 | Lrrtm4 | 0.008229 | 0 | NRXN1_LRRTM4 | Nrxn1 - Lrrtm4 | NRXN | Cell-Cell Contact | PMID:20519524;PMID:20064387;PMID:20064388; |
| GFAP | NeuN | Nrxn2 | Lrrtm4 | 0.002703 | 0 | NRXN2_LRRTM4 | Nrxn2 - Lrrtm4 | NRXN | Cell-Cell Contact | PMID:20519524;PMID:20064387;PMID:20064388; |
| NeuN | NeuN | Nrxn2 | Lrrtm4 | 0.002996 | 0 | NRXN2_LRRTM4 | Nrxn2 - Lrrtm4 | NRXN | Cell-Cell Contact | PMID:20519524;PMID:20064387;PMID:20064388; |
| NeuN | CD31 | Nrxn3 | Lrrtm4 | 0.003616 | 0 | NRXN3_LRRTM4 | Nrxn3 - Lrrtm4 | NRXN | Cell-Cell Contact | PMID:20519524;PMID:20064387;PMID:20064388; |
| NeuN | NeuN | Nrxn3 | Lrrtm4 | 0.004245 | 0 | NRXN3_LRRTM4 | Nrxn3 - Lrrtm4 | NRXN | Cell-Cell Contact | PMID:20519524;PMID:20064387;PMID:20064388; |
| CD31 | NeuN | Lrrtm4 | Ptprs | 0.006214 | 0 | LRRTM4_PTPRS | Lrrtm4 - Ptprs | PTPR | Cell-Cell Contact | PMID:31995730 |
| NeuN | NeuN | Lrrtm4 | Ptprs | 0.007293 | 0 | LRRTM4_PTPRS | Lrrtm4 - Ptprs | PTPR | Cell-Cell Contact | PMID:31995730 |
| CD31 | NeuN | App | Sorl1 | 0.1014 | 0 | APP_SORL1 | App - Sorl1 | APP | Cell-Cell Contact | PMID: 24523320 |
| NeuN | NeuN | App | Sorl1 | 0.145727 | 0 | APP_SORL1 | App - Sorl1 | APP | Cell-Cell Contact | PMID: 24523320 |
| NeuN | CD31 | App | Tnfrsf21 | 0.011679 | 0 | APP_TNFRSF21 | App - Tnfrsf21 | APP | Cell-Cell Contact | PMID: 33075465 |
| NeuN | NeuN | App | Tnfrsf21 | 0.011829 | 0 | APP_TNFRSF21 | App - Tnfrsf21 | APP | Cell-Cell Contact | PMID: 33075465 |
| GFAP | GFAP | App | TREM2_TYROBP | 0.014333 | 0 | APP_TREM2_TYROBP | App - (Trem2+Tyrobp) | APP | Cell-Cell Contact | PMID:27477013 |
| NeuN | GFAP | App | TREM2_TYROBP | 0.027784 | 0 | APP_TREM2_TYROBP | App - (Trem2+Tyrobp) | APP | Cell-Cell Contact | PMID:27477013 |
| CD31 | CD31 | Apoa2 | TREM2_TYROBP | 0.000524 | 0 | APOA2_TREM2_TYROBP | Apoa2 - (Trem2+Tyrobp) | ApoA | Cell-Cell Contact | uniprot;PMID:27477017 |
| CD31 | GFAP | Apoa2 | TREM2_TYROBP | 0.000699 | 0 | APOA2_TREM2_TYROBP | Apoa2 - (Trem2+Tyrobp) | ApoA | Cell-Cell Contact | uniprot;PMID:27477017 |
| GFAP | CD31 | Apoe | TREM2_TYROBP | 0.018176 | 0 | APOE_TREM2_TYROBP | Apoe - (Trem2+Tyrobp) | ApoE | Cell-Cell Contact | uniprot;PMID:27477015 |
| GFAP | GFAP | Apoe | TREM2_TYROBP | 0.024095 | 0 | APOE_TREM2_TYROBP | Apoe - (Trem2+Tyrobp) | ApoE | Cell-Cell Contact | uniprot;PMID:27477015 |
| NeuN | CD31 | Cadm1 | Nectin3 | 0.002987 | 0 | CADM1_NECTIN3 | Cadm1 - Nectin3 | CADM | Cell-Cell Contact | uniprot |
| NeuN | NeuN | Cadm1 | Nectin3 | 0.00209 | 0 | CADM1_NECTIN3 | Cadm1 - Nectin3 | CADM | Cell-Cell Contact | uniprot |
| CD31 | CD31 | Cd99 | Pilrb1 | 0.004618 | 0 | CD99_PILRB1 | Cd99 - Pilrb1 | CD99 | Cell-Cell Contact | uniprot;PMID:18234675 |
| NeuN | CD31 | Cd99 | Pilrb1 | 0.007675 | 0 | CD99_PILRB1 | Cd99 - Pilrb1 | CD99 | Cell-Cell Contact | uniprot;PMID:18234675 |
| CD31 | CD31 | Ceacam2 | Cd209c | 0.000317 | 0 | CEACAM2_CD209C | Ceacam2 - Cd209c | CEACAM | Cell-Cell Contact | PMID:16282604;PMID:16246332 |
| CD31 | GFAP | Ceacam2 | Cd209c | 0.000295 | 0 | CEACAM2_CD209C | Ceacam2 - Cd209c | CEACAM | Cell-Cell Contact | PMID:16282604;PMID:16246332 |
| CD31 | CD31 | Ceacam2 | Cd209a | 0.000473 | 0 | CEACAM2_CD209A | Ceacam2 - Cd209a | CEACAM | Cell-Cell Contact | PMID:16282604;PMID:16246332 |
| CD31 | GFAP | Ceacam2 | Cd209a | 0.000534 | 0 | CEACAM2_CD209A | Ceacam2 - Cd209a | CEACAM | Cell-Cell Contact | PMID:16282604;PMID:16246332 |
| CD31 | GFAP | Ceacam2 | Cd209d | 0.000309 | 0 | CEACAM2_CD209D | Ceacam2 - Cd209d | CEACAM | Cell-Cell Contact | PMID:16282604;PMID:16246332 |
| GFAP | GFAP | Ceacam2 | Cd209d | 0.000256 | 0 | CEACAM2_CD209D | Ceacam2 - Cd209d | CEACAM | Cell-Cell Contact | PMID:16282604;PMID:16246332 |
| CD31 | GFAP | Ceacam2 | Cd209b | 0.000418 | 0 | CEACAM2_CD209B | Ceacam2 - Cd209b | CEACAM | Cell-Cell Contact | PMID:16282604;PMID:16246332 |
| GFAP | GFAP | Ceacam2 | Cd209b | 0.000347 | 0 | CEACAM2_CD209B | Ceacam2 - Cd209b | CEACAM | Cell-Cell Contact | PMID:16282604;PMID:16246332 |
| CD31 | CD31 | Ceacam2 | Ceacam2 | 0.000415 | 0 | CEACAM2_CEACAM2 | Ceacam2 - Ceacam2 | CEACAM | Cell-Cell Contact | PMID:21982860 |
| GFAP | CD31 | Ceacam2 | Ceacam2 | 0.000344 | 0 | CEACAM2_CEACAM2 | Ceacam2 - Ceacam2 | CEACAM | Cell-Cell Contact | PMID:21982860 |
| CD31 | GFAP | Ceacam2 | Ceacam2 | 0.000344 | 0 | CEACAM2_CEACAM2 | Ceacam2 - Ceacam2 | CEACAM | Cell-Cell Contact | PMID:21982860 |
| CD31 | GFAP | Ceacam2 | Sele | 0.000495 | 0 | CEACAM2_SELE | Ceacam2 - Sele | CEACAM | Cell-Cell Contact | PMID:1378450 |
| GFAP | GFAP | Ceacam2 | Sele | 0.00041 | 0 | CEACAM2_SELE | Ceacam2 - Sele | CEACAM | Cell-Cell Contact | PMID:1378450 |
| CD31 | CD31 | Ceacam2 | Cd1d1 | 0.000365 | 0 | CEACAM2_CD1D1 | Ceacam2 - Cd1d1 | CEACAM | Cell-Cell Contact | PMID:24104458 |
| CD31 | GFAP | Ceacam2 | Cd1d1 | 0.000374 | 0 | CEACAM2_CD1D1 | Ceacam2 - Cd1d1 | CEACAM | Cell-Cell Contact | PMID:24104458 |
| NeuN | CD31 | Cntn1 | Notch1 | 0.008829 | 0 | CNTN1_NOTCH1 | Cntn1 - Notch1 | CNTN | Cell-Cell Contact | PMID:14567914 |
| GFAP | GFAP | Cntn1 | Notch1 | 0.004697 | 0 | CNTN1_NOTCH1 | Cntn1 - Notch1 | CNTN | Cell-Cell Contact | PMID:14567914 |
| GFAP | CD31 | Clec2h | Klrb1f | 0.000254 | 0 | CLEC2H_KLRB1F | Klrb1f - Clec2h | CLEC | Cell-Cell Contact | PMID:24223577 |
| GFAP | GFAP | Clec2h | Klrb1f | 0.000306 | 0 | CLEC2H_KLRB1F | Klrb1f - Clec2h | CLEC | Cell-Cell Contact | PMID:24223577 |

| | | | | | | | | | | |
|---|---|---|---|---|---|---|---|---|---|---|
| CD31 | CD31 | Clec2d | Klrb1f | 0.00084 | 0 | CLEC2D_KLRB1F | Klrb1f - Clec2d | CLEC | Cell-Cell Contact | PMID:24223577 |
| CD31 | GFAP | Clec2d | Klrb1f | 0.001011 | 0 | CLEC2D_KLRB1F | Klrb1f - Clec2d | CLEC | Cell-Cell Contact | PMID:24223577 |
| GFAP | CD31 | Mmp2 | MMP14_ITGAV_ITGB3 | 0.000545 | 0 | MMP2_INTEGRIN_AVB3_COMP | Mmp2 - Integrin_avb3 | MMP | Cell-Cell Contact | PMID: 29986882 |
| GFAP | GFAP | Mmp2 | MMP14_ITGAV_ITGB3 | 0.00078 | 0 | MMP2_INTEGRIN_AVB3_COMP | Mmp2 - Integrin_avb3 | MMP | Cell-Cell Contact | PMID: 29986882 |
| GFAP | GFAP | Ntn1 | Dcc | 0.000822 | 0 | NTN1_DCC | Ntn1 - Dcc | Netrin | Cell-Cell Contact | PMID: 22685302 |
| GFAP | NeuN | Ntn1 | Dcc | 0.000909 | 0 | NTN1_DCC | Ntn1 - Dcc | Netrin | Cell-Cell Contact | PMID: 22685302 |
| GFAP | GFAP | Ntn1 | Dscam | 0.000999 | 0 | NTN1_DSCAM | Ntn1 - Dscam | Netrin | Cell-Cell Contact | PMID:18585357;PMID: 22685302 |
| GFAP | NeuN | Ntn1 | Dscam | 0.001374 | 0 | NTN1_DSCAM | Ntn1 - Dscam | Netrin | Cell-Cell Contact | PMID:18585357;PMID: 22685302 |
| CD31 | CD31 | Ntn1 | Unc5a | 0.000479 | 0 | NTN1_UNC5A | Ntn1 - Unc5a | Netrin | Cell-Cell Contact | PMID: 22685302 |
| GFAP | NeuN | Ntn1 | Unc5a | 0.000949 | 0 | NTN1_UNC5A | Ntn1 - Unc5a | Netrin | Cell-Cell Contact | PMID: 22685302 |
| GFAP | CD31 | Ntn1 | Unc5b | 0.000558 | 0 | NTN1_UNC5B | Ntn1 - Unc5b | Netrin | Cell-Cell Contact | PMID: 22685302 |
| GFAP | GFAP | Ntn1 | Unc5b | 0.000512 | 0 | NTN1_UNC5B | Ntn1 - Unc5b | Netrin | Cell-Cell Contact | PMID: 22685302 |
| CD31 | GFAP | Ntn1 | Unc5c | 0.000328 | 0 | NTN1_UNC5C | Ntn1 - Unc5c | Netrin | Cell-Cell Contact | PMID: 22685302 |
| GFAP | GFAP | Ntn1 | Unc5c | 0.000448 | 0 | NTN1_UNC5C | Ntn1 - Unc5c | Netrin | Cell-Cell Contact | PMID: 22685302 |
| CD31 | GFAP | Ntn4 | Ntrk2 | 0.008803 | 0 | NTN4_NTRK2 | Ntn4 - Ntrk2 | Netrin | Cell-Cell Contact | PMID: 34531526 |
| GFAP | GFAP | Ntn4 | Ntrk2 | 0.005347 | 0 | NTN4_NTRK2 | Ntn4 - Ntrk2 | Netrin | Cell-Cell Contact | PMID: 34531526 |
| CD31 | CD31 | Ntn4 | Unc5b | 0.000528 | 0 | NTN4_UNC5B | Ntn4 - Unc5b | Netrin | Cell-Cell Contact | PMID: 31914695 |
| CD31 | GFAP | Ntn4 | Unc5b | 0.000484 | 0 | NTN4_UNC5B | Ntn4 - Unc5b | Netrin | Cell-Cell Contact | PMID: 31914695 |
| CD31 | CD31 | Ntn4 | Unc5d | 0.000254 | 0 | NTN4_UNC5D | Ntn4 - Unc5d | Netrin | Cell-Cell Contact | uniprot |
| CD31 | NeuN | Ntn4 | Unc5d | 0.000383 | 0 | NTN4_UNC5D | Ntn4 - Unc5d | Netrin | Cell-Cell Contact | uniprot |
| CD31 | CD31 | Pecam1 | Cd177 | 0.000602 | 0 | PECAM1_CD177 | Pecam1 - Cd177 | PECAM1 | Cell-Cell Contact | PMID:17580308 |
| CD31 | GFAP | Pecam1 | Cd177 | 0.00111 | 0 | PECAM1_CD177 | Pecam1 - Cd177 | PECAM1 | Cell-Cell Contact | PMID:17580308 |
| CD31 | CD31 | Pecam1 | Cd38 | 0.000866 | 0 | PECAM1_CD38 | Pecam1 - Cd38 | PECAM2 | Cell-Cell Contact | PMID:26407101 |
| CD31 | GFAP | Pecam1 | Cd38 | 0.000678 | 0 | PECAM1_CD38 | Pecam1 - Cd38 | PECAM2 | Cell-Cell Contact | PMID:26407101 |
| CD31 | CD31 | Sirpb1a | Cd47 | 0.000507 | 0 | SIRPB1A_CD47 | Sirpb1a - Cd47 | SIRP | Cell-Cell Contact | PMID:15294972 |
| CD31 | NeuN | Sirpb1a | Cd47 | 0.00131 | 0 | SIRPB1A_CD47 | Sirpb1a - Cd47 | SIRP | Cell-Cell Contact | PMID:15294972 |

**Supplementary Table 14: Cell Chat Interactions at P28 in PAE**

| source | target | ligand | receptor | prob | pval | interaction_name | interaction_name_2 | pathway_name | annotation | evidence |
|---|---|---|---|---|---|---|---|---|---|---|
| CD31 | CD31 | Tgfb1 | ACVR1B_TGFbR2 | 0.000379 | 0 | TGFB1_ACVR1B_TGFBR2 | Tgfb1 - (Acvr1b+Tgfbr2) | TGFb | Secreted Signaling | PMID: 27449815 |
| CD31 | NeuN | Tgfb1 | ACVR1B_TGFbR2 | 0.00038 | 0 | TGFB1_ACVR1B_TGFBR2 | Tgfb1 - (Acvr1b+Tgfbr2) | TGFb | Secreted Signaling | PMID: 27449815 |
| CD31 | CD31 | Tgfb1 | ACVR1C_TGFbR2 | 0.000259 | 0 | TGFB1_ACVR1C_TGFBR2 | Tgfb1 - (Acvr1c+Tgfbr2) | TGFb | Secreted Signaling | PMID: 27449815 |
| GFAP | CD31 | Tgfb1 | ACVR1C_TGFbR2 | 0.000254 | 0 | TGFB1_ACVR1C_TGFBR2 | Tgfb1 - (Acvr1c+Tgfbr2) | TGFb | Secreted Signaling | PMID: 27449815 |
| CD31 | CD31 | Tgfb3 | ACVR1B_TGFbR2 | 0.000486 | 0 | TGFB3_ACVR1B_TGFBR2 | Tgfb3 - (Acvr1b+Tgfbr2) | TGFb | Secreted Signaling | PMID: 27449815 |
| CD31 | NeuN | Tgfb3 | ACVR1B_TGFbR2 | 0.000488 | 0 | TGFB3_ACVR1B_TGFBR2 | Tgfb3 - (Acvr1b+Tgfbr2) | TGFb | Secreted Signaling | PMID: 27449815 |
| CD31 | CD31 | Tgfb3 | ACVR1C_TGFbR2 | 0.000333 | 0 | TGFB3_ACVR1C_TGFBR2 | Tgfb3 - (Acvr1c+Tgfbr2) | TGFb | Secreted Signaling | PMID: 27449815 |
| CD31 | GFAP | Tgfb3 | ACVR1C_TGFbR2 | 0.000241 | 0 | TGFB3_ACVR1C_TGFBR2 | Tgfb3 - (Acvr1c+Tgfbr2) | TGFb | Secreted Signaling | PMID: 27449815 |
| CD31 | GFAP | Bmp2 | BMPR1A_ACVR2A | 0.000525 | 0 | BMP2_BMPR1A_ACVR2A | Bmp2 - (Bmpr1a+Acvr2a) | BMP | Secreted Signaling | KEGG: mmu04350; PMID:26893264 |
| GFAP | GFAP | Bmp2 | BMPR1A_ACVR2A | 0.00048 | 0 | BMP2_BMPR1A_ACVR2A | Bmp2 - (Bmpr1a+Acvr2a) | BMP | Secreted Signaling | KEGG: mmu04350; PMID:26893264 |
| CD31 | GFAP | Bmp2 | BMPR1A_ACVR2B | 0.000475 | 0 | BMP2_BMPR1A_ACVR2B | Bmp2 - (Bmpr1a+Acvr2b) | BMP | Secreted Signaling | KEGG: mmu04350; PMID:26893264 |
| GFAP | GFAP | Bmp2 | BMPR1A_ACVR2B | 0.000434 | 0 | BMP2_BMPR1A_ACVR2B | Bmp2 - (Bmpr1a+Acvr2b) | BMP | Secreted Signaling | KEGG: mmu04350; PMID:26893264 |
| CD31 | CD31 | Bmp2 | BMPR1A_BMPR2 | 0.000799 | 0 | BMP2_BMPR1A_BMPR2 | Bmp2 - (Bmpr1a+Bmpr2) | BMP | Secreted Signaling | KEGG: mmu04350; PMID:26893264 |
| GFAP | CD31 | Bmp2 | BMPR1A_BMPR2 | 0.00073 | 0 | BMP2_BMPR1A_BMPR2 | Bmp2 - (Bmpr1a+Bmpr2) | BMP | Secreted Signaling | KEGG: mmu04350; PMID:26893264 |
| CD31 | GFAP | Bmp2 | BMPR1B_ACVR2A | 0.000516 | 0 | BMP2_BMPR1B_ACVR2A | Bmp2 - (Bmpr1b+Acvr2a) | BMP | Secreted Signaling | KEGG: mmu04350; PMID:26893264 |
| GFAP | GFAP | Bmp2 | BMPR1B_ACVR2A | 0.000472 | 0 | BMP2_BMPR1B_ACVR2A | Bmp2 - (Bmpr1b+Acvr2a) | BMP | Secreted Signaling | KEGG: mmu04350; PMID:26893264 |
| CD31 | GFAP | Bmp2 | BMPR1B_ACVR2B | 0.000467 | 0 | BMP2_BMPR1B_ACVR2B | Bmp2 - (Bmpr1b+Acvr2b) | BMP | Secreted Signaling | KEGG: mmu04350; PMID:26893264 |
| GFAP | GFAP | Bmp2 | BMPR1B_ACVR2B | 0.000427 | 0 | BMP2_BMPR1B_ACVR2B | Bmp2 - (Bmpr1b+Acvr2b) | BMP | Secreted Signaling | KEGG: mmu04350; PMID:26893264 |
| CD31 | CD31 | Bmp2 | BMPR1B_BMPR2 | 0.000517 | 0 | BMP2_BMPR1B_BMPR2 | Bmp2 - (Bmpr1b+Bmpr2) | BMP | Secreted Signaling | KEGG: mmu04350; PMID:26893264 |
| CD31 | GFAP | Bmp2 | BMPR1B_BMPR2 | 0.000548 | 0 | BMP2_BMPR1B_BMPR2 | Bmp2 - (Bmpr1b+Bmpr2) | BMP | Secreted Signaling | KEGG: mmu04350; PMID:26893264 |
| GFAP | CD31 | Gdf6 | BMPR1A_ACVR2A | 0.000525 | 0 | GDF6_BMPR1A_ACVR2A | Gdf6 - (Bmpr1a+Acvr2a) | BMP | Secreted Signaling | KEGG: mmu04350; PMID:26893264 |
| GFAP | GFAP | Gdf6 | BMPR1A_ACVR2A | 0.000638 | 0 | GDF6_BMPR1A_ACVR2A | Gdf6 - (Bmpr1a+Acvr2a) | BMP | Secreted Signaling | KEGG: mmu04350; PMID:26893264 |
| GFAP | GFAP | Gdf6 | BMPR1A_ACVR2B | 0.000577 | 0 | GDF6_BMPR1A_ACVR2B | Gdf6 - (Bmpr1a+Acvr2b) | BMP | Secreted Signaling | KEGG: mmu04350; PMID:26893264 |
| GFAP | NeuN | Gdf6 | BMPR1A_ACVR2B | 0.000312 | 0 | GDF6_BMPR1A_ACVR2B | Gdf6 - (Bmpr1a+Acvr2b) | BMP | Secreted Signaling | KEGG: mmu04350; PMID:26893264 |
| GFAP | CD31 | Gdf6 | BMPR1A_BMPR2 | 0.000971 | 0 | GDF6_BMPR1A_BMPR2 | Gdf6 - (Bmpr1a+Bmpr2) | BMP | Secreted Signaling | KEGG: mmu04350; PMID:26893264 |
| GFAP | GFAP | Gdf6 | BMPR1A_BMPR2 | 0.000677 | 0 | GDF6_BMPR1A_BMPR2 | Gdf6 - (Bmpr1a+Bmpr2) | BMP | Secreted Signaling | KEGG: mmu04350; PMID:26893264 |
| GFAP | CD31 | Gdf6 | BMPR1B_ACVR2A | 0.00034 | 0 | GDF6_BMPR1B_ACVR2A | Gdf6 - (Bmpr1b+Acvr2a) | BMP | Secreted Signaling | KEGG: mmu04350; PMID:26893264 |
| GFAP | GFAP | Gdf6 | BMPR1B_ACVR2A | 0.000627 | 0 | GDF6_BMPR1B_ACVR2A | Gdf6 - (Bmpr1b+Acvr2a) | BMP | Secreted Signaling | KEGG: mmu04350; PMID:26893264 |
| CD31 | GFAP | Gdf6 | BMPR1B_ACVR2B | 0.000294 | 0 | GDF6_BMPR1B_ACVR2B | Gdf6 - (Bmpr1b+Acvr2b) | BMP | Secreted Signaling | KEGG: mmu04350; PMID:26893264 |
| GFAP | GFAP | Gdf6 | BMPR1B_ACVR2B | 0.000567 | 0 | GDF6_BMPR1B_ACVR2B | Gdf6 - (Bmpr1b+Acvr2b) | BMP | Secreted Signaling | KEGG: mmu04350; PMID:26893264 |
| GFAP | CD31 | Gdf6 | BMPR1B_BMPR2 | 0.000628 | 0 | GDF6_BMPR1B_BMPR2 | Gdf6 - (Bmpr1b+Bmpr2) | BMP | Secreted Signaling | KEGG: mmu04350; PMID:26893264 |
| GFAP | GFAP | Gdf6 | BMPR1B_BMPR2 | 0.000665 | 0 | GDF6_BMPR1B_BMPR2 | Gdf6 - (Bmpr1b+Bmpr2) | BMP | Secreted Signaling | KEGG: mmu04350; PMID:26893264 |
| GFAP | GFAP | Bmp5 | ACVR1_ACVR2A | 0.000292 | 0 | BMP5_ACVR1_ACVR2A | Bmp5 - (Acvr1+Acvr2a) | BMP | Secreted Signaling | KEGG: mmu04350; PMID:26893264 |
| GFAP | NeuN | Bmp5 | ACVR1_ACVR2A | 0.000353 | 0 | BMP5_ACVR1_ACVR2A | Bmp5 - (Acvr1+Acvr2a) | BMP | Secreted Signaling | KEGG: mmu04350; PMID:26893264 |
| GFAP | GFAP | Bmp5 | ACVR1_ACVR2B | 0.000265 | 0 | BMP5_ACVR1_ACVR2B | Bmp5 - (Acvr1+Acvr2b) | BMP | Secreted Signaling | KEGG: mmu04350; PMID:26893264 |
| GFAP | NeuN | Bmp5 | ACVR1_ACVR2B | 0.000217 | 0 | BMP5_ACVR1_ACVR2B | Bmp5 - (Acvr1+Acvr2b) | BMP | Secreted Signaling | KEGG: mmu04350; PMID:26893264 |
| CD31 | CD31 | Bmp5 | ACVR1_BMPR2 | 0.00033 | 0 | BMP5_ACVR1_BMPR2 | Bmp5 - (Acvr1+Bmpr2) | BMP | Secreted Signaling | KEGG: mmu04350; PMID:26893264 |
| GFAP | CD31 | Bmp5 | ACVR1_BMPR2 | 0.000528 | 0 | BMP5_ACVR1_BMPR2 | Bmp5 - (Acvr1+Bmpr2) | BMP | Secreted Signaling | KEGG: mmu04350; PMID:26893264 |
| GFAP | CD31 | Bmp5 | BMPR1A_ACVR2A | 0.000355 | 0 | BMP5_BMPR1A_ACVR2A | Bmp5 - (Bmpr1a+Acvr2a) | BMP | Secreted Signaling | KEGG: mmu04350; PMID:26893264 |
| GFAP | GFAP | Bmp5 | BMPR1A_ACVR2A | 0.000431 | 0 | BMP5_BMPR1A_ACVR2A | Bmp5 - (Bmpr1a+Acvr2a) | BMP | Secreted Signaling | KEGG: mmu04350; PMID:26893264 |
| CD31 | GFAP | Bmp5 | BMPR1A_ACVR2B | 0.000244 | 0 | BMP5_BMPR1A_ACVR2B | Bmp5 - (Bmpr1a+Acvr2b) | BMP | Secreted Signaling | KEGG: mmu04350; PMID:26893264 |
| GFAP | GFAP | Bmp5 | BMPR1A_ACVR2B | 0.00039 | 0 | BMP5_BMPR1A_ACVR2B | Bmp5 - (Bmpr1a+Acvr2b) | BMP | Secreted Signaling | KEGG: mmu04350; PMID:26893264 |
| GFAP | CD31 | Bmp5 | BMPR1A_BMPR2 | 0.000656 | 0 | BMP5_BMPR1A_BMPR2 | Bmp5 - (Bmpr1a+Bmpr2) | BMP | Secreted Signaling | KEGG: mmu04350; PMID:26893264 |
| GFAP | GFAP | Bmp5 | BMPR1A_BMPR2 | 0.000458 | 0 | BMP5_BMPR1A_BMPR2 | Bmp5 - (Bmpr1a+Bmpr2) | BMP | Secreted Signaling | KEGG: mmu04350; PMID:26893264 |
| CD31 | GFAP | Bmp5 | BMPR1B_ACVR2A | 0.000264 | 0 | BMP5_BMPR1B_ACVR2A | Bmp5 - (Bmpr1b+Acvr2a) | BMP | Secreted Signaling | KEGG: mmu04350; PMID:26893264 |
| GFAP | GFAP | Bmp5 | BMPR1B_ACVR2A | 0.000424 | 0 | BMP5_BMPR1B_ACVR2A | Bmp5 - (Bmpr1b+Acvr2a) | BMP | Secreted Signaling | KEGG: mmu04350; PMID:26893264 |
| CD31 | GFAP | Bmp5 | BMPR1B_ACVR2B | 0.000239 | 0 | BMP5_BMPR1B_ACVR2B | Bmp5 - (Bmpr1b+Acvr2b) | BMP | Secreted Signaling | KEGG: mmu04350; PMID:26893264 |
| GFAP | GFAP | Bmp5 | BMPR1B_ACVR2B | 0.000384 | 0 | BMP5_BMPR1B_ACVR2B | Bmp5 - (Bmpr1b+Acvr2b) | BMP | Secreted Signaling | KEGG: mmu04350; PMID:26893264 |
| GFAP | CD31 | Bmp5 | BMPR1B_BMPR2 | 0.000425 | 0 | BMP5_BMPR1B_BMPR2 | Bmp5 - (Bmpr1b+Bmpr2) | BMP | Secreted Signaling | KEGG: mmu04350; PMID:26893264 |
| GFAP | GFAP | Bmp5 | BMPR1B_BMPR2 | 0.00045 | 0 | BMP5_BMPR1B_BMPR2 | Bmp5 - (Bmpr1b+Bmpr2) | BMP | Secreted Signaling | KEGG: mmu04350; PMID:26893264 |
| GFAP | GFAP | Bmp6 | ACVR1_ACVR2A | 0.000506 | 0 | BMP6_ACVR1_ACVR2A | Bmp6 - (Acvr1+Acvr2a) | BMP | Secreted Signaling | KEGG: mmu04350; PMID:26893264 |
| GFAP | NeuN | Bmp6 | ACVR1_ACVR2A | 0.000611 | 0 | BMP6_ACVR1_ACVR2A | Bmp6 - (Acvr1+Acvr2a) | BMP | Secreted Signaling | KEGG: mmu04350; PMID:26893264 |
| CD31 | GFAP | Bmp6 | ACVR1_ACVR2B | 0.000409 | 0 | BMP6_ACVR1_ACVR2B | Bmp6 - (Acvr1+Acvr2b) | BMP | Secreted Signaling | KEGG: mmu04350; PMID:26893264 |
| GFAP | GFAP | Bmp6 | ACVR1_ACVR2B | 0.000457 | 0 | BMP6_ACVR1_ACVR2B | Bmp6 - (Acvr1+Acvr2b) | BMP | Secreted Signaling | KEGG: mmu04350; PMID:26893264 |
| CD31 | CD31 | Bmp6 | ACVR1_BMPR2 | 0.000816 | 0 | BMP6_ACVR1_BMPR2 | Bmp6 - (Acvr1+Bmpr2) | BMP | Secreted Signaling | KEGG: mmu04350; PMID:26893264 |
| GFAP | CD31 | Bmp6 | ACVR1_BMPR2 | 0.000913 | 0 | BMP6_ACVR1_BMPR2 | Bmp6 - (Acvr1+Bmpr2) | BMP | Secreted Signaling | KEGG: mmu04350; PMID:26893264 |

| | | | | | | | | | | |
|---|---|---|---|---|---|---|---|---|---|---|
| CD31 | GFAP | Bmp6 | BMPR1A_ACVR2A | 0.000666 | 0 | BMP6_BMPR1A_ACVR2A | Bmp6 - (Bmpr1a+Acvr2a) | BMP | Secreted Signaling | KEGG: mmu04350; PMID:26893264 |
| GFAP | GFAP | Bmp6 | BMPR1A_ACVR2A | 0.000746 | 0 | BMP6_BMPR1A_ACVR2A | Bmp6 - (Bmpr1a+Acvr2a) | BMP | Secreted Signaling | KEGG: mmu04350; PMID:26893264 |
| CD31 | GFAP | Bmp6 | BMPR1A_ACVR2B | 0.000603 | 0 | BMP6_BMPR1A_ACVR2B | Bmp6 - (Bmpr1a+Acvr2b) | BMP | Secreted Signaling | KEGG: mmu04350; PMID:26893264 |
| GFAP | GFAP | Bmp6 | BMPR1A_ACVR2B | 0.000675 | 0 | BMP6_BMPR1A_ACVR2B | Bmp6 - (Bmpr1a+Acvr2b) | BMP | Secreted Signaling | KEGG: mmu04350; PMID:26893264 |
| CD31 | CD31 | Bmp6 | BMPR1A_BMPR2 | 0.001013 | 0 | BMP6_BMPR1A_BMPR2 | Bmp6 - (Bmpr1a+Bmpr2) | BMP | Secreted Signaling | KEGG: mmu04350; PMID:26893264 |
| GFAP | CD31 | Bmp6 | BMPR1A_BMPR2 | 0.001134 | 0 | BMP6_BMPR1A_BMPR2 | Bmp6 - (Bmpr1a+Bmpr2) | BMP | Secreted Signaling | KEGG: mmu04350; PMID:26893264 |
| CD31 | GFAP | Bmp6 | BMPR1B_ACVR2A | 0.000655 | 0 | BMP6_BMPR1B_ACVR2A | Bmp6 - (Bmpr1b+Acvr2a) | BMP | Secreted Signaling | KEGG: mmu04350; PMID:26893264 |
| GFAP | GFAP | Bmp6 | BMPR1B_ACVR2A | 0.000733 | 0 | BMP6_BMPR1B_ACVR2A | Bmp6 - (Bmpr1b+Acvr2a) | BMP | Secreted Signaling | KEGG: mmu04350; PMID:26893264 |
| CD31 | GFAP | Bmp6 | BMPR1B_ACVR2B | 0.000592 | 0 | BMP6_BMPR1B_ACVR2B | Bmp6 - (Bmpr1b+Acvr2b) | BMP | Secreted Signaling | KEGG: mmu04350; PMID:26893264 |
| GFAP | GFAP | Bmp6 | BMPR1B_ACVR2B | 0.000663 | 0 | BMP6_BMPR1B_ACVR2B | Bmp6 - (Bmpr1b+Acvr2b) | BMP | Secreted Signaling | KEGG: mmu04350; PMID:26893264 |
| GFAP | CD31 | Bmp6 | BMPR1B_BMPR2 | 0.000734 | 0 | BMP6_BMPR1B_BMPR2 | Bmp6 - (Bmpr1b+Bmpr2) | BMP | Secreted Signaling | KEGG: mmu04350; PMID:26893264 |
| GFAP | GFAP | Bmp6 | BMPR1B_BMPR2 | 0.000778 | 0 | BMP6_BMPR1B_BMPR2 | Bmp6 - (Bmpr1b+Bmpr2) | BMP | Secreted Signaling | KEGG: mmu04350; PMID:26893264 |
| GFAP | GFAP | Bmp8b | ACVR1_ACVR2A | 0.000238 | 0 | BMP8B_ACVR1_ACVR2A | Bmp8b - (Acvr1+Acvr2a) | BMP | Secreted Signaling | KEGG: mmu04350; PMID:26893264 |
| CD31 | NeuN | Bmp8b | ACVR1_ACVR2A | 0.000293 | 0 | BMP8B_ACVR1_ACVR2A | Bmp8b - (Acvr1+Acvr2a) | BMP | Secreted Signaling | KEGG: mmu04350; PMID:26893264 |
| CD31 | GFAP | Bmp8b | ACVR1_ACVR2B | 0.000219 | 0 | BMP8B_ACVR1_ACVR2B | Bmp8b - (Acvr1+Acvr2b) | BMP | Secreted Signaling | KEGG: mmu04350; PMID:26893264 |
| GFAP | GFAP | Bmp8b | ACVR1_ACVR2B | 0.000215 | 0 | BMP8B_ACVR1_ACVR2B | Bmp8b - (Acvr1+Acvr2b) | BMP | Secreted Signaling | KEGG: mmu04350; PMID:26893264 |
| CD31 | CD31 | Bmp8b | ACVR1_BMPR2 | 0.000438 | 0 | BMP8B_ACVR1_BMPR2 | Bmp8b - (Acvr1+Bmpr2) | BMP | Secreted Signaling | KEGG: mmu04350; PMID:26893264 |
| GFAP | CD31 | Bmp8b | ACVR1_BMPR2 | 0.00043 | 0 | BMP8B_ACVR1_BMPR2 | Bmp8b - (Acvr1+Bmpr2) | BMP | Secreted Signaling | KEGG: mmu04350; PMID:26893264 |
| CD31 | GFAP | Bmp8b | BMPR1A_ACVR2A | 0.000357 | 0 | BMP8B_BMPR1A_ACVR2A | Bmp8b - (Bmpr1a+Acvr2a) | BMP | Secreted Signaling | KEGG: mmu04350; PMID:26893264 |
| GFAP | GFAP | Bmp8b | BMPR1A_ACVR2A | 0.000351 | 0 | BMP8B_BMPR1A_ACVR2A | Bmp8b - (Bmpr1a+Acvr2a) | BMP | Secreted Signaling | KEGG: mmu04350; PMID:26893264 |
| CD31 | GFAP | Bmp8b | BMPR1A_ACVR2B | 0.000323 | 0 | BMP8B_BMPR1A_ACVR2B | Bmp8b - (Bmpr1a+Acvr2b) | BMP | Secreted Signaling | KEGG: mmu04350; PMID:26893264 |
| GFAP | GFAP | Bmp8b | BMPR1A_ACVR2B | 0.000318 | 0 | BMP8B_BMPR1A_ACVR2B | Bmp8b - (Bmpr1a+Acvr2b) | BMP | Secreted Signaling | KEGG: mmu04350; PMID:26893264 |
| CD31 | CD31 | Bmp8b | BMPR1A_BMPR2 | 0.000544 | 0 | BMP8B_BMPR1A_BMPR2 | Bmp8b - (Bmpr1a+Bmpr2) | BMP | Secreted Signaling | KEGG: mmu04350; PMID:26893264 |
| GFAP | CD31 | Bmp8b | BMPR1A_BMPR2 | 0.000534 | 0 | BMP8B_BMPR1A_BMPR2 | Bmp8b - (Bmpr1a+Bmpr2) | BMP | Secreted Signaling | KEGG: mmu04350; PMID:26893264 |
| CD31 | GFAP | Bmp8b | BMPR1B_ACVR2A | 0.000351 | 0 | BMP8B_BMPR1B_ACVR2A | Bmp8b - (Bmpr1b+Acvr2a) | BMP | Secreted Signaling | KEGG: mmu04350; PMID:26893264 |
| GFAP | GFAP | Bmp8b | BMPR1B_ACVR2A | 0.000345 | 0 | BMP8B_BMPR1B_ACVR2A | Bmp8b - (Bmpr1b+Acvr2a) | BMP | Secreted Signaling | KEGG: mmu04350; PMID:26893264 |
| CD31 | GFAP | Bmp8b | BMPR1B_ACVR2B | 0.000318 | 0 | BMP8B_BMPR1B_ACVR2B | Bmp8b - (Bmpr1b+Acvr2b) | BMP | Secreted Signaling | KEGG: mmu04350; PMID:26893264 |
| GFAP | GFAP | Bmp8b | BMPR1B_ACVR2B | 0.000312 | 0 | BMP8B_BMPR1B_ACVR2B | Bmp8b - (Bmpr1b+Acvr2b) | BMP | Secreted Signaling | KEGG: mmu04350; PMID:26893264 |
| CD31 | GFAP | Bmp8b | BMPR1B_BMPR2 | 0.000373 | 0 | BMP8B_BMPR1B_BMPR2 | Bmp8b - (Bmpr1b+Bmpr2) | BMP | Secreted Signaling | KEGG: mmu04350; PMID:26893264 |
| GFAP | GFAP | Bmp8b | BMPR1B_BMPR2 | 0.000366 | 0 | BMP8B_BMPR1B_BMPR2 | Bmp8b - (Bmpr1b+Bmpr2) | BMP | Secreted Signaling | KEGG: mmu04350; PMID:26893264 |
| CD31 | CD31 | Gdf9 | ACVR1B_BMPR2 | 0.000903 | 0 | GDF9_ACVR1B_BMPR2 | Gdf9 - (Acvr1b+Bmpr2) | GDF | Secreted Signaling | KEGG: mmu04350 |
| GFAP | NeuN | Gdf9 | ACVR1B_BMPR2 | 0.001492 | 0 | GDF9_ACVR1B_BMPR2 | Gdf9 - (Acvr1b+Bmpr2) | GDF | Secreted Signaling | KEGG: mmu04350 |
| GFAP | GFAP | Gdf10 | ACVR1B_ACVR2A | 0.000972 | 0 | GDF10_ACVR1B_ACVR2A | Gdf10 - (Acvr1b+Acvr2a) | GDF | Secreted Signaling | KEGG: mmu04350 |
| GFAP | NeuN | Gdf10 | ACVR1B_ACVR2A | 0.001863 | 0 | GDF10_ACVR1B_ACVR2A | Gdf10 - (Acvr1b+Acvr2a) | GDF | Secreted Signaling | KEGG: mmu04350 |
| GFAP | CD31 | Gdf11 | TGFBR1_ACVR2A | 0.000835 | 0 | GDF11_TGFBR1_ACVR2A | Gdf11 - (Tgfbr1+Acvr2a) | GDF | Secreted Signaling | KEGG: mmu04350 |
| GFAP | GFAP | Gdf11 | TGFBR1_ACVR2A | 0.000914 | 0 | GDF11_TGFBR1_ACVR2A | Gdf11 - (Tgfbr1+Acvr2a) | GDF | Secreted Signaling | KEGG: mmu04350 |
| CD31 | GFAP | Gdf11 | TGFBR1_ACVR2B | 0.000571 | 0 | GDF11_TGFBR1_ACVR2B | Gdf11 - (Tgfbr1+Acvr2b) | GDF | Secreted Signaling | KEGG: mmu04350 |
| GFAP | GFAP | Gdf11 | TGFBR1_ACVR2B | 0.000827 | 0 | GDF11_TGFBR1_ACVR2B | Gdf11 - (Tgfbr1+Acvr2b) | GDF | Secreted Signaling | KEGG: mmu04350 |
| GFAP | GFAP | Gdf11 | ACVR1B_ACVR2B | 0.000804 | 0 | GDF11_ACVR1B_ACVR2B | Gdf11 - (Acvr1b+Acvr2b) | GDF | Secreted Signaling | KEGG: mmu04350 |
| GFAP | NeuN | Gdf11 | ACVR1B_ACVR2B | 0.001049 | 0 | GDF11_ACVR1B_ACVR2B | Gdf11 - (Acvr1b+Acvr2b) | GDF | Secreted Signaling | KEGG: mmu04350 |
| GFAP | CD31 | Gdnf | GFRA1_RET | 0.000626 | 0 | GDNF_GFRA1 | Gdnf - Gfra1 | GDNF | Secreted Signaling | PMID: 11544105 |
| GFAP | GFAP | Gdnf | GFRA1_RET | 0.0004 | 0 | GDNF_GFRA1 | Gdnf - Gfra1 | GDNF | Secreted Signaling | PMID: 11544105 |
| CD31 | CD31 | Nrtn | GFRA2_RET | 0.000643 | 0 | NRTN_GFRA2 | Nrtn - Gfra2 | GDNF | Secreted Signaling | PMID: 11544105 |
| CD31 | NeuN | Nrtn | GFRA2_RET | 0.000576 | 0 | NRTN_GFRA2 | Nrtn - Gfra2 | GDNF | Secreted Signaling | PMID: 11544105 |
| GFAP | NeuN | Inhba | ACVR1B_ACVR2A | 0.000808 | 0 | INHBA_ACVR1B_ACVR2A | Inhba - (Acvr1b+Acvr2a) | ACTIVIN | Secreted Signaling | KEGG: mmu04350 |
| NeuN | NeuN | Inhba | ACVR1B_ACVR2A | 0.000518 | 0 | INHBA_ACVR1B_ACVR2A | Inhba - (Acvr1b+Acvr2a) | ACTIVIN | Secreted Signaling | KEGG: mmu04350 |
| GFAP | GFAP | Inhba | ACVR1B_ACVR2B | 0.000371 | 0 | INHBA_ACVR1B_ACVR2B | Inhba - (Acvr1b+Acvr2b) | ACTIVIN | Secreted Signaling | KEGG: mmu04350 |
| GFAP | NeuN | Inhba | ACVR1B_ACVR2B | 0.000497 | 0 | INHBA_ACVR1B_ACVR2B | Inhba - (Acvr1b+Acvr2b) | ACTIVIN | Secreted Signaling | KEGG: mmu04350 |
| GFAP | GFAP | Inhbb | ACVR1B_ACVR2A | 0.000786 | 0 | INHBB_ACVR1B_ACVR2A | Inhbb - (Acvr1b+Acvr2a) | ACTIVIN | Secreted Signaling | KEGG: mmu04350 |
| GFAP | NeuN | Inhbb | ACVR1B_ACVR2A | 0.001546 | 0 | INHBB_ACVR1B_ACVR2A | Inhbb - (Acvr1b+Acvr2a) | ACTIVIN | Secreted Signaling | KEGG: mmu04350 |
| GFAP | GFAP | Inhbb | ACVR1B_ACVR2B | 0.000711 | 0 | INHBB_ACVR1B_ACVR2B | Inhbb - (Acvr1b+Acvr2b) | ACTIVIN | Secreted Signaling | KEGG: mmu04350 |
| GFAP | NeuN | Inhbb | ACVR1B_ACVR2B | 0.000951 | 0 | INHBB_ACVR1B_ACVR2B | Inhbb - (Acvr1b+Acvr2b) | ACTIVIN | Secreted Signaling | KEGG: mmu04350 |
| GFAP | CD31 | Inhbb | ACVR1C_ACVR2A | 0.000517 | 0 | INHBB_ACVR1C_ACVR2A | Inhbb - (Acvr1c+Acvr2a) | ACTIVIN | Secreted Signaling | KEGG: mmu04350 |
| GFAP | GFAP | Inhbb | ACVR1C_ACVR2A | 0.000644 | 0 | INHBB_ACVR1C_ACVR2A | Inhbb - (Acvr1c+Acvr2a) | ACTIVIN | Secreted Signaling | KEGG: mmu04350 |
| CD31 | GFAP | Inhbb | ACVR1C_ACVR2B | 0.000302 | 0 | INHBB_ACVR1C_ACVR2B | Inhbb - (Acvr1c+Acvr2b) | ACTIVIN | Secreted Signaling | KEGG: mmu04350 |
| GFAP | GFAP | Inhbb | ACVR1C_ACVR2B | 0.000583 | 0 | INHBB_ACVR1C_ACVR2B | Inhbb - (Acvr1c+Acvr2b) | ACTIVIN | Secreted Signaling | KEGG: mmu04350 |
| CD31 | GFAP | Wnt10b | FZD1_LRP5 | 0.000265 | 0 | WNT10B_FZD1_LRP5 | Wnt10b - (Fzd1+Lrp5) | WNT | Secreted Signaling | KEGG: mmu04310; PMID: 23209147 |
| GFAP | GFAP | Wnt10b | FZD1_LRP5 | 0.000235 | 0 | WNT10B_FZD1_LRP5 | Wnt10b - (Fzd1+Lrp5) | WNT | Secreted Signaling | KEGG: mmu04310; PMID: 23209147 |
| CD31 | GFAP | Wnt10b | FZD2_LRP5 | 0.000443 | 0 | WNT10B_FZD2_LRP5 | Wnt10b - (Fzd2+Lrp5) | WNT | Secreted Signaling | KEGG: mmu04310; PMID: 23209147 |

| | | | | | | | | | | |
|---|---|---|---|---|---|---|---|---|---|---|
| GFAP | GFAP | Wnt10b | FZD2_LRP5 | 0.000393 | 0 | WNT10B_FZD2_LRP5 | Wnt10b - (Fzd2+Lrp5) | WNT | Secreted Signaling | KEGG: mmu04310; PMID: 23209147 |
| CD31 | GFAP | Wnt10b | FZD3_LRP5 | 0.000387 | 0 | WNT10B_FZD3_LRP5 | Wnt10b - (Fzd3+Lrp5) | WNT | Secreted Signaling | KEGG: mmu04310; PMID: 23209147 |
| GFAP | GFAP | Wnt10b | FZD3_LRP5 | 0.000343 | 0 | WNT10B_FZD3_LRP5 | Wnt10b - (Fzd3+Lrp5) | WNT | Secreted Signaling | KEGG: mmu04310; PMID: 23209147 |
| CD31 | CD31 | Wnt10b | FZD4_LRP5 | 0.0004 | 0 | WNT10B_FZD4_LRP5 | Wnt10b - (Fzd4+Lrp5) | WNT | Secreted Signaling | KEGG: mmu04310; PMID: 23209147 |
| GFAP | CD31 | Wnt10b | FZD4_LRP5 | 0.000355 | 0 | WNT10B_FZD4_LRP5 | Wnt10b - (Fzd4+Lrp5) | WNT | Secreted Signaling | KEGG: mmu04310; PMID: 23209147 |
| CD31 | GFAP | Wnt10b | FZD5_LRP5 | 0.000308 | 0 | WNT10B_FZD5_LRP5 | Wnt10b - (Fzd5+Lrp5) | WNT | Secreted Signaling | KEGG: mmu04310; PMID: 23209147 |
| GFAP | GFAP | Wnt10b | FZD5_LRP5 | 0.000274 | 0 | WNT10B_FZD5_LRP5 | Wnt10b - (Fzd5+Lrp5) | WNT | Secreted Signaling | KEGG: mmu04310; PMID: 23209147 |
| CD31 | CD31 | Wnt10b | FZD6_LRP5 | 0.00055 | 0 | WNT10B_FZD6_LRP5 | Wnt10b - (Fzd6+Lrp5) | WNT | Secreted Signaling | KEGG: mmu04310; PMID: 23209147 |
| GFAP | CD31 | Wnt10b | FZD6_LRP5 | 0.000488 | 0 | WNT10B_FZD6_LRP5 | Wnt10b - (Fzd6+Lrp5) | WNT | Secreted Signaling | KEGG: mmu04310; PMID: 23209147 |
| CD31 | GFAP | Wnt10b | FZD8_LRP5 | 0.00038 | 0 | WNT10B_FZD8_LRP5 | Wnt10b - (Fzd8+Lrp5) | WNT | Secreted Signaling | KEGG: mmu04310; PMID: 23209147 |
| GFAP | GFAP | Wnt10b | FZD8_LRP5 | 0.000338 | 0 | WNT10B_FZD8_LRP5 | Wnt10b - (Fzd8+Lrp5) | WNT | Secreted Signaling | KEGG: mmu04310; PMID: 23209147 |
| CD31 | CD31 | Wnt10b | FZD9_LRP5 | 0.000265 | 0 | WNT10B_FZD9_LRP5 | Wnt10b - (Fzd9+Lrp5) | WNT | Secreted Signaling | KEGG: mmu04310; PMID: 23209147 |
| CD31 | GFAP | Wnt10b | FZD9_LRP5 | 0.000255 | 0 | WNT10B_FZD9_LRP5 | Wnt10b - (Fzd9+Lrp5) | WNT | Secreted Signaling | KEGG: mmu04310; PMID: 23209147 |
| GFAP | GFAP | Wnt4 | FZD1_LRP5 | 0.000224 | 0 | WNT4_FZD1_LRP5 | Wnt4 - (Fzd1+Lrp5) | WNT | Secreted Signaling | KEGG: mmu04310; PMID: 23209147 |
| NeuN | GFAP | Wnt4 | FZD1_LRP5 | 0.000244 | 0 | WNT4_FZD1_LRP5 | Wnt4 - (Fzd1+Lrp5) | WNT | Secreted Signaling | KEGG: mmu04310; PMID: 23209147 |
| GFAP | GFAP | Wnt4 | FZD2_LRP5 | 0.000375 | 0 | WNT4_FZD2_LRP5 | Wnt4 - (Fzd2+Lrp5) | WNT | Secreted Signaling | KEGG: mmu04310; PMID: 23209147 |
| NeuN | GFAP | Wnt4 | FZD2_LRP5 | 0.000408 | 0 | WNT4_FZD2_LRP5 | Wnt4 - (Fzd2+Lrp5) | WNT | Secreted Signaling | KEGG: mmu04310; PMID: 23209147 |
| GFAP | GFAP | Wnt4 | FZD3_LRP5 | 0.000328 | 0 | WNT4_FZD3_LRP5 | Wnt4 - (Fzd3+Lrp5) | WNT | Secreted Signaling | KEGG: mmu04310; PMID: 23209147 |
| NeuN | GFAP | Wnt4 | FZD3_LRP5 | 0.000356 | 0 | WNT4_FZD3_LRP5 | Wnt4 - (Fzd3+Lrp5) | WNT | Secreted Signaling | KEGG: mmu04310; PMID: 23209147 |
| NeuN | CD31 | Wnt4 | FZD4_LRP5 | 0.000368 | 0 | WNT4_FZD4_LRP5 | Wnt4 - (Fzd4+Lrp5) | WNT | Secreted Signaling | KEGG: mmu04310; PMID: 23209147 |
| GFAP | GFAP | Wnt4 | FZD4_LRP5 | 0.000287 | 0 | WNT4_FZD4_LRP5 | Wnt4 - (Fzd4+Lrp5) | WNT | Secreted Signaling | KEGG: mmu04310; PMID: 23209147 |
| GFAP | GFAP | Wnt4 | FZD5_LRP5 | 0.000261 | 0 | WNT4_FZD5_LRP5 | Wnt4 - (Fzd5+Lrp5) | WNT | Secreted Signaling | KEGG: mmu04310; PMID: 23209147 |
| NeuN | GFAP | Wnt4 | FZD5_LRP5 | 0.000284 | 0 | WNT4_FZD5_LRP5 | Wnt4 - (Fzd5+Lrp5) | WNT | Secreted Signaling | KEGG: mmu04310; PMID: 23209147 |
| CD31 | CD31 | Wnt4 | FZD6_LRP5 | 0.000394 | 0 | WNT4_FZD6_LRP5 | Wnt4 - (Fzd6+Lrp5) | WNT | Secreted Signaling | KEGG: mmu04310; PMID: 23209147 |
| NeuN | CD31 | Wnt4 | FZD6_LRP5 | 0.000506 | 0 | WNT4_FZD6_LRP5 | Wnt4 - (Fzd6+Lrp5) | WNT | Secreted Signaling | KEGG: mmu04310; PMID: 23209147 |
| GFAP | GFAP | Wnt4 | FZD8_LRP5 | 0.000322 | 0 | WNT4_FZD8_LRP5 | Wnt4 - (Fzd8+Lrp5) | WNT | Secreted Signaling | KEGG: mmu04310; PMID: 23209147 |
| NeuN | GFAP | Wnt4 | FZD8_LRP5 | 0.00035 | 0 | WNT4_FZD8_LRP5 | Wnt4 - (Fzd8+Lrp5) | WNT | Secreted Signaling | KEGG: mmu04310; PMID: 23209147 |
| NeuN | CD31 | Wnt4 | FZD9_LRP5 | 0.000244 | 0 | WNT4_FZD9_LRP5 | Wnt4 - (Fzd9+Lrp5) | WNT | Secreted Signaling | KEGG: mmu04310; PMID: 23209147 |
| GFAP | GFAP | Wnt4 | FZD9_LRP5 | 0.000216 | 0 | WNT4_FZD9_LRP5 | Wnt4 - (Fzd9+Lrp5) | WNT | Secreted Signaling | KEGG: mmu04310; PMID: 23209147 |
| CD31 | GFAP | Wnt7a | FZD1_LRP5 | 0.000197 | 0 | WNT7A_FZD1_LRP5 | Wnt7a - (Fzd1+Lrp5) | WNT | Secreted Signaling | KEGG: mmu04310; PMID: 23209147 |
| GFAP | GFAP | Wnt7a | FZD1_LRP5 | 0.000236 | 0 | WNT7A_FZD1_LRP5 | Wnt7a - (Fzd1+Lrp5) | WNT | Secreted Signaling | KEGG: mmu04310; PMID: 23209147 |
| CD31 | GFAP | Wnt7a | FZD2_LRP5 | 0.000329 | 0 | WNT7A_FZD2_LRP5 | Wnt7a - (Fzd2+Lrp5) | WNT | Secreted Signaling | KEGG: mmu04310; PMID: 23209147 |
| GFAP | GFAP | Wnt7a | FZD2_LRP5 | 0.000394 | 0 | WNT7A_FZD2_LRP5 | Wnt7a - (Fzd2+Lrp5) | WNT | Secreted Signaling | KEGG: mmu04310; PMID: 23209147 |
| CD31 | GFAP | Wnt7a | FZD3_LRP5 | 0.000287 | 0 | WNT7A_FZD3_LRP5 | Wnt7a - (Fzd3+Lrp5) | WNT | Secreted Signaling | KEGG: mmu04310; PMID: 23209147 |
| GFAP | GFAP | Wnt7a | FZD3_LRP5 | 0.000344 | 0 | WNT7A_FZD3_LRP5 | Wnt7a - (Fzd3+Lrp5) | WNT | Secreted Signaling | KEGG: mmu04310; PMID: 23209147 |
| GFAP | CD31 | Wnt7a | FZD4_LRP5 | 0.000356 | 0 | WNT7A_FZD4_LRP5 | Wnt7a - (Fzd4+Lrp5) | WNT | Secreted Signaling | KEGG: mmu04310; PMID: 23209147 |
| GFAP | GFAP | Wnt7a | FZD4_LRP5 | 0.000302 | 0 | WNT7A_FZD4_LRP5 | Wnt7a - (Fzd4+Lrp5) | WNT | Secreted Signaling | KEGG: mmu04310; PMID: 23209147 |
| CD31 | GFAP | Wnt7a | FZD5_LRP5 | 0.000229 | 0 | WNT7A_FZD5_LRP5 | Wnt7a - (Fzd5+Lrp5) | WNT | Secreted Signaling | KEGG: mmu04310; PMID: 23209147 |
| GFAP | GFAP | Wnt7a | FZD5_LRP5 | 0.000274 | 0 | WNT7A_FZD5_LRP5 | Wnt7a - (Fzd5+Lrp5) | WNT | Secreted Signaling | KEGG: mmu04310; PMID: 23209147 |
| CD31 | CD31 | Wnt7a | FZD6_LRP5 | 0.000408 | 0 | WNT7A_FZD6_LRP5 | Wnt7a - (Fzd6+Lrp5) | WNT | Secreted Signaling | KEGG: mmu04310; PMID: 23209147 |
| GFAP | CD31 | Wnt7a | FZD6_LRP5 | 0.000489 | 0 | WNT7A_FZD6_LRP5 | Wnt7a - (Fzd6+Lrp5) | WNT | Secreted Signaling | KEGG: mmu04310; PMID: 23209147 |
| CD31 | GFAP | Wnt7a | FZD8_LRP5 | 0.000283 | 0 | WNT7A_FZD8_LRP5 | Wnt7a - (Fzd8+Lrp5) | WNT | Secreted Signaling | KEGG: mmu04310; PMID: 23209147 |
| GFAP | GFAP | Wnt7a | FZD8_LRP5 | 0.000338 | 0 | WNT7A_FZD8_LRP5 | Wnt7a - (Fzd8+Lrp5) | WNT | Secreted Signaling | KEGG: mmu04310; PMID: 23209147 |
| GFAP | CD31 | Wnt7a | FZD9_LRP5 | 0.000236 | 0 | WNT7A_FZD9_LRP5 | Wnt7a - (Fzd9+Lrp5) | WNT | Secreted Signaling | KEGG: mmu04310; PMID: 23209147 |
| GFAP | GFAP | Wnt7a | FZD9_LRP5 | 0.000227 | 0 | WNT7A_FZD9_LRP5 | Wnt7a - (Fzd9+Lrp5) | WNT | Secreted Signaling | KEGG: mmu04310; PMID: 23209147 |
| GFAP | CD31 | Wnt7b | FZD1_LRP5 | 0.000222 | 0 | WNT7B_FZD1_LRP5 | Wnt7b - (Fzd1+Lrp5) | WNT | Secreted Signaling | KEGG: mmu04310; PMID: 23209147 |
| GFAP | GFAP | Wnt7b | FZD1_LRP5 | 0.000272 | 0 | WNT7B_FZD1_LRP5 | Wnt7b - (Fzd1+Lrp5) | WNT | Secreted Signaling | KEGG: mmu04310; PMID: 23209147 |
| CD31 | GFAP | Wnt7b | FZD2_LRP5 | 0.000301 | 0 | WNT7B_FZD2_LRP5 | Wnt7b - (Fzd2+Lrp5) | WNT | Secreted Signaling | KEGG: mmu04310; PMID: 23209147 |
| GFAP | GFAP | Wnt7b | FZD2_LRP5 | 0.000455 | 0 | WNT7B_FZD2_LRP5 | Wnt7b - (Fzd2+Lrp5) | WNT | Secreted Signaling | KEGG: mmu04310; PMID: 23209147 |
| GFAP | CD31 | Wnt7b | FZD3_LRP5 | 0.000321 | 0 | WNT7B_FZD3_LRP5 | Wnt7b - (Fzd3+Lrp5) | WNT | Secreted Signaling | KEGG: mmu04310; PMID: 23209147 |
| GFAP | GFAP | Wnt7b | FZD3_LRP5 | 0.000397 | 0 | WNT7B_FZD3_LRP5 | Wnt7b - (Fzd3+Lrp5) | WNT | Secreted Signaling | KEGG: mmu04310; PMID: 23209147 |
| GFAP | CD31 | Wnt7b | FZD4_LRP5 | 0.000411 | 0 | WNT7B_FZD4_LRP5 | Wnt7b - (Fzd4+Lrp5) | WNT | Secreted Signaling | KEGG: mmu04310; PMID: 23209147 |
| GFAP | GFAP | Wnt7b | FZD4_LRP5 | 0.000348 | 0 | WNT7B_FZD4_LRP5 | Wnt7b - (Fzd4+Lrp5) | WNT | Secreted Signaling | KEGG: mmu04310; PMID: 23209147 |
| GFAP | CD31 | Wnt7b | FZD5_LRP5 | 0.000232 | 0 | WNT7B_FZD5_LRP5 | Wnt7b - (Fzd5+Lrp5) | WNT | Secreted Signaling | KEGG: mmu04310; PMID: 23209147 |
| GFAP | GFAP | Wnt7b | FZD5_LRP5 | 0.000316 | 0 | WNT7B_FZD5_LRP5 | Wnt7b - (Fzd5+Lrp5) | WNT | Secreted Signaling | KEGG: mmu04310; PMID: 23209147 |
| CD31 | CD31 | Wnt7b | FZD6_LRP5 | 0.000373 | 0 | WNT7B_FZD6_LRP5 | Wnt7b - (Fzd6+Lrp5) | WNT | Secreted Signaling | KEGG: mmu04310; PMID: 23209147 |
| GFAP | CD31 | Wnt7b | FZD6_LRP5 | 0.000564 | 0 | WNT7B_FZD6_LRP5 | Wnt7b - (Fzd6+Lrp5) | WNT | Secreted Signaling | KEGG: mmu04310; PMID: 23209147 |
| GFAP | CD31 | Wnt7b | FZD8_LRP5 | 0.000286 | 0 | WNT7B_FZD8_LRP5 | Wnt7b - (Fzd8+Lrp5) | WNT | Secreted Signaling | KEGG: mmu04310; PMID: 23209147 |
| GFAP | GFAP | Wnt7b | FZD8_LRP5 | 0.00039 | 0 | WNT7B_FZD8_LRP5 | Wnt7b - (Fzd8+Lrp5) | WNT | Secreted Signaling | KEGG: mmu04310; PMID: 23209147 |

| | | | | | | | | | | |
|---|---|---|---|---|---|---|---|---|---|---|
| GFAP | CD31 | Wnt7b | FZD9_LRP5 | 0.000272 | 0 | WNT7B_FZD9_LRP5 | Wnt7b - (Fzd9+Lrp5) | WNT | Secreted Signaling | KEGG: mmu04310; PMID: 23209147 |
| GFAP | GFAP | Wnt7b | FZD9_LRP5 | 0.000262 | 0 | WNT7B_FZD9_LRP5 | Wnt7b - (Fzd9+Lrp5) | WNT | Secreted Signaling | KEGG: mmu04310; PMID: 23209147 |
| GFAP | CD31 | Wnt9a | FZD1_LRP5 | 0.000207 | 0 | WNT9A_FZD1_LRP5 | Wnt9a - (Fzd1+Lrp5) | WNT | Secreted Signaling | KEGG: mmu04310; PMID: 23209147 |
| GFAP | GFAP | Wnt9a | FZD1_LRP5 | 0.000253 | 0 | WNT9A_FZD1_LRP5 | Wnt9a - (Fzd1+Lrp5) | WNT | Secreted Signaling | KEGG: mmu04310; PMID: 23209147 |
| GFAP | GFAP | Wnt9a | FZD2_LRP5 | 0.000424 | 0 | WNT9A_FZD2_LRP5 | Wnt9a - (Fzd2+Lrp5) | WNT | Secreted Signaling | KEGG: mmu04310; PMID: 23209147 |
| NeuN | GFAP | Wnt9a | FZD2_LRP5 | 0.000283 | 0 | WNT9A_FZD2_LRP5 | Wnt9a - (Fzd2+Lrp5) | WNT | Secreted Signaling | KEGG: mmu04310; PMID: 23209147 |
| GFAP | CD31 | Wnt9a | FZD3_LRP5 | 0.000299 | 0 | WNT9A_FZD3_LRP5 | Wnt9a - (Fzd3+Lrp5) | WNT | Secreted Signaling | KEGG: mmu04310; PMID: 23209147 |
| GFAP | GFAP | Wnt9a | FZD3_LRP5 | 0.00037 | 0 | WNT9A_FZD3_LRP5 | Wnt9a - (Fzd3+Lrp5) | WNT | Secreted Signaling | KEGG: mmu04310; PMID: 23209147 |
| GFAP | CD31 | Wnt9a | FZD4_LRP5 | 0.000382 | 0 | WNT9A_FZD4_LRP5 | Wnt9a - (Fzd4+Lrp5) | WNT | Secreted Signaling | KEGG: mmu04310; PMID: 23209147 |
| GFAP | GFAP | Wnt9a | FZD4_LRP5 | 0.000324 | 0 | WNT9A_FZD4_LRP5 | Wnt9a - (Fzd4+Lrp5) | WNT | Secreted Signaling | KEGG: mmu04310; PMID: 23209147 |
| GFAP | CD31 | Wnt9a | FZD5_LRP5 | 0.000216 | 0 | WNT9A_FZD5_LRP5 | Wnt9a - (Fzd5+Lrp5) | WNT | Secreted Signaling | KEGG: mmu04310; PMID: 23209147 |
| GFAP | GFAP | Wnt9a | FZD5_LRP5 | 0.000295 | 0 | WNT9A_FZD5_LRP5 | Wnt9a - (Fzd5+Lrp5) | WNT | Secreted Signaling | KEGG: mmu04310; PMID: 23209147 |
| CD31 | CD31 | Wnt9a | FZD6_LRP5 | 0.000323 | 0 | WNT9A_FZD6_LRP5 | Wnt9a - (Fzd6+Lrp5) | WNT | Secreted Signaling | KEGG: mmu04310; PMID: 23209147 |
| GFAP | CD31 | Wnt9a | FZD6_LRP5 | 0.000526 | 0 | WNT9A_FZD6_LRP5 | Wnt9a - (Fzd6+Lrp5) | WNT | Secreted Signaling | KEGG: mmu04310; PMID: 23209147 |
| GFAP | CD31 | Wnt9a | FZD8_LRP5 | 0.000267 | 0 | WNT9A_FZD8_LRP5 | Wnt9a - (Fzd8+Lrp5) | WNT | Secreted Signaling | KEGG: mmu04310; PMID: 23209147 |
| GFAP | GFAP | Wnt9a | FZD8_LRP5 | 0.000364 | 0 | WNT9A_FZD8_LRP5 | Wnt9a - (Fzd8+Lrp5) | WNT | Secreted Signaling | KEGG: mmu04310; PMID: 23209147 |
| GFAP | CD31 | Wnt9a | FZD9_LRP5 | 0.000253 | 0 | WNT9A_FZD9_LRP5 | Wnt9a - (Fzd9+Lrp5) | WNT | Secreted Signaling | KEGG: mmu04310; PMID: 23209147 |
| GFAP | GFAP | Wnt9a | FZD9_LRP5 | 0.000244 | 0 | WNT9A_FZD9_LRP5 | Wnt9a - (Fzd9+Lrp5) | WNT | Secreted Signaling | KEGG: mmu04310; PMID: 23209147 |
| CD31 | GFAP | Wnt10b | FZD1_LRP6 | 0.000239 | 0 | WNT10B_FZD1_LRP6 | Wnt10b - (Fzd1+Lrp6) | WNT | Secreted Signaling | KEGG: mmu04310; PMID: 23209147 |
| GFAP | GFAP | Wnt10b | FZD1_LRP6 | 0.000212 | 0 | WNT10B_FZD1_LRP6 | Wnt10b - (Fzd1+Lrp6) | WNT | Secreted Signaling | KEGG: mmu04310; PMID: 23209147 |
| CD31 | GFAP | Wnt10b | FZD2_LRP6 | 0.000399 | 0 | WNT10B_FZD2_LRP6 | Wnt10b - (Fzd2+Lrp6) | WNT | Secreted Signaling | KEGG: mmu04310; PMID: 23209147 |
| GFAP | GFAP | Wnt10b | FZD2_LRP6 | 0.000354 | 0 | WNT10B_FZD2_LRP6 | Wnt10b - (Fzd2+Lrp6) | WNT | Secreted Signaling | KEGG: mmu04310; PMID: 23209147 |
| GFAP | GFAP | Wnt10b | FZD3_LRP6 | 0.000309 | 0 | WNT10B_FZD3_LRP6 | Wnt10b - (Fzd3+Lrp6) | WNT | Secreted Signaling | KEGG: mmu04310; PMID: 23209147 |
| CD31 | NeuN | Wnt10b | FZD3_LRP6 | 0.000537 | 0 | WNT10B_FZD3_LRP6 | Wnt10b - (Fzd3+Lrp6) | WNT | Secreted Signaling | KEGG: mmu04310; PMID: 23209147 |
| CD31 | CD31 | Wnt10b | FZD4_LRP6 | 0.000358 | 0 | WNT10B_FZD4_LRP6 | Wnt10b - (Fzd4+Lrp6) | WNT | Secreted Signaling | KEGG: mmu04310; PMID: 23209147 |
| GFAP | CD31 | Wnt10b | FZD4_LRP6 | 0.000318 | 0 | WNT10B_FZD4_LRP6 | Wnt10b - (Fzd4+Lrp6) | WNT | Secreted Signaling | KEGG: mmu04310; PMID: 23209147 |
| CD31 | GFAP | Wnt10b | FZD5_LRP6 | 0.000278 | 0 | WNT10B_FZD5_LRP6 | Wnt10b - (Fzd5+Lrp6) | WNT | Secreted Signaling | KEGG: mmu04310; PMID: 23209147 |
| GFAP | GFAP | Wnt10b | FZD5_LRP6 | 0.000247 | 0 | WNT10B_FZD5_LRP6 | Wnt10b - (Fzd5+Lrp6) | WNT | Secreted Signaling | KEGG: mmu04310; PMID: 23209147 |
| CD31 | CD31 | Wnt10b | FZD6_LRP6 | 0.000492 | 0 | WNT10B_FZD6_LRP6 | Wnt10b - (Fzd6+Lrp6) | WNT | Secreted Signaling | KEGG: mmu04310; PMID: 23209147 |
| GFAP | CD31 | Wnt10b | FZD6_LRP6 | 0.000437 | 0 | WNT10B_FZD6_LRP6 | Wnt10b - (Fzd6+Lrp6) | WNT | Secreted Signaling | KEGG: mmu04310; PMID: 23209147 |
| CD31 | GFAP | Wnt10b | FZD8_LRP6 | 0.000343 | 0 | WNT10B_FZD8_LRP6 | Wnt10b - (Fzd8+Lrp6) | WNT | Secreted Signaling | KEGG: mmu04310; PMID: 23209147 |
| GFAP | GFAP | Wnt10b | FZD8_LRP6 | 0.000304 | 0 | WNT10B_FZD8_LRP6 | Wnt10b - (Fzd8+Lrp6) | WNT | Secreted Signaling | KEGG: mmu04310; PMID: 23209147 |
| CD31 | CD31 | Wnt10b | FZD9_LRP6 | 0.000237 | 0 | WNT10B_FZD9_LRP6 | Wnt10b - (Fzd9+Lrp6) | WNT | Secreted Signaling | KEGG: mmu04310; PMID: 23209147 |
| CD31 | GFAP | Wnt10b | FZD9_LRP6 | 0.00023 | 0 | WNT10B_FZD9_LRP6 | Wnt10b - (Fzd9+Lrp6) | WNT | Secreted Signaling | KEGG: mmu04310; PMID: 23209147 |
| NeuN | GFAP | Wnt4 | FZD1_LRP6 | 0.00022 | 0 | WNT4_FZD1_LRP6 | Wnt4 - (Fzd1+Lrp6) | WNT | Secreted Signaling | KEGG: mmu04310; PMID: 23209147 |
| NeuN | NeuN | Wnt4 | FZD1_LRP6 | 0.000202 | 0 | WNT4_FZD1_LRP6 | Wnt4 - (Fzd1+Lrp6) | WNT | Secreted Signaling | KEGG: mmu04310; PMID: 23209147 |
| GFAP | GFAP | Wnt4 | FZD2_LRP6 | 0.000338 | 0 | WNT4_FZD2_LRP6 | Wnt4 - (Fzd2+Lrp6) | WNT | Secreted Signaling | KEGG: mmu04310; PMID: 23209147 |
| NeuN | GFAP | Wnt4 | FZD2_LRP6 | 0.000368 | 0 | WNT4_FZD2_LRP6 | Wnt4 - (Fzd2+Lrp6) | WNT | Secreted Signaling | KEGG: mmu04310; PMID: 23209147 |
| GFAP | NeuN | Wnt4 | FZD3_LRP6 | 0.000455 | 0 | WNT4_FZD3_LRP6 | Wnt4 - (Fzd3+Lrp6) | WNT | Secreted Signaling | KEGG: mmu04310; PMID: 23209147 |
| NeuN | NeuN | Wnt4 | FZD3_LRP6 | 0.000495 | 0 | WNT4_FZD3_LRP6 | Wnt4 - (Fzd3+Lrp6) | WNT | Secreted Signaling | KEGG: mmu04310; PMID: 23209147 |
| NeuN | CD31 | Wnt4 | FZD4_LRP6 | 0.00033 | 0 | WNT4_FZD4_LRP6 | Wnt4 - (Fzd4+Lrp6) | WNT | Secreted Signaling | KEGG: mmu04310; PMID: 23209147 |
| GFAP | GFAP | Wnt4 | FZD4_LRP6 | 0.000259 | 0 | WNT4_FZD4_LRP6 | Wnt4 - (Fzd4+Lrp6) | WNT | Secreted Signaling | KEGG: mmu04310; PMID: 23209147 |
| GFAP | GFAP | Wnt4 | FZD5_LRP6 | 0.000235 | 0 | WNT4_FZD5_LRP6 | Wnt4 - (Fzd5+Lrp6) | WNT | Secreted Signaling | KEGG: mmu04310; PMID: 23209147 |
| NeuN | GFAP | Wnt4 | FZD5_LRP6 | 0.000256 | 0 | WNT4_FZD5_LRP6 | Wnt4 - (Fzd5+Lrp6) | WNT | Secreted Signaling | KEGG: mmu04310; PMID: 23209147 |
| CD31 | CD31 | Wnt4 | FZD6_LRP6 | 0.000352 | 0 | WNT4_FZD6_LRP6 | Wnt4 - (Fzd6+Lrp6) | WNT | Secreted Signaling | KEGG: mmu04310; PMID: 23209147 |
| NeuN | CD31 | Wnt4 | FZD6_LRP6 | 0.000453 | 0 | WNT4_FZD6_LRP6 | Wnt4 - (Fzd6+Lrp6) | WNT | Secreted Signaling | KEGG: mmu04310; PMID: 23209147 |
| GFAP | GFAP | Wnt4 | FZD8_LRP6 | 0.00029 | 0 | WNT4_FZD8_LRP6 | Wnt4 - (Fzd8+Lrp6) | WNT | Secreted Signaling | KEGG: mmu04310; PMID: 23209147 |
| NeuN | GFAP | Wnt4 | FZD8_LRP6 | 0.000316 | 0 | WNT4_FZD8_LRP6 | Wnt4 - (Fzd8+Lrp6) | WNT | Secreted Signaling | KEGG: mmu04310; PMID: 23209147 |
| NeuN | CD31 | Wnt4 | FZD9_LRP6 | 0.000219 | 0 | WNT4_FZD9_LRP6 | Wnt4 - (Fzd9+Lrp6) | WNT | Secreted Signaling | KEGG: mmu04310; PMID: 23209147 |
| GFAP | GFAP | Wnt4 | FZD9_LRP6 | 0.000195 | 0 | WNT4_FZD9_LRP6 | Wnt4 - (Fzd9+Lrp6) | WNT | Secreted Signaling | KEGG: mmu04310; PMID: 23209147 |
| GFAP | GFAP | Wnt7a | FZD1_LRP6 | 0.000212 | 0 | WNT7A_FZD1_LRP6 | Wnt7a - (Fzd1+Lrp6) | WNT | Secreted Signaling | KEGG: mmu04310; PMID: 23209147 |
| GFAP | NeuN | Wnt7a | FZD1_LRP6 | 0.000195 | 0 | WNT7A_FZD1_LRP6 | Wnt7a - (Fzd1+Lrp6) | WNT | Secreted Signaling | KEGG: mmu04310; PMID: 23209147 |
| CD31 | GFAP | Wnt7a | FZD2_LRP6 | 0.000297 | 0 | WNT7A_FZD2_LRP6 | Wnt7a - (Fzd2+Lrp6) | WNT | Secreted Signaling | KEGG: mmu04310; PMID: 23209147 |
| GFAP | GFAP | Wnt7a | FZD2_LRP6 | 0.000355 | 0 | WNT7A_FZD2_LRP6 | Wnt7a - (Fzd2+Lrp6) | WNT | Secreted Signaling | KEGG: mmu04310; PMID: 23209147 |
| GFAP | GFAP | Wnt7a | FZD3_LRP6 | 0.00031 | 0 | WNT7A_FZD3_LRP6 | Wnt7a - (Fzd3+Lrp6) | WNT | Secreted Signaling | KEGG: mmu04310; PMID: 23209147 |
| GFAP | NeuN | Wnt7a | FZD3_LRP6 | 0.000478 | 0 | WNT7A_FZD3_LRP6 | Wnt7a - (Fzd3+Lrp6) | WNT | Secreted Signaling | KEGG: mmu04310; PMID: 23209147 |
| GFAP | CD31 | Wnt7a | FZD4_LRP6 | 0.000319 | 0 | WNT7A_FZD4_LRP6 | Wnt7a - (Fzd4+Lrp6) | WNT | Secreted Signaling | KEGG: mmu04310; PMID: 23209147 |
| GFAP | GFAP | Wnt7a | FZD4_LRP6 | 0.000272 | 0 | WNT7A_FZD4_LRP6 | Wnt7a - (Fzd4+Lrp6) | WNT | Secreted Signaling | KEGG: mmu04310; PMID: 23209147 |
| CD31 | GFAP | Wnt7a | FZD5_LRP6 | 0.000206 | 0 | WNT7A_FZD5_LRP6 | Wnt7a - (Fzd5+Lrp6) | WNT | Secreted Signaling | KEGG: mmu04310; PMID: 23209147 |

| | | | | | | | | | | |
|---|---|---|---|---|---|---|---|---|---|---|
| GFAP | GFAP | Wnt7a | FZD5_LRP6 | 0.000247 | 0 | WNT7A_FZD5_LRP6 | Wnt7a - (Fzd5+Lrp6) | WNT | Secreted Signaling | KEGG: mmu04310; PMID: 23209147 |
| CD31 | CD31 | Wnt7a | FZD6_LRP6 | 0.000366 | 0 | WNT7A_FZD6_LRP6 | Wnt7a - (Fzd6+Lrp6) | WNT | Secreted Signaling | KEGG: mmu04310; PMID: 23209147 |
| GFAP | CD31 | Wnt7a | FZD6_LRP6 | 0.000438 | 0 | WNT7A_FZD6_LRP6 | Wnt7a - (Fzd6+Lrp6) | WNT | Secreted Signaling | KEGG: mmu04310; PMID: 23209147 |
| CD31 | GFAP | Wnt7a | FZD8_LRP6 | 0.000255 | 0 | WNT7A_FZD8_LRP6 | Wnt7a - (Fzd8+Lrp6) | WNT | Secreted Signaling | KEGG: mmu04310; PMID: 23209147 |
| GFAP | GFAP | Wnt7a | FZD8_LRP6 | 0.000305 | 0 | WNT7A_FZD8_LRP6 | Wnt7a - (Fzd8+Lrp6) | WNT | Secreted Signaling | KEGG: mmu04310; PMID: 23209147 |
| GFAP | CD31 | Wnt7a | FZD9_LRP6 | 0.000211 | 0 | WNT7A_FZD9_LRP6 | Wnt7a - (Fzd9+Lrp6) | WNT | Secreted Signaling | KEGG: mmu04310; PMID: 23209147 |
| GFAP | GFAP | Wnt7a | FZD9_LRP6 | 0.000204 | 0 | WNT7A_FZD9_LRP6 | Wnt7a - (Fzd9+Lrp6) | WNT | Secreted Signaling | KEGG: mmu04310; PMID: 23209147 |
| GFAP | GFAP | Wnt7b | FZD1_LRP6 | 0.000245 | 0 | WNT7B_FZD1_LRP6 | Wnt7b - (Fzd1+Lrp6) | WNT | Secreted Signaling | KEGG: mmu04310; PMID: 23209147 |
| GFAP | NeuN | Wnt7b | FZD1_LRP6 | 0.000225 | 0 | WNT7B_FZD1_LRP6 | Wnt7b - (Fzd1+Lrp6) | WNT | Secreted Signaling | KEGG: mmu04310; PMID: 23209147 |
| CD31 | GFAP | Wnt7b | FZD2_LRP6 | 0.000271 | 0 | WNT7B_FZD2_LRP6 | Wnt7b - (Fzd2+Lrp6) | WNT | Secreted Signaling | KEGG: mmu04310; PMID: 23209147 |
| GFAP | GFAP | Wnt7b | FZD2_LRP6 | 0.00041 | 0 | WNT7B_FZD2_LRP6 | Wnt7b - (Fzd2+Lrp6) | WNT | Secreted Signaling | KEGG: mmu04310; PMID: 23209147 |
| GFAP | GFAP | Wnt7b | FZD3_LRP6 | 0.000358 | 0 | WNT7B_FZD3_LRP6 | Wnt7b - (Fzd3+Lrp6) | WNT | Secreted Signaling | KEGG: mmu04310; PMID: 23209147 |
| GFAP | NeuN | Wnt7b | FZD3_LRP6 | 0.000551 | 0 | WNT7B_FZD3_LRP6 | Wnt7b - (Fzd3+Lrp6) | WNT | Secreted Signaling | KEGG: mmu04310; PMID: 23209147 |
| GFAP | CD31 | Wnt7b | FZD4_LRP6 | 0.000368 | 0 | WNT7B_FZD4_LRP6 | Wnt7b - (Fzd4+Lrp6) | WNT | Secreted Signaling | KEGG: mmu04310; PMID: 23209147 |
| GFAP | GFAP | Wnt7b | FZD4_LRP6 | 0.000314 | 0 | WNT7B_FZD4_LRP6 | Wnt7b - (Fzd4+Lrp6) | WNT | Secreted Signaling | KEGG: mmu04310; PMID: 23209147 |
| GFAP | GFAP | Wnt7b | FZD5_LRP6 | 0.000285 | 0 | WNT7B_FZD5_LRP6 | Wnt7b - (Fzd5+Lrp6) | WNT | Secreted Signaling | KEGG: mmu04310; PMID: 23209147 |
| GFAP | NeuN | Wnt7b | FZD5_LRP6 | 0.000222 | 0 | WNT7B_FZD5_LRP6 | Wnt7b - (Fzd5+Lrp6) | WNT | Secreted Signaling | KEGG: mmu04310; PMID: 23209147 |
| CD31 | CD31 | Wnt7b | FZD6_LRP6 | 0.000334 | 0 | WNT7B_FZD6_LRP6 | Wnt7b - (Fzd6+Lrp6) | WNT | Secreted Signaling | KEGG: mmu04310; PMID: 23209147 |
| GFAP | CD31 | Wnt7b | FZD6_LRP6 | 0.000505 | 0 | WNT7B_FZD6_LRP6 | Wnt7b - (Fzd6+Lrp6) | WNT | Secreted Signaling | KEGG: mmu04310; PMID: 23209147 |
| GFAP | CD31 | Wnt7b | FZD8_LRP6 | 0.000256 | 0 | WNT7B_FZD8_LRP6 | Wnt7b - (Fzd8+Lrp6) | WNT | Secreted Signaling | KEGG: mmu04310; PMID: 23209147 |
| GFAP | GFAP | Wnt7b | FZD8_LRP6 | 0.000352 | 0 | WNT7B_FZD8_LRP6 | Wnt7b - (Fzd8+Lrp6) | WNT | Secreted Signaling | KEGG: mmu04310; PMID: 23209147 |
| GFAP | CD31 | Wnt7b | FZD9_LRP6 | 0.000244 | 0 | WNT7B_FZD9_LRP6 | Wnt7b - (Fzd9+Lrp6) | WNT | Secreted Signaling | KEGG: mmu04310; PMID: 23209147 |
| GFAP | GFAP | Wnt7b | FZD9_LRP6 | 0.000236 | 0 | WNT7B_FZD9_LRP6 | Wnt7b - (Fzd9+Lrp6) | WNT | Secreted Signaling | KEGG: mmu04310; PMID: 23209147 |
| GFAP | GFAP | Wnt9a | FZD1_LRP6 | 0.000228 | 0 | WNT9A_FZD1_LRP6 | Wnt9a - (Fzd1+Lrp6) | WNT | Secreted Signaling | KEGG: mmu04310; PMID: 23209147 |
| GFAP | NeuN | Wnt9a | FZD1_LRP6 | 0.00021 | 0 | WNT9A_FZD1_LRP6 | Wnt9a - (Fzd1+Lrp6) | WNT | Secreted Signaling | KEGG: mmu04310; PMID: 23209147 |
| GFAP | GFAP | Wnt9a | FZD2_LRP6 | 0.000382 | 0 | WNT9A_FZD2_LRP6 | Wnt9a - (Fzd2+Lrp6) | WNT | Secreted Signaling | KEGG: mmu04310; PMID: 23209147 |
| NeuN | GFAP | Wnt9a | FZD2_LRP6 | 0.000255 | 0 | WNT9A_FZD2_LRP6 | Wnt9a - (Fzd2+Lrp6) | WNT | Secreted Signaling | KEGG: mmu04310; PMID: 23209147 |
| GFAP | NeuN | Wnt9a | FZD3_LRP6 | 0.000514 | 0 | WNT9A_FZD3_LRP6 | Wnt9a - (Fzd3+Lrp6) | WNT | Secreted Signaling | KEGG: mmu04310; PMID: 23209147 |
| NeuN | NeuN | Wnt9a | FZD3_LRP6 | 0.000343 | 0 | WNT9A_FZD3_LRP6 | Wnt9a - (Fzd3+Lrp6) | WNT | Secreted Signaling | KEGG: mmu04310; PMID: 23209147 |
| GFAP | CD31 | Wnt9a | FZD4_LRP6 | 0.000342 | 0 | WNT9A_FZD4_LRP6 | Wnt9a - (Fzd4+Lrp6) | WNT | Secreted Signaling | KEGG: mmu04310; PMID: 23209147 |
| GFAP | GFAP | Wnt9a | FZD4_LRP6 | 0.000292 | 0 | WNT9A_FZD4_LRP6 | Wnt9a - (Fzd4+Lrp6) | WNT | Secreted Signaling | KEGG: mmu04310; PMID: 23209147 |
| GFAP | GFAP | Wnt9a | FZD5_LRP6 | 0.000266 | 0 | WNT9A_FZD5_LRP6 | Wnt9a - (Fzd5+Lrp6) | WNT | Secreted Signaling | KEGG: mmu04310; PMID: 23209147 |
| GFAP | NeuN | Wnt9a | FZD5_LRP6 | 0.000206 | 0 | WNT9A_FZD5_LRP6 | Wnt9a - (Fzd5+Lrp6) | WNT | Secreted Signaling | KEGG: mmu04310; PMID: 23209147 |
| CD31 | CD31 | Wnt9a | FZD6_LRP6 | 0.000289 | 0 | WNT9A_FZD6_LRP6 | Wnt9a - (Fzd6+Lrp6) | WNT | Secreted Signaling | KEGG: mmu04310; PMID: 23209147 |
| GFAP | CD31 | Wnt9a | FZD6_LRP6 | 0.000471 | 0 | WNT9A_FZD6_LRP6 | Wnt9a - (Fzd6+Lrp6) | WNT | Secreted Signaling | KEGG: mmu04310; PMID: 23209147 |
| GFAP | CD31 | Wnt9a | FZD8_LRP6 | 0.000239 | 0 | WNT9A_FZD8_LRP6 | Wnt9a - (Fzd8+Lrp6) | WNT | Secreted Signaling | KEGG: mmu04310; PMID: 23209147 |
| GFAP | GFAP | Wnt9a | FZD8_LRP6 | 0.000328 | 0 | WNT9A_FZD8_LRP6 | Wnt9a - (Fzd8+Lrp6) | WNT | Secreted Signaling | KEGG: mmu04310; PMID: 23209147 |
| GFAP | CD31 | Wnt9a | FZD9_LRP6 | 0.000227 | 0 | WNT9A_FZD9_LRP6 | Wnt9a - (Fzd9+Lrp6) | WNT | Secreted Signaling | KEGG: mmu04310; PMID: 23209147 |
| GFAP | GFAP | Wnt9a | FZD9_LRP6 | 0.00022 | 0 | WNT9A_FZD9_LRP6 | Wnt9a - (Fzd9+Lrp6) | WNT | Secreted Signaling | KEGG: mmu04310; PMID: 23209147 |
| CD31 | CD31 | Wnt5a | Fzd1 | 0.000498 | 0 | WNT5A_FZD1 | Wnt5a - Fzd1 | ncWNT | Secreted Signaling | KEGG: mmu04310 |
| CD31 | GFAP | Wnt5a | Fzd1 | 0.000602 | 0 | WNT5A_FZD1 | Wnt5a - Fzd1 | ncWNT | Secreted Signaling | KEGG: mmu04310 |
| CD31 | GFAP | Wnt5a | Fzd2 | 0.001686 | 0 | WNT5A_FZD2 | Wnt5a - Fzd2 | ncWNT | Secreted Signaling | KEGG: mmu04310 |
| GFAP | GFAP | Wnt5a | Fzd2 | 0.001098 | 0 | WNT5A_FZD2 | Wnt5a - Fzd2 | ncWNT | Secreted Signaling | KEGG: mmu04310 |
| CD31 | CD31 | Wnt5a | Fzd3 | 0.001043 | 0 | WNT5A_FZD3 | Wnt5a - Fzd3 | ncWNT | Secreted Signaling | KEGG: mmu04310 |
| CD31 | NeuN | Wnt5a | Fzd3 | 0.001802 | 0 | WNT5A_FZD3 | Wnt5a - Fzd3 | ncWNT | Secreted Signaling | KEGG: mmu04310 |
| CD31 | CD31 | Wnt5a | Fzd4 | 0.001702 | 0 | WNT5A_FZD4 | Wnt5a - Fzd4 | ncWNT | Secreted Signaling | KEGG: mmu04310 |
| GFAP | CD31 | Wnt5a | Fzd4 | 0.001109 | 0 | WNT5A_FZD4 | Wnt5a - Fzd4 | ncWNT | Secreted Signaling | KEGG: mmu04310 |
| CD31 | CD31 | Wnt5a | Fzd5 | 0.000545 | 0 | WNT5A_FZD5 | Wnt5a - Fzd5 | ncWNT | Secreted Signaling | KEGG: mmu04310 |
| CD31 | GFAP | Wnt5a | Fzd5 | 0.000817 | 0 | WNT5A_FZD5 | Wnt5a - Fzd5 | ncWNT | Secreted Signaling | KEGG: mmu04310 |
| CD31 | CD31 | Wnt5a | Fzd6 | 0.00321 | 0 | WNT5A_FZD6 | Wnt5a - Fzd6 | ncWNT | Secreted Signaling | KEGG: mmu04310 |
| GFAP | CD31 | Wnt5a | Fzd6 | 0.002092 | 0 | WNT5A_FZD6 | Wnt5a - Fzd6 | ncWNT | Secreted Signaling | KEGG: mmu04310 |
| CD31 | CD31 | Wnt5a | Fzd8 | 0.000828 | 0 | WNT5A_FZD8 | Wnt5a - Fzd8 | ncWNT | Secreted Signaling | KEGG: mmu04310 |
| CD31 | GFAP | Wnt5a | Fzd8 | 0.001242 | 0 | WNT5A_FZD8 | Wnt5a - Fzd8 | ncWNT | Secreted Signaling | KEGG: mmu04310 |
| CD31 | CD31 | Wnt5a | Fzd9 | 0.000747 | 0 | WNT5A_FZD9 | Wnt5a - Fzd9 | ncWNT | Secreted Signaling | KEGG: mmu04310 |
| CD31 | GFAP | Wnt5a | Fzd9 | 0.000559 | 0 | WNT5A_FZD9 | Wnt5a - Fzd9 | ncWNT | Secreted Signaling | KEGG: mmu04310 |
| CD31 | CD31 | Wnt5a | Mcam | 0.00193 | 0 | WNT5A_MCAM | Wnt5a - Mcam | ncWNT | Secreted Signaling | PMID: 24335906 |
| CD31 | GFAP | Wnt5a | Mcam | 0.002035 | 0 | WNT5A_MCAM | Wnt5a - Mcam | ncWNT | Secreted Signaling | PMID: 24335906 |
| CD31 | CD31 | Hbegf | Erbb4 | 0.00136 | 0 | HBEGF_ERBB4 | Hbegf - Erbb4 | EGF | Secreted Signaling | KEGG: mmu04012 |
| CD31 | GFAP | Hbegf | Erbb4 | 0.001313 | 0 | HBEGF_ERBB4 | Hbegf - Erbb4 | EGF | Secreted Signaling | KEGG: mmu04012 |

| | | | | | | | | | | |
|---|---|---|---|---|---|---|---|---|---|---|
| CD31 | CD31 | Hbegf | ERBB2_ERBB4 | 0.0009 | 0 | HBEGF_ERBB2_ERBB4 | Hbegf - (Erbb2+Erbb4) | EGF | Secreted Signaling | KEGG: mmu04012 |
| CD31 | GFAP | Hbegf | ERBB2_ERBB4 | 0.000881 | 0 | HBEGF_ERBB2_ERBB4 | Hbegf - (Erbb2+Erbb4) | EGF | Secreted Signaling | KEGG: mmu04012 |
| GFAP | CD31 | Ereg | Erbb4 | 0.001273 | 0 | EREG_ERBB4 | Ereg - Erbb4 | EGF | Secreted Signaling | KEGG: mmu04012 |
| GFAP | GFAP | Ereg | Erbb4 | 0.001229 | 0 | EREG_ERBB4 | Ereg - Erbb4 | EGF | Secreted Signaling | KEGG: mmu04012 |
| GFAP | CD31 | Ereg | ERBB2_ERBB4 | 0.000842 | 0 | EREG_ERBB2_ERBB4 | Ereg - (Erbb2+Erbb4) | EGF | Secreted Signaling | KEGG: mmu04012 |
| GFAP | GFAP | Ereg | ERBB2_ERBB4 | 0.000825 | 0 | EREG_ERBB2_ERBB4 | Ereg - (Erbb2+Erbb4) | EGF | Secreted Signaling | KEGG: mmu04012 |
| GFAP | GFAP | Nrg1 | Erbb3 | 0.000703 | 0 | NRG1_ERBB3 | Nrg1 - Erbb3 | NRG | Secreted Signaling | KEGG: mmu04012 |
| NeuN | GFAP | Nrg1 | Erbb3 | 0.000713 | 0 | NRG1_ERBB3 | Nrg1 - Erbb3 | NRG | Secreted Signaling | KEGG: mmu04012 |
| GFAP | GFAP | Nrg1 | ERBB2_ERBB3 | 0.000595 | 0 | NRG1_ERBB2_ERBB3 | Nrg1 - (Erbb2+Erbb3) | NRG | Secreted Signaling | KEGG: mmu04012 |
| NeuN | GFAP | Nrg1 | ERBB2_ERBB3 | 0.000603 | 0 | NRG1_ERBB2_ERBB3 | Nrg1 - (Erbb2+Erbb3) | NRG | Secreted Signaling | KEGG: mmu04012 |
| NeuN | CD31 | Nrg1 | Erbb4 | 0.001174 | 0 | NRG1_ERBB4 | Nrg1 - Erbb4 | NRG | Secreted Signaling | KEGG: mmu04012 |
| GFAP | GFAP | Nrg1 | Erbb4 | 0.001118 | 0 | NRG1_ERBB4 | Nrg1 - Erbb4 | NRG | Secreted Signaling | KEGG: mmu04012 |
| NeuN | CD31 | Nrg1 | ERBB2_ERBB4 | 0.000777 | 0 | NRG1_ERBB2_ERBB4 | Nrg1 - (Erbb2+Erbb4) | NRG | Secreted Signaling | KEGG: mmu04012 |
| GFAP | GFAP | Nrg1 | ERBB2_ERBB4 | 0.00075 | 0 | NRG1_ERBB2_ERBB4 | Nrg1 - (Erbb2+Erbb4) | NRG | Secreted Signaling | KEGG: mmu04012 |
| GFAP | GFAP | Nrg2 | Erbb3 | 0.000716 | 0 | NRG2_ERBB3 | Nrg2 - Erbb3 | NRG | Secreted Signaling | KEGG: mmu04012 |
| NeuN | GFAP | Nrg2 | Erbb3 | 0.000501 | 0 | NRG2_ERBB3 | Nrg2 - Erbb3 | NRG | Secreted Signaling | KEGG: mmu04012 |
| GFAP | GFAP | Nrg2 | ERBB2_ERBB3 | 0.000606 | 0 | NRG2_ERBB2_ERBB3 | Nrg2 - (Erbb2+Erbb3) | NRG | Secreted Signaling | KEGG: mmu04012 |
| NeuN | GFAP | Nrg2 | ERBB2_ERBB3 | 0.000424 | 0 | NRG2_ERBB2_ERBB3 | Nrg2 - (Erbb2+Erbb3) | NRG | Secreted Signaling | KEGG: mmu04012 |
| GFAP | CD31 | Nrg2 | Erbb4 | 0.001179 | 0 | NRG2_ERBB4 | Nrg2 - Erbb4 | NRG | Secreted Signaling | KEGG: mmu04012 |
| GFAP | GFAP | Nrg2 | Erbb4 | 0.001138 | 0 | NRG2_ERBB4 | Nrg2 - Erbb4 | NRG | Secreted Signaling | KEGG: mmu04012 |
| GFAP | CD31 | Nrg2 | ERBB2_ERBB4 | 0.00078 | 0 | NRG2_ERBB2_ERBB4 | Nrg2 - (Erbb2+Erbb4) | NRG | Secreted Signaling | KEGG: mmu04012 |
| GFAP | GFAP | Nrg2 | ERBB2_ERBB4 | 0.000764 | 0 | NRG2_ERBB2_ERBB4 | Nrg2 - (Erbb2+Erbb4) | NRG | Secreted Signaling | KEGG: mmu04012 |
| NeuN | CD31 | Nrg3 | Erbb4 | 0.004825 | 0 | NRG3_ERBB4 | Nrg3 - Erbb4 | NRG | Secreted Signaling | KEGG: mmu04012 |
| NeuN | NeuN | Nrg3 | Erbb4 | 0.002839 | 0 | NRG3_ERBB4 | Nrg3 - Erbb4 | NRG | Secreted Signaling | KEGG: mmu04012 |
| NeuN | CD31 | Nrg3 | ERBB2_ERBB4 | 0.003196 | 0 | NRG3_ERBB2_ERBB4 | Nrg3 - (Erbb2+Erbb4) | NRG | Secreted Signaling | KEGG: mmu04012 |
| NeuN | NeuN | Nrg3 | ERBB2_ERBB4 | 0.001566 | 0 | NRG3_ERBB2_ERBB4 | Nrg3 - (Erbb2+Erbb4) | NRG | Secreted Signaling | KEGG: mmu04012 |
| GFAP | GFAP | Nrg1 | ITGAV_ITGB3 | 0.000681 | 0 | NRG1_ITGAV_ITGB3 | Nrg1 - (Itgav+Itgb3) | NRG | Secreted Signaling | PMID: 20682778 |
| NeuN | GFAP | Nrg1 | ITGAV_ITGB3 | 0.000691 | 0 | NRG1_ITGAV_ITGB3 | Nrg1 - (Itgav+Itgb3) | NRG | Secreted Signaling | PMID: 20682778 |
| CD31 | GFAP | Fgf2 | Fgfr1 | 0.00119 | 0 | FGF2_FGFR1 | Fgf2 - Fgfr1 | FGF | Secreted Signaling | PMC: 4393358 |
| GFAP | GFAP | Fgf2 | Fgfr1 | 0.001927 | 0 | FGF2_FGFR1 | Fgf2 - Fgfr1 | FGF | Secreted Signaling | PMC: 4393358 |
| CD31 | GFAP | Fgf2 | Fgfr2 | 0.001112 | 0 | FGF2_FGFR2 | Fgf2 - Fgfr2 | FGF | Secreted Signaling | PMC: 4393358 |
| GFAP | GFAP | Fgf2 | Fgfr2 | 0.0018 | 0 | FGF2_FGFR2 | Fgf2 - Fgfr2 | FGF | Secreted Signaling | PMC: 4393358 |
| CD31 | GFAP | Fgf2 | Fgfr3 | 0.001168 | 0 | FGF2_FGFR3 | Fgf2 - Fgfr3 | FGF | Secreted Signaling | PMC: 4393358 |
| GFAP | GFAP | Fgf2 | Fgfr3 | 0.001892 | 0 | FGF2_FGFR3 | Fgf2 - Fgfr3 | FGF | Secreted Signaling | PMC: 4393358 |
| GFAP | GFAP | Fgf5 | Fgfr1 | 0.001111 | 0 | FGF5_FGFR1 | Fgf5 - Fgfr1 | FGF | Secreted Signaling | PMC: 4393358 |
| NeuN | GFAP | Fgf5 | Fgfr1 | 0.001107 | 0 | FGF5_FGFR1 | Fgf5 - Fgfr1 | FGF | Secreted Signaling | PMC: 4393358 |
| GFAP | GFAP | Fgf5 | Fgfr2 | 0.001038 | 0 | FGF5_FGFR2 | Fgf5 - Fgfr2 | FGF | Secreted Signaling | PMC: 4393358 |
| NeuN | GFAP | Fgf5 | Fgfr2 | 0.001034 | 0 | FGF5_FGFR2 | Fgf5 - Fgfr2 | FGF | Secreted Signaling | PMC: 4393358 |
| GFAP | GFAP | Fgf5 | Fgfr3 | 0.00109 | 0 | FGF5_FGFR3 | Fgf5 - Fgfr3 | FGF | Secreted Signaling | PMC: 4393358 |
| NeuN | GFAP | Fgf5 | Fgfr3 | 0.001086 | 0 | FGF5_FGFR3 | Fgf5 - Fgfr3 | FGF | Secreted Signaling | PMC: 4393358 |
| CD31 | GFAP | Fgf3 | Fgfr1 | 0.000897 | 0 | FGF3_FGFR1 | Fgf3 - Fgfr1 | FGF | Secreted Signaling | PMC: 4393358 |
| GFAP | GFAP | Fgf3 | Fgfr1 | 0.001383 | 0 | FGF3_FGFR1 | Fgf3 - Fgfr1 | FGF | Secreted Signaling | PMC: 4393358 |
| CD31 | GFAP | Fgf3 | Fgfr2 | 0.000838 | 0 | FGF3_FGFR2 | Fgf3 - Fgfr2 | FGF | Secreted Signaling | PMC: 4393358 |
| GFAP | GFAP | Fgf3 | Fgfr2 | 0.001292 | 0 | FGF3_FGFR2 | Fgf3 - Fgfr2 | FGF | Secreted Signaling | PMC: 4393358 |
| CD31 | GFAP | Fgf10 | Fgfr1 | 0.001791 | 0 | FGF10_FGFR1 | Fgf10 - Fgfr1 | FGF | Secreted Signaling | PMC: 4393358 |
| GFAP | GFAP | Fgf10 | Fgfr1 | 0.001671 | 0 | FGF10_FGFR1 | Fgf10 - Fgfr1 | FGF | Secreted Signaling | PMC: 4393358 |
| CD31 | GFAP | Fgf10 | Fgfr2 | 0.001673 | 0 | FGF10_FGFR2 | Fgf10 - Fgfr2 | FGF | Secreted Signaling | PMC: 4393358 |
| GFAP | GFAP | Fgf10 | Fgfr2 | 0.001561 | 0 | FGF10_FGFR2 | Fgf10 - Fgfr2 | FGF | Secreted Signaling | PMC: 4393358 |
| CD31 | GFAP | Fgf22 | Fgfr2 | 0.001304 | 0 | FGF22_FGFR2 | Fgf22 - Fgfr2 | FGF | Secreted Signaling | PMC: 4393358 |
| GFAP | GFAP | Fgf22 | Fgfr2 | 0.0016 | 0 | FGF22_FGFR2 | Fgf22 - Fgfr2 | FGF | Secreted Signaling | PMC: 4393358 |
| CD31 | GFAP | Fgf18 | Fgfr1 | 0.001043 | 0 | FGF18_FGFR1 | Fgf18 - Fgfr1 | FGF | Secreted Signaling | PMC: 4393358 |
| GFAP | GFAP | Fgf18 | Fgfr1 | 0.001587 | 0 | FGF18_FGFR1 | Fgf18 - Fgfr1 | FGF | Secreted Signaling | PMC: 4393358 |
| CD31 | GFAP | Fgf18 | Fgfr2 | 0.000975 | 0 | FGF18_FGFR2 | Fgf18 - Fgfr2 | FGF | Secreted Signaling | PMC: 4393358 |
| GFAP | GFAP | Fgf18 | Fgfr2 | 0.001483 | 0 | FGF18_FGFR2 | Fgf18 - Fgfr2 | FGF | Secreted Signaling | PMC: 4393358 |
| CD31 | GFAP | Fgf18 | Fgfr3 | 0.001024 | 0 | FGF18_FGFR3 | Fgf18 - Fgfr3 | FGF | Secreted Signaling | PMC: 4393358 |
| GFAP | GFAP | Fgf18 | Fgfr3 | 0.001558 | 0 | FGF18_FGFR3 | Fgf18 - Fgfr3 | FGF | Secreted Signaling | PMC: 4393358 |
| GFAP | GFAP | Fgf9 | Fgfr1 | 0.001456 | 0 | FGF9_FGFR1 | Fgf9 - Fgfr1 | FGF | Secreted Signaling | PMC: 4393358 |
| NeuN | GFAP | Fgf9 | Fgfr1 | 0.00139 | 0 | FGF9_FGFR1 | Fgf9 - Fgfr1 | FGF | Secreted Signaling | PMC: 4393358 |
| GFAP | GFAP | Fgf9 | Fgfr2 | 0.00136 | 0 | FGF9_FGFR2 | Fgf9 - Fgfr2 | FGF | Secreted Signaling | PMC: 4393358 |

| | | | | | | | | | | |
|---|---|---|---|---|---|---|---|---|---|---|
| NeuN | GFAP | Fgf9 | Fgfr2 | 0.001299 | 0 | FGF9_FGFR2 | Fgf9 - Fgfr2 | FGF | Secreted Signaling | PMC: 4393358 |
| GFAP | GFAP | Fgf9 | Fgfr3 | 0.00143 | 0 | FGF9_FGFR3 | Fgf9 - Fgfr3 | FGF | Secreted Signaling | PMC: 4393358 |
| NeuN | GFAP | Fgf9 | Fgfr3 | 0.001365 | 0 | FGF9_FGFR3 | Fgf9 - Fgfr3 | FGF | Secreted Signaling | PMC: 4393358 |
| CD31 | GFAP | Fgf15 | Fgfr1 | 0.000908 | 0 | FGF15_FGFR1 | Fgf15 - Fgfr1 | FGF | Secreted Signaling | PMC: 4393358 |
| GFAP | GFAP | Fgf15 | Fgfr1 | 0.001233 | 0 | FGF15_FGFR1 | Fgf15 - Fgfr1 | FGF | Secreted Signaling | PMC: 4393358 |
| CD31 | GFAP | Fgf15 | Fgfr2 | 0.000848 | 0 | FGF15_FGFR2 | Fgf15 - Fgfr2 | FGF | Secreted Signaling | PMC: 4393358 |
| GFAP | GFAP | Fgf15 | Fgfr2 | 0.001152 | 0 | FGF15_FGFR2 | Fgf15 - Fgfr2 | FGF | Secreted Signaling | PMC: 4393358 |
| CD31 | GFAP | Fgf15 | Fgfr3 | 0.000891 | 0 | FGF15_FGFR3 | Fgf15 - Fgfr3 | FGF | Secreted Signaling | PMC: 4393358 |
| GFAP | GFAP | Fgf15 | Fgfr3 | 0.001211 | 0 | FGF15_FGFR3 | Fgf15 - Fgfr3 | FGF | Secreted Signaling | PMC: 4393358 |
| CD31 | CD31 | Fgf21 | Fgfr1 | 0.001374 | 0 | FGF21_FGFR1 | Fgf21 - Fgfr1 | FGF | Secreted Signaling | PMC: 4393358 |
| CD31 | GFAP | Fgf21 | Fgfr1 | 0.0026 | 0 | FGF21_FGFR1 | Fgf21 - Fgfr1 | FGF | Secreted Signaling | PMC: 4393358 |
| CD31 | GFAP | Fgf21 | Fgfr3 | 0.002552 | 0 | FGF21_FGFR3 | Fgf21 - Fgfr3 | FGF | Secreted Signaling | PMC: 4393358 |
| GFAP | GFAP | Fgf21 | Fgfr3 | 0.001347 | 0 | FGF21_FGFR3 | Fgf21 - Fgfr3 | FGF | Secreted Signaling | PMC: 4393358 |
| GFAP | GFAP | Pdgfa | Pdgfra | 0.001104 | 0 | PDGFA_PDGFRA | Pdgfa - Pdgfra | PDGF | Secreted Signaling | PMID: 15207812 |
| NeuN | GFAP | Pdgfa | Pdgfra | 0.001115 | 0 | PDGFA_PDGFRA | Pdgfa - Pdgfra | PDGF | Secreted Signaling | PMID: 15207812 |
| CD31 | CD31 | Pdgfa | Pdgfrb | 0.000881 | 0 | PDGFA_PDGFRB | Pdgfa - Pdgfrb | PDGF | Secreted Signaling | PMID: 15207812 |
| NeuN | CD31 | Pdgfa | Pdgfrb | 0.001394 | 0 | PDGFA_PDGFRB | Pdgfa - Pdgfrb | PDGF | Secreted Signaling | PMID: 15207812 |
| CD31 | CD31 | Pdgfb | Pdgfra | 0.000813 | 0 | PDGFB_PDGFRA | Pdgfb - Pdgfra | PDGF | Secreted Signaling | PMID: 15207812 |
| CD31 | GFAP | Pdgfb | Pdgfra | 0.001724 | 0 | PDGFB_PDGFRA | Pdgfb - Pdgfra | PDGF | Secreted Signaling | PMID: 15207812 |
| CD31 | CD31 | Pdgfb | Pdgfrb | 0.002154 | 0 | PDGFB_PDGFRB | Pdgfb - Pdgfrb | PDGF | Secreted Signaling | PMID: 15207812 |
| NeuN | CD31 | Pdgfb | Pdgfrb | 0.001168 | 0 | PDGFB_PDGFRB | Pdgfb - Pdgfrb | PDGF | Secreted Signaling | PMID: 15207812 |
| GFAP | CD31 | Pdgfc | Pdgfra | 0.000616 | 0 | PDGFC_PDGFRA | Pdgfc - Pdgfra | PDGF | Secreted Signaling | PMID: 15207812 |
| GFAP | GFAP | Pdgfc | Pdgfra | 0.001307 | 0 | PDGFC_PDGFRA | Pdgfc - Pdgfra | PDGF | Secreted Signaling | PMID: 15207812 |
| CD31 | CD31 | Pdgfd | Pdgfrb | 0.001114 | 0 | PDGFD_PDGFRB | Pdgfd - Pdgfrb | PDGF | Secreted Signaling | PMID: 15207812 |
| GFAP | CD31 | Pdgfd | Pdgfrb | 0.000953 | 0 | PDGFD_PDGFRB | Pdgfd - Pdgfrb | PDGF | Secreted Signaling | PMID: 15207812 |
| CD31 | CD31 | Vegfa | Flt1 | 0.007118 | 0 | VEGFA_VEGFR1 | Vegfa - Vegfr1 | VEGF | Secreted Signaling | KEGG: mmu04370; PMID: 16633338 |
| GFAP | CD31 | Vegfa | Flt1 | 0.018246 | 0 | VEGFA_VEGFR1 | Vegfa - Vegfr1 | VEGF | Secreted Signaling | KEGG: mmu04370; PMID: 16633338 |
| CD31 | CD31 | Vegfa | Kdr | 0.001903 | 0 | VEGFA_VEGFR2 | Vegfa - Vegfr2 | VEGF | Secreted Signaling | KEGG: mmu04370; PMID: 16633338 |
| GFAP | CD31 | Vegfa | Kdr | 0.00492 | 0 | VEGFA_VEGFR2 | Vegfa - Vegfr2 | VEGF | Secreted Signaling | KEGG: mmu04370; PMID: 16633338 |
| CD31 | CD31 | Vegfb | Flt1 | 0.011159 | 0 | VEGFB_VEGFR1 | Vegfb - Vegfr1 | VEGF | Secreted Signaling | KEGG: mmu04370; PMID: 16633338 |
| GFAP | CD31 | Vegfb | Flt1 | 0.010493 | 0 | VEGFB_VEGFR1 | Vegfb - Vegfr1 | VEGF | Secreted Signaling | KEGG: mmu04370; PMID: 16633338 |
| CD31 | CD31 | Pgf | Flt1 | 0.007968 | 0 | PGF_VEGFR1 | Pgf - Vegfr1 | VEGF | Secreted Signaling | KEGG: mmu04370; PMID: 16633338 |
| GFAP | CD31 | Pgf | Flt1 | 0.005914 | 0 | PGF_VEGFR1 | Pgf - Vegfr1 | VEGF | Secreted Signaling | KEGG: mmu04370; PMID: 16633338 |
| CD31 | CD31 | Vegfa | FLT1_KDR | 0.003767 | 0 | VEGFA_VEGFR1R2 | Vegfa - Vegfr1r2 | VEGF | Secreted Signaling | PMID: 16633338 |
| GFAP | CD31 | Vegfa | FLT1_KDR | 0.009707 | 0 | VEGFA_VEGFR1R2 | Vegfa - Vegfr1r2 | VEGF | Secreted Signaling | PMID: 16633338 |
| CD31 | CD31 | Igf1 | Igf1r | 0.000596 | 0 | IGF1_IGF1R | Igf1 - Igf1r | IGF | Secreted Signaling | PMID: 14604834 |
| NeuN | CD31 | Igf1 | Igf1r | 0.000601 | 0 | IGF1_IGF1R | Igf1 - Igf1r | IGF | Secreted Signaling | PMID: 14604834 |
| CD31 | CD31 | Igf2 | Igf1r | 0.00833 | 0 | IGF2_IGF1R | Igf2 - Igf1r | IGF | Secreted Signaling | PMID: 14604834 |
| CD31 | GFAP | Igf2 | Igf1r | 0.002817 | 0 | IGF2_IGF1R | Igf2 - Igf1r | IGF | Secreted Signaling | PMID: 14604834 |
| CD31 | CD31 | Igf2 | Igf2r | 0.00221 | 0 | IGF2_IGF2R | Igf2 - Igf2r | IGF | Secreted Signaling | PMID: 14604834 |
| CD31 | GFAP | Igf2 | Igf2r | 0.00224 | 0 | IGF2_IGF2R | Igf2 - Igf2r | IGF | Secreted Signaling | PMID: 14604834 |
| NeuN | GFAP | Igf1 | ITGAV_ITGB3 | 0.000155 | 0 | IGF1_ITGAV_ITGB3 | Igf1 - (Itgav+Itgb3) | IGF | Secreted Signaling | PMID: 28190785; PMID: 28873464 |
| NeuN | NeuN | Igf1 | ITGAV_ITGB3 | 0.000153 | 0 | IGF1_ITGAV_ITGB3 | Igf1 - (Itgav+Itgb3) | IGF | Secreted Signaling | PMID: 28190785; PMID: 28873464 |
| CD31 | CD31 | Igf2 | ITGAV_ITGB3 | 0.001497 | 0 | IGF2_ITGAV_ITGB3 | Igf2 - (Itgav+Itgb3) | IGF | Secreted Signaling | PMID: 28190785; PMID: 28873464 |
| CD31 | GFAP | Igf2 | ITGAV_ITGB3 | 0.002178 | 0 | IGF2_ITGAV_ITGB3 | Igf2 - (Itgav+Itgb3) | IGF | Secreted Signaling | PMID: 28190785; PMID: 28873464 |
| CD31 | CD31 | Ins1 | Insr | 0.001656 | 0 | INS1_INSSR | Ins1 - Inssr | INSULIN | Secreted Signaling | KEGG: mmu04910; PMID: 24633815 |
| GFAP | CD31 | Ins1 | Insr | 0.001241 | 0 | INS1_INSSR | Ins1 - Inssr | INSULIN | Secreted Signaling | KEGG: mmu04910; PMID: 24633815 |
| CD31 | CD31 | Apln | Aplnr | 0.000271 | 0 | APLN_APLNR | Apln - Aplnr | APELIN | Secreted Signaling | KEGG: mmu04371 |
| GFAP | CD31 | Apln | Aplnr | 0.00043 | 0 | APLN_APLNR | Apln - Aplnr | APELIN | Secreted Signaling | KEGG: mmu04371 |
| CD31 | GFAP | Ihh | PTCH1_SMO | 0.000896 | 0 | IHH_PTCH1 | Ihh - Ptch1 | HH | Secreted Signaling | KEGG: mmu04340; PMID: 25701396 |
| GFAP | GFAP | Ihh | PTCH1_SMO | 0.000652 | 0 | IHH_PTCH1 | Ihh - Ptch1 | HH | Secreted Signaling | KEGG: mmu04340; PMID: 25701396 |
| CD31 | GFAP | Shh | PTCH1_SMO | 0.000949 | 0 | SHH_PTCH1 | Shh - Ptch1 | HH | Secreted Signaling | KEGG: mmu04340; PMID: 25701396 |
| GFAP | GFAP | Shh | PTCH1_SMO | 0.000819 | 0 | SHH_PTCH1 | Shh - Ptch1 | HH | Secreted Signaling | KEGG: mmu04340; PMID: 25701396 |
| CD31 | GFAP | Ihh | PTCH2_SMO | 0.000687 | 0 | IHH_PTCH2 | Ihh - Ptch2 | HH | Secreted Signaling | KEGG: mmu04340; PMID: 25701396 |
| GFAP | GFAP | Ihh | PTCH2_SMO | 0.0005 | 0 | IHH_PTCH2 | Ihh - Ptch2 | HH | Secreted Signaling | KEGG: mmu04340; PMID: 25701396 |
| CD31 | GFAP | Shh | PTCH2_SMO | 0.000728 | 0 | SHH_PTCH2 | Shh - Ptch2 | HH | Secreted Signaling | KEGG: mmu04340; PMID: 25701396 |
| GFAP | GFAP | Shh | PTCH2_SMO | 0.000628 | 0 | SHH_PTCH2 | Shh - Ptch2 | HH | Secreted Signaling | KEGG: mmu04340; PMID: 25701396 |
| GFAP | CD31 | Ccl8 | Ccr1 | 0.000391 | 0 | CCL8_CCR1 | Ccl8 - Ccr1 | CCL | Secreted Signaling | KEGG: mmu04060 |
| GFAP | GFAP | Ccl8 | Ccr1 | 0.000378 | 0 | CCL8_CCR1 | Ccl8 - Ccr1 | CCL | Secreted Signaling | KEGG: mmu04060 |

| | | | | | | | | | | |
|---|---|---|---|---|---|---|---|---|---|---|
| GFAP | CD31 | Ccl6 | Ccr1 | 0.000418 | 0 | CCL6_CCR1 | Ccl6 - Ccr1 | CCL | Secreted Signaling | KEGG: mmu04060 |
| GFAP | GFAP | Ccl6 | Ccr1 | 0.000405 | 0 | CCL6_CCR1 | Ccl6 - Ccr1 | CCL | Secreted Signaling | KEGG: mmu04060 |
| GFAP | CD31 | Ccl8 | Ccr1l1 | 0.000366 | 0 | CCL8_CCR1L1 | Ccl8 - Ccr1l1 | CCL | Secreted Signaling | KEGG: mmu04060 |
| GFAP | GFAP | Ccl8 | Ccr1l1 | 0.000291 | 0 | CCL8_CCR1L1 | Ccl8 - Ccr1l1 | CCL | Secreted Signaling | KEGG: mmu04060 |
| GFAP | CD31 | Ccl6 | Ccr1l1 | 0.000392 | 0 | CCL6_CCR1L1 | Ccl6 - Ccr1l1 | CCL | Secreted Signaling | KEGG: mmu04060 |
| GFAP | GFAP | Ccl6 | Ccr1l1 | 0.000312 | 0 | CCL6_CCR1L1 | Ccl6 - Ccr1l1 | CCL | Secreted Signaling | KEGG: mmu04060 |
| CD31 | GFAP | Ccl6 | Ccr3 | 0.000318 | 0 | CCL6_CCR3 | Ccl6 - Ccr3 | CCL | Secreted Signaling | KEGG: mmu04060 |
| GFAP | GFAP | Ccl6 | Ccr3 | 0.000452 | 0 | CCL6_CCR3 | Ccl6 - Ccr3 | CCL | Secreted Signaling | KEGG: mmu04060 |
| CD31 | GFAP | Ccl11 | Ccr3 | 0.000499 | 0 | CCL11_CCR3 | Ccl11 - Ccr3 | CCL | Secreted Signaling | KEGG: mmu04060 |
| GFAP | GFAP | Ccl11 | Ccr3 | 0.000362 | 0 | CCL11_CCR3 | Ccl11 - Ccr3 | CCL | Secreted Signaling | KEGG: mmu04060 |
| NeuN | GFAP | Ccl27a | Ccr3 | 0.011456 | 0 | CCL27A_CCR3 | Ccl27a - Ccr3 | CCL | Secreted Signaling | KEGG: mmu04060 |
| NeuN | NeuN | Ccl27a | Ccr3 | 0.003627 | 0 | CCL27A_CCR3 | Ccl27a - Ccr3 | CCL | Secreted Signaling | KEGG: mmu04060 |
| NeuN | GFAP | Ccl27b | Ccr3 | 0.006431 | 0 | CCL27B_CCR3 | Ccl27b - Ccr3 | CCL | Secreted Signaling | KEGG: mmu04060 |
| NeuN | NeuN | Ccl27b | Ccr3 | 0.002029 | 0 | CCL27B_CCR3 | Ccl27b - Ccr3 | CCL | Secreted Signaling | KEGG: mmu04060 |
| CD31 | CD31 | Ccl19 | Ccr7 | 0.000671 | 0 | CCL19_CCR7 | Ccl19 - Ccr7 | CCL | Secreted Signaling | KEGG: mmu04060 |
| CD31 | GFAP | Ccl19 | Ccr7 | 0.0008 | 0 | CCL19_CCR7 | Ccl19 - Ccr7 | CCL | Secreted Signaling | KEGG: mmu04060 |
| GFAP | GFAP | Ccl21a | Ccr7 | 0.000715 | 0 | CCL21A_CCR7 | Ccl21a - Ccr7 | CCL | Secreted Signaling | KEGG: mmu04060 |
| NeuN | GFAP | Ccl21a | Ccr7 | 0.000629 | 0 | CCL21A_CCR7 | Ccl21a - Ccr7 | CCL | Secreted Signaling | KEGG: mmu04060 |
| CD31 | GFAP | Ccl25 | Ccr9 | 0.000255 | 0 | CCL25_CCR9 | Ccl25 - Ccr9 | CCL | Secreted Signaling | KEGG: mmu04060 |
| GFAP | GFAP | Ccl25 | Ccr9 | 0.000248 | 0 | CCL25_CCR9 | Ccl25 - Ccr9 | CCL | Secreted Signaling | KEGG: mmu04060 |
| GFAP | CD31 | Ccl4 | Ackr2 | 0.0003 | 0 | CCL4_ACKR2 | Ccl4 - Ackr2 | CCL | Secreted Signaling | PMID: 26740381 |
| GFAP | GFAP | Ccl4 | Ackr2 | 0.00031 | 0 | CCL4_ACKR2 | Ccl4 - Ackr2 | CCL | Secreted Signaling | PMID: 26740381 |
| GFAP | GFAP | Ccl8 | Ackr1 | 0.000628 | 0 | CCL8_ACKR1 | Ccl8 - Ackr1 | CCL | Secreted Signaling | PMID: 26740381 |
| GFAP | NeuN | Ccl8 | Ackr1 | 0.000856 | 0 | CCL8_ACKR1 | Ccl8 - Ackr1 | CCL | Secreted Signaling | PMID: 26740381 |
| CD31 | CD31 | Ccl11 | Ackr1 | 0.000935 | 0 | CCL11_ACKR1 | Ccl11 - Ackr1 | CCL | Secreted Signaling | PMID: 26740381 |
| CD31 | NeuN | Ccl11 | Ackr1 | 0.00101 | 0 | CCL11_ACKR1 | Ccl11 - Ackr1 | CCL | Secreted Signaling | PMID: 26740381 |
| CD31 | CD31 | Ccl11 | Ackr2 | 0.000264 | 0 | CCL11_ACKR2 | Ccl11 - Ackr2 | CCL | Secreted Signaling | PMID: 26740381 |
| CD31 | GFAP | Ccl11 | Ackr2 | 0.000272 | 0 | CCL11_ACKR2 | Ccl11 - Ackr2 | CCL | Secreted Signaling | PMID: 26740381 |
| CD31 | CD31 | Pf4 | Cxcr3 | 0.000651 | 0 | CXCL4_CXCR3 | Cxcl4 - Cxcr3 | CXCL | Secreted Signaling | KEGG: mmu04060 |
| CD31 | GFAP | Pf4 | Cxcr3 | 0.000884 | 0 | CXCL4_CXCR3 | Cxcl4 - Cxcr3 | CXCL | Secreted Signaling | KEGG: mmu04060 |
| GFAP | CD31 | Cxcl10 | Cxcr3 | 0.000329 | 0 | CXCL10_CXCR3 | Cxcl10 - Cxcr3 | CXCL | Secreted Signaling | KEGG: mmu04060 |
| GFAP | GFAP | Cxcl10 | Cxcr3 | 0.000447 | 0 | CXCL10_CXCR3 | Cxcl10 - Cxcr3 | CXCL | Secreted Signaling | KEGG: mmu04060 |
| CD31 | CD31 | Cxcl12 | Cxcr4 | 0.006821 | 0 | CXCL12_CXCR4 | Cxcl12 - Cxcr4 | CXCL | Secreted Signaling | KEGG: mmu04060 |
| CD31 | GFAP | Cxcl12 | Cxcr4 | 0.007126 | 0 | CXCL12_CXCR4 | Cxcl12 - Cxcr4 | CXCL | Secreted Signaling | KEGG: mmu04060 |
| CD31 | CD31 | Cxcl12 | Ackr3 | 0.016279 | 0 | CXCL12_ACKR3 | Cxcl12 - Ackr3 | CXCL | Secreted Signaling | KEGG: mmu04060 |
| CD31 | GFAP | Cxcl12 | Ackr3 | 0.005764 | 0 | CXCL12_ACKR3 | Cxcl12 - Ackr3 | CXCL | Secreted Signaling | KEGG: mmu04060 |
| CD31 | CD31 | Cxcl1 | Ackr1 | 0.000769 | 0 | CXCL1_ACKR1 | Cxcl1 - Ackr1 | CXCL | Secreted Signaling | KEGG: mmu04060 |
| GFAP | NeuN | Cxcl1 | Ackr1 | 0.000859 | 0 | CXCL1_ACKR1 | Cxcl1 - Ackr1 | CXCL | Secreted Signaling | KEGG: mmu04060 |
| GFAP | GFAP | Cxcl10 | Ackr1 | 0.00046 | 0 | CXCL10_ACKR1 | Cxcl10 - Ackr1 | CXCL | Secreted Signaling | KEGG: mmu04060 |
| GFAP | NeuN | Cxcl10 | Ackr1 | 0.000627 | 0 | CXCL10_ACKR1 | Cxcl10 - Ackr1 | CXCL | Secreted Signaling | KEGG: mmu04060 |
| CD31 | CD31 | Mif | Ackr3 | 0.006539 | 0 | MIF_ACKR3 | Mif - Ackr3 | MIF | Secreted Signaling | PMID: 26175090 |
| NeuN | CD31 | Mif | Ackr3 | 0.012114 | 0 | MIF_ACKR3 | Mif - Ackr3 | MIF | Secreted Signaling | PMID: 26175090 |
| NeuN | GFAP | Cx3cl1 | Cx3cr1 | 0.016704 | 0 | CX3CL1_CX3CR1 | Cx3cl1 - Cx3cr1 | CX3C | Secreted Signaling | KEGG: mmu04060 |
| NeuN | NeuN | Cx3cl1 | Cx3cr1 | 0.005215 | 0 | CX3CL1_CX3CR1 | Cx3cl1 - Cx3cr1 | CX3C | Secreted Signaling | KEGG: mmu04060 |
| CD31 | GFAP | Clcf1 | CNTFR_LIFR | 0.000626 | 0 | CLCF1_CNTFR_LIFR | Clcf1 - (Cntfr+Lifr) | LIFR | Secreted Signaling | KEGG: mmu04060 |
| GFAP | GFAP | Clcf1 | CNTFR_LIFR | 0.000605 | 0 | CLCF1_CNTFR_LIFR | Clcf1 - (Cntfr+Lifr) | LIFR | Secreted Signaling | KEGG: mmu04060 |
| CD31 | CD31 | Ctf1 | LIFR_IL6ST | 0.000767 | 0 | CTF1_LIFR_IL6ST | Ctf1 - (Lifr+Il6st) | LIFR | Secreted Signaling | KEGG: mmu04060 |
| CD31 | GFAP | Ctf1 | LIFR_IL6ST | 0.000648 | 0 | CTF1_LIFR_IL6ST | Ctf1 - (Lifr+Il6st) | LIFR | Secreted Signaling | KEGG: mmu04060 |
| CD31 | CD31 | Il34 | Csf1r | 0.002391 | 0 | IL34_CSF1R | Il34 - Csf1r | CSF | Secreted Signaling | KEGG: mmu04060 |
| NeuN | CD31 | Il34 | Csf1r | 0.003494 | 0 | IL34_CSF1R | Il34 - Csf1r | CSF | Secreted Signaling | KEGG: mmu04060 |
| GFAP | CD31 | Csf1 | Csf1r | 0.002034 | 0 | CSF1_CSF1R | Csf1 - Csf1r | CSF | Secreted Signaling | KEGG: mmu04060 |
| GFAP | GFAP | Csf1 | Csf1r | 0.002031 | 0 | CSF1_CSF1R | Csf1 - Csf1r | CSF | Secreted Signaling | KEGG: mmu04060 |
| CD31 | CD31 | Tnfsf10 | Tnfrsf10b | 0.000608 | 0 | TNFSF10_TNFRSF10B | Tnfsf10 - Tnfrsf10b | TRAIL | Secreted Signaling | KEGG: mmu04060 |
| CD31 | GFAP | Tnfsf10 | Tnfrsf10b | 0.000568 | 0 | TNFSF10_TNFRSF10B | Tnfsf10 - Tnfrsf10b | TRAIL | Secreted Signaling | KEGG: mmu04060 |
| GFAP | GFAP | Retn | Cap1 | 0.002964 | 0 | RETN_CAP1 | Retn - Cap1 | RESISTIN | Secreted Signaling | PMID: 30809105 |
| GFAP | NeuN | Retn | Cap1 | 0.003534 | 0 | RETN_CAP1 | Retn - Cap1 | RESISTIN | Secreted Signaling | PMID: 30809105 |
| CD31 | CD31 | Nampt | ITGA5_ITGB1 | 0.005562 | 0 | NAMPT_ITGA5_ITGB1 | Nampt - (Itga5+Itgb1) | VISFATIN | Secreted Signaling | PMID: 28490838 |
| CD31 | GFAP | Nampt | ITGA5_ITGB1 | 0.003606 | 0 | NAMPT_ITGA5_ITGB1 | Nampt - (Itga5+Itgb1) | VISFATIN | Secreted Signaling | PMID: 28490838 |
| CD31 | CD31 | Angptl4 | ITGA5_ITGB1 | 0.001078 | 0 | ANGPTL4_ITGA5_ITGB1 | Angptl4 - (Itga5+Itgb1) | ANGPTL | Secreted Signaling | PMID: 30049845 |

| | | | | | | | | | | |
|---|---|---|---|---|---|---|---|---|---|---|
| GFAP | CD31 | Angptl4 | ITGA5_ITGB1 | 0.001012 | 0 | ANGPTL4_ITGA5_ITGB1 | Angptl4 - (Itga5+Itgb1) | ANGPTL | Secreted Signaling | PMID: 30049845 |
| CD31 | GFAP | Angptl4 | ITGAV_ITGB3 | 0.000515 | 0 | ANGPTL4_ITGAV_ITGB3 | Angptl4 - (Itgav+Itgb3) | ANGPTL | Secreted Signaling | PMID: 30049845 |
| GFAP | GFAP | Angptl4 | ITGAV_ITGB3 | 0.000484 | 0 | ANGPTL4_ITGAV_ITGB3 | Angptl4 - (Itgav+Itgb3) | ANGPTL | Secreted Signaling | PMID: 30049845 |
| CD31 | CD31 | Angptl4 | Cdh5 | 0.002212 | 0 | ANGPTL4_CDH5 | Angptl4 - Cdh5 | ANGPTL | Secreted Signaling | PMID: 30049845 |
| GFAP | CD31 | Angptl4 | Cdh5 | 0.002077 | 0 | ANGPTL4_CDH5 | Angptl4 - Cdh5 | ANGPTL | Secreted Signaling | PMID: 30049845 |
| GFAP | GFAP | Angptl4 | Cdh11 | 0.000872 | 0 | ANGPTL4_CDH11 | Angptl4 - Cdh11 | ANGPTL | Secreted Signaling | PMID: 30049845 |
| CD31 | NeuN | Angptl4 | Cdh11 | 0.001911 | 0 | ANGPTL4_CDH11 | Angptl4 - Cdh11 | ANGPTL | Secreted Signaling | PMID: 30049845 |
| CD31 | GFAP | Angptl4 | Sdc2 | 0.000713 | 0 | ANGPTL4_SDC2 | Angptl4 - Sdc2 | ANGPTL | Secreted Signaling | PMID: 29017031 |
| GFAP | GFAP | Angptl4 | Sdc2 | 0.000669 | 0 | ANGPTL4_SDC2 | Angptl4 - Sdc2 | ANGPTL | Secreted Signaling | PMID: 29017031 |
| CD31 | CD31 | Angptl4 | Sdc3 | 0.001883 | 0 | ANGPTL4_SDC3 | Angptl4 - Sdc3 | ANGPTL | Secreted Signaling | PMID: 29017031 |
| GFAP | CD31 | Angptl4 | Sdc3 | 0.001768 | 0 | ANGPTL4_SDC3 | Angptl4 - Sdc3 | ANGPTL | Secreted Signaling | PMID: 29017031 |
| CD31 | GFAP | Angptl4 | Sdc4 | 0.00235 | 0 | ANGPTL4_SDC4 | Angptl4 - Sdc4 | ANGPTL | Secreted Signaling | PMID: 29017031 |
| GFAP | GFAP | Angptl4 | Sdc4 | 0.002207 | 0 | ANGPTL4_SDC4 | Angptl4 - Sdc4 | ANGPTL | Secreted Signaling | PMID: 29017031 |
| CD31 | CD31 | Angptl3 | ITGAV_ITGB3 | 0.000543 | 0 | ANGPTL3_ITGAV_ITGB3 | Angptl3 - (Itgav+Itgb3) | ANGPTL | Secreted Signaling | PMID: 11877390 |
| CD31 | GFAP | Angptl3 | ITGAV_ITGB3 | 0.000779 | 0 | ANGPTL3_ITGAV_ITGB3 | Angptl3 - (Itgav+Itgb3) | ANGPTL | Secreted Signaling | PMID: 11877390 |
| GFAP | CD31 | Mdk | Sdc2 | 0.001131 | 0 | MDK_SDC2 | Mdk - Sdc2 | MK | Secreted Signaling | PMID: 28356350 |
| GFAP | GFAP | Mdk | Sdc2 | 0.001214 | 0 | MDK_SDC2 | Mdk - Sdc2 | MK | Secreted Signaling | PMID: 28356350 |
| GFAP | CD31 | Mdk | Sdc4 | 0.001809 | 0 | MDK_SDC4 | Mdk - Sdc4 | MK | Secreted Signaling | PMID: 28356350 |
| GFAP | GFAP | Mdk | Sdc4 | 0.004 | 0 | MDK_SDC4 | Mdk - Sdc4 | MK | Secreted Signaling | PMID: 28356350 |
| CD31 | GFAP | Mdk | Ptprz1 | 0.0027 | 0 | MDK_PTPRZ1 | Mdk - Ptprz1 | MK | Secreted Signaling | PMID: 28356350 |
| GFAP | GFAP | Mdk | Ptprz1 | 0.006333 | 0 | MDK_PTPRZ1 | Mdk - Ptprz1 | MK | Secreted Signaling | PMID: 28356350 |
| GFAP | CD31 | Mdk | ITGA4_ITGB1 | 0.001956 | 0 | MDK_ITGA4_ITGB1 | Mdk - (Itga4+Itgb1) | MK | Secreted Signaling | PMID: 28356350 |
| GFAP | GFAP | Mdk | ITGA4_ITGB1 | 0.000963 | 0 | MDK_ITGA4_ITGB1 | Mdk - (Itga4+Itgb1) | MK | Secreted Signaling | PMID: 28356350 |
| GFAP | CD31 | Mdk | ITGA6_ITGB1 | 0.002648 | 0 | MDK_ITGA6_ITGB1 | Mdk - (Itga6+Itgb1) | MK | Secreted Signaling | PMID: 28356350 |
| GFAP | GFAP | Mdk | ITGA6_ITGB1 | 0.00124 | 0 | MDK_ITGA6_ITGB1 | Mdk - (Itga6+Itgb1) | MK | Secreted Signaling | PMID: 28356350 |
| GFAP | GFAP | Mdk | Lrp1 | 0.001509 | 0 | MDK_LRP1 | Mdk - Lrp1 | MK | Secreted Signaling | PMID: 28356350 |
| GFAP | NeuN | Mdk | Lrp1 | 0.001896 | 0 | MDK_LRP1 | Mdk - Lrp1 | MK | Secreted Signaling | PMID: 28356350 |
| GFAP | GFAP | Mdk | Ncl | 0.002933 | 0 | MDK_NCL | Mdk - Ncl | MK | Secreted Signaling | PMID: 28356350 |
| GFAP | NeuN | Mdk | Ncl | 0.004042 | 0 | MDK_NCL | Mdk - Ncl | MK | Secreted Signaling | PMID: 28356350 |
| CD31 | GFAP | Ptn | Ptprz1 | 0.08423 | 0 | PTN_PTPRZ1 | Ptn - Ptprz1 | PTN | Secreted Signaling | PMID: 28356350; PMID: 25620911 |
| GFAP | GFAP | Ptn | Ptprz1 | 0.084622 | 0 | PTN_PTPRZ1 | Ptn - Ptprz1 | PTN | Secreted Signaling | PMID: 28356350; PMID: 25620911 |
| CD31 | GFAP | Ptn | Sdc2 | 0.017245 | 0 | PTN_SDC2 | Ptn - Sdc2 | PTN | Secreted Signaling | PMID: 28356350; PMID: 25620911 |
| GFAP | GFAP | Ptn | Sdc2 | 0.017331 | 0 | PTN_SDC2 | Ptn - Sdc2 | PTN | Secreted Signaling | PMID: 28356350; PMID: 25620911 |
| CD31 | CD31 | Ptn | Sdc3 | 0.044368 | 0 | PTN_SDC3 | Ptn - Sdc3 | PTN | Secreted Signaling | PMID: 28356350; PMID: 25620911 |
| GFAP | CD31 | Ptn | Sdc3 | 0.044583 | 0 | PTN_SDC3 | Ptn - Sdc3 | PTN | Secreted Signaling | PMID: 28356350; PMID: 25620911 |
| CD31 | GFAP | Ptn | Sdc4 | 0.054781 | 0 | PTN_SDC4 | Ptn - Sdc4 | PTN | Secreted Signaling | PMID: 28356350; PMID: 25620911 |
| GFAP | GFAP | Ptn | Sdc4 | 0.055044 | 0 | PTN_SDC4 | Ptn - Sdc4 | PTN | Secreted Signaling | PMID: 28356350; PMID: 25620911 |
| CD31 | GFAP | Ptn | ITGAV_ITGB3 | 0.012528 | 0 | PTN_ITGAV_ITGB3 | Ptn - (Itgav+Itgb3) | PTN | Secreted Signaling | PMID: 28356350; PMID: 25620911 |
| GFAP | GFAP | Ptn | ITGAV_ITGB3 | 0.01259 | 0 | PTN_ITGAV_ITGB3 | Ptn - (Itgav+Itgb3) | PTN | Secreted Signaling | PMID: 28356350; PMID: 25620911 |
| CD31 | CD31 | Ptn | Ncl | 0.052657 | 0 | PTN_NCL | Ptn - Ncl | PTN | Secreted Signaling | PMID: 28356350; PMID: 25620911 |
| GFAP | NeuN | Ptn | Ncl | 0.055597 | 0 | PTN_NCL | Ptn - Ncl | PTN | Secreted Signaling | PMID: 28356350; PMID: 25620911 |
| GFAP | GFAP | Postn | ITGAV_ITGB3 | 0.000568 | 0 | POSTN_ITGAV_ITGB3 | Postn - (Itgav+Itgb3) | PERIOSTIN | Secreted Signaling | PMID: 26721738 |
| GFAP | NeuN | Postn | ITGAV_ITGB3 | 0.000465 | 0 | POSTN_ITGAV_ITGB3 | Postn - (Itgav+Itgb3) | PERIOSTIN | Secreted Signaling | PMID: 26721738 |
| GFAP | CD31 | Postn | ITGAV_ITGB5 | 0.000864 | 0 | POSTN_ITGAV_ITGB5 | Postn - (Itgav+Itgb5) | PERIOSTIN | Secreted Signaling | PMID: 26721738 |
| GFAP | GFAP | Postn | ITGAV_ITGB5 | 0.000797 | 0 | POSTN_ITGAV_ITGB5 | Postn - (Itgav+Itgb5) | PERIOSTIN | Secreted Signaling | PMID: 26721738 |
| GFAP | GFAP | Nmb | Nmbr | 0.000505 | 0 | NMB_NMBR | Nmb - Nmbr | GPR | Secreted Signaling | KEGG: mmu04080 |
| NeuN | GFAP | Nmb | Nmbr | 0.000302 | 0 | NMB_NMBR | Nmb - Nmbr | GPR | Secreted Signaling | KEGG: mmu04080 |
| NeuN | CD31 | Cck | Cckbr | 0.013098 | 0 | CCK_CCKBR | Cck - Cckbr | CCK | Secreted Signaling | KEGG: mmu04080 |
| NeuN | NeuN | Cck | Cckbr | 0.02724 | 0 | CCK_CCKBR | Cck - Cckbr | CCK | Secreted Signaling | KEGG: mmu04080 |
| CD31 | CD31 | Edn1 | Ednra | 0.001507 | 0 | EDN1_EDNRA | Edn1 - Ednra | EDN | Secreted Signaling | KEGG: mmu04080 |
| CD31 | GFAP | Edn1 | Ednra | 0.001417 | 0 | EDN1_EDNRA | Edn1 - Ednra | EDN | Secreted Signaling | KEGG: mmu04080 |
| CD31 | CD31 | Edn3 | Ednra | 0.001877 | 0 | EDN3_EDNRA | Edn3 - Ednra | EDN | Secreted Signaling | KEGG: mmu04080 |
| CD31 | GFAP | Edn3 | Ednra | 0.001764 | 0 | EDN3_EDNRA | Edn3 - Ednra | EDN | Secreted Signaling | KEGG: mmu04080 |
| CD31 | CD31 | Edn1 | Ednrb | 0.001089 | 0 | EDN1_EDNRB | Edn1 - Ednrb | EDN | Secreted Signaling | KEGG: mmu04080 |
| CD31 | GFAP | Edn1 | Ednrb | 0.002501 | 0 | EDN1_EDNRB | Edn1 - Ednrb | EDN | Secreted Signaling | KEGG: mmu04080 |
| CD31 | GFAP | Edn3 | Ednrb | 0.003113 | 0 | EDN3_EDNRB | Edn3 - Ednrb | EDN | Secreted Signaling | KEGG: mmu04080 |
| GFAP | GFAP | Edn3 | Ednrb | 0.001443 | 0 | EDN3_EDNRB | Edn3 - Ednrb | EDN | Secreted Signaling | KEGG: mmu04080 |
| CD31 | CD31 | Nms | Nmur1 | 0.000416 | 0 | NMS_NMUR1 | Nms - Nmur1 | NMU | Secreted Signaling | PMID: 29288685 |
| GFAP | CD31 | Nms | Nmur1 | 0.000427 | 0 | NMS_NMUR1 | Nms - Nmur1 | NMU | Secreted Signaling | PMID: 29288685 |

| | | | | | | | | | | |
|---|---|---|---|---|---|---|---|---|---|---|
| NeuN | GFAP | Npy | Npy1r | 0.00076 | 0 | NPY_NPY1R | Npy - Npy1r | NPY | Secreted Signaling | KEGG: mmu04080 |
| NeuN | NeuN | Npy | Npy1r | 0.000956 | 0 | NPY_NPY1R | Npy - Npy1r | NPY | Secreted Signaling | KEGG: mmu04080 |
| GFAP | GFAP | Npy | Npy2r | 0.00073 | 0 | NPY_NPY2R | Npy - Npy2r | NPY | Secreted Signaling | KEGG: mmu04080 |
| NeuN | GFAP | Npy | Npy2r | 0.001034 | 0 | NPY_NPY2R | Npy - Npy2r | NPY | Secreted Signaling | KEGG: mmu04080 |
| GFAP | GFAP | Npy | Npy5r | 0.000665 | 0 | NPY_NPY5R | Npy - Npy5r | NPY | Secreted Signaling | KEGG: mmu04080 |
| NeuN | GFAP | Npy | Npy5r | 0.000943 | 0 | NPY_NPY5R | Npy - Npy5r | NPY | Secreted Signaling | KEGG: mmu04080 |
| GFAP | GFAP | Npy | Npy6r | 0.000411 | 0 | NPY_NPY6R | Npy - Npy6r | NPY | Secreted Signaling | KEGG: mmu04080 |
| NeuN | GFAP | Npy | Npy6r | 0.000582 | 0 | NPY_NPY6R | Npy - Npy6r | NPY | Secreted Signaling | KEGG: mmu04080 |
| CD31 | GFAP | Npb | Npbwr1 | 0.000414 | 0 | NPB_NPBWR1 | Npb - Npbwr1 | NPW-B | Secreted Signaling | KEGG: mmu04080 |
| GFAP | GFAP | Npb | Npbwr1 | 0.0006 | 0 | NPB_NPBWR1 | Npb - Npbwr1 | NPW-B | Secreted Signaling | KEGG: mmu04080 |
| CD31 | CD31 | Nts | Ntsr1 | 0.000284 | 0 | NTS_NTSR1 | Nts - Ntsr1 | NTS | Secreted Signaling | KEGG: mmu04080 |
| GFAP | CD31 | Nts | Ntsr1 | 0.000305 | 0 | NTS_NTSR1 | Nts - Ntsr1 | NTS | Secreted Signaling | KEGG: mmu04080 |
| CD31 | GFAP | Nts | Ntsr2 | 0.000789 | 0 | NTS_NTSR2 | Nts - Ntsr2 | NTS | Secreted Signaling | KEGG: mmu04080 |
| GFAP | GFAP | Nts | Ntsr2 | 0.000847 | 0 | NTS_NTSR2 | Nts - Ntsr2 | NTS | Secreted Signaling | KEGG: mmu04080 |
| NeuN | GFAP | Pdyn | Oprd1 | 0.000792 | 0 | PDYN_OPRD1 | Pdyn - Oprd1 | OPIOID | Secreted Signaling | KEGG: mmu04080 |
| NeuN | NeuN | Pdyn | Oprd1 | 0.000441 | 0 | PDYN_OPRD1 | Pdyn - Oprd1 | OPIOID | Secreted Signaling | KEGG: mmu04080 |
| CD31 | CD31 | Pdyn | Oprk1 | 0.000466 | 0 | PDYN_OPRK1 | Pdyn - Oprk1 | OPIOID | Secreted Signaling | KEGG: mmu04080 |
| NeuN | CD31 | Pdyn | Oprk1 | 0.00075 | 0 | PDYN_OPRK1 | Pdyn - Oprk1 | OPIOID | Secreted Signaling | KEGG: mmu04080 |
| CD31 | NeuN | Pdyn | Oprl1 | 0.000489 | 0 | PDYN_OPRL1 | Pdyn - Oprl1 | OPIOID | Secreted Signaling | KEGG: mmu04080 |
| NeuN | NeuN | Pdyn | Oprl1 | 0.000786 | 0 | PDYN_OPRL1 | Pdyn - Oprl1 | OPIOID | Secreted Signaling | KEGG: mmu04080 |
| CD31 | CD31 | Pdyn | Oprm1 | 0.000416 | 0 | PDYN_OPRM1 | Pdyn - Oprm1 | OPIOID | Secreted Signaling | KEGG: mmu04080 |
| NeuN | CD31 | Pdyn | Oprm1 | 0.000669 | 0 | PDYN_OPRM1 | Pdyn - Oprm1 | OPIOID | Secreted Signaling | KEGG: mmu04080 |
| CD31 | CD31 | Penk | Oprd1 | 0.00055 | 0 | PENK_OPRD1 | Penk - Oprd1 | OPIOID | Secreted Signaling | KEGG: mmu04080 |
| NeuN | GFAP | Penk | Oprd1 | 0.000952 | 0 | PENK_OPRD1 | Penk - Oprd1 | OPIOID | Secreted Signaling | KEGG: mmu04080 |
| CD31 | CD31 | Penk | Oprk1 | 0.000652 | 0 | PENK_OPRK1 | Penk - Oprk1 | OPIOID | Secreted Signaling | KEGG: mmu04080 |
| NeuN | CD31 | Penk | Oprk1 | 0.000901 | 0 | PENK_OPRK1 | Penk - Oprk1 | OPIOID | Secreted Signaling | KEGG: mmu04080 |
| CD31 | NeuN | Penk | Oprl1 | 0.000683 | 0 | PENK_OPRL1 | Penk - Oprl1 | OPIOID | Secreted Signaling | KEGG: mmu04080 |
| NeuN | NeuN | Penk | Oprl1 | 0.000945 | 0 | PENK_OPRL1 | Penk - Oprl1 | OPIOID | Secreted Signaling | KEGG: mmu04080 |
| CD31 | CD31 | Penk | Oprm1 | 0.000581 | 0 | PENK_OPRM1 | Penk - Oprm1 | OPIOID | Secreted Signaling | KEGG: mmu04080 |
| NeuN | CD31 | Penk | Oprm1 | 0.000804 | 0 | PENK_OPRM1 | Penk - Oprm1 | OPIOID | Secreted Signaling | KEGG: mmu04080 |
| CD31 | GFAP | Oxt | Oxtr | 0.000388 | 0 | OXT_OXTR | Oxt - Oxtr | OXT | Secreted Signaling | KEGG: mmu04080 |
| GFAP | GFAP | Oxt | Oxtr | 0.00047 | 0 | OXT_OXTR | Oxt - Oxtr | OXT | Secreted Signaling | KEGG: mmu04080 |
| NeuN | GFAP | Sst | Sstr1 | 0.004714 | 0 | SST_SSTR1 | Sst - Sstr1 | SOMATOSTATIN | Secreted Signaling | KEGG: mmu04080 |
| NeuN | NeuN | Sst | Sstr1 | 0.005456 | 0 | SST_SSTR1 | Sst - Sstr1 | SOMATOSTATIN | Secreted Signaling | KEGG: mmu04080 |
| NeuN | CD31 | Sst | Sstr2 | 0.004274 | 0 | SST_SSTR2 | Sst - Sstr2 | SOMATOSTATIN | Secreted Signaling | KEGG: mmu04080 |
| NeuN | NeuN | Sst | Sstr2 | 0.007984 | 0 | SST_SSTR2 | Sst - Sstr2 | SOMATOSTATIN | Secreted Signaling | KEGG: mmu04080 |
| NeuN | CD31 | Sst | Sstr3 | 0.004142 | 0 | SST_SSTR3 | Sst - Sstr3 | SOMATOSTATIN | Secreted Signaling | KEGG: mmu04080 |
| NeuN | NeuN | Sst | Sstr3 | 0.003395 | 0 | SST_SSTR3 | Sst - Sstr3 | SOMATOSTATIN | Secreted Signaling | KEGG: mmu04080 |
| NeuN | CD31 | Sst | Sstr4 | 0.007662 | 0 | SST_SSTR4 | Sst - Sstr4 | SOMATOSTATIN | Secreted Signaling | KEGG: mmu04080 |
| NeuN | NeuN | Sst | Sstr4 | 0.014323 | 0 | SST_SSTR4 | Sst - Sstr4 | SOMATOSTATIN | Secreted Signaling | KEGG: mmu04080 |
| GFAP | GFAP | Cort | Sstr1 | 0.000785 | 0 | CORT_SSTR1 | Cort - Sstr1 | SOMATOSTATIN | Secreted Signaling | KEGG: mmu04080 |
| CD31 | NeuN | Cort | Sstr1 | 0.001018 | 0 | CORT_SSTR1 | Cort - Sstr1 | SOMATOSTATIN | Secreted Signaling | KEGG: mmu04080 |
| CD31 | NeuN | Cort | Sstr2 | 0.001492 | 0 | CORT_SSTR2 | Cort - Sstr2 | SOMATOSTATIN | Secreted Signaling | KEGG: mmu04080 |
| NeuN | NeuN | Cort | Sstr2 | 0.00084 | 0 | CORT_SSTR2 | Cort - Sstr2 | SOMATOSTATIN | Secreted Signaling | KEGG: mmu04080 |
| CD31 | CD31 | Cort | Sstr3 | 0.000772 | 0 | CORT_SSTR3 | Cort - Sstr3 | SOMATOSTATIN | Secreted Signaling | KEGG: mmu04080 |
| CD31 | GFAP | Cort | Sstr3 | 0.000759 | 0 | CORT_SSTR3 | Cort - Sstr3 | SOMATOSTATIN | Secreted Signaling | KEGG: mmu04080 |
| CD31 | NeuN | Cort | Sstr4 | 0.002692 | 0 | CORT_SSTR4 | Cort - Sstr4 | SOMATOSTATIN | Secreted Signaling | KEGG: mmu04080 |
| NeuN | NeuN | Cort | Sstr4 | 0.001516 | 0 | CORT_SSTR4 | Cort - Sstr4 | SOMATOSTATIN | Secreted Signaling | KEGG: mmu04080 |
| CD31 | CD31 | Tac1 | Tacr1 | 0.000526 | 0 | TAC1_TACR1 | Tac1 - Tacr1 | TAC | Secreted Signaling | KEGG: mmu04080 |
| CD31 | GFAP | Tac1 | Tacr1 | 0.000501 | 0 | TAC1_TACR1 | Tac1 - Tacr1 | TAC | Secreted Signaling | KEGG: mmu04080 |
| GFAP | GFAP | Tac2 | Tacr3 | 0.000293 | 0 | TAC2_TACR3 | Tac2 - Tacr3 | TAC | Secreted Signaling | KEGG: mmu04080 |
| GFAP | NeuN | Tac2 | Tacr3 | 0.00027 | 0 | TAC2_TACR3 | Tac2 - Tacr3 | TAC | Secreted Signaling | KEGG: mmu04080 |
| GFAP | CD31 | Ctsg | F2r | 0.00061 | 0 | CTSG_F2R | Ctsg - F2r | PARs | Secreted Signaling | KEGG: mmu04080 |
| GFAP | GFAP | Ctsg | F2r | 0.000629 | 0 | CTSG_F2R | Ctsg - F2r | PARs | Secreted Signaling | KEGG: mmu04080 |
| CD31 | CD31 | F2 | F2r | 0.000714 | 0 | F2_F2R | F2 - F2r | PARs | Secreted Signaling | KEGG: mmu04080 |
| CD31 | GFAP | F2 | F2r | 0.000737 | 0 | F2_F2R | F2 - F2r | PARs | Secreted Signaling | KEGG: mmu04080 |
| GFAP | CD31 | Ctsg | F2rl2 | 0.000482 | 0 | CTSG_F2RL2 | Ctsg - F2rl2 | PARs | Secreted Signaling | KEGG: mmu04080 |
| GFAP | GFAP | Ctsg | F2rl2 | 0.000425 | 0 | CTSG_F2RL2 | Ctsg - F2rl2 | PARs | Secreted Signaling | KEGG: mmu04080 |
| CD31 | CD31 | F2 | F2rl2 | 0.000565 | 0 | F2_F2RL2 | F2 - F2rl2 | PARs | Secreted Signaling | KEGG: mmu04080 |

| | | | | | | | | | |
|---|---|---|---|---|---|---|---|---|---|
| CD31 | GFAP | F2 | F2rl2 | 0.000497 | 0 | F2_F2RL2 | F2 - F2rl2 | PARs | Secreted Signaling | KEGG: mmu04080 |
| GFAP | CD31 | Ctsg | Pard3 | 0.000652 | 0 | CTSG_PARD3 | Ctsg - Pard3 | PARs | Secreted Signaling | KEGG: mmu04080 |
| GFAP | GFAP | Ctsg | Pard3 | 0.000685 | 0 | CTSG_PARD3 | Ctsg - Pard3 | PARs | Secreted Signaling | KEGG: mmu04080 |
| CD31 | CD31 | F2 | Pard3 | 0.000764 | 0 | F2_PARD3 | F2 - Pard3 | PARs | Secreted Signaling | KEGG: mmu04080 |
| CD31 | GFAP | F2 | Pard3 | 0.000802 | 0 | F2_PARD3 | F2 - Pard3 | PARs | Secreted Signaling | KEGG: mmu04080 |
| CD31 | CD31 | Plg | Pard3 | 0.000962 | 0 | PLG_PARD3 | Plg - Pard3 | PARs | Secreted Signaling | KEGG: mmu04080 |
| CD31 | GFAP | Plg | Pard3 | 0.00101 | 0 | PLG_PARD3 | Plg - Pard3 | PARs | Secreted Signaling | KEGG: mmu04080 |
| GFAP | GFAP | Pmch | Mchr1 | 0.000722 | 0 | PMCH_MCHR1 | Pmch - Mchr1 | PMCH | Secreted Signaling | KEGG: mmu04080 |
| GFAP | NeuN | Pmch | Mchr1 | 0.000759 | 0 | PMCH_MCHR1 | Pmch - Mchr1 | PMCH | Secreted Signaling | KEGG: mmu04080 |
| CD31 | GFAP | Prok2 | Prokr1 | 0.000431 | 0 | PROK2_PROKR1 | Prok2 - Prokr1 | PROK | Secreted Signaling | PMID: 24496457 |
| GFAP | GFAP | Prok2 | Prokr1 | 0.000493 | 0 | PROK2_PROKR1 | Prok2 - Prokr1 | PROK | Secreted Signaling | PMID: 24496457 |
| GFAP | CD31 | Prok2 | Prokr2 | 0.000279 | 0 | PROK2_PROKR2 | Prok2 - Prokr2 | PROK | Secreted Signaling | PMID: 24496457 |
| GFAP | GFAP | Prok2 | Prokr2 | 0.000292 | 0 | PROK2_PROKR2 | Prok2 - Prokr2 | PROK | Secreted Signaling | PMID: 24496457 |
| NeuN | GFAP | Adcyap1 | Adcyap1r1 | 0.001819 | 0 | ADCYAP1_ADCYAP1R1 | Adcyap1 - Adcyap1r1 | PACAP | Secreted Signaling | PMID: 21350482 |
| NeuN | NeuN | Adcyap1 | Adcyap1r1 | 0.000958 | 0 | ADCYAP1_ADCYAP1R1 | Adcyap1 - Adcyap1r1 | PACAP | Secreted Signaling | PMID: 21350482 |
| CD31 | NeuN | Vip | Vipr1 | 0.000244 | 0 | VIP_VIPR1 | Vip - Vipr1 | VIP | Secreted Signaling | PMID: 10655512 |
| NeuN | NeuN | Vip | Vipr1 | 0.000226 | 0 | VIP_VIPR1 | Vip - Vipr1 | VIP | Secreted Signaling | PMID: 10655512 |
| CD31 | CD31 | Nppa | Npr1 | 0.001326 | 0 | NPPA_NPR1 | Nppa - Npr1 | NPR1 | Secreted Signaling | KEGG: mmu04080; PMID:29934640 |
| NeuN | CD31 | Nppa | Npr1 | 0.001577 | 0 | NPPA_NPR1 | Nppa - Npr1 | NPR1 | Secreted Signaling | KEGG: mmu04080; PMID:29934640 |
| CD31 | CD31 | Nppb | Npr1 | 0.000323 | 0 | NPPB_NPR1 | Nppb - Npr1 | NPR1 | Secreted Signaling | KEGG: mmu04080; PMID:29934640 |
| GFAP | CD31 | Nppb | Npr1 | 0.000316 | 0 | NPPB_NPR1 | Nppb - Npr1 | NPR1 | Secreted Signaling | KEGG: mmu04080; PMID:29934640 |
| CD31 | NeuN | Nppc | Npr2 | 0.000944 | 0 | NPPC_NPR2 | Nppc - Npr2 | NPR2 | Secreted Signaling | KEGG: mmu04080; PMID:29934640 |
| NeuN | NeuN | Nppc | Npr2 | 0.000948 | 0 | NPPC_NPR2 | Nppc - Npr2 | NPR2 | Secreted Signaling | KEGG: mmu04080; PMID:29934640 |
| CD31 | CD31 | Kitl | Kit | 0.002128 | 0 | KITL_KIT | Kitl - Kit | KIT | Secreted Signaling | KEGG: mmu04080 |
| CD31 | NeuN | Kitl | Kit | 0.001455 | 0 | KITL_KIT | Kitl - Kit | KIT | Secreted Signaling | KEGG: mmu04080 |
| CD31 | GFAP | Gip | Gipr | 0.00037 | 0 | GIP_GIPR | Gip - Gipr | GIPR | Secreted Signaling | KEGG: mmu04080 |
| GFAP | GFAP | Gip | Gipr | 0.000353 | 0 | GIP_GIPR | Gip - Gipr | GIPR | Secreted Signaling | KEGG: mmu04080 |
| NeuN | GFAP | Bdnf | Ntrk2 | 0.011117 | 0 | BDNF_NTRK2 | Bdnf - Ntrk2 | NT | Secreted Signaling | PMID: 27908981 |
| NeuN | NeuN | Bdnf | Ntrk2 | 0.010369 | 0 | BDNF_NTRK2 | Bdnf - Ntrk2 | NT | Secreted Signaling | PMID: 27908981 |
| CD31 | GFAP | Flt3l | Flt3 | 0.000919 | 0 | FLT3L_FLT3 | Flt3l - Flt3 | FLT3 | Secreted Signaling | PMID: 25992210 |
| GFAP | GFAP | Flt3l | Flt3 | 0.000436 | 0 | FLT3L_FLT3 | Flt3l - Flt3 | FLT3 | Secreted Signaling | PMID: 25992210 |
| CD31 | CD31 | Hgf | Met | 0.000341 | 0 | HGF_MET | Hgf - Met | HGF | Secreted Signaling | PMID: 28475121 |
| GFAP | CD31 | Hgf | Met | 0.000536 | 0 | HGF_MET | Hgf - Met | HGF | Secreted Signaling | PMID: 28475121 |
| GFAP | CD31 | Ntf3 | Ntrk1 | 0.0004 | 0 | NTF3_NTRK1 | Ntf3 - Ntrk1 | NT | Secreted Signaling | PMID: 27908981 |
| GFAP | GFAP | Ntf3 | Ntrk1 | 0.00036 | 0 | NTF3_NTRK1 | Ntf3 - Ntrk1 | NT | Secreted Signaling | PMID: 27908981 |
| GFAP | GFAP | Ntf3 | Ntrk2 | 0.008117 | 0 | NTF3_NTRK2 | Ntf3 - Ntrk2 | NT | Secreted Signaling | PMID: 27908981 |
| GFAP | NeuN | Ntf3 | Ntrk2 | 0.007569 | 0 | NTF3_NTRK2 | Ntf3 - Ntrk2 | NT | Secreted Signaling | PMID: 27908981 |
| GFAP | NeuN | Ntf3 | Ntrk3 | 0.004255 | 0 | NTF3_NTRK3 | Ntf3 - Ntrk3 | NT | Secreted Signaling | PMID: 27908981 |
| NeuN | NeuN | Ntf3 | Ntrk3 | 0.001188 | 0 | NTF3_NTRK3 | Ntf3 - Ntrk3 | NT | Secreted Signaling | PMID: 27908981 |
| GFAP | NeuN | Sema3a | NRP1_PLXNA1 | 0.001144 | 0 | SEMA3A_NRP1_PLXNA1 | Sema3a - (Nrp1+Plxna1) | SEMA3 | Secreted Signaling | PMID: 27533782 |
| NeuN | NeuN | Sema3a | NRP1_PLXNA1 | 0.001239 | 0 | SEMA3A_NRP1_PLXNA1 | Sema3a - (Nrp1+Plxna1) | SEMA3 | Secreted Signaling | PMID: 27533782 |
| GFAP | NeuN | Sema3a | NRP1_PLXNA2 | 0.001344 | 0 | SEMA3A_NRP1_PLXNA2 | Sema3a - (Nrp1+Plxna2) | SEMA3 | Secreted Signaling | PMID: 27533782 |
| NeuN | NeuN | Sema3a | NRP1_PLXNA2 | 0.001456 | 0 | SEMA3A_NRP1_PLXNA2 | Sema3a - (Nrp1+Plxna2) | SEMA3 | Secreted Signaling | PMID: 27533782 |
| NeuN | CD31 | Sema3a | NRP1_PLXNA3 | 0.000685 | 0 | SEMA3A_NRP1_PLXNA3 | Sema3a - (Nrp1+Plxna3) | SEMA3 | Secreted Signaling | PMID: 27533782 |
| NeuN | NeuN | Sema3a | NRP1_PLXNA3 | 0.000703 | 0 | SEMA3A_NRP1_PLXNA3 | Sema3a - (Nrp1+Plxna3) | SEMA3 | Secreted Signaling | PMID: 27533782 |
| GFAP | NeuN | Sema3a | NRP1_PLXNA4 | 0.00141 | 0 | SEMA3A_NRP1_PLXNA4 | Sema3a - (Nrp1+Plxna4) | SEMA3 | Secreted Signaling | PMID: 27533782 |
| NeuN | NeuN | Sema3a | NRP1_PLXNA4 | 0.001529 | 0 | SEMA3A_NRP1_PLXNA4 | Sema3a - (Nrp1+Plxna4) | SEMA3 | Secreted Signaling | PMID: 27533782 |
| CD31 | CD31 | Sema3b | NRP1_PLXNA1 | 0.000367 | 0 | SEMA3B_NRP1_PLXNA1 | Sema3b - (Nrp1+Plxna1) | SEMA3 | Secreted Signaling | PMID: 27533782 |
| GFAP | NeuN | Sema3b | NRP1_PLXNA1 | 0.000797 | 0 | SEMA3B_NRP1_PLXNA1 | Sema3b - (Nrp1+Plxna1) | SEMA3 | Secreted Signaling | PMID: 27533782 |
| GFAP | GFAP | Sema3b | NRP1_PLXNA2 | 0.000529 | 0 | SEMA3B_NRP1_PLXNA2 | Sema3b - (Nrp1+Plxna2) | SEMA3 | Secreted Signaling | PMID: 27533782 |
| GFAP | NeuN | Sema3b | NRP1_PLXNA2 | 0.000936 | 0 | SEMA3B_NRP1_PLXNA2 | Sema3b - (Nrp1+Plxna2) | SEMA3 | Secreted Signaling | PMID: 27533782 |
| GFAP | GFAP | Sema3b | NRP1_PLXNA3 | 0.000375 | 0 | SEMA3B_NRP1_PLXNA3 | Sema3b - (Nrp1+Plxna3) | SEMA3 | Secreted Signaling | PMID: 27533782 |
| GFAP | NeuN | Sema3b | NRP1_PLXNA3 | 0.000452 | 0 | SEMA3B_NRP1_PLXNA3 | Sema3b - (Nrp1+Plxna3) | SEMA3 | Secreted Signaling | PMID: 27533782 |
| GFAP | GFAP | Sema3b | NRP1_PLXNA4 | 0.000432 | 0 | SEMA3B_NRP1_PLXNA4 | Sema3b - (Nrp1+Plxna4) | SEMA3 | Secreted Signaling | PMID: 27533782 |
| GFAP | NeuN | Sema3b | NRP1_PLXNA4 | 0.000982 | 0 | SEMA3B_NRP1_PLXNA4 | Sema3b - (Nrp1+Plxna4) | SEMA3 | Secreted Signaling | PMID: 27533782 |
| CD31 | CD31 | Sema3c | NRP1_PLXNA1 | 0.001802 | 0 | SEMA3C_NRP1_PLXNA1 | Sema3c - (Nrp1+Plxna1) | SEMA3 | Secreted Signaling | PMID: 27533782 |
| CD31 | NeuN | Sema3c | NRP1_PLXNA1 | 0.002477 | 0 | SEMA3C_NRP1_PLXNA1 | Sema3c - (Nrp1+Plxna1) | SEMA3 | Secreted Signaling | PMID: 27533782 |
| CD31 | CD31 | Sema3c | NRP1_PLXNA2 | 0.002033 | 0 | SEMA3C_NRP1_PLXNA2 | Sema3c - (Nrp1+Plxna2) | SEMA3 | Secreted Signaling | PMID: 27533782 |
| CD31 | NeuN | Sema3c | NRP1_PLXNA2 | 0.00291 | 0 | SEMA3C_NRP1_PLXNA2 | Sema3c - (Nrp1+Plxna2) | SEMA3 | Secreted Signaling | PMID: 27533782 |

| | | | | | | | | | | |
|---|---|---|---|---|---|---|---|---|---|---|
| CD31 | CD31 | Sema3c | NRP1_PLXNA3 | 0.00137 | 0 | SEMA3C_NRP1_PLXNA3 | Sema3c - (Nrp1+Plxna3) | SEMA3 | Secreted Signaling | PMID: 27533782 |
| CD31 | NeuN | Sema3c | NRP1_PLXNA3 | 0.001406 | 0 | SEMA3C_NRP1_PLXNA3 | Sema3c - (Nrp1+Plxna3) | SEMA3 | Secreted Signaling | PMID: 27533782 |
| CD31 | CD31 | Sema3c | NRP1_PLXNA4 | 0.001961 | 0 | SEMA3C_NRP1_PLXNA4 | Sema3c - (Nrp1+Plxna4) | SEMA3 | Secreted Signaling | PMID: 27533782 |
| CD31 | NeuN | Sema3c | NRP1_PLXNA4 | 0.003054 | 0 | SEMA3C_NRP1_PLXNA4 | Sema3c - (Nrp1+Plxna4) | SEMA3 | Secreted Signaling | PMID: 27533782 |
| GFAP | GFAP | Sema3d | NRP1_PLXNA1 | 0.000605 | 0 | SEMA3D_NRP1_PLXNA1 | Sema3d - (Nrp1+Plxna1) | SEMA3 | Secreted Signaling | PMID: 27533782 |
| GFAP | NeuN | Sema3d | NRP1_PLXNA1 | 0.001415 | 0 | SEMA3D_NRP1_PLXNA1 | Sema3d - (Nrp1+Plxna1) | SEMA3 | Secreted Signaling | PMID: 27533782 |
| GFAP | GFAP | Sema3d | NRP1_PLXNA2 | 0.00094 | 0 | SEMA3D_NRP1_PLXNA2 | Sema3d - (Nrp1+Plxna2) | SEMA3 | Secreted Signaling | PMID: 27533782 |
| GFAP | NeuN | Sema3d | NRP1_PLXNA2 | 0.001663 | 0 | SEMA3D_NRP1_PLXNA2 | Sema3d - (Nrp1+Plxna2) | SEMA3 | Secreted Signaling | PMID: 27533782 |
| GFAP | GFAP | Sema3d | NRP1_PLXNA3 | 0.000667 | 0 | SEMA3D_NRP1_PLXNA3 | Sema3d - (Nrp1+Plxna3) | SEMA3 | Secreted Signaling | PMID: 27533782 |
| GFAP | NeuN | Sema3d | NRP1_PLXNA3 | 0.000803 | 0 | SEMA3D_NRP1_PLXNA3 | Sema3d - (Nrp1+Plxna3) | SEMA3 | Secreted Signaling | PMID: 27533782 |
| GFAP | GFAP | Sema3d | NRP1_PLXNA4 | 0.000767 | 0 | SEMA3D_NRP1_PLXNA4 | Sema3d - (Nrp1+Plxna4) | SEMA3 | Secreted Signaling | PMID: 27533782 |
| GFAP | NeuN | Sema3d | NRP1_PLXNA4 | 0.001745 | 0 | SEMA3D_NRP1_PLXNA4 | Sema3d - (Nrp1+Plxna4) | SEMA3 | Secreted Signaling | PMID: 27533782 |
| GFAP | GFAP | Sema3b | NRP2_PLXNA1 | 0.000451 | 0 | SEMA3B_NRP2_PLXNA1 | Sema3b - (Nrp2+Plxna1) | SEMA3 | Secreted Signaling | PMID: 27533782 |
| GFAP | NeuN | Sema3b | NRP2_PLXNA1 | 0.000597 | 0 | SEMA3B_NRP2_PLXNA1 | Sema3b - (Nrp2+Plxna1) | SEMA3 | Secreted Signaling | PMID: 27533782 |
| GFAP | GFAP | Sema3b | NRP2_PLXNA2 | 0.0007 | 0 | SEMA3B_NRP2_PLXNA2 | Sema3b - (Nrp2+Plxna2) | SEMA3 | Secreted Signaling | PMID: 27533782 |
| GFAP | NeuN | Sema3b | NRP2_PLXNA2 | 0.000701 | 0 | SEMA3B_NRP2_PLXNA2 | Sema3b - (Nrp2+Plxna2) | SEMA3 | Secreted Signaling | PMID: 27533782 |
| GFAP | CD31 | Sema3b | NRP2_PLXNA3 | 0.000381 | 0 | SEMA3B_NRP2_PLXNA3 | Sema3b - (Nrp2+Plxna3) | SEMA3 | Secreted Signaling | PMID: 27533782 |
| GFAP | GFAP | Sema3b | NRP2_PLXNA3 | 0.000497 | 0 | SEMA3B_NRP2_PLXNA3 | Sema3b - (Nrp2+Plxna3) | SEMA3 | Secreted Signaling | PMID: 27533782 |
| GFAP | GFAP | Sema3b | NRP2_PLXNA4 | 0.000572 | 0 | SEMA3B_NRP2_PLXNA4 | Sema3b - (Nrp2+Plxna4) | SEMA3 | Secreted Signaling | PMID: 27533782 |
| GFAP | NeuN | Sema3b | NRP2_PLXNA4 | 0.000736 | 0 | SEMA3B_NRP2_PLXNA4 | Sema3b - (Nrp2+Plxna4) | SEMA3 | Secreted Signaling | PMID: 27533782 |
| CD31 | CD31 | Sema3c | NRP2_PLXNA1 | 0.001558 | 0 | SEMA3C_NRP2_PLXNA1 | Sema3c - (Nrp2+Plxna1) | SEMA3 | Secreted Signaling | PMID: 27533782 |
| CD31 | NeuN | Sema3c | NRP2_PLXNA1 | 0.001857 | 0 | SEMA3C_NRP2_PLXNA1 | Sema3c - (Nrp2+Plxna1) | SEMA3 | Secreted Signaling | PMID: 27533782 |
| CD31 | CD31 | Sema3c | NRP2_PLXNA2 | 0.001758 | 0 | SEMA3C_NRP2_PLXNA2 | Sema3c - (Nrp2+Plxna2) | SEMA3 | Secreted Signaling | PMID: 27533782 |
| CD31 | NeuN | Sema3c | NRP2_PLXNA2 | 0.002182 | 0 | SEMA3C_NRP2_PLXNA2 | Sema3c - (Nrp2+Plxna2) | SEMA3 | Secreted Signaling | PMID: 27533782 |
| CD31 | CD31 | Sema3c | NRP2_PLXNA3 | 0.001185 | 0 | SEMA3C_NRP2_PLXNA3 | Sema3c - (Nrp2+Plxna3) | SEMA3 | Secreted Signaling | PMID: 27533782 |
| CD31 | GFAP | Sema3c | NRP2_PLXNA3 | 0.001546 | 0 | SEMA3C_NRP2_PLXNA3 | Sema3c - (Nrp2+Plxna3) | SEMA3 | Secreted Signaling | PMID: 27533782 |
| CD31 | CD31 | Sema3c | NRP2_PLXNA4 | 0.001696 | 0 | SEMA3C_NRP2_PLXNA4 | Sema3c - (Nrp2+Plxna4) | SEMA3 | Secreted Signaling | PMID: 27533782 |
| CD31 | NeuN | Sema3c | NRP2_PLXNA4 | 0.00229 | 0 | SEMA3C_NRP2_PLXNA4 | Sema3c - (Nrp2+Plxna4) | SEMA3 | Secreted Signaling | PMID: 27533782 |
| GFAP | GFAP | Sema3d | NRP2_PLXNA1 | 0.000801 | 0 | SEMA3D_NRP2_PLXNA1 | Sema3d - (Nrp2+Plxna1) | SEMA3 | Secreted Signaling | PMID: 27533782 |
| GFAP | NeuN | Sema3d | NRP2_PLXNA1 | 0.001061 | 0 | SEMA3D_NRP2_PLXNA1 | Sema3d - (Nrp2+Plxna1) | SEMA3 | Secreted Signaling | PMID: 27533782 |
| GFAP | GFAP | Sema3d | NRP2_PLXNA2 | 0.001244 | 0 | SEMA3D_NRP2_PLXNA2 | Sema3d - (Nrp2+Plxna2) | SEMA3 | Secreted Signaling | PMID: 27533782 |
| GFAP | NeuN | Sema3d | NRP2_PLXNA2 | 0.001246 | 0 | SEMA3D_NRP2_PLXNA2 | Sema3d - (Nrp2+Plxna2) | SEMA3 | Secreted Signaling | PMID: 27533782 |
| GFAP | CD31 | Sema3d | NRP2_PLXNA3 | 0.000676 | 0 | SEMA3D_NRP2_PLXNA3 | Sema3d - (Nrp2+Plxna3) | SEMA3 | Secreted Signaling | PMID: 27533782 |
| GFAP | GFAP | Sema3d | NRP2_PLXNA3 | 0.000883 | 0 | SEMA3D_NRP2_PLXNA3 | Sema3d - (Nrp2+Plxna3) | SEMA3 | Secreted Signaling | PMID: 27533782 |
| GFAP | GFAP | Sema3d | NRP2_PLXNA4 | 0.001016 | 0 | SEMA3D_NRP2_PLXNA4 | Sema3d - (Nrp2+Plxna4) | SEMA3 | Secreted Signaling | PMID: 27533782 |
| GFAP | NeuN | Sema3d | NRP2_PLXNA4 | 0.001308 | 0 | SEMA3D_NRP2_PLXNA4 | Sema3d - (Nrp2+Plxna4) | SEMA3 | Secreted Signaling | PMID: 27533782 |
| GFAP | GFAP | Sema3f | NRP2_PLXNA1 | 0.000655 | 0 | SEMA3F_NRP2_PLXNA1 | Sema3f - (Nrp2+Plxna1) | SEMA3 | Secreted Signaling | PMID: 27533782 |
| GFAP | NeuN | Sema3f | NRP2_PLXNA1 | 0.000866 | 0 | SEMA3F_NRP2_PLXNA1 | Sema3f - (Nrp2+Plxna1) | SEMA3 | Secreted Signaling | PMID: 27533782 |
| GFAP | GFAP | Sema3f | NRP2_PLXNA2 | 0.001016 | 0 | SEMA3F_NRP2_PLXNA2 | Sema3f - (Nrp2+Plxna2) | SEMA3 | Secreted Signaling | PMID: 27533782 |
| GFAP | NeuN | Sema3f | NRP2_PLXNA2 | 0.001018 | 0 | SEMA3F_NRP2_PLXNA2 | Sema3f - (Nrp2+Plxna2) | SEMA3 | Secreted Signaling | PMID: 27533782 |
| GFAP | CD31 | Sema3f | NRP2_PLXNA3 | 0.000553 | 0 | SEMA3F_NRP2_PLXNA3 | Sema3f - (Nrp2+Plxna3) | SEMA3 | Secreted Signaling | PMID: 27533782 |
| GFAP | GFAP | Sema3f | NRP2_PLXNA3 | 0.000721 | 0 | SEMA3F_NRP2_PLXNA3 | Sema3f - (Nrp2+Plxna3) | SEMA3 | Secreted Signaling | PMID: 27533782 |
| GFAP | GFAP | Sema3f | NRP2_PLXNA4 | 0.00083 | 0 | SEMA3F_NRP2_PLXNA4 | Sema3f - (Nrp2+Plxna4) | SEMA3 | Secreted Signaling | PMID: 27533782 |
| GFAP | NeuN | Sema3f | NRP2_PLXNA4 | 0.001069 | 0 | SEMA3F_NRP2_PLXNA4 | Sema3f - (Nrp2+Plxna4) | SEMA3 | Secreted Signaling | PMID: 27533782 |
| CD31 | CD31 | Sema3g | NRP2_PLXNA1 | 0.001509 | 0 | SEMA3G_NRP2_PLXNA1 | Sema3g - (Nrp2+Plxna1) | SEMA3 | Secreted Signaling | PMID: 27533782 |
| CD31 | NeuN | Sema3g | NRP2_PLXNA1 | 0.001798 | 0 | SEMA3G_NRP2_PLXNA1 | Sema3g - (Nrp2+Plxna1) | SEMA3 | Secreted Signaling | PMID: 27533782 |
| CD31 | CD31 | Sema3g | NRP2_PLXNA2 | 0.001703 | 0 | SEMA3G_NRP2_PLXNA2 | Sema3g - (Nrp2+Plxna2) | SEMA3 | Secreted Signaling | PMID: 27533782 |
| CD31 | NeuN | Sema3g | NRP2_PLXNA2 | 0.002113 | 0 | SEMA3G_NRP2_PLXNA2 | Sema3g - (Nrp2+Plxna2) | SEMA3 | Secreted Signaling | PMID: 27533782 |
| CD31 | CD31 | Sema3g | NRP2_PLXNA3 | 0.001147 | 0 | SEMA3G_NRP2_PLXNA3 | Sema3g - (Nrp2+Plxna3) | SEMA3 | Secreted Signaling | PMID: 27533782 |
| CD31 | GFAP | Sema3g | NRP2_PLXNA3 | 0.001497 | 0 | SEMA3G_NRP2_PLXNA3 | Sema3g - (Nrp2+Plxna3) | SEMA3 | Secreted Signaling | PMID: 27533782 |
| CD31 | CD31 | Sema3g | NRP2_PLXNA4 | 0.001642 | 0 | SEMA3G_NRP2_PLXNA4 | Sema3g - (Nrp2+Plxna4) | SEMA3 | Secreted Signaling | PMID: 27533782 |
| CD31 | NeuN | Sema3g | NRP2_PLXNA4 | 0.002218 | 0 | SEMA3G_NRP2_PLXNA4 | Sema3g - (Nrp2+Plxna4) | SEMA3 | Secreted Signaling | PMID: 27533782 |
| CD31 | CD31 | Sema3c | NRP1_NRP2 | 0.002159 | 0 | SEMA3C_NRP1_NRP2 | Sema3c - (Nrp1+Nrp2) | SEMA3 | Secreted Signaling | PMID: 27533782 |
| CD31 | NeuN | Sema3c | NRP1_NRP2 | 0.001388 | 0 | SEMA3C_NRP1_NRP2 | Sema3c - (Nrp1+Nrp2) | SEMA3 | Secreted Signaling | PMID: 27533782 |
| CD31 | CD31 | Sema3c | Plxnd1 | 0.003076 | 0 | SEMA3C_PLXND1 | Sema3c - Plxnd1 | SEMA3 | Secreted Signaling | PMID: 27533782 |
| CD31 | GFAP | Sema3c | Plxnd1 | 0.00094 | 0 | SEMA3C_PLXND1 | Sema3c - Plxnd1 | SEMA3 | Secreted Signaling | PMID: 27533782 |
| CD31 | CD31 | Sema3e | Plxnd1 | 0.001708 | 0 | SEMA3E_PLXND1 | Sema3e - Plxnd1 | SEMA3 | Secreted Signaling | PMID: 22325954 |
| NeuN | CD31 | Sema3e | Plxnd1 | 0.003199 | 0 | SEMA3E_PLXND1 | Sema3e - Plxnd1 | SEMA3 | Secreted Signaling | PMID: 22325954 |
| GFAP | NeuN | Crh | Crhr1 | 0.00086 | 0 | CRH_CRHR1 | Crh - Crhr1 | CRH | Secreted Signaling | KEGG: mmu04080 |

| | | | | | | | | | | |
|---|---|---|---|---|---|---|---|---|---|---|
| NeuN | NeuN | Crh | Crhr1 | 0.000419 | 0 | CRH_CRHR1 | Crh - Crhr1 | CRH | Secreted Signaling | KEGG: mmu04080 |
| CD31 | GFAP | Crh | Crhr2 | 0.000368 | 0 | CRH_CRHR2 | Crh - Crhr2 | CRH | Secreted Signaling | KEGG: mmu04080 |
| GFAP | GFAP | Crh | Crhr2 | 0.000431 | 0 | CRH_CRHR2 | Crh - Crhr2 | CRH | Secreted Signaling | KEGG: mmu04080 |
| GFAP | GFAP | Enho | Gpr19 | 0.001698 | 0 | ENHO_GPR19 | Enho - Gpr19 | ENHO | Secreted Signaling | PMID: 28476646 |
| GFAP | NeuN | Enho | Gpr19 | 0.001911 | 0 | ENHO_GPR19 | Enho - Gpr19 | ENHO | Secreted Signaling | PMID: 28476646 |
| CD31 | CD31 | Gas6 | Axl | 0.002702 | 0 | GAS6_AXL | Gas6 - Axl | GAS | Secreted Signaling | PMID: 27801848 |
| CD31 | GFAP | Gas6 | Axl | 0.003164 | 0 | GAS6_AXL | Gas6 - Axl | GAS | Secreted Signaling | PMID: 27801848 |
| CD31 | CD31 | Gas6 | Mertk | 0.005617 | 0 | GAS6_MERTK | Gas6 - Mertk | GAS | Secreted Signaling | PMID: 27801848 |
| NeuN | CD31 | Gas6 | Mertk | 0.00508 | 0 | GAS6_MERTK | Gas6 - Mertk | GAS | Secreted Signaling | PMID: 27801848 |
| CD31 | NeuN | Gas6 | Tyro3 | 0.002869 | 0 | GAS6_TYRO3 | Gas6 - Tyro3 | GAS | Secreted Signaling | PMID: 27801848 |
| NeuN | NeuN | Gas6 | Tyro3 | 0.002595 | 0 | GAS6_TYRO3 | Gas6 - Tyro3 | GAS | Secreted Signaling | PMID: 27801848 |
| GFAP | GFAP | Grn | Sort1 | 0.006141 | 0 | GRN_SORT1 | Grn - Sort1 | GRN | Secreted Signaling | PMID: 29555433 |
| GFAP | NeuN | Grn | Sort1 | 0.009274 | 0 | GRN_SORT1 | Grn - Sort1 | GRN | Secreted Signaling | PMID: 29555433 |
| CD31 | CD31 | Insl5 | Rxfp4 | 0.000502 | 0 | INSL5_RXFP4 | Insl5 - Rxfp4 | RLN | Secreted Signaling | KEGG: mmu04080 |
| GFAP | CD31 | Insl5 | Rxfp4 | 0.000492 | 0 | INSL5_RXFP4 | Insl5 - Rxfp4 | RLN | Secreted Signaling | KEGG: mmu04080 |
| CD31 | CD31 | Insl5 | Rxfp3 | 0.000395 | 0 | INSL5_RXFP3 | Insl5 - Rxfp3 | RLN | Secreted Signaling | KEGG: mmu04080 |
| CD31 | GFAP | Insl5 | Rxfp3 | 0.0004 | 0 | INSL5_RXFP3 | Insl5 - Rxfp3 | RLN | Secreted Signaling | KEGG: mmu04080 |
| CD31 | GFAP | Insl5 | Rxfp2 | 0.000597 | 0 | INSL5_RXFP2 | Insl5 - Rxfp2 | RLN | Secreted Signaling | KEGG: mmu04080 |
| GFAP | GFAP | Insl5 | Rxfp2 | 0.000586 | 0 | INSL5_RXFP2 | Insl5 - Rxfp2 | RLN | Secreted Signaling | KEGG: mmu04080 |
| CD31 | GFAP | Insl5 | Rxfp1 | 0.000688 | 0 | INSL5_RXFP1 | Insl5 - Rxfp1 | RLN | Secreted Signaling | KEGG: mmu04080 |
| GFAP | GFAP | Insl5 | Rxfp1 | 0.000675 | 0 | INSL5_RXFP1 | Insl5 - Rxfp1 | RLN | Secreted Signaling | KEGG: mmu04080 |
| CD31 | CD31 | Lgals9 | Ighm | 0.000395 | 0 | LGALS9_IGHM | Lgals9 - Ighm | GALECTIN | Secreted Signaling | PMID: 30120235 |
| CD31 | NeuN | Lgals9 | Ighm | 0.000824 | 0 | LGALS9_IGHM | Lgals9 - Ighm | GALECTIN | Secreted Signaling | PMID: 30120235 |
| GFAP | CD31 | Pmch | Mertk | 0.002267 | 0 | PMCH_MERTK | Pmch - Mertk | PMCH | Secreted Signaling | PMID: 30501104 |
| GFAP | GFAP | Pmch | Mertk | 0.001307 | 0 | PMCH_MERTK | Pmch - Mertk | PMCH | Secreted Signaling | PMID: 30501104 |
| CD31 | GFAP | Pros1 | Axl | 0.000983 | 0 | PROS1_AXL | Pros1 - Axl | PROS | Secreted Signaling | PMID: 29531161 |
| GFAP | GFAP | Pros1 | Axl | 0.000947 | 0 | PROS1_AXL | Pros1 - Axl | PROS | Secreted Signaling | PMID: 29531161 |
| GFAP | GFAP | Pros1 | Tyro3 | 0.000577 | 0 | PROS1_TYRO3 | Pros1 - Tyro3 | PROS | Secreted Signaling | PMID: 30501104 |
| CD31 | NeuN | Pros1 | Tyro3 | 0.000891 | 0 | PROS1_TYRO3 | Pros1 - Tyro3 | PROS | Secreted Signaling | PMID: 30501104 |
| GFAP | GFAP | Psap | Gpr37l1 | 0.138584 | 0 | PSAP_GPR37L1 | Psap - Gpr37l1 | PSAP | Secreted Signaling | PMID: 23690594 |
| NeuN | GFAP | Psap | Gpr37l1 | 0.208698 | 0 | PSAP_GPR37L1 | Psap - Gpr37l1 | PSAP | Secreted Signaling | PMID: 23690594 |
| GFAP | GFAP | Psap | Gpr37 | 0.049409 | 0 | PSAP_GPR37 | Psap - Gpr37 | PSAP | Secreted Signaling | PMID: 23690594 |
| NeuN | GFAP | Psap | Gpr37 | 0.078519 | 0 | PSAP_GPR37 | Psap - Gpr37 | PSAP | Secreted Signaling | PMID: 23690594 |
| CD31 | CD31 | Qrfp | Qrfpr | 0.001085 | 0 | QRFP_QRFPR | Qrfp - Qrfpr | QRFP | Secreted Signaling | PMID: 25386115 |
| CD31 | GFAP | Qrfp | Qrfpr | 0.000905 | 0 | QRFP_QRFPR | Qrfp - Qrfpr | QRFP | Secreted Signaling | PMID: 25386115 |
| CD31 | CD31 | Rarres2 | Cmklr1 | 0.000951 | 0 | RARRES2_CMKLR1 | Rarres2 - Cmklr1 | CHEMERIN | Secreted Signaling | PMID: 24671882 |
| CD31 | GFAP | Rarres2 | Cmklr1 | 0.00052 | 0 | RARRES2_CMKLR1 | Rarres2 - Cmklr1 | CHEMERIN | Secreted Signaling | PMID: 24671882 |
| CD31 | CD31 | Rln3 | Rxfp3 | 0.000312 | 0 | RLN3_RXFP3 | Rln3 - Rxfp3 | RLN | Secreted Signaling | KEGG: mmu04080 |
| CD31 | GFAP | Rln3 | Rxfp3 | 0.000316 | 0 | RLN3_RXFP3 | Rln3 - Rxfp3 | RLN | Secreted Signaling | KEGG: mmu04080 |
| CD31 | GFAP | Rln3 | Rxfp1 | 0.000545 | 0 | RLN3_RXFP1 | Rln3 - Rxfp1 | RLN | Secreted Signaling | KEGG: mmu04080 |
| GFAP | GFAP | Rln3 | Rxfp1 | 0.000497 | 0 | RLN3_RXFP1 | Rln3 - Rxfp1 | RLN | Secreted Signaling | KEGG: mmu04080 |
| CD31 | CD31 | Rln3 | Rxfp4 | 0.000397 | 0 | RLN3_RXFP4 | Rln3 - Rxfp4 | RLN | Secreted Signaling | KEGG: mmu04080 |
| GFAP | CD31 | Rln3 | Rxfp4 | 0.000362 | 0 | RLN3_RXFP4 | Rln3 - Rxfp4 | RLN | Secreted Signaling | KEGG: mmu04080 |
| CD31 | GFAP | Rln3 | Rxfp2 | 0.000473 | 0 | RLN3_RXFP2 | Rln3 - Rxfp2 | RLN | Secreted Signaling | KEGG: mmu04080 |
| GFAP | GFAP | Rln3 | Rxfp2 | 0.000431 | 0 | RLN3_RXFP2 | Rln3 - Rxfp2 | RLN | Secreted Signaling | KEGG: mmu04080 |
| GFAP | CD31 | Slurp2 | Chrnb2 | 0.000618 | 0 | SLURP2_CHRNB2 | Slurp2 - Chrnb2 | SLURP | Secreted Signaling | KEGG: mmu04080 |
| GFAP | GFAP | Slurp2 | Chrnb2 | 0.000616 | 0 | SLURP2_CHRNB2 | Slurp2 - Chrnb2 | SLURP | Secreted Signaling | KEGG: mmu04080 |
| CD31 | GFAP | Slurp2 | Chrnb3 | 0.000404 | 0 | SLURP2_CHRNB3 | Slurp2 - Chrnb3 | SLURP | Secreted Signaling | KEGG: mmu04080 |
| GFAP | GFAP | Slurp2 | Chrnb3 | 0.000904 | 0 | SLURP2_CHRNB3 | Slurp2 - Chrnb3 | SLURP | Secreted Signaling | KEGG: mmu04080 |
| GFAP | CD31 | Slurp2 | Chrnb4 | 0.000406 | 0 | SLURP2_CHRNB4 | Slurp2 - Chrnb4 | SLURP | Secreted Signaling | KEGG: mmu04080 |
| GFAP | GFAP | Slurp2 | Chrnb4 | 0.00073 | 0 | SLURP2_CHRNB4 | Slurp2 - Chrnb4 | SLURP | Secreted Signaling | KEGG: mmu04080 |
| GFAP | CD31 | Slurp2 | Chrna1 | 0.0003 | 0 | SLURP2_CHRNA1 | Slurp2 - Chrna1 | SLURP | Secreted Signaling | KEGG: mmu04080 |
| GFAP | GFAP | Slurp2 | Chrna1 | 0.00053 | 0 | SLURP2_CHRNA1 | Slurp2 - Chrna1 | SLURP | Secreted Signaling | KEGG: mmu04080 |
| GFAP | GFAP | Slurp2 | Chrna4 | 0.000682 | 0 | SLURP2_CHRNA4 | Slurp2 - Chrna4 | SLURP | Secreted Signaling | KEGG: mmu04080 |
| GFAP | NeuN | Slurp2 | Chrna4 | 0.0012 | 0 | SLURP2_CHRNA4 | Slurp2 - Chrna4 | SLURP | Secreted Signaling | KEGG: mmu04080 |
| GFAP | CD31 | Slurp2 | Chrna5 | 0.000282 | 0 | SLURP2_CHRNA5 | Slurp2 - Chrna5 | SLURP | Secreted Signaling | KEGG: mmu04080 |
| GFAP | GFAP | Slurp2 | Chrna5 | 0.000312 | 0 | SLURP2_CHRNA5 | Slurp2 - Chrna5 | SLURP | Secreted Signaling | KEGG: mmu04080 |
| GFAP | CD31 | Slurp2 | Chrna7 | 0.000388 | 0 | SLURP2_CHRNA7 | Slurp2 - Chrna7 | SLURP | Secreted Signaling | KEGG: mmu04080 |
| GFAP | GFAP | Slurp2 | Chrna7 | 0.000505 | 0 | SLURP2_CHRNA7 | Slurp2 - Chrna7 | SLURP | Secreted Signaling | KEGG: mmu04080 |

| | | | | | | | | | | |
|---|---|---|---|---|---|---|---|---|---|---|
| GFAP | CD31 | Slurp2 | Chrna9 | 0.000314 | 0 | SLURP2_CHRNA9 | Slurp2 - Chrna9 | SLURP | Secreted Signaling | KEGG: mmu04080 |
| GFAP | GFAP | Slurp2 | Chrna9 | 0.000537 | 0 | SLURP2_CHRNA9 | Slurp2 - Chrna9 | SLURP | Secreted Signaling | KEGG: mmu04080 |
| GFAP | CD31 | Slurp2 | Chrnd | 0.000341 | 0 | SLURP2_CHRND | Slurp2 - Chrnd | SLURP | Secreted Signaling | KEGG: mmu04080 |
| GFAP | GFAP | Slurp2 | Chrnd | 0.000283 | 0 | SLURP2_CHRND | Slurp2 - Chrnd | SLURP | Secreted Signaling | KEGG: mmu04080 |
| CD31 | NeuN | Ucn | Crhr1 | 0.001561 | 0 | UCN_CRHR1 | Ucn - Crhr1 | UCN | Secreted Signaling | KEGG: mmu04080 |
| NeuN | NeuN | Ucn | Crhr1 | 0.000949 | 0 | UCN_CRHR1 | Ucn - Crhr1 | UCN | Secreted Signaling | KEGG: mmu04080 |
| CD31 | GFAP | Ucn | Crhr2 | 0.000783 | 0 | UCN_CRHR2 | Ucn - Crhr2 | UCN | Secreted Signaling | KEGG: mmu04080 |
| GFAP | GFAP | Ucn | Crhr2 | 0.000556 | 0 | UCN_CRHR2 | Ucn - Crhr2 | UCN | Secreted Signaling | KEGG: mmu04080 |
| GFAP | CD31 | Klk6 | F2r | 0.0012 | 0 | KLK6_F2R | Klk6 - F2r | KLK | Secreted Signaling | PMID: 35102281 |
| GFAP | GFAP | Klk6 | F2r | 0.001238 | 0 | KLK6_F2R | Klk6 - F2r | KLK | Secreted Signaling | PMID: 35102281 |
| GFAP | CD31 | Klk8 | F2r | 0.000667 | 0 | KLK8_F2R | Klk8 - F2r | KLK | Secreted Signaling | PMID: 34552064 |
| GFAP | GFAP | Klk8 | F2r | 0.000688 | 0 | KLK8_F2R | Klk8 - F2r | KLK | Secreted Signaling | PMID: 34552064 |
| GFAP | CD31 | Adm | CALCR_RAMP1 | 0.001059 | 0 | ADM_CALCR_RAMP1 | Adm - (Calcr+Ramp1) | ADM | Secreted Signaling | PMID: 33462236 |
| GFAP | GFAP | Adm | CALCR_RAMP1 | 0.00094 | 0 | ADM_CALCR_RAMP1 | Adm - (Calcr+Ramp1) | ADM | Secreted Signaling | PMID: 33462236 |
| CD31 | CD31 | Adm | CALCR_RAMP2 | 0.00144 | 0 | ADM_CALCR_RAMP2 | Adm - (Calcr+Ramp2) | ADM | Secreted Signaling | PMID: 33462236 |
| GFAP | CD31 | Adm | CALCR_RAMP2 | 0.002073 | 0 | ADM_CALCR_RAMP2 | Adm - (Calcr+Ramp2) | ADM | Secreted Signaling | PMID: 33462236 |
| GFAP | CD31 | Adm | CALCR_RAMP3 | 0.000615 | 0 | ADM_CALCR_RAMP3 | Adm - (Calcr+Ramp3) | ADM | Secreted Signaling | PMID: 33462236 |
| GFAP | GFAP | Adm | CALCR_RAMP3 | 0.000575 | 0 | ADM_CALCR_RAMP3 | Adm - (Calcr+Ramp3) | ADM | Secreted Signaling | PMID: 33462236 |
| CD31 | CD31 | Calca | CALCR_RAMP1 | 0.001056 | 0 | CALCA_CALCR_RAMP1 | Calca - (Calcr+Ramp1) | CALC | Secreted Signaling | PMID: 30741933;PMID: 18599553 |
| CD31 | GFAP | Calca | CALCR_RAMP1 | 0.000937 | 0 | CALCA_CALCR_RAMP1 | Calca - (Calcr+Ramp1) | CALC | Secreted Signaling | PMID: 30741933;PMID: 18599553 |
| CD31 | CD31 | Calca | CALCR_RAMP2 | 0.002068 | 0 | CALCA_CALCR_RAMP2 | Calca - (Calcr+Ramp2) | CALC | Secreted Signaling | PMID: 30741933;PMID: 18599553 |
| GFAP | CD31 | Calca | CALCR_RAMP2 | 0.001291 | 0 | CALCA_CALCR_RAMP2 | Calca - (Calcr+Ramp2) | CALC | Secreted Signaling | PMID: 30741933;PMID: 18599553 |
| CD31 | CD31 | Calca | CALCR_RAMP3 | 0.000613 | 0 | CALCA_CALCR_RAMP3 | Calca - (Calcr+Ramp3) | CALC | Secreted Signaling | PMID: 30741933;PMID: 18599553 |
| CD31 | GFAP | Calca | CALCR_RAMP3 | 0.000573 | 0 | CALCA_CALCR_RAMP3 | Calca - (Calcr+Ramp3) | CALC | Secreted Signaling | PMID: 30741933;PMID: 18599553 |
| GFAP | CD31 | Calcb | CALCR_RAMP1 | 0.001391 | 0 | CALCB_CALCR_RAMP1 | Calcb - (Calcr+Ramp1) | CALC | Secreted Signaling | PMID: 30741933;PMID: 18599553 |
| GFAP | GFAP | Calcb | CALCR_RAMP1 | 0.001234 | 0 | CALCB_CALCR_RAMP1 | Calcb - (Calcr+Ramp1) | CALC | Secreted Signaling | PMID: 30741933;PMID: 18599553 |
| CD31 | CD31 | Calcb | CALCR_RAMP2 | 0.00227 | 0 | CALCB_CALCR_RAMP2 | Calcb - (Calcr+Ramp2) | CALC | Secreted Signaling | PMID: 30741933;PMID: 18599553 |
| GFAP | CD31 | Calcb | CALCR_RAMP2 | 0.002722 | 0 | CALCB_CALCR_RAMP2 | Calcb - (Calcr+Ramp2) | CALC | Secreted Signaling | PMID: 30741933;PMID: 18599553 |
| GFAP | CD31 | Calcb | CALCR_RAMP3 | 0.000808 | 0 | CALCB_CALCR_RAMP3 | Calcb - (Calcr+Ramp3) | CALC | Secreted Signaling | PMID: 30741933;PMID: 18599553 |
| GFAP | GFAP | Calcb | CALCR_RAMP3 | 0.000755 | 0 | CALCB_CALCR_RAMP3 | Calcb - (Calcr+Ramp3) | CALC | Secreted Signaling | PMID: 30741933;PMID: 18599553 |
| GFAP | CD31 | Ctsg | Fpr1 | 0.00022 | 0 | CTSG_FPR1 | Ctsg - Fpr1 | CTSG | Secreted Signaling | PMID: 26516201 |
| GFAP | GFAP | Ctsg | Fpr1 | 0.000242 | 0 | CTSG_FPR1 | Ctsg - Fpr1 | CTSG | Secreted Signaling | PMID: 26516201 |
| GFAP | GFAP | Grp | Nmbr | 0.000419 | 0 | GRP_NMBR | Grp - Nmbr | GRP | Secreted Signaling | PMID:25209280;PMID:29133874 |
| NeuN | GFAP | Grp | Nmbr | 0.000903 | 0 | GRP_NMBR | Grp - Nmbr | GRP | Secreted Signaling | PMID:25209280;PMID:29133874 |
| GFAP | NeuN | Ptprs | Ntrk3 | 0.018379 | 0 | PTPRS_NTRK3 | Ptprs - Ntrk3 | PTPR | Secreted Signaling | PMID:21262467;PMID:25385546; |
| NeuN | NeuN | Ptprs | Ntrk3 | 0.041247 | 0 | PTPRS_NTRK3 | Ptprs - Ntrk3 | PTPR | Secreted Signaling | PMID:21262467;PMID:25385546; |
| NeuN | GFAP | Slitrk1 | Ptprd | 0.012221 | 0 | SLITRK1_PTPRD | Slitrk1 - Ptprd | SLITRK | Secreted Signaling | PMID: 30648269;PMID: 30822649 |
| NeuN | NeuN | Slitrk1 | Ptprd | 0.013103 | 0 | SLITRK1_PTPRD | Slitrk1 - Ptprd | SLITRK | Secreted Signaling | PMID: 30648269;PMID: 30822649 |
| CD31 | NeuN | Slitrk1 | Ptprs | 0.010024 | 0 | SLITRK1_PTPRS | Slitrk1 - Ptprs | SLITRK | Secreted Signaling | PMID: 30648269;PMID: 30822649 |
| NeuN | NeuN | Slitrk1 | Ptprs | 0.019385 | 0 | SLITRK1_PTPRS | Slitrk1 - Ptprs | SLITRK | Secreted Signaling | PMID: 30648269;PMID: 30822649 |
| GFAP | GFAP | Slitrk2 | Ptprd | 0.002757 | 0 | SLITRK2_PTPRD | Slitrk2 - Ptprd | SLITRK | Secreted Signaling | PMID: 30648269;PMID: 30822649 |
| CD31 | NeuN | Slitrk2 | Ptprd | 0.004415 | 0 | SLITRK2_PTPRD | Slitrk2 - Ptprd | SLITRK | Secreted Signaling | PMID: 30648269;PMID: 30822649 |
| CD31 | NeuN | Slitrk2 | Ptprs | 0.006559 | 0 | SLITRK2_PTPRS | Slitrk2 - Ptprs | SLITRK | Secreted Signaling | PMID: 30648269;PMID: 30822649 |
| NeuN | NeuN | Slitrk2 | Ptprs | 0.004028 | 0 | SLITRK2_PTPRS | Slitrk2 - Ptprs | SLITRK | Secreted Signaling | PMID: 30648269;PMID: 30822649 |
| NeuN | GFAP | Slitrk3 | Ptprd | 0.005641 | 0 | SLITRK3_PTPRD | Slitrk3 - Ptprd | SLITRK | Secreted Signaling | PMID: 30648269;PMID: 30822649 |
| NeuN | NeuN | Slitrk3 | Ptprd | 0.00605 | 0 | SLITRK3_PTPRD | Slitrk3 - Ptprd | SLITRK | Secreted Signaling | PMID: 30648269;PMID: 30822649 |
| GFAP | NeuN | Slitrk4 | Ptprs | 0.004533 | 0 | SLITRK4_PTPRS | Slitrk4 - Ptprs | SLITRK | Secreted Signaling | PMID: 30648269;PMID: 30822649 |
| NeuN | NeuN | Slitrk4 | Ptprs | 0.006402 | 0 | SLITRK4_PTPRS | Slitrk4 - Ptprs | SLITRK | Secreted Signaling | PMID: 30648269;PMID: 30822649 |
| NeuN | GFAP | Slitrk5 | Ptprs | 0.004725 | 0 | SLITRK5_PTPRS | Slitrk5 - Ptprs | SLITRK | Secreted Signaling | PMID: 30648269;PMID: 30822649 |
| NeuN | NeuN | Slitrk5 | Ptprs | 0.010792 | 0 | SLITRK5_PTPRS | Slitrk5 - Ptprs | SLITRK | Secreted Signaling | PMID: 30648269;PMID: 30822649 |
| NeuN | GFAP | Bdnf | Sort1 | 0.002896 | 0 | BDNF_SORT1 | Bdnf - Sort1 | NT | Secreted Signaling | uniprot |
| NeuN | NeuN | Bdnf | Sort1 | 0.00438 | 0 | BDNF_SORT1 | Bdnf - Sort1 | NT | Secreted Signaling | uniprot |
| GFAP | GFAP | Bdnf | Ngfr | 0.000333 | 0 | BDNF_NGFR | Bdnf - Ngfr | NT | Secreted Signaling | uniprot |
| NeuN | GFAP | Bdnf | Ngfr | 0.000681 | 0 | BDNF_NGFR | Bdnf - Ngfr | NT | Secreted Signaling | uniprot |
| CD31 | CD31 | Igfbp3 | Tmem219 | 0.002486 | 0 | IGFBP3_TMEM219 | Igfbp3 - Tmem219 | IGFBP | Secreted Signaling | PMID: 35115561;PMID:20353938 |
| CD31 | GFAP | Igfbp3 | Tmem219 | 0.003374 | 0 | IGFBP3_TMEM219 | Igfbp3 - Tmem219 | IGFBP | Secreted Signaling | PMID: 35115561;PMID:20353938 |
| CD31 | CD31 | Lgals9 | P4hb | 0.001722 | 0 | LGALS9_P4HB | Lgals9 - P4hb | GALECTIN | Secreted Signaling | PMID:21670307;uniprot |
| CD31 | GFAP | Lgals9 | P4hb | 0.001866 | 0 | LGALS9_P4HB | Lgals9 - P4hb | GALECTIN | Secreted Signaling | PMID:21670307;uniprot |
| GFAP | GFAP | Nts | Sort1 | 0.001401 | 0 | NTS_SORT1 | Nts - Sort1 | NTS | Secreted Signaling | PMID: 34038712 |

| | | | | | | | | | | |
|---|---|---|---|---|---|---|---|---|---|---|
| GFAP | NeuN | Nts | Sort1 | 0.002121 | 0 | NTS_SORT1 | Nts - Sort1 | NTS | Secreted Signaling | PMID: 34038712 |
| GFAP | CD31 | Ntf3 | Ngfr | 0.000343 | 0 | NTF3_NGFR | Ntf3 - Ngfr | NT | Secreted Signaling | uniprot |
| GFAP | GFAP | Ntf3 | Ngfr | 0.000496 | 0 | NTF3_NGFR | Ntf3 - Ngfr | NT | Secreted Signaling | uniprot |
| CD31 | CD31 | Rbp4 | Stra6 | 0.001472 | 0 | RBP4_STRA6 | Rbp4 - Stra6 | RBP4 | Secreted Signaling | PMID: 33199363 |
| NeuN | CD31 | Rbp4 | Stra6 | 0.002186 | 0 | RBP4_STRA6 | Rbp4 - Stra6 | RBP4 | Secreted Signaling | PMID: 33199363 |
| NeuN | GFAP | Slit1 | Robo1 | 0.001431 | 0 | SLIT1_ROBO1 | Slit1 - Robo1 | SLIT | Secreted Signaling | PMID:10102268 |
| NeuN | NeuN | Slit1 | Robo1 | 0.001574 | 0 | SLIT1_ROBO1 | Slit1 - Robo1 | SLIT | Secreted Signaling | PMID:10102268 |
| GFAP | NeuN | Slit2 | Robo1 | 0.000554 | 0 | SLIT2_ROBO1 | Slit2 - Robo1 | SLIT | Secreted Signaling | PMID:10102268 |
| NeuN | NeuN | Slit2 | Robo1 | 0.0006 | 0 | SLIT2_ROBO1 | Slit2 - Robo1 | SLIT | Secreted Signaling | PMID:10102268 |
| GFAP | NeuN | Slit2 | Robo2 | 0.001613 | 0 | SLIT2_ROBO2 | Slit2 - Robo2 | SLIT | Secreted Signaling | PMID:10102268 |
| NeuN | NeuN | Slit2 | Robo2 | 0.001747 | 0 | SLIT2_ROBO2 | Slit2 - Robo2 | SLIT | Secreted Signaling | PMID:10102268 |
| CD31 | CD31 | Tub | Mertk | 0.001554 | 0 | TUB_MERTK | Tub - Mertk | TULP | Secreted Signaling | PMID: 20978475 |
| NeuN | CD31 | Tub | Mertk | 0.005711 | 0 | TUB_MERTK | Tub - Mertk | TULP | Secreted Signaling | PMID: 20978475 |
| CD31 | CD31 | Pros1 | Mertk | 0.001747 | 0 | PROS1_MERTK | Pros1 - Mertk | PROS | Secreted Signaling | PMID: 34631419 |
| GFAP | CD31 | Pros1 | Mertk | 0.001684 | 0 | PROS1_MERTK | Pros1 - Mertk | PROS | Secreted Signaling | PMID: 34631419 |
| CD31 | CD31 | Col1a1 | ITGA1_ITGB1 | 0.008527 | 0 | COL1A1_ITGA1_ITGB1 | Col1a1 - (Itga1+Itgb1) | COLLAGEN | ECM-Receptor | KEGG: mmu04512 |
| CD31 | GFAP | Col1a1 | ITGA1_ITGB1 | 0.003719 | 0 | COL1A1_ITGA1_ITGB1 | Col1a1 - (Itga1+Itgb1) | COLLAGEN | ECM-Receptor | KEGG: mmu04512 |
| CD31 | CD31 | Col1a2 | ITGA1_ITGB1 | 0.005164 | 0 | COL1A2_ITGA1_ITGB1 | Col1a2 - (Itga1+Itgb1) | COLLAGEN | ECM-Receptor | KEGG: mmu04512 |
| CD31 | GFAP | Col1a2 | ITGA1_ITGB1 | 0.002248 | 0 | COL1A2_ITGA1_ITGB1 | Col1a2 - (Itga1+Itgb1) | COLLAGEN | ECM-Receptor | KEGG: mmu04512 |
| CD31 | CD31 | Col4a1 | ITGA1_ITGB1 | 0.012276 | 0 | COL4A1_ITGA1_ITGB1 | Col4a1 - (Itga1+Itgb1) | COLLAGEN | ECM-Receptor | KEGG: mmu04512 |
| CD31 | GFAP | Col4a1 | ITGA1_ITGB1 | 0.005366 | 0 | COL4A1_ITGA1_ITGB1 | Col4a1 - (Itga1+Itgb1) | COLLAGEN | ECM-Receptor | KEGG: mmu04512 |
| CD31 | CD31 | Col4a2 | ITGA1_ITGB1 | 0.009213 | 0 | COL4A2_ITGA1_ITGB1 | Col4a2 - (Itga1+Itgb1) | COLLAGEN | ECM-Receptor | KEGG: mmu04512 |
| CD31 | GFAP | Col4a2 | ITGA1_ITGB1 | 0.00402 | 0 | COL4A2_ITGA1_ITGB1 | Col4a2 - (Itga1+Itgb1) | COLLAGEN | ECM-Receptor | KEGG: mmu04512 |
| CD31 | CD31 | Col6a1 | ITGA1_ITGB1 | 0.002041 | 0 | COL6A1_ITGA1_ITGB1 | Col6a1 - (Itga1+Itgb1) | COLLAGEN | ECM-Receptor | KEGG: mmu04512 |
| NeuN | CD31 | Col6a1 | ITGA1_ITGB1 | 0.004122 | 0 | COL6A1_ITGA1_ITGB1 | Col6a1 - (Itga1+Itgb1) | COLLAGEN | ECM-Receptor | KEGG: mmu04512 |
| CD31 | CD31 | Col6a2 | ITGA1_ITGB1 | 0.001778 | 0 | COL6A2_ITGA1_ITGB1 | Col6a2 - (Itga1+Itgb1) | COLLAGEN | ECM-Receptor | KEGG: mmu04512 |
| GFAP | CD31 | Col6a2 | ITGA1_ITGB1 | 0.002417 | 0 | COL6A2_ITGA1_ITGB1 | Col6a2 - (Itga1+Itgb1) | COLLAGEN | ECM-Receptor | KEGG: mmu04512 |
| GFAP | CD31 | Col6a4 | ITGA1_ITGB1 | 0.001998 | 0 | COL6A4_ITGA1_ITGB1 | Col6a4 - (Itga1+Itgb1) | COLLAGEN | ECM-Receptor | KEGG: mmu04512 |
| GFAP | GFAP | Col6a4 | ITGA1_ITGB1 | 0.000868 | 0 | COL6A4_ITGA1_ITGB1 | Col6a4 - (Itga1+Itgb1) | COLLAGEN | ECM-Receptor | KEGG: mmu04512 |
| CD31 | CD31 | Col6a6 | ITGA1_ITGB1 | 0.00079 | 0 | COL6A6_ITGA1_ITGB1 | Col6a6 - (Itga1+Itgb1) | COLLAGEN | ECM-Receptor | KEGG: mmu04512 |
| GFAP | CD31 | Col6a6 | ITGA1_ITGB1 | 0.001141 | 0 | COL6A6_ITGA1_ITGB1 | Col6a6 - (Itga1+Itgb1) | COLLAGEN | ECM-Receptor | KEGG: mmu04512 |
| CD31 | CD31 | Col9a2 | ITGA1_ITGB1 | 0.001517 | 0 | COL9A2_ITGA1_ITGB1 | Col9a2 - (Itga1+Itgb1) | COLLAGEN | ECM-Receptor | KEGG: mmu04512 |
| GFAP | CD31 | Col9a2 | ITGA1_ITGB1 | 0.001441 | 0 | COL9A2_ITGA1_ITGB1 | Col9a2 - (Itga1+Itgb1) | COLLAGEN | ECM-Receptor | KEGG: mmu04512 |
| CD31 | CD31 | Col9a3 | ITGA1_ITGB1 | 0.001145 | 0 | COL9A3_ITGA1_ITGB1 | Col9a3 - (Itga1+Itgb1) | COLLAGEN | ECM-Receptor | KEGG: mmu04512 |
| GFAP | CD31 | Col9a3 | ITGA1_ITGB1 | 0.002372 | 0 | COL9A3_ITGA1_ITGB1 | Col9a3 - (Itga1+Itgb1) | COLLAGEN | ECM-Receptor | KEGG: mmu04512 |
| CD31 | CD31 | Fn1 | ITGA3_ITGB1 | 0.010361 | 0 | FN1_ITGA3_ITGB1 | Fn1 - (Itga3+Itgb1) | FN1 | ECM-Receptor | KEGG: mmu04512 |
| CD31 | GFAP | Fn1 | ITGA3_ITGB1 | 0.007223 | 0 | FN1_ITGA3_ITGB1 | Fn1 - (Itga3+Itgb1) | FN1 | ECM-Receptor | KEGG: mmu04512 |
| CD31 | CD31 | Fn1 | ITGA4_ITGB1 | 0.011415 | 0 | FN1_ITGA4_ITGB1 | Fn1 - (Itga4+Itgb1) | FN1 | ECM-Receptor | KEGG: mmu04512 |
| CD31 | GFAP | Fn1 | ITGA4_ITGB1 | 0.00565 | 0 | FN1_ITGA4_ITGB1 | Fn1 - (Itga4+Itgb1) | FN1 | ECM-Receptor | KEGG: mmu04512 |
| CD31 | CD31 | Fn1 | ITGA5_ITGB1 | 0.010725 | 0 | FN1_ITGA5_ITGB1 | Fn1 - (Itga5+Itgb1) | FN1 | ECM-Receptor | KEGG: mmu04512 |
| CD31 | GFAP | Fn1 | ITGA5_ITGB1 | 0.006966 | 0 | FN1_ITGA5_ITGB1 | Fn1 - (Itga5+Itgb1) | FN1 | ECM-Receptor | KEGG: mmu04512 |
| CD31 | CD31 | Fn1 | ITGA8_ITGB1 | 0.007609 | 0 | FN1_ITGA8_ITGB1 | Fn1 - (Itga8+Itgb1) | FN1 | ECM-Receptor | KEGG: mmu04512 |
| CD31 | GFAP | Fn1 | ITGA8_ITGB1 | 0.005172 | 0 | FN1_ITGA8_ITGB1 | Fn1 - (Itga8+Itgb1) | FN1 | ECM-Receptor | KEGG: mmu04512 |
| CD31 | CD31 | Lama1 | ITGA1_ITGB1 | 0.001408 | 0 | LAMA1_ITGA1_ITGB1 | Lama1 - (Itga1+Itgb1) | LAMININ | ECM-Receptor | KEGG: mmu04512 |
| GFAP | CD31 | Lama1 | ITGA1_ITGB1 | 0.00172 | 0 | LAMA1_ITGA1_ITGB1 | Lama1 - (Itga1+Itgb1) | LAMININ | ECM-Receptor | KEGG: mmu04512 |
| CD31 | CD31 | Lama4 | ITGA1_ITGB1 | 0.003613 | 0 | LAMA4_ITGA1_ITGB1 | Lama4 - (Itga1+Itgb1) | LAMININ | ECM-Receptor | KEGG: mmu04512 |
| GFAP | CD31 | Lama4 | ITGA1_ITGB1 | 0.002327 | 0 | LAMA4_ITGA1_ITGB1 | Lama4 - (Itga1+Itgb1) | LAMININ | ECM-Receptor | KEGG: mmu04512 |
| CD31 | CD31 | Lama5 | ITGA1_ITGB1 | 0.005778 | 0 | LAMA5_ITGA1_ITGB1 | Lama5 - (Itga1+Itgb1) | LAMININ | ECM-Receptor | KEGG: mmu04512 |
| CD31 | GFAP | Lama5 | ITGA1_ITGB1 | 0.002516 | 0 | LAMA5_ITGA1_ITGB1 | Lama5 - (Itga1+Itgb1) | LAMININ | ECM-Receptor | KEGG: mmu04512 |
| CD31 | CD31 | Lamb1 | ITGA1_ITGB1 | 0.001609 | 0 | LAMB1_ITGA1_ITGB1 | Lamb1 - (Itga1+Itgb1) | LAMININ | ECM-Receptor | KEGG: mmu04512 |
| GFAP | CD31 | Lamb1 | ITGA1_ITGB1 | 0.002214 | 0 | LAMB1_ITGA1_ITGB1 | Lamb1 - (Itga1+Itgb1) | LAMININ | ECM-Receptor | KEGG: mmu04512 |
| CD31 | CD31 | Lamb2 | ITGA1_ITGB1 | 0.008421 | 0 | LAMB2_ITGA1_ITGB1 | Lamb2 - (Itga1+Itgb1) | LAMININ | ECM-Receptor | KEGG: mmu04512 |
| CD31 | GFAP | Lamb2 | ITGA1_ITGB1 | 0.003673 | 0 | LAMB2_ITGA1_ITGB1 | Lamb2 - (Itga1+Itgb1) | LAMININ | ECM-Receptor | KEGG: mmu04512 |
| CD31 | CD31 | Lamc1 | ITGA1_ITGB1 | 0.005456 | 0 | LAMC1_ITGA1_ITGB1 | Lamc1 - (Itga1+Itgb1) | LAMININ | ECM-Receptor | KEGG: mmu04512 |
| CD31 | GFAP | Lamc1 | ITGA1_ITGB1 | 0.002376 | 0 | LAMC1_ITGA1_ITGB1 | Lamc1 - (Itga1+Itgb1) | LAMININ | ECM-Receptor | KEGG: mmu04512 |
| CD31 | CD31 | Lamc2 | ITGA1_ITGB1 | 0.001609 | 0 | LAMC2_ITGA1_ITGB1 | Lamc2 - (Itga1+Itgb1) | LAMININ | ECM-Receptor | KEGG: mmu04512 |
| NeuN | CD31 | Lamc2 | ITGA1_ITGB1 | 0.001146 | 0 | LAMC2_ITGA1_ITGB1 | Lamc2 - (Itga1+Itgb1) | LAMININ | ECM-Receptor | KEGG: mmu04512 |
| CD31 | CD31 | Lamc3 | ITGA1_ITGB1 | 0.004413 | 0 | LAMC3_ITGA1_ITGB1 | Lamc3 - (Itga1+Itgb1) | LAMININ | ECM-Receptor | KEGG: mmu04512 |
| GFAP | CD31 | Lamc3 | ITGA1_ITGB1 | 0.00255 | 0 | LAMC3_ITGA1_ITGB1 | Lamc3 - (Itga1+Itgb1) | LAMININ | ECM-Receptor | KEGG: mmu04512 |

| | | | | | | | | | | |
|---|---|---|---|---|---|---|---|---|---|---|
| CD31 | CD31 | Lama1 | ITGA3_ITGB1 | 0.001008 | 0 | LAMA1_ITGA3_ITGB1 | Lama1 - (Itga3+Itgb1) | LAMININ | ECM-Receptor | KEGG: mmu04512 |
| GFAP | CD31 | Lama1 | ITGA3_ITGB1 | 0.001231 | 0 | LAMA1_ITGA3_ITGB1 | Lama1 - (Itga3+Itgb1) | LAMININ | ECM-Receptor | KEGG: mmu04512 |
| CD31 | CD31 | Lama4 | ITGA3_ITGB1 | 0.002588 | 0 | LAMA4_ITGA3_ITGB1 | Lama4 - (Itga3+Itgb1) | LAMININ | ECM-Receptor | KEGG: mmu04512 |
| CD31 | GFAP | Lama4 | ITGA3_ITGB1 | 0.0018 | 0 | LAMA4_ITGA3_ITGB1 | Lama4 - (Itga3+Itgb1) | LAMININ | ECM-Receptor | KEGG: mmu04512 |
| CD31 | CD31 | Lama5 | ITGA3_ITGB1 | 0.004142 | 0 | LAMA5_ITGA3_ITGB1 | Lama5 - (Itga3+Itgb1) | LAMININ | ECM-Receptor | KEGG: mmu04512 |
| CD31 | GFAP | Lama5 | ITGA3_ITGB1 | 0.002882 | 0 | LAMA5_ITGA3_ITGB1 | Lama5 - (Itga3+Itgb1) | LAMININ | ECM-Receptor | KEGG: mmu04512 |
| CD31 | CD31 | Lamb1 | ITGA3_ITGB1 | 0.001152 | 0 | LAMB1_ITGA3_ITGB1 | Lamb1 - (Itga3+Itgb1) | LAMININ | ECM-Receptor | KEGG: mmu04512 |
| GFAP | CD31 | Lamb1 | ITGA3_ITGB1 | 0.001586 | 0 | LAMB1_ITGA3_ITGB1 | Lamb1 - (Itga3+Itgb1) | LAMININ | ECM-Receptor | KEGG: mmu04512 |
| CD31 | CD31 | Lamb2 | ITGA3_ITGB1 | 0.006041 | 0 | LAMB2_ITGA3_ITGB1 | Lamb2 - (Itga3+Itgb1) | LAMININ | ECM-Receptor | KEGG: mmu04512 |
| CD31 | GFAP | Lamb2 | ITGA3_ITGB1 | 0.004206 | 0 | LAMB2_ITGA3_ITGB1 | Lamb2 - (Itga3+Itgb1) | LAMININ | ECM-Receptor | KEGG: mmu04512 |
| CD31 | CD31 | Lamc1 | ITGA3_ITGB1 | 0.003911 | 0 | LAMC1_ITGA3_ITGB1 | Lamc1 - (Itga3+Itgb1) | LAMININ | ECM-Receptor | KEGG: mmu04512 |
| CD31 | GFAP | Lamc1 | ITGA3_ITGB1 | 0.002721 | 0 | LAMC1_ITGA3_ITGB1 | Lamc1 - (Itga3+Itgb1) | LAMININ | ECM-Receptor | KEGG: mmu04512 |
| CD31 | CD31 | Lamc2 | ITGA3_ITGB1 | 0.001152 | 0 | LAMC2_ITGA3_ITGB1 | Lamc2 - (Itga3+Itgb1) | LAMININ | ECM-Receptor | KEGG: mmu04512 |
| NeuN | CD31 | Lamc2 | ITGA3_ITGB1 | 0.00082 | 0 | LAMC2_ITGA3_ITGB1 | Lamc2 - (Itga3+Itgb1) | LAMININ | ECM-Receptor | KEGG: mmu04512 |
| CD31 | CD31 | Lamc3 | ITGA3_ITGB1 | 0.003162 | 0 | LAMC3_ITGA3_ITGB1 | Lamc3 - (Itga3+Itgb1) | LAMININ | ECM-Receptor | KEGG: mmu04512 |
| CD31 | GFAP | Lamc3 | ITGA3_ITGB1 | 0.002199 | 0 | LAMC3_ITGA3_ITGB1 | Lamc3 - (Itga3+Itgb1) | LAMININ | ECM-Receptor | KEGG: mmu04512 |
| NeuN | CD31 | Reln | ITGA3_ITGB1 | 0.001791 | 0 | RELN_ITGA3_ITGB1 | Reln - (Itga3+Itgb1) | RELN | ECM-Receptor | KEGG: mmu04512 |
| GFAP | GFAP | Reln | ITGA3_ITGB1 | 0.000756 | 0 | RELN_ITGA3_ITGB1 | Reln - (Itga3+Itgb1) | RELN | ECM-Receptor | KEGG: mmu04512 |
| GFAP | CD31 | Thbs1 | ITGA3_ITGB1 | 0.001488 | 0 | THBS1_ITGA3_ITGB1 | Thbs1 - (Itga3+Itgb1) | THBS | ECM-Receptor | KEGG: mmu04512 |
| GFAP | GFAP | Thbs1 | ITGA3_ITGB1 | 0.001034 | 0 | THBS1_ITGA3_ITGB1 | Thbs1 - (Itga3+Itgb1) | THBS | ECM-Receptor | KEGG: mmu04512 |
| CD31 | CD31 | Thbs3 | ITGA3_ITGB1 | 0.001197 | 0 | THBS3_ITGA3_ITGB1 | Thbs3 - (Itga3+Itgb1) | THBS | ECM-Receptor | KEGG: mmu04512 |
| GFAP | CD31 | Thbs3 | ITGA3_ITGB1 | 0.000981 | 0 | THBS3_ITGA3_ITGB1 | Thbs3 - (Itga3+Itgb1) | THBS | ECM-Receptor | KEGG: mmu04512 |
| GFAP | CD31 | Thbs4 | ITGA3_ITGB1 | 0.014734 | 0 | THBS4_ITGA3_ITGB1 | Thbs4 - (Itga3+Itgb1) | THBS | ECM-Receptor | KEGG: mmu04512 |
| GFAP | GFAP | Thbs4 | ITGA3_ITGB1 | 0.010285 | 0 | THBS4_ITGA3_ITGB1 | Thbs4 - (Itga3+Itgb1) | THBS | ECM-Receptor | KEGG: mmu04512 |
| CD31 | CD31 | Col1a1 | ITGA3_ITGB1 | 0.006118 | 0 | COL1A1_ITGA3_ITGB1 | Col1a1 - (Itga3+Itgb1) | COLLAGEN | ECM-Receptor | KEGG: mmu04512 |
| CD31 | GFAP | Col1a1 | ITGA3_ITGB1 | 0.004259 | 0 | COL1A1_ITGA3_ITGB1 | Col1a1 - (Itga3+Itgb1) | COLLAGEN | ECM-Receptor | KEGG: mmu04512 |
| CD31 | CD31 | Col1a2 | ITGA3_ITGB1 | 0.003701 | 0 | COL1A2_ITGA3_ITGB1 | Col1a2 - (Itga3+Itgb1) | COLLAGEN | ECM-Receptor | KEGG: mmu04512 |
| CD31 | GFAP | Col1a2 | ITGA3_ITGB1 | 0.002575 | 0 | COL1A2_ITGA3_ITGB1 | Col1a2 - (Itga3+Itgb1) | COLLAGEN | ECM-Receptor | KEGG: mmu04512 |
| CD31 | CD31 | Col4a1 | ITGA3_ITGB1 | 0.008816 | 0 | COL4A1_ITGA3_ITGB1 | Col4a1 - (Itga3+Itgb1) | COLLAGEN | ECM-Receptor | KEGG: mmu04512 |
| CD31 | GFAP | Col4a1 | ITGA3_ITGB1 | 0.006143 | 0 | COL4A1_ITGA3_ITGB1 | Col4a1 - (Itga3+Itgb1) | COLLAGEN | ECM-Receptor | KEGG: mmu04512 |
| CD31 | CD31 | Col4a2 | ITGA3_ITGB1 | 0.006611 | 0 | COL4A2_ITGA3_ITGB1 | Col4a2 - (Itga3+Itgb1) | COLLAGEN | ECM-Receptor | KEGG: mmu04512 |
| CD31 | GFAP | Col4a2 | ITGA3_ITGB1 | 0.004603 | 0 | COL4A2_ITGA3_ITGB1 | Col4a2 - (Itga3+Itgb1) | COLLAGEN | ECM-Receptor | KEGG: mmu04512 |
| CD31 | CD31 | Col6a1 | ITGA3_ITGB1 | 0.001462 | 0 | COL6A1_ITGA3_ITGB1 | Col6a1 - (Itga3+Itgb1) | COLLAGEN | ECM-Receptor | KEGG: mmu04512 |
| NeuN | CD31 | Col6a1 | ITGA3_ITGB1 | 0.002954 | 0 | COL6A1_ITGA3_ITGB1 | Col6a1 - (Itga3+Itgb1) | COLLAGEN | ECM-Receptor | KEGG: mmu04512 |
| CD31 | CD31 | Col6a2 | ITGA3_ITGB1 | 0.001273 | 0 | COL6A2_ITGA3_ITGB1 | Col6a2 - (Itga3+Itgb1) | COLLAGEN | ECM-Receptor | KEGG: mmu04512 |
| GFAP | CD31 | Col6a2 | ITGA3_ITGB1 | 0.001731 | 0 | COL6A2_ITGA3_ITGB1 | Col6a2 - (Itga3+Itgb1) | COLLAGEN | ECM-Receptor | KEGG: mmu04512 |
| GFAP | CD31 | Col6a4 | ITGA3_ITGB1 | 0.001431 | 0 | COL6A4_ITGA3_ITGB1 | Col6a4 - (Itga3+Itgb1) | COLLAGEN | ECM-Receptor | KEGG: mmu04512 |
| GFAP | GFAP | Col6a4 | ITGA3_ITGB1 | 0.000995 | 0 | COL6A4_ITGA3_ITGB1 | Col6a4 - (Itga3+Itgb1) | COLLAGEN | ECM-Receptor | KEGG: mmu04512 |
| GFAP | CD31 | Col6a6 | ITGA3_ITGB1 | 0.000817 | 0 | COL6A6_ITGA3_ITGB1 | Col6a6 - (Itga3+Itgb1) | COLLAGEN | ECM-Receptor | KEGG: mmu04512 |
| GFAP | GFAP | Col6a6 | ITGA3_ITGB1 | 0.000568 | 0 | COL6A6_ITGA3_ITGB1 | Col6a6 - (Itga3+Itgb1) | COLLAGEN | ECM-Receptor | KEGG: mmu04512 |
| CD31 | CD31 | Col9a2 | ITGA3_ITGB1 | 0.001086 | 0 | COL9A2_ITGA3_ITGB1 | Col9a2 - (Itga3+Itgb1) | COLLAGEN | ECM-Receptor | KEGG: mmu04512 |
| GFAP | CD31 | Col9a2 | ITGA3_ITGB1 | 0.001032 | 0 | COL9A2_ITGA3_ITGB1 | Col9a2 - (Itga3+Itgb1) | COLLAGEN | ECM-Receptor | KEGG: mmu04512 |
| GFAP | CD31 | Col9a3 | ITGA3_ITGB1 | 0.001698 | 0 | COL9A3_ITGA3_ITGB1 | Col9a3 - (Itga3+Itgb1) | COLLAGEN | ECM-Receptor | KEGG: mmu04512 |
| GFAP | GFAP | Col9a3 | ITGA3_ITGB1 | 0.001181 | 0 | COL9A3_ITGA3_ITGB1 | Col9a3 - (Itga3+Itgb1) | COLLAGEN | ECM-Receptor | KEGG: mmu04512 |
| CD31 | CD31 | Lama1 | ITGA6_ITGB1 | 0.001505 | 0 | LAMA1_ITGA6_ITGB1 | Lama1 - (Itga6+Itgb1) | LAMININ | ECM-Receptor | KEGG: mmu04512 |
| GFAP | CD31 | Lama1 | ITGA6_ITGB1 | 0.001839 | 0 | LAMA1_ITGA6_ITGB1 | Lama1 - (Itga6+Itgb1) | LAMININ | ECM-Receptor | KEGG: mmu04512 |
| CD31 | CD31 | Lama4 | ITGA6_ITGB1 | 0.003863 | 0 | LAMA4_ITGA6_ITGB1 | Lama4 - (Itga6+Itgb1) | LAMININ | ECM-Receptor | KEGG: mmu04512 |
| GFAP | CD31 | Lama4 | ITGA6_ITGB1 | 0.002487 | 0 | LAMA4_ITGA6_ITGB1 | Lama4 - (Itga6+Itgb1) | LAMININ | ECM-Receptor | KEGG: mmu04512 |
| CD31 | CD31 | Lama5 | ITGA6_ITGB1 | 0.006176 | 0 | LAMA5_ITGA6_ITGB1 | Lama5 - (Itga6+Itgb1) | LAMININ | ECM-Receptor | KEGG: mmu04512 |
| CD31 | GFAP | Lama5 | ITGA6_ITGB1 | 0.002897 | 0 | LAMA5_ITGA6_ITGB1 | Lama5 - (Itga6+Itgb1) | LAMININ | ECM-Receptor | KEGG: mmu04512 |
| CD31 | CD31 | Lamb1 | ITGA6_ITGB1 | 0.00172 | 0 | LAMB1_ITGA6_ITGB1 | Lamb1 - (Itga6+Itgb1) | LAMININ | ECM-Receptor | KEGG: mmu04512 |
| GFAP | CD31 | Lamb1 | ITGA6_ITGB1 | 0.002367 | 0 | LAMB1_ITGA6_ITGB1 | Lamb1 - (Itga6+Itgb1) | LAMININ | ECM-Receptor | KEGG: mmu04512 |
| CD31 | CD31 | Lamb2 | ITGA6_ITGB1 | 0.008999 | 0 | LAMB2_ITGA6_ITGB1 | Lamb2 - (Itga6+Itgb1) | LAMININ | ECM-Receptor | KEGG: mmu04512 |
| CD31 | GFAP | Lamb2 | ITGA6_ITGB1 | 0.004227 | 0 | LAMB2_ITGA6_ITGB1 | Lamb2 - (Itga6+Itgb1) | LAMININ | ECM-Receptor | KEGG: mmu04512 |
| CD31 | CD31 | Lamc1 | ITGA6_ITGB1 | 0.005832 | 0 | LAMC1_ITGA6_ITGB1 | Lamc1 - (Itga6+Itgb1) | LAMININ | ECM-Receptor | KEGG: mmu04512 |
| CD31 | GFAP | Lamc1 | ITGA6_ITGB1 | 0.002735 | 0 | LAMC1_ITGA6_ITGB1 | Lamc1 - (Itga6+Itgb1) | LAMININ | ECM-Receptor | KEGG: mmu04512 |
| CD31 | CD31 | Lamc2 | ITGA6_ITGB1 | 0.00172 | 0 | LAMC2_ITGA6_ITGB1 | Lamc2 - (Itga6+Itgb1) | LAMININ | ECM-Receptor | KEGG: mmu04512 |
| NeuN | CD31 | Lamc2 | ITGA6_ITGB1 | 0.001225 | 0 | LAMC2_ITGA6_ITGB1 | Lamc2 - (Itga6+Itgb1) | LAMININ | ECM-Receptor | KEGG: mmu04512 |
| CD31 | CD31 | Lamc3 | ITGA6_ITGB1 | 0.004717 | 0 | LAMC3_ITGA6_ITGB1 | Lamc3 - (Itga6+Itgb1) | LAMININ | ECM-Receptor | KEGG: mmu04512 |

| | | | | | | | | | | |
|---|---|---|---|---|---|---|---|---|---|---|
| GFAP | CD31 | Lamc3 | ITGA6_ITGB1 | 0.002726 | 0 | LAMC3_ITGA6_ITGB1 | Lamc3 - (Itga6+Itgb1) | LAMININ | ECM-Receptor | KEGG: mmu04512 |
| CD31 | CD31 | Lama1 | ITGA7_ITGB1 | 0.00102 | 0 | LAMA1_ITGA7_ITGB1 | Lama1 - (Itga7+Itgb1) | LAMININ | ECM-Receptor | KEGG: mmu04512 |
| GFAP | CD31 | Lama1 | ITGA7_ITGB1 | 0.001246 | 0 | LAMA1_ITGA7_ITGB1 | Lama1 - (Itga7+Itgb1) | LAMININ | ECM-Receptor | KEGG: mmu04512 |
| CD31 | CD31 | Lama4 | ITGA7_ITGB1 | 0.00262 | 0 | LAMA4_ITGA7_ITGB1 | Lama4 - (Itga7+Itgb1) | LAMININ | ECM-Receptor | KEGG: mmu04512 |
| GFAP | CD31 | Lama4 | ITGA7_ITGB1 | 0.001686 | 0 | LAMA4_ITGA7_ITGB1 | Lama4 - (Itga7+Itgb1) | LAMININ | ECM-Receptor | KEGG: mmu04512 |
| CD31 | CD31 | Lama5 | ITGA7_ITGB1 | 0.004192 | 0 | LAMA5_ITGA7_ITGB1 | Lama5 - (Itga7+Itgb1) | LAMININ | ECM-Receptor | KEGG: mmu04512 |
| CD31 | GFAP | Lama5 | ITGA7_ITGB1 | 0.002617 | 0 | LAMA5_ITGA7_ITGB1 | Lama5 - (Itga7+Itgb1) | LAMININ | ECM-Receptor | KEGG: mmu04512 |
| CD31 | CD31 | Lamb1 | ITGA7_ITGB1 | 0.001166 | 0 | LAMB1_ITGA7_ITGB1 | Lamb1 - (Itga7+Itgb1) | LAMININ | ECM-Receptor | KEGG: mmu04512 |
| GFAP | CD31 | Lamb1 | ITGA7_ITGB1 | 0.001605 | 0 | LAMB1_ITGA7_ITGB1 | Lamb1 - (Itga7+Itgb1) | LAMININ | ECM-Receptor | KEGG: mmu04512 |
| CD31 | CD31 | Lamb2 | ITGA7_ITGB1 | 0.006114 | 0 | LAMB2_ITGA7_ITGB1 | Lamb2 - (Itga7+Itgb1) | LAMININ | ECM-Receptor | KEGG: mmu04512 |
| CD31 | GFAP | Lamb2 | ITGA7_ITGB1 | 0.00382 | 0 | LAMB2_ITGA7_ITGB1 | Lamb2 - (Itga7+Itgb1) | LAMININ | ECM-Receptor | KEGG: mmu04512 |
| CD31 | CD31 | Lamc1 | ITGA7_ITGB1 | 0.003958 | 0 | LAMC1_ITGA7_ITGB1 | Lamc1 - (Itga7+Itgb1) | LAMININ | ECM-Receptor | KEGG: mmu04512 |
| CD31 | GFAP | Lamc1 | ITGA7_ITGB1 | 0.002471 | 0 | LAMC1_ITGA7_ITGB1 | Lamc1 - (Itga7+Itgb1) | LAMININ | ECM-Receptor | KEGG: mmu04512 |
| CD31 | CD31 | Lamc2 | ITGA7_ITGB1 | 0.001166 | 0 | LAMC2_ITGA7_ITGB1 | Lamc2 - (Itga7+Itgb1) | LAMININ | ECM-Receptor | KEGG: mmu04512 |
| NeuN | CD31 | Lamc2 | ITGA7_ITGB1 | 0.00083 | 0 | LAMC2_ITGA7_ITGB1 | Lamc2 - (Itga7+Itgb1) | LAMININ | ECM-Receptor | KEGG: mmu04512 |
| CD31 | CD31 | Lamc3 | ITGA7_ITGB1 | 0.0032 | 0 | LAMC3_ITGA7_ITGB1 | Lamc3 - (Itga7+Itgb1) | LAMININ | ECM-Receptor | KEGG: mmu04512 |
| CD31 | GFAP | Lamc3 | ITGA7_ITGB1 | 0.001997 | 0 | LAMC3_ITGA7_ITGB1 | Lamc3 - (Itga7+Itgb1) | LAMININ | ECM-Receptor | KEGG: mmu04512 |
| CD31 | CD31 | Vtn | ITGA8_ITGB1 | 0.003932 | 0 | VTN_ITGA8_ITGB1 | Vtn - (Itga8+Itgb1) | VTN | ECM-Receptor | KEGG: mmu04512 |
| CD31 | GFAP | Vtn | ITGA8_ITGB1 | 0.00267 | 0 | VTN_ITGA8_ITGB1 | Vtn - (Itga8+Itgb1) | VTN | ECM-Receptor | KEGG: mmu04512 |
| GFAP | CD31 | Tnc | ITGA8_ITGB1 | 0.00106 | 0 | TNC_ITGA8_ITGB1 | Tnc - (Itga8+Itgb1) | TENASCIN | ECM-Receptor | KEGG: mmu04512 |
| GFAP | GFAP | Tnc | ITGA8_ITGB1 | 0.000719 | 0 | TNC_ITGA8_ITGB1 | Tnc - (Itga8+Itgb1) | TENASCIN | ECM-Receptor | KEGG: mmu04512 |
| GFAP | CD31 | Tnr | ITGA8_ITGB1 | 0.001366 | 0 | TNR_ITGA8_ITGB1 | Tnr - (Itga8+Itgb1) | TENASCIN | ECM-Receptor | KEGG: mmu04512 |
| GFAP | GFAP | Tnr | ITGA8_ITGB1 | 0.000927 | 0 | TNR_ITGA8_ITGB1 | Tnr - (Itga8+Itgb1) | TENASCIN | ECM-Receptor | KEGG: mmu04512 |
| CD31 | CD31 | Col1a1 | ITGA11_ITGB1 | 0.005689 | 0 | COL1A1_ITGA11_ITGB1 | Col1a1 - (Itga11+Itgb1) | COLLAGEN | ECM-Receptor | KEGG: mmu04512 |
| CD31 | GFAP | Col1a1 | ITGA11_ITGB1 | 0.003845 | 0 | COL1A1_ITGA11_ITGB1 | Col1a1 - (Itga11+Itgb1) | COLLAGEN | ECM-Receptor | KEGG: mmu04512 |
| CD31 | CD31 | Col1a2 | ITGA11_ITGB1 | 0.003442 | 0 | COL1A2_ITGA11_ITGB1 | Col1a2 - (Itga11+Itgb1) | COLLAGEN | ECM-Receptor | KEGG: mmu04512 |
| CD31 | GFAP | Col1a2 | ITGA11_ITGB1 | 0.002325 | 0 | COL1A2_ITGA11_ITGB1 | Col1a2 - (Itga11+Itgb1) | COLLAGEN | ECM-Receptor | KEGG: mmu04512 |
| CD31 | CD31 | Col4a1 | ITGA11_ITGB1 | 0.0082 | 0 | COL4A1_ITGA11_ITGB1 | Col4a1 - (Itga11+Itgb1) | COLLAGEN | ECM-Receptor | KEGG: mmu04512 |
| CD31 | GFAP | Col4a1 | ITGA11_ITGB1 | 0.005547 | 0 | COL4A1_ITGA11_ITGB1 | Col4a1 - (Itga11+Itgb1) | COLLAGEN | ECM-Receptor | KEGG: mmu04512 |
| CD31 | CD31 | Col4a2 | ITGA11_ITGB1 | 0.006148 | 0 | COL4A2_ITGA11_ITGB1 | Col4a2 - (Itga11+Itgb1) | COLLAGEN | ECM-Receptor | KEGG: mmu04512 |
| CD31 | GFAP | Col4a2 | ITGA11_ITGB1 | 0.004156 | 0 | COL4A2_ITGA11_ITGB1 | Col4a2 - (Itga11+Itgb1) | COLLAGEN | ECM-Receptor | KEGG: mmu04512 |
| CD31 | CD31 | Col6a1 | ITGA11_ITGB1 | 0.001359 | 0 | COL6A1_ITGA11_ITGB1 | Col6a1 - (Itga11+Itgb1) | COLLAGEN | ECM-Receptor | KEGG: mmu04512 |
| NeuN | CD31 | Col6a1 | ITGA11_ITGB1 | 0.002746 | 0 | COL6A1_ITGA11_ITGB1 | Col6a1 - (Itga11+Itgb1) | COLLAGEN | ECM-Receptor | KEGG: mmu04512 |
| CD31 | CD31 | Col6a2 | ITGA11_ITGB1 | 0.001184 | 0 | COL6A2_ITGA11_ITGB1 | Col6a2 - (Itga11+Itgb1) | COLLAGEN | ECM-Receptor | KEGG: mmu04512 |
| GFAP | CD31 | Col6a2 | ITGA11_ITGB1 | 0.001609 | 0 | COL6A2_ITGA11_ITGB1 | Col6a2 - (Itga11+Itgb1) | COLLAGEN | ECM-Receptor | KEGG: mmu04512 |
| GFAP | CD31 | Col6a4 | ITGA11_ITGB1 | 0.00133 | 0 | COL6A4_ITGA11_ITGB1 | Col6a4 - (Itga11+Itgb1) | COLLAGEN | ECM-Receptor | KEGG: mmu04512 |
| GFAP | GFAP | Col6a4 | ITGA11_ITGB1 | 0.000898 | 0 | COL6A4_ITGA11_ITGB1 | Col6a4 - (Itga11+Itgb1) | COLLAGEN | ECM-Receptor | KEGG: mmu04512 |
| CD31 | CD31 | Col6a6 | ITGA11_ITGB1 | 0.000526 | 0 | COL6A6_ITGA11_ITGB1 | Col6a6 - (Itga11+Itgb1) | COLLAGEN | ECM-Receptor | KEGG: mmu04512 |
| GFAP | CD31 | Col6a6 | ITGA11_ITGB1 | 0.000759 | 0 | COL6A6_ITGA11_ITGB1 | Col6a6 - (Itga11+Itgb1) | COLLAGEN | ECM-Receptor | KEGG: mmu04512 |
| CD31 | CD31 | Col9a2 | ITGA11_ITGB1 | 0.00101 | 0 | COL9A2_ITGA11_ITGB1 | Col9a2 - (Itga11+Itgb1) | COLLAGEN | ECM-Receptor | KEGG: mmu04512 |
| GFAP | CD31 | Col9a2 | ITGA11_ITGB1 | 0.000959 | 0 | COL9A2_ITGA11_ITGB1 | Col9a2 - (Itga11+Itgb1) | COLLAGEN | ECM-Receptor | KEGG: mmu04512 |
| GFAP | CD31 | Col9a3 | ITGA11_ITGB1 | 0.001579 | 0 | COL9A3_ITGA11_ITGB1 | Col9a3 - (Itga11+Itgb1) | COLLAGEN | ECM-Receptor | KEGG: mmu04512 |
| GFAP | GFAP | Col9a3 | ITGA11_ITGB1 | 0.001066 | 0 | COL9A3_ITGA11_ITGB1 | Col9a3 - (Itga11+Itgb1) | COLLAGEN | ECM-Receptor | KEGG: mmu04512 |
| CD31 | CD31 | Fn1 | ITGAV_ITGB1 | 0.011816 | 0 | FN1_ITGAV_ITGB1 | Fn1 - (Itgav+Itgb1) | FN1 | ECM-Receptor | KEGG: mmu04512 |
| CD31 | GFAP | Fn1 | ITGAV_ITGB1 | 0.00911 | 0 | FN1_ITGAV_ITGB1 | Fn1 - (Itgav+Itgb1) | FN1 | ECM-Receptor | KEGG: mmu04512 |
| CD31 | CD31 | Vtn | ITGAV_ITGB1 | 0.006118 | 0 | VTN_ITGAV_ITGB1 | Vtn - (Itgav+Itgb1) | VTN | ECM-Receptor | KEGG: mmu04512 |
| CD31 | GFAP | Vtn | ITGAV_ITGB1 | 0.004711 | 0 | VTN_ITGAV_ITGB1 | Vtn - (Itgav+Itgb1) | VTN | ECM-Receptor | KEGG: mmu04512 |
| CD31 | CD31 | Fn1 | ITGA4_ITGB7 | 0.004056 | 0 | FN1_ITGA4_ITGB7 | Fn1 - (Itga4+Itgb7) | FN1 | ECM-Receptor | KEGG: mmu04512 |
| CD31 | GFAP | Fn1 | ITGA4_ITGB7 | 0.003355 | 0 | FN1_ITGA4_ITGB7 | Fn1 - (Itga4+Itgb7) | FN1 | ECM-Receptor | KEGG: mmu04512 |
| CD31 | CD31 | Fn1 | ITGAV_ITGB3 | 0.003598 | 0 | FN1_ITGAV_ITGB3 | Fn1 - (Itgav+Itgb3) | FN1 | ECM-Receptor | KEGG: mmu04512 |
| CD31 | GFAP | Fn1 | ITGAV_ITGB3 | 0.005153 | 0 | FN1_ITGAV_ITGB3 | Fn1 - (Itgav+Itgb3) | FN1 | ECM-Receptor | KEGG: mmu04512 |
| CD31 | CD31 | Vtn | ITGAV_ITGB3 | 0.001856 | 0 | VTN_ITGAV_ITGB3 | Vtn - (Itgav+Itgb3) | VTN | ECM-Receptor | KEGG: mmu04512 |
| CD31 | GFAP | Vtn | ITGAV_ITGB3 | 0.00266 | 0 | VTN_ITGAV_ITGB3 | Vtn - (Itgav+Itgb3) | VTN | ECM-Receptor | KEGG: mmu04512 |
| CD31 | CD31 | Vwf | ITGAV_ITGB3 | 0.000933 | 0 | VWF_ITGAV_ITGB3 | Vwf - (Itgav+Itgb3) | VWF | ECM-Receptor | KEGG: mmu04512 |
| CD31 | GFAP | Vwf | ITGAV_ITGB3 | 0.001337 | 0 | VWF_ITGAV_ITGB3 | Vwf - (Itgav+Itgb3) | VWF | ECM-Receptor | KEGG: mmu04512 |
| GFAP | GFAP | Tnc | ITGAV_ITGB3 | 0.000716 | 0 | TNC_ITGAV_ITGB3 | Tnc - (Itgav+Itgb3) | TENASCIN | ECM-Receptor | KEGG: mmu04512 |
| GFAP | NeuN | Tnc | ITGAV_ITGB3 | 0.000586 | 0 | TNC_ITGAV_ITGB3 | Tnc - (Itgav+Itgb3) | TENASCIN | ECM-Receptor | KEGG: mmu04512 |
| GFAP | GFAP | Tnr | ITGAV_ITGB3 | 0.000923 | 0 | TNR_ITGAV_ITGB3 | Tnr - (Itgav+Itgb3) | TENASCIN | ECM-Receptor | KEGG: mmu04512 |
| GFAP | NeuN | Tnr | ITGAV_ITGB3 | 0.000755 | 0 | TNR_ITGAV_ITGB3 | Tnr - (Itgav+Itgb3) | TENASCIN | ECM-Receptor | KEGG: mmu04512 |

| | | | | | | | | | | |
|---|---|---|---|---|---|---|---|---|---|---|
| GFAP | GFAP | Thbs1 | ITGAV_ITGB3 | 0.000737 | 0 | THBS1_ITGAV_ITGB3 | Thbs1 - (Itgav+Itgb3) | THBS | ECM-Receptor | KEGG: mmu04512 |
| GFAP | NeuN | Thbs1 | ITGAV_ITGB3 | 0.000603 | 0 | THBS1_ITGAV_ITGB3 | Thbs1 - (Itgav+Itgb3) | THBS | ECM-Receptor | KEGG: mmu04512 |
| CD31 | GFAP | Thbs3 | ITGAV_ITGB3 | 0.000592 | 0 | THBS3_ITGAV_ITGB3 | Thbs3 - (Itgav+Itgb3) | THBS | ECM-Receptor | KEGG: mmu04512 |
| GFAP | GFAP | Thbs3 | ITGAV_ITGB3 | 0.000486 | 0 | THBS3_ITGAV_ITGB3 | Thbs3 - (Itgav+Itgb3) | THBS | ECM-Receptor | KEGG: mmu04512 |
| GFAP | GFAP | Thbs4 | ITGAV_ITGB3 | 0.007344 | 0 | THBS4_ITGAV_ITGB3 | Thbs4 - (Itgav+Itgb3) | THBS | ECM-Receptor | KEGG: mmu04512 |
| GFAP | NeuN | Thbs4 | ITGAV_ITGB3 | 0.006017 | 0 | THBS4_ITGAV_ITGB3 | Thbs4 - (Itgav+Itgb3) | THBS | ECM-Receptor | KEGG: mmu04512 |
| CD31 | CD31 | Fn1 | ITGA2B_ITGB3 | 0.002427 | 0 | FN1_ITGA2B_ITGB3 | Fn1 - (Itga2b+Itgb3) | FN1 | ECM-Receptor | KEGG: mmu04512 |
| CD31 | GFAP | Fn1 | ITGA2B_ITGB3 | 0.003087 | 0 | FN1_ITGA2B_ITGB3 | Fn1 - (Itga2b+Itgb3) | FN1 | ECM-Receptor | KEGG: mmu04512 |
| CD31 | CD31 | Vtn | ITGA2B_ITGB3 | 0.001251 | 0 | VTN_ITGA2B_ITGB3 | Vtn - (Itga2b+Itgb3) | VTN | ECM-Receptor | KEGG: mmu04512 |
| CD31 | GFAP | Vtn | ITGA2B_ITGB3 | 0.001592 | 0 | VTN_ITGA2B_ITGB3 | Vtn - (Itga2b+Itgb3) | VTN | ECM-Receptor | KEGG: mmu04512 |
| CD31 | CD31 | Vwf | ITGA2B_ITGB3 | 0.000628 | 0 | VWF_ITGA2B_ITGB3 | Vwf - (Itga2b+Itgb3) | VWF | ECM-Receptor | KEGG: mmu04512 |
| CD31 | GFAP | Vwf | ITGA2B_ITGB3 | 0.0008 | 0 | VWF_ITGA2B_ITGB3 | Vwf - (Itga2b+Itgb3) | VWF | ECM-Receptor | KEGG: mmu04512 |
| GFAP | CD31 | Tnc | ITGA2B_ITGB3 | 0.000336 | 0 | TNC_ITGA2B_ITGB3 | Tnc - (Itga2b+Itgb3) | TENASCIN | ECM-Receptor | KEGG: mmu04512 |
| GFAP | GFAP | Tnc | ITGA2B_ITGB3 | 0.000428 | 0 | TNC_ITGA2B_ITGB3 | Tnc - (Itga2b+Itgb3) | TENASCIN | ECM-Receptor | KEGG: mmu04512 |
| GFAP | CD31 | Tnr | ITGA2B_ITGB3 | 0.000434 | 0 | TNR_ITGA2B_ITGB3 | Tnr - (Itga2b+Itgb3) | TENASCIN | ECM-Receptor | KEGG: mmu04512 |
| GFAP | GFAP | Tnr | ITGA2B_ITGB3 | 0.000552 | 0 | TNR_ITGA2B_ITGB3 | Tnr - (Itga2b+Itgb3) | TENASCIN | ECM-Receptor | KEGG: mmu04512 |
| CD31 | CD31 | Vtn | ITGAV_ITGB5 | 0.004042 | 0 | VTN_ITGAV_ITGB5 | Vtn - (Itgav+Itgb5) | VTN | ECM-Receptor | KEGG: mmu04512 |
| CD31 | GFAP | Vtn | ITGAV_ITGB5 | 0.003732 | 0 | VTN_ITGAV_ITGB5 | Vtn - (Itgav+Itgb5) | VTN | ECM-Receptor | KEGG: mmu04512 |
| CD31 | CD31 | Vtn | ITGAV_ITGB8 | 0.002761 | 0 | VTN_ITGAV_ITGB8 | Vtn - (Itgav+Itgb8) | VTN | ECM-Receptor | KEGG: mmu04512 |
| CD31 | GFAP | Vtn | ITGAV_ITGB8 | 0.004222 | 0 | VTN_ITGAV_ITGB8 | Vtn - (Itgav+Itgb8) | VTN | ECM-Receptor | KEGG: mmu04512 |
| CD31 | CD31 | Fn1 | ITGAV_ITGB8 | 0.005349 | 0 | FN1_ITGAV_ITGB8 | Fn1 - (Itgav+Itgb8) | FN1 | ECM-Receptor | KEGG: mmu04512 |
| CD31 | GFAP | Fn1 | ITGAV_ITGB8 | 0.008168 | 0 | FN1_ITGAV_ITGB8 | Fn1 - (Itgav+Itgb8) | FN1 | ECM-Receptor | KEGG: mmu04512 |
| CD31 | GFAP | Lama1 | ITGAV_ITGB8 | 0.000793 | 0 | LAMA1_ITGAV_ITGB8 | Lama1 - (Itgav+Itgb8) | LAMININ | ECM-Receptor | KEGG: mmu04512 |
| GFAP | GFAP | Lama1 | ITGAV_ITGB8 | 0.000969 | 0 | LAMA1_ITGAV_ITGB8 | Lama1 - (Itgav+Itgb8) | LAMININ | ECM-Receptor | KEGG: mmu04512 |
| CD31 | CD31 | Lama4 | ITGAV_ITGB8 | 0.001331 | 0 | LAMA4_ITGAV_ITGB8 | Lama4 - (Itgav+Itgb8) | LAMININ | ECM-Receptor | KEGG: mmu04512 |
| CD31 | GFAP | Lama4 | ITGAV_ITGB8 | 0.002037 | 0 | LAMA4_ITGAV_ITGB8 | Lama4 - (Itgav+Itgb8) | LAMININ | ECM-Receptor | KEGG: mmu04512 |
| CD31 | CD31 | Lama5 | ITGAV_ITGB8 | 0.002132 | 0 | LAMA5_ITGAV_ITGB8 | Lama5 - (Itgav+Itgb8) | LAMININ | ECM-Receptor | KEGG: mmu04512 |
| CD31 | GFAP | Lama5 | ITGAV_ITGB8 | 0.003261 | 0 | LAMA5_ITGAV_ITGB8 | Lama5 - (Itgav+Itgb8) | LAMININ | ECM-Receptor | KEGG: mmu04512 |
| CD31 | GFAP | Lamb1 | ITGAV_ITGB8 | 0.000906 | 0 | LAMB1_ITGAV_ITGB8 | Lamb1 - (Itgav+Itgb8) | LAMININ | ECM-Receptor | KEGG: mmu04512 |
| GFAP | GFAP | Lamb1 | ITGAV_ITGB8 | 0.001247 | 0 | LAMB1_ITGAV_ITGB8 | Lamb1 - (Itgav+Itgb8) | LAMININ | ECM-Receptor | KEGG: mmu04512 |
| CD31 | CD31 | Lamb2 | ITGAV_ITGB8 | 0.003112 | 0 | LAMB2_ITGAV_ITGB8 | Lamb2 - (Itgav+Itgb8) | LAMININ | ECM-Receptor | KEGG: mmu04512 |
| CD31 | GFAP | Lamb2 | ITGAV_ITGB8 | 0.004758 | 0 | LAMB2_ITGAV_ITGB8 | Lamb2 - (Itgav+Itgb8) | LAMININ | ECM-Receptor | KEGG: mmu04512 |
| CD31 | CD31 | Lamc1 | ITGAV_ITGB8 | 0.002013 | 0 | LAMC1_ITGAV_ITGB8 | Lamc1 - (Itgav+Itgb8) | LAMININ | ECM-Receptor | KEGG: mmu04512 |
| CD31 | GFAP | Lamc1 | ITGAV_ITGB8 | 0.003079 | 0 | LAMC1_ITGAV_ITGB8 | Lamc1 - (Itgav+Itgb8) | LAMININ | ECM-Receptor | KEGG: mmu04512 |
| CD31 | GFAP | Lamc2 | ITGAV_ITGB8 | 0.000906 | 0 | LAMC2_ITGAV_ITGB8 | Lamc2 - (Itgav+Itgb8) | LAMININ | ECM-Receptor | KEGG: mmu04512 |
| GFAP | GFAP | Lamc2 | ITGAV_ITGB8 | 0.000607 | 0 | LAMC2_ITGAV_ITGB8 | Lamc2 - (Itgav+Itgb8) | LAMININ | ECM-Receptor | KEGG: mmu04512 |
| CD31 | CD31 | Lamc3 | ITGAV_ITGB8 | 0.001627 | 0 | LAMC3_ITGAV_ITGB8 | Lamc3 - (Itgav+Itgb8) | LAMININ | ECM-Receptor | KEGG: mmu04512 |
| CD31 | GFAP | Lamc3 | ITGAV_ITGB8 | 0.002489 | 0 | LAMC3_ITGAV_ITGB8 | Lamc3 - (Itgav+Itgb8) | LAMININ | ECM-Receptor | KEGG: mmu04512 |
| CD31 | CD31 | Col1a1 | ITGAV_ITGB8 | 0.003152 | 0 | COL1A1_ITGAV_ITGB8 | Col1a1 - (Itgav+Itgb8) | COLLAGEN | ECM-Receptor | KEGG: mmu04512 |
| CD31 | GFAP | Col1a1 | ITGAV_ITGB8 | 0.004818 | 0 | COL1A1_ITGAV_ITGB8 | Col1a1 - (Itgav+Itgb8) | COLLAGEN | ECM-Receptor | KEGG: mmu04512 |
| CD31 | CD31 | Col1a2 | ITGAV_ITGB8 | 0.001905 | 0 | COL1A2_ITGAV_ITGB8 | Col1a2 - (Itgav+Itgb8) | COLLAGEN | ECM-Receptor | KEGG: mmu04512 |
| CD31 | GFAP | Col1a2 | ITGAV_ITGB8 | 0.002914 | 0 | COL1A2_ITGAV_ITGB8 | Col1a2 - (Itgav+Itgb8) | COLLAGEN | ECM-Receptor | KEGG: mmu04512 |
| CD31 | CD31 | Col4a1 | ITGAV_ITGB8 | 0.004548 | 0 | COL4A1_ITGAV_ITGB8 | Col4a1 - (Itgav+Itgb8) | COLLAGEN | ECM-Receptor | KEGG: mmu04512 |
| CD31 | GFAP | Col4a1 | ITGAV_ITGB8 | 0.006947 | 0 | COL4A1_ITGAV_ITGB8 | Col4a1 - (Itgav+Itgb8) | COLLAGEN | ECM-Receptor | KEGG: mmu04512 |
| CD31 | CD31 | Col4a2 | ITGAV_ITGB8 | 0.003406 | 0 | COL4A2_ITGAV_ITGB8 | Col4a2 - (Itgav+Itgb8) | COLLAGEN | ECM-Receptor | KEGG: mmu04512 |
| CD31 | GFAP | Col4a2 | ITGAV_ITGB8 | 0.005207 | 0 | COL4A2_ITGAV_ITGB8 | Col4a2 - (Itgav+Itgb8) | COLLAGEN | ECM-Receptor | KEGG: mmu04512 |
| GFAP | GFAP | Col6a1 | ITGAV_ITGB8 | 0.001237 | 0 | COL6A1_ITGAV_ITGB8 | Col6a1 - (Itgav+Itgb8) | COLLAGEN | ECM-Receptor | KEGG: mmu04512 |
| NeuN | GFAP | Col6a1 | ITGAV_ITGB8 | 0.002325 | 0 | COL6A1_ITGAV_ITGB8 | Col6a1 - (Itgav+Itgb8) | COLLAGEN | ECM-Receptor | KEGG: mmu04512 |
| CD31 | GFAP | Col6a2 | ITGAV_ITGB8 | 0.001002 | 0 | COL6A2_ITGAV_ITGB8 | Col6a2 - (Itgav+Itgb8) | COLLAGEN | ECM-Receptor | KEGG: mmu04512 |
| GFAP | GFAP | Col6a2 | ITGAV_ITGB8 | 0.001362 | 0 | COL6A2_ITGAV_ITGB8 | Col6a2 - (Itgav+Itgb8) | COLLAGEN | ECM-Receptor | KEGG: mmu04512 |
| GFAP | CD31 | Col6a4 | ITGAV_ITGB8 | 0.000735 | 0 | COL6A4_ITGAV_ITGB8 | Col6a4 - (Itgav+Itgb8) | COLLAGEN | ECM-Receptor | KEGG: mmu04512 |
| GFAP | GFAP | Col6a4 | ITGAV_ITGB8 | 0.001126 | 0 | COL6A4_ITGAV_ITGB8 | Col6a4 - (Itgav+Itgb8) | COLLAGEN | ECM-Receptor | KEGG: mmu04512 |
| CD31 | GFAP | Col6a6 | ITGAV_ITGB8 | 0.000445 | 0 | COL6A6_ITGAV_ITGB8 | Col6a6 - (Itgav+Itgb8) | COLLAGEN | ECM-Receptor | KEGG: mmu04512 |
| GFAP | GFAP | Col6a6 | ITGAV_ITGB8 | 0.000643 | 0 | COL6A6_ITGAV_ITGB8 | Col6a6 - (Itgav+Itgb8) | COLLAGEN | ECM-Receptor | KEGG: mmu04512 |
| CD31 | GFAP | Col9a2 | ITGAV_ITGB8 | 0.000854 | 0 | COL9A2_ITGAV_ITGB8 | Col9a2 - (Itgav+Itgb8) | COLLAGEN | ECM-Receptor | KEGG: mmu04512 |
| GFAP | GFAP | Col9a2 | ITGAV_ITGB8 | 0.000812 | 0 | COL9A2_ITGAV_ITGB8 | Col9a2 - (Itgav+Itgb8) | COLLAGEN | ECM-Receptor | KEGG: mmu04512 |
| GFAP | CD31 | Col9a3 | ITGAV_ITGB8 | 0.000873 | 0 | COL9A3_ITGAV_ITGB8 | Col9a3 - (Itgav+Itgb8) | COLLAGEN | ECM-Receptor | KEGG: mmu04512 |
| GFAP | GFAP | Col9a3 | ITGAV_ITGB8 | 0.001336 | 0 | COL9A3_ITGAV_ITGB8 | Col9a3 - (Itgav+Itgb8) | COLLAGEN | ECM-Receptor | KEGG: mmu04512 |
| CD31 | CD31 | Col1a1 | Sdc4 | 0.006239 | 0 | COL1A1_SDC4 | Col1a1 - Sdc4 | COLLAGEN | ECM-Receptor | KEGG: mmu04512 |

| | | | | | | | | | | |
|---|---|---|---|---|---|---|---|---|---|---|
| CD31 | GFAP | Col1a1 | Sdc4 | 0.013721 | 0 | COL1A1_SDC4 | Col1a1 - Sdc4 | COLLAGEN | ECM-Receptor | KEGG: mmu04512 |
| CD31 | CD31 | Col1a2 | Sdc4 | 0.003775 | 0 | COL1A2_SDC4 | Col1a2 - Sdc4 | COLLAGEN | ECM-Receptor | KEGG: mmu04512 |
| CD31 | GFAP | Col1a2 | Sdc4 | 0.008327 | 0 | COL1A2_SDC4 | Col1a2 - Sdc4 | COLLAGEN | ECM-Receptor | KEGG: mmu04512 |
| CD31 | CD31 | Col4a1 | Sdc4 | 0.008991 | 0 | COL4A1_SDC4 | Col4a1 - Sdc4 | COLLAGEN | ECM-Receptor | KEGG: mmu04512 |
| CD31 | GFAP | Col4a1 | Sdc4 | 0.019707 | 0 | COL4A1_SDC4 | Col4a1 - Sdc4 | COLLAGEN | ECM-Receptor | KEGG: mmu04512 |
| CD31 | CD31 | Col4a2 | Sdc4 | 0.006742 | 0 | COL4A2_SDC4 | Col4a2 - Sdc4 | COLLAGEN | ECM-Receptor | KEGG: mmu04512 |
| CD31 | GFAP | Col4a2 | Sdc4 | 0.014817 | 0 | COL4A2_SDC4 | Col4a2 - Sdc4 | COLLAGEN | ECM-Receptor | KEGG: mmu04512 |
| GFAP | GFAP | Col6a1 | Sdc4 | 0.003548 | 0 | COL6A1_SDC4 | Col6a1 - Sdc4 | COLLAGEN | ECM-Receptor | KEGG: mmu04512 |
| NeuN | GFAP | Col6a1 | Sdc4 | 0.006651 | 0 | COL6A1_SDC4 | Col6a1 - Sdc4 | COLLAGEN | ECM-Receptor | KEGG: mmu04512 |
| CD31 | GFAP | Col6a2 | Sdc4 | 0.002874 | 0 | COL6A2_SDC4 | Col6a2 - Sdc4 | COLLAGEN | ECM-Receptor | KEGG: mmu04512 |
| GFAP | GFAP | Col6a2 | Sdc4 | 0.003903 | 0 | COL6A2_SDC4 | Col6a2 - Sdc4 | COLLAGEN | ECM-Receptor | KEGG: mmu04512 |
| GFAP | CD31 | Col6a4 | Sdc4 | 0.001459 | 0 | COL6A4_SDC4 | Col6a4 - Sdc4 | COLLAGEN | ECM-Receptor | KEGG: mmu04512 |
| GFAP | GFAP | Col6a4 | Sdc4 | 0.003228 | 0 | COL6A4_SDC4 | Col6a4 - Sdc4 | COLLAGEN | ECM-Receptor | KEGG: mmu04512 |
| CD31 | GFAP | Col6a6 | Sdc4 | 0.001277 | 0 | COL6A6_SDC4 | Col6a6 - Sdc4 | COLLAGEN | ECM-Receptor | KEGG: mmu04512 |
| GFAP | GFAP | Col6a6 | Sdc4 | 0.001844 | 0 | COL6A6_SDC4 | Col6a6 - Sdc4 | COLLAGEN | ECM-Receptor | KEGG: mmu04512 |
| CD31 | GFAP | Col9a2 | Sdc4 | 0.002451 | 0 | COL9A2_SDC4 | Col9a2 - Sdc4 | COLLAGEN | ECM-Receptor | KEGG: mmu04512 |
| GFAP | GFAP | Col9a2 | Sdc4 | 0.002328 | 0 | COL9A2_SDC4 | Col9a2 - Sdc4 | COLLAGEN | ECM-Receptor | KEGG: mmu04512 |
| CD31 | GFAP | Col9a3 | Sdc4 | 0.00185 | 0 | COL9A3_SDC4 | Col9a3 - Sdc4 | COLLAGEN | ECM-Receptor | KEGG: mmu04512 |
| GFAP | GFAP | Col9a3 | Sdc4 | 0.00383 | 0 | COL9A3_SDC4 | Col9a3 - Sdc4 | COLLAGEN | ECM-Receptor | KEGG: mmu04512 |
| CD31 | CD31 | Fn1 | Sdc4 | 0.010567 | 0 | FN1_SDC4 | Fn1 - Sdc4 | FN1 | ECM-Receptor | KEGG: mmu04512 |
| CD31 | GFAP | Fn1 | Sdc4 | 0.023116 | 0 | FN1_SDC4 | Fn1 - Sdc4 | FN1 | ECM-Receptor | KEGG: mmu04512 |
| CD31 | GFAP | Tnc | Sdc4 | 0.001627 | 0 | TNC_SDC4 | Tnc - Sdc4 | TENASCIN | ECM-Receptor | KEGG: mmu04512 |
| GFAP | GFAP | Tnc | Sdc4 | 0.003263 | 0 | TNC_SDC4 | Tnc - Sdc4 | TENASCIN | ECM-Receptor | KEGG: mmu04512 |
| GFAP | GFAP | Tnr | Sdc4 | 0.004204 | 0 | TNR_SDC4 | Tnr - Sdc4 | TENASCIN | ECM-Receptor | KEGG: mmu04512 |
| NeuN | GFAP | Tnr | Sdc4 | 0.002947 | 0 | TNR_SDC4 | Tnr - Sdc4 | TENASCIN | ECM-Receptor | KEGG: mmu04512 |
| CD31 | GFAP | Thbs1 | Sdc4 | 0.001527 | 0 | THBS1_SDC4 | Thbs1 - Sdc4 | THBS | ECM-Receptor | KEGG: mmu04512 |
| GFAP | GFAP | Thbs1 | Sdc4 | 0.003357 | 0 | THBS1_SDC4 | Thbs1 - Sdc4 | THBS | ECM-Receptor | KEGG: mmu04512 |
| CD31 | GFAP | Thbs3 | Sdc4 | 0.002701 | 0 | THBS3_SDC4 | Thbs3 - Sdc4 | THBS | ECM-Receptor | KEGG: mmu04512 |
| GFAP | GFAP | Thbs3 | Sdc4 | 0.002215 | 0 | THBS3_SDC4 | Thbs3 - Sdc4 | THBS | ECM-Receptor | KEGG: mmu04512 |
| GFAP | CD31 | Thbs4 | Sdc4 | 0.015025 | 0 | THBS4_SDC4 | Thbs4 - Sdc4 | THBS | ECM-Receptor | KEGG: mmu04512 |
| GFAP | GFAP | Thbs4 | Sdc4 | 0.032694 | 0 | THBS4_SDC4 | Thbs4 - Sdc4 | THBS | ECM-Receptor | KEGG: mmu04512 |
| GFAP | GFAP | Lama1 | Sv2a | 0.001967 | 0 | LAMA1_SV2A | Lama1 - Sv2a | LAMININ | ECM-Receptor | KEGG: mmu04512 |
| GFAP | NeuN | Lama1 | Sv2a | 0.006178 | 0 | LAMA1_SV2A | Lama1 - Sv2a | LAMININ | ECM-Receptor | KEGG: mmu04512 |
| CD31 | CD31 | Lama4 | Sv2a | 0.004255 | 0 | LAMA4_SV2A | Lama4 - Sv2a | LAMININ | ECM-Receptor | KEGG: mmu04512 |
| CD31 | NeuN | Lama4 | Sv2a | 0.012918 | 0 | LAMA4_SV2A | Lama4 - Sv2a | LAMININ | ECM-Receptor | KEGG: mmu04512 |
| CD31 | CD31 | Lama5 | Sv2a | 0.006801 | 0 | LAMA5_SV2A | Lama5 - Sv2a | LAMININ | ECM-Receptor | KEGG: mmu04512 |
| CD31 | NeuN | Lama5 | Sv2a | 0.020543 | 0 | LAMA5_SV2A | Lama5 - Sv2a | LAMININ | ECM-Receptor | KEGG: mmu04512 |
| GFAP | GFAP | Lamb1 | Sv2a | 0.002533 | 0 | LAMB1_SV2A | Lamb1 - Sv2a | LAMININ | ECM-Receptor | KEGG: mmu04512 |
| GFAP | NeuN | Lamb1 | Sv2a | 0.007945 | 0 | LAMB1_SV2A | Lamb1 - Sv2a | LAMININ | ECM-Receptor | KEGG: mmu04512 |
| CD31 | CD31 | Lamb2 | Sv2a | 0.009908 | 0 | LAMB2_SV2A | Lamb2 - Sv2a | LAMININ | ECM-Receptor | KEGG: mmu04512 |
| CD31 | NeuN | Lamb2 | Sv2a | 0.029738 | 0 | LAMB2_SV2A | Lamb2 - Sv2a | LAMININ | ECM-Receptor | KEGG: mmu04512 |
| CD31 | CD31 | Lamc1 | Sv2a | 0.006423 | 0 | LAMC1_SV2A | Lamc1 - Sv2a | LAMININ | ECM-Receptor | KEGG: mmu04512 |
| CD31 | NeuN | Lamc1 | Sv2a | 0.019415 | 0 | LAMC1_SV2A | Lamc1 - Sv2a | LAMININ | ECM-Receptor | KEGG: mmu04512 |
| CD31 | NeuN | Lamc2 | Sv2a | 0.005782 | 0 | LAMC2_SV2A | Lamc2 - Sv2a | LAMININ | ECM-Receptor | KEGG: mmu04512 |
| NeuN | NeuN | Lamc2 | Sv2a | 0.004124 | 0 | LAMC2_SV2A | Lamc2 - Sv2a | LAMININ | ECM-Receptor | KEGG: mmu04512 |
| CD31 | CD31 | Lamc3 | Sv2a | 0.005196 | 0 | LAMC3_SV2A | Lamc3 - Sv2a | LAMININ | ECM-Receptor | KEGG: mmu04512 |
| CD31 | NeuN | Lamc3 | Sv2a | 0.015744 | 0 | LAMC3_SV2A | Lamc3 - Sv2a | LAMININ | ECM-Receptor | KEGG: mmu04512 |
| GFAP | NeuN | Lama1 | Sv2b | 0.007276 | 0 | LAMA1_SV2B | Lama1 - Sv2b | LAMININ | ECM-Receptor | KEGG: mmu04512 |
| NeuN | NeuN | Lama1 | Sv2b | 0.002251 | 0 | LAMA1_SV2B | Lama1 - Sv2b | LAMININ | ECM-Receptor | KEGG: mmu04512 |
| CD31 | CD31 | Lama4 | Sv2b | 0.003458 | 0 | LAMA4_SV2B | Lama4 - Sv2b | LAMININ | ECM-Receptor | KEGG: mmu04512 |
| CD31 | NeuN | Lama4 | Sv2b | 0.015195 | 0 | LAMA4_SV2B | Lama4 - Sv2b | LAMININ | ECM-Receptor | KEGG: mmu04512 |
| CD31 | CD31 | Lama5 | Sv2b | 0.00553 | 0 | LAMA5_SV2B | Lama5 - Sv2b | LAMININ | ECM-Receptor | KEGG: mmu04512 |
| CD31 | NeuN | Lama5 | Sv2b | 0.024131 | 0 | LAMA5_SV2B | Lama5 - Sv2b | LAMININ | ECM-Receptor | KEGG: mmu04512 |
| GFAP | NeuN | Lamb1 | Sv2b | 0.009353 | 0 | LAMB1_SV2B | Lamb1 - Sv2b | LAMININ | ECM-Receptor | KEGG: mmu04512 |
| NeuN | NeuN | Lamb1 | Sv2b | 0.002541 | 0 | LAMB1_SV2B | Lamb1 - Sv2b | LAMININ | ECM-Receptor | KEGG: mmu04512 |
| CD31 | CD31 | Lamb2 | Sv2b | 0.008061 | 0 | LAMB2_SV2B | Lamb2 - Sv2b | LAMININ | ECM-Receptor | KEGG: mmu04512 |
| CD31 | NeuN | Lamb2 | Sv2b | 0.034875 | 0 | LAMB2_SV2B | Lamb2 - Sv2b | LAMININ | ECM-Receptor | KEGG: mmu04512 |
| CD31 | CD31 | Lamc1 | Sv2b | 0.005222 | 0 | LAMC1_SV2B | Lamc1 - Sv2b | LAMININ | ECM-Receptor | KEGG: mmu04512 |
| CD31 | NeuN | Lamc1 | Sv2b | 0.022811 | 0 | LAMC1_SV2B | Lamc1 - Sv2b | LAMININ | ECM-Receptor | KEGG: mmu04512 |

| | | | | | | | | | | |
|---|---|---|---|---|---|---|---|---|---|---|
| CD31 | NeuN | Lamc2 | Sv2b | 0.006809 | 0 | LAMC2_SV2B | Lamc2 - Sv2b | LAMININ | ECM-Receptor | KEGG: mmu04512 |
| NeuN | NeuN | Lamc2 | Sv2b | 0.004858 | 0 | LAMC2_SV2B | Lamc2 - Sv2b | LAMININ | ECM-Receptor | KEGG: mmu04512 |
| CD31 | CD31 | Lamc3 | Sv2b | 0.004223 | 0 | LAMC3_SV2B | Lamc3 - Sv2b | LAMININ | ECM-Receptor | KEGG: mmu04512 |
| CD31 | NeuN | Lamc3 | Sv2b | 0.01851 | 0 | LAMC3_SV2B | Lamc3 - Sv2b | LAMININ | ECM-Receptor | KEGG: mmu04512 |
| GFAP | CD31 | Lama1 | Sv2c | 0.000489 | 0 | LAMA1_SV2C | Lama1 - Sv2c | LAMININ | ECM-Receptor | KEGG: mmu04512 |
| GFAP | GFAP | Lama1 | Sv2c | 0.000402 | 0 | LAMA1_SV2C | Lama1 - Sv2c | LAMININ | ECM-Receptor | KEGG: mmu04512 |
| CD31 | CD31 | Lama4 | Sv2c | 0.001029 | 0 | LAMA4_SV2C | Lama4 - Sv2c | LAMININ | ECM-Receptor | KEGG: mmu04512 |
| CD31 | GFAP | Lama4 | Sv2c | 0.000846 | 0 | LAMA4_SV2C | Lama4 - Sv2c | LAMININ | ECM-Receptor | KEGG: mmu04512 |
| CD31 | CD31 | Lama5 | Sv2c | 0.001648 | 0 | LAMA5_SV2C | Lama5 - Sv2c | LAMININ | ECM-Receptor | KEGG: mmu04512 |
| CD31 | GFAP | Lama5 | Sv2c | 0.001355 | 0 | LAMA5_SV2C | Lama5 - Sv2c | LAMININ | ECM-Receptor | KEGG: mmu04512 |
| GFAP | CD31 | Lamb1 | Sv2c | 0.00063 | 0 | LAMB1_SV2C | Lamb1 - Sv2c | LAMININ | ECM-Receptor | KEGG: mmu04512 |
| GFAP | GFAP | Lamb1 | Sv2c | 0.000518 | 0 | LAMB1_SV2C | Lamb1 - Sv2c | LAMININ | ECM-Receptor | KEGG: mmu04512 |
| CD31 | CD31 | Lamb2 | Sv2c | 0.002407 | 0 | LAMB2_SV2C | Lamb2 - Sv2c | LAMININ | ECM-Receptor | KEGG: mmu04512 |
| CD31 | GFAP | Lamb2 | Sv2c | 0.001978 | 0 | LAMB2_SV2C | Lamb2 - Sv2c | LAMININ | ECM-Receptor | KEGG: mmu04512 |
| CD31 | CD31 | Lamc1 | Sv2c | 0.001556 | 0 | LAMC1_SV2C | Lamc1 - Sv2c | LAMININ | ECM-Receptor | KEGG: mmu04512 |
| CD31 | GFAP | Lamc1 | Sv2c | 0.001279 | 0 | LAMC1_SV2C | Lamc1 - Sv2c | LAMININ | ECM-Receptor | KEGG: mmu04512 |
| CD31 | CD31 | Lamc2 | Sv2c | 0.000458 | 0 | LAMC2_SV2C | Lamc2 - Sv2c | LAMININ | ECM-Receptor | KEGG: mmu04512 |
| CD31 | GFAP | Lamc2 | Sv2c | 0.000376 | 0 | LAMC2_SV2C | Lamc2 - Sv2c | LAMININ | ECM-Receptor | KEGG: mmu04512 |
| CD31 | CD31 | Lamc3 | Sv2c | 0.001258 | 0 | LAMC3_SV2C | Lamc3 - Sv2c | LAMININ | ECM-Receptor | KEGG: mmu04512 |
| CD31 | GFAP | Lamc3 | Sv2c | 0.001034 | 0 | LAMC3_SV2C | Lamc3 - Sv2c | LAMININ | ECM-Receptor | KEGG: mmu04512 |
| GFAP | CD31 | Thbs1 | Cd36 | 0.000277 | 0 | THBS1_CD36 | Thbs1 - Cd36 | THBS | ECM-Receptor | KEGG: mmu04512 |
| GFAP | GFAP | Thbs1 | Cd36 | 0.000533 | 0 | THBS1_CD36 | Thbs1 - Cd36 | THBS | ECM-Receptor | KEGG: mmu04512 |
| CD31 | GFAP | Thbs3 | Cd36 | 0.000428 | 0 | THBS3_CD36 | Thbs3 - Cd36 | THBS | ECM-Receptor | KEGG: mmu04512 |
| GFAP | GFAP | Thbs3 | Cd36 | 0.000351 | 0 | THBS3_CD36 | Thbs3 - Cd36 | THBS | ECM-Receptor | KEGG: mmu04512 |
| GFAP | CD31 | Thbs4 | Cd36 | 0.002772 | 0 | THBS4_CD36 | Thbs4 - Cd36 | THBS | ECM-Receptor | KEGG: mmu04512 |
| GFAP | GFAP | Thbs4 | Cd36 | 0.005318 | 0 | THBS4_CD36 | Thbs4 - Cd36 | THBS | ECM-Receptor | KEGG: mmu04512 |
| GFAP | GFAP | Thbs1 | Cd47 | 0.00121 | 0 | THBS1_CD47 | Thbs1 - Cd47 | THBS | ECM-Receptor | KEGG: mmu04512 |
| GFAP | NeuN | Thbs1 | Cd47 | 0.001677 | 0 | THBS1_CD47 | Thbs1 - Cd47 | THBS | ECM-Receptor | KEGG: mmu04512 |
| GFAP | GFAP | Thbs3 | Cd47 | 0.000798 | 0 | THBS3_CD47 | Thbs3 - Cd47 | THBS | ECM-Receptor | KEGG: mmu04512 |
| CD31 | NeuN | Thbs3 | Cd47 | 0.001349 | 0 | THBS3_CD47 | Thbs3 - Cd47 | THBS | ECM-Receptor | KEGG: mmu04512 |
| GFAP | GFAP | Thbs4 | Cd47 | 0.012013 | 0 | THBS4_CD47 | Thbs4 - Cd47 | THBS | ECM-Receptor | KEGG: mmu04512 |
| GFAP | NeuN | Thbs4 | Cd47 | 0.016574 | 0 | THBS4_CD47 | Thbs4 - Cd47 | THBS | ECM-Receptor | KEGG: mmu04512 |
| CD31 | CD31 | Agrn | Dag1 | 0.003714 | 0 | AGRN_DAG1 | Agrn - Dag1 | AGRN | ECM-Receptor | KEGG: mmu04512 |
| CD31 | GFAP | Agrn | Dag1 | 0.002855 | 0 | AGRN_DAG1 | Agrn - Dag1 | AGRN | ECM-Receptor | KEGG: mmu04512 |
| CD31 | CD31 | Hspg2 | Dag1 | 0.001735 | 0 | HSPG2_DAG1 | Hspg2 - Dag1 | HSPG | ECM-Receptor | KEGG: mmu04512 |
| CD31 | GFAP | Hspg2 | Dag1 | 0.001333 | 0 | HSPG2_DAG1 | Hspg2 - Dag1 | HSPG | ECM-Receptor | KEGG: mmu04512 |
| CD31 | CD31 | Lama1 | Dag1 | 0.000738 | 0 | LAMA1_DAG1 | Lama1 - Dag1 | LAMININ | ECM-Receptor | KEGG: mmu04512 |
| GFAP | CD31 | Lama1 | Dag1 | 0.000902 | 0 | LAMA1_DAG1 | Lama1 - Dag1 | LAMININ | ECM-Receptor | KEGG: mmu04512 |
| CD31 | CD31 | Lama4 | Dag1 | 0.001897 | 0 | LAMA4_DAG1 | Lama4 - Dag1 | LAMININ | ECM-Receptor | KEGG: mmu04512 |
| CD31 | GFAP | Lama4 | Dag1 | 0.001458 | 0 | LAMA4_DAG1 | Lama4 - Dag1 | LAMININ | ECM-Receptor | KEGG: mmu04512 |
| CD31 | CD31 | Lama5 | Dag1 | 0.003036 | 0 | LAMA5_DAG1 | Lama5 - Dag1 | LAMININ | ECM-Receptor | KEGG: mmu04512 |
| CD31 | GFAP | Lama5 | Dag1 | 0.002334 | 0 | LAMA5_DAG1 | Lama5 - Dag1 | LAMININ | ECM-Receptor | KEGG: mmu04512 |
| GFAP | CD31 | Lamb1 | Dag1 | 0.001161 | 0 | LAMB1_DAG1 | Lamb1 - Dag1 | LAMININ | ECM-Receptor | KEGG: mmu04512 |
| GFAP | GFAP | Lamb1 | Dag1 | 0.000893 | 0 | LAMB1_DAG1 | Lamb1 - Dag1 | LAMININ | ECM-Receptor | KEGG: mmu04512 |
| CD31 | CD31 | Lamb2 | Dag1 | 0.004431 | 0 | LAMB2_DAG1 | Lamb2 - Dag1 | LAMININ | ECM-Receptor | KEGG: mmu04512 |
| CD31 | GFAP | Lamb2 | Dag1 | 0.003407 | 0 | LAMB2_DAG1 | Lamb2 - Dag1 | LAMININ | ECM-Receptor | KEGG: mmu04512 |
| CD31 | CD31 | Lamc1 | Dag1 | 0.002867 | 0 | LAMC1_DAG1 | Lamc1 - Dag1 | LAMININ | ECM-Receptor | KEGG: mmu04512 |
| CD31 | GFAP | Lamc1 | Dag1 | 0.002204 | 0 | LAMC1_DAG1 | Lamc1 - Dag1 | LAMININ | ECM-Receptor | KEGG: mmu04512 |
| CD31 | CD31 | Lamc2 | Dag1 | 0.000844 | 0 | LAMC2_DAG1 | Lamc2 - Dag1 | LAMININ | ECM-Receptor | KEGG: mmu04512 |
| CD31 | GFAP | Lamc2 | Dag1 | 0.000648 | 0 | LAMC2_DAG1 | Lamc2 - Dag1 | LAMININ | ECM-Receptor | KEGG: mmu04512 |
| CD31 | CD31 | Lamc3 | Dag1 | 0.002317 | 0 | LAMC3_DAG1 | Lamc3 - Dag1 | LAMININ | ECM-Receptor | KEGG: mmu04512 |
| CD31 | GFAP | Lamc3 | Dag1 | 0.001781 | 0 | LAMC3_DAG1 | Lamc3 - Dag1 | LAMININ | ECM-Receptor | KEGG: mmu04512 |
| NeuN | CD31 | Reln | Lrp8 | 0.00066 | 0 | RELN_LRP8 | Reln - Lrp8 | RELN | ECM-Receptor | PMID: 14578885 |
| NeuN | NeuN | Reln | Lrp8 | 0.000844 | 0 | RELN_LRP8 | Reln - Lrp8 | RELN | ECM-Receptor | PMID: 14578885 |
| GFAP | GFAP | Reln | Vldlr | 0.000591 | 0 | RELN_VLDLR | Reln - Vldlr | RELN | ECM-Receptor | PMID: 14578885 |
| NeuN | GFAP | Reln | Vldlr | 0.000973 | 0 | RELN_VLDLR | Reln - Vldlr | RELN | ECM-Receptor | PMID: 14578885 |
| GFAP | NeuN | Pnoc | Oprl1 | 0.000544 | 0 | PNOC_OPRL1 | Pnoc - Oprl1 | OPIOID | Non-protein Signaling | PMID: 29934561;PMID: 34724150 |
| NeuN | NeuN | Pnoc | Oprl1 | 0.000368 | 0 | PNOC_OPRL1 | Pnoc - Oprl1 | OPIOID | Non-protein Signaling | PMID: 29934561;PMID: 34724150 |
| CD31 | CD31 | Ppia | Bsg | 0.456259 | 0 | PPIA_BSG | Ppia - Bsg | CypA | Non-protein Signaling | PMID: 36012604 |

| | | | | | | | | | | |
|---|---|---|---|---|---|---|---|---|---|---|
| NeuN | CD31 | Ppia | Bsg | 0.543083 | 0 | PPIA_BSG | Ppia - Bsg | CypA | Non-protein Signaling | PMID: 36012604 |
| NeuN | CD31 | Cadm1 | Cadm1 | 0.004608 | 0 | CADM1_CADM1 | Cadm1 - Cadm1 | CADM | Cell-Cell Contact | PMID: 24503895 |
| CD31 | NeuN | Cadm1 | Cadm1 | 0.004608 | 0 | CADM1_CADM1 | Cadm1 - Cadm1 | CADM | Cell-Cell Contact | PMID: 24503895 |
| NeuN | NeuN | Cadm1 | Cadm1 | 0.009049 | 0 | CADM1_CADM1 | Cadm1 - Cadm1 | CADM | Cell-Cell Contact | PMID: 24503895 |
| CD31 | CD31 | Cd200 | Cd200r1 | 0.001056 | 0 | CD200_CD200R1 | Cd200 - Cd200r1 | CD200 | Cell-Cell Contact | PMID: 15187158 |
| CD31 | GFAP | Cd200 | Cd200r1 | 0.001456 | 0 | CD200_CD200R1 | Cd200 - Cd200r1 | CD200 | Cell-Cell Contact | PMID: 15187158 |
| CD31 | GFAP | Cd200 | Cd200r2 | 0.00186 | 0 | CD200_CD200R2 | Cd200 - Cd200r2 | CD200 | Cell-Cell Contact | KEGG: mmu04514 |
| NeuN | NeuN | Cd200 | Cd200r2 | 0.001235 | 0 | CD200_CD200R2 | Cd200 - Cd200r2 | CD200 | Cell-Cell Contact | KEGG: mmu04514 |
| CD31 | CD31 | Cd200 | Cd200r3 | 0.001238 | 0 | CD200_CD200R3 | Cd200 - Cd200r3 | CD200 | Cell-Cell Contact | PMID: 22020332 |
| NeuN | CD31 | Cd200 | Cd200r3 | 0.001155 | 0 | CD200_CD200R3 | Cd200 - Cd200r3 | CD200 | Cell-Cell Contact | PMID: 22020332 |
| CD31 | CD31 | Entpd1 | Adora1 | 0.003171 | 0 | ENTPD1_ADORA1 | Entpd1 - Adora1 | CD39 | Cell-Cell Contact | PMID: 21677139 |
| CD31 | NeuN | Entpd1 | Adora1 | 0.005718 | 0 | ENTPD1_ADORA1 | Entpd1 - Adora1 | CD39 | Cell-Cell Contact | PMID: 21677139 |
| CD31 | CD31 | Entpd1 | Adora2a | 0.001501 | 0 | ENTPD1_ADORA2A | Entpd1 - Adora2a | CD39 | Cell-Cell Contact | PMID: 21677139 |
| CD31 | GFAP | Entpd1 | Adora2a | 0.001175 | 0 | ENTPD1_ADORA2A | Entpd1 - Adora2a | CD39 | Cell-Cell Contact | PMID: 21677139 |
| CD31 | CD31 | Entpd1 | Adora2b | 0.00151 | 0 | ENTPD1_ADORA2B | Entpd1 - Adora2b | CD39 | Cell-Cell Contact | PMID: 21677139 |
| CD31 | GFAP | Entpd1 | Adora2b | 0.001732 | 0 | ENTPD1_ADORA2B | Entpd1 - Adora2b | CD39 | Cell-Cell Contact | PMID: 21677139 |
| CD31 | CD31 | Cd46 | Jag1 | 0.000755 | 0 | CD46_JAG1 | Cd46 - Jag1 | CD46 | Cell-Cell Contact | PMID: 23086448 |
| GFAP | CD31 | Cd46 | Jag1 | 0.001411 | 0 | CD46_JAG1 | Cd46 - Jag1 | CD46 | Cell-Cell Contact | PMID: 23086448 |
| CD31 | GFAP | Cd6 | Alcam | 0.000687 | 0 | CD6_ALCAM | Cd6 - Alcam | CD6 | Cell-Cell Contact | PMID: 23602662 |
| GFAP | GFAP | Cd6 | Alcam | 0.00058 | 0 | CD6_ALCAM | Cd6 - Alcam | CD6 | Cell-Cell Contact | PMID: 23602662 |
| GFAP | CD31 | Cd96 | Pvr | 0.000279 | 0 | CD96_PVR | Cd96 - Pvr | CD96 | Cell-Cell Contact | PMID: 15034010 |
| GFAP | GFAP | Cd96 | Pvr | 0.000247 | 0 | CD96_PVR | Cd96 - Pvr | CD96 | Cell-Cell Contact | PMID: 15034010 |
| NeuN | CD31 | Cd99 | Cd99 | 0.097586 | 0 | CD99_CD99 | Cd99 - Cd99 | CD99 | Cell-Cell Contact | KEGG: mmu04514 |
| CD31 | NeuN | Cd99 | Cd99 | 0.097586 | 0 | CD99_CD99 | Cd99 - Cd99 | CD99 | Cell-Cell Contact | KEGG: mmu04514 |
| NeuN | NeuN | Cd99 | Cd99 | 0.135997 | 0 | CD99_CD99 | Cd99 - Cd99 | CD99 | Cell-Cell Contact | KEGG: mmu04514 |
| GFAP | CD31 | Cdh4 | Cdh4 | 0.000528 | 0 | CDH4_CDH4 | Cdh4 - Cdh4 | CDH | Cell-Cell Contact | KEGG: mmu04514 |
| CD31 | GFAP | Cdh4 | Cdh4 | 0.000528 | 0 | CDH4_CDH4 | Cdh4 - Cdh4 | CDH | Cell-Cell Contact | KEGG: mmu04514 |
| GFAP | GFAP | Cdh4 | Cdh4 | 0.000781 | 0 | CDH4_CDH4 | Cdh4 - Cdh4 | CDH | Cell-Cell Contact | KEGG: mmu04514 |
| GFAP | GFAP | Cdh2 | Cdh2 | 0.006244 | 0 | CDH2_CDH2 | Cdh2 - Cdh2 | CDH | Cell-Cell Contact | KEGG: mmu04514 |
| NeuN | GFAP | Cdh2 | Cdh2 | 0.005213 | 0 | CDH2_CDH2 | Cdh2 - Cdh2 | CDH | Cell-Cell Contact | KEGG: mmu04514 |
| GFAP | NeuN | Cdh2 | Cdh2 | 0.005213 | 0 | CDH2_CDH2 | Cdh2 - Cdh2 | CDH | Cell-Cell Contact | KEGG: mmu04514 |
| CD31 | CD31 | Cdh5 | Cdh5 | 0.013593 | 0 | CDH5_CDH5 | Cdh5 - Cdh5 | CDH5 | Cell-Cell Contact | KEGG: mmu04514 |
| GFAP | CD31 | Cdh5 | Cdh5 | 0.002296 | 0 | CDH5_CDH5 | Cdh5 - Cdh5 | CDH5 | Cell-Cell Contact | KEGG: mmu04514 |
| CD31 | GFAP | Cdh5 | Cdh5 | 0.002296 | 0 | CDH5_CDH5 | Cdh5 - Cdh5 | CDH5 | Cell-Cell Contact | KEGG: mmu04514 |
| GFAP | CD31 | Cldn11 | Cldn11 | 0.087007 | 0 | CLDN11_CLDN11 | Cldn11 - Cldn11 | CLDN | Cell-Cell Contact | KEGG: mmu04514 |
| CD31 | GFAP | Cldn11 | Cldn11 | 0.087007 | 0 | CLDN11_CLDN11 | Cldn11 - Cldn11 | CLDN | Cell-Cell Contact | KEGG: mmu04514 |
| GFAP | GFAP | Cldn11 | Cldn11 | 0.36747 | 0 | CLDN11_CLDN11 | Cldn11 - Cldn11 | CLDN | Cell-Cell Contact | KEGG: mmu04514 |
| CD31 | GFAP | Clec2f | Klrb1f | 0.000499 | 0 | CLEC2F_KLRB1F | Clec2f - Klrb1f | CLEC | Cell-Cell Contact | PMID: 24223577 |
| GFAP | GFAP | Clec2f | Klrb1f | 0.000669 | 0 | CLEC2F_KLRB1F | Clec2f - Klrb1f | CLEC | Cell-Cell Contact | PMID: 24223577 |
| GFAP | NeuN | Nfasc | CNTN1_CNTNAP1 | 0.04709 | 0 | NFASC_CNTN1_CNTNAP1 | Nfasc - (Cntn1+Cntnap1) | CNTN | Cell-Cell Contact | KEGG: mmu04514 |
| NeuN | NeuN | Nfasc | CNTN1_CNTNAP1 | 0.024757 | 0 | NFASC_CNTN1_CNTNAP1 | Nfasc - (Cntn1+Cntnap1) | CNTN | Cell-Cell Contact | KEGG: mmu04514 |
| GFAP | GFAP | Cntn2 | CNTN2_CNTNAP2 | 0.002235 | 0 | CNTN2_CNTN2_CNTNAP2 | Cntn2 - (Cntn2+Cntnap2) | CNTN | Cell-Cell Contact | KEGG: mmu04514 |
| GFAP | NeuN | Cntn2 | CNTN2_CNTNAP2 | 0.001443 | 0 | CNTN2_CNTN2_CNTNAP2 | Cntn2 - (Cntn2+Cntnap2) | CNTN | Cell-Cell Contact | KEGG: mmu04514 |
| GFAP | NeuN | Cntn2 | L1cam | 0.003639 | 0 | CNTN2_L1CAM | Cntn2 - L1cam | CNTN | Cell-Cell Contact | KEGG: mmu04514 |
| NeuN | NeuN | Cntn2 | L1cam | 0.001676 | 0 | CNTN2_L1CAM | Cntn2 - L1cam | CNTN | Cell-Cell Contact | KEGG: mmu04514 |
| GFAP | GFAP | Cntn2 | Cntn2 | 0.002357 | 0 | CNTN2_CNTN2 | Cntn2 - Cntn2 | CNTN | Cell-Cell Contact | KEGG: mmu04514 |
| NeuN | GFAP | Cntn2 | Cntn2 | 0.001085 | 0 | CNTN2_CNTN2 | Cntn2 - Cntn2 | CNTN | Cell-Cell Contact | KEGG: mmu04514 |
| GFAP | NeuN | Cntn2 | Cntn2 | 0.001085 | 0 | CNTN2_CNTN2 | Cntn2 - Cntn2 | CNTN | Cell-Cell Contact | KEGG: mmu04514 |
| GFAP | NeuN | Cntn1 | Nrcam | 0.02264 | 0 | CNTN1_NRCAM | Cntn1 - Nrcam | CNTN | Cell-Cell Contact | KEGG: mmu04514 |
| NeuN | NeuN | Cntn1 | Nrcam | 0.055605 | 0 | CNTN1_NRCAM | Cntn1 - Nrcam | CNTN | Cell-Cell Contact | KEGG: mmu04514 |
| CD31 | CD31 | Cspg4 | ITGA3_ITGB1 | 0.001241 | 0 | CSPG4_ITGA2_ITGB1 | Cspg4 - (Itga2+Itgb1) | CSPG4 | Cell-Cell Contact | PMID: 30213051 |
| GFAP | CD31 | Cspg4 | ITGA3_ITGB1 | 0.001306 | 0 | CSPG4_ITGA2_ITGB1 | Cspg4 - (Itga2+Itgb1) | CSPG4 | Cell-Cell Contact | PMID: 30213051 |
| GFAP | CD31 | Efna1 | Epha3 | 0.000498 | 0 | EFNA1_EPHA3 | Efna1 - Epha3 | EPHA | Cell-Cell Contact | PMID: 15114347 |
| GFAP | GFAP | Efna1 | Epha3 | 0.000477 | 0 | EFNA1_EPHA3 | Efna1 - Epha3 | EPHA | Cell-Cell Contact | PMID: 15114347 |
| GFAP | NeuN | Efna1 | Epha4 | 0.004531 | 0 | EFNA1_EPHA4 | Efna1 - Epha4 | EPHA | Cell-Cell Contact | PMID: 15114347 |
| NeuN | NeuN | Efna1 | Epha4 | 0.001816 | 0 | EFNA1_EPHA4 | Efna1 - Epha4 | EPHA | Cell-Cell Contact | PMID: 15114347 |
| GFAP | GFAP | Efna1 | Epha5 | 0.001089 | 0 | EFNA1_EPHA5 | Efna1 - Epha5 | EPHA | Cell-Cell Contact | PMID: 15114347 |
| GFAP | NeuN | Efna1 | Epha5 | 0.001761 | 0 | EFNA1_EPHA5 | Efna1 - Epha5 | EPHA | Cell-Cell Contact | PMID: 15114347 |
| GFAP | GFAP | Efna1 | Epha7 | 0.000782 | 0 | EFNA1_EPHA7 | Efna1 - Epha7 | EPHA | Cell-Cell Contact | PMID: 15114347 |

| | | | | | | | | | | |
|---|---|---|---|---|---|---|---|---|---|---|
| GFAP | NeuN | Efna1 | Epha7 | 0.001458 | 0 | EFNA1_EPHA7 | Efna1 - Epha7 | EPHA | Cell-Cell Contact | PMID: 15114347 |
| GFAP | CD31 | Efna1 | Epha8 | 0.000465 | 0 | EFNA1_EPHA8 | Efna1 - Epha8 | EPHA | Cell-Cell Contact | PMID: 15114347 |
| GFAP | GFAP | Efna1 | Epha8 | 0.000356 | 0 | EFNA1_EPHA8 | Efna1 - Epha8 | EPHA | Cell-Cell Contact | PMID: 15114347 |
| GFAP | CD31 | Efna2 | Epha3 | 0.00039 | 0 | EFNA2_EPHA3 | Efna2 - Epha3 | EPHA | Cell-Cell Contact | PMID: 15114347 |
| GFAP | GFAP | Efna2 | Epha3 | 0.000373 | 0 | EFNA2_EPHA3 | Efna2 - Epha3 | EPHA | Cell-Cell Contact | PMID: 15114347 |
| GFAP | NeuN | Efna2 | Epha4 | 0.003552 | 0 | EFNA2_EPHA4 | Efna2 - Epha4 | EPHA | Cell-Cell Contact | PMID: 15114347 |
| NeuN | NeuN | Efna2 | Epha4 | 0.002813 | 0 | EFNA2_EPHA4 | Efna2 - Epha4 | EPHA | Cell-Cell Contact | PMID: 15114347 |
| GFAP | NeuN | Efna2 | Epha5 | 0.001379 | 0 | EFNA2_EPHA5 | Efna2 - Epha5 | EPHA | Cell-Cell Contact | PMID: 15114347 |
| NeuN | NeuN | Efna2 | Epha5 | 0.001092 | 0 | EFNA2_EPHA5 | Efna2 - Epha5 | EPHA | Cell-Cell Contact | PMID: 15114347 |
| GFAP | NeuN | Efna2 | Epha7 | 0.001142 | 0 | EFNA2_EPHA7 | Efna2 - Epha7 | EPHA | Cell-Cell Contact | PMID: 15114347 |
| NeuN | NeuN | Efna2 | Epha7 | 0.000904 | 0 | EFNA2_EPHA7 | Efna2 - Epha7 | EPHA | Cell-Cell Contact | PMID: 15114347 |
| GFAP | CD31 | Efna2 | Epha8 | 0.000364 | 0 | EFNA2_EPHA8 | Efna2 - Epha8 | EPHA | Cell-Cell Contact | PMID: 15114347 |
| GFAP | GFAP | Efna2 | Epha8 | 0.000279 | 0 | EFNA2_EPHA8 | Efna2 - Epha8 | EPHA | Cell-Cell Contact | PMID: 15114347 |
| NeuN | CD31 | Efna3 | Epha3 | 0.001031 | 0 | EFNA3_EPHA3 | Efna3 - Epha3 | EPHA | Cell-Cell Contact | PMID: 15114347 |
| GFAP | GFAP | Efna3 | Epha3 | 0.000552 | 0 | EFNA3_EPHA3 | Efna3 - Epha3 | EPHA | Cell-Cell Contact | PMID: 15114347 |
| GFAP | NeuN | Efna3 | Epha4 | 0.005243 | 0 | EFNA3_EPHA4 | Efna3 - Epha4 | EPHA | Cell-Cell Contact | PMID: 15114347 |
| NeuN | NeuN | Efna3 | Epha4 | 0.009337 | 0 | EFNA3_EPHA4 | Efna3 - Epha4 | EPHA | Cell-Cell Contact | PMID: 15114347 |
| NeuN | GFAP | Efna3 | Epha5 | 0.002253 | 0 | EFNA3_EPHA5 | Efna3 - Epha5 | EPHA | Cell-Cell Contact | PMID: 15114347 |
| NeuN | NeuN | Efna3 | Epha5 | 0.003639 | 0 | EFNA3_EPHA5 | Efna3 - Epha5 | EPHA | Cell-Cell Contact | PMID: 15114347 |
| NeuN | CD31 | Efna3 | Epha7 | 0.002015 | 0 | EFNA3_EPHA7 | Efna3 - Epha7 | EPHA | Cell-Cell Contact | PMID: 15114347 |
| NeuN | NeuN | Efna3 | Epha7 | 0.003014 | 0 | EFNA3_EPHA7 | Efna3 - Epha7 | EPHA | Cell-Cell Contact | PMID: 15114347 |
| NeuN | CD31 | Efna3 | Epha8 | 0.000963 | 0 | EFNA3_EPHA8 | Efna3 - Epha8 | EPHA | Cell-Cell Contact | PMID: 15114347 |
| NeuN | NeuN | Efna3 | Epha8 | 0.0005 | 0 | EFNA3_EPHA8 | Efna3 - Epha8 | EPHA | Cell-Cell Contact | PMID: 15114347 |
| CD31 | CD31 | Efna4 | Epha3 | 0.00043 | 0 | EFNA4_EPHA3 | Efna4 - Epha3 | EPHA | Cell-Cell Contact | PMID: 15114347 |
| CD31 | GFAP | Efna4 | Epha3 | 0.000411 | 0 | EFNA4_EPHA3 | Efna4 - Epha3 | EPHA | Cell-Cell Contact | PMID: 15114347 |
| CD31 | CD31 | Efna4 | Epha4 | 0.002015 | 0 | EFNA4_EPHA4 | Efna4 - Epha4 | EPHA | Cell-Cell Contact | PMID: 15114347 |
| CD31 | NeuN | Efna4 | Epha4 | 0.003914 | 0 | EFNA4_EPHA4 | Efna4 - Epha4 | EPHA | Cell-Cell Contact | PMID: 15114347 |
| GFAP | GFAP | Efna4 | Epha5 | 0.000838 | 0 | EFNA4_EPHA5 | Efna4 - Epha5 | EPHA | Cell-Cell Contact | PMID: 15114347 |
| CD31 | NeuN | Efna4 | Epha5 | 0.00152 | 0 | EFNA4_EPHA5 | Efna4 - Epha5 | EPHA | Cell-Cell Contact | PMID: 15114347 |
| CD31 | CD31 | Efna4 | Epha7 | 0.000841 | 0 | EFNA4_EPHA7 | Efna4 - Epha7 | EPHA | Cell-Cell Contact | PMID: 15114347 |
| CD31 | NeuN | Efna4 | Epha7 | 0.001259 | 0 | EFNA4_EPHA7 | Efna4 - Epha7 | EPHA | Cell-Cell Contact | PMID: 15114347 |
| CD31 | CD31 | Efna4 | Epha8 | 0.000402 | 0 | EFNA4_EPHA8 | Efna4 - Epha8 | EPHA | Cell-Cell Contact | PMID: 15114347 |
| GFAP | CD31 | Efna4 | Epha8 | 0.000358 | 0 | EFNA4_EPHA8 | Efna4 - Epha8 | EPHA | Cell-Cell Contact | PMID: 15114347 |
| CD31 | CD31 | Efna5 | Epha3 | 0.000606 | 0 | EFNA5_EPHA3 | Efna5 - Epha3 | EPHA | Cell-Cell Contact | PMID: 15114347 |
| NeuN | CD31 | Efna5 | Epha3 | 0.000872 | 0 | EFNA5_EPHA3 | Efna5 - Epha3 | EPHA | Cell-Cell Contact | PMID: 15114347 |
| CD31 | NeuN | Efna5 | Epha4 | 0.00551 | 0 | EFNA5_EPHA4 | Efna5 - Epha4 | EPHA | Cell-Cell Contact | PMID: 15114347 |
| NeuN | NeuN | Efna5 | Epha4 | 0.007911 | 0 | EFNA5_EPHA4 | Efna5 - Epha4 | EPHA | Cell-Cell Contact | PMID: 15114347 |
| CD31 | NeuN | Efna5 | Epha5 | 0.002143 | 0 | EFNA5_EPHA5 | Efna5 - Epha5 | EPHA | Cell-Cell Contact | PMID: 15114347 |
| NeuN | NeuN | Efna5 | Epha5 | 0.003081 | 0 | EFNA5_EPHA5 | Efna5 - Epha5 | EPHA | Cell-Cell Contact | PMID: 15114347 |
| CD31 | NeuN | Efna5 | Epha7 | 0.001774 | 0 | EFNA5_EPHA7 | Efna5 - Epha7 | EPHA | Cell-Cell Contact | PMID: 15114347 |
| NeuN | NeuN | Efna5 | Epha7 | 0.002552 | 0 | EFNA5_EPHA7 | Efna5 - Epha7 | EPHA | Cell-Cell Contact | PMID: 15114347 |
| CD31 | CD31 | Efna5 | Epha8 | 0.000566 | 0 | EFNA5_EPHA8 | Efna5 - Epha8 | EPHA | Cell-Cell Contact | PMID: 15114347 |
| NeuN | CD31 | Efna5 | Epha8 | 0.000815 | 0 | EFNA5_EPHA8 | Efna5 - Epha8 | EPHA | Cell-Cell Contact | PMID: 15114347 |
| CD31 | NeuN | Efna5 | Ephb2 | 0.001444 | 0 | EFNA5_EPHB2 | Efna5 - Ephb2 | EPHA | Cell-Cell Contact | PMID:15107857; PMID: 15114347 |
| NeuN | NeuN | Efna5 | Ephb2 | 0.002077 | 0 | EFNA5_EPHB2 | Efna5 - Ephb2 | EPHA | Cell-Cell Contact | PMID:15107857; PMID: 15114347 |
| CD31 | NeuN | Efnb1 | Epha4 | 0.006407 | 0 | EFNB1_EPHA4 | Efnb1 - Epha4 | EPHB | Cell-Cell Contact | PMID: 15114347 |
| NeuN | NeuN | Efnb1 | Epha4 | 0.004115 | 0 | EFNB1_EPHA4 | Efnb1 - Epha4 | EPHB | Cell-Cell Contact | PMID: 15114347 |
| CD31 | GFAP | Efnb1 | Ephb1 | 0.003947 | 0 | EFNB1_EPHB1 | Efnb1 - Ephb1 | EPHB | Cell-Cell Contact | PMID: 15114347 |
| GFAP | GFAP | Efnb1 | Ephb1 | 0.003293 | 0 | EFNB1_EPHB1 | Efnb1 - Ephb1 | EPHB | Cell-Cell Contact | PMID: 15114347 |
| CD31 | NeuN | Efnb1 | Ephb2 | 0.00168 | 0 | EFNB1_EPHB2 | Efnb1 - Ephb2 | EPHB | Cell-Cell Contact | PMID: 15114347 |
| NeuN | NeuN | Efnb1 | Ephb2 | 0.001077 | 0 | EFNB1_EPHB2 | Efnb1 - Ephb2 | EPHB | Cell-Cell Contact | PMID: 15114347 |
| CD31 | GFAP | Efnb1 | Ephb3 | 0.000916 | 0 | EFNB1_EPHB3 | Efnb1 - Ephb3 | EPHB | Cell-Cell Contact | PMID: 15114347 |
| GFAP | GFAP | Efnb1 | Ephb3 | 0.000764 | 0 | EFNB1_EPHB3 | Efnb1 - Ephb3 | EPHB | Cell-Cell Contact | PMID: 15114347 |
| CD31 | CD31 | Efnb1 | Ephb4 | 0.002955 | 0 | EFNB1_EPHB4 | Efnb1 - Ephb4 | EPHB | Cell-Cell Contact | PMID: 15114347 |
| GFAP | CD31 | Efnb1 | Ephb4 | 0.002465 | 0 | EFNB1_EPHB4 | Efnb1 - Ephb4 | EPHB | Cell-Cell Contact | PMID: 15114347 |
| CD31 | NeuN | Efnb2 | Epha4 | 0.009315 | 0 | EFNB2_EPHA4 | Efnb2 - Epha4 | EPHB | Cell-Cell Contact | PMID: 15114347 |
| NeuN | NeuN | Efnb2 | Epha4 | 0.009894 | 0 | EFNB2_EPHA4 | Efnb2 - Epha4 | EPHB | Cell-Cell Contact | PMID: 15114347 |
| NeuN | GFAP | Efnb2 | Ephb1 | 0.006104 | 0 | EFNB2_EPHB1 | Efnb2 - Ephb1 | EPHB | Cell-Cell Contact | PMID: 15114347 |
| NeuN | NeuN | Efnb2 | Ephb1 | 0.003486 | 0 | EFNB2_EPHB1 | Efnb2 - Ephb1 | EPHB | Cell-Cell Contact | PMID: 15114347 |

| | | | | | | | | | | |
|---|---|---|---|---|---|---|---|---|---|---|
| CD31 | NeuN | Efnb2 | Ephb2 | 0.002448 | 0 | EFNB2_EPHB2 | Efnb2 - Ephb2 | EPHB | Cell-Cell Contact | PMID: 15114347 |
| NeuN | NeuN | Efnb2 | Ephb2 | 0.002601 | 0 | EFNB2_EPHB2 | Efnb2 - Ephb2 | EPHB | Cell-Cell Contact | PMID: 15114347 |
| NeuN | GFAP | Efnb2 | Ephb3 | 0.001418 | 0 | EFNB2_EPHB3 | Efnb2 - Ephb3 | EPHB | Cell-Cell Contact | PMID: 15114347 |
| NeuN | NeuN | Efnb2 | Ephb3 | 0.001234 | 0 | EFNB2_EPHB3 | Efnb2 - Ephb3 | EPHB | Cell-Cell Contact | PMID: 15114347 |
| CD31 | CD31 | Efnb2 | Ephb4 | 0.004303 | 0 | EFNB2_EPHB4 | Efnb2 - Ephb4 | EPHB | Cell-Cell Contact | PMID: 15114347 |
| NeuN | CD31 | Efnb2 | Ephb4 | 0.004572 | 0 | EFNB2_EPHB4 | Efnb2 - Ephb4 | EPHB | Cell-Cell Contact | PMID: 15114347 |
| GFAP | GFAP | Efnb3 | Epha4 | 0.007491 | 0 | EFNB3_EPHA4 | Efnb3 - Epha4 | EPHB | Cell-Cell Contact | PMID: 15114347 |
| GFAP | NeuN | Efnb3 | Epha4 | 0.025714 | 0 | EFNB3_EPHA4 | Efnb3 - Epha4 | EPHB | Cell-Cell Contact | PMID: 15114347 |
| GFAP | GFAP | Efnb3 | Ephb1 | 0.01596 | 0 | EFNB3_EPHB1 | Efnb3 - Ephb1 | EPHB | Cell-Cell Contact | PMID: 15114347 |
| GFAP | NeuN | Efnb3 | Ephb1 | 0.009155 | 0 | EFNB3_EPHB1 | Efnb3 - Ephb1 | EPHB | Cell-Cell Contact | PMID: 15114347 |
| GFAP | GFAP | Efnb3 | Ephb2 | 0.004407 | 0 | EFNB3_EPHB2 | Efnb3 - Ephb2 | EPHB | Cell-Cell Contact | PMID: 15114347 |
| GFAP | NeuN | Efnb3 | Ephb2 | 0.00684 | 0 | EFNB3_EPHB2 | Efnb3 - Ephb2 | EPHB | Cell-Cell Contact | PMID: 15114347 |
| GFAP | GFAP | Efnb3 | Ephb3 | 0.003737 | 0 | EFNB3_EPHB3 | Efnb3 - Ephb3 | EPHB | Cell-Cell Contact | PMID: 15114347 |
| GFAP | NeuN | Efnb3 | Ephb3 | 0.003253 | 0 | EFNB3_EPHB3 | Efnb3 - Ephb3 | EPHB | Cell-Cell Contact | PMID: 15114347 |
| CD31 | CD31 | Efnb3 | Ephb4 | 0.00441 | 0 | EFNB3_EPHB4 | Efnb3 - Ephb4 | EPHB | Cell-Cell Contact | PMID: 15114347 |
| GFAP | CD31 | Efnb3 | Ephb4 | 0.011985 | 0 | EFNB3_EPHB4 | Efnb3 - Ephb4 | EPHB | Cell-Cell Contact | PMID: 15114347 |
| GFAP | NeuN | Efnb3 | Ephb6 | 0.019719 | 0 | EFNB3_EPHB6 | Efnb3 - Ephb6 | EPHB | Cell-Cell Contact | PMID: 15114347 |
| NeuN | NeuN | Efnb3 | Ephb6 | 0.004917 | 0 | EFNB3_EPHB6 | Efnb3 - Ephb6 | EPHB | Cell-Cell Contact | PMID: 15114347 |
| CD31 | CD31 | Esam | Esam | 0.018925 | 0 | ESAM_ESAM | Esam - Esam | ESAM | Cell-Cell Contact | KEGG: mmu04514; PMID: 11279107 |
| GFAP | CD31 | Esam | Esam | 0.002393 | 0 | ESAM_ESAM | Esam - Esam | ESAM | Cell-Cell Contact | KEGG: mmu04514; PMID: 11279107 |
| CD31 | GFAP | Esam | Esam | 0.002393 | 0 | ESAM_ESAM | Esam - Esam | ESAM | Cell-Cell Contact | KEGG: mmu04514; PMID: 11279107 |
| NeuN | CD31 | Icam5 | Cd209a | 0.000858 | 0 | ICAM5_CD209A | Icam5 - Cd209a | ICAM | Cell-Cell Contact | PMID: 15795245 |
| NeuN | NeuN | Icam5 | Cd209a | 0.000524 | 0 | ICAM5_CD209A | Icam5 - Cd209a | ICAM | Cell-Cell Contact | PMID: 15795245 |
| CD31 | CD31 | Jam2 | ITGA3_ITGB1 | 0.002026 | 0 | JAM2_ITGA3_ITGB1 | Jam2 - (Itga3+Itgb1) | JAM | Cell-Cell Contact | PMID: 12070135 |
| GFAP | CD31 | Jam2 | ITGA3_ITGB1 | 0.001665 | 0 | JAM2_ITGA3_ITGB1 | Jam2 - (Itga3+Itgb1) | JAM | Cell-Cell Contact | PMID: 12070135 |
| CD31 | CD31 | Jam2 | ITGAV_ITGB1 | 0.002313 | 0 | JAM2_ITGAV_ITGB1 | Jam2 - (Itgav+Itgb1) | JAM | Cell-Cell Contact | KEGG: mmu04514 |
| GFAP | CD31 | Jam2 | ITGAV_ITGB1 | 0.001901 | 0 | JAM2_ITGAV_ITGB1 | Jam2 - (Itgav+Itgb1) | JAM | Cell-Cell Contact | KEGG: mmu04514 |
| CD31 | GFAP | Jam2 | Jam3 | 0.002725 | 0 | JAM2_JAM3 | Jam2 - Jam3 | JAM | Cell-Cell Contact | PMC1237098; PMID: 11590146 |
| GFAP | GFAP | Jam2 | Jam3 | 0.00224 | 0 | JAM2_JAM3 | Jam2 - Jam3 | JAM | Cell-Cell Contact | PMC1237098; PMID: 11590146 |
| CD31 | CD31 | F11r | F11r | 0.002227 | 0 | F11R_F11R | F11r - F11r | JAM | Cell-Cell Contact | KEGG: mmu04514 |
| GFAP | CD31 | F11r | F11r | 0.000844 | 0 | F11R_F11R | F11r - F11r | JAM | Cell-Cell Contact | KEGG: mmu04514 |
| CD31 | GFAP | F11r | F11r | 0.000844 | 0 | F11R_F11R | F11r - F11r | JAM | Cell-Cell Contact | KEGG: mmu04514 |
| CD31 | CD31 | Jam2 | F11r | 0.001735 | 0 | JAM2_F11R | Jam2 - F11r | JAM | Cell-Cell Contact | KEGG: mmu04514 |
| GFAP | CD31 | Jam2 | F11r | 0.001426 | 0 | JAM2_F11R | Jam2 - F11r | JAM | Cell-Cell Contact | KEGG: mmu04514 |
| CD31 | CD31 | Jam2 | Jam2 | 0.001351 | 0 | JAM2_JAM2 | Jam2 - Jam2 | JAM | Cell-Cell Contact | KEGG: mmu04514 |
| GFAP | CD31 | Jam2 | Jam2 | 0.001111 | 0 | JAM2_JAM2 | Jam2 - Jam2 | JAM | Cell-Cell Contact | KEGG: mmu04514 |
| CD31 | GFAP | Jam2 | Jam2 | 0.001111 | 0 | JAM2_JAM2 | Jam2 - Jam2 | JAM | Cell-Cell Contact | KEGG: mmu04514 |
| CD31 | CD31 | Jam3 | F11r | 0.00164 | 0 | JAM3_F11R | Jam3 - F11r | JAM | Cell-Cell Contact | KEGG: mmu04514 |
| GFAP | CD31 | Jam3 | F11r | 0.003496 | 0 | JAM3_F11R | Jam3 - F11r | JAM | Cell-Cell Contact | KEGG: mmu04514 |
| GFAP | CD31 | Jam3 | Jam3 | 0.002576 | 0 | JAM3_JAM3 | Jam3 - Jam3 | JAM | Cell-Cell Contact | KEGG: mmu04514 |
| CD31 | GFAP | Jam3 | Jam3 | 0.002576 | 0 | JAM3_JAM3 | Jam3 - Jam3 | JAM | Cell-Cell Contact | KEGG: mmu04514 |
| GFAP | GFAP | Jam3 | Jam3 | 0.005486 | 0 | JAM3_JAM3 | Jam3 - Jam3 | JAM | Cell-Cell Contact | KEGG: mmu04514 |
| NeuN | GFAP | L1cam | ITGAV_ITGB3 | 0.002047 | 0 | L1CAM_ITGAV_ITGB3 | L1cam - (Itgav+Itgb3) | L1CAM | Cell-Cell Contact | PMID: 9396761 |
| NeuN | NeuN | L1cam | ITGAV_ITGB3 | 0.001675 | 0 | L1CAM_ITGAV_ITGB3 | L1cam - (Itgav+Itgb3) | L1CAM | Cell-Cell Contact | PMID: 9396761 |
| NeuN | CD31 | L1cam | ITGA4_ITGB7 | 0.00161 | 0 | L1CAM_ITGA4_ITGB7 | L1cam - (Itga4+Itgb7) | L1CAM | Cell-Cell Contact | PMID: 8557754 |
| NeuN | NeuN | L1cam | ITGA4_ITGB7 | 0.000828 | 0 | L1CAM_ITGA4_ITGB7 | L1cam - (Itga4+Itgb7) | L1CAM | Cell-Cell Contact | PMID: 8557754 |
| NeuN | GFAP | L1cam | L1cam | 0.002479 | 0 | L1CAM_L1CAM | L1cam - L1cam | L1CAM | Cell-Cell Contact | PMID: 18701456 |
| GFAP | NeuN | L1cam | L1cam | 0.002479 | 0 | L1CAM_L1CAM | L1cam - L1cam | L1CAM | Cell-Cell Contact | PMID: 18701456 |
| NeuN | NeuN | L1cam | L1cam | 0.005613 | 0 | L1CAM_L1CAM | L1cam - L1cam | L1CAM | Cell-Cell Contact | PMID: 18701456 |
| GFAP | CD31 | Mag | Mag | 0.009595 | 0 | MAG_MAG | Mag - Mag | MAG | Cell-Cell Contact | KEGG: mmu04514 |
| CD31 | GFAP | Mag | Mag | 0.009595 | 0 | MAG_MAG | Mag - Mag | MAG | Cell-Cell Contact | KEGG: mmu04514 |
| GFAP | GFAP | Mag | Mag | 0.042777 | 0 | MAG_MAG | Mag - Mag | MAG | Cell-Cell Contact | KEGG: mmu04514 |
| CD31 | CD31 | Mpzl1 | Mpzl1 | 0.007623 | 0 | MPZL1_MPZL1 | Mpzl1 - Mpzl1 | MPZ | Cell-Cell Contact | KEGG: mmu04514 |
| GFAP | CD31 | Mpzl1 | Mpzl1 | 0.004687 | 0 | MPZL1_MPZL1 | Mpzl1 - Mpzl1 | MPZ | Cell-Cell Contact | KEGG: mmu04514 |
| CD31 | GFAP | Mpzl1 | Mpzl1 | 0.004687 | 0 | MPZL1_MPZL1 | Mpzl1 - Mpzl1 | MPZ | Cell-Cell Contact | KEGG: mmu04514 |
| GFAP | GFAP | Ncam1 | Fgfr1 | 0.01414 | 0 | NCAM1_FGFR1 | Ncam1 - Fgfr1 | NCAM | Cell-Cell Contact | PMID: 12791257 |
| NeuN | GFAP | Ncam1 | Fgfr1 | 0.011855 | 0 | NCAM1_FGFR1 | Ncam1 - Fgfr1 | NCAM | Cell-Cell Contact | PMID: 12791257 |
| GFAP | GFAP | Ncam1 | Ncam1 | 0.036981 | 0 | NCAM1_NCAM1 | Ncam1 - Ncam1 | NCAM | Cell-Cell Contact | KEGG: mmu04514 |
| NeuN | GFAP | Ncam1 | Ncam1 | 0.031122 | 0 | NCAM1_NCAM1 | Ncam1 - Ncam1 | NCAM | Cell-Cell Contact | KEGG: mmu04514 |

| | | | | | | | | | | |
|---|---|---|---|---|---|---|---|---|---|---|
| GFAP | NeuN | Ncam1 | Ncam1 | 0.031122 | 0 | NCAM1_NCAM1 | Ncam1 - Ncam1 | NCAM | Cell-Cell Contact | KEGG: mmu04514 |
| GFAP | NeuN | Ncam1 | Ncam2 | 0.014816 | 0 | NCAM1_NCAM2 | Ncam1 - Ncam2 | NCAM | Cell-Cell Contact | KEGG: mmu04514 |
| NeuN | NeuN | Ncam1 | Ncam2 | 0.012424 | 0 | NCAM1_NCAM2 | Ncam1 - Ncam2 | NCAM | Cell-Cell Contact | KEGG: mmu04514 |
| GFAP | NeuN | Ncam1 | L1cam | 0.014509 | 0 | NCAM1_L1CAM | Ncam1 - L1cam | NCAM | Cell-Cell Contact | KEGG: mmu04514 |
| NeuN | NeuN | Ncam1 | L1cam | 0.012165 | 0 | NCAM1_L1CAM | Ncam1 - L1cam | NCAM | Cell-Cell Contact | KEGG: mmu04514 |
| GFAP | NeuN | Ncam2 | L1cam | 0.00256 | 0 | NCAM2_L1CAM | Ncam2 - L1cam | NCAM | Cell-Cell Contact | KEGG: mmu04514 |
| NeuN | NeuN | Ncam2 | L1cam | 0.005733 | 0 | NCAM2_L1CAM | Ncam2 - L1cam | NCAM | Cell-Cell Contact | KEGG: mmu04514 |
| CD31 | GFAP | Nectin1 | Cd96 | 0.000659 | 0 | NECTIN1_CD96 | Nectin1 - Cd96 | NECTIN | Cell-Cell Contact | PMID: 23027581 |
| GFAP | GFAP | Nectin1 | Cd96 | 0.000571 | 0 | NECTIN1_CD96 | Nectin1 - Cd96 | NECTIN | Cell-Cell Contact | PMID: 23027581 |
| GFAP | CD31 | Nectin3 | Nectin2 | 0.001043 | 0 | NECTIN3_NECTIN2 | Nectin3 - Nectin2 | NECTIN | Cell-Cell Contact | PMID: 23027581 |
| GFAP | GFAP | Nectin3 | Nectin2 | 0.001038 | 0 | NECTIN3_NECTIN2 | Nectin3 - Nectin2 | NECTIN | Cell-Cell Contact | PMID: 23027581 |
| GFAP | CD31 | Nectin3 | Pvr | 0.000626 | 0 | NECTIN3_PVR | Nectin3 - Pvr | NECTIN | Cell-Cell Contact | PMID: 23027581 |
| GFAP | GFAP | Nectin3 | Pvr | 0.000555 | 0 | NECTIN3_PVR | Nectin3 - Pvr | NECTIN | Cell-Cell Contact | PMID: 23027581 |
| CD31 | GFAP | Nectin1 | Nectin3 | 0.001479 | 0 | NECTIN1_NECTIN3 | Nectin1 - Nectin3 | NECTIN | Cell-Cell Contact | KEGG: mmu04514 |
| GFAP | GFAP | Nectin1 | Nectin3 | 0.001281 | 0 | NECTIN1_NECTIN3 | Nectin1 - Nectin3 | NECTIN | Cell-Cell Contact | KEGG: mmu04514 |
| NeuN | CD31 | Negr1 | Negr1 | 0.003902 | 0 | NEGR1_NEGR1 | Negr1 - Negr1 | NEGR | Cell-Cell Contact | KEGG: mmu04514 |
| CD31 | NeuN | Negr1 | Negr1 | 0.003902 | 0 | NEGR1_NEGR1 | Negr1 - Negr1 | NEGR | Cell-Cell Contact | KEGG: mmu04514 |
| NeuN | NeuN | Negr1 | Negr1 | 0.009402 | 0 | NEGR1_NEGR1 | Negr1 - Negr1 | NEGR | Cell-Cell Contact | KEGG: mmu04514 |
| NeuN | GFAP | Lrrc4c | Ntng1 | 0.001796 | 0 | LRRC4C_NTNG1 | Lrrc4c - Ntng1 | NGL | Cell-Cell Contact | KEGG: mmu04514 |
| NeuN | NeuN | Lrrc4c | Ntng1 | 0.001812 | 0 | LRRC4C_NTNG1 | Lrrc4c - Ntng1 | NGL | Cell-Cell Contact | KEGG: mmu04514 |
| NeuN | GFAP | Lrrc4 | Ntng2 | 0.001221 | 0 | LRRC4_NTNG2 | Lrrc4 - Ntng2 | NGL | Cell-Cell Contact | KEGG: mmu04514 |
| NeuN | NeuN | Lrrc4 | Ntng2 | 0.001345 | 0 | LRRC4_NTNG2 | Lrrc4 - Ntng2 | NGL | Cell-Cell Contact | KEGG: mmu04514 |
| NeuN | GFAP | Lrrc4b | Ptprf | 0.005171 | 0 | LRRC4B_PTPRF | Lrrc4b - Ptprf | NGL | Cell-Cell Contact | KEGG: mmu04514 |
| NeuN | NeuN | Lrrc4b | Ptprf | 0.005426 | 0 | LRRC4B_PTPRF | Lrrc4b - Ptprf | NGL | Cell-Cell Contact | KEGG: mmu04514 |
| GFAP | CD31 | Dlk1 | Notch1 | 0.001445 | 0 | DLK1_NOTCH1 | Dlk1 - Notch1 | NOTCH | Cell-Cell Contact | PMID: 22353464 |
| GFAP | GFAP | Dlk1 | Notch1 | 0.001767 | 0 | DLK1_NOTCH1 | Dlk1 - Notch1 | NOTCH | Cell-Cell Contact | PMID: 22353464 |
| CD31 | CD31 | Dlk1 | Notch3 | 0.002146 | 0 | DLK1_NOTCH3 | Dlk1 - Notch3 | NOTCH | Cell-Cell Contact | PMID: 22353464 |
| GFAP | CD31 | Dlk1 | Notch3 | 0.003667 | 0 | DLK1_NOTCH3 | Dlk1 - Notch3 | NOTCH | Cell-Cell Contact | PMID: 22353464 |
| CD31 | CD31 | Dlk1 | Notch4 | 0.000999 | 0 | DLK1_NOTCH4 | Dlk1 - Notch4 | NOTCH | Cell-Cell Contact | PMID: 22353464 |
| GFAP | CD31 | Dlk1 | Notch4 | 0.001709 | 0 | DLK1_NOTCH4 | Dlk1 - Notch4 | NOTCH | Cell-Cell Contact | PMID: 22353464 |
| GFAP | CD31 | Dll1 | Notch1 | 0.001315 | 0 | DLL1_NOTCH1 | Dll1 - Notch1 | NOTCH | Cell-Cell Contact | PMID: 22353464 |
| GFAP | GFAP | Dll1 | Notch1 | 0.001608 | 0 | DLL1_NOTCH1 | Dll1 - Notch1 | NOTCH | Cell-Cell Contact | PMID: 22353464 |
| CD31 | CD31 | Dll1 | Notch3 | 0.002476 | 0 | DLL1_NOTCH3 | Dll1 - Notch3 | NOTCH | Cell-Cell Contact | PMID: 22353464 |
| GFAP | CD31 | Dll1 | Notch3 | 0.003337 | 0 | DLL1_NOTCH3 | Dll1 - Notch3 | NOTCH | Cell-Cell Contact | PMID: 22353464 |
| CD31 | CD31 | Dll1 | Notch4 | 0.001153 | 0 | DLL1_NOTCH4 | Dll1 - Notch4 | NOTCH | Cell-Cell Contact | PMID: 22353464 |
| GFAP | CD31 | Dll1 | Notch4 | 0.001555 | 0 | DLL1_NOTCH4 | Dll1 - Notch4 | NOTCH | Cell-Cell Contact | PMID: 22353464 |
| CD31 | CD31 | Dll3 | Notch1 | 0.00091 | 0 | DLL3_NOTCH1 | Dll3 - Notch1 | NOTCH | Cell-Cell Contact | PMID: 22353464 |
| CD31 | GFAP | Dll3 | Notch1 | 0.001113 | 0 | DLL3_NOTCH1 | Dll3 - Notch1 | NOTCH | Cell-Cell Contact | PMID: 22353464 |
| CD31 | CD31 | Dll3 | Notch3 | 0.002311 | 0 | DLL3_NOTCH3 | Dll3 - Notch3 | NOTCH | Cell-Cell Contact | PMID: 22353464 |
| GFAP | CD31 | Dll3 | Notch3 | 0.001827 | 0 | DLL3_NOTCH3 | Dll3 - Notch3 | NOTCH | Cell-Cell Contact | PMID: 22353464 |
| CD31 | CD31 | Dll3 | Notch4 | 0.001076 | 0 | DLL3_NOTCH4 | Dll3 - Notch4 | NOTCH | Cell-Cell Contact | PMID: 22353464 |
| GFAP | CD31 | Dll3 | Notch4 | 0.00085 | 0 | DLL3_NOTCH4 | Dll3 - Notch4 | NOTCH | Cell-Cell Contact | PMID: 22353464 |
| CD31 | CD31 | Dll4 | Notch1 | 0.001698 | 0 | DLL4_NOTCH1 | Dll4 - Notch1 | NOTCH | Cell-Cell Contact | PMID: 22353464 |
| CD31 | GFAP | Dll4 | Notch1 | 0.002076 | 0 | DLL4_NOTCH1 | Dll4 - Notch1 | NOTCH | Cell-Cell Contact | PMID: 22353464 |
| CD31 | CD31 | Dll4 | Notch3 | 0.004307 | 0 | DLL4_NOTCH3 | Dll4 - Notch3 | NOTCH | Cell-Cell Contact | PMID: 22353464 |
| GFAP | CD31 | Dll4 | Notch3 | 0.001446 | 0 | DLL4_NOTCH3 | Dll4 - Notch3 | NOTCH | Cell-Cell Contact | PMID: 22353464 |
| CD31 | CD31 | Dll4 | Notch4 | 0.002008 | 0 | DLL4_NOTCH4 | Dll4 - Notch4 | NOTCH | Cell-Cell Contact | PMID: 22353464 |
| GFAP | CD31 | Dll4 | Notch4 | 0.000673 | 0 | DLL4_NOTCH4 | Dll4 - Notch4 | NOTCH | Cell-Cell Contact | PMID: 22353464 |
| CD31 | CD31 | Jag1 | Notch1 | 0.001994 | 0 | JAG1_NOTCH1 | Jag1 - Notch1 | NOTCH | Cell-Cell Contact | PMID: 22353464 |
| CD31 | GFAP | Jag1 | Notch1 | 0.002438 | 0 | JAG1_NOTCH1 | Jag1 - Notch1 | NOTCH | Cell-Cell Contact | PMID: 22353464 |
| CD31 | CD31 | Jag1 | Notch3 | 0.005055 | 0 | JAG1_NOTCH3 | Jag1 - Notch3 | NOTCH | Cell-Cell Contact | PMID: 22353464 |
| GFAP | CD31 | Jag1 | Notch3 | 0.002119 | 0 | JAG1_NOTCH3 | Jag1 - Notch3 | NOTCH | Cell-Cell Contact | PMID: 22353464 |
| CD31 | CD31 | Jag1 | Notch4 | 0.002357 | 0 | JAG1_NOTCH4 | Jag1 - Notch4 | NOTCH | Cell-Cell Contact | PMID: 22353464 |
| GFAP | CD31 | Jag1 | Notch4 | 0.000987 | 0 | JAG1_NOTCH4 | Jag1 - Notch4 | NOTCH | Cell-Cell Contact | PMID: 22353464 |
| CD31 | CD31 | Jag2 | Notch1 | 0.002679 | 0 | JAG2_NOTCH1 | Jag2 - Notch1 | NOTCH | Cell-Cell Contact | PMID: 22353464 |
| CD31 | GFAP | Jag2 | Notch1 | 0.003275 | 0 | JAG2_NOTCH1 | Jag2 - Notch1 | NOTCH | Cell-Cell Contact | PMID: 22353464 |
| CD31 | CD31 | Jag2 | Notch3 | 0.006785 | 0 | JAG2_NOTCH3 | Jag2 - Notch3 | NOTCH | Cell-Cell Contact | PMID: 22353464 |
| NeuN | CD31 | Jag2 | Notch3 | 0.002814 | 0 | JAG2_NOTCH3 | Jag2 - Notch3 | NOTCH | Cell-Cell Contact | PMID: 22353464 |
| CD31 | CD31 | Jag2 | Notch4 | 0.003167 | 0 | JAG2_NOTCH4 | Jag2 - Notch4 | NOTCH | Cell-Cell Contact | PMID: 22353464 |

| | | | | | | | | | |
|---|---|---|---|---|---|---|---|---|---|
| NeuN | CD31 | Jag2 | Notch4 | 0.001311 | 0 | JAG2_NOTCH4 | Jag2 - Notch4 | NOTCH | Cell-Cell Contact | PMID: 22353464 |
| CD31 | NeuN | Nrxn1 | Nlgn1 | 0.00816 | 0 | NRXN1_NLGN1 | Nrxn1 - Nlgn1 | NRXN | Cell-Cell Contact | KEGG: mmu04514 |
| NeuN | NeuN | Nrxn1 | Nlgn1 | 0.015889 | 0 | NRXN1_NLGN1 | Nrxn1 - Nlgn1 | NRXN | Cell-Cell Contact | KEGG: mmu04514 |
| CD31 | NeuN | Nrxn1 | Nlgn2 | 0.018177 | 0 | NRXN1_NLGN2 | Nrxn1 - Nlgn2 | NRXN | Cell-Cell Contact | KEGG: mmu04514 |
| NeuN | NeuN | Nrxn1 | Nlgn2 | 0.035059 | 0 | NRXN1_NLGN2 | Nrxn1 - Nlgn2 | NRXN | Cell-Cell Contact | KEGG: mmu04514 |
| NeuN | CD31 | Nrxn1 | Nlgn3 | 0.016597 | 0 | NRXN1_NLGN3 | Nrxn1 - Nlgn3 | NRXN | Cell-Cell Contact | KEGG: mmu04514 |
| NeuN | NeuN | Nrxn1 | Nlgn3 | 0.031503 | 0 | NRXN1_NLGN3 | Nrxn1 - Nlgn3 | NRXN | Cell-Cell Contact | KEGG: mmu04514 |
| GFAP | NeuN | Nrxn2 | Nlgn1 | 0.005484 | 0 | NRXN2_NLGN1 | Nrxn2 - Nlgn1 | NRXN | Cell-Cell Contact | KEGG: mmu04514 |
| NeuN | NeuN | Nrxn2 | Nlgn1 | 0.006956 | 0 | NRXN2_NLGN1 | Nrxn2 - Nlgn1 | NRXN | Cell-Cell Contact | KEGG: mmu04514 |
| GFAP | NeuN | Nrxn2 | Nlgn2 | 0.012256 | 0 | NRXN2_NLGN2 | Nrxn2 - Nlgn2 | NRXN | Cell-Cell Contact | KEGG: mmu04514 |
| NeuN | NeuN | Nrxn2 | Nlgn2 | 0.015517 | 0 | NRXN2_NLGN2 | Nrxn2 - Nlgn2 | NRXN | Cell-Cell Contact | KEGG: mmu04514 |
| GFAP | NeuN | Nrxn2 | Nlgn3 | 0.010986 | 0 | NRXN2_NLGN3 | Nrxn2 - Nlgn3 | NRXN | Cell-Cell Contact | KEGG: mmu04514 |
| NeuN | NeuN | Nrxn2 | Nlgn3 | 0.013915 | 0 | NRXN2_NLGN3 | Nrxn2 - Nlgn3 | NRXN | Cell-Cell Contact | KEGG: mmu04514 |
| NeuN | GFAP | Nrxn3 | Nlgn1 | 0.003336 | 0 | NRXN3_NLGN1 | Nrxn3 - Nlgn1 | NRXN | Cell-Cell Contact | KEGG: mmu04514 |
| NeuN | NeuN | Nrxn3 | Nlgn1 | 0.008108 | 0 | NRXN3_NLGN1 | Nrxn3 - Nlgn1 | NRXN | Cell-Cell Contact | KEGG: mmu04514 |
| NeuN | CD31 | Nrxn3 | Nlgn2 | 0.007845 | 0 | NRXN3_NLGN2 | Nrxn3 - Nlgn2 | NRXN | Cell-Cell Contact | KEGG: mmu04514 |
| NeuN | NeuN | Nrxn3 | Nlgn2 | 0.018062 | 0 | NRXN3_NLGN2 | Nrxn3 - Nlgn2 | NRXN | Cell-Cell Contact | KEGG: mmu04514 |
| NeuN | CD31 | Nrxn3 | Nlgn3 | 0.008472 | 0 | NRXN3_NLGN3 | Nrxn3 - Nlgn3 | NRXN | Cell-Cell Contact | KEGG: mmu04514 |
| NeuN | NeuN | Nrxn3 | Nlgn3 | 0.016201 | 0 | NRXN3_NLGN3 | Nrxn3 - Nlgn3 | NRXN | Cell-Cell Contact | KEGG: mmu04514 |
| CD31 | CD31 | Ocln | Ocln | 0.005861 | 0 | OCLN_OCLN | Ocln - Ocln | OCLN | Cell-Cell Contact | KEGG: mmu04514 |
| GFAP | CD31 | Ocln | Ocln | 0.001189 | 0 | OCLN_OCLN | Ocln - Ocln | OCLN | Cell-Cell Contact | KEGG: mmu04514 |
| CD31 | GFAP | Ocln | Ocln | 0.001189 | 0 | OCLN_OCLN | Ocln - Ocln | OCLN | Cell-Cell Contact | KEGG: mmu04514 |
| CD31 | CD31 | Pecam1 | Pecam1 | 0.004327 | 0 | PECAM1_PECAM1 | Pecam1 - Pecam1 | PECAM1 | Cell-Cell Contact | KEGG: mmu04514 |
| GFAP | CD31 | Pecam1 | Pecam1 | 0.001088 | 0 | PECAM1_PECAM1 | Pecam1 - Pecam1 | PECAM1 | Cell-Cell Contact | KEGG: mmu04514 |
| CD31 | GFAP | Pecam1 | Pecam1 | 0.001088 | 0 | PECAM1_PECAM1 | Pecam1 - Pecam1 | PECAM1 | Cell-Cell Contact | KEGG: mmu04514 |
| CD31 | CD31 | Ptprm | Ptprm | 0.02895 | 0 | PTPRM_PTPRM | Ptprm - Ptprm | PTPRM | Cell-Cell Contact | KEGG: mmu04514 |
| NeuN | CD31 | Ptprm | Ptprm | 0.008966 | 0 | PTPRM_PTPRM | Ptprm - Ptprm | PTPRM | Cell-Cell Contact | KEGG: mmu04514 |
| CD31 | NeuN | Ptprm | Ptprm | 0.008966 | 0 | PTPRM_PTPRM | Ptprm - Ptprm | PTPRM | Cell-Cell Contact | KEGG: mmu04514 |
| CD31 | CD31 | Sele | Glg1 | 0.000542 | 0 | SELE_GLG1 | Sele - Glg1 | SELE | Cell-Cell Contact | PMID: 11404363 |
| GFAP | NeuN | Sele | Glg1 | 0.001076 | 0 | SELE_GLG1 | Sele - Glg1 | SELE | Cell-Cell Contact | PMID: 11404363 |
| CD31 | CD31 | Sell | Cd34 | 0.004009 | 0 | SELL_CD34 | Sell - Cd34 | SELL | Cell-Cell Contact | PMID: 7692600; PMID: 8977216 |
| GFAP | CD31 | Sell | Cd34 | 0.005071 | 0 | SELL_CD34 | Sell - Cd34 | SELL | Cell-Cell Contact | PMID: 7692600; PMID: 8977216 |
| CD31 | CD31 | Sell | Podxl | 0.003608 | 0 | SELL_PODXL | Sell - Podxl | SELL | Cell-Cell Contact | PMID: 22814396 |
| GFAP | CD31 | Sell | Podxl | 0.004565 | 0 | SELL_PODXL | Sell - Podxl | SELL | Cell-Cell Contact | PMID: 22814396 |
| CD31 | GFAP | Selplg | Sele | 0.000972 | 0 | SELPLG_SELE | Selplg - Sele | SELPLG | Cell-Cell Contact | PMC4571854 |
| GFAP | GFAP | Selplg | Sele | 0.000783 | 0 | SELPLG_SELE | Selplg - Sele | SELPLG | Cell-Cell Contact | PMC4571854 |
| CD31 | GFAP | Selplg | Sell | 0.001469 | 0 | SELPLG_SELL | Selplg - Sell | SELPLG | Cell-Cell Contact | PMC2431087; PMID: 8892633,PMID: 8896607 |
| GFAP | GFAP | Selplg | Sell | 0.001183 | 0 | SELPLG_SELL | Selplg - Sell | SELPLG | Cell-Cell Contact | PMC2431087; PMID: 8892633,PMID: 8896607 |
| NeuN | CD31 | Sema4a | NRP1_PLXNA1 | 0.001869 | 0 | SEMA4A_NRP1_PLXNA1 | Sema4a - (Nrp1+Plxna1) | SEMA4 | Cell-Cell Contact | PMID: 27533782 |
| NeuN | NeuN | Sema4a | NRP1_PLXNA1 | 0.00257 | 0 | SEMA4A_NRP1_PLXNA1 | Sema4a - (Nrp1+Plxna1) | SEMA4 | Cell-Cell Contact | PMID: 27533782 |
| NeuN | CD31 | Sema4a | NRP1_PLXNA2 | 0.00211 | 0 | SEMA4A_NRP1_PLXNA2 | Sema4a - (Nrp1+Plxna2) | SEMA4 | Cell-Cell Contact | PMID: 27533782 |
| NeuN | NeuN | Sema4a | NRP1_PLXNA2 | 0.003019 | 0 | SEMA4A_NRP1_PLXNA2 | Sema4a - (Nrp1+Plxna2) | SEMA4 | Cell-Cell Contact | PMID: 27533782 |
| NeuN | CD31 | Sema4a | NRP1_PLXNA3 | 0.001422 | 0 | SEMA4A_NRP1_PLXNA3 | Sema4a - (Nrp1+Plxna3) | SEMA4 | Cell-Cell Contact | PMID: 27533782 |
| NeuN | NeuN | Sema4a | NRP1_PLXNA3 | 0.001459 | 0 | SEMA4A_NRP1_PLXNA3 | Sema4a - (Nrp1+Plxna3) | SEMA4 | Cell-Cell Contact | PMID: 27533782 |
| NeuN | CD31 | Sema4a | NRP1_PLXNA4 | 0.002035 | 0 | SEMA4A_NRP1_PLXNA4 | Sema4a - (Nrp1+Plxna4) | SEMA4 | Cell-Cell Contact | PMID: 27533782 |
| NeuN | NeuN | Sema4a | NRP1_PLXNA4 | 0.003169 | 0 | SEMA4A_NRP1_PLXNA4 | Sema4a - (Nrp1+Plxna4) | SEMA4 | Cell-Cell Contact | PMID: 27533782 |
| GFAP | GFAP | Sema4a | Plxnb1 | 0.001505 | 0 | SEMA4A_PLXNB1 | Sema4a - Plxnb1 | SEMA4 | Cell-Cell Contact | PMID: 27533782 |
| NeuN | GFAP | Sema4a | Plxnb1 | 0.002639 | 0 | SEMA4A_PLXNB1 | Sema4a - Plxnb1 | SEMA4 | Cell-Cell Contact | PMID: 27533782 |
| NeuN | CD31 | Sema4a | Plxnb2 | 0.002586 | 0 | SEMA4A_PLXNB2 | Sema4a - Plxnb2 | SEMA4 | Cell-Cell Contact | PMID: 27533782 |
| GFAP | GFAP | Sema4a | Plxnb2 | 0.001417 | 0 | SEMA4A_PLXNB2 | Sema4a - Plxnb2 | SEMA4 | Cell-Cell Contact | PMID: 27533782 |
| GFAP | GFAP | Sema4a | Plxnb3 | 0.00132 | 0 | SEMA4A_PLXNB3 | Sema4a - Plxnb3 | SEMA4 | Cell-Cell Contact | PMID: 27533782 |
| NeuN | GFAP | Sema4a | Plxnb3 | 0.002315 | 0 | SEMA4A_PLXNB3 | Sema4a - Plxnb3 | SEMA4 | Cell-Cell Contact | PMID: 27533782 |
| GFAP | CD31 | Sema4d | Plxnb1 | 0.004383 | 0 | SEMA4D_PLXNB1 | Sema4d - Plxnb1 | SEMA4 | Cell-Cell Contact | PMID: 27533782 |
| GFAP | GFAP | Sema4d | Plxnb1 | 0.004949 | 0 | SEMA4D_PLXNB1 | Sema4d - Plxnb1 | SEMA4 | Cell-Cell Contact | PMID: 27533782 |
| GFAP | CD31 | Sema4d | Plxnb2 | 0.004851 | 0 | SEMA4D_PLXNB2 | Sema4d - Plxnb2 | SEMA4 | Cell-Cell Contact | PMID: 27533782 |
| GFAP | GFAP | Sema4d | Plxnb2 | 0.004662 | 0 | SEMA4D_PLXNB2 | Sema4d - Plxnb2 | SEMA4 | Cell-Cell Contact | PMID: 27533782 |
| GFAP | CD31 | Sema4d | Plxnb3 | 0.001772 | 0 | SEMA4D_PLXNB3 | Sema4d - Plxnb3 | SEMA4 | Cell-Cell Contact | PMID: 27533782 |
| GFAP | GFAP | Sema4d | Plxnb3 | 0.004344 | 0 | SEMA4D_PLXNB3 | Sema4d - Plxnb3 | SEMA4 | Cell-Cell Contact | PMID: 27533782 |
| CD31 | CD31 | Sema4c | Plxnb2 | 0.001121 | 0 | SEMA4C_PLXNB2 | Sema4c - Plxnb2 | SEMA4 | Cell-Cell Contact | PMID: 27533782 |

| | | | | | | | | | | |
|---|---|---|---|---|---|---|---|---|---|---|
| CD31 | GFAP | Sema4c | Plxnb2 | 0.001077 | 0 | SEMA4C_PLXNB2 | Sema4c - Plxnb2 | SEMA4 | Cell-Cell Contact | PMID: 27533782 |
| GFAP | CD31 | Sema4g | Plxnb2 | 0.000865 | 0 | SEMA4G_PLXNB2 | Sema4g - Plxnb2 | SEMA4 | Cell-Cell Contact | PMID: 27533782 |
| GFAP | GFAP | Sema4g | Plxnb2 | 0.000831 | 0 | SEMA4G_PLXNB2 | Sema4g - Plxnb2 | SEMA4 | Cell-Cell Contact | PMID: 27533782 |
| NeuN | CD31 | Sema4a | Timd2 | 0.001897 | 0 | SEMA4A_TIMD2 | Sema4a - Timd2 | SEMA4 | Cell-Cell Contact | PMID: 22325954 |
| GFAP | GFAP | Sema4a | Timd2 | 0.000631 | 0 | SEMA4A_TIMD2 | Sema4a - Timd2 | SEMA4 | Cell-Cell Contact | PMID: 22325954 |
| GFAP | NeuN | Sema5a | Plxna1 | 0.002132 | 0 | SEMA5A_PLXNA1 | Sema5a - Plxna1 | SEMA5 | Cell-Cell Contact | PMID: 27533782 |
| NeuN | NeuN | Sema5a | Plxna1 | 0.001498 | 0 | SEMA5A_PLXNA1 | Sema5a - Plxna1 | SEMA5 | Cell-Cell Contact | PMID: 27533782 |
| GFAP | NeuN | Sema5b | Plxna1 | 0.001369 | 0 | SEMA5B_PLXNA1 | Sema5b - Plxna1 | SEMA5 | Cell-Cell Contact | PMID: 27533782 |
| NeuN | NeuN | Sema5b | Plxna1 | 0.001597 | 0 | SEMA5B_PLXNA1 | Sema5b - Plxna1 | SEMA5 | Cell-Cell Contact | PMID: 27533782 |
| GFAP | CD31 | Sema5a | Plxna3 | 0.000712 | 0 | SEMA5A_PLXNA3 | Sema5a - Plxna3 | SEMA5 | Cell-Cell Contact | PMID: 27533782 |
| GFAP | GFAP | Sema5a | Plxna3 | 0.000886 | 0 | SEMA5A_PLXNA3 | Sema5a - Plxna3 | SEMA5 | Cell-Cell Contact | PMID: 27533782 |
| GFAP | GFAP | Sema5b | Plxna3 | 0.000569 | 0 | SEMA5B_PLXNA3 | Sema5b - Plxna3 | SEMA5 | Cell-Cell Contact | PMID: 27533782 |
| NeuN | GFAP | Sema5b | Plxna3 | 0.000663 | 0 | SEMA5B_PLXNA3 | Sema5b - Plxna3 | SEMA5 | Cell-Cell Contact | PMID: 27533782 |
| GFAP | GFAP | Sema5a | Plxnb3 | 0.00177 | 0 | SEMA5A_PLXNB3 | Sema5a - Plxnb3 | SEMA5 | Cell-Cell Contact | PMID: 27533782 |
| NeuN | GFAP | Sema5a | Plxnb3 | 0.001243 | 0 | SEMA5A_PLXNB3 | Sema5a - Plxnb3 | SEMA5 | Cell-Cell Contact | PMID: 27533782 |
| CD31 | CD31 | Sema6d | PLXNA1_KDR | 0.002565 | 0 | SEMA6D_PLXNA1_KDR | Sema6d - (Plxna1+Kdr) | SEMA6 | Cell-Cell Contact | PMID: 22325954 |
| GFAP | CD31 | Sema6d | PLXNA1_KDR | 0.002567 | 0 | SEMA6D_PLXNA1_KDR | Sema6d - (Plxna1+Kdr) | SEMA6 | Cell-Cell Contact | PMID: 22325954 |
| CD31 | GFAP | Sema6d | PLXNA1_TREM2 | 0.00134 | 0 | SEMA6D_PLXNA1_TREM2 | Sema6d - (Plxna1+Trem2) | SEMA6 | Cell-Cell Contact | PMID: 22325954 |
| GFAP | GFAP | Sema6d | PLXNA1_TREM2 | 0.001341 | 0 | SEMA6D_PLXNA1_TREM2 | Sema6d - (Plxna1+Trem2) | SEMA6 | Cell-Cell Contact | PMID: 22325954 |
| CD31 | CD31 | Sema6d | Plxna1 | 0.001459 | 0 | SEMA6D_PLXNA1 | Sema6d - Plxna1 | SEMA6 | Cell-Cell Contact | PMID: 27533782 |
| GFAP | NeuN | Sema6d | Plxna1 | 0.00253 | 0 | SEMA6D_PLXNA1 | Sema6d - Plxna1 | SEMA6 | Cell-Cell Contact | PMID: 27533782 |
| GFAP | GFAP | Sema6a | Plxna2 | 0.003687 | 0 | SEMA6A_PLXNA2 | Sema6a - Plxna2 | SEMA6 | Cell-Cell Contact | PMID: 27533782 |
| GFAP | NeuN | Sema6a | Plxna2 | 0.006157 | 0 | SEMA6A_PLXNA2 | Sema6a - Plxna2 | SEMA6 | Cell-Cell Contact | PMID: 27533782 |
| CD31 | NeuN | Sema6b | Plxna2 | 0.002566 | 0 | SEMA6B_PLXNA2 | Sema6b - Plxna2 | SEMA6 | Cell-Cell Contact | PMID: 27533782 |
| NeuN | NeuN | Sema6b | Plxna2 | 0.004251 | 0 | SEMA6B_PLXNA2 | Sema6b - Plxna2 | SEMA6 | Cell-Cell Contact | PMID: 27533782 |
| GFAP | GFAP | Sema6a | Plxna4 | 0.002459 | 0 | SEMA6A_PLXNA4 | Sema6a - Plxna4 | SEMA6 | Cell-Cell Contact | PMID: 27533782 |
| GFAP | NeuN | Sema6a | Plxna4 | 0.006781 | 0 | SEMA6A_PLXNA4 | Sema6a - Plxna4 | SEMA6 | Cell-Cell Contact | PMID: 27533782 |
| CD31 | NeuN | Sema6b | Plxna4 | 0.002827 | 0 | SEMA6B_PLXNA4 | Sema6b - Plxna4 | SEMA6 | Cell-Cell Contact | PMID: 27533782 |
| NeuN | NeuN | Sema6b | Plxna4 | 0.004682 | 0 | SEMA6B_PLXNA4 | Sema6b - Plxna4 | SEMA6 | Cell-Cell Contact | PMID: 27533782 |
| CD31 | CD31 | Sema7a | ITGB1_ITGA1 | 0.021324 | 0 | SEMA7A_ITGB1_ITGA1 | Sema7a - (Itgb1+Itga1) | SEMA7 | Cell-Cell Contact | PMID: 22325954 |
| CD31 | GFAP | Sema7a | ITGB1_ITGA1 | 0.009369 | 0 | SEMA7A_ITGB1_ITGA1 | Sema7a - (Itgb1+Itga1) | SEMA7 | Cell-Cell Contact | PMID: 22325954 |
| CD31 | CD31 | Sema7a | Plxnc1 | 0.003455 | 0 | SEMA7A_PLXNC1 | Sema7a - Plxnc1 | SEMA7 | Cell-Cell Contact | PMID: 27533782 |
| CD31 | GFAP | Sema7a | Plxnc1 | 0.007279 | 0 | SEMA7A_PLXNC1 | Sema7a - Plxnc1 | SEMA7 | Cell-Cell Contact | PMID: 27533782 |
| NeuN | GFAP | Thy1 | ITGAV_ITGB3 | 0.020529 | 0 | THY1_ITGAV_ITGB3 | Thy1 - (Itgav+Itgb3) | THY1 | Cell-Cell Contact | PMID: 29212879; PMID: 18346467 |
| NeuN | NeuN | Thy1 | ITGAV_ITGB3 | 0.016859 | 0 | THY1_ITGAV_ITGB3 | Thy1 - (Itgav+Itgb3) | THY1 | Cell-Cell Contact | PMID: 29212879; PMID: 18346467 |
| CD31 | CD31 | Cdh3 | Cdh3 | 0.000621 | 0 | CDH3_CDH3 | Cdh3 - Cdh3 | CDH | Cell-Cell Contact | KEGG: hsa04514 |
| GFAP | CD31 | Cdh3 | Cdh3 | 0.000476 | 0 | CDH3_CDH3 | Cdh3 - Cdh3 | CDH | Cell-Cell Contact | KEGG: hsa04514 |
| CD31 | GFAP | Cdh3 | Cdh3 | 0.000476 | 0 | CDH3_CDH3 | Cdh3 - Cdh3 | CDH | Cell-Cell Contact | KEGG: hsa04514 |
| CD31 | CD31 | Cdh12 | ITGA1_ITGB1 | 0.001732 | 0 | CDH12_ITGA1_ITGB1 | Cdh12 - ITGA1_ITGB1 | CDH | Cell-Cell Contact | PMID: 34385456 |
| NeuN | CD31 | Cdh12 | ITGA1_ITGB1 | 0.004063 | 0 | CDH12_ITGA1_ITGB1 | Cdh12 - ITGA1_ITGB1 | CDH | Cell-Cell Contact | PMID: 34385456 |
| GFAP | CD31 | Gja1 | Gja1 | 0.037404 | 0 | GJA1_GJA1 | Gja1 - Gja1 | GAP | Cell-Cell Contact | PMID: 32532966;PMID: 15094344 |
| CD31 | GFAP | Gja1 | Gja1 | 0.037404 | 0 | GJA1_GJA1 | Gja1 - Gja1 | GAP | Cell-Cell Contact | PMID: 32532966;PMID: 15094344 |
| GFAP | GFAP | Gja1 | Gja1 | 0.110322 | 0 | GJA1_GJA1 | Gja1 - Gja1 | GAP | Cell-Cell Contact | PMID: 32532966;PMID: 15094344 |
| GFAP | GFAP | Gja1 | Gja6 | 0.005451 | 0 | GJA1_GJA6 | Gja1 - Gja6 | GAP | Cell-Cell Contact | PMID: 32532966;PMID: 15094344 |
| GFAP | NeuN | Gja1 | Gja6 | 0.006756 | 0 | GJA1_GJA6 | Gja1 - Gja6 | GAP | Cell-Cell Contact | PMID: 32532966;PMID: 15094344 |
| NeuN | GFAP | Gja6 | Gja6 | 0.000301 | 0 | GJA6_GJA6 | Gja6 - Gja6 | GAP | Cell-Cell Contact | PMID: 32532966;PMID: 15094344 |
| GFAP | NeuN | Gja6 | Gja6 | 0.000301 | 0 | GJA6_GJA6 | Gja6 - Gja6 | GAP | Cell-Cell Contact | PMID: 32532966;PMID: 15094344 |
| NeuN | NeuN | Gja6 | Gja6 | 0.000373 | 0 | GJA6_GJA6 | Gja6 - Gja6 | GAP | Cell-Cell Contact | PMID: 32532966;PMID: 15094344 |
| CD31 | GFAP | Gjc1 | Gja1 | 0.018344 | 0 | GJC1_GJA1 | Gjc1 - Gja1 | GAP | Cell-Cell Contact | PMID: 32532966;PMID: 15094344 |
| GFAP | GFAP | Gjc1 | Gja1 | 0.007652 | 0 | GJC1_GJA1 | Gjc1 - Gja1 | GAP | Cell-Cell Contact | PMID: 32532966;PMID: 15094344 |
| CD31 | CD31 | Gjc1 | Gja6 | 0.000789 | 0 | GJC1_GJA6 | Gjc1 - Gja6 | GAP | Cell-Cell Contact | PMID: 32532966;PMID: 15094344 |
| CD31 | NeuN | Gjc1 | Gja6 | 0.001024 | 0 | GJC1_GJA6 | Gjc1 - Gja6 | GAP | Cell-Cell Contact | PMID: 32532966;PMID: 15094344 |
| CD31 | CD31 | Gjc1 | Gjc1 | 0.002808 | 0 | GJC1_GJC1 | Gjc1 - Gjc1 | GAP | Cell-Cell Contact | PMID: 32532966;PMID: 15094344 |
| GFAP | CD31 | Gjc1 | Gjc1 | 0.001161 | 0 | GJC1_GJC1 | Gjc1 - Gjc1 | GAP | Cell-Cell Contact | PMID: 32532966;PMID: 15094344 |
| CD31 | GFAP | Gjc1 | Gjc1 | 0.001161 | 0 | GJC1_GJC1 | Gjc1 - Gjc1 | GAP | Cell-Cell Contact | PMID: 32532966;PMID: 15094344 |
| GFAP | CD31 | Cldn1 | Cldn1 | 0.000293 | 0 | CLDN1_CLDN1 | Cldn1 - Cldn1 | CLDN | Cell-Cell Contact | PMID: 31942240 |
| CD31 | GFAP | Cldn1 | Cldn1 | 0.000293 | 0 | CLDN1_CLDN1 | Cldn1 - Cldn1 | CLDN | Cell-Cell Contact | PMID: 31942240 |
| GFAP | GFAP | Cldn1 | Cldn1 | 0.00046 | 0 | CLDN1_CLDN1 | Cldn1 - Cldn1 | CLDN | Cell-Cell Contact | PMID: 31942240 |
| CD31 | CD31 | Cldn3 | Cldn3 | 0.000388 | 0 | CLDN3_CLDN3 | Cldn3 - Cldn3 | CLDN | Cell-Cell Contact | PMID: 31942240 |

| | | | | | | | | | | |
|---|---|---|---|---|---|---|---|---|---|---|
| GFAP | CD31 | Cldn3 | Cldn3 | 0.000354 | 0 | CLDN3_CLDN3 | Cldn3 - Cldn3 | CLDN | Cell-Cell Contact | PMID: 31942240 |
| CD31 | GFAP | Cldn3 | Cldn3 | 0.000354 | 0 | CLDN3_CLDN3 | Cldn3 - Cldn3 | CLDN | Cell-Cell Contact | PMID: 31942240 |
| GFAP | CD31 | Pcdha9 | Pcdha9 | 0.000224 | 0 | PCDHA9_PCDHA9 | Pcdha9 - Pcdha9 | PCDH | Cell-Cell Contact | PMID: 35253643; PMID: 32694982 |
| CD31 | GFAP | Pcdha9 | Pcdha9 | 0.000224 | 0 | PCDHA9_PCDHA9 | Pcdha9 - Pcdha9 | PCDH | Cell-Cell Contact | PMID: 35253643; PMID: 32694982 |
| GFAP | GFAP | Pcdha9 | Pcdha9 | 0.000304 | 0 | PCDHA9_PCDHA9 | Pcdha9 - Pcdha9 | PCDH | Cell-Cell Contact | PMID: 35253643; PMID: 32694982 |
| NeuN | GFAP | Pcdha11 | Pcdha11 | 0.000431 | 0 | PCDHA11_PCDHA11 | Pcdha11 - Pcdha11 | PCDH | Cell-Cell Contact | PMID: 35253643; PMID: 32694982 |
| GFAP | NeuN | Pcdha11 | Pcdha11 | 0.000431 | 0 | PCDHA11_PCDHA11 | Pcdha11 - Pcdha11 | PCDH | Cell-Cell Contact | PMID: 35253643; PMID: 32694982 |
| NeuN | NeuN | Pcdha11 | Pcdha11 | 0.000453 | 0 | PCDHA11_PCDHA11 | Pcdha11 - Pcdha11 | PCDH | Cell-Cell Contact | PMID: 35253643; PMID: 32694982 |
| CD31 | CD31 | Pcdha10 | Pcdha10 | 0.00086 | 0 | PCDHA10_PCDHA10 | Pcdha10 - Pcdha10 | PCDH | Cell-Cell Contact | PMID: 35253643; PMID: 32694982 |
| GFAP | CD31 | Pcdha10 | Pcdha10 | 0.000708 | 0 | PCDHA10_PCDHA10 | Pcdha10 - Pcdha10 | PCDH | Cell-Cell Contact | PMID: 35253643; PMID: 32694982 |
| CD31 | GFAP | Pcdha10 | Pcdha10 | 0.000708 | 0 | PCDHA10_PCDHA10 | Pcdha10 - Pcdha10 | PCDH | Cell-Cell Contact | PMID: 35253643; PMID: 32694982 |
| GFAP | GFAP | Pcdha12 | Pcdha12 | 0.000665 | 0 | PCDHA12_PCDHA12 | Pcdha12 - Pcdha12 | PCDH | Cell-Cell Contact | PMID: 35253643; PMID: 32694982 |
| NeuN | GFAP | Pcdha12 | Pcdha12 | 0.000505 | 0 | PCDHA12_PCDHA12 | Pcdha12 - Pcdha12 | PCDH | Cell-Cell Contact | PMID: 35253643; PMID: 32694982 |
| GFAP | NeuN | Pcdha12 | Pcdha12 | 0.000505 | 0 | PCDHA12_PCDHA12 | Pcdha12 - Pcdha12 | PCDH | Cell-Cell Contact | PMID: 35253643; PMID: 32694982 |
| CD31 | CD31 | Pcdha2 | Pcdha2 | 0.000672 | 0 | PCDHA2_PCDHA2 | Pcdha2 - Pcdha2 | PCDH | Cell-Cell Contact | PMID: 35253643; PMID: 32694982 |
| GFAP | CD31 | Pcdha2 | Pcdha2 | 0.000589 | 0 | PCDHA2_PCDHA2 | Pcdha2 - Pcdha2 | PCDH | Cell-Cell Contact | PMID: 35253643; PMID: 32694982 |
| CD31 | GFAP | Pcdha2 | Pcdha2 | 0.000589 | 0 | PCDHA2_PCDHA2 | Pcdha2 - Pcdha2 | PCDH | Cell-Cell Contact | PMID: 35253643; PMID: 32694982 |
| GFAP | CD31 | Pcdha3 | Pcdha3 | 0.000358 | 0 | PCDHA3_PCDHA3 | Pcdha3 - Pcdha3 | PCDH | Cell-Cell Contact | PMID: 35253643; PMID: 32694982 |
| CD31 | GFAP | Pcdha3 | Pcdha3 | 0.000358 | 0 | PCDHA3_PCDHA3 | Pcdha3 - Pcdha3 | PCDH | Cell-Cell Contact | PMID: 35253643; PMID: 32694982 |
| GFAP | GFAP | Pcdha3 | Pcdha3 | 0.000439 | 0 | PCDHA3_PCDHA3 | Pcdha3 - Pcdha3 | PCDH | Cell-Cell Contact | PMID: 35253643; PMID: 32694982 |
| NeuN | GFAP | Pcdha5 | Pcdha5 | 0.000244 | 0 | PCDHA5_PCDHA5 | Pcdha5 - Pcdha5 | PCDH | Cell-Cell Contact | PMID: 35253643; PMID: 32694982 |
| GFAP | NeuN | Pcdha5 | Pcdha5 | 0.000244 | 0 | PCDHA5_PCDHA5 | Pcdha5 - Pcdha5 | PCDH | Cell-Cell Contact | PMID: 35253643; PMID: 32694982 |
| NeuN | NeuN | Pcdha5 | Pcdha5 | 0.000291 | 0 | PCDHA5_PCDHA5 | Pcdha5 - Pcdha5 | PCDH | Cell-Cell Contact | PMID: 35253643; PMID: 32694982 |
| GFAP | GFAP | Pcdha8 | Pcdha8 | 0.000261 | 0 | PCDHA8_PCDHA8 | Pcdha8 - Pcdha8 | PCDH | Cell-Cell Contact | PMID: 35253643; PMID: 32694982 |
| NeuN | GFAP | Pcdha8 | Pcdha8 | 0.000218 | 0 | PCDHA8_PCDHA8 | Pcdha8 - Pcdha8 | PCDH | Cell-Cell Contact | PMID: 35253643; PMID: 32694982 |
| GFAP | NeuN | Pcdha8 | Pcdha8 | 0.000218 | 0 | PCDHA8_PCDHA8 | Pcdha8 - Pcdha8 | PCDH | Cell-Cell Contact | PMID: 35253643; PMID: 32694982 |
| NeuN | CD31 | Pcdhac1 | Pcdhac1 | 0.000335 | 0 | PCDHAC1_PCDHAC1 | Pcdhac1 - Pcdhac1 | PCDH | Cell-Cell Contact | PMID: 35253643; PMID: 32694982 |
| CD31 | NeuN | Pcdhac1 | Pcdhac1 | 0.000335 | 0 | PCDHAC1_PCDHAC1 | Pcdhac1 - Pcdhac1 | PCDH | Cell-Cell Contact | PMID: 35253643; PMID: 32694982 |
| NeuN | NeuN | Pcdhac1 | Pcdhac1 | 0.000358 | 0 | PCDHAC1_PCDHAC1 | Pcdhac1 - Pcdhac1 | PCDH | Cell-Cell Contact | PMID: 35253643; PMID: 32694982 |
| NeuN | GFAP | Pcdhac2 | Pcdhac2 | 0.001037 | 0 | PCDHAC2_PCDHAC2 | Pcdhac2 - Pcdhac2 | PCDH | Cell-Cell Contact | PMID: 35253643; PMID: 32694982 |
| GFAP | NeuN | Pcdhac2 | Pcdhac2 | 0.001037 | 0 | PCDHAC2_PCDHAC2 | Pcdhac2 - Pcdhac2 | PCDH | Cell-Cell Contact | PMID: 35253643; PMID: 32694982 |
| NeuN | NeuN | Pcdhac2 | Pcdhac2 | 0.002055 | 0 | PCDHAC2_PCDHAC2 | Pcdhac2 - Pcdhac2 | PCDH | Cell-Cell Contact | PMID: 35253643; PMID: 32694982 |
| GFAP | CD31 | Pcdhb18 | Pcdhb18 | 0.00055 | 0 | PCDHB18_PCDHB18 | Pcdhb18 - Pcdhb18 | PCDH | Cell-Cell Contact | PMID: 35253643; PMID: 32694982 |
| CD31 | GFAP | Pcdhb18 | Pcdhb18 | 0.00055 | 0 | PCDHB18_PCDHB18 | Pcdhb18 - Pcdhb18 | PCDH | Cell-Cell Contact | PMID: 35253643; PMID: 32694982 |
| GFAP | GFAP | Pcdhb18 | Pcdhb18 | 0.000598 | 0 | PCDHB18_PCDHB18 | Pcdhb18 - Pcdhb18 | PCDH | Cell-Cell Contact | PMID: 35253643; PMID: 32694982 |
| NeuN | CD31 | Pcdhb19 | Pcdhb19 | 0.000163 | 0 | PCDHB19_PCDHB19 | Pcdhb19 - Pcdhb19 | PCDH | Cell-Cell Contact | PMID: 35253643; PMID: 32694982 |
| CD31 | NeuN | Pcdhb19 | Pcdhb19 | 0.000163 | 0 | PCDHB19_PCDHB19 | Pcdhb19 - Pcdhb19 | PCDH | Cell-Cell Contact | PMID: 35253643; PMID: 32694982 |
| NeuN | NeuN | Pcdhb19 | Pcdhb19 | 0.000177 | 0 | PCDHB19_PCDHB19 | Pcdhb19 - Pcdhb19 | PCDH | Cell-Cell Contact | PMID: 35253643; PMID: 32694982 |
| CD31 | CD31 | Pcdhb19 | Pcdhb21 | 0.000259 | 0 | PCDHB19_PCDHB21 | Pcdhb19 - Pcdhb21 | PCDH | Cell-Cell Contact | PMID: 35253643; PMID: 32694982 |
| NeuN | CD31 | Pcdhb19 | Pcdhb21 | 0.000281 | 0 | PCDHB19_PCDHB21 | Pcdhb19 - Pcdhb21 | PCDH | Cell-Cell Contact | PMID: 35253643; PMID: 32694982 |
| CD31 | CD31 | Pcdhb21 | Pcdhb21 | 0.000445 | 0 | PCDHB21_PCDHB21 | Pcdhb21 - Pcdhb21 | PCDH | Cell-Cell Contact | PMID: 35253643; PMID: 32694982 |
| GFAP | CD31 | Pcdhb21 | Pcdhb21 | 0.000427 | 0 | PCDHB21_PCDHB21 | Pcdhb21 - Pcdhb21 | PCDH | Cell-Cell Contact | PMID: 35253643; PMID: 32694982 |
| CD31 | GFAP | Pcdhb21 | Pcdhb21 | 0.000427 | 0 | PCDHB21_PCDHB21 | Pcdhb21 - Pcdhb21 | PCDH | Cell-Cell Contact | PMID: 35253643; PMID: 32694982 |
| CD31 | CD31 | Pcdhb16 | Pcdhb16 | 0.000292 | 0 | PCDHB16_PCDHB16 | Pcdhb16 - Pcdhb16 | PCDH | Cell-Cell Contact | PMID: 35253643; PMID: 32694982 |
| NeuN | CD31 | Pcdhb16 | Pcdhb16 | 0.000206 | 0 | PCDHB16_PCDHB16 | Pcdhb16 - Pcdhb16 | PCDH | Cell-Cell Contact | PMID: 35253643; PMID: 32694982 |
| CD31 | NeuN | Pcdhb16 | Pcdhb16 | 0.000206 | 0 | PCDHB16_PCDHB16 | Pcdhb16 - Pcdhb16 | PCDH | Cell-Cell Contact | PMID: 35253643; PMID: 32694982 |
| GFAP | GFAP | Pcdhb20 | Pcdhb20 | 0.000807 | 0 | PCDHB20_PCDHB20 | Pcdhb20 - Pcdhb20 | PCDH | Cell-Cell Contact | PMID: 35253643; PMID: 32694982 |
| NeuN | GFAP | Pcdhb20 | Pcdhb20 | 0.000445 | 0 | PCDHB20_PCDHB20 | Pcdhb20 - Pcdhb20 | PCDH | Cell-Cell Contact | PMID: 35253643; PMID: 32694982 |
| GFAP | NeuN | Pcdhb20 | Pcdhb20 | 0.000445 | 0 | PCDHB20_PCDHB20 | Pcdhb20 - Pcdhb20 | PCDH | Cell-Cell Contact | PMID: 35253643; PMID: 32694982 |
| CD31 | CD31 | Pcdhb22 | Pcdhb22 | 0.000672 | 0 | PCDHB22_PCDHB22 | Pcdhb22 - Pcdhb22 | PCDH | Cell-Cell Contact | PMID: 35253643; PMID: 32694982 |
| GFAP | CD31 | Pcdhb22 | Pcdhb22 | 0.000591 | 0 | PCDHB22_PCDHB22 | Pcdhb22 - Pcdhb22 | PCDH | Cell-Cell Contact | PMID: 35253643; PMID: 32694982 |
| CD31 | GFAP | Pcdhb22 | Pcdhb22 | 0.000591 | 0 | PCDHB22_PCDHB22 | Pcdhb22 - Pcdhb22 | PCDH | Cell-Cell Contact | PMID: 35253643; PMID: 32694982 |
| CD31 | CD31 | Pcdhb17 | Pcdhb17 | 0.000278 | 0 | PCDHB17_PCDHB17 | Pcdhb17 - Pcdhb17 | PCDH | Cell-Cell Contact | PMID: 35253643; PMID: 32694982 |
| NeuN | CD31 | Pcdhb17 | Pcdhb17 | 0.00026 | 0 | PCDHB17_PCDHB17 | Pcdhb17 - Pcdhb17 | PCDH | Cell-Cell Contact | PMID: 35253643; PMID: 32694982 |
| CD31 | NeuN | Pcdhb17 | Pcdhb17 | 0.00026 | 0 | PCDHB17_PCDHB17 | Pcdhb17 - Pcdhb17 | PCDH | Cell-Cell Contact | PMID: 35253643; PMID: 32694982 |
| GFAP | CD31 | Pcdhb2 | Pcdhb2 | 0.000182 | 0 | PCDHB2_PCDHB2 | Pcdhb2 - Pcdhb2 | PCDH | Cell-Cell Contact | PMID: 35253643; PMID: 32694982 |
| CD31 | GFAP | Pcdhb2 | Pcdhb2 | 0.000182 | 0 | PCDHB2_PCDHB2 | Pcdhb2 - Pcdhb2 | PCDH | Cell-Cell Contact | PMID: 35253643; PMID: 32694982 |
| GFAP | GFAP | Pcdhb2 | Pcdhb2 | 0.000221 | 0 | PCDHB2_PCDHB2 | Pcdhb2 - Pcdhb2 | PCDH | Cell-Cell Contact | PMID: 35253643; PMID: 32694982 |
| CD31 | CD31 | Pcdhb3 | Pcdhb3 | 0.000313 | 0 | PCDHB3_PCDHB3 | Pcdhb3 - Pcdhb3 | PCDH | Cell-Cell Contact | PMID: 35253643; PMID: 32694982 |

| | | | | | | | | | | |
|---|---|---|---|---|---|---|---|---|---|---|
| GFAP | CD31 | Pcdhb3 | Pcdhb3 | 0.000244 | 0 | PCDHB3_PCDHB3 | Pcdhb3 - Pcdhb3 | PCDH | Cell-Cell Contact | PMID: 35253643; PMID: 32694982 |
| CD31 | GFAP | Pcdhb3 | Pcdhb3 | 0.000244 | 0 | PCDHB3_PCDHB3 | Pcdhb3 - Pcdhb3 | PCDH | Cell-Cell Contact | PMID: 35253643; PMID: 32694982 |
| GFAP | CD31 | Pcdhb7 | Pcdhb7 | 0.000298 | 0 | PCDHB7_PCDHB7 | Pcdhb7 - Pcdhb7 | PCDH | Cell-Cell Contact | PMID: 35253643; PMID: 32694982 |
| CD31 | GFAP | Pcdhb7 | Pcdhb7 | 0.000298 | 0 | PCDHB7_PCDHB7 | Pcdhb7 - Pcdhb7 | PCDH | Cell-Cell Contact | PMID: 35253643; PMID: 32694982 |
| GFAP | GFAP | Pcdhb7 | Pcdhb7 | 0.000305 | 0 | PCDHB7_PCDHB7 | Pcdhb7 - Pcdhb7 | PCDH | Cell-Cell Contact | PMID: 35253643; PMID: 32694982 |
| CD31 | CD31 | Pcdhb7 | Pcdhb5 | 0.000415 | 0 | PCDHB7_PCDHB5 | Pcdhb7 - Pcdhb5 | PCDH | Cell-Cell Contact | PMID: 35253643; PMID: 32694982 |
| GFAP | CD31 | Pcdhb7 | Pcdhb5 | 0.000425 | 0 | PCDHB7_PCDHB5 | Pcdhb7 - Pcdhb5 | PCDH | Cell-Cell Contact | PMID: 35253643; PMID: 32694982 |
| CD31 | CD31 | Pcdhb7 | Pcdhb9 | 0.000391 | 0 | PCDHB7_PCDHB9 | Pcdhb7 - Pcdhb9 | PCDH | Cell-Cell Contact | PMID: 35253643; PMID: 32694982 |
| GFAP | CD31 | Pcdhb7 | Pcdhb9 | 0.0004 | 0 | PCDHB7_PCDHB9 | Pcdhb7 - Pcdhb9 | PCDH | Cell-Cell Contact | PMID: 35253643; PMID: 32694982 |
| CD31 | CD31 | Pcdhb5 | Pcdhb5 | 0.000591 | 0 | PCDHB5_PCDHB5 | Pcdhb5 - Pcdhb5 | PCDH | Cell-Cell Contact | PMID: 35253643; PMID: 32694982 |
| GFAP | CD31 | Pcdhb5 | Pcdhb5 | 0.000394 | 0 | PCDHB5_PCDHB5 | Pcdhb5 - Pcdhb5 | PCDH | Cell-Cell Contact | PMID: 35253643; PMID: 32694982 |
| CD31 | GFAP | Pcdhb5 | Pcdhb5 | 0.000394 | 0 | PCDHB5_PCDHB5 | Pcdhb5 - Pcdhb5 | PCDH | Cell-Cell Contact | PMID: 35253643; PMID: 32694982 |
| CD31 | CD31 | Pcdhb5 | Pcdhb9 | 0.000557 | 0 | PCDHB5_PCDHB9 | Pcdhb5 - Pcdhb9 | PCDH | Cell-Cell Contact | PMID: 35253643; PMID: 32694982 |
| CD31 | GFAP | Pcdhb5 | Pcdhb9 | 0.000433 | 0 | PCDHB5_PCDHB9 | Pcdhb5 - Pcdhb9 | PCDH | Cell-Cell Contact | PMID: 35253643; PMID: 32694982 |
| CD31 | CD31 | Pcdhb9 | Pcdhb9 | 0.000525 | 0 | PCDHB9_PCDHB9 | Pcdhb9 - Pcdhb9 | PCDH | Cell-Cell Contact | PMID: 35253643; PMID: 32694982 |
| GFAP | CD31 | Pcdhb9 | Pcdhb9 | 0.000408 | 0 | PCDHB9_PCDHB9 | Pcdhb9 - Pcdhb9 | PCDH | Cell-Cell Contact | PMID: 35253643; PMID: 32694982 |
| CD31 | GFAP | Pcdhb9 | Pcdhb9 | 0.000408 | 0 | PCDHB9_PCDHB9 | Pcdhb9 - Pcdhb9 | PCDH | Cell-Cell Contact | PMID: 35253643; PMID: 32694982 |
| CD31 | CD31 | Pcdhb12 | Pcdhb12 | 0.000292 | 0 | PCDHB12_PCDHB12 | Pcdhb12 - Pcdhb12 | PCDH | Cell-Cell Contact | PMID: 35253643; PMID: 32694982 |
| GFAP | CD31 | Pcdhb12 | Pcdhb12 | 0.000273 | 0 | PCDHB12_PCDHB12 | Pcdhb12 - Pcdhb12 | PCDH | Cell-Cell Contact | PMID: 35253643; PMID: 32694982 |
| CD31 | GFAP | Pcdhb12 | Pcdhb12 | 0.000273 | 0 | PCDHB12_PCDHB12 | Pcdhb12 - Pcdhb12 | PCDH | Cell-Cell Contact | PMID: 35253643; PMID: 32694982 |
| CD31 | GFAP | Pcdhb12 | Pcdhb10 | 0.000288 | 0 | PCDHB12_PCDHB10 | Pcdhb12 - Pcdhb10 | PCDH | Cell-Cell Contact | PMID: 35253643; PMID: 32694982 |
| GFAP | GFAP | Pcdhb12 | Pcdhb10 | 0.000269 | 0 | PCDHB12_PCDHB10 | Pcdhb12 - Pcdhb10 | PCDH | Cell-Cell Contact | PMID: 35253643; PMID: 32694982 |
| CD31 | GFAP | Pcdhb12 | Pcdhb11 | 0.000413 | 0 | PCDHB12_PCDHB11 | Pcdhb12 - Pcdhb11 | PCDH | Cell-Cell Contact | PMID: 35253643; PMID: 32694982 |
| GFAP | GFAP | Pcdhb12 | Pcdhb11 | 0.000386 | 0 | PCDHB12_PCDHB11 | Pcdhb12 - Pcdhb11 | PCDH | Cell-Cell Contact | PMID: 35253643; PMID: 32694982 |
| CD31 | GFAP | Pcdhb12 | Pcdhb8 | 0.000495 | 0 | PCDHB12_PCDHB8 | Pcdhb12 - Pcdhb8 | PCDH | Cell-Cell Contact | PMID: 35253643; PMID: 32694982 |
| GFAP | GFAP | Pcdhb12 | Pcdhb8 | 0.000462 | 0 | PCDHB12_PCDHB8 | Pcdhb12 - Pcdhb8 | PCDH | Cell-Cell Contact | PMID: 35253643; PMID: 32694982 |
| CD31 | GFAP | Pcdhb12 | Pcdhb6 | 0.000325 | 0 | PCDHB12_PCDHB6 | Pcdhb12 - Pcdhb6 | PCDH | Cell-Cell Contact | PMID: 35253643; PMID: 32694982 |
| GFAP | GFAP | Pcdhb12 | Pcdhb6 | 0.000304 | 0 | PCDHB12_PCDHB6 | Pcdhb12 - Pcdhb6 | PCDH | Cell-Cell Contact | PMID: 35253643; PMID: 32694982 |
| GFAP | GFAP | Pcdhb10 | Pcdhb10 | 0.000285 | 0 | PCDHB10_PCDHB10 | Pcdhb10 - Pcdhb10 | PCDH | Cell-Cell Contact | PMID: 35253643; PMID: 32694982 |
| NeuN | GFAP | Pcdhb10 | Pcdhb10 | 0.000131 | 0 | PCDHB10_PCDHB10 | Pcdhb10 - Pcdhb10 | PCDH | Cell-Cell Contact | PMID: 35253643; PMID: 32694982 |
| GFAP | NeuN | Pcdhb10 | Pcdhb10 | 0.000131 | 0 | PCDHB10_PCDHB10 | Pcdhb10 - Pcdhb10 | PCDH | Cell-Cell Contact | PMID: 35253643; PMID: 32694982 |
| GFAP | GFAP | Pcdhb10 | Pcdhb11 | 0.000408 | 0 | PCDHB10_PCDHB11 | Pcdhb10 - Pcdhb11 | PCDH | Cell-Cell Contact | PMID: 35253643; PMID: 32694982 |
| NeuN | GFAP | Pcdhb10 | Pcdhb11 | 0.000188 | 0 | PCDHB10_PCDHB11 | Pcdhb10 - Pcdhb11 | PCDH | Cell-Cell Contact | PMID: 35253643; PMID: 32694982 |
| GFAP | CD31 | Pcdhb10 | Pcdhb8 | 0.000254 | 0 | PCDHB10_PCDHB8 | Pcdhb10 - Pcdhb8 | PCDH | Cell-Cell Contact | PMID: 35253643; PMID: 32694982 |
| GFAP | GFAP | Pcdhb10 | Pcdhb8 | 0.000489 | 0 | PCDHB10_PCDHB8 | Pcdhb10 - Pcdhb8 | PCDH | Cell-Cell Contact | PMID: 35253643; PMID: 32694982 |
| GFAP | CD31 | Pcdhb10 | Pcdhb6 | 0.000275 | 0 | PCDHB10_PCDHB6 | Pcdhb10 - Pcdhb6 | PCDH | Cell-Cell Contact | PMID: 35253643; PMID: 32694982 |
| GFAP | GFAP | Pcdhb10 | Pcdhb6 | 0.000321 | 0 | PCDHB10_PCDHB6 | Pcdhb10 - Pcdhb6 | PCDH | Cell-Cell Contact | PMID: 35253643; PMID: 32694982 |
| GFAP | GFAP | Pcdhb11 | Pcdhb11 | 0.000586 | 0 | PCDHB11_PCDHB11 | Pcdhb11 - Pcdhb11 | PCDH | Cell-Cell Contact | PMID: 35253643; PMID: 32694982 |
| NeuN | GFAP | Pcdhb11 | Pcdhb11 | 0.000243 | 0 | PCDHB11_PCDHB11 | Pcdhb11 - Pcdhb11 | PCDH | Cell-Cell Contact | PMID: 35253643; PMID: 32694982 |
| GFAP | NeuN | Pcdhb11 | Pcdhb11 | 0.000243 | 0 | PCDHB11_PCDHB11 | Pcdhb11 - Pcdhb11 | PCDH | Cell-Cell Contact | PMID: 35253643; PMID: 32694982 |
| GFAP | CD31 | Pcdhb11 | Pcdhb8 | 0.000365 | 0 | PCDHB11_PCDHB8 | Pcdhb11 - Pcdhb8 | PCDH | Cell-Cell Contact | PMID: 35253643; PMID: 32694982 |
| GFAP | GFAP | Pcdhb11 | Pcdhb8 | 0.000701 | 0 | PCDHB11_PCDHB8 | Pcdhb11 - Pcdhb8 | PCDH | Cell-Cell Contact | PMID: 35253643; PMID: 32694982 |
| GFAP | CD31 | Pcdhb11 | Pcdhb6 | 0.000394 | 0 | PCDHB11_PCDHB6 | Pcdhb11 - Pcdhb6 | PCDH | Cell-Cell Contact | PMID: 35253643; PMID: 32694982 |
| GFAP | GFAP | Pcdhb11 | Pcdhb6 | 0.000461 | 0 | PCDHB11_PCDHB6 | Pcdhb11 - Pcdhb6 | PCDH | Cell-Cell Contact | PMID: 35253643; PMID: 32694982 |
| GFAP | CD31 | Pcdhb8 | Pcdhb8 | 0.000436 | 0 | PCDHB8_PCDHB8 | Pcdhb8 - Pcdhb8 | PCDH | Cell-Cell Contact | PMID: 35253643; PMID: 32694982 |
| CD31 | GFAP | Pcdhb8 | Pcdhb8 | 0.000436 | 0 | PCDHB8_PCDHB8 | Pcdhb8 - Pcdhb8 | PCDH | Cell-Cell Contact | PMID: 35253643; PMID: 32694982 |
| GFAP | GFAP | Pcdhb8 | Pcdhb8 | 0.000839 | 0 | PCDHB8_PCDHB8 | Pcdhb8 - Pcdhb8 | PCDH | Cell-Cell Contact | PMID: 35253643; PMID: 32694982 |
| GFAP | CD31 | Pcdhb8 | Pcdhb6 | 0.000471 | 0 | PCDHB8_PCDHB6 | Pcdhb8 - Pcdhb6 | PCDH | Cell-Cell Contact | PMID: 35253643; PMID: 32694982 |
| GFAP | GFAP | Pcdhb8 | Pcdhb6 | 0.000551 | 0 | PCDHB8_PCDHB6 | Pcdhb8 - Pcdhb6 | PCDH | Cell-Cell Contact | PMID: 35253643; PMID: 32694982 |
| GFAP | CD31 | Pcdhb6 | Pcdhb6 | 0.00031 | 0 | PCDHB6_PCDHB6 | Pcdhb6 - Pcdhb6 | PCDH | Cell-Cell Contact | PMID: 35253643; PMID: 32694982 |
| CD31 | GFAP | Pcdhb6 | Pcdhb6 | 0.00031 | 0 | PCDHB6_PCDHB6 | Pcdhb6 - Pcdhb6 | PCDH | Cell-Cell Contact | PMID: 35253643; PMID: 32694982 |
| GFAP | GFAP | Pcdhb6 | Pcdhb6 | 0.000362 | 0 | PCDHB6_PCDHB6 | Pcdhb6 - Pcdhb6 | PCDH | Cell-Cell Contact | PMID: 35253643; PMID: 32694982 |
| GFAP | GFAP | Pcdhb13 | Pcdhb13 | 0.000485 | 0 | PCDHB13_PCDHB13 | Pcdhb13 - Pcdhb13 | PCDH | Cell-Cell Contact | PMID: 35253643; PMID: 32694982 |
| NeuN | GFAP | Pcdhb13 | Pcdhb13 | 0.00033 | 0 | PCDHB13_PCDHB13 | Pcdhb13 - Pcdhb13 | PCDH | Cell-Cell Contact | PMID: 35253643; PMID: 32694982 |
| GFAP | NeuN | Pcdhb13 | Pcdhb13 | 0.00033 | 0 | PCDHB13_PCDHB13 | Pcdhb13 - Pcdhb13 | PCDH | Cell-Cell Contact | PMID: 35253643; PMID: 32694982 |
| GFAP | GFAP | Pcdhb15 | Pcdhb15 | 0.000306 | 0 | PCDHB15_PCDHB15 | Pcdhb15 - Pcdhb15 | PCDH | Cell-Cell Contact | PMID: 35253643; PMID: 32694982 |
| NeuN | GFAP | Pcdhb15 | Pcdhb15 | 0.000183 | 0 | PCDHB15_PCDHB15 | Pcdhb15 - Pcdhb15 | PCDH | Cell-Cell Contact | PMID: 35253643; PMID: 32694982 |
| GFAP | NeuN | Pcdhb15 | Pcdhb15 | 0.000183 | 0 | PCDHB15_PCDHB15 | Pcdhb15 - Pcdhb15 | PCDH | Cell-Cell Contact | PMID: 35253643; PMID: 32694982 |
| GFAP | CD31 | Pcdhga1 | Pcdhga1 | 0.000221 | 0 | PCDHGA1_PCDHGA1 | Pcdhga1 - Pcdhga1 | PCDH | Cell-Cell Contact | PMID: 35253643; PMID: 32694982 |

| | | | | | | | | | | |
|---|---|---|---|---|---|---|---|---|---|---|
| CD31 | GFAP | Pcdhga1 | Pcdhga1 | 0.000221 | 0 | PCDHGA1_PCDHGA1 | Pcdhga1 - Pcdhga1 | PCDH | Cell-Cell Contact | PMID: 35253643; PMID: 32694982 |
| GFAP | GFAP | Pcdhga1 | Pcdhga1 | 0.000241 | 0 | PCDHGA1_PCDHGA1 | Pcdhga1 - Pcdhga1 | PCDH | Cell-Cell Contact | PMID: 35253643; PMID: 32694982 |
| CD31 | CD31 | Pcdhga10 | Pcdhga10 | 0.000388 | 0 | PCDHGA10_PCDHGA10 | Pcdhga10 - Pcdhga10 | PCDH | Cell-Cell Contact | PMID: 35253643; PMID: 32694982 |
| GFAP | CD31 | Pcdhga10 | Pcdhga10 | 0.000371 | 0 | PCDHGA10_PCDHGA10 | Pcdhga10 - Pcdhga10 | PCDH | Cell-Cell Contact | PMID: 35253643; PMID: 32694982 |
| CD31 | GFAP | Pcdhga10 | Pcdhga10 | 0.000371 | 0 | PCDHGA10_PCDHGA10 | Pcdhga10 - Pcdhga10 | PCDH | Cell-Cell Contact | PMID: 35253643; PMID: 32694982 |
| CD31 | CD31 | Pcdhga11 | Pcdhga11 | 0.001847 | 0 | PCDHGA11_PCDHGA11 | Pcdhga11 - Pcdhga11 | PCDH | Cell-Cell Contact | PMID: 35253643; PMID: 32694982 |
| GFAP | CD31 | Pcdhga11 | Pcdhga11 | 0.001619 | 0 | PCDHGA11_PCDHGA11 | Pcdhga11 - Pcdhga11 | PCDH | Cell-Cell Contact | PMID: 35253643; PMID: 32694982 |
| CD31 | GFAP | Pcdhga11 | Pcdhga11 | 0.001619 | 0 | PCDHGA11_PCDHGA11 | Pcdhga11 - Pcdhga11 | PCDH | Cell-Cell Contact | PMID: 35253643; PMID: 32694982 |
| CD31 | CD31 | Pcdhga12 | Pcdhga12 | 0.000933 | 0 | PCDHGA12_PCDHGA12 | Pcdhga12 - Pcdhga12 | PCDH | Cell-Cell Contact | PMID: 35253643; PMID: 32694982 |
| GFAP | CD31 | Pcdhga12 | Pcdhga12 | 0.000575 | 0 | PCDHGA12_PCDHGA12 | Pcdhga12 - Pcdhga12 | PCDH | Cell-Cell Contact | PMID: 35253643; PMID: 32694982 |
| CD31 | GFAP | Pcdhga12 | Pcdhga12 | 0.000575 | 0 | PCDHGA12_PCDHGA12 | Pcdhga12 - Pcdhga12 | PCDH | Cell-Cell Contact | PMID: 35253643; PMID: 32694982 |
| GFAP | CD31 | Pcdhga2 | Pcdhga2 | 0.000328 | 0 | PCDHGA2_PCDHGA2 | Pcdhga2 - Pcdhga2 | PCDH | Cell-Cell Contact | PMID: 35253643; PMID: 32694982 |
| CD31 | GFAP | Pcdhga2 | Pcdhga2 | 0.000328 | 0 | PCDHGA2_PCDHGA2 | Pcdhga2 - Pcdhga2 | PCDH | Cell-Cell Contact | PMID: 35253643; PMID: 32694982 |
| GFAP | GFAP | Pcdhga2 | Pcdhga2 | 0.000439 | 0 | PCDHGA2_PCDHGA2 | Pcdhga2 - Pcdhga2 | PCDH | Cell-Cell Contact | PMID: 35253643; PMID: 32694982 |
| GFAP | CD31 | Pcdhga3 | Pcdhga3 | 0.000421 | 0 | PCDHGA3_PCDHGA3 | Pcdhga3 - Pcdhga3 | PCDH | Cell-Cell Contact | PMID: 35253643; PMID: 32694982 |
| CD31 | GFAP | Pcdhga3 | Pcdhga3 | 0.000421 | 0 | PCDHGA3_PCDHGA3 | Pcdhga3 - Pcdhga3 | PCDH | Cell-Cell Contact | PMID: 35253643; PMID: 32694982 |
| GFAP | GFAP | Pcdhga3 | Pcdhga3 | 0.00043 | 0 | PCDHGA3_PCDHGA3 | Pcdhga3 - Pcdhga3 | PCDH | Cell-Cell Contact | PMID: 35253643; PMID: 32694982 |
| CD31 | CD31 | Pcdhga4 | Pcdhga4 | 0.000534 | 0 | PCDHGA4_PCDHGA4 | Pcdhga4 - Pcdhga4 | PCDH | Cell-Cell Contact | PMID: 35253643; PMID: 32694982 |
| GFAP | CD31 | Pcdhga4 | Pcdhga4 | 0.000465 | 0 | PCDHGA4_PCDHGA4 | Pcdhga4 - Pcdhga4 | PCDH | Cell-Cell Contact | PMID: 35253643; PMID: 32694982 |
| CD31 | GFAP | Pcdhga4 | Pcdhga4 | 0.000465 | 0 | PCDHGA4_PCDHGA4 | Pcdhga4 - Pcdhga4 | PCDH | Cell-Cell Contact | PMID: 35253643; PMID: 32694982 |
| GFAP | CD31 | Pcdhga5 | Pcdhga5 | 0.000535 | 0 | PCDHGA5_PCDHGA5 | Pcdhga5 - Pcdhga5 | PCDH | Cell-Cell Contact | PMID: 35253643; PMID: 32694982 |
| CD31 | GFAP | Pcdhga5 | Pcdhga5 | 0.000535 | 0 | PCDHGA5_PCDHGA5 | Pcdhga5 - Pcdhga5 | PCDH | Cell-Cell Contact | PMID: 35253643; PMID: 32694982 |
| GFAP | GFAP | Pcdhga5 | Pcdhga5 | 0.000657 | 0 | PCDHGA5_PCDHGA5 | Pcdhga5 - Pcdhga5 | PCDH | Cell-Cell Contact | PMID: 35253643; PMID: 32694982 |
| GFAP | CD31 | Pcdhga6 | Pcdhga6 | 0.000237 | 0 | PCDHGA6_PCDHGA6 | Pcdhga6 - Pcdhga6 | PCDH | Cell-Cell Contact | PMID: 35253643; PMID: 32694982 |
| CD31 | GFAP | Pcdhga6 | Pcdhga6 | 0.000237 | 0 | PCDHGA6_PCDHGA6 | Pcdhga6 - Pcdhga6 | PCDH | Cell-Cell Contact | PMID: 35253643; PMID: 32694982 |
| GFAP | GFAP | Pcdhga6 | Pcdhga6 | 0.000247 | 0 | PCDHGA6_PCDHGA6 | Pcdhga6 - Pcdhga6 | PCDH | Cell-Cell Contact | PMID: 35253643; PMID: 32694982 |
| GFAP | GFAP | Pcdhga7 | Pcdhga7 | 0.000678 | 0 | PCDHGA7_PCDHGA7 | Pcdhga7 - Pcdhga7 | PCDH | Cell-Cell Contact | PMID: 35253643; PMID: 32694982 |
| NeuN | GFAP | Pcdhga7 | Pcdhga7 | 0.000407 | 0 | PCDHGA7_PCDHGA7 | Pcdhga7 - Pcdhga7 | PCDH | Cell-Cell Contact | PMID: 35253643; PMID: 32694982 |
| GFAP | NeuN | Pcdhga7 | Pcdhga7 | 0.000407 | 0 | PCDHGA7_PCDHGA7 | Pcdhga7 - Pcdhga7 | PCDH | Cell-Cell Contact | PMID: 35253643; PMID: 32694982 |
| CD31 | CD31 | Pcdhga8 | Pcdhga8 | 0.000563 | 0 | PCDHGA8_PCDHGA8 | Pcdhga8 - Pcdhga8 | PCDH | Cell-Cell Contact | PMID: 35253643; PMID: 32694982 |
| GFAP | CD31 | Pcdhga8 | Pcdhga8 | 0.000455 | 0 | PCDHGA8_PCDHGA8 | Pcdhga8 - Pcdhga8 | PCDH | Cell-Cell Contact | PMID: 35253643; PMID: 32694982 |
| CD31 | GFAP | Pcdhga8 | Pcdhga8 | 0.000455 | 0 | PCDHGA8_PCDHGA8 | Pcdhga8 - Pcdhga8 | PCDH | Cell-Cell Contact | PMID: 35253643; PMID: 32694982 |
| GFAP | GFAP | Pcdhga9 | Pcdhga9 | 0.000263 | 0 | PCDHGA9_PCDHGA9 | Pcdhga9 - Pcdhga9 | PCDH | Cell-Cell Contact | PMID: 35253643; PMID: 32694982 |
| NeuN | GFAP | Pcdhga9 | Pcdhga9 | 0.000136 | 0 | PCDHGA9_PCDHGA9 | Pcdhga9 - Pcdhga9 | PCDH | Cell-Cell Contact | PMID: 35253643; PMID: 32694982 |
| GFAP | NeuN | Pcdhga9 | Pcdhga9 | 0.000136 | 0 | PCDHGA9_PCDHGA9 | Pcdhga9 - Pcdhga9 | PCDH | Cell-Cell Contact | PMID: 35253643; PMID: 32694982 |
| GFAP | CD31 | Pcdhgb1 | Pcdhgb1 | 0.000479 | 0 | PCDHGB1_PCDHGB1 | Pcdhgb1 - Pcdhgb1 | PCDH | Cell-Cell Contact | PMID: 35253643; PMID: 32694982 |
| CD31 | GFAP | Pcdhgb1 | Pcdhgb1 | 0.000479 | 0 | PCDHGB1_PCDHGB1 | Pcdhgb1 - Pcdhgb1 | PCDH | Cell-Cell Contact | PMID: 35253643; PMID: 32694982 |
| GFAP | GFAP | Pcdhgb1 | Pcdhgb1 | 0.000604 | 0 | PCDHGB1_PCDHGB1 | Pcdhgb1 - Pcdhgb1 | PCDH | Cell-Cell Contact | PMID: 35253643; PMID: 32694982 |
| GFAP | CD31 | Pcdhgb2 | Pcdhgb2 | 0.000931 | 0 | PCDHGB2_PCDHGB2 | Pcdhgb2 - Pcdhgb2 | PCDH | Cell-Cell Contact | PMID: 35253643; PMID: 32694982 |
| CD31 | GFAP | Pcdhgb2 | Pcdhgb2 | 0.000931 | 0 | PCDHGB2_PCDHGB2 | Pcdhgb2 - Pcdhgb2 | PCDH | Cell-Cell Contact | PMID: 35253643; PMID: 32694982 |
| GFAP | GFAP | Pcdhgb2 | Pcdhgb2 | 0.00108 | 0 | PCDHGB2_PCDHGB2 | Pcdhgb2 - Pcdhgb2 | PCDH | Cell-Cell Contact | PMID: 35253643; PMID: 32694982 |
| CD31 | CD31 | Pcdhgb8 | Pcdhgb8 | 0.000131 | 0 | PCDHGB8_PCDHGB8 | Pcdhgb8 - Pcdhgb8 | PCDH | Cell-Cell Contact | PMID: 35253643; PMID: 32694982 |
| GFAP | CD31 | Pcdhgb8 | Pcdhgb8 | 9.73E-05 | 0 | PCDHGB8_PCDHGB8 | Pcdhgb8 - Pcdhgb8 | PCDH | Cell-Cell Contact | PMID: 35253643; PMID: 32694982 |
| CD31 | GFAP | Pcdhgb8 | Pcdhgb8 | 9.73E-05 | 0 | PCDHGB8_PCDHGB8 | Pcdhgb8 - Pcdhgb8 | PCDH | Cell-Cell Contact | PMID: 35253643; PMID: 32694982 |
| GFAP | CD31 | Pcdhgb4 | Pcdhgb4 | 0.000671 | 0 | PCDHGB4_PCDHGB4 | Pcdhgb4 - Pcdhgb4 | PCDH | Cell-Cell Contact | PMID: 35253643; PMID: 32694982 |
| CD31 | GFAP | Pcdhgb4 | Pcdhgb4 | 0.000671 | 0 | PCDHGB4_PCDHGB4 | Pcdhgb4 - Pcdhgb4 | PCDH | Cell-Cell Contact | PMID: 35253643; PMID: 32694982 |
| GFAP | GFAP | Pcdhgb4 | Pcdhgb4 | 0.000844 | 0 | PCDHGB4_PCDHGB4 | Pcdhgb4 - Pcdhgb4 | PCDH | Cell-Cell Contact | PMID: 35253643; PMID: 32694982 |
| GFAP | CD31 | Pcdhgb5 | Pcdhgb5 | 0.000491 | 0 | PCDHGB5_PCDHGB5 | Pcdhgb5 - Pcdhgb5 | PCDH | Cell-Cell Contact | PMID: 35253643; PMID: 32694982 |
| CD31 | GFAP | Pcdhgb5 | Pcdhgb5 | 0.000491 | 0 | PCDHGB5_PCDHGB5 | Pcdhgb5 - Pcdhgb5 | PCDH | Cell-Cell Contact | PMID: 35253643; PMID: 32694982 |
| GFAP | GFAP | Pcdhgb5 | Pcdhgb5 | 0.000585 | 0 | PCDHGB5_PCDHGB5 | Pcdhgb5 - Pcdhgb5 | PCDH | Cell-Cell Contact | PMID: 35253643; PMID: 32694982 |
| CD31 | CD31 | Pcdhgb6 | Pcdhgb6 | 0.000471 | 0 | PCDHGB6_PCDHGB6 | Pcdhgb6 - Pcdhgb6 | PCDH | Cell-Cell Contact | PMID: 35253643; PMID: 32694982 |
| NeuN | CD31 | Pcdhgb6 | Pcdhgb6 | 0.000309 | 0 | PCDHGB6_PCDHGB6 | Pcdhgb6 - Pcdhgb6 | PCDH | Cell-Cell Contact | PMID: 35253643; PMID: 32694982 |
| CD31 | NeuN | Pcdhgb6 | Pcdhgb6 | 0.000309 | 0 | PCDHGB6_PCDHGB6 | Pcdhgb6 - Pcdhgb6 | PCDH | Cell-Cell Contact | PMID: 35253643; PMID: 32694982 |
| CD31 | CD31 | Pcdhgb7 | Pcdhgb7 | 0.001279 | 0 | PCDHGB7_PCDHGB7 | Pcdhgb7 - Pcdhgb7 | PCDH | Cell-Cell Contact | PMID: 35253643; PMID: 32694982 |
| GFAP | CD31 | Pcdhgb7 | Pcdhgb7 | 0.000965 | 0 | PCDHGB7_PCDHGB7 | Pcdhgb7 - Pcdhgb7 | PCDH | Cell-Cell Contact | PMID: 35253643; PMID: 32694982 |
| CD31 | GFAP | Pcdhgb7 | Pcdhgb7 | 0.000965 | 0 | PCDHGB7_PCDHGB7 | Pcdhgb7 - Pcdhgb7 | PCDH | Cell-Cell Contact | PMID: 35253643; PMID: 32694982 |
| GFAP | CD31 | Pcdhgc3 | Pcdhgc3 | 0.006216 | 0 | PCDHGC3_PCDHGC3 | Pcdhgc3 - Pcdhgc3 | PCDH | Cell-Cell Contact | PMID: 35253643; PMID: 32694982 |
| CD31 | GFAP | Pcdhgc3 | Pcdhgc3 | 0.006216 | 0 | PCDHGC3_PCDHGC3 | Pcdhgc3 - Pcdhgc3 | PCDH | Cell-Cell Contact | PMID: 35253643; PMID: 32694982 |
| GFAP | GFAP | Pcdhgc3 | Pcdhgc3 | 0.013604 | 0 | PCDHGC3_PCDHGC3 | Pcdhgc3 - Pcdhgc3 | PCDH | Cell-Cell Contact | PMID: 35253643; PMID: 32694982 |

| | | | | | | | | | | |
|---|---|---|---|---|---|---|---|---|---|---|
| GFAP | GFAP | Pcdhgc4 | Pcdhgc4 | 0.000881 | 0 | PCDHGC4_PCDHGC4 | Pcdhgc4 - Pcdhgc4 | PCDH | Cell-Cell Contact | PMID: 35253643; PMID: 32694982 |
| NeuN | GFAP | Pcdhgc4 | Pcdhgc4 | 0.000512 | 0 | PCDHGC4_PCDHGC4 | Pcdhgc4 - Pcdhgc4 | PCDH | Cell-Cell Contact | PMID: 35253643; PMID: 32694982 |
| GFAP | NeuN | Pcdhgc4 | Pcdhgc4 | 0.000512 | 0 | PCDHGC4_PCDHGC4 | Pcdhgc4 - Pcdhgc4 | PCDH | Cell-Cell Contact | PMID: 35253643; PMID: 32694982 |
| NeuN | CD31 | Pcdhgc5 | Pcdhgc5 | 0.003119 | 0 | PCDHGC5_PCDHGC5 | Pcdhgc5 - Pcdhgc5 | PCDH | Cell-Cell Contact | PMID: 35253643; PMID: 32694982 |
| CD31 | NeuN | Pcdhgc5 | Pcdhgc5 | 0.003119 | 0 | PCDHGC5_PCDHGC5 | Pcdhgc5 - Pcdhgc5 | PCDH | Cell-Cell Contact | PMID: 35253643; PMID: 32694982 |
| NeuN | NeuN | Pcdhgc5 | Pcdhgc5 | 0.005626 | 0 | PCDHGC5_PCDHGC5 | Pcdhgc5 - Pcdhgc5 | PCDH | Cell-Cell Contact | PMID: 35253643; PMID: 32694982 |
| CD31 | CD31 | Sdc2 | Adgra2 | 0.001821 | 0 | SDC2_ADGRA2 | Sdc2 - Adgra2 | ADGRA | Cell-Cell Contact | PMID: 31462748 |
| GFAP | CD31 | Sdc2 | Adgra2 | 0.001954 | 0 | SDC2_ADGRA2 | Sdc2 - Adgra2 | ADGRA | Cell-Cell Contact | PMID: 31462748 |
| CD31 | CD31 | ITGAV_ITG | Adgra2 | 0.000986 | 0 | ITGAV_ITGB3_ADGRA2 | (Itgav+Itgb3) - Adgra2 | ADGRA | Cell-Cell Contact | PMID: 31462748 |
| GFAP | CD31 | ITGAV_ITG | Adgra2 | 0.001414 | 0 | ITGAV_ITGB3_ADGRA2 | (Itgav+Itgb3) - Adgra2 | ADGRA | Cell-Cell Contact | PMID: 31462748 |
| CD31 | CD31 | Tax1bp3 | Adgrb1 | 0.002681 | 0 | TAX1BP3_ADGRB1 | Tax1bp3 - Adgrb1 | ADGRB | Cell-Cell Contact | PMID: 31462748 |
| CD31 | NeuN | Tax1bp3 | Adgrb1 | 0.006939 | 0 | TAX1BP3_ADGRB1 | Tax1bp3 - Adgrb1 | ADGRB | Cell-Cell Contact | PMID: 31462748 |
| NeuN | CD31 | Rtn4r | Adgrb1 | 0.004078 | 0 | RTN4R_ADGRB1 | Rtn4r - Adgrb1 | ADGRB | Cell-Cell Contact | PMID: 31462748 |
| NeuN | NeuN | Rtn4r | Adgrb1 | 0.01053 | 0 | RTN4R_ADGRB1 | Rtn4r - Adgrb1 | ADGRB | Cell-Cell Contact | PMID: 31462748 |
| CD31 | CD31 | ITGAV_ITG | Adgrb1 | 0.001717 | 0 | ITGAV_ITGB5_ADGRB1 | (Itgav+Itgb5) - Adgrb1 | ADGRB | Cell-Cell Contact | PMID: 31462748 |
| CD31 | NeuN | ITGAV_ITG | Adgrb1 | 0.004451 | 0 | ITGAV_ITGB5_ADGRB1 | (Itgav+Itgb5) - Adgrb1 | ADGRB | Cell-Cell Contact | PMID: 31462748 |
| CD31 | CD31 | Tax1bp3 | Adgrb2 | 0.006431 | 0 | TAX1BP3_ADGRB2 | Tax1bp3 - Adgrb2 | ADGRB | Cell-Cell Contact | PMID: 31462748 |
| CD31 | NeuN | Tax1bp3 | Adgrb2 | 0.020302 | 0 | TAX1BP3_ADGRB2 | Tax1bp3 - Adgrb2 | ADGRB | Cell-Cell Contact | PMID: 31462748 |
| GFAP | GFAP | C1ql1 | Adgrb3 | 0.001285 | 0 | C1QL1_ADGRB3 | C1ql1 - Adgrb3 | ADGRB | Cell-Cell Contact | PMID: 31462748 |
| GFAP | NeuN | C1ql1 | Adgrb3 | 0.003297 | 0 | C1QL1_ADGRB3 | C1ql1 - Adgrb3 | ADGRB | Cell-Cell Contact | PMID: 31462748 |
| CD31 | NeuN | C1ql2 | Adgrb3 | 0.002283 | 0 | C1QL2_ADGRB3 | C1ql2 - Adgrb3 | ADGRB | Cell-Cell Contact | PMID: 31462748 |
| NeuN | NeuN | C1ql2 | Adgrb3 | 0.00133 | 0 | C1QL2_ADGRB3 | C1ql2 - Adgrb3 | ADGRB | Cell-Cell Contact | PMID: 31462748 |
| NeuN | GFAP | C1ql3 | Adgrb3 | 0.003961 | 0 | C1QL3_ADGRB3 | C1ql3 - Adgrb3 | ADGRB | Cell-Cell Contact | PMID: 31462748 |
| NeuN | NeuN | C1ql3 | Adgrb3 | 0.010119 | 0 | C1QL3_ADGRB3 | C1ql3 - Adgrb3 | ADGRB | Cell-Cell Contact | PMID: 31462748 |
| CD31 | CD31 | Thy1 | Adgre5 | 0.017716 | 0 | THY1_ADGRE5 | Thy1 - Adgre5 | ADGRE | Cell-Cell Contact | PMID: 31462748 |
| NeuN | CD31 | Thy1 | Adgre5 | 0.065613 | 0 | THY1_ADGRE5 | Thy1 - Adgre5 | ADGRE | Cell-Cell Contact | PMID: 31462748 |
| CD31 | CD31 | ITGAV_ITG | Adgre5 | 0.00573 | 0 | ITGAV_ITGB1_ADGRE5 | (Itgav+Itgb1) - Adgre5 | ADGRE | Cell-Cell Contact | PMID: 31462748 |
| GFAP | CD31 | ITGAV_ITG | Adgre5 | 0.004412 | 0 | ITGAV_ITGB1_ADGRE5 | (Itgav+Itgb1) - Adgre5 | ADGRE | Cell-Cell Contact | PMID: 31462748 |
| CD31 | CD31 | ITGAV_ITG | Adgre5 | 0.001738 | 0 | ITGAV_ITGB3_ADGRE5 | (Itgav+Itgb3) - Adgre5 | ADGRE | Cell-Cell Contact | PMID: 31462748 |
| GFAP | CD31 | ITGAV_ITG | Adgre5 | 0.002491 | 0 | ITGAV_ITGB3_ADGRE5 | (Itgav+Itgb3) - Adgre5 | ADGRE | Cell-Cell Contact | PMID: 31462748 |
| CD31 | CD31 | Cd55 | Adgre5 | 0.001798 | 0 | CD55_ADGRE5 | Cd55 - Adgre5 | ADGRE | Cell-Cell Contact | PMID: 31462748 |
| GFAP | CD31 | Cd55 | Adgre5 | 0.00222 | 0 | CD55_ADGRE5 | Cd55 - Adgre5 | ADGRE | Cell-Cell Contact | PMID: 31462748 |
| CD31 | CD31 | Lpar1 | Adgre5 | 0.002604 | 0 | LPAR1_ADGRE5 | Lpar1 - Adgre5 | ADGRE | Cell-Cell Contact | PMID: 31462748 |
| GFAP | CD31 | Lpar1 | Adgre5 | 0.00726 | 0 | LPAR1_ADGRE5 | Lpar1 - Adgre5 | ADGRE | Cell-Cell Contact | PMID: 31462748 |
| CD31 | CD31 | Lpar2 | Adgre5 | 0.002348 | 0 | LPAR2_ADGRE5 | Lpar2 - Adgre5 | ADGRE | Cell-Cell Contact | PMID: 31462748 |
| GFAP | CD31 | Lpar2 | Adgre5 | 0.001714 | 0 | LPAR2_ADGRE5 | Lpar2 - Adgre5 | ADGRE | Cell-Cell Contact | PMID: 31462748 |
| CD31 | CD31 | Col3a1 | Adgrg1 | 0.002857 | 0 | COL3A1_ADGRG1 | Col3a1 - Adgrg1 | ADGRG | Cell-Cell Contact | PMID: 31462748;UniProtKB |
| CD31 | GFAP | Col3a1 | Adgrg1 | 0.006692 | 0 | COL3A1_ADGRG1 | Col3a1 - Adgrg1 | ADGRG | Cell-Cell Contact | PMID: 31462748;UniProtKB |
| CD31 | CD31 | Tgm2 | Adgrg1 | 0.001645 | 0 | TGM2_ADGRG1 | Tgm2 - Adgrg1 | ADGRG | Cell-Cell Contact | PMID: 31462748 |
| CD31 | GFAP | Tgm2 | Adgrg1 | 0.003859 | 0 | TGM2_ADGRG1 | Tgm2 - Adgrg1 | ADGRG | Cell-Cell Contact | PMID: 31462748 |
| CD31 | CD31 | Col4a1 | Adgrg6 | 0.003035 | 0 | COL4A1_ADGRG6 | Col4a1 - Adgrg6 | ADGRG | Cell-Cell Contact | PMID: 31462748 |
| CD31 | GFAP | Col4a1 | Adgrg6 | 0.003059 | 0 | COL4A1_ADGRG6 | Col4a1 - Adgrg6 | ADGRG | Cell-Cell Contact | PMID: 31462748 |
| CD31 | CD31 | Col4a2 | Adgrg6 | 0.002273 | 0 | COL4A2_ADGRG6 | Col4a2 - Adgrg6 | ADGRG | Cell-Cell Contact | PMID: 31462748 |
| CD31 | GFAP | Col4a2 | Adgrg6 | 0.002291 | 0 | COL4A2_ADGRG6 | Col4a2 - Adgrg6 | ADGRG | Cell-Cell Contact | PMID: 31462748 |
| NeuN | GFAP | Prnp | Adgrg6 | 0.008791 | 0 | PRNP_ADGRG6 | Prnp - Adgrg6 | ADGRG | Cell-Cell Contact | PMID: 31462748 |
| NeuN | NeuN | Prnp | Adgrg6 | 0.005254 | 0 | PRNP_ADGRG6 | Prnp - Adgrg6 | ADGRG | Cell-Cell Contact | PMID: 31462748 |
| NeuN | CD31 | Flrt1 | Adgrl1 | 0.008775 | 0 | FLRT1_ADGRL1 | Flrt1 - Adgrl1 | ADGRL | Cell-Cell Contact | PMID: 25728924;PMID: 34022279;PMID: 27091502;PMID: 35907405 |
| NeuN | NeuN | Flrt1 | Adgrl1 | 0.018098 | 0 | FLRT1_ADGRL1 | Flrt1 - Adgrl1 | ADGRL | Cell-Cell Contact | PMID: 25728924;PMID: 34022279;PMID: 27091502;PMID: 35907405 |
| NeuN | CD31 | Flrt2 | Adgrl1 | 0.008724 | 0 | FLRT2_ADGRL1 | Flrt2 - Adgrl1 | ADGRL | Cell-Cell Contact | PMID: 25728924;PMID: 34022279;PMID: 27091502;PMID: 35907405 |
| NeuN | NeuN | Flrt2 | Adgrl1 | 0.017995 | 0 | FLRT2_ADGRL1 | Flrt2 - Adgrl1 | ADGRL | Cell-Cell Contact | PMID: 25728924;PMID: 34022279;PMID: 27091502;PMID: 35907405 |
| GFAP | NeuN | Flrt3 | Adgrl1 | 0.007063 | 0 | FLRT3_ADGRL1 | Flrt3 - Adgrl1 | ADGRL | Cell-Cell Contact | PMID: 25728924;PMID: 34022279;PMID: 27091502;PMID: 35907405 |
| NeuN | NeuN | Flrt3 | Adgrl1 | 0.006834 | 0 | FLRT3_ADGRL1 | Flrt3 - Adgrl1 | ADGRL | Cell-Cell Contact | PMID: 25728924;PMID: 34022279;PMID: 27091502;PMID: 35907405 |
| CD31 | NeuN | Nrxn1 | Adgrl1 | 0.0433 | 0 | NRXN1_ADGRL1 | Nrxn1 - Adgrl1 | ADGRL | Cell-Cell Contact | PMID: 25728924;PMID: 34022279;PMID: 27091502;PMID: 35907405 |
| NeuN | NeuN | Nrxn1 | Adgrl1 | 0.081578 | 0 | NRXN1_ADGRL1 | Nrxn1 - Adgrl1 | ADGRL | Cell-Cell Contact | PMID: 25728924;PMID: 34022279;PMID: 27091502;PMID: 35907405 |
| GFAP | NeuN | Nrxn2 | Adgrl1 | 0.029441 | 0 | NRXN2_ADGRL1 | Nrxn2 - Adgrl1 | ADGRL | Cell-Cell Contact | PMID: 25728924;PMID: 34022279;PMID: 27091502;PMID: 35907405 |
| NeuN | NeuN | Nrxn2 | Adgrl1 | 0.037104 | 0 | NRXN2_ADGRL1 | Nrxn2 - Adgrl1 | ADGRL | Cell-Cell Contact | PMID: 25728924;PMID: 34022279;PMID: 27091502;PMID: 35907405 |
| NeuN | CD31 | Nrxn3 | Adgrl1 | 0.021141 | 0 | NRXN3_ADGRL1 | Nrxn3 - Adgrl1 | ADGRL | Cell-Cell Contact | PMID: 25728924;PMID: 34022279;PMID: 27091502;PMID: 35907405 |
| NeuN | NeuN | Nrxn3 | Adgrl1 | 0.043033 | 0 | NRXN3_ADGRL1 | Nrxn3 - Adgrl1 | ADGRL | Cell-Cell Contact | PMID: 25728924;PMID: 34022279;PMID: 27091502;PMID: 35907405 |
| GFAP | NeuN | Tenm1 | Adgrl1 | 0.005608 | 0 | TENM1_ADGRL1 | Tenm1 - Adgrl1 | ADGRL | Cell-Cell Contact | PMID: 25728924;PMID: 34022279;PMID: 27091502;PMID: 35907405 |

| | | | | | | | | | |
|---|---|---|---|---|---|---|---|---|---|
| NeuN | NeuN | Tenm1 | Adgrl1 | 0.004612 | 0 TENM1_ADGRL1 | Tenm1 - Adgrl1 | ADGRL | Cell-Cell Contact | PMID: 25728924;PMID: 34022279;PMID: 27091502;PMID: 35907405 |
| GFAP | NeuN | Tenm2 | Adgrl1 | 0.005408 | 0 TENM2_ADGRL1 | Tenm2 - Adgrl1 | ADGRL | Cell-Cell Contact | PMID: 25728924;PMID: 34022279;PMID: 27091502;PMID: 35907405 |
| NeuN | NeuN | Tenm2 | Adgrl1 | 0.006593 | 0 TENM2_ADGRL1 | Tenm2 - Adgrl1 | ADGRL | Cell-Cell Contact | PMID: 25728924;PMID: 34022279;PMID: 27091502;PMID: 35907405 |
| GFAP | NeuN | Tenm3 | Adgrl1 | 0.008995 | 0 TENM3_ADGRL1 | Tenm3 - Adgrl1 | ADGRL | Cell-Cell Contact | PMID: 25728924;PMID: 34022279;PMID: 27091502;PMID: 35907405 |
| NeuN | NeuN | Tenm3 | Adgrl1 | 0.010996 | 0 TENM3_ADGRL1 | Tenm3 - Adgrl1 | ADGRL | Cell-Cell Contact | PMID: 25728924;PMID: 34022279;PMID: 27091502;PMID: 35907405 |
| CD31 | NeuN | Tenm4 | Adgrl1 | 0.00781 | 0 TENM4_ADGRL1 | Tenm4 - Adgrl1 | ADGRL | Cell-Cell Contact | PMID: 25728924;PMID: 34022279;PMID: 27091502;PMID: 35907405 |
| NeuN | NeuN | Tenm4 | Adgrl1 | 0.010834 | 0 TENM4_ADGRL1 | Tenm4 - Adgrl1 | ADGRL | Cell-Cell Contact | PMID: 25728924;PMID: 34022279;PMID: 27091502;PMID: 35907405 |
| GFAP | NeuN | TENM2_FL | Adgrl1 | 0.00671 | 0 TENM2_FLRT1_ADGRL1 | (Tenm2+Flrt1) - Adgrl1 | ADGRL | Cell-Cell Contact | PMID: 25728924;PMID: 34022279;PMID: 27091502;PMID: 35907405 |
| NeuN | NeuN | TENM2_FL | Adgrl1 | 0.010939 | 0 TENM2_FLRT1_ADGRL1 | (Tenm2+Flrt1) - Adgrl1 | ADGRL | Cell-Cell Contact | PMID: 25728924;PMID: 34022279;PMID: 27091502;PMID: 35907405 |
| GFAP | NeuN | TENM2_FL | Adgrl1 | 0.006181 | 0 TENM2_FLRT3_ADGRL1 | (Tenm2+Flrt3) - Adgrl1 | ADGRL | Cell-Cell Contact | PMID: 25728924;PMID: 34022279;PMID: 27091502;PMID: 35907405 |
| NeuN | NeuN | TENM2_FL | Adgrl1 | 0.006712 | 0 TENM2_FLRT3_ADGRL1 | (Tenm2+Flrt3) - Adgrl1 | ADGRL | Cell-Cell Contact | PMID: 25728924;PMID: 34022279;PMID: 27091502;PMID: 35907405 |
| CD31 | NeuN | TENM4_FL | Adgrl1 | 0.007845 | 0 TENM4_FLRT1_ADGRL1 | (Tenm4+Flrt1) - Adgrl1 | ADGRL | Cell-Cell Contact | PMID: 25728924;PMID: 34022279;PMID: 27091502;PMID: 35907405 |
| NeuN | NeuN | TENM4_FL | Adgrl1 | 0.014009 | 0 TENM4_FLRT1_ADGRL1 | (Tenm4+Flrt1) - Adgrl1 | ADGRL | Cell-Cell Contact | PMID: 25728924;PMID: 34022279;PMID: 27091502;PMID: 35907405 |
| CD31 | NeuN | TENM4_FL | Adgrl1 | 0.006839 | 0 TENM4_FLRT3_ADGRL1 | (Tenm4+Flrt3) - Adgrl1 | ADGRL | Cell-Cell Contact | PMID: 25728924;PMID: 34022279;PMID: 27091502;PMID: 35907405 |
| NeuN | NeuN | TENM4_FL | Adgrl1 | 0.008606 | 0 TENM4_FLRT3_ADGRL1 | (Tenm4+Flrt3) - Adgrl1 | ADGRL | Cell-Cell Contact | PMID: 25728924;PMID: 34022279;PMID: 27091502;PMID: 35907405 |
| NeuN | CD31 | Flrt1 | Adgrl2 | 0.003122 | 0 FLRT1_ADGRL2 | Flrt1 - Adgrl2 | ADGRL | Cell-Cell Contact | PMID: 25728924;PMID: 34022279;PMID: 27091502;PMID: 35907405 |
| NeuN | NeuN | Flrt1 | Adgrl2 | 0.003508 | 0 FLRT1_ADGRL2 | Flrt1 - Adgrl2 | ADGRL | Cell-Cell Contact | PMID: 25728924;PMID: 34022279;PMID: 27091502;PMID: 35907405 |
| NeuN | CD31 | Flrt2 | Adgrl2 | 0.003104 | 0 FLRT2_ADGRL2 | Flrt2 - Adgrl2 | ADGRL | Cell-Cell Contact | PMID: 25728924;PMID: 34022279;PMID: 27091502;PMID: 35907405 |
| NeuN | NeuN | Flrt2 | Adgrl2 | 0.003488 | 0 FLRT2_ADGRL2 | Flrt2 - Adgrl2 | ADGRL | Cell-Cell Contact | PMID: 25728924;PMID: 34022279;PMID: 27091502;PMID: 35907405 |
| GFAP | NeuN | Flrt3 | Adgrl2 | 0.001357 | 0 FLRT3_ADGRL2 | Flrt3 - Adgrl2 | ADGRL | Cell-Cell Contact | PMID: 25728924;PMID: 34022279;PMID: 27091502;PMID: 35907405 |
| NeuN | NeuN | Flrt3 | Adgrl2 | 0.001312 | 0 FLRT3_ADGRL2 | Flrt3 - Adgrl2 | ADGRL | Cell-Cell Contact | PMID: 25728924;PMID: 34022279;PMID: 27091502;PMID: 35907405 |
| GFAP | NeuN | Tenm1 | Adgrl2 | 0.001076 | 0 TENM1_ADGRL2 | Tenm1 - Adgrl2 | ADGRL | Cell-Cell Contact | PMID: 25728924;PMID: 34022279;PMID: 27091502;PMID: 35907405 |
| NeuN | NeuN | Tenm1 | Adgrl2 | 0.000884 | 0 TENM1_ADGRL2 | Tenm1 - Adgrl2 | ADGRL | Cell-Cell Contact | PMID: 25728924;PMID: 34022279;PMID: 27091502;PMID: 35907405 |
| NeuN | CD31 | Tenm2 | Adgrl2 | 0.001126 | 0 TENM2_ADGRL2 | Tenm2 - Adgrl2 | ADGRL | Cell-Cell Contact | PMID: 25728924;PMID: 34022279;PMID: 27091502;PMID: 35907405 |
| NeuN | NeuN | Tenm2 | Adgrl2 | 0.001266 | 0 TENM2_ADGRL2 | Tenm2 - Adgrl2 | ADGRL | Cell-Cell Contact | PMID: 25728924;PMID: 34022279;PMID: 27091502;PMID: 35907405 |
| NeuN | CD31 | Tenm3 | Adgrl2 | 0.001886 | 0 TENM3_ADGRL2 | Tenm3 - Adgrl2 | ADGRL | Cell-Cell Contact | PMID: 25728924;PMID: 34022279;PMID: 27091502;PMID: 35907405 |
| NeuN | NeuN | Tenm3 | Adgrl2 | 0.002119 | 0 TENM3_ADGRL2 | Tenm3 - Adgrl2 | ADGRL | Cell-Cell Contact | PMID: 25728924;PMID: 34022279;PMID: 27091502;PMID: 35907405 |
| NeuN | CD31 | Tenm4 | Adgrl2 | 0.001858 | 0 TENM4_ADGRL2 | Tenm4 - Adgrl2 | ADGRL | Cell-Cell Contact | PMID: 25728924;PMID: 34022279;PMID: 27091502;PMID: 35907405 |
| NeuN | NeuN | Tenm4 | Adgrl2 | 0.002088 | 0 TENM4_ADGRL2 | Tenm4 - Adgrl2 | ADGRL | Cell-Cell Contact | PMID: 25728924;PMID: 34022279;PMID: 27091502;PMID: 35907405 |
| GFAP | NeuN | TENM2_FL | Adgrl2 | 0.001186 | 0 TENM2_FLRT3_ADGRL2 | (Tenm2+Flrt3) - Adgrl2 | ADGRL | Cell-Cell Contact | PMID: 25728924;PMID: 34022279;PMID: 27091502;PMID: 35907405 |
| NeuN | NeuN | TENM2_FL | Adgrl2 | 0.001289 | 0 TENM2_FLRT3_ADGRL2 | (Tenm2+Flrt3) - Adgrl2 | ADGRL | Cell-Cell Contact | PMID: 25728924;PMID: 34022279;PMID: 27091502;PMID: 35907405 |
| NeuN | CD31 | Flrt1 | Adgrl3 | 0.003414 | 0 FLRT1_ADGRL3 | Flrt1 - Adgrl3 | ADGRL | Cell-Cell Contact | PMID: 25728924;PMID: 34022279;PMID: 27091502;PMID: 35907405 |
| NeuN | NeuN | Flrt1 | Adgrl3 | 0.004979 | 0 FLRT1_ADGRL3 | Flrt1 - Adgrl3 | ADGRL | Cell-Cell Contact | PMID: 25728924;PMID: 34022279;PMID: 27091502;PMID: 35907405 |
| NeuN | CD31 | Flrt2 | Adgrl3 | 0.003394 | 0 FLRT2_ADGRL3 | Flrt2 - Adgrl3 | ADGRL | Cell-Cell Contact | PMID: 25728924;PMID: 34022279;PMID: 27091502;PMID: 35907405 |
| NeuN | NeuN | Flrt2 | Adgrl3 | 0.004951 | 0 FLRT2_ADGRL3 | Flrt2 - Adgrl3 | ADGRL | Cell-Cell Contact | PMID: 25728924;PMID: 34022279;PMID: 27091502;PMID: 35907405 |
| GFAP | NeuN | Flrt3 | Adgrl3 | 0.001928 | 0 FLRT3_ADGRL3 | Flrt3 - Adgrl3 | ADGRL | Cell-Cell Contact | PMID: 25728924;PMID: 34022279;PMID: 27091502;PMID: 35907405 |
| NeuN | NeuN | Flrt3 | Adgrl3 | 0.001865 | 0 FLRT3_ADGRL3 | Flrt3 - Adgrl3 | ADGRL | Cell-Cell Contact | PMID: 25728924;PMID: 34022279;PMID: 27091502;PMID: 35907405 |
| GFAP | NeuN | Tenm1 | Adgrl3 | 0.001529 | 0 TENM1_ADGRL3 | Tenm1 - Adgrl3 | ADGRL | Cell-Cell Contact | PMID: 25728924;PMID: 34022279;PMID: 27091502;PMID: 35907405 |
| NeuN | NeuN | Tenm1 | Adgrl3 | 0.001257 | 0 TENM1_ADGRL3 | Tenm1 - Adgrl3 | ADGRL | Cell-Cell Contact | PMID: 25728924;PMID: 34022279;PMID: 27091502;PMID: 35907405 |
| GFAP | NeuN | Tenm2 | Adgrl3 | 0.001474 | 0 TENM2_ADGRL3 | Tenm2 - Adgrl3 | ADGRL | Cell-Cell Contact | PMID: 25728924;PMID: 34022279;PMID: 27091502;PMID: 35907405 |
| NeuN | NeuN | Tenm2 | Adgrl3 | 0.001799 | 0 TENM2_ADGRL3 | Tenm2 - Adgrl3 | ADGRL | Cell-Cell Contact | PMID: 25728924;PMID: 34022279;PMID: 27091502;PMID: 35907405 |
| GFAP | NeuN | Tenm3 | Adgrl3 | 0.002458 | 0 TENM3_ADGRL3 | Tenm3 - Adgrl3 | ADGRL | Cell-Cell Contact | PMID: 25728924;PMID: 34022279;PMID: 27091502;PMID: 35907405 |
| NeuN | NeuN | Tenm3 | Adgrl3 | 0.00301 | 0 TENM3_ADGRL3 | Tenm3 - Adgrl3 | ADGRL | Cell-Cell Contact | PMID: 25728924;PMID: 34022279;PMID: 27091502;PMID: 35907405 |
| CD31 | NeuN | Tenm4 | Adgrl3 | 0.002133 | 0 TENM4_ADGRL3 | Tenm4 - Adgrl3 | ADGRL | Cell-Cell Contact | PMID: 25728924;PMID: 34022279;PMID: 27091502;PMID: 35907405 |
| NeuN | NeuN | Tenm4 | Adgrl3 | 0.002965 | 0 TENM4_ADGRL3 | Tenm4 - Adgrl3 | ADGRL | Cell-Cell Contact | PMID: 25728924;PMID: 34022279;PMID: 27091502;PMID: 35907405 |
| NeuN | CD31 | Unc5a | Adgrl3 | 0.002799 | 0 UNC5A_ADGRL3 | Unc5a - Adgrl3 | ADGRL | Cell-Cell Contact | PMID: 25728924;PMID: 34022279;PMID: 27091502;PMID: 35907405 |
| NeuN | NeuN | Unc5a | Adgrl3 | 0.004084 | 0 UNC5A_ADGRL3 | Unc5a - Adgrl3 | ADGRL | Cell-Cell Contact | PMID: 25728924;PMID: 34022279;PMID: 27091502;PMID: 35907405 |
| NeuN | CD31 | TENM3_FL | Adgrl3 | 0.002654 | 0 TENM3_FLRT1_ADGRL3 | (Tenm3+Flrt1) - Adgrl3 | ADGRL | Cell-Cell Contact | PMID: 25728924;PMID: 34022279;PMID: 27091502;PMID: 35907405 |
| NeuN | NeuN | TENM3_FL | Adgrl3 | 0.003872 | 0 TENM3_FLRT1_ADGRL3 | (Tenm3+Flrt1) - Adgrl3 | ADGRL | Cell-Cell Contact | PMID: 25728924;PMID: 34022279;PMID: 27091502;PMID: 35907405 |
| GFAP | NeuN | TENM3_FL | Adgrl3 | 0.002177 | 0 TENM3_FLRT3_ADGRL3 | (Tenm3+Flrt3) - Adgrl3 | ADGRL | Cell-Cell Contact | PMID: 25728924;PMID: 34022279;PMID: 27091502;PMID: 35907405 |
| NeuN | NeuN | TENM3_FL | Adgrl3 | 0.002369 | 0 TENM3_FLRT3_ADGRL3 | (Tenm3+Flrt3) - Adgrl3 | ADGRL | Cell-Cell Contact | PMID: 25728924;PMID: 34022279;PMID: 27091502;PMID: 35907405 |
| NeuN | GFAP | Flrt1 | Flrt1 | 0.001331 | 0 FLRT1_FLRT1 | Flrt1 - Flrt1 | FLRT | Cell-Cell Contact | PMID: 27091502 |
| GFAP | NeuN | Flrt1 | Flrt1 | 0.001331 | 0 FLRT1_FLRT1 | Flrt1 - Flrt1 | FLRT | Cell-Cell Contact | PMID: 27091502 |
| NeuN | NeuN | Flrt1 | Flrt1 | 0.002919 | 0 FLRT1_FLRT1 | Flrt1 - Flrt1 | FLRT | Cell-Cell Contact | PMID: 27091502 |
| NeuN | GFAP | Flrt2 | Flrt2 | 0.001349 | 0 FLRT2_FLRT2 | Flrt2 - Flrt2 | FLRT | Cell-Cell Contact | PMID: 27091502 |
| GFAP | NeuN | Flrt2 | Flrt2 | 0.001349 | 0 FLRT2_FLRT2 | Flrt2 - Flrt2 | FLRT | Cell-Cell Contact | PMID: 27091502 |
| NeuN | NeuN | Flrt2 | Flrt2 | 0.002885 | 0 FLRT2_FLRT2 | Flrt2 - Flrt2 | FLRT | Cell-Cell Contact | PMID: 27091502 |
| GFAP | GFAP | Flrt3 | Flrt3 | 0.000436 | 0 FLRT3_FLRT3 | Flrt3 - Flrt3 | FLRT | Cell-Cell Contact | PMID: 27091502 |
| NeuN | GFAP | Flrt3 | Flrt3 | 0.000422 | 0 FLRT3_FLRT3 | Flrt3 - Flrt3 | FLRT | Cell-Cell Contact | PMID: 27091502 |

| | | | | | | | | | | |
|---|---|---|---|---|---|---|---|---|---|---|
| GFAP | NeuN | Flrt3 | Flrt3 | 0.000422 | 0 | FLRT3_FLRT3 | Flrt3 - Flrt3 | FLRT | Cell-Cell Contact | PMID: 27091502 |
| NeuN | CD31 | Flrt1 | Unc5a | 0.001292 | 0 | FLRT1_UNC5A | Flrt1 - Unc5a | UNC5 | Cell-Cell Contact | PMID: 27091502 |
| NeuN | NeuN | Flrt1 | Unc5a | 0.002393 | 0 | FLRT1_UNC5A | Flrt1 - Unc5a | UNC5 | Cell-Cell Contact | PMID: 27091502 |
| NeuN | CD31 | Flrt2 | Unc5a | 0.001285 | 0 | FLRT2_UNC5A | Flrt2 - Unc5a | UNC5 | Cell-Cell Contact | PMID: 27091502 |
| NeuN | NeuN | Flrt2 | Unc5a | 0.002379 | 0 | FLRT2_UNC5A | Flrt2 - Unc5a | UNC5 | Cell-Cell Contact | PMID: 27091502 |
| GFAP | NeuN | Flrt3 | Unc5a | 0.000925 | 0 | FLRT3_UNC5A | Flrt3 - Unc5a | UNC5 | Cell-Cell Contact | PMID: 27091502 |
| NeuN | NeuN | Flrt3 | Unc5a | 0.000895 | 0 | FLRT3_UNC5A | Flrt3 - Unc5a | UNC5 | Cell-Cell Contact | PMID: 27091502 |
| GFAP | GFAP | Flrt1 | Unc5b | 0.00084 | 0 | FLRT1_UNC5B | Flrt1 - Unc5b | UNC5 | Cell-Cell Contact | PMID: 27091502 |
| NeuN | GFAP | Flrt1 | Unc5b | 0.001842 | 0 | FLRT1_UNC5B | Flrt1 - Unc5b | UNC5 | Cell-Cell Contact | PMID: 27091502 |
| GFAP | GFAP | Flrt2 | Unc5b | 0.000856 | 0 | FLRT2_UNC5B | Flrt2 - Unc5b | UNC5 | Cell-Cell Contact | PMID: 27091502 |
| NeuN | GFAP | Flrt2 | Unc5b | 0.001832 | 0 | FLRT2_UNC5B | Flrt2 - Unc5b | UNC5 | Cell-Cell Contact | PMID: 27091502 |
| GFAP | GFAP | Flrt3 | Unc5b | 0.000712 | 0 | FLRT3_UNC5B | Flrt3 - Unc5b | UNC5 | Cell-Cell Contact | PMID: 27091502 |
| NeuN | GFAP | Flrt3 | Unc5b | 0.000689 | 0 | FLRT3_UNC5B | Flrt3 - Unc5b | UNC5 | Cell-Cell Contact | PMID: 27091502 |
| NeuN | GFAP | Flrt1 | Unc5c | 0.000938 | 0 | FLRT1_UNC5C | Flrt1 - Unc5 | UNC5 | Cell-Cell Contact | PMID: 27091502 |
| NeuN | NeuN | Flrt1 | Unc5c | 0.000699 | 0 | FLRT1_UNC5C | Flrt1 - Unc5 | UNC5 | Cell-Cell Contact | PMID: 27091502 |
| NeuN | GFAP | Flrt2 | Unc5c | 0.000933 | 0 | FLRT2_UNC5C | Flrt2 - Unc5 | UNC5 | Cell-Cell Contact | PMID: 27091502 |
| NeuN | NeuN | Flrt2 | Unc5c | 0.000695 | 0 | FLRT2_UNC5C | Flrt2 - Unc5 | UNC5 | Cell-Cell Contact | PMID: 27091502 |
| GFAP | GFAP | Flrt3 | Unc5c | 0.000362 | 0 | FLRT3_UNC5C | Flrt3 - Unc5 | UNC5 | Cell-Cell Contact | PMID: 27091502 |
| NeuN | GFAP | Flrt3 | Unc5c | 0.00035 | 0 | FLRT3_UNC5C | Flrt3 - Unc5 | UNC5 | Cell-Cell Contact | PMID: 27091502 |
| NeuN | GFAP | Flrt1 | Unc5d | 0.00056 | 0 | FLRT1_UNC5D | Flrt1 - Unc5d | UNC5 | Cell-Cell Contact | PMID: 27091502 |
| NeuN | NeuN | Flrt1 | Unc5d | 0.000645 | 0 | FLRT1_UNC5D | Flrt1 - Unc5d | UNC5 | Cell-Cell Contact | PMID: 27091502 |
| NeuN | GFAP | Flrt2 | Unc5d | 0.000557 | 0 | FLRT2_UNC5D | Flrt2 - Unc5d | UNC5 | Cell-Cell Contact | PMID: 27091502 |
| NeuN | NeuN | Flrt2 | Unc5d | 0.000641 | 0 | FLRT2_UNC5D | Flrt2 - Unc5d | UNC5 | Cell-Cell Contact | PMID: 27091502 |
| GFAP | NeuN | Flrt3 | Unc5d | 0.000249 | 0 | FLRT3_UNC5D | Flrt3 - Unc5d | UNC5 | Cell-Cell Contact | PMID: 27091502 |
| NeuN | NeuN | Flrt3 | Unc5d | 0.000241 | 0 | FLRT3_UNC5D | Flrt3 - Unc5d | UNC5 | Cell-Cell Contact | PMID: 27091502 |
| CD31 | NeuN | Nrxn1 | Clstn1 | 0.034661 | 0 | NRXN1_CLSTN1 | Nrxn1 - Clstn1 | NRXN | Cell-Cell Contact | PMID: 32434929 |
| NeuN | NeuN | Nrxn1 | Clstn1 | 0.065826 | 0 | NRXN1_CLSTN1 | Nrxn1 - Clstn1 | NRXN | Cell-Cell Contact | PMID: 32434929 |
| GFAP | NeuN | Nrxn2 | Clstn1 | 0.023498 | 0 | NRXN2_CLSTN1 | Nrxn2 - Clstn1 | NRXN | Cell-Cell Contact | PMID: 32434929 |
| NeuN | NeuN | Nrxn2 | Clstn1 | 0.029662 | 0 | NRXN2_CLSTN1 | Nrxn2 - Clstn1 | NRXN | Cell-Cell Contact | PMID: 32434929 |
| CD31 | NeuN | Nrxn1 | Clstn2 | 0.010723 | 0 | NRXN1_CLSTN2 | Nrxn1 - Clstn2 | NRXN | Cell-Cell Contact | PMID: 32434929 |
| NeuN | NeuN | Nrxn1 | Clstn2 | 0.020829 | 0 | NRXN1_CLSTN2 | Nrxn1 - Clstn2 | NRXN | Cell-Cell Contact | PMID: 32434929 |
| GFAP | NeuN | Nrxn2 | Clstn2 | 0.007212 | 0 | NRXN2_CLSTN2 | Nrxn2 - Clstn2 | NRXN | Cell-Cell Contact | PMID: 32434929 |
| NeuN | NeuN | Nrxn2 | Clstn2 | 0.009144 | 0 | NRXN2_CLSTN2 | Nrxn2 - Clstn2 | NRXN | Cell-Cell Contact | PMID: 32434929 |
| CD31 | NeuN | Nrxn1 | Clstn3 | 0.036761 | 0 | NRXN1_CLSTN3 | Nrxn1 - Clstn3 | NRXN | Cell-Cell Contact | PMID: 32434929 |
| NeuN | NeuN | Nrxn1 | Clstn3 | 0.069678 | 0 | NRXN1_CLSTN3 | Nrxn1 - Clstn3 | NRXN | Cell-Cell Contact | PMID: 32434929 |
| GFAP | NeuN | Nrxn2 | Clstn3 | 0.024939 | 0 | NRXN2_CLSTN3 | Nrxn2 - Clstn3 | NRXN | Cell-Cell Contact | PMID: 32434929 |
| NeuN | NeuN | Nrxn2 | Clstn3 | 0.03147 | 0 | NRXN2_CLSTN3 | Nrxn2 - Clstn3 | NRXN | Cell-Cell Contact | PMID: 32434929 |
| NeuN | CD31 | Nrxn1 | Dag1 | 0.010425 | 0 | NRXN1_DAG1 | Nrxn1 - Dag1 | NRXN | Cell-Cell Contact | PMID: 21628410 |
| NeuN | NeuN | Nrxn1 | Dag1 | 0.006019 | 0 | NRXN1_DAG1 | Nrxn1 - Dag1 | NRXN | Cell-Cell Contact | PMID: 21628410 |
| NeuN | CD31 | Nrxn2 | Dag1 | 0.00455 | 0 | NRXN2_DAG1 | Nrxn2 - Dag1 | NRXN | Cell-Cell Contact | PMID: 21628410 |
| GFAP | GFAP | Nrxn2 | Dag1 | 0.002756 | 0 | NRXN2_DAG1 | Nrxn2 - Dag1 | NRXN | Cell-Cell Contact | PMID: 21628410 |
| NeuN | GFAP | Lrfn4 | Ptprd | 0.00605 | 0 | LRFN4_PTPRD | Lrfn4 - Ptprd | PTPR | Cell-Cell Contact | PMID: 29674953;PMID: 23916315 |
| NeuN | NeuN | Lrfn4 | Ptprd | 0.00649 | 0 | LRFN4_PTPRD | Lrfn4 - Ptprd | PTPR | Cell-Cell Contact | PMID: 29674953;PMID: 23916315 |
| NeuN | GFAP | Lrfn4 | Ptprf | 0.002343 | 0 | LRFN4_PTPRF | Lrfn4 - Ptprf | PTPR | Cell-Cell Contact | PMID: 29674953;PMID: 23916315 |
| NeuN | NeuN | Lrfn4 | Ptprf | 0.002459 | 0 | LRFN4_PTPRF | Lrfn4 - Ptprf | PTPR | Cell-Cell Contact | PMID: 29674953;PMID: 23916315 |
| CD31 | NeuN | Lrfn4 | Ptprs | 0.005256 | 0 | LRFN4_PTPRS | Lrfn4 - Ptprs | PTPR | Cell-Cell Contact | PMID: 29674953;PMID: 23916315 |
| NeuN | NeuN | Lrfn4 | Ptprs | 0.009632 | 0 | LRFN4_PTPRS | Lrfn4 - Ptprs | PTPR | Cell-Cell Contact | PMID: 29674953;PMID: 23916315 |
| NeuN | GFAP | Lrfn5 | Ptprd | 0.004678 | 0 | LRFN5_PTPRD | Lrfn5 - Ptprd | PTPR | Cell-Cell Contact | PMID: 29674953;PMID: 23916315 |
| NeuN | NeuN | Lrfn5 | Ptprd | 0.005018 | 0 | LRFN5_PTPRD | Lrfn5 - Ptprd | PTPR | Cell-Cell Contact | PMID: 29674953;PMID: 23916315 |
| NeuN | GFAP | Lrfn5 | Ptprf | 0.00181 | 0 | LRFN5_PTPRF | Lrfn5 - Ptprf | PTPR | Cell-Cell Contact | PMID: 29674953;PMID: 23916315 |
| NeuN | NeuN | Lrfn5 | Ptprf | 0.0019 | 0 | LRFN5_PTPRF | Lrfn5 - Ptprf | PTPR | Cell-Cell Contact | PMID: 29674953;PMID: 23916315 |
| CD31 | NeuN | Lrfn5 | Ptprs | 0.004965 | 0 | LRFN5_PTPRS | Lrfn5 - Ptprs | PTPR | Cell-Cell Contact | PMID: 29674953;PMID: 23916315 |
| NeuN | NeuN | Lrfn5 | Ptprs | 0.007453 | 0 | LRFN5_PTPRS | Lrfn5 - Ptprs | PTPR | Cell-Cell Contact | PMID: 29674953;PMID: 23916315 |
| NeuN | GFAP | Lrrc4b | Ptprd | 0.013293 | 0 | LRRC4B_PTPRD | Lrrc4b - Ptprd | PTPR | Cell-Cell Contact | PMID: 29674953;PMID: 23916315 |
| NeuN | NeuN | Lrrc4b | Ptprd | 0.014251 | 0 | LRRC4B_PTPRD | Lrrc4b - Ptprd | PTPR | Cell-Cell Contact | PMID: 29674953;PMID: 23916315 |
| GFAP | NeuN | Lrrc4b | Ptprs | 0.013081 | 0 | LRRC4B_PTPRS | Lrrc4b - Ptprs | PTPR | Cell-Cell Contact | PMID: 29674953;PMID: 23916315 |
| NeuN | NeuN | Lrrc4b | Ptprs | 0.021072 | 0 | LRRC4B_PTPRS | Lrrc4b - Ptprs | PTPR | Cell-Cell Contact | PMID: 29674953;PMID: 23916315 |
| NeuN | GFAP | Lrrc4c | Ptprf | 0.004228 | 0 | LRRC4C_PTPRF | Lrrc4c - Ptprf | PTPR | Cell-Cell Contact | PMID: 29674953;PMID: 23916315 |
| NeuN | NeuN | Lrrc4c | Ptprf | 0.004436 | 0 | LRRC4C_PTPRF | Lrrc4c - Ptprf | PTPR | Cell-Cell Contact | PMID: 29674953;PMID: 23916315 |

| | | | | | | | | | | |
|---|---|---|---|---|---|---|---|---|---|---|
| NeuN | CD31 | Nrxn1 | Lrrtm1 | 0.005318 | 0 | NRXN1_LRRTM1 | Nrxn1 - Lrrtm1 | NRXN | Cell-Cell Contact | PMID:20519524;PMID:20064387;PMID:20064388; |
| NeuN | NeuN | Nrxn1 | Lrrtm1 | 0.009416 | 0 | NRXN1_LRRTM1 | Nrxn1 - Lrrtm1 | NRXN | Cell-Cell Contact | PMID:20519524;PMID:20064387;PMID:20064388; |
| GFAP | NeuN | Nrxn2 | Lrrtm1 | 0.003236 | 0 | NRXN2_LRRTM1 | Nrxn2 - Lrrtm1 | NRXN | Cell-Cell Contact | PMID:20519524;PMID:20064387;PMID:20064388; |
| NeuN | NeuN | Nrxn2 | Lrrtm1 | 0.004107 | 0 | NRXN2_LRRTM1 | Nrxn2 - Lrrtm1 | NRXN | Cell-Cell Contact | PMID:20519524;PMID:20064387;PMID:20064388; |
| NeuN | CD31 | Nrxn3 | Lrrtm1 | 0.002699 | 0 | NRXN3_LRRTM1 | Nrxn3 - Lrrtm1 | NRXN | Cell-Cell Contact | PMID:20519524;PMID:20064387;PMID:20064388; |
| NeuN | NeuN | Nrxn3 | Lrrtm1 | 0.004789 | 0 | NRXN3_LRRTM1 | Nrxn3 - Lrrtm1 | NRXN | Cell-Cell Contact | PMID:20519524;PMID:20064387;PMID:20064388; |
| NeuN | CD31 | Nrxn1 | Lrrtm2 | 0.005264 | 0 | NRXN1_LRRTM2 | Nrxn1 - Lrrtm2 | NRXN | Cell-Cell Contact | PMID:20519524;PMID:20064387;PMID:20064388; |
| NeuN | NeuN | Nrxn1 | Lrrtm2 | 0.009055 | 0 | NRXN1_LRRTM2 | Nrxn1 - Lrrtm2 | NRXN | Cell-Cell Contact | PMID:20519524;PMID:20064387;PMID:20064388; |
| GFAP | NeuN | Nrxn2 | Lrrtm2 | 0.003111 | 0 | NRXN2_LRRTM2 | Nrxn2 - Lrrtm2 | NRXN | Cell-Cell Contact | PMID:20519524;PMID:20064387;PMID:20064388; |
| NeuN | NeuN | Nrxn2 | Lrrtm2 | 0.003949 | 0 | NRXN2_LRRTM2 | Nrxn2 - Lrrtm2 | NRXN | Cell-Cell Contact | PMID:20519524;PMID:20064387;PMID:20064388; |
| NeuN | CD31 | Nrxn3 | Lrrtm2 | 0.002672 | 0 | NRXN3_LRRTM2 | Nrxn3 - Lrrtm2 | NRXN | Cell-Cell Contact | PMID:20519524;PMID:20064387;PMID:20064388; |
| NeuN | NeuN | Nrxn3 | Lrrtm2 | 0.004605 | 0 | NRXN3_LRRTM2 | Nrxn3 - Lrrtm2 | NRXN | Cell-Cell Contact | PMID:20519524;PMID:20064387;PMID:20064388; |
| NeuN | CD31 | Nrxn1 | Lrrtm3 | 0.005318 | 0 | NRXN1_LRRTM3 | Nrxn1 - Lrrtm3 | NRXN | Cell-Cell Contact | PMID:20519524;PMID:20064387;PMID:20064388; |
| NeuN | NeuN | Nrxn1 | Lrrtm3 | 0.010089 | 0 | NRXN1_LRRTM3 | Nrxn1 - Lrrtm3 | NRXN | Cell-Cell Contact | PMID:20519524;PMID:20064387;PMID:20064388; |
| GFAP | NeuN | Nrxn2 | Lrrtm3 | 0.003468 | 0 | NRXN2_LRRTM3 | Nrxn2 - Lrrtm3 | NRXN | Cell-Cell Contact | PMID:20519524;PMID:20064387;PMID:20064388; |
| NeuN | NeuN | Nrxn2 | Lrrtm3 | 0.004402 | 0 | NRXN2_LRRTM3 | Nrxn2 - Lrrtm3 | NRXN | Cell-Cell Contact | PMID:20519524;PMID:20064387;PMID:20064388; |
| NeuN | CD31 | Nrxn3 | Lrrtm3 | 0.002699 | 0 | NRXN3_LRRTM3 | Nrxn3 - Lrrtm3 | NRXN | Cell-Cell Contact | PMID:20519524;PMID:20064387;PMID:20064388; |
| NeuN | NeuN | Nrxn3 | Lrrtm3 | 0.005133 | 0 | NRXN3_LRRTM3 | Nrxn3 - Lrrtm3 | NRXN | Cell-Cell Contact | PMID:20519524;PMID:20064387;PMID:20064388; |
| NeuN | CD31 | Nrxn1 | Lrrtm4 | 0.005111 | 0 | NRXN1_LRRTM4 | Nrxn1 - Lrrtm4 | NRXN | Cell-Cell Contact | PMID:20519524;PMID:20064387;PMID:20064388; |
| NeuN | NeuN | Nrxn1 | Lrrtm4 | 0.008517 | 0 | NRXN1_LRRTM4 | Nrxn1 - Lrrtm4 | NRXN | Cell-Cell Contact | PMID:20519524;PMID:20064387;PMID:20064388; |
| GFAP | NeuN | Nrxn2 | Lrrtm4 | 0.002925 | 0 | NRXN2_LRRTM4 | Nrxn2 - Lrrtm4 | NRXN | Cell-Cell Contact | PMID:20519524;PMID:20064387;PMID:20064388; |
| NeuN | NeuN | Nrxn2 | Lrrtm4 | 0.003713 | 0 | NRXN2_LRRTM4 | Nrxn2 - Lrrtm4 | NRXN | Cell-Cell Contact | PMID:20519524;PMID:20064387;PMID:20064388; |
| NeuN | CD31 | Nrxn3 | Lrrtm4 | 0.002594 | 0 | NRXN3_LRRTM4 | Nrxn3 - Lrrtm4 | NRXN | Cell-Cell Contact | PMID:20519524;PMID:20064387;PMID:20064388; |
| NeuN | NeuN | Nrxn3 | Lrrtm4 | 0.00433 | 0 | NRXN3_LRRTM4 | Nrxn3 - Lrrtm4 | NRXN | Cell-Cell Contact | PMID:20519524;PMID:20064387;PMID:20064388; |
| CD31 | NeuN | Lrrtm4 | Ptprs | 0.004772 | 0 | LRRTM4_PTPRS | Lrrtm4 - Ptprs | PTPR | Cell-Cell Contact | PMID:31995730 |
| NeuN | NeuN | Lrrtm4 | Ptprs | 0.007953 | 0 | LRRTM4_PTPRS | Lrrtm4 - Ptprs | PTPR | Cell-Cell Contact | PMID:31995730 |
| CD31 | NeuN | App | Sorl1 | 0.14143 | 0 | APP_SORL1 | App - Sorl1 | APP | Cell-Cell Contact | PMID: 24523320 |
| NeuN | NeuN | App | Sorl1 | 0.204939 | 0 | APP_SORL1 | App - Sorl1 | APP | Cell-Cell Contact | PMID: 24523320 |
| NeuN | CD31 | App | Tnfrsf21 | 0.016219 | 0 | APP_TNFRSF21 | App - Tnfrsf21 | APP | Cell-Cell Contact | PMID: 33075465 |
| NeuN | NeuN | App | Tnfrsf21 | 0.016044 | 0 | APP_TNFRSF21 | App - Tnfrsf21 | APP | Cell-Cell Contact | PMID: 33075465 |
| GFAP | GFAP | App | TREM2_TYROBP | 0.015593 | 0 | APP_TREM2_TYROBP | App - (Trem2+Tyrobp) | APP | Cell-Cell Contact | PMID:27477013 |
| NeuN | GFAP | App | TREM2_TYROBP | 0.034584 | 0 | APP_TREM2_TYROBP | App - (Trem2+Tyrobp) | APP | Cell-Cell Contact | PMID:27477013 |
| CD31 | GFAP | Apoa2 | TREM2_TYROBP | 0.000504 | 0 | APOA2_TREM2_TYROBP | Apoa2 - (Trem2+Tyrobp) | ApoA | Cell-Cell Contact | uniprot;PMID:27477017 |
| GFAP | GFAP | Apoa2 | TREM2_TYROBP | 0.000724 | 0 | APOA2_TREM2_TYROBP | Apoa2 - (Trem2+Tyrobp) | ApoA | Cell-Cell Contact | uniprot;PMID:27477017 |
| GFAP | CD31 | Apoe | TREM2_TYROBP | 0.019486 | 0 | APOE_TREM2_TYROBP | Apoe - (Trem2+Tyrobp) | ApoE | Cell-Cell Contact | uniprot;PMID:27477015 |
| GFAP | GFAP | Apoe | TREM2_TYROBP | 0.028795 | 0 | APOE_TREM2_TYROBP | Apoe - (Trem2+Tyrobp) | ApoE | Cell-Cell Contact | uniprot;PMID:27477015 |
| GFAP | GFAP | Cadm1 | Nectin3 | 0.001405 | 0 | CADM1_NECTIN3 | Cadm1 - Nectin3 | CADM | Cell-Cell Contact | uniprot |
| NeuN | GFAP | Cadm1 | Nectin3 | 0.003 | 0 | CADM1_NECTIN3 | Cadm1 - Nectin3 | CADM | Cell-Cell Contact | uniprot |
| CD31 | CD31 | Cd99 | Pilrb1 | 0.007179 | 0 | CD99_PILRB1 | Cd99 - Pilrb1 | CD99 | Cell-Cell Contact | uniprot;PMID:18234675 |
| NeuN | GFAP | Cd99 | Pilrb1 | 0.011526 | 0 | CD99_PILRB1 | Cd99 - Pilrb1 | CD99 | Cell-Cell Contact | uniprot;PMID:18234675 |
| GFAP | CD31 | Ceacam2 | Cd209c | 0.000402 | 0 | CEACAM2_CD209C | Ceacam2 - Cd209c | CEACAM | Cell-Cell Contact | PMID:16282604;PMID:16246332 |
| GFAP | GFAP | Ceacam2 | Cd209c | 0.000553 | 0 | CEACAM2_CD209C | Ceacam2 - Cd209c | CEACAM | Cell-Cell Contact | PMID:16282604;PMID:16246332 |
| GFAP | CD31 | Ceacam2 | Cd209a | 0.000687 | 0 | CEACAM2_CD209A | Ceacam2 - Cd209a | CEACAM | Cell-Cell Contact | PMID:16282604;PMID:16246332 |
| GFAP | GFAP | Ceacam2 | Cd209a | 0.000599 | 0 | CEACAM2_CD209A | Ceacam2 - Cd209a | CEACAM | Cell-Cell Contact | PMID:16282604;PMID:16246332 |
| GFAP | CD31 | Ceacam2 | Cd209d | 0.00021 | 0 | CEACAM2_CD209D | Ceacam2 - Cd209d | CEACAM | Cell-Cell Contact | PMID:16282604;PMID:16246332 |
| GFAP | GFAP | Ceacam2 | Cd209d | 0.000374 | 0 | CEACAM2_CD209D | Ceacam2 - Cd209d | CEACAM | Cell-Cell Contact | PMID:16282604;PMID:16246332 |
| GFAP | CD31 | Ceacam2 | Cd209b | 0.000643 | 0 | CEACAM2_CD209B | Ceacam2 - Cd209b | CEACAM | Cell-Cell Contact | PMID:16282604;PMID:16246332 |
| GFAP | GFAP | Ceacam2 | Cd209b | 0.000534 | 0 | CEACAM2_CD209B | Ceacam2 - Cd209b | CEACAM | Cell-Cell Contact | PMID:16282604;PMID:16246332 |
| GFAP | CD31 | Ceacam2 | Ceacam2 | 0.000458 | 0 | CEACAM2_CEACAM2 | Ceacam2 - Ceacam2 | CEACAM | Cell-Cell Contact | PMID:21982860 |
| CD31 | GFAP | Ceacam2 | Ceacam2 | 0.000458 | 0 | CEACAM2_CEACAM2 | Ceacam2 - Ceacam2 | CEACAM | Cell-Cell Contact | PMID:21982860 |
| GFAP | GFAP | Ceacam2 | Ceacam2 | 0.000948 | 0 | CEACAM2_CEACAM2 | Ceacam2 - Ceacam2 | CEACAM | Cell-Cell Contact | PMID:21982860 |
| GFAP | CD31 | Ceacam2 | Sele | 0.000464 | 0 | CEACAM2_SELE | Ceacam2 - Sele | CEACAM | Cell-Cell Contact | PMID:1378450 |
| GFAP | GFAP | Ceacam2 | Sele | 0.000699 | 0 | CEACAM2_SELE | Ceacam2 - Sele | CEACAM | Cell-Cell Contact | PMID:1378450 |
| GFAP | CD31 | Ceacam2 | Cd1d1 | 0.000705 | 0 | CEACAM2_CD1D1 | Ceacam2 - Cd1d1 | CEACAM | Cell-Cell Contact | PMID:24104458 |
| GFAP | GFAP | Ceacam2 | Cd1d1 | 0.000499 | 0 | CEACAM2_CD1D1 | Ceacam2 - Cd1d1 | CEACAM | Cell-Cell Contact | PMID:24104458 |
| GFAP | GFAP | Cntn1 | Notch1 | 0.00495 | 0 | CNTN1_NOTCH1 | Cntn1 - Notch1 | CNTN | Cell-Cell Contact | PMID:14567914 |
| NeuN | GFAP | Cntn1 | Notch1 | 0.012487 | 0 | CNTN1_NOTCH1 | Cntn1 - Notch1 | CNTN | Cell-Cell Contact | PMID:14567914 |
| CD31 | GFAP | Clec2h | Klrb1f | 0.000494 | 0 | CLEC2H_KLRB1F | Klrb1f - Clec2h | CLEC | Cell-Cell Contact | PMID:24223577 |
| GFAP | GFAP | Clec2h | Klrb1f | 0.000403 | 0 | CLEC2H_KLRB1F | Klrb1f - Clec2h | CLEC | Cell-Cell Contact | PMID:24223577 |

| | | | | | | | | | | |
|---|---|---|---|---|---|---|---|---|---|---|
| CD31 | CD31 | Clec2d | Klrb1f | 0.001295 | 0 | CLEC2D_KLRB1F | Klrb1f - Clec2d | CLEC | Cell-Cell Contact | PMID:24223577 |
| CD31 | GFAP | Clec2d | Klrb1f | 0.002624 | 0 | CLEC2D_KLRB1F | Klrb1f - Clec2d | CLEC | Cell-Cell Contact | PMID:24223577 |
| CD31 | GFAP | Mmp2 | MMP14_ITGAV_ITGB3 | 0.000654 | 0 | MMP2_INTEGRIN_AVB3_COMP | Mmp2 - Integrin_avb3 | MMP | Cell-Cell Contact | PMID: 29986882 |
| GFAP | GFAP | Mmp2 | MMP14_ITGAV_ITGB3 | 0.0006 | 0 | MMP2_INTEGRIN_AVB3_COMP | Mmp2 - Integrin_avb3 | MMP | Cell-Cell Contact | PMID: 29986882 |
| CD31 | CD31 | Ntn1 | Dcc | 0.00086 | 0 | NTN1_DCC | Ntn1 - Dcc | Netrin | Cell-Cell Contact | PMID: 22685302 |
| CD31 | NeuN | Ntn1 | Dcc | 0.001158 | 0 | NTN1_DCC | Ntn1 - Dcc | Netrin | Cell-Cell Contact | PMID: 22685302 |
| GFAP | GFAP | Ntn1 | Dscam | 0.001017 | 0 | NTN1_DSCAM | Ntn1 - Dscam | Netrin | Cell-Cell Contact | PMID:18585357;PMID: 22685302 |
| CD31 | NeuN | Ntn1 | Dscam | 0.001655 | 0 | NTN1_DSCAM | Ntn1 - Dscam | Netrin | Cell-Cell Contact | PMID:18585357;PMID: 22685302 |
| CD31 | CD31 | Ntn1 | Unc5a | 0.000721 | 0 | NTN1_UNC5A | Ntn1 - Unc5a | Netrin | Cell-Cell Contact | PMID: 22685302 |
| CD31 | NeuN | Ntn1 | Unc5a | 0.001335 | 0 | NTN1_UNC5A | Ntn1 - Unc5a | Netrin | Cell-Cell Contact | PMID: 22685302 |
| CD31 | GFAP | Ntn1 | Unc5b | 0.001027 | 0 | NTN1_UNC5B | Ntn1 - Unc5b | Netrin | Cell-Cell Contact | PMID: 22685302 |
| GFAP | GFAP | Ntn1 | Unc5b | 0.000849 | 0 | NTN1_UNC5B | Ntn1 - Unc5b | Netrin | Cell-Cell Contact | PMID: 22685302 |
| CD31 | GFAP | Ntn1 | Unc5c | 0.000523 | 0 | NTN1_UNC5C | Ntn1 - Unc5c | Netrin | Cell-Cell Contact | PMID: 22685302 |
| GFAP | GFAP | Ntn1 | Unc5c | 0.000432 | 0 | NTN1_UNC5C | Ntn1 - Unc5c | Netrin | Cell-Cell Contact | PMID: 22685302 |
| CD31 | GFAP | Ntn4 | Ntrk2 | 0.016699 | 0 | NTN4_NTRK2 | Ntn4 - Ntrk2 | Netrin | Cell-Cell Contact | PMID: 34531526 |
| GFAP | GFAP | Ntn4 | Ntrk2 | 0.015189 | 0 | NTN4_NTRK2 | Ntn4 - Ntrk2 | Netrin | Cell-Cell Contact | PMID: 34531526 |
| CD31 | GFAP | Ntn4 | Unc5b | 0.001725 | 0 | NTN4_UNC5B | Ntn4 - Unc5b | Netrin | Cell-Cell Contact | PMID: 31914695 |
| GFAP | GFAP | Ntn4 | Unc5b | 0.001567 | 0 | NTN4_UNC5B | Ntn4 - Unc5b | Netrin | Cell-Cell Contact | PMID: 31914695 |
| GFAP | GFAP | Ntn4 | Unc5d | 0.000476 | 0 | NTN4_UNC5D | Ntn4 - Unc5d | Netrin | Cell-Cell Contact | uniprot |
| CD31 | NeuN | Ntn4 | Unc5d | 0.000604 | 0 | NTN4_UNC5D | Ntn4 - Unc5d | Netrin | Cell-Cell Contact | uniprot |
| CD31 | CD31 | Pecam1 | Cd177 | 0.000979 | 0 | PECAM1_CD177 | Pecam1 - Cd177 | PECAM1 | Cell-Cell Contact | PMID:17580308 |
| CD31 | GFAP | Pecam1 | Cd177 | 0.001857 | 0 | PECAM1_CD177 | Pecam1 - Cd177 | PECAM1 | Cell-Cell Contact | PMID:17580308 |
| CD31 | CD31 | Pecam1 | Cd38 | 0.000755 | 0 | PECAM1_CD38 | Pecam1 - Cd38 | PECAM2 | Cell-Cell Contact | PMID:26407101 |
| CD31 | GFAP | Pecam1 | Cd38 | 0.001059 | 0 | PECAM1_CD38 | Pecam1 - Cd38 | PECAM2 | Cell-Cell Contact | PMID:26407101 |
| GFAP | GFAP | Sirpb1a | Cd47 | 0.001514 | 0 | SIRPB1A_CD47 | Sirpb1a - Cd47 | SIRP | Cell-Cell Contact | PMID:15294972 |
| GFAP | NeuN | Sirpb1a | Cd47 | 0.002098 | 0 | SIRPB1A_CD47 | Sirpb1a - Cd47 | SIRP | Cell-Cell Contact | PMID:15294972 |
